# A robust metaproteomics pipeline for a holistic taxonomic and functional characterization of microbial communities from marine particles

**DOI:** 10.1101/667428

**Authors:** Doreen Schultz, Daniela Zühlke, Jörg Bernhardt, Thomas Ben Francis, Dirk Albrecht, Claudia Hirschfeld, Stephanie Markert, Katharina Riedel

**Affiliations:** Institute of Microbiology, University of Greifswald, Greifswald, Germany; Max-Planck Institute for Marine Microbiology, Bremen, Germany; Institute of Pharmacy, University of Greifswald, Greifswald, Germany

## Abstract

This study aimed to establish a robust, reproducible and reliable metaproteomic pipeline for an in-depth characterization of marine particle-associated (PA) bacteria. To this end, we compared six well-established protein extraction protocols together with different MS-sample preparation techniques using particles sampled during a North Sea spring algae bloom in 2009. In this optimized workflow, proteins are extracted using a combination of SDS-containing lysis buffer and cell disruption by bead-beating, separated by SDS-PAGE, in-gel digested and analysed by LC-MS/MS, before MASCOT search against a metagenome-based database and data processing/visualization with the *in-house*-developed bioinformatics tools *Prophane* and *Paver*.

As proof of principle, free-living (FL) and particulate communities sampled in April 2009 were analysed, resulting in an as yet unprecedented number of 9,354 and 5,034 identified protein groups for FL and PA bacteria, respectively. Our data revealed that FL and PA communities appeared similar in their taxonomic distribution, with notable exceptions: eukaryotic proteins and proteins assigned to *Flavobacteriia*, *Cyanobacteria*, and some proteobacterial genera were found more abundant on particles, whilst overall proteins belonging to *Proteobacteria* were more dominant in the FL fraction. In contrast, significant functional differences including proteins involved in polysaccharide degradation, sugar- and phosphorus uptake, adhesion, motility, and stress response were detected.

**Originality-Significance Statement:** Marine particles consist of organic particulate matter (e.g. phyto- or zooplankton) and particle-associated (PA) microbial communities, which are often embedded in a sugary matrix. A significant fraction of the decaying algal biomass in marine ecosystems is expected to be mineralized by PA heterotrophic communities, which are thus greatly contributing to large-scale carbon fluxes. Whilst numerous studies have investigated the succession of planktonic marine bacteria along phytoplankton blooms, the community structure and functionality of PA bacterial communities remained largely unexplored and knowledge on specific contributions of these microorganisms to carbon cycling is still surprisingly limited. This has been mostly been due to technical problems, i.e. to the difficulty to retrieve genomic DNA and proteins from these polysaccharide-rich entities, their enormous complexity and the high abundance of eukaryotic microorganisms.

Our study presents an innovative, robust, reproducible, and reliable metaproteomics pipeline for marine particles, which will help to address and fill the above-described knowledge gap. Employing the here established workflow enabled us to identify more than 5,000 PA proteins, which is, at least to our knowledge, the largest number of protein groups ever assigned to marine particles. Notably, the novel pipeline has been validated by a first, comparative metaproteome analysis of free-living and PA bacterial communities indicating a significant functional shift enabling surface-associated bacteria to adapt to particle-specific living conditions. In conclusion, our novel metaproteomics pipeline presents a solid and promising methodological groundwork for future culture-independent analyses of seasonal taxonomic and functional successions of PA microbial communities in aquatic habitats.

## Introduction

The oceans provide about one half of the global net primary production (NPP) (Field *et al.*, 1998; Falkowski *et al.*, 1998; Azam and Malfatti, 2007), a large part of which is reprocessed by bacterioplankton in the so-called “microbial loop”, a pathway during which dissolved organic carbon is returned to higher trophic levels via its incorporation into bacterial biomass (Azam, 1998). Notably, about 20% of the bacterioplankton lives attached to algae or marine particles (Azam *et al.*, 1983). These marine particles consist of various kinds of organic matter, i.e. dead/dying zoo- or phytoplankton, bacterioplankton, as well as inorganic small particles held together by a sugary matrix consisting of polysaccharide-composed transparent extracellular particles (TEPs) composed of polysaccharides, which are exuded mostly by phytoplankton but also bacteria (Alldredge *et al.*, 1993). These particle-associated (PA) microbial communities present one of the most ancient forms of “multicellularity” (Hall-Stoodley *et al.*, 2004) and have adapted to strive and survive in marine environments. Whilst some bacteria are only loosely associated with algae, others colonize algal surfaces (Grossart, 1999), where they form commensalistic or symbiotic communities with their host or even predate on algae (Sohn *et al.*, 2004; Amin *et al.*, 2012). Marine particles grow while sinking and thus contribute largely to the “biological pump” by transporting carbon to deeper waters and sediments (Volkman and Tanoue, 2002). These aggregates may reach several centimetres in diameter. They are enzymatically well equipped to metabolize high molecular weight substrates, thus providing nutrition to the attached community as well as leaving nutrients to the surrounding water column community (Simon *et al.*, 2002; Grossart, 2010).

About one decade ago, scientists started to link molecular systems biology of microorganisms to ecosystem level processes (e.g. reviewed in Raes and Bork, 2008). Metagenomic studies were initiated to provide valuable knowledge about diversity and distribution of microorganisms in natural environments. Moreover, metatranscriptomics and metaproteomics approaches were established to investigate, which genes are expressed at a given time point and which proteins are particularly abundant in complex biological systems. Metaproteomics has meanwhile widely proven its potential to revisit microbial ecology concepts by linking genetic and functional diversity in microbial communities and relating taxonomic and functional diversity to ecosystem stability (Schneider and Riedel, 2010). Numerous studies, describing large-scale proteome analyses of acid-mine drainage (AMD) biofilms (Ram *et al.*, 2005), wastewater treatment plants (Wilmes *et al.*, 2008), and fresh-water stream biofilms (Hall *et al.*, 2012) have demonstrated the power of metaproteomics to unveil molecular mechanisms involved in function, physiology, and evolution of surface-associated aquatic microbial communities. Marine metaproteomics has meanwhile been widely applied (Saito *et al.*, 2019; Wang *et al.*, 2014), in particular in habitats such as ocean scale shifts (Morris *et al.*, 2010), the Atlantic (Bergauer *et al.*, 2017) or Antarctic oceans (Williams *et al.*, 2012), e.g., to investigate *Roseobacter* clade (Christie-Oleza and Armengaud 2015) and bacterioplankton (e.g. Wöhlbrand *et al.*, 2017a) physiology. Teeling *et al*. (2012) studied the bacterioplankton response to a diatom bloom in the North Sea by an integrated meta-omics approach employing metagenomics and metaproteomics and provided strong evidence that distinct free-living (FL) populations of *Bacteroidetes*, *Gammaproteobacteria*, and *Alphaproteobacteria* specialized in a successive decomposition of algal-derived organic matter. As mentioned above, a significant fraction of decaying algal biomass is, however, mineralized by heterotrophic bacteria living on particles, which process a large fraction of the biosynthesized organic matter (Azam, 1998) and are thus greatly contributing to large-scale carbon fluxes (Battin *et al.*, 2003; Bauer *et al.*, 2006).

So far, the majority of the published studies focused on FL bacterioplankton, thereby leaving the PA bacterial communities largely unexplored. The particulate fraction of bacterioplankton has proven challenging to comprehensive meta-omics characterization, due to its high complexity, presence of DNA/protein-binding polysaccharides, process-interfering substances and lack of (meta)genomic information on marine particles (e.g. Wöhlbrand *et al.*, 2017b), despite the fact that information on marine metagenomes is constantly growing (reviewed by Mineta and Gojobori 2016; Alma’abadi *et al.*, 2015). Previous experiments also indicate that a high abundance of eukaryotic proteins contributes to these challenges (Smith *et al.*, 2017; Saito *et al.*, 2019).

Our goal was therefore to establish a robust, reproducible and reliable pipeline enabling in-depth metaproteomics analyses of marine particles by testing different established protocols for their applicability for protein extraction from PA bacterioplankton in order to unravel the PA community’s specific contribution to polysaccharide decomposition in marine habitats. As stated above, heterotrophic microbial communities are supposed to be well equipped to metabolize algal high molecular weight substrates (Simon *et al.*, 2002; Grossart, 2010). We thus hypothesize that these communities either differ taxonomically from their FL counterparts (as shown by Bizic-Ionescu *et al*. (2015)) and/or express different genes to adapt to the sessile life style and the availability of specific polysaccharides such as insoluble glycan fibres (as observed by Ganesh *et al*. (2014)). Moreover, we postulated that this adaption will mostly affect the expression of proteins involved in motility, adhesion, stress response as well as CAZymes together with appropriate sugar transporters. In order to test these hypotheses, a metaproteomics pipeline for marine particles was established and tested as a proof of concept on spring bloom samples collected in April 2009, for which the FL bacterial fraction had previously been characterized (Teeling *et al.*, 2012).

## Results and Discussion

### Establishment of a metaproteomics pipeline for PA microbial communities

As stated above, the metaproteomics analysis of PA microbial communities is severely hampered by their high complexity, the presence of a large proportion of eukaryotic proteins, the sugary particle-matrix as well as the lack of (meta)genomic information on PA-specific pro- and eukaryotes (Wöhlbrand *et al.*, 2017b; Saito *et al.*, 2019). Whilst the metaproteomics analyses by Teeing *et al*. (2012) and Kappelmann *et al.* (2019) of FL bacterioplankton (harvested on 0.2 μm filters) sampled during spring blooms from 2009 to 2012 at “Kabeltonne” Helgoland resulted in the identification of several thousand protein groups, the PA microbial communities retained on 3 and 10 μm pore-sized filters emerged as difficult to analyse by the integrated metagenomic/metaproteomic approach employed at that time.

We thus aimed to develop a suitable and effective metaproteomics pipeline for in-depth analyses of taxonomic composition and functionality of marine PA microbial communities. To this end, six different well-established protein extraction protocols for various particulate samples were tested using biomass from 3 μm and 10 μm pore-sized filters (assigned as PA fraction) of the MIMAS bio-archive (www.mimas-project.de) covering different stages of the bloom, which had been collected in 2009 during the spring phytoplankton bloom sampling campaign by Teeling *et al*. (2012). Moreover, various MS sample preparation protocols and different protein sequence databases were evaluated.

**Table 1:** Comparison of the six tested protein extraction protocols

| Protocol Nr. | 1 - Phenol | 2 - SDS-TCA | 3 - TRI-reagent® | 4 - Freeze and thaw | 5 - SDS-acetone | 6 - Bead beating |
| --- | --- | --- | --- | --- | --- | --- |
| Reference | Kuhn *et al.* (2011) | Schneider *et al.* (2012) | Sigma Aldrich | Chourey *et al.* (2010)<br>Thompson *et al.* (2008) | Hall *et al.* (2012) | Moog (2012) |
| Originally used for | sewage sludge from biomembrane reactors | leaf litter | simultaneous extraction of RNA, DNA, and proteins | soil | stream hyporheic biofilms | hypersaline microbial mats |
| Composition of the protein extraction buffer | 0.1 M NaOH | 1% (w/v) SDS, 50 mM Tris/HCl, pH 7 | TRI-Reagent® (guanidine thiocyanate and phenol monophasic solution) | 5% (w/v) SDS, 50 mM Tris/HCl, 0.1 mM EDTA, 0.15 M NaCl, 1 mM MgCl<sub>2</sub>, 50mM DTT, pH 8.5 | 1% (w/v) SDS, 50 mM Tris/HCl, pH 6.8 | 5% (w/v) SDS, 0.05 mM Tris/HCl, 0.1 M DTT, 0.01 M EDTA, 10% (v/v) glycerol, 1.7 mM PMSF, pH 6.8 |
| Cell disruption methodology | sonication<br>3 x 30 s (20% power output) | sonication<br>3 x 40 s (20% power output) | TRI-Reagent® | 2 freeze and thaw cycles (liquid nitrogen, rt), 10 min boiling | sonication<br>5 x 1 min (20% power output),<br>15 min boiling,<br>procedure repeated on the pellet | FastPrep® 6.5 m/s,<br>4 x 30 s |
| Additional protein purification | phenol extraction (2x) | / | chloroform extraction, ethanol extraction | / | / | / |
| Protein precipitation | 0.1 M ammonium-acetate in methanol (1:5) | 10% TCA | 2-propanol (1:1.5) | 25% TCA | acetone (1:5) | acetone (1:4) |
| Mean total protein amount [µg]<br>3 - 10 µm fraction | 8.9 | 22.7 | 16.1 | 25.2 | 38.6 | 27.3 |
| Mean total protein amount [µg]<br>≥ 10 µm fraction | 8.8 | 12.8 | 38.2 | 24.3 | 102.1 | 114.2 |

#### Protein extraction

Efficient protein extraction is a crucial step for successful metaproteomics analyses of the microbial communities. In a first step, we therefore tested five different protein extraction methods that employ different strategies and that were already successfully applied for metaproteome analyses of microbial communities from different environments, i.e. sewage sludge (phenol extraction; Kuhn *et al.*, 2011), leaf litter (SDS-TCA; Schneider *et al.*, 2012), stream hyporheic biofilms (SDS-acetone; Hall *et al.*, 2012), hypersaline microbial mats (bead beating; Moog, 2012), and soil (freezing and thawing; Chourey *et al.*, 2010; Thompson *et al.*, 2008). In addition, the commercially available TRI-Reagent^®^ (Sigma Aldrich) for simultaneous isolation of RNA, DNA and proteins was tested (**Table 1**). Filter samples used for protocol evaluation originated from several sampling dates in the 2009 spring bloom sampling campaign (9^th^ of February, 7^th^ of April, 21^st^ of April, and 16^th^ of June 2009). Filters were cut into pieces and subjected to the respective extraction protocol (**Fig. 1A**). Total protein amounts extracted from the filters by each of the applied methods were quite variable. Highest protein yield as determined by the Pierce™ BCA Protein Assay and 1D SDS-PAGE was obtained using the SDS-acetone or bead beating approach (**Table 1**; **Fig. 1B**). In conclusion, SDS-acetone- and bead beating-based protocols turned out to be most efficient for protein extraction from particles and were therefore used for optimizing the downstream MS sample preparation procedure.

#### MS sample preparation

Total protein was extracted by the SDS-acetone and bead beating method from filters collected on 28^th^ of April 2009 and separated by 1D SDS-PAGE (**Fig. 1B**). Even though MS sample preparation via GeLC MS/MS is more time-consuming compared to 1D or 2D-LC approaches, it has been proven valuable to purify protein extracts and remove polymeric contaminants (e.g. Lassek *et al*. 2015; Keiblinger and Riedel, 2018) and yields comparable results as LC-based peptide fractionation (Hinzke *et al.*, 2019). To determine whether an increase in the total number of individual gel sub-fractions will lead to more protein identifications, gel lanes were cut in either 10 or 20 equal-sized fractions, proteins were *in gel* trypsin-digested and the resulting peptides were subjected to LC-MS/MS analysis. Moreover, we tested whether reduction and alkylation of the proteins prior to tryptic digestion increased protein identification rates (**Fig. 1B**). Searching the acquired spectra in the so far available 0.2 μm 2009 (MIMAS) database (Teeling *et al.*, 2012) revealed that the best results (**Fig. 1B**, **Fig. S1**) were obtained by higher fractionation (20 gel pieces) without reduction and alkylation.

**Figure 1:**
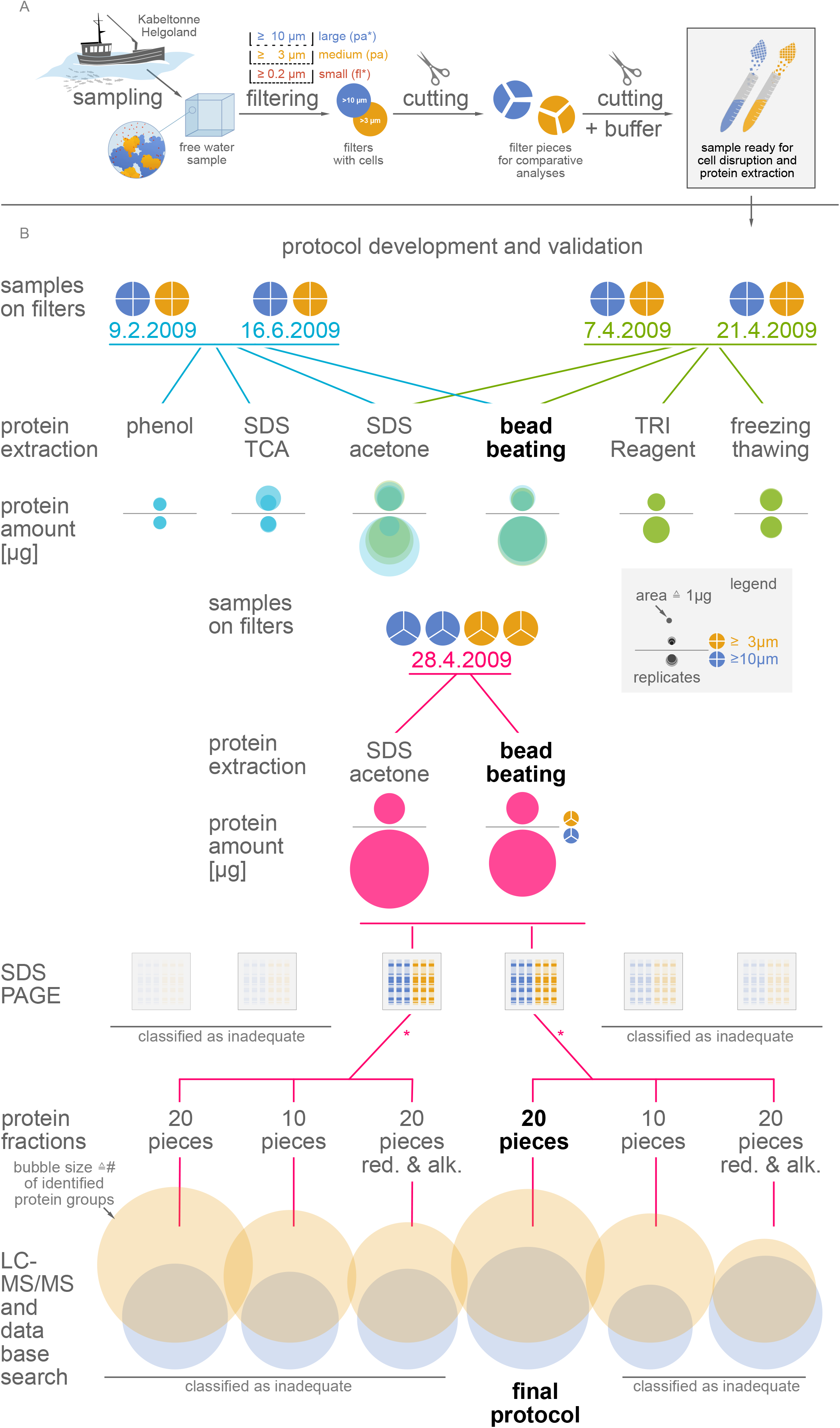
(A) Sampling strategy and (B) evaluation of protein extraction and MS sample preparation protocols. **(A)** Water samples collected at “Kabeltonne” Helgoland during the spring bloom 2009 were sequentially filtered to obtain the 0.2 - 3 μm (FL) and two PA (3 - 10 μm = medium, ≥ 10 μm = large) fractions as described in Teeling *et al.* (2012). Filters were initially cut into three or four pieces, which were subsequently shredded and mixed with the respective extraction buffer. **(B)** Filters (medium particles = yellow; large particles = blue) from different sampling time points (turquoise, green and red) were processed according to the six different protocols describe in the experimental procedure section. With regard to the extracted protein amount the bead beating and SDS-acetone approaches obviously outcompeted the four other protocols. However, the SDS-acetone protocol was less reproducible than the bead beating protocol. Considering bead beating and SDS-acetone as best performing protocols, they were employed to test different MS sample preparation approaches, i.e., different number of SDS gel fractions for tryptic digestion together with protein reduction (red.) and alkylation (alk.) prior to tryptic digestion. The subsequent LC-MS/MS analyses revealed best results for the bead beating protocol followed by GeLC-MS/MS from 20 fractions without protein reduction and alkylation as shown in the bottom line of the figure. Bubble sizes for the large (blue) and medium (yellow) particles correspond to the number of identified protein groups (see also **Fig. S1**).

#### Optimizing databases

Metagenomic sequencing, assembling and annotation of FL (0.2 μm pore-sized filters) and PA (3 and 10 μm pore-sized filters) fractions of water samples collected during the Helgoland spring bloom 2009 was performed in parallel to the optimization of the metaproteomics protocol (for details see experimental procedures). Unfortunately, the coverage and quality of the metagenome sequences of the large particulate fraction (10 μm pore-sized filters) was not sufficient to be correctly assembled, annotated and translated. Thus, the metagenomic database used for subsequent database searches was only composed of sequences of FL bacteria (0.2 μm pore-sized filters) and microbial communities present in the medium particulate fraction (3 μm pore-sized filters).

The LC-MS/MS spectra obtained with the bead beating protocol were searched against four different databases to identify the database that results in the highest number of reliably identified protein groups (**Fig. S2**): (I) the non-redundant NCBI database (NCBInr), (II) a database with Uniprot sequences from abundant bacteria and diatoms identified by Teeling *et al.* (2012) (PABD), (III) the database used by Teeling *et al.* (2012) containing proteins based on translated metagenomes of FL bacteria (0.2 μm pore-sized filters from different sampling time points) of the spring bloom 2009 (MIMAS) and (IV) a database based on the metagenomes of the 0.2 and 3 μm pore-sized filters from samples of the 14^th^ of April 2009 (0.2 + 3 μm 2009). Best results were obtained with the 0.2 + 3 μm 2009 database (**Fig. S2**), which is not surprising as the resolving power of metaproteome analyses relies heavily upon the database used for protein identification (e.g. Schneider and Riedel, 2010; Teeling *et al.*, 2012). It is, moreover, well accepted that metaproteomic data are most informative in combination with complementary omics approaches, i.e. genomics and transcriptomics (e.g. Banfield *et al.*, 2005; Ram *et al.*, 2005).

Based on our results, we finally decided to use the bead beating-based protocol for protein extraction since this method resulted in more reproducible protein yields compared to the SDS-acetone extraction protocol (**Fig. 1B**). Furthermore, this method was less time-intensive compared to the SDS-acetone approach and resulted in the identification of the highest number of unique protein groups, most probably due to the effective disintegration of the particulate matrix by EDTA added to the extraction buffer (Passow, 2002). In summary, the optimized metaproteomic pipeline for marine particles (**Fig. 2**) comprises protein extraction by bead beating, protein fractionation by 1D SDS-PAGE (20 fractions), followed by in-gel tryptic digestion, LC-MS/MS and database search against the matching metagenome database (0.2 + 3 μm 2009).

**Figure 2:**
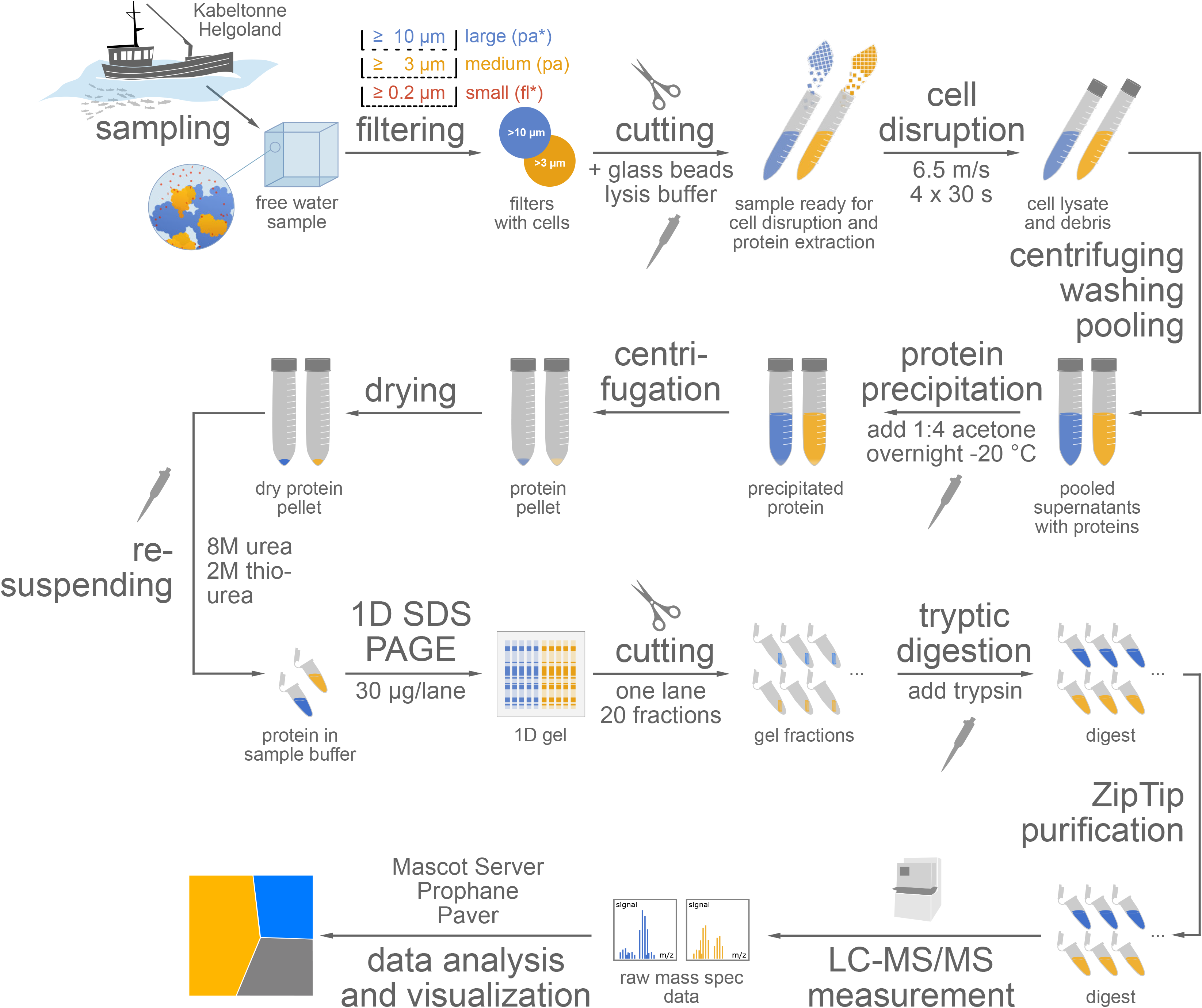
Final metaproteomics pipeline. Protein extraction from filters was conducted using 5% (w/v) SDS containing lysis buffer, cell disruption by FastPrep-mediated bead beating, separation of proteins by 1D-SDS-PAGE, tryptic in-gel digestion, LC-MS/MS analyses on an Orbitrap Velos™ mass spectrometer, MASCOT database search against the metagenome-based database (0.2 + 3 μm 2009) and data-processing and visualization with the *in-house*-developed bioinformatics tools *Prophane 3.1* and *Paver*.

### Proof of principle – comparative metaproteome analyses of FL and PA bacterioplankton

As proof of principle, the newly established protocol was applied for a comparative metaproteomic analysis of FL and PA microbial communities sampled at the 14^th^ of April 2009 in 0.2 μm - 3 μm (= FL), 3 - 10 μm and ≥ 10 μm (= PA) fractions. Five technical replicates of each sample were subjected to the final optimized workflow and the resulting MS/MS-data were searched against the matching metagenome-based database (0.2 + 3 μm 2009). Employing our optimized pipeline, we were able to record 460,000, 360,000, and 440,000 spectra, which subsequently led to the identification of 9,354 protein groups (19.4% of spectral IDs), 2,263 protein groups (10.2% of spectral IDs), and 2,771 protein groups (10.7% of spectral IDs) for the 0.2 - 3 μm (**Table S1**), 3 - 10 μm (**Table S2**), and ≥ 10 μm (**Table S3**) fractions, respectively. This is, at least to our knowledge, the largest number of protein groups ever identified for marine particles. Comparable studies addressing metaproteomic analyses of marine sediments of the Bering Sea (Moore *et al.*, 2012), the coastal North Sea, and the Pacific Ocean (Wöhlbrand *et al.*, 2017b) identified less than 10% of the protein identification numbers resulting from the here presented novel metaproteomic pipeline.

1,956 of the identified protein groups of the two PA fractions were also identified in the FL fraction and only 276 proteins were exclusively found in the PA fractions (**Fig. S3**). This suggests that protein expression profiles of planktonic and particulate bacteria vary less than expected. However, this might also be due to the fact that PA bacteria are known to hop on and off particles and, e.g. as offspring cells searching for a place to settle, may thus only temporarily be part of the planktonic community (Ghiglione *et al.*, 2007; Grossart, 2010; Crespo *et al.*, 2013). Moreover, clogging of filter pores by particles may cause retention of FL bacteria thus contaminating the PA fractions by planktonic bacteria. In addition, the lack of ≥ 10 μm pore-sized filter metagenomic sequences hampers comprehensive protein identifications in this PA fraction that may result in a virtually lower abundance than expected.

#### Taxonomic differences between FL and PA bacterioplankton

Besides the somewhat unexpected similarity of the FL and PA metaproteomic datasets, the phylogenetic assignment of the identified protein groups indicated some notable taxonomic differences between the FL and PA fractions (**Fig. 3, Table S4**).

**Figure 3:**
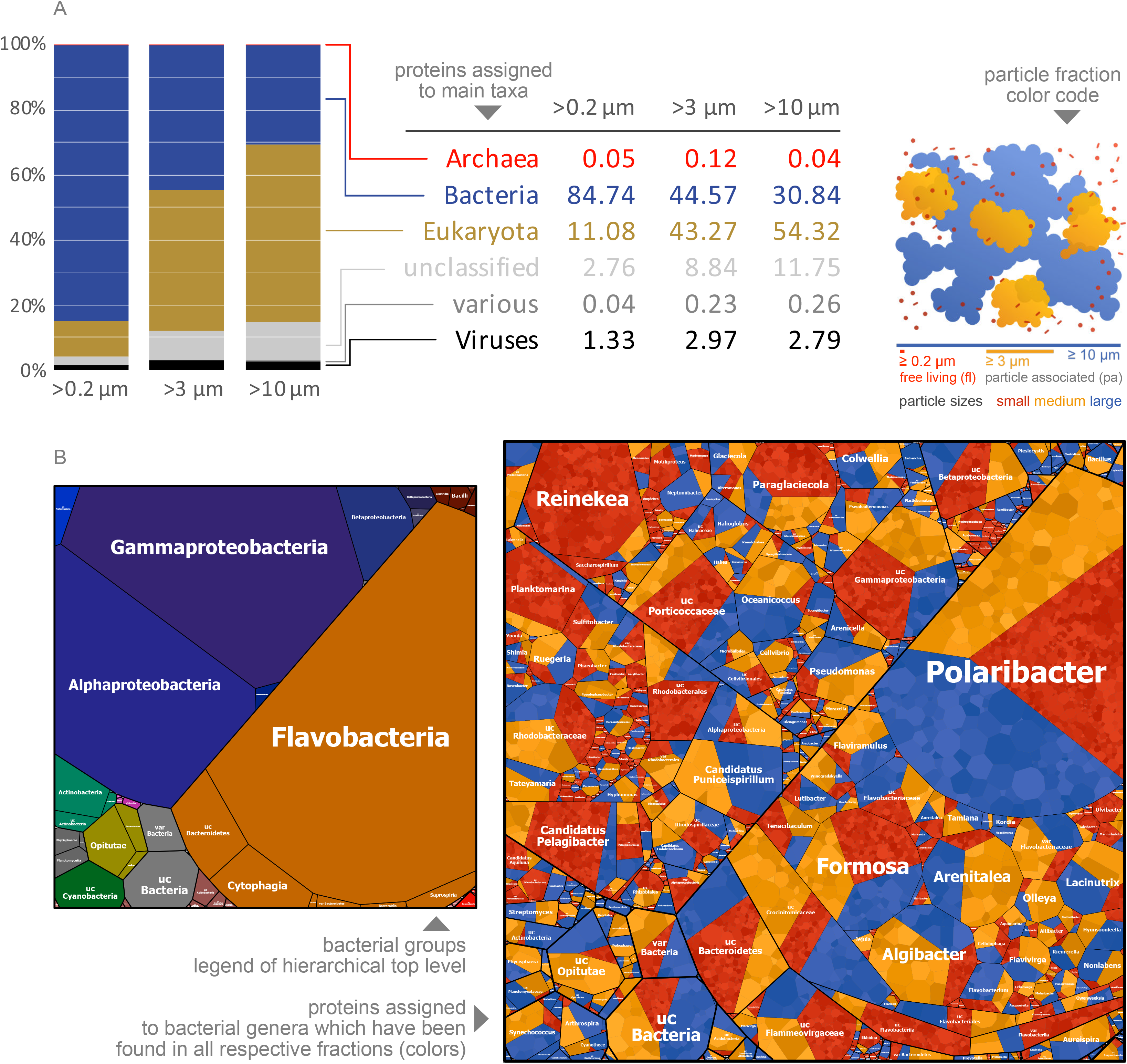
Taxonomic affiliation of proteins of FL and PA metaproteomes during the spring bloom on 14^th^ of April 2009 at “Kabeltonne” Helgoland. **(A)** Distribution of pro- and eukaryotes in the FL (0.2 - 3 μm) and PA (3 - 10 μm, ≥ 10 μm) fractions based on the relative abundance of protein groups assigned to the different phylogenetic groups. **(B)** Voronoi treemaps visualizing the phylogenetic assignment of bacterial protein groups identified in FL (red) and PA (yellow and blue) fractions. Cell size corresponds to the relative abundance of the respective bacterial genus on protein level. Proteins of *Reinekea* for example are most abundant in the FL fraction and are therefore encoded by a large red treemap cell. In the PA fractions they can be detected only in traces resulting in very small cell sizes (coloured in yellow and blue). *Algibacter* protein abundance, on the other hand, was notably higher in the PA fractions, compared to the FL fraction.

The number of eukaryotic protein groups was significantly higher in the PA fractions (**Fig. 3A**) and comprised 43% and 54% of the protein groups identified for the 3 - 10 μm and ≥ 10 μm fractions, respectively, in contrast to only 11% of the protein groups identified for the FL phytoplankton. Moreover, the number of viral protein groups was found to be almost three times higher in the two particulate fractions when compared to their planktonic counterpart (**Fig. 3A**). The most abundant phyla within both, the FL and PA fractions, were *Proteobacteria* (FL 55%; PA 41% and 39%, 3 μm and 10 μm pore-sized filters) and *Bacteroidetes* (FL 40%; PA 48% and 47%, 3 μm and 10 μm pore-sized filters). Proteins expressed by *Alpha*-, *Beta*- and *Gammaproteobacteria* were generally more dominant in the FL bacteria, whilst proteins assigned to *Cyanobacteria* (e.g. *Synechococcus*, *Arthrospira*), *Opitutae*, *Flavobacteriia* (e.g. *Arenitalea*, *Olleya*, *Algibacter*, *Lacinutrix*), and some proteobacterial genera (e.g. *Oceanicoccus*, *Candidatus* Puniceispirillum, *Neptuniibacter*, *Halioglobus*, *Ramlibacter*) were more abundant in the PA fraction (**Fig. 3B**). This is in good accordance to other studies confirming that *Bacteroidetes* have been identified in both, FL and PA, bacterioplankton (DeLong *et al.*, 1993; Eilers *et al.*, 2001; Abell and Bowman, 2005; Alonso *et al.*, 2007). Moreover, *Flavobacteriia* have been found highly abundant during phytoplankton blooms indicating that they play an important role as consumers of algal-derived organic matter (Simon *et al.*, 1999; Riemann *et al.*, 2000; Pinhassi *et al.*, 2004; Grossart *et al.*, 2005; Teeling *et al.*, 2016; Chafee *et al.*, 2018).

#### Functional differences between FL and PA bacterioplankton

Notably, differences in the protein profiles between FL and PA bacteria were more evident on the functional level (**Fig. 4**). Most importantly, the SusC/D utilization system, specific glycoside hydrolases, i.e. GH family 1, 13, and 16 (including beta-glucosidases, alpha-1,4-amylases, and exo- and endo-1,3-beta-glycanases), glycosyl transferases and TonB-dependent transporters were found with higher overall expression levels in the PA fractions compared to the FL fraction (**Fig. 4A**). This is in good accordance with the high substrate availability (Caron *et al.*, 1982; Grossart *et al.*, 2003; Fernández-Gómez *et al.*, 2013), especially the presence of highly abundant microalgae storage polysaccharides, i.e. alpha- and beta-glucans (Kroth *et al.*, 2008), in the particles. Sulfatases, capable of cleaving sulphate sugar ester bonds, are contributing to the degradation of specific sulphated algal polysaccharides such as mannans and fucans (Gómez-Pereira *et al.*, 2012). This is well supported by our finding that sulfatases are strongly expressed by PA *Flavobacteriia*, especially *Formosa* sp. (**Fig. 4A & B**).

**Figure 4:**
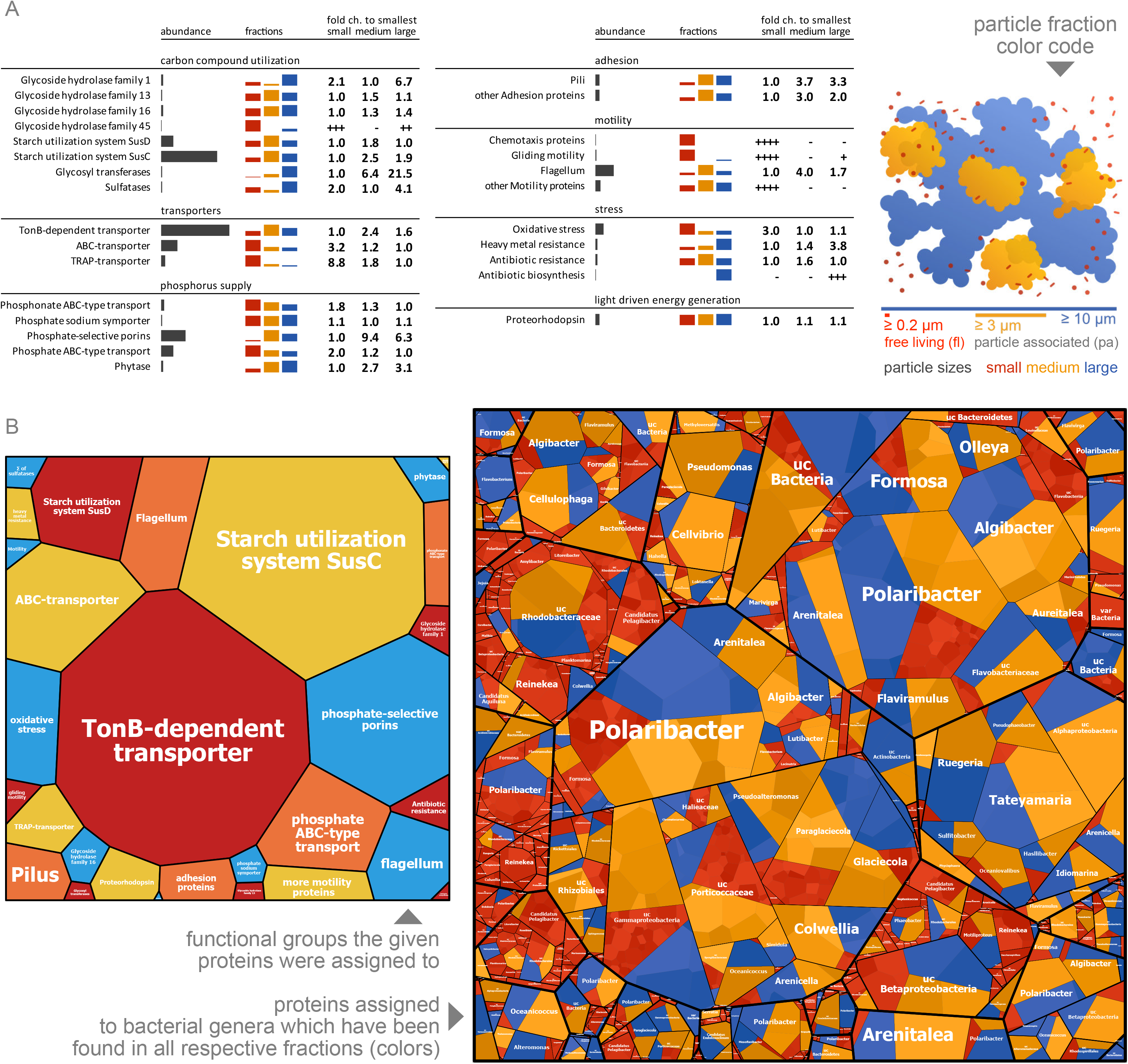
Functional assignment of proteins in FL and PA metaproteomes during the spring bloom on 14^th^ of April 2009 at “Kabeltonne” Helgoland. **(A)** Total abundance of selected protein groups with assigned functions in the FL (0.2 – 3 μm = small) and PA (3 - 10 μm = medium and ≥ 10 μm = large) fractions. **(B)** Voronoi treemaps showing the phylogenetic assignment of selected functional protein groups identified in FL (red) and PA (yellow and blue) fractions. Cell size corresponds to the relative abundance of the respective genus within specific functional categories.

FL and PA bacteria seem to employ different phosphate acquisition strategies: whilst in FL bacteria phosphate and phosphonate ABC-type transporters appeared highly expressed, PA bacteria rather seem to employ phytases and phosphate-selective porins. As expected, various proteins involved in stress response were differentially expressed. Interestingly, functions involved in oxidative stress defense appeared to be less abundant in the PA fraction (maybe due to shading, reducing solar irradiation stress in the particles), whilst proteins for heavy metal and antibiotic resistance were strongly expressed in the ≥ 10 μm fraction, which also contained the highest proportion of eukaryotic proteins (**Fig. 4A**). This might be due to the fact that some algae take up and store heavy metals (Gaudry *et al.*, 2007) and are capable of producing antibiotics (Grossart, 1999). This indicates that close eukaryote-bacterial interactions in particles require such defense strategies of the associated bacteria. As expected, adhesion proteins as well as proteins involved in motility, i.e. flagella and type IV-pili, were more abundant on the particles, emphasizing their importance for biofilm/aggregate formation (O’Toole & Kolter, 1998; Lemon *et al.*, 2007; Houry *et al.*, 2010; Burke *et al.*, 2011). Interestingly, proteorhodopsin, an inner membrane protein involved in light-dependent energy generation, which has been proposed to enable FL bacteria such as *Polaribacter* (Fernández-Gómez *et al.*, 2013) and *Pelagibacter* (Giovannoni *et al.*, 2005) to survive under low nutrient conditions, was also abundantly identified in PA bacteria such as *Polaribacter*, *Paraglaciecola*, and *Marinosulfomonas* in our analyses.

#### Eukaryotes are highly abundant and might contribute to polysaccharide degradation on marine particles

Our metaproteomics data demonstrate a high abundance of eukaryotes on the particles (**Fig. 5**). Preliminary analyses indicate that these include numerous microalgal groups, e.g. diatoms, *Pelagophytes*, *Raphidiophytes*, *Cryptophytes*, *Dinoflagellates* and *Haptophytes*, but also fungi and various protozoa (**Table S2 and S3**). This clearly sets particles apart from the FL fraction and highlights the importance of direct eukaryote-bacterial interactions in particles. Previous work on FL bacteria showed that bacterial succession was largely independent of phytoplankton composition, and instead determined by broad substrate availability (Teeling *et al.*, 2016). PA bacterial composition is more likely to be directly controlled by algal composition due to the intimate nature of their interactions (Grossart *et al.*, 2005), although functional redundancy may be substantial (Burke *et al.*, 2011). Moreover, eukaryotes may also contribute to polysaccharide degradation in concert with bacteria. For example, fungal taxa can be abundant in marine particles and have been shown to utilize algal polysaccharides such as laminarin (Bochdansky *et al.*, 2016; Cunliffe *et al.*, 2017).

**Figure 5:**
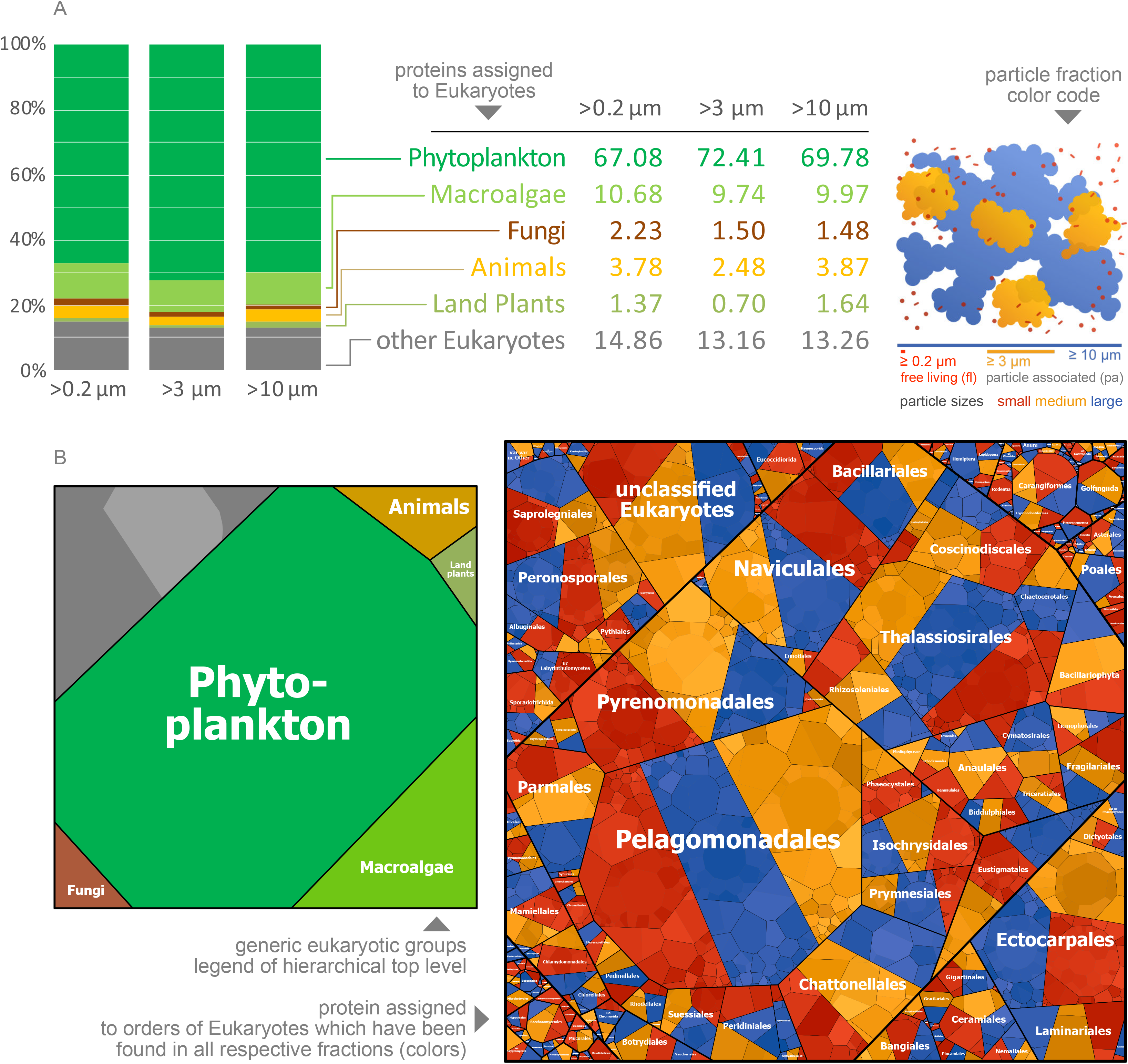
Phylogenetic assignment of eukaryotic proteins present in the FL and PA fractions during the spring bloom on 14^th^ of April 2009 at “Kabeltonne” Helgoland. **(A)** Distribution of different eukaryotes in the FL (0.2 μm) and PA (0.2 - 3 μm and ≥ 10 μm) fractions as shown by the relative protein abundances assigned to the different eukaryotic phylogenetic groups. **(B)** Voronoi treemap visualizing the relative abundance of eukaryotic taxa based on the abundance of assigned proteins extracted from the FL (red) and PA (yellow and blue) fractions. Cell size corresponds to the relative abundance of the respective genus. In this preliminary analysis, protein identification is based on metagenomic (DNA-based) information from the filtered fractions, which suffers limitations for eukaryotic protein identification, probably resulting in incomplete functional and taxonomic profiles.

### Conclusions and outlook

Our comparative metaproteomic analyses of marine microbial communities living either planktonically or attached to particles resulted in an as yet unequalled number of identified protein groups for marine particles. Interestingly, the great overlap between metaproteomes of FL and PA heterotrophic bacterial communities indicates that taxonomic differences between them might be less pronounced than previously thought. This might be due to the fact that (I) FL bacteria can rapidly adapt to the surface-associated life style, as the majority of these bacteria seems to be also present on the particles and proteins important for biofilm-formation, i.e. motility and adhesion proteins, are also expressed when living planktonically, and that (II) FL or PA-specific bacteria are frequently hopping on and off the particles. Notably, there is strong evidence that bacteria, when living on the particles, express life style-specific functions, i.e. special CAZymes, sugar transporters and proteins involved in certain stress responses, which enable them to cope with the unique living conditions on marine particles.

Although our optimized metaproteomic workflow significantly improved the identification rate of PA proteins, the number of protein identifications from the particles is still considerably lower compared to FL bacterial communities. We assume that especially the high abundance of eukaryotic proteins poses problems in protein identification due to the complexity and diversity of microbial eukaryote genomes and the presence of introns and repeats in the metagenomic DNA sequence databases, which hinders peptide identification (Saito *et al.*, 2019). Metaproteome coverage of marine particles could be significantly improved by employing customized databases including eukaryotic metatranscriptomic (RNA-based) sequence data (Keeling *et al.*, 2014). This can be achieved by generating metatranscriptomes from the particular fractions. Alternatively, protein identification could also be substantially improved by extracting already existing metatranscriptomic and metagenomic data from relevant eukaryotic taxa from public databases. Key to the latter approach is reliable information on which eukaryotic organisms make up the particles, which can be attained by 18S rRNA gene amplicon sequencing. Perspectively, we will extend our analyses on eukaryotic taxa and analyse multiple time points during phytoplankton bloom to investigate succession of taxonomical clades and expressed functions of marine particles from pre-bloom to post-bloom conditions.

## Experimental Procedures

### Bacterial biomass samples

Sampling of bacterioplankton was performed as described previously (Teeling *et al.*, 2012). Briefly, surface water samples were taken during spring 2009 at the station “Kabeltonne” (50° 11.3’ N, 7° 54.0’ E) between the main island Helgoland and the minor island Düne about 40 km offshore in the southeastern North Sea in the German Bight. Bacterial biomass for protein extraction was sequentially filtered with peristaltic pumps onto 10 μm, 3 μm, and 0.2 μm pore-sized filters (142 mm diameter) to separate PA and FL bacteria. All filters were stored at −80 °C until further analyses.

### Testing of protein extraction protocols

To test six different existing protein extraction protocols for their applicability on PA bacteria, filters from several time points containing varying amounts of biomass were chosen. Sample preparation for the metaproteomic analysis included cutting the filters into quarters and subsequently into small pieces (1-2 mm in diameter). Pieces of one quarter filter were transferred into 15 ml falcon tubes and treated according to the respective protocol.

#### Protocol 1 - Phenol

Filter pieces were incubated in 2.4 ml of a 0.1 M NaOH solution for 10 min at room temperature and were then sonicated three times for 30 s at 20% power output (Sonopuls HD2200 with microtip MS 73; Bandelin electronic, Germany). Subsequently the sample was centrifuged for 15 min at 12,500 x g at 20 °C to separate the supernatant from the filter pieces. The supernatant was transferred into a new tube and protein extraction using phenol was performed according to the protocol published by Kuhn and colleagues (Kuhn *et al.*, 2011).

#### Protocol 2 - SDS-TCA

Filter pieces were mixed with 5 ml extraction buffer (1% (w/v) SDS, 50 mM Tris/HCl, pH 7.0) and vigorously shaken for 2 min at room temperature. The cell disruption by sonication, boiling and shaking was performed according to the protein extraction protocol published by Schneider and colleagues (Schneider *et al.*, 2012). Subsequently, proteins were precipitated with 10% TCA over night at 4 °C. The precipitated proteins were centrifuged for 20 min at 12,500 x g at 4 °C and the pellet was washed two times in ice-cold acetone.

#### Protocol 3 - TRI-Reagent^®^

The TRI-Reagent^®^ (Sigma-Aldrich, product-number T9424) is used for the simultaneous isolation of RNA, DNA, and proteins. Filter pieces were transferred into 4 ml TRI-Reagent and shaken vigorously for 5 min. Subsequently the proteins were extracted according to the manufacturer’s guidelines.

#### Protocol 4 - Freeze and Thaw

Protein extraction was carried out according to a combination of the extraction protocols of Chourey *et al.* (2010) and Thompson *et al.* (2008). To this end, filter pieces were mixed with 4 ml lysis buffer (5% SDS, 50 mM Tris/HCl, 0.1 mM EDTA, 0.15 M NaCl, 1 mM MgCl_2_, 50 mM DTT, pH 8.5) and vigorously shaken for 3 min. Subsequently, the samples were boiled for 10 min, followed by two freezing and thawing cycles with liquid nitrogen. After cooling at 4 °C the samples were vigorously shaken for 3 min. To remove cell debris, samples were centrifuged for 20 min at 12,500 × g at 4 °C. The proteins in the supernatant were precipitated with 25% TCA over night at 4 °C. Precipitated proteins were centrifuged for 20 min at 12,500 × g at 4 °C and the resulting protein pellet was washed with ice-cold acetone.

#### Protocol 5 - SDS-Acetone

Filter pieces were mixed with 5 ml extraction buffer (50 mM Tris, 1% (w/v) SDS, pH 7.5) and vortexed vigorously. Proteins were extracted by sonication, boiling and shaking as described by Hall and colleagues (Hall *et al.*, 2012). Subsequently, proteins were precipitated with five volumes acetone over night at −20 °C. The precipitated proteins were centrifuged for 20 min at 12,500 × g at 4 °C and the pellet washed two times in ice-cold acetone.

#### Protocol 6 – bead beating

Protein extraction was carried out according to the extraction protocol of Moog (2012), which is based on the protocol of Teeling and colleagues (Teeling *et al.*, 2012). To this end, filter pieces were covered with 4 ml lysis buffer (0.1 M DTT, 0.01 M EDTA, 10% Glycerol (v/v), 1.7 mM PMSF, 5% SDS (w/v), 0.05 M Tris/HCl, pH 6.8) and 2 ml glass beads (0.1 – 0.11 mm diameter) were added. The cells on the filter pieces were subsequently disrupted four times for 30 s with 6.5 m/s via bead beating with a Fast Prep™- 24 (MP Biomedicals, Germany). To remove cell debris and glass beads, samples were centrifuged for 20 min at 12,500 × g at 4 °C and the supernatant was transferred into new tubes. This washing step was repeated 2 to 4 times until the beads were colourless. The glass beads were washed with 3 ml lysis buffer and vigorously shaken. Proteins enriched in the pooled supernatants were precipitated with 1:4 acetone at −20 °C over night. Precipitated proteins were centrifuged for 20 min at 12,500 × g at 4 °C and the resulting protein pellet washed with ice-cold acetone.

All resulting protein pellets were air-dried and resolved in 8 M urea / 2 M thiourea.

### Determination of protein concentrations

Protein concentrations were determined using the Pierce™ BCA Protein Assay Kit (Thermo Fisher Scientific). Protein extracts were prepared with the Compat-Able™ Protein Assay Preparation Reagent Kit (Thermo Fisher Scientific) according to the manufacturer’s guidelines.

### SDS-PAGE protein separation

30 μg protein or 30 μl protein extract was mixed with 4x SDS sample buffer (20% glycerol, 100 mM Tris/HCl, 10% (w/v) SDS, 5% β-mercaptoethanol, 0.8% bromphenol blue, pH 6.8) and loaded on TGX precast 4-20% gels (Biorad, Germany). Samples were separated by electrophoresis at 150 V for 45 min. After fixation (10% acetic acid, 40% ethanol, 30 min) the gels were stained with Brilliant Blue G250 Coomassie and imaged.

### Protein digestion and MS-sample preparation

Three different protocols were tested on proteins extracted from 3 μm and 10 μm filters.

#### Protocol 1 - 10 gel pieces

Protein lanes were cut into 10 equal-sized pieces and washed with a buffer containing 50 mM ammoniumbicarbonate and 30% (v/v) acetonitrile. Prior to tryptic digestion, gel pieces were dried in a vacuum concentrator and re-swollen with 2 ng/μl trypsin solution (sequencing grade trypsin, Promega, USA) followed by overnight digestion at 37 °C. After digestion the gel pieces were covered with water and peptides were eluted from the gel in an ultrasonic bath. The eluted peptides were desalted with C18 Millipore^®^ ZipTip columns (Millipore) according to the manufacturer’s guidelines.

#### Protocol 2 - 20 gel pieces

The protein lanes were cut into 20 equal-sized pieces and treated as described above.

#### Protocol 3 - 20 gel pieces with reduction and alkylation

The protein lanes were cut into 20 equal pieces and washed with a buffer containing 100 mM ammoniumbicarbonate (NH_4_HCO_3_) and 50% (v/v) methanol. Subsequently, proteins were reduced in 50 mM NH_4_HCO_3_ containing 10 mM DTT for 30 min at 60 °C, followed by alkylation in 50 mM NH_4_HCO_3_ containing 50 mM iodoacetamide (IAA) for 60 min in the dark at room temperature. Prior to tryptic digestion, the gel pieces were dehydrated using 100% acetonitrile and dried, re-swollen with 2 ng/μl trypsin solution and incubated at 37 °C over night. Peptides were eluted from the gel pieces by a six-step procedure, using acetonitrile, 1% (v/v) acetic acid in water, acetonitrile, 10% (v/v) acetic acid and two times acetonitrile. Peptide-containing supernatants were pooled and completely dried in a vacuum concentrator. Samples were subsequently resolved in buffer A (5% (v/v) acetonitrile, 0.1% (v/v) formic acid) and desalted with C18 Millipore^®^ ZipTip columns (Millipore) according to the manufacturer’s guidelines.

### Constructions of a protein sequence database from marine particle metagenomes

Environmental DNA was extracted from the 0.2 μm, 3 μm and 10 μm pore-sized filters sampled on the 14^th^ of April 2009 by a modified standard protocol of Zhou *et al.* (1996). In detail one polycarbonate filter was cut into 4 pieces and mixed with 13.5 ml extraction buffer (100 mM Tris-HCl (pH 8.0), 100 mM EDTA (pH 8.0), 100 mM Na-phosphate (pH 8.0), 1.5 M NaCl, 1% CTAB (Hexadecyltrimethylammonium-bromide)). Subsequently 100 μl 10 mg/ml Proteinase K was added and the sample was incubated shaking at 37 °C for 30 min. 1.5 ml 20% SDS was added and the sample was incubated shaking at 65 °C for 2 h. The sample was centrifuged at 6,000 × g for 10 min at room temperature and the supernatants were transferred to fresh tubes. Subsequently, an equal volume of chloroform/isoamylalcohol was added and the sample was mixed carefully by shaking and was centrifuged at 10,000 x g for 10 min at room temperature. Afterwards the aqueous upper phase was transferred into a new tube and the DNA was precipitated by addition of 0.6 volumes isopropanol. The sample was moderately shaken over night at 4°C. After centrifugation at 50,000 x g for 20 min at room temperature, the pellet was washed with 10 ml 80% (v/v) ethanol and dried. The pellet was resuspended in 200 μl TE buffer (10 mM Tris-HCl, 1 mM EDTA, pH 7) and stored at −20 °C until sequencing.

DNA was sequenced at the Max Planck Sequencing Centre (Cologne, Germany), using the Illumina HiSeq 2500 platform and 2 × 250 bp chemistry. Sequences were then trimmed using bbduk v35.14 (http://bbtools.jgi.doe.gov) with the following parameters: ktrim = r k = 28 mink = 12 hdist = 1 tbo = t tpe = t qtrim = rl trimq = 20 minlength = 100. Read quality for each sample was then confirmed using FastQC v0.11.2 (Andrews, 2010). Trimmed and filtered reads from the three metagenomic datasets were then assembled individually. The 0.2 μm pore-sized filter sample was assembled with metaSPAdes v3.10.1 (Nurk *et al.*, 2017) with kmers of length 21, 33, 55, 77, 99, and 127, and error correction mode switched on. Assembly of the larger size fraction was done with MEGAHIT v1.1.3 (Li et al., 2016) with kmers 21, 33, 55, 77, 99, 127, 155, 183, and 211. Assembled contigs longer than 1500 base pairs were kept for gene predictions. Genes were predicted and annotated using Prokka v1.11 (Seeman, 2014), which implements prodigal v2.6.3 (Hyatt *et al.*, 2010) for ORF prediction.

Raw read sequences and assembled contig sequences have been deposited in the European nucleotide archive (ENA) under the project accession number PRJEB2888.

### LC-MS/MS data acquisition and data analysis

Peptides were separated by reversed-phase chromatography on an Easy-nLC 1000 (Thermo Scientific) with self-packed C18 analytical columns (100 μm × 20 cm) and coupled to a LTQ Orbitrap Velos mass spectrometer(Thermo Scientific) using a non-linear binary gradient of 80 minutes from 5 % solvent A (0.1 % (v/v) acetic acid) to 99 % solvent B (0.1 % acetic acid (v/v), 99.9 % acetonitrile (v/v)) and a flow rate of 300 nl/min. Survey scans at a resolution of 30,000 were recorded in the Orbitrap analyser (m/z 300 −1700) and the 20 most intense precursor ions were selected for CID fragmentation in the LTQ. Dynamic exclusion of precursor ions was enabled; single-charged ions and ions with unknown charge state were excluded from fragmentation. Internal lock mass calibration was enabled (lock mass 445.120025).

The mass spectrometry raw data were converted into mgf files using MSConvert (64-bit, Proteowizard 3) and subsequently subjected to database searching via Mascot (Matrix Science; version 2.6.0). Four different protein sequence databases were used for peptide to spectrum matching: I) the non-redundant NCBI database (NCBI nr - NCBIprot_20171030 database (136,216,794 entries)), II) a database containing protein sequences of abundant bacteria and diatoms (PABD) based on the study of Teeling *et al*. (2012), and retrieved from Uniprot KB (Uniprot_DoS_complete_20170829 database (2,638,314 entries)), III) a database containing protein sequences of the free-living fraction from Teeling *et al*. (2012) (0.2 μm 2009 (MIMAS) - MIMAS_forward_reverse_all_contaminants database (1,579,724 entries)), and IV) a database based on translated metagenomes from 0.2 μm and 3 μm filters (see below for details) (0.2 + 3 μm 2009 - 02_plus_3_POMPU_nr97_fw_cont_20181015 database (1,463,571 entries). Mascot was searched with a fragment ion mass tolerance of 0.80 Da and a parent ion tolerance of 10.0 ppm. Oxidation of methionine was specified as a variable modification, trypsin was set as digestion enzyme and a maximum of two missed cleavages was allowed.

Scaffold (version Scaffold_4.8.7, Proteome Software Inc.) was used to validate MS/MS based peptide and protein identifications. Peptide identifications were accepted if they could be established at greater than 95.0% probability by the Peptide Prophet algorithm (Keller *et al.*, 2002) with Scaffold delta-mass correction. Protein identifications were accepted if they could be established at greater than 99.0% probability and contained at least one identified peptide. Protein probabilities were assigned by the Protein Prophet algorithm (Nesvizhskii *et al.*, 2003). Proteins that contained similar peptides and could not be differentiated based on MS/MS analysis alone were grouped to satisfy the principles of parsimony. Peptides that were only found in one of the replicates were excluded from the following data analysis. Mass spectrometry proteomics data have been deposited to the ProteomeXchange Consortium via the PRIDE partner repository (Perez-Riverol *et al.*, 2019) with the data set identifier PXD12699 (reviewer account details: username; password 5CKUi0AF).

For further data analysis, the software *ProPHAnE* (Proteomics result Pruning and Homology group Annotation Engine; version 3.1.1) (Schneider *et al.*, 2011) was used. For the taxonomical classification of the identified protein groups the NCBI NR database (version 2018-08-02; e-value 0.01, query cover 0.9, max-target-seqs 1) and the diamond blastp algorithm (version 0.8.22) were used. For functional classification of the identified protein groups the eggmap database (version 4.5.1, downloaded at 2018-07-31) and the algorithm e-mapper were used.

A list of common contaminants was added to all translated ORF sequences found by metagenome analysis of the 0.2 μm and 3 μm filters from the sampling date 14^th^ of April 2009. Redundant sequences were eliminated (97% redundancy, elimination of shorter sequence) using CD-HIT (www.cdhit.org), a program for clustering and comparing protein or nucleotide sequences, resulting in the database 02_plus_3_POMPU_nr97_fw_cont_20181015 (1,463,571 entries).

## Acknowledgements

We thank the DFG for financial support (RI 969/9-1) in the frame of the Research Unit “POMPU” (FOR 2406). We are grateful to Sabine Kühn for technical assistance and to Thomas Schweder for helpful discussions.

## Supplemental Material

**Figure S1. Protein identifications obtained by different extraction and protein prefractionation protocols**. For the medium particle size fraction (3 - 10 μm, yellow), 20 gel fractions after standard treatment, i.e. without protein reduction (red.) and alkylation (alk.), resulted in the highest number of identified protein groups, no matter which protein extraction protocol (SDS-acetone (red) or bead beating (green)) was applied. For the large particle fraction (≥ 10 μm, blue) the general trend was similar. However, the beat beating protocol performed better compared to the SDS-acetone protocol.

**Figure S2: Number of identified protein groups obtained with different databases: (I)** the non-redundant NCBI database (NCBInr, 136,216,794 entries), **(II)** a database with Uniprot sequences of known abundant bacteria and diatoms identified by the study of Teeling *et al.* (2012) (PABD, 2,638,314 entries), **(III)** a metagenome-based database employed for the FL bacterial fraction within the study of Teeling *et al*. (2012) (MIMAS, 1,579,724 entries) and **(IV)** a database based on translated metagenomes of the FL fraction on the 0.2 μm filters and particles on the 3 μm filters sampled on the 14^th^ of April 2009 (0.2 + 3 μm 2009, 1,463,572 entries).

**Figure S3: Venn diagram of overlapping and fraction-specific protein sets.**

**Table S1:** Prophane output for proteins extracted from 0.2 μm pore-sized filters

**Table S2:** Prophane output for proteins extracted from 3 μm pore-sized filters

**Table S3:** Prophane output for proteins extracted from 10 μm pore-sized filters

**Table S4:** Distribution of phylogenetic groups within proteins extracted from the 0.2 μm, 3 μm and 10 μm pore-sized filters.

