## Supplementary Figures and Tables for "A robust metaproteomics pipeline for a holistic taxonomic and functional characterization of microbial communities from marine particles"

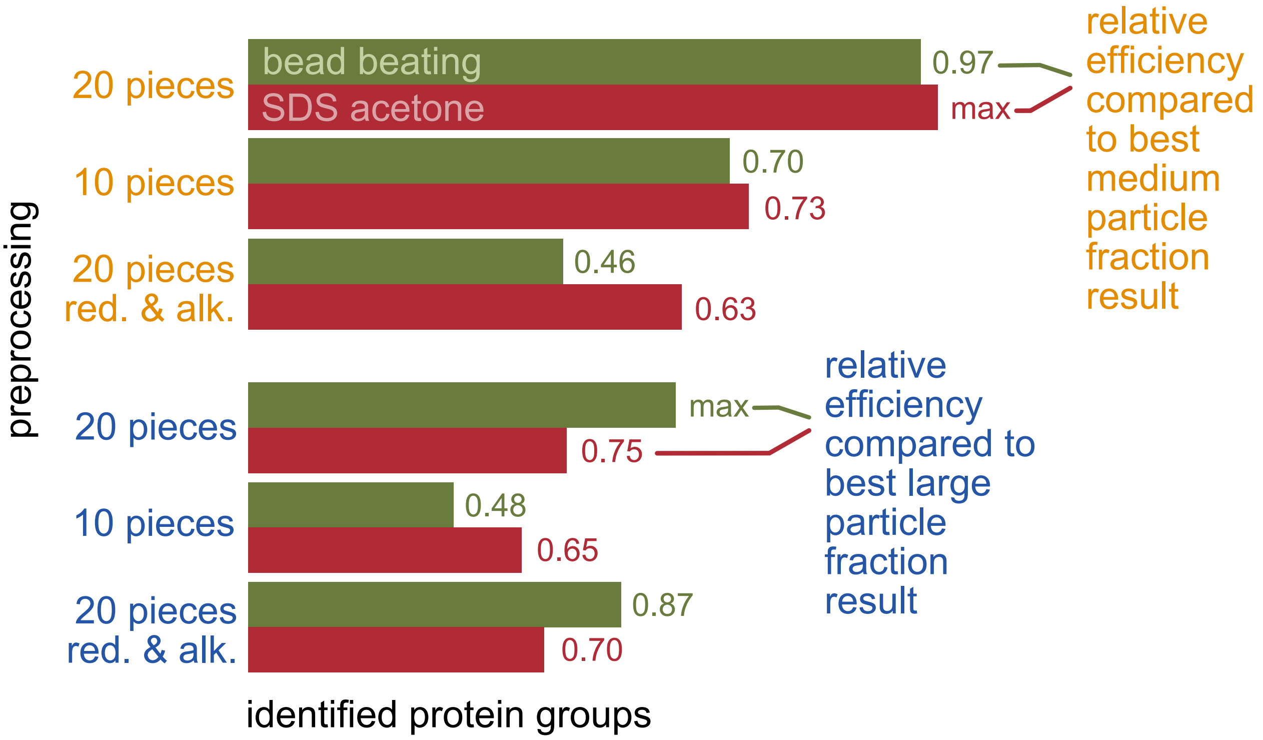
bead beating
SDS acetone
20 pieces
10 pieces
20 pieces red. & alk.
20 pieces
10 pieces
20 pieces red. & alk.
preprocessing
identified protein groups
0.97
max
0.70
0.73
0.46
0.63
max
0.75
0.48
0.65
0.87
0.70
relative efficiency compared to best medium particle fraction result
relative efficiency compared to best large particle fraction result

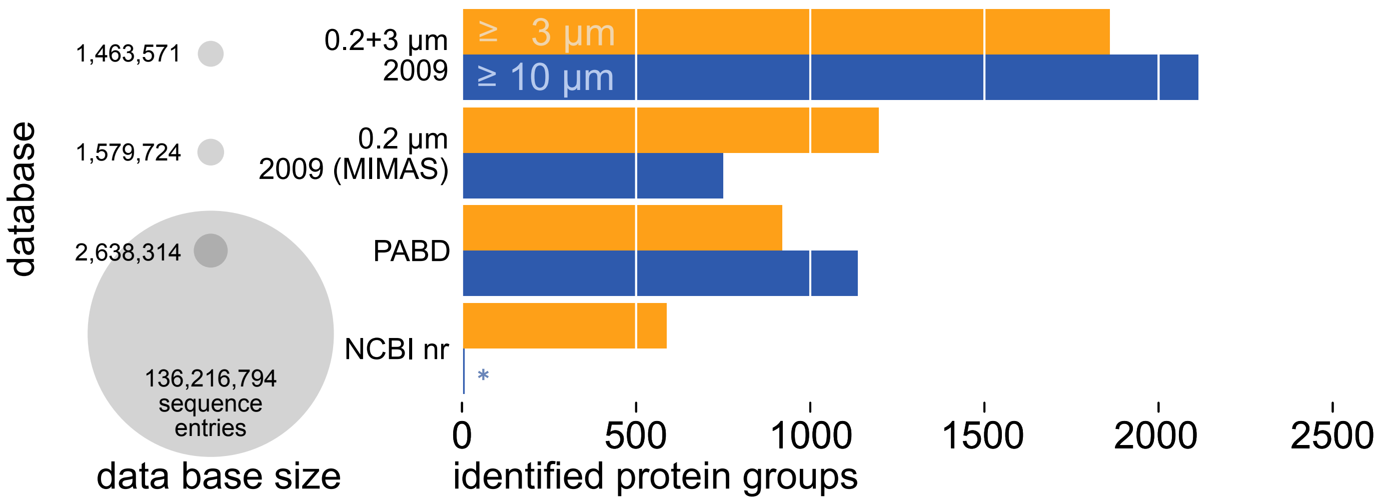
database
1,463,571
1,579,724
2,638,314
136,216,794
sequence
entries
data base size
0.2+3 µm
2009
0.2 µm
2009 (MIMAS)
PABD
NCBI nr
≥ 3 µm
≥ 10 µm
*
0
500
1000
1500
2000
2500
identified protein groups

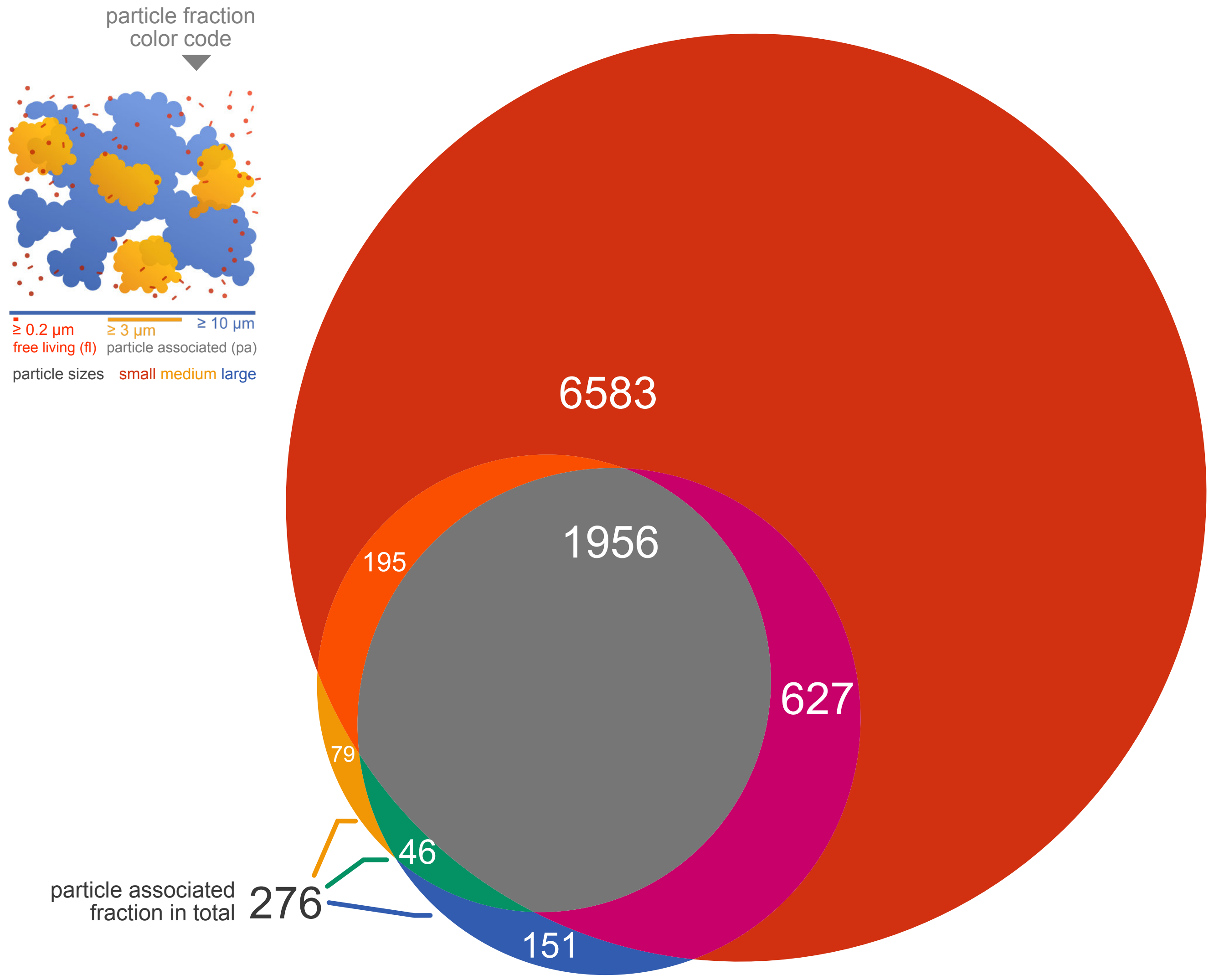

particle fraction
color code
≥ 0.2 µm
free living (fl)
≥ 3 µm
≥ 10 µm
particle associated (pa)
particle sizes
small medium large
6583
1956
195
627
79
46
151
particle associated
fraction in total
276

**PROPHANE JOB INFORMATION**

**program versions**

prophane: 3.1.1
blast+: 2.7.1
diamond: 0.8.22
hmmer: 3.1b2
krona: 2.7

**tasks**

task: *taxonomic assignment*
task comment: tax_from_nr_20180808_qcover90
algorithm: diamond blastp
query: sequences without taxonomic classification (missing_taxa.faa)
database: NCBI NR database
database version: 2018-08-02
database comment: downloaded at 2018-08-02
database scope: tax
parameter:
- evalue 0.01
- query-cover 0.9
- max-target-seqs 1

task: *functional assignment*
task comment: fun_from_eggNog_4.5.1
algorithm: emapper
query: all sequences (all.faa)
database: eggmap database
database version: 4.5.1
database comment: includes euk, viruses, bakt, arch; downloaded at 2018-07-31
database scope: func
parameter:
- m diamond

| | | | taxonomic assignment | | | | | | | functional assignment | | | | | quantitative data mean | | | | quantitative data replicate 1 | | | | quantitative data replicate 2 | | | | quantitative data replicate 3 | | | | quantitative data replicate 4 | | | | quantitative data replicate 5 | | | |
|---|---|---|---|---|---|---|---|---|---|---|---|---|---|---|---|---|---|---|---|---|---|---|---|---|---|---|---|---|---|---|---|---|---|---|---|---|---|---|
| protein group | members_count | protein group: members_identifier | superkingdom | phylum | class | order | family | genus | species | main role | sub role | og | descr | prot | exclusive spetrum count | exclusive spetrum count, sd | NSAF | NSAF, sd | exclusive spetrum count | exclusive spetrum count, sd | NSAF | NSAF, sd | exclusive spetrum count | exclusive spetrum count, sd | NSAF | NSAF, sd | exclusive spetrum count | exclusive spetrum count, sd | NSAF | NSAF, sd | exclusive spetrum count | exclusive spetrum count, sd | NSAF | NSAF, sd | exclusive spetrum count | exclusive spetrum count, sd | NSAF | NSAF, sd |
| 0 | 1 | 0.2um_NOKCPHJN_13538 | Bacteria | Bacteroidetes | Flavobacteriia | Flavobacterial | Flavobacteriac | unclassified | Flavobacteriac | poorly charact | Function unkn | 0P88U | NA | 313603.FB217 | 2,6 | 1,019803903 | 6,35E-05 | 2,51E-05 | 3 | - | 6,25E-05 | - | 4 | - | 0,000103462 | - | 3 | - | 7,38E-05 | - | 2 | - | 4,99E-05 | - | 1 | - | 2,81E-05 | - |
| 1 | 1 | 0.2um_NOKCPHJN_808747 | Bacteria | Bacteroidetes | Flavobacteriia | Flavobacterial | Flavobacteriac | Flavobacteriu | various | metabolism | Energy produc | 0PBIY | subunit C | 706436.HMPR | 2 | 0 | 0,000415234 | 3,94E-05 | 2 | - | 0,000347808 | - | 2 | - | 0,000431875 | - | 2 | - | 0,000410552 | - | 2 | - | 0,000416616 | - | 2 | - | 0,000469317 | - |
| 2 | 1 | 3um_FIJJAFBF_189740 | Bacteria | Bacteroidetes | Flavobacteriia | Flavobacterial | Flavobacteriac | Winogradskye | Winogradskye | unclassified | unclassified | unclassified | unclassified | unclassified | 0,6 | 0,489897949 | 1,59E-05 | 1,31E-05 | 1 | - | 2,35E-05 | - | 0 | - | 0 | - | 1 | - | 2,77E-05 | - | 1 | - | 2,81E-05 | - | 0 | - | 0 | - |
| 3 | 1 | 3um_FIJJAFBF_649060 | Bacteria | Proteobacteri | Gammaprotec | Oceanospirilla | Saccharospirill | Reinekea | Reinekea fors | metabolism | Carbohydrate | 0QK8I | Catalyzes the | 314283.MED2 | 4,8 | 0,748331477 | 0,000127465 | 2,12E-05 | 5 | - | 0,000111651 | - | 6 | - | 0,000166364 | - | 5 | - | 0,000131792 | - | 4 | - | 0,000106991 | - | 4 | - | 0,000120525 | - |
| 4 | 1 | 3um_FIJJAFBF_169953 | Bacteria | Proteobacteri | Alphaproteob | Rhodobactera | Rhodobactera | Roseovarius | Roseovarius a | cellular proces | Posttranslatio | COG0330 | Band 7 protei | 391589.RGAI1 | 8,6 | 1,356465997 | 0,000394808 | 3,70E-05 | 11 | - | 0,000427981 | - | 8 | - | 0,000386492 | - | 9 | - | 0,000413336 | - | 7 | - | 0,000326232 | - | 8 | - | 0,000419999 | - |
| 5 | 1 | 3um_FIJJAFBF_594169 | Eukaryota | unclassified | Synurophycea | Parmales | Triparmaceae | Triparma | Triparma laev | unclassified | unclassified | unclassified | unclassified | unclassified | 1,4 | 1,356465997 | 2,97E-05 | 2,88E-05 | 3 | - | 5,63E-05 | - | 3 | - | 6,99E-05 | - | 0 | - | 0 | - | 1 | - | 2,25E-05 | - | 0 | - | 0 | - |
| 6 | 1 | 0.2um_NOKCPHJN_441222 | Eukaryota | Chlorophyta | Mamiellophyc | Mamiellales | Bathycoccace | Ostreococcus | Ostreococcus | cellular proces | Signal transdu | COG5040 | Tyrosine 3-mo | 436017.A4S8V | 1 | 0 | 7,03E-05 | 6,67E-06 | 1 | - | 5,89E-05 | - | 1 | - | 7,31E-05 | - | 1 | - | 6,95E-05 | - | 1 | - | 7,05E-05 | - | 1 | - | 7,94E-05 | - |
| 7 | 1 | 3um_FIJJAFBF_70343 | Bacteria | Proteobacteri | Betaproteoba | unclassified | unclassified | unclassified | Betaproteoba | information st | Translation, ri | COG0186 | One of the pri | 365046.Rta_0 | 1,4 | 0,489897949 | 0,000215304 | 7,61E-05 | 1 | - | 0,000128963 | - | 1 | - | 0,000160134 | - | 2 | - | 0,000304454 | - | 2 | - | 0,000308951 | - | 1 | - | 0,000174016 | - |
| 8 | 1 | 0.2um_NOKCPHJN_207824 | Bacteria | Proteobacteri | Alphaproteob | Kordiimonada | unclassified | unclassified | Kordiimonada | metabolism | Energy produc | COG0056 | Produces ATP | 70448.Q0P3H | 2,8 | 1,166190379 | 7,07E-05 | 2,34E-05 | 5 | - | 0,000108485 | - | 2 | - | 5,39E-05 | - | 2 | - | 5,12E-05 | - | 2 | - | 5,20E-05 | - | 3 | - | 8,78E-05 | - |
| 9 | 1 | 0.2um_NOKCPHJN_556796 | Bacteria | Proteobacteri | Alphaproteob | Pelagibacteral | Pelagibacterac | Candidatus Pe | Candidatus Pe | unclassified | unclassified | unclassified | unclassified | unclassified | 10 | 1,788854382 | 0,000277556 | 6,21E-05 | 9 | - | 0,000208264 | - | 8 | - | 0,000229869 | - | 9 | - | 0,000245835 | - | 13 | - | 0,000360339 | - | 11 | - | 0,000343472 | - |
| 10 | 1 | 0.2um_NOKCPHJN_837800 | Bacteria | Bacteroidetes | Flavobacteriia | Flavobacterial | Flavobacteriac | unclassified | Flavobacteriac | unclassified | unclassified | unclassified | unclassified | unclassified | 0,4 | 0,489897949 | 2,93E-05 | 3,59E-05 | 0 | - | 0 | - | 1 | - | 7,46E-05 | - | 0 | - | 0 | - | 1 | - | 7,20E-05 | - | 0 | - | 0 | - |
| 11 | 1 | 3um_FIJJAFBF_568797 | Bacteria | Proteobacteri | Alphaproteob | Rhodobactera | Rhodobactera | Marinosulfonc | Marinosulfonc | metabolism | Lipid transport | COG2067 | lipid transport | 391624.OIHEL | 2,2 | 0,979795897 | 9,00E-05 | 4,01E-05 | 2 | - | 6,89E-05 | - | 1 | - | 4,28E-05 | - | 4 | - | 0,000162741 | - | 2 | - | 8,26E-05 | - | 2 | - | 9,30E-05 | - |
| 12 | 1 | 3um_FIJJAFBF_510230 | Bacteria | Bacteroidetes | Flavobacteriia | Flavobacterial | Flavobacteriac | Polaribacter | Polaribacter s | information st | Translation, ri | COG0087 | One of the pri | 313594.PI23P | 10,4 | 0,8 | 0,00069082 | 3,42E-05 | 12 | - | 0,000671864 | - | 10 | - | 0,000695214 | - | 10 | - | 0,000660889 | - | 10 | - | 0,00067065 | - | 10 | - | 0,000755485 | - |
| 13 | 1 | 0.2um_NOKCPHJN_773309 | Bacteria | Proteobacteri | Alphaproteob | Pelagibacteral | Pelagibacterac | Candidatus Pe | Candidatus Pe | information st | Translation, ri | 05C64 | Binds directly | 335992.SAR11 | 1,4 | 0,489897949 | 8,23E-05 | 2,59E-05 | 2 | - | 0,000100242 | - | 2 | - | 0,000124471 | - | 1 | - | 5,92E-05 | - | 1 | - | 6,00E-05 | - | 1 | - | 6,76E-05 | - |
| 14 | 1 | 3um_FIJJAFBF_561880 | Bacteria | Proteobacteri | Gammaprotec | Chromatiales | unclassified | unclassified | Chromatiales | unclassified | unclassified | unclassified | unclassified | unclassified | 1,2 | 0,4 | 3,76E-05 | 1,30E-05 | 1 | - | 2,63E-05 | - | 1 | - | 3,26E-05 | - | 1 | - | 3,10E-05 | - | 2 | - | 6,29E-05 | - | 1 | - | 3,54E-05 | - |
| 15 | 1 | 3um_FIJJAFBF_770106 | Eukaryota | unclassified | Spirotrichea | Sporadotrichi | Oxytrichidae | Oxytricha | Oxytricha trifa | metabolism | Energy produc | 0IFIU | Pyrophosphat | 4577.GRMZM | 0,8 | 0,748331477 | 3,79E-05 | 3,32E-05 | 2 | - | 8,53E-05 | - | 1 | - | 5,30E-05 | - | 0 | - | 0 | - | 1 | - | 5,11E-05 | - | 0 | - | 0 | - |
| 16 | 1 | 0.2um_NOKCPHJN_378545 | Bacteria | Bacteroidetes | Flavobacteriia | Flavobacterial | Flavobacteriac | Altibacter | Altibacter lent | unclassified | unclassified | unclassified | unclassified | unclassified | 2,8 | 0,4 | 0,000287652 | 4,66E-05 | 3 | - | 0,000258895 | - | 2 | - | 0,000214314 | - | 3 | - | 0,000305599 | - | 3 | - | 0,000310113 | - | 3 | - | 0,000349341 | - |
| 17 | 1 | 3um_FIJJAFBF_430473 | Bacteria | Proteobacteri | Betaproteoba | unclassified | unclassified | unclassified | Betaproteoba | metabolism | Energy produc | 05MPD | Cytochrome | 398578.Daci_ | 0,8 | 0,748331477 | 6,38E-05 | 6,14E-05 | 1 | - | 7,00E-05 | - | 0 | - | 0 | - | 2 | - | 0,000165222 | - | 1 | - | 8,38E-05 | - | 0 | - | 0 | - |
| 18 | 1 | 3um_FIJJAFBF_481015 | Bacteria | Bacteroidetes | Flavobacteriia | Flavobacterial | Flavobacteriac | Polaribacter | Polaribacter s | unclassified | unclassified | unclassified | unclassified | unclassified | 0,2 | 0,4 | 3,02E-05 | 6,04E-05 | 0 | - | 0 | - | 0 | - | 0 | - | 0 | - | 0 | - | 1 | - | 0,000151081 | - | 0 | - | 0 | - |
| 19 | 1 | 0.2um_NOKCPHJN_408548 | Bacteria | Proteobacteri | Gammaprotec | Alteromonada | Alteromonada | Bowmanella | Bowmanella d | metabolism | Inorganic ion t | 0QJVD | TonB-depende | 743721.Psesu | 9,4 | 1,356465997 | 0,000144802 | 1,32E-05 | 11 | - | 0,000143634 | - | 10 | - | 0,000162138 | - | 10 | - | 0,000154132 | - | 9 | - | 0,000140768 | - | 7 | - | 0,000123336 | - |
| 20 | 1 | 3um_FIJJAFBF_267368 | Bacteria | Proteobacteri | Gammaprotec | unclassified | unclassified | unclassified | Gammaprotec | information st | Translation, ri | COG0292 | Binds directly | 380703.AHA_ | 0,4 | 0,489897949 | 4,17E-05 | 5,14E-05 | 1 | - | 9,49E-05 | - | 0 | - | 0 | - | 0 | - | 0 | - | 1 | - | 0,000113623 | - | 0 | - | 0 | - |
| 21 | 1 | 3um_FIJJAFBF_229416 | Bacteria | various | various | various | various | various | various | metabolism | Inorganic ion t | COG3119 | Sulfatase | 314230.DSM3 | 2,8 | 1,326649916 | 6,83E-05 | 2,61E-05 | 5 | - | 0,000106868 | - | 2 | - | 5,31E-05 | - | 3 | - | 7,57E-05 | - | 3 | - | 7,68E-05 | - | 1 | - | 2,88E-05 | - |
| 22 | 1 | 0.2um_NOKCPHJN_54371 | Bacteria | Proteobacteri | Alphaproteob | Rhodospirillal | Rhodospirillac | Nisaea | Nisaea denitri | unclassified | unclassified | unclassified | unclassified | unclassified | 1,6 | 0,489897949 | 6,54E-05 | 1,85E-05 | 2 | - | 6,96E-05 | - | 2 | - | 8,64E-05 | - | 1 | - | 4,11E-05 | - | 2 | - | 8,33E-05 | - | 1 | - | 4,69E-05 | - |
| 23 | 1 | 3um_FIJJAFBF_37831 | Bacteria | Proteobacteri | unclassified | unclassified | unclassified | unclassified | Proteobacteri | unclassified | unclassified | unclassified | unclassified | unclassified | 2,4 | 0,489897949 | 0,000101859 | 2,28E-05 | 3 | - | 0,000106935 | - | 2 | - | 8,85E-05 | - | 2 | - | 8,42E-05 | - | 2 | - | 8,54E-05 | - | 3 | - | 0,000144293 | - |
| 24 | 1 | 3um_FIJJAFBF_525361 | Bacteria | Proteobacteri | Gammaprotec | Oceanospirilla | Saccharospirill | Reinekea | Reinekea fors | unclassified | unclassified | unclassified | unclassified | unclassified | 11,8 | 1,326649916 | 0,00044881 | 4,31E-05 | 14 | - | 0,000448848 | - | 11 | - | 0,000437907 | - | 12 | - | 0,00045413 | - | 10 | - | 0,000384032 | - | 12 | - | 0,000519132 | - |
| 25 | 1 | 3um_FIJJAFBF_472985 | Bacteria | Proteobacteri | Gammaprotec | Alteromonada | Alteromonada | Paraglaciecola | Paraglaciecola | cellular proces | Intracellular tr | 0QJ7H | motA TolQ Exl | 342610.Patl_1 | 7 | 1,414213562 | 0,00020094 | 3,25E-05 | 9 | - | 0,000218854 | - | 8 | - | 0,000241557 | - | 5 | - | 0,000143519 | - | 7 | - | 0,000203895 | - | 6 | - | 0,000196874 | - |
| 26 | 1 | 0.2um_NOKCPHJN_554855 | Bacteria | Proteobacteri | Gammaprotec | Enterobactera | Yersiniaceae | Serratia | Serratia marc | cellular proces | Signal transdu | KOG0613 | fibronectin typ | 7165.AGAP00 | 0,4 | 0,489897949 | 7,48E-06 | 9,25E-06 | 1 | - | 1,67E-05 | - | 1 | - | 2,07E-05 | - | 0 | - | 0 | - | 0 | - | 0 | - | 0 | - | 0 | - |
| 27 | 1 | 3um_FIJJAFBF_686400 | Bacteria | Proteobacteri | Alphaproteob | Rhodobactera | Rhodobactera | Loktanella | Loktanella sp. | metabolism | Coenzyme tra | COG2099 | precorrin-6x r | 391593.RCCS2 | 0,2 | 0,4 | 1,13E-05 | 2,26E-05 | 0 | - | 0 | - | 0 | - | 0 | - | 1 | - | 5,65E-05 | - | 0 | - | 0 | - | 0 | - | 0 | - |
| 28 | 1 | 3um_FIJJAFBF_341105 | Viruses | unclassified | unclassified | unclassified | unclassified | unclassified | uncultured vir | unclassified | unclassified | unclassified | unclassified | unclassified | 0,6 | 0,489897949 | 1,79E-05 | 1,49E-05 | 1 | - | 2,53E-05 | - | 0 | - | 0 | - | 1 | - | 2,99E-05 | - | 0 | - | 0 | - | 1 | - | 3,42E-05 | - |
| 29 | 1 | 3um_FIJJAFBF_353321 | Bacteria | Proteobacteri | Alphaproteob | Rhodobactera | Rhodobactera | unclassified | Rhodobactera | unclassified | unclassified | unclassified | unclassified | unclassified | 2,8 | 0,748331477 | 8,55E-05 | 2,15E-05 | 3 | - | 7,76E-05 | - | 3 | - | 9,63E-05 | - | 4 | - | 0,000122056 | - | 2 | - | 6,19E-05 | - | 2 | - | 6,98E-05 | - |
| 30 | 1 | 0.2um_NOKCPHJN_407190 | Bacteria | Proteobacteri | Gammaprotec | unclassified | unclassified | unclassified | Gammaprotec | cellular proces | Posttranslatio | 05C2Z | ATP-depender | 83406.HDN1F | 1 | 0 | 1,60E-05 | 1,52E-06 | 1 | - | 1,34E-05 | - | 1 | - | 1,66E-05 | - | 1 | - | 1,58E-05 | - | 1 | - | 1,60E-05 | - | 1 | - | 1,81E-05 | - |
| 31 | 1 | 3um_FIJJAFBF_449011 | Bacteria | Proteobacteri | Gammaprotec | unclassified | unclassified | unclassified | Gammaprotec | cellular proces | Intracellular tr | 17IXQ | transport prot | 498211.CJA_0 | 0,4 | 0,489897949 | 4,40E-05 | 5,41E-05 | 0 | - | 0 | - | 0 | - | 0 | - | 1 | - | 0,000102638 | - | 0 | - | 0 | - | 1 | - | 0,000117329 | - |
| 32 | 1 | 0.2um_NOKCPHJN_130788 | Bacteria | Proteobacteri | Gammaprotec | Cellvibrionale | Halieaceae | unclassified | marine gamm | metabolism | Inorganic ion t | 16TQC | TONB-depend | 247639.MGP2 | 2,8 | 1,326649916 | 3,98E-05 | 1,96E-05 | 3 | - | 3,62E-05 | - | 5 | - | 7,49E-05 | - | 3 | - | 4,27E-05 | - | 2 | - | 2,89E-05 | - | 1 | - | 1,63E-05 | - |
| 33 | 1 | 0.2um_NOKCPHJN_743723 | Bacteria | Bacteroidetes | Flavobacteriia | Flavobacterial | Flavobacteriac | Polaribacter | Polaribacter s | cellular proces | Cell wall/mem | 0P9R9 | TonB family | 313598.MED1 | 0,8 | 0,4 | 4,49E-05 | 2,28E-05 | 1 | - | 4,86E-05 | - | 1 | - | 6,04E-05 | - | 1 | - | 5,74E-05 | - | 1 | - | 5,83E-05 | - | 0 | - | 0 | - |
| 34 | 1 | 0.2um_NOKCPHJN_190469 | Bacteria | Bacteroidetes | Flavobacteriia | Flavobacterial | Crocinitomica | unclassified | Crocinitomica | unclassified | unclassified | unclassified | unclassified | unclassified | 0,2 | 0,4 | 1,05E-05 | 2,09E-05 | 0 | - | 0 | - | 0 | - | 0 | - | 0 | - | 0 | - | 1 | - | 5,23E-05 | - | 0 | - | 0 | - |
| 35 | 1 | 3um_FIJJAFBF_675676 | Viruses | unclassified | unclassified | unclassified | Phycodnavirid | unclassified | Chrysochromu | unclassified | unclassified | unclassified | unclassified | unclassified | 1 | 0 | 6,92E-05 | 6,57E-06 | 1 | - | 5,80E-05 | - | 1 | - | 7,20E-05 | - | 1 | - | 6,84E-05 | - | 1 | - | 6,94E-05 | - | 1 | - | 7,82E-05 | - |
| 36 | 1 | 0.2um_NOKCPHJN_06727 | Bacteria | Proteobacteri | Gammaprotec | Alteromonada | Pseudoalteror | Pseudoalteror | Pseudoalteror | cellular proces | Posttranslatio | 0QJ3C | Acts as a chap | 1085623.GNIT | 5 | 1,264911064 | 0,000103126 | 2,31E-05 | 6 | - | 0,000104979 | - | 4 | - | 8,69E-05 | - | 4 | - | 8,26E-05 | - | 7 | - | 0,000146705 | - | 4 | - | 9,44E-05 | - |
| 37 | 2 | 0.2um_NOKCPHJN_66260;3um_FIJJAFBF_578514 | Bacteria | Proteobacteri | Gammaprotec | Cellvibrionale | Porticoccacea | unclassified | various | various | various | various | various | various | 0,8 | 0,4 | 6,28E-05 | 3,19E-05 | 1 | - | 6,79E-05 | - | 1 | - | 8,43E-05 | - | 1 | - | 8,02E-05 | - | 1 | - | 8,14E-05 | - | 0 | - | 0 | - |
| 38 | 1 | 3um_FIJJAFBF_200325 | Bacteria | Bacteroidetes | unclassified | unclassified | unclassified | unclassified | Bacteroidetes | poorly charact | Function unkn | 09ARK | Protein of unk | 886377.Murru | 1,6 | 1,019803903 | 7,76E-05 | 5,03E-05 | 2 | - | 8,23E-05 | - | 3 | - | 0,000153246 | - | 2 | - | 9,71E-05 | - | 0 | - | 0 | - | 1 | - | 5,55E-05 | - |
| 39 | 1 | 3um_FIJJAFBF_361750 | Bacteria | unclassified | unclassified | unclassified | unclassified | unclassified | uncultured ba | unclassified | unclassified | unclassified | unclassified | unclassified | 1,6 | 0,8 | 5,70E-05 | 2,48E-05 | 3 | - | 9,18E-05 | - | 1 | - | 3,80E-05 | - | 1 | - | 3,61E-05 | - | 1 | - | 3,67E-05 | - | 2 | - | 8,26E-05 | - |
| 40 | 1 | 3um_FIJJAFBF_417063 | Bacteria | Proteobacteri | Gammaprotec | Alteromonada | Alteromonada | Paraglaciecola | various | information st | Translation, ri | COG0256 | This is one of | 1085623.GNIT | 2 | 0 | 0,000234234 | 2,22E-05 | 2 | - | 0,0001962 | - | 2 | - | 0,000243622 | - | 2 | - | 0,000231594 | - | 2 | - | 0,000235014 | - | 2 | - | 0,000264743 | - |
| 41 | 1 | 0.2um_NOKCPHJN_52646 | Bacteria | Proteobacteri | Gammaprotec | Aeromonadale | Aeromonadac | Oceanimonas | Oceanimonas | poorly charact | Function unkn | 17CZ8 | Lipoprotein | 511062.GU3_ | 2,4 | 1,2 | 0,000173707 | 8,41E-05 | 3 | - | 0,000187136 | - | 1 | - | 7,75E-05 | - | 3 | - | 0,000220895 | - | 4 | - | 0,000298877 | - | 1 | - | 8,42E-05 | - |
| 42 | 1 | 0.2um_NOKCPHJN_176471 | Eukaryota | unclassified | unclassified | Isochrysidales | Noelaerhabda | Emiliania | Emiliania huxl | metabolism | Carbohydrate | COG1850 | RuBisCO catal | 45157.CMV01 | 0,8 | 0,4 | 2,25E-05 | 1,16E-05 | 1 | - | 2,35E-05 | - | 1 | - | 2,92E-05 | - | 0 | - | 0 | - | 1 | - | 2,82E-05 | - | 1 | - | 3,17E-05 | - |
| 43 | 4 | 0.2um_NOKCPHJN_224501;0.2um_NOKCPHJN_5 | Bacteria | Bacteroidetes | Flavobacteriia | Flavobacterial | Flavobacteriac | unclassified | Flavobacteriac | metabolism | Amino acid tra | 09GJ6 | nitrogen regul | 313598.MED1 | 0,2 | 0,4 | 2,05E-05 | 4,10E-05 | 1 | - | 0,000102479 | - | 0 | - | 0 | - | 0 | - | 0 | - | 0 | - | 0 | - | 0 | - | 0 | - |
| 44 | 1 | 3um_FIJJAFBF_40862 | Bacteria | Bacteroidetes | unclassified | unclassified | unclassified | unclassified | Bacteroidetes | cellular proces | Cell motility | COG1360 | Ompa motb d | 926562.Oweh | 0,8 | 0,748331477 | 3,15E-05 | 2,78E-05 | 2 | - | 7,20E-05 | - | 0 | - | 0 | - | 1 | - | 4,25E-05 | - | 1 | - | 4,31E-05 | - | 0 | - | 0 | - |
| 45 | 1 | 0.2um_NOKCPHJN_293118 | Bacteria | Proteobacteri | Gammaprotec | Cellvibrionale | Porticoccacea | unclassified | SAR92 bacteri | information st | Translation, ri | COG0088 | One of the pri | 314287.GB22 | 0,2 | 0,4 | 1,52E-05 | 3,04E-05 | 0 | - | 0 | - | 0 | - | 0 | - | 0 | - | 0 | - | 0 | - | 0 | - | 1 | - | 7,59E-05 | - |
| 46 | 1 | 0.2um_NOKCPHJN_1023367 | Bacteria | Bacteroidetes | Flavobacteriia | Flavobacterial | Flavobacteriac | Marinirhabdu | Marinirhabdu | metabolism | Inorganic ion t | 05C27 | TonB-depende | 983548.Krodi_ | 6,2 | 0,748331477 | 0,000214377 | 2,86E-05 | 6 | - | 0,000174344 | - | 7 | - | 0,000252565 | - | 7 | - | 0,000240095 | - | 6 | - | 0,000208835 | - | 5 | - | 0,000196044 | - |
| 47 | 1 | 0.2um_NOKCPHJN_357602 | Bacteria | Bacteroidetes | Flavobacteriia | Flavobacterial | Flavobacteriac | Polaribacter | Polaribacter a | information st | Translation, ri | 05CAV | thus facilitatin | 391587.KAOT | 1 | 0 | 2,27E-05 | 2,16E-06 | 1 | - | 1,90E-05 | - | 1 | - | 2,36E-05 | - | 1 | - | 2,25E-05 | - | 1 | - | 2,28E-05 | - | 1 | - | 2,57E-05 | - |
| 48 | 1 | 3um_FIJJAFBF_618229 | Bacteria | Bacteroidetes | Chitinophagia | Chitinophagal | Chitinophagac | Hydrobacter | Hydrobacter p | unclassified | unclassified | unclassified | unclassified | unclassified | 0,6 | 0,489897949 | 3,10E-05 | 2,55E-05 | 1 | - | 4,59E-05 | - | 0 | - | 0 | - | 1 | - | 5,42E-05 | - | 1 | - | 5,50E-05 | - | 0 | - | 0 | - |
| 49 | 1 | 3um_FIJJAFBF_250840 | Bacteria | Proteobacteri | Betaproteoba | unclassified | unclassified | unclassified | Betaproteoba | unclassified | unclassified | unclassified | unclassified | unclassified | 2,4 | 2,059126028 | 4,73E-05 | 3,63E-05 | 6 | - | 0,000106769 | - | 0 | - | 0 | - | 3 | - | 6,30E-05 | - | 2 | - | 4,26E-05 | - | 1 | - | 2,40E-05 | - |
| 50 | 1 | 0.2um_NOKCPHJN_483485 | Bacteria | Bacteroidetes | Flavobacteriia | Flavobacterial | Flavobacteriac | Arenitalea | Arenitalea lut | metabolism | Inorganic ion t | 05C27 | TonB-depende | 886377.Murru | 2,2 | 0,979795897 | 3,03E-05 | 1,02E-05 | 4 | - | 4,82E-05 | - | 2 | - | 2,99E-05 | - | 2 | - | 2,84E-05 | - | 2 | - | 2,89E-05 | - | 1 | - | 1,63E-05 | - |
| 51 | 1 | 0.2um_NOKCPHJN_37945 | Bacteria | Bacteroidetes | Flavobacteriia | Flavobacterial | Flavobacteriac | unclassified | Flavobacteriac | unclassified | unclassified | unclassified | unclassified | unclassified | 2,6 | 0,489897949 | 0,000145272 | 2,45E-05 | 3 | - | 0,0001417 | - | 3 | - | 0,000175949 | - | 2 | - | 0,000111508 | - | 3 | - | 0,000169732 | - | 2 | - | 0,000127469 | - |
| 52 | 1 | 3um_FIJJAFBF_581775 | Bacteria | Bacteroidetes | Flavobacteriia | Flavobacterial | Flavobacteriac | Polaribacter | Polaribacter r | unclassified | unclassified | unclassified | unclassified | unclassified | 17,8 | 2,13541565 | 0,000444394 | 4,40E-05 | 21 | - | 0,000442259 | - | 15 | - | 0,000392254 | - | 19 | - | 0,000472323 | - | 16 | - | 0,000403621 | - | 18 | - | 0,000511512 | - |
| 53 | 1 | 3um_FIJJAFBF_176504 | Bacteria | Bacteroidetes | Flavobacteriia | various | various | various | various | information st | Transcription | 09EME | Necessary for | 487797.Flav3 | 0,8 | 0,4 | 6,87E-05 | 3,53E-05 | 1 | - | 7,26E-05 | - | 0 | - | 0 | - | 1 | - | 8,57E-05 | - | 1 | - | 8,70E-05 | - | 1 | - | 9,80E-05 | - |
| 54 | 1 | 0.2um_NOKCPHJN_347515 | Bacteria | unclassified | unclassified | unclassified | unclassified | unclassified | bacterium TM | unclassified | unclassified | unclassified | unclassified | unclassified | 0,4 | 0,489897949 | 6,74E-06 | 8,31E-06 | 1 | - | 1,53E-05 | - | 0 | - | 0 | - | 0 | - | 0 | - | 1 | - | 1,84E-05 | - | 0 | - | 0 | - |
| 55 | 1 | 0.2um_NOKCPHJN_61949 | Bacteria | Bacteroidetes | Flavobacteriia | Flavobacterial | Flavobacteriac | Formosa | Formosa sp. H | unclassified | unclassified | unclassified | unclassified | unclassified | 7,6 | 0,489897949 | 0,000136219 | 2,00E-05 | 7 | - | 0,000104614 | - | 8 | - | 0,000148457 | - | 7 | - | 0,000123486 | - | 8 | - | 0,000143212 | - | 8 | - | 0,000161328 | - |
| 56 | 1 | 3um_FIJJAFBF_51301 | Bacteria | Proteobacteri | Gammaprotec | Oceanospirilla | Saccharospirill | Reinekea | Reinekea blan | metabolism | Amino acid tra | COG0473 | 3-isopropylma | 314283.MED2 | 1,2 | 0,979795897 | 4,12E-05 | 3,39E-05 | 2 | - | 6,11E-05 | - | 0 | - | 0 | - | 2 | - | 7,21E-05 | - | 2 | - | 7,31E-05 | - | 0 | - | 0 | - |
| 57 | 1 | 0.2um_NOKCPHJN_93808 | Bacteria | Bacteroidetes | Flavobacteriia | Flavobacterial | Flavobacteriac | Formosa | Formosa sp. H | metabolism | Energy produc | COG0437 | 4fe-4S ferredc | 983544.Lacal_ | 1 | 0 | 1,32E-05 | 1,25E-06 | 1 | - | 1,10E-05 | - | 1 | - | 1,37E-05 | - | 1 | - | 1,30E-05 | - | 1 | - | 1,32E-05 | - | 1 | - | 1,49E-05 | - |
| 58 | 1 | 0.2um_NOKCPHJN_44940 | Bacteria | Proteobacteri | Oligoflexia | Bacteriovorac | Bacteriovorac | Bacteriovorax | Bacteriovorax | cellular proces | Signal transdu | KOG0613 | fibronectin typ | 7070.TC00296 | 1 | 0,632455532 | 2,99E-05 | 1,91E-05 | 1 | - | 2,50E-05 | - | 1 | - | 3,10E-05 | - | 0 | - | 0 | - | 2 | - | 5,98E-05 | - | 1 | - | 3,37E-05 | - |
| 59 | 1 | 0.2um_NOKCPHJN_309353 | Bacteria | Proteobacteri | Gammaprotec | Alteromonada | Pseudoalteror | Pseudoalteror | Pseudoalteror | information st | Transcription | 05D27 | DNA-depende | 1085623.GNIT | 14,2 | 1,939071943 | 0,000138919 | 1,90E-05 | 16 | - | 0,000131738 | - | 16 | - | 0,00016358 | - | 11 | - | 0,000106909 | - | 15 | - | 0,000147938 | - | 13 | - | 0,000144431 | - |
| 60 | 1 | 3um_FIJJAFBF_151895 | unclassified | unclassified | unclassified | unclassified | unclassified | unclassified | unclassified | unclassified | unclassified | unclassified | unclassified | unclassified | 0,4 | 0,489897949 | 2,51E-05 | 3,10E-05 | 1 | - | 5,60E-05 | - | 1 | - | 6,95E-05 | - | 0 | - | 0 | - | 0 | - | 0 | - | 0 | - | 0 | - |
| 61 | 1 | 0.2um_NOKCPHJN_66080 | Bacteria | Bacteroidetes | Cytophagia | Cytophagales | Cytophagacea | Sporocytopha | Sporocytopha | metabolism | Energy produc | COG0055 | Produces ATP | 880070.Cycma | 3,2 | 1,326649916 | 8,64E-05 | 3,61E-05 | 3 | - | 6,90E-05 | - | 3 | - | 8,57E-05 | - | 4 | - | 0,000108603 | - | 5 | - | 0,000137759 | - | 1 | - | 3,10E-05 | - |
| 62 | 1 | 3um_FIJJAFBF_152174 | Bacteria | Proteobacteri | Alphaproteob | Rhodospirillal | Rhodospirillac | unclassified | Rhodospirillac | unclassified | unclassified | unclassified | unclassified | unclassified | 1 | 0,632455532 | 4,23E-05 | 2,77E-05 | 1 | - | 3,63E-05 | - | 1 | - | 4,51E-05 | - | 1 | - | 4,29E-05 | - | 2 | - | 8,70E-05 | - | 0 | - | 0 | - |
| 63 | 1 | 3um_FIJJAFBF_517420 | Bacteria | Bacteroidetes | Flavobacteriia | Flavobacterial | Flavobacteriac | Polaribacter | various | information st | Transcription | COG0250 | Participates in | 313598.MED1 | 4,2 | 0,4 | 0,00031432 | 3,98E-05 | 4 | - | 0,000250878 | - | 4 | - | 0,000311517 | - | 5 | - | 0,00037017 | - | 4 | - | 0,00030051 | - | 4 | - | 0,000338523 | - |
| 64 | 1 | 3um_FIJJAFBF_216063 | Bacteria | Proteobacteri | Gammaprotec | Cellvibrionale | unclassified | unclassified | Cellvibrionale | information st | Translation, ri | 177SZ | This protein bi | 247633.GP214 | 2,6 | 0,8 | 0,000245867 | 7,13E-05 | 3 | - | 0,000242486 | - | 3 | - | 0,000301096 | - | 3 | - | 0,00028623 | - | 3 | - | 0,000290458 | - | 1 | - | 0,000109067 | - |
| 65 | 2 | 0.2um_NOKCPHJN_311550;3um_FIJJAFBF_10824 | Bacteria | Proteobacteri | Alphaproteob | Rhodobactera | Rhodobactera | Planktomarina | Planktomarina | poorly charact | Function unkn | 01ZSM | NA | 467661.RKLH1 | 0,8 | 0,748331477 | 5,98E-05 | 5,20E-05 | 2 | - | 0,000131927 | - | 0 | - | 0 | - | 1 | - | 7,79E-05 | - | 0 | - | 0 | - | 1 | - | 8,90E-05 | - |
| 66 | 1 | 0.2um_NOKCPHJN_444451 | Bacteria | unclassified | unclassified | unclassified | unclassified | unclassified | uncultured ba | metabolism | Inorganic ion t | 05C27 | TonB-depende | 700598.Niako | 0,8 | 0,4 | 3,78E-05 | 1,94E-05 | 1 | - | 4,00E-05 | - | 0 | - | 0 | - | 1 | - | 4,72E-05 | - | 1 | - | 4,79E-05 | - | 1 | - | 5,40E-05 | - |
| 67 | 2 | 3um_FIJJAFBF_139456;3um_FIJJAFBF_690608 | Bacteria | Proteobacteri | Alphaproteob | Rhodobactera | Rhodobactera | various | various | unclassified | unclassified | unclassified | unclassified | unclassified | 0,6 | 0,8 | 3,18E-05 | 4,09E-05 | 2 | - | 9,94E-05 | - | 0 | - | 0 | - | 0 | - | 0 | - | 1 | - | 5,95E-05 | - | 0 | - | 0 | - |
| 68 | 1 | 3um_FIJJAFBF_537764 | Bacteria | Bacteroidetes | Flavobacteriia | Flavobacterial | Crocinitomica | unclassified | Crocinitomica | unclassified | unclassified | unclassified | unclassified | unclassified | 0,4 | 0,489897949 | 4,22E-05 | 5,22E-05 | 1 | - | 9,41E-05 | - | 1 | - | 0,000116819 | - | 0 | - | 0 | - | 0 | - | 0 | - | 0 | - | 0 | - |
| 69 | 1 | 0.2um_NOKCPHJN_42956 | Bacteria | Bacteroidetes | Flavobacteriia | Flavobacterial | Flavobacteriac | Polaribacter | Polaribacter s | information st | Translation, ri | 05KK9 | This protein bi | 313594.PI23P | 3,2 | 0,4 | 0,000259277 | 3,84E-05 | 3 | - | 0,000203746 | - | 3 | - | 0,000252992 | - | 4 | - | 0,000320668 | - | 3 | - | 0,000244053 | - | 3 | - | 0,000274925 | - |
| 70 | 1 | 3um_FIJJAFBF_292915 | Eukaryota | Bacillariophyt | Coscinodiscop | Coscinodiscale | Coscinodiscac | Coscinodiscus | Coscinodiscus | metabolism | Energy produc | COG0711 | F(1)F(0) ATP s | 35128.Thapsd | 5,8 | 0,748331477 | 0,000438905 | 2,24E-05 | 7 | - | 0,000448848 | - | 5 | - | 0,000398097 | - | 6 | - | 0,00045413 | - | 6 | - | 0,000460838 | - | 5 | - | 0,00043261 | - |
| 71 | 1 | 3um_FIJJAFBF_532873 | Bacteria | Proteobacteri | Alphaproteob | Rhodobactera | Rhodobactera | unclassified | Rhodobactera | information st | Replication, re | COG0178 | The UvrABC re | 314262.MED1 | 0,8 | 0,4 | 1,15E-05 | 5,93E-06 | 1 | - | 1,21E-05 | - | 1 | - | 1,50E-05 | - | 1 | - | 1,43E-05 | - | 0 | - | 0 | - | 1 | - | 1,63E-05 | - |
| 72 | 1 | 0.2um_NOKCPHJN_158780 | Bacteria | Bacteroidetes | Flavobacteriia | Flavobacterial | Flavobacteriac | Sediminicola | Sediminicola s | unclassified | unclassified | unclassified | unclassified | unclassified | 0,6 | 0,489897949 | 9,68E-06 | 8,05E-06 | 1 | - | 1,35E-05 | - | 1 | - | 1,67E-05 | - | 0 | - | 0 | - | 0 | - | 0 | - | 1 | - | 1,82E-05 | - |
| 73 | 1 | 3um_FIJJAFBF_510029 | Bacteria | Proteobacteri | Gammaprotec | Cellvibrionale | Porticoccacea | unclassified | gamma protec | unclassified | unclassified | unclassified | unclassified | unclassified | 2 | 0,632455532 | 0,000182103 | 4,77E-05 | 3 | - | 0,000232656 | - | 2 | - | 0,000192593 | - | 2 | - | 0,000183084 | - | 1 | - | 9,29E-05 | - | 2 | - | 0,00020929 | - |
| 74 | 1 | 0.2um_NOKCPHJN_321483 | Bacteria | Proteobacteri | Gammaprotec | Cellvibrionale | Halieaceae | unclassified | marine gamm | metabolism | Inorganic ion t | 0QJVD | TonB-depende | 247639.MGP2 | 0,6 | 0,489897949 | 9,18E-06 | 7,50E-06 | 0 | - | 0 | - | 1 | - | 1,57E-05 | - | 1 | - | 1,50E-05 | - | 1 | - | 1,52E-05 | - | 0 | - | 0 | - |
| 75 | 1 | 0.2um_NOKCPHJN_200764 | Bacteria | Proteobacteri | Gammaprotec | Oceanospirilla | Saccharospirill | Reinekea | Reinekea fors | unclassified | unclassified | unclassified | unclassified | unclassified | 3,6 | 1,356465997 | 9,07E-05 | 3,68E-05 | 2 | - | 4,15E-05 | - | 3 | - | 7,73E-05 | - | 3 | - | 7,35E-05 | - | 6 | - | 0,000149168 | - | 4 | - | 0,000112025 | - |
| 76 | 1 | 0.2um_NOKCPHJN_639109 | Bacteria | Proteobacteri | Alphaproteob | Pelagibacteral | Pelagibacterac | unclassified | Pelagibactera | cellular proces | Cell wall/mem | 7257 | NA | 439493.PB721 | 1 | 0,632455532 | 4,16E-05 | 2,54E-05 | 0 | - | 0 | - | 1 | - | 4,20E-05 | - | 2 | - | 7,99E-05 | - | 1 | - | 4,06E-05 | - | 1 | - | 4,57E-05 | - |
| 77 | 1 | 0.2um_NOKCPHJN_991962 | unclassified | unclassified | unclassified | unclassified | unclassified | unclassified | unclassified | unclassified | unclassified | unclassified | unclassified | unclassified | 0,4 | 0,489897949 | 3,60E-05 | 4,44E-05 | 1 | - | 8,20E-05 | - | 0 | - | 0 | - | 0 | - | 0 | - | 1 | - | 9,82E-05 | - | 0 | - | 0 | - |
| 78 | 1 | 3um_FIJJAFBF_195995 | Bacteria | Bacteroidetes | Flavobacteriia | Flavobacterial | Flavobacteriac | Polaribacter | Polaribacter s | unclassified | unclassified | unclassified | unclassified | unclassified | 2,6 | 0,489897949 | 0,000139796 | 2,84E-05 | 3 | - | 0,000135563 | - | 2 | - | 0,00011222 | - | 3 | - | 0,000160018 | - | 2 | - | 0,000108255 | - | 3 | - | 0,000182923 | - |
| 79 | 1 | 0.2um_NOKCPHJN_406654 | Bacteria | Proteobacteri | Alphaproteob | Pelagibacteral | Pelagibacterac | Candidatus Pe | Candidatus Pe | information st | Translation, ri | COG0050 | This protein p | 1002672.SAR1 | 0,6 | 0,489897949 | 1,96E-05 | 1,61E-05 | 1 | - | 2,90E-05 | - | 0 | - | 0 | - | 1 | - | 3,42E-05 | - | 1 | - | 3,47E-05 | - | 0 | - | 0 | - |
| 80 | 1 | 0.2um_NOKCPHJN_88734 | Bacteria | Proteobacteri | Gammaprotec | unclassified | unclassified | unclassified | various | unclassified | unclassified | unclassified | unclassified | unclassified | 0,4 | 0,489897949 | 1,25E-05 | 1,54E-05 | 1 | - | 2,79E-05 | - | 1 | - | 3,46E-05 | - | 0 | - | 0 | - | 0 | - | 0 | - | 0 | - | 0 | - |
| 81 | 1 | 0.2um_NOKCPHJN_29228 | Bacteria | Bacteroidetes | Flavobacteriia | Flavobacterial | Flavobacteriac | Arenibacter | Arenibacter tr | unclassified | unclassified | unclassified | unclassified | unclassified | 1,4 | 0,8 | 6,09E-05 | 3,39E-05 | 2 | - | 7,40E-05 | - | 2 | - | 9,19E-05 | - | 0 | - | 0 | - | 2 | - | 8,87E-05 | - | 1 | - | 5,00E-05 | - |
| 82 | 1 | 3um_FIJJAFBF_481951 | Bacteria | Proteobacteri | Alphaproteob | Rhodobactera | Rhodobactera | Aliiroseovariu | Aliiroseovariu | metabolism | Energy produc | 01RNF | Produces ATP | 314262.MED1 | 9,2 | 1,166190379 | 0,000264669 | 3,39E-05 | 11 | - | 0,00026636 | - | 8 | - | 0,000240538 | - | 8 | - | 0,000228662 | - | 9 | - | 0,000261044 | - | 10 | - | 0,000326739 | - |
| 83 | 2 | 0.2um_NOKCPHJN_549863;3um_FIJJAFBF_17664 | Bacteria | Bacteroidetes | various | various | various | various | various | information st | Translation, ri | 0PAI9 | Binds 16S rRN | various | 0,8 | 0,4 | 0,000128171 | 6,45E-05 | 0 | - | 0 | - | 1 | - | 0,000160134 | - | 1 | - | 0,000152227 | - | 1 | - | 0,000154476 | - | 1 | - | 0,000174016 | - |
| 84 | 1 | 3um_FIJJAFBF_420755 | Bacteria | Bacteroidetes | Cytophagia | Cytophagales | Flammeovirga | unclassified | Flammeovirga | poorly charact | Function unkn | 11H69 | NA | 761193.Runsl_ | 1,2 | 0,748331477 | 3,68E-05 | 2,20E-05 | 2 | - | 5,24E-05 | - | 2 | - | 6,51E-05 | - | 0 | - | 0 | - | 1 | - | 3,14E-05 | - | 1 | - | 3,54E-05 | - |
| 85 | 1 | 0.2um_NOKCPHJN_339587 | Bacteria | Proteobacteri | Alphaproteob | Rhodobactera | Rhodobactera | Amylibacter | Amylibacter k | poorly charact | Function unkn | 01QH8 | abc transporte | 501479.CSE45 | 0,8 | 0,748331477 | 1,89E-05 | 1,64E-05 | 2 | - | 4,17E-05 | - | 0 | - | 0 | - | 1 | - | 2,46E-05 | - | 0 | - | 0 | - | 1 | - | 2,81E-05 | - |
| 86 | 1 | 0.2um_NOKCPHJN_207774 | Bacteria | Actinobacteri | Actinobacteri | Micrococcales | Microbacteria | unclassified | Microbacteria | poorly charact | Function unkn | 0ZQ28 | NA | 979556.MTES_ | 0,2 | 0,4 | 2,31E-05 | 4,62E-05 | 0 | - | 0 | - | 0 | - | 0 | - | 0 | - | 0 | - | 1 | - | 0,000115532 | - | 0 | - | 0 | - |
| 87 | 1 | 3um_FIJJAFBF_100818 | Bacteria | Bacteroidetes | Flavobacteriia | Flavobacterial | Flavobacteriac | Polaribacter | various | information st | Translation, ri | 09EM4 | Binds directly | 313598.MED1 | 3,6 | 0,489897949 | 0,000429857 | 5,35E-05 | 4 | - | 0,000402725 | - | 4 | - | 0,000500066 | - | 3 | - | 0,000356532 | - | 4 | - | 0,000482398 | - | 3 | - | 0,000407564 | - |
| 88 | 1 | 0.2um_NOKCPHJN_385913 | Bacteria | Proteobacteri | Alphaproteob | Rickettsiales | unclassified | unclassified | Rickettsiales b | unclassified | unclassified | unclassified | unclassified | unclassified | 30,4 | 5,238320341 | 0,000425348 | 6,50E-05 | 33 | - | 0,000390075 | - | 32 | - | 0,000469681 | - | 38 | - | 0,000530209 | - | 25 | - | 0,000353973 | - | 24 | - | 0,0003828 | - |
| 89 | 1 | 3um_FIJJAFBF_142609 | Bacteria | Proteobacteri | Alphaproteob | Rhodobactera | Rhodobactera | Pseudooctade | Pseudooctade | cellular proces | Defense mech | 0P56F | Type I restricti | 588581.Cpap_ | 0,4 | 0,489897949 | 1,39E-05 | 1,72E-05 | 1 | - | 3,11E-05 | - | 1 | - | 3,86E-05 | - | 0 | - | 0 | - | 0 | - | 0 | - | 0 | - | 0 | - |
| 90 | 3 | 0.2um_NOKCPHJN_329591;3um_FIJJAFBF_02384 | Bacteria | Bacteroidetes | Flavobacteriia | Flavobacterial | Flavobacteriac | various | various | unclassified | unclassified | unclassified | unclassified | unclassified | 3,6 | 1,019803903 | 0,000222356 | 5,64E-05 | 5 | - | 0,000262047 | - | 2 | - | 0,000130154 | - | 3 | - | 0,000185592 | - | 4 | - | 0,000251111 | - | 4 | - | 0,000282876 | - |
| 91 | 1 | 3um_FIJJAFBF_772310 | Bacteria | Bacteroidetes | Flavobacteriia | Flavobacterial | Flavobacteriac | Psychroflexus | Psychroflexus | poorly charact | Function unkn | 09BP5 | RHS repeat-as | 313595.P7007 | 0,6 | 0,489897949 | 4,34E-05 | 3,56E-05 | 0 | - | 0 | - | 0 | - | 0 | - | 1 | - | 6,88E-05 | - | 1 | - | 6,98E-05 | - | 1 | - | 7,86E-05 | - |
| 92 | 1 | 3um_FIJJAFBF_582415 | Bacteria | Proteobacteri | Betaproteoba | Burkholderial | Oxalobacterac | Collimonas | Collimonas sp | unclassified | unclassified | unclassified | unclassified | unclassified | 0,4 | 0,489897949 | 1,47E-05 | 1,80E-05 | 0 | - | 0 | - | 0 | - | 0 | - | 0 | - | 0 | - | 1 | - | 3,45E-05 | - | 1 | - | 3,88E-05 | - |
| 93 | 1 | 3um_FIJJAFBF_269245 | Bacteria | Bacteroidetes | Flavobacteriia | Flavobacterial | Flavobacteriac | Polaribacter | Polaribacter b | unclassified | unclassified | unclassified | unclassified | unclassified | 1,4 | 0,489897949 | 3,58E-05 | 1,34E-05 | 1 | - | 2,13E-05 | - | 2 | - | 5,29E-05 | - | 1 | - | 2,51E-05 | - | 2 | - | 5,10E-05 | - | 1 | - | 2,87E-05 | - |
| 94 | 1 | 3um_FIJJAFBF_348463 | Bacteria | Verrucomicrol | unclassified | unclassified | unclassified | unclassified | Verrucomicro | poorly charact | Function unkn | 05DN3 | alpha-2-macrc | 240016.VspiD | 1 | 0 | 7,75E-06 | 7,36E-07 | 1 | - | 6,49E-06 | - | 1 | - | 8,06E-06 | - | 1 | - | 7,66E-06 | - | 1 | - | 7,78E-06 | - | 1 | - | 8,76E-06 | - |
| 95 | 1 | 3um_FIJJAFBF_360602 | Bacteria | Tenericutes | Mollicutes | Mycoplasmata | Mycoplasmata | Mycoplasma | Mycoplasma I | poorly charact | Function unkn | 03BSV | Inherit from e | 7070.TC01476 | 0,2 | 0,4 | 2,33E-06 | 4,66E-06 | 0 | - | 0 | - | 0 | - | 0 | - | 0 | - | 0 | - | 1 | - | 1,17E-05 | - | 0 | - | 0 | - |
| 96 | 1 | 0.2um_NOKCPHJN_380864 | Bacteria | Bacteroidetes | Flavobacteriia | Flavobacterial | Cryomorphace | unclassified | Cryomorphace | unclassified | unclassified | unclassified | unclassified | unclassified | 0,2 | 0,4 | 2,69E-06 | 5,38E-06 | 0 | - | 0 | - | 0 | - | 0 | - | 0 | - | 0 | - | 1 | - | 1,34E-05 | - | 0 | - | 0 | - |
| 97 | 1 | 3um_FIJJAFBF_235865 | Bacteria | Proteobacteri | Gammaprotec | Cellvibrionale | Spongiibacter | Oceanicoccus | Oceanicoccus | information st | Translation, ri | 0QSAC | Binds togethe | 247633.GP214 | 2,8 | 0,979795897 | 0,000267256 | 7,65E-05 | 4 | - | 0,000327934 | - | 2 | - | 0,000203598 | - | 4 | - | 0,000387092 | - | 2 | - | 0,000196405 | - | 2 | - | 0,000221249 | - |
| 98 | 1 | 0.2um_NOKCPHJN_75523 | Bacteria | Proteobacteri | Gammaprotec | Oceanospirilla | Saccharospirill | Saccharospiril | Saccharospiril | metabolism | Amino acid tra | 0QJ82 | Aminotransfe | 1028805.GGC | 0,2 | 0,4 | 5,67E-06 | 1,13E-05 | 1 | - | 2,83E-05 | - | 0 | - | 0 | - | 0 | - | 0 | - | 0 | - | 0 | - | 0 | - | 0 | - |
| 99 | 1 | 3um_FIJJAFBF_440876 | Bacteria | Proteobacteri | Gammaprotec | unclassified | unclassified | unclassified | Gammaprotec | information st | Translation, ri | 0QSAC | Binds togethe | 247634.GPB2 | 1 | 0,894427191 | 9,08E-05 | 8,56E-05 | 1 | - | 7,45E-05 | - | 0 | - | 0 | - | 0 | - | 0 | - | 2 | - | 0,00017855 | - | 2 | - | 0,000201136 | - |
| 100 | 1 | 3um_FIJJAFBF_584287 | Bacteria | Proteobacteri | Gammaprotec | Oceanospirilla | Saccharospirill | Reinekea | Reinekea fors | cellular proces | Posttranslatio | COG0652 | peptidyl-proly | 314283.MED2 | 1,4 | 0,489897949 | 0,000113381 | 3,24E-05 | 2 | - | 0,000139123 | - | 1 | - | 8,64E-05 | - | 2 | - | 0,000164221 | - | 1 | - | 8,33E-05 | - | 1 | - | 9,39E-05 | - |
| 101 | 1 | 0.2um_NOKCPHJN_230027 | Bacteria | Proteobacteri | Gammaprotec | Cellvibrionale | Porticoccacea | unclassified | Gammaprotec | information st | Translation, ri | 0QTKX | Seems to be t | 314287.GB22 | 0,4 | 0,489897949 | 4,28E-05 | 5,34E-05 | 1 | - | 9,11E-05 | - | 0 | - | 0 | - | 0 | - | 0 | - | 0 | - | 0 | - | 1 | - | 0,000122916 | - |
| 102 | 1 | 0.2um_NOKCPHJN_401629 | Bacteria | Firmicutes | Clostridia | Clostridiales | unclassified | unclassified | Clostridiales b | poorly charact | Function unkn | 07F5I | NA | 760192.Halhy | 8 | 1,095445115 | 9,68E-05 | 4,34E-06 | 10 | - | 0,000102571 | - | 7 | - | 8,92E-05 | - | 8 | - | 9,69E-05 | - | 8 | - | 9,83E-05 | - | 7 | - | 9,69E-05 | - |
| 103 | 1 | 3um_FIJJAFBF_576716 | Bacteria | Proteobacteri | Alphaproteob | Sphingomona | Sphingomona | Sphingomona | various | unclassified | unclassified | unclassified | unclassified | unclassified | 2,8 | 1,166190379 | 3,65E-05 | 1,48E-05 | 3 | - | 3,33E-05 | - | 1 | - | 1,38E-05 | - | 4 | - | 5,24E-05 | - | 4 | - | 5,31E-05 | - | 2 | - | 2,99E-05 | - |
| 104 | 1 | 0.2um_NOKCPHJN_319829 | Bacteria | Proteobacteri | Gammaprotec | Oceanospirilla | Halomonadac | Halomonas | Halomonas sa | information st | Translation, ri | 0QQXI | This protein is | 999141.GME_ | 1 | 0 | 0,000113246 | 1,08E-05 | 1 | - | 9,49E-05 | - | 1 | - | 0,000117784 | - | 1 | - | 0,000111969 | - | 1 | - | 0,000113623 | - | 1 | - | 0,000127995 | - |
| 105 | 1 | 0.2um_NOKCPHJN_1073579 | Eukaryota | Chromerida | unclassified | unclassified | unclassified | Vitrella | Vitrella brassi | metabolism | Lipid transport | 0CXXX | Acyltransferas | 765952.PUV_ | 0,2 | 0,4 | 6,66E-06 | 1,33E-05 | 0 | - | 0 | - | 0 | - | 0 | - | 0 | - | 0 | - | 1 | - | 3,33E-05 | - | 0 | - | 0 | - |
| 106 | 1 | 3um_FIJJAFBF_305373 | Bacteria | Bacteroidetes | Flavobacteriia | Flavobacterial | Flavobacteriac | Polaribacter | Polaribacter s | unclassified | unclassified | unclassified | unclassified | unclassified | 1 | 0,632455532 | 3,51E-05 | 1,98E-05 | 2 | - | 6,12E-05 | - | 0 | - | 0 | - | 1 | - | 3,61E-05 | - | 1 | - | 3,67E-05 | - | 1 | - | 4,13E-05 | - |
| 107 | 1 | 0.2um_NOKCPHJN_118523 | Bacteria | Proteobacteri | Gammaprotec | Oceanospirilla | Saccharospirill | Reinekea | Reinekea blan | metabolism | Secondary me | 16QPI | Amp-depende | 314283.MED2 | 0,2 | 0,4 | 4,25E-06 | 8,50E-06 | 1 | - | 2,13E-05 | - | 0 | - | 0 | - | 0 | - | 0 | - | 0 | - | 0 | - | 0 | - | 0 | - |
| 108 | 1 | 3um_FIJJAFBF_469902 | Bacteria | Proteobacteri | Alphaproteob | Rhodobactera | unclassified | unclassified | Rhodobactera | information st | Translation, ri | 17FUG | Binds directly | 439496.RBY4I | 2 | 0 | 0,000315005 | 2,99E-05 | 2 | - | 0,000263855 | - | 2 | - | 0,00032763 | - | 2 | - | 0,000311453 | - | 2 | - | 0,000316054 | - | 2 | - | 0,000356033 | - |
| 109 | 1 | 3um_FIJJAFBF_192738 | Bacteria | unclassified | unclassified | unclassified | unclassified | unclassified | uncultured ba | unclassified | unclassified | unclassified | unclassified | unclassified | 1 | 0 | 1,79E-05 | 1,70E-06 | 1 | - | 1,50E-05 | - | 1 | - | 1,86E-05 | - | 1 | - | 1,77E-05 | - | 1 | - | 1,79E-05 | - | 1 | - | 2,02E-05 | - |
| 110 | 1 | 0.2um_NOKCPHJN_387553 | Bacteria | Proteobacteri | Gammaprotec | Cellvibrionale | Halieaceae | unclassified | marine gamm | poorly charact | Function unkn | 05ZFT | NA | 247639.MGP2 | 1,4 | 0,489897949 | 3,17E-05 | 9,33E-06 | 2 | - | 3,88E-05 | - | 1 | - | 2,41E-05 | - | 1 | - | 2,29E-05 | - | 2 | - | 4,65E-05 | - | 1 | - | 2,62E-05 | - |
| 111 | 1 | 3um_FIJJAFBF_320754 | Bacteria | Bacteroidetes | Flavobacteriia | Flavobacterial | Flavobacteriac | Formosa | Formosa sp. H | unclassified | unclassified | unclassified | unclassified | unclassified | 1,2 | 1,166190379 | 3,74E-05 | 3,39E-05 | 3 | - | 8,22E-05 | - | 2 | - | 6,80E-05 | - | 0 | - | 0 | - | 0 | - | 0 | - | 1 | - | 3,70E-05 | - |
| 112 | 1 | 3um_FIJJAFBF_106937 | Bacteria | Proteobacteri | Alphaproteob | Rhodobactera | unclassified | unclassified | Rhodobactera | poorly charact | Function unkn | 05EN8 | Extracellular s | 391589.RGAI1 | 7 | 2 | 0,000137057 | 2,81E-05 | 10 | - | 0,000168294 | - | 7 | - | 0,00014628 | - | 8 | - | 0,000158923 | - | 6 | - | 0,000120953 | - | 4 | - | 9,08E-05 | - |
| 113 | 1 | 3um_FIJJAFBF_176420 | Bacteria | Bacteroidetes | Flavobacteriia | Flavobacterial | Crocinitomica | unclassified | Crocinitomica | information st | Translation, ri | COG0184 | Forms an inte | 511995.CFPG_ | 1 | 0 | 0,000153963 | 1,46E-05 | 1 | - | 0,000128963 | - | 1 | - | 0,000160134 | - | 1 | - | 0,000152227 | - | 1 | - | 0,000154476 | - | 1 | - | 0,000174016 | - |
| 114 | 1 | 3um_FIJJAFBF_187829 | Bacteria | Bacteroidetes | Flavobacteriia | Flavobacterial | Flavobacteriac | Polaribacter | Polaribacter s | unclassified | unclassified | unclassified | unclassified | unclassified | 1 | 0,632455532 | 5,51E-05 | 3,56E-05 | 1 | - | 4,65E-05 | - | 0 | - | 0 | - | 1 | - | 5,49E-05 | - | 2 | - | 0,000111323 | - | 1 | - | 6,27E-05 | - |
| 115 | 1 | 3um_FIJJAFBF_131560 | Bacteria | Proteobacteri | Gammaprotec | Oceanospirilla | unclassified | unclassified | Oceanospirilla | metabolism | Amino acid tra | COG0683 | (ABC) transpo | 488538.SAR11 | 7,6 | 0,8 | 0,00026574 | 4,19E-05 | 7 | - | 0,000204437 | - | 7 | - | 0,00025385 | - | 9 | - | 0,000310265 | - | 7 | - | 0,000244881 | - | 8 | - | 0,000315266 | - |
| 116 | 1 | 3um_FIJJAFBF_103163 | Bacteria | Bacteroidetes | Flavobacteriia | Flavobacterial | Flavobacteriac | Polaribacter | Polaribacter s | unclassified | unclassified | unclassified | unclassified | unclassified | 0,2 | 0,4 | 1,10E-05 | 2,20E-05 | 1 | - | 5,49E-05 | - | 0 | - | 0 | - | 0 | - | 0 | - | 0 | - | 0 | - | 0 | - | 0 | - |
| 117 | 1 | 3um_FIJJAFBF_87393 | Bacteria | Bacteroidetes | Flavobacteriia | Flavobacterial | Flavobacteriac | Polaribacter | Polaribacter s | unclassified | unclassified | unclassified | unclassified | unclassified | 1 | 0 | 8,56E-05 | 8,13E-06 | 1 | - | 7,17E-05 | - | 1 | - | 8,91E-05 | - | 1 | - | 8,47E-05 | - | 1 | - | 8,59E-05 | - | 1 | - | 9,68E-05 | - |
| 118 | 1 | 0.2um_NOKCPHJN_388703 | Bacteria | Bacteroidetes | Flavobacteriia | Flavobacterial | Flavobacteriac | Aquimarina | Aquimarina sp | metabolism | Inorganic ion t | 05C27 | TonB-depende | 886377.Murru | 15,2 | 3,310589071 | 0,000190692 | 4,20E-05 | 16 | - | 0,000168945 | - | 18 | - | 0,000236002 | - | 19 | - | 0,000236813 | - | 10 | - | 0,00012648 | - | 13 | - | 0,000185222 | - |
| 119 | 1 | 3um_FIJJAFBF_556534 | Eukaryota | unclassified | unclassified | Choanoflagelli | Salpingoecida | Monosiga | Monosiga bre | unclassified | unclassified | unclassified | unclassified | unclassified | 1 | 0 | 1,13E-05 | 1,08E-06 | 1 | - | 9,49E-06 | - | 1 | - | 1,18E-05 | - | 1 | - | 1,12E-05 | - | 1 | - | 1,14E-05 | - | 1 | - | 1,28E-05 | - |
| 120 | 1 | 3um_FIJJAFBF_489502 | Bacteria | Bacteroidetes | Flavobacteriia | Flavobacterial | Flavobacteriac | Polaribacter | various | unclassified | unclassified | unclassified | unclassified | unclassified | 32,6 | 3,826225294 | 0,00052874 | 8,52E-05 | 32 | - | 0,000434144 | - | 40 | - | 0,000673848 | - | 29 | - | 0,000464419 | - | 31 | - | 0,00050378 | - | 31 | - | 0,000567507 | - |
| 121 | 1 | 3um_FIJJAFBF_254313 | Bacteria | Proteobacteri | Gammaprotec | Cellvibrionale | Spongiibacter | Spongiibacter | Spongiibacter | information st | Replication, re | 16W6D | single-strande | 207954.MED9 | 1,2 | 0,4 | 9,81E-05 | 2,17E-05 | 2 | - | 0,00014083 | - | 1 | - | 8,74E-05 | - | 1 | - | 8,31E-05 | - | 1 | - | 8,43E-05 | - | 1 | - | 9,50E-05 | - |
| 122 | 1 | 3um_FIJJAFBF_279887 | unclassified | unclassified | unclassified | unclassified | unclassified | unclassified | unclassified | poorly charact | Function unkn | 006NJ | Inherit from b | 204669.Acid3 | 1 | 0,894427191 | 2,32E-05 | 2,06E-05 | 0 | - | 0 | - | 1 | - | 2,40E-05 | - | 2 | - | 4,57E-05 | - | 2 | - | 4,64E-05 | - | 0 | - | 0 | - |
| 123 | 1 | 3um_FIJJAFBF_779429 | Bacteria | Proteobacteri | Alphaproteob | Rhizobiales | unclassified | Hartmannibac | Hartmannibac | metabolism | Inorganic ion t | 05C2J | ABC transport | 388401.RB215 | 4,6 | 1,854723699 | 0,000239781 | 7,53E-05 | 8 | - | 0,000358677 | - | 5 | - | 0,000278357 | - | 3 | - | 0,000158768 | - | 3 | - | 0,000161113 | - | 4 | - | 0,000241991 | - |
| 124 | 1 | 0.2um_NOKCPHJN_41616 | Bacteria | Bacteroidetes | Flavobacteriia | Flavobacterial | Flavobacteriac | Polaribacter | Polaribacter s | metabolism | Amino acid tra | 05C6M | Alpha-keto-be | 313594.PI23P | 1,4 | 0,8 | 4,02E-05 | 2,23E-05 | 0 | - | 0 | - | 2 | - | 5,81E-05 | - | 2 | - | 5,52E-05 | - | 2 | - | 5,60E-05 | - | 1 | - | 3,15E-05 | - |
| 125 | 1 | 3um_FIJJAFBF_217664 | Bacteria | Bacteroidetes | Flavobacteriia | Flavobacterial | Flavobacteriac | Formosa | Formosa sp. H | unclassified | unclassified | unclassified | unclassified | unclassified | 1,6 | 0,489897949 | 0,000106724 | 2,96E-05 | 2 | - | 0,00011364 | - | 2 | - | 0,000141108 | - | 2 | - | 0,000134141 | - | 1 | - | 6,81E-05 | - | 1 | - | 7,67E-05 | - |
| 126 | 1 | 3um_FIJJAFBF_556664 | Bacteria | Proteobacteri | Gammaprotec | Oceanospirilla | Saccharospirill | Reinekea | Reinekea fors | metabolism | Nucleotide tra | 0HQGI | orotidine 5'-pl | 577650.Despr | 0,4 | 0,489897949 | 1,88E-05 | 2,32E-05 | 1 | - | 4,31E-05 | - | 0 | - | 0 | - | 1 | - | 5,09E-05 | - | 0 | - | 0 | - | 0 | - | 0 | - |
| 127 | 1 | 3um_FIJJAFBF_545834 | Bacteria | Bacteroidetes | Flavobacteriia | Flavobacterial | Flavobacteriac | Polaribacter | Polaribacter s | metabolism | Carbohydrate | 099YR | Catalyzes the | 313598.MED1 | 14,6 | 2,059126028 | 0,000465753 | 7,96E-05 | 17 | - | 0,000454826 | - | 14 | - | 0,000465097 | - | 13 | - | 0,000410552 | - | 12 | - | 0,000384569 | - | 17 | - | 0,000613722 | - |
| 128 | 1 | 0.2um_NOKCPHJN_642507 | Bacteria | Proteobacteri | Alphaproteob | Pelagibacteral | Pelagibacterac | Candidatus Pe | Candidatus Pe | unclassified | unclassified | unclassified | unclassified | unclassified | 0,2 | 0,4 | 9,47E-06 | 1,89E-05 | 0 | - | 0 | - | 0 | - | 0 | - | 0 | - | 0 | - | 0 | - | 0 | - | 1 | - | 4,74E-05 | - |
| 129 | 1 | 0.2um_NOKCPHJN_26427 | Bacteria | Bacteroidetes | Flavobacteriia | Flavobacterial | Flavobacteriac | unclassified | Flavobacteriac | metabolism | Energy produc | 05CMV | Succinyl-CoA s | 983544.Lacal_ | 2 | 0 | 6,90E-05 | 6,55E-06 | 2 | - | 5,78E-05 | - | 2 | - | 7,18E-05 | - | 2 | - | 6,83E-05 | - | 2 | - | 6,93E-05 | - | 2 | - | 7,80E-05 | - |
| 130 | 1 | 3um_FIJJAFBF_316866 | Bacteria | Bacteroidetes | Flavobacteriia | Flavobacterial | Flavobacteriac | Algibacter | Algibacter alg | information st | Translation, ri | 09EM4 | Binds directly | 1046627.BZAF | 0,4 | 0,489897949 | 4,79E-05 | 5,87E-05 | 0 | - | 0 | - | 0 | - | 0 | - | 1 | - | 0,000118844 | - | 1 | - | 0,000120599 | - | 0 | - | 0 | - |
| 131 | 1 | 3um_FIJJAFBF_217666 | Bacteria | Bacteroidetes | Flavobacteriia | Flavobacterial | Flavobacteriac | Formosa | Formosa sp. H | cellular proces | Cell wall/mem | COG2885 | Ompa motb d | 983544.Lacal_ | 5,2 | 0,748331477 | 0,000150464 | 2,93E-05 | 5 | - | 0,000120818 | - | 4 | - | 0,000120016 | - | 6 | - | 0,000171135 | - | 5 | - | 0,000144719 | - | 6 | - | 0,000195631 | - |
| 132 | 1 | 3um_FIJJAFBF_320300 | Bacteria | Proteobacteri | Gammaprotec | Oceanospirilla | Saccharospirill | Reinekea | Reinekea fors | unclassified | unclassified | unclassified | unclassified | unclassified | 0,4 | 0,489897949 | 2,73E-05 | 3,36E-05 | 1 | - | 6,20E-05 | - | 0 | - | 0 | - | 0 | - | 0 | - | 1 | - | 7,43E-05 | - | 0 | - | 0 | - |
| 133 | 1 | 3um_FIJJAFBF_342789 | Bacteria | Proteobacteri | Gammaprotec | Alteromonada | Alteromonada | Paraglaciecola | Paraglaciecola | cellular proces | Posttranslatio | 05C2Z | ATP-depender | 926562.Oweh | 0,4 | 0,489897949 | 6,24E-06 | 7,64E-06 | 0 | - | 0 | - | 0 | - | 0 | - | 1 | - | 1,55E-05 | - | 1 | - | 1,57E-05 | - | 0 | - | 0 | - |
| 134 | 1 | 3um_FIJJAFBF_634670 | Bacteria | Proteobacteri | Gammaprotec | unclassified | unclassified | unclassified | Gammaprotec | information st | Translation, ri | COG0050 | This protein p | 396588.Tgr7_ | 1,2 | 0,4 | 4,13E-05 | 1,38E-05 | 1 | - | 2,89E-05 | - | 1 | - | 3,59E-05 | - | 2 | - | 6,83E-05 | - | 1 | - | 3,46E-05 | - | 1 | - | 3,90E-05 | - |
| 135 | 1 | 0.2um_NOKCPHJN_45043 | Bacteria | Bacteroidetes | Flavobacteriia | Flavobacterial | Crocinitomica | unclassified | Crocinitomica | unclassified | unclassified | unclassified | unclassified | unclassified | 0,4 | 0,489897949 | 3,07E-05 | 3,76E-05 | 0 | - | 0 | - | 1 | - | 7,35E-05 | - | 0 | - | 0 | - | 0 | - | 0 | - | 1 | - | 7,98E-05 | - |
| 136 | 1 | 3um_FIJJAFBF_437071 | Bacteria | Proteobacteri | Alphaproteob | Rhodospirillal | Rhodospirillac | unclassified | Rhodospirillac | unclassified | unclassified | unclassified | unclassified | unclassified | 10,4 | 1,019803903 | 0,000451469 | 7,47E-05 | 10 | - | 0,000362072 | - | 12 | - | 0,000539504 | - | 9 | - | 0,000398465 | - | 10 | - | 0,000433701 | - | 11 | - | 0,000537419 | - |
| 137 | 1 | 0.2um_NOKCPHJN_15230 | Bacteria | Proteobacteri | Betaproteoba | unclassified | unclassified | unclassified | Betaproteoba | unclassified | unclassified | unclassified | unclassified | unclassified | 4,6 | 0,489897949 | 0,000417105 | 3,31E-05 | 5 | - | 0,000382589 | - | 4 | - | 0,00038005 | - | 5 | - | 0,000451607 | - | 5 | - | 0,000458278 | - | 4 | - | 0,000412999 | - |

| | | | | | | | | | | | | | | | | | | | | | | | | | | | | |
|---|---|---|---|---|---|---|---|---|---|---|---|---|---|---|---|---|---|---|---|---|---|---|---|---|---|---|---|---|
| 138 | 1 | 3um_FIJJAFBF_42231 | Bacteria | Proteobacteri | Alphaproteob | Rhizobiales | Cohaesibacter | Cohaesibacter | Cohaesibacter | metabolism | Energy produc | 16QP8 | isocitrate deh | 391624.OIHEL | 0,4 | 0,489897949 | 1,48E-05 | 1,81E-05 | 0 - | 0 - | 1 - | 3,55E-05 - | 0 - | 0 - | 0 - | 0 - | 1 - | 3,85E-05 - |
| 139 | 1 | 3um_FIJJAFBF_278309 | Eukaryota | unclassified | Florideophyce | Batrachosperr | Batrachosperr | Kumanoa | Kumanoa ame | metabolism | Energy produc | COG0055 | Produces ATP | 7739.JGI1257 | 1,4 | 1,019803903 | 3,81E-05 | 2,57E-05 | 3 - | 7,26E-05 - | 2 - | 6,01E-05 - | 1 - | 2,86E-05 - | 1 - | 2,90E-05 - | 0 - | 0 - |
| 140 | 1 | 0.2um_NOKCPHJN_679396 | Bacteria | Proteobacteri | Gammaprotec | Pasteurellales | Pasteurellacea | Haemophilus | Haemophilus | unclassified | unclassified | unclassified | unclassified | unclassified | 2 | 0,632455532 | 7,11E-05 | 2,46E-05 | 2 - | 5,93E-05 - | 3 - | 0,00011048 - | 2 - | 7,00E-05 - | 1 - | 3,55E-05 - | 2 - | 8,00E-05 - |
| 141 | 1 | 0.2um_NOKCPHJN_263272 | Bacteria | Proteobacteri | Gammaprotec | Cellvibrionales | Porticoccacea | unclassified | gamma protec | metabolism | Inorganic ion t | 16TQC | TONB-depend | 314287.GB22 | 1,4 | 0,489897949 | 1,87E-05 | 5,36E-06 | 2 - | 2,30E-05 - | 1 - | 1,43E-05 - | 2 - | 2,71E-05 - | 1 - | 1,38E-05 - | 1 - | 1,55E-05 - |
| 142 | 1 | 3um_FIJJAFBF_529567 | Bacteria | Bacteroidetes | Flavobacteriia | Flavobacterial | Flavobacteriac | Polaribacter | various | poorly charact | Function unkn | 11FUA | Membrane | 688270.Celal_ | 0,4 | 0,8 | 2,68E-05 | 5,37E-05 | 2 - | 0,000134242 - | 0 - | 0 - | 0 - | 0 - | 0 - | 0 - | 0 - | 0 - |
| 143 | 1 | 0.2um_NOKCPHJN_149316 | Bacteria | Bacteroidetes | Flavobacteriia | Flavobacterial | Flavobacteriac | Ulvibacter | Ulvibacter sp. | unclassified | unclassified | unclassified | unclassified | unclassified | 0,2 | 0,4 | 1,31E-05 | 2,62E-05 | 0 - | 0 - | 0 - | 0 - | 1 - | 6,55E-05 - | 0 - | 0 - | 0 - | 0 - |
| 144 | 1 | 3um_FIJJAFBF_545835 | Bacteria | Bacteroidetes | Flavobacteriia | Flavobacterial | Flavobacteriac | Polaribacter | Polaribacter s | metabolism | Energy produc | COG0372 | citrate syntha | 313598.MED1 | 4,6 | 1,019803903 | 0,000144543 | 2,23E-05 | 6 - | 0,000160902 - | 5 - | 0,000166494 - | 5 - | 0,000158274 - | 4 - | 0,000128489 - | 3 - | 0,000108557 - |
| 145 | 1 | 0.2um_NOKCPHJN_34584 | Eukaryota | unclassified | Bangiophycea | Cyanidiales | Cyanidiaceae | Galdieria | Galdieria sulp | cellular proces | Posttranslatio | KOG0731 | ATP-depender | 59689.scaffol | 0,2 | 0,4 | 4,22E-06 | 8,45E-06 | 0 - | 0 - | 1 - | 2,11E-05 - | 0 - | 0 - | 0 - | 0 - | 0 - | 0 - |
| 146 | 1 | 3um_FIJJAFBF_291816 | Eukaryota | unclassified | Oomycetes | Peronosporale | Peronosporac | Phytophthora | various | cellular proces | Signal transdu | 152IZ | EFh | 67593.Physo1 | 3,4 | 0,8 | 0,000279521 | 6,97E-05 | 4 - | 0,000276571 - | 2 - | 0,000171709 - | 3 - | 0,000244847 - | 4 - | 0,000331285 - | 4 - | 0,000373192 - |
| 147 | 1 | 3um_FIJJAFBF_545833 | Bacteria | Bacteroidetes | Flavobacteriia | Flavobacterial | Flavobacteriac | Polaribacter | Polaribacter a | metabolism | Nucleotide tra | 0P8S0 | carbamoyl-ph | 313598.MED1 | 2,2 | 0,4 | 8,20E-05 | 2,20E-05 | 2 - | 6,17E-05 - | 2 - | 7,66E-05 - | 2 - | 7,28E-05 - | 2 - | 7,39E-05 - | 3 - | 0,000124899 - |
| 148 | 1 | 3um_FIJJAFBF_226282 | Bacteria | Bacteroidetes | Flavobacteriia | Flavobacterial | Flavobacteriac | Mesonia | Mesonia alga | unclassified | unclassified | unclassified | unclassified | unclassified | 2 | 0 | 0,000221011 | 2,10E-05 | 2 - | 0,000185124 - | 2 - | 0,000229869 - | 2 - | 0,00021852 - | 2 - | 0,000221747 - | 2 - | 0,000249798 - |
| 149 | 1 | 3um_FIJJAFBF_213369 | Bacteria | Proteobacteri | Gammaprotec | Oceanospirilla | Saccharospirill | Reinekea | Reinekea fors | metabolism | Coenzyme trai | 08UTT | Catalyzes the | 314283.MED2 | 2,2 | 0,4 | 0,000186764 | 4,17E-05 | 2 - | 0,0001417 - | 3 - | 0,000263924 - | 2 - | 0,000167262 - | 2 - | 0,000169732 - | 2 - | 0,000191203 - |
| 150 | 1 | 0.2um_NOKCPHJN_210705 | Bacteria | Verrucomicrol | Opitutae | Puniceicoccale | Puniceicoccac | unclassified | Puniceicoccac | unclassified | unclassified | unclassified | unclassified | unclassified | 6,6 | 1,2 | 0,000191619 | 1,76E-05 | 9 - | 0,000222149 - | 6 - | 0,000183895 - | 6 - | 0,000174816 - | 6 - | 0,000177398 - | 6 - | 0,000199838 - |
| 151 | 1 | 0.2um_NOKCPHJN_371587 | Bacteria | Bacteroidetes | Flavobacteriia | Flavobacterial | Flavobacteriac | Formosa | Formosa sp. H | unclassified | unclassified | unclassified | unclassified | unclassified | 3,2 | 0,4 | 5,05E-05 | 7,83E-06 | 3 - | 3,97E-05 - | 3 - | 4,93E-05 - | 3 - | 4,68E-05 - | 4 - | 6,34E-05 - | 3 - | 5,35E-05 - |
| 152 | 1 | 0.2um_NOKCPHJN_617508 | Eukaryota | Chytridiomyc | Neocallimastig | Neocallimastig | Neocallimastig | Anaeromyces | Anaeromyces | cellular proces | Posttranslatio | 007DF | Inherit from N | 5833.PFL0275 | 0,2 | 0,4 | 1,87E-06 | 3,73E-06 | 1 - | 9,33E-06 - | 0 - | 0 - | 0 - | 0 - | 0 - | 0 - | 0 - | 0 - |
| 153 | 1 | 0.2um_NOKCPHJN_93334 | Bacteria | Proteobacteri | Betaproteoba | unclassified | unclassified | unclassified | Betaproteoba | metabolism | Lipid transpor | 0B7RE | 3-hydroxybuty | 395495.Lcho_ | 0,2 | 0,4 | 8,11E-06 | 1,62E-05 | 1 - | 4,06E-05 - | 0 - | 0 - | 0 - | 0 - | 0 - | 0 - | 0 - | 0 - |
| 154 | 1 | 0.2um_NOKCPHJN_482344 | Bacteria | Proteobacteri | Betaproteoba | unclassified | unclassified | unclassified | Betaproteoba | information st | Transcription | COG0085 | DNA-depende | 688245.CtCNB | 0,6 | 0,489897949 | 1,51E-05 | 1,24E-05 | 1 - | 2,23E-05 - | 0 - | 0 - | 1 - | 2,63E-05 - | 1 - | 2,67E-05 - | 0 - | 0 - |
| 155 | 1 | 3um_FIJJAFBF_715087 | Eukaryota | Bacillariophyt | Bacillariophyc | Naviculales | Phaeodactylac | Phaeodactylu | Phaeodactylu | unclassified | unclassified | unclassified | unclassified | unclassified | 2 | 0 | 0,000226491 | 2,15E-05 | 2 - | 0,000189714 - | 2 - | 0,000235568 - | 2 - | 0,000223938 - | 2 - | 0,000227245 - | 2 - | 0,000255991 - |
| 156 | 1 | 3um_FIJJAFBF_715674 | Eukaryota | unclassified | Spirotrichea | Sporadotrichi | Oxytrichidae | Stylonychia | Stylonychia le | unclassified | unclassified | unclassified | unclassified | unclassified | 0,8 | 0,4 | 6,20E-05 | 3,15E-05 | 1 - | 6,71E-05 - | 1 - | 8,33E-05 - | 1 - | 7,92E-05 - | 1 - | 8,04E-05 - | 0 - | 0 - |
| 157 | 1 | 3um_FIJJAFBF_285194 | Bacteria | Proteobacteri | Betaproteoba | Nitrosomonad | Thiobacillacea | Thiobacillus | Thiobacillus s | poorly charact | Function unkn | 08D4J | NA | 314607.KB13_ | 1 | 0 | 0,000126877 | 1,20E-05 | 1 - | 0,000106275 - | 1 - | 0,000131962 - | 1 - | 0,000125447 - | 1 - | 0,000127299 - | 1 - | 0,000143402 - |
| 158 | 1 | 0.2um_NOKCPHJN_131051 | Bacteria | Proteobacteri | Alphaproteob | Rhodobactera | Rhodobactera | Actibacterium | Actibacterium | unclassified | unclassified | unclassified | unclassified | unclassified | 0,6 | 0,489897949 | 2,50E-05 | 2,04E-05 | 0 - | 0 - | 1 - | 4,29E-05 - | 1 - | 4,08E-05 - | 1 - | 4,14E-05 - | 0 - | 0 - |
| 159 | 1 | 3um_FIJJAFBF_65230 | Bacteria | Proteobacteri | Alphaproteob | Rhodobactera | Rhodobactera | Roseobacter | Roseobacter c | unclassified | unclassified | unclassified | unclassified | unclassified | 5,8 | 1,720465053 | 0,000144872 | 3,04E-05 | 9 - | 0,000192363 - | 6 - | 0,000159239 - | 5 - | 0,000126147 - | 4 - | 0,000102408 - | 5 - | 0,000144203 - |
| 160 | 1 | 3um_FIJJAFBF_481952 | Bacteria | Proteobacteri | Alphaproteob | Rhodobactera | unclassified | unclassified | Rhodobactera | metabolism | Energy produc | 16W2S | Produces ATP | 246200.SPO3 | 0,6 | 0,489897949 | 5,69E-05 | 4,69E-05 | 1 - | 8,32E-05 - | 1 - | 0,000103275 - | 1 - | 9,82E-05 - | 0 - | 0 - | 0 - | 0 - |
| 161 | 1 | 3um_FIJJAFBF_510237 | Bacteria | Bacteroidetes | Flavobacteriia | Flavobacterial | Flavobacteriac | Polaribacter | Polaribacter s | unclassified | unclassified | unclassified | unclassified | unclassified | 3 | 0,894427191 | 0,000281582 | 6,86E-05 | 4 - | 0,000321054 - | 3 - | 0,000298991 - | 4 - | 0,000378971 - | 2 - | 0,000192284 - | 2 - | 0,000216608 - |
| 162 | 1 | 0.2um_NOKCPHJN_119709 | Bacteria | Proteobacteri | Gammaprotec | unclassified | unclassified | unclassified | Gammaprotec | cellular proces | Intracellular tr | 05E4G | Type IV pilus | 270374.MELB | 0,8 | 1,166190379 | 1,38E-05 | 1,93E-05 | 3 - | 4,94E-05 - | 0 - | 0 - | 1 - | 1,94E-05 - | 0 - | 0 - | 0 - | 0 - |
| 163 | 1 | 3um_FIJJAFBF_346688 | Bacteria | Proteobacteri | Gammaprotec | Oceanospirilla | Oceanospirilla | Neptuniibacte | Neptuniibacte | metabolism | Energy produc | 17C3H | Produces ATP | 207949.RED6 | 1,6 | 0,489897949 | 0,000156116 | 5,03E-05 | 1 - | 8,14E-05 - | 2 - | 0,000202154 - | 2 - | 0,000192173 - | 2 - | 0,000195012 - | 1 - | 0,00010984 - |
| 164 | 1 | 0.2um_NOKCPHJN_139057 | Bacteria | Proteobacteri | Alphaproteob | Rhodobactera | Hyphomonad | unclassified | Hyphomonad | information st | Replication, re | 0PB0H | DNA-binding | 755732.Fluta_ | 0,6 | 0,489897949 | 8,87E-05 | 7,32E-05 | 1 - | 0,000128963 - | 1 - | 0,000160134 - | 0 - | 0 - | 1 - | 0,000154476 - | 0 - | 0 - |
| 165 | 1 | 3um_FIJJAFBF_535171 | Bacteria | Proteobacteri | Alphaproteob | Rhodobactera | unclassified | unclassified | Rhodobactera | poorly charact | Function unkn | 177S4 | Type VI secret | 439496.RBY4 | 1,2 | 0,4 | 9,62E-05 | 3,32E-05 | 1 - | 6,71E-05 - | 1 - | 8,33E-05 - | 1 - | 7,92E-05 - | 2 - | 0,000160799 - | 1 - | 9,06E-05 - |
| 166 | 1 | 0.2um_NOKCPHJN_820822 | Bacteria | Proteobacteri | Alphaproteob | Rickettsiales | unclassified | unclassified | Rickettsiales b | unclassified | unclassified | unclassified | unclassified | unclassified | 3,6 | 0,489897949 | 0,00012738 | 2,67E-05 | 3 - | 8,81E-05 - | 4 - | 0,000145799 - | 3 - | 0,000103951 - | 4 - | 0,000140648 - | 4 - | 0,000158439 - |
| 167 | 1 | 3um_FIJJAFBF_295857 | Bacteria | Proteobacteri | Gammaprotec | Oceanospirilla | Saccharospirill | Reinekea | Reinekea fors | information st | Translation, ri | 0QTKZ | 30S ribosomal | 314283.MED2 | 1,8 | 0,748331477 | 0,000346409 | 0,000147669 | 2 - | 0,000323315 - | 1 - | 0,000200731 - | 1 - | 0,00019082 - | 3 - | 0,000580915 - | 2 - | 0,000436266 - |
| 168 | 1 | 0.2um_NOKCPHJN_205904 | Bacteria | Bacteroidetes | Flavobacteriia | Flavobacterial | Flavobacteriac | Kordia | Kordia sp. SM | metabolism | Lipid transpor | 05CEY | Amp-depende | 1042163.BRL | 0,2 | 0,4 | 5,36E-07 | 1,07E-06 | 1 - | 2,68E-06 - | 0 - | 0 - | 0 - | 0 - | 0 - | 0 - | 0 - | 0 - |
| 169 | 1 | 0.2um_NOKCPHJN_453780 | Bacteria | Bacteroidetes | Flavobacteriia | unclassified | unclassified | unclassified | Flavobacteria | poorly charact | Function unkn | 126RR | cell surface- a | 43179.ENSST | 2,6 | 1,2 | 4,13E-05 | 1,77E-05 | 4 - | 5,49E-05 - | 4 - | 6,81E-05 - | 2 - | 3,24E-05 - | 2 - | 3,29E-05 - | 1 - | 1,85E-05 - |
| 170 | 1 | 3um_FIJJAFBF_405927 | Bacteria | Bacteroidetes | Flavobacteriia | unclassified | unclassified | unclassified | Flavobacteria | poorly charact | Function unkn | 09A4Z | rhodopsin | 487796.Flav2 | 1,2 | 0,748331477 | 6,67E-05 | 4,29E-05 | 1 - | 4,76E-05 - | 1 - | 5,91E-05 - | 2 - | 0,000112433 - | 2 - | 0,000114094 - | 0 - | 0 - |
| 171 | 1 | 3um_FIJJAFBF_321054 | unclassified | unclassified | unclassified | unclassified | unclassified | unclassified | unclassified | unclassified | unclassified | unclassified | unclassified | unclassified | 1 | 0 | 0,000142737 | 1,36E-05 | 1 - | 0,000119559 - | 1 - | 0,000148457 - | 1 - | 0,000141127 - | 1 - | 0,000143212 - | 1 - | 0,000161328 - |
| 172 | 1 | 3um_FIJJAFBF_229532 | Bacteria | Bacteroidetes | Flavobacteriia | Flavobacterial | Flavobacteriac | Marinirhabdu | Marinirhabdu | unclassified | unclassified | unclassified | unclassified | unclassified | 0,8 | 0,748331477 | 2,42E-05 | 2,33E-05 | 1 - | 2,61E-05 - | 1 - | 3,25E-05 - | 0 - | 0 - | 2 - | 6,26E-05 - | 0 - | 0 - |
| 173 | 1 | 3um_FIJJAFBF_285267 | Bacteria | Bacteroidetes | Cytophagia | Cytophagales | Flammeovirga | Marivirga | Marivirga lum | metabolism | Inorganic ion t | 0FW1N | Destroys radic | 643867.Ftrac_ | 2 | 0,894427191 | 0,000131655 | 5,15E-05 | 3 - | 0,000171309 - | 1 - | 7,09E-05 - | 3 - | 0,000202212 - | 2 - | 0,000136799 - | 1 - | 7,71E-05 - |
| 174 | 1 | 0.2um_NOKCPHJN_649835 | Bacteria | Proteobacteri | Gammaprotec | unclassified | unclassified | unclassified | Gammaprotec | unclassified | unclassified | unclassified | unclassified | unclassified | 1,2 | 0,4 | 7,60E-05 | 2,55E-05 | 1 - | 5,31E-05 - | 1 - | 6,60E-05 - | 2 - | 0,000125447 - | 1 - | 6,36E-05 - | 1 - | 7,17E-05 - |
| 175 | 1 | 0.2um_NOKCPHJN_742184 | Eukaryota | unclassified | Raphidophyce | Chattonellales | Chattonellace | Heterosigma | Heterosigma | unclassified | unclassified | unclassified | unclassified | unclassified | 0,4 | 0,489897949 | 2,48E-05 | 3,05E-05 | 1 - | 5,68E-05 - | 0 - | 0 - | 1 - | 6,71E-05 - | 0 - | 0 - | 0 - | 0 - |
| 176 | 1 | 0.2um_NOKCPHJN_445514 | Eukaryota | unclassified | unclassified | Isochrysidales | Noelaerhabda | Emiliania | Emiliania hux | poorly charact | Function unkn | COG2340 | Scp-like extrac | 6085.XP_0021 | 1 | 0 | 2,25E-05 | 2,14E-06 | 1 - | 1,89E-05 - | 1 - | 2,34E-05 - | 1 - | 2,23E-05 - | 1 - | 2,26E-05 - | 1 - | 2,55E-05 - |
| 177 | 1 | 3um_FIJJAFBF_49800 | Bacteria | Proteobacteri | Gammaprotec | Cellvibrionales | Spongiibacter | Dasania | Dasania marin | information st | Translation, ri | 17FIT | 50s ribosomal | 247633.GP214 | 0,2 | 0,4 | 2,94E-05 | 5,89E-05 | 1 - | 0,00014715 - | 0 - | 0 - | 0 - | 0 - | 0 - | 0 - | 0 - | 0 - |
| 178 | 1 | 3um_FIJJAFBF_652495 | Bacteria | Proteobacteri | Alphaproteob | Rhodobactera | Rhodobactera | Nereida | Nereida ignav | information st | Translation, ri | 01RJH | 30S ribosomal | 633131.TR2A | 3,8 | 1,166190379 | 0,000194423 | 3,95E-05 | 6 - | 0,000263855 - | 3 - | 0,000163815 - | 3 - | 0,000155727 - | 4 - | 0,000210702 - | 3 - | 0,000178017 - |
| 179 | 1 | 3um_FIJJAFBF_73682 | unclassified | unclassified | unclassified | unclassified | unclassified | unclassified | unclassified | unclassified | unclassified | unclassified | unclassified | unclassified | 1 | 0 | 2,43E-05 | 2,31E-06 | 1 - | 2,04E-05 - | 1 - | 2,53E-05 - | 1 - | 2,41E-05 - | 1 - | 2,44E-05 - | 1 - | 2,75E-05 - |
| 180 | 2 | 0.2um_NOKCPHJN_127543;3um_FIJJAFBF_13947 | unclassified | unclassified | unclassified | unclassified | unclassified | unclassified | unclassified | unclassified | unclassified | unclassified | unclassified | unclassified | 0,2 | 0,4 | 2,13E-05 | 4,25E-05 | 0 - | 0 - | 1 - | 0,000106357 - | 0 - | 0 - | 0 - | 0 - | 0 - | 0 - |
| 181 | 1 | 3um_FIJJAFBF_694013 | Bacteria | Bacteroidetes | unclassified | Bacteroidetes | Rhodotherma | unclassified | Rhodotherma | unclassified | unclassified | unclassified | unclassified | unclassified | 1 | 0 | 0,000126877 | 1,20E-05 | 1 - | 0,000106275 - | 1 - | 0,000131962 - | 1 - | 0,000125447 - | 1 - | 0,000127299 - | 1 - | 0,000143402 - |
| 182 | 1 | 0.2um_NOKCPHJN_253861 | Bacteria | Proteobacteri | Gammaprotec | Oceanospirilla | Saccharospirill | Reinekea | Reinekea fors | unclassified | unclassified | unclassified | unclassified | unclassified | 0,2 | 0,4 | 8,47E-06 | 1,69E-05 | 0 - | 0 - | 0 - | 0 - | 1 - | 4,23E-05 - | 0 - | 0 - | 0 - | 0 - |
| 183 | 1 | 3um_FIJJAFBF_198476 | Bacteria | Proteobacteri | Gammaprotec | Oceanospirilla | Saccharospirill | Reinekea | Reinekea fors | cellular proces | Cell motility | 0GF3K | ompa motb d | 177437.HRM2 | 0,8 | 0,748331477 | 2,40E-05 | 2,07E-05 | 2 - | 5,23E-05 - | 1 - | 3,25E-05 - | 0 - | 0 - | 0 - | 0 - | 1 - | 3,53E-05 - |
| 184 | 1 | 3um_FIJJAFBF_624840 | Bacteria | Proteobacteri | Gammaprotec | Oceanospirilla | Saccharospirill | Reinekea | Reinekea fors | unclassified | unclassified | unclassified | unclassified | unclassified | 4,2 | 0,4 | 0,000252683 | 1,20E-05 | 5 - | 0,000253931 - | 4 - | 0,000252246 - | 4 - | 0,000239792 - | 4 - | 0,000243333 - | 4 - | 0,000274114 - |
| 185 | 1 | 0.2um_NOKCPHJN_649335 | Bacteria | Bacteroidetes | Flavobacteriia | Flavobacterial | Crocinitomica | Fluviicola | Fluviicola sp. | metabolism | Energy produc | COG0056 | Produces ATP | 755732.Fluta_ | 0,2 | 0,4 | 5,15E-06 | 1,03E-05 | 0 - | 0 - | 0 - | 0 - | 1 - | 2,58E-05 - | 0 - | 0 - | 0 - | 0 - |
| 186 | 1 | 0.2um_NOKCPHJN_150079 | Bacteria | Proteobacteri | Betaproteoba | unclassified | unclassified | unclassified | Betaproteoba | unclassified | unclassified | unclassified | unclassified | unclassified | 13,4 | 1,356465997 | 0,000443841 | 3,58E-05 | 15 - | 0,000418893 - | 13 - | 0,00045079 - | 12 - | 0,000395569 - | 15 - | 0,000501764 - | 12 - | 0,000452188 - |
| 187 | 1 | 3um_FIJJAFBF_491687 | Bacteria | Proteobacteri | Gammaprotec | Cellvibrionales | Porticoccacea | unclassified | various | information st | Translation, ri | 05G6W | One of the pri | 247634.GPB2 | 1,2 | 0,4 | 7,77E-05 | 1,72E-05 | 2 - | 0,000111434 - | 1 - | 6,92E-05 - | 1 - | 6,58E-05 - | 1 - | 6,67E-05 - | 1 - | 7,52E-05 - |
| 188 | 1 | 3um_FIJJAFBF_736125 | Bacteria | Bacteroidetes | Flavobacteriia | various | various | various | various | unclassified | unclassified | unclassified | unclassified | unclassified | 1,4 | 0,489897949 | 0,000104513 | 2,99E-05 | 2 - | 0,000128242 - | 1 - | 7,96E-05 - | 2 - | 0,000151377 - | 1 - | 7,68E-05 - | 1 - | 8,65E-05 - |
| 189 | 1 | 3um_FIJJAFBF_536004 | unclassified | unclassified | unclassified | unclassified | unclassified | unclassified | unclassified | unclassified | unclassified | unclassified | unclassified | unclassified | 1 | 0 | 0,000130502 | 1,24E-05 | 1 - | 0,000109311 - | 1 - | 0,000135732 - | 1 - | 0,000129031 - | 1 - | 0,000130936 - | 1 - | 0,0001475 - |
| 190 | 1 | 3um_FIJJAFBF_439554 | Bacteria | Bacteroidetes | Flavobacteriia | Flavobacterial | Flavobacteriac | Lacinutrix | Lacinutrix him | unclassified | unclassified | unclassified | unclassified | unclassified | 1,4 | 0,489897949 | 0,000151193 | 4,45E-05 | 2 - | 0,000185124 - | 1 - | 0,000114935 - | 1 - | 0,00010926 - | 2 - | 0,000221747 - | 1 - | 0,000124899 - |
| 191 | 1 | 3um_FIJJAFBF_28972 | Bacteria | Proteobacteri | Gammaprotec | unclassified | unclassified | unclassified | Gammaprotec | poorly charact | Function unkn | 070MQ | NA | 425104.Ssed_ | 9,4 | 0,489897949 | 0,000287688 | 3,19E-05 | 9 - | 0,000230578 - | 10 - | 0,000318122 - | 10 - | 0,000302416 - | 9 - | 0,000276194 - | 9 - | 0,000311132 - |
| 192 | 1 | 0.2um_NOKCPHJN_580018 | Bacteria | Bacteroidetes | Flavobacteriia | Flavobacterial | Flavobacteriac | Polaribacter | Polaribacter b | unclassified | unclassified | unclassified | unclassified | unclassified | 0,6 | 0,489897949 | 1,87E-05 | 1,53E-05 | 0 - | 0 - | 1 - | 3,20E-05 - | 1 - | 3,04E-05 - | 1 - | 3,09E-05 - | 0 - | 0 - |
| 193 | 1 | 3um_FIJJAFBF_545826 | Bacteria | Bacteroidetes | Flavobacteriia | Flavobacterial | Flavobacteriac | Polaribacter | Polaribacter s | cellular proces | Intracellular tr | 0P8JN | The central su | 313598.MED1 | 1,8 | 0,4 | 5,51E-05 | 1,09E-05 | 2 - | 5,21E-05 - | 2 - | 6,46E-05 - | 2 - | 6,14E-05 - | 2 - | 6,24E-05 - | 1 - | 3,51E-05 - |
| 194 | 1 | 3um_FIJJAFBF_302477 | Eukaryota | unclassified | Pelagophycea | Pelagomonad | unclassified | Aureococcus | Aureococcus a | unclassified | unclassified | unclassified | unclassified | unclassified | 1 | 0 | 0,000139824 | 1,33E-05 | 1 - | 0,000117119 - | 1 - | 0,000145427 - | 1 - | 0,000138247 - | 1 - | 0,000140289 - | 1 - | 0,000158035 - |
| 195 | 1 | 3um_FIJJAFBF_234031 | Bacteria | Bacteroidetes | Flavobacteriia | Flavobacterial | Flavobacteriac | Formosa | Formosa sp. H | unclassified | unclassified | unclassified | unclassified | unclassified | 0,8 | 0,4 | 7,66E-05 | 3,85E-05 | 0 - | 0 - | 1 - | 9,57E-05 - | 1 - | 9,09E-05 - | 1 - | 9,23E-05 - | 1 - | 0,000103943 - |
| 196 | 1 | 0.2um_NOKCPHJN_1038894 | Eukaryota | Phaeophyceae | unclassified | Ectocarpales | Ectocarpaceae | Ectocarpus | Ectocarpus sil | information st | Transcription | 0VC52 | Chromosome | 7227.FBpp007 | 2,2 | 0,4 | 0,000145593 | 1,34E-05 | 3 - | 0,000168789 - | 2 - | 0,000139724 - | 2 - | 0,000132826 - | 2 - | 0,000134788 - | 2 - | 0,000151838 - |
| 197 | 1 | 3um_FIJJAFBF_463735 | unclassified | unclassified | unclassified | unclassified | unclassified | unclassified | unclassified | unclassified | unclassified | unclassified | unclassified | unclassified | 0,4 | 0,489897949 | 7,25E-05 | 8,96E-05 | 1 - | 0,000161657 - | 1 - | 0,000200731 - | 0 - | 0 - | 0 - | 0 - | 0 - | 0 - |
| 198 | 1 | 0.2um_NOKCPHJN_860507 | Bacteria | Proteobacteri | Betaproteoba | Burkholderial | Comamonada | Acidovorax | Acidovorax sp | unclassified | unclassified | unclassified | unclassified | unclassified | 2,6 | 0,489897949 | 9,19E-05 | 1,55E-05 | 3 - | 8,97E-05 - | 3 - | 0,000111343 - | 2 - | 7,06E-05 - | 3 - | 0,000107409 - | 2 - | 8,07E-05 - |
| 199 | 3 | 0.2um_NOKCPHJN_1021687;0.2um_NOKCPHJN_ | Bacteria | Proteobacteri | Gammaprotec | unclassified | unclassified | unclassified | Gammaprotec | cellular proces | Cell wall/mem | 16QB0 | Part of the ou | various | 1 | 0 | 2,10E-05 | 2,00E-06 | 1 - | 1,76E-05 - | 1 - | 2,19E-05 - | 1 - | 2,08E-05 - | 1 - | 2,11E-05 - | 1 - | 2,38E-05 - |
| 200 | 1 | 3um_FIJJAFBF_172362 | Bacteria | Proteobacteri | Gammaprotec | Oceanospirilla | Saccharospirill | Reinekea | Reinekea fors | information st | Translation, ri | 177SZ | This protein b | 314283.MED2 | 5,8 | 0,748331477 | 0,00054656 | 4,23E-05 | 7 - | 0,000557943 - | 6 - | 0,000593829 - | 5 - | 0,000470424 - | 6 - | 0,000572847 - | 5 - | 0,000537759 - |
| 201 | 2 | 0.2um_NOKCPHJN_137432;3um_FIJJAFBF_6716 | Bacteria | Bacteroidetes | Flavobacteriia | Flavobacterial | Flavobacteriac | Formosa | Formosa sp. H | poorly charact | Function unkn | 099QV | gliding motilit | various | 0,4 | 0,489897949 | 1,59E-05 | 1,94E-05 | 0 - | 0 - | 1 - | 4,04E-05 - | 0 - | 0 - | 1 - | 3,89E-05 - | 0 - | 0 - |
| 202 | 1 | 3um_FIJJAFBF_520093 | Bacteria | Bacteroidetes | Flavobacteriia | Flavobacterial | Flavobacteriac | Polaribacter | Polaribacter s | unclassified | unclassified | unclassified | unclassified | unclassified | 2 | 0,632455532 | 0,000108053 | 3,60E-05 | 2 - | 9,04E-05 - | 2 - | 0,00011222 - | 1 - | 5,33E-05 - | 3 - | 0,000162382 - | 2 - | 0,000121948 - |
| 203 | 1 | 0.2um_NOKCPHJN_914487 | Eukaryota | Chordata | Actinopteri | Beloniformes | Adrianichthyic | Oryzias | Oryzias latipe | cellular proces | Signal transdu | KOG0613 | fibronectin ty | 121225.PHUM | 0,2 | 0,4 | 4,22E-06 | 8,43E-06 | 0 - | 0 - | 0 - | 0 - | 0 - | 0 - | 1 - | 2,11E-05 - | 0 - | 0 - |
| 204 | 1 | 3um_FIJJAFBF_67732 | unclassified | unclassified | unclassified | unclassified | unclassified | unclassified | unclassified | unclassified | unclassified | unclassified | unclassified | unclassified | 2,2 | 1,166190379 | 0,000133912 | 7,11E-05 | 3 - | 0,000155806 - | 4 - | 0,000257953 - | 1 - | 6,13E-05 - | 2 - | 0,000124419 - | 1 - | 7,01E-05 - |
| 205 | 1 | 0.2um_NOKCPHJN_23322 | Eukaryota | unclassified | Pelagophycea | Pelagomonad | unclassified | Pelagomonas | various | cellular proces | Posttranslatio | KOG0730 | cell division cy | 35128.Thapsd | 3,8 | 0,979795897 | 0,000177241 | 4,00E-05 | 5 - | 0,000197891 - | 5 - | 0,000245722 - | 3 - | 0,000140154 - | 3 - | 0,000142224 - | 3 - | 0,000160215 - |
| 206 | 1 | 0.2um_NOKCPHJN_109945 | Eukaryota | unclassified | unclassified | Acytosteliales | Acytosteliacea | Acytostelium | Acytostelium | unclassified | unclassified | unclassified | unclassified | unclassified | 0,4 | 0,489897949 | 3,61E-06 | 4,42E-06 | 0 - | 0 - | 0 - | 0 - | 1 - | 8,95E-06 - | 1 - | 9,08E-06 - | 0 - | 0 - |
| 207 | 1 | 0.2um_NOKCPHJN_476838 | Eukaryota | unclassified | Oomycetes | Saprolegniales | Saprolegniace | Aphanomyces | Aphanomyces | unclassified | unclassified | unclassified | unclassified | unclassified | 1 | 0 | 1,47E-05 | 1,39E-06 | 1 - | 1,23E-05 - | 1 - | 1,53E-05 - | 1 - | 1,45E-05 - | 1 - | 1,47E-05 - | 1 - | 1,66E-05 - |
| 208 | 1 | 3um_FIJJAFBF_12338 | Bacteria | Proteobacteri | Gammaprotec | Cellvibrionales | unclassified | unclassified | Cellvibrionales | information st | Translation, ri | COG0088 | One of the pri | 314287.GB22 | 1 | 0 | 6,72E-05 | 6,38E-06 | 1 - | 5,63E-05 - | 1 - | 6,99E-05 - | 1 - | 6,64E-05 - | 1 - | 6,74E-05 - | 1 - | 7,59E-05 - |
| 209 | 1 | 0.2um_NOKCPHJN_485725 | Bacteria | Proteobacteri | Gammaprotec | Cellvibrionales | Spongiibacter | Oceanicoccus | Oceanicoccus | cellular proces | Cell wall/mem | 0QTSX | Peptidoglycan | 247634.GPB2 | 1,6 | 0,489897949 | 0,00011308 | 3,20E-05 | 2 - | 0,000120185 - | 2 - | 0,000149234 - | 1 - | 7,09E-05 - | 2 - | 0,000143962 - | 1 - | 8,11E-05 - |
| 210 | 2 | 0.2um_NOKCPHJN_718839;0.2um_NOKCPHJN_7 | Bacteria | Proteobacteri | Alphaproteob | Pelagibacteral | Pelagibacterac | Candidatus Pe | Candidatus Pe | poorly charact | Function unkn | 06SN9 | NA | 335992.SAR11 | 2,2 | 0,4 | 0,000140725 | 2,74E-05 | 2 - | 0,000107268 - | 2 - | 0,000133195 - | 3 - | 0,000189928 - | 2 - | 0,000128489 - | 2 - | 0,000144743 - |
| 211 | 1 | 3um_FIJJAFBF_119781 | Eukaryota | Ascomycota | Dothideomyc | Dothideales | Saccotheciace | Aureobasidiur | Aureobasidiur | unclassified | unclassified | unclassified | unclassified | unclassified | 8,4 | 0,8 | 0,000106449 | 1,93E-05 | 8 - | 8,44E-05 - | 8 - | 0,000104793 - | 8 - | 9,96E-05 - | 8 - | 0,000101091 - | 10 - | 0,000142348 - |
| 212 | 1 | 3um_FIJJAFBF_189725 | Bacteria | Bacteroidetes | Flavobacteriia | Flavobacterial | Flavobacteriac | Polaribacter | Polaribacter s | unclassified | unclassified | unclassified | unclassified | unclassified | 1,2 | 0,4 | 0,000179756 | 3,98E-05 | 2 - | 0,000257925 - | 1 - | 0,000160134 - | 1 - | 0,000152227 - | 1 - | 0,000154476 - | 1 - | 0,000174016 - |
| 213 | 1 | 0.2um_NOKCPHJN_69934 | Bacteria | Proteobacteri | Gammaprotec | Cellvibrionales | unclassified | unclassified | Cellvibrionales | metabolism | Energy produc | 16PVJ | Component o | 314287.GB22 | 0,6 | 0,489897949 | 8,96E-06 | 7,39E-06 | 1 - | 1,30E-05 - | 1 - | 1,62E-05 - | 0 - | 0 - | 1 - | 1,56E-05 - | 0 - | 0 - |
| 214 | 1 | 3um_FIJJAFBF_52796 | Bacteria | Proteobacteri | Alphaproteob | Rhodobactera | Rhodobactera | Planktomarina | Planktomarina | unclassified | unclassified | unclassified | unclassified | unclassified | 0,2 | 0,4 | 1,66E-05 | 3,33E-05 | 1 - | 8,32E-05 - | 0 - | 0 - | 0 - | 0 - | 0 - | 0 - | 0 - | 0 - |
| 215 | 1 | 0.2um_NOKCPHJN_213001 | Bacteria | Bacteroidetes | Flavobacteriia | Flavobacterial | Flavobacteriac | Polaribacter | Polaribacter h | unclassified | unclassified | unclassified | unclassified | unclassified | 2,2 | 0,748331477 | 4,60E-05 | 1,63E-05 | 3 - | 5,26E-05 - | 2 - | 4,36E-05 - | 1 - | 2,07E-05 - | 2 - | 4,20E-05 - | 3 - | 7,10E-05 - |
| 216 | 1 | 3um_FIJJAFBF_17218 | Bacteria | Proteobacteri | Alphaproteob | Rhodobactera | unclassified | unclassified | Rhodobactera | cellular proces | Posttranslatio | COG0330 | Band 7 protei | 439496.RBY4 | 6 | 0,894427191 | 0,000212307 | 2,15E-05 | 7 - | 0,000209775 - | 5 - | 0,000186056 - | 6 - | 0,000212244 - | 7 - | 0,000251275 - | 5 - | 0,000202186 - |
| 217 | 1 | 0.2um_NOKCPHJN_109685 | Bacteria | Bacteroidetes | Flavobacteriia | Flavobacterial | Flavobacteriac | Polaribacter | Polaribacter s | unclassified | unclassified | unclassified | unclassified | unclassified | 0,4 | 0,489897949 | 1,02E-05 | 1,27E-05 | 1 - | 2,17E-05 - | 0 - | 0 - | 0 - | 0 - | 0 - | 0 - | 1 - | 2,92E-05 - |
| 218 | 1 | 3um_FIJJAFBF_677888 | Bacteria | Proteobacteri | Alphaproteob | Rhizobiales | unclassified | unclassified | Rhizobiales ba | unclassified | unclassified | unclassified | unclassified | unclassified | 9,4 | 1,019803903 | 0,000355919 | 2,50E-05 | 11 - | 0,000351684 - | 8 - | 0,000317591 - | 10 - | 0,000377388 - | 9 - | 0,000344666 - | 9 - | 0,000388265 - |
| 219 | 1 | 0.2um_NOKCPHJN_128595 | Eukaryota | Basidiomycota | Ustilaginomyc | Ustilaginales | Anthracoideac | Testicularia | Testicularia c | cellular proces | Cytoskeleton | 01GFG | ADF | 578458.XP_00 | 1 | 0 | 0,000126877 | 1,20E-05 | 1 - | 0,000106275 - | 1 - | 0,000131962 - | 1 - | 0,000125447 - | 1 - | 0,000127299 - | 1 - | 0,000143402 - |
| 220 | 1 | 3um_FIJJAFBF_582148 | Bacteria | Bacteroidetes | Flavobacteriia | Flavobacterial | Crocinitomica | unclassified | Crocinitomica | poorly charact | Function unkn | 09AR5 | Inherit from b | 313606.M231 | 1 | 0 | 2,69E-05 | 2,55E-06 | 1 - | 2,25E-05 - | 1 - | 2,79E-05 - | 1 - | 2,66E-05 - | 1 - | 2,70E-05 - | 1 - | 3,04E-05 - |
| 221 | 1 | 0.2um_NOKCPHJN_149987 | Bacteria | Bacteroidetes | Sphingobacter | Sphingobacter | Sphingobacter | Sphingobacter | uncultured Sp | information st | Translation, ri | COG1185 | Involved in m | 926562.Oweh | 3,4 | 1,019803903 | 6,64E-05 | 2,35E-05 | 3 - | 4,84E-05 - | 5 - | 0,000100224 - | 2 - | 3,81E-05 - | 3 - | 5,80E-05 - | 4 - | 8,71E-05 - |
| 222 | 1 | 3um_FIJJAFBF_210720 | Bacteria | unclassified | unclassified | unclassified | unclassified | unclassified | uncultured ba | unclassified | unclassified | unclassified | unclassified | unclassified | 5 | 0,894427191 | 6,80E-05 | 1,11E-05 | 6 - | 6,89E-05 - | 4 - | 5,70E-05 - | 4 - | 5,42E-05 - | 6 - | 8,25E-05 - | 5 - | 7,74E-05 - |
| 223 | 1 | 3um_FIJJAFBF_597915 | Bacteria | Bacteroidetes | Flavobacteriia | Flavobacterial | Flavobacteriac | Polaribacter | Polaribacter s | cellular proces | Posttranslatio | COG0330 | Band 7 protei | 313594.PI23P | 3 | 0 | 0,00013434 | 1,28E-05 | 3 - | 0,000112526 - | 3 - | 0,000139724 - | 3 - | 0,000132826 - | 3 - | 0,000134788 - | 3 - | 0,000151838 - |
| 224 | 1 | 0.2um_NOKCPHJN_131409 | Bacteria | Bacteroidetes | Flavobacteriia | Flavobacterial | Flavobacteriac | Polaribacter | Polaribacter a | unclassified | unclassified | unclassified | unclassified | unclassified | 1,8 | 1,166190379 | 8,50E-05 | 4,50E-05 | 4 - | 0,000167557 - | 1 - | 5,20E-05 - | 2 - | 9,89E-05 - | 1 - | 5,02E-05 - | 1 - | 5,65E-05 - |
| 225 | 1 | 3um_FIJJAFBF_520338 | Viruses | unclassified | unclassified | unclassified | unclassified | unclassified | uncultured M | unclassified | unclassified | unclassified | unclassified | unclassified | 0,2 | 0,4 | 1,93E-06 | 3,86E-06 | 0 - | 0 - | 1 - | 9,65E-06 - | 0 - | 0 - | 0 - | 0 - | 0 - | 0 - |
| 226 | 1 | 3um_FIJJAFBF_594175 | Eukaryota | Phaeophyceae | unclassified | Ectocarpales | various | various | various | metabolism | Energy produc | COG1290 | Component o | 35128.Thapsd | 5 | 0 | 0,000318668 | 3,03E-05 | 5 - | 0,000266923 - | 5 - | 0,000331439 - | 5 - | 0,000315075 - | 5 - | 0,000319729 - | 5 - | 0,000360173 - |
| 227 | 3 | 0.2um_NOKCPHJN_1045591;3um_FIJJAFBF_324 | Eukaryota | various | various | various | various | various | various | metabolism | Energy produc | COG1145 | 4Fe-4S ferred | various | 2 | 0 | 0,000301159 | 2,86E-05 | 2 - | 0,000252257 - | 2 - | 0,000313228 - | 2 - | 0,000297763 - | 2 - | 0,000302161 - | 2 - | 0,000340383 - |
| 228 | 1 | 0.2um_NOKCPHJN_85372 | Bacteria | Proteobacteri | Gammaprotec | Oceanospirilla | Saccharospirill | Reinekea | Reinekea fors | cellular proces | Cell wall/mem | 16U9X | Curli producti | 203122.Sde_3 | 1,4 | 0,489897949 | 5,88E-05 | 1,68E-05 | 2 - | 7,22E-05 - | 1 - | 4,48E-05 - | 2 - | 8,52E-05 - | 1 - | 4,32E-05 - | 1 - | 4,87E-05 - |
| 229 | 1 | 3um_FIJJAFBF_525903 | Bacteria | Bacteroidetes | Flavobacteriia | Flavobacterial | Flavobacteriac | Polaribacter | Polaribacter v | unclassified | unclassified | unclassified | unclassified | unclassified | 1 | 0,632455532 | 1,94E-05 | 1,27E-05 | 1 - | 1,67E-05 - | 1 - | 2,07E-05 - | 1 - | 1,97E-05 - | 2 - | 4,00E-05 - | 0 - | 0 - |
| 230 | 1 | 3um_FIJJAFBF_190224 | Bacteria | Proteobacteri | Gammaprotec | Alteromonada | Alteromonada | Paraglaciecola | Paraglaciecola | information st | Replication, re | 05C68 | Can catalyze t | 167879.CPS_4 | 0,6 | 0,489897949 | 2,26E-05 | 1,86E-05 | 1 - | 3,35E-05 - | 0 - | 0 - | 1 - | 3,95E-05 - | 1 - | 4,01E-05 - | 0 - | 0 - |
| 231 | 1 | 0.2um_NOKCPHJN_75563 | Bacteria | Proteobacteri | Gammaprotec | Oceanospirilla | Saccharospirill | Reinekea | Reinekea blan | information st | Translation, ri | COG0173 | aspartyl-trna s | 314283.MED2 | 8,4 | 0,8 | 0,000193864 | 1,57E-05 | 9 - | 0,000175083 - | 9 - | 0,000217402 - | 9 - | 0,000206668 - | 8 - | 0,000186418 - | 7 - | 0,000183749 - |
| 232 | 1 | 0.2um_NOKCPHJN_66510 | Bacteria | Bacteroidetes | unclassified | unclassified | unclassified | unclassified | Bacteroidetes | unclassified | unclassified | unclassified | unclassified | unclassified | 1,8 | 0,748331477 | 1,95E-05 | 6,35E-06 | 3 - | 2,83E-05 - | 1 - | 1,17E-05 - | 2 - | 2,23E-05 - | 2 - | 2,26E-05 - | 1 - | 1,27E-05 - |
| 233 | 2 | 0.2um_NOKCPHJN_650096;3um_FIJJAFBF_1351 | Bacteria | Planctomycet | unclassified | unclassified | unclassified | unclassified | Planctomycet | unclassified | unclassified | unclassified | unclassified | unclassified | 0,4 | 0,489897949 | 2,02E-05 | 2,49E-05 | 1 - | 4,59E-05 - | 0 - | 0 - | 0 - | 0 - | 1 - | 5,50E-05 - | 0 - | 0 - |
| 234 | 1 | 0.2um_NOKCPHJN_27394 | Bacteria | unclassified | unclassified | unclassified | unclassified | unclassified | uncultured ba | metabolism | Inorganic ion t | COG3119 | Sulfatase | 485917.Phep_ | 1,4 | 0,489897949 | 2,19E-05 | 6,26E-06 | 2 - | 2,68E-05 - | 1 - | 1,67E-05 - | 2 - | 3,17E-05 - | 1 - | 1,61E-05 - | 1 - | 1,81E-05 - |
| 235 | 2 | 0.2um_NOKCPHJN_494936;3um_FIJJAFBF_2988 | Bacteria | Bacteroidetes | Flavobacteriia | Flavobacterial | Flavobacteriac | Polaribacter | Polaribacter s | unclassified | unclassified | unclassified | unclassified | unclassified | 1 | 0 | 7,88E-05 | 7,48E-06 | 1 - | 6,60E-05 - | 1 - | 8,19E-05 - | 1 - | 7,79E-05 - | 1 - | 7,90E-05 - | 1 - | 8,90E-05 - |
| 236 | 1 | 0.2um_NOKCPHJN_403347 | Bacteria | Proteobacteri | Gammaprotec | Cellvibrionales | Halieaceae | unclassified | marine gamm | metabolism | Inorganic ion t | 16PDP | TonB-depend | 247634.GPB2 | 1,8 | 0,748331477 | 2,53E-05 | 8,52E-06 | 3 - | 3,65E-05 - | 2 - | 3,02E-05 - | 1 - | 1,44E-05 - | 2 - | 2,92E-05 - | 1 - | 1,64E-05 - |
| 237 | 1 | 0.2um_NOKCPHJN_936712 | Bacteria | Proteobacteri | Alphaproteob | Rhodobactera | Rhodobactera | Pseudorueger | Pseudorueger | information st | Replication, re | 05C2R | Single-strande | 644107.SL115 | 1 | 0 | 4,64E-05 | 4,41E-06 | 1 - | 3,89E-05 - | 1 - | 4,83E-05 - | 1 - | 4,59E-05 - | 1 - | 4,66E-05 - | 1 - | 5,25E-05 - |
| 238 | 1 | 3um_FIJJAFBF_571391 | Bacteria | Proteobacteri | Gammaprotec | Cellvibrionales | Porticoccacea | unclassified | SAR92 bacteri | metabolism | Energy produc | 05C26 | Dehydrogenas | 314287.GB22 | 0,2 | 0,4 | 7,06E-06 | 1,41E-05 | 0 - | 0 - | 0 - | 0 - | 1 - | 3,53E-05 - | 0 - | 0 - | 0 - | 0 - |
| 239 | 1 | 3um_FIJJAFBF_722063 | unclassified | unclassified | unclassified | unclassified | unclassified | unclassified | unclassified | unclassified | unclassified | unclassified | unclassified | unclassified | 0,6 | 0,8 | 3,04E-05 | 4,25E-05 | 1 - | 4,36E-05 - | 2 - | 0,000108379 - | 0 - | 0 - | 0 - | 0 - | 0 - | 0 - |
| 240 | 1 | 3um_FIJJAFBF_54466 | Bacteria | Proteobacteri | Gammaprotec | Cellvibrionales | various | various | various | metabolism | Inorganic ion t | 1768K | TONB-depend | 247634.GPB2 | 1,2 | 0,4 | 7,21E-05 | 3,07E-05 | 1 - | 4,93E-05 - | 1 - | 6,12E-05 - | 1 - | 5,81E-05 - | 1 - | 5,90E-05 - | 2 - | 0,000132939 - |
| 241 | 1 | 3um_FIJJAFBF_352681 | Bacteria | Proteobacteri | Gammaprotec | Alteromonada | Alteromonada | Paraglaciecola | Paraglaciecola | cellular proces | Posttranslatio | COG0330 | Band 7 protei | 715451.ambt_ | 4,6 | 1,624807681 | 0,000217049 | 8,09E-05 | 3 - | 0,000117519 - | 6 - | 0,000291848 - | 7 - | 0,000323678 - | 3 - | 0,000140768 - | 4 - | 0,000211433 - |
| 242 | 1 | 0.2um_NOKCPHJN_77268 | Bacteria | Proteobacteri | Betaproteoba | unclassified | unclassified | unclassified | Betaproteoba | metabolism | Energy produc | COG2224 | Isocitrate lyas | 640081.Dsui_ | 0,6 | 0,489897949 | 1,88E-05 | 1,57E-05 | 1 - | 2,62E-05 - | 1 - | 3,25E-05 - | 0 - | 0 - | 0 - | 0 - | 1 - | 3,54E-05 - |
| 243 | 1 | 3um_FIJJAFBF_626704 | Bacteria | Bacteroidetes | Flavobacteriia | Flavobacterial | Flavobacteriac | Polaribacter | Polaribacter s | unclassified | unclassified | unclassified | unclassified | unclassified | 1 | 0 | 0,000217503 | 2,07E-05 | 1 - | 0,000182185 - | 1 - | 0,00022622 - | 1 - | 0,000215051 - | 1 - | 0,000218227 - | 1 - | 0,000245833 - |
| 244 | 1 | 3um_FIJJAFBF_19898 | Bacteria | Proteobacteri | Alphaproteob | Rhodobactera | Rhodobactera | unclassified | Rhodobactera | information st | Transcription | 05DEK | Sigma factors | 314270.RB208 | 1,6 | 0,489897949 | 7,20E-05 | 1,89E-05 | 2 - | 7,70E-05 - | 1 - | 4,78E-05 - | 2 - | 9,09E-05 - | 2 - | 9,23E-05 - | 1 - | 5,20E-05 - |
| 245 | 1 | 3um_FIJJAFBF_350119 | Bacteria | Proteobacteri | Alphaproteob | Rhodobactera | unclassified | unclassified | Rhodobactera | information st | Replication, re | 05C68 | Can catalyze t | 246200.SPO2 | 1 | 0,894427191 | 3,63E-05 | 3,19E-05 | 2 - | 6,48E-05 - | 1 - | 4,03E-05 - | 2 - | 7,65E-05 - | 0 - | 0 - | 0 - | 0 - |
| 246 | 1 | 0.2um_NOKCPHJN_160008 | Bacteria | Proteobacteri | Alphaproteob | Rhodobactera | Rhodobactera | Amylibacter | Amylibacter k | metabolism | Energy produc | 05CMV | Succinyl-CoA s | 314256.OG25 | 0,8 | 0,748331477 | 2,65E-05 | 2,57E-05 | 1 - | 2,89E-05 - | 0 - | 0 - | 1 - | 3,41E-05 - | 2 - | 6,93E-05 - | 0 - | 0 - |
| 247 | 1 | 3um_FIJJAFBF_535169 | Bacteria | Proteobacteri | Alphaproteob | Rhodobactera | unclassified | unclassified | Rhodobactera | unclassified | unclassified | unclassified | unclassified | unclassified | 3,6 | 0,8 | 0,000297091 | 6,99E-05 | 3 - | 0,000207428 - | 3 - | 0,000257564 - | 4 - | 0,000326463 - | 5 - | 0,000414106 - | 3 - | 0,000279894 - |
| 248 | 1 | 0.2um_NOKCPHJN_37838 | Bacteria | Proteobacteri | Alphaproteob | Rhodospirillal | Rhodospirillac | unclassified | Rhodospirillac | unclassified | unclassified | unclassified | unclassified | unclassified | 0,6 | 0,489897949 | 5,47E-05 | 4,47E-05 | 0 - | 0 - | 1 - | 9,38E-05 - | 1 - | 8,91E-05 - | 1 - | 9,04E-05 - | 0 - | 0 - |
| 249 | 1 | 0.2um_NOKCPHJN_364214 | Bacteria | Proteobacteri | Alphaproteob | Rhodobactera | unclassified | unclassified | Rhodobactera | information st | Translation, ri | COG0255 | 50s ribosomal | 367336.OM22 | 1,6 | 0,489897949 | 0,000327554 | 0,000108303 | 2 - | 0,000342617 - | 2 - | 0,000425429 - | 1 - | 0,000202212 - | 1 - | 0,000205199 - | 2 - | 0,000462312 - |
| 250 | 1 | 3um_FIJJAFBF_72475 | Bacteria | Bacteroidetes | Cytophagia | Cytophagales | Flammeovirga | unclassified | Flammeovirga | unclassified | unclassified | unclassified | unclassified | unclassified | 1,6 | 0,489897949 | 0,000175882 | 4,61E-05 | 2 - | 0,000188159 - | 1 - | 0,000116819 - | 2 - | 0,000222102 - | 2 - | 0,000225382 - | 1 - | 0,000126946 - |
| 251 | 1 | 3um_FIJJAFBF_174319 | Bacteria | Proteobacteri | Alphaproteob | Pelagibacteral | Pelagibacterac | Candidatus Pe | Candidatus Pe | information st | Translation, ri | COG0051 | tRNA binding | 335992.SAR11 | 1,4 | 0,489897949 | 0,000183803 | 5,41E-05 | 2 - | 0,000225052 - | 1 - | 0,000139724 - | 1 - | 0,000132826 - | 2 - | 0,000269575 - | 1 - | 0,000151838 - |
| 252 | 1 | 0.2um_NOKCPHJN_207188 | Bacteria | Proteobacteri | Gammaprotec | Oceanospirilla | Saccharospirill | Reinekea | Reinekea blan | unclassified | unclassified | unclassified | unclassified | unclassified | 5,8 | 1,166190379 | 0,000185445 | 3,85E-05 | 5 - | 0,000134399 - | 6 - | 0,000200261 - | 7 - | 0,000222102 - | 7 - | 0,000225382 - | 4 - | 0,000145081 - |
| 253 | 1 | 0.2um_NOKCPHJN_242644 | Bacteria | Proteobacteri | Alphaproteob | Rhodobactera | Rhodobactera | Litoreibacter | Litoreibacter j | metabolism | Nucleotide tra | 16QX0 | Catalyzes the | 314256.OG25 | 0,2 | 0,4 | 4,20E-06 | 8,39E-06 | 1 - | 2,10E-05 - | 0 - | 0 - | 0 - | 0 - | 0 - | 0 - | 0 - | 0 - |
| 254 | 1 | 0.2um_NOKCPHJN_935031 | Bacteria | Proteobacteri | Alphaproteob | Pelagibacteral | Pelagibacterac | Candidatus Pe | Candidatus Pe | metabolism | Amino acid tra | 16RER | (ABC) transpo | 335992.SAR11 | 2 | 0,632455532 | 9,49E-05 | 2,42E-05 | 3 - | 0,000121671 - | 1 - | 5,04E-05 - | 2 - | 9,57E-05 - | 2 - | 9,72E-05 - | 2 - | 0,000109452 - |
| 255 | 1 | 3um_FIJJAFBF_75393 | Bacteria | Proteobacteri | Gammaprotec | Oceanospirilla | Saccharospirill | Reinekea | Reinekea fors | metabolism | Amino acid tra | 16R0H | Acetolactate s | 871585.BDGL_ | 2,4 | 0,8 | 0,000201379 | 6,75E-05 | 2 - | 0,00014083 - | 2 - | 0,00017487 - | 4 - | 0,000332472 - | 2 - | 0,000168691 - | 2 - | 0,00019003 - |
| 256 | 1 | 0.2um_NOKCPHJN_91523 | Bacteria | Proteobacteri | Gammaprotec | Cellvibrionales | unclassified | unclassified | Cellvibrionales | information st | Translation, ri | COG0173 | aspartyl-trna s | 314287.GB22 | 0,4 | 0,489897949 | 9,44E-06 | 1,16E-05 | 0 - | 0 - | 1 - | 2,40E-05 - | 0 - | 0 - | 1 - | 2,32E-05 - | 0 - | 0 - |
| 257 | 1 | 3um_FIJJAFBF_154707 | Bacteria | Bacteroidetes | Flavobacteriia | Flavobacterial | Flavobacteriac | Flaviramulus | Flaviramulus b | unclassified | unclassified | unclassified | unclassified | unclassified | 1 | 0 | 0,000150579 | 1,43E-05 | 1 - | 0,000126128 - | 1 - | 0,000156614 - | 1 - | 0,000148882 - | 1 - | 0,000151081 - | 1 - | 0,000170192 - |
| 258 | 1 | 3um_FIJJAFBF_347241 | Bacteria | Bacteroidetes | Flavobacteriia | Flavobacterial | Flavobacteriac | Polaribacter | Polaribacter in | unclassified | unclassified | unclassified | unclassified | unclassified | 1,2 | 0,748331477 | 8,08E-05 | 5,10E-05 | 1 - | 5,68E-05 - | 0 - | 0 - | 2 - | 0,000134141 - | 2 - | 0,000136122 - | 1 - | 7,67E-05 - |
| 259 | 2 | 0.2um_NOKCPHJN_94093;3um_FIJJAFBF_14080 | Bacteria | Bacteroidetes | Flavobacteriia | Flavobacterial | Flavobacteriac | Ulvibacter | Ulvibacter sp. | unclassified | unclassified | unclassified | unclassified | unclassified | 1 | 0,632455532 | 2,50E-05 | 1,76E-05 | 1 - | 2,04E-05 - | 1 - | 2,54E-05 - | 1 - | 2,41E-05 - | 0 - | 0 - | 2 - | 5,51E-05 - |
| 260 | 1 | 3um_FIJJAFBF_80131 | Bacteria | Bacteroidetes | Flavobacteriia | various | various | various | various | cellular proces | Cell wall/mem | 11TPU | outer membra | 487797.Flav3 | 6,8 | 1,6 | 0,000555414 | 0,000163164 | 7 - | 0,000475407 - | 7 - | 0,000590315 - | 7 - | 0,000561169 - | 4 - | 0,000325404 - | 9 - | 0,000824775 - |
| 261 | 1 | 3um_FIJJAFBF_501482 | Bacteria | Proteobacteri | Gammaprotec | unclassified | unclassified | unclassified | various | information st | Translation, ri | COG0051 | tRNA binding | 876044.IMCC | 1 | 0 | 0,000133036 | 1,26E-05 | 1 - | 0,000111434 - | 1 - | 0,000138368 - | 1 - | 0,000131536 - | 1 - | 0,000133479 - | 1 - | 0,000150364 - |
| 262 | 1 | 0.2um_NOKCPHJN_36762 | Viruses | unclassified | unclassified | unclassified | various | various | various | unclassified | unclassified | unclassified | unclassified | unclassified | 2,2 | 1,6 | 5,68E-05 | 4,00E-05 | 4 - | 8,97E-05 - | 4 - | 0,000111343 - | 2 - | 5,29E-05 - | 0 - | 0 - | 1 - | 3,02E-05 - |
| 263 | 1 | 3um_FIJJAFBF_767292 | Bacteria | Proteobacteri | Gammaprotec | unclassified | unclassified | unclassified | Gammaprotec | information st | Translation, ri | 0QSAC | Binds togethe | 314287.GB22 | 0,8 | 0,748331477 | 6,75E-05 | 5,96E-05 | 2 - | 0,000154063 - | 0 - | 0 - | 1 - | 9,09E-05 - | 1 - | 9,23E-05 - | 0 - | 0 - |
| 264 | 1 | 0.2um_NOKCPHJN_43639 | Bacteria | Proteobacteri | Gammaprotec | Oceanospirilla | Saccharospirill | Reinekea | Reinekea fors | unclassified | unclassified | unclassified | unclassified | unclassified | 1,4 | 0,8 | 4,07E-05 | 1,80E-05 | 3 - | 7,65E-05 - | 1 - | 3,17E-05 - | 1 - | 3,01E-05 - | 1 - | 3,06E-05 - | 1 - | 3,44E-05 - |
| 265 | 1 | 0.2um_NOKCPHJN_132364 | Bacteria | Proteobacteri | Gammaprotec | Cellvibrionales | Halieaceae | unclassified | marine gamm | unclassified | unclassified | unclassified | unclassified | unclassified | 0,4 | 0,489897949 | 3,16E-05 | 3,87E-05 | 0 - | 0 - | 0 - | 0 - | 1 - | 7,83E-05 - | 1 - | 7,95E-05 - | 0 - | 0 - |
| 266 | 1 | 3um_FIJJAFBF_634173 | Bacteria | Actinobacteri | Actinobacteri | Micrococcales | Microbacteria | Candidatus Ac | Candidatus Ac | information st | Transcription | 05VEQ | Cold shock pr | 479433.Caci_ | 1 | 0 | 0,000204518 | 1,94E-05 | 1 - | 0,000171309 - | 1 - | 0,000212715 - | 1 - | 0,000202212 - | 1 - | 0,000205199 - | 1 - | 0,000231156 - |
| 267 | 1 | 3um_FIJJAFBF_27521 | Bacteria | Proteobacteri | Alphaproteob | Rhodobactera | unclassified | unclassified | Rhodobactera | metabolism | Amino acid tra | COG0131 | imidazoleglyc | 367336.OM22 | 1,6 | 0,489897949 | 0,000110687 | 3,48E-05 | 2 - | 0,000116525 - | 1 - | 7,23E-05 - | 2 - | 0,000137545 - | 1 - | 6,98E-05 - | 2 - | 0,000157233 - |
| 268 | 1 | 0.2um_NOKCPHJN_266367 | Bacteria | Bacteroidetes | Flavobacteriia | Flavobacterial | Flavobacteriac | Polaribacter | Polaribacter s | unclassified | unclassified | unclassified | unclassified | unclassified | 1 | 0 | 1,50E-05 | 1,42E-06 | 1 - | 1,25E-05 - | 1 - | 1,56E-05 - | 1 - | 1,48E-05 - | 1 - | 1,50E-05 - | 1 - | 1,69E-05 - |
| 269 | 1 | 3um_FIJJAFBF_475146 | Bacteria | Proteobacteri | Gammaprotec | Oceanospirilla | Saccharospirill | Reinekea | Reinekea fors | poorly charact | Function unkn | 0QQAI | Dihydroorotat | 314283.MED2 | 2 | 0 | 0,000111859 | 1,06E-05 | 2 - | 9,37E-05 - | 2 - | 0,000116342 - | 2 - | 0,000110598 - | 2 - | 0,000112231 - | 2 - | 0,000126428 - |
| 270 | 1 | 3um_FIJJAFBF_78388 | Bacteria | Bacteroidetes | Flavobacteriia | Flavobacterial | Flavobacteriac | Polaribacter | Polaribacter a | information st | Translation, ri | COG0216 | Peptide chain | 983544.Lacal_ | 1,2 | 0,4 | 4,47E-05 | 9,90E-06 | 2 - | 6,41E-05 - | 1 - | 3,98E-05 - | 1 - | 3,78E-05 - | 1 - | 3,84E-05 - | 1 - | 4,33E-05 - |
| 271 | 1 | 3um_FIJJAFBF_432135 | Bacteria | Proteobacteri | Alphaproteob | Rhodobactera | unclassified | unclassified | Rhodobactera | poorly charact | Function unkn | COG3181 | extra-cytoplas | 367336.OM22 | 1 | 1,095445115 | 3,97E-05 | 4,48E-05 | 1 - | 3,46E-05 - | 0 - | 0 - | 3 - | 0,000122424 - | 1 - | 4,14E-05 - | 0 - | 0 - |
| 272 | 1 | 3um_FIJJAFBF_304843 | Bacteria | Proteobacteri | Alphaproteob | Rhodobactera | unclassified | unclassified | Rhodobactera | poorly charact | Function unkn | 05DIH | basic membra | 246200.SPO2 | 0,2 | 0,4 | 6,96E-06 | 1,39E-05 | 1 - | 3,48E-05 - | 0 - | 0 - | 0 - | 0 - | 0 - | 0 - | 0 - | 0 - |
| 273 | 1 | 3um_FIJJAFBF_189352 | Bacteria | Proteobacteri | Gammaprotec | Cellvibrionales | Porticoccacea | unclassified | SAR92 bacteri | unclassified | unclassified | unclassified | unclassified | unclassified | 1,8 | 0,4 | 6,43E-05 | 1,60E-05 | 2 - | 5,98E-05 - | 2 - | 7,42E-05 - | 1 - | 3,53E-05 - | 2 - | 7,16E-05 - | 2 - | 8,07E-05 - |
| 274 | 1 | 0.2um_NOKCPHJN_1110943 | Bacteria | Proteobacteri | Alphaproteob | Pelagibacteral | Pelagibacterac | Candidatus Pe | Candidatus Pe | information st | Translation, ri | 08VZM | 50s ribosomal | 335992.SAR11 | 2,2 | 0,4 | 0,000182652 | 4,91E-05 | 2 - | 0,000137457 - | 2 - | 0,000170681 - | 2 - | 0,000162254 - | 2 - | 0,000164651 - | 3 - | 0,000278218 - |
| 275 | 1 | 0.2um_NOKCPHJN_158468 | Bacteria | Bacteroidetes | unclassified | unclassified | unclassified | unclassified | Bacteroidetes | cellular proces | Posttranslatio | 09GE9 | Thioredoxin | 762903.Pedsa | 1,2 | 0,748331477 | 0,000110996 | 6,69E-05 | 2 - | 0,000162804 - | 1 - | 0,000101077 - | 1 - | 9,61E-05 - | 2 - | 0,000195012 - | 0 - | 0 - |
| 276 | 1 | 3um_FIJJAFBF_438692 | Eukaryota | Bacillariophyt | Coscinodiscop | Thalassiosirale | Thalassiosirac | Thalassiosira | Thalassiosira | unclassified | unclassified | unclassified | unclassified | unclassified | 1,6 | 0,8 | 0,000119338 | 4,73E-05 | 3 - | 0,000195642 - | 1 - | 8,10E-05 - | 2 - | 0,000153957 - | 1 - | 7,81E-05 - | 1 - | 8,80E-05 - |
| 277 | 1 | 3um_FIJJAFBF_59204 | Bacteria | unclassified | unclassified | unclassified | unclassified | unclassified | uncultured ba | poorly charact | Function unkn | 11Q9F | Protein of unk | 313594.PI23P | 2,6 | 0,8 | 0,000191914 | 5,37E-05 | 3 - | 0,000188159 - | 2 - | 0,000155758 - | 4 - | 0,000296136 - | 2 - | 0,000150255 - | 2 - | 0,000169262 - |
| 278 | 1 | 3um_FIJJAFBF_130974 | unclassified | unclassified | unclassified | unclassified | unclassified | unclassified | unclassified | unclassified | unclassified | unclassified | unclassified | unclassified | 0,6 | 0,489897949 | 9,55E-05 | 7,80E-05 | 0 - | 0 - | 1 - | 0,000163815 - | 1 - | 0,000155727 - | 1 - | 0,000158027 - | 0 - | 0 - |
| 279 | 1 | 0.2um_NOKCPHJN_820029 | Bacteria | Proteobacteri | Gammaprotec | Alteromonada | Colwelliaceae | Thalassomona | Thalassomona | unclassified | unclassified | unclassified | unclassified | unclassified | 4,4 | 0,489897949 | 0,000490295 | 3,75E-05 | 5 - | 0,000470397 - | 4 - | 0,000467275 - | 5 - | 0,000555255 - | 4 - | 0,000450765 - | 4 - | 0,000507785 - |
| 280 | 1 | 0.2um_NOKCPHJN_27636 | Bacteria | Actinobacteri | Actinobacteri | Micrococcales | Microbacteria | Candidatus Ac | Candidatus Ac | unclassified | unclassified | unclassified | unclassified | unclassified | 0,2 | 0,4 | 1,32E-05 | 2,64E-05 | 0 - | 0 - | 1 - | 6,60E-05 - | 0 - | 0 - | 0 - | 0 - | 0 - | 0 - |

| | | | | | | | | | | | | | | | | | | | | | | | | | | | | |
|---|---|---|---|---|---|---|---|---|---|---|---|---|---|---|---|---|---|---|---|---|---|---|---|---|---|---|---|---|
| 281 | 1 | 0.2um_NOKCPHJN_504803 | Bacteria | Bacteroidetes | Flavobacteriia | Flavobacterial | Flavobacteriac | Polaribacter | Polaribacter s | cellular proces | Posttranslatio | 05C2Z | ATP-depender | 886377.Murru | 0,6 | 0,489897949 | 1,23E-05 | 1,00E-05 | 0 - | 0 - | 1 - | 2,10E-05 - | 1 - | 2,00E-05 - | 1 - | 2,03E-05 - | 0 - | 0 - |
| 282 | 1 | 0.2um_NOKCPHJN_617369 | Eukaryota | Chlorophyta | Ulvophyceae | Ulvales | Ulvaceae | Ulva | Ulva linza | unclassified | unclassified | unclassified | unclassified | unclassified | 2 | 0 | 6,90E-05 | 6,55E-06 | 2 - | 5,78E-05 - | 2 - | 7,18E-05 - | 2 - | 6,83E-05 - | 2 - | 6,93E-05 - | 2 - | 7,80E-05 - |
| 283 | 1 | 3um_FIJJAFBF_43832 | Bacteria | Proteobacteri | Gammaprotec | Cellvibrionales | Porticoccacea | unclassified | SAR92 bacteri | cellular proces | Cell motility | 01W75 | Flagellin | 317655.Sala_ | 3 | 1,095445115 | 0,000143211 | 4,02E-05 | 5 - | 0,000204958 - | 2 - | 0,000101799 - | 3 - | 0,00014516 - | 2 - | 9,82E-05 - | 3 - | 0,000165937 - |
| 284 | 1 | 0.2um_NOKCPHJN_728405 | Bacteria | Proteobacteri | Alphaproteob | Rhizobiales | unclassified | unclassified | Rhizobiales ba | metabolism | Inorganic ion t | 05F28 | tonB-depende | 247634.GPB2 | 3,8 | 0,748331477 | 6,79E-05 | 1,73E-05 | 3 - | 4,44E-05 - | 5 - | 9,19E-05 - | 4 - | 6,99E-05 - | 3 - | 5,32E-05 - | 4 - | 7,99E-05 - |
| 285 | 1 | 3um_FIJJAFBF_355455 | Bacteria | Bacteroidetes | unclassified | unclassified | unclassified | unclassified | Bacteroidetes | cellular proces | Intracellular tr | 0BIEB | domain-conta | 1005048.CFU_ | 9,6 | 1,019803903 | 0,00017669 | 2,30E-05 | 10 - | 0,000154477 - | 11 - | 0,000210997 - | 10 - | 0,000182345 - | 8 - | 0,00014803 - | 9 - | 0,0001876 - |
| 286 | 1 | 0.2um_NOKCPHJN_12206 | Bacteria | Proteobacteri | Gammaprotec | Cellvibrionales | Porticoccacea | unclassified | gamma protec | information st | Translation, ri | 0QQXI | This protein is | 314287.GB22 | 0,2 | 0,4 | 1,95E-05 | 3,89E-05 | 1 - | 9,73E-05 - | 0 - | 0 - | 0 - | 0 - | 0 - | 0 - | 0 - | 0 - |
| 287 | 1 | 0.2um_NOKCPHJN_15304 | Bacteria | Proteobacteri | Betaproteoba | unclassified | unclassified | unclassified | Betaproteoba | unclassified | unclassified | unclassified | unclassified | unclassified | 2,8 | 0,748331477 | 3,92E-05 | 7,39E-06 | 4 - | 4,79E-05 - | 3 - | 4,46E-05 - | 3 - | 4,24E-05 - | 2 - | 2,87E-05 - | 2 - | 3,23E-05 - |
| 288 | 1 | 3um_FIJJAFBF_686422 | Bacteria | Proteobacteri | Alphaproteob | Rhodobactera | Rhodobactera | Amylibacter | Amylibacter k | metabolism | Carbohydrate | COG0057 | glyceraldehyd | 367336.OM22 | 0,6 | 0,489897949 | 2,45E-05 | 2,04E-05 | 1 - | 3,46E-05 - | 0 - | 0 - | 0 - | 0 - | 1 - | 4,14E-05 - | 1 - | 4,66E-05 - |
| 289 | 1 | 3um_FIJJAFBF_03426 | Bacteria | Bacteroidetes | Flavobacteriia | Flavobacterial | Flavobacteriac | Algibacter | Algibacter aqu | information st | Transcription | 05D27 | DNA-depende | 983544.Lacal_ | 8,2 | 1,166190379 | 7,79E-05 | 1,00E-05 | 9 - | 7,21E-05 - | 8 - | 7,96E-05 - | 7 - | 6,62E-05 - | 10 - | 9,59E-05 - | 7 - | 7,57E-05 - |
| 290 | 1 | 0.2um_NOKCPHJN_91499 | Bacteria | Proteobacteri | Gammaprotec | Oceanospirilla | Saccharospirill | Reinekea | Reinekea fors | information st | Translation, ri | 179ZI | One of the pri | 314283.MED2 | 6,4 | 1,854723699 | 0,00052598 | 0,000141659 | 8 - | 0,000556493 - | 5 - | 0,000431875 - | 4 - | 0,000328442 - | 9 - | 0,000749909 - | 6 - | 0,00056318 - |
| 291 | 1 | 3um_FIJJAFBF_289952 | Bacteria | Proteobacteri | Gammaprotec | Alteromonada | Alteromonada | Alteromonas | Alteromonas r | cellular proces | Intracellular tr | 08ZUW | Major pilin sul | 697282.Mettu | 2 | 0 | 0,00019859 | 1,89E-05 | 2 - | 0,000166343 - | 2 - | 0,000206549 - | 2 - | 0,000196351 - | 2 - | 0,000199251 - | 2 - | 0,000224456 - |
| 292 | 1 | 3um_FIJJAFBF_475145 | Bacteria | Proteobacteri | Gammaprotec | Oceanospirilla | Saccharospirill | Reinekea | Reinekea fors | unclassified | unclassified | unclassified | unclassified | unclassified | 2,6 | 1,019803903 | 5,72E-05 | 2,20E-05 | 3 - | 5,64E-05 - | 3 - | 7,00E-05 - | 2 - | 4,43E-05 - | 4 - | 9,00E-05 - | 1 - | 2,53E-05 - |
| 293 | 1 | 0.2um_NOKCPHJN_241609 | Bacteria | Proteobacteri | Alphaproteob | Pelagibacteral | Pelagibacterac | Candidatus Pe | Candidatus Pe | information st | Transcription | 05D27 | DNA-depende | 335992.SAR11 | 10,2 | 1,720465053 | 0,000100584 | 1,99E-05 | 10 - | 8,26E-05 - | 13 - | 0,000133291 - | 8 - | 7,80E-05 - | 11 - | 0,0001088 - | 9 - | 0,000100278 - |
| 294 | 1 | 3um_FIJJAFBF_325820 | Bacteria | Proteobacteri | Alphaproteob | Rhodobactera | Rhodobactera | Thalassococcu | Thalassococcu | metabolism | Carbohydrate | 01RFA | ABC transport | 633131.TR2A | 0,4 | 0,489897949 | 3,48E-05 | 4,28E-05 | 1 - | 7,97E-05 - | 0 - | 0 - | 1 - | 9,41E-05 - | 0 - | 0 - | 0 - | 0 - |
| 295 | 1 | 0.2um_NOKCPHJN_334894 | Bacteria | Bacteroidetes | Flavobacteriia | Flavobacterial | Flavobacteriac | Formosa | Formosa sp. H | unclassified | unclassified | unclassified | unclassified | unclassified | 5 | 0,632455532 | 0,000390567 | 3,74E-05 | 6 - | 0,000395782 - | 4 - | 0,00032763 - | 5 - | 0,000389317 - | 5 - | 0,000395067 - | 5 - | 0,000445042 - |
| 296 | 1 | 3um_FIJJAFBF_473687 | Bacteria | Proteobacteri | Alphaproteob | Rhodobactera | unclassified | unclassified | Rhodobactera | metabolism | Inorganic ion t | 16PII | TRAP transpor | 367336.OM22 | 2,2 | 0,4 | 9,02E-05 | 1,76E-05 | 2 - | 6,87E-05 - | 2 - | 8,53E-05 - | 3 - | 0,000121691 - | 2 - | 8,23E-05 - | 2 - | 9,27E-05 - |
| 297 | 1 | 3um_FIJJAFBF_201377 | Bacteria | Bacteroidetes | Flavobacteriia | Flavobacterial | Flavobacteriac | Polaribacter | Polaribacter s | unclassified | unclassified | unclassified | unclassified | unclassified | 18 | 2,607680962 | 0,000213212 | 3,30E-05 | 18 - | 0,000179339 - | 22 - | 0,000272171 - | 19 - | 0,000223452 - | 17 - | 0,000202883 - | 14 - | 0,000188216 - |
| 298 | 2 | 3um_FIJJAFBF_218671;3um_FIJJAFBF_501599 | Bacteria | Bacteroidetes | Flavobacteriia | Flavobacterial | Flavobacteriac | unclassified | Flavobacteriac | unclassified | unclassified | unclassified | unclassified | unclassified | 0,6 | 0,489897949 | 3,70E-05 | 3,07E-05 | 1 - | 5,15E-05 - | 1 - | 6,39E-05 - | 0 - | 0 - | 0 - | 0 - | 1 - | 6,95E-05 - |
| 299 | 1 | 3um_FIJJAFBF_238496 | Bacteria | Bacteroidetes | Flavobacteriia | Flavobacterial | Flavobacteriac | Flaviramulus | Flaviramulus b | metabolism | Carbohydrate | 099YR | Catalyzes the | 391603.FBALC | 2,8 | 0,748331477 | 8,83E-05 | 2,22E-05 | 3 - | 8,01E-05 - | 3 - | 9,94E-05 - | 4 - | 0,00012603 - | 2 - | 6,39E-05 - | 2 - | 7,20E-05 - |
| 300 | 1 | 0.2um_NOKCPHJN_152947 | Bacteria | Proteobacteri | Alphaproteob | Rhodobactera | Rhodobactera | Planktotalea | Planktotalea f | unclassified | unclassified | unclassified | unclassified | unclassified | 0,2 | 0,4 | 1,20E-05 | 2,39E-05 | 0 - | 0 - | 0 - | 0 - | 0 - | 0 - | 0 - | 0 - | 1 - | 5,98E-05 - |
| 301 | 1 | 3um_FIJJAFBF_53317 | Bacteria | Bacteroidetes | Flavobacteriia | Flavobacterial | Flavobacteriac | Polaribacter | Polaribacter s | unclassified | unclassified | unclassified | unclassified | unclassified | 0,2 | 0,4 | 8,21E-06 | 1,64E-05 | 0 - | 0 - | 0 - | 0 - | 0 - | 0 - | 1 - | 4,10E-05 - | 0 - | 0 - |
| 302 | 1 | 0.2um_NOKCPHJN_11098 | Bacteria | Bacteroidetes | Flavobacteriia | Flavobacterial | Crocinitomica | unclassified | Crocinitomica | unclassified | unclassified | unclassified | unclassified | unclassified | 0,2 | 0,4 | 1,55E-05 | 3,10E-05 | 1 - | 7,76E-05 - | 0 - | 0 - | 0 - | 0 - | 0 - | 0 - | 0 - | 0 - |
| 303 | 1 | 3um_FIJJAFBF_87924 | Bacteria | Bacteroidetes | Flavobacteriia | Flavobacterial | Crocinitomica | unclassified | Crocinitomica | unclassified | unclassified | unclassified | unclassified | unclassified | 0,6 | 0,489897949 | 1,36E-05 | 1,11E-05 | 0 - | 0 - | 1 - | 2,23E-05 - | 0 - | 0 - | 1 - | 2,15E-05 - | 1 - | 2,42E-05 - |
| 304 | 2 | 0.2um_NOKCPHJN_602562;0.2um_NOKCPHJN_6 | Bacteria | Proteobacteri | Alphaproteob | Pelagibacteral | Pelagibacterac | Candidatus Pe | various | metabolism | Amino acid tra | 01QVA | Part of the AB | 439493.PB721 | 1 | 0 | 3,80E-05 | 3,60E-06 | 1 - | 3,18E-05 - | 1 - | 3,95E-05 - | 1 - | 3,75E-05 - | 1 - | 3,81E-05 - | 1 - | 4,29E-05 - |
| 305 | 1 | 3um_FIJJAFBF_563929 | Bacteria | Proteobacteri | Gammaprotec | Oceanospirilla | Saccharospirill | Gynuella | Gynuella suns | metabolism | Amino acid tra | 05EZ7 | amino acid AB | 999141.GME_ | 0,8 | 0,748331477 | 2,58E-05 | 2,26E-05 | 2 - | 5,81E-05 - | 1 - | 3,61E-05 - | 0 - | 0 - | 1 - | 3,48E-05 - | 0 - | 0 - |
| 306 | 1 | 3um_FIJJAFBF_423475 | Eukaryota | Bacillariophyt | Coscinodiscop | Thalassiosirale | Thalassiosirac | Thalassiosira | Thalassiosira | unclassified | unclassified | unclassified | unclassified | unclassified | 2,2 | 0,4 | 4,58E-05 | 4,20E-06 | 3 - | 5,31E-05 - | 2 - | 4,40E-05 - | 2 - | 4,18E-05 - | 2 - | 4,24E-05 - | 2 - | 4,78E-05 - |
| 307 | 1 | 0.2um_NOKCPHJN_198723 | Eukaryota | unclassified | unclassified | Dictyosteliales | Dictyosteliace | Dictyostelium | Dictyostelium | metabolism | Lipid transpor | COG2267 | alpha beta hy | 44689.DDB02 | 1 | 0 | 5,25E-05 | 4,98E-06 | 1 - | 4,40E-05 - | 1 - | 5,46E-05 - | 1 - | 5,19E-05 - | 1 - | 5,27E-05 - | 1 - | 5,93E-05 - |
| 308 | 1 | 0.2um_NOKCPHJN_21340 | Bacteria | Proteobacteri | Alphaproteob | Rhodospirillal | unclassified | unclassified | Rhodospirillal | information st | Translation, ri | COG1185 | Involved in m | 1064539.AZO | 0,2 | 0,4 | 3,11E-06 | 6,22E-06 | 1 - | 1,56E-05 - | 0 - | 0 - | 0 - | 0 - | 0 - | 0 - | 0 - | 0 - |
| 309 | 1 | 3um_FIJJAFBF_663803 | Bacteria | Bacteroidetes | Flavobacteriia | Flavobacterial | Flavobacteriac | Polaribacter | unclassified | information st | Replication, re | 0PB0H | DNA-binding | 313598.MED1 | 1,6 | 0,489897949 | 0,000242282 | 7,62E-05 | 2 - | 0,000255059 - | 1 - | 0,000158354 - | 2 - | 0,000301072 - | 1 - | 0,000152759 - | 2 - | 0,000344166 - |
| 310 | 1 | 3um_FIJJAFBF_257111 | Eukaryota | Bacillariophyt | Coscinodiscop | Leptocylindral | Leptocylindrac | Leptocylindru | Leptocylindru | information st | Translation, ri | COG0087 | One of the pri | 489825.LYNG | 6,2 | 0,4 | 0,000387972 | 4,40E-05 | 6 - | 0,000314457 - | 6 - | 0,000390463 - | 6 - | 0,000371184 - | 7 - | 0,000439444 - | 6 - | 0,000424314 - |
| 311 | 1 | 3um_FIJJAFBF_310912 | Bacteria | Proteobacteri | Gammaprotec | Chromatiales | Chromatiacea | Candidatus Te | Candidatus Te | cellular proces | Cell wall/mem | 0GRGC | Protein of unk | 96561.Dole_2 | 0,8 | 0,4 | 4,47E-05 | 2,27E-05 | 1 - | 4,84E-05 - | 1 - | 6,01E-05 - | 1 - | 5,72E-05 - | 1 - | 5,80E-05 - | 0 - | 0 - |
| 312 | 1 | 3um_FIJJAFBF_09283 | unclassified | unclassified | unclassified | unclassified | unclassified | unclassified | unclassified | unclassified | unclassified | unclassified | unclassified | unclassified | 0,8 | 0,4 | 6,91E-05 | 3,55E-05 | 1 - | 7,22E-05 - | 1 - | 8,96E-05 - | 0 - | 0 - | 1 - | 8,65E-05 - | 1 - | 9,74E-05 - |
| 313 | 1 | 3um_FIJJAFBF_189717 | Bacteria | Bacteroidetes | Flavobacteriia | Flavobacterial | Flavobacteriac | Polaribacter | Polaribacter s | metabolism | Nucleotide tra | 0P87W | Catalyzes the | 313598.MED1 | 0,2 | 0,4 | 1,08E-05 | 2,16E-05 | 1 - | 5,39E-05 - | 0 - | 0 - | 0 - | 0 - | 0 - | 0 - | 0 - | 0 - |
| 314 | 1 | 3um_FIJJAFBF_77806 | Bacteria | Proteobacteri | Alphaproteob | Rhodobactera | unclassified | unclassified | Rhodobactera | unclassified | unclassified | unclassified | unclassified | unclassified | 5,2 | 0,4 | 0,000595034 | 2,95E-05 | 6 - | 0,000578706 - | 5 - | 0,000598819 - | 5 - | 0,000569253 - | 5 - | 0,000577661 - | 5 - | 0,000650733 - |
| 315 | 1 | 0.2um_NOKCPHJN_99876 | Bacteria | Proteobacteri | Gammaprotec | unclassified | unclassified | Thiohalobacte | Thiohalobacte | unclassified | unclassified | unclassified | unclassified | unclassified | 1,6 | 1,2 | 5,73E-05 | 4,13E-05 | 3 - | 9,56E-05 - | 1 - | 3,96E-05 - | 3 - | 0,000112902 - | 1 - | 3,82E-05 - | 0 - | 0 - |
| 316 | 1 | 0.2um_NOKCPHJN_168209 | Bacteria | Bacteroidetes | Flavobacteriia | Flavobacterial | Flavobacteriac | Maribacter | Maribacter fo | unclassified | unclassified | unclassified | unclassified | unclassified | 1 | 0,632455532 | 2,17E-05 | 1,38E-05 | 1 - | 1,82E-05 - | 1 - | 2,27E-05 - | 2 - | 4,31E-05 - | 0 - | 0 - | 1 - | 2,46E-05 - |
| 317 | 1 | 3um_FIJJAFBF_520516 | Bacteria | Bacteroidetes | Flavobacteriia | Flavobacterial | Flavobacteriac | Polaribacter | Polaribacter s | metabolism | Amino acid tra | COG0137 | Citrulline--asp | 313594.PI23P | 1,2 | 0,979795897 | 4,17E-05 | 3,62E-05 | 1 - | 2,94E-05 - | 3 - | 0,00010935 - | 1 - | 3,47E-05 - | 1 - | 3,52E-05 - | 0 - | 0 - |
| 318 | 1 | 3um_FIJJAFBF_105182 | Bacteria | Bacteroidetes | Flavobacteriia | Flavobacterial | Flavobacteriac | Polaribacter | Polaribacter s | unclassified | unclassified | unclassified | unclassified | unclassified | 1 | 0 | 2,57E-05 | 2,44E-06 | 1 - | 2,15E-05 - | 1 - | 2,67E-05 - | 1 - | 2,54E-05 - | 1 - | 2,57E-05 - | 1 - | 2,90E-05 - |
| 319 | 10 | 0.2um_NOKCPHJN_21010;0.2um_NOKCPHJN_26 | Bacteria | various | various | various | various | various | various | poorly charact | Function unkn | 0PAEZ | Inherit from b | 391598.FBBAL | 0,6 | 0,489897949 | 1,27E-05 | 1,05E-05 | 1 - | 1,88E-05 - | 0 - | 0 - | 1 - | 2,22E-05 - | 1 - | 2,25E-05 - | 0 - | 0 - |
| 320 | 1 | 0.2um_NOKCPHJN_489734 | Bacteria | Proteobacteri | Alphaproteob | Pelagibacteral | Pelagibacterac | Candidatus Pe | Candidatus Pe | cellular proces | Cell wall/mem | 7257 | NA | 335992.SAR11 | 0,8 | 0,4 | 3,24E-05 | 1,65E-05 | 1 - | 3,51E-05 - | 1 - | 4,36E-05 - | 1 - | 4,14E-05 - | 1 - | 4,20E-05 - | 0 - | 0 - |
| 321 | 1 | 3um_FIJJAFBF_341042 | Bacteria | Proteobacteri | Gammaprotec | Pseudomonad | Moraxellaceae | unclassified | Moraxellaceae | unclassified | unclassified | unclassified | unclassified | unclassified | 1,6 | 0,8 | 9,77E-05 | 4,25E-05 | 3 - | 0,000157228 - | 1 - | 6,51E-05 - | 1 - | 6,19E-05 - | 1 - | 6,28E-05 - | 2 - | 0,000141438 - |
| 322 | 1 | 3um_FIJJAFBF_227243 | Bacteria | Proteobacteri | Gammaprotec | Oceanospirilla | Oceanospirilla | Thalassolituus | Thalassolituus | unclassified | unclassified | unclassified | unclassified | unclassified | 0,6 | 0,489897949 | 2,75E-05 | 2,25E-05 | 0 - | 0 - | 1 - | 4,51E-05 - | 0 - | 0 - | 1 - | 4,35E-05 - | 1 - | 4,90E-05 - |
| 323 | 1 | 3um_FIJJAFBF_279332 | Bacteria | Bacteroidetes | Flavobacteriia | unclassified | unclassified | unclassified | Flavobacteria | unclassified | unclassified | unclassified | unclassified | unclassified | 0,2 | 0,4 | 2,05E-05 | 4,10E-05 | 0 - | 0 - | 0 - | 0 - | 0 - | 0 - | 0 - | 0 - | 1 - | 0,000102566 - |
| 324 | 1 | 0.2um_NOKCPHJN_194064 | Bacteria | Bacteroidetes | Flavobacteriia | Flavobacterial | Flavobacteriac | Formosa | Formosa sp. H | unclassified | unclassified | unclassified | unclassified | unclassified | 2,4 | 1,356465997 | 0,000171999 | 0,000106368 | 2 - | 0,00011772 - | 1 - | 7,31E-05 - | 1 - | 6,95E-05 - | 4 - | 0,000282017 - | 4 - | 0,000317691 - |
| 325 | 1 | 3um_FIJJAFBF_387539 | Bacteria | Bacteroidetes | unclassified | unclassified | unclassified | unclassified | Bacteroidetes | information st | Translation, ri | 08UHY | One of the pri | 553177.CAPS | 0,6 | 0,489897949 | 5,32E-05 | 4,42E-05 | 1 - | 7,40E-05 - | 1 - | 9,19E-05 - | 0 - | 0 - | 0 - | 0 - | 1 - | 9,99E-05 - |
| 326 | 1 | 3um_FIJJAFBF_713338 | Bacteria | Proteobacteri | Alphaproteob | Rhodobactera | Rhodobactera | Phaeobacter | Phaeobacter g | metabolism | Energy produc | 01R4M | Acetoacetyl-C | 388739.RSK20 | 2,8 | 1,326649916 | 0,000156175 | 7,23E-05 | 3 - | 0,000143471 - | 2 - | 0,000118766 - | 5 - | 0,000282255 - | 3 - | 0,000171854 - | 1 - | 6,45E-05 - |
| 327 | 1 | 0.2um_NOKCPHJN_190079 | Bacteria | Bacteroidetes | Flavobacteriia | Flavobacterial | Flavobacteriac | Polaribacter | Polaribacter r | unclassified | unclassified | unclassified | unclassified | unclassified | 3 | 1,095445115 | 3,87E-05 | 9,92E-06 | 5 - | 5,59E-05 - | 2 - | 2,78E-05 - | 3 - | 3,96E-05 - | 3 - | 4,02E-05 - | 2 - | 3,02E-05 - |
| 328 | 1 | 0.2um_NOKCPHJN_664587 | Eukaryota | Cnidaria | Anthozoa | Scleractinia | Pocilloporidae | Stylophora | Stylophora pis | unclassified | unclassified | unclassified | unclassified | unclassified | 0,6 | 0,8 | 6,96E-05 | 9,55E-05 | 0 - | 0 - | 0 - | 0 - | 1 - | 0,000105845 - | 0 - | 0 - | 2 - | 0,000241991 - |
| 329 | 1 | 3um_FIJJAFBF_131278 | Bacteria | Bacteroidetes | unclassified | unclassified | unclassified | unclassified | Bacteroidetes | metabolism | Carbohydrate | 05CA9 | Pyruvate kinas | 1046627.BZAF | 0,4 | 0,489897949 | 1,05E-05 | 1,29E-05 | 1 - | 2,39E-05 - | 0 - | 0 - | 0 - | 0 - | 1 - | 2,86E-05 - | 0 - | 0 - |
| 330 | 1 | 0.2um_NOKCPHJN_1062283 | Eukaryota | unclassified | Oomycetes | Peronosporale | Peronosporac | Phytophthora | Phytophthora | unclassified | unclassified | unclassified | unclassified | unclassified | 6 | 0,894427191 | 0,000166204 | 1,59E-05 | 7 - | 0,000164302 - | 5 - | 0,000145725 - | 7 - | 0,000193942 - | 6 - | 0,000168691 - | 5 - | 0,000158358 - |
| 331 | 1 | 0.2um_NOKCPHJN_61952 | Bacteria | Bacteroidetes | Flavobacteriia | Flavobacterial | Flavobacteriac | Formosa | Formosa sp. H | poorly charact | Function unkn | 0XRCJ | NA | 391587.KAOT | 4 | 1,095445115 | 5,13E-05 | 9,23E-06 | 6 - | 6,59E-05 - | 4 - | 5,46E-05 - | 3 - | 3,89E-05 - | 4 - | 5,26E-05 - | 3 - | 4,45E-05 - |
| 332 | 1 | 3um_FIJJAFBF_63530 | Bacteria | Bacteroidetes | unclassified | unclassified | unclassified | unclassified | Bacteroidetes | metabolism | Amino acid tra | 09GJ6 | nitrogen regul | 688270.Celal_ | 0,6 | 0,489897949 | 0,000133808 | 0,0001096 | 0 - | 0 - | 1 - | 0,00021926 - | 0 - | 0 - | 1 - | 0,000211513 - | 1 - | 0,000238268 - |
| 333 | 1 | 3um_FIJJAFBF_608458 | Bacteria | Bacteroidetes | Flavobacteriia | Flavobacterial | Flavobacteriac | Tenacibaculur | Tenacibaculur | unclassified | unclassified | unclassified | unclassified | unclassified | 0,6 | 0,8 | 3,17E-05 | 4,05E-05 | 2 - | 9,77E-05 - | 1 - | 6,06E-05 - | 0 - | 0 - | 0 - | 0 - | 0 - | 0 - |
| 334 | 1 | 3um_FIJJAFBF_747819 | Bacteria | Bacteroidetes | unclassified | unclassified | unclassified | unclassified | Bacteroidetes | unclassified | unclassified | unclassified | unclassified | unclassified | 0,2 | 0,4 | 6,62E-06 | 1,32E-05 | 1 - | 3,31E-05 - | 0 - | 0 - | 0 - | 0 - | 0 - | 0 - | 0 - | 0 - |
| 335 | 1 | 0.2um_NOKCPHJN_394976 | Bacteria | Proteobacteri | Gammaprotec | Cellvibrionales | Porticoccacea | unclassified | SAR92 bacteri | metabolism | Inorganic ion t | COG0175 | Reduction of a | 207954.MED9 | 1 | 0 | 2,73E-05 | 2,59E-06 | 1 - | 2,29E-05 - | 1 - | 2,84E-05 - | 1 - | 2,70E-05 - | 1 - | 2,74E-05 - | 1 - | 3,09E-05 - |
| 336 | 1 | 3um_FIJJAFBF_357557 | Bacteria | Bacteroidetes | Flavobacteriia | Flavobacterial | Flavobacteriac | Polaribacter | Polaribacter s | poorly charact | Function unkn | 0XQTX | Chromosome | 313594.PI23P | 2,4 | 0,489897949 | 3,22E-05 | 7,59E-06 | 2 - | 2,24E-05 - | 3 - | 4,17E-05 - | 2 - | 2,64E-05 - | 3 - | 4,02E-05 - | 2 - | 3,02E-05 - |
| 337 | 1 | 3um_FIJJAFBF_45923 | Bacteria | Bacteroidetes | Flavobacteriia | Flavobacterial | Flavobacteriac | Polaribacter | Polaribacter ir | metabolism | Energy produc | 05C26 | Dehydrogenas | 313594.PI23P | 1,2 | 0,4 | 3,02E-05 | 6,69E-06 | 2 - | 4,33E-05 - | 1 - | 2,69E-05 - | 1 - | 2,56E-05 - | 1 - | 2,59E-05 - | 1 - | 2,92E-05 - |
| 338 | 1 | 3um_FIJJAFBF_210816 | Bacteria | Proteobacteri | Gammaprotec | Oceanospirilla | Saccharospirill | Reinekea | Reinekea fors | unclassified | unclassified | unclassified | unclassified | unclassified | 0,8 | 0,748331477 | 7,79E-05 | 6,89E-05 | 2 - | 0,000177948 - | 0 - | 0 - | 1 - | 0,000105025 - | 1 - | 0,000106576 - | 0 - | 0 - |
| 339 | 1 | 0.2um_NOKCPHJN_60394 | Bacteria | Proteobacteri | Gammaprotec | Oceanospirilla | Oceanospirilla | Motiliproteus | Motiliproteus | unclassified | unclassified | unclassified | unclassified | unclassified | 5,6 | 1,019803903 | 0,000424924 | 7,14E-05 | 6 - | 0,000384727 - | 6 - | 0,000477717 - | 5 - | 0,000378442 - | 7 - | 0,000537644 - | 4 - | 0,000346088 - |
| 340 | 1 | 0.2um_NOKCPHJN_198752 | Bacteria | Proteobacteri | Alphaproteob | Rhodobactera | Rhodobactera | unclassified | Rhodobactera | metabolism | Carbohydrate | 01U6R | trap dicarboxy | 488538.SAR11 | 2 | 0,894427191 | 7,87E-05 | 3,89E-05 | 2 - | 6,52E-05 - | 1 - | 4,05E-05 - | 1 - | 3,85E-05 - | 3 - | 0,000117173 - | 3 - | 0,000131995 - |
| 341 | 1 | 3um_FIJJAFBF_357541 | Bacteria | Bacteroidetes | Flavobacteriia | Flavobacterial | Flavobacteriac | Polaribacter | Polaribacter s | metabolism | Amino acid tra | 05C6M | Alpha-keto-be | 313594.PI23P | 12,8 | 0,748331477 | 0,000355362 | 1,71E-05 | 14 - | 0,000327266 - | 12 - | 0,000348315 - | 13 - | 0,000358711 - | 13 - | 0,000364009 - | 12 - | 0,000378512 - |
| 342 | 1 | 0.2um_NOKCPHJN_186703 | Bacteria | Proteobacteri | Alphaproteob | unclassified | unclassified | unclassified | Alphaproteob | metabolism | Inorganic ion t | 05PWN | phosphate ab | 488538.SAR11 | 2,6 | 1,019803903 | 0,000100889 | 3,38E-05 | 4 - | 0,000133075 - | 3 - | 0,000123929 - | 3 - | 0,000117811 - | 1 - | 3,99E-05 - | 2 - | 8,98E-05 - |
| 343 | 1 | 0.2um_NOKCPHJN_765989 | Bacteria | Proteobacteri | Alphaproteob | Rhodobactera | Rhodobactera | unclassified | Rhodobactera | cellular proces | Cell wall/mem | 01XF2 | NA | 391626.OA30 | 0,6 | 0,489897949 | 1,89E-05 | 1,56E-05 | 1 - | 2,75E-05 - | 1 - | 3,41E-05 - | 0 - | 0 - | 1 - | 3,29E-05 - | 0 - | 0 - |
| 344 | 1 | 3um_FIJJAFBF_318795 | Viruses | unclassified | unclassified | Phycodnavirid | unclassified | unclassified | Aureococcus | unclassified | unclassified | unclassified | unclassified | unclassified | 3,4 | 0,8 | 8,59E-05 | 2,15E-05 | 4 - | 8,49E-05 - | 3 - | 7,90E-05 - | 4 - | 0,000100172 - | 2 - | 5,08E-05 - | 4 - | 0,00011451 - |
| 345 | 1 | 3um_FIJJAFBF_612093 | Bacteria | Proteobacteri | Gammaprotec | Oceanospirilla | Saccharospirill | Reinekea | Reinekea blan | cellular proces | Posttranslatio | 0QJ3C | Acts as a chap | 491952.Mar18 | 13,4 | 1,624807681 | 0,000286793 | 4,28E-05 | 15 - | 0,000269429 - | 11 - | 0,000245338 - | 12 - | 0,000254427 - | 14 - | 0,000301215 - | 15 - | 0,000363555 - |
| 346 | 1 | 3um_FIJJAFBF_375383 | Bacteria | Bacteroidetes | Flavobacteriia | Flavobacterial | Flavobacteriac | Formosa | Formosa sp. H | information st | Transcription | 05D27 | DNA-depende | 391603.FBALC | 0,6 | 0,489897949 | 1,01E-05 | 8,31E-06 | 1 - | 1,46E-05 - | 1 - | 1,82E-05 - | 0 - | 0 - | 1 - | 1,75E-05 - | 0 - | 0 - |
| 347 | 2 | 0.2um_NOKCPHJN_565212;3um_FIJJAFBF_82647 | Bacteria | Verrucomicrol | Verrucomicrol | various | various | various | various | poorly charact | Function unkn | 056FM | YitL protein | 585543.HMPR | 0,2 | 0,4 | 1,00E-05 | 2,01E-05 | 0 - | 0 - | 1 - | 5,02E-05 - | 0 - | 0 - | 0 - | 0 - | 0 - | 0 - |
| 348 | 1 | 3um_FIJJAFBF_19382 | Bacteria | Proteobacteri | Alphaproteob | Rhodobactera | Rhodobactera | Ruegeria | Ruegeria med | information st | Replication, re | COG0187 | DNA gyrase ne | 314270.RB208 | 1,4 | 0,489897949 | 2,42E-05 | 9,85E-06 | 1 - | 1,43E-05 - | 1 - | 1,77E-05 - | 2 - | 3,37E-05 - | 1 - | 1,71E-05 - | 2 - | 3,85E-05 - |
| 349 | 1 | 0.2um_NOKCPHJN_207190 | Bacteria | Proteobacteri | Gammaprotec | Oceanospirilla | Saccharospirill | Reinekea | Reinekea fors | unclassified | unclassified | unclassified | unclassified | unclassified | 6,8 | 2,315167381 | 0,000208019 | 5,21E-05 | 11 - | 0,000288912 - | 6 - | 0,000195678 - | 7 - | 0,00021702 - | 4 - | 0,000125843 - | 6 - | 0,000212642 - |
| 350 | 1 | 0.2um_NOKCPHJN_40909 | Bacteria | Proteobacteri | Gammaprotec | Cellvibrionales | Halieaceae | unclassified | marine gamm | metabolism | Inorganic ion t | 16TQC | TONB-depend | 247639.MGP2 | 3,6 | 0,8 | 4,91E-05 | 6,44E-06 | 5 - | 5,83E-05 - | 3 - | 4,34E-05 - | 4 - | 5,50E-05 - | 3 - | 4,19E-05 - | 3 - | 4,72E-05 - |
| 351 | 1 | 0.2um_NOKCPHJN_135858 | Eukaryota | Cnidaria | Anthozoa | Scleractinia | Pocilloporidae | Stylophora | Stylophora pis | unclassified | unclassified | unclassified | unclassified | unclassified | 0,4 | 0,489897949 | 3,76E-06 | 4,61E-06 | 0 - | 0 - | 1 - | 9,64E-06 - | 1 - | 9,17E-06 - | 0 - | 0 - | 0 - | 0 - |
| 352 | 1 | 0.2um_NOKCPHJN_454769 | unclassified | unclassified | unclassified | unclassified | unclassified | unclassified | unclassified | unclassified | unclassified | unclassified | unclassified | unclassified | 0,6 | 0,489897949 | 3,71E-05 | 3,05E-05 | 1 - | 5,49E-05 - | 0 - | 0 - | 1 - | 6,48E-05 - | 1 - | 6,58E-05 - | 0 - | 0 - |
| 353 | 2 | 0.2um_NOKCPHJN_44989;0.2um_NOKCPHJN_62 | Bacteria | Proteobacteri | Gammaprotec | unclassified | unclassified | unclassified | various | cellular proces | Cell wall/mem | 0QTSX | Peptidoglycan | various | 0,6 | 0,489897949 | 4,39E-05 | 3,62E-05 | 1 - | 6,41E-05 - | 1 - | 7,96E-05 - | 1 - | 7,57E-05 - | 0 - | 0 - | 0 - | 0 - |
| 354 | 1 | 0.2um_NOKCPHJN_99558 | Bacteria | Proteobacteri | Gammaprotec | Oceanospirilla | Saccharospirill | Reinekea | Reinekea blan | metabolism | Inorganic ion t | COG0841 | acriflavin resis | 314283.MED2 | 3 | 1,095445115 | 3,86E-05 | 1,31E-05 | 4 - | 4,42E-05 - | 4 - | 5,49E-05 - | 3 - | 3,91E-05 - | 3 - | 3,97E-05 - | 1 - | 1,49E-05 - |
| 355 | 1 | 3um_FIJJAFBF_491676 | Bacteria | Proteobacteri | Gammaprotec | Cellvibrionales | unclassified | unclassified | Cellvibrionale | unclassified | unclassified | unclassified | unclassified | unclassified | 1,2 | 0,4 | 8,94E-05 | 1,98E-05 | 2 - | 0,000128242 - | 1 - | 7,96E-05 - | 1 - | 7,57E-05 - | 1 - | 7,68E-05 - | 1 - | 8,65E-05 - |
| 356 | 1 | 0.2um_NOKCPHJN_07665 | Bacteria | Bacteroidetes | Flavobacteriia | Flavobacterial | Flavobacteriac | Polaribacter | Polaribacter s | metabolism | Energy produc | 099YH | Pyruvate dehy | 50743.SCB49_ | 0,6 | 0,489897949 | 2,39E-05 | 1,96E-05 | 1 - | 3,53E-05 - | 0 - | 0 - | 1 - | 4,17E-05 - | 1 - | 4,23E-05 - | 0 - | 0 - |
| 357 | 1 | 3um_FIJJAFBF_122446 | Bacteria | Bacteroidetes | Flavobacteriia | Flavobacterial | Flavobacteriac | Polaribacter | Polaribacter a | metabolism | Amino acid tra | COG2171 | Catalyzes the | 313598.MED1 | 4,6 | 0,8 | 0,000229194 | 2,05E-05 | 6 - | 0,000254118 - | 4 - | 0,00021036 - | 5 - | 0,000249967 - | 4 - | 0,000202927 - | 4 - | 0,000228597 - |
| 358 | 1 | 0.2um_NOKCPHJN_151232 | Bacteria | Bacteroidetes | Flavobacteriia | Flavobacterial | Flavobacteriac | Formosa | Formosa sp. H | unclassified | unclassified | unclassified | unclassified | unclassified | 12,6 | 1,624807681 | 0,000179377 | 1,83E-05 | 14 - | 0,000168259 - | 12 - | 0,000179081 - | 15 - | 0,000212799 - | 11 - | 0,000158358 - | 11 - | 0,000178389 - |
| 359 | 1 | 3um_FIJJAFBF_259139 | Bacteria | Proteobacteri | Alphaproteob | Rhodobactera | unclassified | unclassified | Rhodobactera | metabolism | Inorganic ion t | COG3639 | Phosphonate | 367336.OM22 | 0,8 | 0,4 | 3,91E-05 | 1,99E-05 | 1 - | 4,24E-05 - | 1 - | 5,26E-05 - | 1 - | 5,00E-05 - | 1 - | 5,07E-05 - | 0 - | 0 - |
| 360 | 1 | 3um_FIJJAFBF_578494 | Bacteria | Proteobacteri | Gammaprotec | Cellvibrionales | Porticoccacea | unclassified | gamma protec | metabolism | Energy produc | 16R80 | pyrophosphat | 314287.GB22 | 1,6 | 0,489897949 | 0,00012507 | 4,03E-05 | 1 - | 6,52E-05 - | 2 - | 0,000161953 - | 2 - | 0,000153957 - | 2 - | 0,000156231 - | 1 - | 8,80E-05 - |
| 361 | 1 | 0.2um_NOKCPHJN_623414 | Eukaryota | Streptophyta | Klebsormidiop | Klebsormidial | Klebsormidiac | Klebsormidiur | Klebsormidiur | unclassified | unclassified | unclassified | unclassified | unclassified | 1 | 0 | 1,87E-05 | 1,78E-06 | 1 - | 1,57E-05 - | 1 - | 1,95E-05 - | 1 - | 1,85E-05 - | 1 - | 1,88E-05 - | 1 - | 2,12E-05 - |
| 362 | 1 | 0.2um_NOKCPHJN_1086528 | Eukaryota | Placozoa | unclassified | unclassified | unclassified | Trichoplax | Trichoplax adh | cellular proces | Cell wall/mem | COG3637 | Membrane | 983545.Glaag | 0,4 | 0,489897949 | 2,21E-05 | 2,73E-05 | 1 - | 5,03E-05 - | 0 - | 0 - | 0 - | 0 - | 1 - | 6,03E-05 - | 0 - | 0 - |
| 363 | 1 | 0.2um_NOKCPHJN_576524 | Bacteria | Bacteroidetes | Flavobacteriia | Flavobacterial | Flavobacteriac | Tenacibaculur | Tenacibaculur | unclassified | unclassified | unclassified | unclassified | unclassified | 5 | 0,894427191 | 0,000123849 | 3,10E-05 | 4 - | 8,20E-05 - | 5 - | 0,000127249 - | 4 - | 9,68E-05 - | 6 - | 0,000147304 - | 6 - | 0,000165937 - |
| 364 | 1 | 3um_FIJJAFBF_514974 | Eukaryota | unclassified | Pelagophycea | Pelagomonad | unclassified | Aureococcus | Aureococcus a | unclassified | unclassified | unclassified | unclassified | unclassified | 0,8 | 0,748331477 | 2,68E-05 | 2,36E-05 | 2 - | 6,06E-05 - | 1 - | 3,76E-05 - | 1 - | 3,57E-05 - | 0 - | 0 - | 0 - | 0 - |
| 365 | 1 | 3um_FIJJAFBF_236816 | Bacteria | Proteobacteri | Alphaproteob | unclassified | unclassified | Candidatus Pu | Candidatus Pu | unclassified | unclassified | unclassified | unclassified | unclassified | 0,4 | 0,489897949 | 6,31E-05 | 7,78E-05 | 1 - | 0,000143471 - | 0 - | 0 - | 0 - | 0 - | 1 - | 0,000171854 - | 0 - | 0 - |
| 366 | 1 | 0.2um_NOKCPHJN_1054690 | Bacteria | Proteobacteri | Alphaproteob | Pelagibacteral | Pelagibacterac | unclassified | Pelagibacterac | cellular proces | Cell wall/mem | 7257 | NA | 439493.PB721 | 3,8 | 0,4 | 0,00015557 | 2,05E-05 | 4 - | 0,000137457 - | 3 - | 0,000128011 - | 4 - | 0,000162254 - | 4 - | 0,000164651 - | 4 - | 0,000185478 - |
| 367 | 1 | 3um_FIJJAFBF_265244 | Bacteria | Bacteroidetes | Flavobacteriia | Flavobacterial | Flavobacteriac | Polaribacter | Polaribacter b | cellular proces | Posttranslatio | 09ADN | alkyl hydroper | 391587.KAOT | 1 | 0 | 6,43E-05 | 6,11E-06 | 1 - | 5,39E-05 - | 1 - | 6,69E-05 - | 1 - | 6,36E-05 - | 1 - | 6,45E-05 - | 1 - | 7,27E-05 - |
| 368 | 1 | 0.2um_NOKCPHJN_177444 | Bacteria | Bacteroidetes | Flavobacteriia | Flavobacterial | Flavobacteriac | Polaribacter | Polaribacter s | metabolism | Inorganic ion t | 05C27 | TonB-depende | 313598.MED1 | 7,4 | 1,2 | 9,18E-05 | 8,62E-06 | 9 - | 9,48E-05 - | 6 - | 7,85E-05 - | 8 - | 9,94E-05 - | 8 - | 0,000100905 - | 6 - | 8,53E-05 - |
| 369 | 1 | 3um_FIJJAFBF_231278 | Bacteria | Proteobacteri | Gammaprotec | Oceanospirilla | Oceanospirilla | Oceanobacter | Oceanobacter | metabolism | Carbohydrate | 0QKP9 | ABC-type suga | 400668.Mmw | 9,6 | 0,8 | 0,000316948 | 3,85E-05 | 9 - | 0,000248913 - | 9 - | 0,000309077 - | 10 - | 0,000326463 - | 11 - | 0,000364414 - | 9 - | 0,000335872 - |
| 370 | 1 | 3um_FIJJAFBF_186687 | Bacteria | Bacteroidetes | Flavobacteriia | Flavobacterial | Flavobacteriac | Polaribacter | Polaribacter fi | metabolism | Carbohydrate | COG0057 | glyceraldehyd | 313594.PI23P | 0,2 | 0,4 | 5,94E-06 | 1,19E-05 | 0 - | 0 - | 1 - | 2,97E-05 - | 0 - | 0 - | 0 - | 0 - | 0 - | 0 - |
| 371 | 1 | 3um_FIJJAFBF_67265 | Bacteria | Proteobacteri | Gammaprotec | Thiotrichales | Piscirickettsia | Methylophaga | Methylophaga | information st | Translation, ri | COG0256 | This is one of | 1637616.MDM | 0,4 | 0,8 | 4,70E-05 | 9,40E-05 | 0 - | 0 - | 0 - | 0 - | 0 - | 0 - | 2 - | 0,000235014 - | 0 - | 0 - |
| 372 | 1 | 0.2um_NOKCPHJN_463228 | Bacteria | Bacteroidetes | unclassified | unclassified | unclassified | unclassified | Bacteroidetes | information st | Translation, ri | COG0050 | This protein p | 487796.Flav2 | 3,4 | 0,8 | 0,000118081 | 3,11E-05 | 4 - | 0,00011623 - | 4 - | 0,000144323 - | 2 - | 6,86E-05 - | 3 - | 0,000104418 - | 4 - | 0,000156835 - |
| 373 | 1 | 0.2um_NOKCPHJN_977312 | Eukaryota | Phaeophyceae | unclassified | Ectocarpales | Ectocarpaceae | Ectocarpus | Ectocarpus sili | information st | Translation, ri | 0TJFM | 40S ribosomal | 4641.GSMUA | 1,2 | 0,4 | 0,000124986 | 2,77E-05 | 2 - | 0,000179339 - | 1 - | 0,000111343 - | 1 - | 0,000105845 - | 1 - | 0,000107409 - | 1 - | 0,000120996 - |
| 374 | 1 | 3um_FIJJAFBF_183640 | Bacteria | Actinobacteria | Actinobacteria | Micrococcales | Microbacteria | various | various | information st | Translation, ri | 05NRA | 50s ribosomal | 312284.A20C1 | 0,6 | 0,489897949 | 9,63E-05 | 7,94E-05 | 1 - | 0,000139972 - | 1 - | 0,000173803 - | 0 - | 0 - | 1 - | 0,000167663 - | 0 - | 0 - |
| 375 | 1 | 0.2um_NOKCPHJN_374274 | Bacteria | Bacteroidetes | unclassified | unclassified | unclassified | unclassified | Bacteroidetes | metabolism | Amino acid tra | 0P7Z9 | aminopeptida | 391603.FBALC | 0,8 | 0,4 | 1,39E-05 | 7,07E-06 | 1 - | 1,51E-05 - | 1 - | 1,87E-05 - | 1 - | 1,78E-05 - | 1 - | 1,80E-05 - | 0 - | 0 - |
| 376 | 1 | 3um_FIJJAFBF_130336 | Bacteria | Proteobacteri | Gammaprotec | Oceanospirilla | Saccharospirill | Saccharospiril | Saccharospiril | cellular proces | Posttranslatio | 16QAM | Protease subu | 345073.VC39 | 0,8 | 0,4 | 6,31E-05 | 3,24E-05 | 1 - | 6,67E-05 - | 0 - | 0 - | 1 - | 7,88E-05 - | 1 - | 7,99E-05 - | 1 - | 9,00E-05 - |
| 377 | 1 | 3um_FIJJAFBF_546283 | Eukaryota | unclassified | Oomycetes | Saprolegniales | Saprolegniace | Thraustotheca | Thraustotheca | information st | Translation, ri | 11UVJ | Ribosomal pro | 29760.VIT_05 | 2,2 | 0,748331477 | 0,000263301 | 0,000103341 | 2 - | 0,000197891 - | 2 - | 0,000245722 - | 1 - | 0,000116795 - | 3 - | 0,00035556 - | 3 - | 0,000400537 - |
| 378 | 2 | 0.2um_NOKCPHJN_767113;0.2um_NOKCPHJN_7 | Bacteria | Proteobacteri | Alphaproteob | Pelagibacteral | Pelagibacterac | Candidatus Pe | Candidatus Pe | unclassified | unclassified | unclassified | unclassified | unclassified | 0,4 | 0,489897949 | 1,77E-05 | 2,19E-05 | 1 - | 3,94E-05 - | 1 - | 4,90E-05 - | 0 - | 0 - | 0 - | 0 - | 0 - | 0 - |
| 379 | 1 | 3um_FIJJAFBF_739116 | Bacteria | Proteobacteri | Alphaproteob | Rhodobactera | Rhodobactera | Sulfitobacter | Sulfitobacter s | metabolism | Carbohydrate | 05CV1 | Transketolase | 52598.EE36_1 | 0,2 | 0,4 | 5,41E-06 | 1,08E-05 | 1 - | 2,71E-05 - | 0 - | 0 - | 0 - | 0 - | 0 - | 0 - | 0 - | 0 - |
| 380 | 2 | 0.2um_NOKCPHJN_126550;0.2um_NOKCPHJN_3 | Bacteria | Verrucomicrol | Opitutae | unclassified | unclassified | unclassified | Opitutae bact | unclassified | unclassified | unclassified | unclassified | unclassified | 0,8 | 0,748331477 | 0,00010496 | 0,00010326 | 1 - | 0,000112526 - | 2 - | 0,000279449 - | 1 - | 0,000132826 - | 0 - | 0 - | 0 - | 0 - |
| 381 | 1 | 3um_FIJJAFBF_05019 | Bacteria | Bacteroidetes | Flavobacteriia | Flavobacterial | Flavobacteriac | Polaribacter | various | unclassified | unclassified | unclassified | unclassified | unclassified | 0,6 | 0,489897949 | 9,82E-05 | 8,08E-05 | 1 - | 0,000145287 - | 0 - | 0 - | 1 - | 0,000171496 - | 1 - | 0,00017403 - | 0 - | 0 - |
| 382 | 1 | 3um_FIJJAFBF_693134 | Bacteria | Proteobacteri | Gammaprotec | unclassified | unclassified | unclassified | various | metabolism | Coenzyme tra | 08UTT | Catalyzes the | 247639.MGP2 | 0,4 | 0,489897949 | 3,68E-05 | 4,52E-05 | 0 - | 0 - | 0 - | 0 - | 1 - | 8,57E-05 - | 0 - | 0 - | 1 - | 9,80E-05 - |
| 383 | 1 | 0.2um_NOKCPHJN_194065 | Bacteria | Bacteroidetes | Flavobacteriia | Flavobacterial | Flavobacteriac | Formosa | Formosa sp. H | poorly charact | Function unkn | 11GNC | SnoaL-like pol | 487796.Flav2 | 1 | 0 | 8,16E-05 | 7,74E-06 | 1 - | 6,83E-05 - | 1 - | 8,48E-05 - | 1 - | 8,06E-05 - | 1 - | 8,18E-05 - | 1 - | 9,22E-05 - |
| 384 | 1 | 0.2um_NOKCPHJN_613243 | Bacteria | Proteobacteri | Gammaprotec | unclassified | unclassified | unclassified | Gammaprotec | unclassified | unclassified | unclassified | unclassified | unclassified | 0,8 | 0,4 | 7,92E-05 | 4,07E-05 | 1 - | 8,38E-05 - | 0 - | 0 - | 1 - | 9,89E-05 - | 1 - | 0,000100353 - | 1 - | 0,000113047 - |
| 385 | 1 | 0.2um_NOKCPHJN_798703 | unclassified | unclassified | unclassified | unclassified | unclassified | unclassified | unclassified | unclassified | unclassified | unclassified | unclassified | unclassified | 0,2 | 0,4 | 3,63E-06 | 7,26E-06 | 0 - | 0 - | 0 - | 0 - | 0 - | 0 - | 1 - | 1,81E-05 - | 0 - | 0 - |
| 386 | 1 | 0.2um_NOKCPHJN_06847 | Bacteria | Bacteroidetes | Cytophagia | Cytophagales | Flammeovirga | unclassified | Flammeovirga | unclassified | unclassified | unclassified | unclassified | unclassified | 0,8 | 0,4 | 8,79E-05 | 4,52E-05 | 1 - | 9,18E-05 - | 1 - | 0,000114015 - | 0 - | 0 - | 1 - | 0,000109987 - | 1 - | 0,0001239 - |
| 387 | 1 | 3um_FIJJAFBF_744568 | Bacteria | unclassified | unclassified | unclassified | unclassified | unclassified | uncultured ba | poorly charact | Function unkn | 0P8NI | Inherit from b | 983544.Lacal_ | 7,2 | 0,4 | 0,000238965 | 1,39E-05 | 8 - | 0,00022341 - | 7 - | 0,000242733 - | 7 - | 0,000230748 - | 7 - | 0,000234156 - | 7 - | 0,000263776 - |
| 388 | 1 | 0.2um_NOKCPHJN_583857 | Bacteria | Bacteroidetes | Flavobacteriia | unclassified | unclassified | unclassified | Flavobacteria | poorly charact | Function unkn | 0NVRZ | NA | 170187.SP_17 | 0,6 | 0,489897949 | 1,70E-05 | 1,39E-05 | 0 - | 0 - | 1 - | 2,79E-05 - | 1 - | 2,66E-05 - | 0 - | 0 - | 1 - | 3,04E-05 - |
| 389 | 1 | 3um_FIJJAFBF_652364 | Bacteria | Bacteroidetes | Flavobacteriia | Flavobacterial | Flavobacteriac | Lutibacter | Lutibacter aga | poorly charact | Function unkn | 0XRCJ | NA | 391587.KAOT | 0,2 | 0,4 | 2,17E-06 | 4,34E-06 | 1 - | 1,09E-05 - | 0 - | 0 - | 0 - | 0 - | 0 - | 0 - | 0 - | 0 - |
| 390 | 1 | 3um_FIJJAFBF_689814 | Bacteria | Bacteroidetes | Flavobacteriia | Flavobacterial | Crocinitomica | unclassified | Crocinitomica | unclassified | unclassified | unclassified | unclassified | unclassified | 1 | 0 | 7,88E-05 | 7,48E-06 | 1 - | 6,60E-05 - | 1 - | 8,19E-05 - | 1 - | 7,79E-05 - | 1 - | 7,90E-05 - | 1 - | 8,90E-05 - |
| 391 | 1 | 3um_FIJJAFBF_335808 | Bacteria | Proteobacteri | Alphaproteob | Rhodobactera | Rhodobactera | Phaeobacter | Phaeobacter g | metabolism | Energy produc | 16QPK | Succinate deh | 644107.SL115 | 3 | 0,632455532 | 6,77E-05 | 1,18E-05 | 4 - | 7,64E-05 - | 3 - | 7,11E-05 - | 2 - | 4,51E-05 - | 3 - | 6,86E-05 - | 3 - | 7,73E-05 - |
| 392 | 1 | 3um_FIJJAFBF_460907 | Bacteria | Bacteroidetes | Flavobacteriia | Flavobacterial | Flavobacteriac | Polaribacter | Polaribacter s | metabolism | Amino acid tra | 08IQK | o-acetylhomo | 313598.MED1 | 8,4 | 1,959591794 | 0,000272095 | 7,60E-05 | 9 - | 0,00024363 - | 5 - | 0,000168065 - | 9 - | 0,00028758 - | 8 - | 0,000259402 - | 11 - | 0,000401797 - |
| 393 | 1 | 3um_FIJJAFBF_391553 | Bacteria | Bacteroidetes | Flavobacteriia | Flavobacterial | Flavobacteriac | Algibacter | Algibacter sp. | metabolism | Inorganic ion t | 0PA08 | Destroys radic | 983544.Lacal_ | 0,4 | 0,8 | 2,43E-05 | 4,86E-05 | 2 - | 0,000121457 - | 0 - | 0 - | 0 - | 0 - | 0 - | 0 - | 0 - | 0 - |
| 394 | 1 | 0.2um_NOKCPHJN_123980 | Bacteria | Bacteroidetes | Flavobacteriia | Flavobacterial | Flavobacteriac | Polaribacter | Polaribacter fi | metabolism | Amino acid tra | 099QE | chorismate m | 867900.Celly_ | 0,2 | 0,4 | 6,38E-06 | 1,28E-05 | 1 - | 3,19E-05 - | 0 - | 0 - | 0 - | 0 - | 0 - | 0 - | 0 - | 0 - |
| 395 | 1 | 3um_FIJJAFBF_371988 | Eukaryota | Bacillariophyt | Coscinodiscop | Thalassiosirale | Thalassiosirac | Thalassiosira | Thalassiosira | metabolism | Energy produc | KOG0369 | Catalyzes a 2- | 35128.Thaps2 | 0,2 | 0,4 | 2,71E-06 | 5,41E-06 | 1 - | 1,35E-05 - | 0 - | 0 - | 0 - | 0 - | 0 - | 0 - | 0 - | 0 - |
| 396 | 1 | 3um_FIJJAFBF_599964 | Bacteria | Proteobacteri | Gammaprotec | Oceanospirilla | Saccharospirill | Reinekea | Reinekea blan | poorly charact | Function unkn | 05E6I | conserved pro | 314283.MED2 | 3,4 | 1,356465997 | 0,000127016 | 4,40E-05 | 5 - | 0,000162114 - | 3 - | 0,000120779 - | 4 - | 0,000153087 - | 4 - | 0,000155348 - | 1 - | 4,37E-05 - |
| 397 | 1 | 3um_FIJJAFBF_135233 | Eukaryota | Eustigmatoph | unclassified | Eustigmatales | Monodopsida | Nannochlorop | Nannochlorop | unclassified | unclassified | unclassified | unclassified | unclassified | 1,8 | 1,166190379 | 6,54E-05 | 4,39E-05 | 0 - | 0 - | 3 - | 0,000107426 - | 2 - | 6,81E-05 - | 1 - | 3,45E-05 - | 3 - | 0,00011674 - |
| 398 | 1 | 3um_FIJJAFBF_246767 | Bacteria | Bacteroidetes | Flavobacteriia | Flavobacterial | Flavobacteriac | Bergeyella | Bergeyella zoo | poorly charact | Function unkn | 0FXH3 | Inherit from b | 643867.Ftrac_ | 3,2 | 1,469693846 | 0,000102847 | 5,09E-05 | 1 - | 2,60E-05 - | 5 - | 0,000161586 - | 2 - | 6,14E-05 - | 4 - | 0,000124701 - | 4 - | 0,000140476 - |
| 399 | 1 | 3um_FIJJAFBF_184050 | Viruses | unclassified | unclassified | Caudovirales | Siphoviridae | unclassified | Idiomarinacea | unclassified | unclassified | unclassified | unclassified | unclassified | 1,2 | 0,748331477 | 9,77E-05 | 6,17E-05 | 1 - | 6,87E-05 - | 0 - | 0 - | 2 - | 0,000162254 - | 2 - | 0,000164651 - | 1 - | 9,27E-05 - |
| 400 | 1 | 3um_FIJJAFBF_628820 | Bacteria | Actinobacteria | Actinobacteria | Micrococcales | Intrasporangia | Janibacter | Janibacter hoy | unclassified | unclassified | unclassified | unclassified | unclassified | 0,4 | 0,489897949 | 3,04E-05 | 3,77E-05 | 1 - | 6,79E-05 - | 1 - | 8,43E-05 - | 0 - | 0 - | 0 - | 0 - | 0 - | 0 - |
| 401 | 1 | 0.2um_NOKCPHJN_249921 | Eukaryota | Chordata | Mammalia | Primates | Cheirogaleida | Microcebus | Microcebus m | cellular proces | Intracellular tr | 0BIEB | domain-conta | 1005048.CFU_ | 0,6 | 0,489897949 | 1,94E-05 | 1,58E-05 | 0 - | 0 - | 1 - | 3,32E-05 - | 1 - | 3,16E-05 - | 1 - | 3,20E-05 - | 0 - | 0 - |
| 402 | 1 | 0.2um_NOKCPHJN_643038 | Viruses | unclassified | unclassified | unclassified | unclassified | unclassified | Pleurochrysis | unclassified | unclassified | unclassified | unclassified | unclassified | 1 | 0,632455532 | 2,74E-05 | 1,85E-05 | 1 - | 2,34E-05 - | 2 - | 5,81E-05 - | 1 - | 2,76E-05 - | 1 - | 2,80E-05 - | 0 - | 0 - |
| 403 | 1 | 0.2um_NOKCPHJN_388003 | Bacteria | Bacteroidetes | Flavobacteriia | Flavobacterial | Flavobacteriac | unclassified | Flavobacteriac | information st | Translation, ri | COG0050 | This protein p | 886377.Murru | 5,6 | 0,489897949 | 0,000194489 | 2,55E-05 | 5 - | 0,000145287 - | 6 - | 0,000216484 - | 6 - | 0,000205796 - | 6 - | 0,000208835 - | 5 - | 0,000196044 - |
| 404 | 1 | 0.2um_NOKCPHJN_172110 | Bacteria | Bacteroidetes | Flavobacteriia | Flavobacterial | Flavobacteriac | Formosa | Formosa sp. H | poorly charact | Function unkn | 11NCV | Protein of unk | 983544.Lacal_ | 2,4 | 0,489897949 | 0,000195486 | 3,04E-05 | 3 - | 0,000207428 - | 2 - | 0,000171709 - | 2 - | 0,000163232 - | 3 - | 0,000248464 - | 2 - | 0,000186596 - |
| 405 | 1 | 0.2um_NOKCPHJN_420835 | Bacteria | unclassified | unclassified | unclassified | unclassified | unclassified | bacterium TM | unclassified | unclassified | unclassified | unclassified | unclassified | 0,2 | 0,4 | 3,83E-06 | 7,67E-06 | 0 - | 0 - | 0 - | 0 - | 0 - | 0 - | 0 - | 0 - | 1 - | 1,92E-05 - |
| 406 | 1 | 0.2um_NOKCPHJN_146801 | Eukaryota | Apicomplexa | Aconoidasida | Haemosporida | Plasmodiidae | Plasmodium | Plasmodium c | unclassified | unclassified | unclassified | unclassified | unclassified | 0,4 | 0,489897949 | 2,40E-06 | 2,96E-06 | 1 - | 5,50E-06 - | 0 - | 0 - | 1 - | 6,49E-06 - | 0 - | 0 - | 0 - | 0 - |
| 407 | 1 | 3um_FIJJAFBF_175109 | Bacteria | Proteobacteri | Alphaproteob | Rhodobactera | Rhodobactera | unclassified | Rhodobactera | unclassified | unclassified | unclassified | unclassified | unclassified | 0,2 | 0,4 | 1,85E-05 | 3,70E-05 | 0 - | 0 - | 1 - | 9,25E-05 - | 0 - | 0 - | 0 - | 0 - | 0 - | 0 - |
| 408 | 1 | 0.2um_NOKCPHJN_634668 | Bacteria | Bacteroidetes | Flavobacteriia | Flavobacterial | Flavobacteriac | Polaribacter | Polaribacter a | metabolism | Amino acid tra | 09AVX | Catalyzes the | 391603.FBALC | 0,4 | 0,489897949 | 1,90E-05 | 2,32E-05 | 0 - | 0 - | 0 - | 0 - | 1 - | 4,70E-05 - | 1 - | 4,77E-05 - | 0 - | 0 - |
| 409 | 1 | 0.2um_NOKCPHJN_260628 | Bacteria | Proteobacteri | Alphaproteob | Pelagibacteral | Pelagibacterac | Candidatus Pe | Candidatus Pe | unclassified | unclassified | unclassified | unclassified | unclassified | 2,2 | 0,4 | 9,38E-05 | 1,83E-05 | 2 - | 7,15E-05 - | 2 - | 8,88E-05 - | 3 - | 0,000126619 - | 2 - | 8,57E-05 - | 2 - | 9,65E-05 - |
| 410 | 1 | 3um_FIJJAFBF_189718 | Bacteria | Bacteroidetes | Flavobacteriia | Flavobacterial | Flavobacteriac | Polaribacter | Polaribacter s | poorly charact | Function unkn | 09FA2 | NA | 313594.PI23P | 1 | 0,632455532 | 8,90E-05 | 5,07E-05 | 2 - | 0,000158313 - | 1 - | 9,83E-05 - | 1 - | 9,34E-05 - | 1 - | 9,48E-05 - | 0 - | 0 - |
| 411 | 1 | 3um_FIJJAFBF_128631 | Eukaryota | Chlorophyta | Mamiellophyc | Mamiellales | Mamiellaceae | Micromonas | Micromonas c | unclassified | unclassified | unclassified | unclassified | unclassified | 2,2 | 0,4 | 0,000260535 | 2,39E-05 | 3 - | 0,000302044 - | 2 - | 0,000250033 - | 2 - | 0,000237688 - | 2 - | 0,000241199 - | 2 - | 0,00027171 - |
| 412 | 1 | 3um_FIJJAFBF_623894 | Bacteria | Proteobacteri | Alphaproteob | Rhodobactera | Rhodobactera | unclassified | Rhodobactera | unclassified | unclassified | unclassified | unclassified | unclassified | 0,6 | 0,489897949 | 2,79E-05 | 2,32E-05 | 1 - | 3,96E-05 - | 0 - | 0 - | 1 - | 4,67E-05 - | 0 - | 0 - | 1 - | 5,34E-05 - |
| 413 | 1 | 3um_FIJJAFBF_154877 | Bacteria | Proteobacteri | Betaproteoba | unclassified | unclassified | unclassified | Betaproteoba | metabolism | Energy produc | COG1290 | Component o | 420662.Mpe_ | 0,2 | 0,4 | 5,81E-06 | 1,16E-05 | 0 - | 0 - | 0 - | 0 - | 1 - | 2,91E-05 - | 0 - | 0 - | 0 - | 0 - |
| 414 | 1 | 0.2um_NOKCPHJN_816138 | Bacteria | Proteobacteri | Alphaproteob | Rhodospirillal | Rhodospirillac | unclassified | Rhodobactera | cellular proces | Cell wall/mem | 16QB0 | Part of the ou | 367336.OM22 | 0,2 | 0,4 | 3,18E-06 | 6,37E-06 | 1 - | 1,59E-05 - | 0 - | 0 - | 0 - | 0 - | 0 - | 0 - | 0 - | 0 - |
| 415 | 2 | 0.2um_NOKCPHJN_457461;3um_FIJJAFBF_60004 | Bacteria | Proteobacteri | Gammaprotec | Cellvibrionales | Porticoccacea | unclassified | various | cellular proces | Intracellular tr | 05VIE | preprotein tra | 314287.GB22 | 1 | 0 | 0,000125713 | 1,19E-05 | 1 - | 0,0001053 - | 1 - | 0,000130751 - | 1 - | 0,000124296 - | 1 - | 0,000126131 - | 1 - | 0,000142087 - |
| 416 | 1 | 0.2um_NOKCPHJN_786539 | Bacteria | Proteobacteri | Gammaprotec | Oceanospirilla | Halomonadac | Halomonas | Halomonas zi | unclassified | unclassified | unclassified | unclassified | unclassified | 2 | 1,095445115 | 4,34E-05 | 2,75E-05 | 2 - | 3,56E-05 - | 1 - | 2,21E-05 - | 2 - | 4,20E-05 - | 1 - | 2,13E-05 - | 4 - | 9,60E-05 - |
| 417 | 1 | 0.2um_NOKCPHJN_652146 | Eukaryota | unclassified | Oomycetes | Albuginales | Albuginaceae | Albugo | Albugo candid | unclassified | unclassified | unclassified | unclassified | unclassified | 1,6 | 0,489897949 | 5,01E-05 | 1,83E-05 | 1 - | 2,57E-05 - | 2 - | 6,38E-05 - | 1 - | 3,03E-05 - | 2 - | 6,15E-05 - | 2 - | 6,93E-05 - |
| 418 | 1 | 3um_FIJJAFBF_99779 | Bacteria | Bacteroidetes | Flavobacteriia | Flavobacterial | Flavobacteriac | Formosa | Formosa sp. H | unclassified | unclassified | unclassified | unclassified | unclassified | 10,2 | 0,748331477 | 0,000283103 | 3,24E-05 | 11 - | 0,000256094 - | 9 - | 0,000260176 - | 10 - | 0,000274812 - | 10 - | 0,000278871 - | 11 - | 0,000345562 - |
| 419 | 1 | 3um_FIJJAFBF_89469 | Bacteria | Bacteroidetes | Flavobacteriia | Flavobacterial | Flavobacteriac | Algibacter | Algibacter pe | unclassified | unclassified | unclassified | unclassified | unclassified | 1,2 | 0,4 | 2,64E-05 | 9,73E-06 | 1 - | 1,83E-05 - | 2 - | 4,54E-05 - | 1 - | 2,16E-05 - | 1 - | 2,19E-05 - | 1 - | 2,47E-05 - |
| 420 | 1 | 0.2um_NOKCPHJN_71343 | Bacteria | Proteobacteri | Gammaprotec | Cellvibrionales | unclassified | unclassified | Cellvibrionale | unclassified | unclassified | unclassified | unclassified | unclassified | 0,8 | 0,4 | 4,89E-05 | 2,51E-05 | 1 - | 5,12E-05 - | 1 - | 6,36E-05 - | 1 - | 6,05E-05 - | 0 - | 0 - | 1 - | 6,91E-05 - |
| 421 | 1 | 3um_FIJJAFBF_13490 | Bacteria | Bacteroidetes | Flavobacteriia | Flavobacterial | Flavobacteriac | Flavivirga | Flavivirga eckl | unclassified | unclassified | unclassified | unclassified | unclassified | 0,4 | 0,489897949 | 2,61E-05 | 3,22E-05 | 1 - | 5,95E-05 - | 0 - | 0 - | 0 - | 0 - | 1 - | 7,12E-05 - | 0 - | 0 - |
| 422 | 1 | 0.2um_NOKCPHJN_505349 | unclassified | unclassified | unclassified | unclassified | unclassified | unclassified | unclassified | unclassified | unclassified | unclassified | unclassified | unclassified | 0,2 | 0,4 | 1,16E-05 | 2,32E-05 | 0 - | 0 - | 0 - | 0 - | 1 - | 5,79E-05 - | 0 - | 0 - | 0 - | 0 - |
| 423 | 1 | 3um_FIJJAFBF_73819 | Bacteria | Bacteroidetes | Flavobacteriia | Flavobacterial | Flavobacteriac | Polaribacter | Polaribacter s | unclassified | unclassified | unclassified | unclassified | unclassified | 13,8 | 1,720465053 | 0,000269977 | 3,09E-05 | 14 - | 0,000230873 - | 13 - | 0,000266199 - | 15 - | 0,000291988 - | 16 - | 0,000316054 - | 11 - | 0,000244773 - |

| | | | | | | | | | | | | | | | | | | | | | | | | | | | | |
|---|---|---|---|---|---|---|---|---|---|---|---|---|---|---|---|---|---|---|---|---|---|---|---|---|---|---|---|---|
| 424 | 1 | 3um_FIJJAFBF_426211 | Eukaryota | Bacillariophyt | Bacillariophyc | Naviculales | Phaeodactylac | Phaeodactylu | Phaeodactylu | cellular proces | Cell wall/mem | KOG0916 | synthase | 35128.Thaps1 | 1,8 | 0,748331477 | 3,96E-05 | 1,63E-05 | 2 - | 3,73E-05 - | 2 - | 4,63E-05 - | 1 - | 2,20E-05 - | 3 - | 6,71E-05 - | 1 - | 2,52E-05 - |
| 425 | 1 | 0.2um_NOKCPHJN_490937 | Bacteria | Proteobacteri | Gammaprotec | unclassified | unclassified | Candidatus Th | Candidatus Th | metabolism | Energy produc | COG0055 | Produces ATP | 412965.COSY_ | 0,4 | 0,489897949 | 1,09E-05 | 1,34E-05 | 1 - | 2,50E-05 - | 0 - | 0 - | 1 - | 2,95E-05 - | 0 - | 0 - | 0 - | 0 - |
| 426 | 1 | 0.2um_NOKCPHJN_735378 | Eukaryota | Mollusca | Bivalvia | Pterioida | Pteriidae | Pinctada | Pinctada fucat | poorly charact | Function unkn | 1ABT4 | RRM | 578455.XP_00 | 0,2 | 0,4 | 1,27E-05 | 2,53E-05 | 0 - | 0 - | 0 - | 0 - | 1 - | 6,33E-05 - | 0 - | 0 - | 0 - | 0 - |
| 427 | 1 | 3um_FIJJAFBF_337913 | Bacteria | unclassified | unclassified | unclassified | unclassified | unclassified | uncultured ba | poorly charact | Function unkn | 0XNQI | domain protei | 313594.PI23P | 0,4 | 0,489897949 | 1,26E-05 | 1,54E-05 | 0 - | 0 - | 1 - | 3,22E-05 - | 1 - | 3,06E-05 - | 0 - | 0 - | 0 - | 0 - |
| 428 | 1 | 3um_FIJJAFBF_434534 | Bacteria | Bacteroidetes | Flavobacteriia | Flavobacterial | Flavobacteriac | Polaribacter | various | unclassified | unclassified | unclassified | unclassified | unclassified | 3,6 | 1,356465997 | 0,00028126 | 8,24E-05 | 6 - | 0,000402725 - | 4 - | 0,000333377 - | 3 - | 0,000237688 - | 2 - | 0,000160799 - | 3 - | 0,00027171 - |
| 429 | 1 | 3um_FIJJAFBF_259339 | Bacteria | Proteobacteri | Gammaprotec | Cellvibrionales | Porticoccacea | unclassified | SAR92 bacteri | cellular proces | Intracellular tr | 175GA | One of the pro | 314287.GB220 | 3 | 1,095445115 | 0,000243208 | 8,79E-05 | 3 - | 0,000207428 - | 3 - | 0,000257564 - | 4 - | 0,000326463 - | 4 - | 0,000331285 - | 1 - | 9,33E-05 - |
| 430 | 1 | 0.2um_NOKCPHJN_243077 | Bacteria | Bacteroidetes | Flavobacteriia | Flavobacterial | Flavobacteriac | Muricauda | Muricauda sp. | cellular proces | Intracellular tr | 18ATJ | Pfam:YadA | 626418.bglu_ | 1,4 | 0,8 | 1,63E-05 | 9,08E-06 | 2 - | 1,98E-05 - | 2 - | 2,46E-05 - | 0 - | 0 - | 2 - | 2,38E-05 - | 1 - | 1,34E-05 - |
| 431 | 1 | 3um_FIJJAFBF_697727 | Bacteria | Proteobacteri | Gammaprotec | Alteromonada | Alteromonada | Paraglaciecola | Paraglaciecola | cellular proces | Cell wall/mem | COG4105 | Part of the ou | 342610.Patl_1 | 0,6 | 0,489897949 | 3,02E-05 | 2,48E-05 | 1 - | 4,47E-05 - | 0 - | 0 - | 1 - | 5,27E-05 - | 1 - | 5,35E-05 - | 0 - | 0 - |
| 432 | 1 | 0.2um_NOKCPHJN_28563 | Bacteria | Bacteroidetes | Flavobacteriia | unclassified | unclassified | unclassified | Flavobacteria | unclassified | unclassified | unclassified | unclassified | unclassified | 14,4 | 1,854723699 | 0,000313237 | 2,34E-05 | 18 - | 0,000331087 - | 13 - | 0,000296914 - | 13 - | 0,000282255 - | 14 - | 0,000308456 - | 14 - | 0,000347475 - |
| 433 | 1 | 0.2um_NOKCPHJN_675062 | Bacteria | Bacteroidetes | Flavobacteriia | Flavobacterial | Flavobacteriac | Tenacibaculur | Tenacibaculur | unclassified | unclassified | unclassified | unclassified | unclassified | 9,6 | 1,959591794 | 0,000136348 | 2,72E-05 | 10 - | 0,000119684 - | 7 - | 0,000104028 - | 13 - | 0,000183657 - | 9 - | 0,000129025 - | 9 - | 0,000145346 - |
| 434 | 1 | 0.2um_NOKCPHJN_255845 | Bacteria | Bacteroidetes | Flavobacteriia | Flavobacterial | Flavobacteriac | Polaribacter | various | information st | Replication, re | 0PB0H | DNA-binding p | 313598.MED1 | 1,4 | 0,489897949 | 0,000216776 | 8,81E-05 | 1 - | 0,00012753 - | 1 - | 0,000158354 - | 2 - | 0,000301072 - | 1 - | 0,000152759 - | 2 - | 0,000344166 - |
| 435 | 1 | 3um_FIJJAFBF_144524 | Bacteria | Bacteroidetes | Flavobacteriia | Flavobacterial | Flavobacteriac | Formosa | Formosa sp. H | metabolism | Inorganic ion t | 09C49 | tonB-depende | 867900.Celly_ | 0,2 | 0,4 | 5,03E-06 | 1,01E-05 | 0 - | 0 - | 0 - | 0 - | 0 - | 0 - | 0 - | 0 - | 1 - | 2,51E-05 - |
| 436 | 1 | 3um_FIJJAFBF_715071 | Eukaryota | unclassified | Florideophyce | Ceramiales | Rhodomelace | Rhodomela | Rhodomela co | information st | Translation, ril | COG0051 | tRNA binding | 41431.PCC880 | 4 | 0,632455532 | 0,000523683 | 8,02E-05 | 4 - | 0,000441449 - | 4 - | 0,000548149 - | 4 - | 0,000521085 - | 5 - | 0,000660977 - | 3 - | 0,000446753 - |
| 437 | 1 | 3um_FIJJAFBF_224307 | Bacteria | Bacteroidetes | Flavobacteriia | Flavobacterial | Flavobacteriac | Polaribacter | Polaribacter s | cellular proces | Cell motility | COG1360 | Ompa motb d | 313594.PI23P | 1,8 | 1,166190379 | 8,36E-05 | 5,54E-05 | 2 - | 8,05E-05 - | 1 - | 5,00E-05 - | 3 - | 0,000142613 - | 3 - | 0,000144719 - | 0 - | 0 - |
| 438 | 1 | 3um_FIJJAFBF_45956 | Eukaryota | Bacillariophyt | Bacillariophyc | Naviculales | Phaeodactylac | Phaeodactylu | Phaeodactylu | unclassified | unclassified | unclassified | unclassified | unclassified | 0,6 | 0,489897949 | 6,85E-05 | 5,61E-05 | 0 - | 0 - | 1 - | 0,00011222 - | 0 - | 0 - | 1 - | 0,000108255 - | 1 - | 0,000121948 - |
| 439 | 1 | 0.2um_NOKCPHJN_24719 | Bacteria | Bacteroidetes | Flavobacteriia | Flavobacterial | Flavobacteriac | unclassified | Flavobacteriac | unclassified | unclassified | unclassified | unclassified | unclassified | 0,2 | 0,4 | 7,70E-06 | 1,54E-05 | 0 - | 0 - | 1 - | 3,85E-05 - | 0 - | 0 - | 0 - | 0 - | 0 - | 0 - |
| 440 | 1 | 3um_FIJJAFBF_475672 | Bacteria | Bacteroidetes | unclassified | unclassified | unclassified | unclassified | Bacteroidetes | information st | Translation, ril | COG0256 | This is one of t | 391603.FBALC | 0,8 | 0,4 | 9,28E-05 | 4,76E-05 | 1 - | 9,81E-05 - | 0 - | 0 - | 1 - | 0,000115797 - | 1 - | 0,000117507 - | 1 - | 0,000132371 - |
| 441 | 1 | 3um_FIJJAFBF_477870 | Eukaryota | unclassified | Cryptophyta | Pyrenomonad | Chroomonada | Chroomonas | Chroomonas r | metabolism | Energy produc | COG0056 | Produces ATP | 41431.PCC880 | 3,4 | 1,019803903 | 9,19E-05 | 2,07E-05 | 5 - | 0,000115702 - | 2 - | 5,75E-05 - | 4 - | 0,00010926 - | 3 - | 8,32E-05 - | 3 - | 9,37E-05 - |
| 442 | 1 | 3um_FIJJAFBF_390409 | Bacteria | Proteobacteri | Alphaproteob | Rhodobactera | Rhodobactera | Jannaschia | Jannaschia aq | metabolism | Lipid transpor | 01SS3 | Dehydrogenas | 314232.SKA53 | 0,6 | 0,8 | 3,19E-05 | 4,21E-05 | 0 - | 0 - | 1 - | 5,50E-05 - | 2 - | 0,000104619 - | 0 - | 0 - | 0 - | 0 - |
| 443 | 1 | 0.2um_NOKCPHJN_123788 | Bacteria | Proteobacteri | Gammaprotec | Oceanospirilla | Saccharospirill | Reinekea | Reinekea fors | metabolism | Carbohydrate | 05ECY | inositol mono | 314283.MED2 | 0,6 | 0,489897949 | 3,11E-05 | 2,56E-05 | 1 - | 4,61E-05 - | 0 - | 0 - | 1 - | 5,44E-05 - | 1 - | 5,52E-05 - | 0 - | 0 - |
| 444 | 1 | 0.2um_NOKCPHJN_256411 | Bacteria | Bacteroidetes | Flavobacteriia | Flavobacterial | Flavobacteriac | Formosa | Formosa sp. H | information st | Translation, ril | 05C22 | threonyL-tRN | 216432.CA255 | 0,4 | 0,489897949 | 8,99E-06 | 1,11E-05 | 0 - | 0 - | 0 - | 0 - | 1 - | 2,10E-05 - | 0 - | 0 - | 1 - | 2,40E-05 - |
| 445 | 1 | 0.2um_NOKCPHJN_990672 | Bacteria | Proteobacteri | Gammaprotec | unclassified | unclassified | unclassified | OM182 bacte | metabolism | Energy produc | 0QIIJ | Succinyl-CoA l | 247633.GP214 | 0,2 | 0,4 | 9,34E-06 | 1,87E-05 | 0 - | 0 - | 0 - | 0 - | 1 - | 4,67E-05 - | 0 - | 0 - | 0 - | 0 - |
| 446 | 1 | 3um_FIJJAFBF_296519 | Viruses | unclassified | unclassified | unclassified | various | various | various | unclassified | unclassified | unclassified | unclassified | unclassified | 0,2 | 0,4 | 1,38E-05 | 2,77E-05 | 1 - | 6,91E-05 - | 0 - | 0 - | 0 - | 0 - | 0 - | 0 - | 0 - | 0 - |
| 447 | 1 | 3um_FIJJAFBF_167714 | Bacteria | Firmicutes | Clostridia | Clostridiales | Ruminococcac | Faecalibacteri | Faecalibacteri | unclassified | unclassified | unclassified | unclassified | unclassified | 0,4 | 0,489897949 | 1,25E-05 | 1,54E-05 | 0 - | 0 - | 1 - | 3,00E-05 - | 0 - | 0 - | 0 - | 0 - | 1 - | 3,26E-05 - |
| 448 | 1 | 0.2um_NOKCPHJN_264062 | Viruses | unclassified | unclassified | unclassified | Phycodnavirid | Prasinovirus | Ostreococcus | unclassified | unclassified | unclassified | unclassified | unclassified | 4 | 0,632455532 | 0,000128807 | 1,66E-05 | 5 - | 0,000135991 - | 4 - | 0,000135089 - | 4 - | 0,000128419 - | 3 - | 9,77E-05 - | 4 - | 0,0001468 - |
| 449 | 1 | 0.2um_NOKCPHJN_85215 | Bacteria | Bacteroidetes | Flavobacteriia | Flavobacterial | Flavobacteriac | Polaribacter | Polaribacter b | metabolism | Energy produc | COG0056 | Produces ATP | 983544.Lacal_ | 1,4 | 0,489897949 | 3,65E-05 | 1,29E-05 | 1 - | 2,19E-05 - | 1 - | 2,71E-05 - | 2 - | 5,16E-05 - | 2 - | 5,24E-05 - | 1 - | 2,95E-05 - |
| 450 | 1 | 0.2um_NOKCPHJN_611665 | Bacteria | Bacteroidetes | Flavobacteriia | Flavobacterial | Schleiferiacea | Schleiferia | Schleiferia the | metabolism | Inorganic ion t | 05C27 | TonB-depende | 694427.Palpr_ | 3,2 | 0,979795897 | 4,27E-05 | 1,12E-05 | 4 - | 4,56E-05 - | 2 - | 2,83E-05 - | 4 - | 5,39E-05 - | 4 - | 5,47E-05 - | 2 - | 3,08E-05 - |
| 451 | 1 | 0.2um_NOKCPHJN_838460 | Bacteria | Proteobacteri | Gammaprotec | unclassified | unclassified | unclassified | gamma prote | metabolism | Lipid transpor | 125CX | Outer membr | 637616.MDM | 0,6 | 0,489897949 | 1,91E-05 | 1,56E-05 | 0 - | 0 - | 1 - | 3,27E-05 - | 1 - | 3,11E-05 - | 1 - | 3,15E-05 - | 0 - | 0 - |
| 452 | 1 | 0.2um_NOKCPHJN_447983 | Bacteria | Proteobacteri | Alphaproteob | Rhodobactera | Rhodobactera | Tropicimonas | Tropicimonas | metabolism | Amino acid tra | 01QH7 | sulfoacetaldel | 89187.ISM_10 | 0,8 | 0,4 | 1,79E-05 | 9,10E-06 | 1 - | 1,94E-05 - | 1 - | 2,41E-05 - | 1 - | 2,29E-05 - | 1 - | 2,32E-05 - | 0 - | 0 - |
| 453 | 1 | 3um_FIJJAFBF_589704 | Eukaryota | unclassified | Cryptophyta | Pyrenomonad | Geminigerace | Teleaulax | Teleaulax amp | metabolism | Energy produc | COG4451 | RuBisCO catal | 45157.CMV01 | 3,8 | 0,4 | 0,000377808 | 6,79E-05 | 3 - | 0,00024772 - | 4 - | 0,000410126 - | 4 - | 0,000389877 - | 4 - | 0,000395635 - | 4 - | 0,000445682 - |
| 454 | 1 | 3um_FIJJAFBF_62158 | Bacteria | Proteobacteri | Deltaproteoba | Myxococcales | Kofleriaceae | Haliangium | Haliangium oc | unclassified | unclassified | unclassified | unclassified | unclassified | 0,4 | 0,489897949 | 4,10E-05 | 5,05E-05 | 1 - | 9,41E-05 - | 0 - | 0 - | 1 - | 0,000111051 - | 0 - | 0 - | 0 - | 0 - |
| 455 | 2 | 0.2um_NOKCPHJN_597208;0.2um_NOKCPHJN_8 | Bacteria | Bacteroidetes | Flavobacteriia | Flavobacterial | Flavobacteriac | Polaribacter | Polaribacter ir | unclassified | unclassified | unclassified | unclassified | unclassified | 1 | 0 | 4,15E-05 | 3,94E-06 | 1 - | 3,48E-05 - | 1 - | 4,32E-05 - | 1 - | 4,11E-05 - | 1 - | 4,17E-05 - | 1 - | 4,69E-05 - |
| 456 | 1 | 3um_FIJJAFBF_231017 | Bacteria | Actinobacteria | Actinobacteria | Micrococcales | Microbacteria | Candidatus Ac | Candidatus Ac | metabolism | Energy produc | COG0055 | Produces ATP | 281090.Lxx07 | 0,6 | 0,489897949 | 1,62E-05 | 1,34E-05 | 1 - | 2,40E-05 - | 0 - | 0 - | 1 - | 2,83E-05 - | 1 - | 2,88E-05 - | 0 - | 0 - |
| 457 | 1 | 3um_FIJJAFBF_452323 | Bacteria | Bacteroidetes | Flavobacteriia | Flavobacterial | unclassified | unclassified | Flavobacterial | unclassified | unclassified | unclassified | unclassified | unclassified | 1 | 0,894427191 | 5,15E-05 | 4,47E-05 | 2 - | 8,97E-05 - | 0 - | 0 - | 0 - | 0 - | 2 - | 0,000107409 - | 1 - | 6,05E-05 - |
| 458 | 1 | 0.2um_NOKCPHJN_810494 | Eukaryota | Chlorophyta | Mamiellophyc | Mamiellales | Bathycoccacea | Ostreococcus | Ostreococcus | information st | Translation, ril | KOG0052 | This protein p | 436017.A4S6E | 2,8 | 0,4 | 8,22E-05 | 1,73E-05 | 2 - | 4,86E-05 - | 3 - | 9,06E-05 - | 3 - | 8,61E-05 - | 3 - | 8,74E-05 - | 3 - | 9,84E-05 - |
| 459 | 1 | 3um_FIJJAFBF_254416 | Bacteria | Cyanobacteria | unclassified | Oscillatoriales | Microcoleacea | Trichodesmiu | Trichodesmiu | information st | Translation, ril | 08UIK | This protein bi | 411154.GFO_ | 0,8 | 0,748331477 | 6,93E-05 | 6,13E-05 | 2 - | 0,000158313 - | 0 - | 0 - | 1 - | 9,34E-05 - | 1 - | 9,48E-05 - | 0 - | 0 - |
| 460 | 3 | 3um_FIJJAFBF_316930;3um_FIJJAFBF_587443;3u | Bacteria | Bacteroidetes | Flavobacteriia | Flavobacterial | Flavobacteriac | various | various | metabolism | Amino acid tra | 0PF77 | Dehydrogenas | 983548.Krodi_ | 2,4 | 0,489897949 | 0,000102565 | 1,51E-05 | 3 - | 0,000108965 - | 2 - | 9,02E-05 - | 3 - | 0,000128622 - | 2 - | 8,70E-05 - | 2 - | 9,80E-05 - |
| 461 | 1 | 3um_FIJJAFBF_291412 | Bacteria | Bacteroidetes | Flavobacteriia | Flavobacterial | Flavobacteriac | Polaribacter | Polaribacter s | unclassified | unclassified | unclassified | unclassified | unclassified | 1 | 0 | 2,53E-05 | 2,40E-06 | 1 - | 2,12E-05 - | 1 - | 2,63E-05 - | 1 - | 2,50E-05 - | 1 - | 2,54E-05 - | 1 - | 2,86E-05 - |
| 462 | 1 | 0.2um_NOKCPHJN_75389 | Bacteria | Bacteroidetes | Flavobacteriia | Flavobacterial | Flavobacteriac | Polaribacter | various | information st | Transcription | 0P88Z | transcriptiona | 688270.Celal_ | 0,6 | 0,489897949 | 4,88E-05 | 4,01E-05 | 1 - | 7,22E-05 - | 0 - | 0 - | 1 - | 8,52E-05 - | 1 - | 8,65E-05 - | 0 - | 0 - |
| 463 | 1 | 0.2um_NOKCPHJN_767795 | Eukaryota | unclassified | unclassified | Longamoebia | Acanthamoeb | Acanthamoeb | Acanthamoeb | unclassified | unclassified | unclassified | unclassified | unclassified | 0,8 | 0,4 | 4,80E-05 | 2,44E-05 | 1 - | 5,19E-05 - | 1 - | 6,45E-05 - | 1 - | 6,13E-05 - | 1 - | 6,22E-05 - | 0 - | 0 - |
| 464 | 1 | 3um_FIJJAFBF_120755 | Bacteria | Bacteroidetes | Flavobacteriia | Flavobacterial | Flavobacteriac | Polaribacter | Polaribacter s | metabolism | Energy produc | 099YH | Pyruvate dehy | 983544.Lacal_ | 0,2 | 0,4 | 7,06E-06 | 1,41E-05 | 1 - | 3,53E-05 - | 0 - | 0 - | 0 - | 0 - | 0 - | 0 - | 0 - | 0 - |
| 465 | 1 | 3um_FIJJAFBF_143872 | Bacteria | Proteobacteri | Gammaprotec | Oceanospirilla | Saccharospirill | Reinekea | Reinekea blan | unclassified | unclassified | unclassified | unclassified | unclassified | 1,2 | 0,4 | 7,67E-05 | 3,26E-05 | 1 - | 5,24E-05 - | 1 - | 6,51E-05 - | 1 - | 6,19E-05 - | 1 - | 6,28E-05 - | 2 - | 0,000141438 - |
| 466 | 1 | 3um_FIJJAFBF_233982 | Bacteria | Bacteroidetes | Flavobacteriia | Flavobacterial | Flavobacteriac | Polaribacter | Polaribacter v | unclassified | unclassified | unclassified | unclassified | unclassified | 1 | 0 | 0,000217503 | 2,07E-05 | 1 - | 0,000182185 - | 1 - | 0,00022622 - | 1 - | 0,000215051 - | 1 - | 0,000218227 - | 1 - | 0,000245833 - |
| 467 | 1 | 0.2um_NOKCPHJN_1044599 | unclassified | unclassified | unclassified | unclassified | unclassified | unclassified | unclassified | unclassified | unclassified | unclassified | unclassified | unclassified | 0,2 | 0,4 | 5,59E-06 | 1,12E-05 | 0 - | 0 - | 1 - | 2,79E-05 - | 0 - | 0 - | 0 - | 0 - | 0 - | 0 - |
| 468 | 1 | 0.2um_NOKCPHJN_378561 | Bacteria | Proteobacteri | Alphaproteob | Pelagibacteral | Pelagibacterac | Candidatus Pe | Candidatus Pe | cellular proces | Cell wall/mem | 7257 | NA | 335992.SAR11 | 1 | 0 | 4,16E-05 | 3,95E-06 | 1 - | 3,49E-05 - | 1 - | 4,33E-05 - | 1 - | 4,12E-05 - | 1 - | 4,18E-05 - | 1 - | 4,71E-05 - |
| 469 | 1 | 3um_FIJJAFBF_383588 | unclassified | unclassified | unclassified | unclassified | unclassified | unclassified | unclassified | unclassified | unclassified | unclassified | unclassified | unclassified | 0,2 | 0,4 | 4,59E-06 | 9,18E-06 | 0 - | 0 - | 0 - | 0 - | 0 - | 0 - | 1 - | 2,30E-05 - | 0 - | 0 - |
| 470 | 1 | 3um_FIJJAFBF_13586 | Bacteria | Proteobacteri | Gammaprotec | Cellvibrionales | Porticoccacea | unclassified | SAR92 bacteri | unclassified | unclassified | unclassified | unclassified | unclassified | 1 | 0 | 0,000161208 | 1,53E-05 | 1 - | 0,000135031 - | 1 - | 0,000167669 - | 1 - | 0,000159391 - | 1 - | 0,000161745 - | 1 - | 0,000182205 - |
| 471 | 1 | 3um_FIJJAFBF_129174 | Bacteria | Bacteroidetes | Flavobacteriia | Flavobacterial | Flavobacteriac | Polaribacter | Polaribacter s | unclassified | unclassified | unclassified | unclassified | unclassified | 1,6 | 0,489897949 | 4,01E-05 | 1,11E-05 | 2 - | 4,27E-05 - | 2 - | 5,31E-05 - | 2 - | 5,05E-05 - | 1 - | 2,56E-05 - | 1 - | 2,88E-05 - |
| 472 | 1 | 3um_FIJJAFBF_242895 | Bacteria | Bacteroidetes | Flavobacteriia | Flavobacterial | Flavobacteriac | Polaribacter | Polaribacter d | poorly charact | Function unkn | 08ME3 | Alpha amylase | 983548.Krodi_ | 2,4 | 0,489897949 | 9,44E-05 | 1,67E-05 | 3 - | 9,98E-05 - | 3 - | 0,000123929 - | 2 - | 7,85E-05 - | 2 - | 7,97E-05 - | 2 - | 8,98E-05 - |
| 473 | 1 | 0.2um_NOKCPHJN_200781 | Bacteria | Proteobacteri | Gammaprotec | Oceanospirilla | Saccharospirill | Reinekea | Reinekea fors | metabolism | Nucleotide tra | 05CAT | Cytidine monc | 314283.MED2 | 1,4 | 0,489897949 | 8,28E-05 | 2,37E-05 | 2 - | 0,000101572 - | 1 - | 6,31E-05 - | 2 - | 0,000119896 - | 1 - | 6,08E-05 - | 1 - | 6,85E-05 - |
| 474 | 1 | 3um_FIJJAFBF_332924 | Bacteria | Bacteroidetes | unclassified | unclassified | unclassified | unclassified | Bacteroidetes | unclassified | unclassified | unclassified | unclassified | unclassified | 3 | 0,632455532 | 0,000272846 | 4,64E-05 | 4 - | 0,000308126 - | 3 - | 0,000286951 - | 3 - | 0,000272783 - | 2 - | 0,000184541 - | 3 - | 0,000311828 - |
| 475 | 1 | 0.2um_NOKCPHJN_434211 | Bacteria | Verrucomicrol | Opitutae | Opitutales | Opitutaceae | Ereboglobus | Ereboglobus l | metabolism | Amino acid tra | COG1505 | prolyl oligope | 2850.Phatr46 | 0,4 | 0,489897949 | 5,77E-06 | 7,10E-06 | 1 - | 1,32E-05 - | 0 - | 0 - | 1 - | 1,56E-05 - | 0 - | 0 - | 0 - | 0 - |
| 476 | 1 | 3um_FIJJAFBF_54816 | Bacteria | Bacteroidetes | Flavobacteriia | Flavobacterial | Flavobacteriac | Polaribacter | Polaribacter s | unclassified | unclassified | unclassified | unclassified | unclassified | 0,8 | 0,748331477 | 3,34E-05 | 2,88E-05 | 2 - | 7,26E-05 - | 1 - | 4,51E-05 - | 0 - | 0 - | 0 - | 0 - | 1 - | 4,90E-05 - |
| 477 | 1 | 3um_FIJJAFBF_505329 | Bacteria | Bacteroidetes | Flavobacteriia | Flavobacterial | unclassified | unclassified | Flavobacterial | metabolism | Inorganic ion t | 05C27 | TonB-depende | 391603.FBALC | 2 | 1,897366596 | 2,44E-05 | 2,42E-05 | 3 - | 3,17E-05 - | 5 - | 6,56E-05 - | 2 - | 2,49E-05 - | 0 - | 0 - | 0 - | 0 - |
| 478 | 1 | 0.2um_NOKCPHJN_28392 | Bacteria | Proteobacteri | Gammaprotec | unclassified | unclassified | unclassified | Gammaprotec | metabolism | Inorganic ion t | 0QK8T | TonB-depende | 247633.GP214 | 4 | 1,095445115 | 0,00031461 | 0,000114209 | 3 - | 0,000193444 - | 4 - | 0,000320267 - | 3 - | 0,000228341 - | 4 - | 0,000308951 - | 6 - | 0,000522049 - |
| 479 | 1 | 3um_FIJJAFBF_57965 | Bacteria | Bacteroidetes | Cytophagia | Cytophagales | Flammeovirga | unclassified | Flammeovirga | metabolism | Inorganic ion t | 0ZVMK | efflux transpo | 411154.GFO_ | 1 | 0,632455532 | 3,55E-05 | 2,00E-05 | 2 - | 6,20E-05 - | 0 - | 0 - | 1 - | 3,66E-05 - | 1 - | 3,72E-05 - | 1 - | 4,19E-05 - |
| 480 | 1 | 3um_FIJJAFBF_216059 | Bacteria | Proteobacteri | Gammaprotec | Xanthomonad | Rhodanobacte | Oleiagrimonas | Oleiagrimonas | information st | Translation, ril | COG0050 | This protein p | 523791.Kkor_ | 0,8 | 0,4 | 0,000187109 | 9,61E-05 | 1 - | 0,000197891 - | 0 - | 0 - | 1 - | 0,00023359 - | 1 - | 0,00023704 - | 1 - | 0,000267025 - |
| 481 | 2 | 3um_FIJJAFBF_02497;3um_FIJJAFBF_45525 | Bacteria | Proteobacteri | Alphaproteob | Rhodobactera | Rhodobactera | various | various | information st | Translation, ril | 17C3B | The globular d | various | 2,6 | 0,489897949 | 0,000285414 | 6,96E-05 | 2 - | 0,000182185 - | 2 - | 0,00022622 - | 3 - | 0,000322577 - | 3 - | 0,000327341 - | 3 - | 0,000368749 - |
| 482 | 1 | 3um_FIJJAFBF_660321 | Bacteria | Proteobacteri | Alphaproteob | Rhodobactera | Rhodobactera | unclassified | Rhodobactera | metabolism | Inorganic ion t | 05EKV | Phosphonate | 246200.SPO04 | 0,4 | 0,489897949 | 1,41E-05 | 1,75E-05 | 1 - | 3,15E-05 - | 1 - | 3,92E-05 - | 0 - | 0 - | 0 - | 0 - | 0 - | 0 - |
| 483 | 1 | 3um_FIJJAFBF_56732 | Bacteria | Proteobacteri | Gammaprotec | Alteromonada | Alteromonada | Paraglaciecola | Paraglaciecola | metabolism | Energy produc | 0QIAU | Electron trans | 314275.MADE | 0,8 | 0,748331477 | 4,19E-05 | 3,64E-05 | 2 - | 9,22E-05 - | 0 - | 0 - | 0 - | 0 - | 1 - | 5,52E-05 - | 1 - | 6,22E-05 - |
| 484 | 1 | 0.2um_NOKCPHJN_103568 | Bacteria | Proteobacteri | Gammaprotec | Alteromonada | Alteromonada | Paraglaciecola | Paraglaciecola | unclassified | unclassified | unclassified | unclassified | unclassified | 2 | 0 | 9,62E-05 | 9,13E-06 | 2 - | 8,05E-05 - | 2 - | 0,000100013 - | 2 - | 9,51E-05 - | 2 - | 9,65E-05 - | 2 - | 0,000108684 - |
| 485 | 1 | 3um_FIJJAFBF_128449 | Bacteria | Bacteroidetes | Flavobacteriia | Flavobacterial | Flavobacteriac | Polaribacter | Polaribacter s | unclassified | unclassified | unclassified | unclassified | unclassified | 4,8 | 0,4 | 0,000161732 | 1,89E-05 | 5 - | 0,000141351 - | 4 - | 0,000140413 - | 5 - | 0,00016685 - | 5 - | 0,000169314 - | 5 - | 0,000190732 - |
| 486 | 2 | 3um_FIJJAFBF_236977;3um_FIJJAFBF_467035 | Bacteria | Bacteroidetes | Flavobacteriia | Flavobacterial | Flavobacteriac | Polaribacter | various | unclassified | unclassified | unclassified | unclassified | unclassified | 1,2 | 0,748331477 | 2,16E-05 | 1,33E-05 | 2 - | 3,15E-05 - | 2 - | 3,91E-05 - | 1 - | 1,86E-05 - | 1 - | 1,89E-05 - | 0 - | 0 - |
| 487 | 1 | 0.2um_NOKCPHJN_212097 | Viruses | unclassified | unclassified | unclassified | Phycodnavirid | Prasinovirus | Micromonas p | unclassified | unclassified | unclassified | unclassified | unclassified | 3 | 0,632455532 | 9,57E-05 | 1,19E-05 | 4 - | 0,000109052 - | 3 - | 0,000101557 - | 3 - | 9,65E-05 - | 3 - | 9,80E-05 - | 2 - | 7,36E-05 - |
| 488 | 1 | 0.2um_NOKCPHJN_694782 | Bacteria | Proteobacteri | Alphaproteob | Pelagibacteral | Pelagibacterac | Candidatus Pe | Candidatus Pe | unclassified | unclassified | unclassified | unclassified | unclassified | 3 | 0,632455532 | 0,00011927 | 2,80E-05 | 3 - | 9,98E-05 - | 3 - | 0,000123929 - | 2 - | 7,85E-05 - | 4 - | 0,000159401 - | 3 - | 0,000134673 - |
| 489 | 1 | 3um_FIJJAFBF_225879 | Bacteria | Bacteroidetes | Flavobacteriia | Flavobacterial | Flavobacteriac | Flaviramulus | Flaviramulus b | cellular proces | Cell wall/mem | 06B97 | phosphate-sel | 391598.FBBAL | 3 | 0,894427191 | 0,000100052 | 3,21E-05 | 2 - | 5,54E-05 - | 2 - | 6,88E-05 - | 4 - | 0,000130901 - | 4 - | 0,000132834 - | 3 - | 0,000112228 - |
| 490 | 2 | 0.2um_NOKCPHJN_206762;3um_FIJJAFBF_7945 | Bacteria | Bacteroidetes | Flavobacteriia | unclassified | unclassified | unclassified | Flavobacteria | information st | Translation, ril | COG0184 | Forms an inte | 487796.Flav2A | 1 | 0 | 0,000153963 | 1,46E-05 | 1 - | 0,000128963 - | 1 - | 0,000160134 - | 1 - | 0,000152227 - | 1 - | 0,000154476 - | 1 - | 0,000174016 - |
| 491 | 1 | 0.2um_NOKCPHJN_170706 | Bacteria | Bacteroidetes | unclassified | unclassified | unclassified | unclassified | Bacteroidetes | unclassified | unclassified | unclassified | unclassified | unclassified | 2,4 | 0,8 | 4,62E-05 | 1,48E-05 | 3 - | 4,88E-05 - | 3 - | 6,06E-05 - | 1 - | 1,92E-05 - | 3 - | 5,84E-05 - | 2 - | 4,39E-05 - |
| 492 | 1 | 3um_FIJJAFBF_556042 | Bacteria | Proteobacteri | Gammaprotec | Cellvibrionales | Halieaceae | unclassified | marine gamm | information st | Translation, ril | 08YY1 | This protein is | 247639.MGP2 | 0,2 | 0,4 | 1,43E-05 | 2,87E-05 | 1 - | 7,17E-05 - | 0 - | 0 - | 0 - | 0 - | 0 - | 0 - | 0 - | 0 - |
| 493 | 1 | 0.2um_NOKCPHJN_67716 | Bacteria | Proteobacteri | Alphaproteob | Rhodobactera | Rhodobactera | Litoreibacter | Litoreibacter a | metabolism | Amino acid tra | 01RBF | Branched-cha | 367336.OM22 | 5 | 0,632455532 | 0,000153092 | 1,84E-05 | 5 - | 0,000128963 - | 5 - | 0,000160134 - | 6 - | 0,000182673 - | 5 - | 0,000154476 - | 4 - | 0,000139213 - |
| 494 | 1 | 3um_FIJJAFBF_163703 | Bacteria | Bacteroidetes | Flavobacteriia | Flavobacterial | Flavobacteriac | Lacinutrix | Lacinutrix sp. | unclassified | unclassified | unclassified | unclassified | unclassified | 3,2 | 1,326649916 | 7,66E-05 | 2,79E-05 | 5 - | 0,000102115 - | 3 - | 7,61E-05 - | 1 - | 2,41E-05 - | 4 - | 9,79E-05 - | 3 - | 8,27E-05 - |
| 495 | 1 | 3um_FIJJAFBF_197932 | Bacteria | Bacteroidetes | Cytophagia | Cytophagales | Flammeovirga | Roseivirga | Roseivirga spo | poorly charact | Function unkn | 0FXQ6 | NA | 643867.Ftrac_ | 0,2 | 0,4 | 1,54E-05 | 3,07E-05 | 0 - | 0 - | 0 - | 0 - | 0 - | 0 - | 1 - | 7,68E-05 - | 0 - | 0 - |
| 496 | 1 | 0.2um_NOKCPHJN_941524 | Eukaryota | unclassified | Spirotrichea | Sporadotrichi | Oxytrichidae | Oxytricha | Oxytricha trifa | unclassified | unclassified | unclassified | unclassified | unclassified | 3 | 0,632455532 | 9,12E-05 | 2,00E-05 | 3 - | 7,69E-05 - | 4 - | 0,000127249 - | 3 - | 9,07E-05 - | 3 - | 9,21E-05 - | 2 - | 6,91E-05 - |
| 497 | 1 | 0.2um_NOKCPHJN_320071 | Eukaryota | unclassified | unclassified | Choanoflagelli | Salpingoecida | Monosiga | Monosiga bre | unclassified | unclassified | unclassified | unclassified | unclassified | 0,2 | 0,4 | 3,28E-06 | 6,57E-06 | 0 - | 0 - | 1 - | 1,64E-05 - | 0 - | 0 - | 0 - | 0 - | 0 - | 0 - |
| 498 | 1 | 3um_FIJJAFBF_58647 | Eukaryota | unclassified | Spirotrichea | Sporadotrichi | Oxytrichidae | Stylonychia | Stylonychia le | unclassified | unclassified | unclassified | unclassified | unclassified | 0,2 | 0,4 | 1,49E-05 | 2,98E-05 | 0 - | 0 - | 0 - | 0 - | 1 - | 7,44E-05 - | 0 - | 0 - | 0 - | 0 - |
| 499 | 1 | 0.2um_NOKCPHJN_174935 | Eukaryota | Chlorophyta | Chlorophycea | Sphaeropleale | Selenastracea | Raphidocelis | Raphidocelis s | unclassified | unclassified | unclassified | unclassified | unclassified | 0,4 | 0,489897949 | 2,42E-06 | 2,97E-06 | 0 - | 0 - | 1 - | 6,21E-06 - | 1 - | 5,90E-06 - | 0 - | 0 - | 0 - | 0 - |
| 500 | 1 | 0.2um_NOKCPHJN_66685 | Bacteria | Bacteroidetes | Flavobacteriia | Flavobacterial | Flavobacteriac | Polaribacter | Polaribacter s | metabolism | Amino acid tra | 08IQK | o-acetylhomo | 1046627.BZAF | 1,2 | 0,748331477 | 3,81E-05 | 2,28E-05 | 2 - | 5,41E-05 - | 2 - | 6,72E-05 - | 0 - | 0 - | 1 - | 3,24E-05 - | 1 - | 3,65E-05 - |
| 501 | 1 | 3um_FIJJAFBF_283713 | Eukaryota | unclassified | Pelagophycea | Pelagomonad | unclassified | Aureococcus | Aureococcus a | information st | Translation, ril | KOG3204 | 60s ribosomal | 65071.PYU1_T | 0,8 | 0,4 | 5,40E-05 | 2,77E-05 | 1 - | 5,65E-05 - | 1 - | 7,02E-05 - | 1 - | 6,67E-05 - | 0 - | 0 - | 1 - | 7,63E-05 - |
| 502 | 1 | 0.2um_NOKCPHJN_1080019 | Bacteria | Proteobacteri | Alphaproteob | Rhodobactera | Rhodobactera | Sulfitobacter | Sulfitobacter r | metabolism | Secondary me | 01Q86 | extracellular s | 367336.OM22 | 1 | 0 | 3,76E-05 | 3,57E-06 | 1 - | 3,15E-05 - | 1 - | 3,92E-05 - | 1 - | 3,72E-05 - | 1 - | 3,78E-05 - | 1 - | 4,25E-05 - |
| 503 | 1 | 3um_FIJJAFBF_487371 | Bacteria | Proteobacteri | Gammaprotec | Oceanospirilla | Saccharospirill | Reinekea | Reinekea fors | poorly charact | Function unkn | 0XW1Y | transmembra | 314287.GB220 | 0,4 | 0,489897949 | 2,81E-05 | 3,46E-05 | 1 - | 6,45E-05 - | 0 - | 0 - | 1 - | 7,61E-05 - | 0 - | 0 - | 0 - | 0 - |
| 504 | 1 | 3um_FIJJAFBF_488138 | Bacteria | Bacteroidetes | Flavobacteriia | Flavobacterial | Flavobacteriac | Algibacter | Algibacter pec | cellular proces | Posttranslatio | 05E6M | Sulphatase-m | 983544.Lacal_ | 0,4 | 0,489897949 | 1,08E-05 | 1,33E-05 | 1 - | 2,46E-05 - | 0 - | 0 - | 0 - | 0 - | 1 - | 2,94E-05 - | 0 - | 0 - |
| 505 | 1 | 3um_FIJJAFBF_241168 | Bacteria | Proteobacteri | Gammaprotec | Oceanospirilla | Saccharospirill | Reinekea | Reinekea fors | metabolism | Carbohydrate | 0QKP5 | TRAP dicarbox | 717774.Marm | 3,8 | 0,4 | 0,000151048 | 1,99E-05 | 4 - | 0,000133461 - | 3 - | 0,00012429 - | 4 - | 0,000157537 - | 4 - | 0,000159864 - | 4 - | 0,000180087 - |
| 506 | 1 | 3um_FIJJAFBF_321858 | Bacteria | Bacteroidetes | Flavobacteriia | Flavobacterial | Flavobacteriac | Polaribacter | Polaribacter ta | cellular proces | Cell wall/mem | COG2885 | Ompa motb d | 313594.PI23P | 2,2 | 0,4 | 6,56E-05 | 6,01E-06 | 3 - | 7,60E-05 - | 2 - | 6,29E-05 - | 2 - | 5,98E-05 - | 2 - | 6,07E-05 - | 2 - | 6,84E-05 - |
| 507 | 1 | 3um_FIJJAFBF_736134 | Bacteria | Bacteroidetes | Flavobacteriia | various | various | various | various | unclassified | unclassified | unclassified | unclassified | unclassified | 2,2 | 0,4 | 0,000175304 | 4,71E-05 | 2 - | 0,000131927 - | 2 - | 0,000163815 - | 2 - | 0,000155727 - | 2 - | 0,000158027 - | 3 - | 0,000267025 - |
| 508 | 1 | 0.2um_NOKCPHJN_311161 | Bacteria | Proteobacteri | Alphaproteob | Rhodobactera | Rhodobactera | Sulfitobacter | Sulfitobacter r | metabolism | Carbohydrate | 0XPV5 | extracellular s | 367336.OM22 | 1 | 0,632455532 | 3,44E-05 | 2,12E-05 | 0 - | 0 - | 1 - | 3,46E-05 - | 1 - | 3,29E-05 - | 2 - | 6,67E-05 - | 1 - | 3,76E-05 - |
| 509 | 1 | 0.2um_NOKCPHJN_96085 | Bacteria | Proteobacteri | Gammaprotec | Alteromonada | Alteromonada | Glaciecola | Glaciecola sp. | unclassified | unclassified | unclassified | unclassified | unclassified | 0,2 | 0,4 | 1,54E-06 | 3,09E-06 | 1 - | 7,71E-06 - | 0 - | 0 - | 0 - | 0 - | 0 - | 0 - | 0 - | 0 - |
| 510 | 1 | 3um_FIJJAFBF_123783 | Bacteria | Bacteroidetes | Flavobacteriia | Flavobacterial | Flavobacteriac | Polaribacter | Polaribacter s | unclassified | unclassified | unclassified | unclassified | unclassified | 0,6 | 0,489897949 | 1,88E-05 | 1,56E-05 | 1 - | 2,61E-05 - | 1 - | 3,25E-05 - | 0 - | 0 - | 0 - | 0 - | 1 - | 3,53E-05 - |
| 511 | 1 | 0.2um_NOKCPHJN_30998 | Bacteria | Proteobacteri | Alphaproteob | Rhodobactera | Rhodobactera | Amylibacter | Amylibacter k | metabolism | Amino acid tra | 16PK4 | phospho-2-de | 375451.RD1_2 | 1 | 0 | 3,00E-05 | 2,85E-06 | 1 - | 2,52E-05 - | 1 - | 3,13E-05 - | 1 - | 2,97E-05 - | 1 - | 3,01E-05 - | 1 - | 3,40E-05 - |
| 512 | 1 | 3um_FIJJAFBF_638896 | Bacteria | Proteobacteri | Alphaproteob | Rhodobactera | Rhodobactera | Roseobacter | Roseobacter s | metabolism | Coenzyme tra | 05DEJ | cobaltochelat | 246200.SPO28 | 1,2 | 0,4 | 4,88E-05 | 1,08E-05 | 2 - | 7,00E-05 - | 1 - | 4,35E-05 - | 1 - | 4,13E-05 - | 1 - | 4,19E-05 - | 1 - | 4,72E-05 - |
| 513 | 1 | 0.2um_NOKCPHJN_60422 | Bacteria | Bacteroidetes | Flavobacteriia | Flavobacterial | Flavobacteriac | unclassified | Flavobacteriac | information st | Translation, ril | 177SZ | This protein bi | 314283.MED2 | 3,4 | 1,019803903 | 0,000316656 | 7,25E-05 | 5 - | 0,00039853 - | 2 - | 0,000197943 - | 3 - | 0,000282255 - | 4 - | 0,000381898 - | 3 - | 0,000322655 - |
| 514 | 3 | 3um_FIJJAFBF_205206;3um_FIJJAFBF_41554;3ur | Eukaryota | Bacillariophyt | various | various | various | various | various | cellular proces | Posttranslatio | 16R8D | CbbX protein | various | 0,2 | 0,4 | 7,92E-06 | 1,58E-05 | 1 - | 3,96E-05 - | 0 - | 0 - | 0 - | 0 - | 0 - | 0 - | 0 - | 0 - |
| 515 | 1 | 3um_FIJJAFBF_363842 | Bacteria | unclassified | unclassified | unclassified | unclassified | unclassified | uncultured ba | unclassified | unclassified | unclassified | unclassified | unclassified | 0,8 | 0,748331477 | 5,65E-05 | 4,92E-05 | 2 - | 0,000124757 - | 0 - | 0 - | 1 - | 7,36E-05 - | 0 - | 0 - | 1 - | 8,42E-05 - |
| 516 | 2 | 0.2um_NOKCPHJN_1053692;3um_FIJJAFBF_5729 | Bacteria | Proteobacteri | Alphaproteob | Rhodobactera | Rhodobactera | various | various | cellular proces | Posttranslatio | 01SC2 | alkyl hydroper | various | 1 | 0 | 6,75E-05 | 6,41E-06 | 1 - | 5,65E-05 - | 1 - | 7,02E-05 - | 1 - | 6,67E-05 - | 1 - | 6,77E-05 - | 1 - | 7,63E-05 - |
| 517 | 1 | 3um_FIJJAFBF_51818 | Bacteria | Proteobacteri | Alphaproteob | Rhodobactera | Rhodobactera | Planktomarina | Planktomarina | metabolism | Amino acid tra | 01R4A | Polyamine AB | 488538.SAR11 | 0,4 | 0,489897949 | 1,61E-05 | 1,98E-05 | 0 - | 0 - | 0 - | 0 - | 0 - | 0 - | 1 - | 3,79E-05 - | 1 - | 4,27E-05 - |
| 518 | 1 | 0.2um_NOKCPHJN_71124 | Eukaryota | unclassified | Cryptophyta | Pyrenomonad | Geminigerace | Guillardia | Guillardia the | unclassified | unclassified | unclassified | unclassified | unclassified | 0,4 | 0,489897949 | 3,17E-06 | 3,89E-06 | 0 - | 0 - | 1 - | 8,13E-06 - | 1 - | 7,73E-06 - | 0 - | 0 - | 0 - | 0 - |
| 519 | 1 | 3um_FIJJAFBF_12518 | unclassified | unclassified | unclassified | unclassified | unclassified | unclassified | unclassified | unclassified | unclassified | unclassified | unclassified | unclassified | 1,2 | 0,4 | 0,00013597 | 4,69E-05 | 1 - | 9,49E-05 - | 1 - | 0,000117784 - | 1 - | 0,000111969 - | 2 - | 0,000227245 - | 1 - | 0,000127995 - |
| 520 | 1 | 3um_FIJJAFBF_58539 | Eukaryota | Phaeophyceae | unclassified | Ectocarpales | Ectocarpaceae | Ectocarpus | Ectocarpus sili | information st | Translation, ril | 1508W | Ribosomal pro | 4787.PITG_09 | 0,8 | 0,4 | 0,000103331 | 5,31E-05 | 1 - | 0,00010828 - | 1 - | 0,000134452 - | 1 - | 0,000127813 - | 0 - | 0 - | 1 - | 0,000146108 - |
| 521 | 2 | 0.2um_NOKCPHJN_374941;3um_FIJJAFBF_45107 | Bacteria | Bacteroidetes | unclassified | unclassified | unclassified | unclassified | Bacteroidetes | cellular proces | Posttranslatio | 09ADN | alkyl hydroper | 926562.Oweh | 0,2 | 0,4 | 1,32E-05 | 2,63E-05 | 0 - | 0 - | 0 - | 0 - | 0 - | 0 - | 1 - | 6,58E-05 - | 0 - | 0 - |
| 522 | 1 | 3um_FIJJAFBF_469778 | Bacteria | Bacteroidetes | Flavobacteriia | Flavobacterial | Flavobacteriac | Hyunsoonleell | Hyunsoonleell | information st | Translation, ril | 0PA05 | 30S ribosomal | 391603.FBALC | 3,4 | 1,624807681 | 0,000244821 | 0,000113099 | 5 - | 0,000305257 - | 2 - | 0,000151616 - | 1 - | 7,21E-05 - | 5 - | 0,000365647 - | 4 - | 0,00032952 - |
| 523 | 1 | 3um_FIJJAFBF_295856 | Bacteria | Proteobacteri | Gammaprotec | Oceanospirilla | Saccharospirill | Reinekea | Reinekea fors | poorly charact | Function unkn | 05KTH | gatB Yqey | 314283.MED2 | 0,2 | 0,4 | 1,55E-05 | 3,10E-05 | 1 - | 7,76E-05 - | 0 - | 0 - | 0 - | 0 - | 0 - | 0 - | 0 - | 0 - |
| 524 | 1 | 0.2um_NOKCPHJN_585129 | Eukaryota | Chordata | unclassified | Petromyzontif | Petromyzontic | Petromyzon | Petromyzon m | cellular proces | Cell cycle cont | 0V9W7 | retinitis pigme | 8128.ENSONIF | 0,6 | 0,489897949 | 7,69E-06 | 6,31E-06 | 0 - | 0 - | 0 - | 0 - | 1 - | 1,22E-05 - | 1 - | 1,24E-05 - | 1 - | 1,39E-05 - |
| 525 | 1 | 3um_FIJJAFBF_111334 | Eukaryota | Apicomplexa | Conoidasida | Eucoccidiorida | Eimeriidae | Eimeria | Eimeria mitis | unclassified | unclassified | unclassified | unclassified | unclassified | 1 | 0,632455532 | 3,81E-05 | 2,32E-05 | 0 - | 0 - | 1 - | 3,84E-05 - | 2 - | 7,30E-05 - | 1 - | 3,71E-05 - | 1 - | 4,17E-05 - |
| 526 | 1 | 3um_FIJJAFBF_679233 | Bacteria | Bacteroidetes | unclassified | unclassified | unclassified | unclassified | unidentified e | unclassified | unclassified | unclassified | unclassified | unclassified | 1 | 0,632455532 | 8,72E-05 | 4,89E-05 | 2 - | 0,000151022 - | 1 - | 9,38E-05 - | 1 - | 8,91E-05 - | 0 - | 0 - | 1 - | 0,000101891 - |
| 527 | 1 | 3um_FIJJAFBF_184101 | Bacteria | Bacteroidetes | Flavobacteriia | Flavobacterial | Flavobacteriac | Algibacter | Algibacter sp. | metabolism | Carbohydrate | 05CV1 | Transketolase | 313603.FB217 | 2,2 | 0,748331477 | 4,30E-05 | 1,23E-05 | 3 - | 5,05E-05 - | 2 - | 4,18E-05 - | 3 - | 5,96E-05 - | 2 - | 4,03E-05 - | 1 - | 2,27E-05 - |
| 528 | 1 | 0.2um_NOKCPHJN_224594 | Bacteria | Proteobacteri | Betaproteoba | unclassified | unclassified | unclassified | Betaproteoba | unclassified | unclassified | unclassified | unclassified | unclassified | 5,2 | 0,979795897 | 6,34E-05 | 8,20E-06 | 7 - | 7,23E-05 - | 5 - | 6,41E-05 - | 4 - | 4,88E-05 - | 5 - | 6,19E-05 - | 5 - | 6,97E-05 - |
| 529 | 1 | 0.2um_NOKCPHJN_402769 | Eukaryota | unclassified | Oomycetes | Peronosporale | Peronosporac | Phytophthora | Phytophthora | unclassified | unclassified | unclassified | unclassified | unclassified | 0,2 | 0,4 | 5,86E-06 | 1,17E-05 | 0 - | 0 - | 1 - | 2,93E-05 - | 0 - | 0 - | 0 - | 0 - | 0 - | 0 - |
| 530 | 1 | 0.2um_NOKCPHJN_197671 | Bacteria | Proteobacteri | Gammaprotec | Cellvibrionales | Halieaceae | unclassified | marine gamm | unclassified | unclassified | unclassified | unclassified | unclassified | 3 | 0,632455532 | 4,62E-05 | 9,08E-06 | 3 - | 3,90E-05 - | 3 - | 4,85E-05 - | 4 - | 6,14E-05 - | 3 - | 4,68E-05 - | 2 - | 3,51E-05 - |
| 531 | 1 | 0.2um_NOKCPHJN_325512 | Bacteria | Proteobacteri | Gammaprotec | Xanthomonad | Xanthomonad | Pseudoxantho | Pseudoxantho | metabolism | Secondary me | 0BCQ5 | DsbA oxidorec | 381666.H16_E | 0,2 | 0,4 | 1,13E-05 | 2,25E-05 | 0 - | 0 - | 0 - | 0 - | 0 - | 0 - | 1 - | 5,63E-05 - | 0 - | 0 - |
| 532 | 1 | 0.2um_NOKCPHJN_217486 | Bacteria | Proteobacteri | Gammaprotec | Oceanospirilla | Saccharospirill | Reinekea | Reinekea fors | metabolism | Inorganic ion t | COG3639 | Phosphonate | 270374.MELB | 1,6 | 0,489897949 | 8,12E-05 | 2,96E-05 | 1 - | 4,16E-05 - | 2 - | 0,000103275 - | 1 - | 4,91E-05 - | 2 - | 9,96E-05 - | 2 - | 0,000112228 - |
| 533 | 1 | 3um_FIJJAFBF_715077 | Bacteria | Cyanobacteria | unclassified | Synechococca | Leptolyngbyac | Leptolyngbya | Leptolyngbya | unclassified | unclassified | unclassified | unclassified | unclassified | 1,6 | 0,489897949 | 0,000149011 | 3,90E-05 | 2 - | 0,000159412 - | 1 - | 9,90E-05 - | 2 - | 0,00018817 - | 2 - | 0,000190949 - | 1 - | 0,000107552 - |
| 534 | 1 | 3um_FIJJAFBF_14772 | Bacteria | Bacteroidetes | Flavobacteriia | unclassified | unclassified | unclassified | Flavobacteria | unclassified | unclassified | unclassified | unclassified | unclassified | 0,4 | 0,489897949 | 2,55E-05 | 3,14E-05 | 1 - | 5,80E-05 - | 0 - | 0 - | 0 - | 0 - | 1 - | 6,94E-05 - | 0 - | 0 - |
| 535 | 1 | 3um_FIJJAFBF_242849 | Bacteria | Bacteroidetes | Flavobacteriia | Flavobacterial | Flavobacteriac | Polaribacter | Polaribacter s | poorly charact | Function unkn | 11PIQ | YCE I like fami | 313598.MED1 | 1,6 | 0,8 | 9,94E-05 | 4,94E-05 | 1 - | 5,22E-05 - | 1 - | 6,48E-05 - | 3 - | 0,000184748 - | 2 - | 0,000124985 - | 1 - | 7,04E-05 - |
| 536 | 1 | 3um_FIJJAFBF_275377 | Bacteria | Proteobacteri | Gammaprotec | Cellvibrionales | Porticoccacea | unclassified | gamma prote | cellular proces | Posttranslatio | 17FI0 | Binds to Cpn6 | 314287.GB220 | 0,8 | 0,4 | 0,000112918 | 5,80E-05 | 1 - | 0,000118327 - | 1 - | 0,000146927 - | 1 - | 0,000139672 - | 0 - | 0 - | 1 - | 0,000159664 - |
| 537 | 8 | 0.2um_NOKCPHJN_260186;0.2um_NOKCPHJN_4 | Bacteria | Proteobacteri | various | various | various | various | various | unclassified | unclassified | unclassified | unclassified | unclassified | 0,6 | 0,489897949 | 6,66E-05 | 5,49E-05 | 1 - | 9,73E-05 - | 1 - | 0,000120779 - | 1 - | 0,000114815 - | 0 - | 0 - | 0 - | 0 - |
| 538 | 1 | 0.2um_NOKCPHJN_462774 | Viruses | unclassified | unclassified | unclassified | unclassified | unclassified | uncultured vir | unclassified | unclassified | unclassified | unclassified | unclassified | 0,2 | 0,4 | 1,20E-06 | 2,40E-06 | 1 - | 6,01E-06 - | 0 - | 0 - | 0 - | 0 - | 0 - | 0 - | 0 - | 0 - |
| 539 | 1 | 3um_FIJJAFBF_410533 | Bacteria | Proteobacteri | Gammaprotec | Alteromonada | Alteromonada | Paraglaciecola | Paraglaciecola | metabolism | Coenzyme tra | 08UTT | Catalyzes the | 983545.Glaag | 0,4 | 0,489897949 | 3,19E-05 | 3,94E-05 | 1 - | 7,26E-05 - | 0 - | 0 - | 0 - | 0 - | 1 - | 8,70E-05 - | 0 - | 0 - |
| 540 | 1 | 0.2um_NOKCPHJN_620130 | Eukaryota | Bacillariophyt | Bacillariophyc | Naviculales | Phaeodactylac | Phaeodactylu | Phaeodactylu | unclassified | unclassified | unclassified | unclassified | unclassified | 1,4 | 0,8 | 0,000135393 | 7,91E-05 | 2 - | 0,000163967 - | 0 - | 0 - | 2 - | 0,000193546 - | 1 - | 9,82E-05 - | 2 - | 0,000221249 - |
| 541 | 1 | 3um_FIJJAFBF_681880 | Eukaryota | Chlorophyta | unclassified | Pyramimonad | unclassified | Cymbomonas | Cymbomonas | metabolism | Energy produc | COG0055 | Produces ATP | 39947.LOC_O | 1 | 0 | 2,85E-05 | 2,70E-06 | 1 - | 2,39E-05 - | 1 - | 2,96E-05 - | 1 - | 2,82E-05 - | 1 - | 2,86E-05 - | 1 - | 3,22E-05 - |
| 542 | 1 | 0.2um_NOKCPHJN_291736 | unclassified | unclassified | unclassified | unclassified | unclassified | unclassified | unclassified | unclassified | unclassified | unclassified | unclassified | unclassified | 0,2 | 0,4 | 5,33E-06 | 1,07E-05 | 0 - | 0 - | 0 - | 0 - | 0 - | 0 - | 1 - | 2,66E-05 - | 0 - | 0 - |
| 543 | 2 | 0.2um_NOKCPHJN_672696;3um_FIJJAFBF_14771 | Bacteria | Bacteroidetes | Flavobacteriia | Flavobacterial | Flavobacteriac | Zobellia | Zobellia uligin | unclassified | unclassified | unclassified | unclassified | unclassified | 0,6 | 0,489897949 | 4,49E-05 | 3,70E-05 | 1 - | 6,52E-05 - | 1 - | 8,10E-05 - | 0 - | 0 - | 1 - | 7,81E-05 - | 0 - | 0 - |
| 544 | 1 | 3um_FIJJAFBF_725016 | Bacteria | Proteobacteri | Gammaprotec | Alteromonada | Alteromonada | Paraglaciecola | Paraglaciecola | cellular proces | Intracellular tr | COG0848 | Biopolymer tr | 342610.Patl_4 | 0,6 | 0,489897949 | 5,85E-05 | 4,83E-05 | 1 - | 8,50E-05 - | 1 - | 0,00010557 - | 0 - | 0 - | 1 - | 0,000101839 - | 0 - | 0 - |
| 545 | 1 | 0.2um_NOKCPHJN_392884 | Eukaryota | Chytridiomyc | Chytridiomyce | Chytridiales | Chytriomyceta | Rhizoclosmati | Rhizoclosmati | metabolism | Energy produc | 0D9WN | Mitochondrial | 436017.A4S9X | 0,8 | 0,4 | 3,19E-05 | 1,62E-05 | 1 - | 3,46E-05 - | 1 - | 4,29E-05 - | 1 - | 4,08E-05 - | 1 - | 4,14E-05 - | 0 - | 0 - |
| 546 | 1 | 0.2um_NOKCPHJN_21673 | Bacteria | Proteobacteri | Alphaproteob | Rhodobactera | Rhodobactera | Nautella | Nautella sp. E | unclassified | unclassified | unclassified | unclassified | unclassified | 0,8 | 0,4 | 3,42E-05 | 1,76E-05 | 1 - | 3,59E-05 - | 1 - | 4,45E-05 - | 1 - | 4,23E-05 - | 0 - | 0 - | 1 - | 4,84E-05 - |
| 547 | 1 | 3um_FIJJAFBF_390935 | Viruses | unclassified | unclassified | unclassified | Phycodnavirid | unclassified | Organic Lake p | unclassified | unclassified | unclassified | unclassified | unclassified | 2,2 | 0,4 | 0,000354295 | 6,91E-05 | 2 - | 0,000270063 - | 2 - | 0,000335339 - | 3 - | 0,000478173 - | 2 - | 0,00032349 - | 2 - | 0,000364411 - |
| 548 | 1 | 0.2um_NOKCPHJN_318849 | Bacteria | Proteobacteri | Gammaprotec | unclassified | unclassified | unclassified | Gammaprotec | poorly charact | Function unkn | 0QRAP | NA | 455436.GHTC | 0,2 | 0,4 | 1,76E-06 | 3,52E-06 | 0 - | 0 - | 0 - | 0 - | 1 - | 8,79E-06 - | 0 - | 0 - | 0 - | 0 - |
| 549 | 1 | 0.2um_NOKCPHJN_159358 | Viruses | unclassified | unclassified | unclassified | unclassified | unclassified | uncultured ph | poorly charact | Function unkn | 17EUU | NA | 204773.HEAR | 0,6 | 0,489897949 | 6,22E-05 | 5,13E-05 | 1 - | 9,04E-05 - | 1 - | 0,00011222 - | 0 - | 0 - | 1 - | 0,000108255 - | 0 - | 0 - |
| 550 | 1 | 3um_FIJJAFBF_573648 | Eukaryota | unclassified | unclassified | Isochrysidales | Noelaerhabda | Emiliania | Emiliania huxl | unclassified | unclassified | unclassified | unclassified | unclassified | 0,2 | 0,4 | 1,73E-05 | 3,45E-05 | 1 - | 8,63E-05 - | 0 - | 0 - | 0 - | 0 - | 0 - | 0 - | 0 - | 0 - |
| 551 | 1 | 3um_FIJJAFBF_190069 | Bacteria | Bacteroidetes | Flavobacteriia | unclassified | unclassified | unclassified | Flavobacteria | unclassified | unclassified | unclassified | unclassified | unclassified | 1,6 | 0,8 | 7,66E-05 | 3,14E-05 | 3 - | 0,000124757 - | 2 - | 0,000103275 - | 1 - | 4,91E-05 - | 1 - | 4,98E-05 - | 1 - | 5,61E-05 - |
| 552 | 1 | 3um_FIJJAFBF_456479 | Eukaryota | unclassified | Glaucocystopl | Gloeochaetale | Gloeochaetac | Gloeochaete | Gloeochaete v | unclassified | unclassified | unclassified | unclassified | unclassified | 2 | 0,632455532 | 0,000206228 | 4,77E-05 | 3 - | 0,000266923 - | 2 - | 0,000220959 - | 2 - | 0,00021005 - | 2 - | 0,000213152 - | 1 - | 0,000120058 - |
| 553 | 1 | 0.2um_NOKCPHJN_321804 | Eukaryota | Eustigmatoph | unclassified | Eustigmatales | Monodopsida | Nannochlorop | Nannochlorop | unclassified | unclassified | unclassified | unclassified | unclassified | 0,6 | 0,489897949 | 4,34E-06 | 3,55E-06 | 0 - | 0 - | 1 - | 7,11E-06 - | 0 - | 0 - | 1 - | 6,86E-06 - | 1 - | 7,72E-06 - |
| 554 | 1 | 3um_FIJJAFBF_42230 | Bacteria | Proteobacteri | Alphaproteob | Rhodobactera | unclassified | unclassified | Rhodobactera | poorly charact | Function unkn | 01Y9B | Protein of unk | 367336.OM22 | 0,2 | 0,4 | 2,39E-05 | 4,78E-05 | 1 - | 0,000119559 - | 0 - | 0 - | 0 - | 0 - | 0 - | 0 - | 0 - | 0 - |
| 555 | 1 | 0.2um_NOKCPHJN_452060 | Eukaryota | unclassified | Labyrinthulom | unclassified | Thraustochytr | Aurantiochytr | Aurantiochytr | unclassified | unclassified | unclassified | unclassified | unclassified | 1 | 0 | 7,49E-05 | 7,11E-06 | 1 - | 6,27E-05 - | 1 - | 7,79E-05 - | 1 - | 7,40E-05 - | 1 - | 7,51E-05 - | 1 - | 8,46E-05 - |
| 556 | 1 | 3um_FIJJAFBF_494165 | Bacteria | Bacteroidetes | Flavobacteriia | Flavobacterial | Flavobacteriac | Polaribacter | various | unclassified | unclassified | unclassified | unclassified | unclassified | 0,8 | 0,4 | 5,86E-05 | 2,98E-05 | 1 - | 6,34E-05 - | 1 - | 7,87E-05 - | 1 - | 7,49E-05 - | 1 - | 7,60E-05 - | 0 - | 0 - |
| 557 | 1 | 0.2um_NOKCPHJN_64570 | Bacteria | Proteobacteri | Gammaprotec | Oceanospirilla | Saccharospirill | Reinekea | Reinekea blan | metabolism | Nucleotide tra | COG0138 | bifunctional p | 314283.MED2 | 1,8 | 0,748331477 | 4,57E-05 | 1,84E-05 | 2 - | 4,32E-05 - | 2 - | 5,37E-05 - | 3 - | 7,65E-05 - | 1 - | 2,59E-05 - | 1 - | 2,92E-05 - |
| 558 | 1 | 3um_FIJJAFBF_451550 | Bacteria | Proteobacteri | Deltaproteoba | Myxococcales | Archangiaceae | Melittangium | Melittangium | unclassified | unclassified | unclassified | unclassified | unclassified | 0,4 | 0,489897949 | 5,50E-05 | 6,77E-05 | 1 - | 0,000126128 - | 0 - | 0 - | 1 - | 0,000148882 - | 0 - | 0 - | 0 - | 0 - |
| 559 | 1 | 0.2um_NOKCPHJN_97671 | Bacteria | Proteobacteri | Betaproteoba | unclassified | unclassified | unclassified | Betaproteoba | unclassified | unclassified | unclassified | unclassified | unclassified | 0,2 | 0,4 | 5,44E-06 | 1,09E-05 | 0 - | 0 - | 0 - | 0 - | 1 - | 2,72E-05 - | 0 - | 0 - | 0 - | 0 - |
| 560 | 1 | 3um_FIJJAFBF_507105 | Eukaryota | Phaeophyceae | unclassified | Ectocarpales | Ectocarpaceae | Ectocarpus | Ectocarpus sili | unclassified | unclassified | unclassified | unclassified | unclassified | 2 | 0,632455532 | 0,000155788 | 4,08E-05 | 3 - | 0,000199035 - | 2 - | 0,000164762 - | 2 - | 0,000156627 - | 1 - | 7,95E-05 - | 2 - | 0,000179046 - |
| 561 | 1 | 3um_FIJJAFBF_220178 | Bacteria | Proteobacteri | Gammaprotec | Cellvibrionales | Porticoccacea | unclassified | Gammaprotec | unclassified | unclassified | unclassified | unclassified | unclassified | 1,4 | 1,019803903 | 4,55E-05 | 3,51E-05 | 1 - | 2,74E-05 - | 3 - | 0,000102042 - | 1 - | 3,23E-05 - | 2 - | 6,56E-05 - | 0 - | 0 - |
| 562 | 1 | 0.2um_NOKCPHJN_204138 | Bacteria | Proteobacteri | Gammaprotec | Alteromonada | Colwelliaceae | Colwellia | Colwellia sp. P | information st | Translation, ril | 0QSAC | Binds togethe | 167879.CPS_0 | 0,4 | 0,489897949 | 4,18E-05 | 5,14E-05 | 0 - | 0 - | 0 - | 0 - | 1 - | 9,75E-05 - | 0 - | 0 - | 1 - | 0,00011142 - |
| 563 | 1 | 3um_FIJJAFBF_808023 | Bacteria | Verrucomicrol | Verrucomicrol | Verrucomicrol | Akkermansiac | Akkermansia | Akkermansia r | unclassified | unclassified | unclassified | unclassified | unclassified | 1 | 0 | 3,50E-05 | 3,33E-06 | 1 - | 2,94E-05 - | 1 - | 3,64E-05 - | 1 - | 3,47E-05 - | 1 - | 3,52E-05 - | 1 - | 3,96E-05 - |
| 564 | 1 | 3um_FIJJAFBF_715094 | Eukaryota | Bacillariophyt | Bacillariophyc | Eunotiales | Eunotiaceae | Eunotia | Eunotia naege | information st | Translation, ril | 0FNT6 | One of the ea | 497965.Cyan7 | 1 | 0,632455532 | 0,000145589 | 8,16E-05 | 2 - | 0,000252257 - | 1 - | 0,000156614 - | 1 - | 0,000148882 - | 0 - | 0 - | 1 - | 0,000170192 - |
| 565 | 1 | 3um_FIJJAFBF_203164 | Bacteria | Proteobacteri | Gammaprotec | Arenicellales | Arenicellaceae | Arenicella | Arenicella xan | metabolism | Energy produc | 17CGT | F(1)F(0) ATP s | 1026882.MAN | 0,8 | 0,4 | 6,83E-05 | 3,51E-05 | 1 - | 7,22E-05 - | 0 - | 0 - | 1 - | 8,52E-05 - | 1 - | 8,65E-05 - | 1 - | 9,74E-05 - |
| 566 | 3 | 0.2um_NOKCPHJN_11819;3um_FIJJAFBF_159892 | Bacteria | Bacteroidetes | Flavobacteriia | Flavobacterial | Flavobacteriac | various | various | poorly charact | Function unkn | various | various | various | 0,4 | 0,489897949 | 3,34E-05 | 4,12E-05 | 1 - | 7,60E-05 - | 0 - | 0 - | 0 - | 0 - | 1 - | 9,10E-05 - | 0 - | 0 - |

| | | | | | | | | | | | | | | | | | | | | | | | | | | | | |
|---|---|---|---|---|---|---|---|---|---|---|---|---|---|---|---|---|---|---|---|---|---|---|---|---|---|---|---|---|
| 567 | 1 | 0.2um_NOKCPHJN_823628 | Viruses | unclassified | unclassified | Herpesvirales | Herpesviridae | Varicellovirus | Equid alphahe | poorly charact | Function unkn | 0YHPQ | DCD | 7739.JGI8903 | 1 | 0 | 3,50E-05 | 3,32E-06 | 1 - | 2,93E-05 - | 1 - | 3,64E-05 - | 1 - | 3,46E-05 - | 1 - | 3,51E-05 - | 1 - | 3,95E-05 - |
| 568 | 1 | 0.2um_NOKCPHJN_80614 | Bacteria | Bacteroidetes | Flavobacteriia | Flavobacterial | Flavobacteriac | Polaribacter | Polaribacter a | unclassified | unclassified | unclassified | unclassified | unclassified | 2,2 | 0,979795897 | 0,000161585 | 5,67E-05 | 4 - | 0,00025365 - | 2 - | 0,000157479 - | 2 - | 0,000149704 - | 1 - | 7,60E-05 - | 2 - | 0,000171132 - |
| 569 | 1 | 3um_FIJJAFBF_810588 | Bacteria | Proteobacteri | Gammaprotec | unclassified | unclassified | unclassified | various | cellular proces | Intracellular tr | 175GA | One of the pro | 443152.MDG8 | 2,4 | 1,019803903 | 0,000188732 | 6,58E-05 | 4 - | 0,000271661 - | 1 - | 8,43E-05 - | 2 - | 0,000160334 - | 3 - | 0,000244053 - | 2 - | 0,000183283 - |
| 570 | 1 | 3um_FIJJAFBF_86167 | Bacteria | Proteobacteri | Alphaproteob | Rickettsiales | unclassified | unclassified | Rickettsiales b | unclassified | unclassified | unclassified | unclassified | unclassified | 0,4 | 0,489897949 | 3,48E-05 | 4,34E-05 | 1 - | 7,40E-05 - | 0 - | 0 - | 0 - | 0 - | 0 - | 0 - | 1 - | 9,99E-05 - |
| 571 | 1 | 3um_FIJJAFBF_649054 | Bacteria | Bacteroidetes | Flavobacteriia | Flavobacterial | Flavobacteriac | Formosa | Formosa sp. H | metabolism | Energy produc | 05CMV | Succinyl-CoA s | 867900.Celly_ | 1,6 | 0,489897949 | 5,45E-05 | 1,54E-05 | 2 - | 5,80E-05 - | 2 - | 7,20E-05 - | 1 - | 3,42E-05 - | 2 - | 6,94E-05 - | 1 - | 3,91E-05 - |
| 572 | 1 | 3um_FIJJAFBF_470534 | Bacteria | Proteobacteri | Alphaproteob | Rhodobactera | unclassified | unclassified | Rhodobactera | metabolism | Amino acid tra | 02DU9 | Lactoylglutath | 367336.OM22 | 1 | 0 | 9,32E-05 | 8,85E-06 | 1 - | 7,81E-05 - | 1 - | 9,70E-05 - | 1 - | 9,22E-05 - | 1 - | 9,35E-05 - | 1 - | 0,000105357 - |
| 573 | 1 | 3um_FIJJAFBF_214617 | Bacteria | Proteobacteri | Alphaproteob | Rhodobactera | Rhodobactera | Ruegeria | Ruegeria litor | information st | Transcription | COG0085 | DNA-depende | 388739.RSK20 | 8 | 0,632455532 | 7,94E-05 | 7,53E-06 | 8 - | 6,67E-05 - | 8 - | 8,28E-05 - | 8 - | 7,87E-05 - | 9 - | 8,99E-05 - | 7 - | 7,87E-05 - |
| 574 | 1 | 3um_FIJJAFBF_467045 | Bacteria | unclassified | unclassified | unclassified | unclassified | unclassified | uncultured ba | cellular proces | Signal transdu | 09BM1 | Two compone | 313594.PI23P | 2,8 | 1,326649916 | 0,000162792 | 7,57E-05 | 3 - | 0,000147781 - | 3 - | 0,000183501 - | 5 - | 0,000290734 - | 1 - | 5,90E-05 - | 2 - | 0,000132939 - |
| 575 | 1 | 3um_FIJJAFBF_744864 | unclassified | unclassified | unclassified | unclassified | unclassified | unclassified | unclassified | unclassified | unclassified | unclassified | unclassified | unclassified | 0,6 | 0,489897949 | 0,00010254 | 8,46E-05 | 1 - | 0,000149061 - | 1 - | 0,000185089 - | 0 - | 0 - | 1 - | 0,00017855 - | 0 - | 0 - |
| 576 | 1 | 3um_FIJJAFBF_559940 | Bacteria | unclassified | unclassified | unclassified | unclassified | unclassified | uncultured ba | poorly charact | Function unkn | 05X9M | NA | 313595.P7007 | 0,4 | 0,489897949 | 1,02E-05 | 1,25E-05 | 0 - | 0 - | 0 - | 0 - | 1 - | 2,37E-05 - | 0 - | 0 - | 1 - | 2,71E-05 - |
| 577 | 1 | 0.2um_NOKCPHJN_84810 | Bacteria | Bacteroidetes | Flavobacteriia | Flavobacterial | Crocinitomica | unclassified | Crocinitomica | poorly charact | Function unkn | 05CZH | peptidase | 926562.Oweh | 1,2 | 0,748331477 | 2,17E-05 | 1,43E-05 | 1 - | 1,53E-05 - | 2 - | 3,81E-05 - | 1 - | 1,81E-05 - | 2 - | 3,68E-05 - | 0 - | 0 - |
| 578 | 2 | 0.2um_NOKCPHJN_56987;3um_FIJJAFBF_32849 | Bacteria | Bacteroidetes | Flavobacteriia | Flavobacterial | Flavobacteriac | Mangrovimon | Mangrovimon | poorly charact | Function unkn | 1015T | domain protei | various | 1,8 | 0,4 | 5,80E-05 | 1,44E-05 | 2 - | 5,39E-05 - | 2 - | 6,69E-05 - | 1 - | 3,18E-05 - | 2 - | 6,45E-05 - | 2 - | 7,27E-05 - |
| 579 | 1 | 0.2um_NOKCPHJN_472850 | Bacteria | Proteobacteri | Alphaproteob | Pelagibacteral | Pelagibacterac | Candidatus Pe | Candidatus Pe | unclassified | unclassified | unclassified | unclassified | unclassified | 1 | 0,632455532 | 3,95E-05 | 2,44E-05 | 0 - | 0 - | 1 - | 3,98E-05 - | 1 - | 3,78E-05 - | 2 - | 7,68E-05 - | 1 - | 4,33E-05 - |
| 580 | 1 | 3um_FIJJAFBF_226594 | Viruses | unclassified | unclassified | unclassified | Phycodnavirid | unclassified | Organic Lake p | unclassified | unclassified | unclassified | unclassified | unclassified | 1,4 | 0,489897949 | 0,000158219 | 5,92E-05 | 1 - | 9,41E-05 - | 2 - | 0,000233637 - | 1 - | 0,000111051 - | 2 - | 0,000225382 - | 1 - | 0,000126946 - |
| 581 | 1 | 0.2um_NOKCPHJN_199220 | unclassified | unclassified | unclassified | unclassified | unclassified | unclassified | unclassified | unclassified | unclassified | unclassified | unclassified | unclassified | 0,2 | 0,4 | 7,34E-06 | 1,47E-05 | 0 - | 0 - | 0 - | 0 - | 1 - | 3,67E-05 - | 0 - | 0 - | 0 - | 0 - |
| 582 | 1 | 0.2um_NOKCPHJN_400251 | Bacteria | Proteobacteri | Gammaprotec | unclassified | unclassified | unclassified | various | unclassified | unclassified | unclassified | unclassified | unclassified | 0,4 | 0,489897949 | 1,50E-05 | 1,85E-05 | 0 - | 0 - | 0 - | 0 - | 0 - | 0 - | 1 - | 3,53E-05 - | 1 - | 3,98E-05 - |
| 583 | 1 | 3um_FIJJAFBF_755962 | Bacteria | Proteobacteri | Gammaprotec | Oceanospirilla | Saccharospirill | Reinekea | Reinekea fors | poorly charact | Function unkn | 1799F | Protein of unk | 314283.MED2 | 0,2 | 0,4 | 9,60E-06 | 1,92E-05 | 1 - | 4,80E-05 - | 0 - | 0 - | 0 - | 0 - | 0 - | 0 - | 0 - | 0 - |
| 584 | 1 | 3um_FIJJAFBF_644353 | Bacteria | Bacteroidetes | Flavobacteriia | Flavobacterial | Flavobacteriac | Ulvibacter | Ulvibacter sp. | unclassified | unclassified | unclassified | unclassified | unclassified | 1 | 0,632455532 | 0,000116129 | 7,61E-05 | 1 - | 9,98E-05 - | 1 - | 0,000123929 - | 1 - | 0,000117811 - | 2 - | 0,000239101 - | 0 - | 0 - |
| 585 | 1 | 3um_FIJJAFBF_660159 | Bacteria | Bacteroidetes | Flavobacteriia | Flavobacterial | Flavobacteriac | Formosa | Formosa sp. H | poorly charact | Function unkn | 11P35 | NA | 867900.Celly_ | 3 | 0 | 0,000111404 | 1,06E-05 | 3 - | 9,33E-05 - | 3 - | 0,000115869 - | 3 - | 0,000110148 - | 3 - | 0,000111775 - | 3 - | 0,000125914 - |
| 586 | 1 | 0.2um_NOKCPHJN_227786 | Eukaryota | unclassified | Pelagophycea | Pelagomonad | unclassified | Aureococcus | Aureococcus a | unclassified | unclassified | unclassified | unclassified | unclassified | 1 | 0,632455532 | 2,92E-05 | 1,91E-05 | 1 - | 2,51E-05 - | 1 - | 3,11E-05 - | 1 - | 2,96E-05 - | 2 - | 6,00E-05 - | 0 - | 0 - |
| 587 | 1 | 0.2um_NOKCPHJN_187384 | Bacteria | Proteobacteri | Alphaproteob | unclassified | unclassified | unclassified | SAR116 cluste | information st | Translation, ri | 16Q16 | Binds the lowe | 331869.BAL19 | 0,8 | 0,4 | 4,74E-05 | 2,43E-05 | 1 - | 4,95E-05 - | 1 - | 6,14E-05 - | 0 - | 0 - | 1 - | 5,93E-05 - | 1 - | 6,68E-05 - |
| 588 | 1 | 3um_FIJJAFBF_45648 | Bacteria | Proteobacteri | Alphaproteob | Rhodobactera | Rhodobactera | Planktomarina | Planktomarina | cellular proces | Posttranslatio | 17FI0 | Binds to Cpn6 | 398580.Dshi_ | 2 | 0 | 0,000288478 | 2,74E-05 | 2 - | 0,000241635 - | 2 - | 0,00030004 - | 2 - | 0,000285226 - | 2 - | 0,000289439 - | 2 - | 0,000326052 - |
| 589 | 1 | 3um_FIJJAFBF_339594 | Bacteria | Proteobacteri | Alphaproteob | Rhodobactera | Rhodobactera | Sulfitobacter | Sulfitobacter r | cellular proces | Posttranslatio | 01QWG | Prevents misf | 398580.Dshi_ | 2,6 | 0,8 | 6,37E-05 | 1,45E-05 | 4 - | 8,39E-05 - | 3 - | 7,82E-05 - | 2 - | 4,95E-05 - | 2 - | 5,03E-05 - | 2 - | 5,66E-05 - |
| 590 | 1 | 3um_FIJJAFBF_382228 | Eukaryota | Apicomplexa | Aconoidasida | Haemosporida | Plasmodiidae | Plasmodium | Plasmodium r | poorly charact | Function unkn | 1AMB2 | NA | 759914.BP951 | 0,2 | 0,4 | 6,75E-07 | 1,35E-06 | 0 - | 0 - | 0 - | 0 - | 1 - | 3,37E-06 - | 0 - | 0 - | 0 - | 0 - |
| 591 | 1 | 3um_FIJJAFBF_94316 | Bacteria | Bacteroidetes | Flavobacteriia | Flavobacterial | Flavobacteriac | unclassified | Flavobacteriac | unclassified | unclassified | unclassified | unclassified | unclassified | 1,4 | 0,489897949 | 3,52E-05 | 1,28E-05 | 2 - | 4,24E-05 - | 1 - | 2,63E-05 - | 1 - | 2,50E-05 - | 1 - | 2,54E-05 - | 2 - | 5,71E-05 - |
| 592 | 1 | 0.2um_NOKCPHJN_592028 | Eukaryota | unclassified | Pelagophycea | Pelagomonad | unclassified | Aureococcus | Aureococcus a | cellular proces | Posttranslatio | KOG0019 | heat shock pro | 67593.Physo1 | 1 | 0,632455532 | 1,87E-05 | 1,04E-05 | 2 - | 3,22E-05 - | 1 - | 2,00E-05 - | 0 - | 0 - | 1 - | 1,93E-05 - | 1 - | 2,18E-05 - |
| 593 | 1 | 3um_FIJJAFBF_12502 | Viruses | unclassified | unclassified | unclassified | Phycodnavirid | unclassified | Chrysochromu | unclassified | unclassified | unclassified | unclassified | unclassified | 1,8 | 1,166190379 | 5,70E-05 | 3,86E-05 | 1 - | 2,62E-05 - | 4 - | 0,000130154 - | 2 - | 6,19E-05 - | 1 - | 3,14E-05 - | 1 - | 3,54E-05 - |
| 594 | 1 | 3um_FIJJAFBF_417060 | Bacteria | Proteobacteri | Gammaprotec | Alteromonada | Alteromonada | Paraglaciecola | various | information st | Translation, ri | 05VBI | Binds 16S rRN | 715451.ambt_ | 1 | 0 | 0,00013567 | 1,29E-05 | 1 - | 0,00011364 - | 1 - | 0,000141108 - | 1 - | 0,000134141 - | 1 - | 0,000136122 - | 1 - | 0,000153341 - |
| 595 | 2 | 0.2um_NOKCPHJN_164027;0.2um_NOKCPHJN_5 | Eukaryota | unclassified | Oomycetes | Peronosporale | Peronosporac | Phytophthora | various | unclassified | unclassified | unclassified | unclassified | unclassified | 0,6 | 0,489897949 | 7,92E-05 | 6,50E-05 | 0 - | 0 - | 0 - | 0 - | 1 - | 0,000125447 - | 1 - | 0,000127299 - | 1 - | 0,000143402 - |
| 596 | 1 | 0.2um_NOKCPHJN_834208 | Eukaryota | Bacillariophyt | Coscinodiscop | Thalassiosirale | Thalassiosirac | Thalassiosira | Thalassiosira p | unclassified | unclassified | unclassified | unclassified | unclassified | 0,6 | 0,489897949 | 1,16E-05 | 9,55E-06 | 1 - | 1,68E-05 - | 1 - | 2,09E-05 - | 0 - | 0 - | 1 - | 2,02E-05 - | 0 - | 0 - |
| 597 | 1 | 0.2um_NOKCPHJN_571617 | Bacteria | Bacteroidetes | unclassified | unclassified | unclassified | unclassified | Bacteroidetes | metabolism | Energy produc | COG0711 | F(1)F(0) ATP s | 487796.Flav2A | 1,8 | 0,4 | 0,000146433 | 2,89E-05 | 2 - | 0,000138285 - | 2 - | 0,000171709 - | 2 - | 0,000163232 - | 2 - | 0,000165643 - | 1 - | 9,33E-05 - |
| 598 | 1 | 0.2um_NOKCPHJN_206819 | Bacteria | Proteobacteri | Gammaprotec | Oceanospirilla | Oceanospirilla | Neptuniibacte | Neptuniibacte | unclassified | unclassified | unclassified | unclassified | unclassified | 1 | 0 | 1,54E-05 | 1,47E-06 | 1 - | 1,29E-05 - | 1 - | 1,61E-05 - | 1 - | 1,53E-05 - | 1 - | 1,55E-05 - | 1 - | 1,75E-05 - |
| 599 | 1 | 0.2um_NOKCPHJN_270964 | Viruses | unclassified | unclassified | unclassified | unclassified | unclassified | uncultured vir | unclassified | unclassified | unclassified | unclassified | unclassified | 0,4 | 0,489897949 | 1,67E-05 | 2,05E-05 | 1 - | 3,79E-05 - | 0 - | 0 - | 0 - | 0 - | 1 - | 4,54E-05 - | 0 - | 0 - |
| 600 | 1 | 0.2um_NOKCPHJN_46802 | Bacteria | Proteobacteri | Alphaproteob | Rhodobactera | Rhodobactera | Confluentimic | Confluentimic | information st | Translation, ri | COG0255 | 50s ribosomal | 314270.RB208 | 1 | 0 | 0,000204518 | 1,94E-05 | 1 - | 0,000171309 - | 1 - | 0,000212715 - | 1 - | 0,000202212 - | 1 - | 0,000205199 - | 1 - | 0,000231156 - |
| 601 | 1 | 0.2um_NOKCPHJN_184685 | Bacteria | Proteobacteri | Gammaprotec | unclassified | unclassified | unclassified | Gammaprotec | information st | Translation, ri | 0QK14 | This protein bi | 351348.Maqu | 3,8 | 0,4 | 0,000292166 | 2,60E-05 | 4 - | 0,000259383 - | 4 - | 0,000322077 - | 4 - | 0,000306175 - | 4 - | 0,000310697 - | 3 - | 0,000262499 - |
| 602 | 1 | 3um_FIJJAFBF_11563 | Bacteria | Bacteroidetes | Flavobacteriia | Flavobacterial | Flavobacteriac | Lutibacter | Lutibacter sp. | unclassified | unclassified | unclassified | unclassified | unclassified | 1 | 0 | 3,33E-05 | 3,17E-06 | 1 - | 2,79E-05 - | 1 - | 3,47E-05 - | 1 - | 3,30E-05 - | 1 - | 3,35E-05 - | 1 - | 3,77E-05 - |
| 603 | 1 | 3um_FIJJAFBF_136304 | unclassified | unclassified | unclassified | unclassified | unclassified | unclassified | unclassified | unclassified | unclassified | unclassified | unclassified | unclassified | 0,2 | 0,4 | 4,46E-06 | 8,91E-06 | 1 - | 2,23E-05 - | 0 - | 0 - | 0 - | 0 - | 0 - | 0 - | 0 - | 0 - |
| 604 | 1 | 3um_FIJJAFBF_274861 | Bacteria | Bacteroidetes | Flavobacteriia | Flavobacterial | Flavobacteriac | unclassified | Flavobacteriac | unclassified | unclassified | unclassified | unclassified | unclassified | 1 | 0 | 8,78E-05 | 8,34E-06 | 1 - | 7,36E-05 - | 1 - | 9,14E-05 - | 1 - | 8,68E-05 - | 1 - | 8,81E-05 - | 1 - | 9,93E-05 - |
| 605 | 1 | 0.2um_NOKCPHJN_236909 | Eukaryota | Chordata | unclassified | unclassified | Branchiostomi | Branchiostom | Branchiostom | poorly charact | Function unkn | 0XR88 | semaphorin re | 4787.PITG_05 | 0,2 | 0,4 | 4,95E-07 | 9,91E-07 | 0 - | 0 - | 0 - | 0 - | 0 - | 0 - | 1 - | 2,48E-06 - | 0 - | 0 - |
| 606 | 1 | 3um_FIJJAFBF_377866 | Bacteria | Bacteroidetes | Flavobacteriia | unclassified | unclassified | unclassified | Flavobacteria | unclassified | unclassified | unclassified | unclassified | unclassified | 10,4 | 0,489897949 | 0,000151248 | 1,68E-05 | 10 - | 0,000121714 - | 11 - | 0,000166247 - | 10 - | 0,000143672 - | 11 - | 0,000160373 - | 10 - | 0,000164236 - |
| 607 | 1 | 0.2um_NOKCPHJN_39143 | Eukaryota | unclassified | Pelagophycea | Pelagomonad | unclassified | Aureococcus | Aureococcus a | metabolism | Carbohydrate | COG1850 | RuBisCO catal | 45157.CMV01 | 16,8 | 0,4 | 0,000471001 | 3,78E-05 | 17 - | 0,000399837 - | 17 - | 0,00049648 - | 17 - | 0,000471967 - | 17 - | 0,000478938 - | 16 - | 0,000507785 - |
| 608 | 1 | 3um_FIJJAFBF_649062 | Bacteria | Proteobacteri | Gammaprotec | Oceanospirilla | Saccharospirill | Reinekea | Reinekea fors | cellular proces | Cell wall/mem | 16QAK | protease | 314283.MED2 | 4,6 | 0,8 | 0,000147807 | 3,15E-05 | 4 - | 0,000107268 - | 6 - | 0,000199793 - | 4 - | 0,000126619 - | 5 - | 0,000160611 - | 4 - | 0,000144743 - |
| 609 | 1 | 0.2um_NOKCPHJN_314333 | Bacteria | Proteobacteri | Gammaprotec | unclassified | unclassified | unclassified | Gammaprotec | metabolism | Inorganic ion t | 16PDP | TonB-depende | 314260.PB250 | 2,8 | 1,166190379 | 4,03E-05 | 1,67E-05 | 2 - | 2,42E-05 - | 2 - | 3,00E-05 - | 5 - | 7,14E-05 - | 3 - | 4,35E-05 - | 2 - | 3,26E-05 - |
| 610 | 1 | 0.2um_NOKCPHJN_227680 | Bacteria | Bacteroidetes | Flavobacteriia | Flavobacterial | Flavobacteriac | Flaviramulus | Flaviramulus b | unclassified | unclassified | unclassified | unclassified | unclassified | 2,4 | 1,019803903 | 4,51E-05 | 1,92E-05 | 2 - | 3,17E-05 - | 1 - | 1,97E-05 - | 4 - | 7,47E-05 - | 3 - | 5,69E-05 - | 2 - | 4,27E-05 - |
| 611 | 1 | 0.2um_NOKCPHJN_724920 | Bacteria | Bacteroidetes | Flavobacteriia | Flavobacterial | Flavobacteriac | Polaribacter | Polaribacter s | metabolism | Inorganic ion t | 0P8Q4 | tonB-depende | 688270.Celal_ | 16,2 | 2,13541565 | 0,000257922 | 3,55E-05 | 18 - | 0,000241071 - | 19 - | 0,000315969 - | 16 - | 0,000252942 - | 13 - | 0,000208551 - | 15 - | 0,000271076 - |
| 612 | 1 | 0.2um_NOKCPHJN_1095134 | Eukaryota | unclassified | unclassified | Euglyphida | Paulinellidae | Paulinella | Paulinella mic | unclassified | unclassified | unclassified | unclassified | unclassified | 1,2 | 0,979795897 | 5,94E-05 | 4,85E-05 | 0 - | 0 - | 3 - | 0,000148457 - | 1 - | 4,70E-05 - | 1 - | 4,77E-05 - | 1 - | 5,38E-05 - |
| 613 | 1 | 3um_FIJJAFBF_292916 | Eukaryota | Bacillariophyt | Coscinodiscop | Coscinodiscale | Coscinodiscac | Coscinodiscus | Coscinodiscus | unclassified | unclassified | unclassified | unclassified | unclassified | 3,6 | 0,489897949 | 0,00031387 | 3,71E-05 | 4 - | 0,000294299 - | 4 - | 0,000365433 - | 4 - | 0,00034739 - | 3 - | 0,000264391 - | 3 - | 0,000297836 - |
| 614 | 1 | 0.2um_NOKCPHJN_46822 | Bacteria | Proteobacteri | Alphaproteob | Rhodobactera | Rhodobactera | Ruegeria | Ruegeria prof | information st | Translation, ri | 05C64 | Binds directly | 467661.RKLH1 | 0,2 | 0,4 | 1,17E-05 | 2,34E-05 | 0 - | 0 - | 0 - | 0 - | 1 - | 5,84E-05 - | 0 - | 0 - | 0 - | 0 - |
| 615 | 1 | 3um_FIJJAFBF_53313 | Bacteria | Bacteroidetes | Flavobacteriia | Flavobacterial | Flavobacteriac | Lutibacter | Lutibacter sp. | unclassified | unclassified | unclassified | unclassified | unclassified | 12 | 1,897366596 | 0,000161559 | 2,43E-05 | 15 - | 0,000170124 - | 12 - | 0,000168995 - | 10 - | 0,000133876 - | 10 - | 0,000135853 - | 13 - | 0,000198949 - |
| 616 | 1 | 0.2um_NOKCPHJN_450475 | Bacteria | Bacteroidetes | Flavobacteriia | Flavobacterial | Flavobacteriac | Maribacter | Maribacter co | unclassified | unclassified | unclassified | unclassified | unclassified | 1,4 | 1,2 | 4,08E-05 | 3,57E-05 | 2 - | 5,09E-05 - | 2 - | 6,32E-05 - | 3 - | 9,01E-05 - | 0 - | 0 - | 0 - | 0 - |
| 617 | 1 | 0.2um_NOKCPHJN_216476 | unclassified | unclassified | unclassified | unclassified | unclassified | unclassified | unclassified | information st | Transcription | KOG2252 | homeobox | 7460.GB13918 | 0,8 | 0,4 | 2,47E-06 | 1,25E-06 | 0 - | 0 - | 1 - | 3,09E-06 - | 1 - | 2,94E-06 - | 1 - | 2,98E-06 - | 1 - | 3,36E-06 - |
| 618 | 1 | 3um_FIJJAFBF_72766 | Bacteria | Proteobacteri | Gammaprotec | unclassified | unclassified | unclassified | gamma protec | unclassified | unclassified | unclassified | unclassified | unclassified | 1 | 0 | 5,25E-05 | 4,98E-06 | 1 - | 4,40E-05 - | 1 - | 5,46E-05 - | 1 - | 5,19E-05 - | 1 - | 5,27E-05 - | 1 - | 5,93E-05 - |
| 619 | 1 | 3um_FIJJAFBF_19225 | Bacteria | Bacteroidetes | Flavobacteriia | unclassified | unclassified | unclassified | Flavobacteria | poorly charact | Function unkn | 0XP53 | Ragb susd dor | 487797.Flav3 | 12,4 | 1,2 | 0,000381076 | 2,91E-05 | 14 - | 0,000362726 - | 13 - | 0,000418227 - | 11 - | 0,000336412 - | 13 - | 0,00040345 - | 11 - | 0,000384564 - |
| 620 | 1 | 3um_FIJJAFBF_753491 | Viruses | unclassified | unclassified | unclassified | Phycodnavirid | unclassified | Aureococcus a | unclassified | unclassified | unclassified | unclassified | unclassified | 0,4 | 0,489897949 | 1,05E-05 | 1,29E-05 | 1 - | 2,39E-05 - | 0 - | 0 - | 0 - | 0 - | 1 - | 2,86E-05 - | 0 - | 0 - |
| 621 | 1 | 3um_FIJJAFBF_88826 | Bacteria | Proteobacteri | Gammaprotec | unclassified | unclassified | unclassified | various | information st | Translation, ri | COG0256 | This is one of | 314283.MED2 | 1 | 0 | 0,000118127 | 1,12E-05 | 1 - | 9,89E-05 - | 1 - | 0,000122861 - | 1 - | 0,000116795 - | 1 - | 0,00011852 - | 1 - | 0,000133512 - |
| 622 | 1 | 0.2um_NOKCPHJN_853396 | Eukaryota | Bacillariophyt | Coscinodiscop | Thalassiosirale | Thalassiosirac | Thalassiosira | Thalassiosira p | unclassified | unclassified | unclassified | unclassified | unclassified | 1 | 0 | 7,33E-05 | 6,96E-06 | 1 - | 6,14E-05 - | 1 - | 7,62E-05 - | 1 - | 7,25E-05 - | 1 - | 7,35E-05 - | 1 - | 8,28E-05 - |
| 623 | 1 | 3um_FIJJAFBF_48986 | Bacteria | Bacteroidetes | Flavobacteriia | Flavobacterial | Flavobacteriac | Pricia | Pricia antarcti | unclassified | unclassified | unclassified | unclassified | unclassified | 1 | 0 | 4,73E-05 | 4,49E-06 | 1 - | 3,96E-05 - | 1 - | 4,91E-05 - | 1 - | 4,67E-05 - | 1 - | 4,74E-05 - | 1 - | 5,34E-05 - |
| 624 | 1 | 3um_FIJJAFBF_691181 | Eukaryota | Phaeophyceae | unclassified | Ectocarpales | Ectocarpaceae | Ectocarpus | Ectocarpus sili | cellular proces | Signal transdu | COG5040 | Tyrosine 3-mo | 162425.CADA | 2,6 | 0,8 | 0,000144758 | 4,72E-05 | 2 - | 9,29E-05 - | 3 - | 0,0001731 - | 2 - | 0,000109702 - | 4 - | 0,000222645 - | 2 - | 0,000125404 - |
| 625 | 1 | 0.2um_NOKCPHJN_241055 | Eukaryota | Ascomycota | Sordariomyce | Hypocreales | Nectriaceae | Fusarium | Fusarium gran | cellular proces | Posttranslatio | 0V7QW | ankyrin repea | 7668.SPU_024 | 0,8 | 0,4 | 7,70E-06 | 3,91E-06 | 1 - | 8,34E-06 - | 1 - | 1,03E-05 - | 1 - | 9,84E-06 - | 1 - | 9,98E-06 - | 0 - | 0 - |
| 626 | 1 | 0.2um_NOKCPHJN_15324 | Bacteria | Proteobacteri | Betaproteoba | unclassified | unclassified | unclassified | Betaproteoba | cellular proces | Posttranslatio | COG0330 | Band 7 protei | 688245.CtCNB | 5 | 1,264911064 | 0,000227172 | 4,90E-05 | 6 - | 0,000231872 - | 3 - | 0,000143958 - | 6 - | 0,000273701 - | 6 - | 0,000277744 - | 4 - | 0,000208585 - |
| 627 | 1 | 3um_FIJJAFBF_421473 | Bacteria | Proteobacteri | Gammaprotec | Alteromonada | Alteromonada | Paraglaciecola | Paraglaciecola | cellular proces | Posttranslatio | 08V1T | peptidylprolyl | 342610.Patl_ | 1,4 | 0,8 | 8,88E-05 | 3,93E-05 | 3 - | 0,000167151 - | 1 - | 6,92E-05 - | 1 - | 6,58E-05 - | 1 - | 6,67E-05 - | 1 - | 7,52E-05 - |
| 628 | 1 | 3um_FIJJAFBF_09089 | Bacteria | Proteobacteri | Alphaproteob | Pelagibacteral | Pelagibacterac | Candidatus Pe | Candidatus Pe | information st | Translation, ri | 177SZ | This protein bi | 335992.SAR11 | 1,4 | 0,8 | 0,000133962 | 7,45E-05 | 2 - | 0,000162804 - | 2 - | 0,000202154 - | 0 - | 0 - | 2 - | 0,000195012 - | 1 - | 0,00010984 - |
| 629 | 1 | 3um_FIJJAFBF_613662 | Bacteria | Bacteroidetes | Flavobacteriia | Flavobacterial | Flavobacteriac | Polaribacter | Polaribacter s | metabolism | Carbohydrate | 05D2N | Aldolase | 313598.MED1 | 4,8 | 0,748331477 | 0,000184084 | 2,60E-05 | 6 - | 0,000193989 - | 5 - | 0,000200731 - | 4 - | 0,000152656 - | 4 - | 0,000154911 - | 5 - | 0,000218133 - |
| 630 | 1 | 3um_FIJJAFBF_132509 | Bacteria | Proteobacteri | Gammaprotec | Cellvibrionales | Porticoccacea | unclassified | Gammaprotec | metabolism | Energy produc | 17CGT | F(1)F(0) ATP s | 314287.GB22 | 1,2 | 0,4 | 0,000102553 | 2,27E-05 | 2 - | 0,00014715 - | 1 - | 9,14E-05 - | 1 - | 8,68E-05 - | 1 - | 8,81E-05 - | 1 - | 9,93E-05 - |
| 631 | 1 | 0.2um_NOKCPHJN_13684 | Bacteria | Bacteroidetes | Flavobacteriia | unclassified | unclassified | unclassified | Flavobacteria | unclassified | unclassified | unclassified | unclassified | unclassified | 0,2 | 0,4 | 2,95E-06 | 5,89E-06 | 1 - | 1,47E-05 - | 0 - | 0 - | 0 - | 0 - | 0 - | 0 - | 0 - | 0 - |
| 632 | 1 | 3um_FIJJAFBF_24248 | Bacteria | Bacteroidetes | Cytophagia | Cytophagales | Cyclobacteriac | Algoriphagus | Algoriphagus | metabolism | Carbohydrate | COG0057 | glyceraldehyd | 388413.ALPR1 | 1,6 | 1,019803903 | 6,42E-05 | 4,04E-05 | 2 - | 6,85E-05 - | 2 - | 8,51E-05 - | 3 - | 0,000121327 - | 0 - | 0 - | 1 - | 4,62E-05 - |
| 633 | 1 | 3um_FIJJAFBF_226197 | Bacteria | Bacteroidetes | Cytophagia | Cytophagales | Flammeovirga | unclassified | Flammeovirga | poorly charact | Function unkn | 08NCC | gliding motilit | 649349.Lbys_ | 1,4 | 1,019803903 | 6,67E-05 | 4,46E-05 | 3 - | 0,000128004 - | 1 - | 5,30E-05 - | 1 - | 5,04E-05 - | 2 - | 0,000102218 - | 0 - | 0 - |
| 634 | 1 | 0.2um_NOKCPHJN_169342 | Bacteria | Bacteroidetes | Flavobacteriia | Flavobacterial | Flavobacteriac | Mesoflavibact | Mesoflavibact | unclassified | unclassified | unclassified | unclassified | unclassified | 1 | 0 | 5,57E-05 | 5,29E-06 | 1 - | 4,67E-05 - | 1 - | 5,79E-05 - | 1 - | 5,51E-05 - | 1 - | 5,59E-05 - | 1 - | 6,30E-05 - |
| 635 | 1 | 0.2um_NOKCPHJN_585767 | Bacteria | Proteobacteri | Betaproteoba | Burkholderial | Comamonada | Variovorax | Variovorax sp. | cellular proces | Intracellular tr | 0BIEB | domain-conta | 1005048.CFU_ | 2 | 0,632455532 | 5,32E-05 | 1,23E-05 | 3 - | 6,89E-05 - | 2 - | 5,70E-05 - | 2 - | 5,42E-05 - | 2 - | 5,50E-05 - | 1 - | 3,10E-05 - |
| 636 | 1 | 0.2um_NOKCPHJN_491959 | Bacteria | Proteobacteri | Alphaproteob | Rhodospirillal | Rhodospirillac | Candidatus En | Candidatus En | unclassified | unclassified | unclassified | unclassified | unclassified | 0,8 | 0,4 | 2,53E-05 | 1,30E-05 | 1 - | 2,68E-05 - | 0 - | 0 - | 1 - | 3,16E-05 - | 1 - | 3,20E-05 - | 1 - | 3,61E-05 - |
| 637 | 1 | 3um_FIJJAFBF_481970 | Bacteria | Proteobacteri | Alphaproteob | Rhodobactera | Rhodobactera | Phaeobacter | Phaeobacter g | cellular proces | Posttranslatio | 01QWG | Prevents misf | 439497.RR11_ | 17 | 0,894427191 | 0,000422666 | 2,77E-05 | 18 - | 0,000376317 - | 16 - | 0,000415356 - | 17 - | 0,000419526 - | 18 - | 0,000450765 - | 16 - | 0,000451365 - |
| 638 | 1 | 3um_FIJJAFBF_180623 | Bacteria | Bacteroidetes | Flavobacteriia | Flavobacterial | Flavobacteriac | Formosa | Formosa sp. H | metabolism | Inorganic ion t | 0PA08 | Destroys radic | 391603.FBALC | 1,6 | 0,489897949 | 0,000112935 | 2,96E-05 | 2 - | 0,000120818 - | 1 - | 7,50E-05 - | 2 - | 0,000142613 - | 2 - | 0,000144719 - | 1 - | 8,15E-05 - |
| 639 | 1 | 3um_FIJJAFBF_773329 | Eukaryota | Bacillariophyt | Coscinodiscop | Thalassiosirale | Stephanodisca | Cyclotella | Cyclotella sp. | cellular proces | Posttranslatio | 0FJCG | Prevents misf | 497965.Cyan7 | 0,2 | 0,4 | 6,19E-06 | 1,24E-05 | 0 - | 0 - | 0 - | 0 - | 0 - | 0 - | 1 - | 3,10E-05 - | 0 - | 0 - |
| 640 | 1 | 3um_FIJJAFBF_326272 | Bacteria | Proteobacteri | Alphaproteob | Rhodobactera | Rhodobactera | Phaeobacter | Phaeobacter s | information st | Translation, ri | COG1185 | Involved in m | 391595.RLO14 | 5,6 | 0,8 | 0,000107527 | 1,28E-05 | 6 - | 9,74E-05 - | 6 - | 0,00012095 - | 6 - | 0,000114978 - | 6 - | 0,000116676 - | 4 - | 8,76E-05 - |
| 641 | 1 | 0.2um_NOKCPHJN_206817 | Bacteria | Proteobacteri | Gammaprotec | Oceanospirilla | Oceanospirilla | Neptuniibacte | Neptuniibacte | unclassified | unclassified | unclassified | unclassified | unclassified | 13 | 1,414213562 | 0,000489565 | 7,36E-05 | 13 - | 0,000409917 - | 11 - | 0,000430689 - | 15 - | 0,000558306 - | 12 - | 0,000453242 - | 14 - | 0,000595671 - |
| 642 | 1 | 3um_FIJJAFBF_776437 | Bacteria | Bacteroidetes | Flavobacteriia | Flavobacterial | Flavobacteriac | Formosa | Formosa sp. H | poorly charact | Function unkn | 09EIV | NA | 983544.Lacal_ | 0,2 | 0,4 | 2,15E-05 | 4,30E-05 | 0 - | 0 - | 0 - | 0 - | 0 - | 0 - | 1 - | 0,000107409 - | 0 - | 0 - |
| 643 | 1 | 0.2um_NOKCPHJN_395040 | Bacteria | Proteobacteri | Deltaproteoba | Myxococcales | Nannocystace | Plesiocystis | Plesiocystis pa | cellular proces | Posttranslatio | COG1225 | alkyl hydroper | 391625.PPSIR | 0,2 | 0,4 | 9,37E-07 | 1,87E-06 | 1 - | 4,68E-06 - | 0 - | 0 - | 0 - | 0 - | 0 - | 0 - | 0 - | 0 - |
| 644 | 1 | 3um_FIJJAFBF_461750 | Bacteria | unclassified | unclassified | unclassified | unclassified | unclassified | uncultured ba | poorly charact | Function unkn | 09BCW | Protein of unk | 313594.PI23P | 0,6 | 0,489897949 | 3,42E-05 | 2,82E-05 | 1 - | 4,99E-05 - | 1 - | 6,20E-05 - | 1 - | 5,89E-05 - | 0 - | 0 - | 0 - | 0 - |
| 645 | 1 | 3um_FIJJAFBF_80670 | Eukaryota | unclassified | Cryptophyta | Pyrenomonad | Geminigerace | Guillardia | Guillardia thet | metabolism | Energy produc | 123BF | photosystem I | 35128.Thapsd | 1 | 0 | 0,000204518 | 1,94E-05 | 1 - | 0,000171309 - | 1 - | 0,000212715 - | 1 - | 0,000202212 - | 1 - | 0,000205199 - | 1 - | 0,000231156 - |
| 646 | 1 | 0.2um_NOKCPHJN_926017 | Bacteria | Proteobacteri | Alphaproteob | Rhodobactera | Rhodobactera | unclassified | various | unclassified | unclassified | unclassified | unclassified | unclassified | 2 | 0,632455532 | 0,000105848 | 2,80E-05 | 3 - | 0,000135031 - | 2 - | 0,00011178 - | 1 - | 5,31E-05 - | 2 - | 0,00010783 - | 2 - | 0,00012147 - |
| 647 | 1 | 3um_FIJJAFBF_226993 | Bacteria | unclassified | unclassified | unclassified | unclassified | unclassified | uncultured ba | metabolism | Inorganic ion t | 09C0C | TonB depende | 742767.HMPR | 1,6 | 0,489897949 | 4,63E-05 | 1,21E-05 | 2 - | 4,96E-05 - | 1 - | 3,08E-05 - | 2 - | 5,85E-05 - | 2 - | 5,94E-05 - | 1 - | 3,35E-05 - |
| 648 | 1 | 0.2um_NOKCPHJN_608299 | Bacteria | Bacteroidetes | unclassified | unclassified | unclassified | unclassified | Bacteroidetes | cellular proces | Signal transdu | 0P8JH | Histidine kina | 926562.Oweh | 0,4 | 0,489897949 | 1,51E-05 | 1,86E-05 | 1 - | 3,46E-05 - | 0 - | 0 - | 1 - | 4,08E-05 - | 0 - | 0 - | 0 - | 0 - |
| 649 | 1 | 3um_FIJJAFBF_307113 | Bacteria | Bacteroidetes | Flavobacteriia | Flavobacterial | Flavobacteriac | Psychroserpe | Psychroserpe | unclassified | unclassified | unclassified | unclassified | unclassified | 0,4 | 0,8 | 1,84E-05 | 3,69E-05 | 0 - | 0 - | 0 - | 0 - | 2 - | 9,22E-05 - | 0 - | 0 - | 0 - | 0 - |
| 650 | 1 | 3um_FIJJAFBF_467908 | Bacteria | Proteobacteri | Betaproteoba | Burkholderial | Comamonada | Limnohabitan | Limnohabitan | unclassified | unclassified | unclassified | unclassified | unclassified | 2,4 | 1,854723699 | 9,75E-05 | 7,75E-05 | 2 - | 7,00E-05 - | 1 - | 4,35E-05 - | 5 - | 0,000206528 - | 4 - | 0,000167663 - | 0 - | 0 - |
| 651 | 1 | 0.2um_NOKCPHJN_178604 | Bacteria | Proteobacteri | Gammaprotec | Oceanospirilla | Oceanospirilla | Marinomonas | various | poorly charact | Function unkn | 05MHK | YceI family | 258594.RPA1 | 4,6 | 1,2 | 0,000283016 | 4,45E-05 | 7 - | 0,000368549 - | 4 - | 0,000261503 - | 4 - | 0,000248591 - | 4 - | 0,000252263 - | 4 - | 0,000284173 - |
| 652 | 1 | 3um_FIJJAFBF_333855 | Bacteria | Bacteroidetes | Saprospiria | Saprospirales | Saprospiracea | Aureispira | Aureispira sp. | cellular proces | Cell wall/mem | 01S9I | epimerase del | 316055.RPE_3 | 0,2 | 0,4 | 6,91E-06 | 1,38E-05 | 1 - | 3,46E-05 - | 0 - | 0 - | 0 - | 0 - | 0 - | 0 - | 0 - | 0 - |
| 653 | 1 | 0.2um_NOKCPHJN_304819 | Bacteria | Proteobacteri | Betaproteoba | Nitrosomonad | unclassified | unclassified | Methylophilal | metabolism | Nucleotide tra | 16QX0 | Catalyzes the | 383631.MB21 | 0,8 | 0,4 | 2,10E-05 | 1,06E-05 | 0 - | 0 - | 1 - | 2,62E-05 - | 1 - | 2,49E-05 - | 1 - | 2,53E-05 - | 1 - | 2,85E-05 - |
| 654 | 3 | 0.2um_NOKCPHJN_26370;0.2um_NOKCPHJN_40 | Bacteria | Bacteroidetes | Flavobacteriia | various | various | various | various | information st | Transcription | COG0250 | Participates in | 487796.Flav2A | 1 | 0 | 7,45E-05 | 7,07E-06 | 1 - | 6,24E-05 - | 1 - | 7,75E-05 - | 1 - | 7,36E-05 - | 1 - | 7,47E-05 - | 1 - | 8,42E-05 - |
| 655 | 1 | 0.2um_NOKCPHJN_611548 | Eukaryota | Phaeophyceae | unclassified | Ectocarpales | Ectocarpaceae | Ectocarpus | Ectocarpus sili | unclassified | unclassified | unclassified | unclassified | unclassified | 3 | 0 | 7,06E-05 | 6,71E-06 | 3 - | 5,92E-05 - | 3 - | 7,35E-05 - | 3 - | 6,98E-05 - | 3 - | 7,09E-05 - | 3 - | 7,98E-05 - |
| 656 | 1 | 3um_FIJJAFBF_501483 | Bacteria | Proteobacteri | Gammaprotec | Pseudomonad | Pseudomonad | Pseudomonas | Pseudomonas | information st | Translation, ri | COG0050 | This protein p | 247633.GP214 | 1,6 | 0,489897949 | 0,000107378 | 3,46E-05 | 1 - | 5,60E-05 - | 2 - | 0,000139043 - | 2 - | 0,000132178 - | 2 - | 0,00013413 - | 1 - | 7,55E-05 - |
| 657 | 1 | 3um_FIJJAFBF_60140 | Viruses | unclassified | unclassified | unclassified | unclassified | unclassified | uncultured M | unclassified | unclassified | unclassified | unclassified | unclassified | 1 | 0 | 2,32E-05 | 2,20E-06 | 1 - | 1,94E-05 - | 1 - | 2,41E-05 - | 1 - | 2,29E-05 - | 1 - | 2,33E-05 - | 1 - | 2,62E-05 - |
| 658 | 1 | 3um_FIJJAFBF_69074 | Bacteria | Proteobacteri | Gammaprotec | Alteromonada | Alteromonada | Paraglaciecola | Paraglaciecola | unclassified | unclassified | unclassified | unclassified | unclassified | 3,2 | 0,979795897 | 0,000412976 | 9,27E-05 | 5 - | 0,000551811 - | 3 - | 0,000411112 - | 3 - | 0,000390814 - | 2 - | 0,000264391 - | 3 - | 0,000446753 - |
| 659 | 1 | 0.2um_NOKCPHJN_143516 | Bacteria | Proteobacteri | Alphaproteob | Rhodobactera | unclassified | unclassified | Rhodobactera | cellular proces | Intracellular tr | COG0552 | Involved in tar | 367336.OM22 | 0,2 | 0,4 | 7,20E-06 | 1,44E-05 | 0 - | 0 - | 1 - | 3,60E-05 - | 0 - | 0 - | 0 - | 0 - | 0 - | 0 - |
| 660 | 2 | 0.2um_NOKCPHJN_577760;3um_FIJJAFBF_2827 | Bacteria | Bacteroidetes | Flavobacteriia | Flavobacterial | Flavobacteriac | Marinirhabdu | Marinirhabdu | unclassified | unclassified | unclassified | unclassified | unclassified | 0,2 | 0,4 | 9,08E-06 | 1,82E-05 | 0 - | 0 - | 1 - | 4,54E-05 - | 0 - | 0 - | 0 - | 0 - | 0 - | 0 - |
| 661 | 1 | 3um_FIJJAFBF_17379 | unclassified | unclassified | unclassified | unclassified | unclassified | unclassified | unclassified | unclassified | unclassified | unclassified | unclassified | unclassified | 0,4 | 0,489897949 | 7,72E-05 | 9,47E-05 | 0 - | 0 - | 1 - | 0,000185089 - | 0 - | 0 - | 0 - | 0 - | 1 - | 0,000201136 - |
| 662 | 1 | 3um_FIJJAFBF_324101 | Bacteria | Proteobacteri | Gammaprotec | Alteromonada | Alteromonada | Paraglaciecola | Paraglaciecola | unclassified | unclassified | unclassified | unclassified | unclassified | 2,2 | 0,4 | 0,000229345 | 6,17E-05 | 2 - | 0,000172597 - | 2 - | 0,000214314 - | 2 - | 0,000203733 - | 2 - | 0,000206742 - | 3 - | 0,000349341 - |
| 663 | 1 | 3um_FIJJAFBF_332804 | Bacteria | Proteobacteri | Alphaproteob | Rhodobactera | Rhodobactera | Litoreibacter | Litoreibacter a | unclassified | unclassified | unclassified | unclassified | unclassified | 0,6 | 0,8 | 3,84E-05 | 4,96E-05 | 2 - | 0,000120818 - | 0 - | 0 - | 1 - | 7,13E-05 - | 0 - | 0 - | 0 - | 0 - |
| 664 | 1 | 0.2um_NOKCPHJN_92216 | Bacteria | Bacteroidetes | Flavobacteriia | Flavobacterial | Flavobacteriac | Polaribacter | Polaribacter s | metabolism | Amino acid tra | 05DX4 | Peptidase M1 | 313603.FB217 | 0,2 | 0,4 | 2,82E-06 | 5,65E-06 | 1 - | 1,41E-05 - | 0 - | 0 - | 0 - | 0 - | 0 - | 0 - | 0 - | 0 - |
| 665 | 1 | 3um_FIJJAFBF_196001 | Bacteria | Bacteroidetes | Flavobacteriia | Flavobacterial | Flavobacteriac | Polaribacter | Polaribacter s | cellular proces | Posttranslatio | 08V1T | peptidylprolyl | 313594.PI23P | 6,8 | 0,979795897 | 0,000400872 | 6,84E-05 | 8 - | 0,000395782 - | 6 - | 0,000368583 - | 6 - | 0,000350385 - | 6 - | 0,00035556 - | 8 - | 0,00053405 - |
| 666 | 1 | 3um_FIJJAFBF_28660 | Bacteria | Proteobacteri | Alphaproteob | Rhodobactera | Rhodobactera | Planktomarina | Planktomarina | unclassified | unclassified | unclassified | unclassified | unclassified | 0,2 | 0,4 | 1,23E-05 | 2,47E-05 | 1 - | 6,17E-05 - | 0 - | 0 - | 0 - | 0 - | 0 - | 0 - | 0 - | 0 - |
| 667 | 1 | 3um_FIJJAFBF_715581 | Bacteria | Proteobacteri | Gammaprotec | Cellvibrionales | Halieaceae | unclassified | marine gamm | information st | Translation, ri | 177XJ | One of the pri | 247639.MGP2 | 1 | 0 | 8,78E-05 | 8,34E-06 | 1 - | 7,36E-05 - | 1 - | 9,14E-05 - | 1 - | 8,68E-05 - | 1 - | 8,81E-05 - | 1 - | 9,93E-05 - |
| 668 | 1 | 3um_FIJJAFBF_405248 | Bacteria | Bacteroidetes | Flavobacteriia | Flavobacterial | Flavobacteriac | Polaribacter | Polaribacter s | metabolism | Amino acid tra | 09GJ6 | nitrogen regul | 313598.MED1 | 4,2 | 0,979795897 | 0,000514241 | 0,000134529 | 5 - | 0,000512396 - | 5 - | 0,000636245 - | 3 - | 0,000362899 - | 3 - | 0,000368259 - | 5 - | 0,000691404 - |
| 669 | 1 | 0.2um_NOKCPHJN_221460 | Bacteria | Proteobacteri | Alphaproteob | Rhodobactera | Rhodobactera | unclassified | Rhodobactera | unclassified | unclassified | unclassified | unclassified | unclassified | 0,2 | 0,4 | 7,68E-06 | 1,54E-05 | 1 - | 3,84E-05 - | 0 - | 0 - | 0 - | 0 - | 0 - | 0 - | 0 - | 0 - |
| 670 | 1 | 0.2um_NOKCPHJN_78687 | Bacteria | Proteobacteri | Gammaprotec | Alteromonada | Alteromonada | Agarivorans | Agarivorans g | poorly charact | Function unkn | 0NVRZ | NA | 170187.SP_17 | 3,4 | 0,8 | 8,31E-05 | 2,18E-05 | 4 - | 8,18E-05 - | 4 - | 0,000101618 - | 3 - | 7,25E-05 - | 2 - | 4,90E-05 - | 4 - | 0,000110427 - |
| 671 | 1 | 3um_FIJJAFBF_711021 | Bacteria | Proteobacteri | Betaproteoba | Burkholderial | Comamonada | Comamonas | Comamonas d | metabolism | Amino acid tra | 16R0H | Acetolactate s | 398578.Daci_ | 0,8 | 0,748331477 | 6,17E-05 | 5,45E-05 | 2 - | 0,00014083 - | 0 - | 0 - | 1 - | 8,31E-05 - | 1 - | 8,43E-05 - | 0 - | 0 - |
| 672 | 1 | 0.2um_NOKCPHJN_88300 | Bacteria | Proteobacteri | Alphaproteob | Rhodobactera | Rhodobactera | Planktomarina | Planktomarina | metabolism | Inorganic ion t | COG0841 | acriflavin resis | 633131.TR2A6 | 0,6 | 0,489897949 | 7,21E-06 | 5,99E-06 | 1 - | 1,02E-05 - | 0 - | 0 - | 0 - | 0 - | 1 - | 1,22E-05 - | 1 - | 1,37E-05 - |
| 673 | 1 | 3um_FIJJAFBF_532821 | Bacteria | Proteobacteri | Alphaproteob | Rhodobactera | Rhodobactera | Leisingera | various | metabolism | Lipid transpor | 16QA8 | reductase | 439496.RBY4I | 0,4 | 0,489897949 | 2,23E-05 | 2,73E-05 | 0 - | 0 - | 0 - | 0 - | 1 - | 5,53E-05 - | 1 - | 5,61E-05 - | 0 - | 0 - |
| 674 | 1 | 0.2um_NOKCPHJN_442338 | Bacteria | Bacteroidetes | Flavobacteriia | Flavobacterial | Cryomorphace | Owenweeksia | Owenweeksia | information st | Transcription | COG0085 | DNA-depende | 926562.Oweh | 0,2 | 0,4 | 2,44E-06 | 4,88E-06 | 0 - | 0 - | 0 - | 0 - | 0 - | 0 - | 0 - | 0 - | 1 - | 1,22E-05 - |
| 675 | 1 | 3um_FIJJAFBF_452509 | Bacteria | Actinobacteri | Actinobacteri | Corynebacteri | Corynebacteri | Corynebacteri | Corynebacteri | poorly charact | Function unkn | 16YV3 | Large repetiti | 272942.RCAP_ | 2,4 | 1,2 | 3,59E-05 | 1,93E-05 | 1 - | 1,21E-05 - | 4 - | 6,03E-05 - | 1 - | 1,43E-05 - | 3 - | 4,36E-05 - | 3 - | 4,91E-05 - |
| 676 | 2 | 0.2um_NOKCPHJN_229463;0.2um_NOKCPHJN_4 | Bacteria | Proteobacteri | Alphaproteob | Pelagibacteral | Pelagibacterac | Candidatus Pe | Candidatus Pe | information st | Translation, ri | 17C3B | The globular d | 335992.SAR11 | 0,2 | 0,4 | 2,08E-05 | 4,17E-05 | 0 - | 0 - | 0 - | 0 - | 0 - | 0 - | 1 - | 0,000104154 - | 0 - | 0 - |
| 677 | 1 | 0.2um_NOKCPHJN_65576 | Bacteria | Proteobacteri | Alphaproteob | Rhodobactera | Rhodobactera | Planktomarina | Planktomarina | information st | Translation, ri | 179R6 | This protein is | 391613.RTM1 | 1,6 | 0,489897949 | 0,00014372 | 4,59E-05 | 2 - | 0,000151022 - | 1 - | 9,38E-05 - | 1 - | 8,91E-05 - | 2 - | 0,000180899 - | 2 - | 0,000203782 - |
| 678 | 1 | 3um_FIJJAFBF_322027 | Bacteria | Proteobacteri | Gammaprotec | Oceanospirilla | Saccharospirill | Reinekea | Reinekea fors | cellular proces | Intracellular tr | 175GA | One of the pro | 314283.MED2 | 2,6 | 0,489897949 | 0,000242988 | 5,93E-05 | 2 - | 0,000155104 - | 2 - | 0,000192593 - | 3 - | 0,000274626 - | 3 - | 0,000278682 - | 3 - | 0,000313935 - |
| 679 | 1 | 0.2um_NOKCPHJN_190348 | Bacteria | Proteobacteri | Gammaprotec | Alteromonada | Alteromonada | Paraglaciecola | Paraglaciecola | poorly charact | Function unkn | 0R65K | Protein of unk | 983545.Glaag | 0,4 | 0,8 | 2,89E-05 | 5,79E-05 | 0 - | 0 - | 2 - | 0,000144689 - | 0 - | 0 - | 0 - | 0 - | 0 - | 0 - |
| 680 | 1 | 0.2um_NOKCPHJN_1021675 | Bacteria | Bacteroidetes | Cytophagia | Cytophagales | Cytophagacea | Ekhidna | Ekhidna lutea | unclassified | unclassified | unclassified | unclassified | unclassified | 0,6 | 0,489897949 | 4,63E-05 | 3,85E-05 | 1 - | 6,52E-05 - | 0 - | 0 - | 0 - | 0 - | 1 - | 7,81E-05 - | 1 - | 8,80E-05 - |
| 681 | 1 | 0.2um_NOKCPHJN_933764 | Bacteria | Proteobacteri | Alphaproteob | Pelagibacteral | Pelagibacterac | Candidatus Pe | Candidatus Pe | unclassified | unclassified | unclassified | unclassified | unclassified | 0,2 | 0,4 | 1,09E-05 | 2,17E-05 | 0 - | 0 - | 0 - | 0 - | 0 - | 0 - | 1 - | 5,43E-05 - | 0 - | 0 - |
| 682 | 1 | 0.2um_NOKCPHJN_963901 | Bacteria | Proteobacteri | Gammaprotec | unclassified | unclassified | unclassified | OM182 bacte | unclassified | unclassified | unclassified | unclassified | unclassified | 0,2 | 0,4 | 3,71E-06 | 7,43E-06 | 1 - | 1,86E-05 - | 0 - | 0 - | 0 - | 0 - | 0 - | 0 - | 0 - | 0 - |
| 683 | 1 | 3um_FIJJAFBF_102286 | Bacteria | Proteobacteri | Alphaproteob | Rhodobactera | Rhodobactera | Amylibacter | Amylibacter k | unclassified | unclassified | unclassified | unclassified | unclassified | 15,4 | 0,489897949 | 0,000621079 | 4,69E-05 | 16 - | 0,000541719 - | 15 - | 0,000630614 - | 16 - | 0,000639444 - | 15 - | 0,000608333 - | 15 - | 0,000685285 - |
| 684 | 1 | 3um_FIJJAFBF_130199 | Bacteria | Proteobacteri | Betaproteoba | Burkholderial | Comamonada | Hydrogenoph | various | unclassified | unclassified | unclassified | unclassified | unclassified | 0,8 | 0,4 | 8,20E-05 | 4,21E-05 | 1 - | 8,57E-05 - | 1 - | 0,000106357 - | 0 - | 0 - | 1 - | 0,000102599 - | 1 - | 0,000115578 - |
| 685 | 1 | 0.2um_NOKCPHJN_15616 | Bacteria | Bacteroidetes | Flavobacteriia | unclassified | unclassified | unclassified | Flavobacteria | unclassified | unclassified | unclassified | unclassified | unclassified | 4,4 | 0,8 | 6,00E-05 | 5,50E-06 | 6 - | 6,96E-05 - | 4 - | 5,76E-05 - | 4 - | 5,47E-05 - | 4 - | 5,55E-05 - | 4 - | 6,26E-05 - |
| 686 | 1 | 3um_FIJJAFBF_526632 | Bacteria | Bacteroidetes | Flavobacteriia | Flavobacterial | Crocinitomica | unclassified | Crocinitomica | cellular proces | Intracellular tr | 09S7I | Preprotein tra | 402612.FP198 | 0,8 | 0,4 | 0,000103331 | 5,31E-05 | 1 - | 0,00010828 - | 1 - | 0,000134452 - | 1 - | 0,000127813 - | 0 - | 0 - | 1 - | 0,000146108 - |
| 687 | 1 | 0.2um_NOKCPHJN_718159 | Eukaryota | Bacillariophyt | Bacillariophyc | Naviculales | Phaeodactylac | Phaeodactylu | Phaeodactylu | information st | Translation, ri | KOG0052 | This protein p | 2850.Phatr18 | 3,6 | 1,019803903 | 0,000110755 | 3,19E-05 | 4 - | 0,000103636 - | 5 - | 0,000160857 - | 4 - | 0,000122332 - | 2 - | 6,21E-05 - | 3 - | 0,000104881 - |
| 688 | 1 | 3um_FIJJAFBF_805819 | Bacteria | Proteobacteri | Gammaprotec | unclassified | unclassified | unclassified | Gammaprotec | unclassified | unclassified | unclassified | unclassified | unclassified | 0,8 | 0,4 | 3,14E-05 | 1,59E-05 | 1 - | 3,40E-05 - | 1 - | 4,22E-05 - | 1 - | 4,01E-05 - | 1 - | 4,07E-05 - | 0 - | 0 - |
| 689 | 1 | 3um_FIJJAFBF_560778 | Bacteria | Proteobacteri | Alphaproteob | Pelagibacteral | Pelagibacterac | unclassified | Pelagibactera | unclassified | unclassified | unclassified | unclassified | unclassified | 0,2 | 0,4 | 1,51E-05 | 3,03E-05 | 0 - | 0 - | 0 - | 0 - | 1 - | 7,57E-05 - | 0 - | 0 - | 0 - | 0 - |
| 690 | 1 | 3um_FIJJAFBF_21475 | Eukaryota | unclassified | Cryptophyta | Pyrenomonad | Geminigerace | Guillardia | Guillardia thet | metabolism | Carbohydrate | 0IVA1 | Photosystem I | 3218.PP1S345 | 0,8 | 0,4 | 6,23E-05 | 3,14E-05 | 0 - | 0 - | 1 - | 7,79E-05 - | 1 - | 7,40E-05 - | 1 - | 7,51E-05 - | 1 - | 8,46E-05 - |
| 691 | 1 | 0.2um_NOKCPHJN_93316 | Bacteria | Proteobacteri | Betaproteoba | unclassified | unclassified | unclassified | Betaproteoba | poorly charact | Function unkn | 0BF8N | signal peptide | 420662.Mpe_ | 0,8 | 0,4 | 5,74E-05 | 2,95E-05 | 1 - | 6,07E-05 - | 0 - | 0 - | 1 - | 7,17E-05 - | 1 - | 7,27E-05 - | 1 - | 8,19E-05 - |
| 692 | 1 | 0.2um_NOKCPHJN_1065032 | Eukaryota | unclassified | Spirotrichea | Sporadotrichi | Oxytrichidae | Stylonychia | Stylonychia le | metabolism | Carbohydrate | COG0057 | glyceraldehyd | 423536.XP_00 | 1,2 | 0,4 | 4,78E-05 | 1,65E-05 | 1 - | 3,34E-05 - | 1 - | 4,14E-05 - | 1 - | 3,94E-05 - | 2 - | 7,99E-05 - | 1 - | 4,50E-05 - |
| 693 | 1 | 3um_FIJJAFBF_216065 | Bacteria | Proteobacteri | Gammaprotec | Cellvibrionales | Cellvibrionace | Candidatus En | Candidatus En | information st | Translation, ri | 08VZM | 50s ribosomal | 314287.GB22 | 0,2 | 0,4 | 1,55E-05 | 3,11E-05 | 0 - | 0 - | 0 - | 0 - | 0 - | 0 - | 1 - | 7,77E-05 - | 0 - | 0 - |
| 694 | 1 | 3um_FIJJAFBF_747826 | Bacteria | Bacteroidetes | Flavobacteriia | Flavobacterial | Flavobacteriac | Polaribacter | Polaribacter a | cellular proces | Posttranslatio | 0P7DX | ATP-depende | 313603.FB217 | 1,4 | 1,019803903 | 4,45E-05 | 2,88E-05 | 3 - | 8,36E-05 - | 1 - | 3,46E-05 - | 0 - | 0 - | 2 - | 6,67E-05 - | 1 - | 3,76E-05 - |
| 695 | 1 | 3um_FIJJAFBF_300355 | Eukaryota | Apicomplexa | Conoidasida | Eucoccidiorida | Eimeriidae | Eimeria | Eimeria acerv | unclassified | unclassified | unclassified | unclassified | unclassified | 4 | 0,632455532 | 0,00019344 | 2,50E-05 | 5 - | 0,000204229 - | 4 - | 0,000202874 - | 4 - | 0,000192857 - | 3 - | 0,000146779 - | 4 - | 0,000220462 - |
| 696 | 1 | 0.2um_NOKCPHJN_538976 | Bacteria | Proteobacteri | Alphaproteob | Pelagibacteral | Pelagibacterac | Candidatus Pe | Candidatus Pe | unclassified | unclassified | unclassified | unclassified | unclassified | 2 | 0,894427191 | 5,46E-05 | 2,13E-05 | 3 - | 7,10E-05 - | 1 - | 2,94E-05 - | 3 - | 8,38E-05 - | 2 - | 5,67E-05 - | 1 - | 3,19E-05 - |
| 697 | 1 | 3um_FIJJAFBF_254284 | Bacteria | Proteobacteri | Gammaprotec | Oceanospirilla | Oceanospirilla | Marinobacteri | various | information st | Translation, ri | COG0051 | tRNA binding | 876044.IMCC | 1,4 | 0,489897949 | 0,000180268 | 5,30E-05 | 2 - | 0,000220725 - | 1 - | 0,000137037 - | 1 - | 0,000130271 - | 2 - | 0,000264391 - | 1 - | 0,000148918 - |
| 698 | 1 | 0.2um_NOKCPHJN_1083488 | unclassified | unclassified | unclassified | unclassified | unclassified | unclassified | unclassified | unclassified | unclassified | unclassified | unclassified | unclassified | 1,4 | 0,489897949 | 3,65E-05 | 1,32E-05 | 2 - | 4,39E-05 - | 1 - | 2,73E-05 - | 1 - | 2,59E-05 - | 1 - | 2,63E-05 - | 2 - | 5,92E-05 - |
| 699 | 1 | 0.2um_NOKCPHJN_1047183 | Bacteria | Bacteroidetes | Flavobacteriia | Flavobacterial | Flavobacteriac | Winogradskye | Winogradskye | poorly charact | Function unkn | 0PAEZ | Inherit from b | 391598.FBBAL | 0,6 | 0,489897949 | 2,92E-05 | 2,43E-05 | 1 - | 4,07E-05 - | 1 - | 5,05E-05 - | 0 - | 0 - | 0 - | 0 - | 1 - | 5,49E-05 - |
| 700 | 1 | 0.2um_NOKCPHJN_683378 | Eukaryota | Phaeophyceae | unclassified | Ectocarpales | Ectocarpaceae | Ectocarpus | Ectocarpus sili | unclassified | unclassified | unclassified | unclassified | unclassified | 0,8 | 0,4 | 1,36E-05 | 7,01E-06 | 1 - | 1,43E-05 - | 1 - | 1,77E-05 - | 1 - | 1,69E-05 - | 0 - | 0 - | 1 - | 1,93E-05 - |
| 701 | 1 | 0.2um_NOKCPHJN_163522 | Bacteria | Proteobacteri | Gammaprotec | Cellvibrionales | Porticoccacea | unclassified | SAR92 bacteri | unclassified | unclassified | unclassified | unclassified | unclassified | 2,2 | 0,748331477 | 4,83E-05 | 1,55E-05 | 3 - | 5,58E-05 - | 3 - | 6,93E-05 - | 2 - | 4,39E-05 - | 1 - | 2,23E-05 - | 2 - | 5,02E-05 - |
| 702 | 1 | 3um_FIJJAFBF_145781 | Eukaryota | Bacillariophyt | Coscinodiscop | Chaetocerotal | Chaetocerotac | Chaetoceros | Chaetoceros g | unclassified | unclassified | unclassified | unclassified | unclassified | 0,2 | 0,4 | 4,17E-05 | 8,35E-05 | 1 - | 0,000208685 - | 0 - | 0 - | 0 - | 0 - | 0 - | 0 - | 0 - | 0 - |
| 703 | 1 | 0.2um_NOKCPHJN_26075 | Bacteria | Proteobacteri | Alphaproteob | unclassified | unclassified | unclassified | Alphaproteob | metabolism | Amino acid tra | 177KB | Extracellular s | 331869.BAL19 | 6,6 | 2,059126028 | 0,000165701 | 4,92E-05 | 8 - | 0,000171309 - | 9 - | 0,000239304 - | 6 - | 0,000151659 - | 7 - | 0,000179549 - | 3 - | 8,67E-05 - |
| 704 | 1 | 3um_FIJJAFBF_492315 | Eukaryota | unclassified | Cryptophyta | Pyrenomonad | Geminigerace | Teleaulax | Teleaulax amp | unclassified | unclassified | unclassified | unclassified | unclassified | 2,8 | 0,4 | 0,000216146 | 3,50E-05 | 3 - | 0,000194537 - | 2 - | 0,000161038 - | 3 - | 0,000229631 - | 3 - | 0,000233023 - | 3 - | 0,000262499 - |
| 705 | 1 | 3um_FIJJAFBF_103116 | Bacteria | Bacteroidetes | Flavobacteriia | Flavobacterial | Flavobacteriac | Polaribacter | Polaribacter s | unclassified | unclassified | unclassified | unclassified | unclassified | 6,8 | 0,748331477 | 9,83E-05 | 3,95E-06 | 8 - | 9,78E-05 - | 6 - | 9,11E-05 - | 7 - | 0,000100998 - | 7 - | 0,00010249 - | 6 - | 9,90E-05 - |
| 706 | 1 | 0.2um_NOKCPHJN_881844 | Bacteria | Proteobacteri | Alphaproteob | Pelagibacteral | Pelagibacterac | Candidatus Pe | Candidatus Pe | metabolism | Amino acid tra | 01R4A | Polyamine AB | 335992.SAR11 | 0,2 | 0,4 | 7,37E-06 | 1,47E-05 | 0 - | 0 - | 0 - | 0 - | 0 - | 0 - | 1 - | 3,69E-05 - | 0 - | 0 - |
| 707 | 1 | 0.2um_NOKCPHJN_498936 | Bacteria | Bacteroidetes | Flavobacteriia | Flavobacterial | Flavobacteriac | Algibacter | Algibacter aqu | poorly charact | Function unkn | 08SSK | NA | 313603.FB217 | 0,2 | 0,4 | 1,41E-05 | 2,82E-05 | 1 - | 7,04E-05 - | 0 - | 0 - | 0 - | 0 - | 0 - | 0 - | 0 - | 0 - |
| 708 | 1 | 0.2um_NOKCPHJN_568002 | Bacteria | Proteobacteri | Alphaproteob | Pelagibacteral | Pelagibacterac | Candidatus Pe | Candidatus Pe | cellular proces | Cell wall/mem | 7257 | NA | 439493.PB721 | 2,4 | 0,489897949 | 0,000105344 | 1,86E-05 | 3 - | 0,000111434 - | 3 - | 0,000138368 - | 2 - | 8,77E-05 - | 2 - | 8,90E-05 - | 2 - | 0,000100242 - |
| 709 | 1 | 0.2um_NOKCPHJN_382024 | Eukaryota | Streptophyta | Klebsormidio | Klebsormidial | Klebsormidiac | Klebsormidiur | Klebsormidiur | unclassified | unclassified | unclassified | unclassified | unclassified | 1,8 | 0,4 | 0,000120862 | 2,96E-05 | 2 - | 0,000112526 - | 2 - | 0,000139724 - | 2 - | 0,000132826 - | 1 - | 6,74E-05 - | 2 - | 0,000151838 - |

| | | | | | | | | | | | | | | | | | | | | | | | | | | | | |
|---|---|---|---|---|---|---|---|---|---|---|---|---|---|---|---|---|---|---|---|---|---|---|---|---|---|---|---|---|
| 710 | 1 | 3um_FIJJAFBF_73809 | Bacteria | Bacteroidetes | Flavobacteriia | Flavobacterial | Flavobacteriac | Polaribacter | Polaribacter s | information st | Replication, re | 09AJH | The RuvA-Ruv | 391603.FBALC | 1,6 | 1,2 | 6,57E-05 | 4,75E-05 | 0 - | 0 - | 1 - | 4,19E-05 - | 3 - | 0,000119543 - | 3 - | 0,000121309 - | 1 - | 4,56E-05 - |
| 711 | 1 | 0.2um_NOKCPHJN_582145 | Bacteria | Proteobacteri | Alphaproteob | Rhodobactera | Rhodobactera | Planktomarina | Planktomarina | metabolism | Energy produc | 01RN9 | Subunits I and | 314270.RB208 | 2 | 0 | 9,72E-05 | 9,23E-06 | 2 - | 8,14E-05 - | 2 - | 0,000101077 - | 2 - | 9,61E-05 - | 2 - | 9,75E-05 - | 2 - | 0,00010984 - |
| 712 | 1 | 3um_FIJJAFBF_728785 | Eukaryota | unclassified | Oomycetes | Saprolegniales | Saprolegniace | Aphanomyces | Aphanomyces | information st | Chromatin str | KOG1744 | Core compone | 65071.PYU1_ | 0,8 | 0,4 | 9,60E-05 | 4,93E-05 | 1 - | 0,000101572 - | 0 - | 0 - | 1 - | 0,000119896 - | 1 - | 0,000121667 - | 1 - | 0,000137057 - |
| 713 | 1 | 3um_FIJJAFBF_131459 | Bacteria | Proteobacteri | Alphaproteob | Rhodobactera | Rhodobactera | unclassified | Rhodobactera | unclassified | unclassified | unclassified | unclassified | unclassified | 0,4 | 0,489897949 | 1,64E-05 | 2,05E-05 | 1 - | 3,50E-05 - | 0 - | 0 - | 0 - | 0 - | 0 - | 0 - | 1 - | 4,72E-05 - |
| 714 | 1 | 3um_FIJJAFBF_350299 | Bacteria | Proteobacteri | Gammaprotec | Oceanospirilla | Oceanospirilla | Balneatrix | Balneatrix alp | unclassified | unclassified | unclassified | unclassified | unclassified | 5,4 | 0,489897949 | 0,000442884 | 3,02E-05 | 6 - | 0,000414856 - | 5 - | 0,000429274 - | 6 - | 0,000489695 - | 5 - | 0,000414106 - | 5 - | 0,000466489 - |
| 715 | 1 | 3um_FIJJAFBF_481950 | Bacteria | Proteobacteri | Alphaproteob | Rhodobactera | Rhodobactera | unclassified | Rhodobactera | metabolism | Energy produc | 16RH6 | Produces ATP | 388739.RSK20 | 0,8 | 0,4 | 3,73E-05 | 1,92E-05 | 1 - | 3,94E-05 - | 0 - | 0 - | 1 - | 4,66E-05 - | 1 - | 4,72E-05 - | 1 - | 5,32E-05 - |
| 716 | 1 | 0.2um_NOKCPHJN_340340 | Eukaryota | Bacillariophyt | Bacillariophyc | Naviculales | Naviculaceae | Fistulifera | Fistulifera sola | cellular proces | Cell cycle cont | 04M7D | Regulator of c | 9103.ENSMGA | 0,8 | 0,4 | 5,46E-06 | 2,80E-06 | 1 - | 5,77E-06 - | 0 - | 0 - | 1 - | 6,81E-06 - | 1 - | 6,91E-06 - | 1 - | 7,79E-06 - |
| 717 | 1 | 0.2um_NOKCPHJN_826353 | unclassified | unclassified | unclassified | unclassified | unclassified | unclassified | unclassified | unclassified | unclassified | unclassified | unclassified | unclassified | 0,8 | 0,4 | 3,48E-05 | 1,79E-05 | 1 - | 3,68E-05 - | 0 - | 0 - | 1 - | 4,34E-05 - | 1 - | 4,41E-05 - | 1 - | 4,96E-05 - |
| 718 | 1 | 0.2um_NOKCPHJN_416852 | Eukaryota | unclassified | unclassified | Choanoflagelli | Salpingoecida | Salpingoeca | Salpingoeca r | unclassified | unclassified | unclassified | unclassified | unclassified | 1 | 0 | 1,75E-05 | 1,66E-06 | 1 - | 1,46E-05 - | 1 - | 1,82E-05 - | 1 - | 1,73E-05 - | 1 - | 1,75E-05 - | 1 - | 1,97E-05 - |
| 719 | 1 | 3um_FIJJAFBF_701041 | Bacteria | Bacteroidetes | unclassified | unclassified | unclassified | unclassified | Bacteroidetes | information st | Replication, re | 0PB0H | DNA-binding p | 313598.MED1 | 1,8 | 0,4 | 0,000278999 | 7,72E-05 | 1 - | 0,00012753 - | 2 - | 0,000316709 - | 2 - | 0,000301072 - | 2 - | 0,000305518 - | 2 - | 0,000344166 - |
| 720 | 1 | 0.2um_NOKCPHJN_264011 | Bacteria | Proteobacteri | Gammaprotec | unclassified | unclassified | unclassified | Gammaprotec | information st | Translation, ri | 177XJ | One of the pri | 207954.MED9 | 1 | 0 | 8,78E-05 | 8,34E-06 | 1 - | 7,36E-05 - | 1 - | 9,14E-05 - | 1 - | 8,68E-05 - | 1 - | 8,81E-05 - | 1 - | 9,93E-05 - |
| 721 | 1 | 3um_FIJJAFBF_346237 | Bacteria | Bacteroidetes | Flavobacteriia | unclassified | unclassified | unclassified | Flavobacteria | unclassified | unclassified | unclassified | unclassified | unclassified | 2 | 0,894427191 | 0,000199338 | 9,92E-05 | 1 - | 8,14E-05 - | 1 - | 0,000101077 - | 2 - | 0,000192173 - | 3 - | 0,000292518 - | 3 - | 0,00032952 - |
| 722 | 1 | 0.2um_NOKCPHJN_470026 | Bacteria | Proteobacteri | Gammaprotec | Thiotrichales | Thiotrichacea | Thiothrix | Thiothrix nive | information st | Translation, ri | 0FK5W | Catalyzes the | 32049.SYNPC | 2,4 | 0,489897949 | 4,72E-05 | 1,02E-05 | 2 - | 3,30E-05 - | 2 - | 4,10E-05 - | 3 - | 5,84E-05 - | 3 - | 5,93E-05 - | 2 - | 4,45E-05 - |
| 723 | 1 | 3um_FIJJAFBF_732537 | Bacteria | Proteobacteri | Alphaproteob | Rhodobactera | Rhodobactera | Sediminimona | Sediminimona | metabolism | Lipid transpor | 08IQE | Amp-depende | 314232.SKA53 | 0,6 | 0,8 | 1,30E-05 | 1,85E-05 | 1 - | 1,76E-05 - | 0 - | 0 - | 0 - | 0 - | 0 - | 0 - | 2 - | 4,74E-05 - |
| 724 | 1 | 3um_FIJJAFBF_280854 | Bacteria | Proteobacteri | Alphaproteob | Rhodobactera | Rhodobactera | Sulfitobacter | various | information st | Transcription | 05VEQ | Cold shock pr | 391624.OIHEL | 3,2 | 0,4 | 0,000638289 | 3,50E-05 | 4 - | 0,000675157 - | 3 - | 0,00062876 - | 3 - | 0,000597716 - | 3 - | 0,000606544 - | 3 - | 0,00068327 - |
| 725 | 1 | 3um_FIJJAFBF_301944 | Eukaryota | unclassified | Oomycetes | Peronosporale | Peronosporac | Phytophthora | various | metabolism | Inorganic ion t | 0IG0G | Calcium-bindi | 2850.Phatr31 | 1 | 0 | 3,95E-05 | 3,75E-06 | 1 - | 3,31E-05 - | 1 - | 4,11E-05 - | 1 - | 3,90E-05 - | 1 - | 3,96E-05 - | 1 - | 4,46E-05 - |
| 726 | 1 | 0.2um_NOKCPHJN_313838 | Eukaryota | Eustigmatoph | unclassified | Eustigmatales | Monodopsida | Nannochlorop | Nannochlorop | cellular proces | Signal transdu | COG5040 | Tyrosine 3-mo | 2850.Phatr56 | 0,2 | 0,4 | 2,15E-05 | 4,29E-05 | 1 - | 0,000107268 - | 0 - | 0 - | 0 - | 0 - | 0 - | 0 - | 0 - | 0 - |
| 727 | 1 | 0.2um_NOKCPHJN_179649 | Bacteria | Bacteroidetes | Flavobacteriia | Flavobacterial | Flavobacteriac | Formosa | Formosa sp. H | unclassified | unclassified | unclassified | unclassified | unclassified | 0,4 | 0,489897949 | 2,74E-05 | 3,36E-05 | 0 - | 0 - | 1 - | 7,02E-05 - | 1 - | 6,67E-05 - | 0 - | 0 - | 0 - | 0 - |
| 728 | 1 | 3um_FIJJAFBF_244228 | Bacteria | Proteobacteri | Alphaproteob | Rhodospirillal | Rhodospirillac | unclassified | Rhodobactera | unclassified | unclassified | unclassified | unclassified | unclassified | 2,2 | 0,4 | 0,000117637 | 2,29E-05 | 2 - | 8,97E-05 - | 2 - | 0,000111343 - | 3 - | 0,000158768 - | 2 - | 0,000107409 - | 2 - | 0,000120996 - |
| 729 | 2 | 0.2um_NOKCPHJN_476676;0.2um_NOKCPHJN_5 | Bacteria | Bacteroidetes | unclassified | unclassified | unclassified | unclassified | Bacteroidetes | poorly charact | Function unkn | 05RNW | Domain-Conta | 926562.Oweh | 2,2 | 0,4 | 1,55E-05 | 1,42E-06 | 3 - | 1,80E-05 - | 2 - | 1,49E-05 - | 2 - | 1,41E-05 - | 2 - | 1,44E-05 - | 2 - | 1,62E-05 - |
| 730 | 1 | 3um_FIJJAFBF_56165 | Bacteria | Proteobacteri | Alphaproteob | Rhodobactera | Rhodobactera | Salipiger | Salipiger muc | cellular proces | Intracellular tr | 175GA | One of the pro | 388399.SSE37 | 0,8 | 0,4 | 6,09E-05 | 3,10E-05 | 1 - | 6,60E-05 - | 1 - | 8,19E-05 - | 1 - | 7,79E-05 - | 1 - | 7,90E-05 - | 0 - | 0 - |
| 731 | 1 | 0.2um_NOKCPHJN_684604 | Bacteria | unclassified | unclassified | unclassified | unclassified | unclassified | uncultured ba | poorly charact | Function unkn | 0ZSK2 | outer membra | 166314.SH810 | 0,2 | 0,4 | 5,74E-06 | 1,15E-05 | 0 - | 0 - | 0 - | 0 - | 1 - | 2,87E-05 - | 0 - | 0 - | 0 - | 0 - |
| 732 | 1 | 3um_FIJJAFBF_56204 | Eukaryota | unclassified | unclassified | Prymnesiales | Chrysochrom | Chrysochrom | various | metabolism | Energy produc | COG0055 | Produces ATP | 329726.AM1_ | 0,6 | 0,8 | 1,57E-05 | 2,00E-05 | 2 - | 4,83E-05 - | 1 - | 3,00E-05 - | 0 - | 0 - | 0 - | 0 - | 0 - | 0 - |
| 733 | 1 | 3um_FIJJAFBF_257128 | Bacteria | Firmicutes | Clostridia | Clostridiales | Clostridiaceae | Clostridium | various | unclassified | unclassified | unclassified | unclassified | unclassified | 1 | 0 | 0,000112317 | 1,07E-05 | 1 - | 9,41E-05 - | 1 - | 0,000116819 - | 1 - | 0,000111051 - | 1 - | 0,000112691 - | 1 - | 0,000126946 - |
| 734 | 1 | 3um_FIJJAFBF_688468 | Bacteria | Proteobacteri | Betaproteoba | unclassified | unclassified | unclassified | Betaproteoba | unclassified | unclassified | unclassified | unclassified | unclassified | 8,8 | 1,326649916 | 0,000330881 | 2,71E-05 | 11 - | 0,000350707 - | 7 - | 0,00027712 - | 9 - | 0,000338706 - | 9 - | 0,000343708 - | 8 - | 0,000344166 - |
| 735 | 1 | 3um_FIJJAFBF_24519 | Bacteria | Proteobacteri | Alphaproteob | Rhodobactera | Rhodobactera | Actibacterium | Actibacterium | metabolism | Carbohydrate | 0XPV5 | extracellular s | 367336.OM22 | 1 | 0 | 3,33E-05 | 3,16E-06 | 1 - | 2,79E-05 - | 1 - | 3,46E-05 - | 1 - | 3,29E-05 - | 1 - | 3,34E-05 - | 1 - | 3,76E-05 - |
| 736 | 1 | 3um_FIJJAFBF_143962 | Bacteria | unclassified | unclassified | unclassified | unclassified | unclassified | uncultured ba | unclassified | unclassified | unclassified | unclassified | unclassified | 0,6 | 0,489897949 | 3,79E-05 | 3,11E-05 | 0 - | 0 - | 0 - | 0 - | 1 - | 5,99E-05 - | 1 - | 6,08E-05 - | 1 - | 6,85E-05 - |
| 737 | 1 | 3um_FIJJAFBF_548868 | Bacteria | Planctomycet | Planctomyceti | Planctomycet | Planctomycet | unclassified | Planctomycet | cellular proces | Cell wall/mem | COG0750 | Membrane-as | 530564.Psta_ | 0,4 | 0,8 | 8,17E-06 | 1,63E-05 | 0 - | 0 - | 0 - | 0 - | 0 - | 0 - | 2 - | 4,09E-05 - | 0 - | 0 - |
| 738 | 1 | 0.2um_NOKCPHJN_1090664 | Bacteria | Proteobacteri | Betaproteoba | Burkholderial | Comamonada | Limnohabitan | Limnohabitan | poorly charact | Function unkn | 0Y9GN | Up-regulator c | 7955.ENSDAR | 0,2 | 0,4 | 2,20E-05 | 4,40E-05 | 0 - | 0 - | 0 - | 0 - | 0 - | 0 - | 1 - | 0,000109987 - | 0 - | 0 - |
| 739 | 1 | 3um_FIJJAFBF_39170 | Eukaryota | Chordata | Actinopteri | Osteoglossifor | Mormyridae | Paramormyro | Paramormyro | unclassified | unclassified | unclassified | unclassified | unclassified | 0,8 | 0,4 | 3,04E-05 | 1,56E-05 | 1 - | 3,22E-05 - | 0 - | 0 - | 1 - | 3,80E-05 - | 1 - | 3,85E-05 - | 1 - | 4,34E-05 - |
| 740 | 1 | 3um_FIJJAFBF_420754 | Bacteria | Bacteroidetes | Cytophagia | Cytophagales | Flammeovirga | unclassified | Flammeovirga | metabolism | Carbohydrate | COG0297 | starch synthas | 643867.Ftrac_ | 0,6 | 0,8 | 2,72E-05 | 3,50E-05 | 2 - | 8,50E-05 - | 0 - | 0 - | 0 - | 0 - | 1 - | 5,09E-05 - | 0 - | 0 - |
| 741 | 1 | 0.2um_NOKCPHJN_12462 | Bacteria | Bacteroidetes | Flavobacteriia | Flavobacterial | Flavobacteriac | Formosa | Formosa sp. H | metabolism | Inorganic ion t | COG3119 | Sulfatase | 313628.LNTAR | 1,2 | 0,4 | 3,43E-05 | 1,15E-05 | 1 - | 2,40E-05 - | 1 - | 2,98E-05 - | 2 - | 5,66E-05 - | 1 - | 2,87E-05 - | 1 - | 3,23E-05 - |
| 742 | 1 | 0.2um_NOKCPHJN_193716 | Bacteria | Bacteroidetes | Flavobacteriia | Flavobacterial | Flavobacteriac | Polaribacter | Polaribacter s | cellular proces | Posttranslatio | 05C6P | ATP-depender | 313594.PI23P | 3 | 0,632455532 | 4,99E-05 | 1,01E-05 | 3 - | 4,21E-05 - | 3 - | 5,23E-05 - | 3 - | 4,97E-05 - | 4 - | 6,73E-05 - | 2 - | 3,79E-05 - |
| 743 | 1 | 3um_FIJJAFBF_800063 | Bacteria | Proteobacteri | Alphaproteob | Rhodobactera | Rhodobactera | Planktotalea | Planktotalea f | poorly charact | Function unkn | 05ERF | Extracellular s | 488538.SAR11 | 2,6 | 1,019803903 | 7,11E-05 | 2,82E-05 | 3 - | 6,98E-05 - | 4 - | 0,000115634 - | 2 - | 5,50E-05 - | 3 - | 8,37E-05 - | 1 - | 3,14E-05 - |
| 744 | 1 | 3um_FIJJAFBF_191134 | Bacteria | Proteobacteri | Alphaproteob | various | various | various | various | information st | Translation, ri | COG0051 | tRNA binding | 367336.OM22 | 3 | 0,894427191 | 0,000397974 | 0,000107254 | 4 - | 0,000450105 - | 2 - | 0,000279449 - | 2 - | 0,000265651 - | 4 - | 0,00053915 - | 3 - | 0,000455513 - |
| 745 | 1 | 3um_FIJJAFBF_768961 | Bacteria | Proteobacteri | Gammaprotec | Oceanospirilla | Saccharospirill | Reinekea | Reinekea blan | unclassified | unclassified | unclassified | unclassified | unclassified | 0,8 | 0,748331477 | 1,90E-05 | 1,65E-05 | 2 - | 4,19E-05 - | 0 - | 0 - | 0 - | 0 - | 1 - | 2,51E-05 - | 1 - | 2,83E-05 - |
| 746 | 2 | 3um_FIJJAFBF_305741;3um_FIJJAFBF_513413 | Eukaryota | Bacillariophyt | Coscinodiscop | Thalassiosirale | Thalassiosirac | Thalassiosira | Thalassiosira | unclassified | unclassified | unclassified | unclassified | unclassified | 0,6 | 0,489897949 | 5,97E-05 | 4,91E-05 | 1 - | 8,83E-05 - | 0 - | 0 - | 1 - | 0,000104217 - | 1 - | 0,000105756 - | 0 - | 0 - |
| 747 | 1 | 3um_FIJJAFBF_222641 | Bacteria | Proteobacteri | Alphaproteob | Rhodobactera | Rhodobactera | Loktanella | Loktanella sp. | unclassified | unclassified | unclassified | unclassified | unclassified | 0,2 | 0,4 | 3,84E-06 | 7,69E-06 | 0 - | 0 - | 0 - | 0 - | 1 - | 1,92E-05 - | 0 - | 0 - | 0 - | 0 - |
| 748 | 1 | 3um_FIJJAFBF_768960 | Bacteria | Proteobacteri | Gammaprotec | Oceanospirilla | Saccharospirill | Reinekea | Reinekea fors | metabolism | Coenzyme tra | 08Z2X | Catalyzes the | 314283.MED2 | 1 | 0 | 0,000107895 | 1,02E-05 | 1 - | 9,04E-05 - | 1 - | 0,00011222 - | 1 - | 0,000106679 - | 1 - | 0,000108255 - | 1 - | 0,000121948 - |
| 749 | 1 | 3um_FIJJAFBF_99521 | Viruses | unclassified | unclassified | Caudovirales | Siphoviridae | Cbastvirus | Cellulophaga | unclassified | unclassified | unclassified | unclassified | unclassified | 5,6 | 0,8 | 0,000235707 | 4,93E-05 | 5 - | 0,000174965 - | 7 - | 0,000304156 - | 5 - | 0,000206528 - | 5 - | 0,000209578 - | 6 - | 0,000283307 - |
| 750 | 1 | 3um_FIJJAFBF_671982 | Bacteria | Proteobacteri | Alphaproteob | Rhodospirillal | Rhodospirillac | Thalassospira | Thalassospira | metabolism | Energy produc | 01QSC | Dihydrolipoyl | 331869.BAL19 | 1 | 0 | 3,80E-05 | 3,60E-06 | 1 - | 3,18E-05 - | 1 - | 3,95E-05 - | 1 - | 3,75E-05 - | 1 - | 3,81E-05 - | 1 - | 4,29E-05 - |
| 751 | 1 | 0.2um_NOKCPHJN_142338 | Bacteria | Proteobacteri | Gammaprotec | Cellvibrionales | Porticoccacea | unclassified | Gammaprotec | metabolism | Lipid transpor | COG1960 | acyl-CoA dehy | 314287.GB22 | 1 | 0 | 2,27E-05 | 2,16E-06 | 1 - | 1,90E-05 - | 1 - | 2,36E-05 - | 1 - | 2,25E-05 - | 1 - | 2,28E-05 - | 1 - | 2,57E-05 - |
| 752 | 1 | 3um_FIJJAFBF_510425 | Bacteria | Bacteroidetes | Flavobacteriia | Flavobacterial | Flavobacteriac | Flavobacteriu | Flavobacteriu | unclassified | unclassified | unclassified | unclassified | unclassified | 0,2 | 0,4 | 7,72E-06 | 1,54E-05 | 0 - | 0 - | 0 - | 0 - | 1 - | 3,86E-05 - | 0 - | 0 - | 0 - | 0 - |
| 753 | 1 | 3um_FIJJAFBF_524479 | Bacteria | Proteobacteri | Alphaproteob | Rhodobactera | Rhodobactera | Sulfitobacter | Sulfitobacter | poorly charact | Function unkn | COG3181 | extra-cytoplas | 367336.OM22 | 0,4 | 0,489897949 | 1,57E-05 | 1,95E-05 | 1 - | 3,51E-05 - | 1 - | 4,36E-05 - | 0 - | 0 - | 0 - | 0 - | 0 - | 0 - |
| 754 | 1 | 3um_FIJJAFBF_56211 | Eukaryota | unclassified | unclassified | Prymnesiales | Chrysochrom | Chrysochrom | various | unclassified | unclassified | unclassified | unclassified | unclassified | 1,2 | 0,748331477 | 0,000108629 | 6,70E-05 | 2 - | 0,000158313 - | 2 - | 0,000196578 - | 1 - | 9,34E-05 - | 1 - | 9,48E-05 - | 0 - | 0 - |
| 755 | 1 | 0.2um_NOKCPHJN_460479 | Bacteria | Proteobacteri | Alphaproteob | Pelagibacteral | Pelagibactera | Candidatus Pe | Candidatus Pe | unclassified | unclassified | unclassified | unclassified | unclassified | 0,4 | 0,489897949 | 3,22E-05 | 3,94E-05 | 0 - | 0 - | 1 - | 7,70E-05 - | 0 - | 0 - | 0 - | 0 - | 1 - | 8,37E-05 - |
| 756 | 1 | 0.2um_NOKCPHJN_585649 | Bacteria | Proteobacteri | Gammaprotec | unclassified | unclassified | unclassified | SAR86 cluster | unclassified | unclassified | unclassified | unclassified | unclassified | 0,2 | 0,4 | 5,31E-06 | 1,06E-05 | 1 - | 2,66E-05 - | 0 - | 0 - | 0 - | 0 - | 0 - | 0 - | 0 - | 0 - |
| 757 | 1 | 0.2um_NOKCPHJN_27392 | Bacteria | unclassified | unclassified | unclassified | unclassified | unclassified | uncultured ba | metabolism | Inorganic ion t | 05C27 | TonB-depende | 313596.RB250 | 42,6 | 2,8 | 0,000555921 | 3,54E-05 | 48 - | 0,000527204 - | 41 - | 0,000559165 - | 42 - | 0,000544522 - | 40 - | 0,000526252 - | 42 - | 0,000622462 - |
| 758 | 1 | 0.2um_NOKCPHJN_708457 | Bacteria | Firmicutes | Bacilli | Lactobacillales | Streptococcac | Streptococcus | Streptococcus | cellular proces | Posttranslatio | 16QU3 | Hemolysin-typ | 744980.TRICH | 1 | 0 | 1,45E-05 | 1,37E-06 | 1 - | 1,21E-05 - | 1 - | 1,50E-05 - | 1 - | 1,43E-05 - | 1 - | 1,45E-05 - | 1 - | 1,63E-05 - |
| 759 | 1 | 0.2um_NOKCPHJN_823005 | Eukaryota | Chromerida | unclassified | unclassified | unclassified | Vitrella | Vitrella brassi | unclassified | unclassified | unclassified | unclassified | unclassified | 0,8 | 0,748331477 | 6,82E-05 | 6,62E-05 | 1 - | 7,45E-05 - | 0 - | 0 - | 1 - | 8,80E-05 - | 2 - | 0,00017855 - | 0 - | 0 - |
| 760 | 1 | 3um_FIJJAFBF_238480 | Bacteria | Proteobacteri | Gammaprotec | Oceanospirilla | unclassified | Terasakiispira | Terasakiispira | information st | Translation, ri | COG0050 | This protein p | 207954.MED9 | 24,6 | 1,019803903 | 0,000849509 | 9,48E-05 | 25 - | 0,000722776 - | 23 - | 0,000825676 - | 24 - | 0,000819036 - | 25 - | 0,000865764 - | 26 - | 0,001014291 - |
| 761 | 1 | 3um_FIJJAFBF_116593 | Bacteria | Cyanobacteria | unclassified | Synechococca | Synechococca | Synechococcu | Synechococcu | unclassified | unclassified | unclassified | unclassified | unclassified | 1,2 | 0,4 | 6,34E-05 | 2,12E-05 | 1 - | 4,43E-05 - | 1 - | 5,50E-05 - | 2 - | 0,000104619 - | 1 - | 5,31E-05 - | 1 - | 5,98E-05 - |
| 762 | 1 | 3um_FIJJAFBF_44700 | Bacteria | Bacteroidetes | Flavobacteriia | Flavobacterial | Flavobacteriac | Polaribacter | Polaribacter s | unclassified | unclassified | unclassified | unclassified | unclassified | 5 | 1,095445115 | 7,92E-05 | 1,46E-05 | 6 - | 8,08E-05 - | 6 - | 0,000100365 - | 5 - | 7,95E-05 - | 5 - | 8,07E-05 - | 3 - | 5,45E-05 - |
| 763 | 1 | 3um_FIJJAFBF_301269 | Bacteria | Bacteroidetes | unclassified | unclassified | unclassified | unclassified | unidentified e | cellular proces | Posttranslatio | COG0652 | peptidyl-proly | 50743.SCB49_ | 0,6 | 0,489897949 | 2,45E-05 | 2,01E-05 | 1 - | 3,62E-05 - | 0 - | 0 - | 1 - | 4,27E-05 - | 1 - | 4,34E-05 - | 0 - | 0 - |
| 764 | 1 | 3um_FIJJAFBF_494162 | Bacteria | Bacteroidetes | Flavobacteriia | Flavobacterial | Flavobacteriac | Polaribacter | Polaribacter s | unclassified | unclassified | unclassified | unclassified | unclassified | 5,8 | 0,748331477 | 0,000435594 | 8,24E-05 | 6 - | 0,000376317 - | 5 - | 0,000389396 - | 6 - | 0,000444204 - | 5 - | 0,000375637 - | 7 - | 0,000592416 - |
| 765 | 1 | 3um_FIJJAFBF_296063 | Bacteria | Proteobacteri | Gammaprotec | unclassified | unclassified | unclassified | Gammaprotec | metabolism | Energy produc | 16RH6 | Produces ATP | 498211.CJA_3 | 0,4 | 0,489897949 | 1,94E-05 | 2,38E-05 | 0 - | 0 - | 1 - | 4,98E-05 - | 1 - | 4,74E-05 - | 0 - | 0 - | 0 - | 0 - |
| 766 | 2 | 3um_FIJJAFBF_582844;3um_FIJJAFBF_652362 | Bacteria | Bacteroidetes | Flavobacteriia | Flavobacterial | Flavobacteriac | various | various | unclassified | unclassified | unclassified | unclassified | unclassified | 1,2 | 0,748331477 | 5,42E-05 | 3,17E-05 | 2 - | 7,76E-05 - | 1 - | 4,81E-05 - | 0 - | 0 - | 2 - | 9,29E-05 - | 1 - | 5,23E-05 - |
| 767 | 1 | 0.2um_NOKCPHJN_202380 | Eukaryota | unclassified | Oomycetes | Peronosporale | Peronosporac | Phytophthora | Phytophthora | unclassified | unclassified | unclassified | unclassified | unclassified | 1 | 0 | 1,83E-05 | 1,74E-06 | 1 - | 1,53E-05 - | 1 - | 1,90E-05 - | 1 - | 1,81E-05 - | 1 - | 1,84E-05 - | 1 - | 2,07E-05 - |
| 768 | 1 | 0.2um_NOKCPHJN_121982 | Bacteria | Bacteroidetes | Flavobacteriia | Flavobacterial | Flavobacteriac | Polaribacter | Polaribacter s | information st | Transcription | 05D27 | DNA-depende | 313594.PI23P | 5,4 | 1,496662955 | 5,09E-05 | 9,95E-06 | 8 - | 6,45E-05 - | 6 - | 6,01E-05 - | 4 - | 3,81E-05 - | 5 - | 4,83E-05 - | 4 - | 4,35E-05 - |
| 769 | 1 | 3um_FIJJAFBF_79208 | Bacteria | Proteobacteri | Alphaproteob | Rhodobactera | Rhodobactera | Shimia | Shimia aestua | information st | Translation, ri | 16Q16 | Binds the low | 367336.OM22 | 2,2 | 0,979795897 | 0,000120916 | 4,08E-05 | 4 - | 0,000192095 - | 2 - | 0,000119263 - | 2 - | 0,000113374 - | 2 - | 0,000115049 - | 1 - | 6,48E-05 - |
| 770 | 1 | 3um_FIJJAFBF_713337 | Bacteria | Proteobacteri | Alphaproteob | Rhodobactera | Rhodobactera | Leisingera | Leisingera aqu | metabolism | Lipid transpor | 01R4W | acetyl-coa ace | 644076.SCH4 | 1 | 0,894427191 | 3,33E-05 | 2,98E-05 | 2 - | 5,87E-05 - | 2 - | 7,29E-05 - | 1 - | 3,47E-05 - | 0 - | 0 - | 0 - | 0 - |
| 771 | 1 | 3um_FIJJAFBF_150219 | Bacteria | Bacteroidetes | Flavobacteriia | Flavobacterial | Flavobacteriac | Formosa | Formosa sp. H | unclassified | unclassified | unclassified | unclassified | unclassified | 1,6 | 0,489897949 | 4,08E-05 | 1,13E-05 | 2 - | 4,34E-05 - | 2 - | 5,39E-05 - | 2 - | 5,12E-05 - | 1 - | 2,60E-05 - | 1 - | 2,93E-05 - |
| 772 | 1 | 3um_FIJJAFBF_344296 | Bacteria | Proteobacteri | Gammaprotec | unclassified | unclassified | unclassified | Gammaprotec | metabolism | Inorganic ion t | 0ZZVV | tonB-depende | 314275.MADE | 22,2 | 1,6 | 0,000386811 | 2,93E-05 | 24 - | 0,000351806 - | 22 - | 0,000400436 - | 24 - | 0,000415271 - | 20 - | 0,000351171 - | 21 - | 0,000415372 - |
| 773 | 1 | 3um_FIJJAFBF_453322 | Bacteria | Proteobacteri | Alphaproteob | Rhodobactera | Rhodobactera | Ruegeria | Ruegeria med | metabolism | Lipid transpor | COG1788 | Key enzyme fo | 391619.RGBS | 0,6 | 0,489897949 | 3,26E-05 | 2,68E-05 | 1 - | 4,82E-05 - | 0 - | 0 - | 1 - | 5,69E-05 - | 1 - | 5,78E-05 - | 0 - | 0 - |
| 774 | 1 | 3um_FIJJAFBF_391433 | Eukaryota | Apicomplexa | Conoidasida | Eucoccidiorida | Sarcocystidae | Besnoitia | Besnoitia besr | unclassified | unclassified | unclassified | unclassified | unclassified | 1 | 0 | 5,09E-05 | 4,84E-06 | 1 - | 4,27E-05 - | 1 - | 5,30E-05 - | 1 - | 5,04E-05 - | 1 - | 5,11E-05 - | 1 - | 5,76E-05 - |
| 775 | 1 | 3um_FIJJAFBF_730452 | Bacteria | Bacteroidetes | Flavobacteriia | Flavobacterial | unclassified | unclassified | Flavobacterial | poorly charact | Function unkn | 09AR5 | Inherit from b | 388413.ALPR1 | 0,8 | 0,4 | 2,81E-05 | 1,41E-05 | 0 - | 0 - | 1 - | 3,51E-05 - | 1 - | 3,34E-05 - | 1 - | 3,39E-05 - | 1 - | 3,81E-05 - |
| 776 | 1 | 0.2um_NOKCPHJN_427809 | Bacteria | Proteobacteri | Gammaprotec | Oceanospirilla | Oceanospirilla | Marinobacteri | Marinobacteri | metabolism | Inorganic ion t | 0QJ3Y | Part of the AB | 207954.MED9 | 1,2 | 0,748331477 | 5,81E-05 | 3,83E-05 | 1 - | 4,11E-05 - | 2 - | 0,000102164 - | 1 - | 4,86E-05 - | 2 - | 9,86E-05 - | 0 - | 0 - |
| 777 | 1 | 3um_FIJJAFBF_41187 | Bacteria | Proteobacteri | Alphaproteob | Rhodobactera | unclassified | unclassified | Rhodobactera | unclassified | unclassified | unclassified | unclassified | unclassified | 4,4 | 1,356465997 | 0,000335852 | 0,000101682 | 4 - | 0,000257925 - | 3 - | 0,0002402 - | 6 - | 0,000456682 - | 6 - | 0,000463427 - | 3 - | 0,000261024 - |
| 778 | 1 | 3um_FIJJAFBF_257131 | Eukaryota | unclassified | Raphidophyce | Chattonellales | Chattonellace | Heterosigma | Heterosigma a | unclassified | unclassified | unclassified | unclassified | unclassified | 0,8 | 0,4 | 7,47E-05 | 3,79E-05 | 1 - | 8,08E-05 - | 1 - | 0,000100365 - | 1 - | 9,54E-05 - | 1 - | 9,68E-05 - | 0 - | 0 - |
| 779 | 1 | 0.2um_NOKCPHJN_448907 | Eukaryota | unclassified | Pelagophycea | Pelagomonad | unclassified | Aureococcus | Aureococcus a | unclassified | unclassified | unclassified | unclassified | unclassified | 0,2 | 0,4 | 5,45E-06 | 1,09E-05 | 1 - | 2,73E-05 - | 0 - | 0 - | 0 - | 0 - | 0 - | 0 - | 0 - | 0 - |
| 780 | 1 | 3um_FIJJAFBF_140338 | unclassified | unclassified | unclassified | unclassified | unclassified | unclassified | unclassified | unclassified | unclassified | unclassified | unclassified | unclassified | 0,8 | 0,4 | 3,03E-05 | 1,54E-05 | 1 - | 3,28E-05 - | 1 - | 4,07E-05 - | 1 - | 3,87E-05 - | 1 - | 3,93E-05 - | 0 - | 0 - |
| 781 | 1 | 3um_FIJJAFBF_17267 | Bacteria | Proteobacteri | Alphaproteob | Rhodobactera | Rhodobactera | Leisingera | Leisingera sp. | metabolism | Energy produc | 07QM5 | aconitate hyd | 391619.RGBS | 3,8 | 1,326649916 | 5,64E-05 | 1,47E-05 | 6 - | 7,69E-05 - | 4 - | 6,37E-05 - | 4 - | 6,06E-05 - | 3 - | 4,61E-05 - | 2 - | 3,46E-05 - |
| 782 | 1 | 0.2um_NOKCPHJN_181896 | Bacteria | Proteobacteri | Gammaprotec | Cellvibrionales | Porticoccacea | unclassified | gamma prote | unclassified | unclassified | unclassified | unclassified | unclassified | 0,2 | 0,4 | 2,76E-06 | 5,52E-06 | 1 - | 1,38E-05 - | 0 - | 0 - | 0 - | 0 - | 0 - | 0 - | 0 - | 0 - |
| 783 | 1 | 0.2um_NOKCPHJN_11367 | Bacteria | Proteobacteri | Alphaproteob | unclassified | unclassified | unclassified | SAR116 cluste | unclassified | unclassified | unclassified | unclassified | unclassified | 1,8 | 0,4 | 9,85E-05 | 2,72E-05 | 1 - | 4,50E-05 - | 2 - | 0,00011178 - | 2 - | 0,000106261 - | 2 - | 0,00010783 - | 2 - | 0,00012147 - |
| 784 | 1 | 0.2um_NOKCPHJN_1075202 | unclassified | unclassified | unclassified | unclassified | unclassified | unclassified | unclassified | unclassified | unclassified | unclassified | unclassified | unclassified | 0,2 | 0,4 | 2,96E-05 | 5,91E-05 | 0 - | 0 - | 0 - | 0 - | 0 - | 0 - | 1 - | 0,000147832 - | 0 - | 0 - |
| 785 | 1 | 3um_FIJJAFBF_75453 | Bacteria | Bacteroidetes | Flavobacteriia | Flavobacterial | Flavobacteriac | Polaribacter | Polaribacter r | unclassified | unclassified | unclassified | unclassified | unclassified | 1,2 | 0,748331477 | 0,000123056 | 7,59E-05 | 2 - | 0,000179339 - | 2 - | 0,000222686 - | 1 - | 0,000105845 - | 1 - | 0,000107409 - | 0 - | 0 - |
| 786 | 1 | 0.2um_NOKCPHJN_251411 | Bacteria | Proteobacteri | Gammaprotec | unclassified | unclassified | unclassified | Gammaprotec | poorly charact | Function unkn | 16R8I | Lipoprotein | 314287.GB22 | 2,2 | 0,748331477 | 5,34E-05 | 1,63E-05 | 3 - | 6,24E-05 - | 3 - | 7,75E-05 - | 2 - | 4,91E-05 - | 2 - | 4,98E-05 - | 1 - | 2,81E-05 - |
| 787 | 1 | 3um_FIJJAFBF_328494 | Bacteria | Bacteroidetes | Flavobacteriia | Flavobacterial | Flavobacteriac | Arenitalea | Arenitalea lut | unclassified | unclassified | unclassified | unclassified | unclassified | 1 | 0 | 4,49E-05 | 4,27E-06 | 1 - | 3,76E-05 - | 1 - | 4,67E-05 - | 1 - | 4,44E-05 - | 1 - | 4,51E-05 - | 1 - | 5,08E-05 - |
| 788 | 1 | 3um_FIJJAFBF_594160 | Eukaryota | unclassified | Bangiophycea | Bangiales | Bangiaceae | Wildemania | Wildemania s | unclassified | unclassified | unclassified | unclassified | unclassified | 7 | 1,414213562 | 0,000625535 | 9,02E-05 | 9 - | 0,0006841 - | 8 - | 0,000755067 - | 6 - | 0,00053834 - | 7 - | 0,00063734 - | 5 - | 0,000512829 - |
| 789 | 1 | 3um_FIJJAFBF_155848 | Bacteria | Proteobacteri | Gammaprotec | Oceanospirilla | Saccharospirill | Reinekea | Reinekea fors | unclassified | unclassified | unclassified | unclassified | unclassified | 0,4 | 0,489897949 | 2,37E-05 | 2,91E-05 | 0 - | 0 - | 1 - | 6,04E-05 - | 0 - | 0 - | 1 - | 5,83E-05 - | 0 - | 0 - |
| 790 | 1 | 0.2um_NOKCPHJN_401292 | unclassified | unclassified | unclassified | unclassified | unclassified | unclassified | unclassified | unclassified | unclassified | unclassified | unclassified | unclassified | 0,4 | 0,489897949 | 1,09E-05 | 1,34E-05 | 1 - | 2,48E-05 - | 0 - | 0 - | 0 - | 0 - | 1 - | 2,97E-05 - | 0 - | 0 - |
| 791 | 1 | 0.2um_NOKCPHJN_154746 | Bacteria | Bacteroidetes | Flavobacteriia | Flavobacterial | Flavobacteriac | Formosa | Formosa sp. H | metabolism | Carbohydrate | COG0149 | triosephospha | 1046627.BZAR | 1,6 | 0,489897949 | 8,66E-05 | 2,40E-05 | 2 - | 9,22E-05 - | 2 - | 0,000114473 - | 2 - | 0,000108821 - | 1 - | 5,52E-05 - | 1 - | 6,22E-05 - |
| 792 | 1 | 3um_FIJJAFBF_371763 | Bacteria | Bacteroidetes | Flavobacteriia | Flavobacterial | Flavobacteriac | Flaviramulus | Flaviramulus i | metabolism | Inorganic ion t | 05C27 | TonB-depende | 688270.Celal_ | 7,4 | 1,356465997 | 0,00010003 | 1,96E-05 | 8 - | 9,09E-05 - | 5 - | 7,06E-05 - | 9 - | 0,000120727 - | 7 - | 9,53E-05 - | 8 - | 0,000122673 - |
| 793 | 1 | 0.2um_NOKCPHJN_316872 | Bacteria | Proteobacteri | Alphaproteob | unclassified | unclassified | unclassified | various | metabolism | Energy produc | 01RNF | Produces ATP | 862719.AZOLI | 1,2 | 0,4 | 3,41E-05 | 7,56E-06 | 2 - | 4,89E-05 - | 1 - | 3,04E-05 - | 1 - | 2,89E-05 - | 1 - | 2,93E-05 - | 1 - | 3,30E-05 - |
| 794 | 1 | 3um_FIJJAFBF_552490 | Bacteria | Proteobacteri | Gammaprotec | Arenicellales | Arenicellaceae | Arenicella | Arenicella xan | unclassified | unclassified | unclassified | unclassified | unclassified | 7,6 | 1,624807681 | 0,000126073 | 2,40E-05 | 10 - | 0,000140143 - | 8 - | 0,000139213 - | 7 - | 0,000115797 - | 5 - | 8,39E-05 - | 8 - | 0,000151282 - |
| 795 | 1 | 3um_FIJJAFBF_12504 | unclassified | unclassified | unclassified | unclassified | unclassified | unclassified | unclassified | unclassified | unclassified | unclassified | unclassified | unclassified | 1 | 0 | 0,000119154 | 1,13E-05 | 1 - | 9,98E-05 - | 1 - | 0,000123929 - | 1 - | 0,000117811 - | 1 - | 0,000119551 - | 1 - | 0,000134673 - |
| 796 | 1 | 3um_FIJJAFBF_319727 | Bacteria | Bacteroidetes | Flavobacteriia | Flavobacterial | Flavobacteriac | Polaribacter | Polaribacter g | metabolism | Inorganic ion t | 05C27 | TonB-depende | 886377.Murru | 6,6 | 2,244994432 | 8,43E-05 | 1,92E-05 | 11 - | 0,000121165 - | 6 - | 8,21E-05 - | 6 - | 7,80E-05 - | 5 - | 6,60E-05 - | 5 - | 7,43E-05 - |
| 797 | 1 | 0.2um_NOKCPHJN_119448 | Bacteria | Proteobacteri | Gammaprotec | Oceanospirilla | Oceanospirilla | Oceanospirillu | Oceanospirillu | metabolism | Lipid transpor | COG1960 | acyl-CoA dehy | 207954.MED9 | 1,2 | 0,748331477 | 2,88E-05 | 1,96E-05 | 1 - | 1,96E-05 - | 2 - | 4,86E-05 - | 1 - | 2,31E-05 - | 0 - | 0 - | 2 - | 5,28E-05 - |
| 798 | 1 | 0.2um_NOKCPHJN_249402 | Bacteria | Bacteroidetes | unclassified | unclassified | unclassified | unclassified | Bacteroidetes | information st | Translation, ri | 05CAV | thus facilitatin | 755732.Fluta_ | 0,6 | 0,489897949 | 1,07E-05 | 8,94E-06 | 1 - | 1,50E-05 - | 1 - | 1,86E-05 - | 0 - | 0 - | 0 - | 0 - | 1 - | 2,02E-05 - |
| 799 | 1 | 3um_FIJJAFBF_296064 | Bacteria | Proteobacteri | Gammaprotec | unclassified | unclassified | unclassified | Gammaprotec | metabolism | Energy produc | COG0056 | Produces ATP | 349521.HCH_ | 6,8 | 1,166190379 | 0,000179222 | 2,45E-05 | 8 - | 0,000178641 - | 8 - | 0,000221819 - | 7 - | 0,000184509 - | 6 - | 0,000160486 - | 5 - | 0,000150656 - |
| 800 | 1 | 3um_FIJJAFBF_493469 | Bacteria | Proteobacteri | Alphaproteob | Rhodobactera | Rhodobactera | various | various | information st | Transcription | 01QD3 | Facilitates tra | 391624.OIHEL | 4 | 0,894427191 | 0,000129733 | 3,08E-05 | 3 - | 8,14E-05 - | 4 - | 0,00013477 - | 5 - | 0,000160144 - | 5 - | 0,00016251 - | 3 - | 0,00010984 - |
| 801 | 1 | 0.2um_NOKCPHJN_454667 | Bacteria | Bacteroidetes | unclassified | unclassified | unclassified | unclassified | Bacteroidetes | unclassified | unclassified | unclassified | unclassified | unclassified | 1,6 | 0,8 | 0,000141915 | 5,62E-05 | 3 - | 0,000232656 - | 1 - | 9,63E-05 - | 2 - | 0,000183084 - | 1 - | 9,29E-05 - | 1 - | 0,000104645 - |
| 802 | 1 | 0.2um_NOKCPHJN_396835 | Eukaryota | unclassified | Pelagophycea | Pelagomonad | unclassified | Aureococcus | Aureococcus a | information st | Translation, ri | KOG3204 | 60s ribosomal | 67593.Physo1 | 0,4 | 0,489897949 | 2,81E-05 | 3,48E-05 | 1 - | 6,27E-05 - | 1 - | 7,79E-05 - | 0 - | 0 - | 0 - | 0 - | 0 - | 0 - |
| 803 | 1 | 0.2um_NOKCPHJN_529800 | Eukaryota | Chordata | unclassified | Squamata | Pythonidae | Python | Python bivitta | information st | Translation, ri | 0STSV | ribosomal pro | 34740.HMEL0 | 0,8 | 0,4 | 6,84E-05 | 3,48E-05 | 1 - | 7,40E-05 - | 1 - | 9,19E-05 - | 1 - | 8,74E-05 - | 1 - | 8,87E-05 - | 0 - | 0 - |
| 804 | 1 | 0.2um_NOKCPHJN_121475 | Bacteria | Proteobacteri | Gammaprotec | Oceanospirilla | Oceanospirilla | Amphritea | Amphritea atl | unclassified | unclassified | unclassified | unclassified | unclassified | 1 | 0,894427191 | 3,25E-05 | 3,01E-05 | 1 - | 2,83E-05 - | 0 - | 0 - | 2 - | 6,67E-05 - | 2 - | 6,77E-05 - | 0 - | 0 - |
| 805 | 1 | 0.2um_NOKCPHJN_83828 | Bacteria | Bacteroidetes | Flavobacteriia | Flavobacterial | unclassified | unclassified | Flavobacterial | metabolism | Amino acid tra | 08IQK | o-acetylhomo | 391603.FBALC | 4 | 0,632455532 | 0,000126706 | 1,63E-05 | 5 - | 0,000133772 - | 4 - | 0,000132885 - | 4 - | 0,000126324 - | 3 - | 9,61E-05 - | 4 - | 0,000144405 - |
| 806 | 1 | 0.2um_NOKCPHJN_179256 | Bacteria | Proteobacteri | Gammaprotec | unclassified | unclassified | unclassified | Gammaprotec | metabolism | Inorganic ion t | 0QK8T | TonB-depende | 247633.GP214 | 1 | 0,632455532 | 1,61E-05 | 1,04E-05 | 1 - | 1,39E-05 - | 1 - | 1,72E-05 - | 2 - | 3,27E-05 - | 1 - | 1,66E-05 - | 0 - | 0 - |
| 807 | 1 | 0.2um_NOKCPHJN_99550 | Bacteria | Proteobacteri | Gammaprotec | Oceanospirilla | Saccharospirill | Reinekea | Reinekea blan | information st | Replication, re | COG0550 | Releases the s | 314283.MED2 | 0,6 | 0,489897949 | 8,66E-06 | 7,14E-06 | 1 - | 1,26E-05 - | 1 - | 1,56E-05 - | 0 - | 0 - | 1 - | 1,51E-05 - | 0 - | 0 - |
| 808 | 1 | 0.2um_NOKCPHJN_237696 | Bacteria | Proteobacteri | Gammaprotec | Oceanospirilla | Saccharospirill | Reinekea | Reinekea fors | poorly charact | Function unkn | 0QNMQ | NA | 314283.MED2 | 0,4 | 0,489897949 | 4,86E-06 | 5,96E-06 | 0 - | 0 - | 1 - | 1,24E-05 - | 0 - | 0 - | 1 - | 1,19E-05 - | 0 - | 0 - |
| 809 | 1 | 3um_FIJJAFBF_430953 | Bacteria | Bacteroidetes | Flavobacteriia | Flavobacterial | Flavobacteriac | Polaribacter | Polaribacter v | unclassified | unclassified | unclassified | unclassified | unclassified | 0,2 | 0,4 | 1,40E-05 | 2,81E-05 | 0 - | 0 - | 0 - | 0 - | 0 - | 0 - | 1 - | 7,01E-05 - | 0 - | 0 - |
| 810 | 1 | 0.2um_NOKCPHJN_121978 | Bacteria | Bacteroidetes | Flavobacteriia | Flavobacterial | Flavobacteriac | Polaribacter | Polaribacter a | information st | Translation, ri | 05C64 | Binds directly | 313594.PI23P | 1 | 0 | 5,98E-05 | 5,68E-06 | 1 - | 5,01E-05 - | 1 - | 6,22E-05 - | 1 - | 5,92E-05 - | 1 - | 6,00E-05 - | 1 - | 6,76E-05 - |
| 811 | 1 | 3um_FIJJAFBF_552358 | Viruses | unclassified | unclassified | unclassified | unclassified | unclassified | uncultured vir | unclassified | unclassified | unclassified | unclassified | unclassified | 5,8 | 1,326649916 | 0,000251642 | 3,71E-05 | 8 - | 0,000296198 - | 4 - | 0,000183895 - | 6 - | 0,000262224 - | 6 - | 0,000266097 - | 5 - | 0,000249798 - |
| 812 | 1 | 3um_FIJJAFBF_552509 | Bacteria | Proteobacteri | Gammaprotec | Arenicellales | Arenicellaceae | Arenicella | Arenicella xan | information st | Translation, ri | 05C64 | Binds directly | 634503.NT01 | 0,2 | 0,4 | 9,94E-06 | 1,99E-05 | 1 - | 4,97E-05 - | 0 - | 0 - | 0 - | 0 - | 0 - | 0 - | 0 - | 0 - |
| 813 | 1 | 0.2um_NOKCPHJN_226136 | Bacteria | Bacteroidetes | Flavobacteriia | Flavobacterial | Crocinitomica | unclassified | Crocinitomica | metabolism | Carbohydrate | arCOG02510 | Cell surface pr | 456442.Mboo | 0,6 | 0,489897949 | 1,37E-05 | 1,13E-05 | 1 - | 2,04E-05 - | 0 - | 0 - | 1 - | 2,40E-05 - | 1 - | 2,44E-05 - | 0 - | 0 - |
| 814 | 1 | 0.2um_NOKCPHJN_152859 | Bacteria | Bacteroidetes | Flavobacteriia | Flavobacterial | Flavobacteriac | Wenyingzhuai | various | unclassified | unclassified | unclassified | unclassified | unclassified | 1,6 | 0,8 | 0,000346735 | 0,000168222 | 2 - | 0,000370248 - | 1 - | 0,000229869 - | 1 - | 0,00021852 - | 3 - | 0,000665242 - | 1 - | 0,000249798 - |
| 815 | 1 | 0.2um_NOKCPHJN_184065 | Bacteria | Bacteroidetes | Flavobacteriia | Flavobacterial | Flavobacteriac | Ulvibacter | Ulvibacter lito | cellular proces | Intracellular tr | COG0653 | Part of the Se | 50743.SCB49_ | 1,4 | 0,8 | 1,69E-05 | 9,40E-06 | 2 - | 2,06E-05 - | 2 - | 2,55E-05 - | 0 - | 0 - | 2 - | 2,46E-05 - | 1 - | 1,39E-05 - |
| 816 | 1 | 3um_FIJJAFBF_611435 | Bacteria | Proteobacteri | Betaproteoba | Burkholderial | unclassified | unclassified | various | metabolism | Amino acid tra | COG0131 | imidazoleglyc | 365046.Rta_0 | 1 | 0 | 6,75E-05 | 6,41E-06 | 1 - | 5,65E-05 - | 1 - | 7,02E-05 - | 1 - | 6,67E-05 - | 1 - | 6,77E-05 - | 1 - | 7,63E-05 - |
| 817 | 1 | 3um_FIJJAFBF_715069 | Eukaryota | unclassified | Compsopogor | Compsopogor | Compsopogor | Compsopogor | Compsopogor | unclassified | unclassified | unclassified | unclassified | unclassified | 0,8 | 0,4 | 0,000246633 | 0,000125267 | 1 - | 0,000266923 - | 1 - | 0,000331439 - | 1 - | 0,000315075 - | 1 - | 0,000319729 - | 0 - | 0 - |
| 818 | 1 | 3um_FIJJAFBF_121131 | Bacteria | Proteobacteri | Gammaprotec | Cellvibrionales | Halieaceae | Halioglobus | Halioglobus sp | metabolism | Lipid transpor | COG1024 | Enoyl-CoA hyc | 208964.PA17 | 1 | 0 | 2,16E-05 | 2,06E-06 | 1 - | 1,81E-05 - | 1 - | 2,25E-05 - | 1 - | 2,14E-05 - | 1 - | 2,17E-05 - | 1 - | 2,45E-05 - |
| 819 | 1 | 3um_FIJJAFBF_338021 | Bacteria | Acidobacteria | unclassified | unclassified | unclassified | unclassified | Acidobacteria | cellular proces | Posttranslatio | COG0652 | peptidyl-proly | 2850.Phatr14 | 1,4 | 0,489897949 | 0,00038261 | 0,000112569 | 2 - | 0,000468477 - | 1 - | 0,000290855 - | 1 - | 0,000276494 - | 2 - | 0,000561156 - | 1 - | 0,00031607 - |
| 820 | 1 | 0.2um_NOKCPHJN_590792 | Bacteria | Bacteroidetes | Flavobacteriia | Flavobacterial | Flavobacteriac | Polaribacter | Polaribacter s | metabolism | Inorganic ion t | 05C27 | TonB-depende | 867900.Celly_ | 1 | 0 | 1,46E-05 | 1,39E-06 | 1 - | 1,22E-05 - | 1 - | 1,52E-05 - | 1 - | 1,45E-05 - | 1 - | 1,47E-05 - | 1 - | 1,65E-05 - |
| 821 | 1 | 3um_FIJJAFBF_189622 | Bacteria | Bacteroidetes | Flavobacteriia | Flavobacterial | Flavobacteriac | Polaribacter | Polaribacter s | unclassified | unclassified | unclassified | unclassified | unclassified | 0,6 | 0,489897949 | 3,06E-05 | 2,51E-05 | 0 - | 0 - | 1 - | 5,04E-05 - | 1 - | 4,79E-05 - | 0 - | 0 - | 1 - | 5,47E-05 - |
| 822 | 1 | 0.2um_NOKCPHJN_406227 | Bacteria | Bacteroidetes | Flavobacteriia | Flavobacterial | Flavobacteriac | Polaribacter | Polaribacter s | unclassified | unclassified | unclassified | unclassified | unclassified | 1,2 | 0,4 | 2,28E-05 | 7,64E-06 | 1 - | 1,59E-05 - | 1 - | 1,98E-05 - | 2 - | 3,76E-05 - | 1 - | 1,91E-05 - | 1 - | 2,15E-05 - |
| 823 | 1 | 3um_FIJJAFBF_453267 | Bacteria | Proteobacteri | Alphaproteob | Rhodobactera | Rhodobactera | unclassified | Rhodobactera | unclassified | unclassified | unclassified | unclassified | unclassified | 0,2 | 0,4 | 2,81E-06 | 5,61E-06 | 0 - | 0 - | 0 - | 0 - | 0 - | 0 - | 1 - | 1,40E-05 - | 0 - | 0 - |
| 824 | 1 | 3um_FIJJAFBF_441623 | Bacteria | Planctomycet | Planctomyceti | Candidatus Br | Candidatus Br | Candidatus Sc | Candidatus Sc | cellular proces | Defense mech | 0BD5S | Type I restricti | 261292.Nit79 | 0,2 | 0,4 | 8,41E-06 | 1,68E-05 | 0 - | 0 - | 0 - | 0 - | 0 - | 0 - | 1 - | 4,20E-05 - | 0 - | 0 - |
| 825 | 1 | 3um_FIJJAFBF_30776 | Bacteria | Bacteroidetes | Cytophagia | Cytophagales | Flammeovirga | unclassified | Flammeovirga | poorly charact | Function unkn | 11NCV | Protein of unk | 643867.Ftrac_ | 3,4 | 0,489897949 | 0,000263794 | 5,64E-05 | 3 - | 0,000193444 - | 3 - | 0,0002402 - | 3 - | 0,000228341 - | 4 - | 0,000308951 - | 4 - | 0,000348033 - |
| 826 | 1 | 3um_FIJJAFBF_256304 | Bacteria | Bacteroidetes | Flavobacteriia | Flavobacterial | Flavobacteriac | Seonamhaeic | Seonamhaeic | unclassified | unclassified | unclassified | unclassified | unclassified | 1,4 | 0,489897949 | 4,39E-05 | 1,86E-05 | 1 - | 2,56E-05 - | 2 - | 6,36E-05 - | 1 - | 3,02E-05 - | 1 - | 3,07E-05 - | 2 - | 6,91E-05 - |
| 827 | 1 | 3um_FIJJAFBF_393744 | Bacteria | Bacteroidetes | Flavobacteriia | unclassified | unclassified | unclassified | Flavobacteria | unclassified | unclassified | unclassified | unclassified | unclassified | 1,4 | 0,8 | 7,61E-05 | 4,45E-05 | 2 - | 9,22E-05 - | 0 - | 0 - | 2 - | 0,000108821 - | 1 - | 5,52E-05 - | 2 - | 0,000124397 - |
| 828 | 1 | 3um_FIJJAFBF_786150 | Bacteria | Proteobacteri | Gammaprotec | Oceanospirilla | Saccharospirill | Saccharospiril | Saccharospiril | unclassified | unclassified | unclassified | unclassified | unclassified | 8 | 1,549193338 | 0,000620216 | 0,000173358 | 7 - | 0,000448848 - | 7 - | 0,000557336 - | 8 - | 0,000605507 - | 7 - | 0,000537644 - | 11 - | 0,000951743 - |
| 829 | 1 | 0.2um_NOKCPHJN_117065 | Eukaryota | Bacillariophyt | Bacillariophyc | Naviculales | Naviculaceae | Fistulifera | Fistulifera sola | metabolism | Energy produc | 0IFIU | Pyrophosphat | 2850.Phatr15 | 4 | 0,632455532 | 6,94E-05 | 1,06E-05 | 4 - | 5,85E-05 - | 4 - | 7,26E-05 - | 4 - | 6,90E-05 - | 5 - | 8,76E-05 - | 3 - | 5,92E-05 - |
| 830 | 1 | 0.2um_NOKCPHJN_69511 | Bacteria | Bacteroidetes | Flavobacteriia | Flavobacterial | unclassified | unclassified | Flavobacterial | metabolism | Inorganic ion t | 05C27 | TonB-depende | 216432.CA25 | 0,6 | 0,489897949 | 8,31E-06 | 6,81E-06 | 0 - | 0 - | 0 - | 0 - | 1 - | 1,32E-05 - | 1 - | 1,33E-05 - | 1 - | 1,50E-05 - |
| 831 | 1 | 0.2um_NOKCPHJN_86901 | Bacteria | Proteobacteri | Gammaprotec | unclassified | unclassified | unclassified | Gammaprotec | information st | Translation, ri | 05G6W | One of the pri | 247634.GPB2 | 1 | 0 | 6,65E-05 | 6,32E-06 | 1 - | 5,57E-05 - | 1 - | 6,92E-05 - | 1 - | 6,58E-05 - | 1 - | 6,67E-05 - | 1 - | 7,52E-05 - |
| 832 | 1 | 3um_FIJJAFBF_244792 | Bacteria | Bacteroidetes | Flavobacteriia | Flavobacterial | Cryomorphace | unclassified | Cryomorphace | information st | Transcription | 05D27 | DNA-depende | 926562.Oweh | 1,6 | 1,2 | 2,38E-05 | 1,91E-05 | 1 - | 1,19E-05 - | 3 - | 4,44E-05 - | 0 - | 0 - | 1 - | 1,43E-05 - | 3 - | 4,82E-05 - |
| 833 | 1 | 0.2um_NOKCPHJN_62911 | Bacteria | Proteobacteri | Gammaprotec | Oceanospirilla | Saccharospirill | Reinekea | Reinekea fors | unclassified | unclassified | unclassified | unclassified | unclassified | 7 | 0,894427191 | 0,000461863 | 6,46E-05 | 8 - | 0,000443582 - | 8 - | 0,000550798 - | 6 - | 0,000392702 - | 6 - | 0,000398502 - | 7 - | 0,00052373 - |
| 834 | 1 | 3um_FIJJAFBF_763426 | Bacteria | Proteobacteri | Gammaprotec | unclassified | unclassified | unclassified | Gammaprotec | metabolism | Inorganic ion t | 16TQC | TONB-depend | 314287.GB22 | 1,4 | 0,489897949 | 7,91E-05 | 2,33E-05 | 2 - | 9,69E-05 - | 1 - | 6,01E-05 - | 1 - | 5,72E-05 - | 2 - | 0,00011602 - | 1 - | 6,53E-05 - |
| 835 | 2 | 3um_FIJJAFBF_165523;3um_FIJJAFBF_233965 | Bacteria | Proteobacteri | Gammaprotec | various | various | various | various | poorly charact | Function unkn | 0QRAP | NA | 393595.ABO_ | 1,8 | 0,4 | 2,68E-05 | 5,30E-06 | 2 - | 2,53E-05 - | 2 - | 3,15E-05 - | 2 - | 2,99E-05 - | 2 - | 3,03E-05 - | 1 - | 1,71E-05 - |
| 836 | 1 | 0.2um_NOKCPHJN_308086 | Bacteria | Actinobacteri | Actinobacteri | Micrococcales | Microbacteria | Candidatus Ac | Candidatus A | unclassified | unclassified | unclassified | unclassified | unclassified | 3,4 | 0,489897949 | 5,34E-05 | 5,19E-06 | 4 - | 5,32E-05 - | 3 - | 4,95E-05 - | 4 - | 6,28E-05 - | 3 - | 4,78E-05 - | 3 - | 5,38E-05 - |
| 837 | 1 | 0.2um_NOKCPHJN_302685 | Eukaryota | Streptophyta | unclassified | Brassicales | Brassicaceae | Camelina | Camelina sati | cellular proces | Cytoskeleton | 1DU1B | Armadillo rep | 3218.PP1S14_ | 0,2 | 0,4 | 2,03E-06 | 4,06E-06 | 0 - | 0 - | 0 - | 0 - | 1 - | 1,02E-05 - | 0 - | 0 - | 0 - | 0 - |
| 838 | 1 | 3um_FIJJAFBF_515059 | Bacteria | Proteobacteri | Gammaprotec | Cellvibrionales | Porticoccacea | unclassified | SAR92 bacteri | metabolism | Energy produc | COG0056 | Produces ATP | 247634.GPB2 | 6,4 | 1,959591794 | 0,000165765 | 3,30E-05 | 10 - | 0,000223301 - | 6 - | 0,000166364 - | 6 - | 0,00015815 - | 6 - | 0,000160486 - | 4 - | 0,000120525 - |
| 839 | 1 | 3um_FIJJAFBF_15027 | Bacteria | Bacteroidetes | Flavobacteriia | Flavobacterial | Flavobacteriac | Polaribacter | Polaribacter s | unclassified | unclassified | unclassified | unclassified | unclassified | 3 | 0,632455532 | 0,000208523 | 5,85E-05 | 3 - | 0,00017303 - | 3 - | 0,000214853 - | 2 - | 0,000136163 - | 3 - | 0,000207261 - | 4 - | 0,000311305 - |
| 840 | 1 | 3um_FIJJAFBF_236209 | Bacteria | Proteobacteri | Gammaprotec | Oceanospirilla | Saccharospirill | Reinekea | Reinekea blan | metabolism | Amino acid tra | 0QIP6 | NAG kinase | 314283.MED2 | 0,2 | 0,4 | 8,91E-06 | 1,78E-05 | 0 - | 0 - | 0 - | 0 - | 1 - | 4,46E-05 - | 0 - | 0 - | 0 - | 0 - |
| 841 | 2 | 0.2um_NOKCPHJN_941271;3um_FIJJAFBF_3965 | Bacteria | Bacteroidetes | Flavobacteriia | Flavobacterial | Flavobacteriac | unclassified | Flavobacteriac | metabolism | Nucleotide tra | various | various | various | 0,2 | 0,4 | 9,60E-06 | 1,92E-05 | 1 - | 4,80E-05 - | 0 - | 0 - | 0 - | 0 - | 0 - | 0 - | 0 - | 0 - |
| 842 | 2 | 0.2um_NOKCPHJN_934022;3um_FIJJAFBF_8145 | Bacteria | Bacteroidetes | Flavobacteriia | Flavobacterial | Flavobacteriac | Formosa | Formosa sp. H | metabolism | Carbohydrate | 16QEQ | hydrolase, fan | various | 1 | 0 | 6,46E-05 | 6,14E-06 | 1 - | 5,41E-05 - | 1 - | 6,72E-05 - | 1 - | 6,39E-05 - | 1 - | 6,49E-05 - | 1 - | 7,31E-05 - |
| 843 | 9 | 0.2um_NOKCPHJN_1036533;0.2um_NOKCPHJN_ | Bacteria | Bacteroidetes | Flavobacteriia | Flavobacterial | Flavobacteriac | Formosa | Formosa sp. H | poorly charact | Function unkn | 0900R | pur-alpha bet | 391603.FBALC | 1 | 0 | 0,000115149 | 1,09E-05 | 1 - | 9,65E-05 - | 1 - | 0,000119764 - | 1 - | 0,000113851 - | 1 - | 0,000115532 - | 1 - | 0,000130147 - |
| 844 | 1 | 0.2um_NOKCPHJN_195327 | Bacteria | Proteobacteri | Gammaprotec | Cellvibrionales | Porticoccacea | unclassified | SAR92 bacteri | cellular proces | Cell wall/mem | 16Q41 | Efflux transpo | 314287.GB22 | 0,6 | 0,489897949 | 2,18E-05 | 1,80E-05 | 1 - | 3,23E-05 - | 0 - | 0 - | 1 - | 3,82E-05 - | 1 - | 3,87E-05 - | 0 - | 0 - |
| 845 | 1 | 3um_FIJJAFBF_288420 | unclassified | unclassified | unclassified | unclassified | unclassified | unclassified | unclassified | unclassified | unclassified | unclassified | unclassified | unclassified | 2,4 | 0,489897949 | 0,000360423 | 8,05E-05 | 3 - | 0,000378385 - | 2 - | 0,000313228 - | 2 - | 0,000297763 - | 2 - | 0,000302161 - | 3 - | 0,000510575 - |
| 846 | 1 | 3um_FIJJAFBF_583511 | Bacteria | Bacteroidetes | unclassified | unclassified | unclassified | unclassified | unidentified e | metabolism | Energy produc | COG0056 | Produces ATP | 398720.MED2 | 3,6 | 0,489897949 | 9,31E-05 | 1,10E-05 | 4 - | 8,73E-05 - | 4 - | 0,000108379 - | 4 - | 0,000103028 - | 3 - | 7,84E-05 - | 3 - | 8,83E-05 - |
| 847 | 1 | 0.2um_NOKCPHJN_62928 | Bacteria | Bacteroidetes | Flavobacteriia | Flavobacterial | Flavobacteriac | Polaribacter | Polaribacter s | information st | Replication, re | COG0178 | The UvrABC re | 313594.PI23P | 3,2 | 0,748331477 | 4,64E-05 | 8,09E-06 | 4 - | 4,95E-05 - | 3 - | 4,61E-05 - | 4 - | 5,84E-05 - | 3 - | 4,44E-05 - | 2 - | 3,34E-05 - |
| 848 | 1 | 3um_FIJJAFBF_630917 | Bacteria | Proteobacteri | Gammaprotec | Oceanospirilla | Saccharospirill | Reinekea | Reinekea blan | metabolism | Energy produc | 05C6K | Malic enzyme | 314283.MED2 | 6,8 | 0,979795897 | 0,000224857 | 4,31E-05 | 7 - | 0,000193599 - | 5 - | 0,000171709 - | 7 - | 0,000228524 - | 7 - | 0,0002319 - | 8 - | 0,000298553 - |
| 849 | 2 | 3um_FIJJAFBF_355622;3um_FIJJAFBF_477127 | Bacteria | various | various | various | various | various | various | cellular proces | Intracellular tr | 175GA | One of the pro | 367336.OM22 | 1,4 | 0,489897949 | 0,000122051 | 4,31E-05 | 1 - | 7,31E-05 - | 1 - | 9,08E-05 - | 2 - | 0,000172589 - | 2 - | 0,000175138 - | 1 - | 9,86E-05 - |
| 850 | 1 | 3um_FIJJAFBF_167961 | Bacteria | Proteobacteri | Alphaproteob | Rhodobactera | Rhodobactera | Planktomarina | Planktomarina | information st | Translation, ri | 08YY1 | This protein is | 375451.RD1_ | 3,4 | 1,019803903 | 0,000372682 | 9,13E-05 | 5 - | 0,000466572 - | 4 - | 0,000463476 - | 2 - | 0,000220296 - | 3 - | 0,000335325 - | 3 - | 0,000377743 - |
| 851 | 1 | 0.2um_NOKCPHJN_1069625 | Bacteria | Bacteroidetes | Flavobacteriia | Flavobacterial | Flavobacteriac | Formosa | Formosa sp. H | poorly charact | Function unkn | 05END | lipolytic prote | 391603.FBALC | 1,2 | 0,748331477 | 3,17E-05 | 1,90E-05 | 2 - | 4,52E-05 - | 2 - | 5,61E-05 - | 1 - | 2,67E-05 - | 0 - | 0 - | 1 - | 3,05E-05 - |
| 852 | 1 | 0.2um_NOKCPHJN_883124 | Bacteria | Cyanobacteria | unclassified | Synechococca | Synechococca | Synechococcu | Synechococcu | unclassified | unclassified | unclassified | unclassified | unclassified | 1,2 | 0,4 | 2,42E-05 | 5,35E-06 | 2 - | 3,47E-05 - | 1 - | 2,15E-05 - | 1 - | 2,05E-05 - | 1 - | 2,08E-05 - | 1 - | 2,34E-05 - |

| | | | | | | | | | | | | | | | | | | | | | | | | | | | | |
|---|---|---|---|---|---|---|---|---|---|---|---|---|---|---|---|---|---|---|---|---|---|---|---|---|---|---|---|---|
| 853 | 1 | 0.2um_NOKCPHJN_586666 | Bacteria | Proteobacteri | Betaproteoba | unclassified | unclassified | unclassified | Betaproteoba | poorly charact | Function unkn | 0XRND | NA | 314607.KB13_ | 1 | 0 | 6,23E-05 | 5,91E-06 | 1 - | 5,22E-05 - | 1 - | 6,48E-05 - | 1 - | 6,16E-05 - | 1 - | 6,25E-05 - | 1 - | 7,04E-05 - |
| 854 | 1 | 0.2um_NOKCPHJN_82821 | Bacteria | Proteobacteri | Alphaproteob | Rhodobactera | Hyphomonada | Robiginitomac | Robiginitomac | unclassified | unclassified | unclassified | unclassified | unclassified | 3,8 | 1,166190379 | 6,36E-05 | 1,35E-05 | 6 - | 8,62E-05 - | 4 - | 7,13E-05 - | 3 - | 5,09E-05 - | 3 - | 5,16E-05 - | 3 - | 5,82E-05 - |
| 855 | 1 | 0.2um_NOKCPHJN_364158 | Bacteria | Proteobacteri | Alphaproteob | Rhodobactera | Rhodobactera | Donghicola | Donghicola ty | unclassified | unclassified | unclassified | unclassified | unclassified | 1 | 0 | 6,65E-05 | 6,32E-06 | 1 - | 5,57E-05 - | 1 - | 6,92E-05 - | 1 - | 6,58E-05 - | 1 - | 6,67E-05 - | 1 - | 7,52E-05 - |
| 856 | 1 | 3um_FIJJAFBF_175546 | Bacteria | Proteobacteri | Gammaprote | Pseudomonad | Pseudomonad | Pseudomonas | Pseudomonas | unclassified | unclassified | unclassified | unclassified | unclassified | 0,2 | 0,4 | 3,59E-06 | 7,18E-06 | 1 - | 1,80E-05 - | 0 - | 0 - | 0 - | 0 - | 0 - | 0 - | 0 - | 0 - |
| 857 | 1 | 3um_FIJJAFBF_615878 | Bacteria | Bacteroidetes | Flavobacteriia | Flavobacterial | Flavobacteria | Polaribacter | Polaribacter s | poorly charact | Function unkn | 0P8NI | Inherit from b | 313594.PI23P | 12,2 | 1,720465053 | 0,000252516 | 5,03E-05 | 13 - | 0,000224714 - | 11 - | 0,000236101 - | 12 - | 0,000244847 - | 10 - | 0,000207053 - | 15 - | 0,000349867 - |
| 858 | 1 | 3um_FIJJAFBF_786146 | Bacteria | Proteobacteri | Gammaprote | Oceanospirilla | Saccharospirill | Reinekea | Reinekea fors | information st | Translation, ri | COG0256 | This is one of | 314283.MED2 | 2 | 0 | 0,000236254 | 2,24E-05 | 2 - | 0,000197891 - | 2 - | 0,000245722 - | 2 - | 0,00023359 - | 2 - | 0,00023704 - | 2 - | 0,000267025 - |
| 859 | 1 | 0.2um_NOKCPHJN_669432 | Bacteria | Proteobacteri | Alphaproteob | Pelagibacteral | Pelagibactera | unclassified | Pelagibactera | information st | Transcription | 05D27 | DNA-depende | 335992.SAR11 | 7,6 | 1,356465997 | 9,91E-05 | 1,83E-05 | 7 - | 7,67E-05 - | 7 - | 9,53E-05 - | 8 - | 0,00010352 - | 10 - | 0,000131312 - | 6 - | 8,88E-05 - |
| 860 | 1 | 3um_FIJJAFBF_417933 | Bacteria | Proteobacteri | Gammaprote | unclassified | unclassified | unclassified | OM182 bacte | poorly charact | Function unkn | 0XZCS | NA | 314345.SPV1_ | 0,8 | 0,4 | 9,60E-05 | 4,93E-05 | 1 - | 0,000101572 - | 0 - | 0 - | 1 - | 0,000119896 - | 1 - | 0,000121667 - | 1 - | 0,000137057 - |
| 861 | 1 | 3um_FIJJAFBF_176633 | Bacteria | Bacteroidetes | unclassified | unclassified | unclassified | unclassified | Bacteroidetes | unclassified | unclassified | unclassified | unclassified | unclassified | 1,8 | 0,4 | 0,000184151 | 3,64E-05 | 2 - | 0,000173904 - | 2 - | 0,000215938 - | 2 - | 0,000205276 - | 2 - | 0,000208308 - | 1 - | 0,000117329 - |
| 862 | 1 | 3um_FIJJAFBF_639264 | Bacteria | Proteobacteri | Gammaprote | Chromatiales | Ectothiorhodo | Thioalkalivibri | unclassified | cellular proces | Posttranslatio | 17FI0 | Binds to Cpn6 | 396595.TK90_ | 0,4 | 0,8 | 4,83E-05 | 9,67E-05 | 2 - | 0,000241635 - | 0 - | 0 - | 0 - | 0 - | 0 - | 0 - | 0 - | 0 - |
| 863 | 1 | 3um_FIJJAFBF_809400 | Bacteria | Proteobacteri | Alphaproteob | various | various | various | various | cellular proces | Posttranslatio | 17FI0 | Binds to Cpn6 | 367336.OM22 | 2,4 | 0,8 | 0,000340388 | 7,54E-05 | 4 - | 0,000488412 - | 2 - | 0,000303232 - | 2 - | 0,00028826 - | 2 - | 0,000292518 - | 2 - | 0,00032952 - |
| 864 | 1 | 0.2um_NOKCPHJN_841638 | Bacteria | Bacteroidetes | Flavobacteriia | various | various | various | various | cellular proces | Cell wall/mem | 09DZB | Fasciclin | 487797.Flav3 | 0,4 | 0,489897949 | 3,17E-05 | 3,96E-05 | 1 - | 6,75E-05 - | 0 - | 0 - | 0 - | 0 - | 0 - | 0 - | 1 - | 9,11E-05 - |
| 865 | 1 | 0.2um_NOKCPHJN_191839 | Bacteria | Proteobacteri | Gammaprote | Oceanospirilla | Saccharospirill | Reinekea | Reinekea fors | metabolism | Coenzyme tra | 08S7T | Catalyzes the | 314283.MED2 | 0,8 | 0,748331477 | 4,92E-05 | 4,28E-05 | 2 - | 0,00010828 - | 0 - | 0 - | 0 - | 0 - | 1 - | 6,49E-05 - | 1 - | 7,31E-05 - |
| 866 | 1 | 3um_FIJJAFBF_243896 | Bacteria | Proteobacteri | Gammaprote | Alteromonada | Alteromonada | Paraglaciecola | Paraglaciecola | metabolism | Energy produc | 16R80 | pyrophosphat | 342610.Patl_ | 1 | 0 | 7,53E-05 | 7,15E-06 | 1 - | 6,31E-05 - | 1 - | 7,83E-05 - | 1 - | 7,44E-05 - | 1 - | 7,55E-05 - | 1 - | 8,51E-05 - |
| 867 | 1 | 0.2um_NOKCPHJN_653231 | Bacteria | Proteobacteri | Alphaproteob | Pelagibacteral | Pelagibactera | Candidatus Pe | Candidatus Pe | unclassified | unclassified | unclassified | unclassified | unclassified | 0,4 | 0,489897949 | 1,22E-05 | 1,50E-05 | 0 - | 0 - | 0 - | 0 - | 1 - | 3,04E-05 - | 1 - | 3,08E-05 - | 0 - | 0 - |
| 868 | 1 | 3um_FIJJAFBF_201375 | Bacteria | Bacteroidetes | Flavobacteriia | Flavobacterial | Flavobacteria | Polaribacter | Polaribacter b | metabolism | Carbohydrate | 064TE | hydrolase fam | 391587.KAOT | 1 | 0 | 2,72E-05 | 2,58E-06 | 1 - | 2,28E-05 - | 1 - | 2,83E-05 - | 1 - | 2,69E-05 - | 1 - | 2,73E-05 - | 1 - | 3,07E-05 - |
| 869 | 1 | 0.2um_NOKCPHJN_85198 | Bacteria | Proteobacteri | Alphaproteob | Rhodobactera | Rhodobactera | Ascidiaceihabi | Ascidiaceihabi | metabolism | Carbohydrate | 07SNC | solute-binding | 367336.OM22 | 2,8 | 0,4 | 9,11E-05 | 1,92E-05 | 2 - | 5,39E-05 - | 3 - | 0,000100365 - | 3 - | 9,54E-05 - | 3 - | 9,68E-05 - | 3 - | 0,000109067 - |
| 870 | 1 | 0.2um_NOKCPHJN_189831 | Bacteria | Bacteroidetes | Flavobacteriia | Flavobacterial | Flavobacteria | Polaribacter | Polaribacter s | unclassified | unclassified | unclassified | unclassified | unclassified | 1 | 0 | 9,12E-06 | 8,66E-07 | 1 - | 7,64E-06 - | 1 - | 9,48E-06 - | 1 - | 9,01E-06 - | 1 - | 9,15E-06 - | 1 - | 1,03E-05 - |
| 871 | 1 | 3um_FIJJAFBF_605381 | Bacteria | Proteobacteri | Gammaprote | Oceanospirilla | Saccharospirill | Reinekea | Reinekea blan | cellular proces | Posttranslatio | 16PM6 | feS assembly | 314283.MED2 | 0,8 | 0,748331477 | 4,50E-05 | 4,57E-05 | 1 - | 4,61E-05 - | 0 - | 0 - | 1 - | 5,44E-05 - | 0 - | 0 - | 2 - | 0,000124397 - |
| 872 | 1 | 0.2um_NOKCPHJN_538357 | Bacteria | Bacteroidetes | Flavobacteriia | Flavobacterial | Cryomorphace | unclassified | Cryomorphace | unclassified | unclassified | unclassified | unclassified | unclassified | 1,2 | 0,4 | 2,44E-05 | 8,42E-06 | 1 - | 1,70E-05 - | 1 - | 2,11E-05 - | 1 - | 2,01E-05 - | 2 - | 4,08E-05 - | 1 - | 2,30E-05 - |
| 873 | 1 | 0.2um_NOKCPHJN_79972 | Bacteria | Proteobacteri | unclassified | unclassified | unclassified | unclassified | Proteobacteri | cellular proces | Intracellular tr | 0QITE | Outer membr | 1001585.MDS | 0,6 | 0,8 | 1,83E-05 | 2,31E-05 | 2 - | 5,47E-05 - | 0 - | 0 - | 0 - | 0 - | 0 - | 0 - | 1 - | 3,69E-05 - |
| 874 | 1 | 3um_FIJJAFBF_644846 | Eukaryota | Bacillariophyt | Bacillariophyc | Naviculales | Naviculaceae | Fistulifera | Fistulifera sola | unclassified | unclassified | unclassified | unclassified | unclassified | 0,4 | 0,489897949 | 1,72E-05 | 2,12E-05 | 1 - | 3,83E-05 - | 1 - | 4,75E-05 - | 0 - | 0 - | 0 - | 0 - | 0 - | 0 - |
| 875 | 1 | 3um_FIJJAFBF_439954 | Bacteria | Proteobacteri | Alphaproteob | Rhodobactera | unclassified | unclassified | Rhodobactera | unclassified | unclassified | unclassified | unclassified | unclassified | 0,6 | 0,489897949 | 7,68E-05 | 6,32E-05 | 1 - | 0,00011364 - | 0 - | 0 - | 1 - | 0,000134141 - | 1 - | 0,000136122 - | 0 - | 0 - |
| 876 | 2 | 0.2um_NOKCPHJN_11097;3um_FIJJAFBF_21287 | Bacteria | Bacteroidetes | Flavobacteriia | Flavobacterial | Crocinitomica | unclassified | Crocinitomica | poorly charact | Function unkn | 0FZ9A | NA | 643867.Ftrac_ | 1,2 | 0,748331477 | 5,13E-05 | 3,04E-05 | 2 - | 7,40E-05 - | 0 - | 0 - | 1 - | 4,37E-05 - | 2 - | 8,87E-05 - | 1 - | 5,00E-05 - |
| 877 | 1 | 3um_FIJJAFBF_150356 | Bacteria | Bacteroidetes | Flavobacteriia | Flavobacterial | Flavobacteria | Formosa | Formosa sp. H | poorly charact | Function unkn | 11NCV | Protein of unk | 983544.Lacal_ | 3,6 | 0,489897949 | 0,000298559 | 3,53E-05 | 4 - | 0,000279943 - | 4 - | 0,000347607 - | 4 - | 0,000330444 - | 3 - | 0,000251494 - | 3 - | 0,000283307 - |
| 878 | 1 | 3um_FIJJAFBF_194075 | Bacteria | Bacteroidetes | Flavobacteriia | Flavobacterial | Flavobacteria | Polaribacter | Polaribacter s | cellular proces | Posttranslatio | 07SFW | Trigger factor | 313598.MED1 | 14,4 | 1,019803903 | 0,00044722 | 4,33E-05 | 16 - | 0,00041737 - | 14 - | 0,000453469 - | 14 - | 0,00043108 - | 13 - | 0,000406201 - | 15 - | 0,000527981 - |
| 879 | 1 | 0.2um_NOKCPHJN_217493 | Bacteria | Proteobacteri | Gammaprote | Oceanospirilla | Saccharospirill | Reinekea | Reinekea fors | metabolism | Inorganic ion t | 0HCH8 | phosphonate | 411464.DESPI | 3,6 | 0,8 | 0,000205028 | 3,79E-05 | 5 - | 0,000241128 - | 3 - | 0,000179646 - | 3 - | 0,000170776 - | 3 - | 0,000173298 - | 4 - | 0,000260293 - |
| 880 | 1 | 3um_FIJJAFBF_37890 | Bacteria | Actinobacteria | Actinobacteria | Micrococcales | Microbacteria | Candidatus Ac | Candidatus Ac | information st | Translation, ri | COG1185 | Involved in m | 288705.RSal3 | 2,2 | 0,748331477 | 4,07E-05 | 1,42E-05 | 2 - | 3,13E-05 - | 3 - | 5,83E-05 - | 3 - | 5,54E-05 - | 2 - | 3,75E-05 - | 1 - | 2,11E-05 - |
| 881 | 1 | 3um_FIJJAFBF_371413 | unclassified | unclassified | unclassified | unclassified | unclassified | unclassified | unclassified | unclassified | unclassified | unclassified | unclassified | unclassified | 0,2 | 0,4 | 2,56E-05 | 5,11E-05 | 0 - | 0 - | 0 - | 0 - | 1 - | 0,000127813 - | 0 - | 0 - | 0 - | 0 - |
| 882 | 1 | 0.2um_NOKCPHJN_613240 | Bacteria | Proteobacteri | Gammaprote | unclassified | unclassified | unclassified | Gammaprote | unclassified | unclassified | unclassified | unclassified | unclassified | 3,6 | 0,489897949 | 0,00040167 | 5,00E-05 | 4 - | 0,000376317 - | 4 - | 0,000467275 - | 3 - | 0,000333153 - | 4 - | 0,000450765 - | 3 - | 0,000380839 - |
| 883 | 2 | 0.2um_NOKCPHJN_95457;3um_FIJJAFBF_662827 | Bacteria | Proteobacteri | Gammaprote | Cellvibrionales | Porticoccacea | unclassified | various | metabolism | Energy produc | COG2224 | Isocitrate lyas | 314287.GB22 | 0,4 | 0,489897949 | 1,04E-05 | 1,27E-05 | 0 - | 0 - | 0 - | 0 - | 1 - | 2,58E-05 - | 1 - | 2,61E-05 - | 0 - | 0 - |
| 884 | 1 | 0.2um_NOKCPHJN_251566 | Eukaryota | Phaeophyceae | unclassified | Ectocarpales | Ectocarpaceae | Ectocarpus | Ectocarpus sili | unclassified | unclassified | unclassified | unclassified | unclassified | 1,4 | 0,489897949 | 0,000106249 | 4,37E-05 | 1 - | 6,24E-05 - | 1 - | 7,75E-05 - | 1 - | 7,36E-05 - | 2 - | 0,000149438 - | 2 - | 0,000168342 - |
| 885 | 1 | 3um_FIJJAFBF_418289 | Bacteria | Proteobacteri | Alphaproteob | Rhodobactera | Rhodobactera | Sulfitobacter | Sulfitobacter | unclassified | unclassified | unclassified | unclassified | unclassified | 1 | 0 | 2,97E-05 | 2,82E-06 | 1 - | 2,48E-05 - | 1 - | 3,08E-05 - | 1 - | 2,93E-05 - | 1 - | 2,98E-05 - | 1 - | 3,35E-05 - |
| 886 | 1 | 3um_FIJJAFBF_713239 | Bacteria | Bacteroidetes | Flavobacteriia | Flavobacterial | Flavobacteria | Hyunsoonleell | Hyunsoonleell | poorly charact | Function unkn | 05YA2 | NA | 313598.MED1 | 9,8 | 1,166190379 | 0,0001173 | 1,48E-05 | 10 - | 0,000100681 - | 11 - | 0,000137518 - | 9 - | 0,00010696 - | 11 - | 0,000132659 - | 8 - | 0,000108684 - |
| 887 | 1 | 3um_FIJJAFBF_806481 | Bacteria | Proteobacteri | Alphaproteob | Rhodobactera | Rhodobactera | Planktomarina | Planktomarina | information st | Replication, re | 08ZUF | This protein is | 391595.RLO14 | 3,8 | 0,4 | 0,000517134 | 4,61E-05 | 4 - | 0,000459107 - | 4 - | 0,000570075 - | 4 - | 0,000541929 - | 4 - | 0,000549933 - | 3 - | 0,000464623 - |
| 888 | 1 | 0.2um_NOKCPHJN_624695 | Bacteria | Bacteroidetes | unclassified | unclassified | unclassified | unclassified | Bacteroidetes | information st | Transcription | 05D27 | DNA-depende | 755732.Fluta_ | 0,6 | 0,489897949 | 6,42E-06 | 5,29E-06 | 1 - | 9,51E-06 - | 0 - | 0 - | 1 - | 1,12E-05 - | 1 - | 1,14E-05 - | 0 - | 0 - |
| 889 | 1 | 3um_FIJJAFBF_422920 | Bacteria | Proteobacteri | Gammaprote | Alteromonada | Pseudoalteror | Pseudoalteror | Pseudoalteror | poorly charact | Function unkn | 070MQ | NA | 425104.Ssed_ | 0,2 | 0,4 | 1,93E-05 | 3,86E-05 | 1 - | 9,65E-05 - | 0 - | 0 - | 0 - | 0 - | 0 - | 0 - | 0 - | 0 - |
| 890 | 1 | 3um_FIJJAFBF_574196 | unclassified | unclassified | unclassified | unclassified | unclassified | unclassified | unclassified | unclassified | unclassified | unclassified | unclassified | unclassified | 2 | 0 | 0,000129884 | 1,23E-05 | 2 - | 0,000108793 - | 2 - | 0,000135089 - | 2 - | 0,000128419 - | 2 - | 0,000130316 - | 2 - | 0,0001468 - |
| 891 | 1 | 0.2um_NOKCPHJN_917557 | Bacteria | Proteobacteri | Alphaproteob | Rhodobactera | Rhodobactera | Shimia | Shimia haliotis | metabolism | Amino acid tra | 01R4A | Polyamine AB | 644107.SL115 | 3,2 | 1,166190379 | 0,000118391 | 3,91E-05 | 4 - | 0,000126476 - | 3 - | 0,000117784 - | 5 - | 0,000186615 - | 2 - | 7,57E-05 - | 2 - | 8,53E-05 - |
| 892 | 1 | 0.2um_NOKCPHJN_970306 | Bacteria | Proteobacteri | Alphaproteob | Rhodobactera | Rhodobactera | Amylibacter | Amylibacter k | information st | Replication, re | 05CZ8 | DNA polymera | 388401.RB215 | 0,2 | 0,4 | 7,39E-06 | 1,48E-05 | 0 - | 0 - | 0 - | 0 - | 0 - | 0 - | 1 - | 3,70E-05 - | 0 - | 0 - |
| 893 | 1 | 3um_FIJJAFBF_715070 | Eukaryota | Bacillariophyt | Mediophycea | Hemiaulales | Hemiaulaceae | Cerataulina | Cerataulina da | unclassified | unclassified | unclassified | unclassified | unclassified | 28,8 | 2,712931993 | 0,000769328 | 4,43E-05 | 32 - | 0,000721583 - | 30 - | 0,000839993 - | 30 - | 0,00079852 - | 28 - | 0,000756293 - | 24 - | 0,000730253 - |
| 894 | 1 | 0.2um_NOKCPHJN_922823 | Bacteria | Proteobacteri | Gammaprote | Cellvibrionales | Cellvibrionace | Saccharophag | Saccharophag | poorly charact | Function unkn | 0ZVFM | outer membra | 203122.Sde_ | 1 | 0 | 2,08E-05 | 1,97E-06 | 1 - | 1,74E-05 - | 1 - | 2,16E-05 - | 1 - | 2,05E-05 - | 1 - | 2,08E-05 - | 1 - | 2,35E-05 - |
| 895 | 1 | 0.2um_NOKCPHJN_69946 | Bacteria | Proteobacteri | Gammaprote | Cellvibrionales | Porticoccacea | unclassified | Gammaprote | metabolism | Inorganic ion t | 0QJ8I | Na( ) H( ) anti | 314287.GB22 | 1,8 | 0,4 | 5,01E-05 | 1,39E-05 | 1 - | 2,29E-05 - | 2 - | 5,69E-05 - | 2 - | 5,41E-05 - | 2 - | 5,49E-05 - | 2 - | 6,18E-05 - |
| 896 | 1 | 3um_FIJJAFBF_410103 | Bacteria | Proteobacteri | Gammaprote | Cellvibrionales | Porticoccacea | unclassified | gamma prote | cellular proces | Posttranslatio | 08V1T | peptidylprolyl | 314287.GB22 | 0,6 | 0,8 | 3,17E-05 | 4,39E-05 | 1 - | 4,67E-05 - | 0 - | 0 - | 0 - | 0 - | 2 - | 0,000111775 - | 0 - | 0 - |
| 897 | 1 | 0.2um_NOKCPHJN_92230 | Bacteria | Bacteroidetes | Flavobacteriia | Flavobacterial | Flavobacteria | Arenibacter | Arenibacter sp | cellular proces | Intracellular tr | COG0653 | Part of the Se | 313603.FB217 | 0,8 | 0,748331477 | 9,51E-06 | 9,15E-06 | 1 - | 1,03E-05 - | 1 - | 1,27E-05 - | 0 - | 0 - | 2 - | 2,46E-05 - | 0 - | 0 - |
| 898 | 2 | 0.2um_NOKCPHJN_195280;3um_FIJJAFBF_63011 | Bacteria | Proteobacteri | Alphaproteob | Rhodobactera | Rhodobactera | unclassified | Rhodobactera | unclassified | unclassified | unclassified | unclassified | unclassified | 0,2 | 0,4 | 1,15E-05 | 2,30E-05 | 1 - | 5,74E-05 - | 0 - | 0 - | 0 - | 0 - | 0 - | 0 - | 0 - | 0 - |
| 899 | 1 | 0.2um_NOKCPHJN_958796 | Bacteria | Proteobacteri | Alphaproteob | Pelagibacteral | Pelagibactera | Candidatus Pe | Candidatus Pe | poorly charact | Function unkn | 02B70 | NA | 439493.PB721 | 0,8 | 0,4 | 5,91E-05 | 2,98E-05 | 0 - | 0 - | 1 - | 7,38E-05 - | 1 - | 7,02E-05 - | 1 - | 7,12E-05 - | 1 - | 8,02E-05 - |
| 900 | 1 | 3um_FIJJAFBF_189664 | Bacteria | Bacteroidetes | Flavobacteriia | Flavobacterial | Flavobacteria | Polaribacter | Polaribacter s | information st | Translation, ri | 0PBIW | 50S ribosomal | 313598.MED1 | 1,2 | 0,4 | 0,000249973 | 5,54E-05 | 2 - | 0,000358677 - | 1 - | 0,000222686 - | 1 - | 0,000211691 - | 1 - | 0,000214818 - | 1 - | 0,000241991 - |
| 901 | 1 | 0.2um_NOKCPHJN_628011 | Bacteria | Proteobacteri | Gammaprote | Oceanospirilla | Saccharospirill | Reinekea | Reinekea blan | unclassified | unclassified | unclassified | unclassified | unclassified | 8 | 1,788854382 | 0,000153905 | 4,20E-05 | 8 - | 0,000128242 - | 6 - | 0,000119429 - | 6 - | 0,000113533 - | 10 - | 0,000192016 - | 10 - | 0,000216305 - |
| 902 | 1 | 0.2um_NOKCPHJN_813912 | Bacteria | Proteobacteri | Alphaproteob | Rhodobactera | Rhodobactera | Labrenzia | unclassified | metabolism | Amino acid tra | 16QXY | (ABC) transpo | 384765.SIAM | 0,6 | 0,8 | 3,82E-05 | 5,18E-05 | 0 - | 0 - | 1 - | 6,01E-05 - | 0 - | 0 - | 0 - | 0 - | 2 - | 0,000130696 - |
| 903 | 1 | 0.2um_NOKCPHJN_719622 | Bacteria | Proteobacteri | Alphaproteob | Rhizobiales | unclassified | unclassified | Rhizobiales ba | cellular proces | Cell wall/mem | 7257 | NA | 335992.SAR11 | 0,2 | 0,4 | 1,15E-05 | 2,30E-05 | 0 - | 0 - | 1 - | 5,75E-05 - | 0 - | 0 - | 0 - | 0 - | 0 - | 0 - |
| 904 | 1 | 3um_FIJJAFBF_569309 | Bacteria | unclassified | unclassified | unclassified | unclassified | unclassified | uncultured ba | cellular proces | Intracellular tr | 0P8JN | The central su | 1046627.BZA | 0,4 | 0,489897949 | 1,14E-05 | 1,41E-05 | 1 - | 2,54E-05 - | 1 - | 3,16E-05 - | 0 - | 0 - | 0 - | 0 - | 0 - | 0 - |
| 905 | 1 | 0.2um_NOKCPHJN_1096866 | Bacteria | Proteobacteri | Gammaprote | Pseudomonad | Pseudomonad | Pseudomonas | various | metabolism | Energy produc | COG0055 | Produces ATP | 717774.Marm | 0,4 | 0,8 | 1,45E-05 | 2,91E-05 | 0 - | 0 - | 0 - | 0 - | 2 - | 7,26E-05 - | 0 - | 0 - | 0 - | 0 - |
| 906 | 1 | 3um_FIJJAFBF_350725 | Bacteria | Bacteroidetes | Flavobacteriia | Flavobacterial | Crocinitomica | unclassified | Crocinitomica | unclassified | unclassified | unclassified | unclassified | unclassified | 2,4 | 0,489897949 | 0,000222059 | 4,79E-05 | 2 - | 0,000155104 - | 2 - | 0,000192593 - | 3 - | 0,000274626 - | 3 - | 0,000278682 - | 2 - | 0,00020929 - |
| 907 | 1 | 0.2um_NOKCPHJN_46820 | Bacteria | Proteobacteri | Alphaproteob | Rhodobactera | Rhodobactera | Roseovarius | Roseovarius n | unclassified | unclassified | unclassified | unclassified | unclassified | 2 | 0,632455532 | 0,000215931 | 5,71E-05 | 3 - | 0,000275464 - | 2 - | 0,00022803 - | 1 - | 0,000108386 - | 2 - | 0,000219973 - | 2 - | 0,000247799 - |
| 908 | 1 | 3um_FIJJAFBF_143433 | Bacteria | Bacteroidetes | Cytophagia | Cytophagales | Flammeovirga | unclassified | Flammeovirga | information st | Translation, ri | 0FXD7 | This protein is | 504472.Slin_1 | 4 | 1,095445115 | 0,000458075 | 8,25E-05 | 6 - | 0,000588599 - | 4 - | 0,000487244 - | 3 - | 0,00034739 - | 4 - | 0,000470028 - | 3 - | 0,000397114 - |
| 909 | 1 | 0.2um_NOKCPHJN_640148 | Eukaryota | unclassified | Pelagophycea | Pelagomonad | unclassified | Aureococcus | Aureococcus | cellular proces | Intracellular tr | KOG3342 | Catalytic comp | 2850.Phatr51 | 1,8 | 0,4 | 0,000101888 | 2,38E-05 | 2 - | 9,53E-05 - | 1 - | 5,91E-05 - | 2 - | 0,000112433 - | 2 - | 0,000114094 - | 2 - | 0,000128527 - |
| 910 | 1 | 0.2um_NOKCPHJN_637138 | Bacteria | Proteobacteri | Alphaproteob | Pelagibacteral | Pelagibactera | Candidatus Pe | Candidatus Pe | cellular proces | Cell wall/mem | 7257 | NA | 335992.SAR11 | 1 | 0 | 3,74E-05 | 3,55E-06 | 1 - | 3,14E-05 - | 1 - | 3,89E-05 - | 1 - | 3,70E-05 - | 1 - | 3,76E-05 - | 1 - | 4,23E-05 - |
| 911 | 1 | 0.2um_NOKCPHJN_740862 | Eukaryota | Arthropoda | Insecta | Hymenoptera | Apidae | Bombus | Bombus impa | information st | Replication, re | 0V9WU | polymerase (D | 12957.ACEP_ | 0,4 | 0,489897949 | 2,15E-05 | 2,63E-05 | 0 - | 0 - | 0 - | 0 - | 1 - | 5,33E-05 - | 1 - | 5,41E-05 - | 0 - | 0 - |
| 912 | 1 | 3um_FIJJAFBF_332470 | Bacteria | Bacteroidetes | Flavobacteriia | Flavobacterial | Flavobacteria | Polaribacter | Polaribacter in | metabolism | Carbohydrate | 08IUS | Transketolase | 487796.Flav2 | 4,6 | 1,2 | 0,000194629 | 3,06E-05 | 7 - | 0,00025345 - | 4 - | 0,000179835 - | 4 - | 0,000170955 - | 4 - | 0,000173481 - | 4 - | 0,000195425 - |
| 913 | 1 | 0.2um_NOKCPHJN_397290 | Eukaryota | unclassified | Pelagophycea | Pelagomonad | unclassified | Aureococcus | Aureococcus | unclassified | unclassified | unclassified | unclassified | unclassified | 0,4 | 0,489897949 | 1,96E-05 | 2,43E-05 | 1 - | 4,38E-05 - | 1 - | 5,44E-05 - | 0 - | 0 - | 0 - | 0 - | 0 - | 0 - |
| 914 | 1 | 3um_FIJJAFBF_39810 | Bacteria | Proteobacteri | Gammaprote | Oceanospirilla | Oceanospirilla | Marinomonas | Marinomonas | poorly charact | Function unkn | 0QP3H | UPF0317 prot | 225937.HP15_ | 0,4 | 0,8 | 1,72E-05 | 3,44E-05 | 2 - | 8,60E-05 - | 0 - | 0 - | 0 - | 0 - | 0 - | 0 - | 0 - | 0 - |
| 915 | 1 | 0.2um_NOKCPHJN_567095 | Bacteria | Bacteroidetes | Flavobacteriia | Flavobacterial | Flavobacteria | Polaribacter | Polaribacter r | information st | Translation, ri | 05CEJ | Catalyzes the | 391603.FBAL | 2,2 | 0,4 | 4,28E-05 | 8,69E-06 | 2 - | 3,26E-05 - | 2 - | 4,04E-05 - | 2 - | 3,84E-05 - | 3 - | 5,85E-05 - | 2 - | 4,39E-05 - |
| 916 | 1 | 0.2um_NOKCPHJN_45477 | Bacteria | Proteobacteri | Gammaprote | Oceanospirilla | Saccharospirill | Reinekea | Reinekea fors | unclassified | unclassified | unclassified | unclassified | unclassified | 1 | 0 | 1,95E-05 | 1,85E-06 | 1 - | 1,63E-05 - | 1 - | 2,02E-05 - | 1 - | 1,92E-05 - | 1 - | 1,95E-05 - | 1 - | 2,20E-05 - |
| 917 | 1 | 0.2um_NOKCPHJN_1017780 | Eukaryota | unclassified | Labyrinthulom | unclassified | Thraustochytr | Aurantiochytr | Aurantiochytr | cellular proces | Signal transdu | 0XNPH | protein serine | 67593.Physo1 | 0,2 | 0,4 | 5,60E-06 | 1,12E-05 | 1 - | 2,80E-05 - | 0 - | 0 - | 0 - | 0 - | 0 - | 0 - | 0 - | 0 - |
| 918 | 1 | 3um_FIJJAFBF_199186 | Bacteria | Actinobacteria | Actinobacteria | unclassified | unclassified | unclassified | actinobacteri | metabolism | Energy produc | COG0056 | Produces ATP | 281090.Lxx07 | 0,6 | 0,489897949 | 1,52E-05 | 1,25E-05 | 0 - | 0 - | 1 - | 2,62E-05 - | 1 - | 2,49E-05 - | 1 - | 2,52E-05 - | 0 - | 0 - |
| 919 | 1 | 3um_FIJJAFBF_211173 | Bacteria | Proteobacteri | Betaproteoba | unclassified | unclassified | unclassified | Betaproteoba | information st | Translation, ri | COG0256 | This is one of | 596154.Alide2 | 2 | 0 | 0,000226491 | 2,15E-05 | 2 - | 0,000189714 - | 2 - | 0,000235568 - | 2 - | 0,000223938 - | 2 - | 0,000227245 - | 2 - | 0,000255991 - |
| 920 | 1 | 3um_FIJJAFBF_09522 | Bacteria | Actinobacteria | Actinobacteria | Micrococcales | Microbacteria | Candidatus Ac | Candidatus Ac | unclassified | unclassified | unclassified | unclassified | unclassified | 0,2 | 0,4 | 9,61E-06 | 1,92E-05 | 0 - | 0 - | 0 - | 0 - | 1 - | 4,80E-05 - | 0 - | 0 - | 0 - | 0 - |
| 921 | 1 | 3um_FIJJAFBF_421411 | Bacteria | Bacteroidetes | unclassified | unclassified | unclassified | unclassified | unidentified e | unclassified | unclassified | unclassified | unclassified | unclassified | 0,6 | 0,489897949 | 8,08E-05 | 6,72E-05 | 1 - | 0,000112526 - | 1 - | 0,000139724 - | 0 - | 0 - | 0 - | 0 - | 1 - | 0,000151838 - |
| 922 | 1 | 0.2um_NOKCPHJN_611594 | Bacteria | Proteobacteri | Alphaproteob | Pelagibacteral | Pelagibactera | Candidatus Pe | Candidatus Pe | cellular proces | Cell wall/mem | 7257 | NA | 335992.SAR11 | 0,2 | 0,4 | 8,59E-06 | 1,72E-05 | 0 - | 0 - | 0 - | 0 - | 0 - | 0 - | 1 - | 4,30E-05 - | 0 - | 0 - |
| 923 | 1 | 0.2um_NOKCPHJN_241601 | Bacteria | Proteobacteri | Alphaproteob | Pelagibacteral | Pelagibactera | Candidatus Pe | Candidatus Pe | information st | Translation, ri | COG0050 | This protein p | 1002672.SAR1 | 8,4 | 0,8 | 0,000289716 | 3,18E-05 | 9 - | 0,000260856 - | 9 - | 0,000323907 - | 9 - | 0,000307914 - | 7 - | 0,000243026 - | 8 - | 0,000312878 - |
| 924 | 1 | 0.2um_NOKCPHJN_562681 | Bacteria | Proteobacteri | Alphaproteob | Pelagibacteral | Pelagibactera | Candidatus Pe | Candidatus Pe | unclassified | unclassified | unclassified | unclassified | unclassified | 1,4 | 0,489897949 | 3,12E-05 | 1,17E-05 | 1 - | 1,86E-05 - | 2 - | 4,61E-05 - | 1 - | 2,19E-05 - | 2 - | 4,45E-05 - | 1 - | 2,51E-05 - |
| 925 | 2 | 0.2um_NOKCPHJN_127082;0.2um_NOKCPHJN_4 | Bacteria | Proteobacteri | Gammaprote | Cellvibrionales | Porticoccacea | unclassified | gamma prote | cellular proces | Intracellular tr | 175GA | One of the pro | 314287.GB22 | 0,2 | 0,4 | 1,36E-05 | 2,72E-05 | 1 - | 6,79E-05 - | 0 - | 0 - | 0 - | 0 - | 0 - | 0 - | 0 - | 0 - |
| 926 | 1 | 3um_FIJJAFBF_690641 | Bacteria | Bacteroidetes | Cytophagia | Cytophagales | unclassified | unclassified | various | unclassified | unclassified | unclassified | unclassified | unclassified | 0,2 | 0,4 | 1,07E-05 | 2,13E-05 | 0 - | 0 - | 0 - | 0 - | 1 - | 5,33E-05 - | 0 - | 0 - | 0 - | 0 - |
| 927 | 1 | 3um_FIJJAFBF_740443 | Bacteria | Proteobacteri | Alphaproteob | Pelagibacteral | Pelagibactera | Candidatus Pe | Candidatus Pe | information st | Transcription | 16PMT | DNA-depende | 439493.PB721 | 5,6 | 0,8 | 0,00022289 | 2,87E-05 | 6 - | 0,000201363 - | 5 - | 0,000208361 - | 7 - | 0,000277303 - | 5 - | 0,000200999 - | 5 - | 0,000226425 - |
| 928 | 1 | 3um_FIJJAFBF_285270 | Bacteria | Bacteroidetes | Cytophagia | Cytophagales | Flammeovirga | unclassified | Flammeovirga | unclassified | unclassified | unclassified | unclassified | unclassified | 0,8 | 0,4 | 5,56E-05 | 2,86E-05 | 1 - | 5,83E-05 - | 1 - | 7,23E-05 - | 1 - | 6,88E-05 - | 0 - | 0 - | 1 - | 7,86E-05 - |
| 929 | 1 | 3um_FIJJAFBF_30607 | Bacteria | Proteobacteri | Alphaproteob | Rhodobactera | Rhodobactera | Ruegeria | Ruegeria kand | unclassified | unclassified | unclassified | unclassified | unclassified | 1,4 | 1,019803903 | 8,76E-05 | 5,75E-05 | 3 - | 0,000163967 - | 2 - | 0,000135732 - | 1 - | 6,45E-05 - | 0 - | 0 - | 1 - | 7,37E-05 - |
| 930 | 1 | 0.2um_NOKCPHJN_869254 | unclassified | unclassified | unclassified | unclassified | unclassified | unclassified | unclassified | unclassified | unclassified | unclassified | unclassified | unclassified | 0,2 | 0,4 | 3,43E-05 | 6,86E-05 | 0 - | 0 - | 0 - | 0 - | 1 - | 0,000171496 - | 0 - | 0 - | 0 - | 0 - |
| 931 | 1 | 3um_FIJJAFBF_122438 | Bacteria | Bacteroidetes | Flavobacteriia | Flavobacterial | Flavobacteria | Wenyingzhua | Wenyingzhua | unclassified | unclassified | unclassified | unclassified | unclassified | 0,4 | 0,8 | 9,35E-06 | 1,87E-05 | 0 - | 0 - | 2 - | 4,67E-05 - | 0 - | 0 - | 0 - | 0 - | 0 - | 0 - |
| 932 | 1 | 3um_FIJJAFBF_580261 | Bacteria | Proteobacteri | Gammaprote | Pseudomonad | Pseudomonad | Pseudomonas | Pseudomonas | poorly charact | Function unkn | 17B58 | Inherit from b | 471874.PROS | 5,4 | 0,489897949 | 0,000220815 | 2,74E-05 | 5 - | 0,000171309 - | 5 - | 0,000212715 - | 6 - | 0,000242655 - | 6 - | 0,000246239 - | 5 - | 0,000231156 - |
| 933 | 2 | 0.2um_NOKCPHJN_71668;3um_FIJJAFBF_44729 | Bacteria | Bacteroidetes | Flavobacteriia | Flavobacterial | Flavobacteria | Polaribacter | various | unclassified | unclassified | unclassified | unclassified | unclassified | 0,4 | 0,489897949 | 1,04E-05 | 1,28E-05 | 1 - | 2,31E-05 - | 1 - | 2,87E-05 - | 0 - | 0 - | 0 - | 0 - | 0 - | 0 - |
| 934 | 3 | 0.2um_NOKCPHJN_232408;3um_FIJJAFBF_52151 | Bacteria | Bacteroidetes | Flavobacteriia | Flavobacterial | Flavobacteria | various | various | unclassified | unclassified | unclassified | unclassified | unclassified | 0,6 | 0,489897949 | 3,95E-05 | 3,26E-05 | 1 - | 5,77E-05 - | 1 - | 7,16E-05 - | 1 - | 6,81E-05 - | 0 - | 0 - | 0 - | 0 - |
| 935 | 1 | 0.2um_NOKCPHJN_260112 | Bacteria | Proteobacteri | Alphaproteob | Rhodobactera | Rhodobactera | Pseudovibrio | Pseudovibrio | unclassified | unclassified | unclassified | unclassified | unclassified | 0,4 | 0,489897949 | 4,14E-05 | 5,10E-05 | 1 - | 9,41E-05 - | 0 - | 0 - | 0 - | 0 - | 1 - | 0,000112691 - | 0 - | 0 - |
| 936 | 1 | 3um_FIJJAFBF_56092 | Eukaryota | Bacillariophyt | Bacillariophyc | Naviculales | Naviculaceae | Fistulifera | Fistulifera sola | metabolism | Energy produc | COG0055 | Produces ATP | 2850.Phatr54 | 2 | 0 | 0,000108322 | 1,03E-05 | 2 - | 9,07E-05 - | 2 - | 0,000112663 - | 2 - | 0,000107101 - | 2 - | 0,000108682 - | 2 - | 0,00012243 - |
| 937 | 1 | 0.2um_NOKCPHJN_06879 | Bacteria | Bacteroidetes | Cytophagia | Cytophagales | Cytophagacea | Ekhidna | Ekhidna lutea | information st | Translation, ri | 05C64 | Binds directly | 880070.Cycm | 2,4 | 1,019803903 | 0,000144657 | 6,52E-05 | 1 - | 4,95E-05 - | 4 - | 0,000245722 - | 2 - | 0,000116795 - | 3 - | 0,00017778 - | 2 - | 0,000133512 - |
| 938 | 1 | 0.2um_NOKCPHJN_852603 | Eukaryota | Bacillariophyt | Bacillariophyc | Naviculales | Naviculaceae | Fistulifera | Fistulifera sola | unclassified | unclassified | unclassified | unclassified | unclassified | 0,4 | 0,489897949 | 3,02E-05 | 3,72E-05 | 1 - | 6,87E-05 - | 0 - | 0 - | 0 - | 0 - | 1 - | 8,23E-05 - | 0 - | 0 - |
| 939 | 1 | 0.2um_NOKCPHJN_44404 | Bacteria | Proteobacteri | Gammaprote | Oceanospirilla | Oceanospirilla | Motiliproteus | Motiliproteus | information st | Translation, ri | 16QAF | thus facilitatin | 247633.GP21 | 0,6 | 0,489897949 | 1,57E-05 | 1,29E-05 | 0 - | 0 - | 1 - | 2,57E-05 - | 0 - | 0 - | 1 - | 2,48E-05 - | 1 - | 2,80E-05 - |
| 940 | 1 | 0.2um_NOKCPHJN_396121 | unclassified | unclassified | unclassified | unclassified | unclassified | unclassified | unclassified | unclassified | unclassified | unclassified | unclassified | unclassified | 1,8 | 0,979795897 | 5,41E-05 | 3,10E-05 | 2 - | 5,16E-05 - | 3 - | 9,61E-05 - | 2 - | 6,09E-05 - | 2 - | 6,18E-05 - | 0 - | 0 - |
| 941 | 1 | 3um_FIJJAFBF_133470 | Bacteria | Actinobacteria | Actinobacteria | Micrococcales | Microbacteria | various | various | unclassified | unclassified | unclassified | unclassified | unclassified | 2,2 | 0,979795897 | 0,000220804 | 7,67E-05 | 4 - | 0,000347808 - | 1 - | 0,000107969 - | 2 - | 0,000205276 - | 2 - | 0,000208308 - | 2 - | 0,000234658 - |
| 942 | 1 | 3um_FIJJAFBF_693879 | Bacteria | Bacteroidetes | Flavobacteriia | Flavobacterial | Flavobacteria | Formosa | Formosa sp. H | metabolism | Energy produc | COG0437 | 4fe-4S ferredo | 983544.Lacal_ | 5 | 0,632455532 | 6,32E-05 | 2,90E-06 | 6 - | 6,42E-05 - | 5 - | 6,65E-05 - | 5 - | 6,32E-05 - | 5 - | 6,41E-05 - | 4 - | 5,78E-05 - |
| 943 | 1 | 0.2um_NOKCPHJN_309750 | Bacteria | Proteobacteri | Alphaproteob | Pelagibacteral | Pelagibactera | Candidatus Pe | Candidatus Pe | poorly charact | Function unkn | 02CU1 | NA | 335992.SAR11 | 0,8 | 0,748331477 | 2,12E-05 | 1,88E-05 | 2 - | 4,85E-05 - | 0 - | 0 - | 1 - | 2,86E-05 - | 1 - | 2,91E-05 - | 0 - | 0 - |
| 944 | 1 | 3um_FIJJAFBF_533992 | Bacteria | Bacteroidetes | Flavobacteriia | Flavobacterial | Crocinitomica | unclassified | Crocinitomica | poorly charact | Function unkn | 0P884 | membrane | 926562.Oweh | 3,2 | 0,748331477 | 0,000124556 | 2,83E-05 | 3 - | 9,87E-05 - | 3 - | 0,000122509 - | 4 - | 0,00015528 - | 4 - | 0,000157574 - | 2 - | 8,88E-05 - |
| 945 | 1 | 0.2um_NOKCPHJN_315471 | unclassified | unclassified | unclassified | unclassified | unclassified | unclassified | unclassified | unclassified | unclassified | unclassified | unclassified | unclassified | 2,4 | 1,356465997 | 7,37E-05 | 4,32E-05 | 3 - | 7,95E-05 - | 4 - | 0,000131657 - | 3 - | 9,39E-05 - | 2 - | 6,35E-05 - | 0 - | 0 - |
| 946 | 1 | 3um_FIJJAFBF_269250 | Bacteria | Bacteroidetes | Flavobacteriia | Flavobacterial | Flavobacteria | Polaribacter | Polaribacter s | unclassified | unclassified | unclassified | unclassified | unclassified | 0,2 | 0,4 | 1,37E-05 | 2,74E-05 | 0 - | 0 - | 1 - | 6,85E-05 - | 0 - | 0 - | 0 - | 0 - | 0 - | 0 - |
| 947 | 1 | 0.2um_NOKCPHJN_978374 | Archaea | Euryarchaeota | Methanobact | Methanobact | Methanobact | Methanobrevi | Methanobrevi | metabolism | Amino acid tra | arCOG02555 | adhesin-like p | 868132.Metb | 0,4 | 0,489897949 | 9,02E-06 | 1,11E-05 | 0 - | 0 - | 0 - | 0 - | 1 - | 2,24E-05 - | 1 - | 2,27E-05 - | 0 - | 0 - |
| 948 | 1 | 3um_FIJJAFBF_325785 | Bacteria | Proteobacteri | Gammaprote | Cellvibrionales | Porticoccacea | unclassified | Gammaprote | unclassified | unclassified | unclassified | unclassified | unclassified | 1 | 0 | 0,000141265 | 1,34E-05 | 1 - | 0,000118327 - | 1 - | 0,000146927 - | 1 - | 0,000139672 - | 1 - | 0,000141735 - | 1 - | 0,000159664 - |
| 949 | 1 | 0.2um_NOKCPHJN_133069 | Bacteria | Bacteroidetes | Cytophagia | Cytophagales | Flammeovirga | Marivirga | Marivirga seri | metabolism | Inorganic ion t | 05C27 | TonB-depende | 643867.Ftrac_ | 0,8 | 1,166190379 | 9,11E-06 | 1,28E-05 | 3 - | 3,27E-05 - | 0 - | 0 - | 1 - | 1,29E-05 - | 0 - | 0 - | 0 - | 0 - |
| 950 | 1 | 0.2um_NOKCPHJN_66509 | Bacteria | Bacteroidetes | unclassified | unclassified | unclassified | unclassified | Bacteroidetes | poorly charact | Function unkn | 08QKU | NA | 926562.Oweh | 1,6 | 1,019803903 | 1,61E-05 | 9,61E-06 | 3 - | 2,57E-05 - | 2 - | 2,13E-05 - | 0 - | 0 - | 1 - | 1,03E-05 - | 2 - | 2,31E-05 - |
| 951 | 1 | 3um_FIJJAFBF_683762 | Bacteria | Proteobacteri | Gammaprote | unclassified | unclassified | unclassified | Gammaprote | cellular proces | Intracellular tr | 0QITE | Outer membr | 207949.RED6 | 0,4 | 0,489897949 | 1,08E-05 | 1,33E-05 | 1 - | 2,41E-05 - | 1 - | 2,99E-05 - | 0 - | 0 - | 0 - | 0 - | 0 - | 0 - |
| 952 | 1 | 3um_FIJJAFBF_745913 | Bacteria | Proteobacteri | Gammaprote | Cellvibrionales | Halieaceae | Halioglobus | Halioglobus pa | unclassified | unclassified | unclassified | unclassified | unclassified | 1 | 0 | 0,000131757 | 1,25E-05 | 1 - | 0,000110362 - | 1 - | 0,000137037 - | 1 - | 0,000130271 - | 1 - | 0,000132195 - | 1 - | 0,000148918 - |
| 953 | 1 | 0.2um_NOKCPHJN_912086 | Viruses | unclassified | unclassified | Caudovirales | Podoviridae | unclassified | Cellulophaga | unclassified | unclassified | unclassified | unclassified | unclassified | 1 | 0 | 2,77E-05 | 2,63E-06 | 1 - | 2,32E-05 - | 1 - | 2,88E-05 - | 1 - | 2,74E-05 - | 1 - | 2,78E-05 - | 1 - | 3,14E-05 - |
| 954 | 1 | 0.2um_NOKCPHJN_175879 | Eukaryota | unclassified | Oomycetes | Peronosporale | Peronosporac | Phytophthora | Phytophthora | unclassified | unclassified | unclassified | unclassified | unclassified | 0,4 | 0,489897949 | 9,34E-07 | 1,14E-06 | 0 - | 0 - | 0 - | 0 - | 1 - | 2,32E-06 - | 1 - | 2,35E-06 - | 0 - | 0 - |
| 955 | 1 | 0.2um_NOKCPHJN_216411 | Eukaryota | unclassified | Dinophyceae | Suessiales | Symbiodiniac | Symbiodinium | Symbiodinium | metabolism | Secondary me | KOG0054 | ATP-binding c | 5786.XP_0032 | 0,6 | 0,489897949 | 4,91E-06 | 4,08E-06 | 1 - | 6,84E-06 - | 1 - | 8,49E-06 - | 0 - | 0 - | 0 - | 0 - | 1 - | 9,22E-06 - |
| 956 | 2 | 0.2um_NOKCPHJN_298096;3um_FIJJAFBF_35365 | Bacteria | Proteobacteri | Alphaproteob | various | various | various | various | information st | Transcription | 05VEQ | Cold shock pro | 367336.OM22 | 1 | 0 | 0,00020151 | 1,91E-05 | 1 - | 0,000168789 - | 1 - | 0,000209587 - | 1 - | 0,000199239 - | 1 - | 0,000202181 - | 1 - | 0,000227757 - |
| 957 | 1 | 0.2um_NOKCPHJN_38981 | Bacteria | Bacteroidetes | Flavobacteriia | Flavobacterial | Flavobacteria | Formosa | Formosa sp. H | metabolism | Carbohydrate | 099Q3 | phosphohexo | 376686.Fjoh_ | 0,6 | 0,489897949 | 2,64E-05 | 2,16E-05 | 0 - | 0 - | 1 - | 4,35E-05 - | 1 - | 4,13E-05 - | 0 - | 0 - | 1 - | 4,72E-05 - |
| 958 | 1 | 3um_FIJJAFBF_729294 | Bacteria | Proteobacteri | Gammaprote | Cellvibrionales | unclassified | unclassified | Cellvibrionales | information st | Translation, ri | 0QTKX | Seems to be t | 314287.GB22 | 5,6 | 1,496662955 | 0,00059677 | 0,000115153 | 8 - | 0,000728741 - | 6 - | 0,000678661 - | 4 - | 0,000430102 - | 6 - | 0,000654682 - | 4 - | 0,000491665 - |
| 959 | 1 | 0.2um_NOKCPHJN_54710 | Eukaryota | Streptophyta | unclassified | Ericales | Primulaceae | Androsace | Androsace lax | cellular proces | Posttranslatio | 0IEKR | One of the co | 59689.fgenes | 0,8 | 0,4 | 2,38E-05 | 1,22E-05 | 1 - | 2,49E-05 - | 1 - | 3,09E-05 - | 1 - | 2,94E-05 - | 0 - | 0 - | 1 - | 3,36E-05 - |
| 960 | 1 | 3um_FIJJAFBF_04413 | Bacteria | Proteobacteri | Alphaproteob | Rickettsiales | unclassified | unclassified | Rickettsiales b | unclassified | unclassified | unclassified | unclassified | unclassified | 2,6 | 0,489897949 | 9,06E-05 | 1,93E-05 | 2 - | 5,83E-05 - | 3 - | 0,000108517 - | 3 - | 0,000103159 - | 3 - | 0,000104683 - | 2 - | 7,86E-05 - |
| 961 | 1 | 0.2um_NOKCPHJN_857829 | Bacteria | Proteobacteri | Alphaproteob | Pelagibacteral | Pelagibactera | Candidatus Pe | Candidatus Pe | unclassified | unclassified | unclassified | unclassified | unclassified | 0,6 | 0,489897949 | 2,51E-05 | 2,07E-05 | 1 - | 3,67E-05 - | 1 - | 4,55E-05 - | 1 - | 4,33E-05 - | 0 - | 0 - | 0 - | 0 - |
| 962 | 1 | 0.2um_NOKCPHJN_93255 | Bacteria | Proteobacteri | Alphaproteob | Pelagibacteral | Pelagibactera | Candidatus Pe | Candidatus Pe | metabolism | Energy produc | COG0711 | F(1)F(0) ATP s | 335992.SAR11 | 1,2 | 0,748331477 | 8,59E-05 | 5,74E-05 | 1 - | 5,86E-05 - | 1 - | 7,27E-05 - | 0 - | 0 - | 2 - | 0,000140289 - | 2 - | 0,000158035 - |
| 963 | 1 | 0.2um_NOKCPHJN_37852 | Bacteria | Proteobacteri | Alphaproteob | unclassified | unclassified | unclassified | Alphaproteob | unclassified | unclassified | unclassified | unclassified | unclassified | 1,2 | 0,979795897 | 4,82E-05 | 4,08E-05 | 1 - | 3,32E-05 - | 3 - | 0,000123571 - | 0 - | 0 - | 1 - | 3,97E-05 - | 1 - | 4,48E-05 - |
| 964 | 1 | 3um_FIJJAFBF_762327 | Bacteria | Proteobacteri | Alphaproteob | Rhodobactera | Rhodobactera | Planktomarina | Planktomarina | metabolism | Amino acid tra | 01R4A | Polyamine AB | 488538.SAR11 | 5 | 0,632455532 | 0,000188355 | 2,75E-05 | 5 - | 0,000158095 - | 4 - | 0,000157046 - | 6 - | 0,000223938 - | 5 - | 0,000189371 - | 5 - | 0,000213326 - |
| 965 | 1 | 0.2um_NOKCPHJN_275421 | Bacteria | Bacteroidetes | Flavobacteriia | Flavobacterial | Flavobacteria | Polaribacter | Polaribacter s | poorly charact | Function unkn | 05ERK | gliding motilit | 313594.PI23P | 0,6 | 0,489897949 | 1,54E-05 | 1,27E-05 | 1 - | 2,28E-05 - | 0 - | 0 - | 1 - | 2,69E-05 - | 1 - | 2,73E-05 - | 0 - | 0 - |
| 966 | 1 | 3um_FIJJAFBF_123938 | Bacteria | Actinobacteria | Actinobacteria | Micrococcales | Microbacteria | various | various | unclassified | unclassified | unclassified | unclassified | unclassified | 2 | 0 | 0,000326255 | 3,10E-05 | 2 - | 0,000273278 - | 2 - | 0,000339331 - | 2 - | 0,000322577 - | 2 - | 0,000327341 - | 2 - | 0,000368749 - |
| 967 | 1 | 0.2um_NOKCPHJN_726696 | Bacteria | Proteobacteri | Alphaproteob | Pelagibacteral | Pelagibactera | Candidatus Pe | Candidatus Pe | cellular proces | Cell wall/mem | COG2885 | Ompa motb d | 335992.SAR11 | 2 | 0 | 0,000172361 | 1,64E-05 | 2 - | 0,000144373 - | 2 - | 0,000179269 - | 2 - | 0,000170418 - | 2 - | 0,000172935 - | 2 - | 0,000194811 - |
| 968 | 1 | 3um_FIJJAFBF_284885 | Bacteria | Proteobacteri | Gammaprote | Oceanospirilla | Saccharospirill | Reinekea | Reinekea fors | metabolism | Amino acid tra | 05DXH | Alcohol dehyd | 177437.HRM2 | 1,8 | 0,748331477 | 4,18E-05 | 1,68E-05 | 2 - | 3,96E-05 - | 1 - | 2,46E-05 - | 2 - | 4,68E-05 - | 3 - | 7,12E-05 - | 1 - | 2,67E-05 - |
| 969 | 1 | 0.2um_NOKCPHJN_230029 | Bacteria | Proteobacteri | Gammaprote | Cellvibrionales | Porticoccacea | unclassified | Gammaprote | information st | Transcription | 05D27 | DNA-depende | 498211.CJA_0 | 2 | 0,632455532 | 1,89E-05 | 4,37E-06 | 3 - | 2,44E-05 - | 2 - | 2,02E-05 - | 2 - | 1,92E-05 - | 2 - | 1,95E-05 - | 1 - | 1,10E-05 - |
| 970 | 1 | 3um_FIJJAFBF_190431 | Bacteria | Bacteroidetes | Flavobacteriia | Flavobacterial | Flavobacteria | Aquimarina | Aquimarina sp | cellular proces | Cell wall/mem | 06B97 | phosphate-sel | 313603.FB217 | 2,8 | 0,979795897 | 9,41E-05 | 3,19E-05 | 3 - | 8,61E-05 - | 3 - | 0,000106889 - | 4 - | 0,000135482 - | 3 - | 0,000103112 - | 1 - | 3,87E-05 - |
| 971 | 1 | 3um_FIJJAFBF_20041 | Bacteria | Bacteroidetes | Flavobacteriia | Flavobacterial | Flavobacteria | Polaribacter | Polaribacter h | unclassified | unclassified | unclassified | unclassified | unclassified | 3,8 | 0,4 | 0,000315522 | 4,55E-05 | 4 - | 0,000278247 - | 4 - | 0,0003455 - | 4 - | 0,000328442 - | 3 - | 0,00024997 - | 4 - | 0,000375453 - |
| 972 | 1 | 0.2um_NOKCPHJN_28562 | Bacteria | Bacteroidetes | Flavobacteriia | unclassified | unclassified | unclassified | Flavobacteria | metabolism | Inorganic ion t | 05C27 | TonB-depende | 487796.Flav2 | 18,2 | 2,925747768 | 0,000233964 | 2,88E-05 | 21 - | 0,000228466 - | 18 - | 0,00024316 - | 22 - | 0,000282522 - | 15 - | 0,000195474 - | 15 - | 0,000220201 - |
| 973 | 1 | 3um_FIJJAFBF_695952 | Eukaryota | unclassified | Oomycetes | Peronosporale | Peronosporac | Phytophthora | various | unclassified | unclassified | unclassified | unclassified | unclassified | 2 | 0,894427191 | 7,85E-05 | 3,59E-05 | 3 - | 9,92E-05 - | 1 - | 4,11E-05 - | 1 - | 3,90E-05 - | 2 - | 7,92E-05 - | 3 - | 0,000133897 - |
| 974 | 1 | 3um_FIJJAFBF_602272 | Bacteria | Proteobacteri | Alphaproteob | Rhodobactera | Rhodobactera | Planktomarina | Planktomarina | cellular proces | Cell wall/mem | COG2913 | Part of the ou | 89187.ISM_16 | 1,6 | 0,489897949 | 0,000144955 | 4,10E-05 | 2 - | 0,000154063 - | 2 - | 0,0001913 - | 1 - | 9,09E-05 - | 2 - | 0,000184541 - | 1 - | 0,000103943 - |
| 975 | 1 | 3um_FIJJAFBF_69659 | Bacteria | Bacteroidetes | Flavobacteriia | unclassified | unclassified | unclassified | Flavobacteria | poorly charact | Function unkn | 05END | lipolytic prote | 487797.Flav3 | 2,2 | 0,748331477 | 6,35E-05 | 2,04E-05 | 3 - | 7,34E-05 - | 3 - | 9,12E-05 - | 2 - | 5,78E-05 - | 1 - | 2,93E-05 - | 2 - | 6,60E-05 - |
| 976 | 1 | 3um_FIJJAFBF_515657 | Bacteria | Proteobacteri | Alphaproteob | Rhodobactera | Rhodobactera | Ruegeria | Ruegeria sp. T | unclassified | unclassified | unclassified | unclassified | unclassified | 5 | 0 | 0,000218196 | 2,07E-05 | 5 - | 0,000182766 - | 5 - | 0,000226941 - | 5 - | 0,000215736 - | 5 - | 0,000218922 - | 5 - | 0,000246615 - |
| 977 | 1 | 3um_FIJJAFBF_264507 | Bacteria | Proteobacteri | Alphaproteob | Kiloniellales | Kiloniellaceae | Kiloniella | Kiloniella spor | metabolism | Energy produc | KOG3180 | Electron trans | 400682.PAC_1 | 1 | 0 | 4,41E-05 | 4,18E-06 | 1 - | 3,69E-05 - | 1 - | 4,58E-05 - | 1 - | 4,36E-05 - | 1 - | 4,42E-05 - | 1 - | 4,98E-05 - |
| 978 | 1 | 3um_FIJJAFBF_675680 | unclassified | unclassified | unclassified | unclassified | unclassified | unclassified | unclassified | unclassified | unclassified | unclassified | unclassified | unclassified | 1,8 | 0,748331477 | 4,35E-05 | 1,79E-05 | 2 - | 4,10E-05 - | 2 - | 5,09E-05 - | 1 - | 2,42E-05 - | 3 - | 7,37E-05 - | 1 - | 2,77E-05 - |
| 979 | 1 | 3um_FIJJAFBF_487948 | Bacteria | Bacteroidetes | Flavobacteriia | Flavobacterial | Flavobacteria | Formosa | Formosa sp. H | information st | Transcription | 05D27 | DNA-depende | 1046627.BZA | 1 | 0 | 3,09E-05 | 2,93E-06 | 1 - | 2,59E-05 - | 1 - | 3,21E-05 - | 1 - | 3,05E-05 - | 1 - | 3,10E-05 - | 1 - | 3,49E-05 - |
| 980 | 1 | 3um_FIJJAFBF_225010 | Bacteria | Bacteroidetes | Flavobacteriia | various | various | various | various | cellular proces | Intracellular tr | 09S7I | Preprotein tra | 487797.Flav3 | 1 | 0 | 0,000111404 | 1,06E-05 | 1 - | 9,33E-05 - | 1 - | 0,000115869 - | 1 - | 0,000110148 - | 1 - | 0,000111775 - | 1 - | 0,000125914 - |
| 981 | 1 | 0.2um_NOKCPHJN_1079624 | Bacteria | Proteobacteri | Alphaproteob | Pelagibacteral | Pelagibactera | unclassified | Pelagibactera | cellular proces | Cell wall/mem | 7257 | NA | 439493.PB721 | 2,4 | 0,489897949 | 9,90E-05 | 2,28E-05 | 2 - | 6,89E-05 - | 3 - | 0,000128395 - | 3 - | 0,000122056 - | 2 - | 8,26E-05 - | 2 - | 9,30E-05 - |
| 982 | 1 | 3um_FIJJAFBF_280306 | Bacteria | Proteobacteri | Betaproteoba | Rhodocyclales | Zoogloeaceae | Thauera | Thauera terpe | unclassified | unclassified | unclassified | unclassified | unclassified | 1,6 | 0,489897949 | 3,51E-05 | 9,91E-06 | 2 - | 3,73E-05 - | 2 - | 4,63E-05 - | 1 - | 2,20E-05 - | 2 - | 4,46E-05 - | 1 - | 2,51E-05 - |
| 983 | 1 | 0.2um_NOKCPHJN_405309 | Bacteria | Bacteroidetes | Flavobacteriia | Flavobacterial | Flavobacteria | Polaribacter | Polaribacter r | unclassified | unclassified | unclassified | unclassified | unclassified | 2,2 | 0,748331477 | 3,21E-05 | 1,11E-05 | 2 - | 2,46E-05 - | 1 - | 1,52E-05 - | 3 - | 4,35E-05 - | 3 - | 4,41E-05 - | 2 - | 3,31E-05 - |
| 984 | 1 | 0.2um_NOKCPHJN_373086 | Bacteria | Proteobacteri | Alphaproteob | Pelagibacteral | Pelagibactera | Candidatus Pe | Candidatus Pe | poorly charact | Function unkn | 0ZQPZ | Phage integra | 335992.SAR11 | 0,2 | 0,4 | 7,53E-06 | 1,51E-05 | 1 - | 3,76E-05 - | 0 - | 0 - | 0 - | 0 - | 0 - | 0 - | 0 - | 0 - |
| 985 | 1 | 0.2um_NOKCPHJN_28389 | Bacteria | Bacteroidetes | Flavobacteriia | Flavobacterial | Flavobacteria | unclassified | Flavobacteria | metabolism | Inorganic ion t | 05C27 | TonB-depende | 487796.Flav2 | 4 | 1,414213562 | 5,03E-05 | 1,48E-05 | 6 - | 6,50E-05 - | 5 - | 6,72E-05 - | 3 - | 3,83E-05 - | 4 - | 5,19E-05 - | 2 - | 2,92E-05 - |
| 986 | 1 | 3um_FIJJAFBF_169112 | Eukaryota | unclassified | Pelagophycea | Pelagomonad | unclassified | Pelagomonas | uncultured Pe | information st | Translation, ri | COG0087 | One of the pri | 489825.LYNG | 1 | 0 | 6,72E-05 | 6,38E-06 | 1 - | 5,63E-05 - | 1 - | 6,99E-05 - | 1 - | 6,64E-05 - | 1 - | 6,74E-05 - | 1 - | 7,59E-05 - |
| 987 | 1 | 3um_FIJJAFBF_142181 | Bacteria | Proteobacteri | Gammaprote | Cellvibrionales | Spongiibacter | Oceanicoccus | Oceanicoccus | metabolism | Energy produc | COG0055 | Produces ATP | 247639.MGP2 | 1,2 | 0,4 | 3,44E-05 | 7,62E-06 | 2 - | 4,94E-05 - | 1 - | 3,06E-05 - | 1 - | 2,91E-05 - | 1 - | 2,96E-05 - | 1 - | 3,33E-05 - |
| 988 | 1 | 0.2um_NOKCPHJN_117132 | Bacteria | Bacteroidetes | Flavobacteriia | Flavobacterial | Flavobacteria | Tenacibaculur | Tenacibaculur | metabolism | Energy produc | 05C6K | Malic enzyme | 313598.MED1 | 0,8 | 0,748331477 | 1,52E-05 | 1,41E-05 | 0 - | 0 - | 2 - | 3,76E-05 - | 0 - | 0 - | 1 - | 1,81E-05 - | 1 - | 2,04E-05 - |
| 989 | 1 | 3um_FIJJAFBF_273274 | Bacteria | Bacteroidetes | Flavobacteriia | Flavobacterial | Flavobacteria | Algibacter | Algibacter sp. | unclassified | unclassified | unclassified | unclassified | unclassified | 2,4 | 0,8 | 4,36E-05 | 1,50E-05 | 2 - | 3,04E-05 - | 2 - | 3,78E-05 - | 2 - | 3,59E-05 - | 4 - | 7,28E-05 - | 2 - | 4,10E-05 - |
| 990 | 1 | 3um_FIJJAFBF_237915 | Bacteria | Proteobacteri | Alphaproteob | Rhizobiales | unclassified | unclassified | Rhizobiales ba | cellular proces | Signal transdu | 0JNCB | NA | 4896.SPBPJ46 | 1 | 0 | 4,62E-06 | 4,38E-07 | 1 - | 3,87E-06 - | 1 - | 4,80E-06 - | 1 - | 4,56E-06 - | 1 - | 4,63E-06 - | 1 - | 5,22E-06 - |
| 991 | 1 | 3um_FIJJAFBF_520473 | Bacteria | Bacteroidetes | Flavobacteriia | Flavobacterial | Flavobacteria | Polaribacter | Polaribacter s | poorly charact | Function unkn | 0900R | pur-alpha bet | 313594.PI23P | 3,8 | 0,748331477 | 0,000405562 | 6,42E-05 | 5 - | 0,000451877 - | 3 - | 0,000336659 - | 4 - | 0,000426716 - | 3 - | 0,000324764 - | 4 - | 0,000487794 - |
| 992 | 1 | 3um_FIJJAFBF_370480 | Bacteria | Bacteroidetes | Flavobacteriia | Flavobacterial | Flavobacteria | Mesoflavibact | various | unclassified | unclassified | unclassified | unclassified | unclassified | 0,8 | 0,4 | 1,99E-05 | 1,01E-05 | 1 - | 2,16E-05 - | 1 - | 2,68E-05 - | 1 - | 2,55E-05 - | 1 - | 2,58E-05 - | 0 - | 0 - |
| 993 | 1 | 3um_FIJJAFBF_56140 | Bacteria | Bacteroidetes | Flavobacteriia | Flavobacterial | Flavobacteria | Polaribacter | Polaribacter s | metabolism | Energy produc | 0P7S4 | Produces ATP | 313598.MED1 | 3,4 | 0,489897949 | 0,000160116 | 1,56E-05 | 4 - | 0,000159412 - | 3 - | 0,000148457 - | 4 - | 0,00018817 - | 3 - | 0,000143212 - | 3 - | 0,000161328 - |
| 994 | 1 | 3um_FIJJAFBF_235465 | Bacteria | Proteobacteri | Alphaproteob | Rhodobactera | Rhodobactera | Sulfitobacter | Sulfitobacter s | unclassified | unclassified | unclassified | unclassified | unclassified | 3 | 1,673320053 | 6,84E-05 | 3,10E-05 | 6 - | 0,000120185 - | 1 - | 2,49E-05 - | 3 - | 7,09E-05 - | 3 - | 7,20E-05 - | 2 - | 5,41E-05 - |
| 995 | 1 | 3um_FIJJAFBF_517424 | Bacteria | Bacteroidetes | Flavobacteriia | Flavobacterial | Flavobacteria | Polaribacter | Polaribacter s | unclassified | unclassified | unclassified | unclassified | unclassified | 5,4 | 0,8 | 0,000592081 | 7,65E-05 | 6 - | 0,000555371 - | 4 - | 0,000459738 - | 6 - | 0,000655559 - | 6 - | 0,000665242 - | 5 - | 0,000624494 - |

| | | | | | | | | | | | | | | | | | | | | | | | | | | | |
|---|---|---|---|---|---|---|---|---|---|---|---|---|---|---|---|---|---|---|---|---|---|---|---|---|---|---|---|
| 996 | 1 | 3um_FIJAFBF_02132 | Bacteria | Proteobacteri | Alphaproteob | various | various | various | various | unclassified | unclassified | unclassified | unclassified | unclassified | 3,4 | 0,489897949 | 0,000379132 | 4,81E-05 | 4 - | 0,000376317 - | 4 - | 0,000467275 - | 3 - | 0,000333153 - | 3 - | 0,000338074 - | 3 - | 0,000380839 - |
| 997 | 1 | 3um_FIJAFBF_388942 | Bacteria | Bacteroidetes | Flavobacteriia | Flavobacterial | Flavobacteriac | Polaribacter | unclassified | unclassified | unclassified | unclassified | unclassified | unclassified | 1 | 0 | 7,92E-05 | 7,52E-06 | 1 - | 6,63E-05 - | 1 - | 8,24E-05 - | 1 - | 7,83E-05 - | 1 - | 7,95E-05 - | 1 - | 8,95E-05 - |
| 998 | 1 | 0.2um_NOKCPHJN_211698 | Bacteria | Bacteroidetes | Flavobacteriia | various | various | various | various | cellular proces | Posttranslatio | 09ADN | alkyl hydroper | 487797.Flav3( | 2 | 0,894427191 | 0,000125677 | 5,05E-05 | 3 - | 0,00016242 - | 2 - | 0,000134452 - | 1 - | 6,39E-05 - | 3 - | 0,000194552 - | 1 - | 7,31E-05 - |
| 999 | 1 | 0.2um_NOKCPHJN_627902 | unclassified | unclassified | unclassified | unclassified | unclassified | unclassified | unclassified | unclassified | unclassified | unclassified | unclassified | unclassified | 0,8 | 0,4 | 3,05E-05 | 1,55E-05 | 1 - | 3,30E-05 - | 1 - | 4,10E-05 - | 1 - | 3,89E-05 - | 1 - | 3,95E-05 - | 0 - | 0 - |
| 1000 | 1 | 0.2um_NOKCPHJN_12988 | Bacteria | Bacteroidetes | Flavobacteriia | Flavobacterial | Flavobacteriac | Formosa | Formosa sp. H | information st | Translation, ril | 0PA05 | 30S ribosomal | 983544.Lacal_ | 1 | 0,894427191 | 5,97E-05 | 5,17E-05 | 2 - | 0,00010387 - | 0 - | 0 - | 0 - | 0 - | 2 - | 0,000124419 - | 1 - | 7,01E-05 - |
| 1001 | 1 | 3um_FIJAFBF_717797 | Bacteria | Proteobacteri | Gammaproteo | Oceanospirilla | Oceanospirilla | Oceanospirillu | Oceanospirillu | information st | Transcription | 17K32 | Transcriptiona | 207954.MED9 | 0,6 | 0,8 | 6,93E-05 | 8,92E-05 | 2 - | 0,00021656 - | 0 - | 0 - | 0 - | 0 - | 1 - | 0,000129701 - | 0 - | 0 - |
| 1002 | 1 | 3um_FIJAFBF_557270 | Bacteria | Proteobacteri | Alphaproteob | unclassified | unclassified | unclassified | Alphaproteob | unclassified | unclassified | unclassified | unclassified | unclassified | 0,2 | 0,4 | 1,17E-05 | 2,33E-05 | 1 - | 5,83E-05 - | 0 - | 0 - | 0 - | 0 - | 0 - | 0 - | 0 - | 0 - |
| 1003 | 1 | 3um_FIJAFBF_581298 | Bacteria | Bacteroidetes | Flavobacteriia | Flavobacterial | Crocinitomica | unclassified | Crocinitomica | information st | Replication, re | 0PB0H | DNA-binding p | 755732.Fluta_ | 6,4 | 1,624807681 | 0,000791864 | 0,000152772 | 9 - | 0,000947698 - | 6 - | 0,000784508 - | 7 - | 0,000870069 - | 4 - | 0,000504526 - | 6 - | 0,00085252 - |
| 1004 | 1 | 3um_FIJAFBF_49007 | Bacteria | Proteobacteri | Gammaproteo | Oceanospirilla | Saccharospirill | Reinekea | Reinekea fors | unclassified | unclassified | unclassified | unclassified | unclassified | 4,8 | 0,748331477 | 0,000277656 | 3,63E-05 | 6 - | 0,000293047 - | 4 - | 0,000242585 - | 4 - | 0,000230608 - | 5 - | 0,000292518 - | 5 - | 0,00032952 - |
| 1005 | 1 | 3um_FIJAFBF_64708 | Bacteria | Proteobacteri | Gammaproteo | unclassified | unclassified | unclassified | various | cellular proces | Intracellular tr | 0QITE | Outer membra | 247633.GP214 | 0,8 | 0,748331477 | 2,37E-05 | 2,19E-05 | 0 - | 0 - | 1 - | 3,06E-05 - | 2 - | 5,83E-05 - | 1 - | 2,96E-05 - | 0 - | 0 - |
| 1006 | 1 | 0.2um_NOKCPHJN_535633 | Bacteria | Proteobacteri | Gammaproteo | Cellvibrionales | Porticoccacea | unclassified | Gammaproteo | unclassified | unclassified | unclassified | unclassified | unclassified | 1,2 | 0,748331477 | 0,000104372 | 7,05E-05 | 1 - | 7,17E-05 - | 0 - | 0 - | 1 - | 8,47E-05 - | 2 - | 0,000171854 - | 2 - | 0,000193593 - |
| 1007 | 1 | 3um_FIJAFBF_30985 | Bacteria | Bacteroidetes | Flavobacteriia | Flavobacterial | Flavobacteriac | Gilvibacter | Gilvibacter sp. | information st | Transcription | COG0250 | Participates in | 50743.SCB49_ | 1,8 | 0,748331477 | 0,000131748 | 4,81E-05 | 3 - | 0,000187136 - | 2 - | 0,000154912 - | 1 - | 7,36E-05 - | 1 - | 7,47E-05 - | 2 - | 0,000168342 - |
| 1008 | 1 | 3um_FIJAFBF_703977 | Bacteria | Proteobacteri | Gammaproteo | Oceanospirilla | Saccharospirill | Reinekea | Reinekea fors | cellular proces | Cell wall/mem | COG2885 | Ompa motb d | 314283.MED2 | 0,6 | 0,489897949 | 2,67E-05 | 2,22E-05 | 1 - | 3,76E-05 - | 0 - | 0 - | 0 - | 0 - | 1 - | 4,51E-05 - | 1 - | 5,08E-05 - |
| 1009 | 1 | 3um_FIJAFBF_214094 | unclassified | unclassified | unclassified | unclassified | unclassified | unclassified | unclassified | unclassified | unclassified | unclassified | unclassified | unclassified | 0,2 | 0,4 | 3,67E-06 | 7,33E-06 | 0 - | 0 - | 0 - | 0 - | 0 - | 0 - | 1 - | 1,83E-05 - | 0 - | 0 - |
| 1010 | 2 | 0.2um_NOKCPHJN_208514;0.2um_NOKCPHJN_8 | Bacteria | Bacteroidetes | Flavobacteriia | Flavobacterial | Flavobacteriac | Formosa | Formosa sp. H | cellular proces | Intracellular tr | 05FGH | MotA TolQ exl | 391603.FBALC | 3 | 0,632455532 | 0,000155916 | 2,71E-05 | 4 - | 0,000175903 - | 3 - | 0,000163815 - | 2 - | 0,000103818 - | 3 - | 0,000158027 - | 3 - | 0,000178017 - |
| 1011 | 1 | 3um_FIJAFBF_818243 | Bacteria | Proteobacteri | Gammaproteo | unclassified | unclassified | unclassified | Gammaproteo | unclassified | unclassified | unclassified | unclassified | unclassified | 0,6 | 0,489897949 | 8,27E-05 | 6,87E-05 | 1 - | 0,000117119 - | 0 - | 0 - | 1 - | 0,000138247 - | 0 - | 0 - | 1 - | 0,000158035 - |
| 1012 | 1 | 3um_FIJAFBF_514040 | Bacteria | Bacteroidetes | Flavobacteriia | Flavobacterial | Flavobacteriac | Tenacibaculur | Tenacibaculur | unclassified | unclassified | unclassified | unclassified | unclassified | 0,6 | 0,489897949 | 4,74E-05 | 3,94E-05 | 1 - | 6,60E-05 - | 1 - | 8,19E-05 - | 0 - | 0 - | 0 - | 0 - | 1 - | 8,90E-05 - |
| 1013 | 1 | 3um_FIJAFBF_355789 | Bacteria | Bacteroidetes | Flavobacteriia | Flavobacterial | Flavobacteriac | Polaribacter | Polaribacter s | poorly charact | Function unkn | 0P8NI | Inherit from b | 313594.PI23P | 10 | 1,264911064 | 0,000208825 | 3,19E-05 | 11 - | 0,000192755 - | 9 - | 0,000195827 - | 11 - | 0,000227527 - | 8 - | 0,000167919 - | 11 - | 0,000260095 - |
| 1014 | 1 | 3um_FIJAFBF_171174 | Bacteria | Proteobacteri | Alphaproteob | Rhodobactera | Rhodobactera | Shimia | Shimia abyssi | cellular proces | Posttranslatio | 179VG | alkyl hydroper | 292414.TM10 | 0,4 | 0,489897949 | 3,07E-05 | 3,76E-05 | 0 - | 0 - | 0 - | 0 - | 1 - | 7,61E-05 - | 1 - | 7,72E-05 - | 0 - | 0 - |
| 1015 | 1 | 3um_FIJAFBF_398124 | Bacteria | Bacteroidetes | Flavobacteriia | Flavobacterial | Flavobacteriac | Lacinutrix | Lacinutrix sp. | metabolism | Energy produc | 099YH | Pyruvate dehy | 983548.Krodi_ | 0,4 | 0,489897949 | 1,54E-05 | 1,90E-05 | 1 - | 3,53E-05 - | 0 - | 0 - | 1 - | 4,17E-05 - | 0 - | 0 - | 0 - | 0 - |
| 1016 | 1 | 3um_FIJAFBF_581779 | Bacteria | Bacteroidetes | Flavobacteriia | Flavobacterial | Flavobacteriac | Polaribacter | Polaribacter a | poorly charact | Function unkn | 11RX2 | NA | 313598.MED1 | 2,6 | 0,8 | 0,000209049 | 5,85E-05 | 3 - | 0,000204958 - | 2 - | 0,000169665 - | 4 - | 0,000322577 - | 2 - | 0,000163671 - | 2 - | 0,000184374 - |
| 1017 | 1 | 3um_FIJAFBF_686532 | Bacteria | Proteobacteri | Gammaproteo | Oceanospirilla | Saccharospirill | Reinekea | Reinekea blan | metabolism | Carbohydrate | COG1109 | Catalyzes the | 314283.MED2 | 1,4 | 0,489897949 | 4,15E-05 | 1,30E-05 | 2 - | 5,06E-05 - | 2 - | 6,28E-05 - | 1 - | 2,98E-05 - | 1 - | 3,03E-05 - | 1 - | 3,41E-05 - |
| 1018 | 1 | 3um_FIJAFBF_793455 | Bacteria | Proteobacteri | Gammaproteo | unclassified | unclassified | unclassified | various | unclassified | unclassified | unclassified | unclassified | unclassified | 1,6 | 0,8 | 0,000231247 | 9,25E-05 | 3 - | 0,000378385 - | 1 - | 0,000156614 - | 1 - | 0,000148882 - | 2 - | 0,000302161 - | 1 - | 0,000170192 - |
| 1019 | 1 | 3um_FIJAFBF_803540 | Bacteria | Bacteroidetes | unclassified | unclassified | unclassified | unclassified | Bacteroidetes | unclassified | unclassified | unclassified | unclassified | unclassified | 0,2 | 0,4 | 6,96E-06 | 1,39E-05 | 1 - | 3,48E-05 - | 0 - | 0 - | 0 - | 0 - | 0 - | 0 - | 0 - | 0 - |
| 1020 | 1 | 0.2um_NOKCPHJN_311210 | Bacteria | Bacteroidetes | Flavobacteriia | Flavobacterial | Cryomorphace | unclassified | Cryomorphace | unclassified | unclassified | unclassified | unclassified | unclassified | 1 | 0 | 1,70E-05 | 1,62E-06 | 1 - | 1,43E-05 - | 1 - | 1,77E-05 - | 1 - | 1,68E-05 - | 1 - | 1,71E-05 - | 1 - | 1,92E-05 - |
| 1021 | 1 | 3um_FIJAFBF_102824 | Bacteria | Bacteroidetes | Flavobacteriia | Flavobacterial | Flavobacteriac | Arenitalea | Arenitalea lut | unclassified | unclassified | unclassified | unclassified | unclassified | 1 | 0,632455532 | 2,65E-05 | 1,72E-05 | 1 - | 2,29E-05 - | 1 - | 2,84E-05 - | 2 - | 5,40E-05 - | 1 - | 2,74E-05 - | 0 - | 0 - |
| 1022 | 1 | 0.2um_NOKCPHJN_176770 | Bacteria | Proteobacteri | Betaproteoba | Burkholderiale | Comamonada | Rhodoferax | Rhodoferax sa | metabolism | Carbohydrate | 08IHS | extracellular s | 395495.Lcho_ | 1,4 | 0,489897949 | 4,31E-05 | 1,27E-05 | 2 - | 5,28E-05 - | 1 - | 3,28E-05 - | 1 - | 3,11E-05 - | 2 - | 6,32E-05 - | 1 - | 3,56E-05 - |
| 1023 | 1 | 3um_FIJAFBF_116569 | Bacteria | Bacteroidetes | Flavobacteriia | Flavobacterial | Flavobacteriac | Polaribacter | Polaribacter ir | information st | Translation, ril | 08YY1 | This protein is | 313594.PI23P | 1,6 | 0,489897949 | 0,000207291 | 5,75E-05 | 2 - | 0,000220725 - | 2 - | 0,000274075 - | 2 - | 0,000260543 - | 1 - | 0,000132195 - | 1 - | 0,000148918 - |
| 1024 | 1 | 3um_FIJAFBF_334796 | Bacteria | Proteobacteri | Gammaproteo | unclassified | unclassified | unclassified | Gammaproteo | metabolism | Amino acid tra | 17C36 | Microcompart | 502025.Hoch_ | 0,4 | 0,489897949 | 5,26E-05 | 6,48E-05 | 1 - | 0,000119559 - | 0 - | 0 - | 0 - | 0 - | 1 - | 0,000143212 - | 0 - | 0 - |
| 1025 | 1 | 0.2um_NOKCPHJN_123808 | Bacteria | Proteobacteri | Gammaproteo | Oceanospirilla | Saccharospirill | Reinekea | Reinekea blan | information st | Translation, ril | 16PEA | histidyl-tRNA | 314283.MED2 | 0,2 | 0,4 | 5,40E-06 | 1,08E-05 | 1 - | 2,70E-05 - | 0 - | 0 - | 0 - | 0 - | 0 - | 0 - | 0 - | 0 - |
| 1026 | 1 | 0.2um_NOKCPHJN_130410 | Bacteria | Proteobacteri | Alphaproteob | Rhodobactera | Rhodobactera | Amylibacter | Amylibacter k | poorly charact | Function unkn | COG1853 | flavin reducta | 207954.MED9 | 0,2 | 0,4 | 7,40E-06 | 1,48E-05 | 1 - | 3,70E-05 - | 0 - | 0 - | 0 - | 0 - | 0 - | 0 - | 0 - | 0 - |
| 1027 | 1 | 0.2um_NOKCPHJN_32623 | Bacteria | Proteobacteri | Gammaproteo | unclassified | unclassified | unclassified | Gammaproteo | metabolism | Inorganic ion t | 1DH50 | phosphate bin | 452637.Oter_ | 6,2 | 1,469693846 | 0,000306701 | 5,56E-05 | 8 - | 0,000337579 - | 7 - | 0,000366776 - | 7 - | 0,000348667 - | 5 - | 0,000252727 - | 4 - | 0,000227757 - |
| 1028 | 1 | 0.2um_NOKCPHJN_384824 | Bacteria | Proteobacteri | Alphaproteob | Pelagibacteral | unclassified | unclassified | Pelagibacteral | cellular proces | Cell wall/mem | 7257 | NA | 439493.PB721 | 1 | 0 | 4,16E-05 | 3,95E-06 | 1 - | 3,49E-05 - | 1 - | 4,33E-05 - | 1 - | 4,12E-05 - | 1 - | 4,18E-05 - | 1 - | 4,71E-05 - |
| 1029 | 1 | 3um_FIJAFBF_12411 | Bacteria | Proteobacteri | Alphaproteob | Rhodobactera | Rhodobactera | Yoonia | Yoonia marico | metabolism | Amino acid tra | COG0136 | Catalyzes the | 391593.RCCS2 | 0,2 | 0,4 | 1,08E-05 | 2,15E-05 | 0 - | 0 - | 0 - | 0 - | 1 - | 5,38E-05 - | 0 - | 0 - | 0 - | 0 - |
| 1030 | 1 | 0.2um_NOKCPHJN_09984 | Bacteria | Bacteroidetes | Flavobacteriia | Flavobacterial | Flavobacteriac | Polaribacter | Polaribacter v | information st | Transcription | COG0085 | DNA-depende | 313594.PI23P | 3,6 | 0,8 | 3,88E-05 | 7,66E-06 | 4 - | 3,66E-05 - | 4 - | 4,55E-05 - | 4 - | 4,33E-05 - | 4 - | 4,39E-05 - | 2 - | 2,47E-05 - |
| 1031 | 1 | 0.2um_NOKCPHJN_460015 | Eukaryota | unclassified | Chrysophycea | Chromulinales | Chromulinace | Ochromonas | Ochromonas | unclassified | unclassified | unclassified | unclassified | unclassified | 1,6 | 0,489897949 | 5,66E-05 | 1,82E-05 | 1 - | 2,95E-05 - | 2 - | 7,33E-05 - | 2 - | 6,97E-05 - | 2 - | 7,07E-05 - | 1 - | 3,98E-05 - |
| 1032 | 1 | 0.2um_NOKCPHJN_271555 | Eukaryota | Arthropoda | Insecta | Hymenoptera | Trichogramma | Trichogramma | Trichogramma | unclassified | unclassified | unclassified | unclassified | unclassified | 0,2 | 0,4 | 9,32E-06 | 1,86E-05 | 0 - | 0 - | 0 - | 0 - | 0 - | 0 - | 1 - | 4,66E-05 - | 0 - | 0 - |
| 1033 | 1 | 0.2um_NOKCPHJN_735038 | Bacteria | Proteobacteri | Alphaproteob | Pelagibacteral | Pelagibacterac | Candidatus Pe | Candidatus Pe | information st | Transcription | 05D27 | DNA-depende | 439493.PB721 | 0,4 | 0,489897949 | 5,91E-06 | 7,37E-06 | 1 - | 1,26E-05 - | 0 - | 0 - | 0 - | 0 - | 0 - | 0 - | 1 - | 1,70E-05 - |
| 1034 | 1 | 3um_FIJAFBF_796498 | Bacteria | Proteobacteri | Gammaproteo | Alteromonada | Alteromonada | various | various | information st | Translation, ril | 16QAF | thus facilitatin | 342610.Patl_2 | 6,2 | 0,748331477 | 0,000153514 | 2,82E-05 | 6 - | 0,00012386 - | 6 - | 0,000153797 - | 5 - | 0,000121837 - | 7 - | 0,00017309 - | 7 - | 0,000194986 - |
| 1035 | 1 | 0.2um_NOKCPHJN_521785 | Eukaryota | Streptophyta | Marchantiops | Marchantiales | Marchantiace | Marchantia | Marchantia p | unclassified | unclassified | unclassified | unclassified | unclassified | 2 | 0,632455532 | 0,000185288 | 7,15E-05 | 2 - | 0,000153036 - | 2 - | 0,000190025 - | 1 - | 9,03E-05 - | 2 - | 0,000183311 - | 3 - | 0,000309749 - |
| 1036 | 1 | 3um_FIJAFBF_360601 | unclassified | unclassified | unclassified | unclassified | unclassified | unclassified | unclassified | unclassified | unclassified | unclassified | unclassified | unclassified | 0,2 | 0,4 | 5,91E-06 | 1,18E-05 | 0 - | 0 - | 1 - | 2,96E-05 - | 0 - | 0 - | 0 - | 0 - | 0 - | 0 - |
| 1037 | 2 | 0.2um_NOKCPHJN_251910;0.2um_NOKCPHJN_6 | Bacteria | Bacteroidetes | Flavobacteriia | Flavobacterial | Crocinitomica | unclassified | various | unclassified | unclassified | unclassified | unclassified | unclassified | 0,6 | 0,489897949 | 1,51E-05 | 1,24E-05 | 1 - | 2,24E-05 - | 0 - | 0 - | 1 - | 2,64E-05 - | 1 - | 2,68E-05 - | 0 - | 0 - |
| 1038 | 1 | 3um_FIJAFBF_277461 | Bacteria | Cyanobacteria | unclassified | Oscillatoriales | unclassified | unclassified | filamentous c | information st | Translation, ril | 08UIK | This protein bi | 1140.Synpcc7 | 0,6 | 0,489897949 | 5,77E-05 | 4,80E-05 | 1 - | 8,14E-05 - | 0 - | 0 - | 0 - | 0 - | 1 - | 9,75E-05 - | 1 - | 0,00010984 - |
| 1039 | 1 | 0.2um_NOKCPHJN_30588 | Bacteria | unclassified | unclassified | unclassified | unclassified | unclassified | uncultured ba | unclassified | unclassified | unclassified | unclassified | unclassified | 1,8 | 0,748331477 | 2,53E-05 | 1,02E-05 | 2 - | 2,40E-05 - | 1 - | 1,49E-05 - | 2 - | 2,83E-05 - | 3 - | 4,31E-05 - | 1 - | 1,62E-05 - |
| 1040 | 1 | 0.2um_NOKCPHJN_409767 | Bacteria | Proteobacteri | Alphaproteob | Pelagibacteral | unclassified | unclassified | Pelagibacteral | cellular proces | Posttranslatio | COG0330 | Band 7 protei | 1002672.SAR1 | 1,4 | 0,489897949 | 6,67E-05 | 2,46E-05 | 1 - | 3,97E-05 - | 2 - | 9,86E-05 - | 2 - | 9,38E-05 - | 1 - | 4,76E-05 - | 1 - | 5,36E-05 - |
| 1041 | 1 | 0.2um_NOKCPHJN_129968 | Bacteria | Proteobacteri | Gammaproteo | Xanthomonad | Rhodanobacte | Dyella | Dyella sp. OK( | poorly charact | Function unkn | 0QJR3 | K13735 adhes | 214092.YPO3 | 4,4 | 1,019803903 | 4,50E-05 | 1,05E-05 | 4 - | 3,44E-05 - | 5 - | 5,34E-05 - | 6 - | 6,09E-05 - | 4 - | 4,12E-05 - | 3 - | 3,48E-05 - |
| 1042 | 1 | 3um_FIJAFBF_140037 | Eukaryota | Bacillariophyt | Bacillariophyc | Naviculales | Phaeodactylac | Phaeodactylu | Phaeodactylu | unclassified | unclassified | unclassified | unclassified | unclassified | 0,6 | 0,489897949 | 3,38E-05 | 2,77E-05 | 0 - | 0 - | 1 - | 5,55E-05 - | 0 - | 0 - | 1 - | 5,35E-05 - | 1 - | 6,03E-05 - |
| 1043 | 1 | 0.2um_NOKCPHJN_74586 | Bacteria | Proteobacteri | Gammaproteo | Oceanospirilla | Saccharospirill | Reinekea | Reinekea blan | information st | Replication, re | COG0178 | The UvrABC re | 314283.MED2 | 2,2 | 0,748331477 | 3,16E-05 | 9,73E-06 | 3 - | 3,66E-05 - | 2 - | 3,03E-05 - | 1 - | 1,44E-05 - | 3 - | 4,38E-05 - | 2 - | 3,29E-05 - |
| 1044 | 1 | 3um_FIJAFBF_709249 | Bacteria | Actinobacteri | Actinobacteri | Micrococcales | Microbacteria | Candidatus Ac | Candidatus Ac | unclassified | unclassified | unclassified | unclassified | unclassified | 0,6 | 0,489897949 | 4,27E-05 | 3,55E-05 | 1 - | 5,95E-05 - | 1 - | 7,38E-05 - | 0 - | 0 - | 0 - | 0 - | 1 - | 8,02E-05 - |
| 1045 | 2 | 0.2um_NOKCPHJN_452346;0.2um_NOKCPHJN_9 | Bacteria | Bacteroidetes | unclassified | unclassified | unclassified | unclassified | Bacteroidetes | unclassified | unclassified | unclassified | unclassified | unclassified | 1,8 | 0,4 | 0,000285976 | 5,65E-05 | 2 - | 0,000270063 - | 2 - | 0,000335339 - | 2 - | 0,000318782 - | 2 - | 0,00032349 - | 1 - | 0,000182205 - |
| 1046 | 1 | 0.2um_NOKCPHJN_139201 | Bacteria | Bacteroidetes | Flavobacteriia | Flavobacterial | Flavobacteriac | Flaviramulus | Flaviramulus i | unclassified | unclassified | unclassified | unclassified | unclassified | 0,6 | 0,489897949 | 1,53E-05 | 1,25E-05 | 0 - | 0 - | 1 - | 2,62E-05 - | 1 - | 2,50E-05 - | 1 - | 2,53E-05 - | 0 - | 0 - |
| 1047 | 1 | 3um_FIJAFBF_438661 | Bacteria | Proteobacteri | Gammaproteo | Oceanospirilla | Saccharospirill | Reinekea | Reinekea fors | unclassified | unclassified | unclassified | unclassified | unclassified | 1 | 0 | 0,000165093 | 1,57E-05 | 1 - | 0,000138285 - | 1 - | 0,000171709 - | 1 - | 0,000163232 - | 1 - | 0,000165643 - | 1 - | 0,000186596 - |
| 1048 | 1 | 0.2um_NOKCPHJN_31660 | Bacteria | Proteobacteri | Betaproteoba | unclassified | unclassified | unclassified | Betaproteoba | unclassified | unclassified | unclassified | unclassified | unclassified | 9 | 0 | 0,000306777 | 2,91E-05 | 9 - | 0,000256963 - | 9 - | 0,000319072 - | 9 - | 0,000303318 - | 9 - | 0,000307798 - | 9 - | 0,000346734 - |
| 1049 | 1 | 3um_FIJAFBF_493300 | Bacteria | Proteobacteri | Gammaproteo | unclassified | unclassified | unclassified | OM182 bacte | unclassified | unclassified | unclassified | unclassified | unclassified | 10,4 | 0,8 | 0,000509418 | 2,53E-05 | 12 - | 0,000495439 - | 10 - | 0,000512658 - | 10 - | 0,000487346 - | 10 - | 0,000494544 - | 10 - | 0,000557102 - |
| 1050 | 1 | 3um_FIJAFBF_597255 | Bacteria | Bacteroidetes | Flavobacteriia | Flavobacterial | Crocinitomica | unclassified | Crocinitomica | cellular proces | Posttranslatio | COG0330 | Band 7 protei | 313596.RB25( | 0,2 | 0,4 | 8,41E-06 | 1,68E-05 | 1 - | 4,20E-05 - | 0 - | 0 - | 0 - | 0 - | 0 - | 0 - | 0 - | 0 - |
| 1051 | 1 | 3um_FIJAFBF_282298 | unclassified | unclassified | unclassified | unclassified | unclassified | unclassified | unclassified | unclassified | unclassified | unclassified | unclassified | unclassified | 2 | 1,264911064 | 0,000260618 | 0,000172741 | 1 - | 0,000106275 - | 1 - | 0,000131962 - | 1 - | 0,000125447 - | 4 - | 0,000509197 - | 3 - | 0,000430207 - |
| 1052 | 1 | 3um_FIJAFBF_777839 | Bacteria | Bacteroidetes | Flavobacteriia | Flavobacterial | Flavobacteriac | Polaribacter | Polaribacter s | information st | Translation, ril | 0P7KA | Involved in pe | 313594.PI23P | 1,2 | 0,979795897 | 8,21E-05 | 5,85E-05 | 3 - | 0,000183154 - | 0 - | 0 - | 1 - | 7,21E-05 - | 1 - | 7,31E-05 - | 1 - | 8,24E-05 - |
| 1053 | 1 | 0.2um_NOKCPHJN_86884 | Bacteria | Proteobacteri | Gammaproteo | Pseudomonad | Pseudomonad | Pseudomonas | various | information st | Translation, ril | 16Q16 | Binds the low | 743720.Psefu | 0,8 | 0,748331477 | 4,77E-05 | 4,63E-05 | 1 - | 5,22E-05 - | 0 - | 0 - | 1 - | 6,16E-05 - | 2 - | 0,000124985 - | 0 - | 0 - |
| 1054 | 1 | 3um_FIJAFBF_596874 | Bacteria | Bacteroidetes | Flavobacteriia | Flavobacterial | Flavobacteriac | Moheibacter | Moheibacter s | poorly charact | Function unkn | 06XI0 | Protein of unk | 313606.M231 | 2 | 0 | 0,000141265 | 1,34E-05 | 2 - | 0,000118327 - | 2 - | 0,000146927 - | 2 - | 0,000139672 - | 2 - | 0,000141735 - | 2 - | 0,000159664 - |
| 1055 | 1 | 3um_FIJAFBF_469802 | Bacteria | Bacteroidetes | Flavobacteriia | Flavobacterial | Flavobacteriac | Formosa | Formosa sp. H | unclassified | unclassified | unclassified | unclassified | unclassified | 0,8 | 0,748331477 | 5,27E-05 | 4,98E-05 | 1 - | 5,60E-05 - | 0 - | 0 - | 2 - | 0,000132178 - | 0 - | 0 - | 1 - | 7,55E-05 - |
| 1056 | 1 | 3um_FIJAFBF_68633 | Eukaryota | Chordata | Actinopteri | Cyprinodontif | unclassified | Austrofundulu | Austrofundulu | unclassified | unclassified | unclassified | unclassified | unclassified | 0,2 | 0,4 | 4,27E-06 | 8,54E-06 | 0 - | 0 - | 0 - | 0 - | 0 - | 0 - | 1 - | 2,13E-05 - | 0 - | 0 - |
| 1057 | 1 | 0.2um_NOKCPHJN_613238 | Bacteria | Proteobacteri | Gammaproteo | unclassified | unclassified | unclassified | Gammaproteo | unclassified | unclassified | unclassified | unclassified | unclassified | 3,8 | 0,979795897 | 0,000301251 | 5,94E-05 | 5 - | 0,000339576 - | 4 - | 0,000337323 - | 4 - | 0,000320668 - | 4 - | 0,000325404 - | 2 - | 0,000183283 - |
| 1058 | 1 | 3um_FIJAFBF_817805 | Bacteria | Bacteroidetes | Flavobacteriia | Flavobacterial | Flavobacteriac | Formosa | Formosa sp. H | unclassified | unclassified | unclassified | unclassified | unclassified | 0,6 | 0,8 | 2,12E-05 | 2,73E-05 | 2 - | 6,63E-05 - | 0 - | 0 - | 0 - | 0 - | 1 - | 3,97E-05 - | 0 - | 0 - |
| 1059 | 1 | 0.2um_NOKCPHJN_221139 | Bacteria | Proteobacteri | Alphaproteob | Rhodobactera | Rhodobactera | Planktomarina | Planktomarina | unclassified | unclassified | unclassified | unclassified | unclassified | 4,4 | 1,356465997 | 0,000107569 | 2,32E-05 | 7 - | 0,000146613 - | 4 - | 0,000104028 - | 4 - | 9,89E-05 - | 3 - | 7,53E-05 - | 4 - | 0,000113047 - |
| 1060 | 1 | 3um_FIJAFBF_780948 | Bacteria | Bacteroidetes | Flavobacteriia | Flavobacterial | Flavobacteriac | Polaribacter | Polaribacter ir | cellular proces | Cell motility | COG1360 | Ompa motb d | 313594.PI23P | 3,2 | 0,748331477 | 0,000154532 | 4,63E-05 | 2 - | 7,94E-05 - | 3 - | 0,000147943 - | 3 - | 0,000140639 - | 4 - | 0,000190288 - | 4 - | 0,000214359 - |
| 1061 | 1 | 0.2um_NOKCPHJN_466945 | Bacteria | Proteobacteri | Alphaproteob | Pelagibacteral | Pelagibacterac | Candidatus Pe | Candidatus Pe | unclassified | unclassified | unclassified | unclassified | unclassified | 0,8 | 0,4 | 4,42E-05 | 2,23E-05 | 0 - | 0 - | 1 - | 5,52E-05 - | 1 - | 5,25E-05 - | 1 - | 5,33E-05 - | 1 - | 6,00E-05 - |
| 1062 | 1 | 0.2um_NOKCPHJN_1071158 | Bacteria | Bacteroidetes | Flavobacteriia | Flavobacterial | Flavobacteriac | Flavobacteriu | Flavobacteriu | metabolism | Energy produc | COG0056 | Produces ATP | 376686.Fjoh_ | 0,8 | 0,4 | 6,59E-05 | 3,32E-05 | 0 - | 0 - | 1 - | 8,24E-05 - | 1 - | 7,83E-05 - | 1 - | 7,95E-05 - | 1 - | 8,95E-05 - |
| 1063 | 1 | 3um_FIJAFBF_298836 | Bacteria | Bacteroidetes | Flavobacteriia | Flavobacterial | Flavobacteriac | Polaribacter | Polaribacter h | unclassified | unclassified | unclassified | unclassified | unclassified | 3,4 | 1,019803903 | 0,000464882 | 0,000104539 | 5 - | 0,000585596 - | 2 - | 0,000290855 - | 4 - | 0,000552989 - | 3 - | 0,000420867 - | 3 - | 0,000474106 - |
| 1064 | 1 | 0.2um_NOKCPHJN_583464 | Bacteria | Proteobacteri | Alphaproteob | Rhodospirillal | Rhodospirillac | unclassified | Rhodospirillac | unclassified | unclassified | unclassified | unclassified | unclassified | 0,2 | 0,4 | 1,14E-05 | 2,28E-05 | 0 - | 0 - | 0 - | 0 - | 1 - | 5,69E-05 - | 0 - | 0 - | 0 - | 0 - |
| 1065 | 1 | 3um_FIJAFBF_355093 | Eukaryota | unclassified | Cryptophyta | Pyrenomonad | Geminigerace | Guillardia | Guillardia thet | unclassified | unclassified | unclassified | unclassified | unclassified | 1,4 | 0,489897949 | 0,00033407 | 9,56E-05 | 2 - | 0,000409917 - | 1 - | 0,000254498 - | 2 - | 0,000483865 - | 1 - | 0,000245506 - | 1 - | 0,000276562 - |
| 1066 | 1 | 3um_FIJAFBF_116533 | Eukaryota | Bacillariophyt | Coscinodiscop | Thalassiosirale | Stephanodisca | Cyclotella | Cyclotella pse | metabolism | Energy produc | 11IIK | Low-potential | 35128.Thapsd | 6 | 0,632455532 | 0,000510384 | 4,31E-05 | 7 - | 0,000502148 - | 5 - | 0,000445371 - | 6 - | 0,000508058 - | 6 - | 0,000515562 - | 6 - | 0,000580779 - |
| 1067 | 1 | 0.2um_NOKCPHJN_458167 | Eukaryota | Arthropoda | Insecta | Hymenoptera | Halictidae | Dufourea | Dufourea nova | poorly charact | Function unkn | 0ANG0 | NA | 8364.ENSXETF | 0,8 | 0,4 | 1,95E-05 | 1,00E-05 | 1 - | 2,05E-05 - | 1 - | 2,54E-05 - | 1 - | 2,42E-05 - | 0 - | 0 - | 1 - | 2,76E-05 - |
| 1068 | 1 | 0.2um_NOKCPHJN_283623 | Bacteria | Proteobacteri | Gammaproteo | Cellvibrionales | Halieaceae | Haliea | Haliea salexig | metabolism | Inorganic ion t | COG0841 | acriflavin resis | 876044.IMCC3 | 0,2 | 0,4 | 2,69E-06 | 5,38E-06 | 0 - | 0 - | 1 - | 1,34E-05 - | 0 - | 0 - | 0 - | 0 - | 0 - | 0 - |
| 1069 | 1 | 0.2um_NOKCPHJN_815268 | Bacteria | Proteobacteri | Gammaproteo | Pseudomonad | Pseudomonad | Pseudomonas | Pseudomonas | information st | Translation, ril | 05C64 | Binds directly | 216595.PFLU5 | 0,2 | 0,4 | 1,50E-05 | 2,99E-05 | 0 - | 0 - | 0 - | 0 - | 1 - | 7,49E-05 - | 0 - | 0 - | 0 - | 0 - |
| 1070 | 1 | 3um_FIJAFBF_38223 | Bacteria | Proteobacteri | Gammaproteo | Alteromonada | Alteromonada | unclassified | Alteromonada | metabolism | Energy produc | 05CMV | Succinyl-CoA s | 715451.ambt_ | 5,8 | 0,979795897 | 0,000204722 | 3,78E-05 | 5 - | 0,000147908 - | 5 - | 0,000183658 - | 7 - | 0,000244427 - | 7 - | 0,000248037 - | 5 - | 0,000199581 - |
| 1071 | 3 | 0.2um_NOKCPHJN_349825;3um_FIJAFBF_3902( | Bacteria | Bacteroidetes | Flavobacteriia | Flavobacterial | unclassified | unclassified | Flavobacterial | unclassified | unclassified | unclassified | unclassified | unclassified | 0,4 | 0,489897949 | 2,31E-05 | 2,85E-05 | 1 - | 5,26E-05 - | 0 - | 0 - | 0 - | 0 - | 1 - | 6,31E-05 - | 0 - | 0 - |
| 1072 | 1 | 3um_FIJAFBF_13028 | Bacteria | Bacteroidetes | Flavobacteriia | Flavobacterial | Flavobacteriac | Aequorivita | Aequorivita sp | poorly charact | Function unkn | 0XT33 | NA | 537011.PREV( | 0,2 | 0,4 | 4,24E-06 | 8,47E-06 | 1 - | 2,12E-05 - | 0 - | 0 - | 0 - | 0 - | 0 - | 0 - | 0 - | 0 - |
| 1073 | 1 | 0.2um_NOKCPHJN_428533 | unclassified | unclassified | unclassified | unclassified | unclassified | unclassified | unclassified | unclassified | unclassified | unclassified | unclassified | unclassified | 0,4 | 0,489897949 | 8,18E-06 | 1,01E-05 | 0 - | 0 - | 0 - | 0 - | 1 - | 1,91E-05 - | 0 - | 0 - | 1 - | 2,18E-05 - |
| 1074 | 1 | 3um_FIJAFBF_475014 | Bacteria | Proteobacteri | Gammaproteo | Cellvibrionales | Cellvibrionace | Cellvibrio | Cellvibrio sp. I | cellular proces | Cell wall/mem | 0QTSX | Peptidoglycan | 314287.GB22( | 3,6 | 0,489897949 | 0,000291425 | 4,75E-05 | 4 - | 0,000271661 - | 3 - | 0,000252992 - | 3 - | 0,000240501 - | 4 - | 0,000325404 - | 4 - | 0,000366567 - |
| 1075 | 1 | 3um_FIJAFBF_44066 | unclassified | unclassified | unclassified | unclassified | unclassified | unclassified | unclassified | unclassified | unclassified | unclassified | unclassified | unclassified | 1 | 0 | 0,000240398 | 2,28E-05 | 1 - | 0,000201363 - | 1 - | 0,000250033 - | 1 - | 0,000237688 - | 1 - | 0,000241199 - | 1 - | 0,00027171 - |
| 1076 | 1 | 3um_FIJAFBF_545830 | Bacteria | Bacteroidetes | Flavobacteriia | Flavobacterial | Flavobacteriac | Polaribacter | Polaribacter h | information st | Translation, ril | 05G6W | One of the pri | 313594.PI23P | 1,6 | 0,489897949 | 0,000109514 | 3,53E-05 | 1 - | 5,71E-05 - | 2 - | 0,00014181 - | 2 - | 0,000134808 - | 2 - | 0,000136799 - | 1 - | 7,71E-05 - |
| 1077 | 1 | 3um_FIJAFBF_673146 | Eukaryota | Bacillariophyt | Coscinodiscop | Thalassiosirale | Thalassiosirac | Thalassiosira | Thalassiosira | unclassified | unclassified | unclassified | unclassified | unclassified | 2,6 | 0,489897949 | 0,000340594 | 8,77E-05 | 2 - | 0,00021656 - | 3 - | 0,000403355 - | 2 - | 0,000255627 - | 3 - | 0,000389104 - | 3 - | 0,000438324 - |
| 1078 | 1 | 3um_FIJAFBF_785885 | Bacteria | Bacteroidetes | Cytophagia | Cytophagales | Flammeovirga | unclassified | Flammeovirga | unclassified | unclassified | unclassified | unclassified | unclassified | 2,6 | 0,489897949 | 7,69E-05 | 1,11E-05 | 3 - | 7,53E-05 - | 2 - | 6,24E-05 - | 3 - | 8,89E-05 - | 3 - | 9,03E-05 - | 2 - | 6,78E-05 - |
| 1079 | 1 | 0.2um_NOKCPHJN_33908 | Bacteria | Proteobacteri | Gammaproteo | Alteromonada | Alteromonada | Paraglaciecola | Paraglaciecola | unclassified | unclassified | unclassified | unclassified | unclassified | 1,4 | 0,489897949 | 3,65E-05 | 1,32E-05 | 2 - | 4,39E-05 - | 1 - | 2,73E-05 - | 1 - | 2,59E-05 - | 1 - | 2,63E-05 - | 2 - | 5,92E-05 - |
| 1080 | 1 | 0.2um_NOKCPHJN_164788 | Bacteria | Proteobacteri | Deltaproteoba | Desulfobacter | Desulfobacter | Candidatus M | Candidatus M | unclassified | unclassified | unclassified | unclassified | unclassified | 3,4 | 2,244994432 | 5,39E-06 | 3,69E-06 | 4 - | 5,38E-06 - | 7 - | 1,17E-05 - | 1 - | 1,59E-06 - | 4 - | 6,45E-06 - | 1 - | 1,82E-06 - |
| 1081 | 1 | 0.2um_NOKCPHJN_510147 | Bacteria | Proteobacteri | Alphaproteob | Rhodobactera | Rhodobactera | various | various | information st | Translation, ril | 16Q16 | Binds the low | 644076.SCH4 | 5,2 | 1,469693846 | 0,000293673 | 7,08E-05 | 7 - | 0,000336166 - | 6 - | 0,000357788 - | 3 - | 0,000170061 - | 6 - | 0,000345146 - | 4 - | 0,000259204 - |
| 1082 | 1 | 0.2um_NOKCPHJN_524231 | Bacteria | Proteobacteri | Betaproteoba | unclassified | unclassified | unclassified | Betaproteoba | unclassified | unclassified | unclassified | unclassified | unclassified | 1 | 0 | 0,000129271 | 1,23E-05 | 1 - | 0,00010828 - | 1 - | 0,000134452 - | 1 - | 0,000127813 - | 1 - | 0,000129701 - | 1 - | 0,000146108 - |
| 1083 | 1 | 0.2um_NOKCPHJN_47181 | Bacteria | Proteobacteri | Gammaproteo | Cellvibrionales | Porticoccacea | unclassified | SAR92 bacteri | unclassified | unclassified | unclassified | unclassified | unclassified | 1 | 0 | 6,59E-05 | 6,26E-06 | 1 - | 5,52E-05 - | 1 - | 6,85E-05 - | 1 - | 6,51E-05 - | 1 - | 6,61E-05 - | 1 - | 7,45E-05 - |
| 1084 | 1 | 0.2um_NOKCPHJN_261542 | Bacteria | Bacteroidetes | Flavobacteriia | Flavobacterial | Flavobacteriac | Lutibacter | Lutibacter pro | poorly charact | Function unkn | 0P88U | NA | 313596.RB25( | 6,2 | 1,166190379 | 0,000147223 | 2,36E-05 | 7 - | 0,000140954 - | 7 - | 0,000175023 - | 6 - | 0,000142613 - | 7 - | 0,000168839 - | 4 - | 0,000108684 - |
| 1085 | 1 | 3um_FIJAFBF_95766 | Bacteria | unclassified | unclassified | unclassified | unclassified | unclassified | uncultured ba | metabolism | Energy produc | 05ERI | cytochrome b | 391603.FBALC | 1 | 0 | 6,04E-05 | 5,73E-06 | 1 - | 5,06E-05 - | 1 - | 6,28E-05 - | 1 - | 5,97E-05 - | 1 - | 6,06E-05 - | 1 - | 6,82E-05 - |
| 1086 | 1 | 0.2um_NOKCPHJN_392043 | Bacteria | Proteobacteri | Gammaproteo | Cellvibrionales | Spongiibacter | Oceanicoccus | Oceanicoccus | information st | Transcription | 05D27 | DNA-depende | 498211.CJA_0 | 1 | 0 | 9,74E-06 | 9,25E-07 | 1 - | 8,16E-06 - | 1 - | 1,01E-05 - | 1 - | 9,63E-06 - | 1 - | 9,77E-06 - | 1 - | 1,10E-05 - |
| 1087 | 1 | 0.2um_NOKCPHJN_514009 | Bacteria | Proteobacteri | Alphaproteob | Rhodobactera | Rhodobactera | Octadecabact | Octadecabact | information st | Replication, re | 08R4N | DNA primase | 391616.OA23 | 0,6 | 0,489897949 | 1,21E-05 | 9,95E-06 | 1 - | 1,76E-05 - | 1 - | 2,19E-05 - | 1 - | 2,08E-05 - | 0 - | 0 - | 0 - | 0 - |
| 1088 | 1 | 3um_FIJAFBF_189654 | Bacteria | Bacteroidetes | Flavobacteriia | Flavobacterial | Flavobacteriac | Polaribacter | Polaribacter s | metabolism | Inorganic ion t | COG0841 | acriflavin resis | 313594.PI23P | 1,6 | 1,356465997 | 1,80E-05 | 1,53E-05 | 3 - | 2,97E-05 - | 3 - | 3,69E-05 - | 2 - | 2,34E-05 - | 0 - | 0 - | 0 - | 0 - |
| 1089 | 1 | 0.2um_NOKCPHJN_303268 | Bacteria | Proteobacteri | Betaproteoba | Burkholderiale | unclassified | unclassified | various | metabolism | Secondary me | 17KHV | Inherit from C | 709032.Sulku_ | 0,8 | 1,6 | 4,33E-05 | 8,66E-05 | 4 - | 0,00021656 - | 0 - | 0 - | 0 - | 0 - | 0 - | 0 - | 0 - | 0 - |
| 1090 | 1 | 3um_FIJAFBF_627986 | Bacteria | Proteobacteri | Gammaproteo | Cellvibrionales | Porticoccacea | unclassified | SAR92 bacteri | information st | Translation, ril | 177XJ | One of the pri | 314287.GB22( | 4,2 | 0,748331477 | 0,000368076 | 7,04E-05 | 4 - | 0,000294299 - | 3 - | 0,000274075 - | 5 - | 0,000434238 - | 5 - | 0,000440652 - | 4 - | 0,000397114 - |
| 1091 | 1 | 3um_FIJAFBF_73184 | Bacteria | Proteobacteri | Alphaproteob | Rhodobactera | Rhodobactera | Pseudopelagic | Pseudopelagic | information st | Translation, ril | COG0088 | One of the pri | 375451.RD1_ | 1 | 0 | 6,68E-05 | 6,35E-06 | 1 - | 5,60E-05 - | 1 - | 6,95E-05 - | 1 - | 6,61E-05 - | 1 - | 6,71E-05 - | 1 - | 7,55E-05 - |
| 1092 | 1 | 3um_FIJAFBF_777831 | Bacteria | Bacteroidetes | Flavobacteriia | Flavobacterial | Flavobacteriac | Polaribacter | Polaribacter s | unclassified | unclassified | unclassified | unclassified | unclassified | 1,2 | 0,4 | 0,000134856 | 4,65E-05 | 1 - | 9,41E-05 - | 1 - | 0,000116819 - | 1 - | 0,000111051 - | 2 - | 0,000225382 - | 1 - | 0,000126946 - |
| 1093 | 1 | 3um_FIJAFBF_50670 | unclassified | unclassified | unclassified | unclassified | unclassified | unclassified | unclassified | unclassified | unclassified | unclassified | unclassified | unclassified | 0,2 | 0,4 | 8,33E-06 | 1,67E-05 | 0 - | 0 - | 0 - | 0 - | 0 - | 0 - | 1 - | 4,17E-05 - | 0 - | 0 - |
| 1094 | 1 | 3um_FIJAFBF_733399 | Eukaryota | unclassified | Pelagophycea | Pelagomonad | unclassified | Aureococcus | Aureococcus a | cellular proces | Signal transdu | KOG0613 | fibronectin ty | 67593.Physo1 | 0,6 | 0,8 | 1,09E-05 | 1,52E-05 | 1 - | 1,56E-05 - | 2 - | 3,87E-05 - | 0 - | 0 - | 0 - | 0 - | 0 - | 0 - |
| 1095 | 1 | 3um_FIJAFBF_206034 | Bacteria | Bacteroidetes | unclassified | unclassified | unclassified | unclassified | unidentified e | poorly charact | Function unkn | 05ERK | gliding motilit | 50743.SCB49_ | 1 | 0 | 2,64E-05 | 2,51E-06 | 1 - | 2,21E-05 - | 1 - | 2,75E-05 - | 1 - | 2,61E-05 - | 1 - | 2,65E-05 - | 1 - | 2,98E-05 - |
| 1096 | 1 | 3um_FIJAFBF_759036 | Eukaryota | Phaeophyceae | unclassified | Ectocarpales | Ectocarpaceae | Ectocarpus | Ectocarpus sili | poorly charact | Function unkn | 11N70 | syndrome nuc | 4787.PITG_13 | 1,8 | 0,4 | 0,000221217 | 5,18E-05 | 2 - | 0,000206805 - | 1 - | 0,000128395 - | 2 - | 0,000244112 - | 2 - | 0,000247718 - | 2 - | 0,000279053 - |
| 1097 | 1 | 3um_FIJAFBF_54806 | Bacteria | Bacteroidetes | Flavobacteriia | Flavobacterial | Flavobacteriac | Polaribacter | Polaribacter b | cellular proces | Posttranslatio | 09ADN | alkyl hydroper | 391587.KAOT | 5,4 | 0,489897949 | 0,000345346 | 2,62E-05 | 6 - | 0,000323315 - | 5 - | 0,000334551 - | 5 - | 0,000318033 - | 6 - | 0,000387277 - | 5 - | 0,000363555 - |
| 1098 | 1 | 3um_FIJAFBF_520323 | Bacteria | Bacteroidetes | Flavobacteriia | Flavobacterial | Flavobacteriac | Polaribacter | Polaribacter s | unclassified | unclassified | unclassified | unclassified | unclassified | 1 | 0 | 6,78E-05 | 6,44E-06 | 1 - | 5,68E-05 - | 1 - | 7,06E-05 - | 1 - | 6,71E-05 - | 1 - | 6,81E-05 - | 1 - | 7,67E-05 - |
| 1099 | 1 | 0.2um_NOKCPHJN_796646 | Eukaryota | Chlorophyta | Mamiellophyc | Mamiellales | Bathycoccace | Ostreococcus | Ostreococcus | unclassified | unclassified | unclassified | unclassified | unclassified | 0,4 | 0,489897949 | 5,69E-05 | 7,00E-05 | 0 - | 0 - | 0 - | 0 - | 1 - | 0,000132826 - | 0 - | 0 - | 1 - | 0,000151838 - |
| 1100 | 1 | 0.2um_NOKCPHJN_72690 | Bacteria | Proteobacteri | Gammaproteo | Cellvibrionales | Porticoccacea | unclassified | SAR92 bacteri | information st | Translation, ril | COG1185 | Involved in mF | 314287.GB22( | 7,8 | 0,4 | 0,000149527 | 1,68E-05 | 8 - | 0,000128422 - | 8 - | 0,000159462 - | 7 - | 0,00013264 - | 8 - | 0,000153827 - | 8 - | 0,000173286 - |
| 1101 | 1 | 3um_FIJAFBF_715060 | Eukaryota | Bacillariophyt | Coscinodiscop | Coscinodiscale | Coscinodiscac | Coscinodiscus | Coscinodiscus | metabolism | Energy produc | 11HAV | photosystem i | 35128.Thapsd | 7,2 | 0,748331477 | 0,000708889 | 4,32E-05 | 8 - | 0,000665373 - | 7 - | 0,000722922 - | 8 - | 0,000785404 - | 7 - | 0,000697379 - | 6 - | 0,000673367 - |
| 1102 | 1 | 3um_FIJAFBF_472987 | Bacteria | Proteobacteri | Gammaproteo | Alteromonada | Alteromonada | Paraglaciecola | Paraglaciecola | cellular proces | Intracellular tr | COG0848 | Biopolymer tr | 983545.Glaag | 1,2 | 0,748331477 | 0,000120336 | 7,69E-05 | 2 - | 0,00017004 - | 0 - | 0 - | 1 - | 0,000100357 - | 1 - | 0,000101839 - | 2 - | 0,000229444 - |
| 1103 | 1 | 0.2um_NOKCPHJN_240988 | Bacteria | Proteobacteri | Gammaproteo | Oceanospirilla | Saccharospirill | Reinekea | Reinekea blan | metabolism | Energy produc | COG1290 | Component of | 314277.MED1 | 0,6 | 1,2 | 1,67E-05 | 3,33E-05 | 3 - | 8,34E-05 - | 0 - | 0 - | 0 - | 0 - | 0 - | 0 - | 0 - | 0 - |
| 1104 | 2 | 3um_FIJAFBF_354716;3um_FIJAFBF_728300 | Bacteria | Proteobacteri | Alphaproteob | Rhodobactera | Rhodobactera | Planktomarina | Planktomarina | poorly charact | Function unkn | 11PM7 | NA | 633131.TR2A | 0,2 | 0,4 | 2,49E-05 | 4,97E-05 | 0 - | 0 - | 0 - | 0 - | 1 - | 0,000124296 - | 0 - | 0 - | 0 - | 0 - |
| 1105 | 1 | 3um_FIJAFBF_199368 | Bacteria | Proteobacteri | Gammaproteo | Cellvibrionales | Spongiibacter | Oceanicoccus | Oceanicoccus | unclassified | unclassified | unclassified | unclassified | unclassified | 1 | 0 | 4,98E-05 | 4,73E-06 | 1 - | 4,17E-05 - | 1 - | 5,18E-05 - | 1 - | 4,93E-05 - | 1 - | 5,00E-05 - | 1 - | 5,63E-05 - |
| 1106 | 1 | 3um_FIJAFBF_762092 | Eukaryota | Bacillariophyt | Coscinodiscop | Thalassiosirale | Thalassiosirac | Thalassiosira | Thalassiosira p | unclassified | unclassified | unclassified | unclassified | unclassified | 0,4 | 0,489897949 | 1,33E-05 | 1,63E-05 | 0 - | 0 - | 1 - | 3,40E-05 - | 1 - | 3,23E-05 - | 0 - | 0 - | 0 - | 0 - |
| 1107 | 1 | 3um_FIJAFBF_311350 | unclassified | unclassified | unclassified | unclassified | unclassified | unclassified | unclassified | unclassified | unclassified | unclassified | unclassified | unclassified | 1 | 0 | 7,49E-05 | 7,11E-06 | 1 - | 6,27E-05 - | 1 - | 7,79E-05 - | 1 - | 7,40E-05 - | 1 - | 7,51E-05 - | 1 - | 8,46E-05 - |
| 1108 | 1 | 3um_FIJAFBF_128630 | Eukaryota | unclassified | Raphidophyce | Chattonellales | Chattonellace | Heterosigma | Heterosigma a | unclassified | unclassified | unclassified | unclassified | unclassified | 1,4 | 0,489897949 | 7,60E-05 | 2,39E-05 | 2 - | 9,26E-05 - | 2 - | 0,000114935 - | 1 - | 5,46E-05 - | 1 - | 5,54E-05 - | 1 - | 6,24E-05 - |
| 1109 | 1 | 3um_FIJAFBF_23703 | Bacteria | Bacteroidetes | unclassified | unclassified | unclassified | unclassified | Bacteroidetes | unclassified | unclassified | unclassified | unclassified | unclassified | 0,6 | 0,489897949 | 7,25E-05 | 5,96E-05 | 1 - | 0,000107268 - | 0 - | 0 - | 1 - | 0,000126619 - | 1 - | 0,000128489 - | 0 - | 0 - |
| 1110 | 1 | 3um_FIJAFBF_346929 | Bacteria | Bacteroidetes | Flavobacteriia | Flavobacterial | Flavobacteriac | Elizabethkingi | various | information st | Translation, ril | 05C64 | Binds directly | 926562.Oweh | 1,2 | 0,748331477 | 6,89E-05 | 4,04E-05 | 2 - | 9,98E-05 - | 0 - | 0 - | 2 - | 0,000117811 - | 1 - | 5,98E-05 - | 1 - | 6,73E-05 - |
| 1111 | 1 | 3um_FIJAFBF_99475 | Bacteria | Proteobacteri | Alphaproteob | Rhodobactera | Rhodobactera | Planktomarina | Planktomarina | unclassified | unclassified | unclassified | unclassified | unclassified | 1,6 | 0,489897949 | 0,000164413 | 5,95E-05 | 1 - | 8,44E-05 - | 2 - | 0,000209587 - | 2 - | 0,000199239 - | 1 - | 0,000101091 - | 2 - | 0,000227757 - |
| 1112 | 1 | 0.2um_NOKCPHJN_118174 | Bacteria | Proteobacteri | Alphaproteob | unclassified | unclassified | unclassified | alpha proteob | unclassified | unclassified | unclassified | unclassified | unclassified | 1,4 | 0,489897949 | 7,88E-05 | 3,24E-05 | 1 - | 4,63E-05 - | 1 - | 5,75E-05 - | 1 - | 5,46E-05 - | 2 - | 0,000110874 - | 2 - | 0,000124899 - |
| 1113 | 1 | 0.2um_NOKCPHJN_441212 | Bacteria | Proteobacteri | Alphaproteob | Pelagibacteral | unclassified | unclassified | Pelagibacteral | unclassified | unclassified | unclassified | unclassified | unclassified | 3,4 | 0,8 | 0,00018405 | 3,81E-05 | 4 - | 0,000183643 - | 2 - | 0,000114015 - | 4 - | 0,000216772 - | 4 - | 0,000219973 - | 3 - | 0,000185849 - |
| 1114 | 1 | 3um_FIJAFBF_155161 | Bacteria | Proteobacteri | Gammaproteo | Cellvibrionales | Porticoccacea | unclassified | SAR92 bacteri | information st | Translation, ril | 179R6 | This protein is | 314287.GB22( | 1,6 | 0,489897949 | 0,000120216 | 4,35E-05 | 1 - | 6,17E-05 - | 2 - | 0,000153246 - | 2 - | 0,00014568 - | 1 - | 7,39E-05 - | 2 - | 0,000166532 - |
| 1115 | 1 | 3um_FIJAFBF_541542 | Bacteria | Bacteroidetes | Flavobacteriia | Flavobacterial | Schleiferiacea | Schleiferia | Schleiferia the | unclassified | unclassified | unclassified | unclassified | unclassified | 0,6 | 0,8 | 1,83E-05 | 2,43E-05 | 0 - | 0 - | 0 - | 0 - | 2 - | 6,06E-05 - | 1 - | 3,08E-05 - | 0 - | 0 - |
| 1116 | 1 | 3um_FIJAFBF_146228 | Bacteria | unclassified | unclassified | unclassified | unclassified | unclassified | uncultured ba | metabolism | Carbohydrate | COG0149 | triosephospha | 1046627.BZAF | 0,6 | 0,8 | 2,93E-05 | 3,78E-05 | 2 - | 9,22E-05 - | 0 - | 0 - | 1 - | 5,44E-05 - | 0 - | 0 - | 0 - | 0 - |
| 1117 | 1 | 3um_FIJAFBF_385889 | Bacteria | Proteobacteri | Alphaproteob | Rhodospirillal | Rhodospirillac | Oceanibaculu | Oceanibaculu | metabolism | Energy produc | COG0056 | Produces ATP | 414684.RC1_2 | 0,8 | 0,4 | 2,09E-05 | 1,06E-05 | 1 - | 2,26E-05 - | 1 - | 2,81E-05 - | 1 - | 2,67E-05 - | 1 - | 2,71E-05 - | 0 - | 0 - |
| 1118 | 1 | 3um_FIJAFBF_711074 | Bacteria | Bacteroidetes | Flavobacteriia | Flavobacterial | Flavobacteriac | Flaviramulus | Flaviramulus i | unclassified | unclassified | unclassified | unclassified | unclassified | 16,6 | 1,019803903 | 0,000238392 | 1,89E-05 | 17 - | 0,000205174 - | 17 - | 0,000254766 - | 18 - | 0,000256433 - | 16 - | 0,000231307 - | 15 - | 0,000244282 - |
| 1119 | 1 | 3um_FIJAFBF_245156 | Bacteria | Proteobacteri | Gammaproteo | Alteromonada | Alteromonada | Paraglaciecola | various | information st | Translation, ril | COG0333 | 50s ribosomal | 342610.Patl_2 | 0,4 | 0,489897949 | 8,14E-05 | 0,000100333 | 1 - | 0,000185124 - | 0 - | 0 - | 0 - | 0 - | 1 - | 0,000221747 - | 0 - | 0 - |
| 1120 | 1 | 0.2um_NOKCPHJN_482343 | Bacteria | Proteobacteri | Betaproteoba | Burkholderiale | Comamonada | Lampropedia | Lampropedia | unclassified | unclassified | unclassified | unclassified | unclassified | 0,4 | 0,489897949 | 4,04E-05 | 4,98E-05 | 1 - | 9,18E-05 - | 0 - | 0 - | 0 - | 0 - | 1 - | 0,000109987 - | 0 - | 0 - |
| 1121 | 1 | 0.2um_NOKCPHJN_80529 | Bacteria | Bacteroidetes | Flavobacteriia | Flavobacterial | Flavobacteriac | Winogradskye | Winogradskye | metabolism | Amino acid tra | 08IQK | o-acetylhomo | 391603.FBALC | 0,8 | 0,4 | 2,55E-05 | 1,31E-05 | 1 - | 2,68E-05 - | 1 - | 3,32E-05 - | 1 - | 3,16E-05 - | 0 - | 0 - | 1 - | 3,61E-05 - |
| 1122 | 1 | 0.2um_NOKCPHJN_73490 | Bacteria | Proteobacteri | Alphaproteob | Rhodobactera | Rhodobactera | Sulfitobacter | Sulfitobacter g | metabolism | Coenzyme tra | 05DEJ | cobaltochelat | 351016.RAZW | 5,8 | 1,720465053 | 0,000238544 | 6,65E-05 | 6 - | 0,000209957 - | 5 - | 0,000217254 - | 8 - | 0,000330444 - | 7 - | 0,000293409 - | 3 - | 0,000141653 - |
| 1123 | 1 | 0.2um_NOKCPHJN_1057723 | Eukaryota | Bacillariophyt | Bacillariophyc | Naviculales | Phaeodactylac | Phaeodactylu | Phaeodactylu | unclassified | unclassified | unclassified | unclassified | unclassified | 0,4 | 0,489897949 | 1,27E-05 | 1,59E-05 | 1 - | 2,71E-05 - | 0 - | 0 - | 0 - | 0 - | 0 - | 0 - | 1 - | 3,66E-05 - |
| 1124 | 1 | 3um_FIJAFBF_524529 | Bacteria | Proteobacteri | Alphaproteob | Rhodobactera | Rhodobactera | Rhodobacter | Rhodobacter | metabolism | Energy produc | 01R4M | Acetoacetyl-C | 272942.RCAP_ | 4 | 0,632455532 | 0,000226763 | 3,33E-05 | 4 - | 0,000191295 - | 4 - | 0,000237531 - | 5 - | 0,000282255 - | 4 - | 0,000229139 - | 3 - | 0,000193593 - |
| 1125 | 1 | 3um_FIJAFBF_449325 | Bacteria | Bacteroidetes | Flavobacteriia | Flavobacterial | Flavobacteriac | Formosa | Formosa sp. H | unclassified | unclassified | unclassified | unclassified | unclassified | 1 | 0,894427191 | 3,33E-05 | 2,99E-05 | 0 - | 0 - | 2 - | 6,85E-05 - | 2 - | 6,51E-05 - | 1 - | 3,30E-05 - | 0 - | 0 - |
| 1126 | 1 | 0.2um_NOKCPHJN_238585 | Bacteria | Proteobacteri | Gammaproteo | Oceanospirilla | Oceanospirilla | Motiliproteus | Motiliproteus | metabolism | Energy produc | 16PVJ | Component of | 207954.MED9 | 1,6 | 0,489897949 | 2,42E-05 | 6,35E-06 | 2 - | 2,59E-05 - | 1 - | 1,61E-05 - | 2 - | 3,06E-05 - | 2 - | 3,10E-05 - | 1 - | 1,75E-05 - |
| 1127 | 1 | 3um_FIJAFBF_146233 | Bacteria | Proteobacteri | Alphaproteob | unclassified | unclassified | Emcibacter | Emcibacter co | information st | Translation, ril | COG0050 | This protein p | 394221.Mmar | 0,4 | 0,489897949 | 1,36E-05 | 1,66E-05 | 0 - | 0 - | 1 - | 3,45E-05 - | 0 - | 0 - | 1 - | 3,33E-05 - | 0 - | 0 - |
| 1128 | 1 | 3um_FIJAFBF_155847 | Bacteria | Proteobacteri | Gammaproteo | Oceanospirilla | Saccharospirill | Reinekea | Reinekea fors | unclassified | unclassified | unclassified | unclassified | unclassified | 1,2 | 0,4 | 6,07E-05 | 2,09E-05 | 1 - | 4,24E-05 - | 1 - | 5,26E-05 - | 1 - | 5,00E-05 - | 2 - | 0,000101464 - | 1 - | 5,71E-05 - |
| 1129 | 1 | 3um_FIJAFBF_690484 | Bacteria | Proteobacteri | Gammaproteo | Alteromonada | Alteromonada | Paraglaciecola | Paraglaciecola | cellular proces | Cell wall/mem | 0QTSX | Peptidoglycan | 983545.Glaag | 5,4 | 0,8 | 0,000391603 | 6,81E-05 | 5 - | 0,000303642 - | 5 - | 0,000377034 - | 5 - | 0,000358419 - | 7 - | 0,000509197 - | 5 - | 0,000409721 - |
| 1130 | 1 | 0.2um_NOKCPHJN_362721 | Bacteria | Proteobacteri | Alphaproteob | Rhodobactera | Rhodobactera | Loktanella | Loktanella sp. | information st | Translation, ril | 05C6A | phenylalanyl-t | 391593.RCCS2 | 0,2 | 0,4 | 3,40E-06 | 6,80E-06 | 0 - | 0 - | 0 - | 0 - | 1 - | 1,70E-05 - | 0 - | 0 - | 0 - | 0 - |
| 1131 | 3 | 0.2um_NOKCPHJN_1056505;0.2um_NOKCPHJN_ | Bacteria | Proteobacteri | Alphaproteob | Pelagibacteral | Pelagibacterac | Candidatus Pe | Candidatus Pe | unclassified | unclassified | unclassified | unclassified | unclassified | 0,2 | 0,4 | 3,86E-06 | 7,71E-06 | 0 - | 0 - | 1 - | 1,93E-05 - | 0 - | 0 - | 0 - | 0 - | 0 - | 0 - |
| 1132 | 1 | 0.2um_NOKCPHJN_177152 | Bacteria | Bacteroidetes | Flavobacteriia | Flavobacterial | Flavobacteriac | Tenacibaculur | Tenacibaculur | cellular proces | Cell wall/mem | COG2885 | Ompa motb d | 313598.MED1 | 1,2 | 0,4 | 7,03E-05 | 2,42E-05 | 1 - | 4,90E-05 - | 1 - | 6,09E-05 - | 1 - | 5,79E-05 - | 2 - | 0,000117507 - | 1 - | 6,62E-05 - |
| 1133 | 1 | 3um_FIJAFBF_681217 | Bacteria | Proteobacteri | Betaproteoba | Burkholderiale | Comamonada | Rhodoferax | Rhodoferax sp | metabolism | Carbohydrate | 0B8CD | Phosphoribois | 543728.Vapar | 0,2 | 0,4 | 1,03E-05 | 2,06E-05 | 1 - | 5,15E-05 - | 0 - | 0 - | 0 - | 0 - | 0 - | 0 - | 0 - | 0 - |
| 1134 | 1 | 3um_FIJAFBF_273272 | Bacteria | Bacteroidetes | Flavobacteriia | Flavobacterial | Flavobacteriac | Algibacter | Algibacter pec | cellular proces | Posttranslatio | 07RXU | peptidylprolyl | 1046627.BZAF | 0,6 | 0,8 | 1,51E-05 | 1,95E-05 | 2 - | 4,75E-05 - | 0 - | 0 - | 1 - | 2,81E-05 - | 0 - | 0 - | 0 - | 0 - |
| 1135 | 1 | 3um_FIJAFBF_303593 | Bacteria | Proteobacteri | Alphaproteob | Rhodobactera | Rhodobactera | Pelagicola | Pelagicola lito | unclassified | unclassified | unclassified | unclassified | unclassified | 0,8 | 0,4 | 3,36E-05 | 1,69E-05 | 0 - | 0 - | 1 - | 4,20E-05 - | 1 - | 4,00E-05 - | 1 - | 4,06E-05 - | 1 - | 4,57E-05 - |
| 1136 | 1 | 3um_FIJAFBF_715097 | Eukaryota | unclassified | Pelagophycea | Pelagomonad | unclassified | Aureococcus | Aureococcus a | metabolism | Energy produc | 1DYB8 | PsaA and PsaB | 3055.DAA014 | 12,2 | 0,748331477 | 0,000222504 | 2,66E-05 | 12 - | 0,000183154 - | 13 - | 0,000246376 - | 11 - | 0,000198179 - | 13 - | 0,000237671 - | 12 - | 0,00024714 - |
| 1137 | 1 | 3um_FIJAFBF_122930 | Bacteria | Proteobacteri | Gammaproteo | unclassified | unclassified | unclassified | OM182 bacte | information st | Transcription | 05VEQ | Cold shock pro | 697282.Mettu | 1,2 | 0,4 | 0,000243481 | 0,000103628 | 1 - | 0,000166343 - | 1 - | 0,000206549 - | 1 - | 0,000196351 - | 1 - | 0,000199251 - | 2 - | 0,000448912 - |
| 1138 | 1 | 0.2um_NOKCPHJN_420926 | Bacteria | Bacteroidetes | Flavobacteriia | Flavobacterial | Flavobacteriac | Polaribacter | Polaribacter a | information st | Transcription | 0P8JP | Facilitates tra | 391603.FBALC | 6,4 | 1,496662955 | 0,000152202 | 2,38E-05 | 9 - | 0,000182507 - | 7 - | 0,00017626 - | 5 - | 0,000119684 - | 6 - | 0,000145742 - | 5 - | 0,000136815 - |

| | | | | | | | | | | | | | | | | | | | | | | | | | | | | |
|---|---|---|---|---|---|---|---|---|---|---|---|---|---|---|---|---|---|---|---|---|---|---|---|---|---|---|---|---|
| 1139 | 1 | 0.2um_NOKCPHJN_450502 | Eukaryota | unclassified | Labyrinthulon | unclassified | Thraustochytr | Aurantiochytr | Aurantiochytr | unclassified | unclassified | unclassified | unclassified | unclassified | 4,2 | 0,748331477 | 0,000151849 | 2,65E-05 | 5 - | 0,000152224 - | 5 - | 0,000189017 - | 3 - | 0,000107811 - | 4 - | 0,000145871 - | 4 - | 0,000164323 - |
| 1140 | 1 | 0.2um_NOKCPHJN_779939 | Viruses | unclassified | unclassified | unclassified | Mimiviridae | Catovirus | Catovirus CTV | unclassified | unclassified | unclassified | unclassified | unclassified | 0,4 | 0,489897949 | 2,53E-05 | 3,13E-05 | 1 - | 5,65E-05 - | 1 - | 7,02E-05 - | 0 - | 0 - | 0 - | 0 - | 0 - | 0 - |
| 1141 | 1 | 3um_FIJJAFBF_779053 | Bacteria | Proteobacteri | Alphaproteob | Rhodobactera | Rhodobactera | Yoonia | Yoonia vestfol | poorly charact | Function unkn | COG3517 | type VI secreti | 314270.RB208 | 0,6 | 0,8 | 1,58E-05 | 2,19E-05 | 1 - | 2,33E-05 - | 0 - | 0 - | 0 - | 0 - | 2 - | 5,58E-05 - | 0 - | 0 - |
| 1142 | 1 | 3um_FIJJAFBF_30146 | Bacteria | Bacteroidetes | unclassified | unclassified | unclassified | unclassified | Bacteroidetes | unclassified | unclassified | unclassified | unclassified | unclassified | 0,4 | 0,489897949 | 2,03E-05 | 2,52E-05 | 1 - | 4,54E-05 - | 1 - | 5,63E-05 - | 0 - | 0 - | 0 - | 0 - | 0 - | 0 - |
| 1143 | 1 | 0.2um_NOKCPHJN_417240 | Eukaryota | unclassified | Oomycetes | Albuginales | Albuginaceae | Albugo | Albugo candid | unclassified | unclassified | unclassified | unclassified | unclassified | 0,2 | 0,4 | 2,23E-06 | 4,46E-06 | 0 - | 0 - | 0 - | 0 - | 0 - | 0 - | 0 - | 0 - | 1 - | 1,12E-05 - |
| 1144 | 1 | 3um_FIJJAFBF_18717 | Bacteria | Bacteroidetes | Flavobacteriia | Flavobacterial | Flavobacteria | Aureitalea | Aureitalea ma | poorly charact | Function unkn | 11NCV | Protein of unk | 50743.SCB49_ | 1,8 | 0,4 | 0,000156049 | 3,82E-05 | 2 - | 0,000145287 - | 2 - | 0,000180404 - | 2 - | 0,000171496 - | 1 - | 8,70E-05 - | 2 - | 0,000196044 - |
| 1145 | 1 | 3um_FIJJAFBF_582313 | Eukaryota | Eustigmatoph | unclassified | Eustigmatales | Monodopsida | Nannochlorop | Nannochlorop | cellular proces | Signal transdu | COG5040 | Tyrosine 3-mo | 35128.Thaps2 | 2,4 | 0,8 | 0,000233142 | 8,60E-05 | 2 - | 0,000161657 - | 4 - | 0,000401462 - | 2 - | 0,00019082 - | 2 - | 0,000193638 - | 2 - | 0,000218133 - |
| 1146 | 1 | 3um_FIJJAFBF_449851 | Bacteria | Proteobacteri | Gammaprote | Oceanospirilla | Saccharospirill | Reinekea | Reinekea fors | metabolism | Carbohydrate | 0XPC7 | Periplasmic bi | 367336.OM22 | 2,6 | 1,019803903 | 0,000104364 | 4,21E-05 | 3 - | 0,000100977 - | 2 - | 8,36E-05 - | 1 - | 3,97E-05 - | 4 - | 0,000161271 - | 3 - | 0,000136253 - |
| 1147 | 1 | 3um_FIJJAFBF_671478 | Bacteria | Bacteroidetes | Cytophagia | Cytophagales | Flammeovirga | unclassified | Flammeovirga | unclassified | unclassified | unclassified | unclassified | unclassified | 1,2 | 0,4 | 0,000111649 | 3,74E-05 | 1 - | 7,81E-05 - | 1 - | 9,70E-05 - | 2 - | 0,00018433 - | 1 - | 9,35E-05 - | 1 - | 0,000105357 - |
| 1148 | 1 | 0.2um_NOKCPHJN_72834 | Bacteria | Proteobacteri | Gammaprote | Oceanospirilla | Saccharospirill | Reinekea | Reinekea blan | unclassified | unclassified | unclassified | unclassified | unclassified | 2,8 | 0,979795897 | 9,08E-05 | 2,69E-05 | 4 - | 0,000111977 - | 3 - | 0,000104282 - | 3 - | 9,91E-05 - | 3 - | 0,000100598 - | 1 - | 3,78E-05 - |
| 1149 | 1 | 3um_FIJJAFBF_346703 | Bacteria | Proteobacteri | Gammaprote | Cellvibrionale | Cellvibrionace | Teredinibacte | unclassified | unclassified | unclassified | unclassified | unclassified | unclassified | 2,2 | 0,4 | 0,000134442 | 2,62E-05 | 2 - | 0,000102479 - | 2 - | 0,000127249 - | 3 - | 0,000181449 - | 2 - | 0,000122753 - | 2 - | 0,000138281 - |
| 1150 | 1 | 0.2um_NOKCPHJN_193472 | Bacteria | Proteobacteri | Gammaprote | Methylococca | Methylococca | unclassified | Methylococca | cellular proces | Cell motility | COG1291 | MotA TolQ exl | 314287.GB22 | 0,4 | 0,489897949 | 1,99E-05 | 2,45E-05 | 1 - | 4,52E-05 - | 0 - | 0 - | 0 - | 0 - | 1 - | 5,41E-05 - | 0 - | 0 - |
| 1151 | 1 | 3um_FIJJAFBF_71348 | Bacteria | Bacteroidetes | Cytophagia | Cytophagales | Flammeovirga | Marivirga | Marivirga trac | poorly charact | Function unkn | 09AR5 | Inherit from b | 643867.Ftrac_ | 2 | 0 | 6,27E-05 | 5,95E-06 | 2 - | 5,25E-05 - | 2 - | 6,52E-05 - | 2 - | 6,20E-05 - | 2 - | 6,29E-05 - | 2 - | 7,09E-05 - |
| 1152 | 1 | 0.2um_NOKCPHJN_110550 | Bacteria | Proteobacteri | Gammaprote | Cellvibrionale | Porticoccacea | unclassified | various | information st | Translation, ri | COG0185 | Protein S19 fo | 314287.GB22 | 2 | 1,095445115 | 0,000294621 | 0,000169982 | 2 - | 0,000252257 - | 3 - | 0,000469842 - | 2 - | 0,000297763 - | 3 - | 0,000453242 - | 0 - | 0 - |
| 1153 | 1 | 3um_FIJJAFBF_43214 | Bacteria | Bacteroidetes | Cytophagia | Cytophagales | Cytophagacea | Candidatus Ne | Candidatus Ne | unclassified | unclassified | unclassified | unclassified | unclassified | 0,4 | 0,489897949 | 2,71E-05 | 3,34E-05 | 1 - | 6,17E-05 - | 0 - | 0 - | 0 - | 0 - | 1 - | 7,39E-05 - | 0 - | 0 - |
| 1154 | 1 | 3um_FIJJAFBF_52145 | Bacteria | Proteobacteri | Gammaprote | Alteromonada | Ferrimonadac | Ferrimonas | Ferrimonas ba | cellular proces | Posttranslatio | 0QSE5 | membrane | 550540.Fbal_ | 0,4 | 0,489897949 | 3,34E-05 | 4,13E-05 | 1 - | 7,45E-05 - | 1 - | 9,25E-05 - | 0 - | 0 - | 0 - | 0 - | 0 - | 0 - |
| 1155 | 1 | 3um_FIJJAFBF_481956 | Bacteria | Proteobacteri | Alphaproteob | Rhodobactera | Rhodobactera | unclassified | Rhodobactera | unclassified | unclassified | unclassified | unclassified | unclassified | 0,6 | 0,8 | 2,15E-05 | 2,77E-05 | 2 - | 6,75E-05 - | 0 - | 0 - | 1 - | 3,98E-05 - | 0 - | 0 - | 0 - | 0 - |
| 1156 | 1 | 0.2um_NOKCPHJN_305282 | Eukaryota | unclassified | unclassified | Dictyosteliales | Dictyosteliace | Dictyostelium | Dictyostelium | unclassified | unclassified | unclassified | unclassified | unclassified | 0,2 | 0,4 | 3,91E-06 | 7,82E-06 | 1 - | 1,96E-05 - | 0 - | 0 - | 0 - | 0 - | 0 - | 0 - | 0 - | 0 - |
| 1157 | 1 | 0.2um_NOKCPHJN_247175 | unclassified | unclassified | unclassified | unclassified | unclassified | unclassified | unclassified | unclassified | unclassified | unclassified | unclassified | unclassified | 0,2 | 0,4 | 2,62E-05 | 5,25E-05 | 0 - | 0 - | 0 - | 0 - | 0 - | 0 - | 0 - | 0 - | 1 - | 0,00013125 - |
| 1158 | 1 | 3um_FIJJAFBF_162960 | Bacteria | Bacteroidetes | Flavobacteriia | Flavobacterial | Flavobacteria | Aequorivita | Aequorivita sp | unclassified | unclassified | unclassified | unclassified | unclassified | 0,8 | 0,4 | 3,13E-05 | 1,59E-05 | 1 - | 3,39E-05 - | 1 - | 4,20E-05 - | 1 - | 4,00E-05 - | 1 - | 4,06E-05 - | 0 - | 0 - |
| 1159 | 1 | 3um_FIJJAFBF_165053 | Bacteria | Bacteroidetes | unclassified | unclassified | unclassified | unclassified | unidentified e | poorly charact | Function unkn | 08PIQ | CHU large pro | 50743.SCB49_ | 0,6 | 0,8 | 6,31E-06 | 8,07E-06 | 2 - | 1,95E-05 - | 1 - | 1,21E-05 - | 0 - | 0 - | 0 - | 0 - | 0 - | 0 - |
| 1160 | 1 | 3um_FIJJAFBF_149820 | Bacteria | Proteobacteri | Betaproteoba | unclassified | unclassified | unclassified | Betaproteoba | metabolism | Energy produc | 17CGT | F(1)F(0) ATP s | 296591.Bpro_ | 4,4 | 1,019803903 | 0,000380136 | 6,33E-05 | 6 - | 0,000441449 - | 3 - | 0,000274075 - | 4 - | 0,00034739 - | 5 - | 0,000440652 - | 4 - | 0,000397114 - |
| 1161 | 1 | 3um_FIJJAFBF_378479 | Bacteria | Candidatus Ha | unclassified | unclassified | unclassified | unclassified | Candidatus Ha | unclassified | unclassified | unclassified | unclassified | unclassified | 0,6 | 0,489897949 | 3,57E-05 | 2,94E-05 | 1 - | 5,22E-05 - | 1 - | 6,48E-05 - | 1 - | 6,16E-05 - | 0 - | 0 - | 0 - | 0 - |
| 1162 | 1 | 3um_FIJJAFBF_21304 | Bacteria | Bacteroidetes | Flavobacteriia | Flavobacterial | Flavobacteria | Polaribacter | Polaribacter b | poorly charact | Function unkn | 09EIH | NA | 867900.Celly_ | 0,4 | 0,8 | 2,72E-05 | 5,43E-05 | 2 - | 0,00013583 - | 0 - | 0 - | 0 - | 0 - | 0 - | 0 - | 0 - | 0 - |
| 1163 | 1 | 0.2um_NOKCPHJN_40847 | Bacteria | Proteobacteri | Gammaprote | unclassified | unclassified | unclassified | Gammaprote | metabolism | Lipid transpor | 16QA8 | reductase | 393595.ABO_ | 0,2 | 0,4 | 1,12E-05 | 2,23E-05 | 0 - | 0 - | 0 - | 0 - | 1 - | 5,58E-05 - | 0 - | 0 - | 0 - | 0 - |
| 1164 | 1 | 3um_FIJJAFBF_374867 | Bacteria | Proteobacteri | Alphaproteob | Pelagibacteral | Pelagibactera | Candidatus Pe | Candidatus Pe | unclassified | unclassified | unclassified | unclassified | unclassified | 0,4 | 0,489897949 | 2,56E-05 | 3,14E-05 | 0 - | 0 - | 0 - | 0 - | 1 - | 6,36E-05 - | 1 - | 6,45E-05 - | 0 - | 0 - |
| 1165 | 1 | 3um_FIJJAFBF_31222 | Bacteria | Bacteroidetes | Flavobacteriia | Flavobacterial | Flavobacteria | Algibacter | Algibacter pe | metabolism | Lipid transpor | COG0439 | acetyl-CoA ca | 391587.KAOT | 1 | 0 | 3,05E-05 | 2,89E-06 | 1 - | 2,55E-05 - | 1 - | 3,17E-05 - | 1 - | 3,01E-05 - | 1 - | 3,06E-05 - | 1 - | 3,44E-05 - |
| 1166 | 1 | 0.2um_NOKCPHJN_434228 | Bacteria | Bacteroidetes | Flavobacteriia | unclassified | unclassified | unclassified | Flavobacteria | metabolism | Inorganic ion t | 05C27 | TonB-depende | 487797.Flav3 | 4,6 | 1,496662955 | 5,49E-05 | 1,52E-05 | 6 - | 6,17E-05 - | 4 - | 5,10E-05 - | 6 - | 7,28E-05 - | 5 - | 6,15E-05 - | 2 - | 2,77E-05 - |
| 1167 | 1 | 3um_FIJJAFBF_620964 | Bacteria | Bacteroidetes | Flavobacteriia | Flavobacterial | Flavobacteria | Hyunsoonleel | Hyunsoonleel | metabolism | Amino acid tra | COG0137 | Citrulline--asp | 313598.MED1 | 1 | 0,632455532 | 3,30E-05 | 1,88E-05 | 2 - | 5,87E-05 - | 1 - | 3,64E-05 - | 1 - | 3,47E-05 - | 1 - | 3,52E-05 - | 0 - | 0 - |
| 1168 | 1 | 0.2um_NOKCPHJN_553017 | Bacteria | Bacteroidetes | Flavobacteriia | Flavobacterial | Flavobacteria | Tenacibaculur | Tenacibaculur | unclassified | unclassified | unclassified | unclassified | unclassified | 0,4 | 0,489897949 | 5,65E-06 | 6,99E-06 | 1 - | 1,26E-05 - | 1 - | 1,57E-05 - | 0 - | 0 - | 0 - | 0 - | 0 - | 0 - |
| 1169 | 1 | 3um_FIJJAFBF_112131 | Bacteria | Bacteroidetes | Flavobacteriia | Flavobacterial | Crocinitomica | unclassified | Crocinitomica | unclassified | unclassified | unclassified | unclassified | unclassified | 0,6 | 0,8 | 2,13E-05 | 2,75E-05 | 2 - | 6,69E-05 - | 0 - | 0 - | 1 - | 3,95E-05 - | 0 - | 0 - | 0 - | 0 - |
| 1170 | 1 | 3um_FIJJAFBF_119746 | Bacteria | Actinobacteri | Actinobacteri | Micrococcales | Microbacteria | Candidatus Ac | Candidatus Ac | unclassified | unclassified | unclassified | unclassified | unclassified | 1,4 | 0,489897949 | 4,11E-05 | 1,29E-05 | 2 - | 5,00E-05 - | 2 - | 6,21E-05 - | 1 - | 2,95E-05 - | 1 - | 3,00E-05 - | 1 - | 3,37E-05 - |
| 1171 | 1 | 0.2um_NOKCPHJN_636078 | Bacteria | Proteobacteri | Gammaprote | Cellvibrionale | Spongiibacter | Oceanicoccus | Oceanicoccus | metabolism | Nucleotide tra | COG0015 | adenylosuccin | 207949.RED6 | 0,2 | 0,4 | 1,27E-05 | 2,55E-05 | 0 - | 0 - | 0 - | 0 - | 0 - | 0 - | 1 - | 6,36E-05 - | 0 - | 0 - |
| 1172 | 1 | 3um_FIJJAFBF_189724 | Bacteria | Bacteroidetes | Flavobacteriia | Flavobacterial | Flavobacteria | Polaribacter | Polaribacter a | metabolism | Lipid transpor | 0P83H | phosphatidyls | 313598.MED1 | 4,4 | 0,8 | 0,000272486 | 2,50E-05 | 6 - | 0,000315899 - | 4 - | 0,000261503 - | 4 - | 0,000248591 - | 4 - | 0,000252263 - | 4 - | 0,000284173 - |
| 1173 | 1 | 0.2um_NOKCPHJN_103501 | Bacteria | Proteobacteri | Alphaproteob | Rhodobactera | Rhodobactera | unclassified | Rhodobactera | unclassified | unclassified | unclassified | unclassified | unclassified | 2,8 | 0,748331477 | 8,56E-05 | 1,90E-05 | 4 - | 0,00010387 - | 2 - | 6,45E-05 - | 2 - | 6,13E-05 - | 3 - | 9,33E-05 - | 3 - | 0,000105118 - |
| 1174 | 1 | 0.2um_NOKCPHJN_495443 | Bacteria | Proteobacteri | Alphaproteob | unclassified | unclassified | Candidatus Pu | Candidatus Pu | metabolism | Inorganic ion t | 05PWN | phosphate ab | 488538.SAR11 | 1,2 | 0,4 | 4,92E-05 | 1,09E-05 | 2 - | 7,06E-05 - | 1 - | 4,39E-05 - | 1 - | 4,17E-05 - | 1 - | 4,23E-05 - | 1 - | 4,77E-05 - |
| 1175 | 1 | 3um_FIJJAFBF_184996 | Eukaryota | unclassified | Oomycetes | Albuginales | Albuginaceae | Albugo | Albugo laibac | information st | Translation, ri | KOG1749 | Ribosomal pro | 653948.CCA1 | 0,2 | 0,4 | 1,90E-05 | 3,79E-05 | 0 - | 0 - | 0 - | 0 - | 0 - | 0 - | 1 - | 9,48E-05 - | 0 - | 0 - |
| 1176 | 1 | 0.2um_NOKCPHJN_84326 | Bacteria | Bacteroidetes | Flavobacteriia | Flavobacterial | Flavobacteria | Algibacter | Algibacter sp. | poorly charact | Function unkn | 11NCV | Protein of unk | 1046627.BZA | 5,4 | 0,489897949 | 0,000464504 | 7,51E-05 | 5 - | 0,000358677 - | 5 - | 0,000445371 - | 6 - | 0,000508058 - | 5 - | 0,000429635 - | 6 - | 0,000580779 - |
| 1177 | 1 | 3um_FIJJAFBF_292460 | Bacteria | Proteobacteri | Gammaprote | Cellvibrionale | Porticoccacea | unclassified | SAR92 bacteri | unclassified | unclassified | unclassified | unclassified | unclassified | 7 | 1,095445115 | 0,001065475 | 0,000199293 | 8 - | 0,001020238 - | 7 - | 0,00110848 - | 5 - | 0,000752679 - | 7 - | 0,001069315 - | 8 - | 0,001376662 - |
| 1178 | 1 | 0.2um_NOKCPHJN_778275 | Bacteria | Proteobacteri | Alphaproteob | Rhodospirillal | Rhodospirillac | unclassified | Rhodobactera | unclassified | unclassified | unclassified | unclassified | unclassified | 1,2 | 0,4 | 5,05E-05 | 1,12E-05 | 2 - | 7,24E-05 - | 1 - | 4,50E-05 - | 1 - | 4,27E-05 - | 1 - | 4,34E-05 - | 1 - | 4,89E-05 - |
| 1179 | 1 | 3um_FIJJAFBF_119801 | unclassified | unclassified | unclassified | unclassified | unclassified | unclassified | unclassified | unclassified | unclassified | unclassified | unclassified | unclassified | 0,6 | 0,489897949 | 0,00011963 | 9,87E-05 | 1 - | 0,000173904 - | 1 - | 0,000215938 - | 0 - | 0 - | 1 - | 0,000208308 - | 0 - | 0 - |
| 1180 | 1 | 0.2um_NOKCPHJN_363833 | Bacteria | Bacteroidetes | Flavobacteriia | Flavobacterial | Flavobacteria | Formosa | Formosa sp. H | information st | Translation, ri | COG0256 | This is one of | 983544.Lacal_ | 4,4 | 1,019803903 | 0,000500458 | 7,35E-05 | 6 - | 0,000583611 - | 4 - | 0,000483115 - | 4 - | 0,000459262 - | 5 - | 0,000582556 - | 3 - | 0,000393749 - |
| 1181 | 1 | 3um_FIJJAFBF_731934 | Bacteria | Proteobacteri | Gammaprote | Oceanospirilla | Saccharospirill | Reinekea | Reinekea fors | poorly charact | Function unkn | 078AD | Bacterial SH3 | 653733.Selin_ | 0,6 | 0,8 | 5,32E-05 | 7,28E-05 | 0 - | 0 - | 0 - | 0 - | 0 - | 0 - | 1 - | 8,18E-05 - | 2 - | 0,000184374 - |
| 1182 | 1 | 3um_FIJJAFBF_737020 | Bacteria | Proteobacteri | Alphaproteob | Rhodospirillal | unclassified | unclassified | Rhodospirillal | cellular proces | Cell wall/mem | COG2885 | Ompa motb d | 331869.BAL1 | 4 | 0,894427191 | 0,000311377 | 5,07E-05 | 5 - | 0,000331725 - | 3 - | 0,000247143 - | 5 - | 0,000391567 - | 4 - | 0,00031788 - | 3 - | 0,000268568 - |
| 1183 | 1 | 3um_FIJJAFBF_85384 | Bacteria | Proteobacteri | Gammaprote | unclassified | unclassified | unclassified | OM182 bacte | cellular proces | Posttranslatio | COG0330 | Band 7 protei | 247634.GPB2 | 1,4 | 0,8 | 4,73E-05 | 2,84E-05 | 1 - | 2,87E-05 - | 2 - | 7,13E-05 - | 2 - | 6,77E-05 - | 2 - | 6,87E-05 - | 0 - | 0 - |
| 1184 | 1 | 3um_FIJJAFBF_142831 | Bacteria | Proteobacteri | Alphaproteob | Rhodobactera | unclassified | unclassified | Rhodobactera | unclassified | unclassified | unclassified | unclassified | unclassified | 1,4 | 0,489897949 | 0,00011038 | 4,00E-05 | 2 - | 0,00013269 - | 1 - | 8,24E-05 - | 1 - | 7,83E-05 - | 1 - | 7,95E-05 - | 2 - | 0,000179046 - |
| 1185 | 1 | 3um_FIJJAFBF_322771 | Bacteria | Proteobacteri | Gammaprote | Oceanospirilla | Oceanospirilla | Marinobacteri | Marinobacteri | poorly charact | Function unkn | 0QITB | UPF0070 prot | 207954.MED9 | 1,2 | 0,748331477 | 7,54E-05 | 4,86E-05 | 1 - | 5,39E-05 - | 1 - | 6,69E-05 - | 2 - | 0,000127213 - | 2 - | 0,000129092 - | 0 - | 0 - |
| 1186 | 1 | 3um_FIJJAFBF_91794 | Bacteria | Proteobacteri | Alphaproteob | Rhizobiales | unclassified | unclassified | Rhizobiales ba | unclassified | unclassified | unclassified | unclassified | unclassified | 0,6 | 0,489897949 | 9,23E-05 | 7,60E-05 | 1 - | 0,000136639 - | 0 - | 0 - | 1 - | 0,000161288 - | 1 - | 0,000163671 - | 0 - | 0 - |
| 1187 | 1 | 3um_FIJJAFBF_328867 | Bacteria | Bacteroidetes | unclassified | unclassified | unclassified | unclassified | Bacteroidetes | unclassified | unclassified | unclassified | unclassified | unclassified | 2,2 | 0,4 | 0,000137504 | 1,26E-05 | 3 - | 0,000159412 - | 2 - | 0,000131962 - | 2 - | 0,000125447 - | 2 - | 0,000127299 - | 2 - | 0,000143402 - |
| 1188 | 1 | 0.2um_NOKCPHJN_560476 | Bacteria | Proteobacteri | Alphaproteob | Pelagibacteral | Pelagibactera | Candidatus Pe | Candidatus Pe | cellular proces | Intracellular tr | 17HSD | domain protei | 96561.Dole_2 | 3,8 | 0,748331477 | 4,91E-05 | 7,78E-06 | 5 - | 5,47E-05 - | 3 - | 4,08E-05 - | 4 - | 5,17E-05 - | 3 - | 3,93E-05 - | 4 - | 5,91E-05 - |
| 1189 | 1 | 3um_FIJJAFBF_788050 | Bacteria | Proteobacteri | Alphaproteob | Rhodobactera | Rhodobactera | unclassified | Rhodobactera | information st | Translation, ri | 17FIH | One of the ea | 467661.RKLH | 2,8 | 0,4 | 0,000391413 | 6,82E-05 | 3 - | 0,000351357 - | 3 - | 0,000436282 - | 3 - | 0,000414742 - | 2 - | 0,000280578 - | 3 - | 0,000474106 - |
| 1190 | 1 | 3um_FIJJAFBF_214623 | Bacteria | Proteobacteri | Alphaproteob | Rhodobactera | unclassified | unclassified | Rhodobactera | information st | Translation, ri | 177XJ | One of the pri | 89187.ISM_01 | 0,4 | 0,489897949 | 3,30E-05 | 4,08E-05 | 1 - | 7,36E-05 - | 1 - | 9,14E-05 - | 0 - | 0 - | 0 - | 0 - | 0 - | 0 - |
| 1191 | 1 | 0.2um_NOKCPHJN_903323 | unclassified | unclassified | unclassified | unclassified | unclassified | unclassified | unclassified | unclassified | unclassified | unclassified | unclassified | unclassified | 0,2 | 0,4 | 4,66E-06 | 9,32E-06 | 0 - | 0 - | 0 - | 0 - | 0 - | 0 - | 0 - | 0 - | 1 - | 2,33E-05 - |
| 1192 | 1 | 3um_FIJJAFBF_448400 | Eukaryota | Bacillariophyt | Bacillariophyc | Naviculales | Phaeodactyla | Phaeodactylu | Phaeodactylu | unclassified | unclassified | unclassified | unclassified | unclassified | 2 | 0,632455532 | 0,000783538 | 0,000308059 | 1 - | 0,000318824 - | 2 - | 0,000791771 - | 2 - | 0,000752679 - | 2 - | 0,000763796 - | 3 - | 0,001290621 - |
| 1193 | 1 | 3um_FIJJAFBF_593553 | Bacteria | Proteobacteri | Betaproteoba | Burkholderial | Comamonada | Polaromonas | Polaromonas | information st | Translation, ri | 16QAF | thus facilitatin | 887062.HGR_ | 6 | 0 | 0,000146292 | 1,39E-05 | 6 - | 0,000122537 - | 6 - | 0,000152155 - | 6 - | 0,000144643 - | 6 - | 0,000146779 - | 6 - | 0,000165346 - |
| 1194 | 1 | 3um_FIJJAFBF_175647 | Bacteria | Proteobacteri | Betaproteoba | unclassified | unclassified | unclassified | Betaproteoba | metabolism | Lipid transpor | 05VNB | Carrier of the | 296591.Bpro_ | 0,2 | 0,4 | 2,91E-05 | 5,81E-05 | 1 - | 0,000145287 - | 0 - | 0 - | 0 - | 0 - | 0 - | 0 - | 0 - | 0 - |
| 1195 | 1 | 3um_FIJJAFBF_72480 | Bacteria | Bacteroidetes | Cytophagia | Cytophagales | Flammeovirga | unclassified | Flammeovirga | unclassified | unclassified | unclassified | unclassified | unclassified | 1,8 | 0,748331477 | 0,000132468 | 4,84E-05 | 3 - | 0,000188159 - | 2 - | 0,000155758 - | 1 - | 7,40E-05 - | 1 - | 7,51E-05 - | 2 - | 0,000169262 - |
| 1196 | 1 | 3um_FIJJAFBF_08764 | Eukaryota | Bacillariophyt | various | various | various | various | various | metabolism | Energy produc | 0J0AS | This b-type cy | 35128.Thapsd | 1 | 0 | 0,000163128 | 1,55E-05 | 1 - | 0,000136639 - | 1 - | 0,000169665 - | 1 - | 0,000161288 - | 1 - | 0,000163671 - | 1 - | 0,000184374 - |
| 1197 | 1 | 0.2um_NOKCPHJN_174369 | Bacteria | Proteobacteri | Betaproteoba | unclassified | unclassified | unclassified | Betaproteoba | unclassified | unclassified | unclassified | unclassified | unclassified | 1 | 0,632455532 | 3,02E-05 | 1,70E-05 | 2 - | 5,26E-05 - | 0 - | 0 - | 1 - | 3,11E-05 - | 1 - | 3,15E-05 - | 1 - | 3,55E-05 - |
| 1198 | 2 | 0.2um_NOKCPHJN_105241;0.2um_NOKCPHJN_2 | Bacteria | Proteobacteri | Alphaproteob | Rhodobactera | Rhodobactera | various | various | metabolism | Inorganic ion t | COG0841 | acriflavin resis | 314270.RB208 | 0,2 | 0,4 | 2,29E-06 | 4,58E-06 | 0 - | 0 - | 0 - | 0 - | 0 - | 0 - | 1 - | 1,15E-05 - | 0 - | 0 - |
| 1199 | 1 | 3um_FIJJAFBF_63911 | Bacteria | Bacteroidetes | Flavobacteriia | Flavobacterial | Flavobacteria | Polaribacter | various | poorly charact | Function unkn | 0P7D8 | NA | 313598.MED1 | 0,4 | 0,489897949 | 2,29E-05 | 2,82E-05 | 1 - | 5,24E-05 - | 0 - | 0 - | 1 - | 6,19E-05 - | 0 - | 0 - | 0 - | 0 - |
| 1200 | 1 | 3um_FIJJAFBF_363664 | unclassified | unclassified | unclassified | unclassified | unclassified | unclassified | unclassified | unclassified | unclassified | unclassified | unclassified | unclassified | 0,2 | 0,4 | 5,63E-06 | 1,13E-05 | 0 - | 0 - | 1 - | 2,82E-05 - | 0 - | 0 - | 0 - | 0 - | 0 - | 0 - |
| 1201 | 1 | 0.2um_NOKCPHJN_364412 | Bacteria | Bacteroidetes | unclassified | unclassified | unclassified | unclassified | Bacteroidetes | information st | Translation, ri | 05C20 | Catalyzes the | 487796.Flav2 | 0,6 | 0,489897949 | 1,60E-05 | 1,33E-05 | 1 - | 2,27E-05 - | 0 - | 0 - | 1 - | 2,68E-05 - | 0 - | 0 - | 1 - | 3,06E-05 - |
| 1202 | 1 | 0.2um_NOKCPHJN_171998 | Bacteria | Proteobacteri | Alphaproteob | Rhodospirillal | Rhodospirillac | unclassified | Rhodobactera | cellular proces | Posttranslatio | COG0330 | Band 7 protei | 367336.OM22 | 6,8 | 1,166190379 | 0,000242391 | 5,12E-05 | 6 - | 0,000178409 - | 6 - | 0,000221532 - | 6 - | 0,000210594 - | 9 - | 0,000320557 - | 7 - | 0,00028086 - |
| 1203 | 1 | 0.2um_NOKCPHJN_88769 | Bacteria | Bacteroidetes | Flavobacteriia | Flavobacterial | Flavobacteria | Flavobacteriu | Flavobacteriu | unclassified | unclassified | unclassified | unclassified | unclassified | 0,4 | 0,489897949 | 1,93E-06 | 2,37E-06 | 0 - | 0 - | 1 - | 4,95E-06 - | 1 - | 4,71E-06 - | 0 - | 0 - | 0 - | 0 - |
| 1204 | 1 | 3um_FIJJAFBF_390778 | Bacteria | Bacteroidetes | Flavobacteriia | Flavobacterial | Flavobacteria | Hanstruepera | Hanstruepera | information st | Transcription | COG0085 | DNA-depende | 391603.FBAL | 1,2 | 0,4 | 2,49E-05 | 9,19E-06 | 1 - | 1,73E-05 - | 2 - | 4,29E-05 - | 1 - | 2,04E-05 - | 1 - | 2,07E-05 - | 1 - | 2,33E-05 - |
| 1205 | 1 | 0.2um_NOKCPHJN_864502 | Bacteria | unclassified | unclassified | unclassified | unclassified | unclassified | uncultured ba | unclassified | unclassified | unclassified | unclassified | unclassified | 2,8 | 0,748331477 | 0,000123084 | 3,07E-05 | 3 - | 0,000111796 - | 2 - | 9,25E-05 - | 3 - | 0,000131963 - | 4 - | 0,00017855 - | 2 - | 0,000100568 - |
| 1206 | 1 | 0.2um_NOKCPHJN_922961 | Eukaryota | unclassified | Chrysophycea | Chromulinales | Chromulinace | Spumella | Spumella sp. | information st | Translation, ri | KOG0052 | This protein p | 4081.Solyc06 | 0,2 | 0,4 | 6,60E-06 | 1,32E-05 | 1 - | 3,30E-05 - | 0 - | 0 - | 0 - | 0 - | 0 - | 0 - | 0 - | 0 - |
| 1207 | 1 | 0.2um_NOKCPHJN_489476 | Bacteria | Proteobacteri | Gammaprote | Alteromonada | Alteromonada | Paraglaciecola | Paraglaciecola | information st | Translation, ri | 177SZ | This protein bi | 455436.GHTC | 2,4 | 0,489897949 | 0,000228526 | 3,56E-05 | 3 - | 0,000242486 - | 2 - | 0,000200731 - | 2 - | 0,00019082 - | 3 - | 0,000290458 - | 2 - | 0,000218133 - |
| 1208 | 1 | 3um_FIJJAFBF_588465 | Bacteria | Bacteroidetes | Flavobacteriia | Flavobacterial | Flavobacteria | Zobellia | Zobellia galac | cellular proces | Posttranslatio | 05DIW | FeS assembly | 886377.Murru | 1 | 0,632455532 | 2,68E-05 | 1,53E-05 | 2 - | 4,77E-05 - | 1 - | 2,96E-05 - | 1 - | 2,82E-05 - | 1 - | 2,86E-05 - | 0 - | 0 - |
| 1209 | 1 | 0.2um_NOKCPHJN_412358 | Bacteria | Bacteroidetes | unclassified | unclassified | unclassified | unclassified | unidentified e | poorly charact | Function unkn | 099QV | gliding motilit | 50743.SCB49_ | 1,4 | 0,8 | 6,08E-05 | 2,69E-05 | 3 - | 0,000114395 - | 1 - | 4,73E-05 - | 1 - | 4,50E-05 - | 1 - | 4,57E-05 - | 1 - | 5,15E-05 - |
| 1210 | 1 | 3um_FIJJAFBF_175150 | Bacteria | Proteobacteri | Alphaproteob | Rhodobactera | Rhodobactera | unclassified | Rhodobactera | unclassified | unclassified | unclassified | unclassified | unclassified | 1,2 | 0,4 | 0,00014028 | 5,18E-05 | 1 - | 9,73E-05 - | 2 - | 0,000241557 - | 1 - | 0,000114815 - | 1 - | 0,000116511 - | 1 - | 0,00013125 - |
| 1211 | 1 | 0.2um_NOKCPHJN_137064 | Bacteria | Bacteroidetes | Flavobacteriia | Flavobacterial | Crocinitomica | unclassified | Crocinitomica | unclassified | unclassified | unclassified | unclassified | unclassified | 0,8 | 0,4 | 5,99E-05 | 3,04E-05 | 1 - | 6,48E-05 - | 1 - | 8,05E-05 - | 1 - | 7,65E-05 - | 1 - | 7,77E-05 - | 0 - | 0 - |
| 1212 | 1 | 3um_FIJJAFBF_638555 | Bacteria | Bacteroidetes | Cytophagia | Cytophagales | Cyclobacteria | Algoriphagus | Algoriphagus | unclassified | unclassified | unclassified | unclassified | unclassified | 0,6 | 0,489897949 | 3,37E-05 | 2,78E-05 | 1 - | 4,90E-05 - | 1 - | 6,09E-05 - | 0 - | 0 - | 1 - | 5,88E-05 - | 0 - | 0 - |
| 1213 | 1 | 0.2um_NOKCPHJN_323682 | Bacteria | Proteobacteri | Alphaproteob | Rhodobactera | Rhodobactera | Shimia | Shimia aestua | information st | Translation, ri | COG0051 | tRNA binding | 398580.Dshi_ | 1,4 | 0,489897949 | 0,000151239 | 6,15E-05 | 1 - | 8,90E-05 - | 1 - | 0,00011048 - | 2 - | 0,00021005 - | 1 - | 0,000106576 - | 2 - | 0,000240115 - |
| 1214 | 1 | 0.2um_NOKCPHJN_27396 | Bacteria | Bacteroidetes | Flavobacteriia | Flavobacterial | Flavobacteria | Formosa | Formosa sp. H | metabolism | Inorganic ion t | 153M9 | iduronate-2-s | 756272.Plabr_ | 0,8 | 0,4 | 2,27E-05 | 1,15E-05 | 1 - | 2,46E-05 - | 1 - | 3,05E-05 - | 1 - | 2,90E-05 - | 1 - | 2,94E-05 - | 0 - | 0 - |
| 1215 | 1 | 3um_FIJJAFBF_749538 | Bacteria | Bacteroidetes | Flavobacteriia | Flavobacterial | Flavobacteria | Lutibacter | Lutibacter sp. | unclassified | unclassified | unclassified | unclassified | unclassified | 1,6 | 0,489897949 | 5,93E-05 | 1,91E-05 | 1 - | 3,09E-05 - | 2 - | 7,68E-05 - | 2 - | 7,30E-05 - | 2 - | 7,41E-05 - | 1 - | 4,17E-05 - |
| 1216 | 1 | 0.2um_NOKCPHJN_574534 | Bacteria | Proteobacteri | Gammaprote | Chromatiales | unclassified | unclassified | Chromatiales | metabolism | Carbohydrate | 07U2E | extracellular s | 367336.OM22 | 12,4 | 1,019803903 | 0,000391132 | 4,53E-05 | 12 - | 0,000317355 - | 11 - | 0,000361223 - | 14 - | 0,000437039 - | 13 - | 0,000411816 - | 12 - | 0,000428224 - |
| 1217 | 1 | 0.2um_NOKCPHJN_359261 | Bacteria | Bacteroidetes | Flavobacteriia | Flavobacterial | Flavobacteria | Formosa | Formosa sp. H | metabolism | Inorganic ion t | 05C27 | TonB-depende | 742766.HMPR | 0,8 | 0,4 | 1,03E-05 | 5,29E-06 | 1 - | 1,09E-05 - | 0 - | 0 - | 1 - | 1,29E-05 - | 1 - | 1,31E-05 - | 1 - | 1,47E-05 - |
| 1218 | 1 | 3um_FIJJAFBF_73701 | Bacteria | Proteobacteri | Gammaprote | Oceanospirilla | Oceanospirilla | Bermanella | Bermanella sp | poorly charact | Function unkn | 17FMV | dNA-binding p | 207949.RED6 | 0,2 | 0,4 | 2,04E-05 | 4,07E-05 | 0 - | 0 - | 0 - | 0 - | 0 - | 0 - | 1 - | 0,000101839 - | 0 - | 0 - |
| 1219 | 1 | 0.2um_NOKCPHJN_489458 | unclassified | unclassified | unclassified | unclassified | unclassified | unclassified | unclassified | unclassified | unclassified | unclassified | unclassified | unclassified | 0,2 | 0,4 | 6,85E-06 | 1,37E-05 | 1 - | 3,43E-05 - | 0 - | 0 - | 0 - | 0 - | 0 - | 0 - | 0 - | 0 - |
| 1220 | 2 | 0.2um_NOKCPHJN_530254;0.2um_NOKCPHJN_5 | Bacteria | Bacteroidetes | Flavobacteriia | Flavobacterial | Flavobacteria | Winogradskye | Winogradskye | poorly charact | Function unkn | 05END | lipolytic prote | various | 0,4 | 0,489897949 | 1,01E-05 | 1,26E-05 | 1 - | 2,26E-05 - | 1 - | 2,81E-05 - | 0 - | 0 - | 0 - | 0 - | 0 - | 0 - |
| 1221 | 1 | 0.2um_NOKCPHJN_1002461 | Bacteria | Proteobacteri | Alphaproteob | Pelagibacteral | Pelagibactera | Candidatus Pe | Candidatus Pe | metabolism | Carbohydrate | 16T1E | Periplasmic bi | 488538.SAR11 | 3 | 0 | 0,000225869 | 2,14E-05 | 3 - | 0,000189192 - | 3 - | 0,000234921 - | 3 - | 0,000223322 - | 3 - | 0,000226621 - | 3 - | 0,000255288 - |
| 1222 | 1 | 0.2um_NOKCPHJN_717324 | Bacteria | Proteobacteri | Gammaprote | Cellvibrionale | Porticoccacea | unclassified | SAR92 bacteri | cellular proces | Cell wall/mem | COG2913 | Part of the ou | 314287.GB22 | 1 | 0,632455532 | 0,000113236 | 6,35E-05 | 2 - | 0,0001962 - | 1 - | 0,000121811 - | 1 - | 0,000115797 - | 0 - | 0 - | 1 - | 0,000132371 - |
| 1223 | 1 | 0.2um_NOKCPHJN_926215 | Eukaryota | unclassified | unclassified | Isochrysidales | Isochrysidace | Chrysotila | Chrysotila car | unclassified | unclassified | unclassified | unclassified | unclassified | 3,2 | 0,4 | 0,000834686 | 4,58E-05 | 4 - | 0,000882898 - | 3 - | 0,000822224 - | 3 - | 0,000781628 - | 3 - | 0,000793173 - | 3 - | 0,000893507 - |
| 1224 | 1 | 0.2um_NOKCPHJN_64582 | Bacteria | Proteobacteri | Gammaprote | Oceanospirilla | Saccharospirill | Reinekea | Reinekea fors | cellular proces | Posttranslatio | 16Q0W | Has an import | 314283.MED2 | 0,6 | 0,8 | 4,01E-05 | 5,30E-05 | 0 - | 0 - | 1 - | 6,92E-05 - | 2 - | 0,000131536 - | 0 - | 0 - | 0 - | 0 - |
| 1225 | 1 | 3um_FIJJAFBF_191342 | Bacteria | Proteobacteri | Gammaprote | Alteromonada | Shewanellace | Shewanella | Shewanella se | unclassified | unclassified | unclassified | unclassified | unclassified | 0,6 | 0,489897949 | 2,22E-05 | 1,82E-05 | 0 - | 0 - | 1 - | 3,64E-05 - | 0 - | 0 - | 1 - | 3,51E-05 - | 1 - | 3,95E-05 - |
| 1226 | 1 | 0.2um_NOKCPHJN_82358 | Bacteria | Bacteroidetes | Flavobacteriia | Flavobacterial | Flavobacteria | Lutibacter | Lutibacter flav | unclassified | unclassified | unclassified | unclassified | unclassified | 1,4 | 1,019803903 | 1,70E-05 | 1,20E-05 | 0 - | 0 - | 1 - | 1,24E-05 - | 2 - | 2,35E-05 - | 3 - | 3,58E-05 - | 1 - | 1,34E-05 - |
| 1227 | 1 | 3um_FIJJAFBF_798846 | Bacteria | Proteobacteri | Gammaprote | Oceanospirilla | Oceanospirilla | Motiliproteus | Motiliproteus | unclassified | unclassified | unclassified | unclassified | unclassified | 12,8 | 1,166190379 | 0,000544002 | 8,08E-05 | 13 - | 0,00046195 - | 12 - | 0,000529482 - | 11 - | 0,000461395 - | 14 - | 0,000595903 - | 14 - | 0,000671283 - |
| 1228 | 1 | 0.2um_NOKCPHJN_364159 | Bacteria | Proteobacteri | Alphaproteob | Rhodobactera | Rhodobactera | Confluentimic | Confluentimic | unclassified | unclassified | unclassified | unclassified | unclassified | 2,2 | 0,4 | 7,05E-05 | 6,47E-06 | 3 - | 8,18E-05 - | 2 - | 6,77E-05 - | 2 - | 6,44E-05 - | 2 - | 6,53E-05 - | 2 - | 7,36E-05 - |
| 1229 | 1 | 0.2um_NOKCPHJN_129011 | Bacteria | Proteobacteri | Alphaproteob | Rhodobactera | unclassified | unclassified | Rhodobactera | metabolism | Amino acid tra | 05TIB | (ABC) transpo | 367336.OM22 | 2,8 | 0,4 | 0,000114622 | 2,05E-05 | 3 - | 0,000102785 - | 3 - | 0,000127629 - | 2 - | 8,09E-05 - | 3 - | 0,000123119 - | 3 - | 0,000138694 - |
| 1230 | 1 | 0.2um_NOKCPHJN_551609 | Bacteria | Bacteroidetes | Flavobacteriia | Flavobacterial | Flavobacteria | Formosa | Formosa alga | poorly charact | Function unkn | 09SDX | Inherit from N | 391598.FBBA | 0,8 | 0,4 | 8,38E-06 | 4,30E-06 | 1 - | 8,75E-06 - | 1 - | 1,09E-05 - | 0 - | 0 - | 1 - | 1,05E-05 - | 1 - | 1,18E-05 - |
| 1231 | 1 | 3um_FIJJAFBF_123266 | Bacteria | Proteobacteri | Gammaprote | Oceanospirilla | Saccharospirill | Reinekea | Reinekea fors | cellular proces | Cell motility | 0QSPW | flagellar basal | 314283.MED2 | 1,8 | 0,4 | 0,0001417 | 3,47E-05 | 2 - | 0,000131927 - | 2 - | 0,000163815 - | 2 - | 0,000155727 - | 1 - | 7,90E-05 - | 2 - | 0,000178017 - |
| 1232 | 1 | 3um_FIJJAFBF_111317 | Viruses | unclassified | unclassified | unclassified | Mimiviridae | unclassified | Yellowstone la | unclassified | unclassified | unclassified | unclassified | unclassified | 4,6 | 1,356465997 | 0,000302215 | 5,86E-05 | 7 - | 0,000395782 - | 4 - | 0,000280825 - | 4 - | 0,00026696 - | 5 - | 0,000338629 - | 3 - | 0,000228879 - |
| 1233 | 1 | 0.2um_NOKCPHJN_358348 | Bacteria | Proteobacteri | Gammaprote | unclassified | unclassified | unclassified | Gammaprote | unclassified | unclassified | unclassified | unclassified | unclassified | 5 | 0,632455532 | 8,48E-05 | 8,11E-06 | 6 - | 8,60E-05 - | 4 - | 7,12E-05 - | 5 - | 8,46E-05 - | 5 - | 8,58E-05 - | 5 - | 9,67E-05 - |
| 1234 | 1 | 3um_FIJJAFBF_102395 | Bacteria | Bacteroidetes | Cytophagia | Cytophagales | Flammeovirga | unclassified | Flammeovirga | poorly charact | Function unkn | 0G01E | NA | 269798.CHU_ | 0,4 | 0,8 | 3,48E-05 | 6,96E-05 | 2 - | 0,000173904 - | 0 - | 0 - | 0 - | 0 - | 0 - | 0 - | 0 - | 0 - |
| 1235 | 1 | 3um_FIJJAFBF_15029 | Bacteria | Bacteroidetes | Flavobacteriia | Flavobacterial | Flavobacteria | Polaribacter | Polaribacter s | metabolism | Amino acid tra | COG0473 | 3-isopropylma | 313598.MED1 | 5,6 | 2,059126028 | 0,000218036 | 8,26E-05 | 6 - | 0,0001962 - | 8 - | 0,000324829 - | 7 - | 0,000270192 - | 2 - | 7,83E-05 - | 5 - | 0,000220619 - |
| 1236 | 1 | 0.2um_NOKCPHJN_187293 | Bacteria | Proteobacteri | Alphaproteob | Rhodobactera | Rhodobactera | unclassified | Rhodobactera | information st | Translation, ri | 01QWB | One of the pri | 314270.RB208 | 0,8 | 0,748331477 | 4,40E-05 | 4,27E-05 | 1 - | 4,80E-05 - | 0 - | 0 - | 1 - | 5,67E-05 - | 2 - | 0,000115049 - | 0 - | 0 - |
| 1237 | 1 | 3um_FIJJAFBF_147295 | Bacteria | Verrucomicrol | Verrucomicrol | Verrucomicrol | Verrucomicrol | Pedosphaera | Pedosphaera | unclassified | unclassified | unclassified | unclassified | unclassified | 0,8 | 0,4 | 4,28E-05 | 2,20E-05 | 1 - | 4,48E-05 - | 1 - | 5,57E-05 - | 1 - | 5,29E-05 - | 0 - | 0 - | 1 - | 6,05E-05 - |
| 1238 | 2 | 3um_FIJJAFBF_185037;3um_FIJJAFBF_293348 | Bacteria | Bacteroidetes | Flavobacteriia | Flavobacterial | Flavobacteria | Formosa | Formosa sp. H | unclassified | unclassified | unclassified | unclassified | unclassified | 3,6 | 1,019803903 | 0,000350767 | 9,84E-05 | 4 - | 0,000330293 - | 5 - | 0,000512658 - | 3 - | 0,000292408 - | 4 - | 0,000395635 - | 2 - | 0,000222841 - |
| 1239 | 1 | 0.2um_NOKCPHJN_1076948 | Bacteria | Bacteroidetes | Cytophagia | Cytophagales | Flammeovirga | unclassified | Flammeovirga | unclassified | unclassified | unclassified | unclassified | unclassified | 0,2 | 0,4 | 7,75E-06 | 1,55E-05 | 0 - | 0 - | 1 - | 3,87E-05 - | 0 - | 0 - | 0 - | 0 - | 0 - | 0 - |
| 1240 | 1 | 0.2um_NOKCPHJN_261463 | Bacteria | Bacteroidetes | unclassified | unclassified | unclassified | unclassified | Bacteroidetes | poorly charact | Function unkn | 099QV | gliding motilit | 926562.Oweh | 1 | 0,894427191 | 4,56E-05 | 4,05E-05 | 2 - | 8,17E-05 - | 0 - | 0 - | 1 - | 4,82E-05 - | 2 - | 9,79E-05 - | 0 - | 0 - |
| 1241 | 1 | 3um_FIJJAFBF_711375 | Bacteria | Bacteroidetes | Flavobacteriia | Flavobacterial | Flavobacteria | Aquimarina | Aquimarina sp | poorly charact | Function unkn | 0PAPS | NA | 873517.HMPR | 1,6 | 0,8 | 0,000169153 | 6,93E-05 | 3 - | 0,000275464 - | 2 - | 0,00022803 - | 1 - | 0,000108386 - | 1 - | 0,000109987 - | 1 - | 0,0001239 - |
| 1242 | 1 | 3um_FIJJAFBF_396481 | Bacteria | Proteobacteri | Gammaprote | Oceanospirilla | Saccharospirill | Reinekea | Reinekea fors | unclassified | unclassified | unclassified | unclassified | unclassified | 1,8 | 0,4 | 0,000142519 | 3,49E-05 | 2 - | 0,00013269 - | 2 - | 0,000164762 - | 2 - | 0,000156627 - | 1 - | 7,95E-05 - | 2 - | 0,000179046 - |
| 1243 | 1 | 3um_FIJJAFBF_295734 | Bacteria | Bacteroidetes | Flavobacteriia | Flavobacterial | Flavobacteria | Algibacter | Algibacter sp. | cellular proces | Posttranslatio | COG1404 | peptidase (S8 | 391603.FBAL | 1 | 0 | 2,53E-05 | 2,40E-06 | 1 - | 2,12E-05 - | 1 - | 2,63E-05 - | 1 - | 2,50E-05 - | 1 - | 2,54E-05 - | 1 - | 2,86E-05 - |
| 1244 | 1 | 0.2um_NOKCPHJN_47947 | Bacteria | Bacteroidetes | Flavobacteriia | Flavobacterial | Flavobacteria | Seonamhaeic | Seonamhaeic | cellular proces | Intracellular tr | 05FGH | MotA TolQ ex | 216432.CA25 | 7,2 | 1,166190379 | 0,000392659 | 8,45E-05 | 8 - | 0,000364371 - | 7 - | 0,000395886 - | 6 - | 0,000322577 - | 6 - | 0,000327341 - | 9 - | 0,000553123 - |
| 1245 | 1 | 0.2um_NOKCPHJN_136729 | Bacteria | Bacteroidetes | Flavobacteriia | unclassified | unclassified | unclassified | Flavobacteria | poorly charact | Function unkn | 0XR4Y | Ragb susd do | 487796.Flav2 | 8,8 | 0,748331477 | 0,000204918 | 7,48E-06 | 10 - | 0,000196536 - | 8 - | 0,000195231 - | 9 - | 0,000208791 - | 9 - | 0,000211875 - | 8 - | 0,000212157 - |
| 1246 | 1 | 0.2um_NOKCPHJN_54240 | Bacteria | Proteobacteri | Gammaprote | Oceanospirilla | Oceanospirilla | Balneatrix | Balneatrix alp | unclassified | unclassified | unclassified | unclassified | unclassified | 0,8 | 0,748331477 | 7,09E-06 | 6,43E-06 | 0 - | 0 - | 1 - | 8,89E-06 - | 2 - | 1,69E-05 - | 0 - | 0 - | 1 - | 9,66E-06 - |
| 1247 | 1 | 0.2um_NOKCPHJN_59488 | Bacteria | Bacteroidetes | Flavobacteriia | Flavobacterial | Flavobacteria | unclassified | Flavobacteria | unclassified | unclassified | unclassified | unclassified | unclassified | 0,8 | 0,748331477 | 1,93E-05 | 1,67E-05 | 2 - | 4,21E-05 - | 1 - | 2,62E-05 - | 0 - | 0 - | 0 - | 0 - | 1 - | 2,84E-05 - |
| 1248 | 1 | 3um_FIJJAFBF_89867 | Eukaryota | unclassified | Oligohymeno | Peniculida | Parameciidae | Paramecium | Paramecium t | unclassified | unclassified | unclassified | unclassified | unclassified | 0,4 | 0,489897949 | 5,70E-06 | 7,00E-06 | 0 - | 0 - | 0 - | 0 - | 0 - | 0 - | 1 - | 1,34E-05 - | 1 - | 1,51E-05 - |
| 1249 | 1 | 0.2um_NOKCPHJN_692479 | Bacteria | Proteobacteri | Alphaproteob | Pelagibacteral | Pelagibactera | Candidatus Pe | Candidatus Pe | metabolism | Secondary me | 01Q86 | extracellular s | 335992.SAR11 | 1 | 0 | 9,39E-05 | 8,91E-06 | 1 - | 7,86E-05 - | 1 - | 9,76E-05 - | 1 - | 9,28E-05 - | 1 - | 9,42E-05 - | 1 - | 0,000106078 - |
| 1250 | 1 | 3um_FIJJAFBF_353348 | Bacteria | Proteobacteri | Gammaprote | unclassified | unclassified | unclassified | various | information st | Translation, ri | 0QQXI | This protein is | 247634.GPB2 | 2,6 | 0,489897949 | 0,000294175 | 4,97E-05 | 3 - | 0,000286942 - | 3 - | 0,000356297 - | 2 - | 0,000225804 - | 3 - | 0,000343708 - | 2 - | 0,000258124 - |
| 1251 | 1 | 0.2um_NOKCPHJN_12869 | Bacteria | Bacteroidetes | Flavobacteriia | Flavobacterial | Flavobacteria | Arenitalea | Arenitalea lut | information st | Translation, ri | 05C64 | Binds directly | 983544.Lacal_ | 1 | 0 | 5,96E-05 | 5,66E-06 | 1 - | 4,99E-05 - | 1 - | 6,20E-05 - | 1 - | 5,89E-05 - | 1 - | 5,98E-05 - | 1 - | 6,73E-05 - |
| 1252 | 2 | 3um_FIJJAFBF_232873;3um_FIJJAFBF_469700 | Eukaryota | various | various | various | various | various | various | poorly charact | Function unkn | COG1100 | GTP-binding P | various | 2 | 0 | 0,000129271 | 1,23E-05 | 2 - | 0,00010828 - | 2 - | 0,000134452 - | 2 - | 0,000127813 - | 2 - | 0,000129701 - | 2 - | 0,000146108 - |
| 1253 | 1 | 3um_FIJJAFBF_611292 | Bacteria | Proteobacteri | Gammaprote | Oceanospirilla | Saccharospirill | Reinekea | Reinekea fors | unclassified | unclassified | unclassified | unclassified | unclassified | 3,6 | 0,8 | 0,000325381 | 8,01E-05 | 3 - | 0,000226533 - | 4 - | 0,00037505 - | 3 - | 0,000267399 - | 5 - | 0,000452248 - | 3 - | 0,000305673 - |
| 1254 | 1 | 0.2um_NOKCPHJN_202533 | Bacteria | Bacteroidetes | Flavobacteriia | Flavobacterial | Crocinitomica | unclassified | Crocinitomica | unclassified | unclassified | unclassified | unclassified | unclassified | 1,2 | 0,748331477 | 0,000254993 | 0,00016416 | 1 - | 0,000182185 - | 1 - | 0,00022622 - | 2 - | 0,000430102 - | 2 - | 0,000436455 - | 0 - | 0 - |
| 1255 | 1 | 0.2um_NOKCPHJN_599362 | Bacteria | Proteobacteri | Alphaproteob | Pelagibacteral | Pelagibactera | Candidatus Pe | Candidatus Pe | unclassified | unclassified | unclassified | unclassified | unclassified | 3,2 | 0,748331477 | 0,000222292 | 5,65E-05 | 4 - | 0,000233049 - | 3 - | 0,000217034 - | 2 - | 0,000137545 - | 3 - | 0,000209365 - | 4 - | 0,000314466 - |
| 1256 | 1 | 0.2um_NOKCPHJN_922925 | unclassified | unclassified | unclassified | unclassified | unclassified | unclassified | unclassified | unclassified | unclassified | unclassified | unclassified | unclassified | 0,2 | 0,4 | 1,82E-05 | 3,63E-05 | 0 - | 0 - | 1 - | 9,08E-05 - | 0 - | 0 - | 0 - | 0 - | 0 - | 0 - |
| 1257 | 1 | 3um_FIJJAFBF_353111 | Bacteria | Bacteroidetes | Cytophagia | Cytophagales | Flammeovirga | unclassified | Flammeovirga | metabolism | Nucleotide tra | 0FX2V | Major role in t | 313606.M231 | 0,4 | 0,489897949 | 3,58E-05 | 4,40E-05 | 1 - | 8,20E-05 - | 0 - | 0 - | 1 - | 9,68E-05 - | 0 - | 0 - | 0 - | 0 - |
| 1258 | 1 | 0.2um_NOKCPHJN_70860 | Bacteria | Proteobacteri | Alphaproteob | unclassified | unclassified | Candidatus Pu | Candidatus Pu | unclassified | unclassified | unclassified | unclassified | unclassified | 0,4 | 0,489897949 | 1,41E-05 | 1,73E-05 | 1 - | 3,20E-05 - | 0 - | 0 - | 0 - | 0 - | 1 - | 3,83E-05 - | 0 - | 0 - |
| 1259 | 1 | 3um_FIJJAFBF_159348 | Bacteria | Bacteroidetes | Flavobacteriia | Flavobacterial | Flavobacteria | Formosa | Formosa sp. H | unclassified | unclassified | unclassified | unclassified | unclassified | 1,8 | 0,4 | 7,56E-05 | 1,77E-05 | 2 - | 7,06E-05 - | 1 - | 4,39E-05 - | 2 - | 8,34E-05 - | 2 - | 8,46E-05 - | 2 - | 9,53E-05 - |
| 1260 | 1 | 3um_FIJJAFBF_148331 | Bacteria | Bacteroidetes | Flavobacteriia | Flavobacterial | Flavobacteria | Tenacibaculur | Tenacibaculur | metabolism | Energy produc | COG2838 | Isocitrate deh | 983544.Lacal_ | 0,6 | 0,489897949 | 1,10E-05 | 9,13E-06 | 1 - | 1,56E-05 - | 0 - | 0 - | 1 - | 1,84E-05 - | 0 - | 0 - | 1 - | 2,10E-05 - |
| 1261 | 1 | 0.2um_NOKCPHJN_92435 | Bacteria | Proteobacteri | Gammaprote | Oceanospirilla | Saccharospirill | Reinekea | Reinekea fors | cellular proces | Cell motility | COG1360 | Ompa motb d | 314283.MED2 | 1,4 | 0,489897949 | 4,39E-05 | 1,29E-05 | 2 - | 5,38E-05 - | 1 - | 3,34E-05 - | 1 - | 3,17E-05 - | 2 - | 6,44E-05 - | 1 - | 3,63E-05 - |
| 1262 | 1 | 3um_FIJJAFBF_390495 | Bacteria | Proteobacteri | Gammaprote | Alteromonada | Alteromonada | Paraglaciecola | Paraglaciecola | unclassified | unclassified | unclassified | unclassified | unclassified | 0,4 | 0,489897949 | 3,77E-05 | 4,61E-05 | 0 - | 0 - | 0 - | 0 - | 1 - | 9,34E-05 - | 1 - | 9,48E-05 - | 0 - | 0 - |
| 1263 | 1 | 0.2um_NOKCPHJN_686117 | Bacteria | Proteobacteri | Gammaprote | Vibrionales | Vibrionaceae | Photobacteriu | Photobacteriu | information st | Translation, ri | 05C64 | Binds directly | 458817.Shal_ | 0,2 | 0,4 | 1,19E-05 | 2,38E-05 | 0 - | 0 - | 0 - | 0 - | 0 - | 0 - | 1 - | 5,95E-05 - | 0 - | 0 - |
| 1264 | 1 | 3um_FIJJAFBF_661300 | Eukaryota | unclassified | unclassified | Prymnesiales | Chrysochrom | Chrysochrom | various | unclassified | unclassified | unclassified | unclassified | unclassified | 0,2 | 0,4 | 7,36E-06 | 1,47E-05 | 1 - | 3,68E-05 - | 0 - | 0 - | 0 - | 0 - | 0 - | 0 - | 0 - | 0 - |
| 1265 | 1 | 0.2um_NOKCPHJN_245528 | Eukaryota | Ascomycota | Saccharomyce | Saccharomyce | Debaryomyce | Candida | Candida parap | unclassified | unclassified | unclassified | unclassified | unclassified | 0,4 | 0,489897949 | 1,87E-06 | 2,30E-06 | 1 - | 4,29E-06 - | 0 - | 0 - | 1 - | 5,06E-06 - | 0 - | 0 - | 0 - | 0 - |
| 1266 | 1 | 3um_FIJJAFBF_734716 | Bacteria | Bacteroidetes | Cytophagia | Cytophagales | Cytophagacea | Ekhidna | Ekhidna lutea | unclassified | unclassified | unclassified | unclassified | unclassified | 3 | 1,095445115 | 6,47E-05 | 2,30E-05 | 3 - | 5,47E-05 - | 3 - | 6,80E-05 - | 5 - | 0,000107697 - | 2 - | 4,37E-05 - | 2 - | 4,92E-05 - |
| 1267 | 1 | 0.2um_NOKCPHJN_39631 | Bacteria | Bacteroidetes | Flavobacteriia | Flavobacterial | Flavobacteria | Polaribacter | Polaribacter s | unclassified | unclassified | unclassified | unclassified | unclassified | 29,6 | 2,576819745 | 0,000449627 | 4,02E-05 | 33 - | 0,000421785 - | 32 - | 0,000507862 - | 29 - | 0,000437526 - | 26 - | 0,000398059 - | 28 - | 0,000482905 - |
| 1268 | 1 | 3um_FIJJAFBF_305842 | Eukaryota | Chlorophyta | unclassified | Pyramimonad | unclassified | Pyramimonas | Pyramimonas | metabolism | Energy produc | 0DCW7 | Produces ATP | 3055.DAA009 | 5 | 0,632455532 | 0,000142147 | 2,08E-05 | 5 - | 0,000119311 - | 4 - | 0,000118519 - | 6 - | 0,000169001 - | 5 - | 0,000142914 - | 5 - | 0,000160992 - |
| 1269 | 1 | 3um_FIJJAFBF_561591 | Bacteria | Bacteroidetes | Flavobacteriia | unclassified | unclassified | unclassified | Flavobacteria | poorly charact | Function unkn | 09JH9 | NA | 487796.Flav2 | 0,2 | 0,4 | 9,53E-06 | 1,91E-05 | 1 - | 4,76E-05 - | 0 - | 0 - | 0 - | 0 - | 0 - | 0 - | 0 - | 0 - |
| 1270 | 1 | 3um_FIJJAFBF_226545 | Bacteria | Proteobacteri | Alphaproteob | Rhodospirillal | Rhodospirillac | Candidatus En | Candidatus En | unclassified | unclassified | unclassified | unclassified | unclassified | 0,6 | 0,489897949 | 1,63E-05 | 1,35E-05 | 1 - | 2,37E-05 - | 1 - | 2,94E-05 - | 0 - | 0 - | 1 - | 2,84E-05 - | 0 - | 0 - |
| 1271 | 1 | 0.2um_NOKCPHJN_344349 | Bacteria | Proteobacteri | Gammaprote | Alteromonada | Colwelliaceae | Colwellia | Colwellia psyc | poorly charact | Function unkn | 0XNQ8 | alkaline phosp | 167879.CPS_1 | 0,4 | 0,8 | 1,17E-05 | 2,33E-05 | 2 - | 5,83E-05 - | 0 - | 0 - | 0 - | 0 - | 0 - | 0 - | 0 - | 0 - |
| 1272 | 1 | 0.2um_NOKCPHJN_479380 | Eukaryota | Ascomycota | Dothideomyce | Pleosporales | Leptosphaeria | Leptosphaeria | Leptosphaeria | information st | Translation, ri | 03M18 | 40S ribosomal | 5022.CBX995 | 0,6 | 0,489897949 | 5,73E-05 | 4,77E-05 | 1 - | 8,08E-05 - | 0 - | 0 - | 0 - | 0 - | 1 - | 9,68E-05 - | 1 - | 0,000109067 - |
| 1273 | 1 | 3um_FIJJAFBF_305344 | Eukaryota | unclassified | Dinophyceae | Gonyaulacales | unclassified | Lingulodinium | Lingulodinium | poorly charact | Function unkn | 0XRVT | NA | 35128.Thaps3 | 1 | 0 | 3,77E-05 | 3,58E-06 | 1 - | 3,16E-05 - | 1 - | 3,93E-05 - | 1 - | 3,73E-05 - | 1 - | 3,79E-05 - | 1 - | 4,27E-05 - |
| 1274 | 1 | 3um_FIJJAFBF_208699 | Bacteria | Bacteroidetes | unclassified | unclassified | unclassified | unclassified | unidentified e | unclassified | unclassified | unclassified | unclassified | unclassified | 2,2 | 0,748331477 | 0,000214877 | 7,49E-05 | 2 - | 0,000165146 - | 3 - | 0,000307595 - | 3 - | 0,000292408 - | 2 - | 0,000197818 - | 1 - | 0,00011142 - |
| 1275 | 1 | 3um_FIJJAFBF_89938 | Bacteria | Bacteroidetes | Flavobacteriia | Flavobacterial | Flavobacteria | unclassified | Flavobacteria | unclassified | unclassified | unclassified | unclassified | unclassified | 0,4 | 0,489897949 | 2,88E-05 | 3,55E-05 | 1 - | 6,56E-05 - | 0 - | 0 - | 0 - | 0 - | 1 - | 7,86E-05 - | 0 - | 0 - |
| 1276 | 1 | 3um_FIJJAFBF_402484 | Bacteria | Bacteroidetes | Flavobacteriia | various | various | various | various | information st | Translation, ri | 09GNN | Protein S19 fo | 487797.Flav3 | 2,6 | 0,489897949 | 0,000383707 | 6,48E-05 | 3 - | 0,000374272 - | 3 - | 0,000464735 - | 2 - | 0,000294527 - | 3 - | 0,000448315 - | 2 - | 0,000336684 - |
| 1277 | 1 | 3um_FIJJAFBF_402490 | Bacteria | Bacteroidetes | Flavobacteriia | Flavobacterial | Flavobacteria | Muricauda | Muricauda ru | information st | Translation, ri | 05CEJ | Catalyzes the | 886377.Murru | 11,8 | 1,166190379 | 0,000228186 | 3,13E-05 | 13 - | 0,000210748 - | 11 - | 0,000221428 - | 10 - | 0,000191359 - | 12 - | 0,000233023 - | 13 - | 0,000284374 - |
| 1278 | 1 | 3um_FIJJAFBF_08084 | Bacteria | Bacteroidetes | Flavobacteriia | Flavobacterial | Flavobacteria | Polaribacter | Polaribacter s | metabolism | Amino acid tra | 05C65 | Catalyzes the | 313598.MED1 | 6,2 | 0,979795897 | 0,000200785 | 3,74E-05 | 5 - | 0,00013535 - | 7 - | 0,000235291 - | 7 - | 0,000223673 - | 7 - | 0,000226977 - | 5 - | 0,000182635 - |
| 1279 | 1 | 0.2um_NOKCPHJN_221328 | Bacteria | Bacteroidetes | unclassified | unclassified | unclassified | unclassified | Bacteroidetes | cellular proces | Cell wall/mem | 09B44 | efflux transpo | 50743.SCB49_ | 0,6 | 0,8 | 1,89E-05 | 2,43E-05 | 2 - | 5,93E-05 - | 0 - | 0 - | 1 - | 3,50E-05 - | 0 - | 0 - | 0 - | 0 - |
| 1280 | 1 | 3um_FIJJAFBF_559087 | Bacteria | Proteobacteri | Gammaprote | Alteromonada | Alteromonada | Paraglaciecola | Paraglaciecola | unclassified | unclassified | unclassified | unclassified | unclassified | 16 | 0,894427191 | 0,000265048 | 2,63E-05 | 17 - | 0,000236223 - | 17 - | 0,00029332 - | 15 - | 0,000246033 - | 15 - | 0,000249667 - | 16 - | 0,000299999 - |
| 1281 | 1 | 3um_FIJJAFBF_669336 | Bacteria | Proteobacteri | Gammaprote | unclassified | unclassified | unclassified | Gammaprote | unclassified | unclassified | unclassified | unclassified | unclassified | 5,4 | 0,8 | 8,09E-05 | 5,47E-06 | 7 - | 8,89E-05 - | 5 - | 7,88E-05 - | 5 - | 7,49E-05 - | 5 - | 7,60E-05 - | 5 - | 8,57E-05 - |

| | | | | | | | | | | | | | | | | | | | | | | | | | | | | |
|---|---|---|---|---|---|---|---|---|---|---|---|---|---|---|---|---|---|---|---|---|---|---|---|---|---|---|---|---|
| 1282 | 1 | 0.2um_NOKCPHJN_97282 | Bacteria | Proteobacteri | Gammaprotec | Alteromonada | Alteromonada | Paraglaciecola | Paraglaciecola | metabolism | Energy produc | COG2838 | Isocitrate deh | 983545.Glaag | 4,6 | 0,8 | 7,92E-05 | 1,17E-05 | 5 - | 7,29E-05 - | 5 - | 9,05E-05 - | 5 - | 8,61E-05 - | 5 - | 8,73E-05 - | 3 - | 5,90E-05 - |
| 1283 | 1 | 3um_FIJJAFBF_37026 | Bacteria | Proteobacteri | Alphaproteob | Rhodobactera | unclassified | unclassified | Rhodobactera | cellular proce | Intracellular tr | 05VIE | preprotein tra | 367336.OM22 | 3,2 | 0,979795897 | 0,0003706 | 8,40E-05 | 5 - | 0,000494727 - | 3 - | 0,000368583 - | 2 - | 0,00023359 - | 3 - | 0,00035556 - | 3 - | 0,000400537 - |
| 1284 | 1 | 3um_FIJJAFBF_370790 | Bacteria | Proteobacteri | Gammaprotec | Oceanospirilla | Saccharospirill | Reinekea | Reinekea fors | metabolism | Inorganic ion t | 08UTW | Plays a role in | 314283.MED2 | 5,4 | 1,356465997 | 0,000309678 | 8,91E-05 | 6 - | 0,000288142 - | 3 - | 0,000178894 - | 6 - | 0,000340123 - | 5 - | 0,000287622 - | 7 - | 0,000453607 - |
| 1285 | 1 | 3um_FIJJAFBF_388074 | Bacteria | Proteobacteri | Alphaproteob | Rhodobactera | Rhodobactera | Planktomarina | Planktomarina | unclassified | unclassified | unclassified | unclassified | unclassified | 1 | 0,632455532 | 3,94E-05 | 2,20E-05 | 2 - | 6,81E-05 - | 1 - | 4,23E-05 - | 0 - | 0 - | 1 - | 4,08E-05 - | 1 - | 4,60E-05 - |
| 1286 | 1 | 3um_FIJJAFBF_607840 | Eukaryota | unclassified | Dinophyceae | Suessiales | Symbiodiniace | Symbiodinium | Symbiodinium | unclassified | unclassified | unclassified | unclassified | unclassified | 4,2 | 0,748331477 | 0,000147352 | 2,54E-05 | 4 - | 0,000118327 - | 4 - | 0,000146927 - | 5 - | 0,00017459 - | 5 - | 0,000177169 - | 3 - | 0,000119748 - |
| 1287 | 1 | 3um_FIJJAFBF_472184 | Bacteria | Bacteroidetes | Cytophagia | Cytophagales | Flammeovirga | unclassified | Flammeovirga | unclassified | unclassified | unclassified | unclassified | unclassified | 0,6 | 0,489897949 | 3,32E-05 | 2,72E-05 | 0 - | 0 - | 1 - | 5,70E-05 - | 1 - | 5,42E-05 - | 1 - | 5,50E-05 - | 0 - | 0 - |
| 1288 | 2 | 0.2um_NOKCPHJN_57663;3um_FIJJAFBF_18332 | Bacteria | Bacteroidetes | Flavobacteriia | Flavobacterial | Flavobacteriac | various | various | information st | Translation, ri | 09G6S | 50S ribosomal | various | 2,6 | 0,489897949 | 0,00040884 | 5,92E-05 | 3 - | 0,000400384 - | 2 - | 0,000331439 - | 3 - | 0,000472612 - | 3 - | 0,000479593 - | 2 - | 0,000360173 - |
| 1289 | 1 | 0.2um_NOKCPHJN_792385 | Viruses | unclassified | unclassified | unclassified | unclassified | unclassified | uncultured M | unclassified | unclassified | unclassified | unclassified | unclassified | 0,2 | 0,4 | 1,47E-05 | 2,95E-05 | 0 - | 0 - | 0 - | 0 - | 1 - | 7,36E-05 - | 0 - | 0 - | 0 - | 0 - |
| 1290 | 1 | 0.2um_NOKCPHJN_58149 | Bacteria | Proteobacteri | Gammaprotec | Oceanospirilla | Saccharospirill | Reinekea | Reinekea fors | unclassified | unclassified | unclassified | unclassified | unclassified | 0,8 | 0,4 | 4,56E-05 | 2,30E-05 | 0 - | 0 - | 1 - | 5,70E-05 - | 1 - | 5,42E-05 - | 1 - | 5,50E-05 - | 1 - | 6,19E-05 - |
| 1291 | 1 | 0.2um_NOKCPHJN_657159 | Bacteria | Proteobacteri | Alphaproteob | Pelagibacteral | Pelagibacterac | Candidatus Pe | Candidatus Pe | cellular proce | Intracellular tr | 17HSD | domain protei | 96561.Dole_2 | 2,2 | 0,979795897 | 2,73E-05 | 1,16E-05 | 3 - | 3,18E-05 - | 3 - | 3,95E-05 - | 1 - | 1,25E-05 - | 3 - | 3,81E-05 - | 1 - | 1,43E-05 - |
| 1292 | 1 | 0.2um_NOKCPHJN_122149 | Bacteria | Bacteroidetes | Flavobacteriia | Flavobacterial | Flavobacteriac | Formosa | Formosa sp. H | poorly charact | Function unkn | 0XRCJ | NA | 391587.KAOT | 1 | 0,632455532 | 3,54E-05 | 2,32E-05 | 1 - | 3,04E-05 - | 1 - | 3,78E-05 - | 1 - | 3,59E-05 - | 2 - | 7,29E-05 - | 0 - | 0 - |
| 1293 | 1 | 0.2um_NOKCPHJN_550676 | Bacteria | Firmicutes | Clostridia | Clostridiales | Eubacteriacea | Eubacterium | Eubacterium | metabolism | Nucleotide tra | arCOG00093 | glutamine phc | 880724.Metig | 0,6 | 0,489897949 | 1,89E-05 | 1,57E-05 | 1 - | 2,67E-05 - | 0 - | 0 - | 0 - | 0 - | 1 - | 3,20E-05 - | 1 - | 3,60E-05 - |
| 1294 | 1 | 0.2um_NOKCPHJN_438098 | Bacteria | Bacteroidetes | unclassified | unclassified | unclassified | unclassified | Bacteroidetes | cellular proce | Intracellular tr | 0P8JN | The central su | 487796.Flav2 | 1 | 0 | 3,05E-05 | 2,89E-06 | 1 - | 2,55E-05 - | 1 - | 3,17E-05 - | 1 - | 3,01E-05 - | 1 - | 3,06E-05 - | 1 - | 3,44E-05 - |
| 1295 | 1 | 0.2um_NOKCPHJN_76326 | Bacteria | Bacteroidetes | Flavobacteriia | Flavobacterial | Flavobacteriac | Formosa | Formosa sp. H | unclassified | unclassified | unclassified | unclassified | unclassified | 0,8 | 0,4 | 8,91E-05 | 4,49E-05 | 0 - | 0 - | 1 - | 0,000111343 - | 1 - | 0,000105845 - | 1 - | 0,000107409 - | 1 - | 0,000120996 - |
| 1296 | 1 | 3um_FIJJAFBF_353345 | Bacteria | Proteobacteri | Gammaprotec | unclassified | unclassified | unclassified | various | unclassified | unclassified | unclassified | unclassified | unclassified | 1,6 | 0,489897949 | 0,000251143 | 7,10E-05 | 2 - | 0,000266923 - | 2 - | 0,000331439 - | 1 - | 0,000157537 - | 2 - | 0,000319729 - | 1 - | 0,000180087 - |
| 1297 | 1 | 3um_FIJJAFBF_476392 | Bacteria | Proteobacteri | Gammaprotec | Oceanospirilla | Halomonadac | Halomonas | Halomonas sp | metabolism | Coenzyme tra | 0CZE3 | Together with | 517418.Ctha_ | 0,4 | 0,489897949 | 3,69E-05 | 4,53E-05 | 0 - | 0 - | 1 - | 8,85E-05 - | 0 - | 0 - | 0 - | 0 - | 1 - | 9,62E-05 - |
| 1298 | 1 | 3um_FIJJAFBF_759391 | Bacteria | Bacteroidetes | Flavobacteriia | unclassified | unclassified | unclassified | Flavobacteria | information st | Translation, ri | 08UHY | One of the pri | 487796.Flav2 | 1 | 0 | 8,67E-05 | 8,23E-06 | 1 - | 7,26E-05 - | 1 - | 9,02E-05 - | 1 - | 8,57E-05 - | 1 - | 8,70E-05 - | 1 - | 9,80E-05 - |
| 1299 | 1 | 0.2um_NOKCPHJN_240674 | Bacteria | Proteobacteri | Gammaprotec | unclassified | unclassified | unclassified | Gammaprotec | unclassified | unclassified | unclassified | unclassified | unclassified | 0,6 | 0,489897949 | 4,04E-05 | 3,32E-05 | 1 - | 5,98E-05 - | 0 - | 0 - | 1 - | 7,06E-05 - | 1 - | 7,16E-05 - | 0 - | 0 - |
| 1300 | 1 | 0.2um_NOKCPHJN_116120 | Bacteria | Bacteroidetes | Flavobacteriia | Flavobacterial | Flavobacteriac | Psychroserpen | Psychroserpen | information st | Translation, ri | 05G6W | One of the pri | 391603.FBALC | 1 | 0 | 6,82E-05 | 6,47E-06 | 1 - | 5,71E-05 - | 1 - | 7,09E-05 - | 1 - | 6,74E-05 - | 1 - | 6,84E-05 - | 1 - | 7,71E-05 - |
| 1301 | 1 | 0.2um_NOKCPHJN_162928 | Bacteria | Proteobacteri | Gammaprotec | Oceanospirilla | Saccharospirill | Reinekea | Reinekea fors | metabolism | Nucleotide tra | 08UHA | Essential for r | 314283.MED2 | 0,6 | 0,8 | 4,37E-05 | 5,93E-05 | 0 - | 0 - | 1 - | 6,88E-05 - | 0 - | 0 - | 0 - | 0 - | 2 - | 0,000149637 - |
| 1302 | 1 | 0.2um_NOKCPHJN_09371 | Bacteria | Bacteroidetes | Flavobacteriia | Flavobacterial | Flavobacteriac | Polaribacter | Polaribacter v | metabolism | Energy produc | COG0055 | Produces ATP | 411154.GFO_ | 6,2 | 0,4 | 0,000168016 | 9,05E-06 | 7 - | 0,000159729 - | 6 - | 0,000170003 - | 6 - | 0,000161609 - | 6 - | 0,000163996 - | 6 - | 0,000184741 - |
| 1303 | 1 | 0.2um_NOKCPHJN_416512 | Bacteria | Bacteroidetes | Flavobacteriia | Flavobacterial | Flavobacteriac | Formosa | Formosa agari | metabolism | Energy produc | 099YH | Pyruvate dehy | 983548.Krodi_ | 1 | 0 | 4,10E-05 | 3,90E-06 | 1 - | 3,44E-05 - | 1 - | 4,27E-05 - | 1 - | 4,06E-05 - | 1 - | 4,12E-05 - | 1 - | 4,64E-05 - |
| 1304 | 1 | 3um_FIJJAFBF_393909 | unclassified | unclassified | unclassified | unclassified | unclassified | unclassified | unclassified | unclassified | unclassified | unclassified | unclassified | unclassified | 0,2 | 0,4 | 6,58E-06 | 1,32E-05 | 0 - | 0 - | 0 - | 0 - | 0 - | 0 - | 1 - | 3,29E-05 - | 0 - | 0 - |
| 1305 | 2 | 0.2um_NOKCPHJN_397156;3um_FIJJAFBF_52275 | Bacteria | Bacteroidetes | Flavobacteriia | Flavobacterial | Flavobacteriac | Formosa | Formosa sp. H | metabolism | Nucleotide tra | 0P7DG | Catalyzes the | 983544.Lacal_ | 0,4 | 0,8 | 2,39E-05 | 4,78E-05 | 2 - | 0,000119559 - | 0 - | 0 - | 0 - | 0 - | 0 - | 0 - | 0 - | 0 - |
| 1306 | 1 | 3um_FIJJAFBF_567396 | Bacteria | Proteobacteri | Alphaproteob | Rhodobactera | unclassified | unclassified | Rhodobactera | unclassified | unclassified | unclassified | unclassified | unclassified | 3 | 1,095445115 | 0,000155279 | 3,97E-05 | 5 - | 0,000224173 - | 2 - | 0,000111343 - | 3 - | 0,000158768 - | 3 - | 0,000161113 - | 2 - | 0,000120996 - |
| 1307 | 1 | 0.2um_NOKCPHJN_817417 | Bacteria | Verrucomicrol | Verrucomicrol | Verrucomicrol | Rubritaleacea | Rubritalea | Rubritalea pro | metabolism | Inorganic ion t | 1DH50 | phosphate bin | 240016.VspiD | 1 | 0 | 6,04E-05 | 5,73E-06 | 1 - | 5,06E-05 - | 1 - | 6,28E-05 - | 1 - | 5,97E-05 - | 1 - | 6,06E-05 - | 1 - | 6,82E-05 - |
| 1308 | 1 | 3um_FIJJAFBF_774175 | Eukaryota | unclassified | Pelagophycea | Pelagomonad | unclassified | Aureococcus | Aureococcus a | unclassified | unclassified | unclassified | unclassified | unclassified | 1,8 | 0,4 | 4,92E-05 | 9,72E-06 | 2 - | 4,65E-05 - | 2 - | 5,77E-05 - | 2 - | 5,49E-05 - | 2 - | 5,57E-05 - | 1 - | 3,14E-05 - |
| 1309 | 1 | 3um_FIJJAFBF_711075 | Bacteria | Bacteroidetes | Flavobacteriia | Flavobacterial | Flavobacteriac | Flaviramulus | Flaviramulus i | poorly charact | Function unkn | 0P8NI | Inherit from b | 391603.FBALC | 1,8 | 0,4 | 6,23E-05 | 1,72E-05 | 1 - | 2,85E-05 - | 2 - | 7,07E-05 - | 2 - | 6,72E-05 - | 2 - | 6,82E-05 - | 2 - | 7,69E-05 - |
| 1310 | 1 | 3um_FIJJAFBF_510246 | Bacteria | Bacteroidetes | Flavobacteriia | Flavobacterial | Flavobacteriac | Polaribacter | Polaribacter s | information st | Translation, ri | COG0256 | This is one of | 313598.MED1 | 3,4 | 1,356465997 | 0,0003924 | 0,000136301 | 5 - | 0,000494727 - | 2 - | 0,000245722 - | 5 - | 0,000583975 - | 2 - | 0,00023704 - | 3 - | 0,000400537 - |
| 1311 | 1 | 3um_FIJJAFBF_246033 | Bacteria | Proteobacteri | Alphaproteob | Rhodobactera | unclassified | unclassified | Rhodobactera | unclassified | unclassified | unclassified | unclassified | unclassified | 1,8 | 0,4 | 0,000151574 | 3,55E-05 | 2 - | 0,0001417 - | 1 - | 8,80E-05 - | 2 - | 0,000167262 - | 2 - | 0,000169732 - | 2 - | 0,000191203 - |
| 1312 | 1 | 3um_FIJJAFBF_485317 | Eukaryota | Phaeophyceae | unclassified | Ectocarpales | Ectocarpaceae | Ectocarpus | Ectocarpus sili | information st | Translation, ri | 151B4 | Ribosomal fan | 67593.Physo1 | 0,8 | 0,4 | 4,69E-05 | 2,36E-05 | 0 - | 0 - | 1 - | 5,86E-05 - | 1 - | 5,58E-05 - | 1 - | 5,66E-05 - | 1 - | 6,37E-05 - |
| 1313 | 1 | 3um_FIJJAFBF_403603 | unclassified | unclassified | unclassified | unclassified | unclassified | unclassified | unclassified | unclassified | unclassified | unclassified | unclassified | unclassified | 0,2 | 0,4 | 1,08E-05 | 2,17E-05 | 0 - | 0 - | 0 - | 0 - | 0 - | 0 - | 0 - | 0 - | 1 - | 5,42E-05 - |
| 1314 | 1 | 3um_FIJJAFBF_199360 | Bacteria | Proteobacteri | Gammaprotec | Oceanospirilla | Oceanospirilla | Motiliproteus | Motiliproteus | unclassified | unclassified | unclassified | unclassified | unclassified | 1 | 0,632455532 | 0,000124046 | 7,07E-05 | 2 - | 0,000220725 - | 1 - | 0,000137037 - | 1 - | 0,000130271 - | 1 - | 0,000132195 - | 0 - | 0 - |
| 1315 | 1 | 3um_FIJJAFBF_660317 | Bacteria | Proteobacteri | Alphaproteob | Rhodobactera | Rhodobactera | Ruegeria | Ruegeria sp. A | unclassified | unclassified | unclassified | unclassified | unclassified | 1,8 | 0,748331477 | 0,000105548 | 3,52E-05 | 3 - | 0,000152359 - | 2 - | 0,000126123 - | 2 - | 0,000119896 - | 1 - | 6,08E-05 - | 1 - | 6,85E-05 - |
| 1316 | 1 | 3um_FIJJAFBF_163319 | Bacteria | Proteobacteri | Gammaprotec | Cellvibrionales | Porticoccacea | unclassified | various | information st | Translation, ri | 179R6 | This protein is | 314287.GB22 | 1,4 | 0,8 | 0,000132455 | 7,35E-05 | 2 - | 0,000161657 - | 2 - | 0,000200731 - | 2 - | 0,00019082 - | 0 - | 0 - | 1 - | 0,000109067 - |
| 1317 | 1 | 3um_FIJJAFBF_223947 | Eukaryota | unclassified | Pelagophycea | Pelagomonad | unclassified | Aureococcus | Aureococcus a | cellular proce | Signal transdu | KOG0613 | fibronectin typ | 65071.PYU1_T | 0,4 | 0,489897949 | 1,56E-05 | 1,92E-05 | 1 - | 3,54E-05 - | 0 - | 0 - | 0 - | 0 - | 1 - | 4,24E-05 - | 0 - | 0 - |
| 1318 | 1 | 3um_FIJJAFBF_108249 | Bacteria | Proteobacteri | Gammaprotec | Oceanospirilla | Saccharospirill | Reinekea | Reinekea fors | cellular proce | Cell wall/mem | 16Q41 | Efflux transpo | 314283.MED2 | 5,4 | 0,489897949 | 0,000204059 | 2,76E-05 | 5 - | 0,000158095 - | 6 - | 0,000235568 - | 6 - | 0,000223938 - | 5 - | 0,000189371 - | 5 - | 0,000213326 - |
| 1319 | 1 | 3um_FIJJAFBF_325339 | Bacteria | Proteobacteri | Gammaprotec | unclassified | unclassified | unclassified | various | metabolism | Lipid transpor | 01YK3 | nodulation pro | 394221.Mmar | 0,8 | 0,979795897 | 6,50E-05 | 8,01E-05 | 2 - | 0,000149061 - | 0 - | 0 - | 2 - | 0,000175951 - | 0 - | 0 - | 0 - | 0 - |
| 1320 | 1 | 0.2um_NOKCPHJN_431475 | Bacteria | Proteobacteri | Alphaproteob | Rhodospirillal | Rhodospirillac | unclassified | Rhodospirillac | unclassified | unclassified | unclassified | unclassified | unclassified | 0,4 | 0,489897949 | 2,39E-06 | 2,93E-06 | 0 - | 0 - | 0 - | 0 - | 1 - | 5,93E-06 - | 1 - | 6,02E-06 - | 0 - | 0 - |
| 1321 | 4 | 0.2um_NOKCPHJN_124548;0.2um_NOKCPHJN_1 | Bacteria | various | various | various | various | various | various | metabolism | Inorganic ion t | various | various | various | 0,4 | 0,489897949 | 1,12E-05 | 1,39E-05 | 1 - | 2,50E-05 - | 1 - | 3,10E-05 - | 0 - | 0 - | 0 - | 0 - | 0 - | 0 - |
| 1322 | 1 | 3um_FIJJAFBF_49902 | Bacteria | Bacteroidetes | Flavobacteriia | Flavobacterial | Flavobacteriac | Polaribacter | Polaribacter ir | metabolism | Nucleotide tra | 09B0C | Provides the p | 313594.PI23P | 0,2 | 0,4 | 3,88E-06 | 7,76E-06 | 1 - | 1,94E-05 - | 0 - | 0 - | 0 - | 0 - | 0 - | 0 - | 0 - | 0 - |
| 1323 | 1 | 3um_FIJJAFBF_806085 | Bacteria | Proteobacteri | Gammaprotec | Oceanospirilla | Saccharospirill | Reinekea | Reinekea fors | unclassified | unclassified | unclassified | unclassified | unclassified | 2 | 0,894427191 | 0,000170194 | 7,03E-05 | 3 - | 0,000217931 - | 1 - | 9,02E-05 - | 1 - | 8,57E-05 - | 3 - | 0,000261044 - | 2 - | 0,000196044 - |
| 1324 | 1 | 0.2um_NOKCPHJN_925765 | Bacteria | Proteobacteri | Gammaprotec | Cellvibrionales | Porticoccacea | unclassified | gamma protec | unclassified | unclassified | unclassified | unclassified | unclassified | 0,4 | 0,489897949 | 1,21E-05 | 1,51E-05 | 1 - | 2,58E-05 - | 0 - | 0 - | 0 - | 0 - | 0 - | 0 - | 1 - | 3,48E-05 - |
| 1325 | 1 | 0.2um_NOKCPHJN_296260 | Bacteria | Proteobacteri | Gammaprotec | Cellvibrionales | Porticoccacea | unclassified | Gammaprotec | unclassified | unclassified | unclassified | unclassified | unclassified | 2 | 0,632455532 | 4,52E-05 | 1,74E-05 | 2 - | 3,73E-05 - | 2 - | 4,63E-05 - | 1 - | 2,20E-05 - | 2 - | 4,47E-05 - | 3 - | 7,55E-05 - |
| 1326 | 1 | 0.2um_NOKCPHJN_111514 | Bacteria | Proteobacteri | Alphaproteob | Rhodobactera | Rhodobactera | Puniceibacteri | Puniceibacteri | cellular proce | Posttranslatio | 17FI0 | Binds to Cpn6 | 89187.ISM_05 | 1,6 | 0,8 | 0,000221089 | 8,76E-05 | 3 - | 0,000362453 - | 1 - | 0,00015002 - | 2 - | 0,000285226 - | 1 - | 0,000144719 - | 1 - | 0,000163026 - |
| 1327 | 1 | 0.2um_NOKCPHJN_1035812 | Bacteria | Proteobacteri | Gammaprotec | Oceanospirilla | Oceanospirilla | Neptuniibacte | Neptuniibacte | metabolism | Carbohydrate | 16PI5 | ribulose-phos | 207954.MED9 | 0,6 | 1,2 | 8,40E-05 | 0,000167966 | 3 - | 0,000419915 - | 0 - | 0 - | 0 - | 0 - | 0 - | 0 - | 0 - | 0 - |
| 1328 | 1 | 3um_FIJJAFBF_513270 | Bacteria | Proteobacteri | Gammaprotec | Oceanospirilla | Oceanospirilla | Marinobacteri | Marinobacteri | metabolism | Energy produc | 0QJ3X | Aconitate hyd | 290339.ESA_C | 1,6 | 0,489897949 | 2,49E-05 | 7,05E-06 | 2 - | 2,65E-05 - | 2 - | 3,29E-05 - | 1 - | 1,56E-05 - | 2 - | 3,18E-05 - | 1 - | 1,79E-05 - |
| 1329 | 1 | 0.2um_NOKCPHJN_20558 | Bacteria | Proteobacteri | Gammaprotec | unclassified | unclassified | unclassified | various | metabolism | Secondary me | 05E7V | fad dependen | 190650.CC_31 | 0,8 | 0,4 | 2,00E-05 | 1,02E-05 | 1 - | 2,17E-05 - | 1 - | 2,69E-05 - | 1 - | 2,56E-05 - | 1 - | 2,60E-05 - | 0 - | 0 - |
| 1330 | 7 | 0.2um_NOKCPHJN_220766;0.2um_NOKCPHJN_3 | Bacteria | Proteobacteri | Alphaproteob | Pelagibacteral | various | various | various | metabolism | Coenzyme tra | 08ZQD | Catalyzes the | 335992.SAR11 | 0,2 | 0,4 | 1,51E-05 | 3,02E-05 | 0 - | 0 - | 1 - | 7,54E-05 - | 0 - | 0 - | 0 - | 0 - | 0 - | 0 - |
| 1331 | 1 | 3um_FIJJAFBF_449669 | Bacteria | Proteobacteri | Alphaproteob | Rhodobactera | Rhodobactera | Planktomarina | Planktomarina | cellular proce | Intracellular tr | 175GA | One of the pro | 314270.RB208 | 2,8 | 0,748331477 | 0,000223789 | 4,32E-05 | 4 - | 0,000273278 - | 3 - | 0,000254498 - | 2 - | 0,000161288 - | 3 - | 0,000245506 - | 2 - | 0,000184374 - |
| 1332 | 1 | 3um_FIJJAFBF_619966 | Bacteria | Proteobacteri | Gammaprotec | unclassified | unclassified | unclassified | Gammaprotec | information st | Translation, ri | 0QVRF | 50S ribosomal | 314285.KT71_ | 2 | 0 | 0,000435007 | 4,13E-05 | 2 - | 0,000364371 - | 2 - | 0,000452441 - | 2 - | 0,000430102 - | 2 - | 0,000436455 - | 2 - | 0,000491665 - |
| 1333 | 1 | 0.2um_NOKCPHJN_530415 | Bacteria | Proteobacteri | Gammaprotec | unclassified | unclassified | unclassified | Gammaprotec | information st | Translation, ri | COG0088 | One of the pri | 1036674.A28L | 6,4 | 1,019803903 | 0,000421884 | 5,26E-05 | 8 - | 0,000445735 - | 7 - | 0,000484287 - | 6 - | 0,000394608 - | 5 - | 0,000333697 - | 6 - | 0,000451091 - |
| 1334 | 1 | 3um_FIJJAFBF_579257 | Bacteria | Proteobacteri | Gammaprotec | Oceanospirilla | Oceanospirilla | Marinomonas | Marinomonas | metabolism | Amino acid tra | COG0174 | glutamine syn | 876044.IMCC | 0,8 | 0,4 | 2,34E-05 | 1,20E-05 | 1 - | 2,45E-05 - | 1 - | 3,05E-05 - | 1 - | 2,89E-05 - | 0 - | 0 - | 1 - | 3,31E-05 - |
| 1335 | 1 | 3um_FIJJAFBF_231373 | unclassified | unclassified | unclassified | unclassified | unclassified | unclassified | unclassified | unclassified | unclassified | unclassified | unclassified | unclassified | 1 | 0 | 9,72E-05 | 9,23E-06 | 1 - | 8,14E-05 - | 1 - | 0,000101077 - | 1 - | 9,61E-05 - | 1 - | 9,75E-05 - | 1 - | 0,00010984 - |
| 1336 | 1 | 0.2um_NOKCPHJN_258750 | Bacteria | Proteobacteri | Gammaprotec | Cellvibrionales | Porticoccacea | unclassified | SAR92 bacteri | metabolism | Amino acid tra | 05C6M | Alpha-keto-be | 167879.CPS_4 | 1 | 0 | 2,78E-05 | 2,64E-06 | 1 - | 2,33E-05 - | 1 - | 2,89E-05 - | 1 - | 2,75E-05 - | 1 - | 2,79E-05 - | 1 - | 3,14E-05 - |
| 1337 | 1 | 3um_FIJJAFBF_82366 | Bacteria | Bacteroidetes | Flavobacteriia | Flavobacterial | Flavobacteriac | Polaribacter | Polaribacter p | metabolism | Energy produc | COG0372 | citrate syntha | 313598.MED1 | 1 | 0 | 3,20E-05 | 3,04E-06 | 1 - | 2,68E-05 - | 1 - | 3,33E-05 - | 1 - | 3,17E-05 - | 1 - | 3,21E-05 - | 1 - | 3,62E-05 - |
| 1338 | 1 | 3um_FIJJAFBF_625747 | Bacteria | Bacteroidetes | Cytophagia | Cytophagales | Cytophagacea | Ekhidna | Ekhidna lutea | unclassified | unclassified | unclassified | unclassified | unclassified | 0,8 | 0,4 | 3,15E-05 | 1,62E-05 | 1 - | 3,30E-05 - | 1 - | 4,10E-05 - | 1 - | 3,89E-05 - | 0 - | 0 - | 1 - | 4,45E-05 - |
| 1339 | 1 | 0.2um_NOKCPHJN_523104 | Eukaryota | Phaeophyceae | unclassified | Ectocarpales | Ectocarpaceae | Ectocarpus | Ectocarpus sili | cellular proce | Intracellular tr | KOG2514 | Anoctamin | 65071.PYU1_T | 0,6 | 0,489897949 | 2,44E-05 | 2,00E-05 | 0 - | 0 - | 1 - | 4,00E-05 - | 0 - | 0 - | 1 - | 3,86E-05 - | 1 - | 4,35E-05 - |
| 1340 | 1 | 3um_FIJJAFBF_523891 | Bacteria | Proteobacteri | Gammaprotec | Pseudomonad | Moraxellaceae | Perlucidibaca | Perlucidibaca | cellular proce | Cell wall/mem | COG2885 | Ompa motb d | 259536.Psyc_ | 2,4 | 0,489897949 | 0,00011194 | 2,50E-05 | 3 - | 0,000117519 - | 2 - | 9,73E-05 - | 2 - | 9,25E-05 - | 2 - | 9,38E-05 - | 3 - | 0,000158575 - |
| 1341 | 1 | 3um_FIJJAFBF_374670 | Bacteria | Proteobacteri | Alphaproteob | Rhodobactera | Rhodobactera | Planktomarina | Planktomarina | cellular proce | Posttranslatio | 08U78 | Glutathione S- | 391626.OA30 | 0,6 | 0,489897949 | 3,86E-05 | 3,17E-05 | 1 - | 5,71E-05 - | 0 - | 0 - | 1 - | 6,74E-05 - | 1 - | 6,84E-05 - | 0 - | 0 - |
| 1342 | 1 | 3um_FIJJAFBF_461081 | Eukaryota | unclassified | Bangiophycea | Bangiales | Bangiaceae | Porphyra | Porphyra umb | metabolism | Inorganic ion t | 1E1DU | Cation transpo | 70448.Q01GZ | 0,6 | 0,489897949 | 3,90E-05 | 3,19E-05 | 0 - | 0 - | 1 - | 6,39E-05 - | 0 - | 0 - | 1 - | 6,17E-05 - | 1 - | 6,95E-05 - |
| 1343 | 1 | 0.2um_NOKCPHJN_912697 | Eukaryota | Basidiomycota | Agaricomycet | Auriculariales | Auriculariacea | Auricularia | Auricularia su | unclassified | unclassified | unclassified | unclassified | unclassified | 0,6 | 0,489897949 | 1,83E-05 | 1,50E-05 | 0 - | 0 - | 1 - | 3,14E-05 - | 1 - | 2,98E-05 - | 1 - | 3,03E-05 - | 0 - | 0 - |
| 1344 | 1 | 3um_FIJJAFBF_311757 | Bacteria | Bacteroidetes | Flavobacteriia | Flavobacterial | Flavobacteriac | Polaribacter | Polaribacter a | information st | Translation, ri | COG0050 | This protein p | 313603.FB217 | 28 | 2,28035085 | 0,000965751 | 6,60E-05 | 31 - | 0,00090078 - | 30 - | 0,001082422 - | 28 - | 0,00096038 - | 26 - | 0,000904953 - | 25 - | 0,000980218 - |
| 1345 | 1 | 3um_FIJJAFBF_587936 | Bacteria | Proteobacteri | Alphaproteob | Rhodobactera | Rhodobactera | Loktanella | Loktanella sp. | metabolism | Amino acid tra | 01RBF | Branched-chai | 375451.RD1_ | 0,4 | 0,489897949 | 1,25E-05 | 1,53E-05 | 0 - | 0 - | 1 - | 3,18E-05 - | 0 - | 0 - | 1 - | 3,07E-05 - | 0 - | 0 - |
| 1346 | 1 | 3um_FIJJAFBF_344729 | Eukaryota | Bacillariophyt | Coscinodiscop | Coscinodiscale | Coscinodiscac | Coscinodiscus | Coscinodiscus | unclassified | unclassified | unclassified | unclassified | unclassified | 1 | 0 | 0,000141265 | 1,34E-05 | 1 - | 0,000118327 - | 1 - | 0,000146927 - | 1 - | 0,000139672 - | 1 - | 0,000141735 - | 1 - | 0,000159664 - |
| 1347 | 1 | 3um_FIJJAFBF_26467 | Bacteria | Proteobacteri | Gammaprotec | Cellvibrionales | Porticoccacea | unclassified | various | information st | Translation, ri | COG0087 | One of the pri | 314287.GB22 | 0,8 | 0,748331477 | 4,95E-05 | 4,30E-05 | 2 - | 0,000108793 - | 0 - | 0 - | 0 - | 0 - | 1 - | 6,52E-05 - | 1 - | 7,34E-05 - |
| 1348 | 1 | 3um_FIJJAFBF_120908 | Bacteria | Bacteroidetes | Cytophagia | Cytophagales | Cytophagacea | Ekhidna | Ekhidna lutea | unclassified | unclassified | unclassified | unclassified | unclassified | 4 | 0,894427191 | 0,000165969 | 3,96E-05 | 5 - | 0,000174433 - | 3 - | 0,000129956 - | 4 - | 0,00016472 - | 3 - | 0,000125365 - | 5 - | 0,000235372 - |
| 1349 | 1 | 3um_FIJJAFBF_247267 | Eukaryota | Apicomplexa | Conoidasida | Eucoccidiorida | Sarcocystidae | Toxoplasma | Toxoplasma g | unclassified | unclassified | unclassified | unclassified | unclassified | 0,6 | 0,489897949 | 5,58E-05 | 4,59E-05 | 1 - | 8,26E-05 - | 0 - | 0 - | 1 - | 9,75E-05 - | 1 - | 9,89E-05 - | 0 - | 0 - |
| 1350 | 2 | 0.2um_NOKCPHJN_123116;3um_FIJJAFBF_4255 | Bacteria | Proteobacteri | Alphaproteob | Rhodobactera | Rhodobactera | various | various | metabolism | Energy produc | COG0056 | Produces ATP | various | 1,6 | 0,489897949 | 4,23E-05 | 1,20E-05 | 2 - | 4,49E-05 - | 2 - | 5,58E-05 - | 1 - | 2,65E-05 - | 2 - | 5,38E-05 - | 1 - | 3,03E-05 - |
| 1351 | 1 | 3um_FIJJAFBF_203160 | Bacteria | Proteobacteri | Gammaprotec | Thiotrichales | Thiotrichacea | Thiothrix | Thiothrix caldi | metabolism | Energy produc | COG0055 | Produces ATP | 572477.Alvin_ | 0,4 | 0,489897949 | 1,29E-05 | 1,58E-05 | 0 - | 0 - | 1 - | 3,09E-05 - | 0 - | 0 - | 0 - | 0 - | 1 - | 3,36E-05 - |
| 1352 | 2 | 0.2um_NOKCPHJN_223180;3um_FIJJAFBF_4412 | Bacteria | Bacteroidetes | Flavobacteriia | Flavobacterial | Flavobacteriac | Tenacibaculum | Tenacibaculum | information st | Translation, ri | 05G6W | One of the pri | 313598.MED1 | 1 | 0,894427191 | 6,90E-05 | 6,19E-05 | 0 - | 0 - | 2 - | 0,00014181 - | 2 - | 0,000134808 - | 1 - | 6,84E-05 - | 0 - | 0 - |
| 1353 | 2 | 0.2um_NOKCPHJN_241607;3um_FIJJAFBF_7371 | Bacteria | Proteobacteri | Alphaproteob | Pelagibacteral | Pelagibacterac | Candidatus Pe | Candidatus Pe | unclassified | unclassified | unclassified | unclassified | unclassified | 0,2 | 0,4 | 2,25E-05 | 4,51E-05 | 0 - | 0 - | 0 - | 0 - | 0 - | 0 - | 1 - | 0,000112691 - | 0 - | 0 - |
| 1354 | 1 | 0.2um_NOKCPHJN_129304 | Bacteria | Bacteroidetes | Flavobacteriia | Flavobacterial | Flavobacteriac | Spongiibacteri | Spongiibacteri | metabolism | Nucleotide tra | 09B0C | Provides the p | 688270.Celal_ | 1,4 | 0,489897949 | 2,31E-05 | 8,36E-06 | 2 - | 2,78E-05 - | 1 - | 1,72E-05 - | 1 - | 1,64E-05 - | 1 - | 1,66E-05 - | 2 - | 3,75E-05 - |
| 1355 | 1 | 3um_FIJJAFBF_580278 | Bacteria | Proteobacteri | Alphaproteob | Rhodobactera | Rhodobactera | Amylibacter | Amylibacter k | information st | Transcription | 16PMT | DNA-depende | 290400.Jann_ | 3,8 | 1,166190379 | 0,000150426 | 4,31E-05 | 5 - | 0,000167802 - | 2 - | 8,33E-05 - | 3 - | 0,000118844 - | 5 - | 0,000200999 - | 4 - | 0,00018114 - |
| 1356 | 1 | 0.2um_NOKCPHJN_307515 | Bacteria | Proteobacteri | Alphaproteob | Rhodobactera | Rhodobactera | Planktomarina | Planktomarina | metabolism | Carbohydrate | 0XPV5 | extracellular s | 391595.RLO14 | 1,4 | 0,489897949 | 4,49E-05 | 1,32E-05 | 2 - | 5,49E-05 - | 1 - | 3,41E-05 - | 1 - | 3,24E-05 - | 2 - | 6,58E-05 - | 1 - | 3,71E-05 - |
| 1357 | 1 | 0.2um_NOKCPHJN_914464 | Bacteria | Bacteroidetes | Bacteroidia | Marinilabiliale | unclassified | unclassified | Marinilabiliale | unclassified | unclassified | unclassified | unclassified | unclassified | 1 | 0 | 0,00010002 | 9,50E-06 | 1 - | 8,38E-05 - | 1 - | 0,000104028 - | 1 - | 9,89E-05 - | 1 - | 0,000100353 - | 1 - | 0,000113047 - |
| 1358 | 1 | 3um_FIJJAFBF_267219 | Bacteria | Proteobacteri | Alphaproteob | Pelagibacteral | Pelagibacterac | unclassified | Pelagibacterac | unclassified | unclassified | unclassified | unclassified | unclassified | 1 | 0 | 5,23E-05 | 4,97E-06 | 1 - | 4,38E-05 - | 1 - | 5,44E-05 - | 1 - | 5,17E-05 - | 1 - | 5,25E-05 - | 1 - | 5,91E-05 - |
| 1359 | 1 | 3um_FIJJAFBF_363802 | Bacteria | Bacteroidetes | Flavobacteriia | Flavobacterial | Crocinitomica | unclassified | Crocinitomica | information st | Translation, ri | 08YY1 | This protein is | 755732.Fluta_ | 2,8 | 0,748331477 | 0,000312971 | 5,90E-05 | 4 - | 0,000382589 - | 3 - | 0,000356297 - | 3 - | 0,000338706 - | 2 - | 0,000229139 - | 2 - | 0,000258124 - |
| 1360 | 1 | 0.2um_NOKCPHJN_166476 | Bacteria | Verrucomicrol | Verrucomicrol | Verrucomicrol | Verrucomicrol | unclassified | Verrucomicrol | unclassified | unclassified | unclassified | unclassified | unclassified | 0,2 | 0,4 | 4,28E-06 | 8,57E-06 | 0 - | 0 - | 0 - | 0 - | 0 - | 0 - | 1 - | 2,14E-05 - | 0 - | 0 - |
| 1361 | 1 | 0.2um_NOKCPHJN_495471 | Bacteria | Proteobacteri | Gammaprotec | Alteromonada | Colwelliaceae | Thalassotalea | Thalassotalea | information st | Transcription | COG0250 | Participates in | 167879.CPS_4 | 0,8 | 0,748331477 | 6,15E-05 | 6,25E-05 | 1 - | 6,31E-05 - | 0 - | 0 - | 1 - | 7,44E-05 - | 0 - | 0 - | 2 - | 0,000170192 - |
| 1362 | 1 | 3um_FIJJAFBF_354715 | Bacteria | Proteobacteri | Alphaproteob | Rhodobactera | Rhodobactera | Planktomarina | Planktomarina | cellular proce | Cell wall/mem | COG2885 | Ompa motb d | 633131.TR2A | 0,2 | 0,4 | 4,54E-06 | 9,08E-06 | 0 - | 0 - | 1 - | 2,27E-05 - | 0 - | 0 - | 0 - | 0 - | 0 - | 0 - |
| 1363 | 1 | 3um_FIJJAFBF_540150 | Bacteria | Bacteroidetes | Flavobacteriia | Flavobacterial | Flavobacteriac | Formosa | Formosa sp. H | unclassified | unclassified | unclassified | unclassified | unclassified | 2 | 0 | 0,000171284 | 1,63E-05 | 2 - | 0,000143471 - | 2 - | 0,000178149 - | 2 - | 0,000169353 - | 2 - | 0,000171854 - | 2 - | 0,000193593 - |
| 1364 | 2 | 3um_FIJJAFBF_254418;3um_FIJJAFBF_581111 | Bacteria | Bacteroidetes | Flavobacteriia | Flavobacterial | Crocinitomica | unclassified | Crocinitomica | cellular proce | Intracellular tr | 0683Z | Preprotein tra | 755732.Fluta_ | 1,2 | 0,979795897 | 0,000216317 | 0,000178469 | 2 - | 0,000314457 - | 2 - | 0,000390463 - | 0 - | 0 - | 2 - | 0,000376667 - | 0 - | 0 - |
| 1365 | 1 | 0.2um_NOKCPHJN_57906 | Bacteria | Proteobacteri | Alphaproteob | Rhodobactera | Rhodobactera | Yoonia | Yoonia litorea | unclassified | unclassified | unclassified | unclassified | unclassified | 3,8 | 0,4 | 8,12E-05 | 1,07E-05 | 4 - | 7,17E-05 - | 3 - | 6,68E-05 - | 4 - | 8,47E-05 - | 4 - | 8,59E-05 - | 4 - | 9,68E-05 - |
| 1366 | 1 | 3um_FIJJAFBF_545652 | Bacteria | Proteobacteri | Gammaprotec | Cellvibrionales | Spongiibacter | Zhongshania | Zhongshania s | information st | Replication, re | COG0513 | purine NTP-de | 342610.Patl_2 | 0,8 | 0,4 | 2,60E-05 | 1,31E-05 | 0 - | 0 - | 1 - | 3,25E-05 - | 1 - | 3,09E-05 - | 1 - | 3,13E-05 - | 1 - | 3,53E-05 - |
| 1367 | 3 | 0.2um_NOKCPHJN_947894;3um_FIJJAFBF_1938 | Viruses | unclassified | unclassified | unclassified | various | various | various | various | various | various | various | various | 0,8 | 0,4 | 4,81E-05 | 2,42E-05 | 0 - | 0 - | 1 - | 6,01E-05 - | 1 - | 5,72E-05 - | 1 - | 5,80E-05 - | 1 - | 6,53E-05 - |
| 1368 | 1 | 3um_FIJJAFBF_604544 | Bacteria | Proteobacteri | Gammaprotec | Pseudomonad | Pseudomonad | Pseudomonas | Pseudomonas | unclassified | unclassified | unclassified | unclassified | unclassified | 0,8 | 0,4 | 6,09E-05 | 3,13E-05 | 1 - | 6,38E-05 - | 1 - | 7,92E-05 - | 1 - | 7,53E-05 - | 0 - | 0 - | 1 - | 8,60E-05 - |
| 1369 | 1 | 0.2um_NOKCPHJN_22823 | Bacteria | Bacteroidetes | Flavobacteriia | Flavobacterial | Flavobacteriac | Flavivirga | Flavivirga eckl | poorly charact | Function unkn | 05YA2 | NA | 391603.FBALC | 0,4 | 0,489897949 | 7,70E-06 | 9,44E-06 | 0 - | 0 - | 1 - | 1,97E-05 - | 1 - | 1,88E-05 - | 0 - | 0 - | 0 - | 0 - |
| 1370 | 1 | 3um_FIJJAFBF_490187 | Bacteria | Proteobacteri | Alphaproteob | Rhodobactera | Rhodobactera | Planktomarina | Planktomarina | metabolism | Carbohydrate | 01RFA | ABC transport | 367336.OM22 | 0,8 | 0,4 | 7,16E-05 | 3,68E-05 | 1 - | 7,50E-05 - | 1 - | 9,31E-05 - | 1 - | 8,86E-05 - | 0 - | 0 - | 1 - | 0,000101225 - |
| 1371 | 1 | 3um_FIJJAFBF_279280 | Eukaryota | Cnidaria | Anthozoa | Scleractinia | Pocilloporidae | Stylophora | Stylophora pis | unclassified | unclassified | unclassified | unclassified | unclassified | 0,4 | 0,489897949 | 1,56E-05 | 1,92E-05 | 0 - | 0 - | 1 - | 3,98E-05 - | 0 - | 0 - | 1 - | 3,84E-05 - | 0 - | 0 - |
| 1372 | 1 | 0.2um_NOKCPHJN_159994 | Bacteria | Proteobacteri | Gammaprotec | Oceanospirilla | Oceanospirilla | Marinobacteri | Marinobacteri | unclassified | unclassified | unclassified | unclassified | unclassified | 0,8 | 0,748331477 | 1,70E-05 | 1,61E-05 | 1 - | 1,79E-05 - | 0 - | 0 - | 0 - | 0 - | 2 - | 4,28E-05 - | 1 - | 2,41E-05 - |
| 1373 | 1 | 0.2um_NOKCPHJN_211939 | Bacteria | Proteobacteri | Deltaproteob | Desulfuromon | Desulfuromon | Desulfuromon | Desulfuromon | cellular proce | Cell wall/mem | 12RRB | ankyrin 3, nod | 400682.PAC_1 | 0,2 | 0,4 | 3,02E-06 | 6,05E-06 | 0 - | 0 - | 0 - | 0 - | 0 - | 0 - | 1 - | 1,51E-05 - | 0 - | 0 - |
| 1374 | 1 | 0.2um_NOKCPHJN_379102 | Bacteria | Proteobacteri | Betaproteoba | unclassified | unclassified | unclassified | Betaproteoba | poorly charact | Function unkn | 0235A | Protein of unk | 331869.BAL19 | 0,2 | 0,4 | 3,06E-05 | 6,12E-05 | 1 - | 0,000153036 - | 0 - | 0 - | 0 - | 0 - | 0 - | 0 - | 0 - | 0 - |
| 1375 | 1 | 0.2um_NOKCPHJN_853067 | Eukaryota | unclassified | Chrysophycea | Chromulinales | Chromulinace | Spumella | Spumella unig | cellular proce | Posttranslatio | KOG0019 | heat shock pro | 67593.Physo1 | 1 | 0,632455532 | 2,35E-05 | 1,34E-05 | 2 - | 4,17E-05 - | 1 - | 2,59E-05 - | 1 - | 2,46E-05 - | 1 - | 2,50E-05 - | 0 - | 0 - |
| 1376 | 1 | 0.2um_NOKCPHJN_899472 | Bacteria | Proteobacteri | Alphaproteob | Kordiimonada | unclassified | unclassified | Kordiimonada | metabolism | Energy produc | COG0056 | Produces ATP | 59729.ENSTG | 3,4 | 0,489897949 | 9,83E-05 | 9,56E-06 | 4 - | 9,79E-05 - | 3 - | 9,12E-05 - | 4 - | 0,00011555 - | 3 - | 8,79E-05 - | 3 - | 9,91E-05 - |
| 1377 | 1 | 0.2um_NOKCPHJN_204056 | Bacteria | Proteobacteri | Gammaprotec | Alteromonada | Colwelliaceae | Colwellia | Colwellia sp. P | unclassified | unclassified | unclassified | unclassified | unclassified | 0,2 | 0,4 | 1,05E-05 | 2,10E-05 | 0 - | 0 - | 1 - | 5,26E-05 - | 0 - | 0 - | 0 - | 0 - | 0 - | 0 - |
| 1378 | 1 | 0.2um_NOKCPHJN_278478 | Bacteria | Proteobacteri | Gammaprotec | Alteromonada | Colwelliaceae | Thalassomona | various | unclassified | unclassified | unclassified | unclassified | unclassified | 1,2 | 0,4 | 0,000189108 | 6,52E-05 | 1 - | 0,000131927 - | 1 - | 0,000163815 - | 1 - | 0,000155727 - | 2 - | 0,000316054 - | 1 - | 0,000178017 - |
| 1379 | 1 | 3um_FIJJAFBF_778060 | Bacteria | Bacteroidetes | Flavobacteriia | Flavobacterial | Flavobacteriac | Polaribacter | Polaribacter s | unclassified | unclassified | unclassified | unclassified | unclassified | 1,2 | 0,4 | 4,94E-05 | 1,82E-05 | 1 - | 3,43E-05 - | 2 - | 8,51E-05 - | 1 - | 4,04E-05 - | 1 - | 4,10E-05 - | 1 - | 4,62E-05 - |
| 1380 | 1 | 3um_FIJJAFBF_62870 | Bacteria | Proteobacteri | Gammaprotec | Oceanospirilla | Oceanospirilla | Amphritea | Amphritea atl | cellular proce | Posttranslatio | 17FI0 | Binds to Cpn6 | 400668.Mmw | 3,4 | 0,489897949 | 0,000481528 | 8,77E-05 | 3 - | 0,00035498 - | 4 - | 0,000587707 - | 3 - | 0,000419017 - | 4 - | 0,000566941 - | 3 - | 0,000478993 - |
| 1381 | 1 | 0.2um_NOKCPHJN_490866 | Bacteria | Proteobacteri | Gammaprotec | unclassified | unclassified | unclassified | Gammaprotec | unclassified | unclassified | unclassified | unclassified | unclassified | 0,6 | 0,8 | 3,88E-05 | 4,90E-05 | 2 - | 0,000115936 - | 0 - | 0 - | 0 - | 0 - | 0 - | 0 - | 1 - | 7,82E-05 - |
| 1382 | 1 | 0.2um_NOKCPHJN_528757 | Bacteria | Proteobacteri | Alphaproteob | Pelagibacteral | Pelagibacterac | unclassified | Pelagibacterac | cellular proce | Cell wall/mem | 7257 NA | | 439493.PB721 | 0,6 | 0,489897949 | 2,50E-05 | 2,08E-05 | 1 - | 3,48E-05 - | 1 - | 4,32E-05 - | 0 - | 0 - | 0 - | 0 - | 1 - | 4,69E-05 - |
| 1383 | 1 | 0.2um_NOKCPHJN_307535 | Bacteria | Bacteroidetes | Flavobacteriia | Flavobacterial | Flavobacteriac | Polaribacter | Polaribacter b | cellular proce | Posttranslatio | 08V1T | peptidylprolyl | 313594.PI23P | 0,6 | 0,489897949 | 3,75E-05 | 3,07E-05 | 0 - | 0 - | 1 - | 6,14E-05 - | 0 - | 0 - | 1 - | 5,93E-05 - | 1 - | 6,68E-05 - |
| 1384 | 1 | 0.2um_NOKCPHJN_413066 | Bacteria | Proteobacteri | Alphaproteob | Rhodobactera | Rhodobactera | Planktomarina | Planktomarina | poorly charact | Function unkn | 173PK | NA | 391589.RGAI1 | 4 | 1,414213562 | 0,000145601 | 4,64E-05 | 5 - | 0,000155947 - | 4 - | 0,000154912 - | 6 - | 0,000220895 - | 3 - | 0,000112079 - | 2 - | 8,42E-05 - |
| 1385 | 1 | 3um_FIJJAFBF_06900 | Bacteria | Bacteroidetes | unclassified | unclassified | unclassified | unclassified | Bacteroidetes | unclassified | unclassified | unclassified | unclassified | unclassified | 11 | 1,897366596 | 0,000334821 | 5,81E-05 | 11 - | 0,000281818 - | 8 - | 0,000254498 - | 13 - | 0,00039314 - | 13 - | 0,000398947 - | 10 - | 0,000345702 - |
| 1386 | 1 | 0.2um_NOKCPHJN_718468 | Viruses | unclassified | unclassified | Caudovirales | Myoviridae | unclassified | Lake Baikal ph | unclassified | unclassified | unclassified | unclassified | unclassified | 1,2 | 0,4 | 0,000186504 | 6,25E-05 | 1 - | 0,000130428 - | 1 - | 0,000161953 - | 2 - | 0,000307914 - | 1 - | 0,000156231 - | 1 - | 0,000175994 - |
| 1387 | 1 | 3um_FIJJAFBF_66683 | Bacteria | Proteobacteri | Betaproteoba | unclassified | unclassified | unclassified | Betaproteoba | metabolism | Energy produc | 05DTD | Dehydrogenas | 395495.Lcho_ | 1 | 0 | 5,08E-05 | 4,82E-06 | 1 - | 4,25E-05 - | 1 - | 5,28E-05 - | 1 - | 5,02E-05 - | 1 - | 5,09E-05 - | 1 - | 5,74E-05 - |
| 1388 | 1 | 0.2um_NOKCPHJN_294415 | Bacteria | Proteobacteri | Gammaprotec | unclassified | unclassified | unclassified | Gammaprotec | metabolism | Inorganic ion t | 16TQC | TONB-depend | 314287.GB22 | 11,4 | 1,019803903 | 0,000156918 | 1,90E-05 | 12 - | 0,000138563 - | 13 - | 0,000186393 - | 11 - | 0,00014993 - | 10 - | 0,000138313 - | 11 - | 0,00017139 - |
| 1389 | 1 | 3um_FIJJAFBF_391739 | Bacteria | Proteobacteri | Gammaprotec | Oceanospirilla | Oceanospirilla | Oceanobacter | Oceanobacter | cellular proce | Posttranslatio | COG0652 | peptidyl-proly | 314277.MED1 | 1 | 0,632455532 | 8,00E-05 | 5,24E-05 | 1 - | 6,87E-05 - | 1 - | 8,53E-05 - | 1 - | 8,11E-05 - | 2 - | 0,000164651 - | 0 - | 0 - |
| 1390 | 1 | 0.2um_NOKCPHJN_123129 | Bacteria | Proteobacteri | Alphaproteob | Rhodobactera | Rhodobactera | Nioella | Nioella sedimi | cellular proce | Posttranslatio | 01QWG | Prevents misf | 633131.TR2A | 1,2 | 0,4 | 3,01E-05 | 1,11E-05 | 1 - | 2,09E-05 - | 2 - | 5,18E-05 - | 1 - | 2,46E-05 - | 1 - | 2,50E-05 - | 1 - | 2,82E-05 - |
| 1391 | 1 | 0.2um_NOKCPHJN_73777 | Bacteria | Proteobacteri | Gammaprotec | Cellvibrionales | Porticoccacea | unclassified | various | unclassified | unclassified | unclassified | unclassified | unclassified | 0,6 | 0,489897949 | 1,01E-05 | 8,31E-06 | 0 - | 0 - | 1 - | 1,67E-05 - | 1 - | 1,59E-05 - | 0 - | 0 - | 1 - | 1,81E-05 - |
| 1392 | 1 | 0.2um_NOKCPHJN_08633 | Bacteria | Proteobacteri | Gammaprotec | Alteromonada | Alteromonada | Paraglaciecola | Paraglaciecola | unclassified | unclassified | unclassified | unclassified | unclassified | 6,6 | 1,019803903 | 0,00035128 | 6,13E-05 | 7 - | 0,000312622 - | 6 - | 0,000332729 - | 8 - | 0,000421735 - | 5 - | 0,000267477 - | 7 - | 0,000421837 - |
| 1393 | 1 | 3um_FIJJAFBF_355378 | Eukaryota | unclassified | Cryptophyta | Cryptomonad | Hemiselmidac | Hemiselmis | Hemiselmis ar | information st | Transcription | COG1405 | Stabilizes TBP | 45157.CML07 | 0,4 | 0,489897949 | 1,46E-05 | 1,80E-05 | 1 - | 3,33E-05 - | 0 - | 0 - | 0 - | 0 - | 1 - | 3,99E-05 - | 0 - | 0 - |
| 1394 | 1 | 0.2um_NOKCPHJN_1065485 | unclassified | unclassified | unclassified | unclassified | unclassified | unclassified | unclassified | unclassified | unclassified | unclassified | unclassified | unclassified | 0,6 | 0,489897949 | 6,79E-05 | 5,57E-05 | 0 - | 0 - | 1 - | 0,000111343 - | 0 - | 0 - | 1 - | 0,000107409 - | 1 - | 0,000120996 - |
| 1395 | 1 | 3um_FIJJAFBF_135487 | Bacteria | Proteobacteri | Deltaproteob | Syntrophobac | Syntrophobac | Syntrophobac | Syntrophobac | cellular proce | Posttranslatio | 0HC06 | Prevents misf | 246197.MXAN | 1,2 | 0,4 | 2,83E-05 | 6,27E-06 | 2 - | 4,06E-05 - | 1 - | 2,52E-05 - | 1 - | 2,40E-05 - | 1 - | 2,43E-05 - | 1 - | 2,74E-05 - |
| 1396 | 1 | 3um_FIJJAFBF_655083 | Bacteria | Proteobacteri | Alphaproteob | Rhodobactera | Rhodobactera | Thalassobium | Thalassobium | metabolism | Amino acid tra | COG0174 | glutamine syn | 391595.RLO14 | 4 | 0 | 0,000117368 | 1,11E-05 | 4 - | 9,83E-05 - | 4 - | 0,000122072 - | 4 - | 0,000116045 - | 4 - | 0,000117759 - | 4 - | 0,000132655 - |
| 1397 | 1 | 0.2um_NOKCPHJN_503367 | Viruses | unclassified | unclassified | unclassified | unclassified | unclassified | uncultured M | poorly charact | Function unkn | 028X2 | NA | 438753.AZC_3 | 0,4 | 0,489897949 | 4,77E-06 | 5,85E-06 | 0 - | 0 - | 1 - | 1,14E-05 - | 0 - | 0 - | 0 - | 0 - | 1 - | 1,24E-05 - |
| 1398 | 1 | 3um_FIJJAFBF_306824 | Bacteria | Proteobacteri | Alphaproteob | Rhodobactera | unclassified | unclassified | Rhodobactera | unclassified | unclassified | unclassified | unclassified | unclassified | 4,6 | 1,624807681 | 0,000116516 | 3,96E-05 | 5 - | 0,000108076 - | 4 - | 0,000107359 - | 5 - | 0,000127573 - | 7 - | 0,00018124 - | 2 - | 5,83E-05 - |
| 1399 | 1 | 3um_FIJJAFBF_49903 | Bacteria | Bacteroidetes | Flavobacteriia | Flavobacterial | Flavobacteriac | Polaribacter | Polaribacter ir | metabolism | Nucleotide tra | 09B0H | reductase | 313594.PI23P | 0,8 | 0,4 | 2,59E-05 | 1,33E-05 | 1 - | 2,71E-05 - | 1 - | 3,36E-05 - | 0 - | 0 - | 1 - | 3,24E-05 - | 1 - | 3,65E-05 - |
| 1400 | 1 | 0.2um_NOKCPHJN_55610 | Bacteria | Bacteroidetes | Flavobacteriia | Flavobacterial | Flavobacteriac | Polaribacter | Polaribacter s | unclassified | unclassified | unclassified | unclassified | unclassified | 0,2 | 0,4 | 3,76E-06 | 7,52E-06 | 0 - | 0 - | 0 - | 0 - | 0 - | 0 - | 0 - | 0 - | 1 - | 1,88E-05 - |
| 1401 | 1 | 0.2um_NOKCPHJN_1010845 | Bacteria | Proteobacteri | Alphaproteob | Rhodobactera | Rhodobactera | unclassified | Rhodobactera | cellular proce | Cell wall/mem | 16RAR | cytidylyl-trans | 272942.RCAP_ | 0,4 | 0,489897949 | 2,59E-05 | 3,17E-05 | 0 - | 0 - | 1 - | 6,20E-05 - | 0 - | 0 - | 0 - | 0 - | 1 - | 6,73E-05 - |
| 1402 | 1 | 3um_FIJJAFBF_788031 | Bacteria | Proteobacteri | Alphaproteob | Rhodobactera | Rhodobactera | Sulfitobacter | Sulfitobacter r | unclassified | unclassified | unclassified | unclassified | unclassified | 2,4 | 0,489897949 | 0,000183337 | 2,86E-05 | 3 - | 0,000194537 - | 2 - | 0,000161038 - | 2 - | 0,000153087 - | 3 - | 0,000233023 - | 2 - | 0,000174999 - |
| 1403 | 1 | 0.2um_NOKCPHJN_249222 | Bacteria | Proteobacteri | Gammaprotec | unclassified | unclassified | unclassified | SAR86 cluster | poorly charact | Function unkn | 05C2A | tonB-depende | 314276.OS14 | 1 | 0,894427191 | 1,65E-05 | 1,47E-05 | 2 - | 2,91E-05 - | 2 - | 3,61E-05 - | 1 - | 1,71E-05 - | 0 - | 0 - | 0 - | 0 - |
| 1404 | 1 | 3um_FIJJAFBF_180449 | Bacteria | Proteobacteri | Alphaproteob | Rhodobactera | Rhodobactera | unclassified | Rhodobactera | metabolism | Amino acid tra | 01QH7 | sulfoacetaldeh | 744980.TRICH | 1 | 0 | 2,31E-05 | 2,20E-06 | 1 - | 1,94E-05 - | 1 - | 2,41E-05 - | 1 - | 2,29E-05 - | 1 - | 2,32E-05 - | 1 - | 2,62E-05 - |
| 1405 | 1 | 0.2um_NOKCPHJN_70815 | Bacteria | Proteobacteri | Alphaproteob | unclassified | unclassified | unclassified | SAR116 cluste | information st | Replication, re | COG0178 | The UvrABC re | 488538.SAR11 | 0,8 | 0,4 | 1,11E-05 | 5,63E-06 | 1 - | 1,20E-05 - | 1 - | 1,49E-05 - | 1 - | 1,42E-05 - | 1 - | 1,44E-05 - | 0 - | 0 - |
| 1406 | 1 | 0.2um_NOKCPHJN_387999 | Bacteria | Bacteroidetes | unclassified | unclassified | unclassified | unclassified | Bacteroidetes | information st | Translation, ri | 08UIK | This protein bi | 391603.FBALC | 0,4 | 0,489897949 | 3,52E-05 | 4,36E-05 | 1 - | 7,86E-05 - | 1 - | 9,76E-05 - | 0 - | 0 - | 0 - | 0 - | 0 - | 0 - |
| 1407 | 1 | 3um_FIJJAFBF_21312 | Bacteria | Bacteroidetes | Flavobacteriia | Flavobacterial | Flavobacteriac | Polaribacter | Polaribacter a | information st | Translation, ri | 0P7DE | amino acids su | 313598.MED1 | 1,8 | 0,748331477 | 2,62E-05 | 8,53E-06 | 3 - | 3,80E-05 - | 1 - | 1,57E-05 - | 2 - | 2,99E-05 - | 2 - | 3,04E-05 - | 1 - | 1,71E-05 - |
| 1408 | 1 | 3um_FIJJAFBF_777843 | Bacteria | Bacteroidetes | Flavobacteriia | Flavobacterial | Flavobacteriac | Polaribacter | Polaribacter s | metabolism | Lipid transpor | 0P7W4 | reductase | 1046627.BZAR | 6,4 | 1,2 | 0,000350542 | 5,95E-05 | 7 - | 0,000323967 - | 7 - | 0,000402271 - | 8 - | 0,000437039 - | 5 - | 0,000277184 - | 5 - | 0,000312247 - |
| 1409 | 1 | 0.2um_NOKCPHJN_102699 | Bacteria | Proteobacteri | Gammaprotec | Cellvibrionales | Porticoccacea | unclassified | gamma protec | unclassified | unclassified | unclassified | unclassified | unclassified | 0,2 | 0,4 | 1,10E-05 | 2,20E-05 | 0 - | 0 - | 0 - | 0 - | 1 - | 5,51E-05 - | 0 - | 0 - | 0 - | 0 - |
| 1410 | 1 | 0.2um_NOKCPHJN_495144 | unclassified | unclassified | unclassified | unclassified | unclassified | unclassified | unclassified | unclassified | unclassified | unclassified | unclassified | unclassified | 0,2 | 0,4 | 2,82E-06 | 5,64E-06 | 0 - | 0 - | 0 - | 0 - | 1 - | 1,41E-05 - | 0 - | 0 - | 0 - | 0 - |
| 1411 | 1 | 3um_FIJJAFBF_15026 | Bacteria | Bacteroidetes | Flavobacteriia | Flavobacterial | Flavobacteriac | Polaribacter | Polaribacter s | unclassified | unclassified | unclassified | unclassified | unclassified | 3,8 | 0,748331477 | 0,000109614 | 1,42E-05 | 5 - | 0,000122625 - | 4 - | 0,000121811 - | 3 - | 8,68E-05 - | 4 - | 0,000117507 - | 3 - | 9,93E-05 - |
| 1412 | 1 | 3um_FIJJAFBF_255961 | Bacteria | Bacteroidetes | Flavobacteriia | unclassified | unclassified | unclassified | Flavobacteria | information st | Translation, ri | 05CAV | thus facilitatin | 402612.FP179 | 0,4 | 0,489897949 | 8,68E-06 | 1,07E-05 | 1 - | 1,94E-05 - | 1 - | 2,40E-05 - | 0 - | 0 - | 0 - | 0 - | 0 - | 0 - |
| 1413 | 1 | 0.2um_NOKCPHJN_284453 | Bacteria | Verrucomicrol | unclassified | unclassified | unclassified | unclassified | Verrucomicrol | unclassified | unclassified | unclassified | unclassified | unclassified | 0,8 | 0,4 | 3,19E-05 | 1,64E-05 | 1 - | 3,35E-05 - | 1 - | 4,16E-05 - | 1 - | 3,95E-05 - | 0 - | 0 - | 1 - | 4,52E-05 - |
| 1414 | 1 | 3um_FIJJAFBF_159538 | Bacteria | Bacteroidetes | Flavobacteriia | Flavobacterial | Crocinitomica | unclassified | Crocinitomica | unclassified | unclassified | unclassified | unclassified | unclassified | 1,6 | 0,489897949 | 3,39E-05 | 9,57E-06 | 2 - | 3,60E-05 - | 2 - | 4,47E-05 - | 1 - | 2,12E-05 - | 2 - | 4,31E-05 - | 1 - | 2,43E-05 - |
| 1415 | 1 | 0.2um_NOKCPHJN_60420 | Bacteria | Proteobacteri | Gammaprotec | Oceanospirilla | Oceanospirilla | Marinomonas | Marinomonas | information st | Translation, ri | 08VZM | 50s ribosomal | 314277.MED1 | 0,8 | 0,4 | 6,87E-05 | 3,46E-05 | 0 - | 0 - | 1 - | 8,59E-05 - | 1 - | 8,16E-05 - | 1 - | 8,28E-05 - | 1 - | 9,33E-05 - |
| 1416 | 1 | 3um_FIJJAFBF_467050 | Bacteria | Bacteroidetes | Flavobacteriia | Flavobacterial | Flavobacteriac | Polaribacter | Polaribacter s | poorly charact | Function unkn | 11NCV | Protein of unk | 313594.PI23P | 2,4 | 1,019803903 | 0,000212357 | 8,71E-05 | 3 - | 0,000225052 - | 2 - | 0,000186299 - | 1 - | 8,86E-05 - | 4 - | 0,000359433 - | 2 - | 0,00020245 - |
| 1417 | 1 | 0.2um_NOKCPHJN_71602 | Bacteria | Proteobacteri | Alphaproteob | Rhodobactera | Rhodobactera | Primorskyibac | Primorskyibac | metabolism | Amino acid tra | 01QVA | Part of the AB | 439496.RBY4 | 0,2 | 0,4 | 6,25E-06 | 1,25E-05 | 1 - | 3,13E-05 - | 0 - | 0 - | 0 - | 0 - | 0 - | 0 - | 0 - | 0 - |
| 1418 | 1 | 0.2um_NOKCPHJN_468547 | Eukaryota | unclassified | Pelagophycea | Pelagomonad | unclassified | Aureoumbra | Aureoumbra l | metabolism | Energy produc | COG0055 | Produces ATP | 329726.AM1_ | 0,4 | 0,489897949 | 1,75E-05 | 2,15E-05 | 1 - | 3,97E-05 - | 0 - | 0 - | 0 - | 0 - | 1 - | 4,76E-05 - | 0 - | 0 - |
| 1419 | 1 | 0.2um_NOKCPHJN_170612 | Bacteria | Bacteroidetes | unclassified | unclassified | unclassified | unclassified | Bacteroidetes | cellular proce | Intracellular tr | 05FGH | MotA TolQ ex | 926562.Oweh | 1,4 | 0,8 | 6,31E-05 | 2,79E-05 | 3 - | 0,000118735 - | 1 - | 4,91E-05 - | 1 - | 4,67E-05 - | 1 - | 4,74E-05 - | 1 - | 5,34E-05 - |
| 1420 | 1 | 3um_FIJJAFBF_302783 | Bacteria | Bacteroidetes | Flavobacteriia | Flavobacterial | Flavobacteriac | Polaribacter | various | information st | Translation, ri | COG0254 | 50S ribosomal | 867900.Celly_ | 2 | 0 | 0,000330186 | 3,14E-05 | 2 - | 0,000276571 - | 2 - | 0,000343419 - | 2 - | 0,000326463 - | 2 - | 0,000331285 - | 2 - | 0,000373192 - |
| 1421 | 1 | 0.2um_NOKCPHJN_127362 | Bacteria | Proteobacteri | Alphaproteob | Rhodobactera | unclassified | unclassified | Rhodobactera | unclassified | unclassified | unclassified | unclassified | unclassified | 2,6 | 1,496662955 | 5,57E-05 | 2,87E-05 | 5 - | 9,20E-05 - | 3 - | 6,85E-05 - | 1 - | 2,17E-05 - | 1 - | 2,20E-05 - | 3 - | 7,45E-05 - |
| 1422 | 1 | 0.2um_NOKCPHJN_523103 | Eukaryota | Phaeophyceae | unclassified | Ectocarpales | Ectocarpaceae | Ectocarpus | Ectocarpus sili | cellular proce | Intracellular tr | KOG2514 | Anoctamin | 4787.PITG_09 | 0,8 | 0,4 | 4,45E-05 | 2,29E-05 | 1 - | 4,65E-05 - | 1 - | 5,77E-05 - | 0 - | 0 - | 1 - | 5,57E-05 - | 1 - | 6,27E-05 - |
| 1423 | 1 | 3um_FIJJAFBF_246180 | Bacteria | Proteobacteri | Gammaprotec | Oceanospirilla | Saccharospirill | Reinekea | Reinekea fors | unclassified | unclassified | unclassified | unclassified | unclassified | 1 | 0 | 9,26E-05 | 8,79E-06 | 1 - | 7,76E-05 - | 1 - | 9,63E-05 - | 1 - | 9,15E-05 - | 1 - | 9,29E-05 - | 1 - | 0,000104645 - |
| 1424 | 1 | 3um_FIJJAFBF_245647 | Bacteria | Bacteroidetes | Flavobacteriia | Flavobacterial | Flavobacteriac | Ulvibacter | Ulvibacter lito | unclassified | unclassified | unclassified | unclassified | unclassified | 1,4 | 0,489897949 | 0,000199604 | 7,05E-05 | 1 - | 0,000119559 - | 1 - | 0,000148457 - | 2 - | 0,000282255 - | 2 - | 0,000286424 - | 1 - | 0,000161328 - |

| | | | | | | | | | | | | | | | | | | | | | | | | | | | | |
|---|---|---|---|---|---|---|---|---|---|---|---|---|---|---|---|---|---|---|---|---|---|---|---|---|---|---|---|---|
| 1425 | 1 | 3um_FIJJAFBF_615992 | Bacteria | Bacteroidetes | Flavobacteriia | Flavobacterial | Flavobacteriac | Polaribacter | Polaribacter s | unclassified | unclassified | unclassified | unclassified | unclassified | 1 | 0 | 7,61E-05 | 7,23E-06 | 1 - | 6,38E-05 - | 1 - | 7,92E-05 - | 1 - | 7,53E-05 - | 1 - | 7,64E-05 - | 1 - | 8,60E-05 - |
| 1426 | 1 | 0.2um_NOKCPHJN_234708 | Eukaryota | unclassified | Cryptophyta | Pyrenomonad | Geminigerace | Guillardia | Guillardia thet | unclassified | unclassified | unclassified | unclassified | unclassified | 0,2 | 0,4 | 1,02E-06 | 2,04E-06 | 0 - | 0 - | 0 - | 0 - | 0 - | 0 - | 1 - | 5,09E-06 - | 0 - | 0 - |
| 1427 | 1 | 0.2um_NOKCPHJN_378532 | Bacteria | Bacteroidetes | Flavobacteriia | Flavobacterial | Flavobacteriac | Marinirhabdu | Marinirhabdu | information st | Translation, ri | COG0088 | One of the pri | 50743.SCB49_ | 2,8 | 0,4 | 0,000183533 | 3,20E-05 | 3 - | 0,000164751 - | 3 - | 0,000204573 - | 3 - | 0,000194472 - | 2 - | 0,000131563 - | 3 - | 0,000222308 - |
| 1428 | 1 | 3um_FIJJAFBF_375099 | Bacteria | Proteobacteri | Alphaproteob | Rhodobactera | Rhodobactera | unclassified | Rhodobactera | metabolism | Carbohydrate | 01U6R | trap dicarboxy | 488538.SAR11 | 0,4 | 0,489897949 | 1,46E-05 | 1,81E-05 | 1 - | 3,26E-05 - | 1 - | 4,05E-05 - | 0 - | 0 - | 0 - | 0 - | 0 - | 0 - |
| 1429 | 1 | 3um_FIJJAFBF_779837 | Bacteria | Proteobacteri | Gammaprote | Oceanospirilla | Oceanospirilla | Marinomonas | Marinomonas | unclassified | unclassified | unclassified | unclassified | unclassified | 0,8 | 0,4 | 2,54E-05 | 1,29E-05 | 1 - | 2,75E-05 - | 1 - | 3,42E-05 - | 1 - | 3,25E-05 - | 1 - | 3,30E-05 - | 0 - | 0 - |
| 1430 | 1 | 0.2um_NOKCPHJN_582287 | Bacteria | Proteobacteri | Deltaproteoba | Desulfobacter | Desulfobulbac | Desulfopila | Desulfopila ae | cellular proces | Posttranslatio | 16QU3 | Hemolysin-typ | 744980.TRICH | 2 | 0 | 6,49E-05 | 6,17E-06 | 2 - | 5,44E-05 - | 2 - | 6,75E-05 - | 2 - | 6,42E-05 - | 2 - | 6,52E-05 - | 2 - | 7,34E-05 - |
| 1431 | 1 | 3um_FIJJAFBF_525062 | Eukaryota | Bacillariophyt | Coscinodiscop | Thalassiosiral | Thalassiosirac | Thalassiosira | Thalassiosira p | cellular proces | Cell wall/mem | 17CVN | Cell envelope | 706191.PANA | 0,2 | 0,4 | 3,75E-06 | 7,49E-06 | 0 - | 0 - | 1 - | 1,87E-05 - | 0 - | 0 - | 0 - | 0 - | 0 - | 0 - |
| 1432 | 1 | 3um_FIJJAFBF_262225 | Bacteria | Candidatus M | unclassified | unclassified | unclassified | unclassified | Candidatus M | information st | Transcription | COG0557 | 3'-5' exoribon | 331113.SNE_A | 0,6 | 0,489897949 | 1,75E-05 | 1,43E-05 | 0 - | 0 - | 1 - | 3,00E-05 - | 1 - | 2,85E-05 - | 1 - | 2,89E-05 - | 0 - | 0 - |
| 1433 | 1 | 3um_FIJJAFBF_18181 | Bacteria | Bacteroidetes | Flavobacteriia | Flavobacterial | Crocinitomica | unclassified | Crocinitomica | unclassified | unclassified | unclassified | unclassified | unclassified | 0,4 | 0,489897949 | 7,01E-05 | 8,64E-05 | 1 - | 0,000159412 - | 0 - | 0 - | 0 - | 0 - | 1 - | 0,000190949 - | 0 - | 0 - |
| 1434 | 1 | 3um_FIJJAFBF_706642 | Bacteria | Actinobacteri | Actinobacteri | unclassified | unclassified | unclassified | Actinobacteri | unclassified | unclassified | unclassified | unclassified | unclassified | 0,2 | 0,4 | 4,58E-06 | 9,17E-06 | 0 - | 0 - | 1 - | 2,29E-05 - | 0 - | 0 - | 0 - | 0 - | 0 - | 0 - |
| 1435 | 1 | 3um_FIJJAFBF_302356 | Bacteria | Bacteroidetes | Flavobacteriia | Flavobacterial | Crocinitomica | Crocinitomix | Crocinitomix s | poorly charact | Function unkn | 0R4M0 | NA | 443152.MDG8 | 0,8 | 0,4 | 5,47E-05 | 2,81E-05 | 1 - | 5,71E-05 - | 1 - | 7,09E-05 - | 0 - | 0 - | 1 - | 6,84E-05 - | 1 - | 7,71E-05 - |
| 1436 | 1 | 0.2um_NOKCPHJN_589037 | Eukaryota | Streptophyta | Liliopsida | Poales | Poaceae | Hordeum | Hordeum vulg | unclassified | unclassified | unclassified | unclassified | unclassified | 2,4 | 0,8 | 0,000107622 | 3,61E-05 | 2 - | 7,53E-05 - | 2 - | 9,35E-05 - | 4 - | 0,000177682 - | 2 - | 9,02E-05 - | 2 - | 0,000101557 - |
| 1437 | 1 | 3um_FIJJAFBF_183316 | Bacteria | Bacteroidetes | Flavobacteriia | Flavobacterial | Flavobacteriac | Polaribacter | Polaribacter ir | unclassified | unclassified | unclassified | unclassified | unclassified | 0,8 | 0,4 | 7,80E-05 | 3,96E-05 | 1 - | 8,44E-05 - | 1 - | 0,000104793 - | 1 - | 9,96E-05 - | 1 - | 0,000101091 - | 0 - | 0 - |
| 1438 | 1 | 3um_FIJJAFBF_44063 | Viruses | unclassified | unclassified | unclassified | Phycodnavirid | unclassified | Chrysochromu | unclassified | unclassified | unclassified | unclassified | unclassified | 2,2 | 0,748331477 | 0,000187905 | 5,36E-05 | 3 - | 0,000220725 - | 2 - | 0,000182716 - | 3 - | 0,000260543 - | 2 - | 0,000176261 - | 1 - | 9,93E-05 - |
| 1439 | 1 | 0.2um_NOKCPHJN_622740 | Bacteria | Proteobacteri | Alphaproteob | Pelagibacteral | Pelagibacterac | unclassified | Pelagibacterac | poorly charact | Function unkn | COG3181 | extra-cytoplas | 384765.SIAM6 | 4,6 | 0,8 | 0,000193662 | 4,18E-05 | 4 - | 0,0001404 - | 4 - | 0,000174335 - | 6 - | 0,000248591 - | 4 - | 0,000168175 - | 5 - | 0,000236811 - |
| 1440 | 1 | 0.2um_NOKCPHJN_27397 | Bacteria | unclassified | unclassified | unclassified | unclassified | unclassified | uncultured ba | metabolism | Inorganic ion t | COG3119 | Sulfatase | 655815.ZPR_3 | 9 | 1,673320053 | 0,000221905 | 3,33E-05 | 11 - | 0,000229554 - | 8 - | 0,0002073 - | 7 - | 0,000172432 - | 11 - | 0,000274967 - | 8 - | 0,000225272 - |
| 1441 | 1 | 3um_FIJJAFBF_55725 | Bacteria | Proteobacteri | Gammaprote | Cellvibrionales | Porticoccacea | unclassified | various | metabolism | Energy produc | 0QJAP | Catalyzes the | 393595.ABO_ | 0,4 | 0,489897949 | 1,78E-05 | 2,19E-05 | 0 - | 0 - | 0 - | 0 - | 1 - | 4,16E-05 - | 0 - | 0 - | 1 - | 4,75E-05 - |
| 1442 | 1 | 3um_FIJJAFBF_444720 | Bacteria | Bacteroidetes | Flavobacteriia | Flavobacterial | Flavobacteriac | Formosa | Formosa sp. H | cellular proces | Intracellular tr | 08SHB | Biopolymer tr | 411154.GFO_ | 0,8 | 0,4 | 6,67E-05 | 3,39E-05 | 1 - | 7,22E-05 - | 1 - | 8,96E-05 - | 1 - | 8,52E-05 - | 1 - | 8,65E-05 - | 0 - | 0 - |
| 1443 | 1 | 0.2um_NOKCPHJN_310259 | Bacteria | Proteobacteri | Gammaprote | unclassified | unclassified | unclassified | Gammaprote | unclassified | unclassified | unclassified | unclassified | unclassified | 0,2 | 0,4 | 9,97E-06 | 1,99E-05 | 0 - | 0 - | 1 - | 4,98E-05 - | 0 - | 0 - | 0 - | 0 - | 0 - | 0 - |
| 1444 | 1 | 0.2um_NOKCPHJN_864847 | Bacteria | Proteobacteri | Gammaprote | unclassified | unclassified | unclassified | Gammaprote | unclassified | unclassified | unclassified | unclassified | unclassified | 0,2 | 0,4 | 1,86E-05 | 3,73E-05 | 0 - | 0 - | 1 - | 9,31E-05 - | 0 - | 0 - | 0 - | 0 - | 0 - | 0 - |
| 1445 | 1 | 3um_FIJJAFBF_251385 | Bacteria | Bacteroidetes | Flavobacteriia | Flavobacterial | Flavobacteriac | Polaribacter | Polaribacter s | unclassified | unclassified | unclassified | unclassified | unclassified | 5,4 | 0,489897949 | 0,000155728 | 1,46E-05 | 6 - | 0,000145594 - | 6 - | 0,000180785 - | 5 - | 0,000143216 - | 5 - | 0,000145331 - | 5 - | 0,000163715 - |
| 1446 | 1 | 3um_FIJJAFBF_88823 | Bacteria | Proteobacteri | Gammaprote | unclassified | unclassified | unclassified | various | information st | Translation, ri | 05VBI | Binds 16S rRN | 876044.IMCC3 | 0,8 | 0,4 | 0,000108842 | 5,59E-05 | 1 - | 0,00011364 - | 1 - | 0,000141108 - | 0 - | 0 - | 1 - | 0,000136122 - | 1 - | 0,000153341 - |
| 1447 | 1 | 3um_FIJJAFBF_213247 | Bacteria | Proteobacteri | Gammaprote | Oceanospirilla | Saccharospirill | Reinekea | Reinekea fors | unclassified | unclassified | unclassified | unclassified | unclassified | 2 | 0,632455532 | 0,00015203 | 4,94E-05 | 2 - | 0,00012753 - | 2 - | 0,000158354 - | 3 - | 0,000225804 - | 1 - | 7,64E-05 - | 2 - | 0,000172083 - |
| 1448 | 1 | 3um_FIJJAFBF_69368 | Bacteria | Proteobacteri | Gammaprote | Alteromonada | Alteromonada | Paraglaciecola | various | unclassified | unclassified | unclassified | unclassified | unclassified | 3,8 | 0,979795897 | 0,000420755 | 8,30E-05 | 5 - | 0,000474284 - | 4 - | 0,000471137 - | 4 - | 0,000447875 - | 4 - | 0,00045449 - | 2 - | 0,000255991 - |
| 1449 | 1 | 3um_FIJJAFBF_165682 | Bacteria | Proteobacteri | Gammaprote | Pseudomonad | Pseudomonad | Pseudomonas | Pseudomonas | unclassified | unclassified | unclassified | unclassified | unclassified | 1 | 0 | 0,000118127 | 1,12E-05 | 1 - | 9,89E-05 - | 1 - | 0,000122861 - | 1 - | 0,000116795 - | 1 - | 0,00011852 - | 1 - | 0,000133512 - |
| 1450 | 2 | 0.2um_NOKCPHJN_156696;3um_FIJJAFBF_55735 | Bacteria | Proteobacteri | Alphaproteob | Rhodobactera | unclassified | unclassified | Rhodobactera | various | various | various | various | various | 3 | 2,097617696 | 0,000338029 | 0,00019685 | 7 - | 0,00070477 - | 2 - | 0,000250033 - | 2 - | 0,000237688 - | 3 - | 0,000361798 - | 1 - | 0,000135855 - |
| 1451 | 1 | 0.2um_NOKCPHJN_888401 | Bacteria | Bacteroidetes | Flavobacteriia | Flavobacterial | Flavobacteriac | Polaribacter | Polaribacter fi | cellular proces | Cell wall/mem | 0P7S9 | Dehydrogenas | 983548.Krodi_ | 0,8 | 0,4 | 2,57E-05 | 1,32E-05 | 1 - | 2,69E-05 - | 1 - | 3,34E-05 - | 1 - | 3,17E-05 - | 0 - | 0 - | 1 - | 3,63E-05 - |
| 1452 | 1 | 3um_FIJJAFBF_442115 | unclassified | unclassified | unclassified | unclassified | unclassified | unclassified | unclassified | unclassified | unclassified | unclassified | unclassified | unclassified | 1,8 | 0,979795897 | 0,000150083 | 7,26E-05 | 3 - | 0,000217931 - | 1 - | 9,02E-05 - | 3 - | 0,000257245 - | 1 - | 8,70E-05 - | 1 - | 9,80E-05 - |
| 1453 | 1 | 3um_FIJJAFBF_388953 | Bacteria | Bacteroidetes | Flavobacteriia | Flavobacterial | Flavobacteriac | Polaribacter | Polaribacter a | poorly charact | Function unkn | 102Q4 | NA | 313598.MED1 | 0,6 | 1,2 | 5,64E-05 | 0,000112895 | 3 - | 0,000282238 - | 0 - | 0 - | 0 - | 0 - | 0 - | 0 - | 0 - | 0 - |
| 1454 | 1 | 3um_FIJJAFBF_610702 | Bacteria | Bacteroidetes | Cytophagia | Cytophagales | Flammeovirga | unclassified | Flammeovirga | cellular proces | Posttranslatio | 0QJZQ | Peptidyl-proly | 318161.Sden_ | 2,8 | 1,166190379 | 0,000166106 | 5,07E-05 | 5 - | 0,000257347 - | 2 - | 0,00012782 - | 2 - | 0,000121509 - | 3 - | 0,000184955 - | 2 - | 0,000138901 - |
| 1455 | 1 | 3um_FIJJAFBF_604541 | Bacteria | Proteobacteri | Gammaprote | Arenicellales | Arenicellaceae | Arenicella | Arenicella xan | information st | Translation, ri | 0QK14 | This protein bi | 1026882.MAN | 0,6 | 0,8 | 4,12E-05 | 5,32E-05 | 2 - | 0,000129691 - | 0 - | 0 - | 1 - | 7,65E-05 - | 0 - | 0 - | 0 - | 0 - |
| 1456 | 1 | 3um_FIJJAFBF_477871 | Eukaryota | unclassified | unclassified | Isochrysidales | Noelaerhabda | Emiliania | Emiliania huxl | metabolism | Energy produc | 0FN1Q | F(1)F(0) ATP s | 696747.NIES3 | 0,8 | 0,4 | 5,86E-05 | 3,01E-05 | 1 - | 6,14E-05 - | 1 - | 7,62E-05 - | 1 - | 7,25E-05 - | 0 - | 0 - | 1 - | 8,28E-05 - |
| 1457 | 1 | 0.2um_NOKCPHJN_261060 | Bacteria | Bacteroidetes | Flavobacteriia | Flavobacterial | Flavobacteriac | Formosa | Formosa sp. H | poorly charact | Function unkn | 09HPY | NA | 1046627.BZAF | 2 | 0 | 0,000221011 | 2,10E-05 | 2 - | 0,000185124 - | 2 - | 0,000229869 - | 2 - | 0,00021852 - | 2 - | 0,000221747 - | 2 - | 0,000249798 - |
| 1458 | 1 | 3um_FIJJAFBF_62752 | Bacteria | Bacteroidetes | Flavobacteriia | Flavobacterial | Flavobacteriac | Formosa | Formosa sp. H | metabolism | Carbohydrate | 05D2N | Aldolase | 391603.FBALC | 1,6 | 0,8 | 5,97E-05 | 3,03E-05 | 2 - | 6,47E-05 - | 2 - | 8,03E-05 - | 2 - | 7,63E-05 - | 2 - | 7,75E-05 - | 0 - | 0 - |
| 1459 | 1 | 0.2um_NOKCPHJN_554191 | Bacteria | Bacteroidetes | Flavobacteriia | Flavobacterial | Flavobacteriac | Formosa | Formosa sp. H | unclassified | unclassified | unclassified | unclassified | unclassified | 0,4 | 0,489897949 | 1,20E-05 | 1,49E-05 | 1 - | 2,55E-05 - | 0 - | 0 - | 0 - | 0 - | 0 - | 0 - | 1 - | 3,44E-05 - |
| 1460 | 1 | 0.2um_NOKCPHJN_289511 | Viruses | unclassified | unclassified | unclassified | unclassified | unclassified | uncultured M | unclassified | unclassified | unclassified | unclassified | unclassified | 0,2 | 0,4 | 4,06E-06 | 8,13E-06 | 1 - | 2,03E-05 - | 0 - | 0 - | 0 - | 0 - | 0 - | 0 - | 0 - | 0 - |
| 1461 | 1 | 0.2um_NOKCPHJN_771062 | Bacteria | unclassified | unclassified | unclassified | unclassified | unclassified | uncultured ba | unclassified | unclassified | unclassified | unclassified | unclassified | 3,8 | 0,748331477 | 0,000129962 | 3,04E-05 | 3 - | 8,54E-05 - | 3 - | 0,000106093 - | 5 - | 0,000168092 - | 4 - | 0,00013646 - | 4 - | 0,000153722 - |
| 1462 | 1 | 3um_FIJJAFBF_605118 | Bacteria | Proteobacteri | Gammaprote | Oceanospirilla | Saccharospirill | Reinekea | Reinekea fors | poorly charact | Function unkn | 05C21 | Displays ATPa | 314283.MED2 | 0,2 | 0,4 | 9,34E-06 | 1,87E-05 | 0 - | 0 - | 0 - | 0 - | 1 - | 4,67E-05 - | 0 - | 0 - | 0 - | 0 - |
| 1463 | 1 | 0.2um_NOKCPHJN_72787 | Bacteria | Proteobacteri | Gammaprote | Oceanospirilla | Saccharospirill | Reinekea | Reinekea fors | metabolism | Amino acid tra | 07RDV | Dehydrogenas | 391591.VSAK1 | 0,4 | 0,489897949 | 1,41E-05 | 1,72E-05 | 0 - | 0 - | 0 - | 0 - | 1 - | 3,49E-05 - | 1 - | 3,54E-05 - | 0 - | 0 - |
| 1464 | 1 | 3um_FIJJAFBF_520476 | Bacteria | Bacteroidetes | Flavobacteriia | Flavobacterial | Flavobacteriac | Polaribacter | Polaribacter s | poorly charact | Function unkn | 11NCV | Protein of unk | 313594.PI23P | 6,6 | 1,019803903 | 0,000578695 | 7,31E-05 | 8 - | 0,000592396 - | 7 - | 0,000643634 - | 5 - | 0,000437039 - | 7 - | 0,000620892 - | 6 - | 0,000599514 - |
| 1465 | 1 | 0.2um_NOKCPHJN_1091763 | Bacteria | Proteobacteri | Alphaproteob | Pelagibacteral | Pelagibacterac | Candidatus Pe | Candidatus Pe | cellular proces | Cell wall/mem | 16QB0 | Part of the ou | 439493.PB721 | 0,2 | 0,4 | 6,77E-06 | 1,35E-05 | 1 - | 3,39E-05 - | 0 - | 0 - | 0 - | 0 - | 0 - | 0 - | 0 - | 0 - |
| 1466 | 1 | 3um_FIJJAFBF_129180 | Bacteria | Bacteroidetes | Flavobacteriia | Flavobacterial | Flavobacteriac | Polaribacter | Polaribacter ir | unclassified | unclassified | unclassified | unclassified | unclassified | 3,2 | 2,13541565 | 8,99E-05 | 5,05E-05 | 7 - | 0,000173904 - | 2 - | 6,17E-05 - | 4 - | 0,000117301 - | 1 - | 2,98E-05 - | 2 - | 6,70E-05 - |
| 1467 | 1 | 0.2um_NOKCPHJN_108660 | Bacteria | Proteobacteri | Alphaproteob | Rhodobactera | Rhodobactera | Planktotalea | Planktotalea a | poorly charact | Function unkn | 17CVT | NA | 314270.RB208 | 0,4 | 0,489897949 | 1,67E-05 | 2,06E-05 | 1 - | 3,84E-05 - | 0 - | 0 - | 1 - | 4,53E-05 - | 0 - | 0 - | 0 - | 0 - |
| 1468 | 1 | 3um_FIJJAFBF_173307 | Bacteria | Bacteroidetes | Flavobacteriia | Flavobacterial | Flavobacteriac | Polaribacter | various | unclassified | unclassified | unclassified | unclassified | unclassified | 2 | 0 | 3,61E-05 | 3,42E-06 | 2 - | 3,02E-05 - | 2 - | 3,75E-05 - | 2 - | 3,57E-05 - | 2 - | 3,62E-05 - | 2 - | 4,08E-05 - |
| 1469 | 2 | 3um_FIJJAFBF_201851;3um_FIJJAFBF_744759 | Bacteria | Proteobacteri | Alphaproteob | Rhodobactera | Rhodobactera | various | various | unclassified | unclassified | unclassified | unclassified | unclassified | 0,2 | 0,4 | 1,07E-05 | 2,15E-05 | 0 - | 0 - | 0 - | 0 - | 0 - | 0 - | 1 - | 5,37E-05 - | 0 - | 0 - |
| 1470 | 1 | 3um_FIJJAFBF_751564 | Bacteria | Bacteroidetes | Cytophagia | Cytophagales | Flammeovirga | unclassified | Flammeovirga | unclassified | unclassified | unclassified | unclassified | unclassified | 0,4 | 0,8 | 9,41E-06 | 1,88E-05 | 2 - | 4,70E-05 - | 0 - | 0 - | 0 - | 0 - | 0 - | 0 - | 0 - | 0 - |
| 1471 | 1 | 3um_FIJJAFBF_305834 | Eukaryota | Chlorophyta | unclassified | Pyramimonad | unclassified | Cymbomonas | Cymbomonas | metabolism | Energy produc | COG1290 | Component of | 70448.Q0P3P | 1 | 0 | 6,37E-05 | 6,05E-06 | 1 - | 5,34E-05 - | 1 - | 6,63E-05 - | 1 - | 6,30E-05 - | 1 - | 6,39E-05 - | 1 - | 7,20E-05 - |
| 1472 | 1 | 3um_FIJJAFBF_477872 | Eukaryota | unclassified | unclassified | Isochrysidales | Noelaerhabda | Emiliania | Emiliania huxl | metabolism | Energy produc | COG0711 | F(1)F(0) ATP s | 32049.SYNPC | 1,2 | 0,4 | 9,57E-05 | 3,53E-05 | 1 - | 6,63E-05 - | 2 - | 0,000164762 - | 1 - | 7,83E-05 - | 1 - | 7,95E-05 - | 1 - | 8,95E-05 - |
| 1473 | 2 | 0.2um_NOKCPHJN_332638;3um_FIJJAFBF_1113C | Viruses | unclassified | unclassified | unclassified | various | various | various | unclassified | unclassified | unclassified | unclassified | unclassified | 1 | 0,632455532 | 6,79E-05 | 4,40E-05 | 1 - | 5,86E-05 - | 1 - | 7,27E-05 - | 2 - | 0,000138247 - | 1 - | 7,01E-05 - | 0 - | 0 - |
| 1474 | 1 | 3um_FIJJAFBF_449836 | Eukaryota | Bacillariophyt | Bacillariophyc | Naviculales | Phaeodactylac | Phaeodactylu | Phaeodactylu | unclassified | unclassified | unclassified | unclassified | unclassified | 3,4 | 1,019803903 | 0,00054592 | 0,000114722 | 5 - | 0,000691426 - | 3 - | 0,000515128 - | 4 - | 0,000652926 - | 3 - | 0,000496928 - | 2 - | 0,000373192 - |
| 1475 | 1 | 3um_FIJJAFBF_359537 | Bacteria | Bacteroidetes | Flavobacteriia | Flavobacterial | Flavobacteriac | Lutibacter | Lutibacter flav | poorly charact | Function unkn | 11P35 | NA | 867900.Celly_ | 8 | 0,894427191 | 0,000297597 | 2,91E-05 | 9 - | 0,000282238 - | 9 - | 0,000350456 - | 8 - | 0,000296136 - | 7 - | 0,000262946 - | 7 - | 0,000296208 - |
| 1476 | 1 | 0.2um_NOKCPHJN_482342 | Bacteria | Proteobacteri | Betaproteoba | unclassified | unclassified | unclassified | Betaproteoba | information st | Translation, ri | 0BBM3 | 50s ribosomal | 595537.Varpa | 3,2 | 0,748331477 | 0,000264569 | 6,83E-05 | 3 - | 0,000207428 - | 4 - | 0,000343419 - | 4 - | 0,000326463 - | 2 - | 0,000165643 - | 3 - | 0,000279894 - |
| 1477 | 1 | 3um_FIJJAFBF_470068 | Bacteria | Bacteroidetes | Flavobacteriia | Flavobacterial | Flavobacteriac | Algibacter | Algibacter sp. | unclassified | unclassified | unclassified | unclassified | unclassified | 11,4 | 2,332380758 | 0,000216573 | 2,33E-05 | 16 - | 0,000259017 - | 10 - | 0,000201014 - | 10 - | 0,000191089 - | 11 - | 0,000213303 - | 10 - | 0,000218441 - |
| 1478 | 1 | 0.2um_NOKCPHJN_25593 | Bacteria | Proteobacteri | Gammaprote | Cellvibrionales | unclassified | unclassified | Cellvibrionales | information st | Translation, ri | 16QAF | thus facilitatin | 314287.GB22 | 0,6 | 0,489897949 | 1,56E-05 | 1,27E-05 | 0 - | 0 - | 1 - | 2,55E-05 - | 0 - | 0 - | 1 - | 2,46E-05 - | 1 - | 2,77E-05 - |
| 1479 | 1 | 0.2um_NOKCPHJN_410090 | Bacteria | Proteobacteri | Gammaprote | unclassified | unclassified | unclassified | Gammaprote | unclassified | unclassified | unclassified | unclassified | unclassified | 2 | 0 | 4,05E-05 | 3,85E-06 | 2 - | 3,40E-05 - | 2 - | 4,22E-05 - | 2 - | 4,01E-05 - | 2 - | 4,07E-05 - | 2 - | 4,58E-05 - |
| 1480 | 1 | 3um_FIJJAFBF_617389 | Bacteria | Proteobacteri | Alphaproteob | Rhodobactera | Rhodobactera | various | various | information st | Translation, ri | 17FIT | 50s ribosomal | 644076.SCH4 | 0,2 | 0,4 | 2,82E-05 | 5,65E-05 | 0 - | 0 - | 0 - | 0 - | 1 - | 0,000141127 - | 0 - | 0 - | 0 - | 0 - |
| 1481 | 1 | 3um_FIJJAFBF_350116 | Bacteria | Proteobacteri | Alphaproteob | Rhodobactera | unclassified | unclassified | Rhodobactera | cellular proces | Signal transdu | 01Q8R | gtp-binding pr | 375451.RD1_ | 1 | 0,632455532 | 2,33E-05 | 1,42E-05 | 0 - | 0 - | 1 - | 2,35E-05 - | 2 - | 4,47E-05 - | 1 - | 2,27E-05 - | 1 - | 2,56E-05 - |
| 1482 | 1 | 3um_FIJJAFBF_464797 | Bacteria | Bacteroidetes | Flavobacteriia | Flavobacterial | unclassified | unclassified | Flavobacterial | unclassified | unclassified | unclassified | unclassified | unclassified | 2,8 | 0,979795897 | 0,00012253 | 3,99E-05 | 4 - | 0,000148099 - | 3 - | 0,000137921 - | 1 - | 4,37E-05 - | 3 - | 0,000133048 - | 3 - | 0,000149879 - |
| 1483 | 1 | 3um_FIJJAFBF_628210 | Bacteria | Bacteroidetes | Cytophagia | Cytophagales | Flammeovirga | Roseivirga | Roseivirga sec | metabolism | Coenzyme tra | 09A4R | May play a ke | 525373.HMPR | 1,6 | 0,8 | 5,18E-05 | 2,84E-05 | 1 - | 2,64E-05 - | 3 - | 9,85E-05 - | 1 - | 3,12E-05 - | 1 - | 3,17E-05 - | 2 - | 7,14E-05 - |
| 1484 | 1 | 3um_FIJJAFBF_709646 | Bacteria | Proteobacteri | Alphaproteob | Rhodobactera | Rhodobactera | Planktotalea | Planktotalea f | unclassified | unclassified | unclassified | unclassified | unclassified | 1 | 0 | 0,000279647 | 2,66E-05 | 1 - | 0,000234238 - | 1 - | 0,000290855 - | 1 - | 0,000276494 - | 1 - | 0,000280578 - | 1 - | 0,00031607 - |
| 1485 | 1 | 0.2um_NOKCPHJN_262932 | Bacteria | Proteobacteri | Alphaproteob | Rhodobactera | Rhodobactera | Planktomarina | Planktomarina | metabolism | Carbohydrate | 07SNC | solute-binding | 367336.OM22 | 6 | 1,264911064 | 0,000189967 | 3,12E-05 | 7 - | 0,0001886 - | 5 - | 0,000167276 - | 7 - | 0,000222623 - | 7 - | 0,000225912 - | 4 - | 0,000145422 - |
| 1486 | 1 | 0.2um_NOKCPHJN_358063 | Bacteria | Cyanobacteria | unclassified | unclassified | unclassified | unclassified | unicellular cya | unclassified | unclassified | unclassified | unclassified | unclassified | 0,2 | 0,4 | 3,60E-06 | 7,21E-06 | 0 - | 0 - | 0 - | 0 - | 0 - | 0 - | 1 - | 1,80E-05 - | 0 - | 0 - |
| 1487 | 1 | 3um_FIJJAFBF_62760 | Bacteria | Proteobacteri | Alphaproteob | Rhodobactera | Rhodobactera | Amylibacter | Amylibacter c | metabolism | Energy produc | COG0056 | Produces ATP | 633131.TR2A6 | 3,2 | 0,4 | 8,65E-05 | 1,82E-05 | 3 - | 6,74E-05 - | 3 - | 8,37E-05 - | 3 - | 7,95E-05 - | 3 - | 8,07E-05 - | 4 - | 0,000121232 - |
| 1488 | 1 | 0.2um_NOKCPHJN_182803 | Bacteria | Bacteroidetes | Flavobacteriia | Flavobacterial | Flavobacteriac | Marinirhabdu | Marinirhabdu | cellular proces | Intracellular tr | 05FGH | MotA TolQ ex | 50743.SCB49_ | 0,2 | 0,4 | 1,12E-05 | 2,24E-05 | 0 - | 0 - | 1 - | 5,59E-05 - | 0 - | 0 - | 0 - | 0 - | 0 - | 0 - |
| 1489 | 1 | 0.2um_NOKCPHJN_993424 | Bacteria | Proteobacteri | Alphaproteob | Pelagibacteral | Pelagibacterac | Candidatus Pe | Candidatus Pe | cellular proces | Posttranslatio | COG0330 | Band 7 protei | 335992.SAR11 | 2,2 | 0,4 | 0,000104343 | 2,12E-05 | 2 - | 7,94E-05 - | 2 - | 9,86E-05 - | 2 - | 9,38E-05 - | 3 - | 0,000142716 - | 2 - | 0,00010718 - |
| 1490 | 1 | 3um_FIJJAFBF_332397 | Bacteria | Proteobacteri | Alphaproteob | Rhodobactera | Rhodobactera | Sulfitobacter | Sulfitobacter s | metabolism | Energy produc | COG0711 | F(1)F(0) ATP s | 391624.OIHEL | 1,6 | 0,489897949 | 0,000118628 | 3,92E-05 | 2 - | 0,000124083 - | 2 - | 0,000154074 - | 1 - | 7,32E-05 - | 1 - | 7,43E-05 - | 2 - | 0,000167432 - |
| 1491 | 1 | 0.2um_NOKCPHJN_323332 | Bacteria | Bacteroidetes | Flavobacteriia | unclassified | unclassified | unclassified | Flavobacteria | unclassified | unclassified | unclassified | unclassified | unclassified | 15,2 | 1,720465053 | 0,000222395 | 2,75E-05 | 16 - | 0,00019683 - | 12 - | 0,000183304 - | 16 - | 0,000232338 - | 17 - | 0,000250505 - | 15 - | 0,000248994 - |
| 1492 | 1 | 0.2um_NOKCPHJN_167490 | Bacteria | Bacteroidetes | unclassified | unclassified | unclassified | unclassified | Bacteroidetes | metabolism | Energy produc | COG0056 | Produces ATP | 216432.CA25 | 2,2 | 0,979795897 | 5,54E-05 | 1,93E-05 | 4 - | 8,73E-05 - | 1 - | 2,71E-05 - | 2 - | 5,15E-05 - | 2 - | 5,23E-05 - | 2 - | 5,89E-05 - |
| 1493 | 1 | 3um_FIJJAFBF_12134 | Bacteria | Proteobacteri | Alphaproteob | Rhodobactera | Rhodobactera | Pelagimonas | Pelagimonas v | unclassified | unclassified | unclassified | unclassified | unclassified | 1 | 0,632455532 | 7,51E-05 | 5,30E-05 | 1 - | 6,14E-05 - | 1 - | 7,62E-05 - | 1 - | 7,25E-05 - | 0 - | 0 - | 2 - | 0,000165641 - |
| 1494 | 1 | 3um_FIJJAFBF_274324 | Viruses | unclassified | unclassified | Caudovirales | Siphoviridae | unclassified | Tenacibaculur | unclassified | unclassified | unclassified | unclassified | unclassified | 0,4 | 0,489897949 | 1,19E-05 | 1,46E-05 | 0 - | 0 - | 1 - | 3,05E-05 - | 1 - | 2,90E-05 - | 0 - | 0 - | 0 - | 0 - |
| 1495 | 1 | 3um_FIJJAFBF_17287 | Bacteria | Proteobacteri | Alphaproteob | Rhodobactera | unclassified | unclassified | Rhodobactera | metabolism | Amino acid tra | 01Y25 | Nitrogen regu | 89187.ISM_05 | 0,6 | 0,489897949 | 6,92E-05 | 5,70E-05 | 1 - | 0,000102479 - | 0 - | 0 - | 1 - | 0,000120966 - | 1 - | 0,000122753 - | 0 - | 0 - |
| 1496 | 1 | 3um_FIJJAFBF_363020 | Bacteria | Proteobacteri | Betaproteoba | unclassified | unclassified | unclassified | Betaproteoba | metabolism | Amino acid tra | 16QAZ | extracellular s | 296591.Bpro_ | 5 | 0,632455532 | 0,000178808 | 2,15E-05 | 5 - | 0,000150626 - | 5 - | 0,000187033 - | 6 - | 0,000213358 - | 5 - | 0,000180424 - | 4 - | 0,000162598 - |
| 1497 | 1 | 3um_FIJJAFBF_346830 | Bacteria | Bacteroidetes | unclassified | unclassified | unclassified | unclassified | Bacteroidetes | unclassified | unclassified | unclassified | unclassified | unclassified | 0,4 | 0,489897949 | 3,94E-05 | 4,91E-05 | 1 - | 8,38E-05 - | 0 - | 0 - | 0 - | 0 - | 0 - | 0 - | 1 - | 0,000113047 - |
| 1498 | 1 | 0.2um_NOKCPHJN_106555 | Bacteria | Proteobacteri | Gammaprote | Oceanospirilla | Saccharospirill | Reinekea | Reinekea fors | unclassified | unclassified | unclassified | unclassified | unclassified | 1 | 0,632455532 | 6,75E-05 | 4,57E-05 | 0 - | 0 - | 1 - | 6,63E-05 - | 1 - | 6,30E-05 - | 1 - | 6,39E-05 - | 2 - | 0,000144069 - |
| 1499 | 1 | 0.2um_NOKCPHJN_138709 | Bacteria | unclassified | unclassified | unclassified | unclassified | unclassified | uncultured ba | unclassified | unclassified | unclassified | unclassified | unclassified | 4,4 | 0,489897949 | 0,000110215 | 1,15E-05 | 5 - | 0,000105493 - | 5 - | 0,000130992 - | 4 - | 9,96E-05 - | 4 - | 0,000101091 - | 4 - | 0,000113878 - |
| 1500 | 1 | 0.2um_NOKCPHJN_128597 | Eukaryota | unclassified | Labyrinthulom | unclassified | Thraustochytr | Aurantiochytr | Aurantiochytr | cellular proces | Posttranslatio | KOG0417 | ubiquitin-conj | 28377.ENSAC | 0,6 | 0,489897949 | 5,82E-05 | 4,77E-05 | 0 - | 0 - | 0 - | 0 - | 1 - | 9,22E-05 - | 1 - | 9,35E-05 - | 1 - | 0,000105357 - |
| 1501 | 1 | 3um_FIJJAFBF_12816 | Bacteria | Bacteroidetes | Flavobacteriia | Flavobacterial | Flavobacteriac | Flavivirga | Flavivirga eckl | metabolism | Energy produc | 099Q7 | Catalyzes the | 391603.FBALC | 0,4 | 0,489897949 | 1,77E-05 | 2,17E-05 | 0 - | 0 - | 0 - | 0 - | 1 - | 4,40E-05 - | 1 - | 4,46E-05 - | 0 - | 0 - |
| 1502 | 1 | 3um_FIJJAFBF_306863 | Bacteria | Proteobacteri | Alphaproteob | Rhodobactera | unclassified | unclassified | Rhodobactera | metabolism | Amino acid tra | COG0683 | (ABC) transpo | 439496.RBY4 | 4,6 | 0,489897949 | 0,000158827 | 2,39E-05 | 5 - | 0,000144555 - | 5 - | 0,000179495 - | 4 - | 0,000136506 - | 4 - | 0,000138522 - | 5 - | 0,000195056 - |
| 1503 | 1 | 0.2um_NOKCPHJN_998892 | Bacteria | Proteobacteri | Alphaproteob | Rhodobactera | Rhodobactera | Sulfitobacter | Sulfitobacter t | unclassified | unclassified | unclassified | unclassified | unclassified | 1,6 | 0,489897949 | 5,64E-05 | 1,57E-05 | 2 - | 6,01E-05 - | 2 - | 7,46E-05 - | 2 - | 7,09E-05 - | 1 - | 3,60E-05 - | 1 - | 4,05E-05 - |
| 1504 | 1 | 3um_FIJJAFBF_69380 | Eukaryota | Chlorophyta | Mamiellophyc | Mamiellales | Bathycoccace | Ostreococcus | Ostreococcus | metabolism | Energy produc | COG0056 | Produces ATP | 70448.Q0P3H | 1 | 0 | 3,14E-05 | 2,98E-06 | 1 - | 2,63E-05 - | 1 - | 3,26E-05 - | 1 - | 3,10E-05 - | 1 - | 3,15E-05 - | 1 - | 3,54E-05 - |
| 1505 | 1 | 0.2um_NOKCPHJN_121696 | Bacteria | Bacteroidetes | Flavobacteriia | Flavobacterial | Flavobacteriac | Polaribacter | various | information st | Replication, re | 05C24 | DNA gyrase ne | 983544.Lacal_ | 2,8 | 0,748331477 | 4,47E-05 | 1,09E-05 | 3 - | 4,06E-05 - | 2 - | 3,36E-05 - | 4 - | 6,39E-05 - | 3 - | 4,86E-05 - | 2 - | 3,65E-05 - |
| 1506 | 1 | 0.2um_NOKCPHJN_503473 | Bacteria | Proteobacteri | Gammaprote | Oceanospirilla | Oceanospirilla | Marinobacteri | Marinobacteri | metabolism | Energy produc | 05C26 | Dehydrogenas | 511062.GU3_ | 1 | 0 | 2,88E-05 | 2,74E-06 | 1 - | 2,42E-05 - | 1 - | 3,00E-05 - | 1 - | 2,85E-05 - | 1 - | 2,89E-05 - | 1 - | 3,26E-05 - |
| 1507 | 1 | 0.2um_NOKCPHJN_256507 | Bacteria | Bacteroidetes | Flavobacteriia | Flavobacterial | Flavobacteriac | Polaribacter | Polaribacter s | metabolism | Inorganic ion t | 05C27 | TonB-depende | 471870.BACIN | 1,6 | 0,8 | 1,95E-05 | 7,73E-06 | 3 - | 3,20E-05 - | 1 - | 1,32E-05 - | 2 - | 2,52E-05 - | 1 - | 1,28E-05 - | 1 - | 1,44E-05 - |
| 1508 | 1 | 3um_FIJJAFBF_21310 | Bacteria | Bacteroidetes | Flavobacteriia | Flavobacterial | Flavobacteriac | Polaribacter | Polaribacter b | metabolism | Energy produc | 0P815 | FAD linked oxi | 313594.PI23P | 0,6 | 1,2 | 8,86E-06 | 1,77E-05 | 0 - | 0 - | 3 - | 4,43E-05 - | 0 - | 0 - | 0 - | 0 - | 0 - | 0 - |
| 1509 | 1 | 0.2um_NOKCPHJN_82690 | Bacteria | Gemmatimon | Gemmatimon | Gemmatimon | Gemmatimon | Gemmatirosa | Gemmatirosa | unclassified | unclassified | unclassified | unclassified | unclassified | 0,4 | 0,489897949 | 6,09E-05 | 7,48E-05 | 0 - | 0 - | 0 - | 0 - | 0 - | 0 - | 1 - | 0,000143212 - | 1 - | 0,000161328 - |
| 1510 | 1 | 3um_FIJJAFBF_235861 | Bacteria | Proteobacteri | Gammaprote | Cellvibrionales | Microbulbifera | Microbulbifer | Microbulbifer | unclassified | unclassified | unclassified | unclassified | unclassified | 2 | 0 | 0,000183929 | 1,75E-05 | 2 - | 0,000154063 - | 2 - | 0,0001913 - | 2 - | 0,000181855 - | 2 - | 0,000184541 - | 2 - | 0,000207885 - |
| 1511 | 1 | 3um_FIJJAFBF_344571 | Bacteria | various | various | various | various | various | various | metabolism | Amino acid tra | 0FNCB | carbon dioxide | 84588.SYNW1 | 4,2 | 0,748331477 | 0,000558196 | 0,000113898 | 5 - | 0,000557169 - | 4 - | 0,000553471 - | 3 - | 0,000394608 - | 4 - | 0,000533916 - | 5 - | 0,000751818 - |
| 1512 | 1 | 3um_FIJJAFBF_462301 | Eukaryota | Bacillariophyt | Coscinodiscop | Rhizosoleniale | Rhizosoleniac | Rhizosolenia | Rhizosolenia i | unclassified | unclassified | unclassified | unclassified | unclassified | 0,4 | 0,489897949 | 2,05E-05 | 2,51E-05 | 0 - | 0 - | 0 - | 0 - | 1 - | 5,09E-05 - | 1 - | 5,17E-05 - | 0 - | 0 - |
| 1513 | 1 | 0.2um_NOKCPHJN_397879 | Bacteria | Proteobacteri | Alphaproteob | Pelagibacteral | unclassified | unclassified | Pelagibacteral | cellular proces | Cell wall/mem | 7257 | NA | 335992.SAR11 | 4 | 1,095445115 | 0,000170388 | 6,01E-05 | 4 - | 0,00014083 - | 4 - | 0,00017487 - | 3 - | 0,000124677 - | 3 - | 0,000126518 - | 6 - | 0,000285045 - |
| 1514 | 1 | 3um_FIJJAFBF_123244 | Viruses | unclassified | unclassified | unclassified | various | various | various | unclassified | unclassified | unclassified | unclassified | unclassified | 3,8 | 0,748331477 | 9,92E-05 | 2,33E-05 | 4 - | 8,73E-05 - | 5 - | 0,000135474 - | 3 - | 7,73E-05 - | 3 - | 7,84E-05 - | 4 - | 0,000117775 - |
| 1515 | 1 | 0.2um_NOKCPHJN_610762 | Bacteria | Bacteroidetes | Flavobacteriia | Flavobacterial | Flavobacteriac | Tenacibaculur | Tenacibaculur | unclassified | unclassified | unclassified | unclassified | unclassified | 4 | 0 | 0,000140181 | 1,33E-05 | 4 - | 0,000117419 - | 4 - | 0,000145799 - | 4 - | 0,000138601 - | 4 - | 0,000140648 - | 4 - | 0,000158439 - |
| 1516 | 1 | 3um_FIJJAFBF_604511 | Bacteria | Proteobacteri | Alphaproteob | Rhodobactera | Rhodobactera | Phaeobacter | Phaeobacter g | unclassified | unclassified | unclassified | unclassified | unclassified | 1 | 0 | 4,08E-05 | 3,87E-06 | 1 - | 3,42E-05 - | 1 - | 4,24E-05 - | 1 - | 4,03E-05 - | 1 - | 4,09E-05 - | 1 - | 4,61E-05 - |
| 1517 | 1 | 3um_FIJJAFBF_611863 | Bacteria | Bacteroidetes | Flavobacteriia | Flavobacterial | Flavobacteriac | Muricauda | Muricauda lut | poorly charact | Function unkn | 0XR4Y | Ragb susd dor | 886377.Murru | 0,2 | 0,4 | 4,86E-06 | 9,73E-06 | 0 - | 0 - | 0 - | 0 - | 1 - | 2,43E-05 - | 0 - | 0 - | 0 - | 0 - |
| 1518 | 1 | 0.2um_NOKCPHJN_129623 | Bacteria | Bacteroidetes | Cytophagia | Cytophagales | Flammeovirga | unclassified | Flammeovirga | poorly charact | Function unkn | 0FZ9D | FG-GAP repea | 643867.Ftrac_ | 1,8 | 1,166190379 | 6,31E-06 | 4,06E-06 | 3 - | 9,19E-06 - | 3 - | 1,14E-05 - | 1 - | 3,61E-06 - | 2 - | 7,34E-06 - | 0 - | 0 - |
| 1519 | 1 | 3um_FIJJAFBF_525959 | Bacteria | Bacteroidetes | Cytophagia | Cytophagales | Flammeovirga | unclassified | Flammeovirga | information st | Translation, ri | 0FWTQ | This protein is | 880070.Cycma | 1,2 | 0,4 | 0,000111921 | 3,86E-05 | 1 - | 7,81E-05 - | 1 - | 9,70E-05 - | 1 - | 9,22E-05 - | 2 - | 0,000187052 - | 1 - | 0,000105357 - |
| 1520 | 1 | 3um_FIJJAFBF_357535 | Bacteria | Bacteroidetes | Flavobacteriia | Flavobacterial | Flavobacteriac | Polaribacter | Polaribacter h | information st | Translation, ri | COG0173 | aspartyl-trna s | 313594.PI23P | 0,8 | 0,748331477 | 1,74E-05 | 1,53E-05 | 2 - | 3,94E-05 - | 1 - | 2,45E-05 - | 1 - | 2,33E-05 - | 0 - | 0 - | 0 - | 0 - |
| 1521 | 1 | 0.2um_NOKCPHJN_183034 | Bacteria | Proteobacteri | Gammaprote | Cellvibrionales | Porticoccacea | unclassified | various | unclassified | unclassified | unclassified | unclassified | unclassified | 2 | 0 | 0,000194365 | 1,85E-05 | 2 - | 0,000162804 - | 2 - | 0,000202154 - | 2 - | 0,000192173 - | 2 - | 0,000195012 - | 2 - | 0,00021968 - |
| 1522 | 1 | 3um_FIJJAFBF_03394 | Bacteria | Proteobacteri | Alphaproteob | Rhodobactera | Rhodobactera | Oceanicola | Oceanicola sp | unclassified | unclassified | unclassified | unclassified | unclassified | 0,4 | 0,489897949 | 9,34E-06 | 1,15E-05 | 1 - | 2,13E-05 - | 0 - | 0 - | 0 - | 0 - | 1 - | 2,55E-05 - | 0 - | 0 - |
| 1523 | 1 | 3um_FIJJAFBF_327780 | Bacteria | Bacteroidetes | Cytophagia | Cytophagales | Flammeovirga | unclassified | Flammeovirga | cellular proces | Posttranslatio | 07T41 | NifU domain p | 388413.ALPR1 | 0,2 | 0,4 | 1,14E-05 | 2,28E-05 | 1 - | 5,71E-05 - | 0 - | 0 - | 0 - | 0 - | 0 - | 0 - | 0 - | 0 - |
| 1524 | 1 | 0.2um_NOKCPHJN_678020 | Bacteria | Proteobacteri | Gammaprote | Pseudomonad | Pseudomonad | Pseudomonas | Pseudomonas | metabolism | Energy produc | 0QIAU | Electron trans | 743720.Psefu | 0,4 | 0,489897949 | 2,64E-05 | 3,24E-05 | 0 - | 0 - | 1 - | 6,72E-05 - | 0 - | 0 - | 1 - | 6,49E-05 - | 0 - | 0 - |
| 1525 | 1 | 0.2um_NOKCPHJN_224348 | Bacteria | Proteobacteri | Gammaprote | Cellvibrionales | Porticoccacea | unclassified | Gammaprote | poorly charact | Function unkn | 05EK9 | Dehydrogenas | 314287.GB22 | 0,4 | 0,489897949 | 2,12E-05 | 2,60E-05 | 0 - | 0 - | 0 - | 0 - | 1 - | 5,27E-05 - | 1 - | 5,35E-05 - | 0 - | 0 - |
| 1526 | 1 | 3um_FIJJAFBF_717041 | Bacteria | Proteobacteri | Gammaprote | Cellvibrionales | Halieaceae | unclassified | marine gamm | unclassified | unclassified | unclassified | unclassified | unclassified | 0,2 | 0,4 | 5,84E-06 | 1,17E-05 | 1 - | 2,92E-05 - | 0 - | 0 - | 0 - | 0 - | 0 - | 0 - | 0 - | 0 - |
| 1527 | 1 | 3um_FIJJAFBF_465306 | Bacteria | Bacteroidetes | Cytophagia | Cytophagales | Flammeovirga | unclassified | Flammeovirga | cellular proces | Cell wall/mem | 09B44 | efflux transpo | 388413.ALPR1 | 1,4 | 0,489897949 | 5,11E-05 | 2,08E-05 | 1 - | 3,00E-05 - | 1 - | 3,73E-05 - | 2 - | 7,09E-05 - | 1 - | 3,60E-05 - | 2 - | 8,11E-05 - |
| 1528 | 1 | 3um_FIJJAFBF_417840 | Bacteria | Proteobacteri | Betaproteoba | unclassified | unclassified | unclassified | Betaproteoba | information st | Translation, ri | COG0184 | Forms an inte | 398578.Daci_ | 2 | 0 | 0,000315005 | 2,99E-05 | 2 - | 0,000263855 - | 2 - | 0,00032763 - | 2 - | 0,000311453 - | 2 - | 0,000316054 - | 2 - | 0,000356033 - |
| 1529 | 1 | 3um_FIJJAFBF_92058 | Bacteria | Bacteroidetes | Flavobacteriia | unclassified | unclassified | unclassified | Flavobacteria | unclassified | unclassified | unclassified | unclassified | unclassified | 1 | 0 | 0,00012457 | 1,18E-05 | 1 - | 0,000104343 - | 1 - | 0,000129563 - | 1 - | 0,000123166 - | 1 - | 0,000124985 - | 1 - | 0,000140795 - |
| 1530 | 1 | 3um_FIJJAFBF_818266 | Bacteria | Bacteroidetes | Flavobacteriia | Flavobacterial | Flavobacteriac | Tenacibaculur | Tenacibaculur | metabolism | Lipid transpor | 05CRW | Short chain fa | 391587.KAOT | 0,2 | 0,4 | 5,01E-06 | 1,00E-05 | 1 - | 2,51E-05 - | 0 - | 0 - | 0 - | 0 - | 0 - | 0 - | 0 - | 0 - |
| 1531 | 1 | 3um_FIJJAFBF_652762 | Archaea | Euryarchaeota | unclassified | unclassified | unclassified | unclassified | Euryarchaeota | unclassified | unclassified | unclassified | unclassified | unclassified | 1,8 | 1,166190379 | 9,83E-05 | 5,20E-05 | 4 - | 0,000193716 - | 1 - | 6,01E-05 - | 2 - | 0,000114331 - | 1 - | 5,80E-05 - | 1 - | 6,53E-05 - |
| 1532 | 1 | 3um_FIJJAFBF_466853 | Bacteria | Proteobacteri | Alphaproteob | Rhizobiales | unclassified | Pseudorhodop | Pseudorhodop | information st | Transcription | 05VEQ | Cold shock pr | 323097.Nham | 0,4 | 0,489897949 | 8,60E-05 | 0,000105623 | 0 - | 0 - | 0 - | 0 - | 0 - | 0 - | 1 - | 0,000202181 - | 1 - | 0,000227757 - |
| 1533 | 1 | 0.2um_NOKCPHJN_392575 | Bacteria | Proteobacteri | Gammaprote | Alteromonada | Alteromonada | Catenovulum | Catenovulum | poorly charact | Function unkn | 11HMD | NA | 377629.TERTU | 1 | 0,632455532 | 2,50E-05 | 1,68E-05 | 1 - | 2,13E-05 - | 2 - | 5,29E-05 - | 1 - | 2,51E-05 - | 1 - | 2,55E-05 - | 0 - | 0 - |
| 1534 | 1 | 3um_FIJJAFBF_407949 | Bacteria | Proteobacteri | Gammaprote | Alteromonada | Alteromonada | Paraglaciecola | Paraglaciecola | poorly charact | Function unkn | 0XNQ8 | alkaline phosp | 314276.OS145 | 8,4 | 2,059126028 | 0,00018511 | 4,40E-05 | 11 - | 0,000204626 - | 6 - | 0,000138592 - | 9 - | 0,000197624 - | 6 - | 0,000133695 - | 10 - | 0,000251012 - |
| 1535 | 1 | 0.2um_NOKCPHJN_563768 | Bacteria | Proteobacteri | Alphaproteob | Pelagibacteral | Pelagibacterac | Candidatus Pe | Candidatus Pe | unclassified | unclassified | unclassified | unclassified | unclassified | 1 | 0 | 1,86E-05 | 1,77E-06 | 1 - | 1,56E-05 - | 1 - | 1,94E-05 - | 1 - | 1,84E-05 - | 1 - | 1,87E-05 - | 1 - | 2,10E-05 - |
| 1536 | 2 | 0.2um_NOKCPHJN_184467;0.2um_NOKCPHJN_7 | Bacteria | Proteobacteri | Gammaprote | Oceanospirilla | various | various | various | metabolism | Amino acid tra | 16Q0N | Phosphoribos | various | 0,6 | 0,489897949 | 3,44E-05 | 2,82E-05 | 0 - | 0 - | 0 - | 0 - | 1 - | 5,44E-05 - | 1 - | 5,52E-05 - | 1 - | 6,22E-05 - |
| 1537 | 7 | 0.2um_NOKCPHJN_279845;0.2um_NOKCPHJN_9 | Bacteria | various | various | various | various | various | various | unclassified | unclassified | unclassified | unclassified | unclassified | 1 | 0 | 0,000125713 | 1,19E-05 | 1 - | 0,0001053 - | 1 - | 0,000130751 - | 1 - | 0,000124296 - | 1 - | 0,000126131 - | 1 - | 0,000142087 - |
| 1538 | 1 | 3um_FIJJAFBF_43065 | Bacteria | Bacteroidetes | Flavobacteriia | Flavobacterial | Flavobacteriac | Formosa | Formosa sp. H | cellular proces | Posttranslatio | 09GE9 | Thioredoxin | 156586.BBFL7 | 3,2 | 0,4 | 0,000417312 | 6,18E-05 | 3 - | 0,000327934 - | 3 - | 0,000407197 - | 4 - | 0,000516123 - | 3 - | 0,000392809 - | 3 - | 0,000442499 - |
| 1539 | 1 | 3um_FIJJAFBF_591462 | Bacteria | Proteobacteri | Gammaprote | Alteromonada | Alteromonada | Paraglaciecola | Paraglaciecola | unclassified | unclassified | unclassified | unclassified | unclassified | 4,2 | 1,166190379 | 7,07E-05 | 1,79E-05 | 5 - | 7,15E-05 - | 4 - | 7,10E-05 - | 3 - | 5,06E-05 - | 6 - | 0,000102727 - | 3 - | 5,79E-05 - |
| 1540 | 1 | 0.2um_NOKCPHJN_483238 | Eukaryota | Arthropoda | Insecta | Lepidoptera | Pyralidae | Amyelois | Amyelois tran | unclassified | unclassified | unclassified | unclassified | unclassified | 0,2 | 0,4 | 1,66E-06 | 3,32E-06 | 0 - | 0 - | 0 - | 0 - | 0 - | 0 - | 0 - | 0 - | 1 - | 8,29E-06 - |
| 1541 | 1 | 3um_FIJJAFBF_658130 | Bacteria | Bacteroidetes | Flavobacteriia | Flavobacterial | Flavobacteriac | Ulvibacter | Ulvibacter lito | unclassified | unclassified | unclassified | unclassified | unclassified | 1,2 | 0,748331477 | 0,000162839 | 9,47E-05 | 2 - | 0,000234238 - | 1 - | 0,000145427 - | 2 - | 0,000276494 - | 0 - | 0 - | 1 - | 0,000158035 - |
| 1542 | 1 | 0.2um_NOKCPHJN_95271 | Bacteria | Proteobacteri | Alphaproteob | Rhodobactera | Rhodobactera | Roseovarius | Roseovarius n | poorly charact | Function unkn | 05EN8 | Extracellular s | 89187.ISM_11 | 1,4 | 1,019803903 | 2,64E-05 | 1,75E-05 | 3 - | 5,07E-05 - | 1 - | 2,10E-05 - | 2 - | 3,99E-05 - | 1 - | 2,02E-05 - | 0 - | 0 - |
| 1543 | 1 | 0.2um_NOKCPHJN_244300 | Bacteria | Proteobacteri | Gammaprote | Cellvibrionales | Porticoccacea | unclassified | SAR92 bacteri | unclassified | unclassified | unclassified | unclassified | unclassified | 4,4 | 1,019803903 | 0,000128325 | 2,38E-05 | 6 - | 0,000148418 - | 5 - | 0,000153576 - | 3 - | 8,76E-05 - | 4 - | 0,00011852 - | 4 - | 0,000133512 - |
| 1544 | 1 | 3um_FIJJAFBF_99477 | Bacteria | Proteobacteri | Alphaproteob | Rhodobactera | Rhodobactera | Planktomarina | Planktomarina | information st | Translation, ri | 08UHH | Located on the | 388401.RB215 | 4 | 0,632455532 | 0,000420589 | 4,92E-05 | 5 - | 0,000444871 - | 3 - | 0,000331439 - | 4 - | 0,0004201 - | 4 - | 0,000426305 - | 4 - | 0,000480231 - |
| 1545 | 1 | 0.2um_NOKCPHJN_956251 | Bacteria | Proteobacteri | Alphaproteob | Pelagibacteral | Pelagibacterac | Candidatus Pe | Candidatus Pe | cellular proces | Cell wall/mem | 7257 | NA | 439493.PB721 | 2,2 | 0,748331477 | 0,000149757 | 4,80E-05 | 3 - | 0,00017303 - | 3 - | 0,000214853 - | 2 - | 0,000136163 - | 1 - | 6,91E-05 - | 2 - | 0,000155653 - |
| 1546 | 1 | 3um_FIJJAFBF_434161 | Bacteria | Proteobacteri | Gammaprote | unclassified | unclassified | unclassified | various | unclassified | unclassified | unclassified | unclassified | unclassified | 1,2 | 0,748331477 | 3,84E-05 | 2,47E-05 | 1 - | 2,75E-05 - | 1 - | 3,41E-05 - | 2 - | 6,48E-05 - | 2 - | 6,58E-05 - | 0 - | 0 - |
| 1547 | 6 | 0.2um_NOKCPHJN_11686;0.2um_NOKCPHJN_14 | Bacteria | various | various | various | various | various | various | metabolism | Energy produc | COG0712 | F(1)F(0) ATP s | 1046627.BZAF | 1 | 0 | 7,70E-05 | 7,31E-06 | 1 - | 6,45E-05 - | 1 - | 8,01E-05 - | 1 - | 7,61E-05 - | 1 - | 7,72E-05 - | 1 - | 8,70E-05 - |
| 1548 | 1 | 3um_FIJJAFBF_287676 | Bacteria | Bacteroidetes | Flavobacteriia | Flavobacterial | Flavobacteriac | various | various | information st | Translation, ri | 0PBIW | 50S ribosomal | 391603.FBALC | 0,8 | 0,4 | 0,000158287 | 8,04E-05 | 1 - | 0,000171309 - | 1 - | 0,000212715 - | 1 - | 0,000202212 - | 1 - | 0,000205199 - | 0 - | 0 - |
| 1549 | 2 | 0.2um_NOKCPHJN_186389;3um_FIJJAFBF_74039 | Bacteria | Bacteroidetes | various | various | various | various | various | unclassified | unclassified | unclassified | unclassified | unclassified | 0,4 | 0,489897949 | 2,53E-05 | 3,11E-05 | 1 - | 5,80E-05 - | 0 - | 0 - | 1 - | 6,84E-05 - | 0 - | 0 - | 0 - | 0 - |
| 1550 | 1 | 3um_FIJJAFBF_14178 | Bacteria | Proteobacteri | Gammaprote | Cellvibrionales | Porticoccacea | unclassified | SAR92 bacteri | metabolism | Amino acid tra | 16Q4S | oligopeptide A | 314287.GB22 | 0,8 | 0,748331477 | 1,85E-05 | 1,62E-05 | 2 - | 4,17E-05 - | 1 - | 2,59E-05 - | 1 - | 2,46E-05 - | 0 - | 0 - | 0 - | 0 - |
| 1551 | 1 | 3um_FIJJAFBF_601350 | Bacteria | Bacteroidetes | unclassified | unclassified | unclassified | unclassified | Bacteroidetes | cellular proces | Cell motility | COG1360 | Ompa motb d | 398720.MED2 | 2,8 | 0,4 | 0,000141652 | 2,99E-05 | 2 - | 8,38E-05 - | 3 - | 0,000156043 - | 3 - | 0,000148338 - | 3 - | 0,000150529 - | 3 - | 0,000169571 - |
| 1552 | 1 | 3um_FIJJAFBF_449354 | Bacteria | Bacteroidetes | Chitinophagia | Chitinophagal | Chitinophagac | Flavihumibact | Flavihumibact | metabolism | Energy produc | 099Q7 | Catalyzes the | 700598.Niako | 0,2 | 0,4 | 8,68E-06 | 1,74E-05 | 0 - | 0 - | 0 - | 0 - | 1 - | 4,34E-05 - | 0 - | 0 - | 0 - | 0 - |
| 1553 | 1 | 0.2um_NOKCPHJN_707513 | Eukaryota | unclassified | unclassified | Longamoebia | Acanthamoeb | Acanthamoeb | Acanthamoeb | unclassified | unclassified | unclassified | unclassified | unclassified | 0,4 | 0,489897949 | 1,02E-05 | 1,26E-05 | 1 - | 2,32E-05 - | 0 - | 0 - | 0 - | 0 - | 1 - | 2,78E-05 - | 0 - | 0 - |
| 1554 | 1 | 3um_FIJJAFBF_52802 | Bacteria | Proteobacteri | Alphaproteob | Rhodobactera | Rhodobactera | Planktomarina | Planktomarina | cellular proces | Posttranslatio | 17FI0 | Binds to Cpn6 | 398580.Dshi_ | 2,8 | 0,748331477 | 0,000398631 | 9,70E-05 | 3 - | 0,000362453 - | 2 - | 0,00030004 - | 4 - | 0,000570451 - | 3 - | 0,000434158 - | 2 - | 0,000326052 - |
| 1555 | 1 | 3um_FIJJAFBF_461253 | Bacteria | Proteobacteri | Alphaproteob | Rhodobactera | Rhodobactera | unclassified | Rhodobactera | unclassified | unclassified | unclassified | unclassified | unclassified | 4,4 | 0,489897949 | 0,000183046 | 1,56E-05 | 5 - | 0,0001755 - | 4 - | 0,000174335 - | 4 - | 0,000165727 - | 5 - | 0,000210219 - | 4 - | 0,000189449 - |
| 1556 | 1 | 0.2um_NOKCPHJN_503409 | Eukaryota | unclassified | Oomycetes | Saprolegniales | Saprolegniace | Saprolegnia | Saprolegnia p | unclassified | unclassified | unclassified | unclassified | unclassified | 0,6 | 0,489897949 | 7,47E-06 | 6,15E-06 | 1 - | 1,11E-05 - | 0 - | 0 - | 1 - | 1,31E-05 - | 1 - | 1,32E-05 - | 0 - | 0 - |
| 1557 | 1 | 0.2um_NOKCPHJN_140679 | Bacteria | Bacteroidetes | Flavobacteriia | Flavobacterial | Flavobacteriac | unclassified | Flavobacteria | poorly charact | Function unkn | 11W7M | NA | 983544.Lacal_ | 0,4 | 0,489897949 | 3,90E-05 | 4,79E-05 | 0 - | 0 - | 0 - | 0 - | 1 - | 9,09E-05 - | 0 - | 0 - | 1 - | 0,000103943 - |
| 1558 | 1 | 3um_FIJJAFBF_345884 | Bacteria | Proteobacteri | Betaproteoba | unclassified | unclassified | unclassified | Betaproteoba | unclassified | unclassified | unclassified | unclassified | unclassified | 0,4 | 0,489897949 | 1,00E-05 | 1,23E-05 | 0 - | 0 - | 1 - | 2,55E-05 - | 0 - | 0 - | 1 - | 2,46E-05 - | 0 - | 0 - |
| 1559 | 1 | 0.2um_NOKCPHJN_315149 | Bacteria | Proteobacteri | Gammaprote | unclassified | unclassified | unclassified | OM182 bacte | unclassified | unclassified | unclassified | unclassified | unclassified | 3 | 1,095445115 | 8,84E-05 | 3,01E-05 | 4 - | 0,000101348 - | 4 - | 0,000125844 - | 3 - | 8,97E-05 - | 3 - | 9,10E-05 - | 1 - | 3,42E-05 - |
| 1560 | 1 | 3um_FIJJAFBF_287088 | Bacteria | Bacteroidetes | Flavobacteriia | Flavobacterial | Flavobacteriac | Ulvibacter | Ulvibacter sp. | unclassified | unclassified | unclassified | unclassified | unclassified | 0,2 | 0,4 | 4,52E-06 | 9,04E-06 | 1 - | 2,26E-05 - | 0 - | 0 - | 0 - | 0 - | 0 - | 0 - | 0 - | 0 - |
| 1561 | 1 | 3um_FIJJAFBF_215516 | Eukaryota | Bacillariophyt | Bacillariophyc | Bacillariales | Bacillariaceae | Fragilariopsis | Fragilariopsis | metabolism | Inorganic ion t | COG0659 | sulfate transp | 4787.PITG_20 | 0,6 | 0,489897949 | 1,21E-05 | 1,01E-05 | 1 - | 1,69E-05 - | 1 - | 2,10E-05 - | 0 - | 0 - | 0 - | 0 - | 1 - | 2,28E-05 - |
| 1562 | 1 | 0.2um_NOKCPHJN_470841 | Bacteria | Proteobacteri | Alphaproteob | Rhodobactera | Rhodobactera | Planktomarina | Planktomarina | unclassified | unclassified | unclassified | unclassified | unclassified | 6,8 | 0,4 | 0,000453993 | 4,72E-05 | 7 - | 0,000391921 - | 6 - | 0,000417128 - | 7 - | 0,000462622 - | 7 - | 0,000469455 - | 7 - | 0,00052884 - |
| 1563 | 1 | 0.2um_NOKCPHJN_450134 | Bacteria | Bacteroidetes | Flavobacteriia | Flavobacterial | Cryomorphace | Owenweeksia | Owenweeksia | unclassified | unclassified | unclassified | unclassified | unclassified | 0,2 | 0,4 | 2,19E-06 | 4,38E-06 | 1 - | 1,09E-05 - | 0 - | 0 - | 0 - | 0 - | 0 - | 0 - | 0 - | 0 - |
| 1564 | 1 | 3um_FIJJAFBF_364498 | Bacteria | unclassified | unclassified | unclassified | unclassified | unclassified | uncultured ba | cellular proces | Posttranslatio | 05E6M | Sulphatase-m | 398720.MED2 | 1 | 0,632455532 | 3,57E-05 | 2,04E-05 | 2 - | 6,36E-05 - | 1 - | 3,95E-05 - | 1 - | 3,75E-05 - | 1 - | 3,81E-05 - | 0 - | 0 - |
| 1565 | 1 | 0.2um_NOKCPHJN_667144 | Bacteria | Proteobacteri | Alphaproteob | Pelagibacteral | Pelagibacterac | Candidatus Pe | Candidatus Pe | metabolism | Amino acid tra | 05TIB | (ABC) transpo | 869210.Marky | 1 | 0 | 4,05E-05 | 3,85E-06 | 1 - | 3,40E-05 - | 1 - | 4,22E-05 - | 1 - | 4,01E-05 - | 1 - | 4,07E-05 - | 1 - | 4,58E-05 - |
| 1566 | 1 | 3um_FIJJAFBF_120611 | Bacteria | Bacteroidetes | Flavobacteriia | Flavobacterial | Flavobacteriac | unclassified | Flavobacteria | cellular proces | Cell wall/mem | 0P88B | Outer membr | 313603.FB217 | 0,8 | 0,748331477 | 2,33E-05 | 2,26E-05 | 1 - | 2,54E-05 - | 0 - | 0 - | 1 - | 3,00E-05 - | 2 - | 6,10E-05 - | 0 - | 0 - |
| 1567 | 1 | 0.2um_NOKCPHJN_24344 | Bacteria | Bacteroidetes | Flavobacteriia | Flavobacterial | Flavobacteriac | Kordia | Kordia sp. SM | unclassified | unclassified | unclassified | unclassified | unclassified | 0,4 | 0,489897949 | 6,45E-06 | 7,94E-06 | 1 - | 1,48E-05 - | 0 - | 0 - | 1 - | 1,75E-05 - | 0 - | 0 - | 0 - | 0 - |

| | | | | | | | | | | | | | | | | | | | | | | | | | | | | |
|---|---|---|---|---|---|---|---|---|---|---|---|---|---|---|---|---|---|---|---|---|---|---|---|---|---|---|---|---|
| 1568 | 1 | 0.2um_NOKCPHJN_656225 | Bacteria | Proteobacteri | Gammaprotec | various | various | various | various | metabolism | Energy produc | 179RI | May be involv | 247634.GPB2 | 0,2 | 0,4 | 1,81E-05 | 3,62E-05 | 1 - | 9,04E-05 - | 0 - | 0 - | 0 - | 0 - | 0 - | 0 - | 0 - | 0 - |
| 1569 | 1 | 3um_FIJJAFBF_292379 | Bacteria | Bacteroidetes | Cytophagia | Cytophagales | Flammeovirga | unclassified | Flammeovirga | poorly charact | Function unkn | 16R8I | Lipoprotein | 314260.PB250 | 10,2 | 1,166190379 | 0,000287234 | 4,54E-05 | 10 - | 0,000235681 - | 8 - | 0,000234117 - | 11 - | 0,000306017 - | 11 - | 0,000310537 - | 11 - | 0,000349819 - |
| 1570 | 1 | 3um_FIJJAFBF_786080 | Bacteria | Proteobacteri | Gammaprotec | Oceanospirilla | Saccharospirill | Reinekea | Reinekea fors | unclassified | unclassified | unclassified | unclassified | unclassified | 0,6 | 0,489897949 | 2,37E-05 | 1,93E-05 | 0 - | 0 - | 1 - | 4,06E-05 - | 1 - | 3,86E-05 - | 1 - | 3,92E-05 - | 0 - | 0 - |
| 1571 | 1 | 3um_FIJJAFBF_254020 | Bacteria | Bacteroidetes | Flavobacteriia | Flavobacterial | Flavobacteriac | Arenitalea | Arenitalea lut | unclassified | unclassified | unclassified | unclassified | unclassified | 0,2 | 0,4 | 6,77E-06 | 1,35E-05 | 0 - | 0 - | 0 - | 0 - | 1 - | 3,39E-05 - | 0 - | 0 - | 0 - | 0 - |
| 1572 | 1 | 3um_FIJJAFBF_418831 | Eukaryota | Bacillariophyt | Bacillariophyc | Naviculales | Naviculaceae | Fistulifera | Fistulifera sola | unclassified | unclassified | unclassified | unclassified | unclassified | 0,6 | 0,489897949 | 1,46E-05 | 1,19E-05 | 0 - | 0 - | 1 - | 2,40E-05 - | 1 - | 2,28E-05 - | 0 - | 0 - | 1 - | 2,61E-05 - |
| 1573 | 1 | 0.2um_NOKCPHJN_1011682 | unclassified | unclassified | unclassified | unclassified | unclassified | unclassified | unclassified | unclassified | unclassified | unclassified | unclassified | unclassified | 0,4 | 0,489897949 | 5,56E-05 | 6,93E-05 | 1 - | 0,000118327 - | 0 - | 0 - | 0 - | 0 - | 0 - | 0 - | 1 - | 0,000159664 - |
| 1574 | 1 | 0.2um_NOKCPHJN_114705 | Bacteria | Proteobacteri | Gammaprotec | Cellvibrionales | Porticoccacea | unclassified | gamma protec | poorly charact | Function unkn | 0QNMQ | NA | 314287.GB220 | 4 | 1,264911064 | 4,68E-05 | 1,42E-05 | 4 - | 3,98E-05 - | 4 - | 4,94E-05 - | 6 - | 7,04E-05 - | 4 - | 4,77E-05 - | 2 - | 2,68E-05 - |
| 1575 | 1 | 0.2um_NOKCPHJN_622970 | Bacteria | Verrucomicrol | unclassified | unclassified | unclassified | unclassified | Verrucomicrol | poorly charact | Function unkn | 1DIXS | Inherit from b | 382464.VDG1 | 0,8 | 0,748331477 | 2,96E-05 | 2,61E-05 | 2 - | 6,75E-05 - | 0 - | 0 - | 1 - | 3,98E-05 - | 1 - | 4,04E-05 - | 0 - | 0 - |
| 1576 | 2 | 0.2um_NOKCPHJN_663424;3um_FIJJAFBF_83991 | Bacteria | Bacteroidetes | Flavobacteriia | Flavobacterial | various | various | various | metabolism | Lipid transpor | COG1960 | acyl-CoA dehy | 983544.Lacal_ | 0,2 | 0,4 | 7,24E-06 | 1,45E-05 | 0 - | 0 - | 0 - | 0 - | 0 - | 0 - | 1 - | 3,62E-05 - | 0 - | 0 - |
| 1577 | 1 | 3um_FIJJAFBF_183675 | Bacteria | Bacteroidetes | Flavobacteriia | Flavobacterial | Flavobacteriac | Arenitalea | Arenitalea lut | unclassified | unclassified | unclassified | unclassified | unclassified | 14,2 | 1,720465053 | 0,00023003 | 2,93E-05 | 16 - | 0,000218103 - | 11 - | 0,000186189 - | 15 - | 0,000241358 - | 14 - | 0,000228595 - | 15 - | 0,000275905 - |
| 1578 | 1 | 0.2um_NOKCPHJN_424838 | Bacteria | Proteobacteri | Alphaproteob | unclassified | unclassified | unclassified | Alphaproteob | unclassified | unclassified | unclassified | unclassified | unclassified | 0,4 | 0,489897949 | 7,01E-06 | 8,64E-06 | 1 - | 1,59E-05 - | 0 - | 0 - | 0 - | 0 - | 1 - | 1,91E-05 - | 0 - | 0 - |
| 1579 | 1 | 0.2um_NOKCPHJN_768533 | Bacteria | Proteobacteri | Gammaprotec | Cellvibrionales | Spongiibacter | unclassified | marine gamm | unclassified | unclassified | unclassified | unclassified | unclassified | 0,2 | 0,4 | 2,01E-05 | 4,01E-05 | 0 - | 0 - | 1 - | 0,000100365 - | 0 - | 0 - | 0 - | 0 - | 0 - | 0 - |
| 1580 | 1 | 3um_FIJJAFBF_76679 | Bacteria | Bacteroidetes | Flavobacteriia | Flavobacterial | Flavobacteriac | Lutibacter | Lutibacter sp. | cellular proces | Signal transdu | 09A4D | gtp-binding pr | 880070.Cycma | 0,8 | 0,4 | 1,84E-05 | 9,45E-06 | 1 - | 1,95E-05 - | 0 - | 0 - | 1 - | 2,30E-05 - | 1 - | 2,33E-05 - | 1 - | 2,62E-05 - |
| 1581 | 1 | 0.2um_NOKCPHJN_114708 | Bacteria | Proteobacteri | Gammaprotec | Cellvibrionales | Porticoccacea | unclassified | gamma protec | metabolism | Carbohydrate | 177G7 | Glycosyl hydro | 314287.GB220 | 0,2 | 0,4 | 4,66E-06 | 9,31E-06 | 0 - | 0 - | 1 - | 2,33E-05 - | 0 - | 0 - | 0 - | 0 - | 0 - | 0 - |
| 1582 | 1 | 0.2um_NOKCPHJN_13356 | Bacteria | Proteobacteri | Gammaprotec | Alteromonada | Alteromonada | Paraglaciecola | Paraglaciecola | unclassified | unclassified | unclassified | unclassified | unclassified | 0,6 | 0,489897949 | 1,84E-05 | 1,52E-05 | 1 - | 2,73E-05 - | 0 - | 0 - | 1 - | 3,22E-05 - | 1 - | 3,27E-05 - | 0 - | 0 - |
| 1583 | 1 | 0.2um_NOKCPHJN_145730 | Bacteria | Proteobacteri | Betaproteoba | Burkholderiale | Comamonada | Hydrogenopha | various | information st | Transcription | COG0086 | DNA-depende | 596154.Alide2 | 15,6 | 2,727636339 | 0,000149676 | 1,93E-05 | 20 - | 0,000162458 - | 17 - | 0,000171466 - | 12 - | 0,000115059 - | 15 - | 0,000145948 - | 14 - | 0,00015345 - |
| 1584 | 1 | 3um_FIJJAFBF_62637 | Bacteria | Proteobacteri | Alphaproteob | Pelagibacteral | Pelagibacterac | Candidatus Pe | Candidatus Pe | metabolism | Inorganic ion t | 05PWN | phosphate ab | 335992.SAR11 | 5,8 | 0,4 | 0,000373461 | 5,59E-05 | 5 - | 0,00026817 - | 6 - | 0,000399586 - | 6 - | 0,000379857 - | 6 - | 0,000385467 - | 6 - | 0,000434228 - |
| 1585 | 1 | 3um_FIJJAFBF_38938 | Bacteria | Proteobacteri | Gammaprotec | Oceanospirilla | Oceanospirilla | Motiliproteus | Motiliproteus | cellular proces | Posttranslatio | 0QHXH | Molecular cha | 377629.TERTU | 6,2 | 1,6 | 0,000131534 | 2,63E-05 | 8 - | 0,000145058 - | 5 - | 0,000112574 - | 6 - | 0,000128419 - | 8 - | 0,000173755 - | 4 - | 9,79E-05 - |
| 1586 | 1 | 3um_FIJJAFBF_152478 | Bacteria | Bacteroidetes | Flavobacteriia | Flavobacterial | Flavobacteriac | Polaribacter | various | unclassified | unclassified | unclassified | unclassified | unclassified | 2,8 | 0,748331477 | 0,00025543 | 5,58E-05 | 4 - | 0,000310207 - | 2 - | 0,000192593 - | 3 - | 0,000274626 - | 2 - | 0,000185788 - | 3 - | 0,000313935 - |
| 1587 | 1 | 0.2um_NOKCPHJN_63178 | Bacteria | Bacteroidetes | Flavobacteriia | Flavobacterial | Flavobacteriac | Gaetbulibacte | Gaetbulibacte | information st | Translation, ri | 05CEJ | Catalyzes the | 313598.MED1 | 1 | 0 | 2,46E-05 | 2,33E-06 | 1 - | 2,06E-05 - | 1 - | 2,55E-05 - | 1 - | 2,43E-05 - | 1 - | 2,46E-05 - | 1 - | 2,78E-05 - |
| 1588 | 1 | 3um_FIJJAFBF_685788 | Bacteria | Proteobacteri | Gammaprotec | Pseudomonad | Moraxellaceae | Psychrobacter | various | information st | Replication, re | COG0178 | The UvrABC re | 335284.Pcryo_ | 0,2 | 0,4 | 4,78E-06 | 9,56E-06 | 0 - | 0 - | 0 - | 0 - | 1 - | 2,39E-05 - | 0 - | 0 - | 0 - | 0 - |
| 1589 | 1 | 3um_FIJJAFBF_123940 | Bacteria | Actinobacteria | Actinobacteria | Micrococcales | Microbacteria | Candidatus Ac | Candidatus Ac | unclassified | unclassified | unclassified | unclassified | unclassified | 0,8 | 0,4 | 8,28E-05 | 4,25E-05 | 1 - | 8,76E-05 - | 0 - | 0 - | 1 - | 0,000103422 - | 1 - | 0,000104949 - | 1 - | 0,000118225 - |
| 1590 | 1 | 0.2um_NOKCPHJN_62771 | Bacteria | Bacteroidetes | Flavobacteriia | Flavobacterial | Flavobacteriac | Zobellia | Zobellia uligin | metabolism | Amino acid tra | 05C65 | Catalyzes the | 313603.FB217 | 1,8 | 1,326649916 | 5,66E-05 | 4,31E-05 | 2 - | 5,41E-05 - | 2 - | 6,72E-05 - | 1 - | 3,20E-05 - | 4 - | 0,000129701 - | 0 - | 0 - |
| 1591 | 1 | 3um_FIJJAFBF_355505 | Bacteria | Proteobacteri | Gammaprotec | Alteromonada | Alteromonada | Paraglaciecola | Paraglaciecola | information st | Translation, ri | COG1185 | Involved in m | 1085623.GNIT | 1 | 0 | 0,000171284 | 1,63E-05 | 1 - | 0,000143471 - | 1 - | 0,000178149 - | 1 - | 0,000169353 - | 1 - | 0,000171854 - | 1 - | 0,000193593 - |
| 1592 | 1 | 3um_FIJJAFBF_296717 | Bacteria | Proteobacteri | Alphaproteob | Rhodobactera | unclassified | unclassified | Rhodobactera | cellular proces | Cell wall/mem | 16Q41 | Efflux transpo | 367336.OM22 | 0,4 | 0,489897949 | 1,27E-05 | 1,56E-05 | 1 - | 2,91E-05 - | 0 - | 0 - | 1 - | 3,43E-05 - | 0 - | 0 - | 0 - | 0 - |
| 1593 | 1 | 3um_FIJJAFBF_63542 | Bacteria | Bacteroidetes | Flavobacteriia | Flavobacterial | Flavobacteriac | Marinirhabdu | Marinirhabdu | cellular proces | Intracellular tr | 05FGH | MotA TolQ ex | 50743.SCB49_ | 4 | 0,632455532 | 0,00021455 | 3,80E-05 | 4 - | 0,000180042 - | 3 - | 0,000167669 - | 4 - | 0,000212521 - | 5 - | 0,000269575 - | 4 - | 0,00024294 - |
| 1594 | 1 | 3um_FIJJAFBF_585135 | Bacteria | Proteobacteri | Alphaproteob | Pelagibacteral | Pelagibacterac | unclassified | Pelagibacterac | unclassified | unclassified | unclassified | unclassified | unclassified | 4,6 | 0,489897949 | 0,00019129 | 3,51E-05 | 4 - | 0,000138285 - | 5 - | 0,000214637 - | 4 - | 0,000163232 - | 5 - | 0,000207053 - | 5 - | 0,000233245 - |
| 1595 | 1 | 0.2um_NOKCPHJN_142270 | Bacteria | Bacteroidetes | Flavobacteriia | Flavobacterial | Flavobacteriac | Tenacibaculur | Tenacibaculur | unclassified | unclassified | unclassified | unclassified | unclassified | 4,6 | 0,489897949 | 0,000121532 | 1,70E-05 | 5 - | 0,000110788 - | 4 - | 0,000110053 - | 4 - | 0,000104619 - | 5 - | 0,000132706 - | 5 - | 0,000149493 - |
| 1596 | 1 | 0.2um_NOKCPHJN_26783 | Bacteria | Proteobacteri | Betaproteoba | unclassified | unclassified | unclassified | Betaproteoba | metabolism | Energy produc | 05CV4 | Subunits I and | 296591.Bpro_ | 2 | 1,414213562 | 6,62E-05 | 4,30E-05 | 4 - | 0,000115354 - | 3 - | 0,000107426 - | 0 - | 0 - | 2 - | 6,91E-05 - | 1 - | 3,89E-05 - |
| 1597 | 1 | 3um_FIJJAFBF_97406 | Bacteria | Bacteroidetes | Flavobacteriia | Flavobacterial | Flavobacteriac | unclassified | Flavobacteriac | unclassified | unclassified | unclassified | unclassified | unclassified | 0,2 | 0,4 | 3,64E-06 | 7,28E-06 | 0 - | 0 - | 0 - | 0 - | 0 - | 0 - | 1 - | 1,82E-05 - | 0 - | 0 - |
| 1598 | 1 | 3um_FIJJAFBF_629960 | Bacteria | Proteobacteri | Alphaproteob | Pelagibacteral | various | various | various | information st | Transcription | 05VEQ | Cold shock pro | 439493.PB721 | 4,6 | 0,489897949 | 0,000933358 | 0,000168212 | 4 - | 0,000675157 - | 5 - | 0,001047933 - | 5 - | 0,000996193 - | 4 - | 0,000808725 - | 5 - | 0,001138783 - |
| 1599 | 1 | 0.2um_NOKCPHJN_206999 | Bacteria | Bacteroidetes | Flavobacteriia | unclassified | unclassified | unclassified | Flavobacteria | unclassified | unclassified | unclassified | unclassified | unclassified | 1 | 0 | 0,000103028 | 9,78E-06 | 1 - | 8,63E-05 - | 1 - | 0,000107157 - | 1 - | 0,000101866 - | 1 - | 0,000103371 - | 1 - | 0,000116447 - |
| 1600 | 1 | 0.2um_NOKCPHJN_481976 | Bacteria | Bacteroidetes | Saprospiria | Saprospirales | Saprospiracea | Aureispira | Aureispira sp. | metabolism | Coenzyme tra | 09A4R | May play a ke | 313606.M231 | 0,4 | 0,489897949 | 1,28E-05 | 1,57E-05 | 0 - | 0 - | 1 - | 3,25E-05 - | 0 - | 0 - | 1 - | 3,14E-05 - | 0 - | 0 - |
| 1601 | 1 | 3um_FIJJAFBF_715072 | Eukaryota | Bacillariophyt | Mediophycea | Triceratiales | Triceratiaceae | Triceratium | Triceratium d | information st | Translation, ri | COG0050 | This protein p | 32049.SYNPC | 3,4 | 1,019803903 | 0,000114609 | 3,80E-05 | 3 - | 8,42E-05 - | 3 - | 0,000104537 - | 5 - | 0,000165626 - | 2 - | 6,72E-05 - | 4 - | 0,000151466 - |
| 1602 | 1 | 3um_FIJJAFBF_273584 | Bacteria | Bacteroidetes | Flavobacteriia | Flavobacterial | Flavobacteriac | Flavobacteriu | Flavobacteriu | unclassified | unclassified | unclassified | unclassified | unclassified | 0,2 | 0,4 | 7,60E-06 | 1,52E-05 | 0 - | 0 - | 1 - | 3,80E-05 - | 0 - | 0 - | 0 - | 0 - | 0 - | 0 - |
| 1603 | 1 | 0.2um_NOKCPHJN_562296 | Bacteria | Proteobacteri | Alphaproteob | Rhodobactera | Rhodobactera | Planktomarina | Planktomarina | information st | Translation, ri | 082FS | 50S ribosomal | 290400.Jann_ | 1,4 | 0,489897949 | 0,000256821 | 7,56E-05 | 2 - | 0,000314457 - | 1 - | 0,000195231 - | 1 - | 0,000185592 - | 2 - | 0,000376667 - | 1 - | 0,000212157 - |
| 1604 | 1 | 0.2um_NOKCPHJN_544661 | Bacteria | unclassified | unclassified | unclassified | unclassified | unclassified | uncultured ba | metabolism | Inorganic ion t | 05C27 | TonB-depende | 688270.Celal_ | 0,6 | 0,8 | 9,47E-06 | 1,22E-05 | 2 - | 2,98E-05 - | 0 - | 0 - | 1 - | 1,76E-05 - | 0 - | 0 - | 0 - | 0 - |
| 1605 | 4 | 0.2um_NOKCPHJN_302225;0.2um_NOKCPHJN_4 | Bacteria | Proteobacteri | Alphaproteob | various | various | various | various | unclassified | unclassified | unclassified | unclassified | unclassified | 0,4 | 0,489897949 | 5,08E-05 | 6,25E-05 | 0 - | 0 - | 0 - | 0 - | 0 - | 0 - | 1 - | 0,000119551 - | 1 - | 0,000134673 - |
| 1606 | 1 | 0.2um_NOKCPHJN_1007807 | Bacteria | Lentisphaerae | unclassified | unclassified | unclassified | unclassified | various | cellular proces | Cell wall/mem | 0NG0S | Methyltransfe | 985665.HPL0 | 0,6 | 0,489897949 | 2,43E-05 | 1,99E-05 | 0 - | 0 - | 1 - | 4,17E-05 - | 1 - | 3,96E-05 - | 1 - | 4,02E-05 - | 0 - | 0 - |
| 1607 | 1 | 0.2um_NOKCPHJN_689763 | Bacteria | Actinobacteria | Actinobacteria | Micrococcales | Microbacteria | Rhodoluna | Candidatus Rh | unclassified | unclassified | unclassified | unclassified | unclassified | 2 | 0,632455532 | 2,91E-05 | 7,63E-06 | 3 - | 3,72E-05 - | 2 - | 3,08E-05 - | 2 - | 2,93E-05 - | 1 - | 1,49E-05 - | 2 - | 3,35E-05 - |
| 1608 | 1 | 3um_FIJJAFBF_177322 | Bacteria | Actinobacteria | Actinobacteria | Micrococcales | Microbacteria | unclassified | Microbacteria | unclassified | unclassified | unclassified | unclassified | unclassified | 0,6 | 0,489897949 | 0,000108159 | 8,92E-05 | 1 - | 0,000157228 - | 1 - | 0,000195231 - | 0 - | 0 - | 1 - | 0,000188333 - | 0 - | 0 - |
| 1609 | 1 | 3um_FIJJAFBF_346928 | Bacteria | Bacteroidetes | unclassified | unclassified | unclassified | unclassified | Bacteroidetes | information st | Translation, ri | 08UIK | This protein bi | 867900.Celly_ | 0,6 | 0,489897949 | 5,42E-05 | 4,47E-05 | 1 - | 7,92E-05 - | 1 - | 9,83E-05 - | 1 - | 9,34E-05 - | 0 - | 0 - | 0 - | 0 - |
| 1610 | 1 | 3um_FIJJAFBF_58538 | Eukaryota | Phaeophyceae | unclassified | Ectocarpales | Ectocarpaceae | Ectocarpus | Ectocarpus sili | information st | Translation, ri | 1508W | Ribosomal pro | 272952.HpaP | 1 | 0 | 0,000130502 | 1,24E-05 | 1 - | 0,000109311 - | 1 - | 0,000135732 - | 1 - | 0,000129031 - | 1 - | 0,000130936 - | 1 - | 0,0001475 - |
| 1611 | 1 | 3um_FIJJAFBF_85799 | Viruses | unclassified | unclassified | unclassified | unclassified | unclassified | uncultured M | unclassified | unclassified | unclassified | unclassified | unclassified | 2,8 | 0,979795897 | 0,000269628 | 7,98E-05 | 4 - | 0,000332686 - | 3 - | 0,000309824 - | 3 - | 0,000294527 - | 3 - | 0,000298877 - | 1 - | 0,000112228 - |
| 1612 | 1 | 0.2um_NOKCPHJN_744618 | Bacteria | Bacteroidetes | Flavobacteriia | Flavobacterial | Flavobacteriac | Polaribacter | various | poorly charact | Function unkn | 0PBMV | NA | 313594.PI23P | 1 | 0 | 7,92E-05 | 7,52E-06 | 1 - | 6,63E-05 - | 1 - | 8,24E-05 - | 1 - | 7,83E-05 - | 1 - | 7,95E-05 - | 1 - | 8,95E-05 - |
| 1613 | 1 | 0.2um_NOKCPHJN_56829 | Bacteria | Proteobacteri | Gammaprotec | Oceanospirilla | Saccharospirill | Reinekea | Reinekea blan | unclassified | unclassified | unclassified | unclassified | unclassified | 2,6 | 0,489897949 | 9,20E-05 | 2,34E-05 | 2 - | 5,86E-05 - | 3 - | 0,000109071 - | 3 - | 0,000103685 - | 2 - | 7,01E-05 - | 3 - | 0,000118526 - |
| 1614 | 1 | 3um_FIJJAFBF_333455 | unclassified | unclassified | unclassified | unclassified | unclassified | unclassified | unclassified | unclassified | unclassified | unclassified | unclassified | unclassified | 1,8 | 0,4 | 0,000279402 | 5,52E-05 | 2 - | 0,000263855 - | 2 - | 0,00032763 - | 2 - | 0,000311453 - | 2 - | 0,000316054 - | 1 - | 0,000178017 - |
| 1615 | 4 | 0.2um_NOKCPHJN_1111222;0.2um_NOKCPHJN_ | Bacteria | Proteobacteri | Alphaproteob | Pelagibacteral | Pelagibacterac | Candidatus Pe | various | information st | Transcription | 05DEK | Sigma factors | 439493.PB721 | 0,2 | 0,4 | 7,73E-06 | 1,55E-05 | 1 - | 3,86E-05 - | 0 - | 0 - | 0 - | 0 - | 0 - | 0 - | 0 - | 0 - |
| 1616 | 1 | 3um_FIJJAFBF_91013 | Eukaryota | unclassified | Florideophyce | Gigartinales | Gigartinaceae | Chondrus | Chondrus crisp | information st | Translation, ri | KOG3280 | ribosomal pro | 35128.Thaps3 | 0,6 | 0,489897949 | 7,25E-05 | 5,96E-05 | 1 - | 0,000107268 - | 0 - | 0 - | 1 - | 0,000126619 - | 1 - | 0,000128489 - | 0 - | 0 - |
| 1617 | 1 | 0.2um_NOKCPHJN_512789 | Bacteria | Proteobacteri | Gammaprotec | Cellvibrionales | Porticoccacea | unclassified | SAR92 bacteri | cellular proces | Cell wall/mem | 0QKXZ | Porin Gram-ne | 314287.GB220 | 4,2 | 0,979795897 | 0,000180483 | 4,50E-05 | 5 - | 0,000180467 - | 3 - | 0,000134452 - | 3 - | 0,000127813 - | 5 - | 0,000216169 - | 5 - | 0,000243513 - |
| 1618 | 1 | 0.2um_NOKCPHJN_790600 | Bacteria | Firmicutes | Bacilli | Lactobacillales | Streptococcac | Streptococcus | Streptococcus | poorly charact | Function unkn | 0GHXM | NA | 439483.CBGD | 3,2 | 0,748331477 | 5,37E-05 | 1,61E-05 | 2 - | 2,76E-05 - | 3 - | 5,15E-05 - | 3 - | 4,89E-05 - | 4 - | 6,62E-05 - | 4 - | 7,45E-05 - |
| 1619 | 1 | 3um_FIJJAFBF_462347 | unclassified | unclassified | unclassified | unclassified | unclassified | unclassified | unclassified | unclassified | unclassified | unclassified | unclassified | unclassified | 0,2 | 0,4 | 1,18E-05 | 2,36E-05 | 0 - | 0 - | 0 - | 0 - | 0 - | 0 - | 1 - | 5,90E-05 - | 0 - | 0 - |
| 1620 | 1 | 0.2um_NOKCPHJN_438106 | Bacteria | Bacteroidetes | unclassified | unclassified | unclassified | unclassified | Bacteroidetes | unclassified | unclassified | unclassified | unclassified | unclassified | 0,8 | 0,4 | 5,80E-05 | 2,94E-05 | 1 - | 6,27E-05 - | 1 - | 7,79E-05 - | 1 - | 7,40E-05 - | 1 - | 7,51E-05 - | 0 - | 0 - |
| 1621 | 1 | 3um_FIJJAFBF_250358 | Bacteria | Bacteroidetes | Flavobacteriia | Flavobacterial | Flavobacteriac | Lacinutrix | Lacinutrix him | poorly charact | Function unkn | 06E9D | NA | 391598.FBBAL | 0,8 | 0,748331477 | 1,69E-05 | 1,49E-05 | 2 - | 3,82E-05 - | 1 - | 2,37E-05 - | 1 - | 2,25E-05 - | 0 - | 0 - | 0 - | 0 - |
| 1622 | 1 | 0.2um_NOKCPHJN_501191 | Bacteria | Bacteroidetes | Flavobacteriia | Flavobacterial | Flavobacteriac | Joostella | Joostella mari | metabolism | Inorganic ion t | 05C27 | TonB-depende | 655815.ZPR_3 | 0,2 | 0,4 | 2,19E-06 | 4,38E-06 | 1 - | 1,09E-05 - | 0 - | 0 - | 0 - | 0 - | 0 - | 0 - | 0 - | 0 - |
| 1623 | 1 | 3um_FIJJAFBF_209215 | Bacteria | Bacteroidetes | unclassified | unclassified | unclassified | unclassified | Bacteroidetes | unclassified | unclassified | unclassified | unclassified | unclassified | 0,2 | 0,4 | 2,85E-06 | 5,70E-06 | 0 - | 0 - | 0 - | 0 - | 1 - | 1,42E-05 - | 0 - | 0 - | 0 - | 0 - |
| 1624 | 1 | 0.2um_NOKCPHJN_760362 | Bacteria | Proteobacteri | Alphaproteob | Pelagibacteral | Pelagibacterac | Candidatus Pe | Candidatus Pe | metabolism | Lipid transpor | 01SHD | Short-chain de | 335992.SAR11 | 0,6 | 0,489897949 | 3,37E-05 | 2,77E-05 | 1 - | 4,99E-05 - | 0 - | 0 - | 1 - | 5,89E-05 - | 1 - | 5,98E-05 - | 0 - | 0 - |
| 1625 | 1 | 0.2um_NOKCPHJN_92348 | Bacteria | Bacteroidetes | Flavobacteriia | Flavobacterial | Flavobacteriac | Polaribacter | Polaribacter s | cellular proces | Intracellular tr | 0P8JN | The central su | 313594.PI23P | 1 | 0 | 3,10E-05 | 2,94E-06 | 1 - | 2,60E-05 - | 1 - | 3,22E-05 - | 1 - | 3,07E-05 - | 1 - | 3,11E-05 - | 1 - | 3,50E-05 - |
| 1626 | 1 | 0.2um_NOKCPHJN_643786 | Bacteria | Proteobacteri | Alphaproteob | Pelagibacteral | Pelagibacterac | Candidatus Pe | Candidatus Pe | unclassified | unclassified | unclassified | unclassified | unclassified | 1 | 0,632455532 | 4,61E-05 | 2,98E-05 | 1 - | 3,97E-05 - | 1 - | 4,93E-05 - | 2 - | 9,38E-05 - | 1 - | 4,76E-05 - | 0 - | 0 - |
| 1627 | 1 | 3um_FIJJAFBF_189700 | Bacteria | Bacteroidetes | Flavobacteriia | Flavobacterial | Flavobacteriac | Polaribacter | Polaribacter s | unclassified | unclassified | unclassified | unclassified | unclassified | 35 | 4,049691346 | 0,000751856 | 7,53E-05 | 42 - | 0,000760351 - | 36 - | 0,000809255 - | 31 - | 0,000662453 - | 31 - | 0,000672237 - | 35 - | 0,000854985 - |
| 1628 | 1 | 0.2um_NOKCPHJN_240980 | Bacteria | Proteobacteri | Gammaprotec | Oceanospirilla | Saccharospirill | Saccharospiril | Saccharospiril | unclassified | unclassified | unclassified | unclassified | unclassified | 9 | 0,894427191 | 0,000273103 | 2,48E-05 | 10 - | 0,000255628 - | 10 - | 0,000317414 - | 9 - | 0,000271568 - | 8 - | 0,000244959 - | 8 - | 0,000275946 - |
| 1629 | 1 | 3um_FIJJAFBF_438717 | Bacteria | Bacteroidetes | Sphingobacte | Sphingobacte | Sphingobacte | Sphingobacte | Sphingobacte | information st | Replication, re | 07RYU | helicase | 755732.Fluta_ | 0,8 | 0,4 | 9,66E-06 | 4,91E-06 | 1 - | 1,05E-05 - | 1 - | 1,30E-05 - | 1 - | 1,23E-05 - | 1 - | 1,25E-05 - | 0 - | 0 - |
| 1630 | 1 | 0.2um_NOKCPHJN_893330 | Bacteria | Bacteroidetes | Flavobacteriia | Flavobacterial | unclassified | unclassified | Flavobacterial | poorly charact | Function unkn | 0644X | Domain-Conta | 1002672.SAR1 | 3 | 0 | 5,94E-05 | 5,64E-06 | 3 - | 4,98E-05 - | 3 - | 6,18E-05 - | 3 - | 5,87E-05 - | 3 - | 5,96E-05 - | 3 - | 6,71E-05 - |
| 1631 | 1 | 0.2um_NOKCPHJN_907916 | Viruses | unclassified | unclassified | unclassified | Circoviridae | unclassified | Avon-Heathco | unclassified | unclassified | unclassified | unclassified | unclassified | 1 | 0 | 5,17E-05 | 4,91E-06 | 1 - | 4,33E-05 - | 1 - | 5,38E-05 - | 1 - | 5,11E-05 - | 1 - | 5,19E-05 - | 1 - | 5,84E-05 - |
| 1632 | 1 | 3um_FIJJAFBF_46521 | Bacteria | Proteobacteri | Gammaprotec | Oceanospirilla | Oceanospirilla | Motiliproteus | Motiliproteus | information st | Translation, ri | COG0088 | One of the pri | 575588.HMPR | 4 | 1,673320053 | 0,000266435 | 8,90E-05 | 7 - | 0,000401719 - | 2 - | 0,000142519 - | 3 - | 0,000203223 - | 4 - | 0,000274967 - | 4 - | 0,000309749 - |
| 1633 | 1 | 3um_FIJJAFBF_08116 | Bacteria | Bacteroidetes | Flavobacteriia | Flavobacterial | Flavobacteriac | Polaribacter | Polaribacter s | poorly charact | Function unkn | 0P8G1 | peptidylprolyl | 313598.MED1 | 1,8 | 1,166190379 | 9,08E-05 | 5,99E-05 | 1 - | 4,16E-05 - | 1 - | 5,16E-05 - | 1 - | 4,91E-05 - | 4 - | 0,000199251 - | 2 - | 0,000112228 - |
| 1634 | 1 | 0.2um_NOKCPHJN_554771 | Bacteria | Proteobacteri | Alphaproteob | Rhodobactera | unclassified | unclassified | Rhodobactera | unclassified | unclassified | unclassified | unclassified | unclassified | 0,8 | 0,4 | 3,19E-05 | 1,64E-05 | 1 - | 3,35E-05 - | 1 - | 4,16E-05 - | 1 - | 3,95E-05 - | 0 - | 0 - | 1 - | 4,52E-05 - |
| 1635 | 1 | 3um_FIJJAFBF_111307 | Viruses | unclassified | unclassified | unclassified | Phycodnavirid | unclassified | Chrysochromu | unclassified | unclassified | unclassified | unclassified | unclassified | 0,2 | 0,4 | 1,05E-05 | 2,09E-05 | 0 - | 0 - | 0 - | 0 - | 0 - | 0 - | 1 - | 5,23E-05 - | 0 - | 0 - |
| 1636 | 1 | 3um_FIJJAFBF_232689 | Bacteria | Proteobacteri | Gammaprotec | Oceanospirilla | Saccharospirill | Reinekea | Reinekea fors | unclassified | unclassified | unclassified | unclassified | unclassified | 0,6 | 0,489897949 | 3,33E-05 | 2,74E-05 | 1 - | 4,93E-05 - | 0 - | 0 - | 1 - | 5,81E-05 - | 1 - | 5,90E-05 - | 0 - | 0 - |
| 1637 | 1 | 0.2um_NOKCPHJN_282157 | Eukaryota | Bacillariophyt | Fragilariophyc | Fragilariales | Fragilariaceae | Ardissonea | Ardissonea cr | unclassified | unclassified | unclassified | unclassified | unclassified | 3,8 | 0,4 | 6,88E-05 | 6,13E-06 | 4 - | 6,11E-05 - | 4 - | 7,58E-05 - | 4 - | 7,21E-05 - | 4 - | 7,31E-05 - | 3 - | 6,18E-05 - |
| 1638 | 1 | 0.2um_NOKCPHJN_167820 | Eukaryota | Phaeophyceae | unclassified | Ectocarpales | Ectocarpaceae | Ectocarpus | Ectocarpus sili | poorly charact | Function unkn | 0J1P2 | NA | 412030.XP_0 | 0,2 | 0,4 | 1,47E-05 | 2,94E-05 | 1 - | 7,36E-05 - | 0 - | 0 - | 0 - | 0 - | 0 - | 0 - | 0 - | 0 - |
| 1639 | 1 | 0.2um_NOKCPHJN_521289 | Eukaryota | unclassified | Bangiophycea | Cyanidiales | Cyanidiaceae | Galdieria | Galdieria sulp | unclassified | unclassified | unclassified | unclassified | unclassified | 0,2 | 0,4 | 1,85E-06 | 3,70E-06 | 0 - | 0 - | 0 - | 0 - | 0 - | 0 - | 1 - | 9,25E-06 - | 0 - | 0 - |
| 1640 | 1 | 3um_FIJJAFBF_388554 | Bacteria | Bacteroidetes | Flavobacteriia | Flavobacterial | Flavobacteriac | Spongiibacteri | Spongiibacteri | poorly charact | Function unkn | 11QQ6 | NA | 313603.FB217 | 2 | 0 | 9,52E-05 | 9,04E-06 | 2 - | 7,97E-05 - | 2 - | 9,90E-05 - | 2 - | 9,41E-05 - | 2 - | 9,55E-05 - | 2 - | 0,000107552 - |
| 1641 | 1 | 3um_FIJJAFBF_508158 | Bacteria | Proteobacteri | Gammaprotec | Cellvibrionales | Porticoccacea | unclassified | SAR92 bacteri | unclassified | unclassified | unclassified | unclassified | unclassified | 6 | 1,095445115 | 0,000544746 | 8,04E-05 | 7 - | 0,00053922 - | 7 - | 0,000669552 - | 6 - | 0,000545566 - | 6 - | 0,000553624 - | 4 - | 0,00041577 - |
| 1642 | 1 | 3um_FIJJAFBF_07325 | Viruses | unclassified | unclassified | Caudovirales | Siphoviridae | unclassified | Tenacibaculur | unclassified | unclassified | unclassified | unclassified | unclassified | 3,4 | 0,489897949 | 8,07E-05 | 1,43E-05 | 3 - | 5,96E-05 - | 4 - | 9,86E-05 - | 4 - | 9,38E-05 - | 3 - | 7,14E-05 - | 3 - | 8,04E-05 - |
| 1643 | 3 | 0.2um_NOKCPHJN_248751;0.2um_NOKCPHJN_9 | Bacteria | Bacteroidetes | various | various | various | various | various | unclassified | unclassified | unclassified | unclassified | unclassified | 0,4 | 0,489897949 | 3,30E-05 | 4,06E-05 | 0 - | 0 - | 0 - | 0 - | 0 - | 0 - | 1 - | 7,77E-05 - | 1 - | 8,75E-05 - |
| 1644 | 1 | 3um_FIJJAFBF_787168 | Bacteria | Proteobacteri | Gammaprotec | Oceanospirilla | Saccharospirill | Saccharospiril | Saccharospiril | information st | Translation, ri | 16REB | Catalyzes the | 314283.MED2 | 1,6 | 0,8 | 5,30E-05 | 2,12E-05 | 3 - | 8,67E-05 - | 1 - | 3,59E-05 - | 1 - | 3,41E-05 - | 2 - | 6,93E-05 - | 1 - | 3,90E-05 - |
| 1645 | 2 | 3um_FIJJAFBF_190681;3um_FIJJAFBF_688633 | Bacteria | Bacteroidetes | Flavobacteriia | Flavobacterial | Flavobacteriac | Marinirhabdu | Marinirhabdu | information st | Translation, ri | COG0251 | Endoribonucle | 50743.SCB49_ | 0,8 | 0,748331477 | 7,71E-05 | 7,38E-05 | 1 - | 8,38E-05 - | 1 - | 0,000104028 - | 2 - | 0,000197784 - | 0 - | 0 - | 0 - | 0 - |
| 1646 | 1 | 0.2um_NOKCPHJN_117608 | Bacteria | Proteobacteri | Betaproteoba | Burkholderiale | Comamonada | Hydrogenopha | Hydrogenopha | unclassified | unclassified | unclassified | unclassified | unclassified | 3,2 | 0,4 | 4,50E-05 | 6,98E-06 | 3 - | 3,54E-05 - | 3 - | 4,39E-05 - | 3 - | 4,17E-05 - | 4 - | 5,65E-05 - | 3 - | 4,77E-05 - |
| 1647 | 1 | 3um_FIJJAFBF_786153 | Bacteria | Proteobacteri | Gammaprotec | Oceanospirilla | Saccharospirill | Reinekea | Reinekea fors | information st | Translation, ri | 0QTNF | One of the pri | 314283.MED2 | 3,2 | 0,4 | 0,000504694 | 2,77E-05 | 4 - | 0,000533845 - | 3 - | 0,000497159 - | 3 - | 0,000472612 - | 3 - | 0,000479593 - | 3 - | 0,00054026 - |
| 1648 | 3 | 0.2um_NOKCPHJN_149315;0.2um_NOKCPHJN_3 | Bacteria | Bacteroidetes | various | various | various | various | various | unclassified | unclassified | unclassified | unclassified | unclassified | 0,2 | 0,4 | 8,47E-06 | 1,69E-05 | 0 - | 0 - | 0 - | 0 - | 1 - | 4,23E-05 - | 0 - | 0 - | 0 - | 0 - |
| 1649 | 1 | 3um_FIJJAFBF_370788 | Bacteria | Proteobacteri | Gammaprotec | Oceanospirilla | Saccharospirill | Reinekea | Reinekea fors | metabolism | Inorganic ion t | 05E2Z | phosphate AB | 314283.MED2 | 3,2 | 0,4 | 0,000100361 | 1,56E-05 | 3 - | 7,88E-05 - | 3 - | 9,78E-05 - | 3 - | 9,30E-05 - | 4 - | 0,000125843 - | 3 - | 0,000106321 - |
| 1650 | 1 | 3um_FIJJAFBF_56370 | Bacteria | Bacteroidetes | unclassified | unclassified | unclassified | unclassified | Bacteroidetes | information st | Translation, ri | 0PA05 | 30S ribosomal | 487796.Flav2 | 2 | 0 | 0,000115635 | 1,10E-05 | 2 - | 9,69E-05 - | 2 - | 0,000120269 - | 2 - | 0,000114331 - | 2 - | 0,00011602 - | 2 - | 0,000130696 - |
| 1651 | 1 | 3um_FIJJAFBF_05421 | Bacteria | Bacteroidetes | Flavobacteriia | Flavobacterial | Flavobacteriac | Polaribacter | Polaribacter s | unclassified | unclassified | unclassified | unclassified | unclassified | 1,2 | 0,4 | 4,28E-05 | 9,47E-06 | 2 - | 6,14E-05 - | 1 - | 3,81E-05 - | 1 - | 3,62E-05 - | 1 - | 3,68E-05 - | 1 - | 4,14E-05 - |
| 1652 | 1 | 3um_FIJJAFBF_321849 | Viruses | unclassified | unclassified | unclassified | unclassified | unclassified | uncultured vir | unclassified | unclassified | unclassified | unclassified | unclassified | 1,4 | 0,8 | 3,91E-05 | 2,62E-05 | 1 - | 2,25E-05 - | 1 - | 2,80E-05 - | 1 - | 2,66E-05 - | 1 - | 2,70E-05 - | 3 - | 9,13E-05 - |
| 1653 | 1 | 3um_FIJJAFBF_566757 | Eukaryota | Bacillariophyt | Bacillariophyc | Naviculales | Naviculaceae | Fistulifera | Fistulifera sola | unclassified | unclassified | unclassified | unclassified | unclassified | 1 | 0,632455532 | 4,29E-05 | 2,81E-05 | 1 - | 3,69E-05 - | 1 - | 4,58E-05 - | 1 - | 4,36E-05 - | 2 - | 8,84E-05 - | 0 - | 0 - |
| 1654 | 1 | 0.2um_NOKCPHJN_15221 | Bacteria | Proteobacteri | Betaproteoba | unclassified | unclassified | unclassified | Betaproteoba | poorly charact | Function unkn | COG4147 | solute sympor | 391597.LMED | 2,8 | 0,748331477 | 6,33E-05 | 1,59E-05 | 3 - | 5,74E-05 - | 3 - | 7,13E-05 - | 4 - | 9,03E-05 - | 2 - | 4,58E-05 - | 2 - | 5,16E-05 - |
| 1655 | 1 | 3um_FIJJAFBF_454663 | Bacteria | Bacteroidetes | Flavobacteriia | Flavobacterial | Flavobacteriac | Polaribacter | Polaribacter s | information st | Replication, re | 05C68 | Can catalyze t | 402612.FP224 | 0,8 | 0,4 | 3,28E-05 | 1,69E-05 | 1 - | 3,44E-05 - | 1 - | 4,27E-05 - | 1 - | 4,06E-05 - | 0 - | 0 - | 1 - | 4,64E-05 - |
| 1656 | 1 | 3um_FIJJAFBF_518709 | Bacteria | Bacteroidetes | Flavobacteriia | Flavobacterial | Flavobacteriac | Polaribacter | Polaribacter b | metabolism | Carbohydrate | COG0297 | starch synthas | 313594.PI23P | 1,2 | 0,4 | 6,25E-05 | 2,66E-05 | 1 - | 4,27E-05 - | 1 - | 5,30E-05 - | 1 - | 5,04E-05 - | 1 - | 5,11E-05 - | 2 - | 0,000115148 - |
| 1657 | 1 | 0.2um_NOKCPHJN_144887 | Bacteria | Proteobacteri | Alphaproteob | unclassified | unclassified | unclassified | SAR116 cluste | unclassified | unclassified | unclassified | unclassified | unclassified | 0,4 | 0,489897949 | 1,37E-05 | 1,68E-05 | 0 - | 0 - | 1 - | 3,48E-05 - | 0 - | 0 - | 1 - | 3,36E-05 - | 0 - | 0 - |
| 1658 | 1 | 0.2um_NOKCPHJN_913764 | Bacteria | Proteobacteri | Alphaproteob | Pelagibacteral | Pelagibacterac | unclassified | Pelagibacterac | unclassified | unclassified | unclassified | unclassified | unclassified | 0,8 | 0,4 | 0,000150094 | 7,56E-05 | 0 - | 0 - | 1 - | 0,000187525 - | 1 - | 0,000178266 - | 1 - | 0,000180899 - | 1 - | 0,000203782 - |
| 1659 | 1 | 0.2um_NOKCPHJN_43981 | Bacteria | Proteobacteri | Gammaprotec | Cellvibrionales | Porticoccacea | unclassified | gamma protec | unclassified | unclassified | unclassified | unclassified | unclassified | 9,4 | 0,8 | 0,000144466 | 1,50E-05 | 10 - | 0,000129108 - | 10 - | 0,000160314 - | 10 - | 0,000152398 - | 8 - | 0,00012372 - | 9 - | 0,000156791 - |
| 1660 | 1 | 0.2um_NOKCPHJN_68925 | Bacteria | Bacteroidetes | various | various | various | various | various | metabolism | Energy produc | 0PBIY | subunit C | 487796.Flav2 | 1,8 | 0,4 | 0,000391361 | 9,59E-05 | 2 - | 0,000364371 - | 2 - | 0,000452441 - | 2 - | 0,000430102 - | 1 - | 0,000218227 - | 2 - | 0,000491665 - |
| 1661 | 1 | 3um_FIJJAFBF_513822 | Viruses | unclassified | unclassified | unclassified | unclassified | unclassified | uncultured M | unclassified | unclassified | unclassified | unclassified | unclassified | 0,6 | 0,8 | 4,95E-05 | 6,61E-05 | 0 - | 0 - | 0 - | 0 - | 1 - | 8,16E-05 - | 2 - | 0,000165643 - | 0 - | 0 - |
| 1662 | 1 | 0.2um_NOKCPHJN_635596 | unclassified | unclassified | unclassified | unclassified | unclassified | unclassified | unclassified | unclassified | unclassified | unclassified | unclassified | unclassified | 1 | 0 | 2,48E-05 | 2,35E-06 | 1 - | 2,08E-05 - | 1 - | 2,58E-05 - | 1 - | 2,45E-05 - | 1 - | 2,49E-05 - | 1 - | 2,80E-05 - |
| 1663 | 1 | 3um_FIJJAFBF_209603 | Bacteria | Bacteroidetes | Flavobacteriia | Flavobacterial | Flavobacteriac | Arenitalea | Arenitalea lut | unclassified | unclassified | unclassified | unclassified | unclassified | 3,6 | 0,489897949 | 0,00026913 | 4,38E-05 | 4 - | 0,000250878 - | 3 - | 0,000233637 - | 3 - | 0,000222102 - | 4 - | 0,00030051 - | 4 - | 0,000338523 - |
| 1664 | 1 | 0.2um_NOKCPHJN_74284 | Bacteria | Bacteroidetes | Flavobacteriia | Flavobacterial | Flavobacteriac | Polaribacter | Polaribacter s | information st | Transcription | 0P7SF | DNA-depende | 313598.MED1 | 0,2 | 0,4 | 1,83E-06 | 3,66E-06 | 1 - | 9,16E-06 - | 0 - | 0 - | 0 - | 0 - | 0 - | 0 - | 0 - | 0 - |
| 1665 | 1 | 3um_FIJJAFBF_177199 | Bacteria | Bacteroidetes | Flavobacteriia | Flavobacterial | Flavobacteriac | Polaribacter | various | unclassified | unclassified | unclassified | unclassified | unclassified | 0,4 | 0,489897949 | 5,31E-05 | 6,55E-05 | 1 - | 0,000120818 - | 0 - | 0 - | 0 - | 0 - | 1 - | 0,000144719 - | 0 - | 0 - |
| 1666 | 1 | 3um_FIJJAFBF_105309 | Bacteria | Bacteroidetes | Flavobacteriia | Flavobacterial | Flavobacteriac | Formosa | Formosa sp. H | information st | Translation, ri | COG0251 | Endoribonucle | 1046627.BZAF | 0,8 | 0,748331477 | 8,45E-05 | 8,13E-05 | 1 - | 9,11E-05 - | 1 - | 0,00011311 - | 0 - | 0 - | 2 - | 0,000218227 - | 0 - | 0 - |
| 1667 | 1 | 3um_FIJJAFBF_331481 | Bacteria | Proteobacteri | Alphaproteob | Rhodobactera | Rhodobactera | Planktomarina | Planktomarina | cellular proces | Cell wall/mem | 08ZG0 | Ompa motb d | 89187.ISM_10 | 0,6 | 0,489897949 | 3,54E-05 | 2,91E-05 | 1 - | 5,24E-05 - | 0 - | 0 - | 1 - | 6,19E-05 - | 1 - | 6,28E-05 - | 0 - | 0 - |
| 1668 | 1 | 0.2um_NOKCPHJN_184907 | Bacteria | Proteobacteri | Gammaprotec | unclassified | unclassified | unclassified | various | cellular proces | Posttranslatio | COG0330 | Band 7 protein | 247634.GPB2 | 1 | 1,095445115 | 4,59E-05 | 5,22E-05 | 1 - | 3,97E-05 - | 0 - | 0 - | 1 - | 4,69E-05 - | 3 - | 0,000142716 - | 0 - | 0 - |
| 1669 | 1 | 3um_FIJJAFBF_402492 | Bacteria | Bacteroidetes | Flavobacteriia | various | various | various | various | unclassified | unclassified | unclassified | unclassified | unclassified | 1 | 0 | 0,000102259 | 9,71E-06 | 1 - | 8,57E-05 - | 1 - | 0,000106357 - | 1 - | 0,000101106 - | 1 - | 0,000102599 - | 1 - | 0,000115578 - |
| 1670 | 2 | 0.2um_NOKCPHJN_577236;3um_FIJJAFBF_7817 | Bacteria | Bacteroidetes | Flavobacteriia | Flavobacterial | Flavobacteriac | Formosa | Formosa sp. H | unclassified | unclassified | unclassified | unclassified | unclassified | 0,2 | 0,4 | 8,11E-06 | 1,62E-05 | 0 - | 0 - | 0 - | 0 - | 1 - | 4,06E-05 - | 0 - | 0 - | 0 - | 0 - |
| 1671 | 1 | 0.2um_NOKCPHJN_264390 | Viruses | unclassified | unclassified | unclassified | unclassified | unclassified | uncultured M | poorly charact | Function unkn | 0R2P9 | NA | 768066.HELO_ | 1 | 0 | 2,07E-05 | 1,97E-06 | 1 - | 1,73E-05 - | 1 - | 2,15E-05 - | 1 - | 2,05E-05 - | 1 - | 2,08E-05 - | 1 - | 2,34E-05 - |
| 1672 | 1 | 0.2um_NOKCPHJN_652382 | Bacteria | Proteobacteri | Gammaprotec | Oceanospirilla | Saccharospirill | Reinekea | Reinekea fors | metabolism | Nucleotide tra | 0QHTA | Catalyzes the | 314283.MED2 | 2,4 | 0,8 | 0,000150914 | 5,36E-05 | 3 - | 0,00015795 - | 3 - | 0,000196127 - | 2 - | 0,000124296 - | 1 - | 6,31E-05 - | 3 - | 0,00021313 - |
| 1673 | 1 | 3um_FIJJAFBF_201374 | Bacteria | Bacteroidetes | Flavobacteriia | Flavobacterial | Flavobacteriac | Lacinutrix | Lacinutrix jang | metabolism | Carbohydrate | 09CFZ | Transporter | 391587.KAOT | 0,4 | 0,489897949 | 6,46E-06 | 7,96E-06 | 1 - | 1,47E-05 - | 0 - | 0 - | 0 - | 0 - | 1 - | 1,76E-05 - | 0 - | 0 - |
| 1674 | 1 | 3um_FIJJAFBF_665678 | Bacteria | Bacteroidetes | Flavobacteriia | Flavobacterial | Flavobacteriac | Polaribacter | Polaribacter s | information st | Translation, ri | COG0013 | Catalyzes the | 313598.MED1 | 0,4 | 0,489897949 | 6,01E-06 | 7,37E-06 | 0 - | 0 - | 1 - | 1,54E-05 - | 1 - | 1,46E-05 - | 0 - | 0 - | 0 - | 0 - |
| 1675 | 1 | 0.2um_NOKCPHJN_347404 | Bacteria | Proteobacteri | Alphaproteob | Pelagibacteral | Pelagibacterac | Candidatus Pe | Candidatus Pe | unclassified | unclassified | unclassified | unclassified | unclassified | 0,2 | 0,4 | 2,00E-06 | 4,01E-06 | 1 - | 1,00E-05 - | 0 - | 0 - | 0 - | 0 - | 0 - | 0 - | 0 - | 0 - |
| 1676 | 1 | 0.2um_NOKCPHJN_60418 | Bacteria | Proteobacteri | Gammaprotec | Oceanospirilla | Oceanospirilla | Amphritea | Amphritea atl | information st | Transcription | COG0085 | DNA-depende | 207954.MED9 | 13,6 | 1,2 | 0,000136881 | 1,64E-05 | 13 - | 0,000109713 - | 15 - | 0,00015719 - | 15 - | 0,000149429 - | 13 - | 0,000131418 - | 12 - | 0,000136654 - |
| 1677 | 1 | 3um_FIJJAFBF_607317 | Bacteria | Proteobacteri | Gammaprotec | unclassified | unclassified | unclassified | SAR86 cluster | information st | Replication, re | 05C68 | Can catalyze t | 207954.MED9 | 0,4 | 0,489897949 | 1,62E-05 | 1,98E-05 | 0 - | 0 - | 0 - | 0 - | 0 - | 0 - | 1 - | 3,80E-05 - | 1 - | 4,28E-05 - |
| 1678 | 1 | 3um_FIJJAFBF_708117 | Eukaryota | unclassified | Cryptophyta | Pyrenomonad | Geminigerace | Guillardia | Guillardia thet | unclassified | unclassified | unclassified | unclassified | unclassified | 0,2 | 0,4 | 1,39E-05 | 2,78E-05 | 1 - | 6,96E-05 - | 0 - | 0 - | 0 - | 0 - | 0 - | 0 - | 0 - | 0 - |
| 1679 | 1 | 3um_FIJJAFBF_603089 | Bacteria | Proteobacteri | Alphaproteob | Rhodospirillale | Rhodospirillac | unclassified | Rhodobactera | metabolism | Secondary me | 01Q86 | extracellular s | 388401.RB215 | 1,2 | 0,748331477 | 4,30E-05 | 2,54E-05 | 2 - | 6,20E-05 - | 0 - | 0 - | 1 - | 3,66E-05 - | 2 - | 7,43E-05 - | 1 - | 4,19E-05 - |
| 1680 | 1 | 0.2um_NOKCPHJN_785802 | Bacteria | Bacteroidetes | unclassified | unclassified | unclassified | unclassified | Bacteroidetes | unclassified | unclassified | unclassified | unclassified | unclassified | 2 | 0 | 0,000111404 | 1,06E-05 | 2 - | 9,33E-05 - | 2 - | 0,000115869 - | 2 - | 0,000110148 - | 2 - | 0,000111775 - | 2 - | 0,000125914 - |
| 1681 | 1 | 3um_FIJJAFBF_211368 | Bacteria | Proteobacteri | Gammaprotec | Arenicellales | Arenicellaceae | Arenicella | Arenicella xan | unclassified | unclassified | unclassified | unclassified | unclassified | 2,6 | 1,019803903 | 0,000390378 | 0,000139816 | 4 - | 0,000510119 - | 2 - | 0,000316709 - | 1 - | 0,000150536 - | 3 - | 0,000458278 - | 3 - | 0,000516248 - |
| 1682 | 1 | 3um_FIJJAFBF_448215 | Bacteria | Actinobacteria | various | various | various | various | various | information st | Translation, ri | COG0050 | This protein p | 58051.PE36_2 | 0,4 | 0,489897949 | 2,03E-05 | 2,51E-05 | 1 - | 4,67E-05 - | 0 - | 0 - | 1 - | 5,51E-05 - | 0 - | 0 - | 0 - | 0 - |
| 1683 | 1 | 0.2um_NOKCPHJN_1001140 | Bacteria | Proteobacteri | Gammaprotec | Pseudomonad | Pseudomonad | Pseudomonas | various | information st | Translation, ri | COG0087 | One of the pri | 629265.PMA4 | 0,6 | 0,8 | 3,66E-05 | 5,05E-05 | 1 - | 5,44E-05 - | 0 - | 0 - | 2 - | 0,000128419 - | 0 - | 0 - | 0 - | 0 - |
| 1684 | 1 | 3um_FIJJAFBF_549861 | Bacteria | Bacteroidetes | Flavobacteriia | Flavobacterial | Crocinitomica | unclassified | Crocinitomica | information st | Translation, ri | COG0186 | One of the pri | 575590.HMPR | 1,6 | 0,489897949 | 0,000227556 | 7,26E-05 | 2 - | 0,000239118 - | 1 - | 0,000148457 - | 1 - | 0,000141127 - | 2 - | 0,000286424 - | 2 - | 0,000322655 - |
| 1685 | 1 | 3um_FIJJAFBF_567038 | Bacteria | Proteobacteri | Alphaproteob | Pelagibacteral | Pelagibacterac | Candidatus Pe | Candidatus Pe | unclassified | unclassified | unclassified | unclassified | unclassified | 0,2 | 0,4 | 1,79E-05 | 3,57E-05 | 0 - | 0 - | 0 - | 0 - | 0 - | 0 - | 1 - | 8,93E-05 - | 0 - | 0 - |
| 1686 | 1 | 0.2um_NOKCPHJN_07998 | Bacteria | Bacteroidetes | Flavobacteriia | Flavobacterial | unclassified | unclassified | Flavobacterial | unclassified | unclassified | unclassified | unclassified | unclassified | 1,8 | 0,4 | 5,47E-05 | 1,08E-05 | 2 - | 5,17E-05 - | 2 - | 6,42E-05 - | 2 - | 6,10E-05 - | 2 - | 6,19E-05 - | 1 - | 3,49E-05 - |
| 1687 | 1 | 0.2um_NOKCPHJN_324789 | Eukaryota | unclassified | unclassified | Acytosteliales | Acytosteliacea | Heterostelium | Heterostelium | poorly charact | Function unkn | 0XXUP | (ABC) transpo | 88036.EFJ371 | 0,4 | 0,489897949 | 2,59E-06 | 3,18E-06 | 0 - | 0 - | 1 - | 6,65E-06 - | 1 - | 6,32E-06 - | 0 - | 0 - | 0 - | 0 - |
| 1688 | 1 | 0.2um_NOKCPHJN_728863 | Eukaryota | unclassified | Labyrinthulom | unclassified | Thraustochytr | Aurantiochytr | Aurantiochytr | poorly charact | Function unkn | 0A4HT | Coiled-coil dor | 7668.SPU_011 | 0,4 | 0,489897949 | 8,14E-06 | 1,00E-05 | 1 - | 1,85E-05 - | 0 - | 0 - | 0 - | 0 - | 1 - | 2,22E-05 - | 0 - | 0 - |
| 1689 | 1 | 3um_FIJJAFBF_122388 | Bacteria | Bacteroidetes | Flavobacteriia | Flavobacterial | Flavobacteriac | Polaribacter | Polaribacter r | cellular proces | Posttranslatio | COG0652 | peptidyl-proly | 313598.MED1 | 2,4 | 0,489897949 | 0,000105018 | 1,64E-05 | 3 - | 0,000111434 - | 2 - | 9,22E-05 - | 2 - | 8,77E-05 - | 3 - | 0,000133479 - | 2 - | 0,000100242 - |
| 1690 | 1 | 3um_FIJJAFBF_44709 | Bacteria | Bacteroidetes | Flavobacteriia | Flavobacterial | Flavobacteriac | Polaribacter | Polaribacter s | unclassified | unclassified | unclassified | unclassified | unclassified | 2,2 | 0,748331477 | 5,98E-05 | 1,92E-05 | 3 - | 6,91E-05 - | 3 - | 8,59E-05 - | 2 - | 5,44E-05 - | 1 - | 2,76E-05 - | 2 - | 6,22E-05 - |
| 1691 | 1 | 3um_FIJJAFBF_197145 | Viruses | unclassified | unclassified | unclassified | various | various | various | unclassified | unclassified | unclassified | unclassified | unclassified | 1 | 0 | 2,06E-05 | 1,96E-06 | 1 - | 1,73E-05 - | 1 - | 2,14E-05 - | 1 - | 2,04E-05 - | 1 - | 2,07E-05 - | 1 - | 2,33E-05 - |
| 1692 | 1 | 3um_FIJJAFBF_337116 | Bacteria | Bacteroidetes | Flavobacteriia | Flavobacterial | Flavobacteriac | Polaribacter | various | information st | Translation, ri | 05CEJ | Catalyzes the | 983548.Krodi_ | 26,8 | 2,993325909 | 0,000532461 | 6,53E-05 | 31 - | 0,000517916 - | 24 - | 0,000497882 - | 27 - | 0,000532463 - | 23 - | 0,000460279 - | 29 - | 0,000653764 - |
| 1693 | 1 | 3um_FIJJAFBF_355903 | Bacteria | Bacteroidetes | Flavobacteriia | Flavobacterial | Flavobacteriac | Polaribacter | Polaribacter ir | metabolism | Inorganic ion t | 05C27 | TonB-depende | 313594.PI23P | 0,6 | 0,8 | 7,03E-06 | 9,00E-06 | 2 - | 2,17E-05 - | 1 - | 1,35E-05 - | 0 - | 0 - | 0 - | 0 - | 0 - | 0 - |
| 1694 | 1 | 3um_FIJJAFBF_67793 | Eukaryota | unclassified | unclassified | Phaeocystales | Phaeocystace | Phaeocystis | Phaeocystis ar | metabolism | Energy produc | 11HAV | photosystem i | 35128.Thapsd | 1,4 | 0,489897949 | 0,000132964 | 3,91E-05 | 2 - | 0,000162804 - | 1 - | 0,000101077 - | 1 - | 9,61E-05 - | 2 - | 0,000195012 - | 1 - | 0,00010984 - |
| 1695 | 1 | 3um_FIJJAFBF_52337 | Bacteria | Proteobacteri | Gammaprotec | Arenicellales | Arenicellaceae | Arenicella | Arenicella xan | unclassified | unclassified | unclassified | unclassified | unclassified | 1 | 0 | 0,000133036 | 1,26E-05 | 1 - | 0,000111434 - | 1 - | 0,000138368 - | 1 - | 0,000131536 - | 1 - | 0,000133479 - | 1 - | 0,000150364 - |
| 1696 | 1 | 3um_FIJJAFBF_611049 | Bacteria | Bacteroidetes | Flavobacteriia | Flavobacterial | Flavobacteriac | Polaribacter | Polaribacter s | cellular proces | Posttranslatio | 05C2Z | ATP-depender | 313598.MED1 | 3,6 | 0,8 | 5,69E-05 | 1,12E-05 | 4 - | 5,37E-05 - | 4 - | 6,67E-05 - | 4 - | 6,34E-05 - | 4 - | 6,43E-05 - | 2 - | 3,62E-05 - |
| 1697 | 2 | 3um_FIJJAFBF_236131;3um_FIJJAFBF_288908 | Viruses | unclassified | unclassified | unclassified | unclassified | unclassified | uncultured M | unclassified | unclassified | unclassified | unclassified | unclassified | 0,8 | 0,4 | 6,20E-05 | 3,18E-05 | 1 - | 6,56E-05 - | 0 - | 0 - | 1 - | 7,74E-05 - | 1 - | 7,86E-05 - | 1 - | 8,85E-05 - |
| 1698 | 1 | 3um_FIJJAFBF_252812 | Bacteria | Proteobacteri | Alphaproteob | Rhodobactera | Rhodobactera | Neptunicoccu | Neptunicoccu | information st | Transcription | 01QD3 | Facilitates trar | 467661.RKLH1 | 1,4 | 0,489897949 | 4,53E-05 | 1,60E-05 | 1 - | 2,71E-05 - | 1 - | 3,37E-05 - | 2 - | 6,41E-05 - | 2 - | 6,50E-05 - | 1 - | 3,66E-05 - |
| 1699 | 1 | 3um_FIJJAFBF_606585 | Eukaryota | Basidiomycota | Microbotryon | Microbotryale | Microbotryac | Microbotryum | Microbotryum | information st | Chromatin stru | KOG3467 | Core compone | 3218.PP1S133 | 1 | 0 | 0,000103808 | 9,86E-06 | 1 - | 8,70E-05 - | 1 - | 0,000107969 - | 1 - | 0,000102638 - | 1 - | 0,000104154 - | 1 - | 0,000117329 - |
| 1700 | 1 | 3um_FIJJAFBF_715091 | Eukaryota | unclassified | Pelagophycea | Pelagomonad | unclassified | Aureococcus | Aureococcus a | unclassified | unclassified | unclassified | unclassified | unclassified | 1 | 0 | 0,000103028 | 9,78E-06 | 1 - | 8,63E-05 - | 1 - | 0,000107157 - | 1 - | 0,000101866 - | 1 - | 0,000103371 - | 1 - | 0,000116447 - |
| 1701 | 1 | 3um_FIJJAFBF_786162 | Bacteria | Proteobacteri | Gammaprotec | Oceanospirilla | Saccharospirill | Reinekea | Reinekea fors | information st | Translation, ri | COG0087 | One of the pri | 314283.MED2 | 10 | 0,632455532 | 0,00065299 | 7,92E-05 | 10 - | 0,000546556 - | 11 - | 0,000746527 - | 10 - | 0,000645153 - | 9 - | 0,000589214 - | 10 - | 0,000737498 - |
| 1702 | 1 | 3um_FIJJAFBF_208701 | Bacteria | Bacteroidetes | Flavobacteriia | Flavobacterial | Flavobacteriac | Ulvibacter | Ulvibacter lito | unclassified | unclassified | unclassified | unclassified | unclassified | 1,2 | 0,4 | 0,000118506 | 2,62E-05 | 2 - | 0,00017004 - | 1 - | 0,00010557 - | 1 - | 0,000100357 - | 1 - | 0,000101839 - | 1 - | 0,000114722 - |
| 1703 | 1 | 0.2um_NOKCPHJN_163369 | Bacteria | Proteobacteri | Gammaprotec | Cellvibrionales | Porticoccacea | unclassified | various | unclassified | unclassified | unclassified | unclassified | unclassified | 1,4 | 0,489897949 | 0,000110127 | 3,89E-05 | 1 - | 6,60E-05 - | 1 - | 8,19E-05 - | 2 - | 0,000155727 - | 2 - | 0,000158027 - | 1 - | 8,90E-05 - |
| 1704 | 1 | 3um_FIJJAFBF_382457 | Eukaryota | unclassified | Ichthyosporea | unclassified | unclassified | Capsaspora | Capsaspora ov | cellular proces | Cell wall/mem | 05D20 | Nad-depender | 502025.Hoch_ | 0,2 | 0,4 | 4,75E-06 | 9,49E-06 | 0 - | 0 - | 0 - | 0 - | 1 - | 2,37E-05 - | 0 - | 0 - | 0 - | 0 - |
| 1705 | 1 | 0.2um_NOKCPHJN_14987 | Bacteria | Bacteroidetes | Flavobacteriia | Flavobacterial | Flavobacteriac | Kordia | Kordia sp. SM | cellular proces | Intracellular tr | COG0653 | Part of the Se | 391587.KAOT | 1 | 0 | 1,22E-05 | 1,16E-06 | 1 - | 1,02E-05 - | 1 - | 1,27E-05 - | 1 - | 1,21E-05 - | 1 - | 1,23E-05 - | 1 - | 1,38E-05 - |
| 1706 | 1 | 0.2um_NOKCPHJN_782226 | Bacteria | Proteobacteri | Gammaprotec | Alteromonada | Pseudoalteror | Pseudoalteror | Pseudoalteror | metabolism | Inorganic ion t | 16PDP | TonB-depende | 637905.SVI_1 | 2,8 | 0,748331477 | 8,57E-05 | 2,41E-05 | 3 - | 7,70E-05 - | 2 - | 6,38E-05 - | 2 - | 6,06E-05 - | 4 - | 0,000123028 - | 3 - | 0,000103943 - |
| 1707 | 1 | 0.2um_NOKCPHJN_328479 | Bacteria | Bacteroidetes | unclassified | unclassified | unclassified | unclassified | Bacteroidetes | unclassified | unclassified | unclassified | unclassified | unclassified | 0,4 | 0,489897949 | 1,53E-06 | 1,87E-06 | 0 - | 0 - | 1 - | 3,92E-06 - | 1 - | 3,72E-06 - | 0 - | 0 - | 0 - | 0 - |
| 1708 | 1 | 0.2um_NOKCPHJN_1052509 | Bacteria | Proteobacteri | Alphaproteob | Rhodobactera | Rhodobactera | Planktotalea | various | unclassified | unclassified | unclassified | unclassified | unclassified | 0,6 | 0,489897949 | 6,44E-05 | 5,31E-05 | 1 - | 9,41E-05 - | 1 - | 0,000116819 - | 1 - | 0,000111051 - | 0 - | 0 - | 0 - | 0 - |
| 1709 | 1 | 3um_FIJJAFBF_748549 | Bacteria | Proteobacteri | Gammaprotec | unclassified | unclassified | unclassified | OM182 bacte | information st | Translation, ri | COG0050 | This protein p | 393595.ABO_ | 3,2 | 0,748331477 | 0,000176131 | 3,89E-05 | 4 - | 0,000185873 - | 4 - | 0,0002308 - | 3 - | 0,000164553 - | 2 - | 0,000111323 - | 3 - | 0,000188107 - |
| 1710 | 1 | 0.2um_NOKCPHJN_910564 | Bacteria | Proteobacteri | Alphaproteob | Pelagibacteral | Pelagibacterac | Candidatus Pe | Candidatus Pe | metabolism | Inorganic ion t | 16PII | TRAP transpor | 744979.R2A13 | 2,2 | 0,4 | 0,00010495 | 9,62E-06 | 3 - | 0,000121671 - | 2 - | 0,00010072 - | 2 - | 9,57E-05 - | 2 - | 9,72E-05 - | 2 - | 0,000109452 - |

| | | | | | | | | | | | | | | | | | | | | | | | | | | | | |
|---|---|---|---|---|---|---|---|---|---|---|---|---|---|---|---|---|---|---|---|---|---|---|---|---|---|---|---|---|
| 1711 | 1 | 0.2um_NOKCPHJN_889559 | Bacteria | Proteobacteri | Alphaproteob | Pelagibacteral | Pelagibacterac | Candidatus Pe | Candidatus Pe | information st | Transcription | 01R4Z | DNA-depende | 439493.PB721 | 1 | 0 | 2,89E-05 | 2,74E-06 | 1 - | 2,42E-05 - | 1 - | 3,01E-05 - | 1 - | 2,86E-05 - | 1 - | 2,90E-05 - | 1 - | 3,27E-05 - |
| 1712 | 1 | 3um_FIJJAFBF_35364 | Bacteria | Bacteroidetes | Flavobacteriia | Flavobacterial | Flavobacteriac | Polaribacter | Polaribacter s | metabolism | Nucleotide tra | COG0138 | bifunctional p | 313598.MED1 | 6,8 | 0,748331477 | 0,000182589 | 2,03E-05 | 8 - | 0,000180751 - | 7 - | 0,000196384 - | 6 - | 0,000160018 - | 6 - | 0,000162382 - | 7 - | 0,00021341 - |
| 1713 | 1 | 0.2um_NOKCPHJN_991107 | Bacteria | Proteobacteri | Alphaproteob | Pelagibacteral | unclassified | unclassified | Pelagibacteral | cellular proces | Cell wall/mem | 7257 | NA | 335992.SAR11 | 2,6 | 0,8 | 0,000104854 | 3,42E-05 | 2 - | 6,73E-05 - | 3 - | 0,000125383 - | 2 - | 7,95E-05 - | 4 - | 0,000161271 - | 2 - | 9,08E-05 - |
| 1714 | 1 | 3um_FIJJAFBF_235863 | Bacteria | Proteobacteri | Gammaprote | Alteromonada | unclassified | unclassified | Alteromonada | unclassified | unclassified | unclassified | unclassified | unclassified | 0,6 | 0,8 | 0,000111329 | 0,000147343 | 0 - | 0 - | 1 - | 0,000190025 - | 0 - | 0 - | 2 - | 0,000366622 - | 0 - | 0 - |
| 1715 | 1 | 0.2um_NOKCPHJN_137195 | Bacteria | Bacteroidetes | unclassified | unclassified | unclassified | unclassified | Bacteroidetes | cellular proces | Posttranslatio | COG0652 | peptidyl-proly | 926562.Oweh | 0,8 | 0,4 | 2,87E-05 | 1,46E-05 | 1 - | 3,11E-05 - | 1 - | 3,86E-05 - | 1 - | 3,67E-05 - | 1 - | 3,73E-05 - | 0 - | 0 - |
| 1716 | 1 | 3um_FIJJAFBF_510915 | Bacteria | Actinobacteri | Actinobacteri | Micrococcales | Microbacteria | unclassified | Microbacteria | unclassified | unclassified | unclassified | unclassified | unclassified | 0,8 | 0,4 | 0,000140937 | 7,24E-05 | 1 - | 0,00014715 - | 1 - | 0,000182716 - | 0 - | 0 - | 1 - | 0,000176261 - | 1 - | 0,000198557 - |
| 1717 | 1 | 0.2um_NOKCPHJN_16172 | Bacteria | Proteobacteri | Gammaprote | Oceanospirilla | Oceanospirilla | Balneatrix | Balneatrix alp | unclassified | unclassified | unclassified | unclassified | unclassified | 2,2 | 0,748331477 | 6,79E-05 | 2,65E-05 | 2 - | 5,12E-05 - | 1 - | 3,18E-05 - | 2 - | 6,05E-05 - | 3 - | 9,21E-05 - | 3 - | 0,000103711 - |
| 1718 | 1 | 3um_FIJJAFBF_652999 | Eukaryota | unclassified | unclassified | Isochrysidales | Noelaerhabda | Emiliania | Emiliania hux | unclassified | unclassified | unclassified | unclassified | unclassified | 0,8 | 0,4 | 6,17E-05 | 3,13E-05 | 1 - | 6,67E-05 - | 1 - | 8,29E-05 - | 1 - | 7,88E-05 - | 1 - | 7,99E-05 - | 0 - | 0 - |
| 1719 | 1 | 3um_FIJJAFBF_336935 | Bacteria | Bacteroidetes | Flavobacteriia | Flavobacterial | Flavobacteriac | Polaribacter | Polaribacter g | metabolism | Inorganic ion t | 09C49 | tonB-depende | 313598.MED1 | 0,6 | 0,489897949 | 1,34E-05 | 1,10E-05 | 0 - | 0 - | 1 - | 2,20E-05 - | 0 - | 0 - | 1 - | 2,12E-05 - | 1 - | 2,39E-05 - |
| 1720 | 1 | 0.2um_NOKCPHJN_1080170 | Bacteria | Proteobacteri | Alphaproteob | Rhodobactera | Rhodobactera | unclassified | Rhodobactera | cellular proces | Cell wall/mem | 01XF2 | NA | 391626.OA30 | 1,2 | 0,4 | 4,09E-05 | 1,51E-05 | 1 - | 2,83E-05 - | 2 - | 7,04E-05 - | 1 - | 3,35E-05 - | 1 - | 3,39E-05 - | 1 - | 3,82E-05 - |
| 1721 | 1 | 0.2um_NOKCPHJN_417758 | Viruses | unclassified | unclassified | unclassified | unclassified | unclassified | Pleurochrysis | unclassified | unclassified | unclassified | unclassified | unclassified | 1 | 0 | 2,79E-05 | 2,65E-06 | 1 - | 2,34E-05 - | 1 - | 2,90E-05 - | 1 - | 2,76E-05 - | 1 - | 2,80E-05 - | 1 - | 3,15E-05 - |
| 1722 | 1 | 0.2um_NOKCPHJN_236302 | Viruses | unclassified | unclassified | unclassified | unclassified | unclassified | uncultured M | poorly charact | Function unkn | 0NVRZ | NA | 170187.SP_17 | 0,8 | 0,748331477 | 8,89E-06 | 8,63E-06 | 0 - | 0 - | 1 - | 1,07E-05 - | 0 - | 0 - | 1 - | 1,04E-05 - | 2 - | 2,33E-05 - |
| 1723 | 2 | 3um_FIJJAFBF_482985;3um_FIJJAFBF_776552 | Bacteria | Bacteroidetes | Flavobacteriia | Flavobacterial | Crocinitomica | unclassified | Crocinitomica | unclassified | unclassified | unclassified | unclassified | unclassified | 0,4 | 0,489897949 | 1,01E-05 | 1,24E-05 | 0 - | 0 - | 0 - | 0 - | 1 - | 2,50E-05 - | 1 - | 2,54E-05 - | 0 - | 0 - |
| 1724 | 1 | 3um_FIJJAFBF_119808 | unclassified | unclassified | unclassified | unclassified | unclassified | unclassified | unclassified | unclassified | unclassified | unclassified | unclassified | unclassified | 4,6 | 0,8 | 0,000530868 | 4,74E-05 | 6 - | 0,000588599 - | 4 - | 0,000487244 - | 5 - | 0,000578984 - | 4 - | 0,000470028 - | 4 - | 0,000529485 - |
| 1725 | 1 | 0.2um_NOKCPHJN_751207 | unclassified | unclassified | unclassified | unclassified | unclassified | unclassified | unclassified | unclassified | unclassified | unclassified | unclassified | unclassified | 0,6 | 0,489897949 | 2,19E-05 | 1,79E-05 | 0 - | 0 - | 1 - | 3,76E-05 - | 1 - | 3,57E-05 - | 1 - | 3,63E-05 - | 0 - | 0 - |
| 1726 | 1 | 3um_FIJJAFBF_08028 | Bacteria | Proteobacteri | Gammaprote | Cellvibrionale | Porticoccacea | unclassified | SAR92 bacteri | cellular proces | Cell wall/mem | 127CM | Membrane | 314287.GB22 | 2,4 | 0,489897949 | 0,000183906 | 3,25E-05 | 3 - | 0,000194537 - | 3 - | 0,000241557 - | 2 - | 0,000153087 - | 2 - | 0,000155348 - | 2 - | 0,000174999 - |
| 1727 | 1 | 3um_FIJJAFBF_552505 | Bacteria | Proteobacteri | Gammaprote | Arenicellales | Arenicellaceae | Arenicella | Arenicella xan | information st | Transcription | 05D27 | DNA-depende | 472759.Nhal_ | 0,4 | 0,489897949 | 3,64E-06 | 4,50E-06 | 1 - | 8,12E-06 - | 1 - | 1,01E-05 - | 0 - | 0 - | 0 - | 0 - | 0 - | 0 - |
| 1728 | 1 | 3um_FIJJAFBF_211629 | Bacteria | Proteobacteri | Gammaprote | Oceanospirilla | Saccharospirill | Reinekea | Reinekea fors | unclassified | unclassified | unclassified | unclassified | unclassified | 2,2 | 0,979795897 | 3,25E-05 | 1,14E-05 | 4 - | 5,11E-05 - | 2 - | 3,17E-05 - | 2 - | 3,01E-05 - | 1 - | 1,53E-05 - | 2 - | 3,45E-05 - |
| 1729 | 1 | 3um_FIJJAFBF_156143 | Bacteria | Actinobacteri | Actinobacteri | various | various | various | various | unclassified | unclassified | unclassified | unclassified | unclassified | 0,6 | 0,489897949 | 8,94E-05 | 7,30E-05 | 0 - | 0 - | 1 - | 0,000153246 - | 1 - | 0,00014568 - | 1 - | 0,000147832 - | 0 - | 0 - |
| 1730 | 1 | 3um_FIJJAFBF_152480 | Bacteria | Bacteroidetes | Flavobacteriia | Flavobacterial | Flavobacteriac | Polaribacter | Polaribacter fi | unclassified | unclassified | unclassified | unclassified | unclassified | 5,6 | 1,356465997 | 0,000669445 | 0,000126257 | 8 - | 0,000812579 - | 4 - | 0,000504492 - | 5 - | 0,000599479 - | 5 - | 0,000608333 - | 6 - | 0,000822342 - |
| 1731 | 1 | 3um_FIJJAFBF_41553 | Eukaryota | Bacillariophyt | Coscinodiscop | Chaetocerotal | Chaetocerotac | Chaetoceros | Chaetoceros s | unclassified | unclassified | unclassified | unclassified | unclassified | 2 | 0 | 0,000181493 | 1,72E-05 | 2 - | 0,000152022 - | 2 - | 0,000188767 - | 2 - | 0,000179447 - | 2 - | 0,000182097 - | 2 - | 0,000205132 - |
| 1732 | 1 | 0.2um_NOKCPHJN_219025 | Bacteria | Bacteroidetes | Cytophagia | Cytophagales | Flammeovirga | unclassified | Flammeovirga | cellular proces | Posttranslatio | 16QU3 | Hemolysin-typ | 744980.TRICH | 2,6 | 0,8 | 2,34E-05 | 7,94E-06 | 2 - | 1,50E-05 - | 4 - | 3,71E-05 - | 2 - | 1,77E-05 - | 3 - | 2,69E-05 - | 2 - | 2,02E-05 - |
| 1733 | 1 | 0.2um_NOKCPHJN_99556 | Bacteria | Proteobacteri | Gammaprote | Oceanospirilla | Saccharospirill | Reinekea | Reinekea fors | unclassified | unclassified | unclassified | unclassified | unclassified | 0,4 | 0,489897949 | 9,08E-06 | 1,11E-05 | 0 - | 0 - | 1 - | 2,31E-05 - | 0 - | 0 - | 1 - | 2,23E-05 - | 0 - | 0 - |
| 1734 | 1 | 0.2um_NOKCPHJN_805964 | Bacteria | Proteobacteri | Alphaproteob | Rhodobactera | Rhodobactera | unclassified | Rhodobactera | poorly charact | Function unkn | 17P8K | NA | 856793.MICA_ | 3,2 | 0,4 | 0,000158187 | 2,34E-05 | 3 - | 0,000124307 - | 3 - | 0,000154353 - | 4 - | 0,000195642 - | 3 - | 0,000148899 - | 3 - | 0,000167734 - |
| 1735 | 1 | 3um_FIJJAFBF_494406 | Bacteria | Proteobacteri | Alphaproteob | Rhizobiales | unclassified | unclassified | Rhizobiales ba | unclassified | unclassified | unclassified | unclassified | unclassified | 0,6 | 0,489897949 | 3,14E-05 | 2,61E-05 | 1 - | 4,45E-05 - | 0 - | 0 - | 1 - | 5,25E-05 - | 0 - | 0 - | 1 - | 6,00E-05 - |
| 1736 | 1 | 3um_FIJJAFBF_258951 | Bacteria | Bacteroidetes | Flavobacteriia | Flavobacterial | Crocinitomica | unclassified | Crocinitomica | unclassified | unclassified | unclassified | unclassified | unclassified | 0,8 | 0,4 | 7,40E-05 | 3,80E-05 | 1 - | 7,76E-05 - | 1 - | 9,63E-05 - | 1 - | 9,15E-05 - | 0 - | 0 - | 1 - | 0,000104645 - |
| 1737 | 1 | 0.2um_NOKCPHJN_81665 | Bacteria | Proteobacteri | Gammaprote | Cellvibrionale | unclassified | unclassified | Cellvibrionale | information st | Translation, ri | 16QAP | Associates wit | 314287.GB22 | 1,8 | 0,4 | 8,47E-05 | 1,67E-05 | 2 - | 8,00E-05 - | 2 - | 9,93E-05 - | 2 - | 9,44E-05 - | 2 - | 9,58E-05 - | 1 - | 5,40E-05 - |
| 1738 | 1 | 3um_FIJJAFBF_176637 | Bacteria | Bacteroidetes | unclassified | unclassified | unclassified | unclassified | Bacteroidetes | information st | Translation, ri | 0PAE2 | One of the pri | 926562.Oweh | 0,2 | 0,4 | 3,15E-05 | 6,30E-05 | 0 - | 0 - | 0 - | 0 - | 1 - | 0,000157537 - | 0 - | 0 - | 0 - | 0 - |
| 1739 | 1 | 3um_FIJJAFBF_737827 | unclassified | unclassified | unclassified | unclassified | unclassified | unclassified | unclassified | unclassified | unclassified | unclassified | unclassified | unclassified | 0,2 | 0,4 | 1,30E-05 | 2,60E-05 | 0 - | 0 - | 1 - | 6,51E-05 - | 0 - | 0 - | 0 - | 0 - | 0 - | 0 - |
| 1740 | 1 | 3um_FIJJAFBF_163754 | Bacteria | Bacteroidetes | Flavobacteriia | Flavobacterial | Flavobacteriac | Polaribacter | various | unclassified | unclassified | unclassified | unclassified | unclassified | 4,6 | 1,019803903 | 0,000420404 | 8,87E-05 | 6 - | 0,000462188 - | 5 - | 0,000478251 - | 3 - | 0,000272783 - | 4 - | 0,000369083 - | 5 - | 0,000519713 - |
| 1741 | 1 | 0.2um_NOKCPHJN_339141 | Bacteria | Verrucomicrol | unclassified | unclassified | unclassified | unclassified | Verrucomicrol | poorly charact | Function unkn | 08NRM | NA | 530564.Psta_ | 0,2 | 0,4 | 4,12E-06 | 8,24E-06 | 0 - | 0 - | 0 - | 0 - | 1 - | 2,06E-05 - | 0 - | 0 - | 0 - | 0 - |
| 1742 | 1 | 3um_FIJJAFBF_464083 | Bacteria | Proteobacteri | Gammaprote | unclassified | unclassified | unclassified | Gammaprote | cellular proces | Cell wall/mem | 0QTZI | Integral meml | 498211.CJA_0 | 0,6 | 0,8 | 4,45E-05 | 5,73E-05 | 2 - | 0,000139123 - | 0 - | 0 - | 0 - | 0 - | 1 - | 8,33E-05 - | 0 - | 0 - |
| 1743 | 1 | 0.2um_NOKCPHJN_994281 | Eukaryota | Chlorophyta | unclassified | Pyramimonad | unclassified | Pyramimonas | Pyramimonas | unclassified | unclassified | unclassified | unclassified | unclassified | 0,8 | 0,748331477 | 3,39E-05 | 3,29E-05 | 1 - | 3,70E-05 - | 0 - | 0 - | 1 - | 4,37E-05 - | 2 - | 8,87E-05 - | 0 - | 0 - |
| 1744 | 1 | 3um_FIJJAFBF_257132 | Eukaryota | unclassified | Florideophyce | Plocamiales | Plocamiaceae | Plocamium | Plocamium ca | unclassified | unclassified | unclassified | unclassified | unclassified | 2 | 0 | 0,000206056 | 1,96E-05 | 2 - | 0,000172597 - | 2 - | 0,000214314 - | 2 - | 0,000203733 - | 2 - | 0,000206742 - | 2 - | 0,000232894 - |
| 1745 | 1 | 3um_FIJJAFBF_380531 | unclassified | unclassified | unclassified | unclassified | unclassified | unclassified | unclassified | unclassified | unclassified | unclassified | unclassified | unclassified | 1 | 0 | 6,99E-05 | 6,64E-06 | 1 - | 5,86E-05 - | 1 - | 7,27E-05 - | 1 - | 6,91E-05 - | 1 - | 7,01E-05 - | 1 - | 7,90E-05 - |
| 1746 | 1 | 3um_FIJJAFBF_436955 | Bacteria | Proteobacteri | Gammaprote | Cellvibrionale | Porticoccacea | Porticoccus | various | metabolism | Energy produc | COG0055 | Produces ATP | 314287.GB22 | 5,6 | 0,8 | 0,000165155 | 2,95E-05 | 6 - | 0,000148099 - | 6 - | 0,000183895 - | 4 - | 0,000116544 - | 6 - | 0,000177398 - | 6 - | 0,000199838 - |
| 1747 | 1 | 3um_FIJJAFBF_788036 | Bacteria | Proteobacteri | Alphaproteob | Rhodobactera | Rhodobactera | Pseudophaeol | Pseudophaeol | unclassified | unclassified | unclassified | unclassified | unclassified | 1,8 | 0,748331477 | 0,000195851 | 6,59E-05 | 3 - | 0,000282238 - | 2 - | 0,000233637 - | 1 - | 0,000111051 - | 2 - | 0,000225382 - | 1 - | 0,000126946 - |
| 1748 | 1 | 0.2um_NOKCPHJN_936620 | Eukaryota | unclassified | Oomycetes | Peronosporale | Peronosporac | Phytophthora | Phytophthora | cellular proces | Signal transdu | 152IZ | EFh | 67593.Physo1 | 1 | 0 | 8,21E-05 | 7,79E-06 | 1 - | 6,87E-05 - | 1 - | 8,53E-05 - | 1 - | 8,11E-05 - | 1 - | 8,23E-05 - | 1 - | 9,27E-05 - |
| 1749 | 1 | 0.2um_NOKCPHJN_1001222 | unclassified | unclassified | unclassified | unclassified | unclassified | unclassified | unclassified | unclassified | unclassified | unclassified | unclassified | unclassified | 0,2 | 0,4 | 8,59E-05 | 0,000171854 | 0 - | 0 - | 0 - | 0 - | 0 - | 0 - | 1 - | 0,000429635 - | 0 - | 0 - |
| 1750 | 1 | 3um_FIJJAFBF_472313 | Bacteria | Proteobacteri | Alphaproteob | Rhodobactera | Rhodobactera | Planktomarina | Planktomarina | unclassified | unclassified | unclassified | unclassified | unclassified | 1,2 | 0,748331477 | 6,03E-05 | 3,60E-05 | 2 - | 8,86E-05 - | 1 - | 5,50E-05 - | 2 - | 0,000104619 - | 1 - | 5,31E-05 - | 0 - | 0 - |
| 1751 | 1 | 0.2um_NOKCPHJN_60421 | Bacteria | Proteobacteri | Gammaprote | Oceanospirilla | Oceanospirilla | Nitrincola | Nitrincola sp. | information st | Translation, ri | 05C64 | Binds directly | 314277.MED1 | 2,4 | 0,489897949 | 0,000141985 | 3,17E-05 | 3 - | 0,000149061 - | 2 - | 0,000123393 - | 2 - | 0,000117301 - | 2 - | 0,000119033 - | 3 - | 0,000201136 - |
| 1752 | 1 | 3um_FIJJAFBF_406905 | Bacteria | Bacteroidetes | Cytophagia | Cytophagales | Cytophagacea | Ekhidna | Ekhidna lutea | metabolism | Energy produc | 0FVN8 | Dihydrolipoyl | 313606.M231 | 1 | 0 | 2,95E-05 | 2,80E-06 | 1 - | 2,47E-05 - | 1 - | 3,07E-05 - | 1 - | 2,92E-05 - | 1 - | 2,96E-05 - | 1 - | 3,34E-05 - |
| 1753 | 1 | 0.2um_NOKCPHJN_505891 | Bacteria | Proteobacteri | Gammaprote | Cellvibrionale | Spongiibacter | unclassified | marine gamm | cellular proces | Cell wall/mem | 0QKXZ | Porin Gram-ne | 247633.GP21 | 1,8 | 0,748331477 | 7,20E-05 | 2,34E-05 | 3 - | 0,000104343 - | 1 - | 4,32E-05 - | 2 - | 8,21E-05 - | 2 - | 8,33E-05 - | 1 - | 4,69E-05 - |
| 1754 | 1 | 3um_FIJJAFBF_717710 | Eukaryota | Eustigmatoph | unclassified | Eustigmatales | Monodopsida | Nannochlorop | Nannochlorop | unclassified | unclassified | unclassified | unclassified | unclassified | 2,4 | 0,489897949 | 0,00019918 | 4,30E-05 | 2 - | 0,000139123 - | 2 - | 0,00017275 - | 3 - | 0,000246331 - | 3 - | 0,00024997 - | 2 - | 0,000187727 - |
| 1755 | 1 | 0.2um_NOKCPHJN_144817 | Bacteria | Bacteroidetes | Flavobacteriia | Flavobacterial | Flavobacteriac | Polaribacter | Polaribacter g | poorly charact | Function unkn | 06HJQ | Hyalin repeat | 696747.NIES3 | 4,4 | 0,8 | 0,000124272 | 1,14E-05 | 6 - | 0,000144071 - | 4 - | 0,000119263 - | 4 - | 0,000113374 - | 4 - | 0,000115049 - | 4 - | 0,000129602 - |
| 1756 | 1 | 3um_FIJJAFBF_103123 | Bacteria | Bacteroidetes | Flavobacteriia | Flavobacterial | Flavobacteriac | Polaribacter | Polaribacter s | metabolism | Carbohydrate | 099QD | pfkb domain p | 313598.MED1 | 1 | 0 | 4,45E-05 | 4,22E-06 | 1 - | 3,73E-05 - | 1 - | 4,63E-05 - | 1 - | 4,40E-05 - | 1 - | 4,46E-05 - | 1 - | 5,03E-05 - |
| 1757 | 1 | 3um_FIJJAFBF_590240 | Bacteria | Proteobacteri | Gammaprote | Oceanospirilla | Saccharospirill | Reinekea | Reinekea fors | metabolism | Inorganic ion t | COG3638 | (ABC) transpo | 1045856.EcW | 11,6 | 1,356465997 | 0,000528013 | 9,81E-05 | 10 - | 0,000378801 - | 10 - | 0,000470359 - | 12 - | 0,000536563 - | 13 - | 0,000589862 - | 13 - | 0,000664478 - |
| 1758 | 1 | 3um_FIJJAFBF_245736 | Bacteria | Bacteroidetes | Flavobacteriia | Flavobacterial | Flavobacteriac | unclassified | Flavobacteriac | information st | Translation, ri | 05CRK | lysyL-tRNA syr | 400668.Mmw | 0,2 | 0,4 | 5,52E-06 | 1,10E-05 | 0 - | 0 - | 0 - | 0 - | 0 - | 0 - | 1 - | 2,76E-05 - | 0 - | 0 - |
| 1759 | 1 | 0.2um_NOKCPHJN_629933 | Bacteria | Proteobacteri | Alphaproteob | Pelagibacteral | Pelagibacterac | Candidatus Pe | Candidatus Pe | information st | Translation, ri | 16QAF | thus facilitatin | 335992.SAR11 | 6 | 0,894427191 | 0,000142447 | 2,48E-05 | 7 - | 0,000139486 - | 5 - | 0,000123714 - | 5 - | 0,000117606 - | 6 - | 0,000143212 - | 7 - | 0,000188216 - |
| 1760 | 1 | 0.2um_NOKCPHJN_341236 | Eukaryota | unclassified | Pelagophycea | Pelagomonad | unclassified | Aureococcus | Aureococcus a | unclassified | unclassified | unclassified | unclassified | unclassified | 4,2 | 0,4 | 8,76E-05 | 1,29E-05 | 4 - | 6,98E-05 - | 5 - | 0,000108297 - | 4 - | 8,24E-05 - | 4 - | 8,36E-05 - | 4 - | 9,41E-05 - |
| 1761 | 1 | 3um_FIJJAFBF_351593 | Bacteria | Proteobacteri | Gammaprote | Oceanospirilla | Oceanospirilla | Neptunomona | Neptunomona | poorly charact | Function unkn | COG3181 | extra-cytoplas | 675812.VHA_ | 6,8 | 0,748331477 | 0,000284297 | 2,95E-05 | 7 - | 0,000246453 - | 6 - | 0,000262305 - | 8 - | 0,000332472 - | 7 - | 0,00029521 - | 6 - | 0,000285045 - |
| 1762 | 1 | 0.2um_NOKCPHJN_43648 | Bacteria | Proteobacteri | Gammaprote | Oceanospirilla | Saccharospirill | Reinekea | Reinekea fors | poorly charact | Function unkn | 17G7E | Protein of unk | 314283.MED2 | 0,2 | 0,4 | 2,15E-05 | 4,30E-05 | 0 - | 0 - | 0 - | 0 - | 1 - | 0,000107526 - | 0 - | 0 - | 0 - | 0 - |
| 1763 | 1 | 3um_FIJJAFBF_724142 | Eukaryota | Chordata | Actinopteri | Osteoglossifor | Osteoglossida | Scleropages | Scleropages fo | unclassified | unclassified | unclassified | unclassified | unclassified | 0,8 | 0,748331477 | 1,02E-05 | 9,28E-06 | 0 - | 0 - | 1 - | 1,27E-05 - | 0 - | 0 - | 2 - | 2,45E-05 - | 1 - | 1,38E-05 - |
| 1764 | 1 | 0.2um_NOKCPHJN_127444 | Bacteria | Proteobacteri | Alphaproteob | Rhizobiales | Aurantimonad | Martelella | Martelella me | unclassified | unclassified | unclassified | unclassified | unclassified | 0,8 | 0,4 | 5,11E-06 | 2,62E-06 | 1 - | 5,40E-06 - | 0 - | 0 - | 1 - | 6,38E-06 - | 1 - | 6,47E-06 - | 1 - | 7,29E-06 - |
| 1765 | 1 | 3um_FIJJAFBF_674253 | Bacteria | Bacteroidetes | Flavobacteriia | Flavobacterial | Flavobacteriac | Tamlana | various | metabolism | Amino acid tra | 0PF77 | Dehydrogenas | 391603.FBALC | 3,4 | 1,2 | 0,000144046 | 4,90E-05 | 4 - | 0,000145287 - | 4 - | 0,000180404 - | 4 - | 0,000171496 - | 4 - | 0,00017403 - | 1 - | 4,90E-05 - |
| 1766 | 1 | 3um_FIJJAFBF_802260 | unclassified | unclassified | unclassified | unclassified | unclassified | unclassified | unclassified | unclassified | unclassified | unclassified | unclassified | unclassified | 0,4 | 0,489897949 | 2,75E-05 | 3,39E-05 | 1 - | 6,31E-05 - | 0 - | 0 - | 1 - | 7,44E-05 - | 0 - | 0 - | 0 - | 0 - |
| 1767 | 1 | 0.2um_NOKCPHJN_145944 | Eukaryota | Phaeophyceae | unclassified | Ectocarpales | Ectocarpaceae | Ectocarpus | Ectocarpus sili | unclassified | unclassified | unclassified | unclassified | unclassified | 1 | 0 | 2,06E-05 | 1,96E-06 | 1 - | 1,73E-05 - | 1 - | 2,15E-05 - | 1 - | 2,04E-05 - | 1 - | 2,07E-05 - | 1 - | 2,33E-05 - |
| 1768 | 1 | 3um_FIJJAFBF_49652 | Bacteria | Proteobacteri | Gammaprote | Cellvibrionale | Porticoccacea | unclassified | Porticoccacea | unclassified | unclassified | unclassified | unclassified | unclassified | 2,4 | 0,8 | 9,79E-05 | 3,72E-05 | 1 - | 3,33E-05 - | 3 - | 0,000123929 - | 3 - | 0,000117811 - | 2 - | 7,97E-05 - | 3 - | 0,000134673 - |
| 1769 | 1 | 3um_FIJJAFBF_245820 | Bacteria | Bacteroidetes | Flavobacteriia | Flavobacterial | Flavobacteriac | unclassified | Flavobacteriac | cellular proces | Cell wall/mem | 09B44 | efflux transpo | 867900.Celly_ | 0,6 | 0,489897949 | 2,08E-05 | 1,73E-05 | 1 - | 2,95E-05 - | 0 - | 0 - | 1 - | 3,48E-05 - | 0 - | 0 - | 1 - | 3,98E-05 - |
| 1770 | 1 | 0.2um_NOKCPHJN_506779 | Bacteria | Proteobacteri | Alphaproteob | Rhodobactera | Rhodobactera | Planktomarina | Planktomarina | metabolism | Lipid transpor | 071I4 | Phosphatidyls | 398580.Dshi_ | 0,6 | 0,489897949 | 3,42E-05 | 2,81E-05 | 1 - | 5,06E-05 - | 0 - | 0 - | 1 - | 5,97E-05 - | 1 - | 6,06E-05 - | 0 - | 0 - |
| 1771 | 1 | 3um_FIJJAFBF_15517 | Bacteria | Bacteroidetes | Flavobacteriia | Flavobacterial | Flavobacteriac | Algibacter | Algibacter pec | cellular proces | Signal transdu | 05WH1 | Two compone | 1046627.BZAF | 1 | 1,095445115 | 5,91E-05 | 6,04E-05 | 3 - | 0,000164751 - | 0 - | 0 - | 1 - | 6,48E-05 - | 1 - | 6,58E-05 - | 0 - | 0 - |
| 1772 | 1 | 3um_FIJJAFBF_137499 | Bacteria | Proteobacteri | Gammaprote | unclassified | unclassified | unclassified | various | information st | Translation, ri | 05KK9 | This protein bi | 247633.GP21 | 2,2 | 0,4 | 0,000292768 | 5,95E-05 | 2 - | 0,000222867 - | 2 - | 0,000276736 - | 2 - | 0,000263072 - | 3 - | 0,000400437 - | 2 - | 0,000300727 - |
| 1773 | 1 | 3um_FIJJAFBF_150221 | Bacteria | Bacteroidetes | Flavobacteriia | Flavobacterial | Flavobacteriac | Formosa | Formosa sp. H | unclassified | unclassified | unclassified | unclassified | unclassified | 1 | 1,095445115 | 0,000114314 | 0,000116835 | 3 - | 0,000318824 - | 0 - | 0 - | 1 - | 0,000125447 - | 1 - | 0,000127299 - | 0 - | 0 - |
| 1774 | 4 | 0.2um_NOKCPHJN_342350;0.2um_NOKCPHJN_4 | Bacteria | various | various | various | various | various | various | various | various | various | various | various | 0,2 | 0,4 | 7,28E-06 | 1,46E-05 | 0 - | 0 - | 0 - | 0 - | 1 - | 3,64E-05 - | 0 - | 0 - | 0 - | 0 - |
| 1775 | 1 | 3um_FIJJAFBF_556069 | unclassified | unclassified | unclassified | unclassified | unclassified | unclassified | unclassified | unclassified | unclassified | unclassified | unclassified | unclassified | 1,8 | 0,748331477 | 0,000219681 | 0,000106341 | 2 - | 0,000201363 - | 2 - | 0,000250033 - | 1 - | 0,000118844 - | 1 - | 0,000120599 - | 3 - | 0,000407564 - |
| 1776 | 1 | 0.2um_NOKCPHJN_28605 | Bacteria | Bacteroidetes | unclassified | unclassified | unclassified | unclassified | Bacteroidetes | information st | Transcription | 05D27 | DNA-depende | 655815.ZPR_1 | 8 | 1,264911064 | 7,59E-05 | 9,79E-06 | 10 - | 8,01E-05 - | 8 - | 7,96E-05 - | 8 - | 7,56E-05 - | 6 - | 5,76E-05 - | 8 - | 8,65E-05 - |
| 1777 | 1 | 3um_FIJJAFBF_298835 | Bacteria | Bacteroidetes | Flavobacteriia | Flavobacterial | Flavobacteriac | Polaribacter | Polaribacter ir | unclassified | unclassified | unclassified | unclassified | unclassified | 2 | 0 | 0,000242526 | 2,30E-05 | 2 - | 0,000203145 - | 2 - | 0,000252246 - | 2 - | 0,000239792 - | 2 - | 0,000243333 - | 2 - | 0,000274114 - |
| 1778 | 1 | 0.2um_NOKCPHJN_98006 | Bacteria | Bacteroidetes | Cytophagia | Cytophagales | Cyclobacteriac | Mongoliibacte | Mongoliibacte | unclassified | unclassified | unclassified | unclassified | unclassified | 0,6 | 0,489897949 | 5,04E-05 | 4,15E-05 | 1 - | 7,36E-05 - | 1 - | 9,14E-05 - | 1 - | 8,68E-05 - | 0 - | 0 - | 0 - | 0 - |
| 1779 | 1 | 3um_FIJJAFBF_455826 | Bacteria | Proteobacteri | Alphaproteob | Rhodobactera | unclassified | unclassified | Rhodobactera | metabolism | Energy produc | 01R8W | NDH-1 shuttle | 367336.OM22 | 3 | 1,095445115 | 0,000201756 | 7,18E-05 | 4 - | 0,000227281 - | 4 - | 0,000282216 - | 3 - | 0,000201211 - | 1 - | 6,81E-05 - | 3 - | 0,000230012 - |
| 1780 | 2 | 0.2um_NOKCPHJN_22624;3um_FIJJAFBF_71813 | Bacteria | Actinobacteri | Actinobacteri | various | various | various | various | unclassified | unclassified | unclassified | unclassified | unclassified | 0,4 | 0,489897949 | 2,08E-05 | 2,57E-05 | 1 - | 4,74E-05 - | 0 - | 0 - | 0 - | 0 - | 1 - | 5,68E-05 - | 0 - | 0 - |
| 1781 | 1 | 0.2um_NOKCPHJN_564048 | Bacteria | Bacteroidetes | Flavobacteriia | Flavobacterial | unclassified | unclassified | Flavobacterial | metabolism | Inorganic ion t | 0XP5Y | tonB-depende | 471854.Dfer_ | 1 | 0,632455532 | 1,62E-05 | 9,26E-06 | 2 - | 2,89E-05 - | 1 - | 1,79E-05 - | 1 - | 1,71E-05 - | 1 - | 1,73E-05 - | 0 - | 0 - |
| 1782 | 1 | 3um_FIJJAFBF_105697 | Bacteria | Bacteroidetes | Saprospiria | Saprospirales | Haliscomenob | Phaeodactylib | Phaeodactylib | cellular proces | Posttranslatio | 05TMR | Functions as a | 465817.ETA_3 | 0,4 | 0,489897949 | 5,26E-05 | 6,48E-05 | 1 - | 0,000119559 - | 0 - | 0 - | 0 - | 0 - | 1 - | 0,000143212 - | 0 - | 0 - |
| 1783 | 1 | 0.2um_NOKCPHJN_180663 | Bacteria | Proteobacteri | Alphaproteob | Rhodobactera | Rhodobactera | Planktomarina | Planktomarina | poorly charact | Function unkn | 01QH8 | abc transporte | 439496.RBY4I | 1,2 | 0,4 | 2,99E-05 | 1,03E-05 | 1 - | 2,08E-05 - | 1 - | 2,59E-05 - | 1 - | 2,46E-05 - | 2 - | 4,99E-05 - | 1 - | 2,81E-05 - |
| 1784 | 1 | 0.2um_NOKCPHJN_106716 | Bacteria | Proteobacteri | Gammaprote | Cellvibrionale | Spongiibacter | Oceanicoccus | Oceanicoccus | poorly charact | Function unkn | 0R9NA | TPR | 857087.Metm | 4 | 0,632455532 | 6,89E-05 | 1,06E-05 | 4 - | 5,81E-05 - | 4 - | 7,22E-05 - | 4 - | 6,86E-05 - | 5 - | 8,70E-05 - | 3 - | 5,88E-05 - |
| 1785 | 1 | 0.2um_NOKCPHJN_875147 | Bacteria | Bacteroidetes | Flavobacteriia | Flavobacterial | Flavobacteriac | Aquimarina | Aquimarina la | cellular proces | Cell wall/mem | 0PENK | NA | 313598.MED1 | 1 | 0 | 2,25E-05 | 2,14E-06 | 1 - | 1,89E-05 - | 1 - | 2,34E-05 - | 1 - | 2,23E-05 - | 1 - | 2,26E-05 - | 1 - | 2,55E-05 - |
| 1786 | 1 | 0.2um_NOKCPHJN_511038 | Bacteria | Bacteroidetes | Flavobacteriia | Flavobacterial | Flavobacteriac | Polaribacter | Polaribacter s | cellular proces | Cell wall/mem | 09B44 | efflux transpo | 487797.Flav3 | 0,8 | 0,4 | 4,19E-05 | 2,11E-05 | 0 - | 0 - | 1 - | 5,24E-05 - | 1 - | 4,98E-05 - | 1 - | 5,05E-05 - | 1 - | 5,69E-05 - |
| 1787 | 1 | 0.2um_NOKCPHJN_356927 | Bacteria | Bacteroidetes | Flavobacteriia | Flavobacterial | Crocinitomica | unclassified | Crocinitomica | unclassified | unclassified | unclassified | unclassified | unclassified | 0,8 | 0,4 | 3,18E-05 | 1,63E-05 | 1 - | 3,32E-05 - | 1 - | 4,12E-05 - | 0 - | 0 - | 1 - | 3,97E-05 - | 1 - | 4,48E-05 - |
| 1788 | 1 | 3um_FIJJAFBF_177316 | Bacteria | Actinobacteri | Actinobacteri | Micrococcales | Microbacteria | Candidatus Ac | Candidatus Ac | unclassified | unclassified | unclassified | unclassified | unclassified | 2 | 0 | 0,000186431 | 1,77E-05 | 2 - | 0,000156159 - | 2 - | 0,000193903 - | 2 - | 0,00018433 - | 2 - | 0,000187052 - | 2 - | 0,000210714 - |
| 1789 | 1 | 0.2um_NOKCPHJN_1058556 | Eukaryota | unclassified | Oomycetes | Pythiales | Pythiaceae | Pythium | Pythium insidi | unclassified | unclassified | unclassified | unclassified | unclassified | 0,6 | 0,489897949 | 2,23E-05 | 1,83E-05 | 0 - | 0 - | 0 - | 0 - | 1 - | 3,54E-05 - | 1 - | 3,59E-05 - | 1 - | 4,04E-05 - |
| 1790 | 1 | 3um_FIJJAFBF_606367 | Bacteria | Proteobacteri | Zetaproteoba | unclassified | unclassified | unclassified | Zetaproteoba | information st | Translation, ri | COG0050 | This protein p | 314345.SPV1_ | 0,8 | 0,4 | 2,68E-05 | 1,36E-05 | 1 - | 2,90E-05 - | 1 - | 3,60E-05 - | 1 - | 3,42E-05 - | 1 - | 3,47E-05 - | 0 - | 0 - |
| 1791 | 1 | 3um_FIJJAFBF_86539 | Bacteria | Bacteroidetes | Flavobacteriia | Flavobacterial | Flavobacteriac | Kordia | Kordia algicida | metabolism | Inorganic ion t | 05C27 | TonB-depende | 391587.KAOT | 2 | 0,632455532 | 2,63E-05 | 8,62E-06 | 2 - | 2,22E-05 - | 3 - | 4,13E-05 - | 2 - | 2,62E-05 - | 2 - | 2,66E-05 - | 1 - | 1,50E-05 - |
| 1792 | 1 | 0.2um_NOKCPHJN_249646 | Eukaryota | unclassified | Labyrinthulon | unclassified | Thraustochytr | Aurantiochytr | Aurantiochytr | poorly charact | Function unkn | 0ZUGC | NA | 7165.AGAP00 | 0,2 | 0,4 | 4,14E-06 | 8,27E-06 | 1 - | 2,07E-05 - | 0 - | 0 - | 0 - | 0 - | 0 - | 0 - | 0 - | 0 - |
| 1793 | 1 | 3um_FIJJAFBF_37676 | Bacteria | Bacteroidetes | Cytophagia | Cytophagales | Flammeovirga | unclassified | Flammeovirga | unclassified | unclassified | unclassified | unclassified | unclassified | 2,4 | 0,8 | 0,000151642 | 3,36E-05 | 4 - | 0,000217586 - | 2 - | 0,000135089 - | 2 - | 0,000128419 - | 2 - | 0,000130316 - | 2 - | 0,0001468 - |
| 1794 | 1 | 3um_FIJJAFBF_537784 | Eukaryota | Chlorophyta | Chlorophycea | Chlamydomor | Tetrabaenace | Tetrabaena | Tetrabaena so | unclassified | unclassified | unclassified | unclassified | unclassified | 13 | 1,414213562 | 0,000413691 | 3,90E-05 | 15 - | 0,000402255 - | 14 - | 0,000466183 - | 11 - | 0,000348202 - | 13 - | 0,000417589 - | 12 - | 0,000434228 - |
| 1795 | 1 | 3um_FIJJAFBF_81022 | Bacteria | Actinobacteri | Actinobacteri | Micrococcales | Microbacteria | Candidatus Ac | Candidatus Ac | metabolism | Amino acid tra | COG0174 | glutamine syn | 281090.Lxx10 | 0,8 | 0,748331477 | 2,26E-05 | 2,22E-05 | 1 - | 2,42E-05 - | 2 - | 6,01E-05 - | 1 - | 2,86E-05 - | 0 - | 0 - | 0 - | 0 - |
| 1796 | 1 | 0.2um_NOKCPHJN_15234 | Bacteria | Proteobacteri | Betaproteoba | unclassified | unclassified | unclassified | Betaproteoba | cellular proces | Posttranslatio | COG1225 | alkyl hydroper | 887062.HGR_ | 1,6 | 0,489897949 | 0,000136311 | 4,51E-05 | 2 - | 0,00014258 - | 2 - | 0,000177042 - | 1 - | 8,42E-05 - | 1 - | 8,54E-05 - | 2 - | 0,000192391 - |
| 1797 | 1 | 3um_FIJJAFBF_53329 | Bacteria | Bacteroidetes | Flavobacteriia | Flavobacterial | Flavobacteriac | Polaribacter | Polaribacter t | metabolism | Nucleotide tra | 09B0H | reductase | 313594.PI23P | 4 | 0,632455532 | 0,000128688 | 2,16E-05 | 4 - | 0,00010828 - | 5 - | 0,000168065 - | 4 - | 0,000127813 - | 4 - | 0,000129701 - | 3 - | 0,000109581 - |
| 1798 | 1 | 0.2um_NOKCPHJN_1068141 | Bacteria | Proteobacteri | Alphaproteob | Rhodobactera | Rhodobactera | Jannaschia | Jannaschia sp. | information st | Translation, ri | COG0050 | This protein p | 314262.MED1 | 1,2 | 0,748331477 | 4,13E-05 | 2,71E-05 | 1 - | 2,94E-05 - | 2 - | 7,29E-05 - | 2 - | 6,93E-05 - | 1 - | 3,52E-05 - | 0 - | 0 - |
| 1799 | 1 | 0.2um_NOKCPHJN_1026860 | Eukaryota | unclassified | unclassified | Protosteliales | Cavosteliacea | Planoprotoste | Planoprotoste | unclassified | unclassified | unclassified | unclassified | unclassified | 0,2 | 0,4 | 8,32E-06 | 1,66E-05 | 1 - | 4,16E-05 - | 0 - | 0 - | 0 - | 0 - | 0 - | 0 - | 0 - | 0 - |
| 1800 | 1 | 3um_FIJJAFBF_788009 | Bacteria | Proteobacteri | Alphaproteob | Rhodobactera | Rhodobactera | unclassified | Rhodobactera | unclassified | unclassified | unclassified | unclassified | unclassified | 11,2 | 1,469693846 | 0,000452541 | 6,46E-05 | 12 - | 0,000407491 - | 13 - | 0,000548149 - | 9 - | 0,000360751 - | 12 - | 0,000488106 - | 10 - | 0,000458209 - |
| 1801 | 2 | 3um_FIJJAFBF_264646;3um_FIJJAFBF_59048 | Bacteria | various | various | various | various | various | various | cellular proces | Posttranslatio | 05DIW | FeS assembly | various | 5,4 | 0,489897949 | 0,00015282 | 1,43E-05 | 6 - | 0,000142876 - | 6 - | 0,000177409 - | 5 - | 0,000140542 - | 5 - | 0,000142618 - | 5 - | 0,000160658 - |
| 1802 | 1 | 0.2um_NOKCPHJN_225231 | Eukaryota | unclassified | Oomycetes | Albuginales | Albuginaceae | Albugo | Albugo candid | cellular proces | Posttranslatio | KOG0734 | YME1-like 1 (S | 65071.PYU1_ | 1 | 0 | 2,12E-05 | 2,01E-06 | 1 - | 1,78E-05 - | 1 - | 2,21E-05 - | 1 - | 2,10E-05 - | 1 - | 2,13E-05 - | 1 - | 2,40E-05 - |
| 1803 | 1 | 3um_FIJJAFBF_620217 | Eukaryota | Streptophyta | Charophyceae | Charales | Characeae | Chara | Chara braunii | cellular proces | Posttranslatio | COG1225 | alkyl hydroper | 391625.PPSIR | 0,4 | 0,489897949 | 1,87E-05 | 2,31E-05 | 1 - | 4,17E-05 - | 1 - | 5,18E-05 - | 0 - | 0 - | 0 - | 0 - | 0 - | 0 - |
| 1804 | 1 | 3um_FIJJAFBF_213471 | Bacteria | Bacteroidetes | Flavobacteriia | Flavobacterial | Flavobacteriac | Algibacter | Algibacter sp. | poorly charact | Function unkn | 05FCH | NA | 1046627.BZAF | 0,8 | 1,166190379 | 4,07E-05 | 5,71E-05 | 3 - | 0,000145903 - | 0 - | 0 - | 1 - | 5,74E-05 - | 0 - | 0 - | 0 - | 0 - |
| 1805 | 1 | 0.2um_NOKCPHJN_184676 | Bacteria | Proteobacteri | Gammaprote | unclassified | unclassified | unclassified | Gammaprote | information st | Transcription | 16PMT | DNA-depende | 351348.Maqu | 2,8 | 0,4 | 0,000114146 | 1,36E-05 | 3 - | 0,000103402 - | 3 - | 0,000128395 - | 3 - | 0,000122056 - | 3 - | 0,000123859 - | 2 - | 9,30E-05 - |
| 1806 | 1 | 0.2um_NOKCPHJN_356192 | Bacteria | Proteobacteri | Gammaprote | Cellvibrionale | Porticoccacea | unclassified | SAR92 bacteri | metabolism | Inorganic ion t | 16TQC | TONB-depend | 314287.GB22 | 5,2 | 0,979795897 | 7,34E-05 | 9,25E-06 | 7 - | 8,38E-05 - | 5 - | 7,43E-05 - | 5 - | 7,06E-05 - | 4 - | 5,73E-05 - | 5 - | 8,07E-05 - |
| 1807 | 1 | 3um_FIJJAFBF_177877 | Bacteria | Bacteroidetes | Cytophagia | Cytophagales | Flammeovirga | Fabibacter | Fabibacter pa | metabolism | Carbohydrate | 0FVR1 | phosphomann | 313606.M231 | 0,6 | 0,489897949 | 1,79E-05 | 1,49E-05 | 1 - | 2,50E-05 - | 1 - | 3,10E-05 - | 0 - | 0 - | 0 - | 0 - | 1 - | 3,37E-05 - |
| 1808 | 1 | 0.2um_NOKCPHJN_900408 | Bacteria | Proteobacteri | Gammaprote | Pseudomonad | various | various | various | information st | Translation, ri | 05VBI | Binds 16S rRN | 216595.PFLU5 | 0,2 | 0,4 | 2,68E-05 | 5,37E-05 | 0 - | 0 - | 0 - | 0 - | 1 - | 0,000134141 - | 0 - | 0 - | 0 - | 0 - |
| 1809 | 1 | 0.2um_NOKCPHJN_184678 | Bacteria | Proteobacteri | Gammaprote | unclassified | unclassified | unclassified | Gammaprote | information st | Translation, ri | 08UHH | Located on th | 87626.PTD2_ | 2 | 0,632455532 | 0,000205755 | 6,69E-05 | 2 - | 0,000172597 - | 2 - | 0,000214314 - | 3 - | 0,000305599 - | 1 - | 0,000103371 - | 2 - | 0,000232894 - |
| 1810 | 1 | 0.2um_NOKCPHJN_07408 | Bacteria | Bacteroidetes | Flavobacteriia | Flavobacterial | Flavobacteriac | Polaribacter | Polaribacter s | information st | Transcription | 0P7SF | DNA-depende | 313598.MED1 | 45 | 3,286335345 | 0,000483789 | 3,73E-05 | 49 - | 0,000443188 - | 42 - | 0,000471694 - | 42 - | 0,000448405 - | 49 - | 0,000530865 - | 43 - | 0,000524791 - |
| 1811 | 1 | 0.2um_NOKCPHJN_61045 | Bacteria | Proteobacteri | Betaproteoba | unclassified | unclassified | unclassified | Betaproteoba | poorly charact | Function unkn | COG5501 | Sulfur oxidatic | 264198.Reut_ | 1 | 0 | 9,07E-05 | 8,62E-06 | 1 - | 7,60E-05 - | 1 - | 9,44E-05 - | 1 - | 8,97E-05 - | 1 - | 9,10E-05 - | 1 - | 0,000102566 - |
| 1812 | 1 | 0.2um_NOKCPHJN_376047 | Bacteria | Proteobacteri | Gammaprote | unclassified | unclassified | unclassified | SAR86 cluster | metabolism | Inorganic ion t | 16PDP | TonB-depende | 342610.Patl_ | 0,4 | 0,489897949 | 5,07E-06 | 6,25E-06 | 1 - | 1,15E-05 - | 0 - | 0 - | 0 - | 0 - | 1 - | 1,38E-05 - | 0 - | 0 - |
| 1813 | 1 | 3um_FIJJAFBF_267067 | Bacteria | Bacteroidetes | Saprospiria | Saprospirales | Lewinellaceae | Lewinella | Lewinella nigr | unclassified | unclassified | unclassified | unclassified | unclassified | 0,6 | 0,489897949 | 7,99E-05 | 6,53E-05 | 0 - | 0 - | 1 - | 0,000137037 - | 1 - | 0,000130271 - | 1 - | 0,000132195 - | 0 - | 0 - |
| 1814 | 1 | 0.2um_NOKCPHJN_108970 | Bacteria | Proteobacteri | Gammaprote | Oceanospirilla | Saccharospirill | Reinekea | Reinekea blan | unclassified | unclassified | unclassified | unclassified | unclassified | 2,8 | 1,166190379 | 2,42E-05 | 9,18E-06 | 4 - | 2,99E-05 - | 3 - | 2,78E-05 - | 2 - | 1,76E-05 - | 4 - | 3,58E-05 - | 1 - | 1,01E-05 - |
| 1815 | 1 | 3um_FIJJAFBF_73801 | Bacteria | Bacteroidetes | Flavobacteriia | Flavobacterial | Flavobacteriac | Polaribacter | Polaribacter s | information st | Translation, ri | 09A0K | glutaminyL-tR | 313594.PI23P | 0,8 | 0,4 | 1,95E-05 | 1,00E-05 | 1 - | 2,05E-05 - | 1 - | 2,54E-05 - | 1 - | 2,42E-05 - | 0 - | 0 - | 1 - | 2,76E-05 - |
| 1816 | 1 | 3um_FIJJAFBF_282908 | Bacteria | Proteobacteri | Alphaproteob | Rhodobactera | Rhodobactera | unclassified | Rhodobactera | unclassified | unclassified | unclassified | unclassified | unclassified | 1,4 | 0,489897949 | 7,52E-05 | 2,77E-05 | 1 - | 4,48E-05 - | 2 - | 0,000111343 - | 2 - | 0,000105845 - | 1 - | 5,37E-05 - | 1 - | 6,05E-05 - |
| 1817 | 1 | 3um_FIJJAFBF_318710 | Bacteria | Proteobacteri | Alphaproteob | Rhodobactera | Rhodobactera | Planktotalea | Planktotalea a | unclassified | unclassified | unclassified | unclassified | unclassified | 4,4 | 0,8 | 0,000180171 | 3,26E-05 | 5 - | 0,000172337 - | 5 - | 0,000213992 - | 3 - | 0,000122056 - | 5 - | 0,000206431 - | 4 - | 0,000186035 - |
| 1818 | 1 | 0.2um_NOKCPHJN_598238 | Eukaryota | Arthropoda | Insecta | Lepidoptera | Noctuidae | Helicoverpa | Helicoverpa a | unclassified | unclassified | unclassified | unclassified | unclassified | 0,2 | 0,4 | 1,09E-05 | 2,18E-05 | 0 - | 0 - | 0 - | 0 - | 0 - | 0 - | 1 - | 5,46E-05 - | 0 - | 0 - |
| 1819 | 1 | 3um_FIJJAFBF_448564 | unclassified | unclassified | unclassified | unclassified | unclassified | unclassified | unclassified | unclassified | unclassified | unclassified | unclassified | unclassified | 0,6 | 0,8 | 5,88E-05 | 8,22E-05 | 1 - | 8,44E-05 - | 2 - | 0,000209587 - | 0 - | 0 - | 0 - | 0 - | 0 - | 0 - |
| 1820 | 1 | 0.2um_NOKCPHJN_39345 | Bacteria | Proteobacteri | Gammaprote | Cellvibrionale | Porticoccacea | unclassified | gamma prote | information st | Translation, ri | 05CRK | lysyL-tRNA syr | 314287.GB22 | 0,2 | 0,4 | 4,62E-06 | 9,24E-06 | 1 - | 2,31E-05 - | 0 - | 0 - | 0 - | 0 - | 0 - | 0 - | 0 - | 0 - |
| 1821 | 1 | 0.2um_NOKCPHJN_192191 | Bacteria | Proteobacteri | Gammaprote | unclassified | unclassified | unclassified | Gammaprote | information st | Translation, ri | 179R6 | This protein is | 247634.GPB2 | 1 | 0 | 9,65E-05 | 9,16E-06 | 1 - | 8,08E-05 - | 1 - | 0,000100365 - | 1 - | 9,54E-05 - | 1 - | 9,68E-05 - | 1 - | 0,000109067 - |
| 1822 | 1 | 3um_FIJJAFBF_516487 | Bacteria | Proteobacteri | Gammaprote | Chromatiales | Chromatiacea | unclassified | Chromatiacea | unclassified | unclassified | unclassified | unclassified | unclassified | 8,4 | 1,743559577 | 0,000199103 | 3,63E-05 | 11 - | 0,000220339 - | 7 - | 0,000174107 - | 6 - | 0,000141866 - | 9 - | 0,000215942 - | 9 - | 0,000243258 - |
| 1823 | 1 | 0.2um_NOKCPHJN_965640 | Viruses | unclassified | unclassified | unclassified | unclassified | unclassified | uncultured vir | unclassified | unclassified | unclassified | unclassified | unclassified | 0,6 | 0,489897949 | 1,75E-05 | 1,44E-05 | 1 - | 2,59E-05 - | 0 - | 0 - | 1 - | 3,06E-05 - | 1 - | 3,10E-05 - | 0 - | 0 - |
| 1824 | 1 | 3um_FIJJAFBF_172363 | Bacteria | Proteobacteri | Gammaprote | Oceanospirilla | Saccharospirill | Reinekea | Reinekea blan | information st | Transcription | COG0250 | Participates in | 207949.RED6 | 2,2 | 0,4 | 0,000167802 | 1,54E-05 | 3 - | 0,000194537 - | 2 - | 0,000161038 - | 2 - | 0,000153087 - | 2 - | 0,000155348 - | 2 - | 0,000174999 - |
| 1825 | 1 | 3um_FIJJAFBF_506191 | Bacteria | Bacteroidetes | Flavobacteriia | Flavobacterial | Flavobacteriac | Polaribacter | Polaribacter h | information st | Translation, ri | 0P8SN | lysyl-tRNA syn | 313594.PI23P | 3 | 1,095445115 | 7,16E-05 | 2,65E-05 | 3 - | 6,09E-05 - | 4 - | 0,000100898 - | 4 - | 9,59E-05 - | 3 - | 7,30E-05 - | 1 - | 2,74E-05 - |
| 1826 | 1 | 0.2um_NOKCPHJN_155861 | Bacteria | Proteobacteri | Alphaproteob | Rhodobactera | Rhodobactera | Planktomarina | Planktomarina | poorly charact | Function unkn | 05DIH | basic membra | 633131.TR2A | 0,6 | 0,489897949 | 2,44E-05 | 2,03E-05 | 1 - | 3,46E-05 - | 0 - | 0 - | 1 - | 4,08E-05 - | 0 - | 0 - | 1 - | 4,66E-05 - |
| 1827 | 1 | 3um_FIJJAFBF_747510 | Bacteria | Proteobacteri | Gammaprote | Alteromonada | Colwelliaceae | Colwellia | Colwellia myti | metabolism | Inorganic ion t | COG0175 | Reduction of a | 167879.CPS_2 | 0,4 | 0,8 | 1,46E-05 | 2,92E-05 | 2 - | 7,31E-05 - | 0 - | 0 - | 0 - | 0 - | 0 - | 0 - | 0 - | 0 - |
| 1828 | 1 | 3um_FIJJAFBF_43427 | Bacteria | Bacteroidetes | Flavobacteriia | Flavobacterial | Flavobacteriac | Polaribacter | Polaribacter s | poorly charact | Function unkn | 11QQ6 | NA | 487797.Flav3 | 0,2 | 0,4 | 8,00E-06 | 1,60E-05 | 1 - | 4,00E-05 - | 0 - | 0 - | 0 - | 0 - | 0 - | 0 - | 0 - | 0 - |
| 1829 | 1 | 3um_FIJJAFBF_191789 | Eukaryota | unclassified | Synurophycea | Parmales | Triparmaceae | Triparma | Triparma laev | metabolism | Energy produc | COG0712 | F(1)F(0) ATP s | 35128.Thapsd | 16 | 0,632455532 | 0,001169996 | 9,71E-05 | 17 - | 0,001043425 - | 16 - | 0,001219413 - | 16 - | 0,001159206 - | 15 - | 0,001102807 - | 16 - | 0,001325129 - |
| 1830 | 1 | 0.2um_NOKCPHJN_218204 | Bacteria | Proteobacteri | Alphaproteob | Rhodobactera | Rhodobactera | Pseudooctade | Pseudooctade | metabolism | Nucleotide tra | 16UID | reductase | 644107.SL115 | 2,4 | 1,356465997 | 2,62E-05 | 1,52E-05 | 3 - | 2,83E-05 - | 3 - | 3,52E-05 - | 2 - | 2,23E-05 - | 4 - | 4,53E-05 - | 0 - | 0 - |
| 1831 | 1 | 3um_FIJJAFBF_403033 | Bacteria | Proteobacteri | Alphaproteob | Rhodobactera | Rhodobactera | Maribius | Maribius salin | metabolism | Amino acid tra | 01R85 | Catalyzes the | 314265.R2601 | 1 | 0 | 6,82E-05 | 6,47E-06 | 1 - | 5,71E-05 - | 1 - | 7,09E-05 - | 1 - | 6,74E-05 - | 1 - | 6,84E-05 - | 1 - | 7,71E-05 - |
| 1832 | 1 | 0.2um_NOKCPHJN_782778 | Bacteria | Proteobacteri | Gammaprote | Cellvibrionale | Porticoccacea | unclassified | SAR92 bacteri | metabolism | Energy produc | 0QJ3X | Aconitate hyd | 203122.Sde_1 | 2,6 | 0,489897949 | 4,37E-05 | 7,11E-06 | 3 - | 4,27E-05 - | 3 - | 5,30E-05 - | 3 - | 5,04E-05 - | 2 - | 3,41E-05 - | 2 - | 3,84E-05 - |
| 1833 | 1 | 3um_FIJJAFBF_550181 | Eukaryota | Bacillariophyt | Bacillariophyc | Bacillariales | Bacillariaceae | Fragilariopsis | Fragilariopsis | poorly charact | Function unkn | 0890F | Dynamin fami | 765420.OSCT_ | 0,2 | 0,4 | 3,44E-06 | 6,88E-06 | 1 - | 1,72E-05 - | 0 - | 0 - | 0 - | 0 - | 0 - | 0 - | 0 - | 0 - |
| 1834 | 1 | 3um_FIJJAFBF_507104 | Eukaryota | unclassified | Cryptophyta | Pyrenomonad | Geminigerace | Guillardia | Guillardia thet | unclassified | unclassified | unclassified | unclassified | unclassified | 8,6 | 0,489897949 | 0,000981036 | 0,000103327 | 9 - | 0,000860826 - | 8 - | 0,000950126 - | 9 - | 0,001016117 - | 8 - | 0,000916555 - | 9 - | 0,001161559 - |
| 1835 | 1 | 3um_FIJJAFBF_84668 | Bacteria | Proteobacteri | Gammaprote | Oceanospirilla | Saccharospirill | Reinekea | Reinekea fors | poorly charact | Function unkn | 0QSE4 | Pfam:DUF400 | 314283.MED2 | 6 | 1,264911064 | 0,000409601 | 6,42E-05 | 8 - | 0,000463744 - | 7 - | 0,000503855 - | 5 - | 0,000342127 - | 5 - | 0,00034718 - | 5 - | 0,000391097 - |
| 1836 | 1 | 3um_FIJJAFBF_545831 | Bacteria | Bacteroidetes | Flavobacteriia | Flavobacterial | Flavobacteriac | Tenacibaculur | Tenacibaculur | unclassified | unclassified | unclassified | unclassified | unclassified | 18 | 1,095445115 | 0,000746124 | 7,09E-05 | 18 - | 0,000626055 - | 18 - | 0,000777376 - | 20 - | 0,000821104 - | 17 - | 0,000708247 - | 17 - | 0,000797838 - |
| 1837 | 1 | 0.2um_NOKCPHJN_46813 | Bacteria | Proteobacteri | Alphaproteob | Rhodobactera | Rhodobactera | Antarctobacte | Antarctobacte | information st | Translation, ri | 05CEJ | Catalyzes the | 89187.ISM_01 | 1,6 | 1,019803903 | 3,05E-05 | 1,83E-05 | 3 - | 4,88E-05 - | 2 - | 4,04E-05 - | 1 - | 1,92E-05 - | 0 - | 0 - | 2 - | 4,39E-05 - |
| 1838 | 1 | 3um_FIJJAFBF_328499 | Bacteria | Bacteroidetes | Flavobacteriia | Flavobacterial | Flavobacteriac | Arenitalea | Arenitalea lut | information st | Translation, ri | 08UHY | One of the pri | 1046627.BZAF | 2,4 | 0,489897949 | 0,00020513 | 3,01E-05 | 3 - | 0,000217931 - | 2 - | 0,000180404 - | 3 - | 0,000257245 - | 2 - | 0,00017403 - | 2 - | 0,000196044 - |
| 1839 | 1 | 0.2um_NOKCPHJN_45726 | Bacteria | Bacteroidetes | Flavobacteriia | Flavobacterial | Flavobacteriac | Winogradskye | Winogradskye | metabolism | Inorganic ion t | 05C27 | TonB-depende | 391598.FBBAL | 1 | 0,632455532 | 1,24E-05 | 6,93E-06 | 2 - | 2,14E-05 - | 1 - | 1,33E-05 - | 0 - | 0 - | 1 - | 1,28E-05 - | 1 - | 1,44E-05 - |
| 1840 | 3 | 0.2um_NOKCPHJN_270961;0.2um_NOKCPHJN_9 | various | various | various | various | various | various | various | unclassified | unclassified | unclassified | unclassified | unclassified | 0,4 | 0,489897949 | 3,74E-05 | 4,61E-05 | 1 - | 8,50E-05 - | 0 - | 0 - | 0 - | 0 - | 1 - | 0,000101839 - | 0 - | 0 - |
| 1841 | 1 | 3um_FIJJAFBF_405471 | Bacteria | Proteobacteri | Gammaprote | Alteromonada | Alteromonada | Paraglaciecola | Paraglaciecola | poorly charact | Function unkn | 0NMWQ | Collagen triple | 324057.Pjdr2_ | 11,8 | 1,6 | 0,000251619 | 2,92E-05 | 14 - | 0,000251861 - | 10 - | 0,000223384 - | 13 - | 0,000276061 - | 10 - | 0,000215491 - | 12 - | 0,0002913 - |
| 1842 | 1 | 3um_FIJJAFBF_449012 | Bacteria | Proteobacteri | Gammaprote | unclassified | unclassified | unclassified | Gammaprote | unclassified | unclassified | unclassified | unclassified | unclassified | 1,8 | 0,4 | 0,0001131 | 2,77E-05 | 2 - | 0,0001053 - | 2 - | 0,000130751 - | 2 - | 0,000124296 - | 1 - | 6,31E-05 - | 2 - | 0,000142087 - |
| 1843 | 1 | 3um_FIJJAFBF_91702 | Bacteria | Bacteroidetes | unclassified | unclassified | unclassified | unclassified | Bacteroidetes | unclassified | unclassified | unclassified | unclassified | unclassified | 0,4 | 0,489897949 | 8,52E-06 | 1,05E-05 | 1 - | 1,90E-05 - | 1 - | 2,36E-05 - | 0 - | 0 - | 0 - | 0 - | 0 - | 0 - |
| 1844 | 1 | 3um_FIJJAFBF_297063 | Bacteria | Cyanobacteria | unclassified | Synechococca | Synechococca | Synechococcu | Synechococcu | poorly charact | Function unkn | 08SDR | phycocyanin, a | 316278.SynR | 0,8 | 0,4 | 6,76E-05 | 3,47E-05 | 1 - | 7,08E-05 - | 1 - | 8,80E-05 - | 1 - | 8,36E-05 - | 0 - | 0 - | 1 - | 9,56E-05 - |
| 1845 | 1 | 3um_FIJJAFBF_380757 | Bacteria | Proteobacteri | Gammaprote | Thiotrichales | Piscirickettsia | Methylophaga | Methylophaga | information st | Translation, ri | 05CRK | lysyL-tRNA syr | 637616.MDM | 0,6 | 0,489897949 | 1,59E-05 | 1,31E-05 | 1 - | 2,31E-05 - | 1 - | 2,87E-05 - | 0 - | 0 - | 1 - | 2,77E-05 - | 0 - | 0 - |
| 1846 | 1 | 0.2um_NOKCPHJN_851899 | Eukaryota | Bacillariophyt | Bacillariophyc | Bacillariales | Bacillariaceae | Fragilariopsis | Fragilariopsis | unclassified | unclassified | unclassified | unclassified | unclassified | 4,6 | 0,489897949 | 0,000245356 | 1,95E-05 | 5 - | 0,000225052 - | 4 - | 0,000223559 - | 5 - | 0,000265651 - | 5 - | 0,000269575 - | 4 - | 0,00024294 - |
| 1847 | 1 | 0.2um_NOKCPHJN_273768 | Bacteria | Proteobacteri | Alphaproteob | Rhodobactera | Rhodobactera | Thalassobacte | Thalassobacte | unclassified | unclassified | unclassified | unclassified | unclassified | 1 | 0 | 4,81E-05 | 4,57E-06 | 1 - | 4,03E-05 - | 1 - | 5,00E-05 - | 1 - | 4,75E-05 - | 1 - | 4,82E-05 - | 1 - | 5,43E-05 - |
| 1848 | 1 | 3um_FIJJAFBF_694667 | Bacteria | Bacteroidetes | unclassified | unclassified | unclassified | unclassified | Bacteroidetes | information st | Translation, ri | COG0088 | One of the pri | 391598.FBBAL | 2 | 0 | 0,000131757 | 1,25E-05 | 2 - | 0,000110362 - | 2 - | 0,000137037 - | 2 - | 0,000130271 - | 2 - | 0,000132195 - | 2 - | 0,000148918 - |
| 1849 | 1 | 3um_FIJJAFBF_734480 | Eukaryota | Platyhelminth | Trematoda | Opisthorchiid | Opisthorchiid | Opisthorchis | Opisthorchis v | unclassified | unclassified | unclassified | unclassified | unclassified | 0,6 | 0,489897949 | 3,01E-05 | 2,46E-05 | 0 - | 0 - | 1 - | 5,16E-05 - | 1 - | 4,91E-05 - | 1 - | 4,98E-05 - | 0 - | 0 - |
| 1850 | 1 | 3um_FIJJAFBF_420046 | unclassified | unclassified | unclassified | unclassified | unclassified | unclassified | unclassified | unclassified | unclassified | unclassified | unclassified | unclassified | 1 | 0 | 0,000169169 | 1,61E-05 | 1 - | 0,0001417 - | 1 - | 0,000175949 - | 1 - | 0,000167262 - | 1 - | 0,000169732 - | 1 - | 0,000191203 - |
| 1851 | 1 | 3um_FIJJAFBF_549855 | Bacteria | Bacteroidetes | Flavobacteriia | Flavobacterial | Crocinitomica | unclassified | Crocinitomica | unclassified | unclassified | unclassified | unclassified | unclassified | 0,4 | 0,489897949 | 2,98E-05 | 3,65E-05 | 0 - | 0 - | 0 - | 0 - | 1 - | 7,40E-05 - | 1 - | 7,51E-05 - | 0 - | 0 - |
| 1852 | 1 | 0.2um_NOKCPHJN_1115842 | Bacteria | Bacteroidetes | Flavobacteriia | Flavobacterial | Flavobacteriac | Flavobacteriu | Flavobacteriu | cellular proces | Cell wall/mem | 09ARE | ompa motb d | 376686.Fjoh_ | 1 | 0,632455532 | 3,18E-05 | 2,06E-05 | 1 - | 2,74E-05 - | 1 - | 3,40E-05 - | 2 - | 6,47E-05 - | 1 - | 3,28E-05 - | 0 - | 0 - |
| 1853 | 1 | 3um_FIJJAFBF_457311 | Bacteria | Proteobacteri | Gammaprote | Cellvibrionale | Spongiibacter | unclassified | marine gamm | unclassified | unclassified | unclassified | unclassified | unclassified | 0,6 | 0,489897949 | 1,31E-05 | 1,08E-05 | 1 - | 1,91E-05 - | 1 - | 2,37E-05 - | 1 - | 2,25E-05 - | 0 - | 0 - | 0 - | 0 - |

| | | | | | | | | | | | | | | | | | | | | | | | | | | | | |
|---|---|---|---|---|---|---|---|---|---|---|---|---|---|---|---|---|---|---|---|---|---|---|---|---|---|---|---|---|
| 1854 | 1 | 3um_FIJJAFBF_688308 | Bacteria | Proteobacteri | Alphaproteob | Rhodobactera | Rhodobactera | Marivita | Marivita hally | information st | Translation, ri | 01WSM | This is one of t | 391619.RGBS | 1 | 0 | 6,46E-05 | 6,14E-06 | 1 - | 5,41E-05 - | 1 - | 6,72E-05 - | 1 - | 6,39E-05 - | 1 - | 6,49E-05 - | 1 - | 7,31E-05 - |
| 1855 | 1 | 3um_FIJJAFBF_164365 | Bacteria | Proteobacteri | Alphaproteob | Rhodobactera | Rhodobactera | Maritimibacte | Maritimibacte | metabolism | Nucleotide tra | COG0015 | adenylosuccin | 644107.SL115 | 5,6 | 0,489897949 | 0,000143013 | 1,25E-05 | 6 - | 0,000128963 - | 6 - | 0,000160134 - | 6 - | 0,000152227 - | 5 - | 0,00012873 - | 5 - | 0,000145014 - |
| 1856 | 1 | 3um_FIJJAFBF_674227 | Bacteria | Proteobacteri | Alphaproteob | Rhodobactera | Rhodobactera | Ruegeria | various | metabolism | Secondary me | 01Q86 | extracellular s | 391595.RLO14 | 0,2 | 0,4 | 6,36E-06 | 1,27E-05 | 1 - | 3,18E-05 - | 0 - | 0 - | 0 - | 0 - | 0 - | 0 - | 0 - | 0 - |
| 1857 | 1 | 3um_FIJJAFBF_131607 | Bacteria | Proteobacteri | Gammaprotec | Cellvibrionale | Porticoccacea | unclassified | SAR92 bacteri | poorly charact | Function unkn | 16R8I | Lipoprotein | 377629.TERTU | 9 | 1,095445115 | 0,000131866 | 7,63E-06 | 11 - | 0,000136344 - | 9 - | 0,000138517 - | 8 - | 0,000117047 - | 9 - | 0,000133623 - | 8 - | 0,000133801 - |
| 1858 | 2 | 3um_FIJJAFBF_14126;3um_FIJJAFBF_425975 | Bacteria | Proteobacteri | Alphaproteob | Rhodobactera | Rhodobactera | Planktomarina | Planktomarina | unclassified | unclassified | unclassified | unclassified | unclassified | 0,2 | 0,4 | 4,97E-06 | 9,94E-06 | 1 - | 2,48E-05 - | 0 - | 0 - | 0 - | 0 - | 0 - | 0 - | 0 - | 0 - |
| 1859 | 1 | 0.2um_NOKCPHJN_804113 | Bacteria | Cyanobacteria | unclassified | unclassified | unclassified | unclassified | filamentous c | poorly charact | Function unkn | COG1100 | GTP-binding P | 756067.Micva | 0,6 | 0,489897949 | 1,25E-05 | 1,03E-05 | 0 - | 0 - | 1 - | 2,07E-05 - | 1 - | 1,96E-05 - | 0 - | 0 - | 1 - | 2,24E-05 - |
| 1860 | 1 | 0.2um_NOKCPHJN_97372 | Bacteria | Proteobacteri | Gammaprotec | unclassified | unclassified | unclassified | uncultured ga | unclassified | unclassified | unclassified | unclassified | unclassified | 1,2 | 0,4 | 3,53E-05 | 7,82E-06 | 2 - | 5,07E-05 - | 1 - | 3,15E-05 - | 1 - | 2,99E-05 - | 1 - | 3,03E-05 - | 1 - | 3,42E-05 - |
| 1861 | 2 | 0.2um_NOKCPHJN_604214;3um_FIJJAFBF_0069 | Bacteria | Proteobacteri | Alphaproteob | Pelagibacteral | Pelagibacterac | Candidatus Pe | Candidatus Pe | metabolism | Coenzyme tra | 08ZQD | Catalyzes the | 335992.SAR11 | 0 | 0 | 0 | 0 | 0 - | 0 - | 0 - | 0 - | 0 - | 0 - | 0 - | 0 - | 0 - | 0 - |
| 1862 | 2 | 0.2um_NOKCPHJN_683043;3um_FIJJAFBF_3240 | Bacteria | unclassified | unclassified | unclassified | unclassified | unclassified | uncultured ba | unclassified | unclassified | unclassified | unclassified | unclassified | 0,4 | 0,489897949 | 1,04E-05 | 1,27E-05 | 0 - | 0 - | 0 - | 0 - | 1 - | 2,58E-05 - | 1 - | 2,62E-05 - | 0 - | 0 - |
| 1863 | 1 | 0.2um_NOKCPHJN_460038 | Bacteria | Bacteroidetes | Flavobacteriia | Flavobacterial | Flavobacteriac | Lacinutrix | Lacinutrix him | poorly charact | Function unkn | 06E9D | NA | 391598.FBBAL | 1,2 | 0,4 | 2,82E-05 | 1,04E-05 | 1 - | 1,96E-05 - | 2 - | 4,86E-05 - | 1 - | 2,31E-05 - | 1 - | 2,34E-05 - | 1 - | 2,64E-05 - |
| 1864 | 1 | 0.2um_NOKCPHJN_106091 | Bacteria | Bacteroidetes | Flavobacteriia | Flavobacterial | Flavobacteriac | Lutibacter | Lutibacter sp. | poorly charact | Function unkn | 0XR4Y | Ragb susd dor | 886377.Murru | 1,8 | 0,748331477 | 4,23E-05 | 1,38E-05 | 3 - | 6,14E-05 - | 1 - | 2,54E-05 - | 2 - | 4,83E-05 - | 2 - | 4,90E-05 - | 1 - | 2,76E-05 - |
| 1865 | 1 | 0.2um_NOKCPHJN_174959 | Bacteria | Bacteroidetes | Flavobacteriia | Flavobacterial | Crocinitomica | unclassified | Crocinitomica | metabolism | Energy produc | COG0055 | Produces ATP | 313596.RB250 | 0,4 | 0,489897949 | 1,17E-05 | 1,44E-05 | 0 - | 0 - | 0 - | 0 - | 0 - | 0 - | 1 - | 2,76E-05 - | 1 - | 3,10E-05 - |
| 1866 | 1 | 3um_FIJJAFBF_744825 | Bacteria | Proteobacteri | Gammaprotec | Cellvibrionale | Halieaceae | Chromatocurv | Chromatocurv | unclassified | unclassified | unclassified | unclassified | unclassified | 0,4 | 0,489897949 | 2,96E-05 | 3,65E-05 | 1 - | 6,79E-05 - | 0 - | 0 - | 1 - | 8,02E-05 - | 0 - | 0 - | 0 - | 0 - |
| 1867 | 1 | 3um_FIJJAFBF_350137 | Bacteria | Proteobacteri | Gammaprotec | Alteromonada | Colwelliaceae | Colwellia | Colwellia sp. F | cellular proces | Cell wall/mem | 0QTSX | Peptidoglycan | 167879.CPS_1 | 1,6 | 0,489897949 | 0,000121923 | 4,03E-05 | 2 - | 0,00012753 - | 2 - | 0,000158354 - | 1 - | 7,53E-05 - | 1 - | 7,64E-05 - | 2 - | 0,000172083 - |
| 1868 | 1 | 0.2um_NOKCPHJN_383541 | Bacteria | Bacteroidetes | Flavobacteriia | unclassified | unclassified | unclassified | Flavobacteria | poorly charact | Function unkn | 11QQ6 | NA | 487796.Flav2A | 2,6 | 1,019803903 | 0,000114443 | 4,10E-05 | 4 - | 0,000149546 - | 2 - | 9,28E-05 - | 1 - | 4,41E-05 - | 3 - | 0,000134349 - | 3 - | 0,000151343 - |
| 1869 | 1 | 0.2um_NOKCPHJN_666584 | Eukaryota | Bacillariophyta | Bacillariophyc | Bacillariales | Bacillariaceae | Nitzschia | Nitzschia sp. I | cellular proces | Posttranslatio | KOG0730 | cell division cy | 35128.Thaps2 | 0,6 | 0,489897949 | 1,06E-05 | 8,82E-06 | 1 - | 1,48E-05 - | 1 - | 1,83E-05 - | 0 - | 0 - | 0 - | 0 - | 1 - | 1,99E-05 - |
| 1870 | 1 | 3um_FIJJAFBF_494166 | Bacteria | Bacteroidetes | Flavobacteriia | Flavobacterial | Flavobacteriac | Polaribacter | various | information st | Translation, ri | COG0256 | This is one of t | 313598.MED1 | 1,4 | 0,489897949 | 0,000161275 | 4,61E-05 | 2 - | 0,000197891 - | 1 - | 0,000122861 - | 2 - | 0,00023359 - | 1 - | 0,00011852 - | 1 - | 0,000133512 - |
| 1871 | 2 | 0.2um_NOKCPHJN_527245;3um_FIJJAFBF_7410 | Bacteria | Proteobacteri | Gammaprotec | Thiotrichales | Piscirickettsiac | Methylophaga | Methylophaga | metabolism | Lipid transpor | 05CEY | Amp-depende | 572477.Alvin_ | 0,6 | 0,489897949 | 7,72E-06 | 6,32E-06 | 0 - | 0 - | 1 - | 1,26E-05 - | 0 - | 0 - | 1 - | 1,22E-05 - | 1 - | 1,37E-05 - |
| 1872 | 1 | 3um_FIJJAFBF_452134 | Bacteria | Proteobacteri | Alphaproteob | Rhodobactera | Rhodobactera | unclassified | Rhodobactera | metabolism | Carbohydrate | 16THB | transporter, d | 744979.R2A13 | 0,6 | 0,489897949 | 0,000160324 | 0,000131408 | 0 - | 0 - | 1 - | 0,000263924 - | 1 - | 0,000250893 - | 0 - | 0 - | 1 - | 0,000286805 - |
| 1873 | 1 | 3um_FIJJAFBF_618298 | Bacteria | Bacteroidetes | Cytophagia | Cytophagales | Flammeovirga | unclassified | Flammeovirga | unclassified | unclassified | unclassified | unclassified | unclassified | 0,4 | 0,489897949 | 2,03E-05 | 2,50E-05 | 1 - | 4,65E-05 - | 0 - | 0 - | 1 - | 5,49E-05 - | 0 - | 0 - | 0 - | 0 - |
| 1874 | 1 | 0.2um_NOKCPHJN_972342 | unclassified | unclassified | unclassified | unclassified | unclassified | unclassified | unclassified | unclassified | unclassified | unclassified | unclassified | unclassified | 0,2 | 0,4 | 1,58E-05 | 3,16E-05 | 0 - | 0 - | 0 - | 0 - | 0 - | 0 - | 1 - | 7,90E-05 - | 0 - | 0 - |
| 1875 | 1 | 3um_FIJJAFBF_53314 | Bacteria | Bacteroidetes | Flavobacteriia | Flavobacterial | Flavobacteriac | Polaribacter | Polaribacter s | poorly charact | Function unkn | COG0714 | ATPase activit | 313598.MED1 | 6,8 | 1,469693846 | 0,00027951 | 5,04E-05 | 8 - | 0,000279092 - | 6 - | 0,000259913 - | 6 - | 0,00024708 - | 9 - | 0,000376094 - | 5 - | 0,000235372 - |
| 1876 | 1 | 3um_FIJJAFBF_92871 | Eukaryota | unclassified | Cryptophyta | Pyrenomonad | Geminigerace | Guillardia | Guillardia thet | metabolism | Energy produc | COG0055 | Produces ATP | 4081.Solyc01g | 1 | 0 | 2,88E-05 | 2,74E-06 | 1 - | 2,42E-05 - | 1 - | 3,00E-05 - | 1 - | 2,85E-05 - | 1 - | 2,89E-05 - | 1 - | 3,26E-05 - |
| 1877 | 1 | 0.2um_NOKCPHJN_488045 | Bacteria | Bacteroidetes | Flavobacteriia | Flavobacterial | Flavobacteriac | Imtechella | Imtechella hal | unclassified | unclassified | unclassified | unclassified | unclassified | 0,2 | 0,4 | 1,14E-05 | 2,27E-05 | 1 - | 5,68E-05 - | 0 - | 0 - | 0 - | 0 - | 0 - | 0 - | 0 - | 0 - |
| 1878 | 1 | 0.2um_NOKCPHJN_712928 | Bacteria | Bacteroidetes | Flavobacteriia | Flavobacterial | Flavobacteriac | Polaribacter | Polaribacter ir | unclassified | unclassified | unclassified | unclassified | unclassified | 0,2 | 0,4 | 5,26E-06 | 1,05E-05 | 1 - | 2,63E-05 - | 0 - | 0 - | 0 - | 0 - | 0 - | 0 - | 0 - | 0 - |
| 1879 | 1 | 3um_FIJJAFBF_194216 | Bacteria | Proteobacteri | Gammaprotec | Alteromonada | Pseudoalteror | Pseudoalteror | Pseudoalteror | metabolism | Carbohydrate | 0QX81 | Inherit from p | 94122.Shewar | 0,2 | 0,4 | 2,03E-06 | 4,06E-06 | 0 - | 0 - | 0 - | 0 - | 1 - | 1,01E-05 - | 0 - | 0 - | 0 - | 0 - |
| 1880 | 1 | 3um_FIJJAFBF_624839 | Bacteria | Proteobacteri | Gammaprotec | Oceanospirilla | Saccharospirill | Reinekea | Reinekea fors | unclassified | unclassified | unclassified | unclassified | unclassified | 0,8 | 0,4 | 0,000114072 | 5,74E-05 | 0 - | 0 - | 1 - | 0,000142519 - | 1 - | 0,000135482 - | 1 - | 0,000137483 - | 1 - | 0,000154874 - |
| 1881 | 1 | 3um_FIJJAFBF_508692 | Bacteria | Proteobacteri | Gammaprotec | Cellvibrionale | unclassified | unclassified | Cellvibrionale | unclassified | unclassified | unclassified | unclassified | unclassified | 1 | 0 | 3,29E-05 | 3,13E-06 | 1 - | 2,76E-05 - | 1 - | 3,43E-05 - | 1 - | 3,26E-05 - | 1 - | 3,30E-05 - | 1 - | 3,72E-05 - |
| 1882 | 1 | 0.2um_NOKCPHJN_119453 | Bacteria | Proteobacteri | Gammaprotec | Oceanospirilla | Oceanospirilla | Oceanospirillu | Oceanospirillu | unclassified | unclassified | unclassified | unclassified | unclassified | 0,2 | 0,4 | 3,04E-06 | 6,07E-06 | 1 - | 1,52E-05 - | 0 - | 0 - | 0 - | 0 - | 0 - | 0 - | 0 - | 0 - |
| 1883 | 1 | 3um_FIJJAFBF_238401 | Bacteria | Firmicutes | Bacilli | Lactobacillales | Streptococcac | Streptococcus | Streptococcus | unclassified | unclassified | unclassified | unclassified | unclassified | 0,2 | 0,4 | 3,50E-06 | 7,00E-06 | 0 - | 0 - | 0 - | 0 - | 0 - | 0 - | 0 - | 0 - | 1 - | 1,75E-05 - |
| 1884 | 1 | 3um_FIJJAFBF_320826 | Bacteria | Proteobacteri | Gammaprotec | unclassified | unclassified | unclassified | Gammaprotec | information st | Replication, re | 06H78 | methyltransfe | 1026882.MAN | 1 | 0 | 7,06E-05 | 6,71E-06 | 1 - | 5,92E-05 - | 1 - | 7,35E-05 - | 1 - | 6,98E-05 - | 1 - | 7,09E-05 - | 1 - | 7,98E-05 - |
| 1885 | 1 | 3um_FIJJAFBF_153772 | Bacteria | Proteobacteri | Deltaproteoba | unclassified | unclassified | unclassified | Deltaproteoba | metabolism | Amino acid tra | 05TIB | (ABC) transpo | 367336.OM22 | 5,4 | 1,2 | 0,000203624 | 3,48E-05 | 7 - | 0,000224424 - | 6 - | 0,000238858 - | 4 - | 0,000151377 - | 6 - | 0,000230419 - | 4 - | 0,000173044 - |
| 1886 | 1 | 0.2um_NOKCPHJN_377107 | Bacteria | Bacteroidetes | Flavobacteriia | Flavobacterial | Flavobacteriac | Oceanihabitar | Oceanihabitar | metabolism | Energy produc | COG0055 | Produces ATP | 391587.KAOT | 0,8 | 0,4 | 2,11E-05 | 1,07E-05 | 1 - | 2,29E-05 - | 1 - | 2,84E-05 - | 1 - | 2,70E-05 - | 1 - | 2,74E-05 - | 0 - | 0 - |
| 1887 | 1 | 0.2um_NOKCPHJN_334042 | Bacteria | Firmicutes | Bacilli | Lactobacillales | Lactobacillace | Lactobacillus | Lactobacillus a | poorly charact | Function unkn | 0KN5V | NA | 140110.Necha | 0,2 | 0,4 | 1,12E-06 | 2,24E-06 | 1 - | 5,60E-06 - | 0 - | 0 - | 0 - | 0 - | 0 - | 0 - | 0 - | 0 - |
| 1888 | 1 | 3um_FIJJAFBF_236176 | Eukaryota | unclassified | unclassified | unclassified | unclassified | Blastocystis | Blastocystis h | cellular proces | Cell cycle cont | 0V9W7 | retinitis pigme | 8128.ENSONI | 0,8 | 0,4 | 1,77E-05 | 9,09E-06 | 1 - | 1,87E-05 - | 0 - | 0 - | 1 - | 2,21E-05 - | 1 - | 2,24E-05 - | 1 - | 2,53E-05 - |
| 1889 | 10 | 0.2um_NOKCPHJN_147933;0.2um_NOKCPHJN_2 | Bacteria | various | various | various | various | various | various | metabolism | Energy produc | 0P7WF | aldo keto redu | various | 0,2 | 0,4 | 7,85E-06 | 1,57E-05 | 0 - | 0 - | 0 - | 0 - | 1 - | 3,93E-05 - | 0 - | 0 - | 0 - | 0 - |
| 1890 | 1 | 3um_FIJJAFBF_12511 | Viruses | unclassified | unclassified | unclassified | Phycodnavirid | unclassified | Organic Lake | poorly charact | Function unkn | 0QZS3 | FN3 | 349521.HCH_ | 0,6 | 0,489897949 | 1,51E-05 | 1,24E-05 | 1 - | 2,23E-05 - | 0 - | 0 - | 1 - | 2,64E-05 - | 1 - | 2,67E-05 - | 0 - | 0 - |
| 1891 | 1 | 0.2um_NOKCPHJN_239349 | Bacteria | Proteobacteri | Gammaprotec | Cellvibrionale | Porticoccacea | unclassified | Gammaprotec | poorly charact | Function unkn | 0QJ8Y | phosphate-sel | 314287.GB22 | 1,2 | 0,4 | 4,18E-05 | 1,44E-05 | 1 - | 2,91E-05 - | 1 - | 3,62E-05 - | 1 - | 3,44E-05 - | 2 - | 6,98E-05 - | 1 - | 3,93E-05 - |
| 1892 | 1 | 0.2um_NOKCPHJN_192192 | Bacteria | Proteobacteri | Gammaprotec | unclassified | unclassified | unclassified | Gammaprotec | information st | Translation, ri | 179RU | 30S ribosomal | 876044.IMCC | 2,6 | 0,489897949 | 0,000279024 | 5,95E-05 | 2 - | 0,000179339 - | 3 - | 0,000334029 - | 3 - | 0,000317536 - | 3 - | 0,000322226 - | 2 - | 0,000241991 - |
| 1893 | 1 | 3um_FIJJAFBF_115673 | Bacteria | Proteobacteri | Gammaprotec | Alteromonada | Colwelliaceae | Colwellia | various | unclassified | unclassified | unclassified | unclassified | unclassified | 1,4 | 0,489897949 | 9,71E-05 | 3,94E-05 | 1 - | 5,71E-05 - | 1 - | 7,09E-05 - | 2 - | 0,000134808 - | 1 - | 6,84E-05 - | 2 - | 0,000154104 - |
| 1894 | 1 | 3um_FIJJAFBF_51581 | Bacteria | Bacteroidetes | Flavobacteriia | Flavobacterial | Flavobacteriac | unclassified | Flavobacteriac | metabolism | Nucleotide tra | COG0138 | bifunctional p | 487797.Flav3 | 1,4 | 0,489897949 | 3,67E-05 | 1,05E-05 | 2 - | 4,50E-05 - | 1 - | 2,79E-05 - | 2 - | 5,31E-05 - | 1 - | 2,70E-05 - | 1 - | 3,04E-05 - |
| 1895 | 1 | 0.2um_NOKCPHJN_66855 | Bacteria | Proteobacteri | Gammaprotec | Arenicellales | Arenicellaceae | Arenicella | Arenicella xan | unclassified | unclassified | unclassified | unclassified | unclassified | 0,4 | 0,489897949 | 2,44E-05 | 2,99E-05 | 0 - | 0 - | 0 - | 0 - | 0 - | 0 - | 1 - | 5,73E-05 - | 1 - | 6,45E-05 - |
| 1896 | 1 | 3um_FIJJAFBF_746098 | Bacteria | Bacteroidetes | Cytophagia | Cytophagales | Flammeovirga | unclassified | Flammeovirga | information st | Translation, ri | COG0254 | 50S ribosomal | 452471.Aasi_ | 1,4 | 0,489897949 | 0,000235608 | 7,41E-05 | 2 - | 0,000286942 - | 2 - | 0,000356297 - | 1 - | 0,000169353 - | 1 - | 0,000171854 - | 1 - | 0,000193593 - |
| 1897 | 1 | 0.2um_NOKCPHJN_154378 | Bacteria | Proteobacteri | Gammaprotec | Cellvibrionale | Cellvibrionace | Candidatus En | Candidatus En | metabolism | Energy produc | COG0055 | Produces ATP | 314285.KT71_ | 1 | 0 | 2,95E-05 | 2,80E-06 | 1 - | 2,47E-05 - | 1 - | 3,06E-05 - | 1 - | 2,91E-05 - | 1 - | 2,96E-05 - | 1 - | 3,33E-05 - |
| 1898 | 1 | 3um_FIJJAFBF_417062 | Bacteria | Proteobacteri | Gammaprotec | Alteromonada | Alteromonada | Paraglaciecola | Paraglaciecola | information st | Translation, ri | 0QK14 | This protein bi | 455436.GHTC | 3,2 | 0,748331477 | 0,00024741 | 6,29E-05 | 4 - | 0,000259383 - | 3 - | 0,000241557 - | 2 - | 0,000153087 - | 3 - | 0,000233023 - | 4 - | 0,000349999 - |
| 1899 | 1 | 3um_FIJJAFBF_188134 | Bacteria | Proteobacteri | Gammaprotec | Cellvibrionale | Porticoccacea | unclassified | various | information st | Translation, ri | 05C64 | Binds directly | 314287.GB22 | 0,2 | 0,4 | 1,17E-05 | 2,35E-05 | 0 - | 0 - | 0 - | 0 - | 1 - | 5,87E-05 - | 0 - | 0 - | 0 - | 0 - |
| 1900 | 1 | 0.2um_NOKCPHJN_110583 | Bacteria | Bacteroidetes | Cytophagia | Cytophagales | Cytophagacea | Ekhidna | Ekhidna lutea | metabolism | Carbohydrate | COG0176 | Transaldolase | 388413.ALPR1 | 0,2 | 0,4 | 1,25E-05 | 2,50E-05 | 0 - | 0 - | 0 - | 0 - | 1 - | 6,24E-05 - | 0 - | 0 - | 0 - | 0 - |
| 1901 | 1 | 0.2um_NOKCPHJN_1059394 | Eukaryota | unclassified | Oomycetes | Saprolegniales | Saprolegniace | Saprolegnia | Saprolegnia di | unclassified | unclassified | unclassified | unclassified | unclassified | 0,8 | 0,4 | 2,41E-05 | 1,21E-05 | 0 - | 0 - | 1 - | 3,01E-05 - | 1 - | 2,86E-05 - | 1 - | 2,91E-05 - | 1 - | 3,27E-05 - |
| 1902 | 1 | 3um_FIJJAFBF_469903 | Bacteria | Proteobacteri | Alphaproteob | Rhodobactera | Rhodobactera | unclassified | Rhodobactera | metabolism | Lipid transpor | COG1024 | Enoyl-CoA hyc | 388739.RSK20 | 0,2 | 0,4 | 8,90E-06 | 1,78E-05 | 1 - | 4,45E-05 - | 0 - | 0 - | 0 - | 0 - | 0 - | 0 - | 0 - | 0 - |
| 1903 | 1 | 3um_FIJJAFBF_50495 | Bacteria | Bacteroidetes | Flavobacteriia | Flavobacterial | Flavobacteriac | Polaribacter | various | unclassified | unclassified | unclassified | unclassified | unclassified | 1 | 0 | 7,97E-05 | 7,56E-06 | 1 - | 6,67E-05 - | 1 - | 8,29E-05 - | 1 - | 7,88E-05 - | 1 - | 7,99E-05 - | 1 - | 9,00E-05 - |
| 1904 | 1 | 3um_FIJJAFBF_304515 | Bacteria | Proteobacteri | Alphaproteob | Rhodobactera | unclassified | unclassified | Rhodobactera | poorly charact | Function unkn | COG1744 | basic membra | 391624.OIHEL | 2 | 0,894427191 | 7,72E-05 | 3,56E-05 | 1 - | 3,22E-05 - | 3 - | 0,000119764 - | 3 - | 0,000113851 - | 2 - | 7,70E-05 - | 1 - | 4,34E-05 - |
| 1905 | 1 | 0.2um_NOKCPHJN_436704 | Bacteria | Bacteroidetes | Flavobacteriia | Flavobacterial | Cryomorphace | unclassified | Cryomorphace | metabolism | Inorganic ion t | 05C27 | TonB-depende | 655815.ZPR_3 | 15,6 | 2,576819745 | 0,000211338 | 2,79E-05 | 18 - | 0,000206598 - | 18 - | 0,000256534 - | 15 - | 0,000203223 - | 16 - | 0,000219973 - | 11 - | 0,000170362 - |
| 1906 | 1 | 0.2um_NOKCPHJN_245109 | Bacteria | Bacteroidetes | Flavobacteriia | Flavobacterial | unclassified | unclassified | Flavobacterial | metabolism | Inorganic ion t | 0YT3X | TonB-depende | 518766.Rmar_ | 2 | 1,414213562 | 2,40E-05 | 1,58E-05 | 4 - | 4,27E-05 - | 2 - | 2,65E-05 - | 3 - | 3,78E-05 - | 1 - | 1,28E-05 - | 0 - | 0 - |
| 1907 | 1 | 3um_FIJJAFBF_254273 | Bacteria | Bacteroidetes | Flavobacteriia | Flavobacterial | Flavobacteriac | Formosa | Formosa sp. H | poorly charact | Function unkn | 0P9WP | NA | 1046627.BZA | 0,8 | 0,4 | 6,43E-05 | 3,30E-05 | 1 - | 6,71E-05 - | 1 - | 8,33E-05 - | 0 - | 0 - | 1 - | 8,04E-05 - | 1 - | 9,06E-05 - |
| 1908 | 1 | 0.2um_NOKCPHJN_641797 | Bacteria | Proteobacteri | Alphaproteob | Pelagibacteral | Pelagibacterac | Candidatus Pe | Candidatus Pe | information st | Translation, ri | 179RU | 30S ribosomal | 335992.SAR11 | 0,6 | 0,489897949 | 5,30E-05 | 4,37E-05 | 1 - | 7,70E-05 - | 1 - | 9,57E-05 - | 0 - | 0 - | 1 - | 9,23E-05 - | 0 - | 0 - |
| 1909 | 1 | 3um_FIJJAFBF_91415 | Bacteria | Proteobacteri | Alphaproteob | Rhodobactera | Rhodobactera | Planktomarina | Planktomarina | metabolism | Energy produc | COG0711 | F(1)F(0) ATP s | 439497.RR11_ | 7,2 | 0,4 | 0,000553165 | 7,88E-05 | 7 - | 0,000448848 - | 7 - | 0,000557336 - | 7 - | 0,000529819 - | 7 - | 0,000537644 - | 8 - | 0,000692176 - |
| 1910 | 1 | 3um_FIJJAFBF_32479 | Eukaryota | Bacillariophyta | Bacillariophyc | unclassified | Entomoneidac | Entomoneis | Entomoneis s | unclassified | unclassified | unclassified | unclassified | unclassified | 1,8 | 0,748331477 | 4,91E-05 | 2,13E-05 | 1 - | 2,25E-05 - | 1 - | 2,80E-05 - | 3 - | 7,99E-05 - | 2 - | 5,40E-05 - | 2 - | 6,09E-05 - |
| 1911 | 1 | 3um_FIJJAFBF_402488 | Bacteria | Bacteroidetes | Flavobacteriia | various | various | various | various | information st | Translation, ri | COG0087 | One of the pri | 487797.Flav3 | 0,8 | 0,4 | 5,17E-05 | 2,63E-05 | 1 - | 5,60E-05 - | 1 - | 6,95E-05 - | 1 - | 6,61E-05 - | 1 - | 6,71E-05 - | 0 - | 0 - |
| 1912 | 2 | 0.2um_NOKCPHJN_418156;3um_FIJJAFBF_5399 | Bacteria | Proteobacteri | Alphaproteob | various | various | various | various | poorly charact | Function unkn | 05N8B | NA | 367336.OM22 | 0,2 | 0,4 | 1,69E-05 | 3,37E-05 | 0 - | 0 - | 1 - | 8,43E-05 - | 0 - | 0 - | 0 - | 0 - | 0 - | 0 - |
| 1913 | 1 | 0.2um_NOKCPHJN_17065 | Bacteria | Bacteroidetes | Flavobacteriia | Flavobacterial | Flavobacteriac | Polaribacter | Polaribacter t | poorly charact | Function unkn | 0ZVFQ | Pkd domain c | 269798.CHU_ | 3 | 0,894427191 | 5,89E-06 | 1,89E-06 | 2 - | 3,27E-06 - | 4 - | 8,12E-06 - | 3 - | 5,79E-06 - | 4 - | 7,84E-06 - | 2 - | 4,41E-06 - |
| 1914 | 1 | 3um_FIJJAFBF_254290 | Bacteria | Proteobacteri | Gammaprotec | Cellvibrionale | Spongiibacter | Oceanicoccus | Oceanicoccus | information st | Translation, ri | 17C3B | The globular d | 247633.GP21 | 3,6 | 0,489897949 | 0,000451842 | 7,36E-05 | 4 - | 0,000421199 - | 3 - | 0,000392254 - | 3 - | 0,000372887 - | 4 - | 0,000504526 - | 4 - | 0,000568347 - |
| 1915 | 1 | 3um_FIJJAFBF_216067 | Bacteria | Proteobacteri | Gammaprotec | Cellvibrionale | Microbulbifer | Microbulbifer | Microbulbifer | information st | Transcription | COG0085 | DNA-depende | 247633.GP21 | 4,4 | 1,743559577 | 4,30E-05 | 1,58E-05 | 6 - | 5,06E-05 - | 5 - | 5,24E-05 - | 5 - | 4,98E-05 - | 5 - | 5,05E-05 - | 1 - | 1,14E-05 - |
| 1916 | 1 | 3um_FIJJAFBF_672155 | Bacteria | Proteobacteri | Gammaprotec | Oceanospirilla | Saccharospirill | Reinekea | Reinekea blan | information st | Replication, re | 05CZ8 | DNA polymera | 379731.PST_0 | 1,4 | 0,489897949 | 5,22E-05 | 1,92E-05 | 1 - | 3,11E-05 - | 2 - | 7,72E-05 - | 2 - | 7,34E-05 - | 1 - | 3,73E-05 - | 1 - | 4,20E-05 - |
| 1917 | 1 | 0.2um_NOKCPHJN_76749 | unclassified | unclassified | unclassified | unclassified | unclassified | unclassified | unclassified | unclassified | unclassified | unclassified | unclassified | unclassified | 1,2 | 0,748331477 | 4,06E-05 | 2,40E-05 | 2 - | 5,86E-05 - | 0 - | 0 - | 1 - | 3,46E-05 - | 2 - | 7,01E-05 - | 1 - | 3,95E-05 - |
| 1918 | 1 | 0.2um_NOKCPHJN_626446 | Bacteria | Proteobacteri | Alphaproteob | Pelagibacteral | Pelagibacterac | Candidatus Pe | Candidatus Pe | poorly charact | Function unkn | 16Q8K | Catalyzes the | 335992.SAR11 | 0,4 | 0,489897949 | 2,86E-05 | 3,50E-05 | 0 - | 0 - | 0 - | 0 - | 1 - | 7,09E-05 - | 1 - | 7,20E-05 - | 0 - | 0 - |
| 1919 | 1 | 3um_FIJJAFBF_14773 | Bacteria | Bacteroidetes | Flavobacteriia | unclassified | unclassified | unclassified | Flavobacteria | information st | Replication, re | 0PB0H | DNA-binding p | 487796.Flav2A | 7,8 | 0,979795897 | 0,001180183 | 0,00013778 | 9 - | 0,001147768 - | 6 - | 0,000950126 - | 8 - | 0,001204286 - | 8 - | 0,001222074 - | 8 - | 0,001376662 - |
| 1920 | 1 | 0.2um_NOKCPHJN_250106 | Bacteria | Proteobacteri | Alphaproteob | Pelagibacteral | Pelagibacterac | Candidatus Pe | Candidatus Pe | metabolism | Inorganic ion t | 01Q7S | magnesium tr | 335992.SAR11 | 0,2 | 0,4 | 6,76E-06 | 1,35E-05 | 0 - | 0 - | 0 - | 0 - | 0 - | 0 - | 0 - | 0 - | 1 - | 3,38E-05 - |
| 1921 | 1 | 0.2um_NOKCPHJN_20493 | Bacteria | Proteobacteri | Gammaprotec | Cellvibrionale | Spongiibacter | unclassified | marine gamm | metabolism | Amino acid tra | COG0174 | glutamine syn | 247633.GP21 | 0,4 | 0,489897949 | 1,08E-05 | 1,33E-05 | 1 - | 2,45E-05 - | 0 - | 0 - | 0 - | 0 - | 1 - | 2,94E-05 - | 0 - | 0 - |
| 1922 | 1 | 0.2um_NOKCPHJN_06848 | Bacteria | Bacteroidetes | Cytophagia | Cytophagales | Flammeovirga | unclassified | Flammeovirga | unclassified | unclassified | unclassified | unclassified | unclassified | 0,8 | 0,4 | 0,000152682 | 7,84E-05 | 1 - | 0,000159412 - | 1 - | 0,000197943 - | 0 - | 0 - | 1 - | 0,000190949 - | 1 - | 0,000215103 - |
| 1923 | 1 | 3um_FIJJAFBF_426981 | Bacteria | Bacteroidetes | Flavobacteriia | Flavobacterial | Flavobacteriac | unclassified | Flavobacteriac | unclassified | unclassified | unclassified | unclassified | unclassified | 0,6 | 0,8 | 6,72E-05 | 9,32E-05 | 1 - | 9,89E-05 - | 0 - | 0 - | 0 - | 0 - | 2 - | 0,00023704 - | 0 - | 0 - |
| 1924 | 1 | 0.2um_NOKCPHJN_645530 | Bacteria | Proteobacteri | Gammaprotec | Pseudomonad | Pseudomonad | Pseudomonas | Pseudomonas | information st | Transcription | 05D27 | DNA-depende | 247634.GPB2 | 0,4 | 0,8 | 4,74E-06 | 9,47E-06 | 0 - | 0 - | 0 - | 0 - | 2 - | 2,37E-05 - | 0 - | 0 - | 0 - | 0 - |
| 1925 | 1 | 0.2um_NOKCPHJN_591591 | Eukaryota | unclassified | Oomycetes | Peronosporale | Peronosporac | Plasmopara | Plasmopara h | information st | Translation, ri | 14XM1 | EFG_IV | 67593.Physo1 | 0,6 | 0,8 | 9,34E-06 | 1,31E-05 | 1 - | 1,34E-05 - | 2 - | 3,33E-05 - | 0 - | 0 - | 0 - | 0 - | 0 - | 0 - |
| 1926 | 1 | 0.2um_NOKCPHJN_220959 | Bacteria | unclassified | unclassified | unclassified | unclassified | unclassified | uncultured ba | metabolism | Inorganic ion t | COG4206 | tonB-depende | 1046627.BZA | 1 | 0,632455532 | 2,41E-05 | 1,56E-05 | 1 - | 2,04E-05 - | 0 - | 0 - | 1 - | 2,40E-05 - | 2 - | 4,88E-05 - | 1 - | 2,75E-05 - |
| 1927 | 1 | 3um_FIJJAFBF_34372 | Bacteria | Proteobacteri | Gammaprotec | Oceanospirilla | Saccharospirill | Reinekea | Reinekea fors | unclassified | unclassified | unclassified | unclassified | unclassified | 0,2 | 0,4 | 8,41E-06 | 1,68E-05 | 1 - | 4,20E-05 - | 0 - | 0 - | 0 - | 0 - | 0 - | 0 - | 0 - | 0 - |
| 1928 | 1 | 0.2um_NOKCPHJN_114165 | Bacteria | Bacteroidetes | Flavobacteriia | Flavobacterial | Flavobacteriac | Formosa | Formosa sp. H | poorly charact | Function unkn | 06E9D | NA | 391598.FBBAL | 5,8 | 1,166190379 | 0,000137282 | 1,64E-05 | 8 - | 0,00016109 - | 6 - | 0,00015002 - | 5 - | 0,000118844 - | 5 - | 0,000120599 - | 5 - | 0,000135855 - |
| 1929 | 1 | 0.2um_NOKCPHJN_138294 | Bacteria | Bacteroidetes | Flavobacteriia | Flavobacterial | Flavobacteriac | unclassified | Flavobacteriac | unclassified | unclassified | unclassified | unclassified | unclassified | 2,4 | 0,489897949 | 0,000141934 | 3,78E-05 | 2 - | 9,81E-05 - | 2 - | 0,000121811 - | 3 - | 0,000173695 - | 2 - | 0,000117507 - | 3 - | 0,000198557 - |
| 1930 | 1 | 0.2um_NOKCPHJN_28609 | Bacteria | Bacteroidetes | Flavobacteriia | unclassified | unclassified | unclassified | Flavobacteria | information st | Translation, ri | 05C64 | Binds directly | 1046627.BZA | 0,4 | 0,489897949 | 2,55E-05 | 3,14E-05 | 0 - | 0 - | 0 - | 0 - | 0 - | 0 - | 1 - | 6,00E-05 - | 1 - | 6,76E-05 - |
| 1931 | 1 | 0.2um_NOKCPHJN_318481 | Bacteria | Proteobacteri | Alphaproteob | Rickettsiales | unclassified | unclassified | Rickettsiales b | information st | Transcription | 01R4Z | DNA-depende | 488538.SAR11 | 0,4 | 0,489897949 | 3,70E-06 | 4,58E-06 | 1 - | 8,26E-06 - | 1 - | 1,03E-05 - | 0 - | 0 - | 0 - | 0 - | 0 - | 0 - |
| 1932 | 1 | 0.2um_NOKCPHJN_67379 | Bacteria | Proteobacteri | Betaproteoba | unclassified | unclassified | unclassified | Betaproteoba | metabolism | Energy produc | COG1053 | Succinate DeH | 572477.Alvin_ | 1,4 | 0,489897949 | 3,07E-05 | 1,11E-05 | 2 - | 3,69E-05 - | 1 - | 2,29E-05 - | 1 - | 2,18E-05 - | 1 - | 2,21E-05 - | 2 - | 4,98E-05 - |
| 1933 | 1 | 3um_FIJJAFBF_06804 | Bacteria | Bacteroidetes | Flavobacteriia | Flavobacterial | Flavobacteriac | Cellulophaga | Cellulophaga | unclassified | unclassified | unclassified | unclassified | unclassified | 2 | 0 | 0,000101502 | 9,64E-06 | 2 - | 8,50E-05 - | 2 - | 0,00010557 - | 2 - | 0,000100357 - | 2 - | 0,000101839 - | 2 - | 0,000114722 - |
| 1934 | 1 | 3um_FIJJAFBF_667351 | Bacteria | Proteobacteri | Gammaprotec | Oceanospirilla | Saccharospirill | Saccharospiril | Saccharospiril | unclassified | unclassified | unclassified | unclassified | unclassified | 2 | 0 | 0,000141997 | 1,35E-05 | 2 - | 0,00011894 - | 2 - | 0,000147688 - | 2 - | 0,000140396 - | 2 - | 0,00014247 - | 2 - | 0,000160492 - |
| 1935 | 1 | 3um_FIJJAFBF_56137 | Bacteria | Bacteroidetes | Flavobacteriia | Flavobacterial | Flavobacteriac | Polaribacter | Polaribacter s | metabolism | Energy produc | COG0711 | F(1)F(0) ATP s | 313598.MED1 | 13,6 | 1,2 | 0,001122073 | 0,000137071 | 13 - | 0,000898854 - | 13 - | 0,001116112 - | 16 - | 0,001305853 - | 13 - | 0,001076676 - | 13 - | 0,001212872 - |
| 1936 | 1 | 0.2um_NOKCPHJN_521860 | unclassified | unclassified | unclassified | unclassified | unclassified | unclassified | unclassified | unclassified | unclassified | unclassified | unclassified | unclassified | 0,4 | 0,489897949 | 4,22E-05 | 5,17E-05 | 0 - | 0 - | 1 - | 0,000101077 - | 0 - | 0 - | 0 - | 0 - | 1 - | 0,00010984 - |
| 1937 | 1 | 0.2um_NOKCPHJN_122841 | Bacteria | Bacteroidetes | Flavobacteriia | unclassified | unclassified | unclassified | Flavobacteria | unclassified | unclassified | unclassified | unclassified | unclassified | 1,2 | 0,748331477 | 3,88E-05 | 2,45E-05 | 1 - | 2,73E-05 - | 0 - | 0 - | 2 - | 6,45E-05 - | 2 - | 6,55E-05 - | 1 - | 3,69E-05 - |
| 1938 | 1 | 3um_FIJJAFBF_662769 | Eukaryota | Bacillariophyta | Coscinodiscop | Thalassiosirale | Thalassiosirac | Thalassiosira | Thalassiosira | unclassified | unclassified | unclassified | unclassified | unclassified | 0,4 | 0,8 | 3,35E-05 | 6,70E-05 | 2 - | 0,000167557 - | 0 - | 0 - | 0 - | 0 - | 0 - | 0 - | 0 - | 0 - |
| 1939 | 1 | 3um_FIJJAFBF_287052 | Bacteria | Proteobacteri | Gammaprotec | Cellvibrionale | Porticoccacea | unclassified | Gammaprotec | unclassified | unclassified | unclassified | unclassified | unclassified | 0,4 | 0,489897949 | 7,03E-06 | 8,61E-06 | 0 - | 0 - | 0 - | 0 - | 1 - | 1,74E-05 - | 1 - | 1,77E-05 - | 0 - | 0 - |
| 1940 | 1 | 0.2um_NOKCPHJN_646842 | Eukaryota | Eustigmatoph | unclassified | Eustigmatales | Monodopsida | Nannochlorop | Nannochlorop | unclassified | unclassified | unclassified | unclassified | unclassified | 0,6 | 0,489897949 | 6,07E-05 | 4,96E-05 | 0 - | 0 - | 1 - | 0,000104028 - | 1 - | 9,89E-05 - | 1 - | 0,000100353 - | 0 - | 0 - |
| 1941 | 1 | 0.2um_NOKCPHJN_278438 | Bacteria | Proteobacteri | Alphaproteob | unclassified | unclassified | unclassified | Alphaproteob | unclassified | unclassified | unclassified | unclassified | unclassified | 0,4 | 0,489897949 | 5,54E-05 | 6,84E-05 | 1 - | 0,000126128 - | 0 - | 0 - | 0 - | 0 - | 1 - | 0,000151081 - | 0 - | 0 - |
| 1942 | 1 | 0.2um_NOKCPHJN_63758 | Bacteria | Bacteroidetes | Flavobacteriia | Flavobacterial | Crocinitomica | unclassified | Crocinitomica | metabolism | Inorganic ion t | 05C27 | TonB-depende | 470145.BACC | 1,8 | 0,748331477 | 2,35E-05 | 7,90E-06 | 3 - | 3,39E-05 - | 2 - | 2,80E-05 - | 1 - | 1,33E-05 - | 2 - | 2,70E-05 - | 1 - | 1,52E-05 - |
| 1943 | 1 | 0.2um_NOKCPHJN_167891 | Bacteria | Bacteroidetes | Flavobacteriia | Flavobacterial | Flavobacteriac | Formosa | Formosa sp. H | unclassified | unclassified | unclassified | unclassified | unclassified | 0,8 | 0,748331477 | 6,07E-05 | 5,27E-05 | 2 - | 0,000133461 - | 0 - | 0 - | 0 - | 0 - | 1 - | 7,99E-05 - | 1 - | 9,00E-05 - |
| 1944 | 1 | 3um_FIJJAFBF_43033 | unclassified | unclassified | unclassified | unclassified | unclassified | unclassified | unclassified | unclassified | unclassified | unclassified | unclassified | unclassified | 0,2 | 0,4 | 1,69E-05 | 3,38E-05 | 1 - | 8,44E-05 - | 0 - | 0 - | 0 - | 0 - | 0 - | 0 - | 0 - | 0 - |
| 1945 | 1 | 0.2um_NOKCPHJN_15732 | Bacteria | Proteobacteri | Gammaprotec | Alteromonada | Alteromonada | Paraglaciecola | Paraglaciecola | metabolism | Energy produc | 0QIAX | dihydrolipoyl | 983545.Glaag | 0,4 | 0,489897949 | 1,12E-05 | 1,39E-05 | 1 - | 2,38E-05 - | 0 - | 0 - | 0 - | 0 - | 0 - | 0 - | 1 - | 3,21E-05 - |
| 1946 | 1 | 0.2um_NOKCPHJN_539312 | unclassified | unclassified | unclassified | unclassified | unclassified | unclassified | unclassified | unclassified | unclassified | unclassified | unclassified | unclassified | 3 | 1,264911064 | 0,00017312 | 6,72E-05 | 4 - | 0,000199612 - | 2 - | 0,000123929 - | 4 - | 0,000235621 - | 4 - | 0,000239101 - | 1 - | 6,73E-05 - |
| 1947 | 1 | 3um_FIJJAFBF_720178 | Bacteria | Proteobacteri | Alphaproteob | Sphingomona | Erythrobacter | Porphyrobact | Porphyrobact | information st | Translation, ri | COG0050 | This protein p | 237727.NAP1_ | 1 | 0,632455532 | 3,57E-05 | 2,20E-05 | 0 - | 0 - | 1 - | 3,60E-05 - | 1 - | 3,42E-05 - | 2 - | 6,94E-05 - | 1 - | 3,91E-05 - |
| 1948 | 1 | 3um_FIJJAFBF_550914 | Eukaryota | Bacillariophyta | Coscinodiscop | Thalassiosirale | Thalassiosirac | Thalassiosira | Thalassiosira | unclassified | unclassified | unclassified | unclassified | unclassified | 4,2 | 0,4 | 0,000714918 | 0,000129298 | 4 - | 0,000566799 - | 4 - | 0,000703797 - | 4 - | 0,000669048 - | 4 - | 0,00067893 - | 5 - | 0,000956015 - |
| 1949 | 1 | 0.2um_NOKCPHJN_481276 | Bacteria | Proteobacteri | Alphaproteob | Pelagibacteral | Pelagibacterac | unclassified | SAR11 cluster | unclassified | unclassified | unclassified | unclassified | unclassified | 3 | 1,549193338 | 0,000116953 | 6,05E-05 | 3 - | 9,89E-05 - | 2 - | 8,19E-05 - | 2 - | 7,79E-05 - | 6 - | 0,00023704 - | 2 - | 8,90E-05 - |
| 1950 | 2 | 0.2um_NOKCPHJN_12932;0.2um_NOKCPHJN_21 | Bacteria | Bacteroidetes | Flavobacteriia | Flavobacterial | Flavobacteriac | various | various | metabolism | Amino acid tra | COG0174 | glutamine syn | various | 0,6 | 0,8 | 2,43E-05 | 3,21E-05 | 0 - | 0 - | 1 - | 4,19E-05 - | 2 - | 7,97E-05 - | 0 - | 0 - | 0 - | 0 - |
| 1951 | 1 | 0.2um_NOKCPHJN_1090264 | Eukaryota | Phaeophyceae | unclassified | Ectocarpales | Ectocarpaceae | Ectocarpus | Ectocarpus sil | cellular proces | Cytoskeleton | 0DFD2 | Kinesin family | 31033.ENSTR | 0,6 | 0,8 | 2,16E-05 | 2,94E-05 | 0 - | 0 - | 1 - | 3,41E-05 - | 0 - | 0 - | 0 - | 0 - | 2 - | 7,41E-05 - |
| 1952 | 1 | 3um_FIJJAFBF_804851 | Bacteria | Proteobacteri | Betaproteoba | unclassified | unclassified | unclassified | Betaproteoba | metabolism | Inorganic ion t | 01V15 | trap transport | 1064539.AZO | 6,2 | 1,469693846 | 0,000262763 | 3,68E-05 | 9 - | 0,000325865 - | 6 - | 0,000269752 - | 5 - | 0,000213694 - | 6 - | 0,000260221 - | 5 - | 0,000244282 - |
| 1953 | 1 | 3um_FIJJAFBF_115637 | Eukaryota | unclassified | Oomycetes | Saprolegniales | Saprolegniace | Aphanomyces | Aphanomyces | poorly charact | Function unkn | 0XQGZ | guanine nucle | 436017.A4S6 | 0,6 | 0,489897949 | 2,22E-05 | 1,85E-05 | 1 - | 3,14E-05 - | 0 - | 0 - | 1 - | 3,71E-05 - | 0 - | 0 - | 1 - | 4,24E-05 - |
| 1954 | 4 | 3um_FIJJAFBF_215759;3um_FIJJAFBF_274939;3u | Bacteria | Bacteroidetes | Flavobacteriia | Flavobacterial | Flavobacteriac | Polaribacter | various | poorly charact | Function unkn | 08WMU | Domain of unl | 313594.PI23P | 2 | 0 | 0,000206056 | 1,96E-05 | 2 - | 0,000172597 - | 2 - | 0,000214314 - | 2 - | 0,000203733 - | 2 - | 0,000206742 - | 2 - | 0,000232894 - |
| 1955 | 1 | 3um_FIJJAFBF_682052 | Bacteria | Bacteroidetes | Flavobacteriia | Flavobacterial | Flavobacteriac | Polaribacter | Polaribacter s | metabolism | Lipid transpor | 0P83H | phosphatidyls | 313594.PI23P | 1,2 | 1,166190379 | 7,10E-05 | 7,02E-05 | 2 - | 0,0001053 - | 0 - | 0 - | 3 - | 0,000186443 - | 1 - | 6,31E-05 - | 0 - | 0 - |
| 1956 | 1 | 3um_FIJJAFBF_24509 | Bacteria | Proteobacteri | Alphaproteob | Rhodobactera | unclassified | unclassified | Rhodobactera | unclassified | unclassified | unclassified | unclassified | unclassified | 5,2 | 1,166190379 | 0,000213726 | 6,51E-05 | 4 - | 0,00013543 - | 5 - | 0,000210205 - | 4 - | 0,000159861 - | 6 - | 0,000243333 - | 7 - | 0,0003198 - |
| 1957 | 1 | 3um_FIJJAFBF_741500 | Bacteria | Bacteroidetes | unclassified | unclassified | unclassified | unclassified | Bacteroidetes | cellular proces | Intracellular tr | 0PBE9 | Preprotein tra | 926562.Oweh | 1,4 | 0,8 | 0,000180001 | 0,000101402 | 2 - | 0,000225052 - | 1 - | 0,000139724 - | 2 - | 0,000265651 - | 2 - | 0,000269575 - | 0 - | 0 - |
| 1958 | 1 | 0.2um_NOKCPHJN_900264 | Bacteria | Proteobacteri | Gammaprotec | unclassified | unclassified | unclassified | OM182 bacter | poorly charact | Function unkn | 16R8I | Lipoprotein | 394221.Mmar | 4 | 1,095445115 | 0,000111363 | 1,87E-05 | 6 - | 0,000143471 - | 3 - | 8,91E-05 - | 4 - | 0,000112902 - | 4 - | 0,000114569 - | 3 - | 9,68E-05 - |
| 1959 | 1 | 0.2um_NOKCPHJN_91466 | Bacteria | Proteobacteri | Gammaprotec | Oceanospirilla | Saccharospirill | Reinekea | Reinekea fors | information st | Translation, ri | COG0251 | Endoribonucle | 314283.MED2 | 3 | 0 | 0,000323686 | 3,07E-05 | 3 - | 0,000271126 - | 3 - | 0,000336659 - | 3 - | 0,000320037 - | 3 - | 0,000324764 - | 3 - | 0,000365845 - |
| 1960 | 1 | 3um_FIJJAFBF_452605 | Bacteria | Proteobacteri | Alphaproteob | Sphingomona | Erythrobacter | Altererythrob | Altererythrob | cellular proces | Signal transdu | 01WWM | Histidine kina | 314225.ELI_1 | 0,2 | 0,4 | 6,12E-06 | 1,22E-05 | 0 - | 0 - | 0 - | 0 - | 0 - | 0 - | 0 - | 0 - | 1 - | 3,06E-05 - |
| 1961 | 1 | 0.2um_NOKCPHJN_04208 | Bacteria | Bacteroidetes | Cytophagia | Cytophagales | Cytophagacea | Ekhidna | Ekhidna lutea | information st | Replication, re | 05C24 | DNA gyrase n | 643867.Ftrac_ | 0,4 | 0,489897949 | 6,90E-06 | 8,48E-06 | 0 - | 0 - | 0 - | 0 - | 0 - | 0 - | 1 - | 1,62E-05 - | 1 - | 1,83E-05 - |
| 1962 | 1 | 3um_FIJJAFBF_241619 | Bacteria | Proteobacteri | Alphaproteob | Rhodospirillal | Rhodospirillac | Candidatus En | Candidatus En | unclassified | unclassified | unclassified | unclassified | unclassified | 0,2 | 0,4 | 2,45E-06 | 4,91E-06 | 0 - | 0 - | 0 - | 0 - | 0 - | 0 - | 0 - | 0 - | 1 - | 1,23E-05 - |
| 1963 | 1 | 3um_FIJJAFBF_182748 | Viruses | unclassified | unclassified | unclassified | various | various | various | unclassified | unclassified | unclassified | unclassified | unclassified | 0,2 | 0,4 | 1,23E-05 | 2,47E-05 | 1 - | 6,17E-05 - | 0 - | 0 - | 0 - | 0 - | 0 - | 0 - | 0 - | 0 - |
| 1964 | 1 | 0.2um_NOKCPHJN_964069 | unclassified | unclassified | unclassified | unclassified | unclassified | unclassified | unclassified | unclassified | unclassified | unclassified | unclassified | unclassified | 2,6 | 0,489897949 | 0,000137202 | 2,23E-05 | 3 - | 0,000133981 - | 3 - | 0,000166364 - | 3 - | 0,00015815 - | 2 - | 0,000106991 - | 2 - | 0,000120525 - |
| 1965 | 1 | 0.2um_NOKCPHJN_156216 | Bacteria | Proteobacteri | Gammaprotec | Cellvibrionale | Porticoccacea | unclassified | various | unclassified | unclassified | unclassified | unclassified | unclassified | 0,4 | 0,489897949 | 7,80E-05 | 9,58E-05 | 0 - | 0 - | 0 - | 0 - | 0 - | 0 - | 1 - | 0,000183311 - | 1 - | 0,000206499 - |
| 1966 | 1 | 3um_FIJJAFBF_650081 | Bacteria | Proteobacteri | Alphaproteob | Rhodobactera | Rhodobactera | Planktomarina | Planktomarina | unclassified | unclassified | unclassified | unclassified | unclassified | 1,2 | 0,4 | 0,000130339 | 4,81E-05 | 1 - | 9,04E-05 - | 2 - | 0,000224439 - | 1 - | 0,000106679 - | 1 - | 0,000108255 - | 1 - | 0,000121948 - |
| 1967 | 1 | 0.2um_NOKCPHJN_449708 | Bacteria | Proteobacteri | Alphaproteob | Rhodobactera | Rhodobactera | Yoonia | Yoonia tamler | metabolism | Lipid transpor | 01R4W | acetyl-coa ace | 314232.SKA53 | 0,2 | 0,4 | 6,93E-06 | 1,39E-05 | 0 - | 0 - | 0 - | 0 - | 1 - | 3,47E-05 - | 0 - | 0 - | 0 - | 0 - |
| 1968 | 2 | 0.2um_NOKCPHJN_649050;0.2um_NOKCPHJN_6 | unclassified | unclassified | unclassified | unclassified | unclassified | unclassified | unclassified | unclassified | unclassified | unclassified | unclassified | unclassified | 0,2 | 0,4 | 1,24E-05 | 2,47E-05 | 0 - | 0 - | 0 - | 0 - | 1 - | 6,19E-05 - | 0 - | 0 - | 0 - | 0 - |
| 1969 | 2 | 0.2um_NOKCPHJN_148411;3um_FIJJAFBF_5854 | Bacteria | Proteobacteri | various | various | various | various | various | unclassified | unclassified | unclassified | unclassified | unclassified | 0,2 | 0,4 | 1,10E-05 | 2,21E-05 | 1 - | 5,52E-05 - | 0 - | 0 - | 0 - | 0 - | 0 - | 0 - | 0 - | 0 - |
| 1970 | 1 | 0.2um_NOKCPHJN_245227 | Bacteria | unclassified | unclassified | unclassified | unclassified | unclassified | uncultured ba | poorly charact | Function unkn | 16YV3 | Large repetiti | 640513.Entas_ | 1 | 0,894427191 | 3,40E-05 | 2,99E-05 | 2 - | 6,07E-05 - | 1 - | 3,77E-05 - | 2 - | 7,17E-05 - | 0 - | 0 - | 0 - | 0 - |
| 1971 | 1 | 0.2um_NOKCPHJN_64539 | Bacteria | Proteobacteri | Gammaprotec | Oceanospirilla | Saccharospirill | Saccharospiril | Saccharospiril | unclassified | unclassified | unclassified | unclassified | unclassified | 1,4 | 0,8 | 4,76E-05 | 2,70E-05 | 1 - | 2,86E-05 - | 1 - | 3,55E-05 - | 3 - | 0,000101106 - | 1 - | 3,42E-05 - | 1 - | 3,85E-05 - |
| 1972 | 1 | 3um_FIJJAFBF_260209 | Bacteria | Bacteroidetes | Flavobacteriia | Flavobacterial | Flavobacteriac | Ulvibacter | Ulvibacter sp. | unclassified | unclassified | unclassified | unclassified | unclassified | 0,4 | 0,489897949 | 4,44E-05 | 5,44E-05 | 0 - | 0 - | 0 - | 0 - | 1 - | 0,000110148 - | 1 - | 0,000111775 - | 0 - | 0 - |
| 1973 | 1 | 3um_FIJJAFBF_91795 | Bacteria | Candidatus Sa | unclassified | unclassified | unclassified | unclassified | Candidatus Sa | poorly charact | Function unkn | 0ZMXK | NA | 292414.TM10 | 0,4 | 0,489897949 | 7,16E-06 | 8,78E-06 | 0 - | 0 - | 1 - | 1,72E-05 - | 0 - | 0 - | 0 - | 0 - | 1 - | 1,86E-05 - |
| 1974 | 1 | 0.2um_NOKCPHJN_522906 | Bacteria | Proteobacteri | Alphaproteob | Pelagibacteral | Pelagibacterac | Candidatus Pe | Candidatus Pe | metabolism | Secondary me | 01Q86 | extracellular s | 335992.SAR11 | 3,2 | 0,748331477 | 0,000122393 | 3,64E-05 | 3 - | 9,51E-05 - | 4 - | 0,000157479 - | 3 - | 0,000112278 - | 2 - | 7,60E-05 - | 4 - | 0,000171132 - |
| 1975 | 1 | 0.2um_NOKCPHJN_699107 | Bacteria | Proteobacteri | Alphaproteob | Pelagibacteral | Pelagibacterac | Candidatus Pe | Candidatus Pe | metabolism | Amino acid tra | 01R4A | Polyamine AB | 439493.PB721 | 1,6 | 0,489897949 | 5,94E-05 | 2,09E-05 | 1 - | 3,06E-05 - | 1 - | 3,80E-05 - | 2 - | 7,23E-05 - | 2 - | 7,33E-05 - | 2 - | 8,26E-05 - |
| 1976 | 1 | 3um_FIJJAFBF_31273 | Bacteria | Proteobacteri | Gammaprotec | Alteromonada | Alteromonada | Paraglaciecola | Paraglaciecola | unclassified | unclassified | unclassified | unclassified | unclassified | 0,2 | 0,4 | 1,41E-05 | 2,82E-05 | 0 - | 0 - | 1 - | 7,06E-05 - | 0 - | 0 - | 0 - | 0 - | 0 - | 0 - |
| 1977 | 1 | 3um_FIJJAFBF_165020 | Bacteria | Proteobacteri | Deltaproteoba | Desulfuromon | Geobacterace | Geothermoba | Geothermoba | information st | Translation, ri | COG0052 | 30S ribosomal | 338963.Pcar_ | 0,2 | 0,4 | 8,35E-06 | 1,67E-05 | 1 - | 4,17E-05 - | 0 - | 0 - | 0 - | 0 - | 0 - | 0 - | 0 - | 0 - |
| 1978 | 1 | 3um_FIJJAFBF_213857 | Bacteria | Proteobacteri | Gammaprotec | Oceanospirilla | Saccharospirill | Reinekea | Reinekea fors | unclassified | unclassified | unclassified | unclassified | unclassified | 1 | 0 | 0,000133036 | 1,26E-05 | 1 - | 0,000111434 - | 1 - | 0,000138368 - | 1 - | 0,000131536 - | 1 - | 0,000133479 - | 1 - | 0,000150364 - |
| 1979 | 1 | 0.2um_NOKCPHJN_172111 | Bacteria | Bacteroidetes | Flavobacteriia | Flavobacterial | Flavobacteriac | Formosa | Formosa sp. H | cellular proces | Intracellular tr | 0PBE9 | Preprotein tra | 1046627.BZA | 0,8 | 0,4 | 0,000114511 | 5,88E-05 | 1 - | 0,000119559 - | 1 - | 0,000148457 - | 0 - | 0 - | 1 - | 0,000143212 - | 1 - | 0,000161328 - |
| 1980 | 1 | 0.2um_NOKCPHJN_95021 | Bacteria | Bacteroidetes | Flavobacteriia | Flavobacterial | Flavobacteriac | Psychroserper | Psychroserper | information st | Translation, ri | COG0050 | This protein p | 688270.Celal_ | 5,8 | 0,748331477 | 0,000200046 | 2,45E-05 | 6 - | 0,000174344 - | 5 - | 0,000180404 - | 6 - | 0,000205796 - | 7 - | 0,000243641 - | 5 - | 0,000196044 - |
| 1981 | 1 | 0.2um_NOKCPHJN_806856 | Bacteria | Proteobacteri | Alphaproteob | Rhodobactera | unclassified | unclassified | Rhodobactera | metabolism | Amino acid tra | 16PKZ | extracellular li | 367336.OM22 | 0,4 | 0,489897949 | 1,57E-05 | 1,92E-05 | 0 - | 0 - | 1 - | 3,75E-05 - | 0 - | 0 - | 0 - | 0 - | 1 - | 4,08E-05 - |
| 1982 | 3 | 0.2um_NOKCPHJN_287646;0.2um_NOKCPHJN_6 | Bacteria | Proteobacteri | Alphaproteob | Rhodobactera | Rhodobactera | Planktomarina | Planktomarina | unclassified | unclassified | unclassified | unclassified | unclassified | 1,4 | 0,489897949 | 0,000111407 | 3,94E-05 | 1 - | 6,67E-05 - | 1 - | 8,29E-05 - | 2 - | 0,000157537 - | 2 - | 0,000159864 - | 1 - | 9,00E-05 - |
| 1983 | 1 | 3um_FIJJAFBF_119804 | Viruses | unclassified | unclassified | unclassified | Phycodnavirid | unclassified | Chrysochromu | information st | Transcription | 12NXR | DNA-depende | 486041.XP_00 | 0,8 | 0,4 | 5,43E-05 | 2,73E-05 | 0 - | 0 - | 1 - | 6,79E-05 - | 1 - | 6,45E-05 - | 1 - | 6,55E-05 - | 1 - | 7,37E-05 - |
| 1984 | 1 | 0.2um_NOKCPHJN_1093443 | Bacteria | Proteobacteri | Alphaproteob | Pelagibacteral | Pelagibacterac | Candidatus Pe | Candidatus Pe | unclassified | unclassified | unclassified | unclassified | unclassified | 0,4 | 0,489897949 | 1,15E-05 | 1,41E-05 | 0 - | 0 - | 1 - | 2,76E-05 - | 0 - | 0 - | 0 - | 0 - | 1 - | 3,00E-05 - |
| 1985 | 1 | 3um_FIJJAFBF_165290 | Bacteria | Proteobacteri | Gammaprotec | Oceanospirilla | Oceanospirilla | Motiliproteus | Motiliproteus | unclassified | unclassified | unclassified | unclassified | unclassified | 0,8 | 0,4 | 1,62E-05 | 8,21E-06 | 1 - | 1,75E-05 - | 1 - | 2,17E-05 - | 1 - | 2,07E-05 - | 1 - | 2,10E-05 - | 0 - | 0 - |
| 1986 | 1 | 3um_FIJJAFBF_387408 | Bacteria | Bacteroidetes | Flavobacteriia | Flavobacterial | Flavobacteriac | Polaribacter | Polaribacter s | unclassified | unclassified | unclassified | unclassified | unclassified | 1 | 0,632455532 | 1,99E-05 | 1,14E-05 | 2 - | 3,55E-05 - | 1 - | 2,20E-05 - | 1 - | 2,09E-05 - | 1 - | 2,12E-05 - | 0 - | 0 - |
| 1987 | 1 | 3um_FIJJAFBF_549918 | Bacteria | Proteobacteri | Gammaprotec | Alteromonada | Alteromonada | Paraglaciecola | Paraglaciecola | poorly charact | Function unkn | 061D9 | Protein with u | 983545.Glaag | 0,6 | 0,489897949 | 7,27E-05 | 6,04E-05 | 1 - | 0,000102479 - | 0 - | 0 - | 0 - | 0 - | 1 - | 0,000122753 - | 1 - | 0,000138281 - |
| 1988 | 1 | 3um_FIJJAFBF_769961 | Bacteria | Bacteroidetes | Flavobacteriia | Flavobacterial | Flavobacteriac | Polaribacter | Polaribacter h | metabolism | Amino acid tra | COG2171 | Catalyzes the | 313598.MED1 | 1 | 0,894427191 | 4,88E-05 | 4,52E-05 | 1 - | 4,24E-05 - | 0 - | 0 - | 2 - | 1,00E-04 - | 2 - | 0,000101464 - | 0 - | 0 - |
| 1989 | 1 | 3um_FIJJAFBF_27717 | Bacteria | Actinobacteria | Actinobacteria | Micrococcales | Microbacteria | unclassified | Microbacteria | information st | Translation, ri | 08VZM | 50s ribosomal | 31964.CMS_0 | 5,8 | 1,326649916 | 0,000457423 | 6,03E-05 | 8 - | 0,000540126 - | 5 - | 0,000419173 - | 6 - | 0,000478173 - | 6 - | 0,000485235 - | 4 - | 0,000364411 - |
| 1990 | 1 | 3um_FIJJAFBF_607174 | Bacteria | Proteobacteri | Gammaprotec | Oceanospirilla | Oceanospirilla | Motiliproteus | Motiliproteus | information st | Translation, ri | 179R6 | This protein is | 290398.Csal_ | 1,6 | 0,8 | 0,00015285 | 7,85E-05 | 2 - | 0,000161657 - | 0 - | 0 - | 2 - | 0,00019082 - | 2 - | 0,000193638 - | 2 - | 0,000218133 - |
| 1991 | 1 | 0.2um_NOKCPHJN_11270 | Bacteria | Proteobacteri | Alphaproteob | unclassified | unclassified | unclassified | SAR116 cluste | unclassified | unclassified | unclassified | unclassified | unclassified | 1 | 0 | 7,66E-05 | 7,27E-06 | 1 - | 6,41E-05 - | 1 - | 7,96E-05 - | 1 - | 7,57E-05 - | 1 - | 7,68E-05 - | 1 - | 8,65E-05 - |
| 1992 | 1 | 3um_FIJJAFBF_39072 | Bacteria | Proteobacteri | Alphaproteob | Rhodobactera | Rhodobactera | Planktomarina | Planktomarina | unclassified | unclassified | unclassified | unclassified | unclassified | 2,6 | 1,019803903 | 0,000251578 | 9,56E-05 | 3 - | 0,00024595 - | 3 - | 0,000305398 - | 4 - | 0,000387092 - | 1 - | 9,82E-05 - | 2 - | 0,000221249 - |
| 1993 | 1 | 0.2um_NOKCPHJN_343613 | Bacteria | Proteobacteri | Gammaprotec | Alteromonada | Colwelliaceae | Colwellia | Colwellia myti | unclassified | unclassified | unclassified | unclassified | unclassified | 0,2 | 0,4 | 5,03E-06 | 1,01E-05 | 0 - | 0 - | 0 - | 0 - | 0 - | 0 - | 1 - | 2,51E-05 - | 0 - | 0 - |
| 1994 | 1 | 3um_FIJJAFBF_291694 | Bacteria | Proteobacteri | Alphaproteob | Pelagibacteral | Pelagibacterac | Candidatus Pe | Candidatus Pe | unclassified | unclassified | unclassified | unclassified | unclassified | 5 | 1,095445115 | 0,000220216 | 5,34E-05 | 6 - | 0,000221434 - | 5 - | 0,00022913 - | 3 - | 0,00013069 - | 5 - | 0,000221034 - | 6 - | 0,000298793 - |
| 1995 | 1 | 3um_FIJJAFBF_269230 | Bacteria | Bacteroidetes | Flavobacteriia | Flavobacterial | Flavobacteriac | Polaribacter | Polaribacter s | unclassified | unclassified | unclassified | unclassified | unclassified | 1,6 | 0,489897949 | 2,69E-05 | 8,88E-06 | 2 - | 2,81E-05 - | 2 - | 3,49E-05 - | 1 - | 1,66E-05 - | 1 - | 1,68E-05 - | 2 - | 3,79E-05 - |
| 1996 | 1 | 3um_FIJJAFBF_405813 | unclassified | unclassified | unclassified | unclassified | unclassified | unclassified | unclassified | unclassified | unclassified | unclassified | unclassified | unclassified | 1,8 | 0,4 | 0,000194881 | 4,56E-05 | 2 - | 0,000182185 - | 1 - | 0,00011311 - | 2 - | 0,000215051 - | 2 - | 0,000218227 - | 2 - | 0,000245833 - |

| | | | | | | | | | | | | | | | | | | | | | | | | | | | | |
|---|---|---|---|---|---|---|---|---|---|---|---|---|---|---|---|---|---|---|---|---|---|---|---|---|---|---|---|---|
| 1997 | 1 | 0.2um_NOKCPHJN_665338 | Bacteria | Proteobacteri | Gammaprote | Cellvibrionale | Spongiibacter | unclassified | marine gamm | metabolism | Inorganic ion t | 05F28 | tonB-depende | 582402.Hbal_ | 0,6 | 0,489897949 | 1,21E-05 | 9,94E-06 | 1 - | 1,79E-05 - | 0 - | 0 - | 1 - | 2,11E-05 - | 1 - | 2,14E-05 - | 0 - | 0 - |
| 1998 | 1 | 0.2um_NOKCPHJN_1058181 | Bacteria | Proteobacteri | Alphaproteob | Rhodospirillal | Rhodospirillac | Magnetovibri | Magnetovibri | metabolism | Carbohydrate | 05CV1 | Transketolase | 342108.amb1 | 1 | 0 | 2,60E-05 | 2,46E-06 | 1 - | 2,17E-05 - | 1 - | 2,70E-05 - | 1 - | 2,57E-05 - | 1 - | 2,60E-05 - | 1 - | 2,93E-05 - |
| 1999 | 1 | 0.2um_NOKCPHJN_56878 | Bacteria | Proteobacteri | Gammaprote | Oceanospirilla | Saccharospirill | Reinekea | Reinekea blan | metabolism | Energy produc | 0QIAU | Electron trans | 314283.MED2 | 0,6 | 0,489897949 | 3,14E-05 | 2,59E-05 | 1 - | 4,59E-05 - | 1 - | 5,70E-05 - | 1 - | 5,42E-05 - | 0 - | 0 - | 0 - | 0 - |
| 2000 | 1 | 3um_FIJJAFBF_350105 | Bacteria | Proteobacteri | Alphaproteob | Rhodobactera | unclassified | unclassified | Rhodobactera | cellular proces | Posttranslatio | 16PM6 | feS assembly | 391619.RGBS | 0,8 | 0,4 | 4,32E-05 | 2,22E-05 | 1 - | 4,57E-05 - | 0 - | 0 - | 1 - | 5,40E-05 - | 1 - | 5,48E-05 - | 1 - | 6,17E-05 - |
| 2001 | 1 | 3um_FIJJAFBF_15435 | Bacteria | Proteobacteri | Gammaprote | Pseudomonad | Moraxellaceae | Psychrobacter | various | unclassified | unclassified | unclassified | unclassified | unclassified | 0,2 | 0,4 | 2,55E-05 | 5,09E-05 | 0 - | 0 - | 0 - | 0 - | 0 - | 0 - | 1 - | 0,000127299 - | 0 - | 0 - |
| 2002 | 1 | 3um_FIJJAFBF_673183 | Bacteria | Proteobacteri | Gammaprote | Alteromonada | Colwelliaceae | Colwellia | Colwellia sp. F | cellular proces | Posttranslatio | COG0330 | Band 7 protei | 575788.VS_II | 0,2 | 0,4 | 7,60E-06 | 1,52E-05 | 1 - | 3,80E-05 - | 0 - | 0 - | 0 - | 0 - | 0 - | 0 - | 0 - | 0 - |
| 2003 | 1 | 0.2um_NOKCPHJN_174554 | Bacteria | Bacteroidetes | Flavobacteriia | Flavobacterial | Crocinitomica | unclassified | Crocinitomica | poorly charact | Function unkn | 09AR5 | Inherit from b | 313606.M231 | 0,6 | 0,489897949 | 1,41E-05 | 1,16E-05 | 1 - | 2,09E-05 - | 0 - | 0 - | 1 - | 2,47E-05 - | 1 - | 2,50E-05 - | 0 - | 0 - |
| 2004 | 1 | 3um_FIJJAFBF_436413 | Bacteria | Proteobacteri | Alphaproteob | Rhodobactera | Rhodobactera | Yoonia | Yoonia sedimi | poorly charact | Function unkn | 173PW | Pfam:DUF88 | 391593.RCCS | 2,4 | 0,489897949 | 0,000169899 | 2,65E-05 | 3 - | 0,000180278 - | 2 - | 0,000149234 - | 2 - | 0,000141866 - | 3 - | 0,000215942 - | 2 - | 0,000162172 - |
| 2005 | 1 | 3um_FIJJAFBF_294668 | Bacteria | Bacteroidetes | Flavobacteriia | Flavobacterial | Flavobacteriac | Algibacter | Algibacter sp. | unclassified | unclassified | unclassified | unclassified | unclassified | 5,8 | 0,748331477 | 0,000460158 | 8,00E-05 | 6 - | 0,00039807 - | 7 - | 0,000576666 - | 5 - | 0,000391567 - | 5 - | 0,000397351 - | 6 - | 0,000537137 - |
| 2006 | 1 | 0.2um_NOKCPHJN_700347 | Eukaryota | Eustigmatoph | unclassified | Eustigmatales | Monodopsida | Nannochlorop | Nannochlorop | metabolism | Carbohydrate | KOG2638 | UTP-glucose-1 | 653948.CCA1 | 10 | 1,673320053 | 0,000149052 | 2,44E-05 | 10 - | 0,000125714 - | 11 - | 0,000171709 - | 10 - | 0,000148392 - | 12 - | 0,000180701 - | 7 - | 0,000118743 - |
| 2007 | 1 | 3um_FIJJAFBF_222380 | Bacteria | Proteobacteri | Alphaproteob | Rhodobactera | unclassified | unclassified | Rhodobactera | unclassified | unclassified | unclassified | unclassified | unclassified | 0,2 | 0,4 | 1,25E-05 | 2,50E-05 | 1 - | 6,24E-05 - | 0 - | 0 - | 0 - | 0 - | 0 - | 0 - | 0 - | 0 - |
| 2008 | 1 | 3um_FIJJAFBF_656627 | Bacteria | Proteobacteri | Alphaproteob | Rhodobactera | Rhodobactera | Loktanella | Loktanella sp. | information st | Transcription | COG0250 | Participates in | 391593.RCCS | 1,4 | 0,489897949 | 0,000108499 | 3,93E-05 | 2 - | 0,000130428 - | 1 - | 8,10E-05 - | 1 - | 7,70E-05 - | 1 - | 7,81E-05 - | 2 - | 0,000175994 - |
| 2009 | 1 | 0.2um_NOKCPHJN_739339 | Bacteria | Proteobacteri | Alphaproteob | Pelagibacteral | Pelagibacterac | Candidatus Pe | Candidatus Pe | unclassified | unclassified | unclassified | unclassified | unclassified | 0,8 | 0,4 | 3,79E-05 | 1,95E-05 | 1 - | 4,01E-05 - | 0 - | 0 - | 1 - | 4,74E-05 - | 1 - | 4,81E-05 - | 1 - | 5,42E-05 - |
| 2010 | 1 | 0.2um_NOKCPHJN_257178 | Bacteria | Bacteroidetes | unclassified | unclassified | unclassified | unclassified | Bacteroidetes | unclassified | unclassified | unclassified | unclassified | unclassified | 1 | 0 | 2,16E-05 | 2,05E-06 | 1 - | 1,81E-05 - | 1 - | 2,24E-05 - | 1 - | 2,13E-05 - | 1 - | 2,17E-05 - | 1 - | 2,44E-05 - |
| 2011 | 1 | 0.2um_NOKCPHJN_83677 | Bacteria | Bacteroidetes | Flavobacteriia | various | various | various | various | unclassified | unclassified | unclassified | unclassified | unclassified | 0,6 | 0,489897949 | 2,99E-05 | 2,46E-05 | 1 - | 4,43E-05 - | 0 - | 0 - | 1 - | 5,23E-05 - | 1 - | 5,31E-05 - | 0 - | 0 - |
| 2012 | 1 | 3um_FIJJAFBF_46933 | Bacteria | Proteobacteri | Gammaprote | Alteromonada | Alteromonada | Paraglaciecola | Paraglaciecola | information st | Translation, ri | 179R6 | This protein is | 342610.Patl_3 | 1,8 | 0,748331477 | 0,000167985 | 5,60E-05 | 3 - | 0,000242486 - | 2 - | 0,000200731 - | 2 - | 0,00019082 - | 1 - | 9,68E-05 - | 1 - | 0,000109067 - |
| 2013 | 1 | 0.2um_NOKCPHJN_1028364 | Bacteria | Bacteroidetes | Flavobacteriia | Flavobacterial | Flavobacteriac | Marinirhabdu | Marinirhabdu | unclassified | unclassified | unclassified | unclassified | unclassified | 1,6 | 0,489897949 | 0,000113465 | 3,15E-05 | 2 - | 0,000120818 - | 2 - | 0,00015002 - | 2 - | 0,000142613 - | 1 - | 7,24E-05 - | 1 - | 8,15E-05 - |
| 2014 | 1 | 3um_FIJJAFBF_86508 | Bacteria | Bacteroidetes | Flavobacteriia | Flavobacterial | Flavobacteriac | Polaribacter | Polaribacter s | poorly charact | Function unkn | 05ERK | gliding motilit | 313594.PI23P | 1 | 0,632455532 | 2,64E-05 | 1,78E-05 | 1 - | 2,25E-05 - | 2 - | 5,60E-05 - | 1 - | 2,66E-05 - | 1 - | 2,70E-05 - | 0 - | 0 - |
| 2015 | 1 | 0.2um_NOKCPHJN_229977 | Bacteria | Proteobacteri | Alphaproteob | unclassified | unclassified | unclassified | Alphaproteob | unclassified | unclassified | unclassified | unclassified | unclassified | 3,2 | 0,4 | 0,00012343 | 1,83E-05 | 3 - | 9,70E-05 - | 3 - | 0,000120438 - | 4 - | 0,000152656 - | 3 - | 0,000116183 - | 3 - | 0,00013088 - |
| 2016 | 2 | 0.2um_NOKCPHJN_125931;0.2um_NOKCPHJN_3 | Bacteria | Proteobacteri | Alphaproteob | various | various | various | various | information st | Translation, ri | 05CA4 | amino acids s | 367336.OM22 | 0,2 | 0,4 | 2,40E-06 | 4,80E-06 | 1 - | 1,20E-05 - | 0 - | 0 - | 0 - | 0 - | 0 - | 0 - | 0 - | 0 - |
| 2017 | 1 | 0.2um_NOKCPHJN_378529 | Bacteria | Bacteroidetes | Flavobacteriia | Flavobacterial | Flavobacteriac | Aquimarina | Aquimarina m | information st | Translation, ri | 05CEJ | Catalyzes the | 983544.Lacal_ | 1 | 0,632455532 | 7,34E-05 | 5,16E-05 | 1 - | 5,98E-05 - | 1 - | 7,42E-05 - | 0 - | 0 - | 1 - | 7,16E-05 - | 2 - | 0,000161328 - |
| 2018 | 1 | 0.2um_NOKCPHJN_427649 | Bacteria | Bacteroidetes | Flavobacteriia | unclassified | unclassified | unclassified | Flavobacteria | cellular proces | Intracellular tr | 0BIEB | domain-conta | 1005048.CFU_ | 1,2 | 0,4 | 1,78E-05 | 6,58E-06 | 1 - | 1,24E-05 - | 2 - | 3,07E-05 - | 1 - | 1,46E-05 - | 1 - | 1,48E-05 - | 1 - | 1,67E-05 - |
| 2019 | 1 | 0.2um_NOKCPHJN_430032 | unclassified | unclassified | unclassified | unclassified | unclassified | unclassified | unclassified | unclassified | unclassified | unclassified | unclassified | unclassified | 0,4 | 0,489897949 | 1,20E-05 | 1,47E-05 | 0 - | 0 - | 1 - | 2,88E-05 - | 0 - | 0 - | 0 - | 0 - | 1 - | 3,13E-05 - |
| 2020 | 1 | 0.2um_NOKCPHJN_881016 | Bacteria | Proteobacteri | Gammaprote | Cellvibrionale | Porticoccacea | unclassified | gamma prote | metabolism | Inorganic ion t | 0QHT5 | TonB-depende | 314287.GB22 | 14,6 | 1,356465997 | 0,000330058 | 4,53E-05 | 16 - | 0,000303041 - | 15 - | 0,000352769 - | 13 - | 0,000290638 - | 13 - | 0,000294931 - | 16 - | 0,00040891 - |
| 2021 | 1 | 3um_FIJJAFBF_103104 | Bacteria | Bacteroidetes | Flavobacteriia | Flavobacterial | Flavobacteriac | Polaribacter | Polaribacter s | metabolism | Energy produc | COG0055 | Produces ATP | 313594.PI23P | 2 | 0 | 0,000114189 | 1,08E-05 | 2 - | 9,56E-05 - | 2 - | 0,000118766 - | 2 - | 0,000112902 - | 2 - | 0,000114569 - | 2 - | 0,000129062 - |
| 2022 | 1 | 3um_FIJJAFBF_191135 | Bacteria | Proteobacteri | Alphaproteob | Rhodobactera | unclassified | unclassified | Rhodobactera | information st | Translation, ri | 01QWB | One of the pri | 314270.RB208 | 5 | 1,414213562 | 0,000272728 | 6,33E-05 | 7 - | 0,000325278 - | 3 - | 0,0001731 - | 6 - | 0,000329107 - | 4 - | 0,000222645 - | 5 - | 0,000313511 - |
| 2023 | 1 | 3um_FIJJAFBF_655084 | Bacteria | Proteobacteri | Alphaproteob | Rhodobactera | Rhodobactera | Planktomarina | Planktomarina | metabolism | Amino acid tra | 01Y25 | Nitrogen regu | 246200.SPO2 | 1 | 0 | 0,000122346 | 1,16E-05 | 1 - | 0,000102479 - | 1 - | 0,000127249 - | 1 - | 0,000120966 - | 1 - | 0,000122753 - | 1 - | 0,000138281 - |
| 2024 | 1 | 0.2um_NOKCPHJN_234809 | Bacteria | Bacteroidetes | Flavobacteriia | Flavobacterial | Flavobacteriac | Polaribacter | Polaribacter h | information st | Translation, ri | 08RWN | rrna methyltr | 313594.PI23P | 0,8 | 0,4 | 4,33E-05 | 2,20E-05 | 1 - | 4,68E-05 - | 1 - | 5,82E-05 - | 1 - | 5,53E-05 - | 1 - | 5,61E-05 - | 0 - | 0 - |
| 2025 | 1 | 3um_FIJJAFBF_760120 | Bacteria | Bacteroidetes | Flavobacteriia | Flavobacterial | Flavobacteriac | Dokdonia | Dokdonia paci | poorly charact | Function unkn | 101IS | NA | 50743.SCB49_ | 1 | 0 | 1,40E-05 | 1,33E-06 | 1 - | 1,18E-05 - | 1 - | 1,46E-05 - | 1 - | 1,39E-05 - | 1 - | 1,41E-05 - | 1 - | 1,59E-05 - |
| 2026 | 1 | 0.2um_NOKCPHJN_254615 | Bacteria | Bacteroidetes | Flavobacteriia | Flavobacterial | Crocinitomica | unclassified | Crocinitomica | unclassified | unclassified | unclassified | unclassified | unclassified | 0,4 | 0,8 | 2,40E-05 | 4,81E-05 | 2 - | 0,000120185 - | 0 - | 0 - | 0 - | 0 - | 0 - | 0 - | 0 - | 0 - |
| 2027 | 1 | 0.2um_NOKCPHJN_71634 | Bacteria | Proteobacteri | Alphaproteob | unclassified | unclassified | Candidatus Pu | Candidatus Pu | metabolism | Carbohydrate | 01U6R | trap dicarboxy | 488538.SAR1 | 2,4 | 1,356465997 | 8,86E-05 | 3,99E-05 | 5 - | 0,000161657 - | 1 - | 4,01E-05 - | 2 - | 7,63E-05 - | 2 - | 7,75E-05 - | 2 - | 8,73E-05 - |
| 2028 | 1 | 0.2um_NOKCPHJN_491324 | Bacteria | Proteobacteri | Alphaproteob | unclassified | unclassified | unclassified | alpha proteob | poorly charact | Function unkn | 16SEZ | PTS system m | 156889.Mmc1 | 0,4 | 0,489897949 | 1,00E-05 | 1,23E-05 | 0 - | 0 - | 1 - | 2,41E-05 - | 0 - | 0 - | 0 - | 0 - | 1 - | 2,62E-05 - |
| 2029 | 1 | 3um_FIJJAFBF_510243 | Bacteria | Bacteroidetes | Flavobacteriia | Flavobacterial | Flavobacteriac | Tenacibaculur | Tenacibaculur | information st | Translation, ri | 0PAI9 | Binds 16S rRN | 313594.PI23P | 2,2 | 0,748331477 | 0,000340403 | 0,000124101 | 2 - | 0,000257925 - | 3 - | 0,000480401 - | 1 - | 0,000152227 - | 3 - | 0,000463427 - | 2 - | 0,000348033 - |
| 2030 | 1 | 3um_FIJJAFBF_527414 | Bacteria | Proteobacteri | Alphaproteob | Rhodobactera | Rhodobactera | Jhaorihella | Jhaorihella the | metabolism | Amino acid tra | 01QVA | Part of the AB | 439496.RBY4I | 1,6 | 0,489897949 | 5,74E-05 | 1,50E-05 | 2 - | 6,14E-05 - | 1 - | 3,81E-05 - | 2 - | 7,25E-05 - | 2 - | 7,35E-05 - | 1 - | 4,14E-05 - |
| 2031 | 1 | 0.2um_NOKCPHJN_396742 | Eukaryota | unclassified | Labyrinthulor | unclassified | Thraustochytr | Aurantiochytr | Aurantiochytr | metabolism | Lipid transpor | KOG1202 | polyketide syr | 572307.XP_0 | 1 | 0 | 8,69E-06 | 8,26E-07 | 1 - | 7,28E-06 - | 1 - | 9,04E-06 - | 1 - | 8,60E-06 - | 1 - | 8,72E-06 - | 1 - | 9,83E-06 - |
| 2032 | 1 | 0.2um_NOKCPHJN_50537 | Bacteria | Proteobacteri | Betaproteoba | Burkholderial | Comamonada | Hylemonella | Hylemonella g | information st | Translation, ri | COG1185 | Involved in m | 338969.Rfer_ | 6,2 | 1,166190379 | 0,00011668 | 1,73E-05 | 7 - | 0,000111899 - | 6 - | 0,000119097 - | 7 - | 0,000132086 - | 7 - | 0,000134037 - | 4 - | 8,63E-05 - |
| 2033 | 1 | 3um_FIJJAFBF_91113 | Eukaryota | unclassified | Florideophyce | Ceramiales | Rhodomelace | Polysiphonia | Polysiphonia s | unclassified | unclassified | unclassified | unclassified | unclassified | 6 | 1,414213562 | 0,000615813 | 0,000122371 | 8 - | 0,000695617 - | 5 - | 0,000539844 - | 4 - | 0,000410552 - | 7 - | 0,000729078 - | 6 - | 0,000703975 - |
| 2034 | 1 | 0.2um_NOKCPHJN_360926 | Bacteria | Proteobacteri | Alphaproteob | Pelagibacteral | Pelagibacterac | Candidatus Pe | Candidatus Pe | cellular proces | Cell wall/mem | 16QB0 | Part of the ou | 335992.SAR1 | 1,2 | 0,4 | 2,21E-05 | 7,60E-06 | 1 - | 1,54E-05 - | 1 - | 1,91E-05 - | 1 - | 1,82E-05 - | 2 - | 3,69E-05 - | 1 - | 2,08E-05 - |
| 2035 | 1 | 3um_FIJJAFBF_250885 | Bacteria | Proteobacteri | Alphaproteob | Pelagibacteral | Pelagibacterac | unclassified | Pelagibactera | unclassified | unclassified | unclassified | unclassified | unclassified | 1,8 | 0,748331477 | 5,86E-05 | 1,91E-05 | 3 - | 8,50E-05 - | 1 - | 3,52E-05 - | 2 - | 6,69E-05 - | 2 - | 6,79E-05 - | 1 - | 3,82E-05 - |
| 2036 | 1 | 0.2um_NOKCPHJN_520515 | Bacteria | Bacteroidetes | Flavobacteriia | Flavobacterial | Flavobacteriac | Marinirhabdu | Marinirhabdu | unclassified | unclassified | unclassified | unclassified | unclassified | 0,6 | 0,8 | 4,29E-05 | 5,54E-05 | 2 - | 0,000135031 - | 0 - | 0 - | 1 - | 7,97E-05 - | 0 - | 0 - | 0 - | 0 - |
| 2037 | 1 | 0.2um_NOKCPHJN_882166 | Bacteria | Bacteroidetes | Flavobacteriia | Flavobacterial | Flavobacteriac | Aquimarina | Aquimarina s | metabolism | Secondary me | COG1228 | amidohydrola | 643867.Ftrac_ | 0,8 | 0,4 | 2,71E-05 | 1,36E-05 | 0 - | 0 - | 1 - | 3,39E-05 - | 1 - | 3,22E-05 - | 1 - | 3,27E-05 - | 1 - | 3,68E-05 - |
| 2038 | 1 | 3um_FIJJAFBF_459237 | Bacteria | Proteobacteri | Alphaproteob | Rhodospirillal | Rhodospirillac | unclassified | Rhodobactera | unclassified | unclassified | unclassified | unclassified | unclassified | 1 | 0 | 0,000131757 | 1,25E-05 | 1 - | 0,000110362 - | 1 - | 0,000137037 - | 1 - | 0,000130271 - | 1 - | 0,000132195 - | 1 - | 0,000148918 - |
| 2039 | 1 | 0.2um_NOKCPHJN_110537 | Bacteria | Proteobacteri | Gammaprote | Cellvibrionale | Porticoccacea | unclassified | various | unclassified | unclassified | unclassified | unclassified | unclassified | 5,6 | 1,624807681 | 0,000447976 | 0,000112105 | 8 - | 0,000543322 - | 5 - | 0,000421653 - | 6 - | 0,000481002 - | 3 - | 0,000244053 - | 6 - | 0,00054985 - |
| 2040 | 2 | 0.2um_NOKCPHJN_152866;3um_FIJJAFBF_7771 | Bacteria | Bacteroidetes | various | various | various | various | various | poorly charact | Function unkn | 09A4Z | rhodopsin | various | 0,2 | 0,4 | 9,26E-06 | 1,85E-05 | 1 - | 4,63E-05 - | 0 - | 0 - | 0 - | 0 - | 0 - | 0 - | 0 - | 0 - |
| 2041 | 2 | 0.2um_NOKCPHJN_188518;3um_FIJJAFBF_1089 | Bacteria | Bacteroidetes | Saprospiria | Saprospirales | Haliscomenob | Phaeodactylib | Phaeodactylib | poorly charact | Function unkn | 0FX0D | NA | 643867.Ftrac_ | 2 | 0 | 5,47E-05 | 5,19E-06 | 2 - | 4,58E-05 - | 2 - | 5,69E-05 - | 2 - | 5,41E-05 - | 2 - | 5,49E-05 - | 2 - | 6,18E-05 - |
| 2042 | 1 | 0.2um_NOKCPHJN_123799 | Bacteria | Proteobacteri | Gammaprote | Oceanospirilla | Saccharospirill | Reinekea | Reinekea fors | cellular proces | Posttranslatio | 05JRK | feS assembly | 314283.MED2 | 0,2 | 0,4 | 6,66E-06 | 1,33E-05 | 0 - | 0 - | 1 - | 3,33E-05 - | 0 - | 0 - | 0 - | 0 - | 0 - | 0 - |
| 2043 | 1 | 3um_FIJJAFBF_50289 | Bacteria | Bacteroidetes | Flavobacteriia | Flavobacterial | Flavobacteriac | Arenitalea | Arenitalea lut | metabolism | Carbohydrate | 05D2N | Aldolase | 313603.FB217 | 6 | 0,632455532 | 0,000231311 | 3,11E-05 | 6 - | 0,000193989 - | 5 - | 0,000200731 - | 6 - | 0,000228984 - | 7 - | 0,000271094 - | 6 - | 0,00026176 - |
| 2044 | 1 | 0.2um_NOKCPHJN_502609 | Bacteria | Proteobacteri | Gammaprote | unclassified | unclassified | unclassified | Gammaprote | metabolism | Lipid transpor | 05C9H | acyl-CoA dehy | 247633.GP21 | 0,4 | 0,489897949 | 1,33E-05 | 1,63E-05 | 0 - | 0 - | 1 - | 3,41E-05 - | 1 - | 3,24E-05 - | 0 - | 0 - | 0 - | 0 - |
| 2045 | 3 | 0.2um_NOKCPHJN_1034081;0.2um_NOKCPHJN_ | Bacteria | various | various | various | various | various | various | unclassified | unclassified | unclassified | unclassified | unclassified | 0,2 | 0,4 | 1,43E-05 | 2,87E-05 | 1 - | 7,17E-05 - | 0 - | 0 - | 0 - | 0 - | 0 - | 0 - | 0 - | 0 - |
| 2046 | 1 | 0.2um_NOKCPHJN_264230 | Viruses | unclassified | unclassified | unclassified | Phycodnavirid | Prasinovirus | Bathycoccus s | unclassified | unclassified | unclassified | unclassified | unclassified | 0,2 | 0,4 | 4,45E-06 | 8,90E-06 | 0 - | 0 - | 0 - | 0 - | 0 - | 0 - | 0 - | 0 - | 1 - | 2,23E-05 - |
| 2047 | 1 | 0.2um_NOKCPHJN_79504 | Bacteria | Bacteroidetes | Flavobacteriia | Flavobacterial | Flavobacteriac | unclassified | Flavobacteria | information st | Translation, ri | 05CAV | thus facilitatin | 487796.Flav2 | 1 | 0 | 1,95E-05 | 1,85E-06 | 1 - | 1,63E-05 - | 1 - | 2,03E-05 - | 1 - | 1,93E-05 - | 1 - | 1,96E-05 - | 1 - | 2,20E-05 - |
| 2048 | 1 | 0.2um_NOKCPHJN_230052 | Bacteria | Bacteroidetes | Flavobacteriia | Flavobacterial | Flavobacteriac | Aureicoccus | Aureicoccus m | unclassified | unclassified | unclassified | unclassified | unclassified | 5,6 | 1,356465997 | 0,000165956 | 2,31E-05 | 8 - | 0,000203145 - | 5 - | 0,000157654 - | 6 - | 0,000179844 - | 5 - | 0,000152083 - | 4 - | 0,000137057 - |
| 2049 | 1 | 3um_FIJJAFBF_176995 | Eukaryota | unclassified | Bangiophycea | Porphyridiales | Porphyridiace | Porphyridium | Porphyridium | unclassified | unclassified | unclassified | unclassified | unclassified | 4,2 | 0,4 | 0,000112193 | 5,31E-06 | 5 - | 0,000112747 - | 4 - | 0,000111999 - | 4 - | 0,000106469 - | 4 - | 0,000108042 - | 4 - | 0,000121709 - |
| 2050 | 1 | 0.2um_NOKCPHJN_194701 | Eukaryota | unclassified | Oomycetes | Saprolegniales | Saprolegniace | Aphanomyces | Aphanomyces | information st | Chromatin str | KOG0889 | Transformatio | 4787.PITG_03 | 0,2 | 0,4 | 6,74E-07 | 1,35E-06 | 0 - | 0 - | 1 - | 3,37E-06 - | 0 - | 0 - | 0 - | 0 - | 0 - | 0 - |
| 2051 | 1 | 0.2um_NOKCPHJN_601806 | Bacteria | Firmicutes | Bacilli | Lactobacillales | Enterococcace | Enterococcus | Enterococcus | poorly charact | Function unkn | 0XQTW | domain protei | 999141.GME_ | 0,2 | 0,4 | 2,34E-06 | 4,67E-06 | 0 - | 0 - | 0 - | 0 - | 1 - | 1,17E-05 - | 0 - | 0 - | 0 - | 0 - |
| 2052 | 1 | 3um_FIJJAFBF_02801 | Bacteria | Proteobacteri | Gammaprote | Oceanospirilla | Saccharospirill | Reinekea | Reinekea fors | unclassified | unclassified | unclassified | unclassified | unclassified | 0,2 | 0,4 | 1,74E-05 | 3,47E-05 | 0 - | 0 - | 0 - | 0 - | 1 - | 8,68E-05 - | 0 - | 0 - | 0 - | 0 - |
| 2053 | 1 | 3um_FIJJAFBF_122458 | Bacteria | Bacteroidetes | Flavobacteriia | Flavobacterial | Flavobacteriac | Polaribacter | Polaribacter s | cellular proces | Cell wall/mem | 0ZZRH | outer membra | 313598.MED1 | 1,4 | 0,8 | 6,73E-05 | 3,95E-05 | 2 - | 8,03E-05 - | 2 - | 9,97E-05 - | 0 - | 0 - | 1 - | 4,81E-05 - | 2 - | 0,000108304 - |
| 2054 | 1 | 3um_FIJJAFBF_386703 | Bacteria | Proteobacteri | Gammaprote | Oceanospirilla | Saccharospirill | Reinekea | Reinekea fors | poorly charact | Function unkn | 0QKWJ | NA | 314283.MED2 | 0,4 | 0,489897949 | 1,25E-05 | 1,54E-05 | 1 - | 2,87E-05 - | 0 - | 0 - | 1 - | 3,39E-05 - | 0 - | 0 - | 0 - | 0 - |
| 2055 | 1 | 0.2um_NOKCPHJN_93902 | Bacteria | Bacteroidetes | Flavobacteriia | Flavobacterial | Flavobacteriac | Arenibacter | Arenibacter e | information st | Translation, ri | COG0052 | 30S ribosomal | 487797.Flav3 | 3,4 | 1,019803903 | 0,00018123 | 5,57E-05 | 4 - | 0,000179339 - | 5 - | 0,000278357 - | 2 - | 0,000105845 - | 3 - | 0,000161113 - | 3 - | 0,000181494 - |
| 2056 | 1 | 0.2um_NOKCPHJN_247996 | Bacteria | Proteobacteri | Gammaprote | Alteromonada | Idiomarinacea | Idiomarina | various | metabolism | Inorganic ion t | 177TK | phosphate-sel | 455436.GHTC | 1,6 | 0,489897949 | 7,08E-05 | 1,86E-05 | 2 - | 7,58E-05 - | 1 - | 4,70E-05 - | 2 - | 8,94E-05 - | 2 - | 9,07E-05 - | 1 - | 5,11E-05 - |
| 2057 | 1 | 0.2um_NOKCPHJN_482245 | Bacteria | Proteobacteri | Alphaproteob | Pelagibacteral | Pelagibacterac | unclassified | Pelagibactera | unclassified | unclassified | unclassified | unclassified | unclassified | 0,4 | 0,489897949 | 1,19E-05 | 1,46E-05 | 0 - | 0 - | 1 - | 3,03E-05 - | 0 - | 0 - | 1 - | 2,93E-05 - | 0 - | 0 - |
| 2058 | 1 | 3um_FIJJAFBF_410570 | Bacteria | Proteobacteri | Gammaprote | Cellvibrionale | Halieaceae | unclassified | marine gamm | unclassified | unclassified | unclassified | unclassified | unclassified | 0,8 | 0,4 | 1,37E-05 | 7,05E-06 | 1 - | 1,45E-05 - | 0 - | 0 - | 1 - | 1,71E-05 - | 1 - | 1,74E-05 - | 1 - | 1,96E-05 - |
| 2059 | 1 | 0.2um_NOKCPHJN_348703 | Bacteria | Proteobacteri | Gammaprote | Cellvibrionale | Halieaceae | unclassified | marine gamm | information st | Transcription | 05D27 | DNA-depende | 314285.KT71_ | 0,4 | 0,489897949 | 3,53E-06 | 4,36E-06 | 1 - | 8,04E-06 - | 0 - | 0 - | 0 - | 0 - | 1 - | 9,63E-06 - | 0 - | 0 - |
| 2060 | 1 | 3um_FIJJAFBF_350584 | Bacteria | Bacteroidetes | Cytophagia | Cytophagales | Cytophagacea | Ekhidna | Ekhidna lutea | metabolism | Amino acid tra | 05C6M | Alpha-keto-be | 643867.Ftrac_ | 0,4 | 0,489897949 | 1,57E-05 | 1,92E-05 | 0 - | 0 - | 0 - | 0 - | 1 - | 3,89E-05 - | 1 - | 3,95E-05 - | 0 - | 0 - |
| 2061 | 1 | 0.2um_NOKCPHJN_94471 | Bacteria | Proteobacteri | Betaproteoba | unclassified | unclassified | unclassified | Betaproteoba | metabolism | Lipid transpor | 0B93T | Dehydrogenas | 296591.Bpro_ | 0,4 | 0,489897949 | 1,62E-05 | 2,00E-05 | 1 - | 3,71E-05 - | 0 - | 0 - | 1 - | 4,38E-05 - | 0 - | 0 - | 0 - | 0 - |
| 2062 | 1 | 3um_FIJJAFBF_795314 | Bacteria | Bacteroidetes | unclassified | unclassified | unclassified | unclassified | Bacteroidetes | cellular proces | Cell wall/mem | 09ARE | ompa motb d | 926562.Oweh | 1,2 | 0,4 | 3,26E-05 | 7,23E-06 | 2 - | 4,68E-05 - | 1 - | 2,91E-05 - | 1 - | 2,76E-05 - | 1 - | 2,81E-05 - | 1 - | 3,16E-05 - |
| 2063 | 1 | 3um_FIJJAFBF_111372 | Viruses | unclassified | unclassified | unclassified | Phycodnavirid | unclassified | Chrysochrom | unclassified | unclassified | unclassified | unclassified | unclassified | 0,6 | 0,489897949 | 5,62E-05 | 4,67E-05 | 1 - | 7,92E-05 - | 0 - | 0 - | 0 - | 0 - | 1 - | 9,48E-05 - | 1 - | 0,00010681 - |
| 2064 | 1 | 0.2um_NOKCPHJN_383255 | Bacteria | Bacteroidetes | Flavobacteriia | Flavobacterial | Flavobacteriac | Mangrovimon | Mangrovimon | metabolism | Energy produc | COG0056 | Produces ATP | 398720.MED2 | 1,2 | 0,748331477 | 3,00E-05 | 1,85E-05 | 2 - | 4,37E-05 - | 2 - | 5,43E-05 - | 1 - | 2,58E-05 - | 1 - | 2,62E-05 - | 0 - | 0 - |
| 2065 | 1 | 0.2um_NOKCPHJN_227778 | Eukaryota | unclassified | Oomycetes | Pythiales | Pythiaceae | Pythium | Pythium insidi | unclassified | unclassified | unclassified | unclassified | unclassified | 0,2 | 0,4 | 2,43E-06 | 4,86E-06 | 0 - | 0 - | 0 - | 0 - | 0 - | 0 - | 1 - | 1,22E-05 - | 0 - | 0 - |
| 2066 | 1 | 3um_FIJJAFBF_370976 | Bacteria | Proteobacteri | Gammaprote | Alteromonada | Alteromonada | Paraglaciecola | various | unclassified | unclassified | unclassified | unclassified | unclassified | 0,4 | 0,489897949 | 5,05E-05 | 6,19E-05 | 0 - | 0 - | 0 - | 0 - | 1 - | 0,000125447 - | 1 - | 0,000127299 - | 0 - | 0 - |
| 2067 | 1 | 0.2um_NOKCPHJN_99578 | Bacteria | Proteobacteri | Gammaprote | Oceanospirilla | Saccharospirill | Reinekea | Reinekea fors | unclassified | unclassified | unclassified | unclassified | unclassified | 1,6 | 0,489897949 | 0,000148678 | 4,12E-05 | 2 - | 0,000158313 - | 2 - | 0,000196578 - | 2 - | 0,000186872 - | 1 - | 9,48E-05 - | 1 - | 0,00010681 - |
| 2068 | 1 | 0.2um_NOKCPHJN_597511 | Eukaryota | unclassified | Chrysophycea | Chromulinales | Chromulinace | Ochromonas | Ochromonas d | unclassified | unclassified | unclassified | unclassified | unclassified | 1,6 | 0,489897949 | 0,000106889 | 3,36E-05 | 2 - | 0,000112526 - | 1 - | 6,99E-05 - | 2 - | 0,000132826 - | 1 - | 6,74E-05 - | 2 - | 0,000151838 - |
| 2069 | 1 | 3um_FIJJAFBF_520504 | Bacteria | Bacteroidetes | Flavobacteriia | Flavobacterial | Flavobacteriac | Algibacter | Algibacter sp. | unclassified | unclassified | unclassified | unclassified | unclassified | 5,6 | 0,489897949 | 0,000208659 | 1,83E-05 | 6 - | 0,000188159 - | 6 - | 0,000233637 - | 6 - | 0,000222102 - | 5 - | 0,000187819 - | 5 - | 0,000211577 - |
| 2070 | 1 | 0.2um_NOKCPHJN_52094 | Bacteria | Proteobacteri | Gammaprote | Pseudomonad | Pseudomonad | Pseudomonas | various | metabolism | Inorganic ion t | 17FMU | ApaG protein | 322710.Avin_ | 1 | 0 | 0,000107052 | 1,02E-05 | 1 - | 8,97E-05 - | 1 - | 0,000111343 - | 1 - | 0,000105845 - | 1 - | 0,000107409 - | 1 - | 0,000120996 - |
| 2071 | 1 | 0.2um_NOKCPHJN_192196 | Bacteria | Proteobacteri | Gammaprote | unclassified | unclassified | unclassified | Gammaprote | cellular proces | Posttranslatio | 0QIPK | Stringent star | 393595.ABO_ | 0,4 | 0,489897949 | 2,73E-05 | 3,36E-05 | 1 - | 6,20E-05 - | 0 - | 0 - | 0 - | 0 - | 1 - | 7,43E-05 - | 0 - | 0 - |
| 2072 | 1 | 3um_FIJJAFBF_744837 | Bacteria | Proteobacteri | Gammaprote | Cellvibrionale | Spongiibacter | Oceanicoccus | Oceanicoccus | information st | Translation, ri | 17C3B | The globular d | 247633.GP21 | 1,2 | 0,4 | 0,000150572 | 5,04E-05 | 1 - | 0,0001053 - | 1 - | 0,000130751 - | 2 - | 0,000248591 - | 1 - | 0,000126131 - | 1 - | 0,000142087 - |
| 2073 | 1 | 0.2um_NOKCPHJN_83411 | Bacteria | Bacteroidetes | Flavobacteriia | Flavobacterial | Flavobacteriac | Polaribacter | Polaribacter s | metabolism | Inorganic ion t | 05C27 | TonB-depende | 313598.MED1 | 0,4 | 0,489897949 | 4,73E-06 | 5,82E-06 | 1 - | 1,08E-05 - | 0 - | 0 - | 1 - | 1,28E-05 - | 0 - | 0 - | 0 - | 0 - |
| 2074 | 1 | 0.2um_NOKCPHJN_409498 | Bacteria | Proteobacteri | Alphaproteob | Rhodospirillal | Rhodospirillac | unclassified | Rhodobactera | unclassified | unclassified | unclassified | unclassified | unclassified | 0,6 | 0,489897949 | 9,45E-05 | 7,74E-05 | 0 - | 0 - | 1 - | 0,000154912 - | 0 - | 0 - | 1 - | 0,000149438 - | 1 - | 0,000168342 - |
| 2075 | 1 | 3um_FIJJAFBF_78392 | Bacteria | Bacteroidetes | Flavobacteriia | Flavobacterial | Flavobacteriac | Winogradskye | Winogradskye | metabolism | Amino acid tra | COG0174 | glutamine syn | 313598.MED1 | 0,8 | 0,4 | 3,21E-05 | 1,65E-05 | 1 - | 3,37E-05 - | 1 - | 4,18E-05 - | 1 - | 3,97E-05 - | 0 - | 0 - | 1 - | 4,54E-05 - |
| 2076 | 1 | 3um_FIJJAFBF_187827 | Bacteria | Bacteroidetes | Flavobacteriia | Flavobacterial | Flavobacteriac | Polaribacter | Polaribacter s | information st | Translation, ri | 1226R | 50S ribosomal | 313594.PI23P | 0,4 | 0,8 | 7,29E-05 | 0,000145748 | 2 - | 0,000364371 - | 0 - | 0 - | 0 - | 0 - | 0 - | 0 - | 0 - | 0 - |
| 2077 | 2 | 3um_FIJJAFBF_01992;3um_FIJJAFBF_120839 | unclassified | unclassified | unclassified | unclassified | unclassified | unclassified | unclassified | unclassified | unclassified | unclassified | unclassified | unclassified | 1 | 0 | 0,000130502 | 1,24E-05 | 1 - | 0,000109311 - | 1 - | 0,000135732 - | 1 - | 0,000129031 - | 1 - | 0,000130936 - | 1 - | 0,0001475 - |
| 2078 | 1 | 3um_FIJJAFBF_40039 | Bacteria | Bacteroidetes | Flavobacteriia | various | various | various | various | unclassified | unclassified | unclassified | unclassified | unclassified | 13,8 | 1,166190379 | 0,00019292 | 2,53E-05 | 15 - | 0,000175679 - | 14 - | 0,000203598 - | 12 - | 0,000165897 - | 13 - | 0,000182376 - | 15 - | 0,000237053 - |
| 2079 | 1 | 0.2um_NOKCPHJN_375834 | Bacteria | Proteobacteri | Gammaprote | unclassified | unclassified | unclassified | Gammaprote | metabolism | Inorganic ion t | 16PDP | TonB-depende | 502025.Hoch_ | 1,4 | 1,2 | 1,98E-05 | 1,75E-05 | 2 - | 2,49E-05 - | 0 - | 0 - | 2 - | 2,94E-05 - | 3 - | 4,48E-05 - | 0 - | 0 - |
| 2080 | 1 | 3um_FIJJAFBF_75029 | Bacteria | Actinobacteri | Actinobacteri | Micrococcales | Microbacteria | Candidatus Ac | Candidatus Ac | unclassified | unclassified | unclassified | unclassified | unclassified | 0,6 | 0,489897949 | 3,85E-05 | 3,18E-05 | 1 - | 5,60E-05 - | 1 - | 6,95E-05 - | 0 - | 0 - | 1 - | 6,71E-05 - | 0 - | 0 - |
| 2081 | 1 | 3um_FIJJAFBF_336133 | Bacteria | unclassified | unclassified | unclassified | unclassified | unclassified | uncultured ba | unclassified | unclassified | unclassified | unclassified | unclassified | 1 | 0 | 5,42E-05 | 5,14E-06 | 1 - | 4,54E-05 - | 1 - | 5,63E-05 - | 1 - | 5,36E-05 - | 1 - | 5,43E-05 - | 1 - | 6,12E-05 - |
| 2082 | 1 | 3um_FIJJAFBF_788023 | Bacteria | Proteobacteri | Alphaproteob | Rhodobactera | Rhodobactera | unclassified | Rhodobactera | unclassified | unclassified | unclassified | unclassified | unclassified | 1,8 | 1,326649916 | 0,000160103 | 0,000113445 | 0 - | 0 - | 2 - | 0,000181553 - | 4 - | 0,000345178 - | 2 - | 0,000175138 - | 1 - | 9,86E-05 - |
| 2083 | 1 | 0.2um_NOKCPHJN_149429 | Bacteria | Bacteroidetes | unclassified | unclassified | unclassified | unclassified | Bacteroidetes | metabolism | Inorganic ion t | 05C27 | TonB-depende | 886377.Murr | 0,6 | 0,489897949 | 8,55E-06 | 7,00E-06 | 0 - | 0 - | 1 - | 1,40E-05 - | 0 - | 0 - | 1 - | 1,35E-05 - | 1 - | 1,52E-05 - |
| 2084 | 1 | 3um_FIJJAFBF_124661 | Bacteria | Proteobacteri | Alphaproteob | Pelagibacteral | Pelagibacterac | Candidatus Pe | Candidatus Pe | unclassified | unclassified | unclassified | unclassified | unclassified | 0,8 | 0,4 | 1,78E-05 | 9,12E-06 | 1 - | 1,85E-05 - | 1 - | 2,30E-05 - | 0 - | 0 - | 1 - | 2,22E-05 - | 1 - | 2,50E-05 - |
| 2085 | 1 | 3um_FIJJAFBF_605316 | Bacteria | Bacteroidetes | Flavobacteriia | Flavobacterial | Flavobacteriac | Confluentibac | Confluentibac | metabolism | Carbohydrate | 08RWB | NmrA family | 219305.MCAG | 0,2 | 0,4 | 1,95E-05 | 3,90E-05 | 0 - | 0 - | 0 - | 0 - | 1 - | 9,75E-05 - | 0 - | 0 - | 0 - | 0 - |
| 2086 | 1 | 0.2um_NOKCPHJN_161604 | unclassified | unclassified | unclassified | unclassified | unclassified | unclassified | unclassified | unclassified | unclassified | unclassified | unclassified | unclassified | 1 | 0 | 3,36E-05 | 3,19E-06 | 1 - | 2,81E-05 - | 1 - | 3,49E-05 - | 1 - | 3,32E-05 - | 1 - | 3,37E-05 - | 1 - | 3,80E-05 - |
| 2087 | 1 | 3um_FIJJAFBF_660889 | Bacteria | Proteobacteri | Gammaprote | Cellvibrionale | Porticoccacea | unclassified | gamma prote | cellular proces | Signal transdu | 16Q0Y | phosphate reg | 314287.GB22 | 1,2 | 0,4 | 7,00E-05 | 2,98E-05 | 1 - | 4,78E-05 - | 1 - | 5,94E-05 - | 1 - | 5,65E-05 - | 1 - | 5,73E-05 - | 2 - | 0,000129062 - |
| 2088 | 1 | 0.2um_NOKCPHJN_205010 | Bacteria | Proteobacteri | Gammaprote | Alteromonada | Alteromonada | Paraglaciecola | Paraglaciecola | metabolism | Lipid transpor | COG1960 | acyl-CoA dehy | 342610.Patl_ | 0,4 | 0,489897949 | 9,00E-06 | 1,12E-05 | 1 - | 1,92E-05 - | 0 - | 0 - | 0 - | 0 - | 0 - | 0 - | 1 - | 2,59E-05 - |
| 2089 | 1 | 3um_FIJJAFBF_91103 | Bacteria | Proteobacteri | Gammaprote | Oceanospirilla | Oceanospirilla | Marinobacteri | Marinobacteri | unclassified | unclassified | unclassified | unclassified | unclassified | 3,8 | 0,748331477 | 0,000352213 | 7,53E-05 | 3 - | 0,000232656 - | 4 - | 0,000385186 - | 5 - | 0,00045771 - | 4 - | 0,000371577 - | 3 - | 0,000313935 - |
| 2090 | 1 | 0.2um_NOKCPHJN_880073 | Bacteria | Cyanobacteria | unclassified | Synechococca | Synechococca | Synechococcu | Synechococcu | unclassified | unclassified | unclassified | unclassified | unclassified | 1,2 | 0,748331477 | 3,54E-05 | 2,39E-05 | 1 - | 2,40E-05 - | 2 - | 5,95E-05 - | 0 - | 0 - | 1 - | 2,87E-05 - | 2 - | 6,47E-05 - |
| 2091 | 1 | 3um_FIJJAFBF_740188 | Bacteria | Proteobacteri | Alphaproteob | Rhodobactera | Rhodobactera | Planktomarina | Planktomarina | unclassified | unclassified | unclassified | unclassified | unclassified | 4,4 | 1,019803903 | 0,000158106 | 3,65E-05 | 4 - | 0,000121136 - | 4 - | 0,000150416 - | 5 - | 0,000178736 - | 6 - | 0,000217652 - | 3 - | 0,000122592 - |
| 2092 | 2 | 0.2um_NOKCPHJN_877796;3um_FIJJAFBF_1836 | Bacteria | Bacteroidetes | Flavobacteriia | Flavobacterial | Flavobacteriac | various | various | metabolism | Carbohydrate | COG0297 | starch synthas | 391603.FBAL | 1 | 0,632455532 | 4,94E-05 | 2,76E-05 | 2 - | 8,53E-05 - | 1 - | 5,30E-05 - | 0 - | 0 - | 1 - | 5,11E-05 - | 1 - | 5,76E-05 - |
| 2093 | 1 | 3um_FIJJAFBF_532450 | Eukaryota | unclassified | Cryptophyta | Pyrenomonad | Geminigerace | Teleaulax | Teleaulax amp | metabolism | Energy produc | COG4451 | RuBisCO catal | 45157.CMV01 | 5,2 | 1,326649916 | 0,000503837 | 0,000110303 | 6 - | 0,000495439 - | 5 - | 0,000512658 - | 7 - | 0,000682285 - | 5 - | 0,000494544 - | 3 - | 0,000334261 - |
| 2094 | 1 | 3um_FIJJAFBF_243088 | Eukaryota | Bacillariophyt | Bacillariophyc | Naviculales | Phaeodactylac | Phaeodactylu | Phaeodactylu | unclassified | unclassified | unclassified | unclassified | unclassified | 0,6 | 0,489897949 | 2,63E-05 | 2,19E-05 | 1 - | 3,73E-05 - | 0 - | 0 - | 1 - | 4,40E-05 - | 0 - | 0 - | 1 - | 5,03E-05 - |
| 2095 | 1 | 3um_FIJJAFBF_788029 | Bacteria | Proteobacteri | Alphaproteob | Rhodobactera | unclassified | unclassified | Rhodobactera | unclassified | unclassified | unclassified | unclassified | unclassified | 2,6 | 0,8 | 0,000190315 | 7,43E-05 | 2 - | 0,000120185 - | 2 - | 0,000149234 - | 2 - | 0,000141866 - | 3 - | 0,000215942 - | 4 - | 0,000324344 - |
| 2096 | 1 | 3um_FIJJAFBF_592540 | Bacteria | Proteobacteri | Gammaprote | Alteromonada | Alteromonada | Glaciecola | Glaciecola sp. | unclassified | unclassified | unclassified | unclassified | unclassified | 10 | 1,414213562 | 0,000142194 | 2,63E-05 | 9 - | 0,000106824 - | 11 - | 0,000162121 - | 12 - | 0,000168127 - | 8 - | 0,00011374 - | 10 - | 0,00016016 - |
| 2097 | 1 | 3um_FIJJAFBF_594167 | Eukaryota | unclassified | Pelagophycea | Pelagomonad | unclassified | Aureococcus | Aureococcus a | metabolism | Energy produc | 0FN1Q | F(1)F(0) ATP s | 497965.Cyan7 | 9,2 | 0,748331477 | 0,000673624 | 8,41E-05 | 10 - | 0,000613779 - | 9 - | 0,00068592 - | 9 - | 0,000652054 - | 8 - | 0,000588164 - | 10 - | 0,000828206 - |
| 2098 | 1 | 3um_FIJJAFBF_503982 | Bacteria | Proteobacteri | Alphaproteob | Rhodobactera | Rhodobactera | Sulfitobacter | Sulfitobacter d | metabolism | Inorganic ion t | 05PWN | phosphate ab | 391624.OIHEL | 2,6 | 0,489897949 | 0,000104236 | 2,30E-05 | 3 - | 0,000100681 - | 3 - | 0,000125017 - | 2 - | 7,92E-05 - | 2 - | 8,04E-05 - | 3 - | 0,000135855 - |
| 2099 | 1 | 3um_FIJJAFBF_306744 | Bacteria | Proteobacteri | Alphaproteob | Rhodobactera | Rhodobactera | Ruegeria | Ruegeria sp. T | unclassified | unclassified | unclassified | unclassified | unclassified | 1,2 | 0,4 | 6,18E-05 | 2,28E-05 | 1 - | 4,28E-05 - | 2 - | 0,000106357 - | 1 - | 5,06E-05 - | 1 - | 5,13E-05 - | 1 - | 5,78E-05 - |
| 2100 | 1 | 0.2um_NOKCPHJN_397201 | Bacteria | Proteobacteri | Gammaprote | Oceanospirilla | Saccharospirill | Reinekea | Reinekea fors | metabolism | Energy produc | 0QIPC | FAD linked oxi | 314283.MED2 | 1 | 0 | 1,35E-05 | 1,28E-06 | 1 - | 1,13E-05 - | 1 - | 1,41E-05 - | 1 - | 1,34E-05 - | 1 - | 1,36E-05 - | 1 - | 1,53E-05 - |
| 2101 | 1 | 3um_FIJJAFBF_590407 | Bacteria | Proteobacteri | Gammaprote | Alteromonada | Alteromonada | Paraglaciecola | Paraglaciecola | information st | Translation, ri | COG0255 | 50s ribosomal | 342610.Patl_ | 0,6 | 0,489897949 | 0,000125327 | 0,000103399 | 1 - | 0,000182185 - | 1 - | 0,00022622 - | 0 - | 0 - | 1 - | 0,000218227 - | 0 - | 0 - |
| 2102 | 1 | 0.2um_NOKCPHJN_1086062 | Bacteria | Proteobacteri | Alphaproteob | Pelagibacteral | Pelagibacterac | Candidatus Pe | Candidatus Pe | unclassified | unclassified | unclassified | unclassified | unclassified | 0,6 | 0,489897949 | 3,54E-05 | 2,91E-05 | 1 - | 5,24E-05 - | 0 - | 0 - | 1 - | 6,19E-05 - | 1 - | 6,28E-05 - | 0 - | 0 - |
| 2103 | 1 | 0.2um_NOKCPHJN_131261 | Bacteria | Proteobacteri | Gammaprote | Oceanospirilla | Alcanivoraca | Ketobacter | Ketobacter alk | unclassified | unclassified | unclassified | unclassified | unclassified | 6,2 | 0,979795897 | 0,000181245 | 1,95E-05 | 8 - | 0,000197891 - | 6 - | 0,000184292 - | 5 - | 0,000145994 - | 6 - | 0,00017778 - | 6 - | 0,000200269 - |
| 2104 | 1 | 0.2um_NOKCPHJN_505357 | Bacteria | Chlamydiae | Chlamydiia | Parachlamydi | Parachlamydi | Neochlamydia | Neochlamydia | poorly charact | Function unkn | COG1100 | GTP-binding P | 91464.S7335_ | 0,4 | 0,489897949 | 8,95E-06 | 1,10E-05 | 0 - | 0 - | 0 - | 0 - | 1 - | 2,22E-05 - | 1 - | 2,25E-05 - | 0 - | 0 - |
| 2105 | 1 | 0.2um_NOKCPHJN_36287 | Bacteria | Bacteroidetes | unclassified | unclassified | unclassified | unclassified | Bacteroidetes | metabolism | Energy produc | COG0056 | Produces ATP | 926562.Oweh | 0,4 | 0,489897949 | 1,06E-05 | 1,30E-05 | 0 - | 0 - | 1 - | 2,72E-05 - | 1 - | 2,59E-05 - | 0 - | 0 - | 0 - | 0 - |
| 2106 | 1 | 3um_FIJJAFBF_453294 | Bacteria | Proteobacteri | Alphaproteob | Rhodobactera | unclassified | unclassified | Rhodobactera | cellular proces | Cell wall/mem | COG2885 | Ompa motb d | 439497.RR11_ | 0,6 | 0,489897949 | 4,76E-05 | 3,93E-05 | 1 - | 6,96E-05 - | 1 - | 8,64E-05 - | 1 - | 8,21E-05 - | 0 - | 0 - | 0 - | 0 - |
| 2107 | 1 | 0.2um_NOKCPHJN_292740 | Bacteria | Proteobacteri | Alphaproteob | Rhodobactera | Rhodobactera | Planktomarina | Planktomarina | metabolism | Carbohydrate | 07SNC | solute-binding | 367336.OM22 | 0,6 | 0,489897949 | 2,04E-05 | 1,67E-05 | 0 - | 0 - | 1 - | 3,35E-05 - | 0 - | 0 - | 1 - | 3,23E-05 - | 1 - | 3,64E-05 - |
| 2108 | 1 | 3um_FIJJAFBF_565795 | Bacteria | Bacteroidetes | Flavobacteriia | Flavobacterial | Flavobacteriac | Hyunsoonleel | Hyunsoonleel | unclassified | unclassified | unclassified | unclassified | unclassified | 1 | 0,632455532 | 3,38E-05 | 2,21E-05 | 1 - | 2,91E-05 - | 1 - | 3,61E-05 - | 1 - | 3,43E-05 - | 2 - | 6,96E-05 - | 0 - | 0 - |
| 2109 | 2 | 0.2um_NOKCPHJN_229446;0.2um_NOKCPHJN_2 | Bacteria | Proteobacteri | Alphaproteob | Pelagibacteral | Pelagibacterac | Candidatus Pe | Candidatus Pe | information st | Translation, ri | 08UHH | Located on th | 335992.SAR1 | 1,2 | 0,4 | 0,000109416 | 3,67E-05 | 1 - | 7,65E-05 - | 1 - | 9,50E-05 - | 2 - | 0,000180643 - | 1 - | 9,17E-05 - | 1 - | 0,00010325 - |
| 2110 | 1 | 3um_FIJJAFBF_406349 | Bacteria | Bacteroidetes | Cytophagia | Cytophagales | Flammeovirga | unclassified | Flammeovirga | metabolism | Energy produc | 090JD | Blue (Type 1) | 643867.Ftrac_ | 0,4 | 0,8 | 2,68E-05 | 5,37E-05 | 2 - | 0,000134242 - | 0 - | 0 - | 0 - | 0 - | 0 - | 0 - | 0 - | 0 - |
| 2111 | 1 | 0.2um_NOKCPHJN_59661 | Bacteria | Bacteroidetes | Flavobacteriia | Flavobacterial | Crocinitomica | unclassified | Crocinitomica | metabolism | Energy produc | COG0437 | 4fe-4S ferred | 755732.Fluta_ | 0,6 | 0,8 | 7,29E-06 | 1,01E-05 | 1 - | 1,08E-05 - | 0 - | 0 - | 2 - | 2,56E-05 - | 0 - | 0 - | 0 - | 0 - |
| 2112 | 1 | 0.2um_NOKCPHJN_409618 | Bacteria | Proteobacteri | Gammaprote | unclassified | unclassified | unclassified | various | metabolism | Inorganic ion t | 16PDP | TonB-depende | 314276.OS14 | 1,4 | 0,489897949 | 4,91E-05 | 1,54E-05 | 2 - | 5,98E-05 - | 2 - | 7,42E-05 - | 1 - | 3,53E-05 - | 1 - | 3,58E-05 - | 1 - | 4,03E-05 - |
| 2113 | 1 | 0.2um_NOKCPHJN_11426 | Bacteria | Proteobacteri | Gammaprote | Cellvibrionale | unclassified | unclassified | Cellvibrionale | metabolism | Amino acid tra | COG0174 | glutamine syn | 377629.TERT | 2,6 | 1,854723699 | 7,19E-05 | 4,37E-05 | 6 - | 0,00014715 - | 3 - | 9,14E-05 - | 1 - | 2,89E-05 - | 2 - | 5,88E-05 - | 1 - | 3,31E-05 - |
| 2114 | 1 | 0.2um_NOKCPHJN_571084 | unclassified | unclassified | unclassified | unclassified | unclassified | unclassified | unclassified | unclassified | unclassified | unclassified | unclassified | unclassified | 0,6 | 0,489897949 | 4,62E-05 | 3,77E-05 | 0 - | 0 - | 1 - | 7,92E-05 - | 1 - | 7,53E-05 - | 1 - | 7,64E-05 - | 0 - | 0 - |
| 2115 | 1 | 3um_FIJJAFBF_301239 | Bacteria | Proteobacteri | Gammaprote | Oceanospirilla | Oceanospirilla | Marinobacteri | Marinobacteri | metabolism | Energy produc | 179RI | May be involv | 207954.MED9 | 0,4 | 0,8 | 3,67E-05 | 7,35E-05 | 2 - | 0,000183643 - | 0 - | 0 - | 0 - | 0 - | 0 - | 0 - | 0 - | 0 - |
| 2116 | 1 | 0.2um_NOKCPHJN_97620 | Bacteria | Bacteroidetes | Flavobacteriia | Flavobacterial | Flavobacteriac | Polaribacter | various | metabolism | Inorganic ion t | COG0659 | sulfate transp | 313598.MED1 | 2 | 0 | 5,20E-05 | 4,94E-06 | 2 - | 4,36E-05 - | 2 - | 5,41E-05 - | 2 - | 5,14E-05 - | 2 - | 5,22E-05 - | 2 - | 5,88E-05 - |
| 2117 | 1 | 3um_FIJJAFBF_458294 | Bacteria | Bacteroidetes | Flavobacteriia | Flavobacterial | Cryomorphace | Owenweeksia | Owenweeksia | metabolism | Amino acid tra | 0P7DH | glutamine syn | 926562.Oweh | 1,2 | 0,979795897 | 2,13E-05 | 1,51E-05 | 3 - | 4,72E-05 - | 1 - | 1,95E-05 - | 1 - | 1,86E-05 - | 0 - | 0 - | 1 - | 2,12E-05 - |
| 2118 | 1 | 0.2um_NOKCPHJN_14312 | Bacteria | Bacteroidetes | Cytophagia | Cytophagales | Flammeovirga | unclassified | Flammeovirga | poorly charact | Function unkn | 07EF7 | domain protei | 649349.Lbys_ | 0,2 | 0,4 | 5,99E-07 | 1,20E-06 | 0 - | 0 - | 0 - | 0 - | 1 - | 3,00E-06 - | 0 - | 0 - | 0 - | 0 - |
| 2119 | 1 | 0.2um_NOKCPHJN_56666 | Bacteria | Proteobacteri | Gammaprote | Alteromonada | Alteromonada | Alteromonas | Alteromonas s | metabolism | Carbohydrate | 16QEQ | hydrolase, fan | 722419.PH50 | 16,8 | 1,720465053 | 0,000213264 | 1,73E-05 | 18 - | 0,000192722 - | 17 - | 0,000226009 - | 19 - | 0,000240127 - | 16 - | 0,000205199 - | 14 - | 0,000202261 - |
| 2120 | 1 | 3um_FIJJAFBF_86507 | Bacteria | Bacteroidetes | Flavobacteriia | Flavobacterial | Flavobacteriac | Polaribacter | Polaribacter t | unclassified | unclassified | unclassified | unclassified | unclassified | 1,4 | 0,489897949 | 8,95E-05 | 3,16E-05 | 1 - | 5,36E-05 - | 1 - | 6,66E-05 - | 2 - | 0,000126619 - | 2 - | 0,000128489 - | 1 - | 7,24E-05 - |
| 2121 | 1 | 0.2um_NOKCPHJN_352637 | Bacteria | Proteobacteri | Gammaprote | Cellvibrionale | Cellvibrionace | Simiduia | Simiduia agari | metabolism | Inorganic ion t | 16TQC | TONB-depend | 190485.XCC3 | 1,6 | 1,019803903 | 2,12E-05 | 1,25E-05 | 3 - | 3,54E-05 - | 1 - | 1,46E-05 - | 2 - | 2,78E-05 - | 2 - | 2,82E-05 - | 0 - | 0 - |
| 2122 | 1 | 0.2um_NOKCPHJN_1087615 | Bacteria | Bacteroidetes | Flavobacteriia | unclassified | unclassified | unclassified | Flavobacteria | unclassified | unclassified | unclassified | unclassified | unclassified | 1,8 | 1,166190379 | 4,78E-05 | 3,11E-05 | 1 - | 2,20E-05 - | 1 - | 2,74E-05 - | 4 - | 0,000104017 - | 1 - | 2,64E-05 - | 2 - | 5,95E-05 - |
| 2123 | 1 | 3um_FIJJAFBF_636516 | Bacteria | Bacteroidetes | Flavobacteriia | Flavobacterial | Flavobacteriac | Polaribacter | Polaribacter s | metabolism | Inorganic ion t | 0P8Q4 | tonB-depende | 867900.Celly_ | 2 | 0 | 3,85E-05 | 3,66E-06 | 2 - | 3,23E-05 - | 2 - | 4,01E-05 - | 2 - | 3,81E-05 - | 2 - | 3,87E-05 - | 2 - | 4,36E-05 - |
| 2124 | 1 | 3um_FIJJAFBF_660319 | Bacteria | Proteobacteri | Alphaproteob | Rhodobactera | Rhodobactera | Ruegeria | Ruegeria sp. P | unclassified | unclassified | unclassified | unclassified | unclassified | 0,8 | 0,4 | 3,63E-05 | 1,86E-05 | 1 - | 3,79E-05 - | 1 - | 4,70E-05 - | 0 - | 0 - | 1 - | 4,54E-05 - | 1 - | 5,11E-05 - |
| 2125 | 1 | 0.2um_NOKCPHJN_248044 | Bacteria | Proteobacteri | Alphaproteob | Rhodospirillal | Rhodospirillac | unclassified | Rhodobactera | poorly charact | Function unkn | 05DIH | basic membra | 246200.SPO2 | 1 | 0 | 4,08E-05 | 3,87E-06 | 1 - | 3,42E-05 - | 1 - | 4,24E-05 - | 1 - | 4,03E-05 - | 1 - | 4,09E-05 - | 1 - | 4,61E-05 - |
| 2126 | 1 | 0.2um_NOKCPHJN_308647 | Eukaryota | Chordata | Mammalia | Sirenia | Trichechidae | Trichechus | Trichechus m | metabolism | Inorganic ion t | 0V7QC | Calcium chan | 7955.ENSDAR | 0,6 | 0,489897949 | 5,82E-06 | 4,80E-06 | 1 - | 8,50E-06 - | 1 - | 1,06E-05 - | 1 - | 1,00E-05 - | 0 - | 0 - | 0 - | 0 - |
| 2127 | 1 | 3um_FIJJAFBF_120907 | Bacteria | Bacteroidetes | Cytophagia | Cytophagales | Cytophagacea | Ekhidna | Ekhidna lutea | information st | Translation, ri | 05G6W | One of the pri | 643867.Ftrac_ | 0,8 | 0,748331477 | 5,17E-05 | 4,50E-05 | 2 - | 0,000114206 - | 0 - | 0 - | 1 - | 6,74E-05 - | 0 - | 0 - | 1 - | 7,71E-05 - |
| 2128 | 1 | 3um_FIJJAFBF_652392 | Bacteria | Proteobacteri | Gammaprote | Alteromonada | Colwelliaceae | Colwellia | various | information st | Translation, ri | 17FIH | One of the ea | 167879.CPS_ | 1,6 | 1,019803903 | 0,00021739 | 0,00014057 | 2 - | 0,000229554 - | 3 - | 0,000427557 - | 0 - | 0 - | 2 - | 0,000274967 - | 1 - | 0,000154874 - |
| 2129 | 1 | 0.2um_NOKCPHJN_66074 | Bacteria | Bacteroidetes | Cytophagia | Cytophagales | Flammeovirga | unclassified | Flammeovirga | unclassified | unclassified | unclassified | unclassified | unclassified | 3,8 | 1,166190379 | 8,08E-05 | 2,44E-05 | 4 - | 7,18E-05 - | 3 - | 6,69E-05 - | 3 - | 6,36E-05 - | 6 - | 0,000129092 - | 3 - | 7,27E-05 - |
| 2130 | 1 | 3um_FIJJAFBF_83546 | Bacteria | Bacteroidetes | Flavobacteriia | Flavobacterial | Flavobacteriac | Polaribacter | Polaribacter b | information st | Translation, ri | COG0087 | One of the pri | 313594.PI23P | 1 | 0 | 6,68E-05 | 6,35E-06 | 1 - | 5,60E-05 - | 1 - | 6,95E-05 - | 1 - | 6,61E-05 - | 1 - | 6,71E-05 - | 1 - | 7,55E-05 - |
| 2131 | 1 | 3um_FIJJAFBF_44702 | Bacteria | Bacteroidetes | Flavobacteriia | Flavobacterial | Flavobacteriac | Polaribacter | Polaribacter s | cellular proces | Cell wall/mem | 11TPU | outer membra | 313598.MED1 | 11,8 | 0,979795897 | 0,000957263 | 7,05E-05 | 13 - | 0,000888153 - | 11 - | 0,000933159 - | 11 - | 0,000887086 - | 13 - | 0,001063859 - | 11 - | 0,001014059 - |
| 2132 | 1 | 3um_FIJJAFBF_85780 | unclassified | unclassified | unclassified | unclassified | unclassified | unclassified | unclassified | unclassified | unclassified | unclassified | unclassified | unclassified | 11 | 1,264911064 | 0,000395342 | 3,13E-05 | 12 - | 0,000364371 - | 10 - | 0,000377034 - | 12 - | 0,000430102 - | 12 - | 0,000436455 - | 9 - | 0,000368749 - |
| 2133 | 1 | 3um_FIJJAFBF_254285 | Bacteria | Proteobacteri | Gammaprote | Cellvibrionale | Spongiibacter | Oceanicoccus | Oceanicoccus | information st | Translation, ri | COG0087 | One of the pri | 247633.GP21 | 1,8 | 1,166190379 | 0,000109696 | 6,86E-05 | 3 - | 0,000161657 - | 1 - | 6,69E-05 - | 3 - | 0,00019082 - | 2 - | 0,000129092 - | 0 - | 0 - |
| 2134 | 1 | 0.2um_NOKCPHJN_217494 | Bacteria | Proteobacteri | Gammaprote | Pseudomonad | Pseudomonad | Pseudomonas | Pseudomonas | metabolism | Inorganic ion t | 16QE8 | phosphonate | 399739.Pmen | 4,4 | 0,8 | 0,000155532 | 2,34E-05 | 5 - | 0,000149839 - | 5 - | 0,000186056 - | 5 - | 0,00017687 - | 4 - | 0,000143586 - | 3 - | 0,000121312 - |
| 2135 | 1 | 3um_FIJJAFBF_624005 | Bacteria | Bacteroidetes | Flavobacteriia | Flavobacterial | Flavobacteriac | Marixanthom | Marixanthom | poorly charact | Function unkn | 0P8NI | Inherit from b | 313594.PI23P | 1,8 | 0,748331477 | 3,86E-05 | 1,69E-05 | 1 - | 1,77E-05 - | 1 - | 2,20E-05 - | 2 - | 4,18E-05 - | 3 - | 6,36E-05 - | 2 - | 4,78E-05 - |
| 2136 | 1 | 3um_FIJJAFBF_180758 | Bacteria | Bacteroidetes | Flavobacteriia | unclassified | unclassified | unclassified | Flavobacteria | information st | Translation, ri | COG0052 | 30S ribosomal | 487796.Flav2 | 1 | 0 | 4,77E-05 | 4,53E-06 | 1 - | 4,00E-05 - | 1 - | 4,97E-05 - | 1 - | 4,72E-05 - | 1 - | 4,79E-05 - | 1 - | 5,40E-05 - |
| 2137 | 1 | 0.2um_NOKCPHJN_329715 | Eukaryota | unclassified | Oomycetes | Pythiales | Pythiaceae | Pythium | Pythium insidi | metabolism | Energy produc | COG0055 | Produces ATP | 65071.PYU1_ | 0,8 | 0,4 | 2,16E-05 | 1,10E-05 | 1 - | 2,34E-05 - | 1 - | 2,91E-05 - | 1 - | 2,76E-05 - | 1 - | 2,81E-05 - | 0 - | 0 - |
| 2138 | 1 | 0.2um_NOKCPHJN_370661 | Bacteria | Firmicutes | Bacilli | Lactobacillales | Streptococcac | Streptococcus | Streptococcus | unclassified | unclassified | unclassified | unclassified | unclassified | 0,2 | 0,4 | 1,71E-06 | 3,42E-06 | 1 - | 8,56E-06 - | 0 - | 0 - | 0 - | 0 - | 0 - | 0 - | 0 - | 0 - |
| 2139 | 1 | 0.2um_NOKCPHJN_987129 | unclassified | unclassified | unclassified | unclassified | unclassified | unclassified | unclassified | unclassified | unclassified | unclassified | unclassified | unclassified | 1 | 0 | 0,000167106 | 1,59E-05 | 1 - | 0,000139972 - | 1 - | 0,000173803 - | 1 - | 0,000165222 - | 1 - | 0,000167663 - | 1 - | 0,000188871 - |

| | | | | | | | | | | | | | | | | | | | | | | | | | | | | |
|---|---|---|---|---|---|---|---|---|---|---|---|---|---|---|---|---|---|---|---|---|---|---|---|---|---|---|---|---|
| 2140 | 1 | 3um_FIJJAFBF_298838 | Bacteria | Bacteroidetes | Flavobacteriia | Flavobacterial | Flavobacteriac | Polaribacter | Polaribacter ir | poorly charact | Function unkn | 05C2A | tonB-depende | 313594.PI23P | 11,4 | 1,959591794 | 0,000148485 | 2,46E-05 | 14 - | 0,000153621 - | 12 - | 0,000163502 - | 11 - | 0,000142477 - | 8 - | 0,00010515 - | 12 - | 0,000177676 - |
| 2141 | 1 | 3um_FIJJAFBF_269706 | Bacteria | Proteobacteri | Gammaprotec | Oceanospirilla | Saccharospirill | Reinekea | Reinekea fors | metabolism | Amino acid tra | COG2988 | Transforms N( | 314283.MED2 | 0,2 | 0,4 | 9,00E-06 | 1,80E-05 | 0 - | 0 - | 0 - | 0 - | 1 - | 4,50E-05 - | 0 - | 0 - | 0 - | 0 - |
| 2142 | 1 | 0.2um_NOKCPHJN_401480 | Eukaryota | Echinodermat | Holothuroidea | Aspidochirotic | Stichopodidae | Apostichopus | Apostichopus | unclassified | unclassified | unclassified | unclassified | unclassified | 0,6 | 0,489897949 | 1,03E-05 | 8,52E-06 | 1 - | 1,51E-05 - | 1 - | 1,88E-05 - | 1 - | 1,78E-05 - | 0 - | 0 - | 0 - | 0 - |
| 2143 | 1 | 0.2um_NOKCPHJN_800127 | Bacteria | Bacteroidetes | Flavobacteriia | Flavobacterial | Flavobacteriac | Formosa | Formosa sp. H | poorly charact | Function unkn | 05C2A | tonB-depende | 867900.Celly_ | 5,6 | 1,2 | 0,000115291 | 1,36E-05 | 8 - | 0,0001404 - | 5 - | 0,000108959 - | 5 - | 0,00010358 - | 5 - | 0,00010511 - | 5 - | 0,000118406 - |
| 2144 | 1 | 3um_FIJJAFBF_147393 | Bacteria | Proteobacteri | Alphaproteob | Rhodobactera | Rhodobactera | Planktomarina | Planktomarina | information st | Translation, ri | 01WSM | This is one of t | 314270.RB208 | 1,8 | 1,166190379 | 0,00011786 | 7,79E-05 | 1 - | 5,44E-05 - | 0 - | 0 - | 3 - | 0,000192629 - | 3 - | 0,000195474 - | 2 - | 0,0001468 - |
| 2145 | 1 | 3um_FIJJAFBF_254852 | Bacteria | Proteobacteri | Alphaproteob | Rhodobactera | Rhodobactera | unclassified | Rhodobactera | information st | Translation, ri | COG0184 | Forms an inte | 89187.ISM_06 | 1 | 0 | 0,000153963 | 1,46E-05 | 1 - | 0,000128963 - | 1 - | 0,000160134 - | 1 - | 0,000152227 - | 1 - | 0,000154476 - | 1 - | 0,000174016 - |
| 2146 | 1 | 0.2um_NOKCPHJN_118667 | Bacteria | Proteobacteri | Gammaprotec | Oceanospirilla | Halomonadac | Halomonas | Halomonas an | metabolism | Carbohydrate | 01S87 | ABC transport | 467661.RKLH1 | 2,6 | 1,356465997 | 0,000108991 | 4,61E-05 | 5 - | 0,000182185 - | 2 - | 9,05E-05 - | 1 - | 4,30E-05 - | 3 - | 0,000130936 - | 2 - | 9,83E-05 - |
| 2147 | 1 | 3um_FIJJAFBF_173758 | Bacteria | Bacteroidetes | Flavobacteriia | Flavobacterial | Flavobacteriac | Lutibacter | Lutibacter sp. | poorly charact | Function unkn | 0ZVFQ | Pkd domain cc | 269798.CHU_ | 5,6 | 0,489897949 | 1,34E-05 | 1,83E-06 | 6 - | 1,20E-05 - | 6 - | 1,49E-05 - | 5 - | 1,18E-05 - | 5 - | 1,20E-05 - | 6 - | 1,62E-05 - |
| 2148 | 1 | 0.2um_NOKCPHJN_857061 | Bacteria | Proteobacteri | Alphaproteob | Pelagibacteral | Pelagibacterac | Candidatus Pe | Candidatus Pe | unclassified | unclassified | unclassified | unclassified | unclassified | 3,6 | 0,8 | 0,000218238 | 2,99E-05 | 5 - | 0,000258506 - | 3 - | 0,000192593 - | 3 - | 0,000183084 - | 4 - | 0,000247718 - | 3 - | 0,00020929 - |
| 2149 | 1 | 3um_FIJJAFBF_100702 | Bacteria | Bacteroidetes | Flavobacteriia | Flavobacterial | Flavobacteriac | Formosa | Formosa sp. H | unclassified | unclassified | unclassified | unclassified | unclassified | 11,8 | 0,748331477 | 0,000190148 | 1,91E-05 | 12 - | 0,000162229 - | 13 - | 0,000218227 - | 12 - | 0,000191494 - | 11 - | 0,000178129 - | 11 - | 0,000200662 - |
| 2150 | 1 | 3um_FIJJAFBF_70542 | Bacteria | Bacteroidetes | Cytophagia | Cytophagales | Catalimonada | Catalinimonas | Catalinimonas | unclassified | unclassified | unclassified | unclassified | unclassified | 6,6 | 0,8 | 0,000148214 | 2,73E-05 | 7 - | 0,000131281 - | 6 - | 0,000139724 - | 6 - | 0,000132826 - | 6 - | 0,000134788 - | 8 - | 0,00020245 - |
| 2151 | 1 | 3um_FIJJAFBF_16158 | Bacteria | Bacteroidetes | Flavobacteriia | Flavobacterial | Flavobacteriac | Polaribacter | Polaribacter s | poorly charact | Function unkn | COG0714 | ATPase activit | 313594.PI23P | 1 | 0 | 4,16E-05 | 3,95E-06 | 1 - | 3,49E-05 - | 1 - | 4,33E-05 - | 1 - | 4,12E-05 - | 1 - | 4,18E-05 - | 1 - | 4,71E-05 - |
| 2152 | 2 | 0.2um_NOKCPHJN_159357;0.2um_NOKCPHJN_2 | Bacteria | unclassified | unclassified | unclassified | unclassified | unclassified | uncultured ba | unclassified | unclassified | unclassified | unclassified | unclassified | 0,6 | 0,489897949 | 1,39E-05 | 1,15E-05 | 1 - | 1,93E-05 - | 1 - | 2,40E-05 - | 0 - | 0 - | 0 - | 0 - | 1 - | 2,61E-05 - |
| 2153 | 1 | 0.2um_NOKCPHJN_304674 | Bacteria | Proteobacteri | Gammaprotec | Cellvibrionales | Microbulbifer | Microbulbifer | Microbulbifer | metabolism | Energy produc | 0QJAP | Catalyzes the | 393595.ABO_ | 0,6 | 0,8 | 2,25E-05 | 2,90E-05 | 2 - | 7,06E-05 - | 0 - | 0 - | 1 - | 4,17E-05 - | 0 - | 0 - | 0 - | 0 - |
| 2154 | 1 | 0.2um_NOKCPHJN_745823 | Bacteria | Proteobacteri | Alphaproteob | Rhodobactera | Rhodobactera | Planktomarina | Planktomarina | metabolism | Energy produc | COG0711 | F(1)F(0) ATP s | 439497.RR11_ | 2,4 | 1,019803903 | 0,000182445 | 7,98E-05 | 2 - | 0,000128242 - | 3 - | 0,000238858 - | 2 - | 0,000151377 - | 4 - | 0,000307225 - | 1 - | 8,65E-05 - |
| 2155 | 2 | 0.2um_NOKCPHJN_688653;3um_FIJJAFBF_4331 | Bacteria | various | various | various | various | various | various | unclassified | unclassified | unclassified | unclassified | unclassified | 0,2 | 0,4 | 2,33E-06 | 4,65E-06 | 0 - | 0 - | 0 - | 0 - | 0 - | 0 - | 0 - | 0 - | 1 - | 1,16E-05 - |
| 2156 | 1 | 0.2um_NOKCPHJN_333747 | Bacteria | Proteobacteri | Alphaproteob | unclassified | unclassified | unclassified | Alphaproteob | unclassified | unclassified | unclassified | unclassified | unclassified | 2,8 | 1,326649916 | 0,000105106 | 4,08E-05 | 5 - | 0,000162573 - | 3 - | 0,000121121 - | 3 - | 0,000115141 - | 1 - | 3,89E-05 - | 2 - | 8,77E-05 - |
| 2157 | 1 | 3um_FIJJAFBF_375635 | Bacteria | Proteobacteri | Alphaproteob | Rhodobactera | Rhodobactera | Pseudaestuari | Pseudaestuari | unclassified | unclassified | unclassified | unclassified | unclassified | 0,4 | 0,489897949 | 2,90E-05 | 3,62E-05 | 1 - | 6,17E-05 - | 0 - | 0 - | 0 - | 0 - | 0 - | 0 - | 1 - | 8,33E-05 - |
| 2158 | 1 | 3um_FIJJAFBF_41143 | Bacteria | Bacteroidetes | Cytophagia | Cytophagales | Flammeovirga | unclassified | Flammeovirga | poorly charact | Function unkn | 09EIV | NA | 643867.Ftrac_ | 1,8 | 0,748331477 | 0,000193662 | 8,26E-05 | 2 - | 0,000180751 - | 1 - | 0,00011222 - | 1 - | 0,000106679 - | 3 - | 0,000324764 - | 2 - | 0,000243897 - |
| 2159 | 1 | 3um_FIJJAFBF_110775 | Bacteria | Proteobacteri | Gammaprotec | Cellvibrionales | Halieaceae | Halioglobus | Halioglobus s | information st | Replication, re | COG0513 | purine NTP-de | 247634.GPB2 | 0,4 | 0,489897949 | 1,04E-05 | 1,28E-05 | 1 - | 2,38E-05 - | 0 - | 0 - | 1 - | 2,81E-05 - | 0 - | 0 - | 0 - | 0 - |
| 2160 | 1 | 0.2um_NOKCPHJN_932636 | unclassified | unclassified | unclassified | unclassified | unclassified | unclassified | unclassified | unclassified | unclassified | unclassified | unclassified | unclassified | 0,2 | 0,4 | 1,40E-05 | 2,79E-05 | 0 - | 0 - | 0 - | 0 - | 1 - | 6,98E-05 - | 0 - | 0 - | 0 - | 0 - |
| 2161 | 1 | 0.2um_NOKCPHJN_109747 | Bacteria | Bacteroidetes | Flavobacteriia | Flavobacterial | Flavobacteriac | Formosa | Formosa sp. H | information st | Translation, ri | COG0256 | This is one of t | 983544.Lacal_ | 2 | 0 | 0,000232249 | 2,21E-05 | 2 - | 0,000194537 - | 2 - | 0,000241557 - | 2 - | 0,000229631 - | 2 - | 0,000233023 - | 2 - | 0,000262499 - |
| 2162 | 3 | 0.2um_NOKCPHJN_173921;0.2um_NOKCPHJN_2 | Bacteria | Bacteroidetes | Flavobacteriia | Flavobacterial | Flavobacteriac | Polaribacter | Polaribacter s | information st | Replication, re | 05C6E | dna polymera | various | 0,6 | 0,489897949 | 1,37E-05 | 1,13E-05 | 1 - | 2,03E-05 - | 0 - | 0 - | 1 - | 2,39E-05 - | 1 - | 2,43E-05 - | 0 - | 0 - |
| 2163 | 1 | 3um_FIJJAFBF_39657 | Bacteria | Bacteroidetes | Flavobacteriia | Flavobacterial | Flavobacteriac | Polaribacter | Polaribacter s | unclassified | unclassified | unclassified | unclassified | unclassified | 0,6 | 0,489897949 | 8,22E-05 | 6,79E-05 | 1 - | 0,000119559 - | 1 - | 0,000148457 - | 0 - | 0 - | 1 - | 0,000143212 - | 0 - | 0 - |
| 2164 | 1 | 0.2um_NOKCPHJN_168484 | Bacteria | Proteobacteri | Alphaproteob | unclassified | unclassified | unclassified | various | information st | Transcription | 05VEQ | Cold shock pro | 945681.APO_ | 1,4 | 0,489897949 | 0,000271129 | 7,76E-05 | 2 - | 0,000332686 - | 1 - | 0,000206549 - | 2 - | 0,000392702 - | 1 - | 0,000199251 - | 1 - | 0,000224456 - |
| 2165 | 1 | 3um_FIJJAFBF_758945 | Bacteria | Bacteroidetes | Flavobacteriia | Flavobacterial | Crocinitomica | unclassified | Crocinitomica | information st | Translation, ri | COG0087 | One of the pri | 755732.Fluta_ | 5,4 | 1,496662955 | 0,00034732 | 7,01E-05 | 8 - | 0,000439337 - | 4 - | 0,000272763 - | 4 - | 0,000259296 - | 6 - | 0,000394689 - | 5 - | 0,000370513 - |
| 2166 | 1 | 3um_FIJJAFBF_57576 | Viruses | unclassified | unclassified | unclassified | Phycodnavirid | unclassified | Chrysochrom | unclassified | unclassified | unclassified | unclassified | unclassified | 1,2 | 0,4 | 7,85E-05 | 2,63E-05 | 1 - | 5,49E-05 - | 1 - | 6,82E-05 - | 2 - | 0,000129648 - | 1 - | 6,58E-05 - | 1 - | 7,41E-05 - |
| 2167 | 1 | 3um_FIJJAFBF_119040 | Bacteria | Bacteroidetes | Flavobacteriia | Flavobacterial | Crocinitomica | unclassified | Crocinitomica | unclassified | unclassified | unclassified | unclassified | unclassified | 0,8 | 0,748331477 | 7,53E-05 | 6,50E-05 | 2 - | 0,000163967 - | 1 - | 0,000101799 - | 0 - | 0 - | 0 - | 0 - | 1 - | 0,000110625 - |
| 2168 | 1 | 0.2um_NOKCPHJN_326585 | Bacteria | Proteobacteri | Gammaprotec | Oceanospirilla | Saccharospirill | Saccharospiril | Saccharospiril | unclassified | unclassified | unclassified | unclassified | unclassified | 0,8 | 0,4 | 1,74E-05 | 8,84E-06 | 1 - | 1,88E-05 - | 1 - | 2,34E-05 - | 1 - | 2,22E-05 - | 1 - | 2,26E-05 - | 0 - | 0 - |
| 2169 | 1 | 3um_FIJJAFBF_714940 | Bacteria | Proteobacteri | Alphaproteob | unclassified | unclassified | Candidatus Pu | Candidatus Pu | metabolism | Carbohydrate | 0IVA1 | Photosystem I | 35128.Thapsd | 1,4 | 0,489897949 | 0,000103579 | 3,66E-05 | 1 - | 6,20E-05 - | 1 - | 7,70E-05 - | 2 - | 0,000146467 - | 2 - | 0,000148631 - | 1 - | 8,37E-05 - |
| 2170 | 1 | 0.2um_NOKCPHJN_223184 | Bacteria | Bacteroidetes | Flavobacteriia | Flavobacterial | Flavobacteriac | Polaribacter | Polaribacter s | cellular proces | Intracellular tr | 0P8JN | The central su | 313598.MED1 | 0,8 | 0,4 | 2,40E-05 | 1,22E-05 | 1 - | 2,60E-05 - | 1 - | 3,23E-05 - | 1 - | 3,07E-05 - | 1 - | 3,12E-05 - | 0 - | 0 - |
| 2171 | 1 | 3um_FIJJAFBF_355787 | Bacteria | Bacteroidetes | Flavobacteriia | Flavobacterial | Flavobacteriac | Polaribacter | Polaribacter ir | poorly charact | Function unkn | 0PBET | NA | 313594.PI23P | 1 | 0 | 8,25E-05 | 7,84E-06 | 1 - | 6,91E-05 - | 1 - | 8,59E-05 - | 1 - | 8,16E-05 - | 1 - | 8,28E-05 - | 1 - | 9,33E-05 - |
| 2172 | 1 | 0.2um_NOKCPHJN_651657 | unclassified | unclassified | unclassified | unclassified | unclassified | unclassified | unclassified | unclassified | unclassified | unclassified | unclassified | unclassified | 0,2 | 0,4 | 6,89E-06 | 1,38E-05 | 0 - | 0 - | 0 - | 0 - | 0 - | 0 - | 1 - | 3,45E-05 - | 0 - | 0 - |
| 2173 | 1 | 3um_FIJJAFBF_230047 | Bacteria | Bacteroidetes | Flavobacteriia | various | various | various | various | unclassified | unclassified | unclassified | unclassified | unclassified | 3,4 | 0,8 | 0,000310183 | 7,53E-05 | 3 - | 0,000229554 - | 3 - | 0,000285038 - | 5 - | 0,000451607 - | 3 - | 0,000274967 - | 3 - | 0,000309749 - |
| 2174 | 1 | 3um_FIJJAFBF_437252 | Bacteria | Bacteroidetes | Flavobacteriia | Flavobacterial | Flavobacteriac | Polaribacter | Polaribacter t | poorly charact | Function unkn | 0XSVM | NA | 867900.Celly_ | 13,8 | 2,13541565 | 0,000396359 | 7,56E-05 | 11 - | 0,000263579 - | 15 - | 0,000446301 - | 14 - | 0,000395981 - | 17 - | 0,000487937 - | 12 - | 0,000387995 - |
| 2175 | 1 | 0.2um_NOKCPHJN_75555 | Bacteria | Proteobacteri | Gammaprotec | Oceanospirilla | Saccharospirill | Reinekea | Reinekea fors | unclassified | unclassified | unclassified | unclassified | unclassified | 0,8 | 0,748331477 | 2,35E-05 | 2,06E-05 | 2 - | 5,29E-05 - | 1 - | 3,28E-05 - | 0 - | 0 - | 1 - | 3,17E-05 - | 0 - | 0 - |
| 2176 | 1 | 3um_FIJJAFBF_391936 | Bacteria | Proteobacteri | Gammaprotec | Oceanospirilla | Oceanospirilla | Motiliproteus | Motiliproteus | information st | Replication, re | 05C24 | DNA gyrase ne | 207954.MED9 | 0,2 | 0,4 | 2,60E-06 | 5,21E-06 | 1 - | 1,30E-05 - | 0 - | 0 - | 0 - | 0 - | 0 - | 0 - | 0 - | 0 - |
| 2177 | 2 | 0.2um_NOKCPHJN_785823;3um_FIJJAFBF_3080 | Bacteria | Proteobacteri | Gammaprotec | Cellvibrionales | Porticoccacea | unclassified | SAR92 bacteri | metabolism | Energy produc | 0QIIJ | Succinyl-CoA l | various | 0,4 | 0,489897949 | 1,73E-05 | 2,13E-05 | 1 - | 3,96E-05 - | 0 - | 0 - | 1 - | 4,67E-05 - | 0 - | 0 - | 0 - | 0 - |
| 2178 | 1 | 0.2um_NOKCPHJN_1003783 | Viruses | unclassified | unclassified | Caudovirales | Siphoviridae | T5virus | Escherichia ph | unclassified | unclassified | unclassified | unclassified | unclassified | 0,2 | 0,4 | 1,73E-05 | 3,46E-05 | 0 - | 0 - | 1 - | 8,64E-05 - | 0 - | 0 - | 0 - | 0 - | 0 - | 0 - |
| 2179 | 1 | 3um_FIJJAFBF_594166 | Eukaryota | unclassified | Pelagophycea | Pelagomonad | unclassified | Aureococcus | Aureococcus | metabolism | Energy produc | COG0711 | F(1)F(0) ATP s | 35128.Thapsd | 10,2 | 0,979795897 | 0,000791755 | 8,64E-05 | 10 - | 0,000652141 - | 10 - | 0,000809766 - | 12 - | 0,000923742 - | 10 - | 0,000781155 - | 9 - | 0,000791972 - |
| 2180 | 1 | 3um_FIJJAFBF_575487 | Bacteria | Proteobacteri | Gammaprotec | Cellvibrionales | Porticoccacea | unclassified | various | unclassified | unclassified | unclassified | unclassified | unclassified | 0,8 | 0,4 | 0,000139153 | 7,15E-05 | 1 - | 0,000145287 - | 1 - | 0,000180404 - | 0 - | 0 - | 1 - | 0,00017403 - | 1 - | 0,000196044 - |
| 2181 | 1 | 0.2um_NOKCPHJN_424906 | Bacteria | Bacteroidetes | Flavobacteriia | Flavobacterial | Flavobacteriac | Altibacter | Altibacter lent | unclassified | unclassified | unclassified | unclassified | unclassified | 1,4 | 0,489897949 | 5,31E-05 | 1,67E-05 | 2 - | 6,47E-05 - | 2 - | 8,03E-05 - | 1 - | 3,82E-05 - | 1 - | 3,87E-05 - | 1 - | 4,36E-05 - |
| 2182 | 1 | 3um_FIJJAFBF_153771 | Bacteria | Proteobacteri | Deltaproteoba | unclassified | unclassified | unclassified | Deltaproteoba | unclassified | unclassified | unclassified | unclassified | unclassified | 0,6 | 0,489897949 | 2,74E-05 | 2,28E-05 | 1 - | 3,88E-05 - | 0 - | 0 - | 1 - | 4,58E-05 - | 0 - | 0 - | 1 - | 5,23E-05 - |
| 2183 | 1 | 0.2um_NOKCPHJN_1068277 | Eukaryota | unclassified | Chrysophycea | Chromulinales | Chromulinace | Spumella | Spumella sp. ( | information st | Translation, ri | KOG0052 | This protein p | 164328.Phyra | 1,2 | 0,4 | 6,30E-05 | 2,17E-05 | 1 - | 4,40E-05 - | 1 - | 5,46E-05 - | 1 - | 5,19E-05 - | 2 - | 0,000105351 - | 1 - | 5,93E-05 - |
| 2184 | 1 | 3um_FIJJAFBF_736127 | Bacteria | Bacteroidetes | Flavobacteriia | Flavobacterial | Flavobacteriac | Arenibacter | Arenibacter h | information st | Translation, ri | 05G6W | One of the pri | 867900.Celly_ | 2,6 | 0,489897949 | 0,000177687 | 3,79E-05 | 2 - | 0,000114206 - | 3 - | 0,000212715 - | 3 - | 0,000202212 - | 3 - | 0,000205199 - | 2 - | 0,000154104 - |
| 2185 | 1 | 0.2um_NOKCPHJN_15245 | Bacteria | Proteobacteri | Betaproteoba | unclassified | unclassified | unclassified | Betaproteoba | unclassified | unclassified | unclassified | unclassified | unclassified | 0,4 | 0,8 | 5,91E-05 | 0,000118265 | 0 - | 0 - | 0 - | 0 - | 0 - | 0 - | 2 - | 0,000295663 - | 0 - | 0 - |
| 2186 | 1 | 0.2um_NOKCPHJN_620349 | Bacteria | Proteobacteri | Alphaproteob | Rhodospirillal | Rhodospirillac | Candidatus En | Candidatus En | cellular proces | Cell motility | 01YGF | Flagellin | 488538.SAR11 | 1 | 0 | 1,28E-05 | 1,21E-06 | 1 - | 1,07E-05 - | 1 - | 1,33E-05 - | 1 - | 1,27E-05 - | 1 - | 1,28E-05 - | 1 - | 1,45E-05 - |
| 2187 | 1 | 3um_FIJJAFBF_588184 | Bacteria | Proteobacteri | Alphaproteob | Rhodobactera | Rhodobactera | Sulfitobacter | Sulfitobacter g | unclassified | unclassified | unclassified | unclassified | unclassified | 0,6 | 0,489897949 | 2,17E-05 | 1,78E-05 | 1 - | 3,21E-05 - | 0 - | 0 - | 1 - | 3,78E-05 - | 1 - | 3,84E-05 - | 0 - | 0 - |
| 2188 | 1 | 3um_FIJJAFBF_27716 | Bacteria | Actinobacteri | Actinobacteri | Micrococcales | Microbacteria | unclassified | Microbacteria | unclassified | unclassified | unclassified | unclassified | unclassified | 2,6 | 0,8 | 0,000284595 | 8,83E-05 | 2 - | 0,000183643 - | 2 - | 0,00022803 - | 4 - | 0,000433543 - | 3 - | 0,00032996 - | 2 - | 0,000247799 - |
| 2189 | 1 | 0.2um_NOKCPHJN_29615 | Bacteria | Proteobacteri | Gammaprotec | Alteromonada | Colwelliaceae | Thalassotalea | Thalassotalea | metabolism | Carbohydrate | 16QEQ | hydrolase, fan | 392500.Swoo_ | 3,8 | 0,4 | 5,46E-05 | 4,87E-06 | 4 - | 4,85E-05 - | 4 - | 6,02E-05 - | 4 - | 5,72E-05 - | 4 - | 5,81E-05 - | 3 - | 4,91E-05 - |
| 2190 | 1 | 3um_FIJJAFBF_224019 | Bacteria | Bacteroidetes | Flavobacteriia | Flavobacterial | Flavobacteriac | Polaribacter | Polaribacter s | poorly charact | Function unkn | 11P35 | NA | 398720.MED2 | 2,8 | 1,166190379 | 6,97E-05 | 2,71E-05 | 4 - | 8,57E-05 - | 4 - | 0,000106357 - | 2 - | 5,06E-05 - | 3 - | 7,69E-05 - | 1 - | 2,89E-05 - |
| 2191 | 1 | 0.2um_NOKCPHJN_54289 | Bacteria | Proteobacteri | Gammaprotec | Oceanospirilla | Oceanospirilla | Marinobacteri | Marinobacteri | metabolism | Carbohydrate | 0QHXA | transketolase | 207954.MED9 | 0,2 | 0,4 | 4,29E-06 | 8,59E-06 | 0 - | 0 - | 1 - | 2,15E-05 - | 0 - | 0 - | 0 - | 0 - | 0 - | 0 - |
| 2192 | 1 | 3um_FIJJAFBF_513820 | Viruses | unclassified | unclassified | unclassified | unclassified | unclassified | uncultured M | unclassified | unclassified | unclassified | unclassified | unclassified | 1,2 | 0,748331477 | 3,76E-05 | 2,25E-05 | 2 - | 5,53E-05 - | 1 - | 3,43E-05 - | 2 - | 6,53E-05 - | 1 - | 3,31E-05 - | 0 - | 0 - |
| 2193 | 1 | 3um_FIJJAFBF_578579 | Bacteria | Proteobacteri | Gammaprotec | Alteromonada | Ferrimonadac | Ferrimonas | Ferrimonas fu | unclassified | unclassified | unclassified | unclassified | unclassified | 0,4 | 0,489897949 | 2,30E-05 | 2,83E-05 | 1 - | 5,26E-05 - | 0 - | 0 - | 1 - | 6,21E-05 - | 0 - | 0 - | 0 - | 0 - |
| 2194 | 1 | 0.2um_NOKCPHJN_389109 | Eukaryota | Mollusca | Bivalvia | Pectinoida | Pectinidae | Azumapecten | Azumapecten | cellular proces | Posttranslatio | COG0652 | peptidyl-proly | 7739.JGI1148 | 0,4 | 0,489897949 | 3,12E-05 | 3,86E-05 | 1 - | 6,96E-05 - | 1 - | 8,64E-05 - | 0 - | 0 - | 0 - | 0 - | 0 - | 0 - |
| 2195 | 1 | 3um_FIJJAFBF_05128 | Bacteria | Proteobacteri | Gammaprotec | Oceanospirilla | Endozoicomor | Endozoicomor | Endozoicomor | unclassified | unclassified | unclassified | unclassified | unclassified | 1 | 0,632455532 | 3,02E-05 | 1,72E-05 | 2 - | 5,38E-05 - | 1 - | 3,34E-05 - | 1 - | 3,17E-05 - | 1 - | 3,22E-05 - | 0 - | 0 - |
| 2196 | 1 | 0.2um_NOKCPHJN_420700 | Eukaryota | Bacillariophyt | Coscinodiscop | Thalassiosirale | Thalassiosirac | Thalassiosira | Thalassiosira p | metabolism | Energy produc | COG0056 | Produces ATP | 35128.Thaps2 | 2,2 | 0,748331477 | 5,71E-05 | 1,75E-05 | 3 - | 6,67E-05 - | 3 - | 8,29E-05 - | 2 - | 5,25E-05 - | 2 - | 5,33E-05 - | 1 - | 3,00E-05 - |
| 2197 | 1 | 3um_FIJJAFBF_788032 | Bacteria | Proteobacteri | Alphaproteob | Rhodobactera | Rhodobactera | unclassified | Rhodobactera | information st | Translation, ri | 179ZI | One of the pri | 52598.EE36_0 | 6,2 | 0,979795897 | 0,000646905 | 6,96E-05 | 8 - | 0,000706319 - | 6 - | 0,000657779 - | 5 - | 0,000521085 - | 6 - | 0,000634538 - | 6 - | 0,000714805 - |
| 2198 | 1 | 0.2um_NOKCPHJN_704317 | unclassified | unclassified | unclassified | unclassified | unclassified | unclassified | unclassified | unclassified | unclassified | unclassified | unclassified | unclassified | 0,2 | 0,4 | 2,96E-06 | 5,91E-06 | 0 - | 0 - | 1 - | 1,48E-05 - | 0 - | 0 - | 0 - | 0 - | 0 - | 0 - |
| 2199 | 1 | 3um_FIJJAFBF_459209 | Eukaryota | unclassified | unclassified | Isochrysidales | Noelaerhabda | Emiliania | Emiliania huxl | cellular proces | Posttranslatio | KOG0730 | cell division cy | 35128.Thapsd | 0,2 | 0,4 | 7,92E-06 | 1,58E-05 | 1 - | 3,96E-05 - | 0 - | 0 - | 0 - | 0 - | 0 - | 0 - | 0 - | 0 - |
| 2200 | 2 | 0.2um_NOKCPHJN_141387;3um_FIJJAFBF_3112 | Bacteria | Proteobacteri | Alphaproteob | various | various | various | various | cellular proces | Cell wall/mem | COG2885 | Ompa motb d | 367336.OM22 | 3,4 | 0,489897949 | 0,000282904 | 2,75E-05 | 4 - | 0,000281661 - | 3 - | 0,000262305 - | 4 - | 0,000332472 - | 3 - | 0,000253037 - | 3 - | 0,000285045 - |
| 2201 | 1 | 0.2um_NOKCPHJN_559361 | Bacteria | Spirochaetes | Spirochaetia | Brachyspirales | Brachyspirace | Brachyspira | Brachyspira in | unclassified | unclassified | unclassified | unclassified | unclassified | 0,4 | 0,489897949 | 4,00E-06 | 4,93E-06 | 1 - | 9,18E-06 - | 0 - | 0 - | 1 - | 1,08E-05 - | 0 - | 0 - | 0 - | 0 - |
| 2202 | 1 | 0.2um_NOKCPHJN_260804 | Bacteria | Bacteroidetes | Flavobacteriia | Flavobacterial | Flavobacteriac | Aureitalea | Aureitalea sp. | cellular proces | Cell wall/mem | COG2885 | Ompa motb d | 50743.SCB49_ | 4,8 | 0,4 | 0,000141593 | 1,09E-05 | 5 - | 0,000124217 - | 5 - | 0,000154241 - | 5 - | 0,000146626 - | 5 - | 0,000148791 - | 4 - | 0,00013409 - |
| 2203 | 1 | 3um_FIJJAFBF_155850 | Bacteria | Proteobacteri | Gammaprotec | Oceanospirilla | Saccharospirill | Reinekea | Reinekea fors | unclassified | unclassified | unclassified | unclassified | unclassified | 1,4 | 0,8 | 9,68E-05 | 4,28E-05 | 3 - | 0,000182185 - | 1 - | 7,54E-05 - | 1 - | 7,17E-05 - | 1 - | 7,27E-05 - | 1 - | 8,19E-05 - |
| 2204 | 1 | 3um_FIJJAFBF_814758 | Bacteria | Bacteroidetes | Cytophagia | Cytophagales | Flammeovirga | unclassified | Flammeovirga | unclassified | unclassified | unclassified | unclassified | unclassified | 1,4 | 0,489897949 | 6,16E-05 | 2,23E-05 | 2 - | 7,40E-05 - | 1 - | 4,60E-05 - | 1 - | 4,37E-05 - | 1 - | 4,43E-05 - | 2 - | 9,99E-05 - |
| 2205 | 1 | 3um_FIJJAFBF_62550 | Bacteria | Bacteroidetes | Flavobacteriia | Flavobacterial | Flavobacteriac | Gilvibacter | Gilvibacter sp. | poorly charact | Function unkn | 09EHV | Inherit from b | 50743.SCB49_ | 1,6 | 0,489897949 | 0,000113778 | 3,63E-05 | 2 - | 0,000119559 - | 1 - | 7,42E-05 - | 1 - | 7,06E-05 - | 2 - | 0,000143212 - | 2 - | 0,000161328 - |
| 2206 | 1 | 0.2um_NOKCPHJN_562377 | Bacteria | Proteobacteri | Gammaprotec | unclassified | unclassified | unclassified | Gammaprotec | unclassified | unclassified | unclassified | unclassified | unclassified | 3 | 0,632455532 | 0,000186285 | 5,09E-05 | 3 - | 0,000155104 - | 2 - | 0,000128395 - | 3 - | 0,000183084 - | 3 - | 0,000185788 - | 4 - | 0,000279053 - |
| 2207 | 1 | 0.2um_NOKCPHJN_264014 | Bacteria | Proteobacteri | Gammaprotec | unclassified | unclassified | unclassified | Gammaprotec | information st | Transcription | COG0085 | DNA-depende | 377629.TERTU | 10 | 1,414213562 | 0,000100367 | 1,24E-05 | 12 - | 0,000101572 - | 9 - | 9,46E-05 - | 8 - | 7,99E-05 - | 11 - | 0,000111528 - | 10 - | 0,000114214 - |
| 2208 | 1 | 3um_FIJJAFBF_786159 | Bacteria | Proteobacteri | Gammaprotec | Oceanospirilla | Saccharospirill | Reinekea | Reinekea blan | unclassified | unclassified | unclassified | unclassified | unclassified | 3 | 1,673320053 | 0,000147587 | 8,13E-05 | 3 - | 0,000125668 - | 1 - | 5,20E-05 - | 6 - | 0,000296676 - | 3 - | 0,000150529 - | 2 - | 0,000113047 - |
| 2209 | 1 | 0.2um_NOKCPHJN_305735 | Eukaryota | unclassified | Oomycetes | Saprolegniales | Saprolegniace | Saprolegnia | Saprolegnia di | unclassified | unclassified | unclassified | unclassified | unclassified | 3 | 1,414213562 | 0,000145241 | 6,77E-05 | 3 - | 0,000124307 - | 2 - | 0,000102902 - | 5 - | 0,000244553 - | 4 - | 0,000198532 - | 1 - | 5,59E-05 - |
| 2210 | 1 | 0.2um_NOKCPHJN_291978 | Bacteria | Proteobacteri | Gammaprotec | unclassified | unclassified | unclassified | SAR86 cluster | unclassified | unclassified | unclassified | unclassified | unclassified | 4 | 0,632455532 | 4,86E-05 | 6,47E-06 | 5 - | 5,12E-05 - | 4 - | 5,09E-05 - | 3 - | 3,63E-05 - | 4 - | 4,91E-05 - | 4 - | 5,53E-05 - |
| 2211 | 1 | 0.2um_NOKCPHJN_60742 | Bacteria | Proteobacteri | Alphaproteob | Rhodobactera | Rhodobactera | Amylibacter | Amylibacter k | information st | Translation, ri | 05CEJ | Catalyzes the | 314262.MED1 | 6,8 | 0,748331477 | 0,000131861 | 1,74E-05 | 7 - | 0,000113963 - | 8 - | 0,000161724 - | 6 - | 0,000115304 - | 7 - | 0,000136508 - | 6 - | 0,000131808 - |
| 2212 | 1 | 3um_FIJJAFBF_81225 | Bacteria | Proteobacteri | Gammaprotec | Oceanospirilla | Oceanospirilla | Marinomonas | Marinomonas | information st | Translation, ri | COG0050 | This protein p | 314277.MED1 | 0,6 | 0,489897949 | 0,00012425 | 0,000101772 | 0 - | 0 - | 1 - | 0,000203598 - | 0 - | 0 - | 1 - | 0,000196405 - | 1 - | 0,000221249 - |
| 2213 | 1 | 3um_FIJJAFBF_611848 | Bacteria | Proteobacteri | Gammaprotec | Alteromonada | Alteromonada | Paraglaciecola | various | unclassified | unclassified | unclassified | unclassified | unclassified | 3,2 | 0,4 | 0,000387533 | 2,13E-05 | 4 - | 0,000409917 - | 3 - | 0,000381747 - | 3 - | 0,000362899 - | 3 - | 0,000368259 - | 3 - | 0,000414842 - |
| 2214 | 1 | 0.2um_NOKCPHJN_270941 | Bacteria | Proteobacteri | Betaproteoba | Burkholderial | Burkholderiac | Cupriavidus | Cupriavidus m | poorly charact | Function unkn | 020MJ | NA | 114615.BRAD | 0,2 | 0,4 | 4,74E-06 | 9,49E-06 | 0 - | 0 - | 1 - | 2,37E-05 - | 0 - | 0 - | 0 - | 0 - | 0 - | 0 - |
| 2215 | 1 | 3um_FIJJAFBF_72571 | Bacteria | Bacteroidetes | Flavobacteriia | Flavobacterial | Flavobacteriac | Psychroserper | Psychroserper | unclassified | unclassified | unclassified | unclassified | unclassified | 4 | 0,894427191 | 5,81E-05 | 1,05E-05 | 5 - | 6,17E-05 - | 4 - | 6,13E-05 - | 3 - | 4,37E-05 - | 5 - | 7,39E-05 - | 3 - | 5,00E-05 - |
| 2216 | 1 | 3um_FIJJAFBF_189643 | Bacteria | Bacteroidetes | Flavobacteriia | Flavobacterial | Flavobacteriac | Polaribacter | Polaribacter s | unclassified | unclassified | unclassified | unclassified | unclassified | 1 | 0,632455532 | 6,36E-05 | 3,62E-05 | 2 - | 0,000113081 - | 1 - | 7,02E-05 - | 1 - | 6,67E-05 - | 1 - | 6,77E-05 - | 0 - | 0 - |
| 2217 | 1 | 0.2um_NOKCPHJN_370963 | Bacteria | Bacteroidetes | Flavobacteriia | Flavobacterial | Flavobacteriac | Formosa | Formosa sp. H | unclassified | unclassified | unclassified | unclassified | unclassified | 1 | 0 | 6,49E-05 | 6,17E-06 | 1 - | 5,44E-05 - | 1 - | 6,75E-05 - | 1 - | 6,42E-05 - | 1 - | 6,52E-05 - | 1 - | 7,34E-05 - |
| 2218 | 1 | 3um_FIJJAFBF_715065 | Eukaryota | Bacillariophyt | Mediophyceae | Anaulales | Anaulaceae | Eunotogramm | Eunotogramm | unclassified | unclassified | unclassified | unclassified | unclassified | 2,6 | 0,8 | 0,00054074 | 0,000177817 | 3 - | 0,000521713 - | 3 - | 0,000647813 - | 1 - | 0,000205276 - | 3 - | 0,000624924 - | 3 - | 0,000703975 - |
| 2219 | 1 | 3um_FIJJAFBF_788028 | Bacteria | Proteobacteri | Alphaproteob | Rhodobactera | Rhodobactera | Sulfitobacter | Sulfitobacter r | unclassified | unclassified | unclassified | unclassified | unclassified | 1 | 0 | 0,000221011 | 2,10E-05 | 1 - | 0,000185124 - | 1 - | 0,000229869 - | 1 - | 0,00021852 - | 1 - | 0,000221747 - | 1 - | 0,000249798 - |
| 2220 | 1 | 3um_FIJJAFBF_528250 | Bacteria | Proteobacteri | Betaproteoba | unclassified | unclassified | unclassified | Betaproteoba | poorly charact | Function unkn | 16ZZU | DsrE/DsrF-like | 580332.Slit_1 | 2,6 | 1,2 | 0,000243206 | 8,25E-05 | 5 - | 0,00040701 - | 2 - | 0,000202154 - | 2 - | 0,000192173 - | 2 - | 0,000195012 - | 2 - | 0,00021968 - |
| 2221 | 1 | 3um_FIJJAFBF_300579 | Bacteria | Proteobacteri | Gammaprotec | Arenicellales | Arenicellaceae | Arenicella | Arenicella xan | information st | Transcription | 05VEQ | Cold shock pro | 717774.Marm | 0,8 | 0,4 | 0,00015728 | 8,08E-05 | 1 - | 0,000166343 - | 0 - | 0 - | 1 - | 0,000196351 - | 1 - | 0,000199251 - | 1 - | 0,000224456 - |
| 2222 | 1 | 0.2um_NOKCPHJN_540891 | Bacteria | Proteobacteri | Alphaproteob | Pelagibacteral | Pelagibacterac | Candidatus Pe | Candidatus Pe | unclassified | unclassified | unclassified | unclassified | unclassified | 1,2 | 0,4 | 3,22E-05 | 7,13E-06 | 2 - | 4,62E-05 - | 1 - | 2,87E-05 - | 1 - | 2,73E-05 - | 1 - | 2,77E-05 - | 1 - | 3,12E-05 - |
| 2223 | 1 | 3um_FIJJAFBF_518929 | Bacteria | Proteobacteri | Gammaprotec | Alteromonada | Alteromonada | Paraglaciecola | Paraglaciecola | information st | Transcription | 173HH | transcriptiona | 342610.Patl_2 | 1 | 0 | 2,64E-05 | 2,51E-06 | 1 - | 2,21E-05 - | 1 - | 2,75E-05 - | 1 - | 2,61E-05 - | 1 - | 2,65E-05 - | 1 - | 2,98E-05 - |
| 2224 | 1 | 3um_FIJJAFBF_103105 | Bacteria | Bacteroidetes | Flavobacteriia | Flavobacterial | Flavobacteriac | Polaribacter | Polaribacter s | cellular proces | Cell wall/mem | COG2885 | Ompa motb d | 313594.PI23P | 12,4 | 0,489897949 | 0,000362875 | 3,70E-05 | 13 - | 0,000318824 - | 12 - | 0,000365433 - | 12 - | 0,00034739 - | 12 - | 0,000352521 - | 13 - | 0,000430207 - |
| 2225 | 1 | 0.2um_NOKCPHJN_253984 | Eukaryota | Streptophyta | Lycopodiopsic | Selaginellales | Selaginellacea | Selaginella | Selaginella mo | information st | Transcription | 1BPB5 | nascent polyp | 3694.POPTR_ | 1 | 0 | 6,14E-05 | 5,83E-06 | 1 - | 5,15E-05 - | 1 - | 6,39E-05 - | 1 - | 6,08E-05 - | 1 - | 6,17E-05 - | 1 - | 6,95E-05 - |
| 2226 | 1 | 0.2um_NOKCPHJN_371627 | unclassified | unclassified | unclassified | unclassified | unclassified | unclassified | unclassified | unclassified | unclassified | unclassified | unclassified | unclassified | 0,2 | 0,4 | 2,99E-05 | 5,98E-05 | 0 - | 0 - | 0 - | 0 - | 0 - | 0 - | 1 - | 0,000149438 - | 0 - | 0 - |
| 2227 | 1 | 3um_FIJJAFBF_393845 | Bacteria | Bacteroidetes | Cytophagia | Cytophagales | Flammeovirga | unclassified | Flammeovirga | information st | Translation, ri | 09EM4 | Binds directly | 643867.Ftrac_ | 2 | 0,632455532 | 0,000240047 | 7,80E-05 | 2 - | 0,000201363 - | 2 - | 0,000250033 - | 3 - | 0,000356532 - | 1 - | 0,000120599 - | 2 - | 0,00027171 - |
| 2228 | 1 | 0.2um_NOKCPHJN_296663 | Bacteria | Proteobacteri | Alphaproteob | Rhodobactera | Rhodobactera | Planktomarina | Planktomarina | unclassified | unclassified | unclassified | unclassified | unclassified | 5 | 0,894427191 | 0,000137987 | 1,93E-05 | 6 - | 0,000140257 - | 5 - | 0,000145131 - | 4 - | 0,000110372 - | 6 - | 0,000168004 - | 4 - | 0,000126171 - |
| 2229 | 1 | 0.2um_NOKCPHJN_112737 | Bacteria | Bacteroidetes | unclassified | unclassified | unclassified | unclassified | Bacteroidetes | unclassified | unclassified | unclassified | unclassified | unclassified | 3,4 | 0,489897949 | 0,000142594 | 3,19E-05 | 3 - | 0,000104343 - | 4 - | 0,00017275 - | 3 - | 0,000123166 - | 3 - | 0,000124985 - | 4 - | 0,000187727 - |
| 2230 | 1 | 0.2um_NOKCPHJN_40481 | Bacteria | Bacteroidetes | Flavobacteriia | Flavobacterial | Flavobacteriac | Algibacter | Algibacter sp. | cellular proces | Cell wall/mem | 0ZZRH | outer membra | 983544.Lacal_ | 0,2 | 0,4 | 9,71E-06 | 1,94E-05 | 0 - | 0 - | 0 - | 0 - | 1 - | 4,86E-05 - | 0 - | 0 - | 0 - | 0 - |
| 2231 | 1 | 3um_FIJJAFBF_652494 | Bacteria | Proteobacteri | Alphaproteob | Rhodobactera | Rhodobactera | Planktomarina | Planktomarina | information st | Transcription | 05VEQ | Cold shock pro | 375451.RD1_ | 5 | 1,549193338 | 0,00098151 | 0,000198195 | 8 - | 0,001350315 - | 4 - | 0,000838346 - | 4 - | 0,000796954 - | 5 - | 0,001010907 - | 4 - | 0,000911026 - |
| 2232 | 1 | 0.2um_NOKCPHJN_590542 | Bacteria | Proteobacteri | Alphaproteob | Rhodobactera | Rhodobactera | Planktomarina | Planktomarina | cellular proces | Defense mech | 08JJA | ABC, transpor | 351016.RAZW | 1,6 | 0,489897949 | 3,03E-05 | 9,66E-06 | 2 - | 3,18E-05 - | 1 - | 1,97E-05 - | 1 - | 1,88E-05 - | 2 - | 3,81E-05 - | 2 - | 4,29E-05 - |
| 2233 | 1 | 3um_FIJJAFBF_210525 | Bacteria | Proteobacteri | Betaproteoba | unclassified | unclassified | unclassified | Betaproteoba | unclassified | unclassified | unclassified | unclassified | unclassified | 3,4 | 1,019803903 | 0,00032629 | 9,59E-05 | 4 - | 0,000325608 - | 5 - | 0,000505386 - | 3 - | 0,00028826 - | 3 - | 0,000292518 - | 2 - | 0,00021968 - |
| 2234 | 1 | 3um_FIJJAFBF_17238 | Bacteria | Proteobacteri | Alphaproteob | Rhodobactera | Rhodobactera | Phaeobacter | Phaeobacter g | metabolism | Energy produc | 05C26 | Dehydrogenas | 314262.MED1 | 3,4 | 0,8 | 9,34E-05 | 2,35E-05 | 3 - | 6,90E-05 - | 3 - | 8,57E-05 - | 3 - | 8,15E-05 - | 5 - | 0,000137759 - | 3 - | 9,31E-05 - |
| 2235 | 1 | 0.2um_NOKCPHJN_189459 | Bacteria | Proteobacteri | Alphaproteob | Rhodospirillal | Rhodospirillac | unclassified | Rhodospirillac | unclassified | unclassified | unclassified | unclassified | unclassified | 1 | 0 | 0,000107052 | 1,02E-05 | 1 - | 8,97E-05 - | 1 - | 0,000111343 - | 1 - | 0,000105845 - | 1 - | 0,000107409 - | 1 - | 0,000120996 - |
| 2236 | 1 | 3um_FIJJAFBF_494279 | Bacteria | Proteobacteri | Gammaprotec | Oceanospirilla | Saccharospirill | Reinekea | Reinekea fors | unclassified | unclassified | unclassified | unclassified | unclassified | 2,4 | 0,489897949 | 0,000206021 | 3,64E-05 | 3 - | 0,000217931 - | 3 - | 0,000270605 - | 2 - | 0,000171496 - | 2 - | 0,00017403 - | 2 - | 0,000196044 - |
| 2237 | 1 | 3um_FIJJAFBF_320776 | Bacteria | Proteobacteri | Gammaprotec | Oceanospirilla | Saccharospirill | Reinekea | Reinekea blan | metabolism | Lipid transpor | 16PEQ | Amp-depende | 314283.MED2 | 0,8 | 0,4 | 1,98E-05 | 1,02E-05 | 1 - | 2,08E-05 - | 1 - | 2,58E-05 - | 1 - | 2,45E-05 - | 0 - | 0 - | 1 - | 2,80E-05 - |
| 2238 | 1 | 0.2um_NOKCPHJN_323681 | Bacteria | Proteobacteri | Alphaproteob | Rhodobactera | Rhodobactera | Sedimentitale | Sedimentitale | information st | Translation, ri | 01QWB | One of the pri | 467661.RKLH1 | 4,4 | 0,489897949 | 0,000246321 | 2,10E-05 | 5 - | 0,000236166 - | 4 - | 0,000234599 - | 4 - | 0,000223016 - | 5 - | 0,000282887 - | 4 - | 0,000254937 - |
| 2239 | 2 | 0.2um_NOKCPHJN_350088;3um_FIJJAFBF_5672 | Bacteria | Proteobacteri | Alphaproteob | Pelagibacteral | Pelagibacterac | unclassified | Pelagibacterac | information st | Translation, ri | 179R6 | This protein is | 335992.SAR11 | 0,8 | 0,4 | 6,26E-05 | 3,22E-05 | 1 - | 6,56E-05 - | 1 - | 8,14E-05 - | 1 - | 7,74E-05 - | 0 - | 0 - | 1 - | 8,85E-05 - |
| 2240 | 1 | 3um_FIJJAFBF_02134 | Bacteria | Proteobacteri | Alphaproteob | Rhodospirillal | Rhodospirillac | unclassified | Rhodobactera | information st | Translation, ri | COG0255 | 50s ribosomal | 367336.OM22 | 1 | 0 | 0,000204518 | 1,94E-05 | 1 - | 0,000171309 - | 1 - | 0,000212715 - | 1 - | 0,000202212 - | 1 - | 0,000205199 - | 1 - | 0,000231156 - |
| 2241 | 1 | 3um_FIJJAFBF_736136 | Bacteria | Bacteroidetes | Flavobacteriia | various | various | various | various | unclassified | unclassified | unclassified | unclassified | unclassified | 2,4 | 0,489897949 | 0,000180059 | 2,65E-05 | 3 - | 0,000191295 - | 2 - | 0,000158354 - | 3 - | 0,000225804 - | 2 - | 0,000152759 - | 2 - | 0,000172083 - |
| 2242 | 1 | 0.2um_NOKCPHJN_95012 | Bacteria | Bacteroidetes | Flavobacteriia | Flavobacterial | Flavobacteriac | Ulvibacter | Ulvibacter lito | information st | Transcription | 05D27 | DNA-depende | 50743.SCB49_ | 12,6 | 3,072458299 | 0,000119073 | 2,54E-05 | 16 - | 0,000128153 - | 15 - | 0,000149182 - | 8 - | 7,56E-05 - | 14 - | 0,000134317 - | 10 - | 0,000108077 - |
| 2243 | 1 | 0.2um_NOKCPHJN_510008 | Bacteria | Proteobacteri | Alphaproteob | Pelagibacteral | Pelagibacterac | Candidatus Pe | Candidatus Pe | information st | Translation, ri | 05G6W | One of the pri | 439493.PB721 | 2 | 0,632455532 | 0,000131793 | 4,01E-05 | 2 - | 0,000111977 - | 2 - | 0,000139043 - | 3 - | 0,000198267 - | 2 - | 0,00013413 - | 1 - | 7,55E-05 - |
| 2244 | 1 | 3um_FIJJAFBF_306877 | Bacteria | Proteobacteri | Alphaproteob | Rhodobactera | unclassified | unclassified | Rhodobactera | metabolism | Inorganic ion t | 08UTW | Plays a role in | 391619.RGBS | 1,8 | 1,166190379 | 0,000109088 | 8,04E-05 | 1 - | 4,86E-05 - | 1 - | 6,04E-05 - | 1 - | 5,74E-05 - | 2 - | 0,000116511 - | 4 - | 0,000262499 - |
| 2245 | 1 | 0.2um_NOKCPHJN_12466 | Bacteria | Bacteroidetes | Flavobacteriia | Flavobacterial | Flavobacteriac | Formosa | Formosa sp. H | metabolism | Inorganic ion t | COG3119 | Sulfatase | 320771.Cflav_ | 1,4 | 0,489897949 | 3,77E-05 | 1,33E-05 | 1 - | 2,26E-05 - | 1 - | 2,81E-05 - | 2 - | 5,33E-05 - | 2 - | 5,41E-05 - | 1 - | 3,05E-05 - |
| 2246 | 1 | 0.2um_NOKCPHJN_832206 | Bacteria | Proteobacteri | Alphaproteob | Pelagibacteral | Pelagibacterac | Candidatus Pe | Candidatus Pe | cellular proces | Cell wall/mem | 7257 | NA | 335992.SAR11 | 2 | 0 | 8,48E-05 | 8,06E-06 | 2 - | 7,11E-05 - | 2 - | 8,82E-05 - | 2 - | 8,39E-05 - | 2 - | 8,51E-05 - | 2 - | 9,59E-05 - |
| 2247 | 1 | 0.2um_NOKCPHJN_159881 | Bacteria | Bacteroidetes | Flavobacteriia | Flavobacterial | Flavobacteriac | Algibacter | Algibacter pec | cellular proces | Intracellular tr | COG0653 | Part of the Se | 1046627.BZAF | 3,2 | 1,166190379 | 3,83E-05 | 1,25E-05 | 4 - | 4,11E-05 - | 2 - | 2,55E-05 - | 5 - | 6,06E-05 - | 3 - | 3,69E-05 - | 2 - | 2,77E-05 - |
| 2248 | 1 | 3um_FIJJAFBF_596752 | Bacteria | unclassified | unclassified | unclassified | unclassified | unclassified | uncultured ba | unclassified | unclassified | unclassified | unclassified | unclassified | 0,6 | 0,489897949 | 1,44E-05 | 1,19E-05 | 1 - | 2,10E-05 - | 1 - | 2,61E-05 - | 0 - | 0 - | 1 - | 2,51E-05 - | 0 - | 0 - |
| 2249 | 1 | 3um_FIJJAFBF_213457 | Bacteria | Bacteroidetes | Flavobacteriia | Flavobacterial | Flavobacteriac | Algibacter | Algibacter pec | metabolism | Nucleotide tra | 09B0C | Provides the p | 391603.FBALC | 0,4 | 0,489897949 | 8,58E-06 | 1,06E-05 | 1 - | 1,91E-05 - | 1 - | 2,38E-05 - | 0 - | 0 - | 0 - | 0 - | 0 - | 0 - |
| 2250 | 1 | 3um_FIJJAFBF_72471 | Bacteria | Bacteroidetes | Cytophagia | Cytophagales | Flammeovirga | Reichenbachie | Reichenbachie | unclassified | unclassified | unclassified | unclassified | unclassified | 2,6 | 0,8 | 0,000144009 | 3,02E-05 | 4 - | 0,000190501 - | 2 - | 0,000118273 - | 3 - | 0,00016865 - | 2 - | 0,000114094 - | 2 - | 0,000128527 - |
| 2251 | 1 | 0.2um_NOKCPHJN_237428 | Bacteria | Proteobacteri | Gammaprotec | unclassified | unclassified | unclassified | Gammaprotec | metabolism | Energy produc | COG0055 | Produces ATP | 377629.TERTU | 2,8 | 0,979795897 | 8,47E-05 | 3,58E-05 | 2 - | 4,95E-05 - | 4 - | 0,000122861 - | 2 - | 5,84E-05 - | 2 - | 5,93E-05 - | 4 - | 0,000133512 - |
| 2252 | 1 | 3um_FIJJAFBF_337087 | Bacteria | Proteobacteri | Gammaprotec | Cellvibrionales | Cellvibrionace | Saccharophag | Saccharophag | unclassified | unclassified | unclassified | unclassified | unclassified | 1,8 | 0,4 | 0,000198025 | 4,63E-05 | 2 - | 0,000185124 - | 1 - | 0,000114935 - | 2 - | 0,00021852 - | 2 - | 0,000221747 - | 2 - | 0,000249798 - |
| 2253 | 1 | 3um_FIJJAFBF_660316 | Bacteria | Proteobacteri | Alphaproteob | Rhodobactera | Rhodobactera | Sulfitobacter | Sulfitobacter s | unclassified | unclassified | unclassified | unclassified | unclassified | 1,6 | 0,489897949 | 0,000121363 | 3,87E-05 | 2 - | 0,00012753 - | 1 - | 7,92E-05 - | 1 - | 7,53E-05 - | 2 - | 0,000152759 - | 2 - | 0,000172083 - |
| 2254 | 1 | 3um_FIJJAFBF_177319 | Bacteria | Actinobacteri | Actinobacteri | Micrococcales | Microbacteria | Rhodoluna | Candidatus Rh | unclassified | unclassified | unclassified | unclassified | unclassified | 3,4 | 1,2 | 0,000144592 | 5,80E-05 | 2 - | 6,98E-05 - | 5 - | 0,000216594 - | 4 - | 0,00016472 - | 2 - | 8,36E-05 - | 4 - | 0,000188297 - |
| 2255 | 2 | 3um_FIJJAFBF_128491;3um_FIJJAFBF_494667 | Bacteria | unclassified | unclassified | unclassified | unclassified | unclassified | uncultured ba | metabolism | Nucleotide tra | 0P81Z | Catalyzes the | 313596.RB250 | 2,6 | 1,2 | 0,000146547 | 4,97E-05 | 5 - | 0,000245249 - | 2 - | 0,000121811 - | 2 - | 0,000115797 - | 2 - | 0,000117507 - | 2 - | 0,000132371 - |
| 2256 | 1 | 0.2um_NOKCPHJN_497672 | Bacteria | Proteobacteri | Alphaproteob | Pelagibacteral | Pelagibacterac | Candidatus Pe | Candidatus Pe | unclassified | unclassified | unclassified | unclassified | unclassified | 2,4 | 0,8 | 0,00021961 | 9,35E-05 | 2 - | 0,000150035 - | 2 - | 0,000186299 - | 2 - | 0,000177101 - | 2 - | 0,000179717 - | 4 - | 0,000404901 - |
| 2257 | 1 | 0.2um_NOKCPHJN_191674 | Eukaryota | Chlorophyta | Chlorophyceae | Chlamydomor | Chlamydomor | Chlamydomor | Chlamydomor | poorly charact | Function unkn | 0ZUA4 | chromosome | 65071.PYU1_ | 0,2 | 0,4 | 8,54E-07 | 1,71E-06 | 0 - | 0 - | 0 - | 0 - | 1 - | 4,27E-06 - | 0 - | 0 - | 0 - | 0 - |
| 2258 | 1 | 0.2um_NOKCPHJN_1004726 | Bacteria | Bacteroidetes | Flavobacteriia | Flavobacterial | Flavobacteriac | Polaribacter | Polaribacter s | poorly charact | Function unkn | 0P8NI | Inherit from b | 983548.Krodi_ | 9,4 | 0,8 | 0,000277625 | 3,69E-05 | 10 - | 0,000247364 - | 10 - | 0,000307153 - | 9 - | 0,000262789 - | 8 - | 0,00023704 - | 10 - | 0,000333781 - |
| 2259 | 1 | 0.2um_NOKCPHJN_06932 | Bacteria | Bacteroidetes | Cytophagia | Cytophagales | Flammeovirga | unclassified | Flammeovirga | unclassified | unclassified | unclassified | unclassified | unclassified | 2 | 0,632455532 | 8,55E-05 | 2,60E-05 | 2 - | 7,26E-05 - | 2 - | 9,02E-05 - | 3 - | 0,000128622 - | 2 - | 8,70E-05 - | 1 - | 4,90E-05 - |
| 2260 | 1 | 3um_FIJJAFBF_476734 | Bacteria | Bacteroidetes | Flavobacteriia | Flavobacterial | Flavobacteriac | Tenacibaculur | Tenacibaculur | unclassified | unclassified | unclassified | unclassified | unclassified | 1,4 | 0,489897949 | 6,92E-05 | 2,04E-05 | 2 - | 8,47E-05 - | 1 - | 5,26E-05 - | 1 - | 5,00E-05 - | 2 - | 0,000101464 - | 1 - | 5,71E-05 - |
| 2261 | 1 | 3um_FIJJAFBF_720082 | Bacteria | Planctomycet | Planctomyceti | Planctomycet | Planctomycet | unclassified | Planctomycet | unclassified | unclassified | unclassified | unclassified | unclassified | 0,2 | 0,4 | 1,11E-05 | 2,22E-05 | 0 - | 0 - | 1 - | 5,55E-05 - | 0 - | 0 - | 0 - | 0 - | 0 - | 0 - |
| 2262 | 1 | 0.2um_NOKCPHJN_450181 | Bacteria | Proteobacteri | Alphaproteob | Rhodobactera | Rhodobactera | Celeribacter | Celeribacter b | metabolism | Inorganic ion t | COG0376 | Bifunctional e | 314283.MED2 | 4 | 0,894427191 | 7,42E-05 | 1,41E-05 | 5 - | 7,87E-05 - | 5 - | 9,77E-05 - | 4 - | 7,43E-05 - | 3 - | 5,66E-05 - | 3 - | 6,37E-05 - |
| 2263 | 1 | 3um_FIJJAFBF_674421 | Viruses | unclassified | unclassified | unclassified | unclassified | unclassified | uncultured M | unclassified | unclassified | unclassified | unclassified | unclassified | 1 | 1,095445115 | 6,25E-05 | 6,24E-05 | 3 - | 0,000169621 - | 0 - | 0 - | 1 - | 6,67E-05 - | 0 - | 0 - | 1 - | 7,63E-05 - |
| 2264 | 1 | 3um_FIJJAFBF_501239 | Eukaryota | unclassified | Spirotrichea | Sporadotrichi | Oxytrichidae | Stylonychia | Stylonychia le | metabolism | Energy produc | 0IFIU | Pyrophosphat | 59689.scaffol | 1,8 | 0,748331477 | 4,22E-05 | 1,37E-05 | 3 - | 6,12E-05 - | 1 - | 2,53E-05 - | 2 - | 4,81E-05 - | 2 - | 4,88E-05 - | 1 - | 2,75E-05 - |
| 2265 | 1 | 3um_FIJJAFBF_508732 | Bacteria | Proteobacteri | Gammaprotec | Alteromonada | Alteromonada | Paraglaciecola | Paraglaciecola | unclassified | unclassified | unclassified | unclassified | unclassified | 1,2 | 0,4 | 7,92E-05 | 1,75E-05 | 2 - | 0,00011364 - | 1 - | 7,06E-05 - | 1 - | 6,71E-05 - | 1 - | 6,81E-05 - | 1 - | 7,67E-05 - |
| 2266 | 1 | 0.2um_NOKCPHJN_166875 | Bacteria | Proteobacteri | Alphaproteob | Rhodospirillal | Rhodospirillac | unclassified | Rhodobactera | poorly charact | Function unkn | 05ERF | Extracellular s | 367336.OM22 | 1 | 0,632455532 | 2,67E-05 | 1,70E-05 | 0 - | 0 - | 2 - | 5,35E-05 - | 1 - | 2,54E-05 - | 1 - | 2,58E-05 - | 1 - | 2,91E-05 - |
| 2267 | 1 | 0.2um_NOKCPHJN_424093 | Bacteria | Proteobacteri | Gammaprotec | unclassified | unclassified | unclassified | various | information st | Transcription | 05VEQ | Cold shock pro | 1036674.A28L | 1,2 | 0,4 | 0,000210503 | 4,66E-05 | 2 - | 0,000302044 - | 1 - | 0,000187525 - | 1 - | 0,000178266 - | 1 - | 0,000180899 - | 1 - | 0,000203782 - |
| 2268 | 1 | 3um_FIJJAFBF_659907 | Bacteria | Proteobacteri | Gammaprotec | Oceanospirilla | Saccharospirill | Reinekea | Reinekea fors | unclassified | unclassified | unclassified | unclassified | unclassified | 0,2 | 0,4 | 7,50E-06 | 1,50E-05 | 1 - | 3,75E-05 - | 0 - | 0 - | 0 - | 0 - | 0 - | 0 - | 0 - | 0 - |
| 2269 | 1 | 0.2um_NOKCPHJN_309398 | Bacteria | Proteobacteri | Alphaproteob | Rhodobactera | Rhodobactera | unclassified | Rhodobactera | metabolism | Energy produc | 01RNF | Produces ATP | 314264.ROS2 | 1 | 0 | 2,89E-05 | 2,74E-06 | 1 - | 2,42E-05 - | 1 - | 3,01E-05 - | 1 - | 2,86E-05 - | 1 - | 2,90E-05 - | 1 - | 3,27E-05 - |
| 2270 | 1 | 3um_FIJJAFBF_496264 | Bacteria | Bacteroidetes | Flavobacteriia | Flavobacterial | Flavobacteriac | Polaribacter | Polaribacter s | unclassified | unclassified | unclassified | unclassified | unclassified | 0,8 | 0,4 | 0,000112918 | 5,80E-05 | 1 - | 0,000118327 - | 1 - | 0,000146927 - | 1 - | 0,000139672 - | 0 - | 0 - | 1 - | 0,000159664 - |
| 2271 | 1 | 3um_FIJJAFBF_554715 | Bacteria | Proteobacteri | Gammaprotec | Alteromonada | Alteromonada | Paraglaciecola | Paraglaciecola | unclassified | unclassified | unclassified | unclassified | unclassified | 1,4 | 0,489897949 | 0,000131104 | 3,86E-05 | 2 - | 0,000160527 - | 1 - | 9,97E-05 - | 1 - | 9,47E-05 - | 2 - | 0,000192284 - | 1 - | 0,000108304 - |
| 2272 | 1 | 0.2um_NOKCPHJN_918711 | Eukaryota | unclassified | unclassified | Isochrysidales | Noelaerhabda | Emiliania | Emiliania huxl | cellular proces | Posttranslatio | KOG0731 | ATP-depender | 45157.CMQ2 | 3,4 | 1,019803903 | 7,72E-05 | 2,47E-05 | 3 - | 5,72E-05 - | 5 - | 0,000118371 - | 3 - | 6,75E-05 - | 4 - | 9,14E-05 - | 2 - | 5,15E-05 - |
| 2273 | 1 | 3um_FIJJAFBF_163672 | Bacteria | Proteobacteri | Betaproteoba | unclassified | unclassified | unclassified | Betaproteoba | unclassified | unclassified | unclassified | unclassified | unclassified | 0,4 | 0,489897949 | 9,62E-06 | 1,18E-05 | 0 - | 0 - | 0 - | 0 - | 0 - | 0 - | 1 - | 2,26E-05 - | 1 - | 2,55E-05 - |
| 2274 | 1 | 0.2um_NOKCPHJN_64589 | Bacteria | Proteobacteri | Gammaprotec | Oceanospirilla | Saccharospirill | Reinekea | Reinekea blan | metabolism | Amino acid tra | 16PQJ | Dihydroxy-aci | 314283.MED2 | 3 | 1,549193338 | 6,41E-05 | 2,56E-05 | 6 - | 0,000111977 - | 3 - | 6,95E-05 - | 2 - | 4,41E-05 - | 2 - | 4,47E-05 - | 2 - | 5,04E-05 - |
| 2275 | 1 | 3um_FIJJAFBF_72470 | Bacteria | Bacteroidetes | Cytophagia | Cytophagales | Flammeovirga | unclassified | Flammeovirga | unclassified | unclassified | unclassified | unclassified | unclassified | 2 | 0,632455532 | 0,000169757 | 4,49E-05 | 3 - | 0,00021656 - | 2 - | 0,000179269 - | 1 - | 8,52E-05 - | 2 - | 0,000172935 - | 2 - | 0,000194811 - |
| 2276 | 1 | 3um_FIJJAFBF_266861 | Bacteria | Bacteroidetes | Flavobacteriia | unclassified | unclassified | unclassified | Flavobacteria | metabolism | Energy produc | COG0711 | F(1)F(0) ATP s | 487797.Flav3 | 5,2 | 1,166190379 | 0,000429455 | 0,000101208 | 4 - | 0,000276571 - | 5 - | 0,000429274 - | 7 - | 0,000571311 - | 6 - | 0,000496928 - | 4 - | 0,000373192 - |
| 2277 | 1 | 0.2um_NOKCPHJN_12866 | Bacteria | Bacteroidetes | Flavobacteriia | Flavobacterial | Flavobacteriac | Formosa | Formosa sp. H | information st | Transcription | COG0085 | DNA-depende | 391603.FBALC | 7,4 | 1,2 | 7,94E-05 | 1,30E-05 | 8 - | 7,23E-05 - | 9 - | 0,000100998 - | 6 - | 6,40E-05 - | 8 - | 8,66E-05 - | 6 - | 7,32E-05 - |
| 2278 | 1 | 3um_FIJJAFBF_532809 | Bacteria | Proteobacteri | Alphaproteob | Rhodobactera | Rhodobactera | unclassified | Rhodobactera | metabolism | Nucleotide tra | 01RWV | 2,3-cyclic nucl | 439497.RR11_ | 0,2 | 0,4 | 3,52E-06 | 7,04E-06 | 1 - | 1,76E-05 - | 0 - | 0 - | 0 - | 0 - | 0 - | 0 - | 0 - | 0 - |
| 2279 | 1 | 3um_FIJJAFBF_194067 | Bacteria | Bacteroidetes | Flavobacteriia | Flavobacterial | Flavobacteriac | Polaribacter | Polaribacter h | information st | Translation, ri | COG1185 | Involved in mF | 688270.Celal_ | 7,2 | 1,166190379 | 0,000133041 | 2,81E-05 | 8 - | 0,000123582 - | 6 - | 0,000115089 - | 6 - | 0,000109407 - | 7 - | 0,000129527 - | 9 - | 0,0001876 - |
| 2280 | 1 | 0.2um_NOKCPHJN_487914 | Eukaryota | unclassified | Ichthyosporea | unclassified | unclassified | Capsaspora | Capsaspora ov | unclassified | unclassified | unclassified | unclassified | unclassified | 1,4 | 0,489897949 | 3,71E-05 | 1,09E-05 | 2 - | 4,54E-05 - | 1 - | 2,82E-05 - | 1 - | 2,68E-05 - | 2 - | 5,43E-05 - | 1 - | 3,06E-05 - |
| 2281 | 1 | 3um_FIJJAFBF_162021 | Bacteria | Bacteroidetes | Flavobacteriia | Flavobacterial | Flavobacteriac | Cellulophaga | Cellulophaga a | metabolism | Energy produc | 0P8GN | aconitate hydr | 688270.Celal_ | 2,2 | 0,4 | 3,21E-05 | 2,94E-06 | 3 - | 3,72E-05 - | 2 - | 3,08E-05 - | 2 - | 2,93E-05 - | 2 - | 2,97E-05 - | 2 - | 3,35E-05 - |
| 2282 | 1 | 3um_FIJJAFBF_608990 | Bacteria | Bacteroidetes | unclassified | unclassified | unclassified | unclassified | Bacteroidetes | unclassified | unclassified | unclassified | unclassified | unclassified | 2,4 | 0,489897949 | 4,88E-05 | 1,15E-05 | 2 - | 3,40E-05 - | 3 - | 6,32E-05 - | 2 - | 4,01E-05 - | 3 - | 6,10E-05 - | 2 - | 4,58E-05 - |

| | | | | | | | | | | | | | | | | | | | | | | | | | | | | |
|---|---|---|---|---|---|---|---|---|---|---|---|---|---|---|---|---|---|---|---|---|---|---|---|---|---|---|---|---|
| 2283 | 1 | 3um_FIJJAFBF_147734 | Bacteria | Bacteroidetes | Flavobacteriia | Flavobacterial | Crocinitomica | unclassified | Crocinitomica | unclassified | unclassified | unclassified | unclassified | unclassified | 0,6 | 0,489897949 | 1,23E-05 | 1,02E-05 | 1 - | 1,80E-05 - | 1 - | 2,24E-05 - | 1 - | 2,13E-05 - | 0 - | 0 - | 0 - | 0 - |
| 2284 | 1 | 3um_FIJJAFBF_54653 | Bacteria | Bacteroidetes | unclassified | unclassified | unclassified | unclassified | Bacteroidetes | information st | Translation, ri | COG1185 | Involved in m | 688270.Celal_ | 3,2 | 0,979795897 | 5,89E-05 | 1,66E-05 | 4 - | 6,25E-05 - | 4 - | 7,77E-05 - | 2 - | 3,69E-05 - | 4 - | 7,49E-05 - | 2 - | 4,22E-05 - |
| 2285 | 1 | 3um_FIJJAFBF_433420 | Bacteria | Proteobacteri | Alphaproteob | Rhodobactera | Rhodobactera | unclassified | Rhodobactera | cellular proces | Signal transdu | 1CP7D | hepatocyte gr | 13616.ENSMO | 0,6 | 0,489897949 | 1,93E-05 | 1,59E-05 | 1 - | 2,86E-05 - | 0 - | 0 - | 1 - | 3,38E-05 - | 1 - | 3,43E-05 - | 0 - | 0 - |
| 2286 | 1 | 0.2um_NOKCPHJN_165213 | Bacteria | Bacteroidetes | Flavobacteriia | Flavobacterial | Flavobacteriac | Flaviramulus | Flaviramulus i | unclassified | unclassified | unclassified | unclassified | unclassified | 4,8 | 1,6 | 0,000130587 | 4,20E-05 | 5 - | 0,000115936 - | 5 - | 0,000143958 - | 7 - | 0,000191591 - | 5 - | 0,000138872 - | 2 - | 6,26E-05 - |
| 2287 | 2 | 3um_FIJJAFBF_374488;3um_FIJJAFBF_385008 | Bacteria | Proteobacteri | Alphaproteob | Rhodobactera | Rhodobactera | various | various | unclassified | unclassified | unclassified | unclassified | unclassified | 0,2 | 0,4 | 1,25E-05 | 2,50E-05 | 0 - | 0 - | 0 - | 0 - | 1 - | 6,24E-05 - | 0 - | 0 - | 0 - | 0 - |
| 2288 | 2 | 0.2um_NOKCPHJN_326347;3um_FIJJAFBF_4656 | Eukaryota | unclassified | Oomycetes | Saprolegniales | Saprolegniace | Aphanomyces | Aphanomyces | cellular proces | Intracellular tr | various | various | various | 1,4 | 0,489897949 | 9,40E-05 | 2,69E-05 | 2 - | 0,000115354 - | 1 - | 7,16E-05 - | 2 - | 0,000136163 - | 1 - | 6,91E-05 - | 1 - | 7,78E-05 - |
| 2289 | 1 | 3um_FIJJAFBF_377020 | Eukaryota | Bacillariophyt | Coscinodiscop | Thalassiosirale | Thalassiosirac | Thalassiosira | Thalassiosira p | unclassified | unclassified | unclassified | unclassified | unclassified | 0,4 | 0,489897949 | 4,75E-05 | 5,84E-05 | 0 - | 0 - | 0 - | 0 - | 0 - | 0 - | 1 - | 0,000111775 - | 1 - | 0,000125914 - |
| 2290 | 1 | 3um_FIJJAFBF_22061 | Bacteria | Proteobacteri | Alphaproteob | Rhodobactera | Rhodobactera | unclassified | Rhodobactera | poorly charact | Function unkn | 01TVT | photosyntheti | 314270.RB208 | 0,2 | 0,4 | 7,48E-06 | 1,50E-05 | 1 - | 3,74E-05 - | 0 - | 0 - | 0 - | 0 - | 0 - | 0 - | 0 - | 0 - |
| 2291 | 4 | 0.2um_NOKCPHJN_154729;0.2um_NOKCPHJN_9 | Bacteria | Bacteroidetes | Flavobacteriia | Flavobacterial | Flavobacteriac | various | various | unclassified | unclassified | unclassified | unclassified | unclassified | 1 | 0 | 5,73E-05 | 5,44E-06 | 1 - | 4,80E-05 - | 1 - | 5,96E-05 - | 1 - | 5,67E-05 - | 1 - | 5,75E-05 - | 1 - | 6,48E-05 - |
| 2292 | 1 | 0.2um_NOKCPHJN_93809 | Bacteria | Bacteroidetes | Flavobacteriia | Flavobacterial | Flavobacteriac | Formosa | Formosa sp. H | unclassified | unclassified | unclassified | unclassified | unclassified | 1,8 | 0,748331477 | 5,60E-05 | 2,47E-05 | 1 - | 2,59E-05 - | 3 - | 9,63E-05 - | 2 - | 6,10E-05 - | 2 - | 6,19E-05 - | 1 - | 3,49E-05 - |
| 2293 | 1 | 0.2um_NOKCPHJN_587146 | Bacteria | Proteobacteri | Alphaproteob | Pelagibacteral | Pelagibacterac | Candidatus Pe | Candidatus Pe | unclassified | unclassified | unclassified | unclassified | unclassified | 1 | 0 | 5,64E-05 | 5,35E-06 | 1 - | 4,72E-05 - | 1 - | 5,86E-05 - | 1 - | 5,58E-05 - | 1 - | 5,66E-05 - | 1 - | 6,37E-05 - |
| 2294 | 3 | 0.2um_NOKCPHJN_259503;0.2um_NOKCPHJN_4 | Bacteria | Proteobacteri | Gammaprotec | Cellvibrionales | Porticoccacea | unclassified | various | unclassified | unclassified | unclassified | unclassified | unclassified | 0,6 | 0,489897949 | 9,08E-05 | 7,49E-05 | 1 - | 0,000131927 - | 1 - | 0,000163815 - | 0 - | 0 - | 1 - | 0,000158027 - | 0 - | 0 - |
| 2295 | 1 | 3um_FIJJAFBF_675307 | Bacteria | Proteobacteri | Alphaproteob | unclassified | unclassified | unclassified | Alphaproteob | cellular proces | Posttranslatio | 01QWG | Prevents misf | 331869.BAL19 | 0,2 | 0,4 | 5,10E-06 | 1,02E-05 | 0 - | 0 - | 1 - | 2,55E-05 - | 0 - | 0 - | 0 - | 0 - | 0 - | 0 - |
| 2296 | 1 | 0.2um_NOKCPHJN_537639 | Bacteria | Proteobacteri | Gammaprotec | Alteromonada | Alteromonada | Marinobacter | Marinobacter | cellular proces | Intracellular tr | 175GA | One of the pr | 207954.MED9 | 0,2 | 0,4 | 1,94E-05 | 3,87E-05 | 0 - | 0 - | 0 - | 0 - | 0 - | 0 - | 0 - | 0 - | 1 - | 9,68E-05 - |
| 2297 | 2 | 0.2um_NOKCPHJN_17831;0.2um_NOKCPHJN_76 | Bacteria | Bacteroidetes | Cytophagia | Cytophagales | Flammeovirga | unclassified | Flammeovirga | unclassified | unclassified | unclassified | unclassified | unclassified | 1,8 | 0,4 | 5,68E-05 | 1,39E-05 | 2 - | 5,29E-05 - | 2 - | 6,57E-05 - | 2 - | 6,24E-05 - | 1 - | 3,17E-05 - | 2 - | 7,14E-05 - |
| 2298 | 1 | 0.2um_NOKCPHJN_846338 | Bacteria | Proteobacteri | Alphaproteob | Pelagibacteral | unclassified | Candidatus Fo | Candidatus Fo | unclassified | unclassified | unclassified | unclassified | unclassified | 1 | 0 | 3,75E-05 | 3,56E-06 | 1 - | 3,14E-05 - | 1 - | 3,90E-05 - | 1 - | 3,71E-05 - | 1 - | 3,77E-05 - | 1 - | 4,24E-05 - |
| 2299 | 1 | 0.2um_NOKCPHJN_390928 | Bacteria | Bacteroidetes | Flavobacteriia | Flavobacterial | Flavobacteriac | Polaribacter | Polaribacter g | information st | Translation, ri | COG1185 | Involved in m | 313594.PI23P | 0,2 | 0,4 | 3,08E-06 | 6,15E-06 | 1 - | 1,54E-05 - | 0 - | 0 - | 0 - | 0 - | 0 - | 0 - | 0 - | 0 - |
| 2300 | 1 | 3um_FIJJAFBF_729551 | Bacteria | Bacteroidetes | Flavobacteriia | Flavobacterial | Flavobacteriac | Arenibacter | Arenibacter la | metabolism | Amino acid tra | COG2171 | Catalyzes the | 487797.Flav3 | 2,4 | 0,489897949 | 0,000120116 | 2,12E-05 | 3 - | 0,000127059 - | 3 - | 0,00015777 - | 2 - | 1,00E-04 - | 2 - | 0,000101464 - | 2 - | 0,000114299 - |
| 2301 | 1 | 3um_FIJJAFBF_424139 | Bacteria | Bacteroidetes | Flavobacteriia | Flavobacterial | Flavobacteriac | Polaribacter | Polaribacter s | unclassified | unclassified | unclassified | unclassified | unclassified | 1 | 0 | 8,78E-05 | 8,34E-06 | 1 - | 7,36E-05 - | 1 - | 9,14E-05 - | 1 - | 8,68E-05 - | 1 - | 8,81E-05 - | 1 - | 9,93E-05 - |
| 2302 | 1 | 3um_FIJJAFBF_391576 | Bacteria | Bacteroidetes | Flavobacteriia | Flavobacterial | Flavobacteriac | Aquimarina | Aquimarina p | metabolism | Amino acid tra | 05C6M | Alpha-keto-be | 313603.FB217 | 2,6 | 1,019803903 | 7,19E-05 | 2,87E-05 | 2 - | 4,68E-05 - | 3 - | 8,71E-05 - | 4 - | 0,000110372 - | 3 - | 8,40E-05 - | 1 - | 3,15E-05 - |
| 2303 | 1 | 0.2um_NOKCPHJN_405887 | Bacteria | Proteobacteri | Gammaprotec | Oceanospirilla | Saccharospirill | Reinekea | Reinekea fors | cellular proces | Signal transdu | COG0784 | response regu | 380703.AHA_ | 0,8 | 0,4 | 3,27E-05 | 1,68E-05 | 1 - | 3,46E-05 - | 0 - | 0 - | 1 - | 4,08E-05 - | 1 - | 4,14E-05 - | 1 - | 4,66E-05 - |
| 2304 | 1 | 0.2um_NOKCPHJN_69353 | Bacteria | Bacteroidetes | Flavobacteriia | Flavobacterial | Flavobacteriac | Formosa | Formosa sp. H | unclassified | unclassified | unclassified | unclassified | unclassified | 0,2 | 0,4 | 8,94E-06 | 1,79E-05 | 0 - | 0 - | 1 - | 4,47E-05 - | 0 - | 0 - | 0 - | 0 - | 0 - | 0 - |
| 2305 | 1 | 0.2um_NOKCPHJN_172835 | Bacteria | Proteobacteri | Gammaprotec | Oceanospirilla | Saccharospirill | Reinekea | Reinekea blan | unclassified | unclassified | unclassified | unclassified | unclassified | 2,8 | 0,4 | 0,000164096 | 2,93E-05 | 3 - | 0,00014715 - | 3 - | 0,000182716 - | 2 - | 0,000115797 - | 3 - | 0,000176261 - | 3 - | 0,000198557 - |
| 2306 | 1 | 3um_FIJJAFBF_563613 | Bacteria | Proteobacteri | Gammaprotec | Cellvibrionales | Cellvibrionace | Cellvibrio | Cellvibrio sp. F | unclassified | unclassified | unclassified | unclassified | unclassified | 1,8 | 0,4 | 0,000125281 | 2,93E-05 | 2 - | 0,000117119 - | 1 - | 7,27E-05 - | 2 - | 0,000138247 - | 2 - | 0,000140289 - | 2 - | 0,000158035 - |
| 2307 | 1 | 3um_FIJJAFBF_638942 | Bacteria | Proteobacteri | Alphaproteob | Rhodobactera | Rhodobactera | Ruegeria | Ruegeria sp. A | unclassified | unclassified | unclassified | unclassified | unclassified | 0,2 | 0,4 | 5,93E-06 | 1,19E-05 | 1 - | 2,97E-05 - | 0 - | 0 - | 0 - | 0 - | 0 - | 0 - | 0 - | 0 - |
| 2308 | 1 | 0.2um_NOKCPHJN_426052 | Bacteria | Bacteroidetes | Cytophagia | Cytophagales | Cyclobacteriac | Cecembia | Cecembia lon | unclassified | unclassified | unclassified | unclassified | unclassified | 1 | 0 | 3,99E-05 | 3,79E-06 | 1 - | 3,35E-05 - | 1 - | 4,16E-05 - | 1 - | 3,95E-05 - | 1 - | 4,01E-05 - | 1 - | 4,52E-05 - |
| 2309 | 1 | 0.2um_NOKCPHJN_174378 | Bacteria | Proteobacteri | Betaproteoba | Burkholderial | Comamonada | Malikia | Malikia spinos | metabolism | Amino acid tra | 05TIB | (ABC) transpo | 411684.HPDF | 5,8 | 1,326649916 | 0,000231392 | 3,71E-05 | 8 - | 0,000271661 - | 6 - | 0,000252992 - | 6 - | 0,000240501 - | 4 - | 0,000162702 - | 5 - | 0,000229104 - |
| 2310 | 1 | 3um_FIJJAFBF_123939 | Bacteria | Actinobacteri | Actinobacteri | Micrococcales | Microbacteria | various | various | information st | Replication, re | 07YH6 | single-strande | 312284.A20C1 | 4,4 | 1,019803903 | 0,00036205 | 5,14E-05 | 6 - | 0,000422491 - | 4 - | 0,00034974 - | 5 - | 0,00041559 - | 4 - | 0,000337382 - | 3 - | 0,000285045 - |
| 2311 | 1 | 0.2um_NOKCPHJN_142704 | Bacteria | Bacteroidetes | Flavobacteriia | Flavobacterial | Flavobacteriac | Formosa | Formosa sp. H | poorly charact | Function unkn | 07BR5 | NA | 481448.Minf_ | 0,6 | 0,489897949 | 0,000119305 | 9,82E-05 | 1 - | 0,00017658 - | 0 - | 0 - | 1 - | 0,000208434 - | 1 - | 0,000211513 - | 0 - | 0 - |
| 2312 | 1 | 3um_FIJJAFBF_581776 | Bacteria | Bacteroidetes | Flavobacteriia | Flavobacterial | Flavobacteriac | Polaribacter | Polaribacter s | unclassified | unclassified | unclassified | unclassified | unclassified | 1 | 0 | 0,000150579 | 1,43E-05 | 1 - | 0,000126128 - | 1 - | 0,000156614 - | 1 - | 0,000148882 - | 1 - | 0,000151081 - | 1 - | 0,000170192 - |
| 2313 | 1 | 3um_FIJJAFBF_85328 | Bacteria | Proteobacteri | Gammaprotec | Alteromonada | Alteromonada | various | various | metabolism | Energy produc | COG0056 | Produces ATP | 1085623.GNIT | 11,6 | 1,959591794 | 0,000307099 | 4,74E-05 | 12 - | 0,000268484 - | 12 - | 0,000333377 - | 12 - | 0,000316917 - | 14 - | 0,000375198 - | 8 - | 0,00024152 - |
| 2314 | 1 | 3um_FIJJAFBF_111320 | Viruses | unclassified | unclassified | unclassified | Mimiviridae | unclassified | Yellowstone la | unclassified | unclassified | unclassified | unclassified | unclassified | 0,6 | 0,489897949 | 6,74E-05 | 5,55E-05 | 1 - | 9,98E-05 - | 0 - | 0 - | 1 - | 0,000117811 - | 1 - | 0,000119551 - | 0 - | 0 - |
| 2315 | 1 | 3um_FIJJAFBF_200671 | Bacteria | Verrucomicrol | Verrucomicrol | unclassified | unclassified | unclassified | Verrucomicro | unclassified | unclassified | unclassified | unclassified | unclassified | 0,6 | 0,489897949 | 2,72E-05 | 2,22E-05 | 0 - | 0 - | 1 - | 4,66E-05 - | 1 - | 4,43E-05 - | 1 - | 4,49E-05 - | 0 - | 0 - |
| 2316 | 1 | 0.2um_NOKCPHJN_143372 | Bacteria | Proteobacteri | Gammaprotec | unclassified | unclassified | unclassified | SAR86 cluster | poorly charact | Function unkn | 05C2A | tonB-depende | 391008.Smal_ | 0,2 | 0,4 | 7,76E-06 | 1,55E-05 | 1 - | 3,88E-05 - | 0 - | 0 - | 0 - | 0 - | 0 - | 0 - | 0 - | 0 - |
| 2317 | 1 | 3um_FIJJAFBF_90373 | Bacteria | Bacteroidetes | Flavobacteriia | Flavobacterial | Crocinitomica | unclassified | Crocinitomica | unclassified | unclassified | unclassified | unclassified | unclassified | 1,2 | 0,4 | 0,000149516 | 3,31E-05 | 2 - | 0,000214536 - | 1 - | 0,000133195 - | 1 - | 0,000126619 - | 1 - | 0,000128489 - | 1 - | 0,000144743 - |
| 2318 | 1 | 3um_FIJJAFBF_221603 | Eukaryota | unclassified | Pelagophycea | Pelagomonad | unclassified | Aureococcus | Aureococcus a | information st | Translation, ri | 1520E | S25 ribosomal | 4787.PITG_13 | 1,2 | 0,4 | 0,000138255 | 4,77E-05 | 1 - | 9,65E-05 - | 1 - | 0,000119764 - | 1 - | 0,000113851 - | 2 - | 0,000231064 - | 1 - | 0,000130147 - |
| 2319 | 2 | 0.2um_NOKCPHJN_512306;3um_FIJJAFBF_18964 | Bacteria | Bacteroidetes | Flavobacteriia | Flavobacterial | Flavobacteriac | Polaribacter | Polaribacter s | poorly charact | Function unkn | 05WQR | glycoside hydr | 313598.MED1 | 1 | 0 | 0,000139824 | 1,33E-05 | 1 - | 0,000117119 - | 1 - | 0,000145427 - | 1 - | 0,000138247 - | 1 - | 0,000140289 - | 1 - | 0,000158035 - |
| 2320 | 1 | 0.2um_NOKCPHJN_47355 | Bacteria | Bacteroidetes | Flavobacteriia | Flavobacterial | Flavobacteriac | unclassified | Flavobacteriac | information st | Replication, re | 05C68 | Can catalyze t | 926562.Oweh | 1 | 0 | 4,10E-05 | 3,90E-06 | 1 - | 3,44E-05 - | 1 - | 4,27E-05 - | 1 - | 4,06E-05 - | 1 - | 4,12E-05 - | 1 - | 4,64E-05 - |
| 2321 | 1 | 3um_FIJJAFBF_451976 | Bacteria | Bacteroidetes | Flavobacteriia | Flavobacterial | Flavobacteriac | Polaribacter | Polaribacter s | metabolism | Nucleotide tra | 09B0H | reductase | 313594.PI23P | 0,2 | 0,4 | 6,39E-06 | 1,28E-05 | 0 - | 0 - | 0 - | 0 - | 1 - | 3,20E-05 - | 0 - | 0 - | 0 - | 0 - |
| 2322 | 1 | 0.2um_NOKCPHJN_23157 | Bacteria | Bacteroidetes | Flavobacteriia | Flavobacterial | Flavobacteriac | Polaribacter | Polaribacter s | unclassified | unclassified | unclassified | unclassified | unclassified | 0,2 | 0,4 | 3,67E-06 | 7,35E-06 | 1 - | 1,84E-05 - | 0 - | 0 - | 0 - | 0 - | 0 - | 0 - | 0 - | 0 - |
| 2323 | 1 | 0.2um_NOKCPHJN_113633 | Bacteria | Bacteroidetes | Flavobacteriia | Flavobacterial | Flavobacteriac | Kordia | Kordia sp. SM | information st | Translation, ri | 05C6A | phenylalanyl-t | 391603.FBALC | 0,6 | 0,8 | 1,09E-05 | 1,42E-05 | 0 - | 0 - | 2 - | 3,52E-05 - | 0 - | 0 - | 0 - | 0 - | 1 - | 1,91E-05 - |
| 2324 | 1 | 3um_FIJJAFBF_448200 | Bacteria | Proteobacteri | Gammaprotec | Cellvibrionales | Porticoccacea | unclassified | various | information st | Translation, ri | 08UHH | Located on th | 314287.GB22 | 2,8 | 0,748331477 | 0,000291615 | 7,28E-05 | 3 - | 0,000264869 - | 2 - | 0,00021926 - | 3 - | 0,000312651 - | 4 - | 0,000423026 - | 2 - | 0,000238268 - |
| 2325 | 1 | 3um_FIJJAFBF_08073 | Bacteria | Bacteroidetes | Flavobacteriia | Flavobacterial | Flavobacteriac | Polaribacter | Polaribacter g | metabolism | Nucleotide tra | 0P85S | Plays an impo | 411154.GFO_ | 5,2 | 0,748331477 | 0,000166481 | 1,48E-05 | 6 - | 0,000162804 - | 5 - | 0,000168462 - | 6 - | 0,000192173 - | 5 - | 0,00016251 - | 4 - | 0,000146453 - |
| 2326 | 1 | 3um_FIJJAFBF_25488 | Bacteria | Proteobacteri | Gammaprotec | unclassified | unclassified | unclassified | SAR86 cluster | unclassified | unclassified | unclassified | unclassified | unclassified | 0,6 | 0,489897949 | 1,88E-05 | 1,56E-05 | 1 - | 2,66E-05 - | 0 - | 0 - | 1 - | 3,14E-05 - | 0 - | 0 - | 1 - | 3,59E-05 - |
| 2327 | 1 | 0.2um_NOKCPHJN_537008 | Bacteria | Bacteroidetes | Flavobacteriia | Flavobacterial | Flavobacteriac | Formosa | Formosa sp. H | unclassified | unclassified | unclassified | unclassified | unclassified | 2,2 | 0,4 | 9,54E-05 | 1,94E-05 | 2 - | 7,26E-05 - | 2 - | 9,02E-05 - | 2 - | 8,57E-05 - | 3 - | 0,000130522 - | 2 - | 9,80E-05 - |
| 2328 | 1 | 3um_FIJJAFBF_353891 | Bacteria | Bacteroidetes | Flavobacteriia | Flavobacterial | Flavobacteriac | Polaribacter | Polaribacter a | poorly charact | Function unkn | 05ERK | gliding motilit | 313598.MED1 | 0,8 | 0,4 | 2,53E-05 | 1,28E-05 | 1 - | 2,73E-05 - | 1 - | 3,39E-05 - | 1 - | 3,23E-05 - | 1 - | 3,27E-05 - | 0 - | 0 - |
| 2329 | 1 | 0.2um_NOKCPHJN_764423 | Bacteria | Proteobacteri | Alphaproteob | Rickettsiales | unclassified | unclassified | Rickettsiales b | unclassified | unclassified | unclassified | unclassified | unclassified | 0,6 | 0,489897949 | 5,47E-05 | 4,47E-05 | 0 - | 0 - | 1 - | 9,38E-05 - | 1 - | 8,91E-05 - | 1 - | 9,04E-05 - | 0 - | 0 - |
| 2330 | 1 | 0.2um_NOKCPHJN_344835 | Eukaryota | Bacillariophyt | Coscinodiscop | Thalassiosirale | Thalassiosirac | Thalassiosira | Thalassiosira p | unclassified | unclassified | unclassified | unclassified | unclassified | 0,2 | 0,4 | 4,28E-06 | 8,56E-06 | 0 - | 0 - | 0 - | 0 - | 1 - | 2,14E-05 - | 0 - | 0 - | 0 - | 0 - |
| 2331 | 1 | 0.2um_NOKCPHJN_387586 | Bacteria | Chlorobi | Chlorobia | Chlorobiales | Chlorobiaceae | Chloroherpet | Chloroherpet | unclassified | unclassified | unclassified | unclassified | unclassified | 0,2 | 0,4 | 3,86E-06 | 7,73E-06 | 1 - | 1,93E-05 - | 0 - | 0 - | 0 - | 0 - | 0 - | 0 - | 0 - | 0 - |
| 2332 | 1 | 3um_FIJJAFBF_704591 | Bacteria | Proteobacteri | Gammaprotec | Thiotrichales | Thiolinaceae | Thiolinea | Thiolinea disc | information st | Transcription | 05S3D | Transcription | 52598.EE36_1 | 0,2 | 0,4 | 3,65E-05 | 7,31E-05 | 0 - | 0 - | 1 - | 0,000182716 - | 0 - | 0 - | 0 - | 0 - | 0 - | 0 - |
| 2333 | 2 | 0.2um_NOKCPHJN_123861;3um_FIJJAFBF_5609 | Eukaryota | unclassified | Florideophyce | Ceramiales | Rhodomelace | Polysiphonia | Polysiphonia t | metabolism | Energy produc | 0FJ7X | This is one of | various | 1,2 | 0,4 | 4,67E-05 | 1,99E-05 | 1 - | 3,19E-05 - | 1 - | 3,96E-05 - | 1 - | 3,76E-05 - | 1 - | 3,82E-05 - | 2 - | 8,60E-05 - |
| 2334 | 1 | 3um_FIJJAFBF_33501 | Bacteria | Proteobacteri | Alphaproteob | Rhodobactera | Rhodobactera | Planktomarin | Planktomarin | poorly charact | Function unkn | 01Y9B | Protein of unk | 391593.RCCS2 | 1,2 | 1,166190379 | 0,000164492 | 0,000153222 | 3 - | 0,000358677 - | 0 - | 0 - | 1 - | 0,000141127 - | 0 - | 0 - | 2 - | 0,000322655 - |
| 2335 | 1 | 3um_FIJJAFBF_191799 | Bacteria | Bacteroidetes | Flavobacteriia | Flavobacterial | Flavobacteriac | Psychroserper | Psychroserper | poorly charact | Function unkn | 05YA2 | NA | 391603.FBALC | 1 | 0,632455532 | 2,20E-05 | 1,23E-05 | 2 - | 3,81E-05 - | 1 - | 2,36E-05 - | 1 - | 2,25E-05 - | 0 - | 0 - | 1 - | 2,57E-05 - |
| 2336 | 1 | 3um_FIJJAFBF_629418 | Bacteria | Bacteroidetes | Flavobacteriia | Flavobacterial | Flavobacteriac | Polaribacter | Polaribacter t | unclassified | unclassified | unclassified | unclassified | unclassified | 5,8 | 0,748331477 | 0,000159702 | 1,96E-05 | 7 - | 0,000162311 - | 6 - | 0,00017275 - | 5 - | 0,000136851 - | 5 - | 0,000138872 - | 6 - | 0,000187727 - |
| 2337 | 1 | 3um_FIJJAFBF_719256 | Bacteria | Bacteroidetes | Flavobacteriia | Flavobacterial | Flavobacteriac | Polaribacter | Polaribacter ir | unclassified | unclassified | unclassified | unclassified | unclassified | 0,6 | 0,489897949 | 1,49E-05 | 1,23E-05 | 1 - | 2,18E-05 - | 1 - | 2,70E-05 - | 1 - | 2,57E-05 - | 0 - | 0 - | 0 - | 0 - |
| 2338 | 1 | 3um_FIJJAFBF_713309 | Bacteria | Proteobacteri | Alphaproteob | Rhodobactera | Rhodobactera | Phaeobacter | Phaeobacter g | metabolism | Energy produc | 05CMV | Succinyl-CoA s | 52598.EE36_0 | 1 | 0,894427191 | 3,22E-05 | 2,87E-05 | 2 - | 5,78E-05 - | 0 - | 0 - | 1 - | 3,41E-05 - | 2 - | 6,93E-05 - | 0 - | 0 - |
| 2339 | 1 | 0.2um_NOKCPHJN_203022 | Bacteria | Bacteroidetes | Flavobacteriia | Flavobacterial | unclassified | unclassified | Flavobacterial | unclassified | unclassified | unclassified | unclassified | unclassified | 0,8 | 0,748331477 | 7,44E-05 | 6,58E-05 | 2 - | 0,00017004 - | 0 - | 0 - | 1 - | 0,000100357 - | 1 - | 0,000101839 - | 0 - | 0 - |
| 2340 | 1 | 3um_FIJJAFBF_331284 | Bacteria | Proteobacteri | Alphaproteob | Rhodobactera | Rhodobactera | Pseudooceani | Pseudooceani | metabolism | Amino acid tra | 16PKZ | extracellular li | 314271.RB265 | 0,8 | 0,4 | 2,77E-05 | 1,42E-05 | 1 - | 2,93E-05 - | 0 - | 0 - | 1 - | 3,46E-05 - | 1 - | 3,51E-05 - | 1 - | 3,95E-05 - |
| 2341 | 1 | 0.2um_NOKCPHJN_1095133 | Eukaryota | Platyhelminth | Trematoda | Plagiorchiida | Fasciolidae | Fasciola | Fasciola hepat | cellular proces | Cytoskeleton | 0VQU4 | ACTIN | 7897.ENSLACF | 1 | 0 | 9,01E-05 | 8,56E-06 | 1 - | 7,55E-05 - | 1 - | 9,38E-05 - | 1 - | 8,91E-05 - | 1 - | 9,04E-05 - | 1 - | 0,000101891 - |
| 2342 | 1 | 3um_FIJJAFBF_46932 | Bacteria | Proteobacteri | Gammaprotec | Alteromonada | Alteromonada | Paraglaciecola | various | information st | Translation, ri | 179RU | 30S ribosomal | 342610.Patl_3 | 2 | 0 | 0,000210811 | 2,00E-05 | 2 - | 0,00017658 - | 2 - | 0,00021926 - | 2 - | 0,000208434 - | 2 - | 0,000211513 - | 2 - | 0,000238268 - |
| 2343 | 1 | 3um_FIJJAFBF_612092 | Bacteria | Proteobacteri | Gammaprotec | Oceanospirilla | Saccharospirill | Reinekea | Reinekea blan | unclassified | unclassified | unclassified | unclassified | unclassified | 0,4 | 0,8 | 1,24E-05 | 2,47E-05 | 2 - | 6,19E-05 - | 0 - | 0 - | 0 - | 0 - | 0 - | 0 - | 0 - | 0 - |
| 2344 | 1 | 3um_FIJJAFBF_205215 | Eukaryota | unclassified | Raphidophyce | Chattonellales | Chattonellace | Heterosigma | Heterosigma a | unclassified | unclassified | unclassified | unclassified | unclassified | 1,4 | 0,489897949 | 3,70E-05 | 1,16E-05 | 2 - | 4,51E-05 - | 2 - | 5,60E-05 - | 1 - | 2,66E-05 - | 1 - | 2,70E-05 - | 1 - | 3,04E-05 - |
| 2345 | 1 | 0.2um_NOKCPHJN_515759 | Bacteria | Bacteroidetes | Flavobacteriia | Flavobacterial | Flavobacteriac | Gaetbulibacte | Gaetbulibacte | metabolism | Energy produc | 05C26 | Dehydrogenas | 983544.Lacal_ | 0,6 | 0,489897949 | 1,90E-05 | 1,56E-05 | 0 - | 0 - | 1 - | 3,11E-05 - | 0 - | 0 - | 1 - | 3,00E-05 - | 1 - | 3,38E-05 - |
| 2346 | 1 | 3um_FIJJAFBF_21369 | Bacteria | Proteobacteri | Alphaproteob | Rhizobiales | Phyllobacteria | Mesorhizobiu | Mesorhizobiu | unclassified | unclassified | unclassified | unclassified | unclassified | 1 | 0,894427191 | 0,000130715 | 0,000115064 | 2 - | 0,000231872 - | 1 - | 0,000143958 - | 0 - | 0 - | 2 - | 0,000277744 - | 0 - | 0 - |
| 2347 | 1 | 3um_FIJJAFBF_34875 | Bacteria | Proteobacteri | Gammaprotec | Oceanospirilla | Saccharospirill | Reinekea | Reinekea blan | metabolism | Amino acid tra | COG0339 | oligopeptidas | 314283.MED2 | 1 | 1,095445115 | 1,82E-05 | 1,86E-05 | 3 - | 5,06E-05 - | 0 - | 0 - | 1 - | 1,99E-05 - | 1 - | 2,02E-05 - | 0 - | 0 - |
| 2348 | 1 | 0.2um_NOKCPHJN_163868 | Bacteria | Bacteroidetes | Flavobacteriia | unclassified | unclassified | unclassified | Flavobacteria | poorly charact | Function unkn | 04X2K | Mediates bind | 176279.SERP2 | 15,4 | 0,489897949 | 0,00028887 | 2,25E-05 | 16 - | 0,000251911 - | 15 - | 0,000293249 - | 15 - | 0,00027877 - | 16 - | 0,000301747 - | 15 - | 0,000318672 - |
| 2349 | 1 | 3um_FIJJAFBF_673120 | Bacteria | Proteobacteri | Gammaprotec | Pseudomonad | Moraxellaceae | Psychrobacter | Psychrobacter | unclassified | unclassified | unclassified | unclassified | unclassified | 1,2 | 0,4 | 0,000121199 | 2,68E-05 | 2 - | 0,000173904 - | 1 - | 0,000107969 - | 1 - | 0,000102638 - | 1 - | 0,000104154 - | 1 - | 0,000117329 - |
| 2350 | 1 | 0.2um_NOKCPHJN_753785 | Bacteria | Bacteroidetes | Flavobacteriia | Flavobacterial | Flavobacteriac | unclassified | Flavobacteriac | unclassified | unclassified | unclassified | unclassified | unclassified | 2 | 0 | 0,000224635 | 2,13E-05 | 2 - | 0,000188159 - | 2 - | 0,000233637 - | 2 - | 0,000222102 - | 2 - | 0,000225382 - | 2 - | 0,000253893 - |
| 2351 | 1 | 0.2um_NOKCPHJN_30621 | Bacteria | Bacteroidetes | Flavobacteriia | Flavobacterial | Flavobacteriac | Formosa | Formosa sp. H | unclassified | unclassified | unclassified | unclassified | unclassified | 0,6 | 0,489897949 | 2,18E-05 | 1,79E-05 | 1 - | 3,22E-05 - | 0 - | 0 - | 1 - | 3,81E-05 - | 1 - | 3,86E-05 - | 0 - | 0 - |
| 2352 | 1 | 0.2um_NOKCPHJN_910811 | Bacteria | Proteobacteri | Alphaproteob | various | various | various | various | information st | Translation, ri | 17FUG | Binds directly | 367336.OM22 | 2,8 | 0,748331477 | 0,000349993 | 0,000105516 | 3 - | 0,000313028 - | 4 - | 0,00051825 - | 2 - | 0,000246331 - | 2 - | 0,00024997 - | 3 - | 0,000422385 - |
| 2353 | 1 | 3um_FIJJAFBF_148353 | Bacteria | Proteobacteri | Alphaproteob | Pelagibacteral | Pelagibacterac | unclassified | alpha proteob | cellular proces | Cell wall/mem | 7257 | NA | 335992.SAR11 | 0,8 | 0,4 | 5,77E-05 | 2,96E-05 | 1 - | 6,11E-05 - | 0 - | 0 - | 1 - | 7,21E-05 - | 1 - | 7,31E-05 - | 1 - | 8,24E-05 - |
| 2354 | 1 | 3um_FIJJAFBF_465307 | Bacteria | Bacteroidetes | Cytophagia | Cytophagales | Flammeovirga | unclassified | Flammeovirga | metabolism | Inorganic ion t | COG0841 | acriflavin resis | 643867.Ftrac_ | 0,4 | 0,489897949 | 4,89E-06 | 5,99E-06 | 0 - | 0 - | 1 - | 1,25E-05 - | 1 - | 1,19E-05 - | 0 - | 0 - | 0 - | 0 - |
| 2355 | 1 | 3um_FIJJAFBF_33869 | Bacteria | Bacteroidetes | Flavobacteriia | Flavobacterial | unclassified | unclassified | Flavobacteriac | metabolism | Carbohydrate | 099YR | Catalyzes the | 867900.Celly_ | 4,2 | 0,4 | 0,000133769 | 1,69E-05 | 4 - | 0,000106769 - | 4 - | 0,000132576 - | 5 - | 0,000157537 - | 4 - | 0,000127891 - | 4 - | 0,000144069 - |
| 2356 | 1 | 3um_FIJJAFBF_116351 | Bacteria | Bacteroidetes | Cytophagia | Cytophagales | Flammeovirga | unclassified | Flammeovirga | cellular proces | Posttranslatio | 09GE9 | Thioredoxin | 643867.Ftrac_ | 1,2 | 0,4 | 0,000140002 | 5,96E-05 | 1 - | 9,56E-05 - | 1 - | 0,000118766 - | 1 - | 0,000112902 - | 1 - | 0,000114569 - | 2 - | 0,000258124 - |
| 2357 | 1 | 0.2um_NOKCPHJN_202274 | Bacteria | Bacteroidetes | Flavobacteriia | Flavobacterial | Flavobacteriac | Formosa | Formosa sp. H | information st | Transcription | 05D27 | DNA-depende | 391603.FBALC | 13,4 | 2,059126028 | 0,000126492 | 9,87E-06 | 17 - | 0,000136162 - | 14 - | 0,000139237 - | 12 - | 0,000113453 - | 13 - | 0,000124723 - | 11 - | 0,000118885 - |
| 2358 | 2 | 0.2um_NOKCPHJN_59698;3um_FIJJAFBF_75371 | Bacteria | Bacteroidetes | unclassified | unclassified | unclassified | unclassified | Bacteroidetes | cellular proces | Posttranslatio | 05E6M | Sulphatase-m | various | 0,2 | 0,4 | 5,92E-06 | 1,18E-05 | 0 - | 0 - | 0 - | 0 - | 1 - | 2,96E-05 - | 0 - | 0 - | 0 - | 0 - |
| 2359 | 1 | 3um_FIJJAFBF_718677 | Eukaryota | unclassified | Cryptophyta | Pyrenomonad | Pyrenomonad | Rhodomonas | Rhodomonas | unclassified | unclassified | unclassified | unclassified | unclassified | 0,4 | 0,8 | 3,06E-05 | 6,12E-05 | 2 - | 0,000153036 - | 0 - | 0 - | 0 - | 0 - | 0 - | 0 - | 0 - | 0 - |
| 2360 | 1 | 3um_FIJJAFBF_39854 | Bacteria | Proteobacteri | Gammaprotec | Cellvibrionales | unclassified | unclassified | Cellvibrionales | information st | Translation, ri | 177SZ | This protein b | 566466.NOR5 | 1 | 0 | 9,65E-05 | 9,16E-06 | 1 - | 8,08E-05 - | 1 - | 0,000100365 - | 1 - | 9,54E-05 - | 1 - | 9,68E-05 - | 1 - | 0,000109067 - |
| 2361 | 1 | 0.2um_NOKCPHJN_56676 | Bacteria | Bacteroidetes | unclassified | unclassified | unclassified | unclassified | Bacteroidetes | information st | Replication, re | 0PB0H | DNA-binding p | 487796.Flav2 | 1 | 0 | 0,000153963 | 1,46E-05 | 1 - | 0,000128963 - | 1 - | 0,000160134 - | 1 - | 0,000152227 - | 1 - | 0,000154476 - | 1 - | 0,000174016 - |
| 2362 | 1 | 0.2um_NOKCPHJN_278290 | Eukaryota | Phaeophyceae | unclassified | Ectocarpales | Ectocarpaceae | Ectocarpus | Ectocarpus sili | metabolism | Energy produc | COG0055 | Produces ATP | 35128.Thaps4 | 8,4 | 1,019803903 | 0,000225774 | 2,89E-05 | 8 - | 0,000180751 - | 8 - | 0,000224439 - | 10 - | 0,000266697 - | 9 - | 0,000243573 - | 7 - | 0,00021341 - |
| 2363 | 1 | 3um_FIJJAFBF_410383 | Bacteria | Bacteroidetes | unclassified | unclassified | unclassified | unclassified | unidentified e | cellular proces | Posttranslatio | COG1404 | peptidase (S8 | 50743.SCB49_ | 1 | 0 | 2,47E-05 | 2,35E-06 | 1 - | 2,07E-05 - | 1 - | 2,57E-05 - | 1 - | 2,45E-05 - | 1 - | 2,48E-05 - | 1 - | 2,80E-05 - |
| 2364 | 1 | 0.2um_NOKCPHJN_463002 | Bacteria | Proteobacteri | Alphaproteob | Rhodobactera | Rhodobactera | unclassified | Rhodobactera | unclassified | unclassified | unclassified | unclassified | unclassified | 1,4 | 1,019803903 | 2,33E-05 | 1,57E-05 | 3 - | 4,45E-05 - | 2 - | 3,68E-05 - | 1 - | 1,75E-05 - | 1 - | 1,78E-05 - | 0 - | 0 - |
| 2365 | 1 | 3um_FIJJAFBF_786160 | Bacteria | Proteobacteri | Gammaprotec | Oceanospirilla | Saccharospirill | Reinekea | Reinekea fors | information st | Translation, ri | 17FIH | One of the ea | 314283.MED2 | 7,4 | 1,019803903 | 0,001021655 | 5,50E-05 | 9 - | 0,001054072 - | 7 - | 0,001017992 - | 8 - | 0,001105977 - | 7 - | 0,000982024 - | 6 - | 0,000948211 - |
| 2366 | 1 | 3um_FIJJAFBF_328496 | Bacteria | Bacteroidetes | Flavobacteriia | Flavobacterial | Flavobacteriac | Arenitalea | Arenitalea lut | metabolism | Inorganic ion t | 05C27 | TonB-depend | 886377.Murru | 29,2 | 2,13541565 | 0,000382649 | 2,91E-05 | 32 - | 0,00035282 - | 29 - | 0,000397027 - | 31 - | 0,000403453 - | 26 - | 0,000343378 - | 28 - | 0,000416569 - |
| 2367 | 1 | 0.2um_NOKCPHJN_619394 | Bacteria | Proteobacteri | Alphaproteob | Pelagibacteral | unclassified | unclassified | Pelagibacteral | metabolism | Secondary me | 01Q86 | extracellular s | 439493.PB721 | 0,2 | 0,4 | 5,83E-06 | 1,17E-05 | 1 - | 2,91E-05 - | 0 - | 0 - | 0 - | 0 - | 0 - | 0 - | 0 - | 0 - |
| 2368 | 1 | 3um_FIJJAFBF_122423 | Bacteria | Bacteroidetes | Flavobacteriia | Flavobacterial | Flavobacteriac | Polaribacter | various | unclassified | unclassified | unclassified | unclassified | unclassified | 3 | 0,894427191 | 0,000513839 | 0,000138479 | 4 - | 0,000581148 - | 2 - | 0,000360807 - | 2 - | 0,000342993 - | 4 - | 0,000696118 - | 3 - | 0,000588131 - |
| 2369 | 1 | 3um_FIJJAFBF_26346 | Bacteria | Bacteroidetes | Flavobacteriia | Flavobacterial | Flavobacteriac | Seonamhaeic | Seonamhaeic | poorly charact | Function unkn | 09AVF | Protein of unk | 391587.KAOT | 1,8 | 1,166190379 | 7,79E-05 | 4,81E-05 | 3 - | 0,000111434 - | 1 - | 4,61E-05 - | 3 - | 0,000131536 - | 0 - | 0 - | 2 - | 0,000100242 - |
| 2370 | 1 | 3um_FIJJAFBF_159469 | Bacteria | Proteobacteri | Gammaprotec | Oceanospirilla | Saccharospiril | Saccharospiril | Saccharospiril | metabolism | Energy produc | COG0843 | Cytochrome C | 208964.PA01 | 0,8 | 0,748331477 | 2,11E-05 | 2,01E-05 | 0 - | 0 - | 2 - | 5,40E-05 - | 1 - | 2,57E-05 - | 1 - | 2,60E-05 - | 0 - | 0 - |
| 2371 | 1 | 0.2um_NOKCPHJN_324343 | Eukaryota | unclassified | Oomycetes | Peronosporale | Peronosporac | Phytophthora | Phytophthora | poorly charact | Function unkn | 0IG8C | kiaa1009 | 7897.ENSLACF | 1,4 | 0,489897949 | 1,42E-05 | 5,83E-06 | 1 - | 8,32E-06 - | 1 - | 1,03E-05 - | 1 - | 9,82E-06 - | 2 - | 1,99E-05 - | 2 - | 2,25E-05 - |
| 2372 | 1 | 3um_FIJJAFBF_123889 | Bacteria | Bacteroidetes | Flavobacteriia | Flavobacterial | Flavobacteriac | Ochrovirga | Ochrovirga pa | metabolism | Inorganic ion t | 05C27 | TonB-depend | 313594.PI23P | 0,4 | 0,489897949 | 4,87E-06 | 6,00E-06 | 1 - | 1,11E-05 - | 0 - | 0 - | 0 - | 0 - | 1 - | 1,33E-05 - | 0 - | 0 - |
| 2373 | 1 | 3um_FIJJAFBF_128442 | Bacteria | Bacteroidetes | Flavobacteriia | Flavobacterial | Flavobacteriac | Polaribacter | Polaribacter s | cellular proces | Cell wall/mem | 06B97 | phosphate-sel | 313594.PI23P | 1,8 | 0,979795897 | 5,59E-05 | 2,71E-05 | 3 - | 8,12E-05 - | 1 - | 3,36E-05 - | 3 - | 9,59E-05 - | 1 - | 3,24E-05 - | 1 - | 3,65E-05 - |
| 2374 | 1 | 0.2um_NOKCPHJN_79748 | Bacteria | Proteobacteri | Betaproteoba | unclassified | unclassified | unclassified | Betaproteoba | information st | Replication, re | 05CZ8 | DNA polymera | 543728.Vapar | 0,8 | 0,4 | 2,87E-05 | 1,46E-05 | 1 - | 3,11E-05 - | 1 - | 3,86E-05 - | 1 - | 3,67E-05 - | 1 - | 3,73E-05 - | 0 - | 0 - |
| 2375 | 1 | 3um_FIJJAFBF_718806 | unclassified | unclassified | unclassified | unclassified | unclassified | unclassified | unclassified | unclassified | unclassified | unclassified | unclassified | unclassified | 0,2 | 0,4 | 2,44E-06 | 4,88E-06 | 0 - | 0 - | 0 - | 0 - | 1 - | 1,22E-05 - | 0 - | 0 - | 0 - | 0 - |
| 2376 | 1 | 0.2um_NOKCPHJN_395768 | Bacteria | Proteobacteri | Gammaprotec | Cellvibrionales | Porticoccacea | unclassified | various | unclassified | unclassified | unclassified | unclassified | unclassified | 5 | 0,632455532 | 8,06E-05 | 1,30E-05 | 5 - | 6,74E-05 - | 5 - | 8,37E-05 - | 4 - | 6,37E-05 - | 6 - | 9,69E-05 - | 5 - | 9,10E-05 - |
| 2377 | 1 | 3um_FIJJAFBF_682946 | Bacteria | Proteobacteri | Alphaproteob | Rhodobactera | Rhodobactera | various | various | information st | Translation, ri | COG0185 | Protein S19 fo | 391593.RCCS2 | 1,2 | 0,4 | 0,000173894 | 3,85E-05 | 2 - | 0,000249515 - | 1 - | 0,000154912 - | 1 - | 0,000147263 - | 1 - | 0,000149438 - | 1 - | 0,000168342 - |
| 2378 | 1 | 3um_FIJJAFBF_292927 | Eukaryota | Bacillariophyt | Coscinodiscop | Coscinodiscale | Coscinodiscac | Coscinodiscus | Coscinodiscus | cellular proces | Posttranslatio | KOG0731 | ATP-depende | 35128.Thapsd | 3 | 0,894427191 | 6,32E-05 | 1,54E-05 | 4 - | 7,21E-05 - | 3 - | 6,71E-05 - | 4 - | 8,51E-05 - | 2 - | 4,32E-05 - | 2 - | 4,86E-05 - |
| 2379 | 2 | 0.2um_NOKCPHJN_587130;3um_FIJJAFBF_30992 | Bacteria | Actinobacteri | Actinobacteri | Micrococcales | Microbacteria | various | various | unclassified | unclassified | unclassified | unclassified | unclassified | 1 | 0 | 0,000152252 | 1,45E-05 | 1 - | 0,00012753 - | 1 - | 0,000158354 - | 1 - | 0,000150536 - | 1 - | 0,000152759 - | 1 - | 0,000172083 - |
| 2380 | 1 | 3um_FIJJAFBF_605114 | Bacteria | Proteobacteri | Gammaprotec | Oceanospirilla | Saccharospirill | Reinekea | Reinekea fors | information st | Transcription | 17DYV | RNA polymera | 314283.MED2 | 0,8 | 0,4 | 5,19E-05 | 2,67E-05 | 1 - | 5,49E-05 - | 0 - | 0 - | 1 - | 6,48E-05 - | 1 - | 6,58E-05 - | 1 - | 7,41E-05 - |
| 2381 | 1 | 0.2um_NOKCPHJN_357119 | Bacteria | Proteobacteri | Alphaproteob | Pelagibacteral | Pelagibacterac | Candidatus Pe | Candidatus Pe | metabolism | Amino acid tra | 16RER | (ABC) transpo | 439493.PB721 | 0,2 | 0,4 | 1,09E-05 | 2,19E-05 | 0 - | 0 - | 0 - | 0 - | 0 - | 0 - | 0 - | 0 - | 1 - | 5,47E-05 - |
| 2382 | 1 | 3um_FIJJAFBF_382755 | Viruses | unclassified | unclassified | unclassified | various | various | various | unclassified | unclassified | unclassified | unclassified | unclassified | 1,8 | 1,166190379 | 7,29E-05 | 4,94E-05 | 1 - | 3,35E-05 - | 4 - | 0,000166203 - | 1 - | 3,95E-05 - | 2 - | 8,02E-05 - | 1 - | 4,52E-05 - |
| 2383 | 1 | 3um_FIJJAFBF_192348 | Bacteria | Proteobacteri | Gammaprotec | Oceanospirilla | Saccharospirill | Reinekea | Reinekea fors | poorly charact | Function unkn | 17IX4 | NA | 207954.MED9 | 0,4 | 0,489897949 | 2,77E-05 | 3,40E-05 | 0 - | 0 - | 0 - | 0 - | 1 - | 6,45E-05 - | 0 - | 0 - | 1 - | 7,37E-05 - |
| 2384 | 1 | 3um_FIJJAFBF_633437 | Bacteria | Proteobacteri | Betaproteoba | unclassified | unclassified | unclassified | Betaproteoba | metabolism | Energy produc | 0BBMN | Rubrerythrin | 395495.Lcho_ | 4 | 0,632455532 | 0,000384794 | 4,51E-05 | 5 - | 0,00040701 - | 3 - | 0,000303232 - | 4 - | 0,000384347 - | 4 - | 0,000390024 - | 4 - | 0,00043936 - |
| 2385 | 2 | 0.2um_NOKCPHJN_44534;3um_FIJJAFBF_434864 | Bacteria | Bacteroidetes | Flavobacteriia | Flavobacterial | Flavobacteriac | Formosa | Formosa sp. H | unclassified | unclassified | unclassified | unclassified | unclassified | 2,2 | 0,748331477 | 0,000135009 | 5,28E-05 | 1 - | 5,01E-05 - | 2 - | 0,000124471 - | 3 - | 0,000177488 - | 2 - | 0,000120073 - | 3 - | 0,000202892 - |
| 2386 | 1 | 3um_FIJJAFBF_124745 | Bacteria | Proteobacteri | Alphaproteob | Rhodobactera | Rhodobactera | Thalassobium | Thalassobium | unclassified | unclassified | unclassified | unclassified | unclassified | 0,8 | 0,4 | 2,43E-05 | 1,25E-05 | 1 - | 2,54E-05 - | 1 - | 3,15E-05 - | 0 - | 0 - | 1 - | 3,04E-05 - | 1 - | 3,43E-05 - |
| 2387 | 1 | 0.2um_NOKCPHJN_33623 | Bacteria | Bacteroidetes | Flavobacteriia | Flavobacterial | Flavobacteriac | Algibacter | Algibacter sp. | unclassified | unclassified | unclassified | unclassified | unclassified | 0,4 | 0,489897949 | 1,23E-05 | 1,51E-05 | 0 - | 0 - | 1 - | 3,16E-05 - | 1 - | 3,00E-05 - | 0 - | 0 - | 0 - | 0 - |
| 2388 | 1 | 0.2um_NOKCPHJN_400970 | Viruses | unclassified | unclassified | unclassified | unclassified | unclassified | uncultured M | unclassified | unclassified | unclassified | unclassified | unclassified | 0,2 | 0,4 | 7,68E-06 | 1,54E-05 | 1 - | 3,84E-05 - | 0 - | 0 - | 0 - | 0 - | 0 - | 0 - | 0 - | 0 - |
| 2389 | 2 | 3um_FIJJAFBF_115677;3um_FIJJAFBF_313955 | Bacteria | Bacteroidetes | Flavobacteriia | Flavobacterial | Cryomorphace | unclassified | Cryomorphace | cellular proces | Cell wall/mem | 0PAE7 | biosynthesis p | 313594.PI23P | 0,4 | 0,489897949 | 1,54E-05 | 1,89E-05 | 0 - | 0 - | 1 - | 3,95E-05 - | 1 - | 3,75E-05 - | 0 - | 0 - | 0 - | 0 - |
| 2390 | 1 | 3um_FIJJAFBF_20002 | Bacteria | Proteobacteri | Gammaprotec | Alteromonada | Colwelliaceae | Colwellia | Colwellia sp. C | metabolism | Inorganic ion t | 16TQC | TONB-depend | 314287.GB22 | 6 | 1,414213562 | 0,000152698 | 2,35E-05 | 8 - | 0,000174234 - | 5 - | 0,000135217 - | 7 - | 0,000179957 - | 6 - | 0,000156527 - | 4 - | 0,000117552 - |
| 2391 | 1 | 3um_FIJJAFBF_266308 | Bacteria | Proteobacteri | Betaproteoba | Burkholderial | Comamonada | Rhodoferax | Rhodoferax fe | unclassified | unclassified | unclassified | unclassified | unclassified | 0,2 | 0,4 | 3,62E-05 | 7,24E-05 | 0 - | 0 - | 0 - | 0 - | 0 - | 0 - | 1 - | 0,000180899 - | 0 - | 0 - |
| 2392 | 1 | 3um_FIJJAFBF_343472 | Bacteria | Proteobacteri | Alphaproteob | Pelagibacteral | Pelagibacterac | Candidatus Pe | Candidatus Pe | information st | Translation, ri | 177XJ | One of the pri | 335992.SAR11 | 2 | 0,632455532 | 0,000173446 | 5,39E-05 | 2 - | 0,00014715 - | 2 - | 0,000182716 - | 2 - | 0,000173695 - | 3 - | 0,000264391 - | 1 - | 9,93E-05 - |
| 2393 | 1 | 0.2um_NOKCPHJN_330840 | Eukaryota | unclassified | Cryptophyta | Pyrenomonad | Geminigerace | Guillardia | Guillardia thet | metabolism | Carbohydrate | COG1850 | RuBisCO catal | 35128.Thapsd | 2,2 | 0,748331477 | 0,000114606 | 3,51E-05 | 3 - | 0,000133981 - | 3 - | 0,000166364 - | 2 - | 0,000105434 - | 2 - | 0,000106991 - | 1 - | 6,03E-05 - |
| 2394 | 1 | 0.2um_NOKCPHJN_149220 | Bacteria | Bacteroidetes | unclassified | unclassified | unclassified | unclassified | Bacteroidetes | unclassified | unclassified | unclassified | unclassified | unclassified | 0,2 | 0,4 | 7,48E-06 | 1,50E-05 | 0 - | 0 - | 1 - | 3,74E-05 - | 0 - | 0 - | 0 - | 0 - | 0 - | 0 - |
| 2395 | 1 | 3um_FIJJAFBF_713387 | Bacteria | Proteobacteri | Alphaproteob | Rhodobactera | Rhodobactera | unclassified | Rhodobactera | metabolism | Coenzyme tra | 01Q80 | Glutamate--cy | 314262.MED1 | 0,4 | 0,489897949 | 1,22E-05 | 1,49E-05 | 0 - | 0 - | 1 - | 3,13E-05 - | 1 - | 2,97E-05 - | 0 - | 0 - | 0 - | 0 - |
| 2396 | 1 | 0.2um_NOKCPHJN_386655 | Bacteria | Proteobacteri | Alphaproteob | Pelagibacteral | Pelagibacterac | Candidatus Pe | Candidatus Pe | metabolism | Amino acid tra | 01R4A | Polyamine AB | 335992.SAR11 | 1 | 0 | 3,67E-05 | 3,49E-06 | 1 - | 3,08E-05 - | 1 - | 3,82E-05 - | 1 - | 3,63E-05 - | 1 - | 3,69E-05 - | 1 - | 4,15E-05 - |
| 2397 | 1 | 0.2um_NOKCPHJN_906787 | Bacteria | unclassified | unclassified | unclassified | unclassified | unclassified | uncultured ba | unclassified | unclassified | unclassified | unclassified | unclassified | 0,2 | 0,4 | 4,21E-06 | 8,42E-06 | 0 - | 0 - | 1 - | 2,11E-05 - | 0 - | 0 - | 0 - | 0 - | 0 - | 0 - |
| 2398 | 1 | 3um_FIJJAFBF_43556 | Bacteria | Proteobacteri | Alphaproteob | Rhodobactera | Rhodobactera | unclassified | Rhodobactera | cellular proces | Posttranslatio | 05C6P | ATP-depende | 375451.RD1_ | 0,2 | 0,4 | 3,43E-06 | 6,86E-06 | 0 - | 0 - | 0 - | 0 - | 0 - | 0 - | 1 - | 1,71E-05 - | 0 - | 0 - |
| 2399 | 1 | 3um_FIJJAFBF_681711 | unclassified | unclassified | unclassified | unclassified | unclassified | unclassified | unclassified | unclassified | unclassified | unclassified | unclassified | unclassified | 0,2 | 0,4 | 3,45E-06 | 6,90E-06 | 1 - | 1,73E-05 - | 0 - | 0 - | 0 - | 0 - | 0 - | 0 - | 0 - | 0 - |
| 2400 | 1 | 0.2um_NOKCPHJN_144312 | Bacteria | Proteobacteri | Alphaproteob | Rhodobactera | Rhodobactera | Planktomarin | Planktomarin | cellular proces | Intracellular tr | 125AN | Part of the tw | 388739.RSK20 | 1,2 | 0,4 | 0,000249973 | 5,54E-05 | 2 - | 0,000358677 - | 1 - | 0,000222686 - | 1 - | 0,000211691 - | 1 - | 0,000214818 - | 1 - | 0,000241991 - |
| 2401 | 1 | 0.2um_NOKCPHJN_864503 | Bacteria | Bacteroidetes | Flavobacteriia | Flavobacterial | Flavobacteriac | Formosa | Formosa sp. H | metabolism | Inorganic ion t | 153M9 | iduronate-2-s | 756272.Plabr_ | 3,2 | 0,4 | 0,000106009 | 2,23E-05 | 3 - | 8,26E-05 - | 3 - | 0,000102532 - | 3 - | 9,75E-05 - | 3 - | 9,89E-05 - | 4 - | 0,000148561 - |
| 2402 | 1 | 3um_FIJJAFBF_788043 | Bacteria | Proteobacteri | Alphaproteob | Rhodobactera | unclassified | unclassified | Rhodobactera | unclassified | unclassified | unclassified | unclassified | unclassified | 0,8 | 0,4 | 8,33E-05 | 4,19E-05 | 0 - | 0 - | 1 - | 0,000104028 - | 1 - | 9,89E-05 - | 1 - | 0,000100353 - | 1 - | 0,000113047 - |
| 2403 | 1 | 3um_FIJJAFBF_302476 | Eukaryota | unclassified | Cryptophyta | Pyrenomonad | Geminigerace | Guillardia | Guillardia thet | unclassified | unclassified | unclassified | unclassified | unclassified | 0,8 | 0,4 | 9,91E-05 | 5,03E-05 | 1 - | 0,000107268 - | 1 - | 0,000133195 - | 1 - | 0,000126619 - | 1 - | 0,000128489 - | 0 - | 0 - |
| 2404 | 1 | 3um_FIJJAFBF_132513 | Bacteria | Proteobacteri | Gammaprotec | Cellvibrionales | Porticoccacea | Porticoccus | various | metabolism | Energy produc | COG0055 | Produces ATP | 314285.KT71_ | 1,2 | 0,4 | 3,44E-05 | 7,62E-06 | 2 - | 4,94E-05 - | 1 - | 3,06E-05 - | 1 - | 2,91E-05 - | 1 - | 2,96E-05 - | 1 - | 3,33E-05 - |
| 2405 | 1 | 3um_FIJJAFBF_647278 | Bacteria | Bacteroidetes | Flavobacteriia | Flavobacterial | Flavobacteriac | unclassified | Flavobacteriac | information st | Translation, ri | 05G6W | One of the pri | 391587.KAOT | 1,8 | 0,748331477 | 0,000123695 | 5,45E-05 | 1 - | 5,71E-05 - | 3 - | 0,000212715 - | 2 - | 0,000134808 - | 2 - | 0,000136799 - | 1 - | 7,71E-05 - |
| 2406 | 1 | 0.2um_NOKCPHJN_33105 | Archaea | Euryarchaeota | unclassified | unclassified | unclassified | unclassified | Euryarchaeota | poorly charact | Function unkn | 099QK | NA | 983544.Lacal_ | 0,6 | 0,489897949 | 2,08E-05 | 1,72E-05 | 1 - | 3,04E-05 - | 1 - | 3,78E-05 - | 1 - | 3,59E-05 - | 0 - | 0 - | 0 - | 0 - |
| 2407 | 1 | 3um_FIJJAFBF_505330 | Bacteria | Bacteroidetes | Flavobacteriia | Flavobacterial | unclassified | unclassified | Flavobacterial | unclassified | unclassified | unclassified | unclassified | unclassified | 2,2 | 0,4 | 6,83E-05 | 1,33E-05 | 2 - | 5,21E-05 - | 2 - | 6,46E-05 - | 3 - | 9,22E-05 - | 2 - | 6,24E-05 - | 2 - | 7,02E-05 - |
| 2408 | 1 | 3um_FIJJAFBF_44604 | Bacteria | Bacteroidetes | Flavobacteriia | Flavobacterial | Flavobacteriac | Polaribacter | various | cellular proces | Cell wall/mem | 0P9R9 | TonB family | 313594.PI23P | 2 | 1,095445115 | 0,000109689 | 5,83E-05 | 3 - | 0,000144071 - | 2 - | 0,000119263 - | 3 - | 0,000170061 - | 2 - | 0,000115049 - | 0 - | 0 - |
| 2409 | 1 | 3um_FIJJAFBF_616012 | Bacteria | Bacteroidetes | Flavobacteriia | Flavobacterial | Flavobacteriac | Polaribacter | Polaribacter s | unclassified | unclassified | unclassified | unclassified | unclassified | 1 | 0 | 5,91E-05 | 5,61E-06 | 1 - | 4,95E-05 - | 1 - | 6,14E-05 - | 1 - | 5,84E-05 - | 1 - | 5,93E-05 - | 1 - | 6,68E-05 - |
| 2410 | 1 | 3um_FIJJAFBF_596611 | Eukaryota | unclassified | unclassified | Phaeocystales | Phaeocystace | Phaeocystis | Phaeocystis g | unclassified | unclassified | unclassified | unclassified | unclassified | 1 | 0 | 9,45E-05 | 8,97E-06 | 1 - | 7,92E-05 - | 1 - | 9,83E-05 - | 1 - | 9,34E-05 - | 1 - | 9,48E-05 - | 1 - | 0,00010681 - |
| 2411 | 1 | 0.2um_NOKCPHJN_125830 | Bacteria | Bacteroidetes | Flavobacteriia | Flavobacterial | Flavobacteriac | Formosa | Formosa sp. H | cellular proces | Posttranslatio | 09GE9 | Thioredoxin | 156586.BBFL7 | 1 | 0 | 0,000130502 | 1,24E-05 | 1 - | 0,000109311 - | 1 - | 0,000135732 - | 1 - | 0,000129031 - | 1 - | 0,000130936 - | 1 - | 0,0001475 - |
| 2412 | 1 | 0.2um_NOKCPHJN_141391 | Bacteria | Proteobacteri | Alphaproteob | unclassified | unclassified | Candidatus Pu | Candidatus Pu | unclassified | unclassified | unclassified | unclassified | unclassified | 0,4 | 0,489897949 | 9,04E-06 | 1,11E-05 | 1 - | 2,06E-05 - | 0 - | 0 - | 0 - | 0 - | 1 - | 2,46E-05 - | 0 - | 0 - |
| 2413 | 1 | 3um_FIJJAFBF_468046 | Bacteria | Proteobacteri | Alphaproteob | Rhodobactera | Rhodobactera | Octadecabact | Octadecabact | metabolism | Carbohydrate | 16THB | transporter, d | 391626.OA30 | 1,2 | 0,4 | 4,87E-05 | 1,63E-05 | 1 - | 3,41E-05 - | 1 - | 4,23E-05 - | 2 - | 8,04E-05 - | 1 - | 4,08E-05 - | 1 - | 4,60E-05 - |
| 2414 | 1 | 3um_FIJJAFBF_214612 | Bacteria | Proteobacteri | Alphaproteob | Rhodobactera | Rhodobactera | Phaeobacter | various | information st | Translation, ri | 05C64 | Binds directly | 467661.RKLH1 | 2,2 | 0,748331477 | 0,000126956 | 3,88E-05 | 3 - | 0,000148418 - | 3 - | 0,000184292 - | 2 - | 0,000116795 - | 2 - | 0,00011852 - | 1 - | 6,68E-05 - |
| 2415 | 1 | 3um_FIJJAFBF_607280 | Bacteria | Proteobacteri | Gammaprotec | unclassified | unclassified | unclassified | Gammaprotec | unclassified | unclassified | unclassified | unclassified | unclassified | 1 | 0 | 3,52E-05 | 3,34E-06 | 1 - | 2,95E-05 - | 1 - | 3,66E-05 - | 1 - | 3,48E-05 - | 1 - | 3,53E-05 - | 1 - | 3,98E-05 - |
| 2416 | 1 | 0.2um_NOKCPHJN_224151 | Bacteria | Bacteroidetes | Flavobacteriia | unclassified | unclassified | unclassified | Flavobacteria | unclassified | unclassified | unclassified | unclassified | unclassified | 0,8 | 0,4 | 4,03E-05 | 2,07E-05 | 1 - | 4,20E-05 - | 1 - | 5,22E-05 - | 0 - | 0 - | 1 - | 5,04E-05 - | 1 - | 5,67E-05 - |
| 2417 | 1 | 3um_FIJJAFBF_526759 | Bacteria | Proteobacteri | Gammaprotec | Cellvibrionales | Halieaceae | unclassified | gamma prote | metabolism | Inorganic ion t | 16TQC | TONB-depend | 326297.Sama_ | 8,6 | 1,854723699 | 0,000109691 | 1,67E-05 | 11 - | 0,000119559 - | 9 - | 0,000121465 - | 10 - | 0,000128298 - | 7 - | 9,11E-05 - | 6 - | 8,80E-05 - |
| 2418 | 1 | 3um_FIJJAFBF_327843 | Bacteria | Bacteroidetes | Flavobacteriia | Flavobacterial | Flavobacteriac | Polaribacter | various | information st | Translation, ri | COG0051 | tRNA binding | 313598.MED1 | 5,8 | 0,748331477 | 0,000778858 | 5,54E-05 | 7 - | 0,000795482 - | 6 - | 0,000846647 - | 6 - | 0,000804845 - | 5 - | 0,00068061 - | 5 - | 0,000766705 - |
| 2419 | 1 | 0.2um_NOKCPHJN_55557 | Bacteria | Proteobacteri | Gammaprotec | Oceanospirilla | Saccharospirill | Reinekea | Reinekea blan | unclassified | unclassified | unclassified | unclassified | unclassified | 0,4 | 0,489897949 | 7,23E-06 | 8,91E-06 | 1 - | 1,64E-05 - | 0 - | 0 - | 0 - | 0 - | 1 - | 1,97E-05 - | 0 - | 0 - |
| 2420 | 1 | 0.2um_NOKCPHJN_582029 | Bacteria | Proteobacteri | Betaproteoba | Burkholderial | Comamonada | Rhodoferax | Rhodoferax s | unclassified | unclassified | unclassified | unclassified | unclassified | 2,8 | 0,748331477 | 0,000151822 | 3,32E-05 | 4 - | 0,00018438 - | 2 - | 0,000114473 - | 3 - | 0,000163232 - | 2 - | 0,000110428 - | 3 - | 0,000186596 - |
| 2421 | 1 | 3um_FIJJAFBF_206808 | Bacteria | Proteobacteri | Gammaprotec | Oceanospirilla | Saccharospirill | Reinekea | Reinekea blan | metabolism | Amino acid tra | COG0174 | glutamine syn | 314287.GB22 | 8,2 | 1,166190379 | 0,00023802 | 2,04E-05 | 10 - | 0,000245775 - | 8 - | 0,000244144 - | 7 - | 0,000203078 - | 9 - | 0,000264957 - | 7 - | 0,000232146 - |
| 2422 | 1 | 3um_FIJJAFBF_301936 | Bacteria | Bacteroidetes | Flavobacteriia | Flavobacterial | various | various | various | information st | Translation, ri | COG0051 | tRNA binding | 216432.CA25 | 3,2 | 0,748331477 | 0,000375274 | 7,36E-05 | 4 - | 0,000399224 - | 4 - | 0,000495718 - | 3 - | 0,000353432 - | 3 - | 0,000358652 - | 2 - | 0,000269347 - |
| 2423 | 1 | 3um_FIJJAFBF_814056 | Bacteria | Bacteroidetes | Flavobacteriia | Flavobacterial | unclassified | unclassified | Flavobacterial | unclassified | unclassified | unclassified | unclassified | unclassified | 0,8 | 0,4 | 2,48E-05 | 1,27E-05 | 1 - | 2,60E-05 - | 1 - | 3,22E-05 - | 1 - | 3,07E-05 - | 0 - | 0 - | 1 - | 3,50E-05 - |
| 2424 | 1 | 3um_FIJJAFBF_348325 | Bacteria | Bacteroidetes | Flavobacteriia | Flavobacterial | Flavobacteriac | Algibacter | Algibacter lect | unclassified | unclassified | unclassified | unclassified | unclassified | 11,8 | 0,748331477 | 0,000177738 | 2,32E-05 | 11 - | 0,000138437 - | 12 - | 0,000187525 - | 13 - | 0,000193122 - | 11 - | 0,000165824 - | 12 - | 0,000203782 - |
| 2425 | 1 | 3um_FIJJAFBF_495044 | Bacteria | Bacteroidetes | Flavobacteriia | Flavobacterial | Flavobacteriac | Cellulophaga | Cellulophaga t | poorly charact | Function unkn | 0XP53 | Ragb susd dor | 313598.MED1 | 0,6 | 0,489897949 | 5,64E-05 | 4,65E-05 | 1 - | 8,20E-05 - | 1 - | 0,000101799 - | 0 - | 0 - | 1 - | 9,82E-05 - | 0 - | 0 - |

| | | | | | | | | | | | | | | | | | | | | | | | | | | | | |
|---|---|---|---|---|---|---|---|---|---|---|---|---|---|---|---|---|---|---|---|---|---|---|---|---|---|---|---|---|
| 2426 | 1 | 0.2um_NOKCPHJN_719205 | Eukaryota | Phaeophyceae | unclassified | Ectocarpales | Ectocarpaceae | Ectocarpus | Ectocarpus sili | unclassified | unclassified | unclassified | unclassified | unclassified | 0,2 | 0,4 | 3,24E-06 | 6,48E-06 | 1 - | 1,62E-05 - | 0 - | 0 - | 0 - | 0 - | 0 - | 0 - | 0 - | 0 - |
| 2427 | 1 | 0.2um_NOKCPHJN_196537 | Bacteria | Proteobacteri | Alphaproteob | Rhodobactera | Rhodobactera | Roseovarius | Roseovarius s | metabolism | Amino acid tra | 01QH7 | sulfoacetaldel | 314264.ROS2 | 1 | 0 | 2,33E-05 | 2,21E-06 | 1 - | 1,95E-05 - | 1 - | 2,42E-05 - | 1 - | 2,30E-05 - | 1 - | 2,33E-05 - | 1 - | 2,63E-05 - |
| 2428 | 1 | 3um_FIJJAFBF_191137 | Bacteria | Proteobacteri | Alphaproteob | Rhodobactera | unclassified | unclassified | Rhodobactera | information st | Translation, ri | 17FIH | One of the ea | 367336.OM22 | 1 | 0 | 0,000139824 | 1,33E-05 | 1 - | 0,000117119 - | 1 - | 0,000145427 - | 1 - | 0,000138247 - | 1 - | 0,000140289 - | 1 - | 0,000158035 - |
| 2429 | 1 | 3um_FIJJAFBF_07064 | Bacteria | Bacteroidetes | Flavobacteriia | Flavobacterial | Flavobacteriac | Kordia | Kordia periser | unclassified | unclassified | unclassified | unclassified | unclassified | 3,4 | 1,019803903 | 9,62E-05 | 3,61E-05 | 2 - | 4,63E-05 - | 3 - | 8,62E-05 - | 4 - | 0,00010926 - | 3 - | 8,32E-05 - | 5 - | 0,000156123 - |
| 2430 | 1 | 3um_FIJJAFBF_206047 | Bacteria | Proteobacteri | Alphaproteob | Rhodobactera | Rhodobactera | Planktomarina | Planktomarina | information st | Translation, ri | 177XJ | One of the pri | 314265.R2601 | 1,2 | 0,4 | 0,000102553 | 2,27E-05 | 2 - | 0,00014715 - | 1 - | 9,14E-05 - | 1 - | 8,68E-05 - | 1 - | 8,81E-05 - | 1 - | 9,93E-05 - |
| 2431 | 1 | 0.2um_NOKCPHJN_239171 | Bacteria | Bacteroidetes | unclassified | unclassified | unclassified | unclassified | Bacteroidetes | poorly charact | Function unkn | 0P8NI | Inherit from b | 313590.MED1 | 5 | 1,264911064 | 0,00016168 | 5,28E-05 | 4 - | 0,000106769 - | 4 - | 0,000132576 - | 6 - | 0,000189045 - | 4 - | 0,000127891 - | 7 - | 0,000252121 - |
| 2432 | 1 | 3um_FIJJAFBF_788019 | Bacteria | Proteobacteri | Alphaproteob | Rhodobactera | unclassified | unclassified | Rhodobactera | unclassified | unclassified | unclassified | unclassified | unclassified | 1,4 | 0,489897949 | 0,000153343 | 4,39E-05 | 2 - | 0,000188159 - | 1 - | 0,000116819 - | 2 - | 0,000222102 - | 1 - | 0,000112691 - | 1 - | 0,000126946 - |
| 2433 | 1 | 0.2um_NOKCPHJN_813287 | Eukaryota | Streptophyta | unclassified | Solanales | Solanaceae | Solanum | Solanum melo | unclassified | unclassified | unclassified | unclassified | unclassified | 0,2 | 0,4 | 5,93E-06 | 1,19E-05 | 0 - | 0 - | 0 - | 0 - | 0 - | 0 - | 1 - | 2,96E-05 - | 0 - | 0 - |
| 2434 | 1 | 3um_FIJJAFBF_693935 | Bacteria | Bacteroidetes | Flavobacteriia | Flavobacterial | Flavobacteriac | Gramella | Gramella sp. S | unclassified | unclassified | unclassified | unclassified | unclassified | 1,4 | 0,8 | 5,47E-05 | 3,03E-05 | 2 - | 6,67E-05 - | 2 - | 8,29E-05 - | 2 - | 7,88E-05 - | 0 - | 0 - | 1 - | 4,50E-05 - |
| 2435 | 1 | 0.2um_NOKCPHJN_76394 | Bacteria | Proteobacteri | Betaproteoba | unclassified | unclassified | unclassified | Betaproteoba | cellular proces | Cell wall/mem | COG2980 | Rare lipoprote | 62928.azo321 | 0,2 | 0,4 | 1,08E-05 | 2,16E-05 | 1 - | 5,39E-05 - | 0 - | 0 - | 0 - | 0 - | 0 - | 0 - | 0 - | 0 - |
| 2436 | 1 | 0.2um_NOKCPHJN_1032143 | Bacteria | Proteobacteri | Alphaproteob | Pelagibacteral | Pelagibacterac | Candidatus Pe | Candidatus Pe | metabolism | Amino acid tra | COG0683 | (ABC) transpo | 335992.SAR11 | 9 | 0,632455532 | 0,000312123 | 2,84E-05 | 9 - | 0,00026218 - | 9 - | 0,000325551 - | 9 - | 0,000309477 - | 10 - | 0,000348942 - | 8 - | 0,000314466 - |
| 2437 | 1 | 3um_FIJJAFBF_667558 | Bacteria | Bacteroidetes | Cytophagia | Cytophagales | Flammeovirga | unclassified | Flammeovirga | unclassified | unclassified | unclassified | unclassified | unclassified | 0,8 | 0,4 | 5,12E-05 | 2,58E-05 | 0 - | 0 - | 1 - | 6,39E-05 - | 1 - | 6,08E-05 - | 1 - | 6,17E-05 - | 1 - | 6,95E-05 - |
| 2438 | 2 | 0.2um_NOKCPHJN_673522;0.2um_NOKCPHJN_8 | Bacteria | Proteobacteri | Alphaproteob | Pelagibacteral | Pelagibacterac | unclassified | Pelagibacterac | unclassified | unclassified | unclassified | unclassified | unclassified | 0,2 | 0,4 | 7,76E-06 | 1,55E-05 | 1 - | 3,88E-05 - | 0 - | 0 - | 0 - | 0 - | 0 - | 0 - | 0 - | 0 - |
| 2439 | 1 | 3um_FIJJAFBF_444705 | Bacteria | Proteobacteri | Gammaprotec | Oceanospirilla | Saccharospirill | Reinekea | Reinekea blan | unclassified | unclassified | unclassified | unclassified | unclassified | 13,8 | 1,720465053 | 0,000352498 | 2,07E-05 | 17 - | 0,000367459 - | 12 - | 0,000322077 - | 14 - | 0,000357204 - | 13 - | 0,000336588 - | 13 - | 0,000379165 - |
| 2440 | 1 | 3um_FIJJAFBF_306742 | Bacteria | Proteobacteri | Alphaproteob | Rhodobactera | unclassified | unclassified | Rhodobactera | metabolism | Nucleotide tra | 01SSM | Acts on guanir | 439496.RBY4I | 0,6 | 0,489897949 | 4,62E-05 | 3,80E-05 | 1 - | 6,83E-05 - | 0 - | 0 - | 1 - | 8,06E-05 - | 1 - | 8,18E-05 - | 0 - | 0 - |
| 2441 | 1 | 0.2um_NOKCPHJN_63177 | Bacteria | Bacteroidetes | Flavobacteriia | Flavobacterial | Flavobacteriac | Marinirhabdus | Marinirhabdus | information st | Translation, ri | 08UHY | One of the pri | 50743.SCB49_ | 2,6 | 0,489897949 | 0,0002285 | 5,88E-05 | 2 - | 0,000145287 - | 3 - | 0,000270605 - | 2 - | 0,000171496 - | 3 - | 0,000261044 - | 3 - | 0,000294065 - |
| 2442 | 1 | 0.2um_NOKCPHJN_433505 | Bacteria | Bacteroidetes | Flavobacteriia | Flavobacterial | unclassified | unclassified | Flavobacterial | poorly charact | Function unkn | 0NVRZ | NA | 170187.SP_17 | 5,2 | 0,748331477 | 5,67E-05 | 5,03E-06 | 6 - | 5,54E-05 - | 5 - | 5,73E-05 - | 6 - | 6,54E-05 - | 5 - | 5,53E-05 - | 4 - | 4,98E-05 - |
| 2443 | 1 | 3um_FIJJAFBF_154390 | Bacteria | unclassified | unclassified | unclassified | unclassified | unclassified | bacterium B17 | unclassified | unclassified | unclassified | unclassified | unclassified | 5,2 | 1,166190379 | 0,000134011 | 3,54E-05 | 3 - | 6,39E-05 - | 6 - | 0,000158648 - | 6 - | 0,000150815 - | 6 - | 0,000153043 - | 5 - | 0,000143668 - |
| 2444 | 1 | 0.2um_NOKCPHJN_93297 | Bacteria | Proteobacteri | Alphaproteob | Pelagibacteral | Pelagibacterac | Candidatus Pe | Candidatus Pe | cellular proces | Signal transdu | 01Q8R | gtp-binding pr | 1002672.SAR1 | 1,4 | 0,489897949 | 3,10E-05 | 9,74E-06 | 2 - | 3,78E-05 - | 2 - | 4,69E-05 - | 1 - | 2,23E-05 - | 1 - | 2,26E-05 - | 1 - | 2,55E-05 - |
| 2445 | 1 | 3um_FIJJAFBF_280044 | Bacteria | Proteobacteri | Gammaprotec | unclassified | unclassified | unclassified | Gammaprotec | metabolism | Inorganic ion t | 08UTW | Plays a role in | 338963.Pcar_ | 0,4 | 0,489897949 | 2,32E-05 | 2,85E-05 | 1 - | 5,31E-05 - | 0 - | 0 - | 1 - | 6,27E-05 - | 0 - | 0 - | 0 - | 0 - |
| 2446 | 1 | 0.2um_NOKCPHJN_39938 | Bacteria | unclassified | unclassified | unclassified | unclassified | unclassified | uncultured ba | unclassified | unclassified | unclassified | unclassified | unclassified | 2,2 | 0,979795897 | 0,000166086 | 5,60E-05 | 4 - | 0,000263855 - | 2 - | 0,000163815 - | 2 - | 0,000155727 - | 2 - | 0,000158027 - | 1 - | 8,90E-05 - |
| 2447 | 1 | 3um_FIJJAFBF_16521 | Bacteria | Proteobacteri | Gammaprotec | Alteromonada | Alteromonada | Aliiglaciecola | Aliiglaciecola l | cellular proces | Cell wall/mem | 05DAY | polysaccharid | 715451.ambt_ | 0,4 | 0,489897949 | 2,45E-05 | 3,03E-05 | 1 - | 5,47E-05 - | 1 - | 6,79E-05 - | 0 - | 0 - | 0 - | 0 - | 0 - | 0 - |
| 2448 | 1 | 3um_FIJJAFBF_126834 | Eukaryota | unclassified | unclassified | Phaeocystales | Phaeocystace | Phaeocystis | Phaeocystis gl | metabolism | Energy produc | COG0711 | F(1)F(0) ATP s | 63737.Npun_ | 1 | 0 | 7,70E-05 | 7,31E-06 | 1 - | 6,45E-05 - | 1 - | 8,01E-05 - | 1 - | 7,61E-05 - | 1 - | 7,72E-05 - | 1 - | 8,70E-05 - |
| 2449 | 1 | 3um_FIJJAFBF_784994 | Bacteria | Proteobacteri | Alphaproteob | Rhodobactera | Rhodobactera | Roseovarius | Roseovarius s | information st | Translation, ri | 01S8S | This is 1 of the | 759362.KVU_ | 0,2 | 0,4 | 1,23E-05 | 2,46E-05 | 1 - | 6,14E-05 - | 0 - | 0 - | 0 - | 0 - | 0 - | 0 - | 0 - | 0 - |
| 2450 | 1 | 0.2um_NOKCPHJN_442404 | Eukaryota | unclassified | Pelagophycea | Pelagomonad | unclassified | Aureococcus | Aureococcus a | unclassified | unclassified | unclassified | unclassified | unclassified | 0,2 | 0,4 | 1,97E-06 | 3,95E-06 | 0 - | 0 - | 0 - | 0 - | 1 - | 9,87E-06 - | 0 - | 0 - | 0 - | 0 - |
| 2451 | 1 | 3um_FIJJAFBF_35344 | Bacteria | Bacteroidetes | Cytophagia | Cytophagales | Flammeovirga | unclassified | Flammeovirga | cellular proces | Posttranslatio | 0QHXH | Molecular cha | 247633.GP214 | 0,8 | 0,748331477 | 1,61E-05 | 1,54E-05 | 1 - | 1,74E-05 - | 1 - | 2,17E-05 - | 2 - | 4,12E-05 - | 0 - | 0 - | 0 - | 0 - |
| 2452 | 1 | 3um_FIJJAFBF_71722 | Bacteria | Proteobacteri | Gammaprotec | Alteromonada | Alteromonada | Paraglaciecola | Paraglaciecola | poorly charact | Function unkn | 0R5I6 | late embryoge | 314275.MADE | 1,6 | 1,019803903 | 0,000156437 | 0,000102094 | 2 - | 0,000165146 - | 0 - | 0 - | 1 - | 9,75E-05 - | 3 - | 0,000296727 - | 2 - | 0,000222841 - |
| 2453 | 1 | 3um_FIJJAFBF_186153 | Bacteria | Bacteroidetes | unclassified | unclassified | unclassified | unclassified | Bacteroidetes | unclassified | unclassified | unclassified | unclassified | unclassified | 2 | 0,632455532 | 0,00010848 | 3,61E-05 | 2 - | 9,07E-05 - | 2 - | 0,000112663 - | 1 - | 5,36E-05 - | 3 - | 0,000163024 - | 2 - | 0,00012243 - |
| 2454 | 1 | 0.2um_NOKCPHJN_229866 | Bacteria | Proteobacteri | Alphaproteob | unclassified | unclassified | unclassified | Alphaproteob | information st | Translation, ri | 179RU | 30S ribosomal | 398580.Dshi_ | 0,8 | 0,4 | 6,70E-05 | 3,44E-05 | 1 - | 7,08E-05 - | 0 - | 0 - | 1 - | 8,36E-05 - | 1 - | 8,49E-05 - | 1 - | 9,56E-05 - |
| 2455 | 1 | 0.2um_NOKCPHJN_267312 | Bacteria | Bacteroidetes | Flavobacteriia | Flavobacterial | Flavobacteriac | Formosa | Formosa sp. H | metabolism | Inorganic ion t | 05C27 | TonB-depende | 398720.MED2 | 0,6 | 0,489897949 | 7,72E-06 | 6,36E-06 | 1 - | 1,14E-05 - | 0 - | 0 - | 1 - | 1,35E-05 - | 1 - | 1,37E-05 - | 0 - | 0 - |
| 2456 | 1 | 3um_FIJJAFBF_800979 | Eukaryota | Bacillariophyt | Mediophyceae | Triceratiales | Triceratiaceae | Trieres | Trieres chinen | unclassified | unclassified | unclassified | unclassified | unclassified | 2 | 0,632455532 | 0,000134537 | 4,48E-05 | 2 - | 0,000112526 - | 2 - | 0,000139724 - | 1 - | 6,64E-05 - | 3 - | 0,000202181 - | 2 - | 0,000151838 - |
| 2457 | 1 | 3um_FIJJAFBF_801828 | Bacteria | Proteobacteri | Alphaproteob | Rhodobactera | Rhodobactera | Loktanella | Loktanella sp. | metabolism | Carbohydrate | 01QKD | xylose isomer | 314232.SKA53 | 1,8 | 0,979795897 | 5,52E-05 | 3,08E-05 | 2 - | 5,30E-05 - | 2 - | 6,58E-05 - | 3 - | 9,39E-05 - | 2 - | 6,35E-05 - | 0 - | 0 - |
| 2458 | 2 | 3um_FIJJAFBF_653239;3um_FIJJAFBF_704910 | Bacteria | Proteobacteri | Alphaproteob | Rhodobactera | Rhodobactera | various | various | metabolism | Amino acid tra | 05CM2 | brancheD-cha | various | 0,2 | 0,4 | 7,94E-06 | 1,59E-05 | 1 - | 3,97E-05 - | 0 - | 0 - | 0 - | 0 - | 0 - | 0 - | 0 - | 0 - |
| 2459 | 1 | 0.2um_NOKCPHJN_421642 | Bacteria | Proteobacteri | Alphaproteob | Rickettsiales | unclassified | unclassified | Rickettsiales b | unclassified | unclassified | unclassified | unclassified | unclassified | 0,6 | 0,489897949 | 1,50E-05 | 1,23E-05 | 0 - | 0 - | 1 - | 2,46E-05 - | 0 - | 0 - | 1 - | 2,37E-05 - | 1 - | 2,67E-05 - |
| 2460 | 1 | 3um_FIJJAFBF_560869 | Bacteria | Bacteroidetes | unclassified | unclassified | unclassified | unclassified | Bacteroidetes | unclassified | unclassified | unclassified | unclassified | unclassified | 2,4 | 0,489897949 | 0,000257543 | 5,94E-05 | 2 - | 0,000179339 - | 3 - | 0,000334029 - | 3 - | 0,000317536 - | 2 - | 0,000214818 - | 2 - | 0,000241991 - |
| 2461 | 1 | 0.2um_NOKCPHJN_55567 | Bacteria | Bacteroidetes | Flavobacteriia | Flavobacterial | Flavobacteriac | Polaribacter | Polaribacter t | unclassified | unclassified | unclassified | unclassified | unclassified | 1,4 | 0,489897949 | 3,86E-05 | 1,63E-05 | 1 - | 2,25E-05 - | 2 - | 5,60E-05 - | 1 - | 2,66E-05 - | 1 - | 2,70E-05 - | 2 - | 6,09E-05 - |
| 2462 | 1 | 3um_FIJJAFBF_780740 | Bacteria | Proteobacteri | Gammaprotec | Cellvibrionales | Porticoccacea | unclassified | SAR92 bacteri | unclassified | unclassified | unclassified | unclassified | unclassified | 1,6 | 0,489897949 | 0,00014401 | 3,77E-05 | 2 - | 0,000154063 - | 1 - | 9,57E-05 - | 2 - | 0,000181855 - | 2 - | 0,000184541 - | 1 - | 0,000103943 - |
| 2463 | 1 | 0.2um_NOKCPHJN_319512 | Eukaryota | Apicomplexa | Conoidasida | Eucoccidiorida | Eimeriidae | Eimeria | Eimeria mitis | cellular proces | Intracellular tr | COG0552 | Involved in tar | 637905.SVI_0 | 0,4 | 0,489897949 | 2,97E-06 | 3,64E-06 | 0 - | 0 - | 1 - | 7,56E-06 - | 0 - | 0 - | 1 - | 7,30E-06 - | 0 - | 0 - |
| 2464 | 1 | 3um_FIJJAFBF_454092 | Bacteria | Proteobacteri | Gammaprotec | Alteromonada | Alteromonada | Marinobacter | Marinobacter | metabolism | Amino acid tra | 16R0H | Acetolactate s | 270374.MELB | 0,2 | 0,4 | 1,41E-05 | 2,82E-05 | 1 - | 7,04E-05 - | 0 - | 0 - | 0 - | 0 - | 0 - | 0 - | 0 - | 0 - |
| 2465 | 1 | 0.2um_NOKCPHJN_79980 | Bacteria | Proteobacteri | Gammaprotec | Oceanospirilla | Oceanospirilla | Motiliproteus | Motiliproteus | metabolism | Energy produc | 0QIAX | dihydrolipoyl | 314277.MED1 | 0,8 | 1,166190379 | 2,01E-05 | 2,81E-05 | 3 - | 7,17E-05 - | 0 - | 0 - | 0 - | 0 - | 1 - | 2,86E-05 - | 0 - | 0 - |
| 2466 | 1 | 3um_FIJJAFBF_390442 | Bacteria | Bacteroidetes | Flavobacteriia | Flavobacterial | Flavobacteriac | Algibacter | Algibacter pec | poorly charact | Function unkn | 09F62 | Protein of unk | 391587.KAOT | 0,6 | 0,489897949 | 7,27E-05 | 6,04E-05 | 1 - | 0,000102479 - | 0 - | 0 - | 0 - | 0 - | 1 - | 0,000122753 - | 1 - | 0,000138281 - |
| 2467 | 1 | 3um_FIJJAFBF_72570 | Bacteria | Bacteroidetes | Flavobacteriia | Flavobacterial | Flavobacteriac | unclassified | Flavobacteriac | poorly charact | Function unkn | 05END | lipolytic prote | 313596.RB250 | 0,6 | 0,489897949 | 1,58E-05 | 1,30E-05 | 1 - | 2,34E-05 - | 0 - | 0 - | 1 - | 2,76E-05 - | 1 - | 2,80E-05 - | 0 - | 0 - |
| 2468 | 1 | 3um_FIJJAFBF_135245 | Bacteria | Cyanobacteria | unclassified | Synechococca | Synechococca | Synechococcu | Synechococcu | unclassified | unclassified | unclassified | unclassified | unclassified | 0,8 | 0,4 | 1,59E-05 | 8,06E-06 | 1 - | 1,72E-05 - | 1 - | 2,13E-05 - | 1 - | 2,03E-05 - | 1 - | 2,06E-05 - | 0 - | 0 - |
| 2469 | 1 | 0.2um_NOKCPHJN_163581 | Eukaryota | unclassified | Oomycetes | Pythiales | Pythiaceae | Pythium | Pythium insidi | unclassified | unclassified | unclassified | unclassified | unclassified | 1 | 0 | 2,33E-05 | 2,21E-06 | 1 - | 1,95E-05 - | 1 - | 2,42E-05 - | 1 - | 2,30E-05 - | 1 - | 2,34E-05 - | 1 - | 2,63E-05 - |
| 2470 | 1 | 3um_FIJJAFBF_159751 | Bacteria | Proteobacteri | Gammaprotec | Alteromonada | Alteromonada | Paraglaciecola | Paraglaciecola | metabolism | Amino acid tra | 05CM2 | brancheD-cha | 983545.Glaag | 0,8 | 0,4 | 3,32E-05 | 1,71E-05 | 1 - | 3,47E-05 - | 1 - | 4,31E-05 - | 0 - | 0 - | 1 - | 4,15E-05 - | 1 - | 4,68E-05 - |
| 2471 | 1 | 0.2um_NOKCPHJN_210477 | Bacteria | Proteobacteri | Alphaproteob | Pelagibacteral | Pelagibacterac | unclassified | alpha proteob | unclassified | unclassified | unclassified | unclassified | unclassified | 1,6 | 0,8 | 0,000106395 | 5,46E-05 | 2 - | 0,000112526 - | 0 - | 0 - | 2 - | 0,000132826 - | 2 - | 0,000134788 - | 2 - | 0,000151838 - |
| 2472 | 1 | 0.2um_NOKCPHJN_192568 | Bacteria | Proteobacteri | Gammaprotec | Alteromonada | Colwelliaceae | Colwellia | Colwellia sp. E | metabolism | Inorganic ion t | 16TQC | TONB-depend | 87626.PTD2_ | 3,2 | 0,979795897 | 4,32E-05 | 1,20E-05 | 4 - | 4,60E-05 - | 4 - | 5,71E-05 - | 4 - | 5,42E-05 - | 2 - | 2,75E-05 - | 2 - | 3,10E-05 - |
| 2473 | 1 | 3um_FIJJAFBF_361450 | Eukaryota | unclassified | Pelagophycea | Pelagomonad | unclassified | Aureococcus | Aureococcus a | unclassified | unclassified | unclassified | unclassified | unclassified | 0,6 | 0,489897949 | 5,24E-05 | 4,32E-05 | 1 - | 7,65E-05 - | 1 - | 9,50E-05 - | 1 - | 9,03E-05 - | 0 - | 0 - | 0 - | 0 - |
| 2474 | 5 | 0.2um_NOKCPHJN_1106292;0.2um_NOKCPHJN_ | Bacteria | Bacteroidetes | unclassified | unclassified | unclassified | unclassified | Bacteroidetes | various | various | various | various | various | 0,8 | 0,4 | 1,69E-05 | 8,58E-06 | 1 - | 1,83E-05 - | 1 - | 2,27E-05 - | 1 - | 2,16E-05 - | 1 - | 2,19E-05 - | 0 - | 0 - |
| 2475 | 1 | 3um_FIJJAFBF_520514 | Bacteria | Bacteroidetes | Flavobacteriia | Flavobacterial | Flavobacteriac | Polaribacter | Polaribacter s | unclassified | unclassified | unclassified | unclassified | unclassified | 0,4 | 0,8 | 2,06E-05 | 4,12E-05 | 0 - | 0 - | 0 - | 0 - | 2 - | 0,000103028 - | 0 - | 0 - | 0 - | 0 - |
| 2476 | 1 | 3um_FIJJAFBF_757368 | Bacteria | Proteobacteri | Gammaprotec | Oceanospirilla | Saccharospirill | Reinekea | Reinekea fors | cellular proces | Cell wall/mem | 0QTZI | Integral memb | 314283.MED2 | 0,8 | 0,4 | 7,36E-05 | 3,74E-05 | 1 - | 7,97E-05 - | 1 - | 9,90E-05 - | 1 - | 9,41E-05 - | 1 - | 9,55E-05 - | 0 - | 0 - |
| 2477 | 1 | 3um_FIJJAFBF_53412 | Bacteria | Proteobacteri | Alphaproteob | Rhodobactera | unclassified | unclassified | Rhodobactera | unclassified | unclassified | unclassified | unclassified | unclassified | 2 | 0 | 8,36E-05 | 7,93E-06 | 2 - | 7,00E-05 - | 2 - | 8,69E-05 - | 2 - | 8,26E-05 - | 2 - | 8,38E-05 - | 2 - | 9,44E-05 - |
| 2478 | 1 | 3um_FIJJAFBF_398175 | Bacteria | Proteobacteri | Gammaprotec | Oceanospirilla | Saccharospirill | Reinekea | Reinekea fors | unclassified | unclassified | unclassified | unclassified | unclassified | 1,4 | 0,489897949 | 3,91E-05 | 1,41E-05 | 2 - | 4,69E-05 - | 1 - | 2,91E-05 - | 1 - | 2,77E-05 - | 1 - | 2,81E-05 - | 2 - | 6,33E-05 - |
| 2479 | 1 | 3um_FIJJAFBF_86540 | Bacteria | Bacteroidetes | Flavobacteriia | Flavobacterial | Flavobacteriac | Kordia | Kordia zhangz | poorly charact | Function unkn | 08PAV | RagB SusD dor | 391587.KAOT | 1,4 | 0,489897949 | 3,98E-05 | 1,25E-05 | 2 - | 4,84E-05 - | 2 - | 6,01E-05 - | 1 - | 2,86E-05 - | 1 - | 2,90E-05 - | 1 - | 3,27E-05 - |
| 2480 | 1 | 0.2um_NOKCPHJN_100729 | Bacteria | Bacteroidetes | Flavobacteriia | Flavobacterial | Flavobacteriac | Altibacter | Altibacter lent | unclassified | unclassified | unclassified | unclassified | unclassified | 9,6 | 1,743559577 | 0,000141399 | 3,03E-05 | 11 - | 0,000135612 - | 11 - | 0,00016839 - | 7 - | 0,000101866 - | 8 - | 0,000118138 - | 11 - | 0,000182988 - |
| 2481 | 1 | 0.2um_NOKCPHJN_14938 | Bacteria | Bacteroidetes | Flavobacteriia | Flavobacterial | Flavobacteriac | Kordia | Kordia jejudor | unclassified | unclassified | unclassified | unclassified | unclassified | 1 | 0 | 2,14E-05 | 2,03E-06 | 1 - | 1,79E-05 - | 1 - | 2,23E-05 - | 1 - | 2,12E-05 - | 1 - | 2,15E-05 - | 1 - | 2,42E-05 - |
| 2482 | 1 | 3um_FIJJAFBF_416695 | unclassified | unclassified | unclassified | unclassified | unclassified | unclassified | unclassified | unclassified | unclassified | unclassified | unclassified | unclassified | 0,8 | 0,4 | 0,000150042 | 7,71E-05 | 1 - | 0,000157228 - | 1 - | 0,000195231 - | 1 - | 0,000185592 - | 0 - | 0 - | 1 - | 0,000212157 - |
| 2483 | 1 | 0.2um_NOKCPHJN_654971 | Bacteria | unclassified | unclassified | unclassified | unclassified | unclassified | uncultured ba | cellular proces | Posttranslatio | 0XNQ9 | peptidase (S8 | 391598.FBBAL | 0,2 | 0,4 | 3,28E-06 | 6,56E-06 | 0 - | 0 - | 0 - | 0 - | 1 - | 1,64E-05 - | 0 - | 0 - | 0 - | 0 - |
| 2484 | 1 | 0.2um_NOKCPHJN_862435 | Eukaryota | unclassified | unclassified | Prymnesiales | Chrysochrom | Chrysochrom | Chrysochrom | unclassified | unclassified | unclassified | unclassified | unclassified | 1,4 | 0,8 | 5,59E-05 | 3,28E-05 | 2 - | 6,75E-05 - | 0 - | 0 - | 1 - | 3,98E-05 - | 2 - | 8,09E-05 - | 2 - | 9,11E-05 - |
| 2485 | 1 | 0.2um_NOKCPHJN_167546 | Bacteria | Bacteroidetes | Flavobacteriia | Flavobacterial | Flavobacteriac | Polaribacter | Polaribacter t | unclassified | unclassified | unclassified | unclassified | unclassified | 1,2 | 0,4 | 4,41E-05 | 1,52E-05 | 1 - | 3,08E-05 - | 1 - | 3,82E-05 - | 1 - | 3,63E-05 - | 2 - | 7,37E-05 - | 1 - | 4,15E-05 - |
| 2486 | 1 | 3um_FIJJAFBF_13035 | Bacteria | Bacteroidetes | Flavobacteriia | Flavobacterial | Flavobacteriac | Aequorivita | Aequorivita sp | cellular proces | Signal transdu | 06FN3 | tellurium resis | 313612.L8106 | 0,4 | 0,489897949 | 2,29E-05 | 2,82E-05 | 1 - | 5,24E-05 - | 0 - | 0 - | 1 - | 6,19E-05 - | 0 - | 0 - | 0 - | 0 - |
| 2487 | 1 | 3um_FIJJAFBF_88376 | Bacteria | Bacteroidetes | Flavobacteriia | Flavobacterial | Flavobacteriac | Lutibacter | Lutibacter flav | poorly charact | Function unkn | 11P35 | NA | 867900.Celly_ | 2,8 | 0,748331477 | 0,000106376 | 2,59E-05 | 3 - | 9,67E-05 - | 2 - | 8,01E-05 - | 4 - | 0,000152227 - | 3 - | 0,000115857 - | 2 - | 8,70E-05 - |
| 2488 | 1 | 3um_FIJJAFBF_371767 | Bacteria | Bacteroidetes | Flavobacteriia | Flavobacterial | Flavobacteriac | Flaviramulus | Flaviramulus i | information st | Transcription | COG0557 | 3'-5' exoribon | 391603.FBALC | 1 | 0 | 1,87E-05 | 1,78E-06 | 1 - | 1,57E-05 - | 1 - | 1,95E-05 - | 1 - | 1,85E-05 - | 1 - | 1,88E-05 - | 1 - | 2,12E-05 - |
| 2489 | 1 | 3um_FIJJAFBF_154705 | Bacteria | Bacteroidetes | Flavobacteriia | Flavobacterial | Flavobacteriac | Arenitalea | Arenitalea lut | unclassified | unclassified | unclassified | unclassified | unclassified | 3,6 | 0,8 | 0,000311215 | 4,71E-05 | 5 - | 0,000367874 - | 4 - | 0,000365433 - | 3 - | 0,000260543 - | 3 - | 0,000264391 - | 3 - | 0,000297836 - |
| 2490 | 1 | 0.2um_NOKCPHJN_685253 | Bacteria | Proteobacteri | Alphaproteob | Rhodobactera | Rhodobactera | Planktomarina | Planktomarina | unclassified | unclassified | unclassified | unclassified | unclassified | 1 | 0 | 4,11E-05 | 3,91E-06 | 1 - | 3,45E-05 - | 1 - | 4,28E-05 - | 1 - | 4,07E-05 - | 1 - | 4,13E-05 - | 1 - | 4,65E-05 - |
| 2491 | 1 | 3um_FIJJAFBF_122453 | Bacteria | Bacteroidetes | Flavobacteriia | Flavobacterial | Flavobacteriac | Polaribacter | Polaribacter ir | unclassified | unclassified | unclassified | unclassified | unclassified | 1,2 | 0,748331477 | 5,82E-05 | 3,48E-05 | 2 - | 8,57E-05 - | 1 - | 5,32E-05 - | 2 - | 0,000101106 - | 1 - | 5,13E-05 - | 0 - | 0 - |
| 2492 | 1 | 3um_FIJJAFBF_274235 | Bacteria | Bacteroidetes | Flavobacteriia | Flavobacterial | Flavobacteriac | Lutibacter | Lutibacter flav | cellular proces | Cell wall/mem | 09G63 | Channel that c | 688270.Celal_ | 0,8 | 0,4 | 8,51E-05 | 4,29E-05 | 0 - | 0 - | 1 - | 0,000106357 - | 1 - | 0,000101106 - | 1 - | 0,000102599 - | 1 - | 0,000115578 - |
| 2493 | 1 | 3um_FIJJAFBF_283180 | Bacteria | Bacteroidetes | Flavobacteriia | Flavobacterial | Flavobacteriac | Flavobacteriu | various | unclassified | unclassified | unclassified | unclassified | unclassified | 1 | 0 | 0,000112317 | 1,07E-05 | 1 - | 9,41E-05 - | 1 - | 0,000116819 - | 1 - | 0,000111051 - | 1 - | 0,000112691 - | 1 - | 0,000126946 - |
| 2494 | 1 | 0.2um_NOKCPHJN_265832 | Bacteria | Bacteroidetes | unclassified | unclassified | unclassified | unclassified | Bacteroidetes | poorly charact | Function unkn | 11PIQ | YCE I like fami | 926562.Oweh | 2 | 0 | 0,000124006 | 1,18E-05 | 2 - | 0,00010387 - | 2 - | 0,000128976 - | 2 - | 0,000122608 - | 2 - | 0,000124419 - | 2 - | 0,000140158 - |
| 2495 | 1 | 0.2um_NOKCPHJN_778065 | Bacteria | Proteobacteri | Betaproteoba | Burkholderiale | unclassified | unclassified | Burkholderiale | unclassified | unclassified | unclassified | unclassified | unclassified | 3,6 | 0,8 | 7,74E-05 | 2,40E-05 | 3 - | 5,31E-05 - | 4 - | 8,80E-05 - | 3 - | 6,27E-05 - | 3 - | 6,36E-05 - | 5 - | 0,000119502 - |
| 2496 | 1 | 0.2um_NOKCPHJN_818796 | Bacteria | Proteobacteri | Alphaproteob | Pelagibacteral | Pelagibacterac | unclassified | alpha proteob | metabolism | Energy produc | 01RNF | Produces ATP | 331869.BAL19 | 0,4 | 0,489897949 | 1,09E-05 | 1,34E-05 | 1 - | 2,42E-05 - | 1 - | 3,01E-05 - | 0 - | 0 - | 0 - | 0 - | 0 - | 0 - |
| 2497 | 1 | 3um_FIJJAFBF_660323 | Bacteria | Proteobacteri | Alphaproteob | Rhodobactera | Rhodobactera | Pseudodonghi | Pseudodonghi | unclassified | unclassified | unclassified | unclassified | unclassified | 3,8 | 0,748331477 | 0,000350656 | 5,69E-05 | 5 - | 0,000390397 - | 3 - | 0,000290855 - | 3 - | 0,000276494 - | 4 - | 0,000374104 - | 4 - | 0,000421427 - |
| 2498 | 1 | 3um_FIJJAFBF_41764 | Bacteria | Bacteroidetes | Flavobacteriia | Flavobacterial | Crocinitomica | unclassified | Crocinitomica | metabolism | Energy produc | COG0711 | F(1)F(0) ATP s | 926562.Oweh | 1,2 | 0,4 | 8,89E-05 | 1,97E-05 | 2 - | 0,00012753 - | 1 - | 7,92E-05 - | 1 - | 7,53E-05 - | 1 - | 7,64E-05 - | 1 - | 8,60E-05 - |
| 2499 | 1 | 0.2um_NOKCPHJN_684776 | Eukaryota | unclassified | unclassified | Choanoflagelli | Salpingoecida | Monosiga | Monosiga bre | metabolism | Carbohydrate | COG1501 | hydrolase, fan | 431895.XP_00 | 0,2 | 0,4 | 3,08E-06 | 6,16E-06 | 0 - | 0 - | 1 - | 1,54E-05 - | 0 - | 0 - | 0 - | 0 - | 0 - | 0 - |
| 2500 | 1 | 3um_FIJJAFBF_683488 | Bacteria | Proteobacteri | Alphaproteob | Rhodobactera | Rhodobactera | Amylibacter | Amylibacter k | metabolism | Amino acid tra | COG0683 | (ABC) transpo | 367336.OM22 | 2,2 | 0,4 | 7,76E-05 | 2,09E-05 | 2 - | 5,84E-05 - | 2 - | 7,25E-05 - | 2 - | 6,89E-05 - | 2 - | 7,00E-05 - | 3 - | 0,000118225 - |
| 2501 | 1 | 3um_FIJJAFBF_755754 | Bacteria | Bacteroidetes | Flavobacteriia | Flavobacterial | Flavobacteriac | Polaribacter | Polaribacter s | unclassified | unclassified | unclassified | unclassified | unclassified | 10,4 | 1,496662955 | 0,000480945 | 3,88E-05 | 13 - | 0,000509248 - | 11 - | 0,000535054 - | 10 - | 0,000462397 - | 9 - | 0,000422304 - | 9 - | 0,000475724 - |
| 2502 | 2 | 0.2um_NOKCPHJN_595439;3um_FIJJAFBF_80437 | Bacteria | Bacteroidetes | various | various | various | various | various | unclassified | unclassified | unclassified | unclassified | unclassified | 1 | 0 | 9,14E-05 | 8,67E-06 | 1 - | 7,65E-05 - | 1 - | 9,50E-05 - | 1 - | 9,03E-05 - | 1 - | 9,17E-05 - | 1 - | 0,00010325 - |
| 2503 | 1 | 0.2um_NOKCPHJN_15319 | Bacteria | Proteobacteri | Betaproteoba | unclassified | unclassified | unclassified | Betaproteoba | cellular proces | Cell wall/mem | 0B897 | Part of the ou | 365044.Pnap_ | 0,4 | 0,489897949 | 1,58E-05 | 1,94E-05 | 0 - | 0 - | 1 - | 3,78E-05 - | 0 - | 0 - | 0 - | 0 - | 1 - | 4,11E-05 - |
| 2504 | 1 | 0.2um_NOKCPHJN_67468 | Bacteria | Bacteroidetes | Flavobacteriia | Flavobacterial | Flavobacteriac | Gaetbulibacte | Gaetbulibacte | information st | Translation, ri | 05CEJ | Catalyzes the | 398720.MED2 | 0,4 | 0,489897949 | 8,26E-06 | 1,01E-05 | 0 - | 0 - | 0 - | 0 - | 0 - | 0 - | 1 - | 1,94E-05 - | 1 - | 2,19E-05 - |
| 2505 | 1 | 3um_FIJJAFBF_588324 | Bacteria | Proteobacteri | Betaproteoba | unclassified | unclassified | unclassified | Betaproteoba | information st | Transcription | COG0085 | DNA-depende | 296591.Bpro_ | 16,4 | 2,154065923 | 0,000157293 | 1,46E-05 | 18 - | 0,000146006 - | 15 - | 0,00015108 - | 19 - | 0,00018192 - | 17 - | 0,000165174 - | 13 - | 0,000142288 - |
| 2506 | 1 | 0.2um_NOKCPHJN_138902 | Bacteria | Proteobacteri | Gammaprotec | Cellvibrionales | Porticoccacea | unclassified | SAR92 bacteri | unclassified | unclassified | unclassified | unclassified | unclassified | 6 | 1,414213562 | 9,73E-05 | 1,86E-05 | 8 - | 0,000110098 - | 7 - | 0,00011962 - | 4 - | 6,50E-05 - | 6 - | 9,89E-05 - | 5 - | 9,29E-05 - |
| 2507 | 1 | 3um_FIJJAFBF_339918 | Bacteria | Proteobacteri | Gammaprotec | unclassified | unclassified | unclassified | Gammaprotec | information st | Transcription | 0QWTY | 'Cold-shock' D | 225937.HP15_ | 1 | 0,632455532 | 8,66E-05 | 4,94E-05 | 2 - | 0,000154063 - | 1 - | 9,57E-05 - | 1 - | 9,09E-05 - | 1 - | 9,23E-05 - | 0 - | 0 - |
| 2508 | 1 | 3um_FIJJAFBF_245503 | Bacteria | Bacteroidetes | Flavobacteriia | Flavobacterial | Flavobacteriac | Lutibacter | Lutibacter flav | unclassified | unclassified | unclassified | unclassified | unclassified | 0,2 | 0,4 | 1,39E-05 | 2,78E-05 | 1 - | 6,96E-05 - | 0 - | 0 - | 0 - | 0 - | 0 - | 0 - | 0 - | 0 - |
| 2509 | 1 | 3um_FIJJAFBF_494212 | Bacteria | Proteobacteri | Gammaprotec | Oceanospirilla | Saccharospirill | Saccharospiril | Saccharospiril | metabolism | Lipid transpor | COG0511 | Acetyl-CoA ca | 314283.MED2 | 4,8 | 1,6 | 0,00011215 | 4,22E-05 | 3 - | 5,77E-05 - | 7 - | 0,000167108 - | 3 - | 6,81E-05 - | 6 - | 0,000138174 - | 5 - | 0,000129711 - |
| 2510 | 1 | 3um_FIJJAFBF_212147 | Bacteria | Proteobacteri | Alphaproteob | Rhodobactera | Rhodobactera | Loktanella | various | information st | Transcription | 05VEQ | Cold shock pr | 391593.RCCS2 | 0,8 | 0,748331477 | 0,000168341 | 0,000153372 | 0 - | 0 - | 1 - | 0,000209587 - | 0 - | 0 - | 2 - | 0,000404363 - | 1 - | 0,000227757 - |
| 2511 | 1 | 3um_FIJJAFBF_292964 | Bacteria | Proteobacteri | Alphaproteob | Rhodobactera | Rhodobactera | Tateyamaria | Tateyamaria s | unclassified | unclassified | unclassified | unclassified | unclassified | 5 | 0,632455532 | 0,000225029 | 2,15E-05 | 6 - | 0,000228033 - | 4 - | 0,000188767 - | 5 - | 0,000224308 - | 5 - | 0,000227621 - | 5 - | 0,000256415 - |
| 2512 | 1 | 3um_FIJJAFBF_238677 | Bacteria | Proteobacteri | Alphaproteob | Rhodobactera | Rhodobactera | unclassified | Rhodobactera | unclassified | unclassified | unclassified | unclassified | unclassified | 1,6 | 0,489897949 | 0,000119209 | 3,12E-05 | 2 - | 0,00012753 - | 1 - | 7,92E-05 - | 2 - | 0,000150536 - | 2 - | 0,000152759 - | 1 - | 8,60E-05 - |
| 2513 | 1 | 3um_FIJJAFBF_82196 | Bacteria | unclassified | unclassified | unclassified | unclassified | unclassified | uncultured ba | poorly charact | Function unkn | 0PAEN | NA | 867900.Celly_ | 2,4 | 0,489897949 | 6,77E-05 | 1,91E-05 | 2 - | 4,66E-05 - | 3 - | 8,67E-05 - | 2 - | 5,50E-05 - | 2 - | 5,58E-05 - | 3 - | 9,42E-05 - |
| 2514 | 1 | 0.2um_NOKCPHJN_157663 | Bacteria | Proteobacteri | Gammaprotec | Cellvibrionales | Porticoccacea | unclassified | SAR92 bacteri | metabolism | Inorganic ion t | 1768K | TONB-depend | 247634.GPB2 | 0,8 | 0,748331477 | 1,20E-05 | 1,06E-05 | 2 - | 2,73E-05 - | 0 - | 0 - | 1 - | 1,61E-05 - | 1 - | 1,63E-05 - | 0 - | 0 - |
| 2515 | 1 | 3um_FIJJAFBF_176634 | Bacteria | Bacteroidetes | Flavobacteriia | Flavobacterial | Cryomorphace | Owenweeksia | Owenweeksia | unclassified | unclassified | unclassified | unclassified | unclassified | 2 | 1,414213562 | 0,000113348 | 7,78E-05 | 3 - | 0,00014715 - | 2 - | 0,000121811 - | 4 - | 0,000231594 - | 0 - | 0 - | 1 - | 6,62E-05 - |
| 2516 | 1 | 3um_FIJJAFBF_160914 | Bacteria | Actinobacteri | Actinobacteri | Micrococcales | Microbacteria | Candidatus Ac | Candidatus Ac | poorly charact | Function unkn | 00JIV | LemA family | 979556.MTES_ | 0,4 | 0,489897949 | 2,65E-05 | 3,26E-05 | 1 - | 6,07E-05 - | 0 - | 0 - | 1 - | 7,17E-05 - | 0 - | 0 - | 0 - | 0 - |
| 2517 | 1 | 0.2um_NOKCPHJN_530228 | Bacteria | Proteobacteri | Gammaprotec | unclassified | unclassified | unclassified | Gammaprotec | unclassified | unclassified | unclassified | unclassified | unclassified | 0,6 | 0,489897949 | 9,44E-06 | 7,79E-06 | 1 - | 1,38E-05 - | 1 - | 1,71E-05 - | 1 - | 1,63E-05 - | 0 - | 0 - | 0 - | 0 - |
| 2518 | 1 | 0.2um_NOKCPHJN_183925 | Eukaryota | unclassified | Oomycetes | Saprolegniales | Saprolegniace | Saprolegnia | Saprolegnia p | unclassified | unclassified | unclassified | unclassified | unclassified | 0,6 | 0,489897949 | 2,70E-05 | 2,24E-05 | 1 - | 3,80E-05 - | 0 - | 0 - | 0 - | 0 - | 1 - | 4,55E-05 - | 1 - | 5,13E-05 - |
| 2519 | 1 | 3um_FIJJAFBF_118447 | Bacteria | Proteobacteri | Alphaproteob | Rhodobactera | Rhodobactera | unclassified | Rhodobactera | unclassified | unclassified | unclassified | unclassified | unclassified | 7,2 | 0,748331477 | 0,000218074 | 3,62E-05 | 6 - | 0,000151354 - | 8 - | 0,000250583 - | 7 - | 0,000208434 - | 8 - | 0,000241729 - | 7 - | 0,000238268 - |
| 2520 | 1 | 3um_FIJJAFBF_664332 | Bacteria | Proteobacteri | Gammaprotec | Pseudomonad | Moraxellaceae | Moraxella | Moraxella boe | unclassified | unclassified | unclassified | unclassified | unclassified | 1 | 0 | 0,000138411 | 1,31E-05 | 1 - | 0,000115936 - | 1 - | 0,000143958 - | 1 - | 0,000136851 - | 1 - | 0,000138872 - | 1 - | 0,000156439 - |
| 2521 | 1 | 3um_FIJJAFBF_552508 | Bacteria | Proteobacteri | Gammaprotec | Arenicellales | Arenicellaceae | Arenicella | Arenicella xan | information st | Translation, ri | 0BBM3 | 50s ribosomal | 314607.KB13_ | 0,6 | 0,489897949 | 4,95E-05 | 4,06E-05 | 0 - | 0 - | 0 - | 0 - | 1 - | 7,83E-05 - | 1 - | 7,95E-05 - | 1 - | 8,95E-05 - |
| 2522 | 1 | 3um_FIJJAFBF_671477 | Bacteria | Bacteroidetes | Cytophagia | Cytophagales | Flammeovirga | unclassified | Flammeovirga | unclassified | unclassified | unclassified | unclassified | unclassified | 1,2 | 0,4 | 0,00019275 | 4,27E-05 | 2 - | 0,000276571 - | 1 - | 0,000171709 - | 1 - | 0,000163232 - | 1 - | 0,000165643 - | 1 - | 0,000186596 - |
| 2523 | 1 | 3um_FIJJAFBF_189702 | Bacteria | Bacteroidetes | Flavobacteriia | Flavobacterial | Flavobacteriac | Polaribacter | Polaribacter fi | poorly charact | Function unkn | 09A4Z | rhodopsin | 313598.MED1 | 2 | 0 | 0,000114189 | 1,08E-05 | 2 - | 9,56E-05 - | 2 - | 0,000118766 - | 2 - | 0,000112902 - | 2 - | 0,000114569 - | 2 - | 0,000129062 - |
| 2524 | 1 | 3um_FIJJAFBF_370622 | Bacteria | Proteobacteri | Alphaproteob | unclassified | unclassified | unclassified | Alphaproteob | unclassified | unclassified | unclassified | unclassified | unclassified | 0,4 | 0,489897949 | 1,43E-05 | 1,77E-05 | 1 - | 3,29E-05 - | 0 - | 0 - | 1 - | 3,88E-05 - | 0 - | 0 - | 0 - | 0 - |
| 2525 | 1 | 3um_FIJJAFBF_388371 | Bacteria | Bacteroidetes | Flavobacteriia | Flavobacterial | Flavobacteriac | Formosa | Formosa sp. H | unclassified | unclassified | unclassified | unclassified | unclassified | 0,6 | 0,489897949 | 1,24E-05 | 1,02E-05 | 1 - | 1,80E-05 - | 1 - | 2,24E-05 - | 0 - | 0 - | 1 - | 2,16E-05 - | 0 - | 0 - |
| 2526 | 1 | 0.2um_NOKCPHJN_786267 | Bacteria | Proteobacteri | Alphaproteob | Rhodospirillal | Rhodospirillac | Candidatus En | Candidatus En | unclassified | unclassified | unclassified | unclassified | unclassified | 0,4 | 0,489897949 | 9,56E-06 | 1,18E-05 | 1 - | 2,17E-05 - | 0 - | 0 - | 0 - | 0 - | 1 - | 2,60E-05 - | 0 - | 0 - |
| 2527 | 1 | 3um_FIJJAFBF_95703 | Eukaryota | unclassified | Pelagophycea | Pelagomonad | unclassified | Pelagomonas | uncultured Pe | information st | Translation, ri | COG0051 | tRNA binding | 123214.PERM | 1 | 0 | 0,000123448 | 1,17E-05 | 1 - | 0,000103402 - | 1 - | 0,000128395 - | 1 - | 0,000122056 - | 1 - | 0,000123859 - | 1 - | 0,000139527 - |
| 2528 | 1 | 3um_FIJJAFBF_339692 | Bacteria | Proteobacteri | Gammaprotec | Alteromonada | Alteromonada | Paraglaciecola | Paraglaciecola | poorly charact | Function unkn | 05EK9 | Dehydrogenas | 983545.Glaag | 0,2 | 0,4 | 9,07E-06 | 1,81E-05 | 1 - | 4,54E-05 - | 0 - | 0 - | 0 - | 0 - | 0 - | 0 - | 0 - | 0 - |
| 2529 | 3 | 0.2um_NOKCPHJN_110536;0.2um_NOKCPHJN_2 | Bacteria | Proteobacteri | Gammaprotec | Cellvibrionales | Porticoccacea | unclassified | Gammaprotec | information st | Translation, ri | 0QVRF | 50S ribosomal | 314287.GB22 | 1 | 0,632455532 | 0,00020867 | 0,0001367 | 1 - | 0,000179339 - | 1 - | 0,000222686 - | 1 - | 0,000211691 - | 2 - | 0,000429635 - | 0 - | 0 - |
| 2530 | 4 | 3um_FIJJAFBF_201254;3um_FIJJAFBF_351376;3u | Bacteria | Bacteroidetes | Flavobacteriia | Flavobacterial | Flavobacteriac | Formosa | Formosa sp. H | poorly charact | Function unkn | 11FUA | Membrane | 391603.FBALC | 1 | 0 | 7,25E-05 | 6,88E-06 | 1 - | 6,07E-05 - | 1 - | 7,54E-05 - | 1 - | 7,17E-05 - | 1 - | 7,27E-05 - | 1 - | 8,19E-05 - |
| 2531 | 1 | 3um_FIJJAFBF_72477 | Bacteria | Bacteroidetes | Cytophagia | Cytophagales | Flammeovirga | unclassified | Flammeovirga | unclassified | unclassified | unclassified | unclassified | unclassified | 3,6 | 0,8 | 0,000239648 | 3,14E-05 | 5 - | 0,000284101 - | 3 - | 0,000211662 - | 4 - | 0,000268282 - | 3 - | 0,000204183 - | 3 - | 0,000230012 - |
| 2532 | 1 | 3um_FIJJAFBF_40656 | Eukaryota | unclassified | Pelagophycea | Pelagomonad | unclassified | Aureococcus | Aureococcus a | unclassified | unclassified | unclassified | unclassified | unclassified | 1 | 0 | 0,000285473 | 2,71E-05 | 1 - | 0,000239118 - | 1 - | 0,000296914 - | 1 - | 0,000282255 - | 1 - | 0,000286424 - | 1 - | 0,000322655 - |
| 2533 | 1 | 3um_FIJJAFBF_583670 | Bacteria | Cyanobacteria | unclassified | Synechococca | Synechococca | Synechococcu | Synechococcu | unclassified | unclassified | unclassified | unclassified | unclassified | 1,4 | 0,489897949 | 0,000101926 | 3,60E-05 | 1 - | 6,11E-05 - | 1 - | 7,58E-05 - | 2 - | 0,00014413 - | 2 - | 0,000146259 - | 1 - | 8,24E-05 - |
| 2534 | 1 | 3um_FIJJAFBF_520490 | Bacteria | Bacteroidetes | Flavobacteriia | Flavobacterial | Flavobacteriac | Polaribacter | Polaribacter s | poorly charact | Function unkn | 0P8S3 | adenosyltrans | 313594.PI23P | 0,6 | 0,8 | 3,82E-05 | 4,93E-05 | 2 - | 0,000120185 - | 0 - | 0 - | 1 - | 7,09E-05 - | 0 - | 0 - | 0 - | 0 - |
| 2535 | 1 | 3um_FIJJAFBF_304381 | Bacteria | Bacteroidetes | Flavobacteriia | Flavobacterial | Flavobacteriac | Polaribacter | various | unclassified | unclassified | unclassified | unclassified | unclassified | 2 | 0 | 3,63E-05 | 3,45E-06 | 2 - | 3,04E-05 - | 2 - | 3,78E-05 - | 2 - | 3,59E-05 - | 2 - | 3,64E-05 - | 2 - | 4,10E-05 - |
| 2536 | 1 | 3um_FIJJAFBF_121417 | Bacteria | Proteobacteri | Gammaprotec | Cellvibrionales | Spongiibacter | various | various | information st | Transcription | 05VEQ | Cold shock pr | 247633.GP214 | 0,8 | 0,748331477 | 0,000139589 | 0,000123364 | 2 - | 0,000318824 - | 0 - | 0 - | 1 - | 0,00018817 - | 1 - | 0,000190949 - | 0 - | 0 - |
| 2537 | 1 | 3um_FIJJAFBF_345861 | Bacteria | Bacteroidetes | Flavobacteriia | Flavobacterial | Flavobacteriac | Formosa | Formosa sp. H | cellular proces | Posttranslatio | COG0330 | Band 7 protei | 391603.FBALC | 2 | 0,632455532 | 8,74E-05 | 2,31E-05 | 3 - | 0,000111434 - | 2 - | 9,22E-05 - | 1 - | 4,38E-05 - | 2 - | 8,90E-05 - | 2 - | 0,000100242 - |
| 2538 | 1 | 0.2um_NOKCPHJN_159989 | Bacteria | Proteobacteri | Gammaprotec | Oceanospirilla | Saccharospirill | Reinekea | Reinekea fors | metabolism | Carbohydrate | 16RY1 | ABC transport | 536019.Meso | 1,2 | 0,4 | 4,47E-05 | 9,90E-06 | 2 - | 6,41E-05 - | 1 - | 3,98E-05 - | 1 - | 3,78E-05 - | 1 - | 3,84E-05 - | 1 - | 4,33E-05 - |
| 2539 | 1 | 3um_FIJJAFBF_71076 | Bacteria | Bacteroidetes | Flavobacteriia | Flavobacterial | Flavobacteriac | Polaribacter | Polaribacter s | metabolism | Coenzyme tra | 09GJQ | dihydroneopt | 313594.PI23P | 0,6 | 0,489897949 | 7,37E-05 | 6,04E-05 | 0 - | 0 - | 1 - | 0,000120779 - | 0 - | 0 - | 1 - | 0,000116511 - | 1 - | 0,00013125 - |
| 2540 | 1 | 3um_FIJJAFBF_280836 | Bacteria | Bacteroidetes | Flavobacteriia | Flavobacterial | Crocinitomica | unclassified | Crocinitomica | information st | Translation, ri | COG1185 | Involved in m | 743722.Sph21 | 4,6 | 0,8 | 8,99E-05 | 2,35E-05 | 4 - | 6,47E-05 - | 5 - | 0,000100365 - | 4 - | 7,63E-05 - | 4 - | 7,75E-05 - | 6 - | 0,00013088 - |
| 2541 | 1 | 0.2um_NOKCPHJN_111478 | Bacteria | Bacteroidetes | Cytophagia | Cytophagales | Flammeovirga | unclassified | Flammeovirga | unclassified | unclassified | unclassified | unclassified | unclassified | 0,6 | 0,489897949 | 6,51E-06 | 5,37E-06 | 1 - | 9,46E-06 - | 1 - | 1,17E-05 - | 0 - | 0 - | 1 - | 1,13E-05 - | 0 - | 0 - |
| 2542 | 1 | 3um_FIJJAFBF_369614 | Bacteria | Bacteroidetes | Flavobacteriia | Flavobacterial | Flavobacteriac | Winogradskye | Winogradskye | unclassified | unclassified | unclassified | unclassified | unclassified | 2 | 0 | 0,00013567 | 1,29E-05 | 2 - | 0,00011364 - | 2 - | 0,000141108 - | 2 - | 0,000134141 - | 2 - | 0,000136122 - | 2 - | 0,000153341 - |
| 2543 | 1 | 0.2um_NOKCPHJN_148359 | Bacteria | Proteobacteri | Alphaproteob | Rhodobactera | Rhodobactera | Amylibacter | Amylibacter c | unclassified | unclassified | unclassified | unclassified | unclassified | 1,2 | 0,748331477 | 2,97E-05 | 1,78E-05 | 2 - | 4,24E-05 - | 2 - | 5,26E-05 - | 1 - | 2,50E-05 - | 0 - | 0 - | 1 - | 2,86E-05 - |
| 2544 | 3 | 0.2um_NOKCPHJN_335948;0.2um_NOKCPHJN_6 | Bacteria | Proteobacteri | Alphaproteob | Rhodobactera | Rhodobactera | Planktomarina | Planktomarina | metabolism | Coenzyme tra | 08S7T | Catalyzes the | various | 0,4 | 0,489897949 | 2,29E-05 | 2,82E-05 | 1 - | 5,24E-05 - | 0 - | 0 - | 1 - | 6,19E-05 - | 0 - | 0 - | 0 - | 0 - |
| 2545 | 1 | 3um_FIJJAFBF_714668 | Bacteria | Proteobacteri | Alphaproteob | Pelagibacteral | Pelagibacterac | Candidatus Pe | Candidatus Pe | metabolism | Inorganic ion t | 05PWN | phosphate ab | 335992.SAR11 | 4,4 | 0,489897949 | 0,000176035 | 2,54E-05 | 5 - | 0,000167802 - | 4 - | 0,000166689 - | 4 - | 0,000158459 - | 4 - | 0,000160799 - | 5 - | 0,000226425 - |
| 2546 | 1 | 3um_FIJJAFBF_744827 | Bacteria | Proteobacteri | Gammaprotec | Alteromonada | unclassified | unclassified | Alteromonada | information st | Translation, ri | 0QK14 | This protein bi | 377629.TERTU | 1,4 | 0,489897949 | 0,000110225 | 4,48E-05 | 1 - | 6,48E-05 - | 1 - | 8,05E-05 - | 2 - | 0,000153087 - | 1 - | 7,77E-05 - | 2 - | 0,000174999 - |
| 2547 | 1 | 0.2um_NOKCPHJN_216840 | Bacteria | Bacteroidetes | Flavobacteriia | Flavobacterial | Flavobacteriac | Marixanthom | Marixanthom | metabolism | Amino acid tra | COG2171 | Catalyzes the | 50743.SCB49_ | 0,8 | 0,4 | 4,04E-05 | 2,08E-05 | 1 - | 4,24E-05 - | 1 - | 5,26E-05 - | 1 - | 5,00E-05 - | 0 - | 0 - | 1 - | 5,71E-05 - |
| 2548 | 1 | 3um_FIJJAFBF_294670 | Bacteria | Bacteroidetes | Flavobacteriia | Flavobacterial | Flavobacteriac | Algibacter | Algibacter aqu | information st | Translation, ri | 08UIK | This protein bi | 983544.Lacal_ | 0,2 | 0,4 | 1,58E-05 | 3,17E-05 | 1 - | 7,92E-05 - | 0 - | 0 - | 0 - | 0 - | 0 - | 0 - | 0 - | 0 - |
| 2549 | 1 | 3um_FIJJAFBF_279094 | unclassified | unclassified | unclassified | unclassified | unclassified | unclassified | unclassified | unclassified | unclassified | unclassified | unclassified | unclassified | 0,8 | 0,4 | 7,66E-05 | 3,94E-05 | 1 - | 8,03E-05 - | 1 - | 9,97E-05 - | 1 - | 9,47E-05 - | 0 - | 0 - | 1 - | 0,000108304 - |
| 2550 | 1 | 0.2um_NOKCPHJN_265603 | Bacteria | Proteobacteri | Alphaproteob | Rhodobactera | Rhodobactera | Sediminimona | Sediminimona | unclassified | unclassified | unclassified | unclassified | unclassified | 0,2 | 0,4 | 5,86E-07 | 1,17E-06 | 1 - | 2,93E-06 - | 0 - | 0 - | 0 - | 0 - | 0 - | 0 - | 0 - | 0 - |
| 2551 | 1 | 3um_FIJJAFBF_56138 | Bacteria | Bacteroidetes | Flavobacteriia | Flavobacterial | Flavobacteriac | Polaribacter | various | metabolism | Energy produc | COG0712 | F(1)F(0) ATP s | 313598.MED1 | 2,6 | 0,8 | 0,000190272 | 5,52E-05 | 3 - | 0,000186124 - | 2 - | 0,000154074 - | 2 - | 0,000146467 - | 4 - | 0,000297261 - | 2 - | 0,000167432 - |
| 2552 | 1 | 3um_FIJJAFBF_179665 | Bacteria | Proteobacteri | Gammaprotec | Cellvibrionales | Porticoccacea | unclassified | Gammaprotec | unclassified | unclassified | unclassified | unclassified | unclassified | 2 | 0 | 0,000104203 | 9,89E-06 | 2 - | 8,73E-05 - | 2 - | 0,000108379 - | 2 - | 0,000103028 - | 2 - | 0,00010455 - | 2 - | 0,000117775 - |
| 2553 | 1 | 3um_FIJJAFBF_355792 | Bacteria | Bacteroidetes | Flavobacteriia | Flavobacterial | Flavobacteriac | Polaribacter | Polaribacter b | unclassified | unclassified | unclassified | unclassified | unclassified | 3,2 | 0,4 | 5,85E-05 | 9,07E-06 | 3 - | 4,59E-05 - | 3 - | 5,70E-05 - | 3 - | 5,42E-05 - | 4 - | 7,33E-05 - | 3 - | 6,19E-05 - |
| 2554 | 1 | 3um_FIJJAFBF_564084 | Viruses | unclassified | unclassified | unclassified | Mimiviridae | Tupanvirus | Tupanvirus so | cellular proces | Posttranslatio | KOG1971 | Procollagen-ly | 7070.TC01339 | 1 | 0 | 2,15E-05 | 2,04E-06 | 1 - | 1,80E-05 - | 1 - | 2,24E-05 - | 1 - | 2,13E-05 - | 1 - | 2,16E-05 - | 1 - | 2,43E-05 - |
| 2555 | 1 | 0.2um_NOKCPHJN_510387 | Bacteria | Proteobacteri | Gammaprotec | unclassified | unclassified | unclassified | Gammaprotec | cellular proces | Posttranslatio | 05C6P | ATP-depender | 208964.PA180 | 0,8 | 0,4 | 1,67E-05 | 8,50E-06 | 1 - | 1,81E-05 - | 1 - | 2,25E-05 - | 1 - | 2,14E-05 - | 1 - | 2,17E-05 - | 0 - | 0 - |
| 2556 | 1 | 3um_FIJJAFBF_795423 | Bacteria | Verrucomicrol | Opitutae | unclassified | unclassified | unclassified | Opitutae bact | unclassified | unclassified | unclassified | unclassified | unclassified | 1,8 | 0,748331477 | 0,000257524 | 0,000103727 | 2 - | 0,000244206 - | 1 - | 0,000151616 - | 2 - | 0,00028826 - | 3 - | 0,000438777 - | 1 - | 0,00016476 - |
| 2557 | 1 | 3um_FIJJAFBF_737461 | Bacteria | Proteobacteri | Gammaprotec | Oceanospirilla | Saccharospirill | Saccharospiril | Saccharospiril | information st | Transcription | 16PMT | DNA-depende | 207949.RED65 | 8,2 | 0,748331477 | 0,000334923 | 1,87E-05 | 9 - | 0,000310207 - | 8 - | 0,000342388 - | 9 - | 0,000366168 - | 8 - | 0,00033029 - | 7 - | 0,000325562 - |
| 2558 | 1 | 3um_FIJJAFBF_604550 | Bacteria | Proteobacteri | Gammaprotec | Arenicellales | Arenicellaceae | Arenicella | Arenicella xan | information st | Translation, ri | 16Q16 | Binds the lowe | 290398.Csal_ | 0,2 | 0,4 | 1,03E-05 | 2,06E-05 | 1 - | 5,15E-05 - | 0 - | 0 - | 0 - | 0 - | 0 - | 0 - | 0 - | 0 - |
| 2559 | 1 | 0.2um_NOKCPHJN_206596 | unclassified | unclassified | unclassified | unclassified | unclassified | unclassified | unclassified | unclassified | unclassified | unclassified | unclassified | unclassified | 0,2 | 0,4 | 1,90E-06 | 3,81E-06 | 0 - | 0 - | 0 - | 0 - | 0 - | 0 - | 1 - | 9,52E-06 - | 0 - | 0 - |
| 2560 | 1 | 0.2um_NOKCPHJN_147529 | Bacteria | Proteobacteri | Alphaproteob | unclassified | unclassified | unclassified | Alphaproteob | poorly charact | Function unkn | 067BE | NA | 270374.MELB | 7 | 0,894427191 | 0,000233326 | 2,92E-05 | 7 - | 0,000196439 - | 6 - | 0,000209074 - | 8 - | 0,000265002 - | 8 - | 0,000268916 - | 6 - | 0,0002272 - |
| 2561 | 1 | 0.2um_NOKCPHJN_178388 | Bacteria | Bacteroidetes | Flavobacteriia | Flavobacterial | Flavobacteriac | Polaribacter | Polaribacter s | unclassified | unclassified | unclassified | unclassified | unclassified | 4 | 1,897366596 | 0,000109591 | 5,31E-05 | 2 - | 4,56E-05 - | 5 - | 0,000141669 - | 4 - | 0,000107739 - | 7 - | 0,000191329 - | 2 - | 6,16E-05 - |
| 2562 | 1 | 3um_FIJJAFBF_362463 | Bacteria | Bacteroidetes | Flavobacteriia | Flavobacterial | Flavobacteriac | Polaribacter | Polaribacter s | unclassified | unclassified | unclassified | unclassified | unclassified | 1 | 0 | 9,20E-05 | 8,73E-06 | 1 - | 7,70E-05 - | 1 - | 9,57E-05 - | 1 - | 9,09E-05 - | 1 - | 9,23E-05 - | 1 - | 0,000103943 - |
| 2563 | 1 | 3um_FIJJAFBF_531311 | Bacteria | Bacteroidetes | Flavobacteriia | Flavobacterial | Flavobacteriac | Polaribacter | Polaribacter s | cellular proces | Cell wall/mem | 05DAF | Involved in th | 391603.FBALC | 1 | 0 | 5,23E-05 | 4,97E-06 | 1 - | 4,38E-05 - | 1 - | 5,44E-05 - | 1 - | 5,17E-05 - | 1 - | 5,25E-05 - | 1 - | 5,91E-05 - |
| 2564 | 1 | 0.2um_NOKCPHJN_1006750 | unclassified | unclassified | unclassified | unclassified | unclassified | unclassified | unclassified | unclassified | unclassified | unclassified | unclassified | unclassified | 0,2 | 0,4 | 4,73E-06 | 9,47E-06 | 0 - | 0 - | 0 - | 0 - | 0 - | 0 - | 1 - | 2,37E-05 - | 0 - | 0 - |
| 2565 | 1 | 3um_FIJJAFBF_71624 | Bacteria | Bacteroidetes | Flavobacteriia | Flavobacterial | Flavobacteriac | Formosa | Formosa sp. H | metabolism | Carbohydrate | COG0176 | Transaldolase | 983544.Lacal_ | 0,8 | 0,4 | 5,00E-05 | 2,57E-05 | 1 - | 5,29E-05 - | 0 - | 0 - | 1 - | 6,24E-05 - | 1 - | 6,34E-05 - | 1 - | 7,14E-05 - |
| 2566 | 1 | 0.2um_NOKCPHJN_236630 | Bacteria | Proteobacteri | Gammaprotec | Cellvibrionales | Porticoccacea | unclassified | Gammaprotec | unclassified | unclassified | unclassified | unclassified | unclassified | 0,4 | 0,489897949 | 1,17E-05 | 1,46E-05 | 1 - | 2,48E-05 - | 0 - | 0 - | 0 - | 0 - | 0 - | 0 - | 1 - | 3,35E-05 - |
| 2567 | 1 | 3um_FIJJAFBF_01147 | Bacteria | Bacteroidetes | Flavobacteriia | Flavobacterial | Flavobacteriac | Polaribacter | Polaribacter s | unclassified | unclassified | unclassified | unclassified | unclassified | 0,6 | 0,489897949 | 5,20E-05 | 4,29E-05 | 1 - | 7,60E-05 - | 1 - | 9,44E-05 - | 1 - | 8,97E-05 - | 0 - | 0 - | 0 - | 0 - |
| 2568 | 1 | 0.2um_NOKCPHJN_28606 | Bacteria | Bacteroidetes | Flavobacteriia | Flavobacterial | Flavobacteriac | unclassified | Flavobacteriac | information st | Transcription | 0P7SF | DNA-depende | 391587.KAOT | 0,4 | 0,489897949 | 4,06E-06 | 5,01E-06 | 1 - | 9,04E-06 - | 1 - | 1,12E-05 - | 0 - | 0 - | 0 - | 0 - | 0 - | 0 - |

| | | | | | | | | | | | | | | | | | | | | | | | | | | | | |
|---|---|---|---|---|---|---|---|---|---|---|---|---|---|---|---|---|---|---|---|---|---|---|---|---|---|---|---|---|
| 2569 | 1 | 3um_FIJJAFBF_343504 | Bacteria | Proteobacteri | Alphaproteob | Rhodobactera | Rhodobactera | Nioella | Nioella sedimi | poorly charact | Function unkn | 05ERF | Extracellular s | 488538.SAR11 | 0,8 | 0,4 | 2,23E-05 | 1,15E-05 | 1 - | 2,33E-05 - | 1 - | 2,89E-05 - | 0 - | 0 - | 1 - | 2,79E-05 - | 1 - | 3,14E-05 - |
| 2570 | 1 | 0.2um_NOKCPHJN_165645 | Bacteria | Proteobacteri | Gammaprote | Oceanospirilla | Saccharospirill | Reinekea | Reinekea fors | metabolism | Coenzyme tra | 05ECF | c-methyltrans | 314283.MED2 | 1,8 | 0,748331477 | 6,34E-05 | 2,66E-05 | 1 - | 2,94E-05 - | 2 - | 7,31E-05 - | 3 - | 0,000104217 - | 2 - | 7,05E-05 - | 1 - | 3,97E-05 - |
| 2571 | 1 | 3um_FIJJAFBF_555396 | Bacteria | Proteobacteri | Alphaproteob | Rhizobiales | Rhizobiaceae | Ensifer | various | cellular proces | Cell wall/mem | COG3637 | Membrane | 444612.REIS_ | 0,6 | 0,489897949 | 3,03E-05 | 2,49E-05 | 1 - | 4,48E-05 - | 0 - | 0 - | 1 - | 5,29E-05 - | 1 - | 5,37E-05 - | 0 - | 0 - |
| 2572 | 1 | 0.2um_NOKCPHJN_429314 | Viruses | unclassified | unclassified | unclassified | unclassified | unclassified | uncultured vir | unclassified | unclassified | unclassified | unclassified | unclassified | 0,6 | 0,489897949 | 2,63E-05 | 2,17E-05 | 1 - | 3,83E-05 - | 1 - | 4,75E-05 - | 0 - | 0 - | 1 - | 4,58E-05 - | 0 - | 0 - |
| 2573 | 1 | 0.2um_NOKCPHJN_387115 | Bacteria | Proteobacteri | Alphaproteob | Rhodobactera | Rhodobactera | Pseudooceani | Pseudooceani | metabolism | Lipid transpor | COG0439 | acetyl-CoA car | 52598.EE36_1 | 1,6 | 0,8 | 4,71E-05 | 2,39E-05 | 2 - | 5,10E-05 - | 2 - | 6,33E-05 - | 2 - | 6,02E-05 - | 2 - | 6,11E-05 - | 0 - | 0 - |
| 2574 | 1 | 0.2um_NOKCPHJN_859046 | Bacteria | Proteobacteri | Alphaproteob | Pelagibacteral | Pelagibactera | Candidatus Pe | Candidatus Pe | cellular proces | Cell wall/mem | 7257 | NA | 335992.SAR11 | 1 | 0 | 4,31E-05 | 4,09E-06 | 1 - | 3,61E-05 - | 1 - | 4,48E-05 - | 1 - | 4,26E-05 - | 1 - | 4,32E-05 - | 1 - | 4,87E-05 - |
| 2575 | 1 | 3um_FIJJAFBF_676570 | Bacteria | Proteobacteri | Alphaproteob | Rhodobactera | Rhodobactera | Amylibacter | Amylibacter k | unclassified | unclassified | unclassified | unclassified | unclassified | 1 | 0 | 5,13E-05 | 4,87E-06 | 1 - | 4,30E-05 - | 1 - | 5,34E-05 - | 1 - | 5,07E-05 - | 1 - | 5,15E-05 - | 1 - | 5,80E-05 - |
| 2576 | 1 | 0.2um_NOKCPHJN_855027 | Eukaryota | unclassified | unclassified | Acytosteliales | Acytosteliacea | Heterostelium | Heterostelium | poorly charact | Function unkn | 0644X | Domain-Conta | 398578.Daci_ | 0,4 | 0,489897949 | 1,07E-05 | 1,32E-05 | 1 - | 2,46E-05 - | 0 - | 0 - | 1 - | 2,91E-05 - | 0 - | 0 - | 0 - | 0 - |
| 2577 | 1 | 3um_FIJJAFBF_647537 | Bacteria | Proteobacteri | Gammaprote | Cellvibrionale | Porticoccacea | unclassified | various | metabolism | Energy produc | 05MPD | Cytochrome | 314287.GB22 | 1 | 0 | 9,72E-05 | 9,23E-06 | 1 - | 8,14E-05 - | 1 - | 0,000101077 - | 1 - | 9,61E-05 - | 1 - | 9,75E-05 - | 1 - | 0,00010984 - |
| 2578 | 1 | 0.2um_NOKCPHJN_965417 | Bacteria | Proteobacteri | Alphaproteob | Pelagibacteral | Pelagibactera | Candidatus Pe | Candidatus Pe | metabolism | Carbohydrate | COG2721 | Dehydratase | 335992.SAR11 | 1 | 0 | 0,000145774 | 1,38E-05 | 1 - | 0,000122103 - | 1 - | 0,000151616 - | 1 - | 0,00014413 - | 1 - | 0,000146259 - | 1 - | 0,00016476 - |
| 2579 | 1 | 0.2um_NOKCPHJN_91493 | Bacteria | Proteobacteri | Gammaprote | Oceanospirilla | Saccharospirill | Saccharospiril | Saccharospiril | unclassified | unclassified | unclassified | unclassified | unclassified | 0,6 | 0,8 | 1,67E-05 | 2,15E-05 | 2 - | 5,22E-05 - | 0 - | 0 - | 0 - | 0 - | 1 - | 3,12E-05 - | 0 - | 0 - |
| 2580 | 1 | 3um_FIJJAFBF_638102 | Bacteria | Proteobacteri | Alphaproteob | Rhodobactera | Rhodobactera | Pseudophaeol | Pseudophaeo | metabolism | Coenzyme tra | 01R34 | Catalyzes the | 391619.RGBS | 1 | 0 | 3,49E-05 | 3,31E-06 | 1 - | 2,92E-05 - | 1 - | 3,63E-05 - | 1 - | 3,45E-05 - | 1 - | 3,50E-05 - | 1 - | 3,94E-05 - |
| 2581 | 1 | 3um_FIJJAFBF_362905 | Bacteria | Proteobacteri | Alphaproteob | Rhodobactera | Rhodobactera | Thalassococcu | Thalassococcu | information st | Translation, ri | 01RJH | 30S ribosomal | 314270.RB208 | 0,2 | 0,4 | 9,68E-06 | 1,94E-05 | 0 - | 0 - | 0 - | 0 - | 1 - | 4,84E-05 - | 0 - | 0 - | 0 - | 0 - |
| 2582 | 1 | 3um_FIJJAFBF_72463 | Bacteria | Bacteroidetes | Cytophagia | Cytophagales | Cytophagacea | Ekhidna | Ekhidna lutea | information st | Translation, ri | 05CEJ | Catalyzes the | 269798.CHU_ | 2,8 | 1,166190379 | 5,38E-05 | 2,04E-05 | 4 - | 6,57E-05 - | 3 - | 6,12E-05 - | 1 - | 1,94E-05 - | 4 - | 7,87E-05 - | 2 - | 4,43E-05 - |
| 2583 | 1 | 0.2um_NOKCPHJN_144469 | Bacteria | Proteobacteri | Gammaprote | Oceanospirilla | Saccharospirill | Reinekea | Reinekea blan | metabolism | Energy produc | COG0055 | Produces ATP | 314283.MED2 | 21,2 | 1,166190379 | 0,000625558 | 8,53E-05 | 20 - | 0,000492604 - | 23 - | 0,000703419 - | 21 - | 0,000610542 - | 20 - | 0,000590057 - | 22 - | 0,000731167 - |
| 2584 | 1 | 0.2um_NOKCPHJN_447104 | Bacteria | Bacteroidetes | Flavobacteriia | Flavobacterial | Flavobacteriac | Formosa | Formosa sp. H | poorly charact | Function unkn | 0XRCJ | NA | 391603.FBALC | 1 | 0 | 1,93E-05 | 1,83E-06 | 1 - | 1,62E-05 - | 1 - | 2,01E-05 - | 1 - | 1,91E-05 - | 1 - | 1,94E-05 - | 1 - | 2,18E-05 - |
| 2585 | 1 | 3um_FIJJAFBF_642735 | Eukaryota | Bacillariophyt | Bacillariophyc | Naviculales | Phaeodactylac | Phaeodactylu | Phaeodactylu | unclassified | unclassified | unclassified | unclassified | unclassified | 0,6 | 0,489897949 | 4,53E-05 | 3,71E-05 | 0 - | 0 - | 0 - | 0 - | 1 - | 7,17E-05 - | 1 - | 7,27E-05 - | 1 - | 8,19E-05 - |
| 2586 | 1 | 0.2um_NOKCPHJN_190080 | Bacteria | Bacteroidetes | Flavobacteriia | Flavobacterial | Flavobacteriac | Polaribacter | Polaribacter r | unclassified | unclassified | unclassified | unclassified | unclassified | 0,8 | 0,4 | 2,27E-05 | 1,16E-05 | 1 - | 2,37E-05 - | 1 - | 2,94E-05 - | 0 - | 0 - | 1 - | 2,83E-05 - | 1 - | 3,19E-05 - |
| 2587 | 1 | 3um_FIJJAFBF_455827 | Bacteria | Proteobacteri | Alphaproteob | Rhodobactera | Rhodobactera | Amylibacter | Amylibacter k | unclassified | unclassified | unclassified | unclassified | unclassified | 0,6 | 0,489897949 | 2,06E-05 | 1,68E-05 | 0 - | 0 - | 1 - | 3,54E-05 - | 1 - | 3,36E-05 - | 1 - | 3,41E-05 - | 0 - | 0 - |
| 2588 | 1 | 0.2um_NOKCPHJN_38279 | Bacteria | Bacteroidetes | Flavobacteriia | Flavobacterial | Crocinitomica | unclassified | Crocinitomica | unclassified | unclassified | unclassified | unclassified | unclassified | 0,2 | 0,4 | 3,36E-06 | 6,71E-06 | 1 - | 1,68E-05 - | 0 - | 0 - | 0 - | 0 - | 0 - | 0 - | 0 - | 0 - |
| 2589 | 1 | 3um_FIJJAFBF_796050 | Bacteria | Bacteroidetes | Flavobacteriia | Flavobacterial | various | various | various | metabolism | Energy produc | COG0055 | Produces ATP | 398720.MED2 | 2,2 | 0,748331477 | 8,85E-05 | 2,67E-05 | 3 - | 0,000102785 - | 2 - | 8,51E-05 - | 3 - | 0,000121327 - | 1 - | 4,10E-05 - | 2 - | 9,25E-05 - |
| 2590 | 1 | 0.2um_NOKCPHJN_100824 | Bacteria | Proteobacteri | Betaproteoba | Burkholderial | Comamonada | Polaromonas | Polaromonas | unclassified | unclassified | unclassified | unclassified | unclassified | 0,6 | 0,489897949 | 2,09E-05 | 1,71E-05 | 0 - | 0 - | 1 - | 3,59E-05 - | 1 - | 3,41E-05 - | 1 - | 3,46E-05 - | 0 - | 0 - |
| 2591 | 1 | 0.2um_NOKCPHJN_518944 | Bacteria | Proteobacteri | Gammaprote | Cellvibrionale | Porticoccacea | unclassified | gamma prote | unclassified | unclassified | unclassified | unclassified | unclassified | 0,6 | 0,489897949 | 2,36E-05 | 1,95E-05 | 1 - | 3,45E-05 - | 1 - | 4,28E-05 - | 1 - | 4,07E-05 - | 0 - | 0 - | 0 - | 0 - |
| 2592 | 1 | 0.2um_NOKCPHJN_1102021 | Bacteria | Verrucomicrol | Verrucomicrol | Verrucomicrol | Rubritaleacea | Rubritalea | Rubritalea squ | cellular proces | Posttranslatio | 1DME7 | Molecular cha | 240016.VspiD | 1 | 0 | 2,75E-05 | 2,61E-06 | 1 - | 2,30E-05 - | 1 - | 2,86E-05 - | 1 - | 2,72E-05 - | 1 - | 2,76E-05 - | 1 - | 3,11E-05 - |
| 2593 | 1 | 0.2um_NOKCPHJN_61248 | Viruses | unclassified | unclassified | unclassified | Phycodnavirid | Prasinovirus | Ostreococcus | unclassified | unclassified | unclassified | unclassified | unclassified | 0,4 | 0,489897949 | 1,33E-05 | 1,63E-05 | 0 - | 0 - | 0 - | 0 - | 1 - | 3,31E-05 - | 1 - | 3,36E-05 - | 0 - | 0 - |
| 2594 | 1 | 0.2um_NOKCPHJN_426944 | Bacteria | Proteobacteri | Gammaprote | unclassified | unclassified | unclassified | SAR86 cluster | information st | Transcription | COG0085 | DNA-depende | 247633.GP214 | 1 | 0,894427191 | 9,84E-06 | 8,59E-06 | 2 - | 1,69E-05 - | 2 - | 2,09E-05 - | 0 - | 0 - | 0 - | 0 - | 1 - | 1,14E-05 - |
| 2595 | 1 | 0.2um_NOKCPHJN_260393 | Eukaryota | unclassified | Dinophyceae | Suessiales | Symbiodiniace | Symbiodinium | Symbiodinium | unclassified | unclassified | unclassified | unclassified | unclassified | 0,2 | 0,4 | 2,78E-06 | 5,57E-06 | 0 - | 0 - | 0 - | 0 - | 1 - | 1,39E-05 - | 0 - | 0 - | 0 - | 0 - |
| 2596 | 1 | 3um_FIJJAFBF_344758 | Eukaryota | Bacillariophyt | Coscinodiscop | Coscinodiscale | Coscinodiscac | Coscinodiscus | Coscinodiscus | information st | Translation, ri | 0FJK5 | Binds the lowe | 197221.tlr008 | 0,2 | 0,4 | 1,27E-05 | 2,53E-05 | 0 - | 0 - | 0 - | 0 - | 1 - | 6,33E-05 - | 0 - | 0 - | 0 - | 0 - |
| 2597 | 1 | 0.2um_NOKCPHJN_696831 | Eukaryota | Bacillariophyt | Coscinodiscop | Thalassiosirale | Thalassiosirac | Thalassiosira | Thalassiosira | cellular proces | Intracellular tr | 11J6D | SEC16 homolo | 2850.Phatr49 | 1 | 0 | 1,80E-05 | 1,71E-06 | 1 - | 1,51E-05 - | 1 - | 1,87E-05 - | 1 - | 1,78E-05 - | 1 - | 1,80E-05 - | 1 - | 2,03E-05 - |
| 2598 | 1 | 0.2um_NOKCPHJN_702618 | Bacteria | Proteobacteri | Alphaproteob | Rhodobactera | Rhodobactera | Planktomarina | Planktomarina | metabolism | Energy produc | 05FG6 | Dehydrogenas | 246200.SPO2 | 0,4 | 0,489897949 | 1,93E-05 | 2,37E-05 | 1 - | 4,41E-05 - | 0 - | 0 - | 1 - | 5,21E-05 - | 0 - | 0 - | 0 - | 0 - |
| 2599 | 1 | 0.2um_NOKCPHJN_768355 | Bacteria | Proteobacteri | Gammaprote | Pseudomonad | Pseudomonad | Pseudomonas | Pseudomonas | poorly charact | Function unkn | 0XNRY | Methyl-accep | 62928.azo058 | 0,6 | 0,489897949 | 1,62E-05 | 1,34E-05 | 1 - | 2,36E-05 - | 1 - | 2,93E-05 - | 0 - | 0 - | 1 - | 2,82E-05 - | 0 - | 0 - |
| 2600 | 1 | 3um_FIJJAFBF_597256 | Bacteria | Bacteroidetes | Bacteroidia | various | various | various | various | unclassified | unclassified | unclassified | unclassified | unclassified | 0,4 | 0,489897949 | 6,31E-05 | 7,78E-05 | 1 - | 0,000143471 - | 0 - | 0 - | 0 - | 0 - | 1 - | 0,000171854 - | 0 - | 0 - |
| 2601 | 1 | 3um_FIJJAFBF_656820 | Bacteria | Bacteroidetes | unclassified | unclassified | unclassified | unclassified | unidentified e | unclassified | unclassified | unclassified | unclassified | unclassified | 0,6 | 0,489897949 | 0,000238314 | 0,000198125 | 1 - | 0,000337579 - | 0 - | 0 - | 1 - | 0,000398477 - | 0 - | 0 - | 1 - | 0,000455513 - |
| 2602 | 1 | 0.2um_NOKCPHJN_88322 | Bacteria | Proteobacteri | Alphaproteob | Rhodobactera | Rhodobactera | Thalassobius | Thalassobius a | metabolism | Lipid transpor | 01R4W | acetyl-coa ace | 439496.RBY4 | 2 | 1,095445115 | 6,72E-05 | 3,66E-05 | 3 - | 8,78E-05 - | 3 - | 0,000109071 - | 2 - | 6,91E-05 - | 2 - | 7,01E-05 - | 0 - | 0 - |
| 2603 | 1 | 3um_FIJJAFBF_518449 | Bacteria | Bacteroidetes | Flavobacteriia | Flavobacterial | Flavobacteriac | Ochrovirga | Ochrovirga pa | information st | Translation, ri | COG0254 | 50S ribosomal | 313598.MED1 | 1 | 0 | 0,000165093 | 1,57E-05 | 1 - | 0,000138285 - | 1 - | 0,000171709 - | 1 - | 0,000163232 - | 1 - | 0,000165643 - | 1 - | 0,000186596 - |
| 2604 | 1 | 3um_FIJJAFBF_757375 | Eukaryota | Bacillariophyt | Coscinodiscop | Coscinodiscale | Coscinodiscac | Coscinodiscus | Coscinodiscus | unclassified | unclassified | unclassified | unclassified | unclassified | 0,6 | 0,489897949 | 0,000117844 | 9,72E-05 | 1 - | 0,000171309 - | 1 - | 0,000212715 - | 0 - | 0 - | 1 - | 0,000205199 - | 0 - | 0 - |
| 2605 | 1 | 0.2um_NOKCPHJN_111935 | Bacteria | Proteobacteri | Gammaprote | Oceanospirilla | Saccharospirill | Reinekea | Reinekea blan | metabolism | Amino acid tra | 08JEH | abc transporte | 349521.HCH_ | 0,4 | 0,489897949 | 1,96E-05 | 2,42E-05 | 1 - | 4,47E-05 - | 0 - | 0 - | 0 - | 0 - | 1 - | 5,35E-05 - | 0 - | 0 - |
| 2606 | 1 | 3um_FIJJAFBF_564204 | Bacteria | Bacteroidetes | Cytophagia | Cytophagales | Flammeovirga | unclassified | Flammeovirga | information st | Translation, ri | 09G6S | 50S ribosomal | 643867.Ftrac_ | 0,6 | 0,489897949 | 9,24E-05 | 7,62E-05 | 1 - | 0,000135031 - | 1 - | 0,000167669 - | 1 - | 0,000159391 - | 0 - | 0 - | 0 - | 0 - |
| 2607 | 1 | 0.2um_NOKCPHJN_86593 | Bacteria | Bacteroidetes | Flavobacteriia | unclassified | unclassified | unclassified | Flavobacteria | unclassified | unclassified | unclassified | unclassified | unclassified | 0,6 | 0,489897949 | 1,72E-05 | 1,40E-05 | 0 - | 0 - | 1 - | 2,94E-05 - | 1 - | 2,80E-05 - | 1 - | 2,84E-05 - | 0 - | 0 - |
| 2608 | 2 | 0.2um_NOKCPHJN_18114;3um_FIJJAFBF_722753 | Eukaryota | Mucoromycot | Glomeromyce | Glomerales | Glomeraceae | Rhizophagus | Rhizophagus | unclassified | unclassified | unclassified | unclassified | unclassified | 2,4 | 0,489897949 | 0,000137503 | 2,14E-05 | 3 - | 0,000145903 - | 2 - | 0,000120779 - | 2 - | 0,000114815 - | 3 - | 0,000174767 - | 2 - | 0,00013125 - |
| 2609 | 1 | 0.2um_NOKCPHJN_163367 | Bacteria | Proteobacteri | Gammaprote | Cellvibrionale | Porticoccacea | unclassified | gamma prote | metabolism | Inorganic ion t | 0R7AT | Pfam:TonB | 314287.GB22 | 1 | 0 | 6,82E-05 | 6,47E-06 | 1 - | 5,71E-05 - | 1 - | 7,09E-05 - | 1 - | 6,74E-05 - | 1 - | 6,84E-05 - | 1 - | 7,71E-05 - |
| 2610 | 1 | 3um_FIJJAFBF_641265 | Bacteria | Proteobacteri | Gammaprote | Oceanospirilla | Saccharospirill | Reinekea | Reinekea blan | metabolism | Nucleotide tra | 0QIPM | dihydroorotas | 314283.MED2 | 0,8 | 0,4 | 3,08E-05 | 1,57E-05 | 1 - | 3,34E-05 - | 1 - | 4,14E-05 - | 1 - | 3,94E-05 - | 1 - | 4,00E-05 - | 0 - | 0 - |
| 2611 | 1 | 0.2um_NOKCPHJN_409097 | Eukaryota | Bacillariophyt | Coscinodiscop | Thalassiosirale | Thalassiosirac | Thalassiosira | Thalassiosira | cellular proces | Posttranslatio | KOG0019 | heat shock pr | 67593.Physo1 | 1,4 | 0,489897949 | 2,67E-05 | 8,38E-06 | 2 - | 3,25E-05 - | 2 - | 4,03E-05 - | 1 - | 1,92E-05 - | 1 - | 1,94E-05 - | 1 - | 2,19E-05 - |
| 2612 | 2 | 0.2um_NOKCPHJN_898522;3um_FIJJAFBF_67938 | Bacteria | Bacteroidetes | Flavobacteriia | Flavobacterial | Flavobacteriac | Polaribacter | Polaribacter s | poorly charact | Function unkn | 099QV | gliding motilit | 313598.MED1 | 0,4 | 0,489897949 | 1,76E-05 | 2,17E-05 | 1 - | 4,04E-05 - | 0 - | 0 - | 1 - | 4,77E-05 - | 0 - | 0 - | 0 - | 0 - |
| 2613 | 1 | 3um_FIJJAFBF_151694 | Bacteria | Bacteroidetes | Flavobacteriia | Flavobacterial | unclassified | unclassified | Flavobacterial | metabolism | Coenzyme tra | 09A4R | May play a ke | 487797.Flav3 | 0,4 | 0,489897949 | 1,17E-05 | 1,45E-05 | 1 - | 2,61E-05 - | 1 - | 3,25E-05 - | 0 - | 0 - | 0 - | 0 - | 0 - | 0 - |
| 2614 | 1 | 3um_FIJJAFBF_320476 | Bacteria | Bacteroidetes | Flavobacteriia | various | various | various | various | unclassified | unclassified | unclassified | unclassified | unclassified | 2,2 | 0,4 | 0,000478652 | 9,72E-05 | 2 - | 0,000364371 - | 2 - | 0,000452441 - | 2 - | 0,000430102 - | 3 - | 0,000654682 - | 2 - | 0,000491665 - |
| 2615 | 1 | 3um_FIJJAFBF_694668 | Bacteria | Bacteroidetes | unclassified | unclassified | unclassified | unclassified | Bacteroidetes | information st | Translation, ri | COG0087 | One of the pri | 391596.PBAL3 | 4 | 1,095445115 | 0,000265283 | 7,03E-05 | 4 - | 0,000223955 - | 4 - | 0,000278086 - | 6 - | 0,000396533 - | 3 - | 0,000201195 - | 3 - | 0,000226646 - |
| 2616 | 1 | 3um_FIJJAFBF_412461 | Bacteria | Bacteroidetes | Flavobacteriia | Flavobacterial | unclassified | unclassified | Flavobacterial | poorly charact | Function unkn | 08QKU | NA | 755732.Fluta_ | 0,2 | 0,4 | 2,03E-06 | 4,07E-06 | 0 - | 0 - | 0 - | 0 - | 1 - | 1,02E-05 - | 0 - | 0 - | 0 - | 0 - |
| 2617 | 1 | 0.2um_NOKCPHJN_143029 | Bacteria | Bacteroidetes | Flavobacteriia | Flavobacterial | Flavobacteriac | Flavivirga | Flavivirga eckl | metabolism | Carbohydrate | COG0057 | glyceraldehyd | 313598.MED1 | 4,2 | 0,4 | 0,000172217 | 2,18E-05 | 4 - | 0,000137457 - | 4 - | 0,000170681 - | 5 - | 0,000202818 - | 4 - | 0,000164651 - | 4 - | 0,000185478 - |
| 2618 | 1 | 3um_FIJJAFBF_164018 | Bacteria | Proteobacteri | Gammaprote | Cellvibrionale | Halieaceae | unclassified | marine gamm | information st | Translation, ri | COG0052 | 30S ribosomal | 566466.NOR5 | 1,4 | 0,489897949 | 7,09E-05 | 2,03E-05 | 2 - | 8,70E-05 - | 1 - | 5,40E-05 - | 2 - | 0,000102638 - | 1 - | 5,21E-05 - | 1 - | 5,87E-05 - |
| 2619 | 1 | 3um_FIJJAFBF_226619 | Bacteria | Bacteroidetes | Flavobacteriia | Flavobacterial | Flavobacteriac | Formosa | Formosa sp. H | metabolism | Inorganic ion t | COG3119 | Sulfatase | 655815.ZPR_3 | 9 | 1,414213562 | 0,000233311 | 3,44E-05 | 9 - | 0,00019676 - | 8 - | 0,000217172 - | 10 - | 0,000258061 - | 11 - | 0,00028806 - | 7 - | 0,000206499 - |
| 2620 | 1 | 0.2um_NOKCPHJN_257142 | Bacteria | Bacteroidetes | Flavobacteriia | Flavobacterial | Flavobacteriac | Jejuia | Jejuia pallidilu | poorly charact | Function unkn | 11P35 | NA | 398720.MED2 | 5 | 0,632455532 | 0,000122003 | 5,59E-06 | 6 - | 0,000124083 - | 5 - | 0,000128395 - | 5 - | 0,000122056 - | 5 - | 0,000123859 - | 4 - | 0,000111621 - |
| 2621 | 1 | 0.2um_NOKCPHJN_45917 | Bacteria | Bacteroidetes | Flavobacteriia | Flavobacterial | Flavobacteriac | Algibacter | Algibacter sp. | metabolism | Energy produc | COG0437 | 4fe-4S ferred | 983544.Lacal_ | 1,8 | 1,166190379 | 2,39E-05 | 1,56E-05 | 1 - | 1,11E-05 - | 1 - | 1,38E-05 - | 2 - | 2,63E-05 - | 4 - | 5,34E-05 - | 1 - | 1,50E-05 - |
| 2622 | 1 | 0.2um_NOKCPHJN_27572 | Bacteria | Actinobacteri | Actinobacteri | Micrococcales | Microbacteria | unclassified | Microbacteria | poorly charact | Function unkn | 07R4F | basic membra | 312284.A20C1 | 1,4 | 0,489897949 | 5,25E-05 | 1,85E-05 | 1 - | 3,14E-05 - | 1 - | 3,90E-05 - | 2 - | 7,42E-05 - | 2 - | 7,53E-05 - | 1 - | 4,24E-05 - |
| 2623 | 1 | 0.2um_NOKCPHJN_42962 | Bacteria | Bacteroidetes | Flavobacteriia | Flavobacterial | Flavobacteriac | Polaribacter | Polaribacter s | unclassified | unclassified | unclassified | unclassified | unclassified | 12,6 | 0,489897949 | 0,000188853 | 2,23E-05 | 12 - | 0,000150363 - | 12 - | 0,000186706 - | 13 - | 0,000192278 - | 13 - | 0,000195118 - | 13 - | 0,0002198 - |
| 2624 | 1 | 0.2um_NOKCPHJN_227986 | Bacteria | Bacteroidetes | Flavobacteriia | Flavobacterial | Crocinitomica | unclassified | Crocinitomica | unclassified | unclassified | unclassified | unclassified | unclassified | 3,8 | 0,748331477 | 5,37E-05 | 1,17E-05 | 3 - | 3,54E-05 - | 4 - | 5,86E-05 - | 4 - | 5,58E-05 - | 5 - | 7,07E-05 - | 3 - | 4,78E-05 - |
| 2625 | 1 | 0.2um_NOKCPHJN_702864 | Eukaryota | unclassified | Oomycetes | Pythiales | Pythiaceae | Pythium | Pythium insidi | unclassified | unclassified | unclassified | unclassified | unclassified | 1,8 | 0,748331477 | 7,04E-05 | 2,35E-05 | 3 - | 0,000101572 - | 2 - | 8,41E-05 - | 2 - | 7,99E-05 - | 1 - | 4,06E-05 - | 1 - | 4,57E-05 - |
| 2626 | 1 | 0.2um_NOKCPHJN_04159 | Bacteria | Bacteroidetes | Cytophagia | Cytophagales | Flammeovirga | unclassified | Flammeovirga | cellular proces | Posttranslatio | COG0330 | Band 7 protei | 926562.Oweh | 7 | 1,673320053 | 0,000333213 | 8,55E-05 | 5 - | 0,000198576 - | 6 - | 0,000295887 - | 9 - | 0,000421917 - | 9 - | 0,000428149 - | 6 - | 0,000321539 - |
| 2627 | 2 | 0.2um_NOKCPHJN_968216;3um_FIJJAFBF_40305 | Eukaryota | various | various | various | various | various | various | information st | Chromatin str | various | various | various | 1 | 0 | 8,90E-05 | 8,45E-06 | 1 - | 7,45E-05 - | 1 - | 9,25E-05 - | 1 - | 8,80E-05 - | 1 - | 8,93E-05 - | 1 - | 0,000100568 - |
| 2628 | 1 | 3um_FIJJAFBF_786158 | Bacteria | Proteobacteri | Gammaprote | Oceanospirilla | Saccharospirill | Reinekea | Reinekea fors | information st | Translation, ri | COG0185 | Protein S19 fo | 314283.MED2 | 1,2 | 0,4 | 0,000180795 | 6,23E-05 | 1 - | 0,000126128 - | 1 - | 0,000156614 - | 1 - | 0,000148882 - | 2 - | 0,000302161 - | 1 - | 0,000170192 - |
| 2629 | 1 | 3um_FIJJAFBF_58623 | unclassified | unclassified | unclassified | unclassified | unclassified | unclassified | unclassified | unclassified | unclassified | unclassified | unclassified | unclassified | 2 | 0,632455532 | 0,000166272 | 3,85E-05 | 3 - | 0,000215206 - | 2 - | 0,000178149 - | 2 - | 0,000169353 - | 2 - | 0,000171854 - | 1 - | 9,68E-05 - |
| 2630 | 1 | 0.2um_NOKCPHJN_622716 | Bacteria | Proteobacteri | Betaproteoba | Burkholderial | Burkholderiac | Burkholderia | Burkholderia | poorly charact | Function unkn | 11D80 | mucin 21, cell | 9483.ENSCJAF | 0,2 | 0,4 | 7,75E-06 | 1,55E-05 | 0 - | 0 - | 0 - | 0 - | 0 - | 0 - | 1 - | 3,87E-05 - | 0 - | 0 - |
| 2631 | 1 | 0.2um_NOKCPHJN_326608 | Bacteria | Bacteroidetes | Flavobacteriia | Flavobacterial | Flavobacteriac | Polaribacter | Polaribacter d | metabolism | Energy produc | COG0056 | Produces ATP | 983544.Lacal_ | 0,8 | 0,4 | 2,09E-05 | 1,07E-05 | 1 - | 2,19E-05 - | 1 - | 2,71E-05 - | 1 - | 2,58E-05 - | 0 - | 0 - | 1 - | 2,95E-05 - |
| 2632 | 1 | 3um_FIJJAFBF_225137 | Bacteria | Proteobacteri | Alphaproteob | Rhodobactera | Rhodobactera | Planktomarina | Planktomarina | unclassified | unclassified | unclassified | unclassified | unclassified | 2,8 | 0,4 | 0,000314742 | 5,63E-05 | 3 - | 0,000282238 - | 3 - | 0,000350456 - | 2 - | 0,000222102 - | 3 - | 0,000338074 - | 3 - | 0,000380839 - |
| 2633 | 1 | 0.2um_NOKCPHJN_179937 | Bacteria | Proteobacteri | Alphaproteob | Rhodobactera | Rhodobactera | Ruegeria | Ruegeria med | metabolism | Energy produc | 05C26 | Dehydrogenas | 246200.SPO2 | 0,4 | 0,489897949 | 1,09E-05 | 1,34E-05 | 0 - | 0 - | 0 - | 0 - | 1 - | 2,72E-05 - | 1 - | 2,76E-05 - | 0 - | 0 - |
| 2634 | 1 | 0.2um_NOKCPHJN_12461 | Bacteria | Bacteroidetes | Flavobacteriia | Flavobacterial | Flavobacteriac | Formosa | Formosa sp. H | unclassified | unclassified | unclassified | unclassified | unclassified | 6 | 0,632455532 | 0,000745174 | 9,18E-05 | 6 - | 0,000626055 - | 7 - | 0,000906938 - | 6 - | 0,000738994 - | 6 - | 0,000749909 - | 5 - | 0,000703975 - |
| 2635 | 1 | 0.2um_NOKCPHJN_689103 | Viruses | unclassified | unclassified | unclassified | unclassified | unclassified | uncultured M | unclassified | unclassified | unclassified | unclassified | unclassified | 0,8 | 0,748331477 | 3,09E-05 | 2,73E-05 | 2 - | 7,06E-05 - | 0 - | 0 - | 1 - | 4,17E-05 - | 1 - | 4,23E-05 - | 0 - | 0 - |
| 2636 | 1 | 3um_FIJJAFBF_315886 | Bacteria | Bacteroidetes | Flavobacteriia | Flavobacterial | Flavobacteriac | Marinirhabdu | Marinirhabdu | unclassified | unclassified | unclassified | unclassified | unclassified | 0,2 | 0,4 | 5,48E-06 | 1,10E-05 | 0 - | 0 - | 0 - | 0 - | 0 - | 0 - | 0 - | 0 - | 1 - | 2,74E-05 - |
| 2637 | 1 | 0.2um_NOKCPHJN_773732 | Eukaryota | Ascomycota | Sordariomyce | Sordariales | Lasiosphaeria | Podospora | Podospora an | cellular proces | Posttranslatio | 0VPU6 | TSP1 | 6238.CBG018 | 0,8 | 0,4 | 1,38E-05 | 7,00E-06 | 1 - | 1,49E-05 - | 1 - | 1,85E-05 - | 1 - | 1,76E-05 - | 1 - | 1,79E-05 - | 0 - | 0 - |
| 2638 | 2 | 0.2um_NOKCPHJN_229467;0.2um_NOKCPHJN_7 | Bacteria | Proteobacteri | Alphaproteob | Pelagibacteral | Pelagibactera | Candidatus Pe | Candidatus Pe | information st | Translation, ri | COG0088 | One of the pri | 335992.SAR11 | 0,8 | 0,4 | 5,43E-05 | 2,73E-05 | 0 - | 0 - | 1 - | 6,79E-05 - | 1 - | 6,45E-05 - | 1 - | 6,55E-05 - | 1 - | 7,37E-05 - |
| 2639 | 1 | 3um_FIJJAFBF_358760 | Eukaryota | unclassified | Pelagophycea | Pelagomonad | unclassified | Aureococcus | Aureococcus a | unclassified | unclassified | unclassified | unclassified | unclassified | 2,4 | 0,489897949 | 0,000449359 | 0,0001216 | 2 - | 0,000310207 - | 2 - | 0,000385186 - | 2 - | 0,000366168 - | 3 - | 0,000557365 - | 3 - | 0,00062787 - |
| 2640 | 1 | 0.2um_NOKCPHJN_133096 | Bacteria | Bacteroidetes | Flavobacteriia | Flavobacterial | Crocinitomica | unclassified | Crocinitomica | cellular proces | Cell wall/mem | 09B44 | efflux transpo | 313596.RB25 | 0,8 | 0,4 | 3,13E-05 | 1,61E-05 | 1 - | 3,31E-05 - | 0 - | 0 - | 1 - | 3,90E-05 - | 1 - | 3,96E-05 - | 1 - | 4,46E-05 - |
| 2641 | 1 | 0.2um_NOKCPHJN_552431 | Bacteria | Proteobacteri | Betaproteoba | Nitrosomonad | Sterolibacteri | Methyloversa | Methyloversa | unclassified | unclassified | unclassified | unclassified | unclassified | 1,2 | 1,166190379 | 3,12E-05 | 3,11E-05 | 2 - | 4,61E-05 - | 0 - | 0 - | 1 - | 2,72E-05 - | 3 - | 8,28E-05 - | 0 - | 0 - |
| 2642 | 1 | 0.2um_NOKCPHJN_785139 | Bacteria | unclassified | unclassified | unclassified | unclassified | unclassified | uncultured ba | metabolism | Inorganic ion t | 05C27 | TonB-depende | 313596.RB25 | 2,6 | 0,8 | 4,46E-05 | 1,41E-05 | 2 - | 2,87E-05 - | 2 - | 3,57E-05 - | 3 - | 5,09E-05 - | 4 - | 6,88E-05 - | 2 - | 3,88E-05 - |
| 2643 | 1 | 3um_FIJJAFBF_123778 | Bacteria | Bacteroidetes | Flavobacteriia | Flavobacterial | Flavobacteriac | Polaribacter | Polaribacter s | unclassified | unclassified | unclassified | unclassified | unclassified | 1,2 | 0,4 | 2,36E-05 | 8,15E-06 | 1 - | 1,65E-05 - | 1 - | 2,05E-05 - | 1 - | 1,95E-05 - | 2 - | 3,95E-05 - | 1 - | 2,23E-05 - |
| 2644 | 1 | 3um_FIJJAFBF_211162 | Bacteria | Bacteroidetes | Flavobacteriia | various | various | various | various | cellular proces | Posttranslatio | 0900B | Thioredoxin | 487797.Flav3 | 0,8 | 0,4 | 0,000101973 | 5,18E-05 | 1 - | 0,000110362 - | 1 - | 0,000137037 - | 1 - | 0,000130271 - | 1 - | 0,000132195 - | 0 - | 0 - |
| 2645 | 1 | 3um_FIJJAFBF_742569 | Bacteria | Proteobacteri | Gammaprote | Alteromonada | Alteromonada | Paraglaciecola | Paraglaciecola | unclassified | unclassified | unclassified | unclassified | unclassified | 3,6 | 0,8 | 0,000187722 | 4,85E-05 | 3 - | 0,000130428 - | 5 - | 0,000269922 - | 3 - | 0,000153957 - | 4 - | 0,000208308 - | 3 - | 0,000175994 - |
| 2646 | 1 | 0.2um_NOKCPHJN_30243 | Bacteria | Proteobacteri | Gammaprote | Oceanospirilla | Oceanospirilla | Motiliproteus | Motiliproteus | unclassified | unclassified | unclassified | unclassified | unclassified | 4 | 0 | 0,000272691 | 2,59E-05 | 4 - | 0,000228411 - | 4 - | 0,00028362 - | 4 - | 0,000269616 - | 4 - | 0,000273599 - | 4 - | 0,000308208 - |
| 2647 | 1 | 3um_FIJJAFBF_533416 | Bacteria | Proteobacteri | Alphaproteob | Rhodobactera | Rhodobactera | Planktomarina | Planktomarina | poorly charact | Function unkn | 02A72 | NA | 314232.SKA53 | 2 | 0 | 0,000330186 | 3,14E-05 | 2 - | 0,000276571 - | 2 - | 0,000343419 - | 2 - | 0,000326463 - | 2 - | 0,000331285 - | 2 - | 0,000373192 - |
| 2648 | 1 | 3um_FIJJAFBF_214624 | Bacteria | Proteobacteri | Alphaproteob | Rhodobactera | Rhodobactera | Shimia | Shimia sp. SK | information st | Translation, ri | 05CEJ | Catalyzes the | 391593.RCCS2 | 3 | 1,095445115 | 5,64E-05 | 1,44E-05 | 5 - | 8,14E-05 - | 2 - | 4,04E-05 - | 3 - | 5,77E-05 - | 3 - | 5,85E-05 - | 2 - | 4,39E-05 - |
| 2649 | 1 | 0.2um_NOKCPHJN_255309 | Bacteria | Proteobacteri | Alphaproteob | Rhodobactera | Rhodobactera | unclassified | Rhodobactera | unclassified | unclassified | unclassified | unclassified | unclassified | 0,6 | 0,489897949 | 8,04E-06 | 6,62E-06 | 1 - | 1,19E-05 - | 0 - | 0 - | 1 - | 1,41E-05 - | 1 - | 1,43E-05 - | 0 - | 0 - |
| 2650 | 1 | 3um_FIJJAFBF_175203 | Bacteria | Proteobacteri | Alphaproteob | Rhodobactera | Rhodobactera | unclassified | Rhodobactera | unclassified | unclassified | unclassified | unclassified | unclassified | 0,8 | 0,4 | 9,40E-05 | 4,83E-05 | 1 - | 9,81E-05 - | 1 - | 0,000121811 - | 0 - | 0 - | 1 - | 0,000117507 - | 1 - | 0,000132371 - |
| 2651 | 1 | 3um_FIJJAFBF_209926 | Bacteria | Bacteroidetes | Flavobacteriia | Flavobacterial | Crocinitomica | Brumimicrobi | Brumimicrobi | poorly charact | Function unkn | 068WQ | NA | 706436.HMPR | 1 | 0 | 0,000145774 | 1,38E-05 | 1 - | 0,000122103 - | 1 - | 0,000151616 - | 1 - | 0,00014413 - | 1 - | 0,000146259 - | 1 - | 0,00016476 - |
| 2652 | 1 | 3um_FIJJAFBF_654925 | Bacteria | Bacteroidetes | Flavobacteriia | various | various | various | various | information st | Replication, re | 0PB0H | DNA-binding p | 487797.Flav3 | 19,6 | 1,356465997 | 0,002968194 | 0,000173415 | 22 - | 0,002805654 - | 18 - | 0,002850377 - | 19 - | 0,00286018 - | 20 - | 0,003055185 - | 19 - | 0,003269572 - |
| 2653 | 1 | 3um_FIJJAFBF_281258 | Bacteria | Proteobacteri | Gammaprote | Oceanospirilla | Saccharospirill | Reinekea | Reinekea fors | unclassified | unclassified | unclassified | unclassified | unclassified | 2,4 | 0,8 | 0,000301772 | 9,68E-05 | 3 - | 0,000318824 - | 3 - | 0,000395886 - | 1 - | 0,000125447 - | 3 - | 0,000381898 - | 2 - | 0,000286805 - |
| 2654 | 2 | 0.2um_NOKCPHJN_217327;0.2um_NOKCPHJN_2 | Bacteria | Proteobacteri | Alphaproteob | Rhodobactera | Rhodobactera | various | various | metabolism | Amino acid tra | 01SW7 | Tetrahydrodip | 89187.ISM_02 | 1,4 | 1,019803903 | 6,85E-05 | 5,11E-05 | 1 - | 4,19E-05 - | 1 - | 5,20E-05 - | 3 - | 0,000148338 - | 2 - | 0,000100353 - | 0 - | 0 - |
| 2655 | 1 | 0.2um_NOKCPHJN_275425 | Bacteria | Bacteroidetes | Flavobacteriia | Flavobacterial | Flavobacteriac | Polaribacter | Polaribacter ir | information st | Replication, re | COG0550 | Releases the s | 313594.PI23P | 0,2 | 0,4 | 2,92E-06 | 5,84E-06 | 1 - | 1,46E-05 - | 0 - | 0 - | 0 - | 0 - | 0 - | 0 - | 0 - | 0 - |
| 2656 | 1 | 3um_FIJJAFBF_203195 | Bacteria | Proteobacteri | Alphaproteob | Sphingomona | Sphingomona | Sphingorhabd | Sphingorhabd | poorly charact | Function unkn | 0YKK5 | NA | 167879.CPS_2 | 4,6 | 1,019803903 | 0,000424643 | 9,89E-05 | 5 - | 0,000387759 - | 6 - | 0,000577779 - | 3 - | 0,000274626 - | 5 - | 0,000464471 - | 4 - | 0,00041858 - |
| 2657 | 1 | 3um_FIJJAFBF_08771 | Eukaryota | unclassified | Raphidophyce | Chattonellales | Chattonellace | Heterosigma | Heterosigma a | metabolism | Energy produc | 0IFBF | One of the tw | 45157.CMV08 | 1 | 0 | 3,90E-05 | 3,71E-06 | 1 - | 3,27E-05 - | 1 - | 4,06E-05 - | 1 - | 3,86E-05 - | 1 - | 3,92E-05 - | 1 - | 4,41E-05 - |
| 2658 | 1 | 3um_FIJJAFBF_628719 | Bacteria | Proteobacteri | Alphaproteob | Rhodobactera | Rhodobactera | Roseovarius | Roseovarius a | metabolism | Energy produc | 16RMM | alcohol dehyd | 89187.ISM_11 | 1 | 0 | 6,68E-05 | 6,35E-06 | 1 - | 5,60E-05 - | 1 - | 6,95E-05 - | 1 - | 6,61E-05 - | 1 - | 6,71E-05 - | 1 - | 7,55E-05 - |
| 2659 | 1 | 3um_FIJJAFBF_396485 | Bacteria | Proteobacteri | Gammaprote | Oceanospirilla | Saccharospirill | Reinekea | Reinekea fors | unclassified | unclassified | unclassified | unclassified | unclassified | 4 | 1,095445115 | 0,000261063 | 7,68E-05 | 3 - | 0,00016319 - | 3 - | 0,000202633 - | 6 - | 0,000385258 - | 4 - | 0,000260632 - | 4 - | 0,000293601 - |
| 2660 | 1 | 0.2um_NOKCPHJN_739786 | Bacteria | Proteobacteri | Gammaprote | Alteromonada | Colwelliaceae | Thalassotalea | Thalassotalea | unclassified | unclassified | unclassified | unclassified | unclassified | 0,2 | 0,4 | 8,74E-06 | 1,75E-05 | 0 - | 0 - | 0 - | 0 - | 1 - | 4,37E-05 - | 0 - | 0 - | 0 - | 0 - |
| 2661 | 1 | 3um_FIJJAFBF_66142 | Bacteria | Bacteroidetes | Flavobacteriia | Flavobacterial | Flavobacteriac | Arenitalea | Arenitalea lut | unclassified | unclassified | unclassified | unclassified | unclassified | 1,8 | 0,748331477 | 0,000163952 | 5,83E-05 | 3 - | 0,000234238 - | 1 - | 9,70E-05 - | 2 - | 0,00018433 - | 1 - | 9,35E-05 - | 2 - | 0,000210714 - |
| 2662 | 1 | 3um_FIJJAFBF_652733 | Bacteria | Bacteroidetes | Flavobacteriia | Flavobacterial | Flavobacteriac | Ulvibacter | Ulvibacter lito | poorly charact | Function unkn | 11W7M | NA | 411154.GFO_ | 0,6 | 0,489897949 | 5,69E-05 | 4,73E-05 | 1 - | 8,03E-05 - | 0 - | 0 - | 0 - | 0 - | 1 - | 9,61E-05 - | 1 - | 0,000108304 - |
| 2663 | 2 | 0.2um_NOKCPHJN_259099;3um_FIJJAFBF_67299 | Viruses | unclassified | unclassified | unclassified | Mimiviridae | Hokovirus | Hokovirus HK | cellular proces | Cell cycle cont | KOG0580 | serine threoni | 352914.XP_72 | 1 | 0 | 1,26E-05 | 1,20E-06 | 1 - | 1,06E-05 - | 1 - | 1,31E-05 - | 1 - | 1,25E-05 - | 1 - | 1,27E-05 - | 1 - | 1,43E-05 - |
| 2664 | 1 | 0.2um_NOKCPHJN_95447 | Bacteria | Proteobacteri | Gammaprote | Cellvibrionale | Porticoccacea | unclassified | SAR92 bacteri | unclassified | unclassified | unclassified | unclassified | unclassified | 0,8 | 0,4 | 2,63E-05 | 1,35E-05 | 1 - | 2,75E-05 - | 1 - | 3,42E-05 - | 1 - | 3,25E-05 - | 0 - | 0 - | 1 - | 3,71E-05 - |
| 2665 | 1 | 0.2um_NOKCPHJN_530412 | Bacteria | Proteobacteri | Gammaprote | unclassified | unclassified | unclassified | Gammaprote | information st | Translation, ri | COG0050 | This protein p | 393595.ABO_ | 6 | 1,095445115 | 0,000207348 | 4,26E-05 | 6 - | 0,000173904 - | 8 - | 0,000287917 - | 5 - | 0,000171063 - | 6 - | 0,000208308 - | 5 - | 0,000195549 - |
| 2666 | 2 | 0.2um_NOKCPHJN_108637;3um_FIJJAFBF_77866 | Bacteria | Proteobacteri | Alphaproteob | Rhodobactera | Rhodobactera | unclassified | Rhodobactera | unclassified | unclassified | unclassified | unclassified | unclassified | 4,6 | 1,019803903 | 0,000187123 | 4,47E-05 | 4 - | 0,000136639 - | 6 - | 0,000254498 - | 5 - | 0,00020161 - | 5 - | 0,000204588 - | 3 - | 0,000138281 - |
| 2667 | 1 | 3um_FIJJAFBF_30298 | Bacteria | Bacteroidetes | Flavobacteriia | Flavobacterial | Flavobacteriac | Polaribacter | Polaribacter s | unclassified | unclassified | unclassified | unclassified | unclassified | 0,8 | 0,4 | 6,24E-05 | 3,20E-05 | 1 - | 6,60E-05 - | 0 - | 0 - | 1 - | 7,79E-05 - | 1 - | 7,90E-05 - | 1 - | 8,90E-05 - |
| 2668 | 1 | 0.2um_NOKCPHJN_149062 | Bacteria | Bacteroidetes | Flavobacteriia | Flavobacterial | Flavobacteriac | Polaribacter | Polaribacter b | information st | Translation, ri | COG0089 | One of the ea | 313598.MED1 | 1,2 | 0,748331477 | 0,000169913 | 0,000107401 | 1 - | 0,000119559 - | 0 - | 0 - | 2 - | 0,000282255 - | 2 - | 0,000286424 - | 1 - | 0,000161328 - |
| 2669 | 1 | 3um_FIJJAFBF_662596 | Bacteria | Bacteroidetes | Flavobacteriia | Flavobacterial | Flavobacteriac | Formosa | Formosa sp. H | unclassified | unclassified | unclassified | unclassified | unclassified | 0,2 | 0,4 | 8,73E-06 | 1,75E-05 | 1 - | 4,36E-05 - | 0 - | 0 - | 0 - | 0 - | 0 - | 0 - | 0 - | 0 - |
| 2670 | 1 | 0.2um_NOKCPHJN_166533 | Eukaryota | unclassified | Oomycetes | Peronosporale | Peronosporac | Phytophthora | Phytophthora | cellular proces | Signal transdu | 0XNRX | calcium calmo | 4787.PITG_17 | 0,6 | 0,489897949 | 2,07E-05 | 1,70E-05 | 1 - | 3,02E-05 - | 1 - | 3,75E-05 - | 1 - | 3,57E-05 - | 0 - | 0 - | 0 - | 0 - |
| 2671 | 1 | 0.2um_NOKCPHJN_384144 | Bacteria | Bacteroidetes | Flavobacteriia | unclassified | unclassified | unclassified | Flavobacteria | metabolism | Nucleotide tra | 0P7KB | dihydroorotas | 487796.Flav2 | 0,2 | 0,4 | 5,16E-06 | 1,03E-05 | 1 - | 2,58E-05 - | 0 - | 0 - | 0 - | 0 - | 0 - | 0 - | 0 - | 0 - |
| 2672 | 1 | 0.2um_NOKCPHJN_119309 | Bacteria | Proteobacteri | Gammaprote | Cellvibrionale | Porticoccacea | unclassified | gamma prote | unclassified | unclassified | unclassified | unclassified | unclassified | 1,8 | 1,326649916 | 2,66E-05 | 1,75E-05 | 4 - | 5,13E-05 - | 2 - | 3,18E-05 - | 0 - | 0 - | 1 - | 1,54E-05 - | 2 - | 3,46E-05 - |
| 2673 | 1 | 0.2um_NOKCPHJN_753040 | Eukaryota | Apicomplexa | Conoidasida | Eucoccidiorida | Eimeriidae | Eimeria | Eimeria necat | unclassified | unclassified | unclassified | unclassified | unclassified | 0,6 | 0,489897949 | 2,71E-05 | 2,25E-05 | 1 - | 3,84E-05 - | 0 - | 0 - | 1 - | 4,53E-05 - | 0 - | 0 - | 1 - | 5,18E-05 - |
| 2674 | 1 | 3um_FIJJAFBF_321707 | unclassified | unclassified | unclassified | unclassified | unclassified | unclassified | unclassified | unclassified | unclassified | unclassified | unclassified | unclassified | 0,2 | 0,4 | 1,54E-05 | 3,07E-05 | 0 - | 0 - | 0 - | 0 - | 0 - | 0 - | 1 - | 7,68E-05 - | 0 - | 0 - |
| 2675 | 1 | 0.2um_NOKCPHJN_444498 | Bacteria | Proteobacteri | Alphaproteob | Rhodospirillal | Rhodospirillac | Candidatus En | Candidatus En | unclassified | unclassified | unclassified | unclassified | unclassified | 0,8 | 0,748331477 | 9,32E-06 | 8,57E-06 | 0 - | 0 - | 0 - | 0 - | 1 - | 1,12E-05 - | 2 - | 2,27E-05 - | 1 - | 1,28E-05 - |
| 2676 | 1 | 3um_FIJJAFBF_409565 | Bacteria | Proteobacteri | Alphaproteob | Pelagibacteral | Pelagibactera | Candidatus Pe | Candidatus Pe | metabolism | Inorganic ion t | 05PWN | phosphate ab | 335992.SAR11 | 10,4 | 1,496662955 | 0,00045532 | 6,02E-05 | 11 - | 0,000405963 - | 10 - | 0,00045826 - | 13 - | 0,000566324 - | 9 - | 0,000397862 - | 9 - | 0,00044819 - |
| 2677 | 2 | 0.2um_NOKCPHJN_319863;3um_FIJJAFBF_20857 | Bacteria | Proteobacteri | Alphaproteob | Rhodobactera | Rhodobactera | unclassified | Rhodobactera | unclassified | unclassified | unclassified | unclassified | unclassified | 0,2 | 0,4 | 1,20E-05 | 2,40E-05 | 1 - | 6,01E-05 - | 0 - | 0 - | 0 - | 0 - | 0 - | 0 - | 0 - | 0 - |
| 2678 | 1 | 3um_FIJJAFBF_419664 | Bacteria | Proteobacteri | Gammaprote | Oceanospirilla | Saccharospirill | Reinekea | Reinekea fors | unclassified | unclassified | unclassified | unclassified | unclassified | 8,8 | 1,469693846 | 0,001346037 | 0,000294316 | 9 - | 0,001147768 - | 10 - | 0,001583543 - | 6 - | 0,000903215 - | 9 - | 0,001374833 - | 10 - | 0,001720828 - |
| 2679 | 1 | 3um_FIJJAFBF_218763 | Bacteria | Proteobacteri | Alphaproteob | Pelagibacteral | Pelagibactera | Candidatus Pe | Candidatus Pe | metabolism | Inorganic ion t | 08UTW | Plays a role in | 335992.SAR11 | 0,8 | 0,4 | 4,72E-05 | 2,43E-05 | 1 - | 4,95E-05 - | 1 - | 6,14E-05 - | 1 - | 5,84E-05 - | 0 - | 0 - | 1 - | 6,68E-05 - |
| 2680 | 1 | 0.2um_NOKCPHJN_494513 | Bacteria | Bacteroidetes | Flavobacteriia | Flavobacterial | Flavobacteriac | Cellulophaga | Cellulophaga | poorly charact | Function unkn | 0P8NI | Inherit from b | 867900.Celly_ | 9,2 | 0,4 | 0,00016827 | 1,68E-05 | 9 - | 0,000137916 - | 9 - | 0,000171251 - | 10 - | 0,000180884 - | 9 - | 0,0001652 - | 9 - | 0,000186098 - |
| 2681 | 1 | 3um_FIJJAFBF_150582 | Bacteria | Proteobacteri | Gammaprote | Oceanospirilla | Saccharospirill | Reinekea | Reinekea fors | cellular proces | Posttranslatio | 17C3T | Peptidyl-proly | 87626.PTD2_ | 0,2 | 0,4 | 1,43E-05 | 2,87E-05 | 1 - | 7,17E-05 - | 0 - | 0 - | 0 - | 0 - | 0 - | 0 - | 0 - | 0 - |
| 2682 | 1 | 0.2um_NOKCPHJN_983192 | Bacteria | Bacteroidetes | Flavobacteriia | Flavobacterial | Flavobacteriac | Tamlana | Tamlana sedin | information st | Translation, ri | 05C64 | Binds directly | 983544.Lacal_ | 4,6 | 1,624807681 | 0,000267553 | 7,31E-05 | 7 - | 0,000350846 - | 4 - | 0,000248941 - | 6 - | 0,000354975 - | 3 - | 0,000180109 - | 3 - | 0,000202892 - |
| 2683 | 1 | 3um_FIJJAFBF_74442 | Bacteria | Bacteroidetes | unclassified | unclassified | unclassified | unclassified | Bacteroidetes | information st | Translation, ri | 05CAV | thus facilitatin | 313594.PI23P | 1 | 0,632455532 | 2,21E-05 | 1,43E-05 | 1 - | 1,86E-05 - | 0 - | 0 - | 1 - | 2,20E-05 - | 2 - | 4,46E-05 - | 1 - | 2,51E-05 - |
| 2684 | 1 | 0.2um_NOKCPHJN_165716 | Bacteria | Proteobacteri | Alphaproteob | Rhodospirillal | Rhodospirillac | Candidatus En | Candidatus En | cellular proces | Cell wall/mem | 05C6Y | Udp-glucose 6 | 313598.MED1 | 0,8 | 0,4 | 2,40E-05 | 1,22E-05 | 1 - | 2,60E-05 - | 1 - | 3,23E-05 - | 1 - | 3,07E-05 - | 1 - | 3,12E-05 - | 0 - | 0 - |
| 2685 | 1 | 0.2um_NOKCPHJN_11860 | Bacteria | Actinobacteri | Actinobacteri | Micrococcales | Microbacteria | Rhodoluna | Rhodoluna la | metabolism | Coenzyme tra | COG0214 | Involved in th | 312284.A20C1 | 0,2 | 0,4 | 7,76E-06 | 1,55E-05 | 1 - | 3,88E-05 - | 0 - | 0 - | 0 - | 0 - | 0 - | 0 - | 0 - | 0 - |
| 2686 | 1 | 3um_FIJJAFBF_46929 | Bacteria | Proteobacteri | Gammaprote | Alteromonada | Alteromonada | Paraglaciecola | Paraglaciecola | unclassified | unclassified | unclassified | unclassified | unclassified | 1,2 | 0,4 | 6,53E-05 | 1,45E-05 | 2 - | 9,37E-05 - | 1 - | 5,82E-05 - | 1 - | 5,53E-05 - | 1 - | 5,61E-05 - | 1 - | 6,32E-05 - |
| 2687 | 1 | 0.2um_NOKCPHJN_411250 | Bacteria | Bacteroidetes | Flavobacteriia | Flavobacterial | Cryomorphace | Owenweeksia | Owenweeksia | unclassified | unclassified | unclassified | unclassified | unclassified | 0,2 | 0,4 | 8,03E-06 | 1,61E-05 | 1 - | 4,01E-05 - | 0 - | 0 - | 0 - | 0 - | 0 - | 0 - | 0 - | 0 - |
| 2688 | 1 | 0.2um_NOKCPHJN_46819 | Bacteria | Proteobacteri | Alphaproteob | Rhodobactera | Rhodobactera | Ruegeria | Ruegeria med | information st | Transcription | COG0085 | DNA-depende | 314270.RB208 | 2,8 | 1,326649916 | 2,72E-05 | 1,11E-05 | 5 - | 4,16E-05 - | 3 - | 3,10E-05 - | 1 - | 9,83E-06 - | 2 - | 2,00E-05 - | 3 - | 3,37E-05 - |
| 2689 | 1 | 3um_FIJJAFBF_544756 | Bacteria | Bacteroidetes | Flavobacteriia | Flavobacterial | Flavobacteriac | Polaribacter | Polaribacter s | cellular proces | Intracellular tr | 08SHB | Biopolymer tra | 313594.PI23P | 0,6 | 0,489897949 | 5,00E-05 | 4,12E-05 | 1 - | 7,40E-05 - | 0 - | 0 - | 1 - | 8,74E-05 - | 1 - | 8,87E-05 - | 0 - | 0 - |
| 2690 | 1 | 3um_FIJJAFBF_194577 | Bacteria | Proteobacteri | Gammaprote | Oceanospirilla | Saccharospirill | Reinekea | Reinekea fors | unclassified | unclassified | unclassified | unclassified | unclassified | 1,8 | 0,748331477 | 4,73E-05 | 1,68E-05 | 3 - | 6,76E-05 - | 1 - | 2,80E-05 - | 2 - | 5,32E-05 - | 1 - | 2,70E-05 - | 2 - | 6,09E-05 - |
| 2691 | 1 | 3um_FIJJAFBF_175876 | Bacteria | Proteobacteri | Gammaprote | unclassified | unclassified | unclassified | Gammaprote | information st | Translation, ri | 0QTKX | Seems to be t | 565045.NOR5 | 6,2 | 1,833030278 | 0,000665416 | 0,000191474 | 6 - | 0,000546556 - | 6 - | 0,000678661 - | 8 - | 0,000860205 - | 8 - | 0,00087291 - | 3 - | 0,000368749 - |
| 2692 | 1 | 0.2um_NOKCPHJN_235596 | Bacteria | Proteobacteri | Gammaprote | Cellvibrionale | Porticoccacea | unclassified | SAR92 bacteri | information st | Transcription | 05D27 | DNA-depende | 498211.CJA_0 | 14,6 | 1,854723699 | 0,000140817 | 1,27E-05 | 17 - | 0,00013858 - | 16 - | 0,000161953 - | 13 - | 0,00012509 - | 15 - | 0,000146467 - | 12 - | 0,000131995 - |
| 2693 | 1 | 0.2um_NOKCPHJN_595673 | Bacteria | Bacteroidetes | Cytophagia | Cytophagales | Microscillacea | Eisenibacter | Eisenibacter e | unclassified | unclassified | unclassified | unclassified | unclassified | 0,6 | 0,489897949 | 4,05E-05 | 3,37E-05 | 1 - | 5,71E-05 - | 0 - | 0 - | 0 - | 0 - | 1 - | 6,84E-05 - | 1 - | 7,71E-05 - |
| 2694 | 1 | 0.2um_NOKCPHJN_969243 | unclassified | unclassified | unclassified | unclassified | unclassified | unclassified | unclassified | unclassified | unclassified | unclassified | unclassified | unclassified | 0,2 | 0,4 | 6,31E-06 | 1,26E-05 | 0 - | 0 - | 0 - | 0 - | 0 - | 0 - | 0 - | 0 - | 1 - | 3,15E-05 - |
| 2695 | 1 | 3um_FIJJAFBF_126542 | Eukaryota | Streptophyta | unclassified | Solanales | Solanaceae | Nicotiana | Nicotiana atte | information st | Translation, ri | KOG3181 | ribosomal pro | 88036.EFJ310 | 1,8 | 0,748331477 | 0,000111969 | 4,85E-05 | 1 - | 5,15E-05 - | 1 - | 6,39E-05 - | 3 - | 0,000182263 - | 2 - | 0,000123303 - | 2 - | 0,000138901 - |
| 2696 | 1 | 3um_FIJJAFBF_308457 | Bacteria | Proteobacteri | Alphaproteob | Rhodobactera | Rhodobactera | Pseudopelagic | Pseudopelagic | information st | Translation, ri | COG0088 | One of the pri | 52598.EE36_ | 5 | 1,095445115 | 0,000329291 | 5,02E-05 | 7 - | 0,000391921 - | 4 - | 0,000278086 - | 5 - | 0,000330444 - | 4 - | 0,00026826 - | 5 - | 0,000377743 - |
| 2697 | 1 | 3um_FIJJAFBF_406146 | Bacteria | Proteobacteri | Alphaproteob | Pelagibacteral | Pelagibactera | Candidatus Pe | Candidatus Pe | unclassified | unclassified | unclassified | unclassified | unclassified | 0,2 | 0,4 | 1,83E-05 | 3,65E-05 | 0 - | 0 - | 1 - | 9,14E-05 - | 0 - | 0 - | 0 - | 0 - | 0 - | 0 - |
| 2698 | 1 | 0.2um_NOKCPHJN_286635 | Bacteria | Proteobacteri | Gammaprote | Cellvibrionale | Porticoccacea | unclassified | SAR92 bacteri | metabolism | Inorganic ion t | 0QHT5 | TonB-depende | 314287.GB22 | 5,2 | 0,979795897 | 8,59E-05 | 1,40E-05 | 6 - | 8,39E-05 - | 6 - | 0,000104155 - | 6 - | 9,90E-05 - | 4 - | 6,70E-05 - | 4 - | 7,55E-05 - |
| 2699 | 1 | 3um_FIJJAFBF_581665 | Bacteria | Proteobacteri | Gammaprote | Oceanospirilla | Saccharospirill | Reinekea | Reinekea fors | unclassified | unclassified | unclassified | unclassified | unclassified | 12,2 | 0,748331477 | 0,000586231 | 6,62E-05 | 13 - | 0,000523543 - | 12 - | 0,000600079 - | 12 - | 0,000570451 - | 11 - | 0,000530637 - | 13 - | 0,000706445 - |
| 2700 | 1 | 3um_FIJJAFBF_350084 | Bacteria | Proteobacteri | Alphaproteob | Rhodobactera | Rhodobactera | Pseudophaeol | Pseudophaeo | information st | Translation, ri | 16QAP | Associates wit | 388739.RSK2 | 6 | 1,095445115 | 0,000278351 | 2,78E-05 | 8 - | 0,000315537 - | 6 - | 0,000293853 - | 5 - | 0,000232787 - | 6 - | 0,000283471 - | 5 - | 0,000266107 - |
| 2701 | 1 | 0.2um_NOKCPHJN_99542 | Bacteria | Proteobacteri | Gammaprote | Oceanospirilla | Saccharospirill | Reinekea | Reinekea fors | metabolism | Energy produc | 0QIAX | dihydrolipoyl | 314283.MED2 | 3 | 0,632455532 | 8,57E-05 | 1,94E-05 | 3 - | 7,19E-05 - | 3 - | 8,93E-05 - | 4 - | 0,000113138 - | 2 - | 5,74E-05 - | 3 - | 9,70E-05 - |
| 2702 | 1 | 3um_FIJJAFBF_370786 | Bacteria | Proteobacteri | Gammaprote | Oceanospirilla | Saccharospirill | Reinekea | Reinekea fors | metabolism | Inorganic ion t | 05PWN | phosphate ab | 314283.MED2 | 14,6 | 1,2 | 0,00058099 | 6,72E-05 | 14 - | 0,000467115 - | 16 - | 0,000662878 - | 16 - | 0,00063015 - | 14 - | 0,000559525 - | 13 - | 0,000585281 - |
| 2703 | 1 | 3um_FIJJAFBF_450132 | Eukaryota | unclassified | Cryptophyta | Pyrenomonad | Chroomonada | Chroomonas | various | metabolism | Energy produc | 11HAV | photosystem i | 35128.Thapsd | 1 | 0 | 9,52E-05 | 9,04E-06 | 1 - | 7,97E-05 - | 1 - | 9,90E-05 - | 1 - | 9,41E-05 - | 1 - | 9,55E-05 - | 1 - | 0,000107552 - |
| 2704 | 1 | 3um_FIJJAFBF_461082 | Eukaryota | Arthropoda | Arachnida | Trombidiform | Tetranychidae | Tetranychus | Tetranychus u | metabolism | Inorganic ion t | 0DGSY | ATPase Na K t | 9305.ENSSHA | 1,2 | 0,4 | 6,24E-05 | 2,09E-05 | 1 - | 4,36E-05 - | 1 - | 5,42E-05 - | 2 - | 0,000103028 - | 1 - | 5,23E-05 - | 1 - | 5,89E-05 - |
| 2705 | 1 | 3um_FIJJAFBF_555835 | Bacteria | Proteobacteri | Gammaprote | Pseudomonad | Moraxellacea | Psychrobacter | Psychrobacter | unclassified | unclassified | unclassified | unclassified | unclassified | 0,2 | 0,4 | 6,19E-06 | 1,24E-05 | 0 - | 0 - | 0 - | 0 - | 1 - | 3,09E-05 - | 0 - | 0 - | 0 - | 0 - |
| 2706 | 1 | 3um_FIJJAFBF_84669 | Bacteria | Proteobacteri | Gammaprote | Oceanospirilla | Saccharospirill | Reinekea | Reinekea fors | poorly charact | Function unkn | 065FR | NA | 314283.MED2 | 1,6 | 0,489897949 | 3,14E-05 | 8,89E-06 | 2 - | 3,34E-05 - | 2 - | 4,15E-05 - | 1 - | 1,97E-05 - | 2 - | 4,00E-05 - | 1 - | 2,25E-05 - |
| 2707 | 1 | 0.2um_NOKCPHJN_58141 | Bacteria | Proteobacteri | Gammaprote | Oceanospirilla | Saccharospirill | Reinekea | Reinekea blan | metabolism | Energy produc | 05D26 | NQR complex | 314283.MED2 | 4,6 | 1,356465997 | 0,000155573 | 4,88E-05 | 3 - | 8,44E-05 - | 4 - | 0,000139724 - | 5 - | 0,000166032 - | 7 - | 0,000235878 - | 4 - | 0,000151838 - |
| 2708 | 1 | 0.2um_NOKCPHJN_232579 | Bacteria | Bacteroidetes | unclassified | unclassified | unclassified | unclassified | unidentified e | metabolism | Energy produc | COG0055 | Produces ATP | 398720.MED2 | 22,2 | 2,638181192 | 0,000601564 | 3,81E-05 | 27 - | 0,000618557 - | 23 - | 0,000654278 - | 21 - | 0,00056789 - | 20 - | 0,000548836 - | 20 - | 0,000618261 - |
| 2709 | 2 | 0.2um_NOKCPHJN_174385;3um_FIJJAFBF_36499 | Bacteria | Proteobacteri | Betaproteoba | unclassified | unclassified | unclassified | Betaproteoba | metabolism | Inorganic ion t | 07QMV | Part of the AB | various | 2 | 0 | 0,000342568 | 3,25E-05 | 2 - | 0,000286942 - | 2 - | 0,000356297 - | 2 - | 0,000338706 - | 2 - | 0,000343708 - | 2 - | 0,000387186 - |
| 2710 | 1 | 0.2um_NOKCPHJN_159457 | Bacteria | Actinobacteri | Actinobacteri | Actinomycetal | unclassified | unclassified | Actinomyceta | information st | Transcription | 05VEQ | Cold shock pr | 313589.JNB_1 | 1,8 | 0,4 | 0,000362805 | 7,16E-05 | 2 - | 0,000342617 - | 2 - | 0,000425429 - | 2 - | 0,000404425 - | 2 - | 0,000410398 - | 1 - | 0,000231156 - |
| 2711 | 1 | 0.2um_NOKCPHJN_06114 | Bacteria | Proteobacteri | Alphaproteob | Rhodobactera | Rhodobactera | Epibacterium | Epibacterium | unclassified | unclassified | unclassified | unclassified | unclassified | 0,2 | 0,4 | 1,99E-05 | 3,99E-05 | 0 - | 0 - | 0 - | 0 - | 0 - | 0 - | 1 - | 9,96E-05 - | 0 - | 0 - |

| | | | | | | | | | | | | | | | | | | | | | | | | | | | | |
|---|---|---|---|---|---|---|---|---|---|---|---|---|---|---|---|---|---|---|---|---|---|---|---|---|---|---|---|---|
| 2712 | 1 | 0.2um_NOKCPHJN_277659 | Bacteria | Bacteroidetes | unclassified | unclassified | unclassified | unclassified | Bacteroidetes | poorly charact | Function unkn | 11NCV | Protein of unk | 487796.Flav2 | 5,2 | 0,748331477 | 0,000411095 | 5,10E-05 | 6 - | 0,000400384 - | 4 - | 0,000331439 - | 5 - | 0,000393844 - | 6 - | 0,000479593 - | 5 - | 0,000450217 - |
| 2713 | 1 | 0.2um_NOKCPHJN_62391 | Bacteria | Bacteroidetes | Flavobacteriia | Flavobacterial | Cryomorphace | unclassified | Cryomorphace | metabolism | Carbohydrate | 16QEQ | hydrolase, fan | 455436.GHTC | 2 | 0,632455532 | 7,30E-05 | 2,37E-05 | 2 - | 6,12E-05 - | 2 - | 7,60E-05 - | 3 - | 0,000108386 - | 1 - | 3,67E-05 - | 2 - | 8,26E-05 - |
| 2714 | 1 | 3um_FIJJAFBF_699297 | unclassified | unclassified | unclassified | unclassified | unclassified | unclassified | unclassified | unclassified | unclassified | unclassified | unclassified | unclassified | 1 | 0 | 0,000456757 | 4,34E-05 | 1 - | 0,000382589 - | 1 - | 0,000475063 - | 1 - | 0,000451607 - | 1 - | 0,000458278 - | 1 - | 0,000516248 - |
| 2715 | 1 | 0.2um_NOKCPHJN_888436 | Bacteria | Bacteroidetes | Flavobacteriia | unclassified | unclassified | unclassified | Flavobacteria | unclassified | unclassified | unclassified | unclassified | unclassified | 27,2 | 1,166190379 | 0,000541439 | 5,44E-05 | 28 - | 0,000467115 - | 29 - | 0,000600734 - | 26 - | 0,000511997 - | 26 - | 0,000519559 - | 27 - | 0,000607792 - |
| 2716 | 1 | 3um_FIJJAFBF_50504 | Bacteria | Bacteroidetes | Flavobacteriia | Flavobacterial | Flavobacteriac | Polaribacter | Polaribacter h | unclassified | unclassified | unclassified | unclassified | unclassified | 1,6 | 1,019803903 | 5,33E-05 | 3,43E-05 | 2 - | 5,81E-05 - | 1 - | 3,61E-05 - | 3 - | 0,000102898 - | 2 - | 6,96E-05 - | 0 - | 0 - |
| 2717 | 1 | 0.2um_NOKCPHJN_285187 | Bacteria | Proteobacteri | Gammaprotec | unclassified | unclassified | unclassified | SAR86 cluster | information st | Translation, ril | COG0087 | One of the pri | 396588.Tgr7_ | 5,2 | 0,4 | 0,000330884 | 1,64E-05 | 6 - | 0,000321804 - | 5 - | 0,000332988 - | 5 - | 0,000316547 - | 5 - | 0,000321223 - | 5 - | 0,000361856 - |
| 2718 | 1 | 3um_FIJJAFBF_383850 | Bacteria | Proteobacteri | Gammaprotec | Cellvibrionales | Halieaceae | Luminiphilus | Luminiphilus s | unclassified | unclassified | unclassified | unclassified | unclassified | 0,6 | 0,489897949 | 1,56E-05 | 1,28E-05 | 0 - | 0 - | 0 - | 0 - | 1 - | 2,48E-05 - | 1 - | 2,51E-05 - | 1 - | 2,83E-05 - |
| 2719 | 1 | 0.2um_NOKCPHJN_628012 | Bacteria | Proteobacteri | Gammaprotec | Oceanospirilla | Saccharospirill | Reinekea | Reinekea fors | unclassified | unclassified | unclassified | unclassified | unclassified | 0,6 | 0,489897949 | 3,33E-05 | 2,74E-05 | 1 - | 4,93E-05 - | 0 - | 0 - | 1 - | 5,81E-05 - | 1 - | 5,90E-05 - | 0 - | 0 - |
| 2720 | 1 | 0.2um_NOKCPHJN_15786 | Bacteria | Proteobacteri | Gammaprotec | Alteromonada | Alteromonada | various | various | unclassified | unclassified | unclassified | unclassified | unclassified | 1,8 | 0,748331477 | 9,31E-05 | 3,03E-05 | 3 - | 0,000135031 - | 1 - | 5,59E-05 - | 2 - | 0,000106261 - | 2 - | 0,00010783 - | 1 - | 6,07E-05 - |
| 2721 | 1 | 0.2um_NOKCPHJN_28455 | Bacteria | Proteobacteri | Gammaprotec | unclassified | unclassified | unclassified | Gammaprotec | unclassified | unclassified | unclassified | unclassified | unclassified | 1 | 0 | 0,000152252 | 1,45E-05 | 1 - | 0,00012753 - | 1 - | 0,000158354 - | 1 - | 0,000150536 - | 1 - | 0,000152759 - | 1 - | 0,000172083 - |
| 2722 | 1 | 3um_FIJJAFBF_151691 | Bacteria | Bacteroidetes | Flavobacteriia | Flavobacterial | Flavobacteriac | unclassified | Flavobacteriac | information st | Translation, ril | 05KK9 | This protein bi | 886377.Murru | 2 | 1,095445115 | 0,000156343 | 9,87E-05 | 1 - | 6,31E-05 - | 1 - | 7,83E-05 - | 2 - | 0,000148882 - | 2 - | 0,000151081 - | 4 - | 0,000340383 - |
| 2723 | 1 | 0.2um_NOKCPHJN_225855 | Eukaryota | unclassified | Oomycetes | Albuginales | Albuginaceae | Albugo | Albugo laibach | unclassified | unclassified | unclassified | unclassified | unclassified | 0,2 | 0,4 | 3,85E-06 | 7,70E-06 | 1 - | 1,93E-05 - | 0 - | 0 - | 0 - | 0 - | 0 - | 0 - | 0 - | 0 - |
| 2724 | 2 | 0.2um_NOKCPHJN_134820;3um_FIJJAFBF_1245 | Bacteria | Bacteroidetes | unclassified | unclassified | unclassified | unclassified | Bacteroidetes | unclassified | unclassified | unclassified | unclassified | unclassified | 0,8 | 0,4 | 8,62E-05 | 4,38E-05 | 1 - | 9,33E-05 - | 1 - | 0,000115869 - | 1 - | 0,000110148 - | 1 - | 0,000111775 - | 0 - | 0 - |
| 2725 | 1 | 3um_FIJJAFBF_176595 | Eukaryota | Phaeophyceae | unclassified | Ectocarpales | Ectocarpaceae | Ectocarpus | Ectocarpus sili | cellular proces | Posttranslatio | 0XRR7 | Calreticulin | 67593.Physo1 | 1 | 0 | 0,000103028 | 9,78E-06 | 1 - | 8,63E-05 - | 1 - | 0,000107157 - | 1 - | 0,000101866 - | 1 - | 0,000103371 - | 1 - | 0,000116447 - |
| 2726 | 1 | 0.2um_NOKCPHJN_326595 | Eukaryota | unclassified | Oomycetes | Peronosporale | Peronosporac | Phytophthora | Phytophthora | unclassified | unclassified | unclassified | unclassified | unclassified | 4,4 | 1,743559577 | 3,17E-05 | 1,26E-05 | 5 - | 3,08E-05 - | 6 - | 4,59E-05 - | 5 - | 3,64E-05 - | 5 - | 3,69E-05 - | 1 - | 8,31E-06 - |
| 2727 | 1 | 3um_FIJJAFBF_424816 | Bacteria | Proteobacteri | Alphaproteob | Rhodobactera | Rhodobactera | unclassified | Rhodobactera | unclassified | unclassified | unclassified | unclassified | unclassified | 6 | 1,095445115 | 0,000127982 | 2,41E-05 | 6 - | 0,000107603 - | 6 - | 0,000133611 - | 5 - | 0,000105845 - | 8 - | 0,000171854 - | 5 - | 0,000120996 - |
| 2728 | 1 | 3um_FIJJAFBF_144021 | Bacteria | Bacteroidetes | Flavobacteriia | Flavobacterial | Flavobacteriac | Polaribacter | Polaribacter s | unclassified | unclassified | unclassified | unclassified | unclassified | 1,6 | 0,489897949 | 0,000149293 | 4,94E-05 | 2 - | 0,000156159 - | 2 - | 0,000193903 - | 1 - | 9,22E-05 - | 1 - | 9,35E-05 - | 2 - | 0,000210714 - |
| 2729 | 1 | 0.2um_NOKCPHJN_03796 | Bacteria | Proteobacteri | Gammaprotec | Alteromonada | Alteromonada | Paraglaciecola | Paraglaciecola | metabolism | Energy produc | 17CGT | F(1)F(0) ATP s | 983545.Glaag | 7 | 0,632455532 | 0,000615122 | 8,24E-05 | 7 - | 0,000515024 - | 7 - | 0,000639508 - | 6 - | 0,000521085 - | 8 - | 0,000705043 - | 7 - | 0,00069495 - |
| 2730 | 1 | 3um_FIJJAFBF_264644 | Bacteria | Bacteroidetes | Flavobacteriia | Flavobacterial | Flavobacteriac | Polaribacter | Polaribacter ir | cellular proces | Posttranslatio | 05JRK | feS assembly | 313594.PI23P | 3,2 | 0,748331477 | 9,99E-05 | 2,47E-05 | 4 - | 0,000105059 - | 2 - | 6,52E-05 - | 3 - | 9,30E-05 - | 3 - | 9,44E-05 - | 4 - | 0,000141762 - |
| 2731 | 1 | 3um_FIJJAFBF_13282 | Bacteria | Proteobacteri | Gammaprotec | Cellvibrionales | Halieaceae | unclassified | marine gamm | poorly charact | Function unkn | 16R8I | Lipoprotein | 247639.MGP2 | 0,6 | 0,489897949 | 3,00E-05 | 2,50E-05 | 1 - | 4,24E-05 - | 0 - | 0 - | 0 - | 0 - | 1 - | 5,07E-05 - | 1 - | 5,71E-05 - |
| 2732 | 1 | 3um_FIJJAFBF_03427 | Bacteria | Bacteroidetes | Flavobacteriia | Flavobacterial | Flavobacteriac | Flavivirga | Flavivirga eckl | information st | Transcription | COG0085 | DNA-depende | 391603.FBALC | 6,2 | 1,166190379 | 6,58E-05 | 8,74E-06 | 8 - | 7,21E-05 - | 7 - | 7,83E-05 - | 5 - | 5,32E-05 - | 6 - | 6,47E-05 - | 5 - | 6,08E-05 - |
| 2733 | 1 | 3um_FIJJAFBF_604570 | Bacteria | Proteobacteri | Alphaproteob | Rhodobactera | unclassified | unclassified | Rhodobactera | metabolism | Energy produc | 01RN9 | Subunits I and | 367336.OM22 | 0,2 | 0,4 | 9,20E-06 | 1,84E-05 | 0 - | 0 - | 0 - | 0 - | 0 - | 0 - | 1 - | 4,60E-05 - | 0 - | 0 - |
| 2734 | 1 | 0.2um_NOKCPHJN_369117 | Bacteria | Bacteroidetes | Flavobacteriia | Flavobacterial | Flavobacteriac | Mangrovimon | Mangrovimon | unclassified | unclassified | unclassified | unclassified | unclassified | 0,4 | 0,489897949 | 5,05E-05 | 6,19E-05 | 0 - | 0 - | 0 - | 0 - | 1 - | 0,000125447 - | 1 - | 0,000127299 - | 0 - | 0 - |
| 2735 | 1 | 0.2um_NOKCPHJN_92016 | Bacteria | Bacteroidetes | Flavobacteriia | Flavobacterial | Flavobacteriac | Polaribacter | Polaribacter b | poorly charact | Function unkn | 0900R | pur-alpha bet | 313598.MED1 | 0,2 | 0,4 | 2,24E-05 | 4,48E-05 | 0 - | 0 - | 0 - | 0 - | 1 - | 0,000111969 - | 0 - | 0 - | 0 - | 0 - |
| 2736 | 1 | 3um_FIJJAFBF_713652 | Eukaryota | Bacillariophyt | Bacillariophyc | Bacillariales | Bacillariaceae | Fragilariopsis | Fragilariopsis | unclassified | unclassified | unclassified | unclassified | unclassified | 0,2 | 0,4 | 7,73E-06 | 1,55E-05 | 1 - | 3,86E-05 - | 0 - | 0 - | 0 - | 0 - | 0 - | 0 - | 0 - | 0 - |
| 2737 | 1 | 0.2um_NOKCPHJN_411672 | Bacteria | Bacteroidetes | unclassified | unclassified | unclassified | unclassified | Bacteroidetes | poorly charact | Function unkn | 0P8NI | Inherit from b | 487796.Flav2 | 0,4 | 0,489897949 | 1,48E-05 | 1,83E-05 | 1 - | 3,30E-05 - | 1 - | 4,10E-05 - | 0 - | 0 - | 0 - | 0 - | 0 - | 0 - |
| 2738 | 1 | 0.2um_NOKCPHJN_228342 | Bacteria | Proteobacteri | Gammaprotec | Cellvibrionales | Porticoccacea | unclassified | gamma protec | information st | Translation, ril | 179ZI | One of the pri | 314287.GB22 | 0,6 | 0,489897949 | 6,04E-05 | 4,98E-05 | 1 - | 8,83E-05 - | 1 - | 0,00010963 - | 1 - | 0,000104217 - | 0 - | 0 - | 0 - | 0 - |
| 2739 | 1 | 3um_FIJJAFBF_130194 | Bacteria | Proteobacteri | Betaproteoba | unclassified | unclassified | unclassified | Betaproteoba | unclassified | unclassified | unclassified | unclassified | unclassified | 0,4 | 0,489897949 | 7,47E-06 | 9,24E-06 | 1 - | 1,67E-05 - | 1 - | 2,07E-05 - | 0 - | 0 - | 0 - | 0 - | 0 - | 0 - |
| 2740 | 1 | 3um_FIJJAFBF_406308 | Bacteria | various | various | various | various | various | various | information st | Translation, ril | 05KK9 | This protein bi | 411154.GFO_ | 0,8 | 0,4 | 6,71E-05 | 3,38E-05 | 0 - | 0 - | 1 - | 8,38E-05 - | 1 - | 7,97E-05 - | 1 - | 8,09E-05 - | 1 - | 9,11E-05 - |
| 2741 | 1 | 3um_FIJJAFBF_154012 | Bacteria | Proteobacteri | Gammaprotec | Oceanospirilla | Saccharospirill | Reinekea | Reinekea fors | unclassified | unclassified | unclassified | unclassified | unclassified | 1,4 | 0,8 | 9,40E-05 | 5,18E-05 | 2 - | 0,000115936 - | 0 - | 0 - | 2 - | 0,000136851 - | 2 - | 0,000138872 - | 1 - | 7,82E-05 - |
| 2742 | 1 | 0.2um_NOKCPHJN_644966 | Bacteria | Actinobacteria | Actinobacteria | Micrococcales | Microbacteria | Candidatus Ac | Candidatus Ac | unclassified | unclassified | unclassified | unclassified | unclassified | 1,8 | 0,748331477 | 2,19E-05 | 7,37E-06 | 3 - | 3,16E-05 - | 2 - | 2,62E-05 - | 1 - | 1,24E-05 - | 2 - | 2,52E-05 - | 1 - | 1,42E-05 - |
| 2743 | 1 | 0.2um_NOKCPHJN_41537 | Bacteria | Bacteroidetes | Cytophagia | Cytophagales | Persicobacter | Persicobacter | Persicobacter | poorly charact | Function unkn | 11TE0 | Domain-Conta | 7668.SPU_002 | 1 | 0 | 3,81E-06 | 3,62E-07 | 1 - | 3,19E-06 - | 1 - | 3,97E-06 - | 1 - | 3,77E-06 - | 1 - | 3,83E-06 - | 1 - | 4,31E-06 - |
| 2744 | 2 | 0.2um_NOKCPHJN_187474;0.2um_NOKCPHJN_2 | Bacteria | Proteobacteri | Gammaprotec | Cellvibrionales | Porticoccacea | unclassified | various | cellular proces | Cell cycle cont | 0QJVR | Required for c | 314287.GB22 | 0,2 | 0,4 | 1,97E-06 | 3,93E-06 | 1 - | 9,83E-06 - | 0 - | 0 - | 0 - | 0 - | 0 - | 0 - | 0 - | 0 - |
| 2745 | 1 | 0.2um_NOKCPHJN_914906 | Eukaryota | Phaeophyceae | unclassified | Ectocarpales | Ectocarpaceae | Ectocarpus | Ectocarpus sili | unclassified | unclassified | unclassified | unclassified | unclassified | 1,2 | 0,4 | 4,31E-05 | 1,59E-05 | 1 - | 2,99E-05 - | 2 - | 7,42E-05 - | 1 - | 3,53E-05 - | 1 - | 3,58E-05 - | 1 - | 4,03E-05 - |
| 2746 | 1 | 0.2um_NOKCPHJN_152857 | Bacteria | Bacteroidetes | Flavobacteriia | Flavobacterial | Crocinitomica | unclassified | Crocinitomica | unclassified | unclassified | unclassified | unclassified | unclassified | 0,4 | 0,489897949 | 2,99E-05 | 3,70E-05 | 1 - | 6,67E-05 - | 1 - | 8,29E-05 - | 0 - | 0 - | 0 - | 0 - | 0 - | 0 - |
| 2747 | 1 | 0.2um_NOKCPHJN_225369 | Bacteria | Bacteroidetes | Flavobacteriia | Flavobacterial | Flavobacteriac | Polaribacter | Polaribacter s | cellular proces | Posttranslatio | COG0330 | Band 7 protei | 1046627.BZAR | 1,6 | 0,489897949 | 7,17E-05 | 2,31E-05 | 1 - | 3,74E-05 - | 2 - | 9,28E-05 - | 2 - | 8,83E-05 - | 2 - | 8,96E-05 - | 1 - | 5,04E-05 - |
| 2748 | 1 | 0.2um_NOKCPHJN_105414 | Bacteria | Proteobacteri | Gammaprotec | Oceanospirilla | Saccharospirill | Reinekea | Reinekea fors | cellular proces | Cell wall/mem | COG4105 | Part of the ou | 314283.MED2 | 0,2 | 0,4 | 9,86E-06 | 1,97E-05 | 0 - | 0 - | 0 - | 0 - | 0 - | 0 - | 1 - | 4,93E-05 - | 0 - | 0 - |
| 2749 | 1 | 3um_FIJJAFBF_112542 | Bacteria | Bacteroidetes | unclassified | unclassified | unclassified | unclassified | Bacteroidetes | unclassified | unclassified | unclassified | unclassified | unclassified | 1 | 1,095445115 | 5,48E-05 | 5,56E-05 | 3 - | 0,000151687 - | 1 - | 6,28E-05 - | 1 - | 5,97E-05 - | 0 - | 0 - | 0 - | 0 - |
| 2750 | 2 | 3um_FIJJAFBF_277229;3um_FIJJAFBF_405814 | Bacteria | Proteobacteri | Alphaproteob | unclassified | unclassified | unclassified | Alphaproteob | information st | Transcription | 05VEQ | Cold shock pro | 323097.Nham | 0,6 | 0,489897949 | 0,000112222 | 9,26E-05 | 1 - | 0,000163967 - | 1 - | 0,000203598 - | 1 - | 0,000193546 - | 0 - | 0 - | 0 - | 0 - |
| 2751 | 1 | 3um_FIJJAFBF_587813 | Bacteria | Proteobacteri | Gammaprotec | Pseudomonad | Pseudomonad | Pseudomonas | Pseudomonas | poorly charact | Function unkn | 05W4M | Transport-ass | 264730.PSPPH | 1 | 0,632455532 | 6,83E-05 | 3,89E-05 | 2 - | 0,000121457 - | 1 - | 7,54E-05 - | 1 - | 7,17E-05 - | 1 - | 7,27E-05 - | 0 - | 0 - |
| 2752 | 1 | 3um_FIJJAFBF_721316 | Bacteria | Proteobacteri | Gammaprotec | unclassified | unclassified | unclassified | Gammaprotec | information st | Transcription | 1CGWD | cold-shock DN | 416591.Tlet_1 | 3,2 | 0,748331477 | 0,000633174 | 0,000114386 | 4 - | 0,000675157 - | 3 - | 0,00062876 - | 3 - | 0,000597716 - | 4 - | 0,000808725 - | 2 - | 0,000455513 - |
| 2753 | 1 | 3um_FIJJAFBF_92884 | Eukaryota | unclassified | Cryptophyta | Pyrenomonad | Chroomonada | Chroomonas | Chroomonas | metabolism | Energy produc | COG0711 | F(1)F(0) ATP s | 118168.MC74 | 0,4 | 0,489897949 | 3,41E-05 | 4,18E-05 | 0 - | 0 - | 1 - | 8,74E-05 - | 1 - | 8,31E-05 - | 0 - | 0 - | 0 - | 0 - |
| 2754 | 1 | 0.2um_NOKCPHJN_190758 | Bacteria | Proteobacteri | Gammaprotec | Alteromonada | Alteromonada | Paraglaciecola | Paraglaciecola | cellular proces | Posttranslatio | 0QHXH | Molecular cha | 342610.Patl_2 | 2,8 | 0,748331477 | 5,85E-05 | 1,03E-05 | 4 - | 7,17E-05 - | 2 - | 4,45E-05 - | 3 - | 6,35E-05 - | 3 - | 6,44E-05 - | 2 - | 4,84E-05 - |
| 2755 | 1 | 3um_FIJJAFBF_569845 | Bacteria | Actinobacteria | Actinobacteria | Micrococcales | Microbacteria | Candidatus Ac | Candidatus Ac | unclassified | unclassified | unclassified | unclassified | unclassified | 0,8 | 0,748331477 | 3,30E-05 | 3,32E-05 | 1 - | 3,34E-05 - | 1 - | 4,14E-05 - | 0 - | 0 - | 0 - | 0 - | 2 - | 9,00E-05 - |
| 2756 | 1 | 3um_FIJJAFBF_153260 | Bacteria | Bacteroidetes | Flavobacteriia | Flavobacterial | Flavobacteriac | Leeuwenhoek | Leeuwenhoek | information st | Replication, re | 16TU1 | Transposase (I | 426114.THI_2 | 0,2 | 0,4 | 9,53E-06 | 1,91E-05 | 0 - | 0 - | 1 - | 4,77E-05 - | 0 - | 0 - | 0 - | 0 - | 0 - | 0 - |
| 2757 | 1 | 3um_FIJJAFBF_141770 | Bacteria | Proteobacteri | Gammaprotec | Oceanospirilla | Saccharospirill | Reinekea | Reinekea fors | unclassified | unclassified | unclassified | unclassified | unclassified | 0,2 | 0,4 | 2,73E-05 | 5,47E-05 | 1 - | 0,000136639 - | 0 - | 0 - | 0 - | 0 - | 0 - | 0 - | 0 - | 0 - |
| 2758 | 1 | 0.2um_NOKCPHJN_1066228 | Eukaryota | Chlorophyta | Mamiellophyc | Mamiellales | Mamiellaceae | Micromonas | Micromonas p | unclassified | unclassified | unclassified | unclassified | unclassified | 1,2 | 0,4 | 6,11E-05 | 1,35E-05 | 2 - | 8,76E-05 - | 1 - | 5,44E-05 - | 1 - | 5,17E-05 - | 1 - | 5,25E-05 - | 1 - | 5,91E-05 - |
| 2759 | 1 | 0.2um_NOKCPHJN_857856 | Eukaryota | unclassified | Oomycetes | Saprolegniale | Saprolegniace | Thraustotheca | Thraustotheca | unclassified | unclassified | unclassified | unclassified | unclassified | 1 | 0 | 3,74E-05 | 3,55E-06 | 1 - | 3,14E-05 - | 1 - | 3,89E-05 - | 1 - | 3,70E-05 - | 1 - | 3,76E-05 - | 1 - | 4,23E-05 - |
| 2760 | 1 | 0.2um_NOKCPHJN_891066 | Bacteria | Bacteroidetes | Flavobacteriia | unclassified | unclassified | unclassified | Flavobacteria | metabolism | Inorganic ion t | 05C27 | TonB-depende | 487797.Flav3 | 25,6 | 2,497999199 | 0,000659911 | 6,37E-05 | 28 - | 0,000607514 - | 23 - | 0,000619647 - | 29 - | 0,000742719 - | 23 - | 0,000597754 - | 25 - | 0,000731921 - |
| 2761 | 1 | 0.2um_NOKCPHJN_733232 | Bacteria | Proteobacteri | Gammaprotec | Alteromonada | Colwelliaceae | Colwellia | Colwellia sp. F | metabolism | Inorganic ion t | 173T4 | TONB-depend | 565045.NOR5 | 1 | 0 | 1,61E-05 | 1,53E-06 | 1 - | 1,35E-05 - | 1 - | 1,68E-05 - | 1 - | 1,59E-05 - | 1 - | 1,62E-05 - | 1 - | 1,82E-05 - |
| 2762 | 1 | 3um_FIJJAFBF_689421 | Bacteria | Bacteroidetes | Flavobacteriia | unclassified | unclassified | unclassified | Flavobacteria | unclassified | unclassified | unclassified | unclassified | unclassified | 0,2 | 0,4 | 8,77E-06 | 1,75E-05 | 0 - | 0 - | 0 - | 0 - | 1 - | 4,38E-05 - | 0 - | 0 - | 0 - | 0 - |
| 2763 | 1 | 3um_FIJJAFBF_397469 | Eukaryota | unclassified | unclassified | Choanoflagelli | Salpingoecida | Monosiga | Monosiga bre | unclassified | unclassified | unclassified | unclassified | unclassified | 0,4 | 0,489897949 | 2,31E-05 | 2,84E-05 | 0 - | 0 - | 0 - | 0 - | 1 - | 5,40E-05 - | 0 - | 0 - | 1 - | 6,17E-05 - |
| 2764 | 1 | 0.2um_NOKCPHJN_71220 | Bacteria | Proteobacteri | Alphaproteob | Pelagibacteral | Pelagibacterac | Candidatus Pe | Candidatus Pe | metabolism | Energy produc | 01RNF | Produces ATP | 1002672.SAR1 | 13 | 1,414213562 | 0,000374957 | 3,38E-05 | 15 - | 0,000364757 - | 14 - | 0,000422725 - | 13 - | 0,00037315 - | 11 - | 0,000320406 - | 12 - | 0,000393749 - |
| 2765 | 1 | 3um_FIJJAFBF_177936 | Bacteria | Proteobacteri | Gammaprotec | Thiotrichales | Piscirickettsiac | Methylophaga | Methylophaga | unclassified | unclassified | unclassified | unclassified | unclassified | 1 | 0 | 3,33E-05 | 3,16E-06 | 1 - | 2,79E-05 - | 1 - | 3,46E-05 - | 1 - | 3,29E-05 - | 1 - | 3,34E-05 - | 1 - | 3,76E-05 - |
| 2766 | 1 | 0.2um_NOKCPHJN_155154 | Eukaryota | Streptophyta | Charophyceae | Charales | Characeae | Chara | Chara braunii | cellular proces | Signal transdu | 0B4QA | Diaphanous h | 59729.ENSTG | 1,2 | 0,4 | 3,29E-05 | 7,29E-06 | 2 - | 4,72E-05 - | 1 - | 2,93E-05 - | 1 - | 2,79E-05 - | 1 - | 2,83E-05 - | 1 - | 3,19E-05 - |
| 2767 | 1 | 0.2um_NOKCPHJN_938882 | Viruses | unclassified | unclassified | unclassified | Phycodnavirid | Prasinovirus | Bathycoccus s | unclassified | unclassified | unclassified | unclassified | unclassified | 0,8 | 0,4 | 5,92E-05 | 3,01E-05 | 1 - | 6,41E-05 - | 1 - | 7,96E-05 - | 1 - | 7,57E-05 - | 1 - | 7,68E-05 - | 0 - | 0 - |
| 2768 | 1 | 0.2um_NOKCPHJN_998134 | Eukaryota | Apicomplexa | Aconoidasida | Haemosporida | Plasmodiidae | Plasmodium | Plasmodium k | unclassified | unclassified | unclassified | unclassified | unclassified | 0,6 | 0,489897949 | 1,65E-05 | 1,37E-05 | 1 - | 2,33E-05 - | 0 - | 0 - | 1 - | 2,75E-05 - | 0 - | 0 - | 1 - | 3,15E-05 - |
| 2769 | 1 | 0.2um_NOKCPHJN_737146 | Eukaryota | unclassified | Labyrinthulom | unclassified | Thraustochytr | Aurantiochytr | Aurantiochytr | metabolism | Energy produc | COG0056 | Produces ATP | 622637.Met4 | 0,2 | 0,4 | 5,28E-06 | 1,06E-05 | 0 - | 0 - | 1 - | 2,64E-05 - | 0 - | 0 - | 0 - | 0 - | 0 - | 0 - |
| 2770 | 1 | 0.2um_NOKCPHJN_149428 | Bacteria | Bacteroidetes | unclassified | unclassified | unclassified | unclassified | Bacteroidetes | poorly charact | Function unkn | 0XR4Y | Ragb susd dor | 886377.Murru | 1,2 | 0,4 | 4,17E-05 | 1,77E-05 | 1 - | 2,85E-05 - | 1 - | 3,54E-05 - | 1 - | 3,36E-05 - | 1 - | 3,41E-05 - | 2 - | 7,69E-05 - |
| 2771 | 1 | 0.2um_NOKCPHJN_170897 | Bacteria | Proteobacteri | Alphaproteob | Rhodospirillale | Rhodospirillac | unclassified | Rhodobactera | metabolism | Energy produc | COG0711 | F(1)F(0) ATP s | 367336.OM22 | 1,2 | 0,4 | 8,65E-05 | 1,92E-05 | 2 - | 0,000124083 - | 1 - | 7,70E-05 - | 1 - | 7,32E-05 - | 1 - | 7,43E-05 - | 1 - | 8,37E-05 - |
| 2772 | 1 | 0.2um_NOKCPHJN_368036 | Bacteria | Bacteroidetes | Cytophagia | Cytophagales | Flammeovirga | unclassified | Flammeovirga | unclassified | unclassified | unclassified | unclassified | unclassified | 5,4 | 1,356465997 | 9,22E-05 | 2,49E-05 | 7 - | 0,000100304 - | 5 - | 8,90E-05 - | 4 - | 6,77E-05 - | 4 - | 6,87E-05 - | 7 - | 0,000135346 - |
| 2773 | 1 | 3um_FIJJAFBF_310291 | Bacteria | Proteobacteri | Betaproteoba | unclassified | unclassified | unclassified | Betaproteoba | unclassified | unclassified | unclassified | unclassified | unclassified | 0,8 | 0,748331477 | 5,08E-05 | 4,49E-05 | 2 - | 0,000115936 - | 0 - | 0 - | 1 - | 6,84E-05 - | 1 - | 6,94E-05 - | 0 - | 0 - |
| 2774 | 1 | 3um_FIJJAFBF_129185 | Bacteria | Bacteroidetes | Flavobacteriia | Flavobacterial | Flavobacteriac | Polaribacter | Polaribacter s | metabolism | Energy produc | 0P7D1 | Dehydrogenas | 313598.MED1 | 1,2 | 0,748331477 | 3,56E-05 | 2,14E-05 | 2 - | 5,22E-05 - | 1 - | 3,24E-05 - | 1 - | 3,08E-05 - | 2 - | 6,25E-05 - | 0 - | 0 - |
| 2775 | 1 | 3um_FIJJAFBF_536896 | Eukaryota | unclassified | Dinophyceae | Gymnodiniale | Gymnodiniace | Amphidinium | Amphidinium | poorly charact | Function unkn | COG1100 | GTP-binding P | 423536.XP_0 | 1,2 | 0,979795897 | 0,000106467 | 7,53E-05 | 3 - | 0,000235843 - | 1 - | 9,76E-05 - | 1 - | 9,28E-05 - | 0 - | 0 - | 1 - | 0,000106078 - |
| 2776 | 1 | 0.2um_NOKCPHJN_58105 | Bacteria | Proteobacteri | Gammaprotec | Oceanospirilla | Saccharospirill | Reinekea | Reinekea blan | metabolism | Inorganic ion t | 16REX | Component of | 314283.MED2 | 1,8 | 0,748331477 | 4,32E-05 | 1,45E-05 | 3 - | 6,23E-05 - | 2 - | 5,15E-05 - | 1 - | 2,45E-05 - | 2 - | 4,97E-05 - | 1 - | 2,80E-05 - |
| 2777 | 1 | 3um_FIJJAFBF_201364 | Bacteria | Bacteroidetes | Flavobacteriia | Flavobacterial | Flavobacteriac | Polaribacter | Polaribacter s | metabolism | Carbohydrate | 064TE | hydrolase fam | 391603.FBALC | 1,2 | 0,4 | 3,20E-05 | 7,09E-06 | 2 - | 4,59E-05 - | 1 - | 2,85E-05 - | 1 - | 2,71E-05 - | 1 - | 2,75E-05 - | 1 - | 3,10E-05 - |
| 2778 | 1 | 3um_FIJJAFBF_402481 | Bacteria | Bacteroidetes | Flavobacteriia | various | various | various | various | unclassified | unclassified | unclassified | unclassified | unclassified | 1 | 0,632455532 | 9,28E-05 | 5,29E-05 | 2 - | 0,000165146 - | 1 - | 0,000102532 - | 1 - | 9,75E-05 - | 1 - | 9,89E-05 - | 0 - | 0 - |
| 2779 | 1 | 0.2um_NOKCPHJN_1108596 | unclassified | unclassified | unclassified | unclassified | unclassified | unclassified | unclassified | unclassified | unclassified | unclassified | unclassified | unclassified | 0,2 | 0,4 | 1,43E-05 | 2,85E-05 | 0 - | 0 - | 1 - | 7,13E-05 - | 0 - | 0 - | 0 - | 0 - | 0 - | 0 - |
| 2780 | 1 | 3um_FIJJAFBF_180757 | Bacteria | Bacteroidetes | Flavobacteriia | unclassified | unclassified | unclassified | Flavobacteria | unclassified | unclassified | unclassified | unclassified | unclassified | 1 | 0 | 0,000107052 | 1,02E-05 | 1 - | 8,97E-05 - | 1 - | 0,000111343 - | 1 - | 0,000105845 - | 1 - | 0,000107409 - | 1 - | 0,000120996 - |
| 2781 | 1 | 0.2um_NOKCPHJN_996328 | Bacteria | Bacteroidetes | Flavobacteriia | Flavobacterial | Cryomorphace | unclassified | Cryomorphace | information st | Translation, ril | COG0052 | 30S ribosomal | 376686.Fjoh_ | 0,4 | 0,489897949 | 2,01E-05 | 2,48E-05 | 1 - | 4,57E-05 - | 0 - | 0 - | 0 - | 0 - | 1 - | 5,48E-05 - | 0 - | 0 - |
| 2782 | 1 | 0.2um_NOKCPHJN_151352 | Bacteria | Bacteroidetes | Flavobacteriia | Flavobacterial | Crocinitomica | unclassified | Crocinitomica | poorly charact | Function unkn | 09AR5 | Inherit from b | 313606.M231 | 1,6 | 0,489897949 | 4,04E-05 | 1,42E-05 | 1 - | 2,08E-05 - | 1 - | 2,59E-05 - | 2 - | 4,92E-05 - | 2 - | 4,99E-05 - | 2 - | 5,62E-05 - |
| 2783 | 1 | 3um_FIJJAFBF_312563 | Bacteria | Bacteroidetes | Flavobacteriia | unclassified | unclassified | unclassified | Flavobacteria | metabolism | Energy produc | COG0437 | 4fe-4S ferredo | 487796.Flav2 | 0,4 | 0,489897949 | 4,86E-06 | 5,99E-06 | 1 - | 1,11E-05 - | 0 - | 0 - | 0 - | 0 - | 1 - | 1,32E-05 - | 0 - | 0 - |
| 2784 | 1 | 0.2um_NOKCPHJN_230045 | Bacteria | Proteobacteri | Alphaproteob | Rhodobactera | Rhodobactera | various | various | metabolism | Carbohydrate | 01RFA | ABC transport | 367336.OM22 | 5,4 | 1,2 | 0,000215573 | 5,01E-05 | 6 - | 0,000201363 - | 4 - | 0,000166689 - | 7 - | 0,000277303 - | 4 - | 0,000160799 - | 6 - | 0,00027171 - |
| 2785 | 1 | 3um_FIJJAFBF_347377 | Bacteria | Proteobacteri | Gammaprotec | Enterobactera | Enterobacteri | Escherichia | Escherichia co | unclassified | unclassified | unclassified | unclassified | unclassified | 2 | 1,095445115 | 5,92E-05 | 3,14E-05 | 3 - | 7,77E-05 - | 2 - | 6,43E-05 - | 3 - | 9,17E-05 - | 2 - | 6,21E-05 - | 0 - | 0 - |
| 2786 | 1 | 3um_FIJJAFBF_167874 | Bacteria | Bacteroidetes | Flavobacteriia | Flavobacterial | Flavobacteriac | Polaribacter | various | unclassified | unclassified | unclassified | unclassified | unclassified | 1 | 0 | 9,52E-05 | 9,04E-06 | 1 - | 7,97E-05 - | 1 - | 9,90E-05 - | 1 - | 9,41E-05 - | 1 - | 9,55E-05 - | 1 - | 0,000107552 - |
| 2787 | 1 | 0.2um_NOKCPHJN_342412 | Bacteria | Bacteroidetes | Flavobacteriia | Flavobacterial | Flavobacteriac | Polaribacter | Polaribacter s | information st | Transcription | 0P7SF | DNA-depende | 391587.KAOT | 2,4 | 1,496662955 | 2,54E-05 | 1,52E-05 | 4 - | 3,61E-05 - | 4 - | 4,49E-05 - | 0 - | 0 - | 2 - | 2,16E-05 - | 2 - | 2,44E-05 - |
| 2788 | 1 | 0.2um_NOKCPHJN_293383 | Bacteria | Proteobacteri | Alphaproteob | Pelagibacteral | Pelagibacterac | Candidatus Pe | Candidatus Pe | metabolism | Secondary me | 01Q86 | extracellular s | 335992.SAR11 | 1 | 0,632455532 | 3,46E-05 | 2,24E-05 | 1 - | 2,98E-05 - | 1 - | 3,70E-05 - | 2 - | 7,04E-05 - | 1 - | 3,57E-05 - | 0 - | 0 - |
| 2789 | 1 | 3um_FIJJAFBF_122461 | Bacteria | Bacteroidetes | Flavobacteriia | Flavobacterial | Flavobacteriac | Oceanihabitar | Oceanihabitar | metabolism | Amino acid tra | COG1363 | Peptidase m4 | 313594.PI23P | 4,4 | 0,489897949 | 0,000166309 | 2,40E-05 | 5 - | 0,000158531 - | 4 - | 0,000157479 - | 4 - | 0,000149704 - | 4 - | 0,000151915 - | 5 - | 0,000213915 - |
| 2790 | 1 | 3um_FIJJAFBF_715058 | Eukaryota | Bacillariophyt | Mediophycea | Cymatosirales | Cymatosiracea | Plagiogrammo | Plagiogrammo | information st | Translation, ril | COG0052 | 30S ribosomal | 35128.Thapsd | 1,2 | 0,4 | 7,30E-05 | 3,11E-05 | 1 - | 4,99E-05 - | 1 - | 6,20E-05 - | 1 - | 5,89E-05 - | 1 - | 5,98E-05 - | 2 - | 0,000134673 - |
| 2791 | 2 | 0.2um_NOKCPHJN_123567;3um_FIJJAFBF_4447 | Bacteria | Bacteroidetes | Flavobacteriia | Flavobacterial | Flavobacteriac | Formosa | Formosa sp. H | unclassified | unclassified | unclassified | unclassified | unclassified | 0,4 | 0,489897949 | 2,87E-05 | 3,53E-05 | 0 - | 0 - | 0 - | 0 - | 1 - | 6,71E-05 - | 0 - | 0 - | 1 - | 7,67E-05 - |
| 2792 | 1 | 0.2um_NOKCPHJN_699050 | Bacteria | Proteobacteri | Gammaprotec | unclassified | unclassified | unclassified | Gammaprotec | unclassified | unclassified | unclassified | unclassified | unclassified | 1,6 | 0,8 | 9,77E-05 | 4,25E-05 | 3 - | 0,000157228 - | 1 - | 6,51E-05 - | 1 - | 6,19E-05 - | 1 - | 6,28E-05 - | 2 - | 0,000141438 - |
| 2793 | 1 | 3um_FIJJAFBF_661807 | Bacteria | Proteobacteri | Gammaprotec | unclassified | unclassified | unclassified | various | unclassified | unclassified | unclassified | unclassified | unclassified | 0,4 | 0,489897949 | 3,72E-05 | 4,56E-05 | 0 - | 0 - | 1 - | 8,91E-05 - | 0 - | 0 - | 0 - | 0 - | 1 - | 9,68E-05 - |
| 2794 | 1 | 0.2um_NOKCPHJN_942569 | Bacteria | Proteobacteri | Gammaprotec | Pseudomonad | Pseudomonad | Pseudomonas | Pseudomonas | metabolism | Energy produc | 0QHX5 | phosphoenolp | 390235.PputW | 0,2 | 0,4 | 5,16E-06 | 1,03E-05 | 0 - | 0 - | 0 - | 0 - | 1 - | 2,58E-05 - | 0 - | 0 - | 0 - | 0 - |
| 2795 | 1 | 3um_FIJJAFBF_192887 | Bacteria | Proteobacteri | Alphaproteob | Rhizobiales | unclassified | unclassified | Rhizobiales ba | poorly charact | Function unkn | 05C2A | tonB-depende | 1085623.GNIT | 9,2 | 0,748331477 | 0,000353463 | 2,93E-05 | 10 - | 0,000323315 - | 8 - | 0,000321169 - | 10 - | 0,00038164 - | 9 - | 0,000348549 - | 9 - | 0,00039264 - |
| 2796 | 1 | 3um_FIJJAFBF_356540 | Bacteria | Proteobacteri | Alphaproteob | Rhodobactera | various | various | various | information st | Transcription | 05VEQ | Cold shock pro | 391619.RGBS | 4 | 0 | 0,000806042 | 7,65E-05 | 4 - | 0,000675157 - | 4 - | 0,000838346 - | 4 - | 0,000796954 - | 4 - | 0,000808725 - | 4 - | 0,000911026 - |
| 2797 | 1 | 0.2um_NOKCPHJN_1094039 | Bacteria | Bacteroidetes | Flavobacteriia | Flavobacterial | Flavobacteriac | Formosa | Formosa sp. H | unclassified | unclassified | unclassified | unclassified | unclassified | 1,4 | 0,489897949 | 5,00E-05 | 1,57E-05 | 2 - | 6,09E-05 - | 2 - | 7,56E-05 - | 1 - | 3,59E-05 - | 1 - | 3,65E-05 - | 1 - | 4,11E-05 - |
| 2798 | 1 | 0.2um_NOKCPHJN_24010 | Eukaryota | Arthropoda | Insecta | Lepidoptera | Noctuidae | Helicoverpa | Helicoverpa a | unclassified | unclassified | unclassified | unclassified | unclassified | 1,8 | 0,4 | 9,53E-05 | 1,88E-05 | 2 - | 9,00E-05 - | 2 - | 0,00011178 - | 2 - | 0,000106261 - | 2 - | 0,00010783 - | 1 - | 6,07E-05 - |
| 2799 | 1 | 0.2um_NOKCPHJN_211205 | Eukaryota | Bacillariophyt | Coscinodiscop | Thalassiosirale | Thalassiosirac | Thalassiosira | Thalassiosira | cellular proces | Posttranslatio | KOG0179 | The proteasor | 35128.Thaps2 | 0,4 | 0,489897949 | 3,02E-05 | 3,70E-05 | 0 - | 0 - | 1 - | 7,75E-05 - | 1 - | 7,36E-05 - | 0 - | 0 - | 0 - | 0 - |
| 2800 | 1 | 0.2um_NOKCPHJN_216573 | Bacteria | Bacteroidetes | Flavobacteriia | Flavobacterial | Flavobacteriac | Lutibacter | Lutibacter sp. | unclassified | unclassified | unclassified | unclassified | unclassified | 1 | 0 | 1,38E-05 | 1,31E-06 | 1 - | 1,16E-05 - | 1 - | 1,44E-05 - | 1 - | 1,36E-05 - | 1 - | 1,38E-05 - | 1 - | 1,56E-05 - |
| 2801 | 1 | 3um_FIJJAFBF_160782 | Bacteria | Bacteroidetes | Flavobacteriia | unclassified | unclassified | unclassified | Flavobacteria | information st | Translation, ril | 09FE5 | One of the pro | 487796.Flav2 | 1,6 | 0,8 | 0,000203947 | 0,000103586 | 2 - | 0,000220725 - | 2 - | 0,000274075 - | 2 - | 0,000260543 - | 2 - | 0,000264391 - | 0 - | 0 - |
| 2802 | 1 | 0.2um_NOKCPHJN_620960 | Eukaryota | Apicomplexa | Conoidasida | Eucoccidiorida | Eimeriidae | Eimeria | Eimeria praec | unclassified | unclassified | unclassified | unclassified | unclassified | 0,2 | 0,4 | 3,80E-06 | 7,61E-06 | 0 - | 0 - | 0 - | 0 - | 0 - | 0 - | 1 - | 1,90E-05 - | 0 - | 0 - |
| 2803 | 2 | 0.2um_NOKCPHJN_269193;0.2um_NOKCPHJN_6 | Bacteria | Bacteroidetes | Flavobacteriia | Flavobacterial | Flavobacteriac | Formosa | Formosa sp. H | various | various | various | various | various | 1 | 0 | 4,11E-05 | 3,91E-06 | 1 - | 3,45E-05 - | 1 - | 4,28E-05 - | 1 - | 4,07E-05 - | 1 - | 4,13E-05 - | 1 - | 4,65E-05 - |
| 2804 | 1 | 3um_FIJJAFBF_250215 | Bacteria | Bacteroidetes | Saprospiria | Saprospirales | Saprospiracea | Aureispira | Aureispira sp. | unclassified | unclassified | unclassified | unclassified | unclassified | 0,6 | 0,8 | 2,91E-05 | 3,75E-05 | 2 - | 9,15E-05 - | 0 - | 0 - | 1 - | 5,40E-05 - | 0 - | 0 - | 0 - | 0 - |
| 2805 | 1 | 3um_FIJJAFBF_554470 | Bacteria | Bacteroidetes | Flavobacteriia | Flavobacterial | Flavobacteriac | Polaribacter | Polaribacter g | unclassified | unclassified | unclassified | unclassified | unclassified | 0,4 | 0,489897949 | 1,30E-05 | 1,61E-05 | 1 - | 2,97E-05 - | 0 - | 0 - | 0 - | 0 - | 1 - | 3,55E-05 - | 0 - | 0 - |
| 2806 | 1 | 3um_FIJJAFBF_779430 | Bacteria | Proteobacteri | Alphaproteob | Rhodobactera | Rhodobactera | unclassified | Rhodobactera | poorly charact | Function unkn | 16Q8M | Oxidoreducta | 388399.SSE37 | 3,2 | 0,4 | 0,000131311 | 2,04E-05 | 3 - | 0,000103093 - | 3 - | 0,000128011 - | 3 - | 0,000121691 - | 4 - | 0,000164651 - | 3 - | 0,000139109 - |
| 2807 | 1 | 3um_FIJJAFBF_407841 | Bacteria | Proteobacteri | Alphaproteob | Rhodobactera | Rhodobactera | Puniceibacteri | Puniceibacteri | unclassified | unclassified | unclassified | unclassified | unclassified | 0,2 | 0,4 | 3,59E-06 | 7,17E-06 | 1 - | 1,79E-05 - | 0 - | 0 - | 0 - | 0 - | 0 - | 0 - | 0 - | 0 - |
| 2808 | 1 | 0.2um_NOKCPHJN_576042 | Eukaryota | unclassified | unclassified | Prymnesiales | Chrysochromu | Chrysochromu | Chrysochromu | cellular proces | Cytoskeleton | 0TX0K | KISc | 4533.OB02G1 | 0,4 | 0,489897949 | 8,44E-06 | 1,04E-05 | 0 - | 0 - | 0 - | 0 - | 1 - | 1,97E-05 - | 0 - | 0 - | 1 - | 2,25E-05 - |
| 2809 | 1 | 0.2um_NOKCPHJN_391542 | Bacteria | Proteobacteri | Alphaproteob | Rhodobactera | Rhodobactera | Planktomarina | Planktomarina | unclassified | unclassified | unclassified | unclassified | unclassified | 0,2 | 0,4 | 1,24E-05 | 2,47E-05 | 0 - | 0 - | 0 - | 0 - | 1 - | 6,19E-05 - | 0 - | 0 - | 0 - | 0 - |
| 2810 | 1 | 3um_FIJJAFBF_281680 | Bacteria | Proteobacteri | Alphaproteob | Rhodobactera | Rhodobactera | Amylibacter | Amylibacter k | metabolism | Amino acid tra | COG0174 | glutamine syn | 633131.TR2A | 0,6 | 0,489897949 | 1,69E-05 | 1,39E-05 | 1 - | 2,45E-05 - | 1 - | 3,05E-05 - | 0 - | 0 - | 1 - | 2,94E-05 - | 0 - | 0 - |
| 2811 | 1 | 3um_FIJJAFBF_292921 | Eukaryota | Bacillariophyt | Mediophycea | Biddulphiales | Biddulphiacea | Biddulphia | Biddulphia bid | metabolism | Carbohydrate | COG1850 | RuBisCO catal | 261292.Nit79 | 5,2 | 0,748331477 | 0,000146227 | 3,00E-05 | 5 - | 0,000117119 - | 5 - | 0,000145427 - | 4 - | 0,000110598 - | 6 - | 0,000168347 - | 6 - | 0,000189642 - |
| 2812 | 1 | 0.2um_NOKCPHJN_1100440 | Bacteria | Bacteroidetes | Flavobacteriia | Flavobacterial | Flavobacteriac | Polaribacter | various | information st | Translation, ril | COG0050 | This protein p | 313603.FB217 | 4,2 | 1,166190379 | 0,000172838 | 5,20E-05 | 5 - | 0,000172337 - | 5 - | 0,000213992 - | 4 - | 0,000162741 - | 2 - | 8,26E-05 - | 5 - | 0,000232544 - |
| 2813 | 1 | 0.2um_NOKCPHJN_961644 | Eukaryota | unclassified | Oomycetes | Saprolegniale | Saprolegniace | Thraustotheca | Thraustotheca | cellular proces | Cytoskeleton | KOG0443 | capping prote | 65071.PYU1_ | 0,2 | 0,4 | 7,56E-06 | 1,51E-05 | 0 - | 0 - | 1 - | 3,78E-05 - | 0 - | 0 - | 0 - | 0 - | 0 - | 0 - |
| 2814 | 1 | 0.2um_NOKCPHJN_302339 | Bacteria | Proteobacteri | Betaproteoba | Burkholderiale | Comamonada | Polaromonas | Polaromonas | metabolism | Energy produc | 05CMV | Succinyl-CoA | 365046.Rta_3 | 1,6 | 0,489897949 | 5,30E-05 | 1,92E-05 | 1 - | 2,72E-05 - | 2 - | 6,75E-05 - | 2 - | 6,42E-05 - | 1 - | 3,26E-05 - | 2 - | 7,34E-05 - |
| 2815 | 1 | 3um_FIJJAFBF_739161 | Bacteria | Proteobacteri | Gammaprotec | Cellvibrionales | Porticoccacea | unclassified | gamma protec | metabolism | Inorganic ion t | 16TQC | TONB-depend | 314287.GB22 | 19 | 3,898717738 | 0,000249497 | 3,31E-05 | 26 - | 0,000290291 - | 16 - | 0,000221819 - | 20 - | 0,000263584 - | 15 - | 0,000200608 - | 18 - | 0,000271181 - |
| 2816 | 1 | 3um_FIJJAFBF_518684 | Bacteria | Bacteroidetes | Flavobacteriia | Flavobacterial | Flavobacteriac | Polaribacter | Polaribacter s | metabolism | Carbohydrate | 05CA9 | Pyruvate kina | 313594.PI23P | 8,8 | 0,979795897 | 0,000251421 | 1,48E-05 | 10 - | 0,000241635 - | 9 - | 0,000270036 - | 9 - | 0,000256703 - | 9 - | 0,000260495 - | 7 - | 0,000228236 - |
| 2817 | 1 | 3um_FIJJAFBF_604896 | Bacteria | Bacteroidetes | Flavobacteriia | unclassified | unclassified | unclassified | Flavobacteria | information st | Translation, ril | COG0256 | This is one of | 487796.Flav2 | 0,4 | 0,489897949 | 4,21E-05 | 5,18E-05 | 1 - | 9,65E-05 - | 0 - | 0 - | 1 - | 0,000113851 - | 0 - | 0 - | 0 - | 0 - |
| 2818 | 1 | 3um_FIJJAFBF_308267 | Bacteria | Bacteroidetes | Flavobacteriia | Flavobacterial | Flavobacteriac | Polaribacter | Polaribacter h | unclassified | unclassified | unclassified | unclassified | unclassified | 0,8 | 0,4 | 1,94E-05 | 9,87E-06 | 1 - | 2,10E-05 - | 1 - | 2,61E-05 - | 1 - | 2,48E-05 - | 1 - | 2,52E-05 - | 0 - | 0 - |
| 2819 | 1 | 3um_FIJJAFBF_08158 | Bacteria | Bacteroidetes | Flavobacteriia | Flavobacterial | Flavobacteriac | Polaribacter | Polaribacter s | metabolism | Lipid transpor | COG4247 | Phytase | 688270.Celal_ | 8,4 | 2,416609195 | 0,000332729 | 7,31E-05 | 12 - | 0,000407491 - | 10 - | 0,000421653 - | 8 - | 0,000320668 - | 7 - | 0,000284729 - | 5 - | 0,000229104 - |
| 2820 | 1 | 0.2um_NOKCPHJN_197364 | Bacteria | Proteobacteri | Alphaproteob | various | various | various | various | information st | Translation, ril | 01RJH | 30S ribosomal | 367336.OM22 | 5 | 0 | 0,000199167 | 1,89E-05 | 5 - | 0,000166827 - | 5 - | 0,00020715 - | 5 - | 0,000196922 - | 5 - | 0,00019983 - | 5 - | 0,000225108 - |
| 2821 | 1 | 3um_FIJJAFBF_196055 | Bacteria | Bacteroidetes | Flavobacteriia | Flavobacterial | Flavobacteriac | Polaribacter | Polaribacter s | information st | Transcription | 0P88Z | transcriptiona | 313598.MED1 | 3,8 | 1,166190379 | 0,000321118 | 8,40E-05 | 5 - | 0,000360933 - | 2 - | 0,000179269 - | 4 - | 0,000340836 - | 5 - | 0,000432337 - | 3 - | 0,000292216 - |
| 2822 | 1 | 3um_FIJJAFBF_132821 | Bacteria | Proteobacteri | Alphaproteob | Rhodobactera | Rhodobactera | Nereida | Nereida ignav | unclassified | unclassified | unclassified | unclassified | unclassified | 1,4 | 0,489897949 | 4,18E-05 | 1,48E-05 | 1 - | 2,51E-05 - | 1 - | 3,11E-05 - | 2 - | 5,92E-05 - | 2 - | 6,00E-05 - | 1 - | 3,38E-05 - |
| 2823 | 1 | 3um_FIJJAFBF_762517 | Bacteria | Proteobacteri | Gammaprotec | Oceanospirilla | Saccharospirill | Reinekea | Reinekea fors | information st | Transcription | 05VEQ | Cold shock pro | 314283.MED2 | 5 | 0 | 0,00099295 | 9,43E-05 | 5 - | 0,000831716 - | 5 - | 0,001032745 - | 5 - | 0,000981755 - | 5 - | 0,000996256 - | 5 - | 0,001122279 - |
| 2824 | 1 | 0.2um_NOKCPHJN_140071 | Bacteria | Proteobacteri | Gammaprotec | Oceanospirilla | Saccharospirill | Reinekea | Reinekea blan | information st | Transcription | COG0085 | DNA-depende | 314283.MED2 | 52,6 | 5,083306011 | 0,000526116 | 3,23E-05 | 61 - | 0,000514808 - | 49 - | 0,000513487 - | 48 - | 0,000478173 - | 56 - | 0,000566108 - | 49 - | 0,000558004 - |
| 2825 | 1 | 3um_FIJJAFBF_111241 | Viruses | unclassified | unclassified | unclassified | Phycodnavirid | unclassified | Chrysochromu | unclassified | unclassified | unclassified | unclassified | unclassified | 1,2 | 0,979795897 | 4,39E-05 | 3,62E-05 | 2 - | 6,50E-05 - | 0 - | 0 - | 2 - | 7,68E-05 - | 2 - | 7,79E-05 - | 0 - | 0 - |
| 2826 | 1 | 3um_FIJJAFBF_619427 | Bacteria | Proteobacteri | Gammaprotec | Alteromonada | Colwelliaceae | Thalassomona | Thalassomona | cellular proces | Intracellular tr | 08ZUW | Major pilin sul | 259536.Psyc_ | 1 | 0 | 8,90E-05 | 8,45E-06 | 1 - | 7,45E-05 - | 1 - | 9,25E-05 - | 1 - | 8,80E-05 - | 1 - | 8,93E-05 - | 1 - | 0,000100568 - |
| 2827 | 1 | 0.2um_NOKCPHJN_58168 | Bacteria | Proteobacteri | Gammaprotec | Oceanospirilla | Saccharospirill | Reinekea | Reinekea fors | unclassified | unclassified | unclassified | unclassified | unclassified | 0,8 | 1,166190379 | 2,93E-05 | 4,26E-05 | 0 - | 0 - | 0 - | 0 - | 3 - | 0,000109554 - | 1 - | 3,71E-05 - | 0 - | 0 - |
| 2828 | 1 | 3um_FIJJAFBF_31371 | Bacteria | Proteobacteri | Alphaproteob | Rhodobactera | Rhodobactera | Planktomarina | Planktomarina | unclassified | unclassified | unclassified | unclassified | unclassified | 3,8 | 0,4 | 0,000371146 | 4,90E-05 | 4 - | 0,000327934 - | 3 - | 0,000305398 - | 4 - | 0,000387092 - | 4 - | 0,000392809 - | 4 - | 0,000442499 - |
| 2829 | 1 | 0.2um_NOKCPHJN_141605 | Eukaryota | unclassified | Bangiophycea | Cyanidiales | Cyanidiaceae | Galdieria | Galdieria sulp | unclassified | unclassified | unclassified | unclassified | unclassified | 0,2 | 0,4 | 1,10E-05 | 2,20E-05 | 1 - | 5,49E-05 - | 0 - | 0 - | 0 - | 0 - | 0 - | 0 - | 0 - | 0 - |
| 2830 | 1 | 3um_FIJJAFBF_15942 | Eukaryota | unclassified | Pelagophycea | Pelagomonad | unclassified | Aureococcus | Aureococcus a | unclassified | unclassified | unclassified | unclassified | unclassified | 1 | 0 | 4,32E-05 | 4,10E-06 | 1 - | 3,62E-05 - | 1 - | 4,50E-05 - | 1 - | 4,27E-05 - | 1 - | 4,34E-05 - | 1 - | 4,89E-05 - |
| 2831 | 3 | 0.2um_NOKCPHJN_109746;0.2um_NOKCPHJN_4 | Bacteria | Bacteroidetes | Flavobacteriia | Flavobacterial | Flavobacteriac | Aequorivita | Aequorivita sc | unclassified | unclassified | unclassified | unclassified | unclassified | 1 | 0 | 7,61E-05 | 7,23E-06 | 1 - | 6,38E-05 - | 1 - | 7,92E-05 - | 1 - | 7,53E-05 - | 1 - | 7,64E-05 - | 1 - | 8,60E-05 - |
| 2832 | 1 | 3um_FIJJAFBF_708222 | Viruses | unclassified | unclassified | Caudovirales | Podoviridae | unclassified | Puniceispirillu | unclassified | unclassified | unclassified | unclassified | unclassified | 0,4 | 0,8 | 1,54E-05 | 3,09E-05 | 0 - | 0 - | 0 - | 0 - | 2 - | 7,72E-05 - | 0 - | 0 - | 0 - | 0 - |
| 2833 | 1 | 0.2um_NOKCPHJN_1032920 | Bacteria | Proteobacteri | Alphaproteob | Rhodobactera | Rhodobactera | Yoonia | Yoonia vestfol | metabolism | Secondary me | 08EVA | Inherit from N | 224911.bll356 | 1 | 0 | 2,55E-05 | 2,42E-06 | 1 - | 2,14E-05 - | 1 - | 2,65E-05 - | 1 - | 2,52E-05 - | 1 - | 2,56E-05 - | 1 - | 2,88E-05 - |
| 2834 | 2 | 0.2um_NOKCPHJN_833583;3um_FIJJAFBF_5843 | Bacteria | unclassified | unclassified | unclassified | unclassified | unclassified | uncultured ba | various | various | various | various | various | 1 | 0 | 3,23E-05 | 3,07E-06 | 1 - | 2,71E-05 - | 1 - | 3,36E-05 - | 1 - | 3,20E-05 - | 1 - | 3,24E-05 - | 1 - | 3,65E-05 - |
| 2835 | 1 | 0.2um_NOKCPHJN_451573 | Bacteria | Proteobacteri | Gammaprotec | unclassified | unclassified | unclassified | OM182 bacte | cellular proces | Intracellular tr | 08ZUW | Major pilin sul | 267608.RSc05 | 1,2 | 0,4 | 0,000104792 | 3,61E-05 | 1 - | 7,31E-05 - | 1 - | 9,08E-05 - | 1 - | 8,63E-05 - | 2 - | 0,000175138 - | 1 - | 9,86E-05 - |
| 2836 | 1 | 0.2um_NOKCPHJN_315127 | Bacteria | Proteobacteri | Gammaprotec | Oceanospirilla | Oceanospirilla | Motiliproteus | Motiliproteus | metabolism | Inorganic ion t | 0QIW4 | transporter | 207954.MED9 | 1 | 0 | 2,63E-05 | 2,50E-06 | 1 - | 2,20E-05 - | 1 - | 2,74E-05 - | 1 - | 2,60E-05 - | 1 - | 2,64E-05 - | 1 - | 2,97E-05 - |
| 2837 | 1 | 0.2um_NOKCPHJN_06883 | Bacteria | Bacteroidetes | Cytophagia | Cytophagales | Cytophagacea | Ekhidna | Ekhidna lutea | information st | Translation, ril | 0FVR5 | This protein p | 643867.Ftrac_ | 5 | 1,788854382 | 0,000176922 | 7,42E-05 | 4 - | 0,00011623 - | 6 - | 0,000216484 - | 2 - | 6,86E-05 - | 6 - | 0,000208835 - | 7 - | 0,000274461 - |
| 2838 | 1 | 3um_FIJJAFBF_347511 | Bacteria | Proteobacteri | Gammaprotec | Alteromonada | Colwelliaceae | Colwellia | Colwellia sp. F | information st | Transcription | 05VEQ | Cold shock pro | 167879.CPS_ | 0,2 | 0,4 | 3,93E-05 | 7,85E-05 | 0 - | 0 - | 0 - | 0 - | 1 - | 0,000196351 - | 0 - | 0 - | 0 - | 0 - |
| 2839 | 1 | 3um_FIJJAFBF_101746 | Bacteria | Bacteroidetes | Flavobacteriia | various | various | various | various | unclassified | unclassified | unclassified | unclassified | unclassified | 0,2 | 0,4 | 1,27E-05 | 2,54E-05 | 1 - | 6,34E-05 - | 0 - | 0 - | 0 - | 0 - | 0 - | 0 - | 0 - | 0 - |
| 2840 | 1 | 0.2um_NOKCPHJN_492097 | Bacteria | Proteobacteri | Alphaproteob | Rhodobactera | Rhodobactera | Pelagimonas | Pelagimonas v | metabolism | Carbohydrate | 07U2E | extracellular s | 633131.TR2A | 2,4 | 1,019803903 | 7,66E-05 | 3,80E-05 | 2 - | 5,25E-05 - | 1 - | 3,26E-05 - | 3 - | 9,30E-05 - | 2 - | 6,29E-05 - | 4 - | 0,000141762 - |
| 2841 | 1 | 3um_FIJJAFBF_269831 | Eukaryota | unclassified | Oomycetes | Saprolegniale | Saprolegniace | various | various | unclassified | unclassified | unclassified | unclassified | unclassified | 2,8 | 0,979795897 | 9,87E-05 | 2,92E-05 | 4 - | 0,000121779 - | 3 - | 0,00011341 - | 3 - | 0,000107811 - | 3 - | 0,000109403 - | 1 - | 4,11E-05 - |
| 2842 | 1 | 3um_FIJJAFBF_481949 | Bacteria | Proteobacteri | Alphaproteob | Rhodobactera | Rhodobactera | Pseudooceani | Pseudooceani | metabolism | Energy produc | COG0056 | Produces ATP | 398580.Dshi_ | 3,2 | 0,4 | 8,56E-05 | 1,27E-05 | 3 - | 6,73E-05 - | 3 - | 8,35E-05 - | 4 - | 0,000105845 - | 3 - | 8,06E-05 - | 3 - | 9,07E-05 - |
| 2843 | 1 | 0.2um_NOKCPHJN_533826 | Bacteria | Proteobacteri | Alphaproteob | Pelagibacteral | Pelagibacterac | unclassified | Pelagibacterac | cellular proces | Cell wall/mem | 7257 | NA | 439493.PB721 | 0,2 | 0,4 | 7,58E-06 | 1,52E-05 | 1 - | 3,79E-05 - | 0 - | 0 - | 0 - | 0 - | 0 - | 0 - | 0 - | 0 - |
| 2844 | 1 | 3um_FIJJAFBF_294698 | Bacteria | Proteobacteri | Alphaproteob | Rhodobactera | Rhodobactera | Marinibacteri | Marinibacteri | metabolism | Lipid transpor | COG1788 | Key enzyme fc | 252305.OB25 | 0,4 | 0,489897949 | 2,33E-05 | 2,86E-05 | 0 - | 0 - | 0 - | 0 - | 1 - | 5,79E-05 - | 1 - | 5,88E-05 - | 0 - | 0 - |
| 2845 | 1 | 3um_FIJJAFBF_115233 | Bacteria | Proteobacteri | Gammaprotec | Alteromonada | Alteromonada | Paraglaciecola | Paraglaciecola | unclassified | unclassified | unclassified | unclassified | unclassified | 2,4 | 0,8 | 0,000210939 | 6,25E-05 | 3 - | 0,000226533 - | 2 - | 0,000187525 - | 3 - | 0,000267399 - | 3 - | 0,000271349 - | 1 - | 0,000101891 - |
| 2846 | 1 | 0.2um_NOKCPHJN_54716 | Eukaryota | Streptophyta | unclassified | Ericales | Primulaceae | Androsace | Androsace lax | metabolism | Energy produc | 1DQDB | PsaA and PsaE | 59689.fgenes | 1 | 0 | 1,87E-05 | 1,77E-06 | 1 - | 1,56E-05 - | 1 - | 1,94E-05 - | 1 - | 1,85E-05 - | 1 - | 1,87E-05 - | 1 - | 2,11E-05 - |
| 2847 | 1 | 0.2um_NOKCPHJN_667152 | Eukaryota | unclassified | Oomycetes | Peronosporale | Peronosporac | Phytophthora | various | unclassified | unclassified | unclassified | unclassified | unclassified | 0,4 | 0,489897949 | 9,29E-05 | 0,000113986 | 0 - | 0 - | 1 - | 0,000222686 - | 0 - | 0 - | 0 - | 0 - | 1 - | 0,000241991 - |
| 2848 | 1 | 3um_FIJJAFBF_32131 | Bacteria | Bacteroidetes | Flavobacteriia | Flavobacterial | Flavobacteriac | Algibacter | Algibacter sp. | poorly charact | Function unkn | 09GJP | Protein of unk | 391603.FBALC | 1,2 | 0,748331477 | 0,00012024 | 7,69E-05 | 1 - | 8,32E-05 - | 2 - | 0,000206549 - | 0 - | 0 - | 2 - | 0,000199251 - | 1 - | 0,000112228 - |
| 2849 | 1 | 0.2um_NOKCPHJN_923663 | Eukaryota | Ascomycota | Schizosacchar | Schizosacchar | Schizosacchar | Schizosacchar | Schizosacchar | information st | Translation, ril | KOG0886 | Ribosomal pro | 4896.SPCC31H | 2,6 | 0,489897949 | 0,000229806 | 3,33E-05 | 3 - | 0,000225052 - | 2 - | 0,000186299 - | 3 - | 0,000265651 - | 3 - | 0,000269575 - | 2 - | 0,00020245 - |
| 2850 | 1 | 3um_FIJJAFBF_430380 | Bacteria | Proteobacteri | Alphaproteob | Rhodobactera | unclassified | unclassified | Rhodobactera | cellular proces | Cell wall/mem | COG2885 | Ompa motb d | 388739.RSK20 | 1,2 | 0,4 | 8,80E-05 | 3,25E-05 | 1 - | 6,11E-05 - | 2 - | 0,000151616 - | 1 - | 7,21E-05 - | 1 - | 7,31E-05 - | 1 - | 8,24E-05 - |
| 2851 | 1 | 0.2um_NOKCPHJN_153589 | Bacteria | Bacteroidetes | Flavobacteriia | unclassified | unclassified | unclassified | Flavobacteria | unclassified | unclassified | unclassified | unclassified | unclassified | 0,4 | 0,489897949 | 4,40E-05 | 5,39E-05 | 0 - | 0 - | 0 - | 0 - | 1 - | 0,00010926 - | 1 - | 0,000110874 - | 0 - | 0 - |
| 2852 | 1 | 3um_FIJJAFBF_76674 | Bacteria | Bacteroidetes | Flavobacteriia | Flavobacterial | Flavobacteriac | Polaribacter | Polaribacter s | unclassified | unclassified | unclassified | unclassified | unclassified | 0,2 | 0,4 | 1,53E-05 | 3,06E-05 | 1 - | 7,65E-05 - | 0 - | 0 - | 0 - | 0 - | 0 - | 0 - | 0 - | 0 - |
| 2853 | 1 | 3um_FIJJAFBF_80027 | Bacteria | Bacteroidetes | Flavobacteriia | various | various | various | various | poorly charact | Function unkn | 09ARW | Zinc ribbon do | 867900.Celly_ | 0,6 | 0,489897949 | 3,36E-05 | 2,75E-05 | 0 - | 0 - | 1 - | 5,50E-05 - | 0 - | 0 - | 1 - | 5,31E-05 - | 1 - | 5,98E-05 - |
| 2854 | 1 | 3um_FIJJAFBF_62816 | Bacteria | Bacteroidetes | Flavobacteriia | Flavobacterial | Flavobacteriac | Polaribacter | Polaribacter s | unclassified | unclassified | unclassified | unclassified | unclassified | 5,8 | 0,748331477 | 0,00010494 | 8,11E-06 | 7 - | 0,000107125 - | 6 - | 0,000114015 - | 5 - | 9,03E-05 - | 6 - | 0,000109987 - | 5 - | 0,00010325 - |

| | | | | | | | | | | | | | | | | | | | | | | | | | | | | |
|---|---|---|---|---|---|---|---|---|---|---|---|---|---|---|---|---|---|---|---|---|---|---|---|---|---|---|---|---|
| 2855 | 1 | 0.2um_NOKCPHJN_132225 | Bacteria | Proteobacteri | Gammaprotec | Oceanospirilla | Saccharospirill | Reinekea | Reinekea blan | metabolism | Lipid transpor | 0RAT4 | Catalyzes the | 207949.RED65 | 0,4 | 0,489897949 | 7,95E-06 | 9,83E-06 | 1 - | 1,77E-05 - | 1 - | 2,20E-05 - | 0 - | 0 - | 0 - | 0 - | 0 - | 0 - |
| 2856 | 2 | 0.2um_NOKCPHJN_93629;3um_FIJJAFBF_125373 | Eukaryota | Xanthophycea | unclassified | Botrydiales | Botrydiaceae | Botrydium | Botrydium sto | metabolism | Carbohydrate | COG1850 | RuBisCO catal | various | 5 | 0,632455532 | 0,000138614 | 1,50E-05 | 6 - | 0,000140257 - | 5 - | 0,000145131 - | 5 - | 0,000137966 - | 4 - | 0,000112003 - | 5 - | 0,000157713 - |
| 2857 | 2 | 0.2um_NOKCPHJN_280031;0.2um_NOKCPHJN_6 | Bacteria | Bacteroidetes | Flavobacteriia | Flavobacterial | Flavobacteriac | Gilvibacter | Gilvibacter sp. | unclassified | unclassified | unclassified | unclassified | unclassified | 0,6 | 0,489897949 | 2,69E-05 | 2,22E-05 | 1 - | 3,90E-05 - | 1 - | 4,85E-05 - | 0 - | 0 - | 1 - | 4,68E-05 - | 0 - | 0 - |
| 2858 | 1 | 0.2um_NOKCPHJN_349772 | Bacteria | unclassified | unclassified | unclassified | unclassified | unclassified | uncultured ba | poorly charact | Function unkn | 08PIQ | CHU large pro | 50743.SCB49_ | 3 | 0 | 3,15E-05 | 3,00E-06 | 3 - | 2,64E-05 - | 3 - | 3,28E-05 - | 3 - | 3,12E-05 - | 3 - | 3,17E-05 - | 3 - | 3,57E-05 - |
| 2859 | 1 | 3um_FIJJAFBF_449010 | Bacteria | Proteobacteri | Gammaprotec | unclassified | unclassified | unclassified | Gammaprotec | unclassified | unclassified | unclassified | unclassified | unclassified | 0,4 | 0,489897949 | 3,22E-05 | 3,98E-05 | 1 - | 7,17E-05 - | 1 - | 8,91E-05 - | 0 - | 0 - | 0 - | 0 - | 0 - | 0 - |
| 2860 | 1 | 0.2um_NOKCPHJN_545211 | Bacteria | Bacteroidetes | unclassified | unclassified | unclassified | unclassified | Bacteroidetes | unclassified | unclassified | unclassified | unclassified | unclassified | 0,4 | 0,489897949 | 2,19E-05 | 2,71E-05 | 1 - | 4,88E-05 - | 1 - | 6,06E-05 - | 0 - | 0 - | 0 - | 0 - | 0 - | 0 - |
| 2861 | 1 | 3um_FIJJAFBF_115059 | Eukaryota | unclassified | unclassified | Phaeocystales | Phaeocystace | Phaeocystis | Phaeocystis gl | metabolism | Energy produc | COG0711 | F(1)F(0) ATP s | 91464.S7335_ | 0,2 | 0,4 | 1,41E-05 | 2,82E-05 | 1 - | 7,04E-05 - | 0 - | 0 - | 0 - | 0 - | 0 - | 0 - | 0 - | 0 - |
| 2862 | 1 | 0.2um_NOKCPHJN_791181 | Bacteria | Proteobacteri | Alphaproteob | Rhizobiales | unclassified | unclassified | Rhizobiales ba | unclassified | unclassified | unclassified | unclassified | unclassified | 3,2 | 0,748331477 | 6,11E-05 | 1,74E-05 | 3 - | 4,76E-05 - | 3 - | 5,91E-05 - | 4 - | 7,50E-05 - | 2 - | 3,80E-05 - | 4 - | 8,57E-05 - |
| 2863 | 1 | 0.2um_NOKCPHJN_590883 | Bacteria | Proteobacteri | Gammaprotec | unclassified | unclassified | unclassified | Gammaprotec | unclassified | unclassified | unclassified | unclassified | unclassified | 2,2 | 0,748331477 | 3,47E-05 | 1,17E-05 | 2 - | 2,68E-05 - | 2 - | 3,32E-05 - | 3 - | 4,74E-05 - | 3 - | 4,81E-05 - | 1 - | 1,81E-05 - |
| 2864 | 1 | 0.2um_NOKCPHJN_1089757 | Bacteria | Proteobacteri | Gammaprotec | Cellvibrionales | Spongiibacter | Dasania | Dasania marin | unclassified | unclassified | unclassified | unclassified | unclassified | 1 | 0 | 3,13E-05 | 2,97E-06 | 1 - | 2,62E-05 - | 1 - | 3,25E-05 - | 1 - | 3,09E-05 - | 1 - | 3,14E-05 - | 1 - | 3,54E-05 - |
| 2865 | 1 | 3um_FIJJAFBF_62871 | Bacteria | Proteobacteri | Gammaprotec | Oceanospirilla | Oceanospirilla | Motiliproteus | Motiliproteus | unclassified | unclassified | unclassified | unclassified | unclassified | 13,6 | 0,8 | 0,000337248 | 2,92E-05 | 15 - | 0,000312459 - | 13 - | 0,000336251 - | 13 - | 0,00031965 - | 13 - | 0,000324371 - | 14 - | 0,00039351 - |
| 2866 | 1 | 3um_FIJJAFBF_524930 | Bacteria | Bacteroidetes | Cytophagia | Cytophagales | Flammeovirga | Reichenbachie | Reichenbachie | unclassified | unclassified | unclassified | unclassified | unclassified | 1,4 | 0,8 | 4,51E-05 | 2,48E-05 | 2 - | 5,56E-05 - | 0 - | 0 - | 2 - | 6,56E-05 - | 2 - | 6,66E-05 - | 1 - | 3,75E-05 - |
| 2867 | 1 | 0.2um_NOKCPHJN_511530 | Bacteria | Bacteroidetes | Flavobacteriia | unclassified | unclassified | unclassified | Flavobacteria | cellular proces | Intracellular tr | 0BIEB | domain-conta | 1005048.CFU_ | 2,6 | 1,356465997 | 3,95E-05 | 2,00E-05 | 3 - | 3,89E-05 - | 1 - | 1,61E-05 - | 5 - | 7,65E-05 - | 2 - | 3,10E-05 - | 2 - | 3,50E-05 - |
| 2868 | 1 | 0.2um_NOKCPHJN_107386 | Bacteria | Proteobacteri | Gammaprotec | Cellvibrionales | Porticoccacea | unclassified | Gammaprotec | unclassified | unclassified | unclassified | unclassified | unclassified | 0,2 | 0,4 | 1,96E-06 | 3,92E-06 | 0 - | 0 - | 0 - | 0 - | 1 - | 9,79E-06 - | 0 - | 0 - | 0 - | 0 - |
| 2869 | 1 | 0.2um_NOKCPHJN_886977 | Bacteria | Proteobacteri | Alphaproteob | Pelagibacteral | Pelagibacterac | Candidatus Pe | Candidatus Pe | metabolism | Nucleotide tra | 16UID | reductase | 335992.SAR11 | 0,6 | 0,8 | 3,17E-05 | 4,34E-05 | 0 - | 0 - | 0 - | 0 - | 0 - | 0 - | 1 - | 4,88E-05 - | 2 - | 0,00010984 - |
| 2870 | 1 | 0.2um_NOKCPHJN_186420 | Bacteria | Bacteroidetes | unclassified | unclassified | unclassified | unclassified | Bacteroidetes | metabolism | Inorganic ion t | COG0659 | sulfate transp | 313596.RB250 | 0,8 | 0,4 | 2,16E-05 | 1,11E-05 | 1 - | 2,26E-05 - | 1 - | 2,81E-05 - | 0 - | 0 - | 1 - | 2,71E-05 - | 1 - | 3,05E-05 - |
| 2871 | 1 | 3um_FIJJAFBF_507802 | Bacteria | Proteobacteri | Gammaprotec | Chromatiales | Ectothiorhodo | Ectothiorhodo | Ectothiorhodo | unclassified | unclassified | unclassified | unclassified | unclassified | 1,8 | 0,4 | 0,000220498 | 5,49E-05 | 2 - | 0,000204958 - | 2 - | 0,000254498 - | 1 - | 0,000120966 - | 2 - | 0,000245506 - | 2 - | 0,000276562 - |
| 2872 | 1 | 0.2um_NOKCPHJN_15572 | Bacteria | Bacteroidetes | Flavobacteriia | unclassified | unclassified | unclassified | Flavobacteria | poorly charact | Function unkn | 0XNQI | domain protei | 487796.Flav2A | 1,4 | 0,489897949 | 4,14E-05 | 1,46E-05 | 1 - | 2,48E-05 - | 1 - | 3,08E-05 - | 2 - | 5,85E-05 - | 2 - | 5,94E-05 - | 1 - | 3,35E-05 - |
| 2873 | 1 | 3um_FIJJAFBF_628699 | Bacteria | Bacteroidetes | Flavobacteriia | Flavobacterial | Flavobacteriac | unclassified | Flavobacteriac | poorly charact | Function unkn | 09JH9 | NA | 487796.Flav2A | 2 | 0,632455532 | 0,000110848 | 2,57E-05 | 3 - | 0,000143471 - | 2 - | 0,000118766 - | 2 - | 0,000112902 - | 2 - | 0,000114569 - | 1 - | 6,45E-05 - |
| 2874 | 1 | 0.2um_NOKCPHJN_343386 | Bacteria | Proteobacteri | Gammaprotec | unclassified | unclassified | unclassified | Gammaprotec | metabolism | Inorganic ion t | 16TQC | TONB-depend | 314287.GB220 | 2,8 | 0,4 | 4,96E-05 | 1,05E-05 | 2 - | 2,94E-05 - | 3 - | 5,47E-05 - | 3 - | 5,20E-05 - | 3 - | 5,27E-05 - | 3 - | 5,94E-05 - |
| 2875 | 1 | 0.2um_NOKCPHJN_698824 | Bacteria | Proteobacteri | Alphaproteob | Pelagibacteral | Pelagibacterac | Candidatus Pe | Candidatus Pe | unclassified | unclassified | unclassified | unclassified | unclassified | 0,6 | 0,489897949 | 5,78E-05 | 4,76E-05 | 1 - | 8,44E-05 - | 1 - | 0,000104793 - | 1 - | 9,96E-05 - | 0 - | 0 - | 0 - | 0 - |
| 2876 | 1 | 3um_FIJJAFBF_774340 | Bacteria | Proteobacteri | Gammaprotec | Oceanospirilla | Saccharospirill | Reinekea | Reinekea fors | unclassified | unclassified | unclassified | unclassified | unclassified | 7,6 | 1,019803903 | 0,000697725 | 0,000110669 | 8 - | 0,000616251 - | 6 - | 0,000573901 - | 7 - | 0,000636494 - | 9 - | 0,000830436 - | 8 - | 0,000831541 - |
| 2877 | 1 | 3um_FIJJAFBF_492261 | Bacteria | Bacteroidetes | Flavobacteriia | Flavobacterial | Flavobacteriac | Polaribacter | various | unclassified | unclassified | unclassified | unclassified | unclassified | 0,2 | 0,4 | 1,12E-05 | 2,24E-05 | 1 - | 5,60E-05 - | 0 - | 0 - | 0 - | 0 - | 0 - | 0 - | 0 - | 0 - |
| 2878 | 1 | 3um_FIJJAFBF_133330 | Bacteria | Proteobacteri | Gammaprotec | Alteromonada | Alteromonada | Paraglaciecola | Paraglaciecola | unclassified | unclassified | unclassified | unclassified | unclassified | 0,8 | 0,748331477 | 2,06E-05 | 1,95E-05 | 0 - | 0 - | 2 - | 5,25E-05 - | 1 - | 2,50E-05 - | 1 - | 2,53E-05 - | 0 - | 0 - |
| 2879 | 1 | 3um_FIJJAFBF_220494 | Bacteria | Proteobacteri | Gammaprotec | Alteromonada | Alteromonada | Paraglaciecola | Paraglaciecola | information st | Translation, ri | 16Q16 | Binds the lowe | 342610.Patl_0 | 2 | 1,095445115 | 0,000116014 | 6,09E-05 | 3 - | 0,000147781 - | 2 - | 0,000122334 - | 0 - | 0 - | 3 - | 0,000177017 - | 2 - | 0,000132939 - |
| 2880 | 1 | 3um_FIJJAFBF_292772 | Bacteria | Bacteroidetes | Flavobacteriia | Flavobacterial | Flavobacteriac | Formosa | Formosa sp. H | information st | Translation, ri | 08UIK | This protein bi | 391603.FBALC | 2,4 | 0,489897949 | 0,00022199 | 3,26E-05 | 3 - | 0,000235843 - | 2 - | 0,000195231 - | 3 - | 0,000278388 - | 2 - | 0,000188333 - | 2 - | 0,000212157 - |
| 2881 | 1 | 3um_FIJJAFBF_217613 | Bacteria | unclassified | unclassified | unclassified | unclassified | unclassified | uncultured ba | unclassified | unclassified | unclassified | unclassified | unclassified | 0,8 | 0,748331477 | 5,30E-05 | 4,58E-05 | 2 - | 0,000115354 - | 1 - | 7,16E-05 - | 0 - | 0 - | 0 - | 0 - | 1 - | 7,78E-05 - |
| 2882 | 1 | 0.2um_NOKCPHJN_1068768 | Bacteria | Proteobacteri | Alphaproteob | Pelagibacteral | Pelagibacterac | Candidatus Pe | Candidatus Pe | poorly charact | Function unkn | 01SDU | solute sympor | 684719.HIMB | 1 | 0,632455532 | 2,89E-05 | 1,95E-05 | 1 - | 2,46E-05 - | 2 - | 6,12E-05 - | 1 - | 2,91E-05 - | 1 - | 2,95E-05 - | 0 - | 0 - |
| 2883 | 2 | 0.2um_NOKCPHJN_556777;0.2um_NOKCPHJN_7 | Bacteria | Bacteroidetes | Flavobacteriia | Flavobacterial | unclassified | unclassified | Flavobacterial | unclassified | unclassified | unclassified | unclassified | unclassified | 0,6 | 0,489897949 | 3,57E-05 | 2,94E-05 | 1 - | 5,22E-05 - | 1 - | 6,48E-05 - | 1 - | 6,16E-05 - | 0 - | 0 - | 0 - | 0 - |
| 2884 | 1 | 3um_FIJJAFBF_748496 | Bacteria | Proteobacteri | Alphaproteob | Rhodobactera | Rhodobactera | Planktomarina | Planktomarina | unclassified | unclassified | unclassified | unclassified | unclassified | 1 | 0 | 1,64E-05 | 1,55E-06 | 1 - | 1,37E-05 - | 1 - | 1,70E-05 - | 1 - | 1,62E-05 - | 1 - | 1,64E-05 - | 1 - | 1,85E-05 - |
| 2885 | 1 | 3um_FIJJAFBF_528404 | Bacteria | Bacteroidetes | Flavobacteriia | Flavobacterial | Flavobacteriac | Polaribacter | Polaribacter s | unclassified | unclassified | unclassified | unclassified | unclassified | 2,2 | 0,4 | 8,66E-05 | 7,94E-06 | 3 - | 0,000100388 - | 2 - | 8,31E-05 - | 2 - | 7,90E-05 - | 2 - | 8,02E-05 - | 2 - | 9,03E-05 - |
| 2886 | 1 | 3um_FIJJAFBF_508358 | Bacteria | Proteobacteri | Gammaprotec | Oceanospirilla | Oceanospirilla | Motiliproteus | Motiliproteus | cellular proces | Cell wall/mem | COG2913 | Part of the ou | 637616.MDM | 0,6 | 0,489897949 | 7,45E-05 | 6,15E-05 | 1 - | 0,00010828 - | 1 - | 0,000134452 - | 0 - | 0 - | 1 - | 0,000129701 - | 0 - | 0 - |
| 2887 | 1 | 3um_FIJJAFBF_11581 | Bacteria | Proteobacteri | Gammaprotec | Oceanospirilla | Saccharospirill | Reinekea | Reinekea fors | unclassified | unclassified | unclassified | unclassified | unclassified | 6 | 1,549193338 | 0,000498723 | 8,99E-05 | 9 - | 0,000637649 - | 5 - | 0,000439873 - | 5 - | 0,000418155 - | 5 - | 0,000424331 - | 6 - | 0,000573609 - |
| 2888 | 1 | 3um_FIJJAFBF_240262 | Bacteria | Bacteroidetes | Flavobacteriia | unclassified | unclassified | unclassified | Flavobacteria | metabolism | Inorganic ion t | 08QIB | TonB depende | 435590.BVU_ | 0,4 | 0,489897949 | 3,74E-05 | 4,60E-05 | 1 - | 8,57E-05 - | 0 - | 0 - | 1 - | 0,000101106 - | 0 - | 0 - | 0 - | 0 - |
| 2889 | 1 | 3um_FIJJAFBF_720357 | Bacteria | Proteobacteri | Alphaproteob | Rhodobactera | Rhodobactera | Pseudopelagic | Pseudopelagic | cellular proces | Signal transdu | 05F2E | Two compone | 391593.RCCS2 | 1 | 0,632455532 | 5,81E-05 | 3,26E-05 | 2 - | 0,000100681 - | 1 - | 6,25E-05 - | 1 - | 5,94E-05 - | 0 - | 0 - | 1 - | 6,79E-05 - |
| 2890 | 1 | 0.2um_NOKCPHJN_215620 | Eukaryota | Chromerida | unclassified | unclassified | unclassified | Vitrella | Vitrella brassi | unclassified | unclassified | unclassified | unclassified | unclassified | 1 | 0 | 2,60E-05 | 2,46E-06 | 1 - | 2,17E-05 - | 1 - | 2,70E-05 - | 1 - | 2,57E-05 - | 1 - | 2,60E-05 - | 1 - | 2,93E-05 - |
| 2891 | 1 | 0.2um_NOKCPHJN_677832 | Bacteria | Proteobacteri | Gammaprotec | unclassified | unclassified | unclassified | Gammaprotec | information st | Translation, ri | 0QJSM | Responsible fo | 357804.Ping_ | 0,2 | 0,4 | 5,94E-06 | 1,19E-05 | 0 - | 0 - | 0 - | 0 - | 1 - | 2,97E-05 - | 0 - | 0 - | 0 - | 0 - |
| 2892 | 1 | 3um_FIJJAFBF_149823 | Bacteria | Proteobacteri | Betaproteoba | unclassified | unclassified | unclassified | Betaproteoba | metabolism | Energy produc | 16RH6 | Produces ATP | 397945.Aave_ | 0,8 | 0,4 | 3,67E-05 | 1,86E-05 | 1 - | 3,97E-05 - | 1 - | 4,93E-05 - | 1 - | 4,69E-05 - | 1 - | 4,76E-05 - | 0 - | 0 - |
| 2893 | 1 | 3um_FIJJAFBF_232085 | Bacteria | Bacteroidetes | Flavobacteriia | Flavobacterial | Flavobacteriac | Gilvibacter | Gilvibacter sp. | metabolism | Lipid transpor | 0P7K8 | carboxylase | 983548.Krodi_ | 0,6 | 0,489897949 | 1,56E-05 | 1,28E-05 | 0 - | 0 - | 1 - | 2,68E-05 - | 1 - | 2,55E-05 - | 1 - | 2,59E-05 - | 0 - | 0 - |
| 2894 | 1 | 0.2um_NOKCPHJN_293115 | Bacteria | Proteobacteri | Gammaprotec | Cellvibrionales | Porticoccacea | unclassified | SAR92 bacteri | information st | Translation, ri | COG0050 | This protein p | 203122.Sde_0 | 8 | 0,894427191 | 0,000267618 | 2,62E-05 | 9 - | 0,000253806 - | 9 - | 0,000315152 - | 8 - | 0,000266304 - | 7 - | 0,000236458 - | 7 - | 0,000266369 - |
| 2895 | 2 | 0.2um_NOKCPHJN_87265;3um_FIJJAFBF_361125 | Bacteria | Proteobacteri | Alphaproteob | Rhodobactera | Rhodobactera | Ruegeria | Ruegeria litor | unclassified | unclassified | unclassified | unclassified | unclassified | 0,4 | 0,489897949 | 1,10E-05 | 1,35E-05 | 0 - | 0 - | 0 - | 0 - | 1 - | 2,74E-05 - | 1 - | 2,78E-05 - | 0 - | 0 - |
| 2896 | 1 | 3um_FIJJAFBF_437595 | Bacteria | Bacteroidetes | Flavobacteriia | Flavobacterial | Flavobacteriac | Polaribacter | Polaribacter s | unclassified | unclassified | unclassified | unclassified | unclassified | 1 | 0 | 7,49E-05 | 7,11E-06 | 1 - | 6,27E-05 - | 1 - | 7,79E-05 - | 1 - | 7,40E-05 - | 1 - | 7,51E-05 - | 1 - | 8,46E-05 - |
| 2897 | 1 | 0.2um_NOKCPHJN_170598 | Bacteria | Bacteroidetes | Flavobacteriia | Flavobacterial | Cryomorphace | unclassified | Cryomorphace | metabolism | Nucleotide tra | 0P81Z | Catalyzes the | 926562.Oweh | 0,4 | 0,489897949 | 2,18E-05 | 2,70E-05 | 1 - | 4,86E-05 - | 1 - | 6,04E-05 - | 0 - | 0 - | 0 - | 0 - | 0 - | 0 - |
| 2898 | 1 | 0.2um_NOKCPHJN_541100 | Bacteria | Bacteroidetes | Bacteroidia | Marinilabiliale | Marinilabiliac | Marinilabilia | Marinilabilia s | unclassified | unclassified | unclassified | unclassified | unclassified | 0,2 | 0,4 | 3,47E-06 | 6,93E-06 | 0 - | 0 - | 0 - | 0 - | 0 - | 0 - | 1 - | 1,73E-05 - | 0 - | 0 - |
| 2899 | 1 | 0.2um_NOKCPHJN_816587 | Bacteria | Bacteroidetes | Flavobacteriia | Flavobacterial | Flavobacteriac | unclassified | Flavobacteriac | unclassified | unclassified | unclassified | unclassified | unclassified | 1,2 | 0,4 | 3,90E-05 | 1,31E-05 | 1 - | 2,73E-05 - | 1 - | 3,39E-05 - | 2 - | 6,44E-05 - | 1 - | 3,27E-05 - | 1 - | 3,68E-05 - |
| 2900 | 1 | 0.2um_NOKCPHJN_95495 | Bacteria | Bacteroidetes | Cytophagia | Cytophagales | Flammeovirga | unclassified | Flammeovirga | cellular proces | Cell wall/mem | 09A4Y | export protein | 880070.Cycma | 0,8 | 0,4 | 1,26E-05 | 6,49E-06 | 1 - | 1,32E-05 - | 1 - | 1,64E-05 - | 0 - | 0 - | 1 - | 1,58E-05 - | 1 - | 1,78E-05 - |
| 2901 | 1 | 3um_FIJJAFBF_773342 | Bacteria | Bacteroidetes | Flavobacteriia | Flavobacterial | Flavobacteriac | Polaribacter | various | cellular proces | Intracellular tr | 08SHB | Biopolymer tr | 313594.PI23P | 2,2 | 0,4 | 0,000196466 | 4,38E-05 | 2 - | 0,000149061 - | 3 - | 0,000277634 - | 2 - | 0,000175951 - | 2 - | 0,00017855 - | 2 - | 0,000201136 - |
| 2902 | 1 | 3um_FIJJAFBF_343897 | Eukaryota | unclassified | unclassified | Phaeocystales | Phaeocystace | Phaeocystis | Phaeocystis an | metabolism | Energy produc | 0FN1Q | F(1)F(0) ATP s | 197221.tlr043 | 3 | 0,632455532 | 0,000225985 | 6,17E-05 | 3 - | 0,000188159 - | 2 - | 0,000155758 - | 3 - | 0,000222102 - | 3 - | 0,000225382 - | 4 - | 0,000338523 - |
| 2903 | 1 | 0.2um_NOKCPHJN_208938 | Bacteria | Proteobacteri | Betaproteoba | Burkholderiale | Comamonada | unclassified | Comamonada | unclassified | unclassified | unclassified | unclassified | unclassified | 4,2 | 0,748331477 | 0,000116205 | 2,01E-05 | 4 - | 9,33E-05 - | 4 - | 0,000115869 - | 5 - | 0,000137685 - | 5 - | 0,000139719 - | 3 - | 9,44E-05 - |
| 2904 | 4 | 0.2um_NOKCPHJN_485777;0.2um_NOKCPHJN_8 | Bacteria | Proteobacteri | Alphaproteob | Rhodobactera | Rhodobactera | unclassified | Rhodobactera | unclassified | unclassified | unclassified | unclassified | unclassified | 0,2 | 0,4 | 1,18E-05 | 2,35E-05 | 1 - | 5,89E-05 - | 0 - | 0 - | 0 - | 0 - | 0 - | 0 - | 0 - | 0 - |
| 2905 | 1 | 0.2um_NOKCPHJN_839859 | Bacteria | Proteobacteri | Alphaproteob | Rhizobiales | unclassified | unclassified | Rhizobiales ba | unclassified | unclassified | unclassified | unclassified | unclassified | 1 | 0,632455532 | 1,80E-05 | 1,01E-05 | 2 - | 3,11E-05 - | 1 - | 1,93E-05 - | 1 - | 1,84E-05 - | 0 - | 0 - | 1 - | 2,10E-05 - |
| 2906 | 1 | 3um_FIJJAFBF_114132 | unclassified | unclassified | unclassified | unclassified | unclassified | unclassified | unclassified | unclassified | unclassified | unclassified | unclassified | unclassified | 2 | 0 | 0,000185172 | 1,76E-05 | 2 - | 0,000155104 - | 2 - | 0,000192593 - | 2 - | 0,000183084 - | 2 - | 0,000185788 - | 2 - | 0,00020929 - |
| 2907 | 2 | 0.2um_NOKCPHJN_345482;0.2um_NOKCPHJN_4 | Bacteria | Bacteroidetes | Flavobacteriia | Flavobacterial | Flavobacteriac | Polaribacter | Polaribacter s | unclassified | unclassified | unclassified | unclassified | unclassified | 0,2 | 0,4 | 1,10E-05 | 2,21E-05 | 1 - | 5,52E-05 - | 0 - | 0 - | 0 - | 0 - | 0 - | 0 - | 0 - | 0 - |
| 2908 | 1 | 0.2um_NOKCPHJN_476940 | Bacteria | Proteobacteri | Alphaproteob | Pelagibacteral | unclassified | unclassified | Pelagibacteral | cellular proces | Cell wall/mem | 7257 | NA | 335992.SAR11 | 1 | 0 | 3,85E-05 | 3,65E-06 | 1 - | 3,22E-05 - | 1 - | 4,00E-05 - | 1 - | 3,81E-05 - | 1 - | 3,86E-05 - | 1 - | 4,35E-05 - |
| 2909 | 1 | 0.2um_NOKCPHJN_241612 | Bacteria | Proteobacteri | Alphaproteob | Pelagibacteral | Pelagibacterac | Candidatus Pe | Candidatus Pe | information st | Translation, ri | 05CEJ | Catalyzes the | 684719.HIMB | 2 | 0 | 3,96E-05 | 3,76E-06 | 2 - | 3,32E-05 - | 2 - | 4,12E-05 - | 2 - | 3,92E-05 - | 2 - | 3,97E-05 - | 2 - | 4,48E-05 - |
| 2910 | 1 | 0.2um_NOKCPHJN_323663 | Bacteria | Proteobacteri | Alphaproteob | Rhodobactera | Rhodobactera | Planktomarina | Planktomarina | unclassified | unclassified | unclassified | unclassified | unclassified | 1,2 | 0,4 | 0,000134856 | 4,65E-05 | 1 - | 9,41E-05 - | 1 - | 0,000116819 - | 1 - | 0,000111051 - | 2 - | 0,000225382 - | 1 - | 0,000126946 - |
| 2911 | 1 | 0.2um_NOKCPHJN_391802 | Bacteria | Proteobacteri | Alphaproteob | Pelagibacteral | Pelagibacterac | Candidatus Pe | Candidatus Pe | unclassified | unclassified | unclassified | unclassified | unclassified | 3,2 | 0,748331477 | 9,22E-05 | 1,67E-05 | 4 - | 9,83E-05 - | 3 - | 9,16E-05 - | 3 - | 8,70E-05 - | 4 - | 0,000117759 - | 2 - | 6,63E-05 - |
| 2912 | 1 | 3um_FIJJAFBF_79234 | Bacteria | Bacteroidetes | Flavobacteriia | Flavobacterial | Flavobacteriac | Polaribacter | Polaribacter s | poorly charact | Function unkn | 09EUF | NA | 313598.MED1 | 4,6 | 1,019803903 | 0,000255981 | 5,10E-05 | 6 - | 0,000282238 - | 4 - | 0,000233637 - | 5 - | 0,000277628 - | 3 - | 0,000169037 - | 5 - | 0,000317366 - |
| 2913 | 1 | 0.2um_NOKCPHJN_305910 | Bacteria | Proteobacteri | Gammaprotec | Alteromonada | Colwelliaceae | Colwellia | Colwellia aest | metabolism | Nucleotide tra | 0QIPY | Glutamine ph | 167879.CPS_3 | 1 | 0 | 2,71E-05 | 2,58E-06 | 1 - | 2,27E-05 - | 1 - | 2,82E-05 - | 1 - | 2,68E-05 - | 1 - | 2,72E-05 - | 1 - | 3,07E-05 - |
| 2914 | 1 | 3um_FIJJAFBF_449188 | Bacteria | Bacteroidetes | Cytophagia | Cytophagales | Flammeovirga | Flexithrix | Flexithrix dor | metabolism | Carbohydrate | COG0057 | glyceraldehyd | 643867.Ftrac_ | 4,4 | 1,356465997 | 0,000180125 | 5,29E-05 | 5 - | 0,000174433 - | 5 - | 0,000216594 - | 4 - | 0,00016472 - | 6 - | 0,000250729 - | 2 - | 9,41E-05 - |
| 2915 | 1 | 0.2um_NOKCPHJN_365882 | unclassified | unclassified | unclassified | unclassified | unclassified | unclassified | unclassified | unclassified | unclassified | unclassified | unclassified | unclassified | 0,6 | 0,489897949 | 0,000114091 | 9,36E-05 | 0 - | 0 - | 0 - | 0 - | 1 - | 0,000180643 - | 1 - | 0,000183311 - | 1 - | 0,000206499 - |
| 2916 | 1 | 3um_FIJJAFBF_111294 | Viruses | unclassified | unclassified | unclassified | Mimiviridae | unclassified | Yellowstone la | unclassified | unclassified | unclassified | unclassified | unclassified | 5,8 | 1,469693846 | 0,00048835 | 0,000101453 | 7 - | 0,000502148 - | 5 - | 0,000445371 - | 8 - | 0,000677411 - | 5 - | 0,000429635 - | 4 - | 0,000387186 - |
| 2917 | 1 | 3um_FIJJAFBF_753294 | Bacteria | Proteobacteri | Alphaproteob | Rhizobiales | Hyphomicrobi | Maritalea | Maritalea myr | unclassified | unclassified | unclassified | unclassified | unclassified | 0,6 | 0,489897949 | 1,91E-05 | 1,57E-05 | 1 - | 2,83E-05 - | 0 - | 0 - | 1 - | 3,34E-05 - | 1 - | 3,39E-05 - | 0 - | 0 - |
| 2918 | 1 | 0.2um_NOKCPHJN_576145 | Eukaryota | Zoopagomyco | Harpellomyce | Harpellales | Legeriomycet | Smittium | Smittium culi | information st | RNA processin | 0PHZJ | RNA recogniti | 425265.XP_00 | 0,2 | 0,4 | 4,33E-06 | 8,65E-06 | 0 - | 0 - | 1 - | 2,16E-05 - | 0 - | 0 - | 0 - | 0 - | 0 - | 0 - |
| 2919 | 1 | 3um_FIJJAFBF_422354 | Bacteria | Proteobacteri | Gammaprotec | Oceanospirilla | Saccharospirill | Reinekea | Reinekea fors | cellular proces | Intracellular tr | 0R8TK | general secret | 314283.MED2 | 0,6 | 0,489897949 | 5,00E-05 | 4,16E-05 | 1 - | 7,08E-05 - | 0 - | 0 - | 1 - | 8,36E-05 - | 0 - | 0 - | 1 - | 9,56E-05 - |
| 2920 | 1 | 3um_FIJJAFBF_55330 | Bacteria | Proteobacteri | Gammaprotec | Cellvibrionales | Porticoccacea | unclassified | Gammaprotec | information st | Translation, ri | 0QQXI | This protein is | 314287.GB220 | 7,2 | 0,4 | 0,000839123 | 0,000119463 | 7 - | 0,000680879 - | 7 - | 0,000845451 - | 7 - | 0,000803708 - | 7 - | 0,000815579 - | 8 - | 0,001049996 - |
| 2921 | 1 | 3um_FIJJAFBF_15048 | Bacteria | Bacteroidetes | Flavobacteriia | Flavobacterial | Flavobacteriac | Ochrovirga | Ochrovirga pa | cellular proces | Signal transdu | 09A4D | gtp-binding pr | 880070.Cycma | 9,4 | 1,019803903 | 0,000216149 | 7,31E-06 | 11 - | 0,000213991 - | 9 - | 0,000217402 - | 10 - | 0,000229631 - | 9 - | 0,00020972 - | 8 - | 0,000209999 - |
| 2922 | 1 | 0.2um_NOKCPHJN_445402 | Eukaryota | unclassified | Synurophycea | Parmales | Triparmaceae | Triparma | Triparma laevi | metabolism | Energy produc | 0SSA5 | NADH dehydr | 7460.GB1535 | 1 | 0 | 7,03E-05 | 6,67E-06 | 1 - | 5,89E-05 - | 1 - | 7,31E-05 - | 1 - | 6,95E-05 - | 1 - | 7,05E-05 - | 1 - | 7,94E-05 - |
| 2923 | 1 | 3um_FIJJAFBF_109115 | Bacteria | Proteobacteri | Betaproteoba | unclassified | unclassified | unclassified | Betaproteoba | information st | Transcription | 0B77K | Transcription | 397945.Aave_ | 0,2 | 0,4 | 4,55E-06 | 9,09E-06 | 1 - | 2,27E-05 - | 0 - | 0 - | 0 - | 0 - | 0 - | 0 - | 0 - | 0 - |
| 2924 | 1 | 3um_FIJJAFBF_272637 | Bacteria | Bacteroidetes | Flavobacteriia | Flavobacterial | unclassified | unclassified | Flavobacterial | unclassified | unclassified | unclassified | unclassified | unclassified | 0,4 | 0,8 | 2,00E-05 | 4,01E-05 | 2 - | 0,000100242 - | 0 - | 0 - | 0 - | 0 - | 0 - | 0 - | 0 - | 0 - |
| 2925 | 2 | 3um_FIJJAFBF_260747;3um_FIJJAFBF_694195 | Bacteria | Proteobacteri | Alphaproteob | Rhodobactera | unclassified | unclassified | Rhodobactera | cellular proces | Posttranslatio | COG0330 | Band 7 protei | 367336.OM22 | 4,2 | 0,4 | 0,000198149 | 2,92E-05 | 4 - | 0,000157769 - | 5 - | 0,000244878 - | 4 - | 0,00018623 - | 4 - | 0,00018898 - | 4 - | 0,000212886 - |
| 2926 | 1 | 3um_FIJJAFBF_725646 | Bacteria | Firmicutes | Bacilli | Bacillales | Thermoactino | Thermoactino | Thermoactino | unclassified | unclassified | unclassified | unclassified | unclassified | 0,2 | 0,4 | 7,68E-06 | 1,54E-05 | 0 - | 0 - | 0 - | 0 - | 1 - | 3,84E-05 - | 0 - | 0 - | 0 - | 0 - |
| 2927 | 1 | 3um_FIJJAFBF_540252 | Bacteria | Bacteroidetes | Cytophagia | Cytophagales | Flammeovirga | unclassified | Flammeovirga | unclassified | unclassified | unclassified | unclassified | unclassified | 0,4 | 0,489897949 | 2,96E-05 | 3,65E-05 | 1 - | 6,79E-05 - | 0 - | 0 - | 1 - | 8,02E-05 - | 0 - | 0 - | 0 - | 0 - |
| 2928 | 1 | 3um_FIJJAFBF_27138 | Bacteria | Bacteroidetes | Flavobacteriia | Flavobacterial | Flavobacteriac | Polaribacter | Polaribacter s | poorly charact | Function unkn | 08S7Z | NA | 688270.Celal_ | 0,4 | 0,489897949 | 1,37E-05 | 1,68E-05 | 0 - | 0 - | 0 - | 0 - | 1 - | 3,20E-05 - | 0 - | 0 - | 1 - | 3,65E-05 - |
| 2929 | 1 | 0.2um_NOKCPHJN_183236 | unclassified | unclassified | unclassified | unclassified | unclassified | unclassified | unclassified | unclassified | unclassified | unclassified | unclassified | unclassified | 0,2 | 0,4 | 1,11E-05 | 2,22E-05 | 1 - | 5,54E-05 - | 0 - | 0 - | 0 - | 0 - | 0 - | 0 - | 0 - | 0 - |
| 2930 | 1 | 0.2um_NOKCPHJN_126571 | Bacteria | Proteobacteri | Gammaprotec | Alteromonada | Alteromonada | unclassified | Alteromonada | unclassified | unclassified | unclassified | unclassified | unclassified | 1 | 0 | 5,69E-06 | 5,40E-07 | 1 - | 4,76E-06 - | 1 - | 5,91E-06 - | 1 - | 5,62E-06 - | 1 - | 5,70E-06 - | 1 - | 6,43E-06 - |
| 2931 | 1 | 0.2um_NOKCPHJN_13656 | Bacteria | Bacteroidetes | Flavobacteriia | unclassified | unclassified | unclassified | Flavobacteria | unclassified | unclassified | unclassified | unclassified | unclassified | 0,2 | 0,4 | 9,45E-06 | 1,89E-05 | 1 - | 4,72E-05 - | 0 - | 0 - | 0 - | 0 - | 0 - | 0 - | 0 - | 0 - |
| 2932 | 1 | 3um_FIJJAFBF_71464 | Bacteria | Proteobacteri | Alphaproteob | Rhodobactera | Rhodobactera | various | various | poorly charact | Function unkn | 173PW | Pfam:DUF88 | 314262.MED1 | 1 | 0 | 7,21E-05 | 6,85E-06 | 1 - | 6,04E-05 - | 1 - | 7,50E-05 - | 1 - | 7,13E-05 - | 1 - | 7,24E-05 - | 1 - | 8,15E-05 - |
| 2933 | 1 | 3um_FIJJAFBF_451424 | Bacteria | Proteobacteri | Alphaproteob | Pelagibacteral | Pelagibacterac | Candidatus Pe | Candidatus Pe | metabolism | Amino acid tra | 01RN4 | Para-aminobe | 439493.PB721 | 0,4 | 0,489897949 | 1,45E-05 | 1,77E-05 | 0 - | 0 - | 1 - | 3,68E-05 - | 0 - | 0 - | 1 - | 3,55E-05 - | 0 - | 0 - |
| 2934 | 1 | 3um_FIJJAFBF_191790 | Eukaryota | Bacillariophyt | Fragilariophyc | Fragilariales | Fragilariaceae | Asterionellops | Asterionellops | metabolism | Energy produc | COG0711 | F(1)F(0) ATP s | 35128.Thapsd | 12,4 | 0,489897949 | 0,000896496 | 6,46E-05 | 13 - | 0,00078947 - | 12 - | 0,000904882 - | 13 - | 0,000931888 - | 12 - | 0,00087291 - | 12 - | 0,00098333 - |
| 2935 | 1 | 0.2um_NOKCPHJN_920400 | Bacteria | Proteobacteri | Gammaprotec | unclassified | unclassified | unclassified | Gammaprotec | unclassified | unclassified | unclassified | unclassified | unclassified | 0,8 | 0,4 | 2,52E-05 | 1,29E-05 | 1 - | 2,66E-05 - | 0 - | 0 - | 1 - | 3,14E-05 - | 1 - | 3,19E-05 - | 1 - | 3,59E-05 - |
| 2936 | 1 | 0.2um_NOKCPHJN_472922 | Viruses | unclassified | unclassified | unclassified | unclassified | unclassified | uncultured vir | cellular proces | Signal transdu | KOG0613 | fibronectin typ | 7070.TC00296 | 0,2 | 0,4 | 3,07E-06 | 6,15E-06 | 1 - | 1,54E-05 - | 0 - | 0 - | 0 - | 0 - | 0 - | 0 - | 0 - | 0 - |
| 2937 | 1 | 3um_FIJJAFBF_194946 | Bacteria | Proteobacteri | Gammaprotec | Alteromonada | Colwelliaceae | Colwellia | Colwellia sp. P | information st | Translation, ri | 05C64 | Binds directly | 234831.PSM_ | 0,4 | 0,489897949 | 2,22E-05 | 2,74E-05 | 1 - | 4,95E-05 - | 1 - | 6,14E-05 - | 0 - | 0 - | 0 - | 0 - | 0 - | 0 - |
| 2938 | 2 | 0.2um_NOKCPHJN_659662;3um_FIJJAFBF_72476 | Bacteria | Bacteroidetes | Cytophagia | Cytophagales | Flammeovirga | unclassified | Flammeovirga | information st | Translation, ri | 09FE5 | One of the pro | various | 1,6 | 0,489897949 | 0,000199984 | 5,65E-05 | 2 - | 0,00021255 - | 2 - | 0,000263924 - | 1 - | 0,000125447 - | 2 - | 0,000254599 - | 1 - | 0,000143402 - |
| 2939 | 1 | 0.2um_NOKCPHJN_27665 | Bacteria | Actinobacteri | Actinobacteri | Micrococcales | Microbacteria | unclassified | Microbacteria | information st | Translation, ri | 05CEJ | Catalyzes the | 281090.Lxx20 | 1 | 0 | 1,96E-05 | 1,86E-06 | 1 - | 1,64E-05 - | 1 - | 2,04E-05 - | 1 - | 1,94E-05 - | 1 - | 1,96E-05 - | 1 - | 2,21E-05 - |
| 2940 | 2 | 3um_FIJJAFBF_227445;3um_FIJJAFBF_418154 | Bacteria | Proteobacteri | Gammaprotec | Cellvibrionales | Porticoccacea | unclassified | SAR92 bacteri | cellular proces | Intracellular tr | COG0848 | Biopolymer tr | 314287.GB220 | 1 | 0 | 0,000101502 | 9,64E-06 | 1 - | 8,50E-05 - | 1 - | 0,00010557 - | 1 - | 0,000100357 - | 1 - | 0,000101839 - | 1 - | 0,000114722 - |
| 2941 | 1 | 0.2um_NOKCPHJN_76449 | Bacteria | Bacteroidetes | Flavobacteriia | Flavobacterial | Flavobacteriac | unclassified | Flavobacteriac | metabolism | Amino acid tra | 0P7DW | synthase | 487796.Flav2A | 0,6 | 0,489897949 | 1,63E-05 | 1,36E-05 | 1 - | 2,27E-05 - | 1 - | 2,82E-05 - | 0 - | 0 - | 0 - | 0 - | 1 - | 3,07E-05 - |
| 2942 | 1 | 0.2um_NOKCPHJN_821735 | Bacteria | Proteobacteri | Alphaproteob | Rhodobactera | Rhodobactera | unclassified | Rhodobactera | poorly charact | Function unkn | 0NVRZ | NA | 170187.SP_17 | 1 | 0 | 1,75E-05 | 1,66E-06 | 1 - | 1,46E-05 - | 1 - | 1,82E-05 - | 1 - | 1,73E-05 - | 1 - | 1,75E-05 - | 1 - | 1,97E-05 - |
| 2943 | 1 | 0.2um_NOKCPHJN_106570 | Bacteria | Proteobacteri | Gammaprotec | Oceanospirilla | Saccharospirill | Reinekea | Reinekea fors | cellular proces | Posttranslatio | 0QI83 | Cytochrome c | 314283.MED2 | 0,4 | 0,489897949 | 7,75E-06 | 9,56E-06 | 1 - | 1,76E-05 - | 0 - | 0 - | 0 - | 0 - | 1 - | 2,11E-05 - | 0 - | 0 - |
| 2944 | 1 | 3um_FIJJAFBF_267805 | Bacteria | Bacteroidetes | Flavobacteriia | Flavobacterial | Flavobacteriac | Polaribacter | various | metabolism | Energy produc | 0P9HY | cytochrome C | 487797.Flav30 | 0,4 | 0,489897949 | 2,69E-05 | 3,31E-05 | 1 - | 6,17E-05 - | 0 - | 0 - | 1 - | 7,28E-05 - | 0 - | 0 - | 0 - | 0 - |
| 2945 | 1 | 0.2um_NOKCPHJN_148310 | Bacteria | Bacteroidetes | Flavobacteriia | Flavobacterial | Flavobacteriac | Flavobacteriu | Flavobacteriu | poorly charact | Function unkn | 0XZ2H | Membrane as | 353153.XP_80 | 6,6 | 0,489897949 | 0,000158857 | 1,31E-05 | 7 - | 0,0001417 - | 7 - | 0,000175949 - | 7 - | 0,000167262 - | 6 - | 0,000145485 - | 6 - | 0,000163888 - |
| 2946 | 1 | 3um_FIJJAFBF_378153 | Bacteria | various | various | various | various | various | various | unclassified | unclassified | unclassified | unclassified | unclassified | 2 | 0,632455532 | 0,000116682 | 2,70E-05 | 3 - | 0,000151022 - | 2 - | 0,000125017 - | 2 - | 0,000118844 - | 2 - | 0,000120599 - | 1 - | 6,79E-05 - |
| 2947 | 1 | 3um_FIJJAFBF_477008 | Bacteria | Bacteroidetes | Flavobacteriia | Flavobacterial | Flavobacteriac | Formosa | Formosa sp. H | information st | Transcription | 05D27 | DNA-depende | 983544.Lacal_ | 2,2 | 0,4 | 3,03E-05 | 5,91E-06 | 2 - | 2,31E-05 - | 2 - | 2,87E-05 - | 3 - | 4,09E-05 - | 2 - | 2,77E-05 - | 2 - | 3,12E-05 - |
| 2948 | 1 | 3um_FIJJAFBF_37756 | Bacteria | Proteobacteri | Gammaprotec | Oceanospirilla | Oceanospirilla | Marinospirillu | Marinospirillu | metabolism | Energy produc | 05CMV | Succinyl-CoA s | 999141.GME_ | 6,2 | 0,748331477 | 0,000216257 | 1,54E-05 | 7 - | 0,000206539 - | 6 - | 0,000219823 - | 7 - | 0,000243798 - | 6 - | 0,000212057 - | 5 - | 0,000199067 - |
| 2949 | 1 | 3um_FIJJAFBF_743643 | Bacteria | Bacteroidetes | Flavobacteriia | Flavobacterial | Flavobacteriac | Muricauda | Muricauda ru | metabolism | Energy produc | 099YH | Pyruvate dehy | 886377.Murru | 0,8 | 0,748331477 | 3,53E-05 | 3,28E-05 | 0 - | 0 - | 2 - | 8,72E-05 - | 0 - | 0 - | 1 - | 4,20E-05 - | 1 - | 4,74E-05 - |
| 2950 | 1 | 3um_FIJJAFBF_103762 | Bacteria | Actinobacteri | Actinobacteri | Micrococcales | Microbacteria | unclassified | Microbacteria | unclassified | unclassified | unclassified | unclassified | unclassified | 11,4 | 1,019803903 | 0,000432368 | 2,95E-05 | 13 - | 0,000415626 - | 10 - | 0,000396988 - | 11 - | 0,000415127 - | 12 - | 0,000459554 - | 11 - | 0,000474546 - |
| 2951 | 1 | 0.2um_NOKCPHJN_212384 | Eukaryota | unclassified | Oomycetes | Peronosporale | Peronosporac | Phytophthora | Phytophthora | poorly charact | Function unkn | COG5307 | and Sec7 dom | 65071.PYU1_ | 0,4 | 0,489897949 | 3,15E-06 | 3,87E-06 | 0 - | 0 - | 1 - | 8,03E-06 - | 0 - | 0 - | 1 - | 7,75E-06 - | 0 - | 0 - |
| 2952 | 1 | 0.2um_NOKCPHJN_178394 | Bacteria | Bacteroidetes | Flavobacteriia | Flavobacterial | Flavobacteriac | Arenibacter | Arenibacter s | unclassified | unclassified | unclassified | unclassified | unclassified | 1,4 | 0,489897949 | 2,03E-05 | 7,47E-06 | 1 - | 1,21E-05 - | 2 - | 3,00E-05 - | 2 - | 2,85E-05 - | 1 - | 1,45E-05 - | 1 - | 1,63E-05 - |
| 2953 | 1 | 3um_FIJJAFBF_20043 | Bacteria | Bacteroidetes | Flavobacteriia | Flavobacterial | Flavobacteriac | Tenacibaculur | various | information st | Replication, re | COG0187 | DNA gyrase ne | 313590.MED1 | 7,8 | 0,748331477 | 0,000164294 | 5,95E-06 | 9 - | 0,000160154 - | 7 - | 0,000154672 - | 8 - | 0,00016804 - | 8 - | 0,000170522 - | 7 - | 0,000168081 - |
| 2954 | 1 | 3um_FIJJAFBF_258847 | Bacteria | Proteobacteri | Gammaprotec | Alteromonada | Colwelliaceae | Colwellia | Colwellia sp. P | unclassified | unclassified | unclassified | unclassified | unclassified | 2,6 | 0,489897949 | 0,000213627 | 5,50E-05 | 2 - | 0,00013583 - | 3 - | 0,000252992 - | 2 - | 0,000160334 - | 3 - | 0,000244053 - | 3 - | 0,000274925 - |
| 2955 | 1 | 3um_FIJJAFBF_467910 | Bacteria | Proteobacteri | Betaproteoba | unclassified | unclassified | unclassified | Betaproteoba | metabolism | Energy produc | 08UTY | succinate deh | 232721.Ajs_2 | 0,8 | 0,4 | 7,42E-05 | 3,77E-05 | 1 - | 8,03E-05 - | 1 - | 9,97E-05 - | 1 - | 9,47E-05 - | 1 - | 9,61E-05 - | 0 - | 0 - |
| 2956 | 1 | 3um_FIJJAFBF_715089 | Eukaryota | Phaeophyceae | unclassified | Dictyotales | Dictyotaceae | Dictyopteris | Dictyopteris d | unclassified | unclassified | unclassified | unclassified | unclassified | 0,6 | 0,489897949 | 5,90E-05 | 4,91E-05 | 1 - | 8,32E-05 - | 0 - | 0 - | 0 - | 0 - | 1 - | 9,96E-05 - | 1 - | 0,000112228 - |
| 2957 | 1 | 3um_FIJJAFBF_755414 | Bacteria | Bacteroidetes | Flavobacteriia | Flavobacterial | Flavobacteriac | Psychroflexus | Psychroflexus | poorly charact | Function unkn | 05X9M | NA | 313595.P7007 | 0,8 | 0,4 | 1,90E-05 | 9,78E-06 | 1 - | 2,01E-05 - | 0 - | 0 - | 1 - | 2,38E-05 - | 1 - | 2,41E-05 - | 1 - | 2,72E-05 - |
| 2958 | 1 | 0.2um_NOKCPHJN_237702 | Bacteria | Proteobacteri | Gammaprotec | Oceanospirilla | Saccharospirill | Reinekea | Reinekea fors | metabolism | Amino acid tra | 07RUI | Extracellular s | 314283.MED2 | 14 | 0,894427191 | 0,000348676 | 4,07E-05 | 15 - | 0,000313028 - | 14 - | 0,000362775 - | 13 - | 0,000320231 - | 13 - | 0,000324961 - | 15 - | 0,000422385 - |
| 2959 | 1 | 0.2um_NOKCPHJN_405308 | Bacteria | Bacteroidetes | Flavobacteriia | Flavobacterial | Flavobacteriac | Polaribacter | Polaribacter r | unclassified | unclassified | unclassified | unclassified | unclassified | 0,8 | 0,4 | 2,21E-05 | 1,12E-05 | 1 - | 2,39E-05 - | 1 - | 2,97E-05 - | 1 - | 2,82E-05 - | 1 - | 2,86E-05 - | 0 - | 0 - |
| 2960 | 1 | 0.2um_NOKCPHJN_75552 | Bacteria | Proteobacteri | Gammaprotec | Oceanospirilla | Saccharospirill | Reinekea | Reinekea fors | unclassified | unclassified | unclassified | unclassified | unclassified | 0,6 | 0,8 | 3,73E-05 | 4,93E-05 | 0 - | 0 - | 1 - | 6,36E-05 - | 0 - | 0 - | 2 - | 0,000122753 - | 0 - | 0 - |
| 2961 | 1 | 3um_FIJJAFBF_565197 | unclassified | unclassified | unclassified | unclassified | unclassified | unclassified | unclassified | unclassified | unclassified | unclassified | unclassified | unclassified | 0,6 | 0,489897949 | 0,000149957 | 0,000122828 | 0 - | 0 - | 1 - | 0,000245722 - | 0 - | 0 - | 1 - | 0,00023704 - | 1 - | 0,000267025 - |
| 2962 | 1 | 0.2um_NOKCPHJN_1064122 | Eukaryota | Chordata | Mammalia | Rodentia | Muridae | Rattus | Rattus norveg | unclassified | unclassified | unclassified | unclassified | unclassified | 8,8 | 0,4 | 0,000411655 | 4,50E-05 | 9 - | 0,000352557 - | 9 - | 0,000437771 - | 8 - | 0,000369917 - | 9 - | 0,000422304 - | 9 - | 0,000475724 - |
| 2963 | 1 | 0.2um_NOKCPHJN_260113 | Bacteria | Proteobacteri | Alphaproteob | Rhodospirillal | unclassified | unclassified | Rhodospirillal | information st | Translation, ri | 08VZM | 50s ribosomal | 402881.Plav_ | 0,6 | 0,8 | 4,50E-05 | 5,68E-05 | 2 - | 0,000134242 - | 0 - | 0 - | 0 - | 0 - | 0 - | 0 - | 1 - | 9,06E-05 - |
| 2964 | 1 | 3um_FIJJAFBF_779814 | Eukaryota | unclassified | unclassified | Kinetoplastida | Trypanosomat | Trypanosoma | Trypanosoma | poorly charact | Function unkn | 0T9G1 | kinetoplast-as | 353153.XP_80 | 0,6 | 0,489897949 | 5,61E-05 | 4,63E-05 | 1 - | 8,20E-05 - | 1 - | 0,000101799 - | 1 - | 9,68E-05 - | 0 - | 0 - | 0 - | 0 - |
| 2965 | 1 | 0.2um_NOKCPHJN_146552 | Bacteria | Proteobacteri | Betaproteoba | unclassified | unclassified | unclassified | Betaproteoba | unclassified | unclassified | unclassified | unclassified | unclassified | 0,2 | 0,4 | 5,56E-06 | 1,11E-05 | 0 - | 0 - | 1 - | 2,78E-05 - | 0 - | 0 - | 0 - | 0 - | 0 - | 0 - |
| 2966 | 1 | 0.2um_NOKCPHJN_522739 | Bacteria | Proteobacteri | Alphaproteob | Rhodobactera | Rhodobactera | Planktomarina | Planktomarina | information st | Translation, ri | COG0292 | Binds directly | 252305.OB259 | 1,8 | 0,4 | 0,000205798 | 5,12E-05 | 2 - | 0,000191295 - | 2 - | 0,000237531 - | 1 - | 0,000112902 - | 2 - | 0,000229139 - | 2 - | 0,000258124 - |
| 2967 | 1 | 0.2um_NOKCPHJN_435650 | Bacteria | Proteobacteri | Alphaproteob | Rhodobactera | Rhodobactera | Litoreibacter | Litoreibacter j | unclassified | unclassified | unclassified | unclassified | unclassified | 2 | 0,894427191 | 4,52E-05 | 1,87E-05 | 3 - | 5,83E-05 - | 3 - | 7,23E-05 - | 2 - | 4,58E-05 - | 1 - | 2,33E-05 - | 1 - | 2,62E-05 - |
| 2968 | 1 | 3um_FIJJAFBF_305583 | Bacteria | Proteobacteri | Gammaprotec | Alteromonada | Alteromonada | Paraglaciecola | Paraglaciecola | poorly charact | Function unkn | 0QNMQ | NA | 167879.CPS_3 | 2,6 | 2,416609195 | 2,92E-05 | 2,64E-05 | 6 - | 6,00E-05 - | 5 - | 6,21E-05 - | 1 - | 1,18E-05 - | 1 - | 1,20E-05 - | 0 - | 0 - |
| 2969 | 1 | 3um_FIJJAFBF_80729 | Bacteria | Proteobacteri | Gammaprotec | Cellvibrionales | Microbulbifer | Microbulbifer | Microbulbifer | information st | Translation, ri | 0QTWN | Binds directly | 377629.TERTU | 1,2 | 0,4 | 0,000183888 | 4,07E-05 | 2 - | 0,000263855 - | 1 - | 0,000163815 - | 1 - | 0,000155727 - | 1 - | 0,000158027 - | 1 - | 0,000178017 - |
| 2970 | 1 | 3um_FIJJAFBF_294765 | Bacteria | Bacteroidetes | Flavobacteriia | Flavobacterial | Flavobacteriac | Formosa | Formosa sp. H | information st | Replication, re | 05C68 | Can catalyze t | 983544.Lacal_ | 5,4 | 0,489897949 | 0,000220537 | 2,06E-05 | 6 - | 0,000206186 - | 6 - | 0,000256022 - | 5 - | 0,000202818 - | 5 - | 0,000205813 - | 5 - | 0,000231848 - |
| 2971 | 1 | 0.2um_NOKCPHJN_435510 | Bacteria | Bacteroidetes | Flavobacteriia | Flavobacterial | Flavobacteriac | Formosa | Formosa sp. H | unclassified | unclassified | unclassified | unclassified | unclassified | 3,4 | 0,8 | 0,000169818 | 4,18E-05 | 4 - | 0,000168171 - | 2 - | 0,000104409 - | 4 - | 0,000198509 - | 3 - | 0,000151081 - | 4 - | 0,000226922 - |
| 2972 | 1 | 3um_FIJJAFBF_773345 | Bacteria | Bacteroidetes | Flavobacteriia | Flavobacterial | Flavobacteriac | Polaribacter | Polaribacter s | cellular proces | Intracellular tr | 05FGH | MotA TolQ ex | 313594.PI23P | 9,4 | 1,496662955 | 0,000532445 | 5,03E-05 | 12 - | 0,000576285 - | 10 - | 0,000596313 - | 8 - | 0,000453497 - | 9 - | 0,00051772 - | 8 - | 0,000518408 - |
| 2973 | 1 | 3um_FIJJAFBF_543375 | Eukaryota | Bacillariophyt | Bacillariophyc | Naviculales | Naviculaceae | Fistulifera | Fistulifera sola | information st | Replication, re | COG0513 | purine NTP-de | 2850.Phatr25 | 0,8 | 0,4 | 0,000124767 | 6,34E-05 | 1 - | 0,000135031 - | 1 - | 0,000167669 - | 1 - | 0,000159391 - | 1 - | 0,000161745 - | 0 - | 0 - |
| 2974 | 1 | 3um_FIJJAFBF_315982 | Bacteria | Bacteroidetes | Flavobacteriia | Flavobacterial | Flavobacteriac | Flaviramulus | Flaviramulus b | poorly charact | Function unkn | 0P8G1 | peptidylprolyl | 391603.FBALC | 1 | 0,632455532 | 4,23E-05 | 2,74E-05 | 1 - | 3,64E-05 - | 1 - | 4,52E-05 - | 2 - | 8,60E-05 - | 1 - | 4,36E-05 - | 0 - | 0 - |
| 2975 | 1 | 3um_FIJJAFBF_90638 | Bacteria | Proteobacteri | Gammaprotec | Alteromonada | Alteromonada | Paraglaciecola | Paraglaciecola | unclassified | unclassified | unclassified | unclassified | unclassified | 1,2 | 0,4 | 2,66E-05 | 9,82E-06 | 1 - | 1,85E-05 - | 2 - | 4,58E-05 - | 1 - | 2,18E-05 - | 1 - | 2,21E-05 - | 1 - | 2,49E-05 - |
| 2976 | 1 | 0.2um_NOKCPHJN_335418 | Bacteria | Proteobacteri | Alphaproteob | unclassified | unclassified | unclassified | alpha proteob | cellular proces | Posttranslatio | 16QU3 | Hemolysin-typ | 744980.TRICH | 6,2 | 0,748331477 | 7,71E-05 | 7,25E-06 | 7 - | 7,35E-05 - | 7 - | 9,13E-05 - | 6 - | 7,44E-05 - | 6 - | 7,55E-05 - | 5 - | 7,08E-05 - |
| 2977 | 1 | 3um_FIJJAFBF_391577 | Bacteria | Bacteroidetes | Flavobacteriia | Flavobacterial | Flavobacteriac | Polaribacter | various | unclassified | unclassified | unclassified | unclassified | unclassified | 1,2 | 0,979795897 | 9,02E-05 | 7,42E-05 | 2 - | 0,000133461 - | 0 - | 0 - | 2 - | 0,000157537 - | 2 - | 0,000159864 - | 0 - | 0 - |
| 2978 | 1 | 3um_FIJJAFBF_247032 | Bacteria | Proteobacteri | Gammaprotec | unclassified | unclassified | unclassified | Gammaprotec | metabolism | Energy produc | 05CV4 | Subunits I and | 349521.HCH_ | 1,8 | 0,4 | 6,20E-05 | 1,22E-05 | 2 - | 5,86E-05 - | 2 - | 7,27E-05 - | 2 - | 6,91E-05 - | 2 - | 7,01E-05 - | 1 - | 3,95E-05 - |
| 2979 | 1 | 0.2um_NOKCPHJN_49200 | Bacteria | Proteobacteri | Gammaprotec | Cellvibrionales | unclassified | unclassified | Cellvibrionales | cellular proces | Intracellular tr | COG0848 | Biopolymer tr | 314287.GB220 | 0,4 | 0,489897949 | 3,84E-05 | 4,75E-05 | 1 - | 8,57E-05 - | 1 - | 0,000106357 - | 0 - | 0 - | 0 - | 0 - | 0 - | 0 - |
| 2980 | 1 | 3um_FIJJAFBF_19897 | Bacteria | Proteobacteri | Alphaproteob | Rhodobactera | Rhodobactera | Planktomarina | Planktomarina | information st | Translation, ri | 01Q7J | Pseudouridine | 644076.SCH4 | 0,8 | 0,4 | 3,44E-05 | 1,73E-05 | 0 - | 0 - | 1 - | 4,29E-05 - | 1 - | 4,08E-05 - | 1 - | 4,14E-05 - | 1 - | 4,66E-05 - |
| 2981 | 1 | 3um_FIJJAFBF_20039 | Bacteria | Bacteroidetes | Flavobacteriia | Flavobacterial | Flavobacteriac | Polaribacter | Polaribacter s | unclassified | unclassified | unclassified | unclassified | unclassified | 15,6 | 1,356465997 | 0,000215863 | 2,77E-05 | 16 - | 0,000185498 - | 16 - | 0,000230334 - | 17 - | 0,000232646 - | 13 - | 0,000180534 - | 16 - | 0,000250302 - |
| 2982 | 1 | 0.2um_NOKCPHJN_512123 | Bacteria | Bacteroidetes | Flavobacteriia | Flavobacterial | Flavobacteriac | Polaribacter | Polaribacter s | poorly charact | Function unkn | 05DPC | Ragb susd dor | 487797.Flav30 | 5,6 | 1,854723699 | 0,000149978 | 4,80E-05 | 7 - | 0,000158782 - | 8 - | 0,000225326 - | 3 - | 8,03E-05 - | 6 - | 0,000163024 - | 4 - | 0,00012243 - |
| 2983 | 1 | 0.2um_NOKCPHJN_725707 | Bacteria | Bacteroidetes | Flavobacteriia | Flavobacterial | Flavobacteriac | Formosa | Formosa sp. H | unclassified | unclassified | unclassified | unclassified | unclassified | 1 | 0 | 2,25E-05 | 2,14E-06 | 1 - | 1,89E-05 - | 1 - | 2,34E-05 - | 1 - | 2,23E-05 - | 1 - | 2,26E-05 - | 1 - | 2,55E-05 - |
| 2984 | 1 | 3um_FIJJAFBF_641886 | Bacteria | Bacteroidetes | Flavobacteriia | Flavobacterial | Flavobacteriac | Formosa | Formosa sp. H | cellular proces | Cell wall/mem | 11TPU | outer membra | 1046627.BZAR | 0,6 | 0,8 | 5,02E-05 | 7,15E-05 | 1 - | 6,79E-05 - | 0 - | 0 - | 0 - | 0 - | 0 - | 0 - | 2 - | 0,000183283 - |
| 2985 | 2 | 0.2um_NOKCPHJN_282908;0.2um_NOKCPHJN_7 | Bacteria | Proteobacteri | Alphaproteob | Rhodobactera | Rhodobactera | Planktomarina | Planktomarina | unclassified | unclassified | unclassified | unclassified | unclassified | 0,4 | 0,489897949 | 4,63E-05 | 5,67E-05 | 0 - | 0 - | 1 - | 0,000117784 - | 0 - | 0 - | 1 - | 0,000113623 - | 0 - | 0 - |
| 2986 | 1 | 3um_FIJJAFBF_490374 | Bacteria | various | various | various | various | various | various | information st | Translation, ri | 0PBIW | 50S ribosomal | 313594.PI23P | 1 | 0,632455532 | 0,00020867 | 0,0001367 | 1 - | 0,000179339 - | 1 - | 0,000222686 - | 1 - | 0,000211691 - | 2 - | 0,000429635 - | 0 - | 0 - |
| 2987 | 2 | 0.2um_NOKCPHJN_239800;0.2um_NOKCPHJN_6 | Bacteria | Proteobacteri | Alphaproteob | Rhodobactera | Planktomarina | Planktomarina | Planktomarina | metabolism | Energy produc | 16RMF | aldo keto redu | 314232.SKA53 | 0,4 | 0,489897949 | 2,02E-05 | 2,48E-05 | 0 - | 0 - | 1 - | 4,85E-05 - | 0 - | 0 - | 0 - | 0 - | 1 - | 5,27E-05 - |
| 2988 | 1 | 0.2um_NOKCPHJN_968394 | Eukaryota | unclassified | Cryptophyta | Pyrenomonad | various | various | various | unclassified | unclassified | unclassified | unclassified | unclassified | 2 | 0 | 0,000330186 | 3,14E-05 | 2 - | 0,000276571 - | 2 - | 0,000343419 - | 2 - | 0,000326463 - | 2 - | 0,000331285 - | 2 - | 0,000373192 - |
| 2989 | 1 | 0.2um_NOKCPHJN_746047 | Bacteria | Bacteroidetes | Flavobacteriia | Flavobacterial | Flavobacteriac | Polaribacter | Polaribacter s | unclassified | unclassified | unclassified | unclassified | unclassified | 9,8 | 1,166190379 | 0,000244465 | 3,80E-05 | 11 - | 0,000229972 - | 10 - | 0,000259597 - | 9 - | 0,000222102 - | 8 - | 0,00020034 - | 11 - | 0,000310313 - |
| 2990 | 1 | 3um_FIJJAFBF_222295 | Bacteria | Proteobacteri | Alphaproteob | Rhodobactera | Rhodobactera | unclassified | Rhodobactera | unclassified | unclassified | unclassified | unclassified | unclassified | 3,4 | 0,489897949 | 7,27E-05 | 7,72E-06 | 4 - | 7,23E-05 - | 3 - | 6,73E-05 - | 3 - | 6,40E-05 - | 4 - | 8,66E-05 - | 3 - | 7,32E-05 - |
| 2991 | 1 | 3um_FIJJAFBF_19833 | Bacteria | Bacteroidetes | Flavobacteriia | Flavobacterial | Flavobacteriac | Polaribacter | Polaribacter v | metabolism | Energy produc | 09A0I | reductase | 313598.MED1 | 3,2 | 1,166190379 | 6,16E-05 | 1,83E-05 | 5 - | 8,27E-05 - | 4 - | 8,21E-05 - | 2 - | 3,90E-05 - | 3 - | 5,94E-05 - | 2 - | 4,46E-05 - |
| 2992 | 1 | 3um_FIJJAFBF_775490 | Bacteria | Bacteroidetes | Flavobacteriia | Flavobacterial | Flavobacteriac | Mangrovimon | Mangrovimon | unclassified | unclassified | unclassified | unclassified | unclassified | 1,2 | 0,4 | 4,06E-05 | 1,36E-05 | 1 - | 2,84E-05 - | 1 - | 3,53E-05 - | 2 - | 6,71E-05 - | 1 - | 3,40E-05 - | 1 - | 3,83E-05 - |
| 2993 | 1 | 3um_FIJJAFBF_92593 | Eukaryota | unclassified | Chrysophycea | Chromulinales | Chrysocapsac | Chrysocapsa | Chrysocapsa v | metabolism | Carbohydrate | COG1850 | RuBisCO catal | 45157.CMV01 | 2,2 | 0,979795897 | 6,15E-05 | 2,90E-05 | 2 - | 4,70E-05 - | 4 - | 0,000116819 - | 2 - | 5,55E-05 - | 2 - | 5,63E-05 - | 1 - | 3,17E-05 - |
| 2994 | 1 | 0.2um_NOKCPHJN_306769 | Bacteria | Proteobacteri | Alphaproteob | Rhodobactera | Rhodobactera | Planktotalea | Planktotalea a | unclassified | unclassified | unclassified | unclassified | unclassified | 0,4 | 0,8 | 9,29E-06 | 1,86E-05 | 2 - | 4,65E-05 - | 0 - | 0 - | 0 - | 0 - | 0 - | 0 - | 0 - | 0 - |
| 2995 | 1 | 3um_FIJJAFBF_532815 | Bacteria | Proteobacteri | Alphaproteob | Rhodobactera | unclassified | unclassified | Rhodobactera | unclassified | unclassified | unclassified | unclassified | unclassified | 2,6 | 0,8 | 0,000297837 | 6,77E-05 | 4 - | 0,000392399 - | 3 - | 0,000365433 - | 2 - | 0,000231594 - | 2 - | 0,000235014 - | 2 - | 0,000264743 - |
| 2996 | 1 | 3um_FIJJAFBF_194912 | Bacteria | Bacteroidetes | Flavobacteriia | Flavobacterial | Flavobacteriac | Xanthomarina | Xanthomarina | metabolism | Coenzyme tra | 05VQ8 | uroporphyrinc | 391603.FBALC | 1 | 0 | 5,53E-05 | 5,25E-06 | 1 - | 4,63E-05 - | 1 - | 5,75E-05 - | 1 - | 5,46E-05 - | 1 - | 5,54E-05 - | 1 - | 6,24E-05 - |
| 2997 | 1 | 0.2um_NOKCPHJN_425101 | Bacteria | Proteobacteri | Gammaprotec | unclassified | unclassified | unclassified | Gammaprotec | unclassified | unclassified | unclassified | unclassified | unclassified | 4,8 | 0,748331477 | 0,000427123 | 6,03E-05 | 6 - | 0,000450105 - | 5 - | 0,000465748 - | 4 - | 0,000354202 - | 4 - | 0,000359433 - | 5 - | 0,000506126 - |

| | | | | | | | | | | | | | | | | | | | | | | | | | | | | |
|---|---|---|---|---|---|---|---|---|---|---|---|---|---|---|---|---|---|---|---|---|---|---|---|---|---|---|---|---|
| 2998 | 1 | 0.2um_NOKCPHJN_139202 | Bacteria | Bacteroidetes | Flavobacteriia | Flavobacterial | Flavobacteriac | Polaribacter | Polaribacter ir | unclassified | unclassified | unclassified | unclassified | unclassified | 1,8 | 0,4 | 0,000271382 | 6,76E-05 | 2 - | 0,000252257 - | 2 - | 0,000313228 - | 1 - | 0,000148882 - | 2 - | 0,000302161 - | 2 - | 0,000340383 - |
| 2999 | 1 | 3um_FIJJAFBF_04342 | Bacteria | Proteobacteri | Gammaprote | Alteromonada | Alteromonada | Paraglaciecola | Paraglaciecola | metabolism | Inorganic ion t | 0QK8T | TonB-depende | 342610.Patl_0 | 9,8 | 1,326649916 | 0,000153511 | 1,88E-05 | 10 - | 0,000132079 - | 9 - | 0,000147603 - | 12 - | 0,000187087 - | 10 - | 0,000158209 - | 8 - | 0,000142577 - |
| 3000 | 2 | 0.2um_NOKCPHJN_1099008;0.2um_NOKCPHJN_ | Bacteria | Proteobacteri | Gammaprote | unclassified | unclassified | unclassified | SAR86 cluster | unclassified | unclassified | unclassified | unclassified | unclassified | 0,6 | 0,489897949 | 4,53E-05 | 3,73E-05 | 1 - | 6,71E-05 - | 0 - | 0 - | 1 - | 7,92E-05 - | 1 - | 8,04E-05 - | 0 - | 0 - |
| 3001 | 1 | 0.2um_NOKCPHJN_184137 | Bacteria | Bacteroidetes | Flavobacteriia | Flavobacterial | Flavobacteriac | Algibacter | Algibacter pec | unclassified | unclassified | unclassified | unclassified | unclassified | 0,4 | 0,8 | 5,24E-06 | 1,05E-05 | 2 - | 2,62E-05 - | 0 - | 0 - | 0 - | 0 - | 0 - | 0 - | 0 - | 0 - |
| 3002 | 1 | 0.2um_NOKCPHJN_679195 | Bacteria | Proteobacteri | Alphaproteob | Pelagibacteral | Pelagibacterac | Candidatus Pe | Candidatus Pe | poorly charact | Function unkn | 07QSV | RmuC domain | 335992.SAR11 | 0,2 | 0,4 | 7,44E-06 | 1,49E-05 | 0 - | 0 - | 0 - | 0 - | 1 - | 3,72E-05 - | 0 - | 0 - | 0 - | 0 - |
| 3003 | 1 | 3um_FIJJAFBF_120182 | Bacteria | unclassified | unclassified | unclassified | unclassified | unclassified | uncultured ba | unclassified | unclassified | unclassified | unclassified | unclassified | 0,8 | 0,4 | 4,31E-05 | 2,22E-05 | 1 - | 4,52E-05 - | 1 - | 5,61E-05 - | 1 - | 5,33E-05 - | 0 - | 0 - | 1 - | 6,10E-05 - |
| 3004 | 1 | 3um_FIJJAFBF_773343 | Bacteria | Bacteroidetes | Flavobacteriia | Flavobacterial | Flavobacteriac | Polaribacter | Polaribacter s | unclassified | unclassified | unclassified | unclassified | unclassified | 0,8 | 0,4 | 5,10E-05 | 2,59E-05 | 1 - | 5,52E-05 - | 1 - | 6,85E-05 - | 1 - | 6,51E-05 - | 1 - | 6,61E-05 - | 0 - | 0 - |
| 3005 | 1 | 3um_FIJJAFBF_543824 | Bacteria | Proteobacteri | Alphaproteob | Rhodobactera | Rhodobactera | various | various | poorly charact | Function unkn | 11QQ4 | NA | 622637.Met4 | 0,4 | 0,489897949 | 2,73E-05 | 3,36E-05 | 1 - | 6,20E-05 - | 0 - | 0 - | 0 - | 0 - | 1 - | 7,43E-05 - | 0 - | 0 - |
| 3006 | 1 | 3um_FIJJAFBF_360756 | unclassified | unclassified | unclassified | unclassified | unclassified | unclassified | unclassified | unclassified | unclassified | unclassified | unclassified | unclassified | 0,8 | 0,4 | 0,000174493 | 8,96E-05 | 1 - | 0,000182185 - | 1 - | 0,00022622 - | 0 - | 0 - | 1 - | 0,000218227 - | 1 - | 0,000245833 - |
| 3007 | 1 | 3um_FIJJAFBF_138237 | Viruses | unclassified | unclassified | unclassified | unclassified | unclassified | uncultured vir | unclassified | unclassified | unclassified | unclassified | unclassified | 0,2 | 0,4 | 1,04E-06 | 2,08E-06 | 1 - | 5,20E-06 - | 0 - | 0 - | 0 - | 0 - | 0 - | 0 - | 0 - | 0 - |
| 3008 | 1 | 3um_FIJJAFBF_599965 | Bacteria | Proteobacteri | Gammaprote | Oceanospirilla | Saccharospirill | Reinekea | Reinekea fors | metabolism | Energy produc | 16R80 | pyrophosphat | 314283.MED2 | 1,4 | 0,489897949 | 0,000109118 | 3,95E-05 | 2 - | 0,000131173 - | 1 - | 8,14E-05 - | 1 - | 7,74E-05 - | 1 - | 7,86E-05 - | 2 - | 0,000176999 - |
| 3009 | 1 | 0.2um_NOKCPHJN_789774 | unclassified | unclassified | unclassified | unclassified | unclassified | unclassified | unclassified | unclassified | unclassified | unclassified | unclassified | unclassified | 1 | 0 | 2,97E-05 | 2,82E-06 | 1 - | 2,48E-05 - | 1 - | 3,08E-05 - | 1 - | 2,93E-05 - | 1 - | 2,98E-05 - | 1 - | 3,35E-05 - |
| 3010 | 1 | 0.2um_NOKCPHJN_884241 | Bacteria | Proteobacteri | Alphaproteob | Rhodobactera | Rhodobactera | Planktomarina | Planktomarina | metabolism | Coenzyme tra | 05C66 | Catalyzes the | 391593.RCCS2 | 1,4 | 0,489897949 | 4,99E-05 | 1,57E-05 | 2 - | 6,07E-05 - | 2 - | 7,54E-05 - | 1 - | 3,58E-05 - | 1 - | 3,64E-05 - | 1 - | 4,10E-05 - |
| 3011 | 1 | 0.2um_NOKCPHJN_317189 | Eukaryota | Chordata | Actinopteri | unclassified | Pomacentrida | Acanthochron | Acanthochron | unclassified | unclassified | unclassified | unclassified | unclassified | 0,2 | 0,4 | 1,28E-06 | 2,55E-06 | 0 - | 0 - | 0 - | 0 - | 0 - | 0 - | 0 - | 0 - | 1 - | 6,38E-06 - |
| 3012 | 1 | 0.2um_NOKCPHJN_698633 | Bacteria | unclassified | unclassified | unclassified | unclassified | unclassified | uncultured ba | unclassified | unclassified | unclassified | unclassified | unclassified | 0,8 | 0,748331477 | 2,34E-05 | 2,24E-05 | 1 - | 2,54E-05 - | 1 - | 3,16E-05 - | 2 - | 6,01E-05 - | 0 - | 0 - | 0 - | 0 - |
| 3013 | 1 | 3um_FIJJAFBF_132244 | Bacteria | Bacteroidetes | Flavobacteriia | Flavobacterial | Flavobacteriac | Polaribacter | Polaribacter s | unclassified | unclassified | unclassified | unclassified | unclassified | 2,2 | 1,326649916 | 3,25E-05 | 2,03E-05 | 2 - | 2,45E-05 - | 2 - | 3,04E-05 - | 4 - | 5,78E-05 - | 0 - | 0 - | 3 - | 4,96E-05 - |
| 3014 | 1 | 0.2um_NOKCPHJN_587110 | Eukaryota | Basidiomycota | Pucciniomyce | Pucciniales | Pucciniaceae | Puccinia | Puccinia coror | unclassified | unclassified | unclassified | unclassified | unclassified | 0,8 | 0,4 | 4,57E-05 | 2,32E-05 | 1 - | 4,95E-05 - | 1 - | 6,14E-05 - | 1 - | 5,84E-05 - | 1 - | 5,93E-05 - | 0 - | 0 - |
| 3015 | 1 | 3um_FIJJAFBF_616741 | Bacteria | Bacteroidetes | Flavobacteriia | Flavobacterial | Flavobacteriac | Spongiibacteri | Spongiibacteri | information st | Replication, re | 05CZ8 | DNA polymera | 313603.FB217 | 0,6 | 0,489897949 | 2,18E-05 | 1,81E-05 | 1 - | 3,09E-05 - | 0 - | 0 - | 1 - | 3,64E-05 - | 0 - | 0 - | 1 - | 4,16E-05 - |
| 3016 | 1 | 3um_FIJJAFBF_651819 | Bacteria | Bacteroidetes | Flavobacteriia | Flavobacterial | Flavobacteriac | Polaribacter | Polaribacter s | metabolism | Energy produc | COG0712 | F(1)F(0) ATP s | 313594.PI23P | 0,4 | 0,8 | 2,93E-05 | 5,86E-05 | 0 - | 0 - | 0 - | 0 - | 2 - | 0,000146467 - | 0 - | 0 - | 0 - | 0 - |
| 3017 | 1 | 0.2um_NOKCPHJN_247021 | Bacteria | Proteobacteri | Alphaproteob | Rhodobactera | Rhodobactera | Planktotalea | Planktotalea f | unclassified | unclassified | unclassified | unclassified | unclassified | 0,2 | 0,4 | 4,68E-06 | 9,37E-06 | 0 - | 0 - | 0 - | 0 - | 0 - | 0 - | 1 - | 2,34E-05 - | 0 - | 0 - |
| 3018 | 1 | 3um_FIJJAFBF_167847 | Bacteria | Proteobacteri | Gammaprote | Alteromonada | Alteromonada | Paraglaciecola | Paraglaciecola | cellular proces | Intracellular tr | 08ZUW | Major pilin sul | 455436.GHTC | 1 | 0,894427191 | 9,11E-05 | 8,02E-05 | 2 - | 0,000161657 - | 1 - | 0,000100365 - | 0 - | 0 - | 2 - | 0,000193638 - | 0 - | 0 - |
| 3019 | 1 | 0.2um_NOKCPHJN_483965 | Bacteria | Proteobacteri | Alphaproteob | Pelagibacteral | Pelagibacterac | Candidatus Pe | Candidatus Pe | cellular proces | Cell wall/mem | 7257 | NA | 335992.SAR11 | 0,8 | 0,4 | 3,39E-05 | 1,74E-05 | 1 - | 3,54E-05 - | 1 - | 4,40E-05 - | 0 - | 0 - | 1 - | 4,24E-05 - | 1 - | 4,78E-05 - |
| 3020 | 1 | 3um_FIJJAFBF_628028 | Bacteria | Proteobacteri | Alphaproteob | Rhodobactera | Rhodobactera | Thalassobius | Thalassobius s | information st | Translation, ri | 05CEJ | Catalyzes the | 391593.RCCS2 | 13,2 | 0,979795897 | 0,000257798 | 3,92E-05 | 12 - | 0,000195365 - | 14 - | 0,000283016 - | 14 - | 0,000269043 - | 12 - | 0,000234014 - | 14 - | 0,000307552 - |
| 3021 | 1 | 3um_FIJJAFBF_47498 | Bacteria | Bacteroidetes | unclassified | unclassified | unclassified | unclassified | unidentified e | information st | Replication, re | 09GNB | DNA-binding | 50743.SCB49_ | 15,2 | 2,039607805 | 0,002286869 | 0,000110494 | 19 - | 0,002423065 - | 15 - | 0,002375314 - | 14 - | 0,002107501 - | 15 - | 0,002291388 - | 13 - | 0,002237076 - |
| 3022 | 1 | 0.2um_NOKCPHJN_57013 | Bacteria | Bacteroidetes | Flavobacteriia | Flavobacterial | Flavobacteriac | Algibacter | Algibacter pec | unclassified | unclassified | unclassified | unclassified | unclassified | 0,8 | 0,748331477 | 6,52E-05 | 6,56E-05 | 1 - | 6,60E-05 - | 1 - | 8,19E-05 - | 0 - | 0 - | 0 - | 0 - | 2 - | 0,000178017 - |
| 3023 | 1 | 0.2um_NOKCPHJN_243992 | Eukaryota | unclassified | Labyrinthulon | unclassified | Thraustochytr | Aurantiochytr | Aurantiochytr | unclassified | unclassified | unclassified | unclassified | unclassified | 0,4 | 0,489897949 | 5,66E-06 | 6,93E-06 | 0 - | 0 - | 0 - | 0 - | 1 - | 1,40E-05 - | 1 - | 1,42E-05 - | 0 - | 0 - |
| 3024 | 1 | 3um_FIJJAFBF_345764 | Bacteria | Bacteroidetes | Flavobacteriia | Flavobacterial | Flavobacteriac | unclassified | Flavobacteriac | unclassified | unclassified | unclassified | unclassified | unclassified | 1,2 | 0,4 | 7,20E-05 | 2,41E-05 | 1 - | 5,03E-05 - | 1 - | 6,25E-05 - | 2 - | 0,000118844 - | 1 - | 6,03E-05 - | 1 - | 6,79E-05 - |
| 3025 | 1 | 3um_FIJJAFBF_14571 | Bacteria | Proteobacteri | Gammaprote | Alteromonada | Pseudoalteror | Pseudoalteror | Pseudoalteror | cellular proces | Posttranslatio | 07RUK | Involved in iro | 342610.Patl_4 | 0,2 | 0,4 | 1,48E-05 | 2,97E-05 | 0 - | 0 - | 1 - | 7,42E-05 - | 0 - | 0 - | 0 - | 0 - | 0 - | 0 - |
| 3026 | 1 | 0.2um_NOKCPHJN_328413 | Bacteria | Proteobacteri | Alphaproteob | Pelagibacteral | Pelagibacterac | unclassified | Pelagibacterac | unclassified | unclassified | unclassified | unclassified | unclassified | 0,8 | 0,4 | 3,86E-05 | 1,96E-05 | 1 - | 4,17E-05 - | 1 - | 5,18E-05 - | 1 - | 4,93E-05 - | 1 - | 5,00E-05 - | 0 - | 0 - |
| 3027 | 1 | 3um_FIJJAFBF_337496 | Bacteria | Proteobacteri | Gammaprote | Oceanospirilla | Saccharospirill | Reinekea | Reinekea blan | metabolism | Amino acid tra | COG0137 | Citrulline--asp | 314283.MED2 | 2,8 | 0,4 | 9,46E-05 | 1,69E-05 | 3 - | 8,48E-05 - | 3 - | 0,00010531 - | 2 - | 6,67E-05 - | 3 - | 0,000101589 - | 3 - | 0,000114439 - |
| 3028 | 1 | 3um_FIJJAFBF_491680 | Bacteria | Proteobacteri | Gammaprote | Cellvibrionales | Porticoccacea | unclassified | SAR92 bacteri | information st | Translation, ri | COG0256 | This is one of | 314287.GB22 | 1,6 | 0,8 | 0,000182277 | 7,47E-05 | 3 - | 0,000296836 - | 2 - | 0,000245722 - | 1 - | 0,000116795 - | 1 - | 0,00011852 - | 1 - | 0,000133512 - |
| 3029 | 1 | 0.2um_NOKCPHJN_168883 | Bacteria | Bacteroidetes | Flavobacteriia | Flavobacterial | Flavobacteriac | Gilvibacter | Gilvibacter sp. | cellular proces | Posttranslatio | 05C2Z | ATP-depender | 50743.SCB49_ | 0,8 | 0,748331477 | 1,23E-05 | 1,20E-05 | 1 - | 1,35E-05 - | 0 - | 0 - | 1 - | 1,59E-05 - | 2 - | 3,23E-05 - | 0 - | 0 - |
| 3030 | 1 | 0.2um_NOKCPHJN_245216 | Bacteria | Bacteroidetes | Flavobacteriia | Flavobacterial | Flavobacteriac | Formosa | Formosa sp. H | cellular proces | Posttranslatio | 07SFW | Trigger factor | 391603.FBALC | 0,6 | 0,489897949 | 1,79E-05 | 1,48E-05 | 1 - | 2,60E-05 - | 1 - | 3,23E-05 - | 0 - | 0 - | 1 - | 3,12E-05 - | 0 - | 0 - |
| 3031 | 1 | 3um_FIJJAFBF_16488 | Bacteria | Proteobacteri | Gammaprote | various | various | various | various | information st | Translation, ri | 05C64 | Binds directly | 715451.ambt_ | 2,6 | 0,489897949 | 0,00015501 | 3,78E-05 | 2 - | 9,89E-05 - | 2 - | 0,000122861 - | 3 - | 0,000175193 - | 3 - | 0,00017778 - | 3 - | 0,000200269 - |
| 3032 | 1 | 0.2um_NOKCPHJN_655411 | Bacteria | Proteobacteri | Alphaproteob | Pelagibacteral | Pelagibacterac | Candidatus Pe | Candidatus Pe | cellular proces | Cell wall/mem | 7257 | NA | 335992.SAR11 | 3,8 | 0,4 | 0,000158362 | 2,35E-05 | 4 - | 0,000139546 - | 4 - | 0,000173275 - | 3 - | 0,00012354 - | 4 - | 0,000167153 - | 4 - | 0,000188297 - |
| 3033 | 1 | 3um_FIJJAFBF_416829 | Bacteria | Proteobacteri | Gammaprote | Oceanospirilla | Saccharospirill | Reinekea | Reinekea blan | metabolism | Energy produc | 0QJ3X | Aconitate hyd | 314283.MED2 | 8,2 | 1,326649916 | 0,000128767 | 1,32E-05 | 10 - | 0,000133306 - | 9 - | 0,000148974 - | 8 - | 0,000125884 - | 8 - | 0,000127743 - | 6 - | 0,000107926 - |
| 3034 | 1 | 0.2um_NOKCPHJN_640395 | Bacteria | Proteobacteri | Betaproteoba | Burkholderiale | Comamonada | Limnohabitan | Limnohabitan | poorly charact | Function unkn | 0ZK5F | NA | 240292.Ava_4 | 0,8 | 0,4 | 1,15E-05 | 5,91E-06 | 1 - | 1,20E-05 - | 1 - | 1,49E-05 - | 0 - | 0 - | 1 - | 1,44E-05 - | 1 - | 1,62E-05 - |
| 3035 | 1 | 3um_FIJJAFBF_52924 | Bacteria | Proteobacteri | Betaproteoba | unclassified | unclassified | unclassified | Betaproteoba | metabolism | Lipid transpor | 16PEQ | Amp-depende | 296591.Bpro_ | 0,4 | 0,8 | 1,02E-05 | 2,04E-05 | 0 - | 0 - | 2 - | 5,09E-05 - | 0 - | 0 - | 0 - | 0 - | 0 - | 0 - |
| 3036 | 1 | 0.2um_NOKCPHJN_900395 | Bacteria | Proteobacteri | Gammaprote | Thiotrichales | Piscirickettsiac | Cycloclasticus | Cycloclasticus | cellular proces | Posttranslatio | 16QU3 | Hemolysin-typ | 744980.TRICH | 0,4 | 0,489897949 | 7,51E-06 | 9,29E-06 | 1 - | 1,68E-05 - | 1 - | 2,08E-05 - | 0 - | 0 - | 0 - | 0 - | 0 - | 0 - |
| 3037 | 1 | 0.2um_NOKCPHJN_43219 | Bacteria | Bacteroidetes | Flavobacteriia | Flavobacterial | Flavobacteriac | Algibacter | Algibacter sp. | poorly charact | Function unkn | 05WQR | glycoside hydr | 313598.MED1 | 0,2 | 0,4 | 1,96E-05 | 3,92E-05 | 1 - | 9,81E-05 - | 0 - | 0 - | 0 - | 0 - | 0 - | 0 - | 0 - | 0 - |
| 3038 | 1 | 3um_FIJJAFBF_420115 | Bacteria | Proteobacteri | Gammaprote | Cellvibrionales | Halieaceae | unclassified | marine gamm | unclassified | unclassified | unclassified | unclassified | unclassified | 1,6 | 0,8 | 2,51E-05 | 1,03E-05 | 3 - | 4,09E-05 - | 2 - | 3,39E-05 - | 1 - | 1,61E-05 - | 1 - | 1,63E-05 - | 1 - | 1,84E-05 - |
| 3039 | 1 | 0.2um_NOKCPHJN_249844 | Eukaryota | unclassified | Pelagophycea | Pelagomonad | unclassified | Aureococcus | Aureococcus a | poorly charact | Function unkn | 0XT7J | HYDIN, axone | 67593.Physo1 | 1 | 0 | 2,99E-06 | 2,83E-07 | 1 - | 2,50E-06 - | 1 - | 3,10E-06 - | 1 - | 2,95E-06 - | 1 - | 3,00E-06 - | 1 - | 3,37E-06 - |
| 3040 | 1 | 0.2um_NOKCPHJN_74285 | Bacteria | Bacteroidetes | Flavobacteriia | Flavobacterial | Flavobacteriac | Polaribacter | Polaribacter s | unclassified | unclassified | unclassified | unclassified | unclassified | 0,6 | 0,489897949 | 6,96E-05 | 5,71E-05 | 0 - | 0 - | 0 - | 0 - | 1 - | 0,000110148 - | 1 - | 0,000111775 - | 1 - | 0,000125914 - |
| 3041 | 1 | 0.2um_NOKCPHJN_33511 | Bacteria | unclassified | unclassified | unclassified | unclassified | unclassified | bacterium B17 | unclassified | unclassified | unclassified | unclassified | unclassified | 1 | 0 | 2,51E-05 | 2,38E-06 | 1 - | 2,10E-05 - | 1 - | 2,61E-05 - | 1 - | 2,48E-05 - | 1 - | 2,52E-05 - | 1 - | 2,84E-05 - |
| 3042 | 1 | 3um_FIJJAFBF_79207 | Bacteria | Proteobacteri | Alphaproteob | Rhodobactera | unclassified | unclassified | Rhodobactera | information st | Translation, ri | 17C3B | The globular d | 367336.OM22 | 4,8 | 1,720465053 | 0,000493063 | 0,000121329 | 8 - | 0,000711794 - | 4 - | 0,000441919 - | 5 - | 0,000525125 - | 4 - | 0,000426305 - | 3 - | 0,000360173 - |
| 3043 | 4 | 0.2um_NOKCPHJN_22695;0.2um_NOKCPHJN_84 | Bacteria | various | various | various | various | various | various | metabolism | Lipid transpor | 05VNB | Carrier of the | various | 0,8 | 0,748331477 | 0,000124953 | 0,0001226 | 1 - | 0,000133461 - | 2 - | 0,000331439 - | 0 - | 0 - | 1 - | 0,000159864 - | 0 - | 0 - |
| 3044 | 1 | 3um_FIJJAFBF_216671 | Bacteria | Proteobacteri | Gammaprote | Cellvibrionales | Porticoccacea | unclassified | Gammaprote | metabolism | Energy produc | 17CGT | F(1)F(0) ATP s | 314287.GB22 | 7,4 | 1,019803903 | 0,000645197 | 7,25E-05 | 9 - | 0,000662174 - | 8 - | 0,000730866 - | 6 - | 0,000521085 - | 7 - | 0,000616912 - | 7 - | 0,00069495 - |
| 3045 | 1 | 0.2um_NOKCPHJN_530413 | Bacteria | Proteobacteri | Gammaprote | unclassified | unclassified | unclassified | Gammaprote | information st | Translation, ri | COG0051 | tRNA binding | 876044.IMCC3 | 1,4 | 0,489897949 | 0,000140661 | 4,42E-05 | 2 - | 0,000171309 - | 2 - | 0,000212715 - | 1 - | 0,000101106 - | 1 - | 0,000102599 - | 1 - | 0,000115578 - |
| 3046 | 1 | 3um_FIJJAFBF_122415 | Bacteria | Bacteroidetes | Flavobacteriia | Flavobacterial | Flavobacteriac | Polaribacter | Polaribacter h | metabolism | Coenzyme tra | 05C66 | Catalyzes the | 313594.PI23P | 2 | 0,894427191 | 7,06E-05 | 3,11E-05 | 2 - | 6,03E-05 - | 1 - | 3,74E-05 - | 3 - | 0,000106679 - | 3 - | 0,000108255 - | 1 - | 4,06E-05 - |
| 3047 | 1 | 3um_FIJJAFBF_580111 | Bacteria | Bacteroidetes | Saprospiria | Saprospirales | Saprospiracea | Aureispira | Aureispira sp. | unclassified | unclassified | unclassified | unclassified | unclassified | 0,4 | 0,8 | 2,50E-05 | 4,99E-05 | 2 - | 0,000124757 - | 0 - | 0 - | 0 - | 0 - | 0 - | 0 - | 0 - | 0 - |
| 3048 | 1 | 0.2um_NOKCPHJN_37737 | Bacteria | Bacteroidetes | Flavobacteriia | Flavobacterial | unclassified | unclassified | Flavobacterial | poorly charact | Function unkn | 0XSVM | NA | 313603.FB217 | 1 | 0 | 2,61E-05 | 2,47E-06 | 1 - | 2,18E-05 - | 1 - | 2,71E-05 - | 1 - | 2,58E-05 - | 1 - | 2,61E-05 - | 1 - | 2,94E-05 - |
| 3049 | 1 | 3um_FIJJAFBF_244869 | Bacteria | Bacteroidetes | Flavobacteriia | Flavobacterial | Flavobacteriac | Oceanihabitar | Oceanihabitar | cellular proces | Signal transdu | 09A4D | gtp-binding pr | 1046627.BZAF | 3,6 | 1,019803903 | 8,11E-05 | 1,91E-05 | 5 - | 9,58E-05 - | 4 - | 9,52E-05 - | 2 - | 4,52E-05 - | 4 - | 9,18E-05 - | 3 - | 7,76E-05 - |
| 3050 | 1 | 0.2um_NOKCPHJN_80602 | Bacteria | Bacteroidetes | Flavobacteriia | Flavobacterial | Flavobacteriac | Polaribacter | Polaribacter ir | unclassified | unclassified | unclassified | unclassified | unclassified | 1 | 0 | 5,00E-05 | 4,75E-06 | 1 - | 4,19E-05 - | 1 - | 5,20E-05 - | 1 - | 4,94E-05 - | 1 - | 5,02E-05 - | 1 - | 5,65E-05 - |
| 3051 | 1 | 3um_FIJJAFBF_337909 | Bacteria | Bacteroidetes | Flavobacteriia | Flavobacterial | Flavobacteriac | Polaribacter | Polaribacter s | unclassified | unclassified | unclassified | unclassified | unclassified | 7,8 | 0,748331477 | 0,000149886 | 5,43E-06 | 9 - | 0,000146109 - | 7 - | 0,000141108 - | 8 - | 0,000153304 - | 8 - | 0,000155568 - | 7 - | 0,000153341 - |
| 3052 | 1 | 3um_FIJJAFBF_513823 | Viruses | unclassified | unclassified | unclassified | unclassified | unclassified | uncultured M | poorly charact | Function unkn | 05BE3 | Phage portal | 742767.HMPR | 0,4 | 0,489897949 | 8,31E-06 | 1,02E-05 | 1 - | 1,89E-05 - | 0 - | 0 - | 0 - | 0 - | 1 - | 2,26E-05 - | 0 - | 0 - |
| 3053 | 1 | 0.2um_NOKCPHJN_101791 | Bacteria | Bacteroidetes | Flavobacteriia | Flavobacterial | Flavobacteriac | Polaribacter | Polaribacter s | unclassified | unclassified | unclassified | unclassified | unclassified | 2,4 | 1,854723699 | 0,000122552 | 7,96E-05 | 6 - | 0,000271126 - | 2 - | 0,00011222 - | 1 - | 5,33E-05 - | 1 - | 5,41E-05 - | 2 - | 0,000121948 - |
| 3054 | 2 | 0.2um_NOKCPHJN_245985;3um_FIJJAFBF_1488 | Bacteria | Proteobacteri | Alphaproteob | Rhodobactera | Rhodobactera | Planktomarina | Planktomarina | unclassified | unclassified | unclassified | unclassified | unclassified | 1,4 | 1,019803903 | 8,15E-05 | 5,27E-05 | 3 - | 0,000153036 - | 1 - | 6,33E-05 - | 0 - | 0 - | 2 - | 0,000122207 - | 1 - | 6,88E-05 - |
| 3055 | 1 | 0.2um_NOKCPHJN_483058 | unclassified | unclassified | unclassified | unclassified | unclassified | unclassified | unclassified | unclassified | unclassified | unclassified | unclassified | unclassified | 0,2 | 0,4 | 6,31E-06 | 1,26E-05 | 0 - | 0 - | 0 - | 0 - | 0 - | 0 - | 1 - | 3,15E-05 - | 0 - | 0 - |
| 3056 | 1 | 0.2um_NOKCPHJN_427680 | Bacteria | Bacteroidetes | unclassified | unclassified | unclassified | unclassified | Bacteroidetes | unclassified | unclassified | unclassified | unclassified | unclassified | 0,2 | 0,4 | 1,09E-05 | 2,17E-05 | 0 - | 0 - | 0 - | 0 - | 0 - | 0 - | 0 - | 0 - | 1 - | 5,43E-05 - |
| 3057 | 1 | 0.2um_NOKCPHJN_172847 | Bacteria | Proteobacteri | Gammaprote | Oceanospirilla | Saccharospirill | Reinekea | Reinekea blan | unclassified | unclassified | unclassified | unclassified | unclassified | 3,4 | 0,8 | 0,000107506 | 2,23E-05 | 4 - | 0,000107268 - | 2 - | 6,66E-05 - | 4 - | 0,000126619 - | 4 - | 0,000128489 - | 3 - | 0,000108557 - |
| 3058 | 2 | 0.2um_NOKCPHJN_559204;3um_FIJJAFBF_47567 | Bacteria | Bacteroidetes | unclassified | unclassified | unclassified | unclassified | Bacteroidetes | information st | Translation, ri | 09FEM | Binds to the 2 | various | 0,4 | 0,489897949 | 3,60E-05 | 4,48E-05 | 1 - | 7,65E-05 - | 0 - | 0 - | 0 - | 0 - | 0 - | 0 - | 1 - | 0,00010325 - |
| 3059 | 1 | 3um_FIJJAFBF_135237 | Bacteria | Cyanobacteria | unclassified | Synechococca | Synechococca | Synechococcu | Synechococcu | metabolism | Energy produc | COG1290 | Component of | 70448.Q0P3F3 | 1 | 0 | 3,59E-05 | 3,41E-06 | 1 - | 3,00E-05 - | 1 - | 3,73E-05 - | 1 - | 3,55E-05 - | 1 - | 3,60E-05 - | 1 - | 4,05E-05 - |
| 3060 | 1 | 0.2um_NOKCPHJN_72300 | Bacteria | Proteobacteri | Gammaprote | Oceanospirilla | Saccharospirill | Reinekea | Reinekea fors | unclassified | unclassified | unclassified | unclassified | unclassified | 22,4 | 3,136877428 | 0,000892612 | 9,50E-05 | 28 - | 0,000942448 - | 20 - | 0,000835888 - | 22 - | 0,000874079 - | 19 - | 0,000766036 - | 23 - | 0,001044608 - |
| 3061 | 1 | 0.2um_NOKCPHJN_417714 | Bacteria | Proteobacteri | Alphaproteob | Rhodobactera | Rhodobactera | unclassified | Rhodobactera | unclassified | unclassified | unclassified | unclassified | unclassified | 1 | 0 | 4,03E-05 | 3,83E-06 | 1 - | 3,38E-05 - | 1 - | 4,19E-05 - | 1 - | 3,98E-05 - | 1 - | 4,04E-05 - | 1 - | 4,56E-05 - |
| 3062 | 1 | 0.2um_NOKCPHJN_28642 | Bacteria | Proteobacteri | Gammaprote | Alteromonada | Alteromonada | Paraglaciecola | Paraglaciecola | cellular proces | Posttranslatio | 0QHTY | protease | 983545.Glaag | 1,4 | 1,019803903 | 4,23E-05 | 3,10E-05 | 1 - | 2,53E-05 - | 2 - | 6,28E-05 - | 3 - | 8,95E-05 - | 0 - | 0 - | 1 - | 3,41E-05 - |
| 3063 | 1 | 3um_FIJJAFBF_70346 | Bacteria | Proteobacteri | Betaproteoba | unclassified | unclassified | unclassified | Betaproteoba | information st | Translation, ri | 16Q16 | Binds the low | 887062.HGR_ | 4,8 | 1,326649916 | 0,000223203 | 4,00E-05 | 7 - | 0,000279943 - | 4 - | 0,000198633 - | 5 - | 0,000236032 - | 5 - | 0,000239518 - | 3 - | 0,00016189 - |
| 3064 | 1 | 0.2um_NOKCPHJN_169493 | Bacteria | Proteobacteri | Gammaprote | Cellvibrionales | Porticoccacea | unclassified | SAR92 bacteri | cellular proces | Posttranslatio | 0QJ3C | Acts as a chap | 203122.Sde_2 | 0,6 | 0,489897949 | 1,19E-05 | 9,83E-06 | 1 - | 1,77E-05 - | 0 - | 0 - | 1 - | 2,09E-05 - | 1 - | 2,12E-05 - | 0 - | 0 - |
| 3065 | 1 | 3um_FIJJAFBF_45325 | Bacteria | Bacteroidetes | Flavobacteriia | Flavobacterial | Flavobacteriac | Formosa | Formosa sp. H | information st | Transcription | 0P8JP | Facilitates trar | 983544.Lacal_ | 2,2 | 0,748331477 | 5,13E-05 | 1,57E-05 | 3 - | 6,00E-05 - | 3 - | 7,45E-05 - | 2 - | 4,72E-05 - | 2 - | 4,79E-05 - | 1 - | 2,70E-05 - |
| 3066 | 1 | 3um_FIJJAFBF_479269 | Bacteria | Proteobacteri | Alphaproteob | Rhodobactera | Rhodobactera | Lentibacter | Lentibacter al | information st | Translation, ri | 16QAF | thus facilitatin | 292414.TM10 | 9,4 | 2,059126028 | 0,000232448 | 6,05E-05 | 8 - | 0,000164554 - | 9 - | 0,000229869 - | 7 - | 0,00016996 - | 13 - | 0,000320302 - | 10 - | 0,000277553 - |
| 3067 | 1 | 3um_FIJJAFBF_233041 | Bacteria | Actinobacteri | Actinobacteri | Micrococcales | Microbacteria | Candidatus Ac | Candidatus Ac | information st | Translation, ri | 05C64 | Binds directly | 861360.AARI_ | 0,4 | 0,489897949 | 2,20E-05 | 2,72E-05 | 1 - | 5,01E-05 - | 0 - | 0 - | 0 - | 0 - | 1 - | 6,00E-05 - | 0 - | 0 - |
| 3068 | 1 | 3um_FIJJAFBF_128441 | Bacteria | Bacteroidetes | Flavobacteriia | Flavobacterial | Flavobacteriac | Polaribacter | Polaribacter s | unclassified | unclassified | unclassified | unclassified | unclassified | 50,2 | 3,709447398 | 0,000728595 | 5,74E-05 | 56 - | 0,000683776 - | 48 - | 0,000727756 - | 52 - | 0,000749476 - | 45 - | 0,000658165 - | 50 - | 0,0008238 - |
| 3069 | 1 | 3um_FIJJAFBF_690483 | Bacteria | Proteobacteri | Gammaprote | Alteromonada | Alteromonada | Paraglaciecola | Paraglaciecola | unclassified | unclassified | unclassified | unclassified | unclassified | 0,4 | 0,489897949 | 1,21E-05 | 1,48E-05 | 0 - | 0 - | 0 - | 0 - | 1 - | 3,00E-05 - | 1 - | 3,04E-05 - | 0 - | 0 - |
| 3070 | 1 | 3um_FIJJAFBF_157376 | Bacteria | Bacteroidetes | Flavobacteriia | Flavobacterial | Flavobacteriac | Algibacter | Algibacter aqu | information st | Translation, ri | COG0052 | 30S ribosomal | 487797.Flav3 | 2,8 | 0,4 | 0,00013384 | 1,59E-05 | 3 - | 0,000121243 - | 3 - | 0,000150548 - | 3 - | 0,000143115 - | 3 - | 0,000145229 - | 2 - | 0,000109067 - |
| 3071 | 1 | 0.2um_NOKCPHJN_130420 | Bacteria | Proteobacteri | Gammaprote | Oceanospirilla | Saccharospirill | Reinekea | Reinekea fors | metabolism | Amino acid tra | 01RBF | Branched-cha | 911045.PSE_0 | 1,2 | 0,748331477 | 3,44E-05 | 2,12E-05 | 2 - | 5,01E-05 - | 2 - | 6,22E-05 - | 1 - | 2,96E-05 - | 1 - | 3,00E-05 - | 0 - | 0 - |
| 3072 | 1 | 3um_FIJJAFBF_183439 | Bacteria | Bacteroidetes | Cytophagia | Cytophagales | Flammeovirga | unclassified | Flammeovirga | metabolism | Inorganic ion t | 05C27 | TonB-depende | 688270.Celal_ | 26,2 | 0,748331477 | 0,00034527 | 3,47E-05 | 26 - | 0,000286942 - | 27 - | 0,000370001 - | 27 - | 0,000351733 - | 25 - | 0,000330489 - | 26 - | 0,000387186 - |
| 3073 | 1 | 3um_FIJJAFBF_489486 | Bacteria | Bacteroidetes | Flavobacteriia | Flavobacterial | Flavobacteriac | Formosa | Formosa sp. H | metabolism | Carbohydrate | COG3979 | chitinase | 867900.Celly_ | 1,6 | 0,489897949 | 6,01E-05 | 1,57E-05 | 2 - | 6,43E-05 - | 1 - | 3,99E-05 - | 2 - | 7,59E-05 - | 2 - | 7,70E-05 - | 1 - | 4,34E-05 - |
| 3074 | 1 | 3um_FIJJAFBF_587442 | Bacteria | Bacteroidetes | Flavobacteriia | Flavobacterial | Flavobacteriac | Winogradskye | Winogradskye | metabolism | Amino acid tra | COG1932 | Catalyzes the | 983544.Lacal_ | 2,4 | 1,019803903 | 8,99E-05 | 3,17E-05 | 4 - | 0,000130059 - | 3 - | 0,000121121 - | 2 - | 7,68E-05 - | 2 - | 7,79E-05 - | 1 - | 4,39E-05 - |
| 3075 | 1 | 3um_FIJJAFBF_499621 | Eukaryota | Arthropoda | Arachnida | Sarcoptiforme | Sarcoptidae | Sarcoptes | Sarcoptes scal | unclassified | unclassified | unclassified | unclassified | unclassified | 0,8 | 0,4 | 2,51E-05 | 1,29E-05 | 1 - | 2,62E-05 - | 1 - | 3,25E-05 - | 0 - | 0 - | 1 - | 3,14E-05 - | 1 - | 3,54E-05 - |
| 3076 | 1 | 0.2um_NOKCPHJN_549494 | Bacteria | Proteobacteri | Gammaprote | unclassified | unclassified | unclassified | OM182 bacter | information st | Translation, ri | COG0052 | 30S ribosomal | 247634.GPB2 | 1,8 | 0,748331477 | 0,000101737 | 4,26E-05 | 1 - | 4,72E-05 - | 2 - | 0,000117299 - | 3 - | 0,000167262 - | 2 - | 0,000113155 - | 1 - | 6,37E-05 - |
| 3077 | 1 | 0.2um_NOKCPHJN_28695 | Bacteria | Proteobacteri | Gammaprote | Alteromonada | Alteromonada | Alteromonas | Alteromonas s | unclassified | unclassified | unclassified | unclassified | unclassified | 6 | 0,632455532 | 0,000148746 | 1,26E-05 | 7 - | 0,000146346 - | 5 - | 0,000129799 - | 6 - | 0,000148068 - | 6 - | 0,000150255 - | 6 - | 0,000169262 - |
| 3078 | 1 | 0.2um_NOKCPHJN_266402 | Bacteria | Proteobacteri | Alphaproteob | Rhodobactera | unclassified | unclassified | Rhodobactera | unclassified | unclassified | unclassified | unclassified | unclassified | 2 | 0 | 7,19E-05 | 6,83E-06 | 2 - | 6,03E-05 - | 2 - | 7,48E-05 - | 2 - | 7,11E-05 - | 2 - | 7,22E-05 - | 2 - | 8,13E-05 - |
| 3079 | 1 | 3um_FIJJAFBF_318783 | Eukaryota | Chlorophyta | Chlorophycea | Chlamydomor | Chlamydomor | Chlamydomor | Chlamydomor | unclassified | unclassified | unclassified | unclassified | unclassified | 6,4 | 1,624807681 | 0,000163129 | 3,39E-05 | 8 - | 0,000173904 - | 7 - | 0,000188945 - | 8 - | 0,000205276 - | 5 - | 0,000130193 - | 4 - | 0,000117329 - |
| 3080 | 1 | 0.2um_NOKCPHJN_418995 | Bacteria | Proteobacteri | Gammaprote | Alteromonada | Alteromonada | unclassified | Alteromonada | unclassified | unclassified | unclassified | unclassified | unclassified | 0,4 | 0,489897949 | 1,44E-05 | 1,78E-05 | 1 - | 3,21E-05 - | 1 - | 3,98E-05 - | 0 - | 0 - | 0 - | 0 - | 0 - | 0 - |
| 3081 | 1 | 0.2um_NOKCPHJN_30655 | Bacteria | Bacteroidetes | Flavobacteriia | Flavobacterial | unclassified | unclassified | Flavobacterial | poorly charact | Function unkn | 11P35 | NA | 391603.FBALC | 1 | 0 | 2,48E-05 | 2,36E-06 | 1 - | 2,08E-05 - | 1 - | 2,58E-05 - | 1 - | 2,45E-05 - | 1 - | 2,49E-05 - | 1 - | 2,81E-05 - |
| 3082 | 1 | 3um_FIJJAFBF_371960 | Bacteria | Proteobacteri | Gammaprote | Alteromonada | Alteromonada | Paraglaciecola | Paraglaciecola | metabolism | Nucleotide tra | COG0138 | bifunctional p | 715451.ambt_ | 0,6 | 0,489897949 | 1,48E-05 | 1,22E-05 | 1 - | 2,15E-05 - | 1 - | 2,66E-05 - | 0 - | 0 - | 1 - | 2,57E-05 - | 0 - | 0 - |
| 3083 | 1 | 3um_FIJJAFBF_164611 | Bacteria | Proteobacteri | Alphaproteob | Pelagibacteral | Pelagibacterac | Candidatus Pe | Candidatus Pe | unclassified | unclassified | unclassified | unclassified | unclassified | 1,8 | 0,4 | 9,71E-05 | 2,38E-05 | 2 - | 9,04E-05 - | 2 - | 0,00011222 - | 2 - | 0,000106679 - | 1 - | 5,41E-05 - | 2 - | 0,000121948 - |
| 3084 | 1 | 3um_FIJJAFBF_719649 | Bacteria | Proteobacteri | Gammaprote | Chromatiales | Granulosicocc | Granulosicocc | Granulosicocc | cellular proces | Cell wall/mem | 0QKXZ | Porin Gram-ne | 396588.Tgr7_ | 0,8 | 0,748331477 | 3,35E-05 | 2,92E-05 | 2 - | 7,40E-05 - | 0 - | 0 - | 1 - | 4,37E-05 - | 0 - | 0 - | 1 - | 5,00E-05 - |
| 3085 | 1 | 3um_FIJJAFBF_95363 | Bacteria | Proteobacteri | Alphaproteob | Rickettsiales | unclassified | unclassified | Rickettsiales b | unclassified | unclassified | unclassified | unclassified | unclassified | 0,2 | 0,4 | 7,11E-06 | 1,42E-05 | 1 - | 3,55E-05 - | 0 - | 0 - | 0 - | 0 - | 0 - | 0 - | 0 - | 0 - |
| 3086 | 1 | 3um_FIJJAFBF_337088 | Bacteria | Proteobacteri | Gammaprote | Oceanospirilla | Oceanospirilla | Motiliproteus | Motiliproteus | information st | Translation, ri | 177XJ | One of the pri | 205918.Psyr_ | 2 | 0,632455532 | 0,000175419 | 5,70E-05 | 2 - | 0,00014715 - | 2 - | 0,000182716 - | 3 - | 0,000260543 - | 1 - | 8,81E-05 - | 2 - | 0,000198557 - |
| 3087 | 1 | 3um_FIJJAFBF_73095 | Bacteria | Proteobacteri | Gammaprote | Alteromonada | Alteromonada | Paraglaciecola | Paraglaciecola | cellular proces | Intracellular tr | COG0848 | Biopolymer tr | 342610.Patl_2 | 1,4 | 0,489897949 | 0,000144833 | 5,34E-05 | 1 - | 8,63E-05 - | 2 - | 0,000214314 - | 2 - | 0,000203733 - | 1 - | 0,000103371 - | 1 - | 0,000116447 - |
| 3088 | 1 | 3um_FIJJAFBF_48984 | Bacteria | Bacteroidetes | Flavobacteriia | Flavobacterial | Flavobacteriac | Polaribacter | various | cellular proces | Posttranslatio | COG0330 | Band 7 protei | 313603.FB217 | 2 | 0,632455532 | 0,000100608 | 3,22E-05 | 2 - | 8,47E-05 - | 1 - | 5,26E-05 - | 3 - | 0,00014998 - | 2 - | 0,000101464 - | 2 - | 0,000114299 - |
| 3089 | 1 | 3um_FIJJAFBF_175144 | Bacteria | Proteobacteri | Alphaproteob | Rhodobactera | Rhodobactera | Aliiroseovariu | Aliiroseovariu | information st | Translation, ri | 08YY1 | This protein is | 388739.RSK20 | 0,8 | 0,748331477 | 8,44E-05 | 8,12E-05 | 1 - | 9,26E-05 - | 0 - | 0 - | 2 - | 0,00021852 - | 1 - | 0,000110874 - | 0 - | 0 - |
| 3090 | 1 | 3um_FIJJAFBF_72464 | Bacteria | Bacteroidetes | Cytophagia | Cytophagales | Flammeovirga | unclassified | Flammeovirga | information st | Translation, ri | COG0051 | tRNA binding | 643867.Ftrac_ | 2 | 0 | 0,000271341 | 2,58E-05 | 2 - | 0,000227281 - | 2 - | 0,000282216 - | 2 - | 0,000268282 - | 2 - | 0,000272244 - | 2 - | 0,000306682 - |
| 3091 | 2 | 0.2um_NOKCPHJN_257105;3um_FIJJAFBF_1301 | various | various | various | various | various | various | various | unclassified | unclassified | unclassified | unclassified | unclassified | 0,4 | 0,489897949 | 4,52E-05 | 5,54E-05 | 0 - | 0 - | 1 - | 0,000115869 - | 1 - | 0,000110148 - | 0 - | 0 - | 0 - | 0 - |
| 3092 | 1 | 3um_FIJJAFBF_413381 | Bacteria | Proteobacteri | Gammaprote | Cellvibrionales | Porticoccacea | unclassified | gamma prote | metabolism | Inorganic ion t | 05PWN | phosphate ab | 314287.GB22 | 0,2 | 0,4 | 6,52E-06 | 1,30E-05 | 1 - | 3,26E-05 - | 0 - | 0 - | 0 - | 0 - | 0 - | 0 - | 0 - | 0 - |
| 3093 | 1 | 3um_FIJJAFBF_602715 | Bacteria | Proteobacteri | Alphaproteob | Rhodobactera | Rhodobactera | Planktomarina | Planktomarina | poorly charact | Function unkn | 01ZZB | NA | 314256.OG25 | 0,6 | 0,8 | 4,66E-05 | 5,88E-05 | 2 - | 0,000139123 - | 0 - | 0 - | 0 - | 0 - | 0 - | 0 - | 1 - | 9,39E-05 - |
| 3094 | 1 | 3um_FIJJAFBF_132905 | Bacteria | Proteobacteri | Alphaproteob | Rhodospirilla | Rhodospirillac | unclassified | Rhodobactera | unclassified | unclassified | unclassified | unclassified | unclassified | 0,2 | 0,4 | 5,95E-06 | 1,19E-05 | 1 - | 2,97E-05 - | 0 - | 0 - | 0 - | 0 - | 0 - | 0 - | 0 - | 0 - |
| 3095 | 1 | 0.2um_NOKCPHJN_1038484 | Bacteria | Bacteroidetes | Cytophagia | Cytophagales | Hymenobacte | Hymenobacte | Hymenobacte | unclassified | unclassified | unclassified | unclassified | unclassified | 0,8 | 0,4 | 3,77E-05 | 1,92E-05 | 1 - | 4,08E-05 - | 1 - | 5,07E-05 - | 1 - | 4,82E-05 - | 1 - | 4,89E-05 - | 0 - | 0 - |
| 3096 | 1 | 3um_FIJJAFBF_93536 | Bacteria | Bacteroidetes | unclassified | unclassified | unclassified | unclassified | Bacteroidetes | unclassified | unclassified | unclassified | unclassified | unclassified | 0,2 | 0,4 | 1,10E-05 | 2,21E-05 | 1 - | 5,52E-05 - | 0 - | 0 - | 0 - | 0 - | 0 - | 0 - | 0 - | 0 - |
| 3097 | 1 | 3um_FIJJAFBF_113461 | Bacteria | Proteobacteri | Alphaproteob | Rhodobactera | unclassified | unclassified | Rhodobactera | metabolism | Amino acid tra | 08KMW | Glycine betain | 367336.OM22 | 1,4 | 0,489897949 | 6,02E-05 | 1,89E-05 | 2 - | 7,33E-05 - | 2 - | 9,11E-05 - | 1 - | 4,33E-05 - | 1 - | 4,39E-05 - | 1 - | 4,95E-05 - |
| 3098 | 1 | 3um_FIJJAFBF_255534 | Bacteria | Proteobacteri | Alphaproteob | Rhodobactera | Rhodobactera | unclassified | Rhodobactera | poorly charact | Function unkn | COG1744 | basic membra | 391613.RTM1 | 1,4 | 0,489897949 | 5,24E-05 | 1,50E-05 | 2 - | 6,43E-05 - | 1 - | 3,99E-05 - | 2 - | 7,59E-05 - | 1 - | 3,85E-05 - | 1 - | 4,34E-05 - |
| 3099 | 1 | 0.2um_NOKCPHJN_191832 | Bacteria | Proteobacteri | Gammaprote | Oceanospirilla | Saccharospirill | Reinekea | Reinekea blan | cellular proces | Posttranslatio | COG0330 | Band 7 protei | 314283.MED2 | 3,2 | 1,6 | 0,000143017 | 6,83E-05 | 5 - | 0,000190028 - | 5 - | 0,000235958 - | 2 - | 8,97E-05 - | 1 - | 4,55E-05 - | 3 - | 0,000153849 - |
| 3100 | 1 | 3um_FIJJAFBF_793080 | Bacteria | Proteobacteri | Alphaproteob | Rhodospirilla | Rhodospirillac | unclassified | Rhodospirillac | cellular proces | Posttranslatio | COG0330 | Band 7 protei | 414684.RC1_1 | 0,6 | 0,8 | 2,57E-05 | 3,29E-05 | 2 - | 7,94E-05 - | 1 - | 4,93E-05 - | 0 - | 0 - | 0 - | 0 - | 0 - | 0 - |
| 3101 | 1 | 0.2um_NOKCPHJN_151766 | Bacteria | Proteobacteri | Alphaproteob | Rhodobactera | Rhodobactera | Planktomarina | Planktomarina | information st | Replication, re | COG0513 | purine NTP-de | 633131.TR2A | 0,6 | 0,489897949 | 1,43E-05 | 1,18E-05 | 1 - | 2,09E-05 - | 1 - | 2,60E-05 - | 1 - | 2,47E-05 - | 0 - | 0 - | 0 - | 0 - |
| 3102 | 1 | 3um_FIJJAFBF_72462 | Bacteria | Bacteroidetes | Cytophagia | Cytophagales | Flammeovirga | Reichenbachie | Reichenbachie | information st | Translation, ri | 08UHY | One of the pri | 643867.Ftrac_ | 2,6 | 0,8 | 0,000224819 | 5,11E-05 | 4 - | 0,000296198 - | 3 - | 0,000275843 - | 2 - | 0,000174816 - | 2 - | 0,000177398 - | 2 - | 0,000199838 - |
| 3103 | 1 | 0.2um_NOKCPHJN_925180 | Eukaryota | Arthropoda | Insecta | Hymenoptera | Formicidae | Cyphomyrmex | Cyphomyrmex | unclassified | unclassified | unclassified | unclassified | unclassified | 0,4 | 0,489897949 | 5,00E-05 | 6,16E-05 | 1 - | 0,00011364 - | 0 - | 0 - | 0 - | 0 - | 1 - | 0,000136122 - | 0 - | 0 - |
| 3104 | 2 | 0.2um_NOKCPHJN_1088933;0.2um_NOKCPHJN_ | Bacteria | Proteobacteri | Alphaproteob | Rhodobactera | Rhodobactera | Planktomarina | Planktomarina | unclassified | unclassified | unclassified | unclassified | unclassified | 0,2 | 0,4 | 7,70E-06 | 1,54E-05 | 1 - | 3,85E-05 - | 0 - | 0 - | 0 - | 0 - | 0 - | 0 - | 0 - | 0 - |
| 3105 | 1 | 0.2um_NOKCPHJN_403265 | Bacteria | Proteobacteri | Alphaproteob | Rhodobactera | Rhodobactera | Marivivens | Marivivens niv | cellular proces | Cell wall/mem | 01XF2 | NA | 391626.OA30 | 2 | 0,632455532 | 7,67E-05 | 3,01E-05 | 1 - | 3,12E-05 - | 2 - | 7,75E-05 - | 2 - | 7,36E-05 - | 2 - | 7,47E-05 - | 3 - | 0,000126256 - |
| 3106 | 1 | 3um_FIJJAFBF_119125 | Bacteria | Bacteroidetes | Flavobacteriia | Flavobacterial | Flavobacteriac | Polaribacter | various | information st | Translation, ri | 08YY1 | This protein is | 313594.PI23P | 5,8 | 0,4 | 0,000613324 | 4,45E-05 | 6 - | 0,000533845 - | 6 - | 0,000662878 - | 6 - | 0,00063015 - | 6 - | 0,000639457 - | 5 - | 0,000600289 - |
| 3107 | 1 | 0.2um_NOKCPHJN_101831 | Bacteria | Bacteroidetes | Flavobacteriia | various | various | various | various | cellular proces | Intracellular tr | 05FGH | MotA TolQ ex | 487797.Flav3 | 2,8 | 0,4 | 0,000160383 | 3,38E-05 | 2 - | 9,49E-05 - | 3 - | 0,000176676 - | 3 - | 0,000167953 - | 3 - | 0,000170434 - | 3 - | 0,000191993 - |
| 3108 | 1 | 0.2um_NOKCPHJN_481501 | Bacteria | Bacteroidetes | Cytophagia | Cytophagales | Bernardetiace | Hugenholtzia | Hugenholtzia | unclassified | unclassified | unclassified | unclassified | unclassified | 0,2 | 0,4 | 8,19E-06 | 1,64E-05 | 0 - | 0 - | 0 - | 0 - | 1 - | 4,09E-05 - | 0 - | 0 - | 0 - | 0 - |
| 3109 | 1 | 3um_FIJJAFBF_271633 | Bacteria | Bacteroidetes | Flavobacteriia | Flavobacterial | Flavobacteriac | Formosa | Formosa sp. H | unclassified | unclassified | unclassified | unclassified | unclassified | 2,6 | 1,019803903 | 3,69E-05 | 1,39E-05 | 3 - | 3,65E-05 - | 2 - | 3,02E-05 - | 3 - | 4,31E-05 - | 4 - | 5,83E-05 - | 1 - | 1,64E-05 - |
| 3110 | 1 | 3um_FIJJAFBF_641954 | Bacteria | Proteobacteri | Gammaprote | Cellvibrionales | Spongiibacter | Oceanicoccus | Oceanicoccus | metabolism | Inorganic ion t | 01UP7 | TonB-depende | 394221.Mmar | 2,4 | 0,489897949 | 4,10E-05 | 6,39E-06 | 3 - | 4,35E-05 - | 2 - | 3,60E-05 - | 2 - | 3,43E-05 - | 3 - | 5,21E-05 - | 2 - | 3,92E-05 - |
| 3111 | 1 | 3um_FIJJAFBF_643991 | Bacteria | Bacteroidetes | Cytophagia | Cytophagales | Cytophagacea | Ekhidna | Ekhidna lutea | metabolism | Nucleotide tra | 09B0C | Provides the p | 643867.Ftrac_ | 0,4 | 0,489897949 | 6,48E-06 | 8,01E-06 | 1 - | 1,45E-05 - | 1 - | 1,79E-05 - | 0 - | 0 - | 0 - | 0 - | 0 - | 0 - |
| 3112 | 1 | 0.2um_NOKCPHJN_163255 | Bacteria | Bacteroidetes | Flavobacteriia | Flavobacterial | Flavobacteriac | Arenibacter | Arenibacter sp | information st | Translation, ri | COG1185 | Involved in m | 1046627.BZAF | 6,6 | 0,8 | 0,000123188 | 2,06E-05 | 6 - | 9,36E-05 - | 8 - | 0,000154912 - | 6 - | 0,000110447 - | 7 - | 0,000130759 - | 6 - | 0,000126256 - |
| 3113 | 2 | 0.2um_NOKCPHJN_39280;0.2um_NOKCPHJN_47 | Bacteria | Proteobacteri | Gammaprote | Cellvibrionales | Porticoccacea | unclassified | various | cellular proces | Posttranslatio | COG0330 | Band 7 protei | 314287.GB22 | 0,6 | 0,489897949 | 2,80E-05 | 2,33E-05 | 1 - | 3,97E-05 - | 0 - | 0 - | 1 - | 4,69E-05 - | 0 - | 0 - | 1 - | 5,36E-05 - |
| 3114 | 1 | 0.2um_NOKCPHJN_892634 | unclassified | unclassified | unclassified | unclassified | unclassified | unclassified | unclassified | unclassified | unclassified | unclassified | unclassified | unclassified | 0,8 | 0,4 | 0,000204761 | 0,000105155 | 1 - | 0,00021656 - | 0 - | 0 - | 1 - | 0,000255627 - | 1 - | 0,000259402 - | 1 - | 0,000292216 - |
| 3115 | 1 | 0.2um_NOKCPHJN_07553 | Bacteria | Bacteroidetes | unclassified | unclassified | unclassified | unclassified | Bacteroidetes | poorly charact | Function unkn | 09CIG | Ragb susd dor | 655815.ZPR_1 | 5 | 2 | 0,000131106 | 4,34E-05 | 8 - | 0,000181825 - | 6 - | 0,000169329 - | 5 - | 0,000134141 - | 4 - | 0,000108898 - | 2 - | 6,13E-05 - |
| 3116 | 1 | 0.2um_NOKCPHJN_72306 | Bacteria | Proteobacteri | Gammaprote | Oceanospirilla | Saccharospirill | Saccharospiril | Saccharospiril | unclassified | unclassified | unclassified | unclassified | unclassified | 1,2 | 0,748331477 | 1,92E-05 | 1,23E-05 | 1 - | 1,34E-05 - | 2 - | 3,32E-05 - | 2 - | 3,15E-05 - | 0 - | 0 - | 1 - | 1,80E-05 - |
| 3117 | 1 | 3um_FIJJAFBF_266945 | Bacteria | Bacteroidetes | Flavobacteriia | unclassified | unclassified | unclassified | Flavobacteria | metabolism | Inorganic ion t | 05C27 | TonB-depende | 487797.Flav3 | 1,2 | 0,748331477 | 3,70E-05 | 2,37E-05 | 2 - | 5,23E-05 - | 0 - | 0 - | 1 - | 3,09E-05 - | 1 - | 3,13E-05 - | 2 - | 7,06E-05 - |
| 3118 | 1 | 3um_FIJJAFBF_537760 | Bacteria | Bacteroidetes | Flavobacteriia | Flavobacterial | Crocinitomica | unclassified | Crocinitomica | unclassified | unclassified | unclassified | unclassified | unclassified | 1,2 | 0,748331477 | 6,60E-05 | 4,32E-05 | 1 - | 4,68E-05 - | 2 - | 0,000116342 - | 2 - | 0,000110598 - | 1 - | 5,61E-05 - | 0 - | 0 - |
| 3119 | 1 | 3um_FIJJAFBF_103165 | Bacteria | Bacteroidetes | Flavobacteriia | Flavobacterial | Flavobacteriac | Polaribacter | Polaribacter a | unclassified | unclassified | unclassified | unclassified | unclassified | 4,6 | 1,019803903 | 0,000199357 | 3,94E-05 | 6 - | 0,000220019 - | 3 - | 0,0001366 - | 4 - | 0,00017314 - | 5 - | 0,000219622 - | 5 - | 0,000247403 - |
| 3120 | 1 | 3um_FIJJAFBF_734139 | Eukaryota | Streptophyta | Liliopsida | Poales | Poaceae | Dichanthelium | Dichanthelium | unclassified | unclassified | unclassified | unclassified | unclassified | 1,2 | 0,4 | 7,96E-05 | 1,76E-05 | 2 - | 0,000114206 - | 1 - | 7,09E-05 - | 1 - | 6,74E-05 - | 1 - | 6,84E-05 - | 1 - | 7,71E-05 - |
| 3121 | 1 | 0.2um_NOKCPHJN_495421 | Bacteria | Proteobacteri | Alphaproteob | Pelagibacteral | Pelagibacterac | Candidatus Pe | Candidatus Pe | information st | Translation, ri | 01S8S | This is 1 of the | 335992.SAR11 | 3 | 0,632455532 | 0,000219055 | 2,72E-05 | 4 - | 0,000249515 - | 3 - | 0,000232368 - | 3 - | 0,000220895 - | 3 - | 0,000224158 - | 2 - | 0,000168342 - |
| 3122 | 1 | 0.2um_NOKCPHJN_207885 | Bacteria | Bacteroidetes | Flavobacteriia | Flavobacterial | Flavobacteriac | Algibacter | Algibacter sp. | cellular proces | Posttranslatio | COG0330 | Band 7 protei | 983548.Krodi_ | 1,4 | 0,489897949 | 7,21E-05 | 2,97E-05 | 1 - | 4,24E-05 - | 1 - | 5,26E-05 - | 1 - | 5,00E-05 - | 2 - | 0,000101464 - | 2 - | 0,000114299 - |
| 3123 | 1 | 3um_FIJJAFBF_86535 | Bacteria | Bacteroidetes | Flavobacteriia | Flavobacterial | Flavobacteriac | Kordia | Kordia algicida | metabolism | Inorganic ion t | 05C27 | TonB-depende | 391587.KAOT | 0,4 | 0,8 | 4,25E-06 | 8,51E-06 | 2 - | 2,13E-05 - | 0 - | 0 - | 0 - | 0 - | 0 - | 0 - | 0 - | 0 - |
| 3124 | 1 | 3um_FIJJAFBF_326474 | Bacteria | Proteobacteri | Alphaproteob | unclassified | unclassified | unclassified | SAR116 cluste | metabolism | Energy produc | COG0056 | Produces ATP | 331869.BAL19 | 0,8 | 0,4 | 2,79E-05 | 1,42E-05 | 1 - | 3,02E-05 - | 1 - | 3,75E-05 - | 1 - | 3,57E-05 - | 1 - | 3,62E-05 - | 0 - | 0 - |
| 3125 | 1 | 3um_FIJJAFBF_72205 | Bacteria | Proteobacteri | Alphaproteob | Rhizobiales | unclassified | unclassified | Rhizobiales ba | unclassified | unclassified | unclassified | unclassified | unclassified | 1 | 0,632455532 | 4,03E-05 | 2,61E-05 | 1 - | 3,48E-05 - | 1 - | 4,32E-05 - | 2 - | 8,21E-05 - | 1 - | 4,17E-05 - | 0 - | 0 - |
| 3126 | 1 | 3um_FIJJAFBF_324099 | Bacteria | Proteobacteri | Gammaprote | Alteromonada | Alteromonada | Alteromonas | Alteromonas s | information st | Translation, ri | 05G6W | One of the pri | 87626.PTD2_ | 2,2 | 0,4 | 0,000144179 | 1,32E-05 | 3 - | 0,000167151 - | 2 - | 0,000138368 - | 2 - | 0,000131536 - | 2 - | 0,000133479 - | 2 - | 0,000150364 - |
| 3127 | 1 | 3um_FIJJAFBF_161285 | Bacteria | Proteobacteri | Alphaproteob | Rhodobactera | Rhodobactera | Roseobacter | Roseobacter li | unclassified | unclassified | unclassified | unclassified | unclassified | 1,8 | 0,748331477 | 0,000160048 | 6,34E-05 | 2 - | 0,000152022 - | 1 - | 9,44E-05 - | 3 - | 0,00026917 - | 2 - | 0,000182097 - | 1 - | 0,000102566 - |
| 3128 | 1 | 0.2um_NOKCPHJN_213917 | Bacteria | Proteobacteri | Gammaprote | Thiotrichales | Thiotrichacea | Leucothrix | Leucothrix mu | metabolism | Amino acid tra | COG0683 | (ABC) transpo | 744979.R2A13 | 5,2 | 1,6 | 0,000176408 | 5,24E-05 | 5 - | 0,000143831 - | 4 - | 0,000142876 - | 7 - | 0,000237688 - | 7 - | 0,000241199 - | 3 - | 0,000116447 - |
| 3129 | 1 | 3um_FIJJAFBF_788044 | Bacteria | Proteobacteri | Alphaproteob | Rhodobactera | Rhodobactera | Pelagicola | Pelagicola sp. | information st | Translation, ri | 16Q16 | Binds the low | 633131.TR2A | 5,4 | 1,356465997 | 0,000312904 | 7,59E-05 | 7 - | 0,000341888 - | 5 - | 0,000303232 - | 3 - | 0,000172956 - | 6 - | 0,000351021 - | 6 - | 0,000395424 - |
| 3130 | 1 | 0.2um_NOKCPHJN_149047 | Bacteria | Bacteroidetes | Flavobacteriia | Flavobacterial | Flavobacteriac | Polaribacter | Polaribacter b | unclassified | unclassified | unclassified | unclassified | unclassified | 0,2 | 0,4 | 1,57E-05 | 3,13E-05 | 0 - | 0 - | 0 - | 0 - | 1 - | 7,83E-05 - | 0 - | 0 - | 0 - | 0 - |
| 3131 | 1 | 3um_FIJJAFBF_346694 | Bacteria | Proteobacteri | Gammaprote | Oceanospirilla | Oceanospirilla | Amphritea | Amphritea jap | unclassified | unclassified | unclassified | unclassified | unclassified | 0,4 | 0,489897949 | 5,82E-05 | 7,17E-05 | 1 - | 0,000133461 - | 0 - | 0 - | 1 - | 0,000157537 - | 0 - | 0 - | 0 - | 0 - |
| 3132 | 1 | 3um_FIJJAFBF_216124 | Bacteria | Bacteroidetes | Flavobacteriia | Flavobacterial | Crocinitomica | unclassified | Crocinitomica | cellular proces | Cell wall/mem | 09J6U | biosynthesis p | 926562.Oweh | 0,2 | 0,4 | 6,96E-06 | 1,39E-05 | 1 - | 3,48E-05 - | 0 - | 0 - | 0 - | 0 - | 0 - | 0 - | 0 - | 0 - |
| 3133 | 1 | 3um_FIJJAFBF_400352 | Bacteria | Proteobacteri | Alphaproteob | Pelagibacteral | Pelagibacterac | unclassified | Pelagibacterac | metabolism | Energy produc | 05CMV | Succinyl-CoA | 439493.PB721 | 2,4 | 0,8 | 8,45E-05 | 2,90E-05 | 2 - | 5,95E-05 - | 3 - | 0,000110766 - | 3 - | 0,000105297 - | 3 - | 0,000106852 - | 1 - | 4,01E-05 - |
| 3134 | 1 | 0.2um_NOKCPHJN_533272 | Bacteria | Bacteroidetes | unclassified | unclassified | unclassified | unclassified | Bacteroidetes | unclassified | unclassified | unclassified | unclassified | unclassified | 2,4 | 1,019803903 | 6,73E-05 | 2,90E-05 | 3 - | 7,10E-05 - | 4 - | 0,000117541 - | 2 - | 5,59E-05 - | 1 - | 2,83E-05 - | 2 - | 6,39E-05 - |
| 3135 | 1 | 0.2um_NOKCPHJN_179223 | Eukaryota | Phaeophyceae | unclassified | Ectocarpales | Ectocarpaceae | Ectocarpus | Ectocarpus sil | information st | Translation, ri | KOG3204 | 60s ribosomal | 2850.Phatr25 | 0,2 | 0,4 | 1,37E-05 | 2,74E-05 | 0 - | 0 - | 0 - | 0 - | 1 - | 6,84E-05 - | 0 - | 0 - | 0 - | 0 - |
| 3136 | 1 | 0.2um_NOKCPHJN_114104 | Bacteria | Proteobacteri | Gammaprote | Cellvibrionales | Porticoccacea | unclassified | Gammaprote | unclassified | unclassified | unclassified | unclassified | unclassified | 1,8 | 0,748331477 | 0,000269352 | 9,84E-05 | 3 - | 0,000382589 - | 2 - | 0,000316709 - | 1 - | 0,000150536 - | 1 - | 0,000152759 - | 2 - | 0,000344166 - |
| 3137 | 1 | 0.2um_NOKCPHJN_493889 | Bacteria | Proteobacteri | Alphaproteob | Pelagibacteral | Pelagibacterac | Candidatus Pe | Candidatus Pe | unclassified | unclassified | unclassified | unclassified | unclassified | 5 | 0 | 0,000267631 | 2,54E-05 | 5 - | 0,000224173 - | 5 - | 0,000278357 - | 5 - | 0,000264614 - | 5 - | 0,000268522 - | 5 - | 0,000302489 - |
| 3138 | 1 | 3um_FIJJAFBF_609869 | Eukaryota | Bacillariophyt | Coscinodiscop | Thalassiosirale | Thalassiosirac | Thalassiosira | Thalassiosira | unclassified | unclassified | unclassified | unclassified | unclassified | 0,6 | 0,489897949 | 3,18E-05 | 2,60E-05 | 0 - | 0 - | 1 - | 5,46E-05 - | 1 - | 5,19E-05 - | 1 - | 5,27E-05 - | 0 - | 0 - |
| 3139 | 1 | 3um_FIJJAFBF_26279 | Bacteria | Bacteroidetes | Flavobacteriia | Flavobacterial | Flavobacteriac | Flavobacteriu | Flavobacteriu | information st | Replication, re | 08IYB | Dead deah bo | 313628.LNTAI | 1 | 0 | 1,62E-05 | 1,54E-06 | 1 - | 1,36E-05 - | 1 - | 1,69E-05 - | 1 - | 1,60E-05 - | 1 - | 1,63E-05 - | 1 - | 1,83E-05 - |
| 3140 | 1 | 3um_FIJJAFBF_782934 | Bacteria | Bacteroidetes | unclassified | unclassified | unclassified | unclassified | unidentified e | information st | Replication, re | COG0187 | DNA gyrase ne | 50743.SCB49_ | 1,2 | 0,4 | 3,96E-05 | 1,33E-05 | 1 - | 2,77E-05 - | 1 - | 3,44E-05 - | 2 - | 6,55E-05 - | 1 - | 3,32E-05 - | 1 - | 3,74E-05 - |

| | | | | | | | | | | | | | | | | | | | | | | | | | | | | |
|---|---|---|---|---|---|---|---|---|---|---|---|---|---|---|---|---|---|---|---|---|---|---|---|---|---|---|---|---|
| 3141 | 1 | 3um_FIJJAFBF_151044 | Bacteria | Bacteroidetes | Flavobacteriia | Flavobacterial | Flavobacteriac | unclassified | Flavobacteriac | unclassified | unclassified | unclassified | unclassified | unclassified | 2,6 | 0,489897949 | 0,00015704 | 3,83E-05 | 2 - | 0,000100242 - | 2 - | 0,000124471 - | 3 - | 0,000177488 - | 3 - | 0,000180109 - | 3 - | 0,000202892 - |
| 3142 | 1 | 0.2um_NOKCPHJN_836715 | Viruses | unclassified | unclassified | unclassified | unclassified | unclassified | uncultured M | unclassified | unclassified | unclassified | unclassified | unclassified | 1,6 | 0,489897949 | 7,01E-05 | 1,84E-05 | 2 - | 7,50E-05 - | 1 - | 4,66E-05 - | 2 - | 8,86E-05 - | 2 - | 8,99E-05 - | 1 - | 5,06E-05 - |
| 3143 | 1 | 0.2um_NOKCPHJN_143993 | Bacteria | Proteobacteri | Gammaprotec | Oceanospirilla | Saccharospirill | Reinekea | Reinekea blan | unclassified | unclassified | unclassified | unclassified | unclassified | 1,6 | 0,489897949 | 5,94E-05 | 1,65E-05 | 2 - | 6,32E-05 - | 2 - | 7,85E-05 - | 2 - | 7,46E-05 - | 1 - | 3,79E-05 - | 1 - | 4,27E-05 - |
| 3144 | 1 | 3um_FIJJAFBF_682252 | Bacteria | Proteobacteri | Alphaproteob | Pelagibacteral | unclassified | unclassified | Pelagibacteral | cellular proces | Cell wall/mem | COG2885 | Ompa motb d | 335992.SAR11 | 1 | 0 | 8,67E-05 | 8,23E-06 | 1 - | 7,26E-05 - | 1 - | 9,02E-05 - | 1 - | 8,57E-05 - | 1 - | 8,70E-05 - | 1 - | 9,80E-05 - |
| 3145 | 1 | 0.2um_NOKCPHJN_168960 | Bacteria | Bacteroidetes | Flavobacteriia | Flavobacterial | Crocinitomica | unclassified | Crocinitomica | metabolism | Lipid transpor | 0P85Z | acyl-CoA dehy | 755732.Fluta_ | 1 | 0 | 2,30E-05 | 2,18E-06 | 1 - | 1,93E-05 - | 1 - | 2,39E-05 - | 1 - | 2,27E-05 - | 1 - | 2,31E-05 - | 1 - | 2,60E-05 - |
| 3146 | 1 | 3um_FIJJAFBF_366631 | Bacteria | Proteobacteri | Alphaproteob | Rhodobactera | Rhodobactera | Amylibacter | Amylibacter k | information st | Transcription | COG0085 | DNA-depende | 644107.SL115 | 18 | 1,264911064 | 0,0001782 | 1,33E-05 | 19 - | 0,000158255 - | 19 - | 0,000196506 - | 19 - | 0,000186804 - | 17 - | 0,000169609 - | 16 - | 0,000179825 - |
| 3147 | 1 | 0.2um_NOKCPHJN_355685 | Bacteria | Proteobacteri | Alphaproteob | Rhizobiales | unclassified | unclassified | Rhizobiales ba | metabolism | Amino acid tra | 01VSB | Ectoine hydro | 367336.OM22 | 6,6 | 1,624807681 | 0,000309833 | 5,66E-05 | 9 - | 0,000359927 - | 5 - | 0,000248291 - | 8 - | 0,000377651 - | 5 - | 0,000239518 - | 6 - | 0,000323779 - |
| 3148 | 2 | 0.2um_NOKCPHJN_12254;3um_FIJJAFBF_119805 | Viruses | unclassified | unclassified | unclassified | various | various | various | information st | Transcription | 12N88 | dna-directed r | various | 1,6 | 0,8 | 1,40E-05 | 5,62E-06 | 3 - | 2,30E-05 - | 1 - | 9,51E-06 - | 1 - | 9,04E-06 - | 2 - | 1,84E-05 - | 1 - | 1,03E-05 - |
| 3149 | 1 | 3um_FIJJAFBF_399377 | Bacteria | Proteobacteri | Gammaprotec | Cellvibrionale | unclassified | unclassified | Cellvibrionale | poorly charact | Function unkn | 0QHT3 | Catalyzes the | 743720.Psefu_ | 1 | 0 | 4,36E-05 | 4,14E-06 | 1 - | 3,66E-05 - | 1 - | 4,54E-05 - | 1 - | 4,31E-05 - | 1 - | 4,38E-05 - | 1 - | 4,93E-05 - |
| 3150 | 1 | 3um_FIJJAFBF_37154 | Bacteria | Proteobacteri | Gammaprotec | Alteromonada | Alteromonada | Paraglaciecola | Paraglaciecola | unclassified | unclassified | unclassified | unclassified | unclassified | 1 | 0 | 9,52E-05 | 9,04E-06 | 1 - | 7,97E-05 - | 1 - | 9,90E-05 - | 1 - | 9,41E-05 - | 1 - | 9,55E-05 - | 1 - | 0,000107552 - |
| 3151 | 1 | 0.2um_NOKCPHJN_154905 | Eukaryota | unclassified | Oomycetes | Peronosporale | Peronosporac | Phytophthora | Phytophthora | unclassified | unclassified | unclassified | unclassified | unclassified | 4,6 | 0,489897949 | 0,000138336 | 1,35E-05 | 5 - | 0,000126685 - | 5 - | 0,000157306 - | 5 - | 0,000149539 - | 4 - | 0,000121398 - | 4 - | 0,000136755 - |
| 3152 | 1 | 0.2um_NOKCPHJN_293112 | Bacteria | Proteobacteri | Gammaprotec | Cellvibrionale | Porticoccacea | unclassified | Gammaprotec | information st | Translation, ri | 177XJ | One of the pri | 314287.GB22 | 2 | 0 | 0,000175676 | 1,67E-05 | 2 - | 0,00014715 - | 2 - | 0,000182716 - | 2 - | 0,000173695 - | 2 - | 0,000176261 - | 2 - | 0,000198557 - |
| 3153 | 1 | 3um_FIJJAFBF_239813 | Bacteria | Bacteroidetes | Flavobacteriia | Flavobacterial | Flavobacteriac | Formosa | Formosa sp. H | unclassified | unclassified | unclassified | unclassified | unclassified | 4,2 | 0,748331477 | 0,000311838 | 4,82E-05 | 5 - | 0,000313598 - | 4 - | 0,000311517 - | 5 - | 0,00037017 - | 3 - | 0,000225382 - | 4 - | 0,000338523 - |
| 3154 | 1 | 0.2um_NOKCPHJN_242649 | Bacteria | Proteobacteri | Alphaproteob | Rhodobactera | Rhodobactera | Planktomarina | Planktomarina | metabolism | Amino acid tra | 16RER | (ABC) transpo | 314270.RB208 | 0,4 | 0,489897949 | 2,13E-05 | 2,62E-05 | 1 - | 4,84E-05 - | 0 - | 0 - | 0 - | 0 - | 1 - | 5,80E-05 - | 0 - | 0 - |
| 3155 | 1 | 3um_FIJJAFBF_343463 | Bacteria | Bacteroidetes | Flavobacteriia | Flavobacterial | Flavobacteriac | Muricauda | Muricauda sp. | unclassified | unclassified | unclassified | unclassified | unclassified | 0,8 | 0,4 | 2,71E-05 | 1,37E-05 | 1 - | 2,93E-05 - | 1 - | 3,64E-05 - | 1 - | 3,46E-05 - | 1 - | 3,51E-05 - | 0 - | 0 - |
| 3156 | 1 | 3um_FIJJAFBF_343895 | Eukaryota | unclassified | unclassified | Phaeocystales | Phaeocystace | Phaeocystis | Phaeocystis gl | metabolism | Energy produc | COG0711 | F(1)F(0) ATP s | 32049.SYNPC | 4,8 | 0,979795897 | 0,000403147 | 7,71E-05 | 6 - | 0,000425099 - | 5 - | 0,000439873 - | 5 - | 0,000418155 - | 3 - | 0,000254599 - | 5 - | 0,000478008 - |
| 3157 | 1 | 3um_FIJJAFBF_110577 | Bacteria | Bacteroidetes | Flavobacteriia | Flavobacterial | Flavobacteriac | Polaribacter | various | unclassified | unclassified | unclassified | unclassified | unclassified | 1,6 | 0,8 | 0,000112532 | 4,50E-05 | 3 - | 0,000184134 - | 1 - | 7,62E-05 - | 1 - | 7,25E-05 - | 2 - | 0,000147041 - | 1 - | 8,28E-05 - |
| 3158 | 1 | 3um_FIJJAFBF_10571 | unclassified | unclassified | unclassified | unclassified | unclassified | unclassified | unclassified | unclassified | unclassified | unclassified | unclassified | unclassified | 0,4 | 0,489897949 | 6,44E-05 | 7,89E-05 | 0 - | 0 - | 1 - | 0,000163815 - | 0 - | 0 - | 1 - | 0,000158027 - | 0 - | 0 - |
| 3159 | 1 | 0.2um_NOKCPHJN_402249 | Bacteria | Proteobacteri | Alphaproteob | Pelagibacteral | Pelagibacterac | unclassified | Pelagibacterac | cellular proces | Cell wall/mem | 7257 | NA | 439493.PB721 | 1,8 | 0,4 | 7,22E-05 | 1,69E-05 | 2 - | 6,75E-05 - | 1 - | 4,19E-05 - | 2 - | 7,97E-05 - | 2 - | 8,09E-05 - | 2 - | 9,11E-05 - |
| 3160 | 1 | 3um_FIJJAFBF_282151 | Bacteria | Proteobacteri | Gammaprotec | Alteromonada | Pseudoalteror | Pseudoalteror | Pseudoalteror | metabolism | Energy produc | 05CV4 | Subunits I and | 342610.Patl_4 | 5 | 0,632455532 | 0,000177238 | 1,91E-05 | 6 - | 0,000179339 - | 5 - | 0,000185571 - | 5 - | 0,000176409 - | 4 - | 0,000143212 - | 5 - | 0,000201659 - |
| 3161 | 1 | 3um_FIJJAFBF_69470 | Bacteria | Proteobacteri | Betaproteoba | unclassified | unclassified | unclassified | Betaproteoba | unclassified | unclassified | unclassified | unclassified | unclassified | 0,6 | 0,489897949 | 4,75E-05 | 3,88E-05 | 0 - | 0 - | 1 - | 8,14E-05 - | 1 - | 7,74E-05 - | 1 - | 7,86E-05 - | 0 - | 0 - |
| 3162 | 1 | 3um_FIJJAFBF_566955 | Bacteria | Bacteroidetes | Flavobacteriia | Flavobacterial | Flavobacteriac | unclassified | Flavobacteriac | metabolism | Carbohydrate | 07SNC | solute-binding | 411684.HPDFI | 4,8 | 0,748331477 | 0,000150251 | 1,90E-05 | 6 - | 0,000158678 - | 4 - | 0,000131354 - | 5 - | 0,000156086 - | 4 - | 0,000126713 - | 5 - | 0,000178427 - |
| 3163 | 1 | 3um_FIJJAFBF_791666 | Bacteria | Proteobacteri | Gammaprotec | Oceanospirilla | Oceanospirilla | Motiliproteus | Motiliproteus | unclassified | unclassified | unclassified | unclassified | unclassified | 0,4 | 0,489897949 | 1,16E-05 | 1,43E-05 | 1 - | 2,66E-05 - | 0 - | 0 - | 1 - | 3,14E-05 - | 0 - | 0 - | 0 - | 0 - |
| 3164 | 1 | 0.2um_NOKCPHJN_40861 | Bacteria | Proteobacteri | Gammaprotec | unclassified | unclassified | unclassified | Gammaprotec | unclassified | unclassified | unclassified | unclassified | unclassified | 0,8 | 0,748331477 | 3,90E-05 | 3,44E-05 | 2 - | 8,90E-05 - | 0 - | 0 - | 1 - | 5,25E-05 - | 1 - | 5,33E-05 - | 0 - | 0 - |
| 3165 | 2 | 3um_FIJJAFBF_283093;3um_FIJJAFBF_559732 | Bacteria | Bacteroidetes | Flavobacteriia | Flavobacterial | Flavobacteriac | various | various | metabolism | Inorganic ion t | 09BXD | ABC transport | 983544.Lacal_ | 0,8 | 0,4 | 4,33E-05 | 2,20E-05 | 1 - | 4,68E-05 - | 1 - | 5,82E-05 - | 1 - | 5,53E-05 - | 1 - | 5,61E-05 - | 0 - | 0 - |
| 3166 | 1 | 3um_FIJJAFBF_44072 | Viruses | unclassified | unclassified | unclassified | Phycodnavirid | unclassified | Chrysochromu | information st | Transcription | COG0086 | DNA-depende | 88036.EFJ195 | 3 | 0,894427191 | 2,74E-05 | 8,41E-06 | 3 - | 2,31E-05 - | 4 - | 3,82E-05 - | 2 - | 1,81E-05 - | 4 - | 3,68E-05 - | 2 - | 2,07E-05 - |
| 3167 | 1 | 0.2um_NOKCPHJN_1003070 | Bacteria | Proteobacteri | Alphaproteob | Rhodobactera | Rhodobactera | unclassified | Rhodobactera | unclassified | unclassified | unclassified | unclassified | unclassified | 1 | 0 | 2,35E-05 | 2,23E-06 | 1 - | 1,97E-05 - | 1 - | 2,44E-05 - | 1 - | 2,32E-05 - | 1 - | 2,35E-05 - | 1 - | 2,65E-05 - |
| 3168 | 1 | 3um_FIJJAFBF_378361 | Bacteria | Bacteroidetes | Flavobacteriia | Flavobacterial | Flavobacteriac | Formosa | Formosa sp. H | unclassified | unclassified | unclassified | unclassified | unclassified | 1 | 0 | 6,49E-05 | 6,17E-06 | 1 - | 5,44E-05 - | 1 - | 6,75E-05 - | 1 - | 6,42E-05 - | 1 - | 6,52E-05 - | 1 - | 7,34E-05 - |
| 3169 | 1 | 0.2um_NOKCPHJN_504894 | Bacteria | Proteobacteri | Alphaproteob | Rhodobactera | Rhodobactera | Oceaniovalibu | Oceaniovalibu | unclassified | unclassified | unclassified | unclassified | unclassified | 17,6 | 1,019803903 | 0,000551592 | 4,79E-05 | 19 - | 0,000500174 - | 16 - | 0,000523005 - | 18 - | 0,00055933 - | 17 - | 0,000536059 - | 18 - | 0,00063939 - |
| 3170 | 1 | 3um_FIJJAFBF_188038 | Bacteria | Proteobacteri | Gammaprotec | Cellvibrionale | Porticoccacea | unclassified | SAR92 bacteri | unclassified | unclassified | unclassified | unclassified | unclassified | 2,2 | 0,4 | 7,56E-05 | 1,54E-05 | 2 - | 5,75E-05 - | 2 - | 7,14E-05 - | 2 - | 6,79E-05 - | 3 - | 0,000103371 - | 2 - | 7,76E-05 - |
| 3171 | 1 | 0.2um_NOKCPHJN_478111 | Eukaryota | Streptophyta | unclassified | Fabales | Fabaceae | Glycine | Glycine max | cellular proces | Posttranslatio | KOG0019 | heat shock pr | 3847.GLYMA0 | 1 | 0 | 1,90E-05 | 1,80E-06 | 1 - | 1,59E-05 - | 1 - | 1,97E-05 - | 1 - | 1,88E-05 - | 1 - | 1,90E-05 - | 1 - | 2,15E-05 - |
| 3172 | 1 | 3um_FIJJAFBF_552802 | Bacteria | Proteobacteri | Alphaproteob | Rhodospirillal | Rhodospirillac | Candidatus En | Candidatus En | unclassified | unclassified | unclassified | unclassified | unclassified | 2,4 | 0,8 | 0,000167868 | 4,97E-05 | 3 - | 0,000180278 - | 2 - | 0,000149234 - | 3 - | 0,000212799 - | 3 - | 0,000215942 - | 1 - | 8,11E-05 - |
| 3173 | 1 | 3um_FIJJAFBF_90137 | Bacteria | Bacteroidetes | Flavobacteriia | Flavobacterial | Cryomorphace | unclassified | Cryomorphace | poorly charact | Function unkn | 0XSVM | NA | 865938.Weev | 2,6 | 0,489897949 | 7,62E-05 | 1,62E-05 | 2 - | 4,89E-05 - | 3 - | 9,12E-05 - | 3 - | 8,67E-05 - | 3 - | 8,79E-05 - | 2 - | 6,60E-05 - |
| 3174 | 2 | 3um_FIJJAFBF_537765;3um_FIJJAFBF_549859 | Bacteria | Bacteroidetes | Flavobacteriia | Flavobacterial | Crocinitomica | unclassified | Crocinitomica | information st | Translation, ri | 09FE5 | One of the pr | 643867.Ftrac_ | 0,4 | 0,489897949 | 5,15E-05 | 6,31E-05 | 0 - | 0 - | 0 - | 0 - | 1 - | 0,000127813 - | 1 - | 0,000129701 - | 0 - | 0 - |
| 3175 | 1 | 0.2um_NOKCPHJN_93918 | Bacteria | Bacteroidetes | Flavobacteriia | unclassified | unclassified | unclassified | Flavobacteria | metabolism | Energy produc | COG0437 | 4fe-4S ferredc | 487797.Flav3 | 3,2 | 0,748331477 | 4,25E-05 | 1,02E-05 | 3 - | 3,36E-05 - | 4 - | 5,57E-05 - | 4 - | 5,29E-05 - | 3 - | 4,03E-05 - | 2 - | 3,02E-05 - |
| 3176 | 1 | 3um_FIJJAFBF_119764 | Bacteria | Bacteroidetes | Flavobacteriia | Flavobacterial | Flavobacteriac | Winogradskye | Winogradskye | poorly charact | Function unkn | 0XY6Y | Pkd domain c | 867900.Celly_ | 1,8 | 0,979795897 | 3,39E-05 | 1,74E-05 | 3 - | 4,97E-05 - | 2 - | 4,11E-05 - | 2 - | 3,91E-05 - | 2 - | 3,97E-05 - | 0 - | 0 - |
| 3177 | 1 | 0.2um_NOKCPHJN_140937 | Bacteria | Bacteroidetes | unclassified | unclassified | unclassified | unclassified | Bacteroidetes | metabolism | Inorganic ion t | COG0841 | acriflavin resis | 643867.Ftrac_ | 0,8 | 0,748331477 | 8,74E-06 | 7,72E-06 | 2 - | 2,00E-05 - | 0 - | 0 - | 1 - | 1,18E-05 - | 1 - | 1,20E-05 - | 0 - | 0 - |
| 3178 | 1 | 3um_FIJJAFBF_449813 | Bacteria | Proteobacteri | Gammaprotec | Alteromonada | Colwelliaceae | Colwellia | Colwellia sp. P | unclassified | unclassified | unclassified | unclassified | unclassified | 0,8 | 0,4 | 2,43E-05 | 1,24E-05 | 1 - | 2,63E-05 - | 1 - | 3,27E-05 - | 1 - | 3,11E-05 - | 1 - | 3,15E-05 - | 0 - | 0 - |
| 3179 | 1 | 3um_FIJJAFBF_780957 | Bacteria | Bacteroidetes | unclassified | unclassified | unclassified | unclassified | unidentified e | unclassified | unclassified | unclassified | unclassified | unclassified | 1 | 0 | 3,83E-05 | 3,63E-06 | 1 - | 3,21E-05 - | 1 - | 3,98E-05 - | 1 - | 3,78E-05 - | 1 - | 3,84E-05 - | 1 - | 4,33E-05 - |
| 3180 | 1 | 0.2um_NOKCPHJN_275444 | Eukaryota | Bacillariophyt | Bacillariophyc | Bacillariales | Bacillariaceae | Fragilariopsis | Fragilariopsis | unclassified | unclassified | unclassified | unclassified | unclassified | 0,8 | 0,748331477 | 7,03E-05 | 6,21E-05 | 2 - | 0,000160527 - | 0 - | 0 - | 1 - | 9,47E-05 - | 1 - | 9,61E-05 - | 0 - | 0 - |
| 3181 | 1 | 3um_FIJJAFBF_255599 | Bacteria | unclassified | unclassified | unclassified | unclassified | unclassified | uncultured ba | poorly charact | Function unkn | 0XNPG | NA | 768066.HELO_ | 0,4 | 0,489897949 | 3,78E-05 | 4,68E-05 | 1 - | 8,44E-05 - | 1 - | 0,000104793 - | 0 - | 0 - | 0 - | 0 - | 0 - | 0 - |
| 3182 | 1 | 0.2um_NOKCPHJN_269101 | Viruses | unclassified | unclassified | unclassified | unclassified | unclassified | uncultured M | unclassified | unclassified | unclassified | unclassified | unclassified | 0,2 | 0,4 | 6,91E-06 | 1,38E-05 | 1 - | 3,46E-05 - | 0 - | 0 - | 0 - | 0 - | 0 - | 0 - | 0 - | 0 - |
| 3183 | 1 | 3um_FIJJAFBF_158908 | Bacteria | Proteobacteri | Alphaproteob | Rhodobactera | Rhodobactera | Planktotalea | Planktotalea a | unclassified | unclassified | unclassified | unclassified | unclassified | 2,4 | 0,489897949 | 6,62E-05 | 1,03E-05 | 3 - | 7,03E-05 - | 2 - | 5,82E-05 - | 2 - | 5,53E-05 - | 3 - | 8,42E-05 - | 2 - | 6,32E-05 - |
| 3184 | 1 | 0.2um_NOKCPHJN_485418 | Eukaryota | Streptophyta | Marchantiops | Marchantiales | Marchantiace | Marchantia | Marchantia p | information st | Translation, ri | 1DTV3 | 60S ribosomal | 3218.PP1S459 | 1 | 0 | 8,41E-05 | 7,98E-06 | 1 - | 7,04E-05 - | 1 - | 8,74E-05 - | 1 - | 8,31E-05 - | 1 - | 8,43E-05 - | 1 - | 9,50E-05 - |
| 3185 | 1 | 0.2um_NOKCPHJN_140037 | Bacteria | Proteobacteri | Gammaprotec | unclassified | unclassified | unclassified | Gammaprotec | unclassified | unclassified | unclassified | unclassified | unclassified | 0,4 | 0,489897949 | 9,86E-06 | 1,23E-05 | 1 - | 2,10E-05 - | 0 - | 0 - | 0 - | 0 - | 0 - | 0 - | 1 - | 2,83E-05 - |
| 3186 | 1 | 3um_FIJJAFBF_504258 | Eukaryota | unclassified | Pelagophycea | Pelagomonad | unclassified | Aureoumbra | Aureoumbra l | metabolism | Energy produc | COG0056 | Produces ATP | 35128.Thapsd | 0,2 | 0,4 | 5,48E-06 | 1,10E-05 | 0 - | 0 - | 0 - | 0 - | 0 - | 0 - | 1 - | 2,74E-05 - | 0 - | 0 - |
| 3187 | 1 | 0.2um_NOKCPHJN_266228 | Bacteria | Bacteroidetes | Flavobacteriia | Flavobacterial | Flavobacteriac | Ulvibacter | Ulvibacter lito | metabolism | Energy produc | 0P7ZB | cytochrome c | 983548.Krodi_ | 0,8 | 0,4 | 2,32E-05 | 1,19E-05 | 1 - | 2,43E-05 - | 1 - | 3,01E-05 - | 0 - | 0 - | 1 - | 2,91E-05 - | 1 - | 3,27E-05 - |
| 3188 | 1 | 3um_FIJJAFBF_543671 | Bacteria | unclassified | unclassified | unclassified | unclassified | unclassified | uncultured ba | poorly charact | Function unkn | 0P9SU | NA | 1046627.BZAF | 0,2 | 0,4 | 6,43E-06 | 1,29E-05 | 1 - | 3,22E-05 - | 0 - | 0 - | 0 - | 0 - | 0 - | 0 - | 0 - | 0 - |
| 3189 | 1 | 3um_FIJJAFBF_274002 | Bacteria | Proteobacteri | Alphaproteob | Rhodobactera | Rhodobactera | Roseovarius | Roseovarius n | poorly charact | Function unkn | COG3181 | extra-cytoplas | 89187.ISM_13 | 1,2 | 0,4 | 5,19E-05 | 1,79E-05 | 1 - | 3,62E-05 - | 1 - | 4,50E-05 - | 1 - | 4,27E-05 - | 2 - | 8,67E-05 - | 1 - | 4,89E-05 - |
| 3190 | 1 | 0.2um_NOKCPHJN_250494 | Bacteria | Proteobacteri | Gammaprotec | unclassified | unclassified | unclassified | various | unclassified | unclassified | unclassified | unclassified | unclassified | 1 | 0,894427191 | 4,43E-05 | 3,89E-05 | 2 - | 7,92E-05 - | 1 - | 4,91E-05 - | 2 - | 9,34E-05 - | 0 - | 0 - | 0 - | 0 - |
| 3191 | 1 | 0.2um_NOKCPHJN_1112807 | Eukaryota | unclassified | Spirotrichea | Sporadotrichi | Oxytrichidae | Sterkiella | Sterkiella hist | information st | Translation, ri | 0T4PB | Elongation fac | 353153.XP_81 | 0,8 | 0,4 | 2,68E-05 | 1,35E-05 | 0 - | 0 - | 1 - | 3,35E-05 - | 1 - | 3,19E-05 - | 1 - | 3,23E-05 - | 1 - | 3,64E-05 - |
| 3192 | 1 | 3um_FIJJAFBF_677318 | Bacteria | various | various | various | various | various | various | unclassified | unclassified | unclassified | unclassified | unclassified | 1,8 | 1,166190379 | 6,13E-05 | 3,25E-05 | 4 - | 0,000120501 - | 1 - | 3,74E-05 - | 1 - | 3,56E-05 - | 2 - | 7,22E-05 - | 1 - | 4,06E-05 - |
| 3193 | 4 | 0.2um_NOKCPHJN_229464;0.2um_NOKCPHJN_2 | Bacteria | Proteobacteri | Alphaproteob | Pelagibacteral | Pelagibacterac | various | various | information st | Translation, ri | COG0185 | Protein S19 fo | 335992.SAR11 | 0,4 | 0,489897949 | 5,05E-05 | 6,22E-05 | 1 - | 0,000114777 - | 0 - | 0 - | 0 - | 0 - | 1 - | 0,000137483 - | 0 - | 0 - |
| 3194 | 1 | 3um_FIJJAFBF_551491 | Eukaryota | unclassified | Pelagophycea | Pelagomonad | unclassified | Aureococcus | Aureococcus a | unclassified | unclassified | unclassified | unclassified | unclassified | 1 | 0,632455532 | 7,35E-05 | 4,14E-05 | 2 - | 0,000128242 - | 0 - | 0 - | 1 - | 7,57E-05 - | 1 - | 7,68E-05 - | 1 - | 8,65E-05 - |
| 3195 | 1 | 0.2um_NOKCPHJN_1044061 | Bacteria | Proteobacteri | Gammaprotec | Cellvibrionale | Spongiibacter | unclassified | marine gamm | cellular proces | Cell wall/mem | 0QTSX | Peptidoglycan | 247633.GP214 | 0,4 | 0,8 | 3,10E-05 | 6,19E-05 | 0 - | 0 - | 0 - | 0 - | 2 - | 0,000154837 - | 0 - | 0 - | 0 - | 0 - |
| 3196 | 1 | 0.2um_NOKCPHJN_84217 | Eukaryota | unclassified | Oomycetes | Saprolegniales | Saprolegniace | Achlya | Achlya hypog | information st | Translation, ri | COG0052 | 30S ribosomal | 653948.CCA1 | 1,6 | 0,489897949 | 6,43E-05 | 2,32E-05 | 1 - | 3,30E-05 - | 2 - | 8,19E-05 - | 2 - | 7,79E-05 - | 1 - | 3,95E-05 - | 2 - | 8,90E-05 - |
| 3197 | 1 | 0.2um_NOKCPHJN_596794 | Bacteria | Bacteroidetes | Flavobacteriia | Flavobacterial | Flavobacteriac | unclassified | Flavobacteriac | unclassified | unclassified | unclassified | unclassified | unclassified | 0,6 | 0,489897949 | 7,29E-05 | 5,96E-05 | 0 - | 0 - | 1 - | 0,000125017 - | 1 - | 0,000118844 - | 1 - | 0,000120599 - | 0 - | 0 - |
| 3198 | 1 | 0.2um_NOKCPHJN_190076 | Bacteria | Bacteroidetes | Flavobacteriia | Flavobacterial | Flavobacteriac | Polaribacter | Polaribacter s | information st | Translation, ri | 08UHY | One of the pri | 313594.PI23P | 1,8 | 0,748331477 | 0,000155665 | 6,64E-05 | 2 - | 0,000145287 - | 1 - | 9,02E-05 - | 1 - | 8,57E-05 - | 3 - | 0,000261044 - | 2 - | 0,000196044 - |
| 3199 | 1 | 0.2um_NOKCPHJN_1004411 | Bacteria | Proteobacteri | Alphaproteob | Pelagibacteral | Pelagibacterac | Candidatus Pe | Candidatus Pe | information st | Translation, ri | COG0088 | One of the pri | 335992.SAR11 | 0,4 | 0,489897949 | 5,77E-05 | 7,07E-05 | 0 - | 0 - | 1 - | 0,000146927 - | 0 - | 0 - | 1 - | 0,000141735 - | 0 - | 0 - |
| 3200 | 1 | 3um_FIJJAFBF_66140 | Bacteria | Bacteroidetes | Flavobacteriia | Flavobacterial | Flavobacteriac | Polaribacter | various | unclassified | unclassified | unclassified | unclassified | unclassified | 1,8 | 0,4 | 0,000218547 | 5,44E-05 | 2 - | 0,000203145 - | 2 - | 0,000252246 - | 1 - | 0,000119896 - | 2 - | 0,000243333 - | 2 - | 0,000274114 - |
| 3201 | 1 | 3um_FIJJAFBF_815021 | Bacteria | Proteobacteri | Alphaproteob | Rhodobactera | Rhodobactera | Aquimixticola | Aquimixticola | information st | Translation, ri | COG0173 | aspartyl-trna s | 314232.SKA53 | 2,8 | 1,166190379 | 6,40E-05 | 2,72E-05 | 3 - | 5,83E-05 - | 4 - | 9,65E-05 - | 2 - | 4,58E-05 - | 4 - | 9,31E-05 - | 1 - | 2,62E-05 - |
| 3202 | 1 | 3um_FIJJAFBF_72465 | Bacteria | Bacteroidetes | Cytophagia | Cytophagales | Cytophagacea | Ekhidna | Ekhidna lutea | information st | Translation, ri | COG0087 | One of the pri | 880070.Cycma | 8,4 | 1,019803903 | 0,000547525 | 7,97E-05 | 8 - | 0,000437245 - | 7 - | 0,000475063 - | 9 - | 0,000580638 - | 10 - | 0,000654682 - | 8 - | 0,000589998 - |
| 3203 | 1 | 3um_FIJJAFBF_257124 | Eukaryota | unclassified | Florideophyce | Ceramiales | Delesseriacea | Membranopte | Membranopte | unclassified | unclassified | unclassified | unclassified | unclassified | 1 | 0 | 0,00010002 | 9,50E-06 | 1 - | 8,38E-05 - | 1 - | 0,000104028 - | 1 - | 9,89E-05 - | 1 - | 0,000100353 - | 1 - | 0,000113047 - |
| 3204 | 1 | 0.2um_NOKCPHJN_930464 | unclassified | unclassified | unclassified | unclassified | unclassified | unclassified | unclassified | unclassified | unclassified | unclassified | unclassified | unclassified | 0,8 | 0,4 | 4,66E-05 | 2,40E-05 | 1 - | 4,88E-05 - | 1 - | 6,06E-05 - | 1 - | 5,77E-05 - | 0 - | 0 - | 1 - | 6,59E-05 - |
| 3205 | 1 | 3um_FIJJAFBF_112632 | Bacteria | Proteobacteri | Gammaprotec | unclassified | unclassified | unclassified | SAR86 cluster | information st | Transcription | 05D27 | DNA-depende | 247634.GPB2 | 1,8 | 0,748331477 | 2,85E-05 | 1,21E-05 | 2 - | 2,68E-05 - | 3 - | 4,99E-05 - | 2 - | 3,17E-05 - | 1 - | 1,61E-05 - | 1 - | 1,81E-05 - |
| 3206 | 1 | 3um_FIJJAFBF_183674 | Bacteria | Bacteroidetes | Flavobacteriia | Flavobacterial | Flavobacteriac | Jejuia | Jejuia pallidilu | poorly charact | Function unkn | 11P35 | NA | 398720.MED2 | 3,8 | 1,326649916 | 9,13E-05 | 2,38E-05 | 6 - | 0,000124307 - | 2 - | 5,15E-05 - | 4 - | 9,78E-05 - | 4 - | 9,93E-05 - | 3 - | 8,39E-05 - |
| 3207 | 1 | 3um_FIJJAFBF_225248 | Eukaryota | Chlorophyta | Trebouxiophy | Chlorellales | Chlorellaceae | Pseudochlorel | Pseudochlorel | metabolism | Carbohydrate | COG1850 | RuBisCO catal | 3055.DAA009 | 1,4 | 0,489897949 | 3,95E-05 | 1,16E-05 | 2 - | 4,83E-05 - | 1 - | 3,00E-05 - | 1 - | 2,85E-05 - | 2 - | 5,79E-05 - | 1 - | 3,26E-05 - |
| 3208 | 1 | 3um_FIJJAFBF_07079 | Bacteria | Proteobacteri | Gammaprotec | Oceanospirilla | Oceanospirilla | Neptuniibacte | Neptuniibacte | unclassified | unclassified | unclassified | unclassified | unclassified | 0,4 | 0,489897949 | 1,21E-05 | 1,50E-05 | 1 - | 2,57E-05 - | 0 - | 0 - | 0 - | 0 - | 0 - | 0 - | 1 - | 3,46E-05 - |
| 3209 | 1 | 3um_FIJJAFBF_17186 | Bacteria | Proteobacteri | Alphaproteob | Rhodobactera | unclassified | unclassified | Rhodobactera | unclassified | unclassified | unclassified | unclassified | unclassified | 1,4 | 0,489897949 | 4,08E-05 | 1,50E-05 | 1 - | 2,43E-05 - | 2 - | 6,04E-05 - | 2 - | 5,74E-05 - | 1 - | 2,91E-05 - | 1 - | 3,28E-05 - |
| 3210 | 1 | 3um_FIJJAFBF_488701 | Bacteria | Proteobacteri | Alphaproteob | Rhizobiales | Methylocystac | Terasakiella | Terasakiella p | unclassified | unclassified | unclassified | unclassified | unclassified | 1 | 0 | 6,47E-06 | 6,15E-07 | 1 - | 5,42E-06 - | 1 - | 6,73E-06 - | 1 - | 6,40E-06 - | 1 - | 6,49E-06 - | 1 - | 7,32E-06 - |
| 3211 | 1 | 0.2um_NOKCPHJN_106487 | Bacteria | Bacteroidetes | Flavobacteriia | Flavobacterial | Flavobacteriac | Aquimarina | Aquimarina m | information st | Translation, ri | 05CEJ | Catalyzes the | 50743.SCB49_ | 7 | 3,033150178 | 0,00013111 | 4,21E-05 | 13 - | 0,000210748 - | 6 - | 0,000120779 - | 5 - | 9,57E-05 - | 5 - | 9,71E-05 - | 6 - | 0,00013125 - |
| 3212 | 1 | 0.2um_NOKCPHJN_93347 | Bacteria | Proteobacteri | Betaproteoba | unclassified | unclassified | unclassified | Betaproteoba | unclassified | unclassified | unclassified | unclassified | unclassified | 0,2 | 0,4 | 1,04E-05 | 2,08E-05 | 0 - | 0 - | 0 - | 0 - | 1 - | 5,19E-05 - | 0 - | 0 - | 0 - | 0 - |
| 3213 | 1 | 3um_FIJJAFBF_111902 | Bacteria | Proteobacteri | Gammaprotec | unclassified | unclassified | unclassified | Gammaprotec | poorly charact | Function unkn | 17FMV | dNA-binding p | 472759.Nhal_ | 1,2 | 0,4 | 0,00014822 | 5,11E-05 | 1 - | 0,000103402 - | 1 - | 0,000128395 - | 1 - | 0,000122056 - | 2 - | 0,000247718 - | 1 - | 0,000139527 - |
| 3214 | 1 | 3um_FIJJAFBF_85330 | Bacteria | Proteobacteri | Gammaprotec | Alteromonada | Alteromonada | Paraglaciecola | Paraglaciecola | metabolism | Energy produc | 17CGT | F(1)F(0) ATP s | 983545.Glaag | 1 | 0,632455532 | 8,85E-05 | 5,84E-05 | 1 - | 7,36E-05 - | 2 - | 0,000182716 - | 1 - | 8,68E-05 - | 0 - | 0 - | 1 - | 9,93E-05 - |
| 3215 | 1 | 3um_FIJJAFBF_481969 | Bacteria | Proteobacteri | Alphaproteob | Rhodobactera | unclassified | unclassified | Rhodobactera | cellular proces | Posttranslatio | 17FI0 | Binds to Cpn6 | 314262.MED1 | 5,2 | 0,748331477 | 0,000749442 | 0,000130857 | 6 - | 0,000724906 - | 5 - | 0,000750099 - | 4 - | 0,000570451 - | 5 - | 0,000723596 - | 6 - | 0,000978155 - |
| 3216 | 2 | 0.2um_NOKCPHJN_225555;3um_FIJJAFBF_3414 | Bacteria | Bacteroidetes | Flavobacteriia | Flavobacterial | Flavobacteriac | Formosa | Formosa sp. H | unclassified | unclassified | unclassified | unclassified | unclassified | 0,8 | 0,4 | 2,27E-05 | 1,15E-05 | 1 - | 2,46E-05 - | 1 - | 3,05E-05 - | 1 - | 2,90E-05 - | 1 - | 2,94E-05 - | 0 - | 0 - |
| 3217 | 1 | 0.2um_NOKCPHJN_55584 | Bacteria | Bacteroidetes | Flavobacteriia | Flavobacterial | Flavobacteriac | Polaribacter | Polaribacter g | poorly charact | Function unkn | 06E9D | NA | 983544.Lacal_ | 3,6 | 1,019803903 | 0,00018874 | 6,30E-05 | 4 - | 0,000174565 - | 3 - | 0,000162569 - | 2 - | 0,000103028 - | 4 - | 0,0002091 - | 5 - | 0,000294438 - |
| 3218 | 1 | 3um_FIJJAFBF_08089 | Bacteria | Bacteroidetes | Flavobacteriia | Flavobacterial | Flavobacteriac | Polaribacter | Polaribacter s | unclassified | unclassified | unclassified | unclassified | unclassified | 0,6 | 0,489897949 | 1,60E-05 | 1,31E-05 | 1 - | 2,36E-05 - | 0 - | 0 - | 1 - | 2,79E-05 - | 1 - | 2,83E-05 - | 0 - | 0 - |
| 3219 | 1 | 3um_FIJJAFBF_346309 | Bacteria | Bacteroidetes | unclassified | unclassified | unclassified | unclassified | unidentified e | unclassified | unclassified | unclassified | unclassified | unclassified | 0,2 | 0,4 | 3,16E-06 | 6,32E-06 | 1 - | 1,58E-05 - | 0 - | 0 - | 0 - | 0 - | 0 - | 0 - | 0 - | 0 - |
| 3220 | 1 | 3um_FIJJAFBF_429762 | Bacteria | Proteobacteri | Gammaprotec | Oceanospirilla | Saccharospirill | Reinekea | Reinekea fors | unclassified | unclassified | unclassified | unclassified | unclassified | 0,6 | 0,489897949 | 3,91E-05 | 3,22E-05 | 1 - | 5,71E-05 - | 1 - | 7,09E-05 - | 1 - | 6,74E-05 - | 0 - | 0 - | 0 - | 0 - |
| 3221 | 1 | 3um_FIJJAFBF_137143 | Bacteria | Proteobacteri | Betaproteoba | unclassified | unclassified | unclassified | Betaproteoba | information st | Translation, ri | 179R6 | This protein is | 232721.Ajs_0 | 2,8 | 0,979795897 | 0,00027187 | 0,000101702 | 2 - | 0,000161657 - | 4 - | 0,000401462 - | 2 - | 0,00019082 - | 4 - | 0,000387277 - | 2 - | 0,000218133 - |
| 3222 | 1 | 3um_FIJJAFBF_751139 | Bacteria | Proteobacteri | Gammaprotec | unclassified | unclassified | unclassified | SAR86 cluster | unclassified | unclassified | unclassified | unclassified | unclassified | 5,8 | 0,748331477 | 0,000442791 | 6,49E-05 | 6 - | 0,000384727 - | 7 - | 0,000557336 - | 5 - | 0,000378442 - | 6 - | 0,000460838 - | 5 - | 0,00043261 - |
| 3223 | 1 | 3um_FIJJAFBF_132845 | Bacteria | Proteobacteri | Gammaprotec | Cellvibrionale | Porticoccacea | unclassified | Gammaprotec | information st | Translation, ri | 05G6W | One of the pri | 203122.Sde_0 | 3,4 | 0,489897949 | 0,000224045 | 2,38E-05 | 4 - | 0,000222867 - | 3 - | 0,000207552 - | 3 - | 0,000197304 - | 4 - | 0,000266958 - | 3 - | 0,000225545 - |
| 3224 | 1 | 0.2um_NOKCPHJN_369026 | Bacteria | Proteobacteri | Alphaproteob | Rhodobactera | Rhodobactera | unclassified | Rhodobactera | unclassified | unclassified | unclassified | unclassified | unclassified | 3,2 | 0,748331477 | 0,000109567 | 2,66E-05 | 3 - | 8,63E-05 - | 2 - | 7,14E-05 - | 4 - | 0,000135822 - | 4 - | 0,000137828 - | 3 - | 0,000116447 - |
| 3225 | 1 | 0.2um_NOKCPHJN_571549 | Bacteria | Proteobacteri | Alphaproteob | Pelagibacteral | Pelagibacterac | Candidatus Pe | Candidatus Pe | cellular proces | Posttranslatio | COG0330 | Band 7 protei | 439493.PB721 | 2,6 | 0,489897949 | 9,76E-05 | 2,08E-05 | 2 - | 6,27E-05 - | 3 - | 0,000116819 - | 3 - | 0,000111051 - | 3 - | 0,000112691 - | 2 - | 8,46E-05 - |
| 3226 | 1 | 3um_FIJJAFBF_33603 | Bacteria | Proteobacteri | Alphaproteob | Rhodospirillal | Rhodospirillac | unclassified | Rhodobactera | metabolism | Carbohydrate | 01QS3 | extracellular s | 388401.RB215 | 0,2 | 0,4 | 6,17E-06 | 1,23E-05 | 0 - | 0 - | 0 - | 0 - | 0 - | 0 - | 1 - | 3,08E-05 - | 0 - | 0 - |
| 3227 | 1 | 3um_FIJJAFBF_663789 | Bacteria | Bacteroidetes | Flavobacteriia | Flavobacterial | Flavobacteriac | Polaribacter | Polaribacter s | information st | Translation, ri | 05CAV | thus facilitatin | 391587.KAOT | 3,6 | 0,8 | 8,11E-05 | 2,47E-05 | 3 - | 5,58E-05 - | 3 - | 6,93E-05 - | 3 - | 6,59E-05 - | 4 - | 8,91E-05 - | 5 - | 0,000125506 - |
| 3228 | 2 | 0.2um_NOKCPHJN_95248;3um_FIJJAFBF_691251 | Eukaryota | unclassified | Chrysophycea | Chromulinales | Chromulinace | Ochromonas | Ochromonas s | unclassified | unclassified | unclassified | unclassified | unclassified | 0,2 | 0,4 | 9,82E-06 | 1,96E-05 | 0 - | 0 - | 0 - | 0 - | 1 - | 4,91E-05 - | 0 - | 0 - | 0 - | 0 - |
| 3229 | 1 | 0.2um_NOKCPHJN_513503 | Bacteria | Proteobacteri | Alphaproteob | Rhodobactera | Rhodobactera | Planktomarina | Planktomarina | unclassified | unclassified | unclassified | unclassified | unclassified | 10,2 | 0,979795897 | 0,001123459 | 0,000180893 | 9 - | 0,000826393 - | 12 - | 0,001368181 - | 10 - | 0,001083858 - | 10 - | 0,001099866 - | 10 - | 0,001238996 - |
| 3230 | 1 | 3um_FIJJAFBF_384698 | Bacteria | Bacteroidetes | Flavobacteriia | Flavobacterial | Flavobacteriac | Polaribacter | Polaribacter s | information st | Translation, ri | 09FEM | Binds to the 2 | 313594.PI23P | 1,8 | 0,748331477 | 0,000161751 | 5,75E-05 | 3 - | 0,000231094 - | 1 - | 9,57E-05 - | 2 - | 0,000181855 - | 1 - | 9,23E-05 - | 2 - | 0,000207885 - |
| 3231 | 1 | 3um_FIJJAFBF_269588 | Bacteria | Proteobacteri | Alphaproteob | Rhodobactera | Rhodobactera | unclassified | Rhodobactera | cellular proces | Defense mech | COG0610 | Type I site-spe | 388401.RB215 | 1 | 0 | 1,26E-05 | 1,19E-06 | 1 - | 1,05E-05 - | 1 - | 1,31E-05 - | 1 - | 1,24E-05 - | 1 - | 1,26E-05 - | 1 - | 1,42E-05 - |
| 3232 | 2 | 0.2um_NOKCPHJN_340055;0.2um_NOKCPHJN_5 | Bacteria | Bacteroidetes | Flavobacteriia | Flavobacterial | Flavobacteriac | various | various | unclassified | unclassified | unclassified | unclassified | unclassified | 5,6 | 1,019803903 | 0,000616692 | 0,000118842 | 6 - | 0,000555371 - | 7 - | 0,000804542 - | 6 - | 0,000655559 - | 4 - | 0,000443495 - | 5 - | 0,000624494 - |
| 3233 | 1 | 0.2um_NOKCPHJN_333008 | Bacteria | Proteobacteri | Gammaprotec | Alteromonada | Alteromonada | Glaciecola | Glaciecola sp. | cellular proces | Posttranslatio | 16QU3 | Hemolysin-typ | 455436.GHTC | 3 | 0,632455532 | 2,26E-05 | 5,54E-06 | 3 - | 1,89E-05 - | 4 - | 3,13E-05 - | 3 - | 2,23E-05 - | 2 - | 1,51E-05 - | 3 - | 2,55E-05 - |
| 3234 | 1 | 0.2um_NOKCPHJN_30024 | Bacteria | Bacteroidetes | Flavobacteriia | Flavobacterial | Flavobacteriac | Polaribacter | Polaribacter a | information st | Translation, ri | 0P97G | This is one of | 313594.PI23P | 1,8 | 0,748331477 | 0,000120564 | 5,14E-05 | 2 - | 0,000112526 - | 1 - | 6,99E-05 - | 1 - | 6,64E-05 - | 3 - | 0,000202181 - | 2 - | 0,000151838 - |
| 3235 | 1 | 0.2um_NOKCPHJN_1095963 | Bacteria | Proteobacteri | Alphaproteob | unclassified | unclassified | unclassified | Alphaproteob | metabolism | Amino acid tra | 05TIB | (ABC) transpo | 684719.HIMB | 0,8 | 0,4 | 3,07E-05 | 1,56E-05 | 1 - | 3,33E-05 - | 1 - | 4,13E-05 - | 1 - | 3,93E-05 - | 1 - | 3,99E-05 - | 0 - | 0 - |
| 3236 | 1 | 3um_FIJJAFBF_175877 | Bacteria | Proteobacteri | Gammaprotec | unclassified | unclassified | unclassified | Gammaprotec | information st | Translation, ri | 08VZM | 50s ribosomal | 876044.IMCC | 2 | 0 | 0,000156602 | 1,49E-05 | 2 - | 0,000131173 - | 2 - | 0,000162879 - | 2 - | 0,000154837 - | 2 - | 0,000157124 - | 2 - | 0,000176999 - |
| 3237 | 2 | 0.2um_NOKCPHJN_146469;0.2um_NOKCPHJN_3 | Bacteria | Bacteroidetes | unclassified | unclassified | unclassified | unclassified | Bacteroidetes | metabolism | Inorganic ion t | 099UQ | tonB-depende | 313606.M231 | 0,2 | 0,4 | 2,88E-06 | 5,77E-06 | 1 - | 1,44E-05 - | 0 - | 0 - | 0 - | 0 - | 0 - | 0 - | 0 - | 0 - |
| 3238 | 1 | 3um_FIJJAFBF_95716 | Eukaryota | unclassified | Raphidophyce | Chattonellales | Chattonellace | Heterosigma | Heterosigma a | cellular proces | Posttranslatio | 0IEKR | One of the co | 35128.Thapsd | 13,8 | 0,748331477 | 0,000410886 | 2,09E-05 | 15 - | 0,000375906 - | 13 - | 0,00040453 - | 14 - | 0,000414138 - | 14 - | 0,000420255 - | 13 - | 0,0004396 - |
| 3239 | 1 | 3um_FIJJAFBF_521164 | Bacteria | Proteobacteri | Betaproteoba | unclassified | unclassified | unclassified | Betaproteoba | metabolism | Carbohydrate | 0B8CV | Transketolase | 296591.Bpro_ | 1,2 | 0,748331477 | 2,33E-05 | 1,44E-05 | 2 - | 3,40E-05 - | 2 - | 4,22E-05 - | 1 - | 2,00E-05 - | 1 - | 2,03E-05 - | 0 - | 0 - |
| 3240 | 2 | 0.2um_NOKCPHJN_567918;3um_FIJJAFBF_41518 | Eukaryota | unclassified | Pelagophycea | Pelagomonad | unclassified | Aureococcus | Aureococcus a | cellular proces | Signal transdu | KOG0613 | fibronectin typ | various | 1,2 | 0,4 | 1,50E-05 | 5,18E-06 | 1 - | 1,05E-05 - | 1 - | 1,30E-05 - | 1 - | 1,24E-05 - | 2 - | 2,51E-05 - | 1 - | 1,41E-05 - |
| 3241 | 1 | 3um_FIJJAFBF_596486 | Bacteria | Proteobacteri | Gammaprotec | Alteromonada | Alteromonada | Paraglaciecola | Paraglaciecola | unclassified | unclassified | unclassified | unclassified | unclassified | 0,2 | 0,4 | 3,29E-06 | 6,58E-06 | 1 - | 1,64E-05 - | 0 - | 0 - | 0 - | 0 - | 0 - | 0 - | 0 - | 0 - |
| 3242 | 1 | 0.2um_NOKCPHJN_805291 | Eukaryota | unclassified | Oomycetes | Pythiales | Pythiaceae | Pythium | Pythium insidi | unclassified | unclassified | unclassified | unclassified | unclassified | 0,4 | 0,8 | 2,50E-05 | 4,99E-05 | 2 - | 0,000124757 - | 0 - | 0 - | 0 - | 0 - | 0 - | 0 - | 0 - | 0 - |
| 3243 | 1 | 3um_FIJJAFBF_154878 | Bacteria | Proteobacteri | Betaproteoba | unclassified | unclassified | unclassified | Betaproteoba | unclassified | unclassified | unclassified | unclassified | unclassified | 1,4 | 1,019803903 | 7,12E-05 | 4,76E-05 | 3 - | 0,000136639 - | 1 - | 5,66E-05 - | 1 - | 5,38E-05 - | 2 - | 0,000109114 - | 0 - | 0 - |
| 3244 | 1 | 0.2um_NOKCPHJN_562106 | Bacteria | Proteobacteri | Gammaprotec | Cellvibrionale | Porticoccacea | unclassified | Gammaprotec | metabolism | Inorganic ion t | 0QHT5 | TonB-depende | 314287.GB22 | 0,4 | 0,489897949 | 6,51E-06 | 8,12E-06 | 1 - | 1,39E-05 - | 0 - | 0 - | 0 - | 0 - | 0 - | 0 - | 1 - | 1,87E-05 - |
| 3245 | 1 | 0.2um_NOKCPHJN_110539 | Bacteria | Proteobacteri | Gammaprotec | Cellvibrionale | Porticoccacea | unclassified | SAR92 bacteri | information st | Translation, ri | 0QK14 | This protein bi | 314287.GB22 | 5,4 | 1,019803903 | 0,000415017 | 7,41E-05 | 6 - | 0,000389074 - | 4 - | 0,000322077 - | 5 - | 0,000382718 - | 7 - | 0,000543719 - | 5 - | 0,000437499 - |
| 3246 | 2 | 0.2um_NOKCPHJN_460311;3um_FIJJAFBF_5060 | Bacteria | Proteobacteri | Alphaproteob | Rhodobactera | Rhodobactera | Planktomarina | Planktomarina | unclassified | unclassified | unclassified | unclassified | unclassified | 0,6 | 0,489897949 | 5,15E-05 | 4,28E-05 | 1 - | 7,17E-05 - | 1 - | 8,91E-05 - | 0 - | 0 - | 0 - | 0 - | 1 - | 9,68E-05 - |
| 3247 | 1 | 3um_FIJJAFBF_218232 | Bacteria | Proteobacteri | Gammaprotec | Alteromonada | Alteromonada | Glaciecola | Glaciecola pal | unclassified | unclassified | unclassified | unclassified | unclassified | 3,4 | 0,489897949 | 5,87E-05 | 1,32E-05 | 3 - | 4,30E-05 - | 4 - | 7,12E-05 - | 3 - | 5,07E-05 - | 3 - | 5,15E-05 - | 4 - | 7,73E-05 - |
| 3248 | 1 | 0.2um_NOKCPHJN_56834 | Bacteria | Proteobacteri | Gammaprotec | Oceanospirilla | Saccharospirill | Reinekea | Reinekea blan | cellular proces | Posttranslatio | 05C2Z | ATP-depender | 225937.HP15_ | 3,2 | 0,979795897 | 5,87E-05 | 2,26E-05 | 3 - | 4,55E-05 - | 2 - | 3,77E-05 - | 3 - | 5,37E-05 - | 3 - | 5,45E-05 - | 5 - | 0,000102295 - |
| 3249 | 1 | 3um_FIJJAFBF_292922 | Eukaryota | Bacillariophyt | Coscinodiscop | Coscinodiscale | Coscinodiscac | Coscinodiscus | Coscinodiscus | metabolism | Energy produc | COG4451 | RuBisCO catal | 977880.RALTA | 8 | 1,095445115 | 0,000781674 | 7,49E-05 | 10 - | 0,000825732 - | 7 - | 0,000717721 - | 7 - | 0,000682285 - | 8 - | 0,000791271 - | 8 - | 0,000891364 - |
| 3250 | 1 | 3um_FIJJAFBF_227245 | Bacteria | Proteobacteri | Gammaprotec | Chromatiales | Granulosicocc | Granulosicocc | Granulosicocc | cellular proces | Posttranslatio | 08V1T | peptidylprolyl | 357804.Ping_ | 2 | 0 | 0,000122346 | 1,16E-05 | 2 - | 0,000102479 - | 2 - | 0,000127249 - | 2 - | 0,000120966 - | 2 - | 0,000122753 - | 2 - | 0,000138281 - |
| 3251 | 1 | 0.2um_NOKCPHJN_1025680 | unclassified | unclassified | unclassified | unclassified | unclassified | unclassified | unclassified | unclassified | unclassified | unclassified | unclassified | unclassified | 0,2 | 0,4 | 1,82E-05 | 3,64E-05 | 0 - | 0 - | 0 - | 0 - | 1 - | 9,09E-05 - | 0 - | 0 - | 0 - | 0 - |
| 3252 | 2 | 0.2um_NOKCPHJN_18165;3um_FIJJAFBF_38273 | Bacteria | Proteobacteri | Epsilonproteo | Nautiliales | Nautiliaceae | Nautilia | Nautilia profu | various | various | various | various | various | 1 | 0 | 0,000102259 | 9,71E-06 | 1 - | 8,57E-05 - | 1 - | 0,000106357 - | 1 - | 0,000101106 - | 1 - | 0,000102599 - | 1 - | 0,000115578 - |
| 3253 | 1 | 3um_FIJJAFBF_328402 | Bacteria | Proteobacteri | Gammaprotec | Cellvibrionale | Porticoccacea | unclassified | various | unclassified | unclassified | unclassified | unclassified | unclassified | 0,4 | 0,8 | 4,10E-05 | 8,20E-05 | 2 - | 0,000204958 - | 0 - | 0 - | 0 - | 0 - | 0 - | 0 - | 0 - | 0 - |
| 3254 | 1 | 3um_FIJJAFBF_00233 | Bacteria | Bacteroidetes | Flavobacteriia | Flavobacterial | Flavobacteriac | various | various | unclassified | unclassified | unclassified | unclassified | unclassified | 0,4 | 0,489897949 | 5,26E-05 | 6,48E-05 | 1 - | 0,000119559 - | 0 - | 0 - | 0 - | 0 - | 1 - | 0,000143212 - | 0 - | 0 - |
| 3255 | 1 | 3um_FIJJAFBF_196746 | Bacteria | Proteobacteri | Alphaproteob | unclassified | unclassified | unclassified | Alphaproteob | unclassified | unclassified | unclassified | unclassified | unclassified | 3 | 0 | 0,000173452 | 1,65E-05 | 3 - | 0,000145287 - | 3 - | 0,000180404 - | 3 - | 0,000171496 - | 3 - | 0,00017403 - | 3 - | 0,000196044 - |
| 3256 | 1 | 0.2um_NOKCPHJN_374320 | Bacteria | Proteobacteri | Alphaproteob | Rhodospirillal | unclassified | unclassified | Rhodospirillal | metabolism | Amino acid tra | COG0683 | (ABC) transpo | 504832.OCA5 | 1 | 0 | 3,14E-05 | 2,98E-06 | 1 - | 2,63E-05 - | 1 - | 3,27E-05 - | 1 - | 3,11E-05 - | 1 - | 3,15E-05 - | 1 - | 3,55E-05 - |
| 3257 | 1 | 3um_FIJJAFBF_594171 | Eukaryota | unclassified | Dinophyceae | Peridiniales | Kryptoperidini | Kryptoperidini | Kryptoperidin | metabolism | Energy produc | 11HAV | photosystem i | 35128.Thapsd | 8 | 0,894427191 | 0,000749786 | 5,65E-05 | 9 - | 0,000712407 - | 7 - | 0,000688022 - | 8 - | 0,000747488 - | 9 - | 0,000853345 - | 7 - | 0,00074767 - |
| 3258 | 1 | 3um_FIJJAFBF_800346 | Bacteria | Proteobacteri | Gammaprotec | Oceanospirilla | Saccharospirill | Reinekea | Reinekea fors | information st | Translation, ri | 16QAP | Associates wit | 314283.MED2 | 12 | 0,632455532 | 0,000571724 | 4,92E-05 | 12 - | 0,000479903 - | 12 - | 0,000595898 - | 12 - | 0,000566476 - | 13 - | 0,000622747 - | 11 - | 0,000593596 - |
| 3259 | 1 | 3um_FIJJAFBF_394456 | Bacteria | Bacteroidetes | Flavobacteriia | Flavobacterial | Flavobacteriac | Altibacter | Altibacter lent | poorly charact | Function unkn | 0XQTX | Chromosome | 50743.SCB49_ | 0,2 | 0,4 | 2,51E-06 | 5,03E-06 | 0 - | 0 - | 0 - | 0 - | 0 - | 0 - | 1 - | 1,26E-05 - | 0 - | 0 - |
| 3260 | 1 | 3um_FIJJAFBF_347374 | Bacteria | Proteobacteri | Alphaproteob | Rhizobiales | unclassified | unclassified | Rhizobiales ba | cellular proces | Intracellular tr | 18ATJ | Pfam:YadA | 1005048.CFU_ | 0,6 | 0,8 | 5,04E-06 | 6,45E-06 | 2 - | 1,55E-05 - | 1 - | 9,65E-06 - | 0 - | 0 - | 0 - | 0 - | 0 - | 0 - |
| 3261 | 1 | 3um_FIJJAFBF_654574 | Bacteria | Proteobacteri | Alphaproteob | Rhodobactera | Rhodobactera | Tateyamaria | Tateyamaria s | information st | Translation, ri | 177SZ | This protein bi | 314262.MED1 | 0,2 | 0,4 | 1,63E-05 | 3,26E-05 | 1 - | 8,14E-05 - | 0 - | 0 - | 0 - | 0 - | 0 - | 0 - | 0 - | 0 - |
| 3262 | 7 | 0.2um_NOKCPHJN_189631;0.2um_NOKCPHJN_3 | Bacteria | various | various | various | various | various | various | unclassified | unclassified | unclassified | unclassified | unclassified | 0,2 | 0,4 | 1,36E-05 | 2,72E-05 | 0 - | 0 - | 0 - | 0 - | 1 - | 6,81E-05 - | 0 - | 0 - | 0 - | 0 - |
| 3263 | 1 | 3um_FIJJAFBF_420069 | Bacteria | Proteobacteri | Alphaproteob | Rhodobactera | Rhodobactera | unclassified | Rhodobactera | unclassified | unclassified | unclassified | unclassified | unclassified | 1,8 | 0,748331477 | 4,57E-05 | 2,01E-05 | 1 - | 2,11E-05 - | 3 - | 7,86E-05 - | 2 - | 4,98E-05 - | 2 - | 5,05E-05 - | 1 - | 2,85E-05 - |
| 3264 | 1 | 0.2um_NOKCPHJN_873954 | Bacteria | Bacteroidetes | Flavobacteriia | Flavobacterial | Flavobacteriac | Polaribacter | Polaribacter r | unclassified | unclassified | unclassified | unclassified | unclassified | 0,6 | 0,489897949 | 1,19E-05 | 9,86E-06 | 1 - | 1,65E-05 - | 1 - | 2,05E-05 - | 0 - | 0 - | 0 - | 0 - | 1 - | 2,23E-05 - |
| 3265 | 1 | 0.2um_NOKCPHJN_571787 | Bacteria | Bacteroidetes | Flavobacteriia | Flavobacterial | Flavobacteriac | unclassified | Flavobacteriac | unclassified | unclassified | unclassified | unclassified | unclassified | 0,6 | 0,489897949 | 2,69E-05 | 2,20E-05 | 0 - | 0 - | 1 - | 4,61E-05 - | 1 - | 4,38E-05 - | 1 - | 4,45E-05 - | 0 - | 0 - |
| 3266 | 1 | 3um_FIJJAFBF_386560 | Bacteria | Bacteroidetes | Flavobacteriia | Flavobacterial | Flavobacteriac | Tenacibaculur | Tenacibaculur | metabolism | Lipid transpor | COG1960 | acyl-CoA dehy | 313598.MED1 | 0,4 | 0,489897949 | 8,29E-06 | 1,02E-05 | 1 - | 1,90E-05 - | 0 - | 0 - | 1 - | 2,24E-05 - | 0 - | 0 - | 0 - | 0 - |
| 3267 | 1 | 3um_FIJJAFBF_731933 | Bacteria | Proteobacteri | Gammaprotec | Oceanospirilla | Saccharospirill | Reinekea | Reinekea fors | cellular proces | Posttranslatio | COG4783 | peptidase m4 | 653733.Selin_ | 1 | 0,632455532 | 4,71E-05 | 2,68E-05 | 2 - | 8,38E-05 - | 1 - | 5,20E-05 - | 1 - | 4,94E-05 - | 1 - | 5,02E-05 - | 0 - | 0 - |
| 3268 | 1 | 0.2um_NOKCPHJN_721458 | Bacteria | Proteobacteri | Alphaproteob | Pelagibacteral | Pelagibacterac | Candidatus Pe | Candidatus Pe | metabolism | Amino acid tra | 16R0H | Acetolactate s | 335992.SAR11 | 0,2 | 0,4 | 1,26E-05 | 2,52E-05 | 1 - | 6,31E-05 - | 0 - | 0 - | 0 - | 0 - | 0 - | 0 - | 0 - | 0 - |
| 3269 | 1 | 3um_FIJJAFBF_134913 | Bacteria | Bacteroidetes | Cytophagia | Cytophagales | Flammeovirga | unclassified | Flammeovirga | poorly charact | Function unkn | 0XY6Y | Pkd domain c | 391587.KAOT | 4,2 | 1,326649916 | 7,99E-05 | 2,41E-05 | 5 - | 8,04E-05 - | 4 - | 7,98E-05 - | 6 - | 0,000113851 - | 2 - | 3,85E-05 - | 4 - | 8,68E-05 - |
| 3270 | 1 | 0.2um_NOKCPHJN_52520 | Bacteria | Proteobacteri | Gammaprotec | Cellvibrionale | Porticoccacea | Porticoccus | Porticoccus sp | information st | Translation, ri | 16QAF | thus facilitatin | 314287.GB22 | 0,2 | 0,4 | 4,94E-06 | 9,87E-06 | 0 - | 0 - | 0 - | 0 - | 0 - | 0 - | 1 - | 2,47E-05 - | 0 - | 0 - |
| 3271 | 1 | 0.2um_NOKCPHJN_264013 | Bacteria | Proteobacteri | Gammaprotec | unclassified | unclassified | unclassified | Gammaprotec | information st | Transcription | 05D27 | DNA-depende | 314285.KT71_ | 7,8 | 1,166190379 | 7,64E-05 | 1,59E-05 | 7 - | 5,71E-05 - | 10 - | 0,000101221 - | 7 - | 6,74E-05 - | 7 - | 6,84E-05 - | 8 - | 8,80E-05 - |
| 3272 | 1 | 3um_FIJJAFBF_793459 | Bacteria | Bacteroidetes | Flavobacteriia | unclassified | unclassified | unclassified | Flavobacteria | poorly charact | Function unkn | 0XR4Y | Ragb susd dor | 487796.Flav2 | 11,8 | 0,748331477 | 0,0002765 | 2,84E-05 | 12 - | 0,000235843 - | 13 - | 0,000317251 - | 11 - | 0,000255189 - | 12 - | 0,0002825 - | 11 - | 0,000291716 - |
| 3273 | 1 | 0.2um_NOKCPHJN_431176 | Bacteria | Proteobacteri | Alphaproteob | Rhodobactera | Rhodobactera | Planktomarina | Planktomarina | cellular proces | Posttranslatio | 16QU3 | Hemolysin-typ | 744980.TRICH | 4 | 0,894427191 | 4,34E-05 | 9,50E-06 | 5 - | 4,57E-05 - | 5 - | 5,67E-05 - | 3 - | 3,24E-05 - | 3 - | 3,28E-05 - | 4 - | 4,93E-05 - |
| 3274 | 1 | 3um_FIJJAFBF_397633 | Bacteria | Bacteroidetes | Flavobacteriia | Flavobacterial | Flavobacteriac | Marixanthom | Marixanthom | unclassified | unclassified | unclassified | unclassified | unclassified | 2,4 | 1,019803903 | 8,45E-05 | 3,62E-05 | 2 - | 5,96E-05 - | 2 - | 7,40E-05 - | 3 - | 0,000105571 - | 4 - | 0,00014284 - | 1 - | 4,02E-05 - |
| 3275 | 1 | 3um_FIJJAFBF_422340 | Bacteria | Bacteroidetes | Flavobacteriia | Flavobacterial | Flavobacteriac | Polaribacter | Polaribacter s | poorly charact | Function unkn | 11RX2 | NA | 313594.PI23P | 0,8 | 0,4 | 6,31E-05 | 3,21E-05 | 1 - | 6,83E-05 - | 1 - | 8,48E-05 - | 1 - | 8,06E-05 - | 1 - | 8,18E-05 - | 0 - | 0 - |
| 3276 | 2 | 0.2um_NOKCPHJN_190470;3um_FIJJAFBF_1642 | Bacteria | Bacteroidetes | Flavobacteriia | Flavobacterial | Crocinitomica | unclassified | Crocinitomica | cellular proces | Posttranslatio | 05E6M | Sulphatase-m | 760192.Halhy | 0,2 | 0,4 | 6,04E-06 | 1,21E-05 | 0 - | 0 - | 0 - | 0 - | 0 - | 0 - | 1 - | 3,02E-05 - | 0 - | 0 - |
| 3277 | 1 | 3um_FIJJAFBF_283091 | Bacteria | Bacteroidetes | Flavobacteriia | Flavobacterial | Flavobacteriac | Algibacter | Algibacter sp. | poorly charact | Function unkn | 08KQQ | NA | 391603.FBALC | 1 | 0,632455532 | 5,30E-05 | 3,72E-05 | 1 - | 4,31E-05 - | 1 - | 5,36E-05 - | 0 - | 0 - | 1 - | 5,17E-05 - | 2 - | 0,000116447 - |
| 3278 | 1 | 3um_FIJJAFBF_545216 | Bacteria | Bacteroidetes | Flavobacteriia | Flavobacterial | Flavobacteriac | Flavivirga | Flavivirga eckl | unclassified | unclassified | unclassified | unclassified | unclassified | 4,8 | 0,979795897 | 0,000366168 | 9,13E-05 | 5 - | 0,000318824 - | 3 - | 0,000237531 - | 5 - | 0,00037634 - | 5 - | 0,000381898 - | 6 - | 0,000516248 - |
| 3279 | 1 | 3um_FIJJAFBF_512335 | Bacteria | Proteobacteri | Alphaproteob | Rhodobactera | Rhodobactera | Tropicibacter | Tropicibacter | metabolism | Energy produc | 16QM4 | alcohol dehyd | 391595.RLO14 | 0,4 | 0,489897949 | 1,52E-05 | 1,87E-05 | 1 - | 3,48E-05 - | 0 - | 0 - | 1 - | 4,11E-05 - | 0 - | 0 - | 0 - | 0 - |
| 3280 | 1 | 3um_FIJJAFBF_246178 | Bacteria | Proteobacteri | Gammaprotec | Oceanospirilla | Saccharospirill | Reinekea | Reinekea fors | poorly charact | Function unkn | 11NWN | Predicted peri | 351348.Maqu | 1,6 | 0,489897949 | 0,000137548 | 3,60E-05 | 2 - | 0,00014715 - | 1 - | 9,14E-05 - | 2 - | 0,000173695 - | 2 - | 0,000176261 - | 1 - | 9,93E-05 - |
| 3281 | 1 | 3um_FIJJAFBF_320586 | Bacteria | Bacteroidetes | Flavobacteriia | Flavobacterial | Flavobacteriac | Polaribacter | Polaribacter s | metabolism | Energy produc | COG0055 | Produces ATP | 411154.GFO_ | 0,2 | 0,4 | 4,57E-06 | 9,15E-06 | 1 - | 2,29E-05 - | 0 - | 0 - | 0 - | 0 - | 0 - | 0 - | 0 - | 0 - |
| 3282 | 1 | 0.2um_NOKCPHJN_274158 | Bacteria | Proteobacteri | Betaproteoba | Nitrosomonad | unclassified | unclassified | various | unclassified | unclassified | unclassified | unclassified | unclassified | 1 | 0 | 6,56E-05 | 6,23E-06 | 1 - | 5,49E-05 - | 1 - | 6,82E-05 - | 1 - | 6,48E-05 - | 1 - | 6,58E-05 - | 1 - | 7,41E-05 - |
| 3283 | 1 | 0.2um_NOKCPHJN_48075 | Bacteria | Bacteroidetes | Flavobacteriia | unclassified | unclassified | unclassified | Flavobacteria | information st | Transcription | 05D27 | DNA-depende | 487796.Flav2 | 2,8 | 1,469693846 | 2,57E-05 | 1,19E-05 | 5 - | 4,00E-05 - | 4 - | 3,98E-05 - | 2 - | 1,89E-05 - | 2 - | 1,92E-05 - | 1 - | 1,08E-05 - |

| | | | | | | | | | | | | | | | | | | | | | | | | | | | | |
|---|---|---|---|---|---|---|---|---|---|---|---|---|---|---|---|---|---|---|---|---|---|---|---|---|---|---|---|---|
| 3284 | 1 | 0.2um_NOKCPHJN_46917 | Bacteria | Bacteroidetes | Flavobacteriia | Flavobacterial | Flavobacteriac | Formosa | Formosa sp. H | poorly charact | Function unkn | 0PAEZ | Inherit from b | 391598.FBBAL | 0,8 | 0,4 | 4,08E-05 | 2,07E-05 | 1 - | 4,41E-05 - | 1 - | 5,48E-05 - | 1 - | 5,21E-05 - | 1 - | 5,29E-05 - | 0 - | 0 - |
| 3285 | 1 | 3um_FIJJAFBF_346215 | Bacteria | Bacteroidetes | Flavobacteriia | Flavobacterial | Crocinitomica | unclassified | Crocinitomica | unclassified | unclassified | unclassified | unclassified | unclassified | 2,4 | 0,489897949 | 0,000432431 | 9,33E-05 | 2 - | 0,000302044 - | 2 - | 0,00037505 - | 3 - | 0,000534798 - | 3 - | 0,000542697 - | 2 - | 0,000407564 - |
| 3286 | 1 | 3um_FIJJAFBF_275255 | Bacteria | Proteobacteri | Gammaprotec | unclassified | unclassified | unclassified | OM182 bacte | metabolism | Inorganic ion t | COG4206 | tonB-depende | 314285.KT71_ | 0,4 | 0,489897949 | 8,71E-06 | 1,07E-05 | 0 - | 0 - | 0 - | 0 - | 0 - | 0 - | 1 - | 2,05E-05 - | 1 - | 2,31E-05 - |
| 3287 | 1 | 0.2um_NOKCPHJN_799008 | Bacteria | Firmicutes | Negativicutes | Veillonellales | Veillonellacea | Megasphaera | Megasphaera | cellular proces | Intracellular tr | 17HSD | domain protei | 96561.Dole_2 | 1 | 0,632455532 | 1,77E-05 | 1,20E-05 | 0 - | 0 - | 1 - | 1,74E-05 - | 1 - | 1,66E-05 - | 1 - | 1,68E-05 - | 2 - | 3,79E-05 - |
| 3288 | 1 | 3um_FIJJAFBF_233543 | Bacteria | Proteobacteri | Alphaproteob | Rhodobactera | unclassified | unclassified | Rhodobactera | cellular proces | Posttranslatio | 01ZS8 | Thioredoxin | 367336.OM22 | 1,8 | 0,4 | 0,000232979 | 5,80E-05 | 2 - | 0,00021656 - | 2 - | 0,000268904 - | 1 - | 0,000127813 - | 2 - | 0,000259402 - | 2 - | 0,000292216 - |
| 3289 | 1 | 0.2um_NOKCPHJN_50314 | Bacteria | Bacteroidetes | Flavobacteriia | Flavobacterial | Flavobacteriac | Polaribacter | Polaribacter h | information st | Translation, ril | COG0017 | asparaginyl-tR | 313598.MED1 | 0,8 | 0,748331477 | 2,24E-05 | 2,21E-05 | 1 - | 2,41E-05 - | 2 - | 5,98E-05 - | 1 - | 2,84E-05 - | 0 - | 0 - | 0 - | 0 - |
| 3290 | 1 | 3um_FIJJAFBF_465011 | Bacteria | Proteobacteri | Gammaprotec | Legionellales | Legionellacea | Legionella | Legionella fee | unclassified | unclassified | unclassified | unclassified | unclassified | 0,2 | 0,4 | 9,64E-06 | 1,93E-05 | 0 - | 0 - | 0 - | 0 - | 1 - | 4,82E-05 - | 0 - | 0 - | 0 - | 0 - |
| 3291 | 1 | 0.2um_NOKCPHJN_27668 | Bacteria | Actinobacteria | Actinobacteria | Micrococcales | Microbacteria | Candidatus Ac | Candidatus Ad | information st | Transcription | 05D27 | DNA-depende | 312284.A20C1 | 1 | 0,894427191 | 9,37E-06 | 8,37E-06 | 2 - | 1,65E-05 - | 2 - | 2,05E-05 - | 0 - | 0 - | 1 - | 9,88E-06 - | 0 - | 0 - |
| 3292 | 1 | 3um_FIJJAFBF_388951 | Bacteria | Bacteroidetes | Flavobacteriia | Flavobacterial | Flavobacteriac | Polaribacter | Polaribacter s | metabolism | Energy produc | COG0437 | 4fe-4S ferredo | 313598.MED1 | 17,2 | 1,469693846 | 0,000230583 | 1,53E-05 | 20 - | 0,000225939 - | 16 - | 0,000224439 - | 16 - | 0,000213358 - | 17 - | 0,000230041 - | 17 - | 0,00025914 - |
| 3293 | 1 | 3um_FIJJAFBF_674254 | Bacteria | Bacteroidetes | Flavobacteriia | Flavobacterial | Flavobacteriac | Flaviramulus | Flaviramulus i | metabolism | Amino acid tra | COG1932 | Catalyzes the | 983544.Lacal_ | 0,8 | 0,4 | 3,07E-05 | 1,57E-05 | 1 - | 3,24E-05 - | 0 - | 0 - | 1 - | 3,83E-05 - | 1 - | 3,88E-05 - | 1 - | 4,37E-05 - |
| 3294 | 1 | 3um_FIJJAFBF_510235 | Bacteria | Bacteroidetes | Flavobacteriia | Flavobacterial | Flavobacteriac | Polaribacter | Polaribacter s | unclassified | unclassified | unclassified | unclassified | unclassified | 2,2 | 0,4 | 0,000220845 | 4,93E-05 | 2 - | 0,000167557 - | 3 - | 0,000312085 - | 2 - | 0,000197784 - | 2 - | 0,000200706 - | 2 - | 0,000226094 - |
| 3295 | 1 | 0.2um_NOKCPHJN_815267 | Bacteria | Proteobacteri | Gammaprotec | Pseudomonad | Pseudomonad | Pseudomonas | various | information st | Translation, ril | 08VZM | 50s ribosomal | 746360.PFWH | 0,4 | 0,8 | 3,26E-05 | 6,53E-05 | 0 - | 0 - | 0 - | 0 - | 2 - | 0,000163232 - | 0 - | 0 - | 0 - | 0 - |
| 3296 | 1 | 3um_FIJJAFBF_450895 | Bacteria | Bacteroidetes | Flavobacteriia | Flavobacterial | Flavobacteriac | Flaviramulus | Flaviramulus b | unclassified | unclassified | unclassified | unclassified | unclassified | 2,8 | 0,4 | 0,000111507 | 1,94E-05 | 3 - | 0,000100096 - | 3 - | 0,00012429 - | 3 - | 0,000118153 - | 2 - | 7,99E-05 - | 3 - | 0,000135065 - |
| 3297 | 1 | 3um_FIJJAFBF_106687 | Bacteria | Proteobacteri | Alphaproteob | Rhizobiales | unclassified | unclassified | Rhizobiales ba | metabolism | Energy produc | COG1146 | metal cluster | 396595.TK90_ | 0,6 | 0,489897949 | 4,88E-05 | 4,01E-05 | 1 - | 7,22E-05 - | 0 - | 0 - | 1 - | 8,52E-05 - | 1 - | 8,65E-05 - | 0 - | 0 - |
| 3298 | 1 | 0.2um_NOKCPHJN_220621 | Bacteria | Bacteroidetes | unclassified | unclassified | unclassified | unclassified | Bacteroidetes | metabolism | Inorganic ion t | 05C27 | TonB-depende | 886377.Murru | 21,8 | 1,720465053 | 0,000292075 | 1,87E-05 | 25 - | 0,000282145 - | 22 - | 0,0003083 - | 21 - | 0,000279757 - | 20 - | 0,00027037 - | 21 - | 0,0003198 - |
| 3299 | 1 | 0.2um_NOKCPHJN_938013 | Bacteria | Proteobacteri | Alphaproteob | Rhodobactera | Rhodobactera | Amylibacter | Amylibacter k | unclassified | unclassified | unclassified | unclassified | unclassified | 0,6 | 0,489897949 | 3,66E-05 | 3,00E-05 | 0 - | 0 - | 0 - | 0 - | 1 - | 5,79E-05 - | 1 - | 5,88E-05 - | 1 - | 6,62E-05 - |
| 3300 | 1 | 3um_FIJJAFBF_50170 | Bacteria | Bacteroidetes | Cytophagia | Cytophagales | Flammeovirga | unclassified | Flammeovirga | unclassified | unclassified | unclassified | unclassified | unclassified | 5 | 1,414213562 | 0,000385394 | 9,13E-05 | 7 - | 0,000459107 - | 3 - | 0,000244318 - | 4 - | 0,000309674 - | 6 - | 0,000471371 - | 5 - | 0,000442499 - |
| 3301 | 2 | 0.2um_NOKCPHJN_1042131;3um_FIJJAFBF_129 | Eukaryota | unclassified | Oomycetes | Peronosporale | Peronosporac | Plasmopara | Plasmopara h | information st | Replication, re | COG0513 | purine NTP-de | 65071.PYU1_ | 4,8 | 1,469693846 | 0,000160265 | 4,35E-05 | 6 - | 0,000171736 - | 5 - | 0,000177704 - | 5 - | 0,00016893 - | 6 - | 0,000205711 - | 2 - | 7,72E-05 - |
| 3302 | 1 | 0.2um_NOKCPHJN_779672 | Bacteria | Bacteroidetes | Flavobacteriia | unclassified | unclassified | unclassified | Flavobacteria | cellular proces | Intracellular tr | 0BIEB | domain-conta | 1005048.CFU_ | 0,6 | 0,8 | 9,62E-06 | 1,28E-05 | 0 - | 0 - | 0 - | 0 - | 2 - | 3,19E-05 - | 1 - | 1,62E-05 - | 0 - | 0 - |
| 3303 | 1 | 3um_FIJJAFBF_353994 | Bacteria | Bacteroidetes | Flavobacteriia | Flavobacterial | Crocinitomica | unclassified | Crocinitomica | metabolism | Carbohydrate | COG0057 | glyceraldehyd | 643867.Ftrac_ | 0,4 | 0,489897949 | 1,81E-05 | 2,22E-05 | 0 - | 0 - | 1 - | 4,35E-05 - | 0 - | 0 - | 0 - | 0 - | 1 - | 4,72E-05 - |
| 3304 | 1 | 3um_FIJJAFBF_486460 | Eukaryota | Streptophyta | Liliopsida | Arecales | Arecaceae | various | various | information st | Translation, ril | KOG0887 | Ribosomal pro | 3847.GLYMA0 | 0,8 | 0,4 | 7,77E-05 | 3,99E-05 | 1 - | 8,14E-05 - | 1 - | 0,000101077 - | 1 - | 9,61E-05 - | 0 - | 0 - | 1 - | 0,00010984 - |
| 3305 | 1 | 3um_FIJJAFBF_15046 | Bacteria | Bacteroidetes | Flavobacteriia | Flavobacterial | Flavobacteriac | Polaribacter | Polaribacter s | metabolism | Amino acid tra | COG0337 | 3-dehydroquir | 313598.MED1 | 0,6 | 0,489897949 | 2,34E-05 | 1,95E-05 | 1 - | 3,26E-05 - | 1 - | 4,05E-05 - | 0 - | 0 - | 0 - | 0 - | 1 - | 4,40E-05 - |
| 3306 | 1 | 3um_FIJJAFBF_396421 | Bacteria | Proteobacteri | Betaproteoba | unclassified | unclassified | unclassified | Betaproteoba | information st | Translation, ril | 0BCDE | Binds to the 2 | 381666.H16_A | 1,2 | 0,4 | 0,000112688 | 3,89E-05 | 1 - | 7,86E-05 - | 1 - | 9,76E-05 - | 1 - | 9,28E-05 - | 2 - | 0,000188333 - | 1 - | 0,000106078 - |
| 3307 | 1 | 0.2um_NOKCPHJN_348678 | Bacteria | Proteobacteri | Alphaproteob | Rhodobactera | Rhodobactera | unclassified | Rhodobactera | metabolism | Lipid transpor | COG2067 | lipid transport | 371731.Rsw2 | 2,6 | 0,8 | 0,000110137 | 3,42E-05 | 2 - | 7,11E-05 - | 2 - | 8,82E-05 - | 4 - | 0,00016778 - | 3 - | 0,000127693 - | 2 - | 9,59E-05 - |
| 3308 | 1 | 3um_FIJJAFBF_211950 | Eukaryota | Bacillariophyt | Coscinodiscop | Thalassiosirale | Thalassiosirac | Thalassiosira | Thalassiosira c | unclassified | unclassified | unclassified | unclassified | unclassified | 1 | 0 | 0,000103808 | 9,86E-06 | 1 - | 8,70E-05 - | 1 - | 0,000107969 - | 1 - | 0,000102638 - | 1 - | 0,000104154 - | 1 - | 0,000117329 - |
| 3309 | 1 | 0.2um_NOKCPHJN_333506 | Eukaryota | unclassified | Labyrinthulom | unclassified | Thraustochytr | Aurantiochytr | Aurantiochytr | poorly charact | Function unkn | 0NVRZ | NA | 170187.SP_17 | 0,4 | 0,489897949 | 1,42E-06 | 1,76E-06 | 1 - | 3,24E-06 - | 0 - | 0 - | 0 - | 0 - | 1 - | 3,88E-06 - | 0 - | 0 - |
| 3310 | 1 | 3um_FIJJAFBF_593664 | Bacteria | Proteobacteri | Alphaproteob | Rhodobactera | unclassified | unclassified | Rhodobactera | metabolism | Carbohydrate | 16QKM | Fructose-1,6-b | 439497.RR11_ | 1 | 0,632455532 | 4,11E-05 | 2,30E-05 | 2 - | 7,11E-05 - | 1 - | 4,41E-05 - | 0 - | 0 - | 1 - | 4,26E-05 - | 1 - | 4,79E-05 - |
| 3311 | 1 | 0.2um_NOKCPHJN_21383 | Bacteria | Proteobacteri | Alphaproteob | Rhodospirillal | unclassified | unclassified | Rhodospirillal | unclassified | unclassified | unclassified | unclassified | unclassified | 1 | 0,632455532 | 1,97E-05 | 1,12E-05 | 2 - | 3,50E-05 - | 1 - | 2,17E-05 - | 1 - | 2,07E-05 - | 1 - | 2,10E-05 - | 0 - | 0 - |
| 3312 | 1 | 0.2um_NOKCPHJN_945535 | Bacteria | Proteobacteri | Alphaproteob | Rhodobactera | Rhodobactera | unclassified | Rhodobactera | poorly charact | Function unkn | 05DIH | basic membra | 314270.RB208 | 0,6 | 0,489897949 | 2,63E-05 | 2,15E-05 | 0 - | 0 - | 1 - | 4,31E-05 - | 0 - | 0 - | 1 - | 4,15E-05 - | 1 - | 4,68E-05 - |
| 3313 | 1 | 3um_FIJJAFBF_736139 | Bacteria | Bacteroidetes | Flavobacteriia | various | various | various | various | unclassified | unclassified | unclassified | unclassified | unclassified | 1,2 | 0,4 | 8,74E-05 | 1,94E-05 | 2 - | 0,000125439 - | 1 - | 7,79E-05 - | 1 - | 7,40E-05 - | 1 - | 7,51E-05 - | 1 - | 8,46E-05 - |
| 3314 | 2 | 0.2um_NOKCPHJN_287598;0.2um_NOKCPHJN_6 | Bacteria | Proteobacteri | Alphaproteob | Pelagibacteral | Pelagibacterac | various | various | unclassified | unclassified | unclassified | unclassified | unclassified | 1,4 | 0,489897949 | 2,92E-05 | 9,19E-06 | 2 - | 3,56E-05 - | 2 - | 4,42E-05 - | 1 - | 2,10E-05 - | 1 - | 2,13E-05 - | 1 - | 2,40E-05 - |
| 3315 | 1 | 3um_FIJJAFBF_62150 | Bacteria | Proteobacteri | Gammaprotec | Nevskiales | Sinobacterace | Sinimarinibact | Sinimarinibact | information st | Translation, ril | 17FIH | One of the ea | 694569.D7S_1 | 1,2 | 0,4 | 0,000167881 | 5,79E-05 | 1 - | 0,000117119 - | 1 - | 0,000145427 - | 1 - | 0,000138247 - | 2 - | 0,000280578 - | 1 - | 0,000158035 - |
| 3316 | 1 | 3um_FIJJAFBF_136469 | Bacteria | Proteobacteri | Alphaproteob | Rhodobactera | Rhodobactera | Thalassobium | Thalassobium | metabolism | Amino acid tra | COG0683 | (ABC) transpo | 633131.TR2A | 2,2 | 0,748331477 | 7,57E-05 | 2,68E-05 | 3 - | 8,67E-05 - | 2 - | 7,18E-05 - | 2 - | 6,83E-05 - | 1 - | 3,46E-05 - | 3 - | 0,000117034 - |
| 3317 | 1 | 3um_FIJJAFBF_350304 | Bacteria | Proteobacteri | Gammaprotec | Oceanospirilla | Oceanospirilla | Motiliproteus | Motiliproteus | unclassified | unclassified | unclassified | unclassified | unclassified | 0,2 | 0,4 | 2,62E-05 | 5,25E-05 | 0 - | 0 - | 0 - | 0 - | 0 - | 0 - | 0 - | 0 - | 1 - | 0,00013125 - |
| 3318 | 1 | 0.2um_NOKCPHJN_92319 | Bacteria | Bacteroidetes | Flavobacteriia | Flavobacterial | Flavobacteriac | Polaribacter | Polaribacter s | information st | Translation, ril | COG0254 | 50S ribosomal | 313594.PI23P | 0,8 | 0,4 | 0,000130751 | 6,71E-05 | 1 - | 0,000138285 - | 0 - | 0 - | 1 - | 0,000163232 - | 1 - | 0,000165643 - | 1 - | 0,000186596 - |
| 3319 | 1 | 3um_FIJJAFBF_786161 | Bacteria | Proteobacteri | Gammaprotec | Oceanospirilla | Saccharospirill | Reinekea | Reinekea blan | information st | Translation, ril | COG0088 | One of the pri | 314283.MED2 | 10,8 | 1,166190379 | 0,000726555 | 4,35E-05 | 13 - | 0,000738662 - | 11 - | 0,000776093 - | 10 - | 0,000670704 - | 10 - | 0,00068061 - | 10 - | 0,000766705 - |
| 3320 | 1 | 3um_FIJJAFBF_69933 | Bacteria | Bacteroidetes | Cytophagia | Cytophagales | Flammeovirga | unclassified | Flammeovirga | metabolism | Nucleotide tra | COG0015 | adenylosuccin | 207949.RED6 | 0,2 | 0,4 | 5,96E-06 | 1,19E-05 | 0 - | 0 - | 1 - | 2,98E-05 - | 0 - | 0 - | 0 - | 0 - | 0 - | 0 - |
| 3321 | 1 | 3um_FIJJAFBF_225012 | Bacteria | Bacteroidetes | Flavobacteriia | various | various | various | various | unclassified | unclassified | unclassified | unclassified | unclassified | 1,2 | 1,166190379 | 0,000169243 | 0,000166263 | 2 - | 0,000249515 - | 1 - | 0,000154912 - | 3 - | 0,00044179 - | 0 - | 0 - | 0 - | 0 - |
| 3322 | 1 | 3um_FIJJAFBF_580394 | Bacteria | Proteobacteri | Gammaprotec | Oceanospirilla | Saccharospirill | Reinekea | Reinekea fors | unclassified | unclassified | unclassified | unclassified | unclassified | 1 | 0 | 0,000106223 | 1,01E-05 | 1 - | 8,90E-05 - | 1 - | 0,00011048 - | 1 - | 0,000105025 - | 1 - | 0,000106576 - | 1 - | 0,000120058 - |
| 3323 | 1 | 0.2um_NOKCPHJN_154975 | Bacteria | Proteobacteri | Gammaprotec | Oceanospirilla | Saccharospirill | Reinekea | Reinekea blan | unclassified | unclassified | unclassified | unclassified | unclassified | 1,4 | 1,019803903 | 1,40E-05 | 9,20E-06 | 3 - | 2,62E-05 - | 2 - | 2,17E-05 - | 1 - | 1,03E-05 - | 0 - | 0 - | 1 - | 1,18E-05 - |
| 3324 | 1 | 0.2um_NOKCPHJN_267389 | Bacteria | Proteobacteri | Gammaprotec | Cellvibrionales | Porticoccacea | unclassified | SAR92 bacteri | poorly charact | Function unkn | 16R8I | Lipoprotein | 377629.TERTU | 4 | 0 | 5,85E-05 | 5,55E-06 | 4 - | 4,90E-05 - | 4 - | 6,08E-05 - | 4 - | 5,78E-05 - | 4 - | 5,87E-05 - | 4 - | 6,61E-05 - |
| 3325 | 1 | 3um_FIJJAFBF_581243 | Bacteria | Proteobacteri | Gammaprotec | Oceanospirilla | Saccharospirill | Reinekea | Reinekea blan | metabolism | Amino acid tra | 16PEK | Catalyzes two | 314283.MED2 | 1,2 | 0,748331477 | 3,87E-05 | 2,39E-05 | 2 - | 5,64E-05 - | 2 - | 7,00E-05 - | 1 - | 3,33E-05 - | 1 - | 3,38E-05 - | 0 - | 0 - |
| 3326 | 2 | 3um_FIJJAFBF_339019;3um_FIJJAFBF_520505 | Bacteria | Bacteroidetes | Flavobacteriia | Flavobacterial | Flavobacteriac | Polaribacter | various | unclassified | unclassified | unclassified | unclassified | unclassified | 0,2 | 0,4 | 6,38E-06 | 1,28E-05 | 0 - | 0 - | 0 - | 0 - | 1 - | 3,19E-05 - | 0 - | 0 - | 0 - | 0 - |
| 3327 | 1 | 3um_FIJJAFBF_165470 | Bacteria | Bacteroidetes | Flavobacteriia | Flavobacterial | Flavobacteriac | Algibacter | Algibacter sp. | metabolism | Energy produc | 090JD | Blue (Type 1) | 313598.MED1 | 0,8 | 0,748331477 | 6,81E-05 | 6,21E-05 | 0 - | 0 - | 1 - | 8,48E-05 - | 0 - | 0 - | 2 - | 0,000163671 - | 1 - | 9,22E-05 - |
| 3328 | 1 | 3um_FIJJAFBF_777883 | Bacteria | Bacteroidetes | Flavobacteriia | Flavobacterial | Flavobacteriac | Polaribacter | various | unclassified | unclassified | unclassified | unclassified | unclassified | 0,8 | 0,4 | 4,51E-05 | 2,32E-05 | 1 - | 4,72E-05 - | 1 - | 5,86E-05 - | 1 - | 5,58E-05 - | 0 - | 0 - | 1 - | 6,37E-05 - |
| 3329 | 1 | 0.2um_NOKCPHJN_1116524 | unclassified | unclassified | unclassified | unclassified | unclassified | unclassified | unclassified | unclassified | unclassified | unclassified | unclassified | unclassified | 1 | 0 | 0,000150579 | 1,43E-05 | 1 - | 0,000126128 - | 1 - | 0,000156614 - | 1 - | 0,000148882 - | 1 - | 0,000151081 - | 1 - | 0,000170192 - |
| 3330 | 1 | 3um_FIJJAFBF_178104 | Bacteria | unclassified | unclassified | unclassified | unclassified | unclassified | bacterium B1 | unclassified | unclassified | unclassified | unclassified | unclassified | 0,8 | 0,748331477 | 1,98E-05 | 1,90E-05 | 1 - | 2,05E-05 - | 2 - | 5,09E-05 - | 0 - | 0 - | 0 - | 0 - | 1 - | 2,77E-05 - |
| 3331 | 1 | 0.2um_NOKCPHJN_1052819 | Eukaryota | Basidiomycota | Agaricomycet | Agaricales | Omphalotacea | Lentinula | Lentinula edo | poorly charact | Function unkn | 04DV9 | retrotranspos | 339724.XP_0 | 0,2 | 0,4 | 1,09E-05 | 2,18E-05 | 0 - | 0 - | 0 - | 0 - | 0 - | 0 - | 0 - | 0 - | 1 - | 5,45E-05 - |
| 3332 | 1 | 0.2um_NOKCPHJN_738440 | Bacteria | Bacteroidetes | unclassified | unclassified | unclassified | unclassified | Bacteroidetes | unclassified | unclassified | unclassified | unclassified | unclassified | 5,6 | 1,356465997 | 0,000209349 | 3,95E-05 | 7 - | 0,000223177 - | 4 - | 0,000158354 - | 6 - | 0,000225804 - | 7 - | 0,000267329 - | 4 - | 0,000172083 - |
| 3333 | 1 | 3um_FIJJAFBF_67727 | Bacteria | Proteobacteri | Gammaprotec | Cellvibrionales | unclassified | unclassified | Cellvibrionales | unclassified | unclassified | unclassified | unclassified | unclassified | 0,8 | 0,4 | 6,44E-05 | 3,31E-05 | 1 - | 6,75E-05 - | 1 - | 8,38E-05 - | 1 - | 7,97E-05 - | 0 - | 0 - | 1 - | 9,11E-05 - |
| 3334 | 1 | 3um_FIJJAFBF_67737 | unclassified | unclassified | unclassified | unclassified | unclassified | unclassified | unclassified | unclassified | unclassified | unclassified | unclassified | unclassified | 1,6 | 0,489897949 | 4,59E-05 | 1,48E-05 | 1 - | 2,39E-05 - | 2 - | 5,94E-05 - | 2 - | 5,65E-05 - | 2 - | 5,73E-05 - | 1 - | 3,23E-05 - |
| 3335 | 1 | 3um_FIJJAFBF_444587 | Bacteria | Proteobacteri | Alphaproteob | Rhodobactera | Rhodobactera | Planktomarina | Planktomarina | cellular proces | Signal transdu | 01WFG | universal stres | 89187.ISM_02 | 0,4 | 0,489897949 | 3,41E-05 | 4,21E-05 | 1 - | 7,60E-05 - | 1 - | 9,44E-05 - | 0 - | 0 - | 0 - | 0 - | 0 - | 0 - |
| 3336 | 1 | 3um_FIJJAFBF_133415 | Bacteria | Bacteroidetes | Flavobacteriia | Flavobacterial | Flavobacteriac | Formosa | Formosa spon | unclassified | unclassified | unclassified | unclassified | unclassified | 0,2 | 0,4 | 8,43E-06 | 1,69E-05 | 0 - | 0 - | 0 - | 0 - | 0 - | 0 - | 1 - | 4,22E-05 - | 0 - | 0 - |
| 3337 | 1 | 3um_FIJJAFBF_397767 | Bacteria | Proteobacteri | Alphaproteob | Rhodobactera | unclassified | unclassified | Rhodobactera | unclassified | unclassified | unclassified | unclassified | unclassified | 4 | 0 | 0,000388729 | 3,69E-05 | 4 - | 0,000325608 - | 4 - | 0,000404309 - | 4 - | 0,000384347 - | 4 - | 0,000390024 - | 4 - | 0,00043936 - |
| 3338 | 1 | 3um_FIJJAFBF_150012 | Bacteria | Proteobacteri | Betaproteoba | Burkholderial | Comamonada | Limnohabitan | Limnohabitan | information st | Translation, ril | COG0050 | This protein p | 339670.Bamb | 10 | 0,632455532 | 0,000351872 | 4,92E-05 | 9 - | 0,000264192 - | 11 - | 0,000400948 - | 10 - | 0,000346502 - | 10 - | 0,00035162 - | 10 - | 0,000396098 - |
| 3339 | 1 | 0.2um_NOKCPHJN_03800 | Bacteria | Proteobacteri | Gammaprotec | Alteromonada | Alteromonada | Paraglaciecola | Paraglaciecola | metabolism | Energy produc | COG0055 | Produces ATP | 234831.PSM_ | 7,2 | 0,748331477 | 0,000215685 | 3,91E-05 | 6 - | 0,000149384 - | 7 - | 0,000216406 - | 8 - | 0,00023511 - | 7 - | 0,00020876 - | 8 - | 0,000268763 - |
| 3340 | 1 | 0.2um_NOKCPHJN_297617 | Bacteria | Proteobacteri | Alphaproteob | Rhodobactera | Rhodobactera | Planktomarina | Planktomarina | unclassified | unclassified | unclassified | unclassified | unclassified | 1 | 0,632455532 | 2,64E-05 | 1,74E-05 | 1 - | 2,19E-05 - | 2 - | 5,45E-05 - | 1 - | 2,59E-05 - | 0 - | 0 - | 1 - | 2,96E-05 - |
| 3341 | 1 | 3um_FIJJAFBF_18183 | Bacteria | Bacteroidetes | Flavobacteriia | Flavobacterial | Crocinitomica | unclassified | Crocinitomica | cellular proces | Intracellular tr | 0P8JN | The central su | 926562.Oweh | 1,6 | 0,489897949 | 4,80E-05 | 1,36E-05 | 2 - | 5,10E-05 - | 2 - | 6,33E-05 - | 1 - | 3,01E-05 - | 2 - | 6,11E-05 - | 1 - | 3,44E-05 - |
| 3342 | 2 | 0.2um_NOKCPHJN_426945;0.2um_NOKCPHJN_8 | Bacteria | Proteobacteri | Gammaprotec | unclassified | unclassified | unclassified | various | information st | Translation, ril | 0QTKX | Seems to be tl | various | 1 | 0 | 0,000109622 | 1,04E-05 | 1 - | 9,18E-05 - | 1 - | 0,000114015 - | 1 - | 0,000108386 - | 1 - | 0,000109987 - | 1 - | 0,0001239 - |
| 3343 | 1 | 3um_FIJJAFBF_56974 | Bacteria | Actinobacteria | Actinobacteria | Micrococcales | Microbacteria | Candidatus Ac | Candidatus Ad | unclassified | unclassified | unclassified | unclassified | unclassified | 0,6 | 0,489897949 | 2,31E-05 | 1,91E-05 | 1 - | 3,38E-05 - | 1 - | 4,19E-05 - | 1 - | 3,98E-05 - | 0 - | 0 - | 0 - | 0 - |
| 3344 | 1 | 3um_FIJJAFBF_388988 | Viruses | unclassified | unclassified | unclassified | Mimiviridae | Mimivirus | Acanthamoeb | metabolism | Amino acid tra | 12TQ9 | proline iminop | 431895.XP_0 | 0,2 | 0,4 | 3,13E-06 | 6,25E-06 | 1 - | 1,56E-05 - | 0 - | 0 - | 0 - | 0 - | 0 - | 0 - | 0 - | 0 - |
| 3345 | 1 | 3um_FIJJAFBF_96040 | Bacteria | Proteobacteri | Gammaprotec | Enterobactera | Enterobacteri | Citrobacter | Citrobacter w | unclassified | unclassified | unclassified | unclassified | unclassified | 1 | 0 | 1,88E-05 | 1,78E-06 | 1 - | 1,57E-05 - | 1 - | 1,95E-05 - | 1 - | 1,86E-05 - | 1 - | 1,89E-05 - | 1 - | 2,12E-05 - |
| 3346 | 1 | 0.2um_NOKCPHJN_646950 | Bacteria | Proteobacteri | Gammaprotec | Oceanospirilla | Saccharospirill | Reinekea | Reinekea fors | poorly charact | Function unkn | 11U8M | neurotransmit | 765910.Marp | 0,4 | 0,489897949 | 1,49E-05 | 1,84E-05 | 1 - | 3,40E-05 - | 0 - | 0 - | 0 - | 0 - | 1 - | 4,07E-05 - | 0 - | 0 - |
| 3347 | 1 | 0.2um_NOKCPHJN_426924 | Bacteria | Proteobacteri | Alphaproteob | Pelagibacteral | unclassified | unclassified | Pelagibacteral | cellular proces | Cell wall/mem | 7257 | NA | 335992.SAR11 | 2 | 0,632455532 | 8,49E-05 | 2,94E-05 | 2 - | 7,08E-05 - | 3 - | 0,000131962 - | 2 - | 8,36E-05 - | 1 - | 4,24E-05 - | 2 - | 9,56E-05 - |
| 3348 | 1 | 0.2um_NOKCPHJN_248048 | Bacteria | Proteobacteri | Alphaproteob | Rhodobactera | unclassified | unclassified | Rhodobactera | metabolism | Carbohydrate | 16Q4I | Abc transport | 367336.OM22 | 0,8 | 0,4 | 2,42E-05 | 1,23E-05 | 1 - | 2,62E-05 - | 1 - | 3,25E-05 - | 1 - | 3,09E-05 - | 1 - | 3,14E-05 - | 0 - | 0 - |
| 3349 | 1 | 0.2um_NOKCPHJN_630073 | Bacteria | Firmicutes | Clostridia | Clostridiales | Clostridiaceae | Butyricicoccus | Butyricicoccus | unclassified | unclassified | unclassified | unclassified | unclassified | 0,2 | 0,4 | 5,57E-06 | 1,11E-05 | 0 - | 0 - | 0 - | 0 - | 0 - | 0 - | 0 - | 0 - | 1 - | 2,79E-05 - |
| 3350 | 1 | 3um_FIJJAFBF_601456 | Bacteria | Proteobacteri | Gammaprotec | Alteromonada | Alteromonada | Paraglaciecola | various | metabolism | Lipid transpor | 05VNB | Carrier of the | 715451.ambt_ | 2,4 | 0,489897949 | 0,000410767 | 6,40E-05 | 3 - | 0,000435861 - | 2 - | 0,000360807 - | 2 - | 0,000342993 - | 3 - | 0,000522089 - | 2 - | 0,000392087 - |
| 3351 | 1 | 3um_FIJJAFBF_465439 | Bacteria | Proteobacteri | Gammaprotec | Cellvibrionales | Porticoccacea | unclassified | gamma prote | metabolism | Energy produc | 05D26 | NQR complex | 314287.GB22 | 0,6 | 0,489897949 | 1,93E-05 | 1,59E-05 | 1 - | 2,82E-05 - | 1 - | 3,50E-05 - | 1 - | 3,33E-05 - | 0 - | 0 - | 0 - | 0 - |
| 3352 | 2 | 0.2um_NOKCPHJN_348581;3um_FIJJAFBF_7100 | Bacteria | Bacteroidetes | Flavobacteriia | unclassified | unclassified | unclassified | Flavobacteria | unclassified | unclassified | unclassified | unclassified | unclassified | 0,6 | 0,489897949 | 2,45E-05 | 2,00E-05 | 0 - | 0 - | 1 - | 4,03E-05 - | 1 - | 3,83E-05 - | 0 - | 0 - | 1 - | 4,37E-05 - |
| 3353 | 1 | 0.2um_NOKCPHJN_457100 | Bacteria | Bacteroidetes | unclassified | unclassified | unclassified | unclassified | Bacteroidetes | information st | Translation, ril | COG0087 | One of the pri | 487796.Flav2 | 1,6 | 0,489897949 | 0,000105162 | 2,92E-05 | 2 - | 0,000111977 - | 2 - | 0,000139043 - | 2 - | 0,000132178 - | 1 - | 6,71E-05 - | 1 - | 7,55E-05 - |
| 3354 | 1 | 0.2um_NOKCPHJN_458223 | Bacteria | Bacteroidetes | Flavobacteriia | Flavobacterial | Cryomorphace | unclassified | Cryomorphace | metabolism | Inorganic ion t | 05C27 | TonB-depende | 1046627.BZA | 9 | 2,607680962 | 0,000107893 | 2,61E-05 | 12 - | 0,000123085 - | 11 - | 0,000140099 - | 10 - | 0,000121074 - | 7 - | 8,60E-05 - | 5 - | 6,92E-05 - |
| 3355 | 1 | 3um_FIJJAFBF_627510 | Bacteria | Bacteroidetes | unclassified | unclassified | unclassified | unclassified | Bacteroidetes | unclassified | unclassified | unclassified | unclassified | unclassified | 0,6 | 0,489897949 | 4,89E-05 | 4,01E-05 | 0 - | 0 - | 1 - | 8,05E-05 - | 1 - | 7,65E-05 - | 0 - | 0 - | 1 - | 8,75E-05 - |
| 3356 | 1 | 3um_FIJJAFBF_669258 | Eukaryota | Bacillariophyt | various | various | various | various | various | metabolism | Energy produc | 0J0AS | This b-type cy | 35128.Thapsd | 6,2 | 0,748331477 | 0,001004418 | 0,000103014 | 7 - | 0,000956473 - | 5 - | 0,000848327 - | 7 - | 0,001129019 - | 6 - | 0,000982024 - | 6 - | 0,001106246 - |
| 3357 | 1 | 0.2um_NOKCPHJN_310980 | Eukaryota | unclassified | unclassified | unclassified | unclassified | Gymnochlora | Gymnochlora | information st | Chromatin stru | KOG3467 | Core compone | 2850.Phatr34 | 2,4 | 0,489897949 | 0,000348931 | 5,43E-05 | 3 - | 0,000370248 - | 2 - | 0,000306492 - | 2 - | 0,00029136 - | 3 - | 0,000443495 - | 2 - | 0,000333063 - |
| 3358 | 1 | 3um_FIJJAFBF_724305 | Bacteria | Proteobacteri | Gammaprotec | Oceanospirilla | Saccharospirill | Reinekea | Reinekea fors | cellular proces | Intracellular tr | 16R4P | MSHA biogen | 314283.MED2 | 0,6 | 0,489897949 | 2,80E-05 | 2,29E-05 | 0 - | 0 - | 0 - | 0 - | 1 - | 4,43E-05 - | 1 - | 4,49E-05 - | 1 - | 5,06E-05 - |
| 3359 | 1 | 0.2um_NOKCPHJN_21172 | Bacteria | Proteobacteri | Alphaproteob | Rhodobactera | Rhodobactera | Roseovarius | Roseovarius n | information st | Replication, re | 05CZ8 | DNA polymera | 89187.ISM_05 | 2,6 | 1,2 | 9,67E-05 | 4,79E-05 | 2 - | 6,17E-05 - | 5 - | 0,000191558 - | 2 - | 7,28E-05 - | 2 - | 7,39E-05 - | 2 - | 8,33E-05 - |
| 3360 | 1 | 0.2um_NOKCPHJN_529648 | Bacteria | Proteobacteri | Alphaproteob | Pelagibacteral | Pelagibacterac | unclassified | Pelagibacterac | cellular proces | Cell wall/mem | 7257 | NA | 335992.SAR11 | 0,4 | 0,489897949 | 1,95E-05 | 2,39E-05 | 0 - | 0 - | 0 - | 0 - | 1 - | 4,84E-05 - | 1 - | 4,91E-05 - | 0 - | 0 - |
| 3361 | 1 | 3um_FIJJAFBF_99130 | Bacteria | Bacteroidetes | Flavobacteriia | Flavobacterial | Flavobacteriac | Gaetbulibacte | Gaetbulibacte | unclassified | unclassified | unclassified | unclassified | unclassified | 2,4 | 1,019803903 | 0,000102353 | 3,59E-05 | 4 - | 0,00014668 - | 2 - | 9,11E-05 - | 1 - | 4,33E-05 - | 3 - | 0,000131773 - | 2 - | 9,90E-05 - |
| 3362 | 1 | 0.2um_NOKCPHJN_216465 | Bacteria | Bacteroidetes | Cytophagia | Cytophagales | Hymenobacte | Hymenobacte | Hymenobacte | metabolism | Coenzyme tra | 09A4R | May play a ke | 388413.ALPR1 | 0,8 | 0,4 | 2,42E-05 | 1,23E-05 | 1 - | 2,61E-05 - | 1 - | 3,25E-05 - | 1 - | 3,09E-05 - | 1 - | 3,13E-05 - | 0 - | 0 - |
| 3363 | 1 | 0.2um_NOKCPHJN_199548 | Bacteria | Proteobacteri | Alphaproteob | Rhodobactera | Rhodobactera | Amylibacter | Amylibacter k | metabolism | Carbohydrate | 01U6R | trap dicarboxy | 367336.OM22 | 1,4 | 0,489897949 | 5,43E-05 | 1,92E-05 | 1 - | 3,25E-05 - | 1 - | 4,04E-05 - | 2 - | 7,68E-05 - | 2 - | 7,79E-05 - | 1 - | 4,39E-05 - |
| 3364 | 1 | 3um_FIJJAFBF_560542 | Eukaryota | Chlorophyta | Chlorophycea | Chlamydomor | Volvocaceae | Volvox | Volvox carteri | unclassified | unclassified | unclassified | unclassified | unclassified | 1 | 0 | 6,46E-05 | 6,14E-06 | 1 - | 5,41E-05 - | 1 - | 6,72E-05 - | 1 - | 6,39E-05 - | 1 - | 6,49E-05 - | 1 - | 7,31E-05 - |
| 3365 | 1 | 3um_FIJJAFBF_199366 | Bacteria | Proteobacteri | Gammaprotec | Oceanospirilla | Oceanospirilla | Neptuniibacte | Neptuniibacte | information st | Translation, ril | 17C3B | The globular d | 207954.MED9 | 1,4 | 0,8 | 0,000166307 | 7,36E-05 | 3 - | 0,000313028 - | 1 - | 0,000129563 - | 1 - | 0,000123166 - | 1 - | 0,000124985 - | 1 - | 0,000140795 - |
| 3366 | 1 | 0.2um_NOKCPHJN_629992 | Bacteria | Bacteroidetes | Flavobacteriia | Flavobacterial | Cryomorphace | unclassified | Cryomorphace | unclassified | unclassified | unclassified | unclassified | unclassified | 0,2 | 0,4 | 9,71E-06 | 1,94E-05 | 0 - | 0 - | 0 - | 0 - | 1 - | 4,86E-05 - | 0 - | 0 - | 0 - | 0 - |
| 3367 | 1 | 3um_FIJJAFBF_348593 | Bacteria | Acidobacteria | unclassified | unclassified | unclassified | unclassified | Acidobacteria | metabolism | Energy produc | 07EQ1 | H -transportin | 682795.AciX8 | 1 | 0 | 0,000130502 | 1,24E-05 | 1 - | 0,000109311 - | 1 - | 0,000135732 - | 1 - | 0,000129031 - | 1 - | 0,000130936 - | 1 - | 0,0001475 - |
| 3368 | 1 | 0.2um_NOKCPHJN_725518 | unclassified | unclassified | unclassified | unclassified | unclassified | unclassified | unclassified | unclassified | unclassified | unclassified | unclassified | unclassified | 1 | 0 | 4,54E-05 | 4,31E-06 | 1 - | 3,80E-05 - | 1 - | 4,72E-05 - | 1 - | 4,49E-05 - | 1 - | 4,55E-05 - | 1 - | 5,13E-05 - |
| 3369 | 1 | 0.2um_NOKCPHJN_31016 | Bacteria | Proteobacteri | Alphaproteob | Rhodobactera | Rhodobactera | Amylibacter | Amylibacter k | metabolism | Energy produc | 01RNF | Produces ATP | 314270.RB208 | 2,2 | 0,4 | 6,34E-05 | 1,41E-05 | 2 - | 4,81E-05 - | 3 - | 8,96E-05 - | 2 - | 5,68E-05 - | 2 - | 5,76E-05 - | 2 - | 6,49E-05 - |
| 3370 | 1 | 0.2um_NOKCPHJN_371016 | Bacteria | Proteobacteri | Betaproteoba | unclassified | unclassified | unclassified | Betaproteoba | unclassified | unclassified | unclassified | unclassified | unclassified | 0,2 | 0,4 | 5,58E-06 | 1,12E-05 | 0 - | 0 - | 0 - | 0 - | 0 - | 0 - | 1 - | 2,79E-05 - | 0 - | 0 - |
| 3371 | 2 | 0.2um_NOKCPHJN_230826;3um_FIJJAFBF_1482 | Bacteria | Bacteroidetes | Flavobacteriia | Flavobacterial | Flavobacteriac | Polaribacter | Polaribacter h | poorly charact | Function unkn | 0PAA9 | Inherit from b | 313598.MED1 | 0,8 | 0,4 | 5,34E-05 | 2,74E-05 | 1 - | 5,57E-05 - | 1 - | 6,92E-05 - | 0 - | 0 - | 1 - | 6,67E-05 - | 1 - | 7,52E-05 - |
| 3372 | 1 | 3um_FIJJAFBF_384596 | Bacteria | Proteobacteri | Alphaproteob | Rhodobactera | Rhodobactera | Planktotalea | Planktotalea f | cellular proces | Posttranslatio | COG0330 | Band 7 protei | 314270.RB208 | 1,4 | 0,8 | 6,33E-05 | 3,57E-05 | 2 - | 7,92E-05 - | 1 - | 4,91E-05 - | 2 - | 9,34E-05 - | 2 - | 9,48E-05 - | 0 - | 0 - |
| 3373 | 1 | 0.2um_NOKCPHJN_105859 | Bacteria | Proteobacteri | Alphaproteob | Rhodobactera | Rhodobactera | Actibacterium | Actibacterium | metabolism | Amino acid tra | 16PQJ | Dihydroxy-aci | 388399.SSE37 | 1 | 1,095445115 | 2,16E-05 | 2,19E-05 | 3 - | 5,97E-05 - | 1 - | 2,47E-05 - | 0 - | 0 - | 1 - | 2,38E-05 - | 0 - | 0 - |
| 3374 | 1 | 3um_FIJJAFBF_19624 | Bacteria | Proteobacteri | Gammaprotec | Cellvibrionales | Porticoccacea | unclassified | gamma prote | metabolism | Inorganic ion t | 16TQC | TONB-depend | 314287.GB22 | 18,4 | 1,356465997 | 0,000244431 | 1,54E-05 | 21 - | 0,000234923 - | 18 - | 0,000250033 - | 18 - | 0,000237688 - | 17 - | 0,000227799 - | 18 - | 0,00027171 - |
| 3375 | 1 | 3um_FIJJAFBF_225461 | Bacteria | Bacteroidetes | Flavobacteriia | Flavobacterial | Flavobacteriac | Flavivirga | Flavivirga eckl | cellular proces | Posttranslatio | COG0330 | Band 7 protei | 983548.Krodi_ | 0,8 | 0,4 | 4,04E-05 | 2,08E-05 | 1 - | 4,24E-05 - | 1 - | 5,26E-05 - | 1 - | 5,00E-05 - | 0 - | 0 - | 1 - | 5,71E-05 - |
| 3376 | 1 | 0.2um_NOKCPHJN_12281 | unclassified | unclassified | unclassified | unclassified | unclassified | unclassified | unclassified | unclassified | unclassified | unclassified | unclassified | unclassified | 8,6 | 1,743559577 | 0,000488602 | 0,00010033 | 10 - | 0,000478236 - | 11 - | 0,000653211 - | 8 - | 0,000451607 - | 6 - | 0,000343708 - | 8 - | 0,000516248 - |
| 3377 | 1 | 0.2um_NOKCPHJN_378546 | Bacteria | Bacteroidetes | unclassified | unclassified | unclassified | unclassified | unidentified e | unclassified | unclassified | unclassified | unclassified | unclassified | 2 | 1,095445115 | 0,000146869 | 6,65E-05 | 4 - | 0,000255059 - | 1 - | 7,92E-05 - | 1 - | 7,53E-05 - | 2 - | 0,000152759 - | 2 - | 0,000172083 - |
| 3378 | 1 | 0.2um_NOKCPHJN_845630 | Bacteria | Actinobacteria | Actinobacteria | Corynebacteri | Mycobacteria | Mycobacteriu | Mycobacteriu | poorly charact | Function unkn | 0IQEP | NA | 5057.CADACL | 1 | 0 | 2,72E-05 | 2,58E-06 | 1 - | 2,28E-05 - | 1 - | 2,83E-05 - | 1 - | 2,69E-05 - | 1 - | 2,73E-05 - | 1 - | 3,07E-05 - |
| 3379 | 1 | 0.2um_NOKCPHJN_933150 | Eukaryota | unclassified | Pelagophycea | Pelagomonad | unclassified | Aureococcus | Aureococcus a | unclassified | unclassified | unclassified | unclassified | unclassified | 1 | 0 | 4,91E-05 | 4,66E-06 | 1 - | 4,11E-05 - | 1 - | 5,11E-05 - | 1 - | 4,86E-05 - | 1 - | 4,93E-05 - | 1 - | 5,55E-05 - |
| 3380 | 1 | 3um_FIJJAFBF_284937 | Bacteria | Proteobacteri | Alphaproteob | Rhodospirillal | Rhodospirillac | unclassified | Rhodobactera | metabolism | Energy produc | 16RH6 | Produces ATP | 398580.Dshi_ | 1,4 | 1,019803903 | 6,17E-05 | 4,11E-05 | 3 - | 0,000118735 - | 1 - | 4,91E-05 - | 2 - | 9,34E-05 - | 1 - | 4,74E-05 - | 0 - | 0 - |
| 3381 | 1 | 3um_FIJJAFBF_211494 | Bacteria | Proteobacteri | Gammaprotec | Alteromonada | Alteromonada | Paraglaciecola | Paraglaciecola | metabolism | Inorganic ion t | 16TQC | TONB-depend | 87626.PTD2_ | 15,2 | 0,748331477 | 0,000211543 | 1,72E-05 | 16 - | 0,000187009 - | 14 - | 0,000203184 - | 15 - | 0,000206948 - | 16 - | 0,000224005 - | 15 - | 0,00023657 - |
| 3382 | 1 | 3um_FIJJAFBF_794580 | Bacteria | Bacteroidetes | Flavobacteriia | Flavobacterial | Flavobacteriac | unclassified | Flavobacteriac | information st | Translation, ril | COG1185 | Involved in mF | 313603.FB217 | 1,2 | 0,748331477 | 2,39E-05 | 1,46E-05 | 0 - | 0 - | 2 - | 4,02E-05 - | 2 - | 3,82E-05 - | 1 - | 1,94E-05 - | 1 - | 2,18E-05 - |
| 3383 | 1 | 3um_FIJJAFBF_605438 | Bacteria | Bacteroidetes | Flavobacteriia | Flavobacterial | Crocinitomica | unclassified | Crocinitomica | cellular proces | Signal transdu | 1AKRM | anti-anti-sigm | 714943.Mucp | 0,2 | 0,4 | 1,91E-05 | 3,83E-05 | 1 - | 9,56E-05 - | 0 - | 0 - | 0 - | 0 - | 0 - | 0 - | 0 - | 0 - |
| 3384 | 1 | 0.2um_NOKCPHJN_371014 | Bacteria | Proteobacteri | Betaproteoba | unclassified | unclassified | unclassified | Betaproteoba | metabolism | Inorganic ion t | 07QMV | Part of the AB | 261292.Nit79 | 1,2 | 0,4 | 0,000202509 | 4,49E-05 | 2 - | 0,000290574 - | 1 - | 0,000180404 - | 1 - | 0,000171496 - | 1 - | 0,00017403 - | 1 - | 0,000196044 - |
| 3385 | 1 | 3um_FIJJAFBF_101610 | Bacteria | Proteobacteri | Gammaprotec | Pseudomonad | Moraxellaceae | Psychrobacter | various | information st | Translation, ril | 0QTKX | Seems to be tl | 335284.Pcryo_ | 1 | 0,894427191 | 0,000180937 | 0,00015793 | 2 - | 0,000310207 - | 2 - | 0,000385186 - | 0 - | 0 - | 0 - | 0 - | 1 - | 0,00020929 - |
| 3386 | 1 | 0.2um_NOKCPHJN_200775 | Bacteria | Proteobacteri | Gammaprotec | Oceanospirilla | Saccharospirill | Reinekea | Reinekea fors | metabolism | Coenzyme tra | COG2227 | Non-specific C | 314283.MED2 | 0,2 | 0,4 | 1,16E-05 | 2,32E-05 | 0 - | 0 - | 0 - | 0 - | 1 - | 5,79E-05 - | 0 - | 0 - | 0 - | 0 - |
| 3387 | 1 | 0.2um_NOKCPHJN_262967 | Bacteria | Proteobacteri | Gammaprotec | unclassified | unclassified | unclassified | Gammaprotec | unclassified | unclassified | unclassified | unclassified | unclassified | 1 | 0,632455532 | 9,75E-05 | 6,39E-05 | 1 - | 8,38E-05 - | 1 - | 0,000104028 - | 1 - | 9,89E-05 - | 2 - | 0,000200706 - | 0 - | 0 - |
| 3388 | 1 | 3um_FIJJAFBF_25444 | Bacteria | Bacteroidetes | Flavobacteriia | unclassified | unclassified | unclassified | Flavobacteria | cellular proces | Cell wall/mem | 0P81E | outer membra | 487797.Flav3 | 2,4 | 0,489897949 | 7,40E-05 | 1,75E-05 | 2 - | 5,15E-05 - | 3 - | 9,59E-05 - | 2 - | 6,08E-05 - | 3 - | 9,25E-05 - | 2 - | 6,95E-05 - |
| 3389 | 1 | 0.2um_NOKCPHJN_176004 | Eukaryota | unclassified | unclassified | Prymnesiales | Chrysochromu | Chrysochromu | various | unclassified | unclassified | unclassified | unclassified | unclassified | 1,2 | 0,4 | 2,19E-05 | 7,54E-06 | 1 - | 1,53E-05 - | 1 - | 1,90E-05 - | 1 - | 1,80E-05 - | 2 - | 3,66E-05 - | 1 - | 2,06E-05 - |
| 3390 | 1 | 3um_FIJJAFBF_540151 | Bacteria | Bacteroidetes | Flavobacteriia | Flavobacterial | Flavobacteriac | Formosa | Formosa sp. H | cellular proces | Posttranslatio | 05E6M | Sulphatase-m | 391603.FBALC | 1,8 | 0,4 | 6,12E-05 | 1,43E-05 | 2 - | 5,72E-05 - | 1 - | 3,55E-05 - | 2 - | 6,76E-05 - | 2 - | 6,86E-05 - | 2 - | 7,72E-05 - |
| 3391 | 1 | 0.2um_NOKCPHJN_326367 | Bacteria | Proteobacteri | Alphaproteob | Pelagibacteral | Pelagibacterac | Candidatus Pe | Candidatus Pe | cellular proces | Intracellular tr | 05VIE | preprotein tra | 439493.PB721 | 2 | 0 | 0,000288478 | 2,74E-05 | 2 - | 0,000241635 - | 2 - | 0,00030004 - | 2 - | 0,000285226 - | 2 - | 0,000289439 - | 2 - | 0,000326052 - |
| 3392 | 1 | 0.2um_NOKCPHJN_24736 | Bacteria | Bacteroidetes | Flavobacteriia | Flavobacterial | Flavobacteriac | unclassified | Flavobacteriac | cellular proces | Cell motility | COG1360 | Ompa motb d | 487796.Flav2 | 8,2 | 0,748331477 | 0,000395609 | 4,04E-05 | 9 - | 0,000365014 - | 8 - | 0,00040288 - | 7 - | 0,000335115 - | 9 - | 0,000437226 - | 8 - | 0,000437808 - |
| 3393 | 1 | 3um_FIJJAFBF_715054 | Bacteria | Cyanobacteria | unclassified | Oscillatoriales | Cyanothecace | Cyanothece | various | unclassified | unclassified | unclassified | unclassified | unclassified | 1 | 0 | 0,000145774 | 1,38E-05 | 1 - | 0,000122103 - | 1 - | 0,000151616 - | 1 - | 0,00014413 - | 1 - | 0,000146259 - | 1 - | 0,00016476 - |
| 3394 | 1 | 3um_FIJJAFBF_803372 | Bacteria | Proteobacteri | Betaproteoba | Burkholderial | Comamonada | Hydrogenoph | Hydrogenoph | metabolism | Lipid transpor | 05C9H | acyl-CoA dehy | 296591.Bpro_ | 0,2 | 0,4 | 5,48E-06 | 1,10E-05 | 1 - | 2,74E-05 - | 0 - | 0 - | 0 - | 0 - | 0 - | 0 - | 0 - | 0 - |
| 3395 | 1 | 0.2um_NOKCPHJN_134831 | Bacteria | Bacteroidetes | unclassified | unclassified | unclassified | unclassified | Bacteroidetes | unclassified | unclassified | unclassified | unclassified | unclassified | 3,4 | 0,8 | 0,000248726 | 4,76E-05 | 4 - | 0,000249515 - | 3 - | 0,000232368 - | 4 - | 0,000294527 - | 4 - | 0,000298877 - | 2 - | 0,000168342 - |
| 3396 | 1 | 3um_FIJJAFBF_405475 | Bacteria | Proteobacteri | Gammaprotec | Alteromonada | Alteromonada | Paraglaciecola | Paraglaciecola | cellular proces | Posttranslatio | COG0652 | peptidyl-proly | 529507.PMI2 | 1 | 0 | 8,36E-05 | 7,93E-06 | 1 - | 7,00E-05 - | 1 - | 8,69E-05 - | 1 - | 8,26E-05 - | 1 - | 8,38E-05 - | 1 - | 9,44E-05 - |
| 3397 | 1 | 0.2um_NOKCPHJN_126396 | Bacteria | Bacteroidetes | Flavobacteriia | Flavobacterial | Flavobacteriac | Polaribacter | Polaribacter b | unclassified | unclassified | unclassified | unclassified | unclassified | 5,6 | 1,356465997 | 9,80E-05 | 2,49E-05 | 6 - | 8,83E-05 - | 3 - | 5,48E-05 - | 6 - | 0,000104217 - | 7 - | 0,000123382 - | 6 - | 0,000119134 - |
| 3398 | 1 | 0.2um_NOKCPHJN_342285 | Bacteria | Actinobacteria | Actinobacteria | Micrococcales | Microbacteria | Candidatus Ac | Candidatus Ad | unclassified | unclassified | unclassified | unclassified | unclassified | 1,2 | 1,469693846 | 1,39E-05 | 1,58E-05 | 4 - | 4,32E-05 - | 1 - | 1,34E-05 - | 1 - | 1,27E-05 - | 0 - | 0 - | 0 - | 0 - |
| 3399 | 1 | 0.2um_NOKCPHJN_46052 | Bacteria | Bacteroidetes | Cytophagia | Cytophagales | Flammeovirga | unclassified | Flammeovirga | metabolism | Amino acid tra | 05DX4 | Peptidase M1 | 760192.Halhy | 0,2 | 0,4 | 3,25E-06 | 6,51E-06 | 0 - | 0 - | 1 - | 1,63E-05 - | 0 - | 0 - | 0 - | 0 - | 0 - | 0 - |
| 3400 | 1 | 0.2um_NOKCPHJN_418379 | Bacteria | Proteobacteri | Alphaproteob | Pelagibacteral | Pelagibacterac | Candidatus Pe | Candidatus Pe | metabolism | Coenzyme tra | 179RB | Reversibly tra | 335992.SAR11 | 0,2 | 0,4 | 1,68E-05 | 3,35E-05 | 0 - | 0 - | 0 - | 0 - | 0 - | 0 - | 1 - | 8,38E-05 - | 0 - | 0 - |
| 3401 | 1 | 3um_FIJJAFBF_610640 | Bacteria | Proteobacteri | Gammaprotec | Oceanospirilla | Saccharospirill | Reinekea | Reinekea fors | unclassified | unclassified | unclassified | unclassified | unclassified | 1 | 0,632455532 | 5,45E-05 | 3,04E-05 | 2 - | 9,41E-05 - | 1 - | 5,84E-05 - | 0 - | 0 - | 1 - | 5,63E-05 - | 1 - | 6,35E-05 - |
| 3402 | 1 | 3um_FIJJAFBF_00348 | Bacteria | Bacteroidetes | Flavobacteriia | Flavobacterial | Flavobacteriac | Polaribacter | Polaribacter s | cellular proces | Intracellular tr | 08SHB | Biopolymer tr | 313594.PI23P | 2,2 | 0,748331477 | 0,000192209 | 5,83E-05 | 3 - | 0,000223591 - | 1 - | 9,25E-05 - | 2 - | 0,000175951 - | 3 - | 0,000267825 - | 2 - | 0,000201136 - |
| 3403 | 1 | 3um_FIJJAFBF_123890 | Bacteria | Bacteroidetes | Flavobacteriia | Flavobacterial | Flavobacteriac | Ochrovirga | Ochrovirga pa | unclassified | unclassified | unclassified | unclassified | unclassified | 0,4 | 0,489897949 | 1,04E-05 | 1,27E-05 | 0 - | 0 - | 0 - | 0 - | 1 - | 2,58E-05 - | 1 - | 2,62E-05 - | 0 - | 0 - |
| 3404 | 1 | 0.2um_NOKCPHJN_574317 | unclassified | unclassified | unclassified | unclassified | unclassified | unclassified | unclassified | unclassified | unclassified | unclassified | unclassified | unclassified | 0,2 | 0,4 | 3,70E-05 | 7,40E-05 | 0 - | 0 - | 1 - | 0,000185089 - | 0 - | 0 - | 0 - | 0 - | 0 - | 0 - |
| 3405 | 1 | 0.2um_NOKCPHJN_532535 | Eukaryota | Chordata | Chondrichthye | Orectolobiforr | Rhincodontida | Rhincodon | Rhincodon typ | unclassified | unclassified | unclassified | unclassified | unclassified | 0,2 | 0,4 | 1,27E-05 | 2,53E-05 | 0 - | 0 - | 0 - | 0 - | 1 - | 6,33E-05 - | 0 - | 0 - | 0 - | 0 - |
| 3406 | 1 | 3um_FIJJAFBF_726464 | Bacteria | Proteobacteri | Alphaproteob | Rhodobactera | Rhodobactera | unclassified | Rhodobactera | unclassified | unclassified | unclassified | unclassified | unclassified | 0,8 | 0,979795897 | 3,02E-05 | 3,72E-05 | 2 - | 6,87E-05 - | 0 - | 0 - | 0 - | 0 - | 2 - | 8,23E-05 - | 0 - | 0 - |
| 3407 | 1 | 3um_FIJJAFBF_173250 | unclassified | unclassified | unclassified | unclassified | unclassified | unclassified | unclassified | unclassified | unclassified | unclassified | unclassified | unclassified | 2,2 | 1,166190379 | 0,000100816 | 5,40E-05 | 3 - | 0,000115936 - | 4 - | 0,000191945 - | 1 - | 4,56E-05 - | 1 - | 4,63E-05 - | 2 - | 0,000104293 - |
| 3408 | 1 | 0.2um_NOKCPHJN_49514 | Bacteria | Proteobacteri | Alphaproteob | Rhodospirillal | unclassified | unclassified | Rhodospirillal | cellular proces | Posttranslatio | COG0330 | Band 7 protei | 414684.RC1_1 | 0,4 | 0,489897949 | 1,50E-05 | 1,83E-05 | 0 - | 0 - | 1 - | 3,81E-05 - | 0 - | 0 - | 1 - | 3,68E-05 - | 0 - | 0 - |
| 3409 | 1 | 3um_FIJJAFBF_753148 | Bacteria | Proteobacteri | Alphaproteob | Rhodobactera | Rhodobactera | Planktotalea | Planktotalea f | cellular proces | Posttranslatio | COG0330 | Band 7 protei | 314270.RB208 | 1 | 0,632455532 | 5,43E-05 | 3,68E-05 | 0 - | 0 - | 1 - | 5,34E-05 - | 1 - | 5,07E-05 - | 1 - | 5,15E-05 - | 2 - | 0,000116011 - |
| 3410 | 1 | 3um_FIJJAFBF_360710 | Bacteria | Bacteroidetes | Flavobacteriia | Flavobacterial | Flavobacteriac | Polaribacter | Polaribacter s | unclassified | unclassified | unclassified | unclassified | unclassified | 0,8 | 0,4 | 7,63E-05 | 3,88E-05 | 1 - | 8,26E-05 - | 1 - | 0,000102532 - | 1 - | 9,75E-05 - | 1 - | 9,89E-05 - | 0 - | 0 - |
| 3411 | 1 | 3um_FIJJAFBF_01771 | Eukaryota | unclassified | Dinophyceae | Suessiales | Symbiodiniace | Symbiodinium | Symbiodinium | unclassified | unclassified | unclassified | unclassified | unclassified | 0,2 | 0,4 | 2,56E-06 | 5,13E-06 | 1 - | 1,28E-05 - | 0 - | 0 - | 0 - | 0 - | 0 - | 0 - | 0 - | 0 - |
| 3412 | 2 | 0.2um_NOKCPHJN_1055616;0.2um_NOKCPHJN_ | Bacteria | Proteobacteri | Alphaproteob | Pelagibacteral | Pelagibacterac | Candidatus Pe | Candidatus Pe | metabolism | Lipid transpor | COG0511 | Acetyl-CoA ca | 335992.SAR11 | 0,2 | 0,4 | 1,95E-05 | 3,90E-05 | 0 - | 0 - | 0 - | 0 - | 0 - | 0 - | 1 - | 9,75E-05 - | 0 - | 0 - |
| 3413 | 1 | 0.2um_NOKCPHJN_551427 | Bacteria | Bacteroidetes | Flavobacteriia | Flavobacterial | Flavobacteriac | Pseudozobelli | Pseudozobelli | metabolism | Energy produc | COG0055 | Produces ATP | 886377.Murru | 2,4 | 1,019803903 | 6,30E-05 | 2,13E-05 | 4 - | 9,15E-05 - | 2 - | 5,68E-05 - | 3 - | 8,10E-05 - | 2 - | 5,48E-05 - | 1 - | 3,09E-05 - |
| 3414 | 1 | 3um_FIJJAFBF_233139 | Bacteria | Bacteroidetes | Flavobacteriia | Flavobacterial | Flavobacteriac | Maribacter | Maribacter an | metabolism | Amino acid tra | 099QE | chorismate m | 391603.FBALC | 2,8 | 0,748331477 | 0,000106847 | 3,03E-05 | 2 - | 6,38E-05 - | 3 - | 0,000118766 - | 3 - | 0,000112902 - | 4 - | 0,000152759 - | 2 - | 8,60E-05 - |
| 3415 | 1 | 0.2um_NOKCPHJN_43890 | Bacteria | Proteobacteri | Gammaprotec | Oceanospirilla | Oceanospirilla | Marinobacteri | Marinobacteri | cellular proces | Cell wall/mem | 16QB0 | Part of the ou | 207949.RED6 | 1 | 0,894427191 | 1,90E-05 | 1,73E-05 | 0 - | 0 - | 0 - | 0 - | 1 - | 1,78E-05 - | 2 - | 3,62E-05 - | 2 - | 4,08E-05 - |
| 3416 | 1 | 0.2um_NOKCPHJN_708100 | Eukaryota | unclassified | unclassified | Choanoflagelli | Salpingoecida | Monosiga | Monosiga bre | information st | Chromatin stru | COG5648 | high mobility | 431895.XP_0 | 1 | 0 | 8,11E-05 | 7,70E-06 | 1 - | 6,79E-05 - | 1 - | 8,43E-05 - | 1 - | 8,02E-05 - | 1 - | 8,14E-05 - | 1 - | 9,16E-05 - |
| 3417 | 1 | 0.2um_NOKCPHJN_729736 | Bacteria | Proteobacteri | Gammaprotec | Cellvibrionales | Porticoccacea | unclassified | various | unclassified | unclassified | unclassified | unclassified | unclassified | 0,4 | 0,489897949 | 6,48E-06 | 7,94E-06 | 0 - | 0 - | 0 - | 0 - | 1 - | 1,61E-05 - | 1 - | 1,63E-05 - | 0 - | 0 - |
| 3418 | 1 | 3um_FIJJAFBF_41558 | Eukaryota | Bacillariophyt | Mediophycea | Lithodesmiale | Lithodesmiace | Lithodesmium | Lithodesmium | metabolism | Carbohydrate | 0IVA1 | Photosystem I | 35128.Thapsd | 3,4 | 0,489897949 | 0,000250022 | 3,17E-05 | 4 - | 0,000248166 - | 4 - | 0,000308149 - | 3 - | 0,000219701 - | 3 - | 0,000222946 - | 3 - | 0,000251148 - |
| 3419 | 1 | 3um_FIJJAFBF_752256 | Bacteria | Bacteroidetes | unclassified | unclassified | unclassified | unclassified | Bacteroidetes | cellular proces | Cell wall/mem | COG2885 | Ompa motb d | 487796.Flav2 | 3,2 | 1,166190379 | 9,46E-05 | 3,19E-05 | 4 - | 0,000101798 - | 3 - | 9,48E-05 - | 4 - | 0,000120162 - | 4 - | 0,000121936 - | 1 - | 3,43E-05 - |
| 3420 | 1 | 3um_FIJJAFBF_242749 | Bacteria | Bacteroidetes | Flavobacteriia | unclassified | unclassified | unclassified | Flavobacteria | unclassified | unclassified | unclassified | unclassified | unclassified | 26,4 | 2,727636339 | 0,000475627 | 2,25E-05 | 31 - | 0,000471894 - | 25 - | 0,000472543 - | 28 - | 0,000503117 - | 24 - | 0,000437613 - | 24 - | 0,000492969 - |
| 3421 | 1 | 3um_FIJJAFBF_238483 | Bacteria | Proteobacteri | Gammaprotec | Oceanospirilla | Saccharospirill | Reinekea | Reinekea fors | unclassified | unclassified | unclassified | unclassified | unclassified | 3 | 0 | 0,000331517 | 3,15E-05 | 3 - | 0,000277686 - | 3 - | 0,000344804 - | 3 - | 0,00032778 - | 3 - | 0,000332621 - | 3 - | 0,000374696 - |
| 3422 | 1 | 3um_FIJJAFBF_510236 | Bacteria | Bacteroidetes | Flavobacteriia | Flavobacterial | Flavobacteriac | Polaribacter | various | unclassified | unclassified | unclassified | unclassified | unclassified | 1,8 | 0,4 | 0,000101888 | 2,38E-05 | 2 - | 9,53E-05 - | 1 - | 5,91E-05 - | 2 - | 0,000112433 - | 2 - | 0,000114094 - | 2 - | 0,000128527 - |
| 3423 | 1 | 3um_FIJJAFBF_388958 | Bacteria | Bacteroidetes | Flavobacteriia | Flavobacterial | Flavobacteriac | Polaribacter | Polaribacter s | cellular proces | Cell wall/mem | COG0750 | Membrane-as | 313594.PI23P | 5,4 | 1,2 | 0,000163103 | 3,16E-05 | 6 - | 0,000153719 - | 4 - | 0,000127249 - | 6 - | 0,000181449 - | 7 - | 0,000214818 - | 4 - | 0,000138281 - |
| 3424 | 1 | 3um_FIJJAFBF_491669 | Bacteria | Proteobacteri | Gammaprotec | Cellvibrionales | Porticoccacea | unclassified | Gammaprotec | information st | Translation, ril | 17C3B | The globular d | 314287.GB22 | 0,2 | 0,4 | 2,09E-05 | 4,17E-05 | 1 - | 0,000104343 - | 0 - | 0 - | 0 - | 0 - | 0 - | 0 - | 0 - | 0 - |
| 3425 | 1 | 3um_FIJJAFBF_468775 | Bacteria | Bacteroidetes | Flavobacteriia | Flavobacterial | Flavobacteriac | Polaribacter | various | cellular proces | Posttranslatio | 09GE9 | Thioredoxin | 313594.PI23P | 1,8 | 1,469693846 | 0,000235526 | 0,000195853 | 3 - | 0,000327934 - | 3 - | 0,000407197 - | 0 - | 0 - | 0 - | 0 - | 3 - | 0,000442499 - |
| 3426 | 1 | 3um_FIJJAFBF_674223 | Bacteria | Proteobacteri | Alphaproteob | Rhodobactera | Rhodobactera | Roseovarius | Roseovarius n | information st | Translation, ril | 05KK9 | This protein b | 391595.RLO1 | 0,2 | 0,4 | 1,21E-05 | 2,42E-05 | 0 - | 0 - | 0 - | 0 - | 1 - | 6,05E-05 - | 0 - | 0 - | 0 - | 0 - |

| | | | | | | | | | | | | | | | | | | | | | | | | | | | | |
|---|---|---|---|---|---|---|---|---|---|---|---|---|---|---|---|---|---|---|---|---|---|---|---|---|---|---|---|---|
| 3427 | 1 | 3um_FIJJAFBF_155643 | Bacteria | Bacteroidetes | Flavobacteriia | Flavobacterial | Crocinitomica | Fluviicola | Fluviicola sp. X | unclassified | unclassified | unclassified | unclassified | unclassified | 0,4 | 0,489897949 | 1,62E-05 | 1,98E-05 | 0 - | 0 - | 0 - | 0 - | 1 - | 4,02E-05 - | 1 - | 4,08E-05 - | 0 - | 0 - |
| 3428 | 1 | 3um_FIJJAFBF_758946 | Bacteria | Bacteroidetes | Flavobacteriia | Flavobacterial | Crocinitomica | unclassified | Crocinitomica | information st | Translation, ri | COG0088 | One of the pri | 388413.ALPR1 | 0,6 | 0,489897949 | 3,94E-05 | 3,28E-05 | 1 - | 5,49E-05 - | 1 - | 6,82E-05 - | 0 - | 0 - | 0 - | 0 - | 1 - | 7,41E-05 - |
| 3429 | 1 | 0.2um_NOKCPHJN_25052 | Bacteria | Bacteroidetes | Flavobacteriia | Flavobacterial | Flavobacteriac | Ulvibacter | Ulvibacter lito | unclassified | unclassified | unclassified | unclassified | unclassified | 0,4 | 0,489897949 | 1,22E-05 | 1,50E-05 | 0 - | 0 - | 1 - | 3,11E-05 - | 0 - | 0 - | 1 - | 3,00E-05 - | 0 - | 0 - |
| 3430 | 1 | 3um_FIJJAFBF_439966 | Bacteria | Proteobacteri | Alphaproteob | Rhodobactera | unclassified | unclassified | Rhodobactera | poorly charact | Function unkn | 05DIH | basic membra | 391595.RLO14 | 3,4 | 0,8 | 0,000140143 | 3,40E-05 | 3 - | 0,000103714 - | 3 - | 0,000128782 - | 5 - | 0,000204039 - | 3 - | 0,000124232 - | 3 - | 0,000139947 - |
| 3431 | 1 | 3um_FIJJAFBF_79036 | Bacteria | Bacteroidetes | Flavobacteriia | Flavobacterial | Flavobacteriac | Formosa | Formosa sp. H | cellular proces | Cell motility | COG1360 | Ompa motb d | 391587.KAOT | 0,4 | 0,489897949 | 1,81E-05 | 2,24E-05 | 1 - | 4,04E-05 - | 1 - | 5,02E-05 - | 0 - | 0 - | 0 - | 0 - | 0 - | 0 - |
| 3432 | 1 | 0.2um_NOKCPHJN_757859 | Eukaryota | unclassified | Pelagophycea | Pelagomonad | unclassified | Aureococcus | Aureococcus a | unclassified | unclassified | unclassified | unclassified | unclassified | 1,2 | 0,4 | 2,40E-05 | 5,32E-06 | 2 - | 3,45E-05 - | 1 - | 2,14E-05 - | 1 - | 2,03E-05 - | 1 - | 2,06E-05 - | 1 - | 2,33E-05 - |
| 3433 | 1 | 0.2um_NOKCPHJN_217813 | Bacteria | unclassified | unclassified | unclassified | unclassified | unclassified | bacterium B17 | unclassified | unclassified | unclassified | unclassified | unclassified | 2,8 | 0,748331477 | 6,82E-05 | 1,88E-05 | 3 - | 6,16E-05 - | 4 - | 0,000101981 - | 2 - | 4,85E-05 - | 3 - | 7,38E-05 - | 2 - | 5,54E-05 - |
| 3434 | 1 | 3um_FIJJAFBF_356615 | Bacteria | Actinobacteria | Actinobacteria | Micrococcales | Microbacteria | Candidatus Ac | Candidatus Ac | unclassified | unclassified | unclassified | unclassified | unclassified | 0,2 | 0,4 | 3,68E-06 | 7,37E-06 | 1 - | 1,84E-05 - | 0 - | 0 - | 0 - | 0 - | 0 - | 0 - | 0 - | 0 - |
| 3435 | 1 | 3um_FIJJAFBF_544964 | Bacteria | Proteobacteri | Alphaproteob | Pelagibacteral | Pelagibacterac | unclassified | alpha proteob | metabolism | Amino acid tra | 0QI1W | alanine dehyd | 882102.VAA_ | 0,6 | 0,489897949 | 2,45E-05 | 2,04E-05 | 1 - | 3,42E-05 - | 1 - | 4,24E-05 - | 0 - | 0 - | 0 - | 0 - | 1 - | 4,61E-05 - |
| 3436 | 1 | 0.2um_NOKCPHJN_627168 | Bacteria | Bacteroidetes | Flavobacteriia | unclassified | unclassified | unclassified | Flavobacteria | cellular proces | Intracellular tr | 18ATJ | Pfam:YadA | 626418.bglu_ | 1,8 | 0,748331477 | 2,85E-05 | 1,13E-05 | 2 - | 2,71E-05 - | 1 - | 1,68E-05 - | 3 - | 4,80E-05 - | 2 - | 3,25E-05 - | 1 - | 1,83E-05 - |
| 3437 | 1 | 3um_FIJJAFBF_426353 | Bacteria | Proteobacteri | Alphaproteob | Rhodobactera | Rhodobactera | Sulfitobacter | Sulfitobacter g | unclassified | unclassified | unclassified | unclassified | unclassified | 0,4 | 0,489897949 | 1,99E-05 | 2,46E-05 | 1 - | 4,54E-05 - | 0 - | 0 - | 0 - | 0 - | 1 - | 5,43E-05 - | 0 - | 0 - |
| 3438 | 1 | 0.2um_NOKCPHJN_436986 | Bacteria | Bacteroidetes | Cytophagia | Cytophagales | Flammeovirga | unclassified | Flammeovirga | metabolism | Inorganic ion t | 05C27 | TonB-depende | 688270.Celal_ | 2,2 | 0,4 | 2,89E-05 | 5,63E-06 | 2 - | 2,20E-05 - | 2 - | 2,73E-05 - | 3 - | 3,90E-05 - | 2 - | 2,64E-05 - | 2 - | 2,97E-05 - |
| 3439 | 1 | 3um_FIJJAFBF_379822 | Bacteria | Proteobacteri | Betaproteoba | unclassified | unclassified | unclassified | Betaproteoba | metabolism | Energy produc | 05FG6 | Dehydrogenas | 365046.Rta_1 | 2 | 0,632455532 | 0,000104738 | 3,65E-05 | 2 - | 8,73E-05 - | 3 - | 0,000162569 - | 1 - | 5,15E-05 - | 2 - | 0,00010455 - | 2 - | 0,000117775 - |
| 3440 | 1 | 3um_FIJJAFBF_719685 | Bacteria | Bacteroidetes | Flavobacteriia | Flavobacterial | Flavobacteriac | Formosa | Formosa sp. H | unclassified | unclassified | unclassified | unclassified | unclassified | 0,4 | 0,489897949 | 5,54E-05 | 6,84E-05 | 1 - | 0,000126128 - | 0 - | 0 - | 0 - | 0 - | 1 - | 0,000151081 - | 0 - | 0 - |
| 3441 | 1 | 3um_FIJJAFBF_373563 | Bacteria | Proteobacteri | Gammaprotec | Oceanospirilla | Saccharospirill | Reinekea | Reinekea fors | unclassified | unclassified | unclassified | unclassified | unclassified | 0,4 | 0,489897949 | 1,81E-05 | 2,23E-05 | 1 - | 4,11E-05 - | 0 - | 0 - | 0 - | 0 - | 1 - | 4,93E-05 - | 0 - | 0 - |
| 3442 | 1 | 3um_FIJJAFBF_694116 | Bacteria | Bacteroidetes | Flavobacteriia | Flavobacterial | Flavobacteriac | Polaribacter | Polaribacter d | metabolism | Amino acid tra | COG2171 | Catalyzes the | 313598.MED1 | 0,6 | 0,489897949 | 2,86E-05 | 2,35E-05 | 1 - | 4,24E-05 - | 0 - | 0 - | 1 - | 5,00E-05 - | 1 - | 5,07E-05 - | 0 - | 0 - |
| 3443 | 1 | 3um_FIJJAFBF_764272 | Bacteria | Proteobacteri | Alphaproteob | Rhodobactera | Rhodobactera | various | various | information st | Translation, ri | COG0051 | tRNA binding | 391593.RCCS2 | 0,6 | 0,489897949 | 0,000151068 | 0,000124588 | 1 - | 0,000220725 - | 1 - | 0,000274075 - | 1 - | 0,000260543 - | 0 - | 0 - | 0 - | 0 - |
| 3444 | 1 | 3um_FIJJAFBF_239323 | Eukaryota | Arthropoda | Insecta | Diptera | Culicidae | Anopheles | Anopheles gar | cellular proces | Signal transdu | KOG0613 | fibronectin typ | 7165.AGAP00 | 0,2 | 0,4 | 6,47E-06 | 1,29E-05 | 0 - | 0 - | 0 - | 0 - | 0 - | 0 - | 1 - | 3,23E-05 - | 0 - | 0 - |
| 3445 | 1 | 0.2um_NOKCPHJN_68849 | Bacteria | Proteobacteri | Alphaproteob | Rhodobactera | Rhodobactera | Ruegeria | Ruegeria mari | unclassified | unclassified | unclassified | unclassified | unclassified | 1 | 0,632455532 | 6,05E-05 | 3,39E-05 | 2 - | 0,000104819 - | 1 - | 6,51E-05 - | 1 - | 6,19E-05 - | 0 - | 0 - | 1 - | 7,07E-05 - |
| 3446 | 1 | 3um_FIJJAFBF_337838 | Bacteria | Proteobacteri | Alphaproteob | Rhodobactera | Rhodobactera | various | various | poorly charact | Function unkn | 05DIH | basic membra | 439497.RR11_ | 0,4 | 0,489897949 | 1,68E-05 | 2,06E-05 | 0 - | 0 - | 1 - | 4,31E-05 - | 1 - | 4,09E-05 - | 0 - | 0 - | 0 - | 0 - |
| 3447 | 1 | 3um_FIJJAFBF_718194 | Bacteria | Bacteroidetes | unclassified | unclassified | unclassified | unclassified | Bacteroidetes | cellular proces | Intracellular tr | 0BIEB | domain-conta | 1005048.CFU_ | 1,4 | 0,489897949 | 2,43E-05 | 1,03E-05 | 1 - | 1,42E-05 - | 2 - | 3,53E-05 - | 1 - | 1,68E-05 - | 1 - | 1,70E-05 - | 2 - | 3,83E-05 - |
| 3448 | 1 | 3um_FIJJAFBF_09976 | Bacteria | Bacteroidetes | Flavobacteriia | Flavobacterial | Flavobacteriac | Polaribacter | Polaribacter s | unclassified | unclassified | unclassified | unclassified | unclassified | 1,2 | 0,4 | 5,02E-05 | 1,85E-05 | 1 - | 3,48E-05 - | 2 - | 8,64E-05 - | 1 - | 4,11E-05 - | 1 - | 4,17E-05 - | 1 - | 4,69E-05 - |
| 3449 | 1 | 0.2um_NOKCPHJN_1076217 | Bacteria | Bacteroidetes | unclassified | unclassified | unclassified | unclassified | Bacteroidetes | unclassified | unclassified | unclassified | unclassified | unclassified | 1,4 | 1,2 | 3,73E-05 | 3,31E-05 | 2 - | 4,59E-05 - | 3 - | 8,55E-05 - | 0 - | 0 - | 2 - | 5,50E-05 - | 0 - | 0 - |
| 3450 | 1 | 0.2um_NOKCPHJN_296665 | Bacteria | Proteobacteri | Alphaproteob | Rhodobactera | Rhodobactera | Planktomarina | Planktomarina | unclassified | unclassified | unclassified | unclassified | unclassified | 1 | 0 | 0,000128063 | 1,22E-05 | 1 - | 0,000107268 - | 1 - | 0,000133195 - | 1 - | 0,000126619 - | 1 - | 0,000128489 - | 1 - | 0,000144743 - |
| 3451 | 1 | 3um_FIJJAFBF_08772 | Eukaryota | unclassified | Pelagophycea | Pelagomonad | unclassified | Aureococcus | Aureococcus a | cellular proces | Posttranslatio | 0IEKR | One of the co | 35128.Thapsd | 6,6 | 0,8 | 0,000195759 | 1,51E-05 | 8 - | 0,000200483 - | 7 - | 0,000217824 - | 6 - | 0,000177488 - | 6 - | 0,000180109 - | 6 - | 0,000202892 - |
| 3452 | 2 | 3um_FIJJAFBF_242747;3um_FIJJAFBF_86357 | Bacteria | Proteobacteri | Alphaproteob | Rhodobactera | Rhodobactera | Planktomarina | Planktomarina | unclassified | unclassified | unclassified | unclassified | unclassified | 0,2 | 0,4 | 7,83E-06 | 1,57E-05 | 0 - | 0 - | 0 - | 0 - | 1 - | 3,92E-05 - | 0 - | 0 - | 0 - | 0 - |
| 3453 | 1 | 0.2um_NOKCPHJN_489599 | Eukaryota | unclassified | Pelagophycea | Pelagomonad | unclassified | Aureococcus | Aureococcus a | unclassified | unclassified | unclassified | unclassified | unclassified | 2 | 0 | 0,000181493 | 1,72E-05 | 2 - | 0,000152022 - | 2 - | 0,000188767 - | 2 - | 0,000179447 - | 2 - | 0,000182097 - | 2 - | 0,000205132 - |
| 3454 | 1 | 3um_FIJJAFBF_740209 | Bacteria | Proteobacteri | Gammaprotec | Pseudomonad | Moraxellaceae | Psychrobacter | various | cellular proces | Cell wall/mem | 08ZG0 | Ompa motb d | 335284.Pcryo_ | 3,2 | 1,469693846 | 0,000194313 | 7,80E-05 | 5 - | 0,000263249 - | 1 - | 6,54E-05 - | 4 - | 0,000248591 - | 4 - | 0,000252263 - | 2 - | 0,000142087 - |
| 3455 | 1 | 3um_FIJJAFBF_122400 | Bacteria | Bacteroidetes | Flavobacteriia | Flavobacterial | Flavobacteriac | Polaribacter | Polaribacter s | poorly charact | Function unkn | 0PCT6 | reductase (By | 376686.Fjoh_ | 0,2 | 0,4 | 6,94E-06 | 1,39E-05 | 0 - | 0 - | 0 - | 0 - | 0 - | 0 - | 1 - | 3,47E-05 - | 0 - | 0 - |
| 3456 | 1 | 3um_FIJJAFBF_614741 | Bacteria | Proteobacteri | Gammaprotec | Oceanospirilla | Saccharospirill | Reinekea | Reinekea fors | unclassified | unclassified | unclassified | unclassified | unclassified | 0,4 | 0,8 | 1,66E-05 | 3,31E-05 | 2 - | 8,29E-05 - | 0 - | 0 - | 0 - | 0 - | 0 - | 0 - | 0 - | 0 - |
| 3457 | 1 | 0.2um_NOKCPHJN_94128 | Bacteria | Bacteroidetes | Flavobacteriia | Flavobacterial | Flavobacteriac | Flavobacteriu | Flavobacteriu | metabolism | Inorganic ion t | 05C27 | TonB-depende | 983548.Krodi_ | 0,2 | 0,4 | 3,14E-06 | 6,28E-06 | 1 - | 1,57E-05 - | 0 - | 0 - | 0 - | 0 - | 0 - | 0 - | 0 - | 0 - |
| 3458 | 1 | 3um_FIJJAFBF_56687 | Eukaryota | Platyhelminth | Trematoda | Strigeidida | Schistosomati | Schistosoma | Schistosoma h | unclassified | unclassified | unclassified | unclassified | unclassified | 0,4 | 0,489897949 | 2,89E-05 | 3,56E-05 | 1 - | 6,63E-05 - | 0 - | 0 - | 1 - | 7,83E-05 - | 0 - | 0 - | 0 - | 0 - |
| 3459 | 1 | 3um_FIJJAFBF_214618 | Bacteria | Proteobacteri | Alphaproteob | Rhodobactera | Rhodobactera | Phaeobacter | Phaeobacter g | information st | Transcription | 05D27 | DNA-depende | 391619.RGBS1 | 2,4 | 1,496662955 | 2,23E-05 | 1,18E-05 | 5 - | 4,06E-05 - | 1 - | 1,01E-05 - | 1 - | 9,58E-06 - | 3 - | 2,92E-05 - | 2 - | 2,19E-05 - |
| 3460 | 1 | 3um_FIJJAFBF_179678 | Bacteria | Bacteroidetes | Flavobacteriia | Flavobacterial | Flavobacteriac | Formosa | Formosa sp. H | information st | Translation, ri | 0PBIW | 50S ribosomal | 1046627.BZAF | 1 | 0 | 0,000204518 | 1,94E-05 | 1 - | 0,000171309 - | 1 - | 0,000212715 - | 1 - | 0,000202212 - | 1 - | 0,000205199 - | 1 - | 0,000231156 - |
| 3461 | 1 | 0.2um_NOKCPHJN_373963 | Eukaryota | unclassified | Pelagophycea | Pelagomonad | unclassified | Aureococcus | Aureococcus a | poorly charact | Function unkn | 0J2G0 | C-Myc binding | 67593.Physo1 | 0,8 | 0,748331477 | 0,00011279 | 9,92E-05 | 2 - | 0,000255059 - | 1 - | 0,000158354 - | 1 - | 0,000150536 - | 0 - | 0 - | 0 - | 0 - |
| 3462 | 1 | 3um_FIJJAFBF_479211 | Bacteria | Bacteroidetes | Cytophagia | Cytophagales | Flammeovirga | unclassified | Flammeovirga | unclassified | unclassified | unclassified | unclassified | unclassified | 14,4 | 1,356465997 | 0,000330935 | 3,56E-05 | 16 - | 0,000309163 - | 12 - | 0,000287917 - | 14 - | 0,000319318 - | 15 - | 0,00034718 - | 15 - | 0,000391097 - |
| 3463 | 1 | 3um_FIJJAFBF_421711 | Bacteria | Proteobacteri | Alphaproteob | Pelagibacteral | Pelagibacterac | Candidatus Pe | Candidatus Pe | information st | Translation, ri | 16Y2T | Binds directly | 335992.SAR11 | 1,2 | 0,4 | 0,000154702 | 5,71E-05 | 1 - | 0,000107268 - | 2 - | 0,00026639 - | 1 - | 0,000126619 - | 1 - | 0,000128489 - | 1 - | 0,000144743 - |
| 3464 | 1 | 3um_FIJJAFBF_24564 | Bacteria | Proteobacteri | Alphaproteob | Rhodobactera | Rhodobactera | unclassified | Rhodobactera | metabolism | Carbohydrate | 16T1E | Periplasmic bi | 391619.RGBS1 | 5,8 | 1,326649916 | 0,000248856 | 4,74E-05 | 8 - | 0,000290574 - | 6 - | 0,000270605 - | 4 - | 0,000171496 - | 5 - | 0,000217537 - | 6 - | 0,000294065 - |
| 3465 | 1 | 0.2um_NOKCPHJN_80603 | Bacteria | Bacteroidetes | Flavobacteriia | Flavobacterial | Flavobacteriac | Polaribacter | Polaribacter t | information st | Translation, ri | 09GNN | Protein S19 fo | 313594.PI23P | 1 | 0 | 0,000148943 | 1,41E-05 | 1 - | 0,000124757 - | 1 - | 0,000154912 - | 1 - | 0,000147263 - | 1 - | 0,000149438 - | 1 - | 0,000168342 - |
| 3466 | 1 | 3um_FIJJAFBF_707230 | Bacteria | Proteobacteri | Gammaprotec | Oceanospirilla | Saccharospirill | Reinekea | Reinekea blan | metabolism | Amino acid tra | COG1448 | aminotransfer | 314283.MED2 | 0,2 | 0,4 | 6,98E-06 | 1,40E-05 | 0 - | 0 - | 0 - | 0 - | 0 - | 0 - | 1 - | 3,49E-05 - | 0 - | 0 - |
| 3467 | 1 | 3um_FIJJAFBF_118448 | Bacteria | Proteobacteri | Alphaproteob | Rhodobactera | Rhodobactera | unclassified | Rhodobactera | unclassified | unclassified | unclassified | unclassified | unclassified | 4 | 0,632455532 | 0,00012122 | 2,09E-05 | 4 - | 0,000101798 - | 3 - | 9,48E-05 - | 5 - | 0,000150202 - | 4 - | 0,000121936 - | 4 - | 0,000137361 - |
| 3468 | 1 | 0.2um_NOKCPHJN_442499 | Bacteria | Bacteroidetes | Flavobacteriia | Flavobacterial | Flavobacteriac | Lacinutrix | Lacinutrix sp. | unclassified | unclassified | unclassified | unclassified | unclassified | 19,6 | 0,489897949 | 0,000316901 | 3,60E-05 | 19 - | 0,000256862 - | 20 - | 0,000335733 - | 19 - | 0,000303199 - | 20 - | 0,000323871 - | 20 - | 0,00036484 - |
| 3469 | 1 | 3um_FIJJAFBF_267391 | unclassified | unclassified | unclassified | unclassified | unclassified | unclassified | unclassified | unclassified | unclassified | unclassified | unclassified | unclassified | 0,2 | 0,4 | 4,01E-06 | 8,03E-06 | 1 - | 2,01E-05 - | 0 - | 0 - | 0 - | 0 - | 0 - | 0 - | 0 - | 0 - |
| 3470 | 3 | 3um_FIJJAFBF_210226;3um_FIJJAFBF_471892;3u | Bacteria | Bacteroidetes | unclassified | unclassified | unclassified | unclassified | Bacteroidetes | cellular proces | Posttranslatio | 08JYM | peptidase, m3 | various | 0,4 | 0,489897949 | 1,28E-05 | 1,56E-05 | 0 - | 0 - | 0 - | 0 - | 1 - | 3,17E-05 - | 1 - | 3,21E-05 - | 0 - | 0 - |
| 3471 | 1 | 3um_FIJJAFBF_461592 | Bacteria | Proteobacteri | Gammaprotec | Cellvibrionale | Porticoccacea | unclassified | Gammaprotec | metabolism | Inorganic ion t | 16TQC | TONB-depend | 314287.GB22 | 9 | 0,632455532 | 0,000124741 | 1,05E-05 | 10 - | 0,000116525 - | 9 - | 0,00013022 - | 9 - | 0,000123791 - | 8 - | 0,000111662 - | 9 - | 0,00014151 - |
| 3472 | 1 | 0.2um_NOKCPHJN_356306 | Bacteria | Bacteroidetes | Flavobacteriia | Flavobacterial | unclassified | unclassified | Flavobacterial | unclassified | unclassified | unclassified | unclassified | unclassified | 0,6 | 0,489897949 | 4,50E-05 | 3,74E-05 | 1 - | 6,34E-05 - | 0 - | 0 - | 0 - | 0 - | 1 - | 7,60E-05 - | 1 - | 8,56E-05 - |
| 3473 | 1 | 3um_FIJJAFBF_174092 | Bacteria | Bacteroidetes | Sphingobacter | Sphingobacter | Sphingobacter | Sphingobacter | Sphingobacter | cellular proces | Posttranslatio | COG3590 | endothelin-co | 865938.Weev | 0,6 | 0,489897949 | 1,41E-05 | 1,16E-05 | 0 - | 0 - | 0 - | 0 - | 1 - | 2,23E-05 - | 1 - | 2,26E-05 - | 1 - | 2,55E-05 - |
| 3474 | 1 | 0.2um_NOKCPHJN_113052 | Bacteria | Proteobacteri | Alphaproteob | Rhodobactera | Hyphomonada | unclassified | Hyphomonada | unclassified | unclassified | unclassified | unclassified | unclassified | 0,2 | 0,4 | 9,82E-06 | 1,96E-05 | 0 - | 0 - | 0 - | 0 - | 1 - | 4,91E-05 - | 0 - | 0 - | 0 - | 0 - |
| 3475 | 1 | 0.2um_NOKCPHJN_474302 | Bacteria | Proteobacteri | Gammaprotec | Cellvibrionale | Porticoccacea | unclassified | gamma protec | information st | Replication, re | 16Q42 | DNA protectin | 314287.GB22 | 0,2 | 0,4 | 2,11E-05 | 4,22E-05 | 0 - | 0 - | 1 - | 0,00010557 - | 0 - | 0 - | 0 - | 0 - | 0 - | 0 - |
| 3476 | 1 | 0.2um_NOKCPHJN_728627 | Eukaryota | Bacillariophyt | Coscinodiscop | Thalassiosirale | Thalassiosirac | Thalassiosira | Thalassiosira p | unclassified | unclassified | unclassified | unclassified | unclassified | 1 | 0 | 9,72E-05 | 9,23E-06 | 1 - | 8,14E-05 - | 1 - | 0,000101077 - | 1 - | 9,61E-05 - | 1 - | 9,75E-05 - | 1 - | 0,00010984 - |
| 3477 | 1 | 0.2um_NOKCPHJN_291934 | Eukaryota | Chlorophyta | Chlorophycea | Chlamydomor | Chlamydomor | Chlamydomor | Chlamydomor | cellular proces | Cytoskeleton | 13DKH | CH | 7955.ENSDAR | 0,6 | 0,489897949 | 4,50E-06 | 3,69E-06 | 0 - | 0 - | 0 - | 0 - | 1 - | 7,12E-06 - | 1 - | 7,23E-06 - | 1 - | 8,14E-06 - |
| 3478 | 1 | 3um_FIJJAFBF_211171 | Bacteria | Proteobacteri | Betaproteoba | Burkholderial | unclassified | Tepidimonas | Tepidimonas f | information st | Translation, ri | 179ZI | One of the pri | 887062.HGR_ | 3,8 | 0,748331477 | 0,000394252 | 6,80E-05 | 5 - | 0,000438079 - | 4 - | 0,000435172 - | 3 - | 0,000310265 - | 3 - | 0,000314847 - | 4 - | 0,000472899 - |
| 3479 | 1 | 0.2um_NOKCPHJN_469204 | Bacteria | Proteobacteri | Alphaproteob | Pelagibacteral | Pelagibacterac | Candidatus Pe | Candidatus Pe | poorly charact | Function unkn | 16XWJ | Domain of unl | 335992.SAR11 | 0,2 | 0,4 | 1,60E-05 | 3,20E-05 | 0 - | 0 - | 1 - | 8,01E-05 - | 0 - | 0 - | 0 - | 0 - | 0 - | 0 - |
| 3480 | 1 | 0.2um_NOKCPHJN_23906 | Bacteria | Proteobacteri | Gammaprotec | Alteromonada | Alteromonada | Paraglaciecola | Paraglaciecola | cellular proces | Posttranslatio | 0QIPK | Stringent star | 342610.Patl_3 | 1,6 | 0,489897949 | 0,000102658 | 2,85E-05 | 2 - | 0,000109311 - | 2 - | 0,000135732 - | 2 - | 0,000129031 - | 1 - | 6,55E-05 - | 1 - | 7,37E-05 - |
| 3481 | 1 | 0.2um_NOKCPHJN_90062 | Bacteria | Bacteroidetes | Flavobacteriia | Flavobacterial | Flavobacteriac | Polaribacter | Polaribacter s | poorly charact | Function unkn | 11P35 | NA | 398720.MED2 | 0,2 | 0,4 | 4,37E-06 | 8,74E-06 | 1 - | 2,19E-05 - | 0 - | 0 - | 0 - | 0 - | 0 - | 0 - | 0 - | 0 - |
| 3482 | 1 | 3um_FIJJAFBF_670752 | Bacteria | Bacteroidetes | Flavobacteriia | Flavobacterial | Flavobacteriac | Arenitalea | Arenitalea lut | metabolism | Inorganic ion t | 05C27 | TonB-depende | 886377.Murru | 12,6 | 0,489897949 | 0,000246328 | 2,62E-05 | 13 - | 0,000212853 - | 13 - | 0,0002643 - | 12 - | 0,000231924 - | 12 - | 0,000235349 - | 13 - | 0,000287214 - |
| 3483 | 1 | 3um_FIJJAFBF_687190 | unclassified | unclassified | unclassified | unclassified | unclassified | unclassified | unclassified | unclassified | unclassified | unclassified | unclassified | unclassified | 3,2 | 1,6 | 3,64E-05 | 1,44E-05 | 6 - | 5,97E-05 - | 2 - | 2,47E-05 - | 4 - | 4,70E-05 - | 2 - | 2,38E-05 - | 2 - | 2,68E-05 - |
| 3484 | 1 | 3um_FIJJAFBF_316082 | Bacteria | Proteobacteri | Alphaproteob | Rhodobactera | Rhodobactera | Planktomarina | Planktomarina | poorly charact | Function unkn | COG2351 | 5-hydroxyisou | 388739.RSK20 | 0,8 | 0,4 | 9,36E-05 | 4,81E-05 | 1 - | 9,81E-05 - | 1 - | 0,000121811 - | 1 - | 0,000115797 - | 0 - | 0 - | 1 - | 0,000132371 - |
| 3485 | 1 | 3um_FIJJAFBF_520515 | Bacteria | Bacteroidetes | Flavobacteriia | Flavobacterial | Flavobacteriac | Polaribacter | Polaribacter g | metabolism | Amino acid tra | 09AJ2 | N-acetyl-gluta | 313598.MED1 | 0,8 | 0,4 | 3,24E-05 | 1,65E-05 | 1 - | 3,51E-05 - | 1 - | 4,36E-05 - | 1 - | 4,14E-05 - | 1 - | 4,20E-05 - | 0 - | 0 - |
| 3486 | 1 | 3um_FIJJAFBF_116900 | Bacteria | Bacteroidetes | Flavobacteriia | Flavobacterial | Flavobacteriac | Polaribacter | Polaribacter s | poorly charact | Function unkn | 1262F | NA | 485917.Phep_ | 0,4 | 0,489897949 | 1,11E-05 | 1,35E-05 | 0 - | 0 - | 0 - | 0 - | 1 - | 2,74E-05 - | 1 - | 2,78E-05 - | 0 - | 0 - |
| 3487 | 2 | 0.2um_NOKCPHJN_821529;3um_FIJJAFBF_4622( | Viruses | unclassified | unclassified | unclassified | various | various | various | poorly charact | Function unkn | 0NVRZ | NA | 170187.SP_17 | 0,2 | 0,4 | 3,95E-06 | 7,90E-06 | 0 - | 0 - | 0 - | 0 - | 0 - | 0 - | 0 - | 0 - | 1 - | 1,98E-05 - |
| 3488 | 1 | 3um_FIJJAFBF_202365 | Bacteria | Bacteroidetes | Cytophagia | Cytophagales | Flammeovirga | unclassified | Flammeovirga | cellular proces | Cell wall/mem | COG2885 | Ompa motb d | 649349.Lbys_ | 4 | 1,095445115 | 6,97E-05 | 1,58E-05 | 5 - | 7,46E-05 - | 4 - | 7,41E-05 - | 5 - | 8,81E-05 - | 4 - | 7,15E-05 - | 2 - | 4,03E-05 - |
| 3489 | 1 | 0.2um_NOKCPHJN_274421 | Bacteria | Proteobacteri | Acidithiobacill | Acidithiobacill | unclassified | unclassified | Acidithiobacill | metabolism | Carbohydrate | COG0058 | Phosphorylase | 187272.Mlg_( | 0,8 | 0,4 | 1,57E-05 | 8,09E-06 | 1 - | 1,65E-05 - | 1 - | 2,05E-05 - | 1 - | 1,95E-05 - | 0 - | 0 - | 1 - | 2,23E-05 - |
| 3490 | 1 | 0.2um_NOKCPHJN_643250 | Bacteria | Proteobacteri | Alphaproteob | Pelagibacteral | Pelagibacterac | Candidatus Pe | Candidatus Pe | poorly charact | Function unkn | COG3181 | extra-cytoplas | 89187.ISM_13 | 2,4 | 0,489897949 | 0,000262918 | 5,67E-05 | 2 - | 0,000183643 - | 2 - | 0,00022803 - | 3 - | 0,000325157 - | 3 - | 0,00032996 - | 2 - | 0,000247799 - |
| 3491 | 1 | 0.2um_NOKCPHJN_891697 | Eukaryota | unclassified | Pelagophycea | Pelagomonad | unclassified | Aureococcus | Aureococcus a | unclassified | unclassified | unclassified | unclassified | unclassified | 18,4 | 0,8 | 0,001417726 | 0,000159999 | 18 - | 0,001160664 - | 17 - | 0,001361135 - | 19 - | 0,001446159 - | 19 - | 0,001467518 - | 19 - | 0,001653155 - |
| 3492 | 1 | 0.2um_NOKCPHJN_919228 | Eukaryota | unclassified | Pelagophycea | Pelagomonad | unclassified | Aureococcus | Aureococcus a | metabolism | Energy produc | COG0055 | Produces ATP | 2850.Phatr54( | 1,4 | 0,489897949 | 0,00020831 | 6,13E-05 | 2 - | 0,000255059 - | 1 - | 0,000158354 - | 1 - | 0,000150536 - | 2 - | 0,000305518 - | 1 - | 0,000172083 - |
| 3493 | 1 | 0.2um_NOKCPHJN_128438 | Bacteria | Proteobacteri | Gammaprotec | Cellvibrionale | Porticoccacea | unclassified | various | information st | Translation, ri | 05KK9 | This protein bi | 314287.GB22( | 1 | 0 | 0,000131757 | 1,25E-05 | 1 - | 0,000110362 - | 1 - | 0,000137037 - | 1 - | 0,000130271 - | 1 - | 0,000132195 - | 1 - | 0,000148918 - |
| 3494 | 1 | 0.2um_NOKCPHJN_160746 | unclassified | unclassified | unclassified | unclassified | unclassified | unclassified | unclassified | unclassified | unclassified | unclassified | unclassified | unclassified | 2 | 0 | 0,000148137 | 1,41E-05 | 2 - | 0,000124083 - | 2 - | 0,000154074 - | 2 - | 0,000146467 - | 2 - | 0,000148631 - | 2 - | 0,000167432 - |
| 3495 | 1 | 3um_FIJJAFBF_239849 | Eukaryota | Apicomplexa | Aconoidasida | Haemosporida | Plasmodiidae | Plasmodium | Plasmodium c | unclassified | unclassified | unclassified | unclassified | unclassified | 1 | 0 | 4,23E-05 | 4,02E-06 | 1 - | 3,54E-05 - | 1 - | 4,40E-05 - | 1 - | 4,18E-05 - | 1 - | 4,24E-05 - | 1 - | 4,78E-05 - |
| 3496 | 1 | 3um_FIJJAFBF_714063 | Bacteria | Bacteroidetes | Flavobacteriia | Flavobacterial | Flavobacteriac | Lutibacter | Lutibacter sp. | information st | Translation, ri | COG0256 | This is one of t | 313598.MED1 | 0,4 | 0,489897949 | 4,28E-05 | 5,27E-05 | 1 - | 9,81E-05 - | 0 - | 0 - | 1 - | 0,000115797 - | 0 - | 0 - | 0 - | 0 - |
| 3497 | 1 | 3um_FIJJAFBF_257126 | Eukaryota | unclassified | Florideophyce | Nemaliales | Liagoraceae | Dermonema | Dermonema v | unclassified | unclassified | unclassified | unclassified | unclassified | 2,6 | 1,019803903 | 0,000194106 | 7,74E-05 | 2 - | 0,000126128 - | 3 - | 0,000234921 - | 4 - | 0,000297763 - | 3 - | 0,000226621 - | 1 - | 8,51E-05 - |
| 3498 | 1 | 0.2um_NOKCPHJN_719978 | Bacteria | Verrucomicrol | unclassified | unclassified | unclassified | unclassified | Verrucomicro | poorly charact | Function unkn | 1DIXS | Inherit from b | 382464.VDG1 | 1,6 | 0,489897949 | 6,79E-05 | 2,19E-05 | 1 - | 3,54E-05 - | 2 - | 8,80E-05 - | 2 - | 8,36E-05 - | 2 - | 8,49E-05 - | 1 - | 4,78E-05 - |
| 3499 | 1 | 3um_FIJJAFBF_26686 | Bacteria | Bacteroidetes | Flavobacteriia | Flavobacterial | Flavobacteriac | Polaribacter | Polaribacter s | unclassified | unclassified | unclassified | unclassified | unclassified | 1,6 | 0,8 | 1,44E-05 | 7,37E-06 | 2 - | 1,53E-05 - | 3 - | 2,84E-05 - | 1 - | 9,01E-06 - | 1 - | 9,15E-06 - | 1 - | 1,03E-05 - |
| 3500 | 1 | 0.2um_NOKCPHJN_416514 | unclassified | unclassified | unclassified | unclassified | unclassified | unclassified | unclassified | unclassified | unclassified | unclassified | unclassified | unclassified | 0,4 | 0,489897949 | 1,54E-05 | 1,90E-05 | 0 - | 0 - | 0 - | 0 - | 1 - | 3,60E-05 - | 0 - | 0 - | 1 - | 4,12E-05 - |
| 3501 | 1 | 0.2um_NOKCPHJN_685442 | Bacteria | Proteobacteri | Alphaproteob | Rhodobactera | unclassified | unclassified | Rhodobactera | unclassified | unclassified | unclassified | unclassified | unclassified | 1,4 | 1,019803903 | 4,97E-05 | 3,49E-05 | 0 - | 0 - | 2 - | 7,20E-05 - | 3 - | 0,000102638 - | 1 - | 3,47E-05 - | 1 - | 3,91E-05 - |
| 3502 | 1 | 3um_FIJJAFBF_448540 | Bacteria | Proteobacteri | Gammaprotec | Oceanospirilla | Saccharospirill | Reinekea | Reinekea fors | unclassified | unclassified | unclassified | unclassified | unclassified | 3,8 | 0,979795897 | 0,000141391 | 3,61E-05 | 4 - | 0,000126128 - | 5 - | 0,000195768 - | 4 - | 0,000148882 - | 4 - | 0,000151081 - | 2 - | 8,51E-05 - |
| 3503 | 2 | 0.2um_NOKCPHJN_138231;3um_FIJJAFBF_3740( | Bacteria | Bacteroidetes | Flavobacteriia | Flavobacterial | unclassified | unclassified | Flavobacterial | unclassified | unclassified | unclassified | unclassified | unclassified | 0,6 | 0,489897949 | 2,69E-05 | 2,23E-05 | 1 - | 3,74E-05 - | 1 - | 4,64E-05 - | 0 - | 0 - | 0 - | 0 - | 1 - | 5,04E-05 - |
| 3504 | 1 | 0.2um_NOKCPHJN_525976 | Bacteria | Proteobacteri | Gammaprotec | Alteromonada | Alteromonada | Glaciecola | Glaciecola sp. | metabolism | Carbohydrate | 16QEQ | hydrolase, fan | 455436.GHTC | 1,6 | 0,489897949 | 2,11E-05 | 5,53E-06 | 2 - | 2,26E-05 - | 1 - | 1,40E-05 - | 2 - | 2,66E-05 - | 2 - | 2,70E-05 - | 1 - | 1,52E-05 - |
| 3505 | 1 | 3um_FIJJAFBF_811417 | Eukaryota | Streptophyta | Liliopsida | Arecales | Arecaceae | various | various | information st | Translation, ri | KOG0887 | Ribosomal prc | 3694.POPTR_( | 1,2 | 0,4 | 0,00011558 | 3,87E-05 | 1 - | 8,08E-05 - | 1 - | 0,000100365 - | 2 - | 0,00019082 - | 1 - | 9,68E-05 - | 1 - | 0,000109067 - |
| 3506 | 1 | 0.2um_NOKCPHJN_398540 | Eukaryota | unclassified | Oomycetes | Peronosporale | Peronosporac | Phytophthora | Phytophthora | unclassified | unclassified | unclassified | unclassified | unclassified | 0,2 | 0,4 | 1,21E-05 | 2,42E-05 | 1 - | 6,04E-05 - | 0 - | 0 - | 0 - | 0 - | 0 - | 0 - | 0 - | 0 - |
| 3507 | 1 | 3um_FIJJAFBF_363291 | Bacteria | Bacteroidetes | Flavobacteriia | Flavobacterial | Flavobacteriac | Formosa | Formosa sp. H | metabolism | Carbohydrate | COG0176 | Transaldolase | 391587.KAOT | 0,2 | 0,4 | 1,31E-05 | 2,63E-05 | 0 - | 0 - | 1 - | 6,57E-05 - | 0 - | 0 - | 0 - | 0 - | 0 - | 0 - |
| 3508 | 1 | 0.2um_NOKCPHJN_300129 | Eukaryota | Chlorophyta | Chlorophycea | Sphaeropleale | Selenastracea | Raphidocelis | Raphidocelis s | unclassified | unclassified | unclassified | unclassified | unclassified | 0,6 | 0,489897949 | 3,89E-06 | 3,19E-06 | 0 - | 0 - | 1 - | 6,41E-06 - | 1 - | 6,09E-06 - | 0 - | 0 - | 1 - | 6,97E-06 - |
| 3509 | 1 | 0.2um_NOKCPHJN_111361 | Bacteria | Bacteroidetes | Flavobacteriia | Flavobacterial | Flavobacteriac | Formosa | Formosa sp. H | unclassified | unclassified | unclassified | unclassified | unclassified | 0,2 | 0,4 | 2,50E-06 | 5,00E-06 | 1 - | 1,25E-05 - | 0 - | 0 - | 0 - | 0 - | 0 - | 0 - | 0 - | 0 - |
| 3510 | 1 | 3um_FIJJAFBF_34585 | Bacteria | Bacteroidetes | Flavobacteriia | Flavobacterial | unclassified | unclassified | Flavobacterial | unclassified | unclassified | unclassified | unclassified | unclassified | 0,4 | 0,489897949 | 4,04E-05 | 4,97E-05 | 1 - | 9,26E-05 - | 0 - | 0 - | 1 - | 0,00010926 - | 0 - | 0 - | 0 - | 0 - |
| 3511 | 1 | 3um_FIJJAFBF_173775 | Bacteria | Proteobacteri | Alphaproteob | Rhodobactera | Rhodobactera | Leisingera | unclassified | information st | Replication, re | 05C24 | DNA gyrase ne | 644107.SL115 | 1 | 0,632455532 | 1,41E-05 | 8,01E-06 | 2 - | 2,50E-05 - | 1 - | 1,55E-05 - | 1 - | 1,48E-05 - | 1 - | 1,50E-05 - | 0 - | 0 - |
| 3512 | 1 | 3um_FIJJAFBF_08132 | Bacteria | Bacteroidetes | Flavobacteriia | Flavobacterial | Flavobacteriac | Cellulophaga | Cellulophaga a | metabolism | Energy produc | 0P8GN | aconitate hyd | 688270.Celal_ | 9,6 | 1,743559577 | 0,000140234 | 1,77E-05 | 12 - | 0,0001489 - | 11 - | 0,000169482 - | 9 - | 0,000131821 - | 9 - | 0,000133768 - | 7 - | 0,000117202 - |
| 3513 | 1 | 0.2um_NOKCPHJN_622978 | Eukaryota | unclassified | Labyrinthulom | unclassified | Thraustochytr | Aurantiochytr | Aurantiochytr | cellular proces | Signal transdu | KOG0614 | cgmp-depend | 653948.CCA21 | 1 | 0 | 1,92E-05 | 1,82E-06 | 1 - | 1,61E-05 - | 1 - | 2,00E-05 - | 1 - | 1,90E-05 - | 1 - | 1,93E-05 - | 1 - | 2,17E-05 - |
| 3514 | 1 | 0.2um_NOKCPHJN_137368 | Bacteria | Bacteroidetes | Flavobacteriia | Flavobacterial | Crocinitomica | unclassified | Crocinitomica | unclassified | unclassified | unclassified | unclassified | unclassified | 1,8 | 0,4 | 1,63E-05 | 4,06E-06 | 2 - | 1,52E-05 - | 2 - | 1,88E-05 - | 1 - | 8,94E-06 - | 2 - | 1,81E-05 - | 2 - | 2,04E-05 - |
| 3515 | 1 | 3um_FIJJAFBF_82424 | Bacteria | Actinobacteria | Actinobacteria | Micrococcales | Microbacteria | Candidatus Ac | Candidatus Ac | poorly charact | Function unkn | 11D80 | mucin 21, cell | 10090.ENSMU | 4 | 0,632455532 | 5,40E-05 | 8,26E-06 | 4 - | 4,55E-05 - | 4 - | 5,65E-05 - | 4 - | 5,37E-05 - | 5 - | 6,81E-05 - | 3 - | 4,60E-05 - |
| 3516 | 1 | 0.2um_NOKCPHJN_623380 | Bacteria | Bacteroidetes | Bacteroidia | Marinilabiliale | unclassified | unclassified | Marinilabiliale | metabolism | Carbohydrate | 05D2N | Aldolase | 749222.Nitsa_ | 0,4 | 0,489897949 | 1,53E-05 | 1,88E-05 | 0 - | 0 - | 0 - | 0 - | 1 - | 3,81E-05 - | 1 - | 3,86E-05 - | 0 - | 0 - |
| 3517 | 1 | 0.2um_NOKCPHJN_132724 | Eukaryota | unclassified | Oomycetes | Saprolegniales | Saprolegniace | Achlya | Achlya hypogy | poorly charact | Function unkn | 0XNUF | Extracellular n | 340101.Vvad_ | 0,4 | 0,489897949 | 6,44E-07 | 7,89E-07 | 0 - | 0 - | 1 - | 1,65E-06 - | 1 - | 1,57E-06 - | 0 - | 0 - | 0 - | 0 - |
| 3518 | 1 | 0.2um_NOKCPHJN_245989 | Bacteria | Proteobacteri | Alphaproteob | Rhodobactera | Rhodobactera | Planktotalea | Planktotalea f | unclassified | unclassified | unclassified | unclassified | unclassified | 0,8 | 0,4 | 3,39E-05 | 1,74E-05 | 1 - | 3,59E-05 - | 0 - | 0 - | 1 - | 4,23E-05 - | 1 - | 4,30E-05 - | 1 - | 4,84E-05 - |
| 3519 | 2 | 0.2um_NOKCPHJN_83311;3um_FIJJAFBF_558168 | Bacteria | Bacteroidetes | Flavobacteriia | Flavobacterial | Flavobacteriac | Formosa | Formosa sp. H | unclassified | unclassified | unclassified | unclassified | unclassified | 1 | 0 | 7,74E-05 | 7,35E-06 | 1 - | 6,48E-05 - | 1 - | 8,05E-05 - | 1 - | 7,65E-05 - | 1 - | 7,77E-05 - | 1 - | 8,75E-05 - |
| 3520 | 1 | 3um_FIJJAFBF_32567 | Bacteria | Bacteroidetes | Flavobacteriia | Flavobacterial | Flavobacteriac | Formosa | Formosa sp. H | unclassified | unclassified | unclassified | unclassified | unclassified | 0,2 | 0,4 | 1,03E-05 | 2,06E-05 | 0 - | 0 - | 0 - | 0 - | 1 - | 5,15E-05 - | 0 - | 0 - | 0 - | 0 - |
| 3521 | 1 | 3um_FIJJAFBF_305831 | Eukaryota | Chlorophyta | Nephroselmid | unclassified | unclassified | Nephroselmis | Nephroselmis | unclassified | unclassified | unclassified | unclassified | unclassified | 1,8 | 0,4 | 4,86E-05 | 1,21E-05 | 2 - | 4,52E-05 - | 2 - | 5,61E-05 - | 1 - | 2,67E-05 - | 2 - | 5,41E-05 - | 2 - | 6,10E-05 - |
| 3522 | 1 | 3um_FIJJAFBF_230046 | Bacteria | Bacteroidetes | Flavobacteriia | various | various | various | various | unclassified | unclassified | unclassified | unclassified | unclassified | 2,6 | 0,489897949 | 0,000278506 | 6,14E-05 | 3 - | 0,000269008 - | 3 - | 0,000334029 - | 2 - | 0,000211691 - | 2 - | 0,000214818 - | 3 - | 0,000362987 - |
| 3523 | 1 | 3um_FIJJAFBF_225131 | Bacteria | Proteobacteri | Alphaproteob | Rhodobactera | Rhodobactera | Planktomarina | Planktomarina | information st | Translation, ri | COG0256 | This is one of t | 644107.SL115 | 1,6 | 0,489897949 | 0,000184978 | 5,96E-05 | 1 - | 9,65E-05 - | 2 - | 0,000239528 - | 2 - | 0,000227701 - | 2 - | 0,000231064 - | 1 - | 0,000130147 - |
| 3524 | 1 | 0.2um_NOKCPHJN_359329 | Bacteria | Proteobacteri | Gammaprotec | Oceanospirilla | Saccharospirill | Reinekea | Reinekea fors | metabolism | Amino acid tra | 0QIE5 | Catalyzes the | 314283.MED2 | 1,6 | 0,8 | 4,91E-05 | 2,52E-05 | 2 - | 5,15E-05 - | 2 - | 6,39E-05 - | 2 - | 6,08E-05 - | 0 - | 0 - | 2 - | 6,95E-05 - |
| 3525 | 1 | 3um_FIJJAFBF_242311 | Bacteria | Proteobacteri | Gammaprotec | Oceanospirilla | Saccharospirill | Reinekea | Reinekea fors | unclassified | unclassified | unclassified | unclassified | unclassified | 2 | 0,632455532 | 9,25E-05 | 2,81E-05 | 2 - | 7,86E-05 - | 2 - | 9,76E-05 - | 3 - | 0,000139194 - | 2 - | 9,42E-05 - | 1 - | 5,30E-05 - |
| 3526 | 1 | 3um_FIJJAFBF_161153 | Viruses | unclassified | unclassified | unclassified | unclassified | unclassified | Vibrio phage 1 | unclassified | unclassified | unclassified | unclassified | unclassified | 6 | 1,095445115 | 0,000253045 | 4,29E-05 | 6 - | 0,00021387 - | 6 - | 0,000265563 - | 7 - | 0,000294527 - | 7 - | 0,000298877 - | 4 - | 0,000192391 - |
| 3527 | 1 | 3um_FIJJAFBF_500695 | Eukaryota | Bacillariophyt | Mediophycea | Cymatosirales | Cymatosiracea | Plagiogrammo | Plagiogrammo | metabolism | Energy produc | COG4451 | RuBisCO catal | 323097.Nham | 3,2 | 0,4 | 0,000312257 | 1,71E-05 | 4 - | 0,000330293 - | 3 - | 0,000307595 - | 3 - | 0,000292408 - | 3 - | 0,000296727 - | 3 - | 0,000334261 - |
| 3528 | 1 | 3um_FIJJAFBF_263579 | Eukaryota | Phaeophyceae | unclassified | Ectocarpales | Ectocarpaceae | Ectocarpus | Ectocarpus sil | cellular proces | Posttranslatio | KOG0880 | PPIases accele | 39946.BGIOSG | 0,8 | 0,4 | 3,90E-05 | 2,00E-05 | 1 - | 4,07E-05 - | 1 - | 5,05E-05 - | 0 - | 0 - | 1 - | 4,88E-05 - | 1 - | 5,49E-05 - |
| 3529 | 1 | 0.2um_NOKCPHJN_89225 | Eukaryota | unclassified | Oomycetes | Saprolegniales | Saprolegniace | Saprolegnia | Saprolegnia di | unclassified | unclassified | unclassified | unclassified | unclassified | 1 | 0 | 8,62E-06 | 8,18E-07 | 1 - | 7,22E-06 - | 1 - | 8,96E-06 - | 1 - | 8,52E-06 - | 1 - | 8,65E-06 - | 1 - | 9,74E-06 - |
| 3530 | 1 | 0.2um_NOKCPHJN_30617 | Bacteria | Bacteroidetes | Flavobacteriia | Flavobacterial | Flavobacteriac | Psychroserpen | Psychroserpen | unclassified | unclassified | unclassified | unclassified | unclassified | 0,4 | 0,489897949 | 1,20E-05 | 1,47E-05 | 0 - | 0 - | 0 - | 0 - | 0 - | 0 - | 1 - | 2,81E-05 - | 1 - | 3,17E-05 - |
| 3531 | 1 | 3um_FIJJAFBF_179138 | Bacteria | Bacteroidetes | Cytophagia | Cytophagales | Flammeovirga | unclassified | Flammeovirga | unclassified | unclassified | unclassified | unclassified | unclassified | 0,8 | 0,979795897 | 3,96E-05 | 4,87E-05 | 2 - | 9,07E-05 - | 0 - | 0 - | 2 - | 0,000107101 - | 0 - | 0 - | 0 - | 0 - |
| 3532 | 1 | 0.2um_NOKCPHJN_62561 | Bacteria | Proteobacteri | Alphaproteob | Rhodobactera | unclassified | unclassified | Rhodobactera | unclassified | unclassified | unclassified | unclassified | unclassified | 0,8 | 0,748331477 | 4,27E-05 | 3,75E-05 | 2 - | 9,65E-05 - | 1 - | 5,99E-05 - | 1 - | 5,69E-05 - | 0 - | 0 - | 0 - | 0 - |
| 3533 | 2 | 0.2um_NOKCPHJN_335300;3um_FIJJAFBF_38129 | Bacteria | Bacteroidetes | Flavobacteriia | Flavobacterial | Flavobacteriac | Ulvibacter | Ulvibacter lito | poorly charact | Function unkn | 11U8P | endonuclease | 391587.KAOT | 0,6 | 0,489897949 | 1,91E-05 | 1,58E-05 | 1 - | 2,78E-05 - | 1 - | 3,45E-05 - | 0 - | 0 - | 1 - | 3,33E-05 - | 0 - | 0 - |
| 3534 | 1 | 3um_FIJJAFBF_85805 | Viruses | unclassified | unclassified | unclassified | various | various | various | unclassified | unclassified | unclassified | unclassified | unclassified | 1,6 | 0,489897949 | 3,48E-05 | 9,83E-06 | 2 - | 3,70E-05 - | 2 - | 4,59E-05 - | 1 - | 2,18E-05 - | 2 - | 4,43E-05 - | 1 - | 2,49E-05 - |
| 3535 | 1 | 0.2um_NOKCPHJN_112745 | Bacteria | Bacteroidetes | unclassified | unclassified | unclassified | unclassified | Bacteroidetes | unclassified | unclassified | unclassified | unclassified | unclassified | 1,2 | 1,166190379 | 8,71E-05 | 8,02E-05 | 3 - | 0,000197891 - | 1 - | 8,19E-05 - | 2 - | 0,000155727 - | 0 - | 0 - | 0 - | 0 - |
| 3536 | 1 | 3um_FIJJAFBF_410099 | Bacteria | Bacteroidetes | Flavobacteriia | Flavobacterial | Flavobacteriac | unclassified | Flavobacteriac | unclassified | unclassified | unclassified | unclassified | unclassified | 0,8 | 0,979795897 | 4,07E-05 | 5,01E-05 | 2 - | 9,33E-05 - | 0 - | 0 - | 2 - | 0,000110148 - | 0 - | 0 - | 0 - | 0 - |
| 3537 | 3 | 0.2um_NOKCPHJN_427226;0.2um_NOKCPHJN_5 | Bacteria | Bacteroidetes | various | various | various | various | various | unclassified | unclassified | unclassified | unclassified | unclassified | 1,2 | 0,748331477 | 5,84E-05 | 3,70E-05 | 2 - | 8,20E-05 - | 1 - | 5,09E-05 - | 1 - | 4,84E-05 - | 0 - | 0 - | 2 - | 0,000110625 - |
| 3538 | 1 | 0.2um_NOKCPHJN_393903 | Bacteria | Proteobacteri | Gammaprotec | Alteromonada | Alteromonada | Agarivorans | Agarivorans al | metabolism | Inorganic ion t | COG3221 | phosphonate | 351348.Maqu | 1,8 | 0,4 | 8,48E-05 | 2,35E-05 | 1 - | 3,88E-05 - | 2 - | 9,63E-05 - | 2 - | 9,15E-05 - | 2 - | 9,29E-05 - | 2 - | 0,000104645 - |
| 3539 | 1 | 3um_FIJJAFBF_37824 | Bacteria | Actinobacteria | unclassified | unclassified | unclassified | unclassified | Actinobacteria | unclassified | unclassified | unclassified | unclassified | unclassified | 3,4 | 1,624807681 | 4,63E-05 | 2,10E-05 | 5 - | 5,89E-05 - | 5 - | 7,31E-05 - | 4 - | 5,56E-05 - | 2 - | 2,82E-05 - | 1 - | 1,59E-05 - |
| 3540 | 1 | 0.2um_NOKCPHJN_131026 | Bacteria | Bacteroidetes | unclassified | unclassified | unclassified | unclassified | Bacteroidetes | unclassified | unclassified | unclassified | unclassified | unclassified | 0,2 | 0,4 | 3,64E-06 | 7,28E-06 | 0 - | 0 - | 0 - | 0 - | 0 - | 0 - | 1 - | 1,82E-05 - | 0 - | 0 - |
| 3541 | 1 | 3um_FIJJAFBF_341378 | Bacteria | Bacteroidetes | Flavobacteriia | Flavobacterial | Flavobacteriac | Algibacter | Algibacter sp. | cellular proces | Cell wall/mem | COG2885 | Ompa motb d | 1046627.BZAF | 5,6 | 1,2 | 0,000164629 | 1,94E-05 | 8 - | 0,000200483 - | 5 - | 0,000155588 - | 5 - | 0,000147906 - | 5 - | 0,000150091 - | 5 - | 0,000169077 - |
| 3542 | 1 | 3um_FIJJAFBF_199365 | Bacteria | Proteobacteri | Gammaprotec | Oceanospirilla | Oceanospirilla | unclassified | Osedax symbi | information st | Translation, ri | 16Q16 | Binds the low | 314277.MED1 | 4 | 0 | 0,000242526 | 2,30E-05 | 4 - | 0,000203145 - | 4 - | 0,000252246 - | 4 - | 0,000239792 - | 4 - | 0,000243333 - | 4 - | 0,000274114 - |
| 3543 | 1 | 3um_FIJJAFBF_401346 | Bacteria | Bacteroidetes | unclassified | unclassified | unclassified | unclassified | Bacteroidetes | unclassified | unclassified | unclassified | unclassified | unclassified | 2,4 | 1,019803903 | 0,000225609 | 7,92E-05 | 4 - | 0,000323315 - | 2 - | 0,000200731 - | 1 - | 9,54E-05 - | 3 - | 0,000290458 - | 2 - | 0,000218133 - |
| 3544 | 1 | 3um_FIJJAFBF_742422 | Viruses | unclassified | unclassified | unclassified | unclassified | unclassified | uncultured M | poorly charact | Function unkn | 18A3I | NA | 78245.Xaut_3 | 0,2 | 0,4 | 1,58E-05 | 3,17E-05 | 1 - | 7,92E-05 - | 0 - | 0 - | 0 - | 0 - | 0 - | 0 - | 0 - | 0 - |
| 3545 | 1 | 3um_FIJJAFBF_766095 | Bacteria | Proteobacteri | Gammaprotec | Oceanospirilla | Saccharospirill | Reinekea | Reinekea fors | poorly charact | Function unkn | 05Y75 | tpr repeat pro | 384676.PSEEN | 1 | 0 | 0,000155713 | 1,48E-05 | 1 - | 0,000130428 - | 1 - | 0,000161953 - | 1 - | 0,000153957 - | 1 - | 0,000156231 - | 1 - | 0,000175994 - |
| 3546 | 1 | 3um_FIJJAFBF_12600 | Bacteria | Proteobacteri | Gammaprotec | Cellvibrionale | Spongiibacter | Dasania | Dasania marin | unclassified | unclassified | unclassified | unclassified | unclassified | 0,2 | 0,4 | 2,65E-06 | 5,30E-06 | 0 - | 0 - | 1 - | 1,32E-05 - | 0 - | 0 - | 0 - | 0 - | 0 - | 0 - |
| 3547 | 1 | 3um_FIJJAFBF_206948 | Bacteria | Bacteroidetes | Flavobacteriia | Flavobacterial | Flavobacteriac | Polaribacter | various | poorly charact | Function unkn | 0P88U | NA | 398720.MED2 | 1,8 | 0,748331477 | 4,05E-05 | 1,72E-05 | 2 - | 3,81E-05 - | 3 - | 7,10E-05 - | 2 - | 4,50E-05 - | 1 - | 2,28E-05 - | 1 - | 2,57E-05 - |
| 3548 | 1 | 3um_FIJJAFBF_55088 | Bacteria | Bacteroidetes | Flavobacteriia | Flavobacterial | Flavobacteriac | Polaribacter | Polaribacter s | unclassified | unclassified | unclassified | unclassified | unclassified | 0,4 | 0,489897949 | 2,90E-05 | 3,58E-05 | 1 - | 6,60E-05 - | 0 - | 0 - | 0 - | 0 - | 1 - | 7,90E-05 - | 0 - | 0 - |
| 3549 | 1 | 0.2um_NOKCPHJN_316701 | Bacteria | Proteobacteri | Alphaproteob | Pelagibacteral | unclassified | unclassified | Pelagibacteral | cellular proces | Cell wall/mem | 7257 | NA | 439493.PB721 | 2 | 0 | 8,23E-05 | 7,81E-06 | 2 - | 6,89E-05 - | 2 - | 8,56E-05 - | 2 - | 8,14E-05 - | 2 - | 8,26E-05 - | 2 - | 9,30E-05 - |
| 3550 | 1 | 0.2um_NOKCPHJN_540900 | Bacteria | Bacteroidetes | Cytophagia | Cytophagales | Flammeovirga | unclassified | Flammeovirga | unclassified | unclassified | unclassified | unclassified | unclassified | 6,6 | 0,8 | 9,09E-05 | 1,38E-05 | 6 - | 6,92E-05 - | 7 - | 0,000100265 - | 8 - | 0,00010893 - | 6 - | 8,29E-05 - | 6 - | 9,34E-05 - |
| 3551 | 1 | 0.2um_NOKCPHJN_121627 | Bacteria | Proteobacteri | Gammaprotec | Alteromonada | Alteromonada | Paraglaciecola | Paraglaciecola | metabolism | Energy produc | 0QJAP | Catalyzes the | 342610.Patl_( | 2,4 | 0,489897949 | 0,000106793 | 2,84E-05 | 2 - | 7,38E-05 - | 2 - | 9,17E-05 - | 3 - | 0,00013069 - | 2 - | 8,84E-05 - | 3 - | 0,000149397 - |
| 3552 | 1 | 3um_FIJJAFBF_49815 | Eukaryota | Bacillariophyt | Bacillariophyc | Naviculales | Naviculaceae | Fistulifera | Fistulifera sola | unclassified | unclassified | unclassified | unclassified | unclassified | 6,6 | 1,624807681 | 0,000632366 | 0,000123905 | 9 - | 0,000732618 - | 7 - | 0,00070754 - | 7 - | 0,000672607 - | 4 - | 0,000390024 - | 6 - | 0,00065904 - |
| 3553 | 1 | 0.2um_NOKCPHJN_460196 | Bacteria | Proteobacteri | Gammaprotec | Xanthomonad | Xanthomonad | Stenotrophon | Stenotrophon | cellular proces | Cell wall/mem | 08NJY | Repeat protei | 169963.lmo11 | 0,4 | 0,489897949 | 4,89E-06 | 6,03E-06 | 1 - | 1,11E-05 - | 0 - | 0 - | 0 - | 0 - | 1 - | 1,33E-05 - | 0 - | 0 - |
| 3554 | 1 | 3um_FIJJAFBF_30988 | Bacteria | Bacteroidetes | Flavobacteriia | Flavobacterial | Flavobacteriac | Aureitalea | Aureitalea sp. | unclassified | unclassified | unclassified | unclassified | unclassified | 2,6 | 0,489897949 | 0,000205249 | 4,17E-05 | 3 - | 0,000199035 - | 2 - | 0,000164762 - | 3 - | 0,00023494 - | 2 - | 0,00015894 - | 3 - | 0,000268568 - |
| 3555 | 1 | 3um_FIJJAFBF_62812 | Bacteria | Bacteroidetes | Flavobacteriia | Flavobacterial | Flavobacteriac | Polaribacter | Polaribacter s | unclassified | unclassified | unclassified | unclassified | unclassified | 1,2 | 0,4 | 0,000131619 | 4,54E-05 | 1 - | 9,18E-05 - | 1 - | 0,000114015 - | 1 - | 0,000108386 - | 2 - | 0,000219973 - | 1 - | 0,0001239 - |
| 3556 | 1 | 0.2um_NOKCPHJN_64621 | Bacteria | Bacteroidetes | unclassified | unclassified | unclassified | unclassified | Bacteroidetes | cellular proces | Posttranslatio | 0HFM8 | Peptidylprolyl | 485915.Dret_ | 3,6 | 0,489897949 | 0,000207643 | 2,58E-05 | 4 - | 0,000194537 - | 4 - | 0,000241557 - | 3 - | 0,000172223 - | 4 - | 0,000233023 - | 3 - | 0,000196874 - |
| 3557 | 1 | 3um_FIJJAFBF_93285 | Bacteria | Proteobacteri | Alphaproteob | Rhodobactera | Rhodobactera | Pseudooceani | Pseudooceani | information st | Translation, ri | COG0051 | tRNA binding | 391595.RLO14 | 1,8 | 0,748331477 | 0,000227745 | 8,16E-05 | 3 - | 0,00032484 - | 1 - | 0,000134452 - | 1 - | 0,000127813 - | 2 - | 0,000259402 - | 2 - | 0,000292216 - |
| 3558 | 1 | 3um_FIJJAFBF_102176 | Bacteria | Proteobacteri | Alphaproteob | Rhodobactera | Rhodobactera | Ruegeria | Ruegeria med | metabolism | Inorganic ion t | COG3221 | phosphonate | 388739.RSK20 | 13,4 | 1,356465997 | 0,00060749 | 8,50E-05 | 15 - | 0,000570083 - | 13 - | 0,000613492 - | 12 - | 0,00053834 - | 12 - | 0,000546291 - | 15 - | 0,000769244 - |
| 3559 | 1 | 3um_FIJJAFBF_214949 | Bacteria | Bacteroidetes | Flavobacteriia | Flavobacterial | Flavobacteriac | Flaviramulus | Flaviramulus i | poorly charact | Function unkn | 08WMU | Domain of unl | 391603.FBALC | 0,8 | 0,4 | 8,79E-05 | 4,52E-05 | 1 - | 9,18E-05 - | 1 - | 0,000114015 - | 0 - | 0 - | 1 - | 0,000109987 - | 1 - | 0,0001239 - |
| 3560 | 2 | 0.2um_NOKCPHJN_362674;3um_FIJJAFBF_22425 | Bacteria | Bacteroidetes | Flavobacteriia | Flavobacterial | Flavobacteriac | Formosa | Formosa sp. H | unclassified | unclassified | unclassified | unclassified | unclassified | 0,6 | 0,489897949 | 2,75E-05 | 2,27E-05 | 1 - | 4,01E-05 - | 1 - | 4,98E-05 - | 1 - | 4,74E-05 - | 0 - | 0 - | 0 - | 0 - |
| 3561 | 1 | 0.2um_NOKCPHJN_788328 | Bacteria | Proteobacteri | Gammaprotec | Alteromonada | Colwelliaceae | Colwellia | Colwellia sp. F | metabolism | Carbohydrate | 0QX81 | Inherit from p | 94122.Shewar | 1,2 | 0,4 | 1,92E-05 | 4,24E-06 | 2 - | 2,75E-05 - | 1 - | 1,71E-05 - | 1 - | 1,62E-05 - | 1 - | 1,65E-05 - | 1 - | 1,85E-05 - |
| 3562 | 1 | 3um_FIJJAFBF_538012 | Eukaryota | Bacillariophyt | Bacillariophyc | Naviculales | Naviculaceae | Fistulifera | Fistulifera sola | unclassified | unclassified | unclassified | unclassified | unclassified | 1,2 | 0,748331477 | 4,89E-05 | 3,26E-05 | 1 - | 3,35E-05 - | 1 - | 4,16E-05 - | 2 - | 7,90E-05 - | 0 - | 0 - | 2 - | 9,03E-05 - |
| 3563 | 1 | 3um_FIJJAFBF_176097 | Bacteria | Proteobacteri | Gammaprotec | Cellvibrionale | Spongiibacter | Oceanicoccus | Oceanicoccus | cellular proces | Intracellular tr | 124XR | MSHA pilin pr | 903510.vfu_A | 2 | 0 | 0,000168131 | 1,60E-05 | 2 - | 0,00014083 - | 2 - | 0,00017487 - | 2 - | 0,000166236 - | 2 - | 0,000168691 - | 2 - | 0,00019003 - |
| 3564 | 1 | 3um_FIJJAFBF_257135 | Eukaryota | Bacillariophyt | Mediophycea | Biddulphiales | Biddulphiacea | Biddulphia | Biddulphia tri | information st | Translation, ri | 08UHY | One of the pri | 489825.LYNGE | 4,6 | 0,489897949 | 0,000401062 | 3,18E-05 | 5 - | 0,000367874 - | 4 - | 0,000365433 - | 5 - | 0,000434238 - | 5 - | 0,000440652 - | 4 - | 0,000397114 - |
| 3565 | 1 | 3um_FIJJAFBF_262661 | Bacteria | Proteobacteri | Alphaproteob | Pelagibacteral | Pelagibacterac | Candidatus Pe | Candidatus Pe | unclassified | unclassified | unclassified | unclassified | unclassified | 4,4 | 0,8 | 0,000261083 | 5,30E-05 | 4 - | 0,000198748 - | 4 - | 0,000246786 - | 4 - | 0,000234601 - | 6 - | 0,000357099 - | 4 - | 0,000268181 - |
| 3566 | 1 | 0.2um_NOKCPHJN_125665 | Bacteria | Proteobacteri | Gammaprotec | Cellvibrionale | Porticoccacea | unclassified | various | cellular proces | Posttranslatio | 08V1T | peptidylprolyl | 314287.GB22( | 0,8 | 0,4 | 4,66E-05 | 2,34E-05 | 0 - | 0 - | 1 - | 5,82E-05 - | 1 - | 5,53E-05 - | 1 - | 5,61E-05 - | 1 - | 6,32E-05 - |
| 3567 | 1 | 0.2um_NOKCPHJN_550489 | Eukaryota | Cnidaria | Anthozoa | Actiniaria | Edwardsiidae | Nematostella | Nematostella | information st | Translation, ri | KOG3141 | Ribosomal prc | 45351.NEMVE | 2,2 | 1,166190379 | 0,000141462 | 6,52E-05 | 4 - | 0,000223955 - | 3 - | 0,000208564 - | 2 - | 0,000132178 - | 1 - | 6,71E-05 - | 1 - | 7,55E-05 - |
| 3568 | 1 | 0.2um_NOKCPHJN_951164 | unclassified | unclassified | unclassified | unclassified | unclassified | unclassified | unclassified | unclassified | unclassified | unclassified | unclassified | unclassified | 0,8 | 0,4 | 0,000105703 | 5,43E-05 | 1 - | 0,000110362 - | 1 - | 0,000137037 - | 0 - | 0 - | 1 - | 0,000132195 - | 1 - | 0,000148918 - |
| 3569 | 1 | 0.2um_NOKCPHJN_61027 | Bacteria | Proteobacteri | Betaproteoba | Burkholderial | Comamonada | Hydrogenoph | Hydrogenoph | poorly charact | Function unkn | COG1744 | basic membra | 365044.Pnap_ | 12 | 0,632455532 | 0,000432067 | 4,05E-05 | 12 - | 0,000362453 - | 13 - | 0,000487565 - | 12 - | 0,000427839 - | 12 - | 0,000434158 - | 11 - | 0,000448321 - |

| | | | | | | | | | | | | | | | | | | | | | | | | | | | | |
|---|---|---|---|---|---|---|---|---|---|---|---|---|---|---|---|---|---|---|---|---|---|---|---|---|---|---|---|---|
| 3570 | 1 | 0.2um_NOKCPHJN_925580 | Eukaryota | unclassified | Synurophycea | Synurales | Mallomonada | Mallomonas | Mallomonas r | metabolism | Carbohydrate | COG0149 | triosephospha | 35128.Thaps2 | 1 | 0 | 4,46E-05 | 4,24E-06 | 1 - | 3,74E-05 - | 1 - | 4,64E-05 - | 1 - | 4,41E-05 - | 1 - | 4,48E-05 - | 1 - | 5,04E-05 - |
| 3571 | 1 | 3um_FIJJAFBF_485635 | Bacteria | Bacteroidetes | Flavobacteriia | Flavobacterial | Flavobacteriac | Tenacibaculur | Tenacibaculur | unclassified | unclassified | unclassified | unclassified | unclassified | 0,8 | 0,4 | 9,73E-05 | 5,00E-05 | 1 - | 0,000101572 - | 1 - | 0,000126123 - | 0 - | 0 - | 1 - | 0,000121667 - | 1 - | 0,000137057 - |
| 3572 | 1 | 0.2um_NOKCPHJN_451579 | Bacteria | Proteobacteri | Gammaprotec | unclassified | unclassified | unclassified | OM182 bacte | cellular proces | Cell wall/mem | 16Q41 | Efflux transpo | 247633.GP214 | 0,2 | 0,4 | 6,25E-06 | 1,25E-05 | 1 - | 3,13E-05 - | 0 - | 0 - | 0 - | 0 - | 0 - | 0 - | 0 - | 0 - |
| 3573 | 1 | 3um_FIJJAFBF_758937 | Bacteria | Proteobacteri | Alphaproteob | Holosporales | Holosporacea | unclassified | endosymbiont | information st | Transcription | COG0086 | DNA-depende | 342108.amb3 | 1 | 0 | 2,72E-05 | 2,59E-06 | 1 - | 2,28E-05 - | 1 - | 2,83E-05 - | 1 - | 2,69E-05 - | 1 - | 2,73E-05 - | 1 - | 3,08E-05 - |
| 3574 | 1 | 3um_FIJJAFBF_471535 | Bacteria | Proteobacteri | Gammaprotec | Alteromonada | Alteromonada | Paraglaciecola | Paraglaciecola | unclassified | unclassified | unclassified | unclassified | unclassified | 0,8 | 0,4 | 0,00013087 | 6,72E-05 | 1 - | 0,000136639 - | 1 - | 0,000169665 - | 0 - | 0 - | 1 - | 0,000163671 - | 1 - | 0,000184374 - |
| 3575 | 1 | 3um_FIJJAFBF_187826 | Bacteria | Bacteroidetes | Flavobacteriia | Flavobacterial | Flavobacteriac | Polaribacter | various | information st | Translation, ril | 0PAE2 | One of the pri | 313594.PI23P | 1,2 | 0,748331477 | 0,000183652 | 0,000109718 | 2 - | 0,000270063 - | 1 - | 0,000167669 - | 2 - | 0,000318782 - | 1 - | 0,000161745 - | 0 - | 0 - |
| 3576 | 1 | 3um_FIJJAFBF_810299 | Bacteria | Bacteroidetes | unclassified | unclassified | unclassified | unclassified | Bacteroidetes | cellular proces | Cell wall/mem | 0P88B | Outer membr | 926562.Oweh | 0,8 | 0,4 | 2,43E-05 | 1,23E-05 | 1 - | 2,63E-05 - | 1 - | 3,26E-05 - | 1 - | 3,10E-05 - | 1 - | 3,15E-05 - | 0 - | 0 - |
| 3577 | 1 | 3um_FIJJAFBF_462349 | Eukaryota | unclassified | Oomycetes | Saprolegniales | Saprolegniace | Aphanomyces | Aphanomyces | cellular proces | Intracellular tr | KOG2514 | Anoctamin | 65071.PYU1_ | 0,6 | 0,489897949 | 2,81E-05 | 2,31E-05 | 0 - | 0 - | 1 - | 4,61E-05 - | 0 - | 0 - | 1 - | 4,45E-05 - | 1 - | 5,01E-05 - |
| 3578 | 1 | 0.2um_NOKCPHJN_187025 | Bacteria | Nitrospinae | unclassified | unclassified | unclassified | unclassified | Nitrospinae ba | unclassified | unclassified | unclassified | unclassified | unclassified | 6 | 1,095445115 | 0,000238821 | 2,29E-05 | 8 - | 0,00027086 - | 6 - | 0,000252246 - | 6 - | 0,000239792 - | 5 - | 0,000202778 - | 5 - | 0,000228428 - |
| 3579 | 1 | 3um_FIJJAFBF_80806 | Bacteria | Proteobacteri | Gammaprotec | unclassified | unclassified | unclassified | Gammaprotec | unclassified | unclassified | unclassified | unclassified | unclassified | 3,6 | 1,019803903 | 0,000105456 | 3,32E-05 | 3 - | 7,33E-05 - | 2 - | 6,06E-05 - | 4 - | 0,000115304 - | 5 - | 0,000146259 - | 4 - | 0,000131808 - |
| 3580 | 1 | 0.2um_NOKCPHJN_390882 | Bacteria | Proteobacteri | Alphaproteob | Pelagibacteral | Pelagibacterac | Candidatus Pe | Candidatus Pe | metabolism | Carbohydrate | 16T1E | Periplasmic bi | 391595.RLO14 | 6,6 | 1,019803903 | 0,000290912 | 2,80E-05 | 8 - | 0,000299093 - | 7 - | 0,000324962 - | 6 - | 0,000264786 - | 7 - | 0,00031348 - | 5 - | 0,000252239 - |
| 3581 | 1 | 3um_FIJJAFBF_606954 | Bacteria | Bacteroidetes | Flavobacteriia | Flavobacterial | Flavobacteriac | Tenacibaculur | Tenacibaculur | unclassified | unclassified | unclassified | unclassified | unclassified | 3,2 | 0,4 | 0,000253823 | 1,39E-05 | 4 - | 0,000268484 - | 3 - | 0,000250033 - | 3 - | 0,000237688 - | 3 - | 0,000241199 - | 3 - | 0,00027171 - |
| 3582 | 1 | 0.2um_NOKCPHJN_521674 | unclassified | unclassified | unclassified | unclassified | unclassified | unclassified | unclassified | unclassified | unclassified | unclassified | unclassified | unclassified | 1 | 0 | 2,02E-05 | 1,92E-06 | 1 - | 1,69E-05 - | 1 - | 2,10E-05 - | 1 - | 2,00E-05 - | 1 - | 2,02E-05 - | 1 - | 2,28E-05 - |
| 3583 | 1 | 3um_FIJJAFBF_432583 | Bacteria | Bacteroidetes | Flavobacteriia | Flavobacterial | Flavobacteriac | Kordia | Kordia algicida | information st | Translation, ril | COG0013 | Catalyzes the | 391587.KAOT | 1 | 0,632455532 | 1,55E-05 | 1,04E-05 | 1 - | 1,32E-05 - | 2 - | 3,28E-05 - | 1 - | 1,56E-05 - | 1 - | 1,58E-05 - | 0 - | 0 - |
| 3584 | 1 | 0.2um_NOKCPHJN_651474 | Eukaryota | Chordata | unclassified | Testudines | Emydidae | Chrysemys | Chrysemys pic | unclassified | unclassified | unclassified | unclassified | unclassified | 0,2 | 0,4 | 2,53E-06 | 5,06E-06 | 0 - | 0 - | 1 - | 1,26E-05 - | 0 - | 0 - | 0 - | 0 - | 0 - | 0 - |
| 3585 | 1 | 0.2um_NOKCPHJN_03860 | Bacteria | Proteobacteri | Gammaprotec | Alteromonada | Alteromonada | Paraglaciecola | Paraglaciecola | metabolism | Amino acid tra | 05C6M | Alpha-keto-be | 983545.Glaag | 3,8 | 0,4 | 0,000100311 | 1,45E-05 | 4 - | 8,85E-05 - | 4 - | 0,000109841 - | 4 - | 0,000104418 - | 3 - | 7,95E-05 - | 4 - | 0,000119364 - |
| 3586 | 1 | 3um_FIJJAFBF_267591 | Bacteria | Proteobacteri | Gammaprotec | Oceanospirilla | Oceanospirilla | Marinobacteri | Marinobacteri | poorly charact | Function unkn | 11WAN | NA | 207954.MED9 | 0,2 | 0,4 | 1,55E-05 | 3,10E-05 | 0 - | 0 - | 0 - | 0 - | 1 - | 7,74E-05 - | 0 - | 0 - | 0 - | 0 - |
| 3587 | 1 | 3um_FIJJAFBF_350453 | Eukaryota | Bacillariophyt | Bacillariophyc | Naviculales | Phaeodactylac | Phaeodactylu | Phaeodactylu | unclassified | unclassified | unclassified | unclassified | unclassified | 1 | 0 | 0,00013434 | 1,28E-05 | 1 - | 0,000112526 - | 1 - | 0,000139724 - | 1 - | 0,000132826 - | 1 - | 0,000134788 - | 1 - | 0,000151838 - |
| 3588 | 1 | 0.2um_NOKCPHJN_79895 | Bacteria | Proteobacteri | Alphaproteob | Rhodobactera | Rhodobactera | Planktomarina | Planktomarina | unclassified | unclassified | unclassified | unclassified | unclassified | 1 | 0,894427191 | 8,59E-06 | 7,70E-06 | 2 - | 1,52E-05 - | 2 - | 1,88E-05 - | 1 - | 8,95E-06 - | 0 - | 0 - | 0 - | 0 - |
| 3589 | 1 | 0.2um_NOKCPHJN_106567 | Bacteria | Proteobacteri | Gammaprotec | Oceanospirilla | Saccharospirill | Reinekea | Reinekea fors | cellular proces | Posttranslatio | 16ZEN | Inherit from b | 314283.MED2 | 1,6 | 1,356465997 | 5,23E-05 | 4,57E-05 | 2 - | 5,72E-05 - | 0 - | 0 - | 3 - | 0,000101358 - | 3 - | 0,000102855 - | 0 - | 0 - |
| 3590 | 1 | 3um_FIJJAFBF_335490 | Bacteria | Bacteroidetes | Flavobacteriia | Flavobacterial | Flavobacteriac | Polaribacter | Polaribacter ir | unclassified | unclassified | unclassified | unclassified | unclassified | 0,4 | 0,489897949 | 3,06E-05 | 3,82E-05 | 1 - | 6,52E-05 - | 0 - | 0 - | 0 - | 0 - | 0 - | 0 - | 1 - | 8,80E-05 - |
| 3591 | 1 | 3um_FIJJAFBF_294408 | Bacteria | Bacteroidetes | Flavobacteriia | Flavobacterial | various | various | various | cellular proces | Cell wall/mem | COG2885 | Ompa motb d | 867900.Celly_ | 2,4 | 0,8 | 7,28E-05 | 2,53E-05 | 3 - | 7,63E-05 - | 2 - | 6,32E-05 - | 1 - | 3,00E-05 - | 3 - | 9,15E-05 - | 3 - | 0,000103021 - |
| 3592 | 1 | 0.2um_NOKCPHJN_398033 | Archaea | Euryarchaeota | unclassified | unclassified | unclassified | unclassified | Euryarchaeota | metabolism | Inorganic ion t | 0P8Q4 | tonB-depende | 867900.Celly_ | 1,4 | 0,8 | 2,09E-05 | 1,16E-05 | 2 - | 2,54E-05 - | 2 - | 3,15E-05 - | 0 - | 0 - | 2 - | 3,04E-05 - | 1 - | 1,71E-05 - |
| 3593 | 1 | 3um_FIJJAFBF_70342 | Bacteria | Proteobacteri | Betaproteoba | Burkholderial | Comamonada | Rhodoferax | Rhodoferax kc | cellular proces | Posttranslatio | 0BBUA | Redoxin doma | 365046.Rta_0 | 2 | 0 | 0,000163128 | 1,55E-05 | 2 - | 0,000136639 - | 2 - | 0,000169665 - | 2 - | 0,000161288 - | 2 - | 0,000163671 - | 2 - | 0,000184374 - |
| 3594 | 1 | 0.2um_NOKCPHJN_287456 | Bacteria | Proteobacteri | Gammaprotec | Alteromonada | Colwelliaceae | Colwellia | Colwellia sp. 3 | unclassified | unclassified | unclassified | unclassified | unclassified | 3,8 | 0,748331477 | 6,09E-05 | 1,42E-05 | 3 - | 4,00E-05 - | 3 - | 4,97E-05 - | 5 - | 7,88E-05 - | 4 - | 6,39E-05 - | 4 - | 7,20E-05 - |
| 3595 | 1 | 0.2um_NOKCPHJN_669447 | Eukaryota | Chordata | Actinopteri | Cypriniformes | Cyprinidae | Cyprinus | Cyprinus carpi | unclassified | unclassified | unclassified | unclassified | unclassified | 0,2 | 0,4 | 3,64E-06 | 7,27E-06 | 0 - | 0 - | 0 - | 0 - | 0 - | 0 - | 0 - | 0 - | 1 - | 1,82E-05 - |
| 3596 | 1 | 0.2um_NOKCPHJN_1017248 | Eukaryota | unclassified | Spirotrichea | Sporadotrichi | Oxytrichidae | Stylonychia | Stylonychia le | cellular proces | Signal transdu | COG5040 | Tyrosine 3-mc | 412030.XP_00 | 2,8 | 0,4 | 0,00015904 | 1,89E-05 | 3 - | 0,000144071 - | 3 - | 0,000178894 - | 3 - | 0,000170061 - | 3 - | 0,000172573 - | 2 - | 0,000129602 - |
| 3597 | 1 | 3um_FIJJAFBF_79252 | Bacteria | Bacteroidetes | Flavobacteriia | Flavobacterial | Flavobacteriac | Polaribacter | Polaribacter s | information st | Replication, re | 0PB0H | DNA-binding p | 313598.MED1 | 8,8 | 1,720465053 | 0,001323907 | 0,00018738 | 12 - | 0,001530357 - | 8 - | 0,001266834 - | 8 - | 0,001204286 - | 7 - | 0,001069315 - | 9 - | 0,001548745 - |
| 3598 | 1 | 0.2um_NOKCPHJN_440161 | Bacteria | Proteobacteri | Alphaproteob | Rhodobactera | Rhodobactera | Planktomarina | Planktomarina | unclassified | unclassified | unclassified | unclassified | unclassified | 1,4 | 1,356465997 | 3,08E-05 | 3,10E-05 | 1 - | 1,85E-05 - | 3 - | 6,91E-05 - | 0 - | 0 - | 3 - | 6,66E-05 - | 0 - | 0 - |
| 3599 | 1 | 0.2um_NOKCPHJN_1013197 | Eukaryota | Eustigmatoph | unclassified | Eustigmatales | Monodopsida | Nannochlorop | Nannochlorop | unclassified | unclassified | unclassified | unclassified | unclassified | 2,4 | 1,019803903 | 6,22E-05 | 2,95E-05 | 2 - | 4,27E-05 - | 4 - | 0,000105962 - | 2 - | 5,04E-05 - | 1 - | 2,56E-05 - | 3 - | 8,64E-05 - |
| 3600 | 1 | 0.2um_NOKCPHJN_04008 | Bacteria | Proteobacteri | Gammaprotec | Alteromonada | Alteromonada | Paraglaciecola | Paraglaciecola | cellular proces | Cell wall/mem | COG2885 | Ompa motb d | 1085623.GNIT | 1,8 | 0,748331477 | 6,24E-05 | 2,66E-05 | 2 - | 5,83E-05 - | 1 - | 3,62E-05 - | 1 - | 3,44E-05 - | 3 - | 0,000104683 - | 2 - | 7,86E-05 - |
| 3601 | 1 | 0.2um_NOKCPHJN_26712 | Bacteria | Proteobacteri | Betaproteoba | unclassified | unclassified | unclassified | Betaproteoba | unclassified | unclassified | unclassified | unclassified | unclassified | 4 | 1,095445115 | 6,27E-05 | 1,70E-05 | 5 - | 6,60E-05 - | 5 - | 8,19E-05 - | 2 - | 3,11E-05 - | 4 - | 6,32E-05 - | 4 - | 7,12E-05 - |
| 3602 | 1 | 3um_FIJJAFBF_16760 | Bacteria | Bacteroidetes | Flavobacteriia | Flavobacterial | Flavobacteriac | Algibacter | Algibacter algi | metabolism | Amino acid tra | 05C6M | Alpha-keto-be | 867900.Celly_ | 1,6 | 1,019803903 | 4,23E-05 | 2,52E-05 | 3 - | 7,01E-05 - | 2 - | 5,81E-05 - | 2 - | 5,52E-05 - | 1 - | 2,80E-05 - | 0 - | 0 - |
| 3603 | 1 | 3um_FIJJAFBF_23889 | Bacteria | Proteobacteri | Gammaprotec | Cellvibrionales | unclassified | unclassified | Cellvibrionales | poorly charact | Function unkn | 179IZ | NA | 247633.GP214 | 0,4 | 0,489897949 | 1,22E-05 | 1,50E-05 | 0 - | 0 - | 1 - | 3,13E-05 - | 1 - | 2,98E-05 - | 0 - | 0 - | 0 - | 0 - |
| 3604 | 1 | 3um_FIJJAFBF_436831 | Bacteria | Actinobacteria | Actinobacteria | Micrococcales | Microbacteria | unclassified | Microbacteria | unclassified | unclassified | unclassified | unclassified | unclassified | 0,4 | 0,489897949 | 4,18E-05 | 5,17E-05 | 1 - | 9,33E-05 - | 1 - | 0,000115869 - | 0 - | 0 - | 0 - | 0 - | 0 - | 0 - |
| 3605 | 1 | 3um_FIJJAFBF_660012 | Bacteria | Proteobacteri | Gammaprotec | Arenicellales | Arenicellaceae | Arenicella | Arenicella xan | unclassified | unclassified | unclassified | unclassified | unclassified | 0,4 | 0,489897949 | 2,28E-05 | 2,79E-05 | 0 - | 0 - | 1 - | 5,84E-05 - | 1 - | 5,55E-05 - | 0 - | 0 - | 0 - | 0 - |
| 3606 | 1 | 3um_FIJJAFBF_138523 | Bacteria | Proteobacteri | Gammaprotec | Oceanospirilla | Saccharospirill | Reinekea | Reinekea fors | metabolism | Amino acid tra | 01RBF | Branched-chai | 911045.PSE_0 | 6,2 | 1,166190379 | 0,000184287 | 2,45E-05 | 8 - | 0,000201805 - | 7 - | 0,00021926 - | 5 - | 0,000148882 - | 6 - | 0,000181297 - | 5 - | 0,000170192 - |
| 3607 | 1 | 3um_FIJJAFBF_794866 | unclassified | unclassified | unclassified | unclassified | unclassified | unclassified | unclassified | unclassified | unclassified | unclassified | unclassified | unclassified | 0,8 | 0,748331477 | 0,000154478 | 0,000149918 | 1 - | 0,000168789 - | 0 - | 0 - | 1 - | 0,000199239 - | 2 - | 0,000404363 - | 0 - | 0 - |
| 3608 | 2 | 0.2um_NOKCPHJN_429288;3um_FIJJAFBF_7252 | Bacteria | Proteobacteri | Gammaprotec | Oceanospirilla | Saccharospirill | Reinekea | Reinekea fors | poorly charact | Function unkn | 0R976 | von Willebran | 314283.MED2 | 0,2 | 0,4 | 3,29E-06 | 6,58E-06 | 1 - | 1,64E-05 - | 0 - | 0 - | 0 - | 0 - | 0 - | 0 - | 0 - | 0 - |
| 3609 | 1 | 0.2um_NOKCPHJN_1019565 | Bacteria | Proteobacteri | Alphaproteob | unclassified | unclassified | unclassified | Alphaproteob | unclassified | unclassified | unclassified | unclassified | unclassified | 0,8 | 0,4 | 3,17E-05 | 1,63E-05 | 1 - | 3,32E-05 - | 1 - | 4,12E-05 - | 1 - | 3,92E-05 - | 0 - | 0 - | 1 - | 4,48E-05 - |
| 3610 | 1 | 0.2um_NOKCPHJN_359987 | Bacteria | Proteobacteri | Gammaprotec | Oceanospirilla | unclassified | unclassified | Oceanospirilla | cellular proces | Posttranslatio | 0IEKR | One of the co | 35128.Thapsd | 1 | 0 | 2,97E-05 | 2,82E-06 | 1 - | 2,49E-05 - | 1 - | 3,09E-05 - | 1 - | 2,94E-05 - | 1 - | 2,98E-05 - | 1 - | 3,36E-05 - |
| 3611 | 1 | 3um_FIJJAFBF_88897 | Bacteria | Bacteroidetes | Flavobacteriia | Flavobacterial | Flavobacteriac | Polaribacter | Polaribacter ir | cellular proces | Posttranslatio | 0900B | Thioredoxin | 313594.PI23P | 0,6 | 0,489897949 | 7,74E-05 | 6,39E-05 | 1 - | 0,000112526 - | 1 - | 0,000139724 - | 0 - | 0 - | 1 - | 0,000134788 - | 0 - | 0 - |
| 3612 | 1 | 3um_FIJJAFBF_571786 | Bacteria | Proteobacteri | Gammaprotec | Oceanospirilla | Saccharospirill | Reinekea | Reinekea fors | unclassified | unclassified | unclassified | unclassified | unclassified | 0,8 | 0,748331477 | 8,11E-05 | 7,40E-05 | 0 - | 0 - | 0 - | 0 - | 2 - | 0,000194938 - | 1 - | 9,89E-05 - | 1 - | 0,00011142 - |
| 3613 | 1 | 0.2um_NOKCPHJN_237698 | Bacteria | Proteobacteri | Gammaprotec | Oceanospirilla | Saccharospirill | Reinekea | Reinekea fors | metabolism | Amino acid tra | COG4608 | (ABC) transpo | 314283.MED2 | 0,2 | 0,4 | 8,02E-06 | 1,60E-05 | 0 - | 0 - | 0 - | 0 - | 1 - | 4,01E-05 - | 0 - | 0 - | 0 - | 0 - |
| 3614 | 1 | 3um_FIJJAFBF_54938 | Bacteria | Bacteroidetes | Flavobacteriia | Flavobacterial | Flavobacteriac | Gaetbulibacte | Gaetbulibacte | poorly charact | Function unkn | 11PZJ | NA | 1046627.BZAF | 1,4 | 0,489897949 | 0,000131104 | 3,86E-05 | 2 - | 0,000160527 - | 1 - | 9,97E-05 - | 1 - | 9,47E-05 - | 2 - | 0,000192284 - | 1 - | 0,000108304 - |
| 3615 | 1 | 0.2um_NOKCPHJN_213918 | Bacteria | Proteobacteri | Alphaproteob | Rhodobactera | Rhodobactera | Celeribacter | Celeribacter h | metabolism | Lipid transpor | COG2067 | lipid transport | 644107.SL115 | 0,4 | 0,8 | 5,50E-05 | 0,000109987 | 0 - | 0 - | 0 - | 0 - | 0 - | 0 - | 2 - | 0,000274967 - | 0 - | 0 - |
| 3616 | 1 | 3um_FIJJAFBF_419639 | Bacteria | Bacteroidetes | Flavobacteriia | Flavobacterial | Flavobacteriac | Polaribacter | various | unclassified | unclassified | unclassified | unclassified | unclassified | 3 | 1,414213562 | 0,000154127 | 6,15E-05 | 5 - | 0,000223301 - | 1 - | 5,55E-05 - | 4 - | 0,000210867 - | 3 - | 0,000160486 - | 2 - | 0,000120525 - |
| 3617 | 1 | 3um_FIJJAFBF_239316 | Bacteria | Bacteroidetes | Cytophagia | Cytophagales | Flammeovirga | unclassified | Flammeovirga | unclassified | unclassified | unclassified | unclassified | unclassified | 2,6 | 1,356465997 | 0,000134035 | 5,72E-05 | 5 - | 0,000225939 - | 3 - | 0,000168329 - | 2 - | 0,000106679 - | 2 - | 0,000108255 - | 1 - | 6,10E-05 - |
| 3618 | 2 | 0.2um_NOKCPHJN_486326;0.2um_NOKCPHJN_8 | Eukaryota | various | various | various | various | various | various | unclassified | unclassified | unclassified | unclassified | unclassified | 1,8 | 0,4 | 0,000200779 | 5,00E-05 | 2 - | 0,000186629 - | 2 - | 0,000231738 - | 1 - | 0,000110148 - | 2 - | 0,00022355 - | 2 - | 0,000251828 - |
| 3619 | 1 | 3um_FIJJAFBF_238978 | unclassified | unclassified | unclassified | unclassified | unclassified | unclassified | unclassified | unclassified | unclassified | unclassified | unclassified | unclassified | 1 | 0,632455532 | 8,43E-05 | 5,46E-05 | 1 - | 7,26E-05 - | 1 - | 9,02E-05 - | 2 - | 0,000171496 - | 1 - | 8,70E-05 - | 0 - | 0 - |
| 3620 | 1 | 0.2um_NOKCPHJN_978445 | Bacteria | Proteobacteri | Gammaprotec | Pseudomonad | Pseudomonad | Pseudomonas | various | metabolism | Carbohydrate | 05CA9 | Pyruvate kina | 216595.PFLU4 | 0,2 | 0,4 | 5,61E-06 | 1,12E-05 | 0 - | 0 - | 0 - | 0 - | 1 - | 2,81E-05 - | 0 - | 0 - | 0 - | 0 - |
| 3621 | 1 | 3um_FIJJAFBF_187034 | Bacteria | Proteobacteri | Gammaprotec | Alteromonada | Alteromonada | Paraglaciecola | Paraglaciecola | cellular proces | Cell motility | 0QY5Y | flagellar prote | 342610.Patl_3 | 0,2 | 0,4 | 1,65E-05 | 3,30E-05 | 1 - | 8,26E-05 - | 0 - | 0 - | 0 - | 0 - | 0 - | 0 - | 0 - | 0 - |
| 3622 | 1 | 0.2um_NOKCPHJN_251410 | Bacteria | Proteobacteri | Gammaprotec | unclassified | unclassified | unclassified | Gammaprotec | unclassified | unclassified | unclassified | unclassified | unclassified | 0,8 | 0,748331477 | 1,28E-05 | 1,19E-05 | 0 - | 0 - | 2 - | 3,15E-05 - | 0 - | 0 - | 1 - | 1,52E-05 - | 1 - | 1,71E-05 - |
| 3623 | 1 | 3um_FIJJAFBF_237591 | Bacteria | Proteobacteri | Alphaproteob | Rhodobactera | Rhodobactera | Planktomarina | Planktomarina | unclassified | unclassified | unclassified | unclassified | unclassified | 0,2 | 0,4 | 5,75E-06 | 1,15E-05 | 1 - | 2,88E-05 - | 0 - | 0 - | 0 - | 0 - | 0 - | 0 - | 0 - | 0 - |
| 3624 | 1 | 3um_FIJJAFBF_500137 | Bacteria | Bacteroidetes | Flavobacteriia | unclassified | unclassified | unclassified | Flavobacteria | poorly charact | Function unkn | 05ERK | gliding motilit | 487797.Flav3 | 2,6 | 0,8 | 6,72E-05 | 1,74E-05 | 4 - | 8,80E-05 - | 2 - | 5,46E-05 - | 2 - | 5,19E-05 - | 2 - | 5,27E-05 - | 3 - | 8,90E-05 - |
| 3625 | 1 | 3um_FIJJAFBF_253700 | Bacteria | Proteobacteri | Gammaprotec | Cellvibrionales | Halieaceae | Chromatocurv | Chromatocurv | unclassified | unclassified | unclassified | unclassified | unclassified | 2,2 | 0,979795897 | 3,50E-05 | 1,59E-05 | 1 - | 1,32E-05 - | 3 - | 4,93E-05 - | 3 - | 4,69E-05 - | 3 - | 4,76E-05 - | 1 - | 1,79E-05 - |
| 3626 | 1 | 0.2um_NOKCPHJN_222203 | Bacteria | Firmicutes | Bacilli | Bacillales | Staphylococca | Staphylococcu | Staphylococcu | poorly charact | Function unkn | 0ZPU4 | NA | 5786.XP_0032 | 0,6 | 0,489897949 | 8,58E-06 | 7,01E-06 | 0 - | 0 - | 1 - | 1,47E-05 - | 1 - | 1,40E-05 - | 1 - | 1,42E-05 - | 0 - | 0 - |
| 3627 | 1 | 3um_FIJJAFBF_742004 | Bacteria | Bacteroidetes | Cytophagia | Cytophagales | Flammeovirga | unclassified | Flammeovirga | poorly charact | Function unkn | 05DI5 | UPF0365 prot | 388413.ALPR1 | 2,6 | 1,019803903 | 0,000103789 | 3,92E-05 | 3 - | 0,000102479 - | 3 - | 0,000127249 - | 4 - | 0,000161288 - | 2 - | 8,18E-05 - | 1 - | 4,61E-05 - |
| 3628 | 1 | 0.2um_NOKCPHJN_56819 | Bacteria | Proteobacteri | Gammaprotec | Oceanospirilla | Saccharospirill | Reinekea | Reinekea blan | cellular proces | Posttranslatio | 0QHXH | Molecular cha | 314283.MED2 | 14,6 | 1,2 | 0,000311031 | 5,06E-05 | 13 - | 0,000230618 - | 14 - | 0,000308387 - | 14 - | 0,000293161 - | 16 - | 0,00033999 - | 16 - | 0,000382997 - |
| 3629 | 1 | 0.2um_NOKCPHJN_547497 | Bacteria | unclassified | unclassified | unclassified | unclassified | unclassified | uncultured ba | metabolism | Inorganic ion t | 05C27 | TonB-depende | 700598.Niako | 1 | 0 | 1,54E-05 | 1,46E-06 | 1 - | 1,29E-05 - | 1 - | 1,60E-05 - | 1 - | 1,52E-05 - | 1 - | 1,54E-05 - | 1 - | 1,74E-05 - |
| 3630 | 1 | 3um_FIJJAFBF_796580 | Bacteria | Bacteroidetes | unclassified | unclassified | unclassified | unclassified | Bacteroidetes | poorly charact | Function unkn | 11NCV | Protein of unk | 643867.Ftrac_ | 2 | 0,632455532 | 0,00015594 | 5,19E-05 | 2 - | 0,000130428 - | 2 - | 0,000161953 - | 1 - | 7,70E-05 - | 3 - | 0,000234347 - | 2 - | 0,000175994 - |
| 3631 | 1 | 3um_FIJJAFBF_92946 | Bacteria | Proteobacteri | Gammaprotec | Cellvibrionales | unclassified | unclassified | Cellvibrionales | unclassified | unclassified | unclassified | unclassified | unclassified | 5,2 | 0,748331477 | 0,000129428 | 2,15E-05 | 6 - | 0,000125439 - | 4 - | 0,000103839 - | 5 - | 0,00012339 - | 5 - | 0,000125212 - | 6 - | 0,000169262 - |
| 3632 | 1 | 3um_FIJJAFBF_192709 | Bacteria | Bacteroidetes | Flavobacteriia | Flavobacterial | Flavobacteriac | Formosa | Formosa sp. H | information st | Translation, ril | 0PA05 | 30S ribosomal | 398720.MED2 | 1,4 | 0,489897949 | 8,96E-05 | 3,30E-05 | 1 - | 5,34E-05 - | 2 - | 0,000132576 - | 2 - | 0,00012603 - | 1 - | 6,39E-05 - | 1 - | 7,20E-05 - |
| 3633 | 1 | 3um_FIJJAFBF_39858 | Bacteria | Proteobacteri | Gammaprotec | Cellvibrionales | Spongiibacter | Oceanicoccus | Oceanicoccus | information st | Translation, ril | COG0050 | This protein p | 247633.GP214 | 1,2 | 0,4 | 5,42E-05 | 1,20E-05 | 2 - | 7,78E-05 - | 1 - | 4,83E-05 - | 1 - | 4,59E-05 - | 1 - | 4,66E-05 - | 1 - | 5,25E-05 - |
| 3634 | 1 | 0.2um_NOKCPHJN_316391 | Bacteria | Proteobacteri | Gammaprotec | unclassified | unclassified | unclassified | SAR86 cluster | unclassified | unclassified | unclassified | unclassified | unclassified | 1 | 0 | 1,12E-05 | 1,07E-06 | 1 - | 9,42E-06 - | 1 - | 1,17E-05 - | 1 - | 1,11E-05 - | 1 - | 1,13E-05 - | 1 - | 1,27E-05 - |
| 3635 | 1 | 0.2um_NOKCPHJN_530493 | Eukaryota | unclassified | Labyrinthulom | unclassified | Thraustochytr | Aurantiochytr | Aurantiochytr | unclassified | unclassified | unclassified | unclassified | unclassified | 10,2 | 1,469693846 | 8,79E-05 | 5,74E-06 | 13 - | 9,50E-05 - | 10 - | 9,07E-05 - | 9 - | 7,76E-05 - | 10 - | 8,75E-05 - | 9 - | 8,87E-05 - |
| 3636 | 1 | 3um_FIJJAFBF_282599 | Eukaryota | unclassified | Pelagophycea | Pelagomonad | unclassified | Aureococcus | Aureococcus a | unclassified | unclassified | unclassified | unclassified | unclassified | 1,2 | 0,4 | 8,65E-05 | 1,92E-05 | 2 - | 0,000124083 - | 1 - | 7,70E-05 - | 1 - | 7,32E-05 - | 1 - | 7,43E-05 - | 1 - | 8,37E-05 - |
| 3637 | 1 | 0.2um_NOKCPHJN_812248 | Bacteria | Bacteroidetes | Saprospiria | Saprospirales | Lewinellaceae | Lewinella | Lewinella nigr | unclassified | unclassified | unclassified | unclassified | unclassified | 0,2 | 0,4 | 6,77E-06 | 1,35E-05 | 0 - | 0 - | 1 - | 3,39E-05 - | 0 - | 0 - | 0 - | 0 - | 0 - | 0 - |
| 3638 | 1 | 3um_FIJJAFBF_365652 | Bacteria | Bacteroidetes | Flavobacteriia | Flavobacterial | Flavobacteriac | Aureitalea | Aureitalea ma | unclassified | unclassified | unclassified | unclassified | unclassified | 23,8 | 1,939071943 | 0,000367351 | 2,78E-05 | 25 - | 0,000324963 - | 25 - | 0,000403508 - | 24 - | 0,000368242 - | 25 - | 0,000389251 - | 20 - | 0,000350792 - |
| 3639 | 1 | 3um_FIJJAFBF_437251 | Bacteria | Bacteroidetes | Flavobacteriia | Flavobacterial | Flavobacteriac | Flaviramulus | Flaviramulus i | unclassified | unclassified | unclassified | unclassified | unclassified | 16,6 | 1,019803903 | 0,000227286 | 2,03E-05 | 18 - | 0,000207012 - | 15 - | 0,000214207 - | 16 - | 0,000217206 - | 17 - | 0,00023419 - | 17 - | 0,000263814 - |
| 3640 | 1 | 3um_FIJJAFBF_715282 | Bacteria | Bacteroidetes | Flavobacteriia | unclassified | unclassified | unclassified | Flavobacteria | unclassified | unclassified | unclassified | unclassified | unclassified | 6,4 | 1,019803903 | 0,000160643 | 2,00E-05 | 8 - | 0,000169725 - | 7 - | 0,000184405 - | 6 - | 0,000150258 - | 5 - | 0,000127064 - | 6 - | 0,000171765 - |
| 3641 | 1 | 3um_FIJJAFBF_60991 | Bacteria | Bacteroidetes | Flavobacteriia | various | various | various | various | poorly charact | Function unkn | 0900R | pur-alpha bet | 487797.Flav3 | 1 | 0 | 0,000103808 | 9,86E-06 | 1 - | 8,70E-05 - | 1 - | 0,000107969 - | 1 - | 0,000102638 - | 1 - | 0,000104154 - | 1 - | 0,000117329 - |
| 3642 | 1 | 3um_FIJJAFBF_456845 | Bacteria | Proteobacteri | Gammaprotec | Oceanospirilla | Saccharospirill | Reinekea | Reinekea blan | information st | Translation, ril | COG0018 | arginyL-tRNA | 314283.MED2 | 0,4 | 0,489897949 | 9,17E-06 | 1,13E-05 | 1 - | 2,05E-05 - | 1 - | 2,54E-05 - | 0 - | 0 - | 0 - | 0 - | 0 - | 0 - |
| 3643 | 1 | 3um_FIJJAFBF_748411 | Bacteria | Proteobacteri | Gammaprotec | Cellvibrionales | unclassified | unclassified | Cellvibrionales | cellular proces | Posttranslatio | COG0330 | Band 7 protei | 247633.GP214 | 0,2 | 0,4 | 7,26E-06 | 1,45E-05 | 0 - | 0 - | 0 - | 0 - | 0 - | 0 - | 1 - | 3,63E-05 - | 0 - | 0 - |
| 3644 | 1 | 3um_FIJJAFBF_08081 | Bacteria | Bacteroidetes | Flavobacteriia | Flavobacterial | Flavobacteriac | Polaribacter | Polaribacter ir | metabolism | Carbohydrate | 0PF2Z | Transketolase | 313594.PI23P | 4 | 0,632455532 | 0,000194914 | 3,51E-05 | 4 - | 0,000163383 - | 4 - | 0,000202874 - | 5 - | 0,000241072 - | 3 - | 0,000146779 - | 4 - | 0,000220462 - |
| 3645 | 1 | 0.2um_NOKCPHJN_483486 | Bacteria | Bacteroidetes | Flavobacteriia | Flavobacterial | Flavobacteriac | Arenitalea | Arenitalea lut | poorly charact | Function unkn | 0PAEN | NA | 886377.Murru | 1,6 | 1,2 | 4,60E-05 | 3,51E-05 | 1 - | 2,39E-05 - | 3 - | 8,91E-05 - | 3 - | 8,47E-05 - | 0 - | 0 - | 1 - | 3,23E-05 - |
| 3646 | 1 | 3um_FIJJAFBF_70033 | Bacteria | Proteobacteri | Gammaprotec | Cellvibrionales | Porticoccacea | unclassified | gamma prote | unclassified | unclassified | unclassified | unclassified | unclassified | 1,2 | 0,4 | 3,62E-05 | 1,21E-05 | 1 - | 2,53E-05 - | 1 - | 3,15E-05 - | 2 - | 5,98E-05 - | 1 - | 3,03E-05 - | 1 - | 3,42E-05 - |
| 3647 | 1 | 3um_FIJJAFBF_01802 | Bacteria | Proteobacteri | Gammaprotec | Alteromonada | Alteromonada | Paraglaciecola | Paraglaciecola | metabolism | Inorganic ion t | 0QIT9 | receptor | 983545.Glaag | 0,6 | 0,489897949 | 1,07E-05 | 8,83E-06 | 1 - | 1,56E-05 - | 1 - | 1,93E-05 - | 0 - | 0 - | 1 - | 1,86E-05 - | 0 - | 0 - |
| 3648 | 1 | 3um_FIJJAFBF_116542 | Eukaryota | Chordata | Actinopteri | Carangiformes | Carangidae | Seriola | Seriola lalandi | unclassified | unclassified | unclassified | unclassified | unclassified | 3,6 | 0,489897949 | 0,000730367 | 0,000144022 | 3 - | 0,000506368 - | 3 - | 0,00062876 - | 4 - | 0,000796954 - | 4 - | 0,000808725 - | 4 - | 0,000911026 - |
| 3649 | 1 | 0.2um_NOKCPHJN_467912 | Eukaryota | unclassified | Oomycetes | Saprolegniales | Saprolegniace | various | various | unclassified | unclassified | unclassified | unclassified | unclassified | 2,6 | 0,489897949 | 0,000418575 | 6,06E-05 | 3 - | 0,000409917 - | 2 - | 0,000339331 - | 3 - | 0,000483865 - | 3 - | 0,000491012 - | 2 - | 0,000368749 - |
| 3650 | 1 | 0.2um_NOKCPHJN_393294 | Eukaryota | unclassified | Oomycetes | Saprolegniales | Saprolegniace | Aphanomyces | Aphanomyces | information st | Translation, ril | 1524W | Ribosomal S1 | 164328.Phyra | 0,4 | 0,489897949 | 3,62E-05 | 4,43E-05 | 0 - | 0 - | 0 - | 0 - | 1 - | 8,97E-05 - | 1 - | 9,10E-05 - | 0 - | 0 - |
| 3651 | 1 | 0.2um_NOKCPHJN_61811 | Bacteria | Bacteroidetes | Flavobacteriia | Flavobacterial | Flavobacteriac | Algibacter | Algibacter sp. | metabolism | Inorganic ion t | 05C27 | TonB-depende | 391587.KAOT | 36,2 | 2,039607805 | 0,000477967 | 4,63E-05 | 39 - | 0,000432075 - | 37 - | 0,000508996 - | 35 - | 0,00045771 - | 33 - | 0,000437929 - | 37 - | 0,000553123 - |
| 3652 | 1 | 3um_FIJJAFBF_589299 | Bacteria | Proteobacteri | Gammaprotec | Oceanospirilla | Saccharospirill | Reinekea | Reinekea fors | cellular proces | Posttranslatio | 17FI0 | Binds to Cpn6 | 314283.MED2 | 1,6 | 0,8 | 0,000238393 | 0,000138749 | 1 - | 0,000120818 - | 1 - | 0,00015002 - | 1 - | 0,000142613 - | 2 - | 0,000289439 - | 3 - | 0,000489077 - |
| 3653 | 1 | 0.2um_NOKCPHJN_522831 | Bacteria | Acidobacteria | unclassified | unclassified | unclassified | unclassified | Acidobacteria | poorly charact | Function unkn | 11FY2 | radical SAM d | 862908.BMS_ | 0,2 | 0,4 | 6,75E-06 | 1,35E-05 | 1 - | 3,38E-05 - | 0 - | 0 - | 0 - | 0 - | 0 - | 0 - | 0 - | 0 - |
| 3654 | 1 | 3um_FIJJAFBF_655407 | Bacteria | Proteobacteri | Gammaprotec | Cellvibrionales | Porticoccacea | unclassified | SAR92 bacteri | information st | Replication, re | 05CZ8 | DNA polymera | 314287.GB22 | 0,2 | 0,4 | 7,53E-06 | 1,51E-05 | 0 - | 0 - | 0 - | 0 - | 0 - | 0 - | 1 - | 3,77E-05 - | 0 - | 0 - |
| 3655 | 1 | 0.2um_NOKCPHJN_156754 | Bacteria | Proteobacteri | Betaproteoba | Burkholderial | Comamonada | Limnohabitan | Limnohabitan | unclassified | unclassified | unclassified | unclassified | unclassified | 0,8 | 0,4 | 1,14E-05 | 5,86E-06 | 1 - | 1,21E-05 - | 0 - | 0 - | 1 - | 1,42E-05 - | 1 - | 1,45E-05 - | 1 - | 1,63E-05 - |
| 3656 | 1 | 3um_FIJJAFBF_35365 | Bacteria | Bacteroidetes | Flavobacteriia | Flavobacterial | Flavobacteriac | Polaribacter | Polaribacter s | unclassified | unclassified | unclassified | unclassified | unclassified | 1 | 0,632455532 | 4,11E-05 | 2,90E-05 | 1 - | 3,36E-05 - | 1 - | 4,17E-05 - | 1 - | 3,96E-05 - | 0 - | 0 - | 2 - | 9,06E-05 - |
| 3657 | 1 | 3um_FIJJAFBF_256676 | Bacteria | Bacteroidetes | Flavobacteriia | Flavobacterial | Flavobacteriac | Algibacter | Algibacter sp. | cellular proces | Cell wall/mem | 0P9R9 | TonB family | 1046627.BZAF | 1 | 0,632455532 | 5,40E-05 | 3,08E-05 | 2 - | 9,60E-05 - | 1 - | 5,96E-05 - | 1 - | 5,67E-05 - | 1 - | 5,75E-05 - | 0 - | 0 - |
| 3658 | 1 | 3um_FIJJAFBF_545651 | Bacteria | Proteobacteri | Gammaprotec | Oceanospirilla | Oceanospirilla | Marinobacteri | Marinobacteri | unclassified | unclassified | unclassified | unclassified | unclassified | 1 | 0 | 2,56E-05 | 2,43E-06 | 1 - | 2,14E-05 - | 1 - | 2,66E-05 - | 1 - | 2,53E-05 - | 1 - | 2,56E-05 - | 1 - | 2,89E-05 - |
| 3659 | 1 | 0.2um_NOKCPHJN_343133 | Bacteria | Proteobacteri | Alphaproteob | Rhizobiales | unclassified | unclassified | Rhizobiales ba | unclassified | unclassified | unclassified | unclassified | unclassified | 2 | 0 | 3,26E-05 | 3,09E-06 | 2 - | 2,73E-05 - | 2 - | 3,39E-05 - | 2 - | 3,22E-05 - | 2 - | 3,27E-05 - | 2 - | 3,68E-05 - |
| 3660 | 1 | 3um_FIJJAFBF_367008 | Bacteria | Proteobacteri | Gammaprotec | Alteromonada | Alteromonada | Paraglaciecola | Paraglaciecola | unclassified | unclassified | unclassified | unclassified | unclassified | 0,4 | 0,489897949 | 2,25E-05 | 2,78E-05 | 1 - | 5,17E-05 - | 0 - | 0 - | 1 - | 6,10E-05 - | 0 - | 0 - | 0 - | 0 - |
| 3661 | 1 | 3um_FIJJAFBF_352268 | Bacteria | Proteobacteri | Gammaprotec | Cellvibrionales | Porticoccacea | unclassified | Gammaprotec | metabolism | Amino acid tra | COG1448 | aminotransfer | 314287.GB22 | 1,4 | 0,8 | 4,62E-05 | 2,61E-05 | 2 - | 5,78E-05 - | 1 - | 3,59E-05 - | 2 - | 6,83E-05 - | 2 - | 6,93E-05 - | 0 - | 0 - |
| 3662 | 1 | 0.2um_NOKCPHJN_538171 | Bacteria | Proteobacteri | Alphaproteob | Rhodobactera | Rhodobactera | Tateyamaria | Tateyamaria s | unclassified | unclassified | unclassified | unclassified | unclassified | 0,8 | 0,4 | 3,50E-05 | 1,80E-05 | 1 - | 3,70E-05 - | 0 - | 0 - | 1 - | 4,37E-05 - | 1 - | 4,43E-05 - | 1 - | 5,00E-05 - |
| 3663 | 1 | 3um_FIJJAFBF_308962 | Eukaryota | Chlorophyta | Chlorophycea | Sphaeropleale | Selenastracea | Monoraphidiu | Monoraphidiu | cellular proces | Posttranslatio | COG0652 | peptidyl-proly | 35128.Thaps2 | 1 | 0 | 5,21E-05 | 4,95E-06 | 1 - | 4,36E-05 - | 1 - | 5,42E-05 - | 1 - | 5,15E-05 - | 1 - | 5,23E-05 - | 1 - | 5,89E-05 - |
| 3664 | 1 | 3um_FIJJAFBF_444690 | Eukaryota | unclassified | unclassified | Phaeocystales | Phaeocystace | Phaeocystis | Phaeocystis ar | metabolism | Energy produc | COG4451 | RuBisCO catal | 977880.RALTA | 1,4 | 0,489897949 | 0,000135602 | 4,26E-05 | 2 - | 0,000165146 - | 2 - | 0,000205063 - | 1 - | 9,75E-05 - | 1 - | 9,89E-05 - | 1 - | 0,00011142 - |
| 3665 | 1 | 3um_FIJJAFBF_526650 | Bacteria | Proteobacteri | Alphaproteob | Pelagibacteral | Pelagibacterac | Candidatus Pe | Candidatus Pe | unclassified | unclassified | unclassified | unclassified | unclassified | 0,2 | 0,4 | 4,45E-06 | 8,90E-06 | 1 - | 2,22E-05 - | 0 - | 0 - | 0 - | 0 - | 0 - | 0 - | 0 - | 0 - |
| 3666 | 1 | 3um_FIJJAFBF_380454 | Bacteria | Proteobacteri | Gammaprotec | Cellvibrionales | Porticoccacea | unclassified | gamma prote | metabolism | Energy produc | 16RH6 | Produces ATP | 314287.GB22 | 1 | 0 | 4,79E-05 | 4,55E-06 | 1 - | 4,01E-05 - | 1 - | 4,98E-05 - | 1 - | 4,74E-05 - | 1 - | 4,81E-05 - | 1 - | 5,42E-05 - |
| 3667 | 1 | 3um_FIJJAFBF_490872 | Bacteria | Bacteroidetes | Flavobacteriia | Flavobacterial | Flavobacteriac | Formosa | Formosa sp. H | unclassified | unclassified | unclassified | unclassified | unclassified | 1,6 | 0,489897949 | 0,000168737 | 4,77E-05 | 2 - | 0,000179339 - | 2 - | 0,000222686 - | 1 - | 0,000105845 - | 2 - | 0,000214818 - | 1 - | 0,000120996 - |
| 3668 | 1 | 0.2um_NOKCPHJN_745300 | Bacteria | Proteobacteri | Alphaproteob | Pelagibacteral | Pelagibacterac | Candidatus Pe | Candidatus Pe | metabolism | Inorganic ion t | 05PWN | phosphate ab | 335992.SAR11 | 1,4 | 1,019803903 | 6,49E-05 | 4,57E-05 | 0 - | 0 - | 2 - | 9,41E-05 - | 3 - | 0,000134141 - | 1 - | 4,54E-05 - | 1 - | 5,11E-05 - |
| 3669 | 1 | 0.2um_NOKCPHJN_07641 | Bacteria | Bacteroidetes | Flavobacteriia | Flavobacterial | Flavobacteriac | Polaribacter | Polaribacter s | unclassified | unclassified | unclassified | unclassified | unclassified | 0,8 | 0,748331477 | 2,88E-05 | 2,55E-05 | 2 - | 6,58E-05 - | 0 - | 0 - | 1 - | 3,88E-05 - | 1 - | 3,94E-05 - | 0 - | 0 - |
| 3670 | 1 | 3um_FIJJAFBF_684478 | Bacteria | Proteobacteri | Gammaprotec | Oceanospirilla | Oceanospirilla | Motiliproteus | Motiliproteus | metabolism | Lipid transpor | 0QN5A | Involved in un | 491952.Mar18 | 0,6 | 0,489897949 | 5,34E-05 | 4,41E-05 | 1 - | 7,81E-05 - | 1 - | 9,70E-05 - | 1 - | 9,22E-05 - | 0 - | 0 - | 0 - | 0 - |
| 3671 | 1 | 3um_FIJJAFBF_175174 | Bacteria | Proteobacteri | Alphaproteob | Rhodobactera | unclassified | unclassified | Rhodobactera | metabolism | Energy produc | COG0711 | F(1)F(0) ATP s | 391619.RGBS | 3,2 | 0,748331477 | 0,000235855 | 6,70E-05 | 3 - | 0,000184134 - | 2 - | 0,000152427 - | 3 - | 0,000217351 - | 4 - | 0,000294082 - | 4 - | 0,000331282 - |
| 3672 | 1 | 0.2um_NOKCPHJN_384271 | Bacteria | Proteobacteri | Gammaprotec | unclassified | unclassified | unclassified | OM182 bacte | unclassified | unclassified | unclassified | unclassified | unclassified | 3,8 | 1,166190379 | 5,27E-05 | 1,64E-05 | 3 - | 3,51E-05 - | 4 - | 5,81E-05 - | 5 - | 6,90E-05 - | 5 - | 7,00E-05 - | 2 - | 3,15E-05 - |
| 3673 | 1 | 0.2um_NOKCPHJN_412759 | Bacteria | Verrucomicrol | Opitutae | unclassified | unclassified | unclassified | Opitutae bact | unclassified | unclassified | unclassified | unclassified | unclassified | 0,8 | 0,4 | 2,27E-05 | 1,17E-05 | 1 - | 2,38E-05 - | 1 - | 2,95E-05 - | 1 - | 2,81E-05 - | 0 - | 0 - | 1 - | 3,21E-05 - |
| 3674 | 1 | 0.2um_NOKCPHJN_860838 | Bacteria | Proteobacteri | Alphaproteob | Pelagibacteral | Pelagibacterac | Candidatus Pe | Candidatus Pe | information st | Translation, ril | 07ZGG | Associates wit | 335992.SAR11 | 1 | 0 | 0,00013434 | 1,28E-05 | 1 - | 0,000112526 - | 1 - | 0,000139724 - | 1 - | 0,000132826 - | 1 - | 0,000134788 - | 1 - | 0,000151838 - |
| 3675 | 2 | 0.2um_NOKCPHJN_610964;3um_FIJJAFBF_3658 | Bacteria | Bacteroidetes | Flavobacteriia | Flavobacterial | unclassified | unclassified | Flavobacterial | unclassified | unclassified | unclassified | unclassified | unclassified | 0,2 | 0,4 | 1,65E-05 | 3,29E-05 | 0 - | 0 - | 0 - | 0 - | 0 - | 0 - | 1 - | 8,23E-05 - | 0 - | 0 - |
| 3676 | 1 | 0.2um_NOKCPHJN_403575 | Bacteria | Bacteroidetes | unclassified | unclassified | unclassified | unclassified | Bacteroidetes | information st | Translation, ril | COG0050 | This protein p | 487796.Flav2 | 1,2 | 0,979795897 | 4,00E-05 | 3,30E-05 | 2 - | 5,81E-05 - | 2 - | 7,22E-05 - | 0 - | 0 - | 2 - | 6,96E-05 - | 0 - | 0 - |
| 3677 | 1 | 3um_FIJJAFBF_01139 | Bacteria | Bacteroidetes | Flavobacteriia | Flavobacterial | Flavobacteriac | Polaribacter | Polaribacter s | information st | Translation, ril | 05KK9 | This protein b | 313594.PI23P | 10,8 | 1,833030278 | 0,000876745 | 0,000108041 | 14 - | 0,0009622 - | 9 - | 0,000768066 - | 11 - | 0,000892398 - | 9 - | 0,000740928 - | 11 - | 0,001020131 - |
| 3678 | 1 | 0.2um_NOKCPHJN_1101372 | unclassified | unclassified | unclassified | unclassified | unclassified | unclassified | unclassified | unclassified | unclassified | unclassified | unclassified | unclassified | 0,6 | 0,489897949 | 2,09E-05 | 1,72E-05 | 1 - | 3,09E-05 - | 0 - | 0 - | 1 - | 3,65E-05 - | 1 - | 3,71E-05 - | 0 - | 0 - |
| 3679 | 1 | 3um_FIJJAFBF_662493 | Bacteria | Bacteroidetes | Flavobacteriia | Flavobacterial | Flavobacteriac | Polaribacter | Polaribacter s | cellular proces | Cell wall/mem | 0ZZRH | outer membra | 313594.PI23P | 0,4 | 0,489897949 | 1,76E-05 | 2,17E-05 | 1 - | 4,00E-05 - | 0 - | 0 - | 0 - | 0 - | 1 - | 4,79E-05 - | 0 - | 0 - |
| 3680 | 1 | 3um_FIJJAFBF_782165 | Bacteria | Bacteroidetes | unclassified | unclassified | unclassified | unclassified | Bacteroidetes | cellular proces | Cell wall/mem | 09B44 | efflux transpo | 391587.KAOT | 0,4 | 0,489897949 | 1,53E-05 | 1,88E-05 | 0 - | 0 - | 1 - | 3,66E-05 - | 0 - | 0 - | 0 - | 0 - | 1 - | 3,98E-05 - |
| 3681 | 1 | 3um_FIJJAFBF_188725 | Bacteria | Proteobacteri | Gammaprotec | Cellvibrionales | Porticoccacea | unclassified | Gammaprotec | information st | Translation, ril | COG0088 | One of the pri | 314287.GB22 | 7,4 | 0,489897949 | 0,000498854 | 7,17E-05 | 7 - | 0,000393842 - | 7 - | 0,000489035 - | 7 - | 0,00046489 - | 8 - | 0,00053915 - | 8 - | 0,000607351 - |
| 3682 | 1 | 0.2um_NOKCPHJN_08418 | Bacteria | Bacteroidetes | Flavobacteriia | Flavobacterial | Flavobacteriac | Polaribacter | Polaribacter s | unclassified | unclassified | unclassified | unclassified | unclassified | 0,2 | 0,4 | 1,01E-05 | 2,02E-05 | 0 - | 0 - | 1 - | 5,05E-05 - | 0 - | 0 - | 0 - | 0 - | 0 - | 0 - |
| 3683 | 1 | 0.2um_NOKCPHJN_367244 | Bacteria | Proteobacteri | Alphaproteob | Pelagibacteral | Pelagibacterac | Candidatus Pe | Candidatus Pe | metabolism | Secondary me | 01Q86 | extracellular s | 1002672.SAR1 | 1 | 0 | 3,43E-05 | 3,26E-06 | 1 - | 2,88E-05 - | 1 - | 3,57E-05 - | 1 - | 3,40E-05 - | 1 - | 3,45E-05 - | 1 - | 3,88E-05 - |
| 3684 | 1 | 0.2um_NOKCPHJN_350081 | Bacteria | Proteobacteri | Alphaproteob | Rhodospirillal | Rhodospirillac | Tistrella | Tistrella mobil | unclassified | unclassified | unclassified | unclassified | unclassified | 0,6 | 0,489897949 | 3,88E-05 | 3,19E-05 | 1 - | 5,74E-05 - | 0 - | 0 - | 1 - | 6,77E-05 - | 1 - | 6,87E-05 - | 0 - | 0 - |
| 3685 | 1 | 0.2um_NOKCPHJN_262617 | Bacteria | unclassified | unclassified | unclassified | unclassified | unclassified | uncultured ba | metabolism | Inorganic ion t | 05C27 | TonB-depende | 313596.RB250 | 5,4 | 0,8 | 7,00E-05 | 4,73E-06 | 7 - | 7,69E-05 - | 5 - | 6,82E-05 - | 5 - | 6,48E-05 - | 5 - | 6,58E-05 - | 5 - | 7,41E-05 - |
| 3686 | 1 | 0.2um_NOKCPHJN_635290 | Eukaryota | unclassified | Pelagophycea | Pelagomonad | unclassified | Aureococcus | Aureococcus a | unclassified | unclassified | unclassified | unclassified | unclassified | 2,4 | 0,489897949 | 8,56E-05 | 2,28E-05 | 2 - | 5,92E-05 - | 2 - | 7,35E-05 - | 3 - | 0,000104754 - | 2 - | 7,09E-05 - | 3 - | 0,000119748 - |
| 3687 | 1 | 3um_FIJJAFBF_631539 | Bacteria | Bacteroidetes | Flavobacteriia | Flavobacterial | Flavobacteriac | Ulvibacter | Ulvibacter sp. | unclassified | unclassified | unclassified | unclassified | unclassified | 1 | 0 | 0,000150579 | 1,43E-05 | 1 - | 0,000126128 - | 1 - | 0,000156614 - | 1 - | 0,000148882 - | 1 - | 0,000151081 - | 1 - | 0,000170192 - |
| 3688 | 2 | 0.2um_NOKCPHJN_559208;3um_FIJJAFBF_47567 | Bacteria | Bacteroidetes | unclassified | unclassified | unclassified | unclassified | Bacteroidetes | unclassified | unclassified | unclassified | unclassified | unclassified | 1 | 0,632455532 | 7,32E-05 | 4,10E-05 | 2 - | 0,000126825 - | 1 - | 7,87E-05 - | 1 - | 7,49E-05 - | 0 - | 0 - | 1 - | 8,56E-05 - |
| 3689 | 1 | 3um_FIJJAFBF_109908 | Bacteria | Proteobacteri | Gammaprotec | Alteromonada | Alteromonada | Paraglaciecola | Paraglaciecola | cellular proces | Posttranslatio | 0QHT2 | Peroxiredoxin | 342610.Patl_3 | 4,8 | 0,748331477 | 0,00026877 | 5,42E-05 | 4 - | 0,000186629 - | 4 - | 0,000231738 - | 5 - | 0,00027537 - | 6 - | 0,000335325 - | 5 - | 0,000314786 - |
| 3690 | 1 | 0.2um_NOKCPHJN_243769 | Bacteria | Proteobacteri | Gammaprotec | unclassified | unclassified | unclassified | Gammaprotec | cellular proces | Cell wall/mem | 0QTSX | Peptidoglycan | 207954.MED9 | 2,2 | 0,979795897 | 0,00016748 | 8,06E-05 | 1 - | 6,24E-05 - | 1 - | 7,75E-05 - | 3 - | 0,000220895 - | 3 - | 0,000224158 - | 3 - | 0,000252513 - |
| 3691 | 1 | 3um_FIJJAFBF_715085 | Eukaryota | unclassified | Pelagophycea | Pelagomonad | unclassified | Aureoumbra | Aureoumbra l | information st | Translation, ril | 0FKC1 | This is 1 of the | 449447.MAE_ | 1,8 | 0,979795897 | 0,000136393 | 7,55E-05 | 2 - | 0,000128242 - | 0 - | 0 - | 3 - | 0,000227065 - | 2 - | 0,000153613 - | 2 - | 0,000173044 - |
| 3692 | 1 | 0.2um_NOKCPHJN_337845 | Bacteria | Proteobacteri | Gammaprotec | unclassified | unclassified | unclassified | Gammaprotec | cellular proces | Cell cycle cont | 0QJVR | Required for c | 247633.GP214 | 0,8 | 0,4 | 9,43E-06 | 4,84E-06 | 1 - | 9,84E-06 - | 1 - | 1,22E-05 - | 0 - | 0 - | 1 - | 1,18E-05 - | 1 - | 1,33E-05 - |
| 3693 | 1 | 3um_FIJJAFBF_100817 | Bacteria | Bacteroidetes | Flavobacteriia | Flavobacterial | Flavobacteriac | Polaribacter | Polaribacter s | poorly charact | Function unkn | 0PAIH | NA | 313598.MED1 | 2,8 | 0,4 | 0,000175164 | 2,08E-05 | 3 - | 0,000158678 - | 3 - | 0,000197031 - | 3 - | 0,000187303 - | 3 - | 0,000190069 - | 2 - | 0,000142741 - |
| 3694 | 2 | 0.2um_NOKCPHJN_423545;3um_FIJJAFBF_6638 | Bacteria | Proteobacteri | Gammaprotec | Cellvibrionales | Porticoccacea | unclassified | SAR92 bacteri | cellular proces | Posttranslatio | 0QIPK | Stringent star | 314287.GB22 | 0,4 | 0,489897949 | 2,84E-05 | 3,49E-05 | 0 - | 0 - | 0 - | 0 - | 0 - | 0 - | 1 - | 6,67E-05 - | 1 - | 7,52E-05 - |
| 3695 | 1 | 0.2um_NOKCPHJN_508767 | Eukaryota | unclassified | Dinophyceae | Suessiales | Symbiodiniace | Symbiodinium | Symbiodinium | unclassified | unclassified | unclassified | unclassified | unclassified | 0,2 | 0,4 | 2,23E-06 | 4,45E-06 | 0 - | 0 - | 0 - | 0 - | 0 - | 0 - | 0 - | 0 - | 1 - | 1,11E-05 - |
| 3696 | 1 | 3um_FIJJAFBF_411761 | Bacteria | Bacteroidetes | Saprospiria | Saprospirales | Saprospiracea | Aureispira | Aureispira sp. | poorly charact | Function unkn | 09S0J | NA | 760192.Halhy | 0,4 | 0,489897949 | 7,64E-06 | 9,35E-06 | 0 - | 0 - | 0 - | 0 - | 1 - | 1,89E-05 - | 1 - | 1,92E-05 - | 0 - | 0 - |
| 3697 | 1 | 0.2um_NOKCPHJN_207712 | Bacteria | Proteobacteri | Alphaproteob | Rhodospirillal | Rhodospirillac | unclassified | Rhodobactera | unclassified | unclassified | unclassified | unclassified | unclassified | 1 | 0,894427191 | 7,51E-05 | 6,63E-05 | 2 - | 0,000135031 - | 0 - | 0 - | 2 - | 0,000159391 - | 1 - | 8,09E-05 - | 0 - | 0 - |
| 3698 | 1 | 3um_FIJJAFBF_180276 | Bacteria | Bacteroidetes | Flavobacteriia | Flavobacterial | Flavobacteriac | Lacinutrix | Lacinutrix him | unclassified | unclassified | unclassified | unclassified | unclassified | 1 | 0 | 4,13E-05 | 3,92E-06 | 1 - | 3,46E-05 - | 1 - | 4,29E-05 - | 1 - | 4,08E-05 - | 1 - | 4,14E-05 - | 1 - | 4,66E-05 - |
| 3699 | 1 | 3um_FIJJAFBF_44830 | Bacteria | Bacteroidetes | Flavobacteriia | Flavobacterial | Flavobacteriac | Algibacter | Algibacter sp. | metabolism | Carbohydrate | 05CA9 | Pyruvate kinas | 983544.Lacal_ | 1,4 | 0,489897949 | 3,99E-05 | 1,41E-05 | 1 - | 2,39E-05 - | 1 - | 2,97E-05 - | 2 - | 5,65E-05 - | 2 - | 5,73E-05 - | 1 - | 3,23E-05 - |
| 3700 | 1 | 3um_FIJJAFBF_155408 | Bacteria | Proteobacteri | Alphaproteob | Rhodobactera | Rhodobactera | unclassified | Rhodobactera | information st | Translation, ril | 177XJ | One of the pri | 252305.OB25 | 0,6 | 0,489897949 | 5,04E-05 | 4,15E-05 | 1 - | 7,36E-05 - | 1 - | 9,14E-05 - | 1 - | 8,68E-05 - | 0 - | 0 - | 0 - | 0 - |
| 3701 | 1 | 3um_FIJJAFBF_770097 | Bacteria | Proteobacteri | Alphaproteob | Rhodobactera | Rhodobactera | unclassified | Rhodobactera | metabolism | Carbohydrate | 16THB | transporter, d | 375451.RD1_ | 1,8 | 0,748331477 | 7,34E-05 | 3,18E-05 | 1 - | 3,38E-05 - | 1 - | 4,19E-05 - | 3 - | 0,000119543 - | 2 - | 8,09E-05 - | 2 - | 9,11E-05 - |
| 3702 | 1 | 3um_FIJJAFBF_305946 | Bacteria | Proteobacteri | Alphaproteob | Rhodobactera | Rhodobactera | unclassified | Rhodobactera | metabolism | Carbohydrate | 16THB | transporter, d | 744979.R2A13 | 2,6 | 0,8 | 0,000104659 | 3,44E-05 | 3 - | 0,000100977 - | 3 - | 0,000125383 - | 1 - | 3,97E-05 - | 3 - | 0,000120953 - | 3 - | 0,000136253 - |
| 3703 | 1 | 0.2um_NOKCPHJN_78717 | Bacteria | Proteobacteri | Gammaprotec | Cellvibrionales | Porticoccacea | unclassified | gamma prote | unclassified | unclassified | unclassified | unclassified | unclassified | 0,8 | 1,166190379 | 1,19E-05 | 1,72E-05 | 0 - | 0 - | 0 - | 0 - | 3 - | 4,43E-05 - | 1 - | 1,50E-05 - | 0 - | 0 - |
| 3704 | 1 | 3um_FIJJAFBF_435520 | Bacteria | Proteobacteri | Gammaprotec | Alteromonada | Pseudoalteror | Pseudoalteror | Pseudoalteror | cellular proces | Cell motility | 0QTZY | Msha pilin pro | 87626.PTD2_ | 0,8 | 0,4 | 5,00E-05 | 2,54E-05 | 1 - | 5,41E-05 - | 1 - | 6,72E-05 - | 1 - | 6,39E-05 - | 1 - | 6,49E-05 - | 0 - | 0 - |
| 3705 | 1 | 3um_FIJJAFBF_805383 | Bacteria | Proteobacteri | Gammaprotec | Oceanospirilla | Saccharospirill | Reinekea | Reinekea fors | unclassified | unclassified | unclassified | unclassified | unclassified | 4 | 0,632455532 | 0,000290309 | 6,57E-05 | 4 - | 0,000241635 - | 4 - | 0,00030004 - | 4 - | 0,000285226 - | 3 - | 0,000217079 - | 5 - | 0,000407564 - |
| 3706 | 1 | 0.2um_NOKCPHJN_860971 | Bacteria | Verrucomicrol | Opitutae | Puniceicoccale | Puniceicoccac | Coraliomargar | Coraliomargar | unclassified | unclassified | unclassified | unclassified | unclassified | 0,6 | 0,489897949 | 2,67E-05 | 2,22E-05 | 1 - | 3,76E-05 - | 0 - | 0 - | 0 - | 0 - | 1 - | 4,51E-05 - | 1 - | 5,08E-05 - |
| 3707 | 1 | 0.2um_NOKCPHJN_597266 | Bacteria | Actinobacteria | Actinobacteria | Micrococcales | Microbacteria | Rathayibacter | Rathayibacter | unclassified | unclassified | unclassified | unclassified | unclassified | 0,6 | 0,489897949 | 6,40E-06 | 5,23E-06 | 0 - | 0 - | 1 - | 1,10E-05 - | 1 - | 1,04E-05 - | 1 - | 1,06E-05 - | 0 - | 0 - |
| 3708 | 1 | 3um_FIJJAFBF_189685 | Bacteria | Bacteroidetes | Flavobacteriia | Flavobacterial | Flavobacteriac | Polaribacter | Polaribacter g | poorly charact | Function unkn | COG0596 | Alpha beta hy | 313594.PI23P | 0,2 | 0,4 | 9,04E-06 | 1,81E-05 | 1 - | 4,52E-05 - | 0 - | 0 - | 0 - | 0 - | 0 - | 0 - | 0 - | 0 - |
| 3709 | 1 | 0.2um_NOKCPHJN_1045255 | Bacteria | Proteobacteri | Alphaproteob | Rhodobactera | Hyphomonad | Hyphomonas | Hyphomonas | poorly charact | Function unkn | 11H20 | NA | 483219.LILAB | 0,2 | 0,4 | 9,50E-06 | 1,90E-05 | 0 - | 0 - | 1 - | 4,75E-05 - | 0 - | 0 - | 0 - | 0 - | 0 - | 0 - |
| 3710 | 1 | 3um_FIJJAFBF_306827 | Bacteria | Proteobacteri | Alphaproteob | Rhodobactera | unclassified | unclassified | Rhodobactera | cellular proces | Defense mech | 16Q8Z | efflux transpo | 467661.RKLH1 | 0,2 | 0,4 | 7,02E-06 | 1,40E-05 | 0 - | 0 - | 0 - | 0 - | 1 - | 3,51E-05 - | 0 - | 0 - | 0 - | 0 - |
| 3711 | 2 | 0.2um_NOKCPHJN_246326;0.2um_NOKCPHJN_9 | Bacteria | Bacteroidetes | various | various | various | various | various | unclassified | unclassified | unclassified | unclassified | unclassified | 1,8 | 0,4 | 0,00014275 | 3,55E-05 | 2 - | 0,00013269 - | 2 - | 0,000164762 - | 1 - | 7,83E-05 - | 2 - | 0,00015894 - | 2 - | 0,000179046 - |
| 3712 | 1 | 3um_FIJJAFBF_30304 | Bacteria | Bacteroidetes | Flavobacteriia | Flavobacterial | Flavobacteriac | Polaribacter | Polaribacter s | unclassified | unclassified | unclassified | unclassified | unclassified | 1,4 | 0,489897949 | 5,42E-05 | 1,70E-05 | 2 - | 6,60E-05 - | 2 - | 8,19E-05 - | 1 - | 3,89E-05 - | 1 - | 3,95E-05 - | 1 - | 4,45E-05 - |

| | | | | | | | | | | | | | | | | | | | | | | | | | | | | |
|---|---|---|---|---|---|---|---|---|---|---|---|---|---|---|---|---|---|---|---|---|---|---|---|---|---|---|---|---|
| 3713 | 1 | 3um_FIJJAFBF_00854 | Bacteria | Bacteroidetes | Flavobacteriia | unclassified | unclassified | unclassified | Flavobacteria | information st | Translation, ril | COG0052 | 30S ribosomal | 398720.MED2 | 5 | 1,264911064 | 0,000214644 | 4,16E-05 | 7 - | 0,000255872 - | 6 - | 0,000272329 - | 4 - | 0,000172589 - | 4 - | 0,000175138 - | 4 - | 0,000197292 - |
| 3714 | 1 | 3um_FIJJAFBF_189650 | Bacteria | Bacteroidetes | Flavobacteriia | Flavobacterial | Flavobacteriac | Polaribacter | Polaribacter a | poorly charact | Function unkn | 08V90 | NA | 860228.Ccan_ | 0,6 | 0,489897949 | 4,91E-05 | 4,05E-05 | 1 - | 7,17E-05 - | 1 - | 8,91E-05 - | 1 - | 8,47E-05 - | 0 - | 0 - | 0 - | 0 - |
| 3715 | 2 | 0.2um_NOKCPHJN_644516;3um_FIJJAFBF_6156 | Bacteria | Proteobacteri | Alphaproteob | Rhodobactera | Rhodobactera | Planktomarina | Planktomarina | poorly charact | Function unkn | COG5501 | Sulfur oxidatic | 388401.RB215 | 0,8 | 0,4 | 7,58E-05 | 3,85E-05 | 1 - | 8,20E-05 - | 1 - | 0,000101799 - | 1 - | 9,68E-05 - | 1 - | 9,82E-05 - | 0 - | 0 - |
| 3716 | 1 | 3um_FIJJAFBF_129483 | Bacteria | Proteobacteri | Gammaprotec | unclassified | unclassified | unclassified | OM182 bacte | unclassified | unclassified | unclassified | unclassified | unclassified | 0,2 | 0,4 | 3,66E-06 | 7,32E-06 | 1 - | 1,83E-05 - | 0 - | 0 - | 0 - | 0 - | 0 - | 0 - | 0 - | 0 - |
| 3717 | 1 | 0.2um_NOKCPHJN_360238 | Bacteria | Proteobacteri | Alphaproteob | Rhodobactera | Rhodobactera | Planktomarina | Planktomarina | unclassified | unclassified | unclassified | unclassified | unclassified | 4,4 | 0,489897949 | 0,000396133 | 3,03E-05 | 5 - | 0,000380055 - | 4 - | 0,000377533 - | 5 - | 0,000448617 - | 4 - | 0,000364194 - | 4 - | 0,000410264 - |
| 3718 | 1 | 3um_FIJJAFBF_208856 | Eukaryota | Eustigmatoph | unclassified | Eustigmatales | Monodopsida | Nannochlorop | Nannochlorop | unclassified | unclassified | unclassified | unclassified | unclassified | 1,2 | 0,748331477 | 6,97E-05 | 4,41E-05 | 1 - | 4,90E-05 - | 0 - | 0 - | 2 - | 0,000115797 - | 2 - | 0,000117507 - | 1 - | 6,62E-05 - |
| 3719 | 1 | 0.2um_NOKCPHJN_62623 | Bacteria | Proteobacteri | Alphaproteob | Rhodospirillal | unclassified | unclassified | Rhodospirillal | information st | Translation, ril | COG0051 | tRNA binding | 394221.Mmar | 0,2 | 0,4 | 2,25E-05 | 4,50E-05 | 1 - | 0,000112526 - | 0 - | 0 - | 0 - | 0 - | 0 - | 0 - | 0 - | 0 - |
| 3720 | 1 | 3um_FIJJAFBF_115058 | Eukaryota | unclassified | unclassified | Phaeocystales | Phaeocystace | Phaeocystis | Phaeocystis gl | metabolism | Energy produc | COG0711 | F(1)F(0) ATP s | 63737.Npun_ | 4,6 | 1,019803903 | 0,000356564 | 7,05E-05 | 6 - | 0,00039352 - | 3 - | 0,000244318 - | 4 - | 0,000309674 - | 5 - | 0,000392809 - | 5 - | 0,000442499 - |
| 3721 | 1 | 0.2um_NOKCPHJN_156087 | Bacteria | Bacteroidetes | unclassified | unclassified | unclassified | unclassified | Bacteroidetes | poorly charact | Function unkn | 11NCV | Protein of unk | 487796.Flav2 | 1,6 | 0,8 | 0,000130788 | 6,58E-05 | 1 - | 6,83E-05 - | 2 - | 0,000169665 - | 3 - | 0,000241933 - | 1 - | 8,18E-05 - | 1 - | 9,22E-05 - |
| 3722 | 1 | 0.2um_NOKCPHJN_521950 | Bacteria | Proteobacteri | Gammaprotec | unclassified | unclassified | unclassified | Gammaprotec | metabolism | Inorganic ion t | 05F28 | tonB-depende | 247633.GP214 | 0,8 | 0,748331477 | 2,01E-05 | 1,75E-05 | 2 - | 4,42E-05 - | 0 - | 0 - | 0 - | 0 - | 1 - | 2,65E-05 - | 1 - | 2,98E-05 - |
| 3723 | 1 | 0.2um_NOKCPHJN_124941 | Bacteria | unclassified | unclassified | unclassified | unclassified | unclassified | bacterium TM | unclassified | unclassified | unclassified | unclassified | unclassified | 0,2 | 0,4 | 5,58E-06 | 1,12E-05 | 0 - | 0 - | 0 - | 0 - | 0 - | 0 - | 1 - | 2,79E-05 - | 0 - | 0 - |
| 3724 | 1 | 0.2um_NOKCPHJN_97296 | Bacteria | Proteobacteri | Gammaprotec | Alteromonada | Alteromonada | Paraglaciecola | various | information st | Transcription | 05VEQ | Cold shock pro | 203122.Sde_3 | 2,4 | 0,489897949 | 0,000381377 | 8,52E-05 | 3 - | 0,000400384 - | 2 - | 0,000331439 - | 2 - | 0,000315075 - | 2 - | 0,000319729 - | 3 - | 0,00054026 - |
| 3725 | 1 | 0.2um_NOKCPHJN_1008302 | Bacteria | Proteobacteri | Alphaproteob | Pelagibacteral | Pelagibacterac | Candidatus Pe | Candidatus Pe | cellular proces | Posttranslatio | 16PVM | this subunit ha | 439493.PB721 | 1 | 0,632455532 | 7,69E-05 | 5,18E-05 | 1 - | 6,56E-05 - | 2 - | 0,000162879 - | 1 - | 7,74E-05 - | 1 - | 7,86E-05 - | 0 - | 0 - |
| 3726 | 1 | 3um_FIJJAFBF_550696 | Bacteria | Proteobacteri | Alphaproteob | Rhodobactera | Rhodobactera | Tateyamaria | Tateyamaria s | unclassified | unclassified | unclassified | unclassified | unclassified | 2 | 0,632455532 | 0,00020308 | 4,70E-05 | 3 - | 0,000262848 - | 2 - | 0,000217586 - | 2 - | 0,000206843 - | 2 - | 0,000209898 - | 1 - | 0,000118225 - |
| 3727 | 1 | 0.2um_NOKCPHJN_97695 | Bacteria | Proteobacteri | Betaproteoba | unclassified | unclassified | unclassified | Betaproteoba | information st | Translation, ril | 177XJ | One of the pri | 365046.Rta_0 | 3,2 | 0,4 | 0,000281565 | 5,93E-05 | 3 - | 0,000219319 - | 3 - | 0,000272329 - | 3 - | 0,000258883 - | 3 - | 0,000262707 - | 4 - | 0,000394585 - |
| 3728 | 1 | 0.2um_NOKCPHJN_775128 | Bacteria | Proteobacteri | Gammaprotec | Alteromonada | Colwelliaceae | Colwellia | Colwellia sp. P | poorly charact | Function unkn | 06U2P | ATPase involv | 156578.ATW7 | 0,2 | 0,4 | 3,78E-06 | 7,55E-06 | 1 - | 1,89E-05 - | 0 - | 0 - | 0 - | 0 - | 0 - | 0 - | 0 - | 0 - |
| 3729 | 1 | 0.2um_NOKCPHJN_715171 | Bacteria | Proteobacteri | Alphaproteob | Rhodobactera | Rhodobactera | unclassified | Rhodobactera | metabolism | Inorganic ion t | 16RE3 | may be the G1 | 314266.SKA58 | 0,2 | 0,4 | 6,09E-06 | 1,22E-05 | 0 - | 0 - | 0 - | 0 - | 1 - | 3,04E-05 - | 0 - | 0 - | 0 - | 0 - |
| 3730 | 1 | 3um_FIJJAFBF_52144 | Bacteria | Proteobacteri | Gammaprotec | Pseudomonad | Pseudomonad | Pseudomonas | various | metabolism | Amino acid tra | 16Q4U | Catalyzes the | 555778.Hneap | 1,4 | 0,489897949 | 4,53E-05 | 1,42E-05 | 2 - | 5,52E-05 - | 2 - | 6,85E-05 - | 1 - | 3,26E-05 - | 1 - | 3,30E-05 - | 1 - | 3,72E-05 - |
| 3731 | 1 | 0.2um_NOKCPHJN_388436 | Bacteria | Proteobacteri | Alphaproteob | Pelagibacteral | Pelagibacterac | Candidatus Pe | Candidatus Pe | poorly charact | Function unkn | 084DF | Protein of unk | 335992.SAR11 | 0,2 | 0,4 | 1,08E-05 | 2,16E-05 | 1 - | 5,39E-05 - | 0 - | 0 - | 0 - | 0 - | 0 - | 0 - | 0 - | 0 - |
| 3732 | 1 | 3um_FIJJAFBF_179417 | Bacteria | Bacteroidetes | Flavobacteriia | Flavobacterial | Flavobacteriac | Formosa | Formosa sp. H | poorly charact | Function unkn | 09HPY | NA | 1046627.BZAR | 1,2 | 0,748331477 | 0,000130153 | 7,79E-05 | 2 - | 0,000185124 - | 2 - | 0,000229869 - | 0 - | 0 - | 1 - | 0,000110874 - | 1 - | 0,000124899 - |
| 3733 | 2 | 0.2um_NOKCPHJN_159613;3um_FIJJAFBF_4360 | Eukaryota | Mollusca | Gastropoda | unclassified | various | various | various | poorly charact | Function unkn | 11NK2 | dynein, light c | various | 1,6 | 1,019803903 | 0,000163598 | 9,46E-05 | 3 - | 0,000266923 - | 2 - | 0,000220959 - | 2 - | 0,00021005 - | 0 - | 0 - | 1 - | 0,000120058 - |
| 3734 | 1 | 3um_FIJJAFBF_680258 | Bacteria | Bacteroidetes | Flavobacteriia | Flavobacterial | Flavobacteriac | Polaribacter | various | unclassified | unclassified | unclassified | unclassified | unclassified | 3,2 | 0,4 | 0,00017021 | 9,34E-06 | 4 - | 0,000180042 - | 3 - | 0,000167669 - | 3 - | 0,000159391 - | 3 - | 0,000161745 - | 3 - | 0,000182205 - |
| 3735 | 1 | 3um_FIJJAFBF_565482 | Eukaryota | unclassified | Pelagophycea | Pelagomonad | unclassified | Aureococcus | Aureococcus a | metabolism | Carbohydrate | KOG1203 | Complex I inte | 35128.Thapsd | 1,6 | 1,019803903 | 6,48E-05 | 3,83E-05 | 3 - | 0,000107941 - | 1 - | 4,47E-05 - | 2 - | 8,49E-05 - | 2 - | 8,62E-05 - | 0 - | 0 - |
| 3736 | 1 | 3um_FIJJAFBF_533046 | Bacteria | Proteobacteri | Gammaprotec | Thiotrichales | Thiotrichaceae | Leucothrix | Leucothrix mu | unclassified | unclassified | unclassified | unclassified | unclassified | 8,4 | 0,8 | 0,000158721 | 2,03E-05 | 9 - | 0,000142678 - | 7 - | 0,000137794 - | 8 - | 0,000149704 - | 9 - | 0,000170905 - | 9 - | 0,000192524 - |
| 3737 | 1 | 3um_FIJJAFBF_332461 | Bacteria | Bacteroidetes | Flavobacteriia | Flavobacterial | Flavobacteriac | Polaribacter | Polaribacter s | cellular proces | Posttranslatio | COG0330 | Band 7 protein | 313598.MED1 | 2,2 | 0,748331477 | 0,000108508 | 3,81E-05 | 3 - | 0,000124757 - | 1 - | 5,16E-05 - | 2 - | 9,82E-05 - | 2 - | 9,96E-05 - | 3 - | 0,000168342 - |
| 3738 | 1 | 0.2um_NOKCPHJN_257082 | Bacteria | Bacteroidetes | Flavobacteriia | Flavobacterial | Crocinitomica | unclassified | Crocinitomica | unclassified | unclassified | unclassified | unclassified | unclassified | 2,4 | 0,489897949 | 4,64E-05 | 7,23E-06 | 3 - | 4,93E-05 - | 2 - | 4,08E-05 - | 2 - | 3,88E-05 - | 3 - | 5,90E-05 - | 2 - | 4,43E-05 - |
| 3739 | 1 | 3um_FIJJAFBF_583512 | Bacteria | Bacteroidetes | Flavobacteriia | Flavobacterial | Flavobacteriac | Ulvibacter | Ulvibacter lito | unclassified | unclassified | unclassified | unclassified | unclassified | 2 | 1,095445115 | 9,25E-05 | 4,21E-05 | 4 - | 0,000159968 - | 2 - | 9,93E-05 - | 1 - | 4,72E-05 - | 1 - | 4,79E-05 - | 2 - | 0,000107926 - |
| 3740 | 1 | 3um_FIJJAFBF_195179 | Bacteria | Proteobacteri | Gammaprotec | Cellvibrionales | Porticoccacea | unclassified | gamma prote | cellular proces | Intracellular tr | 0QJ7H | motA TolQ Exl | 314287.GB22 | 1,6 | 0,489897949 | 4,71E-05 | 1,23E-05 | 2 - | 5,03E-05 - | 1 - | 3,13E-05 - | 2 - | 5,94E-05 - | 2 - | 6,03E-05 - | 1 - | 3,40E-05 - |
| 3741 | 1 | 3um_FIJJAFBF_234032 | Bacteria | Bacteroidetes | Flavobacteriia | Flavobacterial | Flavobacteriac | Formosa | Formosa sp. H | poorly charact | Function unkn | 05C2A | tonB-depende | 983544.Lacal_ | 2,4 | 0,8 | 3,09E-05 | 9,50E-06 | 3 - | 3,31E-05 - | 3 - | 4,11E-05 - | 3 - | 3,90E-05 - | 2 - | 2,64E-05 - | 1 - | 1,49E-05 - |
| 3742 | 1 | 0.2um_NOKCPHJN_837545 | Bacteria | Proteobacteri | Alphaproteob | Pelagibacteral | Pelagibacterac | Candidatus Pe | Candidatus Pe | cellular proces | Posttranslatio | COG0330 | Band 7 protein | 335992.SAR11 | 10,6 | 1,624807681 | 0,000392807 | 4,34E-05 | 13 - | 0,000407677 - | 12 - | 0,000467275 - | 10 - | 0,00037017 - | 9 - | 0,000338074 - | 9 - | 0,000380839 - |
| 3743 | 2 | 0.2um_NOKCPHJN_18883;0.2um_NOKCPHJN_53 | Bacteria | Proteobacteri | Gammaprotec | Cellvibrionales | Porticoccacea | unclassified | gamma prote | cellular proces | Posttranslatio | COG1225 | alkyl hydroper | 314287.GB22 | 0,2 | 0,4 | 1,71E-05 | 3,43E-05 | 0 - | 0 - | 0 - | 0 - | 1 - | 8,57E-05 - | 0 - | 0 - | 0 - | 0 - |
| 3744 | 1 | 3um_FIJJAFBF_63073 | Bacteria | Proteobacteri | Gammaprotec | Oceanospirilla | Oceanospirilla | Amphritea | Amphritea jap | information st | Translation, ril | COG0052 | 30S ribosomal | 876044.IMCC | 2,4 | 0,489897949 | 0,000132519 | 2,86E-05 | 2 - | 9,26E-05 - | 2 - | 0,000114935 - | 3 - | 0,00016389 - | 3 - | 0,00016631 - | 2 - | 0,000124899 - |
| 3745 | 1 | 0.2um_NOKCPHJN_790970 | Eukaryota | Ascomycota | Sordariomyce | Hypocreales | Nectriaceae | Fusarium | Fusarium gran | unclassified | unclassified | unclassified | unclassified | unclassified | 0,2 | 0,4 | 4,26E-06 | 8,51E-06 | 0 - | 0 - | 0 - | 0 - | 0 - | 0 - | 1 - | 2,13E-05 - | 0 - | 0 - |
| 3746 | 2 | 0.2um_NOKCPHJN_313863;3um_FIJJAFBF_5063 | Bacteria | Bacteroidetes | Flavobacteriia | Flavobacterial | Flavobacteriac | Formosa | Formosa sp. H | unclassified | unclassified | unclassified | unclassified | unclassified | 1,6 | 1,356465997 | 5,01E-05 | 4,39E-05 | 2 - | 5,28E-05 - | 4 - | 0,000131052 - | 1 - | 3,11E-05 - | 0 - | 0 - | 1 - | 3,56E-05 - |
| 3747 | 1 | 3um_FIJJAFBF_458234 | Bacteria | Proteobacteri | Alphaproteob | Rhodobactera | Rhodobactera | Marinosulfonc | Marinosulfonc | information st | Translation, ril | COG0051 | tRNA binding | 501479.CSE45 | 1 | 0 | 0,00013434 | 1,28E-05 | 1 - | 0,000112526 - | 1 - | 0,000139724 - | 1 - | 0,000132826 - | 1 - | 0,000134788 - | 1 - | 0,000151838 - |
| 3748 | 1 | 0.2um_NOKCPHJN_164120 | Bacteria | Bacteroidetes | Flavobacteriia | Flavobacterial | Flavobacteriac | Polaribacter | Polaribacter a | metabolism | Energy produc | COG0056 | Produces ATP | 391603.FBALC | 1,2 | 1,166190379 | 2,90E-05 | 2,67E-05 | 3 - | 6,56E-05 - | 1 - | 2,71E-05 - | 0 - | 0 - | 2 - | 5,24E-05 - | 0 - | 0 - |
| 3749 | 1 | 0.2um_NOKCPHJN_80615 | Bacteria | Bacteroidetes | Flavobacteriia | Flavobacterial | Flavobacteriac | Polaribacter | Polaribacter s | information st | Translation, ril | COG0256 | This is one of | 313594.PI23P | 2,6 | 0,489897949 | 0,000307317 | 6,77E-05 | 3 - | 0,000296836 - | 3 - | 0,000368583 - | 2 - | 0,00023359 - | 2 - | 0,00023704 - | 3 - | 0,000400537 - |
| 3750 | 1 | 0.2um_NOKCPHJN_72807 | Bacteria | Proteobacteri | Gammaprotec | Oceanospirilla | Saccharospirill | Reinekea | Reinekea fors | metabolism | Lipid transpor | 0QN5A | Involved in un | 314283.MED2 | 2,2 | 0,4 | 0,000188219 | 3,67E-05 | 2 - | 0,000143471 - | 2 - | 0,000178149 - | 3 - | 0,000254029 - | 2 - | 0,000171854 - | 2 - | 0,000193593 - |
| 3751 | 1 | 3um_FIJJAFBF_434533 | Bacteria | Bacteroidetes | Flavobacteriia | Flavobacterial | Flavobacteriac | Cellulophaga | Cellulophaga | information st | Translation, ril | 05C64 | Binds directly | 886377.Murru | 2,8 | 0,4 | 0,000168414 | 3,01E-05 | 3 - | 0,000151022 - | 3 - | 0,000187525 - | 2 - | 0,000118844 - | 3 - | 0,000180899 - | 3 - | 0,000203782 - |
| 3752 | 1 | 0.2um_NOKCPHJN_369084 | Bacteria | Proteobacteri | Betaproteoba | unclassified | unclassified | unclassified | Betaproteoba | poorly charact | Function unkn | COG3034 | ErfK ybiS ycfS | 338969.Rfer_ | 0,8 | 0,4 | 2,23E-05 | 1,13E-05 | 1 - | 2,42E-05 - | 1 - | 3,00E-05 - | 1 - | 2,85E-05 - | 1 - | 2,89E-05 - | 0 - | 0 - |
| 3753 | 1 | 3um_FIJJAFBF_531870 | Bacteria | Proteobacteri | Gammaprotec | unclassified | unclassified | unclassified | Gammaprotec | unclassified | unclassified | unclassified | unclassified | unclassified | 0,6 | 0,489897949 | 2,05E-05 | 1,69E-05 | 1 - | 3,00E-05 - | 1 - | 3,72E-05 - | 1 - | 3,54E-05 - | 0 - | 0 - | 0 - | 0 - |
| 3754 | 1 | 0.2um_NOKCPHJN_316214 | Bacteria | Bacteroidetes | Flavobacteriia | Flavobacterial | Flavobacteriac | Maribacter | Maribacter sp | metabolism | Energy produc | 0P7SC | Delta-1-pyrrol | 867900.Celly_ | 1 | 0 | 2,53E-05 | 2,40E-06 | 1 - | 2,12E-05 - | 1 - | 2,63E-05 - | 1 - | 2,50E-05 - | 1 - | 2,54E-05 - | 1 - | 2,86E-05 - |
| 3755 | 1 | 0.2um_NOKCPHJN_342379 | Bacteria | Bacteroidetes | Flavobacteriia | Flavobacterial | Flavobacteriac | Polaribacter | Polaribacter s | poorly charact | Function unkn | 08KQQ | NA | 983544.Lacal_ | 0,4 | 0,489897949 | 1,87E-05 | 2,31E-05 | 1 - | 4,30E-05 - | 0 - | 0 - | 1 - | 5,07E-05 - | 0 - | 0 - | 0 - | 0 - |
| 3756 | 1 | 0.2um_NOKCPHJN_91496 | Bacteria | Proteobacteri | Gammaprotec | Oceanospirilla | Saccharospirill | Reinekea | Reinekea fors | unclassified | unclassified | unclassified | unclassified | unclassified | 10,6 | 1,743559577 | 0,000848636 | 9,45E-05 | 13 - | 0,000882898 - | 12 - | 0,001011968 - | 10 - | 0,00080167 - | 10 - | 0,000813511 - | 8 - | 0,000733134 - |
| 3757 | 1 | 3um_FIJJAFBF_475533 | Eukaryota | unclassified | Oomycetes | Albuginales | Albuginaceae | Albugo | Albugo laibach | information st | Translation, ril | COG0081 | Binds directly | 65071.PYU1_ | 0,2 | 0,4 | 2,22E-05 | 4,43E-05 | 0 - | 0 - | 0 - | 0 - | 0 - | 0 - | 1 - | 0,000110874 - | 0 - | 0 - |
| 3758 | 1 | 3um_FIJJAFBF_639894 | Bacteria | Proteobacteri | Gammaprotec | Oceanospirilla | Saccharospirill | Reinekea | Reinekea fors | poorly charact | Function unkn | 08VI5 | type VI secreti | 83406.HDN1F | 2,2 | 0,4 | 0,000188219 | 3,67E-05 | 2 - | 0,000143471 - | 2 - | 0,000178149 - | 3 - | 0,000254029 - | 2 - | 0,000171854 - | 2 - | 0,000193593 - |
| 3759 | 1 | 0.2um_NOKCPHJN_140291 | Bacteria | Proteobacteri | Alphaproteob | Rhodobactera | Rhodobactera | Planktomarina | Planktomarina | unclassified | unclassified | unclassified | unclassified | unclassified | 0,6 | 0,489897949 | 1,87E-05 | 1,54E-05 | 1 - | 2,73E-05 - | 1 - | 3,39E-05 - | 1 - | 3,22E-05 - | 0 - | 0 - | 0 - | 0 - |
| 3760 | 1 | 0.2um_NOKCPHJN_393866 | Bacteria | Bacteroidetes | Flavobacteriia | Flavobacterial | Flavobacteriac | Lutibacter | Lutibacter pro | poorly charact | Function unkn | 0P88U | NA | 313596.RB250 | 0,4 | 0,8 | 8,05E-06 | 1,61E-05 | 2 - | 4,03E-05 - | 0 - | 0 - | 0 - | 0 - | 0 - | 0 - | 0 - | 0 - |
| 3761 | 1 | 3um_FIJJAFBF_554090 | Bacteria | Verrucomicrol | Opitutae | unclassified | unclassified | unclassified | Opitutae bact | unclassified | unclassified | unclassified | unclassified | unclassified | 3,4 | 0,8 | 0,000496731 | 6,72E-05 | 5 - | 0,000623787 - | 3 - | 0,000464735 - | 3 - | 0,00044179 - | 3 - | 0,000448315 - | 3 - | 0,000505025 - |
| 3762 | 1 | 0.2um_NOKCPHJN_78024 | Bacteria | Bacteroidetes | Flavobacteriia | Flavobacterial | Crocinitomica | unclassified | Crocinitomica | metabolism | Carbohydrate | 05EVR | Inherit from C | 926562.Oweh | 0,6 | 0,8 | 1,87E-05 | 2,43E-05 | 0 - | 0 - | 0 - | 0 - | 0 - | 0 - | 2 - | 5,98E-05 - | 1 - | 3,37E-05 - |
| 3763 | 1 | 0.2um_NOKCPHJN_631676 | Bacteria | Proteobacteri | Alphaproteob | Rhodobactera | unclassified | unclassified | Rhodobactera | unclassified | unclassified | unclassified | unclassified | unclassified | 1,2 | 0,4 | 4,71E-05 | 1,04E-05 | 2 - | 6,75E-05 - | 1 - | 4,19E-05 - | 1 - | 3,98E-05 - | 1 - | 4,04E-05 - | 1 - | 4,56E-05 - |
| 3764 | 10 | 0.2um_NOKCPHJN_140870;0.2um_NOKCPHJN_2 | Bacteria | various | various | various | various | various | various | cellular proces | Cell wall/mem | 0P9R9 | TonB family | various | 0,4 | 0,489897949 | 2,49E-05 | 3,05E-05 | 0 - | 0 - | 1 - | 5,96E-05 - | 0 - | 0 - | 0 - | 0 - | 1 - | 6,48E-05 - |
| 3765 | 1 | 3um_FIJJAFBF_706694 | Bacteria | Proteobacteri | Gammaprotec | Alteromonada | Alteromonada | Paraglaciecola | Paraglaciecola | unclassified | unclassified | unclassified | unclassified | unclassified | 0,8 | 0,4 | 4,96E-05 | 2,52E-05 | 1 - | 5,36E-05 - | 1 - | 6,66E-05 - | 1 - | 6,33E-05 - | 1 - | 6,42E-05 - | 0 - | 0 - |
| 3766 | 1 | 3um_FIJJAFBF_735612 | Bacteria | Proteobacteri | Gammaprotec | Cellvibrionales | Porticoccacea | unclassified | SAR92 bacteri | cellular proces | Posttranslatio | 17FI0 | Binds to Cpn6 | 377629.TERTU | 0,6 | 0,489897949 | 8,39E-05 | 6,98E-05 | 1 - | 0,000118327 - | 0 - | 0 - | 0 - | 0 - | 1 - | 0,000141735 - | 1 - | 0,000159664 - |
| 3767 | 1 | 3um_FIJJAFBF_265636 | Bacteria | Proteobacteri | Gammaprotec | Alteromonada | Colwelliaceae | Colwellia | Colwellia pola | unclassified | unclassified | unclassified | unclassified | unclassified | 2,8 | 0,748331477 | 0,000188277 | 4,18E-05 | 4 - | 0,000228411 - | 2 - | 0,00014181 - | 2 - | 0,000134808 - | 3 - | 0,000205199 - | 3 - | 0,000231156 - |
| 3768 | 1 | 0.2um_NOKCPHJN_94453 | Bacteria | Proteobacteri | Betaproteoba | Burkholderiale | Comamonada | Acidovorax | various | metabolism | Energy produc | COG0055 | Produces ATP | 420662.Mpe_ | 6,8 | 1,6 | 0,000201763 | 6,06E-05 | 5 - | 0,000122363 - | 9 - | 0,00027349 - | 7 - | 0,000202212 - | 5 - | 0,000146571 - | 8 - | 0,000264178 - |
| 3769 | 1 | 0.2um_NOKCPHJN_125628 | Bacteria | Proteobacteri | Gammaprotec | Cellvibrionales | Porticoccacea | unclassified | gamma prote | information st | Translation, ril | COG0088 | One of the pri | 314287.GB22 | 0,8 | 0,4 | 5,59E-05 | 2,82E-05 | 0 - | 0 - | 1 - | 6,99E-05 - | 1 - | 6,64E-05 - | 1 - | 6,74E-05 - | 1 - | 7,59E-05 - |
| 3770 | 1 | 3um_FIJJAFBF_520521 | Bacteria | Bacteroidetes | Flavobacteriia | Flavobacterial | Flavobacteriac | Polaribacter | Polaribacter s | unclassified | unclassified | unclassified | unclassified | unclassified | 1,2 | 0,979795897 | 3,81E-05 | 2,98E-05 | 0 - | 0 - | 1 - | 3,22E-05 - | 3 - | 9,20E-05 - | 1 - | 3,11E-05 - | 1 - | 3,50E-05 - |
| 3771 | 1 | 3um_FIJJAFBF_266932 | Bacteria | Proteobacteri | Alphaproteob | Rhodobactera | Rhodobactera | Planktomarina | Planktomarina | unclassified | unclassified | unclassified | unclassified | unclassified | 4,6 | 0,489897949 | 0,000189551 | 2,74E-05 | 4 - | 0,00013787 - | 5 - | 0,000213992 - | 5 - | 0,000203427 - | 5 - | 0,000206431 - | 4 - | 0,000186035 - |
| 3772 | 1 | 0.2um_NOKCPHJN_1004243 | Bacteria | Firmicutes | Bacilli | Lactobacillales | Streptococcac | Streptococcus | Streptococcus | poorly charact | Function unkn | 0JRJ6 | NA | 5786.XP_0032 | 0,2 | 0,4 | 4,72E-06 | 9,43E-06 | 0 - | 0 - | 0 - | 0 - | 0 - | 0 - | 1 - | 2,36E-05 - | 0 - | 0 - |
| 3773 | 1 | 0.2um_NOKCPHJN_1024349 | Bacteria | Proteobacteri | Alphaproteob | Pelagibacteral | Pelagibacterac | Candidatus Pe | Candidatus Pe | metabolism | Secondary me | 01Q86 | extracellular s | 439493.PB721 | 20,2 | 1,6 | 0,000714336 | 3,80E-05 | 22 - | 0,000655867 - | 19 - | 0,00070334 - | 22 - | 0,000774184 - | 20 - | 0,000714199 - | 18 - | 0,000724088 - |
| 3774 | 1 | 3um_FIJJAFBF_342192 | Bacteria | Proteobacteri | Gammaprotec | Cellvibrionales | Porticoccacea | unclassified | gamma prote | unclassified | unclassified | unclassified | unclassified | unclassified | 1,4 | 0,489897949 | 3,87E-05 | 1,14E-05 | 2 - | 4,74E-05 - | 1 - | 2,94E-05 - | 1 - | 2,80E-05 - | 2 - | 5,68E-05 - | 1 - | 3,20E-05 - |
| 3775 | 1 | 0.2um_NOKCPHJN_379868 | Bacteria | Proteobacteri | Betaproteoba | unclassified | unclassified | unclassified | Betaproteoba | information st | Transcription | 16PMT | DNA-depende | 596154.Alide2 | 2,6 | 0,489897949 | 0,000106973 | 1,81E-05 | 3 - | 0,000104343 - | 3 - | 0,000129563 - | 2 - | 8,21E-05 - | 3 - | 0,000124985 - | 2 - | 9,39E-05 - |
| 3776 | 1 | 3um_FIJJAFBF_236214 | Bacteria | Proteobacteri | Gammaprotec | Oceanospirilla | Saccharospirill | Reinekea | Reinekea fors | information st | Translation, ril | 0QVRJ | 50S ribosomal | 314283.MED2 | 1 | 0 | 0,000263514 | 2,50E-05 | 1 - | 0,000220725 - | 1 - | 0,000274075 - | 1 - | 0,000260543 - | 1 - | 0,000264391 - | 1 - | 0,000297836 - |
| 3777 | 1 | 3um_FIJJAFBF_11305 | Bacteria | Bacteroidetes | unclassified | unclassified | unclassified | unclassified | Bacteroidetes | cellular proces | Cell cycle cont | 05CR9 | regulator of ch | 391587.KAOT | 2 | 0,632455532 | 6,31E-05 | 2,13E-05 | 1 - | 2,60E-05 - | 2 - | 6,46E-05 - | 3 - | 9,22E-05 - | 2 - | 6,24E-05 - | 2 - | 7,02E-05 - |
| 3778 | 1 | 3um_FIJJAFBF_392953 | Viruses | unclassified | unclassified | unclassified | Phycodnavirid | unclassified | Chrysochromu | information st | Transcription | COG0086 | DNA-depende | 284813.NP_5 | 0,4 | 0,489897949 | 1,07E-05 | 1,31E-05 | 0 - | 0 - | 0 - | 0 - | 1 - | 2,66E-05 - | 1 - | 2,70E-05 - | 0 - | 0 - |
| 3779 | 1 | 3um_FIJJAFBF_260210 | Bacteria | Bacteroidetes | Flavobacteriia | Flavobacterial | Flavobacteriac | Ulvibacter | Ulvibacter sp. | unclassified | unclassified | unclassified | unclassified | unclassified | 0,8 | 0,4 | 1,57E-05 | 7,96E-06 | 1 - | 1,70E-05 - | 1 - | 2,11E-05 - | 1 - | 2,00E-05 - | 1 - | 2,03E-05 - | 0 - | 0 - |
| 3780 | 1 | 0.2um_NOKCPHJN_724390 | Bacteria | Proteobacteri | Gammaprotec | Cellvibrionales | Spongiibacter | unclassified | marine gamm | information st | Translation, ril | 179ZI | One of the pri | 247633.GP214 | 0,4 | 0,489897949 | 4,24E-05 | 5,20E-05 | 0 - | 0 - | 1 - | 0,000108793 - | 1 - | 0,000103422 - | 0 - | 0 - | 0 - | 0 - |
| 3781 | 1 | 3um_FIJJAFBF_269224 | Bacteria | Bacteroidetes | Flavobacteriia | Flavobacterial | Flavobacteriac | Riemerella | Riemerella col | unclassified | unclassified | unclassified | unclassified | unclassified | 5,2 | 0,979795897 | 0,000398967 | 8,11E-05 | 6 - | 0,000386888 - | 4 - | 0,000320267 - | 6 - | 0,000456682 - | 4 - | 0,000308951 - | 6 - | 0,000522049 - |
| 3782 | 1 | 0.2um_NOKCPHJN_309355 | Bacteria | Proteobacteri | Gammaprotec | Alteromonada | Alteromonada | Paraglaciecola | Paraglaciecola | information st | Translation, ril | 177XJ | One of the pri | 983545.Glaag | 3,4 | 1,019803903 | 0,00030258 | 0,000112154 | 3 - | 0,000220725 - | 2 - | 0,000182716 - | 3 - | 0,000260543 - | 4 - | 0,000352521 - | 5 - | 0,000496393 - |
| 3783 | 1 | 3um_FIJJAFBF_222385 | Bacteria | Proteobacteri | Alphaproteob | Rhodobactera | Rhodobactera | unclassified | Rhodobactera | poorly charact | Function unkn | 01WWX | Protein of unk | 314270.RB208 | 1 | 0 | 0,000121263 | 1,15E-05 | 1 - | 0,000101572 - | 1 - | 0,000126123 - | 1 - | 0,000119896 - | 1 - | 0,000121667 - | 1 - | 0,000137057 - |
| 3784 | 1 | 0.2um_NOKCPHJN_286046 | Eukaryota | unclassified | Pelagophycea | Pelagomonad | unclassified | Aureococcus | Aureococcus a | cellular proces | Cytoskeleton | 1514Z | Tubulin is the | 272952.HpaP | 1,8 | 0,4 | 0,000318897 | 7,46E-05 | 2 - | 0,000298121 - | 1 - | 0,000185089 - | 2 - | 0,000351902 - | 2 - | 0,000357099 - | 2 - | 0,000402271 - |
| 3785 | 1 | 0.2um_NOKCPHJN_408187 | Bacteria | Verrucomicrol | unclassified | unclassified | unclassified | unclassified | Verrucomicro | unclassified | unclassified | unclassified | unclassified | unclassified | 0,6 | 0,489897949 | 9,26E-06 | 7,59E-06 | 0 - | 0 - | 1 - | 1,52E-05 - | 0 - | 0 - | 1 - | 1,46E-05 - | 1 - | 1,65E-05 - |
| 3786 | 1 | 3um_FIJJAFBF_169756 | Bacteria | Bacteroidetes | Saprospiria | Saprospirales | Lewinellaceae | Lewinella | Lewinella pers | metabolism | Inorganic ion t | 05C27 | TonB-depende | 391603.FBALC | 14,8 | 2,561249695 | 0,000186578 | 2,39E-05 | 19 - | 0,000202861 - | 11 - | 0,000145833 - | 15 - | 0,000189045 - | 14 - | 0,000179048 - | 15 - | 0,000216104 - |
| 3787 | 1 | 3um_FIJJAFBF_218855 | Bacteria | Proteobacteri | Gammaprotec | Oceanospirilla | Saccharospirill | Reinekea | Reinekea fors | information st | Translation, ril | COG0333 | 50s ribosomal | 314283.MED2 | 1 | 0 | 0,000232249 | 2,21E-05 | 1 - | 0,000194537 - | 1 - | 0,000241557 - | 1 - | 0,000229631 - | 1 - | 0,000233023 - | 1 - | 0,000262499 - |
| 3788 | 1 | 0.2um_NOKCPHJN_657860 | Bacteria | Proteobacteri | Gammaprotec | unclassified | unclassified | unclassified | Gammaprotec | metabolism | Inorganic ion t | 17AJ4 | TonB-depend | 450851.PHZ_ | 1,6 | 0,8 | 2,24E-05 | 1,15E-05 | 2 - | 2,37E-05 - | 0 - | 0 - | 2 - | 2,80E-05 - | 2 - | 2,84E-05 - | 2 - | 3,20E-05 - |
| 3789 | 1 | 0.2um_NOKCPHJN_318631 | Bacteria | Proteobacteri | Betaproteoba | Burkholderiale | Burkholderiac | Paraburkhold | Paraburkhold | unclassified | unclassified | unclassified | unclassified | unclassified | 0,8 | 0,4 | 4,51E-06 | 2,31E-06 | 1 - | 4,70E-06 - | 1 - | 5,84E-06 - | 0 - | 0 - | 1 - | 5,63E-06 - | 1 - | 6,35E-06 - |
| 3790 | 1 | 3um_FIJJAFBF_182957 | Bacteria | Proteobacteri | Gammaprotec | Alteromonada | Alteromonada | Paraglaciecola | Paraglaciecola | metabolism | Inorganic ion t | 16TQC | TONB-depend | 314287.GB22 | 33,4 | 1,854723699 | 0,000441438 | 2,48E-05 | 37 - | 0,000411506 - | 33 - | 0,000455729 - | 32 - | 0,0004201 - | 33 - | 0,000439627 - | 32 - | 0,000480231 - |
| 3791 | 1 | 3um_FIJJAFBF_309177 | unclassified | unclassified | unclassified | unclassified | unclassified | unclassified | unclassified | unclassified | unclassified | unclassified | unclassified | unclassified | 0,2 | 0,4 | 2,27E-05 | 4,54E-05 | 0 - | 0 - | 0 - | 0 - | 0 - | 0 - | 1 - | 0,000113623 - | 0 - | 0 - |
| 3792 | 1 | 3um_FIJJAFBF_132511 | Bacteria | Proteobacteri | Gammaprotec | Cellvibrionales | Porticoccacea | unclassified | gamma prote | metabolism | Energy produc | COG0056 | Produces ATP | 314287.GB22 | 2 | 0,894427191 | 5,24E-05 | 2,30E-05 | 2 - | 4,47E-05 - | 1 - | 2,77E-05 - | 3 - | 7,91E-05 - | 3 - | 8,02E-05 - | 1 - | 3,01E-05 - |
| 3793 | 1 | 0.2um_NOKCPHJN_415539 | Bacteria | Proteobacteri | Alphaproteob | Rhodobactera | Rhodobactera | Planktotalea | Planktotalea a | unclassified | unclassified | unclassified | unclassified | unclassified | 4 | 0,632455532 | 0,000162931 | 1,91E-05 | 5 - | 0,000172337 - | 3 - | 0,000128395 - | 4 - | 0,000162741 - | 4 - | 0,000165145 - | 4 - | 0,000186035 - |
| 3794 | 1 | 3um_FIJJAFBF_49008 | Bacteria | Proteobacteri | Gammaprotec | Oceanospirilla | Saccharospirill | Reinekea | Reinekea blan | metabolism | Energy produc | 16QPK | Succinate deh | 314283.MED2 | 11,8 | 2,039607805 | 0,000271208 | 3,88E-05 | 14 - | 0,000272352 - | 14 - | 0,00033818 - | 10 - | 0,000229631 - | 12 - | 0,000279627 - | 9 - | 0,000236249 - |
| 3795 | 1 | 0.2um_NOKCPHJN_75803 | Bacteria | Bacteroidetes | Flavobacteriia | Flavobacterial | Flavobacteriac | Muricauda | Muricauda sp. | metabolism | Inorganic ion t | 05C27 | TonB-depende | 655815.ZPR_ | 1,2 | 0,4 | 1,64E-05 | 5,50E-06 | 1 - | 1,15E-05 - | 1 - | 1,43E-05 - | 2 - | 2,71E-05 - | 1 - | 1,37E-05 - | 1 - | 1,55E-05 - |
| 3796 | 1 | 0.2um_NOKCPHJN_571776 | Eukaryota | Ascomycota | Eurotiomycete | Chaetothyriale | Herpotrichiell | Exophiala | Exophiala mes | unclassified | unclassified | unclassified | unclassified | unclassified | 1,8 | 0,979795897 | 3,37E-05 | 1,89E-05 | 2 - | 3,22E-05 - | 2 - | 4,00E-05 - | 2 - | 3,81E-05 - | 3 - | 5,79E-05 - | 0 - | 0 - |
| 3797 | 1 | 0.2um_NOKCPHJN_735975 | Eukaryota | unclassified | Oomycetes | Saprolegniales | Saprolegniace | Achlya | Achlya hypogy | cellular proces | Defense mech | KOG1823 | UTP20, small s | 65071.PYU1_ | 0,2 | 0,4 | 2,96E-06 | 5,92E-06 | 0 - | 0 - | 0 - | 0 - | 0 - | 0 - | 1 - | 1,48E-05 - | 0 - | 0 - |
| 3798 | 1 | 0.2um_NOKCPHJN_18132 | Viruses | unclassified | unclassified | unclassified | Phycodnavirid | unclassified | Chrysochromu | information st | Transcription | 1BKE5 | SWIB complex | 3711.Bra0223 | 0,4 | 0,489897949 | 2,30E-05 | 2,84E-05 | 1 - | 5,24E-05 - | 0 - | 0 - | 0 - | 0 - | 1 - | 6,28E-05 - | 0 - | 0 - |
| 3799 | 1 | 0.2um_NOKCPHJN_288026 | Bacteria | Proteobacteri | Alphaproteob | Rhodobactera | Rhodobactera | Aliiroseovariu | Aliiroseovariu | unclassified | unclassified | unclassified | unclassified | unclassified | 0,6 | 0,489897949 | 2,29E-05 | 1,88E-05 | 1 - | 3,39E-05 - | 0 - | 0 - | 1 - | 4,00E-05 - | 1 - | 4,06E-05 - | 0 - | 0 - |
| 3800 | 1 | 0.2um_NOKCPHJN_170714 | Bacteria | Bacteroidetes | unclassified | unclassified | unclassified | unclassified | Bacteroidetes | poorly charact | Function unkn | 05WQ7 | NA | 313595.P7007 | 3,4 | 1,019803903 | 0,000112376 | 3,94E-05 | 4 - | 0,000110362 - | 2 - | 6,85E-05 - | 3 - | 9,77E-05 - | 3 - | 9,91E-05 - | 5 - | 0,000186147 - |
| 3801 | 1 | 3um_FIJJAFBF_40751 | Bacteria | Proteobacteri | Gammaprotec | unclassified | unclassified | unclassified | SAR86 cluster | unclassified | unclassified | unclassified | unclassified | unclassified | 0,4 | 0,489897949 | 1,66E-05 | 2,03E-05 | 0 - | 0 - | 0 - | 0 - | 1 - | 4,12E-05 - | 1 - | 4,18E-05 - | 0 - | 0 - |
| 3802 | 1 | 3um_FIJJAFBF_175086 | Bacteria | Proteobacteri | Alphaproteob | Rhodobactera | Rhodobactera | Phaeobacter | Phaeobacter g | metabolism | Inorganic ion t | 16PAX | extracellular s | 439496.RBY4 | 8,8 | 0,748331477 | 0,000276862 | 4,06E-05 | 8 - | 0,000210118 - | 9 - | 0,000293517 - | 10 - | 0,000310028 - | 8 - | 0,000251686 - | 9 - | 0,000318963 - |
| 3803 | 1 | 3um_FIJJAFBF_176961 | Eukaryota | unclassified | Rhodellophyc | Rhodellales | Rhodellaceae | Corynoplastis | Corynoplastis | unclassified | unclassified | unclassified | unclassified | unclassified | 2,8 | 0,748331477 | 5,13E-05 | 1,67E-05 | 3 - | 4,58E-05 - | 2 - | 3,79E-05 - | 3 - | 5,40E-05 - | 2 - | 3,66E-05 - | 4 - | 8,24E-05 - |
| 3804 | 1 | 0.2um_NOKCPHJN_460017 | Eukaryota | unclassified | Chrysophycea | Chromulinales | Chromulinace | Ochromonas | Ochromonas | unclassified | unclassified | unclassified | unclassified | unclassified | 0,4 | 0,489897949 | 4,55E-05 | 5,57E-05 | 0 - | 0 - | 0 - | 0 - | 1 - | 0,000112902 - | 1 - | 0,000114569 - | 0 - | 0 - |
| 3805 | 1 | 0.2um_NOKCPHJN_212186 | Bacteria | Bacteroidetes | Flavobacteriia | Flavobacterial | Flavobacteriac | Tenacibaculur | Tenacibaculur | unclassified | unclassified | unclassified | unclassified | unclassified | 2,2 | 0,4 | 4,99E-05 | 4,58E-06 | 3 - | 5,79E-05 - | 2 - | 4,79E-05 - | 2 - | 4,55E-05 - | 2 - | 4,62E-05 - | 2 - | 5,21E-05 - |
| 3806 | 1 | 3um_FIJJAFBF_766830 | Bacteria | Proteobacteri | Gammaprotec | Oceanospirilla | Oceanospirilla | Nitrincola | Nitrincola sp. | unclassified | unclassified | unclassified | unclassified | unclassified | 1,2 | 0,4 | 0,000228505 | 7,88E-05 | 1 - | 0,000159412 - | 1 - | 0,000197943 - | 1 - | 0,00018817 - | 2 - | 0,000381898 - | 1 - | 0,000215103 - |
| 3807 | 1 | 0.2um_NOKCPHJN_31647 | Bacteria | Proteobacteri | Betaproteoba | unclassified | unclassified | unclassified | Betaproteoba | cellular proces | Cell wall/mem | COG2885 | Ompa motb d | 365044.Pnap_ | 5,8 | 0,748331477 | 0,00041336 | 4,65E-05 | 7 - | 0,000420648 - | 5 - | 0,000373086 - | 5 - | 0,000354666 - | 6 - | 0,000431885 - | 6 - | 0,000486517 - |
| 3808 | 1 | 0.2um_NOKCPHJN_457862 | Bacteria | Bacteroidetes | Flavobacteriia | Flavobacterial | Flavobacteriac | Psychroflexus | Psychroflexus | unclassified | unclassified | unclassified | unclassified | unclassified | 0,8 | 0,4 | 9,34E-06 | 4,80E-06 | 1 - | 9,75E-06 - | 1 - | 1,21E-05 - | 0 - | 0 - | 1 - | 1,17E-05 - | 1 - | 1,32E-05 - |
| 3809 | 1 | 0.2um_NOKCPHJN_365441 | Bacteria | Proteobacteri | Alphaproteob | Rhodobactera | Rhodobactera | Planktomarina | Planktomarina | unclassified | unclassified | unclassified | unclassified | unclassified | 1,6 | 1,356465997 | 0,000105354 | 9,89E-05 | 2 - | 0,000107772 - | 0 - | 0 - | 1 - | 6,36E-05 - | 1 - | 6,45E-05 - | 4 - | 0,000290844 - |
| 3810 | 1 | 0.2um_NOKCPHJN_133366 | Bacteria | Bacteroidetes | unclassified | unclassified | unclassified | unclassified | Bacteroidetes | information st | Translation, ril | 05KK9 | This protein bi | 313603.FB217 | 0,2 | 0,4 | 1,22E-05 | 2,44E-05 | 1 - | 6,11E-05 - | 0 - | 0 - | 0 - | 0 - | 0 - | 0 - | 0 - | 0 - |
| 3811 | 1 | 0.2um_NOKCPHJN_09068 | Bacteria | Candidatus Ul | unclassified | unclassified | unclassified | unclassified | Candidatus Ul | poorly charact | Function unkn | 1EB8T | NA | 436017.A4S9 | 0,8 | 0,4 | 7,96E-06 | 4,09E-06 | 1 - | 8,41E-06 - | 0 - | 0 - | 1 - | 9,93E-06 - | 1 - | 1,01E-05 - | 1 - | 1,14E-05 - |
| 3812 | 1 | 0.2um_NOKCPHJN_1015207 | Bacteria | Bacteroidetes | Flavobacteriia | Flavobacterial | Crocinitomica | unclassified | Crocinitomica | information st | Replication, re | COG0187 | DNA gyrase ne | 50743.SCB49_ | 1 | 0 | 3,07E-05 | 2,91E-06 | 1 - | 2,57E-05 - | 1 - | 3,19E-05 - | 1 - | 3,03E-05 - | 1 - | 3,08E-05 - | 1 - | 3,46E-05 - |
| 3813 | 1 | 3um_FIJJAFBF_779113 | unclassified | unclassified | unclassified | unclassified | unclassified | unclassified | unclassified | unclassified | unclassified | unclassified | unclassified | unclassified | 0,2 | 0,4 | 2,07E-05 | 4,13E-05 | 0 - | 0 - | 0 - | 0 - | 0 - | 0 - | 1 - | 0,000103371 - | 0 - | 0 - |
| 3814 | 1 | 0.2um_NOKCPHJN_1095303 | Bacteria | Actinobacteria | Actinobacteria | Corynebacteri | Nocardiaceae | Rhodococcus | Rhodococcus | unclassified | unclassified | unclassified | unclassified | unclassified | 0,2 | 0,4 | 7,03E-06 | 1,41E-05 | 0 - | 0 - | 0 - | 0 - | 0 - | 0 - | 1 - | 3,52E-05 - | 0 - | 0 - |
| 3815 | 1 | 3um_FIJJAFBF_659921 | Bacteria | Proteobacteri | Gammaprotec | Cellvibrionales | Porticoccacea | unclassified | SAR92 bacteri | information st | Replication, re | COG0513 | purine NTP-de | 314287.GB22 | 0,4 | 0,489897949 | 8,32E-06 | 1,02E-05 | 0 - | 0 - | 1 - | 2,13E-05 - | 1 - | 2,03E-05 - | 0 - | 0 - | 0 - | 0 - |
| 3816 | 1 | 3um_FIJJAFBF_434552 | Bacteria | Bacteroidetes | Flavobacteriia | Flavobacterial | Flavobacteriac | Polaribacter | Polaribacter s | unclassified | unclassified | unclassified | unclassified | unclassified | 1 | 0,632455532 | 7,43E-05 | 4,19E-05 | 2 - | 0,000129691 - | 0 - | 0 - | 1 - | 7,65E-05 - | 1 - | 7,77E-05 - | 1 - | 8,75E-05 - |
| 3817 | 1 | 3um_FIJJAFBF_25764 | Bacteria | Bacteroidetes | Flavobacteriia | unclassified | unclassified | unclassified | Flavobacteria | unclassified | unclassified | unclassified | unclassified | unclassified | 1,2 | 0,4 | 0,000131619 | 4,54E-05 | 1 - | 9,18E-05 - | 1 - | 0,000114015 - | 1 - | 0,000108386 - | 2 - | 0,000219973 - | 1 - | 0,0001239 - |
| 3818 | 1 | 0.2um_NOKCPHJN_37492 | Bacteria | Bacteroidetes | Flavobacteriia | unclassified | unclassified | unclassified | Flavobacteria | information st | Translation, ril | 09AI8 | Associates wit | 487796.Flav2 | 0,2 | 0,4 | 9,96E-06 | 1,99E-05 | 0 - | 0 - | 0 - | 0 - | 1 - | 4,98E-05 - | 0 - | 0 - | 0 - | 0 - |
| 3819 | 1 | 0.2um_NOKCPHJN_422835 | Eukaryota | Streptophyta | unclassified | Rosales | Rosaceae | Rosa | Rosa chinensis | metabolism | Energy produc | KOG0017 | Retrotranspos | 423536.XP_00 | 0,4 | 0,489897949 | 3,20E-06 | 3,94E-06 | 1 - | 7,34E-06 - | 0 - | 0 - | 1 - | 8,67E-06 - | 0 - | 0 - | 0 - | 0 - |
| 3820 | 1 | 3um_FIJJAFBF_70349 | Bacteria | Proteobacteri | Betaproteoba | unclassified | unclassified | unclassified | Betaproteoba | unclassified | unclassified | unclassified | unclassified | unclassified | 0,2 | 0,4 | 9,89E-06 | 1,98E-05 | 0 - | 0 - | 0 - | 0 - | 1 - | 4,94E-05 - | 0 - | 0 - | 0 - | 0 - |
| 3821 | 1 | 0.2um_NOKCPHJN_1090570 | Bacteria | Proteobacteri | Alphaproteob | Pelagibacteral | unclassified | unclassified | Pelagibacteral | cellular proces | Cell wall/mem | 7257 | NA | 335992.SAR11 | 0,8 | 0,4 | 3,26E-05 | 1,64E-05 | 0 - | 0 - | 1 - | 4,07E-05 - | 1 - | 3,87E-05 - | 1 - | 3,93E-05 - | 1 - | 4,42E-05 - |
| 3822 | 1 | 0.2um_NOKCPHJN_1109835 | Bacteria | Proteobacteri | Alphaproteob | Pelagibacteral | Pelagibacterac | Candidatus Pe | Candidatus Pe | cellular proces | Cell wall/mem | 7257 | NA | 335992.SAR11 | 1,4 | 0,489897949 | 5,24E-05 | 1,65E-05 | 2 - | 6,38E-05 - | 2 - | 7,92E-05 - | 1 - | 3,76E-05 - | 1 - | 3,82E-05 - | 1 - | 4,30E-05 - |
| 3823 | 1 | 0.2um_NOKCPHJN_350974 | Bacteria | Proteobacteri | Gammaprotec | Cellvibrionales | Porticoccacea | unclassified | gamma prote | cellular proces | Signal transdu | 16Q0Y | phosphate reg | 203122.Sde_3 | 0,2 | 0,4 | 9,56E-06 | 1,91E-05 | 1 - | 4,78E-05 - | 0 - | 0 - | 0 - | 0 - | 0 - | 0 - | 0 - | 0 - |
| 3824 | 1 | 0.2um_NOKCPHJN_139211 | Bacteria | Bacteroidetes | Flavobacteriia | Flavobacterial | Flavobacteriac | Polaribacter | Polaribacter t | unclassified | unclassified | unclassified | unclassified | unclassified | 1 | 0 | 6,40E-05 | 6,08E-06 | 1 - | 5,36E-05 - | 1 - | 6,66E-05 - | 1 - | 6,33E-05 - | 1 - | 6,42E-05 - | 1 - | 7,24E-05 - |
| 3825 | 1 | 0.2um_NOKCPHJN_114298 | Bacteria | Proteobacteri | Alphaproteob | Pelagibacteral | Pelagibacterac | Candidatus Pe | Candidatus Pe | information st | Transcription | 178T0 | Transcriptiona | 335992.SAR11 | 1,4 | 0,489897949 | 6,33E-05 | 1,86E-05 | 2 - | 7,76E-05 - | 1 - | 4,81E-05 - | 1 - | 4,58E-05 - | 2 - | 9,29E-05 - | 1 - | 5,23E-05 - |
| 3826 | 1 | 0.2um_NOKCPHJN_170447 | Bacteria | Proteobacteri | Alphaproteob | Rhodobactera | Rhodobactera | Pseudorueger | Pseudorueger | metabolism | Amino acid tra | 07QMX | Aminotransfe | 398580.Dshi_ | 2,6 | 0,489897949 | 9,35E-05 | 1,52E-05 | 3 - | 9,13E-05 - | 3 - | 0,00011341 - | 3 - | 0,000107811 - | 2 - | 7,29E-05 - | 2 - | 8,22E-05 - |
| 3827 | 1 | 3um_FIJJAFBF_193128 | Bacteria | Proteobacteri | Gammaprotec | Oceanospirilla | Oceanospirilla | Marinomonas | Marinomonas | cellular proces | Posttranslatio | 0QW5X | glutaredoxin 3 | 717774.Marm | 1 | 0 | 0,000159334 | 1,51E-05 | 1 - | 0,000133461 - | 1 - | 0,00016572 - | 1 - | 0,000157537 - | 1 - | 0,000159864 - | 1 - | 0,000180087 - |
| 3828 | 1 | 0.2um_NOKCPHJN_721141 | Bacteria | Bacteroidetes | Flavobacteriia | Flavobacterial | Flavobacteriac | Altibacter | Altibacter lent | poorly charact | Function unkn | 0P8NI | Inherit from b | 313594.PI23P | 7,2 | 1,469693846 | 0,000151853 | 3,16E-05 | 9 - | 0,000159659 - | 8 - | 0,000176221 - | 6 - | 0,00012564 - | 5 - | 0,000106247 - | 8 - | 0,000191499 - |
| 3829 | 1 | 3um_FIJJAFBF_211273 | unclassified | unclassified | unclassified | unclassified | unclassified | unclassified | unclassified | unclassified | unclassified | unclassified | unclassified | unclassified | 0,8 | 0,4 | 9,55E-05 | 4,85E-05 | 1 - | 0,000103402 - | 1 - | 0,000128395 - | 1 - | 0,000122056 - | 1 - | 0,000123859 - | 0 - | 0 - |
| 3830 | 1 | 0.2um_NOKCPHJN_474066 | Bacteria | Proteobacteri | Alphaproteob | Pelagibacteral | Pelagibacterac | Candidatus Pe | Candidatus Pe | cellular proces | Cell wall/mem | 7257 | NA | 335992.SAR11 | 1,2 | 0,4 | 4,85E-05 | 1,07E-05 | 2 - | 6,96E-05 - | 1 - | 4,32E-05 - | 1 - | 4,11E-05 - | 1 - | 4,17E-05 - | 1 - | 4,69E-05 - |
| 3831 | 1 | 3um_FIJJAFBF_224533 | Bacteria | Proteobacteri | Betaproteoba | Nitrosomonad | unclassified | unclassified | unclassified | unclassified | unclassified | unclassified | unclassified | unclassified | 0,2 | 0,4 | 1,54E-05 | 3,08E-05 | 1 - | 7,70E-05 - | 0 - | 0 - | 0 - | 0 - | 0 - | 0 - | 0 - | 0 - |
| 3832 | 1 | 3um_FIJJAFBF_301047 | Bacteria | Bacteroidetes | Flavobacteriia | Flavobacterial | Flavobacteriac | Flavobacteriu | Flavobacteriu | information st | Transcription | 0P7SF | DNA-depende | 376686.Fjoh_ | 0,2 | 0,4 | 3,91E-06 | 7,81E-06 | 0 - | 0 - | 0 - | 0 - | 0 - | 0 - | 0 - | 0 - | 1 - | 1,95E-05 - |
| 3833 | 1 | 3um_FIJJAFBF_293518 | Bacteria | Bacteroidetes | Cytophagia | Cytophagales | Flammeovirga | unclassified | Flammeovirga | unclassified | unclassified | unclassified | unclassified | unclassified | 1,4 | 0,489897949 | 7,37E-05 | 2,60E-05 | 1 - | 4,41E-05 - | 1 - | 5,48E-05 - | 2 - | 0,000104217 - | 2 - | 0,000105756 - | 1 - | 5,96E-05 - |
| 3834 | 1 | 0.2um_NOKCPHJN_615820 | Bacteria | Bacteroidetes | Flavobacteriia | Flavobacterial | Flavobacteriac | Polaribacter | Polaribacter s | metabolism | Carbohydrate | COG1109 | Catalyzes the | 313598.MED1 | 1 | 0,632455532 | 2,39E-05 | 1,52E-05 | 1 - | 2,01E-05 - | 1 - | 2,50E-05 - | 2 - | 4,75E-05 - | 0 - | 0 - | 1 - | 2,71E-05 - |
| 3835 | 1 | 3um_FIJJAFBF_118035 | Bacteria | Proteobacteri | Betaproteoba | unclassified | unclassified | unclassified | Betaproteoba | metabolism | Inorganic ion t | 08UTW | Plays a role in | 596154.Alide2 | 3,4 | 1,019803903 | 0,000200121 | 6,27E-05 | 3 - | 0,000147781 - | 4 - | 0,000244668 - | 5 - | 0,000290734 - | 2 - | 0,000118011 - | 3 - | 0,000199409 - |
| 3836 | 1 | 0.2um_NOKCPHJN_178136 | Bacteria | Proteobacteri | Alphaproteob | Rhodobactera | Rhodobactera | Litoreibacter | Litoreibacter p | unclassified | unclassified | unclassified | unclassified | unclassified | 0,4 | 0,489897949 | 1,48E-05 | 1,82E-05 | 1 - | 3,39E-05 - | 0 - | 0 - | 1 - | 4,00E-05 - | 0 - | 0 - | 0 - | 0 - |
| 3837 | 1 | 3um_FIJJAFBF_20736 | Bacteria | Proteobacteri | Gammaprotec | Oceanospirilla | Saccharospirill | Reinekea | Reinekea fors | metabolism | Energy produc | COG0712 | F(1)F(0) ATP s | 314283.MED2 | 1 | 0 | 7,70E-05 | 7,31E-06 | 1 - | 6,45E-05 - | 1 - | 8,01E-05 - | 1 - | 7,61E-05 - | 1 - | 7,72E-05 - | 1 - | 8,70E-05 - |
| 3838 | 1 | 0.2um_NOKCPHJN_72662 | Bacteria | Proteobacteri | unclassified | unclassified | unclassified | unclassified | Proteobacteri | metabolism | Energy produc | 0QJAP | Catalyzes the | 247634.GPB2 | 1,2 | 0,4 | 5,07E-05 | 1,70E-05 | 1 - | 3,54E-05 - | 1 - | 4,40E-05 - | 2 - | 8,36E-05 - | 1 - | 4,24E-05 - | 1 - | 4,78E-05 - |
| 3839 | 1 | 3um_FIJJAFBF_782573 | Bacteria | Proteobacteri | Gammaprotec | Alteromonada | Colwelliaceae | Colwellia | Colwellia sp. P | information st | Translation, ril | 179R6 | This protein is | 167879.CPS_4 | 0,4 | 0,489897949 | 3,62E-05 | 4,48E-05 | 1 - | 8,08E-05 - | 1 - | 0,000100365 - | 0 - | 0 - | 0 - | 0 - | 0 - | 0 - |
| 3840 | 1 | 3um_FIJJAFBF_396107 | Bacteria | Bacteroidetes | Flavobacteriia | Flavobacterial | Flavobacteriac | Zobellia | various | metabolism | Energy produc | 05D26 | NQR complex | 487797.Flav3 | 1 | 0,632455532 | 3,05E-05 | 1,71E-05 | 2 - | 5,28E-05 - | 1 - | 3,28E-05 - | 1 - | 3,11E-05 - | 0 - | 0 - | 1 - | 3,56E-05 - |
| 3841 | 1 | 0.2um_NOKCPHJN_596982 | Bacteria | Proteobacteri | Alphaproteob | Pelagibacteral | Pelagibacterac | Candidatus Pe | Candidatus Pe | poorly charact | Function unkn | COG3181 | extra-cytoplas | 1002672.SAR1 | 0,8 | 0,4 | 3,39E-05 | 1,74E-05 | 1 - | 3,55E-05 - | 1 - | 4,41E-05 - | 1 - | 4,19E-05 - | 0 - | 0 - | 1 - | 4,79E-05 - |
| 3842 | 1 | 0.2um_NOKCPHJN_480630 | Bacteria | Proteobacteri | Alphaproteob | Pelagibacteral | Pelagibacterac | Candidatus Pe | Candidatus Pe | cellular proces | Posttranslatio | COG0652 | peptidyl-proly | 335992.SAR11 | 0,4 | 0,489897949 | 2,74E-05 | 3,38E-05 | 1 - | 6,24E-05 - | 0 - | 0 - | 0 - | 0 - | 1 - | 7,47E-05 - | 0 - | 0 - |
| 3843 | 1 | 3um_FIJJAFBF_149818 | Bacteria | Proteobacteri | Betaproteoba | unclassified | unclassified | unclassified | Betaproteoba | metabolism | Energy produc | 16UMU | it plays a direc | 397945.Aave_ | 1 | 0 | 4,68E-05 | 4,44E-06 | 1 - | 3,92E-05 - | 1 - | 4,86E-05 - | 1 - | 4,62E-05 - | 1 - | 4,69E-05 - | 1 - | 5,29E-05 - |
| 3844 | 1 | 0.2um_NOKCPHJN_970808 | Bacteria | Proteobacteri | Betaproteoba | Rhodocyclales | Zoogloeaceae | Thauera | Thauera terpe | unclassified | unclassified | unclassified | unclassified | unclassified | 0,2 | 0,4 | 3,75E-06 | 7,50E-06 | 1 - | 1,88E-05 - | 0 - | 0 - | 0 - | 0 - | 0 - | 0 - | 0 - | 0 - |
| 3845 | 1 | 3um_FIJJAFBF_575011 | Bacteria | Proteobacteri | Alphaproteob | Rhodobactera | Rhodobactera | Planktomarina | Planktomarina | unclassified | unclassified | unclassified | unclassified | unclassified | 1,8 | 0,748331477 | 4,19E-05 | 1,40E-05 | 3 - | 6,05E-05 - | 2 - | 5,01E-05 - | 2 - | 4,76E-05 - | 1 - | 2,42E-05 - | 1 - | 2,72E-05 - |
| 3846 | 1 | 3um_FIJJAFBF_263212 | unclassified | unclassified | unclassified | unclassified | unclassified | unclassified | unclassified | unclassified | unclassified | unclassified | unclassified | unclassified | 4 | 0,632455532 | 0,000275922 | 3,56E-05 | 5 - | 0,000291312 - | 4 - | 0,000289378 - | 4 - | 0,000275091 - | 3 - | 0,000209365 - | 4 - | 0,000314466 - |
| 3847 | 1 | 0.2um_NOKCPHJN_368552 | Bacteria | Bacteroidetes | Flavobacteriia | Flavobacterial | Flavobacteriac | Polaribacter | Polaribacter s | cellular proces | Intracellular tr | 0PBE9 | Preprotein tra | 313594.PI23P | 0,2 | 0,4 | 2,69E-05 | 5,38E-05 | 0 - | 0 - | 1 - | 0,000134452 - | 0 - | 0 - | 0 - | 0 - | 0 - | 0 - |
| 3848 | 1 | 0.2um_NOKCPHJN_111451 | Bacteria | Bacteroidetes | Cytophagia | Cytophagales | Cytophagacea | Ekhidna | Ekhidna lutea | unclassified | unclassified | unclassified | unclassified | unclassified | 0,8 | 1,166190379 | 1,16E-05 | 1,77E-05 | 1 - | 1,22E-05 - | 3 - | 4,56E-05 - | 0 - | 0 - | 0 - | 0 - | 0 - | 0 - |
| 3849 | 1 | 3um_FIJJAFBF_359622 | Bacteria | Actinobacteria | Actinobacteria | Micrococcales | Microbacteria | Candidatus Ac | Candidatus Ac | unclassified | unclassified | unclassified | unclassified | unclassified | 1,2 | 0,4 | 4,28E-05 | 9,47E-06 | 2 - | 6,14E-05 - | 1 - | 3,81E-05 - | 1 - | 3,62E-05 - | 1 - | 3,68E-05 - | 1 - | 4,14E-05 - |
| 3850 | 1 | 3um_FIJJAFBF_08101 | Bacteria | Bacteroidetes | Flavobacteriia | Flavobacterial | Flavobacteriac | Tenacibaculur | Tenacibaculur | metabolism | Amino acid tra | COG0131 | imidazoleglyce | 313594.PI23P | 0,2 | 0,4 | 7,26E-06 | 1,45E-05 | 0 - | 0 - | 0 - | 0 - | 0 - | 0 - | 1 - | 3,63E-05 - | 0 - | 0 - |
| 3851 | 1 | 3um_FIJJAFBF_123074 | Bacteria | Proteobacteri | Gammaprotec | Alteromonada | Alteromonada | Paraglaciecola | various | unclassified | unclassified | unclassified | unclassified | unclassified | 1,6 | 0,489897949 | 0,000287444 | 7,97E-05 | 2 - | 0,000306071 - | 2 - | 0,00038005 - | 2 - | 0,000361286 - | 1 - | 0,000183311 - | 1 - | 0,000206499 - |
| 3852 | 1 | 0.2um_NOKCPHJN_1097847 | Eukaryota | Phaeophyceae | unclassified | Ectocarpales | Ectocarpaceae | Ectocarpus | Ectocarpus sili | unclassified | unclassified | unclassified | unclassified | unclassified | 0,4 | 0,489897949 | 3,09E-05 | 3,81E-05 | 1 - | 7,08E-05 - | 0 - | 0 - | 1 - | 8,36E-05 - | 0 - | 0 - | 0 - | 0 - |
| 3853 | 2 | 3um_FIJJAFBF_137570;3um_FIJJAFBF_402671 | Bacteria | Bacteroidetes | unclassified | unclassified | unclassified | unclassified | Bacteroidetes | cellular proces | Signal transdu | 0XVSK | REC | 388413.ALPR1 | 2,4 | 0,8 | 0,000201343 | 6,92E-05 | 2 - | 0,0001417 - | 3 - | 0,000263924 - | 3 - | 0,000250893 - | 3 - | 0,000254599 - | 1 - | 9,56E-05 - |
| 3854 | 1 | 3um_FIJJAFBF_79247 | Bacteria | Bacteroidetes | Flavobacteriia | Flavobacterial | Flavobacteriac | Polaribacter | Polaribacter s | cellular proces | Posttranslatio | 07S5S | Peptidyl-proly | 313598.MED1 | 1,2 | 0,979795897 | 2,86E-05 | 2,02E-05 | 3 - | 6,33E-05 - | 1 - | 2,62E-05 - | 1 - | 2,49E-05 - | 0 - | 0 - | 1 - | 2,85E-05 - |
| 3855 | 1 | 3um_FIJJAFBF_10572 | Eukaryota | unclassified | Oomycetes | Albuginales | Albuginaceae | Albugo | Albugo candid | unclassified | unclassified | unclassified | unclassified | unclassified | 2,4 | 0,489897949 | 0,000114912 | 2,48E-05 | 2 - | 8,03E-05 - | 2 - | 9,97E-05 - | 3 - | 0,000142114 - | 3 - | 0,000144213 - | 2 - | 0,000108304 - |

| | | | | | | | | | | | | | | | | | | | | | | | | | | | | |
|---|---|---|---|---|---|---|---|---|---|---|---|---|---|---|---|---|---|---|---|---|---|---|---|---|---|---|---|---|
| 3856 | 1 | 0.2um_NOKCPHJN_673912 | Bacteria | Actinobacteria | Actinobacteria | Micrococcales | Microbacteria | Candidatus Ac | Candidatus Ac | unclassified | unclassified | unclassified | unclassified | unclassified | 0,8 | 0,748331477 | 1,09E-05 | 1,03E-05 | 1 - | 1,15E-05 - | 0 - | 0 - | 2 - | 2,73E-05 - | 0 - | 0 - | 1 - | 1,56E-05 - |
| 3857 | 1 | 0.2um_NOKCPHJN_480841 | Bacteria | Bacteroidetes | Flavobacteriia | Flavobacterial | Flavobacteriac | Flavobacteriu | Flavobacteriu | unclassified | unclassified | unclassified | unclassified | unclassified | 1,8 | 1,326649916 | 4,74E-05 | 3,11E-05 | 4 - | 9,16E-05 - | 1 - | 2,84E-05 - | 0 - | 0 - | 2 - | 5,49E-05 - | 2 - | 6,18E-05 - |
| 3858 | 1 | 3um_FIJJAFBF_566364 | Bacteria | Proteobacteri | Gammaprotec | Pseudomonad | Moraxellaceae | Psychrobacter | various | cellular proces | Cell wall/mem | 07Q6S | Gram-negativ | 335284.Pcryo_ | 1,4 | 0,489897949 | 5,19E-05 | 1,83E-05 | 1 - | 3,11E-05 - | 1 - | 3,86E-05 - | 2 - | 7,34E-05 - | 2 - | 7,45E-05 - | 1 - | 4,20E-05 - |
| 3859 | 1 | 0.2um_NOKCPHJN_443356 | Eukaryota | Phaeophyceae | unclassified | Ectocarpales | Ectocarpaceae | Ectocarpus | Ectocarpus sili | metabolism | Energy produc | 0IFIU | Pyrophosphat | 2850.Phatr21 | 1 | 0,632455532 | 2,07E-05 | 1,34E-05 | 1 - | 1,79E-05 - | 1 - | 2,22E-05 - | 2 - | 4,22E-05 - | 1 - | 2,14E-05 - | 0 - | 0 - |
| 3860 | 1 | 0.2um_NOKCPHJN_147751 | Bacteria | Bacteroidetes | Flavobacteriia | Flavobacterial | unclassified | unclassified | Flavobacterial | information st | Translation, ril | COG0617 | Catalyzes the | 487796.Flav2A | 0,4 | 0,489897949 | 1,05E-05 | 1,30E-05 | 1 - | 2,41E-05 - | 0 - | 0 - | 1 - | 2,85E-05 - | 0 - | 0 - | 0 - | 0 - |
| 3861 | 1 | 3um_FIJJAFBF_230032 | Bacteria | Proteobacteri | Alphaproteob | Rhodobactera | Rhodobactera | Lentibacter | Lentibacter al | unclassified | unclassified | unclassified | unclassified | unclassified | 0,2 | 0,4 | 8,91E-06 | 1,78E-05 | 0 - | 0 - | 1 - | 4,45E-05 - | 0 - | 0 - | 0 - | 0 - | 0 - | 0 - |
| 3862 | 2 | 0.2um_NOKCPHJN_04268;3um_FIJJAFBF_54303 | Bacteria | Bacteroidetes | Cytophagia | Cytophagales | Flammeovirga | various | various | unclassified | unclassified | unclassified | unclassified | unclassified | 0,2 | 0,4 | 6,36E-06 | 1,27E-05 | 1 - | 3,18E-05 - | 0 - | 0 - | 0 - | 0 - | 0 - | 0 - | 0 - | 0 - |
| 3863 | 1 | 0.2um_NOKCPHJN_01097 | Bacteria | Bacteroidetes | Flavobacteriia | Flavobacterial | Flavobacteriac | Lacinutrix | Lacinutrix algi | metabolism | Amino acid tra | 05C65 | Catalyzes the | 983544.Lacal_ | 0,6 | 0,489897949 | 1,86E-05 | 1,53E-05 | 1 - | 2,71E-05 - | 1 - | 3,37E-05 - | 1 - | 3,20E-05 - | 0 - | 0 - | 0 - | 0 - |
| 3864 | 1 | 0.2um_NOKCPHJN_196703 | Bacteria | Bacteroidetes | Flavobacteriia | Flavobacterial | Flavobacteriac | Polaribacter | Polaribacter s | information st | Translation, ril | 05CAV | thus facilitatin | 313594.PI23P_ | 1,2 | 0,4 | 2,64E-05 | 9,09E-06 | 1 - | 1,84E-05 - | 1 - | 2,28E-05 - | 1 - | 2,17E-05 - | 2 - | 4,41E-05 - | 1 - | 2,48E-05 - |
| 3865 | 1 | 3um_FIJJAFBF_644311 | Bacteria | Bacteroidetes | Flavobacteriia | Flavobacterial | Flavobacteriac | Formosa | Formosa sp. H | unclassified | unclassified | unclassified | unclassified | unclassified | 3,4 | 1,019803903 | 7,26E-05 | 1,77E-05 | 5 - | 9,09E-05 - | 4 - | 9,03E-05 - | 3 - | 6,44E-05 - | 2 - | 4,36E-05 - | 3 - | 7,36E-05 - |
| 3866 | 1 | 3um_FIJJAFBF_792408 | Bacteria | Proteobacteri | Gammaprotec | Oceanospirilla | Saccharospirill | Reinekea | Reinekea fors | poorly charact | Function unkn | 178PC | Domain of Un | 314283.MED2 | 1,2 | 0,979795897 | 3,32E-05 | 2,82E-05 | 1 - | 2,29E-05 - | 3 - | 8,53E-05 - | 1 - | 2,70E-05 - | 0 - | 0 - | 1 - | 3,09E-05 - |
| 3867 | 1 | 3um_FIJJAFBF_20059 | Bacteria | Bacteroidetes | Flavobacteriia | Flavobacterial | Flavobacteriac | Polaribacter | Polaribacter s | unclassified | unclassified | unclassified | unclassified | unclassified | 2,2 | 0,4 | 0,000200682 | 1,84E-05 | 3 - | 0,000232656 - | 2 - | 0,000192593 - | 2 - | 0,000183084 - | 2 - | 0,000185788 - | 2 - | 0,00020929 - |
| 3868 | 1 | 3um_FIJJAFBF_95709 | Eukaryota | unclassified | Florideophyce | Gelidiales | Gelidiaceae | Gelidium | Gelidium eleg | unclassified | unclassified | unclassified | unclassified | unclassified | 0,2 | 0,4 | 2,34E-05 | 4,68E-05 | 1 - | 0,000117119 - | 0 - | 0 - | 0 - | 0 - | 0 - | 0 - | 0 - | 0 - |
| 3869 | 1 | 0.2um_NOKCPHJN_114364 | Bacteria | Proteobacteri | Alphaproteob | Rhizobiales | unclassified | unclassified | Rhizobiales ba | cellular proces | Cell wall/mem | 101WM | NA | 626418.bglu_ | 4 | 0,632455532 | 0,000129825 | 1,73E-05 | 5 - | 0,000136965 - | 4 - | 0,000136056 - | 3 - | 9,70E-05 - | 4 - | 0,000131249 - | 4 - | 0,000147852 - |
| 3870 | 1 | 0.2um_NOKCPHJN_144343 | Bacteria | Proteobacteri | Alphaproteob | Rhodobactera | Rhodobactera | Planktomarina | Planktomarina | unclassified | unclassified | unclassified | unclassified | unclassified | 0,4 | 0,489897949 | 1,69E-05 | 2,08E-05 | 1 - | 3,88E-05 - | 0 - | 0 - | 1 - | 4,58E-05 - | 0 - | 0 - | 0 - | 0 - |
| 3871 | 1 | 3um_FIJJAFBF_22423 | Bacteria | Bacteroidetes | Flavobacteriia | Flavobacterial | unclassified | unclassified | Flavobacterial | unclassified | unclassified | unclassified | unclassified | unclassified | 0,8 | 0,4 | 2,65E-05 | 1,36E-05 | 1 - | 2,77E-05 - | 1 - | 3,44E-05 - | 1 - | 3,27E-05 - | 0 - | 0 - | 1 - | 3,74E-05 - |
| 3872 | 1 | 3um_FIJJAFBF_143871 | Bacteria | Proteobacteri | Gammaprotec | Oceanospirilla | Saccharospirill | Reinekea | Reinekea fors | unclassified | unclassified | unclassified | unclassified | unclassified | 4,4 | 0,489897949 | 0,000234256 | 3,39E-05 | 5 - | 0,000223301 - | 4 - | 0,000221819 - | 4 - | 0,000210867 - | 4 - | 0,000213982 - | 5 - | 0,000301312 - |
| 3873 | 1 | 3um_FIJJAFBF_135085 | unclassified | unclassified | unclassified | unclassified | unclassified | unclassified | unclassified | unclassified | unclassified | unclassified | unclassified | unclassified | 8,4 | 1,854723699 | 0,000109468 | 1,94E-05 | 10 - | 0,000111003 - | 9 - | 0,000124049 - | 10 - | 0,000131027 - | 8 - | 0,00010637 - | 5 - | 7,49E-05 - |
| 3874 | 1 | 0.2um_NOKCPHJN_676059 | Bacteria | Proteobacteri | Alphaproteob | Pelagibacteral | Pelagibacterac | unclassified | Pelagibacterac | metabolism | Inorganic ion t | 05PWN | phosphate ab | 439493.PB721 | 0,8 | 0,4 | 3,09E-05 | 1,57E-05 | 1 - | 3,35E-05 - | 1 - | 4,16E-05 - | 1 - | 3,95E-05 - | 1 - | 4,01E-05 - | 0 - | 0 - |
| 3875 | 1 | 3um_FIJJAFBF_393745 | Bacteria | Bacteroidetes | Flavobacteriia | unclassified | unclassified | unclassified | Flavobacteria | unclassified | unclassified | unclassified | unclassified | unclassified | 7,6 | 0,8 | 0,000419848 | 6,05E-05 | 8 - | 0,000370248 - | 8 - | 0,000459738 - | 8 - | 0,000437039 - | 6 - | 0,000332621 - | 8 - | 0,000499595 - |
| 3876 | 1 | 0.2um_NOKCPHJN_209213 | Bacteria | Proteobacteri | Alphaproteob | Rhodobactera | unclassified | unclassified | Rhodobactera | unclassified | unclassified | unclassified | unclassified | unclassified | 1 | 0,632455532 | 4,38E-05 | 3,08E-05 | 1 - | 3,56E-05 - | 1 - | 4,43E-05 - | 0 - | 0 - | 1 - | 4,27E-05 - | 2 - | 9,62E-05 - |
| 3877 | 2 | 0.2um_NOKCPHJN_199426;3um_FIJJAFBF_4517 | Bacteria | Proteobacteri | Gammaprotec | Cellvibrionales | Porticoccacea | unclassified | Gammaprotec | unclassified | unclassified | unclassified | unclassified | unclassified | 0,8 | 0,4 | 4,29E-05 | 2,21E-05 | 1 - | 4,48E-05 - | 1 - | 5,57E-05 - | 0 - | 0 - | 1 - | 5,37E-05 - | 1 - | 6,05E-05 - |
| 3878 | 1 | 3um_FIJJAFBF_498449 | Bacteria | Proteobacteri | Gammaprotec | Arenicellales | Arenicellaceae | Arenicella | Arenicella xan | unclassified | unclassified | unclassified | unclassified | unclassified | 6,8 | 0,748331477 | 0,00011238 | 1,63E-05 | 6 - | 8,30E-05 - | 7 - | 0,000120197 - | 7 - | 0,000114262 - | 8 - | 0,000132514 - | 6 - | 0,000111957 - |
| 3879 | 1 | 3um_FIJJAFBF_209608 | Bacteria | Bacteroidetes | Flavobacteriia | Flavobacterial | Flavobacteriac | Tamlana | various | unclassified | unclassified | unclassified | unclassified | unclassified | 1 | 0 | 9,86E-05 | 9,36E-06 | 1 - | 8,26E-05 - | 1 - | 0,000102532 - | 1 - | 9,75E-05 - | 1 - | 9,89E-05 - | 1 - | 0,00011142 - |
| 3880 | 1 | 3um_FIJJAFBF_129173 | Bacteria | Bacteroidetes | Flavobacteriia | Flavobacterial | Flavobacteriac | Polaribacter | Polaribacter s | unclassified | unclassified | unclassified | unclassified | unclassified | 13,2 | 2,13541565 | 0,000187887 | 2,11E-05 | 16 - | 0,000193105 - | 15 - | 0,000224793 - | 13 - | 0,000185202 - | 12 - | 0,000173481 - | 10 - | 0,000162854 - |
| 3881 | 1 | 0.2um_NOKCPHJN_185874 | Bacteria | Bacteroidetes | Flavobacteriia | Flavobacterial | Flavobacteriac | Formosa | Formosa sp. H | cellular proces | Intracellular tr | 09S7I | Preprotein tra | 313595.P7007 | 0,8 | 0,4 | 8,97E-05 | 4,61E-05 | 1 - | 9,49E-05 - | 0 - | 0 - | 1 - | 0,000111969 - | 1 - | 0,000113623 - | 1 - | 0,000127995 - |
| 3882 | 1 | 3um_FIJJAFBF_118415 | Viruses | unclassified | unclassified | unclassified | Phycodnavirid | unclassified | Aureococcus a | unclassified | unclassified | unclassified | unclassified | unclassified | 5,2 | 0,4 | 0,000148073 | 1,71E-05 | 5 - | 0,000119311 - | 5 - | 0,000148149 - | 6 - | 0,000169001 - | 5 - | 0,000142914 - | 5 - | 0,000160992 - |
| 3883 | 1 | 0.2um_NOKCPHJN_403801 | Bacteria | Proteobacteri | Alphaproteob | Rhodobactera | Rhodobactera | Pseudooceani | Pseudooceani | cellular proces | Intracellular tr | COG0653 | Part of the Se | 314265.R2601 | 0,8 | 0,748331477 | 1,65E-05 | 1,53E-05 | 0 - | 0 - | 2 - | 4,07E-05 - | 0 - | 0 - | 1 - | 1,96E-05 - | 1 - | 2,21E-05 - |
| 3884 | 1 | 3um_FIJJAFBF_460430 | Bacteria | Bacteroidetes | Flavobacteriia | Flavobacterial | Flavobacteriac | Tenacibaculur | Tenacibaculur | information st | Replication, re | 0PB0H | DNA-binding p | 313598.MED1 | 10 | 1,788854382 | 0,001510576 | 0,000237766 | 13 - | 0,001657887 - | 9 - | 0,001425189 - | 9 - | 0,001354822 - | 8 - | 0,001222074 - | 11 - | 0,00189291 - |
| 3885 | 1 | 3um_FIJJAFBF_149168 | Bacteria | Proteobacteri | Alphaproteob | Rhodobactera | Rhodobactera | Lentibacter | Lentibacter al | information st | Translation, ril | COG0050 | This protein p | 314265.R2601 | 12,8 | 1,939071943 | 0,000453343 | 9,99E-05 | 10 - | 0,000293547 - | 15 - | 0,000546748 - | 14 - | 0,000485103 - | 11 - | 0,000386782 - | 14 - | 0,000554538 - |
| 3886 | 1 | 0.2um_NOKCPHJN_105386 | Bacteria | Proteobacteri | Gammaprotec | Oceanospirilla | Saccharospirill | Reinekea | Reinekea blan | metabolism | Amino acid tra | 16Q4U | Catalyzes the | 314283.MED2 | 5,4 | 1,019803903 | 0,000174121 | 2,69E-05 | 6 - | 0,000163967 - | 5 - | 0,000169665 - | 7 - | 0,000225804 - | 5 - | 0,000163671 - | 4 - | 0,0001475 - |
| 3887 | 1 | 0.2um_NOKCPHJN_217330 | Bacteria | Proteobacteri | Alphaproteob | Rhodobactera | Rhodobactera | Amylibacter | Amylibacter k | unclassified | unclassified | unclassified | unclassified | unclassified | 0,4 | 0,489897949 | 9,54E-06 | 1,18E-05 | 1 - | 2,17E-05 - | 0 - | 0 - | 0 - | 0 - | 1 - | 2,60E-05 - | 0 - | 0 - |
| 3888 | 1 | 3um_FIJJAFBF_800451 | Bacteria | Bacteroidetes | Flavobacteriia | Flavobacterial | Flavobacteriac | Polaribacter | Polaribacter s | information st | Translation, ril | 0P97G | This is one of t | 313598.MED1 | 0,2 | 0,4 | 1,12E-05 | 2,24E-05 | 1 - | 5,60E-05 - | 0 - | 0 - | 0 - | 0 - | 0 - | 0 - | 0 - | 0 - |
| 3889 | 1 | 3um_FIJJAFBF_193522 | Eukaryota | unclassified | Cryptophyta | Pyrenomonad | Geminigerace | Teleaulax | Teleaulax amp | metabolism | Energy produc | COG0056 | Produces ATP | 400682.PAC_1 | 0,2 | 0,4 | 5,30E-06 | 1,06E-05 | 0 - | 0 - | 0 - | 0 - | 1 - | 2,65E-05 - | 0 - | 0 - | 0 - | 0 - |
| 3890 | 1 | 3um_FIJJAFBF_310410 | Bacteria | Proteobacteri | Gammaprotec | Alteromonada | Alteromonada | Paraglaciecola | Paraglaciecola | metabolism | Inorganic ion t | 05F28 | tonB-depende | 983545.Glaag_ | 7 | 1,095445115 | 0,000122111 | 1,92E-05 | 7 - | 0,000102741 - | 7 - | 0,000127574 - | 9 - | 0,000155926 - | 6 - | 0,000105486 - | 6 - | 0,00011883 - |
| 3891 | 1 | 0.2um_NOKCPHJN_386012 | Bacteria | Proteobacteri | Alphaproteob | Pelagibacteral | Pelagibacterac | Candidatus Pe | Candidatus Pe | unclassified | unclassified | unclassified | unclassified | unclassified | 0,4 | 0,489897949 | 9,36E-06 | 1,16E-05 | 1 - | 2,09E-05 - | 1 - | 2,59E-05 - | 0 - | 0 - | 0 - | 0 - | 0 - | 0 - |
| 3892 | 1 | 0.2um_NOKCPHJN_469361 | Bacteria | Bacteroidetes | unclassified | unclassified | unclassified | unclassified | unidentified e | unclassified | unclassified | unclassified | unclassified | unclassified | 1 | 0 | 1,07E-05 | 1,02E-06 | 1 - | 8,97E-06 - | 1 - | 1,11E-05 - | 1 - | 1,06E-05 - | 1 - | 1,07E-05 - | 1 - | 1,21E-05 - |
| 3893 | 1 | 3um_FIJJAFBF_25780 | Bacteria | Proteobacteri | Gammaprotec | Alteromonada | Alteromonada | Alteromonas | various | metabolism | Amino acid tra | COG0174 | glutamine syn | 314275.MADE | 2,6 | 0,489897949 | 7,53E-05 | 1,27E-05 | 3 - | 7,34E-05 - | 3 - | 9,12E-05 - | 2 - | 5,78E-05 - | 3 - | 8,79E-05 - | 2 - | 6,60E-05 - |
| 3894 | 1 | 3um_FIJJAFBF_636610 | Bacteria | unclassified | unclassified | unclassified | unclassified | unclassified | uncultured ba | unclassified | unclassified | unclassified | unclassified | unclassified | 0,8 | 0,748331477 | 5,81E-05 | 5,60E-05 | 1 - | 6,38E-05 - | 0 - | 0 - | 2 - | 0,000150536 - | 1 - | 7,64E-05 - | 0 - | 0 - |
| 3895 | 1 | 0.2um_NOKCPHJN_270882 | Bacteria | Proteobacteri | Betaproteoba | unclassified | unclassified | unclassified | Betaproteoba | unclassified | unclassified | unclassified | unclassified | unclassified | 1 | 0 | 0,000165093 | 1,57E-05 | 1 - | 0,000138285 - | 1 - | 0,000171709 - | 1 - | 0,000163232 - | 1 - | 0,000165643 - | 1 - | 0,000186596 - |
| 3896 | 1 | 3um_FIJJAFBF_491681 | Bacteria | Proteobacteri | Gammaprotec | Cellvibrionales | Porticoccacea | unclassified | SAR92 bacteri | unclassified | unclassified | unclassified | unclassified | unclassified | 2 | 0 | 0,000162162 | 1,54E-05 | 2 - | 0,00013583 - | 2 - | 0,000168661 - | 2 - | 0,000160334 - | 2 - | 0,000162702 - | 2 - | 0,000183283 - |
| 3897 | 1 | 0.2um_NOKCPHJN_672660 | Bacteria | Bacteroidetes | Flavobacteriia | Flavobacterial | Flavobacteriac | Ulvibacter | Ulvibacter lito | unclassified | unclassified | unclassified | unclassified | unclassified | 2,2 | 0,4 | 7,09E-05 | 6,50E-06 | 3 - | 8,22E-05 - | 2 - | 6,80E-05 - | 2 - | 6,47E-05 - | 2 - | 6,56E-05 - | 2 - | 7,39E-05 - |
| 3898 | 1 | 3um_FIJJAFBF_199396 | Bacteria | Bacteroidetes | Flavobacteriia | Flavobacterial | Crocinitomica | unclassified | Crocinitomica | metabolism | Nucleotide tra | 0P7DG | Catalyzes the | 398720.MED2 | 0,6 | 0,489897949 | 4,35E-05 | 3,55E-05 | 0 - | 0 - | 1 - | 7,46E-05 - | 1 - | 7,09E-05 - | 1 - | 7,20E-05 - | 0 - | 0 - |
| 3899 | 1 | 0.2um_NOKCPHJN_208663 | Bacteria | Proteobacteri | Alphaproteob | Rhodobactera | Rhodobactera | Sulfitobacter | Sulfitobacter s | poorly charact | Function unkn | COG3181 | extra-cytoplas | 375451.RD1_ | 1,4 | 0,8 | 5,21E-05 | 2,89E-05 | 2 - | 6,36E-05 - | 2 - | 7,90E-05 - | 2 - | 7,51E-05 - | 0 - | 0 - | 1 - | 4,29E-05 - |
| 3900 | 1 | 3um_FIJJAFBF_66407 | Bacteria | Proteobacteri | Alphaproteob | Rhodobactera | unclassified | unclassified | Rhodobactera | unclassified | unclassified | unclassified | unclassified | unclassified | 1,6 | 0,8 | 4,11E-05 | 1,63E-05 | 3 - | 6,74E-05 - | 1 - | 2,79E-05 - | 2 - | 5,30E-05 - | 1 - | 2,69E-05 - | 1 - | 3,03E-05 - |
| 3901 | 1 | 3um_FIJJAFBF_60256 | Eukaryota | unclassified | unclassified | unclassified | unclassified | Ancoracysta | Ancoracysta t | unclassified | unclassified | unclassified | unclassified | unclassified | 2,4 | 0,8 | 0,000243415 | 8,37E-05 | 2 - | 0,000171309 - | 3 - | 0,000319072 - | 3 - | 0,000303318 - | 3 - | 0,000307798 - | 1 - | 0,000115578 - |
| 3902 | 1 | 3um_FIJJAFBF_177511 | Bacteria | Bacteroidetes | Flavobacteriia | Flavobacterial | Flavobacteriac | Aureitalea | Aureitalea ma | metabolism | Inorganic ion t | 05C27 | TonB-depende | 398720.MED2 | 2,6 | 0,489897949 | 3,50E-05 | 5,91E-06 | 3 - | 3,41E-05 - | 3 - | 4,24E-05 - | 2 - | 2,69E-05 - | 3 - | 4,09E-05 - | 2 - | 3,07E-05 - |
| 3903 | 1 | 0.2um_NOKCPHJN_292785 | Eukaryota | Bacillariophyt | Bacillariophyc | Bacillariales | Bacillariaceae | Fragilariopsis | Fragilariopsis | unclassified | unclassified | unclassified | unclassified | unclassified | 0,4 | 0,489897949 | 1,91E-06 | 2,38E-06 | 1 - | 4,06E-06 - | 0 - | 0 - | 0 - | 0 - | 0 - | 0 - | 1 - | 5,47E-06 - |
| 3904 | 1 | 3um_FIJJAFBF_146669 | Bacteria | Bacteroidetes | Flavobacteriia | Flavobacterial | Flavobacteriac | Polaribacter | Polaribacter s | unclassified | unclassified | unclassified | unclassified | unclassified | 1 | 0,632455532 | 3,20E-05 | 2,07E-05 | 1 - | 2,76E-05 - | 1 - | 3,43E-05 - | 2 - | 6,51E-05 - | 1 - | 3,30E-05 - | 0 - | 0 - |
| 3905 | 1 | 0.2um_NOKCPHJN_1042585 | Bacteria | Proteobacteri | Alphaproteob | Rhodospirillal | Rhodospirillac | unclassified | Rhodospirillac | poorly charact | Function unkn | 0XPG5 | KH domain co | 443218.AS9A_ | 0,8 | 0,4 | 1,92E-05 | 9,74E-06 | 1 - | 2,08E-05 - | 1 - | 2,58E-05 - | 1 - | 2,45E-05 - | 1 - | 2,49E-05 - | 0 - | 0 - |
| 3906 | 1 | 3um_FIJJAFBF_194945 | Bacteria | Proteobacteri | Gammaprotec | Alteromonada | Colwelliaceae | Thalassotalea | Thalassotalea | information st | Translation, ril | 177SZ | This protein bi | 167879.CPS_4 | 1,4 | 0,489897949 | 0,000131746 | 3,77E-05 | 2 - | 0,000161657 - | 1 - | 0,000100365 - | 2 - | 0,00019082 - | 1 - | 9,68E-05 - | 1 - | 0,000109067 - |
| 3907 | 1 | 3um_FIJJAFBF_234946 | Eukaryota | Bacillariophyt | Bacillariophyc | Bacillariales | Bacillariaceae | Fragilariopsis | Fragilariopsis | unclassified | unclassified | unclassified | unclassified | unclassified | 0,2 | 0,4 | 1,73E-05 | 3,45E-05 | 1 - | 8,63E-05 - | 0 - | 0 - | 0 - | 0 - | 0 - | 0 - | 0 - | 0 - |
| 3908 | 1 | 0.2um_NOKCPHJN_955130 | Eukaryota | Bacillariophyt | Bacillariophyc | Naviculales | Naviculaceae | Fistulifera | Fistulifera sola | metabolism | Energy produc | COG0056 | Produces ATP | 13037.EHJ650 | 5,6 | 0,8 | 0,000192779 | 1,73E-05 | 7 - | 0,000203918 - | 6 - | 0,000217034 - | 5 - | 0,000171932 - | 5 - | 0,000174471 - | 5 - | 0,000196541 - |
| 3909 | 1 | 3um_FIJJAFBF_332923 | Bacteria | Bacteroidetes | unclassified | unclassified | unclassified | unclassified | Bacteroidetes | unclassified | unclassified | unclassified | unclassified | unclassified | 1 | 0 | 0,000150579 | 1,43E-05 | 1 - | 0,000126128 - | 1 - | 0,000156614 - | 1 - | 0,000148882 - | 1 - | 0,000151081 - | 1 - | 0,000170192 - |
| 3910 | 1 | 3um_FIJJAFBF_167550 | Bacteria | Proteobacteri | Gammaprotec | unclassified | unclassified | unclassified | Gammaprotec | unclassified | unclassified | unclassified | unclassified | unclassified | 0,4 | 0,489897949 | 3,39E-05 | 4,19E-05 | 1 - | 7,55E-05 - | 1 - | 9,38E-05 - | 0 - | 0 - | 0 - | 0 - | 0 - | 0 - |
| 3911 | 1 | 3um_FIJJAFBF_339167 | Bacteria | Bacteroidetes | Flavobacteriia | Flavobacterial | Flavobacteriac | Polaribacter | Polaribacter s | cellular proces | Cell wall/mem | 11TPU | outer membra | 313594.PI23P_ | 10,2 | 2,039607805 | 0,000781317 | 0,00012859 | 13 - | 0,000842993 - | 8 - | 0,000644153 - | 8 - | 0,000612349 - | 12 - | 0,00093209 - | 10 - | 0,000874997 - |
| 3912 | 1 | 3um_FIJJAFBF_652763 | Bacteria | Bacteroidetes | Cytophagia | Cytophagales | Flammeovirga | unclassified | Flammeovirga | unclassified | unclassified | unclassified | unclassified | unclassified | 1,8 | 0,748331477 | 0,00014335 | 6,89E-05 | 2 - | 0,000131927 - | 1 - | 8,19E-05 - | 1 - | 7,79E-05 - | 2 - | 0,000158027 - | 3 - | 0,000267025 - |
| 3913 | 1 | 0.2um_NOKCPHJN_186098 | Bacteria | Bacteroidetes | Cytophagia | Cytophagales | Flammeovirga | Reichenbachie | Reichenbachie | unclassified | unclassified | unclassified | unclassified | unclassified | 16,2 | 1,166190379 | 0,000402327 | 2,71E-05 | 18 - | 0,000376317 - | 17 - | 0,000441315 - | 15 - | 0,00037017 - | 16 - | 0,00040068 - | 15 - | 0,000423154 - |
| 3914 | 1 | 3um_FIJJAFBF_510255 | Bacteria | Bacteroidetes | Flavobacteriia | Flavobacterial | Flavobacteriac | Mangrovimon | Mangrovimon | unclassified | unclassified | unclassified | unclassified | unclassified | 0,4 | 0,489897949 | 1,53E-05 | 1,89E-05 | 1 - | 3,48E-05 - | 0 - | 0 - | 0 - | 0 - | 1 - | 4,17E-05 - | 0 - | 0 - |
| 3915 | 1 | 0.2um_NOKCPHJN_717701 | Bacteria | Proteobacteri | Alphaproteob | Pelagibacteral | unclassified | unclassified | Pelagibacteral | cellular proces | Cell wall/mem | 7257 | NA | 335992.SAR11 | 2 | 0 | 0,000114189 | 1,08E-05 | 2 - | 9,56E-05 - | 2 - | 0,000118766 - | 2 - | 0,000112902 - | 2 - | 0,000114569 - | 2 - | 0,000129062 - |
| 3916 | 1 | 3um_FIJJAFBF_608092 | Bacteria | Cyanobacteria | unclassified | Synechococca | Synechococca | Synechococcu | Synechococcu | unclassified | unclassified | unclassified | unclassified | unclassified | 0,2 | 0,4 | 3,78E-06 | 7,56E-06 | 1 - | 1,89E-05 - | 0 - | 0 - | 0 - | 0 - | 0 - | 0 - | 0 - | 0 - |
| 3917 | 1 | 0.2um_NOKCPHJN_25283 | Bacteria | unclassified | unclassified | unclassified | unclassified | unclassified | uncultured ba | unclassified | unclassified | unclassified | unclassified | unclassified | 11,8 | 0,748331477 | 0,000169348 | 9,70E-06 | 13 - | 0,000157063 - | 12 - | 0,000180024 - | 12 - | 0,000171135 - | 11 - | 0,000159191 - | 11 - | 0,000179328 - |
| 3918 | 1 | 3um_FIJJAFBF_326249 | Bacteria | Proteobacteri | Alphaproteob | Rhodobactera | Rhodobactera | unclassified | Rhodobactera | metabolism | Inorganic ion t | 01QSG | Zinc mangane | 314262.MED1 | 0,6 | 0,8 | 2,40E-05 | 3,31E-05 | 1 - | 3,56E-05 - | 0 - | 0 - | 2 - | 8,42E-05 - | 0 - | 0 - | 0 - | 0 - |
| 3919 | 1 | 0.2um_NOKCPHJN_337648 | Bacteria | Proteobacteri | Alphaproteob | Rhodobactera | Rhodobactera | Amylibacter | Amylibacter k | unclassified | unclassified | unclassified | unclassified | unclassified | 1,2 | 0,748331477 | 5,00E-05 | 2,99E-05 | 2 - | 7,36E-05 - | 1 - | 4,57E-05 - | 2 - | 8,68E-05 - | 1 - | 4,41E-05 - | 0 - | 0 - |
| 3920 | 1 | 3um_FIJJAFBF_385424 | Bacteria | Proteobacteri | Betaproteoba | Nitrosomonac | unclassified | unclassified | various | information st | Translation, ril | COG0051 | tRNA binding | 383631.MB21 | 1,4 | 0,489897949 | 0,000183803 | 5,41E-05 | 2 - | 0,000225052 - | 1 - | 0,000139724 - | 1 - | 0,000132826 - | 2 - | 0,000269575 - | 1 - | 0,000151838 - |
| 3921 | 1 | 0.2um_NOKCPHJN_902273 | Viruses | unclassified | unclassified | Caudovirales | Podoviridae | unclassified | Cellulophaga p | unclassified | unclassified | unclassified | unclassified | unclassified | 0,2 | 0,4 | 3,97E-06 | 7,95E-06 | 0 - | 0 - | 0 - | 0 - | 1 - | 1,99E-05 - | 0 - | 0 - | 0 - | 0 - |
| 3922 | 4 | 0.2um_NOKCPHJN_141739;0.2um_NOKCPHJN_5 | Bacteria | Proteobacteri | Gammaprotec | Cellvibrionales | Porticoccacea | unclassified | various | unclassified | unclassified | unclassified | unclassified | unclassified | 1 | 0 | 7,61E-05 | 7,23E-06 | 1 - | 6,38E-05 - | 1 - | 7,92E-05 - | 1 - | 7,53E-05 - | 1 - | 7,64E-05 - | 1 - | 8,60E-05 - |
| 3923 | 1 | 3um_FIJJAFBF_85182 | Bacteria | Proteobacteri | Gammaprotec | Oceanospirilla | Oceanospirilla | Amphritea | Amphritea jap | information st | Replication, re | 05C68 | Can catalyze t | 207954.MED9 | 0,4 | 0,489897949 | 1,72E-05 | 2,11E-05 | 0 - | 0 - | 1 - | 4,13E-05 - | 0 - | 0 - | 0 - | 0 - | 1 - | 4,49E-05 - |
| 3924 | 1 | 3um_FIJJAFBF_174305 | Bacteria | Proteobacteri | Gammaprotec | Oceanospirilla | Saccharospirill | Reinekea | Reinekea fors | cellular proces | Posttranslatio | 0QHTY | protease | 314283.MED2 | 1,6 | 1,019803903 | 4,60E-05 | 2,98E-05 | 2 - | 4,99E-05 - | 2 - | 6,20E-05 - | 3 - | 8,84E-05 - | 1 - | 2,99E-05 - | 0 - | 0 - |
| 3925 | 1 | 3um_FIJJAFBF_207711 | Bacteria | Proteobacteri | Gammaprotec | Cellvibrionales | Porticoccacea | unclassified | SAR92 bacteri | unclassified | unclassified | unclassified | unclassified | unclassified | 1,4 | 0,8 | 5,04E-05 | 3,07E-05 | 1 - | 2,95E-05 - | 2 - | 7,33E-05 - | 2 - | 6,97E-05 - | 0 - | 0 - | 2 - | 7,96E-05 - |
| 3926 | 1 | 0.2um_NOKCPHJN_195816 | Bacteria | Proteobacteri | Gammaprotec | Oceanospirilla | Saccharospirill | Reinekea | Reinekea fors | metabolism | Energy produc | 16PVJ | Component of | 314283.MED2 | 11 | 2,097617696 | 0,000167961 | 1,95E-05 | 15 - | 0,000194537 - | 11 - | 0,000177142 - | 9 - | 0,000137779 - | 10 - | 0,000155348 - | 10 - | 0,000174999 - |
| 3927 | 1 | 3um_FIJJAFBF_126833 | Eukaryota | unclassified | unclassified | unclassified | unclassified | unclassified | uncultured pr | metabolism | Energy produc | COG0711 | F(1)F(0) ATP s | 32049.SYNPC | 0,4 | 0,489897949 | 3,52E-05 | 4,31E-05 | 0 - | 0 - | 1 - | 9,02E-05 - | 1 - | 8,57E-05 - | 0 - | 0 - | 0 - | 0 - |
| 3928 | 1 | 3um_FIJJAFBF_71177 | Bacteria | Bacteroidetes | unclassified | unclassified | unclassified | unclassified | Bacteroidetes | unclassified | unclassified | unclassified | unclassified | unclassified | 0,4 | 0,489897949 | 6,18E-06 | 7,62E-06 | 1 - | 1,41E-05 - | 0 - | 0 - | 0 - | 0 - | 1 - | 1,68E-05 - | 0 - | 0 - |
| 3929 | 1 | 0.2um_NOKCPHJN_45872 | Bacteria | Bacteroidetes | Flavobacteriia | Flavobacterial | Crocinitomica | unclassified | Crocinitomica | poorly charact | Function unkn | 09AR5 | Inherit from b | 313606.M231 | 1,6 | 0,489897949 | 3,93E-05 | 1,11E-05 | 2 - | 4,18E-05 - | 2 - | 5,19E-05 - | 1 - | 2,47E-05 - | 2 - | 5,01E-05 - | 1 - | 2,82E-05 - |
| 3930 | 1 | 0.2um_NOKCPHJN_1097228 | Eukaryota | Platyhelminth | unclassified | Macrostomida | Macrostomida | Macrostomun | Macrostomun | unclassified | unclassified | unclassified | unclassified | unclassified | 0,4 | 0,489897949 | 1,33E-05 | 1,64E-05 | 1 - | 3,04E-05 - | 0 - | 0 - | 1 - | 3,59E-05 - | 0 - | 0 - | 0 - | 0 - |
| 3931 | 2 | 3um_FIJJAFBF_389124;3um_FIJJAFBF_663790 | Bacteria | Bacteroidetes | Flavobacteriia | Flavobacterial | Flavobacteriac | Polaribacter | Polaribacter h | metabolism | Carbohydrate | COG1109 | Catalyzes the | 313594.PI23P_ | 2 | 0 | 5,92E-05 | 5,62E-06 | 2 - | 4,96E-05 - | 2 - | 6,16E-05 - | 2 - | 5,85E-05 - | 2 - | 5,94E-05 - | 2 - | 6,69E-05 - |
| 3932 | 1 | 3um_FIJJAFBF_36487 | Bacteria | Proteobacteri | Gammaprotec | Alteromonada | Alteromonada | Paraglaciecola | Paraglaciecola | unclassified | unclassified | unclassified | unclassified | unclassified | 1,8 | 0,979795897 | 0,000298022 | 0,000165259 | 2 - | 0,000276571 - | 2 - | 0,000343419 - | 0 - | 0 - | 3 - | 0,000496928 - | 2 - | 0,000373192 - |
| 3933 | 1 | 3um_FIJJAFBF_408398 | Bacteria | Bacteroidetes | Flavobacteriia | Flavobacterial | Flavobacteriac | Kordia | Kordia sp. SM | cellular proces | Intracellular tr | 05FGH | MotA TolQ ex | 391587.KAOT | 0,2 | 0,4 | 9,77E-06 | 1,95E-05 | 1 - | 4,88E-05 - | 0 - | 0 - | 0 - | 0 - | 0 - | 0 - | 0 - | 0 - |
| 3934 | 1 | 3um_FIJJAFBF_697470 | Bacteria | Bacteroidetes | Cytophagia | Cytophagales | Flammeovirga | unclassified | Flammeovirga | metabolism | Lipid transpor | 09EIU | Acetyl-CoA ca | 643867.Ftrac_ | 0,4 | 0,489897949 | 3,50E-05 | 4,37E-05 | 1 - | 7,45E-05 - | 0 - | 0 - | 0 - | 0 - | 0 - | 0 - | 1 - | 0,000100568 - |
| 3935 | 1 | 3um_FIJJAFBF_99324 | Bacteria | Bacteroidetes | Flavobacteriia | Flavobacterial | Flavobacteriac | unclassified | various | metabolism | Inorganic ion t | 05C27 | TonB-depende | 313598.MED1 | 25,4 | 0,8 | 0,000349922 | 2,90E-05 | 26 - | 0,000300523 - | 24 - | 0,000344456 - | 26 - | 0,000354737 - | 26 - | 0,000359976 - | 25 - | 0,000389916 - |
| 3936 | 1 | 3um_FIJJAFBF_122410 | Bacteria | Bacteroidetes | Flavobacteriia | Flavobacterial | Flavobacteriac | Polaribacter | Polaribacter s | cellular proces | Posttranslatio | 08V1J | Has antioxidar | 313594.PI23P_ | 1,8 | 1,166190379 | 0,000142059 | 7,60E-05 | 4 - | 0,000278247 - | 2 - | 0,00017275 - | 1 - | 8,21E-05 - | 1 - | 8,33E-05 - | 1 - | 9,39E-05 - |
| 3937 | 1 | 3um_FIJJAFBF_175115 | Bacteria | Proteobacteri | Alphaproteob | Rhodobactera | Rhodobactera | unclassified | Rhodobactera | unclassified | unclassified | unclassified | unclassified | unclassified | 0,6 | 0,489897949 | 1,93E-05 | 1,59E-05 | 1 - | 2,86E-05 - | 0 - | 0 - | 1 - | 3,37E-05 - | 1 - | 3,42E-05 - | 0 - | 0 - |
| 3938 | 1 | 3um_FIJJAFBF_410100 | Bacteria | Bacteroidetes | unclassified | unclassified | unclassified | unclassified | Bacteroidetes | unclassified | unclassified | unclassified | unclassified | unclassified | 2,4 | 0,489897949 | 0,000134973 | 3,02E-05 | 3 - | 0,0001417 - | 2 - | 0,000117299 - | 2 - | 0,000111508 - | 2 - | 0,000113155 - | 3 - | 0,000191203 - |
| 3939 | 2 | 0.2um_NOKCPHJN_405836;3um_FIJJAFBF_3555 | Bacteria | Proteobacteri | Alphaproteob | Pelagibacteral | Pelagibacterac | Candidatus Pe | Candidatus Pe | poorly charact | Function unkn | 07FW0 | short-chain de | 335992.SAR11 | 0,2 | 0,4 | 1,07E-05 | 2,14E-05 | 0 - | 0 - | 0 - | 0 - | 1 - | 5,36E-05 - | 0 - | 0 - | 0 - | 0 - |
| 3940 | 1 | 3um_FIJJAFBF_352147 | Viruses | unclassified | unclassified | unclassified | unclassified | unclassified | uncultured M | unclassified | unclassified | unclassified | unclassified | unclassified | 0,2 | 0,4 | 1,09E-05 | 2,18E-05 | 1 - | 5,44E-05 - | 0 - | 0 - | 0 - | 0 - | 0 - | 0 - | 0 - | 0 - |
| 3941 | 1 | 3um_FIJJAFBF_316516 | Bacteria | Proteobacteri | Gammaprotec | Alteromonada | Alteromonada | Paraglaciecola | Paraglaciecola | unclassified | unclassified | unclassified | unclassified | unclassified | 5,4 | 1,2 | 8,63E-05 | 2,06E-05 | 4 - | 5,34E-05 - | 6 - | 9,95E-05 - | 7 - | 0,000110405 - | 6 - | 9,60E-05 - | 4 - | 7,21E-05 - |
| 3942 | 1 | 3um_FIJJAFBF_694666 | Bacteria | Bacteroidetes | unclassified | unclassified | unclassified | unclassified | Bacteroidetes | information st | Translation, ril | COG0089 | One of the ea | 926562.Oweh | 3 | 1,095445115 | 0,000415126 | 0,000109056 | 5 - | 0,000597796 - | 3 - | 0,000445371 - | 3 - | 0,000423382 - | 2 - | 0,000286424 - | 2 - | 0,000322655 - |
| 3943 | 1 | 3um_FIJJAFBF_32106 | Bacteria | Bacteroidetes | Flavobacteriia | Flavobacterial | Flavobacteriac | Polaribacter | Polaribacter t | poorly charact | Function unkn | 0P81C | conserved rep | 688270.Celal_ | 1,2 | 0,748331477 | 1,33E-05 | 7,94E-06 | 2 - | 1,96E-05 - | 1 - | 1,21E-05 - | 2 - | 2,31E-05 - | 1 - | 1,17E-05 - | 0 - | 0 - |
| 3944 | 1 | 0.2um_NOKCPHJN_27919 | Bacteria | Bacteroidetes | Flavobacteriia | Flavobacterial | Flavobacteriac | Formosa | Formosa sp. H | unclassified | unclassified | unclassified | unclassified | unclassified | 1,6 | 0,489897949 | 0,000127239 | 4,10E-05 | 1 - | 6,63E-05 - | 2 - | 0,000164762 - | 2 - | 0,000156627 - | 2 - | 0,00015894 - | 1 - | 8,95E-05 - |
| 3945 | 2 | 0.2um_NOKCPHJN_356919;3um_FIJJAFBF_37662 | Bacteria | Bacteroidetes | Flavobacteriia | Flavobacterial | Flavobacteriac | Ulvibacter | Ulvibacter sp. | unclassified | unclassified | unclassified | unclassified | unclassified | 1,4 | 1,019803903 | 0,0001239 | 8,65E-05 | 0 - | 0 - | 1 - | 9,02E-05 - | 3 - | 0,000257245 - | 2 - | 0,00017403 - | 1 - | 9,80E-05 - |
| 3946 | 1 | 3um_FIJJAFBF_509337 | Bacteria | Bacteroidetes | unclassified | unclassified | unclassified | unclassified | Bacteroidetes | unclassified | unclassified | unclassified | unclassified | unclassified | 0,8 | 0,748331477 | 9,32E-05 | 8,10E-05 | 2 - | 0,000204958 - | 0 - | 0 - | 0 - | 0 - | 1 - | 0,000122753 - | 1 - | 0,000138281 - |
| 3947 | 1 | 0.2um_NOKCPHJN_601504 | Bacteria | Proteobacteri | Epsilonproteo | Campylobacte | Helicobactera | Sulfurimonas | Sulfurimonas | cellular proces | Posttranslatio | 16QU3 | Hemolysin-typ | 744980.TRICH | 1 | 0 | 1,65E-05 | 1,56E-06 | 1 - | 1,38E-05 - | 1 - | 1,71E-05 - | 1 - | 1,63E-05 - | 1 - | 1,65E-05 - | 1 - | 1,86E-05 - |
| 3948 | 1 | 3um_FIJJAFBF_100610 | Bacteria | Proteobacteri | Alphaproteob | Rhodobactera | Rhodobactera | Litoreibacter | Litoreibacter a | unclassified | unclassified | unclassified | unclassified | unclassified | 1,6 | 1,2 | 8,22E-05 | 6,32E-05 | 1 - | 4,38E-05 - | 1 - | 5,44E-05 - | 3 - | 0,000155132 - | 3 - | 0,000157424 - | 0 - | 0 - |
| 3949 | 1 | 3um_FIJJAFBF_470658 | Eukaryota | unclassified | Cryptophyta | Pyrenomonad | Chroomonada | Chroomonas | Chroomonas r | metabolism | Energy produc | COG0056 | Produces ATP | 43989.cce_44 | 2,6 | 1,019803903 | 7,14E-05 | 2,97E-05 | 2 - | 4,57E-05 - | 1 - | 2,84E-05 - | 4 - | 0,000107954 - | 3 - | 8,22E-05 - | 3 - | 9,26E-05 - |
| 3950 | 1 | 0.2um_NOKCPHJN_487257 | Bacteria | unclassified | unclassified | unclassified | unclassified | unclassified | uncultured ba | unclassified | unclassified | unclassified | unclassified | unclassified | 1 | 0 | 4,54E-05 | 4,31E-06 | 1 - | 3,80E-05 - | 1 - | 4,72E-05 - | 1 - | 4,49E-05 - | 1 - | 4,55E-05 - | 1 - | 5,13E-05 - |
| 3951 | 2 | 0.2um_NOKCPHJN_601733;3um_FIJJAFBF_4114 | Bacteria | Proteobacteri | Alphaproteob | Rhodobactera | Rhodobactera | various | various | unclassified | unclassified | unclassified | unclassified | unclassified | 1,2 | 0,4 | 7,23E-05 | 2,42E-05 | 1 - | 5,06E-05 - | 1 - | 6,28E-05 - | 2 - | 0,000119368 - | 1 - | 6,06E-05 - | 1 - | 6,82E-05 - |
| 3952 | 1 | 0.2um_NOKCPHJN_521516 | Bacteria | Proteobacteri | Alphaproteob | Rhodobactera | unclassified | unclassified | Rhodobactera | unclassified | unclassified | unclassified | unclassified | unclassified | 1 | 0 | 3,70E-05 | 3,52E-06 | 1 - | 3,10E-05 - | 1 - | 3,85E-05 - | 1 - | 3,66E-05 - | 1 - | 3,72E-05 - | 1 - | 4,19E-05 - |
| 3953 | 1 | 0.2um_NOKCPHJN_15318 | Bacteria | Proteobacteri | Betaproteoba | unclassified | unclassified | unclassified | Betaproteoba | poorly charact | Function unkn | 0BC9X | Pfam:DUF213 | 365044.Pnap_ | 1,2 | 0,979795897 | 6,86E-05 | 4,89E-05 | 3 - | 0,000153036 - | 0 - | 0 - | 1 - | 6,02E-05 - | 1 - | 6,11E-05 - | 1 - | 6,88E-05 - |
| 3954 | 1 | 3um_FIJJAFBF_126100 | Eukaryota | Bacillariophyt | Coscinodiscop | Thalassiosirale | Thalassiosirac | Thalassiosira | Thalassiosira | unclassified | unclassified | unclassified | unclassified | unclassified | 1,2 | 0,4 | 7,44E-05 | 1,65E-05 | 2 - | 0,000106769 - | 1 - | 6,63E-05 - | 1 - | 6,30E-05 - | 1 - | 6,39E-05 - | 1 - | 7,20E-05 - |
| 3955 | 2 | 3um_FIJJAFBF_434826;3um_FIJJAFBF_516009 | Bacteria | Bacteroidetes | Flavobacteriia | Flavobacterial | Flavobacteriac | various | various | unclassified | unclassified | unclassified | unclassified | unclassified | 0,8 | 0,4 | 8,35E-05 | 4,24E-05 | 1 - | 9,04E-05 - | 1 - | 0,00011222 - | 1 - | 0,000106679 - | 1 - | 0,000108255 - | 0 - | 0 - |
| 3956 | 1 | 3um_FIJJAFBF_226829 | Bacteria | Bacteroidetes | Flavobacteriia | Flavobacterial | Flavobacteriac | Polaribacter | Polaribacter ir | unclassified | unclassified | unclassified | unclassified | unclassified | 6 | 1,095445115 | 0,000123395 | 2,14E-05 | 6 - | 0,000104027 - | 5 - | 0,000107643 - | 8 - | 0,000163725 - | 6 - | 0,000124607 - | 5 - | 0,000116975 - |
| 3957 | 2 | 0.2um_NOKCPHJN_454631;3um_FIJJAFBF_27872 | Bacteria | Proteobacteri | Alphaproteob | Pelagibacteral | Pelagibacterac | Candidatus Pe | Candidatus Pe | poorly charact | Function unkn | COG0718 | Binds to DNA | 335992.SAR11 | 0,8 | 0,4 | 0,000104696 | 5,38E-05 | 1 - | 0,000109311 - | 1 - | 0,000135732 - | 0 - | 0 - | 1 - | 0,000130936 - | 1 - | 0,0001475 - |
| 3958 | 2 | 3um_FIJJAFBF_189055;3um_FIJJAFBF_71147 | Bacteria | Proteobacteri | Alphaproteob | Rhodobactera | Rhodobactera | unclassified | Rhodobactera | unclassified | unclassified | unclassified | unclassified | unclassified | 1,2 | 0,4 | 5,19E-05 | 1,15E-05 | 2 - | 7,45E-05 - | 1 - | 4,63E-05 - | 1 - | 4,40E-05 - | 1 - | 4,46E-05 - | 1 - | 5,03E-05 - |
| 3959 | 1 | 0.2um_NOKCPHJN_249144 | Bacteria | Proteobacteri | Gammaprotec | Cellvibrionales | Halieaceae | unclassified | Halieaceae ba | information st | Translation, ril | COG0052 | 30S ribosomal | 208964.PA365 | 2,4 | 1,019803903 | 0,000106832 | 4,52E-05 | 2 - | 7,55E-05 - | 2 - | 9,38E-05 - | 4 - | 0,000178266 - | 3 - | 0,000135674 - | 1 - | 5,09E-05 - |
| 3960 | 1 | 3um_FIJJAFBF_741499 | Bacteria | Bacteroidetes | unclassified | unclassified | unclassified | unclassified | Bacteroidetes | poorly charact | Function unkn | 11NCV | Protein of unk | 926562.Oweh | 1,8 | 0,748331477 | 0,000224746 | 9,41E-05 | 1 - | 0,000104343 - | 2 - | 0,000259125 - | 3 - | 0,000369497 - | 2 - | 0,00024997 - | 1 - | 0,000140795 - |
| 3961 | 1 | 3um_FIJJAFBF_172447 | Bacteria | Proteobacteri | Gammaprotec | Oceanospirilla | Saccharospirill | Reinekea | Reinekea fors | unclassified | unclassified | unclassified | unclassified | unclassified | 1,4 | 0,489897949 | 0,00010322 | 3,74E-05 | 2 - | 0,000124083 - | 1 - | 7,70E-05 - | 1 - | 7,32E-05 - | 1 - | 7,43E-05 - | 2 - | 0,000167432 - |
| 3962 | 2 | 3um_FIJJAFBF_51341;3um_FIJJAFBF_624299 | Bacteria | Bacteroidetes | Flavobacteriia | Flavobacterial | Flavobacteriac | various | various | metabolism | Lipid transpor | COG1960 | acyl-CoA dehy | 50743.SCB49_ | 0,8 | 0,748331477 | 2,67E-05 | 2,57E-05 | 1 - | 2,93E-05 - | 0 - | 0 - | 2 - | 6,91E-05 - | 1 - | 3,51E-05 - | 0 - | 0 - |
| 3963 | 1 | 0.2um_NOKCPHJN_414320 | Bacteria | Bacteroidetes | Flavobacteriia | Flavobacterial | Flavobacteriac | Gaetbulibacte | Gaetbulibacte | cellular proces | Defense mech | 07VZM | 5-methylcytos | 865938.Weev | 0,2 | 0,4 | 6,41E-06 | 1,28E-05 | 0 - | 0 - | 0 - | 0 - | 0 - | 0 - | 1 - | 3,20E-05 - | 0 - | 0 - |
| 3964 | 1 | 3um_FIJJAFBF_117668 | Bacteria | Proteobacteri | Gammaprotec | Cellvibrionales | Microbulbifer | Microbulbifer | Microbulbifer | metabolism | Energy produc | 05CMV | Succinyl-CoA s | 203122.Sde_2 | 0,8 | 0,4 | 2,94E-05 | 1,48E-05 | 0 - | 0 - | 1 - | 3,67E-05 - | 1 - | 3,49E-05 - | 1 - | 3,54E-05 - | 1 - | 3,99E-05 - |
| 3965 | 1 | 0.2um_NOKCPHJN_777552 | Bacteria | Proteobacteri | Alphaproteob | Pelagibacteral | Pelagibacterac | Candidatus Pe | Candidatus Pe | cellular proces | Cell wall/mem | COG2885 | Ompa motb d | 439493.PB721 | 6,4 | 0,489897949 | 0,000533475 | 7,98E-05 | 6 - | 0,00041737 - | 6 - | 0,00051825 - | 7 - | 0,000574773 - | 6 - | 0,000499939 - | 7 - | 0,000657043 - |
| 3966 | 1 | 0.2um_NOKCPHJN_134092 | Eukaryota | unclassified | Oomycetes | Peronosporale | Peronosporac | Phytophthora | Phytophthora | unclassified | unclassified | unclassified | unclassified | unclassified | 0,2 | 0,4 | 2,36E-07 | 4,72E-07 | 0 - | 0 - | 0 - | 0 - | 0 - | 0 - | 1 - | 1,18E-06 - | 0 - | 0 - |
| 3967 | 1 | 3um_FIJJAFBF_272420 | Bacteria | Proteobacteri | Gammaprotec | Alteromonada | Alteromonada | Salinimonas | Salinimonas sp | cellular proces | Signal transdu | 16Q0Y | phosphate reg | 1085623.GNIT | 3,2 | 1,469693846 | 0,000184225 | 6,38E-05 | 6 - | 0,000300725 - | 3 - | 0,000186706 - | 2 - | 0,000118325 - | 3 - | 0,000180109 - | 2 - | 0,000135262 - |
| 3968 | 1 | 0.2um_NOKCPHJN_61048 | Bacteria | Proteobacteri | Betaproteoba | unclassified | unclassified | unclassified | Betaproteoba | unclassified | unclassified | unclassified | unclassified | unclassified | 0,8 | 0,4 | 4,91E-05 | 2,49E-05 | 1 - | 5,31E-05 - | 1 - | 6,60E-05 - | 1 - | 6,27E-05 - | 1 - | 6,36E-05 - | 0 - | 0 - |
| 3969 | 2 | 0.2um_NOKCPHJN_109516;3um_FIJJAFBF_1505 | Bacteria | Bacteroidetes | Flavobacteriia | Flavobacterial | Crocinitomica | unclassified | Crocinitomica | unclassified | unclassified | unclassified | unclassified | unclassified | 1 | 0 | 1,13E-05 | 1,08E-06 | 1 - | 9,49E-06 - | 1 - | 1,18E-05 - | 1 - | 1,12E-05 - | 1 - | 1,14E-05 - | 1 - | 1,28E-05 - |
| 3970 | 1 | 3um_FIJJAFBF_737464 | Bacteria | Proteobacteri | Gammaprotec | Oceanospirilla | Saccharospirill | Reinekea | Reinekea fors | unclassified | unclassified | unclassified | unclassified | unclassified | 3,4 | 0,489897949 | 0,000394078 | 6,73E-05 | 4 - | 0,000389074 - | 3 - | 0,000362336 - | 3 - | 0,000344446 - | 3 - | 0,000349534 - | 4 - | 0,000524998 - |
| 3971 | 1 | 3um_FIJJAFBF_753878 | Bacteria | Proteobacteri | Alphaproteob | Rhodospirillal | Rhodospirillac | unclassified | Rhodospirillac | metabolism | Inorganic ion t | COG0841 | acriflavin resis | 391593.RCCS2 | 0,6 | 0,489897949 | 9,18E-06 | 7,63E-06 | 1 - | 1,28E-05 - | 1 - | 1,59E-05 - | 0 - | 0 - | 0 - | 0 - | 1 - | 1,72E-05 - |
| 3972 | 1 | 0.2um_NOKCPHJN_1033199 | Eukaryota | unclassified | Labyrinthulom | unclassified | Thraustochytr | Aurantiochytr | Aurantiochytr | metabolism | Inorganic ion t | KOG0441 | Destroys radic | 65071.PYU1_T | 0,2 | 0,4 | 1,77E-05 | 3,54E-05 | 0 - | 0 - | 1 - | 8,85E-05 - | 0 - | 0 - | 0 - | 0 - | 0 - | 0 - |
| 3973 | 1 | 0.2um_NOKCPHJN_232178 | Eukaryota | unclassified | Chrysophycea | Chromulinales | Chromulinace | Ochromonas | Ochromonas c | unclassified | unclassified | unclassified | unclassified | unclassified | 0,2 | 0,4 | 1,84E-05 | 3,67E-05 | 1 - | 9,18E-05 - | 0 - | 0 - | 0 - | 0 - | 0 - | 0 - | 0 - | 0 - |
| 3974 | 1 | 3um_FIJJAFBF_217186 | unclassified | unclassified | unclassified | unclassified | unclassified | unclassified | unclassified | unclassified | unclassified | unclassified | unclassified | unclassified | 0,2 | 0,4 | 5,25E-05 | 0,000105 | 0 - | 0 - | 0 - | 0 - | 0 - | 0 - | 0 - | 0 - | 1 - | 0,000262499 - |
| 3975 | 1 | 3um_FIJJAFBF_432329 | Bacteria | Proteobacteri | Betaproteoba | Burkholderial | Comamonada | Limnohabitan | Limnohabitan | unclassified | unclassified | unclassified | unclassified | unclassified | 1,4 | 0,8 | 5,71E-05 | 3,17E-05 | 2 - | 6,94E-05 - | 2 - | 8,61E-05 - | 0 - | 0 - | 2 - | 8,31E-05 - | 1 - | 4,68E-05 - |
| 3976 | 1 | 3um_FIJJAFBF_121956 | Bacteria | Proteobacteri | Alphaproteob | Rhodobactera | Rhodobactera | Yoonia | Yoonia rosea | metabolism | Energy produc | 01QS0 | Aldehyde oxid | 467661.RKLH1 | 0,2 | 0,4 | 3,66E-06 | 7,32E-06 | 0 - | 0 - | 0 - | 0 - | 0 - | 0 - | 1 - | 1,83E-05 - | 0 - | 0 - |
| 3977 | 3 | 0.2um_NOKCPHJN_181446;0.2um_NOKCPHJN_2 | Bacteria | Bacteroidetes | Flavobacteriia | Flavobacterial | Flavobacteriac | various | various | metabolism | Carbohydrate | COG0297 | starch synthas | 50743.SCB49_ | 0,8 | 0,4 | 3,93E-05 | 1,99E-05 | 1 - | 4,25E-05 - | 1 - | 5,28E-05 - | 1 - | 5,02E-05 - | 1 - | 5,09E-05 - | 0 - | 0 - |
| 3978 | 1 | 3um_FIJJAFBF_171399 | Bacteria | Bacteroidetes | Cytophagia | Cytophagales | Flammeovirga | unclassified | Flammeovirga | unclassified | unclassified | unclassified | unclassified | unclassified | 2,4 | 0,489897949 | 0,000143954 | 3,32E-05 | 2 - | 0,000100242 - | 3 - | 0,000186706 - | 3 - | 0,000177488 - | 2 - | 0,000120073 - | 2 - | 0,000135262 - |
| 3979 | 1 | 3um_FIJJAFBF_605115 | Bacteria | Proteobacteri | Gammaprotec | Oceanospirilla | Saccharospirill | Reinekea | Reinekea fors | poorly charact | Function unkn | 05N42 | UPF0307 prot | 314283.MED2 | 0,2 | 0,4 | 1,33E-05 | 2,65E-05 | 1 - | 6,63E-05 - | 0 - | 0 - | 0 - | 0 - | 0 - | 0 - | 0 - | 0 - |
| 3980 | 1 | 3um_FIJJAFBF_521152 | Bacteria | Bacteroidetes | Flavobacteriia | Flavobacterial | Flavobacteriac | Gaetbulibacte | Gaetbulibacte | metabolism | Inorganic ion t | 0P8Q4 | tonB-depende | 867900.Celly_ | 2,4 | 0,8 | 3,44E-05 | 7,62E-06 | 4 - | 4,94E-05 - | 2 - | 3,06E-05 - | 2 - | 2,91E-05 - | 2 - | 2,96E-05 - | 2 - | 3,33E-05 - |
| 3981 | 1 | 3um_FIJJAFBF_436834 | Bacteria | Actinobacteria | Actinobacteria | Micrococcales | Microbacteria | various | various | information st | Translation, ril | COG0089 | One of the ea | 312284.A20C1 | 3,4 | 1,019803903 | 0,000474965 | 0,000151238 | 2 - | 0,000231872 - | 3 - | 0,000431875 - | 4 - | 0,000547403 - | 5 - | 0,00069436 - | 3 - | 0,000469317 - |
| 3982 | 1 | 3um_FIJJAFBF_334777 | Bacteria | Proteobacteri | Alphaproteob | Rhodobactera | Rhodobactera | Sulfitobacter | Sulfitobacter s | unclassified | unclassified | unclassified | unclassified | unclassified | 1 | 1,095445115 | 3,82E-05 | 4,31E-05 | 1 - | 3,33E-05 - | 0 - | 0 - | 3 - | 0,000117811 - | 1 - | 3,99E-05 - | 0 - | 0 - |
| 3983 | 1 | 3um_FIJJAFBF_516119 | Eukaryota | Bacillariophyt | Coscinodiscop | Thalassiosirale | Thalassiosirac | Thalassiosira | Thalassiosira c | cellular proces | Posttranslatio | KOG0730 | cell division cy | 35128.Thaps4 | 1 | 0 | 4,31E-05 | 4,09E-06 | 1 - | 3,61E-05 - | 1 - | 4,48E-05 - | 1 - | 4,26E-05 - | 1 - | 4,32E-05 - | 1 - | 4,87E-05 - |
| 3984 | 1 | 0.2um_NOKCPHJN_480926 | Bacteria | Proteobacteri | Gammaprotec | unclassified | unclassified | unclassified | various | unclassified | unclassified | unclassified | unclassified | unclassified | 0,2 | 0,4 | 2,99E-05 | 5,98E-05 | 0 - | 0 - | 0 - | 0 - | 0 - | 0 - | 1 - | 0,000149438 - | 0 - | 0 - |
| 3985 | 1 | 3um_FIJJAFBF_323070 | Bacteria | Bacteroidetes | Flavobacteriia | Flavobacterial | Flavobacteriac | Psychroflexus | Psychroflexus | unclassified | unclassified | unclassified | unclassified | unclassified | 0,8 | 0,4 | 3,66E-05 | 1,88E-05 | 1 - | 3,84E-05 - | 1 - | 4,77E-05 - | 1 - | 4,53E-05 - | 0 - | 0 - | 1 - | 5,18E-05 - |
| 3986 | 1 | 3um_FIJJAFBF_283140 | Bacteria | Bacteroidetes | Flavobacteriia | Flavobacterial | Flavobacteriac | Polaribacter | Polaribacter b | unclassified | unclassified | unclassified | unclassified | unclassified | 1,6 | 1,019803903 | 9,28E-05 | 5,55E-05 | 3 - | 0,000148418 - | 2 - | 0,000122861 - | 0 - | 0 - | 1 - | 5,93E-05 - | 2 - | 0,000133512 - |
| 3987 | 1 | 3um_FIJJAFBF_333508 | Bacteria | Proteobacteri | Alphaproteob | Rhodobactera | Rhodobactera | Phaeobacter | Phaeobacter s | metabolism | Amino acid tra | 01RBF | Branched-cha | 388739.RSK20 | 2 | 0,632455532 | 6,21E-05 | 2,39E-05 | 2 - | 5,14E-05 - | 2 - | 6,38E-05 - | 2 - | 6,06E-05 - | 1 - | 3,08E-05 - | 3 - | 0,000103943 - |
| 3988 | 1 | 3um_FIJJAFBF_575460 | Bacteria | Bacteroidetes | Flavobacteriia | Flavobacterial | Flavobacteriac | Formosa | Formosa sp. H | metabolism | Carbohydrate | 099Q3 | phosphohexol | 313603.FB217 | 1,6 | 1,2 | 6,66E-05 | 4,95E-05 | 1 - | 3,50E-05 - | 1 - | 4,35E-05 - | 4 - | 0,000165222 - | 1 - | 4,19E-05 - | 1 - | 4,72E-05 - |
| 3989 | 1 | 0.2um_NOKCPHJN_283508 | Bacteria | Proteobacteri | Alphaproteob | Pelagibacteral | Pelagibacterac | Candidatus Pe | Candidatus Pe | metabolism | Amino acid tra | 05C6M | Alpha-keto-be | 439493.PB721 | 0,8 | 0,4 | 3,12E-05 | 1,58E-05 | 1 - | 3,38E-05 - | 1 - | 4,19E-05 - | 1 - | 3,98E-05 - | 1 - | 4,04E-05 - | 0 - | 0 - |
| 3990 | 1 | 3um_FIJJAFBF_600657 | Bacteria | Proteobacteri | Alphaproteob | Rhodobactera | Rhodobactera | Aliiroseovariu | Aliiroseovariu | unclassified | unclassified | unclassified | unclassified | unclassified | 9,2 | 1,469693846 | 0,000371355 | 2,83E-05 | 12 - | 0,000411141 - | 9 - | 0,000382887 - | 8 - | 0,00032354 - | 9 - | 0,000369358 - | 8 - | 0,000369849 - |
| 3991 | 1 | 3um_FIJJAFBF_189715 | Bacteria | Bacteroidetes | Flavobacteriia | Flavobacterial | Flavobacteriac | Polaribacter | Polaribacter r | metabolism | Inorganic ion t | COG0659 | sulfate transp | 313598.MED1 | 0,8 | 0,4 | 1,89E-05 | 9,58E-06 | 1 - | 2,04E-05 - | 1 - | 2,54E-05 - | 1 - | 2,41E-05 - | 1 - | 2,45E-05 - | 0 - | 0 - |
| 3992 | 1 | 0.2um_NOKCPHJN_331514 | Bacteria | Proteobacteri | Alphaproteob | Rhodobactera | Rhodobactera | Nioella | Nioella sedimi | metabolism | Energy produc | 02EEZ | Catalyzes a 2- | 644107.SL115 | 0,8 | 0,748331477 | 9,65E-06 | 8,99E-06 | 0 - | 0 - | 1 - | 1,24E-05 - | 1 - | 1,18E-05 - | 2 - | 2,40E-05 - | 0 - | 0 - |
| 3993 | 1 | 0.2um_NOKCPHJN_108398 | Bacteria | Proteobacteri | Gammaprotec | Oceanospirilla | Saccharospirill | Saccharospiril | Saccharospiril | poorly charact | Function unkn | COG3181 | extra-cytoplas | 270374.MELB | 12,6 | 1,019803903 | 0,000524882 | 5,39E-05 | 14 - | 0,000489901 - | 12 - | 0,00052141 - | 11 - | 0,000454361 - | 13 - | 0,000544903 - | 13 - | 0,000613832 - |
| 3994 | 1 | 3um_FIJJAFBF_552626 | Bacteria | Proteobacteri | Gammaprotec | Oceanospirilla | Saccharospirill | Reinekea | Reinekea fors | unclassified | unclassified | unclassified | unclassified | unclassified | 14,4 | 1,2 | 0,000790878 | 0,000116561 | 15 - | 0,000688661 - | 15 - | 0,000855113 - | 13 - | 0,000704508 - | 13 - | 0,000714913 - | 16 - | 0,000991197 - |
| 3995 | 1 | 0.2um_NOKCPHJN_27544 | Bacteria | Bacteroidetes | Flavobacteriia | Flavobacterial | Flavobacteriac | Polaribacter | Polaribacter s | unclassified | unclassified | unclassified | unclassified | unclassified | 18,6 | 3,440930107 | 0,000168064 | 2,76E-05 | 22 - | 0,000168003 - | 22 - | 0,00020861 - | 14 - | 0,000126198 - | 20 - | 0,000182945 - | 15 - | 0,000154565 - |
| 3996 | 1 | 3um_FIJJAFBF_44438 | Eukaryota | Arthropoda | Insecta | Blattodea | Ectobiidae | Blattella | Blattella germ | unclassified | unclassified | unclassified | unclassified | unclassified | 0,2 | 0,4 | 6,03E-06 | 1,21E-05 | 0 - | 0 - | 1 - | 3,01E-05 - | 0 - | 0 - | 0 - | 0 - | 0 - | 0 - |
| 3997 | 3 | 0.2um_NOKCPHJN_497193;0.2um_NOKCPHJN_5 | Bacteria | Proteobacteri | Alphaproteob | Pelagibacteral | Pelagibacterac | Candidatus Pe | Candidatus Pe | poorly charact | Function unkn | 01QBG | Protein of unk | 335992.SAR11 | 0,8 | 0,4 | 5,85E-05 | 2,95E-05 | 0 - | 0 - | 1 - | 7,31E-05 - | 1 - | 6,95E-05 - | 1 - | 7,05E-05 - | 1 - | 7,94E-05 - |
| 3998 | 1 | 0.2um_NOKCPHJN_107812 | Bacteria | Proteobacteri | Alphaproteob | Rhodospirillal | unclassified | unclassified | Rhodospirillal | poorly charact | Function unkn | COG3181 | extra-cytoplas | 331869.BAL19 | 1,2 | 0,748331477 | 5,02E-05 | 3,17E-05 | 1 - | 3,53E-05 - | 0 - | 0 - | 2 - | 8,34E-05 - | 2 - | 8,46E-05 - | 1 - | 4,77E-05 - |

| | | | | | | | | | | | | | | | | | | | | | | | | | | | | |
|---|---|---|---|---|---|---|---|---|---|---|---|---|---|---|---|---|---|---|---|---|---|---|---|---|---|---|---|---|
| 3999 | 1 | 0.2um_NOKCPHJN_871977 | Bacteria | Bacteroidetes | unclassified | unclassified | unclassified | unclassified | unidentified e | poorly charact | Function unkn | 05ERK | gliding motilit | 50743.SCB49_ | 1,4 | 0,489897949 | 4,51E-05 | 1,42E-05 | 2 - | 5,49E-05 - | 2 - | 6,82E-05 - | 1 - | 3,24E-05 - | 1 - | 3,29E-05 - | 1 - | 3,71E-05 - |
| 4000 | 1 | 0.2um_NOKCPHJN_573978 | Eukaryota | Eustigmatoph | unclassified | Eustigmatales | Monodopsida | Nannochlorop | Nannochlorop | unclassified | unclassified | unclassified | unclassified | unclassified | 1 | 1,095445115 | 1,97E-05 | 2,22E-05 | 1 - | 1,69E-05 - | 1 - | 2,10E-05 - | 0 - | 0 - | 3 - | 6,07E-05 - | 0 - | 0 - |
| 4001 | 1 | 3um_FIJJAFBF_383275 | Eukaryota | unclassified | Oomycetes | Peronosporale | Peronosporac | Phytophthora | Phytophthora | cellular proces | Signal transdu | KOG0613 | fibronectin typ | 4787.PITG_04 | 4,2 | 1,720465053 | 3,40E-05 | 1,52E-05 | 4 - | 2,68E-05 - | 6 - | 4,99E-05 - | 1 - | 7,91E-06 - | 5 - | 4,01E-05 - | 5 - | 4,52E-05 - |
| 4002 | 1 | 3um_FIJJAFBF_21104 | Bacteria | Verrucomicrob | Opitutae | Opitutales | Opitutaceae | unclassified | Opitutaceae b | unclassified | unclassified | unclassified | unclassified | unclassified | 0,4 | 0,8 | 5,16E-05 | 0,00010317 | 2 - | 0,000257925 - | 0 - | 0 - | 0 - | 0 - | 0 - | 0 - | 0 - | 0 - |
| 4003 | 1 | 3um_FIJJAFBF_84926 | Bacteria | Proteobacteri | Alphaproteob | Rhodobactera | unclassified | unclassified | Rhodobactera | information st | Translation, ri | COG0292 | Binds directly | 367336.OM22 | 2,8 | 0,748331477 | 0,000320264 | 8,74E-05 | 3 - | 0,000289353 - | 4 - | 0,000479055 - | 3 - | 0,000341552 - | 2 - | 0,000231064 - | 2 - | 0,000260293 - |
| 4004 | 1 | 0.2um_NOKCPHJN_116081 | Bacteria | Proteobacteri | Alphaproteob | Rhodobactera | Rhodobactera | unclassified | Rhodobactera | unclassified | unclassified | unclassified | unclassified | unclassified | 1,6 | 1,356465997 | 6,45E-05 | 5,56E-05 | 0 - | 0 - | 0 - | 0 - | 2 - | 7,61E-05 - | 3 - | 0,000115857 - | 3 - | 0,000130512 - |
| 4005 | 1 | 3um_FIJJAFBF_743614 | Bacteria | Proteobacteri | Alphaproteob | Rhodospirillale | Rhodospirillac | unclassified | Rhodobactera | unclassified | unclassified | unclassified | unclassified | unclassified | 1 | 0 | 2,99E-05 | 2,84E-06 | 1 - | 2,51E-05 - | 1 - | 3,11E-05 - | 1 - | 2,96E-05 - | 1 - | 3,00E-05 - | 1 - | 3,38E-05 - |
| 4006 | 1 | 3um_FIJJAFBF_248095 | Bacteria | Proteobacteri | Betaproteoba | Burkholderiale | Comamonada | Hydrogenophi | various | unclassified | unclassified | unclassified | unclassified | unclassified | 1 | 0,632455532 | 3,08E-05 | 1,74E-05 | 2 - | 5,38E-05 - | 0 - | 0 - | 1 - | 3,17E-05 - | 1 - | 3,22E-05 - | 1 - | 3,63E-05 - |
| 4007 | 1 | 3um_FIJJAFBF_570475 | Bacteria | Bacteroidetes | Cytophagia | Cytophagales | Flammeovirga | Fabibacter | Fabibacter sp. | metabolism | Nucleotide tra | 0FVCM | Aspartate trar | 269798.CHU_ | 0,2 | 0,4 | 8,80E-06 | 1,76E-05 | 0 - | 0 - | 0 - | 0 - | 1 - | 4,40E-05 - | 0 - | 0 - | 0 - | 0 - |
| 4008 | 1 | 3um_FIJJAFBF_434164 | Bacteria | Bacteroidetes | Flavobacteriia | Flavobacterial | Flavobacteriac | Olleya | Olleya sp. Bg1 | unclassified | unclassified | unclassified | unclassified | unclassified | 2,2 | 0,4 | 0,000112974 | 3,04E-05 | 2 - | 8,50E-05 - | 2 - | 0,00010557 - | 2 - | 0,000100357 - | 2 - | 0,000101839 - | 3 - | 0,000172083 - |
| 4009 | 1 | 3um_FIJJAFBF_284143 | Bacteria | Actinobacteri | Actinobacteri | Actinomycetal | unclassified | unclassified | Actinomycetal | cellular proces | Cell cycle cont | COG3599 | Divisome com | 351607.Acel_ | 0,2 | 0,4 | 1,58E-05 | 3,16E-05 | 0 - | 0 - | 0 - | 0 - | 0 - | 0 - | 1 - | 7,90E-05 - | 0 - | 0 - |
| 4010 | 1 | 0.2um_NOKCPHJN_859297 | Bacteria | Proteobacteri | Gammaprotec | Cellvibrionales | Porticoccacea | unclassified | gamma protec | poorly charact | Function unkn | 0XY6Y | Pkd domain co | 314287.GB22 | 6 | 0,632455532 | 0,000120581 | 1,86E-05 | 6 - | 0,000100829 - | 7 - | 0,000146066 - | 5 - | 9,92E-05 - | 6 - | 0,000120776 - | 6 - | 0,000136054 - |
| 4011 | 1 | 0.2um_NOKCPHJN_535342 | Bacteria | Actinobacteri | Actinobacteri | Micrococcales | Microbacteria | Candidatus Ac | Candidatus Ac | unclassified | unclassified | unclassified | unclassified | unclassified | 7 | 0 | 9,44E-05 | 8,96E-06 | 7 - | 7,91E-05 - | 7 - | 9,82E-05 - | 7 - | 9,33E-05 - | 7 - | 9,47E-05 - | 7 - | 0,000106705 - |
| 4012 | 1 | 0.2um_NOKCPHJN_479275 | Bacteria | Proteobacteri | Alphaproteob | Rhodobactera | Rhodobactera | Planktomarina | Planktomarina | information st | Replication, re | COG0178 | The UvrABC re | 391616.OA23 | 1,8 | 0,4 | 2,57E-05 | 6,40E-06 | 2 - | 2,39E-05 - | 2 - | 2,97E-05 - | 1 - | 1,41E-05 - | 2 - | 2,86E-05 - | 2 - | 3,22E-05 - |
| 4013 | 1 | 3um_FIJJAFBF_150023 | Bacteria | Proteobacteri | Alphaproteob | Rhodobactera | Rhodobactera | Yoonia | Yoonia rosea | unclassified | unclassified | unclassified | unclassified | unclassified | 0,2 | 0,4 | 6,47E-06 | 1,29E-05 | 1 - | 3,23E-05 - | 0 - | 0 - | 0 - | 0 - | 0 - | 0 - | 0 - | 0 - |
| 4014 | 1 | 0.2um_NOKCPHJN_116820 | Bacteria | Proteobacteri | Alphaproteob | Rhodobactera | Rhodobactera | Sulfitobacter | Sulfitobacter g | metabolism | Amino acid tra | 16Q4U | Catalyzes the | 314264.ROS21 | 3 | 0,894427191 | 9,60E-05 | 2,95E-05 | 3 - | 8,08E-05 - | 4 - | 0,000133821 - | 2 - | 6,36E-05 - | 4 - | 0,000129092 - | 2 - | 7,27E-05 - |
| 4015 | 1 | 3um_FIJJAFBF_284332 | Bacteria | Proteobacteri | Alphaproteob | Rhodobactera | Rhodobactera | Roseobacter | Roseobacter s | information st | Transcription | COG0085 | DNA-depende | 391593.RCCS2 | 0,2 | 0,4 | 5,63E-06 | 1,13E-05 | 0 - | 0 - | 1 - | 2,82E-05 - | 0 - | 0 - | 0 - | 0 - | 0 - | 0 - |
| 4016 | 1 | 0.2um_NOKCPHJN_697301 | Bacteria | Bacteroidetes | Flavobacteriia | Flavobacterial | Flavobacteriac | Polaribacter | Polaribacter s | unclassified | unclassified | unclassified | unclassified | unclassified | 0,6 | 0,489897949 | 4,79E-05 | 3,94E-05 | 1 - | 7,08E-05 - | 0 - | 0 - | 1 - | 8,36E-05 - | 1 - | 8,49E-05 - | 0 - | 0 - |
| 4017 | 1 | 0.2um_NOKCPHJN_830367 | Bacteria | Proteobacteri | Alphaproteob | Pelagibacteral | Pelagibacterac | Candidatus Pe | Candidatus Pe | metabolism | Energy produc | COG0711 | F(1)F(0) ATP s | 335992.SAR11 | 0,6 | 0,489897949 | 4,91E-05 | 4,08E-05 | 1 - | 6,96E-05 - | 0 - | 0 - | 1 - | 8,21E-05 - | 0 - | 0 - | 1 - | 9,39E-05 - |
| 4018 | 1 | 3um_FIJJAFBF_15537 | Bacteria | Bacteroidetes | Flavobacteriia | Flavobacterial | Flavobacteriac | Flaviramulus | Flaviramulus i | unclassified | unclassified | unclassified | unclassified | unclassified | 5,2 | 0,979795897 | 0,000128927 | 1,11E-05 | 7 - | 0,000147963 - | 5 - | 0,000131233 - | 5 - | 0,000124753 - | 5 - | 0,000126596 - | 4 - | 0,000114088 - |
| 4019 | 1 | 0.2um_NOKCPHJN_75092 | Eukaryota | unclassified | Oomycetes | Pythiales | Pythiaceae | Pythium | Pythium insidi | metabolism | Energy produc | COG0055 | Produces ATP | 65071.PYU1_T | 2,6 | 0,8 | 7,35E-05 | 2,50E-05 | 2 - | 4,70E-05 - | 4 - | 0,000116819 - | 2 - | 5,55E-05 - | 3 - | 8,45E-05 - | 2 - | 6,35E-05 - |
| 4020 | 1 | 3um_FIJJAFBF_259434 | Bacteria | Proteobacteri | Alphaproteob | Rhodobactera | Rhodobactera | Planktomarina | Planktomarina | unclassified | unclassified | unclassified | unclassified | unclassified | 6,2 | 1,166190379 | 0,000129481 | 2,77E-05 | 7 - | 0,000122475 - | 6 - | 0,000130353 - | 4 - | 8,26E-05 - | 7 - | 0,000146705 - | 7 - | 0,000165262 - |
| 4021 | 1 | 3um_FIJJAFBF_512101 | Bacteria | Proteobacteri | Alphaproteob | Rhizobiales | unclassified | unclassified | Rhizobiales ba | unclassified | unclassified | unclassified | unclassified | unclassified | 0,4 | 0,489897949 | 1,51E-05 | 1,86E-05 | 1 - | 3,47E-05 - | 0 - | 0 - | 1 - | 4,09E-05 - | 0 - | 0 - | 0 - | 0 - |
| 4022 | 1 | 0.2um_NOKCPHJN_141760 | Bacteria | Proteobacteri | Gammaprotec | Cellvibrionales | Porticoccacea | unclassified | SAR92 bacteri | unclassified | unclassified | unclassified | unclassified | unclassified | 0,2 | 0,4 | 1,54E-05 | 3,08E-05 | 1 - | 7,70E-05 - | 0 - | 0 - | 0 - | 0 - | 0 - | 0 - | 0 - | 0 - |
| 4023 | 1 | 0.2um_NOKCPHJN_62024 | Bacteria | Bacteroidetes | Cytophagia | Cytophagales | Flammeovirga | unclassified | Flammeovirga | poorly charact | Function unkn | 0FVGW | gliding motilit | 643867.Ftrac_ | 6,4 | 0,8 | 0,000321509 | 1,76E-05 | 8 - | 0,000340079 - | 6 - | 0,000316709 - | 6 - | 0,000301072 - | 6 - | 0,000305518 - | 6 - | 0,000344166 - |
| 4024 | 2 | 0.2um_NOKCPHJN_71209;0.2um_NOKCPHJN_83 | Bacteria | Proteobacteri | Alphaproteob | Pelagibacteral | various | various | various | metabolism | Energy produc | 05C26 | Dehydrogenas | various | 0,8 | 0,4 | 2,18E-05 | 1,12E-05 | 1 - | 2,31E-05 - | 0 - | 0 - | 1 - | 2,73E-05 - | 1 - | 2,77E-05 - | 1 - | 3,12E-05 - |
| 4025 | 1 | 3um_FIJJAFBF_511859 | Viruses | unclassified | unclassified | unclassified | Phycodnavirid | unclassified | Chrysochromu | information st | Replication, re | COG0513 | purine NTP-de | 5141.EFNCRP | 1 | 0 | 4,48E-05 | 4,25E-06 | 1 - | 3,75E-05 - | 1 - | 4,66E-05 - | 1 - | 4,43E-05 - | 1 - | 4,49E-05 - | 1 - | 5,06E-05 - |
| 4026 | 1 | 0.2um_NOKCPHJN_368946 | Bacteria | Candidatus M | unclassified | unclassified | unclassified | unclassified | Candidatus M | information st | Replication, re | COG0210 | helicase | 523850.TON_ | 1 | 0 | 2,80E-05 | 2,66E-06 | 1 - | 2,35E-05 - | 1 - | 2,91E-05 - | 1 - | 2,77E-05 - | 1 - | 2,81E-05 - | 1 - | 3,17E-05 - |
| 4027 | 1 | 3um_FIJJAFBF_31373 | Bacteria | Proteobacteri | Alphaproteob | Rhodobactera | Rhodobactera | Roseobacter | Roseobacter li | metabolism | Secondary me | 05ERH | Fumarylaceto | 391595.RLO14 | 0,4 | 0,489897949 | 1,77E-05 | 2,19E-05 | 1 - | 4,07E-05 - | 0 - | 0 - | 1 - | 4,80E-05 - | 0 - | 0 - | 0 - | 0 - |
| 4028 | 1 | 3um_FIJJAFBF_337330 | Bacteria | Bacteroidetes | Flavobacteriia | Flavobacterial | Flavobacteriac | Polaribacter | various | cellular proces | Cell wall/mem | 0ZZRH | outer membra | 886377.Murru | 0,4 | 0,489897949 | 1,97E-05 | 2,45E-05 | 1 - | 4,19E-05 - | 0 - | 0 - | 0 - | 0 - | 0 - | 0 - | 1 - | 5,65E-05 - |
| 4029 | 1 | 3um_FIJJAFBF_514490 | Bacteria | Proteobacteri | Betaproteoba | Burkholderiale | Comamonada | Polaromonas | Polaromonas | unclassified | unclassified | unclassified | unclassified | unclassified | 7,8 | 0,748331477 | 0,000210875 | 1,19E-05 | 9 - | 0,000205366 - | 8 - | 0,00022667 - | 8 - | 0,000215479 - | 7 - | 0,000191329 - | 7 - | 0,000215531 - |
| 4030 | 1 | 3um_FIJJAFBF_523727 | Bacteria | Bacteroidetes | unclassified | unclassified | unclassified | unclassified | Bacteroidetes | unclassified | unclassified | unclassified | unclassified | unclassified | 0,6 | 0,489897949 | 5,04E-05 | 4,15E-05 | 1 - | 7,36E-05 - | 1 - | 9,14E-05 - | 1 - | 8,68E-05 - | 0 - | 0 - | 0 - | 0 - |
| 4031 | 1 | 3um_FIJJAFBF_392874 | Bacteria | Proteobacteri | Gammaprotec | Alteromonada | Alteromonada | Paraglaciecola | Paraglaciecola | unclassified | unclassified | unclassified | unclassified | unclassified | 1,8 | 0,4 | 0,000142152 | 2,81E-05 | 2 - | 0,000134242 - | 2 - | 0,000166689 - | 2 - | 0,000158459 - | 2 - | 0,000160799 - | 1 - | 9,06E-05 - |
| 4032 | 1 | 3um_FIJJAFBF_681308 | Bacteria | Proteobacteri | Gammaprotec | unclassified | unclassified | unclassified | Gammaprotec | unclassified | unclassified | unclassified | unclassified | unclassified | 1,2 | 0,4 | 4,47E-05 | 1,90E-05 | 1 - | 3,05E-05 - | 1 - | 3,79E-05 - | 1 - | 3,60E-05 - | 1 - | 3,66E-05 - | 2 - | 8,24E-05 - |
| 4033 | 1 | 3um_FIJJAFBF_02136 | Bacteria | Proteobacteri | Alphaproteob | various | various | various | various | unclassified | unclassified | unclassified | unclassified | unclassified | 1 | 0 | 0,00010002 | 9,50E-06 | 1 - | 8,38E-05 - | 1 - | 0,000104028 - | 1 - | 9,89E-05 - | 1 - | 0,000100353 - | 1 - | 0,000113047 - |
| 4034 | 1 | 3um_FIJJAFBF_532806 | Bacteria | Proteobacteri | Alphaproteob | Rhodobactera | Rhodobactera | unclassified | Rhodobactera | unclassified | unclassified | unclassified | unclassified | unclassified | 5 | 1,095445115 | 0,000156091 | 4,57E-05 | 5 - | 0,000129545 - | 4 - | 0,000128685 - | 4 - | 0,000122332 - | 5 - | 0,000155173 - | 7 - | 0,000244723 - |
| 4035 | 1 | 0.2um_NOKCPHJN_649776 | Viruses | unclassified | unclassified | unclassified | Phycodnavirid | unclassified | Aureococcus a | unclassified | unclassified | unclassified | unclassified | unclassified | 0,2 | 0,4 | 5,26E-06 | 1,05E-05 | 0 - | 0 - | 0 - | 0 - | 1 - | 2,63E-05 - | 0 - | 0 - | 0 - | 0 - |
| 4036 | 3 | 0.2um_NOKCPHJN_475856;3um_FIJJAFBF_7281 | Bacteria | Bacteroidetes | unclassified | unclassified | unclassified | unclassified | unidentified e | unclassified | unclassified | unclassified | unclassified | unclassified | 0,2 | 0,4 | 8,29E-06 | 1,66E-05 | 0 - | 0 - | 0 - | 0 - | 1 - | 4,14E-05 - | 0 - | 0 - | 0 - | 0 - |
| 4037 | 1 | 0.2um_NOKCPHJN_64536 | Bacteria | Proteobacteri | Gammaprotec | Oceanospirilla | Saccharospirill | Reinekea | Reinekea blan | cellular proces | Intracellular tr | COG0653 | Part of the Sec | 314283.MED2 | 6,2 | 1,469693846 | 9,24E-05 | 1,47E-05 | 9 - | 0,000114269 - | 5 - | 7,88E-05 - | 5 - | 7,49E-05 - | 6 - | 9,12E-05 - | 6 - | 0,000102793 - |
| 4038 | 1 | 0.2um_NOKCPHJN_439510 | Bacteria | Proteobacteri | Gammaprotec | Oceanospirilla | Saccharospirill | Reinekea | Reinekea fors | metabolism | Nucleotide tra | 05CM0 | Catalyzes the | 314283.MED2 | 0,2 | 0,4 | 3,03E-05 | 6,06E-05 | 0 - | 0 - | 1 - | 0,000151616 - | 0 - | 0 - | 0 - | 0 - | 0 - | 0 - |
| 4039 | 1 | 0.2um_NOKCPHJN_720301 | Bacteria | Proteobacteri | Gammaprotec | unclassified | unclassified | unclassified | Gammaprotec | unclassified | unclassified | unclassified | unclassified | unclassified | 1,4 | 0,489897949 | 2,11E-05 | 6,63E-06 | 2 - | 2,57E-05 - | 2 - | 3,19E-05 - | 1 - | 1,52E-05 - | 1 - | 1,54E-05 - | 1 - | 1,73E-05 - |
| 4040 | 1 | 3um_FIJJAFBF_474963 | Bacteria | Proteobacteri | Alphaproteob | Rhodobactera | Rhodobactera | unclassified | Rhodobactera | metabolism | Secondary me | 01Q86 | extracellular s | 367336.OM22 | 2 | 0 | 7,53E-05 | 7,15E-06 | 2 - | 6,31E-05 - | 2 - | 7,83E-05 - | 2 - | 7,44E-05 - | 2 - | 7,55E-05 - | 2 - | 8,51E-05 - |
| 4041 | 1 | 0.2um_NOKCPHJN_04273 | Bacteria | Bacteroidetes | Cytophagia | Cytophagales | Flammeovirga | unclassified | Flammeovirga | metabolism | Energy produc | COG0437 | 4fe-4S ferredc | 643867.Ftrac_ | 3,8 | 1,469693846 | 4,92E-05 | 1,53E-05 | 6 - | 6,69E-05 - | 3 - | 4,16E-05 - | 5 - | 6,58E-05 - | 2 - | 2,67E-05 - | 3 - | 4,52E-05 - |
| 4042 | 1 | 3um_FIJJAFBF_518685 | Bacteria | Bacteroidetes | Flavobacteriia | Flavobacterial | Flavobacteriac | Polaribacter | Polaribacter s | unclassified | unclassified | unclassified | unclassified | unclassified | 4,4 | 1,019803903 | 4,89E-05 | 1,19E-05 | 5 - | 4,67E-05 - | 6 - | 6,96E-05 - | 3 - | 3,31E-05 - | 4 - | 4,48E-05 - | 4 - | 5,04E-05 - |
| 4043 | 1 | 0.2um_NOKCPHJN_754069 | Bacteria | Proteobacteri | Alphaproteob | Pelagibacteral | Pelagibacterac | Candidatus Pe | Candidatus Pe | poorly charact | Function unkn | 17AGM | Protein of unk | 335992.SAR11 | 0,6 | 0,489897949 | 4,13E-05 | 3,44E-05 | 1 - | 5,83E-05 - | 0 - | 0 - | 0 - | 0 - | 1 - | 6,98E-05 - | 1 - | 7,86E-05 - |
| 4044 | 1 | 0.2um_NOKCPHJN_239172 | Bacteria | Bacteroidetes | unclassified | unclassified | unclassified | unclassified | Bacteroidetes | unclassified | unclassified | unclassified | unclassified | unclassified | 11,4 | 1,496662955 | 0,000164977 | 8,57E-06 | 14 - | 0,000171675 - | 10 - | 0,000152264 - | 11 - | 0,000159221 - | 12 - | 0,000176261 - | 10 - | 0,000165464 - |
| 4045 | 1 | 0.2um_NOKCPHJN_904622 | Eukaryota | unclassified | Oomycetes | Peronosporale | Peronosporac | Phytophthora | Phytophthora | information st | Replication, re | COG0513 | purine NTP-de | 653948.CCA1 | 1 | 0 | 3,34E-05 | 3,17E-06 | 1 - | 2,80E-05 - | 1 - | 3,48E-05 - | 1 - | 3,30E-05 - | 1 - | 3,35E-05 - | 1 - | 3,78E-05 - |
| 4046 | 1 | 0.2um_NOKCPHJN_560824 | Bacteria | Bacteroidetes | Flavobacteriia | Flavobacterial | Flavobacteriac | Winogradskye | Winogradskye | poorly charact | Function unkn | COG2340 | Scp-like extrac | 6085.XP_0021 | 0,2 | 0,4 | 4,24E-06 | 8,47E-06 | 0 - | 0 - | 0 - | 0 - | 0 - | 0 - | 1 - | 2,12E-05 - | 0 - | 0 - |
| 4047 | 1 | 3um_FIJJAFBF_03574 | Bacteria | Bacteroidetes | Flavobacteriia | unclassified | unclassified | unclassified | Flavobacteria | information st | Translation, ri | 05C64 | Binds directly | 391603.FBALC | 0,6 | 0,489897949 | 3,54E-05 | 2,94E-05 | 1 - | 5,01E-05 - | 0 - | 0 - | 1 - | 5,92E-05 - | 0 - | 0 - | 1 - | 6,76E-05 - |
| 4048 | 1 | 0.2um_NOKCPHJN_705011 | Bacteria | Proteobacteri | Alphaproteob | Pelagibacteral | Pelagibacterac | Candidatus Pe | Candidatus Pe | unclassified | unclassified | unclassified | unclassified | unclassified | 0,2 | 0,4 | 3,35E-06 | 6,70E-06 | 0 - | 0 - | 1 - | 1,67E-05 - | 0 - | 0 - | 0 - | 0 - | 0 - | 0 - |
| 4049 | 1 | 3um_FIJJAFBF_163899 | Bacteria | Proteobacteri | Alphaproteob | Rhodobactera | Rhodobactera | unclassified | Rhodobactera | unclassified | unclassified | unclassified | unclassified | unclassified | 0,6 | 0,489897949 | 1,82E-05 | 1,50E-05 | 1 - | 2,65E-05 - | 1 - | 3,29E-05 - | 0 - | 0 - | 1 - | 3,18E-05 - | 0 - | 0 - |
| 4050 | 1 | 3um_FIJJAFBF_658391 | Bacteria | Proteobacteri | Gammaprotec | Arenicellales | Arenicellaceae | Arenicella | Arenicella xan | cellular proces | Cell wall/mem | COG2885 | Ompa motb d | 159087.Daro_ | 6,4 | 0,8 | 0,000526483 | 9,19E-05 | 6 - | 0,000412372 - | 5 - | 0,000426703 - | 7 - | 0,00056789 - | 7 - | 0,000576277 - | 7 - | 0,000649174 - |
| 4051 | 1 | 0.2um_NOKCPHJN_192139 | Bacteria | Proteobacteri | Alphaproteob | Rhodobactera | Rhodobactera | Planktomarina | Planktomarina | unclassified | unclassified | unclassified | unclassified | unclassified | 1 | 0,894427191 | 1,39E-05 | 1,25E-05 | 2 - | 2,46E-05 - | 2 - | 3,06E-05 - | 1 - | 1,45E-05 - | 0 - | 0 - | 0 - | 0 - |
| 4052 | 1 | 3um_FIJJAFBF_684977 | Bacteria | Proteobacteri | Gammaprotec | Oceanospirilla | Saccharospirill | Reinekea | Reinekea fors | poorly charact | Function unkn | 0QW1W | NA | 314283.MED2 | 1,8 | 0,748331477 | 6,15E-05 | 2,05E-05 | 3 - | 8,87E-05 - | 2 - | 7,35E-05 - | 2 - | 6,98E-05 - | 1 - | 3,54E-05 - | 1 - | 3,99E-05 - |
| 4053 | 1 | 3um_FIJJAFBF_776677 | Bacteria | Bacteroidetes | unclassified | unclassified | unclassified | unclassified | Bacteroidetes | information st | Replication, re | 0PB0H | DNA-binding p | 487796.Flav2 | 6,6 | 1,356465997 | 0,000986167 | 0,000106792 | 9 - | 0,001147768 - | 6 - | 0,000950126 - | 6 - | 0,000903215 - | 7 - | 0,001069315 - | 5 - | 0,000860414 - |
| 4054 | 1 | 0.2um_NOKCPHJN_375707 | Bacteria | Bacteroidetes | Flavobacteriia | Flavobacterial | Flavobacteriac | Nonlabens | Nonlabens sp. | unclassified | unclassified | unclassified | unclassified | unclassified | 0,8 | 0,4 | 7,97E-05 | 4,05E-05 | 1 - | 8,63E-05 - | 1 - | 0,000107157 - | 1 - | 0,000101866 - | 1 - | 0,000103371 - | 0 - | 0 - |
| 4055 | 1 | 0.2um_NOKCPHJN_956641 | Bacteria | Proteobacteri | Alphaproteob | Rhodobactera | Rhodobactera | unclassified | Rhodobactera | unclassified | unclassified | unclassified | unclassified | unclassified | 5,6 | 0,489897949 | 0,000202303 | 1,43E-05 | 6 - | 0,000182669 - | 5 - | 0,000189017 - | 6 - | 0,000215622 - | 6 - | 0,000218806 - | 5 - | 0,000205404 - |
| 4056 | 1 | 0.2um_NOKCPHJN_220431 | Bacteria | Proteobacteri | Gammaprotec | Alteromonada | Pseudoalteror | Pseudoalteror | Pseudoalteror | metabolism | Inorganic ion t | 16TQC | TONB-depend | 87626.PTD2_ | 0,2 | 0,4 | 2,88E-06 | 5,77E-06 | 0 - | 0 - | 1 - | 1,44E-05 - | 0 - | 0 - | 0 - | 0 - | 0 - | 0 - |
| 4057 | 1 | 3um_FIJJAFBF_143299 | Bacteria | Bacteroidetes | Flavobacteriia | Flavobacterial | Flavobacteriac | Marinirhabdu | Marinirhabdu | information st | Translation, ri | 09EM4 | Binds directly | 50743.SCB49_ | 1 | 0 | 0,000120199 | 1,14E-05 | 1 - | 0,000100681 - | 1 - | 0,000125017 - | 1 - | 0,000118844 - | 1 - | 0,000120599 - | 1 - | 0,000135855 - |
| 4058 | 1 | 3um_FIJJAFBF_644662 | Bacteria | Bacteroidetes | Flavobacteriia | Flavobacterial | Flavobacteriac | Formosa | Formosa sp. H | poorly charact | Function unkn | 0P8NI | Inherit from b | 983544.Lacal_ | 1,2 | 0,4 | 3,85E-05 | 8,52E-06 | 2 - | 5,52E-05 - | 1 - | 3,43E-05 - | 1 - | 3,26E-05 - | 1 - | 3,30E-05 - | 1 - | 3,72E-05 - |
| 4059 | 1 | 3um_FIJJAFBF_584729 | Eukaryota | Bacillariophyta | Fragilariophyc | Fragilariales | Fragilariaceae | Asterionella | Asterionella fo | metabolism | Energy produc | 11IIK | Low-potential | 35128.Thapsd | 2,6 | 0,489897949 | 0,000221491 | 5,70E-05 | 2 - | 0,00014083 - | 3 - | 0,000262305 - | 2 - | 0,000166236 - | 3 - | 0,000253037 - | 3 - | 0,000285045 - |
| 4060 | 1 | 0.2um_NOKCPHJN_69725 | Bacteria | Proteobacteri | Betaproteoba | Nitrosomonad | Methylophilac | Candidatus M | Candidatus M | metabolism | Energy produc | COG0055 | Produces ATP | 1007105.PT7_ | 1 | 0,632455532 | 2,97E-05 | 1,81E-05 | 0 - | 0 - | 1 - | 3,00E-05 - | 2 - | 5,70E-05 - | 1 - | 2,89E-05 - | 1 - | 3,26E-05 - |
| 4061 | 1 | 3um_FIJJAFBF_155806 | Bacteria | Bacteroidetes | Flavobacteriia | Flavobacterial | Flavobacteriac | Arenibacter | Arenibacter n | metabolism | Carbohydrate | 05D2N | Aldolase | 313603.FB217 | 4,8 | 0,979795897 | 0,000182595 | 2,68E-05 | 6 - | 0,000193989 - | 4 - | 0,000160585 - | 6 - | 0,000228984 - | 4 - | 0,000154911 - | 4 - | 0,000174506 - |
| 4062 | 2 | 0.2um_NOKCPHJN_49124;3um_FIJJAFBF_66945 | Bacteria | Bacteroidetes | Flavobacteriia | Flavobacterial | unclassified | unclassified | Flavobacterial | cellular proces | Cell wall/mem | 09GQK | Pfam:Hep_Ha | 391603.FBALC | 0,2 | 0,4 | 5,66E-06 | 1,13E-05 | 0 - | 0 - | 0 - | 0 - | 1 - | 2,83E-05 - | 0 - | 0 - | 0 - | 0 - |
| 4063 | 1 | 3um_FIJJAFBF_193084 | Bacteria | Bacteroidetes | Flavobacteriia | Flavobacterial | Crocinitomica | unclassified | Crocinitomica | unclassified | unclassified | unclassified | unclassified | unclassified | 2 | 0,632455532 | 0,000120793 | 3,76E-05 | 2 - | 0,000102479 - | 2 - | 0,000127249 - | 2 - | 0,000120966 - | 3 - | 0,000184129 - | 1 - | 6,91E-05 - |
| 4064 | 1 | 3um_FIJJAFBF_555528 | Bacteria | Proteobacteri | Alphaproteob | Rhodobactera | Hyphomonada | Hyphomonas | Hyphomonas | cellular proces | Cell wall/mem | COG3637 | Membrane | 411684.HPDF | 5,4 | 0,489897949 | 0,000260822 | 3,52E-05 | 5 - | 0,000202072 - | 6 - | 0,000301096 - | 6 - | 0,00028623 - | 5 - | 0,000242048 - | 5 - | 0,000272666 - |
| 4065 | 1 | 0.2um_NOKCPHJN_450338 | Bacteria | Proteobacteri | Betaproteoba | Burkholderiale | Burkholderiac | Paraburkholde | Paraburkholde | poorly charact | Function unkn | 08K5Z | Endo alpha-1, | 448385.sce40 | 0,2 | 0,4 | 2,60E-06 | 5,20E-06 | 0 - | 0 - | 1 - | 1,30E-05 - | 0 - | 0 - | 0 - | 0 - | 0 - | 0 - |
| 4066 | 1 | 0.2um_NOKCPHJN_370453 | Bacteria | Bacteroidetes | Flavobacteriia | Flavobacterial | Crocinitomica | unclassified | Crocinitomica | unclassified | unclassified | unclassified | unclassified | unclassified | 0,2 | 0,4 | 4,96E-06 | 9,93E-06 | 0 - | 0 - | 0 - | 0 - | 1 - | 2,48E-05 - | 0 - | 0 - | 0 - | 0 - |
| 4067 | 1 | 3um_FIJJAFBF_472237 | Bacteria | Proteobacteri | Gammaprotec | Alteromonada | Alteromonada | Aestuariibacte | Aestuariibacte | metabolism | Carbohydrate | 0NF43 | Glycoside hyd | 373903.Hore_ | 0,4 | 0,489897949 | 1,05E-05 | 1,29E-05 | 1 - | 2,41E-05 - | 0 - | 0 - | 1 - | 2,84E-05 - | 0 - | 0 - | 0 - | 0 - |
| 4068 | 1 | 3um_FIJJAFBF_74175 | Bacteria | Bacteroidetes | Flavobacteriia | Flavobacterial | Flavobacteriac | Algibacter | Algibacter alg | unclassified | unclassified | unclassified | unclassified | unclassified | 0,6 | 0,8 | 2,43E-05 | 3,36E-05 | 1 - | 3,62E-05 - | 0 - | 0 - | 2 - | 8,55E-05 - | 0 - | 0 - | 0 - | 0 - |
| 4069 | 1 | 3um_FIJJAFBF_226843 | Bacteria | Bacteroidetes | Flavobacteriia | Flavobacterial | Flavobacteriac | Winogradskye | Winogradskye | metabolism | Inorganic ion t | 05C27 | TonB-depende | 391598.FBBAL | 5,6 | 1,496662955 | 7,00E-05 | 1,51E-05 | 7 - | 7,49E-05 - | 5 - | 6,65E-05 - | 7 - | 8,85E-05 - | 6 - | 7,69E-05 - | 3 - | 4,33E-05 - |
| 4070 | 1 | 0.2um_NOKCPHJN_229451 | Bacteria | Proteobacteri | Alphaproteob | Pelagibacteral | Pelagibacterac | unclassified | Pelagibacterac | unclassified | unclassified | unclassified | unclassified | unclassified | 1 | 0 | 8,06E-05 | 7,65E-06 | 1 - | 6,75E-05 - | 1 - | 8,38E-05 - | 1 - | 7,97E-05 - | 1 - | 8,09E-05 - | 1 - | 9,11E-05 - |
| 4071 | 2 | 0.2um_NOKCPHJN_321190;3um_FIJJAFBF_1198 | unclassified | unclassified | unclassified | unclassified | unclassified | unclassified | unclassified | unclassified | unclassified | unclassified | unclassified | unclassified | 0,4 | 0,489897949 | 3,08E-05 | 3,79E-05 | 1 - | 7,00E-05 - | 0 - | 0 - | 0 - | 0 - | 1 - | 8,38E-05 - | 0 - | 0 - |
| 4072 | 1 | 0.2um_NOKCPHJN_727671 | Bacteria | Proteobacteri | Alphaproteob | Pelagibacteral | Pelagibacterac | Candidatus Pe | Candidatus Pe | unclassified | unclassified | unclassified | unclassified | unclassified | 1,6 | 0,489897949 | 5,10E-05 | 1,63E-05 | 2 - | 5,36E-05 - | 1 - | 3,33E-05 - | 1 - | 3,17E-05 - | 2 - | 6,42E-05 - | 2 - | 7,24E-05 - |
| 4073 | 1 | 3um_FIJJAFBF_40882 | Bacteria | Bacteroidetes | Flavobacteriia | Flavobacterial | Flavobacteriac | Aequorivita | Aequorivita sc | metabolism | Nucleotide tra | 0P7KC | ribonucleoside | 313603.FB217 | 0,8 | 0,4 | 3,33E-05 | 1,71E-05 | 1 - | 3,52E-05 - | 0 - | 0 - | 1 - | 4,16E-05 - | 1 - | 4,22E-05 - | 1 - | 4,75E-05 - |
| 4074 | 1 | 3um_FIJJAFBF_699804 | unclassified | unclassified | unclassified | unclassified | unclassified | unclassified | unclassified | unclassified | unclassified | unclassified | unclassified | unclassified | 0,2 | 0,4 | 3,12E-05 | 6,25E-05 | 0 - | 0 - | 0 - | 0 - | 0 - | 0 - | 1 - | 0,000156231 - | 0 - | 0 - |
| 4075 | 1 | 3um_FIJJAFBF_556488 | Bacteria | Proteobacteri | Alphaproteob | Rhodobactera | Rhodobactera | various | various | metabolism | Amino acid tra | 01RBF | Branched-chai | 388739.RSK20 | 0,8 | 0,4 | 2,44E-05 | 1,26E-05 | 1 - | 2,56E-05 - | 1 - | 3,18E-05 - | 1 - | 3,02E-05 - | 0 - | 0 - | 1 - | 3,46E-05 - |
| 4076 | 1 | 0.2um_NOKCPHJN_152610 | Bacteria | Bacteroidetes | Flavobacteriia | Flavobacterial | Crocinitomica | unclassified | Crocinitomica | metabolism | Coenzyme tra | 09A4R | May play a ke | 388413.ALPR1 | 0,4 | 0,489897949 | 1,27E-05 | 1,55E-05 | 0 - | 0 - | 1 - | 3,25E-05 - | 1 - | 3,09E-05 - | 0 - | 0 - | 0 - | 0 - |
| 4077 | 1 | 3um_FIJJAFBF_149181 | Bacteria | Bacteroidetes | Flavobacteriia | Flavobacterial | Flavobacteriac | Algibacter | Algibacter sp. | unclassified | unclassified | unclassified | unclassified | unclassified | 0,6 | 0,489897949 | 7,11E-05 | 5,85E-05 | 1 - | 0,0001053 - | 0 - | 0 - | 1 - | 0,000124296 - | 1 - | 0,000126131 - | 0 - | 0 - |
| 4078 | 1 | 0.2um_NOKCPHJN_400172 | Eukaryota | unclassified | Oomycetes | Peronosporale | Peronosporac | Plasmopara | Plasmopara h | unclassified | unclassified | unclassified | unclassified | unclassified | 0,8 | 0,4 | 5,26E-06 | 2,67E-06 | 1 - | 5,70E-06 - | 1 - | 7,07E-06 - | 1 - | 6,72E-06 - | 1 - | 6,82E-06 - | 0 - | 0 - |
| 4079 | 1 | 0.2um_NOKCPHJN_50864 | Bacteria | Bacteroidetes | Flavobacteriia | Flavobacterial | Flavobacteriac | Polaribacter | various | poorly charact | Function unkn | 0P8NI | Inherit from b | 313594.PI23P | 1,8 | 0,748331477 | 3,70E-05 | 1,49E-05 | 2 - | 3,51E-05 - | 1 - | 2,18E-05 - | 2 - | 4,14E-05 - | 3 - | 6,31E-05 - | 1 - | 2,37E-05 - |
| 4080 | 1 | 0.2um_NOKCPHJN_602244 | Bacteria | Bacteroidetes | Flavobacteriia | Flavobacterial | Flavobacteriac | unclassified | Flavobacteriac | unclassified | unclassified | unclassified | unclassified | unclassified | 1 | 0 | 1,64E-05 | 1,56E-06 | 1 - | 1,38E-05 - | 1 - | 1,71E-05 - | 1 - | 1,62E-05 - | 1 - | 1,65E-05 - | 1 - | 1,86E-05 - |
| 4081 | 1 | 3um_FIJJAFBF_337085 | Bacteria | Proteobacteri | Gammaprotec | Oceanospirilla | Oceanospirilla | Motiliproteus | Motiliproteus | information st | Transcription | 05D27 | DNA-depende | 207954.MED9 | 14,4 | 1,743559577 | 0,000140828 | 2,61E-05 | 13 - | 0,000105898 - | 13 - | 0,000131494 - | 13 - | 0,000125001 - | 17 - | 0,000165878 - | 16 - | 0,000175869 - |
| 4082 | 1 | 3um_FIJJAFBF_500056 | Bacteria | Bacteroidetes | Flavobacteriia | Flavobacterial | Flavobacteriac | Kordia | Kordia jejudor | unclassified | unclassified | unclassified | unclassified | unclassified | 0,2 | 0,4 | 1,49E-05 | 2,98E-05 | 0 - | 0 - | 0 - | 0 - | 0 - | 0 - | 0 - | 0 - | 1 - | 7,45E-05 - |
| 4083 | 1 | 0.2um_NOKCPHJN_770695 | unclassified | unclassified | unclassified | unclassified | unclassified | unclassified | unclassified | unclassified | unclassified | unclassified | unclassified | unclassified | 1 | 0 | 0,000190315 | 1,81E-05 | 1 - | 0,000159412 - | 1 - | 0,000197943 - | 1 - | 0,00018817 - | 1 - | 0,000190949 - | 1 - | 0,000215103 - |
| 4084 | 1 | 0.2um_NOKCPHJN_166967 | Bacteria | Proteobacteri | Alphaproteob | Rhodobactera | Rhodobactera | Pseudooceani | Pseudooceani | metabolism | Energy produc | 02EEZ | Catalyzes a 2- | 644107.SL115 | 0,4 | 0,489897949 | 4,49E-06 | 5,55E-06 | 1 - | 1,00E-05 - | 1 - | 1,24E-05 - | 0 - | 0 - | 0 - | 0 - | 0 - | 0 - |
| 4085 | 1 | 3um_FIJJAFBF_580417 | Bacteria | Proteobacteri | Gammaprotec | unclassified | unclassified | unclassified | various | information st | Translation, ri | 177SZ | This protein bi | 247633.GP214 | 2 | 0 | 0,000191646 | 1,82E-05 | 2 - | 0,000160527 - | 2 - | 0,000199327 - | 2 - | 0,000189486 - | 2 - | 0,000192284 - | 2 - | 0,000216608 - |
| 4086 | 1 | 0.2um_NOKCPHJN_45606 | Bacteria | Proteobacteri | Alphaproteob | Rhodospirillale | unclassified | unclassified | Rhodospirillale | unclassified | unclassified | unclassified | unclassified | unclassified | 0,4 | 0,489897949 | 7,38E-06 | 9,13E-06 | 1 - | 1,65E-05 - | 1 - | 2,04E-05 - | 0 - | 0 - | 0 - | 0 - | 0 - | 0 - |
| 4087 | 1 | 3um_FIJJAFBF_545106 | Bacteria | Proteobacteri | Alphaproteob | Rhodobactera | Rhodobactera | Marivita | Marivita geoje | unclassified | unclassified | unclassified | unclassified | unclassified | 2,4 | 1,019803903 | 0,000241593 | 8,27E-05 | 4 - | 0,000350463 - | 2 - | 0,000217586 - | 2 - | 0,000206843 - | 3 - | 0,000314847 - | 1 - | 0,000118225 - |
| 4088 | 1 | 0.2um_NOKCPHJN_96930 | Bacteria | Proteobacteri | Alphaproteob | Rhodobactera | Rhodobactera | Litoreibacter | Litoreibacter p | metabolism | Secondary me | 01Q86 | extracellular s | 391595.RLO14 | 8,4 | 0,8 | 0,000316917 | 4,01E-05 | 8 - | 0,000252952 - | 8 - | 0,000314091 - | 10 - | 0,000373229 - | 8 - | 0,000302994 - | 8 - | 0,000341321 - |
| 4089 | 1 | 0.2um_NOKCPHJN_57060 | Bacteria | Proteobacteri | Gammaprotec | Oceanospirilla | Saccharospirill | Reinekea | Reinekea blan | metabolism | Carbohydrate | 0QHXA | transketolase | 314283.MED2 | 6,8 | 0,979795897 | 0,000140475 | 2,30E-05 | 6 - | 0,00010387 - | 6 - | 0,000128976 - | 8 - | 0,000163478 - | 8 - | 0,000165892 - | 6 - | 0,000140158 - |
| 4090 | 1 | 3um_FIJJAFBF_520502 | Bacteria | Bacteroidetes | Flavobacteriia | Flavobacterial | Flavobacteriac | Polaribacter | Polaribacter s | unclassified | unclassified | unclassified | unclassified | unclassified | 2,6 | 0,489897949 | 7,64E-05 | 1,29E-05 | 3 - | 7,45E-05 - | 3 - | 9,25E-05 - | 2 - | 5,87E-05 - | 3 - | 8,93E-05 - | 2 - | 6,70E-05 - |
| 4091 | 1 | 0.2um_NOKCPHJN_138275 | Bacteria | Bacteroidetes | Cytophagia | Cytophagales | Flammeovirga | Reichenbachie | Reichenbachie | cellular proces | Intracellular tr | 0FVR3 | Protein-expor | 643867.Ftrac_ | 7 | 1,095445115 | 9,49E-05 | 1,17E-05 | 8 - | 9,18E-05 - | 8 - | 0,000114015 - | 7 - | 9,48E-05 - | 7 - | 9,62E-05 - | 5 - | 7,74E-05 - |
| 4092 | 1 | 0.2um_NOKCPHJN_54434 | Bacteria | Proteobacteri | Gammaprotec | Cellvibrionales | Spongiibacter | Oceanicoccus | Oceanicoccus | cellular proces | Cell wall/mem | COG2885 | Ompa motb d | 247633.GP214 | 1,2 | 0,748331477 | 4,26E-05 | 2,74E-05 | 1 - | 3,04E-05 - | 1 - | 3,78E-05 - | 2 - | 7,19E-05 - | 2 - | 7,29E-05 - | 0 - | 0 - |
| 4093 | 1 | 3um_FIJJAFBF_103164 | Bacteria | Bacteroidetes | Flavobacteriia | Flavobacterial | Flavobacteriac | Polaribacter | Polaribacter b | information st | Translation, ri | 0P97G | This is one of t | 313598.MED1 | 4,4 | 0,489897949 | 0,000290369 | 2,22E-05 | 5 - | 0,000278584 - | 4 - | 0,000276736 - | 5 - | 0,00032884 - | 4 - | 0,000266958 - | 4 - | 0,000300727 - |
| 4094 | 1 | 3um_FIJJAFBF_420433 | Bacteria | Proteobacteri | Betaproteoba | Burkholderiale | Burkholderiac | Caballeronia | Caballeronia t | unclassified | unclassified | unclassified | unclassified | unclassified | 1,6 | 1,019803903 | 5,04E-05 | 2,92E-05 | 3 - | 8,20E-05 - | 2 - | 6,79E-05 - | 0 - | 0 - | 2 - | 6,55E-05 - | 1 - | 3,69E-05 - |
| 4095 | 1 | 3um_FIJJAFBF_471961 | Bacteria | Proteobacteri | Alphaproteob | Rhodobactera | Rhodobactera | Tateyamaria | Tateyamaria c | metabolism | Amino acid tra | 16QXJ | Sarcosine oxid | 391589.RGAI1 | 0,4 | 0,489897949 | 1,22E-05 | 1,50E-05 | 1 - | 2,77E-05 - | 0 - | 0 - | 0 - | 0 - | 1 - | 3,32E-05 - | 0 - | 0 - |
| 4096 | 1 | 3um_FIJJAFBF_363299 | Bacteria | Proteobacteri | Alphaproteob | Rhodospirillale | Rhodospirillac | unclassified | Rhodobactera | unclassified | unclassified | unclassified | unclassified | unclassified | 2,4 | 1,2 | 0,000177478 | 8,63E-05 | 3 - | 0,000190238 - | 3 - | 0,000236219 - | 4 - | 0,000299408 - | 1 - | 7,60E-05 - | 1 - | 8,56E-05 - |
| 4097 | 1 | 3um_FIJJAFBF_631322 | Bacteria | Bacteroidetes | Cytophagia | Cytophagales | Flammeovirga | unclassified | Flammeovirga | information st | Transcription | 09BX3 | Facilitates trar | 643867.Ftrac_ | 2,4 | 0,8 | 6,14E-05 | 2,27E-05 | 2 - | 4,26E-05 - | 4 - | 0,000105765 - | 2 - | 5,03E-05 - | 2 - | 5,10E-05 - | 2 - | 5,75E-05 - |
| 4098 | 1 | 0.2um_NOKCPHJN_351457 | Bacteria | Proteobacteri | Alphaproteob | Rhodospirillale | Rhodospirillac | unclassified | Rhodospirillac | unclassified | unclassified | unclassified | unclassified | unclassified | 1 | 0 | 4,43E-05 | 4,21E-06 | 1 - | 3,71E-05 - | 1 - | 4,61E-05 - | 1 - | 4,38E-05 - | 1 - | 4,45E-05 - | 1 - | 5,01E-05 - |
| 4099 | 1 | 3um_FIJJAFBF_137497 | Bacteria | Proteobacteri | Gammaprotec | unclassified | unclassified | unclassified | SAR86 cluster | poorly charact | Function unkn | 0QI1P | An essential G | 247634.GPB2 | 0,4 | 0,489897949 | 1,30E-05 | 1,61E-05 | 1 - | 2,90E-05 - | 1 - | 3,60E-05 - | 0 - | 0 - | 0 - | 0 - | 0 - | 0 - |
| 4100 | 1 | 3um_FIJJAFBF_558106 | Bacteria | Bacteroidetes | Flavobacteriia | various | various | various | various | unclassified | unclassified | unclassified | unclassified | unclassified | 0,2 | 0,4 | 1,65E-05 | 3,30E-05 | 1 - | 8,26E-05 - | 0 - | 0 - | 0 - | 0 - | 0 - | 0 - | 0 - | 0 - |
| 4101 | 1 | 3um_FIJJAFBF_227853 | Bacteria | Proteobacteri | Alphaproteob | Pelagibacteral | Pelagibacterac | Candidatus Pe | Candidatus Pe | unclassified | unclassified | unclassified | unclassified | unclassified | 1,4 | 0,489897949 | 0,00021594 | 9,13E-05 | 1 - | 0,000126128 - | 2 - | 0,000313228 - | 1 - | 0,000148882 - | 1 - | 0,000151081 - | 2 - | 0,000340383 - |
| 4102 | 1 | 3um_FIJJAFBF_234498 | Bacteria | Proteobacteri | Betaproteoba | Burkholderiale | Comamonada | Macromonas | Macromonas | information st | Translation, ri | COG0051 | tRNA binding | 391735.Veis_ | 1 | 0 | 0,000133036 | 1,26E-05 | 1 - | 0,000111434 - | 1 - | 0,000138368 - | 1 - | 0,000131536 - | 1 - | 0,000133479 - | 1 - | 0,000150364 - |
| 4103 | 1 | 3um_FIJJAFBF_788051 | Bacteria | Proteobacteri | Alphaproteob | Rhodobactera | unclassified | unclassified | Rhodobactera | information st | Translation, ri | COG0088 | One of the pri | 314262.MED1 | 2,8 | 0,4 | 0,000187114 | 3,26E-05 | 3 - | 0,000167966 - | 3 - | 0,000208564 - | 3 - | 0,000198267 - | 2 - | 0,00013413 - | 3 - | 0,000226646 - |
| 4104 | 1 | 3um_FIJJAFBF_189630 | Bacteria | Bacteroidetes | Flavobacteriia | Flavobacterial | Flavobacteriac | Polaribacter | Polaribacter s | poorly charact | Function unkn | 0P88S | domain protei | 313598.MED1 | 0,2 | 0,4 | 5,43E-06 | 1,09E-05 | 0 - | 0 - | 1 - | 2,71E-05 - | 0 - | 0 - | 0 - | 0 - | 0 - | 0 - |
| 4105 | 1 | 0.2um_NOKCPHJN_824556 | Eukaryota | unclassified | Oomycetes | Peronosporale | Peronosporac | Phytophthora | Phytophthora | unclassified | unclassified | unclassified | unclassified | unclassified | 0,8 | 0,4 | 2,24E-05 | 1,15E-05 | 1 - | 2,34E-05 - | 1 - | 2,91E-05 - | 0 - | 0 - | 1 - | 2,81E-05 - | 1 - | 3,16E-05 - |
| 4106 | 1 | 0.2um_NOKCPHJN_122616 | Bacteria | unclassified | unclassified | unclassified | unclassified | unclassified | uncultured ba | unclassified | unclassified | unclassified | unclassified | unclassified | 0,4 | 0,489897949 | 8,67E-05 | 0,000106136 | 0 - | 0 - | 0 - | 0 - | 1 - | 0,000215051 - | 1 - | 0,000218227 - | 0 - | 0 - |
| 4107 | 1 | 3um_FIJJAFBF_377867 | Bacteria | Bacteroidetes | Flavobacteriia | unclassified | unclassified | unclassified | Flavobacteria | unclassified | unclassified | unclassified | unclassified | unclassified | 0,4 | 0,489897949 | 7,32E-06 | 8,96E-06 | 0 - | 0 - | 0 - | 0 - | 1 - | 1,82E-05 - | 1 - | 1,84E-05 - | 0 - | 0 - |
| 4108 | 1 | 3um_FIJJAFBF_709327 | Bacteria | Proteobacteri | Gammaprotec | Oceanospirilla | Saccharospirill | Reinekea | Reinekea fors | metabolism | Energy produc | 1780T | Rubrerythrin | 342108.amb3 | 3,4 | 0,489897949 | 0,000300435 | 4,92E-05 | 3 - | 0,000222149 - | 3 - | 0,000275843 - | 4 - | 0,000349632 - | 4 - | 0,000354796 - | 3 - | 0,000299757 - |
| 4109 | 1 | 0.2um_NOKCPHJN_22758 | Bacteria | Bacteroidetes | Flavobacteriia | Flavobacterial | Crocinitomica | unclassified | Crocinitomica | cellular proces | Cell wall/mem | 0G0DD | Catalyzes the | 761193.Runsl_ | 0,2 | 0,4 | 7,86E-06 | 1,57E-05 | 1 - | 3,93E-05 - | 0 - | 0 - | 0 - | 0 - | 0 - | 0 - | 0 - | 0 - |
| 4110 | 1 | 0.2um_NOKCPHJN_116297 | Bacteria | Proteobacteri | Gammaprotec | Oceanospirilla | Saccharospirill | Reinekea | Reinekea fors | metabolism | Coenzyme tra | 05DP9 | ABC transport | 314283.MED2 | 0,2 | 0,4 | 7,00E-06 | 1,40E-05 | 1 - | 3,50E-05 - | 0 - | 0 - | 0 - | 0 - | 0 - | 0 - | 0 - | 0 - |
| 4111 | 1 | 3um_FIJJAFBF_163471 | Bacteria | Proteobacteri | Alphaproteob | Rhodobactera | Rhodobactera | Tateyamaria | Tateyamaria c | information st | Replication, re | 08ZUF | This protein is | 391589.RGAI1 | 0,2 | 0,4 | 2,30E-05 | 4,59E-05 | 1 - | 0,000114777 - | 0 - | 0 - | 0 - | 0 - | 0 - | 0 - | 0 - | 0 - |
| 4112 | 1 | 0.2um_NOKCPHJN_92477 | Bacteria | Proteobacteri | Gammaprotec | Oceanospirilla | Saccharospirill | Reinekea | Reinekea blan | unclassified | unclassified | unclassified | unclassified | unclassified | 11 | 1,673320053 | 0,000139585 | 2,06E-05 | 11 - | 0,000117555 - | 11 - | 0,000145969 - | 14 - | 0,000176606 - | 10 - | 0,000128011 - | 9 - | 0,000129783 - |
| 4113 | 1 | 3um_FIJJAFBF_346691 | Bacteria | Proteobacteri | Gammaprotec | Oceanospirilla | Oceanospirilla | Amphritea | Amphritea atl | metabolism | Energy produc | COG0056 | Produces ATP | 207954.MED9 | 11,6 | 0,489897949 | 0,000308532 | 2,46E-05 | 12 - | 0,000267961 - | 12 - | 0,000332729 - | 12 - | 0,000316301 - | 11 - | 0,000294225 - | 11 - | 0,000331443 - |
| 4114 | 1 | 0.2um_NOKCPHJN_344094 | Bacteria | Proteobacteri | Alphaproteob | Pelagibacteral | Pelagibacterac | Candidatus Pe | Candidatus Pe | poorly charact | Function unkn | COG3181 | extra-cytoplas | 335992.SAR11 | 0,4 | 0,489897949 | 1,54E-05 | 1,89E-05 | 1 - | 3,52E-05 - | 0 - | 0 - | 1 - | 4,16E-05 - | 0 - | 0 - | 0 - | 0 - |
| 4115 | 1 | 0.2um_NOKCPHJN_640706 | Eukaryota | Eustigmatoph | unclassified | Eustigmatales | Monodopsida | Nannochlorop | Nannochlorop | unclassified | unclassified | unclassified | unclassified | unclassified | 1,6 | 0,489897949 | 0,000131026 | 4,22E-05 | 1 - | 6,83E-05 - | 2 - | 0,000169665 - | 2 - | 0,000161288 - | 2 - | 0,000163671 - | 1 - | 9,22E-05 - |
| 4116 | 1 | 3um_FIJJAFBF_395548 | Eukaryota | unclassified | unclassified | Isochrysidales | Noelaerhabda | Emiliania | Emiliania huxl | poorly charact | Function unkn | 17F80 | NA | 391597.LMED | 1,8 | 0,748331477 | 0,000259021 | 0,000106368 | 2 - | 0,000244206 - | 2 - | 0,000303232 - | 1 - | 0,00014413 - | 3 - | 0,000438777 - | 1 - | 0,00016476 - |
| 4117 | 1 | 3um_FIJJAFBF_88825 | Bacteria | Proteobacteri | Gammaprotec | unclassified | unclassified | unclassified | various | information st | Translation, ri | 0QK14 | This protein bi | 207949.RED6 | 3 | 0 | 0,000232249 | 2,21E-05 | 3 - | 0,000194537 - | 3 - | 0,000241557 - | 3 - | 0,000229631 - | 3 - | 0,000233023 - | 3 - | 0,000262499 - |
| 4118 | 1 | 3um_FIJJAFBF_769006 | Bacteria | Proteobacteri | Betaproteoba | Nitrosomonad | unclassified | unclassified | Methylophilal | metabolism | Inorganic ion t | 05PWN | phosphate ab | 383631.MB21 | 0,2 | 0,4 | 6,87E-06 | 1,37E-05 | 1 - | 3,44E-05 - | 0 - | 0 - | 0 - | 0 - | 0 - | 0 - | 0 - | 0 - |
| 4119 | 1 | 3um_FIJJAFBF_81132 | Eukaryota | unclassified | unclassified | unclassified | unclassified | unclassified | uncultured pr | metabolism | Energy produc | COG0055 | Produces ATP | 533240.CRC_ | 1,4 | 0,8 | 4,53E-05 | 2,51E-05 | 2 - | 5,53E-05 - | 2 - | 6,87E-05 - | 2 - | 6,53E-05 - | 0 - | 0 - | 1 - | 3,73E-05 - |
| 4120 | 1 | 3um_FIJJAFBF_411737 | Eukaryota | Chlorophyta | unclassified | Pyramimonad | unclassified | Cymbomonas | Cymbomonas | unclassified | unclassified | unclassified | unclassified | unclassified | 0,8 | 0,4 | 8,11E-05 | 4,17E-05 | 1 - | 8,50E-05 - | 1 - | 0,00010557 - | 1 - | 0,000100357 - | 0 - | 0 - | 1 - | 0,000114722 - |
| 4121 | 1 | 3um_FIJJAFBF_204945 | Bacteria | Proteobacteri | Gammaprotec | Oceanospirilla | Saccharospirill | Reinekea | Reinekea fors | cellular proces | Intracellular tr | 16REI | general secret | 314283.MED2 | 1,2 | 0,748331477 | 2,57E-05 | 1,75E-05 | 1 - | 1,75E-05 - | 2 - | 4,34E-05 - | 1 - | 2,06E-05 - | 0 - | 0 - | 2 - | 4,71E-05 - |
| 4122 | 1 | 3um_FIJJAFBF_478439 | Bacteria | unclassified | unclassified | unclassified | unclassified | unclassified | uncultured ba | unclassified | unclassified | unclassified | unclassified | unclassified | 3 | 1,095445115 | 8,03E-05 | 2,32E-05 | 5 - | 0,000114548 - | 3 - | 8,53E-05 - | 2 - | 5,41E-05 - | 2 - | 5,49E-05 - | 3 - | 9,27E-05 - |
| 4123 | 1 | 0.2um_NOKCPHJN_64565 | Bacteria | Proteobacteri | Gammaprotec | Oceanospirilla | Saccharospirill | Reinekea | Reinekea blan | metabolism | Lipid transpor | COG0439 | acetyl-CoA car | 314283.MED2 | 3,6 | 0,489897949 | 0,000109294 | 1,29E-05 | 4 - | 0,000102479 - | 4 - | 0,000127249 - | 4 - | 0,000120966 - | 3 - | 9,21E-05 - | 3 - | 0,000103711 - |
| 4124 | 1 | 0.2um_NOKCPHJN_349796 | Bacteria | Bacteroidetes | unclassified | unclassified | unclassified | unclassified | Bacteroidetes | unclassified | unclassified | unclassified | unclassified | unclassified | 1 | 0 | 0,000210811 | 2,00E-05 | 1 - | 0,00017658 - | 1 - | 0,00021926 - | 1 - | 0,000208434 - | 1 - | 0,000211513 - | 1 - | 0,000238268 - |
| 4125 | 1 | 0.2um_NOKCPHJN_301303 | Eukaryota | unclassified | unclassified | unclassified | Apusomonadi | Thecamonas | Thecamonas t | unclassified | unclassified | unclassified | unclassified | unclassified | 1,6 | 0,489897949 | 6,58E-05 | 1,72E-05 | 2 - | 7,04E-05 - | 1 - | 4,37E-05 - | 2 - | 8,31E-05 - | 2 - | 8,43E-05 - | 1 - | 4,75E-05 - |
| 4126 | 1 | 3um_FIJJAFBF_128440 | Bacteria | Bacteroidetes | Flavobacteriia | Flavobacterial | Flavobacteriac | Polaribacter | Polaribacter s | poorly charact | Function unkn | 0P8NI | Inherit from b | 313594.PI23P | 18,2 | 0,748331477 | 0,000374292 | 3,16E-05 | 19 - | 0,000327934 - | 18 - | 0,000385765 - | 19 - | 0,000387092 - | 17 - | 0,000351461 - | 18 - | 0,000419209 - |
| 4127 | 1 | 3um_FIJJAFBF_46519 | Bacteria | Proteobacteri | Gammaprotec | Oceanospirilla | Oceanospirilla | Motiliproteus | Motiliproteus | information st | Translation, ri | COG0051 | tRNA binding | 314283.MED2 | 1,4 | 0,489897949 | 0,00018163 | 5,20E-05 | 2 - | 0,000222867 - | 1 - | 0,000138368 - | 2 - | 0,000263072 - | 1 - | 0,000133479 - | 1 - | 0,000150364 - |
| 4128 | 1 | 0.2um_NOKCPHJN_358015 | Eukaryota | unclassified | Labyrinthulom | unclassified | Thraustochytr | Aurantiochytr | Aurantiochytr | metabolism | Energy produc | COG0056 | Produces ATP | 70448.Q0P3H | 1 | 0 | 2,55E-05 | 2,42E-06 | 1 - | 2,14E-05 - | 1 - | 2,65E-05 - | 1 - | 2,52E-05 - | 1 - | 2,56E-05 - | 1 - | 2,88E-05 - |
| 4129 | 1 | 0.2um_NOKCPHJN_239918 | Bacteria | Proteobacteri | Gammaprotec | Cellvibrionales | Porticoccacea | unclassified | gamma protec | metabolism | Carbohydrate | 16QEQ | hydrolase, fan | 314287.GB22 | 2,4 | 0,8 | 5,01E-05 | 1,74E-05 | 3 - | 5,26E-05 - | 2 - | 4,35E-05 - | 1 - | 2,07E-05 - | 3 - | 6,30E-05 - | 3 - | 7,09E-05 - |
| 4130 | 1 | 3um_FIJJAFBF_106326 | Bacteria | Proteobacteri | Gammaprotec | Oceanospirilla | Saccharospirill | Reinekea | Reinekea fors | cellular proces | Cell wall/mem | 0R8PW | Cell division ai | 697282.Mettu | 2,4 | 0,489897949 | 0,000132323 | 2,34E-05 | 3 - | 0,000139972 - | 3 - | 0,000173803 - | 2 - | 0,000110148 - | 2 - | 0,000111775 - | 2 - | 0,000125914 - |
| 4131 | 1 | 3um_FIJJAFBF_48212 | unclassified | unclassified | unclassified | unclassified | unclassified | unclassified | unclassified | unclassified | unclassified | unclassified | unclassified | unclassified | 0,2 | 0,4 | 1,53E-05 | 3,07E-05 | 0 - | 0 - | 0 - | 0 - | 0 - | 0 - | 0 - | 0 - | 1 - | 7,67E-05 - |
| 4132 | 1 | 3um_FIJJAFBF_631097 | Bacteria | Proteobacteri | Gammaprotec | Oceanospirilla | Oceanospirilla | Marinobacteri | Marinobacteri | metabolism | Energy produc | 16QA5 | electron trans | 207954.MED9 | 0,2 | 0,4 | 4,93E-06 | 9,85E-06 | 0 - | 0 - | 0 - | 0 - | 1 - | 2,46E-05 - | 0 - | 0 - | 0 - | 0 - |
| 4133 | 1 | 0.2um_NOKCPHJN_158122 | Bacteria | Proteobacteri | Gammaprotec | Cellvibrionales | Porticoccacea | unclassified | gamma protec | information st | Translation, ri | 0QK14 | This protein bi | 314287.GB22 | 0,2 | 0,4 | 1,30E-05 | 2,59E-05 | 1 - | 6,48E-05 - | 0 - | 0 - | 0 - | 0 - | 0 - | 0 - | 0 - | 0 - |
| 4134 | 1 | 0.2um_NOKCPHJN_192436 | Bacteria | Proteobacteri | Gammaprotec | Oceanospirilla | Saccharospirill | Reinekea | Reinekea fors | poorly charact | Function unkn | 0Y0W9 | NA | 314283.MED2 | 0,2 | 0,4 | 7,68E-06 | 1,54E-05 | 0 - | 0 - | 0 - | 0 - | 0 - | 0 - | 1 - | 3,84E-05 - | 0 - | 0 - |
| 4135 | 1 | 3um_FIJJAFBF_788006 | Bacteria | Proteobacteri | Alphaproteob | Rhodobactera | Rhodobactera | Pseudopelagic | Pseudopelagic | unclassified | unclassified | unclassified | unclassified | unclassified | 2,8 | 1,166190379 | 0,000142648 | 5,33E-05 | 4 - | 0,000174565 - | 3 - | 0,000162569 - | 4 - | 0,000206057 - | 1 - | 5,23E-05 - | 2 - | 0,000117775 - |
| 4136 | 1 | 0.2um_NOKCPHJN_133299 | Bacteria | Bacteroidetes | Flavobacteriia | Flavobacterial | Flavobacteriac | Formosa | Formosa sp. H | information st | Translation, ri | 08YY1 | This protein is | 983544.Lacal_ | 0,4 | 0,489897949 | 4,30E-05 | 5,27E-05 | 0 - | 0 - | 0 - | 0 - | 1 - | 0,000106679 - | 1 - | 0,000108255 - | 0 - | 0 - |
| 4137 | 1 | 3um_FIJJAFBF_790068 | Bacteria | Bacteroidetes | Flavobacteriia | Flavobacterial | Flavobacteriac | Hanstruepera | Hanstruepera | unclassified | unclassified | unclassified | unclassified | unclassified | 5 | 1,095445115 | 0,000205509 | 4,07E-05 | 6 - | 0,000208685 - | 3 - | 0,000129563 - | 6 - | 0,000246331 - | 5 - | 0,000208308 - | 5 - | 0,000234658 - |
| 4138 | 1 | 3um_FIJJAFBF_744863 | unclassified | unclassified | unclassified | unclassified | unclassified | unclassified | unclassified | unclassified | unclassified | unclassified | unclassified | unclassified | 0,8 | 0,4 | 0,00010634 | 5,46E-05 | 1 - | 0,000111434 - | 1 - | 0,000138368 - | 1 - | 0,000131536 - | 0 - | 0 - | 1 - | 0,000150364 - |
| 4139 | 1 | 0.2um_NOKCPHJN_107901 | Bacteria | Proteobacteri | Alphaproteob | Rhodobactera | Rhodobactera | Cognatishimia | Cognatishimia | cellular proces | Posttranslatio | 05C6P | ATP-depender | 290400.Jann_ | 4 | 1,673320053 | 6,59E-05 | 2,45E-05 | 6 - | 8,59E-05 - | 4 - | 7,11E-05 - | 4 - | 6,76E-05 - | 5 - | 8,57E-05 - | 1 - | 1,93E-05 - |
| 4140 | 1 | 3um_FIJJAFBF_349618 | Bacteria | Bacteroidetes | Flavobacteriia | Flavobacterial | Flavobacteriac | Marinirhabdu | Marinirhabdu | unclassified | unclassified | unclassified | unclassified | unclassified | 1 | 0 | 5,39E-05 | 5,12E-06 | 1 - | 4,52E-05 - | 1 - | 5,61E-05 - | 1 - | 5,33E-05 - | 1 - | 5,41E-05 - | 1 - | 6,10E-05 - |
| 4141 | 1 | 0.2um_NOKCPHJN_41350 | Viruses | unclassified | unclassified | unclassified | various | various | various | information st | Transcription | 12N88 | dna-directed r | 45351.NEMVE | 0,2 | 0,4 | 1,72E-06 | 3,45E-06 | 0 - | 0 - | 0 - | 0 - | 1 - | 8,62E-06 - | 0 - | 0 - | 0 - | 0 - |

| | | | | | | | | | | | | | | | | | | | | | | | | | | | | |
|---|---|---|---|---|---|---|---|---|---|---|---|---|---|---|---|---|---|---|---|---|---|---|---|---|---|---|---|---|
| 4142 | 1 | 3um_FIJJAFBF_190341 | Bacteria | Bacteroidetes | Flavobacteriia | Flavobacterial | Flavobacteriac | Algibacter | Algibacter sp. | unclassified | unclassified | unclassified | unclassified | unclassified | 0,2 | 0,4 | 3,17E-05 | 6,33E-05 | 0 - | 0 - | 1 - | 0,000158354 - | 0 - | 0 - | 0 - | 0 - | 0 - | 0 - |
| 4143 | 1 | 3um_FIJJAFBF_296780 | Bacteria | Bacteroidetes | Cytophagia | Cytophagales | Flammeovirga | unclassified | Flammeovirga | unclassified | unclassified | unclassified | unclassified | unclassified | 0,2 | 0,4 | 1,09E-05 | 2,19E-05 | 1 - | 5,47E-05 - | 0 - | 0 - | 0 - | 0 - | 0 - | 0 - | 0 - | 0 - |
| 4144 | 1 | 3um_FIJJAFBF_346689 | Bacteria | Proteobacteri | Gammaprotec | Oceanospirilla | Oceanospirilla | Motiliproteus | Motiliproteus | metabolism | Energy produc | COG0055 | Produces ATP | 999141.GME_ | 18,8 | 2,638181192 | 0,000557926 | 6,90E-05 | 20 - | 0,000501209 - | 21 - | 0,000653471 - | 21 - | 0,000621207 - | 18 - | 0,000540327 - | 14 - | 0,000473415 - |
| 4145 | 3 | 0.2um_NOKCPHJN_671360;3um_FIJJAFBF_0305 | Bacteria | Bacteroidetes | Flavobacteriia | Flavobacterial | Flavobacteriac | Polaribacter | Polaribacter s | poorly charact | Function unkn | 09DAV | Protein of unk | 313598.MED1 | 0,2 | 0,4 | 9,96E-06 | 1,99E-05 | 0 - | 0 - | 0 - | 0 - | 0 - | 0 - | 1 - | 4,98E-05 - | 0 - | 0 - |
| 4146 | 1 | 3um_FIJJAFBF_399094 | Bacteria | Bacteroidetes | Flavobacteriia | Flavobacterial | Flavobacteriac | Polaribacter | Polaribacter ir | poorly charact | Function unkn | 09FAE | YqeY family | 313594.PI23P | 1,2 | 0,4 | 0,00011015 | 3,69E-05 | 1 - | 7,70E-05 - | 1 - | 9,57E-05 - | 2 - | 0,000181855 - | 1 - | 9,23E-05 - | 1 - | 0,000103943 - |
| 4147 | 1 | 0.2um_NOKCPHJN_892804 | Eukaryota | Bacillariophyta | Coscinodiscop | Thalassiosirale | Thalassiosirac | Conticribra | Conticribra we | unclassified | unclassified | unclassified | unclassified | unclassified | 12,4 | 1,2 | 0,000322365 | 4,44E-05 | 13 - | 0,000283131 - | 13 - | 0,000351565 - | 13 - | 0,000334207 - | 10 - | 0,000260879 - | 13 - | 0,000382043 - |
| 4148 | 1 | 0.2um_NOKCPHJN_200777 | Bacteria | Proteobacteri | Gammaprotec | Oceanospirilla | Saccharospirill | Reinekea | Reinekea blan | information st | Replication, re | 05C24 | DNA gyrase ne | 314283.MED2 | 4,6 | 1,854723699 | 6,97E-05 | 2,72E-05 | 6 - | 7,77E-05 - | 7 - | 0,0001126 - | 5 - | 7,65E-05 - | 3 - | 4,66E-05 - | 2 - | 3,50E-05 - |
| 4149 | 1 | 3um_FIJJAFBF_308458 | Bacteria | Proteobacteri | Alphaproteob | Rhodobactera | Rhodobactera | Planktomarina | Planktomarina | information st | Translation, ril | 17FIH | One of the ear | 388739.RSK20 | 1 | 0 | 0,000139824 | 1,33E-05 | 1 - | 0,000117119 - | 1 - | 0,000145427 - | 1 - | 0,000138247 - | 1 - | 0,000140289 - | 1 - | 0,000158035 - |
| 4150 | 1 | 3um_FIJJAFBF_306787 | Bacteria | Proteobacteri | Alphaproteob | Rhodobactera | unclassified | unclassified | Rhodobactera | unclassified | unclassified | unclassified | unclassified | unclassified | 0,6 | 0,489897949 | 4,35E-05 | 3,55E-05 | 0 - | 0 - | 1 - | 7,46E-05 - | 1 - | 7,09E-05 - | 1 - | 7,20E-05 - | 0 - | 0 - |
| 4151 | 1 | 0.2um_NOKCPHJN_450286 | Bacteria | Proteobacteri | Alphaproteob | Pelagibacteral | Pelagibacterac | unclassified | Pelagibacterac | cellular proces | Cell wall/mem | 7257 | NA | 439493.PB721 | 1,4 | 0,489897949 | 5,48E-05 | 1,61E-05 | 2 - | 6,71E-05 - | 1 - | 4,17E-05 - | 1 - | 3,96E-05 - | 2 - | 8,04E-05 - | 1 - | 4,53E-05 - |
| 4152 | 1 | 0.2um_NOKCPHJN_311782 | Bacteria | Proteobacteri | Gammaprotec | unclassified | unclassified | unclassified | Gammaprotec | cellular proces | Cell wall/mem | 0QKII | polysaccharide | 765910.Marpu | 0,2 | 0,4 | 2,54E-06 | 5,08E-06 | 0 - | 0 - | 0 - | 0 - | 0 - | 0 - | 0 - | 0 - | 1 - | 1,27E-05 - |
| 4153 | 1 | 3um_FIJJAFBF_602259 | Bacteria | Proteobacteri | Gammaprotec | Cellvibrionales | Porticoccacea | unclassified | gamma protec | metabolism | Amino acid tra | COG0174 | glutamine syn | 377629.TERTU | 1,8 | 0,4 | 5,19E-05 | 1,03E-05 | 2 - | 4,90E-05 - | 2 - | 6,09E-05 - | 2 - | 5,79E-05 - | 2 - | 5,88E-05 - | 1 - | 3,31E-05 - |
| 4154 | 1 | 0.2um_NOKCPHJN_342665 | Bacteria | Proteobacteri | Alphaproteob | Rhodobactera | Rhodobactera | Aquimixticola | Aquimixticola | metabolism | Energy produc | 07QM5 | aconitate hydi | 388401.RB215 | 0,4 | 0,489897949 | 6,01E-06 | 7,50E-06 | 1 - | 1,28E-05 - | 0 - | 0 - | 0 - | 0 - | 0 - | 0 - | 1 - | 1,73E-05 - |
| 4155 | 1 | 0.2um_NOKCPHJN_997351 | Eukaryota | unclassified | Oligohymenop | Philasterida | Philasteridae | Miamiensis | Miamiensis av | cellular proces | Signal transdu | KOG0699 | protein phosp | 3055.EDP0089 | 1 | 0 | 5,31E-05 | 5,04E-06 | 1 - | 4,45E-05 - | 1 - | 5,52E-05 - | 1 - | 5,25E-05 - | 1 - | 5,33E-05 - | 1 - | 6,00E-05 - |
| 4156 | 1 | 3um_FIJJAFBF_495119 | Bacteria | Proteobacteri | Gammaprotec | Oceanospirilla | Saccharospirill | Reinekea | Reinekea fors | unclassified | unclassified | unclassified | unclassified | unclassified | 5,2 | 0,748331477 | 0,000354214 | 6,18E-05 | 6 - | 0,000342617 - | 5 - | 0,000354525 - | 4 - | 0,000269616 - | 5 - | 0,000341998 - | 6 - | 0,000462312 - |
| 4157 | 1 | 3um_FIJJAFBF_07962 | Viruses | unclassified | unclassified | unclassified | unclassified | unclassified | uncultured M | unclassified | unclassified | unclassified | unclassified | unclassified | 0,4 | 0,489897949 | 3,01E-06 | 3,69E-06 | 0 - | 0 - | 0 - | 0 - | 1 - | 7,01E-06 - | 0 - | 0 - | 1 - | 8,02E-06 - |
| 4158 | 1 | 0.2um_NOKCPHJN_707260 | Bacteria | Bacteroidetes | unclassified | unclassified | unclassified | unclassified | unidentified e | information st | Translation, ril | COG1185 | Involved in mf | 50743.SCB49_ | 3,6 | 0,489897949 | 9,26E-05 | 1,94E-05 | 3 - | 6,40E-05 - | 4 - | 0,000105962 - | 3 - | 7,55E-05 - | 4 - | 0,000102218 - | 4 - | 0,000115148 - |
| 4159 | 1 | 3um_FIJJAFBF_736135 | Bacteria | Bacteroidetes | Flavobacteriia | various | various | various | various | information st | Translation, ril | COG0256 | This is one of t | 487797.Flav3 | 3 | 0,894427191 | 0,000342431 | 8,98E-05 | 4 - | 0,000389074 - | 4 - | 0,000483115 - | 3 - | 0,000344446 - | 2 - | 0,000233023 - | 2 - | 0,000262499 - |
| 4160 | 1 | 3um_FIJJAFBF_07065 | Bacteria | Bacteroidetes | Flavobacteriia | Flavobacterial | Flavobacteriac | Polaribacter | Polaribacter s | metabolism | Inorganic ion t | 05C27 | TonB-depende | 688270.Celal_ | 5,4 | 2,059126028 | 6,71E-05 | 1,86E-05 | 9 - | 9,72E-05 - | 4 - | 5,36E-05 - | 6 - | 7,65E-05 - | 5 - | 6,47E-05 - | 3 - | 4,37E-05 - |
| 4161 | 1 | 3um_FIJJAFBF_319584 | Bacteria | Proteobacteri | Gammaprotec | Cellvibrionales | Porticoccacea | unclassified | Gammaprotec | unclassified | unclassified | unclassified | unclassified | unclassified | 1 | 0 | 1,52E-05 | 1,44E-06 | 1 - | 1,27E-05 - | 1 - | 1,58E-05 - | 1 - | 1,50E-05 - | 1 - | 1,53E-05 - | 1 - | 1,72E-05 - |
| 4162 | 1 | 3um_FIJJAFBF_332469 | Bacteria | Bacteroidetes | Flavobacteriia | Flavobacterial | Flavobacteriac | Polaribacter | Polaribacter s | unclassified | unclassified | unclassified | unclassified | unclassified | 1,6 | 0,489897949 | 0,000105184 | 2,76E-05 | 2 - | 0,000112526 - | 1 - | 6,99E-05 - | 2 - | 0,000132826 - | 2 - | 0,000134788 - | 1 - | 7,59E-05 - |
| 4163 | 1 | 0.2um_NOKCPHJN_116666 | Bacteria | Proteobacteri | Gammaprotec | Cellvibrionales | Porticoccacea | unclassified | Gammaprotec | metabolism | Inorganic ion t | 16TQC | TONB-depend | 314287.GB22 | 19,6 | 1,854723699 | 0,000274531 | 8,04E-06 | 23 - | 0,000272151 - | 18 - | 0,000264468 - | 20 - | 0,000279345 - | 19 - | 0,000269297 - | 18 - | 0,000287396 - |
| 4164 | 1 | 0.2um_NOKCPHJN_272121 | Bacteria | Bacteroidetes | unclassified | unclassified | unclassified | unclassified | unidentified e | unclassified | unclassified | unclassified | unclassified | unclassified | 0,4 | 0,489897949 | 1,35E-05 | 1,66E-05 | 0 - | 0 - | 0 - | 0 - | 1 - | 3,35E-05 - | 1 - | 3,40E-05 - | 0 - | 0 - |
| 4165 | 1 | 3um_FIJJAFBF_104824 | Bacteria | Bacteroidetes | Flavobacteriia | Flavobacterial | Flavobacteriac | Marinirhabdus | Marinirhabdus | metabolism | Lipid transpor | 08IQE | Amp-depende | 50743.SCB49_ | 1,2 | 0,4 | 2,77E-05 | 9,27E-06 | 1 - | 1,94E-05 - | 1 - | 2,40E-05 - | 2 - | 4,57E-05 - | 1 - | 2,32E-05 - | 1 - | 2,61E-05 - |
| 4166 | 1 | 0.2um_NOKCPHJN_717043 | Bacteria | Proteobacteri | Alphaproteob | Rhodobactera | Rhodobactera | Dinoroseobac | Dinoroseobac | information st | Translation, ril | COG0018 | arginyL-tRNA s | 398580.Dshi_ | 0,4 | 0,489897949 | 1,02E-05 | 1,26E-05 | 0 - | 0 - | 1 - | 2,45E-05 - | 0 - | 0 - | 0 - | 0 - | 1 - | 2,67E-05 - |
| 4167 | 1 | 0.2um_NOKCPHJN_1036199 | Bacteria | Proteobacteri | Gammaprotec | Cellvibrionales | Porticoccacea | unclassified | Gammaprotec | unclassified | unclassified | unclassified | unclassified | unclassified | 0,2 | 0,4 | 2,01E-05 | 4,01E-05 | 0 - | 0 - | 0 - | 0 - | 1 - | 0,000100357 - | 0 - | 0 - | 0 - | 0 - |
| 4168 | 1 | 3um_FIJJAFBF_305857 | Bacteria | Actinobacteria | Actinobacteria | Micrococcales | Microbacteria | unclassified | Microbacteria | unclassified | unclassified | unclassified | unclassified | unclassified | 2,8 | 0,748331477 | 0,000100115 | 2,22E-05 | 4 - | 0,000121457 - | 2 - | 7,54E-05 - | 2 - | 7,17E-05 - | 3 - | 0,000109114 - | 3 - | 0,000122916 - |
| 4169 | 1 | 0.2um_NOKCPHJN_447989 | Bacteria | Bacteroidetes | unclassified | unclassified | unclassified | unclassified | Bacteroidetes | unclassified | unclassified | unclassified | unclassified | unclassified | 1,8 | 0,4 | 0,000147639 | 3,62E-05 | 2 - | 0,000137457 - | 2 - | 0,000170681 - | 2 - | 0,000162254 - | 1 - | 8,23E-05 - | 2 - | 0,000185478 - |
| 4170 | 1 | 3um_FIJJAFBF_563548 | Bacteria | Proteobacteri | Alphaproteob | Rhodobactera | Rhodobactera | Planktomarina | Planktomarina | unclassified | unclassified | unclassified | unclassified | unclassified | 0,6 | 0,489897949 | 3,15E-05 | 2,59E-05 | 1 - | 4,67E-05 - | 0 - | 0 - | 1 - | 5,51E-05 - | 1 - | 5,59E-05 - | 0 - | 0 - |
| 4171 | 1 | 0.2um_NOKCPHJN_820163 | Bacteria | Proteobacteri | Alphaproteob | Rhizobiales | unclassified | unclassified | Rhizobiales ba | cellular proces | Posttranslatio | COG1404 | peptidase (S8 | 314287.GB22 | 0,6 | 0,489897949 | 2,34E-05 | 1,95E-05 | 1 - | 3,30E-05 - | 0 - | 0 - | 0 - | 0 - | 1 - | 3,95E-05 - | 1 - | 4,45E-05 - |
| 4172 | 1 | 3um_FIJJAFBF_110387 | Bacteria | Bacteroidetes | Flavobacteriia | Flavobacterial | Flavobacteriac | Formosa | Formosa sp. H | cellular proces | Cell wall/mem | COG2885 | Ompa motb d | 983544.Lacal_ | 3 | 0 | 8,62E-05 | 8,18E-06 | 3 - | 7,22E-05 - | 3 - | 8,96E-05 - | 3 - | 8,52E-05 - | 3 - | 8,65E-05 - | 3 - | 9,74E-05 - |
| 4173 | 1 | 3um_FIJJAFBF_402486 | Bacteria | Bacteroidetes | Flavobacteriia | various | various | various | various | information st | Translation, ril | COG0089 | One of the ear | 487797.Flav3 | 6,6 | 0,8 | 0,000932469 | 4,90E-05 | 8 - | 0,000956473 - | 6 - | 0,000890743 - | 7 - | 0,000987891 - | 6 - | 0,000859271 - | 6 - | 0,000967965 - |
| 4174 | 1 | 0.2um_NOKCPHJN_111949 | Bacteria | Proteobacteri | Gammaprotec | Oceanospirilla | Saccharospirill | Reinekea | Reinekea fors | unclassified | unclassified | unclassified | unclassified | unclassified | 1,4 | 0,8 | 6,32E-05 | 3,70E-05 | 0 - | 0 - | 2 - | 8,96E-05 - | 1 - | 4,26E-05 - | 2 - | 8,65E-05 - | 2 - | 9,74E-05 - |
| 4175 | 1 | 0.2um_NOKCPHJN_1115200 | Eukaryota | Chlorophyta | Chlorophycea | Chlamydomor | Chlamydomor | Chlamydomor | Chlamydomor | information st | Translation, ril | KOG3181 | ribosomal pro | 4558.Sb02g03 | 4,4 | 1,356465997 | 0,000269597 | 8,06E-05 | 6 - | 0,000310207 - | 5 - | 0,000320988 - | 4 - | 0,000244112 - | 2 - | 0,000123859 - | 5 - | 0,000348816 - |
| 4176 | 1 | 3um_FIJJAFBF_630150 | Bacteria | Proteobacteri | Alphaproteob | Rhodobactera | Rhodobactera | Pseudorueger | Pseudorueger | unclassified | unclassified | unclassified | unclassified | unclassified | 3,8 | 1,166190379 | 0,000175879 | 5,12E-05 | 4 - | 0,000156692 - | 3 - | 0,000145924 - | 6 - | 0,000277438 - | 3 - | 0,000140768 - | 3 - | 0,000158575 - |
| 4177 | 1 | 3um_FIJJAFBF_357550 | Bacteria | Bacteroidetes | Flavobacteriia | Flavobacterial | Flavobacteriac | Polaribacter | Polaribacter s | unclassified | unclassified | unclassified | unclassified | unclassified | 0,6 | 0,489897949 | 1,95E-05 | 1,59E-05 | 0 - | 0 - | 1 - | 3,34E-05 - | 1 - | 3,17E-05 - | 1 - | 3,22E-05 - | 0 - | 0 - |
| 4178 | 1 | 0.2um_NOKCPHJN_301507 | Bacteria | Bacteroidetes | Flavobacteriia | Flavobacterial | Flavobacteriac | Siansivirga | Siansivirga zea | metabolism | Energy produc | 099Y5 | fumarate hydr | 391603.FBALC | 1 | 0,632455532 | 2,86E-05 | 1,60E-05 | 2 - | 4,94E-05 - | 1 - | 3,06E-05 - | 0 - | 0 - | 1 - | 2,96E-05 - | 1 - | 3,33E-05 - |
| 4179 | 1 | 0.2um_NOKCPHJN_936883 | Bacteria | Proteobacteri | Alphaproteob | Rhizobiales | Aurantimonad | Aureimonas | Aureimonas u | unclassified | unclassified | unclassified | unclassified | unclassified | 1,6 | 0,8 | 4,20E-05 | 2,16E-05 | 2 - | 4,41E-05 - | 2 - | 5,47E-05 - | 2 - | 5,20E-05 - | 0 - | 0 - | 2 - | 5,95E-05 - |
| 4180 | 1 | 3um_FIJJAFBF_397766 | Bacteria | Proteobacteri | Alphaproteob | Rhodobactera | Rhodobactera | Amylibacter | Amylibacter k | information st | Translation, ril | 08VZM | 50s ribosomal | 367336.OM22 | 1,4 | 0,489897949 | 0,000114764 | 4,66E-05 | 1 - | 6,75E-05 - | 1 - | 8,38E-05 - | 2 - | 0,000159391 - | 1 - | 8,09E-05 - | 2 - | 0,000182205 - |
| 4181 | 1 | 3um_FIJJAFBF_42573 | Bacteria | Proteobacteri | Gammaprotec | Oceanospirilla | Saccharospirill | Reinekea | Reinekea fors | information st | Translation, ril | 05KG7 | Associates wit | 314283.MED2 | 0,8 | 0,748331477 | 7,67E-05 | 7,44E-05 | 1 - | 8,38E-05 - | 0 - | 0 - | 1 - | 9,89E-05 - | 2 - | 0,000200706 - | 0 - | 0 - |
| 4182 | 1 | 3um_FIJJAFBF_205803 | unclassified | unclassified | unclassified | unclassified | unclassified | unclassified | unclassified | unclassified | unclassified | unclassified | unclassified | unclassified | 2,4 | 0,489897949 | 5,02E-05 | 8,86E-06 | 3 - | 5,31E-05 - | 3 - | 6,59E-05 - | 2 - | 4,18E-05 - | 2 - | 4,24E-05 - | 2 - | 4,77E-05 - |
| 4183 | 1 | 3um_FIJJAFBF_545824 | Bacteria | Bacteroidetes | Flavobacteriia | Flavobacterial | Flavobacteriac | Polaribacter | Polaribacter g | unclassified | unclassified | unclassified | unclassified | unclassified | 2,6 | 1,019803903 | 0,00057954 | 0,000214783 | 3 - | 0,000573884 - | 2 - | 0,000475063 - | 4 - | 0,000903215 - | 3 - | 0,000687417 - | 1 - | 0,000258124 - |
| 4184 | 1 | 3um_FIJJAFBF_729198 | Bacteria | Proteobacteri | Gammaprotec | Oceanospirilla | Saccharospirill | Reinekea | Reinekea fors | unclassified | unclassified | unclassified | unclassified | unclassified | 8,8 | 0,979795897 | 0,000861826 | 0,000166055 | 8 - | 0,000651216 - | 10 - | 0,001010772 - | 8 - | 0,000768693 - | 8 - | 0,000780047 - | 10 - | 0,001098401 - |
| 4185 | 1 | 3um_FIJJAFBF_117483 | Bacteria | Bacteroidetes | Flavobacteriia | Flavobacterial | Flavobacteriac | Formosa | Formosa sp. H | metabolism | Energy produc | COG0843 | Cytochrome C | 983544.Lacal_ | 0,2 | 0,4 | 4,77E-06 | 9,55E-06 | 0 - | 0 - | 1 - | 2,39E-05 - | 0 - | 0 - | 0 - | 0 - | 0 - | 0 - |
| 4186 | 1 | 3um_FIJJAFBF_361787 | Bacteria | unclassified | unclassified | unclassified | unclassified | unclassified | uncultured ba | metabolism | Inorganic ion t | COG4206 | tonB-depende | 1046627.BZAF | 1,8 | 1,166190379 | 4,11E-05 | 2,78E-05 | 1 - | 1,89E-05 - | 4 - | 9,39E-05 - | 2 - | 4,46E-05 - | 1 - | 2,26E-05 - | 1 - | 2,55E-05 - |
| 4187 | 1 | 3um_FIJJAFBF_393743 | Bacteria | Bacteroidetes | Flavobacteriia | unclassified | unclassified | unclassified | Flavobacteria | poorly charact | Function unkn | 09JH9 | NA | 487796.Flav2A | 1,2 | 0,4 | 6,46E-05 | 2,16E-05 | 1 - | 4,52E-05 - | 1 - | 5,61E-05 - | 2 - | 0,000106679 - | 1 - | 5,41E-05 - | 1 - | 6,10E-05 - |
| 4188 | 1 | 0.2um_NOKCPHJN_530871 | Bacteria | Bacteroidetes | Flavobacteriia | Flavobacterial | Flavobacteriac | Ulvibacter | Ulvibacter lito | unclassified | unclassified | unclassified | unclassified | unclassified | 1 | 0 | 1,44E-05 | 1,37E-06 | 1 - | 1,21E-05 - | 1 - | 1,50E-05 - | 1 - | 1,43E-05 - | 1 - | 1,45E-05 - | 1 - | 1,63E-05 - |
| 4189 | 1 | 3um_FIJJAFBF_510252 | Bacteria | Bacteroidetes | Flavobacteriia | Flavobacterial | Flavobacteriac | Flaviramulus | Flaviramulus i | unclassified | unclassified | unclassified | unclassified | unclassified | 0,6 | 0,489897949 | 6,37E-05 | 5,25E-05 | 1 - | 9,26E-05 - | 1 - | 0,000114935 - | 0 - | 0 - | 1 - | 0,000110874 - | 0 - | 0 - |
| 4190 | 1 | 0.2um_NOKCPHJN_367158 | Bacteria | Proteobacteri | Gammaprotec | unclassified | unclassified | unclassified | Gammaprotec | poorly charact | Function unkn | 0878N | Protein of unk | 566466.NOR5 | 0,4 | 0,489897949 | 5,20E-05 | 6,41E-05 | 1 - | 0,000118327 - | 0 - | 0 - | 0 - | 0 - | 1 - | 0,000141735 - | 0 - | 0 - |
| 4191 | 1 | 3um_FIJJAFBF_544457 | Bacteria | Proteobacteri | Gammaprotec | Oceanospirilla | Saccharospirill | Reinekea | Reinekea blan | cellular proces | Cell cycle cont | 07QMS | site-determini | 351348.Maqu | 0,8 | 0,748331477 | 4,11E-05 | 3,83E-05 | 0 - | 0 - | 1 - | 5,30E-05 - | 1 - | 5,04E-05 - | 2 - | 0,000102218 - | 0 - | 0 - |
| 4192 | 1 | 3um_FIJJAFBF_20711 | Eukaryota | unclassified | unclassified | Prymnesiales | Chrysochromu | Chrysochromu | Chrysochromu | unclassified | unclassified | unclassified | unclassified | unclassified | 1,2 | 0,4 | 9,36E-05 | 2,07E-05 | 2 - | 0,000134242 - | 1 - | 8,33E-05 - | 1 - | 7,92E-05 - | 1 - | 8,04E-05 - | 1 - | 9,06E-05 - |
| 4193 | 1 | 3um_FIJJAFBF_689813 | Bacteria | Bacteroidetes | Flavobacteriia | Flavobacterial | Crocinitomica | unclassified | Crocinitomica | unclassified | unclassified | unclassified | unclassified | unclassified | 0,6 | 0,489897949 | 6,96E-05 | 5,70E-05 | 0 - | 0 - | 1 - | 0,000114015 - | 0 - | 0 - | 1 - | 0,000109987 - | 1 - | 0,0001239 - |
| 4194 | 1 | 0.2um_NOKCPHJN_210688 | Bacteria | Bacteroidetes | Flavobacteriia | Flavobacterial | Flavobacteriac | Flavobacteriu | Flavobacteriu | metabolism | Inorganic ion t | 05C27 | TonB-depende | 983548.Krodi_ | 20,6 | 1,356465997 | 0,000258895 | 1,62E-05 | 23 - | 0,000243306 - | 20 - | 0,000262708 - | 21 - | 0,000262224 - | 19 - | 0,000240754 - | 20 - | 0,000285483 - |
| 4195 | 1 | 0.2um_NOKCPHJN_50569 | Bacteria | Proteobacteri | Betaproteoba | unclassified | unclassified | unclassified | Betaproteoba | metabolism | Coenzyme tra | 05ECF | c-methyltrans | 232721.Ajs_2 | 0,2 | 0,4 | 7,13E-06 | 1,43E-05 | 0 - | 0 - | 1 - | 3,56E-05 - | 0 - | 0 - | 0 - | 0 - | 0 - | 0 - |
| 4196 | 1 | 0.2um_NOKCPHJN_304091 | Bacteria | Proteobacteri | Alphaproteob | Rhodobactera | unclassified | unclassified | Rhodobactera | unclassified | unclassified | unclassified | unclassified | unclassified | 4 | 0 | 0,000346904 | 3,29E-05 | 4 - | 0,000290574 - | 4 - | 0,000360807 - | 4 - | 0,000342993 - | 4 - | 0,000348059 - | 4 - | 0,000392087 - |
| 4197 | 1 | 0.2um_NOKCPHJN_566440 | unclassified | unclassified | unclassified | unclassified | unclassified | unclassified | unclassified | unclassified | unclassified | unclassified | unclassified | unclassified | 0,2 | 0,4 | 2,06E-06 | 4,13E-06 | 0 - | 0 - | 0 - | 0 - | 1 - | 1,03E-05 - | 0 - | 0 - | 0 - | 0 - |
| 4198 | 1 | 3um_FIJJAFBF_459065 | unclassified | unclassified | unclassified | unclassified | unclassified | unclassified | unclassified | unclassified | unclassified | unclassified | unclassified | unclassified | 1 | 0 | 3,07E-05 | 2,91E-06 | 1 - | 2,57E-05 - | 1 - | 3,19E-05 - | 1 - | 3,03E-05 - | 1 - | 3,08E-05 - | 1 - | 3,46E-05 - |
| 4199 | 1 | 0.2um_NOKCPHJN_27322 | Bacteria | Bacteroidetes | Flavobacteriia | Flavobacterial | Flavobacteriac | Formosa | Formosa sp. H | metabolism | Inorganic ion t | 05C27 | TonB-depende | 742766.HMPR | 29,8 | 2,712931993 | 0,000385913 | 3,43E-05 | 32 - | 0,000348799 - | 28 - | 0,000378968 - | 34 - | 0,000437454 - | 27 - | 0,000352521 - | 28 - | 0,000411822 - |
| 4200 | 1 | 3um_FIJJAFBF_11580 | Bacteria | Proteobacteri | Gammaprotec | Cellvibrionales | Cellvibrionace | Marinagarivor | Marinagarivor | metabolism | Energy produc | 0QKED | alcohol dehyd | 1085623.GNIT | 0,4 | 0,489897949 | 1,30E-05 | 1,60E-05 | 1 - | 2,97E-05 - | 0 - | 0 - | 1 - | 3,51E-05 - | 0 - | 0 - | 0 - | 0 - |
| 4201 | 1 | 0.2um_NOKCPHJN_657137 | Eukaryota | unclassified | Cryptophyta | Pyrenomonad | Geminigerace | Guillardia | Guillardia thet | unclassified | unclassified | unclassified | unclassified | unclassified | 6 | 0 | 0,000696748 | 6,62E-05 | 6 - | 0,000583611 - | 6 - | 0,000724672 - | 6 - | 0,000688893 - | 6 - | 0,000699068 - | 6 - | 0,000787497 - |
| 4202 | 1 | 0.2um_NOKCPHJN_565556 | Eukaryota | unclassified | Cryptophyta | Pyrenomonad | Pyrenomonad | Rhodomonas | Rhodomonas | unclassified | unclassified | unclassified | unclassified | unclassified | 0,8 | 0,4 | 9,06E-05 | 4,60E-05 | 1 - | 9,81E-05 - | 1 - | 0,000121811 - | 1 - | 0,000115797 - | 1 - | 0,000117507 - | 0 - | 0 - |
| 4203 | 1 | 0.2um_NOKCPHJN_50584 | Bacteria | Proteobacteri | Betaproteoba | Burkholderiale | Comamonada | Rhodoferax | Rhodoferax k | information st | Translation, ril | 05CA4 | amino acids su | 887062.HGR_ | 0,2 | 0,4 | 2,34E-06 | 4,69E-06 | 1 - | 1,17E-05 - | 0 - | 0 - | 0 - | 0 - | 0 - | 0 - | 0 - | 0 - |
| 4204 | 1 | 0.2um_NOKCPHJN_115345 | Bacteria | Bacteroidetes | Flavobacteriia | Flavobacterial | Flavobacteriac | Olleya | Olleya namha | unclassified | unclassified | unclassified | unclassified | unclassified | 10,2 | 1,166190379 | 0,000152522 | 2,65E-05 | 9 - | 0,00011216 - | 12 - | 0,000185692 - | 9 - | 0,000132393 - | 11 - | 0,000164204 - | 10 - | 0,000168159 - |
| 4205 | 1 | 3um_FIJJAFBF_69734 | Bacteria | Proteobacteri | Gammaprotec | Pseudomonad | Pseudomonad | Pseudomonas | various | unclassified | unclassified | unclassified | unclassified | unclassified | 10,2 | 1,720465053 | 0,000280689 | 4,16E-05 | 11 - | 0,000256094 - | 11 - | 0,000317993 - | 10 - | 0,000274812 - | 12 - | 0,000334645 - | 7 - | 0,000219903 - |
| 4206 | 1 | 3um_FIJJAFBF_99652 | Bacteria | Proteobacteri | Gammaprotec | Oceanospirilla | Saccharospirill | Reinekea | Reinekea fors | unclassified | unclassified | unclassified | unclassified | unclassified | 1,2 | 0,4 | 8,60E-05 | 1,91E-05 | 2 - | 0,000123416 - | 1 - | 7,66E-05 - | 1 - | 7,28E-05 - | 1 - | 7,39E-05 - | 1 - | 8,33E-05 - |
| 4207 | 1 | 3um_FIJJAFBF_635813 | Bacteria | Bacteroidetes | Flavobacteriia | Flavobacterial | Flavobacteriac | Algibacter | Algibacter sp. | metabolism | Inorganic ion t | 09C49 | tonB-depende | 391587.KAOT | 0,4 | 0,489897949 | 9,56E-06 | 1,17E-05 | 0 - | 0 - | 1 - | 2,29E-05 - | 0 - | 0 - | 0 - | 0 - | 1 - | 2,49E-05 - |
| 4208 | 1 | 3um_FIJJAFBF_294188 | Bacteria | Bacteroidetes | Flavobacteriia | Flavobacterial | unclassified | unclassified | Flavobacterial | metabolism | Energy produc | 099Q7 | Catalyzes the | 391598.FBBAL | 0,4 | 0,489897949 | 1,61E-05 | 1,99E-05 | 1 - | 3,70E-05 - | 0 - | 0 - | 1 - | 4,37E-05 - | 0 - | 0 - | 0 - | 0 - |
| 4209 | 1 | 3um_FIJJAFBF_13036 | Bacteria | Bacteroidetes | Flavobacteriia | Flavobacterial | Flavobacteriac | Aequorivita | Aequorivita sp | cellular proces | Signal transdu | 06FN3 | tellurium resis | 203124.Tery_ | 1,4 | 0,489897949 | 0,000101573 | 3,68E-05 | 2 - | 0,000122103 - | 1 - | 7,58E-05 - | 1 - | 7,21E-05 - | 1 - | 7,31E-05 - | 2 - | 0,00016476 - |
| 4210 | 1 | 3um_FIJJAFBF_721247 | Bacteria | Bacteroidetes | unclassified | unclassified | unclassified | unclassified | Bacteroidetes | unclassified | unclassified | unclassified | unclassified | unclassified | 4,8 | 0,4 | 0,00031466 | 4,04E-05 | 5 - | 0,000274586 - | 5 - | 0,000340954 - | 5 - | 0,00032412 - | 4 - | 0,000263126 - | 5 - | 0,000370513 - |
| 4211 | 1 | 0.2um_NOKCPHJN_211237 | Eukaryota | unclassified | unclassified | Prymnesiales | Chrysochromu | Chrysochromu | Chrysochromu | unclassified | unclassified | unclassified | unclassified | unclassified | 0,4 | 0,489897949 | 2,99E-06 | 3,67E-06 | 0 - | 0 - | 0 - | 0 - | 1 - | 7,43E-06 - | 1 - | 7,54E-06 - | 0 - | 0 - |
| 4212 | 1 | 3um_FIJJAFBF_431819 | Bacteria | Proteobacteri | Alphaproteob | Rhodobactera | Rhodobactera | Planktomarina | Planktomarina | unclassified | unclassified | unclassified | unclassified | unclassified | 1 | 0,632455532 | 7,86E-05 | 4,40E-05 | 2 - | 0,00013583 - | 1 - | 8,43E-05 - | 0 - | 0 - | 1 - | 8,14E-05 - | 1 - | 9,16E-05 - |
| 4213 | 1 | 3um_FIJJAFBF_118512 | Bacteria | Proteobacteri | Gammaprotec | Cellvibrionales | Halieaceae | unclassified | marine gamm | unclassified | unclassified | unclassified | unclassified | unclassified | 1,6 | 1,019803903 | 2,49E-05 | 1,49E-05 | 3 - | 4,13E-05 - | 2 - | 3,42E-05 - | 2 - | 3,25E-05 - | 1 - | 1,65E-05 - | 0 - | 0 - |
| 4214 | 1 | 0.2um_NOKCPHJN_107951 | Bacteria | Bacteroidetes | Flavobacteriia | Flavobacterial | Flavobacteriac | Polaribacter | various | poorly charact | Function unkn | 11QQ6 | NA | 398720.MED2 | 0,2 | 0,4 | 9,09E-06 | 1,82E-05 | 0 - | 0 - | 0 - | 0 - | 1 - | 4,55E-05 - | 0 - | 0 - | 0 - | 0 - |
| 4215 | 1 | 0.2um_NOKCPHJN_391782 | Bacteria | Proteobacteri | Alphaproteob | Sphingomona | unclassified | unclassified | Sphingomona | unclassified | unclassified | unclassified | unclassified | unclassified | 1 | 0 | 4,13E-05 | 3,92E-06 | 1 - | 3,46E-05 - | 1 - | 4,29E-05 - | 1 - | 4,08E-05 - | 1 - | 4,14E-05 - | 1 - | 4,66E-05 - |
| 4216 | 1 | 0.2um_NOKCPHJN_226581 | Bacteria | Proteobacteri | Betaproteoba | Burkholderiale | Oxalobacterac | Massilia | Massilia sp. R | poorly charact | Function unkn | 0231D | TolA protein | 426355.Mrad2 | 0,4 | 0,489897949 | 2,13E-06 | 2,62E-06 | 0 - | 0 - | 1 - | 5,11E-06 - | 0 - | 0 - | 0 - | 0 - | 1 - | 5,55E-06 - |
| 4217 | 1 | 3um_FIJJAFBF_500306 | unclassified | unclassified | unclassified | unclassified | unclassified | unclassified | unclassified | unclassified | unclassified | unclassified | unclassified | unclassified | 0,2 | 0,4 | 1,12E-05 | 2,24E-05 | 0 - | 0 - | 0 - | 0 - | 0 - | 0 - | 1 - | 5,61E-05 - | 0 - | 0 - |
| 4218 | 1 | 3um_FIJJAFBF_139380 | Bacteria | Proteobacteri | Gammaprotec | Alteromonada | Alteromonada | Paraglaciecola | Paraglaciecola | metabolism | Energy produc | 16QPK | Succinate deh | 715451.ambt_ | 0,4 | 0,489897949 | 8,50E-06 | 1,05E-05 | 1 - | 1,95E-05 - | 0 - | 0 - | 1 - | 2,30E-05 - | 0 - | 0 - | 0 - | 0 - |
| 4219 | 1 | 0.2um_NOKCPHJN_473607 | Bacteria | Proteobacteri | Alphaproteob | Pelagibacteral | Pelagibacterac | unclassified | Pelagibacterac | cellular proces | Intracellular tr | 18ATJ | Pfam:YadA | 1005048.CFU_ | 1,6 | 0,489897949 | 2,35E-05 | 7,56E-06 | 1 - | 1,22E-05 - | 2 - | 3,04E-05 - | 2 - | 2,89E-05 - | 2 - | 2,93E-05 - | 1 - | 1,65E-05 - |
| 4220 | 1 | 0.2um_NOKCPHJN_184193 | Eukaryota | unclassified | Pelagophycea | Pelagomonad | unclassified | Aureococcus | Aureococcus a | metabolism | Energy produc | COG0056 | Produces ATP | 2850.Phatr14 | 2 | 0 | 6,16E-05 | 5,85E-06 | 2 - | 5,16E-05 - | 2 - | 6,41E-05 - | 2 - | 6,09E-05 - | 2 - | 6,18E-05 - | 2 - | 6,96E-05 - |
| 4221 | 1 | 3um_FIJJAFBF_373412 | Bacteria | Proteobacteri | Betaproteoba | Burkholderiale | Comamonada | Hydrogenoph | Hydrogenoph | metabolism | Inorganic ion t | 0B7CZ | Destroys radic | 596154.Alide2 | 3 | 0,894427191 | 0,000209913 | 5,51E-05 | 4 - | 0,000237879 - | 2 - | 0,000147688 - | 4 - | 0,000280792 - | 2 - | 0,00014247 - | 3 - | 0,000240738 - |
| 4222 | 1 | 3um_FIJJAFBF_189721 | Bacteria | Bacteroidetes | Flavobacteriia | Flavobacterial | Flavobacteriac | Polaribacter | Polaribacter a | unclassified | unclassified | unclassified | unclassified | unclassified | 7,2 | 0,748331477 | 0,000148883 | 1,45E-05 | 8 - | 0,000139334 - | 6 - | 0,000129759 - | 7 - | 0,000143911 - | 8 - | 0,000166899 - | 7 - | 0,00016451 - |
| 4223 | 1 | 0.2um_NOKCPHJN_557638 | Bacteria | Proteobacteri | Alphaproteob | Rhodobactera | Rhodobactera | Planktomarina | Planktomarina | cellular proces | Cell wall/mem | 16Q41 | Efflux transpo | 351016.RAZW | 2,2 | 0,748331477 | 7,17E-05 | 2,19E-05 | 3 - | 8,38E-05 - | 3 - | 0,000104028 - | 2 - | 6,59E-05 - | 2 - | 6,69E-05 - | 1 - | 3,77E-05 - |
| 4224 | 1 | 3um_FIJJAFBF_25781 | Bacteria | Proteobacteri | Gammaprotec | Alteromonada | Alteromonada | various | various | cellular proces | Signal transdu | 0QIPQ | gtp-binding pr | 342610.Patl_1 | 2,2 | 0,748331477 | 4,86E-05 | 1,56E-05 | 3 - | 5,61E-05 - | 3 - | 6,96E-05 - | 1 - | 2,21E-05 - | 2 - | 4,48E-05 - | 2 - | 5,04E-05 - |
| 4225 | 1 | 3um_FIJJAFBF_39791 | Bacteria | Bacteroidetes | Flavobacteriia | Flavobacterial | Cryomorphace | unclassified | Cryomorphace | unclassified | unclassified | unclassified | unclassified | unclassified | 2,6 | 0,8 | 0,000137413 | 4,75E-05 | 2 - | 8,76E-05 - | 2 - | 0,000108793 - | 2 - | 0,000103422 - | 4 - | 0,000209898 - | 3 - | 0,000177337 - |
| 4226 | 1 | 0.2um_NOKCPHJN_132655 | Bacteria | Bacteroidetes | Flavobacteriia | Flavobacterial | Flavobacteriac | Polaribacter | various | metabolism | Amino acid tra | 09GJ6 | nitrogen regul | 487797.Flav3 | 1,2 | 0,4 | 0,000142841 | 3,16E-05 | 2 - | 0,000204958 - | 1 - | 0,000127249 - | 1 - | 0,000120966 - | 1 - | 0,000122753 - | 1 - | 0,000138281 - |
| 4227 | 1 | 0.2um_NOKCPHJN_828264 | Eukaryota | Chlorophyta | unclassified | Pyramimonad | unclassified | Cymbomonas | Cymbomonas | metabolism | Energy produc | 0IHF0 | This is one of t | 70448.Q0P3J1 | 1 | 0 | 3,88E-05 | 3,69E-06 | 1 - | 3,25E-05 - | 1 - | 4,04E-05 - | 1 - | 3,84E-05 - | 1 - | 3,89E-05 - | 1 - | 4,39E-05 - |
| 4228 | 1 | 3um_FIJJAFBF_383341 | Bacteria | Bacteroidetes | Flavobacteriia | Flavobacterial | Flavobacteriac | Formosa | Formosa sp. H | cellular proces | Cell wall/mem | 11TPU | outer membra | 1046627.BZAF | 4,8 | 0,748331477 | 0,000386089 | 5,05E-05 | 6 - | 0,000407491 - | 4 - | 0,000337323 - | 4 - | 0,000320668 - | 5 - | 0,000406755 - | 5 - | 0,000458209 - |
| 4229 | 1 | 0.2um_NOKCPHJN_409617 | Bacteria | Proteobacteri | Gammaprotec | unclassified | unclassified | unclassified | SAR86 cluster | metabolism | Inorganic ion t | 16PDP | TonB-depende | 190486.XAC3 | 0,2 | 0,4 | 2,75E-06 | 5,49E-06 | 0 - | 0 - | 0 - | 0 - | 1 - | 1,37E-05 - | 0 - | 0 - | 0 - | 0 - |
| 4230 | 1 | 3um_FIJJAFBF_129704 | Bacteria | Proteobacteri | Alphaproteob | Rhodobactera | Rhodobactera | Ascidiaceihabi | Ascidiaceihabi | metabolism | Carbohydrate | COG0057 | glyceraldehyd | 375451.RD1_ | 0,6 | 0,489897949 | 2,61E-05 | 2,14E-05 | 0 - | 0 - | 1 - | 4,29E-05 - | 1 - | 4,08E-05 - | 0 - | 0 - | 1 - | 4,66E-05 - |
| 4231 | 1 | 3um_FIJJAFBF_631312 | Bacteria | Proteobacteri | Gammaprotec | Cellvibrionales | Porticoccacea | unclassified | gamma protec | unclassified | unclassified | unclassified | unclassified | unclassified | 0,8 | 0,4 | 6,28E-05 | 3,19E-05 | 1 - | 6,79E-05 - | 1 - | 8,43E-05 - | 1 - | 8,02E-05 - | 1 - | 8,14E-05 - | 0 - | 0 - |
| 4232 | 1 | 3um_FIJJAFBF_128487 | Bacteria | Bacteroidetes | Flavobacteriia | Flavobacterial | Flavobacteriac | Formosa | Formosa sp. H | unclassified | unclassified | unclassified | unclassified | unclassified | 2,4 | 0,489897949 | 0,000220883 | 6,23E-05 | 2 - | 0,000152022 - | 3 - | 0,00028315 - | 2 - | 0,000179447 - | 2 - | 0,000182097 - | 3 - | 0,000307698 - |
| 4233 | 2 | 0.2um_NOKCPHJN_212846;3um_FIJJAFBF_4894 | Bacteria | Bacteroidetes | Flavobacteriia | Flavobacterial | Flavobacteriac | Formosa | Formosa sp. H | poorly charact | Function unkn | 0P8G1 | peptidylprolyl | 391603.FBALC | 0,4 | 0,489897949 | 1,60E-05 | 1,98E-05 | 1 - | 3,56E-05 - | 1 - | 4,43E-05 - | 0 - | 0 - | 0 - | 0 - | 0 - | 0 - |
| 4234 | 1 | 3um_FIJJAFBF_615694 | Bacteria | Proteobacteri | Alphaproteob | Rhodobactera | Rhodobactera | Planktomarina | Planktomarina | unclassified | unclassified | unclassified | unclassified | unclassified | 2,6 | 0,489897949 | 0,000323862 | 5,47E-05 | 3 - | 0,000315899 - | 3 - | 0,000392254 - | 2 - | 0,000248591 - | 3 - | 0,000378394 - | 2 - | 0,000284173 - |
| 4235 | 1 | 0.2um_NOKCPHJN_1101585 | Bacteria | Bacteroidetes | Flavobacteriia | Flavobacterial | Flavobacteriac | Formosa | Formosa sp. H | unclassified | unclassified | unclassified | unclassified | unclassified | 1 | 0,632455532 | 4,18E-05 | 2,34E-05 | 2 - | 7,22E-05 - | 1 - | 4,48E-05 - | 0 - | 0 - | 1 - | 4,32E-05 - | 1 - | 4,87E-05 - |
| 4236 | 1 | 3um_FIJJAFBF_649594 | Bacteria | Proteobacteri | Gammaprotec | Alteromonada | Alteromonada | Paraglaciecola | Paraglaciecola | unclassified | unclassified | unclassified | unclassified | unclassified | 0,4 | 0,489897949 | 6,97E-06 | 8,59E-06 | 1 - | 1,60E-05 - | 0 - | 0 - | 1 - | 1,89E-05 - | 0 - | 0 - | 0 - | 0 - |
| 4237 | 1 | 0.2um_NOKCPHJN_356926 | Bacteria | Bacteroidetes | unclassified | unclassified | unclassified | unclassified | Bacteroidetes | poorly charact | Function unkn | 08QKU | NA | 755732.Fluta_ | 2,2 | 1,166190379 | 2,11E-05 | 1,11E-05 | 2 - | 1,63E-05 - | 1 - | 1,01E-05 - | 4 - | 3,85E-05 - | 3 - | 2,93E-05 - | 1 - | 1,10E-05 - |
| 4238 | 1 | 3um_FIJJAFBF_127427 | Bacteria | Proteobacteri | Gammaprotec | Cellvibrionales | Porticoccacea | unclassified | Gammaprotec | unclassified | unclassified | unclassified | unclassified | unclassified | 1 | 0,894427191 | 2,58E-05 | 2,26E-05 | 0 - | 0 - | 2 - | 5,15E-05 - | 0 - | 0 - | 2 - | 4,96E-05 - | 1 - | 2,80E-05 - |
| 4239 | 1 | 3um_FIJJAFBF_72482 | Bacteria | Bacteroidetes | Cytophagia | Cytophagales | Flammeovirga | unclassified | Flammeovirga | unclassified | unclassified | unclassified | unclassified | unclassified | 2,2 | 0,748331477 | 0,000170658 | 4,97E-05 | 3 - | 0,000200192 - | 2 - | 0,00016572 - | 2 - | 0,000157537 - | 3 - | 0,000239796 - | 1 - | 9,00E-05 - |
| 4240 | 1 | 3um_FIJJAFBF_312459 | Bacteria | Bacteroidetes | Cytophagia | Cytophagales | Flammeovirga | unclassified | Flammeovirga | cellular proces | Posttranslatio | COG1225 | alkyl hydroper | 643867.Ftrac_ | 0,4 | 0,8 | 3,04E-05 | 6,08E-05 | 2 - | 0,000152022 - | 0 - | 0 - | 0 - | 0 - | 0 - | 0 - | 0 - | 0 - |
| 4241 | 1 | 3um_FIJJAFBF_381160 | Bacteria | Proteobacteri | Alphaproteob | Rhizobiales | Cohaesibacter | Breoghania | Breoghania co | metabolism | Amino acid tra | 05CAI | ABC transport | 1082931.KKY_ | 0,6 | 0,489897949 | 3,10E-05 | 2,55E-05 | 1 - | 4,50E-05 - | 1 - | 5,59E-05 - | 0 - | 0 - | 1 - | 5,39E-05 - | 0 - | 0 - |
| 4242 | 1 | 0.2um_NOKCPHJN_100732 | Bacteria | Bacteroidetes | Flavobacteriia | Flavobacterial | Flavobacteriac | Aureitalea | Aureitalea ma | unclassified | unclassified | unclassified | unclassified | unclassified | 10 | 0,632455532 | 0,000155122 | 1,88E-05 | 10 - | 0,000129838 - | 11 - | 0,000177342 - | 10 - | 0,00015326 - | 9 - | 0,000139972 - | 10 - | 0,000175197 - |
| 4243 | 1 | 0.2um_NOKCPHJN_44818 | Eukaryota | unclassified | unclassified | Isochrysidales | Noelaerhabda | Emiliania | Emiliania huxl | metabolism | Energy produc | 0FJ7X | This is one of t | 32049.SYNPC | 1,2 | 0,4 | 4,44E-05 | 9,84E-06 | 2 - | 6,38E-05 - | 1 - | 3,96E-05 - | 1 - | 3,76E-05 - | 1 - | 3,82E-05 - | 1 - | 4,30E-05 - |
| 4244 | 1 | 0.2um_NOKCPHJN_80666 | Bacteria | unclassified | unclassified | unclassified | unclassified | unclassified | bacterium TM | poorly charact | Function unkn | 0NVRZ | NA | 170187.SP_17 | 1,2 | 0,4 | 2,76E-05 | 6,12E-06 | 2 - | 3,96E-05 - | 1 - | 2,46E-05 - | 1 - | 2,34E-05 - | 1 - | 2,37E-05 - | 1 - | 2,67E-05 - |
| 4245 | 1 | 3um_FIJJAFBF_128529 | unclassified | unclassified | unclassified | unclassified | unclassified | unclassified | unclassified | unclassified | unclassified | unclassified | unclassified | unclassified | 0,8 | 0,4 | 5,59E-05 | 2,87E-05 | 1 - | 5,86E-05 - | 1 - | 7,27E-05 - | 1 - | 6,91E-05 - | 0 - | 0 - | 1 - | 7,90E-05 - |
| 4246 | 1 | 0.2um_NOKCPHJN_246325 | Bacteria | Bacteroidetes | unclassified | unclassified | unclassified | unclassified | Bacteroidetes | unclassified | unclassified | unclassified | unclassified | unclassified | 2 | 0,632455532 | 0,000217267 | 7,13E-05 | 2 - | 0,000183643 - | 3 - | 0,000342045 - | 2 - | 0,000216772 - | 2 - | 0,000219973 - | 1 - | 0,0001239 - |
| 4247 | 1 | 3um_FIJJAFBF_30986 | Bacteria | Bacteroidetes | Flavobacteriia | Flavobacterial | Flavobacteriac | Aequorivita | Aequorivita lip | information st | Translation, ril | 08UIK | This protein bi | 50743.SCB49_ | 3,4 | 0,489897949 | 0,000321849 | 5,71E-05 | 3 - | 0,000237469 - | 4 - | 0,000393155 - | 4 - | 0,000373744 - | 3 - | 0,000284448 - | 3 - | 0,00032043 - |
| 4248 | 1 | 3um_FIJJAFBF_248998 | Bacteria | Bacteroidetes | Flavobacteriia | Flavobacterial | Flavobacteriac | Polaribacter | various | information st | Translation, ril | COG0254 | 50S ribosomal | 313598.MED1 | 3,2 | 0,4 | 0,00051671 | 2,83E-05 | 4 - | 0,000546556 - | 3 - | 0,000508996 - | 3 - | 0,000483865 - | 3 - | 0,000491012 - | 3 - | 0,000553123 - |
| 4249 | 1 | 0.2um_NOKCPHJN_15531 | Bacteria | Bacteroidetes | Flavobacteriia | unclassified | unclassified | unclassified | Flavobacteria | unclassified | unclassified | unclassified | unclassified | unclassified | 1 | 0 | 6,34E-05 | 6,02E-06 | 1 - | 5,31E-05 - | 1 - | 6,60E-05 - | 1 - | 6,27E-05 - | 1 - | 6,36E-05 - | 1 - | 7,17E-05 - |
| 4250 | 1 | 0.2um_NOKCPHJN_505075 | unclassified | unclassified | unclassified | unclassified | unclassified | unclassified | unclassified | unclassified | unclassified | unclassified | unclassified | unclassified | 1,6 | 0,489897949 | 8,48E-05 | 3,10E-05 | 1 - | 4,35E-05 - | 2 - | 0,000107969 - | 1 - | 5,13E-05 - | 2 - | 0,000104154 - | 2 - | 0,000117329 - |
| 4251 | 1 | 3um_FIJJAFBF_521278 | Bacteria | Proteobacteri | Alphaproteob | Rhodobactera | Rhodobactera | Cognatiyoonia | Cognatiyoonia | unclassified | unclassified | unclassified | unclassified | unclassified | 2,2 | 0,4 | 0,000117861 | 1,08E-05 | 3 - | 0,000136639 - | 2 - | 0,00011311 - | 2 - | 0,000107526 - | 2 - | 0,000109114 - | 2 - | 0,000122916 - |
| 4252 | 1 | 0.2um_NOKCPHJN_187377 | Bacteria | Proteobacteri | Alphaproteob | unclassified | unclassified | Candidatus Pu | Candidatus Pu | information st | Translation, ril | COG0051 | tRNA binding | 488538.SAR11 | 0,2 | 0,4 | 2,79E-05 | 5,59E-05 | 0 - | 0 - | 1 - | 0,000139724 - | 0 - | 0 - | 0 - | 0 - | 0 - | 0 - |
| 4253 | 1 | 0.2um_NOKCPHJN_586048 | Bacteria | Bacteroidetes | unclassified | unclassified | unclassified | unclassified | Bacteroidetes | unclassified | unclassified | unclassified | unclassified | unclassified | 3 | 0,632455532 | 3,11E-05 | 3,87E-06 | 4 - | 3,55E-05 - | 3 - | 3,30E-05 - | 3 - | 3,14E-05 - | 3 - | 3,19E-05 - | 2 - | 2,39E-05 - |
| 4254 | 1 | 0.2um_NOKCPHJN_355163 | Eukaryota | unclassified | Dinophyceae | Suessiales | Symbiodiniace | Symbiodinium | Symbiodinium | poorly charact | Function unkn | 0ZUGC | NA | 7165.AGAP00 | 0,8 | 0,4 | 1,13E-05 | 5,76E-06 | 1 - | 1,23E-05 - | 1 - | 1,52E-05 - | 1 - | 1,45E-05 - | 1 - | 1,47E-05 - | 0 - | 0 - |
| 4255 | 1 | 3um_FIJJAFBF_129163 | Bacteria | Proteobacteri | Gammaprotec | unclassified | unclassified | unclassified | OM182 bacte | unclassified | unclassified | unclassified | unclassified | unclassified | 16,8 | 2,039607805 | 0,000230996 | 1,87E-05 | 20 - | 0,000232342 - | 14 - | 0,00020195 - | 16 - | 0,000219404 - | 18 - | 0,000250476 - | 16 - | 0,000250809 - |
| 4256 | 1 | 3um_FIJJAFBF_286853 | Bacteria | Proteobacteri | Betaproteoba | Burkholderiale | Comamonada | Hydrogenoph | Hydrogenoph | unclassified | unclassified | unclassified | unclassified | unclassified | 0,6 | 0,489897949 | 3,57E-05 | 2,94E-05 | 1 - | 5,22E-05 - | 1 - | 6,48E-05 - | 1 - | 6,16E-05 - | 0 - | 0 - | 0 - | 0 - |
| 4257 | 1 | 3um_FIJJAFBF_721624 | Eukaryota | unclassified | Oomycetes | Pythiales | Pythiaceae | Pythium | Pythium insidi | unclassified | unclassified | unclassified | unclassified | unclassified | 0,8 | 0,4 | 0,000115295 | 5,93E-05 | 1 - | 0,000120818 - | 1 - | 0,00015002 - | 1 - | 0,000142613 - | 0 - | 0 - | 1 - | 0,000163026 - |
| 4258 | 1 | 0.2um_NOKCPHJN_11466 | Bacteria | Proteobacteri | Gammaprotec | Cellvibrionales | Porticoccacea | unclassified | various | unclassified | unclassified | unclassified | unclassified | unclassified | 1 | 0 | 7,88E-05 | 7,48E-06 | 1 - | 6,60E-05 - | 1 - | 8,19E-05 - | 1 - | 7,79E-05 - | 1 - | 7,90E-05 - | 1 - | 8,90E-05 - |
| 4259 | 1 | 0.2um_NOKCPHJN_366105 | Viruses | unclassified | unclassified | unclassified | unclassified | unclassified | Tetraselmis vi | unclassified | unclassified | unclassified | unclassified | unclassified | 0,4 | 0,489897949 | 9,39E-06 | 1,16E-05 | 1 - | 2,09E-05 - | 1 - | 2,60E-05 - | 0 - | 0 - | 0 - | 0 - | 0 - | 0 - |
| 4260 | 1 | 3um_FIJJAFBF_784243 | Eukaryota | Bacillariophyta | Bacillariophyc | Bacillariales | Bacillariaceae | Fragilariopsis | Fragilariopsis | metabolism | Carbohydrate | 1255K | major facilitat | 2850.Phatr44 | 1 | 0 | 8,62E-05 | 8,18E-06 | 1 - | 7,22E-05 - | 1 - | 8,96E-05 - | 1 - | 8,52E-05 - | 1 - | 8,65E-05 - | 1 - | 9,74E-05 - |
| 4261 | 1 | 3um_FIJJAFBF_455943 | Bacteria | Proteobacteri | Betaproteoba | unclassified | unclassified | unclassified | Betaproteoba | unclassified | unclassified | unclassified | unclassified | unclassified | 0,4 | 0,489897949 | 9,31E-06 | 1,15E-05 | 1 - | 2,08E-05 - | 1 - | 2,58E-05 - | 0 - | 0 - | 0 - | 0 - | 0 - | 0 - |
| 4262 | 1 | 3um_FIJJAFBF_251956 | Bacteria | Planctomycete | Phycisphaerae | Phycisphaeral | Phycisphaerac | Phycisphaera | Phycisphaera | unclassified | unclassified | unclassified | unclassified | unclassified | 0,2 | 0,4 | 6,43E-06 | 1,29E-05 | 1 - | 3,22E-05 - | 0 - | 0 - | 0 - | 0 - | 0 - | 0 - | 0 - | 0 - |
| 4263 | 1 | 3um_FIJJAFBF_212105 | Bacteria | Bacteroidetes | Saprospiria | Saprospirales | Haliscomenob | Phaeodactylib | Phaeodactylib | unclassified | unclassified | unclassified | unclassified | unclassified | 0,6 | 0,8 | 1,52E-05 | 2,16E-05 | 1 - | 2,05E-05 - | 0 - | 0 - | 0 - | 0 - | 0 - | 0 - | 2 - | 5,53E-05 - |
| 4264 | 1 | 0.2um_NOKCPHJN_903644 | Eukaryota | Mollusca | Bivalvia | Pectinoida | Pectinidae | Mizuhopecter | Mizuhopecter | unclassified | unclassified | unclassified | unclassified | unclassified | 1 | 0 | 2,11E-05 | 2,01E-06 | 1 - | 1,77E-05 - | 1 - | 2,20E-05 - | 1 - | 2,09E-05 - | 1 - | 2,12E-05 - | 1 - | 2,39E-05 - |
| 4265 | 1 | 0.2um_NOKCPHJN_15530 | Bacteria | Bacteroidetes | Flavobacteriia | unclassified | unclassified | unclassified | Flavobacteria | poorly charact | Function unkn | 05ERK | gliding motilit | 487796.Flav2A | 0,6 | 0,489897949 | 1,58E-05 | 1,29E-05 | 0 - | 0 - | 1 - | 2,71E-05 - | 1 - | 2,58E-05 - | 1 - | 2,62E-05 - | 0 - | 0 - |
| 4266 | 1 | 3um_FIJJAFBF_88375 | Bacteria | Bacteroidetes | Flavobacteriia | Flavobacterial | Flavobacteriac | Lacinutrix | Lacinutrix sp. | unclassified | unclassified | unclassified | unclassified | unclassified | 8,2 | 0,748331477 | 0,000132232 | 1,74E-05 | 9 - | 0,000121671 - | 8 - | 0,000134293 - | 8 - | 0,000127663 - | 7 - | 0,000113355 - | 9 - | 0,000164178 - |
| 4267 | 1 | 0.2um_NOKCPHJN_10062 | Bacteria | Bacteroidetes | Flavobacteriia | Flavobacterial | Flavobacteriac | Polaribacter | Polaribacter s | unclassified | unclassified | unclassified | unclassified | unclassified | 2,2 | 1,469693846 | 4,55E-05 | 2,52E-05 | 5 - | 9,05E-05 - | 2 - | 4,50E-05 - | 1 - | 2,14E-05 - | 1 - | 2,17E-05 - | 2 - | 4,89E-05 - |
| 4268 | 1 | 3um_FIJJAFBF_390955 | Viruses | unclassified | unclassified | unclassified | unclassified | unclassified | Tetraselmis vi | unclassified | unclassified | unclassified | unclassified | unclassified | 3,2 | 0,748331477 | 0,000184774 | 4,70E-05 | 4 - | 0,000193716 - | 3 - | 0,000180404 - | 2 - | 0,000114331 - | 3 - | 0,00017403 - | 4 - | 0,000261392 - |
| 4269 | 1 | 3um_FIJJAFBF_11582 | Bacteria | Proteobacteri | Gammaprotec | Oceanospirilla | Saccharospirill | Reinekea | Reinekea fors | unclassified | unclassified | unclassified | unclassified | unclassified | 0,6 | 0,489897949 | 2,47E-05 | 2,02E-05 | 0 - | 0 - | 1 - | 4,24E-05 - | 1 - | 4,03E-05 - | 1 - | 4,09E-05 - | 0 - | 0 - |
| 4270 | 1 | 0.2um_NOKCPHJN_487408 | Bacteria | Proteobacteri | Gammaprotec | Alteromonada | Pseudoalteror | Pseudoalteror | Pseudoalteror | unclassified | unclassified | unclassified | unclassified | unclassified | 2,2 | 0,748331477 | 9,04E-05 | 3,19E-05 | 2 - | 6,94E-05 - | 3 - | 0,000129171 - | 2 - | 8,19E-05 - | 3 - | 0,000124607 - | 1 - | 4,68E-05 - |
| 4271 | 1 | 0.2um_NOKCPHJN_412874 | Bacteria | Bacteroidetes | Flavobacteriia | Flavobacterial | Flavobacteriac | Polaribacter | various | poorly charact | Function unkn | 0XP53 | Ragb susd dor | 487797.Flav3 | 0,6 | 0,489897949 | 5,79E-06 | 4,77E-06 | 1 - | 8,57E-06 - | 0 - | 0 - | 1 - | 1,01E-05 - | 1 - | 1,03E-05 - | 0 - | 0 - |
| 4272 | 1 | 3um_FIJJAFBF_775495 | Bacteria | Bacteroidetes | Flavobacteriia | Flavobacterial | Flavobacteriac | Winogradskye | Winogradskye | unclassified | unclassified | unclassified | unclassified | unclassified | 2,6 | 1,356465997 | 7,07E-05 | 3,51E-05 | 4 - | 9,37E-05 - | 4 - | 0,000116342 - | 1 - | 2,76E-05 - | 3 - | 8,42E-05 - | 1 - | 3,16E-05 - |
| 4273 | 1 | 0.2um_NOKCPHJN_291231 | Bacteria | unclassified | unclassified | unclassified | unclassified | unclassified | uncultured ba | poorly charact | Function unkn | 09AIG | NA | 50743.SCB49_ | 0,8 | 0,979795897 | 1,96E-05 | 2,40E-05 | 0 - | 0 - | 0 - | 0 - | 2 - | 4,86E-05 - | 2 - | 4,94E-05 - | 0 - | 0 - |
| 4274 | 1 | 3um_FIJJAFBF_468768 | Bacteria | Proteobacteri | Gammaprotec | Oceanospirilla | Oceanospirilla | Marinomonas | Marinomonas | unclassified | unclassified | unclassified | unclassified | unclassified | 28,6 | 3,382306905 | 0,000677051 | 8,11E-05 | 34 - | 0,000677502 - | 25 - | 0,000618571 - | 27 - | 0,000635073 - | 26 - | 0,000620584 - | 31 - | 0,000833526 - |
| 4275 | 1 | 3um_FIJJAFBF_690413 | Bacteria | Bacteroidetes | Flavobacteriia | unclassified | unclassified | unclassified | Flavobacteria | poorly charact | Function unkn | 11PV5 | Protein of unk | 487796.Flav2A | 0,6 | 0,489897949 | 4,28E-05 | 3,53E-05 | 1 - | 6,34E-05 - | 0 - | 0 - | 1 - | 7,49E-05 - | 1 - | 7,60E-05 - | 0 - | 0 - |
| 4276 | 1 | 0.2um_NOKCPHJN_247591 | Bacteria | Bacteroidetes | Flavobacteriia | Flavobacterial | Flavobacteriac | Flavivirga | Flavivirga eckl | unclassified | unclassified | unclassified | unclassified | unclassified | 0,4 | 0,489897949 | 5,26E-06 | 6,48E-06 | 1 - | 1,21E-05 - | 0 - | 0 - | 1 - | 1,42E-05 - | 0 - | 0 - | 0 - | 0 - |
| 4277 | 1 | 3um_FIJJAFBF_441367 | unclassified | unclassified | unclassified | unclassified | unclassified | unclassified | unclassified | unclassified | unclassified | unclassified | unclassified | unclassified | 0,4 | 0,489897949 | 5,63E-05 | 6,89E-05 | 0 - | 0 - | 0 - | 0 - | 1 - | 0,000139672 - | 1 - | 0,000141735 - | 0 - | 0 - |
| 4278 | 1 | 0.2um_NOKCPHJN_66784 | Bacteria | Proteobacteri | Alphaproteob | Rhodobactera | Rhodobactera | Roseobacter | Roseobacter d | unclassified | unclassified | unclassified | unclassified | unclassified | 0,2 | 0,4 | 5,31E-06 | 1,06E-05 | 0 - | 0 - | 1 - | 2,65E-05 - | 0 - | 0 - | 0 - | 0 - | 0 - | 0 - |
| 4279 | 1 | 3um_FIJJAFBF_786147 | Bacteria | Proteobacteri | Gammaprotec | Oceanospirilla | Saccharospirill | Reinekea | Reinekea fors | information st | Translation, ril | 0QK14 | This protein bi | 314283.MED2 | 7,8 | 2,712931993 | 0,000590282 | 0,000154911 | 13 - | 0,000842993 - | 6 - | 0,000483115 - | 6 - | 0,000459262 - | 6 - | 0,000466045 - | 8 - | 0,000699998 - |
| 4280 | 1 | 0.2um_NOKCPHJN_371316 | Bacteria | Proteobacteri | Gammaprotec | Cellvibrionales | Porticoccacea | unclassified | SAR92 bacteri | unclassified | unclassified | unclassified | unclassified | unclassified | 0,4 | 0,489897949 | 5,47E-06 | 6,77E-06 | 1 - | 1,22E-05 - | 1 - | 1,52E-05 - | 0 - | 0 - | 0 - | 0 - | 0 - | 0 - |
| 4281 | 1 | 3um_FIJJAFBF_308626 | Bacteria | Bacteroidetes | Flavobacteriia | Flavobacterial | Flavobacteriac | Formosa | Formosa sp. H | unclassified | unclassified | unclassified | unclassified | unclassified | 5 | 1,264911064 | 0,000452337 | 8,77E-05 | 7 - | 0,00053922 - | 6 - | 0,000573901 - | 4 - | 0,000363711 - | 4 - | 0,000369083 - | 4 - | 0,00041577 - |
| 4282 | 1 | 3um_FIJJAFBF_402071 | Bacteria | Bacteroidetes | Flavobacteriia | Flavobacterial | Flavobacteriac | Kordia | Kordia periser | unclassified | unclassified | unclassified | unclassified | unclassified | 0,2 | 0,4 | 5,17E-06 | 1,03E-05 | 1 - | 2,59E-05 - | 0 - | 0 - | 0 - | 0 - | 0 - | 0 - | 0 - | 0 - |
| 4283 | 1 | 3um_FIJJAFBF_170064 | Bacteria | Proteobacteri | Gammaprotec | unclassified | unclassified | unclassified | various | information st | Translation, ril | 179RU | 30S ribosomal | 247634.GPB2 | 1,4 | 0,489897949 | 0,000148483 | 5,55E-05 | 1 - | 8,83E-05 - | 2 - | 0,00021926 - | 1 - | 0,000104217 - | 2 - | 0,000211513 - | 1 - | 0,000119134 - |
| 4284 | 1 | 3um_FIJJAFBF_78607 | Bacteria | Bacteroidetes | Flavobacteriia | unclassified | unclassified | unclassified | Flavobacteria | unclassified | unclassified | unclassified | unclassified | unclassified | 0,2 | 0,4 | 1,02E-05 | 2,03E-05 | 1 - | 5,08E-05 - | 0 - | 0 - | 0 - | 0 - | 0 - | 0 - | 0 - | 0 - |

| | | | | | | | | | | | | | | | | | | | | | | | | | | | | |
|---|---|---|---|---|---|---|---|---|---|---|---|---|---|---|---|---|---|---|---|---|---|---|---|---|---|---|---|---|
| 4285 | 1 | 0.2um_NOKCPHJN_85460 | Bacteria | Bacteroidetes | Flavobacteriia | Flavobacterial | Flavobacteriac | unclassified | Flavobacteriac | unclassified | unclassified | unclassified | unclassified | unclassified | 3,2 | 0,4 | 0,000167243 | 2,48E-05 | 3 - | 0,000131424 - | 3 - | 0,00016319 - | 4 - | 0,000206843 - | 3 - | 0,000157424 - | 3 - | 0,000177337 - |
| 4286 | 1 | 0.2um_NOKCPHJN_237255 | Bacteria | Proteobacteri | Alphaproteob | Pelagibacteral | Pelagibacterac | Candidatus Pe | Candidatus Pe | unclassified | unclassified | unclassified | unclassified | unclassified | 5,8 | 1,326649916 | 0,000120406 | 1,74E-05 | 8 - | 0,000141919 - | 6 - | 0,000132166 - | 5 - | 0,0001047 - | 6 - | 0,000127496 - | 4 - | 9,57E-05 - |
| 4287 | 1 | 3um_FIJJAFBF_331193 | Bacteria | unclassified | unclassified | unclassified | unclassified | unclassified | uncultured ba | metabolism | Energy produc | 05CMV | Succinyl-CoA s | 983544.Lacal_ | 5,8 | 0,979795897 | 0,00019777 | 2,11E-05 | 7 - | 0,000202888 - | 6 - | 0,000215938 - | 6 - | 0,000205276 - | 6 - | 0,000208308 - | 4 - | 0,000156439 - |
| 4288 | 1 | 3um_FIJJAFBF_10380 | Eukaryota | Phaeophyceae | unclassified | Ectocarpales | Ectocarpaceae | Ectocarpus | Ectocarpus sili | unclassified | unclassified | unclassified | unclassified | unclassified | 1 | 0 | 3,66E-05 | 3,48E-06 | 1 - | 3,07E-05 - | 1 - | 3,81E-05 - | 1 - | 3,62E-05 - | 1 - | 3,68E-05 - | 1 - | 4,14E-05 - |
| 4289 | 1 | 0.2um_NOKCPHJN_1087152 | Bacteria | Bacteroidetes | Flavobacteriia | Flavobacterial | Crocinitomica | unclassified | Crocinitomica | cellular proces | Intracellular tr | 0PBE9 | Preprotein tra | 755732.Fluta_ | 0,4 | 0,489897949 | 6,05E-05 | 7,44E-05 | 0 - | 0 - | 0 - | 0 - | 1 - | 0,000141127 - | 0 - | 0 - | 1 - | 0,000161328 - |
| 4290 | 1 | 0.2um_NOKCPHJN_1105032 | Bacteria | Proteobacteri | Alphaproteob | Pelagibacteral | Pelagibacterac | Candidatus Pe | Candidatus Pe | metabolism | Amino acid tra | 16V6G | Glycine betain | 335992.SAR11 | 4,6 | 1,356465997 | 0,000191718 | 3,72E-05 | 7 - | 0,000251074 - | 4 - | 0,000178149 - | 4 - | 0,000169353 - | 5 - | 0,000214818 - | 3 - | 0,000145195 - |
| 4291 | 1 | 3um_FIJJAFBF_119138 | Bacteria | Bacteroidetes | Flavobacteriia | Flavobacterial | Flavobacteriac | Polaribacter | Polaribacter s | unclassified | unclassified | unclassified | unclassified | unclassified | 4 | 1,095445115 | 9,72E-05 | 2,83E-05 | 5 - | 0,000101933 - | 4 - | 0,000101257 - | 4 - | 9,63E-05 - | 2 - | 4,88E-05 - | 5 - | 0,000137544 - |
| 4292 | 1 | 3um_FIJJAFBF_212746 | Bacteria | Bacteroidetes | Flavobacteriia | Flavobacterial | Flavobacteriac | Polaribacter | Polaribacter s | poorly charact | Function unkn | 05ERK | gliding motilit | 313594.PI23P_ | 12,6 | 0,489897949 | 0,000332515 | 3,35E-05 | 13 - | 0,000287495 - | 12 - | 0,000329523 - | 12 - | 0,000313254 - | 13 - | 0,000344371 - | 13 - | 0,000387932 - |
| 4293 | 1 | 3um_FIJJAFBF_116618 | Bacteria | Bacteroidetes | unclassified | unclassified | unclassified | unclassified | Bacteroidetes | poorly charact | Function unkn | 09EUF | NA | 487796.Flav2A | 0,2 | 0,4 | 1,03E-05 | 2,06E-05 | 1 - | 5,15E-05 - | 0 - | 0 - | 0 - | 0 - | 0 - | 0 - | 0 - | 0 - |
| 4294 | 1 | 3um_FIJJAFBF_08157 | Bacteria | Bacteroidetes | Flavobacteriia | Flavobacterial | Flavobacteriac | Polaribacter | various | metabolism | Inorganic ion t | 0P8Q4 | tonB-depende | 867900.Celly_ | 51,2 | 3,18747549 | 0,0007168 | 6,37E-05 | 55 - | 0,000646795 - | 47 - | 0,00068631 - | 54 - | 0,000749594 - | 48 - | 0,000676147 - | 52 - | 0,000825151 - |
| 4295 | 1 | 3um_FIJJAFBF_619864 | Eukaryota | unclassified | unclassified | Tritrichomona | Tritrichomona | Tritrichomona | Tritrichomona | information st | Replication, re | 1DRSG | DNA-binding p | 3711.Bra0173 | 0,6 | 0,489897949 | 8,90E-06 | 7,30E-06 | 0 - | 0 - | 0 - | 0 - | 1 - | 1,41E-05 - | 1 - | 1,43E-05 - | 1 - | 1,61E-05 - |
| 4296 | 1 | 3um_FIJJAFBF_50357 | Bacteria | Proteobacteri | Gammaprotec | Oceanospirilla | Oceanospirilla | Marinobacteri | Marinobacteri | metabolism | Energy produc | 16QPK | Succinate deh | 491952.Mar18 | 1 | 0 | 2,32E-05 | 2,20E-06 | 1 - | 1,94E-05 - | 1 - | 2,41E-05 - | 1 - | 2,29E-05 - | 1 - | 2,33E-05 - | 1 - | 2,62E-05 - |
| 4297 | 1 | 0.2um_NOKCPHJN_363042 | Bacteria | Proteobacteri | Betaproteoba | Neisseriales | Neisseriaceae | Neisseria | Neisseria sp. C | poorly charact | Function unkn | 0XSV1 | NA | 522373.Smlt1 | 0,2 | 0,4 | 1,34E-06 | 2,69E-06 | 0 - | 0 - | 1 - | 6,72E-06 - | 0 - | 0 - | 0 - | 0 - | 0 - | 0 - |
| 4298 | 1 | 0.2um_NOKCPHJN_27339 | Bacteria | Bacteroidetes | Flavobacteriia | Flavobacterial | Flavobacteriac | Flavobacteriu | Flavobacteriu | unclassified | unclassified | unclassified | unclassified | unclassified | 5,6 | 1,019803903 | 8,53E-05 | 2,11E-05 | 4 - | 5,03E-05 - | 7 - | 0,000109389 - | 5 - | 7,43E-05 - | 6 - | 9,04E-05 - | 6 - | 0,000101891 - |
| 4299 | 1 | 3um_FIJJAFBF_80079 | Bacteria | Proteobacteri | Gammaprotec | Oceanospirilla | Saccharospiril | Saccharospiril | Saccharospiril | information st | Translation, ril | COG0016 | phenylalanyL- | 322710.Avin_ | 1,2 | 0,748331477 | 4,66E-05 | 2,87E-05 | 2 - | 6,79E-05 - | 2 - | 8,43E-05 - | 1 - | 4,01E-05 - | 1 - | 4,07E-05 - | 0 - | 0 - |
| 4300 | 1 | 3um_FIJJAFBF_297064 | Bacteria | Cyanobacteria | unclassified | Synechococca | Synechococca | Synechococcu | Synechococcu | unclassified | unclassified | unclassified | unclassified | unclassified | 0,2 | 0,4 | 1,66E-05 | 3,31E-05 | 0 - | 0 - | 1 - | 8,29E-05 - | 0 - | 0 - | 0 - | 0 - | 0 - | 0 - |
| 4301 | 1 | 0.2um_NOKCPHJN_547080 | Bacteria | Actinobacteria | Actinobacteria | Micrococcales | Microbacteria | Candidatus Ac | Candidatus Ac | unclassified | unclassified | unclassified | unclassified | unclassified | 3,6 | 0,8 | 5,06E-05 | 1,21E-05 | 3 - | 3,53E-05 - | 4 - | 5,85E-05 - | 5 - | 6,95E-05 - | 3 - | 4,23E-05 - | 3 - | 4,77E-05 - |
| 4302 | 1 | 3um_FIJJAFBF_437253 | Bacteria | Bacteroidetes | Flavobacteriia | Flavobacterial | Flavobacteriac | Polaribacter | Polaribacter s | metabolism | Inorganic ion t | 05C27 | TonB-depende | 867900.Celly_ | 45 | 2,75680975 | 0,000562591 | 5,01E-05 | 49 - | 0,000514553 - | 43 - | 0,000560687 - | 46 - | 0,000570191 - | 41 - | 0,00051572 - | 46 - | 0,000651805 - |
| 4303 | 1 | 3um_FIJJAFBF_306873 | Bacteria | Proteobacteri | Alphaproteob | Rhodobactera | unclassified | unclassified | Rhodobactera | metabolism | Inorganic ion t | 05PWN | phosphate ab | 439497.RR11_ | 1,2 | 0,4 | 4,79E-05 | 2,04E-05 | 1 - | 3,27E-05 - | 1 - | 4,06E-05 - | 1 - | 3,86E-05 - | 1 - | 3,92E-05 - | 2 - | 8,82E-05 - |
| 4304 | 1 | 0.2um_NOKCPHJN_456211 | Archaea | Euryarchaeota | unclassified | unclassified | unclassified | unclassified | Euryarchaeota | information st | Replication, re | 05C2M | dna polymera | 580327.Tthe_ | 0,2 | 0,4 | 1,08E-05 | 2,15E-05 | 0 - | 0 - | 1 - | 5,38E-05 - | 0 - | 0 - | 0 - | 0 - | 0 - | 0 - |
| 4305 | 1 | 3um_FIJJAFBF_458400 | Bacteria | Bacteroidetes | Cytophagia | Cytophagales | Flammeovirga | unclassified | Flammeovirga | cellular proces | Cell wall/mem | 09DZB | Fasciclin | 487797.Flav3C | 1,8 | 0,4 | 0,00012156 | 2,84E-05 | 2 - | 0,00011364 - | 1 - | 7,06E-05 - | 2 - | 0,000134141 - | 2 - | 0,000136122 - | 2 - | 0,000153341 - |
| 4306 | 1 | 3um_FIJJAFBF_748863 | Bacteria | Proteobacteri | Gammaprotec | unclassified | unclassified | unclassified | various | poorly charact | Function unkn | 17FMV | dNA-binding p | 472759.Nhal_ | 1,2 | 1,166190379 | 0,000128833 | 0,000127949 | 1 - | 8,90E-05 - | 0 - | 0 - | 3 - | 0,000315075 - | 0 - | 0 - | 2 - | 0,000240115 - |
| 4307 | 1 | 3um_FIJJAFBF_352680 | Bacteria | Proteobacteri | Gammaprotec | Alteromonada | Alteromonada | Paraglaciecola | Paraglaciecola | cellular proces | Posttranslatio | COG0330 | Band 7 protei | 342610.Patl_3 | 5,2 | 0,979795897 | 0,000188318 | 4,78E-05 | 4 - | 0,000119871 - | 6 - | 0,000223267 - | 6 - | 0,000212244 - | 4 - | 0,000143586 - | 6 - | 0,000242623 - |
| 4308 | 1 | 3um_FIJJAFBF_606953 | Bacteria | unclassified | unclassified | unclassified | unclassified | unclassified | uncultured ba | information st | Translation, ril | 05C64 | Binds directly | 391603.FBALC | 1 | 1,264911064 | 5,35E-05 | 6,63E-05 | 3 - | 0,000149709 - | 0 - | 0 - | 2 - | 0,000117811 - | 0 - | 0 - | 0 - | 0 - |
| 4309 | 1 | 3um_FIJJAFBF_415869 | Bacteria | Bacteroidetes | Flavobacteriia | Flavobacterial | Flavobacteriac | Polaribacter | Polaribacter s | cellular proces | Posttranslatio | 09GE9 | Thioredoxin | 688270.Celal_ | 1,8 | 0,4 | 0,000231504 | 4,57E-05 | 2 - | 0,000218622 - | 2 - | 0,000271465 - | 2 - | 0,000258061 - | 2 - | 0,000261873 - | 1 - | 0,0001475 - |
| 4310 | 2 | 3um_FIJJAFBF_196006;3um_FIJJAFBF_529319 | Bacteria | Bacteroidetes | Flavobacteriia | Flavobacterial | Flavobacteriac | Polaribacter | various | information st | Translation, ril | COG0012 | gtp-binding pr | 313598.MED1 | 2 | 0 | 7,55E-05 | 7,17E-06 | 2 - | 6,32E-05 - | 2 - | 7,85E-05 - | 2 - | 7,46E-05 - | 2 - | 7,57E-05 - | 2 - | 8,53E-05 - |
| 4311 | 1 | 3um_FIJJAFBF_332119 | Bacteria | Bacteroidetes | Flavobacteriia | unclassified | unclassified | unclassified | Flavobacteria | unclassified | unclassified | unclassified | unclassified | unclassified | 3,2 | 1,166190379 | 5,59E-05 | 1,66E-05 | 5 - | 7,51E-05 - | 4 - | 7,46E-05 - | 2 - | 3,55E-05 - | 3 - | 5,40E-05 - | 2 - | 4,05E-05 - |
| 4312 | 1 | 3um_FIJJAFBF_326268 | Bacteria | Proteobacteri | Alphaproteob | Rhodobactera | various | various | various | information st | Translation, ril | COG0184 | Forms an inte | 314262.MED1 | 1,6 | 0,489897949 | 0,00024733 | 7,97E-05 | 1 - | 0,000128963 - | 2 - | 0,000320267 - | 2 - | 0,000304454 - | 2 - | 0,000308951 - | 1 - | 0,000174016 - |
| 4313 | 2 | 3um_FIJJAFBF_339765;3um_FIJJAFBF_668780 | Bacteria | Bacteroidetes | Flavobacteriia | Flavobacterial | Crocinitomica | unclassified | Crocinitomica | various | various | various | various | various | 1 | 0 | 1,87E-05 | 1,78E-06 | 1 - | 1,57E-05 - | 1 - | 1,95E-05 - | 1 - | 1,85E-05 - | 1 - | 1,88E-05 - | 1 - | 2,12E-05 - |
| 4314 | 1 | 3um_FIJJAFBF_195998 | Bacteria | Bacteroidetes | Flavobacteriia | Flavobacterial | Flavobacteriac | Polaribacter | Polaribacter g | unclassified | unclassified | unclassified | unclassified | unclassified | 0,6 | 0,489897949 | 5,68E-05 | 4,69E-05 | 1 - | 8,26E-05 - | 1 - | 0,000102532 - | 0 - | 0 - | 1 - | 9,89E-05 - | 0 - | 0 - |
| 4315 | 1 | 0.2um_NOKCPHJN_526772 | Bacteria | Proteobacteri | Gammaprotec | Cellvibrionales | Spongiibacter | Oceanicoccus | Oceanicoccus | poorly charact | Function unkn | 0R9NA | TPR | 857087.Metm | 0,4 | 0,489897949 | 8,41E-06 | 1,04E-05 | 1 - | 1,88E-05 - | 1 - | 2,33E-05 - | 0 - | 0 - | 0 - | 0 - | 0 - | 0 - |
| 4316 | 1 | 0.2um_NOKCPHJN_1007957 | Eukaryota | unclassified | Labyrinthulom | unclassified | Thraustochytr | Aurantiochytr | Aurantiochytr | metabolism | Inorganic ion t | COG0475 | Sodium hydro | 35128.Thaps2 | 0,8 | 0,4 | 2,38E-05 | 1,21E-05 | 1 - | 2,58E-05 - | 1 - | 3,20E-05 - | 1 - | 3,04E-05 - | 1 - | 3,09E-05 - | 0 - | 0 - |
| 4317 | 1 | 3um_FIJJAFBF_627437 | Eukaryota | unclassified | Dinophyceae | Suessiales | Symbiodiniace | Symbiodinium | Symbiodinium | metabolism | Energy produc | KOG1647 | subunit D | 423536.XP_00 | 0,4 | 0,489897949 | 2,09E-05 | 2,58E-05 | 1 - | 4,80E-05 - | 0 - | 0 - | 1 - | 5,67E-05 - | 0 - | 0 - | 0 - | 0 - |
| 4318 | 1 | 3um_FIJJAFBF_666073 | Bacteria | Proteobacteri | Gammaprotec | Alteromonada | Alteromonada | Paraglaciecola | Paraglaciecola | information st | Translation, ril | 05KK9 | This protein bi | 342610.Patl_3 | 2,4 | 1,019803903 | 0,000308246 | 0,000108622 | 4 - | 0,000445735 - | 3 - | 0,000415103 - | 2 - | 0,000263072 - | 2 - | 0,000266958 - | 1 - | 0,000150364 - |
| 4319 | 1 | 3um_FIJJAFBF_336807 | Bacteria | Bacteroidetes | Flavobacteriia | Flavobacterial | Cryomorphace | unclassified | various | unclassified | unclassified | unclassified | unclassified | unclassified | 0,8 | 0,4 | 1,30E-05 | 6,68E-06 | 1 - | 1,38E-05 - | 0 - | 0 - | 1 - | 1,62E-05 - | 1 - | 1,65E-05 - | 1 - | 1,86E-05 - |
| 4320 | 1 | 0.2um_NOKCPHJN_538165 | Bacteria | unclassified | unclassified | unclassified | unclassified | unclassified | bacterium TM | information st | Replication, re | COG3973 | Helicase | 404589.Anae1 | 1 | 0 | 1,93E-05 | 1,83E-06 | 1 - | 1,62E-05 - | 1 - | 2,01E-05 - | 1 - | 1,91E-05 - | 1 - | 1,94E-05 - | 1 - | 2,18E-05 - |
| 4321 | 1 | 3um_FIJJAFBF_305710 | Bacteria | Bacteroidetes | Flavobacteriia | Flavobacterial | Flavobacteriac | Formosa | Formosa sp. H | unclassified | unclassified | unclassified | unclassified | unclassified | 0,4 | 0,489897949 | 6,03E-06 | 7,38E-06 | 0 - | 0 - | 0 - | 0 - | 1 - | 1,50E-05 - | 1 - | 1,52E-05 - | 0 - | 0 - |
| 4322 | 1 | 0.2um_NOKCPHJN_403365 | Eukaryota | unclassified | Oomycetes | Peronosporale | Peronosporac | Phytophthora | Phytophthora | unclassified | unclassified | unclassified | unclassified | unclassified | 0,4 | 0,489897949 | 1,51E-05 | 1,85E-05 | 0 - | 0 - | 0 - | 0 - | 1 - | 3,74E-05 - | 1 - | 3,80E-05 - | 0 - | 0 - |
| 4323 | 1 | 0.2um_NOKCPHJN_704076 | Bacteria | Bacteroidetes | Flavobacteriia | Flavobacterial | Flavobacteriac | Polaribacter | Polaribacter s | unclassified | unclassified | unclassified | unclassified | unclassified | 0,6 | 0,8 | 2,92E-05 | 3,77E-05 | 2 - | 9,18E-05 - | 0 - | 0 - | 1 - | 5,42E-05 - | 0 - | 0 - | 0 - | 0 - |
| 4324 | 1 | 3um_FIJJAFBF_483811 | Bacteria | Actinobacteria | Actinobacteria | Micrococcales | Microbacteria | Candidatus Ac | Candidatus Ac | unclassified | unclassified | unclassified | unclassified | unclassified | 12,8 | 1,6 | 0,00015456 | 2,61E-05 | 14 - | 0,000141575 - | 11 - | 0,000138124 - | 13 - | 0,000155178 - | 11 - | 0,000133244 - | 15 - | 0,00020468 - |
| 4325 | 1 | 0.2um_NOKCPHJN_767449 | Bacteria | Proteobacteri | Alphaproteob | Rhodobactera | Rhodobactera | Sulfitobacter | Sulfitobacter s | metabolism | Carbohydrate | 01QKM | inositol mono | 292414.TM10 | 0,2 | 0,4 | 1,03E-05 | 2,06E-05 | 0 - | 0 - | 0 - | 0 - | 1 - | 5,15E-05 - | 0 - | 0 - | 0 - | 0 - |
| 4326 | 1 | 3um_FIJJAFBF_179842 | Bacteria | Bacteroidetes | Flavobacteriia | Flavobacterial | Flavobacteriac | Algibacter | Algibacter pec | metabolism | Energy produc | COG0056 | Produces ATP | 391603.FBALC | 4,2 | 0,748331477 | 0,000108097 | 1,54E-05 | 5 - | 0,000109103 - | 5 - | 0,000135474 - | 4 - | 0,000103028 - | 4 - | 0,00010455 - | 3 - | 8,83E-05 - |
| 4327 | 1 | 3um_FIJJAFBF_632800 | Bacteria | Proteobacteri | Deltaproteoba | Myxococcales | unclassified | Enhygromyxa | Enhygromyxa | cellular proces | Signal transdu | KOG0613 | fibronectin typ | 7165.AGAP00 | 0,2 | 0,4 | 7,94E-06 | 1,59E-05 | 1 - | 3,97E-05 - | 0 - | 0 - | 0 - | 0 - | 0 - | 0 - | 0 - | 0 - |
| 4328 | 1 | 3um_FIJJAFBF_39793 | Bacteria | Bacteroidetes | Flavobacteriia | Flavobacterial | Flavobacteriac | Polaribacter | Polaribacter s | unclassified | unclassified | unclassified | unclassified | unclassified | 4,2 | 0,748331477 | 0,000228382 | 5,55E-05 | 4 - | 0,000180751 - | 5 - | 0,000280549 - | 4 - | 0,000213358 - | 3 - | 0,000162382 - | 5 - | 0,000304871 - |
| 4329 | 1 | 0.2um_NOKCPHJN_38882 | Bacteria | Proteobacteri | Gammaprotec | Alteromonada | Alteromonada | Paraglaciecola | Paraglaciecola | unclassified | unclassified | unclassified | unclassified | unclassified | 0,4 | 0,489897949 | 8,66E-06 | 1,07E-05 | 1 - | 1,99E-05 - | 0 - | 0 - | 1 - | 2,34E-05 - | 0 - | 0 - | 0 - | 0 - |
| 4330 | 1 | 3um_FIJJAFBF_628027 | Bacteria | Proteobacteri | Alphaproteob | Rhodobactera | Rhodobactera | Planktomarina | Planktomarina | information st | Translation, ril | 177XJ | One of the pri | 314265.R2601 | 4,8 | 0,4 | 0,000424475 | 6,86E-05 | 4 - | 0,000294299 - | 5 - | 0,000456791 - | 5 - | 0,000434238 - | 5 - | 0,000440652 - | 5 - | 0,000496393 - |
| 4331 | 1 | 0.2um_NOKCPHJN_641798 | Bacteria | Proteobacteri | Alphaproteob | Pelagibacteral | Pelagibacterac | Candidatus Pe | Candidatus Pe | information st | Translation, ril | 179R6 | This protein is | 335992.SAR11 | 1 | 0 | 8,96E-05 | 8,50E-06 | 1 - | 7,50E-05 - | 1 - | 9,31E-05 - | 1 - | 8,86E-05 - | 1 - | 8,99E-05 - | 1 - | 0,000101225 - |
| 4332 | 1 | 0.2um_NOKCPHJN_138361 | Bacteria | Proteobacteri | Gammaprotec | Cellvibrionales | Porticoccacea | unclassified | gamma prote | metabolism | Inorganic ion t | 05PWN | phosphate ab | 314287.GB220 | 3,6 | 1,2 | 0,000144468 | 5,84E-05 | 2 - | 6,54E-05 - | 5 - | 0,000203018 - | 3 - | 0,000115797 - | 3 - | 0,000117507 - | 5 - | 0,000220619 - |
| 4333 | 1 | 0.2um_NOKCPHJN_454143 | Bacteria | Proteobacteri | Alphaproteob | Rhodobactera | Rhodobactera | Planktomarina | Planktomarina | metabolism | Lipid transpor | 08IQF | Catalyzes the | 388401.RB215 | 2,2 | 1,469693846 | 4,28E-05 | 2,37E-05 | 5 - | 8,57E-05 - | 1 - | 2,13E-05 - | 2 - | 4,04E-05 - | 1 - | 2,05E-05 - | 2 - | 4,62E-05 - |
| 4334 | 1 | 0.2um_NOKCPHJN_495423 | Bacteria | Proteobacteri | Alphaproteob | Pelagibacteral | Pelagibacterac | Candidatus Pe | Candidatus Pe | information st | Translation, ril | 179ZI | One of the pri | 335992.SAR11 | 2,4 | 1,019803903 | 0,000245017 | 8,82E-05 | 4 - | 0,000350463 - | 3 - | 0,000326379 - | 2 - | 0,000206843 - | 1 - | 0,000104949 - | 2 - | 0,00023645 - |
| 4335 | 1 | 0.2um_NOKCPHJN_198737 | Eukaryota | Chordata | unclassified | unclassified | Branchiostomi | Branchiostom | Branchiostom | information st | Replication, re | COG1643 | helicase | 70448.Q01C4 | 0,2 | 0,4 | 2,11E-06 | 4,21E-06 | 1 - | 1,05E-05 - | 0 - | 0 - | 0 - | 0 - | 0 - | 0 - | 0 - | 0 - |
| 4336 | 1 | 3um_FIJJAFBF_01111 | Bacteria | Bacteroidetes | Flavobacteriia | Flavobacterial | Flavobacteriac | various | various | unclassified | unclassified | unclassified | unclassified | unclassified | 1 | 0 | 0,000217503 | 2,07E-05 | 1 - | 0,000182185 - | 1 - | 0,00022622 - | 1 - | 0,000215051 - | 1 - | 0,000218227 - | 1 - | 0,000245833 - |
| 4337 | 1 | 0.2um_NOKCPHJN_04036 | Bacteria | Proteobacteri | Gammaprotec | Alteromonada | Alteromonada | Paraglaciecola | Paraglaciecola | cellular proces | Posttranslatio | COG0612 | peptidase' | 582402.Hbal_ | 0,2 | 0,4 | 2,38E-06 | 4,76E-06 | 1 - | 1,19E-05 - | 0 - | 0 - | 0 - | 0 - | 0 - | 0 - | 0 - | 0 - |
| 4338 | 1 | 3um_FIJJAFBF_281830 | Bacteria | Bacteroidetes | unclassified | unclassified | unclassified | unclassified | Bacteroidetes | information st | Replication, re | 0PB0H | DNA-binding p | 487796.Flav2A | 6,6 | 1,019803903 | 0,001009717 | 0,000145435 | 8 - | 0,001031701 - | 6 - | 0,000960801 - | 7 - | 0,001065591 - | 5 - | 0,000772378 - | 7 - | 0,001218114 - |
| 4339 | 1 | 3um_FIJJAFBF_581782 | Bacteria | Bacteroidetes | Flavobacteriia | Flavobacterial | Flavobacteriac | Tenacibaculur | Tenacibaculur | information st | Replication, re | COG0550 | Releases the s | 313598.MED1 | 4,2 | 0,4 | 6,86E-05 | 3,25E-06 | 5 - | 6,90E-05 - | 4 - | 6,85E-05 - | 4 - | 6,51E-05 - | 4 - | 6,61E-05 - | 4 - | 7,45E-05 - |
| 4340 | 1 | 3um_FIJJAFBF_239853 | Bacteria | Bacteroidetes | unclassified | unclassified | unclassified | unclassified | unidentified e | unclassified | unclassified | unclassified | unclassified | unclassified | 4,4 | 1,496662955 | 8,08E-05 | 2,35E-05 | 6 - | 9,47E-05 - | 4 - | 7,84E-05 - | 4 - | 7,45E-05 - | 6 - | 0,000113466 - | 2 - | 4,26E-05 - |
| 4341 | 1 | 3um_FIJJAFBF_225135 | Bacteria | Proteobacteri | Alphaproteob | Rhodobactera | Rhodobactera | Actibacterium | Actibacterium | information st | Translation, ril | 01S8S | This is 1 of the | 1051075.PDI_ | 3,4 | 0,8 | 0,000245695 | 3,33E-05 | 5 - | 0,00030854 - | 3 - | 0,000229869 - | 3 - | 0,00021852 - | 3 - | 0,000221747 - | 3 - | 0,000249798 - |
| 4342 | 1 | 0.2um_NOKCPHJN_557642 | Bacteria | Proteobacteri | Gammaprotec | Xanthomonad | Xanthomonad | Xanthomonas | Xanthomonas | poorly charact | Function unkn | 0NVRZ | NA | 170187.SP_17 | 3 | 0 | 0,000113558 | 1,08E-05 | 3 - | 9,51E-05 - | 3 - | 0,00011811 - | 3 - | 0,000112278 - | 3 - | 0,000113936 - | 3 - | 0,000128349 - |
| 4343 | 1 | 3um_FIJJAFBF_470660 | Eukaryota | unclassified | Cryptophyta | Pyrenomonad | Geminigerace | Guillardia | Guillardia thet | metabolism | Energy produc | COG0711 | F(1)F(0) ATP s | 197221.tlr043 | 1 | 0 | 7,45E-05 | 7,07E-06 | 1 - | 6,24E-05 - | 1 - | 7,75E-05 - | 1 - | 7,36E-05 - | 1 - | 7,47E-05 - | 1 - | 8,42E-05 - |
| 4344 | 1 | 0.2um_NOKCPHJN_329662 | Bacteria | Bacteroidetes | Flavobacteriia | Flavobacterial | Cryomorphace | unclassified | Cryomorphace | unclassified | unclassified | unclassified | unclassified | unclassified | 0,2 | 0,4 | 6,96E-06 | 1,39E-05 | 1 - | 3,48E-05 - | 0 - | 0 - | 0 - | 0 - | 0 - | 0 - | 0 - | 0 - |
| 4345 | 1 | 3um_FIJJAFBF_455318 | Bacteria | Proteobacteri | Gammaprotec | Alteromonada | Alteromonada | Paraglaciecola | Paraglaciecola | information st | Translation, ril | 05KG7 | Associates wit | 983545.Glaag_ | 0,6 | 0,8 | 6,04E-05 | 8,11E-05 | 0 - | 0 - | 2 - | 0,000203598 - | 0 - | 0 - | 1 - | 9,82E-05 - | 0 - | 0 - |
| 4346 | 1 | 0.2um_NOKCPHJN_75187 | Bacteria | Bacteroidetes | Flavobacteriia | Flavobacterial | Flavobacteriac | Formosa | Formosa sp. H | unclassified | unclassified | unclassified | unclassified | unclassified | 0,2 | 0,4 | 1,53E-06 | 3,06E-06 | 1 - | 7,66E-06 - | 0 - | 0 - | 0 - | 0 - | 0 - | 0 - | 0 - | 0 - |
| 4347 | 1 | 0.2um_NOKCPHJN_219779 | Bacteria | Bacteroidetes | Flavobacteriia | Flavobacterial | Flavobacteriac | Polaribacter | Polaribacter s | information st | Translation, ril | COG0050 | This protein p | 313603.FB217 | 2,4 | 0,489897949 | 8,44E-05 | 2,38E-05 | 2 - | 5,81E-05 - | 3 - | 0,000108242 - | 2 - | 6,86E-05 - | 2 - | 6,96E-05 - | 3 - | 0,000117626 - |
| 4348 | 1 | 3um_FIJJAFBF_88551 | Bacteria | Proteobacteri | Gammaprotec | Oceanospirilla | Saccharospiril | Reinekea | Reinekea fors | unclassified | unclassified | unclassified | unclassified | unclassified | 1,8 | 0,748331477 | 0,000198702 | 7,26E-05 | 3 - | 0,000282238 - | 2 - | 0,000233637 - | 1 - | 0,000111051 - | 1 - | 0,000112691 - | 2 - | 0,000253893 - |
| 4349 | 1 | 3um_FIJJAFBF_254286 | Bacteria | Proteobacteri | Gammaprotec | Cellvibrionales | Spongiibacter | Spongiibacter | Spongiibacter | information st | Translation, ril | COG0088 | One of the pri | 876044.IMCC3 | 4 | 1,264911064 | 0,000265674 | 8,62E-05 | 5 - | 0,000279943 - | 2 - | 0,000139043 - | 5 - | 0,000330444 - | 3 - | 0,000201195 - | 5 - | 0,000377743 - |
| 4350 | 1 | 3um_FIJJAFBF_71023 | Bacteria | Bacteroidetes | Cytophagia | Cytophagales | Flammeovirga | unclassified | Flammeovirga | poorly charact | Function unkn | 11NCV | Protein of unk | 643867.Ftrac_ | 0,8 | 0,748331477 | 7,41E-05 | 6,52E-05 | 2 - | 0,000167557 - | 1 - | 0,000104028 - | 1 - | 9,89E-05 - | 0 - | 0 - | 0 - | 0 - |
| 4351 | 1 | 3um_FIJJAFBF_15909 | Bacteria | Bacteroidetes | Flavobacteriia | Flavobacterial | Flavobacteriac | Polaribacter | Polaribacter s | unclassified | unclassified | unclassified | unclassified | unclassified | 1 | 0 | 5,91E-05 | 5,61E-06 | 1 - | 4,95E-05 - | 1 - | 6,14E-05 - | 1 - | 5,84E-05 - | 1 - | 5,93E-05 - | 1 - | 6,68E-05 - |
| 4352 | 1 | 3um_FIJJAFBF_195976 | Bacteria | Bacteroidetes | Flavobacteriia | Flavobacterial | Flavobacteriac | Polaribacter | Polaribacter s | unclassified | unclassified | unclassified | unclassified | unclassified | 0,8 | 0,4 | 7,81E-05 | 4,01E-05 | 1 - | 8,26E-05 - | 0 - | 0 - | 1 - | 9,75E-05 - | 1 - | 9,89E-05 - | 1 - | 0,00011142 - |
| 4353 | 1 | 0.2um_NOKCPHJN_520265 | Bacteria | Bacteroidetes | unclassified | unclassified | unclassified | unclassified | Bacteroidetes | poorly charact | Function unkn | 09CIG | Ragb susd dor | 655815.ZPR_1 | 0,4 | 0,489897949 | 1,01E-05 | 1,25E-05 | 1 - | 2,26E-05 - | 1 - | 2,81E-05 - | 0 - | 0 - | 0 - | 0 - | 0 - | 0 - |
| 4354 | 1 | 0.2um_NOKCPHJN_557695 | Bacteria | Bacteroidetes | Flavobacteriia | Flavobacterial | Cryomorphace | Owenweeksia | Owenweeksia | information st | Translation, ril | 05C22 | threonyL-tRN | 926562.Oweh | 0,4 | 0,489897949 | 8,63E-06 | 1,06E-05 | 0 - | 0 - | 1 - | 2,21E-05 - | 1 - | 2,10E-05 - | 0 - | 0 - | 0 - | 0 - |
| 4355 | 1 | 3um_FIJJAFBF_562674 | Bacteria | Proteobacteri | Gammaprotec | Chromatiales | Chromatiacea | Rheinheimera | Rheinheimera | cellular proces | Cell wall/mem | 16PMX | Catalyzes the | 743720.Psefu_ | 0,2 | 0,4 | 7,83E-06 | 1,57E-05 | 1 - | 3,92E-05 - | 0 - | 0 - | 0 - | 0 - | 0 - | 0 - | 0 - | 0 - |
| 4356 | 1 | 3um_FIJJAFBF_510698 | Bacteria | Ignavibacteria | Ignavibacteria | Ignavibacteria | unclassified | unclassified | Ignavibacteria | unclassified | unclassified | unclassified | unclassified | unclassified | 0,8 | 0,4 | 4,00E-05 | 2,06E-05 | 1 - | 4,24E-05 - | 0 - | 0 - | 1 - | 5,00E-05 - | 1 - | 5,07E-05 - | 1 - | 5,71E-05 - |
| 4357 | 1 | 0.2um_NOKCPHJN_855491 | Eukaryota | Phaeophyceae | unclassified | Ectocarpales | Ectocarpaceae | Ectocarpus | Ectocarpus sili | unclassified | unclassified | unclassified | unclassified | unclassified | 1 | 0 | 2,14E-05 | 2,03E-06 | 1 - | 1,79E-05 - | 1 - | 2,22E-05 - | 1 - | 2,11E-05 - | 1 - | 2,14E-05 - | 1 - | 2,42E-05 - |
| 4358 | 1 | 0.2um_NOKCPHJN_940010 | Bacteria | Proteobacteri | Alphaproteob | Pelagibacteral | Pelagibacterac | unclassified | Pelagibacterac | cellular proces | Cell wall/mem | 7257 | NA | 335992.SAR11 | 1 | 0 | 5,27E-05 | 5,00E-06 | 1 - | 4,41E-05 - | 1 - | 5,48E-05 - | 1 - | 5,21E-05 - | 1 - | 5,29E-05 - | 1 - | 5,96E-05 - |
| 4359 | 1 | 3um_FIJJAFBF_705369 | Bacteria | Proteobacteri | Gammaprotec | Alteromonada | Alteromonada | Paraglaciecola | Paraglaciecola | metabolism | Amino acid tra | 05C6T | cysteine synth | 715451.ambt_ | 0,8 | 0,748331477 | 3,32E-05 | 3,19E-05 | 1 - | 3,58E-05 - | 1 - | 4,44E-05 - | 0 - | 0 - | 2 - | 8,57E-05 - | 0 - | 0 - |
| 4360 | 1 | 0.2um_NOKCPHJN_06849 | Bacteria | Bacteroidetes | Cytophagia | Cytophagales | Flammeovirga | unclassified | Flammeovirga | unclassified | unclassified | unclassified | unclassified | unclassified | 0,8 | 0,4 | 2,43E-05 | 1,23E-05 | 1 - | 2,63E-05 - | 1 - | 3,26E-05 - | 1 - | 3,10E-05 - | 1 - | 3,15E-05 - | 0 - | 0 - |
| 4361 | 1 | 3um_FIJJAFBF_338236 | Bacteria | Proteobacteri | Gammaprotec | Thiotrichales | Piscirickettsiac | Cycloclasticus | Cycloclasticus | information st | Translation, ril | 05CEJ | Catalyzes the | 575788.VS_24 | 0,4 | 0,489897949 | 8,57E-06 | 1,05E-05 | 0 - | 0 - | 1 - | 2,05E-05 - | 0 - | 0 - | 0 - | 0 - | 1 - | 2,23E-05 - |
| 4362 | 1 | 3um_FIJJAFBF_320528 | Bacteria | Proteobacteri | Gammaprotec | Oceanospirilla | Oceanospirilla | Motiliproteus | Motiliproteus | unclassified | unclassified | unclassified | unclassified | unclassified | 1,2 | 0,4 | 3,24E-05 | 7,17E-06 | 2 - | 4,65E-05 - | 1 - | 2,88E-05 - | 1 - | 2,74E-05 - | 1 - | 2,78E-05 - | 1 - | 3,14E-05 - |
| 4363 | 1 | 3um_FIJJAFBF_262967 | Bacteria | unclassified | unclassified | unclassified | unclassified | unclassified | uncultured ba | metabolism | Carbohydrate | 16QEQ | hydrolase, fan | 983545.Glaag_ | 2 | 0 | 6,46E-05 | 6,14E-06 | 2 - | 5,41E-05 - | 2 - | 6,72E-05 - | 2 - | 6,39E-05 - | 2 - | 6,49E-05 - | 2 - | 7,31E-05 - |
| 4364 | 1 | 3um_FIJJAFBF_348505 | Bacteria | Bacteroidetes | Cytophagia | Cytophagales | Flammeovirga | Marivirga | Marivirga seri | metabolism | Inorganic ion t | 05C27 | TonB-depende | 643867.Ftrac_ | 10,2 | 1,720465053 | 0,000130691 | 1,04E-05 | 13 - | 0,0001417 - | 9 - | 0,000121811 - | 10 - | 0,000128663 - | 11 - | 0,00014362 - | 8 - | 0,000117663 - |
| 4365 | 1 | 0.2um_NOKCPHJN_529425 | Bacteria | Proteobacteri | Betaproteoba | Burkholderiale | Comamonada | Acidovorax | Acidovorax an | metabolism | Secondary me | 16URM | outer membra | 323850.Shew_ | 3,8 | 0,748331477 | 3,22E-05 | 4,00E-06 | 5 - | 3,61E-05 - | 4 - | 3,58E-05 - | 4 - | 3,41E-05 - | 3 - | 2,59E-05 - | 3 - | 2,92E-05 - |
| 4366 | 1 | 0.2um_NOKCPHJN_222458 | Bacteria | Bacteroidetes | Flavobacteriia | Flavobacterial | Flavobacteriac | Flavobacteriu | Flavobacteriu | unclassified | unclassified | unclassified | unclassified | unclassified | 1 | 0 | 8,16E-05 | 7,74E-06 | 1 - | 6,83E-05 - | 1 - | 8,48E-05 - | 1 - | 8,06E-05 - | 1 - | 8,18E-05 - | 1 - | 9,22E-05 - |
| 4367 | 3 | 3um_FIJJAFBF_275521;3um_FIJJAFBF_32276;3ur | Bacteria | Proteobacteri | Alphaproteob | Rhodobactera | Rhodobactera | various | various | cellular proces | Posttranslatio | 01ZS8 | Thioredoxin | various | 0,6 | 0,489897949 | 7,45E-05 | 6,15E-05 | 1 - | 0,00010828 - | 1 - | 0,000134452 - | 0 - | 0 - | 1 - | 0,000129701 - | 0 - | 0 - |
| 4368 | 1 | 3um_FIJJAFBF_621109 | Bacteria | Proteobacteri | Gammaprotec | Chromatiales | unclassified | unclassified | Chromatiales | metabolism | Amino acid tra | 08KMW | Glycine betain | 335992.SAR11 | 0,4 | 0,489897949 | 1,73E-05 | 2,16E-05 | 1 - | 3,68E-05 - | 0 - | 0 - | 0 - | 0 - | 0 - | 0 - | 1 - | 4,96E-05 - |
| 4369 | 1 | 3um_FIJJAFBF_04736 | Bacteria | Bacteroidetes | Flavobacteriia | Flavobacterial | Flavobacteriac | Flavobacteriu | Flavobacteriu | poorly charact | Function unkn | 06D3R | NA | 983548.Krodi_ | 1,2 | 0,4 | 5,60E-05 | 1,88E-05 | 1 - | 3,92E-05 - | 1 - | 4,86E-05 - | 2 - | 9,25E-05 - | 1 - | 4,69E-05 - | 1 - | 5,29E-05 - |
| 4370 | 1 | 3um_FIJJAFBF_636785 | Bacteria | Proteobacteri | Gammaprotec | unclassified | unclassified | Sedimenticola | Sedimenticola | unclassified | unclassified | unclassified | unclassified | unclassified | 6,8 | 0,748331477 | 0,000369141 | 3,75E-05 | 8 - | 0,000365822 - | 6 - | 0,000340683 - | 6 - | 0,000323862 - | 7 - | 0,00038342 - | 7 - | 0,000431921 - |
| 4371 | 1 | 3um_FIJJAFBF_197210 | Bacteria | Bacteroidetes | Flavobacteriia | Flavobacterial | Flavobacteriac | Ulvibacter | Ulvibacter lito | unclassified | unclassified | unclassified | unclassified | unclassified | 1 | 0 | 3,50E-05 | 3,32E-06 | 1 - | 2,93E-05 - | 1 - | 3,64E-05 - | 1 - | 3,46E-05 - | 1 - | 3,51E-05 - | 1 - | 3,95E-05 - |
| 4372 | 1 | 3um_FIJJAFBF_78697 | Bacteria | Proteobacteri | Alphaproteob | Rhodobactera | unclassified | unclassified | Rhodobactera | metabolism | Amino acid tra | 0XRCI | His Glu Gln Ar | 367336.OM22 | 3 | 0 | 0,000153963 | 1,46E-05 | 3 - | 0,000128963 - | 3 - | 0,000160134 - | 3 - | 0,000152227 - | 3 - | 0,000154476 - | 3 - | 0,000174016 - |
| 4373 | 1 | 3um_FIJJAFBF_510245 | Bacteria | Bacteroidetes | Flavobacteriia | Flavobacterial | Flavobacteriac | Polaribacter | various | unclassified | unclassified | unclassified | unclassified | unclassified | 5,8 | 0,748331477 | 0,000434832 | 3,36E-05 | 7 - | 0,000443888 - | 6 - | 0,000472438 - | 5 - | 0,00037426 - | 6 - | 0,000455746 - | 5 - | 0,00042783 - |
| 4374 | 1 | 3um_FIJJAFBF_03913 | Bacteria | Bacteroidetes | Flavobacteriia | Flavobacterial | Flavobacteriac | Polaribacter | various | poorly charact | Function unkn | COG0714 | ATPase activit | 313594.PI23P_ | 0,8 | 0,4 | 3,47E-05 | 1,75E-05 | 0 - | 0 - | 1 - | 4,33E-05 - | 1 - | 4,12E-05 - | 1 - | 4,18E-05 - | 1 - | 4,71E-05 - |
| 4375 | 1 | 0.2um_NOKCPHJN_64674 | Bacteria | Bacteroidetes | Cytophagia | Cytophagales | Flammeovirga | unclassified | Flammeovirga | poorly charact | Function unkn | 0XY6Y | Pkd domain cc | 314287.GB220 | 4 | 0 | 9,00E-05 | 8,55E-06 | 4 - | 7,54E-05 - | 4 - | 9,36E-05 - | 4 - | 8,90E-05 - | 4 - | 9,03E-05 - | 4 - | 0,000101724 - |
| 4376 | 1 | 0.2um_NOKCPHJN_534108 | Eukaryota | Phaeophyceae | unclassified | Ectocarpales | Ectocarpaceae | Ectocarpus | Ectocarpus sili | unclassified | unclassified | unclassified | unclassified | unclassified | 3,2 | 1,166190379 | 5,60E-05 | 2,08E-05 | 4 - | 5,89E-05 - | 5 - | 9,14E-05 - | 2 - | 3,47E-05 - | 2 - | 3,53E-05 - | 3 - | 5,96E-05 - |
| 4377 | 1 | 3um_FIJJAFBF_701188 | Bacteria | Proteobacteri | Gammaprotec | Alteromonada | Alteromonada | Paraglaciecola | Paraglaciecola | unclassified | unclassified | unclassified | unclassified | unclassified | 0,2 | 0,4 | 1,02E-05 | 2,04E-05 | 1 - | 5,10E-05 - | 0 - | 0 - | 0 - | 0 - | 0 - | 0 - | 0 - | 0 - |
| 4378 | 1 | 3um_FIJJAFBF_524442 | Bacteria | Bacteroidetes | unclassified | unclassified | unclassified | unclassified | Bacteroidetes | poorly charact | Function unkn | 09PDM | NA | 553178.CAPGI | 0,4 | 0,489897949 | 2,74E-05 | 3,38E-05 | 1 - | 6,24E-05 - | 0 - | 0 - | 0 - | 0 - | 1 - | 7,47E-05 - | 0 - | 0 - |
| 4379 | 1 | 0.2um_NOKCPHJN_297481 | Bacteria | Proteobacteri | Gammaprotec | Alteromonada | Alteromonada | Glaciecola | Glaciecola pur | metabolism | Inorganic ion t | 0R03K | TonB-depende | 203122.Sde_1 | 3 | 0,632455532 | 4,41E-05 | 7,66E-06 | 4 - | 4,97E-05 - | 3 - | 4,63E-05 - | 2 - | 2,94E-05 - | 3 - | 4,47E-05 - | 3 - | 5,03E-05 - |
| 4380 | 1 | 3um_FIJJAFBF_727603 | unclassified | unclassified | unclassified | unclassified | unclassified | unclassified | unclassified | unclassified | unclassified | unclassified | unclassified | unclassified | 0,2 | 0,4 | 2,96E-05 | 5,91E-05 | 0 - | 0 - | 0 - | 0 - | 0 - | 0 - | 1 - | 0,000147832 - | 0 - | 0 - |
| 4381 | 1 | 3um_FIJJAFBF_423338 | Bacteria | Bacteroidetes | Flavobacteriia | Flavobacterial | Flavobacteriac | Polaribacter | Polaribacter p | cellular proces | Intracellular tr | 0P8JN | The central su | 313598.MED1 | 1 | 0 | 3,11E-05 | 2,95E-06 | 1 - | 2,60E-05 - | 1 - | 3,23E-05 - | 1 - | 3,07E-05 - | 1 - | 3,12E-05 - | 1 - | 3,51E-05 - |
| 4382 | 1 | 0.2um_NOKCPHJN_331161 | Viruses | unclassified | unclassified | unclassified | Phycodnavirid | unclassified | Chrysochromu | unclassified | unclassified | unclassified | unclassified | unclassified | 0,2 | 0,4 | 4,24E-06 | 8,49E-06 | 1 - | 2,12E-05 - | 0 - | 0 - | 0 - | 0 - | 0 - | 0 - | 0 - | 0 - |
| 4383 | 3 | 0.2um_NOKCPHJN_1026952;0.2um_NOKCPHJN_ | Bacteria | Bacteroidetes | Flavobacteriia | Flavobacterial | Flavobacteriac | Formosa | Formosa sp. H | unclassified | unclassified | unclassified | unclassified | unclassified | 1 | 0,894427191 | 7,81E-05 | 7,09E-05 | 0 - | 0 - | 0 - | 0 - | 2 - | 0,000147263 - | 1 - | 7,47E-05 - | 2 - | 0,000168342 - |
| 4384 | 1 | 0.2um_NOKCPHJN_132742 | Eukaryota | unclassified | Oomycetes | Albuginales | Albuginaceae | Albugo | Albugo candid | unclassified | unclassified | unclassified | unclassified | unclassified | 19,4 | 1,019803903 | 0,000593491 | 5,46E-05 | 21 - | 0,00053922 - | 19 - | 0,000605785 - | 19 - | 0,000575875 - | 18 - | 0,000553624 - | 20 - | 0,000692951 - |
| 4385 | 1 | 3um_FIJJAFBF_168836 | Bacteria | Proteobacteri | Alphaproteob | Rhodobactera | Hyphomonada | Hyphobacteriu | Hyphobacteriu | cellular proces | Cell wall/mem | 01TJH | Udp-sulfoquin | 394221.Mmar | 0,2 | 0,4 | 7,78E-06 | 1,56E-05 | 1 - | 3,89E-05 - | 0 - | 0 - | 0 - | 0 - | 0 - | 0 - | 0 - | 0 - |
| 4386 | 1 | 3um_FIJJAFBF_346332 | Bacteria | Proteobacteri | Gammaprotec | Oceanospirilla | Saccharospiril | Reinekea | Reinekea fors | cellular proces | Cell wall/mem | 16PZ8 | Involved in the | 314283.MED2 | 0,8 | 0,748331477 | 4,06E-05 | 3,54E-05 | 2 - | 8,97E-05 - | 0 - | 0 - | 1 - | 5,29E-05 - | 0 - | 0 - | 1 - | 6,05E-05 - |
| 4387 | 1 | 3um_FIJJAFBF_742492 | Eukaryota | Bacillariophyta | Bacillariophyc | Naviculales | Naviculaceae | Fistulifera | Fistulifera sola | unclassified | unclassified | unclassified | unclassified | unclassified | 1 | 0 | 6,72E-05 | 6,38E-06 | 1 - | 5,63E-05 - | 1 - | 6,99E-05 - | 1 - | 6,64E-05 - | 1 - | 6,74E-05 - | 1 - | 7,59E-05 - |
| 4388 | 1 | 3um_FIJJAFBF_244316 | Bacteria | Bacteroidetes | Cytophagia | Cytophagales | Flammeovirga | unclassified | Flammeovirga | unclassified | unclassified | unclassified | unclassified | unclassified | 1 | 0 | 4,49E-05 | 4,27E-06 | 1 - | 3,76E-05 - | 1 - | 4,67E-05 - | 1 - | 4,44E-05 - | 1 - | 4,51E-05 - | 1 - | 5,08E-05 - |
| 4389 | 1 | 3um_FIJJAFBF_346409 | Bacteria | Proteobacteri | Gammaprotec | Pseudomonad | Moraxellaceae | Psychrobacter | Psychrobacter | cellular proces | Cell wall/mem | 08ZG0 | Ompa motb d | 349106.PsycP | 1,6 | 0,489897949 | 9,84E-05 | 2,58E-05 | 2 - | 0,0001053 - | 1 - | 6,54E-05 - | 2 - | 0,000124296 - | 2 - | 0,000126131 - | 1 - | 7,10E-05 - |
| 4390 | 1 | 3um_FIJJAFBF_91420 | Bacteria | Proteobacteri | Gammaprotec | Oceanospirilla | Oceanospirilla | Neptunomona | Neptunomona | metabolism | Inorganic ion t | 16PAX | extracellular s | 572480.Arnit_ | 1,4 | 0,8 | 4,61E-05 | 2,56E-05 | 0 - | 0 - | 2 - | 6,66E-05 - | 2 - | 6,33E-05 - | 2 - | 6,42E-05 - | 1 - | 3,62E-05 - |
| 4391 | 1 | 3um_FIJJAFBF_21313 | Bacteria | Bacteroidetes | Flavobacteriia | Flavobacterial | Flavobacteriac | Polaribacter | Polaribacter a | poorly charact | Function unkn | 09GJP | Protein of unk | 313594.PI23P_ | 1,2 | 0,4 | 0,000122779 | 4,23E-05 | 1 - | 8,57E-05 - | 1 - | 0,000106357 - | 1 - | 0,000101106 - | 2 - | 0,000205199 - | 1 - | 0,000115578 - |
| 4392 | 1 | 0.2um_NOKCPHJN_583156 | Bacteria | Proteobacteri | Alphaproteob | Pelagibacteral | unclassified | unclassified | Pelagibacteral | cellular proces | Cell wall/mem | 7257 | NA | 335992.SAR11 | 1,6 | 0,489897949 | 6,70E-05 | 2,36E-05 | 1 - | 3,46E-05 - | 1 - | 4,29E-05 - | 2 - | 8,16E-05 - | 2 - | 8,28E-05 - | 2 - | 9,33E-05 - |
| 4393 | 1 | 3um_FIJJAFBF_282387 | Eukaryota | Bacillariophyta | Coscinodiscop | Thalassiosirale | Thalassiosirac | Thalassiosira | Thalassiosira p | unclassified | unclassified | unclassified | unclassified | unclassified | 2,8 | 0,4 | 8,60E-05 | 1,02E-05 | 3 - | 7,79E-05 - | 3 - | 9,67E-05 - | 3 - | 9,20E-05 - | 3 - | 9,33E-05 - | 2 - | 7,01E-05 - |
| 4394 | 1 | 3um_FIJJAFBF_76222 | Bacteria | Bacteroidetes | Flavobacteriia | Flavobacterial | Flavobacteriac | Maribacter | Maribacter th | metabolism | Amino acid tra | 0P7Z9 | aminopeptida | 487797.Flav3C | 3,4 | 1,743559577 | 5,87E-05 | 2,65E-05 | 6 - | 8,96E-05 - | 5 - | 9,27E-05 - | 2 - | 3,52E-05 - | 2 - | 3,58E-05 - | 2 - | 4,03E-05 - |
| 4395 | 2 | 0.2um_NOKCPHJN_919657;3um_FIJJAFBF_33445 | Bacteria | Proteobacteri | Betaproteoba | Neisseriales | Chromobacter | Aquaspirillum | Aquaspirillum | poorly charact | Function unkn | 123UY | FecR protein | 640511.BC100 | 0,4 | 0,489897949 | 1,33E-05 | 1,63E-05 | 0 - | 0 - | 1 - | 3,41E-05 - | 1 - | 3,24E-05 - | 0 - | 0 - | 0 - | 0 - |
| 4396 | 1 | 3um_FIJJAFBF_317823 | unclassified | unclassified | unclassified | unclassified | unclassified | unclassified | unclassified | unclassified | unclassified | unclassified | unclassified | unclassified | 1 | 0 | 4,20E-05 | 3,99E-06 | 1 - | 3,52E-05 - | 1 - | 4,37E-05 - | 1 - | 4,16E-05 - | 1 - | 4,22E-05 - | 1 - | 4,75E-05 - |
| 4397 | 2 | 0.2um_NOKCPHJN_590543;0.2um_NOKCPHJN_7 | Bacteria | Proteobacteri | Alphaproteob | Rhodobactera | Rhodobactera | Planktomarina | Planktomarina | cellular proces | Intracellular tr | 0XNME | Type I secretic | 314232.SKA53 | 2,8 | 0,748331477 | 8,85E-05 | 2,62E-05 | 2 - | 5,26E-05 - | 4 - | 0,000130751 - | 3 - | 9,32E-05 - | 3 - | 9,46E-05 - | 2 - | 7,10E-05 - |
| 4398 | 1 | 0.2um_NOKCPHJN_878359 | Bacteria | Proteobacteri | Gammaprotec | unclassified | unclassified | unclassified | Gammaprotec | metabolism | Inorganic ion t | 05F28 | tonB-depende | 565045.NOR5 | 1,4 | 1,019803903 | 3,56E-05 | 2,33E-05 | 3 - | 6,74E-05 - | 0 - | 0 - | 1 - | 2,65E-05 - | 2 - | 5,38E-05 - | 1 - | 3,03E-05 - |
| 4399 | 1 | 0.2um_NOKCPHJN_175599 | Bacteria | Proteobacteri | Alphaproteob | Pelagibacteral | Pelagibacterac | Candidatus Pe | Candidatus Pe | unclassified | unclassified | unclassified | unclassified | unclassified | 1 | 0 | 8,11E-05 | 7,70E-06 | 1 - | 6,79E-05 - | 1 - | 8,43E-05 - | 1 - | 8,02E-05 - | 1 - | 8,14E-05 - | 1 - | 9,16E-05 - |
| 4400 | 1 | 3um_FIJJAFBF_18512 | Bacteria | Bacteroidetes | Flavobacteriia | Flavobacterial | Flavobacteriac | unclassified | Flavobacteriac | unclassified | unclassified | unclassified | unclassified | unclassified | 0,4 | 0,489897949 | 5,31E-06 | 6,55E-06 | 1 - | 1,21E-05 - | 0 - | 0 - | 0 - | 0 - | 1 - | 1,45E-05 - | 0 - | 0 - |
| 4401 | 1 | 3um_FIJJAFBF_736129 | Bacteria | Bacteroidetes | Flavobacteriia | unclassified | unclassified | unclassified | Flavobacteria | unclassified | unclassified | unclassified | unclassified | unclassified | 2,4 | 0,489897949 | 0,00026585 | 6,13E-05 | 2 - | 0,000185124 - | 3 - | 0,000344804 - | 3 - | 0,00032778 - | 2 - | 0,000221747 - | 2 - | 0,000249798 - |
| 4402 | 1 | 3um_FIJJAFBF_226198 | Bacteria | Bacteroidetes | Cytophagia | Cytophagales | Flammeovirga | unclassified | Flammeovirga | poorly charact | Function unkn | 05ERK | gliding motilit | 643867.Ftrac_ | 3,4 | 1,2 | 8,43E-05 | 2,96E-05 | 4 - | 8,41E-05 - | 5 - | 0,000130512 - | 2 - | 4,96E-05 - | 4 - | 0,00010072 - | 2 - | 5,67E-05 - |
| 4403 | 1 | 0.2um_NOKCPHJN_978628 | Bacteria | Proteobacteri | Alphaproteob | Pelagibacteral | Pelagibacterac | Candidatus Pe | Candidatus Pe | unclassified | unclassified | unclassified | unclassified | unclassified | 0,4 | 0,8 | 1,47E-05 | 2,93E-05 | 2 - | 7,33E-05 - | 0 - | 0 - | 0 - | 0 - | 0 - | 0 - | 0 - | 0 - |
| 4404 | 1 | 0.2um_NOKCPHJN_121979 | Bacteria | Bacteroidetes | Flavobacteriia | Flavobacterial | Flavobacteriac | Polaribacter | Polaribacter b | unclassified | unclassified | unclassified | unclassified | unclassified | 0,6 | 0,489897949 | 4,57E-05 | 3,77E-05 | 1 - | 6,67E-05 - | 1 - | 8,29E-05 - | 1 - | 7,88E-05 - | 0 - | 0 - | 0 - | 0 - |
| 4405 | 1 | 0.2um_NOKCPHJN_682442 | unclassified | unclassified | unclassified | unclassified | unclassified | unclassified | unclassified | unclassified | unclassified | unclassified | unclassified | unclassified | 1 | 0 | 0,000103808 | 9,86E-06 | 1 - | 8,70E-05 - | 1 - | 0,000107969 - | 1 - | 0,000102638 - | 1 - | 0,000104154 - | 1 - | 0,000117329 - |
| 4406 | 1 | 0.2um_NOKCPHJN_325765 | Eukaryota | Apicomplexa | Conoidasida | Eucoccidiorida | Eimeriidae | Eimeria | Eimeria mitis | information st | Transcription | 12NU4 | Required for t | 5270.UM0232 | 0,2 | 0,4 | 5,59E-06 | 1,12E-05 | 1 - | 2,79E-05 - | 0 - | 0 - | 0 - | 0 - | 0 - | 0 - | 0 - | 0 - |
| 4407 | 1 | 0.2um_NOKCPHJN_50843 | Bacteria | Bacteroidetes | Flavobacteriia | Flavobacterial | Flavobacteriac | Polaribacter | Polaribacter s | unclassified | unclassified | unclassified | unclassified | unclassified | 3,4 | 0,489897949 | 9,07E-05 | 9,63E-06 | 4 - | 9,02E-05 - | 3 - | 8,40E-05 - | 3 - | 7,99E-05 - | 4 - | 0,000108042 - | 3 - | 9,13E-05 - |
| 4408 | 1 | 3um_FIJJAFBF_365990 | Bacteria | Proteobacteri | Gammaprotec | unclassified | unclassified | unclassified | Gammaprotec | metabolism | Carbohydrate | 16SJ5 | Trap dicarbox | 349521.HCH_ | 0,2 | 0,4 | 6,96E-06 | 1,39E-05 | 1 - | 3,48E-05 - | 0 - | 0 - | 0 - | 0 - | 0 - | 0 - | 0 - | 0 - |
| 4409 | 1 | 3um_FIJJAFBF_281259 | Bacteria | Proteobacteri | Gammaprotec | Oceanospirilla | Saccharospiril | Reinekea | Reinekea blan | unclassified | unclassified | unclassified | unclassified | unclassified | 3,8 | 1,326649916 | 0,000127203 | 4,99E-05 | 2 - | 5,49E-05 - | 6 - | 0,000204573 - | 4 - | 0,000129648 - | 3 - | 9,87E-05 - | 4 - | 0,000148205 - |
| 4410 | 1 | 3um_FIJJAFBF_251401 | Bacteria | Bacteroidetes | Flavobacteriia | Flavobacterial | Flavobacteriac | Polaribacter | Polaribacter s | unclassified | unclassified | unclassified | unclassified | unclassified | 0,8 | 0,4 | 6,87E-05 | 3,53E-05 | 1 - | 7,26E-05 - | 0 - | 0 - | 1 - | 8,57E-05 - | 1 - | 8,70E-05 - | 1 - | 9,80E-05 - |
| 4411 | 1 | 0.2um_NOKCPHJN_280290 | Bacteria | Proteobacteri | Alphaproteob | Rhodobactera | Rhodobactera | unclassified | Rhodobactera | unclassified | unclassified | unclassified | unclassified | unclassified | 0,6 | 0,489897949 | 8,25E-06 | 6,81E-06 | 1 - | 1,21E-05 - | 1 - | 1,50E-05 - | 1 - | 1,42E-05 - | 0 - | 0 - | 0 - | 0 - |
| 4412 | 1 | 0.2um_NOKCPHJN_798550 | Bacteria | Proteobacteri | Alphaproteob | Pelagibacteral | Pelagibacterac | Candidatus Pe | Candidatus Pe | cellular proces | Cell wall/mem | 170GF | mechanosensi | 335992.SAR11 | 1 | 0 | 0,000110506 | 1,05E-05 | 1 - | 9,26E-05 - | 1 - | 0,000114935 - | 1 - | 0,00010926 - | 1 - | 0,000110874 - | 1 - | 0,000124899 - |
| 4413 | 1 | 3um_FIJJAFBF_510241 | Bacteria | Bacteroidetes | Flavobacteriia | Flavobacterial | Flavobacteriac | Polaribacter | Polaribacter s | information st | Translation, ril | 09FE5 | One of the prc | 313598.MED1 | 1,4 | 0,489897949 | 0,000187479 | 5,52E-05 | 2 - | 0,000229554 - | 1 - | 0,000142519 - | 1 - | 0,000135482 - | 2 - | 0,000274967 - | 1 - | 0,000154874 - |
| 4414 | 1 | 0.2um_NOKCPHJN_212630 | Archaea | Candidatus Ba | unclassified | unclassified | unclassified | unclassified | Candidatus Ba | information st | Translation, ril | 0DGFE | This protein p | 99883.ENSTNI | 0,8 | 0,4 | 1,46E-05 | 7,34E-06 | 0 - | 0 - | 1 - | 1,82E-05 - | 1 - | 1,73E-05 - | 1 - | 1,76E-05 - | 1 - | 1,98E-05 - |
| 4415 | 1 | 3um_FIJJAFBF_147391 | Bacteria | Bacteroidetes | Flavobacteriia | Flavobacterial | Flavobacteriac | Polaribacter | Polaribacter s | unclassified | unclassified | unclassified | unclassified | unclassified | 8 | 1,095445115 | 0,000822824 | 7,60E-05 | 10 - | 0,000869521 - | 7 - | 0,000755782 - | 8 - | 0,000821104 - | 7 - | 0,000729078 - | 8 - | 0,000938633 - |
| 4416 | 2 | 0.2um_NOKCPHJN_786247;0.2um_NOKCPHJN_9 | Bacteria | Bacteroidetes | Flavobacteriia | Flavobacterial | Flavobacteriac | Formosa | Formosa sp. H | unclassified | unclassified | unclassified | unclassified | unclassified | 3 | 0 | 0,000234904 | 2,23E-05 | 3 - | 0,00019676 - | 3 - | 0,000244318 - | 3 - | 0,000232255 - | 3 - | 0,000235686 - | 3 - | 0,000265499 - |
| 4417 | 1 | 3um_FIJJAFBF_365082 | Bacteria | Proteobacteri | Alphaproteob | Pelagibacteral | Pelagibacterac | Candidatus Pe | Candidatus Pe | unclassified | unclassified | unclassified | unclassified | unclassified | 0,2 | 0,4 | 2,17E-05 | 4,33E-05 | 0 - | 0 - | 0 - | 0 - | 0 - | 0 - | 1 - | 0,000108255 - | 0 - | 0 - |
| 4418 | 1 | 0.2um_NOKCPHJN_263318 | Eukaryota | Phaeophyceae | unclassified | Ectocarpales | Ectocarpaceae | Ectocarpus | Ectocarpus sili | unclassified | unclassified | unclassified | unclassified | unclassified | 1 | 0,632455532 | 2,02E-05 | 1,14E-05 | 2 - | 3,52E-05 - | 0 - | 0 - | 1 - | 2,08E-05 - | 1 - | 2,11E-05 - | 1 - | 2,38E-05 - |
| 4419 | 1 | 0.2um_NOKCPHJN_950738 | Bacteria | Proteobacteri | Alphaproteob | Rhodobactera | Rhodobactera | Neptunicoccu | Neptunicoccu | metabolism | Inorganic ion t | 05PWN | phosphate ab | 765910.Marpu | 3,8 | 0,4 | 0,000150175 | 1,98E-05 | 4 - | 0,00013269 - | 3 - | 0,000123571 - | 4 - | 0,000156627 - | 4 - | 0,00015894 - | 4 - | 0,000179046 - |
| 4420 | 1 | 3um_FIJJAFBF_695142 | Bacteria | Proteobacteri | Alphaproteob | unclassified | unclassified | unclassified | SAR116 cluste | unclassified | unclassified | unclassified | unclassified | unclassified | 0,8 | 0,4 | 3,76E-05 | 1,91E-05 | 1 - | 4,07E-05 - | 1 - | 5,05E-05 - | 1 - | 4,80E-05 - | 1 - | 4,88E-05 - | 0 - | 0 - |
| 4421 | 1 | 0.2um_NOKCPHJN_746717 | Bacteria | Proteobacteri | Alphaproteob | Pelagibacteral | Pelagibacterac | Candidatus Pe | Candidatus Pe | unclassified | unclassified | unclassified | unclassified | unclassified | 0,8 | 0,4 | 0,000121701 | 6,25E-05 | 1 - | 0,00012753 - | 1 - | 0,000158354 - | 1 - | 0,000150536 - | 0 - | 0 - | 1 - | 0,000172083 - |
| 4422 | 1 | 3um_FIJJAFBF_651417 | Bacteria | Bacteroidetes | Flavobacteriia | Flavobacterial | Flavobacteriac | Flavobacteriu | Flavobacteriu | unclassified | unclassified | unclassified | unclassified | unclassified | 0,6 | 0,489897949 | 1,89E-05 | 1,55E-05 | 1 - | 2,79E-05 - | 0 - | 0 - | 1 - | 3,30E-05 - | 1 - | 3,35E-05 - | 0 - | 0 - |
| 4423 | 1 | 3um_FIJJAFBF_592694 | Bacteria | Bacteroidetes | Flavobacteriia | Flavobacterial | Flavobacteriac | Polaribacter | Polaribacter s | unclassified | unclassified | unclassified | unclassified | unclassified | 5,4 | 1,019803903 | 7,81E-05 | 1,15E-05 | 7 - | 8,57E-05 - | 6 - | 9,13E-05 - | 5 - | 7,23E-05 - | 4 - | 5,87E-05 - | 5 - | 8,26E-05 - |
| 4424 | 1 | 0.2um_NOKCPHJN_839084 | Bacteria | Proteobacteri | Alphaproteob | Rhodospirillal | Rhodospirillac | unclassified | Rhodospirillac | poorly charact | Function unkn | 0NDJX | P22 coat prote | 760568.Desku | 1,8 | 0,748331477 | 7,88E-05 | 3,17E-05 | 2 - | 7,45E-05 - | 2 - | 9,25E-05 - | 3 - | 0,000131963 - | 1 - | 4,46E-05 - | 1 - | 5,03E-05 - |
| 4425 | 1 | 3um_FIJJAFBF_199369 | Bacteria | Proteobacteri | Gammaprotec | Oceanospirilla | Oceanospirilla | Motiliproteus | Motiliproteus | information st | Translation, ril | 17FIH | One of the ear | 207949.RED65 | 3,6 | 1,019803903 | 0,000495053 | 0,000128179 | 4 - | 0,000468477 - | 3 - | 0,000436282 - | 4 - | 0,000552989 - | 5 - | 0,000701445 - | 2 - | 0,00031607 - |
| 4426 | 1 | 3um_FIJJAFBF_24823 | unclassified | unclassified | unclassified | unclassified | unclassified | unclassified | unclassified | unclassified | unclassified | unclassified | unclassified | unclassified | 1 | 0 | 0,000147341 | 1,40E-05 | 1 - | 0,000123416 - | 1 - | 0,000153246 - | 1 - | 0,00014568 - | 1 - | 0,000147832 - | 1 - | 0,000166532 - |
| 4427 | 1 | 0.2um_NOKCPHJN_39754 | Bacteria | Proteobacteri | Alphaproteob | unclassified | unclassified | unclassified | Alphaproteob | unclassified | unclassified | unclassified | unclassified | unclassified | 3 | 1,095445115 | 0,000456465 | 0,000173541 | 4 - | 0,000510119 - | 3 - | 0,000475063 - | 1 - | 0,000150536 - | 3 - | 0,000458278 - | 4 - | 0,000688331 - |

| | | | | | | | | | | | | | | | | | | | | | | | | | | | | |
|---|---|---|---|---|---|---|---|---|---|---|---|---|---|---|---|---|---|---|---|---|---|---|---|---|---|---|---|---|
| 4428 | 1 | 3um_FIJJAFBF_149821 | Bacteria | Proteobacteri | Betaproteoba | Burkholderial | Comamonada | Hydrogenoph | Hydrogenoph | metabolism | Energy produc | COG0712 | F(1)F(0) ATP s | 688245.CtCNE | 1,4 | 0,8 | 0,000110172 | 6,72E-05 | 1 - | 6,45E-05 - | 2 - | 0,000160134 - | 2 - | 0,000152227 - | 0 - | 0 - | 2 - | 0,000174016 - |
| 4429 | 1 | 0.2um_NOKCPHJN_326393 | Bacteria | Bacteroidetes | unclassified | unclassified | unclassified | unclassified | Bacteroidetes | metabolism | Inorganic ion t | 05C27 | TonB-depende | 391603.FBALC | 14,4 | 1,624807681 | 0,000181599 | 2,21E-05 | 15 - | 0,000159118 - | 17 - | 0,000223921 - | 14 - | 0,0001753 - | 14 - | 0,00017789 - | 12 - | 0,000171765 - |
| 4430 | 1 | 0.2um_NOKCPHJN_58123 | Bacteria | Proteobacteri | Gammaprotec | Oceanospirilla | Saccharospirill | Reinekea | Reinekea blan | unclassified | unclassified | unclassified | unclassified | unclassified | 2 | 1,095445115 | 6,86E-05 | 3,12E-05 | 4 - | 0,000118632 - | 2 - | 7,37E-05 - | 1 - | 3,50E-05 - | 1 - | 3,55E-05 - | 2 - | 8,00E-05 - |
| 4431 | 1 | 3um_FIJJAFBF_676243 | Bacteria | Proteobacteri | Alphaproteob | Rhodobactera | unclassified | unclassified | Rhodobactera | metabolism | Carbohydrate | 01RFA | ABC transport | 367336.OM22 | 0,6 | 0,489897949 | 2,54E-05 | 2,08E-05 | 0 - | 0 - | 1 - | 4,16E-05 - | 0 - | 0 - | 1 - | 4,01E-05 - | 1 - | 4,52E-05 - |
| 4432 | 1 | 0.2um_NOKCPHJN_560311 | Bacteria | Proteobacteri | Alphaproteob | Rhodobactera | Rhodobactera | Maribius | Maribius salin | poorly charact | Function unkn | 16V6S | Alkaline phosp | 439497.RR11_ | 1,2 | 0,4 | 2,25E-05 | 4,98E-06 | 2 - | 3,22E-05 - | 1 - | 2,00E-05 - | 1 - | 1,90E-05 - | 1 - | 1,93E-05 - | 1 - | 2,18E-05 - |
| 4433 | 1 | 3um_FIJJAFBF_426982 | Bacteria | Bacteroidetes | Flavobacteriia | Flavobacterial | Flavobacteriac | Hyunsoonleell | Hyunsoonleell | unclassified | unclassified | unclassified | unclassified | unclassified | 0,2 | 0,4 | 4,16E-06 | 8,31E-06 | 0 - | 0 - | 1 - | 2,08E-05 - | 0 - | 0 - | 0 - | 0 - | 0 - | 0 - |
| 4434 | 1 | 0.2um_NOKCPHJN_562145 | Bacteria | Bacteroidetes | Flavobacteriia | Flavobacterial | Crocinitomica | unclassified | Crocinitomica | poorly charact | Function unkn | 16R8I | Lipoprotein | 715451.ambt_ | 7,6 | 2,154065923 | 7,88E-05 | 1,92E-05 | 10 - | 8,83E-05 - | 9 - | 9,87E-05 - | 5 - | 5,21E-05 - | 9 - | 9,52E-05 - | 5 - | 5,96E-05 - |
| 4435 | 1 | 0.2um_NOKCPHJN_431087 | Bacteria | Proteobacteri | Gammaprotec | Oceanospirilla | Saccharospirill | Reinekea | Reinekea fors | cellular proces | Intracellular tr | 0QQ57 | motA TolQ Exl | 314283.MED2 | 0,4 | 0,489897949 | 2,84E-05 | 3,49E-05 | 0 - | 0 - | 0 - | 0 - | 0 - | 0 - | 1 - | 6,67E-05 - | 1 - | 7,52E-05 - |
| 4436 | 1 | 3um_FIJJAFBF_751711 | Bacteria | Bacteroidetes | Flavobacteriia | Flavobacterial | Flavobacteriac | Polaribacter | Polaribacter s | metabolism | Energy produc | COG0711 | F(1)F(0) ATP s | 313598.MED1 | 1,2 | 0,4 | 9,64E-05 | 2,13E-05 | 2 - | 0,000138285 - | 1 - | 8,59E-05 - | 1 - | 8,16E-05 - | 1 - | 8,28E-05 - | 1 - | 9,33E-05 - |
| 4437 | 1 | 0.2um_NOKCPHJN_110548 | Bacteria | Proteobacteri | Gammaprotec | Cellvibrionales | Porticoccacea | unclassified | Gammaprotec | information st | Translation, ri | 16Q16 | Binds the lowe | 498211.CJA_0 | 1,6 | 0,489897949 | 9,51E-05 | 2,69E-05 | 2 - | 0,000101125 - | 2 - | 0,000125567 - | 1 - | 5,97E-05 - | 2 - | 0,000121131 - | 1 - | 6,82E-05 - |
| 4438 | 1 | 3um_FIJJAFBF_151367 | Eukaryota | Annelida | Polychaeta | Golfingiida | Sipunculidae | Sipunculus | Sipunculus nu | information st | Chromatin str | KOG1744 | Core compone | 400682.PAC_1 | 2,6 | 0,8 | 0,000285317 | 7,37E-05 | 4 - | 0,000373258 - | 2 - | 0,000231738 - | 2 - | 0,000220296 - | 2 - | 0,00022355 - | 3 - | 0,000377743 - |
| 4439 | 1 | 3um_FIJJAFBF_810984 | Bacteria | Proteobacteri | Alphaproteob | Rhodobactera | Rhodobactera | Planktomarina | Planktomarina | metabolism | Energy produc | 01R8W | NDH-1 shuttle | 314262.MED1 | 1,2 | 0,4 | 7,88E-05 | 1,75E-05 | 2 - | 0,000113081 - | 1 - | 7,02E-05 - | 1 - | 6,67E-05 - | 1 - | 6,77E-05 - | 1 - | 7,63E-05 - |
| 4440 | 1 | 0.2um_NOKCPHJN_64584 | Bacteria | Proteobacteri | Gammaprotec | Oceanospirilla | Saccharospirill | Reinekea | Reinekea fors | unclassified | unclassified | unclassified | unclassified | unclassified | 7,6 | 1,019803903 | 0,000180256 | 2,26E-05 | 8 - | 0,000159968 - | 8 - | 0,000198633 - | 7 - | 0,000165222 - | 9 - | 0,000215566 - | 6 - | 0,00016189 - |
| 4441 | 1 | 0.2um_NOKCPHJN_562126 | Bacteria | Bacteroidetes | Flavobacteriia | Flavobacterial | Flavobacteriac | unclassified | Flavobacteriac | unclassified | unclassified | unclassified | unclassified | unclassified | 0,2 | 0,4 | 2,81E-05 | 5,61E-05 | 0 - | 0 - | 0 - | 0 - | 0 - | 0 - | 1 - | 0,000140289 - | 0 - | 0 - |
| 4442 | 1 | 3um_FIJJAFBF_537788 | Eukaryota | unclassified | Dinophyceae | Suessiales | Symbiodiniace | Symbiodinium | Symbiodinium | unclassified | unclassified | unclassified | unclassified | unclassified | 0,2 | 0,4 | 9,89E-06 | 1,98E-05 | 0 - | 0 - | 0 - | 0 - | 0 - | 0 - | 1 - | 4,95E-05 - | 0 - | 0 - |
| 4443 | 1 | 3um_FIJJAFBF_99463 | Bacteria | Bacteroidetes | Flavobacteriia | Flavobacterial | Flavobacteriac | Gaetbulibacte | Gaetbulibacte | unclassified | unclassified | unclassified | unclassified | unclassified | 0,4 | 0,489897949 | 1,29E-05 | 1,59E-05 | 1 - | 2,94E-05 - | 0 - | 0 - | 0 - | 0 - | 1 - | 3,52E-05 - | 0 - | 0 - |
| 4444 | 1 | 0.2um_NOKCPHJN_647577 | Eukaryota | Chordata | Actinopteri | Cypriniformes | Cyprinidae | Cyprinus | Cyprinus carpi | cellular proces | Signal transdu | KOG0613 | fibronectin typ | 7070.TC00296 | 0,2 | 0,4 | 6,09E-06 | 1,22E-05 | 0 - | 0 - | 0 - | 0 - | 1 - | 3,04E-05 - | 0 - | 0 - | 0 - | 0 - |
| 4445 | 2 | 0.2um_NOKCPHJN_163193;3um_FIJJAFBF_2208( | Bacteria | Bacteroidetes | Flavobacteriia | Flavobacterial | Flavobacteriac | Ulvibacter | Ulvibacter sp. | poorly charact | Function unkn | 09GJP | Protein of unk | 50743.SCB49_ | 1,2 | 0,4 | 0,000121199 | 2,68E-05 | 2 - | 0,000173904 - | 1 - | 0,000107969 - | 1 - | 0,000102638 - | 1 - | 0,000104154 - | 1 - | 0,000117329 - |
| 4446 | 1 | 0.2um_NOKCPHJN_132233 | Bacteria | Proteobacteri | Gammaprotec | Oceanospirilla | Saccharospirill | Gynuella | Gynuella suns | unclassified | unclassified | unclassified | unclassified | unclassified | 3,2 | 0,748331477 | 0,000111152 | 1,94E-05 | 4 - | 0,000118632 - | 3 - | 0,00011048 - | 4 - | 0,000140033 - | 3 - | 0,000106576 - | 2 - | 8,00E-05 - |
| 4447 | 1 | 0.2um_NOKCPHJN_327326 | Bacteria | Bacteroidetes | Flavobacteriia | Flavobacterial | Flavobacteriac | Formosa | Formosa sp. H | cellular proces | Cell wall/mem | 05C6Y | Udp-glucose 6 | 391598.FBBAL | 0,6 | 0,489897949 | 1,75E-05 | 1,45E-05 | 1 - | 2,47E-05 - | 0 - | 0 - | 1 - | 2,92E-05 - | 0 - | 0 - | 1 - | 3,34E-05 - |
| 4448 | 1 | 0.2um_NOKCPHJN_496089 | Eukaryota | Ascomycota | Dothideomyce | Pleosporales | Pleosporaceae | Alternaria | Alternaria alte | cellular proces | Signal transdu | KOG0613 | fibronectin typ | 7165.AGAP00 | 0,2 | 0,4 | 5,57E-06 | 1,11E-05 | 0 - | 0 - | 1 - | 2,78E-05 - | 0 - | 0 - | 0 - | 0 - | 0 - | 0 - |
| 4449 | 1 | 0.2um_NOKCPHJN_472194 | Bacteria | Bacteroidetes | Flavobacteriia | Flavobacterial | Cryomorphace | unclassified | Cryomorphace | unclassified | unclassified | unclassified | unclassified | unclassified | 2 | 0 | 0,000164104 | 1,56E-05 | 2 - | 0,000137457 - | 2 - | 0,000170681 - | 2 - | 0,000162254 - | 2 - | 0,000164651 - | 2 - | 0,000185478 - |
| 4450 | 1 | 3um_FIJJAFBF_536639 | Bacteria | Bacteroidetes | unclassified | unclassified | unclassified | unclassified | unidentified e | unclassified | unclassified | unclassified | unclassified | unclassified | 0,6 | 0,489897949 | 1,08E-05 | 8,91E-06 | 1 - | 1,58E-05 - | 1 - | 1,96E-05 - | 1 - | 1,86E-05 - | 0 - | 0 - | 0 - | 0 - |
| 4451 | 1 | 3um_FIJJAFBF_123172 | Bacteria | Bacteroidetes | Flavobacteriia | Flavobacterial | Crocinitomica | unclassified | Crocinitomica | unclassified | unclassified | unclassified | unclassified | unclassified | 1 | 0 | 6,49E-05 | 6,17E-06 | 1 - | 5,44E-05 - | 1 - | 6,75E-05 - | 1 - | 6,42E-05 - | 1 - | 6,52E-05 - | 1 - | 7,34E-05 - |
| 4452 | 1 | 3um_FIJJAFBF_398908 | Bacteria | Proteobacteri | Alphaproteob | Rhodobactera | Rhodobactera | Planktomarina | Planktomarina | information st | Translation, ri | 17FIT | 50s ribosomal | 391624.OIHEL | 1 | 1,095445115 | 0,0001351 | 0,000133563 | 3 - | 0,000362453 - | 1 - | 0,00015002 - | 0 - | 0 - | 0 - | 0 - | 1 - | 0,000163026 - |
| 4453 | 1 | 3um_FIJJAFBF_176640 | Bacteria | Bacteroidetes | unclassified | unclassified | unclassified | unclassified | Bacteroidetes | unclassified | unclassified | unclassified | unclassified | unclassified | 1,6 | 1,019803903 | 0,000116761 | 7,00E-05 | 3 - | 0,000189192 - | 0 - | 0 - | 2 - | 0,000148882 - | 1 - | 7,55E-05 - | 2 - | 0,000170192 - |
| 4454 | 1 | 3um_FIJJAFBF_10583 | Bacteria | Bacteroidetes | Flavobacteriia | Flavobacterial | Flavobacteriac | Kordia | Kordia algicida | unclassified | unclassified | unclassified | unclassified | unclassified | 0,2 | 0,4 | 4,56E-06 | 9,13E-06 | 1 - | 2,28E-05 - | 0 - | 0 - | 0 - | 0 - | 0 - | 0 - | 0 - | 0 - |
| 4455 | 1 | 3um_FIJJAFBF_604721 | Eukaryota | Bacillariophyt | Coscinodiscop | Thalassiosirale | Thalassiosirac | Conticribra | Conticribra we | unclassified | unclassified | unclassified | unclassified | unclassified | 0,6 | 0,8 | 1,79E-05 | 2,30E-05 | 2 - | 5,59E-05 - | 0 - | 0 - | 0 - | 0 - | 1 - | 3,35E-05 - | 0 - | 0 - |
| 4456 | 1 | 0.2um_NOKCPHJN_265489 | Eukaryota | unclassified | Pelagophycea | Pelagomonad | unclassified | Aureococcus | Aureococcus a | unclassified | unclassified | unclassified | unclassified | unclassified | 0,2 | 0,4 | 2,18E-06 | 4,35E-06 | 1 - | 1,09E-05 - | 0 - | 0 - | 0 - | 0 - | 0 - | 0 - | 0 - | 0 - |
| 4457 | 1 | 3um_FIJJAFBF_496029 | Eukaryota | Phaeophyceae | unclassified | Ectocarpales | Ectocarpaceae | Ectocarpus | Ectocarpus sili | unclassified | unclassified | unclassified | unclassified | unclassified | 0,6 | 0,489897949 | 1,70E-05 | 1,42E-05 | 1 - | 2,41E-05 - | 0 - | 0 - | 1 - | 2,85E-05 - | 0 - | 0 - | 1 - | 3,25E-05 - |
| 4458 | 1 | 0.2um_NOKCPHJN_366743 | Bacteria | Proteobacteri | Gammaprotec | unclassified | unclassified | unclassified | Gammaprotec | information st | Translation, ri | COG1185 | Involved in mF | 203122.Sde_2 | 5,2 | 0,748331477 | 9,33E-05 | 1,04E-05 | 6 - | 9,11E-05 - | 6 - | 0,00011311 - | 5 - | 8,96E-05 - | 5 - | 9,09E-05 - | 4 - | 8,19E-05 - |
| 4459 | 1 | 3um_FIJJAFBF_337331 | Bacteria | Bacteroidetes | Flavobacteriia | Flavobacterial | Flavobacteriac | Polaribacter | various | unclassified | unclassified | unclassified | unclassified | unclassified | 1 | 0 | 0,000126877 | 1,20E-05 | 1 - | 0,000106275 - | 1 - | 0,000131962 - | 1 - | 0,000125447 - | 1 - | 0,000127299 - | 1 - | 0,000143402 - |
| 4460 | 1 | 3um_FIJJAFBF_322025 | Bacteria | Proteobacteri | Gammaprotec | Oceanospirilla | Saccharospirill | Reinekea | Reinekea fors | unclassified | unclassified | unclassified | unclassified | unclassified | 1,8 | 0,748331477 | 0,000170886 | 5,55E-05 | 3 - | 0,00024772 - | 1 - | 0,000102532 - | 2 - | 0,000194938 - | 2 - | 0,000197818 - | 1 - | 0,00011142 - |
| 4461 | 1 | 3um_FIJJAFBF_534854 | Bacteria | Proteobacteri | Alphaproteob | Caulobacteral | unclassified | unclassified | Caulobacteral | unclassified | unclassified | unclassified | unclassified | unclassified | 0,2 | 0,4 | 2,05E-05 | 4,10E-05 | 0 - | 0 - | 1 - | 0,000102532 - | 0 - | 0 - | 0 - | 0 - | 0 - | 0 - |
| 4462 | 1 | 3um_FIJJAFBF_336152 | Bacteria | Bacteroidetes | Flavobacteriia | Flavobacterial | Flavobacteriac | Polaribacter | Polaribacter s | metabolism | Energy produc | 09A0E | phosphoenolp | 313594.PI23P | 0,2 | 0,4 | 2,67E-06 | 5,34E-06 | 1 - | 1,34E-05 - | 0 - | 0 - | 0 - | 0 - | 0 - | 0 - | 0 - | 0 - |
| 4463 | 1 | 0.2um_NOKCPHJN_674145 | Eukaryota | unclassified | Pelagophycea | Pelagomonad | unclassified | Aureococcus | Aureococcus a | unclassified | unclassified | unclassified | unclassified | unclassified | 1 | 0 | 2,55E-05 | 2,42E-06 | 1 - | 2,14E-05 - | 1 - | 2,65E-05 - | 1 - | 2,52E-05 - | 1 - | 2,56E-05 - | 1 - | 2,88E-05 - |
| 4464 | 1 | 3um_FIJJAFBF_308266 | Bacteria | Bacteroidetes | Flavobacteriia | Flavobacterial | Flavobacteriac | Polaribacter | Polaribacter s | unclassified | unclassified | unclassified | unclassified | unclassified | 4 | 0,632455532 | 0,000167656 | 3,80E-05 | 4 - | 0,000139546 - | 4 - | 0,000173275 - | 4 - | 0,00016472 - | 3 - | 0,000125365 - | 5 - | 0,000235372 - |
| 4465 | 1 | 0.2um_NOKCPHJN_217466 | Eukaryota | unclassified | Oomycetes | Peronosporale | Peronosporac | Phytophthora | Phytophthora | unclassified | unclassified | unclassified | unclassified | unclassified | 0,8 | 0,4 | 1,16E-05 | 5,84E-06 | 0 - | 0 - | 1 - | 1,45E-05 - | 1 - | 1,38E-05 - | 1 - | 1,40E-05 - | 1 - | 1,57E-05 - |
| 4466 | 1 | 3um_FIJJAFBF_492177 | Bacteria | Proteobacteri | Gammaprotec | Cellvibrionales | Halieaceae | unclassified | Halieaceae ba | unclassified | unclassified | unclassified | unclassified | unclassified | 0,8 | 0,4 | 3,56E-05 | 1,81E-05 | 1 - | 3,85E-05 - | 1 - | 4,78E-05 - | 1 - | 4,55E-05 - | 1 - | 4,61E-05 - | 0 - | 0 - |
| 4467 | 1 | 3um_FIJJAFBF_474905 | Bacteria | Proteobacteri | Alphaproteob | Rhodobactera | Rhodobactera | Aliiroseovariu | Aliiroseovariu | unclassified | unclassified | unclassified | unclassified | unclassified | 2 | 0 | 0,000156602 | 1,49E-05 | 2 - | 0,000131173 - | 2 - | 0,000162879 - | 2 - | 0,000154837 - | 2 - | 0,000157124 - | 2 - | 0,000176999 - |
| 4468 | 1 | 0.2um_NOKCPHJN_224828 | Bacteria | Proteobacteri | Betaproteoba | Nitrosomonad | unclassified | unclassified | Methylophilal | unclassified | unclassified | unclassified | unclassified | unclassified | 7,4 | 1,356465997 | 0,000160324 | 2,23E-05 | 9 - | 0,000165279 - | 7 - | 0,000159621 - | 6 - | 0,000130063 - | 9 - | 0,000197976 - | 6 - | 0,000148679 - |
| 4469 | 1 | 3um_FIJJAFBF_715082 | Eukaryota | unclassified | Pelagophycea | Pelagomonad | unclassified | Aureoumbra | Aureoumbra l | unclassified | unclassified | unclassified | unclassified | unclassified | 0,6 | 0,489897949 | 4,91E-05 | 4,08E-05 | 1 - | 6,96E-05 - | 0 - | 0 - | 1 - | 8,21E-05 - | 0 - | 0 - | 1 - | 9,39E-05 - |
| 4470 | 1 | 3um_FIJJAFBF_137029 | Bacteria | Bacteroidetes | Flavobacteriia | Flavobacterial | Flavobacteriac | Flaviramulus | Flaviramulus b | unclassified | unclassified | unclassified | unclassified | unclassified | 0,8 | 0,4 | 1,96E-05 | 9,94E-06 | 1 - | 2,12E-05 - | 1 - | 2,63E-05 - | 1 - | 2,50E-05 - | 1 - | 2,54E-05 - | 0 - | 0 - |
| 4471 | 1 | 3um_FIJJAFBF_83106 | Bacteria | Bacteroidetes | Flavobacteriia | Flavobacterial | Flavobacteriac | Polaribacter | Polaribacter s | unclassified | unclassified | unclassified | unclassified | unclassified | 0,8 | 0,4 | 6,19E-05 | 3,18E-05 | 1 - | 6,48E-05 - | 1 - | 8,05E-05 - | 1 - | 7,65E-05 - | 0 - | 0 - | 1 - | 8,75E-05 - |
| 4472 | 2 | 0.2um_NOKCPHJN_294292;3um_FIJJAFBF_67715 | Bacteria | Proteobacteri | Alphaproteob | Rhodobactera | Rhodobactera | various | various | cellular proces | Posttranslatio | 05DIW | FeS assembly | various | 1 | 0 | 2,70E-05 | 2,56E-06 | 1 - | 2,26E-05 - | 1 - | 2,81E-05 - | 1 - | 2,67E-05 - | 1 - | 2,71E-05 - | 1 - | 3,05E-05 - |
| 4473 | 1 | 0.2um_NOKCPHJN_1015904 | Bacteria | Proteobacteri | Alphaproteob | Rhodobactera | Rhodobactera | Shimia | Shimia marina | metabolism | Inorganic ion t | 05PWN | phosphate ab | 439497.RR11_ | 0,2 | 0,4 | 9,80E-06 | 1,96E-05 | 0 - | 0 - | 1 - | 4,90E-05 - | 0 - | 0 - | 0 - | 0 - | 0 - | 0 - |
| 4474 | 2 | 0.2um_NOKCPHJN_249373;3um_FIJJAFBF_17974 | Bacteria | Bacteroidetes | unclassified | unclassified | unclassified | unclassified | unidentified e | metabolism | Secondary me | 09BC1 | fumarylacetoa | 50743.SCB49_ | 0,8 | 0,4 | 2,50E-05 | 1,27E-05 | 1 - | 2,70E-05 - | 1 - | 3,35E-05 - | 1 - | 3,19E-05 - | 1 - | 3,23E-05 - | 0 - | 0 - |
| 4475 | 1 | 3um_FIJJAFBF_326387 | Bacteria | Proteobacteri | Alphaproteob | Rhodobactera | Rhodobactera | unclassified | Rhodobactera | metabolism | Amino acid tra | 0XRCI | His Glu Gln Ar | 351016.RAZW | 2,2 | 0,4 | 0,000126702 | 2,57E-05 | 2 - | 9,65E-05 - | 2 - | 0,000119764 - | 2 - | 0,000113851 - | 3 - | 0,000173298 - | 2 - | 0,000130147 - |
| 4476 | 1 | 0.2um_NOKCPHJN_245797 | Bacteria | Proteobacteri | Alphaproteob | Rhodobactera | Rhodobactera | Planktomarina | Planktomarina | information st | Transcription | 01R4Z | DNA-depende | 633131.TR2A( | 1 | 0 | 9,95E-06 | 9,45E-07 | 1 - | 8,34E-06 - | 1 - | 1,03E-05 - | 1 - | 9,84E-06 - | 1 - | 9,98E-06 - | 1 - | 1,12E-05 - |
| 4477 | 1 | 3um_FIJJAFBF_332698 | Bacteria | Proteobacteri | Gammaprotec | Oceanospirilla | Saccharospirill | Reinekea | Reinekea fors | metabolism | Coenzyme tra | 0QHUT | riboflavin synt | 314283.MED2 | 0,6 | 0,489897949 | 3,87E-05 | 3,19E-05 | 1 - | 5,65E-05 - | 1 - | 7,02E-05 - | 1 - | 6,67E-05 - | 0 - | 0 - | 0 - | 0 - |
| 4478 | 1 | 0.2um_NOKCPHJN_344794 | Bacteria | Bacteroidetes | Flavobacteriia | Flavobacterial | Flavobacteriac | unclassified | Flavobacteriac | information st | Translation, ri | COG0087 | One of the pri | 487796.Flav2A | 2,8 | 0,748331477 | 0,000188515 | 6,19E-05 | 3 - | 0,000167966 - | 2 - | 0,000139043 - | 2 - | 0,000132178 - | 3 - | 0,000201195 - | 4 - | 0,000302194 - |
| 4479 | 1 | 3um_FIJJAFBF_651483 | Bacteria | Bacteroidetes | Flavobacteriia | Flavobacterial | Flavobacteriac | Formosa | Formosa sp. H | unclassified | unclassified | unclassified | unclassified | unclassified | 6,2 | 0,979795897 | 0,00029928 | 3,22E-05 | 8 - | 0,000326767 - | 6 - | 0,000304311 - | 5 - | 0,000241072 - | 6 - | 0,000293559 - | 6 - | 0,000330693 - |
| 4480 | 1 | 0.2um_NOKCPHJN_767833 | Bacteria | Proteobacteri | Alphaproteob | Pelagibacteral | Pelagibacterac | Candidatus Pe | Candidatus Pe | cellular proces | Posttranslatio | 05C2Z | ATP-depender | 751586.BDIM_ | 0,4 | 0,489897949 | 8,10E-06 | 9,93E-06 | 0 - | 0 - | 1 - | 2,06E-05 - | 0 - | 0 - | 1 - | 1,99E-05 - | 0 - | 0 - |
| 4481 | 1 | 3um_FIJJAFBF_624304 | Bacteria | Bacteroidetes | Flavobacteriia | Flavobacterial | Flavobacteriac | Gilvibacter | Gilvibacter sp. | cellular proces | Posttranslatio | 09GE9 | Thioredoxin | 216432.CA255 | 1,6 | 0,8 | 0,000201373 | 8,25E-05 | 3 - | 0,000327934 - | 2 - | 0,000271465 - | 1 - | 0,000129031 - | 1 - | 0,000130936 - | 1 - | 0,0001475 - |
| 4482 | 1 | 0.2um_NOKCPHJN_970282 | unclassified | unclassified | unclassified | unclassified | unclassified | unclassified | unclassified | unclassified | unclassified | unclassified | unclassified | unclassified | 1 | 0 | 6,14E-05 | 5,83E-06 | 1 - | 5,15E-05 - | 1 - | 6,39E-05 - | 1 - | 6,08E-05 - | 1 - | 6,17E-05 - | 1 - | 6,95E-05 - |
| 4483 | 2 | 0.2um_NOKCPHJN_722408;3um_FIJJAFBF_38959 | Bacteria | Proteobacteri | Alphaproteob | Pelagibacteral | Pelagibacterac | various | various | metabolism | Lipid transpor | 071I4 | Phosphatidyls | 335992.SAR11 | 0,4 | 0,489897949 | 2,51E-05 | 3,13E-05 | 1 - | 5,34E-05 - | 0 - | 0 - | 0 - | 0 - | 0 - | 0 - | 1 - | 7,20E-05 - |
| 4484 | 1 | 3um_FIJJAFBF_49006 | Bacteria | Proteobacteri | Gammaprotec | Oceanospirilla | Saccharospirill | Reinekea | Reinekea blan | unclassified | unclassified | unclassified | unclassified | unclassified | 5,2 | 0,979795897 | 7,39E-05 | 6,34E-06 | 7 - | 8,48E-05 - | 5 - | 7,52E-05 - | 5 - | 7,15E-05 - | 5 - | 7,26E-05 - | 4 - | 6,54E-05 - |
| 4485 | 1 | 3um_FIJJAFBF_507106 | Eukaryota | Bacillariophyt | Coscinodiscop | Thalassiosirale | Thalassiosirac | Thalassiosira | Thalassiosira c | unclassified | unclassified | unclassified | unclassified | unclassified | 3,6 | 0,8 | 0,000557583 | 0,000144617 | 3 - | 0,000386888 - | 3 - | 0,000480401 - | 5 - | 0,000761136 - | 3 - | 0,000463427 - | 4 - | 0,000696065 - |
| 4486 | 1 | 3um_FIJJAFBF_513234 | Bacteria | Proteobacteri | Gammaprotec | Oceanospirilla | Saccharospirill | Reinekea | Reinekea blan | poorly charact | Function unkn | 0QI1P | An essential G | 314283.MED2 | 1,6 | 1,019803903 | 5,54E-05 | 3,40E-05 | 0 - | 0 - | 2 - | 7,04E-05 - | 3 - | 0,000100357 - | 2 - | 6,79E-05 - | 1 - | 3,82E-05 - |
| 4487 | 1 | 0.2um_NOKCPHJN_22719 | Bacteria | Bacteroidetes | unclassified | unclassified | unclassified | unclassified | Bacteroidetes | unclassified | unclassified | unclassified | unclassified | unclassified | 0,4 | 0,489897949 | 1,37E-05 | 1,68E-05 | 0 - | 0 - | 0 - | 0 - | 0 - | 0 - | 1 - | 3,21E-05 - | 1 - | 3,62E-05 - |
| 4488 | 1 | 3um_FIJJAFBF_452750 | Bacteria | Bacteroidetes | Flavobacteriia | Flavobacterial | Flavobacteriac | Polaribacter | Polaribacter s | information st | Translation, ri | COG0016 | phenylalanyL- | 313598.MED1 | 0,8 | 0,748331477 | 3,17E-05 | 3,11E-05 | 1 - | 3,39E-05 - | 2 - | 8,41E-05 - | 0 - | 0 - | 1 - | 4,06E-05 - | 0 - | 0 - |
| 4489 | 1 | 0.2um_NOKCPHJN_412094 | Bacteria | Proteobacteri | Gammaprotec | Cellvibrionales | Porticoccacea | unclassified | SAR92 bacteri | metabolism | Coenzyme tra | 16RAU | gamma-glutar | 314287.GB22( | 0,4 | 0,489897949 | 1,08E-05 | 1,32E-05 | 0 - | 0 - | 1 - | 2,75E-05 - | 0 - | 0 - | 1 - | 2,65E-05 - | 0 - | 0 - |
| 4490 | 1 | 3um_FIJJAFBF_521154 | Bacteria | Bacteroidetes | Flavobacteriia | Flavobacterial | Flavobacteriac | Gaetbulibacte | Gaetbulibacte | information st | Translation, ri | 05KK9 | This protein bi | 313603.FB217 | 1,6 | 1,356465997 | 0,000127049 | 0,000109928 | 3 - | 0,000202547 - | 0 - | 0 - | 2 - | 0,000159391 - | 0 - | 0 - | 3 - | 0,000273308 - |
| 4491 | 1 | 3um_FIJJAFBF_551992 | Bacteria | Proteobacteri | Betaproteoba | unclassified | unclassified | unclassified | Betaproteoba | unclassified | unclassified | unclassified | unclassified | unclassified | 1 | 0 | 0,000195753 | 1,86E-05 | 1 - | 0,000163967 - | 1 - | 0,000203598 - | 1 - | 0,000193546 - | 1 - | 0,000196405 - | 1 - | 0,000221249 - |
| 4492 | 1 | 0.2um_NOKCPHJN_1024464 | Bacteria | Actinobacteri | Actinobacteri | Micrococcales | Microbacteria | unclassified | Microbacteria | information st | Transcription | 05VEQ | Cold shock pr | 644283.Micau | 0,6 | 0,8 | 9,87E-05 | 0,000136803 | 1 - | 0,000145287 - | 0 - | 0 - | 0 - | 0 - | 2 - | 0,000348059 - | 0 - | 0 - |
| 4493 | 1 | 0.2um_NOKCPHJN_1082065 | Bacteria | Verrucomicrol | Verrucomicrol | Verrucomicrol | Rubritaleacea | Rubritalea | Rubritalea squ | poorly charact | Function unkn | 1DMX1 | Permease YjgF | 349741.Amuc | 1 | 0 | 3,73E-05 | 3,55E-06 | 1 - | 3,13E-05 - | 1 - | 3,88E-05 - | 1 - | 3,69E-05 - | 1 - | 3,75E-05 - | 1 - | 4,22E-05 - |
| 4494 | 1 | 0.2um_NOKCPHJN_132232 | Bacteria | Proteobacteri | Gammaprotec | Oceanospirilla | Saccharospirill | Reinekea | Reinekea blan | metabolism | Carbohydrate | 05D2N | Aldolase | 314283.MED2 | 6,4 | 1,356465997 | 0,000240929 | 3,06E-05 | 9 - | 0,000288545 - | 5 - | 0,000199049 - | 6 - | 0,000227065 - | 6 - | 0,000230419 - | 6 - | 0,000259566 - |
| 4495 | 5 | 0.2um_NOKCPHJN_473341;0.2um_NOKCPHJN_6 | Bacteria | Bacteroidetes | Flavobacteriia | Flavobacterial | Flavobacteriac | Formosa | various | information st | Translation, ri | COG0172 | Catalyzes the | various | 0,4 | 0,489897949 | 1,22E-05 | 1,50E-05 | 1 - | 2,71E-05 - | 1 - | 3,37E-05 - | 0 - | 0 - | 0 - | 0 - | 0 - | 0 - |
| 4496 | 1 | 0.2um_NOKCPHJN_112759 | Bacteria | Bacteroidetes | Flavobacteriia | Flavobacterial | Cryomorphace | Owenweeksia | Owenweeksia | unclassified | unclassified | unclassified | unclassified | unclassified | 0,4 | 0,489897949 | 1,83E-05 | 2,25E-05 | 1 - | 4,19E-05 - | 0 - | 0 - | 1 - | 4,94E-05 - | 0 - | 0 - | 0 - | 0 - |
| 4497 | 1 | 0.2um_NOKCPHJN_97158 | Bacteria | Proteobacteri | Betaproteoba | Burkholderial | Comamonada | Limnohabitan | Limnohabitan | cellular proces | Posttranslatio | 08V1T | peptidylprolyl | 338969.Rfer_1 | 0,2 | 0,4 | 2,35E-05 | 4,70E-05 | 0 - | 0 - | 0 - | 0 - | 0 - | 0 - | 1 - | 0,000117507 - | 0 - | 0 - |
| 4498 | 1 | 0.2um_NOKCPHJN_112104 | Bacteria | Bacteroidetes | Flavobacteriia | Flavobacterial | Flavobacteriac | unclassified | Flavobacteriac | cellular proces | Cell motility | COG1360 | Ompa motb d | 487796.Flav2A | 5,6 | 0,489897949 | 0,000268905 | 2,35E-05 | 6 - | 0,000242486 - | 6 - | 0,000301096 - | 6 - | 0,00028623 - | 5 - | 0,000242048 - | 5 - | 0,000272666 - |
| 4499 | 1 | 0.2um_NOKCPHJN_854755 | Bacteria | Proteobacteri | Alphaproteob | Rhodobactera | Rhodobactera | Planktotalea | Planktotalea f | metabolism | Amino acid tra | 16R0H | Acetolactate s | 314270.RB208 | 0,6 | 0,8 | 4,73E-05 | 6,20E-05 | 0 - | 0 - | 2 - | 0,000153246 - | 0 - | 0 - | 0 - | 0 - | 1 - | 8,33E-05 - |
| 4500 | 1 | 3um_FIJJAFBF_333511 | Bacteria | Proteobacteri | Alphaproteob | Rhodobactera | Rhodobactera | unclassified | Rhodobactera | metabolism | Amino acid tra | 05CAI | ABC transport | 439497.RR11_ | 2,2 | 1,166190379 | 0,00011761 | 5,95E-05 | 3 - | 0,000137183 - | 1 - | 5,68E-05 - | 4 - | 0,000215908 - | 1 - | 5,48E-05 - | 2 - | 0,000123406 - |
| 4501 | 1 | 0.2um_NOKCPHJN_64676 | Bacteria | Bacteroidetes | Cytophagia | Cytophagales | Flammeovirga | unclassified | Flammeovirga | metabolism | Inorganic ion t | 05C27 | TonB-depende | 435591.BDI_3 | 0,8 | 0,748331477 | 1,01E-05 | 9,68E-06 | 1 - | 1,10E-05 - | 1 - | 1,37E-05 - | 2 - | 2,60E-05 - | 0 - | 0 - | 0 - | 0 - |
| 4502 | 1 | 0.2um_NOKCPHJN_109032 | Bacteria | Proteobacteri | Gammaprotec | Cellvibrionales | Porticoccacea | unclassified | gamma protec | metabolism | Inorganic ion t | 16TQC | TONB-depend | 314287.GB22( | 0,4 | 0,489897949 | 5,48E-06 | 6,84E-06 | 1 - | 1,17E-05 - | 0 - | 0 - | 0 - | 0 - | 0 - | 0 - | 1 - | 1,57E-05 - |
| 4503 | 1 | 3um_FIJJAFBF_645034 | Bacteria | Proteobacteri | Alphaproteob | Rhodobactera | Rhodobactera | unclassified | Rhodobactera | metabolism | Amino acid tra | COG0683 | (ABC) transpo | 391593.RCCS2 | 2,4 | 0,489897949 | 8,30E-05 | 1,91E-05 | 2 - | 5,78E-05 - | 3 - | 0,000107697 - | 3 - | 0,00010238 - | 2 - | 6,93E-05 - | 2 - | 7,80E-05 - |
| 4504 | 1 | 3um_FIJJAFBF_140885 | Bacteria | Proteobacteri | Gammaprotec | unclassified | Competibacte | Plasticicumula | Plasticicumula | unclassified | unclassified | unclassified | unclassified | unclassified | 3,4 | 0,489897949 | 0,000152189 | 1,48E-05 | 4 - | 0,00015152 - | 3 - | 0,000141108 - | 4 - | 0,000178854 - | 3 - | 0,000136122 - | 3 - | 0,000153341 - |
| 4505 | 1 | 0.2um_NOKCPHJN_352260 | Bacteria | Proteobacteri | Alphaproteob | Rhodobactera | Rhodobactera | Cognatiyoonia | Cognatiyoonia | metabolism | Amino acid tra | 01QDI | sarcosine oxid | 314232.SKA53 | 1,2 | 0,4 | 1,74E-05 | 6,44E-06 | 1 - | 1,21E-05 - | 2 - | 3,00E-05 - | 1 - | 1,43E-05 - | 1 - | 1,45E-05 - | 1 - | 1,63E-05 - |
| 4506 | 1 | 0.2um_NOKCPHJN_807950 | Bacteria | Proteobacteri | Alphaproteob | Pelagibacteral | Pelagibacterac | unclassified | Pelagibacterac | information st | Translation, ri | 05CEJ | Catalyzes the | 684719.HIMB | 5 | 1,673320053 | 9,88E-05 | 3,05E-05 | 7 - | 0,00011729 - | 6 - | 0,000124834 - | 5 - | 9,89E-05 - | 2 - | 4,01E-05 - | 5 - | 0,000113047 - |
| 4507 | 2 | 0.2um_NOKCPHJN_97628;3um_FIJJAFBF_53315 | Bacteria | Bacteroidetes | Flavobacteriia | Flavobacterial | Flavobacteriac | Polaribacter | various | poorly charact | Function unkn | 05FCN | von Willebran | 313598.MED1 | 0,6 | 0,489897949 | 2,72E-05 | 2,24E-05 | 1 - | 3,97E-05 - | 1 - | 4,93E-05 - | 1 - | 4,69E-05 - | 0 - | 0 - | 0 - | 0 - |
| 4508 | 1 | 0.2um_NOKCPHJN_917547 | Bacteria | Proteobacteri | Gammaprotec | unclassified | unclassified | unclassified | Gammaprotec | unclassified | unclassified | unclassified | unclassified | unclassified | 1,6 | 0,489897949 | 7,21E-05 | 2,00E-05 | 2 - | 7,68E-05 - | 2 - | 9,53E-05 - | 2 - | 9,06E-05 - | 1 - | 4,60E-05 - | 1 - | 5,18E-05 - |
| 4509 | 1 | 3um_FIJJAFBF_300641 | Bacteria | Bacteroidetes | Flavobacteriia | various | various | various | various | poorly charact | Function unkn | 0XNQI | domain protei | 487797.Flav3( | 3,2 | 1,166190379 | 9,80E-05 | 3,91E-05 | 2 - | 5,06E-05 - | 2 - | 6,28E-05 - | 5 - | 0,00014921 - | 3 - | 9,08E-05 - | 4 - | 0,000136453 - |
| 4510 | 1 | 3um_FIJJAFBF_19824 | Bacteria | Bacteroidetes | Flavobacteriia | Flavobacterial | Flavobacteriac | Polaribacter | Polaribacter s | metabolism | Amino acid tra | 08IQK | o-acetylhomo | 313594.PI23P | 1 | 0 | 3,23E-05 | 3,07E-06 | 1 - | 2,71E-05 - | 1 - | 3,36E-05 - | 1 - | 3,20E-05 - | 1 - | 3,24E-05 - | 1 - | 3,65E-05 - |
| 4511 | 1 | 3um_FIJJAFBF_586309 | Bacteria | Bacteroidetes | Flavobacteriia | Flavobacterial | Flavobacteriac | Formosa | Formosa sp. H | poorly charact | Function unkn | 0ZU63 | NA | 313603.FB217 | 1,4 | 0,8 | 6,90E-05 | 3,98E-05 | 2 - | 8,57E-05 - | 2 - | 0,000106357 - | 1 - | 5,06E-05 - | 2 - | 0,000102599 - | 0 - | 0 - |
| 4512 | 1 | 3um_FIJJAFBF_777836 | Bacteria | Bacteroidetes | Flavobacteriia | Flavobacterial | Flavobacteriac | Polaribacter | Polaribacter s | unclassified | unclassified | unclassified | unclassified | unclassified | 0,2 | 0,4 | 6,60E-06 | 1,32E-05 | 1 - | 3,30E-05 - | 0 - | 0 - | 0 - | 0 - | 0 - | 0 - | 0 - | 0 - |
| 4513 | 1 | 0.2um_NOKCPHJN_195256 | Bacteria | Bacteroidetes | Flavobacteriia | Flavobacterial | Flavobacteriac | Polaribacter | Polaribacter s | unclassified | unclassified | unclassified | unclassified | unclassified | 1,2 | 0,979795897 | 3,00E-05 | 2,14E-05 | 3 - | 6,70E-05 - | 0 - | 0 - | 1 - | 2,64E-05 - | 1 - | 2,67E-05 - | 1 - | 3,01E-05 - |
| 4514 | 1 | 0.2um_NOKCPHJN_110552 | Bacteria | Proteobacteri | Gammaprotec | Cellvibrionales | Porticoccacea | unclassified | SAR92 bacteri | information st | Translation, ri | 17FIH | One of the ea | 314287.GB22( | 2,2 | 0,4 | 0,000303071 | 2,78E-05 | 3 - | 0,000351357 - | 2 - | 0,000290855 - | 2 - | 0,000276494 - | 2 - | 0,000280578 - | 2 - | 0,00031607 - |
| 4515 | 1 | 3um_FIJJAFBF_422712 | unclassified | unclassified | unclassified | unclassified | unclassified | unclassified | unclassified | unclassified | unclassified | unclassified | unclassified | unclassified | 0,4 | 0,489897949 | 5,69E-05 | 6,97E-05 | 0 - | 0 - | 0 - | 0 - | 1 - | 0,000141127 - | 1 - | 0,000143212 - | 0 - | 0 - |
| 4516 | 1 | 3um_FIJJAFBF_27482 | Bacteria | Bacteroidetes | Flavobacteriia | Flavobacterial | Flavobacteriac | Formosa | Formosa sp. H | metabolism | Energy produc | 05C26 | Dehydrogenas | 391587.KAOT | 0,2 | 0,4 | 5,13E-06 | 1,03E-05 | 0 - | 0 - | 0 - | 0 - | 1 - | 2,57E-05 - | 0 - | 0 - | 0 - | 0 - |
| 4517 | 1 | 3um_FIJJAFBF_457484 | Bacteria | Proteobacteri | Alphaproteob | Rhodobactera | Rhodobactera | Tateyamaria | Tateyamaria s | unclassified | unclassified | unclassified | unclassified | unclassified | 1 | 0 | 0,000126877 | 1,20E-05 | 1 - | 0,000106275 - | 1 - | 0,000131962 - | 1 - | 0,000125447 - | 1 - | 0,000127299 - | 1 - | 0,000143402 - |
| 4518 | 1 | 0.2um_NOKCPHJN_351628 | Bacteria | unclassified | unclassified | unclassified | unclassified | unclassified | bacterium TM | metabolism | Carbohydrate | 0B9GB | TRAP dicarbox | 340100.Bpet3 | 0,2 | 0,4 | 8,29E-06 | 1,66E-05 | 1 - | 4,14E-05 - | 0 - | 0 - | 0 - | 0 - | 0 - | 0 - | 0 - | 0 - |
| 4519 | 1 | 0.2um_NOKCPHJN_1080022 | Bacteria | Bacteroidetes | Flavobacteriia | Flavobacterial | Flavobacteriac | Polaribacter | Polaribacter t | cellular proces | Cell motility | COG1360 | Ompa motb d | 313594.PI23P | 3,2 | 0,4 | 0,000152634 | 2,62E-05 | 3 - | 0,000119559 - | 4 - | 0,000197943 - | 3 - | 0,000141127 - | 3 - | 0,000143212 - | 3 - | 0,000161328 - |
| 4520 | 1 | 3um_FIJJAFBF_784795 | Bacteria | Proteobacteri | Betaproteoba | unclassified | unclassified | unclassified | Betaproteoba | metabolism | Carbohydrate | 0BRDP | Catalyzes the | 596154.Alide2 | 0,6 | 0,489897949 | 1,03E-05 | 8,44E-06 | 0 - | 0 - | 1 - | 1,77E-05 - | 1 - | 1,69E-05 - | 1 - | 1,71E-05 - | 0 - | 0 - |
| 4521 | 1 | 0.2um_NOKCPHJN_237246 | Bacteria | Proteobacteri | Alphaproteob | Pelagibacteral | Pelagibacterac | Candidatus Pe | Candidatus Pe | cellular proces | Cell wall/mem | 7257 | NA | 439493.PB721 | 2,6 | 0,489897949 | 0,000108557 | 2,31E-05 | 2 - | 6,98E-05 - | 3 - | 0,000129956 - | 3 - | 0,00012354 - | 3 - | 0,000125365 - | 2 - | 9,41E-05 - |
| 4522 | 1 | 3um_FIJJAFBF_43326 | Bacteria | Proteobacteri | Alphaproteob | Rhodobactera | Rhodobactera | Planktotalea | Planktotalea f | unclassified | unclassified | unclassified | unclassified | unclassified | 0,4 | 0,489897949 | 8,08E-06 | 9,99E-06 | 1 - | 1,80E-05 - | 1 - | 2,24E-05 - | 0 - | 0 - | 0 - | 0 - | 0 - | 0 - |
| 4523 | 1 | 0.2um_NOKCPHJN_99363 | Bacteria | Bacteroidetes | unclassified | unclassified | unclassified | unclassified | Bacteroidetes | information st | Transcription | 05D27 | DNA-depende | 926562.Oweh | 9,2 | 1,469693846 | 8,71E-05 | 9,96E-06 | 11 - | 8,83E-05 - | 10 - | 9,97E-05 - | 8 - | 7,58E-05 - | 10 - | 9,61E-05 - | 7 - | 7,58E-05 - |
| 4524 | 1 | 3um_FIJJAFBF_328849 | Bacteria | Proteobacteri | Betaproteoba | Neisseriales | Neisseriaceae | Neisseria | Neisseria men | unclassified | unclassified | unclassified | unclassified | unclassified | 0,8 | 0,4 | 0,000101417 | 5,21E-05 | 1 - | 0,000106275 - | 1 - | 0,000131962 - | 1 - | 0,000125447 - | 0 - | 0 - | 1 - | 0,000143402 - |
| 4525 | 1 | 3um_FIJJAFBF_238482 | Bacteria | Proteobacteri | Gammaprotec | Oceanospirilla | Saccharospirill | Saccharospiril | various | information st | Translation, ri | 177XJ | One of the pri | 314283.MED2 | 5,4 | 0,489897949 | 0,000471274 | 3,21E-05 | 6 - | 0,000441449 - | 5 - | 0,000456791 - | 6 - | 0,000521085 - | 5 - | 0,000440652 - | 5 - | 0,000496393 - |
| 4526 | 1 | 0.2um_NOKCPHJN_291840 | Bacteria | Proteobacteri | Alphaproteob | Rhodospirillal | Rhodospirillac | unclassified | Rhodobactera | poorly charact | Function unkn | 17FM2 | Domain of unl | 367336.OM22 | 0,2 | 0,4 | 1,66E-05 | 3,31E-05 | 0 - | 0 - | 0 - | 0 - | 0 - | 0 - | 1 - | 8,28E-05 - | 0 - | 0 - |
| 4527 | 1 | 3um_FIJJAFBF_136473 | Bacteria | Proteobacteri | Gammaprotec | Alteromonada | Colwelliaceae | Colwellia | Colwellia sp. F | cellular proces | Intracellular tr | 0QJ7H | motA TolQ Exl | 167879.CPS_3 | 0,2 | 0,4 | 5,80E-06 | 1,16E-05 | 0 - | 0 - | 0 - | 0 - | 1 - | 2,90E-05 - | 0 - | 0 - | 0 - | 0 - |
| 4528 | 1 | 3um_FIJJAFBF_222361 | Bacteria | Proteobacteri | Alphaproteob | Rhodobactera | Rhodobactera | unclassified | Rhodobactera | cellular proces | Intracellular tr | 175GA | One of the pro | 439497.RR11_ | 3,8 | 0,748331477 | 0,000319021 | 6,89E-05 | 4 - | 0,000281661 - | 3 - | 0,000262305 - | 3 - | 0,000249354 - | 5 - | 0,000421728 - | 4 - | 0,00038006 - |
| 4529 | 1 | 3um_FIJJAFBF_216062 | Bacteria | Proteobacteri | Gammaprotec | Cellvibrionales | Cellvibrionace | Teredinibacte | various | information st | Transcription | COG0250 | Participates in | 377629.TERTU | 0,8 | 0,4 | 6,13E-05 | 3,15E-05 | 1 - | 6,48E-05 - | 0 - | 0 - | 1 - | 7,65E-05 - | 1 - | 7,77E-05 - | 1 - | 8,75E-05 - |
| 4530 | 1 | 0.2um_NOKCPHJN_442561 | Bacteria | Bacteroidetes | unclassified | unclassified | unclassified | unclassified | Bacteroidetes | unclassified | unclassified | unclassified | unclassified | unclassified | 0,4 | 0,489897949 | 1,50E-05 | 1,87E-05 | 1 - | 3,19E-05 - | 0 - | 0 - | 0 - | 0 - | 0 - | 0 - | 1 - | 4,30E-05 - |
| 4531 | 1 | 3um_FIJJAFBF_173293 | Bacteria | Proteobacteri | Gammaprotec | unclassified | unclassified | unclassified | various | unclassified | unclassified | unclassified | unclassified | unclassified | 1,6 | 0,489897949 | 0,000156873 | 4,94E-05 | 2 - | 0,000165146 - | 1 - | 0,000102532 - | 2 - | 0,000194938 - | 1 - | 9,89E-05 - | 2 - | 0,000222841 - |
| 4532 | 1 | 3um_FIJJAFBF_238153 | Bacteria | Proteobacteri | Alphaproteob | Rhodobactera | Rhodobactera | Planktomarina | Planktomarina | unclassified | unclassified | unclassified | unclassified | unclassified | 1,2 | 0,4 | 4,90E-05 | 1,64E-05 | 1 - | 3,43E-05 - | 1 - | 4,25E-05 - | 2 - | 8,09E-05 - | 1 - | 4,10E-05 - | 1 - | 4,62E-05 - |
| 4533 | 1 | 3um_FIJJAFBF_111323 | Viruses | unclassified | unclassified | unclassified | Mimiviridae | unclassified | Yellowstone la | unclassified | unclassified | unclassified | unclassified | unclassified | 3,2 | 1,166190379 | 7,55E-05 | 2,70E-05 | 4 - | 8,01E-05 - | 5 - | 0,000124362 - | 3 - | 7,09E-05 - | 2 - | 4,80E-05 - | 2 - | 5,41E-05 - |
| 4534 | 1 | 0.2um_NOKCPHJN_457094 | Bacteria | Bacteroidetes | Flavobacteriia | Flavobacterial | Flavobacteriac | unclassified | Flavobacteriac | unclassified | unclassified | unclassified | unclassified | unclassified | 1 | 0 | 6,17E-05 | 5,86E-06 | 1 - | 5,17E-05 - | 1 - | 6,42E-05 - | 1 - | 6,10E-05 - | 1 - | 6,19E-05 - | 1 - | 6,98E-05 - |
| 4535 | 1 | 0.2um_NOKCPHJN_343391 | Bacteria | Proteobacteri | Gammaprotec | unclassified | unclassified | unclassified | Gammaprotec | poorly charact | Function unkn | 0XY6Y | Pkd domain co | 314287.GB22( | 4 | 0,632455532 | 6,58E-05 | 4,82E-06 | 5 - | 6,99E-05 - | 4 - | 6,94E-05 - | 4 - | 6,60E-05 - | 4 - | 6,70E-05 - | 3 - | 5,66E-05 - |
| 4536 | 1 | 0.2um_NOKCPHJN_664786 | Bacteria | Actinobacteri | Actinobacteri | Micrococcales | Microbacteria | Candidatus Ac | Candidatus Ac | poorly charact | Function unkn | 126RR | cell surface- a | 43179.ENSST( | 10,8 | 1,720465053 | 0,000135049 | 1,28E-05 | 13 - | 0,000138029 - | 11 - | 0,000145024 - | 12 - | 0,000150397 - | 10 - | 0,000127182 - | 8 - | 0,000114616 - |
| 4537 | 1 | 3um_FIJJAFBF_644895 | Bacteria | Proteobacteri | Alphaproteob | Rhodobactera | Rhodobactera | Planktomarina | Planktomarina | unclassified | unclassified | unclassified | unclassified | unclassified | 0,6 | 0,489897949 | 1,86E-05 | 1,54E-05 | 1 - | 2,59E-05 - | 1 - | 3,21E-05 - | 0 - | 0 - | 0 - | 0 - | 1 - | 3,49E-05 - |
| 4538 | 1 | 0.2um_NOKCPHJN_97147 | Bacteria | Proteobacteri | Betaproteoba | Burkholderial | unclassified | unclassified | Burkholderial | unclassified | unclassified | unclassified | unclassified | unclassified | 0,2 | 0,4 | 8,21E-06 | 1,64E-05 | 0 - | 0 - | 0 - | 0 - | 0 - | 0 - | 1 - | 4,10E-05 - | 0 - | 0 - |
| 4539 | 1 | 0.2um_NOKCPHJN_729548 | Bacteria | Proteobacteri | Gammaprotec | Alteromonada | Alteromonada | Paraglaciecola | Paraglaciecola | metabolism | Inorganic ion t | 0QJE8 | receptor | 342610.Patl_2 | 4,6 | 1,496662955 | 7,75E-05 | 2,34E-05 | 6 - | 8,57E-05 - | 5 - | 8,86E-05 - | 2 - | 3,37E-05 - | 6 - | 0,000102599 - | 4 - | 7,71E-05 - |
| 4540 | 1 | 3um_FIJJAFBF_636371 | Bacteria | Bacteroidetes | Cytophagia | Cytophagales | Flammeovirga | unclassified | Flammeovirga | unclassified | unclassified | unclassified | unclassified | unclassified | 2,4 | 0,489897949 | 0,000188935 | 5,11E-05 | 2 - | 0,000130428 - | 2 - | 0,000161953 - | 2 - | 0,000153957 - | 3 - | 0,000234347 - | 3 - | 0,000263991 - |
| 4541 | 1 | 3um_FIJJAFBF_702421 | unclassified | unclassified | unclassified | unclassified | unclassified | unclassified | unclassified | unclassified | unclassified | unclassified | unclassified | unclassified | 0,2 | 0,4 | 1,30E-05 | 2,60E-05 | 0 - | 0 - | 0 - | 0 - | 0 - | 0 - | 0 - | 0 - | 1 - | 6,51E-05 - |
| 4542 | 1 | 3um_FIJJAFBF_112432 | Bacteria | Bacteroidetes | unclassified | unclassified | unclassified | unclassified | Bacteroidetes | unclassified | unclassified | unclassified | unclassified | unclassified | 2,6 | 0,8 | 9,79E-05 | 2,23E-05 | 4 - | 0,000128963 - | 3 - | 0,0001201 - | 2 - | 7,61E-05 - | 2 - | 7,72E-05 - | 2 - | 8,70E-05 - |
| 4543 | 2 | 0.2um_NOKCPHJN_225811;0.2um_NOKCPHJN_5 | Bacteria | Proteobacteri | Gammaprotec | Cellvibrionales | Porticoccacea | unclassified | SAR92 bacteri | metabolism | Carbohydrate | 16QI2 | glycoside hydr | 314287.GB22( | 1,8 | 0,4 | 1,48E-05 | 2,93E-06 | 2 - | 1,40E-05 - | 2 - | 1,74E-05 - | 2 - | 1,65E-05 - | 2 - | 1,68E-05 - | 1 - | 9,46E-06 - |
| 4544 | 1 | 3um_FIJJAFBF_31607 | Bacteria | Proteobacteri | Alphaproteob | Pelagibacteral | Pelagibacterac | Candidatus Pe | Candidatus Pe | metabolism | Inorganic ion t | 16PII | TRAP transpor | 439493.PB721 | 4,8 | 1,326649916 | 0,000193877 | 4,29E-05 | 7 - | 0,00024055 - | 4 - | 0,000170681 - | 5 - | 0,000202818 - | 3 - | 0,000123488 - | 5 - | 0,000231848 - |
| 4545 | 1 | 3um_FIJJAFBF_05935 | Bacteria | Proteobacteri | Gammaprotec | Cellvibrionales | Porticoccacea | unclassified | SAR92 bacteri | metabolism | Lipid transpor | COG4247 | Phytase | 314287.GB22( | 6,4 | 0,8 | 0,000241332 | 2,70E-05 | 7 - | 0,000222559 - | 5 - | 0,000197395 - | 7 - | 0,000262708 - | 7 - | 0,000266588 - | 6 - | 0,000257409 - |
| 4546 | 1 | 0.2um_NOKCPHJN_27323 | Bacteria | Bacteroidetes | unclassified | unclassified | unclassified | unclassified | Bacteroidetes | poorly charact | Function unkn | 0Y63T | Ragb susd dor | 471854.Dfer_ | 20,4 | 2,939387691 | 0,000544641 | 3,34E-05 | 26 - | 0,000588599 - | 20 - | 0,000562205 - | 20 - | 0,000534447 - | 18 - | 0,000488106 - | 18 - | 0,00054985 - |
| 4547 | 1 | 3um_FIJJAFBF_147642 | Bacteria | Proteobacteri | Betaproteoba | Burkholderial | Comamonada | Curvibacter | Curvibacter sp | unclassified | unclassified | unclassified | unclassified | unclassified | 8,4 | 2,244994432 | 0,000363908 | 0,000104695 | 6 - | 0,00021656 - | 10 - | 0,000448173 - | 12 - | 0,000511254 - | 7 - | 0,000302636 - | 7 - | 0,000340919 - |
| 4548 | 1 | 3um_FIJJAFBF_110199 | Bacteria | various | various | various | various | various | various | information st | Translation, ri | 09EM4 | Binds directly | 688270.Celal_ | 0,2 | 0,4 | 2,01E-05 | 4,03E-05 | 1 - | 0,000100681 - | 0 - | 0 - | 0 - | 0 - | 0 - | 0 - | 0 - | 0 - |
| 4549 | 1 | 0.2um_NOKCPHJN_480443 | Bacteria | Bacteroidetes | Cytophagia | Cytophagales | Flammeovirga | unclassified | Flammeovirga | poorly charact | Function unkn | 0FZ9D | FG-GAP repea | 643867.Ftrac_ | 1,4 | 0,489897949 | 1,04E-05 | 4,40E-06 | 1 - | 6,08E-06 - | 2 - | 1,51E-05 - | 1 - | 7,18E-06 - | 1 - | 7,28E-06 - | 2 - | 1,64E-05 - |
| 4550 | 1 | 3um_FIJJAFBF_260030 | Bacteria | Verrucomicrol | Verrucomicrol | unclassified | unclassified | unclassified | Verrucomicrol | poorly charact | Function unkn | 0ZMMU | Prokaryotic N- | 240016.VspiD | 2 | 0,894427191 | 0,000113519 | 4,49E-05 | 3 - | 0,00014715 - | 2 - | 0,000121811 - | 3 - | 0,000173695 - | 1 - | 5,88E-05 - | 1 - | 6,62E-05 - |
| 4551 | 1 | 0.2um_NOKCPHJN_491288 | Bacteria | Proteobacteri | Alphaproteob | Pelagibacteral | Pelagibacterac | Candidatus Pe | Candidatus Pe | metabolism | Secondary me | 01Q86 | extracellular s | 439493.PB721 | 4 | 0 | 0,000143484 | 1,36E-05 | 4 - | 0,000120185 - | 4 - | 0,000149234 - | 4 - | 0,000141866 - | 4 - | 0,000143962 - | 4 - | 0,000162172 - |
| 4552 | 3 | 0.2um_NOKCPHJN_1039336;0.2um_NOKCPHJN_ | Bacteria | Proteobacteri | Alphaproteob | Rhizobiales | unclassified | unclassified | Rhizobiales ba | information st | Replication, re | COG0210 | helicase | various | 0,2 | 0,4 | 5,91E-06 | 1,18E-05 | 0 - | 0 - | 1 - | 2,96E-05 - | 0 - | 0 - | 0 - | 0 - | 0 - | 0 - |
| 4553 | 4 | 0.2um_NOKCPHJN_78574;3um_FIJJAFBF_31293( | Bacteria | Bacteroidetes | Flavobacteriia | Flavobacterial | Flavobacteriac | Formosa | Formosa sp. H | unclassified | unclassified | unclassified | unclassified | unclassified | 3 | 0 | 0,000183518 | 1,74E-05 | 3 - | 0,000153719 - | 3 - | 0,000190873 - | 3 - | 0,000181449 - | 3 - | 0,000184129 - | 3 - | 0,000207421 - |
| 4554 | 1 | 3um_FIJJAFBF_346214 | Bacteria | Bacteroidetes | Flavobacteriia | Flavobacterial | Crocinitomica | unclassified | Crocinitomica | metabolism | Energy produc | COG0711 | F(1)F(0) ATP s | 926562.Oweh | 2,2 | 0,748331477 | 0,000182114 | 6,44E-05 | 3 - | 0,000208685 - | 2 - | 0,00017275 - | 2 - | 0,000164221 - | 1 - | 8,33E-05 - | 3 - | 0,00028159 - |
| 4555 | 1 | 3um_FIJJAFBF_270756 | Bacteria | Bacteroidetes | Flavobacteriia | Flavobacterial | Flavobacteriac | Lacinutrix | Lacinutrix him | information st | Translation, ri | 08YY1 | This protein is | 391603.FBALC | 0,8 | 0,4 | 9,14E-05 | 4,64E-05 | 1 - | 9,89E-05 - | 1 - | 0,000122861 - | 1 - | 0,000116795 - | 1 - | 0,00011852 - | 0 - | 0 - |
| 4556 | 1 | 3um_FIJJAFBF_103136 | Bacteria | Bacteroidetes | Flavobacteriia | Flavobacterial | Flavobacteriac | Polaribacter | Polaribacter s | unclassified | unclassified | unclassified | unclassified | unclassified | 0,8 | 0,4 | 3,14E-05 | 1,61E-05 | 1 - | 3,29E-05 - | 1 - | 4,08E-05 - | 1 - | 3,88E-05 - | 0 - | 0 - | 1 - | 4,44E-05 - |
| 4557 | 1 | 0.2um_NOKCPHJN_912740 | Viruses | unclassified | unclassified | unclassified | unclassified | unclassified | uncultured M | unclassified | unclassified | unclassified | unclassified | unclassified | 0,8 | 0,748331477 | 8,42E-05 | 7,31E-05 | 2 - | 0,000185124 - | 0 - | 0 - | 0 - | 0 - | 1 - | 0,000110874 - | 1 - | 0,000124899 - |
| 4558 | 1 | 0.2um_NOKCPHJN_605258 | Bacteria | Proteobacteri | Gammaprotec | unclassified | unclassified | unclassified | OM182 bacter | unclassified | unclassified | unclassified | unclassified | unclassified | 1 | 0 | 3,22E-05 | 3,05E-06 | 1 - | 2,69E-05 - | 1 - | 3,35E-05 - | 1 - | 3,18E-05 - | 1 - | 3,23E-05 - | 1 - | 3,64E-05 - |
| 4559 | 1 | 0.2um_NOKCPHJN_345091 | Bacteria | Bacteroidetes | Flavobacteriia | Flavobacterial | Flavobacteriac | Polaribacter | various | unclassified | unclassified | unclassified | unclassified | unclassified | 0,8 | 0,4 | 6,35E-05 | 3,23E-05 | 1 - | 6,87E-05 - | 1 - | 8,53E-05 - | 1 - | 8,11E-05 - | 1 - | 8,23E-05 - | 0 - | 0 - |
| 4560 | 1 | 0.2um_NOKCPHJN_261679 | Bacteria | Bacteroidetes | unclassified | unclassified | unclassified | unclassified | Bacteroidetes | unclassified | unclassified | unclassified | unclassified | unclassified | 0,8 | 0,4 | 8,21E-05 | 4,13E-05 | 0 - | 0 - | 1 - | 0,000102532 - | 1 - | 9,75E-05 - | 1 - | 9,89E-05 - | 1 - | 0,00011142 - |
| 4561 | 1 | 3um_FIJJAFBF_142734 | Bacteria | Proteobacteri | Gammaprotec | Cellvibrionales | Porticoccacea | unclassified | SAR92 bacteri | unclassified | unclassified | unclassified | unclassified | unclassified | 0,2 | 0,4 | 2,70E-05 | 5,40E-05 | 1 - | 0,000135031 - | 0 - | 0 - | 0 - | 0 - | 0 - | 0 - | 0 - | 0 - |
| 4562 | 4 | 3um_FIJJAFBF_165805;3um_FIJJAFBF_300267;3u | Bacteria | Proteobacteri | Gammaprotec | various | various | various | various | unclassified | unclassified | unclassified | unclassified | unclassified | 0,2 | 0,4 | 2,26E-05 | 4,52E-05 | 0 - | 0 - | 0 - | 0 - | 1 - | 0,000112902 - | 0 - | 0 - | 0 - | 0 - |
| 4563 | 1 | 0.2um_NOKCPHJN_51590 | Bacteria | Proteobacteri | Gammaprotec | Cellvibrionales | Porticoccacea | unclassified | Gammaprotec | metabolism | Inorganic ion t | 16TQC | TONB-depend | 314287.GB22( | 3,8 | 0,748331477 | 5,31E-05 | 8,63E-06 | 5 - | 5,92E-05 - | 3 - | 4,41E-05 - | 3 - | 4,19E-05 - | 4 - | 5,67E-05 - | 4 - | 6,39E-05 - |
| 4564 | 1 | 0.2um_NOKCPHJN_1103607 | Bacteria | Proteobacteri | Alphaproteob | Rhodobactera | Rhodobactera | Lentibacter | Lentibacter al | metabolism | Secondary me | 07US8 | Inherit from C | 167555.NATL1 | 0,6 | 0,489897949 | 1,78E-05 | 1,47E-05 | 1 - | 2,59E-05 - | 1 - | 3,21E-05 - | 0 - | 0 - | 1 - | 3,10E-05 - | 0 - | 0 - |
| 4565 | 1 | 3um_FIJJAFBF_243019 | Bacteria | Bacteroidetes | Flavobacteriia | Flavobacterial | Flavobacteriac | unclassified | Flavobacteriac | poorly charact | Function unkn | 11PIQ | YCE I like fami | 983544.Lacal_ | 0,4 | 0,8 | 2,39E-05 | 4,78E-05 | 2 - | 0,000119559 - | 0 - | 0 - | 0 - | 0 - | 0 - | 0 - | 0 - | 0 - |
| 4566 | 1 | 0.2um_NOKCPHJN_613241 | Bacteria | Proteobacteri | Gammaprotec | unclassified | unclassified | unclassified | Gammaprotec | information st | Translation, ri | 0QTNF | One of the pri | 322710.Avin_( | 1,8 | 0,979795897 | 0,000267113 | 0,000136905 | 3 - | 0,000391284 - | 2 - | 0,000323907 - | 2 - | 0,000307914 - | 2 - | 0,000312462 - | 0 - | 0 - |
| 4567 | 1 | 3um_FIJJAFBF_297872 | Bacteria | Proteobacteri | unclassified | unclassified | unclassified | unclassified | uncultured ma | unclassified | unclassified | unclassified | unclassified | unclassified | 4,4 | 1,019803903 | 0,000129714 | 3,13E-05 | 5 - | 0,000123949 - | 6 - | 0,00018469 - | 4 - | 0,000117047 - | 3 - | 8,91E-05 - | 4 - | 0,000133801 - |
| 4568 | 1 | 0.2um_NOKCPHJN_775435 | Eukaryota | Bacillariophyt | Bacillariophyc | Naviculales | Phaeodactylac | Phaeodactylu | Phaeodactylu | metabolism | Energy produc | COG0712 | F(1)F(0) ATP s | 2850.Phatr44( | 0,4 | 0,489897949 | 2,09E-05 | 2,57E-05 | 0 - | 0 - | 0 - | 0 - | 1 - | 4,87E-05 - | 0 - | 0 - | 1 - | 5,57E-05 - |
| 4569 | 1 | 0.2um_NOKCPHJN_588988 | Bacteria | Proteobacteri | Alphaproteob | Pelagibacteral | unclassified | unclassified | Pelagibacteral | cellular proces | Cell wall/mem | 7257 | NA | 439493.PB721 | 1 | 0 | 4,09E-05 | 3,88E-06 | 1 - | 3,43E-05 - | 1 - | 4,25E-05 - | 1 - | 4,04E-05 - | 1 - | 4,10E-05 - | 1 - | 4,62E-05 - |
| 4570 | 1 | 3um_FIJJAFBF_425485 | Bacteria | Proteobacteri | Gammaprotec | Oceanospirilla | Saccharospirill | Saccharospiril | Saccharospirill | unclassified | unclassified | unclassified | unclassified | unclassified | 3 | 0,894427191 | 0,000119015 | 2,99E-05 | 4 - | 0,00013543 - | 3 - | 0,000126123 - | 2 - | 7,99E-05 - | 4 - | 0,000162222 - | 2 - | 9,14E-05 - |

| | | | | | | | | | | | | | | | | | | | | | | | | | | | | |
|---|---|---|---|---|---|---|---|---|---|---|---|---|---|---|---|---|---|---|---|---|---|---|---|---|---|---|---|---|
| 4571 | 1 | 3um_FIJJAFBF_56139 | Bacteria | Bacteroidetes | Flavobacteriia | Flavobacterial | Flavobacteriac | Polaribacter | Polaribacter s | metabolism | Energy produc | COG0056 | Produces ATP | 313598.MED1 | 22,6 | 0,489897949 | 0,000589172 | 4,99E-05 | 23 - | 0,000502832 - | 23 - | 0,000624368 - | 23 - | 0,000593541 - | 22 - | 0,000576121 - | 22 - | 0,000648998 - |
| 4572 | 1 | 0.2um_NOKCPHJN_347562 | Bacteria | Proteobacteri | Alphaproteob | Rhodobactera | Rhodobactera | unclassified | Rhodobactera | unclassified | unclassified | unclassified | unclassified | unclassified | 1 | 0 | 4,05E-05 | 3,85E-06 | 1 - | 3,40E-05 - | 1 - | 4,22E-05 - | 1 - | 4,01E-05 - | 1 - | 4,07E-05 - | 1 - | 4,58E-05 - |
| 4573 | 1 | 0.2um_NOKCPHJN_865341 | Bacteria | Proteobacteri | Alphaproteob | Pelagibacteral | Pelagibacterac | Candidatus Pe | Candidatus Pe | metabolism | Secondary me | 01Q86 | extracellular s | 335992.SAR11 | 2 | 0,632455532 | 7,59E-05 | 2,64E-05 | 2 - | 6,32E-05 - | 3 - | 0,000117784 - | 1 - | 3,73E-05 - | 2 - | 7,57E-05 - | 2 - | 8,53E-05 - |
| 4574 | 1 | 3um_FIJJAFBF_205104 | Bacteria | Bacteroidetes | Flavobacteriia | Flavobacterial | Flavobacteriac | Lacinutrix | Lacinutrix him | metabolism | Carbohydrate | 099YR | Catalyzes the | 391587.KAOT | 0,8 | 0,4 | 2,56E-05 | 1,32E-05 | 1 - | 2,68E-05 - | 1 - | 3,33E-05 - | 1 - | 3,17E-05 - | 0 - | 0 - | 1 - | 3,62E-05 - |
| 4575 | 1 | 3um_FIJJAFBF_584717 | Eukaryota | Bacillariophyta | Mediophyceae | Hemiaulales | Hemiaulaceae | Cerataulina | Cerataulina da | metabolism | Energy produc | 11HAV | photosystem i | 35128.Thapsd | 1,6 | 0,489897949 | 9,64E-05 | 3,39E-05 | 1 - | 4,97E-05 - | 1 - | 6,17E-05 - | 2 - | 0,000117301 - | 2 - | 0,000119033 - | 2 - | 0,00013409 - |
| 4576 | 1 | 3um_FIJJAFBF_328504 | Bacteria | Bacteroidetes | Flavobacteriia | Flavobacterial | Flavobacteriac | Bizionia | Bizionia argen | information st | Translation, ri | COG0089 | One of the ea | 983544.Lacal_ | 0,8 | 0,4 | 0,000114094 | 5,86E-05 | 1 - | 0,000119559 - | 1 - | 0,000148457 - | 1 - | 0,000141127 - | 0 - | 0 - | 1 - | 0,000161328 - |
| 4577 | 1 | 3um_FIJJAFBF_584954 | unclassified | unclassified | unclassified | unclassified | unclassified | unclassified | unclassified | unclassified | unclassified | unclassified | unclassified | unclassified | 0,6 | 0,489897949 | 2,11E-05 | 1,72E-05 | 0 - | 0 - | 1 - | 3,62E-05 - | 1 - | 3,44E-05 - | 1 - | 3,49E-05 - | 0 - | 0 - |
| 4578 | 2 | 3um_FIJJAFBF_56256;3um_FIJJAFBF_585687 | Bacteria | Bacteroidetes | Flavobacteriia | Flavobacterial | Flavobacteriac | Polaribacter | various | unclassified | unclassified | unclassified | unclassified | unclassified | 0,8 | 0,4 | 3,75E-05 | 1,93E-05 | 1 - | 3,93E-05 - | 1 - | 4,88E-05 - | 1 - | 4,64E-05 - | 0 - | 0 - | 1 - | 5,30E-05 - |
| 4579 | 1 | 3um_FIJJAFBF_709642 | Bacteria | Proteobacteri | Gammaprote | Cellvibrionales | Porticoccacea | unclassified | Gammaprote | metabolism | Inorganic ion t | 0QKPS | TonB-depende | 506534.Rhein | 3,4 | 1,2 | 6,72E-05 | 2,33E-05 | 4 - | 6,71E-05 - | 5 - | 0,00010418 - | 4 - | 7,92E-05 - | 2 - | 4,02E-05 - | 2 - | 4,53E-05 - |
| 4580 | 1 | 0.2um_NOKCPHJN_490659 | Bacteria | Verrucomicrol | unclassified | unclassified | unclassified | unclassified | Verrucomicro | unclassified | unclassified | unclassified | unclassified | unclassified | 0,2 | 0,4 | 7,31E-06 | 1,46E-05 | 0 - | 0 - | 0 - | 0 - | 0 - | 0 - | 0 - | 0 - | 1 - | 3,65E-05 - |
| 4581 | 1 | 3um_FIJJAFBF_132758 | Bacteria | Proteobacteri | Gammaprote | Oceanospirilla | Saccharospirill | Reinekea | Reinekea fors | unclassified | unclassified | unclassified | unclassified | unclassified | 0,8 | 0,748331477 | 4,00E-05 | 3,51E-05 | 2 - | 9,04E-05 - | 1 - | 5,61E-05 - | 1 - | 5,33E-05 - | 0 - | 0 - | 0 - | 0 - |
| 4582 | 2 | 0.2um_NOKCPHJN_157941;3um_FIJJAFBF_27397 | Bacteria | Bacteroidetes | Flavobacteriia | Flavobacterial | Flavobacteriac | various | various | metabolism | Amino acid tra | 099QE | chorismate m | 487797.Flav3 | 0,2 | 0,4 | 6,38E-06 | 1,28E-05 | 1 - | 3,19E-05 - | 0 - | 0 - | 0 - | 0 - | 0 - | 0 - | 0 - | 0 - |
| 4583 | 1 | 3um_FIJJAFBF_209610 | Bacteria | Bacteroidetes | Flavobacteriia | Flavobacterial | Flavobacteriac | Algibacter | Algibacter lect | unclassified | unclassified | unclassified | unclassified | unclassified | 0,6 | 0,489897949 | 6,03E-05 | 5,01E-05 | 1 - | 8,50E-05 - | 0 - | 0 - | 0 - | 0 - | 1 - | 0,000101839 - | 1 - | 0,000114722 - |
| 4584 | 1 | 3um_FIJJAFBF_556176 | Bacteria | Proteobacteri | Alphaproteob | Rhodobactera | Rhodobactera | Sulfitobacter | Sulfitobacter | poorly charact | Function unkn | 05DIH | basic membra | 633131.TR2A6 | 0,6 | 0,489897949 | 2,45E-05 | 2,04E-05 | 1 - | 3,46E-05 - | 0 - | 0 - | 0 - | 0 - | 1 - | 4,14E-05 - | 1 - | 4,66E-05 - |
| 4585 | 1 | 3um_FIJJAFBF_20710 | Eukaryota | various | various | various | various | various | various | unclassified | unclassified | unclassified | unclassified | unclassified | 5,2 | 0,4 | 0,000895138 | 0,000145762 | 5 - | 0,000717355 - | 5 - | 0,000890743 - | 5 - | 0,000846764 - | 5 - | 0,000859271 - | 6 - | 0,001161559 - |
| 4586 | 1 | 0.2um_NOKCPHJN_99362 | Bacteria | Bacteroidetes | unclassified | unclassified | unclassified | unclassified | Bacteroidetes | information st | Transcription | COG0085 | DNA-depende | 926562.Oweh | 8,8 | 1,166190379 | 9,40E-05 | 9,77E-06 | 10 - | 9,02E-05 - | 10 - | 0,000112043 - | 9 - | 9,59E-05 - | 8 - | 8,65E-05 - | 7 - | 8,52E-05 - |
| 4587 | 1 | 0.2um_NOKCPHJN_252641 | Bacteria | Proteobacteri | Alphaproteob | Rhodobactera | Rhodobactera | unclassified | Rhodobactera | unclassified | unclassified | unclassified | unclassified | unclassified | 15,4 | 1,019803903 | 0,000247916 | 1,97E-05 | 17 - | 0,000230095 - | 16 - | 0,000268904 - | 15 - | 0,00023965 - | 14 - | 0,000226977 - | 15 - | 0,000273952 - |
| 4588 | 1 | 3um_FIJJAFBF_264144 | Bacteria | Proteobacteri | Gammaprote | Alteromonada | Alteromonada | Paraglaciecola | Paraglaciecola | unclassified | unclassified | unclassified | unclassified | unclassified | 9,8 | 1,469693846 | 0,000794849 | 8,54E-05 | 11 - | 0,000756015 - | 10 - | 0,000853406 - | 11 - | 0,000892398 - | 10 - | 0,000823253 - | 7 - | 0,000649174 - |
| 4589 | 1 | 3um_FIJJAFBF_458111 | Bacteria | Bacteroidetes | Flavobacteriia | Flavobacterial | Flavobacteriac | Kordia | various | unclassified | unclassified | unclassified | unclassified | unclassified | 1 | 0 | 9,86E-05 | 9,36E-06 | 1 - | 8,26E-05 - | 1 - | 0,000102532 - | 1 - | 9,75E-05 - | 1 - | 9,89E-05 - | 1 - | 0,00011142 - |
| 4590 | 1 | 3um_FIJJAFBF_510247 | Bacteria | Bacteroidetes | Flavobacteriia | Flavobacterial | Flavobacteriac | Polaribacter | Polaribacter s | unclassified | unclassified | unclassified | unclassified | unclassified | 5,4 | 0,8 | 0,000427358 | 7,11E-05 | 5 - | 0,000331725 - | 5 - | 0,000411904 - | 7 - | 0,000548194 - | 5 - | 0,000397351 - | 5 - | 0,000447614 - |
| 4591 | 1 | 3um_FIJJAFBF_782661 | Eukaryota | unclassified | Dinophyceae | Gonyaulacales | unclassified | Lingulodinium | Lingulodinium | metabolism | Energy produc | 0FJ7X | This is one of t | 513049.Amax | 1 | 0 | 3,90E-05 | 3,71E-06 | 1 - | 3,27E-05 - | 1 - | 4,06E-05 - | 1 - | 3,86E-05 - | 1 - | 3,92E-05 - | 1 - | 4,41E-05 - |
| 4592 | 2 | 0.2um_NOKCPHJN_1002908;0.2um_NOKCPHJN_ | Bacteria | Proteobacteri | Alphaproteob | Pelagibacteral | Pelagibacterac | Candidatus Pe | Candidatus Pe | metabolism | Amino acid tra | 01R8N | serine o-acety | 335992.SAR11 | 0,6 | 0,8 | 3,82E-05 | 4,92E-05 | 2 - | 0,000119559 - | 0 - | 0 - | 0 - | 0 - | 1 - | 7,16E-05 - | 0 - | 0 - |
| 4593 | 1 | 3um_FIJJAFBF_220490 | Bacteria | Proteobacteri | Gammaprote | Alteromonada | Alteromonada | Paraglaciecola | Paraglaciecola | information st | Translation, ri | 17FIH | One of the ea | 715451.ambt_ | 2,6 | 0,489897949 | 0,000351603 | 5,09E-05 | 3 - | 0,00034433 - | 2 - | 0,000285038 - | 3 - | 0,000406447 - | 3 - | 0,00041245 - | 2 - | 0,000309749 - |
| 4594 | 1 | 3um_FIJJAFBF_274268 | Bacteria | Proteobacteri | Gammaprote | Alteromonada | Alteromonada | Paraglaciecola | Paraglaciecola | cellular proces | Signal transdu | 17JQI | Could accelera | 342610.Patl_3 | 0,2 | 0,4 | 3,23E-05 | 6,47E-05 | 1 - | 0,000161657 - | 0 - | 0 - | 0 - | 0 - | 0 - | 0 - | 0 - | 0 - |
| 4595 | 1 | 3um_FIJJAFBF_722163 | Eukaryota | unclassified | Dinophyceae | Suessiales | Symbiodiniace | Symbiodinium | Symbiodinium | unclassified | unclassified | unclassified | unclassified | unclassified | 2 | 0,632455532 | 0,000130393 | 3,45E-05 | 3 - | 0,000166343 - | 2 - | 0,000137699 - | 1 - | 6,55E-05 - | 2 - | 0,000132834 - | 2 - | 0,000149637 - |
| 4596 | 1 | 0.2um_NOKCPHJN_386535 | Bacteria | Proteobacteri | Alphaproteob | Rhodobactera | unclassified | unclassified | Rhodobactera | metabolism | Amino acid tra | 05TIB | (ABC) transpo | 367336.OM22 | 0,8 | 0,4 | 3,17E-05 | 1,61E-05 | 1 - | 3,43E-05 - | 1 - | 4,25E-05 - | 1 - | 4,04E-05 - | 1 - | 4,10E-05 - | 0 - | 0 - |
| 4597 | 1 | 0.2um_NOKCPHJN_47855 | Bacteria | Bacteroidetes | Flavobacteriia | Flavobacterial | Flavobacteriac | Aureitalea | Aureitalea ma | unclassified | unclassified | unclassified | unclassified | unclassified | 7,4 | 0,489897949 | 0,000115205 | 1,62E-05 | 7 - | 9,10E-05 - | 7 - | 0,000112982 - | 8 - | 0,000122747 - | 7 - | 0,00010899 - | 8 - | 0,000140317 - |
| 4598 | 1 | 3um_FIJJAFBF_189632 | Bacteria | Bacteroidetes | Flavobacteriia | Flavobacterial | Flavobacteriac | Polaribacter | Polaribacter s | cellular proces | Posttranslatio | 05CRE | cytochrome C | 313594.PI23P | 0,4 | 0,489897949 | 4,91E-06 | 6,08E-06 | 1 - | 1,10E-05 - | 1 - | 1,36E-05 - | 0 - | 0 - | 0 - | 0 - | 0 - | 0 - |
| 4599 | 1 | 3um_FIJJAFBF_649923 | Bacteria | Bacteroidetes | Flavobacteriia | Flavobacterial | Flavobacteriac | Formosa | Formosa sp. H | poorly charact | Function unkn | 12B3I | NA | 1034807.FBFL | 1 | 0 | 5,44E-05 | 5,16E-06 | 1 - | 4,55E-05 - | 1 - | 5,66E-05 - | 1 - | 5,38E-05 - | 1 - | 5,46E-05 - | 1 - | 6,15E-05 - |
| 4600 | 1 | 3um_FIJJAFBF_428889 | Bacteria | Proteobacteri | Gammaprote | Oceanospirilla | Oceanospirilla | Marinomonas | various | metabolism | Amino acid tra | COG0131 | imidazoleglyc | 400668.Mmw | 0,8 | 0,4 | 5,56E-05 | 2,86E-05 | 1 - | 5,83E-05 - | 1 - | 7,23E-05 - | 1 - | 6,88E-05 - | 0 - | 0 - | 1 - | 7,86E-05 - |
| 4601 | 1 | 3um_FIJJAFBF_163994 | Eukaryota | unclassified | Pelagophycea | Pelagomonad | unclassified | Aureococcus | Aureococcus a | unclassified | unclassified | unclassified | unclassified | unclassified | 0,2 | 0,4 | 3,08E-05 | 6,16E-05 | 0 - | 0 - | 0 - | 0 - | 1 - | 0,000153957 - | 0 - | 0 - | 0 - | 0 - |
| 4602 | 1 | 0.2um_NOKCPHJN_164218 | Bacteria | Proteobacteri | Gammaprote | Cellvibrionales | Halieaceae | unclassified | marine gamm | poorly charact | Function unkn | 05C2A | tonB-depende | 247639.MGP2 | 1,8 | 0,748331477 | 2,35E-05 | 7,83E-06 | 3 - | 3,39E-05 - | 2 - | 2,81E-05 - | 2 - | 2,67E-05 - | 1 - | 1,35E-05 - | 1 - | 1,52E-05 - |
| 4603 | 1 | 0.2um_NOKCPHJN_457437 | Bacteria | Bacteroidetes | unclassified | unclassified | unclassified | unclassified | Bacteroidetes | unclassified | unclassified | unclassified | unclassified | unclassified | 2,6 | 0,8 | 0,000163708 | 3,44E-05 | 4 - | 0,00021656 - | 2 - | 0,000134452 - | 3 - | 0,00019172 - | 2 - | 0,000129701 - | 2 - | 0,000146108 - |
| 4604 | 1 | 3um_FIJJAFBF_129175 | Bacteria | Bacteroidetes | Flavobacteriia | Flavobacterial | Flavobacteriac | Polaribacter | Polaribacter s | unclassified | unclassified | unclassified | unclassified | unclassified | 7,6 | 0,489897949 | 0,000165878 | 1,11E-05 | 8 - | 0,000146914 - | 7 - | 0,000159621 - | 8 - | 0,000173417 - | 8 - | 0,000175979 - | 7 - | 0,000173459 - |
| 4605 | 1 | 3um_FIJJAFBF_205903 | Bacteria | Proteobacteri | Alphaproteob | Rhizobiales | Aurantimonad | Aurantimonas | Aurantimonas | metabolism | Amino acid tra | 05CAI | ABC transport | 287752.SI859 | 0,6 | 0,489897949 | 4,36E-05 | 3,60E-05 | 1 - | 6,34E-05 - | 1 - | 7,87E-05 - | 0 - | 0 - | 1 - | 7,60E-05 - | 0 - | 0 - |
| 4606 | 1 | 3um_FIJJAFBF_292579 | Bacteria | Proteobacteri | Gammaprote | unclassified | unclassified | unclassified | various | information st | Translation, ri | 177XJ | One of the pri | 322710.Avin_ | 1,8 | 0,748331477 | 0,000155564 | 6,25E-05 | 2 - | 0,00014715 - | 2 - | 0,000182716 - | 3 - | 0,000260543 - | 1 - | 8,81E-05 - | 1 - | 9,93E-05 - |
| 4607 | 1 | 3um_FIJJAFBF_112130 | Bacteria | Bacteroidetes | Flavobacteriia | Flavobacterial | Crocinitomica | unclassified | Crocinitomica | unclassified | unclassified | unclassified | unclassified | unclassified | 0,6 | 0,489897949 | 1,32E-05 | 1,09E-05 | 1 - | 1,96E-05 - | 0 - | 0 - | 1 - | 2,31E-05 - | 1 - | 2,35E-05 - | 0 - | 0 - |
| 4608 | 1 | 0.2um_NOKCPHJN_08638 | Bacteria | Proteobacteri | Gammaprote | Alteromonada | Alteromonada | Paraglaciecola | Paraglaciecola | poorly charact | Function unkn | 06NUI | domain protei | 983545.Glaag | 3,4 | 1,356465997 | 0,000109363 | 4,44E-05 | 3 - | 8,20E-05 - | 4 - | 0,000135732 - | 5 - | 0,000161288 - | 4 - | 0,000130936 - | 1 - | 3,69E-05 - |
| 4609 | 1 | 0.2um_NOKCPHJN_612171 | Bacteria | Proteobacteri | Gammaprote | Cellvibrionales | Porticoccacea | unclassified | gamma prote | unclassified | unclassified | unclassified | unclassified | unclassified | 3 | 0,632455532 | 9,83E-05 | 1,71E-05 | 4 - | 0,000110895 - | 3 - | 0,000103275 - | 2 - | 6,55E-05 - | 3 - | 9,96E-05 - | 3 - | 0,000112228 - |
| 4610 | 1 | 3um_FIJJAFBF_314913 | Bacteria | Proteobacteri | Gammaprote | Alteromonada | Alteromonada | Paraglaciecola | Paraglaciecola | unclassified | unclassified | unclassified | unclassified | unclassified | 1 | 0 | 1,98E-05 | 1,88E-06 | 1 - | 1,66E-05 - | 1 - | 2,06E-05 - | 1 - | 1,96E-05 - | 1 - | 1,99E-05 - | 1 - | 2,24E-05 - |
| 4611 | 1 | 3um_FIJJAFBF_289012 | unclassified | unclassified | unclassified | unclassified | unclassified | unclassified | unclassified | unclassified | unclassified | unclassified | unclassified | unclassified | 0,2 | 0,4 | 1,44E-05 | 2,88E-05 | 0 - | 0 - | 0 - | 0 - | 0 - | 0 - | 1 - | 7,20E-05 - | 0 - | 0 - |
| 4612 | 1 | 3um_FIJJAFBF_623285 | Bacteria | Proteobacteri | Gammaprote | Oceanospirilla | Saccharospirill | Reinekea | Reinekea fors | information st | Transcription | 05MPC | Transcriptiona | 326297.Sama_ | 0,6 | 0,489897949 | 5,71E-05 | 4,74E-05 | 1 - | 8,08E-05 - | 0 - | 0 - | 1 - | 9,54E-05 - | 0 - | 0 - | 1 - | 0,000109067 - |
| 4613 | 1 | 0.2um_NOKCPHJN_09983 | Bacteria | Bacteroidetes | Flavobacteriia | Flavobacterial | Flavobacteriac | Polaribacter | Polaribacter s | information st | Transcription | 05D27 | DNA-depende | 391603.FBALC | 10 | 1,095445115 | 9,54E-05 | 5,03E-06 | 12 - | 9,67E-05 - | 10 - | 0,000100083 - | 9 - | 8,56E-05 - | 10 - | 9,65E-05 - | 9 - | 9,79E-05 - |
| 4614 | 1 | 0.2um_NOKCPHJN_275839 | Eukaryota | unclassified | Labyrinthulom | unclassified | Thraustochytr | Aurantiochytr | Aurantiochytr | unclassified | unclassified | unclassified | unclassified | unclassified | 1 | 0 | 1,17E-05 | 1,11E-06 | 1 - | 9,77E-06 - | 1 - | 1,21E-05 - | 1 - | 1,15E-05 - | 1 - | 1,17E-05 - | 1 - | 1,32E-05 - |
| 4615 | 1 | 3um_FIJJAFBF_47485 | Bacteria | Proteobacteri | Gammaprote | Alteromonada | Alteromonada | Aliiglaciecola | Aliiglaciecola l | metabolism | Lipid transpor | 174G5 | reductase | 391597.LMED | 0,6 | 0,489897949 | 2,09E-05 | 1,74E-05 | 1 - | 2,94E-05 - | 0 - | 0 - | 0 - | 0 - | 1 - | 3,53E-05 - | 1 - | 3,97E-05 - |
| 4616 | 1 | 3um_FIJJAFBF_54799 | Bacteria | Bacteroidetes | Flavobacteriia | Flavobacterial | Flavobacteriac | Polaribacter | Polaribacter b | unclassified | unclassified | unclassified | unclassified | unclassified | 8,6 | 1,496662955 | 0,000215682 | 3,05E-05 | 10 - | 0,00021255 - | 10 - | 0,000263924 - | 8 - | 0,000200714 - | 9 - | 0,000229139 - | 6 - | 0,000172083 - |
| 4617 | 1 | 3um_FIJJAFBF_372060 | Bacteria | Proteobacteri | Alphaproteob | Rhodobactera | Rhodobactera | Antarctobacte | Antarctobacte | metabolism | Carbohydrate | COG0057 | glyceraldehyd | 391593.RCCS2 | 1,8 | 1,166190379 | 7,15E-05 | 4,74E-05 | 2 - | 6,89E-05 - | 1 - | 4,28E-05 - | 3 - | 0,000122056 - | 3 - | 0,000123859 - | 0 - | 0 - |
| 4618 | 1 | 0.2um_NOKCPHJN_600843 | Bacteria | Bacteroidetes | Flavobacteriia | Flavobacterial | Flavobacteriac | Wenyingzhuai | Wenyingzhuai | poorly charact | Function unkn | 0XY6Y | Pkd domain c | 313594.PI23P | 1 | 0 | 2,59E-05 | 2,46E-06 | 1 - | 2,17E-05 - | 1 - | 2,69E-05 - | 1 - | 2,56E-05 - | 1 - | 2,60E-05 - | 1 - | 2,93E-05 - |
| 4619 | 1 | 3um_FIJJAFBF_642657 | Bacteria | Proteobacteri | Alphaproteob | Rhodobactera | Rhodobactera | unclassified | Rhodobactera | metabolism | Coenzyme tra | 05CVM | Molybdenum | 314256.OG25 | 1 | 0 | 2,62E-05 | 2,48E-06 | 1 - | 2,19E-05 - | 1 - | 2,72E-05 - | 1 - | 2,59E-05 - | 1 - | 2,62E-05 - | 1 - | 2,96E-05 - |
| 4620 | 1 | 3um_FIJJAFBF_24295 | Bacteria | Proteobacteri | Gammaprote | Oceanospirilla | Saccharospirill | Reinekea | Reinekea blan | cellular proces | Cell motility | COG1291 | MotA TolQ ex | 314283.MED2 | 2 | 1,264911064 | 0,000105726 | 5,78E-05 | 4 - | 0,000186629 - | 1 - | 5,79E-05 - | 3 - | 0,000165222 - | 1 - | 5,59E-05 - | 1 - | 6,30E-05 - |
| 4621 | 1 | 3um_FIJJAFBF_06420 | Bacteria | Proteobacteri | Betaproteoba | Burkholderial | Comamonada | Limnohabitan | Limnohabitan | unclassified | unclassified | unclassified | unclassified | unclassified | 1 | 0 | 0,000210811 | 2,00E-05 | 1 - | 0,00017658 - | 1 - | 0,00021926 - | 1 - | 0,000208434 - | 1 - | 0,000211513 - | 1 - | 0,000238268 - |
| 4622 | 1 | 0.2um_NOKCPHJN_589358 | Bacteria | Proteobacteri | Alphaproteob | Pelagibacteral | Pelagibacterac | Candidatus Pe | Candidatus Pe | cellular proces | Cell wall/mem | 16QB0 | Part of the ou | 335992.SAR11 | 0,4 | 0,489897949 | 8,73E-06 | 1,07E-05 | 0 - | 0 - | 0 - | 0 - | 1 - | 2,17E-05 - | 1 - | 2,20E-05 - | 0 - | 0 - |
| 4623 | 1 | 3um_FIJJAFBF_604542 | Bacteria | Proteobacteri | Gammaprote | Arenicellales | Arenicellaceae | Arenicella | Arenicella xan | information st | Translation, ri | 179ZI | One of the pri | 1026882.MAN | 1,6 | 0,489897949 | 0,000165058 | 4,33E-05 | 2 - | 0,00017658 - | 1 - | 0,00010963 - | 2 - | 0,000208434 - | 2 - | 0,000211513 - | 1 - | 0,000119134 - |
| 4624 | 2 | 3um_FIJJAFBF_151896;3um_FIJJAFBF_356855 | various | various | various | various | various | various | various | unclassified | unclassified | unclassified | unclassified | unclassified | 1 | 0 | 5,50E-05 | 5,23E-06 | 1 - | 4,61E-05 - | 1 - | 5,72E-05 - | 1 - | 5,44E-05 - | 1 - | 5,52E-05 - | 1 - | 6,22E-05 - |
| 4625 | 1 | 0.2um_NOKCPHJN_201892 | Eukaryota | unclassified | Oomycetes | Myzocytiopsid | Eurychasmata | Eurychasma | Eurychasma d | unclassified | unclassified | unclassified | unclassified | unclassified | 1 | 0 | 2,74E-06 | 2,60E-07 | 1 - | 2,29E-06 - | 1 - | 2,85E-06 - | 1 - | 2,71E-06 - | 1 - | 2,75E-06 - | 1 - | 3,09E-06 - |
| 4626 | 1 | 3um_FIJJAFBF_130278 | Bacteria | Bacteroidetes | Flavobacteriia | Flavobacterial | Flavobacteriac | Polaribacter | Polaribacter s | unclassified | unclassified | unclassified | unclassified | unclassified | 1 | 0 | 9,79E-05 | 9,29E-06 | 1 - | 8,20E-05 - | 1 - | 0,000101799 - | 1 - | 9,68E-05 - | 1 - | 9,82E-05 - | 1 - | 0,000110625 - |
| 4627 | 1 | 3um_FIJJAFBF_73880 | Eukaryota | Bacillariophyta | Coscinodiscop | Thalassiosirale | Thalassiosirac | Thalassiosira | Thalassiosira | unclassified | unclassified | unclassified | unclassified | unclassified | 1 | 0 | 0,000130502 | 1,24E-05 | 1 - | 0,000109311 - | 1 - | 0,000135732 - | 1 - | 0,000129031 - | 1 - | 0,000130936 - | 1 - | 0,0001475 - |
| 4628 | 1 | 0.2um_NOKCPHJN_824137 | Bacteria | Proteobacteri | Alphaproteob | Pelagibacteral | Pelagibacterac | Candidatus Pe | Candidatus Pe | cellular proces | Cell wall/mem | 0F70V | Nad-depende | 656519.Halsa_ | 0,4 | 0,489897949 | 1,67E-05 | 2,04E-05 | 0 - | 0 - | 0 - | 0 - | 1 - | 4,14E-05 - | 1 - | 4,20E-05 - | 0 - | 0 - |
| 4629 | 1 | 0.2um_NOKCPHJN_136728 | Bacteria | Bacteroidetes | Flavobacteriia | unclassified | unclassified | unclassified | Flavobacteria | metabolism | Inorganic ion t | 05C27 | TonB-depende | 487796.Flav2 | 19 | 1,897366596 | 0,00024207 | 2,30E-05 | 21 - | 0,000225263 - | 19 - | 0,000253071 - | 21 - | 0,0002659 - | 16 - | 0,000205583 - | 18 - | 0,000260537 - |
| 4630 | 1 | 3um_FIJJAFBF_70350 | Bacteria | Proteobacteri | Betaproteoba | unclassified | unclassified | unclassified | Betaproteoba | information st | Translation, ri | 17FIH | One of the ea | 397945.Aave_ | 2,8 | 1,6 | 0,000355785 | 0,000204902 | 4 - | 0,000441449 - | 4 - | 0,000548149 - | 2 - | 0,000260543 - | 4 - | 0,000528782 - | 0 - | 0 - |
| 4631 | 1 | 0.2um_NOKCPHJN_157068 | Bacteria | unclassified | unclassified | unclassified | unclassified | unclassified | uncultured ba | poorly charact | Function unkn | 0P8NI | Inherit from b | 983544.Lacal_ | 4,4 | 0,489897949 | 0,000144127 | 1,51E-05 | 5 - | 0,000137953 - | 5 - | 0,000171297 - | 4 - | 0,000130271 - | 4 - | 0,000132195 - | 4 - | 0,000148918 - |
| 4632 | 1 | 0.2um_NOKCPHJN_372322 | Bacteria | Proteobacteri | Alphaproteob | Rhodobactera | Rhodobactera | Planktomarina | Planktomarina | poorly charact | Function unkn | COG3181 | extra-cytoplas | 388401.RB215 | 4,4 | 1,356465997 | 0,000188297 | 6,60E-05 | 5 - | 0,000178225 - | 4 - | 0,000177042 - | 2 - | 8,42E-05 - | 5 - | 0,000213483 - | 6 - | 0,000288586 - |
| 4633 | 1 | 0.2um_NOKCPHJN_479664 | Eukaryota | Streptophyta | Bryopsida | Funariales | Funariaceae | Physcomitrell | Physcomitrella | poorly charact | Function unkn | 0NVRZ | NA | 170187.SP_17 | 1 | 0 | 7,23E-06 | 6,87E-07 | 1 - | 6,06E-06 - | 1 - | 7,52E-06 - | 1 - | 7,15E-06 - | 1 - | 7,26E-06 - | 1 - | 8,17E-06 - |
| 4634 | 1 | 0.2um_NOKCPHJN_139372 | Bacteria | Bacteroidetes | unclassified | unclassified | unclassified | unclassified | Bacteroidetes | metabolism | Inorganic ion t | 05C27 | TonB-depende | 391603.FBALC | 0,6 | 0,489897949 | 7,55E-06 | 6,28E-06 | 1 - | 1,05E-05 - | 1 - | 1,31E-05 - | 0 - | 0 - | 0 - | 0 - | 1 - | 1,42E-05 - |
| 4635 | 1 | 0.2um_NOKCPHJN_1060612 | Bacteria | Bacteroidetes | Flavobacteriia | Flavobacterial | Flavobacteriac | Polaribacter | Polaribacter g | unclassified | unclassified | unclassified | unclassified | unclassified | 0,6 | 0,8 | 1,63E-05 | 2,18E-05 | 0 - | 0 - | 0 - | 0 - | 1 - | 2,69E-05 - | 2 - | 5,47E-05 - | 0 - | 0 - |
| 4636 | 1 | 3um_FIJJAFBF_175143 | Bacteria | Proteobacteri | Alphaproteob | Rhodobactera | Rhodobactera | unclassified | Rhodobactera | information st | Translation, ri | 082FS | 50S ribosomal | 314262.MED1 | 0,2 | 0,4 | 3,14E-05 | 6,29E-05 | 1 - | 0,000157228 - | 0 - | 0 - | 0 - | 0 - | 0 - | 0 - | 0 - | 0 - |
| 4637 | 1 | 0.2um_NOKCPHJN_58285 | Bacteria | Bacteroidetes | Flavobacteriia | unclassified | unclassified | unclassified | Flavobacteria | poorly charact | Function unkn | 11PIQ | YCE I like fami | 487797.Flav3 | 1,8 | 0,748331477 | 0,000117506 | 3,92E-05 | 3 - | 0,000169621 - | 2 - | 0,000140413 - | 2 - | 0,00013348 - | 1 - | 6,77E-05 - | 1 - | 7,63E-05 - |
| 4638 | 1 | 0.2um_NOKCPHJN_442394 | Bacteria | Bacteroidetes | Flavobacteriia | Flavobacterial | Flavobacteriac | Polaribacter | Polaribacter s | cellular proces | Intracellular tr | 05FGH | MotA TolQ ex | 313594.PI23P | 1,2 | 0,748331477 | 6,95E-05 | 4,64E-05 | 1 - | 4,76E-05 - | 1 - | 5,91E-05 - | 2 - | 0,000112433 - | 0 - | 0 - | 2 - | 0,000128527 - |
| 4639 | 1 | 3um_FIJJAFBF_154879 | Bacteria | Proteobacteri | Betaproteoba | Burkholderial | Comamonada | Hydrogenoph | various | cellular proces | Posttranslatio | 07MKT | Stringent star | 397945.Aave_ | 1,6 | 0,489897949 | 0,000108435 | 3,49E-05 | 1 - | 5,65E-05 - | 2 - | 0,000140413 - | 2 - | 0,00013348 - | 2 - | 0,000135452 - | 1 - | 7,63E-05 - |
| 4640 | 1 | 3um_FIJJAFBF_188951 | unclassified | unclassified | unclassified | unclassified | unclassified | unclassified | unclassified | unclassified | unclassified | unclassified | unclassified | unclassified | 0,2 | 0,4 | 1,13E-05 | 2,26E-05 | 1 - | 5,65E-05 - | 0 - | 0 - | 0 - | 0 - | 0 - | 0 - | 0 - | 0 - |
| 4641 | 1 | 3um_FIJJAFBF_67736 | Eukaryota | Arthropoda | Insecta | Hymenoptera | Formicidae | Pseudomyrme | Pseudomyrme | information st | Transcription | 09YI4 | paf1, RNA pol | 12957.ACEP_ | 1,8 | 0,748331477 | 6,06E-05 | 2,63E-05 | 1 - | 2,79E-05 - | 1 - | 3,46E-05 - | 3 - | 9,87E-05 - | 2 - | 6,67E-05 - | 2 - | 7,52E-05 - |
| 4642 | 1 | 0.2um_NOKCPHJN_07993 | Bacteria | Bacteroidetes | Flavobacteriia | Flavobacterial | Flavobacteriac | unclassified | Flavobacteriac | information st | Translation, ri | 08UHY | One of the pri | 487796.Flav2 | 0,6 | 0,489897949 | 4,97E-05 | 4,10E-05 | 1 - | 7,26E-05 - | 1 - | 9,02E-05 - | 1 - | 8,57E-05 - | 0 - | 0 - | 0 - | 0 - |
| 4643 | 1 | 0.2um_NOKCPHJN_127388 | Bacteria | Proteobacteri | Alphaproteob | Rhodobactera | unclassified | unclassified | Rhodobactera | information st | Replication, re | 05C68 | Can catalyze t | 290400.Jann_ | 2 | 0,894427191 | 7,63E-05 | 3,74E-05 | 2 - | 6,34E-05 - | 1 - | 3,94E-05 - | 3 - | 0,000112278 - | 1 - | 3,80E-05 - | 3 - | 0,000128349 - |
| 4644 | 1 | 0.2um_NOKCPHJN_873188 | Eukaryota | Phaeophyceae | unclassified | Ectocarpales | Ectocarpaceae | Ectocarpus | Ectocarpus sili | information st | Translation, ri | COG0052 | 30S ribosomal | 2850.Phatr13 | 0,8 | 0,4 | 3,50E-05 | 1,78E-05 | 1 - | 3,79E-05 - | 1 - | 4,70E-05 - | 1 - | 4,47E-05 - | 1 - | 4,54E-05 - | 0 - | 0 - |
| 4645 | 2 | 0.2um_NOKCPHJN_377278;0.2um_NOKCPHJN_5 | Bacteria | unclassified | unclassified | unclassified | unclassified | unclassified | bacterium TM | unclassified | unclassified | unclassified | unclassified | unclassified | 1,8 | 0,4 | 0,000134298 | 2,65E-05 | 2 - | 0,000126825 - | 2 - | 0,000157479 - | 2 - | 0,000149704 - | 2 - | 0,000151915 - | 1 - | 8,56E-05 - |
| 4646 | 1 | 3um_FIJJAFBF_462596 | unclassified | unclassified | unclassified | unclassified | unclassified | unclassified | unclassified | information st | Replication, re | 0PFXX | DNA repair pr | 578458.XP_00 | 0,2 | 0,4 | 4,47E-06 | 8,93E-06 | 1 - | 2,23E-05 - | 0 - | 0 - | 0 - | 0 - | 0 - | 0 - | 0 - | 0 - |
| 4647 | 1 | 0.2um_NOKCPHJN_82403 | Bacteria | Bacteroidetes | Flavobacteriia | Flavobacterial | Flavobacteriac | Polaribacter | Polaribacter a | metabolism | Energy produc | COG0056 | Produces ATP | 391603.FBALC | 3,2 | 0,979795897 | 8,11E-05 | 1,62E-05 | 5 - | 0,000109311 - | 3 - | 8,14E-05 - | 3 - | 7,74E-05 - | 3 - | 7,86E-05 - | 2 - | 5,90E-05 - |
| 4648 | 1 | 3um_FIJJAFBF_177956 | Eukaryota | Streptophyta | Liliopsida | Poales | Poaceae | Sorghum | Sorghum bico | poorly charact | Function unkn | 11CRP | NA | 4533.OB06G2 | 0,2 | 0,4 | 2,97E-05 | 5,94E-05 | 0 - | 0 - | 1 - | 0,000148457 - | 0 - | 0 - | 0 - | 0 - | 0 - | 0 - |
| 4649 | 1 | 3um_FIJJAFBF_73927 | Bacteria | Bacteroidetes | Flavobacteriia | Flavobacterial | Flavobacteriac | Robiginitalea | Robiginitalea | information st | Translation, ri | COG0052 | 30S ribosomal | 313628.LNTAF | 0,4 | 0,489897949 | 4,05E-05 | 4,96E-05 | 0 - | 0 - | 1 - | 9,70E-05 - | 0 - | 0 - | 0 - | 0 - | 1 - | 0,000105357 - |
| 4650 | 1 | 3um_FIJJAFBF_257120 | Bacteria | Cyanobacteria | unclassified | Synechococca | Synechococca | Synechococcu | Candidatus Sy | information st | Translation, ri | COG0186 | One of the pri | 232348.SCB01 | 1 | 0 | 0,000163128 | 1,55E-05 | 1 - | 0,000136639 - | 1 - | 0,000169665 - | 1 - | 0,000161288 - | 1 - | 0,000163671 - | 1 - | 0,000184374 - |
| 4651 | 1 | 3um_FIJJAFBF_377195 | Bacteria | Bacteroidetes | unclassified | unclassified | unclassified | unclassified | Bacteroidetes | information st | Translation, ri | 05CEJ | Catalyzes the | 391603.FBALC | 1,2 | 0,748331477 | 2,27E-05 | 1,36E-05 | 2 - | 3,24E-05 - | 2 - | 4,03E-05 - | 1 - | 1,91E-05 - | 0 - | 0 - | 1 - | 2,19E-05 - |
| 4652 | 1 | 3um_FIJJAFBF_133813 | Bacteria | Bacteroidetes | Flavobacteriia | Flavobacterial | Flavobacteriac | Seonamhaeic | Seonamhaeic | information st | Transcription | 0P7W9 | Transcription | 983544.Lacal_ | 1,8 | 0,748331477 | 5,90E-05 | 2,38E-05 | 2 - | 5,60E-05 - | 1 - | 3,48E-05 - | 2 - | 6,61E-05 - | 3 - | 0,000100598 - | 1 - | 3,78E-05 - |
| 4653 | 1 | 3um_FIJJAFBF_254304 | Bacteria | Proteobacteri | Gammaprote | Cellvibrionales | Halieaceae | Pseudohaliea | Pseudohaliea | unclassified | unclassified | unclassified | unclassified | unclassified | 2,6 | 1,019803903 | 0,000235629 | 8,89E-05 | 3 - | 0,000232656 - | 3 - | 0,00028889 - | 4 - | 0,000366168 - | 2 - | 0,000185788 - | 1 - | 0,000104645 - |
| 4654 | 1 | 3um_FIJJAFBF_167870 | Bacteria | Bacteroidetes | Flavobacteriia | Flavobacterial | Flavobacteriac | Polaribacter | Polaribacter b | poorly charact | Function unkn | 09BCW | Protein of unk | 313594.PI23P | 0,8 | 0,4 | 4,61E-05 | 2,34E-05 | 1 - | 4,99E-05 - | 1 - | 6,20E-05 - | 1 - | 5,89E-05 - | 1 - | 5,98E-05 - | 0 - | 0 - |
| 4655 | 1 | 0.2um_NOKCPHJN_149936 | unclassified | unclassified | unclassified | unclassified | unclassified | unclassified | unclassified | unclassified | unclassified | unclassified | unclassified | unclassified | 0,2 | 0,4 | 5,64E-06 | 1,13E-05 | 1 - | 2,82E-05 - | 0 - | 0 - | 0 - | 0 - | 0 - | 0 - | 0 - | 0 - |
| 4656 | 2 | 3um_FIJJAFBF_114989;3um_FIJJAFBF_27238 | Bacteria | Bacteroidetes | Flavobacteriia | Flavobacterial | Flavobacteriac | Aureitalea | Aureitalea sp. | poorly charact | Function unkn | 09BD6 | Protein of unk | 983548.Krodi_ | 0,2 | 0,4 | 1,01E-05 | 2,01E-05 | 1 - | 5,03E-05 - | 0 - | 0 - | 0 - | 0 - | 0 - | 0 - | 0 - | 0 - |
| 4657 | 1 | 0.2um_NOKCPHJN_188148 | Bacteria | Bacteroidetes | Flavobacteriia | Flavobacterial | Flavobacteriac | Polaribacter | Polaribacter d | metabolism | Inorganic ion t | 0P78U | receptor | 313598.MED1 | 0,6 | 0,489897949 | 1,06E-05 | 8,76E-06 | 1 - | 1,54E-05 - | 1 - | 1,92E-05 - | 0 - | 0 - | 1 - | 1,85E-05 - | 0 - | 0 - |
| 4658 | 1 | 3um_FIJJAFBF_254300 | Bacteria | Proteobacteri | Gammaprote | Oceanospirilla | Saccharospirill | Saccharospiril | Saccharospiril | information st | Translation, ri | 0QK14 | This protein bi | 351348.Maqu | 2,2 | 0,748331477 | 0,000173276 | 6,31E-05 | 1 - | 6,48E-05 - | 3 - | 0,000241557 - | 3 - | 0,000229631 - | 2 - | 0,000155348 - | 2 - | 0,000174999 - |
| 4659 | 1 | 3um_FIJJAFBF_510231 | Bacteria | Bacteroidetes | Flavobacteriia | Flavobacterial | Flavobacteriac | Polaribacter | Polaribacter r | information st | Translation, ri | COG0088 | One of the pri | 313598.MED1 | 9,4 | 2,059126028 | 0,000608391 | 0,000103223 | 13 - | 0,000713922 - | 7 - | 0,000477336 - | 9 - | 0,000583416 - | 8 - | 0,000526252 - | 10 - | 0,000741026 - |
| 4660 | 1 | 0.2um_NOKCPHJN_282185 | Bacteria | Bacteroidetes | Cytophagia | Cytophagales | Cyclobacteriac | Algoriphagus | Algoriphagus | information st | Translation, ri | COG0013 | Catalyzes the | 525373.HMPR | 0,6 | 0,489897949 | 9,00E-06 | 7,42E-06 | 1 - | 1,31E-05 - | 1 - | 1,63E-05 - | 1 - | 1,55E-05 - | 0 - | 0 - | 0 - | 0 - |
| 4661 | 1 | 3um_FIJJAFBF_397370 | Bacteria | Bacteroidetes | Flavobacteriia | Flavobacterial | Flavobacteriac | Algibacter | Algibacter sp. | metabolism | Amino acid tra | COG2171 | Catalyzes the | 983544.Lacal_ | 0,6 | 0,489897949 | 2,86E-05 | 2,35E-05 | 1 - | 4,24E-05 - | 0 - | 0 - | 1 - | 5,00E-05 - | 1 - | 5,07E-05 - | 0 - | 0 - |
| 4662 | 1 | 0.2um_NOKCPHJN_1055624 | Bacteria | Bacteroidetes | Flavobacteriia | Flavobacterial | Flavobacteriac | Formosa | Formosa sp. H | information st | Translation, ri | COG0088 | One of the pri | 983544.Lacal_ | 2,2 | 0,748331477 | 0,000143966 | 5,11E-05 | 3 - | 0,000164751 - | 2 - | 0,000136382 - | 1 - | 6,48E-05 - | 2 - | 0,000131563 - | 3 - | 0,000222308 - |
| 4663 | 1 | 0.2um_NOKCPHJN_06881 | Bacteria | Bacteroidetes | Cytophagia | Cytophagales | Flammeovirga | unclassified | Flammeovirga | information st | Transcription | COG0250 | Participates in | 643867.Ftrac_ | 1,2 | 0,4 | 8,89E-05 | 3,07E-05 | 1 - | 6,20E-05 - | 1 - | 7,70E-05 - | 1 - | 7,32E-05 - | 2 - | 0,000148631 - | 1 - | 8,37E-05 - |
| 4664 | 1 | 0.2um_NOKCPHJN_240115 | Bacteria | Proteobacteri | Alphaproteob | Rhodobactera | Rhodobactera | Boseongicola | Boseongicola | unclassified | unclassified | unclassified | unclassified | unclassified | 0,6 | 0,489897949 | 1,67E-05 | 1,38E-05 | 1 - | 2,48E-05 - | 0 - | 0 - | 1 - | 2,93E-05 - | 1 - | 2,97E-05 - | 0 - | 0 - |
| 4665 | 1 | 0.2um_NOKCPHJN_16780 | Bacteria | Proteobacteri | Gammaprote | Alteromonada | Alteromonada | Paraglaciecola | Paraglaciecola | unclassified | unclassified | unclassified | unclassified | unclassified | 1 | 0,894427191 | 1,99E-05 | 1,74E-05 | 2 - | 3,55E-05 - | 1 - | 2,20E-05 - | 2 - | 4,19E-05 - | 0 - | 0 - | 0 - | 0 - |
| 4666 | 1 | 3um_FIJJAFBF_428065 | Bacteria | Bacteroidetes | Cytophagia | Cytophagales | Flammeovirga | unclassified | Flammeovirga | cellular proces | Cell wall/mem | 0P8SR | outer membra | 391587.KAOT | 0,8 | 0,4 | 2,41E-05 | 1,22E-05 | 1 - | 2,61E-05 - | 1 - | 3,24E-05 - | 1 - | 3,08E-05 - | 1 - | 3,12E-05 - | 0 - | 0 - |
| 4667 | 1 | 0.2um_NOKCPHJN_241707 | Bacteria | Bacteroidetes | Flavobacteriia | Flavobacterial | Flavobacteriac | unclassified | Flavobacteriac | poorly charact | Function unkn | 0P8NI | Inherit from b | 487796.Flav2 | 0,6 | 0,489897949 | 1,79E-05 | 1,48E-05 | 1 - | 2,61E-05 - | 1 - | 3,25E-05 - | 1 - | 3,09E-05 - | 0 - | 0 - | 0 - | 0 - |
| 4668 | 1 | 0.2um_NOKCPHJN_313349 | Bacteria | Bacteroidetes | Flavobacteriia | Flavobacterial | Flavobacteriac | Algibacter | Algibacter sp. | metabolism | Inorganic ion t | 05C27 | TonB-depende | 391587.KAOT | 2,6 | 1,019803903 | 3,34E-05 | 1,08E-05 | 4 - | 4,47E-05 - | 2 - | 2,77E-05 - | 3 - | 3,95E-05 - | 3 - | 4,01E-05 - | 1 - | 1,51E-05 - |
| 4669 | 1 | 0.2um_NOKCPHJN_199899 | Bacteria | Bacteroidetes | Flavobacteriia | Flavobacterial | Flavobacteriac | Arenibacter | various | unclassified | unclassified | unclassified | unclassified | unclassified | 0,2 | 0,4 | 4,30E-06 | 8,60E-06 | 1 - | 2,15E-05 - | 0 - | 0 - | 0 - | 0 - | 0 - | 0 - | 0 - | 0 - |
| 4670 | 1 | 3um_FIJJAFBF_410098 | Bacteria | Bacteroidetes | Flavobacteriia | Flavobacterial | Flavobacteriac | Polaribacter | Polaribacter s | unclassified | unclassified | unclassified | unclassified | unclassified | 6 | 1,095445115 | 0,000323841 | 3,10E-05 | 8 - | 0,000367286 - | 6 - | 0,000342045 - | 6 - | 0,000325157 - | 5 - | 0,000274967 - | 5 - | 0,000309749 - |
| 4671 | 1 | 0.2um_NOKCPHJN_356144 | Bacteria | Bacteroidetes | Flavobacteriia | Flavobacterial | Flavobacteriac | Gilvibacter | Gilvibacter sp. | poorly charact | Function unkn | 09C0E | Ragb susd dor | 411154.GFO_ | 4,4 | 0,489897949 | 0,000116849 | 1,81E-05 | 4 - | 8,88E-05 - | 5 - | 0,000137833 - | 4 - | 0,000104822 - | 5 - | 0,000132963 - | 4 - | 0,000119826 - |
| 4672 | 1 | 3um_FIJJAFBF_532378 | Bacteria | Proteobacteri | Betaproteoba | unclassified | unclassified | unclassified | Betaproteoba | metabolism | Amino acid tra | 16SET | amino acid | 52598.EE36_ | 4 | 1,095445115 | 0,000131611 | 3,79E-05 | 4 - | 0,000110362 - | 3 - | 0,000102778 - | 3 - | 9,77E-05 - | 6 - | 0,000198293 - | 4 - | 0,000148918 - |
| 4673 | 1 | 0.2um_NOKCPHJN_23410 | Eukaryota | unclassified | Pelagophycea | Pelagomonad | unclassified | various | various | unclassified | unclassified | unclassified | unclassified | unclassified | 1 | 0 | 0,000128063 | 1,22E-05 | 1 - | 0,000107268 - | 1 - | 0,000133195 - | 1 - | 0,000126619 - | 1 - | 0,000128489 - | 1 - | 0,000144743 - |
| 4674 | 1 | 0.2um_NOKCPHJN_1042835 | Bacteria | Acidobacteria | Acidobacteriia | Acidobacterial | Acidobacteria | Granulicella | Granulicella r | cellular proces | Signal transdu | 07WV4 | Serine Threon | 656024.Fsym | 1,8 | 0,4 | 6,17E-05 | 1,71E-05 | 1 - | 2,82E-05 - | 2 - | 7,00E-05 - | 2 - | 6,66E-05 - | 2 - | 6,76E-05 - | 2 - | 7,61E-05 - |
| 4675 | 1 | 0.2um_NOKCPHJN_606052 | unclassified | unclassified | unclassified | unclassified | unclassified | unclassified | unclassified | unclassified | unclassified | unclassified | unclassified | unclassified | 0,6 | 0,489897949 | 0,00010696 | 8,77E-05 | 0 - | 0 - | 0 - | 0 - | 1 - | 0,000169353 - | 1 - | 0,000171854 - | 1 - | 0,000193593 - |
| 4676 | 1 | 0.2um_NOKCPHJN_159285 | Bacteria | Bacteroidetes | Flavobacteriia | Flavobacterial | Crocinitomica | unclassified | Crocinitomica | unclassified | unclassified | unclassified | unclassified | unclassified | 0,2 | 0,4 | 2,01E-06 | 4,02E-06 | 1 - | 1,01E-05 - | 0 - | 0 - | 0 - | 0 - | 0 - | 0 - | 0 - | 0 - |
| 4677 | 2 | 0.2um_NOKCPHJN_1107193;0.2um_NOKCPHJN_ | Bacteria | Bacteroidetes | Flavobacteriia | Flavobacterial | unclassified | unclassified | Flavobacterial | various | various | various | various | various | 0,6 | 0,489897949 | 2,39E-05 | 1,97E-05 | 1 - | 3,49E-05 - | 1 - | 4,33E-05 - | 1 - | 4,12E-05 - | 0 - | 0 - | 0 - | 0 - |
| 4678 | 1 | 3um_FIJJAFBF_79248 | Bacteria | Bacteroidetes | Flavobacteriia | Flavobacterial | Flavobacteriac | Polaribacter | Polaribacter s | unclassified | unclassified | unclassified | unclassified | unclassified | 1,4 | 0,489897949 | 0,00020159 | 5,93E-05 | 2 - | 0,000246832 - | 1 - | 0,000153246 - | 1 - | 0,00014568 - | 2 - | 0,000295663 - | 1 - | 0,000166532 - |
| 4679 | 1 | 3um_FIJJAFBF_123837 | Bacteria | Bacteroidetes | Flavobacteriia | Flavobacterial | Flavobacteriac | Ulvibacter | Ulvibacter lito | cellular proces | Cell wall/mem | 11TPU | outer membra | 50743.SCB49_ | 1,6 | 0,489897949 | 0,000130251 | 4,20E-05 | 1 - | 6,79E-05 - | 2 - | 0,000168661 - | 2 - | 0,000160334 - | 2 - | 0,000162702 - | 1 - | 9,16E-05 - |
| 4680 | 1 | 0.2um_NOKCPHJN_11919 | Bacteria | Actinobacteria | Actinobacteria | Micrococcales | Microbacteria | Candidatus Ac | Candidatus Ad | information st | Translation, ri | 05CAV | thus facilitatin | 281090.Lxx11 | 2,2 | 1,6 | 5,79E-05 | 3,74E-05 | 5 - | 0,000118327 - | 2 - | 5,88E-05 - | 2 - | 5,59E-05 - | 2 - | 5,67E-05 - | 0 - | 0 - |
| 4681 | 1 | 3um_FIJJAFBF_103901 | Bacteria | Bacteroidetes | Flavobacteriia | Flavobacterial | Flavobacteriac | Flaviramulus | Flaviramulus b | metabolism | Amino acid tra | COG0334 | oxidoreductas | 313598.MED1 | 2,2 | 1,166190379 | 6,46E-05 | 2,89E-05 | 4 - | 0,000102709 - | 2 - | 6,38E-05 - | 3 - | 9,09E-05 - | 1 - | 3,08E-05 - | 1 - | 3,46E-05 - |
| 4682 | 1 | 0.2um_NOKCPHJN_401250 | Eukaryota | Phaeophyceae | unclassified | Ectocarpales | Ectocarpaceae | Ectocarpus | Ectocarpus sili | metabolism | Carbohydrate | KOG2638 | UTP-glucose-1 | 164328.Phyra | 1 | 0 | 2,12E-05 | 2,01E-06 | 1 - | 1,77E-05 - | 1 - | 2,20E-05 - | 1 - | 2,09E-05 - | 1 - | 2,12E-05 - | 1 - | 2,39E-05 - |
| 4683 | 1 | 3um_FIJJAFBF_107750 | Bacteria | Proteobacteri | Gammaprote | Oceanospirilla | Saccharospirill | Reinekea | Reinekea blan | unclassified | unclassified | unclassified | unclassified | unclassified | 1 | 0,632455532 | 2,93E-05 | 1,97E-05 | 1 - | 2,50E-05 - | 2 - | 6,20E-05 - | 1 - | 2,95E-05 - | 1 - | 2,99E-05 - | 0 - | 0 - |
| 4684 | 1 | 3um_FIJJAFBF_440201 | Bacteria | Bacteroidetes | Flavobacteriia | Flavobacterial | Flavobacteriac | Polaribacter | Polaribacter s | metabolism | Energy produc | 0P7ZB | cytochrome c | 313598.MED1 | 0,8 | 0,4 | 2,32E-05 | 1,19E-05 | 1 - | 2,43E-05 - | 1 - | 3,01E-05 - | 0 - | 0 - | 1 - | 2,91E-05 - | 1 - | 3,27E-05 - |
| 4685 | 1 | 0.2um_NOKCPHJN_134680 | Bacteria | Bacteroidetes | Flavobacteriia | Flavobacterial | Flavobacteriac | Flaviramulus | Flaviramulus i | unclassified | unclassified | unclassified | unclassified | unclassified | 4,2 | 0,748331477 | 0,000282206 | 4,28E-05 | 5 - | 0,000284101 - | 3 - | 0,000211662 - | 4 - | 0,000268282 - | 5 - | 0,000340305 - | 4 - | 0,000306682 - |
| 4686 | 1 | 0.2um_NOKCPHJN_739034 | unclassified | unclassified | unclassified | unclassified | unclassified | unclassified | unclassified | unclassified | unclassified | unclassified | unclassified | unclassified | 0,2 | 0,4 | 4,39E-06 | 8,78E-06 | 0 - | 0 - | 0 - | 0 - | 0 - | 0 - | 1 - | 2,20E-05 - | 0 - | 0 - |
| 4687 | 1 | 3um_FIJJAFBF_98529 | Bacteria | Proteobacteri | Gammaprote | Pseudomonad | Moraxellaceae | Psychrobacter | Psychrobacter | information st | Translation, ri | COG0184 | Forms an inte | 335284.Pcryo | 1 | 0 | 0,000108752 | 1,03E-05 | 1 - | 9,11E-05 - | 1 - | 0,00011311 - | 1 - | 0,000107526 - | 1 - | 0,000109114 - | 1 - | 0,000122916 - |
| 4688 | 1 | 0.2um_NOKCPHJN_15703 | Bacteria | Proteobacteri | Gammaprote | Chromatiales | Chromatiacea | Rheinheimera | Rheinheimera | poorly charact | Function unkn | 0YKK5 | NA | 167879.CPS_2 | 1,2 | 0,4 | 0,000107371 | 2,38E-05 | 2 - | 0,000154063 - | 1 - | 9,57E-05 - | 1 - | 9,09E-05 - | 1 - | 9,23E-05 - | 1 - | 0,000103943 - |
| 4689 | 1 | 3um_FIJJAFBF_220491 | Bacteria | Proteobacteri | Gammaprote | Alteromonada | various | various | various | unclassified | unclassified | unclassified | unclassified | unclassified | 1,2 | 0,748331477 | 5,70E-05 | 3,40E-05 | 2 - | 8,38E-05 - | 1 - | 5,20E-05 - | 2 - | 9,89E-05 - | 1 - | 5,02E-05 - | 0 - | 0 - |
| 4690 | 1 | 0.2um_NOKCPHJN_524560 | Bacteria | Proteobacteri | Gammaprote | Cellvibrionales | Spongiibacter | unclassified | marine gamm | information st | Translation, ri | 08UHH | Located on th | 247633.GP21 | 1 | 0 | 0,000106223 | 1,01E-05 | 1 - | 8,90E-05 - | 1 - | 0,00011048 - | 1 - | 0,000105025 - | 1 - | 0,000106576 - | 1 - | 0,000120058 - |
| 4691 | 1 | 3um_FIJJAFBF_366579 | Viruses | unclassified | unclassified | unclassified | various | various | various | unclassified | unclassified | unclassified | unclassified | unclassified | 1,2 | 0,4 | 8,04E-05 | 2,97E-05 | 1 - | 5,57E-05 - | 2 - | 0,000138368 - | 1 - | 6,58E-05 - | 1 - | 6,67E-05 - | 1 - | 7,52E-05 - |
| 4692 | 1 | 3um_FIJJAFBF_417620 | Bacteria | Proteobacteri | Gammaprote | unclassified | unclassified | unclassified | SAR86 cluster | information st | Translation, ri | 05G6W | One of the pri | 207954.MED9 | 0,6 | 0,489897949 | 3,81E-05 | 3,14E-05 | 1 - | 5,57E-05 - | 1 - | 6,92E-05 - | 1 - | 6,58E-05 - | 0 - | 0 - | 0 - | 0 - |
| 4693 | 1 | 0.2um_NOKCPHJN_154868 | Bacteria | Bacteroidetes | Flavobacteriia | Flavobacterial | Flavobacteriac | Altibacter | Altibacter lent | unclassified | unclassified | unclassified | unclassified | unclassified | 0,8 | 0,4 | 2,99E-05 | 1,52E-05 | 1 - | 3,23E-05 - | 1 - | 4,01E-05 - | 1 - | 3,82E-05 - | 1 - | 3,87E-05 - | 0 - | 0 - |
| 4694 | 1 | 3um_FIJJAFBF_70692 | Bacteria | Bacteroidetes | Cytophagia | Cytophagales | Flammeovirga | Reichenbachie | Reichenbachie | cellular proces | Posttranslatio | COG0652 | peptidyl-proly | 388413.ALPR1 | 1,2 | 0,4 | 0,000114952 | 4,24E-05 | 1 - | 7,97E-05 - | 2 - | 0,000197943 - | 1 - | 9,41E-05 - | 1 - | 9,55E-05 - | 1 - | 0,000107552 - |
| 4695 | 1 | 3um_FIJJAFBF_440220 | Bacteria | Bacteroidetes | Flavobacteriia | Flavobacterial | Flavobacteriac | Polaribacter | Polaribacter s | poorly charact | Function unkn | 09GNW | Late embryog | 313594.PI23P | 1,2 | 0,4 | 0,000105252 | 2,33E-05 | 2 - | 0,000151022 - | 1 - | 9,38E-05 - | 1 - | 8,91E-05 - | 1 - | 9,04E-05 - | 1 - | 0,000101891 - |
| 4696 | 2 | 0.2um_NOKCPHJN_666257;3um_FIJJAFBF_21732 | Bacteria | various | various | various | various | various | various | poorly charact | Function unkn | 08PDX | NA | 137722.AZL_0 | 1,2 | 0,748331477 | 5,84E-05 | 3,74E-05 | 1 - | 4,04E-05 - | 2 - | 0,000100365 - | 0 - | 0 - | 2 - | 9,68E-05 - | 1 - | 5,45E-05 - |
| 4697 | 1 | 3um_FIJJAFBF_191493 | Bacteria | Bacteroidetes | Flavobacteriia | Flavobacterial | Flavobacteriac | Polaribacter | Polaribacter s | unclassified | unclassified | unclassified | unclassified | unclassified | 1,6 | 0,8 | 2,56E-05 | 1,21E-05 | 2 - | 2,74E-05 - | 1 - | 1,70E-05 - | 3 - | 4,85E-05 - | 1 - | 1,64E-05 - | 1 - | 1,85E-05 - |
| 4698 | 1 | 0.2um_NOKCPHJN_01338 | Bacteria | Bacteroidetes | Flavobacteriia | Flavobacterial | Flavobacteriac | Formosa | Formosa sp. H | metabolism | Carbohydrate | COG0057 | glyceraldehyd | 313598.MED1 | 0,8 | 0,4 | 3,18E-05 | 1,62E-05 | 1 - | 3,45E-05 - | 1 - | 4,28E-05 - | 1 - | 4,07E-05 - | 1 - | 4,13E-05 - | 0 - | 0 - |
| 4699 | 1 | 3um_FIJJAFBF_743051 | Bacteria | Proteobacteri | Gammaprote | unclassified | unclassified | unclassified | various | unclassified | unclassified | unclassified | unclassified | unclassified | 2,6 | 1,2 | 8,64E-05 | 3,50E-05 | 4 - | 0,000115644 - | 2 - | 7,18E-05 - | 4 - | 0,000136506 - | 2 - | 6,93E-05 - | 1 - | 3,90E-05 - |
| 4700 | 2 | 0.2um_NOKCPHJN_315802;0.2um_NOKCPHJN_7 | Bacteria | Bacteroidetes | Flavobacteriia | Flavobacterial | Flavobacteriac | Formosa | Formosa sp. H | poorly charact | Function unkn | 08PIQ | CHU large pro | 391587.KAOT | 0,4 | 0,489897949 | 2,41E-06 | 2,98E-06 | 1 - | 5,38E-06 - | 1 - | 6,68E-06 - | 0 - | 0 - | 0 - | 0 - | 0 - | 0 - |
| 4701 | 1 | 3um_FIJJAFBF_359843 | Bacteria | Bacteroidetes | unclassified | unclassified | unclassified | unclassified | Bacteroidetes | unclassified | unclassified | unclassified | unclassified | unclassified | 1,4 | 0,489897949 | 3,46E-05 | 1,09E-05 | 2 - | 4,22E-05 - | 2 - | 5,24E-05 - | 1 - | 2,49E-05 - | 1 - | 2,53E-05 - | 1 - | 2,85E-05 - |
| 4702 | 1 | 3um_FIJJAFBF_673395 | Bacteria | Bacteroidetes | Flavobacteriia | Flavobacterial | Crocinitomica | unclassified | Crocinitomica | poorly charact | Function unkn | 08WMU | Domain of un | 382464.VDG1 | 1 | 0 | 0,000105405 | 1,00E-05 | 1 - | 8,83E-05 - | 1 - | 0,00010963 - | 1 - | 0,000104217 - | 1 - | 0,000105756 - | 1 - | 0,000119134 - |
| 4703 | 1 | 3um_FIJJAFBF_506031 | Bacteria | Bacteroidetes | Flavobacteriia | Flavobacterial | Flavobacteriac | Ulvibacter | Ulvibacter sp. | unclassified | unclassified | unclassified | unclassified | unclassified | 2 | 0 | 0,000101127 | 9,60E-06 | 2 - | 8,47E-05 - | 2 - | 0,00010518 - | 2 - | 1,00E-04 - | 2 - | 0,000101464 - | 2 - | 0,000114299 - |
| 4704 | 1 | 3um_FIJJAFBF_72466 | Bacteria | Bacteroidetes | Cytophagia | Cytophagales | Flammeovirga | unclassified | Flammeovirga | information st | Translation, ri | COG0088 | One of the pri | 391603.FBALC | 1,6 | 0,489897949 | 0,000103161 | 2,70E-05 | 2 - | 0,000110362 - | 1 - | 6,85E-05 - | 2 - | 0,000130271 - | 2 - | 0,000132195 - | 1 - | 7,45E-05 - |
| 4705 | 1 | 0.2um_NOKCPHJN_566512 | Bacteria | Bacteroidetes | Flavobacteriia | unclassified | unclassified | unclassified | Flavobacteria | cellular proces | Intracellular tr | 18ATJ | Pfam:YadA | 1005048.CFU_ | 0,6 | 0,489897949 | 8,59E-06 | 7,14E-06 | 1 - | 1,20E-05 - | 1 - | 1,48E-05 - | 0 - | 0 - | 0 - | 0 - | 1 - | 1,61E-05 - |
| 4706 | 1 | 3um_FIJJAFBF_390925 | Bacteria | Bacteroidetes | Flavobacteriia | Flavobacterial | Flavobacteriac | Formosa | Formosa sp. H | unclassified | unclassified | unclassified | unclassified | unclassified | 4,6 | 0,8 | 6,81E-05 | 1,40E-05 | 5 - | 6,20E-05 - | 5 - | 7,70E-05 - | 3 - | 4,39E-05 - | 5 - | 7,42E-05 - | 5 - | 8,36E-05 - |
| 4707 | 2 | 0.2um_NOKCPHJN_05620;3um_FIJJAFBF_46313 | Bacteria | Bacteroidetes | Cytophagia | Cytophagales | Flammeovirga | unclassified | Flammeovirga | cellular proces | Cell cycle cont | 05CR9 | regulator of c | 391587.KAOT | 1 | 0 | 4,89E-05 | 4,65E-06 | 1 - | 4,10E-05 - | 1 - | 5,09E-05 - | 1 - | 4,84E-05 - | 1 - | 4,91E-05 - | 1 - | 5,53E-05 - |
| 4708 | 1 | 3um_FIJJAFBF_189828 | Bacteria | Bacteroidetes | Flavobacteriia | Flavobacterial | Flavobacteriac | Ulvibacter | Ulvibacter sp. | information st | Translation, ri | 09FE5 | One of the pr | 50743.SCB49_ | 1,4 | 0,489897949 | 0,000181237 | 5,70E-05 | 2 - | 0,000220725 - | 2 - | 0,000274075 - | 1 - | 0,000130271 - | 1 - | 0,000132195 - | 1 - | 0,000148918 - |
| 4709 | 1 | 0.2um_NOKCPHJN_400179 | Bacteria | Proteobacteri | Gammaprote | Oceanospirilla | Saccharospirill | Reinekea | Reinekea blan | unclassified | unclassified | unclassified | unclassified | unclassified | 18 | 1,673320053 | 0,000461466 | 5,17E-05 | 20 - | 0,000430682 - | 19 - | 0,000508041 - | 15 - | 0,000381282 - | 18 - | 0,000464296 - | 18 - | 0,000523028 - |
| 4710 | 1 | 0.2um_NOKCPHJN_461932 | Eukaryota | Zoopagomyco | Basidiobolom | Basidiobolales | Basidiobolace | Basidiobolus | Basidiobolus r | metabolism | Secondary me | KOG0059 | ATP-binding c | 5786.XP_0032 | 0,8 | 0,4 | 5,71E-06 | 2,90E-06 | 1 - | 6,18E-06 - | 1 - | 7,68E-06 - | 1 - | 7,30E-06 - | 1 - | 7,41E-06 - | 0 - | 0 - |
| 4711 | 1 | 3um_FIJJAFBF_376215 | Bacteria | Bacteroidetes | Flavobacteriia | Flavobacterial | Flavobacteriac | Ulvibacter | Ulvibacter lito | unclassified | unclassified | unclassified | unclassified | unclassified | 1,6 | 0,8 | 2,14E-05 | 1,13E-05 | 1 - | 1,10E-05 - | 1 - | 1,37E-05 - | 1 - | 1,30E-05 - | 3 - | 3,95E-05 - | 2 - | 2,97E-05 - |
| 4712 | 1 | 0.2um_NOKCPHJN_607353 | Bacteria | Bacteroidetes | Flavobacteriia | Flavobacterial | Flavobacteriac | Marixanthom | Marixanthom | poorly charact | Function unkn | 0P8NI | Inherit from b | 313596.RB250 | 0,8 | 0,4 | 3,40E-05 | 1,75E-05 | 1 - | 3,60E-05 - | 0 - | 0 - | 1 - | 4,25E-05 - | 1 - | 4,31E-05 - | 1 - | 4,85E-05 - |
| 4713 | 1 | 0.2um_NOKCPHJN_406490 | Eukaryota | Ascomycota | Eurotiomycet | Chaetothyriale | Herpotrichiell | Capronia | Capronia epim | unclassified | unclassified | unclassified | unclassified | unclassified | 0,2 | 0,4 | 2,48E-06 | 4,96E-06 | 0 - | 0 - | 0 - | 0 - | 0 - | 0 - | 1 - | 1,24E-05 - | 0 - | 0 - |

| | | | | | | | | | | | | | | | | | | | | | | | | | | | | |
|---|---|---|---|---|---|---|---|---|---|---|---|---|---|---|---|---|---|---|---|---|---|---|---|---|---|---|---|---|
| 4714 | 1 | 3um_FIJJAFBF_75111 | Bacteria | Proteobacteri | Alphaproteob | Rickettsiales | unclassified | unclassified | Rickettsiales b | unclassified | unclassified | unclassified | unclassified | unclassified | 0,8 | 0,4 | 0,000152682 | 7,84E-05 | 1 - | 0,000159412 - | 1 - | 0,000197943 - | 0 - | 0 - | 1 - | 0,000190949 - | 1 - | 0,000215103 - |
| 4715 | 1 | 3um_FIJJAFBF_225132 | Bacteria | Proteobacteri | Alphaproteob | Rhodobactera | Rhodobactera | Planktomarina | Planktomarina | unclassified | unclassified | unclassified | unclassified | unclassified | 3,8 | 0,4 | 0,000293562 | 3,88E-05 | 4 - | 0,000259383 - | 3 - | 0,000241557 - | 4 - | 0,000306175 - | 4 - | 0,000310697 - | 4 - | 0,000349999 - |
| 4716 | 1 | 0.2um_NOKCPHJN_184686 | Bacteria | Proteobacteri | Gammaprote | unclassified | unclassified | unclassified | Gammaprote | information st | Translation, ri | 179ZI | One of the pri | 582744.Msip3 | 0,4 | 0,8 | 6,04E-05 | 0,000120818 | 2 - | 0,000302044 - | 0 - | 0 - | 0 - | 0 - | 0 - | 0 - | 0 - | 0 - |
| 4717 | 1 | 3um_FIJJAFBF_465353 | Bacteria | Bacteroidetes | Flavobacteriia | Flavobacterial | Crocinitomica | unclassified | Crocinitomica | cellular proces | Posttranslatio | 1AIH3 | Peptidase M1 | 760192.Halhy | 0,6 | 0,489897949 | 1,20E-05 | 9,82E-06 | 0 - | 0 - | 1 - | 2,06E-05 - | 1 - | 1,96E-05 - | 1 - | 1,99E-05 - | 0 - | 0 - |
| 4718 | 1 | 3um_FIJJAFBF_81189 | Bacteria | Proteobacteri | Alphaproteob | Rhizobiales | Methylobacte | Methylobacte | Methylobacte | poorly charact | Function unkn | 0231D | TolA protein | 426117.M446 | 0,2 | 0,4 | 2,67E-06 | 5,35E-06 | 0 - | 0 - | 0 - | 0 - | 0 - | 0 - | 1 - | 1,34E-05 - | 0 - | 0 - |
| 4719 | 1 | 0.2um_NOKCPHJN_22857 | Bacteria | Bacteroidetes | Flavobacteriia | Flavobacterial | Flavobacteria | Formosa | Formosa sp. H | unclassified | unclassified | unclassified | unclassified | unclassified | 3,6 | 1,2 | 0,000326781 | 0,000100075 | 5 - | 0,000385157 - | 5 - | 0,000478251 - | 2 - | 0,000181855 - | 3 - | 0,000276812 - | 3 - | 0,000311828 - |
| 4720 | 1 | 0.2um_NOKCPHJN_60738 | Bacteria | Proteobacteri | Alphaproteob | Rhodobactera | Rhodobactera | Amylibacter | Amylibacter k | information st | Transcription | 05D27 | DNA-depende | 644107.SL115 | 12,4 | 2,059126028 | 0,000118503 | 1,53E-05 | 15 - | 0,000121328 - | 14 - | 0,000140611 - | 10 - | 9,55E-05 - | 13 - | 0,000125954 - | 10 - | 0,000109143 - |
| 4721 | 1 | 0.2um_NOKCPHJN_110549 | Bacteria | Proteobacteri | Gammaprote | Cellvibrionales | Porticoccacea | unclassified | SAR92 bacteri | information st | Translation, ri | 17C3B | The globular d | 314287.GB22 | 3 | 0,632455532 | 0,000377236 | 0,000105802 | 3 - | 0,000313028 - | 3 - | 0,000388688 - | 2 - | 0,000246331 - | 3 - | 0,000374954 - | 4 - | 0,00056318 - |
| 4722 | 1 | 0.2um_NOKCPHJN_915855 | Bacteria | Proteobacteri | Gammaprote | Alteromonada | Colwelliaceae | Thalassotalea | Thalassotalea | cellular proces | Cell wall/mem | 0QKXZ | Porin Gram-ne | 326442.PSHA | 0,8 | 0,748331477 | 3,50E-05 | 3,30E-05 | 1 - | 3,71E-05 - | 0 - | 0 - | 2 - | 8,77E-05 - | 0 - | 0 - | 1 - | 5,01E-05 - |
| 4723 | 1 | 3um_FIJJAFBF_718087 | Bacteria | Bacteroidetes | Cytophagia | Cytophagales | Flammeovirga | unclassified | Flammeovirga | unclassified | unclassified | unclassified | unclassified | unclassified | 1,6 | 0,8 | 0,000120248 | 4,81E-05 | 3 - | 0,00019676 - | 1 - | 8,14E-05 - | 1 - | 7,74E-05 - | 2 - | 0,000157124 - | 1 - | 8,85E-05 - |
| 4724 | 1 | 3um_FIJJAFBF_357543 | Bacteria | unclassified | unclassified | unclassified | unclassified | unclassified | uncultured ba | unclassified | unclassified | unclassified | unclassified | unclassified | 4,2 | 0,748331477 | 9,91E-05 | 1,71E-05 | 4 - | 7,96E-05 - | 4 - | 9,88E-05 - | 5 - | 0,000117402 - | 5 - | 0,000119136 - | 3 - | 8,05E-05 - |
| 4725 | 1 | 3um_FIJJAFBF_83699 | Bacteria | Bacteroidetes | Flavobacteriia | Flavobacterial | Flavobacteria | Polaribacter | Polaribacter s | metabolism | Carbohydrate | COG0149 | triosephospha | 313594.PI23P | 1,4 | 0,489897949 | 7,42E-05 | 2,12E-05 | 2 - | 9,11E-05 - | 1 - | 5,66E-05 - | 2 - | 0,000107526 - | 1 - | 5,46E-05 - | 1 - | 6,15E-05 - |
| 4726 | 1 | 0.2um_NOKCPHJN_549633 | Bacteria | Proteobacteri | Alphaproteob | Pelagibacteral | Pelagibactera | Candidatus Pe | Candidatus Pe | unclassified | unclassified | unclassified | unclassified | unclassified | 1 | 0 | 0,00024914 | 2,37E-05 | 1 - | 0,000208685 - | 1 - | 0,000259125 - | 1 - | 0,000246331 - | 1 - | 0,00024997 - | 1 - | 0,00028159 - |
| 4727 | 1 | 0.2um_NOKCPHJN_792907 | Bacteria | Proteobacteri | Alphaproteob | Pelagibacteral | Pelagibactera | Candidatus Pe | Candidatus Pe | unclassified | unclassified | unclassified | unclassified | unclassified | 1 | 0 | 5,66E-05 | 5,38E-06 | 1 - | 4,74E-05 - | 1 - | 5,89E-05 - | 1 - | 5,60E-05 - | 1 - | 5,68E-05 - | 1 - | 6,40E-05 - |
| 4728 | 1 | 3um_FIJJAFBF_799680 | Bacteria | Bacteroidetes | Flavobacteriia | Flavobacterial | Flavobacteria | Arenibacter | Arenibacter n | unclassified | unclassified | unclassified | unclassified | unclassified | 2,6 | 0,489897949 | 4,24E-05 | 6,89E-06 | 3 - | 4,14E-05 - | 3 - | 5,14E-05 - | 3 - | 4,89E-05 - | 2 - | 3,30E-05 - | 2 - | 3,72E-05 - |
| 4729 | 1 | 0.2um_NOKCPHJN_370983 | Bacteria | Proteobacteri | Alphaproteob | Rhodobactera | Rhodobactera | Loktanella | Loktanella sp. | metabolism | Energy produc | COG2838 | Isocitrate deh | 388401.RB215 | 0,8 | 0,748331477 | 1,43E-05 | 1,24E-05 | 2 - | 3,15E-05 - | 0 - | 0 - | 0 - | 0 - | 1 - | 1,89E-05 - | 1 - | 2,12E-05 - |
| 4730 | 1 | 0.2um_NOKCPHJN_877186 | Bacteria | Bacteroidetes | Flavobacteriia | Flavobacterial | Flavobacteria | Formosa | Formosa sp. H | information st | Translation, ri | 05KK9 | This protein bi | 313603.FB217 | 0,4 | 0,8 | 2,55E-05 | 5,10E-05 | 2 - | 0,00012753 - | 0 - | 0 - | 0 - | 0 - | 0 - | 0 - | 0 - | 0 - |
| 4731 | 1 | 3um_FIJJAFBF_204881 | Bacteria | Bacteroidetes | Flavobacteriia | Flavobacterial | Flavobacteria | Polaribacter | Polaribacter s | unclassified | unclassified | unclassified | unclassified | unclassified | 2,4 | 0,8 | 3,37E-05 | 1,29E-05 | 2 - | 2,32E-05 - | 3 - | 4,31E-05 - | 1 - | 1,37E-05 - | 3 - | 4,16E-05 - | 3 - | 4,69E-05 - |
| 4732 | 1 | 3um_FIJJAFBF_149027 | Bacteria | Proteobacteri | Alphaproteob | Rhodobactera | Rhodobactera | Amylibacter | Amylibacter c | information st | Translation, ri | 16QAP | Associates wit | 391589.RGAI1 | 1,2 | 0,748331477 | 5,62E-05 | 3,56E-05 | 2 - | 7,89E-05 - | 1 - | 4,90E-05 - | 1 - | 4,66E-05 - | 0 - | 0 - | 2 - | 0,000106443 - |
| 4733 | 1 | 3um_FIJJAFBF_214615 | Bacteria | Proteobacteri | Alphaproteob | Rhodobactera | Rhodobactera | Phaeobacter | Phaeobacter g | information st | Translation, ri | 08VZM | 50s ribosomal | 388739.RSK20 | 1 | 0 | 7,97E-05 | 7,56E-06 | 1 - | 6,67E-05 - | 1 - | 8,29E-05 - | 1 - | 7,88E-05 - | 1 - | 7,99E-05 - | 1 - | 9,00E-05 - |
| 4734 | 2 | 0.2um_NOKCPHJN_100694;0.2um_NOKCPHJN_1 | Bacteria | Proteobacteri | Alphaproteob | Rhodobactera | Rhodobactera | Ruegeria | Ruegeria pom | unclassified | unclassified | unclassified | unclassified | unclassified | 1 | 0 | 1,29E-05 | 1,23E-06 | 1 - | 1,08E-05 - | 1 - | 1,34E-05 - | 1 - | 1,28E-05 - | 1 - | 1,29E-05 - | 1 - | 1,46E-05 - |
| 4735 | 1 | 0.2um_NOKCPHJN_267643 | Eukaryota | Bacillariophyt | Bacillariophyc | Naviculales | Phaeodactyla | Phaeodactylu | Phaeodactylu | unclassified | unclassified | unclassified | unclassified | unclassified | 0,6 | 0,489897949 | 3,62E-05 | 2,99E-05 | 1 - | 5,29E-05 - | 1 - | 6,57E-05 - | 1 - | 6,24E-05 - | 0 - | 0 - | 0 - | 0 - |
| 4736 | 1 | 0.2um_NOKCPHJN_1004542 | Bacteria | Proteobacteri | Alphaproteob | Pelagibacteral | unclassified | Candidatus Fo | Candidatus Fo | cellular proces | Cell wall/mem | 7257 | NA | 335992.SAR11 | 0,2 | 0,4 | 9,29E-06 | 1,86E-05 | 1 - | 4,65E-05 - | 0 - | 0 - | 0 - | 0 - | 0 - | 0 - | 0 - | 0 - |
| 4737 | 1 | 0.2um_NOKCPHJN_150646 | Bacteria | Proteobacteri | Gammaprote | Cellvibrionales | Porticoccacea | unclassified | Gammaprote | cellular proces | Intracellular tr | 16Q1A | type II secretio | 314287.GB22 | 1 | 0 | 2,81E-05 | 2,67E-06 | 1 - | 2,36E-05 - | 1 - | 2,93E-05 - | 1 - | 2,78E-05 - | 1 - | 2,82E-05 - | 1 - | 3,18E-05 - |
| 4738 | 1 | 0.2um_NOKCPHJN_609415 | Bacteria | Proteobacteri | Alphaproteob | Pelagibacteral | Pelagibactera | Candidatus Pe | Candidatus Pe | metabolism | Amino acid tra | 01RJV | Glutamate syr | 335992.SAR11 | 0,2 | 0,4 | 6,87E-06 | 1,37E-05 | 1 - | 3,44E-05 - | 0 - | 0 - | 0 - | 0 - | 0 - | 0 - | 0 - | 0 - |
| 4739 | 1 | 3um_FIJJAFBF_46298 | Bacteria | Bacteroidetes | unclassified | unclassified | unclassified | unclassified | Bacteroidetes | information st | Translation, ri | 0P97G | This is one of | 926562.Oweh | 0,8 | 1,166190379 | 4,61E-05 | 6,48E-05 | 3 - | 0,000165543 - | 0 - | 0 - | 1 - | 6,51E-05 - | 0 - | 0 - | 0 - | 0 - |
| 4740 | 1 | 0.2um_NOKCPHJN_282074 | Bacteria | Proteobacteri | Gammaprote | unclassified | unclassified | unclassified | OM182 bacte | unclassified | unclassified | unclassified | unclassified | unclassified | 8 | 1,414213562 | 0,000130784 | 1,59E-05 | 10 - | 0,000138787 - | 9 - | 0,000155099 - | 7 - | 0,000114677 - | 8 - | 0,000132995 - | 6 - | 0,000112364 - |
| 4741 | 1 | 3um_FIJJAFBF_264291 | Bacteria | Proteobacteri | Alphaproteob | Rhodobactera | Rhodobactera | Roseobacter | Roseobacter s | metabolism | Inorganic ion t | 17B5T | ferritin dps fa | 391593.RCCS2 | 0,4 | 0,489897949 | 3,21E-05 | 3,95E-05 | 1 - | 7,36E-05 - | 0 - | 0 - | 1 - | 8,68E-05 - | 0 - | 0 - | 0 - | 0 - |
| 4742 | 1 | 3um_FIJJAFBF_91101 | Bacteria | Proteobacteri | Gammaprote | Oceanospirilla | Oceanospirilla | Motiliproteus | Motiliproteus | unclassified | unclassified | unclassified | unclassified | unclassified | 1,2 | 0,4 | 0,00021331 | 4,72E-05 | 2 - | 0,000306071 - | 1 - | 0,000190025 - | 1 - | 0,000180643 - | 1 - | 0,000183311 - | 1 - | 0,000206499 - |
| 4743 | 1 | 0.2um_NOKCPHJN_769752 | Bacteria | Proteobacteri | Alphaproteob | Pelagibacteral | Pelagibactera | unclassified | Pelagibactera | poorly charact | Function unkn | 02CBJ | NA | 684719.HIMB | 0,6 | 0,489897949 | 0,000100668 | 8,26E-05 | 0 - | 0 - | 0 - | 0 - | 1 - | 0,000159391 - | 1 - | 0,000161745 - | 1 - | 0,000182205 - |
| 4744 | 1 | 0.2um_NOKCPHJN_899935 | Bacteria | Proteobacteri | Alphaproteob | Rhodobactera | Rhodobactera | Roseobacter | Roseobacter s | information st | Translation, ri | COG0050 | This protein p | 391593.RCCS2 | 0,4 | 0,489897949 | 1,32E-05 | 1,63E-05 | 1 - | 2,94E-05 - | 1 - | 3,64E-05 - | 0 - | 0 - | 0 - | 0 - | 0 - | 0 - |
| 4745 | 1 | 3um_FIJJAFBF_250698 | Bacteria | Bacteroidetes | Cytophagia | Cytophagales | Cytophagacea | Ekhidna | Ekhidna lutea | information st | Translation, ri | 05CAV | thus facilitatin | 388413.ALPR1 | 0,8 | 0,748331477 | 1,36E-05 | 1,18E-05 | 2 - | 2,98E-05 - | 0 - | 0 - | 0 - | 0 - | 1 - | 1,79E-05 - | 1 - | 2,01E-05 - |
| 4746 | 1 | 0.2um_NOKCPHJN_118555 | Bacteria | Bacteroidetes | Flavobacteriia | Flavobacterial | Flavobacteria | Polaribacter | Polaribacter s | metabolism | Inorganic ion t | 0P838 | tonB-depende | 313598.MED1 | 0,4 | 0,489897949 | 6,90E-06 | 8,45E-06 | 0 - | 0 - | 0 - | 0 - | 1 - | 1,71E-05 - | 1 - | 1,74E-05 - | 0 - | 0 - |
| 4747 | 1 | 0.2um_NOKCPHJN_179942 | Bacteria | Proteobacteri | Alphaproteob | Rhodospirillal | Rhodospirillac | unclassified | Rhodobactera | unclassified | unclassified | unclassified | unclassified | unclassified | 0,6 | 0,489897949 | 4,36E-05 | 3,59E-05 | 1 - | 6,45E-05 - | 0 - | 0 - | 1 - | 7,61E-05 - | 1 - | 7,72E-05 - | 0 - | 0 - |
| 4748 | 1 | 0.2um_NOKCPHJN_434036 | Bacteria | Bacteroidetes | unclassified | unclassified | unclassified | unclassified | Bacteroidetes | unclassified | unclassified | unclassified | unclassified | unclassified | 1 | 0 | 3,92E-05 | 3,72E-06 | 1 - | 3,28E-05 - | 1 - | 4,07E-05 - | 1 - | 3,87E-05 - | 1 - | 3,93E-05 - | 1 - | 4,42E-05 - |
| 4749 | 1 | 3um_FIJJAFBF_469742 | Bacteria | Proteobacteri | Gammaprote | Pseudomonad | Pseudomonad | Pseudomonas | Pseudomonas | metabolism | Lipid transpor | 05VNB | Carrier of the | 223283.PSPTC | 0,8 | 0,748331477 | 0,000137691 | 0,000132555 | 1 - | 0,000151022 - | 0 - | 0 - | 2 - | 0,000356532 - | 1 - | 0,000180899 - | 0 - | 0 - |
| 4750 | 1 | 0.2um_NOKCPHJN_98865 | Bacteria | Bacteroidetes | Flavobacteriia | Flavobacterial | Flavobacteria | Polaribacter | Polaribacter b | information st | Replication, re | 05C24 | DNA gyrase ne | 983544.Lacal_ | 7,8 | 1,166190379 | 0,000127053 | 1,86E-05 | 8 - | 0,000109703 - | 7 - | 0,000119191 - | 10 - | 0,000161866 - | 7 - | 0,00011498 - | 7 - | 0,000129525 - |
| 4751 | 1 | 0.2um_NOKCPHJN_1016790 | Bacteria | Proteobacteri | Alphaproteob | Rhodobactera | Rhodobactera | unclassified | Rhodobactera | metabolism | Energy produc | 01RNC | Oxidoreducta | 314270.RB208 | 0,4 | 0,489897949 | 9,02E-06 | 1,11E-05 | 1 - | 2,07E-05 - | 0 - | 0 - | 1 - | 2,44E-05 - | 0 - | 0 - | 0 - | 0 - |
| 4752 | 1 | 3um_FIJJAFBF_90637 | Bacteria | Proteobacteri | Gammaprote | Alteromonada | Alteromonada | Paraglaciecola | Paraglaciecola | cellular proces | Intracellular tr | 05VIE | preprotein tra | 983545.Glaag | 2,2 | 0,4 | 0,000267576 | 2,45E-05 | 3 - | 0,000310207 - | 2 - | 0,000256791 - | 2 - | 0,000244112 - | 2 - | 0,000247718 - | 2 - | 0,000279053 - |
| 4753 | 1 | 3um_FIJJAFBF_549238 | Bacteria | Proteobacteri | Betaproteoba | Burkholderial | Comamonada | Limnohabitan | Limnohabitan | unclassified | unclassified | unclassified | unclassified | unclassified | 4,6 | 0,489897949 | 0,000109394 | 1,58E-05 | 4 - | 7,96E-05 - | 5 - | 0,0001235 - | 5 - | 0,000117402 - | 5 - | 0,000119136 - | 4 - | 0,000107365 - |
| 4754 | 1 | 3um_FIJJAFBF_744551 | Bacteria | Proteobacteri | Alphaproteob | Rhodobactera | Rhodobactera | Yoonia | Yoonia maritir | metabolism | Energy produc | COG0056 | Produces ATP | 391593.RCCS2 | 11,2 | 1,166190379 | 0,000290311 | 1,44E-05 | 13 - | 0,000284752 - | 10 - | 0,000271983 - | 11 - | 0,000284409 - | 12 - | 0,000314847 - | 10 - | 0,000295562 - |
| 4755 | 1 | 3um_FIJJAFBF_213248 | Bacteria | Proteobacteri | Gammaprote | Oceanospirilla | Saccharospirill | Reinekea | Reinekea fors | cellular proces | Posttranslatio | 17JNZ | Redoxin doma | 314283.MED2 | 1 | 0 | 8,51E-05 | 8,08E-06 | 1 - | 7,13E-05 - | 1 - | 8,85E-05 - | 1 - | 8,42E-05 - | 1 - | 8,54E-05 - | 1 - | 9,62E-05 - |
| 4756 | 1 | 3um_FIJJAFBF_563869 | Bacteria | Proteobacteri | Gammaprote | Cellvibrionales | Porticoccacea | unclassified | gamma prote | unclassified | unclassified | unclassified | unclassified | unclassified | 0,2 | 0,4 | 5,01E-06 | 1,00E-05 | 0 - | 0 - | 1 - | 2,50E-05 - | 0 - | 0 - | 0 - | 0 - | 0 - | 0 - |
| 4757 | 1 | 3um_FIJJAFBF_246779 | Bacteria | Bacteroidetes | Flavobacteriia | Flavobacterial | Flavobacteria | Flavobacteriu | Flavobacteriu | metabolism | Inorganic ion t | 05C27 | TonB-depende | 867900.Celly_ | 6 | 1,264911064 | 8,09E-05 | 1,49E-05 | 7 - | 7,99E-05 - | 5 - | 7,08E-05 - | 5 - | 6,73E-05 - | 8 - | 0,000109331 - | 5 - | 7,70E-05 - |
| 4758 | 1 | 0.2um_NOKCPHJN_943127 | Bacteria | Bacteroidetes | unclassified | unclassified | unclassified | unclassified | unidentified e | cellular proces | Signal transdu | 09A4D | gtp-binding pr | 391603.FBALC | 0,4 | 0,489897949 | 8,45E-06 | 1,04E-05 | 1 - | 1,92E-05 - | 0 - | 0 - | 0 - | 0 - | 1 - | 2,30E-05 - | 0 - | 0 - |
| 4759 | 1 | 3um_FIJJAFBF_306011 | Bacteria | Bacteroidetes | Flavobacteriia | Flavobacterial | Flavobacteria | Polaribacter | Polaribacter s | unclassified | unclassified | unclassified | unclassified | unclassified | 1,4 | 0,489897949 | 0,000117804 | 3,70E-05 | 2 - | 0,000143471 - | 2 - | 0,000178149 - | 1 - | 8,47E-05 - | 1 - | 8,59E-05 - | 1 - | 9,68E-05 - |
| 4760 | 1 | 3um_FIJJAFBF_17156 | Bacteria | Proteobacteri | Alphaproteob | Rhodobactera | unclassified | unclassified | Rhodobactera | unclassified | unclassified | unclassified | unclassified | unclassified | 2,6 | 1,2 | 0,000123353 | 5,96E-05 | 2 - | 7,89E-05 - | 4 - | 0,000195902 - | 1 - | 4,66E-05 - | 4 - | 0,00018898 - | 2 - | 0,000106443 - |
| 4761 | 1 | 3um_FIJJAFBF_106934 | Bacteria | Proteobacteri | Alphaproteob | Rhodobactera | unclassified | unclassified | Rhodobactera | unclassified | unclassified | unclassified | unclassified | unclassified | 0,4 | 0,8 | 1,65E-05 | 3,30E-05 | 0 - | 0 - | 0 - | 0 - | 0 - | 0 - | 2 - | 8,26E-05 - | 0 - | 0 - |
| 4762 | 1 | 3um_FIJJAFBF_227251 | Bacteria | Proteobacteri | Gammaprote | Oceanospirilla | Oceanospirilla | Marinobacteri | Marinobacteri | information st | Translation, ri | 17FIT | 50s ribosomal | 207954.MED9 | 0,2 | 0,4 | 2,94E-05 | 5,89E-05 | 1 - | 0,00014715 - | 0 - | 0 - | 0 - | 0 - | 0 - | 0 - | 0 - | 0 - |
| 4763 | 1 | 3um_FIJJAFBF_398712 | Bacteria | Bacteroidetes | Flavobacteriia | Flavobacterial | Flavobacteria | various | various | unclassified | unclassified | unclassified | unclassified | unclassified | 1,8 | 0,748331477 | 0,000334551 | 0,000108706 | 3 - | 0,000484972 - | 1 - | 0,000200731 - | 2 - | 0,00038164 - | 2 - | 0,000387277 - | 1 - | 0,000218133 - |
| 4764 | 1 | 3um_FIJJAFBF_139150 | Bacteria | Proteobacteri | Alphaproteob | Rhodobactera | Rhodobactera | Planktomarina | Planktomarina | metabolism | Amino acid tra | 16R0H | Acetolactate s | 52598.EE36_1 | 1,6 | 0,489897949 | 0,000116531 | 3,23E-05 | 2 - | 0,000124083 - | 2 - | 0,000154074 - | 2 - | 0,000146467 - | 1 - | 7,43E-05 - | 1 - | 8,37E-05 - |
| 4765 | 1 | 0.2um_NOKCPHJN_787005 | Bacteria | Proteobacteri | Alphaproteob | Pelagibacteral | unclassified | unclassified | Pelagibacteral | cellular proces | Cell wall/mem | 7257 | NA | 335992.SAR11 | 3,2 | 0,4 | 0,000127124 | 1,97E-05 | 3 - | 9,98E-05 - | 3 - | 0,000123929 - | 3 - | 0,000117811 - | 4 - | 0,000159401 - | 3 - | 0,000134673 - |
| 4766 | 1 | 0.2um_NOKCPHJN_813821 | Bacteria | Proteobacteri | Gammaprote | Methylococca | Methylococca | Methylomagn | Methylomagn | unclassified | unclassified | unclassified | unclassified | unclassified | 3,4 | 0,8 | 5,74E-05 | 7,77E-06 | 5 - | 7,21E-05 - | 3 - | 5,37E-05 - | 3 - | 5,11E-05 - | 3 - | 5,18E-05 - | 3 - | 5,84E-05 - |
| 4767 | 1 | 0.2um_NOKCPHJN_510145 | Bacteria | Proteobacteri | Alphaproteob | Rhodobactera | Rhodobactera | Planktomarina | Planktomarina | information st | Translation, ri | COG0185 | Protein S19 fo | 633131.TR2A | 3,8 | 1,720465053 | 0,000545539 | 0,000188744 | 7 - | 0,000873301 - | 2 - | 0,000309824 - | 3 - | 0,00044179 - | 4 - | 0,000597754 - | 3 - | 0,000505025 - |
| 4768 | 1 | 3um_FIJJAFBF_282388 | Eukaryota | Bacillariophyt | Bacillariophyc | Naviculales | Phaeodactyla | Phaeodactylu | Phaeodactylu | unclassified | unclassified | unclassified | unclassified | unclassified | 4,2 | 0,748331477 | 0,000480177 | 9,95E-05 | 4 - | 0,000382589 - | 5 - | 0,000593829 - | 5 - | 0,000564509 - | 3 - | 0,000343708 - | 4 - | 0,000516248 - |
| 4769 | 1 | 0.2um_NOKCPHJN_473659 | Bacteria | Proteobacteri | Alphaproteob | Pelagibacteral | Pelagibactera | unclassified | alpha proteob | unclassified | unclassified | unclassified | unclassified | unclassified | 1,8 | 0,4 | 7,03E-05 | 1,39E-05 | 2 - | 6,63E-05 - | 2 - | 8,24E-05 - | 2 - | 7,83E-05 - | 2 - | 7,95E-05 - | 1 - | 4,48E-05 - |
| 4770 | 1 | 3um_FIJJAFBF_358493 | Bacteria | Bacteroidetes | Flavobacteriia | Flavobacterial | Flavobacteria | Formosa | Formosa sp. H | poorly charact | Function unkn | 05FCH | NA | 886377.Murru | 0,2 | 0,4 | 8,93E-06 | 1,79E-05 | 1 - | 4,47E-05 - | 0 - | 0 - | 0 - | 0 - | 0 - | 0 - | 0 - | 0 - |
| 4771 | 1 | 3um_FIJJAFBF_212820 | Eukaryota | unclassified | unclassified | Phaeocystales | Phaeocystace | Phaeocystis | Phaeocystis a | metabolism | Carbohydrate | 0IVA1 | Photosystem I | 35128.Thapsd | 1 | 0 | 7,45E-05 | 7,07E-06 | 1 - | 6,24E-05 - | 1 - | 7,75E-05 - | 1 - | 7,36E-05 - | 1 - | 7,47E-05 - | 1 - | 8,42E-05 - |
| 4772 | 1 | 3um_FIJJAFBF_549854 | Bacteria | Bacteroidetes | Flavobacteriia | Flavobacterial | Crocinitomica | unclassified | Crocinitomica | information st | Translation, ri | COG0256 | This is one of | 888832.HMPR | 3,2 | 0,4 | 0,000368217 | 5,45E-05 | 3 - | 0,000289353 - | 3 - | 0,000359291 - | 4 - | 0,000455402 - | 3 - | 0,000346597 - | 3 - | 0,00039044 - |
| 4773 | 1 | 3um_FIJJAFBF_715083 | Eukaryota | unclassified | Pelagophycea | Pelagomonad | unclassified | Aureoumbra | Aureoumbra l | unclassified | unclassified | unclassified | unclassified | unclassified | 0,2 | 0,4 | 1,28E-05 | 2,56E-05 | 1 - | 6,41E-05 - | 0 - | 0 - | 0 - | 0 - | 0 - | 0 - | 0 - | 0 - |
| 4774 | 1 | 3um_FIJJAFBF_498429 | Bacteria | Proteobacteri | Gammaprote | Arenicellales | Arenicellaceae | Arenicella | Arenicella xan | metabolism | Inorganic ion t | 177TK | phosphate-sel | 455436.GHTC | 7,4 | 0,8 | 0,000297414 | 2,99E-05 | 8 - | 0,00027086 - | 6 - | 0,000252246 - | 8 - | 0,000319722 - | 8 - | 0,000324444 - | 7 - | 0,0003198 - |
| 4775 | 1 | 3um_FIJJAFBF_264641 | Bacteria | Bacteroidetes | Flavobacteriia | Flavobacterial | Flavobacteria | Polaribacter | Polaribacter t | poorly charact | Function unkn | 05END | lipolytic prote | 313598.MED1 | 14,4 | 1,356465997 | 0,000391328 | 6,73E-05 | 12 - | 0,000271126 - | 16 - | 0,000448878 - | 14 - | 0,000373376 - | 15 - | 0,000405955 - | 15 - | 0,000457307 - |
| 4776 | 1 | 3um_FIJJAFBF_213931 | Bacteria | Bacteroidetes | Flavobacteriia | Flavobacterial | Flavobacteria | Formosa | Formosa sp. H | poorly charact | Function unkn | 0XPRT | NA | 391598.FBBAL | 6,8 | 2,039607805 | 0,000186662 | 4,21E-05 | 10 - | 0,000235681 - | 8 - | 0,000234117 - | 6 - | 0,000166919 - | 6 - | 0,000169384 - | 4 - | 0,000127207 - |
| 4777 | 1 | 3um_FIJJAFBF_597540 | Eukaryota | Bacillariophyt | Bacillariophyc | Naviculales | Naviculaceae | Fistulifera | Fistulifera sola | unclassified | unclassified | unclassified | unclassified | unclassified | 0,4 | 0,489897949 | 9,15E-06 | 1,13E-05 | 1 - | 2,10E-05 - | 0 - | 0 - | 1 - | 2,48E-05 - | 0 - | 0 - | 0 - | 0 - |
| 4778 | 1 | 0.2um_NOKCPHJN_313796 | Bacteria | Proteobacteri | Betaproteoba | unclassified | unclassified | unclassified | Betaproteoba | unclassified | unclassified | unclassified | unclassified | unclassified | 0,2 | 0,4 | 1,73E-06 | 3,46E-06 | 0 - | 0 - | 0 - | 0 - | 1 - | 8,64E-06 - | 0 - | 0 - | 0 - | 0 - |
| 4779 | 1 | 0.2um_NOKCPHJN_450476 | Bacteria | Bacteroidetes | Flavobacteriia | Flavobacterial | Flavobacteria | Maribacter | Maribacter th | metabolism | Inorganic ion t | 05C27 | TonB-depende | 655815.ZPR_ | 3,4 | 0,8 | 4,53E-05 | 9,22E-06 | 4 - | 4,53E-05 - | 4 - | 5,63E-05 - | 4 - | 5,35E-05 - | 3 - | 4,07E-05 - | 2 - | 3,06E-05 - |
| 4780 | 1 | 0.2um_NOKCPHJN_144460 | Bacteria | Proteobacteri | Gammaprote | Oceanospirilla | Saccharospirill | Reinekea | Reinekea blan | unclassified | unclassified | unclassified | unclassified | unclassified | 0,2 | 0,4 | 1,00E-05 | 2,00E-05 | 0 - | 0 - | 0 - | 0 - | 1 - | 5,00E-05 - | 0 - | 0 - | 0 - | 0 - |
| 4781 | 1 | 3um_FIJJAFBF_103122 | Bacteria | Bacteroidetes | Flavobacteriia | Flavobacterial | Flavobacteria | Polaribacter | Polaribacter a | poorly charact | Function unkn | 0P9S8 | Fe-S metaboli | 313598.MED1 | 0,6 | 0,8 | 5,28E-05 | 6,75E-05 | 2 - | 0,000162804 - | 1 - | 0,000101077 - | 0 - | 0 - | 0 - | 0 - | 0 - | 0 - |
| 4782 | 1 | 3um_FIJJAFBF_103119 | Bacteria | Bacteroidetes | Flavobacteriia | Flavobacterial | Flavobacteria | Polaribacter | Polaribacter s | poorly charact | Function unkn | 09AVF | Protein of unk | 313598.MED1 | 0,8 | 0,748331477 | 3,69E-05 | 3,37E-05 | 0 - | 0 - | 0 - | 0 - | 2 - | 8,88E-05 - | 1 - | 4,51E-05 - | 1 - | 5,08E-05 - |
| 4783 | 1 | 0.2um_NOKCPHJN_107308 | Eukaryota | unclassified | Labyrinthulom | unclassified | Thraustochytr | Aurantiochytr | Aurantiochytr | cellular proces | Signal transdu | KOG0613 | fibronectin ty | 67593.Physo1 | 0,4 | 0,489897949 | 4,23E-07 | 5,19E-07 | 0 - | 0 - | 0 - | 0 - | 1 - | 1,05E-06 - | 1 - | 1,07E-06 - | 0 - | 0 - |
| 4784 | 1 | 3um_FIJJAFBF_431093 | Eukaryota | unclassified | Cryptophyta | Pyrenomonad | Geminigerace | Guillardia | Guillardia thet | poorly charact | Function unkn | 0T9G1 | kinetoplast-as | 353153.XP_80 | 0,2 | 0,4 | 1,21E-05 | 2,41E-05 | 0 - | 0 - | 0 - | 0 - | 0 - | 0 - | 0 - | 0 - | 1 - | 6,03E-05 - |
| 4785 | 1 | 3um_FIJJAFBF_762340 | Bacteria | Bacteroidetes | Cytophagia | Cytophagales | Flammeovirga | unclassified | Flammeovirga | unclassified | unclassified | unclassified | unclassified | unclassified | 0,2 | 0,4 | 1,85E-05 | 3,70E-05 | 1 - | 9,26E-05 - | 0 - | 0 - | 0 - | 0 - | 0 - | 0 - | 0 - | 0 - |
| 4786 | 1 | 0.2um_NOKCPHJN_520859 | Bacteria | Bacteroidetes | Flavobacteriia | unclassified | unclassified | unclassified | Flavobacteria | cellular proces | Intracellular tr | 0BIEB | domain-conta | 1005048.CFU_ | 2,4 | 1,019803903 | 3,47E-05 | 1,19E-05 | 4 - | 5,04E-05 - | 2 - | 3,13E-05 - | 2 - | 2,97E-05 - | 3 - | 4,53E-05 - | 1 - | 1,70E-05 - |
| 4787 | 1 | 3um_FIJJAFBF_240581 | Bacteria | Bacteroidetes | Flavobacteriia | Flavobacterial | Flavobacteria | Flaviramulus | Flaviramulus i | information st | Transcription | 0P8JP | Facilitates tra | 398720.MED2 | 0,6 | 0,489897949 | 1,40E-05 | 1,16E-05 | 1 - | 2,08E-05 - | 0 - | 0 - | 1 - | 2,45E-05 - | 1 - | 2,49E-05 - | 0 - | 0 - |
| 4788 | 1 | 3um_FIJJAFBF_111295 | unclassified | unclassified | unclassified | unclassified | unclassified | unclassified | unclassified | unclassified | unclassified | unclassified | unclassified | unclassified | 0,8 | 0,748331477 | 6,04E-05 | 5,31E-05 | 2 - | 0,000136639 - | 1 - | 8,48E-05 - | 1 - | 8,06E-05 - | 0 - | 0 - | 0 - | 0 - |
| 4789 | 1 | 3um_FIJJAFBF_55089 | Bacteria | Bacteroidetes | Flavobacteriia | Flavobacterial | Flavobacteria | Polaribacter | Polaribacter s | information st | Translation, ri | 0PA05 | 30S ribosomal | 313594.PI23P | 8 | 1,264911064 | 0,000604166 | 0,000108396 | 9 - | 0,000570713 - | 6 - | 0,000472438 - | 7 - | 0,000523964 - | 9 - | 0,000683619 - | 9 - | 0,000770094 - |
| 4790 | 1 | 0.2um_NOKCPHJN_506775 | Bacteria | Proteobacteri | Alphaproteob | Pelagibacteral | Pelagibactera | various | various | information st | Translation, ri | COG0184 | Forms an inte | 335992.SAR11 | 1 | 0 | 0,000167106 | 1,59E-05 | 1 - | 0,000139972 - | 1 - | 0,000173803 - | 1 - | 0,000165222 - | 1 - | 0,000167663 - | 1 - | 0,000188871 - |
| 4791 | 1 | 3um_FIJJAFBF_414322 | Bacteria | Bacteroidetes | Flavobacteriia | Flavobacterial | Flavobacteria | Formosa | Formosa sp. H | poorly charact | Function unkn | 09GJP | Protein of unk | 391603.FBALC | 1 | 0,632455532 | 9,40E-05 | 5,27E-05 | 2 - | 0,000162804 - | 1 - | 0,000101077 - | 1 - | 9,61E-05 - | 0 - | 0 - | 1 - | 0,00010984 - |
| 4792 | 2 | 0.2um_NOKCPHJN_182124;3um_FIJJAFBF_18075 | Bacteria | Bacteroidetes | Flavobacteriia | various | various | various | various | unclassified | unclassified | unclassified | unclassified | unclassified | 0,4 | 0,489897949 | 5,87E-05 | 7,23E-05 | 1 - | 0,000133461 - | 0 - | 0 - | 0 - | 0 - | 1 - | 0,000159864 - | 0 - | 0 - |
| 4793 | 1 | 3um_FIJJAFBF_20314 | Bacteria | Bacteroidetes | Flavobacteriia | Flavobacterial | Flavobacteria | Polaribacter | Polaribacter h | metabolism | Carbohydrate | COG0057 | glyceraldehyd | 762903.Pedsa | 16,6 | 1,496662955 | 0,000680089 | 6,28E-05 | 17 - | 0,000585947 - | 15 - | 0,000641977 - | 19 - | 0,000773022 - | 17 - | 0,000701867 - | 15 - | 0,000697633 - |
| 4794 | 1 | 0.2um_NOKCPHJN_144610 | Bacteria | Proteobacteri | Gammaprote | Cellvibrionales | Porticoccacea | unclassified | SAR92 bacteri | unclassified | unclassified | unclassified | unclassified | unclassified | 2 | 0 | 0,000118127 | 1,12E-05 | 2 - | 9,89E-05 - | 2 - | 0,000122861 - | 2 - | 0,000116795 - | 2 - | 0,00011852 - | 2 - | 0,000133512 - |
| 4795 | 2 | 3um_FIJJAFBF_118495;3um_FIJJAFBF_385377 | Bacteria | Proteobacteri | Gammaprote | various | various | various | various | poorly charact | Function unkn | COG0714 | ATPase activit | 247634.GPB2 | 1,6 | 0,489897949 | 7,83E-05 | 2,86E-05 | 1 - | 4,01E-05 - | 2 - | 9,97E-05 - | 1 - | 4,74E-05 - | 2 - | 9,61E-05 - | 2 - | 0,000108304 - |
| 4796 | 1 | 3um_FIJJAFBF_777326 | Bacteria | Bacteroidetes | Flavobacteriia | Flavobacterial | Flavobacteria | Formosa | Formosa sp. H | poorly charact | Function unkn | 05ERK | gliding motilit | 1046627.BZAF | 0,6 | 0,489897949 | 2,03E-05 | 1,68E-05 | 1 - | 2,96E-05 - | 1 - | 3,67E-05 - | 0 - | 0 - | 1 - | 3,54E-05 - | 0 - | 0 - |
| 4797 | 1 | 3um_FIJJAFBF_62015 | Bacteria | Proteobacteri | Alphaproteob | Rhodobactera | Rhodobactera | Thalassobacte | various | information st | Translation, ri | COG1185 | Involved in mF | 633131.TR2A | 16,2 | 1,469693846 | 0,000313609 | 5,30E-05 | 16 - | 0,000258288 - | 15 - | 0,000300673 - | 15 - | 0,000285827 - | 16 - | 0,000309386 - | 19 - | 0,00041387 - |
| 4798 | 1 | 0.2um_NOKCPHJN_925967 | Eukaryota | Chlorophyta | Mamiellophyc | Mamiellales | Bathycoccace | Bathycoccus | Bathycoccus p | poorly charact | Function unkn | 0JP5U | NA | 2850.Phatrdra | 0,2 | 0,4 | 4,34E-06 | 8,68E-06 | 0 - | 0 - | 1 - | 2,17E-05 - | 0 - | 0 - | 0 - | 0 - | 0 - | 0 - |
| 4799 | 1 | 3um_FIJJAFBF_301825 | Bacteria | Bacteroidetes | Flavobacteriia | Flavobacterial | Flavobacteria | Aureitalea | Aureitalea ma | metabolism | Inorganic ion t | 05C27 | TonB-depende | 398720.MED2 | 9,2 | 1,720465053 | 0,00012346 | 1,67E-05 | 12 - | 0,000136504 - | 8 - | 0,000112998 - | 10 - | 0,000134274 - | 7 - | 9,54E-05 - | 9 - | 0,000138144 - |
| 4800 | 1 | 3um_FIJJAFBF_120906 | Bacteria | Bacteroidetes | Cytophagia | Cytophagales | Flammeovirga | unclassified | Flammeovirga | information st | Translation, ri | 08UHH | Located on th | 643867.Ftrac_ | 1 | 0 | 0,000105405 | 1,00E-05 | 1 - | 8,83E-05 - | 1 - | 0,00010963 - | 1 - | 0,000104217 - | 1 - | 0,000105756 - | 1 - | 0,000119134 - |
| 4801 | 1 | 3um_FIJJAFBF_496816 | Bacteria | Proteobacteri | Gammaprote | Oceanospirilla | Saccharospirill | Reinekea | Reinekea fors | poorly charact | Function unkn | 0R3Z6 | NA | 167879.CPS_2 | 0,4 | 0,489897949 | 5,34E-05 | 6,66E-05 | 1 - | 0,00011364 - | 0 - | 0 - | 0 - | 0 - | 0 - | 0 - | 1 - | 0,000153341 - |
| 4802 | 1 | 3um_FIJJAFBF_605358 | Bacteria | Bacteroidetes | Flavobacteriia | Flavobacterial | Flavobacteria | Gilvibacter | Gilvibacter sp. | metabolism | Lipid transpor | 071I4 | Phosphatidyls | 50743.SCB49_ | 1 | 0 | 6,34E-05 | 6,02E-06 | 1 - | 5,31E-05 - | 1 - | 6,60E-05 - | 1 - | 6,27E-05 - | 1 - | 6,36E-05 - | 1 - | 7,17E-05 - |
| 4803 | 1 | 3um_FIJJAFBF_47109 | Bacteria | Bacteroidetes | Flavobacteriia | Flavobacterial | Flavobacteria | Cellulophaga | Cellulophaga s | metabolism | Amino acid tra | COG0174 | glutamine syn | 867900.Celly_ | 1 | 0 | 4,08E-05 | 3,87E-06 | 1 - | 3,42E-05 - | 1 - | 4,24E-05 - | 1 - | 4,03E-05 - | 1 - | 4,09E-05 - | 1 - | 4,61E-05 - |
| 4804 | 1 | 3um_FIJJAFBF_228149 | Bacteria | Proteobacteri | Gammaprote | unclassified | unclassified | unclassified | various | information st | Translation, ri | 0QTNF | One of the pri | 351348.Maqu | 2 | 0 | 0,000304505 | 2,89E-05 | 2 - | 0,000255059 - | 2 - | 0,000316709 - | 2 - | 0,000301072 - | 2 - | 0,000305518 - | 2 - | 0,000344166 - |
| 4805 | 1 | 0.2um_NOKCPHJN_103497 | Bacteria | Proteobacteri | Alphaproteob | Rhodobactera | unclassified | unclassified | Rhodobactera | unclassified | unclassified | unclassified | unclassified | unclassified | 0,8 | 0,4 | 7,76E-05 | 3,91E-05 | 0 - | 0 - | 1 - | 9,70E-05 - | 1 - | 9,22E-05 - | 1 - | 9,35E-05 - | 1 - | 0,000105357 - |
| 4806 | 2 | 0.2um_NOKCPHJN_64605;3um_FIJJAFBF_72030 | Bacteria | Bacteroidetes | unclassified | unclassified | unclassified | unclassified | Bacteroidetes | unclassified | unclassified | unclassified | unclassified | unclassified | 0,4 | 0,489897949 | 2,43E-05 | 2,99E-05 | 1 - | 5,57E-05 - | 0 - | 0 - | 1 - | 6,58E-05 - | 0 - | 0 - | 0 - | 0 - |
| 4807 | 1 | 3um_FIJJAFBF_736140 | Bacteria | Bacteroidetes | Flavobacteriia | various | various | various | various | information st | Translation, ri | 09FE5 | One of the pr | 487797.Flav3 | 1 | 0,894427191 | 0,000123661 | 0,000108555 | 2 - | 0,000220725 - | 1 - | 0,000137037 - | 2 - | 0,000260543 - | 0 - | 0 - | 0 - | 0 - |
| 4808 | 2 | 0.2um_NOKCPHJN_292903;0.2um_NOKCPHJN_8 | Bacteria | Bacteroidetes | Flavobacteriia | Flavobacterial | Flavobacteria | Polaribacter | various | information st | Translation, ri | 05C22 | threonyL-tRN | 313598.MED1 | 8 | 0,894427191 | 0,000168772 | 1,79E-05 | 9 - | 0,000159906 - | 7 - | 0,000154432 - | 9 - | 0,000188752 - | 7 - | 0,000148976 - | 8 - | 0,000191795 - |
| 4809 | 1 | 0.2um_NOKCPHJN_770232 | Eukaryota | unclassified | Pelagophycea | Pelagomonad | unclassified | Aureococcus | Aureococcus a | cellular proces | Posttranslatio | KOG0019 | heat shock pr | 272952.HpaP | 5,6 | 1,019803903 | 9,98E-05 | 2,31E-05 | 5 - | 7,41E-05 - | 6 - | 0,00011048 - | 4 - | 7,00E-05 - | 7 - | 0,000124339 - | 6 - | 0,000120058 - |
| 4810 | 1 | 0.2um_NOKCPHJN_102310 | Bacteria | Bacteroidetes | Flavobacteriia | various | various | various | various | information st | Translation, ri | 0P97G | This is one of | 487797.Flav3 | 2,8 | 0,4 | 0,000182219 | 3,84E-05 | 2 - | 0,000107772 - | 3 - | 0,000200731 - | 3 - | 0,00019082 - | 3 - | 0,000193638 - | 3 - | 0,000218133 - |
| 4811 | 1 | 0.2um_NOKCPHJN_159990 | Bacteria | Proteobacteri | Gammaprote | Oceanospirilla | Saccharospirill | Reinekea | Reinekea fors | unclassified | unclassified | unclassified | unclassified | unclassified | 14,6 | 2,154065923 | 0,000627336 | 8,78E-05 | 18 - | 0,000651729 - | 13 - | 0,000584462 - | 14 - | 0,000598344 - | 12 - | 0,000520442 - | 16 - | 0,000781701 - |
| 4812 | 1 | 3um_FIJJAFBF_788038 | Bacteria | Proteobacteri | Alphaproteob | Rhodobactera | unclassified | unclassified | Rhodobactera | information st | Translation, ri | COG0255 | 50s ribosomal | 644076.SCH4 | 1 | 0 | 0,000207617 | 1,97E-05 | 1 - | 0,000173904 - | 1 - | 0,000215938 - | 1 - | 0,000205276 - | 1 - | 0,000208308 - | 1 - | 0,000234658 - |
| 4813 | 1 | 3um_FIJJAFBF_254365 | Viruses | unclassified | unclassified | unclassified | Phycodnavirid | unclassified | Pyramimonas | unclassified | unclassified | unclassified | unclassified | unclassified | 5 | 1,264911064 | 0,000179261 | 3,71E-05 | 6 - | 0,000183643 - | 4 - | 0,00015202 - | 6 - | 0,000216772 - | 6 - | 0,000219973 - | 3 - | 0,0001239 - |
| 4814 | 1 | 3um_FIJJAFBF_276043 | Bacteria | Bacteroidetes | Flavobacteriia | unclassified | unclassified | unclassified | Flavobacteria | cellular proces | Cell wall/mem | 0P9R9 | TonB family | 487797.Flav3 | 1,2 | 0,979795897 | 6,38E-05 | 4,63E-05 | 3 - | 0,000144677 - | 1 - | 5,99E-05 - | 1 - | 5,69E-05 - | 1 - | 5,78E-05 - | 0 - | 0 - |
| 4815 | 1 | 0.2um_NOKCPHJN_318027 | Bacteria | Proteobacteri | Alphaproteob | unclassified | unclassified | unclassified | Alphaproteob | metabolism | Energy produc | 16TH8 | malate L-lacta | 488538.SAR11 | 1 | 0 | 4,08E-05 | 3,87E-06 | 1 - | 3,42E-05 - | 1 - | 4,24E-05 - | 1 - | 4,03E-05 - | 1 - | 4,09E-05 - | 1 - | 4,61E-05 - |
| 4816 | 1 | 3um_FIJJAFBF_347891 | Eukaryota | unclassified | Cryptophyta | Pyrenomonad | Geminigerace | Guillardia | Guillardia thet | unclassified | unclassified | unclassified | unclassified | unclassified | 2,8 | 0,748331477 | 0,000159877 | 4,17E-05 | 3 - | 0,000144677 - | 3 - | 0,000179646 - | 2 - | 0,000113851 - | 4 - | 0,000231064 - | 2 - | 0,000130147 - |
| 4817 | 1 | 0.2um_NOKCPHJN_492829 | Bacteria | Bacteroidetes | Flavobacteriia | Flavobacterial | Flavobacteria | Formosa | Formosa sp. H | poorly charact | Function unkn | 0PAEN | NA | 886377.Murru | 7 | 1,095445115 | 0,000200053 | 3,94E-05 | 7 - | 0,000167383 - | 5 - | 0,000148457 - | 8 - | 0,000225804 - | 7 - | 0,000200496 - | 8 - | 0,000258124 - |
| 4818 | 1 | 3um_FIJJAFBF_210878 | Bacteria | Bacteroidetes | Flavobacteriia | unclassified | unclassified | unclassified | Flavobacteria | cellular proces | Posttranslatio | 09EE9 | Participates a | 487797.Flav3 | 1 | 0 | 6,96E-05 | 6,60E-06 | 1 - | 5,83E-05 - | 1 - | 7,23E-05 - | 1 - | 6,88E-05 - | 1 - | 6,98E-05 - | 1 - | 7,86E-05 - |
| 4819 | 1 | 3um_FIJJAFBF_304726 | Bacteria | Bacteroidetes | Flavobacteriia | Flavobacterial | Flavobacteria | Polaribacter | Polaribacter s | poorly charact | Function unkn | 0XQTX | Chromosome | 313594.PI23P | 1,2 | 0,4 | 1,57E-05 | 5,27E-06 | 1 - | 1,10E-05 - | 1 - | 1,37E-05 - | 2 - | 2,60E-05 - | 1 - | 1,32E-05 - | 1 - | 1,48E-05 - |
| 4820 | 2 | 0.2um_NOKCPHJN_544266;3um_FIJJAFBF_0469 | Bacteria | Bacteroidetes | Flavobacteriia | Flavobacterial | Flavobacteria | Ulvibacter | Ulvibacter sp. | cellular proces | Cell cycle cont | 05CAK | Cell division A | various | 0,2 | 0,4 | 1,36E-05 | 2,73E-05 | 0 - | 0 - | 0 - | 0 - | 0 - | 0 - | 0 - | 0 - | 1 - | 6,82E-05 - |
| 4821 | 1 | 3um_FIJJAFBF_73822 | Bacteria | Bacteroidetes | Flavobacteriia | Flavobacterial | Flavobacteria | Polaribacter | Polaribacter s | poorly charact | Function unkn | 0XNQI | domain protei | 313594.PI23P | 4 | 0,632455532 | 0,00012224 | 1,63E-05 | 5 - | 0,000128963 - | 4 - | 0,000128107 - | 3 - | 9,13E-05 - | 4 - | 0,00012358 - | 4 - | 0,000139213 - |
| 4822 | 1 | 0.2um_NOKCPHJN_457287 | Bacteria | Bacteroidetes | Flavobacteriia | unclassified | unclassified | unclassified | Flavobacteria | information st | Replication, re | 0PB0H | DNA-binding | 487796.Flav2 | 0,8 | 0,979795897 | 0,000121318 | 0,00014859 | 0 - | 0 - | 0 - | 0 - | 2 - | 0,000301072 - | 2 - | 0,000305518 - | 0 - | 0 - |
| 4823 | 1 | 3um_FIJJAFBF_373422 | Bacteria | Actinobacteria | Actinobacteria | Micrococcales | Microbacteria | Pontimonas | Pontimonas sa | metabolism | Carbohydrate | 0NE0D | PTS system m | 621372.POTG | 2 | 0 | 6,90E-05 | 6,55E-06 | 2 - | 5,78E-05 - | 2 - | 7,18E-05 - | 2 - | 6,83E-05 - | 2 - | 6,93E-05 - | 2 - | 7,80E-05 - |
| 4824 | 1 | 0.2um_NOKCPHJN_644354 | Bacteria | Proteobacteri | Betaproteoba | Burkholderial | Comamonada | Rhodoferax | Rhodoferax sa | poorly charact | Function unkn | 01XSR | ghmp kinase | 224911.blr597 | 0,4 | 0,489897949 | 1,10E-05 | 1,36E-05 | 1 - | 2,51E-05 - | 0 - | 0 - | 0 - | 0 - | 1 - | 3,01E-05 - | 0 - | 0 - |
| 4825 | 1 | 0.2um_NOKCPHJN_1072722 | unclassified | unclassified | unclassified | unclassified | unclassified | unclassified | unclassified | unclassified | unclassified | unclassified | unclassified | unclassified | 1,8 | 0,4 | 0,000326102 | 9,02E-05 | 1 - | 0,000149061 - | 2 - | 0,000370179 - | 2 - | 0,000351902 - | 2 - | 0,000357099 - | 2 - | 0,000402271 - |
| 4826 | 1 | 3um_FIJJAFBF_142849 | Bacteria | Proteobacteri | Alphaproteob | Rhodobactera | unclassified | unclassified | Rhodobactera | cellular proces | Cell wall/mem | 16QB0 | Part of the ou | 439497.RR11_ | 2,8 | 0,4 | 4,92E-05 | 7,97E-06 | 3 - | 4,43E-05 - | 2 - | 3,67E-05 - | 3 - | 5,23E-05 - | 3 - | 5,31E-05 - | 3 - | 5,98E-05 - |
| 4827 | 1 | 3um_FIJJAFBF_85816 | Viruses | unclassified | unclassified | unclassified | various | various | various | unclassified | unclassified | unclassified | unclassified | unclassified | 1,4 | 0,8 | 9,95E-05 | 5,48E-05 | 2 - | 0,000122756 - | 0 - | 0 - | 2 - | 0,000144901 - | 2 - | 0,000147041 - | 1 - | 8,28E-05 - |
| 4828 | 1 | 3um_FIJJAFBF_260544 | Bacteria | Proteobacteri | Alphaproteob | unclassified | unclassified | unclassified | Alphaproteob | information st | Translation, ri | COG1185 | Involved in mF | 488538.SAR11 | 1,4 | 0,489897949 | 2,74E-05 | 1,01E-05 | 1 - | 1,63E-05 - | 2 - | 4,05E-05 - | 2 - | 3,85E-05 - | 1 - | 1,96E-05 - | 1 - | 2,20E-05 - |
| 4829 | 1 | 0.2um_NOKCPHJN_09370 | Bacteria | Bacteroidetes | Flavobacteriia | Flavobacterial | Flavobacteria | Polaribacter | Polaribacter s | cellular proces | Cell wall/mem | COG2885 | Ompa motb d | 313598.MED1 | 1,8 | 0,4 | 5,43E-05 | 1,27E-05 | 2 - | 5,08E-05 - | 1 - | 3,15E-05 - | 2 - | 5,99E-05 - | 2 - | 6,08E-05 - | 2 - | 6,85E-05 - |
| 4830 | 2 | 3um_FIJJAFBF_231203;3um_FIJJAFBF_657054 | Bacteria | Bacteroidetes | Flavobacteriia | Flavobacterial | Flavobacteria | Polaribacter | various | unclassified | unclassified | unclassified | unclassified | unclassified | 0,2 | 0,4 | 1,01E-05 | 2,01E-05 | 1 - | 5,03E-05 - | 0 - | 0 - | 0 - | 0 - | 0 - | 0 - | 0 - | 0 - |
| 4831 | 1 | 3um_FIJJAFBF_208517 | Bacteria | Proteobacteri | Gammaprote | Oceanospirilla | Saccharospirill | Reinekea | Reinekea fors | unclassified | unclassified | unclassified | unclassified | unclassified | 0,8 | 0,4 | 0,000102365 | 5,26E-05 | 1 - | 0,000107268 - | 1 - | 0,000133195 - | 1 - | 0,000126619 - | 0 - | 0 - | 1 - | 0,000144743 - |
| 4832 | 1 | 3um_FIJJAFBF_43525 | Bacteria | Bacteroidetes | Flavobacteriia | Flavobacterial | Flavobacteria | Cellulophaga | Cellulophaga t | information st | Transcription | 05D27 | DNA-depende | 867900.Celly_ | 19,8 | 1,833030278 | 0,000189725 | 2,75E-05 | 20 - | 0,000160303 - | 23 - | 0,000228906 - | 18 - | 0,000170299 - | 18 - | 0,000172814 - | 20 - | 0,000216305 - |
| 4833 | 1 | 3um_FIJJAFBF_63617 | Bacteria | Proteobacteri | Gammaprote | Cellvibrionales | Spongiibacter | Zhongshania | Zhongshania s | cellular proces | Posttranslatio | 0HFM8 | Peptidylprolyl | 485915.Dret_ | 0,8 | 0,4 | 4,58E-05 | 2,36E-05 | 1 - | 4,80E-05 - | 1 - | 5,96E-05 - | 1 - | 5,67E-05 - | 0 - | 0 - | 1 - | 6,48E-05 - |
| 4834 | 1 | 3um_FIJJAFBF_95715 | Eukaryota | unclassified | Raphidophyce | Chattonellales | Chattonellace | Heterosigma | Heterosigma a | metabolism | Energy produc | 0IFBF | One of the tw | 45157.CMV08 | 3,2 | 0,4 | 0,000117851 | 2,03E-05 | 3 - | 9,23E-05 - | 4 - | 0,000152835 - | 3 - | 0,000108967 - | 3 - | 0,000110576 - | 3 - | 0,000124564 - |
| 4835 | 1 | 0.2um_NOKCPHJN_873473 | Bacteria | Bacteroidetes | Flavobacteriia | Flavobacterial | Flavobacteria | Mangrovimon | Mangrovimon | unclassified | unclassified | unclassified | unclassified | unclassified | 0,2 | 0,4 | 2,23E-05 | 4,45E-05 | 0 - | 0 - | 1 - | 0,000111343 - | 0 - | 0 - | 0 - | 0 - | 0 - | 0 - |
| 4836 | 1 | 3um_FIJJAFBF_164234 | Bacteria | Bacteroidetes | Flavobacteriia | Flavobacterial | Crocinitomica | unclassified | Crocinitomica | unclassified | unclassified | unclassified | unclassified | unclassified | 1,8 | 0,748331477 | 8,98E-05 | 3,02E-05 | 3 - | 0,000129447 - | 2 - | 0,000107157 - | 1 - | 5,09E-05 - | 2 - | 0,000103371 - | 1 - | 5,82E-05 - |
| 4837 | 1 | 0.2um_NOKCPHJN_515151 | Bacteria | Proteobacteri | Alphaproteob | Rickettsiales | unclassified | unclassified | Rickettsiales b | unclassified | unclassified | unclassified | unclassified | unclassified | 5,8 | 0,4 | 0,000203863 | 2,50E-05 | 6 - | 0,00017658 - | 6 - | 0,00021926 - | 5 - | 0,000173695 - | 6 - | 0,000211513 - | 6 - | 0,000238268 - |
| 4838 | 1 | 3um_FIJJAFBF_459236 | Bacteria | Proteobacteri | Alphaproteob | Rhodobactera | unclassified | unclassified | Rhodobactera | information st | Translation, ri | 01S8S | This is 1 of the | 367336.OM22 | 3,8 | 0,748331477 | 0,000283718 | 5,91E-05 | 4 - | 0,000250878 - | 3 - | 0,000233637 - | 5 - | 0,00037017 - | 3 - | 0,000225382 - | 4 - | 0,000338523 - |
| 4839 | 1 | 0.2um_NOKCPHJN_160027 | Bacteria | Proteobacteri | Alphaproteob | Rhodobactera | unclassified | unclassified | Rhodobactera | information st | Translation, ri | 01WSM | This is one of | 367336.OM22 | 1 | 0,632455532 | 6,01E-05 | 3,36E-05 | 2 - | 0,00010387 - | 1 - | 6,45E-05 - | 0 - | 0 - | 1 - | 6,22E-05 - | 1 - | 7,01E-05 - |
| 4840 | 1 | 3um_FIJJAFBF_629416 | Bacteria | Bacteroidetes | Flavobacteriia | Flavobacterial | Flavobacteria | Polaribacter | Polaribacter s | unclassified | unclassified | unclassified | unclassified | unclassified | 3,4 | 0,8 | 8,28E-05 | 2,01E-05 | 3 - | 6,13E-05 - | 3 - | 7,61E-05 - | 5 - | 0,000120536 - | 3 - | 7,34E-05 - | 3 - | 8,27E-05 - |
| 4841 | 1 | 3um_FIJJAFBF_503587 | unclassified | unclassified | unclassified | unclassified | unclassified | unclassified | unclassified | unclassified | unclassified | unclassified | unclassified | unclassified | 0,2 | 0,4 | 1,34E-05 | 2,68E-05 | 1 - | 6,71E-05 - | 0 - | 0 - | 0 - | 0 - | 0 - | 0 - | 0 - | 0 - |
| 4842 | 1 | 3um_FIJJAFBF_61424 | Bacteria | Bacteroidetes | Flavobacteriia | Flavobacterial | Flavobacteria | Algibacter | Algibacter pe | unclassified | unclassified | unclassified | unclassified | unclassified | 0,6 | 0,8 | 1,60E-05 | 2,04E-05 | 2 - | 4,93E-05 - | 1 - | 3,06E-05 - | 0 - | 0 - | 0 - | 0 - | 0 - | 0 - |
| 4843 | 1 | 3um_FIJJAFBF_151113 | Bacteria | Bacteroidetes | Flavobacteriia | Flavobacterial | Flavobacteria | Polaribacter | Polaribacter s | unclassified | unclassified | unclassified | unclassified | unclassified | 0,8 | 0,748331477 | 4,15E-05 | 3,58E-05 | 2 - | 9,04E-05 - | 1 - | 5,61E-05 - | 0 - | 0 - | 0 - | 0 - | 1 - | 6,10E-05 - |
| 4844 | 1 | 0.2um_NOKCPHJN_285541 | Bacteria | Proteobacteri | Alphaproteob | Rhodospirillal | Rhodospirillac | Candidatus En | Candidatus En | unclassified | unclassified | unclassified | unclassified | unclassified | 0,6 | 0,489897949 | 3,98E-06 | 3,27E-06 | 1 - | 5,89E-06 - | 0 - | 0 - | 1 - | 6,95E-06 - | 1 - | 7,05E-06 - | 0 - | 0 - |
| 4845 | 1 | 3um_FIJJAFBF_295399 | Bacteria | Proteobacteri | Betaproteoba | unclassified | unclassified | unclassified | Betaproteoba | unclassified | unclassified | unclassified | unclassified | unclassified | 1 | 0,632455532 | 6,03E-05 | 3,44E-05 | 2 - | 0,000107268 - | 1 - | 6,66E-05 - | 1 - | 6,33E-05 - | 1 - | 6,42E-05 - | 0 - | 0 - |
| 4846 | 1 | 0.2um_NOKCPHJN_639687 | Bacteria | Bacteroidetes | Flavobacteriia | Flavobacterial | Flavobacteria | Winogradskye | Winogradskye | metabolism | Carbohydrate | 08IUS | Transketolase | 313590.MED1 | 2 | 0,632455532 | 8,54E-05 | 2,65E-05 | 2 - | 7,24E-05 - | 2 - | 8,99E-05 - | 2 - | 8,55E-05 - | 3 - | 0,00013011 - | 1 - | 4,89E-05 - |
| 4847 | 1 | 0.2um_NOKCPHJN_62012 | Bacteria | Bacteroidetes | Cytophagia | Cytophagales | Flammeovirga | Roseivirga | Roseivirga spo | metabolism | Energy produc | COG0056 | Produces ATP | 643867.Ftrac_ | 5,2 | 1,469693846 | 0,000133209 | 3,14E-05 | 7 - | 0,000152745 - | 6 - | 0,000162569 - | 6 - | 0,000154542 - | 3 - | 7,84E-05 - | 4 - | 0,000117775 - |
| 4848 | 1 | 0.2um_NOKCPHJN_811890 | Eukaryota | unclassified | Cryptophyta | Pyrenomonad | Geminigerace | Guillardia | Guillardia thet | metabolism | Energy produc | KOG2799 | beta' subunit | 423536.XP_00 | 1 | 0 | 4,51E-05 | 4,28E-06 | 1 - | 3,78E-05 - | 1 - | 4,69E-05 - | 1 - | 4,46E-05 - | 1 - | 4,52E-05 - | 1 - | 5,09E-05 - |
| 4849 | 1 | 3um_FIJJAFBF_783853 | Bacteria | Bacteroidetes | Flavobacteriia | Flavobacterial | Flavobacteria | Polaribacter | Polaribacter s | unclassified | unclassified | unclassified | unclassified | unclassified | 21,6 | 2,059126028 | 0,000347813 | 2,95E-05 | 24 - | 0,000325608 - | 24 - | 0,000404309 - | 21 - | 0,000336303 - | 20 - | 0,00032502 - | 19 - | 0,000347827 - |
| 4850 | 2 | 0.2um_NOKCPHJN_727728;0.2um_NOKCPHJN_8 | Eukaryota | various | various | various | various | various | various | information st | Translation, ri | KOG0886 | Ribosomal pr | 35128.Thaps3 | 1 | 0 | 8,30E-05 | 7,89E-06 | 1 - | 6,96E-05 - | 1 - | 8,64E-05 - | 1 - | 8,21E-05 - | 1 - | 8,33E-05 - | 1 - | 9,39E-05 - |
| 4851 | 1 | 3um_FIJJAFBF_427162 | Bacteria | Proteobacteri | Alphaproteob | Rhodobactera | Rhodobactera | Planktomarina | Planktomarina | metabolism | Amino acid tra | 01SEG | branched-cha | 314262.MED1 | 1 | 0 | 3,22E-05 | 3,05E-06 | 1 - | 2,69E-05 - | 1 - | 3,35E-05 - | 1 - | 3,18E-05 - | 1 - | 3,23E-05 - | 1 - | 3,64E-05 - |
| 4852 | 1 | 3um_FIJJAFBF_72569 | Bacteria | Bacteroidetes | Flavobacteriia | Flavobacterial | Flavobacteria | Formosa | Formosa sp. H | metabolism | Energy produc | COG0055 | Produces ATP | 391603.FBALC | 0,6 | 0,489897949 | 5,65E-05 | 4,70E-05 | 1 - | 7,86E-05 - | 1 - | 9,76E-05 - | 0 - | 0 - | 0 - | 0 - | 1 - | 0,000106078 - |
| 4853 | 1 | 0.2um_NOKCPHJN_516482 | Bacteria | Bacteroidetes | Flavobacteriia | Flavobacterial | Flavobacteria | unclassified | Flavobacteria | information st | Replication, re | 0PB0H | DNA-binding | 487796.Flav2 | 1,8 | 0,748331477 | 0,000268524 | 0,000106309 | 2 - | 0,000255059 - | 1 - | 0,000158354 - | 3 - | 0,000451607 - | 2 - | 0,000305518 - | 1 - | 0,000172083 - |
| 4854 | 1 | 0.2um_NOKCPHJN_813867 | Eukaryota | unclassified | Pelagophycea | Pelagomonad | unclassified | Aureococcus | Aureococcus a | cellular proces | Cytoskeleton | 0DGS2 | Tubulin is the | 7897.ENSLAC | 1,4 | 0,489897949 | 4,16E-05 | 1,31E-05 | 2 - | 5,07E-05 - | 2 - | 6,29E-05 - | 1 - | 2,99E-05 - | 1 - | 3,03E-05 - | 1 - | 3,42E-05 - |
| 4855 | 1 | 3um_FIJJAFBF_588389 | Bacteria | Bacteroidetes | Flavobacteriia | Flavobacterial | Flavobacteria | Algibacter | Algibacter sp. | information st | Translation, ri | 05C22 | threonyL-tRN | 983544.Lacal_ | 1,2 | 0,4 | 2,55E-05 | 8,81E-06 | 1 - | 1,78E-05 - | 1 - | 2,21E-05 - | 1 - | 2,10E-05 - | 2 - | 4,27E-05 - | 1 - | 2,40E-05 - |
| 4856 | 2 | 0.2um_NOKCPHJN_108830;3um_FIJJAFBF_2958 | Bacteria | Proteobacteri | Alphaproteob | Rhodobactera | Rhodobactera | unclassified | Rhodobactera | unclassified | unclassified | unclassified | unclassified | unclassified | 0,4 | 0,489897949 | 4,10E-05 | 5,05E-05 | 1 - | 9,41E-05 - | 0 - | 0 - | 1 - | 0,000111051 - | 0 - | 0 - | 0 - | 0 - |

| | | | | | | | | | | | | | | | | | | | | | | | | | | | | |
|---|---|---|---|---|---|---|---|---|---|---|---|---|---|---|---|---|---|---|---|---|---|---|---|---|---|---|---|---|
| 4857 | 1 | 3um_FIJJAFBF_396471 | Bacteria | Bacteroidetes | Flavobacteriia | Flavobacterial | Flavobacteriac | Formosa | Formosa sp. H | unclassified | unclassified | unclassified | unclassified | unclassified | 1 | 0 | 1,68E-05 | 1,59E-06 | 1 - | 1,40E-05 - | 1 - | 1,74E-05 - | 1 - | 1,66E-05 - | 1 - | 1,68E-05 - | 1 - | 1,90E-05 - |
| 4858 | 2 | 0.2um_NOKCPHJN_530547;3um_FIJJAFBF_38923 | Eukaryota | various | various | various | various | various | various | information st | Chromatin str | KOG3467 | Core compone | various | 8,8 | 1,166190379 | 0,000811964 | 5,74E-05 | 11 - | 0,000858874 - | 9 - | 0,000872564 - | 8 - | 0,000737318 - | 8 - | 0,000748208 - | 8 - | 0,000842854 - |
| 4859 | 1 | 3um_FIJJAFBF_98259 | unclassified | unclassified | unclassified | unclassified | unclassified | unclassified | unclassified | unclassified | unclassified | unclassified | unclassified | unclassified | 17,6 | 2,727636339 | 0,000183407 | 1,26E-05 | 23 - | 0,000203379 - | 16 - | 0,000175678 - | 16 - | 0,000167004 - | 17 - | 0,000180063 - | 16 - | 0,000190908 - |
| 4860 | 1 | 3um_FIJJAFBF_70345 | Bacteria | Proteobacteri | Betaproteoba | unclassified | unclassified | unclassified | Betaproteoba | unclassified | unclassified | unclassified | unclassified | unclassified | 1,2 | 0,748331477 | 0,000115639 | 6,73E-05 | 2 - | 0,000166343 - | 1 - | 0,000103275 - | 2 - | 0,000196351 - | 0 - | 0 - | 1 - | 0,000112228 - |
| 4861 | 1 | 3um_FIJJAFBF_365122 | Bacteria | Bacteroidetes | Flavobacteriia | Flavobacterial | Flavobacteriac | Winogradskye | Winogradskye | unclassified | unclassified | unclassified | unclassified | unclassified | 0,6 | 0,8 | 4,73E-05 | 6,14E-05 | 0 - | 0 - | 0 - | 0 - | 2 - | 0,000150536 - | 0 - | 0 - | 1 - | 8,60E-05 - |
| 4862 | 1 | 0.2um_NOKCPHJN_156663 | Bacteria | Bacteroidetes | Flavobacteriia | unclassified | unclassified | unclassified | Flavobacteria | unclassified | unclassified | unclassified | unclassified | unclassified | 17,2 | 2,13541565 | 0,000237083 | 1,39E-05 | 21 - | 0,00024495 - | 15 - | 0,000217254 - | 18 - | 0,000247833 - | 16 - | 0,00022355 - | 16 - | 0,000251828 - |
| 4863 | 1 | 0.2um_NOKCPHJN_882210 | Bacteria | Proteobacteri | Alphaproteob | Pelagibacteral | Pelagibacterac | Candidatus Pe | Candidatus Pe | unclassified | unclassified | unclassified | unclassified | unclassified | 0,8 | 0,4 | 3,32E-05 | 1,67E-05 | 0 - | 0 - | 1 - | 4,14E-05 - | 1 - | 3,94E-05 - | 1 - | 4,00E-05 - | 1 - | 4,50E-05 - |
| 4864 | 1 | 3um_FIJJAFBF_156011 | Eukaryota | Chlorophyta | Chlorophycea | Chlamydomor | Volvocaceae | Gonium | Gonium pecto | unclassified | unclassified | unclassified | unclassified | unclassified | 1,6 | 1,019803903 | 4,72E-05 | 3,05E-05 | 2 - | 4,98E-05 - | 3 - | 9,27E-05 - | 0 - | 0 - | 2 - | 5,96E-05 - | 1 - | 3,36E-05 - |
| 4865 | 1 | 0.2um_NOKCPHJN_891694 | Bacteria | Proteobacteri | Alphaproteob | Pelagibacteral | unclassified | unclassified | Pelagibacteral | cellular proces | Cell wall/mem | 7257 | NA | 439493.PB721 | 1,2 | 0,4 | 5,71E-05 | 2,43E-05 | 1 - | 3,90E-05 - | 1 - | 4,85E-05 - | 1 - | 4,61E-05 - | 1 - | 4,68E-05 - | 2 - | 0,000105357 - |
| 4866 | 1 | 3um_FIJJAFBF_532050 | Bacteria | Bacteroidetes | Flavobacteriia | Flavobacterial | Flavobacteriac | Formosa | Formosa sp. H | metabolism | Amino acid tra | 09GJ6 | nitrogen regul | 391603.FBALC | 0,6 | 0,489897949 | 7,64E-05 | 6,27E-05 | 0 - | 0 - | 0 - | 0 - | 1 - | 0,000120966 - | 1 - | 0,000122753 - | 1 - | 0,000138281 - |
| 4867 | 1 | 0.2um_NOKCPHJN_390972 | Bacteria | Proteobacteri | Gammaprotec | unclassified | unclassified | unclassified | Gammaprotec | unclassified | unclassified | unclassified | unclassified | unclassified | 1,6 | 0,489897949 | 5,55E-05 | 1,57E-05 | 2 - | 5,90E-05 - | 2 - | 7,33E-05 - | 1 - | 3,48E-05 - | 2 - | 7,07E-05 - | 1 - | 3,98E-05 - |
| 4868 | 1 | 3um_FIJJAFBF_666650 | Bacteria | Bacteroidetes | Flavobacteriia | Flavobacterial | Flavobacteriac | Hyunsoonleell | Hyunsoonleell | metabolism | Energy produc | 05D26 | NQR complex | 391587.KAOT1 | 0,6 | 0,489897949 | 1,78E-05 | 1,46E-05 | 1 - | 2,63E-05 - | 0 - | 0 - | 1 - | 3,11E-05 - | 1 - | 3,15E-05 - | 0 - | 0 - |
| 4869 | 1 | 3um_FIJJAFBF_680340 | Bacteria | Proteobacteri | Gammaprotec | Oceanospirilla | Saccharospirill | Reinekea | Reinekea fors | cellular proces | Cell motility | COG1261 | basal body p-r | 314283.MED2 | 0,2 | 0,4 | 1,00E-05 | 2,00E-05 | 0 - | 0 - | 0 - | 0 - | 1 - | 5,00E-05 - | 0 - | 0 - | 0 - | 0 - |
| 4870 | 1 | 0.2um_NOKCPHJN_31720 | Bacteria | Proteobacteri | Betaproteoba | unclassified | unclassified | unclassified | Betaproteoba | metabolism | Amino acid tra | COG2171 | Catalyzes the | 232721.Ajs_20 | 3,4 | 1,019803903 | 0,000165516 | 3,55E-05 | 5 - | 0,000209447 - | 3 - | 0,000156043 - | 3 - | 0,000148338 - | 4 - | 0,000200706 - | 2 - | 0,000113047 - |
| 4871 | 1 | 0.2um_NOKCPHJN_750086 | Bacteria | Proteobacteri | Alphaproteob | Pelagibacteral | Pelagibacterac | Candidatus Pe | Candidatus Pe | metabolism | Energy produc | 01RNF | Produces ATP | 439493.PB721 | 0,6 | 0,489897949 | 1,73E-05 | 1,43E-05 | 1 - | 2,43E-05 - | 0 - | 0 - | 0 - | 0 - | 1 - | 2,91E-05 - | 1 - | 3,28E-05 - |
| 4872 | 1 | 3um_FIJJAFBF_520466 | Bacteria | Bacteroidetes | Flavobacteriia | Flavobacterial | Flavobacteriac | Polaribacter | Polaribacter s | metabolism | Energy produc | 05CMV | Succinyl-CoA s | 391587.KAOT1 | 3,8 | 1,166190379 | 0,000127345 | 3,24E-05 | 5 - | 0,000143831 - | 3 - | 0,000107157 - | 4 - | 0,000135822 - | 5 - | 0,000172285 - | 2 - | 7,76E-05 - |
| 4873 | 1 | 3um_FIJJAFBF_48889 | Eukaryota | unclassified | Oomycetes | Saprolegniales | Saprolegniace | Aphanomyces | various | unclassified | unclassified | unclassified | unclassified | unclassified | 3,2 | 0,748331477 | 0,000305361 | 5,52E-05 | 4 - | 0,000325608 - | 3 - | 0,000303232 - | 3 - | 0,00028826 - | 4 - | 0,000390024 - | 2 - | 0,00021968 - |
| 4874 | 1 | 3um_FIJJAFBF_172653 | Bacteria | Proteobacteri | Alphaproteob | Pelagibacteral | Pelagibacterac | Candidatus Pe | Candidatus Pe | metabolism | Amino acid tra | 01VSB | Ectoine hydro | 439493.PB721 | 7 | 0,632455532 | 0,000336103 | 9,82E-06 | 8 - | 0,000324457 - | 7 - | 0,00035252 - | 7 - | 0,000335115 - | 7 - | 0,000340065 - | 6 - | 0,000328356 - |
| 4875 | 3 | 0.2um_NOKCPHJN_142296;0.2um_NOKCPHJN_6 | Bacteria | Bacteroidetes | Flavobacteriia | Flavobacterial | Flavobacteriac | various | various | various | various | various | various | various | 0,4 | 0,489897949 | 3,00E-06 | 3,69E-06 | 1 - | 6,82E-06 - | 0 - | 0 - | 0 - | 0 - | 1 - | 8,16E-06 - | 0 - | 0 - |
| 4876 | 1 | 3um_FIJJAFBF_218547 | Bacteria | Proteobacteri | Alphaproteob | Rhodobactera | Rhodobactera | Planktotalea | Planktotalea f | cellular proces | Posttranslatio | COG0330 | Band 7 protei | 314270.RB208 | 0,6 | 0,489897949 | 2,71E-05 | 2,23E-05 | 1 - | 3,96E-05 - | 1 - | 4,91E-05 - | 1 - | 4,67E-05 - | 0 - | 0 - | 0 - | 0 - |
| 4877 | 1 | 0.2um_NOKCPHJN_1021140 | Bacteria | Actinobacteria | Actinobacteria | Micrococcales | Microbacteria | Pontimonas | Pontimonas s | metabolism | Amino acid tra | 00DZF | branched-chai | 312284.A20C1 | 1,4 | 0,489897949 | 5,88E-05 | 2,20E-05 | 1 - | 3,50E-05 - | 2 - | 8,69E-05 - | 1 - | 4,13E-05 - | 2 - | 8,38E-05 - | 1 - | 4,72E-05 - |
| 4878 | 1 | 0.2um_NOKCPHJN_83189 | Bacteria | Bacteroidetes | Flavobacteriia | Flavobacterial | Flavobacteriac | Polaribacter | Polaribacter s | information st | Replication, re | 09B0W | type iii restrict | 313598.MED1 | 0,4 | 0,489897949 | 1,09E-05 | 1,33E-05 | 0 - | 0 - | 1 - | 2,76E-05 - | 0 - | 0 - | 1 - | 2,66E-05 - | 0 - | 0 - |
| 4879 | 1 | 3um_FIJJAFBF_95704 | Eukaryota | unclassified | Raphidophyce | Chattonellales | Chattonellace | Heterosigma | Heterosigma a | unclassified | unclassified | unclassified | unclassified | unclassified | 12 | 0,632455532 | 0,000310912 | 3,44E-05 | 12 - | 0,000260363 - | 12 - | 0,000323294 - | 11 - | 0,000281721 - | 13 - | 0,000337861 - | 12 - | 0,000351322 - |
| 4880 | 1 | 3um_FIJJAFBF_368199 | Bacteria | Actinobacteria | Actinobacteria | unclassified | unclassified | unclassified | Actinobacteria | unclassified | unclassified | unclassified | unclassified | unclassified | 3,4 | 1,624807681 | 0,000124328 | 4,91E-05 | 6 - | 0,000192363 - | 3 - | 0,000119429 - | 4 - | 0,000151377 - | 3 - | 0,000115209 - | 1 - | 4,33E-05 - |
| 4881 | 1 | 0.2um_NOKCPHJN_11685 | Bacteria | Bacteroidetes | Flavobacteriia | Flavobacterial | Flavobacteriac | Siansivirga | Siansivirga zea | metabolism | Energy produc | COG0056 | Produces ATP | 1046627.BZAR | 4,4 | 1,019803903 | 0,000116248 | 3,61E-05 | 4 - | 8,73E-05 - | 5 - | 0,000135474 - | 3 - | 7,73E-05 - | 4 - | 0,00010455 - | 6 - | 0,000176663 - |
| 4882 | 1 | 3um_FIJJAFBF_17173 | Bacteria | Proteobacteri | Alphaproteob | Rhodobactera | Rhodobactera | Phaeobacter | Phaeobacter g | information st | Translation, ri | 17FIT | 50s ribosomal | 314262.MED1 | 3,4 | 0,8 | 0,000466995 | 8,94E-05 | 4 - | 0,000468477 - | 3 - | 0,000436282 - | 4 - | 0,000552989 - | 4 - | 0,000561156 - | 2 - | 0,00031607 - |
| 4883 | 1 | 3um_FIJJAFBF_300053 | Bacteria | Proteobacteri | Gammaprotec | Cellvibrionales | Halieaceae | unclassified | marine gamm | unclassified | unclassified | unclassified | unclassified | unclassified | 0,4 | 0,489897949 | 2,36E-05 | 2,91E-05 | 1 - | 5,41E-05 - | 0 - | 0 - | 1 - | 6,39E-05 - | 0 - | 0 - | 0 - | 0 - |
| 4884 | 1 | 3um_FIJJAFBF_72467 | Bacteria | Bacteroidetes | Cytophagia | Cytophagales | Flammeovirga | unclassified | Flammeovirga | information st | Translation, ri | COG0089 | One of the ea | 643867.Ftrac_ | 1,6 | 0,489897949 | 0,00022735 | 6,43E-05 | 2 - | 0,000241635 - | 2 - | 0,00030004 - | 1 - | 0,000142613 - | 2 - | 0,000289439 - | 1 - | 0,000163026 - |
| 4885 | 1 | 3um_FIJJAFBF_67252 | Eukaryota | unclassified | Florideophyce | Acrosymphyta | Schimmelman | Schimmelman | Schimmelman | information st | Translation, ri | COG0052 | 30S ribosomal | 321332.CYB_1 | 0,4 | 0,489897949 | 2,20E-05 | 2,70E-05 | 1 - | 5,03E-05 - | 0 - | 0 - | 1 - | 5,94E-05 - | 0 - | 0 - | 0 - | 0 - |
| 4886 | 1 | 3um_FIJJAFBF_290323 | Bacteria | Proteobacteri | Gammaprotec | Oceanospirilla | Saccharospirill | Reinekea | Reinekea fors | information st | Replication, re | 0QUGS | DNA-binding p | 314283.MED2 | 3,4 | 1,019803903 | 0,00050346 | 0,000105799 | 5 - | 0,000637649 - | 3 - | 0,000475063 - | 4 - | 0,000602143 - | 3 - | 0,000458278 - | 2 - | 0,000344166 - |
| 4887 | 1 | 0.2um_NOKCPHJN_728308 | Archaea | Euryarchaeota | unclassified | unclassified | unclassified | unclassified | Euryarchaeota | unclassified | unclassified | unclassified | unclassified | unclassified | 0,2 | 0,4 | 6,39E-05 | 0,000127891 | 0 - | 0 - | 0 - | 0 - | 0 - | 0 - | 1 - | 0,000319729 - | 0 - | 0 - |
| 4888 | 1 | 0.2um_NOKCPHJN_406372 | Eukaryota | Chlorophyta | Mamiellophyc | Mamiellales | Bathycoccace | Bathycoccus | Bathycoccus p | metabolism | Secondary me | 1DU19 | ABC transport | 70448.Q01FX7 | 0,4 | 0,489897949 | 3,61E-06 | 4,42E-06 | 0 - | 0 - | 0 - | 0 - | 1 - | 8,95E-06 - | 1 - | 9,09E-06 - | 0 - | 0 - |
| 4889 | 1 | 3um_FIJJAFBF_585398 | Bacteria | Proteobacteri | Gammaprotec | Oceanospirilla | Saccharospirill | Reinekea | Reinekea fors | poorly charact | Function unkn | 16UXU | Rna binding s1 | 314283.MED2 | 1,4 | 1,019803903 | 6,53E-05 | 4,86E-05 | 1 - | 3,85E-05 - | 1 - | 4,78E-05 - | 3 - | 0,000136392 - | 0 - | 0 - | 2 - | 0,000103943 - |
| 4890 | 1 | 3um_FIJJAFBF_541534 | Bacteria | Proteobacteri | Alphaproteob | Rhodobactera | Rhodobactera | Planktomarina | Planktomarina | cellular proces | Posttranslatio | COG0652 | peptidyl-proly | 351016.RAZW | 0,4 | 0,489897949 | 3,17E-05 | 3,96E-05 | 1 - | 6,75E-05 - | 0 - | 0 - | 0 - | 0 - | 0 - | 0 - | 1 - | 9,11E-05 - |
| 4891 | 1 | 0.2um_NOKCPHJN_141365 | Bacteria | Proteobacteri | Alphaproteob | Rhodobactera | Rhodobactera | Amylibacter | Amylibacter k | unclassified | unclassified | unclassified | unclassified | unclassified | 2 | 0 | 8,28E-05 | 7,86E-06 | 2 - | 6,94E-05 - | 2 - | 8,61E-05 - | 2 - | 8,19E-05 - | 2 - | 8,31E-05 - | 2 - | 9,36E-05 - |
| 4892 | 1 | 3um_FIJJAFBF_613559 | Bacteria | Bacteroidetes | Flavobacteriia | Flavobacterial | Flavobacteriac | Tamlana | Tamlana nanh | cellular proces | Posttranslatio | 07SFW | Trigger factor | 983544.Lacal_ | 2 | 0,632455532 | 6,02E-05 | 1,39E-05 | 3 - | 7,79E-05 - | 2 - | 6,45E-05 - | 2 - | 6,13E-05 - | 2 - | 6,22E-05 - | 1 - | 3,50E-05 - |
| 4893 | 1 | 3um_FIJJAFBF_05650 | Bacteria | Proteobacteri | Gammaprotec | unclassified | unclassified | unclassified | OM182 bacte | metabolism | Inorganic ion t | COG4206 | tonB-depende | 722419.PH505 | 3,8 | 0,4 | 7,62E-05 | 1,37E-05 | 3 - | 5,00E-05 - | 4 - | 8,27E-05 - | 4 - | 7,87E-05 - | 4 - | 7,98E-05 - | 4 - | 8,99E-05 - |
| 4894 | 1 | 0.2um_NOKCPHJN_61751 | Bacteria | Proteobacteri | Alphaproteob | Rhodobactera | Hyphomonada | unclassified | Hyphomonada | information st | Replication, re | 0PB0H | DNA-binding p | 755732.Fluta_ | 2,8 | 0,748331477 | 0,00043422 | 0,000142464 | 3 - | 0,000386888 - | 2 - | 0,000320267 - | 2 - | 0,000304454 - | 3 - | 0,000463427 - | 4 - | 0,000696065 - |
| 4895 | 1 | 0.2um_NOKCPHJN_81672 | Bacteria | Proteobacteri | Gammaprotec | Cellvibrionales | Porticoccacea | unclassified | gamma prote | cellular proces | Cell wall/mem | 16QB0 | Part of the ou | 314287.GB220 | 0,2 | 0,4 | 2,94E-06 | 5,87E-06 | 1 - | 1,47E-05 - | 0 - | 0 - | 0 - | 0 - | 0 - | 0 - | 0 - | 0 - |
| 4896 | 1 | 3um_FIJJAFBF_00231 | Bacteria | Bacteroidetes | Flavobacteriia | Flavobacterial | Flavobacteriac | Algibacter | Algibacter sp. | unclassified | unclassified | unclassified | unclassified | unclassified | 0,4 | 0,489897949 | 3,69E-05 | 4,52E-05 | 0 - | 0 - | 0 - | 0 - | 1 - | 9,15E-05 - | 1 - | 9,29E-05 - | 0 - | 0 - |
| 4897 | 1 | 3um_FIJJAFBF_730482 | unclassified | unclassified | unclassified | unclassified | unclassified | unclassified | unclassified | unclassified | unclassified | unclassified | unclassified | unclassified | 1,2 | 0,4 | 0,000102553 | 2,27E-05 | 2 - | 0,00014715 - | 1 - | 9,14E-05 - | 1 - | 8,68E-05 - | 1 - | 8,81E-05 - | 1 - | 9,93E-05 - |
| 4898 | 1 | 0.2um_NOKCPHJN_633702 | Bacteria | Proteobacteri | Alphaproteob | Pelagibacteral | Pelagibacterac | Candidatus Pe | Candidatus Pe | metabolism | Amino acid tra | 01R4A | Polyamine AB | 335992.SAR11 | 3 | 0,632455532 | 0,00010958 | 1,86E-05 | 4 - | 0,000123749 - | 3 - | 0,000115244 - | 3 - | 0,000109554 - | 2 - | 7,41E-05 - | 3 - | 0,000125235 - |
| 4899 | 1 | 3um_FIJJAFBF_588036 | Bacteria | Proteobacteri | Betaproteoba | unclassified | unclassified | unclassified | Betaproteoba | cellular proces | Cell wall/mem | COG3317 | Part of the ou | 338969.Rfer_2 | 1 | 0 | 3,70E-05 | 3,52E-06 | 1 - | 3,10E-05 - | 1 - | 3,85E-05 - | 1 - | 3,66E-05 - | 1 - | 3,72E-05 - | 1 - | 4,19E-05 - |
| 4900 | 1 | 3um_FIJJAFBF_56132 | Bacteria | Bacteroidetes | Flavobacteriia | Flavobacterial | Flavobacteriac | Polaribacter | Polaribacter s | poorly charact | Function unkn | 0PAX5 | Pfam:DUF583 | 313594.PI23P | 1,4 | 0,489897949 | 0,00014449 | 6,11E-05 | 1 - | 8,44E-05 - | 2 - | 0,000209587 - | 1 - | 9,96E-05 - | 1 - | 0,000101091 - | 2 - | 0,000227757 - |
| 4901 | 1 | 3um_FIJJAFBF_183510 | unclassified | unclassified | unclassified | unclassified | unclassified | unclassified | unclassified | unclassified | unclassified | unclassified | unclassified | unclassified | 1 | 0 | 0,000152252 | 1,45E-05 | 1 - | 0,00012753 - | 1 - | 0,000158354 - | 1 - | 0,000150536 - | 1 - | 0,000152759 - | 1 - | 0,000172083 - |
| 4902 | 1 | 3um_FIJJAFBF_163752 | Bacteria | Bacteroidetes | Flavobacteriia | Flavobacterial | Flavobacteriac | Polaribacter | various | unclassified | unclassified | unclassified | unclassified | unclassified | 3,8 | 1,469693846 | 0,000455158 | 0,00017488 | 4 - | 0,000409917 - | 4 - | 0,000508996 - | 5 - | 0,000604831 - | 5 - | 0,000613765 - | 1 - | 0,000138281 - |
| 4903 | 2 | 0.2um_NOKCPHJN_83496;3um_FIJJAFBF_655088 | Bacteria | Bacteroidetes | Flavobacteriia | Flavobacterial | Flavobacteriac | Polaribacter | Polaribacter s | unclassified | unclassified | unclassified | unclassified | unclassified | 2 | 0,894427191 | 8,10E-05 | 3,71E-05 | 1 - | 3,36E-05 - | 1 - | 4,17E-05 - | 3 - | 0,000118844 - | 3 - | 0,000120599 - | 2 - | 9,06E-05 - |
| 4904 | 1 | 3um_FIJJAFBF_212877 | Bacteria | Bacteroidetes | Flavobacteriia | Flavobacterial | Crocinitomica | unclassified | Crocinitomica | unclassified | unclassified | unclassified | unclassified | unclassified | 1,6 | 0,489897949 | 0,000144407 | 4,54E-05 | 2 - | 0,000152022 - | 1 - | 9,44E-05 - | 2 - | 0,000179447 - | 1 - | 9,10E-05 - | 2 - | 0,000205132 - |
| 4905 | 1 | 0.2um_NOKCPHJN_1098614 | Eukaryota | unclassified | Cryptophyta | Pyrenomonad | Geminigerace | Guillardia | Guillardia thet | unclassified | unclassified | unclassified | unclassified | unclassified | 2,4 | 0,8 | 6,51E-05 | 2,29E-05 | 1 - | 2,24E-05 - | 3 - | 8,33E-05 - | 3 - | 7,92E-05 - | 3 - | 8,04E-05 - | 2 - | 6,04E-05 - |
| 4906 | 1 | 3um_FIJJAFBF_60207 | Bacteria | Actinobacteria | unclassified | unclassified | unclassified | unclassified | Actinobacteria | unclassified | unclassified | unclassified | unclassified | unclassified | 4,2 | 0,4 | 0,000214475 | 3,88E-05 | 4 - | 0,00017004 - | 4 - | 0,000211139 - | 4 - | 0,000200714 - | 4 - | 0,000203679 - | 5 - | 0,000286805 - |
| 4907 | 1 | 0.2um_NOKCPHJN_34121 | Eukaryota | unclassified | Oomycetes | Saprolegniales | Saprolegniace | Aphanomyces | Aphanomyces | cellular proces | Signal transdu | KOG0613 | fibronectin typ | 8083.ENSXMA | 0,2 | 0,4 | 1,43E-07 | 2,87E-07 | 0 - | 0 - | 0 - | 0 - | 0 - | 0 - | 0 - | 0 - | 1 - | 7,17E-07 - |
| 4908 | 2 | 0.2um_NOKCPHJN_757420;0.2um_NOKCPHJN_8 | Bacteria | Proteobacteri | Alphaproteob | various | various | various | various | unclassified | unclassified | unclassified | unclassified | unclassified | 1 | 0,632455532 | 3,57E-05 | 2,04E-05 | 2 - | 6,36E-05 - | 1 - | 3,95E-05 - | 1 - | 3,75E-05 - | 1 - | 3,81E-05 - | 0 - | 0 - |
| 4909 | 1 | 3um_FIJJAFBF_158434 | Bacteria | Bacteroidetes | Flavobacteriia | Flavobacterial | Flavobacteriac | Cellulophaga | Cellulophaga s | unclassified | unclassified | unclassified | unclassified | unclassified | 1,6 | 1,624807681 | 2,83E-05 | 2,89E-05 | 3 - | 4,56E-05 - | 4 - | 7,55E-05 - | 0 - | 0 - | 0 - | 0 - | 1 - | 2,05E-05 - |
| 4910 | 1 | 3um_FIJJAFBF_75280 | Bacteria | Bacteroidetes | Flavobacteriia | unclassified | unclassified | unclassified | Flavobacteria | unclassified | unclassified | unclassified | unclassified | unclassified | 3,6 | 1,019803903 | 0,000160969 | 5,45E-05 | 4 - | 0,000148578 - | 4 - | 0,00018449 - | 2 - | 8,77E-05 - | 3 - | 0,000133479 - | 5 - | 0,000250606 - |
| 4911 | 1 | 3um_FIJJAFBF_682765 | Bacteria | Bacteroidetes | Flavobacteriia | Flavobacterial | Flavobacteriac | Formosa | Formosa sp. H | poorly charact | Function unkn | 08PAV | RagB SusD do | 983544.Lacal_ | 1,4 | 0,489897949 | 3,92E-05 | 1,46E-05 | 1 - | 2,33E-05 - | 2 - | 5,78E-05 - | 1 - | 2,75E-05 - | 2 - | 5,58E-05 - | 1 - | 3,14E-05 - |
| 4912 | 1 | 3um_FIJJAFBF_503090 | Bacteria | Verrucomicrol | Opitutae | unclassified | unclassified | unclassified | Opitutae bact | unclassified | unclassified | unclassified | unclassified | unclassified | 1,8 | 0,748331477 | 5,90E-05 | 2,15E-05 | 3 - | 8,38E-05 - | 2 - | 6,94E-05 - | 1 - | 3,30E-05 - | 1 - | 3,35E-05 - | 2 - | 7,54E-05 - |
| 4913 | 1 | 0.2um_NOKCPHJN_772735 | Bacteria | Proteobacteri | Gammaprotec | Cellvibrionales | Porticoccacea | unclassified | SAR92 bacteri | unclassified | unclassified | unclassified | unclassified | unclassified | 0,2 | 0,4 | 2,69E-06 | 5,39E-06 | 1 - | 1,35E-05 - | 0 - | 0 - | 0 - | 0 - | 0 - | 0 - | 0 - | 0 - |
| 4914 | 1 | 0.2um_NOKCPHJN_981081 | Eukaryota | unclassified | unclassified | Prymnesiales | Chrysochromu | Chrysochromu | Chrysochromu | unclassified | unclassified | unclassified | unclassified | unclassified | 0,8 | 0,4 | 1,82E-05 | 9,37E-06 | 1 - | 1,90E-05 - | 1 - | 2,36E-05 - | 0 - | 0 - | 1 - | 2,28E-05 - | 1 - | 2,57E-05 - |
| 4915 | 1 | 3um_FIJJAFBF_155211 | Bacteria | Proteobacteri | Alphaproteob | Rhizobiales | unclassified | unclassified | Rhizobiales ba | unclassified | unclassified | unclassified | unclassified | unclassified | 0,6 | 0,489897949 | 2,73E-05 | 2,27E-05 | 1 - | 3,85E-05 - | 0 - | 0 - | 0 - | 0 - | 1 - | 4,61E-05 - | 1 - | 5,20E-05 - |
| 4916 | 1 | 0.2um_NOKCPHJN_06908 | Bacteria | Bacteroidetes | Cytophagia | Cytophagales | Cytophagacea | Ekhidna | Ekhidna lutea | poorly charact | Function unkn | 0XQTX | Chromosome | 643867.Ftrac_ | 0,8 | 0,748331477 | 1,07E-05 | 1,04E-05 | 1 - | 1,17E-05 - | 0 - | 0 - | 1 - | 1,38E-05 - | 2 - | 2,79E-05 - | 0 - | 0 - |
| 4917 | 1 | 0.2um_NOKCPHJN_483828 | Bacteria | Proteobacteri | Alphaproteob | Pelagibacteral | unclassified | unclassified | Pelagibacteral | metabolism | Nucleotide tra | 01Q8W | Plays an impo | 684719.HIMB | 0,4 | 0,489897949 | 1,30E-05 | 1,60E-05 | 1 - | 2,95E-05 - | 0 - | 0 - | 0 - | 0 - | 1 - | 3,53E-05 - | 0 - | 0 - |
| 4918 | 1 | 0.2um_NOKCPHJN_1119720 | Bacteria | Planctomycete | Planctomyceti | Planctomyceta | Planctomyceta | Rhodopirellula | Rhodopirellula | unclassified | unclassified | unclassified | unclassified | unclassified | 0,6 | 0,489897949 | 1,55E-05 | 1,28E-05 | 1 - | 2,30E-05 - | 0 - | 0 - | 1 - | 2,71E-05 - | 1 - | 2,75E-05 - | 0 - | 0 - |
| 4919 | 1 | 3um_FIJJAFBF_661566 | Bacteria | Proteobacteri | Gammaprotec | Oceanospirilla | Saccharospirill | Saccharospiril | Saccharospirill | information st | Translation, ri | 0QHTE | Catalyzes the | 706191.PANA | 1,2 | 0,4 | 3,64E-05 | 1,34E-05 | 1 - | 2,52E-05 - | 2 - | 6,26E-05 - | 1 - | 2,98E-05 - | 1 - | 3,02E-05 - | 1 - | 3,40E-05 - |
| 4920 | 1 | 3um_FIJJAFBF_191791 | Eukaryota | Bacillariophyta | Coscinodiscop | Rhizosoleniale | Rhizosoleniace | Rhizosolenia | Rhizosolenia i | unclassified | unclassified | unclassified | unclassified | unclassified | 10,8 | 0,4 | 0,000918506 | 8,85E-05 | 11 - | 0,000784189 - | 10 - | 0,00088521 - | 11 - | 0,000925655 - | 11 - | 0,000939327 - | 11 - | 0,001058149 - |
| 4921 | 1 | 3um_FIJJAFBF_749561 | unclassified | unclassified | unclassified | unclassified | unclassified | unclassified | unclassified | unclassified | unclassified | unclassified | unclassified | unclassified | 0,6 | 0,8 | 1,58E-05 | 2,02E-05 | 2 - | 4,86E-05 - | 1 - | 3,02E-05 - | 0 - | 0 - | 0 - | 0 - | 0 - | 0 - |
| 4922 | 2 | 0.2um_NOKCPHJN_1022248;3um_FIJJAFBF_6609 | Viruses | unclassified | unclassified | unclassified | unclassified | unclassified | uncultured M | unclassified | unclassified | unclassified | unclassified | unclassified | 2 | 0,632455532 | 6,27E-05 | 2,16E-05 | 1 - | 2,59E-05 - | 2 - | 6,42E-05 - | 2 - | 6,10E-05 - | 3 - | 9,29E-05 - | 2 - | 6,98E-05 - |
| 4923 | 1 | 0.2um_NOKCPHJN_574440 | Eukaryota | unclassified | Pelagophycea | Pelagomonad | unclassified | Aureococcus | Aureococcus a | cellular proces | Signal transdu | KOG0613 | fibronectin typ | 65071.PYU1_T | 0,8 | 0,748331477 | 1,28E-05 | 1,23E-05 | 1 - | 1,38E-05 - | 1 - | 1,71E-05 - | 0 - | 0 - | 2 - | 3,30E-05 - | 0 - | 0 - |
| 4924 | 1 | 3um_FIJJAFBF_275939 | Bacteria | Proteobacteri | Betaproteoba | unclassified | unclassified | unclassified | Betaproteoba | unclassified | unclassified | unclassified | unclassified | unclassified | 2,4 | 1,019803903 | 9,88E-05 | 3,91E-05 | 3 - | 0,000106275 - | 2 - | 8,80E-05 - | 4 - | 0,000167262 - | 2 - | 8,49E-05 - | 1 - | 4,78E-05 - |
| 4925 | 1 | 0.2um_NOKCPHJN_133802 | Bacteria | Proteobacteri | Gammaprotec | Oceanospirilla | Oceanospirilla | Marinomonas | Marinomonas | poorly charact | Function unkn | 0QQPZ | UPF0325 prot | 314277.MED1 | 1 | 0,632455532 | 0,000110634 | 7,80E-05 | 1 - | 9,04E-05 - | 1 - | 0,00011222 - | 1 - | 0,000106679 - | 0 - | 0 - | 2 - | 0,000243897 - |
| 4926 | 1 | 3um_FIJJAFBF_315830 | Bacteria | Proteobacteri | Alphaproteob | Rhodobactera | Rhodobactera | Planktomarina | Planktomarina | unclassified | unclassified | unclassified | unclassified | unclassified | 2,6 | 1,019803903 | 0,000372287 | 0,000150116 | 2 - | 0,000241635 - | 3 - | 0,00045006 - | 3 - | 0,000427839 - | 4 - | 0,000578877 - | 1 - | 0,000163026 - |
| 4927 | 1 | 3um_FIJJAFBF_105881 | Bacteria | Proteobacteri | Alphaproteob | Rhodobactera | Rhodobactera | Planktomarina | Planktomarina | metabolism | Energy produc | COG0711 | F(1)F(0) ATP s | 644107.SL115 | 2 | 0 | 0,000148137 | 1,41E-05 | 2 - | 0,000124083 - | 2 - | 0,000154074 - | 2 - | 0,000146467 - | 2 - | 0,000148631 - | 2 - | 0,000167432 - |
| 4928 | 1 | 0.2um_NOKCPHJN_401155 | Bacteria | unclassified | unclassified | unclassified | unclassified | unclassified | uncultured ba | unclassified | unclassified | unclassified | unclassified | unclassified | 12,2 | 2,481934729 | 0,000180501 | 2,93E-05 | 16 - | 0,000200483 - | 14 - | 0,000217824 - | 9 - | 0,000133116 - | 11 - | 0,0001651 - | 11 - | 0,000185985 - |
| 4929 | 1 | 0.2um_NOKCPHJN_987990 | unclassified | unclassified | unclassified | unclassified | unclassified | unclassified | unclassified | unclassified | unclassified | unclassified | unclassified | unclassified | 0,2 | 0,4 | 1,41E-05 | 2,82E-05 | 1 - | 7,04E-05 - | 0 - | 0 - | 0 - | 0 - | 0 - | 0 - | 0 - | 0 - |
| 4930 | 1 | 3um_FIJJAFBF_752878 | Bacteria | Proteobacteri | Alphaproteob | Rhodobactera | Rhodobactera | Amylibacter | Amylibacter k | metabolism | Amino acid tra | 01QH7 | sulfoacetaldeh | 488538.SAR11 | 1,2 | 0,979795897 | 2,77E-05 | 2,25E-05 | 1 - | 1,94E-05 - | 1 - | 2,41E-05 - | 3 - | 6,87E-05 - | 0 - | 0 - | 1 - | 2,62E-05 - |
| 4931 | 1 | 0.2um_NOKCPHJN_175750 | Eukaryota | unclassified | Pelagophycea | Pelagomonad | unclassified | Aureococcus | Aureococcus a | unclassified | unclassified | unclassified | unclassified | unclassified | 1 | 0 | 6,89E-05 | 6,54E-06 | 1 - | 5,77E-05 - | 1 - | 7,16E-05 - | 1 - | 6,81E-05 - | 1 - | 6,91E-05 - | 1 - | 7,78E-05 - |
| 4932 | 1 | 3um_FIJJAFBF_72765 | Bacteria | Proteobacteri | Gammaprotec | Oceanospirilla | Halomonadac | Halomonas | Halomonas ve | metabolism | Energy produc | COG1290 | Component o | 999141.GME_ | 1 | 0 | 3,25E-05 | 3,08E-06 | 1 - | 2,72E-05 - | 1 - | 3,38E-05 - | 1 - | 3,21E-05 - | 1 - | 3,26E-05 - | 1 - | 3,67E-05 - |
| 4933 | 1 | 0.2um_NOKCPHJN_481996 | Bacteria | Proteobacteri | Alphaproteob | Pelagibacteral | Pelagibacterac | Candidatus Pe | Candidatus Pe | unclassified | unclassified | unclassified | unclassified | unclassified | 1,6 | 0,489897949 | 0,000178168 | 4,94E-05 | 2 - | 0,000189714 - | 2 - | 0,000235568 - | 2 - | 0,000223938 - | 1 - | 0,000113623 - | 1 - | 0,000127995 - |
| 4934 | 2 | 3um_FIJJAFBF_223942;3um_FIJJAFBF_640360 | Bacteria | Bacteroidetes | Flavobacteriia | Flavobacterial | Flavobacteriac | Formosa | Formosa sp. H | unclassified | unclassified | unclassified | unclassified | unclassified | 1 | 0 | 2,72E-05 | 2,59E-06 | 1 - | 2,28E-05 - | 1 - | 2,83E-05 - | 1 - | 2,69E-05 - | 1 - | 2,73E-05 - | 1 - | 3,08E-05 - |
| 4935 | 1 | 0.2um_NOKCPHJN_700962 | Bacteria | Proteobacteri | Alphaproteob | Rhodobactera | Rhodobactera | Neptunicoccu | Neptunicoccu | metabolism | Inorganic ion t | 05PWN | phosphate ab | 765910.Marp | 12,2 | 1,469693846 | 0,00048429 | 8,50E-05 | 13 - | 0,000431242 - | 13 - | 0,000535476 - | 11 - | 0,000430724 - | 10 - | 0,000397351 - | 14 - | 0,00062666 - |
| 4936 | 1 | 3um_FIJJAFBF_490377 | Bacteria | Bacteroidetes | Flavobacteriia | Flavobacterial | Flavobacteriac | Polaribacter | Polaribacter s | metabolism | Lipid transpor | COG0439 | acetyl-CoA ca | 391587.KAOT1 | 1,2 | 0,4 | 3,72E-05 | 1,37E-05 | 1 - | 2,58E-05 - | 2 - | 6,41E-05 - | 1 - | 3,04E-05 - | 1 - | 3,09E-05 - | 1 - | 3,48E-05 - |
| 4937 | 1 | 3um_FIJJAFBF_192374 | Bacteria | Proteobacteri | Gammaprotec | unclassified | unclassified | unclassified | SAR86 cluster | metabolism | Energy produc | 16QPK | Succinate deh | 876044.IMCC3 | 3,2 | 0,748331477 | 7,31E-05 | 1,43E-05 | 4 - | 7,78E-05 - | 4 - | 9,66E-05 - | 3 - | 6,89E-05 - | 3 - | 6,99E-05 - | 2 - | 5,25E-05 - |
| 4938 | 1 | 3um_FIJJAFBF_24303 | Bacteria | Proteobacteri | Gammaprotec | Oceanospirilla | Saccharospirill | Reinekea | Reinekea fors | cellular proces | Defense mech | 07SUK | Efflux transpo | 314283.MED2 | 9,4 | 1,496662955 | 0,000298875 | 2,97E-05 | 12 - | 0,000323315 - | 8 - | 0,000267641 - | 10 - | 0,000318033 - | 8 - | 0,000258185 - | 9 - | 0,0003272 - |
| 4939 | 1 | 3um_FIJJAFBF_586240 | Bacteria | Bacteroidetes | Flavobacteriia | Flavobacterial | Flavobacteriac | Lacinutrix | Lacinutrix alg | metabolism | Amino acid tra | COG0174 | glutamine syn | 313594.PI23P | 3,6 | 0,489897949 | 0,000146938 | 2,90E-05 | 3 - | 0,000101873 - | 3 - | 0,000126496 - | 4 - | 0,000160334 - | 4 - | 0,000162702 - | 4 - | 0,000183283 - |
| 4940 | 1 | 3um_FIJJAFBF_119786 | unclassified | unclassified | unclassified | unclassified | unclassified | unclassified | unclassified | unclassified | unclassified | unclassified | unclassified | unclassified | 1 | 0,632455532 | 9,02E-05 | 5,91E-05 | 1 - | 7,76E-05 - | 1 - | 9,63E-05 - | 1 - | 9,15E-05 - | 2 - | 0,000185788 - | 0 - | 0 - |
| 4941 | 1 | 0.2um_NOKCPHJN_307745 | Eukaryota | Phaeophyceae | unclassified | Ectocarpales | Ectocarpaceae | Ectocarpus | Ectocarpus sil | unclassified | unclassified | unclassified | unclassified | unclassified | 0,4 | 0,489897949 | 2,20E-05 | 2,70E-05 | 1 - | 5,03E-05 - | 0 - | 0 - | 1 - | 5,94E-05 - | 0 - | 0 - | 0 - | 0 - |
| 4942 | 1 | 0.2um_NOKCPHJN_100705 | Bacteria | Proteobacteri | Alphaproteob | Rhodobactera | Rhodobactera | Thalassobius | Thalassobius a | unclassified | unclassified | unclassified | unclassified | unclassified | 0,4 | 0,489897949 | 3,19E-05 | 3,91E-05 | 0 - | 0 - | 0 - | 0 - | 1 - | 7,92E-05 - | 1 - | 8,04E-05 - | 0 - | 0 - |
| 4943 | 1 | 3um_FIJJAFBF_212747 | Bacteria | Bacteroidetes | Flavobacteriia | Flavobacterial | Flavobacteriac | Polaribacter | Polaribacter s | poorly charact | Function unkn | 099QV | gliding motilit | 313594.PI23P | 3,2 | 0,748331477 | 0,000145721 | 3,48E-05 | 3 - | 0,000115161 - | 4 - | 0,000190661 - | 4 - | 0,000181247 - | 3 - | 0,000137943 - | 2 - | 0,000103595 - |
| 4944 | 1 | 0.2um_NOKCPHJN_39209 | Eukaryota | unclassified | Pelagophycea | Pelagomonad | unclassified | Aureoumbra | Aureoumbra l | metabolism | Energy produc | COG0055 | Produces ATP | 489825.LYNG | 47 | 3,633180425 | 0,001351943 | 0,00020542 | 47 - | 0,001128558 - | 50 - | 0,001490783 - | 43 - | 0,001218773 - | 43 - | 0,001236775 - | 52 - | 0,001684827 - |
| 4945 | 1 | 0.2um_NOKCPHJN_290986 | Bacteria | Proteobacteri | Alphaproteob | Rhodobactera | Rhodobactera | Sulfitobacter | Sulfitobacter s | unclassified | unclassified | unclassified | unclassified | unclassified | 4,4 | 0,489897949 | 0,000316287 | 4,90E-05 | 4 - | 0,00024037 - | 5 - | 0,000373086 - | 4 - | 0,000283732 - | 5 - | 0,000359904 - | 4 - | 0,000324344 - |
| 4946 | 1 | 0.2um_NOKCPHJN_47862 | Bacteria | Bacteroidetes | unclassified | unclassified | unclassified | unclassified | unidentified e | poorly charact | Function unkn | 08PIQ | CHU large pro | 50743.SCB49_ | 1 | 0 | 1,16E-05 | 1,10E-06 | 1 - | 9,74E-06 - | 1 - | 1,21E-05 - | 1 - | 1,15E-05 - | 1 - | 1,17E-05 - | 1 - | 1,31E-05 - |
| 4947 | 1 | 0.2um_NOKCPHJN_765302 | Bacteria | Bacteroidetes | Bacteroidia | Bacteroidales | Porphyromon | Porphyromon | various | information st | Translation, ri | 0559A | This protein p | 435591.BDI_2 | 1 | 0 | 0,000114189 | 1,08E-05 | 1 - | 9,56E-05 - | 1 - | 0,000118766 - | 1 - | 0,000112902 - | 1 - | 0,000114569 - | 1 - | 0,000129062 - |
| 4948 | 1 | 3um_FIJJAFBF_52801 | Bacteria | Proteobacteri | Alphaproteob | Rhodobactera | Rhodobactera | Sulfitobacter | Sulfitobacter p | cellular proces | Posttranslatio | 01QWG | Prevents misf | 633131.TR2A6 | 35,8 | 2,712931993 | 0,000888503 | 7,93E-05 | 40 - | 0,00083474 - | 32 - | 0,000829201 - | 36 - | 0,000886793 - | 34 - | 0,000849897 - | 37 - | 0,001041883 - |
| 4949 | 1 | 3um_FIJJAFBF_520520 | Bacteria | Bacteroidetes | Flavobacteriia | Flavobacterial | Flavobacteriac | Polaribacter | Polaribacter s | unclassified | unclassified | unclassified | unclassified | unclassified | 0,4 | 0,489897949 | 1,72E-05 | 2,12E-05 | 1 - | 3,94E-05 - | 0 - | 0 - | 1 - | 4,66E-05 - | 0 - | 0 - | 0 - | 0 - |
| 4950 | 1 | 0.2um_NOKCPHJN_329673 | Bacteria | Bacteroidetes | Flavobacteriia | Flavobacterial | Cryomorphace | unclassified | Cryomorphace | unclassified | unclassified | unclassified | unclassified | unclassified | 0,4 | 0,489897949 | 4,09E-05 | 5,10E-05 | 1 - | 8,70E-05 - | 0 - | 0 - | 0 - | 0 - | 0 - | 0 - | 1 - | 0,000117329 - |
| 4951 | 1 | 0.2um_NOKCPHJN_314904 | unclassified | unclassified | unclassified | unclassified | unclassified | unclassified | unclassified | unclassified | unclassified | unclassified | unclassified | unclassified | 1,6 | 0,489897949 | 0,000127008 | 4,05E-05 | 2 - | 0,000133461 - | 1 - | 8,29E-05 - | 1 - | 7,88E-05 - | 2 - | 0,000159864 - | 2 - | 0,000180087 - |
| 4952 | 1 | 3um_FIJJAFBF_448245 | Bacteria | Proteobacteri | Gammaprotec | Oceanospirilla | Saccharospirill | Reinekea | Reinekea blan | metabolism | Amino acid tra | COG1448 | aminotransfer | 314283.MED2 | 4,6 | 1,2 | 0,000156492 | 3,27E-05 | 6 - | 0,000173904 - | 3 - | 0,000107969 - | 6 - | 0,000205276 - | 4 - | 0,000138872 - | 4 - | 0,000156439 - |
| 4953 | 1 | 0.2um_NOKCPHJN_06875 | Bacteria | Bacteroidetes | Cytophagia | Cytophagales | Cytophagacea | Ekhidna | Ekhidna lutea | information st | Transcription | 05D27 | DNA-depende | 643867.Ftrac_ | 11 | 1,897366596 | 0,000103658 | 1,19E-05 | 14 - | 0,000111899 - | 12 - | 0,000119097 - | 9 - | 8,49E-05 - | 11 - | 0,000105315 - | 9 - | 9,71E-05 - |
| 4954 | 1 | 0.2um_NOKCPHJN_394002 | Bacteria | Proteobacteri | Alphaproteob | Rhodobactera | Rhodobactera | Planktomarina | Planktomarina | unclassified | unclassified | unclassified | unclassified | unclassified | 3,8 | 0,748331477 | 0,000230685 | 4,26E-05 | 4 - | 0,000204958 - | 4 - | 0,000254498 - | 5 - | 0,000302416 - | 3 - | 0,000184129 - | 3 - | 0,000207421 - |
| 4955 | 1 | 0.2um_NOKCPHJN_473189 | Bacteria | Proteobacteri | Gammaprotec | unclassified | unclassified | unclassified | Gammaprotec | poorly charact | Function unkn | 070MQ | NA | 425104.Ssed_ | 1,6 | 0,489897949 | 7,27E-05 | 2,40E-05 | 2 - | 7,60E-05 - | 2 - | 9,44E-05 - | 1 - | 4,49E-05 - | 1 - | 4,55E-05 - | 2 - | 0,000102566 - |
| 4956 | 1 | 3um_FIJJAFBF_336106 | Bacteria | Proteobacteri | Betaproteoba | unclassified | unclassified | unclassified | Betaproteoba | unclassified | unclassified | unclassified | unclassified | unclassified | 0,2 | 0,4 | 7,46E-06 | 1,49E-05 | 0 - | 0 - | 0 - | 0 - | 1 - | 3,73E-05 - | 0 - | 0 - | 0 - | 0 - |
| 4957 | 1 | 3um_FIJJAFBF_75454 | Bacteria | Bacteroidetes | Flavobacteriia | Flavobacterial | Flavobacteriac | Tenacibaculur | Tenacibaculur | information st | Translation, ri | COG0052 | 30S ribosomal | 886377.Murru | 0,6 | 0,489897949 | 2,72E-05 | 2,24E-05 | 1 - | 4,03E-05 - | 0 - | 0 - | 1 - | 4,75E-05 - | 1 - | 4,82E-05 - | 0 - | 0 - |
| 4958 | 1 | 0.2um_NOKCPHJN_818091 | Eukaryota | Cnidaria | Anthozoa | Scleractinia | Pocilloporidae | Stylophora | Stylophora pis | cellular proces | Intracellular tr | 14PHI | NA | 135651.CBN1 | 0,2 | 0,4 | 4,45E-06 | 8,90E-06 | 0 - | 0 - | 0 - | 0 - | 0 - | 0 - | 0 - | 0 - | 1 - | 2,23E-05 - |
| 4959 | 1 | 0.2um_NOKCPHJN_81098 | Bacteria | Proteobacteri | Gammaprotec | unclassified | unclassified | unclassified | Gammaprotec | metabolism | Energy produc | 17C3H | Produces ATP | 876044.IMCC3 | 1 | 0,632455532 | 9,36E-05 | 5,23E-05 | 2 - | 0,000161657 - | 1 - | 0,000100365 - | 0 - | 0 - | 1 - | 9,68E-05 - | 1 - | 0,000109067 - |
| 4960 | 1 | 3um_FIJJAFBF_367967 | Bacteria | Proteobacteri | Gammaprotec | Enterobactera | Enterobacteri | Kosakonia | Kosakonia ory | unclassified | unclassified | unclassified | unclassified | unclassified | 1 | 0 | 6,78E-05 | 6,44E-06 | 1 - | 5,68E-05 - | 1 - | 7,06E-05 - | 1 - | 6,71E-05 - | 1 - | 6,81E-05 - | 1 - | 7,67E-05 - |
| 4961 | 1 | 3um_FIJJAFBF_253091 | Bacteria | Proteobacteri | Gammaprotec | Chromatiales | Ectothiorhodo | Thioalkalivibri | Thioalkalivibri | metabolism | Amino acid tra | 11KI6 | polar amino a | 488538.SAR11 | 2,6 | 1,356465997 | 0,000126933 | 6,66E-05 | 2 - | 8,26E-05 - | 3 - | 0,000153797 - | 5 - | 0,000243673 - | 2 - | 9,89E-05 - | 1 - | 5,57E-05 - |
| 4962 | 1 | 3um_FIJJAFBF_163753 | Bacteria | Bacteroidetes | Flavobacteriia | Flavobacterial | Flavobacteriac | Polaribacter | various | unclassified | unclassified | unclassified | unclassified | unclassified | 0,6 | 0,489897949 | 7,99E-05 | 6,58E-05 | 1 - | 0,000118327 - | 0 - | 0 - | 1 - | 0,000139672 - | 1 - | 0,000141735 - | 0 - | 0 - |
| 4963 | 1 | 0.2um_NOKCPHJN_440051 | Bacteria | Proteobacteri | Gammaprotec | Cellvibrionales | Halieaceae | unclassified | gamma prote | metabolism | Inorganic ion t | 16PDP | TonB-depende | 342610.Patl_2 | 0,6 | 0,489897949 | 7,83E-06 | 6,46E-06 | 1 - | 1,14E-05 - | 1 - | 1,42E-05 - | 1 - | 1,35E-05 - | 0 - | 0 - | 0 - | 0 - |
| 4964 | 1 | 3um_FIJJAFBF_388113 | Bacteria | Proteobacteri | Gammaprotec | Cellvibrionales | Halieaceae | unclassified | marine gamm | unclassified | unclassified | unclassified | unclassified | unclassified | 1,2 | 0,4 | 0,000180795 | 6,23E-05 | 1 - | 0,000126128 - | 1 - | 0,000156614 - | 1 - | 0,000148882 - | 2 - | 0,000302161 - | 1 - | 0,000170192 - |
| 4965 | 1 | 3um_FIJJAFBF_02149 | Bacteria | Bacteroidetes | Flavobacteriia | Flavobacterial | Flavobacteriac | Arenitalea | Arenitalea lut | poorly charact | Function unkn | 0PA0H | (Lipo)protein | 391603.FBALC | 0,4 | 0,489897949 | 2,71E-05 | 3,34E-05 | 1 - | 6,17E-05 - | 0 - | 0 - | 0 - | 0 - | 1 - | 7,39E-05 - | 0 - | 0 - |
| 4966 | 1 | 0.2um_NOKCPHJN_820442 | Bacteria | Proteobacteri | Alphaproteob | Rhodospirillal | Rhodospirillac | Thalassobacul | Thalassobacul | unclassified | unclassified | unclassified | unclassified | unclassified | 1 | 0 | 1,95E-05 | 1,85E-06 | 1 - | 1,63E-05 - | 1 - | 2,03E-05 - | 1 - | 1,93E-05 - | 1 - | 1,96E-05 - | 1 - | 2,21E-05 - |
| 4967 | 1 | 3um_FIJJAFBF_425559 | Bacteria | Proteobacteri | Alphaproteob | various | various | various | various | information st | Replication, re | 17D5C | DNA-binding p | 367336.OM22 | 4,2 | 0,4 | 0,000599589 | 7,96E-05 | 4 - | 0,000478236 - | 4 - | 0,000593829 - | 4 - | 0,000564509 - | 5 - | 0,000716059 - | 4 - | 0,00064531 - |
| 4968 | 1 | 3um_FIJJAFBF_639893 | Bacteria | Proteobacteri | Gammaprotec | Oceanospirilla | Saccharospirill | Reinekea | Reinekea fors | metabolism | Amino acid tra | 05F6H | The alpha sub | 314283.MED2 | 0,6 | 0,8 | 2,86E-05 | 3,70E-05 | 2 - | 9,00E-05 - | 0 - | 0 - | 1 - | 5,31E-05 - | 0 - | 0 - | 0 - | 0 - |
| 4969 | 1 | 3um_FIJJAFBF_24824 | unclassified | unclassified | unclassified | unclassified | unclassified | unclassified | unclassified | unclassified | unclassified | unclassified | unclassified | unclassified | 3,6 | 0,489897949 | 0,000424487 | 8,37E-05 | 3 - | 0,000294299 - | 3 - | 0,000365433 - | 4 - | 0,000463187 - | 4 - | 0,000470028 - | 4 - | 0,000529485 - |
| 4970 | 1 | 3um_FIJJAFBF_337968 | Bacteria | Proteobacteri | Alphaproteob | various | various | various | various | information st | Translation, ri | 08UHH | Located on th | 367336.OM22 | 1 | 0 | 0,000107052 | 1,02E-05 | 1 - | 8,97E-05 - | 1 - | 0,000111343 - | 1 - | 0,000105845 - | 1 - | 0,000107409 - | 1 - | 0,000120996 - |
| 4971 | 1 | 3um_FIJJAFBF_197153 | Eukaryota | unclassified | Cryptophyta | Pyrenomonad | Geminigerace | Guillardia | Guillardia thet | metabolism | Carbohydrate | 12R22 | fructose-bisph | 400682.PAC_1 | 0,2 | 0,4 | 7,61E-06 | 1,52E-05 | 0 - | 0 - | 0 - | 0 - | 1 - | 3,81E-05 - | 0 - | 0 - | 0 - | 0 - |
| 4972 | 1 | 3um_FIJJAFBF_117798 | Viruses | unclassified | unclassified | unclassified | unclassified | unclassified | uncultured M | cellular proces | Cell wall/mem | 08T5X | Glycosyl trans | 1031711.RSPO | 0,8 | 0,4 | 2,69E-05 | 1,35E-05 | 0 - | 0 - | 1 - | 3,36E-05 - | 1 - | 3,20E-05 - | 1 - | 3,24E-05 - | 1 - | 3,65E-05 - |
| 4973 | 1 | 3um_FIJJAFBF_402491 | Bacteria | Bacteroidetes | Flavobacteriia | unclassified | unclassified | unclassified | Flavobacteria | information st | Translation, ri | 08UHY | One of the pri | 487797.Flav30 | 1 | 0 | 8,67E-05 | 8,23E-06 | 1 - | 7,26E-05 - | 1 - | 9,02E-05 - | 1 - | 8,57E-05 - | 1 - | 8,70E-05 - | 1 - | 9,80E-05 - |
| 4974 | 1 | 3um_FIJJAFBF_105173 | Bacteria | Bacteroidetes | Flavobacteriia | Flavobacterial | Flavobacteriac | Polaribacter | Polaribacter s | unclassified | unclassified | unclassified | unclassified | unclassified | 0,4 | 0,489897949 | 1,58E-05 | 1,95E-05 | 1 - | 3,52E-05 - | 1 - | 4,37E-05 - | 0 - | 0 - | 0 - | 0 - | 0 - | 0 - |
| 4975 | 1 | 3um_FIJJAFBF_414835 | Bacteria | Cyanobacteria | unclassified | Synechococca | Synechococca | Synechococcu | various | unclassified | unclassified | unclassified | unclassified | unclassified | 1 | 0 | 0,000133036 | 1,26E-05 | 1 - | 0,000111434 - | 1 - | 0,000138368 - | 1 - | 0,000131536 - | 1 - | 0,000133479 - | 1 - | 0,000150364 - |
| 4976 | 1 | 3um_FIJJAFBF_745490 | Eukaryota | Bacillariophyta | Coscinodiscop | Thalassiosirale | Thalassiosirac | Thalassiosira | Thalassiosira c | cellular proces | Posttranslatio | KOG0731 | ATP-depender | 2850.Phatr179 | 4,4 | 0,8 | 0,000438007 | 6,58E-05 | 5 - | 0,000421973 - | 5 - | 0,000523966 - | 5 - | 0,000498096 - | 4 - | 0,000404363 - | 3 - | 0,000341635 - |
| 4977 | 1 | 0.2um_NOKCPHJN_549153 | Bacteria | Proteobacteri | Alphaproteob | Rhodobactera | Rhodobactera | Rhodobacter | Rhodobacter s | metabolism | Secondary me | 07US8 | Inherit from C | 533240.CRC_0 | 0,8 | 0,979795897 | 1,60E-05 | 1,96E-05 | 0 - | 0 - | 2 - | 4,07E-05 - | 0 - | 0 - | 2 - | 3,92E-05 - | 0 - | 0 - |
| 4978 | 1 | 0.2um_NOKCPHJN_162314 | Eukaryota | unclassified | Pelagophycea | Pelagomonad | unclassified | Aureococcus | Aureococcus a | information st | Translation, ri | KOG3181 | ribosomal pro | 272952.HpaP8 | 0,6 | 0,489897949 | 5,51E-05 | 4,58E-05 | 1 - | 7,81E-05 - | 0 - | 0 - | 1 - | 9,22E-05 - | 0 - | 0 - | 1 - | 0,000105357 - |
| 4979 | 1 | 3um_FIJJAFBF_395138 | unclassified | unclassified | unclassified | unclassified | unclassified | unclassified | unclassified | unclassified | unclassified | unclassified | unclassified | unclassified | 0,4 | 0,489897949 | 8,43E-06 | 1,05E-05 | 1 - | 1,79E-05 - | 0 - | 0 - | 0 - | 0 - | 0 - | 0 - | 1 - | 2,42E-05 - |
| 4980 | 1 | 0.2um_NOKCPHJN_730373 | Bacteria | Verrucomicrol | Opitutae | Opitutales | Opitutaceae | Opitutus | Opitutus terra | poorly charact | Function unkn | 0NVRZ | NA | 170187.SP_17 | 1 | 0 | 1,57E-05 | 1,49E-06 | 1 - | 1,31E-05 - | 1 - | 1,63E-05 - | 1 - | 1,55E-05 - | 1 - | 1,57E-05 - | 1 - | 1,77E-05 - |
| 4981 | 1 | 3um_FIJJAFBF_98740 | Bacteria | Bacteroidetes | Flavobacteriia | Flavobacterial | Flavobacteriac | Ulvibacter | Ulvibacter sp. | poorly charact | Function unkn | 11TW9 | Pfam:DUF583 | 50743.SCB49_ | 0,8 | 0,748331477 | 6,75E-05 | 5,96E-05 | 2 - | 0,000154063 - | 0 - | 0 - | 1 - | 9,09E-05 - | 1 - | 9,23E-05 - | 0 - | 0 - |
| 4982 | 1 | 3um_FIJJAFBF_199362 | Bacteria | Proteobacteri | Gammaprotec | Oceanospirilla | Oceanospirilla | Marinobacteri | Marinobacteri | information st | Translation, ri | 0QTNF | One of the pri | 247633.GP214 | 1,4 | 0,489897949 | 0,000215033 | 6,15E-05 | 2 - | 0,000263855 - | 1 - | 0,000163815 - | 2 - | 0,000311453 - | 1 - | 0,000158027 - | 1 - | 0,000178017 - |
| 4983 | 1 | 3um_FIJJAFBF_37652 | Bacteria | Proteobacteri | Gammaprotec | unclassified | unclassified | unclassified | various | unclassified | unclassified | unclassified | unclassified | unclassified | 1,8 | 0,4 | 0,000140508 | 2,77E-05 | 2 - | 0,00013269 - | 2 - | 0,000164762 - | 2 - | 0,000156627 - | 2 - | 0,00015894 - | 1 - | 8,95E-05 - |
| 4984 | 1 | 3um_FIJJAFBF_297317 | Bacteria | Bacteroidetes | Flavobacteriia | Flavobacterial | Flavobacteriac | Polaribacter | Polaribacter s | metabolism | Carbohydrate | 05CA9 | Pyruvate kina | 411154.GFO_ | 2,2 | 0,748331477 | 6,41E-05 | 2,49E-05 | 2 - | 4,83E-05 - | 2 - | 6,00E-05 - | 3 - | 8,56E-05 - | 1 - | 2,89E-05 - | 3 - | 9,78E-05 - |
| 4985 | 1 | 0.2um_NOKCPHJN_710137 | Eukaryota | unclassified | Oomycetes | Peronosporale | Peronosporac | Phytophthora | Phytophthora | poorly charact | Function unkn | 0XQGZ | guanine nucle | 67593.Physo1 | 0,6 | 0,489897949 | 3,04E-05 | 2,53E-05 | 1 - | 4,24E-05 - | 1 - | 5,26E-05 - | 0 - | 0 - | 0 - | 0 - | 1 - | 5,71E-05 - |
| 4986 | 1 | 3um_FIJJAFBF_751701 | Bacteria | Proteobacteri | Alphaproteob | Rhodobactera | Rhodobactera | Planktomarina | Planktomarina | metabolism | Carbohydrate | 0XRJK | TRAP dicarbox | 388739.RSK20 | 3,6 | 0,489897949 | 0,000146938 | 2,90E-05 | 3 - | 0,000101873 - | 3 - | 0,000126496 - | 4 - | 0,000160334 - | 4 - | 0,000162702 - | 4 - | 0,000183283 - |
| 4987 | 1 | 0.2um_NOKCPHJN_779798 | Bacteria | Bacteroidetes | Flavobacteriia | Flavobacterial | Flavobacteriac | Siansivirga | Siansivirga zea | cellular proces | Posttranslatio | 0XNQ9 | peptidase (S8 | 391603.FBALC | 1,2 | 0,4 | 3,01E-05 | 6,66E-06 | 2 - | 4,31E-05 - | 1 - | 2,68E-05 - | 1 - | 2,55E-05 - | 1 - | 2,58E-05 - | 1 - | 2,91E-05 - |
| 4988 | 1 | 0.2um_NOKCPHJN_321472 | unclassified | unclassified | unclassified | unclassified | unclassified | unclassified | unclassified | unclassified | unclassified | unclassified | unclassified | unclassified | 1 | 0 | 1,84E-05 | 1,74E-06 | 1 - | 1,54E-05 - | 1 - | 1,91E-05 - | 1 - | 1,82E-05 - | 1 - | 1,84E-05 - | 1 - | 2,08E-05 - |
| 4989 | 1 | 0.2um_NOKCPHJN_470938 | Bacteria | Bacteroidetes | Flavobacteriia | unclassified | unclassified | unclassified | Flavobacteria | metabolism | Secondary me | 07US8 | Inherit from C | 167555.NATL1 | 4,2 | 0,979795897 | 6,19E-05 | 7,95E-06 | 6 - | 7,58E-05 - | 4 - | 6,27E-05 - | 4 - | 5,96E-05 - | 4 - | 6,05E-05 - | 3 - | 5,11E-05 - |
| 4990 | 2 | 0.2um_NOKCPHJN_252074;3um_FIJJAFBF_16432 | Bacteria | Proteobacteri | Alphaproteob | Rhodobactera | unclassified | unclassified | Rhodobactera | unclassified | unclassified | unclassified | unclassified | unclassified | 0,2 | 0,4 | 3,59E-06 | 7,18E-06 | 1 - | 1,80E-05 - | 0 - | 0 - | 0 - | 0 - | 0 - | 0 - | 0 - | 0 - |
| 4991 | 1 | 3um_FIJJAFBF_185357 | Bacteria | Bacteroidetes | Flavobacteriia | Flavobacterial | Flavobacteriac | Formosa | Formosa sp. H | information st | Translation, ri | 0PAE2 | One of the pri | 655815.ZPR_3 | 0,4 | 0,489897949 | 5,89E-05 | 7,25E-05 | 1 - | 0,000135031 - | 0 - | 0 - | 1 - | 0,000159391 - | 0 - | 0 - | 0 - | 0 - |
| 4992 | 1 | 3um_FIJJAFBF_744832 | Bacteria | Proteobacteri | Gammaprotec | Cellvibrionales | Spongiibacter | various | various | unclassified | unclassified | unclassified | unclassified | unclassified | 0,8 | 0,4 | 8,69E-05 | 4,42E-05 | 1 - | 9,41E-05 - | 1 - | 0,000116819 - | 1 - | 0,000111051 - | 1 - | 0,000112691 - | 0 - | 0 - |
| 4993 | 1 | 3um_FIJJAFBF_296483 | Bacteria | Bacteroidetes | Flavobacteriia | Flavobacterial | Flavobacteriac | Polaribacter | various | poorly charact | Function unkn | 11NCV | Protein of unk | 487797.Flav30 | 3 | 0,632455532 | 0,00026193 | 5,73E-05 | 3 - | 0,000220725 - | 4 - | 0,000365433 - | 3 - | 0,000260543 - | 3 - | 0,000264391 - | 2 - | 0,000198557 - |
| 4994 | 1 | 3um_FIJJAFBF_264645 | Bacteria | Bacteroidetes | Flavobacteriia | Flavobacterial | Flavobacteriac | Polaribacter | Polaribacter b | unclassified | unclassified | unclassified | unclassified | unclassified | 2,6 | 0,489897949 | 0,000137274 | 3,53E-05 | 2 - | 8,73E-05 - | 3 - | 0,000162569 - | 2 - | 0,000103028 - | 3 - | 0,000156825 - | 3 - | 0,000176663 - |
| 4995 | 1 | 0.2um_NOKCPHJN_313403 | unclassified | unclassified | unclassified | unclassified | unclassified | unclassified | unclassified | unclassified | unclassified | unclassified | unclassified | unclassified | 0,2 | 0,4 | 1,76E-05 | 3,52E-05 | 0 - | 0 - | 0 - | 0 - | 1 - | 8,80E-05 - | 0 - | 0 - | 0 - | 0 - |
| 4996 | 1 | 0.2um_NOKCPHJN_440914 | Bacteria | Proteobacteri | Gammaprotec | Cellvibrionales | Porticoccacea | unclassified | various | unclassified | unclassified | unclassified | unclassified | unclassified | 13,8 | 1,166190379 | 0,00021541 | 1,25E-05 | 15 - | 0,000197437 - | 13 - | 0,000212471 - | 14 - | 0,000217517 - | 15 - | 0,000236497 - | 12 - | 0,00021313 - |
| 4997 | 1 | 3um_FIJJAFBF_22964 | Bacteria | Actinobacteria | Actinobacteria | Micrococcales | Microbacteria | Candidatus Ac | Candidatus Ac | unclassified | unclassified | unclassified | unclassified | unclassified | 0,4 | 0,489897949 | 8,93E-06 | 1,11E-05 | 1 - | 1,90E-05 - | 0 - | 0 - | 0 - | 0 - | 0 - | 0 - | 1 - | 2,56E-05 - |
| 4998 | 1 | 0.2um_NOKCPHJN_69744 | Bacteria | Bacteroidetes | Flavobacteriia | Flavobacterial | Flavobacteriac | Formosa | Formosa sp. H | unclassified | unclassified | unclassified | unclassified | unclassified | 3,4 | 0,8 | 8,61E-05 | 1,79E-05 | 4 - | 8,61E-05 - | 4 - | 0,000106956 - | 3 - | 7,63E-05 - | 4 - | 0,000103177 - | 2 - | 5,81E-05 - |
| 4999 | 1 | 3um_FIJJAFBF_516704 | Bacteria | Bacteroidetes | Flavobacteriia | Flavobacterial | Flavobacteriac | Polaribacter | Polaribacter a | cellular proces | Intracellular tr | 0PBE9 | Preprotein tra | 313598.MED1 | 1 | 0 | 0,000130502 | 1,24E-05 | 1 - | 0,000109311 - | 1 - | 0,000135732 - | 1 - | 0,000129031 - | 1 - | 0,000130936 - | 1 - | 0,0001475 - |

| | | | | | | | | | | | | | | | | | | | | | | | | | | | | |
|---|---|---|---|---|---|---|---|---|---|---|---|---|---|---|---|---|---|---|---|---|---|---|---|---|---|---|---|---|
| 5000 | 1 | 0.2um_NOKCPHJN_1017195 | Bacteria | Bacteroidetes | Flavobacteriia | unclassified | unclassified | unclassified | Flavobacteria | poorly charact | Function unkn | 0PA0H | (Lipo)protein | 487797.Flav3( | 0,2 | 0,4 | 1,46E-05 | 2,93E-05 | 0 - | 0 - | 0 - | 0 - | 1 - | 7,32E-05 - | 0 - | 0 - | 0 - | 0 - |
| 5001 | 1 | 3um_FIJJAFBF_744826 | Bacteria | Proteobacteri | Gammaprotec | Pseudomonad | Moraxellaceae | Perlucidibaca | Perlucidibaca | information st | Translation, ri | COG0256 | This is one of t | 314287.GB22( | 1 | 0 | 0,000118127 | 1,12E-05 | 1 - | 9,89E-05 - | 1 - | 0,000122861 - | 1 - | 0,000116795 - | 1 - | 0,00011852 - | 1 - | 0,000133512 - |
| 5002 | 1 | 3um_FIJJAFBF_231202 | Bacteria | Bacteroidetes | Flavobacteriia | Flavobacterial | Flavobacteriac | Polaribacter | Polaribacter s | cellular proces | Intracellular tr | COG0653 | Part of the Sec | 313594.PI23P | 1,6 | 0,489897949 | 1,94E-05 | 5,50E-06 | 2 - | 2,07E-05 - | 2 - | 2,57E-05 - | 1 - | 1,22E-05 - | 2 - | 2,47E-05 - | 1 - | 1,39E-05 - |
| 5003 | 1 | 3um_FIJJAFBF_741169 | Eukaryota | Bacillariophyta | Coscinodiscop | Thalassiosirale | Thalassiosirac | Thalassiosira | Thalassiosira c | unclassified | unclassified | unclassified | unclassified | unclassified | 0,8 | 0,4 | 1,00E-05 | 5,10E-06 | 1 - | 1,09E-05 - | 1 - | 1,35E-05 - | 1 - | 1,28E-05 - | 1 - | 1,30E-05 - | 0 - | 0 - |
| 5004 | 1 | 3um_FIJJAFBF_377576 | Bacteria | Bacteroidetes | Flavobacteriia | Flavobacterial | Flavobacteriac | Polaribacter | Polaribacter s | information st | Translation, ri | COG0089 | One of the ear | 313598.MED1 | 2,6 | 0,489897949 | 0,000367302 | 5,97E-05 | 3 - | 0,000358677 - | 3 - | 0,000445371 - | 3 - | 0,000423382 - | 2 - | 0,000286424 - | 2 - | 0,000322655 - |
| 5005 | 1 | 3um_FIJJAFBF_433042 | Bacteria | Proteobacteri | Alphaproteob | Rhodobactera | Rhodobactera | Planktomarina | Planktomarina | metabolism | Amino acid tra | 16PQJ | Dihydroxy-acid | 351016.RAZW | 1 | 0 | 2,28E-05 | 2,16E-06 | 1 - | 1,91E-05 - | 1 - | 2,37E-05 - | 1 - | 2,25E-05 - | 1 - | 2,29E-05 - | 1 - | 2,58E-05 - |
| 5006 | 1 | 0.2um_NOKCPHJN_224384 | Bacteria | Proteobacteri | Alphaproteob | Pelagibacteral | Pelagibacterac | Candidatus Pe | Candidatus Pe | unclassified | unclassified | unclassified | unclassified | unclassified | 0,4 | 0,489897949 | 3,96E-06 | 4,86E-06 | 0 - | 0 - | 1 - | 9,49E-06 - | 0 - | 0 - | 0 - | 0 - | 1 - | 1,03E-05 - |
| 5007 | 1 | 0.2um_NOKCPHJN_09727 | Bacteria | Bacteroidetes | Flavobacteriia | Flavobacterial | Flavobacteriac | Formosa | Formosa sp. H | cellular proces | Intracellular tr | COG0653 | Part of the Sec | 391603.FBALC | 1,2 | 0,4 | 1,43E-05 | 4,92E-06 | 1 - | 9,95E-06 - | 1 - | 1,23E-05 - | 1 - | 1,17E-05 - | 2 - | 2,38E-05 - | 1 - | 1,34E-05 - |
| 5008 | 1 | 0.2um_NOKCPHJN_751739 | Eukaryota | unclassified | Labyrinthulom | unclassified | Thraustochytr | Aurantiochytr | Aurantiochytr | poorly charact | Function unkn | 0XQX6 | Rhophilin, Rhc | 4787.PITG_00 | 0,6 | 0,8 | 1,01E-05 | 1,33E-05 | 0 - | 0 - | 1 - | 1,73E-05 - | 2 - | 3,30E-05 - | 0 - | 0 - | 0 - | 0 - |
| 5009 | 1 | 3um_FIJJAFBF_159918 | Bacteria | Bacteroidetes | Flavobacteriia | Flavobacterial | Flavobacteriac | unclassified | Flavobacteriac | unclassified | unclassified | unclassified | unclassified | unclassified | 0,6 | 0,489897949 | 3,22E-05 | 2,66E-05 | 1 - | 4,70E-05 - | 1 - | 5,84E-05 - | 1 - | 5,55E-05 - | 0 - | 0 - | 0 - | 0 - |
| 5010 | 1 | 3um_FIJJAFBF_228152 | Bacteria | Proteobacteri | Gammaprotec | unclassified | unclassified | unclassified | Gammaprotec | information st | Translation, ri | 16Q16 | Binds the lowe | 203122.Sde_0 | 1,8 | 0,748331477 | 0,000108563 | 3,86E-05 | 3 - | 0,000155104 - | 1 - | 6,42E-05 - | 2 - | 0,000122056 - | 1 - | 6,19E-05 - | 2 - | 0,000139527 - |
| 5011 | 1 | 0.2um_NOKCPHJN_229045 | Eukaryota | Phaeophyceae | unclassified | Ectocarpales | Ectocarpaceae | Ectocarpus | Ectocarpus sili | cellular proces | Signal transdu | KOG0585 | Calcium calmc | 4787.PITG_08 | 0,6 | 0,489897949 | 5,02E-06 | 4,11E-06 | 0 - | 0 - | 1 - | 8,26E-06 - | 1 - | 7,85E-06 - | 0 - | 0 - | 1 - | 8,97E-06 - |
| 5012 | 2 | 0.2um_NOKCPHJN_326825;0.2um_NOKCPHJN_5 | Bacteria | various | various | various | various | various | various | metabolism | Amino acid tra | COG0174 | glutamine syn | 684719.HIMB | 1,2 | 0,748331477 | 3,44E-05 | 2,06E-05 | 2 - | 4,90E-05 - | 2 - | 6,09E-05 - | 1 - | 2,89E-05 - | 0 - | 0 - | 1 - | 3,31E-05 - |
| 5013 | 1 | 3um_FIJJAFBF_190049 | Bacteria | Proteobacteri | Alphaproteob | Rhodobactera | Rhodobactera | Labrenzia | various | unclassified | unclassified | unclassified | unclassified | unclassified | 3 | 1,095445115 | 0,000149441 | 3,83E-05 | 5 - | 0,000215746 - | 2 - | 0,000107157 - | 3 - | 0,0001528 - | 3 - | 0,000155056 - | 2 - | 0,000116447 - |
| 5014 | 1 | 3um_FIJJAFBF_362243 | unclassified | unclassified | unclassified | unclassified | unclassified | unclassified | unclassified | unclassified | unclassified | unclassified | unclassified | unclassified | 2 | 0,632455532 | 0,000350017 | 9,16E-05 | 3 - | 0,000447182 - | 2 - | 0,000370179 - | 2 - | 0,000351902 - | 1 - | 0,00017855 - | 2 - | 0,000402271 - |
| 5015 | 1 | 0.2um_NOKCPHJN_220732 | Bacteria | Bacteroidetes | Flavobacteriia | Flavobacterial | Flavobacteriac | Winogradskye | Winogradskye | information st | Replication, re | 05C24 | DNA gyrase ne | 313590.MED1 | 1,8 | 0,748331477 | 2,80E-05 | 9,11E-06 | 3 - | 4,07E-05 - | 1 - | 1,68E-05 - | 2 - | 3,20E-05 - | 2 - | 3,25E-05 - | 1 - | 1,83E-05 - |
| 5016 | 1 | 3um_FIJJAFBF_269014 | Bacteria | Bacteroidetes | Flavobacteriia | Flavobacterial | Flavobacteriac | Winogradskye | Winogradskye | metabolism | Carbohydrate | COG0149 | triosephospha | 391603.FBALC | 1,4 | 0,489897949 | 7,42E-05 | 2,12E-05 | 2 - | 9,11E-05 - | 1 - | 5,66E-05 - | 2 - | 0,000107526 - | 1 - | 5,46E-05 - | 1 - | 6,15E-05 - |
| 5017 | 2 | 0.2um_NOKCPHJN_100726;3um_FIJJAFBF_55152 | Bacteria | Bacteroidetes | Flavobacteriia | Flavobacterial | Flavobacteriac | Aureitalea | Aureitalea sp. | unclassified | unclassified | unclassified | unclassified | unclassified | 1,4 | 0,489897949 | 6,11E-05 | 1,75E-05 | 2 - | 7,50E-05 - | 1 - | 4,66E-05 - | 2 - | 8,86E-05 - | 1 - | 4,49E-05 - | 1 - | 5,06E-05 - |
| 5018 | 1 | 3um_FIJJAFBF_155127 | Bacteria | Bacteroidetes | Cytophagia | Cytophagales | Flammeovirga | unclassified | Flammeovirga | unclassified | unclassified | unclassified | unclassified | unclassified | 6,6 | 0,489897949 | 0,00016347 | 1,35E-05 | 7 - | 0,000145814 - | 7 - | 0,000181058 - | 7 - | 0,000172119 - | 6 - | 0,00014971 - | 6 - | 0,000168647 - |
| 5019 | 1 | 0.2um_NOKCPHJN_171709 | Bacteria | Bacteroidetes | Flavobacteriia | Flavobacterial | Flavobacteriac | Psychroserpe | Psychroserpe | metabolism | Coenzyme tra | 09A4R | May play a ke | 391587.KAOT | 1 | 0 | 3,09E-05 | 2,93E-06 | 1 - | 2,59E-05 - | 1 - | 3,21E-05 - | 1 - | 3,05E-05 - | 1 - | 3,10E-05 - | 1 - | 3,49E-05 - |
| 5020 | 1 | 0.2um_NOKCPHJN_334091 | Bacteria | Bacteroidetes | Flavobacteriia | Flavobacterial | Flavobacteriac | Polaribacter | Polaribacter s | unclassified | unclassified | unclassified | unclassified | unclassified | 0,6 | 0,489897949 | 3,80E-05 | 3,13E-05 | 1 - | 5,52E-05 - | 1 - | 6,85E-05 - | 0 - | 0 - | 1 - | 6,61E-05 - | 0 - | 0 - |
| 5021 | 1 | 0.2um_NOKCPHJN_95911 | Bacteria | Bacteroidetes | Flavobacteriia | Flavobacterial | Flavobacteriac | Aquimarina | Aquimarina m | unclassified | unclassified | unclassified | unclassified | unclassified | 4,8 | 0,4 | 0,000103343 | 7,96E-06 | 5 - | 9,07E-05 - | 5 - | 0,000112574 - | 5 - | 0,000107016 - | 5 - | 0,000108597 - | 4 - | 9,79E-05 - |
| 5022 | 1 | 3um_FIJJAFBF_280837 | Bacteria | Bacteroidetes | Flavobacteriia | Flavobacterial | Crocinitomica | unclassified | Crocinitomica | information st | Translation, ri | COG0184 | Forms an inte | 398720.MED2 | 1,4 | 0,489897949 | 0,000210201 | 6,01E-05 | 2 - | 0,000257925 - | 1 - | 0,000160134 - | 2 - | 0,000304454 - | 1 - | 0,000154476 - | 1 - | 0,000174016 - |
| 5023 | 1 | 3um_FIJJAFBF_100079 | Bacteria | Bacteroidetes | Flavobacteriia | Flavobacterial | Flavobacteriac | Formosa | Formosa sp. H | unclassified | unclassified | unclassified | unclassified | unclassified | 0,8 | 0,4 | 1,55E-05 | 7,89E-06 | 1 - | 1,68E-05 - | 1 - | 2,09E-05 - | 1 - | 1,98E-05 - | 1 - | 2,01E-05 - | 0 - | 0 - |
| 5024 | 1 | 3um_FIJJAFBF_183329 | Bacteria | Bacteroidetes | Flavobacteriia | Flavobacterial | Flavobacteriac | Polaribacter | various | information st | Translation, ri | 05KK9 | This protein bi | 313594.PI23P | 4 | 0,632455532 | 0,000325729 | 4,34E-05 | 5 - | 0,000343643 - | 4 - | 0,000341363 - | 3 - | 0,000243381 - | 4 - | 0,000329301 - | 4 - | 0,000370957 - |
| 5025 | 1 | 0.2um_NOKCPHJN_233764 | Bacteria | Proteobacteri | Alphaproteob | Rhodobactera | Rhodobactera | Amylibacter | Amylibacter ci | metabolism | Inorganic ion t | COG3221 | phosphonate | 439497.RR11 | 0,4 | 0,489897949 | 1,67E-05 | 2,05E-05 | 1 - | 3,83E-05 - | 0 - | 0 - | 1 - | 4,52E-05 - | 0 - | 0 - | 0 - | 0 - |
| 5026 | 1 | 3um_FIJJAFBF_755990 | Eukaryota | Bacillariophyta | Coscinodiscop | Thalassiosirale | Thalassiosirac | Thalassiosira | Thalassiosira p | unclassified | unclassified | unclassified | unclassified | unclassified | 0,8 | 0,4 | 4,86E-05 | 2,47E-05 | 1 - | 5,26E-05 - | 1 - | 6,54E-05 - | 1 - | 6,21E-05 - | 1 - | 6,31E-05 - | 0 - | 0 - |
| 5027 | 1 | 3um_FIJJAFBF_326228 | Bacteria | Proteobacteri | Alphaproteob | Rhodobactera | Rhodobactera | Ruegeria | Ruegeria med | metabolism | Amino acid tra | 01SEG | branched-chai | 439496.RBY4I | 3 | 1,095445115 | 9,57E-05 | 3,56E-05 | 3 - | 8,06E-05 - | 1 - | 3,34E-05 - | 4 - | 0,000126915 - | 4 - | 0,00012879 - | 3 - | 0,000108811 - |
| 5028 | 1 | 3um_FIJJAFBF_402483 | Bacteria | Bacteroidetes | Flavobacteriia | various | various | various | various | unclassified | unclassified | unclassified | unclassified | unclassified | 3,8 | 0,979795897 | 0,000377122 | 7,44E-05 | 5 - | 0,000425099 - | 4 - | 0,000422278 - | 4 - | 0,000401429 - | 4 - | 0,000407358 - | 2 - | 0,000229444 - |
| 5029 | 1 | 0.2um_NOKCPHJN_404386 | Bacteria | Proteobacteri | Gammaprotec | unclassified | unclassified | unclassified | Gammaprotec | information st | Translation, ri | 05KK9 | This protein bi | 247639.MGP2 | 4,2 | 0,979795897 | 0,000543765 | 8,37E-05 | 6 - | 0,000662174 - | 3 - | 0,000411112 - | 4 - | 0,000521085 - | 4 - | 0,000528782 - | 4 - | 0,000595671 - |
| 5030 | 1 | 3um_FIJJAFBF_263676 | Bacteria | Proteobacteri | Alphaproteob | unclassified | unclassified | Candidatus Pu | Candidatus Pu | metabolism | Carbohydrate | 0IVA1 | Photosystem I | 45157.CMV20 | 8,8 | 0,979795897 | 0,00065628 | 0,000103293 | 9 - | 0,000561408 - | 10 - | 0,000774559 - | 7 - | 0,000515422 - | 9 - | 0,000672473 - | 9 - | 0,000757538 - |
| 5031 | 1 | 3um_FIJJAFBF_706939 | Bacteria | Proteobacteri | Gammaprotec | Oceanospirilla | Saccharospirill | Reinekea | Reinekea fors | metabolism | Energy produc | 17C3H | Produces ATP | 314283.MED2 | 2 | 0 | 0,000194365 | 1,85E-05 | 2 - | 0,000162804 - | 2 - | 0,000202154 - | 2 - | 0,000192173 - | 2 - | 0,000195012 - | 2 - | 0,00021968 - |
| 5032 | 1 | 3um_FIJJAFBF_116291 | Bacteria | Proteobacteri | Betaproteoba | unclassified | unclassified | unclassified | Betaproteoba | cellular proces | Intracellular tr | 05VIE | preprotein tra | 365046.Rta_3 | 3,6 | 1,2 | 0,000390201 | 8,64E-05 | 6 - | 0,000559887 - | 3 - | 0,000347607 - | 3 - | 0,000330444 - | 3 - | 0,000335325 - | 3 - | 0,000377743 - |
| 5033 | 1 | 3um_FIJJAFBF_20008 | Bacteria | Bacteroidetes | Flavobacteriia | Flavobacterial | Flavobacteriac | Formosa | Formosa sp. H | unclassified | unclassified | unclassified | unclassified | unclassified | 1 | 0 | 1,37E-05 | 1,30E-06 | 1 - | 1,15E-05 - | 1 - | 1,43E-05 - | 1 - | 1,36E-05 - | 1 - | 1,38E-05 - | 1 - | 1,55E-05 - |
| 5034 | 1 | 0.2um_NOKCPHJN_316399 | Bacteria | Proteobacteri | Gammaprotec | unclassified | unclassified | unclassified | SAR86 cluster | metabolism | Inorganic ion t | 0QK8T | TonB-depende | 247633.GP214 | 11 | 2 | 0,000133119 | 1,93E-05 | 14 - | 0,000143343 - | 10 - | 0,000127135 - | 12 - | 0,00014503 - | 8 - | 9,81E-05 - | 11 - | 0,000151973 - |
| 5035 | 1 | 0.2um_NOKCPHJN_627845 | Bacteria | Proteobacteri | Alphaproteob | Rhodobactera | unclassified | unclassified | Rhodobactera | unclassified | unclassified | unclassified | unclassified | unclassified | 1,4 | 0,489897949 | 4,91E-05 | 1,74E-05 | 1 - | 2,94E-05 - | 1 - | 3,65E-05 - | 2 - | 6,95E-05 - | 2 - | 7,05E-05 - | 1 - | 3,97E-05 - |
| 5036 | 1 | 0.2um_NOKCPHJN_04158 | Bacteria | Bacteroidetes | Cytophagia | Cytophagales | unclassified | unclassified | various | metabolism | Energy produc | COG0372 | citrate syntha | 388413.ALPR1 | 0,4 | 0,489897949 | 1,28E-05 | 1,56E-05 | 0 - | 0 - | 0 - | 0 - | 1 - | 3,17E-05 - | 1 - | 3,21E-05 - | 0 - | 0 - |
| 5037 | 1 | 3um_FIJJAFBF_532805 | Bacteria | Proteobacteri | Alphaproteob | Rhodobactera | Rhodobactera | unclassified | Rhodobactera | unclassified | unclassified | unclassified | unclassified | unclassified | 10,2 | 1,720465053 | 0,000371102 | 3,12E-05 | 13 - | 0,000402183 - | 10 - | 0,000384148 - | 11 - | 0,000401699 - | 9 - | 0,000333517 - | 8 - | 0,000333961 - |
| 5038 | 1 | 3um_FIJJAFBF_15538 | Bacteria | Bacteroidetes | Flavobacteriia | Flavobacterial | Flavobacteriac | Algibacter | Algibacter pe | unclassified | unclassified | unclassified | unclassified | unclassified | 1 | 0 | 0,000150579 | 1,43E-05 | 1 - | 0,000126128 - | 1 - | 0,000156614 - | 1 - | 0,000148882 - | 1 - | 0,000151081 - | 1 - | 0,000170192 - |
| 5039 | 1 | 3um_FIJJAFBF_163563 | Bacteria | Proteobacteri | Gammaprotec | Oceanospirilla | Saccharospirill | Reinekea | Reinekea fors | unclassified | unclassified | unclassified | unclassified | unclassified | 0,4 | 0,489897949 | 1,49E-05 | 1,82E-05 | 0 - | 0 - | 0 - | 0 - | 1 - | 3,69E-05 - | 1 - | 3,75E-05 - | 0 - | 0 - |
| 5040 | 1 | 0.2um_NOKCPHJN_09839 | Bacteria | Proteobacteri | Gammaprotec | Alteromonada | Psychromonac | Psychromonas | Psychromonas | poorly charact | Function unkn | COG3517 | type VI secreti | 357804.Ping_( | 10 | 2 | 0,000278226 | 3,46E-05 | 13 - | 0,000308285 - | 10 - | 0,00029446 - | 9 - | 0,00025193 - | 11 - | 0,000312462 - | 7 - | 0,000223992 - |
| 5041 | 1 | 0.2um_NOKCPHJN_548584 | Bacteria | Proteobacteri | Gammaprotec | unclassified | unclassified | unclassified | Gammaprotec | information st | Translation, ri | 16QAF | thus facilitatin | 379731.PST_2 | 4,4 | 1,624807681 | 0,000102476 | 3,15E-05 | 7 - | 0,000139244 - | 5 - | 0,0001235 - | 2 - | 4,70E-05 - | 4 - | 9,53E-05 - | 4 - | 0,000107365 - |
| 5042 | 1 | 0.2um_NOKCPHJN_05749 | Bacteria | Bacteroidetes | Sphingobacter | Sphingobacter | Sphingobacter | Pedobacter | Pedobacter ru | metabolism | Inorganic ion t | COG0376 | Bifunctional e | 761193.Runsl | 3,8 | 1,720465053 | 6,82E-05 | 3,05E-05 | 4 - | 6,16E-05 - | 3 - | 5,74E-05 - | 6 - | 0,000109113 - | 5 - | 9,23E-05 - | 1 - | 2,08E-05 - |
| 5043 | 2 | 0.2um_NOKCPHJN_397965;3um_FIJJAFBF_38635 | Bacteria | Proteobacteri | Alphaproteob | Rhodobactera | Rhodobactera | Marinovum | Marinovum al | metabolism | Lipid transpor | 05CMX | Polyhydroxyal | various | 0,2 | 0,4 | 6,80E-06 | 1,36E-05 | 0 - | 0 - | 1 - | 3,40E-05 - | 0 - | 0 - | 0 - | 0 - | 0 - | 0 - |
| 5044 | 1 | 0.2um_NOKCPHJN_715183 | Bacteria | Actinobacteria | Actinobacteria | Nakamurellale | Nakamurellac | Nakamurella | Nakamurella l | unclassified | unclassified | unclassified | unclassified | unclassified | 0,4 | 0,489897949 | 1,25E-05 | 1,53E-05 | 0 - | 0 - | 0 - | 0 - | 1 - | 3,10E-05 - | 1 - | 3,15E-05 - | 0 - | 0 - |
| 5045 | 1 | 3um_FIJJAFBF_125686 | Eukaryota | Bacillariophyta | Bacillariophyc | Naviculales | Phaeodactylac | Phaeodactylu | Phaeodactylu | unclassified | unclassified | unclassified | unclassified | unclassified | 1 | 0 | 5,35E-05 | 5,08E-06 | 1 - | 4,48E-05 - | 1 - | 5,57E-05 - | 1 - | 5,29E-05 - | 1 - | 5,37E-05 - | 1 - | 6,05E-05 - |
| 5046 | 2 | 3um_FIJJAFBF_180937;3um_FIJJAFBF_598590 | Bacteria | Bacteroidetes | Flavobacteriia | Flavobacterial | Flavobacteriac | various | various | poorly charact | Function unkn | COG0714 | ATPase activit | 1046627.BZAF | 1,8 | 0,4 | 7,35E-05 | 1,72E-05 | 2 - | 6,87E-05 - | 1 - | 4,27E-05 - | 2 - | 8,11E-05 - | 2 - | 8,23E-05 - | 2 - | 9,27E-05 - |
| 5047 | 1 | 0.2um_NOKCPHJN_319417 | Bacteria | Bacteroidetes | Flavobacteriia | various | various | various | various | information st | Translation, ri | 05KK9 | This protein bi | 487797.Flav3( | 4,2 | 0,748331477 | 0,000491083 | 7,18E-05 | 5 - | 0,000494727 - | 3 - | 0,000368583 - | 5 - | 0,000583975 - | 4 - | 0,00047408 - | 4 - | 0,00053405 - |
| 5048 | 2 | 0.2um_NOKCPHJN_424280;3um_FIJJAFBF_4828( | Bacteria | Proteobacteri | Alphaproteob | Rhodobactera | Rhodobactera | Ahrensia | Ahrensia sp. R | metabolism | Lipid transpor | COG1024 | Enoyl-CoA hyc | 744979.R2A13 | 0,2 | 0,4 | 1,03E-05 | 2,05E-05 | 0 - | 0 - | 0 - | 0 - | 1 - | 5,13E-05 - | 0 - | 0 - | 0 - | 0 - |
| 5049 | 1 | 0.2um_NOKCPHJN_428865 | Bacteria | Bacteroidetes | Cytophagia | Cytophagales | Cyclobacteriac | Aquiflexum | Aquiflexum ba | unclassified | unclassified | unclassified | unclassified | unclassified | 1,6 | 0,8 | 3,52E-05 | 1,81E-05 | 2 - | 3,68E-05 - | 2 - | 4,57E-05 - | 0 - | 0 - | 2 - | 4,41E-05 - | 2 - | 4,96E-05 - |
| 5050 | 2 | 0.2um_NOKCPHJN_296468;3um_FIJJAFBF_6601( | Bacteria | Bacteroidetes | Flavobacteriia | unclassified | unclassified | unclassified | various | information st | Translation, ri | COG0052 | 30S ribosomal | 487797.Flav3( | 1,4 | 1,019803903 | 6,30E-05 | 4,66E-05 | 2 - | 7,63E-05 - | 3 - | 0,000142045 - | 1 - | 4,50E-05 - | 0 - | 0 - | 1 - | 5,15E-05 - |
| 5051 | 1 | 0.2um_NOKCPHJN_1052566 | Eukaryota | unclassified | Cryptophyta | Pyrenomonad | Geminigerace | Guillardia | Guillardia thet | unclassified | unclassified | unclassified | unclassified | unclassified | 1,2 | 0,979795897 | 2,79E-05 | 2,02E-05 | 3 - | 6,32E-05 - | 1 - | 2,62E-05 - | 1 - | 2,49E-05 - | 1 - | 2,52E-05 - | 0 - | 0 - |
| 5052 | 1 | 3um_FIJJAFBF_228148 | Bacteria | Proteobacteri | Gammaprotec | unclassified | unclassified | unclassified | various | unclassified | unclassified | unclassified | unclassified | unclassified | 1 | 0,632455532 | 0,000105744 | 6,03E-05 | 2 - | 0,000188159 - | 1 - | 0,000116819 - | 1 - | 0,000111051 - | 1 - | 0,000112691 - | 0 - | 0 - |
| 5053 | 1 | 3um_FIJJAFBF_119151 | Bacteria | Bacteroidetes | Flavobacteriia | Flavobacterial | Flavobacteriac | Tenacibaculur | Tenacibaculur | unclassified | unclassified | unclassified | unclassified | unclassified | 4,8 | 1,326649916 | 0,000273021 | 5,11E-05 | 7 - | 0,000341888 - | 5 - | 0,000303232 - | 5 - | 0,00028826 - | 4 - | 0,000234014 - | 3 - | 0,000197712 - |
| 5054 | 1 | 3um_FIJJAFBF_453412 | Bacteria | Proteobacteri | Alphaproteob | Rhodobactera | unclassified | unclassified | Rhodobactera | poorly charact | Function unkn | COG3181 | extra-cytoplas | 388739.RSK20 | 5,2 | 1,326649916 | 0,000214537 | 4,46E-05 | 7 - | 0,000245699 - | 3 - | 0,000130751 - | 5 - | 0,000207159 - | 6 - | 0,000252263 - | 5 - | 0,000236811 - |
| 5055 | 1 | 3um_FIJJAFBF_801365 | Bacteria | Proteobacteri | Alphaproteob | Rhodobactera | Rhodobactera | Sulfitobacter | Sulfitobacter g | metabolism | Energy produc | COG0372 | citrate syntha | 633131.TR2A6 | 0,8 | 0,4 | 2,40E-05 | 1,22E-05 | 1 - | 2,60E-05 - | 1 - | 3,23E-05 - | 1 - | 3,07E-05 - | 1 - | 3,12E-05 - | 0 - | 0 - |
| 5056 | 1 | 3um_FIJJAFBF_254302 | Bacteria | Proteobacteri | Gammaprotec | unclassified | unclassified | unclassified | SAR86 cluster | unclassified | unclassified | unclassified | unclassified | unclassified | 1,2 | 0,4 | 9,49E-05 | 3,18E-05 | 1 - | 6,63E-05 - | 1 - | 8,24E-05 - | 2 - | 0,000156627 - | 1 - | 7,95E-05 - | 1 - | 8,95E-05 - |
| 5057 | 1 | 0.2um_NOKCPHJN_596448 | Bacteria | Bacteroidetes | Bacteroidia | Marinilabiliale | Prolixibactera | Prolixibacter | Prolixibacter d | unclassified | unclassified | unclassified | unclassified | unclassified | 0,8 | 0,4 | 3,24E-05 | 1,65E-05 | 1 - | 3,51E-05 - | 1 - | 4,36E-05 - | 1 - | 4,14E-05 - | 1 - | 4,20E-05 - | 0 - | 0 - |
| 5058 | 1 | 3um_FIJJAFBF_203258 | Bacteria | Proteobacteri | Gammaprotec | Arenicellales | Arenicellaceae | Arenicella | Arenicella xan | metabolism | Inorganic ion t | 16TQC | TONB-depend | 247639.MGP2 | 6,6 | 1,356465997 | 9,19E-05 | 1,09E-05 | 9 - | 0,000106824 - | 7 - | 0,000103168 - | 6 - | 8,41E-05 - | 6 - | 8,53E-05 - | 5 - | 8,01E-05 - |
| 5059 | 1 | 0.2um_NOKCPHJN_889562 | Bacteria | Proteobacteri | Alphaproteob | Pelagibacteral | Pelagibacterac | Candidatus Pe | Candidatus Pe | metabolism | Secondary me | 01Q86 | extracellular s | 439493.PB721 | 3,6 | 0,8 | 0,000353349 | 4,63E-05 | 5 - | 0,000418893 - | 3 - | 0,000312085 - | 4 - | 0,000395569 - | 3 - | 0,000301058 - | 3 - | 0,000339141 - |
| 5060 | 1 | 3um_FIJJAFBF_347373 | Bacteria | Proteobacteri | Gammaprotec | Cellvibrionales | unclassified | unclassified | Cellvibrionales | poorly charact | Function unkn | 1EB8T | NA | 436017.A4S9C | 2 | 0,894427191 | 3,83E-05 | 1,76E-05 | 2 - | 3,23E-05 - | 3 - | 6,01E-05 - | 1 - | 1,91E-05 - | 3 - | 5,80E-05 - | 1 - | 2,18E-05 - |
| 5061 | 1 | 3um_FIJJAFBF_524832 | Bacteria | Proteobacteri | Gammaprotec | Oceanospirilla | Saccharospirill | Reinekea | Reinekea fors | cellular proces | Posttranslatio | COG1225 | alkyl hydroper | 314283.MED2 | 1,4 | 0,489897949 | 0,000119295 | 3,75E-05 | 2 - | 0,000145287 - | 2 - | 0,000180404 - | 1 - | 8,57E-05 - | 1 - | 8,70E-05 - | 1 - | 9,80E-05 - |
| 5062 | 1 | 3um_FIJJAFBF_66474 | Bacteria | unclassified | unclassified | unclassified | unclassified | unclassified | uncultured ba | unclassified | unclassified | unclassified | unclassified | unclassified | 15,8 | 2,561249695 | 0,000225313 | 2,91E-05 | 18 - | 0,000217472 - | 18 - | 0,000270036 - | 16 - | 0,000228181 - | 16 - | 0,000231551 - | 11 - | 0,000179328 - |
| 5063 | 1 | 3um_FIJJAFBF_99476 | Bacteria | Proteobacteri | Alphaproteob | Rhodobactera | Rhodobactera | Yoonia | Yoonia litorea | information st | Transcription | 16PMT | DNA-depende | 391589.RGAI1 | 10,4 | 0,489897949 | 0,000421361 | 4,42E-05 | 11 - | 0,000373534 - | 10 - | 0,000421653 - | 10 - | 0,000400835 - | 10 - | 0,000406755 - | 11 - | 0,000504029 - |
| 5064 | 1 | 0.2um_NOKCPHJN_239339 | Bacteria | Proteobacteri | Gammaprotec | Cellvibrionales | Porticoccacea | unclassified | gamma protec | cellular proces | Posttranslatio | 0QJ3C | Acts as a chap | 203122.Sde_2 | 1,8 | 0,4 | 3,78E-05 | 7,46E-06 | 2 - | 3,57E-05 - | 2 - | 4,43E-05 - | 2 - | 4,21E-05 - | 2 - | 4,28E-05 - | 1 - | 2,41E-05 - |
| 5065 | 1 | 0.2um_NOKCPHJN_113830 | Bacteria | Proteobacteri | Gammaprotec | Oceanospirilla | Oceanospirilla | Neptuniibacte | Neptuniibacte | cellular proces | Posttranslatio | COG0330 | Band 7 protei | 207954.MED9 | 6,2 | 1,166190379 | 0,000288517 | 3,29E-05 | 8 - | 0,000316626 - | 6 - | 0,000294867 - | 7 - | 0,000327026 - | 5 - | 0,00023704 - | 5 - | 0,000267025 - |
| 5066 | 2 | 0.2um_NOKCPHJN_229458;0.2um_NOKCPHJN_5 | Bacteria | Proteobacteri | Alphaproteob | Pelagibacteral | various | various | various | unclassified | unclassified | unclassified | unclassified | unclassified | 1 | 0 | 0,000112317 | 1,07E-05 | 1 - | 9,41E-05 - | 1 - | 0,000116819 - | 1 - | 0,000111051 - | 1 - | 0,000112691 - | 1 - | 0,000126946 - |
| 5067 | 1 | 0.2um_NOKCPHJN_898928 | Eukaryota | Bacillariophyta | Bacillariophyc | Naviculales | Phaeodactylac | Phaeodactylu | Phaeodactylu | metabolism | Energy produc | COG2838 | Isocitrate deh | 2850.Phatr45( | 0,6 | 0,489897949 | 1,51E-05 | 1,24E-05 | 1 - | 2,19E-05 - | 1 - | 2,72E-05 - | 0 - | 0 - | 1 - | 2,62E-05 - | 0 - | 0 - |
| 5068 | 1 | 3um_FIJJAFBF_385769 | Bacteria | Bacteroidetes | Flavobacteriia | Flavobacterial | Flavobacteriac | Ulvibacter | Ulvibacter lito | metabolism | Energy produc | COG0437 | 4fe-4S ferredc | 50743.SCB49 | 6 | 1,673320053 | 8,33E-05 | 1,90E-05 | 8 - | 9,48E-05 - | 7 - | 0,000102955 - | 7 - | 9,79E-05 - | 4 - | 5,68E-05 - | 4 - | 6,39E-05 - |
| 5069 | 1 | 3um_FIJJAFBF_703978 | Bacteria | Proteobacteri | Gammaprotec | Oceanospirilla | Saccharospirill | Reinekea | Reinekea fors | metabolism | Lipid transpor | 0RAT3 | Amp-depende | 314283.MED2 | 6,8 | 0,979795897 | 0,00016783 | 1,78E-05 | 8 - | 0,000166948 - | 6 - | 0,000155475 - | 6 - | 0,000147799 - | 8 - | 0,000199976 - | 6 - | 0,000168954 - |
| 5070 | 1 | 3um_FIJJAFBF_545825 | Bacteria | Bacteroidetes | Flavobacteriia | Flavobacterial | Flavobacteriac | Polaribacter | Polaribacter s | information st | Translation, ri | 09FEM | Binds to the 2 | 313594.PI23P | 4 | 1,788854382 | 0,000357946 | 0,000155895 | 5 - | 0,000385157 - | 5 - | 0,000478251 - | 6 - | 0,000545566 - | 3 - | 0,000276812 - | 1 - | 0,000103943 - |
| 5071 | 10 | 0.2um_NOKCPHJN_130372;0.2um_NOKCPHJN_3 | Bacteria | Proteobacteri | Gammaprotec | Cellvibrionales | Porticoccacea | unclassified | various | information st | Transcription | 17CVY | Involved in th | 314287.GB22( | 0,2 | 0,4 | 1,82E-05 | 3,64E-05 | 0 - | 0 - | 0 - | 0 - | 1 - | 9,09E-05 - | 0 - | 0 - | 0 - | 0 - |
| 5072 | 1 | 0.2um_NOKCPHJN_07596 | Bacteria | Bacteroidetes | Flavobacteriia | Flavobacterial | Flavobacteriac | Tenacibaculur | various | information st | Replication, re | COG0187 | DNA gyrase ne | 391587.KAOT | 2 | 0,632455532 | 4,19E-05 | 1,30E-05 | 2 - | 3,56E-05 - | 2 - | 4,42E-05 - | 2 - | 4,20E-05 - | 3 - | 6,39E-05 - | 1 - | 2,40E-05 - |
| 5073 | 1 | 0.2um_NOKCPHJN_148150 | Bacteria | Bacteroidetes | Flavobacteriia | Flavobacterial | Flavobacteriac | Formosa | Formosa sp. H | metabolism | Energy produc | COG0711 | F(1)F(0) ATP s | 391587.KAOT | 3,8 | 1,469693846 | 0,000320005 | 0,000126029 | 2 - | 0,000139972 - | 3 - | 0,000260705 - | 5 - | 0,000413056 - | 6 - | 0,000502988 - | 3 - | 0,000283307 - |
| 5074 | 1 | 3um_FIJJAFBF_676357 | Eukaryota | Chordata | Actinopteri | Cyprinodontifi | Nothobranchi | Nothobranchi | Nothobranchi | information st | Chromatin stri | KOG3467 | Core compone | 6085.XP_0021 | 2 | 0 | 0,000212445 | 2,02E-05 | 2 - | 0,000177948 - | 2 - | 0,000220959 - | 2 - | 0,00021005 - | 2 - | 0,000213152 - | 2 - | 0,000240115 - |
| 5075 | 1 | 0.2um_NOKCPHJN_263666 | Viruses | unclassified | unclassified | unclassified | unclassified | unclassified | uncultured vir | unclassified | unclassified | unclassified | unclassified | unclassified | 1,8 | 0,4 | 8,13E-05 | 1,90E-05 | 2 - | 7,60E-05 - | 1 - | 4,72E-05 - | 2 - | 8,97E-05 - | 2 - | 9,10E-05 - | 2 - | 0,000102566 - |
| 5076 | 1 | 0.2um_NOKCPHJN_40862 | Bacteria | Proteobacteri | Gammaprotec | unclassified | unclassified | unclassified | Gammaprotec | cellular proces | Intracellular tr | 0QJ7H | motA TolQ Exl | 342610.Patl_1 | 1,2 | 0,4 | 3,40E-05 | 1,17E-05 | 1 - | 2,37E-05 - | 1 - | 2,94E-05 - | 1 - | 2,80E-05 - | 2 - | 5,68E-05 - | 1 - | 3,20E-05 - |
| 5077 | 1 | 3um_FIJJAFBF_203230 | Bacteria | unclassified | unclassified | unclassified | unclassified | unclassified | bacterium BM | unclassified | unclassified | unclassified | unclassified | unclassified | 2,6 | 1,496662955 | 6,23E-05 | 3,10E-05 | 5 - | 0,000104915 - | 3 - | 7,82E-05 - | 1 - | 2,48E-05 - | 3 - | 7,54E-05 - | 1 - | 2,83E-05 - |
| 5078 | 1 | 0.2um_NOKCPHJN_230055 | Bacteria | unclassified | unclassified | unclassified | unclassified | unclassified | uncultured ba | metabolism | Carbohydrate | 0P7K6 | Beta-glucosid | 867900.Celly | 0,4 | 0,489897949 | 6,52E-06 | 8,03E-06 | 1 - | 1,49E-05 - | 0 - | 0 - | 1 - | 1,76E-05 - | 0 - | 0 - | 0 - | 0 - |
| 5079 | 1 | 3um_FIJJAFBF_72901 | Bacteria | Proteobacteri | Alphaproteob | Rhodobactera | Rhodobactera | Planktomarina | Planktomarina | poorly charact | Function unkn | 173PW | Pfam:DUF88 | 644107.SL115 | 2 | 0 | 0,000145774 | 1,38E-05 | 2 - | 0,000122103 - | 2 - | 0,000151616 - | 2 - | 0,00014413 - | 2 - | 0,000146259 - | 2 - | 0,00016476 - |
| 5080 | 1 | 0.2um_NOKCPHJN_210319 | Bacteria | Proteobacteri | Alphaproteob | Rhodobactera | Rhodobactera | Amylibacter | Amylibacter k | metabolism | Nucleotide tra | COG0138 | bifunctional p | 388401.RB215 | 0,2 | 0,4 | 5,38E-06 | 1,08E-05 | 0 - | 0 - | 1 - | 2,69E-05 - | 0 - | 0 - | 0 - | 0 - | 0 - | 0 - |
| 5081 | 1 | 3um_FIJJAFBF_276224 | unclassified | unclassified | unclassified | unclassified | unclassified | unclassified | unclassified | unclassified | unclassified | unclassified | unclassified | unclassified | 0,6 | 0,489897949 | 2,95E-05 | 2,43E-05 | 1 - | 4,28E-05 - | 1 - | 5,32E-05 - | 0 - | 0 - | 1 - | 5,13E-05 - | 0 - | 0 - |
| 5082 | 1 | 3um_FIJJAFBF_347923 | Bacteria | Proteobacteri | various | various | various | various | various | metabolism | Energy produc | 05C26 | Dehydrogenas | 167879.CPS_1 | 2,4 | 0,489897949 | 6,65E-05 | 1,57E-05 | 2 - | 4,63E-05 - | 3 - | 8,62E-05 - | 2 - | 5,46E-05 - | 3 - | 8,32E-05 - | 2 - | 6,24E-05 - |
| 5083 | 1 | 3um_FIJJAFBF_434531 | Bacteria | Bacteroidetes | Flavobacteriia | Flavobacterial | Flavobacteriac | Polaribacter | various | information st | Transcription | COG0250 | Participates in | 313603.FB217 | 0,6 | 0,8 | 4,07E-05 | 5,20E-05 | 2 - | 0,000125439 - | 1 - | 7,79E-05 - | 0 - | 0 - | 0 - | 0 - | 0 - | 0 - |
| 5084 | 1 | 3um_FIJJAFBF_338981 | Bacteria | Bacteroidetes | unclassified | unclassified | unclassified | unclassified | Bacteroidetes | unclassified | unclassified | unclassified | unclassified | unclassified | 2,6 | 0,489897949 | 6,75E-05 | 1,37E-05 | 3 - | 6,55E-05 - | 2 - | 5,42E-05 - | 3 - | 7,73E-05 - | 2 - | 5,23E-05 - | 3 - | 8,83E-05 - |
| 5085 | 1 | 0.2um_NOKCPHJN_64571 | Bacteria | Proteobacteri | Gammaprotec | Oceanospirilla | Saccharospirill | Reinekea | Reinekea blan | metabolism | Nucleotide tra | 16PVE | Phosphoribos | 314283.MED2 | 0,4 | 0,489897949 | 1,28E-05 | 1,56E-05 | 0 - | 0 - | 0 - | 0 - | 1 - | 3,17E-05 - | 1 - | 3,21E-05 - | 0 - | 0 - |
| 5086 | 1 | 0.2um_NOKCPHJN_291529 | Bacteria | Bacteroidetes | Flavobacteriia | Flavobacterial | Flavobacteriac | Formosa | Formosa sp. H | metabolism | Inorganic ion t | 05C27 | TonB-depende | 742766.HMPR | 1,2 | 0,4 | 1,52E-05 | 3,37E-06 | 2 - | 2,18E-05 - | 1 - | 1,35E-05 - | 1 - | 1,29E-05 - | 1 - | 1,31E-05 - | 1 - | 1,47E-05 - |
| 5087 | 1 | 0.2um_NOKCPHJN_45637 | Bacteria | unclassified | unclassified | unclassified | unclassified | unclassified | uncultured ba | metabolism | Energy produc | 05D26 | NQR complex | 983544.Lacal | 0,6 | 0,489897949 | 1,80E-05 | 1,49E-05 | 1 - | 2,63E-05 - | 1 - | 3,27E-05 - | 1 - | 3,11E-05 - | 0 - | 0 - | 0 - | 0 - |
| 5088 | 1 | 0.2um_NOKCPHJN_04207 | Bacteria | Bacteroidetes | Cytophagia | Cytophagales | Flammeovirga | unclassified | Flammeovirga | poorly charact | Function unkn | 08K5A | Tetratricopept | 643867.Ftrac | 0,4 | 0,489897949 | 1,28E-05 | 1,58E-05 | 1 - | 2,85E-05 - | 1 - | 3,54E-05 - | 0 - | 0 - | 0 - | 0 - | 0 - | 0 - |
| 5089 | 1 | 3um_FIJJAFBF_170065 | Bacteria | Proteobacteri | Gammaprotec | unclassified | unclassified | unclassified | various | information st | Translation, ri | 179R6 | This protein is | 876044.IMCC | 1,4 | 0,489897949 | 0,000134477 | 4,87E-05 | 2 - | 0,000161657 - | 1 - | 0,000100365 - | 1 - | 9,54E-05 - | 1 - | 9,68E-05 - | 2 - | 0,000218133 - |
| 5090 | 1 | 3um_FIJJAFBF_467912 | Bacteria | Proteobacteri | Betaproteoba | Burkholderial | unclassified | unclassified | Burkholderial | metabolism | Energy produc | 16QPK | Succinate deh | 365046.Rta_2 | 0,2 | 0,4 | 3,82E-06 | 7,64E-06 | 1 - | 1,91E-05 - | 0 - | 0 - | 0 - | 0 - | 0 - | 0 - | 0 - | 0 - |
| 5091 | 1 | 3um_FIJJAFBF_228685 | Bacteria | Bacteroidetes | Flavobacteriia | Flavobacterial | Flavobacteriac | Polaribacter | Polaribacter s | information st | Translation, ri | 0PBIW | 50S ribosomal | 313598.MED1 | 0,4 | 0,489897949 | 7,88E-05 | 9,72E-05 | 1 - | 0,000179339 - | 0 - | 0 - | 0 - | 0 - | 1 - | 0,000214818 - | 0 - | 0 - |
| 5092 | 1 | 0.2um_NOKCPHJN_58271 | Bacteria | Bacteroidetes | Flavobacteriia | Flavobacterial | Flavobacteriac | Muricauda | Muricauda sp. | information st | Translation, ri | COG0050 | This protein p | 391598.FBBAL | 20 | 1,414213562 | 0,00069391 | 8,78E-05 | 21 - | 0,000610206 - | 18 - | 0,000649453 - | 20 - | 0,000685986 - | 19 - | 0,000661312 - | 22 - | 0,000862592 - |
| 5093 | 1 | 3um_FIJJAFBF_80395 | Bacteria | Proteobacteri | Gammaprotec | Oceanospirilla | Saccharospirill | Reinekea | Reinekea blan | unclassified | unclassified | unclassified | unclassified | unclassified | 2 | 0,894427191 | 0,000143042 | 5,59E-05 | 3 - | 0,000186124 - | 1 - | 7,70E-05 - | 3 - | 0,000219701 - | 2 - | 0,000148631 - | 1 - | 8,37E-05 - |
| 5094 | 1 | 0.2um_NOKCPHJN_172833 | Bacteria | Proteobacteri | Gammaprotec | Cellvibrionales | Cellvibrionace | Gilvimarinus | Gilvimarinus p | cellular proces | Signal transdu | 0QIPQ | gtp-binding pr | 498211.CJA_3 | 7,6 | 2,154065923 | 0,00017685 | 6,35E-05 | 5 - | 9,55E-05 - | 6 - | 0,000142282 - | 9 - | 0,000202885 - | 7 - | 0,00016013 - | 11 - | 0,000283464 - |
| 5095 | 1 | 0.2um_NOKCPHJN_50816 | Bacteria | Bacteroidetes | Flavobacteriia | Flavobacterial | Flavobacteriac | Polaribacter | various | unclassified | unclassified | unclassified | unclassified | unclassified | 2 | 0 | 0,000181493 | 1,72E-05 | 2 - | 0,000152022 - | 2 - | 0,000188767 - | 2 - | 0,000179447 - | 2 - | 0,000182097 - | 2 - | 0,000205132 - |
| 5096 | 3 | 0.2um_NOKCPHJN_1071703;0.2um_NOKCPHJN_ | Bacteria | Proteobacteri | Alphaproteob | Pelagibacteral | Pelagibacterac | Candidatus Pe | Candidatus Pe | information st | Translation, ri | COG0256 | This is one of t | 335992.SAR11 | 0,4 | 0,489897949 | 4,28E-05 | 5,27E-05 | 1 - | 9,73E-05 - | 0 - | 0 - | 0 - | 0 - | 1 - | 0,000116511 - | 0 - | 0 - |
| 5097 | 1 | 3um_FIJJAFBF_667394 | Bacteria | Proteobacteri | Alphaproteob | Rhodobactera | Rhodobactera | Sulfitobacter | Sulfitobacter g | unclassified | unclassified | unclassified | unclassified | unclassified | 2 | 0 | 9,48E-05 | 9,00E-06 | 2 - | 7,94E-05 - | 2 - | 9,86E-05 - | 2 - | 9,38E-05 - | 2 - | 9,51E-05 - | 2 - | 0,00010718 - |
| 5098 | 1 | 0.2um_NOKCPHJN_15490 | Bacteria | Bacteroidetes | Flavobacteriia | Flavobacterial | Flavobacteriac | Polaribacter | various | information st | Translation, ri | 09AI8 | Associates wit | 313594.PI23P | 1,2 | 0,4 | 5,13E-05 | 1,77E-05 | 1 - | 3,58E-05 - | 1 - | 4,44E-05 - | 1 - | 4,22E-05 - | 2 - | 8,57E-05 - | 1 - | 4,82E-05 - |
| 5099 | 1 | 3um_FIJJAFBF_307539 | Bacteria | Bacteroidetes | Flavobacteriia | Flavobacterial | Flavobacteriac | Polaribacter | Polaribacter h | unclassified | unclassified | unclassified | unclassified | unclassified | 4 | 0,632455532 | 0,000190488 | 3,28E-05 | 4 - | 0,000159968 - | 3 - | 0,000148974 - | 5 - | 0,000236032 - | 4 - | 0,000191614 - | 4 - | 0,000215853 - |
| 5100 | 1 | 0.2um_NOKCPHJN_944412 | Bacteria | Bacteroidetes | Cytophagia | Cytophagales | Flammeovirga | unclassified | Flammeovirga | unclassified | unclassified | unclassified | unclassified | unclassified | 1,4 | 0,489897949 | 0,000108325 | 3,40E-05 | 2 - | 0,000131927 - | 2 - | 0,000163815 - | 1 - | 7,79E-05 - | 1 - | 7,90E-05 - | 1 - | 8,90E-05 - |
| 5101 | 1 | 3um_FIJJAFBF_366559 | Viruses | unclassified | unclassified | unclassified | Phycodnavirid | unclassified | Chrysochromu | information st | Transcription | 12NXR | DNA-depende | 4950.XP_0036 | 0,6 | 0,489897949 | 3,96E-05 | 3,30E-05 | 1 - | 5,52E-05 - | 1 - | 6,85E-05 - | 0 - | 0 - | 0 - | 0 - | 1 - | 7,45E-05 - |
| 5102 | 1 | 0.2um_NOKCPHJN_947787 | Eukaryota | unclassified | Oomycetes | Peronosporale | Peronosporac | Phytophthora | Phytophthora | unclassified | unclassified | unclassified | unclassified | unclassified | 0,2 | 0,4 | 5,00E-06 | 1,00E-05 | 1 - | 2,50E-05 - | 0 - | 0 - | 0 - | 0 - | 0 - | 0 - | 0 - | 0 - |
| 5103 | 1 | 0.2um_NOKCPHJN_119224 | Bacteria | Actinobacteria | Actinobacteria | Actinomycetal | unclassified | unclassified | Actinomycetal | metabolism | Carbohydrate | 00BJE | Glyceraldehyd | 469371.Tbis_1 | 1 | 0 | 4,13E-05 | 3,92E-06 | 1 - | 3,46E-05 - | 1 - | 4,29E-05 - | 1 - | 4,08E-05 - | 1 - | 4,14E-05 - | 1 - | 4,66E-05 - |
| 5104 | 2 | 3um_FIJJAFBF_246857;3um_FIJJAFBF_277458 | Eukaryota | Bacillariophyta | various | various | various | various | various | information st | Translation, ri | 082FX | This protein bi | 197221.tll016 | 1 | 0 | 0,000126877 | 1,20E-05 | 1 - | 0,000106275 - | 1 - | 0,000131962 - | 1 - | 0,000125447 - | 1 - | 0,000127299 - | 1 - | 0,000143402 - |
| 5105 | 1 | 3um_FIJJAFBF_235047 | Bacteria | Proteobacteri | Gammaprotec | unclassified | unclassified | unclassified | various | unclassified | unclassified | unclassified | unclassified | unclassified | 0,4 | 0,489897949 | 4,21E-05 | 5,18E-05 | 1 - | 9,65E-05 - | 0 - | 0 - | 1 - | 0,000113851 - | 0 - | 0 - | 0 - | 0 - |
| 5106 | 1 | 3um_FIJJAFBF_379812 | Bacteria | Bacteroidetes | Cytophagia | Cytophagales | Cytophagacea | Ekhidna | Ekhidna lutea | unclassified | unclassified | unclassified | unclassified | unclassified | 0,2 | 0,4 | 7,92E-06 | 1,58E-05 | 1 - | 3,96E-05 - | 0 - | 0 - | 0 - | 0 - | 0 - | 0 - | 0 - | 0 - |
| 5107 | 1 | 0.2um_NOKCPHJN_332733 | Bacteria | Verrucomicrol | Opitutae | Opitutales | Opitutaceae | unclassified | Opitutaceae b | poorly charact | Function unkn | 028X2 | NA | 438753.AZC_3 | 0,6 | 0,8 | 6,93E-06 | 9,09E-06 | 0 - | 0 - | 2 - | 2,25E-05 - | 0 - | 0 - | 0 - | 0 - | 1 - | 1,22E-05 - |
| 5108 | 1 | 0.2um_NOKCPHJN_372005 | Eukaryota | Phaeophyceae | unclassified | Ectocarpales | Ectocarpaceae | Ectocarpus | Ectocarpus sili | cellular proces | Signal transdu | KOG0613 | fibronectin typ | 653948.CCA1 | 2,6 | 1,356465997 | 2,43E-05 | 1,18E-05 | 4 - | 3,22E-05 - | 3 - | 3,00E-05 - | 1 - | 9,51E-06 - | 4 - | 3,86E-05 - | 1 - | 1,09E-05 - |
| 5109 | 1 | 3um_FIJJAFBF_804605 | Bacteria | Bacteroidetes | Flavobacteriia | Flavobacterial | Flavobacteriac | Zobellia | Zobellia uligin | unclassified | unclassified | unclassified | unclassified | unclassified | 0,4 | 0,489897949 | 1,23E-05 | 1,52E-05 | 1 - | 2,74E-05 - | 1 - | 3,40E-05 - | 0 - | 0 - | 0 - | 0 - | 0 - | 0 - |
| 5110 | 1 | 3um_FIJJAFBF_637911 | Eukaryota | Brachiopoda | Lingulata | Lingulida | Lingulidae | Lingula | Lingula anatin | poorly charact | Function unkn | 0J17T | NA | 7668.SPU_00( | 0,2 | 0,4 | 7,53E-06 | 1,51E-05 | 1 - | 3,76E-05 - | 0 - | 0 - | 0 - | 0 - | 0 - | 0 - | 0 - | 0 - |
| 5111 | 1 | 0.2um_NOKCPHJN_94624 | Bacteria | Proteobacteri | Alphaproteob | Rhodobactera | Rhodobactera | Nereida | Nereida ignav | unclassified | unclassified | unclassified | unclassified | unclassified | 1 | 1,264911064 | 1,50E-05 | 1,87E-05 | 0 - | 0 - | 3 - | 4,34E-05 - | 0 - | 0 - | 0 - | 0 - | 2 - | 3,14E-05 - |
| 5112 | 1 | 3um_FIJJAFBF_106733 | Bacteria | Bacteroidetes | Flavobacteriia | Flavobacterial | Flavobacteriac | Flavobacteriu | Flavobacteriu | cellular proces | Intracellular tr | 0BIEB | domain-conta | 1005048.CFU | 3,6 | 0,8 | 8,70E-05 | 1,72E-05 | 4 - | 8,18E-05 - | 3 - | 7,62E-05 - | 5 - | 0,000120751 - | 3 - | 7,35E-05 - | 3 - | 8,28E-05 - |
| 5113 | 1 | 0.2um_NOKCPHJN_229539 | Bacteria | Bacteroidetes | Flavobacteriia | Flavobacterial | Flavobacteriac | Pricia | Pricia antarcti | unclassified | unclassified | unclassified | unclassified | unclassified | 3,2 | 0,4 | 0,000520066 | 0,000109596 | 3 - | 0,000405094 - | 3 - | 0,000503008 - | 3 - | 0,000478173 - | 3 - | 0,000485235 - | 4 - | 0,000728821 - |
| 5114 | 1 | 3um_FIJJAFBF_163921 | Bacteria | Proteobacteri | Alphaproteob | Rhodobactera | Rhodobactera | Phaeobacter | Phaeobacter s | unclassified | unclassified | unclassified | unclassified | unclassified | 3,6 | 1,624807681 | 5,03E-05 | 2,21E-05 | 5 - | 5,95E-05 - | 6 - | 8,87E-05 - | 2 - | 2,81E-05 - | 3 - | 4,28E-05 - | 2 - | 3,21E-05 - |
| 5115 | 1 | 3um_FIJJAFBF_133469 | Bacteria | Actinobacteria | Actinobacteria | Micrococcales | Microbacteria | Candidatus Ac | Candidatus Ac | unclassified | unclassified | unclassified | unclassified | unclassified | 2,4 | 0,489897949 | 0,000173888 | 4,63E-05 | 2 - | 0,000120185 - | 2 - | 0,000149234 - | 3 - | 0,000212799 - | 2 - | 0,000143962 - | 3 - | 0,000243258 - |
| 5116 | 1 | 3um_FIJJAFBF_245819 | Bacteria | Bacteroidetes | Flavobacteriia | Flavobacterial | Flavobacteriac | unclassified | Flavobacteriac | cellular proces | Cell wall/mem | 0P88B | Outer membra | 391603.FBALC | 0,6 | 0,489897949 | 1,76E-05 | 1,45E-05 | 1 - | 2,57E-05 - | 1 - | 3,20E-05 - | 1 - | 3,04E-05 - | 0 - | 0 - | 0 - | 0 - |
| 5117 | 1 | 3um_FIJJAFBF_70280 | Bacteria | Actinobacteria | Actinobacteria | Micrococcales | Microbacteria | Candidatus Ac | Candidatus Ac | unclassified | unclassified | unclassified | unclassified | unclassified | 0,2 | 0,4 | 9,60E-06 | 1,92E-05 | 1 - | 4,80E-05 - | 0 - | 0 - | 0 - | 0 - | 0 - | 0 - | 0 - | 0 - |
| 5118 | 1 | 0.2um_NOKCPHJN_24507 | Bacteria | Bacteroidetes | Flavobacteriia | Flavobacterial | Flavobacteriac | Formosa | Formosa sp. H | unclassified | unclassified | unclassified | unclassified | unclassified | 0,6 | 0,489897949 | 1,01E-05 | 8,43E-06 | 1 - | 1,43E-05 - | 0 - | 0 - | 0 - | 0 - | 1 - | 1,71E-05 - | 1 - | 1,93E-05 - |
| 5119 | 1 | 3um_FIJJAFBF_375489 | Bacteria | Bacteroidetes | Flavobacteriia | Flavobacterial | Flavobacteriac | Tamlana | Tamlana sp. U | metabolism | Carbohydrate | 099YR | Catalyzes the | 391603.FBALC | 6 | 2,097617696 | 0,000187069 | 5,52E-05 | 9 - | 0,00024023 - | 8 - | 0,000265151 - | 4 - | 0,00012603 - | 5 - | 0,000159864 - | 4 - | 0,000144069 - |
| 5120 | 1 | 3um_FIJJAFBF_189624 | Bacteria | Bacteroidetes | Flavobacteriia | Flavobacterial | Flavobacteriac | Polaribacter | Polaribacter s | unclassified | unclassified | unclassified | unclassified | unclassified | 3,8 | 0,4 | 9,50E-05 | 1,71E-05 | 3 - | 6,23E-05 - | 4 - | 0,000103088 - | 4 - | 9,80E-05 - | 4 - | 9,94E-05 - | 4 - | 0,000112025 - |
| 5121 | 1 | 0.2um_NOKCPHJN_524267 | Eukaryota | Arthropoda | Merostomata | Xiphosura | Limulidae | Limulus | Limulus polyp | unclassified | unclassified | unclassified | unclassified | unclassified | 0,2 | 0,4 | 3,27E-06 | 6,54E-06 | 0 - | 0 - | 0 - | 0 - | 0 - | 0 - | 1 - | 1,63E-05 - | 0 - | 0 - |
| 5122 | 1 | 0.2um_NOKCPHJN_360958 | Bacteria | Bacteroidetes | unclassified | unclassified | unclassified | unclassified | Bacteroidetes | unclassified | unclassified | unclassified | unclassified | unclassified | 3 | 0,632455532 | 0,000153013 | 3,44E-05 | 3 - | 0,000128481 - | 2 - | 0,000106357 - | 3 - | 0,000151659 - | 4 - | 0,000205199 - | 3 - | 0,000173367 - |
| 5123 | 1 | 3um_FIJJAFBF_724446 | Bacteria | Proteobacteri | Gammaprotec | Thiotrichales | Fastidiosibact | Fangia | Fangia hongk | unclassified | unclassified | unclassified | unclassified | unclassified | 0,4 | 0,489897949 | 3,30E-05 | 4,08E-05 | 1 - | 7,36E-05 - | 1 - | 9,14E-05 - | 0 - | 0 - | 0 - | 0 - | 0 - | 0 - |
| 5124 | 1 | 0.2um_NOKCPHJN_441502 | Eukaryota | unclassified | unclassified | unclassified | Apusomonadi | Thecamonas | Thecamonas t | unclassified | unclassified | unclassified | unclassified | unclassified | 0,2 | 0,4 | 5,39E-06 | 1,08E-05 | 0 - | 0 - | 0 - | 0 - | 0 - | 0 - | 1 - | 2,70E-05 - | 0 - | 0 - |
| 5125 | 1 | 3um_FIJJAFBF_130481 | Bacteria | Proteobacteri | Alphaproteob | Rhodobactera | Rhodobactera | Sulfitobacter | Sulfitobacter g | metabolism | Inorganic ion t | 16PII | TRAP transpor | 367336.OM22 | 1 | 0,632455532 | 4,06E-05 | 2,59E-05 | 1 - | 3,44E-05 - | 0 - | 0 - | 2 - | 8,11E-05 - | 1 - | 4,12E-05 - | 1 - | 4,64E-05 - |
| 5126 | 1 | 3um_FIJJAFBF_115451 | Bacteria | Proteobacteri | Gammaprotec | Cellvibrionales | Halieaceae | Congregibacte | Congregibacte | information st | Translation, ri | 179R6 | This protein is | 314285.KT71 | 1,4 | 0,489897949 | 0,000134944 | 4,77E-05 | 1 - | 8,08E-05 - | 1 - | 0,000100365 - | 2 - | 0,00019082 - | 2 - | 0,000193638 - | 1 - | 0,000109067 - |
| 5127 | 1 | 3um_FIJJAFBF_74663 | Bacteria | Bacteroidetes | unclassified | unclassified | unclassified | unclassified | Bacteroidetes | cellular proces | Posttranslatio | COG0330 | Band 7 protei | 926562.Oweh | 1,8 | 0,748331477 | 8,97E-05 | 3,21E-05 | 3 - | 0,000128004 - | 1 - | 5,30E-05 - | 1 - | 5,04E-05 - | 2 - | 0,000102218 - | 2 - | 0,000115148 - |
| 5128 | 1 | 0.2um_NOKCPHJN_63174 | Bacteria | Bacteroidetes | Flavobacteriia | Flavobacterial | Flavobacteriac | Flavobacteriu | Flavobacteriu | metabolism | Inorganic ion t | 05C27 | TonB-depende | 983548.Krodi | 4,8 | 0,4 | 6,10E-05 | 9,86E-06 | 4 - | 4,23E-05 - | 5 - | 6,57E-05 - | 5 - | 6,24E-05 - | 5 - | 6,34E-05 - | 5 - | 7,14E-05 - |
| 5129 | 1 | 3um_FIJJAFBF_620070 | Eukaryota | unclassified | unclassified | Pavlovales | Pavlovaceae | Pavlova | Pavlova luther | unclassified | unclassified | unclassified | unclassified | unclassified | 0,2 | 0,4 | 2,34E-05 | 4,68E-05 | 1 - | 0,000117119 - | 0 - | 0 - | 0 - | 0 - | 0 - | 0 - | 0 - | 0 - |
| 5130 | 1 | 3um_FIJJAFBF_480851 | Bacteria | Proteobacteri | Gammaprotec | unclassified | unclassified | unclassified | SAR86 cluster | information st | Translation, ri | COG1185 | Involved in mI | 290398.Csal_3 | 1,8 | 0,748331477 | 3,50E-05 | 1,56E-05 | 2 - | 3,24E-05 - | 3 - | 6,04E-05 - | 1 - | 1,91E-05 - | 1 - | 1,94E-05 - | 2 - | 4,37E-05 - |
| 5131 | 1 | 3um_FIJJAFBF_545215 | Bacteria | Bacteroidetes | Flavobacteriia | Flavobacterial | Flavobacteriac | Flavivirga | Flavivirga eckl | unclassified | unclassified | unclassified | unclassified | unclassified | 3 | 0 | 0,000311425 | 2,96E-05 | 3 - | 0,000260856 - | 3 - | 0,000323907 - | 3 - | 0,000307914 - | 3 - | 0,000312462 - | 3 - | 0,000351987 - |
| 5132 | 1 | 0.2um_NOKCPHJN_434037 | Bacteria | Bacteroidetes | unclassified | unclassified | unclassified | unclassified | Bacteroidetes | metabolism | Inorganic ion t | 05C27 | TonB-depende | 485918.Cpin | 12 | 1,264911064 | 0,000158123 | 1,66E-05 | 12 - | 0,000133075 - | 12 - | 0,000165239 - | 14 - | 0,000183261 - | 12 - | 0,000159401 - | 10 - | 0,000149637 - |
| 5133 | 1 | 0.2um_NOKCPHJN_998347 | Bacteria | Proteobacteri | Gammaprotec | unclassified | unclassified | unclassified | OM182 bacter | unclassified | unclassified | unclassified | unclassified | unclassified | 0,6 | 0,489897949 | 1,64E-05 | 1,36E-05 | 1 - | 2,39E-05 - | 1 - | 2,97E-05 - | 0 - | 0 - | 1 - | 2,86E-05 - | 0 - | 0 - |
| 5134 | 1 | 0.2um_NOKCPHJN_126550 | Bacteria | Verrucomicrol | Opitutae | unclassified | unclassified | unclassified | Opitutae bact | unclassified | unclassified | unclassified | unclassified | unclassified | 0 | 0 | 0 | 0 | 0 - | 0 - | 0 - | 0 - | 0 - | 0 - | 0 - | 0 - | 0 - | 0 - |
| 5135 | 1 | 0.2um_NOKCPHJN_673524 | Bacteria | Proteobacteri | Gammaprotec | unclassified | unclassified | unclassified | Gammaprotec | unclassified | unclassified | unclassified | unclassified | unclassified | 0,6 | 0,8 | 7,88E-06 | 1,01E-05 | 2 - | 2,46E-05 - | 0 - | 0 - | 0 - | 0 - | 1 - | 1,48E-05 - | 0 - | 0 - |
| 5136 | 1 | 3um_FIJJAFBF_639895 | Bacteria | Proteobacteri | Gammaprotec | Oceanospirilla | various | various | various | poorly charact | Function unkn | COG3517 | type VI secreti | 83406.HDN1F | 18,4 | 1,2 | 0,000522985 | 3,65E-05 | 20 - | 0,000478236 - | 19 - | 0,000564137 - | 17 - | 0,000479833 - | 19 - | 0,000544205 - | 17 - | 0,000548514 - |
| 5137 | 1 | 3um_FIJJAFBF_525961 | Bacteria | Bacteroidetes | Cytophagia | Cytophagales | Flammeovirga | Reichenbachie | Reichenbachie | information st | Translation, ri | COG0052 | 30S ribosomal | 880070.Cycma | 1 | 0 | 5,39E-05 | 5,12E-06 | 1 - | 4,52E-05 - | 1 - | 5,61E-05 - | 1 - | 5,33E-05 - | 1 - | 5,41E-05 - | 1 - | 6,10E-05 - |
| 5138 | 1 | 3um_FIJJAFBF_191785 | Eukaryota | unclassified | Raphidophyce | Chattonellales | Chattonellace | Heterosigma | Heterosigma a | unclassified | unclassified | unclassified | unclassified | unclassified | 3,6 | 0,489897949 | 0,000244818 | 2,89E-05 | 4 - | 0,000229554 - | 4 - | 0,000285038 - | 4 - | 0,000270964 - | 3 - | 0,000206225 - | 3 - | 0,000232312 - |
| 5139 | 1 | 3um_FIJJAFBF_308258 | Bacteria | Bacteroidetes | Flavobacteriia | Flavobacterial | Flavobacteriac | Polaribacter | Polaribacter s | unclassified | unclassified | unclassified | unclassified | unclassified | 0,6 | 0,489897949 | 4,39E-05 | 3,62E-05 | 1 - | 6,41E-05 - | 1 - | 7,96E-05 - | 1 - | 7,57E-05 - | 0 - | 0 - | 0 - | 0 - |
| 5140 | 1 | 3um_FIJJAFBF_352780 | Bacteria | Actinobacteria | Actinobacteria | Micrococcales | Microbacteria | Candidatus Ac | Candidatus Ac | unclassified | unclassified | unclassified | unclassified | unclassified | 0,2 | 0,4 | 9,89E-06 | 1,98E-05 | 0 - | 0 - | 0 - | 0 - | 0 - | 0 - | 1 - | 4,95E-05 - | 0 - | 0 - |
| 5141 | 1 | 3um_FIJJAFBF_85824 | Viruses | unclassified | unclassified | unclassified | Phycodnavirid | unclassified | Chrysochromu | unclassified | unclassified | unclassified | unclassified | unclassified | 4,2 | 0,4 | 4,93E-05 | 2,33E-06 | 5 - | 4,95E-05 - | 4 - | 4,92E-05 - | 4 - | 4,68E-05 - | 4 - | 4,74E-05 - | 4 - | 5,35E-05 - |
| 5142 | 1 | 3um_FIJJAFBF_06969 | Bacteria | Proteobacteri | Gammaprotec | Cellvibrionales | unclassified | unclassified | Cellvibrionales | unclassified | unclassified | unclassified | unclassified | unclassified | 3 | 0 | 0,000244691 | 2,32E-05 | 3 - | 0,000204958 - | 3 - | 0,000254498 - | 3 - | 0,000241933 - | 3 - | 0,000245506 - | 3 - | 0,000276562 - |

| | | | | | | | | | | | | | | | | | | | | | | | | | | | | |
|---|---|---|---|---|---|---|---|---|---|---|---|---|---|---|---|---|---|---|---|---|---|---|---|---|---|---|---|---|
| 5143 | 1 | 0.2um_NOKCPHJN_439144 | Bacteria | Proteobacteri | Gammaprotec | Oceanospirilla | Saccharospirill | Reinekea | Reinekea fors | cellular proces | Intracellular tr | 05VIE | preprotein tra | 314283.MED2 | 3,8 | 0,748331477 | 0,000453063 | 0,000111761 | 3 - | 0,000296836 - | 4 - | 0,000491444 - | 3 - | 0,000350385 - | 5 - | 0,0005926 - | 4 - | 0,00053405 - |
| 5144 | 1 | 0.2um_NOKCPHJN_260016 | Bacteria | Bacteroidetes | Flavobacteriia | Flavobacterial | Flavobacteriac | Formosa | Formosa sp. H | unclassified | unclassified | unclassified | unclassified | unclassified | 0,2 | 0,4 | 2,95E-06 | 5,89E-06 | 1 - | 1,47E-05 - | 0 - | 0 - | 0 - | 0 - | 0 - | 0 - | 0 - | 0 - |
| 5145 | 1 | 0.2um_NOKCPHJN_614431 | Bacteria | Proteobacteri | Gammaprotec | Cellvibrionales | Spongiibacter | Oceanicoccus | Oceanicoccus | metabolism | Amino acid tra | 05C6T | cysteine synth | 207954.MED9 | 1 | 0 | 4,55E-05 | 4,32E-06 | 1 - | 3,81E-05 - | 1 - | 4,73E-05 - | 1 - | 4,50E-05 - | 1 - | 4,57E-05 - | 1 - | 5,15E-05 - |
| 5146 | 1 | 3um_FIJJAFBF_618650 | Bacteria | Proteobacteri | Gammaprotec | Thiotrichales | Piscirickettsiac | Methylophaga | Methylophaga | metabolism | Energy produc | COG0056 | Produces ATP | 857087.Metm | 0,2 | 0,4 | 7,51E-06 | 1,50E-05 | 0 - | 0 - | 0 - | 0 - | 1 - | 3,75E-05 - | 0 - | 0 - | 0 - | 0 - |
| 5147 | 1 | 3um_FIJJAFBF_122455 | Bacteria | Bacteroidetes | Flavobacteriia | Flavobacterial | Flavobacteriac | Polaribacter | Polaribacter s | poorly charact | Function unkn | 09C02 | NA | 313594.PI23P | 2 | 0,632455532 | 9,12E-05 | 2,41E-05 | 3 - | 0,000116328 - | 2 - | 9,63E-05 - | 1 - | 4,58E-05 - | 2 - | 9,29E-05 - | 2 - | 0,000104645 - |
| 5148 | 1 | 0.2um_NOKCPHJN_48462 | Bacteria | Bacteroidetes | Flavobacteriia | Flavobacterial | Flavobacteriac | Tenacibaculur | Tenacibaculur | information st | Translation, ril | 05KK9 | This protein bi | 886377.Murru | 2,4 | 0,8 | 0,000189004 | 6,53E-05 | 3 - | 0,000199035 - | 1 - | 8,24E-05 - | 2 - | 0,000156627 - | 3 - | 0,00023841 - | 3 - | 0,000268568 - |
| 5149 | 1 | 3um_FIJJAFBF_421713 | Bacteria | Proteobacteri | Alphaproteob | Pelagibacteral | Pelagibacterac | Candidatus Pe | Candidatus Pe | information st | Replication, re | 05CZ8 | DNA polymera | 335992.SAR11 | 1 | 0 | 3,70E-05 | 3,52E-06 | 1 - | 3,10E-05 - | 1 - | 3,85E-05 - | 1 - | 3,66E-05 - | 1 - | 3,72E-05 - | 1 - | 4,19E-05 - |
| 5150 | 1 | 0.2um_NOKCPHJN_613022 | Bacteria | Proteobacteri | Alphaproteob | Rhodobactera | Rhodobactera | Planktomarina | Planktomarina | unclassified | unclassified | unclassified | unclassified | unclassified | 2 | 0,632455532 | 5,39E-05 | 1,37E-05 | 3 - | 6,91E-05 - | 1 - | 2,86E-05 - | 2 - | 5,44E-05 - | 2 - | 5,52E-05 - | 2 - | 6,22E-05 - |
| 5151 | 1 | 3um_FIJJAFBF_73796 | Bacteria | Bacteroidetes | Flavobacteriia | Flavobacterial | Flavobacteriac | Polaribacter | Polaribacter b | unclassified | unclassified | unclassified | unclassified | unclassified | 3,2 | 0,748331477 | 0,000109526 | 2,14E-05 | 4 - | 0,00011623 - | 2 - | 7,22E-05 - | 4 - | 0,000137197 - | 3 - | 0,000104418 - | 3 - | 0,000117626 - |
| 5152 | 1 | 0.2um_NOKCPHJN_613239 | Bacteria | Proteobacteri | Gammaprotec | unclassified | unclassified | unclassified | Gammaprotec | unclassified | unclassified | unclassified | unclassified | unclassified | 1,6 | 0,489897949 | 0,000204358 | 5,35E-05 | 2 - | 0,000218622 - | 1 - | 0,000135732 - | 2 - | 0,000258061 - | 2 - | 0,000261873 - | 1 - | 0,0001475 - |
| 5153 | 1 | 3um_FIJJAFBF_03923 | Bacteria | Bacteroidetes | unclassified | unclassified | unclassified | unclassified | Bacteroidetes | poorly charact | Function unkn | 17F80 | NA | 314225.ELI_05 | 1,2 | 0,4 | 0,000103222 | 3,46E-05 | 1 - | 7,22E-05 - | 1 - | 8,96E-05 - | 2 - | 0,000170418 - | 1 - | 8,65E-05 - | 1 - | 9,74E-05 - |
| 5154 | 1 | 0.2um_NOKCPHJN_925851 | Bacteria | Proteobacteri | Gammaprotec | Alteromonada | Pseudoalteror | Psychrosphae | Psychrosphae | information st | Translation, ril | COG0087 | One of the pri | 511062.GU3_ | 0,2 | 0,4 | 1,07E-05 | 2,14E-05 | 1 - | 5,34E-05 - | 0 - | 0 - | 0 - | 0 - | 0 - | 0 - | 0 - | 0 - |
| 5155 | 1 | 3um_FIJJAFBF_292914 | Eukaryota | Bacillariophyt | Coscinodiscop | Coscinodiscale | Coscinodiscac | Coscinodiscus | Coscinodiscus | metabolism | Energy produc | COG0712 | F(1)F(0) ATP s | 35128.Thapsd | 0,4 | 0,489897949 | 2,88E-05 | 3,60E-05 | 1 - | 6,14E-05 - | 0 - | 0 - | 0 - | 0 - | 0 - | 0 - | 1 - | 8,28E-05 - |
| 5156 | 1 | 3um_FIJJAFBF_402485 | Bacteria | Bacteroidetes | Flavobacteriia | Flavobacterial | Flavobacteriac | Maribacter | Maribacter th | unclassified | unclassified | unclassified | unclassified | unclassified | 0,4 | 0,489897949 | 1,99E-05 | 2,44E-05 | 0 - | 0 - | 0 - | 0 - | 1 - | 4,94E-05 - | 1 - | 5,02E-05 - | 0 - | 0 - |
| 5157 | 1 | 3um_FIJJAFBF_172360 | Bacteria | Proteobacteri | Gammaprotec | Oceanospirilla | Saccharospirill | Reinekea | Reinekea fors | information st | Translation, ril | 08VZM | 50s ribosomal | 314283.MED2 | 4,6 | 1,356465997 | 0,000383989 | 0,000118961 | 5 - | 0,000349929 - | 5 - | 0,000434509 - | 6 - | 0,000495667 - | 2 - | 0,000167663 - | 5 - | 0,000472178 - |
| 5158 | 1 | 0.2um_NOKCPHJN_39793 | Bacteria | Bacteroidetes | Flavobacteriia | Flavobacterial | Flavobacteriac | Polaribacter | Polaribacter ir | poorly charact | Function unkn | 11W7M | NA | 983544.Lacal_ | 1,4 | 0,489897949 | 0,000125063 | 5,08E-05 | 1 - | 7,36E-05 - | 1 - | 9,14E-05 - | 2 - | 0,000173695 - | 1 - | 8,81E-05 - | 2 - | 0,000198557 - |
| 5159 | 1 | 3um_FIJJAFBF_55327 | Bacteria | Proteobacteri | Gammaprotec | Cellvibrionales | Porticoccacea | unclassified | SAR92 bacteri | unclassified | unclassified | unclassified | unclassified | unclassified | 0,6 | 0,489897949 | 9,46E-05 | 7,81E-05 | 1 - | 0,000138285 - | 1 - | 0,000171709 - | 1 - | 0,000163232 - | 0 - | 0 - | 0 - | 0 - |
| 5160 | 1 | 3um_FIJJAFBF_539571 | Bacteria | Bacteroidetes | Flavobacteriia | Flavobacterial | Flavobacteriac | Lutibacter | Lutibacter sp. | poorly charact | Function unkn | 09F62 | Protein of unk | 391587.KAOT | 0,8 | 0,748331477 | 8,98E-05 | 7,90E-05 | 2 - | 0,000203145 - | 1 - | 0,000126123 - | 1 - | 0,000119896 - | 0 - | 0 - | 0 - | 0 - |
| 5161 | 1 | 3um_FIJJAFBF_67825 | Bacteria | Proteobacteri | Gammaprotec | Alteromonada | Shewanellacea | Shewanella | Shewanella ps | information st | Transcription | 05VEQ | Cold shock pro | 1085623.GNIT | 0,8 | 1,166190379 | 0,000146825 | 0,000200247 | 3 - | 0,000506368 - | 0 - | 0 - | 0 - | 0 - | 0 - | 0 - | 1 - | 0,000227757 - |
| 5162 | 1 | 3um_FIJJAFBF_368916 | Bacteria | Proteobacteri | Alphaproteob | Rhodobactera | Rhodobactera | Planktomarina | Planktomarina | information st | Translation, ril | COG0184 | Forms an inte | 375451.RD1_ | 1,2 | 0,4 | 0,000179756 | 3,98E-05 | 2 - | 0,000257925 - | 1 - | 0,000160134 - | 1 - | 0,000152227 - | 1 - | 0,000154476 - | 1 - | 0,000174016 - |
| 5163 | 1 | 3um_FIJJAFBF_12818 | Bacteria | Bacteroidetes | Flavobacteriia | Flavobacterial | Flavobacteriac | Arenitalea | Arenitalea lut | unclassified | unclassified | unclassified | unclassified | unclassified | 9 | 1,095445115 | 0,000122722 | 7,10E-06 | 11 - | 0,000126889 - | 9 - | 0,000128912 - | 8 - | 0,00010893 - | 9 - | 0,000124357 - | 8 - | 0,000124522 - |
| 5164 | 1 | 3um_FIJJAFBF_532203 | Bacteria | Proteobacteri | Gammaprotec | Cellvibrionales | Halieaceae | Halioglobus | Halioglobus ja | metabolism | Amino acid tra | 16R0H | Acetolactate s | 377629.TERTU | 0,2 | 0,4 | 1,63E-05 | 3,26E-05 | 0 - | 0 - | 0 - | 0 - | 1 - | 8,16E-05 - | 0 - | 0 - | 0 - | 0 - |
| 5165 | 1 | 3um_FIJJAFBF_709599 | Bacteria | Proteobacteri | Gammaprotec | unclassified | unclassified | unclassified | Gammaprotec | metabolism | Energy produc | 17CGT | F(1)F(0) ATP s | 349521.HCH_ | 1 | 0 | 8,78E-05 | 8,34E-06 | 1 - | 7,36E-05 - | 1 - | 9,14E-05 - | 1 - | 8,68E-05 - | 1 - | 8,81E-05 - | 1 - | 9,93E-05 - |
| 5166 | 1 | 3um_FIJJAFBF_207484 | unclassified | unclassified | unclassified | unclassified | unclassified | unclassified | unclassified | unclassified | unclassified | unclassified | unclassified | unclassified | 2,6 | 0,489897949 | 9,57E-05 | 1,62E-05 | 3 - | 9,33E-05 - | 3 - | 0,000115869 - | 2 - | 7,34E-05 - | 3 - | 0,000111775 - | 2 - | 8,39E-05 - |
| 5167 | 1 | 3um_FIJJAFBF_705334 | Bacteria | Proteobacteri | Alphaproteob | Rhodospirillal | Rhodospirillac | unclassified | Rhodobactera | metabolism | Carbohydrate | 07U2E | extracellular s | 367336.OM22 | 1 | 0 | 3,16E-05 | 3,00E-06 | 1 - | 2,64E-05 - | 1 - | 3,28E-05 - | 1 - | 3,12E-05 - | 1 - | 3,17E-05 - | 1 - | 3,57E-05 - |
| 5168 | 1 | 3um_FIJJAFBF_209609 | Bacteria | Bacteroidetes | Flavobacteriia | Flavobacterial | Flavobacteriac | various | various | unclassified | unclassified | unclassified | unclassified | unclassified | 2 | 0 | 0,000114667 | 1,09E-05 | 2 - | 9,60E-05 - | 2 - | 0,000119263 - | 2 - | 0,000113374 - | 2 - | 0,000115049 - | 2 - | 0,000129602 - |
| 5169 | 1 | 3um_FIJJAFBF_594864 | Bacteria | Proteobacteri | Alphaproteob | Rhodobactera | unclassified | unclassified | Rhodobactera | unclassified | unclassified | unclassified | unclassified | unclassified | 0,4 | 0,489897949 | 5,23E-05 | 6,41E-05 | 0 - | 0 - | 1 - | 0,000133195 - | 0 - | 0 - | 1 - | 0,000128489 - | 0 - | 0 - |
| 5170 | 1 | 0.2um_NOKCPHJN_65240 | Bacteria | Proteobacteri | Gammaprotec | Cellvibrionales | Porticoccacea | unclassified | SAR92 bacteri | unclassified | unclassified | unclassified | unclassified | unclassified | 20,2 | 1,720465053 | 0,000331333 | 5,40E-05 | 20 - | 0,000273604 - | 21 - | 0,000356722 - | 19 - | 0,000306813 - | 18 - | 0,000294958 - | 23 - | 0,000424567 - |
| 5171 | 1 | 0.2um_NOKCPHJN_121521 | Bacteria | Bacteroidetes | Flavobacteriia | Flavobacterial | Flavobacteriac | Tenacibaculur | Tenacibaculur | unclassified | unclassified | unclassified | unclassified | unclassified | 0,2 | 0,4 | 8,29E-06 | 1,66E-05 | 0 - | 0 - | 0 - | 0 - | 1 - | 4,14E-05 - | 0 - | 0 - | 0 - | 0 - |
| 5172 | 1 | 0.2um_NOKCPHJN_558947 | Bacteria | Proteobacteri | Alphaproteob | Rhodobactera | Rhodobactera | Planktomarina | Planktomarina | metabolism | Secondary me | 01Q86 | extracellular s | 388401.RB215 | 4,2 | 1,166190379 | 0,000153129 | 3,77E-05 | 5 - | 0,000155104 - | 3 - | 0,000115556 - | 4 - | 0,000146467 - | 6 - | 0,000222946 - | 3 - | 0,000125574 - |
| 5173 | 1 | 3um_FIJJAFBF_658713 | Eukaryota | unclassified | unclassified | Isochrysidales | Noelaerhabda | Emiliania | Emiliania huxl | unclassified | unclassified | unclassified | unclassified | unclassified | 0,4 | 0,489897949 | 2,23E-05 | 2,78E-05 | 1 - | 4,74E-05 - | 0 - | 0 - | 0 - | 0 - | 0 - | 0 - | 1 - | 6,40E-05 - |
| 5174 | 1 | 0.2um_NOKCPHJN_144384 | Bacteria | Bacteroidetes | Flavobacteriia | Flavobacterial | Flavobacteriac | Formosa | Formosa sp. H | unclassified | unclassified | unclassified | unclassified | unclassified | 0,8 | 0,748331477 | 5,23E-05 | 5,00E-05 | 1 - | 5,68E-05 - | 1 - | 7,06E-05 - | 2 - | 0,000134141 - | 0 - | 0 - | 0 - | 0 - |
| 5175 | 1 | 3um_FIJJAFBF_424915 | Eukaryota | Phaeophyceae | unclassified | Ectocarpales | Ectocarpaceae | Ectocarpus | Ectocarpus sil | unclassified | unclassified | unclassified | unclassified | unclassified | 1 | 0 | 0,000169169 | 1,61E-05 | 1 - | 0,0001417 - | 1 - | 0,000175949 - | 1 - | 0,000167262 - | 1 - | 0,000169732 - | 1 - | 0,000191203 - |
| 5176 | 2 | 3um_FIJJAFBF_167965;3um_FIJJAFBF_419580 | Bacteria | Proteobacteri | Alphaproteob | Rhodobactera | Rhodobactera | Planktomarina | Planktomarina | unclassified | unclassified | unclassified | unclassified | unclassified | 1 | 0 | 0,000107052 | 1,02E-05 | 1 - | 8,97E-05 - | 1 - | 0,000111343 - | 1 - | 0,000105845 - | 1 - | 0,000107409 - | 1 - | 0,000120996 - |
| 5177 | 1 | 3um_FIJJAFBF_333738 | Bacteria | Proteobacteri | Gammaprotec | Cellvibrionales | Porticoccacea | unclassified | gamma protec | information st | Translation, ril | COG0292 | Binds directly | 314287.GB22 | 4 | 0,632455532 | 0,000465352 | 9,17E-05 | 4 - | 0,000389074 - | 5 - | 0,000603893 - | 4 - | 0,000459262 - | 3 - | 0,000349534 - | 4 - | 0,000524998 - |
| 5178 | 1 | 0.2um_NOKCPHJN_445039 | Eukaryota | unclassified | unclassified | Prymnesiales | Chrysochromu | Chrysochromu | Chrysochromu | information st | Transcription | 0B4G2 | Neuronal PAS | 7260.FBpp023 | 0,2 | 0,4 | 1,56E-06 | 3,13E-06 | 1 - | 7,82E-06 - | 0 - | 0 - | 0 - | 0 - | 0 - | 0 - | 0 - | 0 - |
| 5179 | 1 | 3um_FIJJAFBF_306693 | Bacteria | Proteobacteri | Alphaproteob | Rhodobactera | unclassified | unclassified | Rhodobactera | unclassified | unclassified | unclassified | unclassified | unclassified | 2,4 | 1,019803903 | 0,000226062 | 7,74E-05 | 4 - | 0,000327934 - | 2 - | 0,000203598 - | 2 - | 0,000193546 - | 3 - | 0,000294607 - | 1 - | 0,000110625 - |
| 5180 | 1 | 0.2um_NOKCPHJN_159985 | Bacteria | Proteobacteri | Gammaprotec | unclassified | unclassified | unclassified | gamma protec | poorly charact | Function unkn | 16Q04 | oxidoreductas | 388401.RB215 | 6 | 1,414213562 | 0,00021629 | 4,98E-05 | 6 - | 0,000182669 - | 7 - | 0,000264624 - | 8 - | 0,000287495 - | 5 - | 0,000182339 - | 4 - | 0,000164323 - |
| 5181 | 1 | 3um_FIJJAFBF_355790 | Bacteria | Bacteroidetes | Flavobacteriia | Flavobacterial | Flavobacteriac | Polaribacter | Polaribacter s | unclassified | unclassified | unclassified | unclassified | unclassified | 15 | 1,095445115 | 0,000217626 | 1,25E-05 | 16 - | 0,000195573 - | 15 - | 0,000227666 - | 16 - | 0,000230854 - | 15 - | 0,000219622 - | 13 - | 0,000214416 - |
| 5182 | 1 | 3um_FIJJAFBF_35692 | Bacteria | Bacteroidetes | Flavobacteriia | Flavobacterial | Crocinitomica | unclassified | Crocinitomica | poorly charact | Function unkn | 055DA | Tetratricopept | 709991.Odosp | 4,4 | 1,854723699 | 0,000102621 | 3,14E-05 | 8 - | 0,000161943 - | 4 - | 0,000100542 - | 3 - | 7,17E-05 - | 4 - | 9,70E-05 - | 3 - | 8,19E-05 - |
| 5183 | 1 | 0.2um_NOKCPHJN_464262 | Bacteria | Proteobacteri | Alphaproteob | Rhodobactera | Rhodobactera | Yoonia | Yoonia vestfol | unclassified | unclassified | unclassified | unclassified | unclassified | 0,8 | 0,4 | 1,12E-05 | 5,66E-06 | 1 - | 1,21E-05 - | 1 - | 1,50E-05 - | 1 - | 1,42E-05 - | 1 - | 1,45E-05 - | 0 - | 0 - |
| 5184 | 1 | 0.2um_NOKCPHJN_283034 | Bacteria | Proteobacteri | Gammaprotec | Cellvibrionales | Halieaceae | unclassified | gamma protec | cellular proces | Defense mech | 16QBT | Type II restrict | 876044.IMCC3 | 0,4 | 0,489897949 | 5,76E-06 | 7,09E-06 | 1 - | 1,32E-05 - | 0 - | 0 - | 1 - | 1,56E-05 - | 0 - | 0 - | 0 - | 0 - |
| 5185 | 1 | 0.2um_NOKCPHJN_359322 | Bacteria | Proteobacteri | Gammaprotec | Oceanospirilla | Saccharospirill | Reinekea | Reinekea fors | cellular proces | Posttranslatio | 11WF3 | Glutaredoxin | 314283.MED2 | 1 | 0 | 0,000111404 | 1,06E-05 | 1 - | 9,33E-05 - | 1 - | 0,000115869 - | 1 - | 0,000110148 - | 1 - | 0,000111775 - | 1 - | 0,000125914 - |
| 5186 | 1 | 3um_FIJJAFBF_20040 | Bacteria | Bacteroidetes | Flavobacteriia | Flavobacterial | Flavobacteriac | Polaribacter | Polaribacter s | metabolism | Energy produc | 099Q7 | Catalyzes the | 313594.PI23P | 4 | 0 | 0,000177957 | 1,69E-05 | 4 - | 0,000149061 - | 4 - | 0,000185089 - | 4 - | 0,000175951 - | 4 - | 0,00017855 - | 1 - | 0,000201136 - |
| 5187 | 1 | 0.2um_NOKCPHJN_900409 | Bacteria | Proteobacteri | Gammaprotec | Pseudomonad | Pseudomonad | Pseudomonas | various | information st | Translation, ril | 179ZI | One of the pri | 220664.PFL_5 | 0,2 | 0,4 | 2,08E-05 | 4,17E-05 | 0 - | 0 - | 0 - | 0 - | 1 - | 0,000104217 - | 0 - | 0 - | 0 - | 0 - |
| 5188 | 1 | 0.2um_NOKCPHJN_62422 | Bacteria | Proteobacteri | Gammaprotec | unclassified | unclassified | unclassified | Gammaprotec | unclassified | unclassified | unclassified | unclassified | unclassified | 6,2 | 1,166190379 | 9,76E-05 | 1,53E-05 | 7 - | 9,35E-05 - | 7 - | 0,000116139 - | 7 - | 0,000110405 - | 6 - | 9,60E-05 - | 4 - | 7,21E-05 - |
| 5189 | 1 | 0.2um_NOKCPHJN_91534 | Bacteria | Proteobacteri | Gammaprotec | Cellvibrionales | Porticoccacea | unclassified | Gammaprotec | cellular proces | Cell wall/mem | 0QTSX | Peptidoglycan | 314287.GB22 | 1 | 0 | 8,51E-05 | 8,08E-06 | 1 - | 7,13E-05 - | 1 - | 8,85E-05 - | 1 - | 8,42E-05 - | 1 - | 8,54E-05 - | 1 - | 9,62E-05 - |
| 5190 | 1 | 3um_FIJJAFBF_05046 | Bacteria | Proteobacteri | Betaproteoba | unclassified | unclassified | unclassified | Betaproteoba | metabolism | Inorganic ion t | 07QMV | Part of the AB | 316058.RPB_ | 4,2 | 0,4 | 0,000752056 | 0,000136015 | 4 - | 0,000596243 - | 4 - | 0,000740358 - | 4 - | 0,000703804 - | 4 - | 0,000714199 - | 5 - | 0,001005678 - |
| 5191 | 1 | 3um_FIJJAFBF_201886 | Bacteria | Proteobacteri | Betaproteoba | unclassified | unclassified | unclassified | Betaproteoba | poorly charact | Function unkn | 0QSE4 | Pfam:DUF400 | 876044.IMCC3 | 0,4 | 0,489897949 | 2,66E-05 | 3,28E-05 | 1 - | 6,11E-05 - | 0 - | 0 - | 1 - | 7,21E-05 - | 0 - | 0 - | 0 - | 0 - |
| 5192 | 1 | 0.2um_NOKCPHJN_337707 | Bacteria | Firmicutes | Bacilli | Bacillales | Bacillaceae | Bacillus | Bacillus sp. es | cellular proces | Intracellular tr | 17HSD | domain protei | 1035188.HMP | 0,4 | 0,489897949 | 1,96E-06 | 2,41E-06 | 0 - | 0 - | 0 - | 0 - | 1 - | 4,57E-06 - | 0 - | 0 - | 1 - | 5,22E-06 - |
| 5193 | 1 | 0.2um_NOKCPHJN_123350 | Bacteria | Proteobacteri | Gammaprotec | Oceanospirilla | Saccharospirill | Reinekea | Reinekea blan | metabolism | Coenzyme tra | 16Q44 | Methyltransfe | 314283.MED2 | 0,4 | 0,489897949 | 2,01E-05 | 2,49E-05 | 1 - | 4,48E-05 - | 1 - | 5,57E-05 - | 0 - | 0 - | 0 - | 0 - | 0 - | 0 - |
| 5194 | 1 | 0.2um_NOKCPHJN_250770 | Eukaryota | unclassified | Labyrinthulom | unclassified | Thraustochytr | Aurantiochytr | Aurantiochytr | unclassified | unclassified | unclassified | unclassified | unclassified | 0,2 | 0,4 | 5,33E-07 | 1,07E-06 | 1 - | 2,66E-06 - | 0 - | 0 - | 0 - | 0 - | 0 - | 0 - | 0 - | 0 - |
| 5195 | 1 | 3um_FIJJAFBF_189829 | Bacteria | Bacteroidetes | Flavobacteriia | Flavobacterial | Flavobacteriac | Gilvibacter | Gilvibacter sp. | unclassified | unclassified | unclassified | unclassified | unclassified | 3,2 | 1,166190379 | 0,000236191 | 8,04E-05 | 4 - | 0,000250878 - | 2 - | 0,000155758 - | 5 - | 0,00037017 - | 2 - | 0,000150255 - | 3 - | 0,000253893 - |
| 5196 | 1 | 0.2um_NOKCPHJN_530548 | Eukaryota | Bacillariophyt | Coscinodiscop | Thalassiosirale | Thalassiosirac | Thalassiosira | Thalassiosira | information st | Chromatin stri | KOG1744 | Core compone | 2850.Phatr54 | 1 | 0 | 0,000118127 | 1,12E-05 | 1 - | 9,89E-05 - | 1 - | 0,000122861 - | 1 - | 0,000116795 - | 1 - | 0,00011852 - | 1 - | 0,000133512 - |
| 5197 | 1 | 0.2um_NOKCPHJN_339031 | Bacteria | Verrucomicrol | Opitutae | unclassified | unclassified | unclassified | Opitutae bact | unclassified | unclassified | unclassified | unclassified | unclassified | 1,6 | 0,489897949 | 0,000131026 | 4,22E-05 | 1 - | 6,83E-05 - | 2 - | 0,000169665 - | 2 - | 0,000161288 - | 2 - | 0,000163671 - | 1 - | 9,22E-05 - |
| 5198 | 1 | 3um_FIJJAFBF_245641 | Bacteria | various | various | various | various | various | various | information st | Translation, ril | 05KK9 | This protein bi | 886377.Murru | 4,4 | 1,019803903 | 0,000349188 | 5,89E-05 | 6 - | 0,000405094 - | 4 - | 0,000335339 - | 5 - | 0,000398477 - | 3 - | 0,000242618 - | 4 - | 0,000364411 - |
| 5199 | 1 | 0.2um_NOKCPHJN_221370 | Bacteria | Proteobacteri | Alphaproteob | Rhodobactera | Hyphomonada | unclassified | Hyphomonada | information st | Replication, re | 0PB0H | DNA-binding p | 755732.Fluta_ | 1 | 0,632455532 | 0,000154413 | 9,88E-05 | 1 - | 0,000128963 - | 1 - | 0,000160134 - | 0 - | 0 - | 2 - | 0,000308951 - | 1 - | 0,000174016 - |
| 5200 | 1 | 3um_FIJJAFBF_118044 | Bacteria | Bacteroidetes | unclassified | unclassified | unclassified | unclassified | Bacteroidetes | unclassified | unclassified | unclassified | unclassified | unclassified | 0,4 | 0,8 | 1,79E-05 | 3,59E-05 | 0 - | 0 - | 0 - | 0 - | 2 - | 8,97E-05 - | 0 - | 0 - | 0 - | 0 - |
| 5201 | 1 | 3um_FIJJAFBF_64213 | Bacteria | unclassified | unclassified | unclassified | unclassified | unclassified | bacterium TM | cellular proces | Cell wall/mem | 09A4Y | export protein | 518766.Rmar_ | 3,4 | 1,496662955 | 5,16E-05 | 2,13E-05 | 5 - | 6,55E-05 - | 5 - | 8,13E-05 - | 3 - | 4,64E-05 - | 3 - | 4,71E-05 - | 1 - | 1,77E-05 - |
| 5202 | 1 | 0.2um_NOKCPHJN_612534 | Eukaryota | Chlorophyta | Mamiellophyc | Mamiellales | Bathycoccace | Ostreococcus | Ostreococcus | information st | Translation, ril | KOG0886 | Ribosomal pro | 436017.A4RV | 2,2 | 0,979795897 | 0,000169454 | 5,89E-05 | 4 - | 0,000266923 - | 1 - | 8,29E-05 - | 2 - | 0,000157537 - | 2 - | 0,000159864 - | 2 - | 0,000180087 - |
| 5203 | 1 | 3um_FIJJAFBF_163900 | Bacteria | Proteobacteri | Alphaproteob | Rhodobactera | Rhodobactera | unclassified | Rhodobactera | metabolism | Carbohydrate | 01RFA | ABC transport | 644076.SCH4 | 10 | 1,414213562 | 0,000401076 | 7,52E-05 | 11 - | 0,000369165 - | 9 - | 0,00037505 - | 10 - | 0,000396147 - | 8 - | 0,000321598 - | 12 - | 0,000543419 - |
| 5204 | 1 | 0.2um_NOKCPHJN_405305 | Bacteria | Bacteroidetes | Flavobacteriia | Flavobacterial | Flavobacteriac | Polaribacter | Polaribacter s | cellular proces | Cell wall/mem | COG2885 | Ompa motb d | 313598.MED1 | 2,6 | 0,489897949 | 0,000131195 | 1,90E-05 | 3 - | 0,000128481 - | 2 - | 0,000106357 - | 3 - | 0,000151659 - | 3 - | 0,000153899 - | 2 - | 0,000115578 - |
| 5205 | 1 | 0.2um_NOKCPHJN_60423 | Bacteria | Proteobacteri | Gammaprotec | Oceanospirilla | Oceanospirilla | Marinobacteri | Marinobacter | information st | Transcription | COG0250 | Participates in | 247634.GPB2 | 2,2 | 0,4 | 0,000167802 | 1,54E-05 | 3 - | 0,000194537 - | 2 - | 0,000161038 - | 2 - | 0,000153087 - | 2 - | 0,000155348 - | 2 - | 0,000174999 - |
| 5206 | 1 | 0.2um_NOKCPHJN_914567 | Bacteria | Proteobacteri | Alphaproteob | Pelagibacteral | Pelagibacterac | unclassified | Pelagibactera | cellular proces | Cell wall/mem | 7257 | NA | 439493.PB721 | 2,2 | 0,4 | 9,19E-05 | 2,47E-05 | 2 - | 6,91E-05 - | 2 - | 8,59E-05 - | 2 - | 8,16E-05 - | 2 - | 8,28E-05 - | 3 - | 0,000139947 - |
| 5207 | 1 | 0.2um_NOKCPHJN_370628 | Bacteria | Proteobacteri | Alphaproteob | Rhodobactera | Rhodobactera | Amylibacter | Amylibacter k | information st | Replication, re | 05C24 | DNA gyrase ne | 351016.RAZW | 0,8 | 0,748331477 | 1,15E-05 | 1,00E-05 | 2 - | 2,54E-05 - | 0 - | 0 - | 0 - | 0 - | 1 - | 1,52E-05 - | 1 - | 1,71E-05 - |
| 5208 | 1 | 0.2um_NOKCPHJN_45424 | Bacteria | Proteobacteri | Gammaprotec | Oceanospirilla | Saccharospirill | Gynuella | Gynuella suns | unclassified | unclassified | unclassified | unclassified | unclassified | 0,8 | 0,4 | 3,34E-05 | 1,68E-05 | 0 - | 0 - | 1 - | 4,17E-05 - | 1 - | 3,96E-05 - | 1 - | 4,02E-05 - | 1 - | 4,53E-05 - |
| 5209 | 1 | 3um_FIJJAFBF_198878 | Bacteria | Bacteroidetes | Flavobacteriia | Flavobacterial | Flavobacteriac | Polaribacter | Polaribacter b | cellular proces | Posttranslatio | 09ADN | alkyl hydroper | 313594.PI23P | 1,4 | 0,8 | 8,51E-05 | 3,77E-05 | 3 - | 0,000160154 - | 1 - | 6,63E-05 - | 1 - | 6,30E-05 - | 1 - | 6,39E-05 - | 1 - | 7,20E-05 - |
| 5210 | 1 | 3um_FIJJAFBF_83896 | unclassified | unclassified | unclassified | unclassified | unclassified | unclassified | unclassified | unclassified | unclassified | unclassified | unclassified | unclassified | 1 | 0,632455532 | 0,000107712 | 6,04E-05 | 2 - | 0,000186629 - | 1 - | 0,000115869 - | 1 - | 0,000110148 - | 0 - | 0 - | 1 - | 0,000125914 - |
| 5211 | 1 | 0.2um_NOKCPHJN_195403 | Bacteria | Proteobacteri | Gammaprotec | unclassified | unclassified | unclassified | SAR86 cluster | unclassified | unclassified | unclassified | unclassified | unclassified | 0,4 | 0,489897949 | 5,16E-06 | 6,36E-06 | 1 - | 1,17E-05 - | 0 - | 0 - | 0 - | 0 - | 1 - | 1,41E-05 - | 0 - | 0 - |
| 5212 | 1 | 3um_FIJJAFBF_150222 | Bacteria | Bacteroidetes | Flavobacteriia | Flavobacterial | Flavobacteriac | Formosa | Formosa sp. H | unclassified | unclassified | unclassified | unclassified | unclassified | 1 | 0 | 4,13E-05 | 3,92E-06 | 1 - | 3,46E-05 - | 1 - | 4,29E-05 - | 1 - | 4,08E-05 - | 1 - | 4,14E-05 - | 1 - | 4,66E-05 - |
| 5213 | 1 | 0.2um_NOKCPHJN_206886 | Bacteria | Proteobacteri | Alphaproteob | Rhodobactera | Rhodobactera | Sulfitobacter | Sulfitobacter g | unclassified | unclassified | unclassified | unclassified | unclassified | 1,2 | 1,166190379 | 2,78E-05 | 2,78E-05 | 1 - | 2,00E-05 - | 0 - | 0 - | 3 - | 7,09E-05 - | 2 - | 4,80E-05 - | 0 - | 0 - |
| 5214 | 2 | 0.2um_NOKCPHJN_690897;0.2um_NOKCPHJN_8 | Bacteria | Proteobacteri | Alphaproteob | Rhodobactera | unclassified | unclassified | Rhodobactera | unclassified | unclassified | unclassified | unclassified | unclassified | 1 | 0,632455532 | 4,08E-05 | 2,68E-05 | 1 - | 3,51E-05 - | 1 - | 4,36E-05 - | 1 - | 4,14E-05 - | 2 - | 8,41E-05 - | 0 - | 0 - |
| 5215 | 1 | 0.2um_NOKCPHJN_518694 | Bacteria | Proteobacteri | Alphaproteob | Pelagibacteral | Pelagibacterac | Candidatus Pe | Candidatus Pe | metabolism | Carbohydrate | 16THB | transporter, d | 388399.SSE37 | 2,2 | 0,748331477 | 8,68E-05 | 2,53E-05 | 3 - | 0,000101873 - | 2 - | 8,43E-05 - | 2 - | 8,02E-05 - | 3 - | 0,000122027 - | 1 - | 4,58E-05 - |
| 5216 | 1 | 0.2um_NOKCPHJN_102309 | Bacteria | Bacteroidetes | Flavobacteriia | various | various | various | various | unclassified | unclassified | unclassified | unclassified | unclassified | 0,4 | 0,489897949 | 1,51E-05 | 1,86E-05 | 1 - | 3,47E-05 - | 0 - | 0 - | 1 - | 4,09E-05 - | 0 - | 0 - | 0 - | 0 - |
| 5217 | 1 | 3um_FIJJAFBF_377575 | Bacteria | Bacteroidetes | Flavobacteriia | Flavobacterial | Flavobacteriac | Tenacibaculur | Tenacibaculur | information st | Translation, ril | COG0088 | One of the pri | 313598.MED1 | 1,4 | 0,489897949 | 9,33E-05 | 3,79E-05 | 1 - | 5,49E-05 - | 1 - | 6,82E-05 - | 2 - | 0,000129648 - | 1 - | 6,58E-05 - | 2 - | 0,000148205 - |
| 5218 | 1 | 0.2um_NOKCPHJN_197048 | Bacteria | Proteobacteri | Betaproteoba | unclassified | unclassified | unclassified | Betaproteoba | metabolism | Lipid transpor | 05CMX | Polyhydroxyal | 397945.Aave_ | 1,6 | 0,489897949 | 4,68E-05 | 1,55E-05 | 2 - | 4,89E-05 - | 2 - | 6,08E-05 - | 1 - | 2,89E-05 - | 1 - | 2,93E-05 - | 2 - | 6,60E-05 - |
| 5219 | 1 | 3um_FIJJAFBF_20076 | Bacteria | Bacteroidetes | Flavobacteriia | Flavobacterial | Flavobacteriac | Polaribacter | Polaribacter d | unclassified | unclassified | unclassified | unclassified | unclassified | 2 | 1,264911064 | 9,56E-05 | 5,22E-05 | 4 - | 0,000168789 - | 1 - | 5,24E-05 - | 3 - | 0,000149429 - | 1 - | 5,05E-05 - | 1 - | 5,69E-05 - |
| 5220 | 1 | 0.2um_NOKCPHJN_92001 | Bacteria | Bacteroidetes | Flavobacteriia | Flavobacterial | Flavobacteriac | Polaribacter | Polaribacter ir | metabolism | Inorganic ion t | 05C27 | TonB-depende | 313594.PI23P | 5,6 | 0,8 | 7,08E-05 | 1,24E-05 | 6 - | 6,36E-05 - | 6 - | 7,90E-05 - | 6 - | 7,51E-05 - | 4 - | 5,08E-05 - | 6 - | 8,58E-05 - |
| 5221 | 2 | 0.2um_NOKCPHJN_1002580;0.2um_NOKCPHJN_ | Bacteria | Proteobacteri | Alphaproteob | Rhodobactera | Rhodobactera | unclassified | Rhodobactera | unclassified | unclassified | unclassified | unclassified | unclassified | 1,8 | 0,4 | 6,14E-05 | 1,53E-05 | 2 - | 5,71E-05 - | 2 - | 7,09E-05 - | 1 - | 3,37E-05 - | 2 - | 6,84E-05 - | 2 - | 7,71E-05 - |
| 5222 | 1 | 3um_FIJJAFBF_301021 | Bacteria | Bacteroidetes | Flavobacteriia | unclassified | unclassified | unclassified | Flavobacteria | poorly charact | Function unkn | 11NCV | Protein of unk | 487796.Flav2 | 1,8 | 0,979795897 | 0,000141994 | 6,87E-05 | 3 - | 0,000206186 - | 1 - | 8,53E-05 - | 3 - | 0,000243381 - | 1 - | 8,23E-05 - | 1 - | 9,27E-05 - |
| 5223 | 1 | 3um_FIJJAFBF_242312 | Bacteria | Proteobacteri | Gammaprotec | Oceanospirilla | Saccharospirill | Saccharospiril | Saccharospiril | metabolism | Carbohydrate | 0QMVS | (ABC) transpo | 314283.MED2 | 2,8 | 0,748331477 | 0,000126969 | 2,24E-05 | 4 - | 0,00015563 - | 2 - | 9,66E-05 - | 3 - | 0,000137779 - | 3 - | 0,000139814 - | 2 - | 0,000105 - |
| 5224 | 1 | 3um_FIJJAFBF_123452 | Bacteria | Proteobacteri | Gammaprotec | unclassified | unclassified | unclassified | various | metabolism | Energy produc | 17CGT | F(1)F(0) ATP s | 349521.HCH_ | 4,6 | 1,019803903 | 0,000399222 | 7,95E-05 | 5 - | 0,000367874 - | 5 - | 0,000456791 - | 6 - | 0,000521085 - | 4 - | 0,000352521 - | 3 - | 0,000297836 - |
| 5225 | 1 | 3um_FIJJAFBF_175173 | Bacteria | Proteobacteri | Alphaproteob | Rhodobactera | unclassified | unclassified | Rhodobactera | metabolism | Energy produc | COG0711 | F(1)F(0) ATP s | 439497.RR11_ | 3,4 | 0,8 | 0,000258188 | 3,49E-05 | 5 - | 0,000324228 - | 3 - | 0,000241557 - | 3 - | 0,000229631 - | 3 - | 0,000233023 - | 3 - | 0,000262499 - |
| 5226 | 1 | 3um_FIJJAFBF_220297 | Eukaryota | Bacillariophyt | Mediophyceae | Toxariales | Toxariaceae | Toxarium | Toxarium und | cellular proces | Posttranslatio | 0IEKR | One of the co | 45157.CMV08 | 1 | 0 | 2,99E-05 | 2,83E-06 | 1 - | 2,50E-05 - | 1 - | 3,10E-05 - | 1 - | 2,95E-05 - | 1 - | 3,00E-05 - | 1 - | 3,37E-05 - |
| 5227 | 1 | 3um_FIJJAFBF_744836 | Bacteria | Proteobacteri | Gammaprotec | Cellvibrionales | Spongiibacter | unclassified | marine gamm | information st | Translation, ril | 16Q16 | Binds the lowe | 247639.MGP2 | 0,6 | 0,489897949 | 3,49E-05 | 2,87E-05 | 1 - | 5,17E-05 - | 0 - | 0 - | 1 - | 6,10E-05 - | 1 - | 6,19E-05 - | 0 - | 0 - |
| 5228 | 1 | 3um_FIJJAFBF_270535 | unclassified | unclassified | unclassified | unclassified | unclassified | unclassified | unclassified | unclassified | unclassified | unclassified | unclassified | unclassified | 1 | 0 | 2,27E-05 | 2,16E-06 | 1 - | 1,90E-05 - | 1 - | 2,36E-05 - | 1 - | 2,25E-05 - | 1 - | 2,28E-05 - | 1 - | 2,57E-05 - |
| 5229 | 1 | 3um_FIJJAFBF_97618 | Bacteria | Bacteroidetes | Flavobacteriia | Flavobacterial | Flavobacteriac | Polaribacter | Polaribacter s | information st | Translation, ril | 05C2N | RNA methyltra | 313594.PI23P | 1,8 | 0,748331477 | 9,68E-05 | 4,19E-05 | 1 - | 4,45E-05 - | 1 - | 5,52E-05 - | 3 - | 0,000157537 - | 2 - | 0,000106576 - | 2 - | 0,000120058 - |
| 5230 | 1 | 0.2um_NOKCPHJN_33790 | Bacteria | Bacteroidetes | Saprospiria | Saprospirales | Saprospiracea | Aureispira | Aureispira sp. | unclassified | unclassified | unclassified | unclassified | unclassified | 1,2 | 0,4 | 1,29E-05 | 2,86E-06 | 2 - | 1,85E-05 - | 1 - | 1,15E-05 - | 1 - | 1,09E-05 - | 1 - | 1,11E-05 - | 1 - | 1,25E-05 - |
| 5231 | 1 | 0.2um_NOKCPHJN_1038955 | Bacteria | Bacteroidetes | Flavobacteriia | Flavobacterial | Flavobacteriac | Marinirhabdu | Marinirhabdu | unclassified | unclassified | unclassified | unclassified | unclassified | 0,2 | 0,4 | 1,23E-05 | 2,47E-05 | 0 - | 0 - | 0 - | 0 - | 0 - | 0 - | 0 - | 0 - | 1 - | 6,17E-05 - |
| 5232 | 1 | 0.2um_NOKCPHJN_446856 | Bacteria | Proteobacteri | Alphaproteob | Pelagibacteral | Pelagibacterac | Candidatus Pe | Candidatus Pe | unclassified | unclassified | unclassified | unclassified | unclassified | 1 | 0 | 7,57E-05 | 7,19E-06 | 1 - | 6,34E-05 - | 1 - | 7,87E-05 - | 1 - | 7,49E-05 - | 1 - | 7,60E-05 - | 1 - | 8,56E-05 - |
| 5233 | 1 | 0.2um_NOKCPHJN_07540 | Bacteria | Bacteroidetes | Flavobacteriia | Flavobacterial | Flavobacteriac | Algibacter | Algibacter sp. | metabolism | Carbohydrate | 0XNQK | Transporter | 391587.KAOT | 1,8 | 0,4 | 5,38E-05 | 1,34E-05 | 2 - | 5,00E-05 - | 2 - | 6,21E-05 - | 1 - | 2,95E-05 - | 2 - | 5,99E-05 - | 2 - | 6,75E-05 - |
| 5234 | 1 | 3um_FIJJAFBF_276071 | Bacteria | Bacteroidetes | Flavobacteriia | Flavobacterial | Flavobacteriac | Winogradskye | Winogradskye | unclassified | unclassified | unclassified | unclassified | unclassified | 1,2 | 0,4 | 7,02E-05 | 1,55E-05 | 2 - | 0,000100681 - | 1 - | 6,25E-05 - | 1 - | 5,94E-05 - | 1 - | 6,03E-05 - | 1 - | 6,79E-05 - |
| 5235 | 1 | 0.2um_NOKCPHJN_355741 | Bacteria | Proteobacteri | Alphaproteob | Pelagibacteral | Pelagibacterac | Candidatus Pe | Candidatus Pe | poorly charact | Function unkn | COG3181 | extra-cytoplas | 439493.PB721 | 5,6 | 0,8 | 0,000233749 | 2,91E-05 | 7 - | 0,000246453 - | 5 - | 0,000218587 - | 5 - | 0,000207795 - | 5 - | 0,000210864 - | 6 - | 0,000285045 - |
| 5236 | 1 | 0.2um_NOKCPHJN_393841 | Bacteria | Bacteroidetes | Flavobacteriia | Flavobacterial | Flavobacteriac | Mesoflavibact | Mesoflavibact | poorly charact | Function unkn | 0PC4P | Inherit from b | 391587.KAOT | 1,2 | 0,4 | 3,44E-05 | 7,62E-06 | 2 - | 4,94E-05 - | 1 - | 3,06E-05 - | 1 - | 2,91E-05 - | 1 - | 2,96E-05 - | 1 - | 3,33E-05 - |
| 5237 | 1 | 3um_FIJJAFBF_14603 | Bacteria | Proteobacteri | Gammaprotec | Oceanospirilla | Saccharospirill | Reinekea | Reinekea fors | unclassified | unclassified | unclassified | unclassified | unclassified | 0,2 | 0,4 | 8,91E-06 | 1,78E-05 | 0 - | 0 - | 0 - | 0 - | 1 - | 4,46E-05 - | 0 - | 0 - | 0 - | 0 - |
| 5238 | 1 | 3um_FIJJAFBF_786157 | Bacteria | Proteobacteri | Gammaprotec | Oceanospirilla | Saccharospirill | Reinekea | Reinekea fors | information st | Translation, ril | 17C3B | The globular d | 314283.MED2 | 2,8 | 0,748331477 | 0,000337532 | 7,38E-05 | 4 - | 0,000409917 - | 2 - | 0,000254498 - | 3 - | 0,000362899 - | 2 - | 0,000245506 - | 3 - | 0,000414842 - |
| 5239 | 1 | 3um_FIJJAFBF_217104 | Bacteria | Bacteroidetes | Flavobacteriia | Flavobacterial | Crocinitomica | unclassified | Crocinitomica | unclassified | unclassified | unclassified | unclassified | unclassified | 2 | 0 | 8,73E-05 | 8,29E-06 | 2 - | 7,31E-05 - | 2 - | 9,08E-05 - | 2 - | 8,63E-05 - | 2 - | 8,76E-05 - | 2 - | 9,86E-05 - |
| 5240 | 1 | 0.2um_NOKCPHJN_799134 | unclassified | unclassified | unclassified | unclassified | unclassified | unclassified | unclassified | unclassified | unclassified | unclassified | unclassified | unclassified | 3,2 | 0,4 | 0,000185669 | 2,75E-05 | 3 - | 0,000145903 - | 3 - | 0,000181168 - | 4 - | 0,000229631 - | 3 - | 0,000174767 - | 3 - | 0,000196874 - |
| 5241 | 1 | 3um_FIJJAFBF_424281 | Eukaryota | Eustigmatoph | unclassified | Eustigmatales | Monodopsida | Nannochlorop | Nannochlorop | cellular proces | Posttranslatio | KOG0731 | ATP-depender | 2850.Phatr17 | 0,8 | 0,4 | 0,000281874 | 0,000144818 | 1 - | 0,000294299 - | 1 - | 0,000365433 - | 0 - | 0 - | 1 - | 0,000352521 - | 1 - | 0,000397114 - |
| 5242 | 1 | 0.2um_NOKCPHJN_978568 | Eukaryota | unclassified | Oligohymeno | Hymenostoma | Tetrahymenid | Tetrahymena | Tetrahymena | unclassified | unclassified | unclassified | unclassified | unclassified | 8,6 | 1,019803903 | 0,000262235 | 4,30E-05 | 9 - | 0,000229554 - | 10 - | 0,000316709 - | 7 - | 0,00021075 - | 8 - | 0,000244415 - | 9 - | 0,000309749 - |
| 5243 | 1 | 3um_FIJJAFBF_08129 | Bacteria | Bacteroidetes | Flavobacteriia | Flavobacterial | Flavobacteriac | Polaribacter | Polaribacter ir | unclassified | unclassified | unclassified | unclassified | unclassified | 1,6 | 0,8 | 5,06E-05 | 2,59E-05 | 1 - | 2,63E-05 - | 2 - | 6,54E-05 - | 1 - | 3,11E-05 - | 3 - | 9,46E-05 - | 1 - | 3,55E-05 - |
| 5244 | 1 | 0.2um_NOKCPHJN_331924 | Eukaryota | unclassified | Oomycetes | Saprolegniales | Saprolegniace | Thraustotheca | Thraustotheca | information st | Translation, ril | 14ZFT | ribosomal L5P | 67593.Physo1 | 0,6 | 0,489897949 | 4,75E-05 | 3,88E-05 | 0 - | 0 - | 1 - | 8,14E-05 - | 1 - | 7,74E-05 - | 1 - | 7,86E-05 - | 0 - | 0 - |
| 5245 | 1 | 3um_FIJJAFBF_672316 | Bacteria | Bacteroidetes | Cytophagia | Cytophagales | Flammeovirga | unclassified | Flammeovirga | unclassified | unclassified | unclassified | unclassified | unclassified | 7,2 | 0,979795897 | 0,000653776 | 2,48E-05 | 9 - | 0,000693282 - | 7 - | 0,000669552 - | 7 - | 0,000636494 - | 7 - | 0,000645895 - | 6 - | 0,000623656 - |
| 5246 | 1 | 3um_FIJJAFBF_417619 | Bacteria | Proteobacteri | Gammaprotec | unclassified | unclassified | unclassified | OM182 bacter | information st | Translation, ril | 08UHH | Located on the | 876044.IMCC3 | 0,2 | 0,4 | 2,10E-05 | 4,20E-05 | 0 - | 0 - | 0 - | 0 - | 0 - | 0 - | 1 - | 0,000104949 - | 0 - | 0 - |
| 5247 | 1 | 3um_FIJJAFBF_592963 | Bacteria | Proteobacteri | Gammaprotec | Oceanospirilla | Saccharospirill | Reinekea | Reinekea fors | metabolism | Carbohydrate | COG0149 | triosephospha | 314283.MED2 | 4,2 | 1,469693846 | 0,000224298 | 5,40E-05 | 7 - | 0,000322666 - | 4 - | 0,000228946 - | 4 - | 0,000217642 - | 3 - | 0,000165643 - | 3 - | 0,000186596 - |
| 5248 | 1 | 3um_FIJJAFBF_113627 | Bacteria | Proteobacteri | Gammaprotec | Alteromonada | Alteromonada | Lacimicrobium | Lacimicrobium | metabolism | Inorganic ion t | 173T4 | TONB-depend | 506534.Rhein | 3,4 | 1,019803903 | 4,70E-05 | 1,44E-05 | 4 - | 4,65E-05 - | 5 - | 7,21E-05 - | 2 - | 2,74E-05 - | 3 - | 4,17E-05 - | 3 - | 4,70E-05 - |
| 5249 | 1 | 0.2um_NOKCPHJN_975818 | Bacteria | Proteobacteri | Gammaprotec | Pseudomonad | Pseudomonad | Pseudomonas | various | unclassified | unclassified | unclassified | unclassified | unclassified | 0,2 | 0,4 | 2,15E-05 | 4,30E-05 | 0 - | 0 - | 0 - | 0 - | 1 - | 0,000107526 - | 0 - | 0 - | 0 - | 0 - |
| 5250 | 1 | 0.2um_NOKCPHJN_622401 | Bacteria | Proteobacteri | Alphaproteob | Pelagibacteral | Pelagibacterac | Candidatus Pe | Candidatus Pe | cellular proces | Intracellular tr | 05VIE | preprotein tra | 335992.SAR11 | 3,2 | 0,4 | 0,000466147 | 6,90E-05 | 3 - | 0,000366309 - | 3 - | 0,000454847 - | 4 - | 0,00057652 - | 3 - | 0,000438777 - | 3 - | 0,00049428 - |
| 5251 | 1 | 3um_FIJJAFBF_196005 | Bacteria | Bacteroidetes | Flavobacteriia | Flavobacterial | Flavobacteriac | Polaribacter | Polaribacter s | information st | Translation, ril | 05C6A | phenylalanyl-t | 313598.MED1 | 0,6 | 0,8 | 9,04E-06 | 1,17E-05 | 2 - | 2,84E-05 - | 0 - | 0 - | 1 - | 1,68E-05 - | 0 - | 0 - | 0 - | 0 - |
| 5252 | 1 | 3um_FIJJAFBF_151131 | Bacteria | Bacteroidetes | Flavobacteriia | Flavobacterial | Flavobacteriac | Polaribacter | Polaribacter s | unclassified | unclassified | unclassified | unclassified | unclassified | 0,6 | 0,489897949 | 4,97E-05 | 4,10E-05 | 1 - | 7,26E-05 - | 1 - | 9,02E-05 - | 1 - | 8,57E-05 - | 0 - | 0 - | 0 - | 0 - |
| 5253 | 1 | 0.2um_NOKCPHJN_289810 | Bacteria | Proteobacteri | Alphaproteob | Rhodobactera | Rhodobactera | Planktomarina | Planktomarina | cellular proces | Posttranslatio | 05JRK | feS assembly p | 375451.RD1_ | 0,6 | 0,8 | 1,74E-05 | 2,24E-05 | 2 - | 5,45E-05 - | 0 - | 0 - | 0 - | 0 - | 1 - | 3,27E-05 - | 0 - | 0 - |
| 5254 | 1 | 0.2um_NOKCPHJN_1087909 | Eukaryota | Phaeophyceae | unclassified | Ectocarpales | Ectocarpaceae | Ectocarpus | Ectocarpus sil | unclassified | unclassified | unclassified | unclassified | unclassified | 0,8 | 0,4 | 2,19E-05 | 1,10E-05 | 0 - | 0 - | 1 - | 2,74E-05 - | 1 - | 2,60E-05 - | 1 - | 2,64E-05 - | 1 - | 2,97E-05 - |
| 5255 | 1 | 3um_FIJJAFBF_221830 | Bacteria | Proteobacteri | Gammaprotec | Oceanospirilla | Oceanospirilla | Amphritea | Amphritea jap | unclassified | unclassified | unclassified | unclassified | unclassified | 2,4 | 0,489897949 | 7,89E-05 | 1,16E-05 | 3 - | 8,38E-05 - | 2 - | 6,94E-05 - | 3 - | 9,89E-05 - | 2 - | 6,69E-05 - | 2 - | 7,54E-05 - |
| 5256 | 1 | 0.2um_NOKCPHJN_928675 | Bacteria | Proteobacteri | Gammaprotec | unclassified | unclassified | unclassified | Gammaprotec | cellular proces | Posttranslatio | 0QHXH | Molecular cha | 203122.Sde_2 | 0,2 | 0,4 | 6,67E-06 | 1,33E-05 | 1 - | 3,34E-05 - | 0 - | 0 - | 0 - | 0 - | 0 - | 0 - | 0 - | 0 - |
| 5257 | 1 | 3um_FIJJAFBF_217624 | Bacteria | Bacteroidetes | Flavobacteriia | Flavobacterial | unclassified | unclassified | Flavobacterial | poorly charact | Function unkn | 099QK | NA | 983544.Lacal_ | 4,4 | 0,489897949 | 0,000151406 | 1,58E-05 | 5 - | 0,00014492 - | 5 - | 0,000179948 - | 4 - | 0,000136851 - | 4 - | 0,000138872 - | 4 - | 0,000156439 - |
| 5258 | 1 | 0.2um_NOKCPHJN_277651 | Bacteria | Bacteroidetes | unclassified | unclassified | unclassified | unclassified | Bacteroidetes | information st | Replication, re | COG0514 | atp-dependen | 655815.ZPR_1 | 0,4 | 0,489897949 | 7,03E-06 | 8,66E-06 | 1 - | 1,61E-05 - | 0 - | 0 - | 1 - | 1,90E-05 - | 0 - | 0 - | 0 - | 0 - |
| 5259 | 1 | 3um_FIJJAFBF_322984 | unclassified | unclassified | unclassified | unclassified | unclassified | unclassified | unclassified | unclassified | unclassified | unclassified | unclassified | unclassified | 0,8 | 0,4 | 1,56E-05 | 8,04E-06 | 1 - | 1,64E-05 - | 1 - | 2,04E-05 - | 1 - | 1,94E-05 - | 0 - | 0 - | 1 - | 2,21E-05 - |
| 5260 | 1 | 3um_FIJJAFBF_40863 | Bacteria | Bacteroidetes | unclassified | unclassified | unclassified | unclassified | Bacteroidetes | unclassified | unclassified | unclassified | unclassified | unclassified | 1 | 0 | 4,64E-05 | 4,41E-06 | 1 - | 3,89E-05 - | 1 - | 4,83E-05 - | 1 - | 4,59E-05 - | 1 - | 4,66E-05 - | 1 - | 5,25E-05 - |
| 5261 | 1 | 3um_FIJJAFBF_02802 | Bacteria | Proteobacteri | Gammaprotec | Oceanospirilla | Saccharospirill | Reinekea | Reinekea fors | cellular proces | Posttranslatio | 0QIT3 | Thiol disulfide | 314283.MED2 | 1 | 0,632455532 | 7,72E-05 | 4,90E-05 | 1 - | 6,48E-05 - | 1 - | 8,05E-05 - | 2 - | 0,000153087 - | 0 - | 0 - | 1 - | 8,75E-05 - |
| 5262 | 1 | 3um_FIJJAFBF_17653 | Bacteria | Proteobacteri | Gammaprotec | Cellvibrionales | Porticoccacea | unclassified | Gammaprotec | unclassified | unclassified | unclassified | unclassified | unclassified | 1 | 0 | 9,72E-05 | 9,23E-06 | 1 - | 8,14E-05 - | 1 - | 0,000101077 - | 1 - | 9,61E-05 - | 1 - | 9,75E-05 - | 1 - | 0,00010984 - |
| 5263 | 1 | 0.2um_NOKCPHJN_327380 | Bacteria | Bacteroidetes | Flavobacteriia | Flavobacterial | Flavobacteriac | Polaribacter | Polaribacter s | information st | Translation, ril | 05CAV | thus facilitatin | 391587.KAOT | 1,2 | 0,4 | 2,72E-05 | 1,16E-05 | 1 - | 1,86E-05 - | 1 - | 2,31E-05 - | 1 - | 2,19E-05 - | 1 - | 2,22E-05 - | 2 - | 5,01E-05 - |
| 5264 | 1 | 0.2um_NOKCPHJN_53869 | Viruses | unclassified | unclassified | unclassified | Mimiviridae | unclassified | Yellowstone la | unclassified | unclassified | unclassified | unclassified | unclassified | 0,4 | 0,489897949 | 5,98E-05 | 7,40E-05 | 1 - | 0,000133461 - | 1 - | 0,00016572 - | 0 - | 0 - | 0 - | 0 - | 0 - | 0 - |
| 5265 | 1 | 3um_FIJJAFBF_62484 | Bacteria | Proteobacteri | Gammaprotec | Oceanospirilla | Oceanospirilla | Marinomonas | Marinomonas | poorly charact | Function unkn | 05DIH | basic membra | 314277.MED1 | 3,4 | 0,8 | 0,000141189 | 4,06E-05 | 3 - | 0,000103402 - | 4 - | 0,000171194 - | 4 - | 0,000162741 - | 2 - | 8,26E-05 - | 4 - | 0,000186035 - |
| 5266 | 1 | 3um_FIJJAFBF_341477 | Bacteria | Bacteroidetes | Flavobacteriia | Flavobacterial | Flavobacteriac | Ulvibacter | Ulvibacter lito | unclassified | unclassified | unclassified | unclassified | unclassified | 0,4 | 0,489897949 | 1,16E-05 | 1,42E-05 | 0 - | 0 - | 1 - | 2,96E-05 - | 1 - | 2,82E-05 - | 0 - | 0 - | 0 - | 0 - |
| 5267 | 1 | 0.2um_NOKCPHJN_119122 | Bacteria | Bacteroidetes | Flavobacteriia | Flavobacterial | Flavobacteriac | Polaribacter | Polaribacter s | unclassified | unclassified | unclassified | unclassified | unclassified | 1,4 | 0,489897949 | 2,07E-05 | 7,76E-06 | 1 - | 1,23E-05 - | 2 - | 3,06E-05 - | 1 - | 1,46E-05 - | 2 - | 2,95E-05 - | 1 - | 1,66E-05 - |
| 5268 | 1 | 0.2um_NOKCPHJN_241566 | Bacteria | Proteobacteri | Gammaprotec | Cellvibrionales | Porticoccacea | unclassified | SAR92 bacteri | metabolism | Amino acid tra | COG0019 | Specifically ca | 228410.NE09 | 1 | 0 | 3,50E-05 | 3,33E-06 | 1 - | 2,94E-05 - | 1 - | 3,64E-05 - | 1 - | 3,47E-05 - | 1 - | 3,52E-05 - | 1 - | 3,96E-05 - |
| 5269 | 1 | 0.2um_NOKCPHJN_14210 | Bacteria | Bacteroidetes | unclassified | unclassified | unclassified | unclassified | Bacteroidetes | poorly charact | Function unkn | 06SH2 | NA | 313603.FB217 | 0,8 | 0,748331477 | 4,07E-05 | 3,60E-05 | 2 - | 9,29E-05 - | 0 - | 0 - | 1 - | 5,49E-05 - | 1 - | 5,57E-05 - | 0 - | 0 - |
| 5270 | 1 | 0.2um_NOKCPHJN_1026862 | Bacteria | Proteobacteri | Gammaprotec | Alteromonada | Alteromonada | Glaciecola | Glaciecola sp. | information st | Translation, ril | COG0050 | This protein p | 345073.VC03 | 3,4 | 0,489897949 | 0,00011714 | 1,24E-05 | 4 - | 0,000116525 - | 3 - | 0,000108517 - | 3 - | 0,000103159 - | 4 - | 0,000139577 - | 3 - | 0,000117925 - |
| 5271 | 2 | 3um_FIJJAFBF_311063;3um_FIJJAFBF_523040 | Bacteria | Bacteroidetes | Flavobacteriia | Flavobacterial | Flavobacteriac | Formosa | Formosa sp. H | poorly charact | Function unkn | 09ARW | Zinc ribbon do | 1046627.BZAF | 1 | 0,894427191 | 5,40E-05 | 5,09E-05 | 1 - | 4,43E-05 - | 0 - | 0 - | 0 - | 0 - | 2 - | 0,000106165 - | 2 - | 0,000119594 - |
| 5272 | 1 | 3um_FIJJAFBF_124365 | Bacteria | Bacteroidetes | Cytophagia | Cytophagales | Cytophagacea | Spirosoma | Spirosoma spi | information st | Replication, re | 11K60 | integrase fami | 880070.Cycma | 0,2 | 0,4 | 5,17E-06 | 1,03E-05 | 1 - | 2,59E-05 - | 0 - | 0 - | 0 - | 0 - | 0 - | 0 - | 0 - | 0 - |
| 5273 | 1 | 0.2um_NOKCPHJN_716887 | Bacteria | Proteobacteri | Gammaprotec | Oceanospirilla | Saccharospirill | Reinekea | Reinekea fors | metabolism | Inorganic ion t | 0QIIZ | Na( ) H( ) antip | 58051.PE36_0 | 0,8 | 0,4 | 3,70E-05 | 1,90E-05 | 1 - | 3,86E-05 - | 1 - | 4,80E-05 - | 0 - | 0 - | 1 - | 4,63E-05 - | 1 - | 5,21E-05 - |
| 5274 | 1 | 3um_FIJJAFBF_348329 | Bacteria | Bacteroidetes | Flavobacteriia | Flavobacterial | Flavobacteriac | Algibacter | Algibacter sp. | unclassified | unclassified | unclassified | unclassified | unclassified | 3,4 | 1,019803903 | 6,08E-05 | 1,95E-05 | 3 - | 4,48E-05 - | 4 - | 7,41E-05 - | 2 - | 3,52E-05 - | 5 - | 8,94E-05 - | 3 - | 6,04E-05 - |
| 5275 | 5 | 0.2um_NOKCPHJN_1021993;0.2um_NOKCPHJN_ | Bacteria | Proteobacteri | Alphaproteob | Pelagibacteral | Pelagibacterac | Candidatus Pe | various | poorly charact | Function unkn | 17CZ8 | Lipoprotein | 439493.PB721 | 0,4 | 0,489897949 | 2,77E-05 | 3,42E-05 | 1 - | 6,31E-05 - | 0 - | 0 - | 0 - | 0 - | 1 - | 7,55E-05 - | 0 - | 0 - |
| 5276 | 1 | 3um_FIJJAFBF_553555 | Bacteria | Proteobacteri | Alphaproteob | Rhodobactera | Rhodobactera | various | various | information st | Translation, ril | COG0291 | 50s ribosomal | 391619.RGBS | 0,8 | 0,4 | 0,000172836 | 8,70E-05 | 0 - | 0 - | 1 - | 0,000215938 - | 1 - | 0,000205276 - | 1 - | 0,000208308 - | 1 - | 0,000234658 - |
| 5277 | 1 | 0.2um_NOKCPHJN_42581 | Bacteria | Proteobacteri | Alphaproteob | Rhodobactera | Rhodobactera | unclassified | Rhodobactera | unclassified | unclassified | unclassified | unclassified | unclassified | 3,2 | 0,979795897 | 0,000364117 | 7,25E-05 | 5 - | 0,000490499 - | 3 - | 0,000365433 - | 3 - | 0,00034739 - | 3 - | 0,000352521 - | 2 - | 0,000264743 - |
| 5278 | 1 | 3um_FIJJAFBF_91116 | Eukaryota | unclassified | Pelagophycea | Pelagomonad | unclassified | Aureococcus | Aureococcus a | metabolism | Carbohydrate | KOG1203 | Complex I inte | 35128.Thapsd | 3,6 | 1,019803903 | 0,000151105 | 4,07E-05 | 4 - | 0,00014258 - | 4 - | 0,000177042 - | 3 - | 0,000126226 - | 5 - | 0,000213483 - | 2 - | 9,62E-05 - |
| 5279 | 1 | 3um_FIJJAFBF_396395 | Bacteria | Proteobacteri | Alphaproteob | Rhodobactera | Rhodobactera | unclassified | Rhodobactera | unclassified | unclassified | unclassified | unclassified | unclassified | 0,4 | 0,489897949 | 1,22E-05 | 1,51E-05 | 1 - | 2,78E-05 - | 0 - | 0 - | 0 - | 0 - | 1 - | 3,33E-05 - | 0 - | 0 - |
| 5280 | 2 | 0.2um_NOKCPHJN_1054807;0.2um_NOKCPHJN_ | Eukaryota | Eustigmatoph | unclassified | Eustigmatales | Monodopsida | Nannochlorop | Nannochlorop | unclassified | unclassified | unclassified | unclassified | unclassified | 0,4 | 0,489897949 | 4,52E-05 | 5,53E-05 | 0 - | 0 - | 1 - | 0,000114935 - | 0 - | 0 - | 1 - | 0,000110874 - | 0 - | 0 - |
| 5281 | 1 | 3um_FIJJAFBF_316515 | Bacteria | Proteobacteri | Gammaprotec | Alteromonada | Alteromonada | Paraglaciecola | Paraglaciecola | unclassified | unclassified | unclassified | unclassified | unclassified | 11,4 | 1,2 | 0,000192047 | 2,56E-05 | 12 - | 0,000169621 - | 10 - | 0,000175516 - | 13 - | 0,000216905 - | 10 - | 0,000169314 - | 12 - | 0,000228879 - |
| 5282 | 1 | 0.2um_NOKCPHJN_648281 | Bacteria | Proteobacteri | Alphaproteob | Pelagibacteral | Pelagibacterac | Candidatus Pe | Candidatus Pe | information st | Translation, ril | 17FIH | One of the ea | 335992.SAR11 | 1 | 0 | 0,000141265 | 1,34E-05 | 1 - | 0,000118327 - | 1 - | 0,000146927 - | 1 - | 0,000139672 - | 1 - | 0,000141735 - | 1 - | 0,000159664 - |
| 5283 | 1 | 3um_FIJJAFBF_673429 | Bacteria | Proteobacteri | various | various | various | various | various | metabolism | Energy produc | COG3474 | Electron carrie | 314270.RB208 | 0,8 | 0,4 | 7,92E-05 | 3,99E-05 | 0 - | 0 - | 1 - | 9,90E-05 - | 1 - | 9,41E-05 - | 1 - | 9,55E-05 - | 1 - | 0,000107552 - |
| 5284 | 1 | 3um_FIJJAFBF_43758 | Bacteria | Proteobacteri | Gammaprotec | Oceanospirilla | Saccharospirill | Reinekea | Reinekea fors | cellular proces | Posttranslatio | COG0330 | Band 7 protein | 331869.BAL19 | 4 | 1,095445115 | 0,000152051 | 3,95E-05 | 4 - | 0,000128601 - | 3 - | 0,000119764 - | 6 - | 0,000227701 - | 4 - | 0,000154043 - | 3 - | 0,000130147 - |
| 5285 | 1 | 0.2um_NOKCPHJN_09985 | Bacteria | Bacteroidetes | Flavobacteriia | Flavobacterial | Flavobacteriac | Polaribacter | Polaribacter s | unclassified | unclassified | unclassified | unclassified | unclassified | 2 | 0 | 0,000221011 | 2,10E-05 | 2 - | 0,000185124 - | 2 - | 0,000229869 - | 2 - | 0,00021852 - | 2 - | 0,000221747 - | 2 - | 0,000249798 - |

| | | | | | | | | | | | | | | | | | | | | | | | | | | | | |
|---|---|---|---|---|---|---|---|---|---|---|---|---|---|---|---|---|---|---|---|---|---|---|---|---|---|---|---|---|
| 5286 | 1 | 3um_FIJJAFBF_715063 | Eukaryota | unclassified | Raphidophyce | Chattonellales | Chattonellace | Heterosigma | Heterosigma a | cellular proces | Posttranslatio | 0IEKR | One of the cor | 35128.Thapsd | 1 | 0 | 2,99E-05 | 2,84E-06 | 1 - | 2,51E-05 - | 1 - | 3,11E-05 - | 1 - | 2,96E-05 - | 1 - | 3,00E-05 - | 1 - | 3,38E-05 - |
| 5287 | 1 | 0.2um_NOKCPHJN_16452 | Bacteria | Bacteroidetes | Flavobacteriia | Flavobacterial | Flavobacteriac | Formosa | Formosa sp. H | poorly charact | Function unkn | 0XQTX | Chromosome | 1046627.BZAF | 1 | 0 | 1,26E-05 | 1,19E-06 | 1 - | 1,05E-05 - | 1 - | 1,31E-05 - | 1 - | 1,24E-05 - | 1 - | 1,26E-05 - | 1 - | 1,42E-05 - |
| 5288 | 1 | 3um_FIJJAFBF_285894 | Bacteria | Proteobacteri | Alphaproteob | Rhodobactera | Rhodobactera | Planktomarina | Planktomarina | metabolism | Amino acid tra | 05TIB | (ABC) transpo | 367336.OM22 | 2 | 1,095445115 | 8,12E-05 | 3,58E-05 | 4 - | 0,00014258 - | 2 - | 8,85E-05 - | 2 - | 8,42E-05 - | 1 - | 4,27E-05 - | 1 - | 4,81E-05 - |
| 5289 | 1 | 3um_FIJJAFBF_325287 | Bacteria | Bacteroidetes | Flavobacteriia | Flavobacterial | Flavobacteriac | Polaribacter | Polaribacter s | cellular proces | Posttranslatio | COG0330 | Band 7 proteir | 313594.PI23P | 8,4 | 1,743559577 | 0,000371455 | 4,53E-05 | 11 - | 0,000415311 - | 7 - | 0,000328168 - | 9 - | 0,000401099 - | 9 - | 0,000407023 - | 6 - | 0,000305673 - |
| 5290 | 1 | 0.2um_NOKCPHJN_852069 | Bacteria | Cyanobacteria | unclassified | Nostocales | Nostocaceae | Nostoc | Nostoc sp. AT | metabolism | Amino acid tra | COG0082 | 5-enolpyruvyl | 513049.Amax | 0,2 | 0,4 | 8,53E-06 | 1,71E-05 | 0 - | 0 - | 0 - | 0 - | 0 - | 0 - | 0 - | 0 - | 1 - | 4,27E-05 - |
| 5291 | 1 | 0.2um_NOKCPHJN_72833 | Bacteria | Proteobacteri | Gammaprotec | Oceanospirilla | Saccharospirill | Reinekea | Reinekea blan | information st | Translation, ri | COG0013 | Catalyzes the | 314283.MED2 | 4 | 0,894427191 | 6,22E-05 | 1,07E-05 | 5 - | 6,61E-05 - | 4 - | 6,57E-05 - | 5 - | 7,80E-05 - | 3 - | 4,75E-05 - | 3 - | 5,35E-05 - |
| 5292 | 1 | 0.2um_NOKCPHJN_94589 | Bacteria | Proteobacteri | Alphaproteob | Rhodobactera | Rhodobactera | Sulfitobacter | Sulfitobacter s | metabolism | Energy produc | 05C26 | Dehydrogenas | 391624.OIHEL | 0,8 | 0,4 | 2,11E-05 | 1,07E-05 | 1 - | 2,29E-05 - | 1 - | 2,84E-05 - | 1 - | 2,70E-05 - | 1 - | 2,74E-05 - | 0 - | 0 - |
| 5293 | 1 | 3um_FIJJAFBF_44458 | Bacteria | Bacteroidetes | Flavobacteriia | Flavobacterial | Flavobacteriac | Flavobacteriu | Flavobacteriu | metabolism | Coenzyme tra | 09A4R | May play a ke | 376686.Fjoh_ | 0,8 | 0,4 | 3,34E-05 | 1,72E-05 | 1 - | 3,50E-05 - | 1 - | 4,35E-05 - | 1 - | 4,13E-05 - | 0 - | 0 - | 1 - | 4,72E-05 - |
| 5294 | 1 | 3um_FIJJAFBF_122381 | Bacteria | Bacteroidetes | Flavobacteriia | Flavobacterial | Flavobacteriac | Polaribacter | Polaribacter s | poorly charact | Function unkn | 05VUE | NA | 313598.MED1 | 0,8 | 0,748331477 | 0,000116865 | 0,000103282 | 2 - | 0,000266923 - | 0 - | 0 - | 1 - | 0,000157537 - | 1 - | 0,000159864 - | 0 - | 0 - |
| 5295 | 1 | 3um_FIJJAFBF_739999 | Eukaryota | unclassified | unclassified | Phaeocystales | Phaeocystace | Phaeocystis | Phaeocystis gl | cellular proces | Posttranslatio | KOG0730 | cell division cy | 35128.Thapsd | 0,4 | 0,489897949 | 1,73E-05 | 2,13E-05 | 1 - | 3,93E-05 - | 0 - | 0 - | 0 - | 0 - | 1 - | 4,71E-05 - | 0 - | 0 - |
| 5296 | 1 | 0.2um_NOKCPHJN_800097 | Bacteria | Proteobacteri | Alphaproteob | Pelagibacteral | Pelagibacterac | Candidatus Pe | Candidatus Pe | metabolism | Amino acid tra | 01V8M | Aminomethylt | 335992.SAR11 | 1,2 | 0,4 | 3,72E-05 | 1,58E-05 | 1 - | 2,54E-05 - | 1 - | 3,15E-05 - | 1 - | 3,00E-05 - | 1 - | 3,04E-05 - | 2 - | 6,85E-05 - |
| 5297 | 1 | 3um_FIJJAFBF_119650 | Bacteria | Bacteroidetes | Flavobacteriia | Flavobacterial | Flavobacteriac | Formosa | Formosa sp. H | unclassified | unclassified | unclassified | unclassified | unclassified | 0,2 | 0,4 | 3,09E-06 | 6,17E-06 | 1 - | 1,54E-05 - | 0 - | 0 - | 0 - | 0 - | 0 - | 0 - | 0 - | 0 - |
| 5298 | 1 | 0.2um_NOKCPHJN_341333 | Bacteria | Proteobacteri | Gammaprotec | Oceanospirilla | Saccharospirill | Reinekea | Reinekea fors | unclassified | unclassified | unclassified | unclassified | unclassified | 0,6 | 0,489897949 | 4,24E-05 | 3,50E-05 | 1 - | 6,17E-05 - | 1 - | 7,66E-05 - | 0 - | 0 - | 1 - | 7,39E-05 - | 0 - | 0 - |
| 5299 | 1 | 0.2um_NOKCPHJN_15491 | Bacteria | Bacteroidetes | Flavobacteriia | Flavobacterial | Flavobacteriac | Polaribacter | Polaribacter a | information st | Translation, ri | COG0052 | 30S ribosomal | 983548.Krodi_ | 2,6 | 0,489897949 | 0,000117576 | 2,39E-05 | 3 - | 0,000114017 - | 2 - | 9,44E-05 - | 3 - | 0,000134585 - | 2 - | 9,10E-05 - | 3 - | 0,000153849 - |
| 5300 | 1 | 3um_FIJJAFBF_575597 | Bacteria | Proteobacteri | Alphaproteob | Rhodobactera | Rhodobactera | Planktotalea | Planktotalea a | metabolism | Inorganic ion t | 08UTW | Plays a role in | 391619.RGBS1 | 1,8 | 0,979795897 | 0,000104349 | 5,34E-05 | 3 - | 0,000149709 - | 0 - | 0 - | 2 - | 0,000117811 - | 2 - | 0,000119551 - | 2 - | 0,000134673 - |
| 5301 | 1 | 0.2um_NOKCPHJN_179177 | Bacteria | Proteobacteri | Gammaprotec | Alteromonada | Colwelliaceae | Colwellia | Colwellia sp. P | information st | Replication, re | 08ZUF | This protein is | 167879.CPS_2 | 0,6 | 0,489897949 | 8,22E-05 | 6,79E-05 | 1 - | 0,000119559 - | 1 - | 0,000148457 - | 0 - | 0 - | 1 - | 0,000143212 - | 0 - | 0 - |
| 5302 | 1 | 0.2um_NOKCPHJN_113026 | unclassified | unclassified | unclassified | unclassified | unclassified | unclassified | unclassified | unclassified | unclassified | unclassified | unclassified | unclassified | 0,6 | 0,489897949 | 5,00E-05 | 4,13E-05 | 1 - | 7,31E-05 - | 1 - | 9,08E-05 - | 1 - | 8,63E-05 - | 0 - | 0 - | 0 - | 0 - |
| 5303 | 1 | 3um_FIJJAFBF_202841 | Bacteria | Bacteroidetes | Flavobacteriia | Flavobacterial | Flavobacteriac | Arenitalea | Arenitalea lut | cellular proces | Posttranslatio | 09ADN | alkyl hydroper | 391587.KAOT1 | 2,8 | 0,4 | 0,000179432 | 3,21E-05 | 3 - | 0,000160902 - | 3 - | 0,000199793 - | 2 - | 0,000126619 - | 3 - | 0,000192734 - | 3 - | 0,000217114 - |
| 5304 | 1 | 3um_FIJJAFBF_374868 | Bacteria | Proteobacteri | Alphaproteob | Pelagibacteral | Pelagibacterac | Candidatus Pe | Candidatus Pe | unclassified | unclassified | unclassified | unclassified | unclassified | 1 | 0,632455532 | 0,000110634 | 7,80E-05 | 1 - | 9,04E-05 - | 1 - | 0,00011222 - | 1 - | 0,000106679 - | 0 - | 0 - | 2 - | 0,000243897 - |
| 5305 | 1 | 3um_FIJJAFBF_598844 | Bacteria | Bacteroidetes | Flavobacteriia | Flavobacterial | Flavobacteriac | Polaribacter | Polaribacter b | metabolism | Nucleotide tra | COG0138 | bifunctional p | 313598.MED1 | 1 | 0 | 2,70E-05 | 2,56E-06 | 1 - | 2,26E-05 - | 1 - | 2,81E-05 - | 1 - | 2,67E-05 - | 1 - | 2,71E-05 - | 1 - | 3,05E-05 - |
| 5306 | 1 | 3um_FIJJAFBF_123073 | Bacteria | Proteobacteri | Gammaprotec | Alteromonada | Alteromonada | Paraglaciecola | Paraglaciecola | unclassified | unclassified | unclassified | unclassified | unclassified | 5,2 | 0,4 | 0,000475088 | 5,73E-05 | 5 - | 0,000382589 - | 5 - | 0,000475063 - | 5 - | 0,000451607 - | 6 - | 0,000549933 - | 5 - | 0,000516248 - |
| 5307 | 1 | 3um_FIJJAFBF_696362 | Bacteria | Proteobacteri | Gammaprotec | Oceanospirilla | Saccharospirill | Reinekea | Reinekea blan | metabolism | Energy produc | 16RH6 | Produces ATP | 314283.MED2 | 4,6 | 0,489897949 | 0,00021845 | 2,96E-05 | 5 - | 0,000199265 - | 4 - | 0,000197943 - | 5 - | 0,000235212 - | 4 - | 0,000190949 - | 5 - | 0,000268879 - |
| 5308 | 1 | 0.2um_NOKCPHJN_372004 | Eukaryota | Phaeophyceae | unclassified | Ectocarpales | Ectocarpaceae | Ectocarpus | Ectocarpus sili | unclassified | unclassified | unclassified | unclassified | unclassified | 1 | 0 | 2,36E-05 | 2,24E-06 | 1 - | 1,98E-05 - | 1 - | 2,46E-05 - | 1 - | 2,34E-05 - | 1 - | 2,37E-05 - | 1 - | 2,67E-05 - |
| 5309 | 1 | 0.2um_NOKCPHJN_112963 | Bacteria | Proteobacteri | Alphaproteob | Rhodobactera | Rhodobactera | Thalassobius | Thalassobius s | cellular proces | Posttranslatio | 01QWG | Prevents misfo | 391619.RGBS1 | 15,6 | 1,356465997 | 0,000385348 | 2,20E-05 | 18 - | 0,000374951 - | 14 - | 0,000362117 - | 15 - | 0,000368826 - | 16 - | 0,000399226 - | 15 - | 0,000421618 - |
| 5310 | 1 | 3um_FIJJAFBF_412007 | Bacteria | Proteobacteri | Gammaprotec | Cellvibrionales | Spongiibacter | Oceanicoccus | Oceanicoccus | unclassified | unclassified | unclassified | unclassified | unclassified | 0,8 | 0,4 | 2,17E-05 | 1,12E-05 | 1 - | 2,30E-05 - | 0 - | 0 - | 1 - | 2,72E-05 - | 1 - | 2,76E-05 - | 1 - | 3,10E-05 - |
| 5311 | 1 | 0.2um_NOKCPHJN_555040 | Bacteria | Proteobacteri | Alphaproteob | Rhodobactera | Rhodobactera | Planktomarina | Planktomarina | metabolism | Energy produc | 178X6 | carbon monox | 388739.RSK20 | 1,4 | 0,489897949 | 0,000155158 | 6,38E-05 | 1 - | 9,11E-05 - | 1 - | 0,00011311 - | 1 - | 0,000107526 - | 2 - | 0,000218227 - | 2 - | 0,000245833 - |
| 5312 | 1 | 3um_FIJJAFBF_26688 | Bacteria | Bacteroidetes | Flavobacteriia | Flavobacterial | Flavobacteriac | Polaribacter | Polaribacter s | metabolism | Amino acid tra | 09GJ6 | nitrogen regul | 313598.MED1 | 2,8 | 0,4 | 0,000341587 | 5,53E-05 | 3 - | 0,000307438 - | 2 - | 0,000254498 - | 3 - | 0,000362899 - | 3 - | 0,000368259 - | 3 - | 0,000414842 - |
| 5313 | 1 | 0.2um_NOKCPHJN_206813 | Bacteria | Proteobacteri | Gammaprotec | Oceanospirilla | Saccharospirill | Reinekea | Reinekea fors | unclassified | unclassified | unclassified | unclassified | unclassified | 0,6 | 0,489897949 | 2,74E-05 | 2,25E-05 | 1 - | 4,06E-05 - | 0 - | 0 - | 1 - | 4,79E-05 - | 1 - | 4,86E-05 - | 0 - | 0 - |
| 5314 | 1 | 0.2um_NOKCPHJN_15568 | Bacteria | Bacteroidetes | Flavobacteriia | unclassified | unclassified | unclassified | Flavobacteria | unclassified | unclassified | unclassified | unclassified | unclassified | 2 | 1,673320053 | 4,00E-05 | 3,44E-05 | 1 - | 1,63E-05 - | 5 - | 0,000101221 - | 2 - | 3,85E-05 - | 0 - | 0 - | 2 - | 4,40E-05 - |
| 5315 | 1 | 3um_FIJJAFBF_361521 | Eukaryota | unclassified | unclassified | Phaeocystales | Phaeocystace | Phaeocystis | Phaeocystis a | unclassified | unclassified | unclassified | unclassified | unclassified | 2 | 0,632455532 | 8,63E-05 | 2,63E-05 | 2 - | 7,33E-05 - | 2 - | 9,11E-05 - | 3 - | 0,000129855 - | 2 - | 8,78E-05 - | 1 - | 4,95E-05 - |
| 5316 | 1 | 3um_FIJJAFBF_765335 | Eukaryota | Bacillariophyt | Bacillariophyc | Naviculales | Phaeodactylac | Phaeodactylu | Phaeodactylu | unclassified | unclassified | unclassified | unclassified | unclassified | 2,4 | 0,489897949 | 5,71E-05 | 1,35E-05 | 2 - | 3,97E-05 - | 3 - | 7,40E-05 - | 2 - | 4,69E-05 - | 3 - | 7,14E-05 - | 2 - | 5,36E-05 - |
| 5317 | 1 | 0.2um_NOKCPHJN_619380 | Bacteria | Proteobacteri | Alphaproteob | Rhodobactera | Rhodobactera | Thalassobium | Thalassobium | information st | Translation, ri | 05C22 | threonyL-tRN/ | 633131.TR2A6 | 1 | 0,632455532 | 2,18E-05 | 1,35E-05 | 0 - | 0 - | 1 - | 2,20E-05 - | 1 - | 2,09E-05 - | 2 - | 4,24E-05 - | 1 - | 2,39E-05 - |
| 5318 | 1 | 3um_FIJJAFBF_629636 | unclassified | unclassified | unclassified | unclassified | unclassified | unclassified | unclassified | unclassified | unclassified | unclassified | unclassified | unclassified | 1 | 0,632455532 | 2,69E-05 | 1,64E-05 | 0 - | 0 - | 1 - | 2,72E-05 - | 2 - | 5,17E-05 - | 1 - | 2,62E-05 - | 1 - | 2,96E-05 - |
| 5319 | 1 | 3um_FIJJAFBF_128453 | Bacteria | Bacteroidetes | Flavobacteriia | Flavobacterial | Flavobacteriac | Polaribacter | Polaribacter s | poorly charact | Function unkn | 11W7M | NA | 398720.MED2 | 1,2 | 0,4 | 0,000102553 | 2,27E-05 | 2 - | 0,00014715 - | 1 - | 9,14E-05 - | 1 - | 8,68E-05 - | 1 - | 8,81E-05 - | 1 - | 9,93E-05 - |
| 5320 | 1 | 0.2um_NOKCPHJN_380182 | Bacteria | Proteobacteri | Deltaproteoba | Myxococcales | unclassified | Minicystis | Minicystis ros | unclassified | unclassified | unclassified | unclassified | unclassified | 0,2 | 0,4 | 1,42E-06 | 2,83E-06 | 0 - | 0 - | 0 - | 0 - | 0 - | 0 - | 0 - | 0 - | 1 - | 7,08E-06 - |
| 5321 | 1 | 0.2um_NOKCPHJN_248160 | Eukaryota | Phaeophyceae | unclassified | Ectocarpales | Ectocarpaceae | Ectocarpus | Ectocarpus sili | unclassified | unclassified | unclassified | unclassified | unclassified | 1 | 0 | 1,52E-05 | 1,44E-06 | 1 - | 1,27E-05 - | 1 - | 1,58E-05 - | 1 - | 1,50E-05 - | 1 - | 1,52E-05 - | 1 - | 1,71E-05 - |
| 5322 | 1 | 3um_FIJJAFBF_539671 | Bacteria | unclassified | unclassified | unclassified | unclassified | unclassified | bacterium (Ca | cellular proces | Posttranslatio | 066HJ | ATP-depender | 445970.ALIPU | 0,2 | 0,4 | 6,53E-06 | 1,31E-05 | 0 - | 0 - | 0 - | 0 - | 0 - | 0 - | 1 - | 3,27E-05 - | 0 - | 0 - |
| 5323 | 1 | 0.2um_NOKCPHJN_343303 | Bacteria | Proteobacteri | Gammaprotec | Oceanospirilla | Saccharospirill | Reinekea | Reinekea fors | poorly charact | Function unkn | 0XTCR | NA | 1051646.VITU | 1,2 | 0,4 | 2,99E-05 | 1,27E-05 | 1 - | 2,05E-05 - | 1 - | 2,54E-05 - | 1 - | 2,42E-05 - | 1 - | 2,45E-05 - | 2 - | 5,52E-05 - |
| 5324 | 1 | 3um_FIJJAFBF_750093 | Eukaryota | Bacillariophyt | Bacillariophyc | Naviculales | Phaeodactylac | Phaeodactylu | Phaeodactylu | unclassified | unclassified | unclassified | unclassified | unclassified | 0,4 | 0,489897949 | 2,96E-05 | 3,65E-05 | 1 - | 6,79E-05 - | 0 - | 0 - | 1 - | 8,02E-05 - | 0 - | 0 - | 0 - | 0 - |
| 5325 | 1 | 3um_FIJJAFBF_234499 | Bacteria | Proteobacteri | Betaproteoba | unclassified | unclassified | unclassified | Betaproteoba | information st | Translation, ri | 0B8JH | One of the pri | 596154.Alide2 | 6,8 | 1,166190379 | 0,000413455 | 5,09E-05 | 9 - | 0,000463225 - | 6 - | 0,000383459 - | 6 - | 0,000364526 - | 6 - | 0,00036991 - | 7 - | 0,000486153 - |
| 5326 | 1 | 3um_FIJJAFBF_732530 | Bacteria | Bacteroidetes | Cytophagia | Cytophagales | Flammeovirga | unclassified | Flammeovirga | cellular proces | Cell wall/mem | 0QKXZ | Porin Gram-ne | 247633.GP214 | 2,6 | 0,489897949 | 0,00011162 | 1,62E-05 | 3 - | 0,000109311 - | 2 - | 9,05E-05 - | 3 - | 0,000129031 - | 3 - | 0,000130936 - | 2 - | 9,83E-05 - |
| 5327 | 1 | 0.2um_NOKCPHJN_790252 | Bacteria | Proteobacteri | Alphaproteob | Rhodobactera | unclassified | unclassified | Rhodobactera | unclassified | unclassified | unclassified | unclassified | unclassified | 1 | 0 | 4,20E-05 | 3,99E-06 | 1 - | 3,52E-05 - | 1 - | 4,37E-05 - | 1 - | 4,16E-05 - | 1 - | 4,22E-05 - | 1 - | 4,75E-05 - |
| 5328 | 1 | 3um_FIJJAFBF_737960 | Bacteria | Bacteroidetes | Flavobacteriia | Flavobacterial | unclassified | unclassified | Flavobacterial | poorly charact | Function unkn | 0XSVM | NA | 865938.Weev | 1 | 0 | 4,08E-05 | 3,87E-06 | 1 - | 3,42E-05 - | 1 - | 4,24E-05 - | 1 - | 4,03E-05 - | 1 - | 4,09E-05 - | 1 - | 4,61E-05 - |
| 5329 | 1 | 0.2um_NOKCPHJN_629100 | Bacteria | Proteobacteri | Alphaproteob | Rickettsiales | unclassified | unclassified | Rickettsiales b | information st | Replication, re | 0R0E7 | RecF/RecN/SN | 220341.HCM2 | 0,4 | 0,489897949 | 1,41E-05 | 1,74E-05 | 0 - | 0 - | 0 - | 0 - | 1 - | 3,30E-05 - | 0 - | 0 - | 1 - | 3,77E-05 - |
| 5330 | 1 | 3um_FIJJAFBF_103117 | Bacteria | Bacteroidetes | Flavobacteriia | Flavobacterial | Flavobacteriac | Polaribacter | Polaribacter s | information st | Transcription | 0P7W9 | Transcription | 313598.MED1 | 0,8 | 0,4 | 2,63E-05 | 1,35E-05 | 1 - | 2,75E-05 - | 1 - | 3,41E-05 - | 0 - | 0 - | 1 - | 3,29E-05 - | 1 - | 3,71E-05 - |
| 5331 | 1 | 3um_FIJJAFBF_609410 | Bacteria | unclassified | unclassified | unclassified | unclassified | unclassified | uncultured ba | unclassified | unclassified | unclassified | unclassified | unclassified | 1 | 0 | 4,69E-05 | 4,46E-06 | 1 - | 3,93E-05 - | 1 - | 4,88E-05 - | 1 - | 4,64E-05 - | 1 - | 4,71E-05 - | 1 - | 5,30E-05 - |
| 5332 | 1 | 0.2um_NOKCPHJN_134937 | Bacteria | Proteobacteri | Gammaprotec | Cellvibrionales | Halieaceae | Halioglobus | various | metabolism | Energy produc | COG0055 | Produces ATP | 247634.GPB2 | 0,8 | 0,4 | 2,19E-05 | 1,12E-05 | 1 - | 2,28E-05 - | 1 - | 2,83E-05 - | 0 - | 0 - | 1 - | 2,73E-05 - | 1 - | 3,08E-05 - |
| 5333 | 1 | 3um_FIJJAFBF_190021 | Bacteria | Proteobacteri | Gammaprotec | Oceanospirilla | Saccharospirill | Reinekea | Reinekea fors | metabolism | Amino acid tra | 0RATG | O-acetylhomo | 400668.Mmw | 2,6 | 1,019803903 | 8,16E-05 | 2,72E-05 | 4 - | 0,000108536 - | 3 - | 0,000101077 - | 2 - | 6,41E-05 - | 3 - | 9,75E-05 - | 1 - | 3,66E-05 - |
| 5334 | 1 | 0.2um_NOKCPHJN_122868 | Bacteria | Bacteroidetes | Flavobacteriia | Flavobacterial | Flavobacteriac | Arenibacter | Arenibacter s | information st | Translation, ri | 05CAV | thus facilitatin | 313603.FB217 | 5,8 | 1,939071943 | 0,000127953 | 3,71E-05 | 8 - | 0,000150774 - | 7 - | 0,000163815 - | 7 - | 0,000155727 - | 3 - | 6,77E-05 - | 4 - | 0,000101724 - |
| 5335 | 1 | 3um_FIJJAFBF_254309 | Bacteria | Proteobacteri | Gammaprotec | Cellvibrionales | Spongiibacter | Spongiibacter | Spongiibacter | information st | Translation, ri | 05G6W | One of the pri | 247634.GPB2 | 1 | 0 | 6,65E-05 | 6,32E-06 | 1 - | 5,57E-05 - | 1 - | 6,92E-05 - | 1 - | 6,58E-05 - | 1 - | 6,67E-05 - | 1 - | 7,52E-05 - |
| 5336 | 1 | 0.2um_NOKCPHJN_67575 | Bacteria | Bacteroidetes | Flavobacteriia | Flavobacterial | Flavobacteriac | unclassified | Flavobacteriac | metabolism | Coenzyme tra | 09A4R | May play a ke | 313603.FB217 | 1,8 | 0,4 | 5,46E-05 | 1,28E-05 | 2 - | 5,10E-05 - | 1 - | 3,17E-05 - | 2 - | 6,02E-05 - | 2 - | 6,11E-05 - | 2 - | 6,88E-05 - |
| 5337 | 1 | 3um_FIJJAFBF_332143 | Bacteria | Proteobacteri | Alphaproteob | Rhodobactera | Rhodobactera | Tropicibacter | Tropicibacter | metabolism | Energy produc | 16QM4 | alcohol dehyd | 439497.RR11_ | 0,2 | 0,4 | 7,00E-06 | 1,40E-05 | 1 - | 3,50E-05 - | 0 - | 0 - | 0 - | 0 - | 0 - | 0 - | 0 - | 0 - |
| 5338 | 1 | 3um_FIJJAFBF_217017 | Bacteria | Proteobacteri | Gammaprotec | Oceanospirilla | Saccharospirill | Reinekea | Reinekea blan | metabolism | Nucleotide tra | 16SJZ | purine nucleo | 314283.MED2 | 0,2 | 0,4 | 9,73E-06 | 1,95E-05 | 1 - | 4,86E-05 - | 0 - | 0 - | 0 - | 0 - | 0 - | 0 - | 0 - | 0 - |
| 5339 | 1 | 3um_FIJJAFBF_168621 | Bacteria | Proteobacteri | Gammaprotec | Cellvibrionales | Spongiibacter | various | various | information st | Translation, ri | 16QAF | thus facilitatin | 247633.GP214 | 0,2 | 0,4 | 5,11E-06 | 1,02E-05 | 0 - | 0 - | 1 - | 2,55E-05 - | 0 - | 0 - | 0 - | 0 - | 0 - | 0 - |
| 5340 | 1 | 3um_FIJJAFBF_563343 | Bacteria | Proteobacteri | Betaproteoba | Burkholderiale | Comamonada | Macromonas | Macromonas | cellular proces | Cell wall/mem | COG2913 | Part of the ou | 535289.Dtpsy | 1,2 | 0,4 | 8,94E-05 | 1,98E-05 | 2 - | 0,000128242 - | 1 - | 7,96E-05 - | 1 - | 7,57E-05 - | 1 - | 7,68E-05 - | 1 - | 8,65E-05 - |
| 5341 | 1 | 3um_FIJJAFBF_296066 | Bacteria | Proteobacteri | Alphaproteob | Rhizobiales | unclassified | unclassified | Rhizobiales ba | metabolism | Energy produc | 17CGT | F(1)F(0) ATP s | 349521.HCH_ | 7,2 | 1,6 | 0,000645002 | 0,000199973 | 5 - | 0,000367874 - | 7 - | 0,000639508 - | 7 - | 0,000607933 - | 7 - | 0,000616912 - | 10 - | 0,000992785 - |
| 5342 | 1 | 3um_FIJJAFBF_95789 | Bacteria | Proteobacteri | Alphaproteob | Rhodobactera | Rhodobactera | Pseudaestuari | Pseudaestuari | unclassified | unclassified | unclassified | unclassified | unclassified | 1,6 | 0,489897949 | 4,08E-05 | 1,48E-05 | 1 - | 2,09E-05 - | 2 - | 5,20E-05 - | 2 - | 4,94E-05 - | 1 - | 2,51E-05 - | 2 - | 5,65E-05 - |
| 5343 | 1 | 3um_FIJJAFBF_03912 | Bacteria | Bacteroidetes | Flavobacteriia | Flavobacterial | Flavobacteriac | Polaribacter | Polaribacter s | unclassified | unclassified | unclassified | unclassified | unclassified | 10 | 1,095445115 | 0,000151021 | 1,69E-05 | 11 - | 0,000139817 - | 8 - | 0,000126263 - | 10 - | 0,000150036 - | 11 - | 0,000167477 - | 10 - | 0,000171511 - |
| 5344 | 1 | 3um_FIJJAFBF_386348 | Bacteria | Proteobacteri | Alphaproteob | Rhodobactera | Rhodobactera | Planktomarina | Planktomarina | unclassified | unclassified | unclassified | unclassified | unclassified | 13,4 | 1,743559577 | 0,001297981 | 9,75E-05 | 16 - | 0,001311734 - | 14 - | 0,001425189 - | 12 - | 0,001161276 - | 14 - | 0,001374833 - | 11 - | 0,001216871 - |
| 5345 | 1 | 3um_FIJJAFBF_607687 | Bacteria | Proteobacteri | Gammaprotec | Alteromonada | Alteromonada | Paraglaciecola | Paraglaciecola | metabolism | Inorganic ion t | 16RE3 | may be the G1 | 983545.Glaag | 0,8 | 0,748331477 | 2,37E-05 | 2,28E-05 | 1 - | 2,45E-05 - | 2 - | 6,09E-05 - | 0 - | 0 - | 0 - | 0 - | 1 - | 3,31E-05 - |
| 5346 | 1 | 3um_FIJJAFBF_399864 | Bacteria | Proteobacteri | Gammaprotec | Oceanospirilla | Halomonadac | Halomonas | Halomonas sp | unclassified | unclassified | unclassified | unclassified | unclassified | 2,6 | 0,8 | 0,000451249 | 9,69E-05 | 4 - | 0,000596243 - | 2 - | 0,000370179 - | 2 - | 0,000351902 - | 3 - | 0,000535649 - | 2 - | 0,000402271 - |
| 5347 | 1 | 0.2um_NOKCPHJN_397510 | Bacteria | Bacteroidetes | unclassified | unclassified | unclassified | unclassified | Bacteroidetes | information st | Translation, ri | 09AI8 | Associates wit | 926562.Oweh | 0,8 | 0,4 | 3,99E-05 | 2,05E-05 | 1 - | 4,22E-05 - | 0 - | 0 - | 1 - | 4,98E-05 - | 1 - | 5,05E-05 - | 1 - | 5,69E-05 - |
| 5348 | 1 | 3um_FIJJAFBF_35814 | Bacteria | Bacteroidetes | Flavobacteriia | Flavobacterial | Flavobacteriac | Hanstruepera | Hanstruepera | information st | Translation, ri | 08YY1 | This protein is | 391603.FBALC | 1 | 0 | 0,000118127 | 1,12E-05 | 1 - | 9,89E-05 - | 1 - | 0,000122861 - | 1 - | 0,000116795 - | 1 - | 0,00011852 - | 1 - | 0,000133512 - |
| 5349 | 1 | 0.2um_NOKCPHJN_962528 | Bacteria | Proteobacteri | Gammaprotec | unclassified | unclassified | unclassified | Gammaprotec | unclassified | unclassified | unclassified | unclassified | unclassified | 2,6 | 0,489897949 | 0,000382177 | 5,53E-05 | 3 - | 0,000374272 - | 2 - | 0,000309824 - | 3 - | 0,00044179 - | 3 - | 0,000448315 - | 2 - | 0,000336684 - |
| 5350 | 1 | 0.2um_NOKCPHJN_97696 | Bacteria | Proteobacteri | Betaproteoba | unclassified | unclassified | unclassified | Betaproteoba | information st | Translation, ri | 05CEJ | Catalyzes the | 887062.HGR_ | 4 | 0,632455532 | 7,89E-05 | 1,80E-05 | 4 - | 6,56E-05 - | 4 - | 8,14E-05 - | 3 - | 5,81E-05 - | 4 - | 7,86E-05 - | 5 - | 0,000110625 - |
| 5351 | 1 | 3um_FIJJAFBF_594165 | Eukaryota | unclassified | Pelagophycea | Pelagomonad | unclassified | Aureococcus | Aureococcus a | unclassified | unclassified | unclassified | unclassified | unclassified | 11,8 | 0,748331477 | 0,001037292 | 4,68E-05 | 13 - | 0,000962644 - | 11 - | 0,001011424 - | 12 - | 0,001048895 - | 12 - | 0,001064387 - | 11 - | 0,001099109 - |
| 5352 | 1 | 0.2um_NOKCPHJN_315401 | Bacteria | Bacteroidetes | Flavobacteriia | Flavobacterial | Flavobacteriac | Lutibacter | Lutibacter pro | poorly charact | Function unkn | 0P88U | NA | 313596.RB250 | 1 | 0,632455532 | 2,51E-05 | 1,54E-05 | 0 - | 0 - | 1 - | 2,52E-05 - | 1 - | 2,40E-05 - | 2 - | 4,87E-05 - | 1 - | 2,74E-05 - |
| 5353 | 1 | 0.2um_NOKCPHJN_384403 | unclassified | unclassified | unclassified | unclassified | unclassified | unclassified | unclassified | unclassified | unclassified | unclassified | unclassified | unclassified | 0,6 | 0,489897949 | 6,26E-06 | 5,13E-06 | 0 - | 0 - | 1 - | 1,03E-05 - | 1 - | 9,79E-06 - | 0 - | 0 - | 1 - | 1,12E-05 - |
| 5354 | 1 | 0.2um_NOKCPHJN_555993 | Bacteria | Proteobacteri | Alphaproteob | Pelagibacteral | Pelagibacterac | Candidatus Pe | Candidatus Pe | metabolism | Amino acid tra | COG0436 | aminotransfer | 335992.SAR11 | 0,4 | 0,489897949 | 1,29E-05 | 1,59E-05 | 1 - | 2,97E-05 - | 0 - | 0 - | 1 - | 3,50E-05 - | 0 - | 0 - | 0 - | 0 - |
| 5355 | 1 | 0.2um_NOKCPHJN_262674 | Bacteria | Bacteroidetes | unclassified | unclassified | unclassified | unclassified | unidentified e | unclassified | unclassified | unclassified | unclassified | unclassified | 8,6 | 0,489897949 | 0,000675387 | 5,63E-05 | 9 - | 0,000593673 - | 9 - | 0,000737167 - | 8 - | 0,000622907 - | 9 - | 0,000711121 - | 8 - | 0,000712067 - |
| 5356 | 1 | 3um_FIJJAFBF_642401 | unclassified | unclassified | unclassified | unclassified | unclassified | unclassified | unclassified | unclassified | unclassified | unclassified | unclassified | unclassified | 0,2 | 0,4 | 9,03E-06 | 1,81E-05 | 0 - | 0 - | 0 - | 0 - | 1 - | 4,52E-05 - | 0 - | 0 - | 0 - | 0 - |
| 5357 | 1 | 0.2um_NOKCPHJN_274013 | Eukaryota | Bacillariophyt | Bacillariophyc | Naviculales | Naviculaceae | Fistulifera | Fistulifera sola | unclassified | unclassified | unclassified | unclassified | unclassified | 0,2 | 0,4 | 1,30E-06 | 2,59E-06 | 1 - | 6,48E-06 - | 0 - | 0 - | 0 - | 0 - | 0 - | 0 - | 0 - | 0 - |
| 5358 | 2 | 0.2um_NOKCPHJN_295829;3um_FIJJAFBF_77215 | Eukaryota | Bacillariophyt | Bacillariophyc | Naviculales | Naviculaceae | Fistulifera | Fistulifera sola | various | various | various | various | various | 0,6 | 0,489897949 | 1,82E-05 | 1,50E-05 | 1 - | 2,66E-05 - | 1 - | 3,30E-05 - | 1 - | 3,14E-05 - | 0 - | 0 - | 0 - | 0 - |
| 5359 | 1 | 3um_FIJJAFBF_20738 | Bacteria | Proteobacteri | Gammaprotec | Oceanospirilla | Saccharospirill | Reinekea | Reinekea fors | unclassified | unclassified | unclassified | unclassified | unclassified | 0,6 | 0,489897949 | 0,000111507 | 9,13E-05 | 0 - | 0 - | 1 - | 0,000182716 - | 0 - | 0 - | 1 - | 0,000176261 - | 1 - | 0,000198557 - |
| 5360 | 1 | 3um_FIJJAFBF_82367 | Bacteria | Bacteroidetes | Flavobacteriia | Flavobacterial | Flavobacteriac | Polaribacter | Polaribacter ir | metabolism | Carbohydrate | 099YR | Catalyzes the | 313598.MED1 | 1,6 | 0,489897949 | 5,22E-05 | 1,91E-05 | 1 - | 2,68E-05 - | 2 - | 6,64E-05 - | 1 - | 3,16E-05 - | 2 - | 6,41E-05 - | 2 - | 7,22E-05 - |
| 5361 | 1 | 3um_FIJJAFBF_557699 | Bacteria | Proteobacteri | Gammaprotec | Chromatiales | Ectothiorhodo | Thioalkalivibri | Thioalkalivibri | cellular proces | Cell wall/mem | 12RRB | ankyrin 3, nod | 7668.SPU_017 | 0,2 | 0,4 | 6,04E-06 | 1,21E-05 | 0 - | 0 - | 0 - | 0 - | 0 - | 0 - | 0 - | 0 - | 1 - | 3,02E-05 - |
| 5362 | 1 | 3um_FIJJAFBF_123597 | Bacteria | Proteobacteri | Gammaprotec | Oceanospirilla | Saccharospirill | Reinekea | Reinekea fors | metabolism | Inorganic ion t | COG0614 | Periplasmic bi | 314283.MED2 | 0,8 | 0,4 | 3,43E-05 | 1,74E-05 | 1 - | 3,71E-05 - | 1 - | 4,61E-05 - | 1 - | 4,38E-05 - | 1 - | 4,45E-05 - | 0 - | 0 - |
| 5363 | 1 | 0.2um_NOKCPHJN_24031 | Viruses | unclassified | unclassified | unclassified | Phycodnavirid | unclassified | Chrysochrom | unclassified | unclassified | unclassified | unclassified | unclassified | 6,4 | 1,019803903 | 0,000300556 | 2,82E-05 | 8 - | 0,000318824 - | 7 - | 0,0003464 - | 6 - | 0,000282255 - | 6 - | 0,000286424 - | 5 - | 0,000268879 - |
| 5364 | 1 | 3um_FIJJAFBF_17221 | Bacteria | Proteobacteri | Alphaproteob | Rhodobactera | unclassified | unclassified | Rhodobactera | unclassified | unclassified | unclassified | unclassified | unclassified | 1,8 | 0,748331477 | 4,87E-05 | 2,23E-05 | 1 - | 2,22E-05 - | 3 - | 8,25E-05 - | 2 - | 5,23E-05 - | 1 - | 2,65E-05 - | 2 - | 5,98E-05 - |
| 5365 | 1 | 0.2um_NOKCPHJN_987128 | unclassified | unclassified | unclassified | unclassified | unclassified | unclassified | unclassified | unclassified | unclassified | unclassified | unclassified | unclassified | 1 | 0 | 8,51E-05 | 8,08E-06 | 1 - | 7,13E-05 - | 1 - | 8,85E-05 - | 1 - | 8,42E-05 - | 1 - | 8,54E-05 - | 1 - | 9,62E-05 - |
| 5366 | 1 | 3um_FIJJAFBF_810298 | Bacteria | Bacteroidetes | unclassified | unclassified | unclassified | unclassified | Bacteroidetes | cellular proces | Cell wall/mem | 09B44 | efflux transpo | 391598.FBBAL | 1,4 | 1,019803903 | 4,85E-05 | 3,14E-05 | 3 - | 9,11E-05 - | 1 - | 3,77E-05 - | 0 - | 0 - | 2 - | 7,27E-05 - | 1 - | 4,10E-05 - |
| 5367 | 1 | 3um_FIJJAFBF_337516 | Bacteria | Bacteroidetes | Flavobacteriia | Flavobacterial | Flavobacteriac | Polaribacter | various | information st | Translation, ri | 08YY1 | This protein is | 313598.MED1 | 1 | 1,095445115 | 0,000110642 | 0,000109384 | 3 - | 0,000296836 - | 1 - | 0,000122861 - | 0 - | 0 - | 0 - | 0 - | 1 - | 0,000133512 - |
| 5368 | 1 | 3um_FIJJAFBF_426980 | Bacteria | Bacteroidetes | Flavobacteriia | Flavobacterial | Flavobacteriac | Mesoflavibact | Mesoflavibact | unclassified | unclassified | unclassified | unclassified | unclassified | 2,4 | 0,489897949 | 7,06E-05 | 1,63E-05 | 2 - | 4,92E-05 - | 3 - | 9,16E-05 - | 3 - | 8,70E-05 - | 2 - | 5,89E-05 - | 2 - | 6,63E-05 - |
| 5369 | 1 | 3um_FIJJAFBF_221205 | Bacteria | Proteobacteri | Gammaprotec | Oceanospirilla | Saccharospirill | Reinekea | Reinekea fors | information st | Translation, ri | 179RU | 30S ribosomal | 234831.PSM_ | 3,4 | 1,019803903 | 0,000348543 | 8,00E-05 | 5 - | 0,000438079 - | 3 - | 0,000326379 - | 4 - | 0,000413686 - | 2 - | 0,000209898 - | 3 - | 0,000354674 - |
| 5370 | 1 | 0.2um_NOKCPHJN_299473 | unclassified | unclassified | unclassified | unclassified | unclassified | unclassified | unclassified | unclassified | unclassified | unclassified | unclassified | unclassified | 0,2 | 0,4 | 3,39E-05 | 6,79E-05 | 0 - | 0 - | 0 - | 0 - | 0 - | 0 - | 1 - | 0,000169732 - | 0 - | 0 - |
| 5371 | 1 | 0.2um_NOKCPHJN_128823 | Bacteria | Bacteroidetes | Flavobacteriia | unclassified | unclassified | unclassified | Flavobacteria | unclassified | unclassified | unclassified | unclassified | unclassified | 2,4 | 0,8 | 3,36E-05 | 1,03E-05 | 3 - | 3,60E-05 - | 3 - | 4,47E-05 - | 3 - | 4,25E-05 - | 2 - | 2,87E-05 - | 1 - | 1,62E-05 - |
| 5372 | 1 | 3um_FIJJAFBF_189826 | Bacteria | Bacteroidetes | Flavobacteriia | Flavobacterial | Flavobacteriac | Aequorivita | various | information st | Translation, ri | COG0186 | One of the pri | 50743.SCB49_ | 1,2 | 0,748331477 | 0,000195025 | 0,000119023 | 0 - | 0 - | 2 - | 0,00032763 - | 2 - | 0,000311453 - | 1 - | 0,000158027 - | 1 - | 0,000178017 - |
| 5373 | 1 | 3um_FIJJAFBF_564131 | Bacteria | Bacteroidetes | Flavobacteriia | Flavobacterial | Flavobacteriac | Mesoflavibact | various | metabolism | Carbohydrate | COG0057 | glyceraldehyd | 391587.KAOT1 | 1 | 0 | 4,14E-05 | 3,93E-06 | 1 - | 3,47E-05 - | 1 - | 4,31E-05 - | 1 - | 4,09E-05 - | 1 - | 4,15E-05 - | 1 - | 4,68E-05 - |
| 5374 | 1 | 3um_FIJJAFBF_264508 | Bacteria | Proteobacteri | Alphaproteob | Rhodospirillal | Rhodospirillac | Thalassospira | Thalassospira | metabolism | Energy produc | 0QIAU | Electron trans | 523791.Kkor_ | 0,8 | 0,748331477 | 4,04E-05 | 3,57E-05 | 2 - | 9,22E-05 - | 0 - | 0 - | 1 - | 5,44E-05 - | 1 - | 5,52E-05 - | 0 - | 0 - |
| 5375 | 2 | 3um_FIJJAFBF_275863;3um_FIJJAFBF_424984 | Bacteria | Bacteroidetes | Flavobacteriia | Flavobacterial | Flavobacteriac | Formosa | Formosa sp. H | unclassified | unclassified | unclassified | unclassified | unclassified | 1 | 0 | 5,64E-05 | 5,35E-06 | 1 - | 4,72E-05 - | 1 - | 5,86E-05 - | 1 - | 5,58E-05 - | 1 - | 5,66E-05 - | 1 - | 6,37E-05 - |
| 5376 | 1 | 0.2um_NOKCPHJN_86820 | Bacteria | Bacteroidetes | unclassified | unclassified | unclassified | unclassified | Bacteroidetes | metabolism | Lipid transpor | 0PA4W | Biotin carboxy | 926562.Oweh | 0,4 | 0,489897949 | 3,39E-05 | 4,15E-05 | 0 - | 0 - | 0 - | 0 - | 1 - | 8,42E-05 - | 1 - | 8,54E-05 - | 0 - | 0 - |
| 5377 | 1 | 0.2um_NOKCPHJN_102627 | Viruses | unclassified | unclassified | unclassified | unclassified | unclassified | uncultured vir | unclassified | unclassified | unclassified | unclassified | unclassified | 1 | 0 | 2,69E-05 | 2,56E-06 | 1 - | 2,25E-05 - | 1 - | 2,80E-05 - | 1 - | 2,66E-05 - | 1 - | 2,70E-05 - | 1 - | 3,04E-05 - |
| 5378 | 1 | 0.2um_NOKCPHJN_247074 | Bacteria | Bacteroidetes | Flavobacteriia | Flavobacterial | Flavobacteriac | Polaribacter | Polaribacter fi | unclassified | unclassified | unclassified | unclassified | unclassified | 11,8 | 1,469693846 | 0,00018835 | 1,56E-05 | 14 - | 0,000188822 - | 11 - | 0,000184219 - | 13 - | 0,000206965 - | 10 - | 0,000161555 - | 11 - | 0,00020019 - |
| 5379 | 1 | 3um_FIJJAFBF_162847 | Bacteria | Bacteroidetes | unclassified | unclassified | unclassified | unclassified | Bacteroidetes | unclassified | unclassified | unclassified | unclassified | unclassified | 4,4 | 1,854723699 | 0,000312461 | 0,000127882 | 6 - | 0,000362453 - | 1 - | 7,50E-05 - | 6 - | 0,000427839 - | 4 - | 0,000289439 - | 5 - | 0,000407564 - |
| 5380 | 1 | 3um_FIJJAFBF_457028 | Bacteria | Bacteroidetes | Flavobacteriia | Flavobacterial | Flavobacteriac | Aureitalea | Aureitalea sp. | unclassified | unclassified | unclassified | unclassified | unclassified | 1 | 0 | 7,70E-05 | 7,31E-06 | 1 - | 6,45E-05 - | 1 - | 8,01E-05 - | 1 - | 7,61E-05 - | 1 - | 7,72E-05 - | 1 - | 8,70E-05 - |
| 5381 | 1 | 3um_FIJJAFBF_177926 | Eukaryota | Bacillariophyt | Coscinodiscop | Coscinodiscale | Coscinodiscac | Coscinodiscus | Coscinodiscus | unclassified | unclassified | unclassified | unclassified | unclassified | 5,2 | 0,748331477 | 0,000225955 | 3,32E-05 | 6 - | 0,000219319 - | 6 - | 0,000272329 - | 4 - | 0,000172589 - | 5 - | 0,000218922 - | 5 - | 0,000246615 - |
| 5382 | 3 | 0.2um_NOKCPHJN_113413;0.2um_NOKCPHJN_1 | Bacteria | Proteobacteri | Gammaprotec | Cellvibrionales | various | various | various | unclassified | unclassified | unclassified | unclassified | unclassified | 0,2 | 0,4 | 1,16E-05 | 2,32E-05 | 0 - | 0 - | 0 - | 0 - | 1 - | 5,79E-05 - | 0 - | 0 - | 0 - | 0 - |
| 5383 | 1 | 3um_FIJJAFBF_788018 | Bacteria | Proteobacteri | Alphaproteob | Rhodobactera | unclassified | unclassified | Rhodobactera | information st | Translation, ri | 08UHH | Located on the | 314262.MED1 | 1,6 | 0,489897949 | 0,000169034 | 5,32E-05 | 2 - | 0,000177948 - | 1 - | 0,00011048 - | 2 - | 0,00021005 - | 1 - | 0,000106576 - | 2 - | 0,000240115 - |
| 5384 | 1 | 3um_FIJJAFBF_105978 | Bacteria | Proteobacteri | Alphaproteob | Pelagibacteral | Pelagibacterac | unclassified | Pelagibacterac | unclassified | unclassified | unclassified | unclassified | unclassified | 3,2 | 0,979795897 | 0,000108376 | 3,58E-05 | 4 - | 0,00011336 - | 4 - | 0,000140759 - | 2 - | 6,69E-05 - | 2 - | 6,79E-05 - | 4 - | 0,000152962 - |
| 5385 | 1 | 0.2um_NOKCPHJN_11896 | Bacteria | Actinobacteria | Actinobacteria | Micrococcales | Microbacteria | unclassified | Microbacteria | unclassified | unclassified | unclassified | unclassified | unclassified | 0,2 | 0,4 | 6,89E-06 | 1,38E-05 | 0 - | 0 - | 0 - | 0 - | 1 - | 3,45E-05 - | 0 - | 0 - | 0 - | 0 - |
| 5386 | 1 | 0.2um_NOKCPHJN_661601 | Bacteria | Proteobacteri | Alphaproteob | Pelagibacteral | Pelagibacterac | Candidatus Pe | Candidatus Pe | metabolism | Amino acid tra | 01R4A | Polyamine AB | 335992.SAR11 | 2,8 | 0,4 | 0,000101905 | 1,21E-05 | 3 - | 9,23E-05 - | 3 - | 0,000114626 - | 3 - | 0,000108967 - | 3 - | 0,000110576 - | 2 - | 8,30E-05 - |
| 5387 | 1 | 0.2um_NOKCPHJN_1089082 | Bacteria | Proteobacteri | Alphaproteob | Rhodobactera | Rhodobactera | Planktomarina | Planktomarina | unclassified | unclassified | unclassified | unclassified | unclassified | 0,2 | 0,4 | 1,89E-05 | 3,78E-05 | 0 - | 0 - | 0 - | 0 - | 0 - | 0 - | 0 - | 0 - | 1 - | 9,44E-05 - |
| 5388 | 1 | 0.2um_NOKCPHJN_390598 | Bacteria | Proteobacteri | Alphaproteob | Rhodobactera | Rhodobactera | Sulfitobacter | Sulfitobacter s | metabolism | Energy produc | 01RNF | Produces ATP | 391595.RLO14 | 7,4 | 1,019803903 | 0,000210868 | 1,29E-05 | 9 - | 0,000217472 - | 7 - | 0,000210028 - | 7 - | 0,000199658 - | 8 - | 0,000231551 - | 6 - | 0,000195631 - |
| 5389 | 1 | 0.2um_NOKCPHJN_187364 | Bacteria | Bacteroidetes | Flavobacteriia | Flavobacterial | Crocinitomica | unclassified | Crocinitomica | information st | Translation, ri | COG0617 | Catalyzes the | 755732.Fluta_ | 0,2 | 0,4 | 6,19E-06 | 1,24E-05 | 0 - | 0 - | 0 - | 0 - | 0 - | 0 - | 0 - | 0 - | 1 - | 3,10E-05 - |
| 5390 | 1 | 3um_FIJJAFBF_263680 | Bacteria | Proteobacteri | Alphaproteob | unclassified | unclassified | Candidatus Pu | Candidatus Pu | cellular proces | Posttranslatio | 05C2Z | ATP-depender | 313624.N9414 | 3,6 | 0,489897949 | 5,70E-05 | 7,09E-06 | 4 - | 5,34E-05 - | 4 - | 6,63E-05 - | 3 - | 4,73E-05 - | 4 - | 6,39E-05 - | 3 - | 5,40E-05 - |
| 5391 | 1 | 3um_FIJJAFBF_203100 | Bacteria | Bacteroidetes | Flavobacteriia | Flavobacterial | Crocinitomica | unclassified | Crocinitomica | cellular proces | Intracellular tr | 05FGH | MotA TolQ exl | 655815.ZPR_4 | 3,2 | 0,4 | 0,00014704 | 2,18E-05 | 3 - | 0,000115547 - | 3 - | 0,000143475 - | 4 - | 0,000181855 - | 3 - | 0,000138406 - | 3 - | 0,000155914 - |
| 5392 | 1 | 0.2um_NOKCPHJN_150740 | Bacteria | Proteobacteri | Alphaproteob | Rhodospirillal | Rhodospirillac | unclassified | Rhodospirillac | unclassified | unclassified | unclassified | unclassified | unclassified | 0,8 | 0,979795897 | 3,76E-05 | 4,62E-05 | 0 - | 0 - | 0 - | 0 - | 2 - | 8,77E-05 - | 0 - | 0 - | 2 - | 0,000100242 - |
| 5393 | 1 | 3um_FIJJAFBF_482466 | Bacteria | Bacteroidetes | Flavobacteriia | Flavobacterial | Flavobacteriac | unclassified | Flavobacteriac | unclassified | unclassified | unclassified | unclassified | unclassified | 1 | 0 | 7,83E-05 | 7,43E-06 | 1 - | 6,56E-05 - | 1 - | 8,14E-05 - | 1 - | 7,74E-05 - | 1 - | 7,86E-05 - | 1 - | 8,85E-05 - |
| 5394 | 1 | 0.2um_NOKCPHJN_770496 | Eukaryota | Bacillariophyt | Coscinodiscop | Thalassiosirale | Thalassiosirac | Thalassiosira | Thalassiosira c | unclassified | unclassified | unclassified | unclassified | unclassified | 1 | 0 | 6,40E-05 | 6,08E-06 | 1 - | 5,36E-05 - | 1 - | 6,66E-05 - | 1 - | 6,33E-05 - | 1 - | 6,42E-05 - | 1 - | 7,24E-05 - |
| 5395 | 1 | 0.2um_NOKCPHJN_608366 | Bacteria | Bacteroidetes | unclassified | Bacteroidetes | Rhodotherma | Rhodothermu | Rhodothermu | unclassified | unclassified | unclassified | unclassified | unclassified | 6 | 1,095445115 | 0,000111866 | 2,31E-05 | 7 - | 0,00010946 - | 6 - | 0,0001165 - | 6 - | 0,000110748 - | 4 - | 7,49E-05 - | 7 - | 0,0001477 - |
| 5396 | 1 | 3um_FIJJAFBF_126406 | Eukaryota | Chordata | Mammalia | Rodentia | Cricetidae | Cricetulus | Cricetulus gris | unclassified | unclassified | unclassified | unclassified | unclassified | 1 | 0 | 1,33E-05 | 1,26E-06 | 1 - | 1,12E-05 - | 1 - | 1,39E-05 - | 1 - | 1,32E-05 - | 1 - | 1,34E-05 - | 1 - | 1,51E-05 - |
| 5397 | 1 | 0.2um_NOKCPHJN_431814 | Bacteria | Bacteroidetes | Flavobacteriia | Flavobacterial | Flavobacteriac | Aquimarina | Aquimarina sp | information st | Translation, ri | 05CEJ | Catalyzes the | 313594.PI23P | 2,8 | 0,4 | 5,37E-05 | 6,39E-06 | 3 - | 4,86E-05 - | 3 - | 6,04E-05 - | 3 - | 5,74E-05 - | 3 - | 5,83E-05 - | 2 - | 4,37E-05 - |
| 5398 | 1 | 3um_FIJJAFBF_206344 | Bacteria | Bacteroidetes | Flavobacteriia | Flavobacterial | Flavobacteriac | Polaribacter | various | poorly charact | Function unkn | 09E2D | NA | 156586.BBFL7 | 0,4 | 0,8 | 2,00E-05 | 4,01E-05 | 2 - | 0,000100242 - | 0 - | 0 - | 0 - | 0 - | 0 - | 0 - | 0 - | 0 - |
| 5399 | 1 | 3um_FIJJAFBF_452765 | Bacteria | Bacteroidetes | Flavobacteriia | Flavobacterial | Flavobacteriac | Polaribacter | Polaribacter fi | metabolism | Energy produc | 05ERI | cytochrome b | 313594.PI23P | 1 | 0 | 6,09E-05 | 5,78E-06 | 1 - | 5,10E-05 - | 1 - | 6,33E-05 - | 1 - | 6,02E-05 - | 1 - | 6,11E-05 - | 1 - | 6,88E-05 - |
| 5400 | 1 | 0.2um_NOKCPHJN_955301 | Bacteria | Proteobacteri | Alphaproteob | Pelagibacteral | Pelagibacterac | Candidatus Pe | Candidatus Pe | unclassified | unclassified | unclassified | unclassified | unclassified | 21,4 | 1,496662955 | 0,000844338 | 7,13E-05 | 24 - | 0,000796139 - | 20 - | 0,000823808 - | 21 - | 0,000822291 - | 20 - | 0,000794701 - | 22 - | 0,000984751 - |
| 5401 | 1 | 3um_FIJJAFBF_325788 | Bacteria | Proteobacteri | Gammaprotec | Cellvibrionales | Porticoccacea | unclassified | SAR92 bacteri | cellular proces | Posttranslatio | COG0330 | Band 7 proteir | 314287.GB220 | 5 | 0,632455532 | 0,000235868 | 2,98E-05 | 5 - | 0,000198576 - | 5 - | 0,000246572 - | 5 - | 0,000234398 - | 6 - | 0,000285432 - | 4 - | 0,000214359 - |
| 5402 | 1 | 0.2um_NOKCPHJN_375561 | Bacteria | Proteobacteri | Alphaproteob | Pelagibacteral | Pelagibacterac | Candidatus Pe | Candidatus Pe | poorly charact | Function unkn | 01SDU | solute sympor | 335992.SAR11 | 4 | 0 | 9,10E-05 | 8,64E-06 | 4 - | 7,63E-05 - | 4 - | 9,47E-05 - | 4 - | 9,00E-05 - | 4 - | 9,14E-05 - | 4 - | 0,000102907 - |
| 5403 | 1 | 0.2um_NOKCPHJN_1104359 | Bacteria | Proteobacteri | Alphaproteob | unclassified | unclassified | Candidatus Pu | Candidatus Pu | unclassified | unclassified | unclassified | unclassified | unclassified | 0,2 | 0,4 | 1,03E-05 | 2,07E-05 | 1 - | 5,17E-05 - | 0 - | 0 - | 0 - | 0 - | 0 - | 0 - | 0 - | 0 - |
| 5404 | 1 | 3um_FIJJAFBF_115305 | Bacteria | Proteobacteri | Gammaprotec | Arenicellales | Arenicellaceae | Arenicella | Arenicella xan | unclassified | unclassified | unclassified | unclassified | unclassified | 1 | 0,632455532 | 9,23E-05 | 5,86E-05 | 1 - | 7,76E-05 - | 1 - | 9,63E-05 - | 2 - | 0,000183084 - | 0 - | 0 - | 1 - | 0,000104645 - |
| 5405 | 1 | 0.2um_NOKCPHJN_314894 | Bacteria | Proteobacteri | Alphaproteob | Rhodobactera | Rhodobactera | Octadecabact | Octadecabact | cellular proces | Cell motility | 01YGF | Flagellin | 391616.OA238 | 1,8 | 1,326649916 | 8,68E-05 | 6,41E-05 | 2 - | 8,14E-05 - | 1 - | 5,05E-05 - | 4 - | 0,000192173 - | 0 - | 0 - | 2 - | 0,00010984 - |
| 5406 | 1 | 3um_FIJJAFBF_258166 | Bacteria | Proteobacteri | Alphaproteob | Rhodobactera | Rhodobactera | Roseovarius | Roseovarius a | unclassified | unclassified | unclassified | unclassified | unclassified | 3,2 | 0,748331477 | 0,000120472 | 2,93E-05 | 3 - | 9,51E-05 - | 4 - | 0,000157479 - | 3 - | 0,000112278 - | 4 - | 0,000151915 - | 2 - | 8,56E-05 - |
| 5407 | 1 | 0.2um_NOKCPHJN_745669 | Bacteria | Bacteroidetes | unclassified | unclassified | unclassified | unclassified | Bacteroidetes | poorly charact | Function unkn | 0P9X5 | Inherit from b | 926562.Oweh | 1 | 0 | 1,66E-05 | 1,58E-06 | 1 - | 1,39E-05 - | 1 - | 1,73E-05 - | 1 - | 1,64E-05 - | 1 - | 1,66E-05 - | 1 - | 1,87E-05 - |
| 5408 | 1 | 3um_FIJJAFBF_216064 | Bacteria | Proteobacteri | Gammaprotec | Cellvibrionales | Microbulbifera | Microbulbifer | Microbulbifer | information st | Translation, ri | 05C64 | Binds directly | 314277.MED1 | 0,8 | 0,748331477 | 4,68E-05 | 4,42E-05 | 1 - | 4,97E-05 - | 0 - | 0 - | 2 - | 0,000117301 - | 0 - | 0 - | 1 - | 6,70E-05 - |
| 5409 | 1 | 3um_FIJJAFBF_164692 | Bacteria | Proteobacteri | Alphaproteob | Rhodobactera | Hyphomonada | Robiginitomac | Robiginitomac | poorly charact | Function unkn | 05WA2 | Putative phosp | 511062.GU3_ | 1 | 0 | 0,000106223 | 1,01E-05 | 1 - | 8,90E-05 - | 1 - | 0,00011048 - | 1 - | 0,000105025 - | 1 - | 0,000106576 - | 1 - | 0,000120058 - |
| 5410 | 1 | 0.2um_NOKCPHJN_613244 | Bacteria | Proteobacteri | Gammaprotec | unclassified | unclassified | unclassified | Gammaprotec | information st | Translation, ri | 16Q16 | Binds the lowe | 208964.PA425 | 2,6 | 1,019803903 | 0,000156599 | 5,72E-05 | 4 - | 0,000204048 - | 3 - | 0,000190025 - | 1 - | 6,02E-05 - | 2 - | 0,000122207 - | 3 - | 0,000206499 - |
| 5411 | 1 | 3um_FIJJAFBF_13649 | Bacteria | Cyanobacteria | unclassified | Synechococca | Prochloraceae | Prochlorococc | Prochlorococc | unclassified | unclassified | unclassified | unclassified | unclassified | 0,4 | 0,489897949 | 2,52E-05 | 3,10E-05 | 1 - | 5,77E-05 - | 0 - | 0 - | 1 - | 6,81E-05 - | 0 - | 0 - | 0 - | 0 - |
| 5412 | 1 | 3um_FIJJAFBF_494842 | Bacteria | Proteobacteri | Gammaprotec | Cellvibrionales | Porticoccacea | unclassified | gamma protec | metabolism | Inorganic ion t | 05PWN | phosphate abc | 314287.GB220 | 2,2 | 0,4 | 8,56E-05 | 1,67E-05 | 2 - | 6,52E-05 - | 2 - | 8,10E-05 - | 3 - | 0,000115468 - | 2 - | 7,81E-05 - | 2 - | 8,80E-05 - |
| 5413 | 1 | 3um_FIJJAFBF_242901 | Bacteria | Bacteroidetes | Flavobacteriia | Flavobacterial | Flavobacteriac | Polaribacter | Polaribacter s | metabolism | Inorganic ion t | 0ZVMK | efflux transpo | 313598.MED1 | 0,4 | 0,489897949 | 1,47E-05 | 1,80E-05 | 0 - | 0 - | 0 - | 0 - | 1 - | 3,65E-05 - | 1 - | 3,71E-05 - | 0 - | 0 - |
| 5414 | 1 | 3um_FIJJAFBF_335154 | Bacteria | Actinobacteria | Actinobacteria | Micrococcales | Microbacteria | Candidatus Ac | Candidatus Ac | metabolism | Amino acid tra | COG0687 | polyamine bin | 312284.A20C1 | 4,2 | 0,748331477 | 0,000137695 | 2,09E-05 | 5 - | 0,000138619 - | 3 - | 0,000103275 - | 4 - | 0,000130901 - | 5 - | 0,000166043 - | 4 - | 0,000149637 - |
| 5415 | 1 | 3um_FIJJAFBF_178961 | Bacteria | Proteobacteri | Alphaproteob | Rhodobactera | unclassified | unclassified | Rhodobactera | information st | Translation, ri | 082FS | 50S ribosomal | 367336.OM22 | 1,4 | 0,489897949 | 0,000256821 | 7,56E-05 | 2 - | 0,000314457 - | 1 - | 0,000195231 - | 1 - | 0,000185592 - | 2 - | 0,000376667 - | 1 - | 0,000212157 - |
| 5416 | 1 | 3um_FIJJAFBF_467969 | Bacteria | Bacteroidetes | Flavobacteriia | Flavobacterial | Flavobacteriac | unclassified | Flavobacteriac | metabolism | Carbohydrate | 099Q3 | phosphohexol | 983544.Lacal_ | 1 | 0,632455532 | 4,07E-05 | 2,67E-05 | 1 - | 3,50E-05 - | 1 - | 4,35E-05 - | 1 - | 4,13E-05 - | 2 - | 8,38E-05 - | 0 - | 0 - |
| 5417 | 1 | 0.2um_NOKCPHJN_190712 | Bacteria | Proteobacteri | Alphaproteob | Rhodobactera | Hyphomonada | unclassified | Hyphomonada | information st | Translation, ri | 05C64 | Binds directly | 679937.Bcop_ | 0,4 | 0,489897949 | 3,55E-05 | 4,39E-05 | 1 - | 7,92E-05 - | 1 - | 9,83E-05 - | 0 - | 0 - | 0 - | 0 - | 0 - | 0 - |
| 5418 | 1 | 0.2um_NOKCPHJN_15282 | Bacteria | Proteobacteri | Betaproteoba | unclassified | unclassified | unclassified | Betaproteoba | cellular proces | Cell wall/mem | 16QB0 | Part of the ou | 232721.Ajs_2 | 1,2 | 1,939071943 | 1,85E-05 | 2,86E-05 | 5 - | 7,40E-05 - | 1 - | 1,84E-05 - | 0 - | 0 - | 0 - | 0 - | 0 - | 0 - |
| 5419 | 1 | 3um_FIJJAFBF_702258 | Bacteria | Proteobacteri | Alphaproteob | Rhodobactera | Rhodobactera | unclassified | Rhodobactera | metabolism | Carbohydrate | 16THB | transporter, d | 375451.RD1_ | 0,8 | 0,4 | 3,23E-05 | 1,66E-05 | 1 - | 3,38E-05 - | 1 - | 4,19E-05 - | 0 - | 0 - | 1 - | 4,04E-05 - | 1 - | 4,56E-05 - |
| 5420 | 1 | 0.2um_NOKCPHJN_705037 | Bacteria | Proteobacteri | Betaproteoba | Nitrosomonad | Methylophilac | Methylotener | Methylotener | unclassified | unclassified | unclassified | unclassified | unclassified | 1,4 | 0,8 | 4,31E-05 | 2,43E-05 | 2 - | 5,39E-05 - | 1 - | 3,35E-05 - | 2 - | 6,36E-05 - | 2 - | 6,45E-05 - | 0 - | 0 - |
| 5421 | 1 | 0.2um_NOKCPHJN_175137 | Bacteria | Bacteroidetes | unclassified | unclassified | unclassified | unclassified | Bacteroidetes | unclassified | unclassified | unclassified | unclassified | unclassified | 6,6 | 1,356465997 | 9,38E-05 | 1,83E-05 | 7 - | 8,40E-05 - | 5 - | 7,45E-05 - | 9 - | 0,000127413 - | 6 - | 8,62E-05 - | 6 - | 9,71E-05 - |
| 5422 | 1 | 3um_FIJJAFBF_20737 | Bacteria | Proteobacteri | Gammaprotec | Oceanospirilla | Saccharospirill | Reinekea | Reinekea fors | metabolism | Energy produc | 17CGT | F(1)F(0) ATP s | 498211.CJA_3 | 9,2 | 1,166190379 | 0,000801443 | 7,39E-05 | 11 - | 0,000809323 - | 8 - | 0,000730866 - | 10 - | 0,000868476 - | 8 - | 0,000705043 - | 9 - | 0,000893507 - |
| 5423 | 1 | 3um_FIJJAFBF_44713 | Bacteria | Bacteroidetes | Flavobacteriia | Flavobacterial | Flavobacteriac | Polaribacter | Polaribacter s | cellular proces | Cell motility | COG1360 | Ompa motb d | 313598.MED1 | 7,8 | 0,748331477 | 0,000381088 | 6,35E-05 | 7 - | 0,000284907 - | 9 - | 0,000454847 - | 7 - | 0,000336303 - | 8 - | 0,000390024 - | 8 - | 0,00043936 - |
| 5424 | 1 | 3um_FIJJAFBF_223365 | Bacteria | Proteobacteri | Gammaprotec | Cellvibrionales | Porticoccacea | unclassified | gamma protec | metabolism | Lipid transpor | COG4247 | Phytase | 314287.GB220 | 1 | 0,632455532 | 3,68E-05 | 2,06E-05 | 2 - | 6,36E-05 - | 1 - | 3,95E-05 - | 0 - | 0 - | 1 - | 3,81E-05 - | 1 - | 4,29E-05 - |
| 5425 | 1 | 0.2um_NOKCPHJN_671461 | Bacteria | Bacteroidetes | Flavobacteriia | Flavobacterial | Flavobacteriac | Leeuwenhoek | Leeuwenhoek | metabolism | Amino acid tra | 0P7SN | Catalyzes the | 398720.MED2 | 0,4 | 0,489897949 | 2,09E-05 | 2,57E-05 | 1 - | 4,78E-05 - | 0 - | 0 - | 1 - | 5,65E-05 - | 0 - | 0 - | 0 - | 0 - |
| 5426 | 1 | 0.2um_NOKCPHJN_393200 | Bacteria | Proteobacteri | Gammaprotec | Cellvibrionales | Microbulbifera | Microbulbifer | Microbulbifer | cellular proces | Signal transdu | COG0515 | Serine Threon | 234267.Acid_ | 0,4 | 0,489897949 | 6,15E-06 | 7,60E-06 | 1 - | 1,37E-05 - | 1 - | 1,70E-05 - | 0 - | 0 - | 0 - | 0 - | 0 - | 0 - |
| 5427 | 1 | 3um_FIJJAFBF_254308 | Bacteria | Proteobacteri | Gammaprotec | Cellvibrionales | Halieaceae | Chromatocurv | Chromatocurv | information st | Translation, ri | 08UHH | Located on the | 566466.NOR5 | 1,8 | 0,4 | 0,000186984 | 3,69E-05 | 2 - | 0,00017658 - | 2 - | 0,00021926 - | 2 - | 0,000208434 - | 2 - | 0,000211513 - | 1 - | 0,000119134 - |
| 5428 | 1 | 3um_FIJJAFBF_297164 | Bacteria | Proteobacteri | Gammaprotec | Oceanospirilla | Oceanospirilla | Amphritea | Amphritea jap | cellular proces | Signal transdu | 0QIPQ | gtp-binding pr | 207954.MED9 | 1 | 0,632455532 | 2,28E-05 | 1,46E-05 | 1 - | 1,91E-05 - | 1 - | 2,37E-05 - | 0 - | 0 - | 2 - | 4,57E-05 - | 1 - | 2,57E-05 - |

| | | | | | | | | | | | | | | | | | | | | | | | | | | | | |
|---|---|---|---|---|---|---|---|---|---|---|---|---|---|---|---|---|---|---|---|---|---|---|---|---|---|---|---|---|
| 5429 | 1 | 3um_FIJJAFBF_32855 | Bacteria | Proteobacteri | Gammaprotec | Alteromonada | Alteromonada | Paraglaciecola | Paraglaciecola | metabolism | Carbohydrate | 08UHU | Aldolase | 983545.Glaag | 0,8 | 0,4 | 5,09E-05 | 2,62E-05 | 1 - | 5,34E-05 - | 1 - | 6,63E-05 - | 1 - | 6,30E-05 - | 0 - | 0 - | 1 - | 7,20E-05 - |
| 5430 | 1 | 0.2um_NOKCPHJN_481465 | Eukaryota | Ascomycota | Saccharomyce | Saccharomyce | Metschnikowi | Clavispora | [Candida] inte | unclassified | unclassified | unclassified | unclassified | unclassified | 2,4 | 0,489897949 | 9,78E-05 | 2,76E-05 | 2 - | 6,73E-05 - | 3 - | 0,000125383 - | 2 - | 7,95E-05 - | 2 - | 8,06E-05 - | 3 - | 0,000136253 - |
| 5431 | 1 | 3um_FIJJAFBF_79935 | Bacteria | Verrucomicrol | Verrucomicrol | Verrucomicrol | Rubritaleacea | Rubritalea | Rubritalea squ | unclassified | unclassified | unclassified | unclassified | unclassified | 0,6 | 0,489897949 | 1,15E-05 | 9,47E-06 | 1 - | 1,67E-05 - | 1 - | 2,07E-05 - | 0 - | 0 - | 1 - | 2,00E-05 - | 0 - | 0 - |
| 5432 | 7 | 0.2um_NOKCPHJN_338692;0.2um_NOKCPHJN_8 | Bacteria | Bacteroidetes | Flavobacteriia | Flavobacterial | Flavobacteriac | various | various | cellular proces | Signal transdu | 09BM1 | Two compone | various | 1 | 0 | 5,76E-05 | 5,47E-06 | 1 - | 4,82E-05 - | 1 - | 5,99E-05 - | 1 - | 5,69E-05 - | 1 - | 5,78E-05 - | 1 - | 6,51E-05 - |
| 5433 | 1 | 3um_FIJJAFBF_273644 | Bacteria | Bacteroidetes | Flavobacteriia | Flavobacterial | Flavobacteriac | Algibacter | Algibacter sp. | poorly charact | Function unkn | 0XNQI | domain protei | 1046627.BZAF | 0,8 | 0,748331477 | 2,22E-05 | 1,94E-05 | 2 - | 4,99E-05 - | 1 - | 3,10E-05 - | 0 - | 0 - | 1 - | 2,99E-05 - | 0 - | 0 - |
| 5434 | 1 | 3um_FIJJAFBF_788034 | Bacteria | Proteobacteri | Alphaproteob | Rhodobactera | Rhodobactera | unclassified | Rhodobactera | information st | Translation, ri | 01S8S | This is 1 of the | 388739.RSK20 | 4,8 | 0,748331477 | 0,00034928 | 5,04E-05 | 5 - | 0,00030689 - | 4 - | 0,000304853 - | 5 - | 0,000362252 - | 6 - | 0,000441123 - | 4 - | 0,000331282 - |
| 5435 | 1 | 3um_FIJJAFBF_189667 | Bacteria | Bacteroidetes | Flavobacteriia | Flavobacterial | Flavobacteriac | Polaribacter | Polaribacter s | metabolism | Lipid transpor | COG0439 | acetyl-CoA car | 983548.Krodi_ | 6,8 | 0,748331477 | 0,000208361 | 2,26E-05 | 7 - | 0,000180548 - | 6 - | 0,00019216 - | 7 - | 0,000213118 - | 8 - | 0,000247161 - | 6 - | 0,00020882 - |
| 5436 | 1 | 3um_FIJJAFBF_38603 | Bacteria | Proteobacteri | Gammaprotec | Chromatiales | unclassified | unclassified | Chromatiales | metabolism | Inorganic ion t | 0BJE9 | ABC-type nitra | 1000565.MET | 4,6 | 1,2 | 0,000162431 | 3,19E-05 | 6 - | 0,000181226 - | 4 - | 0,00015002 - | 6 - | 0,000213919 - | 4 - | 0,000144719 - | 3 - | 0,000122269 - |
| 5437 | 1 | 0.2um_NOKCPHJN_35232 | Bacteria | Proteobacteri | Alphaproteob | unclassified | unclassified | various | various | metabolism | Energy produc | COG0056 | Produces ATP | 488538.SAR11 | 1,6 | 0,489897949 | 4,36E-05 | 1,53E-05 | 1 - | 2,25E-05 - | 1 - | 2,79E-05 - | 2 - | 5,30E-05 - | 2 - | 5,38E-05 - | 2 - | 6,06E-05 - |
| 5438 | 1 | 3um_FIJJAFBF_85786 | Viruses | unclassified | unclassified | unclassified | Phycodnavirid | unclassified | Chrysochromu | unclassified | unclassified | unclassified | unclassified | unclassified | 1,2 | 0,4 | 9,19E-05 | 2,04E-05 | 2 - | 0,000131927 - | 1 - | 8,19E-05 - | 1 - | 7,79E-05 - | 1 - | 7,90E-05 - | 1 - | 8,90E-05 - |
| 5439 | 1 | 3um_FIJJAFBF_800229 | Bacteria | Bacteroidetes | Flavobacteriia | Flavobacterial | Flavobacteriac | Polaribacter | Polaribacter h | unclassified | unclassified | unclassified | unclassified | unclassified | 1 | 0 | 0,000139824 | 1,33E-05 | 1 - | 0,000117119 - | 1 - | 0,000145427 - | 1 - | 0,000138247 - | 1 - | 0,000140289 - | 1 - | 0,000158035 - |
| 5440 | 1 | 3um_FIJJAFBF_436837 | Bacteria | Actinobacteria | Actinobacteria | Micrococcales | Microbacteria | Candidatus Ac | Candidatus Ac | information st | Translation, ri | COG0051 | tRNA binding | 465515.MlutN | 1,6 | 0,8 | 0,000207946 | 0,000105617 | 2 - | 0,000225052 - | 2 - | 0,000279449 - | 2 - | 0,000265651 - | 2 - | 0,000269575 - | 0 - | 0 - |
| 5441 | 2 | 3um_FIJJAFBF_13108;3um_FIJJAFBF_74383 | Eukaryota | unclassified | Cryptophyta | Pyrenomonad | Geminigerace | various | various | metabolism | Energy produc | COG1290 | Component of | various | 1 | 0 | 6,37E-05 | 6,05E-06 | 1 - | 5,34E-05 - | 1 - | 6,63E-05 - | 1 - | 6,30E-05 - | 1 - | 6,39E-05 - | 1 - | 7,20E-05 - |
| 5442 | 1 | 3um_FIJJAFBF_393573 | Bacteria | Proteobacteri | Gammaprotec | Oceanospirilla | Saccharospirill | Reinekea | Reinekea fors | poorly charact | Function unkn | 17FMV | dNA-binding p | 314283.MED2 | 2,6 | 0,489897949 | 0,000330853 | 8,40E-05 | 2 - | 0,0002106 - | 3 - | 0,000392254 - | 3 - | 0,000372887 - | 2 - | 0,000252263 - | 3 - | 0,00042626 - |
| 5443 | 1 | 3um_FIJJAFBF_101550 | Bacteria | Proteobacteri | Alphaproteob | Rhodospirillal | Rhodospirillac | unclassified | Rhodobactera | information st | Translation, ri | 177XJ | One of the pri | 367336.OM22 | 5,8 | 1,166190379 | 0,000502677 | 8,27E-05 | 7 - | 0,000515024 - | 7 - | 0,000639508 - | 6 - | 0,000521085 - | 5 - | 0,000440652 - | 4 - | 0,000397114 - |
| 5444 | 1 | 0.2um_NOKCPHJN_440915 | Bacteria | Proteobacteri | Gammaprotec | Cellvibrionales | Porticoccacea | unclassified | various | unclassified | unclassified | unclassified | unclassified | unclassified | 0,6 | 0,489897949 | 8,77E-06 | 7,23E-06 | 1 - | 1,28E-05 - | 1 - | 1,59E-05 - | 1 - | 1,51E-05 - | 0 - | 0 - | 0 - | 0 - |
| 5445 | 1 | 3um_FIJJAFBF_109825 | Eukaryota | unclassified | unclassified | Isochrysidales | Noelaerhabda | Emiliania | Emiliania huxl | metabolism | Carbohydrate | 0IVA1 | Photosystem I | 45157.CMV20 | 0,4 | 0,489897949 | 2,80E-05 | 3,46E-05 | 1 - | 6,24E-05 - | 1 - | 7,75E-05 - | 0 - | 0 - | 0 - | 0 - | 0 - | 0 - |
| 5446 | 1 | 3um_FIJJAFBF_257117 | Eukaryota | Bacillariophyta | Fragilariophyc | Fragilariales | Fragilariaceae | Asterionellops | Asterionellops | information st | Translation, ri | 0FJK5 | Binds the lowe | 197221.tlr008 | 1 | 0,632455532 | 6,04E-05 | 3,38E-05 | 2 - | 0,000104343 - | 1 - | 6,48E-05 - | 0 - | 0 - | 1 - | 6,25E-05 - | 1 - | 7,04E-05 - |
| 5447 | 1 | 3um_FIJJAFBF_685913 | Bacteria | Proteobacteri | Gammaprotec | Alteromonada | Alteromonada | Paraglaciecola | Paraglaciecola | unclassified | unclassified | unclassified | unclassified | unclassified | 0,2 | 0,4 | 1,32E-05 | 2,64E-05 | 0 - | 0 - | 1 - | 6,60E-05 - | 0 - | 0 - | 0 - | 0 - | 0 - | 0 - |
| 5448 | 1 | 3um_FIJJAFBF_187036 | Bacteria | Proteobacteri | Gammaprotec | Alteromonada | Alteromonada | Paraglaciecola | Paraglaciecola | unclassified | unclassified | unclassified | unclassified | unclassified | 0,6 | 0,489897949 | 3,02E-05 | 2,47E-05 | 0 - | 0 - | 1 - | 5,18E-05 - | 1 - | 4,93E-05 - | 1 - | 5,00E-05 - | 0 - | 0 - |
| 5449 | 1 | 3um_FIJJAFBF_270785 | Bacteria | Bacteroidetes | Flavobacteriia | Flavobacterial | Flavobacteriac | Bizionia | Bizionia psych | metabolism | Nucleotide tra | 0P7KC | ribonucleoside | 983544.Lacal_ | 1,2 | 0,4 | 5,08E-05 | 1,87E-05 | 1 - | 3,52E-05 - | 2 - | 8,74E-05 - | 1 - | 4,16E-05 - | 1 - | 4,22E-05 - | 1 - | 4,75E-05 - |
| 5450 | 1 | 3um_FIJJAFBF_577077 | Bacteria | Bacteroidetes | Flavobacteriia | Flavobacterial | Flavobacteriac | Lacinutrix | Lacinutrix him | poorly charact | Function unkn | 06E9D | NA | 391598.FBBAL | 0,8 | 0,4 | 1,72E-05 | 8,76E-06 | 1 - | 1,87E-05 - | 1 - | 2,32E-05 - | 1 - | 2,20E-05 - | 1 - | 2,24E-05 - | 0 - | 0 - |
| 5451 | 1 | 3um_FIJJAFBF_15040 | Bacteria | Bacteroidetes | Flavobacteriia | Flavobacterial | Flavobacteriac | Polaribacter | Polaribacter s | information st | Replication, re | COG0513 | purine NTP-de | 313598.MED1 | 0,6 | 0,8 | 2,19E-05 | 2,91E-05 | 0 - | 0 - | 0 - | 0 - | 2 - | 7,26E-05 - | 1 - | 3,69E-05 - | 0 - | 0 - |
| 5452 | 1 | 0.2um_NOKCPHJN_405135 | Bacteria | Proteobacteri | various | various | various | various | various | cellular proces | Cell wall/mem | 0B7GS | outer membra | 583345.Mmol | 1 | 0 | 3,23E-05 | 3,07E-06 | 1 - | 2,71E-05 - | 1 - | 3,36E-05 - | 1 - | 3,20E-05 - | 1 - | 3,24E-05 - | 1 - | 3,65E-05 - |
| 5453 | 1 | 0.2um_NOKCPHJN_746599 | Eukaryota | Phaeophyceae | unclassified | Ectocarpales | Ectocarpaceae | Ectocarpus | Ectocarpus sil | unclassified | unclassified | unclassified | unclassified | unclassified | 2 | 0 | 0,000246896 | 2,34E-05 | 2 - | 0,000206805 - | 2 - | 0,000256791 - | 2 - | 0,000244112 - | 2 - | 0,000247718 - | 2 - | 0,000279053 - |
| 5454 | 1 | 3um_FIJJAFBF_157512 | Bacteria | Proteobacteri | Betaproteoba | unclassified | unclassified | unclassified | Betaproteoba | unclassified | unclassified | unclassified | unclassified | unclassified | 0,6 | 0,489897949 | 7,78E-05 | 6,38E-05 | 0 - | 0 - | 0 - | 0 - | 1 - | 0,000123166 - | 1 - | 0,000124985 - | 1 - | 0,000140795 - |
| 5455 | 1 | 3um_FIJJAFBF_105882 | Bacteria | Proteobacteri | Alphaproteob | Rhodobactera | Rhodobactera | Planktomarina | Planktomarina | metabolism | Energy produc | COG0711 | F(1)F(0) ATP s | 439497.RR11_ | 7,6 | 0,8 | 0,000580564 | 7,66E-05 | 8 - | 0,000512969 - | 6 - | 0,000477717 - | 8 - | 0,000605507 - | 8 - | 0,000614451 - | 8 - | 0,000692176 - |
| 5456 | 1 | 0.2um_NOKCPHJN_56931 | Bacteria | Bacteroidetes | Flavobacteriia | Flavobacterial | Flavobacteriac | Formosa | Formosa sp. H | metabolism | Energy produc | 09A0E | phosphoenolp | 983544.Lacal_ | 0,2 | 0,4 | 3,14E-06 | 6,27E-06 | 0 - | 0 - | 1 - | 1,57E-05 - | 0 - | 0 - | 0 - | 0 - | 0 - | 0 - |
| 5457 | 1 | 3um_FIJJAFBF_665397 | Bacteria | Bacteroidetes | Flavobacteriia | Flavobacterial | Flavobacteriac | Polaribacter | Polaribacter s | unclassified | unclassified | unclassified | unclassified | unclassified | 0,4 | 0,8 | 8,00E-06 | 1,60E-05 | 0 - | 0 - | 0 - | 0 - | 0 - | 0 - | 2 - | 4,00E-05 - | 0 - | 0 - |
| 5458 | 1 | 3um_FIJJAFBF_162557 | Bacteria | Bacteroidetes | Flavobacteriia | Flavobacterial | Flavobacteriac | unclassified | Flavobacteriac | unclassified | unclassified | unclassified | unclassified | unclassified | 1,8 | 0,748331477 | 8,00E-05 | 3,35E-05 | 1 - | 3,71E-05 - | 2 - | 9,22E-05 - | 3 - | 0,000131536 - | 2 - | 8,90E-05 - | 1 - | 5,01E-05 - |
| 5459 | 1 | 0.2um_NOKCPHJN_29655 | Bacteria | Proteobacteri | Gammaprotec | Alteromonada | Alteromonada | Paraglaciecola | Paraglaciecola | metabolism | Energy produc | 0QJ3X | Aconitate hyd | 167879.CPS_4 | 1 | 0,632455532 | 1,42E-05 | 7,95E-06 | 2 - | 2,46E-05 - | 1 - | 1,53E-05 - | 1 - | 1,45E-05 - | 0 - | 0 - | 1 - | 1,66E-05 - |
| 5460 | 1 | 0.2um_NOKCPHJN_35610 | Eukaryota | unclassified | Oomycetes | Myzocytiopsic | Eurychasmata | Eurychasma | Eurychasma d | unclassified | unclassified | unclassified | unclassified | unclassified | 0,4 | 0,489897949 | 1,61E-07 | 1,98E-07 | 0 - | 0 - | 1 - | 4,11E-07 - | 0 - | 0 - | 1 - | 3,96E-07 - | 0 - | 0 - |
| 5461 | 1 | 3um_FIJJAFBF_362287 | Bacteria | Proteobacteri | Gammaprotec | Alteromonada | Psychromonac | Corallincola | Corallincola sp | metabolism | Inorganic ion t | 0QJ8G | extracellular s | 207954.MED9 | 0,6 | 0,489897949 | 2,44E-05 | 2,00E-05 | 0 - | 0 - | 1 - | 4,19E-05 - | 1 - | 3,98E-05 - | 1 - | 4,04E-05 - | 0 - | 0 - |
| 5462 | 1 | 3um_FIJJAFBF_00853 | Bacteria | Bacteroidetes | Flavobacteriia | Flavobacterial | Flavobacteriac | Polaribacter | Polaribacter s | unclassified | unclassified | unclassified | unclassified | unclassified | 8,4 | 1,356465997 | 0,000887327 | 9,42E-05 | 10 - | 0,000896693 - | 9 - | 0,001002086 - | 9 - | 0,000952609 - | 8 - | 0,000859271 - | 6 - | 0,000725974 - |
| 5463 | 1 | 3um_FIJJAFBF_263677 | Eukaryota | Bacillariophyta | Coscinodiscop | Rhizosoleniale | Rhizosoleniac | Rhizosolenia | Rhizosolenia s | metabolism | Energy produc | 11IIK | Low-potential | 35128.Thapsd | 7,4 | 0,489897949 | 0,000630304 | 7,49E-05 | 7 - | 0,000499029 - | 8 - | 0,000708168 - | 8 - | 0,000673204 - | 7 - | 0,000597754 - | 7 - | 0,000673367 - |
| 5464 | 1 | 0.2um_NOKCPHJN_844820 | Bacteria | Bacteroidetes | unclassified | unclassified | unclassified | unclassified | Bacteroidetes | unclassified | unclassified | unclassified | unclassified | unclassified | 1 | 0 | 8,73E-05 | 8,29E-06 | 1 - | 7,31E-05 - | 1 - | 9,08E-05 - | 1 - | 8,63E-05 - | 1 - | 8,76E-05 - | 1 - | 9,86E-05 - |
| 5465 | 1 | 0.2um_NOKCPHJN_286943 | Eukaryota | unclassified | Oomycetes | Peronosporale | Peronosporac | Phytophthora | Phytophthora | unclassified | unclassified | unclassified | unclassified | unclassified | 0,8 | 0,4 | 1,35E-05 | 6,91E-06 | 1 - | 1,42E-05 - | 0 - | 0 - | 1 - | 1,68E-05 - | 1 - | 1,71E-05 - | 1 - | 1,92E-05 - |
| 5466 | 1 | 3um_FIJJAFBF_176833 | Bacteria | Proteobacteri | Alphaproteob | Rhodobactera | Rhodobactera | Aliiroseovarius | Aliiroseovarius | cellular proces | Signal transdu | 01Q8R | gtp-binding pr | 388739.RSK20 | 0,8 | 0,748331477 | 1,74E-05 | 1,50E-05 | 2 - | 3,79E-05 - | 1 - | 2,35E-05 - | 0 - | 0 - | 0 - | 0 - | 1 - | 2,56E-05 - |
| 5467 | 1 | 3um_FIJJAFBF_193971 | Bacteria | Proteobacteri | Gammaprotec | Arenicellales | Arenicellaceae | Arenicella | Arenicella xan | unclassified | unclassified | unclassified | unclassified | unclassified | 0,4 | 0,8 | 1,75E-05 | 3,50E-05 | 2 - | 8,76E-05 - | 0 - | 0 - | 0 - | 0 - | 0 - | 0 - | 0 - | 0 - |
| 5468 | 1 | 0.2um_NOKCPHJN_248194 | Bacteria | Proteobacteri | Alphaproteob | Pelagibacteral | Pelagibacterac | Candidatus Pe | Candidatus Pe | metabolism | Energy produc | COG1053 | Succinate DeH | 335992.SAR11 | 3,6 | 1,854723699 | 7,98E-05 | 4,10E-05 | 2 - | 3,71E-05 - | 4 - | 9,21E-05 - | 7 - | 0,000153211 - | 3 - | 6,66E-05 - | 2 - | 5,00E-05 - |
| 5469 | 1 | 3um_FIJJAFBF_163214 | Bacteria | Bacteroidetes | Flavobacteriia | Flavobacterial | Flavobacteriac | Siansivirga | Siansivirga zea | metabolism | Energy produc | 05CMV | Succinyl-CoA s | 391603.FBALC | 2,4 | 0,489897949 | 8,18E-05 | 1,20E-05 | 3 - | 8,70E-05 - | 2 - | 7,20E-05 - | 3 - | 0,000102638 - | 2 - | 6,94E-05 - | 2 - | 7,82E-05 - |
| 5470 | 1 | 3um_FIJJAFBF_489503 | Bacteria | Bacteroidetes | Flavobacteriia | Flavobacterial | Flavobacteriac | Polaribacter | Polaribacter s | poorly charact | Function unkn | 11P35 | NA | 867900.Celly_ | 12,2 | 0,979795897 | 0,000469731 | 4,72E-05 | 12 - | 0,000387978 - | 12 - | 0,000481754 - | 14 - | 0,000534296 - | 12 - | 0,000464732 - | 11 - | 0,000479893 - |
| 5471 | 1 | 0.2um_NOKCPHJN_305649 | Eukaryota | Cnidaria | Anthozoa | Actiniaria | Aiptasiidae | Exaiptasia | Exaiptasia pall | cellular proces | Cell cycle cont | 0V9W7 | retinitis pigme | 7955.ENSDAR | 0,2 | 0,4 | 1,31E-06 | 2,61E-06 | 0 - | 0 - | 0 - | 0 - | 0 - | 0 - | 1 - | 6,53E-06 - | 0 - | 0 - |
| 5472 | 1 | 3um_FIJJAFBF_566201 | unclassified | unclassified | unclassified | unclassified | unclassified | unclassified | unclassified | unclassified | unclassified | unclassified | unclassified | unclassified | 0,8 | 0,4 | 8,42E-05 | 4,27E-05 | 1 - | 9,11E-05 - | 1 - | 0,00011311 - | 1 - | 0,000107526 - | 1 - | 0,000109114 - | 0 - | 0 - |
| 5473 | 1 | 3um_FIJJAFBF_615446 | Bacteria | Proteobacteri | Alphaproteob | Rhodobactera | Rhodobactera | Ascidiaceihabi | Ascidiaceihabi | metabolism | Carbohydrate | 16SJ5 | Trap dicarbox | 314270.RB208 | 0,2 | 0,4 | 6,81E-06 | 1,36E-05 | 1 - | 3,41E-05 - | 0 - | 0 - | 0 - | 0 - | 0 - | 0 - | 0 - | 0 - |
| 5474 | 1 | 0.2um_NOKCPHJN_127058 | Bacteria | Proteobacteri | Gammaprotec | Cellvibrionales | Porticoccacea | unclassified | gamma protec | poorly charact | Function unkn | 16R8I | Lipoprotein | 314287.GB220 | 0,8 | 0,4 | 2,09E-05 | 1,05E-05 | 0 - | 0 - | 1 - | 2,61E-05 - | 1 - | 2,48E-05 - | 1 - | 2,51E-05 - | 1 - | 2,83E-05 - |
| 5475 | 1 | 3um_FIJJAFBF_334376 | Eukaryota | Apicomplexa | Conoidasida | Eucoccidiorida | Eimeriidae | Eimeria | Eimeria mitis | poorly charact | Function unkn | 0HVC9 | SERTA motif | 7237.FBpp027 | 0,2 | 0,4 | 7,28E-06 | 1,46E-05 | 0 - | 0 - | 0 - | 0 - | 1 - | 3,64E-05 - | 0 - | 0 - | 0 - | 0 - |
| 5476 | 1 | 0.2um_NOKCPHJN_655088 | Bacteria | Proteobacteri | Alphaproteob | Pelagibacteral | Pelagibacterac | Candidatus Pe | Candidatus Pe | metabolism | Lipid transpor | 0QJMG | Acyltransferas | 575788.VS_04 | 0,4 | 0,489897949 | 8,14E-06 | 9,97E-06 | 0 - | 0 - | 0 - | 0 - | 1 - | 2,02E-05 - | 1 - | 2,05E-05 - | 0 - | 0 - |
| 5477 | 2 | 3um_FIJJAFBF_239200;3um_FIJJAFBF_267803 | Bacteria | Bacteroidetes | various | various | various | various | various | unclassified | unclassified | unclassified | unclassified | unclassified | 1 | 0 | 3,87E-05 | 3,68E-06 | 1 - | 3,24E-05 - | 1 - | 4,03E-05 - | 1 - | 3,83E-05 - | 1 - | 3,88E-05 - | 1 - | 4,37E-05 - |
| 5478 | 1 | 3um_FIJJAFBF_763145 | Bacteria | Bacteroidetes | Flavobacteriia | Flavobacterial | Flavobacteriac | Polaribacter | Polaribacter s | metabolism | Amino acid tra | COG0436 | aminotransfer | 313598.MED1 | 0,8 | 0,748331477 | 2,79E-05 | 2,58E-05 | 0 - | 0 - | 1 - | 3,61E-05 - | 2 - | 6,86E-05 - | 1 - | 3,48E-05 - | 0 - | 0 - |
| 5479 | 1 | 3um_FIJJAFBF_88824 | Bacteria | Proteobacteri | Gammaprotec | unclassified | unclassified | unclassified | various | information st | Translation, ri | 179ZI | One of the pri | 247633.GP214 | 2,6 | 0,489897949 | 0,000269473 | 4,55E-05 | 3 - | 0,000262848 - | 3 - | 0,000326379 - | 2 - | 0,000206843 - | 3 - | 0,000314847 - | 2 - | 0,00023645 - |
| 5480 | 1 | 0.2um_NOKCPHJN_149786 | Bacteria | unclassified | unclassified | unclassified | unclassified | unclassified | uncultured ba | poorly charact | Function unkn | 0XNQM | proprotein co | 313598.MED1 | 0,8 | 0,4 | 6,83E-06 | 3,47E-06 | 1 - | 7,39E-06 - | 1 - | 9,18E-06 - | 1 - | 8,72E-06 - | 1 - | 8,85E-06 - | 0 - | 0 - |
| 5481 | 1 | 0.2um_NOKCPHJN_221509 | Bacteria | Proteobacteri | Gammaprotec | Oceanospirilla | Saccharospirill | Reinekea | Reinekea blan | metabolism | Carbohydrate | 0QJR5 | PTS system m | 314283.MED2 | 6,4 | 1,356465997 | 0,000188393 | 3,38E-05 | 8 - | 0,000199612 - | 8 - | 0,000247859 - | 6 - | 0,000176716 - | 5 - | 0,000149438 - | 5 - | 0,000168342 - |
| 5482 | 1 | 3um_FIJJAFBF_469932 | Bacteria | Proteobacteri | Alphaproteob | Rhodobactera | Rhodobactera | Roseobacter | Roseobacter s | metabolism | Amino acid tra | 01SW7 | Tetrahydrodip | 439496.RBY4I | 1 | 0,632455532 | 5,11E-05 | 3,60E-05 | 1 - | 4,17E-05 - | 1 - | 5,18E-05 - | 1 - | 4,93E-05 - | 0 - | 0 - | 2 - | 0,000112636 - |
| 5483 | 1 | 3um_FIJJAFBF_128439 | Bacteria | Bacteroidetes | Flavobacteriia | Flavobacterial | Flavobacteriac | Polaribacter | Polaribacter s | poorly charact | Function unkn | 122UE | nicotinic acid | 313598.MED1 | 1,8 | 0,4 | 0,00021255 | 5,21E-05 | 2 - | 0,000197891 - | 2 - | 0,000245722 - | 2 - | 0,00023359 - | 1 - | 0,00011852 - | 2 - | 0,000267025 - |
| 5484 | 1 | 3um_FIJJAFBF_360762 | Bacteria | Bacteroidetes | Flavobacteriia | Flavobacterial | Flavobacteriac | Polaribacter | Polaribacter s | cellular proces | Signal transdu | 0P8JH | Histidine kinas | 313594.PI23P | 0,8 | 0,748331477 | 3,20E-05 | 2,98E-05 | 0 - | 0 - | 1 - | 4,12E-05 - | 1 - | 3,92E-05 - | 2 - | 7,95E-05 - | 0 - | 0 - |
| 5485 | 1 | 3um_FIJJAFBF_331698 | Bacteria | Proteobacteri | Alphaproteob | Rickettsiales | Candidatus Mi | Candidatus Jic | Candidatus Jid | metabolism | Energy produc | 01RNF | Produces ATP | 696127.midi_ | 0,8 | 0,748331477 | 2,18E-05 | 1,91E-05 | 2 - | 4,90E-05 - | 1 - | 3,05E-05 - | 0 - | 0 - | 1 - | 2,94E-05 - | 0 - | 0 - |
| 5486 | 1 | 0.2um_NOKCPHJN_678877 | Bacteria | Proteobacteri | Alphaproteob | Pelagibacteral | Pelagibacterac | Candidatus Pe | Candidatus Pe | metabolism | Nucleotide tra | 16SYS | cytosine purin | 388739.RSK20 | 1 | 0 | 2,84E-05 | 2,70E-06 | 1 - | 2,38E-05 - | 1 - | 2,96E-05 - | 1 - | 2,81E-05 - | 1 - | 2,85E-05 - | 1 - | 3,21E-05 - |
| 5487 | 1 | 0.2um_NOKCPHJN_530307 | Bacteria | Proteobacteri | Gammaprotec | Cellvibrionales | Halieaceae | unclassified | Halieaceae ba | information st | Translation, ri | 16Q16 | Binds the lowe | 247639.MGP2 | 0,6 | 0,489897949 | 3,68E-05 | 3,06E-05 | 1 - | 5,12E-05 - | 1 - | 6,36E-05 - | 0 - | 0 - | 0 - | 0 - | 1 - | 6,91E-05 - |
| 5488 | 1 | 3um_FIJJAFBF_248470 | Bacteria | Proteobacteri | Alphaproteob | Rhodobactera | Rhodobactera | unclassified | Rhodobactera | unclassified | unclassified | unclassified | unclassified | unclassified | 0,6 | 0,8 | 0,000114014 | 0,000152485 | 0 - | 0 - | 0 - | 0 - | 1 - | 0,00018817 - | 2 - | 0,000381898 - | 0 - | 0 - |
| 5489 | 1 | 3um_FIJJAFBF_357382 | Bacteria | Proteobacteri | Gammaprotec | Oceanospirilla | Saccharospirill | Reinekea | Reinekea blan | information st | Translation, ri | 05CRK | lysyL-tRNA syr | 314283.MED2 | 0,2 | 0,4 | 4,55E-06 | 9,09E-06 | 1 - | 2,27E-05 - | 0 - | 0 - | 0 - | 0 - | 0 - | 0 - | 0 - | 0 - |
| 5490 | 1 | 0.2um_NOKCPHJN_07058 | Bacteria | Bacteroidetes | Cytophagia | Cytophagales | unclassified | Chryseolinea | Chryseolinea s | metabolism | Amino acid tra | 08IQK | o-acetylhomo | 643867.Ftrac_ | 1 | 1,264911064 | 2,88E-05 | 3,74E-05 | 2 - | 5,19E-05 - | 0 - | 0 - | 3 - | 9,20E-05 - | 0 - | 0 - | 0 - | 0 - |
| 5491 | 1 | 0.2um_NOKCPHJN_655230 | Bacteria | Proteobacteri | Gammaprotec | Cellvibrionales | Porticoccacea | unclassified | gamma protec | poorly charact | Function unkn | 17GBW | NA | 314287.GB220 | 0,4 | 0,489897949 | 8,16E-06 | 1,01E-05 | 1 - | 1,82E-05 - | 1 - | 2,26E-05 - | 0 - | 0 - | 0 - | 0 - | 0 - | 0 - |
| 5492 | 1 | 0.2um_NOKCPHJN_72797 | Bacteria | Proteobacteri | various | various | various | various | various | information st | Translation, ri | COG0052 | 30S ribosomal | 349521.HCH_ | 10,2 | 1,166190379 | 0,000551899 | 8,10E-05 | 11 - | 0,000499029 - | 10 - | 0,000563316 - | 11 - | 0,000589053 - | 8 - | 0,00043473 - | 11 - | 0,000673367 - |
| 5493 | 1 | 0.2um_NOKCPHJN_632309 | unclassified | unclassified | unclassified | unclassified | unclassified | unclassified | unclassified | unclassified | unclassified | unclassified | unclassified | unclassified | 3 | 1,414213562 | 7,65E-05 | 3,08E-05 | 5 - | 0,000110788 - | 1 - | 2,75E-05 - | 3 - | 7,85E-05 - | 4 - | 0,000106165 - | 2 - | 5,98E-05 - |
| 5494 | 1 | 3um_FIJJAFBF_184051 | Viruses | unclassified | unclassified | Caudovirales | Siphoviridae | unclassified | Idiomarinacea | poorly charact | Function unkn | 08K17 | NA | 342108.amb0 | 1 | 0 | 5,31E-05 | 5,04E-06 | 1 - | 4,45E-05 - | 1 - | 5,52E-05 - | 1 - | 5,25E-05 - | 1 - | 5,33E-05 - | 1 - | 6,00E-05 - |
| 5495 | 1 | 3um_FIJJAFBF_594150 | Eukaryota | unclassified | Pelagophycea | Pelagomonad | unclassified | Aureoumbra | Aureoumbra l | metabolism | Energy produc | COG4451 | RuBisCO catal | 543728.Vapar | 5,8 | 0,748331477 | 0,000567998 | 6,39E-05 | 7 - | 0,000578012 - | 5 - | 0,000512658 - | 5 - | 0,000487346 - | 6 - | 0,000593453 - | 6 - | 0,000668523 - |
| 5496 | 1 | 3um_FIJJAFBF_340084 | Bacteria | Proteobacteri | Alphaproteob | Rhodobactera | Rhodobactera | Actibacterium | Actibacterium | metabolism | Carbohydrate | 16T1E | Periplasmic bi | 391619.RGBS | 0,2 | 0,4 | 8,55E-06 | 1,71E-05 | 0 - | 0 - | 0 - | 0 - | 1 - | 4,27E-05 - | 0 - | 0 - | 0 - | 0 - |
| 5497 | 1 | 3um_FIJJAFBF_426660 | Bacteria | Proteobacteri | Gammaprotec | Cellvibrionales | Porticoccacea | unclassified | SAR92 bacteri | information st | Translation, ri | COG0333 | 50s ribosomal | 314287.GB220 | 0,8 | 0,748331477 | 0,000184616 | 0,000175233 | 1 - | 0,000194537 - | 0 - | 0 - | 0 - | 0 - | 2 - | 0,000466045 - | 1 - | 0,000262499 - |
| 5498 | 1 | 0.2um_NOKCPHJN_495882 | Bacteria | Actinobacteria | Actinobacteria | Micrococcales | Microbacteria | Candidatus Ac | Candidatus Ac | unclassified | unclassified | unclassified | unclassified | unclassified | 1 | 0 | 1,29E-05 | 1,22E-06 | 1 - | 1,08E-05 - | 1 - | 1,34E-05 - | 1 - | 1,27E-05 - | 1 - | 1,29E-05 - | 1 - | 1,46E-05 - |
| 5499 | 1 | 3um_FIJJAFBF_508693 | Bacteria | Proteobacteri | Gammaprotec | Cellvibrionales | Porticoccacea | unclassified | SAR92 bacteri | unclassified | unclassified | unclassified | unclassified | unclassified | 0,2 | 0,4 | 7,55E-06 | 1,51E-05 | 1 - | 3,78E-05 - | 0 - | 0 - | 0 - | 0 - | 0 - | 0 - | 0 - | 0 - |
| 5500 | 1 | 0.2um_NOKCPHJN_559988 | Bacteria | Proteobacteri | Alphaproteob | Pelagibacteral | Pelagibacterac | Candidatus Pe | Candidatus Pe | metabolism | Energy produc | 16QM4 | alcohol dehyd | 439493.PB721 | 0,6 | 0,8 | 2,47E-05 | 3,30E-05 | 0 - | 0 - | 0 - | 0 - | 1 - | 4,07E-05 - | 2 - | 8,26E-05 - | 0 - | 0 - |
| 5501 | 1 | 0.2um_NOKCPHJN_1077861 | Eukaryota | unclassified | Cryptophyta | Pyrenomonad | Geminigerace | Guillardia | Guillardia thet | unclassified | unclassified | unclassified | unclassified | unclassified | 1,4 | 0,8 | 0,000208718 | 0,00012063 | 1 - | 0,000124757 - | 1 - | 0,000154912 - | 1 - | 0,000147263 - | 3 - | 0,000448315 - | 1 - | 0,000168342 - |
| 5502 | 1 | 0.2um_NOKCPHJN_30850 | Bacteria | Firmicutes | Bacilli | Bacillales | Bacillaceae | Calditerricola | Calditerricola | unclassified | unclassified | unclassified | unclassified | unclassified | 0,2 | 0,4 | 1,52E-05 | 3,04E-05 | 1 - | 7,60E-05 - | 0 - | 0 - | 0 - | 0 - | 0 - | 0 - | 0 - | 0 - |
| 5503 | 1 | 3um_FIJJAFBF_526175 | Eukaryota | Mollusca | Bivalvia | Ostreoida | Ostreidae | Crassostrea | Crassostrea vi | unclassified | unclassified | unclassified | unclassified | unclassified | 0,4 | 0,489897949 | 9,37E-06 | 1,15E-05 | 1 - | 2,15E-05 - | 0 - | 0 - | 1 - | 2,54E-05 - | 0 - | 0 - | 0 - | 0 - |
| 5504 | 1 | 0.2um_NOKCPHJN_503156 | Viruses | unclassified | unclassified | unclassified | various | various | various | unclassified | unclassified | unclassified | unclassified | unclassified | 1 | 0 | 2,71E-05 | 2,58E-06 | 1 - | 2,27E-05 - | 1 - | 2,82E-05 - | 1 - | 2,68E-05 - | 1 - | 2,72E-05 - | 1 - | 3,07E-05 - |
| 5505 | 1 | 0.2um_NOKCPHJN_226683 | Bacteria | Proteobacteri | Gammaprotec | Cellvibrionales | Porticoccacea | unclassified | gamma protec | metabolism | Inorganic ion t | 16TQC | TONB-depend | 314287.GB220 | 2,2 | 0,748331477 | 2,86E-05 | 8,75E-06 | 3 - | 3,34E-05 - | 3 - | 4,15E-05 - | 2 - | 2,63E-05 - | 2 - | 2,67E-05 - | 1 - | 1,50E-05 - |
| 5506 | 1 | 0.2um_NOKCPHJN_19170 | Bacteria | Bacteroidetes | Flavobacteriia | Flavobacterial | Flavobacteriac | Cellulophaga | Cellulophaga l | metabolism | Carbohydrate | 07R46 | aggregation fa | 485918.Cpin_ | 1,8 | 0,4 | 5,91E-05 | 1,47E-05 | 2 - | 5,49E-05 - | 2 - | 6,82E-05 - | 1 - | 3,24E-05 - | 2 - | 6,58E-05 - | 2 - | 7,41E-05 - |
| 5507 | 1 | 0.2um_NOKCPHJN_101833 | Bacteria | Bacteroidetes | Flavobacteriia | various | various | various | various | unclassified | unclassified | unclassified | unclassified | unclassified | 1 | 0 | 6,65E-05 | 6,32E-06 | 1 - | 5,57E-05 - | 1 - | 6,92E-05 - | 1 - | 6,58E-05 - | 1 - | 6,67E-05 - | 1 - | 7,52E-05 - |
| 5508 | 1 | 0.2um_NOKCPHJN_430361 | Bacteria | Proteobacteri | Gammaprotec | Cellvibrionales | Halieaceae | Chromatocurv | Chromatocurv | metabolism | Energy produc | COG2225 | malate syntha | 247634.GPB2 | 2 | 0,632455532 | 3,77E-05 | 1,21E-05 | 2 - | 3,17E-05 - | 1 - | 1,97E-05 - | 3 - | 5,61E-05 - | 2 - | 3,80E-05 - | 2 - | 4,28E-05 - |
| 5509 | 1 | 0.2um_NOKCPHJN_71934 | Bacteria | Bacteroidetes | Flavobacteriia | Flavobacterial | Flavobacteriac | Polaribacter | Polaribacter s | metabolism | Amino acid tra | COG0174 | glutamine syn | 313594.PI23P | 1,2 | 0,748331477 | 4,75E-05 | 3,06E-05 | 1 - | 3,40E-05 - | 1 - | 4,22E-05 - | 2 - | 8,02E-05 - | 2 - | 8,14E-05 - | 0 - | 0 - |
| 5510 | 1 | 0.2um_NOKCPHJN_449592 | Bacteria | Proteobacteri | Gammaprotec | Oceanospirilla | Saccharospirill | Reinekea | Reinekea blan | metabolism | Energy produc | 0QIP5 | 2,4-dienoyl-co | 314283.MED2 | 0,4 | 0,489897949 | 8,50E-06 | 1,05E-05 | 0 - | 0 - | 0 - | 0 - | 1 - | 1,98E-05 - | 0 - | 0 - | 1 - | 2,27E-05 - |
| 5511 | 1 | 0.2um_NOKCPHJN_567304 | Eukaryota | unclassified | Pelagophycea | Pelagomonad | unclassified | Aureococcus | Aureococcus a | unclassified | unclassified | unclassified | unclassified | unclassified | 9,4 | 1,2 | 0,000519331 | 0,000104738 | 8 - | 0,000367286 - | 8 - | 0,00045606 - | 10 - | 0,000541929 - | 10 - | 0,000549933 - | 11 - | 0,000681448 - |
| 5512 | 1 | 3um_FIJJAFBF_429663 | Bacteria | Proteobacteri | Deltaproteoba | unclassified | unclassified | unclassified | Deltaproteoba | information st | Translation, ri | 0NDJY | One of the pri | 428125.CLOLE | 0,2 | 0,4 | 1,27E-05 | 2,54E-05 | 0 - | 0 - | 0 - | 0 - | 1 - | 6,36E-05 - | 0 - | 0 - | 0 - | 0 - |
| 5513 | 1 | 3um_FIJJAFBF_370787 | Bacteria | Proteobacteri | Gammaprotec | Oceanospirilla | Saccharospirill | Reinekea | Reinekea fors | unclassified | unclassified | unclassified | unclassified | unclassified | 2 | 0 | 5,93E-05 | 5,63E-06 | 2 - | 4,97E-05 - | 2 - | 6,17E-05 - | 2 - | 5,87E-05 - | 2 - | 5,95E-05 - | 2 - | 6,70E-05 - |
| 5514 | 1 | 3um_FIJJAFBF_540105 | unclassified | unclassified | unclassified | unclassified | unclassified | unclassified | unclassified | unclassified | unclassified | unclassified | unclassified | unclassified | 0,6 | 0,489897949 | 5,48E-05 | 4,52E-05 | 1 - | 7,97E-05 - | 1 - | 9,90E-05 - | 0 - | 0 - | 1 - | 9,55E-05 - | 0 - | 0 - |
| 5515 | 1 | 3um_FIJJAFBF_79255 | Bacteria | Bacteroidetes | Flavobacteriia | Flavobacterial | Flavobacteriac | Polaribacter | Polaribacter s | poorly charact | Function unkn | 09EIV | NA | 313598.MED1 | 0,8 | 0,748331477 | 8,27E-05 | 8,03E-05 | 1 - | 9,04E-05 - | 0 - | 0 - | 1 - | 0,000106679 - | 2 - | 0,000216509 - | 0 - | 0 - |
| 5516 | 3 | 3um_FIJJAFBF_03932;3um_FIJJAFBF_297328;3ur | Bacteria | Bacteroidetes | Flavobacteriia | Flavobacterial | Flavobacteriac | Polaribacter | various | unclassified | unclassified | unclassified | unclassified | unclassified | 1,4 | 0,489897949 | 2,74E-05 | 8,06E-06 | 2 - | 3,36E-05 - | 1 - | 2,08E-05 - | 1 - | 1,98E-05 - | 2 - | 4,02E-05 - | 1 - | 2,26E-05 - |
| 5517 | 1 | 3um_FIJJAFBF_617115 | Bacteria | Proteobacteri | Gammaprotec | Pseudomonad | Pseudomonad | Pseudomonas | Pseudomonas | unclassified | unclassified | unclassified | unclassified | unclassified | 2 | 0,632455532 | 4,24E-05 | 9,82E-06 | 3 - | 5,49E-05 - | 2 - | 4,55E-05 - | 2 - | 4,32E-05 - | 2 - | 4,39E-05 - | 1 - | 2,47E-05 - |
| 5518 | 1 | 0.2um_NOKCPHJN_32984 | Bacteria | Proteobacteri | Alphaproteob | Rhodobactera | Rhodobactera | Planktomarina | Planktomarina | unclassified | unclassified | unclassified | unclassified | unclassified | 2,2 | 0,4 | 0,000215108 | 4,19E-05 | 2 - | 0,000163967 - | 2 - | 0,000203598 - | 3 - | 0,000290319 - | 2 - | 0,000196405 - | 2 - | 0,000221249 - |
| 5519 | 1 | 3um_FIJJAFBF_423716 | Eukaryota | Bacillariophyta | Coscinodiscop | Thalassiosirale | Thalassiosirac | Thalassiosira | Thalassiosira p | unclassified | unclassified | unclassified | unclassified | unclassified | 1 | 0 | 7,92E-05 | 7,52E-06 | 1 - | 6,63E-05 - | 1 - | 8,24E-05 - | 1 - | 7,83E-05 - | 1 - | 7,95E-05 - | 1 - | 8,95E-05 - |
| 5520 | 1 | 3um_FIJJAFBF_96340 | Bacteria | Actinobacteria | Actinobacteria | Micrococcales | Microbacteria | Microbacteriu | Microbacteriu | unclassified | unclassified | unclassified | unclassified | unclassified | 1,8 | 1,166190379 | 4,45E-05 | 2,78E-05 | 3 - | 6,56E-05 - | 1 - | 2,71E-05 - | 3 - | 7,74E-05 - | 2 - | 5,24E-05 - | 0 - | 0 - |
| 5521 | 1 | 3um_FIJJAFBF_08200 | Bacteria | Bacteroidetes | Flavobacteriia | Flavobacterial | Flavobacteriac | Psychroserper | Psychroserper | unclassified | unclassified | unclassified | unclassified | unclassified | 0,2 | 0,4 | 9,96E-06 | 1,99E-05 | 0 - | 0 - | 0 - | 0 - | 1 - | 4,98E-05 - | 0 - | 0 - | 0 - | 0 - |
| 5522 | 1 | 3um_FIJJAFBF_24024 | Viruses | unclassified | unclassified | unclassified | Phycodnavirid | unclassified | Chrysochromu | unclassified | unclassified | unclassified | unclassified | unclassified | 0,4 | 0,489897949 | 7,69E-06 | 9,45E-06 | 0 - | 0 - | 0 - | 0 - | 0 - | 0 - | 1 - | 1,81E-05 - | 1 - | 2,04E-05 - |
| 5523 | 1 | 0.2um_NOKCPHJN_583911 | unclassified | unclassified | unclassified | unclassified | unclassified | unclassified | unclassified | unclassified | unclassified | unclassified | unclassified | unclassified | 0,6 | 0,8 | 9,61E-05 | 0,000127751 | 0 - | 0 - | 0 - | 0 - | 2 - | 0,000318782 - | 1 - | 0,000161745 - | 0 - | 0 - |
| 5524 | 1 | 0.2um_NOKCPHJN_127771 | Bacteria | Bacteroidetes | Flavobacteriia | Flavobacterial | Flavobacteriac | Gilvibacter | Gilvibacter sp. | poorly charact | Function unkn | 09C0E | Ragb susd dor | 411154.GFO_ | 1 | 0,894427191 | 2,57E-05 | 2,39E-05 | 1 - | 2,20E-05 - | 2 - | 5,46E-05 - | 2 - | 5,19E-05 - | 0 - | 0 - | 0 - | 0 - |
| 5525 | 1 | 0.2um_NOKCPHJN_611589 | Bacteria | Bacteroidetes | Flavobacteriia | Flavobacterial | Flavobacteriac | Winogradskye | Winogradskye | unclassified | unclassified | unclassified | unclassified | unclassified | 1 | 0,632455532 | 3,24E-05 | 2,07E-05 | 1 - | 2,75E-05 - | 0 - | 0 - | 2 - | 6,48E-05 - | 1 - | 3,29E-05 - | 1 - | 3,71E-05 - |
| 5526 | 1 | 0.2um_NOKCPHJN_176077 | Bacteria | Proteobacteri | Gammaprotec | Cellvibrionales | Porticoccacea | unclassified | SAR92 bacteri | poorly charact | Function unkn | COG4805 | secreted prote | 247639.MGP2 | 0,8 | 0,748331477 | 1,84E-05 | 1,75E-05 | 0 - | 0 - | 2 - | 4,70E-05 - | 1 - | 2,24E-05 - | 1 - | 2,27E-05 - | 0 - | 0 - |
| 5527 | 1 | 3um_FIJJAFBF_599702 | Bacteria | Bacteroidetes | Flavobacteriia | Flavobacterial | Flavobacteriac | Formosa | Formosa sp. H | unclassified | unclassified | unclassified | unclassified | unclassified | 1,6 | 0,489897949 | 6,89E-05 | 2,52E-05 | 1 - | 3,53E-05 - | 2 - | 8,77E-05 - | 1 - | 4,17E-05 - | 2 - | 8,46E-05 - | 2 - | 9,53E-05 - |
| 5528 | 1 | 3um_FIJJAFBF_263679 | Bacteria | Proteobacteri | Alphaproteob | unclassified | unclassified | Candidatus Pu | Candidatus Pu | information st | Translation, ri | 0FN1J | This protein is | 533247.CRD_ | 1 | 0,632455532 | 0,000107378 | 6,95E-05 | 1 - | 9,26E-05 - | 1 - | 0,000114935 - | 2 - | 0,00021852 - | 1 - | 0,000110874 - | 0 - | 0 - |
| 5529 | 1 | 0.2um_NOKCPHJN_591574 | Bacteria | Proteobacteri | Gammaprotec | Cellvibrionales | Porticoccacea | unclassified | various | unclassified | unclassified | unclassified | unclassified | unclassified | 3,4 | 0,489897949 | 6,57E-05 | 6,98E-06 | 4 - | 6,54E-05 - | 3 - | 6,09E-05 - | 3 - | 5,79E-05 - | 4 - | 7,83E-05 - | 3 - | 6,62E-05 - |
| 5530 | 1 | 0.2um_NOKCPHJN_1085731 | Bacteria | Proteobacteri | Alphaproteob | Pelagibacteral | Pelagibacterac | Candidatus Pe | Candidatus Pe | unclassified | unclassified | unclassified | unclassified | unclassified | 7,4 | 1,019803903 | 0,000514376 | 7,05E-05 | 8 - | 0,000468477 - | 9 - | 0,000654423 - | 7 - | 0,000483865 - | 7 - | 0,000491012 - | 6 - | 0,000474106 - |
| 5531 | 1 | 0.2um_NOKCPHJN_251027 | Bacteria | Bacteroidetes | Flavobacteriia | Flavobacterial | Flavobacteriac | Polaribacter | Polaribacter s | poorly charact | Function unkn | 05EK8 | von Willebran | 313598.MED1 | 1,4 | 0,489897949 | 5,42E-05 | 1,70E-05 | 2 - | 6,60E-05 - | 2 - | 8,19E-05 - | 1 - | 3,89E-05 - | 1 - | 3,95E-05 - | 1 - | 4,45E-05 - |
| 5532 | 1 | 0.2um_NOKCPHJN_490832 | Bacteria | Bacteroidetes | Flavobacteriia | unclassified | unclassified | unclassified | Flavobacteria | unclassified | unclassified | unclassified | unclassified | unclassified | 0,8 | 0,748331477 | 4,11E-05 | 3,58E-05 | 2 - | 9,07E-05 - | 0 - | 0 - | 1 - | 5,36E-05 - | 0 - | 0 - | 1 - | 6,12E-05 - |
| 5533 | 1 | 0.2um_NOKCPHJN_30325 | Bacteria | Proteobacteri | Betaproteoba | Burkholderiale | Comamonada | Limnohabitan | various | poorly charact | Function unkn | 0QSE4 | Pfam:DUF400 | 314287.GB220 | 0,8 | 0,4 | 7,14E-05 | 3,67E-05 | 1 - | 7,45E-05 - | 1 - | 9,25E-05 - | 0 - | 0 - | 1 - | 8,93E-05 - | 1 - | 0,000100568 - |
| 5534 | 1 | 0.2um_NOKCPHJN_358475 | Bacteria | Proteobacteri | Alphaproteob | various | various | various | various | poorly charact | Function unkn | 020IJ | Inherit from p | 367336.OM22 | 0,8 | 0,4 | 9,44E-05 | 4,85E-05 | 1 - | 9,89E-05 - | 1 - | 0,000122861 - | 1 - | 0,000116795 - | 0 - | 0 - | 1 - | 0,000133512 - |
| 5535 | 1 | 0.2um_NOKCPHJN_51563 | Bacteria | Proteobacteri | Gammaprotec | Cellvibrionales | Porticoccacea | unclassified | gamma protec | unclassified | unclassified | unclassified | unclassified | unclassified | 0,2 | 0,4 | 3,74E-06 | 7,48E-06 | 0 - | 0 - | 1 - | 1,87E-05 - | 0 - | 0 - | 0 - | 0 - | 0 - | 0 - |
| 5536 | 1 | 3um_FIJJAFBF_300655 | Eukaryota | Bacillariophyta | Coscinodiscop | Thalassiosirale | Thalassiosirac | Thalassiosira | Thalassiosira c | unclassified | unclassified | unclassified | unclassified | unclassified | 0,6 | 0,489897949 | 2,61E-05 | 2,15E-05 | 1 - | 3,81E-05 - | 1 - | 4,73E-05 - | 1 - | 4,50E-05 - | 0 - | 0 - | 0 - | 0 - |
| 5537 | 1 | 0.2um_NOKCPHJN_217490 | Bacteria | Proteobacteri | Gammaprotec | Oceanospirilla | Saccharospirill | Reinekea | Reinekea fors | metabolism | Inorganic ion t | 05EKV | Phosphonate | 1007105.PT7_ | 13,2 | 0,4 | 0,000457946 | 3,35E-05 | 14 - | 0,000407836 - | 13 - | 0,00047024 - | 13 - | 0,000447023 - | 13 - | 0,000453625 - | 13 - | 0,000511007 - |
| 5538 | 1 | 0.2um_NOKCPHJN_298547 | Bacteria | Bacteroidetes | Flavobacteriia | Flavobacterial | Flavobacteriac | Formosa | Formosa sp. H | information st | Transcription | 0P7SF | DNA-depende | 983544.Lacal_ | 1,8 | 0,4 | 1,91E-05 | 3,78E-06 | 2 - | 1,81E-05 - | 2 - | 2,24E-05 - | 2 - | 2,13E-05 - | 2 - | 2,17E-05 - | 1 - | 1,22E-05 - |
| 5539 | 1 | 0.2um_NOKCPHJN_164465 | Eukaryota | unclassified | Oomycetes | Albuginales | Albuginaceae | Albugo | Albugo laibach | poorly charact | Function unkn | 109SB | MYND finger | 164328.Phyra | 0,2 | 0,4 | 1,33E-06 | 2,66E-06 | 0 - | 0 - | 1 - | 6,65E-06 - | 0 - | 0 - | 0 - | 0 - | 0 - | 0 - |
| 5540 | 1 | 0.2um_NOKCPHJN_59574 | Bacteria | Proteobacteri | Gammaprotec | unclassified | unclassified | unclassified | Gammaprotec | unclassified | unclassified | unclassified | unclassified | unclassified | 0,8 | 0,4 | 8,27E-05 | 4,16E-05 | 0 - | 0 - | 1 - | 0,000103275 - | 1 - | 9,82E-05 - | 1 - | 9,96E-05 - | 1 - | 0,000112228 - |
| 5541 | 1 | 3um_FIJJAFBF_392895 | Bacteria | Proteobacteri | Gammaprotec | unclassified | unclassified | unclassified | various | unclassified | unclassified | unclassified | unclassified | unclassified | 1 | 0 | 9,79E-05 | 9,29E-06 | 1 - | 8,20E-05 - | 1 - | 0,000101799 - | 1 - | 9,68E-05 - | 1 - | 9,82E-05 - | 1 - | 0,000110625 - |
| 5542 | 1 | 3um_FIJJAFBF_596860 | Bacteria | unclassified | unclassified | unclassified | unclassified | unclassified | uncultured ba | unclassified | unclassified | unclassified | unclassified | unclassified | 1 | 0 | 2,68E-05 | 2,55E-06 | 1 - | 2,25E-05 - | 1 - | 2,79E-05 - | 1 - | 2,65E-05 - | 1 - | 2,69E-05 - | 1 - | 3,03E-05 - |
| 5543 | 2 | 3um_FIJJAFBF_33016;3um_FIJJAFBF_714263 | Bacteria | Proteobacteri | Gammaprotec | Alteromonada | Alteromonada | Paraglaciecola | various | cellular proces | Signal transdu | 0QJ37 | two-compone | various | 0,6 | 0,489897949 | 3,51E-05 | 2,92E-05 | 1 - | 4,88E-05 - | 1 - | 6,06E-05 - | 0 - | 0 - | 0 - | 0 - | 1 - | 6,59E-05 - |
| 5544 | 1 | 0.2um_NOKCPHJN_137058 | Bacteria | Bacteroidetes | Flavobacteriia | Flavobacterial | Flavobacteriac | Wenyingzhuai | various | unclassified | unclassified | unclassified | unclassified | unclassified | 0,6 | 0,489897949 | 9,29E-05 | 7,66E-05 | 1 - | 0,000135031 - | 1 - | 0,000167669 - | 0 - | 0 - | 1 - | 0,000161745 - | 0 - | 0 - |
| 5545 | 1 | 3um_FIJJAFBF_27218 | Bacteria | Bacteroidetes | Flavobacteriia | unclassified | unclassified | unclassified | Flavobacteria | unclassified | unclassified | unclassified | unclassified | unclassified | 0,6 | 0,489897949 | 7,04E-05 | 5,77E-05 | 0 - | 0 - | 1 - | 0,000115869 - | 1 - | 0,000110148 - | 0 - | 0 - | 1 - | 0,000125914 - |
| 5546 | 1 | 3um_FIJJAFBF_62795 | Bacteria | Bacteroidetes | Flavobacteriia | Flavobacterial | Flavobacteriac | Polaribacter | Polaribacter s | unclassified | unclassified | unclassified | unclassified | unclassified | 1,2 | 0,748331477 | 8,99E-05 | 5,86E-05 | 0 - | 0 - | 2 - | 0,000147688 - | 1 - | 7,02E-05 - | 1 - | 7,12E-05 - | 2 - | 0,000160492 - |
| 5547 | 1 | 0.2um_NOKCPHJN_678783 | Bacteria | unclassified | unclassified | unclassified | unclassified | unclassified | uncultured ba | metabolism | Inorganic ion t | 05C27 | TonB-depende | 313596.RB250 | 2,8 | 0,4 | 3,82E-05 | 6,19E-06 | 3 - | 3,44E-05 - | 2 - | 2,85E-05 - | 3 - | 4,06E-05 - | 3 - | 4,12E-05 - | 3 - | 4,64E-05 - |
| 5548 | 1 | 0.2um_NOKCPHJN_342411 | Bacteria | Bacteroidetes | Flavobacteriia | Flavobacterial | Flavobacteriac | Polaribacter | Polaribacter s | information st | Transcription | 05D27 | DNA-depende | 313594.PI23P | 4 | 0,632455532 | 3,89E-05 | 8,51E-06 | 3 - | 2,42E-05 - | 5 - | 5,00E-05 - | 4 - | 3,80E-05 - | 4 - | 3,86E-05 - | 4 - | 4,35E-05 - |
| 5549 | 1 | 0.2um_NOKCPHJN_375462 | Bacteria | Proteobacteri | Alphaproteob | Pelagibacteral | Pelagibacterac | Candidatus Pe | Candidatus Pe | unclassified | unclassified | unclassified | unclassified | unclassified | 1,4 | 0,489897949 | 4,49E-05 | 1,59E-05 | 1 - | 2,69E-05 - | 1 - | 3,34E-05 - | 2 - | 6,35E-05 - | 2 - | 6,44E-05 - | 1 - | 3,63E-05 - |
| 5550 | 1 | 0.2um_NOKCPHJN_42232 | Bacteria | Proteobacteri | Gammaprotec | Cellvibrionales | Porticoccacea | unclassified | SAR92 bacteri | metabolism | Carbohydrate | 0QIP7 | Catalyzes the | 314287.GB220 | 1,4 | 0,8 | 5,46E-05 | 3,08E-05 | 2 - | 6,83E-05 - | 1 - | 4,24E-05 - | 2 - | 8,06E-05 - | 2 - | 8,18E-05 - | 0 - | 0 - |
| 5551 | 1 | 3um_FIJJAFBF_382754 | unclassified | unclassified | unclassified | unclassified | unclassified | unclassified | unclassified | unclassified | unclassified | unclassified | unclassified | unclassified | 1,2 | 0,748331477 | 6,41E-05 | 3,87E-05 | 2 - | 9,41E-05 - | 1 - | 5,84E-05 - | 1 - | 5,55E-05 - | 2 - | 0,000112691 - | 0 - | 0 - |
| 5552 | 1 | 3um_FIJJAFBF_157377 | Bacteria | Bacteroidetes | Flavobacteriia | Flavobacterial | Flavobacteriac | Gaetbulibacte | Gaetbulibacte | information st | Translation, ri | 09AI8 | Associates wit | 391603.FBALC | 0,6 | 0,489897949 | 2,64E-05 | 2,17E-05 | 0 - | 0 - | 0 - | 0 - | 1 - | 4,18E-05 - | 1 - | 4,24E-05 - | 1 - | 4,78E-05 - |
| 5553 | 2 | 0.2um_NOKCPHJN_817281;3um_FIJJAFBF_2073§ | unclassified | unclassified | unclassified | unclassified | unclassified | unclassified | unclassified | unclassified | unclassified | unclassified | unclassified | unclassified | 1 | 0,632455532 | 9,35E-05 | 5,33E-05 | 2 - | 0,000166343 - | 1 - | 0,000103275 - | 1 - | 9,82E-05 - | 1 - | 9,96E-05 - | 0 - | 0 - |
| 5554 | 1 | 0.2um_NOKCPHJN_296782 | Bacteria | Bacteroidetes | Flavobacteriia | Flavobacterial | Flavobacteriac | Marinirhabdus | Marinirhabdus | unclassified | unclassified | unclassified | unclassified | unclassified | 6,8 | 0,4 | 0,00017293 | 2,02E-05 | 7 - | 0,000149061 - | 7 - | 0,000185089 - | 6 - | 0,000150815 - | 7 - | 0,00017855 - | 7 - | 0,000201136 - |
| 5555 | 1 | 0.2um_NOKCPHJN_780014 | Bacteria | unclassified | unclassified | unclassified | unclassified | unclassified | uncultured ba | poorly charact | Function unkn | 09DAY | NA | 50743.SCB49_ | 0,8 | 0,748331477 | 1,79E-05 | 1,57E-05 | 2 - | 4,03E-05 - | 1 - | 2,50E-05 - | 0 - | 0 - | 1 - | 2,42E-05 - | 0 - | 0 - |
| 5556 | 1 | 0.2um_NOKCPHJN_321673 | Bacteria | Bacteroidetes | Flavobacteriia | Flavobacterial | Cryomorphace | unclassified | Cryomorphace | information st | Transcription | 05D27 | DNA-depende | 886377.Murru | 1,4 | 0,489897949 | 1,56E-05 | 4,92E-06 | 2 - | 1,91E-05 - | 2 - | 2,37E-05 - | 1 - | 1,12E-05 - | 1 - | 1,14E-05 - | 1 - | 1,29E-05 - |
| 5557 | 1 | 3um_FIJJAFBF_242856 | Bacteria | Bacteroidetes | Flavobacteriia | Flavobacterial | Flavobacteriac | Flaviramulus | Flaviramulus i | unclassified | unclassified | unclassified | unclassified | unclassified | 0,4 | 0,489897949 | 8,05E-06 | 9,86E-06 | 0 - | 0 - | 0 - | 0 - | 1 - | 2,00E-05 - | 1 - | 2,03E-05 - | 0 - | 0 - |
| 5558 | 1 | 0.2um_NOKCPHJN_364496 | Bacteria | Proteobacteri | Gammaprotec | Cellvibrionales | Halieaceae | unclassified | gamma protec | information st | Transcription | 0QKAX | Negative regu | 247639.MGP2 | 0,2 | 0,4 | 6,67E-06 | 1,33E-05 | 1 - | 3,34E-05 - | 0 - | 0 - | 0 - | 0 - | 0 - | 0 - | 0 - | 0 - |
| 5559 | 1 | 3um_FIJJAFBF_284237 | Bacteria | Bacteroidetes | Flavobacteriia | various | various | various | various | unclassified | unclassified | unclassified | unclassified | unclassified | 0,2 | 0,4 | 9,22E-06 | 1,84E-05 | 1 - | 4,61E-05 - | 0 - | 0 - | 0 - | 0 - | 0 - | 0 - | 0 - | 0 - |
| 5560 | 1 | 3um_FIJJAFBF_436835 | Bacteria | Actinobacteria | Actinobacteria | Micrococcales | Microbacteria | Candidatus Ac | Candidatus Ac | information st | Translation, ri | COG0088 | One of the pri | 281090.Lxx20 | 1,2 | 0,4 | 7,51E-05 | 1,66E-05 | 2 - | 0,000107772 - | 1 - | 6,69E-05 - | 1 - | 6,36E-05 - | 1 - | 6,45E-05 - | 1 - | 7,27E-05 - |
| 5561 | 1 | 0.2um_NOKCPHJN_21190 | Bacteria | Proteobacteri | Alphaproteob | Rhodobactera | Rhodobactera | Planktomarina | Planktomarina | poorly charact | Function unkn | 05ERF | Extracellular s | 388401.RB215 | 2,2 | 0,748331477 | 5,79E-05 | 1,95E-05 | 2 - | 4,47E-05 - | 2 - | 5,55E-05 - | 3 - | 7,91E-05 - | 3 - | 8,02E-05 - | 1 - | 3,01E-05 - |
| 5562 | 1 | 0.2um_NOKCPHJN_286790 | Eukaryota | unclassified | Pelagophycea | Pelagomonad | unclassified | Aureococcus | Aureococcus a | information st | Translation, ri | KOG0886 | Ribosomal pro | 54126.PPA171 | 0,2 | 0,4 | 1,72E-05 | 3,43E-05 | 0 - | 0 - | 1 - | 8,59E-05 - | 0 - | 0 - | 0 - | 0 - | 0 - | 0 - |
| 5563 | 1 | 0.2um_NOKCPHJN_94575 | Bacteria | Bacteroidetes | Flavobacteriia | Flavobacterial | Crocinitomica | unclassified | Crocinitomica | metabolism | Amino acid tra | 09AZK | oligopeptidase | 50743.SCB49_ | 0,2 | 0,4 | 3,14E-06 | 6,29E-06 | 1 - | 1,57E-05 - | 0 - | 0 - | 0 - | 0 - | 0 - | 0 - | 0 - | 0 - |
| 5564 | 1 | 3um_FIJJAFBF_785179 | Eukaryota | Euglenida | unclassified | Euglenales | Euglenaceae | Trachelomona | Trachelomona | cellular proces | Posttranslatio | 0IEKR | One of the cor | 3055.DAA009 | 0,8 | 0,4 | 5,65E-05 | 2,90E-05 | 1 - | 5,98E-05 - | 0 - | 0 - | 1 - | 7,06E-05 - | 1 - | 7,16E-05 - | 1 - | 8,07E-05 - |
| 5565 | 2 | 0.2um_NOKCPHJN_132167;0.2um_NOKCPHJN_2 | unclassified | unclassified | unclassified | unclassified | unclassified | unclassified | unclassified | unclassified | unclassified | unclassified | unclassified | unclassified | 0,8 | 0,748331477 | 7,94E-05 | 7,60E-05 | 1 - | 8,63E-05 - | 1 - | 0,000107157 - | 2 - | 0,000203733 - | 0 - | 0 - | 0 - | 0 - |
| 5566 | 1 | 3um_FIJJAFBF_172525 | Bacteria | Proteobacteri | Betaproteoba | Burkholderiale | Comamonada | Hydrogenoph | various | unclassified | unclassified | unclassified | unclassified | unclassified | 21,6 | 1,743559577 | 0,001636464 | 0,000129824 | 24 - | 0,001530357 - | 21 - | 0,00166272 - | 23 - | 0,001731162 - | 19 - | 0,001451213 - | 21 - | 0,001806869 - |
| 5567 | 1 | 3um_FIJJAFBF_604555 | Bacteria | Proteobacteri | Gammaprotec | Arenicellales | Arenicellaceae | Arenicella | Arenicella xan | information st | Translation, ri | COG0088 | One of the pri | 298386.PBPRA | 0,8 | 0,4 | 5,43E-05 | 2,79E-05 | 1 - | 5,74E-05 - | 0 - | 0 - | 1 - | 6,77E-05 - | 1 - | 6,87E-05 - | 1 - | 7,74E-05 - |
| 5568 | 1 | 3um_FIJJAFBF_195017 | Bacteria | Bacteroidetes | Flavobacteriia | Flavobacterial | Flavobacteriac | Aequorivita | Aequorivita ar | unclassified | unclassified | unclassified | unclassified | unclassified | 1,8 | 0,4 | 0,000177667 | 4,42E-05 | 2 - | 0,000165146 - | 2 - | 0,000205063 - | 1 - | 9,75E-05 - | 2 - | 0,000197818 - | 2 - | 0,000222841 - |
| 5569 | 1 | 3um_FIJJAFBF_674299 | Bacteria | Bacteroidetes | Flavobacteriia | Flavobacterial | Crocinitomica | unclassified | Crocinitomica | cellular proces | Cell wall/mem | COG2885 | Ompa motb d | 203122.Sde_ | 0,2 | 0,4 | 4,04E-06 | 8,08E-06 | 0 - | 0 - | 0 - | 0 - | 1 - | 2,02E-05 - | 0 - | 0 - | 0 - | 0 - |
| 5570 | 1 | 0.2um_NOKCPHJN_154824 | Bacteria | Proteobacteri | Gammaprotec | Alteromonada | Psychromonac | Psychromonas | Psychromonas | poorly charact | Function unkn | COG1744 | basic membra | 722419.PH50 | 7,4 | 1,854723699 | 0,000270994 | 6,65E-05 | 7 - | 0,000215978 - | 6 - | 0,000229869 - | 11 - | 0,000400619 - | 7 - | 0,000258705 - | 6 - | 0,000249798 - |
| 5571 | 1 | 0.2um_NOKCPHJN_457438 | Bacteria | Bacteroidetes | unclassified | unclassified | unclassified | unclassified | Bacteroidetes | cellular proces | Posttranslatio | 05E6M | Sulphatase-m | 926562.Oweh | 2,2 | 0,748331477 | 6,22E-05 | 1,78E-05 | 3 - | 7,31E-05 - | 2 - | 6,05E-05 - | 3 - | 8,63E-05 - | 2 - | 5,84E-05 - | 1 - | 3,29E-05 - |

| | | | | | | | | | | | | | | | | | | | | | | | | | | | | |
|---|---|---|---|---|---|---|---|---|---|---|---|---|---|---|---|---|---|---|---|---|---|---|---|---|---|---|---|---|
| 5572 | 1 | 3um_FIJJAFBF_62128 | Bacteria | Bacteroidetes | unclassified | unclassified | unclassified | unclassified | Bacteroidetes | unclassified | unclassified | unclassified | unclassified | unclassified | 0,4 | 0,489897949 | 6,39E-05 | 7,87E-05 | 1 - | 0,000145287 - | 0 - | 0 - | 0 - | 0 - | 1 - | 0,00017403 - | 0 - | 0 - |
| 5573 | 1 | 3um_FIJJAFBF_115388 | Bacteria | Proteobacteri | Alphaproteob | Pelagibacteral | Pelagibacterac | Candidatus Pe | Candidatus Pe | metabolism | Secondary me | 01Q86 | extracellular s | 439493.PB721 | 1,8 | 0,4 | 0,000202096 | 4,95E-05 | 2 - | 0,000188159 - | 2 - | 0,000233637 - | 2 - | 0,000222102 - | 1 - | 0,000112691 - | 2 - | 0,000253893 - |
| 5574 | 1 | 0.2um_NOKCPHJN_280731 | Eukaryota | unclassified | unclassified | Prymnesiales | Chrysochromu | Chrysochromu | Chrysochromu | information st | Translation, ri | 11UVJ | Ribosomal prc | 272952.HpaP8 | 0,4 | 0,489897949 | 3,02E-05 | 3,69E-05 | 0 - | 0 - | 0 - | 0 - | 1 - | 7,49E-05 - | 1 - | 7,60E-05 - | 0 - | 0 - |
| 5575 | 1 | 3um_FIJJAFBF_42273 | Bacteria | Proteobacteri | Alphaproteob | Rhodobactera | Rhodobactera | Thalassobius | Thalassobius a | poorly charact | Function unkn | 05ERF | Extracellular s | 384765.SIAM6 | 1,8 | 0,4 | 4,79E-05 | 1,33E-05 | 1 - | 2,19E-05 - | 2 - | 5,44E-05 - | 2 - | 5,17E-05 - | 2 - | 5,25E-05 - | 2 - | 5,91E-05 - |
| 5576 | 1 | 3um_FIJJAFBF_63160 | Bacteria | Proteobacteri | Betaproteoba | Burkholderiale | Comamonada | Pseudorhodof | Pseudorhodof | metabolism | Energy produc | 07QSN | dihydrolipoam | 365046.Rta_1 | 1,2 | 0,748331477 | 3,06E-05 | 2,08E-05 | 1 - | 2,08E-05 - | 2 - | 5,15E-05 - | 1 - | 2,45E-05 - | 0 - | 0 - | 2 - | 5,60E-05 - |
| 5577 | 1 | 3um_FIJJAFBF_30331 | Bacteria | Bacteroidetes | Flavobacteriia | Flavobacterial | Flavobacteriac | Polaribacter | Polaribacter s | metabolism | Inorganic ion t | 09B4C | tonB-depende | 313594.PI23P | 4,4 | 0,8 | 7,70E-05 | 1,50E-05 | 4 - | 5,87E-05 - | 4 - | 7,29E-05 - | 6 - | 0,000103951 - | 4 - | 7,03E-05 - | 4 - | 7,92E-05 - |
| 5578 | 1 | 3um_FIJJAFBF_175545 | Bacteria | Proteobacteri | Gammaprote | Oceanospirilla | Oceanospirilla | Bacterioplane | Bacterioplane | unclassified | unclassified | unclassified | unclassified | unclassified | 0,4 | 0,489897949 | 5,72E-05 | 7,07E-05 | 1 - | 0,00012753 - | 1 - | 0,000158354 - | 0 - | 0 - | 0 - | 0 - | 0 - | 0 - |
| 5579 | 1 | 3um_FIJJAFBF_242833 | Bacteria | Bacteroidetes | Flavobacteriia | Flavobacterial | Flavobacteriac | Polaribacter | Polaribacter s | unclassified | unclassified | unclassified | unclassified | unclassified | 1,4 | 0,489897949 | 4,32E-05 | 1,36E-05 | 2 - | 5,26E-05 - | 2 - | 6,54E-05 - | 1 - | 3,11E-05 - | 1 - | 3,15E-05 - | 1 - | 3,55E-05 - |
| 5580 | 1 | 3um_FIJJAFBF_317104 | Eukaryota | unclassified | Pelagophycea | Pelagomonad | unclassified | Aureococcus | Aureococcus a | unclassified | unclassified | unclassified | unclassified | unclassified | 0,4 | 0,489897949 | 1,38E-05 | 1,70E-05 | 0 - | 0 - | 0 - | 0 - | 1 - | 3,22E-05 - | 0 - | 0 - | 1 - | 3,68E-05 - |
| 5581 | 1 | 3um_FIJJAFBF_772926 | Bacteria | Proteobacteri | Gammaprote | Cellvibrionales | Porticoccacea | unclassified | gamma prote | unclassified | unclassified | unclassified | unclassified | unclassified | 1 | 0 | 5,98E-05 | 5,68E-06 | 1 - | 5,01E-05 - | 1 - | 6,22E-05 - | 1 - | 5,92E-05 - | 1 - | 6,00E-05 - | 1 - | 6,76E-05 - |
| 5582 | 3 | 0.2um_NOKCPHJN_612909;0.2um_NOKCPHJN_7 | Bacteria | Proteobacteri | Alphaproteob | Pelagibacteral | Pelagibacterac | Candidatus Pe | Candidatus Pe | information st | Translation, ri | 01RJH | 30S ribosomal | 335992.SAR11 | 2,6 | 1,019803903 | 0,000129551 | 5,22E-05 | 2 - | 8,41E-05 - | 3 - | 0,000156614 - | 3 - | 0,000148882 - | 4 - | 0,000201441 - | 1 - | 5,67E-05 - |
| 5583 | 1 | 3um_FIJJAFBF_264684 | Bacteria | Proteobacteri | Gammaprote | Oceanospirilla | Kangiellaceae | Kangiella | Kangiella spor | information st | Translation, ri | 17FIT | 50s ribosomal | 247639.MGP2 | 1 | 0 | 0,000175676 | 1,67E-05 | 1 - | 0,00014715 - | 1 - | 0,000182716 - | 1 - | 0,000173695 - | 1 - | 0,000176261 - | 1 - | 0,000198557 - |
| 5584 | 1 | 3um_FIJJAFBF_594162 | Eukaryota | unclassified | Pelagophycea | Pelagomonad | unclassified | Pelagomonas | uncultured Pe | unclassified | unclassified | unclassified | unclassified | unclassified | 7 | 1,095445115 | 0,000295638 | 4,64E-05 | 7 - | 0,000248742 - | 7 - | 0,000308864 - | 9 - | 0,000377505 - | 6 - | 0,000255387 - | 6 - | 0,000287693 - |
| 5585 | 1 | 3um_FIJJAFBF_545832 | Bacteria | Bacteroidetes | Flavobacteriia | Flavobacterial | Flavobacteriac | Polaribacter | Polaribacter s | unclassified | unclassified | unclassified | unclassified | unclassified | 3 | 1,549193338 | 0,000250301 | 9,87E-05 | 6 - | 0,000438637 - | 2 - | 0,000181553 - | 3 - | 0,000258883 - | 2 - | 0,000175138 - | 2 - | 0,000197292 - |
| 5586 | 1 | 0.2um_NOKCPHJN_953100 | Bacteria | Proteobacteri | Epsilonproteo | Campylobacte | Helicobactera | Sulfurimonas | Sulfurimonas | unclassified | unclassified | unclassified | unclassified | unclassified | 0,6 | 0,489897949 | 1,36E-05 | 1,12E-05 | 0 - | 0 - | 0 - | 0 - | 1 - | 2,15E-05 - | 1 - | 2,19E-05 - | 1 - | 2,46E-05 - |
| 5587 | 1 | 0.2um_NOKCPHJN_754949 | Bacteria | various | various | various | various | various | various | metabolism | Amino acid tra | 099QE | chorismate m | 391603.FBALC | 0,4 | 0,489897949 | 2,59E-05 | 3,17E-05 | 0 - | 0 - | 0 - | 0 - | 1 - | 6,42E-05 - | 1 - | 6,52E-05 - | 0 - | 0 - |
| 5588 | 1 | 3um_FIJJAFBF_83928 | Bacteria | Bacteroidetes | Flavobacteriia | Flavobacterial | Flavobacteriac | Maribacter | Maribacter pc | metabolism | Amino acid tra | 1AFXU | peptidase S9 | 760192.Halhy | 1 | 0 | 3,03E-05 | 2,88E-06 | 1 - | 2,54E-05 - | 1 - | 3,15E-05 - | 1 - | 3,00E-05 - | 1 - | 3,04E-05 - | 1 - | 3,43E-05 - |
| 5589 | 1 | 3um_FIJJAFBF_34663 | Bacteria | Bacteroidetes | Flavobacteriia | Flavobacterial | Flavobacteriac | Arenibacter | Arenibacter e | information st | Replication, re | COG0513 | purine NTP-de | 688270.Celal | 2,2 | 1,6 | 4,47E-05 | 3,13E-05 | 4 - | 7,21E-05 - | 2 - | 4,47E-05 - | 4 - | 8,51E-05 - | 1 - | 2,16E-05 - | 0 - | 0 - |
| 5590 | 1 | 0.2um_NOKCPHJN_181011 | Bacteria | Proteobacteri | Gammaprote | Oceanospirilla | Saccharospirill | Reinekea | Reinekea fors | metabolism | Coenzyme tra | 16T4M | short chain de | 314283.MED2 | 1,2 | 0,4 | 6,90E-05 | 1,53E-05 | 2 - | 9,89E-05 - | 1 - | 6,14E-05 - | 1 - | 5,84E-05 - | 1 - | 5,93E-05 - | 1 - | 6,68E-05 - |
| 5591 | 1 | 3um_FIJJAFBF_167273 | Bacteria | Bacteroidetes | Flavobacteriia | Flavobacterial | Flavobacteriac | Dokdonia | Dokdonia paci | unclassified | unclassified | unclassified | unclassified | unclassified | 5,6 | 0,489897949 | 0,000246823 | 1,74E-05 | 6 - | 0,000222867 - | 5 - | 0,000230613 - | 6 - | 0,000263072 - | 6 - | 0,000266958 - | 5 - | 0,000250606 - |
| 5592 | 1 | 0.2um_NOKCPHJN_470843 | Bacteria | Proteobacteri | Alphaproteob | Rhodobactera | Rhodobactera | Planktomarina | Planktomarina | unclassified | unclassified | unclassified | unclassified | unclassified | 4 | 1,095445115 | 0,000444974 | 8,98E-05 | 6 - | 0,000569141 - | 3 - | 0,000353353 - | 3 - | 0,000335906 - | 4 - | 0,00045449 - | 4 - | 0,000511982 - |
| 5593 | 1 | 3um_FIJJAFBF_71483 | Bacteria | Proteobacteri | Alphaproteob | Rhodobactera | Rhodobactera | unclassified | Rhodobactera | information st | Translation, ri | COG0050 | This protein p | 314265.R2601 | 6,2 | 0,4 | 0,000217562 | 2,71E-05 | 6 - | 0,000176128 - | 7 - | 0,000255149 - | 6 - | 0,000207901 - | 6 - | 0,000210972 - | 6 - | 0,000237659 - |
| 5594 | 1 | 3um_FIJJAFBF_54367 | Bacteria | Proteobacteri | Alphaproteob | Rhodobactera | Rhodobactera | Planktomarina | Planktomarina | metabolism | Amino acid tra | 01QBN | Extracellular s | 633131.TR2A6 | 0,8 | 0,4 | 1,77E-05 | 9,09E-06 | 1 - | 1,85E-05 - | 1 - | 2,29E-05 - | 0 - | 0 - | 1 - | 2,21E-05 - | 1 - | 2,49E-05 - |
| 5595 | 1 | 3um_FIJJAFBF_592957 | Bacteria | Proteobacteri | Gammaprote | Oceanospirilla | Saccharospirill | Reinekea | Reinekea blan | unclassified | unclassified | unclassified | unclassified | unclassified | 4,6 | 0,489897949 | 0,0001275 | 2,19E-05 | 4 - | 9,24E-05 - | 4 - | 0,000114703 - | 5 - | 0,0001363 - | 5 - | 0,000138313 - | 5 - | 0,000155809 - |
| 5596 | 2 | 0.2um_NOKCPHJN_1007598;0.2um_NOKCPHJN_ | Bacteria | Bacteroidetes | Flavobacteriia | Flavobacterial | Flavobacteriac | Polaribacter | various | poorly charact | Function unkn | 0P884 | membrane | 313603.FB217 | 1,4 | 0,489897949 | 6,34E-05 | 1,81E-05 | 2 - | 7,78E-05 - | 1 - | 4,83E-05 - | 2 - | 9,19E-05 - | 1 - | 4,66E-05 - | 1 - | 5,25E-05 - |
| 5597 | 2 | 0.2um_NOKCPHJN_57913;3um_FIJJAFBF_30941 | Bacteria | Proteobacteri | Alphaproteob | Rhodobactera | Rhodobactera | various | various | unclassified | unclassified | unclassified | unclassified | unclassified | 0,6 | 0,489897949 | 3,71E-05 | 3,03E-05 | 0 - | 0 - | 1 - | 6,36E-05 - | 1 - | 6,05E-05 - | 1 - | 6,14E-05 - | 0 - | 0 - |
| 5598 | 1 | 3um_FIJJAFBF_807498 | Bacteria | Bacteroidetes | Flavobacteriia | Flavobacterial | Flavobacteriac | Robiginitalea | Robiginitalea | cellular proces | Posttranslatio | 099Y4 | peptidase, M1 | 487797.Flav3 | 0,8 | 1,166190379 | 2,18E-05 | 3,06E-05 | 3 - | 7,81E-05 - | 0 - | 0 - | 1 - | 3,07E-05 - | 0 - | 0 - | 0 - | 0 - |
| 5599 | 1 | 3um_FIJJAFBF_128963 | Bacteria | Proteobacteri | Gammaprote | Oceanospirilla | Saccharospirill | Reinekea | Reinekea fors | unclassified | unclassified | unclassified | unclassified | unclassified | 0,8 | 0,748331477 | 4,68E-05 | 4,07E-05 | 2 - | 0,000102939 - | 0 - | 0 - | 0 - | 0 - | 1 - | 6,17E-05 - | 1 - | 6,95E-05 - |
| 5600 | 1 | 3um_FIJJAFBF_175180 | Bacteria | Proteobacteri | Alphaproteob | Rhodobactera | unclassified | unclassified | Rhodobactera | poorly charact | Function unkn | 021QX | NA | 439496.RBY4I | 1 | 0 | 8,84E-05 | 8,39E-06 | 1 - | 7,40E-05 - | 1 - | 9,19E-05 - | 1 - | 8,74E-05 - | 1 - | 8,87E-05 - | 1 - | 9,99E-05 - |
| 5601 | 1 | 3um_FIJJAFBF_131604 | Bacteria | Proteobacteri | Gammaprote | Oceanospirilla | Alcanivoraca | Alcanivorax | Alcanivorax sp | metabolism | Inorganic ion t | 0QJ3Y | Part of the AB | 377629.TERTU | 0,2 | 0,4 | 9,34E-06 | 1,87E-05 | 0 - | 0 - | 0 - | 0 - | 1 - | 4,67E-05 - | 0 - | 0 - | 0 - | 0 - |
| 5602 | 1 | 3um_FIJJAFBF_399088 | Bacteria | Bacteroidetes | Flavobacteriia | Flavobacterial | Flavobacteriac | Polaribacter | Polaribacter s | unclassified | unclassified | unclassified | unclassified | unclassified | 3,4 | 0,489897949 | 0,000185941 | 1,81E-05 | 4 - | 0,000185124 - | 3 - | 0,000172402 - | 4 - | 0,00021852 - | 3 - | 0,00016631 - | 3 - | 0,000187348 - |
| 5603 | 1 | 0.2um_NOKCPHJN_126548 | Archaea | Euryarchaeota | unclassified | unclassified | unclassified | unclassified | Euryarchaeota | poorly charact | Function unkn | 11TP8 | Erf family | 992406.RIA_0 | 0,4 | 0,489897949 | 6,03E-05 | 7,43E-05 | 1 - | 0,000138285 - | 0 - | 0 - | 1 - | 0,000163232 - | 0 - | 0 - | 0 - | 0 - |
| 5604 | 1 | 3um_FIJJAFBF_320297 | Bacteria | Proteobacteri | Gammaprote | Oceanospirilla | Saccharospirill | Reinekea | Reinekea fors | unclassified | unclassified | unclassified | unclassified | unclassified | 2,4 | 0,489897949 | 0,000148935 | 3,96E-05 | 2 - | 0,000102939 - | 2 - | 0,00012782 - | 3 - | 0,000182263 - | 2 - | 0,000123303 - | 3 - | 0,000208351 - |
| 5605 | 1 | 3um_FIJJAFBF_285166 | Bacteria | Proteobacteri | Betaproteoba | Burkholderiale | unclassified | Leptothrix | Leptothrix ch | unclassified | unclassified | unclassified | unclassified | unclassified | 1 | 0,632455532 | 2,03E-05 | 1,16E-05 | 2 - | 3,61E-05 - | 1 - | 2,24E-05 - | 1 - | 2,13E-05 - | 1 - | 2,16E-05 - | 0 - | 0 - |
| 5606 | 1 | 3um_FIJJAFBF_111311 | Viruses | unclassified | unclassified | unclassified | unclassified | unclassified | uncultured M | cellular proces | Intracellular tr | 03K5Y | intracellular p | 436907.XP_00 | 7,6 | 1,2 | 5,32E-05 | 7,07E-06 | 9 - | 5,32E-05 - | 7 - | 5,14E-05 - | 6 - | 4,19E-05 - | 9 - | 6,37E-05 - | 7 - | 5,59E-05 - |
| 5607 | 1 | 0.2um_NOKCPHJN_125415 | Bacteria | Proteobacteri | Alphaproteob | Rhodobactera | Rhodobactera | Labrenzia | Labrenzia alex | poorly charact | Function unkn | 11G4W | NA | 244592.SADFL | 0,2 | 0,4 | 1,64E-06 | 3,28E-06 | 1 - | 8,21E-06 - | 0 - | 0 - | 0 - | 0 - | 0 - | 0 - | 0 - | 0 - |
| 5608 | 1 | 0.2um_NOKCPHJN_111274 | Bacteria | Proteobacteri | Gammaprote | Cellvibrionales | Porticoccacea | unclassified | gamma prote | unclassified | unclassified | unclassified | unclassified | unclassified | 11,2 | 1,6 | 0,000173013 | 2,23E-05 | 13 - | 0,000169364 - | 13 - | 0,0002103 - | 11 - | 0,000169161 - | 9 - | 0,000140448 - | 10 - | 0,000175794 - |
| 5609 | 1 | 3um_FIJJAFBF_350307 | Bacteria | Proteobacteri | Gammaprote | Oceanospirilla | Oleiphilaceae | Oleiphilus | various | information st | Transcription | 16PMT | DNA-depende | 491952.Mar18 | 5,6 | 1,2 | 0,000236816 | 6,71E-05 | 4 - | 0,000139546 - | 5 - | 0,000216594 - | 5 - | 0,0002059 - | 7 - | 0,000292518 - | 7 - | 0,00032952 - |
| 5610 | 1 | 0.2um_NOKCPHJN_492864 | Eukaryota | unclassified | Pelagophycea | Pelagomonad | unclassified | Aureococcus | Aureococcus a | metabolism | Nucleotide tra | KOG0888 | nucleoside dip | 2850.Phatr42 | 1 | 0,632455532 | 8,38E-05 | 4,78E-05 | 2 - | 0,000149061 - | 1 - | 9,25E-05 - | 1 - | 8,80E-05 - | 1 - | 8,93E-05 - | 0 - | 0 - |
| 5611 | 1 | 0.2um_NOKCPHJN_74538 | Bacteria | Proteobacteri | Gammaprote | Oceanospirilla | Saccharospirill | Reinekea | Reinekea fors | unclassified | unclassified | unclassified | unclassified | unclassified | 1,2 | 0,4 | 7,31E-05 | 1,62E-05 | 2 - | 0,000104819 - | 1 - | 6,51E-05 - | 1 - | 6,19E-05 - | 1 - | 6,28E-05 - | 1 - | 7,07E-05 - |
| 5612 | 1 | 0.2um_NOKCPHJN_1082627 | Bacteria | Proteobacteri | Alphaproteob | Pelagibacteral | Pelagibacterac | Candidatus Pe | Candidatus Pe | cellular proces | Cell wall/mem | COG2885 | Ompa motb d | 439493.PB721 | 1 | 0 | 8,41E-05 | 7,98E-06 | 1 - | 7,04E-05 - | 1 - | 8,74E-05 - | 1 - | 8,31E-05 - | 1 - | 8,43E-05 - | 1 - | 9,50E-05 - |
| 5613 | 1 | 0.2um_NOKCPHJN_259073 | Eukaryota | Basidiomycota | Ustilaginomyc | Ustilaginales | Ustilaginaceae | Moesziomyce | Moesziomyce | cellular proces | Intracellular tr | 12PEB | vacuolar prote | 5180.EDO032 | 0,2 | 0,4 | 1,07E-06 | 2,14E-06 | 0 - | 0 - | 0 - | 0 - | 0 - | 0 - | 0 - | 0 - | 1 - | 5,36E-06 - |
| 5614 | 1 | 0.2um_NOKCPHJN_1081452 | Bacteria | Proteobacteri | Alphaproteob | Rhodobactera | Rhodobactera | Sulfitobacter | Sulfitobacter | metabolism | Secondary me | 01Q86 | extracellular s | 367336.OM22 | 1 | 0 | 3,76E-05 | 3,57E-06 | 1 - | 3,15E-05 - | 1 - | 3,92E-05 - | 1 - | 3,72E-05 - | 1 - | 3,78E-05 - | 1 - | 4,25E-05 - |
| 5615 | 1 | 3um_FIJJAFBF_40363 | Eukaryota | unclassified | Raphidophyce | Chattonellales | Chattonellace | Heterosigma | Heterosigma a | unclassified | unclassified | unclassified | unclassified | unclassified | 2,4 | 0,8 | 6,46E-05 | 2,30E-05 | 3 - | 6,76E-05 - | 3 - | 8,40E-05 - | 2 - | 5,32E-05 - | 1 - | 2,70E-05 - | 3 - | 9,13E-05 - |
| 5616 | 1 | 3um_FIJJAFBF_667302 | Bacteria | Proteobacteri | Gammaprote | Pseudomonad | Moraxellaceae | Psychrobacter | various | unclassified | unclassified | unclassified | unclassified | unclassified | 0,6 | 0,489897949 | 9,05E-05 | 7,52E-05 | 1 - | 0,00012753 - | 0 - | 0 - | 0 - | 0 - | 1 - | 0,000152759 - | 1 - | 0,000172083 - |
| 5617 | 1 | 3um_FIJJAFBF_225136 | Bacteria | Proteobacteri | Alphaproteob | Rhodobactera | Rhodobactera | Planktomarina | Planktomarina | unclassified | unclassified | unclassified | unclassified | unclassified | 2,4 | 0,489897949 | 0,000321293 | 5,00E-05 | 3 - | 0,000340921 - | 2 - | 0,000282216 - | 2 - | 0,000268282 - | 3 - | 0,000408366 - | 2 - | 0,000306682 - |
| 5618 | 1 | 0.2um_NOKCPHJN_601063 | Viruses | unclassified | unclassified | Herpesvirales | Herpesviridae | Lymphocryptc | Human gamm | poorly charact | Function unkn | 162JK | NA | 4577.GRMZM | 0,2 | 0,4 | 2,81E-06 | 5,63E-06 | 0 - | 0 - | 0 - | 0 - | 1 - | 1,41E-05 - | 0 - | 0 - | 0 - | 0 - |
| 5619 | 1 | 0.2um_NOKCPHJN_756783 | Bacteria | Proteobacteri | Gammaprote | unclassified | unclassified | unclassified | OM182 bacter | unclassified | unclassified | unclassified | unclassified | unclassified | 16 | 1,095445115 | 0,00024525 | 1,29E-05 | 18 - | 0,000232394 - | 16 - | 0,000256502 - | 15 - | 0,000228598 - | 16 - | 0,000247439 - | 15 - | 0,000261318 - |
| 5620 | 1 | 3um_FIJJAFBF_135416 | Bacteria | Proteobacteri | Alphaproteob | Rhodobactera | Rhodobactera | Planktomarina | Planktomarina | metabolism | Energy produc | 07QM5 | aconitate hyd | 633131.TR2A6 | 0,6 | 0,489897949 | 8,40E-06 | 6,93E-06 | 1 - | 1,22E-05 - | 1 - | 1,52E-05 - | 0 - | 0 - | 1 - | 1,46E-05 - | 0 - | 0 - |
| 5621 | 1 | 3um_FIJJAFBF_95945 | Bacteria | Bacteroidetes | Flavobacteriia | Flavobacterial | unclassified | unclassified | Flavobacterial | unclassified | unclassified | unclassified | unclassified | unclassified | 1 | 0 | 0,000129271 | 1,23E-05 | 1 - | 0,00010828 - | 1 - | 0,000134452 - | 1 - | 0,000127813 - | 1 - | 0,000129701 - | 1 - | 0,000146108 - |
| 5622 | 1 | 0.2um_NOKCPHJN_829615 | Bacteria | Proteobacteri | Gammaprote | Pseudomonad | Pseudomonad | Pseudomonas | Pseudomonas | information st | Translation, ri | 16QAF | thus facilitatin | 205922.Pfl01 | 0,4 | 0,8 | 1,33E-05 | 2,65E-05 | 0 - | 0 - | 0 - | 0 - | 2 - | 6,63E-05 - | 0 - | 0 - | 0 - | 0 - |
| 5623 | 1 | 3um_FIJJAFBF_175649 | Bacteria | Proteobacteri | Gammaprote | unclassified | unclassified | unclassified | Gammaprote | poorly charact | Function unkn | 16R8I | Lipoprotein | 314287.GB22 | 0,8 | 0,748331477 | 7,33E-05 | 6,65E-05 | 0 - | 0 - | 1 - | 9,19E-05 - | 2 - | 0,000174816 - | 0 - | 0 - | 1 - | 9,99E-05 - |
| 5624 | 1 | 3um_FIJJAFBF_30127 | Bacteria | Bacteroidetes | Flavobacteriia | Flavobacterial | Flavobacteriac | Formosa | Formosa sp. H | poorly charact | Function unkn | 0PBEY | Pfam:DUF145 | 313603.FB217 | 0,6 | 0,489897949 | 0,000142828 | 0,000117792 | 1 - | 0,000208685 - | 1 - | 0,000259125 - | 1 - | 0,000246331 - | 0 - | 0 - | 0 - | 0 - |
| 5625 | 1 | 3um_FIJJAFBF_123937 | Bacteria | Actinobacteria | Actinobacteria | Micrococcales | Microbacteria | unclassified | Microbacteria | information st | Translation, ri | 00JY2 | Binds to the 2 | 979556.MTES | 3 | 0 | 0,000274054 | 2,60E-05 | 3 - | 0,000229554 - | 3 - | 0,000285038 - | 3 - | 0,000270964 - | 3 - | 0,000274967 - | 3 - | 0,000309749 - |
| 5626 | 1 | 3um_FIJJAFBF_246732 | Bacteria | Bacteroidetes | Cytophagia | Cytophagales | Flammeovirga | unclassified | Flammeovirga | unclassified | unclassified | unclassified | unclassified | unclassified | 4,2 | 0,4 | 0,000620722 | 2,94E-05 | 5 - | 0,000623787 - | 4 - | 0,000619647 - | 4 - | 0,000589053 - | 4 - | 0,000597754 - | 4 - | 0,000673367 - |
| 5627 | 1 | 3um_FIJJAFBF_92189 | Bacteria | Bacteroidetes | Flavobacteriia | Flavobacterial | Flavobacteriac | Flaviramulus | Flaviramulus b | unclassified | unclassified | unclassified | unclassified | unclassified | 0,6 | 0,489897949 | 2,06E-05 | 1,68E-05 | 0 - | 0 - | 1 - | 3,54E-05 - | 1 - | 3,36E-05 - | 1 - | 3,41E-05 - | 0 - | 0 - |
| 5628 | 2 | 0.2um_NOKCPHJN_481754;0.2um_NOKCPHJN_8 | Bacteria | Proteobacteri | Alphaproteob | Rhodobactera | unclassified | unclassified | Rhodobactera | unclassified | unclassified | unclassified | unclassified | unclassified | 0,4 | 0,489897949 | 1,77E-05 | 2,18E-05 | 1 - | 4,06E-05 - | 0 - | 0 - | 1 - | 4,79E-05 - | 0 - | 0 - | 0 - | 0 - |
| 5629 | 1 | 0.2um_NOKCPHJN_283182 | Bacteria | Proteobacteri | Alphaproteob | Rhodobactera | Rhodobactera | unclassified | Rhodobactera | metabolism | Energy produc | 05C26 | Dehydrogenas | 388401.RB215 | 0,2 | 0,4 | 4,78E-06 | 9,56E-06 | 1 - | 2,39E-05 - | 0 - | 0 - | 0 - | 0 - | 0 - | 0 - | 0 - | 0 - |
| 5630 | 1 | 3um_FIJJAFBF_362607 | Bacteria | Proteobacteri | Alphaproteob | unclassified | unclassified | unclassified | Alphaproteob | metabolism | Energy produc | 02CNG | F(1)F(0) ATP s | 342108.amb3 | 0,6 | 0,489897949 | 4,91E-05 | 4,05E-05 | 1 - | 7,17E-05 - | 1 - | 8,91E-05 - | 1 - | 8,47E-05 - | 0 - | 0 - | 0 - | 0 - |
| 5631 | 1 | 3um_FIJJAFBF_128455 | Bacteria | Bacteroidetes | Flavobacteriia | Flavobacterial | Flavobacteriac | Polaribacter | Polaribacter s | cellular proces | Intracellular tr | 08SHB | Biopolymer tr | 313594.PI23P | 6,4 | 1,743559577 | 0,000565962 | 0,0001414 | 7 - | 0,000525122 - | 4 - | 0,000372598 - | 9 - | 0,000796954 - | 7 - | 0,000629009 - | 5 - | 0,000506126 - |
| 5632 | 1 | 3um_FIJJAFBF_25445 | Bacteria | Bacteroidetes | Flavobacteriia | Flavobacterial | Flavobacteriac | Arenibacter | Arenibacter tr | unclassified | unclassified | unclassified | unclassified | unclassified | 5,6 | 0,489897949 | 0,000203873 | 2,57E-05 | 6 - | 0,000183154 - | 5 - | 0,00018952 - | 5 - | 0,000180163 - | 6 - | 0,000219388 - | 6 - | 0,00024714 - |
| 5633 | 1 | 0.2um_NOKCPHJN_331984 | Eukaryota | Phaeophyceae | unclassified | Ectocarpales | Ectocarpaceae | Ectocarpus | Ectocarpus sili | information st | Translation, ri | 036ZM | ribosomal pro | 34740.HMEL0 | 0,6 | 0,489897949 | 7,16E-05 | 5,85E-05 | 0 - | 0 - | 1 - | 0,000122861 - | 1 - | 0,000116795 - | 1 - | 0,00011852 - | 0 - | 0 - |
| 5634 | 3 | 0.2um_NOKCPHJN_123137;3um_FIJJAFBF_27497 | Bacteria | Proteobacteri | Alphaproteob | Rhodobactera | Rhodobactera | various | various | various | various | various | various | various | 0,2 | 0,4 | 8,34E-06 | 1,67E-05 | 0 - | 0 - | 0 - | 0 - | 1 - | 4,17E-05 - | 0 - | 0 - | 0 - | 0 - |
| 5635 | 1 | 0.2um_NOKCPHJN_234273 | Bacteria | Proteobacteri | Gammaprote | Cellvibrionales | Halieaceae | unclassified | marine gamm | metabolism | Inorganic ion t | 16PDP | TonB-depende | 247639.MGP2 | 3,4 | 0,8 | 4,97E-05 | 1,48E-05 | 2 - | 2,40E-05 - | 4 - | 5,96E-05 - | 3 - | 4,25E-05 - | 4 - | 5,75E-05 - | 4 - | 6,48E-05 - |
| 5636 | 1 | 3um_FIJJAFBF_442870 | Eukaryota | unclassified | Cryptophyta | Cryptomonad | Hemiselmidac | Hemiselmis | Hemiselmis ar | unclassified | unclassified | unclassified | unclassified | unclassified | 1 | 0 | 0,000102259 | 9,71E-06 | 1 - | 8,57E-05 - | 1 - | 0,000106357 - | 1 - | 0,000101106 - | 1 - | 0,000102599 - | 1 - | 0,000115578 - |
| 5637 | 1 | 0.2um_NOKCPHJN_62594 | Bacteria | Proteobacteri | Alphaproteob | Rhodobactera | Rhodobactera | Amylibacter | Amylibacter k | unclassified | unclassified | unclassified | unclassified | unclassified | 1 | 0,632455532 | 3,04E-05 | 1,73E-05 | 2 - | 5,40E-05 - | 1 - | 3,35E-05 - | 1 - | 3,19E-05 - | 1 - | 3,23E-05 - | 0 - | 0 - |
| 5638 | 1 | 3um_FIJJAFBF_785962 | Bacteria | Proteobacteri | Alphaproteob | Rhodobactera | Rhodobactera | Jannaschia | Jannaschia fae | metabolism | Energy produc | 01R4M | Acetoacetyl-C | 314232.SKA53 | 0,6 | 0,489897949 | 3,22E-05 | 2,65E-05 | 1 - | 4,76E-05 - | 0 - | 0 - | 1 - | 5,62E-05 - | 1 - | 5,70E-05 - | 0 - | 0 - |
| 5639 | 1 | 3um_FIJJAFBF_807524 | Bacteria | Proteobacteri | Alphaproteob | Rhodobactera | Rhodobactera | Planktomarina | Planktomarina | cellular proces | Cell wall/mem | 16PAV | penicillin-bind | 644107.SL115 | 1 | 0 | 1,93E-05 | 1,83E-06 | 1 - | 1,61E-05 - | 1 - | 2,00E-05 - | 1 - | 1,91E-05 - | 1 - | 1,93E-05 - | 1 - | 2,18E-05 - |
| 5640 | 1 | 0.2um_NOKCPHJN_307776 | Bacteria | Bacteroidetes | Flavobacteriia | Flavobacterial | Flavobacteriac | Tenacibaculur | Tenacibaculur | unclassified | unclassified | unclassified | unclassified | unclassified | 0,6 | 0,489897949 | 5,14E-05 | 4,23E-05 | 1 - | 7,60E-05 - | 0 - | 0 - | 1 - | 8,97E-05 - | 1 - | 9,10E-05 - | 0 - | 0 - |
| 5641 | 1 | 3um_FIJJAFBF_221344 | Bacteria | Planctomycete | unclassified | unclassified | unclassified | unclassified | Planctomycet | unclassified | unclassified | unclassified | unclassified | unclassified | 0,4 | 0,489897949 | 2,03E-05 | 2,50E-05 | 1 - | 4,65E-05 - | 0 - | 0 - | 1 - | 5,49E-05 - | 0 - | 0 - | 0 - | 0 - |
| 5642 | 1 | 3um_FIJJAFBF_26085 | Bacteria | Bacteroidetes | Flavobacteriia | Flavobacterial | Flavobacteriac | Polaribacter | Polaribacter s | metabolism | Carbohydrate | COG0176 | Transaldolase | 156586.BBFL7 | 4,6 | 0,8 | 0,000288137 | 4,45E-05 | 5 - | 0,000264463 - | 4 - | 0,000262708 - | 6 - | 0,000374605 - | 4 - | 0,000253425 - | 4 - | 0,000285483 - |
| 5643 | 1 | 3um_FIJJAFBF_168406 | Bacteria | Bacteroidetes | Flavobacteriia | Flavobacterial | Flavobacteriac | unclassified | Flavobacteriac | poorly charact | Function unkn | 0ZMES | Caspase doma | 272559.BF343 | 1 | 0,632455532 | 1,38E-05 | 7,80E-06 | 2 - | 2,41E-05 - | 0 - | 0 - | 1 - | 1,42E-05 - | 1 - | 1,45E-05 - | 1 - | 1,63E-05 - |
| 5644 | 1 | 0.2um_NOKCPHJN_121623 | Bacteria | Proteobacteri | Gammaprote | Alteromonada | Alteromonada | Paraglaciecola | Paraglaciecola | metabolism | Amino acid tra | COG0339 | oligopeptidas | 1085623.GNIT | 1 | 0,632455532 | 1,89E-05 | 1,20E-05 | 1 - | 1,59E-05 - | 1 - | 1,97E-05 - | 2 - | 3,75E-05 - | 0 - | 0 - | 1 - | 2,15E-05 - |
| 5645 | 1 | 0.2um_NOKCPHJN_499519 | Eukaryota | Basidiomycota | Tremellomyce | Cystofilobasid | Mrakiaceae | Xanthophyllor | Xanthophyllor | unclassified | unclassified | unclassified | unclassified | unclassified | 0,4 | 0,489897949 | 1,04E-05 | 1,29E-05 | 1 - | 2,32E-05 - | 1 - | 2,88E-05 - | 0 - | 0 - | 0 - | 0 - | 0 - | 0 - |
| 5646 | 1 | 3um_FIJJAFBF_237282 | Bacteria | Proteobacteri | Gammaprote | Cellvibrionales | unclassified | unclassified | Cellvibrionales | information st | Transcription | 16PMT | DNA-depende | 314287.GB22 | 6 | 1,788854382 | 0,000242595 | 5,29E-05 | 9 - | 0,000312082 - | 7 - | 0,000301399 - | 5 - | 0,000204656 - | 5 - | 0,000207679 - | 4 - | 0,000187159 - |
| 5647 | 2 | 0.2um_NOKCPHJN_291233;3um_FIJJAFBF_5095 | Bacteria | unclassified | unclassified | unclassified | unclassified | unclassified | uncultured ba | poorly charact | Function unkn | 11MEK | NA | 50743.SCB49 | 0,2 | 0,4 | 8,10E-06 | 1,62E-05 | 0 - | 0 - | 1 - | 4,05E-05 - | 0 - | 0 - | 0 - | 0 - | 0 - | 0 - |
| 5648 | 1 | 0.2um_NOKCPHJN_27656 | Bacteria | Actinobacteria | Actinobacteria | Micrococcales | Microbacteria | unclassified | Microbacteria | unclassified | unclassified | unclassified | unclassified | unclassified | 2 | 0 | 0,000108752 | 1,03E-05 | 2 - | 9,11E-05 - | 2 - | 0,00011311 - | 2 - | 0,000107526 - | 2 - | 0,000109114 - | 2 - | 0,000122916 - |
| 5649 | 1 | 3um_FIJJAFBF_121142 | Bacteria | Proteobacteri | Gammaprote | Cellvibrionales | Porticoccacea | unclassified | SAR92 bacteri | information st | Translation, ri | COG0087 | One of the pri | 314287.GB22 | 0,2 | 0,4 | 1,09E-05 | 2,18E-05 | 1 - | 5,44E-05 - | 0 - | 0 - | 0 - | 0 - | 0 - | 0 - | 0 - | 0 - |
| 5650 | 1 | 3um_FIJJAFBF_209605 | Bacteria | Bacteroidetes | Flavobacteriia | Flavobacterial | Flavobacteriac | various | various | unclassified | unclassified | unclassified | unclassified | unclassified | 1,4 | 0,489897949 | 0,000157891 | 5,82E-05 | 1 - | 9,41E-05 - | 2 - | 0,000233637 - | 2 - | 0,000222102 - | 1 - | 0,000112691 - | 1 - | 0,000126946 - |
| 5651 | 1 | 3um_FIJJAFBF_225013 | Bacteria | Bacteroidetes | Flavobacteriia | Flavobacterial | Flavobacteriac | Arenibacter | various | unclassified | unclassified | unclassified | unclassified | unclassified | 6,8 | 1,326649916 | 0,000167518 | 1,65E-05 | 9 - | 0,000189192 - | 6 - | 0,000156614 - | 7 - | 0,000173695 - | 7 - | 0,000176261 - | 5 - | 0,000141826 - |
| 5652 | 1 | 3um_FIJJAFBF_629750 | Bacteria | Proteobacteri | Gammaprote | Cellvibrionales | Porticoccacea | unclassified | various | cellular proces | Posttranslatio | 08V1T | peptidylprolyl | 314287.GB22 | 1,4 | 0,489897949 | 7,59E-05 | 2,23E-05 | 2 - | 9,29E-05 - | 1 - | 5,77E-05 - | 1 - | 5,49E-05 - | 2 - | 0,000111323 - | 1 - | 6,27E-05 - |
| 5653 | 1 | 0.2um_NOKCPHJN_145298 | Bacteria | Proteobacteri | Alphaproteob | unclassified | unclassified | various | various | unclassified | unclassified | unclassified | unclassified | unclassified | 0,2 | 0,4 | 8,90E-06 | 1,78E-05 | 1 - | 4,45E-05 - | 0 - | 0 - | 0 - | 0 - | 0 - | 0 - | 0 - | 0 - |
| 5654 | 1 | 3um_FIJJAFBF_100701 | Bacteria | Bacteroidetes | Flavobacteriia | Flavobacterial | Flavobacteriac | Formosa | Formosa sp. H | poorly charact | Function unkn | 11P35 | NA | 867900.Celly | 1 | 0,632455532 | 3,70E-05 | 2,35E-05 | 1 - | 3,11E-05 - | 1 - | 3,86E-05 - | 2 - | 7,34E-05 - | 0 - | 0 - | 1 - | 4,20E-05 - |
| 5655 | 1 | 0.2um_NOKCPHJN_235600 | Bacteria | Proteobacteri | Gammaprote | Oceanospirilla | Oceanospirilla | Motiliproteus | Motiliproteus | information st | Translation, ri | 05CEJ | Catalyzes the | 247634.GPB2 | 8,6 | 0,489897949 | 0,000169689 | 1,84E-05 | 9 - | 0,000148846 - | 8 - | 0,000164287 - | 8 - | 0,000156175 - | 9 - | 0,000178292 - | 9 - | 0,000200846 - |
| 5656 | 1 | 3um_FIJJAFBF_364992 | Bacteria | Proteobacteri | Betaproteoba | Burkholderiale | unclassified | Thiomonas | Thiomonas sp | unclassified | unclassified | unclassified | unclassified | unclassified | 1 | 0 | 8,41E-05 | 7,98E-06 | 1 - | 7,04E-05 - | 1 - | 8,74E-05 - | 1 - | 8,31E-05 - | 1 - | 8,43E-05 - | 1 - | 9,50E-05 - |
| 5657 | 1 | 0.2um_NOKCPHJN_602365 | Viruses | unclassified | unclassified | unclassified | Phycodnavirid | unclassified | Chrysochromu | information st | Replication, re | COG0187 | DNA gyrase ne | 436017.A4S64 | 0,4 | 0,489897949 | 7,40E-06 | 9,12E-06 | 1 - | 1,70E-05 - | 0 - | 0 - | 1 - | 2,00E-05 - | 0 - | 0 - | 0 - | 0 - |
| 5658 | 1 | 0.2um_NOKCPHJN_644072 | Bacteria | Bacteroidetes | Flavobacteriia | Flavobacterial | Cryomorphace | unclassified | Cryomorphace | information st | Translation, ri | COG0173 | aspartyl-trna s | 391587.KAOT | 0,2 | 0,4 | 4,18E-06 | 8,36E-06 | 1 - | 2,09E-05 - | 0 - | 0 - | 0 - | 0 - | 0 - | 0 - | 0 - | 0 - |
| 5659 | 1 | 3um_FIJJAFBF_88567 | unclassified | unclassified | unclassified | unclassified | unclassified | unclassified | unclassified | unclassified | unclassified | unclassified | unclassified | unclassified | 0,2 | 0,4 | 3,70E-05 | 7,40E-05 | 0 - | 0 - | 1 - | 0,000185089 - | 0 - | 0 - | 0 - | 0 - | 0 - | 0 - |
| 5660 | 1 | 0.2um_NOKCPHJN_730058 | Bacteria | Proteobacteri | Gammaprote | Alteromonada | Idiomarinacea | Idiomarina | Idiomarina ae | poorly charact | Function unkn | 0PXWZ | ChtBD1 | 318829.MGG | 1,6 | 0,489897949 | 4,72E-05 | 1,24E-05 | 2 - | 5,05E-05 - | 1 - | 3,13E-05 - | 2 - | 5,96E-05 - | 2 - | 6,04E-05 - | 1 - | 3,40E-05 - |
| 5661 | 1 | 0.2um_NOKCPHJN_327204 | Bacteria | Bacteroidetes | Flavobacteriia | Flavobacterial | Flavobacteriac | Arenibacter | Arenibacter c | information st | Translation, ri | 0PA05 | 30S ribosomal | 983544.Lacal | 1,2 | 0,4 | 6,13E-05 | 1,36E-05 | 2 - | 8,80E-05 - | 1 - | 5,46E-05 - | 1 - | 5,19E-05 - | 1 - | 5,27E-05 - | 1 - | 5,93E-05 - |
| 5662 | 1 | 0.2um_NOKCPHJN_76863 | Bacteria | Proteobacteri | Gammaprote | Alteromonada | Colwelliaceae | Colwellia | Colwellia sp. F | poorly charact | Function unkn | 16R8I | Lipoprotein | 167879.CPS_3 | 1 | 0,894427191 | 1,46E-05 | 1,33E-05 | 2 - | 2,47E-05 - | 0 - | 0 - | 0 - | 0 - | 1 - | 1,48E-05 - | 2 - | 3,34E-05 - |
| 5663 | 1 | 0.2um_NOKCPHJN_833884 | Bacteria | Firmicutes | Bacilli | Bacillales | Paenibacillace | Saccharibacill | Saccharibacill | poorly charact | Function unkn | 0XR88 | semaphorin re | 4787.PITG_05 | 0,4 | 0,489897949 | 7,75E-06 | 9,51E-06 | 0 - | 0 - | 1 - | 1,86E-05 - | 0 - | 0 - | 0 - | 0 - | 1 - | 2,02E-05 - |
| 5664 | 1 | 3um_FIJJAFBF_483536 | Bacteria | Actinobacteria | Actinobacteria | Micrococcales | Microbacteria | unclassified | Microbacteria | unclassified | unclassified | unclassified | unclassified | unclassified | 3,4 | 0,489897949 | 0,000107795 | 1,77E-05 | 3 - | 7,97E-05 - | 3 - | 9,90E-05 - | 4 - | 0,000125447 - | 4 - | 0,000127299 - | 3 - | 0,000107552 - |
| 5665 | 1 | 0.2um_NOKCPHJN_168884 | Bacteria | Bacteroidetes | unclassified | unclassified | unclassified | unclassified | unidentified e | information st | Replication, re | 05C24 | DNA gyrase ne | 50743.SCB49 | 0,2 | 0,4 | 3,36E-06 | 6,72E-06 | 0 - | 0 - | 1 - | 1,68E-05 - | 0 - | 0 - | 0 - | 0 - | 0 - | 0 - |
| 5666 | 1 | 3um_FIJJAFBF_01114 | Bacteria | Bacteroidetes | Flavobacteriia | Flavobacterial | Flavobacteriac | Algibacter | Algibacter sp. | unclassified | unclassified | unclassified | unclassified | unclassified | 1 | 0,632455532 | 5,31E-05 | 3,43E-05 | 1 - | 4,48E-05 - | 0 - | 0 - | 1 - | 5,29E-05 - | 2 - | 0,000107409 - | 1 - | 6,05E-05 - |
| 5667 | 1 | 0.2um_NOKCPHJN_296783 | Bacteria | Bacteroidetes | Flavobacteriia | Flavobacterial | Flavobacteriac | Flavobacteriu | Flavobacteriu | metabolism | Inorganic ion t | 05C27 | TonB-depende | 983548.Krodi | 6,4 | 1,019803903 | 0,000119227 | 1,78E-05 | 8 - | 0,000125611 - | 6 - | 0,000116979 - | 6 - | 0,000111203 - | 5 - | 9,40E-05 - | 7 - | 0,000148307 - |
| 5668 | 1 | 0.2um_NOKCPHJN_878070 | Bacteria | Proteobacteri | Gammaprote | unclassified | unclassified | unclassified | Gammaprote | unclassified | unclassified | unclassified | unclassified | unclassified | 0,4 | 0,489897949 | 2,48E-05 | 3,04E-05 | 0 - | 0 - | 0 - | 0 - | 1 - | 6,16E-05 - | 1 - | 6,25E-05 - | 0 - | 0 - |
| 5669 | 1 | 0.2um_NOKCPHJN_298845 | Archaea | Euryarchaeota | unclassified | unclassified | unclassified | unclassified | Euryarchaeota | unclassified | unclassified | unclassified | unclassified | unclassified | 0,4 | 0,489897949 | 9,43E-06 | 1,16E-05 | 0 - | 0 - | 1 - | 2,26E-05 - | 0 - | 0 - | 0 - | 0 - | 1 - | 2,45E-05 - |
| 5670 | 1 | 3um_FIJJAFBF_152184 | Bacteria | Proteobacteri | Alphaproteob | Rickettsiales | unclassified | unclassified | Rickettsiales b | unclassified | unclassified | unclassified | unclassified | unclassified | 1 | 0 | 9,39E-05 | 8,91E-06 | 1 - | 7,86E-05 - | 1 - | 9,76E-05 - | 1 - | 9,28E-05 - | 1 - | 9,42E-05 - | 1 - | 0,000106078 - |
| 5671 | 1 | 3um_FIJJAFBF_791366 | Bacteria | Proteobacteri | Alphaproteob | Rhodobactera | unclassified | unclassified | Rhodobactera | unclassified | unclassified | unclassified | unclassified | unclassified | 0,6 | 0,489897949 | 5,54E-05 | 4,56E-05 | 1 - | 8,20E-05 - | 0 - | 0 - | 1 - | 9,68E-05 - | 1 - | 9,82E-05 - | 0 - | 0 - |
| 5672 | 1 | 3um_FIJJAFBF_119799 | unclassified | unclassified | unclassified | unclassified | unclassified | unclassified | unclassified | unclassified | unclassified | unclassified | unclassified | unclassified | 8,2 | 1,6 | 0,000234262 | 4,28E-05 | 9 - | 0,000217015 - | 7 - | 0,000209587 - | 7 - | 0,000199239 - | 11 - | 0,000317714 - | 7 - | 0,000227757 - |
| 5673 | 1 | 0.2um_NOKCPHJN_865899 | unclassified | unclassified | unclassified | unclassified | unclassified | unclassified | unclassified | unclassified | unclassified | unclassified | unclassified | unclassified | 0,6 | 0,489897949 | 1,28E-05 | 1,06E-05 | 1 - | 1,88E-05 - | 1 - | 2,33E-05 - | 1 - | 2,21E-05 - | 0 - | 0 - | 0 - | 0 - |
| 5674 | 1 | 0.2um_NOKCPHJN_110299 | Bacteria | Bacteroidetes | Flavobacteriia | Flavobacterial | Flavobacteriac | unclassified | Flavobacteriac | information st | Translation, ri | 05CAV | thus facilitatin | 391603.FBALC | 0,8 | 0,4 | 1,59E-05 | 8,06E-06 | 1 - | 1,72E-05 - | 1 - | 2,13E-05 - | 1 - | 2,03E-05 - | 1 - | 2,06E-05 - | 0 - | 0 - |
| 5675 | 1 | 3um_FIJJAFBF_221206 | Bacteria | Proteobacteri | Gammaprote | Oceanospirilla | Saccharospirill | Reinekea | Reinekea fors | unclassified | unclassified | unclassified | unclassified | unclassified | 2,6 | 0,8 | 0,000178142 | 4,60E-05 | 4 - | 0,000233049 - | 2 - | 0,000144689 - | 2 - | 0,000137545 - | 2 - | 0,000139577 - | 3 - | 0,000235849 - |
| 5676 | 1 | 0.2um_NOKCPHJN_884992 | Eukaryota | unclassified | Cryptophyta | Pyrenomonad | Geminigerace | Guillardia | Guillardia thet | unclassified | unclassified | unclassified | unclassified | unclassified | 0,4 | 0,489897949 | 1,10E-05 | 1,34E-05 | 0 - | 0 - | 0 - | 0 - | 1 - | 2,72E-05 - | 1 - | 2,76E-05 - | 0 - | 0 - |
| 5677 | 1 | 0.2um_NOKCPHJN_207185 | Bacteria | Proteobacteri | Gammaprote | Oceanospirilla | Saccharospirill | Reinekea | Reinekea fors | unclassified | unclassified | unclassified | unclassified | unclassified | 7,4 | 1,496662955 | 0,000163621 | 2,07E-05 | 10 - | 0,000188159 - | 8 - | 0,00018691 - | 7 - | 0,000155471 - | 6 - | 0,000135229 - | 6 - | 0,000152336 - |
| 5678 | 1 | 3um_FIJJAFBF_298889 | Bacteria | Proteobacteri | Alphaproteob | Rhodobactera | Rhodobactera | Loktanella | Loktanella sp. | poorly charact | Function unkn | 05W3P | Protein of unk | 391593.RCCS2 | 0,2 | 0,4 | 4,30E-05 | 8,59E-05 | 0 - | 0 - | 0 - | 0 - | 0 - | 0 - | 1 - | 0,000214818 - | 0 - | 0 - |
| 5679 | 1 | 0.2um_NOKCPHJN_160361 | Eukaryota | unclassified | Pelagophycea | Pelagomonad | unclassified | Aureoumbra | Aureoumbra l | metabolism | Energy produc | COG0056 | Produces ATP | 41431.PCC880 | 0,2 | 0,4 | 4,56E-06 | 9,13E-06 | 1 - | 2,28E-05 - | 0 - | 0 - | 0 - | 0 - | 0 - | 0 - | 0 - | 0 - |
| 5680 | 1 | 0.2um_NOKCPHJN_123118 | Bacteria | Proteobacteri | Alphaproteob | Rhodobactera | Rhodobactera | Planktomarina | Planktomarina | metabolism | Energy produc | 01RNF | Produces ATP | 388401.RB215 | 2,8 | 0,4 | 8,10E-05 | 1,45E-05 | 3 - | 7,26E-05 - | 3 - | 9,02E-05 - | 2 - | 5,72E-05 - | 3 - | 8,70E-05 - | 3 - | 9,80E-05 - |
| 5681 | 2 | 0.2um_NOKCPHJN_109749;3um_FIJJAFBF_63661 | Bacteria | Bacteroidetes | Flavobacteriia | Flavobacterial | Flavobacteriac | Formosa | Formosa sp. H | unclassified | unclassified | unclassified | unclassified | unclassified | 0,4 | 0,489897949 | 9,14E-05 | 0,000113995 | 1 - | 0,000194537 - | 0 - | 0 - | 0 - | 0 - | 0 - | 0 - | 1 - | 0,000262499 - |
| 5682 | 1 | 0.2um_NOKCPHJN_36415 | Bacteria | Proteobacteri | Alphaproteob | Rhodobactera | Rhodobactera | Puniceibacteri | Puniceibacteri | metabolism | Energy produc | 01RSD | Isocitrate deh | 375451.RD1_ | 0,4 | 0,489897949 | 6,90E-06 | 8,53E-06 | 1 - | 1,54E-05 - | 1 - | 1,91E-05 - | 0 - | 0 - | 0 - | 0 - | 0 - | 0 - |
| 5683 | 1 | 3um_FIJJAFBF_415633 | Bacteria | Bacteroidetes | Flavobacteriia | unclassified | unclassified | unclassified | Flavobacteria | unclassified | unclassified | unclassified | unclassified | unclassified | 2,4 | 1,019803903 | 4,98E-05 | 1,91E-05 | 4 - | 7,06E-05 - | 2 - | 4,39E-05 - | 2 - | 4,17E-05 - | 1 - | 2,12E-05 - | 3 - | 7,15E-05 - |
| 5684 | 1 | 3um_FIJJAFBF_135023 | Bacteria | Bacteroidetes | Flavobacteriia | various | various | various | various | unclassified | unclassified | unclassified | unclassified | unclassified | 0,4 | 0,489897949 | 2,21E-05 | 2,73E-05 | 1 - | 5,08E-05 - | 0 - | 0 - | 1 - | 5,99E-05 - | 0 - | 0 - | 0 - | 0 - |
| 5685 | 1 | 3um_FIJJAFBF_739375 | Bacteria | Bacteroidetes | Flavobacteriia | Flavobacterial | Flavobacteriac | Polaribacter | Polaribacter s | unclassified | unclassified | unclassified | unclassified | unclassified | 0,4 | 0,8 | 5,72E-06 | 1,14E-05 | 2 - | 2,86E-05 - | 0 - | 0 - | 0 - | 0 - | 0 - | 0 - | 0 - | 0 - |
| 5686 | 1 | 0.2um_NOKCPHJN_984473 | unclassified | unclassified | unclassified | unclassified | unclassified | unclassified | unclassified | unclassified | unclassified | unclassified | unclassified | unclassified | 1 | 0 | 2,28E-05 | 2,17E-06 | 1 - | 1,91E-05 - | 1 - | 2,38E-05 - | 1 - | 2,26E-05 - | 1 - | 2,29E-05 - | 1 - | 2,58E-05 - |
| 5687 | 1 | 0.2um_NOKCPHJN_19271 | Bacteria | Bacteroidetes | Flavobacteriia | Flavobacterial | Flavobacteriac | Formosa | Formosa sp. H | cellular proces | Posttranslatio | COG0330 | Band 7 protei | 983544.Lacal | 0,6 | 0,8 | 2,85E-05 | 3,60E-05 | 2 - | 8,50E-05 - | 0 - | 0 - | 0 - | 0 - | 0 - | 0 - | 1 - | 5,74E-05 - |
| 5688 | 1 | 0.2um_NOKCPHJN_492975 | Bacteria | Proteobacteri | Alphaproteob | Rhodobactera | Rhodobactera | Planktomarina | Planktomarina | unclassified | unclassified | unclassified | unclassified | unclassified | 1,2 | 0,4 | 5,53E-05 | 1,85E-05 | 1 - | 3,86E-05 - | 1 - | 4,80E-05 - | 2 - | 9,12E-05 - | 1 - | 4,63E-05 - | 1 - | 5,21E-05 - |
| 5689 | 1 | 3um_FIJJAFBF_437894 | Bacteria | Bacteroidetes | unclassified | unclassified | unclassified | unclassified | Bacteroidetes | unclassified | unclassified | unclassified | unclassified | unclassified | 10,2 | 1,469693846 | 0,000312598 | 5,45E-05 | 10 - | 0,000256771 - | 13 - | 0,000414484 - | 10 - | 0,000303092 - | 9 - | 0,000276812 - | 9 - | 0,000311828 - |
| 5690 | 1 | 3um_FIJJAFBF_298837 | Bacteria | Bacteroidetes | Flavobacteriia | Flavobacterial | Flavobacteriac | Polaribacter | Polaribacter g | unclassified | unclassified | unclassified | unclassified | unclassified | 4,2 | 0,4 | 0,000385854 | 1,83E-05 | 5 - | 0,000387759 - | 4 - | 0,000385186 - | 4 - | 0,000366168 - | 4 - | 0,000371577 - | 4 - | 0,00041858 - |
| 5691 | 1 | 3um_FIJJAFBF_629417 | Bacteria | Bacteroidetes | Flavobacteriia | Flavobacterial | Flavobacteriac | Polaribacter | Polaribacter s | unclassified | unclassified | unclassified | unclassified | unclassified | 4,8 | 0,4 | 9,84E-05 | 7,58E-06 | 5 - | 8,63E-05 - | 5 - | 0,000107157 - | 5 - | 0,000101866 - | 5 - | 0,000103371 - | 4 - | 9,32E-05 - |
| 5692 | 1 | 3um_FIJJAFBF_715096 | Eukaryota | unclassified | Pelagophycea | Pelagomonad | unclassified | Pelagomonas | uncultured Pe | metabolism | Energy produc | 0IE88 | oxidoreductas | 35128.Thapsd | 3,2 | 0,748331477 | 5,88E-05 | 1,15E-05 | 4 - | 6,25E-05 - | 4 - | 7,77E-05 - | 3 - | 5,54E-05 - | 3 - | 5,62E-05 - | 2 - | 4,22E-05 - |
| 5693 | 2 | 0.2um_NOKCPHJN_947652;3um_FIJJAFBF_5094 | Eukaryota | various | various | various | various | various | various | various | various | various | various | various | 0,2 | 0,4 | 1,70E-05 | 3,39E-05 | 0 - | 0 - | 1 - | 8,48E-05 - | 0 - | 0 - | 0 - | 0 - | 0 - | 0 - |
| 5694 | 1 | 0.2um_NOKCPHJN_621499 | Bacteria | Proteobacteri | Alphaproteob | Pelagibacteral | Pelagibacterac | unclassified | Pelagibacterac | cellular proces | Cell wall/mem | 7257 | NA | 439493.PB721 | 2 | 0,632455532 | 7,84E-05 | 2,65E-05 | 1 - | 3,23E-05 - | 2 - | 8,03E-05 - | 3 - | 0,000114492 - | 2 - | 7,75E-05 - | 2 - | 8,73E-05 - |
| 5695 | 1 | 3um_FIJJAFBF_85577 | Bacteria | Proteobacteri | Gammaprote | Alteromonada | Alteromonada | Paraglaciecola | Paraglaciecola | unclassified | unclassified | unclassified | unclassified | unclassified | 0,2 | 0,4 | 2,20E-05 | 4,41E-05 | 0 - | 0 - | 0 - | 0 - | 1 - | 0,000110148 - | 0 - | 0 - | 0 - | 0 - |
| 5696 | 1 | 0.2um_NOKCPHJN_205270 | Bacteria | Bacteroidetes | Flavobacteriia | Flavobacterial | Flavobacteriac | Polaribacter | Polaribacter s | information st | Translation, ri | COG0089 | One of the ea | 313594.PI23P | 0,4 | 0,489897949 | 5,26E-05 | 6,48E-05 | 1 - | 0,000119559 - | 0 - | 0 - | 0 - | 0 - | 1 - | 0,000143212 - | 0 - | 0 - |
| 5697 | 1 | 3um_FIJJAFBF_417270 | Bacteria | Proteobacteri | Gammaprote | Cellvibrionales | Porticoccacea | unclassified | Gammaprote | cellular proces | Posttranslatio | 11WF3 | Glutaredoxin | 314287.GB22 | 0,6 | 0,489897949 | 6,46E-05 | 5,32E-05 | 1 - | 9,56E-05 - | 0 - | 0 - | 1 - | 0,000112902 - | 1 - | 0,000114569 - | 0 - | 0 - |
| 5698 | 1 | 3um_FIJJAFBF_209216 | Bacteria | Bacteroidetes | unclassified | unclassified | unclassified | unclassified | Bacteroidetes | poorly charact | Function unkn | 0P8NI | Inherit from b | 313590.MED1 | 5,4 | 0,489897949 | 0,000182882 | 1,25E-05 | 6 - | 0,000171309 - | 5 - | 0,000177262 - | 6 - | 0,000202212 - | 5 - | 0,000170999 - | 5 - | 0,00019263 - |
| 5699 | 1 | 3um_FIJJAFBF_13890 | unclassified | unclassified | unclassified | unclassified | unclassified | unclassified | unclassified | unclassified | unclassified | unclassified | unclassified | unclassified | 0,8 | 0,4 | 0,00011916 | 6,05E-05 | 1 - | 0,000128963 - | 1 - | 0,000160134 - | 1 - | 0,000152227 - | 1 - | 0,000154476 - | 0 - | 0 - |
| 5700 | 1 | 0.2um_NOKCPHJN_317314 | Bacteria | Bacteroidetes | Flavobacteriia | Flavobacterial | Flavobacteriac | Formosa | Formosa sp. H | unclassified | unclassified | unclassified | unclassified | unclassified | 2 | 1,095445115 | 0,000100519 | 4,55E-05 | 4 - | 0,000174565 - | 1 - | 5,42E-05 - | 1 - | 5,15E-05 - | 2 - | 0,00010455 - | 2 - | 0,000117775 - |
| 5701 | 1 | 3um_FIJJAFBF_201850 | Bacteria | Proteobacteri | Alphaproteob | Rhodobactera | Rhodobactera | Litorimicrobiu | Litorimicrobiu | poorly charact | Function unkn | 16QP7 | metallo-beta-l | 467661.RKLH1 | 1 | 0 | 2,47E-05 | 2,35E-06 | 1 - | 2,07E-05 - | 1 - | 2,57E-05 - | 1 - | 2,45E-05 - | 1 - | 2,48E-05 - | 1 - | 2,80E-05 - |
| 5702 | 1 | 3um_FIJJAFBF_155808 | Bacteria | Bacteroidetes | Flavobacteriia | Flavobacterial | Flavobacteriac | Polaribacter | various | information st | Translation, ri | COG0184 | Forms an inte | 391587.KAOT | 0,6 | 0,489897949 | 9,15E-05 | 7,61E-05 | 1 - | 0,000128963 - | 0 - | 0 - | 0 - | 0 - | 1 - | 0,000154476 - | 1 - | 0,000174016 - |
| 5703 | 1 | 0.2um_NOKCPHJN_157490 | Bacteria | Proteobacteri | Alphaproteob | Rhodobactera | Hyphomonad | Robiginitomac | Robiginitomac | unclassified | unclassified | unclassified | unclassified | unclassified | 17,4 | 1,496662955 | 0,000296321 | 3,38E-05 | 18 - | 0,000257283 - | 20 - | 0,000354966 - | 17 - | 0,000286824 - | 16 - | 0,000273939 - | 16 - | 0,000308592 - |
| 5704 | 1 | 3um_FIJJAFBF_63415 | Bacteria | Bacteroidetes | Flavobacteriia | Flavobacterial | Flavobacteriac | Ulvibacter | Ulvibacter lito | cellular proces | Cell motility | COG1360 | Ompa motb d | 983544.Lacal | 3,2 | 0,4 | 0,000135364 | 2,10E-05 | 3 - | 0,000106275 - | 3 - | 0,000131962 - | 3 - | 0,000125447 - | 4 - | 0,000169732 - | 3 - | 0,000143402 - |
| 5705 | 1 | 3um_FIJJAFBF_190545 | Viruses | unclassified | unclassified | unclassified | Phycodnavirid | unclassified | Chrysochromu | information st | RNA processin | KOG1975 | mRNA cap gu | 88036.EFJ119 | 1,6 | 0,489897949 | 2,00E-05 | 5,66E-06 | 2 - | 2,13E-05 - | 2 - | 2,64E-05 - | 1 - | 1,26E-05 - | 2 - | 2,55E-05 - | 1 - | 1,44E-05 - |
| 5706 | 1 | 0.2um_NOKCPHJN_508819 | Bacteria | Proteobacteri | Alphaproteob | Pelagibacteral | Pelagibacterac | Candidatus Pe | Candidatus Pe | metabolism | Inorganic ion t | 16PII | TRAP transpor | 439493.PB721 | 1 | 0 | 4,10E-05 | 3,90E-06 | 1 - | 3,44E-05 - | 1 - | 4,27E-05 - | 1 - | 4,06E-05 - | 1 - | 4,12E-05 - | 1 - | 4,64E-05 - |
| 5707 | 1 | 3um_FIJJAFBF_328502 | Bacteria | Bacteroidetes | Flavobacteriia | Flavobacterial | Flavobacteriac | Algibacter | Algibacter lect | information st | Translation, ri | COG0087 | One of the pri | 983544.Lacal | 1 | 0 | 6,68E-05 | 6,35E-06 | 1 - | 5,60E-05 - | 1 - | 6,95E-05 - | 1 - | 6,61E-05 - | 1 - | 6,71E-05 - | 1 - | 7,55E-05 - |
| 5708 | 1 | 3um_FIJJAFBF_352546 | Eukaryota | unclassified | Pelagophycea | Pelagomonad | unclassified | Aureococcus | Aureococcus a | unclassified | unclassified | unclassified | unclassified | unclassified | 1 | 0 | 4,18E-05 | 3,97E-06 | 1 - | 3,50E-05 - | 1 - | 4,35E-05 - | 1 - | 4,13E-05 - | 1 - | 4,19E-05 - | 1 - | 4,72E-05 - |
| 5709 | 1 | 0.2um_NOKCPHJN_121634 | Bacteria | Proteobacteri | Gammaprote | Alteromonada | Alteromonada | Paraglaciecola | Paraglaciecola | unclassified | unclassified | unclassified | unclassified | unclassified | 2 | 0,632455532 | 0,000195614 | 4,53E-05 | 3 - | 0,000253184 - | 2 - | 0,000209587 - | 2 - | 0,000199239 - | 2 - | 0,000202181 - | 1 - | 0,000113878 - |
| 5710 | 1 | 3um_FIJJAFBF_739631 | Eukaryota | unclassified | Chrysophycea | Chromulinales | Chromulinace | Ochromonas | Ochromonas s | unclassified | unclassified | unclassified | unclassified | unclassified | 0,2 | 0,4 | 4,01E-06 | 8,01E-06 | 1 - | 2,00E-05 - | 0 - | 0 - | 0 - | 0 - | 0 - | 0 - | 0 - | 0 - |
| 5711 | 1 | 0.2um_NOKCPHJN_308002 | Bacteria | Actinobacteria | Actinobacteria | Micrococcales | Microbacteria | Candidatus Ac | Candidatus Ac | unclassified | unclassified | unclassified | unclassified | unclassified | 4,6 | 1,356465997 | 7,23E-05 | 1,91E-05 | 6 - | 8,03E-05 - | 6 - | 9,97E-05 - | 3 - | 4,74E-05 - | 5 - | 8,01E-05 - | 3 - | 5,42E-05 - |
| 5712 | 1 | 0.2um_NOKCPHJN_319100 | Bacteria | Proteobacteri | Alphaproteob | Rhodobactera | Rhodobactera | Litorimicrobiu | Litorimicrobiu | metabolism | Energy produc | 01RNF | Produces ATP | 388401.RB215 | 18,4 | 1,854723699 | 0,00053337 | 8,36E-05 | 18 - | 0,000435861 - | 20 - | 0,000601345 - | 15 - | 0,000428741 - | 20 - | 0,000580098 - | 19 - | 0,000620805 - |
| 5713 | 1 | 0.2um_NOKCPHJN_707387 | Bacteria | Proteobacteri | Alphaproteob | Pelagibacteral | Pelagibacterac | Candidatus Pe | Candidatus Pe | unclassified | unclassified | unclassified | unclassified | unclassified | 0,4 | 0,489897949 | 1,79E-05 | 2,20E-05 | 1 - | 4,10E-05 - | 0 - | 0 - | 1 - | 4,84E-05 - | 0 - | 0 - | 0 - | 0 - |
| 5714 | 1 | 0.2um_NOKCPHJN_92459 | Bacteria | Proteobacteri | Gammaprote | Oceanospirilla | Saccharospirill | Reinekea | Reinekea fors | unclassified | unclassified | unclassified | unclassified | unclassified | 2,2 | 0,748331477 | 0,000197001 | 5,74E-05 | 3 - | 0,000231094 - | 2 - | 0,0001913 - | 2 - | 0,000181855 - | 3 - | 0,000276812 - | 1 - | 0,000103943 - |

| | | | | | | | | | | | | | | | | | | | | | | | | | | | | |
|---|---|---|---|---|---|---|---|---|---|---|---|---|---|---|---|---|---|---|---|---|---|---|---|---|---|---|---|---|
| 5715 | 1 | 3um_FIJJAFBF_122294 | Bacteria | Proteobacteri | Gammaprote | Oceanospirilla | Saccharospirill | Reinekea | Reinekea fors | information st | Translation, ri | COG0292 | Binds directly | 314283.MED2 | 2,6 | 0,8 | 0,00029245 | 7,56E-05 | 4 - | 0,000382589 - | 2 - | 0,000237531 - | 2 - | 0,000225804 - | 2 - | 0,000229139 - | 3 - | 0,000387186 - |
| 5716 | 1 | 3um_FIJJAFBF_603886 | Bacteria | Bacteroidetes | Flavobacteriia | Flavobacterial | Flavobacteriac | Lacinutrix | Lacinutrix sp. | unclassified | unclassified | unclassified | unclassified | unclassified | 1,6 | 1,356465997 | 0,00018037 | 0,000137954 | 4 - | 0,000409917 - | 1 - | 0,000127249 - | 2 - | 0,000241933 - | 1 - | 0,000122753 - | 0 - | 0 - |
| 5717 | 1 | 3um_FIJJAFBF_267386 | Eukaryota | Apicomplexa | Aconoidasida | Haemosporida | Plasmodiidae | Plasmodium | Plasmodium y | unclassified | unclassified | unclassified | unclassified | unclassified | 0,6 | 0,8 | 2,75E-05 | 3,66E-05 | 0 - | 0 - | 0 - | 0 - | 2 - | 9,12E-05 - | 1 - | 4,63E-05 - | 0 - | 0 - |
| 5718 | 1 | 3um_FIJJAFBF_166979 | Bacteria | Bacteroidetes | Flavobacteriia | Flavobacterial | Cryomorphace | unclassified | Cryomorphace | unclassified | unclassified | unclassified | unclassified | unclassified | 0,4 | 0,489897949 | 4,46E-05 | 5,51E-05 | 1 - | 0,000101572 - | 0 - | 0 - | 0 - | 0 - | 1 - | 0,000121667 - | 0 - | 0 - |
| 5719 | 1 | 3um_FIJJAFBF_434613 | Bacteria | Bacteroidetes | Cytophagia | Cytophagales | Flammeovirga | unclassified | Flammeovirga | poorly charact | Function unkn | COG0011 | Domain of unl | 880070.Cycma | 0,6 | 0,489897949 | 8,39E-05 | 6,98E-05 | 1 - | 0,000118327 - | 0 - | 0 - | 0 - | 0 - | 1 - | 0,000141735 - | 1 - | 0,000159664 - |
| 5720 | 1 | 0.2um_NOKCPHJN_327891 | Bacteria | Bacteroidetes | Flavobacteriia | Flavobacterial | Flavobacteriac | Polaribacter | various | metabolism | Inorganic ion t | COG1055 | arsenical pum | 487797.Flav3 | 0,6 | 0,489897949 | 1,77E-05 | 1,46E-05 | 1 - | 2,59E-05 - | 1 - | 3,21E-05 - | 1 - | 3,05E-05 - | 0 - | 0 - | 0 - | 0 - |
| 5721 | 1 | 3um_FIJJAFBF_376893 | Bacteria | Proteobacteri | Gammaprote | Alteromonada | Colwelliaceae | Colwellia | Colwellia psyc | unclassified | unclassified | unclassified | unclassified | unclassified | 0,4 | 0,489897949 | 1,32E-05 | 1,62E-05 | 0 - | 0 - | 0 - | 0 - | 1 - | 3,28E-05 - | 1 - | 3,33E-05 - | 0 - | 0 - |
| 5722 | 1 | 3um_FIJJAFBF_322946 | Bacteria | Proteobacteri | Alphaproteob | Rhodobactera | Rhodobactera | unclassified | Rhodobactera | cellular proces | Signal transdu | 01Q8R | gtp-binding pr | 388401.RB215 | 2 | 0,632455532 | 4,44E-05 | 1,16E-05 | 3 - | 5,67E-05 - | 2 - | 4,70E-05 - | 2 - | 4,46E-05 - | 1 - | 2,26E-05 - | 2 - | 5,10E-05 - |
| 5723 | 1 | 0.2um_NOKCPHJN_473242 | Eukaryota | Ascomycota | Sordariomyce | Hypocreales | Nectriaceae | Fusarium | Fusarium oxys | unclassified | unclassified | unclassified | unclassified | unclassified | 3 | 0 | 9,54E-05 | 9,06E-06 | 3 - | 7,99E-05 - | 3 - | 9,92E-05 - | 3 - | 9,43E-05 - | 3 - | 9,57E-05 - | 3 - | 0,000107801 - |
| 5724 | 1 | 0.2um_NOKCPHJN_83663 | Bacteria | Bacteroidetes | Flavobacteriia | Flavobacterial | Flavobacteriac | Polaribacter | Polaribacter s | cellular proces | Posttranslatio | 05E6M | Sulphatase-m | 487797.Flav3 | 0,8 | 0,4 | 2,42E-05 | 1,24E-05 | 1 - | 2,53E-05 - | 1 - | 3,15E-05 - | 1 - | 2,99E-05 - | 0 - | 0 - | 1 - | 3,42E-05 - |
| 5725 | 1 | 3um_FIJJAFBF_578538 | Bacteria | Proteobacteri | Gammaprote | Alteromonada | Alteromonada | Paraglaciecola | Paraglaciecola | information st | Translation, ri | COG0290 | IF-3 binds to t | 1085623.GNIT | 0,6 | 0,8 | 4,39E-05 | 5,55E-05 | 2 - | 0,000131173 - | 0 - | 0 - | 0 - | 0 - | 0 - | 0 - | 1 - | 8,85E-05 - |
| 5726 | 1 | 3um_FIJJAFBF_225139 | Bacteria | Proteobacteri | Alphaproteob | Rhodobactera | Rhodobactera | Planktomarina | Planktomarina | information st | Translation, ri | COG0255 | 50s ribosomal | 290400.Jann_ | 2,8 | 0,4 | 0,00055898 | 6,65E-05 | 3 - | 0,000506368 - | 3 - | 0,00062876 - | 3 - | 0,000597716 - | 3 - | 0,000606544 - | 2 - | 0,000455513 - |
| 5727 | 1 | 3um_FIJJAFBF_450540 | Bacteria | unclassified | unclassified | unclassified | unclassified | unclassified | uncultured ba | cellular proces | Cell wall/mem | 09DZB | Fasciclin | 487797.Flav3 | 1,8 | 0,4 | 0,000145269 | 3,62E-05 | 2 - | 0,000135031 - | 2 - | 0,000167669 - | 1 - | 7,97E-05 - | 2 - | 0,000161745 - | 2 - | 0,000182205 - |
| 5728 | 1 | 3um_FIJJAFBF_359538 | Bacteria | Bacteroidetes | Flavobacteriia | Flavobacterial | Flavobacteriac | Polaribacter | Polaribacter s | unclassified | unclassified | unclassified | unclassified | unclassified | 5,2 | 0,748331477 | 8,45E-05 | 1,91E-05 | 4 - | 5,38E-05 - | 6 - | 0,00010013 - | 5 - | 7,93E-05 - | 5 - | 8,05E-05 - | 6 - | 0,000108811 - |
| 5729 | 1 | 0.2um_NOKCPHJN_481994 | Bacteria | Proteobacteri | Alphaproteob | Pelagibacteral | Pelagibactera | Candidatus Pe | Candidatus Pe | unclassified | unclassified | unclassified | unclassified | unclassified | 2 | 1,414213562 | 0,000176159 | 0,000125519 | 2 - | 0,000151022 - | 0 - | 0 - | 4 - | 0,000356532 - | 3 - | 0,000271349 - | 1 - | 0,000101891 - |
| 5730 | 1 | 0.2um_NOKCPHJN_617451 | Eukaryota | Chlorophyta | Chlorophycea | Sphaeropleale | Selenastracea | Raphidocelis | Raphidocelis s | poorly charact | Function unkn | 0NVRZ | NA | 170187.SP_17 | 0,2 | 0,4 | 2,52E-06 | 5,05E-06 | 0 - | 0 - | 1 - | 1,26E-05 - | 0 - | 0 - | 0 - | 0 - | 0 - | 0 - |
| 5731 | 1 | 0.2um_NOKCPHJN_117150 | Bacteria | Bacteroidetes | Flavobacteriia | Flavobacterial | Cryomorphace | unclassified | Cryomorphace | metabolism | Inorganic ion t | 05C27 | TonB-depende | 313596.RB25 | 1,4 | 0,489897949 | 1,82E-05 | 5,36E-06 | 2 - | 2,23E-05 - | 1 - | 1,38E-05 - | 1 - | 1,32E-05 - | 2 - | 2,67E-05 - | 1 - | 1,50E-05 - |
| 5732 | 1 | 3um_FIJJAFBF_142183 | Bacteria | Proteobacteri | Gammaprote | Cellvibrionales | Halieaceae | Halioglobus | Halioglobus ja | metabolism | Energy produc | COG0056 | Produces ATP | 314285.KT71_ | 2,2 | 0,748331477 | 5,97E-05 | 2,17E-05 | 1 - | 2,23E-05 - | 3 - | 8,32E-05 - | 3 - | 7,91E-05 - | 2 - | 5,35E-05 - | 2 - | 6,03E-05 - |
| 5733 | 1 | 0.2um_NOKCPHJN_704379 | Bacteria | Proteobacteri | Gammaprote | unclassified | unclassified | unclassified | Gammaprote | poorly charact | Function unkn | 17AGM | Protein of unk | 243233.MCA2 | 0,2 | 0,4 | 1,06E-05 | 2,13E-05 | 1 - | 5,31E-05 - | 0 - | 0 - | 0 - | 0 - | 0 - | 0 - | 0 - | 0 - |
| 5734 | 4 | 0.2um_NOKCPHJN_865894;3um_FIJJAFBF_1168 | Eukaryota | various | various | various | various | various | various | information st | various | various | various | various | 1 | 0 | 3,69E-05 | 3,51E-06 | 1 - | 3,09E-05 - | 1 - | 3,84E-05 - | 1 - | 3,65E-05 - | 1 - | 3,71E-05 - | 1 - | 4,17E-05 - |
| 5735 | 1 | 3um_FIJJAFBF_411197 | Bacteria | Proteobacteri | Betaproteoba | unclassified | unclassified | unclassified | Betaproteoba | metabolism | Carbohydrate | COG0176 | Transaldolase | 688245.CtCNB | 1,2 | 1,166190379 | 5,08E-05 | 4,91E-05 | 0 - | 0 - | 1 - | 4,39E-05 - | 2 - | 8,34E-05 - | 3 - | 0,000126908 - | 0 - | 0 - |
| 5736 | 1 | 0.2um_NOKCPHJN_328778 | Bacteria | Proteobacteri | Gammaprote | Nevskiales | Algiphilaceae | Algiphilus | Algiphilus aro | poorly charact | Function unkn | 0XSV1 | NA | 204773.HEAR | 1 | 0 | 4,50E-06 | 4,28E-07 | 1 - | 3,77E-06 - | 1 - | 4,69E-06 - | 1 - | 4,45E-06 - | 1 - | 4,52E-06 - | 1 - | 5,09E-06 - |
| 5737 | 1 | 0.2um_NOKCPHJN_499931 | Bacteria | Bacteroidetes | Flavobacteriia | unclassified | unclassified | unclassified | Flavobacteria | metabolism | Secondary me | 17KHV | Inherit from C | 709032.Sulku_ | 1,6 | 0,8 | 2,77E-05 | 1,41E-05 | 2 - | 3,00E-05 - | 2 - | 3,72E-05 - | 2 - | 3,54E-05 - | 2 - | 3,59E-05 - | 0 - | 0 - |
| 5738 | 1 | 0.2um_NOKCPHJN_265381 | Bacteria | Proteobacteri | Betaproteoba | Burkholderiale | Burkholderiac | Caballeronia | Caballeronia s | cellular proces | Cell wall/mem | 16PZG | Glycosyltransf | 945681.APO_ | 0,6 | 0,489897949 | 5,30E-06 | 4,34E-06 | 0 - | 0 - | 1 - | 8,72E-06 - | 1 - | 8,29E-06 - | 0 - | 0 - | 1 - | 9,48E-06 - |
| 5739 | 1 | 3um_FIJJAFBF_37757 | Bacteria | Proteobacteri | Gammaprote | Oceanospirilla | Saccharospirill | Reinekea | Reinekea blan | metabolism | Energy produc | 0QIIJ | Succinyl-CoA l | 314283.MED2 | 1,6 | 0,489897949 | 7,52E-05 | 2,37E-05 | 2 - | 7,92E-05 - | 1 - | 4,91E-05 - | 2 - | 9,34E-05 - | 1 - | 4,74E-05 - | 2 - | 0,00010681 - |
| 5740 | 1 | 3um_FIJJAFBF_532747 | unclassified | unclassified | unclassified | unclassified | unclassified | unclassified | unclassified | unclassified | unclassified | unclassified | unclassified | unclassified | 0,6 | 0,489897949 | 1,49E-05 | 1,22E-05 | 0 - | 0 - | 1 - | 2,55E-05 - | 1 - | 2,43E-05 - | 1 - | 2,46E-05 - | 0 - | 0 - |
| 5741 | 1 | 0.2um_NOKCPHJN_351106 | unclassified | unclassified | unclassified | unclassified | unclassified | unclassified | unclassified | unclassified | unclassified | unclassified | unclassified | unclassified | 1 | 0 | 3,26E-05 | 3,10E-06 | 1 - | 2,73E-05 - | 1 - | 3,39E-05 - | 1 - | 3,23E-05 - | 1 - | 3,27E-05 - | 1 - | 3,69E-05 - |
| 5742 | 1 | 3um_FIJJAFBF_346693 | Bacteria | Proteobacteri | Gammaprote | Oceanospirilla | Oceanospirilla | Marinobacteri | Marinobacteri | metabolism | Energy produc | 17CGT | F(1)F(0) ATP s | 1001585.MDS | 8 | 0,632455532 | 0,000697562 | 2,21E-05 | 9 - | 0,000662174 - | 8 - | 0,000730866 - | 8 - | 0,000694781 - | 8 - | 0,000705043 - | 7 - | 0,00069495 - |
| 5743 | 1 | 3um_FIJJAFBF_452766 | Bacteria | Bacteroidetes | Flavobacteriia | Flavobacterial | Flavobacteriac | Polaribacter | Polaribacter s | metabolism | Energy produc | COG1053 | Succinate DeH | 313598.MED1 | 13 | 0,894427191 | 0,000266978 | 2,33E-05 | 14 - | 0,000241635 - | 12 - | 0,000257177 - | 14 - | 0,000285226 - | 12 - | 0,00024809 - | 13 - | 0,000302762 - |
| 5744 | 1 | 0.2um_NOKCPHJN_843966 | Eukaryota | Ascomycota | Eurotiomycete | Eurotiales | Aspergillaceae | Penicillium | Penicillium ch | cellular proces | Cytoskeleton | 0M6P0 | TPR domain p | 500485.XP_00 | 0,2 | 0,4 | 4,11E-06 | 8,23E-06 | 0 - | 0 - | 0 - | 0 - | 0 - | 0 - | 0 - | 0 - | 1 - | 2,06E-05 - |
| 5745 | 1 | 0.2um_NOKCPHJN_772040 | Bacteria | Proteobacteri | Alphaproteob | Rhodobactera | Rhodobactera | Planktomarina | Planktomarina | metabolism | Energy produc | 178X6 | carbon mono | 388739.RSK20 | 1 | 0,632455532 | 8,02E-05 | 4,52E-05 | 2 - | 0,000139972 - | 0 - | 0 - | 1 - | 8,26E-05 - | 1 - | 8,38E-05 - | 1 - | 9,44E-05 - |
| 5746 | 1 | 0.2um_NOKCPHJN_1059827 | Bacteria | Bacteroidetes | Flavobacteriia | Flavobacterial | Flavobacteriac | Polaribacter | Polaribacter h | metabolism | Carbohydrate | 099YR | Catalyzes the | 391603.FBALC | 0,6 | 0,489897949 | 2,02E-05 | 1,66E-05 | 1 - | 2,99E-05 - | 0 - | 0 - | 1 - | 3,53E-05 - | 1 - | 3,58E-05 - | 0 - | 0 - |
| 5747 | 1 | 3um_FIJJAFBF_156358 | Eukaryota | Bacillariophyta | Bacillariophyc | Naviculales | Naviculaceae | Fistulifera | Fistulifera sola | unclassified | unclassified | unclassified | unclassified | unclassified | 0,2 | 0,4 | 7,72E-06 | 1,54E-05 | 0 - | 0 - | 0 - | 0 - | 0 - | 0 - | 1 - | 3,86E-05 - | 0 - | 0 - |
| 5748 | 1 | 3um_FIJJAFBF_738572 | Bacteria | Proteobacteri | Alphaproteob | Rhodobactera | Rhodobactera | Yoonia | Yoonia marico | unclassified | unclassified | unclassified | unclassified | unclassified | 0,6 | 0,489897949 | 1,65E-05 | 1,37E-05 | 1 - | 2,33E-05 - | 0 - | 0 - | 1 - | 2,75E-05 - | 0 - | 0 - | 1 - | 3,15E-05 - |
| 5749 | 1 | 3um_FIJJAFBF_525779 | Bacteria | Proteobacteri | Betaproteoba | Burkholderiale | Comamonada | Hydrogenoph | Hydrogenoph | metabolism | Energy produc | 179RI | May be involv | 338969.Rfer_ | 0,6 | 0,489897949 | 6,09E-05 | 5,02E-05 | 1 - | 8,90E-05 - | 1 - | 0,00011048 - | 1 - | 0,000105025 - | 0 - | 0 - | 0 - | 0 - |
| 5750 | 1 | 3um_FIJJAFBF_264702 | Bacteria | Proteobacteri | Gammaprote | Oceanospirilla | Saccharospirill | Reinekea | Reinekea fors | metabolism | Carbohydrate | 0XPV5 | extracellular s | 314283.MED2 | 17 | 0,632455532 | 0,000574054 | 4,78E-05 | 18 - | 0,000510119 - | 17 - | 0,000598227 - | 17 - | 0,000568691 - | 16 - | 0,000543144 - | 17 - | 0,00065009 - |
| 5751 | 1 | 3um_FIJJAFBF_744842 | Bacteria | Proteobacteri | Gammaprote | Cellvibrionales | Cellvibrionace | Gilvimarinus | Gilvimarinus p | information st | Translation, ri | COG0087 | One of the pri | 247633.GP214 | 1 | 0,632455532 | 6,41E-05 | 4,07E-05 | 1 - | 5,39E-05 - | 1 - | 6,69E-05 - | 2 - | 0,000127213 - | 0 - | 0 - | 1 - | 7,27E-05 - |
| 5752 | 1 | 3um_FIJJAFBF_38224 | Bacteria | Proteobacteri | Gammaprote | Alteromonada | Alteromonada | Paraglaciecola | Paraglaciecola | unclassified | unclassified | unclassified | unclassified | unclassified | 0,2 | 0,4 | 5,49E-06 | 1,10E-05 | 0 - | 0 - | 0 - | 0 - | 0 - | 0 - | 1 - | 2,74E-05 - | 0 - | 0 - |
| 5753 | 1 | 0.2um_NOKCPHJN_957389 | Eukaryota | unclassified | Oomycetes | Saprolegniales | Saprolegniace | Aphanomyces | Aphanomyces | information st | Chromatin str | KOG1744 | Core compone | 4787.PITG_03 | 1,2 | 0,4 | 0,000136737 | 3,03E-05 | 2 - | 0,0001962 - | 1 - | 0,000121811 - | 1 - | 0,000115797 - | 1 - | 0,000117507 - | 1 - | 0,000132371 - |
| 5754 | 1 | 3um_FIJJAFBF_532813 | Bacteria | Proteobacteri | Alphaproteob | Rhodobactera | unclassified | unclassified | Rhodobactera | unclassified | unclassified | unclassified | unclassified | unclassified | 6,8 | 0,748331477 | 0,000465399 | 4,54E-05 | 8 - | 0,000461414 - | 6 - | 0,000429705 - | 7 - | 0,000476571 - | 6 - | 0,000414523 - | 7 - | 0,000544785 - |
| 5755 | 1 | 3um_FIJJAFBF_43263 | Bacteria | Bacteroidetes | Flavobacteriia | Flavobacterial | Flavobacteriac | Gaetbulibacte | Gaetbulibacte | unclassified | unclassified | unclassified | unclassified | unclassified | 0,4 | 0,489897949 | 3,62E-05 | 4,43E-05 | 0 - | 0 - | 0 - | 0 - | 1 - | 8,97E-05 - | 1 - | 9,10E-05 - | 0 - | 0 - |
| 5756 | 1 | 3um_FIJJAFBF_222371 | Bacteria | Proteobacteri | Alphaproteob | Rhodobactera | Rhodobactera | Ruegeria | Ruegeria sp. 6 | cellular proces | Posttranslatio | 01ZS8 | Thioredoxin | 439497.RR11_ | 1,8 | 0,4 | 0,00022932 | 4,53E-05 | 2 - | 0,00021656 - | 2 - | 0,000268904 - | 2 - | 0,000255627 - | 2 - | 0,000259402 - | 1 - | 0,000146108 - |
| 5757 | 1 | 3um_FIJJAFBF_356109 | Bacteria | Proteobacteri | Gammaprote | Oceanospirilla | Saccharospirill | Reinekea | Reinekea blan | unclassified | unclassified | unclassified | unclassified | unclassified | 0,4 | 0,489897949 | 2,35E-05 | 2,89E-05 | 1 - | 5,34E-05 - | 0 - | 0 - | 0 - | 0 - | 1 - | 6,39E-05 - | 0 - | 0 - |
| 5758 | 1 | 3um_FIJJAFBF_73387 | unclassified | unclassified | unclassified | unclassified | unclassified | unclassified | unclassified | unclassified | unclassified | unclassified | unclassified | unclassified | 0,8 | 0,4 | 5,36E-05 | 2,76E-05 | 1 - | 5,60E-05 - | 1 - | 6,95E-05 - | 0 - | 0 - | 1 - | 6,71E-05 - | 1 - | 7,55E-05 - |
| 5759 | 1 | 3um_FIJJAFBF_08103 | Bacteria | Bacteroidetes | Flavobacteriia | Flavobacterial | Flavobacteriac | Polaribacter | Polaribacter s | information st | Translation, ri | COG0251 | Endoribonucle | 153948.NAL21 | 2,8 | 0,748331477 | 0,000300873 | 7,51E-05 | 3 - | 0,000273278 - | 2 - | 0,00022622 - | 3 - | 0,000322577 - | 4 - | 0,000436455 - | 2 - | 0,000245833 - |
| 5760 | 2 | 0.2um_NOKCPHJN_201636;0.2um_NOKCPHJN_4 | Bacteria | various | various | various | various | various | various | poorly charact | Function unkn | 0PBET | NA | 313603.FB217 | 0,6 | 0,489897949 | 4,21E-05 | 3,47E-05 | 1 - | 6,24E-05 - | 0 - | 0 - | 1 - | 7,36E-05 - | 1 - | 7,47E-05 - | 0 - | 0 - |
| 5761 | 1 | 3um_FIJJAFBF_33416 | Bacteria | Bacteroidetes | Flavobacteriia | Flavobacterial | Flavobacteriac | Polaribacter | Polaribacter s | metabolism | Amino acid tra | COG0174 | glutamine syn | 313594.PI23P | 1,4 | 0,489897949 | 5,67E-05 | 2,00E-05 | 1 - | 3,40E-05 - | 1 - | 4,22E-05 - | 2 - | 8,02E-05 - | 2 - | 8,14E-05 - | 1 - | 4,58E-05 - |
| 5762 | 1 | 0.2um_NOKCPHJN_21801 | Bacteria | Bacteroidetes | Flavobacteriia | Flavobacterial | Flavobacteriac | Polaribacter | Polaribacter s | poorly charact | Function unkn | 0P9SU | NA | 313598.MED1 | 4,4 | 0,489897949 | 0,00017582 | 3,23E-05 | 4 - | 0,000133075 - | 4 - | 0,000165239 - | 4 - | 0,000157081 - | 5 - | 0,000199251 - | 5 - | 0,000224456 - |
| 5763 | 1 | 3um_FIJJAFBF_559826 | Bacteria | Proteobacteri | Alphaproteob | Rhodobactera | Rhodobactera | Sulfitobacter | unclassified | unclassified | unclassified | unclassified | unclassified | unclassified | 1,2 | 0,748331477 | 3,99E-05 | 2,57E-05 | 1 - | 2,85E-05 - | 1 - | 3,54E-05 - | 2 - | 6,72E-05 - | 2 - | 6,82E-05 - | 0 - | 0 - |
| 5764 | 1 | 0.2um_NOKCPHJN_181513 | Bacteria | Proteobacteri | Betaproteoba | unclassified | unclassified | unclassified | Betaproteoba | information st | Translation, ri | 0B7GH | Ribonuclease | 887062.HGR_ | 0,2 | 0,4 | 3,20E-06 | 6,41E-06 | 0 - | 0 - | 0 - | 0 - | 0 - | 0 - | 1 - | 1,60E-05 - | 0 - | 0 - |
| 5765 | 1 | 0.2um_NOKCPHJN_957709 | Bacteria | Bacteroidetes | unclassified | unclassified | unclassified | unclassified | Bacteroidetes | information st | Translation, ri | 05C64 | Binds directly | 398720.MED2 | 0,4 | 0,489897949 | 2,18E-05 | 2,69E-05 | 1 - | 4,97E-05 - | 0 - | 0 - | 0 - | 0 - | 1 - | 5,95E-05 - | 0 - | 0 - |
| 5766 | 1 | 0.2um_NOKCPHJN_384897 | Bacteria | Bacteroidetes | Flavobacteriia | Flavobacterial | Crocinitomica | unclassified | Crocinitomica | unclassified | unclassified | unclassified | unclassified | unclassified | 0,2 | 0,4 | 7,94E-06 | 1,59E-05 | 1 - | 3,97E-05 - | 0 - | 0 - | 0 - | 0 - | 0 - | 0 - | 0 - | 0 - |
| 5767 | 1 | 0.2um_NOKCPHJN_315936 | Bacteria | Proteobacteri | Alphaproteob | Pelagibacteral | Pelagibactera | Candidatus Pe | Candidatus Pe | unclassified | unclassified | unclassified | unclassified | unclassified | 1,6 | 0,489897949 | 6,49E-05 | 2,04E-05 | 2 - | 6,83E-05 - | 1 - | 4,24E-05 - | 2 - | 8,06E-05 - | 1 - | 4,09E-05 - | 2 - | 9,22E-05 - |
| 5768 | 1 | 3um_FIJJAFBF_107751 | Bacteria | Proteobacteri | Gammaprote | Oceanospirilla | Saccharospirill | Reinekea | Reinekea blan | unclassified | unclassified | unclassified | unclassified | unclassified | 2,8 | 0,748331477 | 9,86E-05 | 2,40E-05 | 3 - | 8,97E-05 - | 2 - | 7,42E-05 - | 4 - | 0,000141127 - | 3 - | 0,000107409 - | 2 - | 8,07E-05 - |
| 5769 | 1 | 0.2um_NOKCPHJN_220129 | Bacteria | Proteobacteri | Gammaprote | unclassified | unclassified | unclassified | OM182 bacter | cellular proces | Posttranslatio | 0QJ3C | Acts as a chap | 314277.MED1 | 0,8 | 0,748331477 | 1,64E-05 | 1,58E-05 | 1 - | 1,80E-05 - | 0 - | 0 - | 2 - | 4,24E-05 - | 1 - | 2,15E-05 - | 0 - | 0 - |
| 5770 | 1 | 0.2um_NOKCPHJN_588087 | Bacteria | Proteobacteri | Alphaproteob | Pelagibacteral | Pelagibactera | Candidatus Pe | Candidatus Pe | cellular proces | Cell wall/mem | 7257 | NA | 335992.SAR11 | 0,8 | 0,4 | 3,41E-05 | 1,75E-05 | 1 - | 3,56E-05 - | 1 - | 4,43E-05 - | 0 - | 0 - | 1 - | 4,27E-05 - | 1 - | 4,81E-05 - |
| 5771 | 2 | 3um_FIJJAFBF_238002;3um_FIJJAFBF_468833 | Bacteria | Bacteroidetes | unclassified | unclassified | unclassified | unclassified | Bacteroidetes | unclassified | unclassified | unclassified | unclassified | unclassified | 0,2 | 0,4 | 9,12E-06 | 1,82E-05 | 0 - | 0 - | 0 - | 0 - | 1 - | 4,56E-05 - | 0 - | 0 - | 0 - | 0 - |
| 5772 | 1 | 3um_FIJJAFBF_306383 | Bacteria | Proteobacteri | Betaproteoba | Burkholderiale | Comamonada | Simplicispira | Simplicispira p | information st | Translation, ri | 177SZ | This protein bi | 426114.THI_3 | 1,6 | 0,489897949 | 0,000150053 | 3,93E-05 | 2 - | 0,000160527 - | 1 - | 9,97E-05 - | 2 - | 0,000189486 - | 2 - | 0,000192284 - | 1 - | 0,000108304 - |
| 5773 | 1 | 0.2um_NOKCPHJN_248049 | Bacteria | Proteobacteri | Alphaproteob | Rhodobactera | Rhodobactera | Litoreibacter | Litoreibacter a | metabolism | Carbohydrate | 01RFA | ABC transport | 367336.OM22 | 14,2 | 2,481934729 | 0,000538145 | 0,000118684 | 16 - | 0,000507301 - | 11 - | 0,000433068 - | 13 - | 0,000486538 - | 13 - | 0,000493725 - | 18 - | 0,000770094 - |
| 5774 | 1 | 0.2um_NOKCPHJN_444557 | Bacteria | Proteobacteri | Alphaproteob | Pelagibacteral | Pelagibactera | Candidatus Pe | Candidatus Pe | metabolism | Amino acid tra | 01R4A | Polyamine AB | 439493.PB721 | 5 | 1,264911064 | 0,00018458 | 4,85E-05 | 4 - | 0,000123749 - | 4 - | 0,000153659 - | 6 - | 0,000219109 - | 7 - | 0,000259402 - | 4 - | 0,000166981 - |
| 5775 | 1 | 3um_FIJJAFBF_189642 | Bacteria | Bacteroidetes | Flavobacteriia | Flavobacterial | Flavobacteriac | Polaribacter | Polaribacter s | unclassified | unclassified | unclassified | unclassified | unclassified | 2,6 | 1,624807681 | 0,000113805 | 7,13E-05 | 1 - | 3,61E-05 - | 4 - | 0,000179269 - | 5 - | 0,000213022 - | 1 - | 4,32E-05 - | 2 - | 9,74E-05 - |
| 5776 | 1 | 3um_FIJJAFBF_102139 | Bacteria | Bacteroidetes | Flavobacteriia | unclassified | unclassified | unclassified | Flavobacteria | metabolism | Amino acid tra | COG0174 | glutamine syn | 487797.Flav3 | 2,6 | 0,8 | 0,0001036 | 3,00E-05 | 3 - | 0,000102175 - | 3 - | 0,000126871 - | 3 - | 0,000120607 - | 3 - | 0,000122389 - | 1 - | 4,60E-05 - |
| 5777 | 1 | 0.2um_NOKCPHJN_281073 | Bacteria | Bacteroidetes | Flavobacteriia | Flavobacterial | unclassified | unclassified | Flavobacterial | unclassified | unclassified | unclassified | unclassified | unclassified | 0,2 | 0,4 | 1,24E-06 | 2,48E-06 | 1 - | 6,20E-06 - | 0 - | 0 - | 0 - | 0 - | 0 - | 0 - | 0 - | 0 - |
| 5778 | 1 | 3um_FIJJAFBF_581444 | Bacteria | Proteobacteri | Gammaprote | Arenicellales | Arenicellaceae | Arenicella | Arenicella xan | unclassified | unclassified | unclassified | unclassified | unclassified | 2,2 | 1,166190379 | 4,93E-05 | 2,99E-05 | 2 - | 3,65E-05 - | 3 - | 6,80E-05 - | 1 - | 2,15E-05 - | 1 - | 2,19E-05 - | 4 - | 9,85E-05 - |
| 5779 | 1 | 0.2um_NOKCPHJN_714302 | Bacteria | Proteobacteri | Alphaproteob | Pelagibacteral | Pelagibactera | Candidatus Pe | Candidatus Pe | unclassified | unclassified | unclassified | unclassified | unclassified | 0,4 | 0,489897949 | 2,03E-05 | 2,49E-05 | 0 - | 0 - | 1 - | 5,20E-05 - | 1 - | 4,94E-05 - | 0 - | 0 - | 0 - | 0 - |
| 5780 | 1 | 3um_FIJJAFBF_564112 | Bacteria | Proteobacteri | Gammaprote | Oceanospirilla | Saccharospirill | Reinekea | Reinekea fors | poorly charact | Function unkn | COG2847 | Secreted prot | 314283.MED2 | 0,8 | 0,4 | 7,60E-05 | 3,83E-05 | 0 - | 0 - | 1 - | 9,50E-05 - | 1 - | 9,03E-05 - | 1 - | 9,17E-05 - | 1 - | 0,00010325 - |
| 5781 | 1 | 0.2um_NOKCPHJN_177445 | Bacteria | Bacteroidetes | Flavobacteriia | Flavobacterial | Flavobacteriac | Polaribacter | Polaribacter s | poorly charact | Function unkn | 05DPC | Ragb susd dor | 313598.MED1 | 1,6 | 0,489897949 | 4,49E-05 | 1,58E-05 | 1 - | 2,31E-05 - | 1 - | 2,87E-05 - | 2 - | 5,46E-05 - | 2 - | 5,54E-05 - | 2 - | 6,24E-05 - |
| 5782 | 1 | 3um_FIJJAFBF_259311 | Bacteria | Planctomycete | Phycisphaerae | Phycisphaeral | Phycisphaerac | Phycisphaera | Phycisphaera | cellular proces | Cell wall/mem | 06B97 | phosphate-sel | 443144.GM21 | 1 | 0 | 2,99E-05 | 2,84E-06 | 1 - | 2,51E-05 - | 1 - | 3,11E-05 - | 1 - | 2,96E-05 - | 1 - | 3,00E-05 - | 1 - | 3,38E-05 - |
| 5783 | 1 | 0.2um_NOKCPHJN_438104 | Bacteria | Bacteroidetes | unclassified | unclassified | unclassified | unclassified | Bacteroidetes | unclassified | unclassified | unclassified | unclassified | unclassified | 1,6 | 0,489897949 | 0,000165008 | 5,46E-05 | 2 - | 0,000172597 - | 2 - | 0,000214314 - | 1 - | 0,000101866 - | 1 - | 0,000103371 - | 2 - | 0,000232894 - |
| 5784 | 1 | 3um_FIJJAFBF_28489 | Bacteria | Proteobacteri | Alphaproteob | Sphingomona | Sphingomona | Sphingopyxis | Sphingopyxis | unclassified | unclassified | unclassified | unclassified | unclassified | 1 | 0 | 0,000118127 | 1,12E-05 | 1 - | 9,89E-05 - | 1 - | 0,000122861 - | 1 - | 0,000116795 - | 1 - | 0,00011852 - | 1 - | 0,000133512 - |
| 5785 | 1 | 3um_FIJJAFBF_374733 | Bacteria | Bacteroidetes | Flavobacteriia | Flavobacterial | Flavobacteriac | Formosa | Formosa sp. H | information st | Translation, ri | COG0254 | 50S ribosomal | 1046627.BZAR | 3 | 0,894427191 | 0,000487312 | 0,000128944 | 4 - | 0,000553141 - | 4 - | 0,000686838 - | 2 - | 0,000326463 - | 3 - | 0,000496928 - | 2 - | 0,000373192 - |
| 5786 | 1 | 3um_FIJJAFBF_189631 | Bacteria | Bacteroidetes | Flavobacteriia | Flavobacterial | Flavobacteriac | Polaribacter | Polaribacter h | poorly charact | Function unkn | 11FUA | Membrane | 313594.PI23P | 2 | 0,632455532 | 0,000160032 | 5,20E-05 | 2 - | 0,000134242 - | 2 - | 0,000166689 - | 3 - | 0,000237688 - | 1 - | 8,04E-05 - | 2 - | 0,00018114 - |
| 5787 | 1 | 3um_FIJJAFBF_135566 | Bacteria | Proteobacteri | Alphaproteob | Rhodobactera | unclassified | unclassified | Rhodobactera | metabolism | Energy produc | COG0711 | F(1)F(0) ATP s | 367336.OM22 | 2 | 0 | 0,000153963 | 1,46E-05 | 2 - | 0,000128963 - | 2 - | 0,000160134 - | 2 - | 0,000152227 - | 2 - | 0,000154476 - | 2 - | 0,000174016 - |
| 5788 | 1 | 3um_FIJJAFBF_719543 | Viruses | unclassified | unclassified | unclassified | unclassified | unclassified | uncultured M | unclassified | unclassified | unclassified | unclassified | unclassified | 1,6 | 1,496662955 | 0,000106294 | 9,35E-05 | 4 - | 0,00024037 - | 2 - | 0,000149234 - | 2 - | 0,000141866 - | 0 - | 0 - | 0 - | 0 - |
| 5789 | 1 | 3um_FIJJAFBF_93576 | Bacteria | various | various | various | various | various | various | metabolism | Inorganic ion t | 0PAA4 | heavy metal | 313598.MED1 | 0,6 | 0,489897949 | 5,74E-05 | 4,73E-05 | 1 - | 8,50E-05 - | 0 - | 0 - | 1 - | 0,000100357 - | 1 - | 0,000101839 - | 0 - | 0 - |
| 5790 | 1 | 3um_FIJJAFBF_481967 | Bacteria | Proteobacteri | Alphaproteob | Rhodobactera | Rhodobactera | unclassified | Rhodobactera | metabolism | Energy produc | 05E2M | Manganese-d | 314262.MED1 | 0,6 | 0,489897949 | 2,84E-05 | 2,33E-05 | 0 - | 0 - | 1 - | 4,66E-05 - | 0 - | 0 - | 1 - | 4,49E-05 - | 1 - | 5,06E-05 - |
| 5791 | 1 | 0.2um_NOKCPHJN_164695 | Bacteria | Proteobacteri | Gammaprote | unclassified | unclassified | unclassified | Gammaprote | unclassified | unclassified | unclassified | unclassified | unclassified | 6 | 1,095445115 | 0,000314211 | 4,30E-05 | 7 - | 0,00031141 - | 6 - | 0,000331439 - | 6 - | 0,000315075 - | 7 - | 0,000373017 - | 4 - | 0,000240115 - |
| 5792 | 1 | 0.2um_NOKCPHJN_221723 | Eukaryota | unclassified | Oomycetes | Peronosporale | Peronosporac | Phytophthora | Phytophthora | unclassified | unclassified | unclassified | unclassified | unclassified | 0,4 | 0,489897949 | 1,16E-06 | 1,42E-06 | 0 - | 0 - | 1 - | 2,78E-06 - | 0 - | 0 - | 0 - | 0 - | 1 - | 3,02E-06 - |
| 5793 | 1 | 3um_FIJJAFBF_358728 | Bacteria | Proteobacteri | Gammaprote | unclassified | unclassified | unclassified | gamma prote | unclassified | unclassified | unclassified | unclassified | unclassified | 0,4 | 0,489897949 | 6,88E-05 | 8,45E-05 | 0 - | 0 - | 0 - | 0 - | 0 - | 0 - | 1 - | 0,000161745 - | 1 - | 0,000182205 - |
| 5794 | 1 | 0.2um_NOKCPHJN_1068205 | Bacteria | Proteobacteri | Alphaproteob | Rhodobactera | Rhodobactera | Planktomarina | Planktomarina | unclassified | unclassified | unclassified | unclassified | unclassified | 1 | 0 | 0,000153963 | 1,46E-05 | 1 - | 0,000128963 - | 1 - | 0,000160134 - | 1 - | 0,000152227 - | 1 - | 0,000154476 - | 1 - | 0,000174016 - |
| 5795 | 1 | 3um_FIJJAFBF_103157 | Bacteria | Bacteroidetes | Flavobacteriia | Flavobacterial | Flavobacteriac | Polaribacter | Polaribacter s | unclassified | unclassified | unclassified | unclassified | unclassified | 0,4 | 0,489897949 | 1,74E-05 | 2,14E-05 | 1 - | 3,99E-05 - | 0 - | 0 - | 1 - | 4,70E-05 - | 0 - | 0 - | 0 - | 0 - |
| 5796 | 1 | 3um_FIJJAFBF_698322 | Bacteria | Proteobacteri | Gammaprote | Oceanospirilla | Saccharospirill | Reinekea | Reinekea fors | poorly charact | Function unkn | 17NH8 | NA | 314283.MED2 | 1,2 | 0,748331477 | 7,68E-05 | 4,69E-05 | 0 - | 0 - | 2 - | 0,000128976 - | 2 - | 0,000122608 - | 1 - | 6,22E-05 - | 1 - | 7,01E-05 - |
| 5797 | 1 | 3um_FIJJAFBF_15907 | Bacteria | Bacteroidetes | Flavobacteriia | Flavobacterial | Flavobacteriac | Polaribacter | Polaribacter s | information st | Translation, ri | 09AI4 | Nol1 Nop2 Sur | 313594.PI23P | 0,4 | 0,489897949 | 1,21E-05 | 1,50E-05 | 1 - | 2,70E-05 - | 1 - | 3,35E-05 - | 0 - | 0 - | 0 - | 0 - | 0 - | 0 - |
| 5798 | 1 | 3um_FIJJAFBF_532861 | Bacteria | Proteobacteri | Alphaproteob | Rhodobactera | unclassified | unclassified | Rhodobactera | cellular proces | Cell wall/mem | 01XSD | Pfam:DUF583 | 391619.RGBS | 3,4 | 0,489897949 | 0,000260153 | 4,26E-05 | 3 - | 0,000192363 - | 3 - | 0,000238858 - | 4 - | 0,000302754 - | 4 - | 0,000307225 - | 3 - | 0,000259566 - |
| 5799 | 1 | 3um_FIJJAFBF_705236 | Bacteria | Proteobacteri | Gammaprote | Cellvibrionales | Halieaceae | Chromatocurv | Chromatocurv | information st | Translation, ri | COG0051 | tRNA binding | 876044.IMCC3 | 2,4 | 0,489897949 | 0,000323818 | 9,13E-05 | 2 - | 0,000222867 - | 3 - | 0,000415103 - | 2 - | 0,000263072 - | 2 - | 0,000266958 - | 3 - | 0,000451091 - |
| 5800 | 1 | 3um_FIJJAFBF_549765 | Bacteria | Proteobacteri | Gammaprote | Alteromonada | Alteromonada | Paraglaciecola | various | information st | Translation, ri | 17FIT | 50s ribosomal | 983545.Glaag | 1,6 | 0,489897949 | 0,000275097 | 7,21E-05 | 2 - | 0,000294299 - | 1 - | 0,000182716 - | 2 - | 0,00034739 - | 2 - | 0,000352521 - | 1 - | 0,000198557 - |
| 5801 | 1 | 0.2um_NOKCPHJN_503146 | Bacteria | Bacteroidetes | Flavobacteriia | Flavobacterial | Cryomorphace | unclassified | Cryomorphace | unclassified | unclassified | unclassified | unclassified | unclassified | 3 | 1,095445115 | 0,000244898 | 7,98E-05 | 4 - | 0,000281661 - | 3 - | 0,000262305 - | 4 - | 0,000332472 - | 3 - | 0,000253037 - | 1 - | 9,50E-05 - |
| 5802 | 1 | 0.2um_NOKCPHJN_302756 | Bacteria | Proteobacteri | Alphaproteob | Rhodobactera | Rhodobactera | Roseovarius | Roseovarius n | unclassified | unclassified | unclassified | unclassified | unclassified | 1,2 | 0,748331477 | 3,44E-05 | 2,21E-05 | 1 - | 2,46E-05 - | 1 - | 3,05E-05 - | 2 - | 5,80E-05 - | 2 - | 5,89E-05 - | 0 - | 0 - |
| 5803 | 1 | 3um_FIJJAFBF_00855 | Bacteria | Bacteroidetes | Flavobacteriia | Flavobacterial | Flavobacteriac | Maribacter | Maribacter sp | information st | Translation, ri | 09AI8 | Associates wit | 313603.FB217 | 6,4 | 1,624807681 | 0,000315621 | 6,84E-05 | 8 - | 0,000335115 - | 7 - | 0,000364099 - | 8 - | 0,000395569 - | 4 - | 0,000200706 - | 5 - | 0,000282618 - |
| 5804 | 1 | 0.2um_NOKCPHJN_476012 | Eukaryota | unclassified | Oomycetes | Saprolegniales | Saprolegniace | Thraustotheca | Thraustotheca | unclassified | unclassified | unclassified | unclassified | unclassified | 0,2 | 0,4 | 3,86E-06 | 7,71E-06 | 0 - | 0 - | 0 - | 0 - | 0 - | 0 - | 1 - | 1,93E-05 - | 0 - | 0 - |
| 5805 | 1 | 3um_FIJJAFBF_652365 | Bacteria | Bacteroidetes | Flavobacteriia | Flavobacterial | Flavobacteriac | Gaetbulibacte | Gaetbulibacte | poorly charact | Function unkn | 0XPRT | NA | 391598.FBBAL | 0,6 | 0,8 | 2,55E-05 | 3,28E-05 | 2 - | 7,97E-05 - | 0 - | 0 - | 0 - | 0 - | 1 - | 4,77E-05 - | 0 - | 0 - |
| 5806 | 1 | 0.2um_NOKCPHJN_280545 | Bacteria | Proteobacteri | Alphaproteob | Pelagibacteral | Pelagibactera | unclassified | Pelagibactera | unclassified | unclassified | unclassified | unclassified | unclassified | 0,6 | 0,489897949 | 1,24E-05 | 1,02E-05 | 1 - | 1,80E-05 - | 1 - | 2,24E-05 - | 1 - | 2,13E-05 - | 0 - | 0 - | 0 - | 0 - |
| 5807 | 1 | 3um_FIJJAFBF_348254 | Bacteria | Bacteroidetes | Flavobacteriia | Flavobacterial | Flavobacteriac | unclassified | Flavobacteriac | metabolism | Energy produc | COG0056 | Produces ATP | 313596.RB25 | 0,2 | 0,4 | 5,43E-06 | 1,09E-05 | 0 - | 0 - | 1 - | 2,71E-05 - | 0 - | 0 - | 0 - | 0 - | 0 - | 0 - |
| 5808 | 1 | 0.2um_NOKCPHJN_314443 | Bacteria | Bacteroidetes | Flavobacteriia | unclassified | unclassified | unclassified | Flavobacteria | unclassified | unclassified | unclassified | unclassified | unclassified | 6,2 | 0,748331477 | 8,22E-05 | 5,84E-06 | 7 - | 7,85E-05 - | 6 - | 8,36E-05 - | 7 - | 9,27E-05 - | 6 - | 8,06E-05 - | 5 - | 7,57E-05 - |
| 5809 | 1 | 3um_FIJJAFBF_46520 | Bacteria | Proteobacteri | Gammaprote | Oceanospirilla | Oceanospirilla | Amphritea | Amphritea jap | information st | Translation, ri | COG0087 | One of the pri | 377629.TERTU | 4,8 | 0,4 | 0,000309732 | 3,63E-05 | 5 - | 0,0002707 - | 4 - | 0,000268904 - | 5 - | 0,000319534 - | 5 - | 0,000324253 - | 5 - | 0,00036527 - |
| 5810 | 1 | 3um_FIJJAFBF_472754 | Bacteria | Bacteroidetes | Flavobacteriia | Flavobacterial | Flavobacteriac | Formosa | Formosa halio | unclassified | unclassified | unclassified | unclassified | unclassified | 0,4 | 0,489897949 | 2,21E-05 | 2,73E-05 | 1 - | 5,03E-05 - | 0 - | 0 - | 0 - | 0 - | 1 - | 6,03E-05 - | 0 - | 0 - |
| 5811 | 1 | 0.2um_NOKCPHJN_703369 | Bacteria | Bacteroidetes | Flavobacteriia | Flavobacterial | Flavobacteriac | Formosa | Formosa sp. H | unclassified | unclassified | unclassified | unclassified | unclassified | 0,8 | 0,4 | 4,74E-05 | 2,43E-05 | 1 - | 5,01E-05 - | 0 - | 0 - | 1 - | 5,92E-05 - | 1 - | 6,00E-05 - | 1 - | 6,76E-05 - |
| 5812 | 1 | 0.2um_NOKCPHJN_427810 | Bacteria | Proteobacteri | Gammaprote | Oceanospirilla | Oceanospirilla | Motiliproteus | Motiliproteus | metabolism | Inorganic ion t | 08UTW | Plays a role in | 207954.MED9 | 2,2 | 1,166190379 | 0,000121482 | 6,76E-05 | 2 - | 9,45E-05 - | 3 - | 0,000175949 - | 3 - | 0,000167262 - | 3 - | 0,000169732 - | 0 - | 0 - |
| 5813 | 1 | 3um_FIJJAFBF_12339 | Bacteria | Proteobacteri | Gammaprote | Cellvibrionales | Porticoccacea | unclassified | various | information st | Translation, ri | COG0087 | One of the pri | 314287.GB22 | 6,4 | 0,489897949 | 0,000413372 | 2,71E-05 | 7 - | 0,000380776 - | 6 - | 0,000405267 - | 7 - | 0,000449467 - | 6 - | 0,000390948 - | 6 - | 0,000440401 - |
| 5814 | 1 | 3um_FIJJAFBF_444552 | Bacteria | Proteobacteri | Gammaprote | Cellvibrionales | Porticoccacea | unclassified | SAR92 bacteri | poorly charact | Function unkn | 16R8I | Lipoprotein | 207949.RED6 | 2 | 0 | 6,40E-05 | 6,08E-06 | 2 - | 5,36E-05 - | 2 - | 6,66E-05 - | 2 - | 6,33E-05 - | 2 - | 6,42E-05 - | 2 - | 7,24E-05 - |
| 5815 | 1 | 3um_FIJJAFBF_217823 | Eukaryota | unclassified | Pelagophycea | Pelagomonad | unclassified | Aureococcus | Aureococcus a | unclassified | unclassified | unclassified | unclassified | unclassified | 0,4 | 0,489897949 | 1,61E-05 | 1,99E-05 | 1 - | 3,60E-05 - | 1 - | 4,47E-05 - | 0 - | 0 - | 0 - | 0 - | 0 - | 0 - |
| 5816 | 1 | 3um_FIJJAFBF_02496 | Bacteria | Proteobacteri | Alphaproteob | Rhodobactera | Rhodobactera | Pelagicola | Pelagicola sp. | information st | Translation, ri | COG0185 | Protein S19 fo | 391595.RLO14 | 1 | 0 | 0,000148943 | 1,41E-05 | 1 - | 0,000124757 - | 1 - | 0,000154912 - | 1 - | 0,000147263 - | 1 - | 0,000149438 - | 1 - | 0,000168342 - |
| 5817 | 1 | 3um_FIJJAFBF_552359 | Bacteria | Proteobacteri | Alphaproteob | Rhodobactera | Rhodobactera | unclassified | Rhodobactera | unclassified | unclassified | unclassified | unclassified | unclassified | 2 | 0,632455532 | 0,000135434 | 3,55E-05 | 3 - | 0,00017303 - | 2 - | 0,000143235 - | 2 - | 0,000136163 - | 1 - | 6,91E-05 - | 2 - | 0,000155653 - |
| 5818 | 1 | 3um_FIJJAFBF_684978 | Bacteria | Proteobacteri | Gammaprote | Oceanospirilla | Saccharospirill | Reinekea | Reinekea fors | poorly charact | Function unkn | 05Q5Y | NA | 983545.Glaag | 0,2 | 0,4 | 2,44E-05 | 4,88E-05 | 1 - | 0,000122103 - | 0 - | 0 - | 0 - | 0 - | 0 - | 0 - | 0 - | 0 - |
| 5819 | 1 | 3um_FIJJAFBF_381511 | Bacteria | Proteobacteri | Gammaprote | Cellvibrionales | Halieaceae | unclassified | gamma prote | cellular proces | Posttranslatio | 0QHT2 | Peroxiredoxin | 876044.IMCC3 | 0,2 | 0,4 | 1,44E-05 | 2,89E-05 | 1 - | 7,22E-05 - | 0 - | 0 - | 0 - | 0 - | 0 - | 0 - | 0 - | 0 - |
| 5820 | 1 | 0.2um_NOKCPHJN_436369 | Bacteria | Verrucomicrob | Opitutae | unclassified | unclassified | unclassified | Opitutae bact | unclassified | unclassified | unclassified | unclassified | unclassified | 1,2 | 0,4 | 3,38E-05 | 1,25E-05 | 1 - | 2,34E-05 - | 2 - | 5,82E-05 - | 1 - | 2,76E-05 - | 1 - | 2,81E-05 - | 1 - | 3,16E-05 - |
| 5821 | 1 | 3um_FIJJAFBF_180326 | unclassified | unclassified | unclassified | unclassified | unclassified | unclassified | unclassified | unclassified | unclassified | unclassified | unclassified | unclassified | 0,8 | 0,4 | 6,10E-05 | 3,13E-05 | 1 - | 6,45E-05 - | 0 - | 0 - | 1 - | 7,61E-05 - | 1 - | 7,72E-05 - | 1 - | 8,70E-05 - |
| 5822 | 1 | 3um_FIJJAFBF_786614 | Bacteria | Proteobacteri | Gammaprote | Oceanospirilla | Oceanospirilla | Marinobacteri | Marinobacteri | unclassified | unclassified | unclassified | unclassified | unclassified | 0,8 | 0,4 | 0,000129627 | 6,53E-05 | 0 - | 0 - | 1 - | 0,000161953 - | 1 - | 0,000153957 - | 1 - | 0,000156231 - | 1 - | 0,000175994 - |
| 5823 | 1 | 3um_FIJJAFBF_245353 | Bacteria | Proteobacteri | Alphaproteob | Rhodobactera | Rhodobactera | Thalassobius | Thalassobius s | information st | Translation, ri | 17FIH | One of the ea | 398580.Dshi_ | 1,8 | 0,4 | 0,000251589 | 6,17E-05 | 2 - | 0,000234238 - | 2 - | 0,000290855 - | 2 - | 0,000276494 - | 1 - | 0,000140289 - | 2 - | 0,00031607 - |
| 5824 | 1 | 3um_FIJJAFBF_815477 | Eukaryota | Bacillariophyta | Coscinodiscop | Thalassiosirale | Stephanodisca | Cyclotella | Cyclotella cryp | metabolism | Lipid transpor | COG0439 | acetyl-CoA ca | 2850.Phatr54 | 1,4 | 0,489897949 | 0,000127833 | 4,78E-05 | 1 - | 7,60E-05 - | 2 - | 0,000188767 - | 1 - | 8,97E-05 - | 2 - | 0,000182097 - | 1 - | 0,000102566 - |
| 5825 | 1 | 0.2um_NOKCPHJN_29259 | Bacteria | Bacteroidetes | Flavobacteriia | Flavobacterial | Flavobacteriac | Formosa | Formosa sp. H | poorly charact | Function unkn | 0XSGQ | NA | 1046627.BZAR | 0,8 | 1,166190379 | 2,18E-05 | 3,15E-05 | 0 - | 0 - | 1 - | 2,80E-05 - | 0 - | 0 - | 3 - | 8,10E-05 - | 0 - | 0 - |
| 5826 | 1 | 3um_FIJJAFBF_384933 | Bacteria | Proteobacteri | Gammaprote | Alteromonada | Alteromonada | unclassified | Alteromonada | information st | Transcription | COG0250 | Participates in | 983545.Glaag | 0,6 | 0,489897949 | 4,86E-05 | 3,99E-05 | 0 - | 0 - | 1 - | 8,01E-05 - | 1 - | 7,61E-05 - | 0 - | 0 - | 1 - | 8,70E-05 - |
| 5827 | 1 | 0.2um_NOKCPHJN_518631 | Bacteria | Proteobacteri | Alphaproteob | Pelagibacteral | Pelagibactera | Candidatus Pe | Candidatus Pe | cellular proces | Cell wall/mem | 7257 | NA | 335992.SAR11 | 1,8 | 0,748331477 | 7,40E-05 | 2,97E-05 | 2 - | 7,00E-05 - | 2 - | 8,69E-05 - | 3 - | 0,000123917 - | 1 - | 4,19E-05 - | 1 - | 4,72E-05 - |
| 5828 | 1 | 0.2um_NOKCPHJN_993771 | Bacteria | Proteobacteri | Alphaproteob | Pelagibacteral | Pelagibactera | Candidatus Pe | Candidatus Pe | unclassified | unclassified | unclassified | unclassified | unclassified | 1,2 | 0,979795897 | 2,82E-05 | 2,32E-05 | 2 - | 4,11E-05 - | 2 - | 5,11E-05 - | 2 - | 4,86E-05 - | 0 - | 0 - | 0 - | 0 - |
| 5829 | 1 | 0.2um_NOKCPHJN_80624 | Bacteria | Bacteroidetes | Flavobacteriia | Flavobacterial | Flavobacteriac | Polaribacter | Polaribacter h | information st | Translation, ri | 05G6W | One of the pri | 313594.PI23P | 6,4 | 1,496662955 | 0,000430847 | 8,14E-05 | 9 - | 0,000513926 - | 5 - | 0,000354525 - | 6 - | 0,000404425 - | 5 - | 0,000341998 - | 7 - | 0,000539364 - |
| 5830 | 1 | 3um_FIJJAFBF_571927 | Bacteria | Proteobacteri | Gammaprote | Oceanospirilla | Saccharospirill | Reinekea | Reinekea fors | unclassified | unclassified | unclassified | unclassified | unclassified | 0,8 | 0,4 | 0,00011796 | 6,06E-05 | 1 - | 0,000124757 - | 0 - | 0 - | 1 - | 0,000147263 - | 1 - | 0,000149438 - | 1 - | 0,000168342 - |
| 5831 | 1 | 0.2um_NOKCPHJN_983493 | Eukaryota | unclassified | Cryptophyta | Pyrenomonad | Geminigerace | Guillardia | Guillardia thet | unclassified | unclassified | unclassified | unclassified | unclassified | 0,4 | 0,489897949 | 3,29E-05 | 4,06E-05 | 1 - | 7,55E-05 - | 0 - | 0 - | 1 - | 8,91E-05 - | 0 - | 0 - | 0 - | 0 - |
| 5832 | 1 | 0.2um_NOKCPHJN_625025 | Bacteria | Bacteroidetes | Flavobacteriia | Flavobacterial | Flavobacteriac | Formosa | Formosa sp. H | unclassified | unclassified | unclassified | unclassified | unclassified | 3,2 | 1,166190379 | 3,65E-05 | 1,39E-05 | 2 - | 1,90E-05 - | 4 - | 4,73E-05 - | 3 - | 3,37E-05 - | 5 - | 5,70E-05 - | 2 - | 2,57E-05 - |
| 5833 | 1 | 3um_FIJJAFBF_532698 | Bacteria | Bacteroidetes | Flavobacteriia | Flavobacterial | Flavobacteriac | Formosa | Formosa sp. H | unclassified | unclassified | unclassified | unclassified | unclassified | 2 | 0,632455532 | 9,49E-05 | 3,21E-05 | 1 - | 3,92E-05 - | 2 - | 9,73E-05 - | 3 - | 0,000138719 - | 2 - | 9,38E-05 - | 2 - | 0,000105716 - |
| 5834 | 1 | 0.2um_NOKCPHJN_395415 | Bacteria | Proteobacteri | Alphaproteob | Pelagibacteral | Pelagibactera | unclassified | various | unclassified | unclassified | unclassified | unclassified | unclassified | 0,8 | 0,748331477 | 1,91E-05 | 1,88E-05 | 1 - | 2,05E-05 - | 2 - | 5,08E-05 - | 1 - | 2,42E-05 - | 0 - | 0 - | 0 - | 0 - |
| 5835 | 1 | 0.2um_NOKCPHJN_127038 | Bacteria | Bacteroidetes | Flavobacteriia | Flavobacterial | Flavobacteriac | Formosa | Formosa sp. H | unclassified | unclassified | unclassified | unclassified | unclassified | 0,2 | 0,4 | 1,06E-05 | 2,13E-05 | 0 - | 0 - | 0 - | 0 - | 1 - | 5,31E-05 - | 0 - | 0 - | 0 - | 0 - |
| 5836 | 1 | 3um_FIJJAFBF_17255 | Bacteria | Proteobacteri | Alphaproteob | Rhodobactera | unclassified | unclassified | Rhodobactera | cellular proces | Intracellular tr | 05VIE | preprotein tra | 391619.RGBS | 1,8 | 0,748331477 | 0,00026165 | 0,000111537 | 2 - | 0,000244206 - | 1 - | 0,000151616 - | 1 - | 0,00014413 - | 3 - | 0,000438777 - | 2 - | 0,00032952 - |
| 5837 | 1 | 0.2um_NOKCPHJN_682593 | Bacteria | Proteobacteri | Alphaproteob | Pelagibacteral | Pelagibactera | Candidatus Pe | Candidatus Pe | unclassified | unclassified | unclassified | unclassified | unclassified | 1 | 0,632455532 | 5,93E-05 | 3,32E-05 | 2 - | 0,000102479 - | 1 - | 6,36E-05 - | 0 - | 0 - | 1 - | 6,14E-05 - | 1 - | 6,91E-05 - |
| 5838 | 1 | 0.2um_NOKCPHJN_101165 | Bacteria | Proteobacteri | Gammaprote | Cellvibrionales | Halieaceae | unclassified | gamma prote | unclassified | unclassified | unclassified | unclassified | unclassified | 1,4 | 0,8 | 2,05E-05 | 1,18E-05 | 1 - | 1,22E-05 - | 1 - | 1,52E-05 - | 1 - | 1,45E-05 - | 3 - | 4,40E-05 - | 1 - | 1,65E-05 - |
| 5839 | 1 | 3um_FIJJAFBF_329993 | Bacteria | Bacteroidetes | Flavobacteriia | Flavobacterial | Flavobacteriac | Polaribacter | Polaribacter s | cellular proces | Cell wall/mem | COG2885 | Ompa motb d | 313594.PI23P | 5,2 | 0,748331477 | 0,00015893 | 2,99E-05 | 4 - | 0,000101798 - | 5 - | 0,000158003 - | 6 - | 0,000180242 - | 6 - | 0,000182905 - | 5 - | 0,000171701 - |
| 5840 | 1 | 0.2um_NOKCPHJN_141754 | Bacteria | Proteobacteri | Gammaprote | Cellvibrionales | Porticoccacea | unclassified | SAR92 bacteri | metabolism | Amino acid tra | 16R0H | Acetolactate s | 314287.GB22 | 0,2 | 0,4 | 1,74E-05 | 3,48E-05 | 0 - | 0 - | 1 - | 8,69E-05 - | 0 - | 0 - | 0 - | 0 - | 0 - | 0 - |
| 5841 | 1 | 0.2um_NOKCPHJN_464767 | Eukaryota | unclassified | Cryptophyta | Pyrenomonad | Geminigerace | Guillardia | Guillardia thet | unclassified | unclassified | unclassified | unclassified | unclassified | 0,8 | 0,4 | 5,26E-05 | 2,70E-05 | 1 - | 5,49E-05 - | 1 - | 6,82E-05 - | 0 - | 0 - | 1 - | 6,58E-05 - | 1 - | 7,41E-05 - |
| 5842 | 1 | 3um_FIJJAFBF_631659 | Eukaryota | Chlorophyta | Mamiellophyc | Mamiellales | Mamiellaceae | Micromonas | Micromonas c | metabolism | Energy produc | 0IHF0 | This is one of | 88036.EFJ050 | 0,2 | 0,4 | 7,68E-06 | 1,54E-05 | 0 - | 0 - | 0 - | 0 - | 1 - | 3,84E-05 - | 0 - | 0 - | 0 - | 0 - |
| 5843 | 1 | 3um_FIJJAFBF_298752 | Bacteria | Bacteroidetes | Flavobacteriia | Flavobacterial | Flavobacteriac | Polaribacter | Polaribacter s | cellular proces | Intracellular tr | 05FGH | MotA TolQ exl | 313594.PI23P | 2,2 | 0,4 | 0,000126066 | 2,81E-05 | 2 - | 9,56E-05 - | 3 - | 0,000178149 - | 2 - | 0,000112902 - | 2 - | 0,000114569 - | 2 - | 0,000129062 - |
| 5844 | 1 | 3um_FIJJAFBF_405441 | Bacteria | Bacteroidetes | Flavobacteriia | Flavobacterial | Flavobacteriac | Polaribacter | Polaribacter h | unclassified | unclassified | unclassified | unclassified | unclassified | 0,8 | 0,4 | 4,96E-05 | 2,54E-05 | 1 - | 5,24E-05 - | 0 - | 0 - | 1 - | 6,19E-05 - | 1 - | 6,28E-05 - | 1 - | 7,07E-05 - |
| 5845 | 1 | 0.2um_NOKCPHJN_331200 | Bacteria | Bacteroidetes | Flavobacteriia | Flavobacterial | Flavobacteriac | Gramella | Gramella gaet | metabolism | Inorganic ion t | COG3119 | Sulfatase | 436308.Nmar_ | 1 | 0 | 1,85E-05 | 1,76E-06 | 1 - | 1,55E-05 - | 1 - | 1,93E-05 - | 1 - | 1,83E-05 - | 1 - | 1,86E-05 - | 1 - | 2,10E-05 - |
| 5846 | 1 | 0.2um_NOKCPHJN_169103 | Bacteria | Bacteroidetes | Bacteroidia | Marinilabiliale | Salinivirgacea | unclassified | Salinivirgacea | cellular proces | Cell wall/mem | 09BX8 | glycosyl transf | 313594.PI23P | 0,8 | 0,4 | 2,73E-05 | 1,39E-05 | 1 - | 2,96E-05 - | 1 - | 3,67E-05 - | 1 - | 3,49E-05 - | 1 - | 3,54E-05 - | 0 - | 0 - |
| 5847 | 1 | 3um_FIJJAFBF_461514 | Bacteria | Proteobacteri | Gammaprote | Oceanospirilla | Saccharospirill | Reinekea | Reinekea fors | unclassified | unclassified | unclassified | unclassified | unclassified | 0,4 | 0,489897949 | 2,08E-05 | 2,58E-05 | 1 - | 4,65E-05 - | 1 - | 5,77E-05 - | 0 - | 0 - | 0 - | 0 - | 0 - | 0 - |
| 5848 | 1 | 0.2um_NOKCPHJN_374154 | Bacteria | Bacteroidetes | Flavobacteriia | Flavobacterial | Flavobacteriac | Robiginitalea | Robiginitalea | unclassified | unclassified | unclassified | unclassified | unclassified | 0,6 | 0,489897949 | 2,46E-05 | 2,05E-05 | 1 - | 3,47E-05 - | 0 - | 0 - | 0 - | 0 - | 1 - | 4,15E-05 - | 1 - | 4,68E-05 - |
| 5849 | 1 | 3um_FIJJAFBF_361813 | Bacteria | Bacteroidetes | Flavobacteriia | Flavobacterial | Flavobacteriac | Polaribacter | various | information st | Translation, ri | 0PAI9 | Binds 16S rRN | 313603.FB217 | 2 | 0,894427191 | 0,000300047 | 0,000124228 | 3 - | 0,000386888 - | 3 - | 0,000480401 - | 2 - | 0,000304454 - | 1 - | 0,000154476 - | 1 - | 0,000174016 - |
| 5850 | 1 | 3um_FIJJAFBF_736142 | Bacteria | Bacteroidetes | Flavobacteriia | various | various | various | various | information st | Translation, ri | 0PAE2 | One of the pri | 487797.Flav3 | 0,8 | 0,4 | 0,00012933 | 6,64E-05 | 1 - | 0,000135031 - | 1 - | 0,000167669 - | 0 - | 0 - | 1 - | 0,000161745 - | 1 - | 0,000182205 - |
| 5851 | 1 | 3um_FIJJAFBF_211403 | Bacteria | Proteobacteri | Alphaproteob | Rhodobactera | Rhodobactera | Roseovarius | Roseovarius a | cellular proces | Cell wall/mem | 177PY | Molecular cha | 420662.Mpe_ | 3,2 | 0,4 | 0,000229649 | 1,26E-05 | 4 - | 0,000242914 - | 3 - | 0,00022622 - | 3 - | 0,000215051 - | 3 - | 0,000218227 - | 3 - | 0,000245833 - |
| 5852 | 1 | 3um_FIJJAFBF_487035 | Bacteria | Bacteroidetes | Flavobacteriia | Flavobacterial | Flavobacteriac | Marixanthom | Marixanthom | cellular proces | Posttranslatio | COG0330 | Band 7 protei | 50743.SCB49_ | 0,4 | 0,489897949 | 1,64E-05 | 2,01E-05 | 0 - | 0 - | 0 - | 0 - | 1 - | 4,07E-05 - | 1 - | 4,13E-05 - | 0 - | 0 - |
| 5853 | 1 | 3um_FIJJAFBF_16182 | Bacteria | Bacteroidetes | Flavobacteriia | Flavobacterial | Flavobacteriac | Formosa | Formosa sp. H | unclassified | unclassified | unclassified | unclassified | unclassified | 0,8 | 0,748331477 | 0,000157037 | 0,000138785 | 2 - | 0,000358677 - | 0 - | 0 - | 1 - | 0,000211691 - | 1 - | 0,000214818 - | 0 - | 0 - |
| 5854 | 1 | 3um_FIJJAFBF_670946 | Bacteria | Proteobacteri | Gammaprote | Alteromonada | Alteromonada | Paraglaciecola | Paraglaciecola | unclassified | unclassified | unclassified | unclassified | unclassified | 0,6 | 0,489897949 | 2,22E-05 | 1,85E-05 | 1 - | 3,09E-05 - | 1 - | 3,84E-05 - | 0 - | 0 - | 0 - | 0 - | 1 - | 4,17E-05 - |
| 5855 | 1 | 0.2um_NOKCPHJN_75522 | Bacteria | Proteobacteri | Gammaprote | Oceanospirilla | Saccharospirill | Reinekea | Reinekea fors | unclassified | unclassified | unclassified | unclassified | unclassified | 1 | 0 | 0,000103028 | 9,78E-06 | 1 - | 8,63E-05 - | 1 - | 0,000107157 - | 1 - | 0,000101866 - | 1 - | 0,000103371 - | 1 - | 0,000116447 - |
| 5856 | 1 | 3um_FIJJAFBF_92868 | Eukaryota | unclassified | Cryptophyta | Pyrenomonad | Pyrenomonad | Rhodomonas | Rhodomonas | unclassified | unclassified | unclassified | unclassified | unclassified | 0,6 | 0,489897949 | 2,45E-05 | 2,01E-05 | 1 - | 3,62E-05 - | 0 - | 0 - | 1 - | 4,27E-05 - | 1 - | 4,34E-05 - | 0 - | 0 - |
| 5857 | 1 | 0.2um_NOKCPHJN_477124 | Eukaryota | Chordata | Mammalia | Dasyuromorp | Dasyuridae | Sarcophilus | Sarcophilus ha | cellular proces | Cytoskeleton | 0DGS2 | Tubulin is the | 9305.ENSSHA | 3,6 | 0,489897949 | 0,000108465 | 1,89E-05 | 4 - | 0,000100903 - | 4 - | 0,000125291 - | 3 - | 8,93E-05 - | 3 - | 9,06E-05 - | 4 - | 0,000136153 - |

| | | | | | | | | | | | | | | | | | | | | | | | | | | | | |
|---|---|---|---|---|---|---|---|---|---|---|---|---|---|---|---|---|---|---|---|---|---|---|---|---|---|---|---|---|
| 5858 | 1 | 0.2um_NOKCPHJN_868123 | Bacteria | Bacteroidetes | unclassified | unclassified | unclassified | unclassified | Bacteroidetes | unclassified | unclassified | unclassified | unclassified | unclassified | 0,6 | 0,8 | 4,56E-05 | 6,24E-05 | 0 - | 0 - | 0 - | 0 - | 0 - | 0 - | 1 - | 7,01E-05 - | 2 - | 0,000158035 - |
| 5859 | 1 | 3um_FIJJAFBF_153480 | Bacteria | Bacteroidetes | Flavobacteriia | Flavobacterial | Flavobacteriac | Polaribacter | Polaribacter g | unclassified | unclassified | unclassified | unclassified | unclassified | 0,4 | 0,489897949 | 2,59E-05 | 3,23E-05 | 1 - | 5,52E-05 - | 0 - | 0 - | 0 - | 0 - | 0 - | 0 - | 1 - | 7,45E-05 - |
| 5860 | 1 | 0.2um_NOKCPHJN_494189 | Bacteria | Proteobacteri | Alphaproteob | Pelagibacteral | Pelagibacterac | Candidatus Pe | Candidatus Pe | metabolism | Energy produc | 01RNF | Produces ATP | 331869.BAL19 | 2 | 0 | 5,94E-05 | 5,64E-06 | 2 - | 4,98E-05 - | 2 - | 6,18E-05 - | 2 - | 5,88E-05 - | 2 - | 5,96E-05 - | 2 - | 6,72E-05 - |
| 5861 | 1 | 0.2um_NOKCPHJN_970842 | Bacteria | Proteobacteri | Betaproteoba | Rhodocyclales | Rhodocyclace | Azovibrio | Azovibrio rest | unclassified | unclassified | unclassified | unclassified | unclassified | 0,2 | 0,4 | 2,52E-05 | 5,04E-05 | 0 - | 0 - | 0 - | 0 - | 0 - | 0 - | 0 - | 0 - | 1 - | 0,000125914 - |
| 5862 | 1 | 0.2um_NOKCPHJN_221840 | Bacteria | Proteobacteri | Gammaprotec | unclassified | unclassified | unclassified | various | poorly charact | Function unkn | 0ZPVF | NA | 247633.GP214 | 1 | 0 | 9,20E-05 | 8,73E-06 | 1 - | 7,70E-05 - | 1 - | 9,57E-05 - | 1 - | 9,09E-05 - | 1 - | 9,23E-05 - | 1 - | 0,000103943 - |
| 5863 | 1 | 0.2um_NOKCPHJN_992556 | Bacteria | Proteobacteri | Alphaproteob | Pelagibacteral | Pelagibacterac | unclassified | Pelagibacterac | information st | Transcription | 01QD3 | Facilitates trar | 439493.PB721 | 8,6 | 1,019803903 | 0,000274293 | 1,32E-05 | 10 - | 0,000270063 - | 8 - | 0,000268271 - | 9 - | 0,000286904 - | 9 - | 0,000291141 - | 7 - | 0,000255087 - |
| 5864 | 1 | 3um_FIJJAFBF_295528 | Bacteria | Bacteroidetes | Flavobacteriia | unclassified | unclassified | unclassified | Flavobacteria | unclassified | unclassified | unclassified | unclassified | unclassified | 3,2 | 0,4 | 0,000330515 | 5,68E-05 | 3 - | 0,000258895 - | 4 - | 0,000428628 - | 3 - | 0,000305599 - | 3 - | 0,000310113 - | 3 - | 0,000349341 - |
| 5865 | 2 | 0.2um_NOKCPHJN_176926;3um_FIJJAFBF_18021 | Bacteria | Bacteroidetes | Flavobacteriia | Flavobacterial | Flavobacteriac | Polaribacter | Polaribacter s | various | various | various | various | various | 1,4 | 0,489897949 | 3,82E-05 | 1,35E-05 | 1 - | 2,29E-05 - | 1 - | 2,84E-05 - | 2 - | 5,41E-05 - | 2 - | 5,49E-05 - | 1 - | 3,09E-05 - |
| 5866 | 1 | 3um_FIJJAFBF_655392 | Bacteria | Bacteroidetes | Cytophagia | Cytophagales | Flammeovirga | unclassified | Flammeovirga | cellular proces | Posttranslatio | 09FAR | peroxiredoxin | 388413.ALPR1 | 0,8 | 0,4 | 8,08E-05 | 4,15E-05 | 1 - | 8,44E-05 - | 1 - | 0,000104793 - | 0 - | 0 - | 1 - | 0,000101091 - | 1 - | 0,000113878 - |
| 5867 | 1 | 3um_FIJJAFBF_545217 | Bacteria | Bacteroidetes | Flavobacteriia | Flavobacterial | Flavobacteriac | Arenitalea | Arenitalea lut | information st | Translation, ri | COG0256 | This is one of t | 983544.Lacal_ | 1,6 | 0,489897949 | 0,000181844 | 4,77E-05 | 2 - | 0,000194537 - | 1 - | 0,000120779 - | 2 - | 0,000229631 - | 2 - | 0,000233023 - | 1 - | 0,00013125 - |
| 5868 | 1 | 3um_FIJJAFBF_257123 | Eukaryota | Phaeophyceae | unclassified | Dictyotales | Dictyotaceae | Dictyopteris | Dictyopteris d | information st | Translation, ri | 0FKC1 | This is 1 of the | 551115.Aazo_ | 4,2 | 0,748331477 | 0,000318185 | 5,86E-05 | 4 - | 0,000255059 - | 5 - | 0,000395886 - | 5 - | 0,00037634 - | 4 - | 0,000305518 - | 3 - | 0,000258124 - |
| 5869 | 1 | 0.2um_NOKCPHJN_265412 | Eukaryota | unclassified | Pelagophycea | Pelagomonad | unclassified | Aureococcus | Aureococcus a | unclassified | unclassified | unclassified | unclassified | unclassified | 1 | 0 | 3,02E-05 | 2,87E-06 | 1 - | 2,53E-05 - | 1 - | 3,15E-05 - | 1 - | 2,99E-05 - | 1 - | 3,03E-05 - | 1 - | 3,42E-05 - |
| 5870 | 1 | 0.2um_NOKCPHJN_323412 | Bacteria | Bacteroidetes | Flavobacteriia | Flavobacterial | Cryomorphace | unclassified | Cryomorphace | metabolism | Energy produc | COG0056 | Produces ATP | 398720.MED2 | 0,4 | 0,489897949 | 1,04E-05 | 1,28E-05 | 0 - | 0 - | 0 - | 0 - | 1 - | 2,59E-05 - | 1 - | 2,62E-05 - | 0 - | 0 - |
| 5871 | 1 | 3um_FIJJAFBF_55506 | Bacteria | Proteobacteri | Betaproteoba | unclassified | unclassified | unclassified | Betaproteoba | information st | Translation, ri | COG0173 | aspartyl-trna s | 887062.HGR_ | 0,6 | 0,489897949 | 1,39E-05 | 1,14E-05 | 0 - | 0 - | 1 - | 2,38E-05 - | 1 - | 2,27E-05 - | 1 - | 2,30E-05 - | 0 - | 0 - |
| 5872 | 1 | 3um_FIJJAFBF_691942 | Bacteria | Bacteroidetes | Flavobacteriia | Flavobacterial | Flavobacteriac | Formosa | Formosa sp. H | unclassified | unclassified | unclassified | unclassified | unclassified | 2 | 0 | 9,07E-05 | 8,62E-06 | 2 - | 7,60E-05 - | 2 - | 9,44E-05 - | 2 - | 8,97E-05 - | 2 - | 9,10E-05 - | 2 - | 0,000102566 - |
| 5873 | 1 | 3um_FIJJAFBF_590106 | Bacteria | Proteobacteri | Gammaprotec | unclassified | unclassified | unclassified | Gammaprotec | unclassified | unclassified | unclassified | unclassified | unclassified | 0,2 | 0,4 | 2,95E-05 | 5,90E-05 | 0 - | 0 - | 0 - | 0 - | 0 - | 0 - | 0 - | 0 - | 1 - | 0,0001475 - |
| 5874 | 1 | 0.2um_NOKCPHJN_522742 | Bacteria | Proteobacteri | Alphaproteob | Rhodobactera | Rhodobactera | Nereida | Nereida ignav | metabolism | Carbohydrate | 05CA9 | Pyruvate kinas | 391593.RCCS2 | 0,4 | 0,489897949 | 1,04E-05 | 1,28E-05 | 1 - | 2,38E-05 - | 0 - | 0 - | 1 - | 2,81E-05 - | 0 - | 0 - | 0 - | 0 - |
| 5875 | 1 | 0.2um_NOKCPHJN_489582 | Bacteria | Bacteroidetes | unclassified | unclassified | unclassified | unclassified | Bacteroidetes | unclassified | unclassified | unclassified | unclassified | unclassified | 2,8 | 0,979795897 | 4,92E-05 | 1,75E-05 | 3 - | 4,48E-05 - | 4 - | 7,42E-05 - | 3 - | 5,29E-05 - | 3 - | 5,37E-05 - | 1 - | 2,02E-05 - |
| 5876 | 1 | 0.2um_NOKCPHJN_710509 | Eukaryota | Streptophyta | unclassified | Asterales | Asteraceae | Lactuca | Lactuca sativa | metabolism | Energy produc | 1BFZ8 | Cytochrome b | 3847.GLYMA1 | 2 | 0 | 0,000217503 | 2,07E-05 | 2 - | 0,000182185 - | 2 - | 0,00022622 - | 2 - | 0,000215051 - | 2 - | 0,000218227 - | 2 - | 0,000245833 - |
| 5877 | 1 | 0.2um_NOKCPHJN_360630 | Bacteria | Proteobacteri | Alphaproteob | Rhodobactera | Rhodobactera | unclassified | Rhodobactera | metabolism | Carbohydrate | 16SJ5 | Trap dicarbox | 367336.OM22 | 1,6 | 0,489897949 | 6,73E-05 | 2,23E-05 | 2 - | 7,04E-05 - | 2 - | 8,74E-05 - | 1 - | 4,16E-05 - | 1 - | 4,22E-05 - | 2 - | 9,50E-05 - |
| 5878 | 1 | 0.2um_NOKCPHJN_438308 | Eukaryota | unclassified | Oomycetes | Saprolegniales | Saprolegniace | Aphanomyces | Aphanomyces | unclassified | unclassified | unclassified | unclassified | unclassified | 1,6 | 0,8 | 0,000113137 | 4,52E-05 | 3 - | 0,000185124 - | 1 - | 7,66E-05 - | 1 - | 7,28E-05 - | 2 - | 0,000147832 - | 1 - | 8,33E-05 - |
| 5879 | 4 | 3um_FIJJAFBF_101303;3um_FIJJAFBF_396337;3u | Bacteria | Bacteroidetes | Flavobacteriia | Flavobacterial | Flavobacteriac | Formosa | Formosa sp. H | cellular proces | Posttranslatio | 0PAEA | Thioredoxin | 391603.FBALC | 0,4 | 0,489897949 | 5,15E-05 | 6,35E-05 | 1 - | 0,000117119 - | 0 - | 0 - | 0 - | 0 - | 1 - | 0,000140289 - | 0 - | 0 - |
| 5880 | 1 | 3um_FIJJAFBF_93363 | Bacteria | Bacteroidetes | Flavobacteriia | various | various | various | various | unclassified | unclassified | unclassified | unclassified | unclassified | 1 | 0 | 9,45E-05 | 8,97E-06 | 1 - | 7,92E-05 - | 1 - | 9,83E-05 - | 1 - | 9,34E-05 - | 1 - | 9,48E-05 - | 1 - | 0,00010681 - |
| 5881 | 1 | 0.2um_NOKCPHJN_470840 | Bacteria | Proteobacteri | Alphaproteob | Rhodobactera | Rhodobactera | Aliiroseovariu | Aliiroseovariu | unclassified | unclassified | unclassified | unclassified | unclassified | 5,6 | 0,8 | 0,000172014 | 2,14E-05 | 7 - | 0,000181363 - | 5 - | 0,000160857 - | 5 - | 0,000152914 - | 5 - | 0,000155173 - | 6 - | 0,000209762 - |
| 5882 | 1 | 3um_FIJJAFBF_94120 | Bacteria | Proteobacteri | Gammaprotec | Cellvibrionales | Cellvibrionace | Simiduia | Simiduia agari | metabolism | Inorganic ion t | 0QJ3Y | Part of the AB | 203122.Sde_3 | 0,2 | 0,4 | 9,93E-06 | 1,99E-05 | 0 - | 0 - | 0 - | 0 - | 1 - | 4,96E-05 - | 0 - | 0 - | 0 - | 0 - |
| 5883 | 1 | 0.2um_NOKCPHJN_818460 | Bacteria | Proteobacteri | Alphaproteob | Pelagibacteral | Pelagibacterac | Candidatus Pe | Candidatus Pe | metabolism | Amino acid tra | 05EZ7 | amino acid AB | 335992.SAR11 | 1 | 0 | 3,60E-05 | 3,41E-06 | 1 - | 3,01E-05 - | 1 - | 3,74E-05 - | 1 - | 3,56E-05 - | 1 - | 3,61E-05 - | 1 - | 4,06E-05 - |
| 5884 | 1 | 0.2um_NOKCPHJN_213543 | Bacteria | Proteobacteri | Alphaproteob | Pelagibacteral | Pelagibacterac | Candidatus Pe | Candidatus Pe | unclassified | unclassified | unclassified | unclassified | unclassified | 3,8 | 0,4 | 0,000203767 | 2,69E-05 | 4 - | 0,000180042 - | 3 - | 0,000167669 - | 4 - | 0,000212521 - | 4 - | 0,00021566 - | 4 - | 0,00024294 - |
| 5885 | 1 | 0.2um_NOKCPHJN_149872 | Bacteria | Bacteroidetes | Flavobacteriia | Flavobacterial | Crocinitomica | unclassified | Crocinitomica | unclassified | unclassified | unclassified | unclassified | unclassified | 10,2 | 1,939071943 | 0,000142081 | 2,00E-05 | 13 - | 0,000153666 - | 12 - | 0,00017613 - | 9 - | 0,000125576 - | 9 - | 0,00012743 - | 8 - | 0,0001276 - |
| 5886 | 1 | 3um_FIJJAFBF_679232 | Bacteria | Bacteroidetes | Flavobacteriia | Flavobacterial | Flavobacteriac | Aureitalea | Aureitalea ma | unclassified | unclassified | unclassified | unclassified | unclassified | 0,6 | 0,489897949 | 6,69E-05 | 5,48E-05 | 0 - | 0 - | 0 - | 0 - | 1 - | 0,000105845 - | 1 - | 0,000107409 - | 1 - | 0,000120996 - |
| 5887 | 1 | 3um_FIJJAFBF_396426 | Bacteria | Proteobacteri | Betaproteoba | unclassified | unclassified | unclassified | Betaproteoba | information st | Translation, ri | 05G6W | One of the pri | 595537.Varpa | 0,4 | 0,489897949 | 2,44E-05 | 3,01E-05 | 1 - | 5,54E-05 - | 0 - | 0 - | 0 - | 0 - | 1 - | 6,64E-05 - | 0 - | 0 - |
| 5888 | 1 | 3um_FIJJAFBF_462227 | Bacteria | Bacteroidetes | Flavobacteriia | Flavobacterial | Flavobacteriac | Maribacter | Maribacter sp | metabolism | Inorganic ion t | 05C27 | TonB-depende | 714943.Mucp | 0,8 | 0,748331477 | 9,14E-06 | 8,87E-06 | 1 - | 9,99E-06 - | 0 - | 0 - | 1 - | 1,18E-05 - | 2 - | 2,39E-05 - | 0 - | 0 - |
| 5889 | 1 | 0.2um_NOKCPHJN_310371 | Bacteria | Bacteroidetes | Flavobacteriia | Flavobacterial | Cryomorphace | unclassified | Cryomorphace | unclassified | unclassified | unclassified | unclassified | unclassified | 0,6 | 0,489897949 | 2,47E-05 | 2,05E-05 | 1 - | 3,48E-05 - | 0 - | 0 - | 0 - | 0 - | 1 - | 4,17E-05 - | 1 - | 4,69E-05 - |
| 5890 | 1 | 3um_FIJJAFBF_552510 | Bacteria | Proteobacteri | Gammaprotec | Arenicellales | Arenicellaceae | Arenicella | Arenicella xan | information st | Translation, ri | 177SZ | This protein bi | 317025.Tcr_0 | 0,2 | 0,4 | 1,62E-05 | 3,23E-05 | 1 - | 8,08E-05 - | 0 - | 0 - | 0 - | 0 - | 0 - | 0 - | 0 - | 0 - |
| 5891 | 1 | 0.2um_NOKCPHJN_21186 | Bacteria | Proteobacteri | Alphaproteob | Rhodobactera | Rhodobactera | Planktomarina | Planktomarina | unclassified | unclassified | unclassified | unclassified | unclassified | 2,4 | 0,489897949 | 3,95E-05 | 5,80E-06 | 3 - | 4,19E-05 - | 2 - | 3,47E-05 - | 3 - | 4,95E-05 - | 2 - | 3,35E-05 - | 2 - | 3,77E-05 - |
| 5892 | 1 | 0.2um_NOKCPHJN_367529 | Eukaryota | Phaeophyceae | unclassified | Ectocarpales | Ectocarpaceae | Ectocarpus | Ectocarpus sili | information st | Chromatin stru | KOG1744 | Core compone | 2850.Phatr118 | 0,8 | 0,4 | 9,69E-05 | 4,98E-05 | 1 - | 0,000101572 - | 1 - | 0,000126123 - | 1 - | 0,000119896 - | 0 - | 0 - | 1 - | 0,000137057 - |
| 5893 | 1 | 3um_FIJJAFBF_16399 | Bacteria | Actinobacteria | Actinobacteria | Micrococcales | Microbacteria | Candidatus Ac | Candidatus Ac | metabolism | Carbohydrate | 00BJE | Glyceraldehyd | 288705.RSal33 | 2,6 | 0,8 | 0,000105707 | 3,06E-05 | 3 - | 0,000103402 - | 2 - | 8,56E-05 - | 2 - | 8,14E-05 - | 4 - | 0,000165145 - | 2 - | 9,30E-05 - |
| 5894 | 1 | 3um_FIJJAFBF_429764 | Bacteria | Proteobacteri | Gammaprotec | Oceanospirilla | Saccharospirill | Reinekea | Reinekea fors | metabolism | Energy produc | 16QAQ | C-type cytochr | 314283.MED2 | 0,4 | 0,489897949 | 1,74E-05 | 2,15E-05 | 1 - | 3,88E-05 - | 1 - | 4,81E-05 - | 0 - | 0 - | 0 - | 0 - | 0 - | 0 - |
| 5895 | 1 | 0.2um_NOKCPHJN_437814 | Bacteria | Proteobacteri | Gammaprotec | unclassified | unclassified | unclassified | OM182 bacter | unclassified | unclassified | unclassified | unclassified | unclassified | 0,4 | 0,489897949 | 3,44E-05 | 4,22E-05 | 0 - | 0 - | 0 - | 0 - | 0 - | 0 - | 1 - | 8,09E-05 - | 1 - | 9,11E-05 - |
| 5896 | 1 | 0.2um_NOKCPHJN_1110489 | Bacteria | Proteobacteri | Gammaprotec | Cellvibrionales | Porticoccacea | unclassified | gamma protec | cellular proces | Cell wall/mem | 0QKXZ | Porin Gram-ne | 314287.GB220 | 0,2 | 0,4 | 1,06E-05 | 2,12E-05 | 0 - | 0 - | 0 - | 0 - | 0 - | 0 - | 1 - | 5,31E-05 - | 0 - | 0 - |
| 5897 | 2 | 0.2um_NOKCPHJN_296781;0.2um_NOKCPHJN_9 | Bacteria | Bacteroidetes | Flavobacteriia | Flavobacterial | Flavobacteriac | Marinirhabdu | Marinirhabdu | unclassified | unclassified | unclassified | unclassified | unclassified | 0,6 | 0,489897949 | 4,48E-05 | 3,67E-05 | 0 - | 0 - | 0 - | 0 - | 1 - | 7,09E-05 - | 1 - | 7,20E-05 - | 1 - | 8,11E-05 - |
| 5898 | 1 | 0.2um_NOKCPHJN_111995 | Bacteria | Actinobacteria | Actinobacteria | Micrococcales | Microbacteria | Candidatus Ac | Candidatus Ac | unclassified | unclassified | unclassified | unclassified | unclassified | 2 | 0 | 0,000107895 | 1,02E-05 | 2 - | 9,04E-05 - | 2 - | 0,00011222 - | 2 - | 0,000106679 - | 2 - | 0,000108255 - | 2 - | 0,000121948 - |
| 5899 | 1 | 3um_FIJJAFBF_92950 | Bacteria | Proteobacteri | Gammaprotec | Cellvibrionales | Porticoccacea | unclassified | various | cellular proces | Cell wall/mem | COG2885 | Ompa motb d | 314287.GB220 | 5 | 0,894427191 | 0,000217198 | 3,92E-05 | 6 - | 0,000219319 - | 6 - | 0,000272329 - | 4 - | 0,000172589 - | 4 - | 0,000175138 - | 5 - | 0,000246615 - |
| 5900 | 1 | 0.2um_NOKCPHJN_325756 | Bacteria | Proteobacteri | Gammaprotec | Pseudomonad | Pseudomonad | Pseudomonas | Pseudomonas | unclassified | unclassified | unclassified | unclassified | unclassified | 1 | 0,632455532 | 4,01E-05 | 2,63E-05 | 1 - | 3,45E-05 - | 1 - | 4,28E-05 - | 1 - | 4,07E-05 - | 2 - | 8,26E-05 - | 0 - | 0 - |
| 5901 | 1 | 0.2um_NOKCPHJN_241608 | Bacteria | Proteobacteri | Alphaproteob | Pelagibacteral | Pelagibacterac | Candidatus Pe | Candidatus Pe | information st | Transcription | 01R4Z | DNA-depende | 439493.PB721 | 9,4 | 1,624807681 | 9,46E-05 | 1,88E-05 | 10 - | 8,43E-05 - | 12 - | 0,00012566 - | 7 - | 6,97E-05 - | 9 - | 9,09E-05 - | 9 - | 0,000102415 - |
| 5902 | 1 | 3um_FIJJAFBF_189350 | Bacteria | Proteobacteri | Gammaprotec | Cellvibrionales | Porticoccacea | unclassified | various | metabolism | Carbohydrate | 0QHXA | transketolase | 314287.GB220 | 0,4 | 0,489897949 | 8,22E-06 | 1,01E-05 | 0 - | 0 - | 0 - | 0 - | 1 - | 2,04E-05 - | 1 - | 2,07E-05 - | 0 - | 0 - |
| 5903 | 1 | 0.2um_NOKCPHJN_684222 | Bacteria | Proteobacteri | Alphaproteob | Pelagibacteral | unclassified | unclassified | Pelagibacteral | metabolism | Amino acid tra | 16Q4U | Catalyzes the | 684719.HIMB | 1 | 0 | 3,15E-05 | 2,99E-06 | 1 - | 2,64E-05 - | 1 - | 3,28E-05 - | 1 - | 3,11E-05 - | 1 - | 3,16E-05 - | 1 - | 3,56E-05 - |
| 5904 | 1 | 0.2um_NOKCPHJN_170765 | Bacteria | Bacteroidetes | Flavobacteriia | Flavobacterial | Flavobacteriac | Polaribacter | Polaribacter s | metabolism | Energy produc | 099Q7 | Catalyzes the | 313598.MED1 | 4,2 | 0,748331477 | 0,000184151 | 2,20E-05 | 5 - | 0,000186326 - | 4 - | 0,000185089 - | 5 - | 0,000219939 - | 4 - | 0,00017855 - | 3 - | 0,000150852 - |
| 5905 | 1 | 3um_FIJJAFBF_128448 | Bacteria | Bacteroidetes | Flavobacteriia | Flavobacterial | Flavobacteriac | Polaribacter | Polaribacter s | unclassified | unclassified | unclassified | unclassified | unclassified | 1,2 | 0,4 | 1,76E-05 | 3,89E-06 | 2 - | 2,52E-05 - | 1 - | 1,57E-05 - | 1 - | 1,49E-05 - | 1 - | 1,51E-05 - | 1 - | 1,70E-05 - |
| 5906 | 1 | 3um_FIJJAFBF_203938 | Bacteria | Proteobacteri | Gammaprotec | Oceanospirilla | Saccharospirill | Reinekea | Reinekea fors | unclassified | unclassified | unclassified | unclassified | unclassified | 1,4 | 0,8 | 0,00012704 | 5,62E-05 | 3 - | 0,000239118 - | 1 - | 9,90E-05 - | 1 - | 9,41E-05 - | 1 - | 9,55E-05 - | 1 - | 0,000107552 - |
| 5907 | 1 | 3um_FIJJAFBF_183699 | Bacteria | Bacteroidetes | Flavobacteriia | Flavobacterial | Flavobacteriac | Algibacter | Algibacter sp. | unclassified | unclassified | unclassified | unclassified | unclassified | 0,4 | 0,489897949 | 8,34E-06 | 1,03E-05 | 1 - | 1,86E-05 - | 1 - | 2,31E-05 - | 0 - | 0 - | 0 - | 0 - | 0 - | 0 - |
| 5908 | 1 | 0.2um_NOKCPHJN_256891 | Bacteria | Proteobacteri | Gammaprotec | Oceanospirilla | Saccharospirill | Reinekea | Reinekea fors | unclassified | unclassified | unclassified | unclassified | unclassified | 0,6 | 0,489897949 | 3,69E-05 | 3,04E-05 | 1 - | 5,47E-05 - | 0 - | 0 - | 1 - | 6,45E-05 - | 1 - | 6,55E-05 - | 0 - | 0 - |
| 5909 | 1 | 0.2um_NOKCPHJN_132520 | Bacteria | Bacteroidetes | unclassified | unclassified | unclassified | unclassified | Bacteroidetes | metabolism | Energy produc | COG0437 | 4fe-4S ferredc | 926562.Oweh | 5,2 | 0,748331477 | 7,07E-05 | 6,78E-06 | 6 - | 6,91E-05 - | 5 - | 7,15E-05 - | 5 - | 6,80E-05 - | 6 - | 8,28E-05 - | 4 - | 6,22E-05 - |
| 5910 | 1 | 3um_FIJJAFBF_363298 | Bacteria | Verrucomicrob | Opitutae | Puniceicoccale | Puniceicoccac | unclassified | Puniceicoccac | poorly charact | Function unkn | 05C2K | tail SHEATH pr | 552811.Dehly | 0,8 | 0,4 | 1,70E-05 | 8,74E-06 | 1 - | 1,80E-05 - | 0 - | 0 - | 1 - | 2,12E-05 - | 1 - | 2,15E-05 - | 1 - | 2,43E-05 - |
| 5911 | 1 | 3um_FIJJAFBF_552506 | Bacteria | Proteobacteri | Gammaprotec | Arenicellales | Arenicellaceae | Arenicella | Arenicella xan | information st | Transcription | COG0085 | DNA-depende | 396588.Tgr7_ | 0,2 | 0,4 | 1,69E-06 | 3,37E-06 | 1 - | 8,43E-06 - | 0 - | 0 - | 0 - | 0 - | 0 - | 0 - | 0 - | 0 - |
| 5912 | 1 | 3um_FIJJAFBF_12251 | Bacteria | Proteobacteri | Gammaprotec | Cellvibrionales | Porticoccacea | unclassified | Gammaprotec | unclassified | unclassified | unclassified | unclassified | unclassified | 23,6 | 1,743559577 | 0,000319188 | 3,68E-05 | 25 - | 0,000283539 - | 22 - | 0,000309824 - | 25 - | 0,000334689 - | 21 - | 0,000285291 - | 25 - | 0,000382595 - |
| 5913 | 1 | 3um_FIJJAFBF_328500 | Bacteria | Bacteroidetes | Flavobacteriia | Flavobacterial | unclassified | unclassified | Flavobacterial | information st | Translation, ri | 05CEJ | Catalyzes the | 50743.SCB49_ | 1,4 | 0,489897949 | 2,70E-05 | 9,76E-06 | 2 - | 3,24E-05 - | 1 - | 2,01E-05 - | 1 - | 1,91E-05 - | 1 - | 1,94E-05 - | 2 - | 4,37E-05 - |
| 5914 | 1 | 3um_FIJJAFBF_46722 | Bacteria | Proteobacteri | Betaproteoba | unclassified | unclassified | unclassified | Betaproteoba | cellular proces | Posttranslatio | COG0652 | peptidyl-proly | 398578.Daci_ | 1 | 0 | 8,16E-05 | 7,74E-06 | 1 - | 6,83E-05 - | 1 - | 8,48E-05 - | 1 - | 8,06E-05 - | 1 - | 8,18E-05 - | 1 - | 9,22E-05 - |
| 5915 | 1 | 0.2um_NOKCPHJN_823488 | Bacteria | Bacteroidetes | Flavobacteriia | Flavobacterial | unclassified | unclassified | Flavobacterial | metabolism | Inorganic ion t | COG1055 | arsenical pum | 926562.Oweh | 0,8 | 0,4 | 2,38E-05 | 1,22E-05 | 1 - | 2,50E-05 - | 1 - | 3,10E-05 - | 1 - | 2,95E-05 - | 0 - | 0 - | 1 - | 3,37E-05 - |
| 5916 | 1 | 3um_FIJJAFBF_531265 | Bacteria | Proteobacteri | Gammaprotec | Thiotrichales | Piscirickettsia | Methylophaga | various | unclassified | unclassified | unclassified | unclassified | unclassified | 1,2 | 0,4 | 5,19E-05 | 1,79E-05 | 1 - | 3,62E-05 - | 1 - | 4,50E-05 - | 1 - | 4,27E-05 - | 2 - | 8,67E-05 - | 1 - | 4,89E-05 - |
| 5917 | 1 | 3um_FIJJAFBF_788215 | Bacteria | Bacteroidetes | unclassified | unclassified | unclassified | unclassified | Bacteroidetes | information st | Transcription | 0P7W9 | Transcription | 487796.Flav2A | 0,6 | 0,489897949 | 1,93E-05 | 1,59E-05 | 1 - | 2,80E-05 - | 1 - | 3,48E-05 - | 0 - | 0 - | 1 - | 3,35E-05 - | 0 - | 0 - |
| 5918 | 1 | 0.2um_NOKCPHJN_1022996 | Bacteria | Proteobacteri | Gammaprotec | Alteromonada | Colwelliaceae | Colwellia | various | information st | Translation, ri | 177XJ | One of the pri | 167879.CPS_4 | 0,4 | 0,8 | 2,92E-05 | 5,85E-05 | 2 - | 0,000146212 - | 0 - | 0 - | 0 - | 0 - | 0 - | 0 - | 0 - | 0 - |
| 5919 | 1 | 0.2um_NOKCPHJN_123049 | Eukaryota | unclassified | Oomycetes | Saprolegniales | Saprolegniace | Thraustotheca | Thraustotheca | unclassified | unclassified | unclassified | unclassified | unclassified | 0,2 | 0,4 | 4,43E-06 | 8,87E-06 | 0 - | 0 - | 0 - | 0 - | 1 - | 2,22E-05 - | 0 - | 0 - | 0 - | 0 - |
| 5920 | 1 | 0.2um_NOKCPHJN_870446 | Bacteria | Proteobacteri | Gammaprotec | unclassified | unclassified | unclassified | Gammaprotec | poorly charact | Function unkn | 16R8I | Lipoprotein | 314287.GB220 | 1 | 0 | 2,84E-05 | 2,69E-06 | 1 - | 2,38E-05 - | 1 - | 2,95E-05 - | 1 - | 2,81E-05 - | 1 - | 2,85E-05 - | 1 - | 3,21E-05 - |
| 5921 | 1 | 3um_FIJJAFBF_390923 | Bacteria | Bacteroidetes | Flavobacteriia | Flavobacterial | Flavobacteriac | Formosa | Formosa sp. H | metabolism | Energy produc | COG0055 | Produces ATP | 1046627.BZAF | 4,6 | 0,8 | 0,000123728 | 1,11E-05 | 6 - | 0,000137183 - | 4 - | 0,000113561 - | 5 - | 0,000134942 - | 4 - | 0,000109548 - | 4 - | 0,000123406 - |
| 5922 | 1 | 3um_FIJJAFBF_128640 | Bacteria | Bacteroidetes | Flavobacteriia | Flavobacterial | Flavobacteriac | Flaviramulus | Flaviramulus b | poorly charact | Function unkn | 09ARK | Protein of unk | 313603.FB217 | 0,8 | 0,748331477 | 3,75E-05 | 3,24E-05 | 2 - | 8,17E-05 - | 1 - | 5,07E-05 - | 0 - | 0 - | 0 - | 0 - | 1 - | 5,51E-05 - |
| 5923 | 1 | 3um_FIJJAFBF_211284 | Bacteria | Proteobacteri | Gammaprotec | Arenicellales | Arenicellaceae | Arenicella | Arenicella xan | information st | Translation, ri | 16QAF | thus facilitatin | 1026882.MAN | 0,2 | 0,4 | 4,12E-06 | 8,24E-06 | 1 - | 2,06E-05 - | 0 - | 0 - | 0 - | 0 - | 0 - | 0 - | 0 - | 0 - |
| 5924 | 1 | 0.2um_NOKCPHJN_230047 | Bacteria | Proteobacteri | Alphaproteob | Rhodobactera | Rhodobactera | Planktomarina | Planktomarina | metabolism | Inorganic ion t | 05C2J | ABC transport | 272942.RCAP_ | 1 | 0 | 5,44E-05 | 5,16E-06 | 1 - | 4,55E-05 - | 1 - | 5,66E-05 - | 1 - | 5,38E-05 - | 1 - | 5,46E-05 - | 1 - | 6,15E-05 - |
| 5925 | 1 | 3um_FIJJAFBF_571511 | Bacteria | Proteobacteri | Betaproteoba | unclassified | unclassified | unclassified | Betaproteoba | metabolism | Amino acid tra | 0B77R | Aspartokinase | 887062.HGR_ | 0,4 | 0,489897949 | 1,31E-05 | 1,60E-05 | 0 - | 0 - | 1 - | 3,35E-05 - | 1 - | 3,18E-05 - | 0 - | 0 - | 0 - | 0 - |
| 5926 | 1 | 3um_FIJJAFBF_727103 | Bacteria | Proteobacteri | Gammaprotec | Oceanospirilla | Oceanospirilla | Marinobacteri | Marinobacter | unclassified | unclassified | unclassified | unclassified | unclassified | 0,2 | 0,4 | 6,91E-06 | 1,38E-05 | 1 - | 3,46E-05 - | 0 - | 0 - | 0 - | 0 - | 0 - | 0 - | 0 - | 0 - |
| 5927 | 1 | 3um_FIJJAFBF_586239 | Bacteria | Bacteroidetes | Flavobacteriia | Flavobacterial | Flavobacteriac | Polaribacter | Polaribacter s | metabolism | Amino acid tra | 0P7DH | glutamine syn | 313598.MED1 | 0,8 | 0,4 | 1,46E-05 | 7,41E-06 | 1 - | 1,58E-05 - | 1 - | 1,96E-05 - | 1 - | 1,86E-05 - | 1 - | 1,89E-05 - | 0 - | 0 - |
| 5928 | 1 | 3um_FIJJAFBF_235815 | Bacteria | Bacteroidetes | Flavobacteriia | unclassified | unclassified | unclassified | Flavobacteria | metabolism | Inorganic ion t | 05C27 | TonB-depende | 487797.Flav3C | 24,2 | 2,4 | 0,000328922 | 1,70E-05 | 28 - | 0,000321375 - | 23 - | 0,000327793 - | 26 - | 0,000352254 - | 22 - | 0,000302463 - | 22 - | 0,000340724 - |
| 5929 | 1 | 0.2um_NOKCPHJN_172844 | Bacteria | Proteobacteri | Gammaprotec | Oceanospirilla | Saccharospirill | Reinekea | Reinekea fors | metabolism | Energy produc | 16RMF | aldo keto redu | 491952.Mar18 | 1,8 | 0,4 | 8,10E-05 | 1,60E-05 | 2 - | 7,65E-05 - | 2 - | 9,50E-05 - | 2 - | 9,03E-05 - | 2 - | 9,17E-05 - | 1 - | 5,16E-05 - |
| 5930 | 1 | 3um_FIJJAFBF_183020 | Bacteria | Bacteroidetes | Flavobacteriia | Flavobacterial | Flavobacteriac | Tenacibaculur | Tenacibaculur | unclassified | unclassified | unclassified | unclassified | unclassified | 0,8 | 0,4 | 3,32E-05 | 1,71E-05 | 1 - | 3,48E-05 - | 1 - | 4,32E-05 - | 1 - | 4,11E-05 - | 0 - | 0 - | 1 - | 4,69E-05 - |
| 5931 | 1 | 3um_FIJJAFBF_448542 | Bacteria | Proteobacteri | Gammaprotec | Oceanospirilla | Saccharospirill | Reinekea | Reinekea fors | metabolism | Carbohydrate | 08UHU | Aldolase | 716541.ECL_0 | 2 | 0,894427191 | 0,000128897 | 5,25E-05 | 3 - | 0,000165543 - | 1 - | 6,85E-05 - | 3 - | 0,000195407 - | 1 - | 6,61E-05 - | 2 - | 0,000148918 - |
| 5932 | 1 | 3um_FIJJAFBF_521113 | Bacteria | Proteobacteri | Gammaprotec | unclassified | unclassified | unclassified | various | unclassified | unclassified | unclassified | unclassified | unclassified | 2,2 | 0,4 | 0,000115383 | 2,25E-05 | 2 - | 8,80E-05 - | 2 - | 0,00010921 - | 3 - | 0,000155727 - | 2 - | 0,000105351 - | 2 - | 0,000118678 - |
| 5933 | 1 | 3um_FIJJAFBF_592964 | Bacteria | Proteobacteri | Gammaprotec | Oceanospirilla | Saccharospirill | Reinekea | Reinekea blan | metabolism | Carbohydrate | 16PZZ | Catalyzes the | 314283.MED2 | 0,8 | 0,748331477 | 2,24E-05 | 1,98E-05 | 2 - | 5,12E-05 - | 0 - | 0 - | 1 - | 3,02E-05 - | 1 - | 3,07E-05 - | 0 - | 0 - |
| 5934 | 1 | 0.2um_NOKCPHJN_526010 | Bacteria | Proteobacteri | Alphaproteob | Pelagibacteral | Pelagibacterac | Candidatus Pe | Candidatus Pe | metabolism | Amino acid tra | 16QU8 | Methylenetetr | 439493.PB721 | 0,6 | 0,489897949 | 3,35E-05 | 2,78E-05 | 1 - | 4,74E-05 - | 0 - | 0 - | 1 - | 5,60E-05 - | 0 - | 0 - | 1 - | 6,40E-05 - |
| 5935 | 2 | 3um_FIJJAFBF_18776;3um_FIJJAFBF_277840 | Bacteria | Bacteroidetes | Flavobacteriia | Flavobacterial | Flavobacteriac | Polaribacter | Polaribacter r | unclassified | unclassified | unclassified | unclassified | unclassified | 0,2 | 0,4 | 8,36E-06 | 1,67E-05 | 0 - | 0 - | 0 - | 0 - | 0 - | 0 - | 1 - | 4,18E-05 - | 0 - | 0 - |
| 5936 | 1 | 3um_FIJJAFBF_629414 | Bacteria | Bacteroidetes | Flavobacteriia | Flavobacterial | Flavobacteriac | Polaribacter | Polaribacter fi | unclassified | unclassified | unclassified | unclassified | unclassified | 3,8 | 1,166190379 | 4,47E-05 | 1,38E-05 | 4 - | 3,99E-05 - | 5 - | 6,19E-05 - | 3 - | 3,53E-05 - | 5 - | 5,97E-05 - | 2 - | 2,69E-05 - |
| 5937 | 3 | 0.2um_NOKCPHJN_609417;0.2um_NOKCPHJN_7 | Bacteria | Proteobacteri | Alphaproteob | Pelagibacteral | Pelagibacterac | Candidatus Pe | Candidatus Pe | unclassified | unclassified | unclassified | unclassified | unclassified | 0,2 | 0,4 | 8,76E-06 | 1,75E-05 | 0 - | 0 - | 0 - | 0 - | 0 - | 0 - | 1 - | 4,38E-05 - | 0 - | 0 - |
| 5938 | 1 | 0.2um_NOKCPHJN_75279 | Bacteria | Bacteroidetes | unclassified | unclassified | unclassified | unclassified | Bacteroidetes | unclassified | unclassified | unclassified | unclassified | unclassified | 3,6 | 0,489897949 | 3,76E-05 | 5,95E-06 | 4 - | 3,51E-05 - | 3 - | 3,27E-05 - | 4 - | 4,14E-05 - | 3 - | 3,15E-05 - | 4 - | 4,73E-05 - |
| 5939 | 1 | 3um_FIJJAFBF_403782 | Bacteria | Bacteroidetes | Flavobacteriia | unclassified | unclassified | unclassified | Flavobacteria | information st | Translation, ri | COG0254 | 50S ribosomal | 487796.Flav2A | 0,4 | 0,489897949 | 5,87E-05 | 7,23E-05 | 1 - | 0,000133461 - | 0 - | 0 - | 0 - | 0 - | 1 - | 0,000159864 - | 0 - | 0 - |
| 5940 | 1 | 0.2um_NOKCPHJN_132653 | Bacteria | Bacteroidetes | Flavobacteriia | various | various | various | various | unclassified | unclassified | unclassified | unclassified | unclassified | 1 | 0 | 4,04E-05 | 3,84E-06 | 1 - | 3,39E-05 - | 1 - | 4,20E-05 - | 1 - | 4,00E-05 - | 1 - | 4,06E-05 - | 1 - | 4,57E-05 - |
| 5941 | 1 | 3um_FIJJAFBF_615123 | Bacteria | Proteobacteri | Gammaprotec | Cellvibrionales | Porticoccacea | unclassified | Gammaprotec | cellular proces | Cell wall/mem | 0QTSX | Peptidoglycan | 314287.GB220 | 5,4 | 0,8 | 0,000456013 | 5,90E-05 | 6 - | 0,000427739 - | 4 - | 0,000354084 - | 6 - | 0,000504903 - | 6 - | 0,00051236 - | 5 - | 0,000480977 - |
| 5942 | 1 | 0.2um_NOKCPHJN_991012 | Eukaryota | unclassified | Cryptophyta | Pyrenomonad | Geminigerace | Guillardia | Guillardia thet | unclassified | unclassified | unclassified | unclassified | unclassified | 1,8 | 0,748331477 | 9,27E-05 | 3,94E-05 | 1 - | 4,30E-05 - | 2 - | 0,000106756 - | 2 - | 0,000101485 - | 3 - | 0,000154476 - | 1 - | 5,80E-05 - |
| 5943 | 2 | 3um_FIJJAFBF_583293;3um_FIJJAFBF_67794 | Eukaryota | unclassified | unclassified | Phaeocystales | Phaeocystace | Phaeocystis | various | metabolism | Energy produc | 11IIK | Low-potential | 35128.Thapsd | 1 | 0 | 8,41E-05 | 7,98E-06 | 1 - | 7,04E-05 - | 1 - | 8,74E-05 - | 1 - | 8,31E-05 - | 1 - | 8,43E-05 - | 1 - | 9,50E-05 - |
| 5944 | 1 | 3um_FIJJAFBF_421412 | Bacteria | Bacteroidetes | unclassified | unclassified | unclassified | unclassified | unidentified e | unclassified | unclassified | unclassified | unclassified | unclassified | 0,2 | 0,4 | 4,23E-06 | 8,46E-06 | 1 - | 2,11E-05 - | 0 - | 0 - | 0 - | 0 - | 0 - | 0 - | 0 - | 0 - |
| 5945 | 1 | 3um_FIJJAFBF_40823 | Bacteria | Proteobacteri | Alphaproteob | Rhodobactera | Rhodobactera | Pseudooceani | Pseudooceani | metabolism | Energy produc | COG0056 | Produces ATP | 375451.RD1_3 | 1,8 | 0,748331477 | 4,77E-05 | 1,92E-05 | 2 - | 4,53E-05 - | 1 - | 2,81E-05 - | 2 - | 5,34E-05 - | 3 - | 8,14E-05 - | 1 - | 3,05E-05 - |
| 5946 | 1 | 3um_FIJJAFBF_73093 | Bacteria | Proteobacteri | Gammaprotec | Alteromonada | Alteromonada | Paraglaciecola | Paraglaciecola | cellular proces | Intracellular tr | 0QJ7H | motA TolQ Exl | 983545.Glaag | 6,6 | 0,8 | 0,00020108 | 3,05E-05 | 6 - | 0,000153036 - | 7 - | 0,000221696 - | 8 - | 0,000240857 - | 6 - | 0,000183311 - | 6 - | 0,000206499 - |
| 5947 | 1 | 3um_FIJJAFBF_157374 | Bacteria | Bacteroidetes | Flavobacteriia | Flavobacterial | Flavobacteriac | Algibacter | Algibacter pec | unclassified | unclassified | unclassified | unclassified | unclassified | 2,8 | 0,4 | 0,000251726 | 2,99E-05 | 3 - | 0,000228033 - | 3 - | 0,00028315 - | 3 - | 0,00026917 - | 3 - | 0,000273146 - | 2 - | 0,000205132 - |
| 5948 | 1 | 0.2um_NOKCPHJN_945420 | various | various | various | various | various | various | various | metabolism | Coenzyme tra | 09A4R | May play a key | 762903.Pedsa | 0,2 | 0,4 | 1,29E-05 | 2,58E-05 | 0 - | 0 - | 1 - | 6,45E-05 - | 0 - | 0 - | 0 - | 0 - | 0 - | 0 - |
| 5949 | 1 | 0.2um_NOKCPHJN_67466 | Bacteria | Bacteroidetes | Flavobacteriia | Flavobacterial | Flavobacteriac | Xanthomarina | Xanthomarina | information st | Translation, ri | COG0087 | One of the pri | 983544.Lacal_ | 0,2 | 0,4 | 1,39E-05 | 2,78E-05 | 0 - | 0 - | 1 - | 6,95E-05 - | 0 - | 0 - | 0 - | 0 - | 0 - | 0 - |
| 5950 | 1 | 3um_FIJJAFBF_604988 | Bacteria | Bacteroidetes | Flavobacteriia | Flavobacterial | Flavobacteriac | Arenibacter | Arenibacter la | metabolism | Amino acid tra | COG0436 | aminotransfer | 867900.Celly_ | 1,4 | 0,489897949 | 4,72E-05 | 1,39E-05 | 2 - | 5,78E-05 - | 1 - | 3,59E-05 - | 1 - | 3,41E-05 - | 2 - | 6,93E-05 - | 1 - | 3,90E-05 - |
| 5951 | 1 | 0.2um_NOKCPHJN_730413 | Bacteria | Proteobacteri | Alphaproteob | Rhizobiales | Aurantimonad | Aureimonas | Aureimonas u | poorly charact | Function unkn | 0NMWQ | Collagen triple | 388400.BB149 | 5,8 | 0,979795897 | 0,000101019 | 1,94E-05 | 7 - | 0,000102349 - | 5 - | 9,08E-05 - | 5 - | 8,63E-05 - | 5 - | 8,76E-05 - | 7 - | 0,000138105 - |
| 5952 | 1 | 0.2um_NOKCPHJN_62912 | Bacteria | Proteobacteri | Gammaprotec | Oceanospirilla | Saccharospirill | Reinekea | Reinekea fors | metabolism | Secondary me | 05CV6 | ABC transport | 314283.MED2 | 0,4 | 0,489897949 | 6,17E-06 | 7,63E-06 | 1 - | 1,38E-05 - | 1 - | 1,71E-05 - | 0 - | 0 - | 0 - | 0 - | 0 - | 0 - |
| 5953 | 1 | 3um_FIJJAFBF_17286 | Bacteria | Proteobacteri | Alphaproteob | Rhodobactera | Rhodobactera | unclassified | Rhodobactera | metabolism | Amino acid tra | COG0174 | glutamine syn | 351016.RAZW | 2,4 | 0,489897949 | 6,93E-05 | 1,02E-05 | 3 - | 7,36E-05 - | 2 - | 6,09E-05 - | 3 - | 8,68E-05 - | 2 - | 5,88E-05 - | 2 - | 6,62E-05 - |
| 5954 | 1 | 3um_FIJJAFBF_208700 | Bacteria | Bacteroidetes | Flavobacteriia | Flavobacterial | Flavobacteriac | Ulvibacter | Ulvibacter sp. | unclassified | unclassified | unclassified | unclassified | unclassified | 1 | 0 | 5,66E-05 | 5,38E-06 | 1 - | 4,74E-05 - | 1 - | 5,89E-05 - | 1 - | 5,60E-05 - | 1 - | 5,68E-05 - | 1 - | 6,40E-05 - |
| 5955 | 1 | 0.2um_NOKCPHJN_200584 | Bacteria | Proteobacteri | Gammaprotec | unclassified | unclassified | unclassified | Gammaprotec | unclassified | unclassified | unclassified | unclassified | unclassified | 2 | 0,894427191 | 0,000152315 | 6,04E-05 | 3 - | 0,000197891 - | 1 - | 8,19E-05 - | 2 - | 0,000155727 - | 3 - | 0,00023704 - | 1 - | 8,90E-05 - |
| 5956 | 1 | 0.2um_NOKCPHJN_23471 | Bacteria | Proteobacteri | Gammaprotec | Arenicellales | Arenicellaceae | Arenicella | Arenicella xan | metabolism | Energy produc | 0QJ3X | Aconitate hyd | 878320.AcavA | 1 | 0 | 1,59E-05 | 1,51E-06 | 1 - | 1,33E-05 - | 1 - | 1,65E-05 - | 1 - | 1,57E-05 - | 1 - | 1,59E-05 - | 1 - | 1,79E-05 - |
| 5957 | 1 | 3um_FIJJAFBF_70315 | unclassified | unclassified | unclassified | unclassified | unclassified | unclassified | unclassified | unclassified | unclassified | unclassified | unclassified | unclassified | 0,2 | 0,4 | 2,39E-05 | 4,78E-05 | 1 - | 0,000119559 - | 0 - | 0 - | 0 - | 0 - | 0 - | 0 - | 0 - | 0 - |
| 5958 | 1 | 0.2um_NOKCPHJN_25096 | Bacteria | Bacteroidetes | Flavobacteriia | Flavobacterial | Flavobacteriac | Ulvibacter | Ulvibacter sp. | unclassified | unclassified | unclassified | unclassified | unclassified | 0,2 | 0,4 | 3,30E-06 | 6,61E-06 | 0 - | 0 - | 0 - | 0 - | 1 - | 1,65E-05 - | 0 - | 0 - | 0 - | 0 - |
| 5959 | 1 | 3um_FIJJAFBF_209254 | Bacteria | Proteobacteri | Alphaproteob | Pelagibacteral | Pelagibacterac | Candidatus Pe | Candidatus Pe | information st | Translation, ri | COG1185 | Involved in mI | 335992.SAR11 | 0,8 | 0,4 | 5,29E-05 | 2,72E-05 | 1 - | 5,60E-05 - | 0 - | 0 - | 1 - | 6,61E-05 - | 1 - | 6,71E-05 - | 1 - | 7,55E-05 - |
| 5960 | 1 | 3um_FIJJAFBF_780709 | unclassified | unclassified | unclassified | unclassified | unclassified | unclassified | unclassified | unclassified | unclassified | unclassified | unclassified | unclassified | 0,2 | 0,4 | 5,63E-06 | 1,13E-05 | 0 - | 0 - | 0 - | 0 - | 0 - | 0 - | 0 - | 0 - | 1 - | 2,82E-05 - |
| 5961 | 1 | 3um_FIJJAFBF_713189 | Bacteria | Bacteroidetes | Flavobacteriia | Flavobacterial | Flavobacteriac | Kordia | Kordia algicida | poorly charact | Function unkn | 08ZIU | NA | 391587.KAOT | 0,8 | 0,748331477 | 6,65E-05 | 5,78E-05 | 2 - | 0,000146212 - | 0 - | 0 - | 0 - | 0 - | 1 - | 8,76E-05 - | 1 - | 9,86E-05 - |
| 5962 | 1 | 3um_FIJJAFBF_554897 | Bacteria | Proteobacteri | Alphaproteob | Pelagibacteral | Pelagibacterac | Candidatus Pe | Candidatus Pe | information st | Transcription | 01R4Z | DNA-depende | 439493.PB721 | 9,8 | 1,6 | 0,000239287 | 2,20E-05 | 12 - | 0,000249063 - | 10 - | 0,000257719 - | 10 - | 0,000244995 - | 10 - | 0,000248614 - | 7 - | 0,000196044 - |
| 5963 | 1 | 3um_FIJJAFBF_122401 | Bacteria | Bacteroidetes | Flavobacteriia | Flavobacterial | Flavobacteriac | Polaribacter | Polaribacter s | unclassified | unclassified | unclassified | unclassified | unclassified | 2 | 0,632455532 | 8,59E-05 | 2,82E-05 | 2 - | 7,26E-05 - | 3 - | 0,000135303 - | 2 - | 8,57E-05 - | 2 - | 8,70E-05 - | 1 - | 4,90E-05 - |
| 5964 | 1 | 0.2um_NOKCPHJN_270820 | unclassified | unclassified | unclassified | unclassified | unclassified | unclassified | unclassified | unclassified | unclassified | unclassified | unclassified | unclassified | 0,6 | 0,489897949 | 7,43E-06 | 6,10E-06 | 0 - | 0 - | 0 - | 0 - | 1 - | 1,18E-05 - | 1 - | 1,19E-05 - | 1 - | 1,35E-05 - |
| 5965 | 2 | 3um_FIJJAFBF_228388;3um_FIJJAFBF_662095 | Bacteria | Proteobacteri | Gammaprotec | Cellvibrionales | Porticoccacea | unclassified | various | information st | Transcription | 05VEQ | Cold shock prc | 314287.GB220 | 3,2 | 0,4 | 0,000620052 | 3,40E-05 | 4 - | 0,000655867 - | 3 - | 0,000610795 - | 3 - | 0,000580638 - | 3 - | 0,000589214 - | 3 - | 0,000663748 - |
| 5966 | 1 | 3um_FIJJAFBF_242843 | Bacteria | Bacteroidetes | Flavobacteriia | Flavobacterial | Flavobacteriac | Polaribacter | Polaribacter s | information st | Translation, ri | COG0013 | Catalyzes the | 313598.MED1 | 2,8 | 0,4 | 4,36E-05 | 5,19E-06 | 3 - | 3,95E-05 - | 3 - | 4,90E-05 - | 3 - | 4,66E-05 - | 3 - | 4,73E-05 - | 2 - | 3,55E-05 - |
| 5967 | 1 | 0.2um_NOKCPHJN_317159 | Bacteria | Proteobacteri | Gammaprotec | Cellvibrionales | Porticoccacea | unclassified | SAR92 bacteri | information st | Translation, ri | 05C22 | threonyL-tRN | 314287.GB220 | 0,6 | 0,8 | 1,21E-05 | 1,68E-05 | 1 - | 1,79E-05 - | 0 - | 0 - | 0 - | 0 - | 2 - | 4,28E-05 - | 0 - | 0 - |
| 5968 | 1 | 3um_FIJJAFBF_242854 | Bacteria | Bacteroidetes | Flavobacteriia | Flavobacterial | Flavobacteriac | Polaribacter | Polaribacter s | metabolism | Amino acid tra | COG0436 | aminotransfer | 313598.MED1 | 1 | 0 | 3,61E-05 | 3,42E-06 | 1 - | 3,02E-05 - | 1 - | 3,75E-05 - | 1 - | 3,57E-05 - | 1 - | 3,62E-05 - | 1 - | 4,08E-05 - |
| 5969 | 1 | 3um_FIJJAFBF_642087 | Bacteria | Proteobacteri | Gammaprotec | Oceanospirilla | Saccharospirill | Reinekea | Reinekea fors | cellular proces | Posttranslatio | 16QA9 | Chaperone inv | 314283.MED2 | 1 | 0,632455532 | 3,29E-05 | 2,15E-05 | 1 - | 2,83E-05 - | 1 - | 3,51E-05 - | 1 - | 3,34E-05 - | 2 - | 6,77E-05 - | 0 - | 0 - |
| 5970 | 1 | 3um_FIJJAFBF_11812 | Bacteria | Bacteroidetes | Flavobacteriia | Flavobacterial | Flavobacteriac | Psychroserper | Psychroserper | unclassified | unclassified | unclassified | unclassified | unclassified | 0,2 | 0,4 | 1,26E-05 | 2,52E-05 | 1 - | 6,31E-05 - | 0 - | 0 - | 0 - | 0 - | 0 - | 0 - | 0 - | 0 - |
| 5971 | 2 | 0.2um_NOKCPHJN_116477;3um_FIJJAFBF_11015 | Bacteria | Proteobacteri | Alphaproteob | Rhodobactera | Rhodobactera | various | various | unclassified | unclassified | unclassified | unclassified | unclassified | 0,2 | 0,4 | 9,09E-06 | 1,82E-05 | 0 - | 0 - | 0 - | 0 - | 1 - | 4,55E-05 - | 0 - | 0 - | 0 - | 0 - |
| 5972 | 1 | 3um_FIJJAFBF_189618 | Bacteria | Bacteroidetes | Flavobacteriia | Flavobacterial | Flavobacteriac | Polaribacter | Polaribacter s | cellular proces | Cell wall/mem | 16PMX | Catalyzes the | 709032.Sulku_ | 0,2 | 0,4 | 7,86E-06 | 1,57E-05 | 1 - | 3,93E-05 - | 0 - | 0 - | 0 - | 0 - | 0 - | 0 - | 0 - | 0 - |
| 5973 | 1 | 3um_FIJJAFBF_254413 | Bacteria | Bacteroidetes | Flavobacteriia | Flavobacterial | Crocinitomica | unclassified | Crocinitomica | unclassified | unclassified | unclassified | unclassified | unclassified | 2,8 | 1,166190379 | 0,000301898 | 0,000117646 | 4 - | 0,000367286 - | 4 - | 0,00045606 - | 1 - | 0,000108386 - | 3 - | 0,00032996 - | 2 - | 0,000247799 - |
| 5974 | 1 | 3um_FIJJAFBF_284683 | Eukaryota | Bacillariophyta | Coscinodiscop | Leptocylindral | Leptocylindrac | Leptocylindrus | Leptocylindrus | metabolism | Energy produc | 1DYB8 | PsaA and PsaB | 3055.DAA014 | 2 | 0 | 3,64E-05 | 3,46E-06 | 2 - | 3,05E-05 - | 2 - | 3,79E-05 - | 2 - | 3,60E-05 - | 2 - | 3,66E-05 - | 2 - | 4,12E-05 - |
| 5975 | 1 | 3um_FIJJAFBF_438660 | Bacteria | Proteobacteri | Gammaprotec | Oceanospirilla | Saccharospirill | Reinekea | Reinekea blan | unclassified | unclassified | unclassified | unclassified | unclassified | 0,4 | 0,489897949 | 1,07E-05 | 1,32E-05 | 1 - | 2,44E-05 - | 0 - | 0 - | 0 - | 0 - | 1 - | 2,92E-05 - | 0 - | 0 - |
| 5976 | 1 | 0.2um_NOKCPHJN_640771 | Viruses | unclassified | unclassified | unclassified | unclassified | unclassified | uncultured vir | unclassified | unclassified | unclassified | unclassified | unclassified | 0,4 | 0,489897949 | 1,99E-05 | 2,46E-05 | 1 - | 4,43E-05 - | 1 - | 5,50E-05 - | 0 - | 0 - | 0 - | 0 - | 0 - | 0 - |
| 5977 | 1 | 3um_FIJJAFBF_343416 | Bacteria | Actinobacteria | Actinobacteria | Jiangellales | Jiangellaceae | Jiangella | Jiangella sp. G | unclassified | unclassified | unclassified | unclassified | unclassified | 1 | 0 | 2,19E-05 | 2,08E-06 | 1 - | 1,84E-05 - | 1 - | 2,28E-05 - | 1 - | 2,17E-05 - | 1 - | 2,20E-05 - | 1 - | 2,48E-05 - |
| 5978 | 1 | 3um_FIJJAFBF_595341 | Bacteria | Proteobacteri | Alphaproteob | Rhodospirillal | Rhodospirillac | Candidatus En | Candidatus En | unclassified | unclassified | unclassified | unclassified | unclassified | 0,2 | 0,4 | 2,61E-06 | 5,22E-06 | 0 - | 0 - | 0 - | 0 - | 0 - | 0 - | 1 - | 1,31E-05 - | 0 - | 0 - |
| 5979 | 1 | 3um_FIJJAFBF_318785 | Bacteria | Proteobacteri | Alphaproteob | Rhodobactera | Rhodobactera | unclassified | Rhodobactera | unclassified | unclassified | unclassified | unclassified | unclassified | 1,6 | 0,8 | 5,26E-05 | 2,09E-05 | 3 - | 8,63E-05 - | 1 - | 3,57E-05 - | 2 - | 6,79E-05 - | 1 - | 3,45E-05 - | 1 - | 3,88E-05 - |
| 5980 | 1 | 3um_FIJJAFBF_19550 | Bacteria | Bacteroidetes | Flavobacteriia | Flavobacterial | Flavobacteriac | Formosa | Formosa sp. H | unclassified | unclassified | unclassified | unclassified | unclassified | 2,2 | 0,4 | 0,000174712 | 1,60E-05 | 3 - | 0,000202547 - | 2 - | 0,000167669 - | 2 - | 0,000159391 - | 2 - | 0,000161745 - | 2 - | 0,000182205 - |
| 5981 | 1 | 3um_FIJJAFBF_336130 | Bacteria | unclassified | unclassified | unclassified | unclassified | unclassified | uncultured ba | poorly charact | Function unkn | 05X9M | NA | 313595.P7007 | 0,4 | 0,489897949 | 9,09E-06 | 1,12E-05 | 1 - | 2,03E-05 - | 1 - | 2,52E-05 - | 0 - | 0 - | 0 - | 0 - | 0 - | 0 - |
| 5982 | 1 | 3um_FIJJAFBF_620947 | Bacteria | Proteobacteri | Gammaprotec | Oceanospirilla | Oceanospirilla | Marinomonas | Marinomonas | metabolism | Amino acid tra | 17HFC | Ribose-Phospl | 400668.Mmw | 0,8 | 0,4 | 2,84E-05 | 1,43E-05 | 0 - | 0 - | 1 - | 3,55E-05 - | 1 - | 3,38E-05 - | 1 - | 3,43E-05 - | 1 - | 3,86E-05 - |
| 5983 | 1 | 3um_FIJJAFBF_251400 | Bacteria | Bacteroidetes | Flavobacteriia | Flavobacterial | Flavobacteriac | Polaribacter | Polaribacter s | cellular proces | Cell wall/mem | 08ZG0 | Ompa motb d | 402612.FP150 | 2 | 0,632455532 | 0,000115167 | 2,67E-05 | 3 - | 0,000149061 - | 2 - | 0,000123393 - | 2 - | 0,000117301 - | 2 - | 0,000119033 - | 1 - | 6,70E-05 - |
| 5984 | 1 | 3um_FIJJAFBF_77002 | Bacteria | Bacteroidetes | Flavobacteriia | Flavobacterial | Flavobacteriac | Polaribacter | Polaribacter ir | unclassified | unclassified | unclassified | unclassified | unclassified | 0,2 | 0,4 | 1,36E-05 | 2,72E-05 | 0 - | 0 - | 0 - | 0 - | 0 - | 0 - | 1 - | 6,81E-05 - | 0 - | 0 - |
| 5985 | 1 | 3um_FIJJAFBF_253585 | Viruses | unclassified | unclassified | unclassified | unclassified | unclassified | uncultured M | unclassified | unclassified | unclassified | unclassified | unclassified | 3 | 0 | 0,000132607 | 1,26E-05 | 3 - | 0,000111074 - | 3 - | 0,000137921 - | 3 - | 0,000131112 - | 3 - | 0,000133048 - | 3 - | 0,000149879 - |
| 5986 | 1 | 0.2um_NOKCPHJN_563536 | Eukaryota | Phaeophyceae | unclassified | Ectocarpales | Ectocarpaceae | Ectocarpus | Ectocarpus sili | metabolism | Carbohydrate | KOG2638 | UTP-glucose-1 | 653948.CCA18 | 0,2 | 0,4 | 2,17E-06 | 4,34E-06 | 1 - | 1,08E-05 - | 0 - | 0 - | 0 - | 0 - | 0 - | 0 - | 0 - | 0 - |
| 5987 | 1 | 0.2um_NOKCPHJN_90569 | Viruses | unclassified | unclassified | unclassified | various | various | various | unclassified | unclassified | unclassified | unclassified | unclassified | 0,8 | 0,4 | 2,10E-05 | 1,07E-05 | 1 - | 2,28E-05 - | 1 - | 2,83E-05 - | 1 - | 2,69E-05 - | 1 - | 2,73E-05 - | 0 - | 0 - |
| 5988 | 1 | 0.2um_NOKCPHJN_282195 | Bacteria | Bacteroidetes | Flavobacteriia | Flavobacterial | Flavobacteriac | Polaribacter | various | unclassified | unclassified | unclassified | unclassified | unclassified | 0,6 | 0,8 | 2,31E-05 | 2,92E-05 | 2 - | 6,89E-05 - | 0 - | 0 - | 0 - | 0 - | 0 - | 0 - | 1 - | 4,65E-05 - |
| 5989 | 1 | 3um_FIJJAFBF_718642 | Bacteria | Verrucomicrob | Verrucomicrob | Verrucomicrob | Verrucomicrob | unclassified | Verrucomicrob | unclassified | unclassified | unclassified | unclassified | unclassified | 0,6 | 0,8 | 2,19E-05 | 2,83E-05 | 2 - | 6,89E-05 - | 0 - | 0 - | 1 - | 4,07E-05 - | 0 - | 0 - | 0 - | 0 - |
| 5990 | 1 | 0.2um_NOKCPHJN_722538 | Bacteria | Proteobacteri | Betaproteoba | Nitrosomonad | Nitrosomonad | Nitrosomonas | Nitrosomonas | cellular proces | Posttranslatio | 16QU3 | Hemolysin-typ | 744980.TRICH | 0,6 | 0,489897949 | 1,08E-05 | 8,87E-06 | 1 - | 1,57E-05 - | 1 - | 1,95E-05 - | 1 - | 1,86E-05 - | 0 - | 0 - | 0 - | 0 - |
| 5991 | 1 | 3um_FIJJAFBF_778064 | Bacteria | Bacteroidetes | Flavobacteriia | Flavobacterial | Flavobacteriac | Ochrovirga | Ochrovirga pa | unclassified | unclassified | unclassified | unclassified | unclassified | 1 | 0 | 2,81E-05 | 2,67E-06 | 1 - | 2,36E-05 - | 1 - | 2,93E-05 - | 1 - | 2,78E-05 - | 1 - | 2,82E-05 - | 1 - | 3,18E-05 - |
| 5992 | 1 | 3um_FIJJAFBF_437249 | Bacteria | Bacteroidetes | Bacteroidia | Marinilabiliale | Prolixibactera | Prolixibacter | Prolixibacter d | unclassified | unclassified | unclassified | unclassified | unclassified | 0,2 | 0,4 | 9,49E-06 | 1,90E-05 | 1 - | 4,74E-05 - | 0 - | 0 - | 0 - | 0 - | 0 - | 0 - | 0 - | 0 - |
| 5993 | 1 | 3um_FIJJAFBF_88485 | Bacteria | Bacteroidetes | unclassified | unclassified | unclassified | unclassified | Bacteroidetes | poorly charact | Function unkn | 09A4Z | rhodopsin | 487796.Flav2A | 2,2 | 0,4 | 0,000122731 | 1,13E-05 | 3 - | 0,000142285 - | 2 - | 0,000117784 - | 2 - | 0,000111969 - | 2 - | 0,000113623 - | 2 - | 0,000127995 - |
| 5994 | 1 | 3um_FIJJAFBF_403236 | Bacteria | Proteobacteri | Gammaprotec | Cellvibrionales | Cellvibrionace | Gilvimarinus | Gilvimarinus p | information st | Replication, re | 16W6D | single-strande | 207954.MED9 | 2 | 0 | 0,000148137 | 1,41E-05 | 2 - | 0,000124083 - | 2 - | 0,000154074 - | 2 - | 0,000146467 - | 2 - | 0,000148631 - | 2 - | 0,000167432 - |
| 5995 | 1 | 3um_FIJJAFBF_410272 | Bacteria | Bacteroidetes | Flavobacteriia | Flavobacterial | Flavobacteriac | Polaribacter | Polaribacter s | cellular proces | Posttranslatio | 07T41 | NifU domain p | 313594.PI23P | 1,6 | 0,8 | 7,02E-05 | 2,88E-05 | 3 - | 0,000114395 - | 2 - | 9,47E-05 - | 1 - | 4,50E-05 - | 1 - | 4,57E-05 - | 1 - | 5,15E-05 - |
| 5996 | 1 | 3um_FIJJAFBF_672183 | Bacteria | Proteobacteri | Alphaproteob | Pelagibacteral | Pelagibacterac | Candidatus Pe | Candidatus Pe | unclassified | unclassified | unclassified | unclassified | unclassified | 0,4 | 0,489897949 | 1,36E-05 | 1,69E-05 | 1 - | 2,89E-05 - | 0 - | 0 - | 0 - | 0 - | 0 - | 0 - | 1 - | 3,90E-05 - |
| 5997 | 1 | 3um_FIJJAFBF_341696 | Bacteria | Bacteroidetes | Flavobacteriia | Flavobacterial | Crocinitomica | unclassified | Crocinitomica | poorly charact | Function unkn | 09N7H | NA | 50743.SCB49_ | 1,2 | 0,4 | 5,35E-05 | 1,19E-05 | 2 - | 7,68E-05 - | 1 - | 4,77E-05 - | 1 - | 4,53E-05 - | 1 - | 4,60E-05 - | 1 - | 5,18E-05 - |
| 5998 | 1 | 3um_FIJJAFBF_520519 | Bacteria | Bacteroidetes | Flavobacteriia | Flavobacterial | Flavobacteriac | Polaribacter | Polaribacter s | unclassified | unclassified | unclassified | unclassified | unclassified | 2,2 | 0,748331477 | 0,000157771 | 4,76E-05 | 3 - | 0,000183154 - | 2 - | 0,000151616 - | 3 - | 0,000216195 - | 1 - | 7,31E-05 - | 2 - | 0,00016476 - |
| 5999 | 1 | 0.2um_NOKCPHJN_858760 | Bacteria | Bacteroidetes | Flavobacteriia | Flavobacterial | Flavobacteriac | Winogradskye | Winogradskye | unclassified | unclassified | unclassified | unclassified | unclassified | 1,2 | 0,979795897 | 6,85E-05 | 4,84E-05 | 3 - | 0,000151687 - | 1 - | 6,28E-05 - | 1 - | 5,97E-05 - | 0 - | 0 - | 1 - | 6,82E-05 - |
| 6000 | 1 | 0.2um_NOKCPHJN_212073 | Eukaryota | Apicomplexa | Conoidasida | Eucoccidiorida | Eimeriidae | Eimeria | Eimeria tenell | poorly charact | Function unkn | 0KN5V | NA | 140110.Necha | 0,2 | 0,4 | 4,81E-06 | 9,61E-06 | 0 - | 0 - | 1 - | 2,40E-05 - | 0 - | 0 - | 0 - | 0 - | 0 - | 0 - |

| | | | | | | | | | | | | | | | | | | | | | | | | | | | | |
|---|---|---|---|---|---|---|---|---|---|---|---|---|---|---|---|---|---|---|---|---|---|---|---|---|---|---|---|---|
| 6001 | 1 | 3um_FIJJAFBF_455735 | Bacteria | Bacteroidetes | unclassified | unclassified | unclassified | unclassified | Bacteroidetes | unclassified | unclassified | unclassified | unclassified | unclassified | 2,6 | 1,356465997 | 0,000370774 | 0,000186444 | 4 - | 0,000499029 - | 3 - | 0,000464735 - | 3 - | 0,00044179 - | 3 - | 0,000448315 - | 0 - | 0 - |
| 6002 | 1 | 3um_FIJJAFBF_84361 | Eukaryota | Bacillariophyta | Mediophyceae | Cymatosirales | Cymatosiracea | Plagiogrammc | Plagiogrammc | metabolism | Energy produc | 0IHF0 | This is one of | 45157.CMV04 | 3,4 | 0,489897949 | 0,000128483 | 1,63E-05 | 4 - | 0,00012753 - | 4 - | 0,000158354 - | 3 - | 0,000112902 - | 3 - | 0,000114569 - | 3 - | 0,000129062 - |
| 6003 | 1 | 3um_FIJJAFBF_180339 | Bacteria | Proteobacteri | Alphaproteob | Rhodobactera | Rhodobactera | Planktomarina | Planktomarina | information st | Translation, ri | 17FUG | Binds directly | 439496.RBY4I | 2,4 | 0,8 | 0,000378216 | 0,000130421 | 2 - | 0,000263855 - | 2 - | 0,00032763 - | 2 - | 0,000311453 - | 4 - | 0,000632107 - | 2 - | 0,000356033 - |
| 6004 | 1 | 3um_FIJJAFBF_316476 | Bacteria | Bacteroidetes | Flavobacteriia | Flavobacterial | Flavobacteriac | unclassified | Flavobacteriac | unclassified | unclassified | unclassified | unclassified | unclassified | 0,8 | 0,4 | 4,40E-05 | 2,24E-05 | 1 - | 4,76E-05 - | 1 - | 5,91E-05 - | 1 - | 5,62E-05 - | 1 - | 5,70E-05 - | 0 - | 0 - |
| 6005 | 1 | 3um_FIJJAFBF_331893 | Bacteria | Proteobacteri | Alphaproteob | Rhodobactera | Rhodobactera | Roseovarius | Roseovarius s | metabolism | Amino acid tra | 01QH7 | sulfoacetaldel | 314264.ROS21 | 7,8 | 0,748331477 | 0,000181866 | 2,89E-05 | 8 - | 0,000155894 - | 7 - | 0,000169377 - | 8 - | 0,000184017 - | 7 - | 0,000163393 - | 9 - | 0,00023665 - |
| 6006 | 1 | 3um_FIJJAFBF_350726 | Bacteria | Bacteroidetes | Flavobacteriia | Flavobacterial | Crocinitomica | unclassified | Crocinitomica | unclassified | unclassified | unclassified | unclassified | unclassified | 1 | 0 | 0,000155713 | 1,48E-05 | 1 - | 0,000130428 - | 1 - | 0,000161953 - | 1 - | 0,000153957 - | 1 - | 0,000156231 - | 1 - | 0,000175994 - |
| 6007 | 1 | 3um_FIJJAFBF_514037 | Bacteria | Bacteroidetes | Flavobacteriia | Flavobacterial | Flavobacteriac | Polaribacter | Polaribacter s | unclassified | unclassified | unclassified | unclassified | unclassified | 2,8 | 0,748331477 | 0,000283756 | 5,00E-05 | 4 - | 0,000347808 - | 2 - | 0,000215938 - | 3 - | 0,000307914 - | 3 - | 0,000312462 - | 2 - | 0,000234658 - |
| 6008 | 1 | 0.2um_NOKCPHJN_537235 | Eukaryota | Zoopagomyco | Basidiobolomy | Basidiobolales | Basidiobolacea | Basidiobolus | Basidiobolus r | unclassified | unclassified | unclassified | unclassified | unclassified | 1,2 | 0,748331477 | 0,000109225 | 7,16E-05 | 1 - | 7,76E-05 - | 2 - | 0,000192593 - | 2 - | 0,000183084 - | 1 - | 9,29E-05 - | 0 - | 0 - |
| 6009 | 1 | 3um_FIJJAFBF_451551 | Bacteria | Firmicutes | Bacilli | Bacillales | Bacillaceae | Bacillus | Bacillus marin | information st | Translation, ri | 05KK9 | This protein bi | 479434.Sthe_ | 1,6 | 0,8 | 0,000206308 | 8,25E-05 | 3 - | 0,000337579 - | 1 - | 0,000139724 - | 1 - | 0,000132826 - | 2 - | 0,000269575 - | 1 - | 0,000151838 - |
| 6010 | 1 | 3um_FIJJAFBF_55604 | Bacteria | Bacteroidetes | unclassified | unclassified | unclassified | unclassified | Bacteroidetes | information st | Translation, ri | COG0050 | This protein p | 983548.Krodi_ | 8,8 | 0,979795897 | 0,000306451 | 4,62E-05 | 8 - | 0,000233049 - | 10 - | 0,000361723 - | 10 - | 0,000343864 - | 8 - | 0,000279154 - | 8 - | 0,000314466 - |
| 6011 | 1 | 3um_FIJJAFBF_542237 | unclassified | unclassified | unclassified | unclassified | unclassified | unclassified | unclassified | unclassified | unclassified | unclassified | unclassified | unclassified | 0,6 | 0,489897949 | 1,27E-05 | 1,04E-05 | 1 - | 1,88E-05 - | 0 - | 0 - | 1 - | 2,22E-05 - | 1 - | 2,25E-05 - | 0 - | 0 - |
| 6012 | 1 | 0.2um_NOKCPHJN_07200 | Viruses | unclassified | unclassified | unclassified | unclassified | unclassified | Pacmanvirus A | unclassified | unclassified | unclassified | unclassified | unclassified | 1,2 | 0,4 | 2,54E-05 | 8,74E-06 | 1 - | 1,77E-05 - | 1 - | 2,20E-05 - | 1 - | 2,09E-05 - | 2 - | 4,24E-05 - | 1 - | 2,39E-05 - |
| 6013 | 1 | 3um_FIJJAFBF_20027 | Bacteria | Bacteroidetes | Flavobacteriia | Flavobacterial | Flavobacteriac | Polaribacter | Polaribacter a | metabolism | Amino acid tra | 0PF76 | Dehydrogenas | 313598.MED1 | 4,8 | 0,4 | 0,000103835 | 8,00E-06 | 5 - | 9,11E-05 - | 5 - | 0,00011311 - | 5 - | 0,000107526 - | 5 - | 0,000109114 - | 4 - | 9,83E-05 - |
| 6014 | 1 | 3um_FIJJAFBF_568354 | Bacteria | Proteobacteri | Gammaprote | Oceanospirilla | Saccharospirill | Reinekea | Reinekea fors | poorly charact | Function unkn | 17H2T | NA | 314283.MED2 | 0,6 | 0,8 | 5,48E-05 | 7,65E-05 | 1 - | 7,86E-05 - | 2 - | 0,000195231 - | 0 - | 0 - | 0 - | 0 - | 0 - | 0 - |
| 6015 | 1 | 0.2um_NOKCPHJN_673420 | Bacteria | Bacteroidetes | Flavobacteriia | Flavobacterial | Cryomorphace | unclassified | Cryomorphace | unclassified | unclassified | unclassified | unclassified | unclassified | 0,2 | 0,4 | 1,05E-05 | 2,09E-05 | 0 - | 0 - | 0 - | 0 - | 1 - | 5,23E-05 - | 0 - | 0 - | 0 - | 0 - |
| 6016 | 1 | 3um_FIJJAFBF_176419 | Bacteria | Bacteroidetes | Flavobacteriia | Flavobacterial | Crocinitomica | unclassified | Crocinitomica | information st | Translation, ri | COG1185 | Involved in mF | 743722.Sph21 | 1 | 0 | 1,90E-05 | 1,80E-06 | 1 - | 1,59E-05 - | 1 - | 1,97E-05 - | 1 - | 1,88E-05 - | 1 - | 1,90E-05 - | 1 - | 2,15E-05 - |
| 6017 | 1 | 3um_FIJJAFBF_27824 | Bacteria | Proteobacteri | Gammaprote | Alteromonada | Colwelliaceae | Colwellia | various | information st | Transcription | 05VEQ | Cold shock pro | 156578.ATW7 | 2,8 | 0,4 | 0,000573112 | 0,000102448 | 3 - | 0,000513926 - | 3 - | 0,000638144 - | 2 - | 0,000404425 - | 3 - | 0,000615597 - | 3 - | 0,000693468 - |
| 6018 | 1 | 3um_FIJJAFBF_698996 | Bacteria | Proteobacteri | Alphaproteob | Rhodobactera | Rhodobactera | Tateyamaria | Tateyamaria s | metabolism | Amino acid tra | COG1932 | Catalyzes the | 391589.RGAI1 | 0,8 | 0,4 | 2,89E-05 | 1,49E-05 | 1 - | 3,03E-05 - | 1 - | 3,76E-05 - | 1 - | 3,57E-05 - | 0 - | 0 - | 1 - | 4,09E-05 - |
| 6019 | 2 | 0.2um_NOKCPHJN_91525;3um_FIJJAFBF_180241 | Bacteria | various | various | various | various | various | various | unclassified | unclassified | unclassified | unclassified | unclassified | 0,8 | 0,4 | 4,82E-05 | 2,45E-05 | 1 - | 5,22E-05 - | 1 - | 6,48E-05 - | 1 - | 6,16E-05 - | 1 - | 6,25E-05 - | 0 - | 0 - |
| 6020 | 1 | 0.2um_NOKCPHJN_16433 | Bacteria | Bacteroidetes | Flavobacteriia | Flavobacterial | Flavobacteriac | Formosa | Formosa sp. H | poorly charact | Function unkn | 0P8NI | Inherit from b | 983544.Lacal_ | 1 | 0,632455532 | 3,10E-05 | 1,77E-05 | 2 - | 5,52E-05 - | 1 - | 3,43E-05 - | 1 - | 3,26E-05 - | 1 - | 3,30E-05 - | 0 - | 0 - |
| 6021 | 1 | 3um_FIJJAFBF_715084 | Eukaryota | Xanthophycea | unclassified | Vaucheriales | Vaucheriaceae | Vaucheria | Vaucheria lito | unclassified | unclassified | unclassified | unclassified | unclassified | 1,2 | 0,4 | 0,000122779 | 4,23E-05 | 1 - | 8,57E-05 - | 1 - | 0,000106357 - | 1 - | 0,000101106 - | 2 - | 0,000205199 - | 1 - | 0,000115578 - |
| 6022 | 1 | 0.2um_NOKCPHJN_488159 | Bacteria | Proteobacteri | Alphaproteob | Sphingomona | various | various | various | unclassified | unclassified | unclassified | unclassified | unclassified | 0,2 | 0,4 | 1,94E-05 | 3,88E-05 | 0 - | 0 - | 1 - | 9,70E-05 - | 0 - | 0 - | 0 - | 0 - | 0 - | 0 - |
| 6023 | 1 | 3um_FIJJAFBF_64747 | Bacteria | Proteobacteri | Alphaproteob | Rhodobactera | Rhodobactera | various | various | metabolism | Energy produc | COG0437 | 4fe-4S ferredc | 391616.OA23 | 0,2 | 0,4 | 1,17E-05 | 2,33E-05 | 1 - | 5,83E-05 - | 0 - | 0 - | 0 - | 0 - | 0 - | 0 - | 0 - | 0 - |
| 6024 | 1 | 3um_FIJJAFBF_249307 | Bacteria | Bacteroidetes | Flavobacteriia | Flavobacterial | Flavobacteriac | Tenacibaculum | Tenacibaculum | metabolism | Energy produc | COG0437 | 4fe-4S ferredc | 313598.MED1 | 0,6 | 0,8 | 7,07E-06 | 9,12E-06 | 2 - | 2,22E-05 - | 0 - | 0 - | 1 - | 1,31E-05 - | 0 - | 0 - | 0 - | 0 - |
| 6025 | 1 | 3um_FIJJAFBF_37823 | Bacteria | Actinobacteri | unclassified | unclassified | unclassified | unclassified | Actinobacteri | unclassified | unclassified | unclassified | unclassified | unclassified | 0,6 | 0,8 | 1,32E-05 | 1,69E-05 | 2 - | 4,08E-05 - | 1 - | 2,53E-05 - | 0 - | 0 - | 0 - | 0 - | 0 - | 0 - |
| 6026 | 1 | 0.2um_NOKCPHJN_953225 | Bacteria | Bacteroidetes | Flavobacteriia | Flavobacterial | Flavobacteriac | Formosa | Formosa sp. H | metabolism | Inorganic ion t | 05C27 | TonB-depende | 886377.Murru | 4,8 | 0,4 | 0,00011981 | 9,23E-06 | 5 - | 0,000105107 - | 5 - | 0,000130512 - | 5 - | 0,000124068 - | 5 - | 0,0001259 - | 4 - | 0,000113461 - |
| 6027 | 1 | 0.2um_NOKCPHJN_369085 | Bacteria | Proteobacteri | Betaproteoba | unclassified | unclassified | unclassified | Betaproteoba | unclassified | unclassified | unclassified | unclassified | unclassified | 0,2 | 0,4 | 6,45E-06 | 1,29E-05 | 1 - | 3,22E-05 - | 0 - | 0 - | 0 - | 0 - | 0 - | 0 - | 0 - | 0 - |
| 6028 | 1 | 0.2um_NOKCPHJN_466259 | unclassified | unclassified | unclassified | unclassified | unclassified | unclassified | unclassified | unclassified | unclassified | unclassified | unclassified | unclassified | 1 | 0 | 5,96E-05 | 5,66E-06 | 1 - | 4,99E-05 - | 1 - | 6,20E-05 - | 1 - | 5,89E-05 - | 1 - | 5,98E-05 - | 1 - | 6,73E-05 - |
| 6029 | 1 | 0.2um_NOKCPHJN_347120 | Bacteria | Bacteroidetes | Flavobacteriia | Flavobacterial | Flavobacteriac | Polaribacter | Polaribacter s | unclassified | unclassified | unclassified | unclassified | unclassified | 0,6 | 0,489897949 | 6,59E-05 | 5,48E-05 | 1 - | 9,33E-05 - | 0 - | 0 - | 1 - | 0,000110148 - | 0 - | 0 - | 1 - | 0,000125914 - |
| 6030 | 1 | 3um_FIJJAFBF_128488 | Bacteria | Bacteroidetes | Flavobacteriia | Flavobacterial | Flavobacteriac | Formosa | Formosa sp. H | unclassified | unclassified | unclassified | unclassified | unclassified | 1 | 0 | 0,000107052 | 1,02E-05 | 1 - | 8,97E-05 - | 1 - | 0,000111343 - | 1 - | 0,000105845 - | 1 - | 0,000107409 - | 1 - | 0,000120996 - |
| 6031 | 1 | 0.2um_NOKCPHJN_250636 | Bacteria | Bacteroidetes | Flavobacteriia | Flavobacterial | Flavobacteriac | Polaribacter | Polaribacter s | unclassified | unclassified | unclassified | unclassified | unclassified | 0,4 | 0,489897949 | 9,03E-06 | 1,11E-05 | 1 - | 2,05E-05 - | 0 - | 0 - | 0 - | 0 - | 1 - | 2,46E-05 - | 0 - | 0 - |
| 6032 | 1 | 0.2um_NOKCPHJN_45697 | Bacteria | various | various | various | various | various | various | information st | Translation, ri | 08UHH | Located on the | 391603.FBALC | 5 | 0 | 0,000539477 | 5,12E-05 | 5 - | 0,000451877 - | 5 - | 0,000561098 - | 5 - | 0,000533395 - | 5 - | 0,000541273 - | 5 - | 0,000609742 - |
| 6033 | 1 | 3um_FIJJAFBF_303369 | Bacteria | Proteobacteri | Alphaproteob | Rhodobactera | Rhodobactera | Planktomarina | Planktomarina | metabolism | Energy produc | 16RH6 | Produces ATP | 388401.RB215 | 5,8 | 1,720465053 | 0,000267501 | 4,98E-05 | 9 - | 0,000357436 - | 5 - | 0,000246572 - | 6 - | 0,000281278 - | 5 - | 0,00023786 - | 4 - | 0,000214359 - |
| 6034 | 1 | 3um_FIJJAFBF_502399 | Bacteria | Bacteroidetes | Flavobacteriia | Flavobacterial | Flavobacteriac | Formosa | Formosa sp. H | poorly charact | Function unkn | 11PIQ | YCE I like fami | 391598.FBBAL | 2,2 | 1,469693846 | 0,000145111 | 8,04E-05 | 5 - | 0,00028984 - | 1 - | 7,20E-05 - | 1 - | 6,84E-05 - | 2 - | 0,000138872 - | 2 - | 0,000156439 - |
| 6035 | 1 | 0.2um_NOKCPHJN_439546 | Bacteria | Proteobacteri | Alphaproteob | Rhodobactera | Rhodobactera | Marivita | Marivita geoje | metabolism | Inorganic ion t | 16PII | TRAP transpor | 388401.RB215 | 0,4 | 0,489897949 | 1,59E-05 | 1,97E-05 | 1 - | 3,55E-05 - | 1 - | 4,41E-05 - | 0 - | 0 - | 0 - | 0 - | 0 - | 0 - |
| 6036 | 1 | 0.2um_NOKCPHJN_352941 | Bacteria | Proteobacteri | Gammaprote | Alteromonada | Alteromonada | Glaciecola | Glaciecola sp. | cellular proces | Posttranslatio | 16QU3 | Hemolysin-typ | 455436.GHTC | 18,2 | 2,13541565 | 0,000136045 | 1,48E-05 | 21 - | 0,000132217 - | 20 - | 0,000156356 - | 15 - | 0,000111477 - | 18 - | 0,000135749 - | 17 - | 0,000144425 - |
| 6037 | 1 | 0.2um_NOKCPHJN_546115 | Bacteria | Proteobacteri | Gammaprote | Cellvibrionales | Spongiibacter | Oceanicoccus | Oceanicoccus | metabolism | Energy produc | COG1290 | Component of | 247633.GP214 | 0,2 | 0,4 | 3,91E-06 | 7,82E-06 | 0 - | 0 - | 0 - | 0 - | 1 - | 1,96E-05 - | 0 - | 0 - | 0 - | 0 - |
| 6038 | 1 | 3um_FIJJAFBF_87014 | Bacteria | Proteobacteri | Alphaproteob | Rhodospirillale | Rhodospirillac | Azospirillum | Azospirillum b | unclassified | unclassified | unclassified | unclassified | unclassified | 1,6 | 0,489897949 | 1,42E-05 | 4,46E-06 | 2 - | 1,49E-05 - | 1 - | 9,26E-06 - | 2 - | 1,76E-05 - | 1 - | 8,93E-06 - | 2 - | 2,01E-05 - |
| 6039 | 6 | 0.2um_NOKCPHJN_758394;0.2um_NOKCPHJN_7 | Bacteria | Bacteroidetes | various | various | various | various | various | unclassified | unclassified | unclassified | unclassified | unclassified | 0,2 | 0,4 | 9,33E-06 | 1,87E-05 | 1 - | 4,67E-05 - | 0 - | 0 - | 0 - | 0 - | 0 - | 0 - | 0 - | 0 - |
| 6040 | 1 | 3um_FIJJAFBF_192507 | Bacteria | Bacteroidetes | unclassified | unclassified | unclassified | unclassified | Bacteroidetes | unclassified | unclassified | unclassified | unclassified | unclassified | 0,2 | 0,4 | 6,85E-06 | 1,37E-05 | 0 - | 0 - | 1 - | 3,43E-05 - | 0 - | 0 - | 0 - | 0 - | 0 - | 0 - |
| 6041 | 1 | 3um_FIJJAFBF_807627 | Bacteria | Bacteroidetes | Flavobacteriia | Flavobacterial | Flavobacteriac | Chryseobacter | Chryseobacter | unclassified | unclassified | unclassified | unclassified | unclassified | 1 | 0 | 5,48E-05 | 5,20E-06 | 1 - | 4,59E-05 - | 1 - | 5,70E-05 - | 1 - | 5,42E-05 - | 1 - | 5,50E-05 - | 1 - | 6,19E-05 - |
| 6042 | 1 | 0.2um_NOKCPHJN_868369 | Bacteria | Bacteroidetes | Flavobacteriia | Flavobacterial | Flavobacteriac | Polaribacter | Polaribacter s | poorly charact | Function unkn | 0P8NI | Inherit from b | 156586.BBFL7 | 2,2 | 0,4 | 7,70E-05 | 1,50E-05 | 2 - | 5,87E-05 - | 2 - | 7,29E-05 - | 3 - | 0,000103951 - | 2 - | 7,03E-05 - | 2 - | 7,92E-05 - |
| 6043 | 1 | 0.2um_NOKCPHJN_265006 | Eukaryota | Apicomplexa | Conoidasida | Eucoccidiorida | Eimeriidae | Eimeria | Eimeria maxim | unclassified | unclassified | unclassified | unclassified | unclassified | 1 | 0 | 1,07E-05 | 1,01E-06 | 1 - | 8,93E-06 - | 1 - | 1,11E-05 - | 1 - | 1,05E-05 - | 1 - | 1,07E-05 - | 1 - | 1,20E-05 - |
| 6044 | 1 | 3um_FIJJAFBF_590082 | Bacteria | Proteobacteri | Gammaprote | Alteromonada | Alteromonada | Paraglaciecola | Paraglaciecola | metabolism | Carbohydrate | 08Q79 | 6-phosphoglu | 342610.Patl_ | 0,2 | 0,4 | 9,98E-06 | 2,00E-05 | 1 - | 4,99E-05 - | 0 - | 0 - | 0 - | 0 - | 0 - | 0 - | 0 - | 0 - |
| 6045 | 1 | 3um_FIJJAFBF_374069 | Bacteria | Bacteroidetes | Flavobacteriia | Flavobacterial | unclassified | unclassified | Flavobacterial | unclassified | unclassified | unclassified | unclassified | unclassified | 2,2 | 0,4 | 6,17E-05 | 1,66E-05 | 2 - | 4,65E-05 - | 2 - | 5,77E-05 - | 2 - | 5,49E-05 - | 2 - | 5,57E-05 - | 3 - | 9,41E-05 - |
| 6046 | 1 | 3um_FIJJAFBF_352150 | Viruses | unclassified | unclassified | unclassified | Phycodnavirid | unclassified | Chrysochromu | unclassified | unclassified | unclassified | unclassified | unclassified | 0,6 | 0,489897949 | 3,34E-05 | 2,74E-05 | 0 - | 0 - | 0 - | 0 - | 1 - | 5,29E-05 - | 1 - | 5,37E-05 - | 1 - | 6,05E-05 - |
| 6047 | 1 | 3um_FIJJAFBF_58928 | Bacteria | Proteobacteri | Gammaprote | Chromatiales | Ectothiorhodo | Thiohalomona | Thiohalomona | cellular proces | Cell wall/mem | 08ZG0 | Ompa motb d | 637616.MDM | 0,4 | 0,489897949 | 2,59E-05 | 3,18E-05 | 0 - | 0 - | 0 - | 0 - | 0 - | 0 - | 1 - | 6,08E-05 - | 1 - | 6,85E-05 - |
| 6048 | 1 | 3um_FIJJAFBF_111300 | unclassified | unclassified | unclassified | unclassified | unclassified | unclassified | unclassified | unclassified | unclassified | unclassified | unclassified | unclassified | 6,4 | 1,019803903 | 0,000110994 | 1,94E-05 | 7 - | 0,000101959 - | 8 - | 0,000144689 - | 6 - | 0,000103159 - | 5 - | 8,72E-05 - | 6 - | 0,000117925 - |
| 6049 | 1 | 0.2um_NOKCPHJN_1023580 | Bacteria | Bacteroidetes | Flavobacteriia | Flavobacterial | Flavobacteriac | Flavobacteriu | Flavobacteriu | metabolism | Carbohydrate | 0P8NX | Glyceraldehyd | 391598.FBBAL | 0,2 | 0,4 | 8,23E-06 | 1,65E-05 | 0 - | 0 - | 0 - | 0 - | 0 - | 0 - | 1 - | 4,12E-05 - | 0 - | 0 - |
| 6050 | 1 | 3um_FIJJAFBF_774536 | Eukaryota | unclassified | Cryptophyta | Pyrenomonad | Geminigeracea | Guillardia | Guillardia thet | unclassified | unclassified | unclassified | unclassified | unclassified | 4,8 | 0,4 | 0,000370978 | 4,35E-05 | 5 - | 0,000324228 - | 4 - | 0,000322077 - | 5 - | 0,000382718 - | 5 - | 0,000388371 - | 5 - | 0,000437499 - |
| 6051 | 1 | 3um_FIJJAFBF_420946 | Bacteria | Proteobacteri | Gammaprote | Cellvibrionales | Porticoccacea | unclassified | SAR92 bacteri | information st | Translation, ri | 0QTNF | One of the pri | 314287.GB22 | 0,2 | 0,4 | 2,61E-05 | 5,22E-05 | 1 - | 0,000130428 - | 0 - | 0 - | 0 - | 0 - | 0 - | 0 - | 0 - | 0 - |
| 6052 | 1 | 3um_FIJJAFBF_295347 | Bacteria | Proteobacteri | Betaproteoba | unclassified | unclassified | unclassified | Betaproteoba | unclassified | unclassified | unclassified | unclassified | unclassified | 5,6 | 0,8 | 0,000623715 | 0,000108742 | 6 - | 0,000559887 - | 6 - | 0,000695214 - | 6 - | 0,000660889 - | 4 - | 0,0004471 - | 6 - | 0,000755485 - |
| 6053 | 1 | 0.2um_NOKCPHJN_180963 | Bacteria | Bacteroidetes | Cytophagia | Cytophagales | Flammeovirga | Fabibacter | Fabibacter mi | poorly charact | Function unkn | 05BEG | Protein of unk | 694427.Palpr_ | 1 | 0 | 3,73E-05 | 3,55E-06 | 1 - | 3,13E-05 - | 1 - | 3,88E-05 - | 1 - | 3,69E-05 - | 1 - | 3,75E-05 - | 1 - | 4,22E-05 - |
| 6054 | 1 | 0.2um_NOKCPHJN_122677 | Bacteria | Bacteroidetes | Flavobacteriia | Flavobacterial | Flavobacteriac | Polaribacter | Polaribacter d | unclassified | unclassified | unclassified | unclassified | unclassified | 1,2 | 0,748331477 | 8,04E-05 | 5,14E-05 | 2 - | 0,00011364 - | 0 - | 0 - | 1 - | 6,71E-05 - | 1 - | 6,81E-05 - | 2 - | 0,000153341 - |
| 6055 | 1 | 3um_FIJJAFBF_676755 | Bacteria | Proteobacteri | Alphaproteob | Rhodospirillale | Rhodospirillac | unclassified | Rhodobactera | cellular proces | Posttranslatio | 01SH5 | Molecular cha | 367336.OM22 | 0,2 | 0,4 | 4,45E-06 | 8,90E-06 | 1 - | 2,22E-05 - | 0 - | 0 - | 0 - | 0 - | 0 - | 0 - | 0 - | 0 - |
| 6056 | 1 | 3um_FIJJAFBF_73737 | Bacteria | Proteobacteri | Gammaprote | Alteromonada | Alteromonada | Alteromonas | Alteromonas s | unclassified | unclassified | unclassified | unclassified | unclassified | 1 | 0 | 0,000129271 | 1,23E-05 | 1 - | 0,00010828 - | 1 - | 0,000134452 - | 1 - | 0,000127813 - | 1 - | 0,000129701 - | 1 - | 0,000146108 - |
| 6057 | 1 | 3um_FIJJAFBF_19901 | Bacteria | Proteobacteri | Alphaproteob | Rhodobactera | Rhodobactera | Planktomarina | Planktomarina | metabolism | Amino acid tra | 16QM8 | Oligoendopep | 391593.RCCS2 | 0,6 | 0,489897949 | 1,36E-05 | 1,13E-05 | 1 - | 1,90E-05 - | 1 - | 2,36E-05 - | 0 - | 0 - | 0 - | 0 - | 1 - | 2,56E-05 - |
| 6058 | 1 | 0.2um_NOKCPHJN_357557 | Bacteria | Proteobacteri | Alphaproteob | Pelagibacteral | Pelagibacterac | Candidatus Pe | Candidatus Pe | unclassified | unclassified | unclassified | unclassified | unclassified | 14,6 | 1,743559577 | 0,000637125 | 8,62E-05 | 17 - | 0,000623388 - | 14 - | 0,000637465 - | 14 - | 0,000605991 - | 12 - | 0,000527093 - | 16 - | 0,000791691 - |
| 6059 | 1 | 3um_FIJJAFBF_518710 | Bacteria | Bacteroidetes | Flavobacteriia | Flavobacterial | Flavobacteriac | Polaribacter | Polaribacter s | unclassified | unclassified | unclassified | unclassified | unclassified | 3,6 | 1,019803903 | 9,36E-05 | 2,51E-05 | 4 - | 8,81E-05 - | 2 - | 5,47E-05 - | 4 - | 0,000104017 - | 5 - | 0,000131942 - | 3 - | 8,92E-05 - |
| 6060 | 1 | 3um_FIJJAFBF_113994 | Viruses | unclassified | unclassified | unclassified | unclassified | unclassified | uncultured vir | cellular proces | Cell wall/mem | 01Y9Y | glycosyl transf | 488538.SAR11 | 0,4 | 0,489897949 | 1,23E-05 | 1,51E-05 | 0 - | 0 - | 0 - | 0 - | 0 - | 0 - | 1 - | 2,89E-05 - | 1 - | 3,25E-05 - |
| 6061 | 1 | 0.2um_NOKCPHJN_553807 | unclassified | unclassified | unclassified | unclassified | unclassified | unclassified | unclassified | unclassified | unclassified | unclassified | unclassified | unclassified | 0,6 | 0,489897949 | 1,22E-05 | 1,01E-05 | 1 - | 1,78E-05 - | 1 - | 2,21E-05 - | 0 - | 0 - | 1 - | 2,13E-05 - | 0 - | 0 - |
| 6062 | 1 | 0.2um_NOKCPHJN_594836 | Bacteria | Proteobacteri | Alphaproteob | Rhodobactera | Rhodobactera | Planktomarina | Planktomarina | unclassified | unclassified | unclassified | unclassified | unclassified | 1,6 | 0,489897949 | 8,15E-05 | 2,30E-05 | 2 - | 8,66E-05 - | 2 - | 0,000107561 - | 1 - | 5,11E-05 - | 2 - | 0,000103761 - | 1 - | 5,84E-05 - |
| 6063 | 1 | 3um_FIJJAFBF_256247 | Bacteria | Proteobacteri | Alphaproteob | Rhodobactera | Rhodobactera | Roseibium | Roseibium sp. | metabolism | Carbohydrate | 01U6R | trap dicarboxy | 744980.TRICH | 5,8 | 1,166190379 | 0,000221903 | 2,32E-05 | 8 - | 0,000260856 - | 5 - | 0,000202442 - | 6 - | 0,000230936 - | 5 - | 0,000195289 - | 5 - | 0,000219992 - |
| 6064 | 1 | 0.2um_NOKCPHJN_201765 | Bacteria | Proteobacteri | Gammaprote | Oceanospirilla | Oceanospirilla | Neptuniibacte | Neptuniibacte | poorly charact | Function unkn | 179VZ | UPF0234 prot | 207954.MED9 | 0,8 | 0,4 | 6,63E-05 | 3,37E-05 | 1 - | 7,17E-05 - | 1 - | 8,91E-05 - | 1 - | 8,47E-05 - | 1 - | 8,59E-05 - | 0 - | 0 - |
| 6065 | 1 | 0.2um_NOKCPHJN_202559 | Bacteria | Proteobacteri | Gammaprote | Cellvibrionales | Porticoccacea | unclassified | Gammaprote | cellular proces | Cell wall/mem | 05CET | Activates KDO | 314287.GB22 | 0,2 | 0,4 | 1,13E-05 | 2,25E-05 | 0 - | 0 - | 1 - | 5,63E-05 - | 0 - | 0 - | 0 - | 0 - | 0 - | 0 - |
| 6066 | 1 | 3um_FIJJAFBF_190547 | Archaea | Euryarchaeota | unclassified | unclassified | unclassified | unclassified | candidate divi | unclassified | unclassified | unclassified | unclassified | unclassified | 0,2 | 0,4 | 1,12E-05 | 2,23E-05 | 0 - | 0 - | 0 - | 0 - | 1 - | 5,58E-05 - | 0 - | 0 - | 0 - | 0 - |
| 6067 | 1 | 0.2um_NOKCPHJN_409937 | Eukaryota | unclassified | Oomycetes | Pythiales | Pythiaceae | Pythium | Pythium insidi | unclassified | unclassified | unclassified | unclassified | unclassified | 1 | 0 | 0,000103028 | 9,78E-06 | 1 - | 8,63E-05 - | 1 - | 0,000107157 - | 1 - | 0,000101866 - | 1 - | 0,000103371 - | 1 - | 0,000116447 - |
| 6068 | 1 | 3um_FIJJAFBF_561834 | Bacteria | Bacteroidetes | Flavobacteriia | Flavobacterial | Flavobacteriac | Algibacter | Algibacter sp. | metabolism | Amino acid tra | 05C6M | Alpha-keto-be | 313603.FB217 | 0,2 | 0,4 | 5,79E-06 | 1,16E-05 | 0 - | 0 - | 0 - | 0 - | 0 - | 0 - | 1 - | 2,89E-05 - | 0 - | 0 - |
| 6069 | 1 | 3um_FIJJAFBF_202366 | Bacteria | Bacteroidetes | Cytophagia | Cytophagales | Flammeovirga | unclassified | Flammeovirga | unclassified | unclassified | unclassified | unclassified | unclassified | 1 | 0 | 2,37E-05 | 2,25E-06 | 1 - | 1,99E-05 - | 1 - | 2,47E-05 - | 1 - | 2,34E-05 - | 1 - | 2,38E-05 - | 1 - | 2,68E-05 - |
| 6070 | 2 | 0.2um_NOKCPHJN_349824;0.2um_NOKCPHJN_5 | Bacteria | Bacteroidetes | Flavobacteriia | Flavobacterial | unclassified | unclassified | Flavobacterial | unclassified | unclassified | unclassified | unclassified | unclassified | 2,6 | 0,8 | 0,000116591 | 3,01E-05 | 4 - | 0,000152527 - | 2 - | 9,47E-05 - | 2 - | 9,00E-05 - | 2 - | 9,14E-05 - | 3 - | 0,00015436 - |
| 6071 | 1 | 3um_FIJJAFBF_273674 | Bacteria | Proteobacteri | Alphaproteob | Rhodobactera | Rhodobactera | unclassified | Rhodobactera | information st | Translation, ri | 01ZRG | endoribonucle | 314262.MED1 | 0,8 | 0,4 | 9,44E-05 | 4,85E-05 | 1 - | 9,98E-05 - | 0 - | 0 - | 1 - | 0,000117811 - | 1 - | 0,000119551 - | 1 - | 0,000134673 - |
| 6072 | 1 | 3um_FIJJAFBF_722765 | Bacteria | Actinobacteri | Actinobacteri | Corynebacteri | Corynebacteri | Corynebacteri | Corynebacteri | unclassified | unclassified | unclassified | unclassified | unclassified | 0,4 | 0,8 | 7,31E-06 | 1,46E-05 | 2 - | 3,66E-05 - | 0 - | 0 - | 0 - | 0 - | 0 - | 0 - | 0 - | 0 - |
| 6073 | 1 | 0.2um_NOKCPHJN_607864 | Viruses | unclassified | unclassified | unclassified | Mimiviridae | Cafeteriavirus | Cafeteria roen | unclassified | unclassified | unclassified | unclassified | unclassified | 1,6 | 1,019803903 | 3,91E-05 | 2,34E-05 | 3 - | 6,26E-05 - | 2 - | 5,18E-05 - | 0 - | 0 - | 1 - | 2,50E-05 - | 2 - | 5,63E-05 - |
| 6074 | 1 | 3um_FIJJAFBF_377215 | Bacteria | Bacteroidetes | Flavobacteriia | Flavobacterial | Flavobacteriac | Polaribacter | Polaribacter s | unclassified | unclassified | unclassified | unclassified | unclassified | 1,2 | 0,4 | 0,000114253 | 3,94E-05 | 1 - | 7,97E-05 - | 1 - | 9,90E-05 - | 1 - | 9,41E-05 - | 2 - | 0,000190949 - | 1 - | 0,000107552 - |
| 6075 | 1 | 0.2um_NOKCPHJN_399702 | Eukaryota | Mucoromycot | Mucoromycet | Mucorales | Mucoraceae | Mucor | Mucor ambigu | cellular proces | Intracellular tr | 0PI28 | GTP-binding p | 5022.CBX9689 | 0,2 | 0,4 | 1,21E-05 | 2,42E-05 | 0 - | 0 - | 0 - | 0 - | 1 - | 6,05E-05 - | 0 - | 0 - | 0 - | 0 - |
| 6076 | 1 | 3um_FIJJAFBF_774385 | Bacteria | Bacteroidetes | Cytophagia | Cytophagales | Flammeovirga | unclassified | Flammeovirga | unclassified | unclassified | unclassified | unclassified | unclassified | 0,4 | 0,489897949 | 2,61E-05 | 3,23E-05 | 1 - | 5,83E-05 - | 1 - | 7,23E-05 - | 0 - | 0 - | 0 - | 0 - | 0 - | 0 - |
| 6077 | 1 | 0.2um_NOKCPHJN_154863 | Bacteria | Bacteroidetes | Flavobacteriia | Flavobacterial | Flavobacteriac | Ulvibacter | Ulvibacter lito | metabolism | Energy produc | COG0437 | 4fe-4S ferredc | 50743.SCB49_ | 1 | 0 | 1,34E-05 | 1,27E-06 | 1 - | 1,12E-05 - | 1 - | 1,39E-05 - | 1 - | 1,32E-05 - | 1 - | 1,34E-05 - | 1 - | 1,51E-05 - |
| 6078 | 1 | 0.2um_NOKCPHJN_23137 | Bacteria | Bacteroidetes | Flavobacteriia | Flavobacterial | Flavobacteriac | Polaribacter | Polaribacter s | information st | Replication, re | 0P7SM | DNA topoisom | 313594.PI23P | 0,8 | 1,166190379 | 1,57E-05 | 2,19E-05 | 3 - | 5,56E-05 - | 1 - | 2,30E-05 - | 0 - | 0 - | 0 - | 0 - | 0 - | 0 - |
| 6079 | 1 | 0.2um_NOKCPHJN_09461 | Bacteria | Bacteroidetes | Flavobacteriia | Flavobacterial | Flavobacteriac | Polaribacter | various | metabolism | Inorganic ion t | COG0175 | Reduction of a | 313594.PI23P | 1,6 | 1,019803903 | 6,79E-05 | 4,40E-05 | 2 - | 7,38E-05 - | 1 - | 4,58E-05 - | 2 - | 8,71E-05 - | 3 - | 0,000132621 - | 0 - | 0 - |
| 6080 | 1 | 3um_FIJJAFBF_73823 | Bacteria | Bacteroidetes | Flavobacteriia | Flavobacterial | Flavobacteriac | Polaribacter | Polaribacter s | unclassified | unclassified | unclassified | unclassified | unclassified | 5,2 | 0,748331477 | 0,000164152 | 1,46E-05 | 6 - | 0,000160527 - | 5 - | 0,000166106 - | 6 - | 0,000189486 - | 5 - | 0,000160237 - | 4 - | 0,000144405 - |
| 6081 | 1 | 3um_FIJJAFBF_559090 | Bacteria | Proteobacteri | Gammaprote | Alteromonada | Alteromonada | Paraglaciecola | Paraglaciecola | unclassified | unclassified | unclassified | unclassified | unclassified | 0,4 | 0,8 | 4,10E-05 | 8,20E-05 | 2 - | 0,000204958 - | 0 - | 0 - | 0 - | 0 - | 0 - | 0 - | 0 - | 0 - |
| 6082 | 1 | 0.2um_NOKCPHJN_73064 | Bacteria | Bacteroidetes | unclassified | unclassified | unclassified | unclassified | Bacteroidetes | poorly charact | Function unkn | 09GJP | Protein of unk | 926562.Oweh | 2,2 | 0,748331477 | 0,000213221 | 8,34E-05 | 1 - | 7,92E-05 - | 2 - | 0,000196578 - | 3 - | 0,000280308 - | 2 - | 0,000189632 - | 3 - | 0,00032043 - |
| 6083 | 1 | 3um_FIJJAFBF_372861 | Bacteria | Bacteroidetes | Cytophagia | Cytophagales | Flammeovirga | unclassified | Flammeovirga | unclassified | unclassified | unclassified | unclassified | unclassified | 1 | 0 | 8,06E-05 | 7,65E-06 | 1 - | 6,75E-05 - | 1 - | 8,38E-05 - | 1 - | 7,97E-05 - | 1 - | 8,09E-05 - | 1 - | 9,11E-05 - |
| 6084 | 1 | 0.2um_NOKCPHJN_251837 | Eukaryota | unclassified | Oomycetes | Peronosporale | Peronosporac | Phytophthora | Phytophthora | information st | Translation, ri | 14XM1 | EFG_IV | 4787.PITG_10 | 0,6 | 0,489897949 | 8,59E-06 | 7,08E-06 | 1 - | 1,25E-05 - | 1 - | 1,56E-05 - | 1 - | 1,48E-05 - | 0 - | 0 - | 0 - | 0 - |
| 6085 | 1 | 3um_FIJJAFBF_242832 | Bacteria | Bacteroidetes | Flavobacteriia | Flavobacterial | Flavobacteriac | Polaribacter | Polaribacter a | unclassified | unclassified | unclassified | unclassified | unclassified | 0,4 | 0,489897949 | 1,70E-05 | 2,10E-05 | 1 - | 3,90E-05 - | 0 - | 0 - | 1 - | 4,61E-05 - | 0 - | 0 - | 0 - | 0 - |
| 6086 | 1 | 3um_FIJJAFBF_113804 | Bacteria | Proteobacteri | Alphaproteob | Rhizobiales | unclassified | unclassified | Rhizobiales ba | unclassified | unclassified | unclassified | unclassified | unclassified | 12,4 | 1,2 | 0,000202524 | 2,44E-05 | 14 - | 0,00019198 - | 13 - | 0,000221355 - | 11 - | 0,000178053 - | 11 - | 0,000180683 - | 13 - | 0,000240546 - |
| 6087 | 1 | 3um_FIJJAFBF_581784 | Bacteria | Bacteroidetes | Flavobacteriia | Flavobacterial | Flavobacteriac | Polaribacter | Polaribacter s | cellular proces | Posttranslatio | 05E6M | Sulphatase-m | 313598.MED1 | 0,8 | 0,748331477 | 2,25E-05 | 1,98E-05 | 2 - | 5,08E-05 - | 1 - | 3,15E-05 - | 0 - | 0 - | 1 - | 3,04E-05 - | 0 - | 0 - |
| 6088 | 1 | 3um_FIJJAFBF_128463 | Bacteria | Bacteroidetes | Flavobacteriia | Flavobacterial | Flavobacteriac | Polaribacter | Polaribacter a | information st | Translation, ri | 09AI8 | Associates wit | 313598.MED1 | 6,4 | 1,019803903 | 0,000268442 | 2,85E-05 | 8 - | 0,000284277 - | 5 - | 0,000220617 - | 6 - | 0,00025167 - | 7 - | 0,000297951 - | 6 - | 0,000287693 - |
| 6089 | 1 | 0.2um_NOKCPHJN_461763 | Bacteria | Bacteroidetes | Flavobacteriia | Flavobacterial | Flavobacteriac | Polaribacter | Polaribacter s | unclassified | unclassified | unclassified | unclassified | unclassified | 0,8 | 0,748331477 | 1,68E-05 | 1,65E-05 | 1 - | 1,80E-05 - | 2 - | 4,46E-05 - | 1 - | 2,12E-05 - | 0 - | 0 - | 0 - | 0 - |
| 6090 | 1 | 3um_FIJJAFBF_121248 | Bacteria | Bacteroidetes | Cytophagia | Cytophagales | Flammeovirga | unclassified | Flammeovirga | unclassified | unclassified | unclassified | unclassified | unclassified | 0,4 | 0,489897949 | 2,91E-05 | 3,60E-05 | 1 - | 6,48E-05 - | 1 - | 8,05E-05 - | 0 - | 0 - | 0 - | 0 - | 0 - | 0 - |
| 6091 | 1 | 3um_FIJJAFBF_448546 | Bacteria | Proteobacteri | Gammaprote | Oceanospirilla | Saccharospirill | Reinekea | Reinekea fors | unclassified | unclassified | unclassified | unclassified | unclassified | 1,2 | 0,979795897 | 3,15E-05 | 2,59E-05 | 2 - | 4,66E-05 - | 0 - | 0 - | 2 - | 5,50E-05 - | 2 - | 5,58E-05 - | 0 - | 0 - |
| 6092 | 1 | 0.2um_NOKCPHJN_110535 | Bacteria | Proteobacteri | Gammaprote | Cellvibrionales | Porticoccacea | unclassified | gamma prote | unclassified | unclassified | unclassified | unclassified | unclassified | 1,8 | 0,748331477 | 0,000165349 | 5,92E-05 | 3 - | 0,000235843 - | 1 - | 9,76E-05 - | 1 - | 9,28E-05 - | 2 - | 0,000188333 - | 2 - | 0,000212157 - |
| 6093 | 2 | 0.2um_NOKCPHJN_449241;3um_FIJJAFBF_3852C | Bacteria | Bacteroidetes | Flavobacteriia | Flavobacterial | Cryomorphace | unclassified | Cryomorphace | unclassified | unclassified | unclassified | unclassified | unclassified | 0,4 | 0,489897949 | 3,06E-05 | 3,79E-05 | 1 - | 6,83E-05 - | 1 - | 8,48E-05 - | 0 - | 0 - | 0 - | 0 - | 0 - | 0 - |
| 6094 | 1 | 3um_FIJJAFBF_504256 | Eukaryota | unclassified | Synurophycea | Parmales | Triparmaceae | Triparma | Triparma laevi | metabolism | Energy produc | COG0711 | F(1)F(0) ATP s | 35128.Thapsd | 0,6 | 0,489897949 | 4,55E-05 | 3,78E-05 | 1 - | 6,45E-05 - | 0 - | 0 - | 1 - | 7,61E-05 - | 0 - | 0 - | 1 - | 8,70E-05 - |
| 6095 | 1 | 0.2um_NOKCPHJN_833276 | Eukaryota | unclassified | Oomycetes | Pythiales | Pythiaceae | Pythium | Pythium insidi | poorly charact | Function unkn | 14WW8 | ATP synthase | 4787.PITG_09 | 1 | 0 | 0,000111404 | 1,06E-05 | 1 - | 9,33E-05 - | 1 - | 0,000115869 - | 1 - | 0,000110148 - | 1 - | 0,000111775 - | 1 - | 0,000125914 - |
| 6096 | 1 | 0.2um_NOKCPHJN_525701 | Bacteria | Proteobacteri | Gammaprote | Cellvibrionales | Spongiibacter | Oceanicoccus | Oceanicoccus | metabolism | Amino acid tra | 16Q4U | Catalyzes the | 247634.GPB2 | 0,6 | 0,489897949 | 1,93E-05 | 1,60E-05 | 1 - | 2,73E-05 - | 0 - | 0 - | 1 - | 3,23E-05 - | 0 - | 0 - | 1 - | 3,69E-05 - |
| 6097 | 1 | 3um_FIJJAFBF_437250 | Bacteria | Bacteroidetes | Cytophagia | Cytophagales | Flammeovirga | Fabibacter | Fabibacter sp. | unclassified | unclassified | unclassified | unclassified | unclassified | 8,6 | 0,489897949 | 0,000244828 | 2,04E-05 | 9 - | 0,000215206 - | 9 - | 0,000267223 - | 8 - | 0,000225804 - | 9 - | 0,000257781 - | 8 - | 0,000258124 - |
| 6098 | 1 | 3um_FIJJAFBF_118754 | Bacteria | Actinobacteri | Acidimicrobiia | Acidimicrobial | Acidimicrobiac | unclassified | Acidimicrobiac | information st | Transcription | 05D27 | DNA-depende | 525909.Afer_ | 1 | 0 | 1,05E-05 | 9,94E-07 | 1 - | 8,77E-06 - | 1 - | 1,09E-05 - | 1 - | 1,04E-05 - | 1 - | 1,05E-05 - | 1 - | 1,18E-05 - |
| 6099 | 1 | 3um_FIJJAFBF_115452 | Bacteria | Proteobacteri | Gammaprote | unclassified | unclassified | unclassified | SAR86 cluster | information st | Translation, ri | 179RU | 30S ribosomal | 566466.NOR5 | 1,2 | 1,166190379 | 0,000119103 | 0,00010785 | 3 - | 0,000264869 - | 0 - | 0 - | 0 - | 0 - | 2 - | 0,000211513 - | 1 - | 0,000119134 - |
| 6100 | 1 | 0.2um_NOKCPHJN_222290 | Bacteria | Bacteroidetes | Flavobacteriia | Flavobacterial | Flavobacteriac | Formosa | Formosa sp. H | metabolism | Nucleotide tra | 09B0C | Provides the p | 983544.Lacal_ | 4,4 | 0,489897949 | 7,31E-05 | 1,34E-05 | 4 - | 5,53E-05 - | 4 - | 6,87E-05 - | 4 - | 6,53E-05 - | 5 - | 8,28E-05 - | 5 - | 9,33E-05 - |
| 6101 | 1 | 3um_FIJJAFBF_685940 | Bacteria | Bacteroidetes | Cytophagia | Cytophagales | Flammeovirga | unclassified | Flammeovirga | unclassified | unclassified | unclassified | unclassified | unclassified | 0,2 | 0,4 | 5,41E-06 | 1,08E-05 | 1 - | 2,71E-05 - | 0 - | 0 - | 0 - | 0 - | 0 - | 0 - | 0 - | 0 - |
| 6102 | 1 | 3um_FIJJAFBF_564568 | Bacteria | Proteobacteri | Gammaprote | Cellvibrionales | Porticoccacea | unclassified | gamma prote | unclassified | unclassified | unclassified | unclassified | unclassified | 2,6 | 1,019803903 | 3,87E-05 | 1,56E-05 | 2 - | 2,51E-05 - | 3 - | 4,67E-05 - | 3 - | 4,44E-05 - | 4 - | 6,01E-05 - | 1 - | 1,69E-05 - |
| 6103 | 1 | 3um_FIJJAFBF_183700 | Bacteria | Bacteroidetes | Flavobacteriia | Flavobacterial | Flavobacteriac | Algibacter | Algibacter sp. | unclassified | unclassified | unclassified | unclassified | unclassified | 8,6 | 1,854723699 | 0,0001248 | 2,19E-05 | 10 - | 0,000123549 - | 9 - | 0,00013807 - | 10 - | 0,000145837 - | 9 - | 0,000133192 - | 5 - | 8,34E-05 - |
| 6104 | 1 | 0.2um_NOKCPHJN_313351 | Bacteria | Bacteroidetes | Flavobacteriia | Flavobacterial | Flavobacteriac | Winogradskye | Winogradskye | poorly charact | Function unkn | 0XY6Y | Pkd domain co | 391587.KAOT | 0,6 | 0,489897949 | 1,32E-05 | 1,10E-05 | 1 - | 1,86E-05 - | 0 - | 0 - | 0 - | 0 - | 1 - | 2,23E-05 - | 1 - | 2,51E-05 - |
| 6105 | 1 | 0.2um_NOKCPHJN_137431 | Bacteria | Bacteroidetes | Flavobacteriia | Flavobacterial | Flavobacteriac | Formosa | Formosa sp. H | poorly charact | Function unkn | 05ERK | gliding motilit | 391603.FBALC | 3,6 | 1,624807681 | 9,08E-05 | 3,51E-05 | 6 - | 0,000131927 - | 4 - | 0,00010921 - | 3 - | 7,79E-05 - | 4 - | 0,000105351 - | 1 - | 2,97E-05 - |
| 6106 | 1 | 0.2um_NOKCPHJN_27620 | Bacteria | Actinobacteri | Actinobacteri | Micrococcales | Microbacteria | Candidatus Ac | Candidatus Ac | metabolism | Amino acid tra | 00DZF | branched-chai | 312284.A20C1 | 15 | 0 | 0,000485912 | 4,61E-05 | 15 - | 0,00040701 - | 15 - | 0,000505386 - | 15 - | 0,000480433 - | 15 - | 0,000487529 - | 15 - | 0,0005492 - |
| 6107 | 1 | 3um_FIJJAFBF_765129 | Bacteria | Proteobacteri | Betaproteoba | unclassified | unclassified | unclassified | Betaproteoba | cellular proces | Intracellular tr | 175GA | One of the pro | 365046.Rta_3 | 2 | 0 | 0,000183929 | 1,75E-05 | 2 - | 0,000154063 - | 2 - | 0,0001913 - | 2 - | 0,000181855 - | 2 - | 0,000184541 - | 2 - | 0,000207885 - |
| 6108 | 1 | 3um_FIJJAFBF_02611 | Bacteria | Proteobacteri | Alphaproteob | Rhodobactera | Rhodobactera | Ascidiaceihabi | Ascidiaceihabi | metabolism | Amino acid tra | 01RJV | Glutamate syr | 633131.TR2A6 | 0,2 | 0,4 | 6,55E-06 | 1,31E-05 | 0 - | 0 - | 1 - | 3,28E-05 - | 0 - | 0 - | 0 - | 0 - | 0 - | 0 - |
| 6109 | 1 | 0.2um_NOKCPHJN_123315 | Bacteria | Proteobacteri | Gammaprote | Oceanospirilla | Saccharospirill | Reinekea | Reinekea fors | metabolism | Energy produc | 0B7GK | faD-dependen | 391597.LMED | 2 | 0 | 7,37E-05 | 6,99E-06 | 2 - | 6,17E-05 - | 2 - | 7,66E-05 - | 2 - | 7,28E-05 - | 2 - | 7,39E-05 - | 2 - | 8,33E-05 - |
| 6110 | 1 | 0.2um_NOKCPHJN_63759 | Bacteria | Bacteroidetes | Flavobacteriia | Flavobacterial | Crocinitomica | unclassified | Crocinitomica | poorly charact | Function unkn | 1AGHF | SusD family | 485918.Cpin_ | 1,2 | 0,748331477 | 3,13E-05 | 2,05E-05 | 1 - | 2,22E-05 - | 2 - | 5,51E-05 - | 2 - | 5,24E-05 - | 1 - | 2,66E-05 - | 0 - | 0 - |
| 6111 | 1 | 3um_FIJJAFBF_01032 | Bacteria | Bacteroidetes | Flavobacteriia | Flavobacterial | unclassified | unclassified | Flavobacterial | cellular proces | Cell cycle cont | 05CR9 | regulator of ch | 391587.KAOT | 1,2 | 0,4 | 3,26E-05 | 7,23E-06 | 2 - | 4,68E-05 - | 1 - | 2,91E-05 - | 1 - | 2,76E-05 - | 1 - | 2,81E-05 - | 1 - | 3,16E-05 - |
| 6112 | 1 | 3um_FIJJAFBF_481490 | unclassified | unclassified | unclassified | unclassified | unclassified | unclassified | unclassified | unclassified | unclassified | unclassified | unclassified | unclassified | 0,2 | 0,4 | 1,00E-05 | 2,00E-05 | 1 - | 5,01E-05 - | 0 - | 0 - | 0 - | 0 - | 0 - | 0 - | 0 - | 0 - |
| 6113 | 1 | 0.2um_NOKCPHJN_384275 | Bacteria | Proteobacteri | Gammaprote | unclassified | unclassified | unclassified | OM182 bacter | unclassified | unclassified | unclassified | unclassified | unclassified | 1 | 0 | 2,95E-05 | 2,80E-06 | 1 - | 2,47E-05 - | 1 - | 3,06E-05 - | 1 - | 2,91E-05 - | 1 - | 2,96E-05 - | 1 - | 3,33E-05 - |
| 6114 | 1 | 0.2um_NOKCPHJN_170984 | Bacteria | Proteobacteri | Betaproteoba | unclassified | unclassified | unclassified | Betaproteoba | poorly charact | Function unkn | 1,78E+03 | Type VI secret | 381666.H16_ | 0,2 | 0,4 | 1,28E-05 | 2,55E-05 | 1 - | 6,38E-05 - | 0 - | 0 - | 0 - | 0 - | 0 - | 0 - | 0 - | 0 - |
| 6115 | 1 | 3um_FIJJAFBF_95052 | Bacteria | Candidatus Ri | unclassified | unclassified | unclassified | unclassified | Candidatus Ri | cellular proces | Cell motility | 0EQIK | Flagellin | 357809.Cphy_ | 0,8 | 0,4 | 6,13E-05 | 3,09E-05 | 0 - | 0 - | 1 - | 7,66E-05 - | 1 - | 7,28E-05 - | 1 - | 7,39E-05 - | 1 - | 8,33E-05 - |
| 6116 | 1 | 0.2um_NOKCPHJN_254353 | Bacteria | Proteobacteri | Gammaprote | Cellvibrionales | Porticoccacea | unclassified | SAR92 bacteri | metabolism | Energy produc | 05CMV | Succinyl-CoA s | 203122.Sde_2 | 3,2 | 0,748331477 | 0,000111502 | 2,18E-05 | 4 - | 0,000118327 - | 2 - | 7,35E-05 - | 4 - | 0,000139672 - | 3 - | 0,000106302 - | 3 - | 0,000119748 - |
| 6117 | 1 | 0.2um_NOKCPHJN_331418 | Eukaryota | Chordata | Ascidiacea | Enterogona | Cionidae | Ciona | Ciona intestin | unclassified | unclassified | unclassified | unclassified | unclassified | 0,2 | 0,4 | 7,68E-07 | 1,54E-06 | 1 - | 3,84E-06 - | 0 - | 0 - | 0 - | 0 - | 0 - | 0 - | 0 - | 0 - |
| 6118 | 1 | 3um_FIJJAFBF_222383 | Bacteria | Proteobacteri | Alphaproteob | Rhodobactera | Rhodobactera | unclassified | Rhodobactera | unclassified | unclassified | unclassified | unclassified | unclassified | 2,8 | 0,748331477 | 8,29E-05 | 2,35E-05 | 2 - | 4,95E-05 - | 3 - | 9,21E-05 - | 3 - | 8,76E-05 - | 4 - | 0,00011852 - | 2 - | 6,68E-05 - |
| 6119 | 1 | 0.2um_NOKCPHJN_796355 | unclassified | unclassified | unclassified | unclassified | unclassified | unclassified | unclassified | unclassified | unclassified | unclassified | unclassified | unclassified | 0,4 | 0,489897949 | 6,14E-06 | 7,57E-06 | 1 - | 1,40E-05 - | 0 - | 0 - | 0 - | 0 - | 1 - | 1,67E-05 - | 0 - | 0 - |
| 6120 | 1 | 3um_FIJJAFBF_594156 | Eukaryota | unclassified | Raphidophyce | Chattonellales | Chattonellacea | Heterosigma | Heterosigma a | metabolism | Energy produc | COG0055 | Produces ATP | 35128.Thapsd | 7,8 | 0,4 | 0,000628659 | 6,87E-05 | 8 - | 0,000540126 - | 8 - | 0,000670677 - | 8 - | 0,000637563 - | 7 - | 0,000566108 - | 8 - | 0,000728821 - |
| 6121 | 1 | 3um_FIJJAFBF_44705 | Bacteria | Bacteroidetes | Flavobacteriia | Flavobacterial | Flavobacteriac | Polaribacter | Polaribacter s | unclassified | unclassified | unclassified | unclassified | unclassified | 0,4 | 0,489897949 | 8,36E-06 | 1,02E-05 | 0 - | 0 - | 0 - | 0 - | 1 - | 2,07E-05 - | 1 - | 2,11E-05 - | 0 - | 0 - |
| 6122 | 1 | 0.2um_NOKCPHJN_187276 | Bacteria | Bacteroidetes | Flavobacteriia | Flavobacterial | Flavobacteriac | Polaribacter | Polaribacter h | unclassified | unclassified | unclassified | unclassified | unclassified | 1,2 | 1,6 | 1,80E-05 | 2,51E-05 | 2 - | 2,58E-05 - | 4 - | 6,41E-05 - | 0 - | 0 - | 0 - | 0 - | 0 - | 0 - |
| 6123 | 1 | 3um_FIJJAFBF_38607 | Bacteria | Proteobacteri | Gammaprote | Chromatiales | unclassified | unclassified | Chromatiales | metabolism | Secondary me | 027AN | Ring hydroxyla | 488538.SAR11 | 0,8 | 0,4 | 6,64E-05 | 3,41E-05 | 1 - | 6,96E-05 - | 1 - | 8,64E-05 - | 1 - | 8,21E-05 - | 0 - | 0 - | 1 - | 9,39E-05 - |
| 6124 | 1 | 3um_FIJJAFBF_119807 | unclassified | unclassified | unclassified | unclassified | unclassified | unclassified | unclassified | unclassified | unclassified | unclassified | unclassified | unclassified | 2,2 | 0,4 | 0,000326384 | 2,99E-05 | 3 - | 0,000378385 - | 2 - | 0,000313228 - | 2 - | 0,000297763 - | 2 - | 0,000302161 - | 2 - | 0,000340383 - |
| 6125 | 1 | 3um_FIJJAFBF_232387 | Eukaryota | unclassified | Dinophyceae | Suessiales | Symbiodiniace | Symbiodinium | Symbiodinium | metabolism | Amino acid tra | COG0591 | symporter | 35128.Thaps2 | 1,2 | 0,4 | 3,59E-05 | 1,24E-05 | 1 - | 2,51E-05 - | 1 - | 3,11E-05 - | 1 - | 2,96E-05 - | 2 - | 6,00E-05 - | 1 - | 3,38E-05 - |
| 6126 | 1 | 0.2um_NOKCPHJN_28388 | Bacteria | Bacteroidetes | Flavobacteriia | Flavobacterial | Flavobacteriac | unclassified | Flavobacteriac | poorly charact | Function unkn | 0XR4Y | Ragb susd dor | 487796.Flav2 | 2,6 | 0,8 | 6,17E-05 | 2,01E-05 | 3 - | 5,97E-05 - | 3 - | 7,41E-05 - | 3 - | 7,04E-05 - | 1 - | 2,38E-05 - | 3 - | 8,05E-05 - |
| 6127 | 1 | 3um_FIJJAFBF_694562 | Bacteria | Proteobacteri | Alphaproteob | Rhodobactera | Rhodobactera | Planktomarina | Planktomarina | metabolism | Energy produc | 16W2S | Produces ATP | 314270.RB208 | 0,8 | 0,4 | 7,86E-05 | 3,99E-05 | 1 - | 8,50E-05 - | 1 - | 0,00010557 - | 1 - | 0,000100357 - | 1 - | 0,000101839 - | 0 - | 0 - |
| 6128 | 1 | 0.2um_NOKCPHJN_180550 | Bacteria | Bacteroidetes | Flavobacteriia | Flavobacterial | Flavobacteriac | Polaribacter | Polaribacter s | cellular proces | Cell motility | COG1360 | Ompa motb d | 487797.Flav3 | 10 | 1,264911064 | 0,000490916 | 5,08E-05 | 11 - | 0,000455792 - | 8 - | 0,000411607 - | 11 - | 0,000538016 - | 11 - | 0,000545963 - | 9 - | 0,000503202 - |
| 6129 | 1 | 3um_FIJJAFBF_189661 | Bacteria | Bacteroidetes | Flavobacteriia | Flavobacterial | Flavobacteriac | Polaribacter | Polaribacter s | unclassified | unclassified | unclassified | unclassified | unclassified | 0,8 | 0,979795897 | 5,16E-05 | 6,36E-05 | 2 - | 0,000118327 - | 0 - | 0 - | 2 - | 0,000139672 - | 0 - | 0 - | 0 - | 0 - |
| 6130 | 1 | 3um_FIJJAFBF_733763 | Bacteria | Cyanobacteria | unclassified | Synechococca | Synechococca | Synechococcu | Synechococcu | unclassified | unclassified | unclassified | unclassified | unclassified | 0,4 | 0,489897949 | 4,50E-05 | 5,53E-05 | 0 - | 0 - | 0 - | 0 - | 1 - | 0,000105025 - | 0 - | 0 - | 1 - | 0,000120058 - |
| 6131 | 1 | 3um_FIJJAFBF_796763 | Bacteria | Bacteroidetes | Cytophagia | Cytophagales | Hymenobacte | Rufibacter | Rufibacter sp. | poorly charact | Function unkn | 0QK6M | Endonuclease | 314283.MED2 | 1 | 0 | 2,42E-05 | 2,29E-06 | 1 - | 2,02E-05 - | 1 - | 2,51E-05 - | 1 - | 2,39E-05 - | 1 - | 2,42E-05 - | 1 - | 2,73E-05 - |
| 6132 | 1 | 3um_FIJJAFBF_326277 | Bacteria | Proteobacteri | Alphaproteob | Rhodobactera | Rhodobactera | unclassified | Rhodobactera | cellular proces | Posttranslatio | 01SC2 | alkyl hydroper | 391619.RGBS | 0,6 | 0,489897949 | 3,64E-05 | 3,00E-05 | 1 - | 5,31E-05 - | 1 - | 6,60E-05 - | 1 - | 6,27E-05 - | 0 - | 0 - | 0 - | 0 - |
| 6133 | 1 | 3um_FIJJAFBF_22494 | Bacteria | Proteobacteri | Gammaprote | Alteromonada | Alteromonada | Paraglaciecola | Paraglaciecola | cellular proces | Cell wall/mem | 177PY | Molecular cha | 342610.Patl_1 | 0,8 | 0,748331477 | 6,73E-05 | 6,26E-05 | 0 - | 0 - | 2 - | 0,000166689 - | 1 - | 7,92E-05 - | 0 - | 0 - | 1 - | 9,06E-05 - |
| 6134 | 1 | 3um_FIJJAFBF_112505 | Bacteria | Proteobacteri | Deltaproteoba | unclassified | unclassified | unclassified | various | poorly charact | Function unkn | 0HFRS | MotA TolQ Exl | 391625.PPSIR | 0,4 | 0,489897949 | 2,29E-05 | 2,82E-05 | 1 - | 5,24E-05 - | 0 - | 0 - | 1 - | 6,19E-05 - | 0 - | 0 - | 0 - | 0 - |
| 6135 | 1 | 0.2um_NOKCPHJN_83083 | Viruses | unclassified | unclassified | unclassified | various | various | various | unclassified | unclassified | unclassified | unclassified | unclassified | 1 | 0 | 7,23E-06 | 6,87E-07 | 1 - | 6,06E-06 - | 1 - | 7,52E-06 - | 1 - | 7,15E-06 - | 1 - | 7,26E-06 - | 1 - | 8,17E-06 - |
| 6136 | 1 | 3um_FIJJAFBF_43757 | Bacteria | Proteobacteri | Gammaprote | Oceanospirilla | Alcanivoracac | Marinicella | Marinicella lit | cellular proces | Posttranslatio | COG0330 | Band 7 protei | 357804.Ping_ | 4 | 1,095445115 | 0,000167359 | 2,96E-05 | 6 - | 0,000215206 - | 4 - | 0,000178149 - | 4 - | 0,000169353 - | 3 - | 0,000128891 - | 3 - | 0,000145195 - |
| 6137 | 1 | 3um_FIJJAFBF_566924 | Bacteria | Proteobacteri | Gammaprote | unclassified | unclassified | unclassified | various | information st | Translation, ri | 16QAP | Associates wit | 349521.HCH_ | 1,2 | 0,748331477 | 5,46E-05 | 3,52E-05 | 1 - | 3,90E-05 - | 1 - | 4,85E-05 - | 2 - | 9,22E-05 - | 2 - | 9,35E-05 - | 0 - | 0 - |
| 6138 | 1 | 0.2um_NOKCPHJN_06965 | Bacteria | Bacteroidetes | Cytophagia | Cytophagales | Flammeovirga | unclassified | Flammeovirga | unclassified | unclassified | unclassified | unclassified | unclassified | 4,6 | 1,019803903 | 1,58E-05 | 2,78E-06 | 6 - | 1,75E-05 - | 5 - | 1,81E-05 - | 5 - | 1,72E-05 - | 3 - | 1,05E-05 - | 4 - | 1,58E-05 - |
| 6139 | 1 | 3um_FIJJAFBF_339520 | unclassified | unclassified | unclassified | unclassified | unclassified | unclassified | unclassified | unclassified | unclassified | unclassified | unclassified | unclassified | 0,2 | 0,4 | 9,29E-06 | 1,86E-05 | 0 - | 0 - | 0 - | 0 - | 0 - | 0 - | 1 - | 4,64E-05 - | 0 - | 0 - |
| 6140 | 1 | 3um_FIJJAFBF_352965 | Bacteria | Proteobacteri | Gammaprote | Alteromonada | Alteromonada | Glaciecola | Glaciecola sp. | metabolism | Inorganic ion t | 0QJVD | TonB-depende | 455436.GHTC | 33,8 | 2,13541565 | 0,000527511 | 3,42E-05 | 37 - | 0,000485897 - | 34 - | 0,000554421 - | 35 - | 0,000542549 - | 31 - | 0,000487641 - | 32 - | 0,000567046 - |
| 6141 | 1 | 0.2um_NOKCPHJN_272386 | Bacteria | Bacteroidetes | Flavobacteriia | Flavobacterial | Crocinitomica | unclassified | Crocinitomica | unclassified | unclassified | unclassified | unclassified | unclassified | 0,2 | 0,4 | 9,18E-06 | 1,84E-05 | 1 - | 4,59E-05 - | 0 - | 0 - | 0 - | 0 - | 0 - | 0 - | 0 - | 0 - |
| 6142 | 1 | 0.2um_NOKCPHJN_109750 | Bacteria | Bacteroidetes | Flavobacteriia | Flavobacterial | Flavobacteriac | Formosa | Formosa sp. H | information st | Translation, ri | 09FEM | Binds to the 2 | 983544.Lacal_ | 2,2 | 0,748331477 | 0,000198945 | 6,40E-05 | 3 - | 0,000229554 - | 3 - | 0,000285038 - | 1 - | 9,03E-05 - | 2 - | 0,000183311 - | 2 - | 0,000206499 - |
| 6143 | 1 | 3um_FIJJAFBF_675681 | Viruses | unclassified | unclassified | unclassified | unclassified | unclassified | uncultured M | unclassified | unclassified | unclassified | unclassified | unclassified | 4 | 0,632455532 | 0,000204676 | 4,51E-05 | 4 - | 0,000170672 - | 3 - | 0,000158943 - | 4 - | 0,000201461 - | 4 - | 0,000204436 - | 5 - | 0,000287871 - |

| | | | | | | | | | | | | | | | | | | | | | | | | | | | | |
|---|---|---|---|---|---|---|---|---|---|---|---|---|---|---|---|---|---|---|---|---|---|---|---|---|---|---|---|---|
| 6144 | 1 | 0.2um_NOKCPHJN_530417 | Bacteria | Proteobacteri | Gammaprote | unclassified | unclassified | unclassified | Gammaprote | unclassified | unclassified | unclassified | unclassified | unclassified | 2 | 0 | 9,89E-05 | 9,39E-06 | 2 - | 8,29E-05 - | 2 - | 0,000102902 - | 2 - | 9,78E-05 - | 2 - | 9,93E-05 - | 2 - | 0,000111823 - |
| 6145 | 1 | 3um_FIJJAFBF_56982 | unclassified | unclassified | unclassified | unclassified | unclassified | unclassified | unclassified | unclassified | unclassified | unclassified | unclassified | unclassified | 2 | 1,264911064 | 0,000533412 | 0,000344628 | 2 - | 0,000450105 - | 0 - | 0 - | 2 - | 0,000531303 - | 4 - | 0,0010783 - | 2 - | 0,000607351 - |
| 6146 | 1 | 3um_FIJJAFBF_744518 | Viruses | unclassified | unclassified | unclassified | unclassified | unclassified | uncultured vi | unclassified | unclassified | unclassified | unclassified | unclassified | 2 | 0 | 9,10E-05 | 8,64E-06 | 2 - | 7,63E-05 - | 2 - | 9,47E-05 - | 2 - | 9,00E-05 - | 2 - | 9,14E-05 - | 2 - | 0,000102907 - |
| 6147 | 1 | 3um_FIJJAFBF_75356 | Bacteria | Bacteroidetes | Flavobacteriia | Flavobacterial | Flavobacteria | Formosa | Formosa sp. H | unclassified | unclassified | unclassified | unclassified | unclassified | 1 | 0 | 4,11E-05 | 3,91E-06 | 1 - | 3,45E-05 - | 1 - | 4,28E-05 - | 1 - | 4,07E-05 - | 1 - | 4,13E-05 - | 1 - | 4,65E-05 - |
| 6148 | 1 | 0.2um_NOKCPHJN_432655 | Bacteria | Proteobacteri | Alphaproteob | Pelagibacteral | Pelagibactera | Candidatus Pe | Candidatus Pe | cellular proces | Cell wall/mem | 7257 | NA | 335992.SAR11 | 1 | 0 | 4,16E-05 | 3,95E-06 | 1 - | 3,49E-05 - | 1 - | 4,33E-05 - | 1 - | 4,12E-05 - | 1 - | 4,18E-05 - | 1 - | 4,71E-05 - |
| 6149 | 1 | 0.2um_NOKCPHJN_307737 | Bacteria | Bacteroidetes | Flavobacteriia | Flavobacterial | Flavobacteria | Formosa | Formosa sp. H | unclassified | unclassified | unclassified | unclassified | unclassified | 1,4 | 0,489897949 | 2,59E-05 | 9,71E-06 | 1 - | 1,54E-05 - | 2 - | 3,83E-05 - | 1 - | 1,82E-05 - | 2 - | 3,70E-05 - | 1 - | 2,08E-05 - |
| 6150 | 1 | 0.2um_NOKCPHJN_374224 | Bacteria | Actinobacteri | Actinobacteri | Streptomycet | Streptomycet | Streptomyces | Streptomyces | unclassified | unclassified | unclassified | unclassified | unclassified | 1,2 | 1,166190379 | 3,42E-05 | 3,48E-05 | 1 - | 2,41E-05 - | 3 - | 8,98E-05 - | 2 - | 5,69E-05 - | 0 - | 0 - | 0 - | 0 - |
| 6151 | 1 | 3um_FIJJAFBF_757484 | Bacteria | Proteobacteri | Gammaprote | Oceanospirilla | Saccharospirill | Reinekea | Reinekea blan | unclassified | unclassified | unclassified | unclassified | unclassified | 1 | 0 | 4,13E-05 | 3,92E-06 | 1 - | 3,46E-05 - | 1 - | 4,29E-05 - | 1 - | 4,08E-05 - | 1 - | 4,14E-05 - | 1 - | 4,66E-05 - |
| 6152 | 1 | 0.2um_NOKCPHJN_295626 | unclassified | unclassified | unclassified | unclassified | unclassified | unclassified | unclassified | cellular proces | Intracellular tr | KOG2514 | Anoctamin | 67593.Physo1 | 0,4 | 0,489897949 | 6,02E-06 | 7,41E-06 | 1 - | 1,38E-05 - | 0 - | 0 - | 1 - | 1,63E-05 - | 0 - | 0 - | 0 - | 0 - |
| 6153 | 1 | 3um_FIJJAFBF_374544 | Bacteria | Proteobacteri | Alphaproteob | Rhodobactera | Rhodobactera | Sulfitobacter | Sulfitobacter r | metabolism | Secondary me | 01Q86 | extracellular s | 367336.OM22 | 4,8 | 1,166190379 | 0,000179366 | 4,13E-05 | 6 - | 0,000189192 - | 4 - | 0,000156614 - | 3 - | 0,000111661 - | 6 - | 0,000226621 - | 5 - | 0,00021274 - |
| 6154 | 1 | 3um_FIJJAFBF_337746 | unclassified | unclassified | unclassified | unclassified | unclassified | unclassified | unclassified | unclassified | unclassified | unclassified | unclassified | unclassified | 2,2 | 0,4 | 6,92E-05 | 1,35E-05 | 2 - | 5,28E-05 - | 2 - | 6,55E-05 - | 3 - | 9,34E-05 - | 2 - | 6,32E-05 - | 2 - | 7,12E-05 - |
| 6155 | 1 | 3um_FIJJAFBF_304135 | Bacteria | Bacteroidetes | Cytophagia | Cytophagales | Flammeovirga | unclassified | Flammeovirga | unclassified | unclassified | unclassified | unclassified | unclassified | 0,2 | 0,4 | 6,15E-06 | 1,23E-05 | 1 - | 3,08E-05 - | 0 - | 0 - | 0 - | 0 - | 0 - | 0 - | 0 - | 0 - |
| 6156 | 1 | 0.2um_NOKCPHJN_321034 | unclassified | unclassified | unclassified | unclassified | unclassified | unclassified | unclassified | unclassified | unclassified | unclassified | unclassified | unclassified | 0,4 | 0,489897949 | 2,58E-05 | 3,15E-05 | 0 - | 0 - | 0 - | 0 - | 1 - | 6,39E-05 - | 1 - | 6,49E-05 - | 0 - | 0 - |
| 6157 | 1 | 3um_FIJJAFBF_20305 | Bacteria | Bacteroidetes | Flavobacteriia | Flavobacterial | Flavobacteria | Polaribacter | Polaribacter s | unclassified | unclassified | unclassified | unclassified | unclassified | 0,2 | 0,4 | 1,58E-05 | 3,17E-05 | 0 - | 0 - | 0 - | 0 - | 1 - | 7,92E-05 - | 0 - | 0 - | 0 - | 0 - |
| 6158 | 1 | 0.2um_NOKCPHJN_414609 | Bacteria | Proteobacteri | Alphaproteob | Caulobacteral | Caulobacterac | Phenylobacte | Phenylobacte | cellular proces | Posttranslatio | 16QU3 | Hemolysin-typ | 744980.TRICH | 3,4 | 0,489897949 | 7,38E-05 | 7,17E-06 | 4 - | 7,35E-05 - | 3 - | 6,84E-05 - | 4 - | 8,67E-05 - | 3 - | 6,60E-05 - | 3 - | 7,43E-05 - |
| 6159 | 1 | 0.2um_NOKCPHJN_683847 | Eukaryota | unclassified | unclassified | Acytosteliales | Acytosteliacea | Acytostelium | Acytostelium | unclassified | unclassified | unclassified | unclassified | unclassified | 1,4 | 1,019803903 | 1,91E-05 | 1,38E-05 | 0 - | 0 - | 3 - | 4,10E-05 - | 1 - | 1,30E-05 - | 2 - | 2,64E-05 - | 1 - | 1,49E-05 - |
| 6160 | 1 | 3um_FIJJAFBF_100736 | unclassified | unclassified | unclassified | unclassified | unclassified | unclassified | unclassified | unclassified | unclassified | unclassified | unclassified | unclassified | 0,4 | 0,8 | 1,53E-05 | 3,05E-05 | 2 - | 7,63E-05 - | 0 - | 0 - | 0 - | 0 - | 0 - | 0 - | 0 - | 0 - |
| 6161 | 1 | 3um_FIJJAFBF_399525 | Bacteria | Proteobacteri | Gammaprote | Alteromonada | Alteromonada | Paraglaciecola | Paraglaciecola | unclassified | unclassified | unclassified | unclassified | unclassified | 0,2 | 0,4 | 6,06E-06 | 1,21E-05 | 1 - | 3,03E-05 - | 0 - | 0 - | 0 - | 0 - | 0 - | 0 - | 0 - | 0 - |
| 6162 | 1 | 3um_FIJJAFBF_357544 | Bacteria | Bacteroidetes | Flavobacteriia | Flavobacterial | Flavobacteria | Polaribacter | Polaribacter s | unclassified | unclassified | unclassified | unclassified | unclassified | 3,4 | 0,8 | 8,34E-05 | 2,12E-05 | 4 - | 8,23E-05 - | 3 - | 7,66E-05 - | 2 - | 4,86E-05 - | 4 - | 9,86E-05 - | 4 - | 0,000111021 - |
| 6163 | 1 | 3um_FIJJAFBF_401050 | Viruses | unclassified | unclassified | Caudovirales | Siphoviridae | unclassified | Freshwater ph | unclassified | unclassified | unclassified | unclassified | unclassified | 0,2 | 0,4 | 5,51E-06 | 1,10E-05 | 0 - | 0 - | 1 - | 2,76E-05 - | 0 - | 0 - | 0 - | 0 - | 0 - | 0 - |
| 6164 | 1 | 0.2um_NOKCPHJN_171704 | Bacteria | Bacteroidetes | Flavobacteriia | Flavobacterial | Flavobacteria | Formosa | Formosa sp. H | metabolism | Inorganic ion t | 09B4C | tonB-depende | 391603.FBALC | 2 | 0,894427191 | 3,68E-05 | 1,46E-05 | 3 - | 4,78E-05 - | 1 - | 1,98E-05 - | 2 - | 3,76E-05 - | 3 - | 5,73E-05 - | 1 - | 2,15E-05 - |
| 6165 | 1 | 0.2um_NOKCPHJN_783159 | Bacteria | Proteobacteri | Gammaprote | unclassified | unclassified | unclassified | OM182 bacter | metabolism | Energy produc | 17C3H | Produces ATP | 379731.PST_4 | 1,8 | 0,4 | 0,000179234 | 4,19E-05 | 2 - | 0,000167557 - | 1 - | 0,000104028 - | 2 - | 0,000197784 - | 2 - | 0,000200706 - | 2 - | 0,000226094 - |
| 6166 | 1 | 0.2um_NOKCPHJN_695488 | Bacteria | Proteobacteri | Alphaproteob | Pelagibacteral | Pelagibactera | Candidatus Pe | Candidatus Pe | metabolism | Amino acid tra | 16RER | (ABC) transpo | 335992.SAR11 | 4,6 | 1,019803903 | 0,000221202 | 4,64E-05 | 6 - | 0,000243343 - | 5 - | 0,0002518 - | 4 - | 0,000191494 - | 3 - | 0,000145742 - | 5 - | 0,00027363 - |
| 6167 | 1 | 0.2um_NOKCPHJN_398692 | Bacteria | Bacteroidetes | Flavobacteriia | Flavobacterial | Flavobacteria | Polaribacter | Polaribacter s | unclassified | unclassified | unclassified | unclassified | unclassified | 1 | 0,894427191 | 2,12E-05 | 1,97E-05 | 1 - | 1,82E-05 - | 2 - | 4,51E-05 - | 2 - | 4,29E-05 - | 0 - | 0 - | 0 - | 0 - |
| 6168 | 1 | 0.2um_NOKCPHJN_125977 | Bacteria | Proteobacteri | Gammaprote | Alteromonada | Alteromonada | Catenovulum | Catenovulum | poorly charact | Function unkn | 0P8NE | NA | 867900.Celly_ | 0,4 | 0,489897949 | 4,49E-06 | 5,55E-06 | 1 - | 1,00E-05 - | 1 - | 1,24E-05 - | 0 - | 0 - | 0 - | 0 - | 0 - | 0 - |
| 6169 | 1 | 0.2um_NOKCPHJN_05622 | Bacteria | Bacteroidetes | Cytophagia | Cytophagales | Flammeovirga | unclassified | Flammeovirga | metabolism | Inorganic ion t | 0FW52 | TonB-depende | 880070.Cycma | 0,4 | 0,489897949 | 5,12E-06 | 6,32E-06 | 1 - | 1,17E-05 - | 0 - | 0 - | 0 - | 0 - | 1 - | 1,40E-05 - | 0 - | 0 - |
| 6170 | 5 | 0.2um_NOKCPHJN_449820;0.2um_NOKCPHJN_9 | Bacteria | Proteobacteri | Alphaproteob | various | various | various | various | various | various | various | various | various | 2 | 0,632455532 | 0,000219116 | 5,74E-05 | 3 - | 0,000279943 - | 2 - | 0,000231738 - | 2 - | 0,000220296 - | 1 - | 0,000111775 - | 2 - | 0,000251828 - |
| 6171 | 1 | 3um_FIJJAFBF_180756 | Bacteria | Bacteroidetes | Flavobacteriia | unclassified | unclassified | unclassified | Flavobacteria | unclassified | unclassified | unclassified | unclassified | unclassified | 2,4 | 0,8 | 0,000213003 | 6,55E-05 | 3 - | 0,000228033 - | 3 - | 0,00028315 - | 3 - | 0,00026917 - | 2 - | 0,000182097 - | 1 - | 0,000102566 - |
| 6172 | 1 | 0.2um_NOKCPHJN_122528 | Bacteria | Proteobacteri | Gammaprote | Oceanospirilla | unclassified | unclassified | Oceanospirilla | metabolism | Inorganic ion t | 05PWN | phosphate ab | 488538.SAR11 | 0,4 | 0,489897949 | 1,56E-05 | 1,95E-05 | 1 - | 3,33E-05 - | 0 - | 0 - | 0 - | 0 - | 0 - | 0 - | 1 - | 4,49E-05 - |
| 6173 | 1 | 3um_FIJJAFBF_656407 | Bacteria | Proteobacteri | Gammaprote | Alteromonada | Alteromonada | Alteromonas | Alteromonas s | metabolism | Energy produc | 16PVJ | Component of | 715451.ambt_ | 0,8 | 0,4 | 1,19E-05 | 6,03E-06 | 1 - | 1,28E-05 - | 1 - | 1,59E-05 - | 1 - | 1,52E-05 - | 1 - | 1,54E-05 - | 0 - | 0 - |
| 6174 | 1 | 3um_FIJJAFBF_337948 | Bacteria | Bacteroidetes | Flavobacteriia | Flavobacterial | Flavobacteria | Tenacibacul | Tenacibacul | unclassified | unclassified | unclassified | unclassified | unclassified | 0,2 | 0,4 | 6,55E-06 | 1,31E-05 | 0 - | 0 - | 0 - | 0 - | 1 - | 3,27E-05 - | 0 - | 0 - | 0 - | 0 - |
| 6175 | 1 | 0.2um_NOKCPHJN_276340 | Bacteria | Bacteroidetes | Flavobacteriia | Flavobacterial | Flavobacteria | Altibacter | Altibacter lent | metabolism | Lipid transpor | COG2067 | lipid transport | 50743.SCB49_ | 4,6 | 0,489897949 | 0,000148499 | 1,45E-05 | 5 - | 0,000135991 - | 5 - | 0,000168861 - | 5 - | 0,000160524 - | 4 - | 0,000130316 - | 4 - | 0,0001468 - |
| 6176 | 1 | 0.2um_NOKCPHJN_244384 | Eukaryota | Streptophyta | Liliopsida | Poales | Poaceae | Aegilops | Aegilops taus | unclassified | unclassified | unclassified | unclassified | unclassified | 1 | 0 | 5,32E-06 | 5,05E-07 | 1 - | 4,46E-06 - | 1 - | 5,53E-06 - | 1 - | 5,26E-06 - | 1 - | 5,34E-06 - | 1 - | 6,01E-06 - |
| 6177 | 1 | 3um_FIJJAFBF_708514 | Bacteria | Cyanobacteria | unclassified | Synechococca | Synechococca | Synechococcu | Synechococcu | poorly charact | Function unkn | 06QDP | C-phycoerythr | 221359.RS991 | 3,6 | 0,489897949 | 0,0002991 | 5,22E-05 | 4 - | 0,000278247 - | 4 - | 0,0003455 - | 3 - | 0,000246331 - | 3 - | 0,00024997 - | 4 - | 0,000375453 - |
| 6178 | 1 | 0.2um_NOKCPHJN_11467 | Bacteria | Proteobacteri | Gammaprote | Cellvibrionales | Porticoccacea | unclassified | SAR92 bacteri | cellular proces | Intracellular tr | 0QJ7H | motA TolQ Exl | 314287.GB220 | 6,2 | 1,326649916 | 0,000185564 | 3,85E-05 | 7 - | 0,00017658 - | 6 - | 0,000187937 - | 8 - | 0,000238211 - | 4 - | 0,000120864 - | 6 - | 0,00020423 - |
| 6179 | 1 | 0.2um_NOKCPHJN_728287 | Bacteria | Bacteroidetes | Cytophagia | Cytophagales | Flammeovirga | Reichenbachie | Reichenbachie | metabolism | Carbohydrate | 064TE | hydrolase fam | 391603.FBALC | 0,2 | 0,4 | 5,81E-06 | 1,16E-05 | 0 - | 0 - | 0 - | 0 - | 1 - | 2,91E-05 - | 0 - | 0 - | 0 - | 0 - |
| 6180 | 1 | 3um_FIJJAFBF_85327 | Bacteria | Proteobacteri | Gammaprote | Alteromonada | Alteromonada | Paraglaciecola | Paraglaciecola | metabolism | Energy produc | 16RH6 | Produces ATP | 342610.Patl_4 | 0,6 | 0,489897949 | 2,85E-05 | 2,37E-05 | 1 - | 4,01E-05 - | 0 - | 0 - | 0 - | 0 - | 1 - | 4,81E-05 - | 1 - | 5,42E-05 - |
| 6181 | 1 | 0.2um_NOKCPHJN_315400 | Bacteria | Bacteroidetes | Flavobacteriia | Flavobacterial | Flavobacteria | Lutibacter | Lutibacter pro | metabolism | Inorganic ion t | 05C27 | TonB-depende | 487797.Flav30 | 3,2 | 0,979795897 | 4,07E-05 | 1,28E-05 | 4 - | 4,28E-05 - | 2 - | 2,66E-05 - | 4 - | 5,06E-05 - | 2 - | 2,56E-05 - | 4 - | 5,78E-05 - |
| 6182 | 11 | 0.2um_NOKCPHJN_1011510;0.2um_NOKCPHJN_ | Bacteria | Proteobacteri | Gammaprote | various | various | various | various | unclassified | unclassified | unclassified | unclassified | unclassified | 0,4 | 0,489897949 | 2,65E-05 | 3,26E-05 | 1 - | 6,07E-05 - | 0 - | 0 - | 1 - | 7,17E-05 - | 0 - | 0 - | 0 - | 0 - |
| 6183 | 1 | 0.2um_NOKCPHJN_06743 | Bacteria | Proteobacteri | Gammaprote | Alteromonada | Alteromonada | Paraglaciecola | Paraglaciecola | unclassified | unclassified | unclassified | unclassified | unclassified | 0,2 | 0,4 | 2,65E-06 | 5,31E-06 | 0 - | 0 - | 1 - | 1,33E-05 - | 0 - | 0 - | 0 - | 0 - | 0 - | 0 - |
| 6184 | 1 | 3um_FIJJAFBF_478909 | Bacteria | Bacteroidetes | Flavobacteriia | Flavobacterial | Flavobacteria | Polaribacter | Polaribacter s | unclassified | unclassified | unclassified | unclassified | unclassified | 0,4 | 0,8 | 2,44E-05 | 4,88E-05 | 2 - | 0,000122103 - | 0 - | 0 - | 0 - | 0 - | 0 - | 0 - | 0 - | 0 - |
| 6185 | 1 | 0.2um_NOKCPHJN_129925 | Bacteria | Proteobacteri | Gammaprote | unclassified | unclassified | unclassified | Gammaprote | unclassified | unclassified | unclassified | unclassified | unclassified | 9 | 1,673320053 | 0,000222582 | 4,11E-05 | 11 - | 0,000229137 - | 7 - | 0,000181058 - | 7 - | 0,000172119 - | 10 - | 0,000249516 - | 10 - | 0,000281079 - |
| 6186 | 1 | 3um_FIJJAFBF_214637 | Bacteria | Bacteroidetes | unclassified | unclassified | unclassified | unclassified | Bacteroidetes | poorly charact | Function unkn | 09GJP | Protein of unk | 755732.Fluta_ | 1,2 | 0,4 | 0,000112606 | 4,16E-05 | 1 - | 7,81E-05 - | 2 - | 0,000193903 - | 1 - | 9,22E-05 - | 1 - | 9,35E-05 - | 1 - | 0,000105357 - |
| 6187 | 1 | 3um_FIJJAFBF_49005 | Bacteria | Proteobacteri | Gammaprote | Oceanospirilla | Saccharospirill | Reinekea | Reinekea fors | unclassified | unclassified | unclassified | unclassified | unclassified | 3,6 | 0,489897949 | 0,000117742 | 1,17E-05 | 4 - | 0,000110628 - | 3 - | 0,000103026 - | 4 - | 0,000130585 - | 4 - | 0,000132514 - | 3 - | 0,000111957 - |
| 6188 | 1 | 0.2um_NOKCPHJN_1008844 | unclassified | unclassified | unclassified | unclassified | unclassified | unclassified | unclassified | unclassified | unclassified | unclassified | unclassified | unclassified | 0,8 | 0,4 | 2,19E-05 | 1,10E-05 | 0 - | 0 - | 1 - | 2,74E-05 - | 1 - | 2,61E-05 - | 1 - | 2,64E-05 - | 1 - | 2,98E-05 - |
| 6189 | 1 | 3um_FIJJAFBF_21329 | Bacteria | Bacteroidetes | Flavobacteriia | Flavobacterial | Flavobacteria | Polaribacter | Polaribacter s | cellular proces | Cell wall/mem | 09GJR | outer membra | 313598.MED1 | 1,8 | 0,748331477 | 0,000143363 | 6,00E-05 | 2 - | 0,000134242 - | 1 - | 8,33E-05 - | 3 - | 0,000237688 - | 1 - | 8,04E-05 - | 2 - | 0,00018114 - |
| 6190 | 1 | 0.2um_NOKCPHJN_172977 | Bacteria | Proteobacteri | Gammaprote | Oceanospirilla | Saccharospirill | Reinekea | Reinekea fors | cellular proces | Posttranslatio | 07RUK | Involved in iro | 314283.MED2 | 2,2 | 1,6 | 0,000146902 | 9,28E-05 | 5 - | 0,000292798 - | 2 - | 0,000145427 - | 2 - | 0,000138247 - | 0 - | 0 - | 2 - | 0,000158035 - |
| 6191 | 1 | 3um_FIJJAFBF_106480 | Bacteria | Bacteroidetes | Flavobacteriia | Flavobacterial | Flavobacteria | Algibacter | Algibacter sp. | information st | Translation, ri | 08UIK | This protein bi | 983544.Lacal_ | 2 | 0,632455532 | 0,000186328 | 5,67E-05 | 2 - | 0,000158313 - | 2 - | 0,000196578 - | 3 - | 0,000280308 - | 2 - | 0,000189632 - | 1 - | 0,00010681 - |
| 6192 | 1 | 0.2um_NOKCPHJN_165121 | Bacteria | Proteobacteri | Alphaproteob | Rhodobactera | Rhodobactera | Ahrensia | Ahrensia kiele | unclassified | unclassified | unclassified | unclassified | unclassified | 1,2 | 0,748331477 | 3,37E-05 | 2,01E-05 | 2 - | 4,96E-05 - | 1 - | 3,08E-05 - | 2 - | 5,85E-05 - | 1 - | 2,97E-05 - | 0 - | 0 - |
| 6193 | 2 | 0.2um_NOKCPHJN_822213;3um_FIJJAFBF_64562 | Bacteria | Bacteroidetes | Flavobacteriia | Flavobacterial | Flavobacteria | Eudoraea | Eudoraea adri | unclassified | unclassified | unclassified | unclassified | unclassified | 0,2 | 0,4 | 4,42E-06 | 8,85E-06 | 1 - | 2,21E-05 - | 0 - | 0 - | 0 - | 0 - | 0 - | 0 - | 0 - | 0 - |
| 6194 | 1 | 0.2um_NOKCPHJN_678942 | Eukaryota | unclassified | Labyrinthulom | unclassified | Thraustochytr | Aurantiochytr | Aurantiochytr | unclassified | unclassified | unclassified | unclassified | unclassified | 1,2 | 1,166190379 | 2,00E-05 | 2,02E-05 | 2 - | 2,89E-05 - | 3 - | 5,38E-05 - | 1 - | 1,71E-05 - | 0 - | 0 - | 0 - | 0 - |
| 6195 | 1 | 3um_FIJJAFBF_176649 | Bacteria | Bacteroidetes | unclassified | unclassified | unclassified | unclassified | Bacteroidetes | unclassified | unclassified | unclassified | unclassified | unclassified | 1,2 | 0,4 | 0,000127986 | 2,83E-05 | 2 - | 0,000183643 - | 1 - | 0,000114015 - | 1 - | 0,000108386 - | 1 - | 0,000109987 - | 1 - | 0,0001239 - |
| 6196 | 1 | 3um_FIJJAFBF_407524 | unclassified | unclassified | unclassified | unclassified | unclassified | unclassified | unclassified | unclassified | unclassified | unclassified | unclassified | unclassified | 2 | 0 | 2,05E-05 | 1,95E-06 | 2 - | 1,72E-05 - | 2 - | 2,14E-05 - | 2 - | 2,03E-05 - | 2 - | 2,06E-05 - | 2 - | 2,32E-05 - |
| 6197 | 1 | 3um_FIJJAFBF_479083 | Bacteria | Proteobacteri | Alphaproteob | Rhodospirillal | Rhodospirillac | unclassified | Rhodospirillac | cellular proces | Posttranslatio | 01QWG | Prevents misf | 331869.BAL19 | 3,4 | 1,2 | 0,000129175 | 4,22E-05 | 5 - | 0,000161203 - | 2 - | 8,01E-05 - | 2 - | 7,61E-05 - | 4 - | 0,000154476 - | 4 - | 0,000174016 - |
| 6198 | 1 | 0.2um_NOKCPHJN_143408 | Bacteria | Proteobacteri | Gammaprote | unclassified | unclassified | unclassified | SAR86 cluster | unclassified | unclassified | unclassified | unclassified | unclassified | 0,6 | 0,8 | 1,02E-05 | 1,31E-05 | 2 - | 3,18E-05 - | 0 - | 0 - | 0 - | 0 - | 1 - | 1,91E-05 - | 0 - | 0 - |
| 6199 | 1 | 3um_FIJJAFBF_518970 | Bacteria | Proteobacteri | Gammaprote | unclassified | unclassified | Pseudohongie | Pseudohongie | unclassified | unclassified | unclassified | unclassified | unclassified | 0,4 | 0,489897949 | 8,45E-06 | 1,04E-05 | 1 - | 1,92E-05 - | 0 - | 0 - | 0 - | 0 - | 1 - | 2,30E-05 - | 0 - | 0 - |
| 6200 | 1 | 0.2um_NOKCPHJN_1097286 | Bacteria | Chlorobi | Chlorobia | Chlorobiales | Chlorobiaceae | Prosthecochlo | Prosthecochlo | unclassified | unclassified | unclassified | unclassified | unclassified | 0,2 | 0,4 | 1,46E-05 | 2,92E-05 | 1 - | 7,31E-05 - | 0 - | 0 - | 0 - | 0 - | 0 - | 0 - | 0 - | 0 - |
| 6201 | 1 | 3um_FIJJAFBF_89936 | Bacteria | Bacteroidetes | Flavobacteriia | Flavobacterial | Flavobacteria | unclassified | Flavobacteria | unclassified | unclassified | unclassified | unclassified | unclassified | 1 | 0 | 8,16E-05 | 7,74E-06 | 1 - | 6,83E-05 - | 1 - | 8,48E-05 - | 1 - | 8,06E-05 - | 1 - | 8,18E-05 - | 1 - | 9,22E-05 - |
| 6202 | 4 | 0.2um_NOKCPHJN_263260;0.2um_NOKCPHJN_3 | various | various | various | various | various | various | various | various | various | various | various | various | 0,2 | 0,4 | 2,43E-06 | 4,86E-06 | 0 - | 0 - | 0 - | 0 - | 0 - | 0 - | 0 - | 0 - | 1 - | 1,22E-05 - |
| 6203 | 1 | 0.2um_NOKCPHJN_670080 | Bacteria | Bacteroidetes | Flavobacteriia | Flavobacterial | Flavobacteria | Polaribacter | Polaribacter ir | cellular proces | Cell wall/mem | COG2885 | Ompa motb d | 313594.PI23P_ | 0,2 | 0,4 | 9,47E-06 | 1,89E-05 | 0 - | 0 - | 0 - | 0 - | 0 - | 0 - | 0 - | 0 - | 1 - | 4,74E-05 - |
| 6204 | 1 | 3um_FIJJAFBF_194065 | Bacteria | Bacteroidetes | Flavobacteriia | Flavobacterial | Flavobacteria | Polaribacter | Polaribacter ir | metabolism | Lipid transpor | 07QTG | Component of | 313598.MED1 | 1 | 1,095445115 | 4,66E-05 | 5,22E-05 | 1 - | 4,03E-05 - | 1 - | 5,00E-05 - | 3 - | 0,000142613 - | 0 - | 0 - | 0 - | 0 - |
| 6205 | 1 | 0.2um_NOKCPHJN_142153 | Bacteria | Proteobacteri | Gammaprote | unclassified | unclassified | unclassified | various | poorly charact | Function unkn | 05W4M | Transport-ass | 384676.PSEEN | 2,6 | 0,489897949 | 0,000177577 | 2,57E-05 | 3 - | 0,000173904 - | 2 - | 0,000143958 - | 3 - | 0,000205276 - | 3 - | 0,000208308 - | 2 - | 0,000156439 - |
| 6206 | 1 | 0.2um_NOKCPHJN_351959 | Eukaryota | unclassified | Cryptophyta | Pyrenomonad | Geminigerace | Guillardia | Guillardia thet | information st | Translation, ri | 151Y2 | Ribosomal pro | 272952.HpaP8 | 0,4 | 0,489897949 | 5,36E-05 | 6,63E-05 | 1 - | 0,000119559 - | 1 - | 0,000148457 - | 0 - | 0 - | 0 - | 0 - | 0 - | 0 - |
| 6207 | 1 | 3um_FIJJAFBF_393616 | Bacteria | Proteobacteri | Gammaprote | Oceanospirilla | Saccharospirill | Reinekea | Reinekea fors | metabolism | Amino acid tra | 05TIB | (ABC) transpo | 314283.MED2 | 3,6 | 0,489897949 | 0,000148346 | 2,42E-05 | 4 - | 0,000138285 - | 3 - | 0,000128782 - | 3 - | 0,000122424 - | 4 - | 0,000165643 - | 4 - | 0,000186596 - |
| 6208 | 1 | 3um_FIJJAFBF_788017 | Bacteria | Proteobacteri | Alphaproteob | Rhodobactera | Rhodobactera | Pseudopelagic | Pseudopelagic | information st | Transcription | 16PMT | DNA-depende | 391589.RGAI1 | 2 | 0 | 8,11E-05 | 7,70E-06 | 2 - | 6,79E-05 - | 2 - | 8,43E-05 - | 2 - | 8,02E-05 - | 2 - | 8,14E-05 - | 2 - | 9,16E-05 - |
| 6209 | 1 | 3um_FIJJAFBF_747524 | Bacteria | Proteobacteri | Gammaprote | unclassified | unclassified | unclassified | Gammaprote | unclassified | unclassified | unclassified | unclassified | unclassified | 0,2 | 0,4 | 1,58E-05 | 3,17E-05 | 1 - | 7,92E-05 - | 0 - | 0 - | 0 - | 0 - | 0 - | 0 - | 0 - | 0 - |
| 6210 | 1 | 3um_FIJJAFBF_594149 | Eukaryota | unclassified | Pelagophycea | Pelagomonad | unclassified | Aureoumbra | Aureoumbra l | metabolism | Carbohydrate | COG1850 | RuBisCO catal | 45157.CMV01 | 26,8 | 1,166190379 | 0,000755356 | 9,33E-05 | 27 - | 0,000636339 - | 26 - | 0,000760881 - | 26 - | 0,000723314 - | 26 - | 0,000733997 - | 29 - | 0,00092225 - |
| 6211 | 1 | 3um_FIJJAFBF_520097 | Bacteria | Bacteroidetes | Flavobacteriia | Flavobacterial | Flavobacteria | Polaribacter | Polaribacter s | cellular proces | Intracellular tr | COG0681 | Signal peptida | 313598.MED1 | 2 | 0,894427191 | 4,89E-05 | 2,27E-05 | 1 - | 2,03E-05 - | 3 - | 7,57E-05 - | 2 - | 4,80E-05 - | 3 - | 7,30E-05 - | 1 - | 2,74E-05 - |
| 6212 | 1 | 3um_FIJJAFBF_47161 | Bacteria | Proteobacteri | Gammaprote | Oceanospirilla | Saccharospirill | Reinekea | Reinekea fors | cellular proces | Intracellular tr | 0QITE | Outer membra | 314283.MED2 | 5,6 | 1,019803903 | 0,000174841 | 4,18E-05 | 6 - | 0,000156159 - | 6 - | 0,000193903 - | 5 - | 0,000153608 - | 4 - | 0,000124701 - | 7 - | 0,000245833 - |
| 6213 | 1 | 3um_FIJJAFBF_178960 | Bacteria | Proteobacteri | Alphaproteob | Rhodobactera | unclassified | unclassified | Rhodobactera | information st | Translation, ri | 08YY1 | This protein is | 367336.OM22 | 5,8 | 0,748331477 | 0,000642703 | 7,88E-05 | 7 - | 0,000653201 - | 6 - | 0,000695214 - | 5 - | 0,000550741 - | 5 - | 0,000558875 - | 6 - | 0,000755485 - |
| 6214 | 1 | 0.2um_NOKCPHJN_901365 | Bacteria | Proteobacteri | Gammaprote | Pseudomonad | Pseudomonad | Pseudomonas | Pseudomonas | unclassified | unclassified | unclassified | unclassified | unclassified | 0,2 | 0,4 | 6,47E-06 | 1,29E-05 | 0 - | 0 - | 0 - | 0 - | 1 - | 3,23E-05 - | 0 - | 0 - | 0 - | 0 - |
| 6215 | 1 | 3um_FIJJAFBF_254291 | Bacteria | Proteobacteri | Gammaprote | Cellvibrionales | Spongiibacter | Oceanicoccus | Oceanicoccus | information st | Translation, ri | 16Q16 | Binds the low | 247634.GPB2 | 2,2 | 0,4 | 0,000133788 | 1,23E-05 | 3 - | 0,000155104 - | 2 - | 0,000128395 - | 2 - | 0,000122056 - | 2 - | 0,000123859 - | 2 - | 0,000139527 - |
| 6216 | 1 | 0.2um_NOKCPHJN_771264 | Eukaryota | unclassified | Oomycetes | Peronosporal | Peronosporac | Phytophthora | Phytophthora | unclassified | unclassified | unclassified | unclassified | unclassified | 3,2 | 1,166190379 | 0,00010646 | 3,96E-05 | 3 - | 8,48E-05 - | 4 - | 0,000140413 - | 4 - | 0,00013348 - | 4 - | 0,000135452 - | 1 - | 3,81E-05 - |
| 6217 | 1 | 0.2um_NOKCPHJN_1094406 | Bacteria | Proteobacteri | Alphaproteob | Rhodobactera | Rhodobactera | Shimia | Shimia abyssi | unclassified | unclassified | unclassified | unclassified | unclassified | 1 | 0 | 7,79E-05 | 7,39E-06 | 1 - | 6,52E-05 - | 1 - | 8,10E-05 - | 1 - | 7,70E-05 - | 1 - | 7,81E-05 - | 1 - | 8,80E-05 - |
| 6218 | 1 | 3um_FIJJAFBF_191786 | Eukaryota | unclassified | Florideophyce | Gracilariales | Gracilariaceae | Gracilaria | Gracilaria chil | unclassified | unclassified | unclassified | unclassified | unclassified | 1,6 | 0,8 | 0,000212104 | 0,000107729 | 2 - | 0,000229554 - | 2 - | 0,000285038 - | 2 - | 0,000270964 - | 2 - | 0,000274967 - | 0 - | 0 - |
| 6219 | 1 | 0.2um_NOKCPHJN_481615 | Eukaryota | Basidiomycota | Pucciniomycet | Pucciniales | Pucciniaceae | Puccinia | Puccinia coror | unclassified | unclassified | unclassified | unclassified | unclassified | 1 | 0 | 8,31E-06 | 7,89E-07 | 1 - | 6,96E-06 - | 1 - | 8,65E-06 - | 1 - | 8,22E-06 - | 1 - | 8,34E-06 - | 1 - | 9,40E-06 - |
| 6220 | 2 | 0.2um_NOKCPHJN_314557;3um_FIJJAFBF_64704 | Bacteria | Bacteroidetes | Flavobacteriia | unclassified | unclassified | unclassified | Flavobacteria | unclassified | unclassified | unclassified | unclassified | unclassified | 0,6 | 0,489897949 | 3,57E-05 | 2,95E-05 | 1 - | 5,19E-05 - | 1 - | 6,45E-05 - | 0 - | 0 - | 1 - | 6,22E-05 - | 0 - | 0 - |
| 6221 | 1 | 3um_FIJJAFBF_163782 | Bacteria | Bacteroidetes | Flavobacteriia | Flavobacterial | unclassified | unclassified | Flavobacterial | unclassified | unclassified | unclassified | unclassified | unclassified | 1 | 0 | 4,79E-05 | 4,55E-06 | 1 - | 4,01E-05 - | 1 - | 4,98E-05 - | 1 - | 4,74E-05 - | 1 - | 4,81E-05 - | 1 - | 5,42E-05 - |
| 6222 | 1 | 0.2um_NOKCPHJN_207187 | Bacteria | Proteobacteri | Gammaprote | Oceanospirilla | Saccharospirill | Saccharospiril | Saccharospiril | cellular proces | Posttranslatio | 05C6P | ATP-depender | 377629.TERTU | 6,2 | 1,469693846 | 0,000104492 | 2,43E-05 | 7 - | 9,96E-05 - | 8 - | 0,000141283 - | 7 - | 0,000117519 - | 4 - | 6,81E-05 - | 5 - | 9,60E-05 - |
| 6223 | 1 | 3um_FIJJAFBF_385385 | Bacteria | Bacteroidetes | Flavobacteriia | Flavobacterial | Flavobacteria | Polaribacter | Polaribacter s | unclassified | unclassified | unclassified | unclassified | unclassified | 3,6 | 0,489897949 | 5,14E-05 | 5,10E-06 | 4 - | 4,83E-05 - | 3 - | 4,50E-05 - | 4 - | 5,70E-05 - | 4 - | 5,78E-05 - | 3 - | 4,89E-05 - |
| 6224 | 1 | 3um_FIJJAFBF_243316 | Bacteria | Proteobacteri | Alphaproteob | Pelagibacteral | Pelagibactera | unclassified | Pelagibactera | unclassified | unclassified | unclassified | unclassified | unclassified | 5,2 | 0,979795897 | 0,000773132 | 9,75E-05 | 7 - | 0,000882898 - | 5 - | 0,000783071 - | 5 - | 0,000744408 - | 4 - | 0,000604322 - | 5 - | 0,000850959 - |
| 6225 | 1 | 3um_FIJJAFBF_560797 | Eukaryota | Bacillariophyt | Coscinodiscop | Thalassiosiral | Thalassiosirac | Thalassiosira | Thalassiosira p | poorly charact | Function unkn | 0IHC7 | Thylakoid mer | 35128.Thaps4 | 1,4 | 0,489897949 | 8,85E-05 | 3,64E-05 | 1 - | 5,19E-05 - | 1 - | 6,45E-05 - | 1 - | 6,13E-05 - | 2 - | 0,000124419 - | 2 - | 0,000140158 - |
| 6226 | 1 | 3um_FIJJAFBF_241339 | Bacteria | Proteobacteri | Alphaproteob | Rhodobactera | unclassified | unclassified | Rhodobactera | information st | Transcription | COG0250 | Participates in | 367336.OM22 | 1 | 0 | 7,74E-05 | 7,35E-06 | 1 - | 6,48E-05 - | 1 - | 8,05E-05 - | 1 - | 7,65E-05 - | 1 - | 7,77E-05 - | 1 - | 8,75E-05 - |
| 6227 | 1 | 0.2um_NOKCPHJN_93346 | Bacteria | Proteobacteri | Betaproteoba | unclassified | unclassified | unclassified | Betaproteoba | metabolism | Lipid transpor | 08IQE | Amp-depende | 338969.Rfer_ | 0,6 | 0,8 | 1,13E-05 | 1,45E-05 | 2 - | 3,54E-05 - | 0 - | 0 - | 1 - | 2,09E-05 - | 0 - | 0 - | 0 - | 0 - |
| 6228 | 1 | 0.2um_NOKCPHJN_71894 | Bacteria | Proteobacteri | Gammaprote | Oceanospirilla | Saccharospirill | Saccharospiril | Saccharospiril | unclassified | unclassified | unclassified | unclassified | unclassified | 6,6 | 0,8 | 0,000143719 | 3,00E-05 | 6 - | 0,00010845 - | 7 - | 0,000157107 - | 6 - | 0,000128015 - | 6 - | 0,000129905 - | 8 - | 0,000195117 - |
| 6229 | 1 | 3um_FIJJAFBF_128704 | Bacteria | Bacteroidetes | Flavobacteriia | unclassified | unclassified | unclassified | Flavobacteria | unclassified | unclassified | unclassified | unclassified | unclassified | 0,8 | 0,748331477 | 2,38E-05 | 2,35E-05 | 1 - | 2,56E-05 - | 2 - | 6,35E-05 - | 1 - | 3,02E-05 - | 0 - | 0 - | 0 - | 0 - |
| 6230 | 1 | 3um_FIJJAFBF_123075 | Bacteria | Proteobacteri | Gammaprote | Alteromonada | Alteromonada | Paraglaciecola | Paraglaciecola | information st | Translation, ri | 0QSAC | Binds togethe | 342610.Patl_3 | 3,4 | 0,489897949 | 0,000337445 | 5,53E-05 | 3 - | 0,000249515 - | 3 - | 0,000309824 - | 4 - | 0,000392702 - | 4 - | 0,000398502 - | 3 - | 0,000336684 - |
| 6231 | 1 | 0.2um_NOKCPHJN_202276 | Bacteria | Bacteroidetes | Flavobacteriia | Flavobacterial | Flavobacteria | Lacinutrix | Lacinutrix him | unclassified | unclassified | unclassified | unclassified | unclassified | 0,6 | 0,489897949 | 6,53E-05 | 5,43E-05 | 1 - | 9,26E-05 - | 0 - | 0 - | 1 - | 0,00010926 - | 0 - | 0 - | 1 - | 0,000124899 - |
| 6232 | 1 | 0.2um_NOKCPHJN_299571 | Eukaryota | unclassified | Pelagophycea | Pelagomonad | unclassified | Aureococcus | Aureococcus a | information st | Translation, ri | KOG3184 | Ribosomal pro | 35128.Thaps2 | 0,2 | 0,4 | 9,56E-06 | 1,91E-05 | 1 - | 4,78E-05 - | 0 - | 0 - | 0 - | 0 - | 0 - | 0 - | 0 - | 0 - |
| 6233 | 1 | 0.2um_NOKCPHJN_105420 | Bacteria | Proteobacteri | Gammaprote | Oceanospirilla | Saccharospirill | Reinekea | Reinekea blan | metabolism | Amino acid tra | 05C6M | Alpha-keto-be | 314283.MED2 | 9,6 | 1,624807681 | 0,000267507 | 5,35E-05 | 8 - | 0,00018625 - | 11 - | 0,000317993 - | 9 - | 0,000247331 - | 12 - | 0,000334645 - | 8 - | 0,000251318 - |
| 6234 | 1 | 3um_FIJJAFBF_765494 | Bacteria | Proteobacteri | Gammaprote | Alteromonada | Alteromonada | Paraglaciecola | Paraglaciecola | unclassified | unclassified | unclassified | unclassified | unclassified | 1 | 1,095445115 | 5,04E-05 | 5,03E-05 | 3 - | 0,000136639 - | 0 - | 0 - | 1 - | 5,38E-05 - | 0 - | 0 - | 1 - | 6,15E-05 - |
| 6235 | 1 | 3um_FIJJAFBF_813141 | Bacteria | Proteobacteri | Betaproteoba | unclassified | unclassified | unclassified | Betaproteoba | unclassified | unclassified | unclassified | unclassified | unclassified | 0,6 | 0,489897949 | 1,83E-05 | 1,51E-05 | 1 - | 2,67E-05 - | 1 - | 3,31E-05 - | 1 - | 3,15E-05 - | 0 - | 0 - | 0 - | 0 - |
| 6236 | 1 | 0.2um_NOKCPHJN_203995 | Bacteria | Bacteroidetes | Flavobacteriia | Flavobacterial | Flavobacteria | Lacinutrix | Lacinutrix sp. | poorly charact | Function unkn | 0XQTW | domain protei | 999141.GME_ | 0,6 | 0,489897949 | 7,44E-06 | 6,10E-06 | 0 - | 0 - | 0 - | 0 - | 1 - | 1,18E-05 - | 1 - | 1,20E-05 - | 1 - | 1,35E-05 - |
| 6237 | 1 | 3um_FIJJAFBF_403054 | Bacteria | Proteobacteri | Alphaproteob | Rhodobactera | Rhodobactera | unclassified | Rhodobactera | information st | Translation, ri | 01QWF | Allows the for | 246200.SPO28 | 0,2 | 0,4 | 5,47E-06 | 1,09E-05 | 0 - | 0 - | 0 - | 0 - | 0 - | 0 - | 1 - | 2,73E-05 - | 0 - | 0 - |
| 6238 | 1 | 0.2um_NOKCPHJN_687320 | Bacteria | Proteobacteri | Alphaproteob | Rhodobactera | Rhodobactera | Ascidiaceihabi | Ascidiaceihabi | metabolism | Nucleotide tra | 05FCI | Catalyzes a re | 439496.RBY4I | 0,2 | 0,4 | 8,02E-06 | 1,60E-05 | 0 - | 0 - | 0 - | 0 - | 1 - | 4,01E-05 - | 0 - | 0 - | 0 - | 0 - |
| 6239 | 1 | 0.2um_NOKCPHJN_674047 | Bacteria | Proteobacteri | Gammaprote | unclassified | unclassified | unclassified | Gammaprote | unclassified | unclassified | unclassified | unclassified | unclassified | 1,8 | 0,4 | 0,000112537 | 2,22E-05 | 2 - | 0,000106275 - | 2 - | 0,000131962 - | 2 - | 0,000125447 - | 2 - | 0,000127299 - | 1 - | 7,17E-05 - |
| 6240 | 1 | 3um_FIJJAFBF_09090 | Bacteria | Proteobacteri | Alphaproteob | Pelagibacteral | Pelagibactera | Candidatus Pe | Candidatus Pe | information st | Transcription | COG0250 | Participates in | 335992.SAR11 | 0,4 | 0,489897949 | 3,06E-05 | 3,82E-05 | 1 - | 6,52E-05 - | 0 - | 0 - | 0 - | 0 - | 0 - | 0 - | 1 - | 8,80E-05 - |
| 6241 | 1 | 3um_FIJJAFBF_37832 | Bacteria | Proteobacteri | unclassified | unclassified | unclassified | unclassified | Proteobacteri | unclassified | unclassified | unclassified | unclassified | unclassified | 1,2 | 0,748331477 | 5,00E-05 | 3,01E-05 | 2 - | 7,33E-05 - | 1 - | 4,55E-05 - | 1 - | 4,33E-05 - | 2 - | 8,78E-05 - | 0 - | 0 - |
| 6242 | 1 | 3um_FIJJAFBF_344410 | Bacteria | Proteobacteri | Alphaproteob | Rhodobactera | Rhodobactera | various | various | information st | Transcription | 05VEQ | Cold shock pro | 314262.MED1 | 1 | 0 | 0,00020151 | 1,91E-05 | 1 - | 0,000168789 - | 1 - | 0,000209587 - | 1 - | 0,000199239 - | 1 - | 0,000202181 - | 1 - | 0,000227757 - |
| 6243 | 1 | 0.2um_NOKCPHJN_815346 | Bacteria | Proteobacteri | Alphaproteob | Pelagibacteral | Pelagibactera | Candidatus Pe | Candidatus Pe | cellular proces | Posttranslatio | COG0330 | Band 7 protei | 335992.SAR11 | 12 | 0,632455532 | 0,00056937 | 4,69E-05 | 13 - | 0,00051809 - | 12 - | 0,000593829 - | 12 - | 0,000564509 - | 11 - | 0,00052511 - | 12 - | 0,00064531 - |
| 6244 | 1 | 0.2um_NOKCPHJN_275838 | Eukaryota | unclassified | Labyrinthulom | unclassified | Thraustochytr | Aurantiochytr | Aurantiochytr | unclassified | unclassified | unclassified | unclassified | unclassified | 2 | 0,632455532 | 2,80E-05 | 9,18E-06 | 2 - | 2,36E-05 - | 3 - | 4,40E-05 - | 2 - | 2,79E-05 - | 2 - | 2,83E-05 - | 1 - | 1,59E-05 - |
| 6245 | 1 | 0.2um_NOKCPHJN_346435 | Bacteria | Bacteroidetes | unclassified | unclassified | unclassified | unclassified | Bacteroidetes | unclassified | unclassified | unclassified | unclassified | unclassified | 0,6 | 0,489897949 | 5,42E-05 | 4,44E-05 | 0 - | 0 - | 0 - | 0 - | 1 - | 8,57E-05 - | 1 - | 8,70E-05 - | 1 - | 9,80E-05 - |
| 6246 | 1 | 0.2um_NOKCPHJN_409279 | Eukaryota | Phaeophyceae | unclassified | Ectocarpales | Ectocarpaceae | Ectocarpus | Ectocarpus sili | unclassified | unclassified | unclassified | unclassified | unclassified | 0,4 | 0,489897949 | 5,20E-06 | 6,38E-06 | 0 - | 0 - | 0 - | 0 - | 0 - | 0 - | 1 - | 1,22E-05 - | 1 - | 1,38E-05 - |
| 6247 | 2 | 3um_FIJJAFBF_583280;3um_FIJJAFBF_67779 | Eukaryota | unclassified | unclassified | Phaeocystales | Phaeocystacea | Phaeocystis | various | unclassified | unclassified | unclassified | unclassified | unclassified | 0,2 | 0,4 | 2,05E-05 | 4,11E-05 | 0 - | 0 - | 0 - | 0 - | 1 - | 0,000102638 - | 0 - | 0 - | 0 - | 0 - |
| 6248 | 1 | 3um_FIJJAFBF_25763 | Bacteria | Bacteroidetes | Flavobacteriia | unclassified | unclassified | unclassified | Flavobacteria | unclassified | unclassified | unclassified | unclassified | unclassified | 0,6 | 0,8 | 4,27E-05 | 5,51E-05 | 2 - | 0,000134242 - | 0 - | 0 - | 1 - | 7,92E-05 - | 0 - | 0 - | 0 - | 0 - |
| 6249 | 1 | 3um_FIJJAFBF_556071 | Eukaryota | unclassified | Dinophyceae | Peridiniales | Heterocapsac | Heterocapsa | Heterocapsa t | unclassified | unclassified | unclassified | unclassified | unclassified | 2,4 | 0,489897949 | 0,000405031 | 0,000107821 | 2 - | 0,000279943 - | 2 - | 0,000347607 - | 3 - | 0,000495667 - | 2 - | 0,000335325 - | 3 - | 0,000566614 - |
| 6250 | 1 | 3um_FIJJAFBF_475148 | Bacteria | Proteobacteri | Gammaprote | Oceanospirilla | Saccharospirill | Reinekea | Reinekea fors | unclassified | unclassified | unclassified | unclassified | unclassified | 2 | 0,632455532 | 0,000179004 | 4,56E-05 | 3 - | 0,000229554 - | 1 - | 9,50E-05 - | 2 - | 0,000180643 - | 2 - | 0,000183311 - | 2 - | 0,000206499 - |
| 6251 | 1 | 0.2um_NOKCPHJN_908861 | Bacteria | Proteobacteri | Alphaproteob | Pelagibacteral | Pelagibactera | Candidatus Pe | Candidatus Pe | metabolism | Amino acid tra | 01QH7 | sulfoacetaldel | 439493.PB721 | 1,8 | 0,4 | 4,20E-05 | 1,16E-05 | 1 - | 1,92E-05 - | 2 - | 4,77E-05 - | 2 - | 4,53E-05 - | 2 - | 4,60E-05 - | 2 - | 5,18E-05 - |
| 6252 | 1 | 3um_FIJJAFBF_103579 | Bacteria | Proteobacteri | Gammaprote | Oceanospirilla | Saccharospirill | Reinekea | Reinekea fors | metabolism | Amino acid tra | COG0131 | imidazoleglyc | 743720.Psefu_ | 1,2 | 0,979795897 | 8,27E-05 | 6,87E-05 | 2 - | 0,000117119 - | 0 - | 0 - | 2 - | 0,000138247 - | 0 - | 0 - | 2 - | 0,000158035 - |
| 6253 | 1 | 3um_FIJJAFBF_172446 | Bacteria | Proteobacteri | Gammaprote | Oceanospirilla | Saccharospirill | Reinekea | Reinekea fors | unclassified | unclassified | unclassified | unclassified | unclassified | 3,8 | 0,748331477 | 0,000461369 | 8,52E-05 | 4 - | 0,000409917 - | 4 - | 0,000508996 - | 5 - | 0,000604831 - | 3 - | 0,000368259 - | 3 - | 0,000414842 - |
| 6254 | 1 | 0.2um_NOKCPHJN_144001 | Bacteria | Proteobacteri | Gammaprote | Oceanospirilla | Saccharospirill | Reinekea | Reinekea fors | cellular proces | Cell wall/mem | 0QJAT | cyclopropane- | 314283.MED2 | 2 | 0,632455532 | 6,47E-05 | 1,50E-05 | 3 - | 8,38E-05 - | 2 - | 6,94E-05 - | 2 - | 6,59E-05 - | 2 - | 6,69E-05 - | 1 - | 3,77E-05 - |
| 6255 | 1 | 0.2um_NOKCPHJN_602391 | Bacteria | Proteobacteri | Alphaproteob | Pelagibacteral | Pelagibactera | Candidatus Pe | Candidatus Pe | cellular proces | Cell wall/mem | 7257 | NA | 335992.SAR11 | 3 | 1,414213562 | 0,000127874 | 5,63E-05 | 4 - | 0,00014668 - | 1 - | 4,55E-05 - | 5 - | 0,000216425 - | 3 - | 0,000131773 - | 2 - | 9,90E-05 - |
| 6256 | 1 | 3um_FIJJAFBF_189827 | Bacteria | Bacteroidetes | Flavobacteriia | Flavobacterial | Flavobacteria | Arenibacter | Arenibacter sp | unclassified | unclassified | unclassified | unclassified | unclassified | 1 | 0 | 0,000112317 | 1,07E-05 | 1 - | 9,41E-05 - | 1 - | 0,000116819 - | 1 - | 0,000111051 - | 1 - | 0,000112691 - | 1 - | 0,000126946 - |
| 6257 | 1 | 0.2um_NOKCPHJN_510389 | Bacteria | Proteobacteri | Gammaprote | unclassified | unclassified | unclassified | Gammaprote | unclassified | unclassified | unclassified | unclassified | unclassified | 0,2 | 0,4 | 1,06E-05 | 2,13E-05 | 1 - | 5,31E-05 - | 0 - | 0 - | 0 - | 0 - | 0 - | 0 - | 0 - | 0 - |
| 6258 | 1 | 0.2um_NOKCPHJN_319552 | Eukaryota | Phaeophyceae | unclassified | Ectocarpales | Ectocarpaceae | Ectocarpus | Ectocarpus sili | metabolism | Energy produc | COG0055 | Produces ATP | 35128.Thaps4 | 9,2 | 2,315167381 | 0,00030045 | 7,75E-05 | 12 - | 0,000329503 - | 10 - | 0,000340954 - | 7 - | 0,000226884 - | 6 - | 0,000197344 - | 11 - | 0,000407564 - |
| 6259 | 1 | 0.2um_NOKCPHJN_113762 | Bacteria | Proteobacteri | Gammaprote | Cellvibrionales | Porticoccacea | unclassified | SAR92 bacteri | unclassified | unclassified | unclassified | unclassified | unclassified | 0,2 | 0,4 | 2,59E-06 | 5,18E-06 | 1 - | 1,29E-05 - | 0 - | 0 - | 0 - | 0 - | 0 - | 0 - | 0 - | 0 - |
| 6260 | 1 | 0.2um_NOKCPHJN_45826 | Archaea | Euryarchaeota | unclassified | unclassified | unclassified | unclassified | Euryarchaeota | metabolism | Secondary me | 003N9 | Beta-ketoacyl | 204669.Acid34 | 0,6 | 0,489897949 | 2,25E-06 | 1,85E-06 | 1 - | 3,33E-06 - | 0 - | 0 - | 1 - | 3,93E-06 - | 1 - | 3,99E-06 - | 0 - | 0 - |
| 6261 | 1 | 3um_FIJJAFBF_199367 | Bacteria | Proteobacteri | Gammaprote | Oceanospirilla | Oceanospirilla | Amphritea | Amphritea jap | information st | Translation, ri | COG0185 | Protein S19 fo | 207954.MED9 | 2 | 0,894427191 | 0,000297715 | 0,000127757 | 3 - | 0,000378385 - | 3 - | 0,000469842 - | 1 - | 0,000148882 - | 1 - | 0,000151081 - | 2 - | 0,000340383 - |
| 6262 | 2 | 0.2um_NOKCPHJN_301543;0.2um_NOKCPHJN_4 | Bacteria | Proteobacteri | Alphaproteob | Pelagibacteral | Pelagibactera | Candidatus Pe | Candidatus Pe | metabolism | Amino acid tra | 083Z3 | extracellular li | 335992.SAR11 | 2,4 | 0,489897949 | 7,95E-05 | 2,12E-05 | 2 - | 5,49E-05 - | 2 - | 6,82E-05 - | 3 - | 9,72E-05 - | 2 - | 6,58E-05 - | 3 - | 0,000111154 - |
| 6263 | 1 | 0.2um_NOKCPHJN_423416 | Bacteria | Bacteroidetes | Flavobacteriia | Flavobacterial | Flavobacteria | Tenacibacul | Tenacibacul | unclassified | unclassified | unclassified | unclassified | unclassified | 11,6 | 0,8 | 0,000175853 | 2,70E-05 | 11 - | 0,000139047 - | 11 - | 0,000172655 - | 11 - | 0,00016413 - | 12 - | 0,000181696 - | 13 - | 0,000221737 - |
| 6264 | 1 | 3um_FIJJAFBF_658129 | Bacteria | Bacteroidetes | Flavobacteriia | Flavobacterial | Flavobacteria | Ulvibacter | Ulvibacter sp. | unclassified | unclassified | unclassified | unclassified | unclassified | 2,8 | 0,748331477 | 0,000331467 | 5,85E-05 | 4 - | 0,000406289 - | 2 - | 0,000252246 - | 3 - | 0,000359687 - | 3 - | 0,000365 - | 2 - | 0,000274114 - |
| 6265 | 1 | 3um_FIJJAFBF_32817 | Bacteria | Proteobacteri | Alphaproteob | Rhodobactera | Rhodobactera | Pseudophaeol | Pseudophaeol | unclassified | unclassified | unclassified | unclassified | unclassified | 1,6 | 0,489897949 | 6,50E-05 | 1,70E-05 | 2 - | 6,96E-05 - | 1 - | 4,32E-05 - | 2 - | 8,21E-05 - | 2 - | 8,33E-05 - | 1 - | 4,69E-05 - |
| 6266 | 1 | 3um_FIJJAFBF_782497 | Bacteria | Proteobacteri | Alphaproteob | Rhodobactera | Rhodobactera | unclassified | Rhodobactera | metabolism | Nucleotide tra | 16Q8Q | Catalyzes the | 367336.OM22 | 2 | 0,632455532 | 0,00010599 | 2,45E-05 | 3 - | 0,000137183 - | 2 - | 0,000113561 - | 2 - | 0,000107954 - | 2 - | 0,000109548 - | 1 - | 6,17E-05 - |
| 6267 | 1 | 0.2um_NOKCPHJN_1030708 | Bacteria | unclassified | unclassified | unclassified | unclassified | unclassified | uncultured ma | poorly charact | Function unkn | 16WVI | MORN repeat | 633131.TR2A6 | 0,2 | 0,4 | 6,66E-06 | 1,33E-05 | 0 - | 0 - | 0 - | 0 - | 1 - | 3,33E-05 - | 0 - | 0 - | 0 - | 0 - |
| 6268 | 1 | 0.2um_NOKCPHJN_860493 | Bacteria | Proteobacteri | Alphaproteob | Pelagibacteral | Pelagibactera | Candidatus Pe | Candidatus Pe | unclassified | unclassified | unclassified | unclassified | unclassified | 0,6 | 0,489897949 | 2,56E-05 | 2,10E-05 | 0 - | 0 - | 1 - | 4,40E-05 - | 1 - | 4,18E-05 - | 1 - | 4,24E-05 - | 0 - | 0 - |
| 6269 | 1 | 0.2um_NOKCPHJN_860142 | Viruses | unclassified | unclassified | unclassified | unclassified | unclassified | uncultured M | poorly charact | Function unkn | 0NGTE | Major head pr | 608534.GCWU | 1,4 | 0,489897949 | 6,80E-05 | 2,14E-05 | 2 - | 8,29E-05 - | 2 - | 0,000102902 - | 1 - | 4,89E-05 - | 1 - | 4,96E-05 - | 1 - | 5,59E-05 - |
| 6270 | 2 | 0.2um_NOKCPHJN_469736;3um_FIJJAFBF_28215 | Bacteria | Proteobacteri | Gammaprote | Cellvibrionales | Spongiibacter | Oceanicoccus | Oceanicoccus | cellular proces | Signal transdu | 07RYR | Transcriptiona | 247634.GPB2 | 2,2 | 0,4 | 0,000139607 | 2,84E-05 | 2 - | 0,000106275 - | 2 - | 0,000131962 - | 2 - | 0,000125447 - | 3 - | 0,000190949 - | 2 - | 0,000143402 - |
| 6271 | 1 | 3um_FIJJAFBF_158823 | Viruses | unclassified | unclassified | unclassified | unclassified | unclassified | uncultured vi | unclassified | unclassified | unclassified | unclassified | unclassified | 0,8 | 0,748331477 | 2,51E-05 | 2,34E-05 | 0 - | 0 - | 2 - | 6,22E-05 - | 1 - | 2,96E-05 - | 0 - | 0 - | 1 - | 3,38E-05 - |
| 6272 | 1 | 0.2um_NOKCPHJN_183442 | Eukaryota | Eustigmatoph | unclassified | Eustigmatales | Monodopsida | Nannochlorop | Nannochlorop | cellular proces | Posttranslatio | KOG0738 | katanin p60 | 400682.PAC_1 | 0,8 | 0,4 | 1,83E-05 | 9,23E-06 | 0 - | 0 - | 1 - | 2,29E-05 - | 1 - | 2,18E-05 - | 1 - | 2,21E-05 - | 1 - | 2,49E-05 - |
| 6273 | 1 | 0.2um_NOKCPHJN_214190 | Bacteria | Bacteroidetes | Flavobacteriia | Flavobacterial | Flavobacteria | Mangrovimon | Mangrovimon | unclassified | unclassified | unclassified | unclassified | unclassified | 1 | 0,894427191 | 3,15E-05 | 2,77E-05 | 2 - | 5,63E-05 - | 1 - | 3,49E-05 - | 2 - | 6,64E-05 - | 0 - | 0 - | 0 - | 0 - |
| 6274 | 1 | 0.2um_NOKCPHJN_858636 | Eukaryota | Chordata | Actinopteri | unclassified | Sciaenidae | Larimichthys | Larimichthys c | unclassified | unclassified | unclassified | unclassified | unclassified | 0,8 | 0,4 | 1,56E-05 | 7,84E-06 | 0 - | 0 - | 1 - | 1,94E-05 - | 1 - | 1,85E-05 - | 1 - | 1,88E-05 - | 1 - | 2,11E-05 - |
| 6275 | 1 | 0.2um_NOKCPHJN_507733 | Eukaryota | Phaeophyceae | unclassified | Ectocarpales | Ectocarpaceae | Ectocarpus | Ectocarpus sili | poorly charact | Function unkn | 102D8 | PAP_fibrillin | 2850.Phatr46 | 2 | 0,632455532 | 0,000137597 | 5,41E-05 | 1 - | 5,60E-05 - | 2 - | 0,000139043 - | 2 - | 0,000132178 - | 2 - | 0,00013413 - | 3 - | 0,000226646 - |
| 6276 | 1 | 0.2um_NOKCPHJN_817682 | Bacteria | Proteobacteri | Gammaprote | Alteromonada | Colwelliaceae | Colwellia | Colwellia echi | metabolism | Energy produc | 16SYZ | Forms oxaloac | 715451.ambt_ | 0,6 | 0,489897949 | 1,51E-05 | 1,24E-05 | 1 - | 2,24E-05 - | 0 - | 0 - | 1 - | 2,64E-05 - | 1 - | 2,68E-05 - | 0 - | 0 - |
| 6277 | 2 | 3um_FIJJAFBF_132847;3um_FIJJAFBF_237281 | Bacteria | Proteobacteri | Gammaprote | Cellvibrionales | Porticoccacea | unclassified | SAR92 bacteri | unclassified | unclassified | unclassified | unclassified | unclassified | 1,2 | 0,4 | 0,000122779 | 4,23E-05 | 1 - | 8,57E-05 - | 1 - | 0,000106357 - | 1 - | 0,000101106 - | 2 - | 0,000205199 - | 1 - | 0,000115578 - |
| 6278 | 1 | 0.2um_NOKCPHJN_372597 | Bacteria | Proteobacteri | Gammaprote | Oceanospirilla | Saccharospirill | Saccharospiril | Saccharospiril | unclassified | unclassified | unclassified | unclassified | unclassified | 1,6 | 1,496662955 | 1,58E-05 | 1,39E-05 | 4 - | 3,56E-05 - | 2 - | 2,21E-05 - | 0 - | 0 - | 2 - | 2,13E-05 - | 0 - | 0 - |
| 6279 | 1 | 0.2um_NOKCPHJN_27393 | Bacteria | unclassified | unclassified | unclassified | unclassified | unclassified | uncultured ba | unclassified | unclassified | unclassified | unclassified | unclassified | 11,6 | 1,743559577 | 0,000310212 | 4,17E-05 | 14 - | 0,000315692 - | 13 - | 0,000363997 - | 9 - | 0,000239556 - | 11 - | 0,000297115 - | 11 - | 0,000334699 - |
| 6280 | 1 | 3um_FIJJAFBF_649835 | Bacteria | Proteobacteri | Gammaprote | Cellvibrionales | Spongiibacter | Spongiibacter | Spongiibacter | poorly charact | Function unkn | 0QNIY | Dehydrogenas | 247633.GP214 | 0,8 | 0,748331477 | 8,50E-05 | 8,36E-05 | 1 - | 9,11E-05 - | 2 - | 0,00022622 - | 1 - | 0,000107526 - | 0 - | 0 - | 0 - | 0 - |
| 6281 | 1 | 0.2um_NOKCPHJN_154859 | Bacteria | Bacteroidetes | Flavobacteriia | Flavobacterial | Flavobacteria | Aequorivita | Aequorivita sp | unclassified | unclassified | unclassified | unclassified | unclassified | 2,6 | 0,8 | 8,72E-05 | 2,85E-05 | 2 - | 5,60E-05 - | 3 - | 0,000104282 - | 2 - | 6,61E-05 - | 4 - | 0,00013413 - | 2 - | 7,55E-05 - |
| 6282 | 1 | 0.2um_NOKCPHJN_186611 | Eukaryota | Phaeophyceae | unclassified | Ectocarpales | Ectocarpaceae | Ectocarpus | Ectocarpus sili | information st | Translation, ri | KOG1242 | GCN1 general | 65071.PYU1_ | 0,2 | 0,4 | 1,03E-06 | 2,06E-06 | 0 - | 0 - | 1 - | 5,15E-06 - | 0 - | 0 - | 0 - | 0 - | 0 - | 0 - |
| 6283 | 1 | 3um_FIJJAFBF_193151 | Bacteria | Bacteroidetes | Flavobacteriia | Flavobacterial | Flavobacteria | Polaribacter | Polaribacter d | unclassified | unclassified | unclassified | unclassified | unclassified | 0,2 | 0,4 | 2,45E-06 | 4,89E-06 | 1 - | 1,22E-05 - | 0 - | 0 - | 0 - | 0 - | 0 - | 0 - | 0 - | 0 - |
| 6284 | 1 | 3um_FIJJAFBF_467884 | unclassified | unclassified | unclassified | unclassified | unclassified | unclassified | unclassified | unclassified | unclassified | unclassified | unclassified | unclassified | 0,4 | 0,489897949 | 1,27E-05 | 1,56E-05 | 0 - | 0 - | 1 - | 3,23E-05 - | 0 - | 0 - | 1 - | 3,12E-05 - | 0 - | 0 - |
| 6285 | 1 | 3um_FIJJAFBF_388952 | Bacteria | Bacteroidetes | Flavobacteriia | Flavobacterial | Flavobacteria | Polaribacter | Polaribacter s | unclassified | unclassified | unclassified | unclassified | unclassified | 3,4 | 0,489897949 | 0,000106252 | 1,03E-05 | 4 - | 0,000105785 - | 3 - | 9,85E-05 - | 4 - | 0,000124868 - | 3 - | 9,50E-05 - | 3 - | 0,000107056 - |
| 6286 | 1 | 0.2um_NOKCPHJN_12867 | Bacteria | Bacteroidetes | Flavobacteriia | Flavobacterial | Flavobacteria | Lacinutrix | Lacinutrix him | unclassified | unclassified | unclassified | unclassified | unclassified | 1 | 0 | 0,000110506 | 1,05E-05 | 1 - | 9,26E-05 - | 1 - | 0,000114935 - | 1 - | 0,00010926 - | 1 - | 0,000110874 - | 1 - | 0,000124899 - |

| | | | | | | | | | | | | | | | | | | | | | | | | | | | | |
|---|---|---|---|---|---|---|---|---|---|---|---|---|---|---|---|---|---|---|---|---|---|---|---|---|---|---|---|---|
| 6287 | 1 | 0.2um_NOKCPHJN_407176 | Eukaryota | Mucoromycot | Mucoromycet | Mucorales | Syncephalastr | Syncephalastr | Syncephalastr | unclassified | unclassified | unclassified | unclassified | unclassified | 0,4 | 0,489897949 | 4,56E-06 | 5,61E-06 | 1 - | 1,05E-05 - | 0 - | 0 - | 1 - | 1,23E-05 - | 0 - | 0 - | 0 - | 0 - |
| 6288 | 1 | 3um_FIJJAFBF_262855 | Bacteria | Proteobacteri | Alphaproteob | Rhodobactera | Rhodobactera | unclassified | Rhodobactera | information st | Transcription | 05D27 | DNA-depende | 644076.SCH4E | 2,2 | 0,4 | 4,39E-05 | 4,03E-06 | 3 - | 5,09E-05 - | 2 - | 4,22E-05 - | 2 - | 4,01E-05 - | 2 - | 4,07E-05 - | 2 - | 4,58E-05 - |
| 6289 | 1 | 3um_FIJJAFBF_423694 | Bacteria | Bacteroidetes | Flavobacteriia | Flavobacterial | Crocinitomica | unclassified | Crocinitomica | unclassified | unclassified | unclassified | unclassified | unclassified | 1 | 0 | 9,86E-05 | 9,36E-06 | 1 - | 8,26E-05 - | 1 - | 0,000102532 - | 1 - | 9,75E-05 - | 1 - | 9,89E-05 - | 1 - | 0,00011142 - |
| 6290 | 2 | 3um_FIJJAFBF_218994;3um_FIJJAFBF_306072 | Eukaryota | unclassified | various | various | various | various | various | cellular proces | Posttranslatio | 0IEKR | One of the co | 35128.Thapsd | 1 | 0 | 2,99E-05 | 2,83E-06 | 1 - | 2,50E-05 - | 1 - | 3,10E-05 - | 1 - | 2,95E-05 - | 1 - | 3,00E-05 - | 1 - | 3,37E-05 - |
| 6291 | 1 | 0.2um_NOKCPHJN_1050830 | Eukaryota | unclassified | Oomycetes | Peronosporale | Peronosporac | Phytophthora | Phytophthora | information st | Translation, ri | KOG0900 | Ribosomal prc | 436017.A4RTY | 1,2 | 0,4 | 0,000120288 | 2,66E-05 | 2 - | 0,000172597 - | 1 - | 0,000107157 - | 1 - | 0,000101866 - | 1 - | 0,000103371 - | 1 - | 0,000116447 - |
| 6292 | 1 | 3um_FIJJAFBF_517392 | Bacteria | Verrucomicrol | Verrucomicrol | Verrucomicrol | Verrucomicrol | unclassified | Verrucomicrol | unclassified | unclassified | unclassified | unclassified | unclassified | 0,6 | 0,489897949 | 1,73E-05 | 1,43E-05 | 1 - | 2,53E-05 - | 1 - | 3,15E-05 - | 1 - | 2,99E-05 - | 0 - | 0 - | 0 - | 0 - |
| 6293 | 1 | 3um_FIJJAFBF_350296 | Bacteria | Proteobacteri | Gammaprotec | Oceanospirilla | Halomonadac | Halomonas | Halomonas sp | information st | Translation, ri | 179ZI | One of the pri | 351348.Maqu | 4,4 | 1,019803903 | 0,000457756 | 9,34E-05 | 5 - | 0,000441449 - | 4 - | 0,00043852 - | 4 - | 0,000416868 - | 6 - | 0,000634538 - | 3 - | 0,000357403 - |
| 6294 | 1 | 3um_FIJJAFBF_753257 | Bacteria | Bacteroidetes | Flavobacteriia | Flavobacterial | Flavobacteriac | Aquimarina | Aquimarina m | information st | Replication, re | COG0513 | purine NTP-de | 867900.Celly_ | 0,2 | 0,4 | 6,31E-06 | 1,26E-05 | 0 - | 0 - | 0 - | 0 - | 0 - | 0 - | 1 - | 3,15E-05 - | 0 - | 0 - |
| 6295 | 1 | 0.2um_NOKCPHJN_146524 | Bacteria | Proteobacteri | Gammaprotec | Oceanospirilla | Saccharospirill | Reinekea | Reinekea fors | unclassified | unclassified | unclassified | unclassified | unclassified | 0,4 | 0,489897949 | 1,07E-05 | 1,32E-05 | 1 - | 2,38E-05 - | 1 - | 2,95E-05 - | 0 - | 0 - | 0 - | 0 - | 0 - | 0 - |
| 6296 | 1 | 0.2um_NOKCPHJN_1074739 | Bacteria | Proteobacteri | Alphaproteob | Pelagibacteral | Pelagibacterac | Candidatus Pe | Candidatus Pe | unclassified | unclassified | unclassified | unclassified | unclassified | 0,4 | 0,489897949 | 4,18E-05 | 5,17E-05 | 1 - | 9,33E-05 - | 1 - | 0,000115869 - | 0 - | 0 - | 0 - | 0 - | 0 - | 0 - |
| 6297 | 1 | 3um_FIJJAFBF_510254 | Bacteria | Bacteroidetes | Flavobacteriia | Flavobacterial | Flavobacteriac | Flaviramulus | Flaviramulus i | information st | Translation, ri | 05G6W | One of the pri | 983544.Lacal_ | 3,6 | 0,8 | 0,00024277 | 4,49E-05 | 5 - | 0,000285514 - | 3 - | 0,000212715 - | 3 - | 0,000202212 - | 3 - | 0,000205199 - | 4 - | 0,000308208 - |
| 6298 | 1 | 3um_FIJJAFBF_585224 | Bacteria | unclassified | unclassified | unclassified | unclassified | unclassified | uncultured ma | unclassified | unclassified | unclassified | unclassified | unclassified | 16,4 | 2,497999199 | 0,000651803 | 6,02E-05 | 21 - | 0,000706836 - | 17 - | 0,000710505 - | 15 - | 0,000595963 - | 14 - | 0,000564448 - | 15 - | 0,000681266 - |
| 6299 | 1 | 0.2um_NOKCPHJN_981236 | Bacteria | Proteobacteri | Alphaproteob | Pelagibacteral | Pelagibacterac | Candidatus Pe | Candidatus Pe | metabolism | Amino acid tra | 05CM2 | brancheD-cha | 439493.PB721 | 1 | 0,632455532 | 6,48E-05 | 3,70E-05 | 2 - | 0,000115354 - | 1 - | 7,16E-05 - | 1 - | 6,81E-05 - | 1 - | 6,91E-05 - | 0 - | 0 - |
| 6300 | 1 | 3um_FIJJAFBF_532003 | Bacteria | Bacteroidetes | Flavobacteriia | Flavobacterial | Flavobacteriac | Lacinutrix | Lacinutrix jang | metabolism | Lipid transpor | 07QM9 | Component of | 983544.Lacal_ | 0,8 | 0,4 | 3,35E-05 | 1,70E-05 | 1 - | 3,62E-05 - | 1 - | 4,50E-05 - | 1 - | 4,27E-05 - | 1 - | 4,34E-05 - | 0 - | 0 - |
| 6301 | 1 | 0.2um_NOKCPHJN_209509 | Eukaryota | Streptophyta | Liliopsida | Arecales | Arecaceae | Phoenix | Phoenix dacty | unclassified | unclassified | unclassified | unclassified | unclassified | 0,8 | 0,4 | 6,20E-05 | 3,18E-05 | 1 - | 6,56E-05 - | 0 - | 0 - | 1 - | 7,74E-05 - | 1 - | 7,86E-05 - | 1 - | 8,85E-05 - |
| 6302 | 1 | 3um_FIJJAFBF_648168 | Bacteria | Bacteroidetes | Flavobacteriia | Flavobacterial | Flavobacteriac | Gilvibacter | Gilvibacter sp. | unclassified | unclassified | unclassified | unclassified | unclassified | 7,2 | 0,748331477 | 0,000466395 | 7,45E-05 | 7 - | 0,00037898 - | 6 - | 0,000403355 - | 8 - | 0,000511254 - | 7 - | 0,000453954 - | 8 - | 0,000584432 - |
| 6303 | 1 | 0.2um_NOKCPHJN_148485 | Bacteria | Bacteroidetes | Flavobacteriia | Flavobacterial | Flavobacteriac | unclassified | Flavobacteriac | unclassified | unclassified | unclassified | unclassified | unclassified | 0,4 | 0,489897949 | 9,44E-06 | 1,17E-05 | 1 - | 2,11E-05 - | 1 - | 2,62E-05 - | 0 - | 0 - | 0 - | 0 - | 0 - | 0 - |
| 6304 | 1 | 3um_FIJJAFBF_259313 | Bacteria | Planctomycete | Phycisphaerae | Phycisphaeral | Phycisphaerac | Phycisphaera | Phycisphaera | unclassified | unclassified | unclassified | unclassified | unclassified | 1,8 | 1,166190379 | 8,89E-05 | 5,72E-05 | 3 - | 0,000129447 - | 3 - | 0,000160736 - | 1 - | 5,09E-05 - | 2 - | 0,000103371 - | 0 - | 0 - |
| 6305 | 1 | 0.2um_NOKCPHJN_173895 | Bacteria | Bacteroidetes | Flavobacteriia | Flavobacterial | Flavobacteriac | Formosa | Formosa sp. H | unclassified | unclassified | unclassified | unclassified | unclassified | 1 | 0 | 3,14E-05 | 2,98E-06 | 1 - | 2,63E-05 - | 1 - | 3,27E-05 - | 1 - | 3,11E-05 - | 1 - | 3,15E-05 - | 1 - | 3,55E-05 - |
| 6306 | 1 | 0.2um_NOKCPHJN_742695 | unclassified | unclassified | unclassified | unclassified | unclassified | unclassified | unclassified | unclassified | unclassified | unclassified | unclassified | unclassified | 0,8 | 0,748331477 | 2,63E-05 | 2,50E-05 | 1 - | 2,77E-05 - | 0 - | 0 - | 0 - | 0 - | 2 - | 6,64E-05 - | 1 - | 3,74E-05 - |
| 6307 | 1 | 3um_FIJJAFBF_815479 | Eukaryota | unclassified | Cryptophyta | Pyrenomonad | Geminigerace | Guillardia | Guillardia thet | metabolism | Lipid transpor | COG0439 | acetyl-CoA car | 2850.Phatr549 | 1 | 0,632455532 | 9,56E-05 | 5,45E-05 | 2 - | 0,00017004 - | 1 - | 0,00010557 - | 1 - | 0,000100357 - | 1 - | 0,000101839 - | 0 - | 0 - |
| 6308 | 1 | 3um_FIJJAFBF_371423 | Bacteria | Proteobacteri | Betaproteoba | unclassified | unclassified | unclassified | Betaproteoba | information st | Translation, ri | COG0052 | 30S ribosomal | 6085.XP_0021 | 0,8 | 0,748331477 | 4,17E-05 | 4,01E-05 | 1 - | 4,57E-05 - | 0 - | 0 - | 2 - | 0,000107954 - | 1 - | 5,48E-05 - | 0 - | 0 - |
| 6309 | 1 | 3um_FIJJAFBF_143870 | Bacteria | Proteobacteri | Gammaprotec | Oceanospirilla | Saccharospirill | Reinekea | Reinekea fors | unclassified | unclassified | unclassified | unclassified | unclassified | 1,6 | 0,8 | 5,30E-05 | 2,69E-05 | 2 - | 5,74E-05 - | 2 - | 7,13E-05 - | 2 - | 6,77E-05 - | 2 - | 6,87E-05 - | 0 - | 0 - |
| 6310 | 1 | 3um_FIJJAFBF_36534 | Bacteria | Bacteroidetes | Flavobacteriia | Flavobacterial | Flavobacteriac | Flavobacteriu | Flavobacteriu | cellular proces | Cell cycle cont | 05CAK | Cell division A | 376686.Fjoh_ | 0,2 | 0,4 | 1,26E-05 | 2,51E-05 | 0 - | 0 - | 1 - | 6,28E-05 - | 0 - | 0 - | 0 - | 0 - | 0 - | 0 - |
| 6311 | 1 | 0.2um_NOKCPHJN_258777 | Bacteria | Bacteroidetes | unclassified | unclassified | unclassified | unclassified | unidentified e | unclassified | unclassified | unclassified | unclassified | unclassified | 2 | 0 | 4,35E-05 | 4,13E-06 | 2 - | 3,64E-05 - | 2 - | 4,52E-05 - | 2 - | 4,30E-05 - | 2 - | 4,36E-05 - | 2 - | 4,92E-05 - |
| 6312 | 1 | 0.2um_NOKCPHJN_23889 | Bacteria | Proteobacteri | Gammaprotec | Alteromonada | Alteromonada | Paraglaciecola | Paraglaciecola | unclassified | unclassified | unclassified | unclassified | unclassified | 0,4 | 0,489897949 | 1,04E-05 | 1,28E-05 | 1 - | 2,31E-05 - | 1 - | 2,87E-05 - | 0 - | 0 - | 0 - | 0 - | 0 - | 0 - |
| 6313 | 1 | 3um_FIJJAFBF_707228 | Bacteria | Proteobacteri | Gammaprotec | Oceanospirilla | Saccharospirill | Reinekea | Reinekea blan | metabolism | Energy produc | 0QID8 | fumarate hyd | 314283.MED2 | 0,6 | 0,489897949 | 1,64E-05 | 1,36E-05 | 1 - | 2,28E-05 - | 1 - | 2,83E-05 - | 0 - | 0 - | 0 - | 0 - | 1 - | 3,07E-05 - |
| 6314 | 1 | 0.2um_NOKCPHJN_240799 | Bacteria | Proteobacteri | Alphaproteob | Rhodobactera | Rhodobactera | unclassified | Rhodobactera | metabolism | Secondary me | 01Q86 | extracellular s | 388401.RB21 | 0,6 | 0,489897949 | 2,11E-05 | 1,74E-05 | 1 - | 3,13E-05 - | 0 - | 0 - | 1 - | 3,69E-05 - | 1 - | 3,75E-05 - | 0 - | 0 - |
| 6315 | 1 | 3um_FIJJAFBF_130520 | Bacteria | Bacteroidetes | Flavobacteriia | Flavobacterial | Flavobacteriac | Psychroserpe | Psychroserpen | unclassified | unclassified | unclassified | unclassified | unclassified | 0,8 | 0,748331477 | 1,92E-05 | 1,67E-05 | 2 - | 4,22E-05 - | 0 - | 0 - | 0 - | 0 - | 1 - | 2,53E-05 - | 1 - | 2,85E-05 - |
| 6316 | 1 | 0.2um_NOKCPHJN_744823 | Bacteria | Proteobacteri | Alphaproteob | Pelagibacteral | Pelagibacterac | unclassified | alpha proteob | unclassified | unclassified | unclassified | unclassified | unclassified | 0,6 | 0,489897949 | 1,89E-05 | 1,56E-05 | 1 - | 2,75E-05 - | 1 - | 3,42E-05 - | 0 - | 0 - | 1 - | 3,30E-05 - | 0 - | 0 - |
| 6317 | 1 | 3um_FIJJAFBF_740062 | Bacteria | Bacteroidetes | Flavobacteriia | Flavobacterial | Flavobacteriac | Polaribacter | Polaribacter s | cellular proces | Cell wall/mem | 0P7KR | tyrosine-prote | 313598.MED1 | 1,4 | 0,489897949 | 2,38E-05 | 8,60E-06 | 2 - | 2,86E-05 - | 1 - | 1,77E-05 - | 1 - | 1,69E-05 - | 1 - | 1,71E-05 - | 2 - | 3,85E-05 - |
| 6318 | 1 | 3um_FIJJAFBF_80306 | Bacteria | Bacteroidetes | Flavobacteriia | unclassified | unclassified | unclassified | Flavobacteriia | unclassified | unclassified | unclassified | unclassified | unclassified | 0,4 | 0,489897949 | 3,68E-05 | 4,55E-05 | 1 - | 8,20E-05 - | 1 - | 0,000101799 - | 0 - | 0 - | 0 - | 0 - | 0 - | 0 - |
| 6319 | 1 | 3um_FIJJAFBF_477756 | Bacteria | Nitrospirae | unclassified | unclassified | unclassified | unclassified | Nitrospirae ba | unclassified | unclassified | unclassified | unclassified | unclassified | 0,4 | 0,489897949 | 1,03E-05 | 1,27E-05 | 0 - | 0 - | 0 - | 0 - | 0 - | 0 - | 1 - | 2,43E-05 - | 1 - | 2,74E-05 - |
| 6320 | 1 | 3um_FIJJAFBF_194532 | Bacteria | Proteobacteri | Alphaproteob | Rhodobactera | Rhodobactera | Sulfitobacter | Sulfitobacter s | metabolism | Nucleotide tra | 16Q4Q | Catalyzes the | 391595.RLO14 | 2 | 0,632455532 | 5,58E-05 | 1,46E-05 | 3 - | 7,13E-05 - | 2 - | 5,90E-05 - | 2 - | 5,61E-05 - | 1 - | 2,85E-05 - | 2 - | 6,41E-05 - |
| 6321 | 1 | 3um_FIJJAFBF_348503 | Bacteria | Bacteroidetes | Flavobacteriia | Flavobacterial | Crocinitomica | unclassified | Crocinitomica | metabolism | Energy produc | 0P7S4 | Produces ATP | 755732.Fluta_ | 0,6 | 0,489897949 | 2,69E-05 | 2,22E-05 | 1 - | 3,92E-05 - | 1 - | 4,86E-05 - | 0 - | 0 - | 1 - | 4,69E-05 - | 0 - | 0 - |
| 6322 | 1 | 0.2um_NOKCPHJN_186158 | Bacteria | Proteobacteri | Alphaproteob | Rhodobactera | Rhodobactera | Sulfitobacter | various | unclassified | unclassified | unclassified | unclassified | unclassified | 0,8 | 0,4 | 6,24E-05 | 3,17E-05 | 1 - | 6,75E-05 - | 1 - | 8,38E-05 - | 1 - | 7,97E-05 - | 1 - | 8,09E-05 - | 0 - | 0 - |
| 6323 | 1 | 3um_FIJJAFBF_724165 | Bacteria | Bacteroidetes | Flavobacteriia | Flavobacterial | Flavobacteriac | Polaribacter | Polaribacter ir | metabolism | Inorganic ion t | 05C27 | TonB-depende | 313594.PI23P | 0,2 | 0,4 | 5,96E-06 | 1,19E-05 | 0 - | 0 - | 1 - | 2,98E-05 - | 0 - | 0 - | 0 - | 0 - | 0 - | 0 - |
| 6324 | 1 | 3um_FIJJAFBF_638938 | Bacteria | Proteobacteri | Alphaproteob | Rhodobactera | Rhodobactera | unclassified | Rhodobactera | unclassified | unclassified | unclassified | unclassified | unclassified | 10,4 | 0,489897949 | 0,000251991 | 2,79E-05 | 10 - | 0,000202786 - | 11 - | 0,00027698 - | 10 - | 0,000239368 - | 11 - | 0,000267194 - | 10 - | 0,00027363 - |
| 6325 | 1 | 0.2um_NOKCPHJN_75313 | Bacteria | Proteobacteri | Gammaprotec | Alteromonada | Alteromonada | Glaciecola | Glaciecola sp. | metabolism | Carbohydrate | 16QEQ | hydrolase, fan | 455436.GHTC | 7,2 | 0,979795897 | 0,000112364 | 2,22E-05 | 6 - | 7,79E-05 - | 6 - | 9,67E-05 - | 8 - | 0,000122608 - | 8 - | 0,000124419 - | 8 - | 0,000140158 - |
| 6326 | 1 | 3um_FIJJAFBF_437166 | Bacteria | Proteobacteri | Alphaproteob | Rhodobactera | Rhodobactera | Roseovarius | Roseovarius n | metabolism | Energy produc | 05CMV | Succinyl-CoA s | 89187.ISM_05 | 4,2 | 1,720465053 | 0,000143917 | 5,99E-05 | 3 - | 8,65E-05 - | 4 - | 0,000143235 - | 5 - | 0,000170204 - | 7 - | 0,000241805 - | 2 - | 7,78E-05 - |
| 6327 | 1 | 3um_FIJJAFBF_542441 | Bacteria | unclassified | unclassified | unclassified | unclassified | unclassified | bacterium B1 | unclassified | unclassified | unclassified | unclassified | unclassified | 3,8 | 0,4 | 9,52E-05 | 1,41E-05 | 4 - | 8,39E-05 - | 4 - | 0,000104219 - | 3 - | 7,43E-05 - | 4 - | 0,000100536 - | 4 - | 0,000113254 - |
| 6328 | 1 | 3um_FIJJAFBF_189680 | Bacteria | Bacteroidetes | Flavobacteriia | Flavobacterial | Flavobacteriac | Polaribacter | Polaribacter b | unclassified | unclassified | unclassified | unclassified | unclassified | 0,6 | 0,489897949 | 2,08E-05 | 1,72E-05 | 1 - | 3,04E-05 - | 1 - | 3,78E-05 - | 1 - | 3,59E-05 - | 0 - | 0 - | 0 - | 0 - |
| 6329 | 1 | 3um_FIJJAFBF_35958 | Bacteria | Bacteroidetes | Flavobacteriia | Flavobacterial | Flavobacteriac | Arenitalea | Arenitalea lut | unclassified | unclassified | unclassified | unclassified | unclassified | 2,2 | 0,748331477 | 7,83E-05 | 2,88E-05 | 1 - | 2,93E-05 - | 3 - | 0,000109071 - | 2 - | 6,91E-05 - | 3 - | 0,000105217 - | 2 - | 7,90E-05 - |
| 6330 | 1 | 0.2um_NOKCPHJN_263380 | Bacteria | Bacteroidetes | Flavobacteriia | Flavobacterial | Crocinitomica | unclassified | Crocinitomica | metabolism | Inorganic ion t | COG0841 | acriflavin resis | 388413.ALPR1 | 1,8 | 0,979795897 | 2,24E-05 | 1,22E-05 | 0 - | 0 - | 3 - | 3,72E-05 - | 2 - | 2,36E-05 - | 2 - | 2,40E-05 - | 2 - | 2,70E-05 - |
| 6331 | 1 | 0.2um_NOKCPHJN_308183 | Eukaryota | Chordata | Mammalia | Chiroptera | Vespertilionid | Miniopterus | Miniopterus n | unclassified | unclassified | unclassified | unclassified | unclassified | 0,8 | 0,4 | 8,96E-06 | 4,60E-06 | 1 - | 9,35E-06 - | 1 - | 1,16E-05 - | 0 - | 0 - | 1 - | 1,12E-05 - | 1 - | 1,26E-05 - |
| 6332 | 1 | 3um_FIJJAFBF_44906 | Viruses | unclassified | unclassified | unclassified | Phycodnavirid | unclassified | Chrysochrom | information st | Transcription | 12N88 | dna-directed r | 6334.EFV5348 | 0,2 | 0,4 | 1,70E-06 | 3,39E-06 | 0 - | 0 - | 0 - | 0 - | 0 - | 0 - | 1 - | 8,48E-06 - | 0 - | 0 - |
| 6333 | 1 | 0.2um_NOKCPHJN_724458 | Bacteria | Bacteroidetes | Flavobacteriia | unclassified | unclassified | unclassified | Flavobacteriia | unclassified | unclassified | unclassified | unclassified | unclassified | 0,6 | 0,489897949 | 3,13E-05 | 2,57E-05 | 1 - | 4,63E-05 - | 0 - | 0 - | 1 - | 5,46E-05 - | 1 - | 5,54E-05 - | 0 - | 0 - |
| 6334 | 1 | 3um_FIJJAFBF_479265 | Bacteria | Proteobacteri | Alphaproteob | Rhodobactera | Rhodobactera | Planktomarina | Planktomarina | metabolism | Amino acid tra | COG0133 | The beta subu | 391595.RLO14 | 1,2 | 0,4 | 4,04E-05 | 1,49E-05 | 1 - | 2,80E-05 - | 2 - | 6,95E-05 - | 1 - | 3,30E-05 - | 1 - | 3,35E-05 - | 1 - | 3,78E-05 - |
| 6335 | 1 | 0.2um_NOKCPHJN_1056570 | Bacteria | Proteobacteri | Deltaproteoba | Desulfobacter | Desulfobulbac | Desulfopila | Desulfopila ae | unclassified | unclassified | unclassified | unclassified | unclassified | 0,8 | 0,4 | 2,10E-05 | 1,06E-05 | 0 - | 0 - | 1 - | 2,63E-05 - | 1 - | 2,50E-05 - | 1 - | 2,54E-05 - | 1 - | 2,86E-05 - |
| 6336 | 1 | 3um_FIJJAFBF_430948 | Bacteria | Bacteroidetes | Flavobacteriia | Flavobacterial | Flavobacteriac | Ulvibacter | Ulvibacter lito | cellular proces | Cell motility | COG1360 | Ompa motb d | 983544.Lacal_ | 1 | 0 | 4,23E-05 | 4,02E-06 | 1 - | 3,54E-05 - | 1 - | 4,40E-05 - | 1 - | 4,18E-05 - | 1 - | 4,24E-05 - | 1 - | 4,78E-05 - |
| 6337 | 1 | 3um_FIJJAFBF_91100 | Bacteria | Proteobacteri | Gammaprotec | Oceanospirilla | Oceanospirilla | Amphritea | Amphritea jap | information st | Translation, ri | 0QSAC | Binds togethe | 247633.GP214 | 0,6 | 0,489897949 | 4,93E-05 | 4,07E-05 | 1 - | 7,17E-05 - | 1 - | 8,91E-05 - | 0 - | 0 - | 1 - | 8,59E-05 - | 0 - | 0 - |
| 6338 | 1 | 0.2um_NOKCPHJN_264227 | Viruses | unclassified | unclassified | unclassified | Phycodnavirid | Prasinovirus | various | unclassified | unclassified | unclassified | unclassified | unclassified | 0,4 | 0,489897949 | 1,23E-05 | 1,53E-05 | 1 - | 2,75E-05 - | 1 - | 3,42E-05 - | 0 - | 0 - | 0 - | 0 - | 0 - | 0 - |
| 6339 | 1 | 0.2um_NOKCPHJN_557639 | Bacteria | Proteobacteri | Alphaproteob | Rhodobactera | Rhodobactera | Planktomarina | Planktomarina | unclassified | unclassified | unclassified | unclassified | unclassified | 0,8 | 0,4 | 5,26E-05 | 2,70E-05 | 1 - | 5,49E-05 - | 1 - | 6,82E-05 - | 0 - | 0 - | 1 - | 6,58E-05 - | 1 - | 7,41E-05 - |
| 6340 | 1 | 3um_FIJJAFBF_361814 | Bacteria | Bacteroidetes | Flavobacteriia | Flavobacterial | Flavobacteriac | Polaribacter | various | unclassified | unclassified | unclassified | unclassified | unclassified | 0,8 | 0,4 | 5,93E-05 | 3,05E-05 | 1 - | 6,27E-05 - | 0 - | 0 - | 1 - | 7,40E-05 - | 1 - | 7,51E-05 - | 1 - | 8,46E-05 - |
| 6341 | 1 | 3um_FIJJAFBF_238100 | Bacteria | Bacteroidetes | Flavobacteriia | Flavobacterial | Flavobacteriac | Lutibacter | Lutibacter flav | poorly charact | Function unkn | 05DPC | Ragb susd dor | 313594.PI23P | 4,4 | 0,8 | 0,000121998 | 2,31E-05 | 4 - | 9,33E-05 - | 5 - | 0,000144836 - | 5 - | 0,000137685 - | 5 - | 0,000139719 - | 3 - | 9,44E-05 - |
| 6342 | 1 | 0.2um_NOKCPHJN_536668 | Bacteria | Proteobacteri | Gammaprotec | unclassified | unclassified | unclassified | Gammaprotec | unclassified | unclassified | unclassified | unclassified | unclassified | 3,4 | 0,8 | 0,000247925 | 5,05E-05 | 4 - | 0,000248166 - | 4 - | 0,000308149 - | 4 - | 0,000292935 - | 3 - | 0,000222946 - | 2 - | 0,000167432 - |
| 6343 | 1 | 3um_FIJJAFBF_50637 | Bacteria | Bacteroidetes | Flavobacteriia | Flavobacterial | Flavobacteriac | Formosa | Formosa sp. H | poorly charact | Function unkn | 0XNQI | domain protei | 1046627.BZAR | 0,2 | 0,4 | 6,12E-06 | 1,22E-05 | 0 - | 0 - | 1 - | 3,06E-05 - | 0 - | 0 - | 0 - | 0 - | 0 - | 0 - |
| 6344 | 1 | 0.2um_NOKCPHJN_786900 | unclassified | unclassified | unclassified | unclassified | unclassified | unclassified | unclassified | unclassified | unclassified | unclassified | unclassified | unclassified | 1 | 0 | 1,64E-05 | 1,55E-06 | 1 - | 1,37E-05 - | 1 - | 1,70E-05 - | 1 - | 1,62E-05 - | 1 - | 1,64E-05 - | 1 - | 1,85E-05 - |
| 6345 | 1 | 3um_FIJJAFBF_53316 | Bacteria | Bacteroidetes | Flavobacteriia | Flavobacterial | Flavobacteriac | Polaribacter | Polaribacter s | poorly charact | Function unkn | 0P7WW | NA | 313598.MED1 | 1 | 0,632455532 | 4,16E-05 | 2,74E-05 | 1 - | 3,46E-05 - | 2 - | 8,59E-05 - | 1 - | 4,08E-05 - | 0 - | 0 - | 1 - | 4,66E-05 - |
| 6346 | 1 | 0.2um_NOKCPHJN_341331 | Bacteria | Proteobacteri | Gammaprotec | Oceanospirilla | Saccharospirill | Reinekea | Reinekea fors | cellular proces | Posttranslatio | 179VG | alkyl hydroper | 292414.TM10 | 2,6 | 0,489897949 | 0,000199484 | 4,06E-05 | 3 - | 0,000193444 - | 2 - | 0,000160134 - | 3 - | 0,000228341 - | 2 - | 0,000154476 - | 3 - | 0,000261024 - |
| 6347 | 1 | 3um_FIJJAFBF_609699 | Bacteria | Bacteroidetes | Flavobacteriia | Flavobacterial | Flavobacteriac | Maribacter | Maribacter pc | cellular proces | Posttranslatio | 05C2Z | ATP-depender | 688270.Celal_ | 0,8 | 0,4 | 1,25E-05 | 6,33E-06 | 1 - | 1,35E-05 - | 1 - | 1,67E-05 - | 1 - | 1,59E-05 - | 1 - | 1,62E-05 - | 0 - | 0 - |
| 6348 | 1 | 0.2um_NOKCPHJN_27376 | Bacteria | Bacteroidetes | Flavobacteriia | Flavobacterial | Flavobacteriac | Flavobacteriu | Flavobacteriu | unclassified | unclassified | unclassified | unclassified | unclassified | 2,2 | 1,166190379 | 1,82E-05 | 9,86E-06 | 2 - | 1,39E-05 - | 3 - | 2,60E-05 - | 1 - | 8,23E-06 - | 4 - | 3,34E-05 - | 1 - | 9,41E-06 - |
| 6349 | 1 | 0.2um_NOKCPHJN_64220 | Bacteria | Proteobacteri | Alphaproteob | Rhizobiales | unclassified | unclassified | Rhizobiales ba | metabolism | Carbohydrate | 01RFA | ABC transport | 367336.OM22 | 1,2 | 0,4 | 4,69E-05 | 1,04E-05 | 2 - | 6,73E-05 - | 1 - | 4,18E-05 - | 1 - | 3,97E-05 - | 1 - | 4,03E-05 - | 1 - | 4,54E-05 - |
| 6350 | 1 | 0.2um_NOKCPHJN_233934 | Eukaryota | Phaeophyceae | unclassified | Ectocarpales | Ectocarpaceae | Ectocarpus | Ectocarpus sil | unclassified | unclassified | unclassified | unclassified | unclassified | 0,8 | 0,4 | 2,53E-05 | 1,30E-05 | 1 - | 2,64E-05 - | 1 - | 3,28E-05 - | 0 - | 0 - | 1 - | 3,17E-05 - | 1 - | 3,57E-05 - |
| 6351 | 1 | 3um_FIJJAFBF_411342 | Bacteria | Bacteroidetes | unclassified | unclassified | unclassified | unclassified | Bacteroidetes | cellular proces | Posttranslatio | 09GE9 | Thioredoxin | 391603.FBAL0 | 1,4 | 0,489897949 | 0,000178552 | 5,25E-05 | 2 - | 0,000218622 - | 1 - | 0,000135732 - | 1 - | 0,000129031 - | 2 - | 0,000261873 - | 1 - | 0,0001475 - |
| 6352 | 1 | 3um_FIJJAFBF_681129 | Bacteria | Bacteroidetes | Flavobacteriia | Flavobacterial | Flavobacteriac | Polaribacter | Polaribacter s | unclassified | unclassified | unclassified | unclassified | unclassified | 2 | 0,894427191 | 6,71E-05 | 2,65E-05 | 3 - | 8,70E-05 - | 2 - | 7,20E-05 - | 3 - | 0,000102638 - | 1 - | 3,47E-05 - | 1 - | 3,91E-05 - |
| 6353 | 1 | 0.2um_NOKCPHJN_513747 | Bacteria | Proteobacteri | Alphaproteob | Pelagibacteral | Pelagibacterac | Candidatus Pe | Candidatus Pe | poorly charact | Function unkn | COG3181 | extra-cytoplas | 367336.OM22 | 0,6 | 0,489897949 | 2,74E-05 | 2,25E-05 | 0 - | 0 - | 1 - | 4,51E-05 - | 1 - | 4,29E-05 - | 0 - | 0 - | 1 - | 4,90E-05 - |
| 6354 | 1 | 3um_FIJJAFBF_110528 | Eukaryota | unclassified | Cryptophyta | Pyrenomonad | Pyrenomonad | Rhodomonas | Rhodomonas s | cellular proces | Posttranslatio | KOG0730 | cell division cy | 45157.CMT17 | 1 | 0 | 4,66E-05 | 4,43E-06 | 1 - | 3,90E-05 - | 1 - | 4,85E-05 - | 1 - | 4,61E-05 - | 1 - | 4,68E-05 - | 1 - | 5,27E-05 - |
| 6355 | 1 | 3um_FIJJAFBF_478437 | Bacteria | Bacteroidetes | Flavobacteriia | Flavobacterial | Flavobacteriac | Formosa | Formosa sp. H | unclassified | unclassified | unclassified | unclassified | unclassified | 1 | 0 | 4,34E-05 | 4,12E-06 | 1 - | 3,63E-05 - | 1 - | 4,51E-05 - | 1 - | 4,29E-05 - | 1 - | 4,35E-05 - | 1 - | 4,90E-05 - |
| 6356 | 2 | 0.2um_NOKCPHJN_284335;3um_FIJJAFBF_5000 | Bacteria | Proteobacteri | Alphaproteob | Rhodobactera | Rhodobactera | various | various | unclassified | unclassified | unclassified | unclassified | unclassified | 0,6 | 0,489897949 | 6,75E-05 | 5,57E-05 | 1 - | 9,81E-05 - | 1 - | 0,000121811 - | 0 - | 0 - | 1 - | 0,000117507 - | 0 - | 0 - |
| 6357 | 1 | 0.2um_NOKCPHJN_1035876 | Bacteria | Bacteroidetes | Flavobacteriia | Flavobacterial | Flavobacteriac | Ulvibacter | Ulvibacter sp. | unclassified | unclassified | unclassified | unclassified | unclassified | 2 | 0,894427191 | 4,82E-05 | 1,99E-05 | 3 - | 6,17E-05 - | 1 - | 2,55E-05 - | 1 - | 2,43E-05 - | 3 - | 7,39E-05 - | 2 - | 5,55E-05 - |
| 6358 | 1 | 3um_FIJJAFBF_257406 | Bacteria | Bacteroidetes | Flavobacteriia | Flavobacterial | Flavobacteriac | Algibacter | Algibacter sp. | cellular proces | Intracellular tr | 08SHB | Biopolymer tr | 411154.GFO_ | 3,2 | 0,4 | 0,000274707 | 1,51E-05 | 4 - | 0,000290574 - | 3 - | 0,000270605 - | 3 - | 0,000257245 - | 3 - | 0,000261044 - | 3 - | 0,000294065 - |
| 6359 | 1 | 3um_FIJJAFBF_726456 | Bacteria | Proteobacteri | Alphaproteob | Rhodobactera | Rhodobactera | unclassified | Rhodobactera | cellular proces | Posttranslatio | COG0330 | Band 7 protei | 314262.MED1 | 0,2 | 0,4 | 7,76E-06 | 1,55E-05 | 1 - | 3,88E-05 - | 0 - | 0 - | 0 - | 0 - | 0 - | 0 - | 0 - | 0 - |
| 6360 | 1 | 3um_FIJJAFBF_220489 | Bacteria | Proteobacteri | Gammaprotec | Alteromonada | Alteromonada | Paraglaciecola | Paraglaciecola | information st | Translation, ri | COG0088 | One of the pri | 983545.Glaag | 4,6 | 0,489897949 | 0,000313003 | 4,24E-05 | 5 - | 0,000285514 - | 4 - | 0,00028362 - | 5 - | 0,00033702 - | 4 - | 0,000273599 - | 5 - | 0,00038526 - |
| 6361 | 1 | 3um_FIJJAFBF_452453 | Bacteria | Proteobacteri | Alphaproteob | Rhodobactera | Rhodobactera | Planktomarina | Planktomarina | cellular proces | Posttranslatio | COG0652 | peptidyl-proly | 89187.ISM_04 | 0,8 | 0,748331477 | 5,52E-05 | 4,88E-05 | 2 - | 0,000126128 - | 0 - | 0 - | 1 - | 7,44E-05 - | 1 - | 7,55E-05 - | 0 - | 0 - |
| 6362 | 1 | 3um_FIJJAFBF_811879 | Bacteria | Proteobacteri | Gammaprotec | Oceanospirilla | Saccharospirill | Reinekea | Reinekea fors | unclassified | unclassified | unclassified | unclassified | unclassified | 2 | 0,632455532 | 7,01E-05 | 2,24E-05 | 2 - | 5,90E-05 - | 1 - | 3,66E-05 - | 3 - | 0,000104485 - | 2 - | 7,07E-05 - | 2 - | 7,96E-05 - |
| 6363 | 1 | 3um_FIJJAFBF_81227 | Bacteria | Proteobacteri | Gammaprotec | Cellvibrionale | Cellvibrionace | Marinagarivor | Marinagarivor | information st | Translation, ri | 177XJ | One of the pri | 377629.TERTU | 1,4 | 0,8 | 0,000118338 | 6,78E-05 | 2 - | 0,00014715 - | 2 - | 0,000182716 - | 2 - | 0,000173695 - | 1 - | 8,81E-05 - | 0 - | 0 - |
| 6364 | 1 | 3um_FIJJAFBF_105462 | Bacteria | unclassified | unclassified | unclassified | unclassified | unclassified | bacterium TM | unclassified | unclassified | unclassified | unclassified | unclassified | 2,4 | 0,8 | 6,75E-05 | 2,00E-05 | 3 - | 7,25E-05 - | 2 - | 6,00E-05 - | 3 - | 8,56E-05 - | 3 - | 8,68E-05 - | 1 - | 3,26E-05 - |
| 6365 | 1 | 3um_FIJJAFBF_427380 | Bacteria | Bacteroidetes | Cytophagia | Cytophagales | Flammeovirga | unclassified | Flammeovirga | unclassified | unclassified | unclassified | unclassified | unclassified | 9,8 | 1,720465053 | 0,000188668 | 3,04E-05 | 12 - | 0,000194812 - | 9 - | 0,000181424 - | 7 - | 0,000134141 - | 11 - | 0,000213906 - | 10 - | 0,000219059 - |
| 6366 | 1 | 3um_FIJJAFBF_810935 | Bacteria | Proteobacteri | Gammaprotec | unclassified | unclassified | unclassified | Gammaprotec | cellular proces | Intracellular tr | 0QITE | Outer membr | 384676.PSEEN | 1 | 0,894427191 | 2,92E-05 | 2,72E-05 | 1 - | 2,50E-05 - | 2 - | 6,21E-05 - | 2 - | 5,90E-05 - | 0 - | 0 - | 0 - | 0 - |
| 6367 | 1 | 3um_FIJJAFBF_688496 | Bacteria | Proteobacteri | Betaproteoba | Burkholderial | Comamonada | Rhodoferax | Rhodoferax sa | cellular proces | Posttranslatio | 0BDJ9 | Glutaredoxin | 595537.Varpa | 0,2 | 0,4 | 2,55E-05 | 5,09E-05 | 0 - | 0 - | 0 - | 0 - | 0 - | 0 - | 1 - | 0,000127299 - | 0 - | 0 - |
| 6368 | 1 | 0.2um_NOKCPHJN_194011 | Bacteria | Bacteroidetes | Flavobacteriia | Flavobacterial | Flavobacteriac | Flavobacteriu | Flavobacteriu | unclassified | unclassified | unclassified | unclassified | unclassified | 0,4 | 0,489897949 | 1,96E-05 | 2,42E-05 | 1 - | 4,50E-05 - | 0 - | 0 - | 1 - | 5,31E-05 - | 0 - | 0 - | 0 - | 0 - |
| 6369 | 1 | 3um_FIJJAFBF_573440 | Bacteria | Bacteroidetes | Flavobacteriia | Flavobacterial | Flavobacteriac | Polaribacter | Polaribacter s | cellular proces | Posttranslatio | 09FEV | heat shock pr | 313594.PI23P | 0,4 | 0,489897949 | 3,48E-05 | 4,30E-05 | 1 - | 7,76E-05 - | 1 - | 9,63E-05 - | 0 - | 0 - | 0 - | 0 - | 0 - | 0 - |
| 6370 | 1 | 3um_FIJJAFBF_346742 | Bacteria | Bacteroidetes | Flavobacteriia | various | various | various | various | cellular proces | Intracellular tr | 08SHB | Biopolymer tr | 487797.Flav3 | 4,6 | 0,489897949 | 0,000394503 | 5,71E-05 | 4 - | 0,000286942 - | 5 - | 0,000445371 - | 5 - | 0,000423382 - | 5 - | 0,000429635 - | 4 - | 0,000387186 - |
| 6371 | 1 | 0.2um_NOKCPHJN_60625 | Bacteria | Bacteroidetes | Flavobacteriia | Flavobacterial | Flavobacteriac | Polaribacter | Polaribacter s | cellular proces | Cell wall/mem | COG2885 | Ompa motb d | 313598.MED1 | 1 | 0 | 3,03E-05 | 2,88E-06 | 1 - | 2,54E-05 - | 1 - | 3,15E-05 - | 1 - | 3,00E-05 - | 1 - | 3,04E-05 - | 1 - | 3,43E-05 - |
| 6372 | 1 | 3um_FIJJAFBF_763603 | Bacteria | Actinobacteri | Actinobacteri | Micrococcales | Microbacteria | unclassified | Microbacteria | unclassified | unclassified | unclassified | unclassified | unclassified | 0,8 | 0,4 | 9,82E-05 | 4,99E-05 | 1 - | 0,000106275 - | 1 - | 0,000131962 - | 1 - | 0,000125447 - | 1 - | 0,000127299 - | 0 - | 0 - |
| 6373 | 1 | 0.2um_NOKCPHJN_556208 | Bacteria | Proteobacteri | Alphaproteob | Rhodobactera | Rhodobactera | Planktomarina | Planktomarina | unclassified | unclassified | unclassified | unclassified | unclassified | 0,6 | 1,2 | 3,62E-05 | 7,25E-05 | 3 - | 0,000181226 - | 0 - | 0 - | 0 - | 0 - | 0 - | 0 - | 0 - | 0 - |
| 6374 | 1 | 3um_FIJJAFBF_647164 | Bacteria | Bacteroidetes | Flavobacteriia | Flavobacterial | Crocinitomica | unclassified | Crocinitomica | unclassified | unclassified | unclassified | unclassified | unclassified | 0,2 | 0,4 | 1,37E-05 | 2,75E-05 | 1 - | 6,87E-05 - | 0 - | 0 - | 0 - | 0 - | 0 - | 0 - | 0 - | 0 - |
| 6375 | 1 | 3um_FIJJAFBF_191787 | Eukaryota | unclassified | Dictyochophy | Pedinellales | unclassified | Apedinella | Apedinella rac | cellular proces | Posttranslatio | KOG0730 | cell division cy | 45157.CMV01 | 8 | 1,897366596 | 0,000359891 | 7,72E-05 | 11 - | 0,000418061 - | 8 - | 0,000377533 - | 6 - | 0,00026917 - | 6 - | 0,000273146 - | 9 - | 0,000461546 - |
| 6376 | 1 | 0.2um_NOKCPHJN_65138 | Eukaryota | unclassified | Oomycetes | Peronosporale | Peronosporac | Phytophthora | Phytophthora | poorly charact | Function unkn | 0XNUF | Extracellular n | 340101.Vvad_ | 0,2 | 0,4 | 1,41E-07 | 2,81E-07 | 0 - | 0 - | 0 - | 0 - | 0 - | 0 - | 0 - | 0 - | 1 - | 7,03E-07 - |
| 6377 | 1 | 3um_FIJJAFBF_402482 | Bacteria | Bacteroidetes | Flavobacteriia | unclassified | unclassified | unclassified | Flavobacteriia | unclassified | unclassified | unclassified | unclassified | unclassified | 3,6 | 0,8 | 0,00020399 | 3,09E-05 | 5 - | 0,000241128 - | 4 - | 0,000239528 - | 3 - | 0,000170776 - | 3 - | 0,000173298 - | 3 - | 0,00019522 - |
| 6378 | 1 | 0.2um_NOKCPHJN_432271 | Viruses | unclassified | unclassified | unclassified | Phycodnavirid | unclassified | Pyramimonas | unclassified | unclassified | unclassified | unclassified | unclassified | 0,8 | 0,748331477 | 3,85E-05 | 3,91E-05 | 1 - | 3,94E-05 - | 0 - | 0 - | 1 - | 4,66E-05 - | 0 - | 0 - | 2 - | 0,000106443 - |
| 6379 | 1 | 0.2um_NOKCPHJN_213552 | Bacteria | Proteobacteri | Alphaproteob | Pelagibacteral | Pelagibacterac | Candidatus Pe | various | metabolism | Amino acid tra | 01VSB | Ectoine hydro | 439493.PB721 | 0,4 | 0,489897949 | 1,78E-05 | 2,19E-05 | 1 - | 4,08E-05 - | 0 - | 0 - | 1 - | 4,82E-05 - | 0 - | 0 - | 0 - | 0 - |
| 6380 | 1 | 0.2um_NOKCPHJN_514943 | Bacteria | Proteobacteri | Alphaproteob | Rhodobactera | Rhodobactera | Pseudorueger | Pseudorueger | unclassified | unclassified | unclassified | unclassified | unclassified | 3,8 | 1,6 | 4,87E-05 | 2,07E-05 | 4 - | 4,34E-05 - | 5 - | 6,73E-05 - | 2 - | 2,56E-05 - | 6 - | 7,79E-05 - | 2 - | 2,92E-05 - |
| 6381 | 2 | 0.2um_NOKCPHJN_1045583;3um_FIJJAFBF_3692 | Eukaryota | various | various | various | various | various | various | cellular proces | Intracellular tr | KOG0096 | GTP-binding n | various | 1 | 0 | 6,40E-05 | 6,08E-06 | 1 - | 5,36E-05 - | 1 - | 6,66E-05 - | 1 - | 6,33E-05 - | 1 - | 6,42E-05 - | 1 - | 7,24E-05 - |
| 6382 | 1 | 0.2um_NOKCPHJN_159982 | Bacteria | Proteobacteri | Gammaprotec | unclassified | unclassified | unclassified | gamma proteo | unclassified | unclassified | unclassified | unclassified | unclassified | 5 | 1,264911064 | 0,000108253 | 1,84E-05 | 7 - | 0,000129587 - | 4 - | 9,19E-05 - | 6 - | 0,000131112 - | 4 - | 8,87E-05 - | 4 - | 9,99E-05 - |
| 6383 | 1 | 3um_FIJJAFBF_220782 | Bacteria | Actinobacteri | Actinobacteri | Micrococcales | Microbacteria | unclassified | Microbacteria | metabolism | Amino acid tra | COG0141 | Catalyzes the | 525264.HMPR | 0,2 | 0,4 | 7,54E-06 | 1,51E-05 | 0 - | 0 - | 1 - | 3,77E-05 - | 0 - | 0 - | 0 - | 0 - | 0 - | 0 - |
| 6384 | 1 | 3um_FIJJAFBF_33738 | Bacteria | Proteobacteri | Alphaproteob | Rhodobactera | Rhodobactera | Planktomarina | Planktomarina | unclassified | unclassified | unclassified | unclassified | unclassified | 0,6 | 0,489897949 | 0,000113093 | 9,40E-05 | 1 - | 0,000159412 - | 0 - | 0 - | 0 - | 0 - | 1 - | 0,000190949 - | 1 - | 0,000215103 - |
| 6385 | 1 | 0.2um_NOKCPHJN_06876 | Bacteria | Bacteroidetes | Cytophagia | Cytophagales | Cytophagacea | Ekhidna | Ekhidna lutea | information st | Transcription | 09AJX | DNA-depende | 643867.Ftrac_ | 20,4 | 2,332380758 | 0,000215202 | 1,93E-05 | 24 - | 0,000213538 - | 21 - | 0,000232007 - | 17 - | 0,000178542 - | 21 - | 0,00022381 - | 19 - | 0,00022811 - |
| 6386 | 1 | 3um_FIJJAFBF_735890 | Eukaryota | Bacillariophyta | Coscinodiscop | Thalassiosiral | Thalassiosirac | Thalassiosira | Thalassiosira p | unclassified | unclassified | unclassified | unclassified | unclassified | 0,6 | 0,489897949 | 4,57E-05 | 3,80E-05 | 1 - | 6,45E-05 - | 0 - | 0 - | 0 - | 0 - | 1 - | 7,72E-05 - | 1 - | 8,70E-05 - |
| 6387 | 1 | 0.2um_NOKCPHJN_677898 | Bacteria | Bacteroidetes | unclassified | unclassified | unclassified | unclassified | unidentified e | unclassified | unclassified | unclassified | unclassified | unclassified | 0,2 | 0,4 | 6,43E-06 | 1,29E-05 | 0 - | 0 - | 1 - | 3,22E-05 - | 0 - | 0 - | 0 - | 0 - | 0 - | 0 - |
| 6388 | 1 | 3um_FIJJAFBF_44064 | Viruses | unclassified | unclassified | unclassified | various | various | various | unclassified | unclassified | unclassified | unclassified | unclassified | 1,2 | 0,748331477 | 1,35E-05 | 8,06E-06 | 2 - | 1,98E-05 - | 1 - | 1,23E-05 - | 2 - | 2,34E-05 - | 1 - | 1,19E-05 - | 0 - | 0 - |
| 6389 | 1 | 0.2um_NOKCPHJN_586903 | Bacteria | Proteobacteri | Betaproteoba | Burkholderial | Comamonada | Limnohabitan | Limnohabitan | cellular proces | Posttranslatio | 16QU3 | Hemolysin-typ | 744980.TRICH | 3,2 | 0,4 | 3,94E-05 | 6,10E-06 | 3 - | 3,09E-05 - | 3 - | 3,84E-05 - | 3 - | 3,65E-05 - | 4 - | 4,94E-05 - | 3 - | 4,17E-05 - |
| 6390 | 1 | 3um_FIJJAFBF_356795 | Bacteria | Proteobacteri | Gammaprotec | Cellvibrionale | Spongiibacter | Oceanicoccus | Oceanicoccus | cellular proces | Posttranslatio | 0QJ3C | Acts as a chap | 207949.RED65 | 0,6 | 0,489897949 | 1,35E-05 | 1,11E-05 | 0 - | 0 - | 1 - | 2,22E-05 - | 1 - | 2,11E-05 - | 0 - | 0 - | 1 - | 2,41E-05 - |
| 6391 | 1 | 3um_FIJJAFBF_534779 | Bacteria | Proteobacteri | Gammaprotec | Cellvibrionale | Porticoccacea | unclassified | Gammaprotec | information st | Translation, ri | COG0256 | This is one of | 314287.GB220 | 0,2 | 0,4 | 1,98E-05 | 3,96E-05 | 1 - | 9,89E-05 - | 0 - | 0 - | 0 - | 0 - | 0 - | 0 - | 0 - | 0 - |
| 6392 | 1 | 0.2um_NOKCPHJN_241962 | Bacteria | Bacteroidetes | Flavobacteriia | Flavobacterial | Flavobacteriac | Flaviramulus | Flaviramulus t | information st | Translation, ri | 05CAV | thus facilitatin | 983544.Lacal_ | 3,6 | 1,019803903 | 7,80E-05 | 1,71E-05 | 5 - | 9,27E-05 - | 2 - | 4,60E-05 - | 4 - | 8,75E-05 - | 4 - | 8,88E-05 - | 3 - | 7,51E-05 - |
| 6393 | 1 | 0.2um_NOKCPHJN_867613 | Bacteria | Proteobacteri | Alphaproteob | Pelagibacteral | Pelagibacterac | Candidatus Pe | Candidatus Pe | cellular proces | Posttranslatio | 01QWG | Prevents misfo | 439493.PB721 | 1,4 | 0,8 | 3,46E-05 | 2,01E-05 | 2 - | 4,15E-05 - | 1 - | 2,58E-05 - | 0 - | 0 - | 2 - | 4,97E-05 - | 2 - | 5,60E-05 - |
| 6394 | 1 | 3um_FIJJAFBF_715582 | Bacteria | Proteobacteri | Gammaprotec | Cellvibrionale | Halieaceae | unclassified | marine gamm | unclassified | unclassified | unclassified | unclassified | unclassified | 0,2 | 0,4 | 2,20E-05 | 4,41E-05 | 0 - | 0 - | 0 - | 0 - | 1 - | 0,000110148 - | 0 - | 0 - | 0 - | 0 - |
| 6395 | 1 | 3um_FIJJAFBF_544204 | Bacteria | Proteobacteri | Gammaprotec | Cellvibrionale | Porticoccacea | unclassified | Gammaprotec | unclassified | unclassified | unclassified | unclassified | unclassified | 1,2 | 0,748331477 | 5,03E-05 | 3,10E-05 | 2 - | 7,33E-05 - | 2 - | 9,11E-05 - | 1 - | 4,33E-05 - | 1 - | 4,39E-05 - | 0 - | 0 - |
| 6396 | 1 | 3um_FIJJAFBF_527628 | Bacteria | Proteobacteri | Gammaprotec | Oceanospirilla | Saccharospirill | Reinekea | Reinekea fors | poorly charact | Function unkn | 0SZFV | vi secretion sy | 690850.Desaf | 0,8 | 0,748331477 | 4,82E-05 | 4,62E-05 | 1 - | 5,24E-05 - | 1 - | 6,51E-05 - | 2 - | 0,000123728 - | 0 - | 0 - | 0 - | 0 - |
| 6397 | 1 | 3um_FIJJAFBF_332473 | Bacteria | Bacteroidetes | Flavobacteriia | Flavobacterial | Flavobacteriac | Polaribacter | various | unclassified | unclassified | unclassified | unclassified | unclassified | 1 | 0 | 0,000217503 | 2,07E-05 | 1 - | 0,000182185 - | 1 - | 0,00022622 - | 1 - | 0,000215051 - | 1 - | 0,000218227 - | 1 - | 0,000245833 - |
| 6398 | 1 | 0.2um_NOKCPHJN_171716 | Bacteria | Bacteroidetes | Flavobacteriia | Flavobacterial | Flavobacteriac | Formosa | Formosa sp. H | information st | Translation, ri | 05KK9 | This protein b | 313603.FB217 | 2,6 | 1,356465997 | 0,000188357 | 7,88E-05 | 5 - | 0,000318824 - | 2 - | 0,000158354 - | 3 - | 0,000225804 - | 2 - | 0,000152759 - | 1 - | 8,60E-05 - |
| 6399 | 1 | 3um_FIJJAFBF_217983 | Viruses | unclassified | unclassified | unclassified | unclassified | unclassified | uncultured M | unclassified | unclassified | unclassified | unclassified | unclassified | 1 | 0 | 7,83E-05 | 7,43E-06 | 1 - | 6,56E-05 - | 1 - | 8,14E-05 - | 1 - | 7,74E-05 - | 1 - | 7,86E-05 - | 1 - | 8,85E-05 - |
| 6400 | 1 | 0.2um_NOKCPHJN_134694 | Eukaryota | Chordata | Mammalia | Primates | Hominidae | Gorilla | Gorilla gorilla | cellular proces | Posttranslatio | 0JAJ8 | Dentin sialopl | 61853.ENSNLE | 0,2 | 0,4 | 7,63E-07 | 1,53E-06 | 1 - | 3,81E-06 - | 0 - | 0 - | 0 - | 0 - | 0 - | 0 - | 0 - | 0 - |
| 6401 | 1 | 3um_FIJJAFBF_147737 | Bacteria | Bacteroidetes | Flavobacteriia | Flavobacterial | Crocinitomica | unclassified | Crocinitomica | cellular proces | Posttranslatio | COG0330 | Band 7 protei | 216432.CA259 | 0,8 | 0,4 | 3,87E-05 | 1,97E-05 | 1 - | 4,19E-05 - | 1 - | 5,20E-05 - | 1 - | 4,94E-05 - | 1 - | 5,02E-05 - | 0 - | 0 - |
| 6402 | 1 | 3um_FIJJAFBF_571566 | Eukaryota | Chlorophyta | Chlorophyceae | Chlamydomor | Chlamydomor | Chlamydomor | Chlamydomor | cellular proces | Signal transdu | 09UWE | dual-specificit | 12957.ACEP_0 | 0,2 | 0,4 | 2,31E-06 | 4,62E-06 | 0 - | 0 - | 0 - | 0 - | 0 - | 0 - | 0 - | 0 - | 1 - | 1,15E-05 - |
| 6403 | 1 | 0.2um_NOKCPHJN_362249 | Bacteria | Proteobacteri | Alphaproteob | Pelagibacteral | Pelagibacterac | Candidatus Pe | Candidatus Pe | unclassified | unclassified | unclassified | unclassified | unclassified | 0,8 | 0,4 | 2,50E-05 | 1,27E-05 | 1 - | 2,70E-05 - | 1 - | 3,35E-05 - | 1 - | 3,19E-05 - | 1 - | 3,23E-05 - | 0 - | 0 - |
| 6404 | 1 | 0.2um_NOKCPHJN_450544 | Bacteria | Bacteroidetes | Flavobacteriia | unclassified | unclassified | unclassified | Flavobacteriia | cellular proces | Intracellular tr | 0BIE8 | domain-conta | 1005048.CFU_ | 0,8 | 0,748331477 | 1,16E-05 | 1,06E-05 | 0 - | 0 - | 0 - | 0 - | 2 - | 2,80E-05 - | 1 - | 1,42E-05 - | 1 - | 1,60E-05 - |
| 6405 | 1 | 3um_FIJJAFBF_369156 | unclassified | unclassified | unclassified | unclassified | unclassified | unclassified | unclassified | unclassified | unclassified | unclassified | unclassified | unclassified | 0,8 | 0,4 | 4,83E-05 | 2,43E-05 | 0 - | 0 - | 1 - | 6,04E-05 - | 1 - | 5,74E-05 - | 1 - | 5,83E-05 - | 1 - | 6,56E-05 - |
| 6406 | 1 | 0.2um_NOKCPHJN_960876 | Bacteria | Bacteroidetes | Flavobacteriia | Flavobacterial | Flavobacteriac | Formosa | Formosa sp. H | poorly charact | Function unkn | 05C2A | tonB-depende | 867900.Celly_ | 0,2 | 0,4 | 4,42E-06 | 8,84E-06 | 0 - | 0 - | 0 - | 0 - | 0 - | 0 - | 1 - | 2,21E-05 - | 0 - | 0 - |
| 6407 | 1 | 0.2um_NOKCPHJN_1052751 | Bacteria | Proteobacteri | Alphaproteob | Pelagibacteral | Pelagibacterac | Candidatus Pe | Candidatus Pe | metabolism | Secondary me | 0837G | toluene tolera | 335992.SAR11 | 1 | 0 | 7,33E-05 | 6,96E-06 | 1 - | 6,14E-05 - | 1 - | 7,62E-05 - | 1 - | 7,25E-05 - | 1 - | 7,35E-05 - | 1 - | 8,28E-05 - |
| 6408 | 1 | 3um_FIJJAFBF_31776 | Bacteria | Proteobacteri | Betaproteoba | Burkholderial | Comamonada | Limnohabitan | various | poorly charact | Function unkn | 05E4M | (twin-arginine | 1007105.PT7_ | 2 | 1,095445115 | 4,04E-05 | 1,78E-05 | 4 - | 7,10E-05 - | 2 - | 4,41E-05 - | 2 - | 4,19E-05 - | 1 - | 2,12E-05 - | 1 - | 2,39E-05 - |
| 6409 | 1 | 3um_FIJJAFBF_330466 | unclassified | unclassified | unclassified | unclassified | unclassified | unclassified | unclassified | unclassified | unclassified | unclassified | unclassified | unclassified | 1,6 | 0,489897949 | 0,000282336 | 7,40E-05 | 2 - | 0,000302044 - | 1 - | 0,000187525 - | 2 - | 0,000356532 - | 2 - | 0,000361798 - | 1 - | 0,000203782 - |
| 6410 | 1 | 3um_FIJJAFBF_758943 | Bacteria | Bacteroidetes | Flavobacteriia | Flavobacterial | Crocinitomica | unclassified | Crocinitomica | information st | Translation, ri | 05CEJ | Catalyzes the | 755732.Fluta_ | 2 | 0,632455532 | 3,94E-05 | 1,37E-05 | 2 - | 3,28E-05 - | 3 - | 6,12E-05 - | 1 - | 1,94E-05 - | 2 - | 3,93E-05 - | 2 - | 4,43E-05 - |
| 6411 | 1 | 0.2um_NOKCPHJN_86271 | Bacteria | Proteobacteri | Alphaproteob | Rhodobactera | Rhodobactera | Roseovarius | Roseovarius s | unclassified | unclassified | unclassified | unclassified | unclassified | 0,8 | 0,748331477 | 2,17E-05 | 1,96E-05 | 0 - | 0 - | 1 - | 2,71E-05 - | 2 - | 5,16E-05 - | 0 - | 0 - | 1 - | 2,95E-05 - |
| 6412 | 1 | 3um_FIJJAFBF_144690 | Bacteria | Proteobacteri | Gammaprotec | Oceanospirilla | Saccharospirill | Reinekea | Reinekea fors | unclassified | unclassified | unclassified | unclassified | unclassified | 2,2 | 0,979795897 | 0,000194296 | 8,54E-05 | 2 - | 0,000150035 - | 2 - | 0,000186299 - | 4 - | 0,000354202 - | 2 - | 0,000179717 - | 1 - | 0,000101225 - |
| 6413 | 1 | 0.2um_NOKCPHJN_184934 | Bacteria | Proteobacteri | Gammaprotec | unclassified | unclassified | unclassified | Gammaprotec | metabolism | Inorganic ion t | 0QHT5 | TonB-depende | 314287.GB220 | 4,4 | 1,019803903 | 7,43E-05 | 2,23E-05 | 4 - | 5,61E-05 - | 3 - | 5,22E-05 - | 5 - | 8,27E-05 - | 4 - | 6,71E-05 - | 6 - | 0,000113461 - |
| 6414 | 1 | 0.2um_NOKCPHJN_36508 | Bacteria | Bacteroidetes | Cytophagia | Cytophagales | Flammeovirga | unclassified | Flammeovirga | poorly charact | Function unkn | 1AINX | Phage tail she | 760192.Halhy | 3,8 | 0,748331477 | 0,000104405 | 1,14E-05 | 5 - | 0,000117119 - | 3 - | 8,73E-05 - | 4 - | 0,000110598 - | 4 - | 0,000112231 - | 3 - | 9,48E-05 - |
| 6415 | 1 | 0.2um_NOKCPHJN_611626 | Bacteria | Bacteroidetes | Flavobacteriia | Flavobacterial | Cryomorphac | unclassified | Cryomorphac | information st | Translation, ri | COG0050 | This protein p | 886377.Murru | 0,4 | 0,489897949 | 1,49E-05 | 1,84E-05 | 0 - | 0 - | 0 - | 0 - | 1 - | 3,48E-05 - | 0 - | 0 - | 1 - | 3,98E-05 - |
| 6416 | 1 | 3um_FIJJAFBF_112857 | unclassified | unclassified | unclassified | unclassified | unclassified | unclassified | unclassified | unclassified | unclassified | unclassified | unclassified | unclassified | 1 | 0 | 0,000240398 | 2,28E-05 | 1 - | 0,000201363 - | 1 - | 0,000250033 - | 1 - | 0,000237688 - | 1 - | 0,000241199 - | 1 - | 0,00027171 - |
| 6417 | 1 | 3um_FIJJAFBF_229006 | Bacteria | Proteobacteri | Alphaproteob | Rhodobactera | Rhodobactera | Planktomarina | Planktomarina | metabolism | Carbohydrate | 16Q4I | Abc transport | 367336.OM22 | 0,8 | 0,4 | 2,51E-05 | 1,29E-05 | 1 - | 2,63E-05 - | 1 - | 3,26E-05 - | 1 - | 3,10E-05 - | 0 - | 0 - | 1 - | 3,54E-05 - |
| 6418 | 1 | 3um_FIJJAFBF_510122 | Bacteria | Proteobacteri | Betaproteoba | unclassified | unclassified | unclassified | Betaproteoba | unclassified | unclassified | unclassified | unclassified | unclassified | 0,6 | 0,489897949 | 9,99E-06 | 8,16E-06 | 0 - | 0 - | 1 - | 1,71E-05 - | 1 - | 1,63E-05 - | 1 - | 1,65E-05 - | 0 - | 0 - |
| 6419 | 1 | 0.2um_NOKCPHJN_657738 | Bacteria | Proteobacteri | Alphaproteob | Pelagibacteral | Pelagibacterac | Candidatus Pe | Candidatus Pe | unclassified | unclassified | unclassified | unclassified | unclassified | 4,4 | 1,019803903 | 0,000244463 | 4,98E-05 | 6 - | 0,000282238 - | 3 - | 0,000175228 - | 4 - | 0,000222102 - | 4 - | 0,000225382 - | 5 - | 0,000317366 - |
| 6420 | 1 | 0.2um_NOKCPHJN_16777 | Bacteria | Proteobacteri | Gammaprotec | Alteromonada | Alteromonada | Paraglaciecola | Paraglaciecola | cellular proces | Posttranslatio | 17C3T | Peptidyl-proly | 342610.Patl_1 | 1 | 0,632455532 | 7,87E-05 | 4,44E-05 | 2 - | 0,000137457 - | 0 - | 0 - | 1 - | 8,11E-05 - | 1 - | 8,23E-05 - | 1 - | 9,27E-05 - |
| 6421 | 1 | 3um_FIJJAFBF_158111 | Bacteria | Bacteroidetes | Flavobacteriia | unclassified | unclassified | unclassified | Flavobacteriia | unclassified | unclassified | unclassified | unclassified | unclassified | 1,2 | 0,748331477 | 1,71E-05 | 1,12E-05 | 1 - | 1,22E-05 - | 2 - | 3,02E-05 - | 2 - | 2,87E-05 - | 1 - | 1,46E-05 - | 0 - | 0 - |
| 6422 | 1 | 3um_FIJJAFBF_123868 | Bacteria | Bacteroidetes | Flavobacteriia | Flavobacterial | Flavobacteriac | Lacinutrix | Lacinutrix sp. | information st | Replication, re | 05C24 | DNA gyrase ne | 313603.FB217 | 0,4 | 0,489897949 | 6,80E-06 | 8,36E-06 | 0 - | 0 - | 0 - | 0 - | 1 - | 1,59E-05 - | 0 - | 0 - | 1 - | 1,81E-05 - |
| 6423 | 1 | 0.2um_NOKCPHJN_235644 | Bacteria | Bacteroidetes | Flavobacteriia | Flavobacterial | Flavobacteriac | unclassified | various | cellular proces | Cell wall/mem | COG2885 | Ompa motb d | 313598.MED1 | 0,6 | 0,489897949 | 1,72E-05 | 1,42E-05 | 1 - | 2,51E-05 - | 1 - | 3,12E-05 - | 1 - | 2,96E-05 - | 0 - | 0 - | 0 - | 0 - |
| 6424 | 1 | 3um_FIJJAFBF_786152 | Bacteria | Proteobacteri | Gammaprotec | Oceanospirilla | Saccharospirill | Reinekea | Reinekea fors | unclassified | unclassified | unclassified | unclassified | unclassified | 7 | 1,549193338 | 0,000771706 | 9,50E-05 | 10 - | 0,000940793 - | 6 - | 0,000700912 - | 6 - | 0,000666306 - | 7 - | 0,000788839 - | 6 - | 0,000761678 - |
| 6425 | 1 | 0.2um_NOKCPHJN_258383 | Bacteria | Proteobacteri | Gammaprotec | unclassified | unclassified | unclassified | Gammaprotec | unclassified | unclassified | unclassified | unclassified | unclassified | 0,8 | 1,166190379 | 1,80E-05 | 2,56E-05 | 0 - | 0 - | 0 - | 0 - | 0 - | 0 - | 3 - | 6,55E-05 - | 1 - | 2,46E-05 - |
| 6426 | 1 | 3um_FIJJAFBF_205811 | Bacteria | Bacteroidetes | Flavobacteriia | Flavobacterial | Flavobacteriac | Formosa | Formosa sp. H | poorly charact | Function unkn | 0XPRT | NA | 391598.FBBAL | 0,6 | 0,8 | 1,80E-05 | 2,42E-05 | 0 - | 0 - | 2 - | 6,06E-05 - | 0 - | 0 - | 1 - | 2,93E-05 - | 0 - | 0 - |
| 6427 | 1 | 0.2um_NOKCPHJN_702476 | Bacteria | Proteobacteri | Alphaproteob | Pelagibacteral | Pelagibacterac | unclassified | Pelagibacterac | cellular proces | Cell wall/mem | 7257 | NA | 439493.PB721 | 2,4 | 0,489897949 | 9,01E-05 | 2,44E-05 | 2 - | 6,22E-05 - | 2 - | 7,72E-05 - | 2 - | 7,34E-05 - | 3 - | 0,000111775 - | 3 - | 0,000125914 - |
| 6428 | 1 | 3um_FIJJAFBF_709135 | Bacteria | Bacteroidetes | Flavobacteriia | Flavobacterial | Flavobacteriac | Formosa | Formosa sp. H | unclassified | unclassified | unclassified | unclassified | unclassified | 0,2 | 0,4 | 4,99E-06 | 9,98E-06 | 1 - | 2,50E-05 - | 0 - | 0 - | 0 - | 0 - | 0 - | 0 - | 0 - | 0 - |
| 6429 | 1 | 3um_FIJJAFBF_33859 | Bacteria | Bacteroidetes | Flavobacteriia | Flavobacterial | Flavobacteriac | Polaribacter | Polaribacter t | cellular proces | Cell wall/mem | 11TPU | outer membra | 313594.PI23P | 5,2 | 0,979795897 | 0,000388923 | 5,03E-05 | 7 - | 0,000443888 - | 5 - | 0,000393699 - | 4 - | 0,000299408 - | 5 - | 0,000379788 - | 5 - | 0,00042783 - |

| | | | | | | | | | | | | | | | | | | | | | | | | | | | | |
|---|---|---|---|---|---|---|---|---|---|---|---|---|---|---|---|---|---|---|---|---|---|---|---|---|---|---|---|---|
| 6430 | 1 | 3um_FIJJAFBF_119126 | Bacteria | Bacteroidetes | Flavobacteriia | Flavobacterial | Flavobacteriac | Polaribacter | Polaribacter s | metabolism | Energy produc | COG2838 | Isocitrate deh | 313598.MED1 | 3,2 | 0,748331477 | 5,87E-05 | 1,30E-05 | 4 - | 6,20E-05 - | 4 - | 7,69E-05 - | 3 - | 5,49E-05 - | 2 - | 3,71E-05 - | 3 - | 6,27E-05 - |
| 6431 | 1 | 3um_FIJJAFBF_656940 | Bacteria | Bacteroidetes | Flavobacteriia | Flavobacterial | Flavobacteriac | Formosa | Formosa sp. H | poorly charact | Function unkn | 0P8G1 | peptidylprolyl | 391603.FBALC | 0,8 | 0,4 | 8,99E-05 | 4,56E-05 | 1 - | 9,73E-05 - | 1 - | 0,000120779 - | 1 - | 0,000114815 - | 1 - | 0,000116511 - | 0 - | 0 - |
| 6432 | 1 | 3um_FIJJAFBF_56733 | Bacteria | Proteobacteri | Gammaprotec | Alteromonada | Alteromonada | Paraglaciecola | Paraglaciecola | metabolism | Energy produc | 0QIP8 | Electron trans | 342610.Patl_2 | 0,4 | 0,489897949 | 1,82E-05 | 2,23E-05 | 0 - | 0 - | 1 - | 4,63E-05 - | 0 - | 0 - | 1 - | 4,46E-05 - | 0 - | 0 - |
| 6433 | 1 | 0.2um_NOKCPHJN_850912 | Bacteria | Proteobacteri | Betaproteoba | Nitrosomonad | unclassified | unclassified | beta proteoba | cellular proces | Cell wall/mem | COG2885 | Ompa motb d | 383631.MB21 | 1,4 | 0,8 | 0,000103712 | 5,94E-05 | 2 - | 0,000128963 - | 2 - | 0,000160134 - | 2 - | 0,000152227 - | 1 - | 7,72E-05 - | 0 - | 0 - |
| 6434 | 1 | 3um_FIJJAFBF_437893 | Bacteria | Rhodotherma | unclassified | unclassified | unclassified | unclassified | Rhodotherma | unclassified | unclassified | unclassified | unclassified | unclassified | 2,4 | 0,489897949 | 6,00E-05 | 9,34E-06 | 3 - | 6,36E-05 - | 2 - | 5,27E-05 - | 2 - | 5,01E-05 - | 3 - | 7,62E-05 - | 2 - | 5,73E-05 - |
| 6435 | 1 | 0.2um_NOKCPHJN_106633 | Bacteria | Proteobacteri | Gammaprotec | Cellvibrionales | Porticoccacea | unclassified | SAR92 bacteri | metabolism | Energy produc | 16RH6 | Produces ATP | 247634.GPB2 | 0,4 | 0,8 | 1,61E-05 | 3,21E-05 | 2 - | 8,03E-05 - | 0 - | 0 - | 0 - | 0 - | 0 - | 0 - | 0 - | 0 - |
| 6436 | 1 | 3um_FIJJAFBF_369511 | Bacteria | Bacteroidetes | Flavobacteriia | Flavobacterial | Flavobacteriac | Polaribacter | Polaribacter t | metabolism | Carbohydrate | 099Q3 | phosphohexol | 313594.PI23P | 6,2 | 1,166190379 | 0,000260793 | 5,70E-05 | 5 - | 0,0001755 - | 5 - | 0,000217919 - | 7 - | 0,000290023 - | 8 - | 0,000336351 - | 6 - | 0,000284173 - |
| 6437 | 1 | 0.2um_NOKCPHJN_712155 | Bacteria | unclassified | unclassified | unclassified | unclassified | unclassified | bacterium B1 | unclassified | unclassified | unclassified | unclassified | unclassified | 0,4 | 0,489897949 | 9,34E-06 | 1,15E-05 | 1 - | 2,13E-05 - | 0 - | 0 - | 0 - | 0 - | 1 - | 2,55E-05 - | 0 - | 0 - |
| 6438 | 1 | 3um_FIJJAFBF_786149 | Bacteria | Proteobacteri | Gammaprotec | Oceanospirilla | Saccharospirill | Reinekea | Reinekea fors | information st | Translation, ri | 05VBI | Binds 16S rRN | 314283.MED2 | 1,8 | 0,4 | 0,000240673 | 4,75E-05 | 2 - | 0,000227281 - | 2 - | 0,000282216 - | 2 - | 0,000268282 - | 2 - | 0,000272244 - | 1 - | 0,000153341 - |
| 6439 | 1 | 3um_FIJJAFBF_201709 | Bacteria | Bacteroidetes | Flavobacteriia | unclassified | unclassified | unclassified | Flavobacteria | unclassified | unclassified | unclassified | unclassified | unclassified | 1,4 | 0,8 | 4,94E-05 | 2,80E-05 | 1 - | 2,97E-05 - | 1 - | 3,68E-05 - | 3 - | 0,000105025 - | 1 - | 3,55E-05 - | 1 - | 4,00E-05 - |
| 6440 | 1 | 3um_FIJJAFBF_102544 | Bacteria | Bacteroidetes | Flavobacteriia | Flavobacterial | Flavobacteriac | Polaribacter | various | information st | Translation, ri | 08YY1 | This protein is | 867900.Celly_ | 6 | 0,632455532 | 0,000704846 | 6,79E-05 | 7 - | 0,000692618 - | 6 - | 0,000737167 - | 6 - | 0,00070077 - | 5 - | 0,0005926 - | 6 - | 0,000801075 - |
| 6441 | 1 | 0.2um_NOKCPHJN_12874 | Bacteria | Bacteroidetes | Flavobacteriia | Flavobacterial | Flavobacteriac | Hyunsoonleell | Hyunsoonleell | information st | Translation, ri | COG0050 | This protein p | 688270.Celal_ | 10 | 0,894427191 | 0,000346736 | 4,62E-05 | 11 - | 0,000319631 - | 10 - | 0,000360807 - | 9 - | 0,000308694 - | 9 - | 0,000313253 - | 11 - | 0,000431296 - |
| 6442 | 1 | 3um_FIJJAFBF_404819 | Bacteria | Bacteroidetes | Flavobacteriia | Flavobacterial | Flavobacteriac | Polaribacter | Polaribacter s | metabolism | Nucleotide tra | COG0138 | bifunctional p | 313598.MED1 | 0,6 | 0,489897949 | 1,68E-05 | 1,38E-05 | 0 - | 0 - | 0 - | 0 - | 1 - | 2,66E-05 - | 1 - | 2,70E-05 - | 1 - | 3,04E-05 - |
| 6443 | 1 | 3um_FIJJAFBF_103127 | Bacteria | Bacteroidetes | Flavobacteriia | Flavobacterial | Flavobacteriac | Polaribacter | Polaribacter s | unclassified | unclassified | unclassified | unclassified | unclassified | 1,2 | 0,4 | 0,000107371 | 2,38E-05 | 2 - | 0,000154063 - | 1 - | 9,57E-05 - | 1 - | 9,09E-05 - | 1 - | 9,23E-05 - | 1 - | 0,000103943 - |
| 6444 | 1 | 0.2um_NOKCPHJN_137199 | Bacteria | Bacteroidetes | unclassified | unclassified | unclassified | unclassified | Bacteroidetes | information st | Translation, ri | 08YY1 | This protein is | 216432.CA25 | 2 | 0,632455532 | 0,000234359 | 6,14E-05 | 3 - | 0,000299418 - | 2 - | 0,000247859 - | 2 - | 0,000235621 - | 1 - | 0,000119551 - | 2 - | 0,000269347 - |
| 6445 | 1 | 3um_FIJJAFBF_112180 | Bacteria | Proteobacteri | Gammaprotec | Cellvibrionales | Halieaceae | Haliea | Haliea salexig | unclassified | unclassified | unclassified | unclassified | unclassified | 0,2 | 0,4 | 8,46E-06 | 1,69E-05 | 0 - | 0 - | 1 - | 4,23E-05 - | 0 - | 0 - | 0 - | 0 - | 0 - | 0 - |
| 6446 | 1 | 0.2um_NOKCPHJN_423479 | Bacteria | Bacteroidetes | Flavobacteriia | Flavobacterial | Flavobacteriac | Marinirhabdu | Marinirhabdu | cellular proces | Cell wall/mem | COG2885 | Ompa motb d | 50743.SCB49_ | 2,4 | 1,854723699 | 4,87E-05 | 3,16E-05 | 6 - | 0,000109138 - | 2 - | 4,52E-05 - | 2 - | 4,29E-05 - | 1 - | 2,18E-05 - | 1 - | 2,45E-05 - |
| 6447 | 1 | 3um_FIJJAFBF_158303 | Bacteria | Proteobacteri | Alphaproteob | Pelagibacteral | Pelagibacterac | Candidatus Pe | Candidatus Pe | cellular proces | Cell wall/mem | COG2885 | Ompa motb d | 439493.PB721 | 1 | 0 | 8,30E-05 | 7,89E-06 | 1 - | 6,96E-05 - | 1 - | 8,64E-05 - | 1 - | 8,21E-05 - | 1 - | 8,33E-05 - | 1 - | 9,39E-05 - |
| 6448 | 1 | 0.2um_NOKCPHJN_94629 | Bacteria | Proteobacteri | Alphaproteob | Rhodobactera | Rhodobactera | Planktomarina | Planktomarina | poorly charact | Function unkn | 05W3P | Protein of unk | 314270.RB208 | 0,2 | 0,4 | 2,84E-05 | 5,68E-05 | 0 - | 0 - | 0 - | 0 - | 0 - | 0 - | 0 - | 0 - | 1 - | 0,000142087 - |
| 6449 | 1 | 0.2um_NOKCPHJN_179233 | Eukaryota | Eustigmatoph | unclassified | Eustigmatales | Monodopsida | Nannochlorop | Nannochlorop | metabolism | Amino acid tra | 15H0A | Interconversic | 15368.BRADI1 | 1 | 0,632455532 | 2,69E-05 | 1,74E-05 | 1 - | 2,32E-05 - | 1 - | 2,88E-05 - | 2 - | 5,47E-05 - | 1 - | 2,78E-05 - | 0 - | 0 - |
| 6450 | 1 | 0.2um_NOKCPHJN_177683 | Eukaryota | Streptophyta | unclassified | Gentianales | Rubiaceae | Coffea | Coffea caneph | poorly charact | Function unkn | 0UKBB | HEAT repeat c | 9796.ENSECA | 0,8 | 0,4 | 9,63E-06 | 4,85E-06 | 0 - | 0 - | 1 - | 1,20E-05 - | 1 - | 1,14E-05 - | 1 - | 1,16E-05 - | 1 - | 1,31E-05 - |
| 6451 | 1 | 3um_FIJJAFBF_349745 | Bacteria | Proteobacteri | Gammaprotec | Alteromonada | Ferrimonadac | Ferrimonas | Ferrimonas m | information st | Translation, ri | COG0251 | Endoribonucle | 983545.Glaag_ | 1 | 0 | 0,000119154 | 1,13E-05 | 1 - | 9,98E-05 - | 1 - | 0,000123929 - | 1 - | 0,000117811 - | 1 - | 0,000119551 - | 1 - | 0,000134673 - |
| 6452 | 1 | 3um_FIJJAFBF_399091 | Bacteria | Bacteroidetes | Flavobacteriia | Flavobacterial | Flavobacteriac | Polaribacter | Polaribacter b | metabolism | Energy produc | 05D26 | NQR complex | 313598.MED1 | 5,2 | 0,979795897 | 0,000164306 | 3,62E-05 | 4 - | 0,0001053 - | 5 - | 0,000163439 - | 7 - | 0,000217517 - | 5 - | 0,000157664 - | 5 - | 0,000177608 - |
| 6453 | 1 | 0.2um_NOKCPHJN_221606 | Bacteria | Proteobacteri | Alphaproteob | Pelagibacteral | Pelagibacterac | Candidatus Pe | Candidatus Pe | metabolism | Amino acid tra | 16V6G | Glycine betain | 439493.PB721 | 0,4 | 0,489897949 | 1,58E-05 | 1,94E-05 | 1 - | 3,59E-05 - | 0 - | 0 - | 0 - | 0 - | 1 - | 4,30E-05 - | 0 - | 0 - |
| 6454 | 1 | 3um_FIJJAFBF_175283 | Eukaryota | Apicomplexa | Conoidasida | Eucoccidiorida | Sarcocystidae | Hammondia | Hammondia h | unclassified | unclassified | unclassified | unclassified | unclassified | 0,2 | 0,4 | 4,16E-06 | 8,31E-06 | 0 - | 0 - | 1 - | 2,08E-05 - | 0 - | 0 - | 0 - | 0 - | 0 - | 0 - |
| 6455 | 1 | 3um_FIJJAFBF_290357 | Bacteria | Bacteroidetes | Flavobacteriia | unclassified | unclassified | unclassified | Flavobacteria | unclassified | unclassified | unclassified | unclassified | unclassified | 0,6 | 0,489897949 | 6,04E-05 | 4,98E-05 | 1 - | 8,83E-05 - | 1 - | 0,00010963 - | 1 - | 0,000104217 - | 0 - | 0 - | 0 - | 0 - |
| 6456 | 1 | 3um_FIJJAFBF_241431 | Bacteria | Proteobacteri | Gammaprotec | Oceanospirilla | Saccharospirill | Reinekea | Reinekea fors | poorly charact | Function unkn | 16QXM | short-chain de | 314283.MED2 | 1 | 0,894427191 | 5,08E-05 | 4,39E-05 | 2 - | 8,90E-05 - | 0 - | 0 - | 2 - | 0,000105025 - | 0 - | 0 - | 1 - | 6,00E-05 - |
| 6457 | 1 | 3um_FIJJAFBF_359572 | Eukaryota | unclassified | Synurophycea | Synurales | Mallomonada | Synura | Synura synur | unclassified | unclassified | unclassified | unclassified | unclassified | 0,2 | 0,4 | 2,17E-05 | 4,33E-05 | 0 - | 0 - | 0 - | 0 - | 0 - | 0 - | 1 - | 0,000108255 - | 0 - | 0 - |
| 6458 | 1 | 3um_FIJJAFBF_644593 | Bacteria | Bacteroidetes | Flavobacteriia | Flavobacterial | Flavobacteriac | Formosa | Formosa sp. H | information st | Transcription | 0P88Z | transcriptiona | 391603.FBALC | 2,4 | 0,489897949 | 0,000200076 | 5,33E-05 | 2 - | 0,000138285 - | 2 - | 0,000171709 - | 3 - | 0,000244847 - | 2 - | 0,000165643 - | 3 - | 0,000279894 - |
| 6459 | 1 | 0.2um_NOKCPHJN_182039 | Bacteria | Bacteroidetes | unclassified | unclassified | unclassified | unclassified | Bacteroidetes | information st | Translation, ri | 05CAV | thus facilitatin | 487796.Flav2 | 4,2 | 0,748331477 | 8,86E-05 | 1,81E-05 | 5 - | 8,84E-05 - | 4 - | 8,78E-05 - | 3 - | 6,26E-05 - | 4 - | 8,47E-05 - | 5 - | 0,000119318 - |
| 6460 | 1 | 0.2um_NOKCPHJN_93295 | Bacteria | Proteobacteri | Alphaproteob | Pelagibacteral | Pelagibacterac | unclassified | various | cellular proces | Posttranslatio | 01QWG | Prevents misf | 439493.PB721 | 10,8 | 1,469693846 | 0,000264108 | 1,87E-05 | 13 - | 0,000269332 - | 10 - | 0,000257254 - | 10 - | 0,000244553 - | 12 - | 0,000297798 - | 9 - | 0,000251601 - |
| 6461 | 1 | 0.2um_NOKCPHJN_930179 | unclassified | unclassified | unclassified | unclassified | unclassified | unclassified | unclassified | unclassified | unclassified | unclassified | unclassified | unclassified | 0,6 | 0,489897949 | 2,42E-05 | 1,98E-05 | 0 - | 0 - | 0 - | 0 - | 1 - | 3,83E-05 - | 1 - | 3,88E-05 - | 1 - | 4,37E-05 - |
| 6462 | 1 | 3um_FIJJAFBF_383460 | Bacteria | Proteobacteri | Gammaprotec | Oceanospirilla | Oceanospirilla | Marinobacteri | Marinobacteri | unclassified | unclassified | unclassified | unclassified | unclassified | 1 | 0 | 0,000165093 | 1,57E-05 | 1 - | 0,000138285 - | 1 - | 0,000171709 - | 1 - | 0,000163232 - | 1 - | 0,000165643 - | 1 - | 0,000186596 - |
| 6463 | 1 | 3um_FIJJAFBF_245359 | Bacteria | Bacteroidetes | Flavobacteriia | Flavobacterial | Flavobacteriac | Ulvibacter | Ulvibacter lito | poorly charact | Function unkn | 0XNQI | domain protei | 50743.SCB49_ | 0,4 | 0,489897949 | 1,22E-05 | 1,50E-05 | 0 - | 0 - | 1 - | 3,13E-05 - | 1 - | 2,98E-05 - | 0 - | 0 - | 0 - | 0 - |
| 6464 | 1 | 3um_FIJJAFBF_160985 | Bacteria | Bacteroidetes | Flavobacteriia | Flavobacterial | Flavobacteriac | various | various | unclassified | unclassified | unclassified | unclassified | unclassified | 2,4 | 0,489897949 | 0,000111761 | 1,64E-05 | 3 - | 0,000118735 - | 2 - | 9,83E-05 - | 3 - | 0,000140154 - | 2 - | 9,48E-05 - | 2 - | 0,00010681 - |
| 6465 | 1 | 0.2um_NOKCPHJN_245311 | Bacteria | Proteobacteri | Gammaprotec | Alteromonada | Colwelliaceae | Colwellia | Colwellia psyc | cellular proces | Cell wall/mem | 0QJ86 | Dehydrogenas | 1116375.VEJY | 0,2 | 0,4 | 5,39E-06 | 1,08E-05 | 1 - | 2,69E-05 - | 0 - | 0 - | 0 - | 0 - | 0 - | 0 - | 0 - | 0 - |
| 6466 | 1 | 0.2um_NOKCPHJN_190557 | Bacteria | Bacteroidetes | Flavobacteriia | Flavobacterial | Flavobacteriac | Maribacter | Maribacter po | metabolism | Energy produc | 05CMV | Succinyl-CoA s | 886377.Murru | 6,8 | 1,326649916 | 0,000236008 | 5,64E-05 | 5 - | 0,000144192 - | 9 - | 0,000322279 - | 7 - | 0,000238285 - | 7 - | 0,000241805 - | 6 - | 0,000233479 - |
| 6467 | 1 | 3um_FIJJAFBF_31550 | Bacteria | Proteobacteri | Betaproteoba | Burkholderiale | unclassified | unclassified | various | information st | Translation, ri | 08YY1 | This protein is | 391597.LMED | 2,4 | 0,489897949 | 0,000258256 | 5,77E-05 | 3 - | 0,000271126 - | 2 - | 0,000224439 - | 2 - | 0,000213358 - | 2 - | 0,000216509 - | 3 - | 0,000365845 - |
| 6468 | 1 | 0.2um_NOKCPHJN_474960 | Bacteria | Proteobacteri | Alphaproteob | Rhodobactera | Rhodobactera | unclassified | Rhodobactera | information st | Translation, ri | COG1185 | Involved in m | 89187.ISM_06 | 1,8 | 1,720465053 | 3,20E-05 | 2,71E-05 | 5 - | 8,05E-05 - | 1 - | 2,00E-05 - | 2 - | 3,80E-05 - | 0 - | 0 - | 1 - | 2,17E-05 - |
| 6469 | 1 | 3um_FIJJAFBF_454939 | Bacteria | Bacteroidetes | Flavobacteriia | Flavobacterial | Flavobacteriac | Marinirhabdu | Marinirhabdu | unclassified | unclassified | unclassified | unclassified | unclassified | 1,2 | 0,4 | 2,91E-05 | 1,00E-05 | 1 - | 2,03E-05 - | 1 - | 2,52E-05 - | 1 - | 2,40E-05 - | 2 - | 4,87E-05 - | 1 - | 2,74E-05 - |
| 6470 | 1 | 3um_FIJJAFBF_21981 | Bacteria | Cyanobacteria | unclassified | Synechococca | Prochloraceae | Prochlorococc | Prochlorococc | unclassified | unclassified | unclassified | unclassified | unclassified | 0,2 | 0,4 | 1,71E-05 | 3,43E-05 | 1 - | 8,57E-05 - | 0 - | 0 - | 0 - | 0 - | 0 - | 0 - | 0 - | 0 - |
| 6471 | 1 | 3um_FIJJAFBF_70793 | Bacteria | Bacteroidetes | Flavobacteriia | Flavobacterial | Crocinitomica | unclassified | Crocinitomica | unclassified | unclassified | unclassified | unclassified | unclassified | 2 | 0 | 9,32E-05 | 8,85E-06 | 2 - | 7,81E-05 - | 2 - | 9,70E-05 - | 2 - | 9,22E-05 - | 2 - | 9,35E-05 - | 2 - | 0,000105357 - |
| 6472 | 1 | 3um_FIJJAFBF_208756 | Bacteria | Proteobacteri | Alphaproteob | Rhodobactera | Rhodobactera | Roseobacter | Roseobacter s | metabolism | Carbohydrate | 16ZMP | ABC transport | 351016.RAZW | 4,8 | 2,13541565 | 0,000144079 | 4,77E-05 | 9 - | 0,000234238 - | 4 - | 0,000129269 - | 4 - | 0,000122886 - | 3 - | 9,35E-05 - | 4 - | 0,000140476 - |
| 6473 | 1 | 3um_FIJJAFBF_313318 | Bacteria | Proteobacteri | Gammaprotec | Oceanospirilla | Saccharospirill | Reinekea | Reinekea fors | cellular proces | Posttranslatio | COG0330 | Band 7 protei | 314283.MED2 | 7,2 | 0,979795897 | 0,000315031 | 2,52E-05 | 9 - | 0,000333223 - | 6 - | 0,000275843 - | 7 - | 0,000305928 - | 7 - | 0,000310446 - | 7 - | 0,000349717 - |
| 6474 | 1 | 3um_FIJJAFBF_25653 | Bacteria | Proteobacteri | Gammaprotec | Alteromonada | Alteromonada | Paraglaciecola | Paraglaciecola | unclassified | unclassified | unclassified | unclassified | unclassified | 9,4 | 1,2 | 0,000136871 | 9,11E-06 | 11 - | 0,000135612 - | 8 - | 0,000122465 - | 10 - | 0,000145523 - | 10 - | 0,000147673 - | 8 - | 0,000133082 - |
| 6475 | 1 | 3um_FIJJAFBF_735694 | Bacteria | Cyanobacteria | unclassified | Synechococca | Synechococca | Synechococcu | Synechococcu | unclassified | unclassified | unclassified | unclassified | unclassified | 0,2 | 0,4 | 6,68E-06 | 1,34E-05 | 0 - | 0 - | 1 - | 3,34E-05 - | 0 - | 0 - | 0 - | 0 - | 0 - | 0 - |
| 6476 | 1 | 3um_FIJJAFBF_684627 | Bacteria | Bacteroidetes | Flavobacteriia | Flavobacterial | Flavobacteriac | Polaribacter | Polaribacter s | poorly charact | Function unkn | 0P8NI | Inherit from b | 487797.Flav3 | 3 | 0,632455532 | 0,000126177 | 2,84E-05 | 3 - | 0,000105948 - | 2 - | 8,77E-05 - | 3 - | 0,000125061 - | 4 - | 0,00016921 - | 3 - | 0,000142961 - |
| 6477 | 2 | 0.2um_NOKCPHJN_192520;3um_FIJJAFBF_82612 | various | various | various | various | various | various | various | various | various | various | various | various | 0,4 | 0,489897949 | 4,79E-06 | 5,87E-06 | 0 - | 0 - | 1 - | 1,23E-05 - | 1 - | 1,17E-05 - | 0 - | 0 - | 0 - | 0 - |
| 6478 | 1 | 3um_FIJJAFBF_478440 | Bacteria | unclassified | unclassified | unclassified | unclassified | unclassified | uncultured ba | metabolism | Inorganic ion t | 05C27 | TonB-depende | 867900.Celly_ | 7,4 | 1,854723699 | 9,88E-05 | 2,12E-05 | 9 - | 0,000101973 - | 7 - | 9,85E-05 - | 10 - | 0,000133744 - | 5 - | 6,79E-05 - | 6 - | 9,17E-05 - |
| 6479 | 1 | 3um_FIJJAFBF_352896 | Bacteria | Bacteroidetes | Flavobacteriia | Flavobacterial | Flavobacteriac | Polaribacter | various | information st | Translation, ri | 09AI8 | Associates wit | 313598.MED1 | 1,6 | 0,8 | 6,78E-05 | 3,42E-05 | 1 - | 3,55E-05 - | 1 - | 4,41E-05 - | 2 - | 8,39E-05 - | 3 - | 0,000127693 - | 1 - | 4,79E-05 - |
| 6480 | 1 | 3um_FIJJAFBF_602242 | Eukaryota | unclassified | Oomycetes | Pythiales | Pythiaceae | Pythium | Pythium insidi | poorly charact | Function unkn | 0YPRH | FBG | 6085.XP_0021 | 0,4 | 0,489897949 | 9,68E-06 | 1,19E-05 | 0 - | 0 - | 0 - | 0 - | 1 - | 2,40E-05 - | 1 - | 2,44E-05 - | 0 - | 0 - |
| 6481 | 1 | 3um_FIJJAFBF_608035 | Bacteria | Proteobacteri | Gammaprotec | Cellvibrionales | Halieaceae | unclassified | marine gamm | information st | Transcription | 05VEQ | Cold shock pro | 247639.MGP2 | 0,8 | 0,4 | 0,000155959 | 7,92E-05 | 1 - | 0,000168789 - | 1 - | 0,000209587 - | 1 - | 0,000199239 - | 1 - | 0,000202181 - | 0 - | 0 - |
| 6482 | 1 | 0.2um_NOKCPHJN_328414 | Bacteria | Proteobacteri | Alphaproteob | Pelagibacteral | Pelagibacterac | unclassified | Pelagibacterac | cellular proces | Intracellular tr | 05VIE | preprotein tra | 335992.SAR11 | 1 | 0 | 0,000145774 | 1,38E-05 | 1 - | 0,000122103 - | 1 - | 0,000151616 - | 1 - | 0,00014413 - | 1 - | 0,000146259 - | 1 - | 0,00016476 - |
| 6483 | 1 | 3um_FIJJAFBF_180780 | Bacteria | Verrucomicrol | Opitutae | Opitutales | unclassified | unclassified | Opitutales bac | unclassified | unclassified | unclassified | unclassified | unclassified | 1,8 | 0,4 | 0,000186023 | 4,35E-05 | 2 - | 0,000173904 - | 1 - | 0,000107969 - | 2 - | 0,000205276 - | 2 - | 0,000208308 - | 2 - | 0,000234658 - |
| 6484 | 1 | 0.2um_NOKCPHJN_144457 | Bacteria | Proteobacteri | Gammaprotec | Oceanospirilla | Saccharospirill | Reinekea | Reinekea fors | unclassified | unclassified | unclassified | unclassified | unclassified | 5,8 | 1,326649916 | 0,000171891 | 2,76E-05 | 8 - | 0,000201805 - | 6 - | 0,000187937 - | 6 - | 0,000178658 - | 4 - | 0,000120864 - | 5 - | 0,000170192 - |
| 6485 | 1 | 3um_FIJJAFBF_149321 | Bacteria | Proteobacteri | Gammaprotec | unclassified | unclassified | unclassified | Gammaprotec | cellular proces | Posttranslatio | 0QHT2 | Peroxiredoxin | 523791.Kkor_ | 1,2 | 0,748331477 | 0,000139175 | 9,29E-05 | 1 - | 9,49E-05 - | 1 - | 0,000117784 - | 0 - | 0 - | 2 - | 0,000227245 - | 2 - | 0,000255991 - |
| 6486 | 1 | 3um_FIJJAFBF_278688 | Bacteria | Bacteroidetes | Flavobacteriia | Flavobacterial | Flavobacteriac | Polaribacter | Polaribacter s | metabolism | Carbohydrate | COG0149 | triosephospha | 313594.PI23P | 4,2 | 0,979795897 | 0,00022343 | 2,87E-05 | 6 - | 0,000273278 - | 4 - | 0,00022622 - | 4 - | 0,000215051 - | 4 - | 0,000218227 - | 3 - | 0,000184374 - |
| 6487 | 1 | 0.2um_NOKCPHJN_476350 | Bacteria | Proteobacteri | Gammaprotec | unclassified | unclassified | unclassified | Gammaprotec | metabolism | Inorganic ion t | 05CVA | phosphate | 555079.Toce_ | 4,6 | 0,8 | 0,000188562 | 3,53E-05 | 4 - | 0,000137457 - | 4 - | 0,000170681 - | 6 - | 0,000243381 - | 5 - | 0,000205813 - | 4 - | 0,000185478 - |
| 6488 | 1 | 3um_FIJJAFBF_174797 | Eukaryota | unclassified | Chrysophycea | Chromulinales | Chromulinace | Ochromonas | Ochromonas | unclassified | unclassified | unclassified | unclassified | unclassified | 0,6 | 0,489897949 | 1,92E-05 | 1,58E-05 | 0 - | 0 - | 1 - | 3,15E-05 - | 0 - | 0 - | 1 - | 3,04E-05 - | 1 - | 3,43E-05 - |
| 6489 | 2 | 3um_FIJJAFBF_546848;3um_FIJJAFBF_772047 | Bacteria | Proteobacteri | Alphaproteob | various | various | various | various | unclassified | unclassified | unclassified | unclassified | unclassified | 1 | 0,632455532 | 0,00012694 | 8,56E-05 | 1 - | 0,00010828 - | 2 - | 0,000268904 - | 1 - | 0,000127813 - | 1 - | 0,000129701 - | 0 - | 0 - |
| 6490 | 1 | 3um_FIJJAFBF_579113 | Bacteria | Bacteroidetes | Flavobacteriia | Flavobacterial | unclassified | unclassified | Flavobacterial | metabolism | Energy produc | 0PBIY | subunit C | 926562.Oweh | 2,4 | 0,489897949 | 0,000481921 | 0,000130412 | 2 - | 0,000332686 - | 2 - | 0,000413098 - | 2 - | 0,000392702 - | 3 - | 0,000597754 - | 3 - | 0,000673367 - |
| 6491 | 1 | 3um_FIJJAFBF_491674 | Bacteria | Proteobacteri | Gammaprotec | Cellvibrionales | Porticoccacea | unclassified | various | unclassified | unclassified | unclassified | unclassified | unclassified | 0,2 | 0,4 | 1,88E-05 | 3,76E-05 | 1 - | 9,41E-05 - | 0 - | 0 - | 0 - | 0 - | 0 - | 0 - | 0 - | 0 - |
| 6492 | 1 | 3um_FIJJAFBF_490084 | Bacteria | Proteobacteri | Gammaprotec | unclassified | unclassified | unclassified | OM182 bacte | unclassified | unclassified | unclassified | unclassified | unclassified | 2,8 | 0,979795897 | 4,49E-05 | 1,29E-05 | 4 - | 5,51E-05 - | 2 - | 3,42E-05 - | 4 - | 6,51E-05 - | 2 - | 3,30E-05 - | 2 - | 3,72E-05 - |
| 6493 | 1 | 3um_FIJJAFBF_130129 | Viruses | unclassified | unclassified | unclassified | unclassified | unclassified | uncultured vir | unclassified | unclassified | unclassified | unclassified | unclassified | 0,4 | 0,489897949 | 2,11E-05 | 2,60E-05 | 0 - | 0 - | 0 - | 0 - | 1 - | 4,93E-05 - | 0 - | 0 - | 1 - | 5,63E-05 - |
| 6494 | 1 | 0.2um_NOKCPHJN_09992 | Bacteria | Bacteroidetes | Flavobacteriia | Flavobacterial | Flavobacteriac | Polaribacter | Polaribacter v | information st | Translation, ri | COG0050 | This protein p | 313603.FB217 | 3 | 0 | 0,000104071 | 9,88E-06 | 3 - | 8,72E-05 - | 3 - | 0,000108242 - | 3 - | 0,000102898 - | 3 - | 0,000104418 - | 3 - | 0,000117626 - |
| 6495 | 1 | 0.2um_NOKCPHJN_927062 | Eukaryota | Streptophyta | unclassified | Fagales | Fagaceae | Quercus | Quercus sube | unclassified | unclassified | unclassified | unclassified | unclassified | 0,2 | 0,4 | 4,06E-06 | 8,13E-06 | 1 - | 2,03E-05 - | 0 - | 0 - | 0 - | 0 - | 0 - | 0 - | 0 - | 0 - |
| 6496 | 1 | 3um_FIJJAFBF_199015 | Eukaryota | unclassified | Oomycetes | Pythiales | Pythiaceae | Pythium | Pythium insidi | information st | Translation, ri | COG0089 | One of the ea | 65071.PYU1_ | 2 | 0,632455532 | 0,000184746 | 4,28E-05 | 3 - | 0,000239118 - | 2 - | 0,000197943 - | 2 - | 0,00018817 - | 2 - | 0,000190949 - | 1 - | 0,000107552 - |
| 6497 | 1 | 3um_FIJJAFBF_343898 | Eukaryota | unclassified | unclassified | Isochrysidales | Noelaerhabda | Emiliania | Emiliania huxl | metabolism | Energy produc | COG0056 | Produces ATP | 41431.PCC88 | 4,8 | 0,4 | 0,000131265 | 1,69E-05 | 5 - | 0,000114548 - | 5 - | 0,000142234 - | 5 - | 0,000135212 - | 4 - | 0,000109767 - | 5 - | 0,000154565 - |
| 6498 | 1 | 0.2um_NOKCPHJN_180529 | Bacteria | Proteobacteri | Gammaprotec | unclassified | unclassified | unclassified | Gammaprotec | cellular proces | Posttranslatio | 0QHXH | Molecular cha | 498211.CJA_1 | 1 | 0 | 2,16E-05 | 2,06E-06 | 1 - | 1,81E-05 - | 1 - | 2,25E-05 - | 1 - | 2,14E-05 - | 1 - | 2,17E-05 - | 1 - | 2,45E-05 - |
| 6499 | 1 | 0.2um_NOKCPHJN_62639 | Bacteria | Proteobacteri | Alphaproteob | Rhodospirillal | Rhodospirillac | unclassified | Rhodospirillac | unclassified | unclassified | unclassified | unclassified | unclassified | 0,2 | 0,4 | 1,55E-05 | 3,11E-05 | 0 - | 0 - | 0 - | 0 - | 0 - | 0 - | 1 - | 7,77E-05 - | 0 - | 0 - |
| 6500 | 1 | 3um_FIJJAFBF_747859 | Bacteria | Proteobacteri | Gammaprotec | Cellvibrionales | Spongiibacter | Oceanicoccus | Oceanicoccus | poorly charact | Function unkn | 17SP0 | Inherit from C | 491916.RHECI | 1 | 0 | 8,11E-05 | 7,70E-06 | 1 - | 6,79E-05 - | 1 - | 8,43E-05 - | 1 - | 8,02E-05 - | 1 - | 8,14E-05 - | 1 - | 9,16E-05 - |
| 6501 | 1 | 3um_FIJJAFBF_185076 | Bacteria | Proteobacteri | Alphaproteob | Rhodobactera | unclassified | unclassified | Rhodobactera | poorly charact | Function unkn | 06APV | NA | 367336.OM22 | 1 | 0,632455532 | 0,000119954 | 7,77E-05 | 1 - | 0,000103402 - | 1 - | 0,000128395 - | 2 - | 0,000244112 - | 1 - | 0,000123859 - | 0 - | 0 - |
| 6502 | 1 | 3um_FIJJAFBF_173757 | Bacteria | Bacteroidetes | Flavobacteriia | Flavobacterial | Flavobacteriac | Polaribacter | Polaribacter s | unclassified | unclassified | unclassified | unclassified | unclassified | 1,2 | 0,748331477 | 3,23E-05 | 1,99E-05 | 2 - | 4,71E-05 - | 2 - | 5,85E-05 - | 1 - | 2,78E-05 - | 1 - | 2,82E-05 - | 0 - | 0 - |
| 6503 | 1 | 0.2um_NOKCPHJN_904616 | Eukaryota | unclassified | Pelagophycea | Pelagomonad | unclassified | Aureococcus | Aureococcus a | cellular proces | Posttranslatio | KOG3083 | prohibitin | 294747.XP_00 | 1 | 0,632455532 | 0,000119526 | 6,74E-05 | 2 - | 0,000208685 - | 0 - | 0 - | 1 - | 0,000123166 - | 1 - | 0,000124985 - | 1 - | 0,000140795 - |
| 6504 | 1 | 3um_FIJJAFBF_171932 | Bacteria | Bacteroidetes | Flavobacteriia | Flavobacterial | Flavobacteriac | Polaribacter | Polaribacter s | unclassified | unclassified | unclassified | unclassified | unclassified | 1 | 0,632455532 | 1,94E-05 | 1,31E-05 | 1 - | 1,66E-05 - | 2 - | 4,12E-05 - | 1 - | 1,96E-05 - | 1 - | 1,99E-05 - | 0 - | 0 - |
| 6505 | 1 | 3um_FIJJAFBF_480271 | Bacteria | Proteobacteri | Gammaprotec | Oceanospirilla | Saccharospirill | Reinekea | Reinekea fors | unclassified | unclassified | unclassified | unclassified | unclassified | 0,2 | 0,4 | 1,19E-05 | 2,38E-05 | 0 - | 0 - | 0 - | 0 - | 0 - | 0 - | 0 - | 0 - | 1 - | 5,96E-05 - |
| 6506 | 1 | 3um_FIJJAFBF_409104 | Bacteria | Verrucomicrol | Opitutae | unclassified | unclassified | unclassified | Opitutae bact | poorly charact | Function unkn | 11TP8 | Erf family | 992406.RIA_0 | 1,2 | 0,4 | 9,19E-05 | 2,04E-05 | 2 - | 0,000131927 - | 1 - | 8,19E-05 - | 1 - | 7,79E-05 - | 1 - | 7,90E-05 - | 1 - | 8,90E-05 - |
| 6507 | 1 | 3um_FIJJAFBF_214150 | Bacteria | Bacteroidetes | Flavobacteriia | Flavobacterial | Flavobacteriac | Olleya | various | information st | Translation, ri | COG0050 | This protein p | 1046627.BZAR | 4,2 | 0,748331477 | 0,000144741 | 2,50E-05 | 4 - | 0,00011623 - | 4 - | 0,000144323 - | 5 - | 0,000171496 - | 5 - | 0,00017403 - | 3 - | 0,000117626 - |
| 6508 | 1 | 0.2um_NOKCPHJN_191187 | Eukaryota | Platyhelminth | unclassified | Macrostomida | Macrostomida | Macrostomun | Macrostomun | poorly charact | Function unkn | 0NVRZ | NA | 170187.SP_17 | 1 | 0 | 5,94E-06 | 5,64E-07 | 1 - | 4,97E-06 - | 1 - | 6,17E-06 - | 1 - | 5,87E-06 - | 1 - | 5,96E-06 - | 1 - | 6,71E-06 - |
| 6509 | 1 | 3um_FIJJAFBF_330922 | Bacteria | Proteobacteri | Alphaproteob | Rhodospirillal | Rhodospirillac | unclassified | Rhodobactera | unclassified | unclassified | unclassified | unclassified | unclassified | 1,4 | 0,8 | 0,000108314 | 6,33E-05 | 2 - | 0,000131173 - | 0 - | 0 - | 2 - | 0,000154837 - | 1 - | 7,86E-05 - | 2 - | 0,000176999 - |
| 6510 | 1 | 3um_FIJJAFBF_653259 | Bacteria | Bacteroidetes | Flavobacteriia | Flavobacterial | Flavobacteriac | Polaribacter | Polaribacter t | metabolism | Inorganic ion t | 05C27 | TonB-depende | 867900.Celly_ | 20,4 | 1,624807681 | 0,000259283 | 2,32E-05 | 21 - | 0,000224424 - | 18 - | 0,000238858 - | 22 - | 0,000277524 - | 22 - | 0,000281623 - | 19 - | 0,000273987 - |
| 6511 | 1 | 3um_FIJJAFBF_270780 | Bacteria | Bacteroidetes | Flavobacteriia | Flavobacterial | Flavobacteriac | Tenacibaculur | Tenacibaculur | poorly charact | Function unkn | 0P8NI | Inherit from b | 487797.Flav3 | 2 | 0,632455532 | 7,15E-05 | 1,66E-05 | 3 - | 9,26E-05 - | 2 - | 7,66E-05 - | 2 - | 7,28E-05 - | 2 - | 7,39E-05 - | 1 - | 4,16E-05 - |
| 6512 | 1 | 0.2um_NOKCPHJN_99565 | Bacteria | Proteobacteri | Gammaprotec | Oceanospirilla | Saccharospirill | Reinekea | Reinekea fors | poorly charact | Function unkn | 0QY82 | (LipO)protein | 314283.MED2 | 3,4 | 0,8 | 0,000241669 | 6,03E-05 | 4 - | 0,000239118 - | 2 - | 0,000148457 - | 3 - | 0,000211691 - | 4 - | 0,000286424 - | 4 - | 0,000322655 - |
| 6513 | 1 | 0.2um_NOKCPHJN_158025 | Viruses | unclassified | unclassified | Caudovirales | Siphoviridae | Yuavirus | Pseudomonas | poorly charact | Function unkn | 0G6W0 | Phage capsid f | 311424.DhcV | 1,8 | 1,166190379 | 5,28E-05 | 3,50E-05 | 2 - | 5,09E-05 - | 1 - | 3,16E-05 - | 3 - | 9,01E-05 - | 3 - | 9,15E-05 - | 0 - | 0 - |
| 6514 | 1 | 0.2um_NOKCPHJN_76325 | Bacteria | Bacteroidetes | Flavobacteriia | Flavobacterial | Flavobacteriac | Formosa | Formosa sp. H | unclassified | unclassified | unclassified | unclassified | unclassified | 3,2 | 0,4 | 0,000292753 | 6,17E-05 | 3 - | 0,000228033 - | 3 - | 0,00028315 - | 3 - | 0,00026917 - | 3 - | 0,000273146 - | 4 - | 0,000410264 - |
| 6515 | 1 | 3um_FIJJAFBF_257116 | Eukaryota | unclassified | Cryptophyta | Pyrenomonad | Chroomonada | Chroomonas | various | unclassified | unclassified | unclassified | unclassified | unclassified | 2,2 | 1,166190379 | 0,000250343 | 0,000115581 | 4 - | 0,000395782 - | 3 - | 0,000368583 - | 1 - | 0,000116795 - | 2 - | 0,00023704 - | 1 - | 0,000133512 - |
| 6516 | 1 | 0.2um_NOKCPHJN_28005 | Bacteria | Bacteroidetes | Flavobacteriia | Flavobacterial | Flavobacteriac | Formosa | Formosa sp. H | unclassified | unclassified | unclassified | unclassified | unclassified | 0,4 | 0,489897949 | 7,83E-06 | 9,60E-06 | 0 - | 0 - | 1 - | 1,99E-05 - | 0 - | 0 - | 1 - | 1,92E-05 - | 0 - | 0 - |
| 6517 | 1 | 0.2um_NOKCPHJN_156055 | Bacteria | Bacteroidetes | Flavobacteriia | Flavobacterial | Flavobacteriac | unclassified | Flavobacteriac | metabolism | Inorganic ion t | COG0841 | acriflavin resis | 886377.Murru | 1 | 0 | 1,19E-05 | 1,13E-06 | 1 - | 1,00E-05 - | 1 - | 1,24E-05 - | 1 - | 1,18E-05 - | 1 - | 1,20E-05 - | 1 - | 1,35E-05 - |
| 6518 | 1 | 3um_FIJJAFBF_575437 | Bacteria | Proteobacteri | Alphaproteob | Rhodobactera | Rhodobactera | Planktomarina | Planktomarina | unclassified | unclassified | unclassified | unclassified | unclassified | 0,8 | 0,4 | 6,00E-05 | 3,02E-05 | 0 - | 0 - | 1 - | 7,50E-05 - | 1 - | 7,13E-05 - | 1 - | 7,24E-05 - | 1 - | 8,15E-05 - |
| 6519 | 1 | 0.2um_NOKCPHJN_766227 | Bacteria | Proteobacteri | Gammaprotec | unclassified | unclassified | unclassified | Gammaprotec | cellular proces | Signal transdu | KOG0613 | fibronectin ty | 6239.W06H8. | 0,2 | 0,4 | 3,72E-06 | 7,45E-06 | 0 - | 0 - | 0 - | 0 - | 0 - | 0 - | 0 - | 0 - | 1 - | 1,86E-05 - |
| 6520 | 1 | 3um_FIJJAFBF_766981 | Bacteria | Planctomycet | Planctomyceti | Planctomycet | Planctomycet | unclassified | Planctomycet | unclassified | unclassified | unclassified | unclassified | unclassified | 1 | 0 | 1,64E-05 | 1,56E-06 | 1 - | 1,38E-05 - | 1 - | 1,71E-05 - | 1 - | 1,63E-05 - | 1 - | 1,65E-05 - | 1 - | 1,86E-05 - |
| 6521 | 2 | 0.2um_NOKCPHJN_174775;3um_FIJJAFBF_20051 | Bacteria | Bacteroidetes | Flavobacteriia | Flavobacterial | Flavobacteriac | Polaribacter | various | metabolism | Amino acid tra | 05CRQ | synthase (Con | various | 0,8 | 1,166190379 | 2,34E-05 | 3,58E-05 | 1 - | 2,48E-05 - | 3 - | 9,23E-05 - | 0 - | 0 - | 0 - | 0 - | 0 - | 0 - |
| 6522 | 1 | 0.2um_NOKCPHJN_720387 | Bacteria | Proteobacteri | Alphaproteob | Rhodobactera | Rhodobactera | Ascidiaceihabi | Ascidiaceihabi | metabolism | Carbohydrate | 05CV1 | Transketolase | 644107.SL115 | 1 | 0,632455532 | 1,94E-05 | 1,09E-05 | 2 - | 3,38E-05 - | 0 - | 0 - | 1 - | 2,00E-05 - | 1 - | 2,02E-05 - | 1 - | 2,28E-05 - |
| 6523 | 1 | 3um_FIJJAFBF_56135 | Bacteria | Bacteroidetes | Flavobacteriia | Flavobacterial | Flavobacteriac | Polaribacter | Polaribacter s | cellular proces | Cell wall/mem | 0P8W8 | it plays a direc | 313598.MED1 | 2 | 0 | 7,25E-05 | 6,88E-06 | 2 - | 6,07E-05 - | 2 - | 7,54E-05 - | 2 - | 7,17E-05 - | 2 - | 7,27E-05 - | 2 - | 8,19E-05 - |
| 6524 | 1 | 3um_FIJJAFBF_202614 | Bacteria | Bacteroidetes | Flavobacteriia | Flavobacterial | Flavobacteriac | Gillisia | Gillisia sp. CAL | unclassified | unclassified | unclassified | unclassified | unclassified | 0,2 | 0,4 | 5,89E-06 | 1,18E-05 | 0 - | 0 - | 0 - | 0 - | 1 - | 2,95E-05 - | 0 - | 0 - | 0 - | 0 - |
| 6525 | 1 | 3um_FIJJAFBF_37895 | Bacteria | Actinobacteri | Actinobacteri | Micrococcales | Microbacteria | unclassified | Microbacteria | information st | Translation, ri | COG0184 | Forms an inte | 312284.A20C1 | 1 | 1,095445115 | 0,00013985 | 0,000141921 | 3 - | 0,000386888 - | 1 - | 0,000160134 - | 1 - | 0,000152227 - | 0 - | 0 - | 0 - | 0 - |
| 6526 | 1 | 0.2um_NOKCPHJN_25738 | Bacteria | Proteobacteri | Alphaproteob | Rhodobactera | Hyphomonad | unclassified | Hyphomonad | information st | Translation, ri | 08UIK | This protein bi | 596327.PORU | 2,2 | 0,4 | 0,000204834 | 1,88E-05 | 3 - | 0,000237469 - | 2 - | 0,000196578 - | 2 - | 0,000186872 - | 2 - | 0,000189632 - | 2 - | 0,00021362 - |
| 6527 | 1 | 3um_FIJJAFBF_82767 | Bacteria | Proteobacteri | Alphaproteob | Rhodobactera | Rhodobactera | Sulfitobacter | Sulfitobacter s | metabolism | Carbohydrate | 08ERU | trap dicarboxy | 367336.OM22 | 1 | 0,632455532 | 3,97E-05 | 2,60E-05 | 1 - | 3,42E-05 - | 1 - | 4,24E-05 - | 1 - | 4,03E-05 - | 2 - | 8,18E-05 - | 0 - | 0 - |
| 6528 | 1 | 0.2um_NOKCPHJN_218171 | Bacteria | Proteobacteri | Gammaprotec | unclassified | unclassified | unclassified | Gammaprotec | unclassified | unclassified | unclassified | unclassified | unclassified | 0,6 | 0,8 | 3,46E-05 | 4,60E-05 | 0 - | 0 - | 0 - | 0 - | 2 - | 0,000114815 - | 1 - | 5,83E-05 - | 0 - | 0 - |
| 6529 | 1 | 0.2um_NOKCPHJN_205751 | Eukaryota | Phaeophyceae | unclassified | Ectocarpales | Ectocarpaceae | Ectocarpus | Ectocarpus sil | cellular proces | Posttranslatio | KOG3083 | prohibitin | 3218.PP1S24_ | 1,2 | 0,4 | 7,56E-05 | 2,53E-05 | 1 - | 5,29E-05 - | 1 - | 6,57E-05 - | 2 - | 0,000124868 - | 1 - | 6,34E-05 - | 1 - | 7,14E-05 - |
| 6530 | 1 | 3um_FIJJAFBF_296715 | Bacteria | Proteobacteri | Alphaproteob | Rhodobactera | unclassified | unclassified | Rhodobactera | unclassified | unclassified | unclassified | unclassified | unclassified | 0,4 | 0,8 | 2,04E-05 | 4,08E-05 | 2 - | 0,000102024 - | 0 - | 0 - | 0 - | 0 - | 0 - | 0 - | 0 - | 0 - |
| 6531 | 1 | 0.2um_NOKCPHJN_738214 | Bacteria | Bacteroidetes | Flavobacteriia | unclassified | unclassified | unclassified | Flavobacteria | cellular proces | Intracellular tr | 18ATJ | Pfam:YadA | 626418.bglu_ | 1,4 | 0,489897949 | 2,08E-05 | 7,68E-06 | 1 - | 1,24E-05 - | 2 - | 3,08E-05 - | 2 - | 2,93E-05 - | 1 - | 1,49E-05 - | 1 - | 1,68E-05 - |
| 6532 | 1 | 3um_FIJJAFBF_164074 | Bacteria | Bacteroidetes | Flavobacteriia | Flavobacterial | Flavobacteriac | Formosa | Formosa sp. H | unclassified | unclassified | unclassified | unclassified | unclassified | 1 | 0 | 5,88E-05 | 5,58E-06 | 1 - | 4,93E-05 - | 1 - | 6,12E-05 - | 1 - | 5,81E-05 - | 1 - | 5,90E-05 - | 1 - | 6,65E-05 - |
| 6533 | 1 | 3um_FIJJAFBF_135240 | Bacteria | Cyanobacteria | unclassified | Synechococca | Synechococca | Synechococcu | Synechococcu | unclassified | unclassified | unclassified | unclassified | unclassified | 1,2 | 0,748331477 | 5,81E-05 | 3,41E-05 | 2 - | 8,41E-05 - | 0 - | 0 - | 2 - | 9,93E-05 - | 1 - | 5,04E-05 - | 1 - | 5,67E-05 - |
| 6534 | 1 | 0.2um_NOKCPHJN_258975 | Bacteria | Proteobacteri | Alphaproteob | Pelagibacteral | Pelagibacterac | Candidatus Pe | Candidatus Pe | cellular proces | Cell wall/mem | COG2885 | Ompa motb d | 1002672.SAR1 | 1 | 0 | 8,84E-05 | 8,39E-06 | 1 - | 7,40E-05 - | 1 - | 9,19E-05 - | 1 - | 8,74E-05 - | 1 - | 8,87E-05 - | 1 - | 9,99E-05 - |
| 6535 | 1 | 3um_FIJJAFBF_41184 | Bacteria | Proteobacteri | Alphaproteob | various | various | various | various | information st | Translation, ri | 179ZI | One of the pri | 367336.OM22 | 3,2 | 0,979795897 | 0,000330689 | 7,50E-05 | 5 - | 0,000441449 - | 3 - | 0,00032889 - | 2 - | 0,000208434 - | 3 - | 0,000317269 - | 3 - | 0,000357403 - |
| 6536 | 1 | 0.2um_NOKCPHJN_206382 | Bacteria | Proteobacteri | Alphaproteob | Rhodobactera | Rhodobactera | Planktomarina | Planktomarina | information st | Transcription | 05D27 | DNA-depende | 633131.TR2A | 20,2 | 2,785677655 | 0,000194652 | 2,79E-05 | 21 - | 0,00017022 - | 22 - | 0,000221428 - | 17 - | 0,000162655 - | 24 - | 0,000233023 - | 17 - | 0,000185937 - |
| 6537 | 1 | 3um_FIJJAFBF_732538 | Bacteria | Proteobacteri | Alphaproteob | Rhodobactera | Rhodobactera | Leisingera | Leisingera cae | metabolism | Amino acid tra | 01QS2 | ABC transport | 246200.SPO3 | 0,6 | 0,489897949 | 2,56E-05 | 2,13E-05 | 1 - | 3,62E-05 - | 0 - | 0 - | 1 - | 4,27E-05 - | 0 - | 0 - | 1 - | 4,89E-05 - |
| 6538 | 1 | 3um_FIJJAFBF_520014 | Bacteria | Proteobacteri | Alphaproteob | Rhodobactera | Rhodobactera | Thalassobacte | various | unclassified | unclassified | unclassified | unclassified | unclassified | 1,2 | 0,4 | 0,000146773 | 3,25E-05 | 2 - | 0,0002106 - | 1 - | 0,000130751 - | 1 - | 0,000124296 - | 1 - | 0,000126131 - | 1 - | 0,000142087 - |
| 6539 | 1 | 3um_FIJJAFBF_169755 | Bacteria | Bacteroidetes | unclassified | unclassified | unclassified | unclassified | Bacteroidetes | unclassified | unclassified | unclassified | unclassified | unclassified | 7,4 | 1,496662955 | 0,000224154 | 3,38E-05 | 10 - | 0,000256771 - | 8 - | 0,000255067 - | 6 - | 0,000181855 - | 6 - | 0,000184541 - | 7 - | 0,000242533 - |
| 6540 | 1 | 3um_FIJJAFBF_14212 | Bacteria | Bacteroidetes | unclassified | unclassified | unclassified | unclassified | Bacteroidetes | unclassified | unclassified | unclassified | unclassified | unclassified | 3,2 | 1,166190379 | 0,000178631 | 7,13E-05 | 2 - | 9,22E-05 - | 2 - | 0,000114473 - | 5 - | 0,000272053 - | 3 - | 0,000165643 - | 4 - | 0,000248794 - |
| 6541 | 1 | 3um_FIJJAFBF_434532 | Bacteria | Bacteroidetes | Flavobacteriia | Flavobacterial | Flavobacteriac | Polaribacter | various | information st | Translation, ri | 08UIK | This protein bi | 867900.Celly_ | 1 | 0 | 9,45E-05 | 8,97E-06 | 1 - | 7,92E-05 - | 1 - | 9,83E-05 - | 1 - | 9,34E-05 - | 1 - | 9,48E-05 - | 1 - | 0,00010681 - |
| 6542 | 1 | 3um_FIJJAFBF_374067 | Bacteria | Bacteroidetes | Flavobacteriia | Flavobacterial | unclassified | unclassified | Flavobacterial | unclassified | unclassified | unclassified | unclassified | unclassified | 2,2 | 0,4 | 0,000101192 | 2,06E-05 | 2 - | 7,70E-05 - | 2 - | 9,57E-05 - | 2 - | 9,09E-05 - | 3 - | 0,000138406 - | 2 - | 0,000103943 - |
| 6543 | 1 | 0.2um_NOKCPHJN_760798 | Bacteria | Proteobacteri | Alphaproteob | Pelagibacteral | Pelagibacterac | Candidatus Pe | Candidatus Pe | poorly charact | Function unkn | 05W4U | Zinc finger, CD | 335992.SAR11 | 0,2 | 0,4 | 2,94E-05 | 5,89E-05 | 1 - | 0,00014715 - | 0 - | 0 - | 0 - | 0 - | 0 - | 0 - | 0 - | 0 - |
| 6544 | 1 | 0.2um_NOKCPHJN_74584 | Bacteria | Proteobacteri | Gammaprotec | Oceanospirilla | Saccharospirill | Reinekea | Reinekea fors | information st | Replication, re | 16W6D | single-strande | 203122.Sde_0 | 0,6 | 0,8 | 3,86E-05 | 4,88E-05 | 2 - | 0,000115354 - | 0 - | 0 - | 0 - | 0 - | 0 - | 0 - | 1 - | 7,78E-05 - |
| 6545 | 1 | 3um_FIJJAFBF_721248 | Bacteria | Bacteroidetes | unclassified | unclassified | unclassified | unclassified | Bacteroidetes | poorly charact | Function unkn | 09BD6 | Protein of unk | 983544.Lacal_ | 0,6 | 0,8 | 3,46E-05 | 4,78E-05 | 1 - | 5,15E-05 - | 0 - | 0 - | 2 - | 0,000121509 - | 0 - | 0 - | 0 - | 0 - |
| 6546 | 1 | 0.2um_NOKCPHJN_472881 | Bacteria | Bacteroidetes | Flavobacteriia | Flavobacterial | Flavobacteriac | Maribacter | Maribacter po | information st | Translation, ri | 0PA05 | 30S ribosomal | 867900.Celly_ | 2,6 | 0,489897949 | 0,000210089 | 4,48E-05 | 2 - | 0,000135031 - | 3 - | 0,000251504 - | 3 - | 0,000239086 - | 3 - | 0,000242618 - | 2 - | 0,000182205 - |
| 6547 | 1 | 3um_FIJJAFBF_45541 | Bacteria | Bacteroidetes | Flavobacteriia | various | various | various | various | information st | Translation, ri | 0PBIW | 50S ribosomal | 487797.Flav3 | 0,8 | 0,4 | 0,000163157 | 8,29E-05 | 1 - | 0,00017658 - | 1 - | 0,00021926 - | 1 - | 0,000208434 - | 1 - | 0,000211513 - | 0 - | 0 - |
| 6548 | 1 | 3um_FIJJAFBF_119785 | Eukaryota | Apicomplexa | Aconoidasida | Haemosporida | Plasmodiidae | Plasmodium | Plasmodium s | unclassified | unclassified | unclassified | unclassified | unclassified | 1,8 | 0,979795897 | 5,32E-05 | 2,71E-05 | 3 - | 7,63E-05 - | 3 - | 9,48E-05 - | 1 - | 3,00E-05 - | 1 - | 3,05E-05 - | 1 - | 3,43E-05 - |
| 6549 | 1 | 3um_FIJJAFBF_734730 | Eukaryota | Chromerida | unclassified | unclassified | unclassified | Vitrella | Vitrella brassi | poorly charact | Function unkn | 0XPG5 | KH domain co | 67593.Physo1 | 1,4 | 0,489897949 | 3,42E-05 | 1,01E-05 | 2 - | 4,19E-05 - | 1 - | 2,60E-05 - | 1 - | 2,47E-05 - | 2 - | 5,02E-05 - | 1 - | 2,83E-05 - |
| 6550 | 1 | 3um_FIJJAFBF_41185 | Bacteria | Proteobacteri | Alphaproteob | Rhodobactera | unclassified | unclassified | Rhodobactera | unclassified | unclassified | unclassified | unclassified | unclassified | 0,8 | 0,748331477 | 5,68E-05 | 5,02E-05 | 2 - | 0,000129691 - | 0 - | 0 - | 1 - | 7,65E-05 - | 1 - | 7,77E-05 - | 0 - | 0 - |
| 6551 | 1 | 3um_FIJJAFBF_220487 | Bacteria | Proteobacteri | Gammaprotec | Alteromonada | Alteromonada | various | various | information st | Translation, ri | COG0051 | tRNA binding | 715451.ambt_ | 3 | 0,894427191 | 0,000392689 | 0,000103906 | 4 - | 0,000445735 - | 4 - | 0,000553471 - | 2 - | 0,000263072 - | 3 - | 0,000400437 - | 2 - | 0,000300727 - |
| 6552 | 1 | 0.2um_NOKCPHJN_95013 | Bacteria | Bacteroidetes | Flavobacteriia | Flavobacterial | Flavobacteriac | Marinirhabdu | Marinirhabdu | information st | Transcription | COG0085 | DNA-depende | 50743.SCB49_ | 17,8 | 2,712931993 | 0,000189774 | 1,51E-05 | 23 - | 0,000207863 - | 18 - | 0,000201995 - | 16 - | 0,000170686 - | 16 - | 0,000173207 - | 16 - | 0,000195117 - |
| 6553 | 1 | 3um_FIJJAFBF_610478 | Bacteria | Bacteroidetes | unclassified | unclassified | unclassified | unclassified | Bacteroidetes | information st | Translation, ri | 08YY1 | This protein is | 50743.SCB49_ | 1 | 0 | 0,000117117 | 1,11E-05 | 1 - | 9,81E-05 - | 1 - | 0,000121811 - | 1 - | 0,000115797 - | 1 - | 0,000117507 - | 1 - | 0,000132371 - |
| 6554 | 2 | 3um_FIJJAFBF_452324;3um_FIJJAFBF_754707 | Bacteria | Bacteroidetes | Flavobacteriia | Flavobacterial | Flavobacteriac | Polaribacter | Polaribacter fi | unclassified | unclassified | unclassified | unclassified | unclassified | 0,2 | 0,4 | 1,24E-05 | 2,48E-05 | 1 - | 6,20E-05 - | 0 - | 0 - | 0 - | 0 - | 0 - | 0 - | 0 - | 0 - |
| 6555 | 1 | 0.2um_NOKCPHJN_47150 | Bacteria | Proteobacteri | Gammaprotec | Cellvibrionales | Porticoccacea | unclassified | Gammaprotec | unclassified | unclassified | unclassified | unclassified | unclassified | 1 | 0,632455532 | 0,000118648 | 6,63E-05 | 2 - | 0,000204958 - | 1 - | 0,000127249 - | 0 - | 0 - | 1 - | 0,000122753 - | 1 - | 0,000138281 - |
| 6556 | 1 | 3um_FIJJAFBF_774341 | Bacteria | Proteobacteri | Gammaprotec | Oceanospirilla | Saccharospirill | Reinekea | Reinekea fors | unclassified | unclassified | unclassified | unclassified | unclassified | 1,6 | 0,8 | 0,000289396 | 0,000148619 | 2 - | 0,000306071 - | 0 - | 0 - | 2 - | 0,000361286 - | 2 - | 0,000366622 - | 2 - | 0,000412999 - |
| 6557 | 1 | 3um_FIJJAFBF_433876 | Bacteria | Bacteroidetes | Flavobacteriia | Flavobacterial | Flavobacteriac | Polaribacter | Polaribacter s | poorly charact | Function unkn | 0PCAU | NA | 313598.MED1 | 0,4 | 0,8 | 6,04E-05 | 0,000120818 | 2 - | 0,000302044 - | 0 - | 0 - | 0 - | 0 - | 0 - | 0 - | 0 - | 0 - |
| 6558 | 1 | 3um_FIJJAFBF_504263 | Eukaryota | Phaeophyceae | unclassified | Ectocarpales | Chordariaceae | Pleurocladia | Pleurocladia l | metabolism | Energy produc | COG4451 | RuBisCO catal | 45157.CMV01 | 1 | 0,632455532 | 9,53E-05 | 5,34E-05 | 2 - | 0,000165146 - | 1 - | 0,000102532 - | 1 - | 9,75E-05 - | 0 - | 0 - | 1 - | 0,00011142 - |
| 6559 | 1 | 3um_FIJJAFBF_182831 | Viruses | unclassified | unclassified | unclassified | unclassified | unclassified | uncultured M | unclassified | unclassified | unclassified | unclassified | unclassified | 1 | 0,632455532 | 7,72E-05 | 4,94E-05 | 1 - | 6,45E-05 - | 1 - | 8,01E-05 - | 0 - | 0 - | 2 - | 0,000154476 - | 1 - | 8,70E-05 - |
| 6560 | 1 | 0.2um_NOKCPHJN_625061 | Eukaryota | unclassified | unclassified | Isochrysidales | Noelaerhabda | Emiliania | Emiliania huxl | unclassified | unclassified | unclassified | unclassified | unclassified | 1 | 0 | 1,41E-05 | 1,33E-06 | 1 - | 1,18E-05 - | 1 - | 1,46E-05 - | 1 - | 1,39E-05 - | 1 - | 1,41E-05 - | 1 - | 1,59E-05 - |
| 6561 | 1 | 0.2um_NOKCPHJN_164568 | Eukaryota | Eustigmatoph | unclassified | Eustigmatales | Monodopsida | Nannochlorop | Nannochlorop | unclassified | unclassified | unclassified | unclassified | unclassified | 0,6 | 0,489897949 | 2,73E-05 | 2,23E-05 | 0 - | 0 - | 1 - | 4,47E-05 - | 0 - | 0 - | 1 - | 4,31E-05 - | 1 - | 4,85E-05 - |
| 6562 | 1 | 3um_FIJJAFBF_383463 | Bacteria | Proteobacteri | Gammaprotec | Oceanospirilla | Oceanospirilla | Motiliproteus | Motiliproteus | information st | Translation, ri | 0QQXI | This protein is | 314283.MED2 | 3,4 | 1,019803903 | 0,000393859 | 0,000132858 | 2 - | 0,000191295 - | 5 - | 0,000593829 - | 3 - | 0,000338706 - | 4 - | 0,000458278 - | 3 - | 0,000387186 - |
| 6563 | 1 | 3um_FIJJAFBF_532817 | Bacteria | Proteobacteri | Alphaproteob | Rhodobactera | unclassified | unclassified | Rhodobactera | poorly charact | Function unkn | 08YYC | ycei family | 644076.SCH4 | 1,4 | 1,356465997 | 9,30E-05 | 8,91E-05 | 3 - | 0,000180278 - | 0 - | 0 - | 3 - | 0,000212799 - | 1 - | 7,20E-05 - | 0 - | 0 - |
| 6564 | 2 | 0.2um_NOKCPHJN_12438;3um_FIJJAFBF_11124 | various | various | various | various | various | various | various | poorly charact | Function unkn | various | various | various | 3,2 | 1,469693846 | 5,84E-05 | 2,02E-05 | 6 - | 9,53E-05 - | 3 - | 5,91E-05 - | 2 - | 3,75E-05 - | 3 - | 5,70E-05 - | 2 - | 4,28E-05 - |
| 6565 | 1 | 3um_FIJJAFBF_792634 | Eukaryota | Bacillariophyt | Bacillariophyc | Naviculales | Phaeodactylac | Phaeodactylu | Phaeodactylu | unclassified | unclassified | unclassified | unclassified | unclassified | 2 | 0 | 5,38E-05 | 5,11E-06 | 2 - | 4,51E-05 - | 2 - | 5,60E-05 - | 2 - | 5,32E-05 - | 2 - | 5,40E-05 - | 2 - | 6,09E-05 - |
| 6566 | 1 | 0.2um_NOKCPHJN_130099 | Eukaryota | Arthropoda | Insecta | Hymenoptera | Apidae | Habropoda | Habropoda la | poorly charact | Function unkn | 0F9Q7 | NA | 353151.XP_66 | 0,6 | 0,489897949 | 7,53E-07 | 6,15E-07 | 0 - | 0 - | 1 - | 1,29E-06 - | 1 - | 1,23E-06 - | 1 - | 1,25E-06 - | 0 - | 0 - |
| 6567 | 1 | 3um_FIJJAFBF_438666 | Bacteria | Proteobacteri | Gammaprotec | Oceanospirilla | Saccharospirill | Reinekea | Reinekea blan | metabolism | Amino acid tra | 0QIER | threonine syn | 314283.MED2 | 1 | 0,894427191 | 2,80E-05 | 2,50E-05 | 2 - | 4,94E-05 - | 2 - | 6,13E-05 - | 1 - | 2,91E-05 - | 0 - | 0 - | 0 - | 0 - |
| 6568 | 1 | 0.2um_NOKCPHJN_164694 | Bacteria | Proteobacteri | Gammaprotec | unclassified | unclassified | unclassified | Gammaprotec | cellular proces | Intracellular tr | 0QJ7H | motA TolQ Exl | 234831.PSM_ | 4,6 | 1,019803903 | 0,00013515 | 1,94E-05 | 6 - | 0,000150692 - | 4 - | 0,000124743 - | 5 - | 0,00014823 - | 5 - | 0,000150419 - | 3 - | 0,000101668 - |
| 6569 | 1 | 0.2um_NOKCPHJN_118676 | Bacteria | Proteobacteri | Gammaprotec | Oceanospirilla | Saccharospirill | Reinekea | Reinekea blan | cellular proces | Defense mech | 0QJ8P | (ABC) transpo | 314283.MED2 | 0,2 | 0,4 | 8,80E-06 | 1,76E-05 | 0 - | 0 - | 0 - | 0 - | 1 - | 4,40E-05 - | 0 - | 0 - | 0 - | 0 - |
| 6570 | 1 | 3um_FIJJAFBF_16279 | Bacteria | Proteobacteri | Alphaproteob | Rhodobactera | Rhodobactera | unclassified | Rhodobactera | cellular proces | Cell wall/mem | 172GZ | Catalyzes the | 388401.RB215 | 0,2 | 0,4 | 7,76E-06 | 1,55E-05 | 1 - | 3,88E-05 - | 0 - | 0 - | 0 - | 0 - | 0 - | 0 - | 0 - | 0 - |
| 6571 | 1 | 0.2um_NOKCPHJN_112700 | Bacteria | Bacteroidetes | Flavobacteriia | unclassified | unclassified | unclassified | Flavobacteria | unclassified | unclassified | unclassified | unclassified | unclassified | 0,6 | 0,489897949 | 9,79E-06 | 8,00E-06 | 0 - | 0 - | 1 - | 1,68E-05 - | 1 - | 1,60E-05 - | 1 - | 1,62E-05 - | 0 - | 0 - |
| 6572 | 1 | 3um_FIJJAFBF_386707 | Bacteria | Proteobacteri | Gammaprotec | Oceanospirilla | Saccharospirill | Reinekea | Reinekea fors | unclassified | unclassified | unclassified | unclassified | unclassified | 2,2 | 0,979795897 | 9,28E-05 | 4,16E-05 | 1 - | 3,49E-05 - | 2 - | 8,66E-05 - | 4 - | 0,00016472 - | 2 - | 8,36E-05 - | 2 - | 9,41E-05 - |

| | | | | | | | | | | | | | | | | | | | | | | | | | | | | |
|---|---|---|---|---|---|---|---|---|---|---|---|---|---|---|---|---|---|---|---|---|---|---|---|---|---|---|---|---|
| 6573 | 1 | 0.2um_NOKCPHJN_994909 | Bacteria | Proteobacteri | Alphaproteob | Rhodobactera | Rhodobactera | Planktomarina | Planktomarina | unclassified | unclassified | unclassified | unclassified | unclassified | 2,8 | 0,4 | 0,000129153 | 2,25E-05 | 3 - | 0,000115936 - | 3 - | 0,000143958 - | 3 - | 0,000136851 - | 2 - | 9,26E-05 - | 3 - | 0,000156439 - |
| 6574 | 1 | 3um_FIJJAFBF_712425 | Bacteria | Bacteroidetes | Flavobacteriia | Flavobacterial | Flavobacteriac | Marinirhabdu | Marinirhabdu | metabolism | Energy produc | COG0711 | F(1)F(0) ATP s | 50743.SCB49_ | 2,4 | 1,019803903 | 0,000192509 | 6,68E-05 | 4 - | 0,000276571 - | 2 - | 0,000171709 - | 3 - | 0,000244847 - | 1 - | 8,28E-05 - | 2 - | 0,000186596 - |
| 6575 | 1 | 3um_FIJJAFBF_258583 | Bacteria | Proteobacteri | Gammaprotec | Cellvibrionales | Porticoccacea | unclassified | gamma protec | cellular proces | Posttranslatio | 0QHT2 | Peroxiredoxin | 314287.GB22 | 1 | 0 | 8,67E-05 | 8,23E-06 | 1 - | 7,26E-05 - | 1 - | 9,02E-05 - | 1 - | 8,57E-05 - | 1 - | 8,70E-05 - | 1 - | 9,80E-05 - |
| 6576 | 1 | 0.2um_NOKCPHJN_14311 | Bacteria | Bacteroidetes | Cytophagia | Cytophagales | Flammeovirga | unclassified | Flammeovirga | poorly charact | Function unkn | 07EF7 | domain protei | 649349.Lbys_ | 2 | 0,632455532 | 1,08E-05 | 3,29E-06 | 2 - | 9,19E-06 - | 2 - | 1,14E-05 - | 3 - | 1,63E-05 - | 2 - | 1,10E-05 - | 1 - | 6,20E-06 - |
| 6577 | 1 | 3um_FIJJAFBF_806703 | Eukaryota | unclassified | Oomycetes | Saprolegniales | Saprolegniace | Thraustotheca | Thraustotheca | unclassified | unclassified | unclassified | unclassified | unclassified | 3,4 | 0,8 | 0,000113437 | 3,68E-05 | 3 - | 8,26E-05 - | 3 - | 0,000102532 - | 3 - | 9,75E-05 - | 3 - | 9,89E-05 - | 5 - | 0,000185701 - |
| 6578 | 1 | 0.2um_NOKCPHJN_12072 | Bacteria | Proteobacteri | Gammaprotec | Cellvibrionales | Porticoccacea | unclassified | SAR92 bacteri | information st | Transcription | 16PMT | DNA-depende | 498211.CJA_0 | 0,4 | 0,489897949 | 1,68E-05 | 2,06E-05 | 0 - | 0 - | 1 - | 4,31E-05 - | 1 - | 4,09E-05 - | 0 - | 0 - | 0 - | 0 - |
| 6579 | 1 | 3um_FIJJAFBF_791054 | Eukaryota | unclassified | Pelagophycea | Pelagomonad | unclassified | Aureococcus | Aureococcus a | unclassified | unclassified | unclassified | unclassified | unclassified | 0,6 | 0,489897949 | 7,51E-05 | 6,16E-05 | 0 - | 0 - | 0 - | 0 - | 1 - | 0,000118844 - | 1 - | 0,000120599 - | 1 - | 0,000135855 - |
| 6580 | 1 | 3um_FIJJAFBF_344002 | Bacteria | Proteobacteri | Alphaproteob | Rhodobactera | Rhodobactera | Shimia | Shimia halioti | unclassified | unclassified | unclassified | unclassified | unclassified | 0,8 | 0,4 | 8,69E-05 | 4,42E-05 | 1 - | 9,41E-05 - | 1 - | 0,000116819 - | 1 - | 0,000111051 - | 1 - | 0,000112691 - | 0 - | 0 - |
| 6581 | 1 | 0.2um_NOKCPHJN_66499 | Bacteria | Bacteroidetes | Cytophagia | Cytophagales | Cyclobacteriac | Algoriphagus | Algoriphagus | unclassified | unclassified | unclassified | unclassified | unclassified | 1 | 0,632455532 | 1,53E-05 | 8,75E-06 | 2 - | 2,73E-05 - | 1 - | 1,69E-05 - | 1 - | 1,61E-05 - | 1 - | 1,63E-05 - | 0 - | 0 - |
| 6582 | 1 | 0.2um_NOKCPHJN_41564 | Bacteria | Bacteroidetes | Flavobacteriia | Flavobacterial | Crocinitomica | unclassified | Crocinitomica | information st | Transcription | 0P7SF | DNA-depende | 755732.Fluta_ | 10 | 1,673320053 | 0,000106354 | 1,22E-05 | 12 - | 0,000108365 - | 11 - | 0,000123344 - | 10 - | 0,000106595 - | 10 - | 0,000108169 - | 7 - | 8,53E-05 - |
| 6583 | 1 | 0.2um_NOKCPHJN_200782 | Bacteria | Proteobacteri | Gammaprotec | Oceanospirilla | Saccharospiril | Saccharospiril | Saccharospiril | information st | Translation, ril | 16QAF | thus facilitatin | 349521.HCH_ | 9,2 | 0,748331477 | 0,000224763 | 2,68E-05 | 10 - | 0,000204958 - | 8 - | 0,000203598 - | 9 - | 0,000217739 - | 9 - | 0,000220955 - | 10 - | 0,000276562 - |
| 6584 | 1 | 3um_FIJJAFBF_336232 | Bacteria | Bacteroidetes | Cytophagia | Cytophagales | Flammeovirga | unclassified | Flammeovirga | cellular proces | Posttranslatio | 0FW51 | Trigger factor | 643867.Ftrac_ | 0,2 | 0,4 | 7,06E-06 | 1,41E-05 | 0 - | 0 - | 0 - | 0 - | 0 - | 0 - | 0 - | 0 - | 1 - | 3,53E-05 - |
| 6585 | 1 | 0.2um_NOKCPHJN_491689 | Bacteria | Proteobacteri | Alphaproteob | Rhodobactera | unclassified | unclassified | various | metabolism | Lipid transpor | 16PR9 | acyl-Coa dehy | 536019.Meso | 0,4 | 0,489897949 | 1,50E-05 | 1,85E-05 | 0 - | 0 - | 0 - | 0 - | 1 - | 3,51E-05 - | 0 - | 0 - | 1 - | 4,01E-05 - |
| 6586 | 1 | 0.2um_NOKCPHJN_1105933 | Eukaryota | unclassified | Cryptophyta | Pyrenomonad | Geminigerace | Guillardia | Guillardia thet | poorly charact | Function unkn | arCOG02812 | bacteriorhodo | 272569.rrnAC | 0,8 | 0,4 | 0,000102381 | 5,26E-05 | 1 - | 0,00010828 - | 0 - | 0 - | 1 - | 0,000127813 - | 1 - | 0,000129701 - | 1 - | 0,000146108 - |
| 6587 | 1 | 3um_FIJJAFBF_128452 | Bacteria | Bacteroidetes | Flavobacteriia | Flavobacterial | Flavobacteriac | Polaribacter | Polaribacter s | cellular proces | Intracellular tr | 05FGH | MotA TolQ exl | 313594.PI23P | 12,4 | 1,019803903 | 0,000705824 | 7,03E-05 | 13 - | 0,000621707 - | 12 - | 0,000712594 - | 14 - | 0,000790313 - | 11 - | 0,000630132 - | 12 - | 0,000774372 - |
| 6588 | 1 | 3um_FIJJAFBF_225138 | Bacteria | Proteobacteri | Alphaproteob | Rhodobactera | Rhodobactera | Planktomarina | Planktomarina | information st | Translation, ril | COG0186 | One of the pri | 391593.RCCS2 | 0,2 | 0,4 | 3,57E-05 | 7,13E-05 | 0 - | 0 - | 0 - | 0 - | 1 - | 0,000178266 - | 0 - | 0 - | 0 - | 0 - |
| 6589 | 1 | 3um_FIJJAFBF_189629 | Bacteria | Bacteroidetes | Flavobacteriia | Flavobacterial | Flavobacteriac | Polaribacter | Polaribacter s | metabolism | Nucleotide tra | 05CM0 | Catalyzes the | 313598.MED1 | 2,4 | 0,489897949 | 6,45E-05 | 1,49E-05 | 2 - | 4,49E-05 - | 3 - | 8,37E-05 - | 3 - | 7,95E-05 - | 2 - | 5,38E-05 - | 2 - | 6,06E-05 - |
| 6590 | 1 | 3um_FIJJAFBF_768809 | Bacteria | Bacteroidetes | Flavobacteriia | Flavobacterial | Flavobacteriac | Formosa | Formosa sp. H | information st | Transcription | 0P88Z | transcriptiona | 391603.FBALC | 0,6 | 0,489897949 | 4,88E-05 | 4,01E-05 | 1 - | 7,22E-05 - | 0 - | 0 - | 1 - | 8,52E-05 - | 1 - | 8,65E-05 - | 0 - | 0 - |
| 6591 | 1 | 3um_FIJJAFBF_370311 | Bacteria | Proteobacteri | Alphaproteob | Rhodobactera | Rhodobactera | Planktomarina | Planktomarina | metabolism | Secondary me | 01Q86 | extracellular s | 367336.OM22 | 8 | 2,19089023 | 0,00029574 | 6,41E-05 | 11 - | 0,000346853 - | 5 - | 0,000195768 - | 7 - | 0,000260543 - | 10 - | 0,000377701 - | 7 - | 0,000297836 - |
| 6592 | 1 | 3um_FIJJAFBF_135261 | Bacteria | Cyanobacteria | unclassified | Synechococca | Synechococca | Synechococcu | Synechococcu | unclassified | unclassified | unclassified | unclassified | unclassified | 0,6 | 0,489897949 | 1,66E-05 | 1,38E-05 | 1 - | 2,31E-05 - | 1 - | 2,87E-05 - | 0 - | 0 - | 0 - | 0 - | 1 - | 3,12E-05 - |
| 6593 | 1 | 0.2um_NOKCPHJN_199759 | Eukaryota | unclassified | Labyrinthulom | unclassified | Thraustochytr | Aurantiochytr | Aurantiochytr | unclassified | unclassified | unclassified | unclassified | unclassified | 0,4 | 0,489897949 | 5,02E-06 | 6,20E-06 | 1 - | 1,12E-05 - | 1 - | 1,39E-05 - | 0 - | 0 - | 0 - | 0 - | 0 - | 0 - |
| 6594 | 1 | 3um_FIJJAFBF_255080 | Eukaryota | Bacillariophyt | Bacillariophyc | Bacillariales | Bacillariaceae | Fragilariopsis | Fragilariopsis | unclassified | unclassified | unclassified | unclassified | unclassified | 0,2 | 0,4 | 2,28E-05 | 4,56E-05 | 0 - | 0 - | 1 - | 0,000114015 - | 0 - | 0 - | 0 - | 0 - | 0 - | 0 - |
| 6595 | 1 | 0.2um_NOKCPHJN_256901 | Bacteria | Proteobacteri | Gammaprotec | Oceanospirilla | Saccharospiril | Reinekea | Reinekea fors | metabolism | Carbohydrate | 16Q47 | ec 3.2.1.23 | 314283.MED2 | 0,2 | 0,4 | 3,03E-06 | 6,06E-06 | 0 - | 0 - | 0 - | 0 - | 0 - | 0 - | 0 - | 0 - | 1 - | 1,52E-05 - |
| 6596 | 1 | 3um_FIJJAFBF_172201 | Bacteria | unclassified | unclassified | unclassified | unclassified | unclassified | bacterium TM | unclassified | unclassified | unclassified | unclassified | unclassified | 0,2 | 0,4 | 5,88E-06 | 1,18E-05 | 0 - | 0 - | 1 - | 2,94E-05 - | 0 - | 0 - | 0 - | 0 - | 0 - | 0 - |
| 6597 | 1 | 3um_FIJJAFBF_220493 | Bacteria | Proteobacteri | Gammaprotec | Alteromonada | Alteromonada | Paraglaciecola | various | information st | Translation, ril | 17C3B | The globular d | 342610.Patl_ | 2,2 | 0,4 | 0,000270009 | 2,48E-05 | 3 - | 0,000313028 - | 2 - | 0,000259125 - | 2 - | 0,000246331 - | 2 - | 0,00024997 - | 2 - | 0,00028159 - |
| 6598 | 1 | 3um_FIJJAFBF_145290 | Bacteria | Proteobacteri | Alphaproteob | Rhodobactera | Rhodobactera | unclassified | Rhodobactera | metabolism | Amino acid tra | COG0687 | polyamine bin | 314270.RB208 | 0,6 | 0,489897949 | 2,20E-05 | 1,81E-05 | 1 - | 3,20E-05 - | 1 - | 3,97E-05 - | 0 - | 0 - | 1 - | 3,83E-05 - | 0 - | 0 - |
| 6599 | 1 | 3um_FIJJAFBF_500620 | Bacteria | Bacteroidetes | Flavobacteriia | Flavobacterial | Flavobacteriac | Gaetbulibacte | Gaetbulibacte | poorly charact | Function unkn | 09ARF | SSS sodium so | 411154.GFO_ | 0,8 | 0,4 | 1,94E-05 | 9,83E-06 | 1 - | 2,09E-05 - | 1 - | 2,60E-05 - | 1 - | 2,47E-05 - | 1 - | 2,51E-05 - | 0 - | 0 - |
| 6600 | 2 | 0.2um_NOKCPHJN_261462;0.2um_NOKCPHJN_3 | Bacteria | Bacteroidetes | unclassified | unclassified | unclassified | unclassified | Bacteroidetes | poorly charact | Function unkn | 05ERK | gliding motilit | 926562.Oweh | 2,2 | 1,166190379 | 6,23E-05 | 3,37E-05 | 0 - | 0 - | 3 - | 8,45E-05 - | 2 - | 5,36E-05 - | 3 - | 8,15E-05 - | 3 - | 9,18E-05 - |
| 6601 | 1 | 3um_FIJJAFBF_307116 | Bacteria | Bacteroidetes | Cytophagia | Cytophagales | Flammeovirga | unclassified | Flammeovirga | poorly charact | Function unkn | 0P8NK | Alpha beta hy | 487797.Flav3 | 0,2 | 0,4 | 8,67E-06 | 1,73E-05 | 0 - | 0 - | 0 - | 0 - | 0 - | 0 - | 1 - | 4,34E-05 - | 0 - | 0 - |
| 6602 | 1 | 0.2um_NOKCPHJN_87353 | Bacteria | Proteobacteri | Alphaproteob | Rhodobactera | Rhodobactera | unclassified | Rhodobactera | cellular proces | Cell wall/mem | 16ZEI | MltA-interacti | 398580.Dshi_ | 3,4 | 0,8 | 0,000183531 | 2,48E-05 | 5 - | 0,000230475 - | 3 - | 0,000171709 - | 3 - | 0,000163232 - | 3 - | 0,000165643 - | 3 - | 0,000186596 - |
| 6603 | 1 | 3um_FIJJAFBF_132757 | Bacteria | Proteobacteri | Gammaprotec | Oceanospirilla | Saccharospiril | Reinekea | Reinekea fors | cellular proces | Posttranslatio | 0QIPK | Stringent star | 314283.MED2 | 2 | 0,632455532 | 0,000129145 | 3,41E-05 | 3 - | 0,000164751 - | 2 - | 0,000136382 - | 1 - | 6,48E-05 - | 2 - | 0,000131563 - | 2 - | 0,000148205 - |
| 6604 | 1 | 3um_FIJJAFBF_135258 | Eukaryota | unclassified | Synurophycea | Parmales | Triparmaceae | Triparma | Triparma laev | metabolism | Energy produc | COG0711 | F(1)F(0) ATP s | 314260.PB25 | 6,4 | 0,8 | 0,000658897 | 6,61E-05 | 7 - | 0,000608665 - | 7 - | 0,000755782 - | 7 - | 0,000718466 - | 6 - | 0,000624924 - | 5 - | 0,000586646 - |
| 6605 | 1 | 0.2um_NOKCPHJN_626560 | Eukaryota | Phaeophyceae | unclassified | Ectocarpales | Ectocarpaceae | Ectocarpus | Ectocarpus sil | unclassified | unclassified | unclassified | unclassified | unclassified | 1,2 | 0,4 | 9,41E-05 | 2,08E-05 | 2 - | 0,000135031 - | 1 - | 8,38E-05 - | 1 - | 7,97E-05 - | 1 - | 8,09E-05 - | 1 - | 9,11E-05 - |
| 6606 | 1 | 3um_FIJJAFBF_777828 | Bacteria | Bacteroidetes | Flavobacteriia | Flavobacterial | Flavobacteriac | Polaribacter | various | unclassified | unclassified | unclassified | unclassified | unclassified | 8 | 2,097617696 | 0,000723033 | 0,00010948 | 12 - | 0,000930622 - | 7 - | 0,000674076 - | 8 - | 0,000732336 - | 7 - | 0,000650259 - | 6 - | 0,00062787 - |
| 6607 | 1 | 3um_FIJJAFBF_119155 | Bacteria | Bacteroidetes | Flavobacteriia | Flavobacterial | Flavobacteriac | Polaribacter | Polaribacter s | poorly charact | Function unkn | 05C99 | atpase involve | 313594.PI23P | 0,4 | 0,489897949 | 9,30E-06 | 1,14E-05 | 0 - | 0 - | 0 - | 0 - | 0 - | 0 - | 1 - | 2,19E-05 - | 1 - | 2,46E-05 - |
| 6608 | 1 | 0.2um_NOKCPHJN_339044 | unclassified | unclassified | unclassified | unclassified | unclassified | unclassified | unclassified | unclassified | unclassified | unclassified | unclassified | unclassified | 1 | 0 | 0,00013434 | 1,28E-05 | 1 - | 0,000112526 - | 1 - | 0,000139724 - | 1 - | 0,000132826 - | 1 - | 0,000134788 - | 1 - | 0,000151838 - |
| 6609 | 1 | 3um_FIJJAFBF_137600 | Bacteria | Bacteroidetes | Flavobacteriia | Flavobacterial | Crocinitomica | unclassified | Crocinitomica | unclassified | unclassified | unclassified | unclassified | unclassified | 1,4 | 0,8 | 0,000101175 | 5,74E-05 | 1 - | 6,07E-05 - | 1 - | 7,54E-05 - | 3 - | 0,000215051 - | 1 - | 7,27E-05 - | 1 - | 8,19E-05 - |
| 6610 | 1 | 0.2um_NOKCPHJN_79024 | Bacteria | Bacteroidetes | Flavobacteriia | Flavobacterial | Crocinitomica | unclassified | Crocinitomica | information st | Translation, ril | COG0088 | One of the pri | 388413.ALPR1 | 1 | 0,632455532 | 6,39E-05 | 4,19E-05 | 1 - | 5,49E-05 - | 1 - | 6,82E-05 - | 1 - | 6,48E-05 - | 2 - | 0,000131563 - | 0 - | 0 - |
| 6611 | 1 | 0.2um_NOKCPHJN_311017 | Bacteria | Proteobacteri | Alphaproteob | Rhodobactera | Rhodobactera | Thalassobius | Thalassobius s | unclassified | unclassified | unclassified | unclassified | unclassified | 1,8 | 1,326649916 | 4,05E-05 | 2,63E-05 | 4 - | 7,94E-05 - | 2 - | 4,93E-05 - | 2 - | 4,69E-05 - | 0 - | 0 - | 1 - | 2,68E-05 - |
| 6612 | 1 | 0.2um_NOKCPHJN_232960 | Eukaryota | unclassified | Labyrinthulom | unclassified | Thraustochytr | Aurantiochytr | Aurantiochytr | unclassified | unclassified | unclassified | unclassified | unclassified | 0,6 | 0,489897949 | 4,57E-06 | 3,76E-06 | 1 - | 6,76E-06 - | 0 - | 0 - | 1 - | 7,98E-06 - | 1 - | 8,10E-06 - | 0 - | 0 - |
| 6613 | 1 | 3um_FIJJAFBF_187657 | Bacteria | Bacteroidetes | Flavobacteriia | Flavobacterial | Flavobacteriac | Marinirhabdu | Marinirhabdu | poorly charact | Function unkn | 07WJ4 | Metalloprotea | 313606.M231 | 0,4 | 0,489897949 | 1,93E-05 | 2,38E-05 | 1 - | 4,30E-05 - | 1 - | 5,34E-05 - | 0 - | 0 - | 0 - | 0 - | 0 - | 0 - |
| 6614 | 1 | 0.2um_NOKCPHJN_174720 | Bacteria | Bacteroidetes | Flavobacteriia | Flavobacterial | Flavobacteriac | Formosa | Formosa sp. H | cellular proces | Cell wall/mem | 0P7KR | tyrosine-prote | 391603.FBALC | 0,6 | 0,489897949 | 1,10E-05 | 9,02E-06 | 0 - | 0 - | 0 - | 0 - | 1 - | 1,74E-05 - | 1 - | 1,77E-05 - | 1 - | 1,99E-05 - |
| 6615 | 1 | 0.2um_NOKCPHJN_37991 | Bacteria | Bacteroidetes | Cytophagia | Cytophagales | Flammeovirga | unclassified | Flammeovirga | poorly charact | Function unkn | 09ARW | Zinc ribbon do | 313606.M231 | 0,6 | 0,8 | 2,90E-05 | 3,66E-05 | 2 - | 8,66E-05 - | 0 - | 0 - | 0 - | 0 - | 0 - | 0 - | 1 - | 5,84E-05 - |
| 6616 | 1 | 0.2um_NOKCPHJN_60412 | Bacteria | Proteobacteri | Gammaprotec | Oceanospirilla | Oceanospirilla | Motiliproteus | Motiliproteus | information st | Translation, ril | COG0050 | This protein p | 717774.Marm | 4,6 | 0,8 | 0,000154989 | 3,03E-05 | 4 - | 0,000112803 - | 5 - | 0,000175085 - | 6 - | 0,000199728 - | 4 - | 0,000135119 - | 4 - | 0,000152211 - |
| 6617 | 1 | 3um_FIJJAFBF_625163 | Bacteria | Proteobacteri | Gammaprotec | Cellvibrionales | Porticoccacea | unclassified | various | information st | Transcription | 05VEQ | Cold shock pro | 314287.GB22 | 1,8 | 0,748331477 | 0,000345707 | 0,000115247 | 3 - | 0,000499029 - | 2 - | 0,000413098 - | 2 - | 0,000392702 - | 1 - | 0,000199251 - | 1 - | 0,000224456 - |
| 6618 | 1 | 3um_FIJJAFBF_292684 | Bacteria | Proteobacteri | Gammaprotec | Oceanospirilla | Oceanospirilla | Marinobacteri | Marinobacteri | information st | Translation, ril | 16QAF | thus facilitatin | 207954.MED9 | 6 | 0,894427191 | 0,000147856 | 2,99E-05 | 6 - | 0,000123416 - | 5 - | 0,000127705 - | 5 - | 0,0001214 - | 7 - | 0,00017247 - | 7 - | 0,000194287 - |
| 6619 | 1 | 3um_FIJJAFBF_744567 | Bacteria | unclassified | unclassified | unclassified | unclassified | unclassified | uncultured ba | unclassified | unclassified | unclassified | unclassified | unclassified | 37,2 | 2,227105745 | 0,00053822 | 4,43E-05 | 41 - | 0,000498501 - | 35 - | 0,000528407 - | 37 - | 0,000531021 - | 35 - | 0,000509737 - | 38 - | 0,000623435 - |
| 6620 | 1 | 0.2um_NOKCPHJN_99128 | Eukaryota | unclassified | unclassified | Choanoflagelli | Salpingoecida | Monosiga | Monosiga bre | information st | Chromatin stru | COG5648 | high mobility | 431895.XP_00 | 1,6 | 0,489897949 | 8,92E-05 | 2,85E-05 | 2 - | 9,37E-05 - | 1 - | 5,82E-05 - | 1 - | 5,53E-05 - | 2 - | 0,000112231 - | 2 - | 0,000126428 - |
| 6621 | 1 | 0.2um_NOKCPHJN_482341 | Bacteria | Proteobacteri | Betaproteoba | unclassified | unclassified | unclassified | Betaproteoba | information st | Translation, ril | 05C64 | Binds directly | 596154.Alide2 | 2,6 | 0,8 | 0,000152036 | 4,26E-05 | 3 - | 0,000149061 - | 2 - | 0,000123393 - | 4 - | 0,000234601 - | 2 - | 0,000119033 - | 2 - | 0,00013409 - |
| 6622 | 1 | 3um_FIJJAFBF_673147 | unclassified | unclassified | unclassified | unclassified | unclassified | unclassified | unclassified | unclassified | unclassified | unclassified | unclassified | unclassified | 1,8 | 0,4 | 0,000220262 | 6,10E-05 | 1 - | 0,000100681 - | 2 - | 0,000250033 - | 2 - | 0,000237688 - | 2 - | 0,000241199 - | 2 - | 0,00027171 - |
| 6623 | 1 | 3um_FIJJAFBF_619329 | Bacteria | Proteobacteri | Gammaprotec | Cellvibrionales | Halieaceae | unclassified | gamma protec | unclassified | unclassified | unclassified | unclassified | unclassified | 0,8 | 0,748331477 | 2,83E-05 | 2,62E-05 | 0 - | 0 - | 1 - | 3,66E-05 - | 2 - | 6,97E-05 - | 1 - | 3,53E-05 - | 0 - | 0 - |
| 6624 | 1 | 3um_FIJJAFBF_299316 | Bacteria | Proteobacteri | Gammaprotec | Alteromonada | Alteromonada | Paraglaciecola | Paraglaciecola | metabolism | Energy produc | 0QJV5 | alcohol dehyd | 1085623.GNIT | 1,4 | 0,489897949 | 4,82E-05 | 1,77E-05 | 1 - | 2,87E-05 - | 2 - | 7,13E-05 - | 2 - | 6,77E-05 - | 1 - | 3,44E-05 - | 1 - | 3,87E-05 - |
| 6625 | 1 | 0.2um_NOKCPHJN_302870 | Bacteria | Proteobacteri | Alphaproteob | Rhodobactera | Rhodobactera | Roseobacter | uncultured Ro | cellular proces | Posttranslatio | COG0330 | Band 7 protein | 246200.SPO2 | 1,2 | 0,748331477 | 5,46E-05 | 3,58E-05 | 1 - | 3,88E-05 - | 2 - | 9,63E-05 - | 2 - | 9,15E-05 - | 1 - | 4,64E-05 - | 0 - | 0 - |
| 6626 | 2 | 0.2um_NOKCPHJN_584821;0.2um_NOKCPHJN_7 | Bacteria | Proteobacteri | Alphaproteob | Pelagibacteral | Pelagibacterac | Candidatus Pe | Candidatus Pe | metabolism | Energy produc | 16RH6 | Produces ATP | 335992.SAR11 | 1,6 | 0,8 | 7,36E-05 | 3,50E-05 | 2 - | 7,89E-05 - | 1 - | 4,90E-05 - | 3 - | 0,000139672 - | 1 - | 4,72E-05 - | 1 - | 5,32E-05 - |
| 6627 | 1 | 0.2um_NOKCPHJN_305614 | Bacteria | Proteobacteri | Gammaprotec | Cellvibrionales | Halieaceae | unclassified | marine gamm | unclassified | unclassified | unclassified | unclassified | unclassified | 1 | 0 | 9,86E-05 | 9,36E-06 | 1 - | 8,26E-05 - | 1 - | 0,000102532 - | 1 - | 9,75E-05 - | 1 - | 9,89E-05 - | 1 - | 0,00011142 - |
| 6628 | 1 | 3um_FIJJAFBF_753603 | Bacteria | Proteobacteri | Alphaproteob | Rhodobactera | Rhodobactera | Oceanicola | Oceanicola sp | information st | Translation, ril | 179ZI | One of the pri | 388739.RSK20 | 1 | 0,632455532 | 9,92E-05 | 5,66E-05 | 2 - | 0,00017658 - | 1 - | 0,00010963 - | 1 - | 0,000104217 - | 1 - | 0,000105756 - | 0 - | 0 - |
| 6629 | 1 | 3um_FIJJAFBF_108989 | Bacteria | Proteobacteri | Alphaproteob | Rhodospirillal | Rhodospirillac | unclassified | Rhodobactera | metabolism | Energy produc | COG0711 | F(1)F(0) ATP s | 367336.OM22 | 3,4 | 0,8 | 0,000265748 | 8,61E-05 | 3 - | 0,000193444 - | 3 - | 0,0002402 - | 3 - | 0,000228341 - | 3 - | 0,000231713 - | 5 - | 0,000435041 - |
| 6630 | 1 | 0.2um_NOKCPHJN_631188 | Bacteria | Proteobacteri | Alphaproteob | Pelagibacteral | Pelagibacterac | Candidatus Pe | Candidatus Pe | unclassified | unclassified | unclassified | unclassified | unclassified | 3,8 | 0,4 | 0,000165218 | 1,47E-05 | 4 - | 0,00014668 - | 4 - | 0,000182133 - | 4 - | 0,00017314 - | 4 - | 0,000175698 - | 3 - | 0,000148442 - |
| 6631 | 1 | 0.2um_NOKCPHJN_139371 | Bacteria | Bacteroidetes | unclassified | unclassified | unclassified | unclassified | Bacteroidetes | unclassified | unclassified | unclassified | unclassified | unclassified | 1 | 0,632455532 | 3,00E-05 | 2,03E-05 | 1 - | 2,56E-05 - | 2 - | 6,36E-05 - | 1 - | 3,02E-05 - | 1 - | 3,07E-05 - | 0 - | 0 - |
| 6632 | 1 | 0.2um_NOKCPHJN_891139 | Bacteria | Actinobacteria | Actinobacteria | Corynebacteri | Mycobacteria | Mycobacteroi | Mycobacteroi | cellular proces | Cell wall/mem | 08N67 | choline bindin | 203120.LEUM | 0,8 | 0,4 | 3,49E-05 | 1,77E-05 | 1 - | 3,78E-05 - | 1 - | 4,69E-05 - | 1 - | 4,46E-05 - | 1 - | 4,52E-05 - | 0 - | 0 - |
| 6633 | 1 | 3um_FIJJAFBF_427091 | Bacteria | Bacteroidetes | unclassified | unclassified | unclassified | unclassified | Bacteroidetes | unclassified | unclassified | unclassified | unclassified | unclassified | 0,6 | 0,489897949 | 1,38E-05 | 1,13E-05 | 0 - | 0 - | 1 - | 2,37E-05 - | 1 - | 2,25E-05 - | 1 - | 2,28E-05 - | 0 - | 0 - |
| 6634 | 1 | 3um_FIJJAFBF_206033 | Bacteria | Bacteroidetes | Flavobacteriia | Flavobacterial | Flavobacteriac | Gilvibacter | Gilvibacter sp. | unclassified | unclassified | unclassified | unclassified | unclassified | 1 | 0 | 6,49E-05 | 6,17E-06 | 1 - | 5,44E-05 - | 1 - | 6,75E-05 - | 1 - | 6,42E-05 - | 1 - | 6,52E-05 - | 1 - | 7,34E-05 - |
| 6635 | 1 | 3um_FIJJAFBF_810847 | Bacteria | Bacteroidetes | Flavobacteriia | Flavobacterial | Flavobacteriac | Formosa | Formosa sp. H | metabolism | Carbohydrate | 0PF2Z | Transketolase | 1046627.BZAF | 0,6 | 0,489897949 | 2,82E-05 | 2,33E-05 | 1 - | 4,10E-05 - | 1 - | 5,09E-05 - | 0 - | 0 - | 1 - | 4,91E-05 - | 0 - | 0 - |
| 6636 | 1 | 0.2um_NOKCPHJN_327113 | Bacteria | Bacteroidetes | Flavobacteriia | Flavobacterial | Flavobacteriac | Tenacibaculur | Tenacibaculur | information st | Translation, ril | 05CAV | thus facilitatin | 313598.MED1 | 0,4 | 0,489897949 | 8,26E-06 | 1,02E-05 | 1 - | 1,89E-05 - | 0 - | 0 - | 1 - | 2,24E-05 - | 0 - | 0 - | 0 - | 0 - |
| 6637 | 1 | 3um_FIJJAFBF_75391 | Bacteria | Proteobacteri | Gammaprotec | Oceanospirilla | Saccharospiril | Reinekea | Reinekea blan | cellular proces | Posttranslatio | 05C2Z | ATP-depender | 314283.MED2 | 0,8 | 0,4 | 1,26E-05 | 6,46E-06 | 1 - | 1,33E-05 - | 0 - | 0 - | 1 - | 1,57E-05 - | 1 - | 1,59E-05 - | 1 - | 1,79E-05 - |
| 6638 | 1 | 3um_FIJJAFBF_357548 | Bacteria | Bacteroidetes | Flavobacteriia | Flavobacterial | Flavobacteriac | Polaribacter | Polaribacter s | unclassified | unclassified | unclassified | unclassified | unclassified | 5,8 | 0,748331477 | 9,80E-05 | 1,70E-05 | 6 - | 8,48E-05 - | 7 - | 0,000122861 - | 5 - | 8,34E-05 - | 5 - | 8,47E-05 - | 6 - | 0,000114439 - |
| 6639 | 1 | 0.2um_NOKCPHJN_1077244 | Bacteria | Proteobacteri | Alphaproteob | Pelagibacteral | Pelagibacterac | unclassified | Pelagibacterac | metabolism | Inorganic ion t | COG0841 | acriflavin resis | 335992.SAR11 | 0,6 | 0,489897949 | 1,50E-05 | 1,23E-05 | 1 - | 2,17E-05 - | 1 - | 2,70E-05 - | 0 - | 0 - | 1 - | 2,60E-05 - | 0 - | 0 - |
| 6640 | 1 | 0.2um_NOKCPHJN_121973 | Bacteria | Bacteroidetes | Flavobacteriia | Flavobacterial | Flavobacteriac | Polaribacter | Polaribacter v | information st | Translation, ril | COG0050 | This protein p | 313603.FB217 | 6,4 | 0,489897949 | 0,000220814 | 1,45E-05 | 7 - | 0,000203402 - | 6 - | 0,000216484 - | 7 - | 0,000240095 - | 6 - | 0,000208835 - | 6 - | 0,000235252 - |
| 6641 | 1 | 0.2um_NOKCPHJN_462253 | Bacteria | Planctomycete | Phycisphaerae | Phycisphaeral | Phycisphaerac | Phycisphaera | Phycisphaera | information st | Replication, re | COG0210 | helicase | 626939.HMPR | 0,2 | 0,4 | 3,82E-06 | 7,64E-06 | 0 - | 0 - | 1 - | 1,91E-05 - | 0 - | 0 - | 0 - | 0 - | 0 - | 0 - |
| 6642 | 1 | 0.2um_NOKCPHJN_27648 | Bacteria | Actinobacteria | Actinobacteria | Micrococcales | Microbacteria | Candidatus Ac | Candidatus Ac | information st | Translation, ril | 00DS2 | This protein bi | 979556.MTES_ | 0,4 | 0,489897949 | 2,81E-05 | 3,46E-05 | 1 - | 6,45E-05 - | 0 - | 0 - | 1 - | 7,61E-05 - | 0 - | 0 - | 0 - | 0 - |
| 6643 | 1 | 0.2um_NOKCPHJN_95014 | Bacteria | Bacteroidetes | Flavobacteriia | Flavobacterial | Flavobacteriac | Ulvibacter | Ulvibacter sp. | unclassified | unclassified | unclassified | unclassified | unclassified | 2,6 | 0,8 | 0,000279888 | 5,87E-05 | 4 - | 0,000370248 - | 2 - | 0,000229869 - | 3 - | 0,00032778 - | 2 - | 0,000221747 - | 2 - | 0,000249798 - |
| 6644 | 1 | 3um_FIJJAFBF_296764 | Bacteria | Proteobacteri | Gammaprotec | Arenicellales | Arenicellaceae | Arenicella | Arenicella xan | cellular proces | Posttranslatio | 0QJZQ | Peptidyl-proly | 1026882.MAN | 0,2 | 0,4 | 1,15E-05 | 2,30E-05 | 0 - | 0 - | 0 - | 0 - | 0 - | 0 - | 1 - | 5,75E-05 - | 0 - | 0 - |
| 6645 | 1 | 0.2um_NOKCPHJN_184677 | Bacteria | Proteobacteri | Gammaprotec | unclassified | unclassified | unclassified | Gammaprotec | information st | Translation, ril | 05G6W | One of the pri | 876044.IMCC | 1,8 | 0,748331477 | 0,000115989 | 3,90E-05 | 3 - | 0,000167151 - | 2 - | 0,000138368 - | 1 - | 6,58E-05 - | 2 - | 0,000133479 - | 1 - | 7,52E-05 - |
| 6646 | 1 | 0.2um_NOKCPHJN_220455 | Bacteria | unclassified | unclassified | unclassified | unclassified | unclassified | bacterium B1 | poorly charact | Function unkn | 05C2K | tail SHEATH pr | 443143.GM18 | 0,4 | 0,489897949 | 9,67E-06 | 1,18E-05 | 0 - | 0 - | 1 - | 2,48E-05 - | 1 - | 2,36E-05 - | 0 - | 0 - | 0 - | 0 - |
| 6647 | 1 | 3um_FIJJAFBF_715066 | Eukaryota | unclassified | Florideophyce | Gigartinales | Peyssonneliac | Riquetophycu | Riquetophycu | unclassified | unclassified | unclassified | unclassified | unclassified | 20,2 | 2,712931993 | 0,00041755 | 2,66E-05 | 24 - | 0,000420556 - | 19 - | 0,000413414 - | 20 - | 0,000413686 - | 22 - | 0,000461776 - | 16 - | 0,000378319 - |
| 6648 | 1 | 0.2um_NOKCPHJN_730596 | Eukaryota | unclassified | Oomycetes | Saprolegniales | Saprolegniace | Achlya | Achlya hypog | poorly charact | Function unkn | COG1100 | GTP-binding P | 272952.HpaP | 1,2 | 0,4 | 7,58E-05 | 2,61E-05 | 1 - | 5,29E-05 - | 1 - | 6,57E-05 - | 1 - | 6,24E-05 - | 2 - | 0,000126713 - | 1 - | 7,14E-05 - |
| 6649 | 1 | 0.2um_NOKCPHJN_604311 | Bacteria | Proteobacteri | Alphaproteob | Pelagibacteral | Pelagibacterac | Candidatus Pe | Candidatus Pe | metabolism | Energy produc | COG0056 | Produces ATP | 684719.HIMB | 0,8 | 0,4 | 2,36E-05 | 1,19E-05 | 0 - | 0 - | 1 - | 2,95E-05 - | 1 - | 2,81E-05 - | 1 - | 2,85E-05 - | 1 - | 3,21E-05 - |
| 6650 | 1 | 3um_FIJJAFBF_535108 | Bacteria | Bacteroidetes | unclassified | unclassified | unclassified | unclassified | Bacteroidetes | unclassified | unclassified | unclassified | unclassified | unclassified | 2 | 0 | 7,90E-05 | 7,50E-06 | 2 - | 6,62E-05 - | 2 - | 8,21E-05 - | 2 - | 7,81E-05 - | 2 - | 7,92E-05 - | 2 - | 8,93E-05 - |
| 6651 | 1 | 3um_FIJJAFBF_222255 | Bacteria | Proteobacteri | Alphaproteob | Rhodobactera | unclassified | unclassified | Rhodobactera | metabolism | Nucleotide tra | 01R42 | Amp nucleosi | 439496.RBY4I | 0,8 | 0,748331477 | 2,06E-05 | 1,82E-05 | 2 - | 4,70E-05 - | 0 - | 0 - | 1 - | 2,78E-05 - | 1 - | 2,82E-05 - | 0 - | 0 - |
| 6652 | 1 | 0.2um_NOKCPHJN_506774 | Bacteria | Proteobacteri | Alphaproteob | Pelagibacteral | Pelagibacterac | Candidatus Pe | Candidatus Pe | information st | Translation, ril | COG1185 | Involved in mf | 439493.PB721 | 2 | 0 | 5,01E-05 | 4,76E-06 | 2 - | 4,20E-05 - | 2 - | 5,21E-05 - | 2 - | 4,95E-05 - | 2 - | 5,03E-05 - | 2 - | 5,66E-05 - |
| 6653 | 2 | 0.2um_NOKCPHJN_615790;0.2um_NOKCPHJN_6 | Bacteria | Proteobacteri | Alphaproteob | Pelagibacteral | Pelagibacterac | Candidatus Pe | Candidatus Pe | unclassified | unclassified | unclassified | unclassified | unclassified | 0,6 | 0,489897949 | 4,94E-05 | 4,07E-05 | 1 - | 7,22E-05 - | 1 - | 8,96E-05 - | 1 - | 8,52E-05 - | 0 - | 0 - | 0 - | 0 - |
| 6654 | 1 | 3um_FIJJAFBF_28984 | Bacteria | Proteobacteri | Gammaprotec | Alteromonada | Alteromonada | Paraglaciecola | Paraglaciecola | cellular proces | Cell wall/mem | 0QKII | polysaccharid | 722419.PH50 | 0,6 | 0,8 | 9,05E-06 | 1,20E-05 | 0 - | 0 - | 0 - | 0 - | 2 - | 3,00E-05 - | 1 - | 1,52E-05 - | 0 - | 0 - |
| 6655 | 1 | 0.2um_NOKCPHJN_820335 | Bacteria | Proteobacteri | Alphaproteob | Rhodobactera | Rhodobactera | Roseovarius | Roseovarius a | unclassified | unclassified | unclassified | unclassified | unclassified | 0,4 | 0,489897949 | 8,07E-06 | 9,89E-06 | 0 - | 0 - | 1 - | 2,07E-05 - | 1 - | 1,97E-05 - | 0 - | 0 - | 0 - | 0 - |
| 6656 | 1 | 3um_FIJJAFBF_792632 | Eukaryota | Bacillariophyt | Coscinodiscop | Rhizosoleniale | Rhizosoleniac | Rhizosolenia | Rhizosolenia s | information st | Translation, ril | COG0050 | This protein p | 32049.SYNPC | 2,6 | 0,489897949 | 8,73E-05 | 1,86E-05 | 2 - | 5,61E-05 - | 3 - | 0,000104537 - | 3 - | 9,94E-05 - | 3 - | 0,000100844 - | 2 - | 7,57E-05 - |
| 6657 | 1 | 3um_FIJJAFBF_203752 | unclassified | unclassified | unclassified | unclassified | unclassified | unclassified | unclassified | unclassified | unclassified | unclassified | unclassified | unclassified | 1,4 | 0,489897949 | 3,57E-05 | 1,47E-05 | 1 - | 2,10E-05 - | 1 - | 2,61E-05 - | 1 - | 2,48E-05 - | 2 - | 5,03E-05 - | 2 - | 5,66E-05 - |
| 6658 | 1 | 0.2um_NOKCPHJN_974726 | Bacteria | Proteobacteri | Betaproteoba | Burkholderial | Comamonada | Limnohabitan | Limnohabitan | poorly charact | Function unkn | 0ZK5F | NA | 240292.Ava_4 | 1,2 | 0,4 | 2,76E-05 | 1,17E-05 | 1 - | 1,88E-05 - | 1 - | 2,34E-05 - | 1 - | 2,22E-05 - | 1 - | 2,26E-05 - | 2 - | 5,09E-05 - |
| 6659 | 2 | 0.2um_NOKCPHJN_580410;3um_FIJJAFBF_38057 | Bacteria | Proteobacteri | Alphaproteob | Rhodobactera | Rhodobactera | Planktomarina | Planktomarina | metabolism | Nucleotide tra | 01Y2N | Histidine triad | various | 0,2 | 0,4 | 2,26E-05 | 4,52E-05 | 0 - | 0 - | 1 - | 0,00011311 - | 0 - | 0 - | 0 - | 0 - | 0 - | 0 - |
| 6660 | 2 | 0.2um_NOKCPHJN_76327;0.2um_NOKCPHJN_89 | Bacteria | Bacteroidetes | Flavobacteriia | Flavobacterial | Flavobacteriac | Formosa | Formosa sp. H | information st | Translation, ril | COG0052 | 30S ribosomal | 50743.SCB49_ | 5,4 | 1,019803903 | 0,000227865 | 3,18E-05 | 7 - | 0,000250292 - | 5 - | 0,000221992 - | 4 - | 0,000168825 - | 6 - | 0,000256978 - | 5 - | 0,000241238 - |
| 6661 | 1 | 3um_FIJJAFBF_117655 | Eukaryota | various | various | various | various | various | various | metabolism | Energy produc | COG1145 | 4Fe-4S ferred | 35128.Thapsd | 1 | 0 | 0,000169169 | 1,61E-05 | 1 - | 0,0001417 - | 1 - | 0,000175949 - | 1 - | 0,000167262 - | 1 - | 0,000169732 - | 1 - | 0,000191203 - |
| 6662 | 1 | 3um_FIJJAFBF_77386 | unclassified | unclassified | unclassified | unclassified | unclassified | unclassified | unclassified | unclassified | unclassified | unclassified | unclassified | unclassified | 0,2 | 0,4 | 2,01E-05 | 4,01E-05 | 0 - | 0 - | 1 - | 0,000100365 - | 0 - | 0 - | 0 - | 0 - | 0 - | 0 - |
| 6663 | 1 | 0.2um_NOKCPHJN_73718 | Bacteria | Proteobacteri | Betaproteoba | Burkholderial | Comamonada | Limnohabitan | Limnohabitan | cellular proces | Cell wall/mem | 172GZ | Catalyzes the | 338969.Rfer_ | 0,2 | 0,4 | 7,89E-06 | 1,58E-05 | 1 - | 3,94E-05 - | 0 - | 0 - | 0 - | 0 - | 0 - | 0 - | 0 - | 0 - |
| 6664 | 1 | 3um_FIJJAFBF_21302 | Bacteria | Bacteroidetes | Flavobacteriia | Flavobacterial | Flavobacteriac | Polaribacter | Polaribacter s | unclassified | unclassified | unclassified | unclassified | unclassified | 3,4 | 0,489897949 | 0,000229619 | 2,44E-05 | 4 - | 0,000228411 - | 3 - | 0,000212715 - | 3 - | 0,000202212 - | 4 - | 0,000273599 - | 3 - | 0,000231156 - |
| 6665 | 1 | 0.2um_NOKCPHJN_153564 | Bacteria | Bacteroidetes | Flavobacteriia | Flavobacterial | Flavobacteriac | unclassified | Flavobacteriac | information st | Translation, ril | 05CEJ | Catalyzes the | 886377.Murru | 0,4 | 0,8 | 8,75E-06 | 1,75E-05 | 0 - | 0 - | 0 - | 0 - | 0 - | 0 - | 0 - | 0 - | 2 - | 4,37E-05 - |
| 6666 | 1 | 3um_FIJJAFBF_235910 | Eukaryota | Bacillariophyt | Bacillariophyc | Naviculales | Naviculaceae | Fistulifera | Fistulifera sola | metabolism | Carbohydrate | 09WWU | ER degradatio | 31033.ENSTRU | 0,6 | 0,489897949 | 5,66E-05 | 4,66E-05 | 1 - | 8,38E-05 - | 0 - | 0 - | 1 - | 9,89E-05 - | 1 - | 0,000100353 - | 0 - | 0 - |
| 6667 | 1 | 3um_FIJJAFBF_366248 | Bacteria | Proteobacteri | Alphaproteob | unclassified | unclassified | Candidatus Pu | Candidatus Pu | metabolism | Inorganic ion t | 05PWN | phosphate ab | 488538.SAR11 | 4,8 | 1,166190379 | 0,000188736 | 2,92E-05 | 7 - | 0,000234923 - | 5 - | 0,000208361 - | 4 - | 0,000158459 - | 4 - | 0,000160799 - | 4 - | 0,00018114 - |
| 6668 | 1 | 0.2um_NOKCPHJN_544665 | Bacteria | Bacteroidetes | Flavobacteriia | Flavobacterial | Flavobacteriac | Polaribacter | Polaribacter s | unclassified | unclassified | unclassified | unclassified | unclassified | 1,2 | 0,748331477 | 1,73E-05 | 1,01E-05 | 2 - | 2,48E-05 - | 1 - | 1,54E-05 - | 0 - | 0 - | 2 - | 2,97E-05 - | 1 - | 1,67E-05 - |
| 6669 | 1 | 0.2um_NOKCPHJN_89784 | Bacteria | Bacteroidetes | Saprospiria | Saprospirales | Haliscomenob | Phaeodactylib | Phaeodactylib | unclassified | unclassified | unclassified | unclassified | unclassified | 0,8 | 0,4 | 1,10E-06 | 5,64E-07 | 1 - | 1,15E-06 - | 1 - | 1,43E-06 - | 1 - | 1,36E-06 - | 0 - | 0 - | 1 - | 1,55E-06 - |
| 6670 | 1 | 0.2um_NOKCPHJN_163356 | Bacteria | Proteobacteri | Gammaprotec | Cellvibrionales | Porticoccacea | unclassified | SAR92 bacteri | cellular proces | Intracellular tr | 0QITE | Outer membr | 314287.GB22 | 2,8 | 0,4 | 7,97E-05 | 1,29E-05 | 3 - | 7,17E-05 - | 2 - | 5,94E-05 - | 3 - | 8,47E-05 - | 3 - | 8,59E-05 - | 3 - | 9,68E-05 - |
| 6671 | 1 | 3um_FIJJAFBF_528374 | Bacteria | Proteobacteri | Gammaprotec | Oceanospirilla | Oceanospirilla | unclassified | Osedax symbi | unclassified | unclassified | unclassified | unclassified | unclassified | 2,2 | 0,748331477 | 8,34E-05 | 3,22E-05 | 2 - | 6,31E-05 - | 1 - | 3,92E-05 - | 3 - | 0,000111661 - | 2 - | 7,55E-05 - | 3 - | 0,000127644 - |
| 6672 | 1 | 3um_FIJJAFBF_186024 | Bacteria | Proteobacteri | Betaproteoba | unclassified | unclassified | unclassified | Betaproteoba | poorly charact | Function unkn | 0B89A | peptidase U62 | 595537.Varpa | 0,2 | 0,4 | 5,86E-06 | 1,17E-05 | 0 - | 0 - | 1 - | 2,93E-05 - | 0 - | 0 - | 0 - | 0 - | 0 - | 0 - |
| 6673 | 1 | 3um_FIJJAFBF_194263 | Bacteria | Proteobacteri | Betaproteoba | unclassified | unclassified | unclassified | Betaproteoba | unclassified | unclassified | unclassified | unclassified | unclassified | 2,4 | 0,489897949 | 0,000171171 | 3,69E-05 | 2 - | 0,000119559 - | 2 - | 0,000148457 - | 3 - | 0,000211691 - | 3 - | 0,000214818 - | 2 - | 0,000161328 - |
| 6674 | 1 | 3um_FIJJAFBF_426858 | Bacteria | Proteobacteri | Alphaproteob | Rhodobactera | Rhodobactera | Planktotalea | Planktotalea a | cellular proces | Posttranslatio | 01QWG | Prevents misf | 292414.TM10 | 0,2 | 0,4 | 4,98E-06 | 9,96E-06 | 0 - | 0 - | 0 - | 0 - | 0 - | 0 - | 1 - | 2,49E-05 - | 0 - | 0 - |
| 6675 | 1 | 0.2um_NOKCPHJN_48076 | Bacteria | Bacteroidetes | Flavobacteriia | Flavobacterial | Flavobacteriac | unclassified | Flavobacteriac | information st | Transcription | COG0085 | DNA-depende | 487796.Flav2 | 0,8 | 0,4 | 8,99E-06 | 4,53E-06 | 0 - | 0 - | 1 - | 1,12E-05 - | 1 - | 1,07E-05 - | 1 - | 1,08E-05 - | 1 - | 1,22E-05 - |
| 6676 | 1 | 3um_FIJJAFBF_111252 | Bacteria | Bacteroidetes | Flavobacteriia | Flavobacterial | Flavobacteriac | unclassified | Flavobacteriac | unclassified | unclassified | unclassified | unclassified | unclassified | 0,2 | 0,4 | 8,47E-06 | 1,69E-05 | 0 - | 0 - | 0 - | 0 - | 1 - | 4,23E-05 - | 0 - | 0 - | 0 - | 0 - |
| 6677 | 1 | 3um_FIJJAFBF_619213 | Bacteria | Bacteroidetes | Flavobacteriia | Flavobacterial | Flavobacteriac | unclassified | Flavobacteriac | metabolism | Amino acid tra | 0PF77 | Dehydrogenas | 487796.Flav2 | 0,4 | 0,8 | 1,45E-05 | 2,91E-05 | 2 - | 7,26E-05 - | 0 - | 0 - | 0 - | 0 - | 0 - | 0 - | 0 - | 0 - |
| 6678 | 1 | 3um_FIJJAFBF_666764 | Bacteria | Proteobacteri | Epsilonproteo | Campylobacte | Campylobacte | Arcobacter | Arcobacter sp | metabolism | Carbohydrate | 05D2N | Aldolase | 469613.HMPR | 1 | 0 | 8,30E-05 | 7,89E-06 | 1 - | 6,96E-05 - | 1 - | 8,64E-05 - | 1 - | 8,21E-05 - | 1 - | 8,33E-05 - | 1 - | 9,39E-05 - |
| 6679 | 1 | 0.2um_NOKCPHJN_36475 | Bacteria | Bacteroidetes | Cytophagia | Cytophagales | Flammeovirga | unclassified | Flammeovirga | information st | Translation, ril | COG0184 | Forms an inte | 643867.Ftrac_ | 1,4 | 0,489897949 | 0,000215304 | 7,61E-05 | 1 - | 0,000128963 - | 1 - | 0,000160134 - | 2 - | 0,000304454 - | 2 - | 0,000308951 - | 1 - | 0,000174016 - |
| 6680 | 1 | 3um_FIJJAFBF_415981 | Bacteria | Bacteroidetes | Flavobacteriia | Flavobacterial | Flavobacteriac | Formosa | Formosa sp. H | metabolism | Inorganic ion t | 05C27 | TonB-depende | 983544.Lacal_ | 0,4 | 0,489897949 | 5,01E-06 | 6,17E-06 | 1 - | 1,15E-05 - | 0 - | 0 - | 1 - | 1,36E-05 - | 0 - | 0 - | 0 - | 0 - |
| 6681 | 1 | 3um_FIJJAFBF_331695 | Bacteria | Bacteroidetes | Flavobacteriia | Flavobacterial | Crocinitomica | unclassified | Crocinitomica | poorly charact | Function unkn | 09AR5 | Inherit from b | 313606.M231 | 1 | 0 | 2,64E-05 | 2,51E-06 | 1 - | 2,21E-05 - | 1 - | 2,75E-05 - | 1 - | 2,61E-05 - | 1 - | 2,65E-05 - | 1 - | 2,98E-05 - |
| 6682 | 1 | 3um_FIJJAFBF_611864 | Bacteria | Bacteroidetes | Flavobacteriia | Flavobacterial | Flavobacteriac | Lutibacter | Lutibacter sp. | metabolism | Inorganic ion t | 05C27 | TonB-depende | 688270.Celal_ | 7,6 | 0,8 | 0,00010248 | 1,04E-05 | 9 - | 0,000102175 - | 7 - | 9,87E-05 - | 7 - | 9,38E-05 - | 7 - | 9,52E-05 - | 8 - | 0,000122552 - |
| 6683 | 1 | 0.2um_NOKCPHJN_182731 | Bacteria | Bacteroidetes | unclassified | unclassified | unclassified | unclassified | Bacteroidetes | unclassified | unclassified | unclassified | unclassified | unclassified | 0,4 | 0,489897949 | 2,54E-05 | 3,12E-05 | 0 - | 0 - | 0 - | 0 - | 0 - | 0 - | 1 - | 5,98E-05 - | 1 - | 6,73E-05 - |
| 6684 | 1 | 3um_FIJJAFBF_466831 | Bacteria | Proteobacteri | Alphaproteob | Rhodobactera | Rhodobactera | Amylibacter | Amylibacter k | unclassified | unclassified | unclassified | unclassified | unclassified | 2 | 0,632455532 | 9,96E-05 | 2,63E-05 | 3 - | 0,000127059 - | 2 - | 0,00010518 - | 1 - | 5,00E-05 - | 2 - | 0,000101464 - | 2 - | 0,000114299 - |
| 6685 | 1 | 3um_FIJJAFBF_189666 | Bacteria | Bacteroidetes | Flavobacteriia | Flavobacterial | Flavobacteriac | Polaribacter | Polaribacter b | metabolism | Lipid transpor | 0PA4W | Biotin carboxy | 313594.PI23P | 2,4 | 1,2 | 0,000203313 | 9,89E-05 | 3 - | 0,000217931 - | 3 - | 0,000270605 - | 4 - | 0,000342993 - | 1 - | 8,70E-05 - | 1 - | 9,80E-05 - |
| 6686 | 1 | 3um_FIJJAFBF_462772 | Bacteria | Proteobacteri | Gammaprotec | Oceanospirilla | Saccharospiril | Reinekea | Reinekea fors | poorly charact | Function unkn | 0ZA9F | NA | 448385.sce09 | 0,8 | 0,4 | 5,85E-05 | 3,00E-05 | 1 - | 6,11E-05 - | 1 - | 7,58E-05 - | 0 - | 0 - | 1 - | 7,31E-05 - | 1 - | 8,24E-05 - |
| 6687 | 1 | 0.2um_NOKCPHJN_127271 | Bacteria | Bacteroidetes | Flavobacteriia | Flavobacterial | Flavobacteriac | Arenitalea | Arenitalea lut | cellular proces | Posttranslatio | 08V1T | peptidylprolyl | 313594.PI23P | 1,2 | 0,4 | 7,01E-05 | 2,35E-05 | 1 - | 4,90E-05 - | 1 - | 6,09E-05 - | 2 - | 0,000115797 - | 1 - | 5,88E-05 - | 1 - | 6,62E-05 - |
| 6688 | 1 | 3um_FIJJAFBF_552507 | Bacteria | Proteobacteri | Gammaprotec | Arenicellales | Arenicellaceae | Arenicella | Arenicella xan | information st | Translation, ril | 0QTKX | Seems to be t | 349106.PsycP | 1 | 0,632455532 | 0,00011026 | 7,04E-05 | 1 - | 9,33E-05 - | 0 - | 0 - | 2 - | 0,000220296 - | 1 - | 0,000111775 - | 1 - | 0,000125914 - |
| 6689 | 1 | 3um_FIJJAFBF_284149 | Bacteria | Proteobacteri | Betaproteoba | unclassified | unclassified | unclassified | Betaproteoba | unclassified | unclassified | unclassified | unclassified | unclassified | 0,8 | 0,4 | 2,03E-05 | 1,04E-05 | 1 - | 2,13E-05 - | 1 - | 2,64E-05 - | 1 - | 2,51E-05 - | 0 - | 0 - | 1 - | 2,87E-05 - |
| 6690 | 1 | 3um_FIJJAFBF_545823 | Bacteria | Bacteroidetes | Flavobacteriia | Flavobacterial | Flavobacteriac | Polaribacter | Polaribacter t | unclassified | unclassified | unclassified | unclassified | unclassified | 2,2 | 0,748331477 | 0,0001711 | 5,19E-05 | 3 - | 0,000199035 - | 1 - | 8,24E-05 - | 2 - | 0,000156627 - | 3 - | 0,00023841 - | 2 - | 0,000179046 - |
| 6691 | 1 | 3um_FIJJAFBF_671452 | Bacteria | Proteobacteri | Gammaprotec | Alteromonada | Alteromonada | Paraglaciecola | Paraglaciecola | metabolism | Energy produc | 17C3H | Produces ATP | 1085623.GNIT | 1 | 0 | 0,00010002 | 9,50E-06 | 1 - | 8,38E-05 - | 1 - | 0,000104028 - | 1 - | 9,89E-05 - | 1 - | 0,000100353 - | 1 - | 0,000113047 - |
| 6692 | 1 | 3um_FIJJAFBF_37522 | Bacteria | Bacteroidetes | Flavobacteriia | Flavobacterial | Crocinitomica | unclassified | Crocinitomica | information st | Translation, ril | 08UHY | One of the pri | 755732.Fluta_ | 2,4 | 0,8 | 0,000210436 | 7,23E-05 | 2 - | 0,000148099 - | 3 - | 0,000275843 - | 3 - | 0,000262224 - | 3 - | 0,000266097 - | 1 - | 9,99E-05 - |
| 6693 | 1 | 0.2um_NOKCPHJN_262673 | Bacteria | Bacteroidetes | Flavobacteriia | Flavobacterial | Flavobacteriac | Marinirhabdu | Marinirhabdu | information st | Translation, ril | COG0256 | This is one of | 50743.SCB49_ | 3 | 0,632455532 | 0,000347812 | 6,04E-05 | 4 - | 0,000392399 - | 3 - | 0,000365433 - | 2 - | 0,000231594 - | 3 - | 0,000352521 - | 3 - | 0,000397114 - |
| 6694 | 1 | 0.2um_NOKCPHJN_416693 | Bacteria | Bacteroidetes | Flavobacteriia | unclassified | unclassified | unclassified | Flavobacteria | poorly charact | Function unkn | 11QQ6 | NA | 487797.Flav3 | 0,2 | 0,4 | 8,35E-06 | 1,67E-05 | 1 - | 4,17E-05 - | 0 - | 0 - | 0 - | 0 - | 0 - | 0 - | 0 - | 0 - |
| 6695 | 1 | 3um_FIJJAFBF_271513 | Bacteria | Bacteroidetes | Flavobacteriia | Flavobacterial | Flavobacteriac | Gilvibacter | Gilvibacter sp. | poorly charact | Function unkn | 0P9HI | ASPIC and Unl | 391587.KAOT | 1,2 | 0,748331477 | 3,02E-05 | 1,99E-05 | 1 - | 2,14E-05 - | 2 - | 5,32E-05 - | 1 - | 2,53E-05 - | 2 - | 5,13E-05 - | 0 - | 0 - |
| 6696 | 1 | 0.2um_NOKCPHJN_159997 | Bacteria | Proteobacteri | Gammaprotec | Oceanospirilla | unclassified | unclassified | Oceanospirilla | metabolism | Energy produc | 05C26 | Dehydrogenas | 349521.HCH_ | 0,6 | 0,489897949 | 1,71E-05 | 1,40E-05 | 0 - | 0 - | 0 - | 0 - | 1 - | 2,71E-05 - | 1 - | 2,75E-05 - | 1 - | 3,10E-05 - |
| 6697 | 1 | 0.2um_NOKCPHJN_584820 | Bacteria | Proteobacteri | Alphaproteob | Pelagibacteral | Pelagibacterac | Candidatus Pe | Candidatus Pe | metabolism | Energy produc | COG0056 | Produces ATP | 684719.HIMB | 1,6 | 0,489897949 | 4,20E-05 | 1,10E-05 | 2 - | 4,49E-05 - | 1 - | 2,79E-05 - | 2 - | 5,30E-05 - | 2 - | 5,38E-05 - | 1 - | 3,03E-05 - |
| 6698 | 1 | 0.2um_NOKCPHJN_252739 | Bacteria | Proteobacteri | Gammaprotec | Cellvibrionales | Porticoccacea | unclassified | gamma protec | poorly charact | Function unkn | 0XY6Y | Pkd domain co | 314287.GB22 | 1 | 0 | 1,82E-05 | 1,73E-06 | 1 - | 1,53E-05 - | 1 - | 1,90E-05 - | 1 - | 1,80E-05 - | 1 - | 1,83E-05 - | 1 - | 2,06E-05 - |
| 6699 | 1 | 3um_FIJJAFBF_03894 | Bacteria | Proteobacteri | Gammaprotec | unclassified | unclassified | unclassified | Gammaprotec | unclassified | unclassified | unclassified | unclassified | unclassified | 4,4 | 0,489897949 | 0,000162103 | 1,24E-05 | 5 - | 0,000155524 - | 4 - | 0,000154492 - | 5 - | 0,00018358 - | 4 - | 0,000149033 - | 4 - | 0,000167886 - |
| 6700 | 1 | 3um_FIJJAFBF_178047 | Bacteria | Proteobacteri | Gammaprotec | Alteromonada | Alteromonada | Paraglaciecola | Paraglaciecola | poorly charact | Function unkn | 05W4M | Transport-ass | 342610.Patl_ | 1 | 0 | 7,14E-05 | 6,78E-06 | 1 - | 5,98E-05 - | 1 - | 7,42E-05 - | 1 - | 7,06E-05 - | 1 - | 7,16E-05 - | 1 - | 8,07E-05 - |
| 6701 | 1 | 3um_FIJJAFBF_116531 | Bacteria | Proteobacteri | Alphaproteob | unclassified | unclassified | Candidatus Pu | Candidatus Pu | metabolism | Carbohydrate | 0IVA1 | Photosystem I | 35128.Thapsd | 9 | 1,095445115 | 0,000639562 | 7,89E-05 | 9 - | 0,000538016 - | 8 - | 0,000593829 - | 11 - | 0,0007762 - | 9 - | 0,000644453 - | 8 - | 0,00064531 - |
| 6702 | 1 | 0.2um_NOKCPHJN_72792 | Bacteria | Proteobacteri | Gammaprotec | Oceanospirilla | Saccharospiril | Reinekea | Reinekea blan | metabolism | Amino acid tra | COG2171 | Catalyzes the | 314283.MED2 | 1 | 0,632455532 | 4,13E-05 | 2,54E-05 | 0 - | 0 - | 1 - | 4,16E-05 - | 1 - | 3,95E-05 - | 2 - | 8,02E-05 - | 1 - | 4,52E-05 - |
| 6703 | 1 | 3um_FIJJAFBF_36942 | Bacteria | Proteobacteri | Alphaproteob | Rhodospirillal | Rhodospirillac | Candidatus En | Candidatus En | unclassified | unclassified | unclassified | unclassified | unclassified | 3,4 | 0,489897949 | 0,000115266 | 1,89E-05 | 3 - | 8,52E-05 - | 3 - | 0,000105831 - | 4 - | 0,000134141 - | 4 - | 0,000136122 - | 3 - | 0,000115006 - |
| 6704 | 1 | 3um_FIJJAFBF_122480 | Bacteria | Bacteroidetes | Flavobacteriia | Flavobacterial | Flavobacteriac | Polaribacter | Polaribacter v | information st | Transcription | 05D27 | DNA-depende | 313594.PI23P | 48,4 | 4,92341345 | 0,000462646 | 3,65E-05 | 56 - | 0,000451369 - | 52 - | 0,000520434 - | 43 - | 0,000409111 - | 47 - | 0,000453772 - | 44 - | 0,000478545 - |
| 6705 | 1 | 3um_FIJJAFBF_99501 | Bacteria | Bacteroidetes | Flavobacteriia | Flavobacterial | Flavobacteriac | Polaribacter | Polaribacter v | metabolism | Energy produc | COG0055 | Produces ATP | 313594.PI23P | 0,4 | 0,489897949 | 1,60E-05 | 1,97E-05 | 1 - | 3,63E-05 - | 0 - | 0 - | 0 - | 0 - | 1 - | 4,35E-05 - | 0 - | 0 - |
| 6706 | 1 | 0.2um_NOKCPHJN_124489 | Bacteria | Proteobacteri | Alphaproteob | Rhodobactera | Rhodobactera | Planktomarina | Planktomarina | poorly charact | Function unkn | 05DIH | basic membra | 633131.TR2A | 3 | 0 | 0,000120199 | 1,14E-05 | 3 - | 0,000100681 - | 3 - | 0,000125017 - | 3 - | 0,000118844 - | 3 - | 0,000120599 - | 3 - | 0,000135855 - |
| 6707 | 1 | 0.2um_NOKCPHJN_72584 | Bacteria | Proteobacteri | Gammaprotec | Cellvibrionales | Porticoccacea | unclassified | gamma protec | cellular proces | Posttranslatio | 0QHXH | Molecular cha | 203122.Sde_2 | 2 | 0,632455532 | 4,18E-05 | 1,30E-05 | 2 - | 3,54E-05 - | 2 - | 4,40E-05 - | 2 - | 4,18E-05 - | 3 - | 6,36E-05 - | 1 - | 2,39E-05 - |
| 6708 | 1 | 3um_FIJJAFBF_345763 | Bacteria | Bacteroidetes | unclassified | unclassified | unclassified | unclassified | Bacteroidetes | unclassified | unclassified | unclassified | unclassified | unclassified | 1 | 0 | 7,83E-05 | 7,43E-06 | 1 - | 6,56E-05 - | 1 - | 8,14E-05 - | 1 - | 7,74E-05 - | 1 - | 7,86E-05 - | 1 - | 8,85E-05 - |
| 6709 | 1 | 3um_FIJJAFBF_320477 | Bacteria | Bacteroidetes | Flavobacteriia | Flavobacterial | Flavobacteriac | Sediminicola | Sediminicola s | cellular proces | Posttranslatio | COG0652 | peptidyl-proly | 487797.Flav3 | 0,2 | 0,4 | 9,99E-06 | 2,00E-05 | 0 - | 0 - | 0 - | 0 - | 0 - | 0 - | 0 - | 0 - | 1 - | 5,00E-05 - |
| 6710 | 1 | 0.2um_NOKCPHJN_608862 | Bacteria | Bacteroidetes | Flavobacteriia | Flavobacterial | Crocinitomica | unclassified | Crocinitomica | unclassified | unclassified | unclassified | unclassified | unclassified | 1,4 | 0,489897949 | 9,63E-05 | 3,55E-05 | 1 - | 5,74E-05 - | 2 - | 0,000142519 - | 2 - | 0,000135482 - | 1 - | 6,87E-05 - | 1 - | 7,74E-05 - |
| 6711 | 1 | 0.2um_NOKCPHJN_687008 | Viruses | unclassified | unclassified | unclassified | unclassified | unclassified | uncultured M | unclassified | unclassified | unclassified | unclassified | unclassified | 1 | 0 | 1,51E-05 | 1,43E-06 | 1 - | 1,26E-05 - | 1 - | 1,57E-05 - | 1 - | 1,49E-05 - | 1 - | 1,51E-05 - | 1 - | 1,70E-05 - |
| 6712 | 1 | 3um_FIJJAFBF_222955 | Bacteria | Proteobacteri | Alphaproteob | Rhodobactera | Rhodobactera | Planktomarina | Planktomarina | unclassified | unclassified | unclassified | unclassified | unclassified | 1 | 0,632455532 | 4,93E-05 | 3,13E-05 | 1 - | 4,14E-05 - | 1 - | 5,15E-05 - | 2 - | 9,78E-05 - | 0 - | 0 - | 1 - | 5,59E-05 - |
| 6713 | 1 | 0.2um_NOKCPHJN_991621 | Eukaryota | unclassified | unclassified | Isochrysidales | Noelaerhabda | Emiliania | Emiliania huxl | unclassified | unclassified | unclassified | unclassified | unclassified | 1,6 | 1,2 | 0,000156717 | 0,000124194 | 1 - | 7,97E-05 - | 1 - | 9,90E-05 - | 3 - | 0,000282255 - | 0 - | 0 - | 3 - | 0,000322655 - |
| 6714 | 1 | 0.2um_NOKCPHJN_409446 | Eukaryota | unclassified | Oomycetes | Pythiales | Pythiaceae | Pythium | Pythium insidi | unclassified | unclassified | unclassified | unclassified | unclassified | 0,2 | 0,4 | 2,82E-06 | 5,64E-06 | 0 - | 0 - | 0 - | 0 - | 1 - | 1,41E-05 - | 0 - | 0 - | 0 - | 0 - |
| 6715 | 1 | 3um_FIJJAFBF_228147 | Bacteria | Proteobacteri | Gammaprotec | unclassified | unclassified | unclassified | various | unclassified | unclassified | unclassified | unclassified | unclassified | 0,4 | 0,489897949 | 4,95E-05 | 6,12E-05 | 1 - | 0,000110362 - | 1 - | 0,000137037 - | 0 - | 0 - | 0 - | 0 - | 0 - | 0 - |

| | | | | | | | | | | | | | | | | | | | | | | | | | | | | |
|---|---|---|---|---|---|---|---|---|---|---|---|---|---|---|---|---|---|---|---|---|---|---|---|---|---|---|---|---|
| 6716 | 1 | 0.2um_NOKCPHJN_625898 | Eukaryota | Arthropoda | Arachnida | Mesostigmata | Varroidae | Varroa | various | unclassified | unclassified | unclassified | unclassified | unclassified | 0,8 | 0,4 | 2,58E-05 | 1,31E-05 | 1 - | 2,79E-05 - | 1 - | 3,47E-05 - | 1 - | 3,30E-05 - | 1 - | 3,35E-05 - | 0 - | 0 - |
| 6717 | 1 | 0.2um_NOKCPHJN_538278 | Bacteria | Bacteroidetes | Flavobacteriia | Flavobacterial | Flavobacteria | unclassified | Flavobacteria | unclassified | unclassified | unclassified | unclassified | unclassified | 1 | 0 | 1,48E-05 | 1,41E-06 | 1 - | 1,24E-05 - | 1 - | 1,54E-05 - | 1 - | 1,46E-05 - | 1 - | 1,48E-05 - | 1 - | 1,67E-05 - |
| 6718 | 1 | 3um_FIJJAFBF_798999 | Bacteria | Proteobacteri | Gammaprote | Oceanospirilla | Saccharospirill | Reinekea | Reinekea fors | information st | Replication, re | 0QKMF | The RuvA-Ruv | 314283.MED2 | 0,8 | 0,748331477 | 5,24E-05 | 4,53E-05 | 2 - | 0,000114206 - | 1 - | 7,09E-05 - | 0 - | 0 - | 0 - | 0 - | 1 - | 7,71E-05 - |
| 6719 | 1 | 0.2um_NOKCPHJN_266842 | Bacteria | Bacteroidetes | Flavobacteriia | unclassified | unclassified | unclassified | Flavobacteria | poorly charact | Function unkn | 0P8NI | Inherit from b | 487796.Flav2A | 7,6 | 0,8 | 0,000237722 | 3,43E-05 | 8 - | 0,000209638 - | 8 - | 0,000260308 - | 8 - | 0,000247456 - | 6 - | 0,000188333 - | 8 - | 0,000282876 - |
| 6720 | 1 | 0.2um_NOKCPHJN_208843 | Bacteria | Bacteroidetes | Flavobacteriia | Flavobacterial | Flavobacteria | unclassified | Flavobacteria | unclassified | unclassified | unclassified | unclassified | unclassified | 1,4 | 0,489897949 | 3,59E-05 | 1,03E-05 | 2 - | 4,41E-05 - | 1 - | 2,74E-05 - | 2 - | 5,20E-05 - | 1 - | 2,64E-05 - | 1 - | 2,97E-05 - |
| 6721 | 1 | 0.2um_NOKCPHJN_307670 | Bacteria | Bacteroidetes | Flavobacteriia | Flavobacterial | Flavobacteria | Gelidibacter | Gelidibacter a | unclassified | unclassified | unclassified | unclassified | unclassified | 0,8 | 0,4 | 1,12E-05 | 5,68E-06 | 1 - | 1,21E-05 - | 1 - | 1,50E-05 - | 1 - | 1,43E-05 - | 1 - | 1,45E-05 - | 0 - | 0 - |
| 6722 | 1 | 0.2um_NOKCPHJN_363679 | Bacteria | Proteobacteri | Gammaprote | unclassified | unclassified | unclassified | Gammaprote | poorly charact | Function unkn | 16R8I | Lipoprotein | 314287.GB22 | 1 | 0 | 2,45E-05 | 2,32E-06 | 1 - | 2,05E-05 - | 1 - | 2,54E-05 - | 1 - | 2,42E-05 - | 1 - | 2,46E-05 - | 1 - | 2,77E-05 - |
| 6723 | 1 | 3um_FIJJAFBF_780738 | Bacteria | Proteobacteri | Gammaprote | Cellvibrionales | Porticoccacea | unclassified | SAR92 bacteri | unclassified | unclassified | unclassified | unclassified | unclassified | 1,6 | 0,8 | 0,000280579 | 0,000112206 | 3 - | 0,000459107 - | 1 - | 0,000190025 - | 1 - | 0,000180643 - | 2 - | 0,000366622 - | 1 - | 0,000206499 - |
| 6724 | 1 | 3um_FIJJAFBF_67730 | Bacteria | Proteobacteri | Gammaprote | Cellvibrionales | unclassified | unclassified | Cellvibrionales | unclassified | unclassified | unclassified | unclassified | unclassified | 3 | 0 | 0,000297885 | 2,83E-05 | 3 - | 0,000249515 - | 3 - | 0,000309824 - | 3 - | 0,000294527 - | 3 - | 0,000298877 - | 3 - | 0,000336684 - |
| 6725 | 1 | 3um_FIJJAFBF_775040 | Eukaryota | unclassified | Cryptophyta | Pyrenomonad | Pyrenomonad | Rhodomonas | Rhodomonas | unclassified | unclassified | unclassified | unclassified | unclassified | 1 | 0,632455532 | 4,20E-05 | 2,72E-05 | 1 - | 3,62E-05 - | 1 - | 4,50E-05 - | 2 - | 8,55E-05 - | 1 - | 4,34E-05 - | 0 - | 0 - |
| 6726 | 1 | 0.2um_NOKCPHJN_626508 | Bacteria | Bacteroidetes | Flavobacteriia | Flavobacterial | Flavobacteria | unclassified | Flavobacteria | information st | Translation, ril | COG0254 | 50S ribosomal | 487796.Flav2A | 0,6 | 0,489897949 | 9,46E-05 | 7,81E-05 | 1 - | 0,000138285 - | 1 - | 0,000171709 - | 1 - | 0,000163232 - | 0 - | 0 - | 0 - | 0 - |
| 6727 | 1 | 3um_FIJJAFBF_40841 | Bacteria | Bacteroidetes | Flavobacteriia | Flavobacterial | Flavobacteria | Polaribacter | Polaribacter t | information st | Replication, re | 05CZ8 | DNA polymera | 391587.KAOT | 0,8 | 0,748331477 | 2,73E-05 | 2,40E-05 | 2 - | 6,17E-05 - | 1 - | 3,83E-05 - | 1 - | 3,64E-05 - | 0 - | 0 - | 0 - | 0 - |
| 6728 | 1 | 3um_FIJJAFBF_387035 | Bacteria | Proteobacteri | Alphaproteob | unclassified | unclassified | unclassified | Alphaproteob | unclassified | unclassified | unclassified | unclassified | unclassified | 10,8 | 0,4 | 0,000496578 | 5,11E-05 | 11 - | 0,000423673 - | 11 - | 0,000526076 - | 11 - | 0,000500102 - | 10 - | 0,000461353 - | 11 - | 0,000571684 - |
| 6729 | 1 | 3um_FIJJAFBF_346328 | Bacteria | Proteobacteri | Gammaprote | Oceanospirilla | Saccharospirill | Saccharospiril | Saccharospirill | metabolism | Lipid transpor | 07QM9 | Component o | 314283.MED2 | 2,4 | 0,489897949 | 0,000102686 | 1,81E-05 | 3 - | 0,000108622 - | 3 - | 0,000134876 - | 2 - | 8,55E-05 - | 2 - | 8,67E-05 - | 2 - | 9,77E-05 - |
| 6730 | 1 | 3um_FIJJAFBF_165694 | Bacteria | Proteobacteri | Alphaproteob | Rhodobactera | unclassified | unclassified | Rhodobactera | information st | Translation, ril | 179R6 | This protein is | 391624.OIHEL | 2,8 | 0,748331477 | 0,000251959 | 7,10E-05 | 3 - | 0,000226533 - | 2 - | 0,000187525 - | 2 - | 0,000178266 - | 4 - | 0,000361798 - | 3 - | 0,000305673 - |
| 6731 | 1 | 3um_FIJJAFBF_305655 | Eukaryota | unclassified | Oomycetes | Saprolegniales | Saprolegniace | Aphanomyces | Aphanomyces | unclassified | unclassified | unclassified | unclassified | unclassified | 0,8 | 0,4 | 5,76E-05 | 2,93E-05 | 1 - | 6,24E-05 - | 1 - | 7,75E-05 - | 1 - | 7,36E-05 - | 1 - | 7,47E-05 - | 0 - | 0 - |
| 6732 | 1 | 3um_FIJJAFBF_436836 | Bacteria | Actinobacteria | Actinobacteria | Micrococcales | Microbacteria | Candidatus Ac | Candidatus Ac | information st | Translation, ril | COG0087 | One of the pri | 312284.A20C1 | 0,4 | 0,489897949 | 2,29E-05 | 2,82E-05 | 1 - | 5,24E-05 - | 0 - | 0 - | 1 - | 6,19E-05 - | 0 - | 0 - | 0 - | 0 - |
| 6733 | 2 | 0.2um_NOKCPHJN_295945;3um_FIJJAFBF_40134 | Bacteria | Bacteroidetes | unclassified | unclassified | unclassified | unclassified | Bacteroidetes | unclassified | unclassified | unclassified | unclassified | unclassified | 0,2 | 0,4 | 1,88E-05 | 3,76E-05 | 0 - | 0 - | 0 - | 0 - | 1 - | 9,41E-05 - | 0 - | 0 - | 0 - | 0 - |
| 6734 | 1 | 3um_FIJJAFBF_17656 | Bacteria | Proteobacteri | Alphaproteob | Pelagibacteral | Pelagibactera | Candidatus Pe | Candidatus Pe | unclassified | unclassified | unclassified | unclassified | unclassified | 2,2 | 0,4 | 0,000218515 | 4,44E-05 | 2 - | 0,000166343 - | 2 - | 0,000206549 - | 2 - | 0,000196351 - | 3 - | 0,000298877 - | 2 - | 0,000224456 - |
| 6735 | 1 | 0.2um_NOKCPHJN_160377 | Eukaryota | unclassified | Bangiophycea | Bangiales | Bangiaceae | Porphyra | Porphyra purp | unclassified | unclassified | unclassified | unclassified | unclassified | 1 | 0,894427191 | 2,04E-05 | 1,90E-05 | 1 - | 1,75E-05 - | 2 - | 4,34E-05 - | 2 - | 4,12E-05 - | 0 - | 0 - | 0 - | 0 - |
| 6736 | 1 | 3um_FIJJAFBF_223366 | Bacteria | Proteobacteri | Gammaprote | Cellvibrionales | Porticoccacea | unclassified | gamma prote | metabolism | Inorganic ion t | 0QHT5 | TonB-depend | 314287.GB22 | 2 | 0,632455532 | 5,06E-05 | 1,81E-05 | 1 - | 2,08E-05 - | 3 - | 7,73E-05 - | 2 - | 4,90E-05 - | 2 - | 4,97E-05 - | 2 - | 5,60E-05 - |
| 6737 | 1 | 0.2um_NOKCPHJN_302849 | Bacteria | Proteobacteri | Alphaproteob | Pelagibacteral | Pelagibactera | unclassified | various | cellular proces | Posttranslatio | COG0652 | peptidyl-proly | 439493.PB721 | 0,8 | 0,4 | 7,09E-05 | 3,57E-05 | 0 - | 0 - | 1 - | 8,85E-05 - | 1 - | 8,42E-05 - | 1 - | 8,54E-05 - | 1 - | 9,62E-05 - |
| 6738 | 1 | 0.2um_NOKCPHJN_563993 | Bacteria | Bacteroidetes | Saprospiria | Saprospirales | Lewinellaceae | Lewinella | Lewinella nigr | poorly charact | Function unkn | 09AR5 | Inherit from b | 518766.Rmar_ | 1 | 0 | 3,49E-05 | 3,31E-06 | 1 - | 2,92E-05 - | 1 - | 3,63E-05 - | 1 - | 3,45E-05 - | 1 - | 3,50E-05 - | 1 - | 3,94E-05 - |
| 6739 | 1 | 3um_FIJJAFBF_69370 | Eukaryota | Phaeophyceae | unclassified | Ectocarpales | Ectocarpaceae | Ectocarpus | Ectocarpus sili | cellular proces | Signal transdu | KOG0613 | fibronectin typ | 4787.PITG_01 | 2,6 | 1,356465997 | 3,43E-05 | 1,82E-05 | 3 - | 3,37E-05 - | 4 - | 5,57E-05 - | 1 - | 1,32E-05 - | 4 - | 5,38E-05 - | 1 - | 1,51E-05 - |
| 6740 | 1 | 3um_FIJJAFBF_543287 | Bacteria | Bacteroidetes | Flavobacteriia | Flavobacterial | Flavobacteria | Lacinutrix | Lacinutrix him | information st | Replication, re | 0PB0H | DNA-binding p | 983544.Lacal_ | 3,2 | 0,748331477 | 0,000490055 | 0,000139303 | 3 - | 0,000382589 - | 2 - | 0,000316709 - | 3 - | 0,000451607 - | 4 - | 0,000611037 - | 4 - | 0,000688331 - |
| 6741 | 1 | 0.2um_NOKCPHJN_824762 | Eukaryota | unclassified | Cryptophyta | Pyrenomonad | Geminigerace | Guillardia | Guillardia thet | unclassified | unclassified | unclassified | unclassified | unclassified | 1 | 0 | 0,000175676 | 1,67E-05 | 1 - | 0,00014715 - | 1 - | 0,000182716 - | 1 - | 0,000173695 - | 1 - | 0,000176261 - | 1 - | 0,000198557 - |
| 6742 | 1 | 3um_FIJJAFBF_489815 | Bacteria | Bacteroidetes | Flavobacteriia | Flavobacterial | Flavobacteria | Formosa | Formosa sp. H | unclassified | unclassified | unclassified | unclassified | unclassified | 2 | 0,632455532 | 0,000154314 | 3,93E-05 | 3 - | 0,000197891 - | 1 - | 8,19E-05 - | 2 - | 0,000155727 - | 2 - | 0,000158027 - | 2 - | 0,000178017 - |
| 6743 | 1 | 0.2um_NOKCPHJN_389828 | Bacteria | Proteobacteri | Gammaprote | unclassified | unclassified | unclassified | Gammaprote | unclassified | unclassified | unclassified | unclassified | unclassified | 1,6 | 0,489897949 | 0,000156116 | 5,03E-05 | 1 - | 8,14E-05 - | 2 - | 0,000202154 - | 2 - | 0,000192173 - | 2 - | 0,000195012 - | 1 - | 0,00010984 - |
| 6744 | 1 | 3um_FIJJAFBF_485183 | Bacteria | Bacteroidetes | Flavobacteriia | Flavobacterial | Flavobacteria | Polaribacter | Polaribacter ir | unclassified | unclassified | unclassified | unclassified | unclassified | 1 | 0 | 2,20E-05 | 2,09E-06 | 1 - | 1,84E-05 - | 1 - | 2,28E-05 - | 1 - | 2,17E-05 - | 1 - | 2,20E-05 - | 1 - | 2,48E-05 - |
| 6745 | 1 | 3um_FIJJAFBF_174540 | Bacteria | Bacteroidetes | Flavobacteriia | Flavobacterial | Flavobacteria | Flavobacteriu | Flavobacteriu | unclassified | unclassified | unclassified | unclassified | unclassified | 0,2 | 0,4 | 7,27E-06 | 1,45E-05 | 0 - | 0 - | 1 - | 3,64E-05 - | 0 - | 0 - | 0 - | 0 - | 0 - | 0 - |
| 6746 | 1 | 0.2um_NOKCPHJN_278387 | Eukaryota | unclassified | Oomycetes | Peronosporale | Peronosporac | Phytophthora | Phytophthora | unclassified | unclassified | unclassified | unclassified | unclassified | 0,2 | 0,4 | 1,67E-06 | 3,35E-06 | 1 - | 8,37E-06 - | 0 - | 0 - | 0 - | 0 - | 0 - | 0 - | 0 - | 0 - |
| 6747 | 1 | 0.2um_NOKCPHJN_198494 | Bacteria | Bacteroidetes | Flavobacteriia | Flavobacterial | Flavobacteria | Algibacter | Algibacter sp. | cellular proces | Intracellular tr | 05FGH | MotA TolQ exl | 391603.FBALC | 3,8 | 0,4 | 0,000204318 | 3,03E-05 | 4 - | 0,000180042 - | 4 - | 0,000223559 - | 3 - | 0,000159391 - | 4 - | 0,00021566 - | 4 - | 0,00024294 - |
| 6748 | 1 | 3um_FIJJAFBF_379962 | Eukaryota | unclassified | unclassified | Isochrysidales | Noelaerhabda | Emiliania | Emiliania huxl | information st | Replication, re | 1E8C4 | Inherit from N | 436017.A4S7F | 2 | 1,095445115 | 9,13E-05 | 4,03E-05 | 4 - | 0,000159968 - | 2 - | 9,93E-05 - | 1 - | 4,72E-05 - | 2 - | 9,58E-05 - | 1 - | 5,40E-05 - |
| 6749 | 1 | 0.2um_NOKCPHJN_46936 | Bacteria | Proteobacteri | Alphaproteob | Rhodobactera | various | various | various | metabolism | Amino acid tra | 171HI | PBPb | 388399.SSE37 | 1,4 | 1,356465997 | 5,21E-05 | 4,40E-05 | 4 - | 0,00013385 - | 1 - | 4,16E-05 - | 0 - | 0 - | 1 - | 4,01E-05 - | 1 - | 4,52E-05 - |
| 6750 | 1 | 3um_FIJJAFBF_517422 | Bacteria | Bacteroidetes | Flavobacteriia | Flavobacterial | Flavobacteria | Polaribacter | Polaribacter a | information st | Translation, ril | 05C64 | Binds directly | 313594.PI23P | 10,4 | 1,624807681 | 0,000616262 | 7,01E-05 | 13 - | 0,000651571 - | 10 - | 0,000622353 - | 11 - | 0,000650788 - | 8 - | 0,000480291 - | 10 - | 0,000676308 - |
| 6751 | 1 | 3um_FIJJAFBF_482523 | Bacteria | Bacteroidetes | Flavobacteriia | Flavobacterial | Crocinitomica | unclassified | Crocinitomica | unclassified | unclassified | unclassified | unclassified | unclassified | 1 | 0 | 6,65E-05 | 6,32E-06 | 1 - | 5,57E-05 - | 1 - | 6,92E-05 - | 1 - | 6,58E-05 - | 1 - | 6,67E-05 - | 1 - | 7,52E-05 - |
| 6752 | 1 | 3um_FIJJAFBF_333663 | Bacteria | Bacteroidetes | Flavobacteriia | Flavobacterial | Flavobacteria | Arenibacter | Arenibacter al | poorly charact | Function unkn | 0XQTX | Chromosome | 886377.Murru | 0,8 | 0,748331477 | 9,51E-06 | 8,26E-06 | 2 - | 2,09E-05 - | 0 - | 0 - | 0 - | 0 - | 1 - | 1,25E-05 - | 1 - | 1,41E-05 - |
| 6753 | 1 | 0.2um_NOKCPHJN_892782 | unclassified | unclassified | unclassified | unclassified | unclassified | unclassified | unclassified | unclassified | unclassified | unclassified | unclassified | unclassified | 1,8 | 0,748331477 | 9,24E-05 | 3,00E-05 | 3 - | 0,000133981 - | 1 - | 5,55E-05 - | 2 - | 0,000105434 - | 2 - | 0,000106991 - | 1 - | 6,03E-05 - |
| 6754 | 1 | 0.2um_NOKCPHJN_622086 | Bacteria | Actinobacteria | Actinobacteria | Micrococcales | Microbacteria | Candidatus Ac | Candidatus Ac | unclassified | unclassified | unclassified | unclassified | unclassified | 11,6 | 1,356465997 | 0,000139388 | 6,98E-06 | 14 - | 0,000142327 - | 12 - | 0,000151482 - | 11 - | 0,000132002 - | 11 - | 0,000133952 - | 10 - | 0,000137178 - |
| 6755 | 1 | 3um_FIJJAFBF_128443 | Bacteria | Bacteroidetes | Flavobacteriia | Flavobacterial | Flavobacteria | Polaribacter | Polaribacter s | unclassified | unclassified | unclassified | unclassified | unclassified | 9,4 | 1,019803903 | 0,00017042 | 1,27E-05 | 11 - | 0,000168339 - | 8 - | 0,00015202 - | 9 - | 0,000162579 - | 10 - | 0,000183311 - | 9 - | 0,000185849 - |
| 6756 | 1 | 0.2um_NOKCPHJN_435831 | Bacteria | Proteobacteri | Gammaprote | Alteromonada | Alteromonada | Bowmanella | Bowmanella d | metabolism | Inorganic ion t | 0QJVD | TonB-depende | 223926.VP016 | 1,2 | 0,4 | 1,91E-05 | 8,13E-06 | 1 - | 1,31E-05 - | 1 - | 1,62E-05 - | 1 - | 1,54E-05 - | 1 - | 1,56E-05 - | 2 - | 3,52E-05 - |
| 6757 | 1 | 3um_FIJJAFBF_275151 | Bacteria | Bacteroidetes | Flavobacteriia | Flavobacterial | Flavobacteria | Polaribacter | various | unclassified | unclassified | unclassified | unclassified | unclassified | 1,4 | 0,489897949 | 4,39E-05 | 1,55E-05 | 1 - | 2,63E-05 - | 1 - | 3,27E-05 - | 2 - | 6,21E-05 - | 2 - | 6,31E-05 - | 1 - | 3,55E-05 - |
| 6758 | 1 | 3um_FIJJAFBF_314256 | Bacteria | Proteobacteri | Alphaproteob | various | various | various | various | cellular proces | Defense mech | 07SUK | Efflux transpo | 367336.OM22 | 0,2 | 0,4 | 5,91E-06 | 1,18E-05 | 0 - | 0 - | 1 - | 2,96E-05 - | 0 - | 0 - | 0 - | 0 - | 0 - | 0 - |
| 6759 | 1 | 3um_FIJJAFBF_178374 | Bacteria | Proteobacteri | Gammaprote | unclassified | unclassified | unclassified | various | unclassified | unclassified | unclassified | unclassified | unclassified | 0,4 | 0,8 | 4,25E-05 | 8,50E-05 | 2 - | 0,00021255 - | 0 - | 0 - | 0 - | 0 - | 0 - | 0 - | 0 - | 0 - |
| 6760 | 1 | 0.2um_NOKCPHJN_236392 | Bacteria | Bacteroidetes | unclassified | unclassified | unclassified | unclassified | unidentified e | information st | Translation, ril | COG0184 | Forms an inte | 50743.SCB49_ | 3,2 | 0,748331477 | 0,000483324 | 8,43E-05 | 4 - | 0,000515851 - | 3 - | 0,000480401 - | 4 - | 0,000608909 - | 3 - | 0,000463427 - | 2 - | 0,000348033 - |
| 6761 | 1 | 3um_FIJJAFBF_261973 | Bacteria | Proteobacteri | Gammaprote | unclassified | unclassified | unclassified | Gammaprote | poorly charact | Function unkn | 08MVQ | NADPH-deper | 247634.GPB2 | 0,4 | 0,489897949 | 2,24E-05 | 2,76E-05 | 1 - | 5,15E-05 - | 0 - | 0 - | 1 - | 6,08E-05 - | 0 - | 0 - | 0 - | 0 - |
| 6762 | 1 | 3um_FIJJAFBF_182958 | Bacteria | Proteobacteri | Gammaprote | Alteromonada | Alteromonada | Paraglaciecola | Paraglaciecola | poorly charact | Function unkn | 08XUC | SapC family | 314287.GB22 | 0,8 | 0,4 | 4,36E-05 | 2,22E-05 | 1 - | 4,72E-05 - | 1 - | 5,86E-05 - | 1 - | 5,58E-05 - | 1 - | 5,66E-05 - | 0 - | 0 - |
| 6763 | 1 | 0.2um_NOKCPHJN_103488 | Bacteria | Proteobacteri | Alphaproteob | Rhodobactera | unclassified | unclassified | Rhodobactera | cellular proces | Cell wall/mem | 01XSD | Pfam:DUF583 | 367336.OM22 | 0,4 | 0,489897949 | 3,16E-05 | 3,90E-05 | 1 - | 7,04E-05 - | 1 - | 8,74E-05 - | 0 - | 0 - | 0 - | 0 - | 0 - | 0 - |
| 6764 | 1 | 3um_FIJJAFBF_586277 | Eukaryota | unclassified | Pelagophycea | Pelagomonad | unclassified | Aureococcus | Aureococcus a | unclassified | unclassified | unclassified | unclassified | unclassified | 0,4 | 0,8 | 5,11E-05 | 0,000102251 | 0 - | 0 - | 0 - | 0 - | 2 - | 0,000255627 - | 0 - | 0 - | 0 - | 0 - |
| 6765 | 2 | 0.2um_NOKCPHJN_160385;3um_FIJJAFBF_53768 | Eukaryota | Phaeophyceae | unclassified | Laminariales | Agaraceae | Thalassiophyll | Thalassiophyll | metabolism | Energy produc | 0IE88 | oxidoreductas | 35128.Thapsd | 10,4 | 0,8 | 0,000193367 | 1,65E-05 | 11 - | 0,000172009 - | 9 - | 0,000174751 - | 11 - | 0,000203039 - | 11 - | 0,000206038 - | 10 - | 0,000211001 - |
| 6766 | 1 | 3um_FIJJAFBF_214156 | Bacteria | Proteobacteri | Alphaproteob | Rhodobactera | Rhodobactera | Sulfitobacter | Sulfitobacter s | unclassified | unclassified | unclassified | unclassified | unclassified | 6,2 | 0,979795897 | 0,000145498 | 1,57E-05 | 8 - | 0,000158861 - | 6 - | 0,000147943 - | 5 - | 0,000117199 - | 6 - | 0,000142716 - | 6 - | 0,000160769 - |
| 6767 | 2 | 3um_FIJJAFBF_181049;3um_FIJJAFBF_501896 | Bacteria | Bacteroidetes | Flavobacteriia | Flavobacterial | unclassified | unclassified | Flavobacterial | unclassified | unclassified | unclassified | unclassified | unclassified | 0,2 | 0,4 | 9,07E-06 | 1,81E-05 | 1 - | 4,54E-05 - | 0 - | 0 - | 0 - | 0 - | 0 - | 0 - | 0 - | 0 - |
| 6768 | 1 | 0.2um_NOKCPHJN_555039 | Bacteria | Proteobacteri | Alphaproteob | Rhodobactera | Rhodobactera | Sulfitobacter | Sulfitobacter s | unclassified | unclassified | unclassified | unclassified | unclassified | 2 | 0,632455532 | 3,29E-05 | 7,62E-06 | 3 - | 4,26E-05 - | 2 - | 3,53E-05 - | 2 - | 3,35E-05 - | 2 - | 3,40E-05 - | 1 - | 1,92E-05 - |
| 6769 | 1 | 3um_FIJJAFBF_604897 | Bacteria | Bacteroidetes | Flavobacteriia | unclassified | unclassified | unclassified | Flavobacteria | unclassified | unclassified | unclassified | unclassified | unclassified | 1,6 | 0,8 | 0,000120939 | 4,84E-05 | 3 - | 0,000197891 - | 1 - | 8,19E-05 - | 1 - | 7,79E-05 - | 2 - | 0,000158027 - | 1 - | 8,90E-05 - |
| 6770 | 1 | 3um_FIJJAFBF_791565 | Bacteria | Proteobacteri | Alphaproteob | Rhodospirillal | Rhodospirillac | unclassified | Rhodobactera | metabolism | Inorganic ion t | 05C2J | ABC transport | 272942.RCAP_ | 0,4 | 0,489897949 | 2,30E-05 | 2,83E-05 | 0 - | 0 - | 0 - | 0 - | 1 - | 5,38E-05 - | 0 - | 0 - | 1 - | 6,15E-05 - |
| 6771 | 1 | 0.2um_NOKCPHJN_228575 | Bacteria | Bacteroidetes | Flavobacteriia | Flavobacterial | Crocinitomica | unclassified | Crocinitomica | unclassified | unclassified | unclassified | unclassified | unclassified | 0,2 | 0,4 | 1,08E-05 | 2,16E-05 | 0 - | 0 - | 0 - | 0 - | 1 - | 5,40E-05 - | 0 - | 0 - | 0 - | 0 - |
| 6772 | 1 | 3um_FIJJAFBF_727048 | Bacteria | Proteobacteri | Gammaprote | Cellvibrionales | Porticoccacea | unclassified | various | metabolism | Lipid transpor | 0QKY4 | 2-hydroxy-3-o | 314287.GB22 | 1 | 0,632455532 | 4,49E-05 | 2,51E-05 | 2 - | 7,76E-05 - | 1 - | 4,81E-05 - | 0 - | 0 - | 1 - | 4,64E-05 - | 1 - | 5,23E-05 - |
| 6773 | 1 | 3um_FIJJAFBF_514035 | Bacteria | Bacteroidetes | Flavobacteriia | Flavobacterial | Flavobacteria | Polaribacter | Polaribacter s | unclassified | unclassified | unclassified | unclassified | unclassified | 0,6 | 0,8 | 3,99E-05 | 5,15E-05 | 2 - | 0,000125439 - | 0 - | 0 - | 1 - | 7,40E-05 - | 0 - | 0 - | 0 - | 0 - |
| 6774 | 1 | 3um_FIJJAFBF_537683 | Eukaryota | Bacillariophyt | Bacillariophyc | Naviculales | Naviculaceae | Fistulifera | Fistulifera sola | metabolism | Energy produc | 1DYB8 | PsaA and PsaE | 3055.DAA014 | 12,2 | 0,979795897 | 0,000221109 | 1,55E-05 | 14 - | 0,00021368 - | 12 - | 0,000227424 - | 12 - | 0,000216195 - | 11 - | 0,000201106 - | 12 - | 0,00024714 - |
| 6775 | 1 | 0.2um_NOKCPHJN_733053 | Bacteria | Bacteroidetes | Flavobacteriia | Flavobacterial | Flavobacteria | Polaribacter | Polaribacter s | metabolism | Energy produc | COG0711 | F(1)F(0) ATP s | 313598.MED1 | 1 | 0 | 8,30E-05 | 7,89E-06 | 1 - | 6,96E-05 - | 1 - | 8,64E-05 - | 1 - | 8,21E-05 - | 1 - | 8,33E-05 - | 1 - | 9,39E-05 - |
| 6776 | 1 | 0.2um_NOKCPHJN_895539 | Eukaryota | Eustigmatoph | unclassified | Eustigmatales | Monodopsida | Nannochlorop | Nannochlorop | unclassified | unclassified | unclassified | unclassified | unclassified | 0,6 | 0,489897949 | 5,79E-05 | 4,76E-05 | 1 - | 8,57E-05 - | 0 - | 0 - | 1 - | 0,000101106 - | 1 - | 0,000102599 - | 0 - | 0 - |
| 6777 | 1 | 3um_FIJJAFBF_179843 | Bacteria | Bacteroidetes | Flavobacteriia | Flavobacterial | Flavobacteria | Algibacter | Algibacter sp. | metabolism | Energy produc | COG0712 | F(1)F(0) ATP s | 983544.Lacal_ | 0,8 | 0,748331477 | 6,01E-05 | 5,92E-05 | 1 - | 6,45E-05 - | 2 - | 0,000160134 - | 1 - | 7,61E-05 - | 0 - | 0 - | 0 - | 0 - |
| 6778 | 1 | 0.2um_NOKCPHJN_146555 | Bacteria | Proteobacteri | Betaproteoba | Burkholderiale | Comamonada | Curvibacter | Curvibacter sp | metabolism | Amino acid tra | 0B7ZJ | Alpha-keto-be | 232721.Ajs_1 | 3,2 | 0,979795897 | 0,000128491 | 3,82E-05 | 3 - | 0,000101873 - | 3 - | 0,000126496 - | 5 - | 0,000200417 - | 3 - | 0,000122027 - | 2 - | 9,16E-05 - |
| 6779 | 1 | 3um_FIJJAFBF_106373 | unclassified | unclassified | unclassified | unclassified | unclassified | unclassified | unclassified | unclassified | unclassified | unclassified | unclassified | unclassified | 0,2 | 0,4 | 1,65E-05 | 3,30E-05 | 1 - | 8,26E-05 - | 0 - | 0 - | 0 - | 0 - | 0 - | 0 - | 0 - | 0 - |
| 6780 | 1 | 3um_FIJJAFBF_592170 | Bacteria | Bacteroidetes | Flavobacteriia | Flavobacterial | Crocinitomica | unclassified | Crocinitomica | unclassified | unclassified | unclassified | unclassified | unclassified | 1 | 0 | 9,32E-05 | 8,85E-06 | 1 - | 7,81E-05 - | 1 - | 9,70E-05 - | 1 - | 9,22E-05 - | 1 - | 9,35E-05 - | 1 - | 0,000105357 - |
| 6781 | 1 | 0.2um_NOKCPHJN_393858 | Eukaryota | unclassified | Cryptophyta | Pyrenomonad | Geminigerace | Guillardia | Guillardia thet | metabolism | Energy produc | 0IFIU | Pyrophosphat | 65071.PYU1_ | 2,4 | 0,489897949 | 3,61E-05 | 5,62E-06 | 3 - | 3,83E-05 - | 2 - | 3,17E-05 - | 2 - | 3,01E-05 - | 3 - | 4,59E-05 - | 2 - | 3,45E-05 - |
| 6782 | 1 | 0.2um_NOKCPHJN_105145 | Bacteria | Bacteroidetes | Flavobacteriia | Flavobacterial | Flavobacteria | Polaribacter | Polaribacter s | unclassified | unclassified | unclassified | unclassified | unclassified | 0,6 | 0,489897949 | 4,59E-05 | 3,75E-05 | 0 - | 0 - | 1 - | 7,87E-05 - | 1 - | 7,49E-05 - | 1 - | 7,60E-05 - | 0 - | 0 - |
| 6783 | 1 | 0.2um_NOKCPHJN_133072 | Bacteria | Bacteroidetes | Flavobacteriia | Flavobacterial | Crocinitomica | unclassified | Crocinitomica | unclassified | unclassified | unclassified | unclassified | unclassified | 4 | 0 | 0,000174557 | 1,66E-05 | 4 - | 0,000146212 - | 4 - | 0,000181553 - | 4 - | 0,000172589 - | 4 - | 0,000175138 - | 4 - | 0,000197292 - |
| 6784 | 1 | 0.2um_NOKCPHJN_89944 | Bacteria | Proteobacteri | Alphaproteob | unclassified | unclassified | unclassified | uncultured alp | metabolism | Amino acid tra | COG1448 | aminotransfer | 501479.CSE45 | 0,8 | 0,4 | 2,79E-05 | 1,43E-05 | 1 - | 2,91E-05 - | 1 - | 3,62E-05 - | 0 - | 0 - | 1 - | 3,49E-05 - | 1 - | 3,93E-05 - |
| 6785 | 1 | 3um_FIJJAFBF_378657 | Bacteria | Bacteroidetes | Flavobacteriia | Flavobacterial | Flavobacteria | Polaribacter | Polaribacter d | unclassified | unclassified | unclassified | unclassified | unclassified | 1 | 1,095445115 | 2,70E-05 | 3,11E-05 | 1 - | 2,28E-05 - | 3 - | 8,50E-05 - | 0 - | 0 - | 1 - | 2,73E-05 - | 0 - | 0 - |
| 6786 | 1 | 0.2um_NOKCPHJN_353311 | unclassified | unclassified | unclassified | unclassified | unclassified | unclassified | unclassified | unclassified | unclassified | unclassified | unclassified | unclassified | 0,2 | 0,4 | 1,13E-05 | 2,25E-05 | 1 - | 5,63E-05 - | 0 - | 0 - | 0 - | 0 - | 0 - | 0 - | 0 - | 0 - |
| 6787 | 1 | 0.2um_NOKCPHJN_1028497 | Viruses | unclassified | unclassified | unclassified | Phycodnavirid | unclassified | Chrysochromu | unclassified | unclassified | unclassified | unclassified | unclassified | 0,4 | 0,489897949 | 3,24E-05 | 4,00E-05 | 1 - | 7,22E-05 - | 1 - | 8,96E-05 - | 0 - | 0 - | 0 - | 0 - | 0 - | 0 - |
| 6788 | 1 | 0.2um_NOKCPHJN_323757 | Eukaryota | unclassified | Heterotrichea | Heterotrichida | Stentoridae | Stentor | Stentor coeru | unclassified | unclassified | unclassified | unclassified | unclassified | 0,2 | 0,4 | 1,16E-06 | 2,32E-06 | 1 - | 5,81E-06 - | 0 - | 0 - | 0 - | 0 - | 0 - | 0 - | 0 - | 0 - |
| 6789 | 1 | 0.2um_NOKCPHJN_987138 | unclassified | unclassified | unclassified | unclassified | unclassified | unclassified | unclassified | unclassified | unclassified | unclassified | unclassified | unclassified | 0,2 | 0,4 | 1,87E-05 | 3,74E-05 | 0 - | 0 - | 0 - | 0 - | 1 - | 9,34E-05 - | 0 - | 0 - | 0 - | 0 - |
| 6790 | 1 | 3um_FIJJAFBF_673918 | Eukaryota | unclassified | Oomycetes | Peronosporale | Peronosporac | Phytophthora | various | unclassified | unclassified | unclassified | unclassified | unclassified | 1 | 0 | 3,76E-05 | 3,57E-06 | 1 - | 3,15E-05 - | 1 - | 3,92E-05 - | 1 - | 3,72E-05 - | 1 - | 3,78E-05 - | 1 - | 4,25E-05 - |
| 6791 | 1 | 3um_FIJJAFBF_583737 | Bacteria | Bacteroidetes | Flavobacteriia | Flavobacterial | Flavobacteria | Formosa | Formosa sp. H | metabolism | Amino acid tra | COG1932 | Catalyzes the | 983544.Lacal_ | 0,2 | 0,4 | 6,50E-06 | 1,30E-05 | 1 - | 3,25E-05 - | 0 - | 0 - | 0 - | 0 - | 0 - | 0 - | 0 - | 0 - |
| 6792 | 1 | 0.2um_NOKCPHJN_443059 | Bacteria | Bacteroidetes | Flavobacteriia | Flavobacterial | Flavobacteria | Gramella | Gramella port | poorly charact | Function unkn | 11PIQ | YCE I like fami | 983548.Krodi_ | 0,6 | 0,489897949 | 3,65E-05 | 3,03E-05 | 1 - | 5,17E-05 - | 0 - | 0 - | 1 - | 6,10E-05 - | 0 - | 0 - | 1 - | 6,98E-05 - |
| 6793 | 1 | 0.2um_NOKCPHJN_348144 | Bacteria | Actinobacteria | Actinobacteria | Pseudonocard | Pseudonocard | Amycolatopsis | Amycolatopsis | unclassified | unclassified | unclassified | unclassified | unclassified | 0,2 | 0,4 | 9,15E-07 | 1,83E-06 | 0 - | 0 - | 1 - | 4,57E-06 - | 0 - | 0 - | 0 - | 0 - | 0 - | 0 - |
| 6794 | 1 | 0.2um_NOKCPHJN_41625 | Bacteria | Bacteroidetes | Flavobacteriia | Flavobacterial | Flavobacteria | Polaribacter | Polaribacter s | metabolism | Energy produc | COG0437 | 4fe-4S ferredo | 313594.PI23P | 2,4 | 1,496662955 | 3,06E-05 | 1,60E-05 | 5 - | 5,64E-05 - | 2 - | 2,80E-05 - | 3 - | 3,99E-05 - | 1 - | 1,35E-05 - | 1 - | 1,52E-05 - |
| 6795 | 2 | 0.2um_NOKCPHJN_464680;0.2um_NOKCPHJN_6 | Bacteria | Proteobacteri | Alphaproteob | Pelagibacteral | Pelagibactera | Candidatus Pe | Candidatus Pe | unclassified | unclassified | unclassified | unclassified | unclassified | 0,4 | 0,8 | 2,33E-05 | 4,66E-05 | 2 - | 0,000116525 - | 0 - | 0 - | 0 - | 0 - | 0 - | 0 - | 0 - | 0 - |
| 6796 | 1 | 3um_FIJJAFBF_784992 | Bacteria | Proteobacteri | Alphaproteob | Rhodobactera | Rhodobactera | Pelagicola | Pelagicola sp. | unclassified | unclassified | unclassified | unclassified | unclassified | 1 | 0 | 0,000112317 | 1,07E-05 | 1 - | 9,41E-05 - | 1 - | 0,000116819 - | 1 - | 0,000111051 - | 1 - | 0,000112691 - | 1 - | 0,000126946 - |
| 6797 | 1 | 0.2um_NOKCPHJN_357407 | Bacteria | Proteobacteri | Gammaprote | Pseudomonad | Pseudomonad | Pseudomonas | Pseudomonas | unclassified | unclassified | unclassified | unclassified | unclassified | 2,8 | 0,4 | 7,87E-05 | 9,36E-06 | 3 - | 7,13E-05 - | 3 - | 8,85E-05 - | 3 - | 8,42E-05 - | 3 - | 8,54E-05 - | 2 - | 6,41E-05 - |
| 6798 | 1 | 0.2um_NOKCPHJN_121981 | Bacteria | Bacteroidetes | Flavobacteriia | Flavobacterial | Flavobacteria | Polaribacter | Polaribacter d | information st | Transcription | 0P7SF | DNA-depende | 391587.KAOT | 5,2 | 0,748331477 | 5,58E-05 | 7,56E-06 | 6 - | 5,43E-05 - | 5 - | 5,62E-05 - | 4 - | 4,27E-05 - | 6 - | 6,50E-05 - | 5 - | 6,10E-05 - |
| 6799 | 1 | 3um_FIJJAFBF_242064 | Bacteria | Proteobacteri | Alphaproteob | Rhodobactera | Rhodobactera | Phaeobacter | Phaeobacter s | poorly charact | Function unkn | 02DQA | Corrinoid met | 314262.MED1 | 0,8 | 0,4 | 4,92E-05 | 2,48E-05 | 0 - | 0 - | 1 - | 6,14E-05 - | 1 - | 5,84E-05 - | 1 - | 5,93E-05 - | 1 - | 6,68E-05 - |
| 6800 | 1 | 0.2um_NOKCPHJN_90473 | Bacteria | Proteobacteri | Gammaprote | Alteromonada | Colwelliaceae | Colwellia | Colwellia sp. P | cellular proces | Posttranslatio | 0QI1Q | Mg chelatase | 167879.CPS_4 | 0,2 | 0,4 | 5,62E-06 | 1,12E-05 | 0 - | 0 - | 1 - | 2,81E-05 - | 0 - | 0 - | 0 - | 0 - | 0 - | 0 - |
| 6801 | 1 | 3um_FIJJAFBF_222673 | Bacteria | Proteobacteri | Gammaprote | Alteromonada | Alteromonada | Paraglaciecola | Paraglaciecola | cellular proces | Posttranslatio | COG0330 | Band 7 protein | 1085623.GNIT | 0,6 | 0,489897949 | 2,67E-05 | 2,18E-05 | 0 - | 0 - | 1 - | 4,58E-05 - | 1 - | 4,36E-05 - | 1 - | 4,42E-05 - | 0 - | 0 - |
| 6802 | 1 | 3um_FIJJAFBF_137404 | Bacteria | Proteobacteri | Betaproteoba | Burkholderiale | Comamonada | Acidovorax | Acidovorax sp | unclassified | unclassified | unclassified | unclassified | unclassified | 0,4 | 0,489897949 | 5,27E-06 | 6,46E-06 | 0 - | 0 - | 1 - | 1,26E-05 - | 0 - | 0 - | 0 - | 0 - | 1 - | 1,37E-05 - |
| 6803 | 2 | 0.2um_NOKCPHJN_1087598;3um_FIJJAFBF_858( | various | various | various | various | various | various | various | various | various | various | various | various | 0,6 | 0,489897949 | 3,23E-05 | 2,69E-05 | 1 - | 4,50E-05 - | 1 - | 5,59E-05 - | 0 - | 0 - | 0 - | 0 - | 1 - | 6,07E-05 - |
| 6804 | 4 | 0.2um_NOKCPHJN_769801;3um_FIJJAFBF_2031( | Bacteria | Bacteroidetes | Flavobacteriia | Flavobacterial | Flavobacteria | Formosa | various | metabolism | Amino acid tra | 05CM2 | brancheD-cha | 688270.Celal_ | 1,2 | 0,4 | 4,56E-05 | 1,01E-05 | 2 - | 6,54E-05 - | 1 - | 4,06E-05 - | 1 - | 3,86E-05 - | 1 - | 3,92E-05 - | 1 - | 4,41E-05 - |
| 6805 | 1 | 0.2um_NOKCPHJN_550084 | Bacteria | Proteobacteri | Alphaproteob | Rhodobactera | unclassified | unclassified | Rhodobactera | poorly charact | Function unkn | 05DIH | basic membra | 367336.OM22 | 2,6 | 0,489897949 | 0,000102146 | 2,25E-05 | 3 - | 9,87E-05 - | 3 - | 0,000122509 - | 2 - | 7,76E-05 - | 2 - | 7,88E-05 - | 3 - | 0,00013313 - |
| 6806 | 1 | 3um_FIJJAFBF_401900 | Bacteria | Proteobacteri | Alphaproteob | Rhodobactera | unclassified | unclassified | Rhodobactera | metabolism | Energy produc | COG0711 | F(1)F(0) ATP s | 367336.OM22 | 2,4 | 0,489897949 | 0,000178192 | 4,11E-05 | 2 - | 0,000124083 - | 3 - | 0,000231112 - | 3 - | 0,000219701 - | 2 - | 0,000148631 - | 2 - | 0,000167432 - |
| 6807 | 1 | 3um_FIJJAFBF_41560 | Eukaryota | Bacillariophyt | Coscinodiscop | Chaetocerotal | Chaetocerota | Chaetoceros | Chaetoceros s | metabolism | Energy produc | 0IE88 | oxidoreductas | 45157.CMV13 | 3,4 | 0,489897949 | 6,37E-05 | 1,13E-05 | 3 - | 4,70E-05 - | 4 - | 7,78E-05 - | 4 - | 7,39E-05 - | 3 - | 5,63E-05 - | 3 - | 6,34E-05 - |
| 6808 | 1 | 3um_FIJJAFBF_735065 | Bacteria | Actinobacteria | Actinobacteria | Micrococcales | Microbacteria | unclassified | Microbacteria | unclassified | unclassified | unclassified | unclassified | unclassified | 7,8 | 1,326649916 | 0,000286222 | 3,22E-05 | 10 - | 0,000311048 - | 8 - | 0,000308984 - | 8 - | 0,000293728 - | 6 - | 0,00022355 - | 7 - | 0,0002938 - |
| 6809 | 1 | 3um_FIJJAFBF_87953 | Bacteria | Bacteroidetes | Cytophagia | Cytophagales | Flammeovirga | unclassified | Flammeovirga | poorly charact | Function unkn | 061MS | NA | 742767.HMPR | 1,8 | 0,4 | 5,22E-05 | 1,28E-05 | 2 - | 4,86E-05 - | 2 - | 6,04E-05 - | 2 - | 5,74E-05 - | 1 - | 2,91E-05 - | 2 - | 6,56E-05 - |
| 6810 | 1 | 3um_FIJJAFBF_178759 | Bacteria | Actinobacteria | Actinobacteria | Micrococcales | Microbacteria | Rhodoluna | Candidatus Rh | poorly charact | Function unkn | 0NVRZ | NA | 170187.SP_17 | 1,2 | 0,4 | 1,61E-05 | 3,56E-06 | 2 - | 2,31E-05 - | 1 - | 1,43E-05 - | 1 - | 1,36E-05 - | 1 - | 1,38E-05 - | 1 - | 1,56E-05 - |
| 6811 | 1 | 3um_FIJJAFBF_280181 | Bacteria | Bacteroidetes | Flavobacteriia | Flavobacterial | Flavobacteria | Polaribacter | Polaribacter s | metabolism | Energy produc | 0P7ZB | cytochrome c | 313598.MED1 | 0,8 | 0,4 | 2,32E-05 | 1,19E-05 | 1 - | 2,43E-05 - | 1 - | 3,01E-05 - | 0 - | 0 - | 1 - | 2,91E-05 - | 1 - | 3,27E-05 - |
| 6812 | 2 | 0.2um_NOKCPHJN_314313;3um_FIJJAFBF_11213 | Bacteria | Bacteroidetes | Flavobacteriia | Flavobacterial | Crocinitomica | unclassified | Crocinitomica | unclassified | unclassified | unclassified | unclassified | unclassified | 0,2 | 0,4 | 6,42E-06 | 1,28E-05 | 0 - | 0 - | 0 - | 0 - | 1 - | 3,21E-05 - | 0 - | 0 - | 0 - | 0 - |
| 6813 | 1 | 0.2um_NOKCPHJN_498935 | Bacteria | Bacteroidetes | Flavobacteriia | Flavobacterial | Flavobacteria | Dokdonia | Dokdonia paci | cellular proces | Signal transdu | 0PAET | LMWPc | 391587.KAOT | 0,8 | 0,4 | 8,15E-05 | 4,10E-05 | 0 - | 0 - | 1 - | 0,000101799 - | 1 - | 9,68E-05 - | 1 - | 9,82E-05 - | 1 - | 0,000110625 - |
| 6814 | 1 | 0.2um_NOKCPHJN_306394 | Bacteria | Proteobacteri | Alphaproteob | unclassified | unclassified | unclassified | alpha proteob | unclassified | unclassified | unclassified | unclassified | unclassified | 3,6 | 0,489897949 | 7,69E-05 | 1,29E-05 | 3 - | 5,36E-05 - | 4 - | 8,87E-05 - | 4 - | 8,43E-05 - | 4 - | 8,55E-05 - | 3 - | 7,23E-05 - |
| 6815 | 1 | 0.2um_NOKCPHJN_553781 | Bacteria | Proteobacteri | Gammaprote | Cellvibrionales | Halieaceae | unclassified | gamma prote | unclassified | unclassified | unclassified | unclassified | unclassified | 0,2 | 0,4 | 6,14E-06 | 1,23E-05 | 1 - | 3,07E-05 - | 0 - | 0 - | 0 - | 0 - | 0 - | 0 - | 0 - | 0 - |
| 6816 | 1 | 0.2um_NOKCPHJN_471920 | Viruses | unclassified | unclassified | unclassified | unclassified | unclassified | uncultured vir | unclassified | unclassified | unclassified | unclassified | unclassified | 0,8 | 0,4 | 3,62E-05 | 1,86E-05 | 1 - | 3,83E-05 - | 0 - | 0 - | 1 - | 4,52E-05 - | 1 - | 4,58E-05 - | 1 - | 5,16E-05 - |
| 6817 | 1 | 0.2um_NOKCPHJN_446869 | Bacteria | Bacteroidetes | unclassified | unclassified | unclassified | unclassified | Bacteroidetes | information st | Replication, re | 05C2M | dna polymera | 926562.Oweh | 0,8 | 0,4 | 1,12E-05 | 5,71E-06 | 1 - | 1,22E-05 - | 1 - | 1,51E-05 - | 1 - | 1,44E-05 - | 1 - | 1,46E-05 - | 0 - | 0 - |
| 6818 | 1 | 3um_FIJJAFBF_383772 | Bacteria | Bacteroidetes | Flavobacteriia | Flavobacterial | Flavobacteria | Polaribacter | Polaribacter d | metabolism | Lipid transpor | 0P85Z | acyl-CoA dehy | 313598.MED1 | 0,4 | 0,489897949 | 9,24E-06 | 1,13E-05 | 0 - | 0 - | 1 - | 2,37E-05 - | 1 - | 2,25E-05 - | 0 - | 0 - | 0 - | 0 - |
| 6819 | 1 | 0.2um_NOKCPHJN_71903 | Bacteria | Proteobacteri | Gammaprote | Oceanospirilla | Saccharospirill | Reinekea | Reinekea fors | unclassified | unclassified | unclassified | unclassified | unclassified | 6,4 | 1,019803903 | 0,000101858 | 2,33E-05 | 6 - | 7,94E-05 - | 5 - | 8,22E-05 - | 6 - | 9,38E-05 - | 7 - | 0,000111002 - | 8 - | 0,000142906 - |
| 6820 | 1 | 0.2um_NOKCPHJN_389330 | Archaea | Euryarchaeota | unclassified | unclassified | unclassified | unclassified | Euryarchaeota | metabolism | Inorganic ion t | 05C27 | TonB-depende | 391596.PBAL3 | 1,8 | 0,979795897 | 2,66E-05 | 1,48E-05 | 2 - | 2,55E-05 - | 2 - | 3,17E-05 - | 3 - | 4,52E-05 - | 2 - | 3,06E-05 - | 0 - | 0 - |
| 6821 | 1 | 0.2um_NOKCPHJN_533539 | Eukaryota | unclassified | Oomycetes | Pythiales | Pythiaceae | Pythium | Pythium insidi | cellular proces | Posttranslatio | KOG1424 | Large subunit | 65071.PYU1_ | 0,2 | 0,4 | 3,14E-06 | 6,27E-06 | 1 - | 1,57E-05 - | 0 - | 0 - | 0 - | 0 - | 0 - | 0 - | 0 - | 0 - |
| 6822 | 1 | 0.2um_NOKCPHJN_178778 | Bacteria | Proteobacteri | Alphaproteob | Rhodobactera | Rhodobactera | Roseibium | Roseibium sp. | cellular proces | Posttranslatio | 16QU3 | Hemolysin-typ | 744980.TRICH | 3,4 | 0,8 | 2,93E-05 | 6,69E-06 | 4 - | 2,90E-05 - | 4 - | 3,61E-05 - | 2 - | 1,71E-05 - | 4 - | 3,48E-05 - | 3 - | 2,94E-05 - |
| 6823 | 2 | 3um_FIJJAFBF_122412;3um_FIJJAFBF_547977 | Bacteria | Bacteroidetes | Flavobacteriia | Flavobacterial | Flavobacteria | Polaribacter | various | cellular proces | Cell cycle cont | 0P8N7 | Cell division p | 313598.MED1 | 2 | 0,894427191 | 3,32E-05 | 1,53E-05 | 2 - | 2,80E-05 - | 3 - | 5,22E-05 - | 1 - | 1,65E-05 - | 3 - | 5,04E-05 - | 1 - | 1,89E-05 - |
| 6824 | 1 | 3um_FIJJAFBF_653260 | Bacteria | Bacteroidetes | Flavobacteriia | Flavobacterial | Flavobacteria | Polaribacter | Polaribacter t | poorly charact | Function unkn | 0XSVM | NA | 867900.Celly_ | 6,2 | 1,166190379 | 0,000175359 | 2,50E-05 | 8 - | 0,000191694 - | 5 - | 0,000148767 - | 5 - | 0,000141422 - | 7 - | 0,000200915 - | 6 - | 0,000193997 - |
| 6825 | 1 | 0.2um_NOKCPHJN_735680 | Bacteria | Bacteroidetes | Bacteroidia | Bacteroidales | Bacteroidacea | Bacteroides | Bacteroides t | unclassified | unclassified | unclassified | unclassified | unclassified | 0,2 | 0,4 | 4,99E-06 | 9,98E-06 | 1 - | 2,50E-05 - | 0 - | 0 - | 0 - | 0 - | 0 - | 0 - | 0 - | 0 - |
| 6826 | 1 | 3um_FIJJAFBF_91117 | Eukaryota | Bacillariophyt | Coscinodiscop | Leptocylindral | Leptocylindra | Leptocylindru | Leptocylindru | unclassified | unclassified | unclassified | unclassified | unclassified | 1,4 | 0,489897949 | 0,000183803 | 5,41E-05 | 2 - | 0,000225052 - | 1 - | 0,000139724 - | 1 - | 0,000132826 - | 2 - | 0,000269575 - | 1 - | 0,000151838 - |
| 6827 | 1 | 3um_FIJJAFBF_30987 | Bacteria | Bacteroidetes | Flavobacteriia | Flavobacterial | Flavobacteria | Altibacter | Altibacter lent | information st | Translation, ril | 05C64 | Binds directly | 50743.SCB49_ | 3,6 | 0,489897949 | 0,000213034 | 3,37E-05 | 4 - | 0,000198748 - | 3 - | 0,000185089 - | 4 - | 0,000234601 - | 3 - | 0,00017855 - | 4 - | 0,000268181 - |
| 6828 | 1 | 3um_FIJJAFBF_412012 | Bacteria | Proteobacteri | Gammaprote | Alteromonada | Alteromonada | unclassified | Alteromonada | information st | Translation, ril | COG1185 | Involved in mR | 377629.TERTU | 3,8 | 0,979795897 | 7,32E-05 | 1,50E-05 | 5 - | 8,21E-05 - | 3 - | 6,12E-05 - | 3 - | 5,81E-05 - | 5 - | 9,83E-05 - | 3 - | 6,65E-05 - |
| 6829 | 1 | 0.2um_NOKCPHJN_349986 | Bacteria | Proteobacteri | Gammaprote | Vibrionales | Vibrionaceae | Vibrio | Vibrio rumoie | metabolism | Carbohydrate | 0QIP7 | Catalyzes the | 882102.VAA_ | 11 | 2,097617696 | 0,000447604 | 3,95E-05 | 15 - | 0,000520136 - | 10 - | 0,000430571 - | 10 - | 0,000409312 - | 11 - | 0,000456893 - | 9 - | 0,000421109 - |
| 6830 | 1 | 3um_FIJJAFBF_237981 | Eukaryota | unclassified | Compsopogor | Erythropeltida | Erythrotrichia | Erythrotrichia | Erythrotrichia | information st | Translation, ril | COG0333 | 50s ribosomal | 696747.NIES3 | 0,4 | 0,489897949 | 9,33E-05 | 0,000114341 | 0 - | 0 - | 1 - | 0,000237531 - | 0 - | 0 - | 1 - | 0,000229139 - | 0 - | 0 - |
| 6831 | 1 | 0.2um_NOKCPHJN_42229 | Bacteria | Proteobacteri | Gammaprote | Cellvibrionales | Porticoccacea | unclassified | SAR92 bacteri | unclassified | unclassified | unclassified | unclassified | unclassified | 0,2 | 0,4 | 5,59E-06 | 1,12E-05 | 0 - | 0 - | 0 - | 0 - | 1 - | 2,79E-05 - | 0 - | 0 - | 0 - | 0 - |
| 6832 | 1 | 0.2um_NOKCPHJN_163487 | Bacteria | Bacteroidetes | Flavobacteriia | Flavobacterial | Flavobacteria | Polaribacter | various | poorly charact | Function unkn | 11PIQ | YCE I like fami | 313594.PI23P | 1 | 0,894427191 | 6,31E-05 | 5,65E-05 | 2 - | 0,000111434 - | 2 - | 0,000138368 - | 1 - | 6,58E-05 - | 0 - | 0 - | 0 - | 0 - |
| 6833 | 1 | 0.2um_NOKCPHJN_08635 | Bacteria | Proteobacteri | Gammaprote | Alteromonada | Alteromonada | Paraglaciecola | Paraglaciecola | unclassified | unclassified | unclassified | unclassified | unclassified | 2,2 | 1,166190379 | 0,000158012 | 7,05E-05 | 4 - | 0,000252257 - | 1 - | 7,83E-05 - | 3 - | 0,000223322 - | 2 - | 0,000151081 - | 1 - | 8,51E-05 - |
| 6834 | 1 | 0.2um_NOKCPHJN_56879 | Bacteria | Proteobacteri | Gammaprote | Oceanospirilla | Saccharospirill | Reinekea | Reinekea blan | metabolism | Energy produc | 16QA5 | electron trans | 314283.MED2 | 0,4 | 0,489897949 | 9,08E-06 | 1,12E-05 | 1 - | 2,08E-05 - | 0 - | 0 - | 1 - | 2,46E-05 - | 0 - | 0 - | 0 - | 0 - |
| 6835 | 1 | 0.2um_NOKCPHJN_77261 | Bacteria | Proteobacteri | Betaproteoba | Burkholderiale | Comamonada | Hydrogenoph | Hydrogenoph | metabolism | Lipid transpor | 08IQF | Catalyzes the | 397945.Aave_ | 0,2 | 0,4 | 3,45E-06 | 6,90E-06 | 1 - | 1,73E-05 - | 0 - | 0 - | 0 - | 0 - | 0 - | 0 - | 0 - | 0 - |
| 6836 | 3 | 0.2um_NOKCPHJN_56363;0.2um_NOKCPHJN_67 | Bacteria | Proteobacteri | Alphaproteob | Pelagibacteral | Pelagibactera | various | various | information st | Translation, ril | COG0292 | Binds directly | 335992.SAR11 | 1 | 0 | 0,000116125 | 1,10E-05 | 1 - | 9,73E-05 - | 1 - | 0,000120779 - | 1 - | 0,000114815 - | 1 - | 0,000116511 - | 1 - | 0,00013125 - |
| 6837 | 1 | 3um_FIJJAFBF_375923 | Bacteria | Proteobacteri | Gammaprote | Oceanospirilla | Saccharospirill | Saccharospiril | Saccharospiril | unclassified | unclassified | unclassified | unclassified | unclassified | 0,4 | 0,8 | 1,56E-05 | 3,12E-05 | 2 - | 7,81E-05 - | 0 - | 0 - | 0 - | 0 - | 0 - | 0 - | 0 - | 0 - |
| 6838 | 1 | 3um_FIJJAFBF_804151 | Bacteria | Proteobacteri | Gammaprote | Oceanospirilla | Saccharospirill | Reinekea | Reinekea fors | cellular proces | Posttranslatio | 08V1T | peptidylprolyl | 314283.MED2 | 4,2 | 0,4 | 0,000279909 | 4,12E-05 | 4 - | 0,000222867 - | 5 - | 0,00034592 - | 4 - | 0,000263072 - | 4 - | 0,000266958 - | 4 - | 0,000300727 - |
| 6839 | 1 | 3um_FIJJAFBF_689815 | Bacteria | Bacteroidetes | unclassified | unclassified | unclassified | unclassified | Bacteroidetes | information st | Translation, ril | 05C64 | Binds directly | 575590.HMPR | 1 | 0,894427191 | 5,52E-05 | 4,88E-05 | 2 - | 9,94E-05 - | 0 - | 0 - | 2 - | 0,000117301 - | 1 - | 5,95E-05 - | 0 - | 0 - |
| 6840 | 1 | 3um_FIJJAFBF_263218 | Bacteria | Proteobacteri | Gammaprote | Oceanospirilla | Saccharospirill | Reinekea | Reinekea fors | unclassified | unclassified | unclassified | unclassified | unclassified | 0,8 | 0,4 | 3,53E-05 | 1,82E-05 | 1 - | 3,74E-05 - | 0 - | 0 - | 1 - | 4,41E-05 - | 1 - | 4,48E-05 - | 1 - | 5,04E-05 - |
| 6841 | 1 | 3um_FIJJAFBF_738498 | Bacteria | Proteobacteri | Gammaprote | Alteromonada | Alteromonada | Paraglaciecola | Paraglaciecola | poorly charact | Function unkn | 17CR4 | Pfam:DUF213 | 983545.Glaag | 1,4 | 0,8 | 9,18E-05 | 5,39E-05 | 2 - | 0,000110895 - | 0 - | 0 - | 1 - | 6,55E-05 - | 2 - | 0,000132834 - | 2 - | 0,000149637 - |
| 6842 | 1 | 3um_FIJJAFBF_133398 | Bacteria | Bacteroidetes | Flavobacteriia | Flavobacterial | Flavobacteria | Algibacter | Algibacter sp. | unclassified | unclassified | unclassified | unclassified | unclassified | 0,2 | 0,4 | 1,25E-05 | 2,50E-05 | 0 - | 0 - | 0 - | 0 - | 1 - | 6,24E-05 - | 0 - | 0 - | 0 - | 0 - |
| 6843 | 1 | 0.2um_NOKCPHJN_355136 | Bacteria | Bacteroidetes | unclassified | unclassified | unclassified | unclassified | Bacteroidetes | metabolism | Energy produc | COG0711 | F(1)F(0) ATP s | 313598.MED1 | 1,6 | 0,489897949 | 0,000132206 | 4,37E-05 | 2 - | 0,000138285 - | 2 - | 0,000171709 - | 1 - | 8,16E-05 - | 1 - | 8,28E-05 - | 2 - | 0,000186596 - |
| 6844 | 2 | 0.2um_NOKCPHJN_08185;0.2um_NOKCPHJN_21 | Bacteria | Bacteroidetes | Flavobacteriia | various | various | various | various | unclassified | unclassified | unclassified | unclassified | unclassified | 0,4 | 0,489897949 | 2,43E-05 | 2,97E-05 | 0 - | 0 - | 0 - | 0 - | 1 - | 6,02E-05 - | 1 - | 6,11E-05 - | 0 - | 0 - |
| 6845 | 1 | 3um_FIJJAFBF_663734 | Eukaryota | Bacillariophyt | Bacillariophyc | Naviculales | Naviculaceae | Fistulifera | Fistulifera sola | poorly charact | Function unkn | 110WY | Chlorophyll A- | 2850.Phatr18 | 1,8 | 0,4 | 0,000235198 | 5,86E-05 | 2 - | 0,000218622 - | 2 - | 0,000271465 - | 1 - | 0,000129031 - | 2 - | 0,000261873 - | 2 - | 0,000294999 - |
| 6846 | 1 | 0.2um_NOKCPHJN_15250 | Bacteria | Proteobacteri | Betaproteoba | Burkholderiale | unclassified | Xylophilus | Xylophilus am | metabolism | Inorganic ion t | 0B7ZS | receptor | 397945.Aave_ | 1,4 | 0,489897949 | 3,13E-05 | 1,15E-05 | 1 - | 1,86E-05 - | 2 - | 4,63E-05 - | 2 - | 4,40E-05 - | 1 - | 2,23E-05 - | 1 - | 2,51E-05 - |
| 6847 | 1 | 3um_FIJJAFBF_187995 | unclassified | unclassified | unclassified | unclassified | unclassified | unclassified | unclassified | unclassified | unclassified | unclassified | unclassified | unclassified | 2 | 0,632455532 | 7,69E-05 | 2,68E-05 | 2 - | 6,41E-05 - | 3 - | 0,000119429 - | 1 - | 3,78E-05 - | 2 - | 7,68E-05 - | 2 - | 8,65E-05 - |
| 6848 | 1 | 3um_FIJJAFBF_337119 | Bacteria | Bacteroidetes | Flavobacteriia | Flavobacterial | Flavobacteria | Polaribacter | Polaribacter ir | information st | Translation, ril | COG0088 | One of the pri | 313594.PI23P | 3,4 | 0,489897949 | 0,000222494 | 3,80E-05 | 4 - | 0,000219668 - | 3 - | 0,000204573 - | 3 - | 0,000194472 - | 3 - | 0,000197344 - | 4 - | 0,00029641 - |
| 6849 | 1 | 0.2um_NOKCPHJN_674645 | unclassified | unclassified | unclassified | unclassified | unclassified | unclassified | unclassified | unclassified | unclassified | unclassified | unclassified | unclassified | 0,4 | 0,489897949 | 9,06E-06 | 1,11E-05 | 0 - | 0 - | 1 - | 2,32E-05 - | 1 - | 2,21E-05 - | 0 - | 0 - | 0 - | 0 - |
| 6850 | 1 | 3um_FIJJAFBF_21850 | Bacteria | Bacteroidetes | Flavobacteriia | Flavobacterial | Cryomorphace | Owenweeksia | Owenweeksia | unclassified | unclassified | unclassified | unclassified | unclassified | 3,2 | 0,748331477 | 5,76E-05 | 1,04E-05 | 4 - | 6,15E-05 - | 3 - | 5,72E-05 - | 3 - | 5,44E-05 - | 4 - | 7,36E-05 - | 2 - | 4,15E-05 - |
| 6851 | 1 | 3um_FIJJAFBF_188726 | Bacteria | Proteobacteri | Gammaprote | Cellvibrionales | Porticoccacea | unclassified | various | information st | Translation, ril | COG0087 | One of the pri | 314287.GB22 | 2,6 | 0,8 | 0,000166447 | 4,66E-05 | 3 - | 0,00016319 - | 2 - | 0,000135089 - | 4 - | 0,000256838 - | 2 - | 0,000130316 - | 2 - | 0,0001468 - |
| 6852 | 1 | 3um_FIJJAFBF_179285 | Bacteria | Verrucomicrol | Verrucomicrol | Verrucomicrol | Verrucomicrol | Verrucomicrol | Verrucomicrol | cellular proces | Posttranslatio | 1DGZS | Prevents misf | 240016.VspiD | 1 | 1,095445115 | 2,58E-05 | 2,72E-05 | 0 - | 0 - | 1 - | 2,61E-05 - | 3 - | 7,44E-05 - | 0 - | 0 - | 1 - | 2,84E-05 - |
| 6853 | 1 | 3um_FIJJAFBF_209845 | Bacteria | Chlamydiae | Chlamydiia | Parachlamydia | Parachlamydia | Neochlamydia | Neochlamydia | cellular proces | Signal transdu | 0JEMJ | STYKc | 5762.XP_0026 | 1,2 | 0,4 | 5,33E-05 | 1,78E-05 | 1 - | 3,73E-05 - | 1 - | 4,63E-05 - | 2 - | 8,80E-05 - | 1 - | 4,46E-05 - | 1 - | 5,03E-05 - |
| 6854 | 2 | 0.2um_NOKCPHJN_457098;3um_FIJJAFBF_31132 | Bacteria | Bacteroidetes | various | various | various | various | various | information st | Translation, ril | COG0089 | One of the ea | 391587.KAOT | 0,2 | 0,4 | 2,39E-05 | 4,78E-05 | 1 - | 0,000119559 - | 0 - | 0 - | 0 - | 0 - | 0 - | 0 - | 0 - | 0 - |
| 6855 | 1 | 0.2um_NOKCPHJN_778009 | Bacteria | Proteobacteri | Alphaproteob | Rhodobactera | Rhodobactera | Planktomarina | Planktomarina | unclassified | unclassified | unclassified | unclassified | unclassified | 0,2 | 0,4 | 7,04E-06 | 1,41E-05 | 0 - | 0 - | 0 - | 0 - | 1 - | 3,52E-05 - | 0 - | 0 - | 0 - | 0 - |
| 6856 | 1 | 3um_FIJJAFBF_330297 | Bacteria | Proteobacteri | Betaproteoba | unclassified | unclassified | unclassified | Betaproteoba | unclassified | unclassified | unclassified | unclassified | unclassified | 0,4 | 0,489897949 | 4,56E-05 | 5,58E-05 | 0 - | 0 - | 1 - | 0,000116819 - | 1 - | 0,000111051 - | 0 - | 0 - | 0 - | 0 - |
| 6857 | 1 | 3um_FIJJAFBF_361902 | Bacteria | Proteobacteri | Alphaproteob | Rhodobactera | Rhodobactera | Amylibacter | Amylibacter k | information st | Translation, ril | COG1185 | Involved in mR | 314256.OG25 | 8,6 | 1,624807681 | 0,000164121 | 2,29E-05 | 11 - | 0,000178074 - | 10 - | 0,000201014 - | 8 - | 0,000152871 - | 7 - | 0,000135738 - | 7 - | 0,000152909 - |
| 6858 | 1 | 3um_FIJJAFBF_251216 | Bacteria | Proteobacteri | Alphaproteob | Rhodobactera | Rhodobactera | Amylibacter | Amylibacter k | information st | Translation, ril | 16QAF | thus facilitatin | 398580.Dshi_ | 4,4 | 0,489897949 | 0,000107077 | 9,11E-06 | 5 - | 0,000102663 - | 4 - | 0,000101981 - | 4 - | 9,69E-05 - | 5 - | 0,000122973 - | 4 - | 0,000110823 - |

| | | | | | | | | | | | | | | | | | | | | | | | | | | | | |
|---|---|---|---|---|---|---|---|---|---|---|---|---|---|---|---|---|---|---|---|---|---|---|---|---|---|---|---|---|
| 6859 | 1 | 0.2um_NOKCPHJN_461186 | Bacteria | Proteobacteri | Alphaproteob | Pelagibacteral | Pelagibacterac | unclassified | Pelagibacterac | cellular proces | Intracellular tr | 0BIEB | domain-conta | 1005048.CFU_ | 10,8 | 1,326649916 | 0,00014614 | 1,40E-05 | 13 - | 0,000148467 - | 9 - | 0,000127629 - | 11 - | 0,000148289 - | 10 - | 0,000136799 - | 11 - | 0,000169514 - |
| 6860 | 1 | 3um_FIJJAFBF_746784 | Bacteria | Proteobacteri | Gammaprote | Oceanospirilla | Saccharospirill | Reinekea | Reinekea blan | information st | Translation, ril | 05CA4 | amino acids su | 314283.MED2 | 1,4 | 0,489897949 | 1,96E-05 | 6,16E-06 | 2 - | 2,39E-05 - | 2 - | 2,96E-05 - | 1 - | 1,41E-05 - | 1 - | 1,43E-05 - | 1 - | 1,61E-05 - |
| 6861 | 1 | 0.2um_NOKCPHJN_529682 | Bacteria | Proteobacteri | Alphaproteob | Rhodobactera | Rhodobactera | Tateyamaria | Tateyamaria s | unclassified | unclassified | unclassified | unclassified | unclassified | 4,4 | 0,489897949 | 0,0002064 | 1,76E-05 | 5 - | 0,000197891 - | 4 - | 0,000196578 - | 4 - | 0,000186872 - | 5 - | 0,00023704 - | 4 - | 0,00021362 - |
| 6862 | 1 | 0.2um_NOKCPHJN_32103 | Bacteria | Proteobacteri | Alphaproteob | unclassified | unclassified | unclassified | SAR116 cluste | metabolism | Secondary me | 01Q86 | extracellular s | 367336.OM22 | 0,4 | 0,489897949 | 1,54E-05 | 1,88E-05 | 0 - | 0 - | 1 - | 3,94E-05 - | 1 - | 3,74E-05 - | 0 - | 0 - | 0 - | 0 - |
| 6863 | 1 | 0.2um_NOKCPHJN_383542 | Bacteria | Bacteroidetes | Flavobacteriia | unclassified | unclassified | unclassified | Flavobacteria | poorly charact | Function unkn | 0P88U | NA | 487796.Flav2A | 1,6 | 1,2 | 3,80E-05 | 2,34E-05 | 4 - | 8,47E-05 - | 1 - | 2,63E-05 - | 1 - | 2,50E-05 - | 1 - | 2,54E-05 - | 1 - | 2,86E-05 - |
| 6864 | 1 | 3um_FIJJAFBF_122612 | Bacteria | Bacteroidetes | Flavobacteriia | Flavobacterial | Flavobacteriac | Kordia | Kordia jejudor | metabolism | Amino acid tra | COG0136 | Catalyzes the | 1046627.BZAF | 0,4 | 0,489897949 | 1,56E-05 | 1,93E-05 | 1 - | 3,49E-05 - | 1 - | 4,33E-05 - | 0 - | 0 - | 0 - | 0 - | 0 - | 0 - |
| 6865 | 1 | 3um_FIJJAFBF_772383 | Bacteria | Proteobacteri | Alphaproteob | Rhodobactera | Rhodobactera | Phaeobacter | Phaeobacter g | metabolism | Energy produc | 05C26 | Dehydrogenas | 633131.TR2A6 | 0,4 | 0,489897949 | 1,05E-05 | 1,30E-05 | 1 - | 2,34E-05 - | 1 - | 2,90E-05 - | 0 - | 0 - | 0 - | 0 - | 0 - | 0 - |
| 6866 | 1 | 0.2um_NOKCPHJN_51262 | Bacteria | Bacteroidetes | unclassified | unclassified | unclassified | unclassified | Bacteroidetes | unclassified | unclassified | unclassified | unclassified | unclassified | 0,6 | 1,2 | 5,24E-06 | 1,05E-05 | 3 - | 2,62E-05 - | 0 - | 0 - | 0 - | 0 - | 0 - | 0 - | 0 - | 0 - |
| 6867 | 1 | 0.2um_NOKCPHJN_1000553 | Bacteria | Proteobacteri | Alphaproteob | Pelagibacteral | Pelagibacterac | Candidatus Pe | Candidatus Pe | poorly charact | Function unkn | 05FKW | NA | 335992.SAR11 | 0,2 | 0,4 | 1,98E-05 | 3,96E-05 | 1 - | 9,89E-05 - | 0 - | 0 - | 0 - | 0 - | 0 - | 0 - | 0 - | 0 - |
| 6868 | 1 | 3um_FIJJAFBF_371424 | Bacteria | Proteobacteri | Betaproteoba | unclassified | unclassified | unclassified | Betaproteoba | information st | Translation, ril | 086NF | K02433 aspart | 365046.Rta_1 | 0,4 | 0,489897949 | 1,24E-05 | 1,55E-05 | 1 - | 2,64E-05 - | 0 - | 0 - | 0 - | 0 - | 0 - | 0 - | 1 - | 3,57E-05 - |
| 6869 | 1 | 0.2um_NOKCPHJN_334114 | Bacteria | Proteobacteri | Alphaproteob | Pelagibacteral | Pelagibacterac | Candidatus Pe | Candidatus Pe | cellular proces | Cell wall/mem | 7257 | NA | 439493.PB721 | 1,8 | 0,4 | 7,67E-05 | 1,51E-05 | 2 - | 7,24E-05 - | 2 - | 8,99E-05 - | 2 - | 8,55E-05 - | 2 - | 8,67E-05 - | 1 - | 4,89E-05 - |
| 6870 | 1 | 3um_FIJJAFBF_197234 | Viruses | unclassified | unclassified | unclassified | Phycodnavirid | unclassified | Chrysochromu | unclassified | unclassified | unclassified | unclassified | unclassified | 0,2 | 0,4 | 1,22E-05 | 2,43E-05 | 0 - | 0 - | 0 - | 0 - | 1 - | 6,08E-05 - | 0 - | 0 - | 0 - | 0 - |
| 6871 | 1 | 0.2um_NOKCPHJN_315153 | Bacteria | Proteobacteri | Gammaprote | unclassified | unclassified | unclassified | OM182 bacte | cellular proces | Intracellular tr | 0QJ7H | motA TolQ Exl | 314285.KT71_ | 9,8 | 1,6 | 0,000287534 | 3,82E-05 | 11 - | 0,000273278 - | 11 - | 0,000339331 - | 11 - | 0,000322577 - | 9 - | 0,000267825 - | 7 - | 0,000234658 - |
| 6872 | 1 | 0.2um_NOKCPHJN_663091 | Eukaryota | Bacillariophyt | Bacillariophyc | Bacillariales | Bacillariaceae | Fragilariopsis | Fragilariopsis | metabolism | Inorganic ion t | 0INF7 | Two pore segr | 2850.Phatr15 | 0,8 | 0,4 | 2,30E-05 | 1,18E-05 | 1 - | 2,40E-05 - | 1 - | 2,98E-05 - | 0 - | 0 - | 1 - | 2,88E-05 - | 1 - | 3,24E-05 - |
| 6873 | 1 | 0.2um_NOKCPHJN_624039 | Bacteria | Proteobacteri | Alphaproteob | Pelagibacteral | Pelagibacterac | Candidatus Pe | Candidatus Pe | cellular proces | Cell wall/mem | 7257 | NA | 335992.SAR11 | 2 | 0,632455532 | 8,44E-05 | 2,81E-05 | 2 - | 7,06E-05 - | 2 - | 8,77E-05 - | 1 - | 4,17E-05 - | 3 - | 0,000126908 - | 2 - | 9,53E-05 - |
| 6874 | 1 | 0.2um_NOKCPHJN_1040576 | Bacteria | Proteobacteri | Gammaprote | Cellvibrionales | Cellvibrionace | Agarilytica | Agarilytica rho | metabolism | Carbohydrate | 16QI2 | glycoside hydr | 314287.GB22 | 0,2 | 0,4 | 5,19E-06 | 1,04E-05 | 0 - | 0 - | 0 - | 0 - | 0 - | 0 - | 1 - | 2,59E-05 - | 0 - | 0 - |
| 6875 | 1 | 0.2um_NOKCPHJN_62897 | Bacteria | Proteobacteri | Gammaprote | Oceanospirilla | Saccharospirill | Saccharospiril | Saccharospiril | metabolism | Secondary me | 0QIEH | Catalyzes the | 314283.MED2 | 1 | 1,095445115 | 4,46E-05 | 4,97E-05 | 0 - | 0 - | 3 - | 0,000136165 - | 1 - | 4,31E-05 - | 1 - | 4,38E-05 - | 0 - | 0 - |
| 6876 | 1 | 3um_FIJJAFBF_467729 | unclassified | unclassified | unclassified | unclassified | unclassified | unclassified | unclassified | unclassified | unclassified | unclassified | unclassified | unclassified | 0,6 | 0,489897949 | 7,99E-05 | 6,53E-05 | 0 - | 0 - | 1 - | 0,000137037 - | 1 - | 0,000130271 - | 1 - | 0,000132195 - | 0 - | 0 - |
| 6877 | 1 | 3um_FIJJAFBF_172745 | Bacteria | Proteobacteri | Betaproteoba | Burkholderial | unclassified | Thiomonas | Thiomonas de | unclassified | unclassified | unclassified | unclassified | unclassified | 0,6 | 0,489897949 | 1,81E-05 | 1,49E-05 | 1 - | 2,68E-05 - | 0 - | 0 - | 1 - | 3,17E-05 - | 1 - | 3,21E-05 - | 0 - | 0 - |
| 6878 | 1 | 3um_FIJJAFBF_155742 | Viruses | unclassified | unclassified | unclassified | Phycodnavirid | Prymnesioviru | Prymnesium p | unclassified | unclassified | unclassified | unclassified | unclassified | 2 | 1,095445115 | 5,62E-05 | 2,48E-05 | 4 - | 9,87E-05 - | 2 - | 6,13E-05 - | 2 - | 5,83E-05 - | 1 - | 2,96E-05 - | 1 - | 3,33E-05 - |
| 6879 | 1 | 3um_FIJJAFBF_246733 | Bacteria | Bacteroidetes | Cytophagia | Cytophagales | Flammeovirga | unclassified | Flammeovirga | cellular proces | Intracellular tr | 0ZPEJ | Preprotein tra | 643867.Ftrac_ | 0,6 | 0,489897949 | 6,36E-05 | 5,23E-05 | 1 - | 9,41E-05 - | 0 - | 0 - | 1 - | 0,000111051 - | 1 - | 0,000112691 - | 0 - | 0 - |
| 6880 | 1 | 0.2um_NOKCPHJN_219417 | Eukaryota | unclassified | Oomycetes | Peronosporale | Peronosporac | Phytophthora | Phytophthora | unclassified | unclassified | unclassified | unclassified | unclassified | 0,2 | 0,4 | 4,10E-07 | 8,19E-07 | 0 - | 0 - | 0 - | 0 - | 0 - | 0 - | 1 - | 2,05E-06 - | 0 - | 0 - |
| 6881 | 1 | 3um_FIJJAFBF_434535 | Bacteria | Bacteroidetes | Flavobacteriia | Flavobacterial | Flavobacteriac | Polaribacter | various | unclassified | unclassified | unclassified | unclassified | unclassified | 8,6 | 1,019803903 | 0,00095333 | 0,000170961 | 9 - | 0,000833057 - | 9 - | 0,001034411 - | 8 - | 0,000874079 - | 7 - | 0,000776115 - | 10 - | 0,001248988 - |
| 6882 | 1 | 3um_FIJJAFBF_295529 | Bacteria | Bacteroidetes | Flavobacteriia | unclassified | unclassified | unclassified | Flavobacteria | unclassified | unclassified | unclassified | unclassified | unclassified | 0,4 | 0,489897949 | 2,80E-05 | 3,46E-05 | 1 - | 6,38E-05 - | 0 - | 0 - | 0 - | 0 - | 1 - | 7,64E-05 - | 0 - | 0 - |
| 6883 | 1 | 3um_FIJJAFBF_376284 | Bacteria | Proteobacteri | Alphaproteob | Rhodobactera | Rhodobactera | Roseovarius | Roseovarius s | metabolism | Amino acid tra | 11KI6 | polar amino a | 314277.MED1 | 0,8 | 0,4 | 4,05E-05 | 2,08E-05 | 1 - | 4,28E-05 - | 0 - | 0 - | 1 - | 5,06E-05 - | 1 - | 5,13E-05 - | 1 - | 5,78E-05 - |
| 6884 | 1 | 3um_FIJJAFBF_156244 | Bacteria | unclassified | unclassified | unclassified | unclassified | unclassified | uncultured ba | metabolism | Carbohydrate | COG0057 | glyceraldehyd | 50743.SCB49_ | 0,8 | 0,4 | 3,27E-05 | 1,68E-05 | 1 - | 3,46E-05 - | 0 - | 0 - | 1 - | 4,08E-05 - | 1 - | 4,14E-05 - | 1 - | 4,66E-05 - |
| 6885 | 1 | 0.2um_NOKCPHJN_720323 | Bacteria | Proteobacteri | Gammaprote | Alteromonada | Colwelliaceae | Thalassotalea | Thalassotalea | metabolism | Carbohydrate | 08IUN | Phosphorylase | 983545.Glaag | 0,2 | 0,4 | 3,35E-06 | 6,70E-06 | 0 - | 0 - | 0 - | 0 - | 0 - | 0 - | 1 - | 1,67E-05 - | 0 - | 0 - |
| 6886 | 1 | 3um_FIJJAFBF_216550 | Bacteria | Bacteroidetes | Cytophagia | Cytophagales | Flammeovirga | unclassified | Flammeovirga | poorly charact | Function unkn | 09ARK | Protein of unk | 643867.Ftrac_ | 1,4 | 0,489897949 | 6,70E-05 | 2,37E-05 | 1 - | 4,01E-05 - | 1 - | 4,98E-05 - | 2 - | 9,47E-05 - | 2 - | 9,61E-05 - | 1 - | 5,42E-05 - |
| 6887 | 1 | 3um_FIJJAFBF_277463 | Bacteria | Cyanobacteria | unclassified | Synechococca | Leptolyngbyac | Nodosilinea | Nodosilinea n | information st | Translation, ril | 0FP0S | Seems to be tl | 513049.Amax | 1,4 | 0,8 | 0,000146424 | 8,14E-05 | 2 - | 0,000177948 - | 2 - | 0,000220959 - | 0 - | 0 - | 2 - | 0,000213152 - | 1 - | 0,000120058 - |
| 6888 | 1 | 0.2um_NOKCPHJN_246331 | Bacteria | Bacteroidetes | Flavobacteriia | Flavobacterial | Cryomorphace | Owenweeksia | Owenweeksia | information st | Translation, ril | COG0050 | This protein p | 391587.KAOT | 5,2 | 0,748331477 | 0,000178383 | 1,71E-05 | 6 - | 0,000174344 - | 5 - | 0,000180404 - | 5 - | 0,000171496 - | 6 - | 0,000208835 - | 4 - | 0,000156835 - |
| 6889 | 1 | 0.2um_NOKCPHJN_616355 | Eukaryota | unclassified | Oomycetes | Peronosporale | Peronosporac | Plasmopara | Plasmopara h | unclassified | unclassified | unclassified | unclassified | unclassified | 1 | 0 | 2,91E-05 | 2,76E-06 | 1 - | 2,44E-05 - | 1 - | 3,03E-05 - | 1 - | 2,88E-05 - | 1 - | 2,92E-05 - | 1 - | 3,29E-05 - |
| 6890 | 1 | 3um_FIJJAFBF_184625 | Bacteria | Bacteroidetes | Cytophagia | Cytophagales | Catalimonada | Catalinimonas | Catalinimonas | metabolism | Energy produc | 099Q7 | Catalyzes the | 880070.Cycma | 0,2 | 0,4 | 1,18E-05 | 2,37E-05 | 0 - | 0 - | 0 - | 0 - | 1 - | 5,92E-05 - | 0 - | 0 - | 0 - | 0 - |
| 6891 | 1 | 3um_FIJJAFBF_72479 | Bacteria | Bacteroidetes | Cytophagia | Cytophagales | Flammeovirga | unclassified | Flammeovirga | unclassified | unclassified | unclassified | unclassified | unclassified | 6,8 | 1,6 | 0,000696659 | 0,000137516 | 9 - | 0,000782569 - | 4 - | 0,000431875 - | 7 - | 0,000718466 - | 7 - | 0,000729078 - | 7 - | 0,000821304 - |
| 6892 | 1 | 3um_FIJJAFBF_493420 | Bacteria | Verrucomicrol | Opitutae | unclassified | unclassified | unclassified | Opitutae bact | unclassified | unclassified | unclassified | unclassified | unclassified | 0,4 | 0,489897949 | 5,05E-05 | 6,24E-05 | 1 - | 0,000112526 - | 1 - | 0,000139724 - | 0 - | 0 - | 0 - | 0 - | 0 - | 0 - |
| 6893 | 1 | 0.2um_NOKCPHJN_17904 | Bacteria | Bacteroidetes | Flavobacteriia | Flavobacterial | Crocinitomica | Lishizhenia | Lishizhenia tia | unclassified | unclassified | unclassified | unclassified | unclassified | 0,6 | 0,8 | 1,98E-05 | 2,62E-05 | 0 - | 0 - | 1 - | 3,42E-05 - | 2 - | 6,50E-05 - | 0 - | 0 - | 0 - | 0 - |
| 6894 | 1 | 0.2um_NOKCPHJN_304089 | Bacteria | Proteobacteri | Alphaproteob | Rhodobactera | unclassified | unclassified | Rhodobactera | unclassified | unclassified | unclassified | unclassified | unclassified | 0,6 | 0,489897949 | 0,000132958 | 0,000110562 | 1 - | 0,000185124 - | 1 - | 0,000229869 - | 0 - | 0 - | 0 - | 0 - | 1 - | 0,000249798 - |
| 6895 | 1 | 0.2um_NOKCPHJN_740895 | Bacteria | Proteobacteri | Alphaproteob | Pelagibacteral | Pelagibacterac | unclassified | Pelagibacterac | unclassified | unclassified | unclassified | unclassified | unclassified | 0,2 | 0,4 | 9,12E-06 | 1,82E-05 | 0 - | 0 - | 0 - | 0 - | 1 - | 4,56E-05 - | 0 - | 0 - | 0 - | 0 - |
| 6896 | 1 | 3um_FIJJAFBF_09778 | Bacteria | Bacteroidetes | unclassified | unclassified | unclassified | unclassified | unidentified e | information st | Translation, ril | COG0254 | 50S ribosomal | 50743.SCB49_ | 1 | 0 | 0,000147341 | 1,40E-05 | 1 - | 0,000123416 - | 1 - | 0,000153246 - | 1 - | 0,00014568 - | 1 - | 0,000147832 - | 1 - | 0,000166532 - |
| 6897 | 1 | 3um_FIJJAFBF_520094 | Bacteria | Bacteroidetes | Flavobacteriia | Flavobacterial | Flavobacteriac | Polaribacter | Polaribacter a | unclassified | unclassified | unclassified | unclassified | unclassified | 0,8 | 0,4 | 3,73E-05 | 1,91E-05 | 1 - | 3,90E-05 - | 1 - | 4,85E-05 - | 1 - | 4,61E-05 - | 0 - | 0 - | 1 - | 5,27E-05 - |
| 6898 | 1 | 0.2um_NOKCPHJN_646683 | unclassified | unclassified | unclassified | unclassified | unclassified | unclassified | unclassified | unclassified | unclassified | unclassified | unclassified | unclassified | 0,2 | 0,4 | 5,89E-06 | 1,18E-05 | 0 - | 0 - | 1 - | 2,94E-05 - | 0 - | 0 - | 0 - | 0 - | 0 - | 0 - |
| 6899 | 1 | 3um_FIJJAFBF_38492 | Bacteria | Proteobacteri | Alphaproteob | Rhodobactera | Rhodobactera | Pseudophaeol | Pseudophaeol | metabolism | Amino acid tra | 16PMY | Aminotransfer | 644107.SL115 | 0,4 | 0,489897949 | 1,89E-05 | 2,35E-05 | 1 - | 4,01E-05 - | 0 - | 0 - | 0 - | 0 - | 0 - | 0 - | 1 - | 5,42E-05 - |
| 6900 | 1 | 3um_FIJJAFBF_672108 | Bacteria | Bacteroidetes | Flavobacteriia | Flavobacterial | Flavobacteriac | Polaribacter | Polaribacter s | unclassified | unclassified | unclassified | unclassified | unclassified | 1,2 | 0,4 | 3,25E-05 | 7,20E-06 | 2 - | 4,67E-05 - | 1 - | 2,90E-05 - | 1 - | 2,75E-05 - | 1 - | 2,79E-05 - | 1 - | 3,15E-05 - |
| 6901 | 1 | 3um_FIJJAFBF_128464 | Bacteria | Bacteroidetes | Flavobacteriia | Flavobacterial | Flavobacteriac | Polaribacter | Polaribacter s | information st | Translation, ril | COG0052 | 30S ribosomal | 487797.Flav3 | 17,8 | 2,315167381 | 0,000720257 | 5,39E-05 | 21 - | 0,000719496 - | 19 - | 0,000808316 - | 17 - | 0,000687522 - | 18 - | 0,000738716 - | 14 - | 0,000647237 - |
| 6902 | 1 | 0.2um_NOKCPHJN_856350 | Bacteria | Proteobacteri | Alphaproteob | Pelagibacteral | Pelagibacterac | various | various | unclassified | unclassified | unclassified | unclassified | unclassified | 0,8 | 0,4 | 0,000164429 | 8,44E-05 | 1 - | 0,000173904 - | 0 - | 0 - | 1 - | 0,000205276 - | 1 - | 0,000208308 - | 1 - | 0,000234658 - |
| 6903 | 1 | 3um_FIJJAFBF_91115 | Eukaryota | unclassified | Raphidophyce | Chattonellales | Chattonellace | Heterosigma | Heterosigma a | unclassified | unclassified | unclassified | unclassified | unclassified | 20,8 | 1,469693846 | 0,000894695 | 9,39E-05 | 22 - | 0,000794053 - | 21 - | 0,000941162 - | 22 - | 0,000937298 - | 18 - | 0,000778207 - | 21 - | 0,001022756 - |
| 6904 | 1 | 3um_FIJJAFBF_708523 | Bacteria | Cyanobacteria | unclassified | Synechococca | Synechococca | Synechococcu | Synechococcu | poorly charact | Function unkn | 11FY4 | C-phycoerythr | 64471.sync_0 | 2 | 1,095445115 | 0,000161198 | 7,29E-05 | 4 - | 0,000279943 - | 1 - | 8,69E-05 - | 1 - | 8,26E-05 - | 2 - | 0,000167663 - | 2 - | 0,000188871 - |
| 6905 | 1 | 0.2um_NOKCPHJN_100278 | Bacteria | Bacteroidetes | unclassified | unclassified | unclassified | unclassified | Bacteroidetes | metabolism | Inorganic ion t | 05C27 | TonB-depende | 504472.Slin_4 | 32 | 3,847076812 | 0,0003878 | 2,00E-05 | 38 - | 0,000389769 - | 29 - | 0,000369352 - | 35 - | 0,00042376 - | 30 - | 0,000368588 - | 28 - | 0,000387532 - |
| 6906 | 1 | 0.2um_NOKCPHJN_699912 | Bacteria | Proteobacteri | Gammaprote | unclassified | unclassified | unclassified | Gammaprote | metabolism | Inorganic ion t | 0QK8T | TonB-depende | 247633.GP214 | 0,6 | 0,489897949 | 1,84E-05 | 1,52E-05 | 1 - | 2,68E-05 - | 1 - | 3,33E-05 - | 0 - | 0 - | 1 - | 3,21E-05 - | 0 - | 0 - |
| 6907 | 1 | 3um_FIJJAFBF_581772 | Bacteria | Bacteroidetes | Flavobacteriia | Flavobacterial | Flavobacteriac | Polaribacter | Polaribacter s | unclassified | unclassified | unclassified | unclassified | unclassified | 3,8 | 1,469693846 | 6,75E-05 | 2,48E-05 | 5 - | 7,64E-05 - | 5 - | 9,49E-05 - | 4 - | 7,22E-05 - | 4 - | 7,32E-05 - | 1 - | 2,06E-05 - |
| 6908 | 1 | 0.2um_NOKCPHJN_372789 | Eukaryota | unclassified | Oomycetes | Peronosporale | Peronosporac | Phytophthora | Phytophthora | unclassified | unclassified | unclassified | unclassified | unclassified | 0,8 | 0,4 | 1,89E-05 | 9,73E-06 | 1 - | 2,00E-05 - | 0 - | 0 - | 1 - | 2,36E-05 - | 1 - | 2,40E-05 - | 1 - | 2,70E-05 - |
| 6909 | 1 | 3um_FIJJAFBF_617422 | Eukaryota | unclassified | Dictyochophy | Florenciellales | unclassified | Pseudochatto | Pseudochatto | cellular proces | Cytoskeleton | 1514Z | Tubulin is the | 4787.PITG_00 | 2,8 | 0,4 | 0,000307188 | 5,49E-05 | 3 - | 0,000275464 - | 3 - | 0,000342045 - | 2 - | 0,000216772 - | 3 - | 0,00032996 - | 3 - | 0,000371699 - |
| 6910 | 1 | 3um_FIJJAFBF_59594 | Bacteria | Proteobacteri | Alphaproteob | Rhodospirillal | Rhodospirillac | unclassified | Rhodobactera | unclassified | unclassified | unclassified | unclassified | unclassified | 1,2 | 0,4 | 5,35E-05 | 1,19E-05 | 2 - | 7,68E-05 - | 1 - | 4,77E-05 - | 1 - | 4,53E-05 - | 1 - | 4,60E-05 - | 1 - | 5,18E-05 - |
| 6911 | 1 | 3um_FIJJAFBF_172361 | Bacteria | Proteobacteri | Gammaprote | Oceanospirilla | Saccharospirill | Saccharospiril | Saccharospiril | information st | Translation, ril | 05C64 | Binds directly | 314283.MED2 | 11,2 | 0,979795897 | 0,000661205 | 5,79E-05 | 12 - | 0,000596243 - | 12 - | 0,000740358 - | 12 - | 0,000703804 - | 10 - | 0,000595166 - | 10 - | 0,000670452 - |
| 6912 | 1 | 3um_FIJJAFBF_17247 | Bacteria | Proteobacteri | Alphaproteob | Rhodobactera | Rhodobactera | unclassified | Rhodobactera | cellular proces | Cell wall/mem | 16Q41 | Efflux transpo | 439497.RR11_ | 0,4 | 0,489897949 | 1,24E-05 | 1,53E-05 | 1 - | 2,77E-05 - | 1 - | 3,43E-05 - | 0 - | 0 - | 0 - | 0 - | 0 - | 0 - |
| 6913 | 1 | 0.2um_NOKCPHJN_305566 | Eukaryota | Streptophyta | Bryopsida | Funariales | Funariaceae | Physcomitrell | Physcomitrell | unclassified | unclassified | unclassified | unclassified | unclassified | 0,2 | 0,4 | 2,13E-06 | 4,26E-06 | 0 - | 0 - | 0 - | 0 - | 1 - | 1,06E-05 - | 0 - | 0 - | 0 - | 0 - |
| 6914 | 1 | 0.2um_NOKCPHJN_668827 | Bacteria | Proteobacteri | Alphaproteob | Pelagibacteral | Pelagibacterac | unclassified | Pelagibacterac | cellular proces | Cell wall/mem | 7257 | NA | 439493.PB721 | 2 | 0 | 8,28E-05 | 7,86E-06 | 2 - | 6,94E-05 - | 2 - | 8,61E-05 - | 2 - | 8,19E-05 - | 2 - | 8,31E-05 - | 2 - | 9,36E-05 - |
| 6915 | 1 | 3um_FIJJAFBF_161064 | Bacteria | Proteobacteri | Gammaprote | Oceanospirilla | Saccharospirill | Reinekea | Reinekea fors | cellular proces | Posttranslatio | 0QJ7Y | Converts hem | 314283.MED2 | 0,8 | 0,4 | 3,45E-05 | 1,75E-05 | 1 - | 3,74E-05 - | 1 - | 4,64E-05 - | 1 - | 4,41E-05 - | 1 - | 4,48E-05 - | 0 - | 0 - |
| 6916 | 1 | 3um_FIJJAFBF_53324 | Bacteria | Bacteroidetes | Flavobacteriia | Flavobacterial | Flavobacteriac | Tenacibaculur | Tenacibaculur | unclassified | unclassified | unclassified | unclassified | unclassified | 1,8 | 0,748331477 | 0,000114133 | 3,80E-05 | 3 - | 0,000164751 - | 2 - | 0,000136382 - | 2 - | 0,000129648 - | 1 - | 6,58E-05 - | 1 - | 7,41E-05 - |
| 6917 | 1 | 3um_FIJJAFBF_723597 | Bacteria | Bacteroidetes | Flavobacteriia | Flavobacterial | Flavobacteriac | Formosa | Formosa sp. H | metabolism | Energy produc | 0P9HY | cytochrome C | 983544.Lacal_ | 1 | 0 | 7,49E-05 | 7,11E-06 | 1 - | 6,27E-05 - | 1 - | 7,79E-05 - | 1 - | 7,40E-05 - | 1 - | 7,51E-05 - | 1 - | 8,46E-05 - |
| 6918 | 1 | 0.2um_NOKCPHJN_73377 | Bacteria | Proteobacteri | Gammaprote | Cellvibrionales | Porticoccacea | unclassified | SAR92 bacteri | cellular proces | Posttranslatio | COG0330 | Band 7 protei | 314287.GB22 | 2,8 | 1,469693846 | 0,000130534 | 5,89E-05 | 5 - | 0,000202786 - | 1 - | 5,04E-05 - | 2 - | 9,57E-05 - | 4 - | 0,000194323 - | 2 - | 0,000109452 - |
| 6919 | 1 | 3um_FIJJAFBF_783854 | Bacteria | Bacteroidetes | Flavobacteriia | Flavobacterial | Flavobacteriac | Tenacibaculur | Tenacibaculur | poorly charact | Function unkn | 11P35 | NA | 983544.Lacal_ | 8 | 1,897366596 | 0,00019746 | 4,05E-05 | 11 - | 0,000229972 - | 6 - | 0,000155758 - | 8 - | 0,000197424 - | 6 - | 0,000150255 - | 9 - | 0,000253893 - |
| 6920 | 1 | 0.2um_NOKCPHJN_182741 | Bacteria | Bacteroidetes | unclassified | unclassified | unclassified | unclassified | Bacteroidetes | unclassified | unclassified | unclassified | unclassified | unclassified | 5,6 | 1,019803903 | 7,45E-05 | 1,34E-05 | 6 - | 6,73E-05 - | 6 - | 8,35E-05 - | 7 - | 9,26E-05 - | 4 - | 5,37E-05 - | 5 - | 7,56E-05 - |
| 6921 | 1 | 0.2um_NOKCPHJN_31006 | Bacteria | Proteobacteri | Alphaproteob | Rhodobactera | Rhodobactera | Epibacterium | Epibacterium | cellular proces | Posttranslatio | 01QWG | Prevents misf | 52598.EE36_0 | 9,8 | 1,326649916 | 0,000243077 | 2,92E-05 | 12 - | 0,000250878 - | 10 - | 0,000259597 - | 8 - | 0,000197424 - | 9 - | 0,000225382 - | 10 - | 0,000282103 - |
| 6922 | 1 | 3um_FIJJAFBF_124742 | Bacteria | Proteobacteri | Alphaproteob | Rhodobactera | Rhodobactera | Planktomarina | Planktomarina | unclassified | unclassified | unclassified | unclassified | unclassified | 3 | 0 | 0,000216359 | 2,05E-05 | 3 - | 0,000181226 - | 3 - | 0,00022503 - | 3 - | 0,000213919 - | 3 - | 0,000217079 - | 3 - | 0,000244539 - |
| 6923 | 1 | 3um_FIJJAFBF_264827 | Bacteria | Proteobacteri | Alphaproteob | Sphingomona | Sphingomona | Sphingopyxis | Sphingopyxis | cellular proces | Cell wall/mem | COG2885 | Ompa motb d | 161528.ED21_ | 0,6 | 0,489897949 | 4,91E-05 | 4,05E-05 | 1 - | 7,17E-05 - | 1 - | 8,91E-05 - | 1 - | 8,47E-05 - | 0 - | 0 - | 0 - | 0 - |
| 6924 | 1 | 0.2um_NOKCPHJN_539569 | Bacteria | Bacteroidetes | Flavobacteriia | Flavobacterial | Crocinitomica | unclassified | Crocinitomica | information st | Transcription | 05D27 | DNA-depende | 755732.Fluta_ | 0,2 | 0,4 | 3,90E-06 | 7,79E-06 | 0 - | 0 - | 0 - | 0 - | 0 - | 0 - | 0 - | 0 - | 1 - | 1,95E-05 - |
| 6925 | 1 | 0.2um_NOKCPHJN_564928 | Eukaryota | Phaeophyceae | unclassified | Ectocarpales | Ectocarpaceae | Ectocarpus | Ectocarpus sili | information st | Chromatin str | KOG1744 | Core compone | 2850.Phatr54 | 2,6 | 0,8 | 0,000299281 | 9,46E-05 | 2 - | 0,000192902 - | 2 - | 0,000239528 - | 3 - | 0,000341552 - | 4 - | 0,000462129 - | 2 - | 0,000260293 - |
| 6926 | 1 | 3um_FIJJAFBF_313760 | Bacteria | Bacteroidetes | unclassified | unclassified | unclassified | unclassified | Bacteroidetes | unclassified | unclassified | unclassified | unclassified | unclassified | 12,4 | 0,8 | 0,000176503 | 1,09E-05 | 13 - | 0,000155751 - | 12 - | 0,00017852 - | 13 - | 0,000183849 - | 13 - | 0,000186564 - | 11 - | 0,000177831 - |
| 6927 | 1 | 3um_FIJJAFBF_93514 | Bacteria | Bacteroidetes | Flavobacteriia | Flavobacterial | Crocinitomica | unclassified | Crocinitomica | unclassified | unclassified | unclassified | unclassified | unclassified | 0,8 | 0,4 | 9,43E-05 | 4,75E-05 | 0 - | 0 - | 1 - | 0,000117784 - | 1 - | 0,000111969 - | 1 - | 0,000113623 - | 1 - | 0,000127995 - |
| 6928 | 1 | 3um_FIJJAFBF_90521 | Bacteria | Proteobacteri | Gammaprote | Oceanospirilla | Saccharospirill | Reinekea | Reinekea fors | unclassified | unclassified | unclassified | unclassified | unclassified | 0,8 | 0,4 | 5,91E-05 | 2,98E-05 | 0 - | 0 - | 1 - | 7,38E-05 - | 1 - | 7,02E-05 - | 1 - | 7,12E-05 - | 1 - | 8,02E-05 - |
| 6929 | 1 | 0.2um_NOKCPHJN_661124 | Bacteria | Proteobacteri | Alphaproteob | Pelagibacteral | Pelagibacterac | Candidatus Pe | Candidatus Pe | poorly charact | Function unkn | 0ZNGQ | NA | 335992.SAR11 | 1,8 | 0,4 | 0,000286695 | 7,03E-05 | 2 - | 0,000266923 - | 2 - | 0,000331439 - | 2 - | 0,000315075 - | 1 - | 0,000159864 - | 2 - | 0,000360173 - |
| 6930 | 1 | 3um_FIJJAFBF_38435 | Bacteria | Proteobacteri | Alphaproteob | Rhodobactera | Rhodobactera | Leisingera | Leisingera sp. | poorly charact | Function unkn | 0QM7Q | Dehydrogenas | 393595.ABO_ | 1,4 | 0,8 | 7,26E-05 | 3,21E-05 | 3 - | 0,000136639 - | 1 - | 5,66E-05 - | 1 - | 5,38E-05 - | 1 - | 5,46E-05 - | 1 - | 6,15E-05 - |
| 6931 | 1 | 3um_FIJJAFBF_177265 | Bacteria | Proteobacteri | Gammaprote | Alteromonada | Alteromonada | Marinobacter | Marinobacter | information st | Translation, ril | COG0251 | Endoribonucle | 351348.Maqu | 1,6 | 0,8 | 0,000145857 | 5,78E-05 | 3 - | 0,000239118 - | 1 - | 9,90E-05 - | 2 - | 0,00018817 - | 1 - | 9,55E-05 - | 1 - | 0,000107552 - |
| 6932 | 1 | 3um_FIJJAFBF_91229 | Bacteria | Proteobacteri | Gammaprote | unclassified | unclassified | Candidatus Th | Candidatus Th | unclassified | unclassified | unclassified | unclassified | unclassified | 1,2 | 0,4 | 3,02E-05 | 6,70E-06 | 2 - | 4,34E-05 - | 1 - | 2,69E-05 - | 1 - | 2,56E-05 - | 1 - | 2,60E-05 - | 1 - | 2,93E-05 - |
| 6933 | 1 | 3um_FIJJAFBF_30120 | Bacteria | Proteobacteri | Alphaproteob | Rhodobactera | unclassified | unclassified | Rhodobactera | unclassified | unclassified | unclassified | unclassified | unclassified | 1,4 | 0,489897949 | 0,000164639 | 6,07E-05 | 1 - | 9,81E-05 - | 2 - | 0,000243622 - | 2 - | 0,000231594 - | 1 - | 0,000117507 - | 1 - | 0,000132371 - |
| 6934 | 1 | 0.2um_NOKCPHJN_87161 | Eukaryota | unclassified | Bangiophycea | Cyanidiales | Cyanidiaceae | Cyanidioschyz | Cyanidioschyz | unclassified | unclassified | unclassified | unclassified | unclassified | 0,4 | 0,489897949 | 4,73E-06 | 5,79E-06 | 0 - | 0 - | 0 - | 0 - | 1 - | 1,17E-05 - | 1 - | 1,19E-05 - | 0 - | 0 - |
| 6935 | 1 | 3um_FIJJAFBF_438664 | Bacteria | Proteobacteri | Gammaprote | Oceanospirilla | Saccharospirill | Reinekea | Reinekea fors | information st | Translation, ril | 0QQXI | This protein is | 314283.MED2 | 9,4 | 1,019803903 | 0,001039638 | 8,24E-05 | 11 - | 0,001026459 - | 9 - | 0,001042821 - | 10 - | 0,001101482 - | 8 - | 0,0008942 - | 9 - | 0,001133228 - |
| 6936 | 1 | 0.2um_NOKCPHJN_713307 | Bacteria | Bacteroidetes | Flavobacteriia | Flavobacterial | Flavobacteriac | Tenacibaculur | Tenacibaculur | unclassified | unclassified | unclassified | unclassified | unclassified | 2,2 | 0,748331477 | 3,39E-05 | 1,17E-05 | 2 - | 2,59E-05 - | 1 - | 1,61E-05 - | 3 - | 4,59E-05 - | 3 - | 4,66E-05 - | 2 - | 3,50E-05 - |
| 6937 | 1 | 0.2um_NOKCPHJN_01141 | Bacteria | Bacteroidetes | Flavobacteriia | Flavobacterial | Flavobacteriac | Formosa | Formosa sp. H | information st | Translation, ril | 05CAV | thus facilitatin | 688270.Celal_ | 9 | 1,414213562 | 0,000202477 | 3,98E-05 | 9 - | 0,000169343 - | 7 - | 0,000163546 - | 8 - | 0,000177682 - | 11 - | 0,000247921 - | 10 - | 0,000253893 - |
| 6938 | 1 | 0.2um_NOKCPHJN_272972 | Bacteria | Chloroflexi | unclassified | unclassified | unclassified | unclassified | Chloroflexi ba | unclassified | unclassified | unclassified | unclassified | unclassified | 0,2 | 0,4 | 5,71E-07 | 1,14E-06 | 0 - | 0 - | 0 - | 0 - | 1 - | 2,86E-06 - | 0 - | 0 - | 0 - | 0 - |
| 6939 | 1 | 3um_FIJJAFBF_769734 | Bacteria | Bacteroidetes | Flavobacteriia | Flavobacterial | Flavobacteriac | Lacinutrix | Lacinutrix him | poorly charact | Function unkn | 06E9D | NA | 391598.FBBAL | 2,8 | 0,748331477 | 6,37E-05 | 1,59E-05 | 3 - | 5,79E-05 - | 2 - | 4,79E-05 - | 3 - | 6,83E-05 - | 4 - | 9,24E-05 - | 2 - | 5,21E-05 - |
| 6940 | 1 | 3um_FIJJAFBF_168225 | Bacteria | Bacteroidetes | Flavobacteriia | Flavobacterial | Crocinitomica | unclassified | Crocinitomica | metabolism | Energy produc | 05CMV | Succinyl-CoA s | 755732.Fluta_ | 0,2 | 0,4 | 5,71E-06 | 1,14E-05 | 1 - | 2,86E-05 - | 0 - | 0 - | 0 - | 0 - | 0 - | 0 - | 0 - | 0 - |
| 6941 | 1 | 3um_FIJJAFBF_490335 | Bacteria | Bacteroidetes | Flavobacteriia | Flavobacterial | Flavobacteriac | Polaribacter | Polaribacter s | metabolism | Inorganic ion t | COG0841 | acriflavin resis | 313594.PI23P | 1,8 | 1,6 | 2,05E-05 | 1,87E-05 | 3 - | 2,97E-05 - | 4 - | 4,91E-05 - | 0 - | 0 - | 2 - | 2,37E-05 - | 0 - | 0 - |
| 6942 | 1 | 0.2um_NOKCPHJN_157008 | Bacteria | Proteobacteri | Gammaprote | Cellvibrionales | Halieaceae | Luminiphilus | Luminiphilus s | information st | Translation, ril | 16QAF | thus facilitatin | 247639.MGP2 | 0,2 | 0,4 | 4,94E-06 | 9,87E-06 | 0 - | 0 - | 0 - | 0 - | 0 - | 0 - | 1 - | 2,47E-05 - | 0 - | 0 - |
| 6943 | 1 | 3um_FIJJAFBF_158174 | unclassified | unclassified | unclassified | unclassified | unclassified | unclassified | unclassified | unclassified | unclassified | unclassified | unclassified | unclassified | 0,8 | 0,4 | 9,44E-05 | 4,85E-05 | 1 - | 9,98E-05 - | 0 - | 0 - | 1 - | 0,000117811 - | 1 - | 0,000119551 - | 1 - | 0,000134673 - |
| 6944 | 1 | 3um_FIJJAFBF_622719 | Bacteria | Proteobacteri | Gammaprote | Oceanospirilla | Saccharospirill | Reinekea | Reinekea blan | metabolism | Energy produc | 0QIP8 | Electron trans | 314283.MED2 | 1,4 | 0,489897949 | 6,09E-05 | 1,79E-05 | 2 - | 7,45E-05 - | 1 - | 4,63E-05 - | 1 - | 4,40E-05 - | 2 - | 8,93E-05 - | 1 - | 5,03E-05 - |
| 6945 | 1 | 3um_FIJJAFBF_81388 | Bacteria | Bacteroidetes | Flavobacteriia | Flavobacterial | Flavobacteriac | Formosa | Formosa sp. H | poorly charact | Function unkn | 0P9WP | NA | 1046627.BZAF | 1,4 | 0,489897949 | 0,000130154 | 4,80E-05 | 1 - | 7,76E-05 - | 2 - | 0,000192593 - | 2 - | 0,000183084 - | 1 - | 9,29E-05 - | 1 - | 0,000104645 - |
| 6946 | 1 | 0.2um_NOKCPHJN_640379 | Bacteria | Proteobacteri | Alphaproteob | Pelagibacteral | Pelagibacterac | Candidatus Pe | Candidatus Pe | metabolism | Amino acid tra | 01R4A | Polyamine AB | 335992.SAR11 | 15,6 | 1,356465997 | 0,000571013 | 5,83E-05 | 17 - | 0,000523111 - | 13 - | 0,000496715 - | 16 - | 0,000581157 - | 16 - | 0,000589741 - | 16 - | 0,000664341 - |
| 6947 | 1 | 3um_FIJJAFBF_328495 | Bacteria | Bacteroidetes | Flavobacteriia | Flavobacterial | Flavobacteriac | Arenitalea | Arenitalea lut | poorly charact | Function unkn | 0PAEN | NA | 886377.Murru | 6,2 | 0,748331477 | 0,00017716 | 2,72E-05 | 7 - | 0,000167732 - | 6 - | 0,00017852 - | 6 - | 0,000169706 - | 5 - | 0,000143511 - | 7 - | 0,00022633 - |
| 6948 | 1 | 3um_FIJJAFBF_658505 | Bacteria | Proteobacteri | Gammaprote | Oceanospirilla | Saccharospirill | Reinekea | Reinekea fors | poorly charact | Function unkn | 16XUA | Protein of unk | 314283.MED2 | 0,4 | 0,489897949 | 6,31E-05 | 7,78E-05 | 1 - | 0,000143471 - | 0 - | 0 - | 0 - | 0 - | 1 - | 0,000171854 - | 0 - | 0 - |
| 6949 | 1 | 0.2um_NOKCPHJN_917895 | Bacteria | Bacteroidetes | Flavobacteriia | Flavobacterial | Flavobacteriac | Formosa | Formosa sp. H | cellular proces | Cell wall/mem | 09DZB | Fasciclin | 487797.Flav3 | 1 | 0 | 8,06E-05 | 7,65E-06 | 1 - | 6,75E-05 - | 1 - | 8,38E-05 - | 1 - | 7,97E-05 - | 1 - | 8,09E-05 - | 1 - | 9,11E-05 - |
| 6950 | 1 | 0.2um_NOKCPHJN_168385 | Bacteria | Proteobacteri | Gammaprote | Cellvibrionales | Porticoccacea | unclassified | Gammaprote | information st | Translation, ril | COG0052 | 30S ribosomal | 314287.GB22 | 0,2 | 0,4 | 1,01E-05 | 2,02E-05 | 0 - | 0 - | 0 - | 0 - | 1 - | 5,06E-05 - | 0 - | 0 - | 0 - | 0 - |
| 6951 | 1 | 3um_FIJJAFBF_194625 | Bacteria | Bacteroidetes | Cytophagia | Cytophagales | Cytophagacea | Ekhidna | Ekhidna lutea | metabolism | Inorganic ion t | 0P7K4 | Potassium upt | 391587.KAOT | 0,6 | 0,489897949 | 1,76E-05 | 1,45E-05 | 1 - | 2,57E-05 - | 1 - | 3,20E-05 - | 1 - | 3,04E-05 - | 0 - | 0 - | 0 - | 0 - |
| 6952 | 1 | 0.2um_NOKCPHJN_678021 | Bacteria | Proteobacteri | Gammaprote | Pseudomonad | Pseudomonad | Pseudomonas | various | metabolism | Energy produc | 0QIP8 | Electron trans | 746360.PFWH | 3,4 | 1,624807681 | 0,000146724 | 6,80E-05 | 4 - | 0,000148578 - | 3 - | 0,000138368 - | 6 - | 0,000263072 - | 3 - | 0,000133479 - | 1 - | 5,01E-05 - |
| 6953 | 1 | 3um_FIJJAFBF_302340 | Bacteria | Bacteroidetes | Flavobacteriia | Flavobacterial | Flavobacteriac | Winogradskye | Winogradskye | poorly charact | Function unkn | 0PCAY | NA | 391598.FBBAL | 0,8 | 0,4 | 4,46E-05 | 2,24E-05 | 0 - | 0 - | 1 - | 5,57E-05 - | 1 - | 5,29E-05 - | 1 - | 5,37E-05 - | 1 - | 6,05E-05 - |
| 6954 | 1 | 0.2um_NOKCPHJN_736852 | Bacteria | Proteobacteri | Alphaproteob | Rhodobactera | unclassified | unclassified | Rhodobactera | unclassified | unclassified | unclassified | unclassified | unclassified | 1 | 0 | 3,34E-05 | 3,17E-06 | 1 - | 2,80E-05 - | 1 - | 3,48E-05 - | 1 - | 3,30E-05 - | 1 - | 3,35E-05 - | 1 - | 3,78E-05 - |
| 6955 | 1 | 3um_FIJJAFBF_371765 | Bacteria | Bacteroidetes | Flavobacteriia | Flavobacterial | Flavobacteriac | Arenitalea | Arenitalea lut | poorly charact | Function unkn | 09BCW | Protein of unk | 391603.FBALC | 0,4 | 0,489897949 | 2,60E-05 | 3,20E-05 | 0 - | 0 - | 0 - | 0 - | 1 - | 6,08E-05 - | 0 - | 0 - | 1 - | 6,95E-05 - |
| 6956 | 1 | 0.2um_NOKCPHJN_159892 | Bacteria | Proteobacteri | Alphaproteob | Rhodobactera | Rhodobactera | Thalassobium | Thalassobium | unclassified | unclassified | unclassified | unclassified | unclassified | 1,8 | 0,979795897 | 4,53E-05 | 2,60E-05 | 2 - | 4,32E-05 - | 3 - | 8,05E-05 - | 2 - | 5,10E-05 - | 2 - | 5,18E-05 - | 0 - | 0 - |
| 6957 | 1 | 0.2um_NOKCPHJN_260663 | Bacteria | Proteobacteri | Alphaproteob | Rhodobactera | Rhodobactera | Roseobacter | Roseobacter s | unclassified | unclassified | unclassified | unclassified | unclassified | 0,8 | 0,748331477 | 1,48E-05 | 1,41E-05 | 0 - | 0 - | 2 - | 3,79E-05 - | 1 - | 1,80E-05 - | 1 - | 1,83E-05 - | 0 - | 0 - |
| 6958 | 1 | 0.2um_NOKCPHJN_760411 | Bacteria | Proteobacteri | Gammaprote | unclassified | unclassified | unclassified | Gammaprote | metabolism | Inorganic ion t | 05F28 | tonB-depende | 247634.GPB2 | 4,6 | 0,8 | 8,23E-05 | 1,22E-05 | 5 - | 7,58E-05 - | 5 - | 9,41E-05 - | 5 - | 8,95E-05 - | 5 - | 9,08E-05 - | 3 - | 6,14E-05 - |
| 6959 | 1 | 0.2um_NOKCPHJN_605814 | Bacteria | Bacteroidetes | Flavobacteriia | Flavobacterial | Flavobacteriac | Formosa | Formosa sp. H | unclassified | unclassified | unclassified | unclassified | unclassified | 0,6 | 0,489897949 | 4,48E-05 | 3,72E-05 | 1 - | 6,34E-05 - | 0 - | 0 - | 1 - | 7,49E-05 - | 0 - | 0 - | 1 - | 8,56E-05 - |
| 6960 | 1 | 3um_FIJJAFBF_725107 | Bacteria | unclassified | unclassified | unclassified | unclassified | unclassified | uncultured ba | metabolism | Energy produc | 0P7S4 | Produces ATP | 313603.FB217 | 0,6 | 0,8 | 2,91E-05 | 3,84E-05 | 0 - | 0 - | 1 - | 5,02E-05 - | 2 - | 9,54E-05 - | 0 - | 0 - | 0 - | 0 - |
| 6961 | 1 | 0.2um_NOKCPHJN_358994 | Eukaryota | unclassified | Labyrinthulom | unclassified | Thraustochytr | Aurantiochytr | Aurantiochytr | information st | Translation, ril | COG0081 | Binds directly | 35128.Thaps2 | 1 | 0 | 5,46E-05 | 5,18E-06 | 1 - | 4,57E-05 - | 1 - | 5,68E-05 - | 1 - | 5,40E-05 - | 1 - | 5,48E-05 - | 1 - | 6,17E-05 - |
| 6962 | 1 | 0.2um_NOKCPHJN_509333 | Viruses | unclassified | unclassified | unclassified | unclassified | unclassified | uncultured M | unclassified | unclassified | unclassified | unclassified | unclassified | 0,6 | 0,489897949 | 1,82E-05 | 1,51E-05 | 1 - | 2,57E-05 - | 0 - | 0 - | 0 - | 0 - | 1 - | 3,08E-05 - | 1 - | 3,46E-05 - |
| 6963 | 1 | 0.2um_NOKCPHJN_108379 | Bacteria | Proteobacteri | Gammaprote | Oceanospirilla | Saccharospirill | Reinekea | Reinekea fors | unclassified | unclassified | unclassified | unclassified | unclassified | 0,4 | 0,489897949 | 4,66E-06 | 5,71E-06 | 0 - | 0 - | 0 - | 0 - | 1 - | 1,16E-05 - | 1 - | 1,17E-05 - | 0 - | 0 - |
| 6964 | 1 | 3um_FIJJAFBF_191792 | Eukaryota | unclassified | Raphidophyce | Chattonellales | Chattonellace | Heterosigma | Heterosigma a | unclassified | unclassified | unclassified | unclassified | unclassified | 3,6 | 1,2 | 0,000599323 | 0,000207685 | 5 - | 0,000699858 - | 3 - | 0,00052141 - | 3 - | 0,000495667 - | 2 - | 0,000335325 - | 5 - | 0,000944357 - |
| 6965 | 1 | 3um_FIJJAFBF_281958 | Bacteria | Firmicutes | Bacilli | Bacillales | Paenibacillace | Paenibacillus | Paenibacillus | cellular proces | Cell motility | 174CZ | Pilus assembly | 762376.AXYL_ | 0,2 | 0,4 | 1,10E-05 | 2,21E-05 | 0 - | 0 - | 0 - | 0 - | 0 - | 0 - | 1 - | 5,52E-05 - | 0 - | 0 - |
| 6966 | 1 | 3um_FIJJAFBF_133467 | Bacteria | Actinobacteri | Actinobacteri | Micrococcales | Microbacteria | various | various | unclassified | unclassified | unclassified | unclassified | unclassified | 1,2 | 0,4 | 0,000134856 | 4,65E-05 | 1 - | 9,41E-05 - | 1 - | 0,000116819 - | 1 - | 0,000111051 - | 2 - | 0,000225382 - | 1 - | 0,000126946 - |
| 6967 | 1 | 3um_FIJJAFBF_212745 | Bacteria | Bacteroidetes | Flavobacteriia | Flavobacterial | Flavobacteriac | Polaribacter | Polaribacter t | unclassified | unclassified | unclassified | unclassified | unclassified | 6,4 | 1,624807681 | 0,000404861 | 9,07E-05 | 8 - | 0,000429072 - | 7 - | 0,000466183 - | 4 - | 0,000253238 - | 8 - | 0,000513956 - | 5 - | 0,000361856 - |
| 6968 | 1 | 3um_FIJJAFBF_236213 | Bacteria | Proteobacteri | Gammaprote | Oceanospirilla | Saccharospirill | Reinekea | Reinekea fors | information st | Translation, ril | 17FIT | 50s ribosomal | 314283.MED2 | 1 | 0 | 0,000175676 | 1,67E-05 | 1 - | 0,00014715 - | 1 - | 0,000182716 - | 1 - | 0,000173695 - | 1 - | 0,000176261 - | 1 - | 0,000198557 - |
| 6969 | 1 | 3um_FIJJAFBF_93751 | Bacteria | Proteobacteri | Gammaprote | Alteromonada | Ferrimonadac | Ferrimonas | Ferrimonas m | metabolism | Inorganic ion t | COG0376 | Bifunctional e | 523791.Kkor_ | 1 | 0,632455532 | 1,92E-05 | 1,18E-05 | 0 - | 0 - | 1 - | 1,93E-05 - | 1 - | 1,84E-05 - | 2 - | 3,73E-05 - | 1 - | 2,10E-05 - |
| 6970 | 1 | 3um_FIJJAFBF_05548 | Bacteria | Bacteroidetes | Flavobacteriia | Flavobacterial | Flavobacteriac | Polaribacter | various | unclassified | unclassified | unclassified | unclassified | unclassified | 1 | 0 | 0,000214105 | 2,03E-05 | 1 - | 0,000179339 - | 1 - | 0,000222686 - | 1 - | 0,000211691 - | 1 - | 0,000214818 - | 1 - | 0,000241991 - |
| 6971 | 1 | 3um_FIJJAFBF_662492 | Bacteria | Bacteroidetes | Flavobacteriia | Flavobacterial | Flavobacteriac | Polaribacter | Polaribacter s | metabolism | Amino acid tra | 0P7S1 | Catalyzes the | 313594.PI23P | 0,2 | 0,4 | 9,31E-06 | 1,86E-05 | 0 - | 0 - | 0 - | 0 - | 1 - | 4,66E-05 - | 0 - | 0 - | 0 - | 0 - |
| 6972 | 1 | 0.2um_NOKCPHJN_58847 | Bacteria | Proteobacteri | Betaproteoba | unclassified | unclassified | unclassified | Betaproteoba | metabolism | Energy produc | 0B7RT | Acetoacetyl-C | 232721.Ajs_2 | 2,4 | 1,019803903 | 0,000131009 | 4,72E-05 | 4 - | 0,000187391 - | 3 - | 0,000174513 - | 2 - | 0,000110598 - | 1 - | 5,61E-05 - | 2 - | 0,000126428 - |
| 6973 | 1 | 3um_FIJJAFBF_235372 | Bacteria | Proteobacteri | Gammaprote | Cellvibrionales | Porticoccacea | unclassified | Gammaprote | unclassified | unclassified | unclassified | unclassified | unclassified | 0,2 | 0,4 | 8,83E-06 | 1,77E-05 | 1 - | 4,41E-05 - | 0 - | 0 - | 0 - | 0 - | 0 - | 0 - | 0 - | 0 - |
| 6974 | 2 | 0.2um_NOKCPHJN_270892;3um_FIJJAFBF_2214 | Bacteria | Proteobacteri | Betaproteoba | unclassified | unclassified | unclassified | Betaproteoba | unclassified | unclassified | unclassified | unclassified | unclassified | 1,2 | 0,748331477 | 3,28E-05 | 1,91E-05 | 2 - | 4,72E-05 - | 1 - | 2,93E-05 - | 2 - | 5,58E-05 - | 0 - | 0 - | 1 - | 3,19E-05 - |
| 6975 | 1 | 3um_FIJJAFBF_292535 | Bacteria | Bacteroidetes | Flavobacteriia | Flavobacterial | Flavobacteriac | Gaetbulibacte | Gaetbulibacte | unclassified | unclassified | unclassified | unclassified | unclassified | 0,4 | 0,8 | 1,42E-05 | 2,84E-05 | 2 - | 7,11E-05 - | 0 - | 0 - | 0 - | 0 - | 0 - | 0 - | 0 - | 0 - |
| 6976 | 1 | 0.2um_NOKCPHJN_184459 | Bacteria | Proteobacteri | Gammaprote | Alteromonada | Alteromonada | Marinobacter | various | cellular proces | Intracellular tr | 175GA | One of the pro | 443152.MDG8 | 2,6 | 0,8 | 0,000202127 | 6,27E-05 | 2 - | 0,000130428 - | 2 - | 0,000161953 - | 4 - | 0,000307914 - | 3 - | 0,000234347 - | 2 - | 0,000175994 - |
| 6977 | 1 | 0.2um_NOKCPHJN_45444 | Bacteria | Proteobacteri | Gammaprote | Chromatiales | Ectothiorhodo | Aquisalimonas | Aquisalimonas | metabolism | Amino acid tra | COG0082 | 5-enolpyruvyl | 314283.MED2 | 0,4 | 0,489897949 | 1,37E-05 | 1,70E-05 | 1 - | 3,13E-05 - | 0 - | 0 - | 0 - | 0 - | 1 - | 3,75E-05 - | 0 - | 0 - |
| 6978 | 1 | 0.2um_NOKCPHJN_346026 | Eukaryota | Bacillariophyt | Coscinodiscop | Thalassiosirale | Thalassiosirac | Thalassiosira | Thalassiosira | information st | Chromatin str | KOG1744 | Core compone | 35128.Thaps2 | 1 | 0 | 0,000126877 | 1,20E-05 | 1 - | 0,000106275 - | 1 - | 0,000131962 - | 1 - | 0,000125447 - | 1 - | 0,000127299 - | 1 - | 0,000143402 - |
| 6979 | 1 | 3um_FIJJAFBF_697086 | Eukaryota | unclassified | Heterolobosea | Schizopyrenid | Vahlkampfiida | Naegleria | Naegleria grul | unclassified | unclassified | unclassified | unclassified | unclassified | 2 | 0 | 7,35E-05 | 6,98E-06 | 2 - | 6,15E-05 - | 2 - | 7,64E-05 - | 2 - | 7,26E-05 - | 2 - | 7,37E-05 - | 2 - | 8,30E-05 - |
| 6980 | 4 | 0.2um_NOKCPHJN_475274;0.2um_NOKCPHJN_5 | Bacteria | Proteobacteri | Alphaproteob | Pelagibacteral | Pelagibacterac | various | various | information st | Translation, ril | 082FS | 50S ribosomal | 335992.SAR11 | 0,4 | 0,489897949 | 5,87E-05 | 7,23E-05 | 1 - | 0,000133461 - | 0 - | 0 - | 0 - | 0 - | 1 - | 0,000159864 - | 0 - | 0 - |
| 6981 | 1 | 0.2um_NOKCPHJN_74560 | Bacteria | Proteobacteri | Gammaprote | Oceanospirilla | Saccharospirill | Reinekea | Reinekea blan | unclassified | unclassified | unclassified | unclassified | unclassified | 0,2 | 0,4 | 6,07E-06 | 1,21E-05 | 0 - | 0 - | 0 - | 0 - | 0 - | 0 - | 1 - | 3,03E-05 - | 0 - | 0 - |
| 6982 | 1 | 0.2um_NOKCPHJN_104449 | Bacteria | Proteobacteri | Gammaprote | Cellvibrionales | Porticoccacea | unclassified | SAR92 bacteri | unclassified | unclassified | unclassified | unclassified | unclassified | 0,2 | 0,4 | 8,44E-06 | 1,69E-05 | 1 - | 4,22E-05 - | 0 - | 0 - | 0 - | 0 - | 0 - | 0 - | 0 - | 0 - |
| 6983 | 1 | 0.2um_NOKCPHJN_413903 | Bacteria | Bacteroidetes | Flavobacteriia | Flavobacterial | Flavobacteriac | Formosa | Formosa sp. H | unclassified | unclassified | unclassified | unclassified | unclassified | 3,8 | 0,4 | 3,45E-05 | 3,08E-06 | 4 - | 3,06E-05 - | 4 - | 3,81E-05 - | 4 - | 3,62E-05 - | 4 - | 3,67E-05 - | 3 - | 3,10E-05 - |
| 6984 | 1 | 3um_FIJJAFBF_560706 | Bacteria | Proteobacteri | Alphaproteob | Rhodobactera | unclassified | unclassified | Rhodobactera | information st | Translation, ril | 179R6 | This protein is | 367336.OM22 | 2,4 | 0,489897949 | 0,000196398 | 4,39E-05 | 3 - | 0,000206186 - | 2 - | 0,000170681 - | 2 - | 0,000162254 - | 2 - | 0,000164651 - | 3 - | 0,000278218 - |
| 6985 | 1 | 0.2um_NOKCPHJN_60225 | Bacteria | Bacteroidetes | Flavobacteriia | Flavobacterial | Flavobacteriac | unclassified | Flavobacteriac | metabolism | Nucleotide tra | 0P7KC | ribonucleoside | 1041826.FCOL | 1,4 | 1,019803903 | 5,04E-05 | 3,35E-05 | 3 - | 9,70E-05 - | 1 - | 4,01E-05 - | 2 - | 7,63E-05 - | 1 - | 3,87E-05 - | 0 - | 0 - |
| 6986 | 1 | 3um_FIJJAFBF_30299 | Bacteria | Bacteroidetes | Flavobacteriia | Flavobacterial | Flavobacteriac | Polaribacter | Polaribacter s | metabolism | Energy produc | 0P9HY | cytochrome C | 313598.MED1 | 2,8 | 0,979795897 | 0,000190419 | 6,04E-05 | 4 - | 0,000231872 - | 1 - | 7,20E-05 - | 3 - | 0,000205276 - | 3 - | 0,000208308 - | 3 - | 0,000234658 - |
| 6987 | 1 | 0.2um_NOKCPHJN_817880 | Eukaryota | unclassified | Pelagophycea | Pelagomonad | unclassified | Aureococcus | Aureococcus a | unclassified | unclassified | unclassified | unclassified | unclassified | 1,2 | 0,4 | 2,45E-05 | 5,43E-06 | 2 - | 3,52E-05 - | 1 - | 2,19E-05 - | 1 - | 2,08E-05 - | 1 - | 2,11E-05 - | 1 - | 2,38E-05 - |
| 6988 | 1 | 0.2um_NOKCPHJN_684761 | Bacteria | Bacteroidetes | Flavobacteriia | Flavobacterial | Flavobacteriac | Formosa | Formosa sp. H | poorly charact | Function unkn | 06VZ7 | NA | 688270.Celal_ | 1,6 | 0,489897949 | 6,71E-05 | 1,76E-05 | 2 - | 7,17E-05 - | 1 - | 4,45E-05 - | 2 - | 8,47E-05 - | 2 - | 8,59E-05 - | 1 - | 4,84E-05 - |
| 6989 | 1 | 0.2um_NOKCPHJN_473331 | Bacteria | Proteobacteri | Alphaproteob | Rhodobactera | Rhodobactera | Maritimibacte | Maritimibacte | unclassified | unclassified | unclassified | unclassified | unclassified | 0,2 | 0,4 | 4,11E-06 | 8,23E-06 | 1 - | 2,06E-05 - | 0 - | 0 - | 0 - | 0 - | 0 - | 0 - | 0 - | 0 - |
| 6990 | 1 | 0.2um_NOKCPHJN_1092050 | Bacteria | Proteobacteri | Gammaprote | Cellvibrionales | Porticoccacea | unclassified | gamma prote | unclassified | unclassified | unclassified | unclassified | unclassified | 0,2 | 0,4 | 4,44E-06 | 8,88E-06 | 1 - | 2,22E-05 - | 0 - | 0 - | 0 - | 0 - | 0 - | 0 - | 0 - | 0 - |
| 6991 | 1 | 3um_FIJJAFBF_30600 | Eukaryota | unclassified | Dinophyceae | Suessiales | Symbiodiniace | Symbiodinium | Symbiodinium | unclassified | unclassified | unclassified | unclassified | unclassified | 0,2 | 0,4 | 4,15E-06 | 8,29E-06 | 0 - | 0 - | 0 - | 0 - | 0 - | 0 - | 1 - | 2,07E-05 - | 0 - | 0 - |
| 6992 | 1 | 0.2um_NOKCPHJN_112414 | Bacteria | Bacteroidetes | Flavobacteriia | Flavobacterial | Flavobacteriac | Gilvibacter | Gilvibacter sp. | information st | Translation, ril | COG0052 | 30S ribosomal | 313603.FB217 | 3,8 | 0,748331477 | 0,000169647 | 2,98E-05 | 4 - | 0,000151022 - | 3 - | 0,000140644 - | 5 - | 0,000222833 - | 4 - | 0,000180899 - | 3 - | 0,000152837 - |
| 6993 | 1 | 3um_FIJJAFBF_267726 | Bacteria | Bacteroidetes | Flavobacteriia | Flavobacterial | various | various | various | unclassified | unclassified | unclassified | unclassified | unclassified | 3,4 | 0,8 | 0,00065551 | 0,000166851 | 4 - | 0,00064663 - | 3 - | 0,000602192 - | 2 - | 0,00038164 - | 4 - | 0,000774554 - | 4 - | 0,000872532 - |
| 6994 | 1 | 3um_FIJJAFBF_681369 | Bacteria | Bacteroidetes | Cytophagia | Cytophagales | Flammeovirga | Fabibacter | Fabibacter pa | unclassified | unclassified | unclassified | unclassified | unclassified | 0,2 | 0,4 | 5,64E-06 | 1,13E-05 | 1 - | 2,82E-05 - | 0 - | 0 - | 0 - | 0 - | 0 - | 0 - | 0 - | 0 - |
| 6995 | 1 | 3um_FIJJAFBF_62160 | Bacteria | Nitrospirae | Nitrospira | Nitrospirales | Nitrospiraceae | Nitrospira | various | unclassified | unclassified | unclassified | unclassified | unclassified | 1 | 0,632455532 | 7,92E-05 | 5,02E-05 | 0 - | 0 - | 2 - | 0,000158354 - | 1 - | 7,53E-05 - | 1 - | 7,64E-05 - | 1 - | 8,60E-05 - |
| 6996 | 1 | 3um_FIJJAFBF_513825 | Viruses | unclassified | unclassified | unclassified | unclassified | unclassified | uncultured M | unclassified | unclassified | unclassified | unclassified | unclassified | 1 | 0 | 7,83E-05 | 7,43E-06 | 1 - | 6,56E-05 - | 1 - | 8,14E-05 - | 1 - | 7,74E-05 - | 1 - | 7,86E-05 - | 1 - | 8,85E-05 - |
| 6997 | 1 | 0.2um_NOKCPHJN_96088 | Bacteria | Proteobacteri | Gammaprote | Alteromonada | Alteromonada | Paraglaciecola | Paraglaciecola | poorly charact | Function unkn | 05C2A | tonB-depende | 1085623.GNIT | 20,8 | 1,833030278 | 0,000257241 | 1,72E-05 | 23 - | 0,00023977 - | 20 - | 0,00025889 - | 23 - | 0,000283024 - | 19 - | 0,000237255 - | 19 - | 0,000267267 - |
| 6998 | 1 | 3um_FIJJAFBF_25394 | Bacteria | Bacteroidetes | Flavobacteriia | unclassified | unclassified | unclassified | Flavobacteria | information st | Translation, ril | COG0088 | One of the pri | 391587.KAOT | 0,6 | 0,8 | 3,69E-05 | 5,10E-05 | 1 - | 5,49E-05 - | 0 - | 0 - | 2 - | 0,000129648 - | 0 - | 0 - | 0 - | 0 - |
| 6999 | 1 | 0.2um_NOKCPHJN_115354 | Bacteria | Bacteroidetes | Flavobacteriia | Flavobacterial | Flavobacteriac | Lutibacter | Lutibacter sp. | unclassified | unclassified | unclassified | unclassified | unclassified | 2,2 | 0,979795897 | 2,82E-05 | 1,33E-05 | 2 - | 2,15E-05 - | 4 - | 5,35E-05 - | 2 - | 2,54E-05 - | 2 - | 2,58E-05 - | 1 - | 1,45E-05 - |
| 7000 | 1 | 3um_FIJJAFBF_634172 | Bacteria | Actinobacteri | Actinobacteri | Micrococcales | Microbacteria | Rhodoluna | Rhodoluna lac | unclassified | unclassified | unclassified | unclassified | unclassified | 3,4 | 1,624807681 | 8,60E-05 | 4,18E-05 | 3 - | 6,39E-05 - | 1 - | 2,64E-05 - | 4 - | 0,000100543 - | 6 - | 0,000153043 - | 3 - | 8,62E-05 - |
| 7001 | 1 | 3um_FIJJAFBF_120257 | Eukaryota | Arthropoda | Insecta | Blattodea | Kalotermitida | Cryptotermes | Cryptotermes | cellular proces | Intracellular tr | 036AZ | Trafficking pro | 12957.ACEP_ | 0,2 | 0,4 | 5,47E-06 | 1,09E-05 | 0 - | 0 - | 0 - | 0 - | 1 - | 2,74E-05 - | 0 - | 0 - | 0 - | 0 - |

| | | | | | | | | | | | | | | | | | | | | | | | | | | | | |
|---|---|---|---|---|---|---|---|---|---|---|---|---|---|---|---|---|---|---|---|---|---|---|---|---|---|---|---|---|
| 7002 | 1 | 3um_FIJJAFBF_265695 | unclassified | unclassified | unclassified | unclassified | unclassified | unclassified | unclassified | unclassified | unclassified | unclassified | unclassified | unclassified | 24,6 | 2,244994432 | 0,002619986 | 0,000224843 | 28 - | 0,002510742 - | 25 - | 0,002783572 - | 25 - | 0,002646137 - | 21 - | 0,002255585 - | 24 - | 0,002903896 - |
| 7003 | 1 | 3um_FIJJAFBF_596578 | Bacteria | Bacteroidetes | Flavobacteriia | Flavobacterial | Flavobacteriac | Polaribacter | Polaribacter s | cellular proces | Posttranslatio | 08V1T | peptidylprolyl | 313594.PI23P | 3,2 | 1,166190379 | 0,000183657 | 6,06E-05 | 4 - | 0,0001962 - | 3 - | 0,000182716 - | 5 - | 0,000289492 - | 2 - | 0,000117507 - | 2 - | 0,000132371 - |
| 7004 | 1 | 3um_FIJJAFBF_257112 | Eukaryota | unclassified | Dinophyceae | Peridiniales | Kryptoperidini | Durinskia | Durinskia balt | information st | Translation, ril | COG0088 | One of the pri | 329726.AM1_ | 0,6 | 0,8 | 3,59E-05 | 4,54E-05 | 2 - | 0,000107268 - | 0 - | 0 - | 0 - | 0 - | 0 - | 0 - | 1 - | 7,24E-05 - |
| 7005 | 2 | 0.2um_NOKCPHJN_610694;3um_FIJJAFBF_44683 | Bacteria | Bacteroidetes | Flavobacteriia | Flavobacterial | Flavobacteriac | Polaribacter | various | poorly charact | Function unkn | 07MPW | Short-chain de | 313598.MED1 | 1,2 | 0,4 | 7,23E-05 | 2,42E-05 | 1 - | 5,06E-05 - | 1 - | 6,28E-05 - | 2 - | 0,000119368 - | 1 - | 6,06E-05 - | 1 - | 6,82E-05 - |
| 7006 | 1 | 3um_FIJJAFBF_436838 | Bacteria | Actinobacteria | Actinobacteria | Micrococcales | Microbacteria | Candidatus Ac | Candidatus Ac | information st | Translation, ril | COG0050 | This protein p | 566461.SSFG_ | 4,6 | 0,8 | 0,000158604 | 2,95E-05 | 5 - | 0,00014492 - | 6 - | 0,000215938 - | 4 - | 0,000136851 - | 4 - | 0,000138872 - | 4 - | 0,000156439 - |
| 7007 | 1 | 0.2um_NOKCPHJN_586865 | Bacteria | Bacteroidetes | Flavobacteriia | Flavobacterial | Flavobacteriac | Polaribacter | Polaribacter s | metabolism | Inorganic ion t | COG0376 | Bifunctional e | 391603.FBALC | 1 | 0,632455532 | 1,76E-05 | 1,00E-05 | 2 - | 3,14E-05 - | 1 - | 1,95E-05 - | 1 - | 1,85E-05 - | 1 - | 1,88E-05 - | 0 - | 0 - |
| 7008 | 1 | 3um_FIJJAFBF_70344 | Bacteria | Proteobacteria | Betaproteoba | unclassified | unclassified | unclassified | Betaproteoba | information st | Translation, ril | COG0255 | 50s ribosomal | 365044.Pnap_ | 1 | 0 | 0,000214105 | 2,03E-05 | 1 - | 0,000179339 - | 1 - | 0,000222686 - | 1 - | 0,000211691 - | 1 - | 0,000214818 - | 1 - | 0,000241991 - |
| 7009 | 1 | 3um_FIJJAFBF_242261 | Bacteria | unclassified | unclassified | unclassified | unclassified | unclassified | bacterium TM | metabolism | Energy produc | 11HAV | photosystem i | 35128.Thapsd | 0,4 | 0,489897949 | 3,58E-05 | 4,41E-05 | 1 - | 8,14E-05 - | 0 - | 0 - | 0 - | 0 - | 1 - | 9,75E-05 - | 0 - | 0 - |
| 7010 | 1 | 3um_FIJJAFBF_238481 | Bacteria | Proteobacteria | Gammaprotec | Oceanospirilla | Saccharospirill | Saccharospiril | Saccharospiril | information st | Translation, ril | 05CEJ | Catalyzes the | 314283.MED2 | 20 | 1,414213562 | 0,00039286 | 5,45E-05 | 20 - | 0,000328403 - | 19 - | 0,00038739 - | 21 - | 0,000407028 - | 18 - | 0,000354034 - | 22 - | 0,000487445 - |
| 7011 | 1 | 0.2um_NOKCPHJN_231776 | Bacteria | Proteobacteria | Alphaproteob | Rhodobactera | Rhodobactera | Loktanella | Loktanella sp. | unclassified | unclassified | unclassified | unclassified | unclassified | 0,4 | 0,489897949 | 1,69E-05 | 2,07E-05 | 0 - | 0 - | 0 - | 0 - | 1 - | 4,19E-05 - | 1 - | 4,26E-05 - | 0 - | 0 - |
| 7012 | 1 | 3um_FIJJAFBF_254287 | Bacteria | Proteobacteria | Gammaprotec | Cellvibrionales | Halieaceae | Haliea | Haliea salexig | information st | Translation, ril | 17FIH | One of the ea | 566466.NOR5 | 1 | 0 | 0,000139824 | 1,33E-05 | 1 - | 0,000117119 - | 1 - | 0,000145427 - | 1 - | 0,000138247 - | 1 - | 0,000140289 - | 1 - | 0,000158035 - |
| 7013 | 1 | 0.2um_NOKCPHJN_67773 | Bacteria | Proteobacteria | Alphaproteob | Rhodobactera | Rhodobactera | Amylibacter | Amylibacter k | information st | Translation, ril | COG0173 | aspartyl-trna s | 398580.Dshi_ | 0,8 | 0,748331477 | 1,85E-05 | 1,64E-05 | 2 - | 4,23E-05 - | 0 - | 0 - | 1 - | 2,50E-05 - | 1 - | 2,53E-05 - | 0 - | 0 - |
| 7014 | 1 | 0.2um_NOKCPHJN_273441 | Eukaryota | Chlorophyta | Chlorophycea | Chlamydomor | Chlamydomor | Chlamydomor | Chlamydomor | unclassified | unclassified | unclassified | unclassified | unclassified | 0,2 | 0,4 | 4,19E-06 | 8,39E-06 | 0 - | 0 - | 0 - | 0 - | 1 - | 2,10E-05 - | 0 - | 0 - | 0 - | 0 - |
| 7015 | 1 | 3um_FIJJAFBF_17219 | Bacteria | Proteobacteria | Alphaproteob | Rhodobactera | Rhodobactera | Phaeobacter | Phaeobacter g | cellular proces | Posttranslatio | COG0330 | Band 7 protei | 246200.SPO13 | 6,8 | 0,748331477 | 0,000318964 | 5,54E-05 | 7 - | 0,000274211 - | 6 - | 0,000291848 - | 6 - | 0,000277438 - | 7 - | 0,000328458 - | 8 - | 0,000422865 - |
| 7016 | 1 | 0.2um_NOKCPHJN_515270 | Bacteria | Proteobacteria | Gammaprotec | Cellvibrionales | Spongiibacter | unclassified | marine gamm | unclassified | unclassified | unclassified | unclassified | unclassified | 0,2 | 0,4 | 3,95E-06 | 7,89E-06 | 0 - | 0 - | 0 - | 0 - | 0 - | 0 - | 0 - | 0 - | 1 - | 1,97E-05 - |
| 7017 | 1 | 0.2um_NOKCPHJN_447106 | Bacteria | Bacteroidetes | Flavobacteriia | Flavobacterial | Flavobacteriac | Formosa | Formosa sp. H | unclassified | unclassified | unclassified | unclassified | unclassified | 1,2 | 0,4 | 5,12E-05 | 2,18E-05 | 1 - | 3,50E-05 - | 1 - | 4,35E-05 - | 1 - | 4,13E-05 - | 1 - | 4,19E-05 - | 2 - | 9,44E-05 - |
| 7018 | 1 | 3um_FIJJAFBF_84876 | Bacteria | Proteobacteria | Alphaproteob | Rhodobactera | Rhodobactera | Nioella | Nioella sedimi | metabolism | Coenzyme tra | 01R34 | Catalyzes the | 252305.OB25 | 1 | 0 | 3,49E-05 | 3,31E-06 | 1 - | 2,92E-05 - | 1 - | 3,63E-05 - | 1 - | 3,45E-05 - | 1 - | 3,50E-05 - | 1 - | 3,94E-05 - |
| 7019 | 1 | 0.2um_NOKCPHJN_740335 | Bacteria | Bacteroidetes | Flavobacteriia | Flavobacterial | Flavobacteriac | Polaribacter | Polaribacter s | unclassified | unclassified | unclassified | unclassified | unclassified | 28,4 | 2,576819745 | 0,000490697 | 5,49E-05 | 30 - | 0,00043531 - | 31 - | 0,000558544 - | 30 - | 0,000513839 - | 24 - | 0,000417143 - | 27 - | 0,000528649 - |
| 7020 | 1 | 3um_FIJJAFBF_238099 | Bacteria | Bacteroidetes | Flavobacteriia | Flavobacterial | Flavobacteriac | Polaribacter | Polaribacter ir | metabolism | Inorganic ion t | 05C27 | TonB-depende | 313594.PI23P | 17,4 | 2,059126028 | 0,000224067 | 2,16E-05 | 19 - | 0,000206511 - | 19 - | 0,000256426 - | 19 - | 0,000243765 - | 16 - | 0,000208308 - | 14 - | 0,000205326 - |
| 7021 | 1 | 3um_FIJJAFBF_663675 | Bacteria | Bacteroidetes | Flavobacteriia | Flavobacterial | Flavobacteriac | Maribacter | Maribacter sp | unclassified | unclassified | unclassified | unclassified | unclassified | 4,2 | 0,4 | 5,80E-05 | 1,05E-05 | 4 - | 4,60E-05 - | 4 - | 5,71E-05 - | 4 - | 5,43E-05 - | 4 - | 5,51E-05 - | 5 - | 7,76E-05 - |
| 7022 | 1 | 3um_FIJJAFBF_256288 | Bacteria | Proteobacteria | Alphaproteob | Rhodobactera | Rhodobactera | Roseobacter | Roseobacter s | unclassified | unclassified | unclassified | unclassified | unclassified | 0,8 | 0,748331477 | 6,23E-05 | 5,99E-05 | 1 - | 6,71E-05 - | 1 - | 8,33E-05 - | 0 - | 0 - | 2 - | 0,000160799 - | 0 - | 0 - |
| 7023 | 1 | 0.2um_NOKCPHJN_59639 | Bacteria | Bacteroidetes | Flavobacteriia | Flavobacterial | Crocinitomica | unclassified | Crocinitomica | unclassified | unclassified | unclassified | unclassified | unclassified | 1,4 | 0,489897949 | 2,85E-05 | 1,17E-05 | 1 - | 1,68E-05 - | 1 - | 2,08E-05 - | 1 - | 1,98E-05 - | 2 - | 4,01E-05 - | 2 - | 4,52E-05 - |
| 7024 | 1 | 0.2um_NOKCPHJN_109744 | Bacteria | Bacteroidetes | Flavobacteriia | Flavobacterial | Flavobacteriac | Formosa | Formosa sp. H | information st | Translation, ril | 0PAI9 | Binds 16S rRN | 391587.KAOT | 1,8 | 0,4 | 0,000282134 | 7,81E-05 | 1 - | 0,000128963 - | 2 - | 0,000320267 - | 2 - | 0,000304454 - | 2 - | 0,000308951 - | 2 - | 0,000348033 - |
| 7025 | 1 | 3um_FIJJAFBF_177760 | Viruses | unclassified | unclassified | unclassified | Phycodnavirid | unclassified | Phaeocystis p | unclassified | unclassified | unclassified | unclassified | unclassified | 0,2 | 0,4 | 5,82E-06 | 1,16E-05 | 0 - | 0 - | 0 - | 0 - | 0 - | 0 - | 0 - | 0 - | 1 - | 2,91E-05 - |
| 7026 | 1 | 3um_FIJJAFBF_314032 | Eukaryota | Euglenida | unclassified | Euglenales | Euglenaceae | Monomorphir | Monomorphir | metabolism | Energy produc | COG0056 | Produces ATP | 70448.Q0P3K | 0,6 | 0,489897949 | 1,65E-05 | 1,35E-05 | 0 - | 0 - | 1 - | 2,83E-05 - | 1 - | 2,69E-05 - | 1 - | 2,73E-05 - | 0 - | 0 - |
| 7027 | 1 | 0.2um_NOKCPHJN_254213 | Bacteria | Proteobacteria | Gammaprotec | unclassified | unclassified | unclassified | SAR86 cluster | information st | Translation, ril | 16QAF | thus facilitatin | 247633.GP21 | 3 | 1,414213562 | 7,42E-05 | 3,84E-05 | 4 - | 8,23E-05 - | 3 - | 7,66E-05 - | 1 - | 2,43E-05 - | 2 - | 4,93E-05 - | 5 - | 0,000138776 - |
| 7028 | 1 | 0.2um_NOKCPHJN_835463 | Eukaryota | Apicomplexa | Conoidasida | Eucoccidiorida | Eimeriidae | Eimeria | Eimeria mitis | poorly charact | Function unkn | 0NF4H | copper amine | 479436.Vpar_ | 0,6 | 0,489897949 | 1,33E-05 | 1,09E-05 | 1 - | 1,97E-05 - | 0 - | 0 - | 1 - | 2,32E-05 - | 1 - | 2,36E-05 - | 0 - | 0 - |
| 7029 | 1 | 3um_FIJJAFBF_498718 | Viruses | unclassified | unclassified | unclassified | unclassified | unclassified | uncultured vir | unclassified | unclassified | unclassified | unclassified | unclassified | 0,4 | 0,489897949 | 9,54E-06 | 1,17E-05 | 0 - | 0 - | 1 - | 2,44E-05 - | 1 - | 2,32E-05 - | 0 - | 0 - | 0 - | 0 - |
| 7030 | 1 | 3um_FIJJAFBF_474990 | Bacteria | Bacteroidetes | Flavobacteriia | unclassified | unclassified | unclassified | Flavobacteria | cellular proces | Cell wall/mem | 11TPU | outer membra | 487796.Flav2 | 4,2 | 1,833030278 | 0,000336501 | 0,000144575 | 5 - | 0,000341597 - | 6 - | 0,000508996 - | 6 - | 0,000483865 - | 2 - | 0,000163671 - | 2 - | 0,000184374 - |
| 7031 | 1 | 3um_FIJJAFBF_345765 | Bacteria | Bacteroidetes | Flavobacteriia | Flavobacterial | Flavobacteriac | Polaribacter | Polaribacter s | unclassified | unclassified | unclassified | unclassified | unclassified | 1,2 | 0,748331477 | 9,70E-05 | 5,82E-05 | 2 - | 0,000138285 - | 2 - | 0,000171709 - | 1 - | 8,16E-05 - | 0 - | 0 - | 1 - | 9,33E-05 - |
| 7032 | 1 | 3um_FIJJAFBF_79245 | Bacteria | Bacteroidetes | Flavobacteriia | Flavobacterial | Flavobacteriac | Polaribacter | Polaribacter s | cellular proces | Posttranslatio | 07RXU | peptidylprolyl | 313598.MED1 | 2,8 | 0,979795897 | 8,15E-05 | 2,40E-05 | 4 - | 9,98E-05 - | 2 - | 6,20E-05 - | 2 - | 5,89E-05 - | 4 - | 0,000119551 - | 2 - | 6,73E-05 - |
| 7033 | 1 | 3um_FIJJAFBF_725334 | Bacteria | Proteobacteria | Gammaprotec | Oceanospirilla | Saccharospirill | Reinekea | Reinekea fors | unclassified | unclassified | unclassified | unclassified | unclassified | 0,2 | 0,4 | 1,87E-05 | 3,73E-05 | 1 - | 9,33E-05 - | 0 - | 0 - | 0 - | 0 - | 0 - | 0 - | 0 - | 0 - |
| 7034 | 1 | 0.2um_NOKCPHJN_813774 | Bacteria | Proteobacteria | Gammaprotec | Oceanospirilla | Halomonadac | Halomonas | Halomonas sp | unclassified | unclassified | unclassified | unclassified | unclassified | 0,8 | 0,748331477 | 1,28E-05 | 1,12E-05 | 2 - | 2,88E-05 - | 1 - | 1,79E-05 - | 0 - | 0 - | 1 - | 1,73E-05 - | 0 - | 0 - |
| 7035 | 3 | 0.2um_NOKCPHJN_1107039;0.2um_NOKCPHJN_ | Bacteria | Bacteroidetes | various | various | various | various | various | unclassified | unclassified | unclassified | unclassified | unclassified | 1 | 0 | 6,56E-05 | 6,23E-06 | 1 - | 5,49E-05 - | 1 - | 6,82E-05 - | 1 - | 6,48E-05 - | 1 - | 6,58E-05 - | 1 - | 7,41E-05 - |
| 7036 | 1 | 3um_FIJJAFBF_510232 | Bacteria | Bacteroidetes | Flavobacteriia | Flavobacterial | Flavobacteriac | Polaribacter | Polaribacter g | information st | Translation, ril | COG0089 | One of the ea | 313594.PI23P | 6,6 | 1,496662955 | 0,000931146 | 0,000174279 | 9 - | 0,001076032 - | 5 - | 0,000742286 - | 5 - | 0,000705637 - | 7 - | 0,001002482 - | 7 - | 0,001129293 - |
| 7037 | 1 | 3um_FIJJAFBF_285633 | Bacteria | unclassified | unclassified | unclassified | unclassified | unclassified | uncultured ba | poorly charact | Function unkn | 09A4Z | rhodopsin | 983548.Krodi_ | 0,6 | 0,8 | 3,13E-05 | 4,00E-05 | 2 - | 9,65E-05 - | 1 - | 5,99E-05 - | 0 - | 0 - | 0 - | 0 - | 0 - | 0 - |
| 7038 | 1 | 0.2um_NOKCPHJN_638581 | Bacteria | Proteobacteria | Alphaproteob | Pelagibacteral | Pelagibacterac | Candidatus Pe | Candidatus Pe | unclassified | unclassified | unclassified | unclassified | unclassified | 0,4 | 0,489897949 | 1,04E-05 | 1,29E-05 | 1 - | 2,21E-05 - | 0 - | 0 - | 0 - | 0 - | 0 - | 0 - | 1 - | 2,98E-05 - |
| 7039 | 1 | 0.2um_NOKCPHJN_615591 | Bacteria | Proteobacteria | Alphaproteob | unclassified | unclassified | unclassified | alpha proteob | metabolism | Inorganic ion t | 05F28 | tonB-depende | 247634.GPB2 | 1,4 | 0,489897949 | 2,35E-05 | 8,51E-06 | 2 - | 2,83E-05 - | 1 - | 1,76E-05 - | 1 - | 1,67E-05 - | 1 - | 1,69E-05 - | 2 - | 3,81E-05 - |
| 7040 | 1 | 3um_FIJJAFBF_450894 | Bacteria | Bacteroidetes | Flavobacteriia | Flavobacterial | Flavobacteriac | Hyunsoonleell | Hyunsoonleell | metabolism | Amino acid tra | 09GJ6 | nitrogen regul | 983548.Krodi_ | 1 | 0 | 0,000122346 | 1,16E-05 | 1 - | 0,000102479 - | 1 - | 0,000127249 - | 1 - | 0,000120966 - | 1 - | 0,000122753 - | 1 - | 0,000138281 - |
| 7041 | 1 | 3um_FIJJAFBF_65171 | Bacteria | Bacteroidetes | Flavobacteriia | Flavobacterial | Flavobacteriac | unclassified | Flavobacteriac | unclassified | unclassified | unclassified | unclassified | unclassified | 2,2 | 0,979795897 | 0,000265381 | 0,000121066 | 3 - | 0,000304717 - | 1 - | 0,000126123 - | 1 - | 0,000119896 - | 3 - | 0,000365 - | 3 - | 0,000411171 - |
| 7042 | 1 | 0.2um_NOKCPHJN_434594 | Eukaryota | Mollusca | Bivalvia | Pectinoida | Pectinidae | Mizuhopecter | Mizuhopecter | poorly charact | Function unkn | 0IFVP | chromosome | 67593.Physo1 | 0,2 | 0,4 | 2,14E-06 | 4,28E-06 | 0 - | 0 - | 0 - | 0 - | 1 - | 1,07E-05 - | 0 - | 0 - | 0 - | 0 - |
| 7043 | 1 | 3um_FIJJAFBF_713312 | Bacteria | Proteobacteria | Alphaproteob | Rhodobactera | Rhodobactera | Thalassobius | Thalassobius s | unclassified | unclassified | unclassified | unclassified | unclassified | 2,6 | 0,8 | 0,000118451 | 2,49E-05 | 4 - | 0,000156692 - | 2 - | 9,73E-05 - | 3 - | 0,000138719 - | 2 - | 9,38E-05 - | 2 - | 0,000105716 - |
| 7044 | 1 | 3um_FIJJAFBF_640756 | Bacteria | Bacteroidetes | Cytophagia | Cytophagales | Flammeovirga | unclassified | Flammeovirga | metabolism | Energy produc | COG0711 | F(1)F(0) ATP s | 643867.Ftrac_ | 3 | 0,894427191 | 0,000244911 | 5,79E-05 | 4 - | 0,000279943 - | 2 - | 0,000173803 - | 4 - | 0,000330444 - | 3 - | 0,000251494 - | 2 - | 0,000188871 - |
| 7045 | 1 | 0.2um_NOKCPHJN_439145 | Bacteria | Proteobacteria | Gammaprotec | Oceanospirilla | Saccharospirill | Reinekea | Reinekea fors | unclassified | unclassified | unclassified | unclassified | unclassified | 4,6 | 0,489897949 | 0,000102697 | 1,43E-05 | 5 - | 9,36E-05 - | 4 - | 9,30E-05 - | 4 - | 8,84E-05 - | 5 - | 0,00011214 - | 5 - | 0,000126325 - |
| 7046 | 1 | 3um_FIJJAFBF_141769 | Bacteria | Proteobacteria | Gammaprotec | Oceanospirilla | Saccharospirill | Reinekea | Reinekea fors | information st | Translation, ril | 05KK9 | This protein bi | 205922.Pfl01_ | 1,8 | 0,748331477 | 0,000246184 | 0,000120404 | 1 - | 0,000111434 - | 1 - | 0,000138368 - | 2 - | 0,000263072 - | 2 - | 0,000266958 - | 3 - | 0,000451091 - |
| 7047 | 1 | 3um_FIJJAFBF_45126 | Bacteria | Proteobacteria | Gammaprotec | Pseudomonad | Pseudomonad | Pseudomonas | Pseudomonas | metabolism | Amino acid tra | 0QR3G | Sarcosine oxid | 208964.PA541 | 0,2 | 0,4 | 1,13E-05 | 2,25E-05 | 1 - | 5,63E-05 - | 0 - | 0 - | 0 - | 0 - | 0 - | 0 - | 0 - | 0 - |
| 7048 | 1 | 3um_FIJJAFBF_554727 | Bacteria | Proteobacteria | Betaproteoba | Burkholderiale | Comamonada | Hydrogenoph | Hydrogenoph | poorly charact | Function unkn | COG3181 | extra-cytoplas | 384765.SIAM | 1,8 | 0,748331477 | 7,25E-05 | 2,60E-05 | 3 - | 0,000103402 - | 1 - | 4,28E-05 - | 1 - | 4,07E-05 - | 2 - | 8,26E-05 - | 2 - | 9,30E-05 - |
| 7049 | 1 | 0.2um_NOKCPHJN_159900 | Bacteria | Proteobacteria | Alphaproteob | Rhodobactera | Rhodobactera | Ruegeria | Ruegeria halo | metabolism | Amino acid tra | 01SW7 | Tetrahydrodip | 439496.RBY4I | 1,8 | 0,4 | 8,84E-05 | 1,75E-05 | 2 - | 8,35E-05 - | 2 - | 0,00010365 - | 2 - | 9,85E-05 - | 2 - | 1,00E-04 - | 1 - | 5,63E-05 - |
| 7050 | 1 | 3um_FIJJAFBF_91414 | Bacteria | Proteobacteria | Alphaproteob | Rhodobactera | Rhodobactera | Planktomarina | Planktomarina | metabolism | Energy produc | COG0711 | F(1)F(0) ATP s | 644107.SL115 | 3 | 0,894427191 | 0,000219423 | 5,91E-05 | 4 - | 0,000248166 - | 2 - | 0,000154074 - | 2 - | 0,000146467 - | 4 - | 0,000297261 - | 3 - | 0,000251148 - |
| 7051 | 1 | 3um_FIJJAFBF_143730 | unclassified | unclassified | unclassified | unclassified | unclassified | unclassified | unclassified | unclassified | unclassified | unclassified | unclassified | unclassified | 1 | 0 | 2,74E-05 | 2,60E-06 | 1 - | 2,29E-05 - | 1 - | 2,84E-05 - | 1 - | 2,70E-05 - | 1 - | 2,74E-05 - | 1 - | 3,09E-05 - |
| 7052 | 1 | 0.2um_NOKCPHJN_108377 | Bacteria | Proteobacteria | Gammaprotec | Oceanospirilla | Saccharospirill | Reinekea | Reinekea fors | cellular proces | Intracellular tr | 16Q4J | type VI secreti | 634503.NT01 | 0,6 | 0,489897949 | 1,90E-05 | 1,56E-05 | 0 - | 0 - | 1 - | 3,11E-05 - | 0 - | 0 - | 1 - | 3,00E-05 - | 1 - | 3,38E-05 - |
| 7053 | 1 | 0.2um_NOKCPHJN_516033 | Bacteria | Proteobacteria | Alphaproteob | Pelagibacteral | Pelagibacterac | Candidatus Pe | Candidatus Pe | metabolism | Energy produc | COG3474 | Electron carrie | 335992.SAR11 | 0,2 | 0,4 | 1,43E-05 | 2,87E-05 | 0 - | 0 - | 0 - | 0 - | 1 - | 7,17E-05 - | 0 - | 0 - | 0 - | 0 - |
| 7054 | 1 | 3um_FIJJAFBF_402118 | Viruses | unclassified | unclassified | unclassified | unclassified | unclassified | uncultured M | poorly charact | Function unkn | 08PAS | Phage tail tap | 880070.Cycma | 1 | 0 | 1,87E-05 | 1,77E-06 | 1 - | 1,57E-05 - | 1 - | 1,94E-05 - | 1 - | 1,85E-05 - | 1 - | 1,88E-05 - | 1 - | 2,11E-05 - |
| 7055 | 1 | 3um_FIJJAFBF_116680 | Bacteria | Bacteroidetes | Flavobacteriia | Flavobacterial | Flavobacteriac | Polaribacter | Polaribacter s | unclassified | unclassified | unclassified | unclassified | unclassified | 2,6 | 1,743559577 | 6,54E-05 | 4,61E-05 | 2 - | 4,17E-05 - | 6 - | 0,000155193 - | 2 - | 4,92E-05 - | 1 - | 2,50E-05 - | 2 - | 5,62E-05 - |
| 7056 | 1 | 3um_FIJJAFBF_420945 | Bacteria | Proteobacteria | Gammaprotec | Cellvibrionales | unclassified | unclassified | Cellvibrionales | unclassified | unclassified | unclassified | unclassified | unclassified | 4 | 0,894427191 | 0,000441542 | 7,19E-05 | 5 - | 0,000470397 - | 3 - | 0,000350456 - | 5 - | 0,000555255 - | 4 - | 0,000450765 - | 3 - | 0,000380839 - |
| 7057 | 1 | 3um_FIJJAFBF_16490 | Bacteria | Proteobacteria | Gammaprotec | Alteromonada | Alteromonada | Paraglaciecola | Paraglaciecola | information st | Translation, ril | 0QTKX | Seems to be tl | 1085623.GNIT | 6,4 | 1,959591794 | 0,000706122 | 0,000208957 | 7 - | 0,000653201 - | 5 - | 0,000579345 - | 5 - | 0,000550741 - | 10 - | 0,00111775 - | 5 - | 0,000629571 - |
| 7058 | 1 | 0.2um_NOKCPHJN_782759 | Bacteria | Proteobacteria | Alphaproteob | Pelagibacteral | Pelagibacterac | Candidatus Pe | Candidatus Pe | unclassified | unclassified | unclassified | unclassified | unclassified | 1,8 | 0,4 | 0,000213522 | 5,00E-05 | 2 - | 0,000199612 - | 1 - | 0,000123929 - | 2 - | 0,000235621 - | 2 - | 0,000239101 - | 2 - | 0,000269347 - |
| 7059 | 1 | 3um_FIJJAFBF_236095 | Bacteria | Bacteroidetes | Flavobacteriia | Flavobacterial | Flavobacteriac | Algibacter | Algibacter sp. | unclassified | unclassified | unclassified | unclassified | unclassified | 0,2 | 0,4 | 4,24E-06 | 8,47E-06 | 1 - | 2,12E-05 - | 0 - | 0 - | 0 - | 0 - | 0 - | 0 - | 0 - | 0 - |
| 7060 | 1 | 3um_FIJJAFBF_370854 | Bacteria | Bacteroidetes | Flavobacteriia | Flavobacterial | Flavobacteriac | Polaribacter | Polaribacter s | metabolism | Amino acid tra | 09GJ6 | nitrogen regul | 313598.MED1 | 0,8 | 0,4 | 9,78E-05 | 5,03E-05 | 1 - | 0,000102479 - | 1 - | 0,000127249 - | 1 - | 0,000120966 - | 0 - | 0 - | 1 - | 0,000138281 - |
| 7061 | 1 | 3um_FIJJAFBF_785995 | Bacteria | Proteobacteria | Betaproteoba | Nitrosomonad | Methylophilac | Methylotener | Methylotener | cellular proces | Cell wall/mem | 16PMX | Catalyzes the | 583345.Mmol | 0,2 | 0,4 | 7,97E-06 | 1,59E-05 | 1 - | 3,99E-05 - | 0 - | 0 - | 0 - | 0 - | 0 - | 0 - | 0 - | 0 - |
| 7062 | 1 | 0.2um_NOKCPHJN_737628 | Bacteria | Proteobacteria | Betaproteoba | Burkholderiale | Oxalobacterac | Collimonas | Collimonas sp | metabolism | Carbohydrate | 11GRQ | Xylose Isomer | 640510.BC10 | 0,4 | 0,489897949 | 2,00E-05 | 2,45E-05 | 0 - | 0 - | 1 - | 5,13E-05 - | 1 - | 4,87E-05 - | 0 - | 0 - | 0 - | 0 - |
| 7063 | 1 | 0.2um_NOKCPHJN_538589 | Bacteria | Bacteroidetes | Flavobacteriia | Flavobacterial | Flavobacteriac | Formosa | Formosa sp. H | unclassified | unclassified | unclassified | unclassified | unclassified | 10,2 | 1,469693846 | 0,000152254 | 3,31E-05 | 9 - | 0,000111554 - | 11 - | 0,000169299 - | 11 - | 0,00016094 - | 8 - | 0,000118776 - | 12 - | 0,000200701 - |
| 7064 | 1 | 0.2um_NOKCPHJN_124676 | Bacteria | Proteobacteria | Gammaprotec | Cellvibrionales | Porticoccacea | unclassified | Gammaprotec | unclassified | unclassified | unclassified | unclassified | unclassified | 1 | 0,894427191 | 1,85E-05 | 1,67E-05 | 2 - | 3,11E-05 - | 1 - | 1,93E-05 - | 0 - | 0 - | 0 - | 0 - | 2 - | 4,20E-05 - |
| 7065 | 1 | 3um_FIJJAFBF_29544 | Bacteria | Bacteroidetes | Flavobacteriia | unclassified | unclassified | unclassified | Flavobacteria | unclassified | unclassified | unclassified | unclassified | unclassified | 0,6 | 0,489897949 | 3,11E-05 | 2,56E-05 | 1 - | 4,61E-05 - | 0 - | 0 - | 1 - | 5,44E-05 - | 1 - | 5,52E-05 - | 0 - | 0 - |
| 7066 | 1 | 0.2um_NOKCPHJN_284253 | Bacteria | Proteobacteria | Gammaprotec | unclassified | unclassified | unclassified | Gammaprotec | information st | Translation, ril | COG0087 | One of the pri | 377629.TERTU | 1,2 | 1,166190379 | 6,92E-05 | 6,44E-05 | 3 - | 0,000155806 - | 2 - | 0,000128976 - | 1 - | 6,13E-05 - | 0 - | 0 - | 0 - | 0 - |
| 7067 | 1 | 3um_FIJJAFBF_777837 | Bacteria | Bacteroidetes | Flavobacteriia | Flavobacterial | Flavobacteriac | Polaribacter | Polaribacter s | unclassified | unclassified | unclassified | unclassified | unclassified | 4,2 | 1,720465053 | 0,000121435 | 4,97E-05 | 5 - | 0,000123949 - | 6 - | 0,00018469 - | 5 - | 0,000146309 - | 4 - | 0,000118776 - | 1 - | 3,35E-05 - |
| 7068 | 1 | 0.2um_NOKCPHJN_38455 | Bacteria | Proteobacteria | Gammaprotec | Oceanospirilla | Oceanospirilla | Motiliproteus | Motiliproteus | cellular proces | Posttranslatio | 0QJ3C | Acts as a chap | 491952.Mar18 | 0,4 | 0,8 | 7,15E-06 | 1,43E-05 | 2 - | 3,58E-05 - | 0 - | 0 - | 0 - | 0 - | 0 - | 0 - | 0 - | 0 - |
| 7069 | 1 | 0.2um_NOKCPHJN_180010 | Bacteria | Proteobacteria | Alphaproteob | Rhodospirillale | Rhodospirillac | unclassified | Rhodospirillac | unclassified | unclassified | unclassified | unclassified | unclassified | 1,8 | 0,748331477 | 7,73E-05 | 3,12E-05 | 2 - | 7,33E-05 - | 1 - | 4,55E-05 - | 2 - | 8,66E-05 - | 3 - | 0,000131773 - | 1 - | 4,95E-05 - |
| 7070 | 1 | 3um_FIJJAFBF_525960 | Bacteria | Bacteroidetes | Cytophagia | Cytophagales | Flammeovirga | unclassified | Flammeovirga | unclassified | unclassified | unclassified | unclassified | unclassified | 0,8 | 0,4 | 8,29E-05 | 4,21E-05 | 1 - | 8,97E-05 - | 1 - | 0,000111343 - | 1 - | 0,000105845 - | 1 - | 0,000107409 - | 0 - | 0 - |
| 7071 | 1 | 3um_FIJJAFBF_217623 | Bacteria | Bacteroidetes | Flavobacteriia | Flavobacterial | Flavobacteriac | Psychroserper | Psychroserper | unclassified | unclassified | unclassified | unclassified | unclassified | 1 | 0,632455532 | 3,61E-05 | 2,34E-05 | 1 - | 3,11E-05 - | 1 - | 3,86E-05 - | 2 - | 7,34E-05 - | 1 - | 3,73E-05 - | 0 - | 0 - |
| 7072 | 1 | 3um_FIJJAFBF_121845 | Bacteria | Bacteroidetes | Cytophagia | Cytophagales | Flammeovirga | unclassified | Flammeovirga | cellular proces | Cell wall/mem | 0FX0M | TonB protein | 643867.Ftrac_ | 0,4 | 0,489897949 | 2,30E-05 | 2,84E-05 | 1 - | 5,12E-05 - | 1 - | 6,36E-05 - | 0 - | 0 - | 0 - | 0 - | 0 - | 0 - |
| 7073 | 1 | 0.2um_NOKCPHJN_94098 | Bacteria | Bacteroidetes | Flavobacteriia | Flavobacterial | Flavobacteriac | Aureitalea | Aureitalea ma | unclassified | unclassified | unclassified | unclassified | unclassified | 0,2 | 0,4 | 5,17E-06 | 1,03E-05 | 0 - | 0 - | 0 - | 0 - | 0 - | 0 - | 1 - | 2,58E-05 - | 0 - | 0 - |
| 7074 | 1 | 3um_FIJJAFBF_778062 | Bacteria | Bacteroidetes | Flavobacteriia | Flavobacterial | Flavobacteriac | Polaribacter | Polaribacter s | metabolism | Energy produc | 0P9HY | cytochrome C | 313598.MED1 | 0,8 | 0,748331477 | 5,52E-05 | 4,88E-05 | 2 - | 0,000126128 - | 0 - | 0 - | 1 - | 7,44E-05 - | 1 - | 7,55E-05 - | 0 - | 0 - |
| 7075 | 1 | 3um_FIJJAFBF_106924 | Bacteria | Proteobacteria | Alphaproteob | Rhodobactera | Rhodobactera | Yoonia | Yoonia marico | unclassified | unclassified | unclassified | unclassified | unclassified | 0,2 | 0,4 | 3,86E-06 | 7,72E-06 | 0 - | 0 - | 0 - | 0 - | 1 - | 1,93E-05 - | 0 - | 0 - | 0 - | 0 - |
| 7076 | 1 | 3um_FIJJAFBF_516772 | Bacteria | Bacteroidetes | Flavobacteriia | Flavobacterial | unclassified | unclassified | Flavobacterial | poorly charact | Function unkn | 0XSVM | NA | 398720.MED2 | 6,8 | 1,166190379 | 0,000198008 | 4,51E-05 | 7 - | 0,00016986 - | 8 - | 0,000241047 - | 6 - | 0,000171859 - | 5 - | 0,000145331 - | 8 - | 0,000261944 - |
| 7077 | 1 | 3um_FIJJAFBF_333522 | Bacteria | Proteobacteria | Alphaproteob | Rhodobactera | Rhodobactera | Phaeobacter | various | unclassified | unclassified | unclassified | unclassified | unclassified | 0,4 | 0,489897949 | 1,19E-05 | 1,46E-05 | 1 - | 2,72E-05 - | 0 - | 0 - | 1 - | 3,21E-05 - | 0 - | 0 - | 0 - | 0 - |
| 7078 | 1 | 3um_FIJJAFBF_280809 | Bacteria | Bacteroidetes | Flavobacteriia | Flavobacterial | Flavobacteriac | Arenitalea | Arenitalea lut | cellular proces | Cell motility | COG1360 | Ompa motb d | 391587.KAOT | 8,6 | 1,2 | 0,000426216 | 9,61E-05 | 8 - | 0,000329109 - | 8 - | 0,000408656 - | 8 - | 0,00038848 - | 8 - | 0,000394217 - | 11 - | 0,000610616 - |
| 7079 | 1 | 3um_FIJJAFBF_668122 | unclassified | unclassified | unclassified | unclassified | unclassified | unclassified | unclassified | unclassified | unclassified | unclassified | unclassified | unclassified | 0,2 | 0,4 | 1,91E-05 | 3,82E-05 | 0 - | 0 - | 0 - | 0 - | 1 - | 9,54E-05 - | 0 - | 0 - | 0 - | 0 - |
| 7080 | 1 | 0.2um_NOKCPHJN_842736 | Eukaryota | Basidiomycota | Tremellomyce | Cystofilobasid | Mrakiaceae | Xanthophyllor | Xanthophyllor | information st | Translation, ril | 12P22 | ribosomal pro | 5297.EFP9156 | 0,8 | 0,4 | 6,20E-05 | 3,15E-05 | 1 - | 6,71E-05 - | 1 - | 8,33E-05 - | 1 - | 7,92E-05 - | 1 - | 8,04E-05 - | 0 - | 0 - |
| 7081 | 1 | 0.2um_NOKCPHJN_492828 | Bacteria | Bacteroidetes | Flavobacteriia | Flavobacterial | Flavobacteriac | Formosa | Formosa sp. H | unclassified | unclassified | unclassified | unclassified | unclassified | 1 | 0 | 4,49E-05 | 4,27E-06 | 1 - | 3,76E-05 - | 1 - | 4,67E-05 - | 1 - | 4,44E-05 - | 1 - | 4,51E-05 - | 1 - | 5,08E-05 - |
| 7082 | 1 | 3um_FIJJAFBF_165773 | Bacteria | Proteobacteria | Alphaproteob | Rhodobactera | Rhodobactera | unclassified | Rhodobactera | metabolism | Amino acid tra | 16V6G | Glycine betain | 367336.OM22 | 0,4 | 0,489897949 | 1,83E-05 | 2,25E-05 | 0 - | 0 - | 0 - | 0 - | 0 - | 0 - | 1 - | 4,31E-05 - | 1 - | 4,85E-05 - |
| 7083 | 1 | 0.2um_NOKCPHJN_260532 | Eukaryota | Tardigrada | Eutardigrada | Parachela | Ramazzottiida | Ramazzottius | Ramazzottius | unclassified | unclassified | unclassified | unclassified | unclassified | 1 | 0 | 9,32E-05 | 8,85E-06 | 1 - | 7,81E-05 - | 1 - | 9,70E-05 - | 1 - | 9,22E-05 - | 1 - | 9,35E-05 - | 1 - | 0,000105357 - |
| 7084 | 1 | 0.2um_NOKCPHJN_25109 | Bacteria | Bacteroidetes | Flavobacteriia | Flavobacterial | Flavobacteriac | Ulvibacter | Ulvibacter sp. | information st | Translation, ril | 05CAV | thus facilitatin | 313590.MED1 | 3,6 | 0,8 | 7,71E-05 | 1,78E-05 | 4 - | 7,21E-05 - | 5 - | 0,000111867 - | 3 - | 6,38E-05 - | 3 - | 6,47E-05 - | 3 - | 7,29E-05 - |
| 7085 | 1 | 3um_FIJJAFBF_50494 | Bacteria | Bacteroidetes | Flavobacteriia | Flavobacterial | Flavobacteriac | Polaribacter | Polaribacter h | unclassified | unclassified | unclassified | unclassified | unclassified | 1,6 | 0,489897949 | 0,00017418 | 4,92E-05 | 2 - | 0,000185124 - | 2 - | 0,000229869 - | 1 - | 0,00010926 - | 2 - | 0,000221747 - | 1 - | 0,000124899 - |
| 7086 | 1 | 0.2um_NOKCPHJN_486785 | Bacteria | Proteobacteria | Gammaprotec | unclassified | unclassified | unclassified | various | unclassified | unclassified | unclassified | unclassified | unclassified | 1 | 0 | 3,48E-05 | 3,30E-06 | 1 - | 2,91E-05 - | 1 - | 3,62E-05 - | 1 - | 3,44E-05 - | 1 - | 3,49E-05 - | 1 - | 3,93E-05 - |
| 7087 | 1 | 3um_FIJJAFBF_470659 | Eukaryota | unclassified | Cryptophyta | Pyrenomonad | Geminigerace | Guillardia | Guillardia thet | metabolism | Energy produc | 0FN1Q | F(1)F(0) ATP s | 316278.SynRC | 1 | 1,095445115 | 6,80E-05 | 6,90E-05 | 3 - | 0,000188159 - | 1 - | 7,79E-05 - | 1 - | 7,40E-05 - | 0 - | 0 - | 0 - | 0 - |
| 7088 | 1 | 3um_FIJJAFBF_82350 | Bacteria | Bacteroidetes | Flavobacteriia | Flavobacterial | Flavobacteriac | Polaribacter | Polaribacter s | information st | Translation, ril | COG0254 | 50S ribosomal | 313594.PI23P | 1,8 | 0,4 | 0,000293997 | 7,32E-05 | 2 - | 0,000273278 - | 2 - | 0,000339331 - | 1 - | 0,000161288 - | 2 - | 0,000327341 - | 2 - | 0,000368749 - |
| 7089 | 1 | 3um_FIJJAFBF_578540 | Bacteria | Proteobacteria | Gammaprotec | Alteromonada | Alteromonada | Paraglaciecola | various | information st | Translation, ril | COG0292 | Binds directly | 342610.Patl_2 | 1,8 | 0,4 | 0,000205999 | 4,07E-05 | 2 - | 0,000194537 - | 2 - | 0,000241557 - | 2 - | 0,000229631 - | 2 - | 0,000233023 - | 1 - | 0,00013125 - |
| 7090 | 1 | 3um_FIJJAFBF_629963 | Bacteria | Proteobacteria | Alphaproteob | Rhodobactera | Rhodobactera | Amylibacter | Amylibacter k | metabolism | Inorganic ion t | 05CRM | ABC transport | 367336.OM22 | 1 | 0,894427191 | 3,93E-05 | 3,46E-05 | 2 - | 6,98E-05 - | 1 - | 4,33E-05 - | 0 - | 0 - | 2 - | 8,36E-05 - | 0 - | 0 - |
| 7091 | 1 | 0.2um_NOKCPHJN_221102 | Bacteria | Proteobacteria | Gammaprotec | Cellvibrionales | Halieaceae | Congregibacte | Congregibacte | poorly charact | Function unkn | 0XNQ8 | alkaline phosp | 314285.KT71_ | 8,6 | 1,019803903 | 0,000191206 | 3,93E-05 | 7 - | 0,000128963 - | 9 - | 0,000205886 - | 8 - | 0,000173974 - | 9 - | 0,000198612 - | 10 - | 0,000248595 - |
| 7092 | 1 | 0.2um_NOKCPHJN_898820 | Bacteria | Bacteroidetes | Flavobacteriia | Flavobacterial | Flavobacteriac | unclassified | Flavobacteriac | unclassified | unclassified | unclassified | unclassified | unclassified | 0,8 | 0,4 | 6,96E-05 | 3,57E-05 | 1 - | 7,36E-05 - | 0 - | 0 - | 1 - | 8,68E-05 - | 1 - | 8,81E-05 - | 1 - | 9,93E-05 - |
| 7093 | 1 | 3um_FIJJAFBF_541856 | Bacteria | Verrucomicrol | Opitutae | unclassified | unclassified | unclassified | Opitutae bact | unclassified | unclassified | unclassified | unclassified | unclassified | 1,4 | 0,489897949 | 0,000251315 | 7,90E-05 | 2 - | 0,000306071 - | 2 - | 0,00038005 - | 1 - | 0,000180643 - | 1 - | 0,000183311 - | 1 - | 0,000206499 - |
| 7094 | 1 | 0.2um_NOKCPHJN_04094 | Bacteria | Bacteroidetes | Cytophagia | Cytophagales | Cytophagacea | Ekhidna | Ekhidna lutea | poorly charact | Function unkn | 09S7H | NA | 643867.Ftrac_ | 2,8 | 0,4 | 0,000194713 | 3,39E-05 | 3 - | 0,000174787 - | 3 - | 0,000217034 - | 3 - | 0,000206318 - | 2 - | 0,000139577 - | 3 - | 0,000235849 - |
| 7095 | 1 | 3um_FIJJAFBF_10655 | Bacteria | Bacteroidetes | Flavobacteriia | Flavobacterial | Flavobacteriac | Flavobacteriu | Flavobacteriu | unclassified | unclassified | unclassified | unclassified | unclassified | 3,6 | 1,019803903 | 9,03E-05 | 2,29E-05 | 5 - | 0,000106472 - | 2 - | 5,29E-05 - | 3 - | 7,54E-05 - | 4 - | 0,000102028 - | 4 - | 0,000114935 - |
| 7096 | 1 | 0.2um_NOKCPHJN_93915 | Bacteria | Bacteroidetes | Flavobacteriia | Flavobacterial | Flavobacteriac | Cellulophaga | Cellulophaga l | unclassified | unclassified | unclassified | unclassified | unclassified | 0,2 | 0,4 | 2,98E-06 | 5,97E-06 | 0 - | 0 - | 1 - | 1,49E-05 - | 0 - | 0 - | 0 - | 0 - | 0 - | 0 - |
| 7097 | 1 | 3um_FIJJAFBF_313320 | Bacteria | Proteobacteria | Gammaprotec | Oceanospirilla | Saccharospirill | Reinekea | Reinekea blan | metabolism | Nucleotide tra | 16PIX | Plays an impo | 314283.MED2 | 1,8 | 0,4 | 5,74E-05 | 1,43E-05 | 2 - | 5,34E-05 - | 2 - | 6,63E-05 - | 1 - | 3,15E-05 - | 2 - | 6,39E-05 - | 2 - | 7,20E-05 - |
| 7098 | 1 | 3um_FIJJAFBF_63332 | Bacteria | Proteobacteria | Gammaprotec | Alteromonada | Alteromonada | Glaciecola | Glaciecola sp. | metabolism | Inorganic ion t | 0QJVD | TonB-depende | 455436.GHTC | 18,8 | 1,720465053 | 0,000293322 | 7,93E-06 | 22 - | 0,000289907 - | 18 - | 0,000294528 - | 18 - | 0,000279986 - | 19 - | 0,000299906 - | 17 - | 0,000302281 - |
| 7099 | 1 | 3um_FIJJAFBF_324098 | Bacteria | Proteobacteria | Gammaprotec | Alteromonada | Alteromonada | Paraglaciecola | Paraglaciecola | information st | Translation, ril | 08UHH | Located on th | 342610.Patl_1 | 2,4 | 0,489897949 | 0,000252336 | 4,46E-05 | 3 - | 0,000266923 - | 3 - | 0,000331439 - | 2 - | 0,00021005 - | 2 - | 0,000213152 - | 2 - | 0,000240115 - |
| 7100 | 1 | 3um_FIJJAFBF_48476 | Bacteria | Bacteroidetes | Flavobacteriia | Flavobacterial | Flavobacteriac | Algibacter | Algibacter sp. | poorly charact | Function unkn | 0900R | pur-alpha bet | 1046627.BZAF | 0,4 | 0,489897949 | 4,53E-05 | 5,57E-05 | 0 - | 0 - | 0 - | 0 - | 0 - | 0 - | 1 - | 0,000106576 - | 1 - | 0,000120058 - |
| 7101 | 1 | 0.2um_NOKCPHJN_15493 | Bacteria | Bacteroidetes | Flavobacteriia | Flavobacterial | Flavobacteriac | Polaribacter | Polaribacter ir | unclassified | unclassified | unclassified | unclassified | unclassified | 8,6 | 0,8 | 0,000785208 | 0,000138429 | 8 - | 0,000608089 - | 8 - | 0,000755067 - | 8 - | 0,000717787 - | 9 - | 0,000819437 - | 10 - | 0,001025659 - |
| 7102 | 1 | 0.2um_NOKCPHJN_345611 | Eukaryota | unclassified | Labyrinthulom | unclassified | Thraustochytr | Aurantiochytr | Aurantiochytr | poorly charact | Function unkn | 0XNR8 | NIPA-like dom | 164328.Phyra | 0,2 | 0,4 | 2,17E-06 | 4,34E-06 | 1 - | 1,09E-05 - | 0 - | 0 - | 0 - | 0 - | 0 - | 0 - | 0 - | 0 - |
| 7103 | 1 | 3um_FIJJAFBF_80731 | Bacteria | Proteobacteria | Gammaprotec | Alteromonada | unclassified | unclassified | gamma protec | information st | Translation, ril | 05KK9 | This protein bi | 523791.Kkor_ | 4,4 | 1,356465997 | 0,000309069 | 8,26E-05 | 6 - | 0,000358677 - | 3 - | 0,000222686 - | 3 - | 0,000211691 - | 6 - | 0,000429635 - | 4 - | 0,000322655 - |
| 7104 | 1 | 0.2um_NOKCPHJN_822445 | Bacteria | Bacteroidetes | Flavobacteriia | Flavobacterial | Flavobacteriac | Polaribacter | Polaribacter s | unclassified | unclassified | unclassified | unclassified | unclassified | 0,8 | 0,4 | 1,48E-05 | 7,47E-06 | 0 - | 0 - | 1 - | 1,85E-05 - | 1 - | 1,76E-05 - | 1 - | 1,79E-05 - | 1 - | 2,01E-05 - |
| 7105 | 1 | 3um_FIJJAFBF_379821 | Bacteria | Proteobacteria | Betaproteoba | unclassified | unclassified | unclassified | Betaproteoba | unclassified | unclassified | unclassified | unclassified | unclassified | 4,6 | 1,854723699 | 7,71E-05 | 2,64E-05 | 7 - | 0,000101572 - | 5 - | 9,01E-05 - | 6 - | 0,000102768 - | 3 - | 5,21E-05 - | 2 - | 3,92E-05 - |
| 7106 | 1 | 0.2um_NOKCPHJN_57134 | Bacteria | Proteobacteria | Gammaprotec | Oceanospirilla | Saccharospirill | Reinekea | Reinekea fors | unclassified | unclassified | unclassified | unclassified | unclassified | 4,2 | 0,748331477 | 0,000102345 | 1,93E-05 | 4 - | 8,20E-05 - | 5 - | 0,000127249 - | 4 - | 9,68E-05 - | 5 - | 0,000122753 - | 3 - | 8,30E-05 - |
| 7107 | 1 | 0.2um_NOKCPHJN_449496 | Bacteria | Proteobacteria | Gammaprotec | unclassified | unclassified | unclassified | Gammaprotec | unclassified | unclassified | unclassified | unclassified | unclassified | 1 | 0,894427191 | 4,95E-05 | 4,37E-05 | 2 - | 8,90E-05 - | 0 - | 0 - | 2 - | 0,000105025 - | 1 - | 5,33E-05 - | 0 - | 0 - |
| 7108 | 1 | 0.2um_NOKCPHJN_137623 | Eukaryota | Ascomycota | Saccharomyce | Saccharomyce | Saccharomyce | unclassified | Saccharomyce | unclassified | unclassified | unclassified | unclassified | unclassified | 0,4 | 0,489897949 | 8,26E-06 | 1,01E-05 | 0 - | 0 - | 1 - | 2,12E-05 - | 1 - | 2,01E-05 - | 0 - | 0 - | 0 - | 0 - |
| 7109 | 1 | 3um_FIJJAFBF_387044 | Bacteria | Proteobacteria | Alphaproteob | Rhodobactera | unclassified | unclassified | Rhodobactera | poorly charact | Function unkn | 0XR8Q | NA | 439496.RBY4I | 2 | 0,632455532 | 5,25E-05 | 1,21E-05 | 3 - | 6,79E-05 - | 2 - | 5,62E-05 - | 2 - | 5,34E-05 - | 2 - | 5,42E-05 - | 1 - | 3,05E-05 - |
| 7110 | 1 | 3um_FIJJAFBF_313269 | Bacteria | Bacteroidetes | Flavobacteriia | Flavobacterial | Flavobacteriac | Polaribacter | various | poorly charact | Function unkn | 0XQTX | Chromosome | 313598.MED1 | 1 | 0 | 1,32E-05 | 1,26E-06 | 1 - | 1,11E-05 - | 1 - | 1,38E-05 - | 1 - | 1,31E-05 - | 1 - | 1,33E-05 - | 1 - | 1,49E-05 - |
| 7111 | 1 | 0.2um_NOKCPHJN_34376 | Bacteria | Bacteroidetes | unclassified | unclassified | unclassified | unclassified | Bacteroidetes | unclassified | unclassified | unclassified | unclassified | unclassified | 1,8 | 0,4 | 3,91E-05 | 9,58E-06 | 2 - | 3,64E-05 - | 2 - | 4,52E-05 - | 2 - | 4,29E-05 - | 1 - | 2,18E-05 - | 2 - | 4,91E-05 - |
| 7112 | 1 | 3um_FIJJAFBF_102914 | Bacteria | Bacteroidetes | Flavobacteriia | Flavobacterial | Flavobacteriac | Formosa | Formosa sp. H | cellular proces | Posttranslatio | COG1404 | peptidase (S8 | 391603.FBALC | 0,2 | 0,4 | 4,28E-06 | 8,57E-06 | 1 - | 2,14E-05 - | 0 - | 0 - | 0 - | 0 - | 0 - | 0 - | 0 - | 0 - |
| 7113 | 1 | 0.2um_NOKCPHJN_212464 | unclassified | unclassified | unclassified | unclassified | unclassified | unclassified | unclassified | unclassified | unclassified | unclassified | unclassified | unclassified | 0,6 | 0,8 | 3,78E-05 | 5,23E-05 | 1 - | 5,63E-05 - | 0 - | 0 - | 2 - | 0,000132826 - | 0 - | 0 - | 0 - | 0 - |
| 7114 | 1 | 0.2um_NOKCPHJN_461488 | Bacteria | Bacteroidetes | Flavobacteriia | unclassified | unclassified | unclassified | Flavobacteria | metabolism | Inorganic ion t | 05C27 | TonB-depende | 487797.Flav3 | 14,4 | 3,006659276 | 0,000198075 | 3,99E-05 | 15 - | 0,000173904 - | 12 - | 0,00017275 - | 20 - | 0,000273701 - | 12 - | 0,000166646 - | 13 - | 0,000203371 - |
| 7115 | 1 | 3um_FIJJAFBF_540106 | unclassified | unclassified | unclassified | unclassified | unclassified | unclassified | unclassified | unclassified | unclassified | unclassified | unclassified | unclassified | 0,8 | 0,4 | 0,000151503 | 7,69E-05 | 1 - | 0,000163967 - | 1 - | 0,000203598 - | 1 - | 0,000193546 - | 1 - | 0,000196405 - | 0 - | 0 - |
| 7116 | 1 | 3um_FIJJAFBF_28925 | Bacteria | various | various | various | various | various | various | poorly charact | Function unkn | 11PIQ | YCE I like fami | 313594.PI23P | 4 | 1,095445115 | 0,000255023 | 4,51E-05 | 6 - | 0,000327934 - | 4 - | 0,000271465 - | 4 - | 0,000258061 - | 3 - | 0,000196405 - | 3 - | 0,000221249 - |
| 7117 | 1 | 3um_FIJJAFBF_44695 | Bacteria | Bacteroidetes | Flavobacteriia | Flavobacterial | Flavobacteriac | Polaribacter | Polaribacter s | unclassified | unclassified | unclassified | unclassified | unclassified | 0,4 | 0,489897949 | 2,13E-05 | 2,62E-05 | 1 - | 4,84E-05 - | 0 - | 0 - | 0 - | 0 - | 1 - | 5,80E-05 - | 0 - | 0 - |
| 7118 | 1 | 3um_FIJJAFBF_195324 | unclassified | unclassified | unclassified | unclassified | unclassified | unclassified | unclassified | unclassified | unclassified | unclassified | unclassified | unclassified | 11,2 | 2,4 | 0,000311376 | 7,55E-05 | 13 - | 0,000302657 - | 13 - | 0,00037581 - | 10 - | 0,000274812 - | 7 - | 0,00019521 - | 13 - | 0,000408391 - |
| 7119 | 1 | 3um_FIJJAFBF_374333 | Bacteria | Proteobacteria | unclassified | unclassified | unclassified | unclassified | Proteobacteria | metabolism | Inorganic ion t | 05CRM | ABC transport | 367336.OM22 | 1,6 | 1,2 | 6,35E-05 | 4,46E-05 | 3 - | 0,000104343 - | 1 - | 4,32E-05 - | 3 - | 0,000123166 - | 0 - | 0 - | 1 - | 4,69E-05 - |
| 7120 | 1 | 0.2um_NOKCPHJN_198756 | Bacteria | Proteobacteria | Alphaproteob | Rhodobactera | Rhodobactera | Planktomarina | Planktomarina | metabolism | Carbohydrate | 01QS3 | extracellular s | 388401.RB215 | 6,2 | 0,748331477 | 0,000181053 | 2,16E-05 | 7 - | 0,000172042 - | 6 - | 0,000183108 - | 5 - | 0,000145056 - | 7 - | 0,000206078 - | 6 - | 0,000198982 - |
| 7121 | 1 | 3um_FIJJAFBF_726535 | Bacteria | Proteobacteria | Alphaproteob | Rhodobactera | Rhodobactera | various | various | poorly charact | Function unkn | COG3181 | extra-cytoplas | 744980.TRICH | 0,4 | 0,489897949 | 1,58E-05 | 1,95E-05 | 1 - | 3,63E-05 - | 0 - | 0 - | 1 - | 4,29E-05 - | 0 - | 0 - | 0 - | 0 - |
| 7122 | 1 | 0.2um_NOKCPHJN_53956 | Bacteria | Bacteroidetes | Flavobacteriia | unclassified | unclassified | unclassified | Flavobacteria | unclassified | unclassified | unclassified | unclassified | unclassified | 0,4 | 0,489897949 | 1,18E-05 | 1,45E-05 | 0 - | 0 - | 0 - | 0 - | 1 - | 2,93E-05 - | 1 - | 2,98E-05 - | 0 - | 0 - |
| 7123 | 1 | 3um_FIJJAFBF_684171 | Bacteria | Bacteroidetes | Flavobacteriia | Flavobacterial | Flavobacteriac | Polaribacter | Polaribacter s | unclassified | unclassified | unclassified | unclassified | unclassified | 0,2 | 0,4 | 1,54E-05 | 3,08E-05 | 1 - | 7,70E-05 - | 0 - | 0 - | 0 - | 0 - | 0 - | 0 - | 0 - | 0 - |
| 7124 | 1 | 3um_FIJJAFBF_170599 | Bacteria | Bacteroidetes | Flavobacteriia | Flavobacterial | Flavobacteriac | Formosa | Formosa sp. H | metabolism | Carbohydrate | 05CA9 | Pyruvate kina | 1046627.BZAF | 1,6 | 0,489897949 | 4,53E-05 | 1,45E-05 | 2 - | 4,76E-05 - | 1 - | 2,96E-05 - | 1 - | 2,81E-05 - | 2 - | 5,70E-05 - | 2 - | 6,43E-05 - |
| 7125 | 2 | 3um_FIJJAFBF_139394;3um_FIJJAFBF_171312 | Bacteria | Proteobacteria | Alphaproteob | Rhodobactera | Rhodobactera | Planktomarina | Planktomarina | poorly charact | Function unkn | 0248J | NA | various | 0,2 | 0,4 | 3,64E-05 | 7,29E-05 | 1 - | 0,000182185 - | 0 - | 0 - | 0 - | 0 - | 0 - | 0 - | 0 - | 0 - |
| 7126 | 1 | 3um_FIJJAFBF_266560 | Bacteria | Bacteroidetes | Flavobacteriia | unclassified | unclassified | unclassified | Flavobacteria | unclassified | unclassified | unclassified | unclassified | unclassified | 35,8 | 2,481934729 | 0,000547689 | 5,64E-05 | 38 - | 0,000487865 - | 39 - | 0,000621727 - | 35 - | 0,000530411 - | 32 - | 0,00049211 - | 35 - | 0,000606332 - |
| 7127 | 1 | 0.2um_NOKCPHJN_727290 | Bacteria | unclassified | unclassified | unclassified | unclassified | unclassified | uncultured ba | unclassified | unclassified | unclassified | unclassified | unclassified | 2,8 | 0,4 | 4,85E-05 | 5,77E-06 | 3 - | 4,40E-05 - | 3 - | 5,46E-05 - | 3 - | 5,19E-05 - | 3 - | 5,27E-05 - | 2 - | 3,96E-05 - |
| 7128 | 1 | 0.2um_NOKCPHJN_891067 | Bacteria | Bacteroidetes | Flavobacteriia | unclassified | unclassified | unclassified | Flavobacteria | poorly charact | Function unkn | 0XP53 | Ragb susd dor | 487797.Flav3 | 1,6 | 0,489897949 | 0,000136311 | 4,51E-05 | 2 - | 0,00014258 - | 2 - | 0,000177042 - | 1 - | 8,42E-05 - | 1 - | 8,54E-05 - | 2 - | 0,000192391 - |
| 7129 | 1 | 0.2um_NOKCPHJN_794161 | Bacteria | Actinobacteria | Actinobacteria | Micrococcales | Microbacteria | Rhodoluna | Rhodoluna lac | unclassified | unclassified | unclassified | unclassified | unclassified | 1,2 | 0,748331477 | 1,94E-05 | 1,14E-05 | 2 - | 2,78E-05 - | 1 - | 1,73E-05 - | 0 - | 0 - | 2 - | 3,33E-05 - | 1 - | 1,88E-05 - |
| 7130 | 1 | 3um_FIJJAFBF_257125 | Bacteria | Cyanobacteria | unclassified | Oscillatoriales | Microcoleacea | Arthrospira | various | unclassified | unclassified | unclassified | unclassified | unclassified | 5 | 0,632455532 | 0,000381791 | 3,65E-05 | 6 - | 0,000386888 - | 4 - | 0,000320267 - | 5 - | 0,000380568 - | 5 - | 0,000386189 - | 5 - | 0,000435041 - |
| 7131 | 1 | 3um_FIJJAFBF_136320 | Bacteria | Bacteroidetes | Flavobacteriia | Flavobacterial | Flavobacteriac | Formosa | Formosa sp. H | unclassified | unclassified | unclassified | unclassified | unclassified | 0,2 | 0,4 | 1,10E-05 | 2,20E-05 | 1 - | 5,49E-05 - | 0 - | 0 - | 0 - | 0 - | 0 - | 0 - | 0 - | 0 - |
| 7132 | 1 | 0.2um_NOKCPHJN_64664 | Bacteria | Bacteroidetes | Cytophagia | Cytophagales | Flammeovirga | Reichenbachie | Reichenbachie | metabolism | Inorganic ion t | COG0841 | acriflavin resis | 926562.Oweh | 0,4 | 0,489897949 | 4,81E-06 | 5,94E-06 | 1 - | 1,10E-05 - | 0 - | 0 - | 0 - | 0 - | 1 - | 1,31E-05 - | 0 - | 0 - |
| 7133 | 1 | 3um_FIJJAFBF_332654 | Bacteria | Proteobacteria | Alphaproteob | Pelagibacteral | Pelagibacterac | Candidatus Pe | Candidatus Pe | unclassified | unclassified | unclassified | unclassified | unclassified | 5 | 1,414213562 | 0,000185267 | 5,01E-05 | 7 - | 0,00021892 - | 5 - | 0,000194167 - | 4 - | 0,000147665 - | 3 - | 0,000112384 - | 6 - | 0,000253201 - |
| 7134 | 1 | 0.2um_NOKCPHJN_51693 | Bacteria | Bacteroidetes | Flavobacteriia | Flavobacterial | Flavobacteriac | Tenacibaculur | Tenacibaculur | metabolism | Amino acid tra | 05C6T | cysteine synth | 313594.PI23P | 1,2 | 0,4 | 5,39E-05 | 1,19E-05 | 2 - | 7,73E-05 - | 1 - | 4,80E-05 - | 1 - | 4,56E-05 - | 1 - | 4,63E-05 - | 1 - | 5,21E-05 - |
| 7135 | 1 | 3um_FIJJAFBF_189665 | Bacteria | Bacteroidetes | Flavobacteriia | Flavobacterial | Flavobacteriac | Polaribacter | Polaribacter s | metabolism | Lipid transpor | COG0332 | Catalyzes the | 313594.PI23P | 2,6 | 0,489897949 | 0,000106529 | 1,73E-05 | 3 - | 0,000104027 - | 3 - | 0,000129171 - | 3 - | 0,000122794 - | 2 - | 8,31E-05 - | 2 - | 9,36E-05 - |
| 7136 | 6 | 0.2um_NOKCPHJN_16550;0.2um_NOKCPHJN_54 | Bacteria | Bacteroidetes | Flavobacteriia | Flavobacterial | Flavobacteriac | Formosa | Formosa sp. H | poorly charact | Function unkn | 11RX2 | NA | 983544.Lacal_ | 1 | 0 | 8,06E-05 | 7,65E-06 | 1 - | 6,75E-05 - | 1 - | 8,38E-05 - | 1 - | 7,97E-05 - | 1 - | 8,09E-05 - | 1 - | 9,11E-05 - |
| 7137 | 1 | 0.2um_NOKCPHJN_530416 | Bacteria | Proteobacteria | Gammaprotec | unclassified | unclassified | unclassified | Gammaprotec | information st | Translation, ril | 17FIH | One of the ea | 322710.Avin_ | 1,2 | 0,748331477 | 0,000161194 | 9,38E-05 | 2 - | 0,000231872 - | 1 - | 0,000143958 - | 2 - | 0,000273701 - | 0 - | 0 - | 1 - | 0,000156439 - |
| 7138 | 1 | 0.2um_NOKCPHJN_391827 | Bacteria | Proteobacteria | Alphaproteob | unclassified | unclassified | Candidatus Pu | Candidatus Pu | unclassified | unclassified | unclassified | unclassified | unclassified | 0,8 | 0,4 | 4,37E-05 | 2,20E-05 | 0 - | 0 - | 1 - | 5,46E-05 - | 1 - | 5,19E-05 - | 1 - | 5,27E-05 - | 1 - | 5,93E-05 - |
| 7139 | 1 | 3um_FIJJAFBF_804749 | Eukaryota | unclassified | unclassified | Isochrysidales | Noelaerhabda | Emiliania | Emiliania huxl | unclassified | unclassified | unclassified | unclassified | unclassified | 0,2 | 0,4 | 7,97E-06 | 1,59E-05 | 1 - | 3,99E-05 - | 0 - | 0 - | 0 - | 0 - | 0 - | 0 - | 0 - | 0 - |
| 7140 | 1 | 3um_FIJJAFBF_336984 | Bacteria | Proteobacteria | Gammaprotec | Cellvibrionales | Cellvibrionace | Cellvibrio | Cellvibrio sp. I | poorly charact | Function unkn | 129NV | NA | 62977.ACIAD3 | 0,6 | 0,489897949 | 2,89E-05 | 2,41E-05 | 1 - | 4,03E-05 - | 1 - | 5,00E-05 - | 0 - | 0 - | 0 - | 0 - | 1 - | 5,43E-05 - |
| 7141 | 1 | 3um_FIJJAFBF_527610 | Bacteria | Bacteroidetes | Flavobacteriia | Flavobacterial | Flavobacteriac | Formosa | Formosa sp. H | unclassified | unclassified | unclassified | unclassified | unclassified | 1,6 | 0,489897949 | 0,000134953 | 3,54E-05 | 2 - | 0,000144373 - | 1 - | 8,96E-05 - | 2 - | 0,000170418 - | 2 - | 0,000172935 - | 1 - | 9,74E-05 - |
| 7142 | 1 | 3um_FIJJAFBF_550178 | Bacteria | Proteobacteria | Gammaprotec | Oceanospirilla | Saccharospirill | Reinekea | Reinekea fors | cellular proces | Cell wall/mem | 175GI | peptidase | 314283.MED2 | 1 | 0,632455532 | 4,36E-05 | 2,86E-05 | 1 - | 3,75E-05 - | 1 - | 4,66E-05 - | 1 - | 4,43E-05 - | 2 - | 8,99E-05 - | 0 - | 0 - |
| 7143 | 1 | 0.2um_NOKCPHJN_22849 | Bacteria | Bacteroidetes | Flavobacteriia | Flavobacterial | Flavobacteriac | Formosa | Formosa sp. H | metabolism | Lipid transpor | 0P85Z | acyl-CoA dehy | 391603.FBALC | 2 | 0,632455532 | 4,42E-05 | 1,02E-05 | 3 - | 5,72E-05 - | 2 - | 4,73E-05 - | 2 - | 4,50E-05 - | 2 - | 4,57E-05 - | 1 - | 2,57E-05 - |
| 7144 | 1 | 3um_FIJJAFBF_194576 | Bacteria | Proteobacteria | Gammaprotec | Oceanospirilla | Saccharospirill | Reinekea | Reinekea fors | metabolism | Energy produc | 16Q0J | alcohol dehyd | 633131.TR2A6 | 1 | 0 | 3,92E-05 | 3,72E-06 | 1 - | 3,28E-05 - | 1 - | 4,07E-05 - | 1 - | 3,87E-05 - | 1 - | 3,93E-05 - | 1 - | 4,42E-05 - |

| | | | | | | | | | | | | | | | | | | |
|---|---|---|---|---|---|---|---|---|---|---|---|---|---|---|---|---|---|---|
| 7145 | 2 | 0.2um_NOKCPHJN_954744;3um_FIJJAFBF_3375E | Eukaryota | Eustigmatoph unclassified Eustigmatales Monodopsida Nannochlorop Nannochlorop unclassified unclassified unclassified unclassified unclassified | 1,2 | 0,4 | 4,32E-05 | 9,58E-06 | 2 - | 6,20E-05 - | 1 - | 3,85E-05 - | 1 - | 3,66E-05 - | 1 - | 3,72E-05 - | 1 - | 4,19E-05 - |
| 7146 | 1 | 3um_FIJJAFBF_337914 | Bacteria | unclassified unclassified unclassified unclassified unclassified uncultured ba unclassified unclassified unclassified unclassified unclassified | 0,2 | 0,4 | 5,31E-06 | 1,06E-05 | 1 - | 2,66E-05 - | 0 - | 0 - | 0 - | 0 - | 0 - | 0 - | 0 - | 0 - |
| 7147 | 1 | 3um_FIJJAFBF_299455 | Bacteria | Bacteroidetes Saprospiria Saprospirales Saprospiracea Aureispira Aureispira sp. poorly charact Function unkn 09PDM NA 755732.Fluta | 0,6 | 0,489897949 | 2,76E-05 | 2,27E-05 | 1 - | 4,03E-05 - | 1 - | 5,00E-05 - | 0 - | 0 - | 1 - | 4,75E-05 - | 0 - | 0 - |
| 7148 | 1 | 0.2um_NOKCPHJN_438712 | Bacteria | Bacteroidetes Cytophagia Cytophagales Flammeovirga unclassified Flammeovirga unclassified unclassified unclassified unclassified unclassified | 7,4 | 2,244994432 | 0,000409839 | 0,000113898 | 9 - | 0,000423357 - | 5 - | 0,000292047 - | 6 - | 0,000333153 - | 11 - | 0,000619802 - | 6 - | 0,000380839 - |
| 7149 | 1 | 0.2um_NOKCPHJN_392516 | Eukaryota | Mollusca Gastropoda Architaenioglc Ampullariidae Pomacea Pomacea can unclassified unclassified unclassified unclassified unclassified | 0,2 | 0,4 | 1,29E-06 | 2,57E-06 | 0 - | 0 - | 0 - | 0 - | 0 - | 0 - | 0 - | 0 - | 0 - | 0 - |
| 7150 | 1 | 3um_FIJJAFBF_404304 | unclassified | unclassified unclassified unclassified unclassified unclassified unclassified unclassified unclassified unclassified unclassified unclassified | 1 | 0 | 8,51E-05 | 8,08E-06 | 1 - | 7,13E-05 - | 1 - | 8,85E-05 - | 1 - | 8,42E-05 - | 1 - | 8,54E-05 - | 1 - | 9,62E-05 - |
| 7151 | 1 | 3um_FIJJAFBF_47908 | Bacteria | Proteobacteri Alphaproteob Rhodobactera Rhodobactera Sulfitobacter Sulfitobacter s metabolism Inorganic ion 16P29 Extracellular s 391595.RLO | 2,4 | 1,356465997 | 0,000105744 | 4,72E-05 | 5 - | 0,000194537 - | 2 - | 9,66E-05 - | 2 - | 9,19E-05 - | 2 - | 9,32E-05 - | 1 - | 5,25E-05 - |
| 7152 | 1 | 3um_FIJJAFBF_246711 | Bacteria | Proteobacteri Gammaprotec Oceanospirilla Oceanospirilla Oceanobacter Oceanobacter metabolism Carbohydrate 0QMVS (ABC) transpo 945543.VIBR | 0,8 | 0,4 | 3,66E-05 | 1,86E-05 | 1 - | 3,96E-05 - | 1 - | 4,91E-05 - | 1 - | 4,67E-05 - | 1 - | 4,74E-05 - | 0 - | 0 - |
| 7153 | 1 | 3um_FIJJAFBF_437895 | Bacteria | Bacteroidetes Saprospiria Saprospirales Lewinellaceae unclassified Lewinellaceae metabolism Inorganic ion 05C27 TonB-depende 391603.FBAL | 41,8 | 1,833030278 | 0,000531349 | 4,10E-05 | 45 - | 0,000480461 - | 40 - | 0,000530303 - | 42 - | 0,000529326 - | 40 - | 0,000511566 - | 42 - | 0,000605091 - |
| 7154 | 1 | 0.2um_NOKCPHJN_379855 | Bacteria | Proteobacteri Betaproteoba Burkholderiale unclassified Tepidimonas Tepidimonas information st Translation, ri 0B7CD This is 1 of the 596154.Alide | 1,6 | 0,489897949 | 0,000122604 | 4,05E-05 | 2 - | 0,000128242 - | 2 - | 0,000159239 - | 1 - | 7,57E-05 - | 1 - | 7,68E-05 - | 2 - | 0,000170404 - |
| 7155 | 1 | 3um_FIJJAFBF_425939 | Bacteria | Proteobacteri Gammaprotec Thiotrichales Piscirickettsia Methylophaga Methylophaga cellular proce: Intracellular tr 05VIE preprotein tra 1026882.MAI | 1 | 0 | 0,000128063 | 1,22E-05 | 1 - | 0,000107268 - | 1 - | 0,000133195 - | 1 - | 0,000126619 - | 1 - | 0,000128489 - | 1 - | 0,000144743 - |
| 7156 | 1 | 3um_FIJJAFBF_62156 | Bacteria | Deferribacter Deferribacter Deferribacter Deferribacter Geovibrio Geovibrio sp. unclassified unclassified unclassified unclassified unclassified | 0,2 | 0,4 | 2,15E-05 | 4,29E-05 | 1 - | 0,000107268 - | 0 - | 0 - | 0 - | 0 - | 0 - | 0 - | 0 - | 0 - |
| 7157 | 1 | 3um_FIJJAFBF_237055 | Eukaryota | Bacillariophyt Bacillariophyc Bacillariales Bacillariaceae Cylindrotheca Cylindrotheca metabolism Energy produc 0FJ7X This is one of f 103690.alr37 | 11,8 | 0,4 | 0,000448839 | 4,29E-05 | 12 - | 0,000382589 - | 11 - | 0,000435474 - | 12 - | 0,000451607 - | 12 - | 0,000458278 - | 12 - | 0,000516248 - |
| 7158 | 1 | 3um_FIJJAFBF_175263 | Bacteria | Proteobacteri Gammaprotec unclassified unclassified unclassified OM182 bacte poorly charact Function unkn 1283Z NA 743721.Psesu | 0,4 | 0,489897949 | 2,48E-05 | 3,04E-05 | 0 - | 0 - | 1 - | 6,31E-05 - | 0 - | 0 - | 1 - | 6,08E-05 - | 0 - | 0 - |
| 7159 | 1 | 0.2um_NOKCPHJN_347138 | Bacteria | Bacteroidetes Flavobacteria Flavobacterial Flavobacteriac Lutibacter Lutibacter ma poorly charact Function unkn 0P853 adenosyltrans 313598.MED | 0,2 | 0,4 | 1,23E-05 | 2,46E-05 | 1 - | 6,14E-05 - | 0 - | 0 - | 0 - | 0 - | 0 - | 0 - | 0 - | 0 - |
| 7160 | 1 | 0.2um_NOKCPHJN_68988 | Bacteria | Bacteroidetes Flavobacteria Flavobacterial Flavobacteriac Maribacter Maribacter sp unclassified unclassified unclassified unclassified unclassified | 0,2 | 0,4 | 1,81E-05 | 3,62E-05 | 1 - | 9,04E-05 - | 0 - | 0 - | 0 - | 0 - | 0 - | 0 - | 0 - | 0 - |
| 7161 | 1 | 3um_FIJJAFBF_151034 | Bacteria | Bacteroidetes Cytophagia Cytophagales Flammeovirga unclassified Flammeovirga unclassified unclassified unclassified unclassified unclassified | 2,2 | 1,166190379 | 0,000124747 | 6,64E-05 | 3 - | 0,000147781 - | 3 - | 0,000183501 - | 3 - | 0,000174441 - | 2 - | 0,000118011 - | 0 - | 0 - |
| 7162 | 1 | 3um_FIJJAFBF_170869 | Bacteria | Bacteroidetes Flavobacteria Flavobacterial Crocinitomica unclassified Crocinitomica unclassified unclassified unclassified unclassified unclassified | 4,4 | 0,489897949 | 0,000407869 | 4,26E-05 | 5 - | 0,000390397 - | 5 - | 0,000484758 - | 4 - | 0,000368659 - | 4 - | 0,000374104 - | 4 - | 0,000421427 - |
| 7163 | 1 | 3um_FIJJAFBF_398710 | Bacteria | Bacteroidetes unclassified unclassified unclassified unclassified unidentified e information st Translation, ri 0BUHH Located on th 50743.SCB49 | 1 | 0 | 0,000103808 | 9,86E-06 | 1 - | 8,70E-05 - | 1 - | 0,000107969 - | 1 - | 0,000102638 - | 1 - | 0,000104154 - | 1 - | 0,000117329 - |
| 7164 | 1 | 0.2um_NOKCPHJN_31684 | Bacteria | Proteobacteri Betaproteoba unclassified unclassified unclassified Betaproteoba poorly charact Function unkn 0BA6J Short-chain de 338969.Rfer | 0,4 | 0,489897949 | 2,21E-05 | 2,71E-05 | 0 - | 0 - | 0 - | 0 - | 1 - | 5,15E-05 - | 0 - | 0 - | 1 - | 5,89E-05 - |
| 7165 | 4 | 3um_FIJJAFBF_00823;3um_FIJJAFBF_442511;3ur | Bacteria | Bacteroidetes Flavobacteria Flavobacterial Flavobacteriac various various unclassified unclassified unclassified unclassified unclassified | 0,4 | 0,489897949 | 1,88E-05 | 2,31E-05 | 0 - | 0 - | 0 - | 0 - | 1 - | 4,38E-05 - | 0 - | 0 - | 1 - | 5,01E-05 - |
| 7166 | 1 | 0.2um_NOKCPHJN_494952 | Bacteria | Bacteroidetes Flavobacteria Flavobacterial Flavobacteriac Polaribacter Polaribacter s poorly charact Function unkn 09EUF NA 313594.PI23P | 2,6 | 1,356465997 | 0,000142551 | 6,34E-05 | 5 - | 0,000236166 - | 2 - | 0,000117299 - | 2 - | 0,000111508 - | 1 - | 5,66E-05 - | 3 - | 0,000191203 - |
| 7167 | 1 | 3um_FIJJAFBF_113561 | Bacteria | Proteobacteri Gammaprotec Alteromonada Alteromonada Paraglaciecola Paraglaciecola unclassified unclassified unclassified unclassified unclassified | 0,2 | 0,4 | 1,15E-05 | 2,30E-05 | 0 - | 0 - | 0 - | 0 - | 1 - | 5,74E-05 - | 0 - | 0 - | 0 - | 0 - |
| 7168 | 1 | 0.2um_NOKCPHJN_241494 | Bacteria | Proteobacteri Alphaproteob Pelagibacteral Pelagibacterac unclassified alpha proteob metabolism Inorganic ion COG0376 Bifunctional e 684719.HIMB | 3 | 0,632455532 | 5,61E-05 | 1,23E-05 | 3 - | 4,73E-05 - | 4 - | 7,83E-05 - | 3 - | 5,58E-05 - | 3 - | 5,67E-05 - | 2 - | 4,25E-05 - |
| 7169 | 1 | 0.2um_NOKCPHJN_615947 | Eukaryota | Eustigmatoph unclassified Eustigmatales Monodopsida Nannochlorop Nannochlorop unclassified unclassified unclassified unclassified unclassified | 0,8 | 0,4 | 6,70E-05 | 3,44E-05 | 1 - | 7,08E-05 - | 0 - | 0 - | 1 - | 8,36E-05 - | 1 - | 8,49E-05 - | 1 - | 9,56E-05 - |
| 7170 | 1 | 0.2um_NOKCPHJN_113831 | Bacteria | Proteobacteri Gammaprotec Oceanospirilla Oceanospirilla Neptunomona Neptunomona cellular proce: Posttranslatio COG0330 Band 7 protein 207954.MED | 7,4 | 1,019803903 | 0,000257331 | 3,04E-05 | 8 - | 0,000234837 - | 7 - | 0,000255149 - | 7 - | 0,000242551 - | 9 - | 0,000316458 - | 6 - | 0,000237659 - |
| 7171 | 1 | 0.2um_NOKCPHJN_290426 | Bacteria | Proteobacteri Gammaprotec Alteromonada Colwelliaceae Colwellia Colwellia sp. P information st Translation, ri 179ZI One of the pri 167879.CPS | 1,2 | 0,748331477 | 0,000121321 | 7,31E-05 | 2 - | 0,000177948 - | 1 - | 0,000110048 - | 1 - | 0,000105025 - | 2 - | 0,000213152 - | 0 - | 0 - |
| 7172 | 2 | 3um_FIJJAFBF_261452;3um_FIJJAFBF_768793 | Bacteria | Proteobacteri Alphaproteob Pelagibacteral Pelagibacterac Candidatus Pe Candidatus Pe information st Translation, ri 05KK9 This protein b 335992.SAR1 | 1,4 | 0,489897949 | 0,000143 | 5,05E-05 | 1 - | 8,57E-05 - | 1 - | 0,000106357 - | 2 - | 0,000202212 - | 2 - | 0,000205199 - | 1 - | 0,000115578 - |
| 7173 | 1 | 0.2um_NOKCPHJN_246126 | Viruses | unclassified unclassified unclassified unclassified unclassified uncultured M unclassified unclassified unclassified unclassified unclassified | 0,4 | 0,489897949 | 1,05E-05 | 1,30E-05 | 1 - | 2,40E-05 - | 0 - | 0 - | 1 - | 2,87E-05 - | 0 - | 0 - | 0 - | 0 - |
| 7174 | 1 | 3um_FIJJAFBF_189626 | Bacteria | Bacteroidetes Flavobacteria Flavobacterial Flavobacteriac Polaribacter Polaribacter s unclassified unclassified unclassified unclassified unclassified | 5,6 | 1,356465997 | 0,000197796 | 3,20E-05 | 8 - | 0,000241001 - | 5 - | 0,000187033 - | 6 - | 0,000213358 - | 4 - | 0,000144339 - | 5 - | 0,000203247 - |
| 7175 | 1 | 3um_FIJJAFBF_228261 | Bacteria | Bacteroidetes Flavobacteria unclassified unclassified unclassified Flavobacteria unclassified unclassified unclassified unclassified unclassified | 0,6 | 0,489897949 | 2,84E-05 | 2,36E-05 | 1 - | 4,00E-05 - | 0 - | 0 - | 0 - | 0 - | 1 - | 4,79E-05 - | 1 - | 5,40E-05 - |
| 7176 | 1 | 3um_FIJJAFBF_761111 | Eukaryota | Phaeophycea unclassified Ectocarpales Ectocarpaceae Ectocarpus Ectocarpus sil information st Translation, ri 1E2KA 40S ribosomal 29760.VIT_01 | 1,4 | 0,8 | 9,31E-05 | 5,17E-05 | 2 - | 0,00011364 - | 2 - | 0,000141108 - | 2 - | 0,000134141 - | 0 - | 0 - | 1 - | 7,67E-05 - |
| 7177 | 1 | 0.2um_NOKCPHJN_420038 | Eukaryota | Chlorophyta Chlorophycea Chlamydomor Chlamydomor Chlamydomor Chlamydomor cellular proce: Posttranslatio KOG0543 Peptidyl-proly 35128.Thaps | 1 | 0 | 5,37E-05 | 5,10E-06 | 1 - | 4,50E-05 - | 1 - | 5,59E-05 - | 1 - | 5,31E-05 - | 1 - | 5,39E-05 - | 1 - | 6,07E-05 - |
| 7178 | 1 | 0.2um_NOKCPHJN_670374 | Bacteria | Bacteroidetes Flavobacteria Flavobacterial Flavobacteriac Algibacter Algibacter sp. metabolism Carbohydrate 0XNQK Transporter 391603.FBAL | 1 | 0 | 2,87E-05 | 2,72E-06 | 1 - | 2,40E-05 - | 1 - | 2,98E-05 - | 1 - | 2,83E-05 - | 1 - | 2,88E-05 - | 1 - | 3,24E-05 - |
| 7179 | 1 | 3um_FIJJAFBF_23716 | Bacteria | Proteobacteri Gammaprotec unclassified unclassified unclassified various cellular proce: Cell wall/mem 0QT5X Peptidoglycan 270374.MELB | 2 | 0 | 0,000151411 | 1,44E-05 | 2 - | 0,000126825 - | 2 - | 0,000157479 - | 2 - | 0,000149704 - | 2 - | 0,000151915 - | 2 - | 0,000171132 - |
| 7180 | 1 | 0.2um_NOKCPHJN_773405 | Bacteria | Proteobacteri Betaproteoba Burkholderiale Comamonada Limnohabitan Limnohabitan cellular proce: Posttranslatio 16QU3 Hemolysin-typ 311402.Avi_0 | 0,8 | 0,979795897 | 1,77E-05 | 2,21E-05 | 2 - | 3,77E-05 - | 0 - | 0 - | 0 - | 0 - | 0 - | 0 - | 2 - | 5,09E-05 - |
| 7181 | 1 | 0.2um_NOKCPHJN_879656 | Bacteria | Proteobacteri Alphaproteob Rhodobactera Rhodobactera Planktomarin Planktomarin unclassified unclassified unclassified unclassified unclassified | 0,2 | 0,4 | 1,66E-05 | 3,33E-05 | 1 - | 8,32E-05 - | 0 - | 0 - | 0 - | 0 - | 0 - | 0 - | 0 - | 0 - |
| 7182 | 1 | 3um_FIJJAFBF_308402 | Bacteria | Proteobacteri Gammaprotec Alteromonada Colwelliaceae Colwellia Colwellia sp. P unclassified unclassified unclassified unclassified unclassified | 1,2 | 0,4 | 0,000193557 | 6,67E-05 | 1 - | 0,000135031 - | 1 - | 0,000167669 - | 1 - | 0,000159391 - | 2 - | 0,00032349 - | 1 - | 0,000182205 - |
| 7183 | 1 | 3um_FIJJAFBF_736059 | Bacteria | Bacteroidetes Flavobacteria unclassified unclassified unclassified Flavobacteria poorly charact Function unkn 0XSVM NA 487797.Flav | 1,6 | 1,356465997 | 4,64E-05 | 4,08E-05 | 1 - | 2,36E-05 - | 4 - | 0,000117059 - | 1 - | 2,78E-05 - | 0 - | 0 - | 2 - | 6,36E-05 - |
| 7184 | 1 | 0.2um_NOKCPHJN_39166 | Eukaryota | unclassified Synurophycea Parmales Triparmaceae Triparma Triparma lae unclassified unclassified unclassified unclassified unclassified | 5,2 | 0,4 | 0,000498064 | 5,75E-05 | 5 - | 0,000401317 - | 5 - | 0,000498318 - | 6 - | 0,000568457 - | 5 - | 0,000480711 - | 5 - | 0,000541519 - |
| 7185 | 1 | 3um_FIJJAFBF_185959 | Bacteria | Bacteroidetes Flavobacteria Flavobacterial Flavobacteriac Formosa Formosa sp. H poorly charact Function unkn 08PAV RagB SusD dor 983544.Lacal | 6,4 | 1,019803903 | 0,000178496 | 3,13E-05 | 7 - | 0,000163967 - | 8 - | 0,000232684 - | 6 - | 0,000165897 - | 5 - | 0,000140289 - | 6 - | 0,000189642 - |
| 7186 | 1 | 3um_FIJJAFBF_264142 | Bacteria | Proteobacteri Gammaprotec Alteromonada Alteromonada Paraglaciecola Paraglaciecola unclassified unclassified unclassified unclassified unclassified | 1,8 | 0,4 | 0,000168805 | 3,38E-05 | 2 - | 0,000159412 - | 2 - | 0,000197943 - | 2 - | 0,00018817 - | 2 - | 0,000190949 - | 1 - | 0,000107552 - |
| 7187 | 1 | 3um_FIJJAFBF_188724 | Bacteria | Proteobacteri Gammaprotec Cellvibrionale: Porticoccacea unclassified various information st Translation, ri 17FIH One of the ear 314287.GB22 | 1,2 | 0,748331477 | 0,000160726 | 9,91E-05 | 2 - | 0,000234238 - | 2 - | 0,000290855 - | 1 - | 0,000138247 - | 1 - | 0,000140289 - | 0 - | 0 - |
| 7188 | 1 | 0.2um_NOKCPHJN_186648 | Bacteria | Proteobacteri Betaproteoba Burkholderiale unclassified various various unclassified unclassified unclassified unclassified unclassified | 1 | 0,632455532 | 9,84E-05 | 5,50E-05 | 2 - | 0,00017004 - | 1 - | 0,00010557 - | 0 - | 0 - | 1 - | 0,000101839 - | 1 - | 0,000114722 - |
| 7189 | 1 | 3um_FIJJAFBF_651818 | Bacteria | Bacteroidetes Flavobacteria Flavobacterial Flavobacteriac Polaribacter Polaribacter metabolism Energy produc COG0711 F(1)F(0) ATP s 313594.PI23P | 5 | 0,894427191 | 0,000407053 | 4,93E-05 | 6 - | 0,000414856 - | 4 - | 0,000343419 - | 6 - | 0,000489695 - | 5 - | 0,000414106 - | 4 - | 0,000373192 - |
| 7190 | 1 | 3um_FIJJAFBF_538287 | Bacteria | Bacteroidetes Flavobacteria Flavobacterial unclassified unclassified Flavobacteria cellular proce: Intracellular tr 0BIEB domain-conta 1005048.CFU | 1,6 | 0,489897949 | 4,24E-05 | 1,18E-05 | 2 - | 4,52E-05 - | 2 - | 5,61E-05 - | 2 - | 5,33E-05 - | 1 - | 2,71E-05 - | 1 - | 3,05E-05 - |
| 7191 | 1 | 3um_FIJJAFBF_163199 | Bacteria | Proteobacteri Gammaprotec Cellvibrionale: Halieaceae unclassified marine gamm information st Translation, ri 17FIH One of the ear 247639.MGP | 0,4 | 0,489897949 | 5,15E-05 | 6,35E-05 | 1 - | 0,000117119 - | 0 - | 0 - | 0 - | 0 - | 1 - | 0,000140289 - | 0 - | 0 - |
| 7192 | 1 | 3um_FIJJAFBF_22823 | Eukaryota | Bacillariophyt Bacillariophyc Naviculales Phaeodactylac Phaeodactylu Phaeodactylu unclassified unclassified unclassified unclassified unclassified | 1 | 0 | 3,42E-05 | 3,24E-06 | 1 - | 2,86E-05 - | 1 - | 3,55E-05 - | 1 - | 3,38E-05 - | 1 - | 3,43E-05 - | 1 - | 3,86E-05 - |
| 7193 | 1 | 0.2um_NOKCPHJN_397832 | unclassified | unclassified unclassified unclassified unclassified unclassified unclassified unclassified unclassified unclassified unclassified unclassified | 0,2 | 0,4 | 1,82E-06 | 3,63E-06 | 0 - | 0 - | 1 - | 9,08E-06 - | 0 - | 0 - | 0 - | 0 - | 0 - | 0 - |
| 7194 | 1 | 3um_FIJJAFBF_313317 | Bacteria | Proteobacteri Gammaprotec Oceanospirilla Saccharospiril Reinekea Reinekea fors cellular proce: Posttranslatio COG0330 Band 7 protein 314283.MED | 11,6 | 1,854723699 | 0,000397817 | 3,52E-05 | 14 - | 0,000407836 - | 10 - | 0,000361723 - | 12 - | 0,000447023 - | 12 - | 0,000418731 - | 9 - | 0,000353774 - |
| 7195 | 1 | 3um_FIJJAFBF_203162 | Bacteria | Proteobacteri Gammaprotec Oceanospirilla Arenicellales Arenicellacea Arenicella Arenicella xan metabolism Energy produc COG0056 Produces ATP 396588.Tgr7 | 1,4 | 1,019803903 | 3,55E-05 | 2,32E-05 | 3 - | 6,71E-05 - | 0 - | 0 - | 1 - | 2,64E-05 - | 2 - | 5,36E-05 - | 1 - | 3,02E-05 - |
| 7196 | 1 | 0.2um_NOKCPHJN_240238 | Eukaryota | unclassified Oomycetes Saprolegniale: Saprolegniace Aphanomyces Aphanomyces poorly charact Function unkn 0XNUF Extracellular n 340101.Vvad | 0,2 | 0,4 | 4,74E-07 | 9,49E-07 | 1 - | 2,37E-06 - | 0 - | 0 - | 0 - | 0 - | 0 - | 0 - | 0 - | 0 - |
| 7197 | 1 | 3um_FIJJAFBF_636077 | Eukaryota | Bacillariophyt Coscinodiscop Thalassiosirale Thalassiosira Thalassiosira Thalassiosira unclassified unclassified unclassified unclassified unclassified | 0,4 | 0,489897949 | 2,56E-05 | 3,14E-05 | 0 - | 0 - | 1 - | 6,57E-05 - | 1 - | 6,24E-05 - | 0 - | 0 - | 0 - | 0 - |
| 7198 | 1 | 0.2um_NOKCPHJN_530776 | Bacteria | Proteobacteri Alphaproteob Pelagibacteral Pelagibacterac Candidatus Pe Candidatus Pe metabolism Inorganic ion 05PWN phosphate abc 335992.SAR1 | 0,2 | 0,4 | 8,08E-06 | 1,62E-05 | 1 - | 4,04E-05 - | 0 - | 0 - | 0 - | 0 - | 0 - | 0 - | 0 - | 0 - |
| 7199 | 1 | 0.2um_NOKCPHJN_14356 | Bacteria | Bacteroidetes Cytophagia Cytophagales Flammeovirga Reichenbachie Reichenbachie poorly charact Function unkn 05XJE NA 269798.CHU | 2,8 | 0,748331477 | 0,000184712 | 4,82E-05 | 3 - | 0,000167151 - | 3 - | 0,000207552 - | 2 - | 0,000131536 - | 4 - | 0,000266958 - | 2 - | 0,000150364 - |
| 7200 | 1 | 3um_FIJJAFBF_157067 | Bacteria | Bacteroidetes Flavobacteria unclassified unclassified unclassified Flavobacteria poorly charact Intracellular tr 18ATJ Pfam:YadA 1005048.CFU | 9,2 | 0,4 | 0,000131127 | 8,50E-06 | 10 - | 0,000119809 - | 9 - | 0,00013389 - | 9 - | 0,00012728 - | 9 - | 0,00012916 - | 9 - | 0,000145498 - |
| 7201 | 1 | 0.2um_NOKCPHJN_229751 | Bacteria | Bacteroidetes Cytophagia Cytophagales Cyclobacteriac Algoriphagus Algoriphagus s metabolism Inorganic ion 105C27 TonB-depende 388413.ALPR | 9,8 | 4,118252056 | 0,000132022 | 4,59E-05 | 16 - | 0,000187774 - | 10 - | 0,000145725 - | 10 - | 0,000138853 - | 10 - | 0,000140576 - | 3 - | 4,75E-05 - |
| 7202 | 1 | 0.2um_NOKCPHJN_352204 | Eukaryota | Chordata unclassified unclassified Branchiostom Branchiostom Branchiostom unclassified unclassified unclassified unclassified unclassified | 0,8 | 0,4 | 5,47E-06 | 2,75E-06 | 1 - | 6,83E-06 - | 1 - | 6,49E-06 - | 1 - | 6,59E-06 - | 1 - | 7,42E-06 - | 0 - | 0 - |
| 7203 | 1 | 0.2um_NOKCPHJN_227678 | Bacteria | Bacteroidetes Flavobacteria Flavobacterial Flavobacteriac Formosa Formosa sp. H metabolism Inorganic ion 05C27 TonB-depende 867900.Celly | 15,4 | 1,356465997 | 0,000208662 | 1,16E-05 | 17 - | 0,000194343 - | 16 - | 0,000227122 - | 16 - | 0,000215908 - | 15 - | 0,000205403 - | 13 - | 0,000200535 - |
| 7204 | 1 | 3um_FIJJAFBF_490337 | Bacteria | Bacteroidetes Flavobacteria Flavobacterial Flavobacteriac Polaribacter Polaribacter s cellular proce: Cell wall/mem 0P8B8 Outer membr: 313594.PI23P | 0,2 | 0,4 | 4,90E-06 | 9,81E-06 | 1 - | 2,45E-05 - | 0 - | 0 - | 0 - | 0 - | 0 - | 0 - | 0 - | 0 - |
| 7205 | 1 | 3um_FIJJAFBF_675593 | Bacteria | Bacteroidetes Flavobacteria Flavobacterial Flavobacteriac Polaribacter Polaribacter s unclassified unclassified unclassified unclassified unclassified | 3,2 | 0,4 | 0,000481954 | 7,47E-05 | 3 - | 0,000378385 - | 3 - | 0,000469842 - | 3 - | 0,000446645 - | 4 - | 0,000604322 - | 3 - | 0,000510575 - |
| 7206 | 1 | 3um_FIJJAFBF_222363 | Bacteria | Proteobacteri Alphaproteob Rhodobactera Rhodobactera unclassified Rhodobactera unclassified unclassified unclassified unclassified unclassified | 1 | 0 | 6,29E-05 | 5,97E-06 | 1 - | 5,26E-05 - | 1 - | 6,54E-05 - | 1 - | 6,21E-05 - | 1 - | 6,31E-05 - | 1 - | 7,10E-05 - |
| 7207 | 1 | 0.2um_NOKCPHJN_1003141 | Eukaryota | Nemertea Pilidiophora Heteronemert Cerebratulida Cerebratulus Cerebratulus unclassified unclassified unclassified unclassified unclassified | 2,8 | 0,748331477 | 0,000338347 | 6,38E-05 | 4 - | 0,00041361 - | 3 - | 0,000385186 - | 3 - | 0,000366168 - | 2 - | 0,000247718 - | 2 - | 0,000279053 - |
| 7208 | 1 | 3um_FIJJAFBF_222261 | Bacteria | Proteobacteri Alphaproteob Rhodobactera unclassified unclassified Rhodobactera cellular proce: Posttranslatio 16QDW Has an import 314262.MED | 1 | 0 | 8,06E-05 | 7,65E-06 | 1 - | 6,75E-05 - | 1 - | 8,38E-05 - | 1 - | 7,97E-05 - | 1 - | 8,09E-05 - | 1 - | 9,11E-05 - |
| 7209 | 1 | 0.2um_NOKCPHJN_269921 | Eukaryota | unclassified Kinetoplastida Trypanosomat Leishmania Leishmania m poorly charact Function unkn 0NVRZ NA 170187.SP_1 | 0,2 | 0,4 | 2,59E-06 | 5,17E-06 | 0 - | 0 - | 0 - | 0 - | 0 - | 0 - | 1 - | 1,29E-05 - | 0 - | 0 - |
| 7210 | 1 | 0.2um_NOKCPHJN_061362 | Bacteria | Proteobacteri Alphaproteob Pelagibacteral Pelagibacterac Candidatus Pe Candidatus Pe metabolism Amino acid tra 01QDI sarcosine oxid 335992.SAR1 | 1 | 0 | 1,54E-05 | 1,46E-06 | 1 - | 1,29E-05 - | 1 - | 1,60E-05 - | 1 - | 1,52E-05 - | 1 - | 1,55E-05 - | 1 - | 1,74E-05 - |
| 7211 | 1 | 3um_FIJJAFBF_172359 | Bacteria | Proteobacteri Gammaprotec Oceanospirilla Saccharospiril Reinekea Reinekea fors information st Translation, ri 0QTKX Seems to be t 314283.MED | 11,8 | 1,166190379 | 0,000137224 | 7,69E-05 | 14 - | 0,000133906 - | 12 - | 0,001425189 - | 11 - | 0,00124192 - | 11 - | 0,001260264 - | 11 - | 0,001419683 - |
| 7212 | 1 | 3um_FIJJAFBF_395209 | Bacteria | Bacteroidetes Flavobacteria Flavobacterial Crocinitomica unclassified Crocinitomica information st Translation, ri COG0089 One of the ear 313595.P700 | 2,2 | 0,748331477 | 0,000318732 | 0,000124712 | 1 - | 0,000118327 - | 2 - | 0,000293853 - | 3 - | 0,000419017 - | 2 - | 0,000283471 - | 3 - | 0,000478993 - |
| 7213 | 1 | 0.2um_NOKCPHJN_215352 | Bacteria | Bacteroidetes Flavobacteria Flavobacterial Flavobacteriac Formosa Formosa sp. H unclassified unclassified unclassified unclassified unclassified | 12 | 0,632455532 | 0,000260813 | 3,04E-05 | 12 - | 0,000218276 - | 13 - | 0,00029362 - | 11 - | 0,000236181 - | 12 - | 0,000261458 - | 12 - | 0,000294531 - |
| 7214 | 1 | 0.2um_NOKCPHJN_27403 | Bacteria | Bacteroidetes Flavobacteria Flavobacterial Cryomorphace unclassified Cryomorphac unclassified unclassified unclassified unclassified unclassified | 0,8 | 0,748331477 | 7,49E-06 | 7,20E-06 | 1 - | 8,07E-06 - | 2 - | 1,00E-05 - | 0 - | 0 - | 2 - | 1,93E-05 - | 0 - | 0 - |
| 7215 | 1 | 0.2um_NOKCPHJN_625482 | Bacteria | Proteobacteri Gammaprotec Oceanospirilla Saccharospiril Reinekea Reinekea fors unclassified unclassified unclassified unclassified unclassified | 0,4 | 0,8 | 5,96E-05 | 0,000119249 | 2 - | 0,000298121 - | 0 - | 0 - | 0 - | 0 - | 0 - | 0 - | 0 - | 0 - |
| 7216 | 1 | 0.2um_NOKCPHJN_366565 | unclassified | unclassified unclassified unclassified unclassified unclassified unclassified unclassified unclassified unclassified unclassified unclassified | 1,6 | 0,8 | 0,000156263 | 7,87E-05 | 0 - | 0 - | 2 - | 0,000195231 - | 2 - | 0,000185592 - | 2 - | 0,000188333 - | 2 - | 0,000212157 - |
| 7217 | 1 | 0.2um_NOKCPHJN_266251 | Bacteria | Proteobacteri Alphaproteob Caulobacteral Caulobacterac Phenylobacte Phenylobacte unclassified unclassified unclassified unclassified unclassified | 0,2 | 0,4 | 1,65E-05 | 3,30E-05 | 1 - | 8,26E-05 - | 0 - | 0 - | 0 - | 0 - | 0 - | 0 - | 0 - | 0 - |
| 7218 | 1 | 0.2um_NOKCPHJN_529854 | Eukaryota | unclassified unclassified Acytosteliales Acytosteliace Acytostelium Acytostelium poorly charact Function unkn 0PXWZ ChtBD1 318829.MGG | 0,6 | 0,8 | 1,13E-05 | 1,50E-05 | 0 - | 0 - | 0 - | 0 - | 2 - | 3,73E-05 - | 1 - | 1,89E-05 - | 0 - | 0 - |
| 7219 | 1 | 0.2um_NOKCPHJN_57116 | Bacteria | Proteobacteri Gammaprotec Oceanospirilla Saccharospiril Reinekea Reinekea fors metabolism Energy produc 05CV4 Subunits I and 314283.MED | 2,4 | 0,489897949 | 8,70E-05 | 2,01E-05 | 2 - | 6,06E-05 - | 3 - | 0,000112812 - | 3 - | 0,000107242 - | 2 - | 7,26E-05 - | 2 - | 8,17E-05 - |
| 7220 | 1 | 3um_FIJJAFBF_258667 | Bacteria | Proteobacteri Gammaprotec Alteromonada Alteromonada Aliiglaciecola Aliiglaciecola unclassified unclassified unclassified unclassified unclassified | 0,6 | 0,489897949 | 1,79E-05 | 1,49E-05 | 1 - | 2,49E-05 - | 1 - | 3,09E-05 - | 0 - | 0 - | 0 - | 0 - | 1 - | 3,36E-05 - |
| 7221 | 1 | 0.2um_NOKCPHJN_594929 | Bacteria | Proteobacteri Alphaproteob Pelagibacteral Pelagibacterac Candidatus Pe Candidatus Pe unclassified unclassified unclassified unclassified unclassified | 0,8 | 0,4 | 5,43E-05 | 2,79E-05 | 1 - | 5,74E-05 - | 0 - | 0 - | 1 - | 6,77E-05 - | 1 - | 6,87E-05 - | 1 - | 7,74E-05 - |
| 7222 | 1 | 3um_FIJJAFBF_300487 | Bacteria | Proteobacteri Gammaprotec Alteromonada Psychromonac Psychromonas Psychromonas unclassified unclassified unclassified unclassified unclassified | 3,8 | 0,4 | 0,000305322 | 5,49E-05 | 3 - | 0,000200192 - | 4 - | 0,000331439 - | 4 - | 0,000315075 - | 4 - | 0,000319729 - | 4 - | 0,000360173 - |
| 7223 | 1 | 0.2um_NOKCPHJN_606421 | Eukaryota | Basidiomycota Agaricomycet Hymenochaet Hymenochaet Pyrrhoderma Pyrrhoderma cellular proce: Intracellular tr 0BIEB domain-conta 1005048.CFU | 0,4 | 0,489897949 | 1,15E-05 | 1,41E-05 | 1 - | 2,63E-05 - | 0 - | 0 - | 1 - | 3,10E-05 - | 0 - | 0 - | 0 - | 0 - |
| 7224 | 1 | 3um_FIJJAFBF_110579 | Bacteria | Bacteroidetes Flavobacteria Flavobacterial Flavobacteriac Polaribacter various metabolism Carbohydrate 08K9H protein TIM b 886377.Muri | 0,4 | 0,489897949 | 1,71E-05 | 2,10E-05 | 0 - | 0 - | 1 - | 4,36E-05 - | 0 - | 0 - | 1 - | 4,20E-05 - | 0 - | 0 - |
| 7225 | 1 | 0.2um_NOKCPHJN_78018 | Bacteria | Bacteroidetes Flavobacteria Flavobacterial Crocinitomica unclassified Crocinitomica unclassified unclassified unclassified unclassified unclassified | 0,8 | 0,4 | 1,58E-05 | 8,11E-06 | 1 - | 1,65E-05 - | 1 - | 2,05E-05 - | 0 - | 0 - | 1 - | 1,98E-05 - | 1 - | 2,23E-05 - |
| 7226 | 1 | 3um_FIJJAFBF_737884 | Bacteria | Proteobacteri Betaproteoba Burkholderiale unclassified unclassified Betaproteoba unclassified unclassified unclassified unclassified unclassified | 4,4 | 0,489897949 | 0,000443082 | 3,38E-05 | 5 - | 0,000425099 - | 4 - | 0,000422278 - | 5 - | 0,000501786 - | 4 - | 0,000407358 - | 4 - | 0,000458887 - |
| 7227 | 1 | 0.2um_NOKCPHJN_99117 | Bacteria | Bacteroidetes Flavobacteria Flavobacterial Flavobacteriac Polaribacter Polaribacter s metabolism Lipid transpor 0F7W4 reductase 391587.KAO | 1,2 | 0,4 | 6,63E-05 | 2,29E-05 | 1 - | 4,63E-05 - | 1 - | 5,75E-05 - | 1 - | 5,46E-05 - | 2 - | 0,000110874 - | 1 - | 6,24E-05 - |
| 7228 | 1 | 0.2um_NOKCPHJN_225074 | Eukaryota | Phaeophycea unclassified Ectocarpales Ectocarpaceae Ectocarpus Ectocarpus sil cellular proce: Cytoskeleton 13DKH CH 7955.ENSDA | 0,2 | 0,4 | 1,88E-06 | 3,77E-06 | 0 - | 0 - | 1 - | 9,42E-06 - | 0 - | 0 - | 0 - | 0 - | 0 - | 0 - |
| 7229 | 1 | 0.2um_NOKCPHJN_804742 | Eukaryota | unclassified Oomycetes Peronosporale Peronosporac Phytophthora Phytophthora information st Translation, ri COG0052 30S ribosomal 67593.Physo | 2,6 | 0,489897949 | 0,000132064 | 2,15E-05 | 3 - | 0,000128963 - | 3 - | 0,000160134 - | 3 - | 0,000152227 - | 2 - | 0,000102984 - | 2 - | 0,000116011 - |
| 7230 | 1 | 3um_FIJJAFBF_208626 | Bacteria | Bacteroidetes Flavobacteria Flavobacterial Flavobacteriac Ulvibacter Ulvibacter sp. information st Translation, ri COG0089 One of the ear 50743.SCB49 | 2,4 | 0,489897949 | 0,000335581 | 5,93E-05 | 3 - | 0,00035498 - | 3 - | 0,0004078 - | 2 - | 0,000279345 - | 2 - | 0,000283471 - | 2 - | 0,000319329 - |
| 7231 | 1 | 3um_FIJJAFBF_488074 | Bacteria | Proteobacteri Alphaproteob Rhodobactera Rhodobactera Sulfitobacter Sulfitobacter information st Transcription 16PMT DNA-depende 391589.RGAI | 2,6 | 0,8 | 0,000102691 | 2,87E-05 | 3 - | 0,000100681 - | 2 - | 8,33E-05 - | 4 - | 0,000158459 - | 2 - | 8,04E-05 - | 2 - | 9,06E-05 - |
| 7232 | 1 | 3um_FIJJAFBF_640407 | unclassified | unclassified unclassified unclassified unclassified unclassified unclassified unclassified unclassified unclassified unclassified unclassified | 1 | 0 | 5,73E-05 | 5,44E-06 | 1 - | 4,80E-05 - | 1 - | 5,96E-05 - | 1 - | 5,67E-05 - | 1 - | 5,75E-05 - | 1 - | 6,48E-05 - |
| 7233 | 1 | 3um_FIJJAFBF_552795 | Bacteria | Proteobacteri Gammaprotec Cellvibrionale: Porticoccacea unclassified SAR92 bacteri metabolism Coenzyme tra 05HJX Biotin- acetyl- 314287.GB22 | 0,2 | 0,4 | 8,74E-06 | 1,75E-05 | 0 - | 0 - | 1 - | 4,37E-05 - | 0 - | 0 - | 0 - | 0 - | 0 - | 0 - |
| 7234 | 1 | 3um_FIJJAFBF_145237 | Bacteria | Proteobacteri Gammaprotec Oceanospirilla Hyunsoonleell Hyunsoonleell Hyunsoonleell information st Translation, ri 08YY1 This protein is 391603.FBAL | 1,4 | 0,8 | 0,000112638 | 6,35E-05 | 2 - | 0,00014083 - | 1 - | 8,74E-05 - | 2 - | 0,000166236 - | 2 - | 0,000168691 - | 0 - | 0 - |
| 7235 | 1 | 0.2um_NOKCPHJN_905000 | Bacteria | Bacteroidetes unclassified unclassified unclassified unclassified Bacteroidetes poorly charact Function unkn 08HPR NA 269798.CHU | 1,6 | 0,489897949 | 3,18E-05 | 8,80E-06 | 2 - | 3,38E-05 - | 2 - | 4,20E-05 - | 2 - | 3,99E-05 - | 1 - | 2,02E-05 - | 1 - | 2,28E-05 - |
| 7236 | 1 | 3um_FIJJAFBF_272767 | Bacteria | Proteobacteri Alphaproteob Rhodobactera unclassified unclassified Rhodobactera metabolism Amino acid tra COG0683 (ABC) transpo 246200.SPO | 1,8 | 0,4 | 6,25E-05 | 1,46E-05 | 2 - | 3,63E-05 - | 2 - | 6,89E-05 - | 2 - | 7,00E-05 - | 2 - | 7,88E-05 - | 2 - | 7,88E-05 - |
| 7237 | 1 | 0.2um_NOKCPHJN_487599 | Bacteria | Proteobacteri Betaproteoba Burkholderiale Burkholderiac Polynucleoba Polynucleoba information st Replication, re 16QPP helicase 702113.PP1Y | 0,2 | 0,4 | 3,86E-06 | 7,72E-06 | 0 - | 0 - | 0 - | 0 - | 0 - | 0 - | 0 - | 0 - | 1 - | 1,93E-05 - |
| 7238 | 1 | 3um_FIJJAFBF_35010 | Bacteria | Proteobacteri Alphaproteob Rhodobactera Rhodobactera Planktomarin Planktomarin unclassified unclassified unclassified unclassified unclassified | 1 | 0,632455532 | 2,32E-05 | 1,53E-05 | 1 - | 1,93E-05 - | 2 - | 4,80E-05 - | 1 - | 2,28E-05 - | 0 - | 0 - | 1 - | 2,61E-05 - |
| 7239 | 1 | 3um_FIJJAFBF_30305 | Bacteria | Bacteroidetes Flavobacteria Flavobacterial Flavobacteriac Polaribacter Polaribacter s metabolism Amino acid tra COG0136 Catalyzes the 886377.Muri | 1,8 | 0,4 | 7,49E-05 | 1,84E-05 | 2 - | 6,98E-05 - | 2 - | 8,66E-05 - | 2 - | 8,24E-05 - | 1 - | 4,18E-05 - | 2 - | 9,41E-05 - |
| 7240 | 1 | 0.2um_NOKCPHJN_177183 | Eukaryota | unclassified Prymnesiales Chrysochrom Chrysochrom Chrysochrom unclassified unclassified unclassified unclassified unclassified unclassified | 0,6 | 0,489897949 | 1,71E-05 | 1,40E-05 | 0 - | 0 - | 0 - | 0 - | 1 - | 2,70E-05 - | 1 - | 2,74E-05 - | 1 - | 3,09E-05 - |
| 7241 | 1 | 3um_FIJJAFBF_94592 | Bacteria | Proteobacteri Gammaprotec Alteromonada Alteromonada Paraglaciecola Paraglaciecola cellular proce: Intracellular tr 16REI general secret 342610.Patl | 0,2 | 0,4 | 3,32E-06 | 6,64E-06 | 1 - | 1,66E-05 - | 0 - | 0 - | 0 - | 0 - | 0 - | 0 - | 0 - | 0 - |
| 7242 | 1 | 3um_FIJJAFBF_371442 | Bacteria | Proteobacteri Gammaprotec Oceanospirilla Saccharospiril Reinekea Reinekea fors unclassified unclassified unclassified unclassified unclassified | 7,6 | 0,489897949 | 0,000119589 | 1,44E-05 | 8 - | 0,000105421 - | 8 - | 0,000130901 - | 7 - | 0,000108884 - | 7 - | 0,000110492 - | 8 - | 0,00014225 - |
| 7243 | 1 | 3um_FIJJAFBF_468296 | Bacteria | Proteobacteri Gammaprotec Thiotrichales Piscirickettsia Methylophaga Methylophaga unclassified unclassified unclassified unclassified unclassified | 0,4 | 0,489897949 | 1,49E-05 | 1,83E-05 | 0 - | 0 - | 1 - | 3,80E-05 - | 0 - | 0 - | 1 - | 3,67E-05 - | 0 - | 0 - |
| 7244 | 1 | 3um_FIJJAFBF_309166 | Bacteria | Proteobacteri Alphaproteob Rhodobactera Rhodobactera unclassified Rhodobactera metabolism Nucleotide tra 01Q8W Plays an impo 644107.SIL1 | 0,2 | 0,4 | 6,63E-06 | 1,33E-05 | 0 - | 0 - | 1 - | 3,31E-05 - | 0 - | 0 - | 0 - | 0 - | 0 - | 0 - |
| 7245 | 1 | 3um_FIJJAFBF_419416 | Bacteria | Proteobacteri Gammaprotec unclassified unclassified unclassified Gammaprotec metabolism Inorganic ion 16PDP TonB-depende 414684.RC1 | 8,8 | 1,326649916 | 0,000117209 | 1,15E-05 | 11 - | 0,000124022 - | 9 - | 0,000125999 - | 9 - | 0,000119778 - | 7 - | 9,45E-05 - | 8 - | 0,000121709 - |
| 7246 | 1 | 3um_FIJJAFBF_161277 | Bacteria | Proteobacteri Alphaproteob Rhodobactera Rhodobactera Roseovarius Roseovarius s metabolism Inorganic ion COG3638 (ABC) transpo 375451.RD1 | 2,2 | 1,166190379 | 0,000106883 | 5,61E-05 | 3 - | 0,000127059 - | 2 - | 0,00010518 - | 3 - | 0,0001499 - | 3 - | 0,000152196 - | 0 - | 0 - |
| 7247 | 1 | 3um_FIJJAFBF_44672 | Bacteria | Bacteroidetes Flavobacteria Flavobacterial Flavobacteriac Polaribacter Polaribacter s metabolism Amino acid tra 09E2D NA 313594.PI23P | 2,6 | 0,8 | 0,000164781 | 5,58E-05 | 2 - | 0,0001053 - | 2 - | 0,000130751 - | 4 - | 0,000248591 - | 2 - | 0,000126131 - | 3 - | 0,00021313 - |
| 7248 | 1 | 3um_FIJJAFBF_120654 | Bacteria | Proteobacteri Gammaprotec Arenicellales Arenicellacea Arenicella Arenicella xan cellular proce: Posttranslatio COG0330 Band 7 protein 395494.Galf | 0,6 | 0,489897949 | 2,73E-05 | 2,25E-05 | 1 - | 3,99E-05 - | 1 - | 4,95E-05 - | 1 - | 4,70E-05 - | 0 - | 0 - | 0 - | 0 - |
| 7249 | 1 | 0.2um_NOKCPHJN_115245 | Bacteria | Actinobacteri Actinobacteri Actinomycetal unclassified unclassified Actinomycetal unclassified unclassified unclassified unclassified unclassified | 0,4 | 0,489897949 | 2,27E-05 | 2,80E-05 | 1 - | 5,06E-05 - | 1 - | 6,28E-05 - | 0 - | 0 - | 0 - | 0 - | 0 - | 0 - |
| 7250 | 1 | 0.2um_NOKCPHJN_267388 | Bacteria | Proteobacteri Gammaprotec Cellvibrionale: Porticoccacea unclassified SAR92 bacteri unclassified unclassified unclassified unclassified unclassified | 0,4 | 0,489897949 | 5,08E-06 | 6,29E-06 | 1 - | 1,13E-05 - | 1 - | 1,41E-05 - | 0 - | 0 - | 0 - | 0 - | 0 - | 0 - |
| 7251 | 1 | 3um_FIJJAFBF_465006 | Bacteria | Proteobacteri Gammaprotec Oceanospirilla Saccharospiril Reinekea Reinekea fors unclassified unclassified unclassified unclassified unclassified | 1,6 | 0,489897949 | 0,000127008 | 4,05E-05 | 2 - | 0,000133461 - | 1 - | 8,29E-05 - | 1 - | 7,88E-05 - | 2 - | 0,000159864 - | 2 - | 0,000180087 - |
| 7252 | 1 | 3um_FIJJAFBF_62761 | Bacteria | Proteobacteri Alphaproteob Rhodobactera Rhodobactera unclassified Rhodobactera metabolism Energy produc COG0712 F(1)F(0) ATP s 367336.OM2 | 1 | 0 | 7,37E-05 | 6,99E-06 | 1 - | 6,17E-05 - | 1 - | 7,66E-05 - | 1 - | 7,28E-05 - | 1 - | 7,39E-05 - | 1 - | 8,33E-05 - |
| 7253 | 1 | 0.2um_NOKCPHJN_88267 | Bacteria | Proteobacteri Alphaproteob Rhodobactera Rhodobactera Planktomarin Planktomarin information st Translation, ri COG0050 This protein p 388401.RB2 | 1,6 | 0,8 | 0,000534295 | 0,00027458 | 2 - | 0,000559887 - | 2 - | 0,000695214 - | 2 - | 0,000660889 - | 0 - | 0 - | 2 - | 0,000755485 - |
| 7254 | 1 | 0.2um_NOKCPHJN_140070 | Bacteria | Proteobacteri Gammaprotec Oceanospirilla Saccharospiril Reinekea Reinekea blan information st Transcription 05D27 DNA-depende 314283.MED | 35,2 | 3,370459909 | 0,000342816 | 2,62E-05 | 39 - | 0,000320193 - | 35 - | 0,000356807 - | 32 - | 0,000310117 - | 39 - | 0,000383537 - | 31 - | 0,000343427 - |
| 7255 | 1 | 3um_FIJJAFBF_269222 | Bacteria | Bacteroidetes Flavobacteria Flavobacterial Flavobacteriac Polaribacter Polaribacter s unclassified unclassified unclassified unclassified unclassified | 1,2 | 0,4 | 8,57E-05 | 3,65E-05 | 1 - | 5,86E-05 - | 1 - | 7,27E-05 - | 1 - | 6,91E-05 - | 1 - | 7,01E-05 - | 2 - | 0,000158035 - |
| 7256 | 1 | 0.2um_NOKCPHJN_194779 | Bacteria | Proteobacteri Alphaproteob Rhodobactera Amylibacter Amylibacter s metabolism Carbohydrate 05CA9 Pyruvate kinas 388401.RB2 | 0,8 | 0,4 | 2,38E-05 | 1,20E-05 | 0 - | 0 - | 1 - | 2,97E-05 - | 1 - | 2,86E-05 - | 1 - | 3,23E-05 - | 1 - | 3,23E-05 - |
| 7257 | 1 | 3um_FIJJAFBF_52276 | Bacteria | Proteobacteri Alphaproteob Rhodobactera unclassified unclassified Rhodobactera metabolism Secondary me 01Q86 extracellular s 644107.SIL1 | 10 | 1,549193338 | 0,00037636 | 4,48E-05 | 11 - | 0,000350707 - | 10 - | 0,000395886 - | 11 - | 0,000413973 - | 11 - | 0,000420088 - | 7 - | 0,000301145 - |
| 7258 | 1 | 0.2um_NOKCPHJN_915793 | unclassified | unclassified unclassified unclassified unclassified unclassified unclassified unclassified unclassified unclassified unclassified unclassified | 1 | 0 | 0,00013567 | 1,29E-05 | 1 - | 0,00011364 - | 1 - | 0,000141108 - | 1 - | 0,000134141 - | 1 - | 0,000136122 - | 1 - | 0,000153341 - |
| 7259 | 1 | 3um_FIJJAFBF_184933 | Bacteria | Proteobacteri Alphaproteob Rhodobactera Rhodobactera Yoonia Yoonia marin unclassified unclassified unclassified unclassified unclassified | 1,8 | 0,748331477 | 6,48E-05 | 2,18E-05 | 3 - | 9,33E-05 - | 2 - | 7,72E-05 - | 1 - | 3,67E-05 - | 2 - | 7,45E-05 - | 1 - | 4,20E-05 - |
| 7260 | 2 | 0.2um_NOKCPHJN_475432;0.2um_NOKCPHJN_6 | Bacteria | Proteobacteri Betaproteoba Nitrosomonac unclassified unclassified Gallionellales unclassified unclassified unclassified unclassified unclassified | 0,4 | 0,489897949 | 1,02E-05 | 1,26E-05 | 1 - | 2,32E-05 - | 0 - | 0 - | 0 - | 0 - | 1 - | 2,78E-05 - | 0 - | 0 - |
| 7261 | 1 | 3um_FIJJAFBF_176386 | Bacteria | Proteobacteri Gammaprotec Chromatiales unclassified unclassified Chromatiales metabolism Inorganic ion 116STV Binding-protei 1000565.MET | 0,2 | 0,4 | 9,93E-06 | 1,99E-05 | 0 - | 0 - | 0 - | 0 - | 1 - | 4,96E-05 - | 0 - | 0 - | 0 - | 0 - |
| 7262 | 1 | 0.2um_NOKCPHJN_540896 | Bacteria | Proteobacteri Alphaproteob Pelagibacteral Pelagibacterac Candidatus Pe Candidatus Pe unclassified unclassified unclassified unclassified unclassified | 4 | 1,095445115 | 0,000127816 | 2,26E-05 | 6 - | 0,000164358 - | 4 - | 0,000136056 - | 3 - | 0,000129339 - | 3 - | 9,84E-05 - | 3 - | 0,000110889 - |
| 7263 | 1 | 3um_FIJJAFBF_451386 | Bacteria | Proteobacteri Gammaprotec Oceanospirilla Saccharospiril Reinekea Reinekea fors poorly charact Function unkn 075W9 High frequenc 314283.MED | 0,2 | 0,4 | 1,10E-05 | 2,21E-05 | 1 - | 5,52E-05 - | 0 - | 0 - | 0 - | 0 - | 0 - | 0 - | 0 - | 0 - |
| 7264 | 1 | 3um_FIJJAFBF_85783 | Viruses | unclassified unclassified unclassified unclassified unclassified uncultured M unclassified unclassified unclassified unclassified unclassified | 2,6 | 0,8 | 0,000174415 | 5,69E-05 | 2 - | 0,000111977 - | 3 - | 0,000208564 - | 2 - | 0,000132178 - | 4 - | 0,00026826 - | 2 - | 0,000151097 - |
| 7265 | 1 | 3um_FIJJAFBF_711547 | Bacteria | Bacteroidetes unclassified unclassified unclassified unclassified Bacteroidetes unclassified unclassified unclassified unclassified unclassified | 0,2 | 0,4 | 9,07E-06 | 1,81E-05 | 1 - | 4,54E-05 - | 0 - | 0 - | 0 - | 0 - | 0 - | 0 - | 0 - | 0 - |
| 7266 | 1 | 3um_FIJJAFBF_279337 | Bacteria | Actinobacteri Actinobacteri Micrococcales Microbacteria Candidatus Ac Candidatus Ac cellular proce: Posttranslatio 00MJH Peptidyl-proly 281090.Lxx1 | 0,2 | 0,4 | 8,36E-06 | 1,67E-05 | 0 - | 0 - | 0 - | 0 - | 1 - | 4,18E-05 - | 0 - | 0 - | 0 - | 0 - |
| 7267 | 1 | 3um_FIJJAFBF_100642 | Bacteria | Proteobacteri Gammaprotec Cellvibrionale: Porticoccacea unclassified gamma prote information st Translation, ri 16QAF thus facilitati 314287.GB22 | 2,8 | 1,166190379 | 6,84E-05 | 2,93E-05 | 4 - | 8,24E-05 - | 1 - | 2,56E-05 - | 2 - | 4,86E-05 - | 3 - | 7,40E-05 - | 4 - | 0,00011122 - |
| 7268 | 1 | 3um_FIJJAFBF_464954 | Bacteria | Bacteroidetes Flavobacteria Flavobacterial Flavobacteriac Ulvibacter Ulvibacter sp. metabolism Energy produc 0P7SC Delta-1-pyrrol 391603.FBAL | 2,2 | 1,166190379 | 2,86E-05 | 2,83E-05 | 2 - | 4,24E-05 - | 0 - | 0 - | 3 - | 7,51E-05 - | 1 - | 2,54E-05 - | 0 - | 0 - |
| 7269 | 1 | 0.2um_NOKCPHJN_376481 | Bacteria | Bacteroidetes Cytophagia Cytophagales Flammeovirga unclassified Flammeovirga poorly charact Function unkn 0XR4Y Ragb susd dor 886377.Muri | 19,6 | 2,059126028 | 0,000477272 | 4,93E-05 | 23 - | 0,000471405 - | 20 - | 0,000508996 - | 18 - | 0,000435479 - | 17 - | 0,000417736 - | 20 - | 0,000553123 - |
| 7270 | 1 | 0.2um_NOKCPHJN_72842 | Bacteria | Proteobacteri Gammaprotec Oceanospirilla Saccharospiril Reinekea Reinekea blan metabolism Energy produc 05D9Q decarboxylase 314283.MED | 0,4 | 0,8 | 1,22E-05 | 2,44E-05 | 2 - | 6,11E-05 - | 0 - | 0 - | 0 - | 0 - | 0 - | 0 - | 0 - | 0 - |
| 7271 | 1 | 3um_FIJJAFBF_105537 | Eukaryota | Ascomycota Saccharomyce Saccharomyce Saccharomyce Saccharomyce Saccharomyce poorly charact Function unkn 0NVRZ NA 170187.SP_1 | 6,6 | 0,8 | 0,000247395 | 1,49E-05 | 8 - | 0,000253665 - | 6 - | 0,000236219 - | 6 - | 0,000224556 - | 7 - | 0,000265852 - | 6 - | 0,000256698 - |
| 7272 | 1 | 0.2um_NOKCPHJN_09238 | Viruses | unclassified unclassified unclassified Phycodnavirid Aureococcus Aureococcus unclassified unclassified unclassified unclassified unclassified | 2 | 0 | 5,71E-05 | 5,42E-06 | 2 - | 4,78E-05 - | 2 - | 5,94E-05 - | 2 - | 5,65E-05 - | 2 - | 5,73E-05 - | 2 - | 6,45E-05 - |
| 7273 | 1 | 0.2um_NOKCPHJN_717968 | Bacteria | Proteobacteri Betaproteoba Burkholderiale unclassified unclassified Burkholderiale unclassified unclassified unclassified unclassified unclassified | 0,8 | 0,4 | 1,24E-05 | 6,31E-06 | 1 - | 1,35E-05 - | 1 - | 1,67E-05 - | 1 - | 1,59E-05 - | 1 - | 1,61E-05 - | 0 - | 0 - |
| 7274 | 1 | 0.2um_NOKCPHJN_132140 | Eukaryota | unclassified Oomycetes Saprolegniale: Saprolegniace Aphanomyces Aphanomyces cellular proce: Cytoskeleton 1E1TK Dynein heavy 3055.EDO96 | 0,4 | 0,489897949 | 1,20E-06 | 1,47E-06 | 0 - | 0 - | 1 - | 2,98E-06 - | 1 - | 3,03E-06 - | 0 - | 0 - | 0 - | 0 - |
| 7275 | 1 | 0.2um_NOKCPHJN_196006 | Bacteria | Proteobacteri Alphaproteob Pelagibacteral Pelagibacterac Candidatus Pe Candidatus Pe metabolism Secondary me 01Q86 extracellular s 439493.PB72 | 12 | 1,095445115 | 0,000447736 | 3,25E-05 | 14 - | 0,000440024 - | 11 - | 0,000429509 - | 12 - | 0,000445421 - | 11 - | 0,000414333 - | 12 - | 0,000509176 - |
| 7276 | 1 | 3um_FIJJAFBF_67378 | Bacteria | Bacteroidetes Flavobacteria Flavobacterial Flavobacteriac Polaribacter Polaribacter s metabolism Carbohydrate COG0149 triosephospha 313603.FB21 | 0,4 | 0,489897949 | 2,18E-05 | 2,67E-05 | 0 - | 0 - | 1 - | 5,42E-05 - | 1 - | 5,50E-05 - | 0 - | 0 - | 0 - | 0 - |
| 7277 | 2 | 0.2um_NOKCPHJN_194770;3um_FIJJAFBF_1089 | Bacteria | Proteobacteri Alphaproteob Rhodobactera Rhodobactera Pseudaestuari Pseudaestuari metabolism Energy produc 01RS8 it plays a direc 367336.OM2 | 0,6 | 0,489897949 | 3,11E-05 | 2,56E-05 | 1 - | 4,61E-05 - | 0 - | 0 - | 1 - | 5,44E-05 - | 1 - | 5,52E-05 - | 0 - | 0 - |
| 7278 | 3 | 0.2um_NOKCPHJN_585560;0.2um_NOKCPHJN_6 | Bacteria | Proteobacteri Alphaproteob Rhodobactera Rhodobactera Planktomarin Planktomarin cellular proce: Function unkn 01RSK Required for r various | 0,2 | 0,4 | 1,21E-05 | 2,42E-05 | 1 - | 6,04E-05 - | 0 - | 0 - | 0 - | 0 - | 0 - | 0 - | 0 - | 0 - |
| 7279 | 1 | 3um_FIJJAFBF_08159 | Bacteria | Proteobacteri Alphaproteob Pelagibacteral Pelagibacterac Polaribacter various poorly charact Function unkn 05E4M (twin-arginine 313606.M23 | 1,4 | 0,489897949 | 3,45E-05 | 1,27E-05 | 1 - | 2,05E-05 - | 2 - | 5,10E-05 - | 2 - | 4,85E-05 - | 1 - | 2,46E-05 - | 1 - | 2,77E-05 - |
| 7280 | 1 | 3um_FIJJAFBF_176110 | Bacteria | Proteobacteri Gammaprotec Alteromonada Ferrimonadac Paraferrimon Paraferrimon cellular proce: Intracellular tr 17SGA One of the pri 323850.Shew | 1,6 | 0,489897949 | 0,000129869 | 3,60E-05 | 2 - | 0,000138285 - | 2 - | 0,000171709 - | 2 - | 0,000163232 - | 1 - | 8,28E-05 - | 1 - | 9,33E-05 - |
| 7281 | 1 | 3um_FIJJAFBF_442690 | Bacteria | Proteobacteri Gammaprotec Cellvibrionale: Halieaceae unclassified marine gamm unclassified unclassified unclassified unclassified unclassified | 0,2 | 0,4 | 1,35E-05 | 2,71E-05 | 0 - | 0 - | 0 - | 0 - | 1 - | 6,77E-05 - | 0 - | 0 - | 0 - | 0 - |
| 7282 | 1 | 3um_FIJJAFBF_469846 | Bacteria | Proteobacteri Alphaproteob Rhodobactera Leisingera Leisingera aqu cellular proce: Cell wall/mem 16QAK protease 314262.MED | 3,4 | 1,2 | 0,000105258 | 3,81E-05 | 2 - | 5,17E-05 - | 4 - | 0,000128395 - | 5 - | 0,00015257 - | 4 - | 0,000123859 - | 2 - | 6,98E-05 - |
| 7283 | 1 | 0.2um_NOKCPHJN_353236 | Bacteria | Proteobacteri Alphaproteob Pelagibacteral unclassified unclassified Pelagibacteral cellular proce: Cell wall/mem 7257 NA 439493.PB72 | 0,6 | 0,489897949 | 2,52E-05 | 2,07E-05 | 0 - | 0 - | 1 - | 4,00E-05 - | 1 - | 4,06E-05 - | 1 - | 4,06E-05 - | 1 - | 4,57E-05 - |
| 7284 | 1 | 3um_FIJJAFBF_430367 | Bacteria | Proteobacteri Alphaproteob Rhodobactera unclassified unclassified Rhodobactera information st Translation, ri 16QAF thus facilitatin 439496.RB2 | 4,4 | 1,624807681 | 0,000107761 | 4,17E-05 | 6 - | 0,000123195 - | 3 - | 7,27E-05 - | 2 - | 4,92E-05 - | 5 - | 0,000127477 - | 6 - | 0,000166234 - |
| 7285 | 1 | 3um_FIJJAFBF_28898 | Bacteria | Bacteroidetes Flavobacteria Flavobacterial Flavobacteriac unclassified Flavobacteria unclassified unclassified unclassified unclassified unclassified | 1 | 0 | 6,40E-05 | 6,08E-06 | 1 - | 5,36E-05 - | 1 - | 6,66E-05 - | 1 - | 6,33E-05 - | 1 - | 6,42E-05 - | 1 - | 7,24E-05 - |
| 7286 | 2 | 0.2um_NOKCPHJN_845165;3um_FIJJAFBF_3859 | Bacteria | Proteobacteri Gammaprotec Cellvibrionale: Porticoccacea unclassified Porticoccacea unclassified unclassified unclassified unclassified unclassified | 0,2 | 0,4 | 1,93E-05 | 3,86E-05 | 1 - | 9,65E-05 - | 0 - | 0 - | 0 - | 0 - | 0 - | 0 - | 0 - | 0 - |
| 7287 | 1 | 0.2um_NOKCPHJN_1088455 | Eukaryota | Chlorophyta Mamiellophyc Mamiellales Bathycoccace Bathycoccus Bathycoccus metabolism Energy produc COG0055 Produces ATP 436017.A4RS | 1,2 | 0,4 | 0,000150927 | 3,34E-05 | 2 - | 0,00021656 - | 1 - | 0,000134452 - | 1 - | 0,000127813 - | 1 - | 0,000129701 - | 1 - | 0,000146108 - |

| | | | | | | | | | | | | | | | | | | | | | | | | | |
|---|---|---|---|---|---|---|---|---|---|---|---|---|---|---|---|---|---|---|---|---|---|---|---|---|---|
| 7288 | 1 | 3um_FIJJAFBF_536779 | Bacteria | Proteobacteri | Gammaprotec | Oceanospirilla | Saccharospirill | Reinekea | Reinekea fors | cellular proces | Signal transdu | 07EW6 | Cyclic nucleoti | 314283.MED2 | 0,4 | 0,489897949 | 3,78E-05 | 4,63E-05 | 0 - | 0 - | 1 - | 9,70E-05 - | 1 - | 9,22E-05 - | 0 - | 0 - | 0 - | 0 - |
| 7289 | 1 | 3um_FIJJAFBF_369394 | Bacteria | Proteobacteri | Gammaprotec | Alteromonada | Alteromonada | Paraglaciecola | Paraglaciecola | cellular proces | Intracellular tr | 175GA | One of the prc | 983545.Glaag | 6,2 | 0,748331477 | 0,000493987 | 5,51E-05 | 7 - | 0,000469846 - | 6 - | 0,000500066 - | 7 - | 0,000554606 - | 5 - | 0,000401998 - | 6 - | 0,000543419 - |
| 7290 | 1 | 3um_FIJJAFBF_189726 | Bacteria | Bacteroidetes | Flavobacteriia | Flavobacterial | Flavobacteriac | Polaribacter | Polaribacter s | poorly charact | Function unkn | 0PAA9 | Inherit from b | 313598.MED1 | 5,8 | 2,638181192 | 0,000365258 | 0,000121119 | 11 - | 0,000604088 - | 4 - | 0,000272763 - | 5 - | 0,00032412 - | 5 - | 0,000328907 - | 4 - | 0,00029641 - |
| 7291 | 1 | 3um_FIJJAFBF_494164 | Bacteria | Bacteroidetes | Flavobacteriia | Flavobacterial | Flavobacteriac | Polaribacter | Polaribacter t | unclassified | unclassified | unclassified | unclassified | unclassified | 1,4 | 0,8 | 0,000140158 | 8,07E-05 | 2 - | 0,000173904 - | 2 - | 0,000215938 - | 1 - | 0,000102638 - | 2 - | 0,000208308 - | 0 - | 0 - |
| 7292 | 2 | 0.2um_NOKCPHJN_261113;3um_FIJJAFBF_76555 | Bacteria | Proteobacteri | Alphaproteob | Rhodobactera | Rhodobactera | Planktomarina | Planktomarina | poorly charact | Function unkn | 10QGR | glyoxalase fan | 765698.Mesci | 0,6 | 0,489897949 | 4,08E-05 | 3,36E-05 | 1 - | 6,04E-05 - | 0 - | 0 - | 1 - | 7,13E-05 - | 1 - | 7,24E-05 - | 0 - | 0 - |
| 7293 | 1 | 3um_FIJJAFBF_606922 | Bacteria | Proteobacteri | Alphaproteob | Rhodobactera | Rhodobactera | Sulfitobacter | Sulfitobacter g | metabolism | Amino acid tra | COG1448 | aminotransfer | 52598.EE36_0 | 0,4 | 0,489897949 | 1,28E-05 | 1,58E-05 | 1 - | 2,91E-05 - | 0 - | 0 - | 0 - | 0 - | 1 - | 3,49E-05 - | 0 - | 0 - |
| 7294 | 1 | 0.2um_NOKCPHJN_510401 | Bacteria | Proteobacteri | Alphaproteob | Rhodobactera | Rhodobactera | Ruegeria | Ruegeria litor | cellular proces | Posttranslatio | 05C2Z | ATP-depender | 644107.SL115 | 2,6 | 1,356465997 | 4,47E-05 | 2,43E-05 | 3 - | 4,44E-05 - | 4 - | 7,35E-05 - | 3 - | 5,24E-05 - | 3 - | 5,32E-05 - | 0 - | 0 - |
| 7295 | 1 | 3um_FIJJAFBF_502375 | Bacteria | Bacteroidetes | Flavobacteriia | Flavobacterial | Flavobacteriac | Gramella | Gramella sale | poorly charact | Function unkn | 0P88U | NA | 313603.FB217 | 6 | 0,632455532 | 0,000144389 | 1,59E-05 | 6 - | 0,000121457 - | 6 - | 0,000150814 - | 6 - | 0,000143367 - | 7 - | 0,000169732 - | 5 - | 0,000136574 - |
| 7296 | 1 | 3um_FIJJAFBF_321055 | unclassified | unclassified | unclassified | unclassified | unclassified | unclassified | unclassified | unclassified | unclassified | unclassified | unclassified | unclassified | 0,2 | 0,4 | 6,55E-05 | 0,000130936 | 0 - | 0 - | 0 - | 0 - | 0 - | 0 - | 1 - | 0,000327341 - | 0 - | 0 - |
| 7297 | 1 | 3um_FIJJAFBF_586299 | Eukaryota | unclassified | Dinophyceae | Suessiales | Symbiodiniace | Symbiodinium | Symbiodinium | unclassified | unclassified | unclassified | unclassified | unclassified | 2 | 0 | 0,000147341 | 1,40E-05 | 2 - | 0,000123416 - | 2 - | 0,000153246 - | 2 - | 0,00014568 - | 2 - | 0,000147832 - | 2 - | 0,000166532 - |
| 7298 | 1 | 3um_FIJJAFBF_619968 | Bacteria | Proteobacteri | Gammaprotec | unclassified | unclassified | unclassified | Gammaprotec | information st | Translation, ril | COG0256 | This is one of | 999141.GME_ | 2,6 | 0,489897949 | 0,000310887 | 7,89E-05 | 2 - | 0,000197891 - | 3 - | 0,000368583 - | 3 - | 0,000350385 - | 2 - | 0,00023704 - | 3 - | 0,000400537 - |
| 7299 | 1 | 3um_FIJJAFBF_213368 | Bacteria | Proteobacteri | Gammaprotec | Oceanospirilla | Saccharospirill | Reinekea | Reinekea fors | information st | Transcription | 17CVY | Involved in the | 314283.MED2 | 0,4 | 0,489897949 | 3,62E-05 | 4,43E-05 | 0 - | 0 - | 0 - | 0 - | 1 - | 8,97E-05 - | 1 - | 9,10E-05 - | 0 - | 0 - |
| 7300 | 1 | 3um_FIJJAFBF_231021 | Bacteria | Actinobacteri | Actinobacteri | Micrococcales | Microbacteria | Candidatus Ac | Candidatus Ac | metabolism | Energy produc | COG0711 | F(1)F(0) ATP s | 312284.A20C1 | 0,8 | 0,748331477 | 5,70E-05 | 4,95E-05 | 2 - | 0,000125439 - | 0 - | 0 - | 0 - | 0 - | 1 - | 7,51E-05 - | 1 - | 8,46E-05 - |
| 7301 | 1 | 3um_FIJJAFBF_417059 | Bacteria | Proteobacteri | Gammaprotec | Alteromonada | Alteromonada | Paraglaciecola | Paraglaciecola | unclassified | unclassified | unclassified | unclassified | unclassified | 4,6 | 0,489897949 | 0,000350315 | 3,63E-05 | 5 - | 0,000320605 - | 5 - | 0,000398097 - | 4 - | 0,000302754 - | 5 - | 0,000384032 - | 4 - | 0,000346088 - |
| 7302 | 1 | 3um_FIJJAFBF_370789 | Bacteria | Proteobacteri | Gammaprotec | Oceanospirilla | Saccharospirill | Saccharospiril | Saccharospiril | metabolism | Inorganic ion t | 0QJ3Y | Part of the AB | 377629.TERTU | 6,6 | 0,8 | 0,000320849 | 1,69E-05 | 8 - | 0,000329109 - | 6 - | 0,000306492 - | 7 - | 0,00033992 - | 6 - | 0,000295663 - | 6 - | 0,000333063 - |
| 7303 | 1 | 3um_FIJJAFBF_323002 | Bacteria | Proteobacteri | Alphaproteob | Rhodobactera | Rhodobactera | Roseobacter | Roseobacter s | poorly charact | Function unkn | 01QWY | Bacterial prot | 314262.MED1 | 0,4 | 0,489897949 | 1,24E-05 | 1,52E-05 | 0 - | 0 - | 1 - | 3,17E-05 - | 1 - | 3,02E-05 - | 0 - | 0 - | 0 - | 0 - |
| 7304 | 1 | 3um_FIJJAFBF_601585 | Bacteria | Proteobacteri | Gammaprotec | unclassified | unclassified | unclassified | Gammaprotec | unclassified | unclassified | unclassified | unclassified | unclassified | 0,6 | 0,489897949 | 2,82E-05 | 2,32E-05 | 1 - | 4,17E-05 - | 0 - | 0 - | 1 - | 4,93E-05 - | 1 - | 5,00E-05 - | 0 - | 0 - |
| 7305 | 1 | 3um_FIJJAFBF_511983 | Eukaryota | unclassified | Cryptophyta | Pyrenomonad | Geminigerace | Proteomonas | Proteomonas | metabolism | Energy produc | COG0056 | Produces ATP | 70448.Q0P3H | 0,6 | 0,489897949 | 1,89E-05 | 1,55E-05 | 0 - | 0 - | 1 - | 3,10E-05 - | 0 - | 0 - | 1 - | 2,99E-05 - | 1 - | 3,37E-05 - |
| 7306 | 1 | 3um_FIJJAFBF_398711 | Bacteria | Bacteroidetes | Flavobacteriia | Flavobacterial | Flavobacteriac | Ulvibacter | Ulvibacter sp. | unclassified | unclassified | unclassified | unclassified | unclassified | 2,4 | 0,489897949 | 0,000262511 | 4,64E-05 | 3 - | 0,000277686 - | 3 - | 0,000344804 - | 2 - | 0,00021852 - | 2 - | 0,000221747 - | 2 - | 0,000249798 - |
| 7307 | 1 | 3um_FIJJAFBF_778443 | Bacteria | Bacteroidetes | Flavobacteriia | Flavobacterial | Flavobacteriac | Polaribacter | Polaribacter ir | unclassified | unclassified | unclassified | unclassified | unclassified | 0,4 | 0,489897949 | 4,12E-05 | 5,09E-05 | 1 - | 9,18E-05 - | 1 - | 0,000114015 - | 0 - | 0 - | 0 - | 0 - | 0 - | 0 - |
| 7308 | 1 | 3um_FIJJAFBF_348326 | Bacteria | Bacteroidetes | Flavobacteriia | Flavobacterial | Flavobacteriac | Algibacter | Algibacter lect | poorly charact | Function unkn | 0P8NI | Inherit from b | 313594.PI23P | 1,2 | 0,4 | 2,38E-05 | 5,27E-06 | 2 - | 3,42E-05 - | 1 - | 2,12E-05 - | 1 - | 2,02E-05 - | 1 - | 2,05E-05 - | 1 - | 2,30E-05 - |
| 7309 | 1 | 3um_FIJJAFBF_37678 | Bacteria | Bacteroidetes | Cytophagia | Cytophagales | Flammeovirga | unclassified | Flammeovirga | cellular proces | Intracellular tr | 05FGH | MotA TolQ exl | 643867.Ftrac_ | 4,2 | 0,748331477 | 0,000204136 | 4,40E-05 | 3 - | 0,000121243 - | 4 - | 0,000200731 - | 5 - | 0,000238525 - | 5 - | 0,000242048 - | 4 - | 0,000218133 - |
| 7310 | 1 | 0.2um_NOKCPHJN_65176 | Bacteria | Proteobacteri | Gammaprotec | Cellvibrionale | Porticoccacea | unclassified | Gammaprotec | metabolism | Energy produc | 16PVJ | Component of | 377629.TERTU | 0,8 | 0,4 | 1,20E-05 | 6,11E-06 | 1 - | 1,30E-05 - | 1 - | 1,62E-05 - | 1 - | 1,54E-05 - | 1 - | 1,56E-05 - | 0 - | 0 - |
| 7311 | 1 | 0.2um_NOKCPHJN_405787 | Bacteria | Verrucomicrol | unclassified | unclassified | unclassified | unclassified | Verrucomicrol | poorly charact | Function unkn | 05EV9 | Alpha Beta Hy | 583355.Caka_ | 0,2 | 0,4 | 1,05E-05 | 2,10E-05 | 0 - | 0 - | 0 - | 0 - | 1 - | 5,25E-05 - | 0 - | 0 - | 0 - | 0 - |
| 7312 | 1 | 0.2um_NOKCPHJN_129290 | Bacteria | Bacteroidetes | unclassified | unclassified | unclassified | unclassified | Bacteroidetes | unclassified | unclassified | unclassified | unclassified | unclassified | 0,4 | 0,489897949 | 4,82E-06 | 5,94E-06 | 1 - | 1,10E-05 - | 0 - | 0 - | 0 - | 0 - | 1 - | 1,31E-05 - | 0 - | 0 - |
| 7313 | 1 | 3um_FIJJAFBF_103155 | Bacteria | Bacteroidetes | Flavobacteriia | Flavobacterial | Flavobacteriac | Polaribacter | Polaribacter h | unclassified | unclassified | unclassified | unclassified | unclassified | 0,4 | 0,8 | 6,29E-05 | 0,000125783 | 2 - | 0,000314457 - | 0 - | 0 - | 0 - | 0 - | 0 - | 0 - | 0 - | 0 - |
| 7314 | 1 | 3um_FIJJAFBF_365316 | Bacteria | Bacteroidetes | Flavobacteriia | unclassified | unclassified | unclassified | Flavobacteria | unclassified | unclassified | unclassified | unclassified | unclassified | 5,4 | 1,2 | 0,000320308 | 6,08E-05 | 6 - | 0,000302044 - | 4 - | 0,000250033 - | 7 - | 0,000415954 - | 6 - | 0,000361798 - | 4 - | 0,00027171 - |
| 7315 | 1 | 3um_FIJJAFBF_111404 | unclassified | unclassified | unclassified | unclassified | unclassified | unclassified | unclassified | unclassified | unclassified | unclassified | unclassified | unclassified | 0,6 | 0,8 | 6,59E-05 | 8,43E-05 | 2 - | 0,000203145 - | 1 - | 0,000126123 - | 0 - | 0 - | 0 - | 0 - | 0 - | 0 - |
| 7316 | 3 | 0.2um_NOKCPHJN_189206;3um_FIJJAFBF_63645 | Bacteria | Bacteroidetes | Flavobacteriia | Flavobacterial | Flavobacteriac | Formosa | Formosa sp. H | cellular proces | Cell motility | COG1360 | Ompa motb d | various | 0,8 | 0,4 | 3,45E-05 | 1,77E-05 | 1 - | 3,60E-05 - | 1 - | 4,47E-05 - | 0 - | 0 - | 1 - | 4,31E-05 - | 1 - | 4,85E-05 - |
| 7317 | 1 | 0.2um_NOKCPHJN_972643 | Bacteria | Bacteroidetes | Flavobacteriia | Flavobacterial | Flavobacteriac | Chryseobacte | various | information st | Translation, ril | 05C64 | Binds directly | 525257.HMPR | 0,6 | 0,489897949 | 3,66E-05 | 3,02E-05 | 1 - | 5,31E-05 - | 1 - | 6,60E-05 - | 0 - | 0 - | 1 - | 6,36E-05 - | 0 - | 0 - |
| 7318 | 1 | 3um_FIJJAFBF_263771 | Bacteria | Actinobacteri | unclassified | unclassified | unclassified | unclassified | Actinobacteri | unclassified | unclassified | unclassified | unclassified | unclassified | 1,2 | 0,748331477 | 0,000130392 | 7,63E-05 | 2 - | 0,000186629 - | 1 - | 0,000115869 - | 0 - | 0 - | 2 - | 0,00022355 - | 1 - | 0,000125914 - |
| 7319 | 1 | 3um_FIJJAFBF_690639 | Bacteria | Bacteroidetes | Cytophagia | Cytophagales | Flammeovirga | unclassified | Flammeovirga | unclassified | unclassified | unclassified | unclassified | unclassified | 1,2 | 0,979795897 | 0,000104575 | 8,82E-05 | 1 - | 7,45E-05 - | 1 - | 9,25E-05 - | 1 - | 8,80E-05 - | 3 - | 0,000267825 - | 0 - | 0 - |
| 7320 | 1 | 0.2um_NOKCPHJN_93294 | Bacteria | Proteobacteri | Alphaproteob | Pelagibacteral | Pelagibacterac | Candidatus Pe | Candidatus Pe | cellular proces | Posttranslatio | 17FI0 | Binds to Cpn6 | 335992.SAR11 | 2,2 | 0,748331477 | 0,000279554 | 8,14E-05 | 3 - | 0,000327934 - | 2 - | 0,000271465 - | 2 - | 0,000258061 - | 3 - | 0,000392809 - | 1 - | 0,0001475 - |
| 7321 | 1 | 3um_FIJJAFBF_317844 | Bacteria | Bacteroidetes | Flavobacteriia | Flavobacterial | Crocinitomica | unclassified | Crocinitomica | unclassified | unclassified | unclassified | unclassified | unclassified | 3,4 | 0,8 | 6,07E-05 | 1,16E-05 | 4 - | 6,09E-05 - | 3 - | 5,67E-05 - | 4 - | 7,19E-05 - | 4 - | 7,29E-05 - | 2 - | 4,11E-05 - |
| 7322 | 1 | 3um_FIJJAFBF_36156 | Bacteria | Bacteroidetes | Flavobacteriia | Flavobacterial | Crocinitomica | unclassified | Crocinitomica | poorly charact | Function unkn | 099QV | gliding motilit | 743722.Sph21 | 1,6 | 0,489897949 | 7,54E-05 | 2,09E-05 | 2 - | 8,03E-05 - | 2 - | 9,97E-05 - | 2 - | 9,47E-05 - | 1 - | 4,81E-05 - | 1 - | 5,42E-05 - |
| 7323 | 1 | 3um_FIJJAFBF_556412 | Bacteria | Proteobacteri | Gammaprotec | Alteromonada | Alteromonada | Paraglaciecola | Paraglaciecola | metabolism | Lipid transpor | 0QN5A | Involved in un | 342610.Patl_1 | 0,8 | 0,4 | 7,33E-05 | 3,77E-05 | 1 - | 7,76E-05 - | 0 - | 0 - | 1 - | 9,15E-05 - | 1 - | 9,29E-05 - | 1 - | 0,000104645 - |
| 7324 | 1 | 3um_FIJJAFBF_155642 | Bacteria | Bacteroidetes | Flavobacteriia | Flavobacterial | Crocinitomica | unclassified | Crocinitomica | unclassified | unclassified | unclassified | unclassified | unclassified | 4 | 0,894427191 | 0,000420845 | 6,85E-05 | 5 - | 0,000448347 - | 3 - | 0,000334029 - | 5 - | 0,000529227 - | 4 - | 0,000429635 - | 3 - | 0,000362987 - |
| 7325 | 1 | 3um_FIJJAFBF_208625 | Bacteria | Bacteroidetes | Flavobacteriia | Flavobacterial | Flavobacteriac | Ulvibacter | Ulvibacter lito | unclassified | unclassified | unclassified | unclassified | unclassified | 0,8 | 0,4 | 4,01E-05 | 2,06E-05 | 1 - | 4,19E-05 - | 1 - | 5,20E-05 - | 0 - | 0 - | 1 - | 5,02E-05 - | 1 - | 5,65E-05 - |
| 7326 | 1 | 3um_FIJJAFBF_445116 | Bacteria | Bacteroidetes | Flavobacteriia | Flavobacterial | Flavobacteriac | Polaribacter | various | poorly charact | Function unkn | 0PAEG | NA | 688270.Celal_ | 0,8 | 0,4 | 6,20E-05 | 3,12E-05 | 0 - | 0 - | 1 - | 7,75E-05 - | 1 - | 7,36E-05 - | 1 - | 7,47E-05 - | 1 - | 8,42E-05 - |
| 7327 | 1 | 0.2um_NOKCPHJN_329634 | Eukaryota | Chordata | Actinopteri | Cyprinodontif | Poeciliidae | Poecilia | Poecilia latipi | unclassified | unclassified | unclassified | unclassified | unclassified | 1 | 0 | 4,54E-05 | 4,31E-06 | 1 - | 3,80E-05 - | 1 - | 4,72E-05 - | 1 - | 4,49E-05 - | 1 - | 4,55E-05 - | 1 - | 5,13E-05 - |
| 7328 | 1 | 0.2um_NOKCPHJN_861812 | Eukaryota | Bacillariophyt | Bacillariophyc | Naviculales | Naviculaceae | Fistulifera | Fistulifera sola | poorly charact | Function unkn | 110WY | Chlorophyll A- | 2850.Phatr18 | 1 | 0 | 0,000152252 | 1,45E-05 | 1 - | 0,00012753 - | 1 - | 0,000158354 - | 1 - | 0,000150536 - | 1 - | 0,000152759 - | 1 - | 0,000172083 - |
| 7329 | 1 | 0.2um_NOKCPHJN_548160 | Eukaryota | Phaeophyceae | unclassified | Ectocarpales | Ectocarpaceae | Ectocarpus | Ectocarpus sili | cellular proces | Signal transdu | 151NR | Pfam:efhand | 67593.Physo1 | 1,8 | 0,748331477 | 0,000153024 | 7,35E-05 | 2 - | 0,00014083 - | 1 - | 8,74E-05 - | 1 - | 8,31E-05 - | 2 - | 0,000168691 - | 3 - | 0,000285045 - |
| 7330 | 1 | 3um_FIJJAFBF_200767 | Bacteria | Proteobacteri | Gammaprotec | Alteromonada | Alteromonada | Paraglaciecola | Paraglaciecola | poorly charact | Function unkn | 05C2A | tonB-depende | 1085623.GNIT | 0,4 | 0,489897949 | 1,11E-05 | 1,36E-05 | 0 - | 0 - | 0 - | 0 - | 0 - | 0 - | 1 - | 2,61E-05 - | 1 - | 2,94E-05 - |
| 7331 | 1 | 3um_FIJJAFBF_382743 | Viruses | unclassified | unclassified | unclassified | Phycodnavirid | unclassified | Chrysochrom | unclassified | unclassified | unclassified | unclassified | unclassified | 1 | 0 | 7,10E-05 | 6,74E-06 | 1 - | 5,95E-05 - | 1 - | 7,38E-05 - | 1 - | 7,02E-05 - | 1 - | 7,12E-05 - | 1 - | 8,02E-05 - |
| 7332 | 1 | 0.2um_NOKCPHJN_876272 | Eukaryota | unclassified | Pelagophycea | Pelagomonad | unclassified | Aureococcus | Aureococcus a | cellular proces | Posttranslatio | 08V1T | peptidylprolyl | 513049.Amax | 1,8 | 0,4 | 0,000141119 | 3,51E-05 | 2 - | 0,000131173 - | 2 - | 0,000162879 - | 1 - | 7,74E-05 - | 2 - | 0,000157124 - | 2 - | 0,000176999 - |
| 7333 | 1 | 3um_FIJJAFBF_777827 | Bacteria | Bacteroidetes | Flavobacteriia | Flavobacterial | Flavobacteriac | Algibacter | Algibacter sp. | unclassified | unclassified | unclassified | unclassified | unclassified | 0,6 | 0,489897949 | 8,06E-05 | 6,65E-05 | 1 - | 0,000117119 - | 1 - | 0,000145427 - | 0 - | 0 - | 1 - | 0,000140289 - | 0 - | 0 - |
| 7334 | 1 | 3um_FIJJAFBF_305657 | Bacteria | Bacteroidetes | Flavobacteriia | Flavobacterial | Flavobacteriac | Formosa | Formosa sp. H | unclassified | unclassified | unclassified | unclassified | unclassified | 1,2 | 0,748331477 | 6,39E-05 | 3,83E-05 | 2 - | 9,11E-05 - | 2 - | 0,00011311 - | 1 - | 5,38E-05 - | 0 - | 0 - | 1 - | 6,15E-05 - |
| 7335 | 1 | 3um_FIJJAFBF_163209 | Bacteria | Proteobacteri | Gammaprotec | Cellvibrionale | Porticoccacea | unclassified | gamma prote | unclassified | unclassified | unclassified | unclassified | unclassified | 0,2 | 0,4 | 9,94E-06 | 1,99E-05 | 1 - | 4,97E-05 - | 0 - | 0 - | 0 - | 0 - | 0 - | 0 - | 0 - | 0 - |
| 7336 | 1 | 3um_FIJJAFBF_81228 | Bacteria | Proteobacteri | Gammaprotec | Oceanospirilla | Endozoicomor | Endozoicomor | various | unclassified | unclassified | unclassified | unclassified | unclassified | 0,8 | 0,4 | 9,27E-05 | 4,67E-05 | 0 - | 0 - | 1 - | 0,000115869 - | 1 - | 0,000110148 - | 1 - | 0,000111775 - | 1 - | 0,000125914 - |
| 7337 | 1 | 0.2um_NOKCPHJN_355727 | Bacteria | Bacteroidetes | Flavobacteriia | Flavobacterial | Flavobacteriac | Formosa | Formosa sp. H | metabolism | Inorganic ion t | 05C27 | TonB-depende | 983544.Lacal_ | 16,8 | 1,326649916 | 0,000229409 | 5,57E-06 | 19 - | 0,000218952 - | 16 - | 0,000228946 - | 17 - | 0,000231245 - | 17 - | 0,00023466 - | 15 - | 0,000233245 - |
| 7338 | 1 | 3um_FIJJAFBF_574535 | unclassified | unclassified | unclassified | unclassified | unclassified | unclassified | unclassified | unclassified | unclassified | unclassified | unclassified | unclassified | 1 | 0 | 0,000171284 | 1,63E-05 | 1 - | 0,000143471 - | 1 - | 0,000178149 - | 1 - | 0,000169353 - | 1 - | 0,000171854 - | 1 - | 0,000193593 - |
| 7339 | 1 | 0.2um_NOKCPHJN_990892 | Eukaryota | unclassified | unclassified | Prymnesiales | Chrysochrom | Chrysochrom | Chrysochrom | unclassified | unclassified | unclassified | unclassified | unclassified | 1 | 0 | 6,82E-05 | 6,47E-06 | 1 - | 5,71E-05 - | 1 - | 7,09E-05 - | 1 - | 6,74E-05 - | 1 - | 6,84E-05 - | 1 - | 7,71E-05 - |
| 7340 | 1 | 0.2um_NOKCPHJN_893335 | Bacteria | Bacteroidetes | Flavobacteriia | Flavobacterial | Flavobacteriac | Formosa | Formosa sp. H | unclassified | unclassified | unclassified | unclassified | unclassified | 2,6 | 1,356465997 | 0,000129405 | 5,41E-05 | 5 - | 0,00021904 - | 2 - | 0,000108793 - | 3 - | 0,000155132 - | 2 - | 0,000104949 - | 1 - | 5,91E-05 - |
| 7341 | 1 | 3um_FIJJAFBF_176638 | Bacteria | Bacteroidetes | unclassified | unclassified | unclassified | unclassified | Bacteroidetes | unclassified | unclassified | unclassified | unclassified | unclassified | 0,2 | 0,4 | 1,88E-05 | 3,76E-05 | 1 - | 9,41E-05 - | 0 - | 0 - | 0 - | 0 - | 0 - | 0 - | 0 - | 0 - |
| 7342 | 1 | 3um_FIJJAFBF_228661 | Bacteria | Bacteroidetes | Flavobacteriia | Flavobacterial | Flavobacteriac | Formosa | Formosa sp. H | poorly charact | Function unkn | 0P8NI | Inherit from b | 983544.Lacal_ | 4,8 | 0,4 | 0,000158086 | 2,03E-05 | 5 - | 0,000137953 - | 5 - | 0,000171297 - | 5 - | 0,000162839 - | 4 - | 0,000132195 - | 5 - | 0,000186147 - |
| 7343 | 1 | 0.2um_NOKCPHJN_530414 | Bacteria | Proteobacteri | Gammaprotec | unclassified | unclassified | unclassified | Gammaprotec | information st | Translation, ril | COG0087 | One of the pri | 203122.Sde_0 | 3 | 0,894427191 | 0,000191811 | 6,41E-05 | 2 - | 0,000105785 - | 4 - | 0,000262708 - | 4 - | 0,000249737 - | 2 - | 0,000126713 - | 3 - | 0,000214112 - |
| 7344 | 1 | 0.2um_NOKCPHJN_792193 | unclassified | unclassified | unclassified | unclassified | unclassified | unclassified | unclassified | unclassified | unclassified | unclassified | unclassified | unclassified | 7 | 0,632455532 | 0,000420522 | 5,37E-05 | 7 - | 0,000352385 - | 7 - | 0,000437558 - | 8 - | 0,000475376 - | 6 - | 0,000361798 - | 7 - | 0,000475492 - |
| 7345 | 1 | 0.2um_NOKCPHJN_295066 | Bacteria | Proteobacteri | Alphaproteob | Rhodobactera | unclassified | unclassified | Rhodobactera | unclassified | unclassified | unclassified | unclassified | unclassified | 1 | 0 | 2,67E-05 | 2,53E-06 | 1 - | 2,23E-05 - | 1 - | 2,77E-05 - | 1 - | 2,64E-05 - | 1 - | 2,67E-05 - | 1 - | 3,01E-05 - |
| 7346 | 1 | 0.2um_NOKCPHJN_21288 | Bacteria | Proteobacteri | unclassified | unclassified | unclassified | unclassified | uncultured ma | poorly charact | Function unkn | 177TV | Photosyntheti | 633131.TR2A6 | 0,4 | 0,489897949 | 2,00E-05 | 2,48E-05 | 1 - | 4,47E-05 - | 1 - | 5,55E-05 - | 0 - | 0 - | 0 - | 0 - | 0 - | 0 - |
| 7347 | 3 | 0.2um_NOKCPHJN_363828;0.2um_NOKCPHJN_4 | Bacteria | various | various | various | various | various | various | information st | Translation, ril | 09FE5 | One of the prc | 983544.Lacal_ | 1,4 | 0,489897949 | 0,000186039 | 6,57E-05 | 1 - | 0,000111434 - | 1 - | 0,000138368 - | 2 - | 0,000263072 - | 2 - | 0,000266958 - | 1 - | 0,000150364 - |
| 7348 | 1 | 0.2um_NOKCPHJN_938763 | Eukaryota | unclassified | Oomycetes | Peronosporale | Peronosporac | Phytophthora | various | unclassified | unclassified | unclassified | unclassified | unclassified | 1 | 0 | 6,40E-05 | 6,08E-06 | 1 - | 5,36E-05 - | 1 - | 6,66E-05 - | 1 - | 6,33E-05 - | 1 - | 6,42E-05 - | 1 - | 7,24E-05 - |
| 7349 | 1 | 0.2um_NOKCPHJN_41565 | Bacteria | Bacteroidetes | Flavobacteriia | Flavobacterial | Crocinitomica | unclassified | Crocinitomica | information st | Transcription | 05D27 | DNA-depende | 755732.Fluta_ | 7,8 | 1,166190379 | 7,38E-05 | 8,90E-06 | 9 - | 7,20E-05 - | 9 - | 8,94E-05 - | 7 - | 6,61E-05 - | 8 - | 7,67E-05 - | 6 - | 6,48E-05 - |
| 7350 | 1 | 0.2um_NOKCPHJN_785122 | Eukaryota | Arthropoda | Insecta | Hymenoptera | Braconidae | Microplitis | Microplitis de | cellular proces | Posttranslatio | 0SRIS | plasma glutan | 7176.CPIJ0011 | 0,4 | 0,489897949 | 1,76E-05 | 2,19E-05 | 1 - | 3,74E-05 - | 0 - | 0 - | 0 - | 0 - | 0 - | 0 - | 1 - | 5,04E-05 - |
| 7351 | 1 | 3um_FIJJAFBF_290205 | Bacteria | Bacteroidetes | Flavobacteriia | Flavobacterial | Flavobacteriac | Marinirhabdu | Marinirhabdu | cellular proces | Intracellular tr | 05FGH | MotA TolQ exl | 50743.SCB49_ | 1 | 0 | 7,29E-05 | 6,92E-06 | 1 - | 6,11E-05 - | 1 - | 7,58E-05 - | 1 - | 7,21E-05 - | 1 - | 7,31E-05 - | 1 - | 8,24E-05 - |
| 7352 | 1 | 0.2um_NOKCPHJN_543480 | Bacteria | Proteobacteri | Gammaprotec | Cellvibrionale | Halieaceae | unclassified | gamma prote | information st | Translation, ril | COG0087 | One of the pri | 565045.NOR5 | 0,2 | 0,4 | 1,27E-05 | 2,54E-05 | 0 - | 0 - | 0 - | 0 - | 1 - | 6,36E-05 - | 0 - | 0 - | 0 - | 0 - |
| 7353 | 1 | 3um_FIJJAFBF_43944 | Bacteria | Bacteroidetes | Flavobacteriia | Flavobacterial | Flavobacteriac | Formosa | Formosa sp. H | metabolism | Carbohydrate | COG0057 | glyceraldehyd | 313598.MED1 | 4 | 0,632455532 | 0,000163432 | 2,40E-05 | 4 - | 0,00013787 - | 4 - | 0,000171194 - | 5 - | 0,000203427 - | 4 - | 0,000165145 - | 3 - | 0,000139527 - |
| 7354 | 1 | 3um_FIJJAFBF_170538 | Bacteria | Proteobacteri | Alphaproteob | Pelagibacteral | Pelagibacterac | Candidatus Pe | Candidatus Pe | information st | Replication, re | 05CEC | DNA methylas | 325240.Sbal_ | 0,2 | 0,4 | 5,97E-06 | 1,19E-05 | 0 - | 0 - | 0 - | 0 - | 1 - | 2,98E-05 - | 0 - | 0 - | 0 - | 0 - |
| 7355 | 1 | 0.2um_NOKCPHJN_494280 | Eukaryota | Streptophyta | Charophyceae | Charales | Characeae | Chara | Chara braunii | information st | Replication, re | COG0513 | purine NTP-de | 148305.A4RJ4 | 0,8 | 0,4 | 5,50E-05 | 2,82E-05 | 1 - | 5,74E-05 - | 1 - | 7,13E-05 - | 0 - | 0 - | 1 - | 6,87E-05 - | 1 - | 7,74E-05 - |
| 7356 | 1 | 3um_FIJJAFBF_153634 | Bacteria | Bacteroidetes | Flavobacteriia | Flavobacterial | Flavobacteriac | Marinirhabdu | Marinirhabdu | metabolism | Energy produc | COG0711 | F(1)F(0) ATP s | 50743.SCB49_ | 0,6 | 0,489897949 | 4,97E-05 | 4,13E-05 | 1 - | 6,91E-05 - | 1 - | 8,59E-05 - | 0 - | 0 - | 0 - | 0 - | 1 - | 9,33E-05 - |
| 7357 | 1 | 3um_FIJJAFBF_704354 | Bacteria | Proteobacteri | Gammaprotec | unclassified | unclassified | unclassified | Gammaprotec | unclassified | unclassified | unclassified | unclassified | unclassified | 0,4 | 0,489897949 | 3,29E-05 | 4,10E-05 | 1 - | 7,00E-05 - | 0 - | 0 - | 0 - | 0 - | 0 - | 0 - | 1 - | 9,44E-05 - |
| 7358 | 1 | 0.2um_NOKCPHJN_252102 | Eukaryota | Apicomplexa | Conoidasida | Eucoccidiorida | Eimeriidae | Eimeria | Eimeria acerv | unclassified | unclassified | unclassified | unclassified | unclassified | 0,8 | 0,4 | 1,24E-05 | 6,22E-06 | 0 - | 0 - | 1 - | 1,54E-05 - | 1 - | 1,47E-05 - | 1 - | 1,49E-05 - | 1 - | 1,68E-05 - |
| 7359 | 1 | 3um_FIJJAFBF_141658 | Bacteria | Proteobacteri | Gammaprotec | Oceanospirilla | Saccharospirill | Reinekea | Reinekea blan | metabolism | Nucleotide tra | 16QX0 | Catalyzes the | 314283.MED2 | 4,6 | 1,496662955 | 0,000113425 | 3,21E-05 | 6 - | 0,000127059 - | 5 - | 0,000131475 - | 6 - | 0,00014998 - | 4 - | 0,000101464 - | 2 - | 5,71E-05 - |
| 7360 | 1 | 0.2um_NOKCPHJN_72249 | Bacteria | Bacteroidetes | Flavobacteriia | Flavobacterial | Flavobacteriac | Polaribacter | Polaribacter s | cellular proces | Posttranslatio | 07SFW | Trigger factor | 313598.MED1 | 1 | 1,095445115 | 3,08E-05 | 3,56E-05 | 1 - | 2,61E-05 - | 3 - | 9,72E-05 - | 1 - | 3,08E-05 - | 0 - | 0 - | 0 - | 0 - |
| 7361 | 1 | 0.2um_NOKCPHJN_112734 | Bacteria | Bacteroidetes | Cytophagia | Cytophagales | unclassified | unclassified | Cytophagales | metabolism | Carbohydrate | 099YR | Catalyzes the | 926562.Oweh | 1,4 | 0,489897949 | 4,50E-05 | 1,68E-05 | 1 - | 2,68E-05 - | 2 - | 6,64E-05 - | 1 - | 3,16E-05 - | 2 - | 6,41E-05 - | 1 - | 3,61E-05 - |
| 7362 | 1 | 0.2um_NOKCPHJN_56831 | Bacteria | Proteobacteri | Gammaprotec | Oceanospirilla | Saccharospirill | Reinekea | Reinekea blan | metabolism | Energy produc | COG2838 | Isocitrate deh | 314283.MED2 | 8,8 | 0,748331477 | 0,000161255 | 9,70E-06 | 10 - | 0,000154477 - | 9 - | 0,000172634 - | 8 - | 0,000145876 - | 9 - | 0,000166534 - | 8 - | 0,000166756 - |
| 7363 | 1 | 3um_FIJJAFBF_142115 | Bacteria | Proteobacteri | Gammaprotec | Cellvibrionale | Porticoccacea | unclassified | gamma prote | cellular proces | Posttranslatio | 16QA9 | Chaperone in | 314287.GB22 | 0,6 | 0,8 | 1,67E-05 | 2,15E-05 | 2 - | 5,24E-05 - | 0 - | 0 - | 1 - | 3,09E-05 - | 0 - | 0 - | 0 - | 0 - |
| 7364 | 1 | 0.2um_NOKCPHJN_236282 | Bacteria | unclassified | unclassified | unclassified | unclassified | unclassified | uncultured ba | metabolism | Inorganic ion t | 05C27 | TonB-depende | 867900.Celly_ | 7,4 | 1,854723699 | 9,78E-05 | 2,46E-05 | 9 - | 0,000100193 - | 10 - | 0,000138234 - | 6 - | 7,88E-05 - | 5 - | 6,67E-05 - | 7 - | 0,000105152 - |
| 7365 | 1 | 3um_FIJJAFBF_588338 | Bacteria | unclassified | unclassified | unclassified | unclassified | unclassified | uncultured ba | unclassified | unclassified | unclassified | unclassified | unclassified | 1 | 0 | 1,75E-05 | 1,66E-06 | 1 - | 1,47E-05 - | 1 - | 1,82E-05 - | 1 - | 1,73E-05 - | 1 - | 1,76E-05 - | 1 - | 1,98E-05 - |
| 7366 | 1 | 3um_FIJJAFBF_164685 | Bacteria | Bacteroidetes | Flavobacteriia | Flavobacterial | Flavobacteriac | Polaribacter | Polaribacter ir | unclassified | unclassified | unclassified | unclassified | unclassified | 8,2 | 1,326649916 | 0,000210508 | 1,84E-05 | 10 - | 0,000218207 - | 8 - | 0,000216759 - | 9 - | 0,000231814 - | 8 - | 0,0002091 - | 6 - | 0,000176663 - |
| 7367 | 1 | 0.2um_NOKCPHJN_72905 | Bacteria | Proteobacteri | Alphaproteob | Rhodobactera | unclassified | unclassified | Rhodobactera | unclassified | unclassified | unclassified | unclassified | unclassified | 1,4 | 0,8 | 1,97E-05 | 1,18E-05 | 1 - | 1,19E-05 - | 2 - | 2,96E-05 - | 2 - | 2,82E-05 - | 2 - | 2,86E-05 - | 0 - | 0 - |
| 7368 | 1 | 3um_FIJJAFBF_56129 | Bacteria | Bacteroidetes | Flavobacteriia | Flavobacterial | Flavobacteriac | Polaribacter | Polaribacter s | information st | Replication, re | COG0513 | purine NTP-de | 313598.MED1 | 1 | 0 | 3,15E-05 | 2,99E-06 | 1 - | 2,64E-05 - | 1 - | 3,28E-05 - | 1 - | 3,11E-05 - | 1 - | 3,16E-05 - | 1 - | 3,56E-05 - |
| 7369 | 1 | 3um_FIJJAFBF_207478 | unclassified | unclassified | unclassified | unclassified | unclassified | unclassified | unclassified | unclassified | unclassified | unclassified | unclassified | unclassified | 0,6 | 0,489897949 | 0,000105685 | 8,79E-05 | 1 - | 0,00014715 - | 1 - | 0,000182716 - | 0 - | 0 - | 0 - | 0 - | 1 - | 0,000198557 - |
| 7370 | 1 | 3um_FIJJAFBF_463910 | Bacteria | Proteobacteri | Alphaproteob | Rhodobactera | Rhodobactera | Amylibacter | Amylibacter k | metabolism | Carbohydrate | 01RFA | ABC transport | 411684.HPDF | 5,4 | 0,8 | 0,000215781 | 3,75E-05 | 5 - | 0,000167313 - | 5 - | 0,000207753 - | 5 - | 0,000197496 - | 7 - | 0,000280578 - | 5 - | 0,000225765 - |
| 7371 | 1 | 3um_FIJJAFBF_461258 | Bacteria | Proteobacteri | Alphaproteob | Rhodobactera | Rhodobactera | unclassified | Rhodobactera | metabolism | Carbohydrate | 0XRJK | TRAP dicarbox | 439497.RR11_ | 0,4 | 0,489897949 | 1,49E-05 | 1,83E-05 | 1 - | 3,42E-05 - | 0 - | 0 - | 1 - | 4,03E-05 - | 0 - | 0 - | 0 - | 0 - |
| 7372 | 1 | 0.2um_NOKCPHJN_315402 | Bacteria | Bacteroidetes | Flavobacteriia | Flavobacterial | Flavobacteriac | Polaribacter | Polaribacter s | poorly charact | Function unkn | 11QQ6 | NA | 487797.Flav3 | 1,2 | 0,4 | 5,54E-05 | 1,23E-05 | 2 - | 7,94E-05 - | 1 - | 4,93E-05 - | 1 - | 4,69E-05 - | 1 - | 4,76E-05 - | 1 - | 5,36E-05 - |
| 7373 | 1 | 3um_FIJJAFBF_594159 | Eukaryota | unclassified | Pelagophycea | Pelagomonad | unclassified | Aureoumbra | Aureoumbra l | metabolism | Energy produc | COG0055 | Produces ATP | 7739.JGI1257 | 6,6 | 0,489897949 | 0,000325395 | 4,13E-05 | 7 - | 0,000289006 - | 7 - | 0,00035886 - | 6 - | 0,000292408 - | 6 - | 0,000296727 - | 7 - | 0,000389972 - |
| 7374 | 1 | 3um_FIJJAFBF_604557 | Bacteria | Proteobacteri | Gammaprotec | Arenicellales | Arenicellaceae | Arenicella | Arenicella xan | information st | Translation, ril | COG0051 | tRNA binding | 259536.Psyc_ | 0,4 | 0,489897949 | 4,77E-05 | 5,87E-05 | 1 - | 0,000109311 - | 0 - | 0 - | 1 - | 0,000129031 - | 0 - | 0 - | 0 - | 0 - |
| 7375 | 1 | 0.2um_NOKCPHJN_583914 | unclassified | unclassified | unclassified | unclassified | unclassified | unclassified | unclassified | unclassified | unclassified | unclassified | unclassified | unclassified | 1 | 0 | 0,000139824 | 1,33E-05 | 1 - | 0,000117119 - | 1 - | 0,000145427 - | 1 - | 0,000138247 - | 1 - | 0,000140289 - | 1 - | 0,000158035 - |
| 7376 | 1 | 3um_FIJJAFBF_736126 | Bacteria | Bacteroidetes | Flavobacteriia | Flavobacterial | Flavobacteriac | Cellulophaga | Cellulophaga s | unclassified | unclassified | unclassified | unclassified | unclassified | 8,8 | 1,939071943 | 0,000361371 | 6,50E-05 | 12 - | 0,00041737 - | 9 - | 0,000388688 - | 8 - | 0,000328442 - | 6 - | 0,00024997 - | 9 - | 0,000422385 - |
| 7377 | 1 | 3um_FIJJAFBF_350301 | Bacteria | Proteobacteri | Gammaprotec | Oceanospirilla | Oceanospirilla | Marinobacteri | Marinobacter | unclassified | unclassified | unclassified | unclassified | unclassified | 0,2 | 0,4 | 1,59E-05 | 3,19E-05 | 1 - | 7,97E-05 - | 0 - | 0 - | 0 - | 0 - | 0 - | 0 - | 0 - | 0 - |
| 7378 | 1 | 0.2um_NOKCPHJN_359127 | Bacteria | Proteobacteri | Betaproteoba | various | various | various | various | unclassified | unclassified | unclassified | unclassified | unclassified | 20,2 | 2,561249695 | 0,000496617 | 1,82E-05 | 25 - | 0,000520766 - | 18 - | 0,000465579 - | 20 - | 0,000491769 - | 20 - | 0,000499032 - | 18 - | 0,000505942 - |
| 7379 | 1 | 3um_FIJJAFBF_254296 | Bacteria | Proteobacteri | Gammaprotec | Cellvibrionale | Halieaceae | Halioglobus | various | unclassified | unclassified | unclassified | unclassified | unclassified | 1,4 | 0,489897949 | 0,000180268 | 5,30E-05 | 2 - | 0,000220725 - | 1 - | 0,000137037 - | 1 - | 0,000130271 - | 2 - | 0,000264391 - | 1 - | 0,000148918 - |
| 7380 | 1 | 0.2um_NOKCPHJN_356143 | Bacteria | Bacteroidetes | Flavobacteriia | Flavobacterial | Cryomorphac | unclassified | Cryomorphac | metabolism | Inorganic ion t | 05C27 | TonB-depende | 313596.RB25 | 18,2 | 1,720465053 | 0,000242451 | 3,84E-05 | 19 - | 0,000211519 - | 17 - | 0,000234997 - | 16 - | 0,000210254 - | 18 - | 0,000240029 - | 21 - | 0,000315457 - |
| 7381 | 1 | 0.2um_NOKCPHJN_52378 | Bacteria | Proteobacteri | Gammaprotec | unclassified | unclassified | unclassified | various | information st | Transcription | 05D27 | DNA-depende | 314285.KT71_ | 0,2 | 0,4 | 1,95E-06 | 3,89E-06 | 0 - | 0 - | 0 - | 0 - | 1 - | 9,73E-06 - | 0 - | 0 - | 0 - | 0 - |
| 7382 | 1 | 3um_FIJJAFBF_509346 | Bacteria | Proteobacteri | Gammaprotec | unclassified | unclassified | unclassified | Gammaprotec | cellular proces | Intracellular tr | 05VIE | preprotein tra | 76869.PputGB | 0,6 | 0,489897949 | 7,57E-05 | 6,30E-05 | 1 - | 0,000107268 - | 0 - | 0 - | 1 - | 0,000126619 - | 0 - | 0 - | 1 - | 0,000144743 - |
| 7383 | 1 | 0.2um_NOKCPHJN_170431 | Bacteria | Proteobacteri | Gammaprotec | unclassified | unclassified | unclassified | Gammaprotec | cellular proces | Posttranslatio | COG0330 | Band 7 protei | 205922.Pfl01_ | 2,4 | 0,489897949 | 0,000111514 | 1,74E-05 | 3 - | 0,000118327 - | 2 - | 9,80E-05 - | 2 - | 9,31E-05 - | 3 - | 0,000141735 - | 2 - | 0,000106443 - |
| 7384 | 1 | 3um_FIJJAFBF_44189 | Bacteria | Bacteroidetes | unclassified | unclassified | unclassified | unclassified | Bacteroidetes | information st | Replication, re | 05CZ8 | DNA polymera | 983544.Lacal_ | 1,6 | 1,019803903 | 5,66E-05 | 3,64E-05 | 2 - | 6,17E-05 - | 1 - | 3,83E-05 - | 3 - | 0,00010926 - | 2 - | 7,39E-05 - | 0 - | 0 - |
| 7385 | 1 | 3um_FIJJAFBF_164706 | Bacteria | Actinobacteri | Actinobacteri | Micrococcales | Microbacteria | Candidatus Ac | Candidatus Ac | unclassified | unclassified | unclassified | unclassified | unclassified | 1 | 0,894427191 | 9,52E-05 | 8,50E-05 | 2 - | 0,000167557 - | 2 - | 0,000208057 - | 0 - | 0 - | 1 - | 0,000100353 - | 0 - | 0 - |
| 7386 | 1 | 3um_FIJJAFBF_189640 | Bacteria | Bacteroidetes | Flavobacteriia | Flavobacterial | Flavobacteriac | Tenacibaculur | Tenacibaculur | metabolism | Energy produc | 099YH | Pyruvate dehy | 983544.Lacal_ | 7,6 | 0,489897949 | 0,000321464 | 4,41E-05 | 7 - | 0,000247211 - | 7 - | 0,000306964 - | 8 - | 0,000333495 - | 8 - | 0,00033842 - | 8 - | 0,000381229 - |
| 7387 | 1 | 3um_FIJJAFBF_69075 | Bacteria | Proteobacteri | Gammaprotec | Alteromonada | Alteromonada | Paraglaciecola | Paraglaciecola | unclassified | unclassified | unclassified | unclassified | unclassified | 1,2 | 1,166190379 | 2,62E-05 | 2,64E-05 | 1 - | 1,77E-05 - | 3 - | 6,58E-05 - | 0 - | 0 - | 0 - | 0 - | 2 - | 4,77E-05 - |
| 7388 | 1 | 0.2um_NOKCPHJN_944186 | Eukaryota | unclassified | Heterotrichea | Heterotrichida | Stentoridae | Stentor | Stentor coeru | information st | Chromatin str | KOG2130 | jumonji doma | 572307.XP_00 | 0,4 | 0,489897949 | 1,35E-05 | 1,66E-05 | 0 - | 0 - | 0 - | 0 - | 0 - | 0 - | 1 - | 3,18E-05 - | 1 - | 3,59E-05 - |
| 7389 | 1 | 3um_FIJJAFBF_60527 | Bacteria | Proteobacteri | Gammaprotec | Oceanospirilla | Oceanospirilla | Bermanella | Bermanella sp | information st | Replication, re | 0QUGS | DNA-binding p | 207949.RED6 | 0,2 | 0,4 | 2,55E-05 | 5,10E-05 | 1 - | 0,00012753 - | 0 - | 0 - | 0 - | 0 - | 0 - | 0 - | 0 - | 0 - |
| 7390 | 1 | 0.2um_NOKCPHJN_742772 | Bacteria | Proteobacteri | Alphaproteob | unclassified | unclassified | Candidatus Pu | Candidatus Pu | unclassified | unclassified | unclassified | unclassified | unclassified | 1 | 0 | 0,000109622 | 1,04E-05 | 1 - | 9,18E-05 - | 1 - | 0,000114015 - | 1 - | 0,000108386 - | 1 - | 0,000109987 - | 1 - | 0,0001239 - |
| 7391 | 1 | 0.2um_NOKCPHJN_64113 | Bacteria | Bacteroidetes | Flavobacteriia | Flavobacterial | Flavobacteriac | Psychroflexus | Psychroflexus | unclassified | unclassified | unclassified | unclassified | unclassified | 0,2 | 0,4 | 1,56E-06 | 3,11E-06 | 1 - | 7,78E-06 - | 0 - | 0 - | 0 - | 0 - | 0 - | 0 - | 0 - | 0 - |
| 7392 | 1 | 3um_FIJJAFBF_131217 | Bacteria | Proteobacteri | Gammaprotec | Oceanospirilla | Saccharospirill | Reinekea | Reinekea fors | metabolism | Carbohydrate | COG2723 | beta-glucosid | 314283.MED2 | 2 | 0,632455532 | 6,09E-05 | 1,85E-05 | 2 - | 5,17E-05 - | 2 - | 6,42E-05 - | 3 - | 9,15E-05 - | 2 - | 6,19E-05 - | 1 - | 3,49E-05 - |
| 7393 | 1 | 0.2um_NOKCPHJN_211707 | Bacteria | Bacteroidetes | Flavobacteriia | Flavobacterial | Flavobacteriac | Polaribacter | Polaribacter s | poorly charact | Function unkn | 0P8NM | penicillin | 487797.Flav3 | 0,2 | 0,4 | 2,90E-06 | 5,80E-06 | 1 - | 1,45E-05 - | 0 - | 0 - | 0 - | 0 - | 0 - | 0 - | 0 - | 0 - |
| 7394 | 1 | 3um_FIJJAFBF_169089 | Bacteria | Bacteroidetes | Cytophagia | Cytophagales | Flammeovirga | Reichenbachie | Reichenbachie | information st | Translation, ril | COG1185 | Involved in mI | 643867.Ftrac_ | 6 | 0,632455532 | 0,000116031 | 1,17E-05 | 7 - | 0,000113963 - | 6 - | 0,000121293 - | 5 - | 9,61E-05 - | 6 - | 0,000117007 - | 6 - | 0,000131808 - |
| 7395 | 1 | 3um_FIJJAFBF_452290 | Bacteria | Proteobacteri | Gammaprotec | Oceanospirilla | Saccharospirill | Reinekea | Reinekea fors | unclassified | unclassified | unclassified | unclassified | unclassified | 1,2 | 0,4 | 1,30E-05 | 2,89E-06 | 2 - | 1,87E-05 - | 1 - | 1,16E-05 - | 1 - | 1,11E-05 - | 1 - | 1,12E-05 - | 1 - | 1,26E-05 - |
| 7396 | 1 | 0.2um_NOKCPHJN_282656 | Bacteria | Proteobacteri | Gammaprotec | Cellvibrionale | Porticoccacea | unclassified | various | metabolism | Inorganic ion t | COG4206 | tonB-depende | 314287.GB22 | 4,6 | 1,356465997 | 9,66E-05 | 2,30E-05 | 6 - | 0,000107941 - | 3 - | 6,70E-05 - | 6 - | 0,000127413 - | 5 - | 0,000107746 - | 3 - | 7,28E-05 - |
| 7397 | 1 | 0.2um_NOKCPHJN_549634 | Bacteria | Proteobacteri | Alphaproteob | Pelagibacteral | Pelagibacterac | unclassified | alpha proteob | unclassified | unclassified | unclassified | unclassified | unclassified | 1,2 | 0,4 | 7,51E-05 | 1,66E-05 | 2 - | 0,000107772 - | 1 - | 6,69E-05 - | 1 - | 6,36E-05 - | 1 - | 6,45E-05 - | 1 - | 7,27E-05 - |
| 7398 | 1 | 0.2um_NOKCPHJN_328979 | Bacteria | Proteobacteri | Alphaproteob | Rhodobactera | Rhodobactera | Roseobacter | uncultured Ro | cellular proces | Posttranslatio | 05C2Z | ATP-depender | 439497.RR11_ | 4,8 | 0,748331477 | 7,54E-05 | 1,25E-05 | 5 - | 6,60E-05 - | 6 - | 9,84E-05 - | 5 - | 7,80E-05 - | 4 - | 6,33E-05 - | 4 - | 7,13E-05 - |
| 7399 | 1 | 3um_FIJJAFBF_152493 | Bacteria | Bacteroidetes | Flavobacteriia | Flavobacterial | Flavobacteriac | Polaribacter | Polaribacter s | unclassified | unclassified | unclassified | unclassified | unclassified | 0,8 | 0,4 | 5,17E-05 | 2,63E-05 | 1 - | 5,60E-05 - | 1 - | 6,95E-05 - | 1 - | 6,61E-05 - | 1 - | 6,71E-05 - | 0 - | 0 - |
| 7400 | 1 | 3um_FIJJAFBF_16489 | Bacteria | Proteobacteri | Gammaprotec | Alteromonada | Alteromonada | Paraglaciecola | Paraglaciecola | information st | Translation, ril | 08VZM | 50s ribosomal | 715451.ambt_ | 9 | 0,894427191 | 0,000712925 | 6,47E-05 | 10 - | 0,000667307 - | 10 - | 0,000828598 - | 9 - | 0,000708919 - | 8 - | 0,000639457 - | 8 - | 0,000720346 - |
| 7401 | 1 | 3um_FIJJAFBF_385183 | Bacteria | Proteobacteri | Gammaprotec | Cellvibrionale | Cellvibrionace | Gilvimarinus | Gilvimarinus c | unclassified | unclassified | unclassified | unclassified | unclassified | 0,6 | 0,489897949 | 1,27E-05 | 1,05E-05 | 1 - | 1,84E-05 - | 1 - | 2,29E-05 - | 0 - | 0 - | 1 - | 2,21E-05 - | 0 - | 0 - |
| 7402 | 1 | 0.2um_NOKCPHJN_331769 | Eukaryota | Chlorophyta | Chlorophycea | Chlamydomor | Chlamydomor | Chlamydomor | Chlamydomor | unclassified | unclassified | unclassified | unclassified | unclassified | 0,4 | 0,489897949 | 4,76E-06 | 5,89E-06 | 1 - | 1,06E-05 - | 1 - | 1,32E-05 - | 0 - | 0 - | 0 - | 0 - | 0 - | 0 - |
| 7403 | 1 | 3um_FIJJAFBF_391486 | Eukaryota | Bacillariophyt | Bacillariophyc | Naviculales | Naviculaceae | Fistulifera | Fistulifera sola | unclassified | unclassified | unclassified | unclassified | unclassified | 0,2 | 0,4 | 1,09E-05 | 2,18E-05 | 0 - | 0 - | 0 - | 0 - | 1 - | 5,44E-05 - | 0 - | 0 - | 0 - | 0 - |
| 7404 | 1 | 3um_FIJJAFBF_225008 | Bacteria | Bacteroidetes | Flavobacteriia | unclassified | unclassified | unclassified | Flavobacteria | poorly charact | Function unkn | 11RX2 | NA | 487797.Flav3 | 0,4 | 0,489897949 | 2,80E-05 | 3,46E-05 | 1 - | 6,38E-05 - | 0 - | 0 - | 0 - | 0 - | 1 - | 7,64E-05 - | 0 - | 0 - |
| 7405 | 1 | 3um_FIJJAFBF_121184 | Bacteria | Proteobacteri | Gammaprotec | Alteromonada | Alteromonada | Paraglaciecola | Paraglaciecola | unclassified | unclassified | unclassified | unclassified | unclassified | 15,8 | 2,13541565 | 0,000240751 | 2,01E-05 | 19 - | 0,000245029 - | 17 - | 0,000272227 - | 14 - | 0,000213118 - | 16 - | 0,000247161 - | 13 - | 0,000226221 - |
| 7406 | 1 | 3um_FIJJAFBF_609627 | Bacteria | Proteobacteri | Gammaprotec | unclassified | unclassified | unclassified | enodsymbiont | cellular proces | Posttranslatio | 01YC9 | Peptidyl-proly | 460265.Mnod | 0,6 | 0,8 | 4,80E-05 | 6,20E-05 | 2 - | 0,000151022 - | 0 - | 0 - | 1 - | 8,91E-05 - | 0 - | 0 - | 0 - | 0 - |
| 7407 | 1 | 0.2um_NOKCPHJN_133348 | Eukaryota | Zoopagomyco | Basidiobolom | Basidiobolales | Basidiobolace | Basidiobolus | Basidiobolus r | information st | Translation, ril | KOG0901 | ribosomal pro | 65071.PYU1_ | 0,4 | 0,489897949 | 3,68E-05 | 4,55E-05 | 1 - | 8,20E-05 - | 1 - | 0,000101799 - | 0 - | 0 - | 0 - | 0 - | 0 - | 0 - |
| 7408 | 1 | 0.2um_NOKCPHJN_288409 | Bacteria | Bacteroidetes | unclassified | unclassified | unclassified | unclassified | Bacteroidetes | poorly charact | Function unkn | 09A4Z | rhodopsin | 313595.P7007 | 1 | 0 | 5,59E-05 | 5,31E-06 | 1 - | 4,68E-05 - | 1 - | 5,82E-05 - | 1 - | 5,53E-05 - | 1 - | 5,61E-05 - | 1 - | 6,32E-05 - |
| 7409 | 1 | 0.2um_NOKCPHJN_1068757 | Bacteria | Proteobacteri | Alphaproteob | Pelagibacteral | Pelagibacterac | Candidatus Pe | Candidatus Pe | unclassified | unclassified | unclassified | unclassified | unclassified | 2,4 | 0,489897949 | 0,000131459 | 2,84E-05 | 2 - | 9,18E-05 - | 2 - | 0,000114015 - | 3 - | 0,000162579 - | 3 - | 0,00016498 - | 2 - | 0,0001239 - |
| 7410 | 1 | 0.2um_NOKCPHJN_73746 | Bacteria | Proteobacteri | Betaproteoba | Burkholderiale | Comamonada | unclassified | various | metabolism | Secondary me | 01Q86 | extracellular s | 991905.SL003 | 9,8 | 2,227105745 | 0,000343796 | 7,20E-05 | 10 - | 0,000297349 - | 9 - | 0,000332298 - | 12 - | 0,000421188 - | 12 - | 0,000427409 - | 6 - | 0,000240738 - |
| 7411 | 1 | 3um_FIJJAFBF_453302 | Bacteria | Proteobacteri | Alphaproteob | Rhodobactera | unclassified | unclassified | Rhodobactera | unclassified | unclassified | unclassified | unclassified | unclassified | 2 | 0 | 9,17E-05 | 8,70E-06 | 2 - | 7,68E-05 - | 2 - | 9,53E-05 - | 2 - | 9,06E-05 - | 2 - | 9,20E-05 - | 2 - | 0,000103595 - |
| 7412 | 1 | 3um_FIJJAFBF_70034 | Bacteria | Proteobacteri | Gammaprotec | Cellvibrionale | Porticoccacea | unclassified | Gammaprotec | metabolism | Inorganic ion t | 0R7AT | Pfam:TonB | 314287.GB22 | 0,4 | 0,489897949 | 2,56E-05 | 3,17E-05 | 1 - | 5,71E-05 - | 1 - | 7,09E-05 - | 0 - | 0 - | 0 - | 0 - | 0 - | 0 - |
| 7413 | 1 | 3um_FIJJAFBF_788052 | Bacteria | Proteobacteri | Alphaproteob | Rhodobactera | Rhodobactera | unclassified | Rhodobactera | information st | Translation, ril | 01QWB | One of the pri | 391619.RGBS | 5,4 | 0,489897949 | 0,000305223 | 2,32E-05 | 6 - | 0,000285751 - | 5 - | 0,000295682 - | 5 - | 0,000281083 - | 6 - | 0,000342282 - | 5 - | 0,000321316 - |
| 7414 | 1 | 3um_FIJJAFBF_519389 | Bacteria | Proteobacteri | Gammaprotec | Cellvibrionale | Porticoccacea | unclassified | SAR92 bacteri | unclassified | unclassified | unclassified | unclassified | unclassified | 0,2 | 0,4 | 5,37E-06 | 1,07E-05 | 0 - | 0 - | 0 - | 0 - | 0 - | 0 - | 1 - | 2,69E-05 - | 0 - | 0 - |
| 7415 | 1 | 0.2um_NOKCPHJN_112764 | Bacteria | Bacteroidetes | unclassified | unclassified | unclassified | unclassified | Bacteroidetes | information st | Translation, ril | 05CEJ | Catalyzes the | 411154.GFO_ | 1,2 | 0,4 | 2,36E-05 | 8,71E-06 | 1 - | 1,64E-05 - | 2 - | 4,07E-05 - | 1 - | 1,93E-05 - | 1 - | 1,96E-05 - | 1 - | 2,21E-05 - |
| 7416 | 1 | 3um_FIJJAFBF_15541 | Bacteria | Bacteroidetes | Flavobacteriia | Flavobacterial | Flavobacteriac | Algibacter | Algibacter sp. | poorly charact | Function unkn | 11RX2 | NA | 983544.Lacal_ | 1 | 0 | 8,11E-05 | 7,70E-06 | 1 - | 6,79E-05 - | 1 - | 8,43E-05 - | 1 - | 8,02E-05 - | 1 - | 8,14E-05 - | 1 - | 9,16E-05 - |
| 7417 | 1 | 0.2um_NOKCPHJN_960349 | Bacteria | Proteobacteri | Gammaprotec | Pseudomonad | Pseudomonad | Pseudomonas | various | metabolism | Lipid transpor | COG0439 | acetyl-CoA car | 205918.Psyr_ | 0,2 | 0,4 | 1,27E-05 | 2,53E-05 | 0 - | 0 - | 0 - | 0 - | 1 - | 6,33E-05 - | 0 - | 0 - | 0 - | 0 - |
| 7418 | 1 | 3um_FIJJAFBF_75469 | Bacteria | Proteobacteri | Gammaprotec | Pseudomonad | Moraxellaceae | Psychrobacter | Psychrobacter | cellular proces | Cell wall/mem | COG2885 | Ompa motb d | 259536.Psyc_ | 2,6 | 0,8 | 0,000265779 | 8,40E-05 | 2 - | 0,000171309 - | 2 - | 0,000212715 - | 3 - | 0,000303318 - | 4 - | 0,000410398 - | 2 - | 0,000231156 - |
| 7419 | 1 | 3um_FIJJAFBF_33572 | Bacteria | Proteobacteri | Gammaprotec | Oceanospirilla | Oceanospirilla | Motiliproteus | Motiliproteus | unclassified | unclassified | unclassified | unclassified | unclassified | 6 | 0,894427191 | 0,000146933 | 2,53E-05 | 5 - | 0,000102479 - | 6 - | 0,000152699 - | 7 - | 0,000169353 - | 7 - | 0,000171854 - | 5 - | 0,000138281 - |
| 7420 | 1 | 3um_FIJJAFBF_417061 | Bacteria | Proteobacteri | Gammaprotec | Alteromonada | Alteromonada | Paraglaciecola | various | information st | Translation, ril | 179ZI | One of the pri | 342610.Patl_1 | 5,8 | 1,166190379 | 0,000603828 | 8,76E-05 | 8 - | 0,000706319 - | 5 - | 0,000548149 - | 5 - | 0,000521085 - | 5 - | 0,000528782 - | 6 - | 0,000714805 - |
| 7421 | 1 | 0.2um_NOKCPHJN_403043 | Eukaryota | unclassified | Labyrinthulom | unclassified | Thraustochytr | Aurantiochytr | Aurantiochytr | unclassified | unclassified | unclassified | unclassified | unclassified | 0,6 | 0,489897949 | 3,40E-05 | 2,78E-05 | 0 - | 0 - | 1 - | 5,57E-05 - | 0 - | 0 - | 1 - | 5,37E-05 - | 1 - | 6,05E-05 - |
| 7422 | 1 | 0.2um_NOKCPHJN_182375 | Bacteria | Proteobacteri | Gammaprotec | Oceanospirilla | Saccharospirill | Reinekea | Reinekea fors | metabolism | Carbohydrate | 0QIP7 | Catalyzes the | 94122.Shewar | 2,6 | 0,489897949 | 0,000106529 | 1,73E-05 | 3 - | 0,000104027 - | 3 - | 0,000129171 - | 3 - | 0,000122794 - | 2 - | 8,31E-05 - | 2 - | 9,36E-05 - |
| 7423 | 1 | 3um_FIJJAFBF_213370 | Bacteria | Proteobacteri | Gammaprotec | Oceanospirilla | Saccharospirill | Reinekea | Reinekea fors | metabolism | Coenzyme tra | 05C66 | Catalyzes the | 314283.MED2 | 1,6 | 0,8 | 5,79E-05 | 2,81E-05 | 2 - | 6,19E-05 - | 1 - | 3,84E-05 - | 1 - | 3,65E-05 - | 3 - | 0,000111172 - | 1 - | 4,17E-05 - |
| 7424 | 1 | 3um_FIJJAFBF_398891 | Bacteria | Proteobacteri | Gammaprotec | Oceanospirilla | Saccharospirill | Reinekea | Reinekea fors | information st | Replication, re | 16Q01 | hydrolase Tat | 314283.MED2 | 0,2 | 0,4 | 1,02E-05 | 2,04E-05 | 0 - | 0 - | 0 - | 0 - | 1 - | 5,09E-05 - | 0 - | 0 - | 0 - | 0 - |
| 7425 | 1 | 0.2um_NOKCPHJN_714430 | Bacteria | Proteobacteri | Alphaproteob | Pelagibacteral | unclassified | unclassified | Pelagibacteral | unclassified | unclassified | unclassified | unclassified | unclassified | 0,6 | 0,489897949 | 1,31E-05 | 1,09E-05 | 1 - | 1,82E-05 - | 1 - | 2,26E-05 - | 0 - | 0 - | 0 - | 0 - | 1 - | 2,46E-05 - |
| 7426 | 1 | 3um_FIJJAFBF_543312 | Eukaryota | Ascomycota | Sordariomyce | Sordariales | Chaetomiacea | Chaetomium | Chaetomium g | unclassified | unclassified | unclassified | unclassified | unclassified | 0,2 | 0,4 | 1,52E-05 | 3,04E-05 | 1 - | 7,60E-05 - | 0 - | 0 - | 0 - | 0 - | 0 - | 0 - | 0 - | 0 - |
| 7427 | 1 | 3um_FIJJAFBF_737875 | Bacteria | Bacteroidetes | unclassified | unclassified | unclassified | unclassified | Bacteroidetes | unclassified | unclassified | unclassified | unclassified | unclassified | 0,8 | 0,748331477 | 6,44E-05 | 5,87E-05 | 0 - | 0 - | 0 - | 0 - | 2 - | 0,000154837 - | 1 - | 7,86E-05 - | 1 - | 8,85E-05 - |
| 7428 | 1 | 0.2um_NOKCPHJN_1016981 | Eukaryota | unclassified | Ichthyosporea | unclassified | unclassified | Capsaspora | Capsaspora ov | poorly charact | Function unkn | 0Q26Z | NA | 441959.XP_00 | 0,2 | 0,4 | 5,74E-06 | 1,15E-05 | 0 - | 0 - | 0 - | 0 - | 0 - | 0 - | 1 - | 2,87E-05 - | 0 - | 0 - |
| 7429 | 1 | 0.2um_NOKCPHJN_654066 | Bacteria | Proteobacteri | Alphaproteob | Rhodobactera | unclassified | unclassified | Rhodobactera | unclassified | unclassified | unclassified | unclassified | unclassified | 1,8 | 0,4 | 8,16E-05 | 1,91E-05 | 2 - | 7,63E-05 - | 1 - | 4,73E-05 - | 2 - | 9,00E-05 - | 2 - | 9,14E-05 - | 2 - | 0,000102907 - |
| 7430 | 1 | 3um_FIJJAFBF_254414 | Bacteria | Bacteroidetes | Flavobacteriia | Flavobacterial | Crocinitomica | unclassified | Crocinitomica | unclassified | unclassified | unclassified | unclassified | unclassified | 1,4 | 0,489897949 | 0,000107747 | 3,17E-05 | 2 - | 0,000131927 - | 1 - | 8,19E-05 - | 1 - | 7,79E-05 - | 2 - | 0,000158027 - | 1 - | 8,90E-05 - |

| | | | | | | | | | | | | | | | | | | | | | | | | | | | | |
|---|---|---|---|---|---|---|---|---|---|---|---|---|---|---|---|---|---|---|---|---|---|---|---|---|---|---|---|---|
| 7431 | 1 | 0.2um_NOKCPHJN_251145 | Bacteria | Firmicutes | Bacilli | Bacillales | Staphylococca | Staphylococcu | Staphylococcu | cellular proces | Signal transdu | KOG0613 | fibronectin typ | 7070.TC00296 | 0,2 | 0,4 | 1,19E-06 | 2,38E-06 | 0 - | 0 - | 1 - | 5,94E-06 - | 0 - | 0 - | 0 - | 0 - | 0 - | 0 - |
| 7432 | 1 | 3um_FIJJAFBF_570016 | Bacteria | Proteobacteri | Alphaproteob | Rhodobactera | unclassified | unclassified | Rhodobactera | unclassified | unclassified | unclassified | unclassified | unclassified | 0,2 | 0,4 | 5,00E-05 | 1,00E-04 | 0 - | 0 - | 0 - | 0 - | 0 - | 0 - | 1 - | 0,00024997 - | 0 - | 0 - |
| 7433 | 1 | 3um_FIJJAFBF_457406 | Eukaryota | Bacillariophyt | Bacillariophyc | Naviculales | Naviculaceae | Fistulifera | Fistulifera sola | unclassified | unclassified | unclassified | unclassified | unclassified | 2,2 | 1,166190379 | 0,000165794 | 8,93E-05 | 3 - | 0,000190238 - | 2 - | 0,000157479 - | 3 - | 0,000224556 - | 0 - | 0 - | 3 - | 0,000256698 - |
| 7434 | 1 | 3um_FIJJAFBF_170963 | Bacteria | Bacteroidetes | Flavobacteriia | Flavobacterial | Flavobacteriac | Polaribacter | Polaribacter r | unclassified | unclassified | unclassified | unclassified | unclassified | 0,2 | 0,4 | 1,36E-05 | 2,72E-05 | 0 - | 0 - | 0 - | 0 - | 1 - | 6,81E-05 - | 0 - | 0 - | 0 - | 0 - |
| 7435 | 1 | 0.2um_NOKCPHJN_798315 | Bacteria | Proteobacteri | Gammaprote | Oceanospirilla | Saccharospirill | Reinekea | Reinekea fors | metabolism | Amino acid tra | 0QJVS | branched-chai | 999141.GME_ | 6 | 0,894427191 | 0,000206825 | 3,23E-05 | 7 - | 0,000202888 - | 7 - | 0,000251927 - | 5 - | 0,000171063 - | 5 - | 0,00017359 - | 6 - | 0,000234658 - |
| 7436 | 1 | 3um_FIJJAFBF_150333 | Bacteria | Proteobacteri | Alphaproteob | Rhodobactera | Rhodobactera | Ahrensia | Ahrensia sp. R | metabolism | Amino acid tra | 16V6G | Glycine betain | 744979.R2A13 | 0,6 | 0,489897949 | 2,46E-05 | 2,03E-05 | 1 - | 3,58E-05 - | 1 - | 4,44E-05 - | 0 - | 0 - | 1 - | 4,28E-05 - | 0 - | 0 - |
| 7437 | 1 | 3um_FIJJAFBF_25139 | Bacteria | Bacteroidetes | Flavobacteriia | Flavobacterial | Flavobacteriac | unclassified | Flavobacteriac | unclassified | unclassified | unclassified | unclassified | unclassified | 3,4 | 1,019803903 | 0,000196153 | 4,12E-05 | 5 - | 0,000248435 - | 3 - | 0,000185089 - | 4 - | 0,000234601 - | 3 - | 0,00017855 - | 2 - | 0,00013409 - |
| 7438 | 1 | 3um_FIJJAFBF_98338 | Bacteria | Bacteroidetes | Flavobacteriia | Flavobacterial | Flavobacteriac | Formosa | Formosa sp. H | cellular proces | Cell wall/mem | COG2885 | Ompa motb d | 983544.Lacal_ | 0,8 | 0,4 | 2,30E-05 | 1,18E-05 | 1 - | 2,41E-05 - | 1 - | 2,99E-05 - | 0 - | 0 - | 1 - | 2,88E-05 - | 1 - | 3,25E-05 - |
| 7439 | 1 | 0.2um_NOKCPHJN_370876 | Bacteria | Proteobacteri | Alphaproteob | Pelagibacteral | Pelagibacterac | Candidatus Pe | Candidatus Pe | unclassified | unclassified | unclassified | unclassified | unclassified | 1,4 | 1,019803903 | 3,68E-05 | 2,84E-05 | 1 - | 2,22E-05 - | 3 - | 8,27E-05 - | 2 - | 5,24E-05 - | 1 - | 2,66E-05 - | 0 - | 0 - |
| 7440 | 1 | 3um_FIJJAFBF_636517 | Bacteria | Bacteroidetes | Flavobacteriia | Flavobacterial | Flavobacteriac | Flavivirga | Flavivirga eckl | metabolism | Lipid transpor | COG4247 | Phytase | 867900.Celly_ | 2 | 0 | 8,23E-05 | 7,81E-06 | 2 - | 6,89E-05 - | 2 - | 8,56E-05 - | 2 - | 8,14E-05 - | 2 - | 8,26E-05 - | 2 - | 9,30E-05 - |
| 7441 | 1 | 3um_FIJJAFBF_24076 | Bacteria | unclassified | unclassified | unclassified | unclassified | unclassified | uncultured ba | unclassified | unclassified | unclassified | unclassified | unclassified | 0,4 | 0,489897949 | 2,07E-05 | 2,53E-05 | 0 - | 0 - | 1 - | 5,30E-05 - | 1 - | 5,04E-05 - | 0 - | 0 - | 0 - | 0 - |
| 7442 | 1 | 0.2um_NOKCPHJN_626152 | Bacteria | Proteobacteri | Gammaprote | unclassified | unclassified | unclassified | various | poorly charact | Function unkn | 05C2A | tonB-depende | 566466.NOR5 | 3,2 | 1,326649916 | 4,08E-05 | 1,43E-05 | 5 - | 5,54E-05 - | 3 - | 4,13E-05 - | 4 - | 5,24E-05 - | 3 - | 3,99E-05 - | 1 - | 1,50E-05 - |
| 7443 | 1 | 0.2um_NOKCPHJN_397202 | Bacteria | Proteobacteri | Gammaprote | Oceanospirilla | Saccharospirill | Reinekea | Reinekea blan | unclassified | unclassified | unclassified | unclassified | unclassified | 0,8 | 0,4 | 1,90E-05 | 9,77E-06 | 1 - | 1,99E-05 - | 1 - | 2,47E-05 - | 0 - | 0 - | 1 - | 2,38E-05 - | 1 - | 2,68E-05 - |
| 7444 | 1 | 3um_FIJJAFBF_647279 | Bacteria | Bacteroidetes | Flavobacteriia | unclassified | unclassified | unclassified | Flavobacteria | unclassified | unclassified | unclassified | unclassified | unclassified | 2,8 | 0,979795897 | 0,00011899 | 4,46E-05 | 1 - | 3,48E-05 - | 3 - | 0,000129563 - | 3 - | 0,000123166 - | 4 - | 0,000166646 - | 3 - | 0,000140795 - |
| 7445 | 1 | 3um_FIJJAFBF_277270 | Bacteria | Proteobacteri | Gammaprote | Cellvibrionales | Spongiibacter | Oceanicoccus | Oceanicoccus | unclassified | unclassified | unclassified | unclassified | unclassified | 0,6 | 0,489897949 | 5,06E-05 | 4,14E-05 | 0 - | 0 - | 1 - | 8,29E-05 - | 0 - | 0 - | 1 - | 7,99E-05 - | 1 - | 9,00E-05 - |
| 7446 | 1 | 3um_FIJJAFBF_743884 | Bacteria | Proteobacteri | Alphaproteob | Rhizobiales | unclassified | unclassified | Rhizobiales ba | poorly charact | Function unkn | 05C2A | tonB-depende | 1085623.GNIT | 10 | 1,095445115 | 0,000257734 | 3,11E-05 | 10 - | 0,00021656 - | 10 - | 0,000268904 - | 12 - | 0,000306752 - | 9 - | 0,000233462 - | 9 - | 0,000262994 - |
| 7447 | 1 | 0.2um_NOKCPHJN_18329 | Bacteria | Bacteroidetes | Flavobacteriia | Flavobacterial | Flavobacteriac | Ulvibacter | Ulvibacter lito | metabolism | Energy produc | COG0712 | F(1)F(0) ATP s | 50743.SCB49_ | 0,8 | 0,748331477 | 6,09E-05 | 5,78E-05 | 1 - | 6,41E-05 - | 0 - | 0 - | 0 - | 0 - | 2 - | 0,000153613 - | 1 - | 8,65E-05 - |
| 7448 | 1 | 3um_FIJJAFBF_741923 | Bacteria | Proteobacteri | Gammaprote | Cellvibrionales | Halieaceae | unclassified | marine gamm | unclassified | unclassified | unclassified | unclassified | unclassified | 2 | 1,264911064 | 0,000187786 | 0,000105581 | 4 - | 0,000327934 - | 3 - | 0,000305398 - | 1 - | 9,68E-05 - | 1 - | 9,82E-05 - | 1 - | 0,000110625 - |
| 7449 | 1 | 3um_FIJJAFBF_43756 | Bacteria | Proteobacteri | Alphaproteob | Pelagibacteral | Pelagibacterac | Candidatus Pe | Candidatus Pe | unclassified | unclassified | unclassified | unclassified | unclassified | 1 | 0 | 5,39E-05 | 5,12E-06 | 1 - | 4,52E-05 - | 1 - | 5,61E-05 - | 1 - | 5,33E-05 - | 1 - | 5,41E-05 - | 1 - | 6,10E-05 - |
| 7450 | 1 | 0.2um_NOKCPHJN_167124 | Bacteria | Proteobacteri | Alphaproteob | unclassified | unclassified | unclassified | SAR116 cluste | unclassified | unclassified | unclassified | unclassified | unclassified | 1,6 | 0,489897949 | 0,000235797 | 6,18E-05 | 2 - | 0,000252257 - | 1 - | 0,000156614 - | 2 - | 0,000297763 - | 2 - | 0,000302161 - | 1 - | 0,000170192 - |
| 7451 | 1 | 3um_FIJJAFBF_396425 | Bacteria | Proteobacteri | Betaproteoba | unclassified | unclassified | unclassified | Betaproteoba | information st | Translation, ri | 08UHH | Located on th | 365044.Pnap_ | 2,2 | 1,166190379 | 0,000216159 | 0,000113487 | 3 - | 0,000256963 - | 2 - | 0,000212715 - | 3 - | 0,000303318 - | 3 - | 0,000307798 - | 0 - | 0 - |
| 7452 | 1 | 3um_FIJJAFBF_92594 | Bacteria | Fusobacteria | Fusobacteriia | Fusobacteriale | Fusobacteriac | Fusobacterium | Fusobacterium | metabolism | Energy produc | COG4451 | RuBisCO catal | 381666.H16_E | 1 | 0 | 9,79E-05 | 9,29E-06 | 1 - | 8,20E-05 - | 1 - | 0,000101799 - | 1 - | 9,68E-05 - | 1 - | 9,82E-05 - | 1 - | 0,000110625 - |
| 7453 | 1 | 0.2um_NOKCPHJN_750541 | Bacteria | Proteobacteri | Gammaprote | Alteromonada | Alteromonada | Alteromonas | Alteromonas s | unclassified | unclassified | unclassified | unclassified | unclassified | 0,4 | 0,489897949 | 2,49E-05 | 3,07E-05 | 1 - | 5,71E-05 - | 0 - | 0 - | 1 - | 6,74E-05 - | 0 - | 0 - | 0 - | 0 - |
| 7454 | 1 | 3um_FIJJAFBF_53211 | Bacteria | Bacteroidetes | Flavobacteriia | Flavobacterial | Flavobacteriac | Polaribacter | various | unclassified | unclassified | unclassified | unclassified | unclassified | 1,4 | 0,8 | 0,00031541 | 0,000139617 | 3 - | 0,000593673 - | 1 - | 0,000245722 - | 1 - | 0,00023359 - | 1 - | 0,00023704 - | 1 - | 0,000267025 - |
| 7455 | 1 | 0.2um_NOKCPHJN_74926 | Bacteria | Bacteroidetes | Flavobacteriia | Flavobacterial | Flavobacteriac | unclassified | Flavobacteriac | cellular proces | Intracellular tr | 18ATJ | Pfam:YadA | 595537.Varpa | 3,8 | 0,748331477 | 7,18E-05 | 1,14E-05 | 5 - | 8,00E-05 - | 3 - | 5,96E-05 - | 4 - | 7,56E-05 - | 3 - | 5,75E-05 - | 4 - | 8,64E-05 - |
| 7456 | 1 | 3um_FIJJAFBF_111336 | Viruses | unclassified | unclassified | unclassified | Phycodnavirid | unclassified | Organic Lake p | unclassified | unclassified | unclassified | unclassified | unclassified | 6,6 | 0,489897949 | 0,001116786 | 0,000141711 | 7 - | 0,000991898 - | 7 - | 0,001231645 - | 6 - | 0,001003572 - | 6 - | 0,001018395 - | 7 - | 0,001338421 - |
| 7457 | 1 | 3um_FIJJAFBF_271634 | Bacteria | Bacteroidetes | Flavobacteriia | Flavobacterial | Flavobacteriac | Formosa | Formosa sp. H | metabolism | Energy produc | COG0055 | Produces ATP | 983544.Lacal_ | 0,6 | 0,489897949 | 3,66E-05 | 2,99E-05 | 0 - | 0 - | 1 - | 6,28E-05 - | 1 - | 5,97E-05 - | 1 - | 6,06E-05 - | 0 - | 0 - |
| 7458 | 1 | 0.2um_NOKCPHJN_81664 | Bacteria | Proteobacteri | Gammaprote | Cellvibrionales | Porticoccacea | unclassified | Gammaprote | information st | Translation, ri | COG0052 | 30S ribosomal | 565045.NOR5 | 2,8 | 0,748331477 | 0,000135709 | 2,39E-05 | 4 - | 0,000166343 - | 2 - | 0,000103275 - | 3 - | 0,000147263 - | 3 - | 0,000149438 - | 2 - | 0,000112228 - |
| 7459 | 1 | 3um_FIJJAFBF_40376 | Eukaryota | Bacillariophyt | Mediophycea | Triceratiales | Triceratiaceae | Trieres | Trieres chinen | metabolism | Energy produc | COG1290 | Component of | 35128.Thapsd | 2 | 0 | 0,000127467 | 1,21E-05 | 2 - | 0,000106769 - | 2 - | 0,000132576 - | 2 - | 0,00012603 - | 2 - | 0,000127891 - | 2 - | 0,000144069 - |
| 7460 | 1 | 0.2um_NOKCPHJN_497502 | Bacteria | Proteobacteri | Alphaproteob | Rhodobactera | Rhodobactera | Thalassobacte | Thalassobacte | metabolism | Nucleotide tra | 05CM0 | Catalyzes the | 272943.RSP_0 | 0,6 | 0,489897949 | 1,52E-05 | 1,26E-05 | 1 - | 2,22E-05 - | 1 - | 2,75E-05 - | 0 - | 0 - | 1 - | 2,65E-05 - | 0 - | 0 - |
| 7461 | 1 | 0.2um_NOKCPHJN_398196 | Bacteria | Bacteroidetes | Flavobacteriia | Flavobacterial | Crocinitomica | unclassified | Crocinitomica | unclassified | unclassified | unclassified | unclassified | unclassified | 1 | 0,632455532 | 7,64E-05 | 4,27E-05 | 2 - | 0,000131927 - | 1 - | 8,19E-05 - | 0 - | 0 - | 1 - | 7,90E-05 - | 1 - | 8,90E-05 - |
| 7462 | 1 | 3um_FIJJAFBF_350452 | Eukaryota | Bacillariophyt | Coscinodiscop | Thalassiosirale | Thalassiosirac | Thalassiosira | Thalassiosira p | unclassified | unclassified | unclassified | unclassified | unclassified | 4 | 1,095445115 | 0,000143053 | 3,93E-05 | 4 - | 0,000120501 - | 4 - | 0,000149626 - | 3 - | 0,000106679 - | 6 - | 0,000216509 - | 3 - | 0,000121948 - |
| 7463 | 1 | 0.2um_NOKCPHJN_369201 | Bacteria | Proteobacteri | Alphaproteob | Rhodobactera | Rhodobactera | Yoonia | Yoonia vestfol | information st | Transcription | 16QIX | transcriptiona | 391626.OA307 | 0,6 | 0,489897949 | 2,24E-05 | 1,84E-05 | 1 - | 3,32E-05 - | 0 - | 0 - | 1 - | 3,92E-05 - | 1 - | 3,97E-05 - | 0 - | 0 - |
| 7464 | 1 | 0.2um_NOKCPHJN_710508 | Eukaryota | Eustigmatoph | unclassified | Eustigmatales | Monodopsida | Nannochlorop | Nannochlorop | metabolism | Energy produc | 1BFZ8 | Cytochrome b | 3218.PP1S35_ | 4,6 | 0,8 | 0,000509111 | 4,55E-05 | 6 - | 0,000564476 - | 4 - | 0,000467275 - | 5 - | 0,000555255 - | 4 - | 0,000450765 - | 4 - | 0,000507785 - |
| 7465 | 1 | 3um_FIJJAFBF_30335 | Bacteria | Bacteroidetes | Flavobacteriia | Flavobacterial | Flavobacteriac | Tenacibaculum | Tenacibaculum | metabolism | Coenzyme tra | 09A4R | May play a key | 391603.FBALC | 3,6 | 1,2 | 0,00010798 | 3,73E-05 | 3 - | 7,57E-05 - | 5 - | 0,000156614 - | 5 - | 0,000148882 - | 3 - | 9,06E-05 - | 2 - | 6,81E-05 - |
| 7466 | 1 | 0.2um_NOKCPHJN_235007 | Bacteria | Bacteroidetes | Flavobacteriia | Flavobacterial | Flavobacteriac | Flaviramulus | Flaviramulus i | metabolism | Energy produc | 05CMV | Succinyl-CoA s | 1046627.BZAR | 0,4 | 0,489897949 | 1,26E-05 | 1,56E-05 | 1 - | 2,90E-05 - | 0 - | 0 - | 1 - | 3,42E-05 - | 0 - | 0 - | 0 - | 0 - |
| 7467 | 1 | 0.2um_NOKCPHJN_39203 | Eukaryota | Bacillariophyt | Fragilariophyc | Licmophorales | Licmophorace | Licmophora | Licmophora sp | metabolism | Energy produc | COG1290 | Component of | 35128.Thapsd | 6,4 | 0,489897949 | 0,000405867 | 2,94E-05 | 7 - | 0,000373692 - | 6 - | 0,000397727 - | 6 - | 0,00037809 - | 7 - | 0,00044762 - | 6 - | 0,000432208 - |
| 7468 | 1 | 0.2um_NOKCPHJN_350434 | Bacteria | Bacteroidetes | Flavobacteriia | Flavobacterial | unclassified | unclassified | Flavobacterial | information st | Replication, re | 0EQ45 | snf2 family | 720554.Clocl_ | 0,4 | 0,489897949 | 4,90E-05 | 6,11E-05 | 1 - | 0,000104343 - | 0 - | 0 - | 0 - | 0 - | 0 - | 0 - | 1 - | 0,000140795 - |
| 7469 | 1 | 3um_FIJJAFBF_654569 | Bacteria | Proteobacteri | Gammaprote | unclassified | unclassified | unclassified | various | metabolism | Nucleotide tra | 05DIP | Molybdopteri | 644968.DFW1 | 0,2 | 0,4 | 6,65E-06 | 1,33E-05 | 1 - | 3,33E-05 - | 0 - | 0 - | 0 - | 0 - | 0 - | 0 - | 0 - | 0 - |
| 7470 | 1 | 0.2um_NOKCPHJN_109642 | Bacteria | Bacteroidetes | Flavobacteriia | Flavobacterial | Flavobacteriac | Jejuia | Jejuia pallidilu | unclassified | unclassified | unclassified | unclassified | unclassified | 1,2 | 0,4 | 8,12E-05 | 2,72E-05 | 1 - | 5,68E-05 - | 1 - | 7,06E-05 - | 2 - | 0,000134141 - | 1 - | 6,81E-05 - | 1 - | 7,67E-05 - |
| 7471 | 1 | 0.2um_NOKCPHJN_138235 | Eukaryota | Chordata | Actinopteri | Siluriformes | Ictaluridae | Ictalurus | Ictalurus punc | unclassified | unclassified | unclassified | unclassified | unclassified | 0,2 | 0,4 | 6,98E-07 | 1,40E-06 | 0 - | 0 - | 0 - | 0 - | 1 - | 3,49E-06 - | 0 - | 0 - | 0 - | 0 - |
| 7472 | 1 | 3um_FIJJAFBF_524441 | Bacteria | Bacteroidetes | Flavobacteriia | Flavobacterial | Flavobacteriac | Formosa | Formosa sp. H | unclassified | unclassified | unclassified | unclassified | unclassified | 2 | 0 | 0,000156602 | 1,49E-05 | 2 - | 0,000131173 - | 2 - | 0,000162879 - | 2 - | 0,000154837 - | 2 - | 0,000157124 - | 2 - | 0,000176999 - |
| 7473 | 1 | 3um_FIJJAFBF_348252 | Bacteria | Bacteroidetes | Flavobacteriia | unclassified | unclassified | unclassified | Flavobacteria | metabolism | Energy produc | COG0711 | F(1)F(0) ATP s | 487796.Flav2A | 2,4 | 0,8 | 0,00019275 | 4,27E-05 | 4 - | 0,000276571 - | 2 - | 0,000171709 - | 2 - | 0,000163232 - | 2 - | 0,000165643 - | 2 - | 0,000186596 - |
| 7474 | 1 | 3um_FIJJAFBF_293073 | Bacteria | Proteobacteri | Gammaprote | unclassified | unclassified | unclassified | Gammaprote | unclassified | unclassified | unclassified | unclassified | unclassified | 0,4 | 0,489897949 | 1,96E-05 | 2,41E-05 | 1 - | 4,48E-05 - | 0 - | 0 - | 1 - | 5,29E-05 - | 0 - | 0 - | 0 - | 0 - |
| 7475 | 1 | 0.2um_NOKCPHJN_604203 | Bacteria | Proteobacteri | Alphaproteob | Pelagibacteral | Pelagibacterac | Candidatus Pe | Candidatus Pe | unclassified | unclassified | unclassified | unclassified | unclassified | 3,8 | 0,4 | 0,000380245 | 3,39E-05 | 4 - | 0,000337579 - | 4 - | 0,000419173 - | 4 - | 0,000398477 - | 4 - | 0,000404363 - | 3 - | 0,000341635 - |
| 7476 | 1 | 3um_FIJJAFBF_634956 | Bacteria | Proteobacteri | Gammaprote | Oceanospirilla | Oceanospirilla | Motiliproteus | Motiliproteus | information st | Translation, ri | COG1185 | Involved in mF | 207954.MED9 | 8,2 | 0,748331477 | 0,000160858 | 1,64E-05 | 9 - | 0,000148418 - | 8 - | 0,000163815 - | 7 - | 0,000136261 - | 9 - | 0,00017778 - | 8 - | 0,000178017 - |
| 7477 | 1 | 3um_FIJJAFBF_571325 | Bacteria | Bacteroidetes | Flavobacteriia | Flavobacterial | Flavobacteriac | Formosa | Formosa sp. H | unclassified | unclassified | unclassified | unclassified | unclassified | 2,4 | 0,489897949 | 7,77E-05 | 1,37E-05 | 3 - | 8,22E-05 - | 3 - | 0,000102042 - | 2 - | 6,47E-05 - | 2 - | 6,56E-05 - | 2 - | 7,39E-05 - |
| 7478 | 1 | 0.2um_NOKCPHJN_950064 | Bacteria | Proteobacteri | Gammaprote | unclassified | unclassified | unclassified | Gammaprote | metabolism | Inorganic ion t | 0QK8T | TonB-depende | 247633.GP214 | 3,8 | 1,166190379 | 8,14E-05 | 2,33E-05 | 5 - | 9,08E-05 - | 2 - | 4,51E-05 - | 3 - | 6,43E-05 - | 5 - | 0,000108768 - | 4 - | 9,80E-05 - |
| 7479 | 1 | 3um_FIJJAFBF_56609 | Bacteria | Proteobacteri | Alphaproteob | Rhodobactera | Rhodobactera | various | various | unclassified | unclassified | unclassified | unclassified | unclassified | 6,6 | 0,489897949 | 0,000267632 | 3,40E-05 | 7 - | 0,000237703 - | 7 - | 0,000295157 - | 6 - | 0,000240501 - | 6 - | 0,000244053 - | 7 - | 0,000320746 - |
| 7480 | 1 | 3um_FIJJAFBF_203489 | Bacteria | Proteobacteri | Gammaprote | Alteromonada | Alteromonada | Paraglaciecola | Paraglaciecola | information st | Translation, ri | COG0052 | 30S ribosomal | 715451.ambt_ | 2,4 | 0,8 | 0,000135138 | 4,69E-05 | 3 - | 0,0001417 - | 2 - | 0,000117299 - | 1 - | 5,58E-05 - | 3 - | 0,000169732 - | 3 - | 0,000191203 - |
| 7481 | 1 | 3um_FIJJAFBF_142185 | Bacteria | Proteobacteri | Gammaprote | Oceanospirilla | Oceanospirilla | Marinomonas | Marinomonas | metabolism | Energy produc | 17CGT | F(1)F(0) ATP s | 717774.Marm | 4 | 1,264911064 | 0,000346043 | 9,15E-05 | 6 - | 0,000441449 - | 4 - | 0,000365433 - | 2 - | 0,000173695 - | 4 - | 0,000352521 - | 4 - | 0,000397114 - |
| 7482 | 1 | 3um_FIJJAFBF_03328 | Bacteria | Bacteroidetes | Flavobacteriia | Flavobacterial | Flavobacteriac | Formosa | Formosa sp. H | unclassified | unclassified | unclassified | unclassified | unclassified | 0,6 | 0,8 | 1,39E-05 | 1,84E-05 | 0 - | 0 - | 1 - | 2,38E-05 - | 0 - | 0 - | 2 - | 4,58E-05 - | 0 - | 0 - |
| 7483 | 2 | 0.2um_NOKCPHJN_85097;3um_FIJJAFBF_15798 | Bacteria | Bacteroidetes | Flavobacteriia | Flavobacterial | Crocinitomica | unclassified | Crocinitomica | cellular proces | Signal transdu | COG0784 | response regu | 313606.M231 | 0,2 | 0,4 | 1,04E-05 | 2,08E-05 | 1 - | 5,19E-05 - | 0 - | 0 - | 0 - | 0 - | 0 - | 0 - | 0 - | 0 - |
| 7484 | 1 | 3um_FIJJAFBF_452325 | Bacteria | Bacteroidetes | Flavobacteriia | Flavobacterial | Flavobacteriac | Formosa | Formosa sp. H | unclassified | unclassified | unclassified | unclassified | unclassified | 1 | 0,632455532 | 6,28E-05 | 4,07E-05 | 1 - | 5,41E-05 - | 1 - | 6,72E-05 - | 2 - | 0,000127813 - | 1 - | 6,49E-05 - | 0 - | 0 - |
| 7485 | 1 | 0.2um_NOKCPHJN_219472 | Bacteria | Proteobacteri | Gammaprote | Alteromonada | Pseudoalteror | Pseudoalteror | Pseudoalteror | metabolism | Inorganic ion t | 16PDP | TonB-depende | 983545.Glaag | 4,2 | 0,4 | 0,000104399 | 4,94E-06 | 5 - | 0,000104915 - | 4 - | 0,000104219 - | 4 - | 9,91E-05 - | 4 - | 0,000100536 - | 4 - | 0,000113254 - |
| 7486 | 2 | 0.2um_NOKCPHJN_700241;0.2um_NOKCPHJN_7 | Bacteria | Proteobacteri | Alphaproteob | Pelagibacteral | Pelagibacterac | Candidatus Pe | Candidatus Pe | information st | Translation, ri | 16QAP | Associates wit | 335992.SAR11 | 1 | 0 | 4,84E-05 | 4,60E-06 | 1 - | 4,06E-05 - | 1 - | 5,04E-05 - | 1 - | 4,79E-05 - | 1 - | 4,86E-05 - | 1 - | 5,47E-05 - |
| 7487 | 1 | 0.2um_NOKCPHJN_375631 | Bacteria | Proteobacteri | Alphaproteob | Pelagibacteral | Pelagibacterac | Candidatus Pe | Candidatus Pe | metabolism | Inorganic ion t | 01Q8E | flavin-contain | 335992.SAR11 | 1 | 0,632455532 | 3,18E-05 | 2,02E-05 | 1 - | 2,68E-05 - | 1 - | 3,32E-05 - | 2 - | 6,32E-05 - | 0 - | 0 - | 1 - | 3,61E-05 - |
| 7488 | 1 | 0.2um_NOKCPHJN_553565 | Eukaryota | Arthropoda | Insecta | Neuroptera | Chrysopidae | Mallada | Mallada signa | unclassified | unclassified | unclassified | unclassified | unclassified | 0,2 | 0,4 | 2,14E-06 | 4,29E-06 | 0 - | 0 - | 0 - | 0 - | 0 - | 0 - | 0 - | 0 - | 1 - | 1,07E-05 - |
| 7489 | 1 | 3um_FIJJAFBF_419267 | Bacteria | Proteobacteri | Gammaprote | Cellvibrionales | Porticoccacea | unclassified | SAR92 bacteri | poorly charact | Function unkn | 0QKIS | short-chain de | 314287.GB220 | 0,2 | 0,4 | 9,00E-06 | 1,80E-05 | 1 - | 4,50E-05 - | 0 - | 0 - | 0 - | 0 - | 0 - | 0 - | 0 - | 0 - |
| 7490 | 1 | 0.2um_NOKCPHJN_596098 | Eukaryota | unclassified | Oomycetes | Peronosporale | Peronosporac | Phytophthora | Phytophthora | information st | Translation, ri | COG0052 | 30S ribosomal | 65071.PYU1_T | 1,2 | 0,4 | 5,33E-05 | 1,78E-05 | 1 - | 3,73E-05 - | 1 - | 4,63E-05 - | 2 - | 8,80E-05 - | 1 - | 4,46E-05 - | 1 - | 5,03E-05 - |
| 7491 | 1 | 0.2um_NOKCPHJN_397509 | Bacteria | Bacteroidetes | Flavobacteriia | Flavobacterial | Cryomorphace | Owenweeksia | Owenweeksia | information st | Translation, ri | COG0052 | 30S ribosomal | 926562.Oweh | 4,4 | 1,019803903 | 0,000220974 | 3,87E-05 | 6 - | 0,000256008 - | 4 - | 0,000211924 - | 3 - | 0,000151095 - | 5 - | 0,000255545 - | 4 - | 0,000230297 - |
| 7492 | 1 | 3um_FIJJAFBF_142851 | Bacteria | Proteobacteri | Alphaproteob | Rhodobactera | unclassified | unclassified | Rhodobactera | unclassified | unclassified | unclassified | unclassified | unclassified | 0,6 | 0,8 | 5,06E-05 | 7,02E-05 | 1 - | 7,45E-05 - | 0 - | 0 - | 0 - | 0 - | 2 - | 0,00017855 - | 0 - | 0 - |
| 7493 | 2 | 0.2um_NOKCPHJN_105152;3um_FIJJAFBF_26772 | Bacteria | Bacteroidetes | Flavobacteriia | Flavobacterial | Flavobacteriac | Polaribacter | various | unclassified | unclassified | unclassified | unclassified | unclassified | 5,8 | 0,4 | 0,000656749 | 7,83E-05 | 6 - | 0,000569141 - | 6 - | 0,000706705 - | 6 - | 0,000671813 - | 5 - | 0,000568113 - | 6 - | 0,000767973 - |
| 7494 | 1 | 3um_FIJJAFBF_514630 | Bacteria | Proteobacteri | Gammaprote | Oceanospirilla | Saccharospirill | Reinekea | Reinekea blan | metabolism | Lipid transpor | 16QA8 | reductase | 314283.MED2 | 2,4 | 0,8 | 0,000132055 | 4,54E-05 | 2 - | 9,29E-05 - | 3 - | 0,0001731 - | 3 - | 0,000164553 - | 3 - | 0,000166984 - | 1 - | 6,27E-05 - |
| 7495 | 1 | 3um_FIJJAFBF_660192 | Bacteria | Bacteroidetes | unclassified | unclassified | unclassified | unclassified | Bacteroidetes | unclassified | unclassified | unclassified | unclassified | unclassified | 1 | 0,632455532 | 0,000100787 | 5,75E-05 | 2 - | 0,000179339 - | 1 - | 0,000111343 - | 1 - | 0,000105845 - | 1 - | 0,000107409 - | 0 - | 0 - |
| 7496 | 1 | 3um_FIJJAFBF_92885 | Eukaryota | unclassified | Cryptophyta | Pyrenomonad | Pyrenomonad | Storeatula | Storeatula sp. | metabolism | Energy produc | COG0711 | F(1)F(0) ATP s | 91464.S7335_ | 1 | 0 | 7,61E-05 | 7,23E-06 | 1 - | 6,38E-05 - | 1 - | 7,92E-05 - | 1 - | 7,53E-05 - | 1 - | 7,64E-05 - | 1 - | 8,60E-05 - |
| 7497 | 1 | 3um_FIJJAFBF_232081 | Bacteria | Actinobacteri | Actinobacteri | Streptosporan | Streptosporan | Nonomuraea | Nonomuraea j | unclassified | unclassified | unclassified | unclassified | unclassified | 2,4 | 0,8 | 0,000222093 | 6,92E-05 | 3 - | 0,000237469 - | 3 - | 0,000294867 - | 2 - | 0,000186872 - | 3 - | 0,000284448 - | 1 - | 0,00010681 - |
| 7498 | 1 | 3um_FIJJAFBF_619967 | Bacteria | Proteobacteri | Gammaprote | unclassified | unclassified | unclassified | Gammaprote | unclassified | unclassified | unclassified | unclassified | unclassified | 1,4 | 0,489897949 | 0,00011406 | 4,03E-05 | 1 - | 6,83E-05 - | 1 - | 8,48E-05 - | 2 - | 0,000161288 - | 2 - | 0,000163671 - | 1 - | 9,22E-05 - |
| 7499 | 1 | 3um_FIJJAFBF_26690 | Bacteria | Bacteroidetes | Flavobacteriia | Flavobacterial | Flavobacteriac | Polaribacter | Polaribacter s | unclassified | unclassified | unclassified | unclassified | unclassified | 2,2 | 0,4 | 9,08E-05 | 8,33E-06 | 3 - | 0,0001053 - | 2 - | 8,72E-05 - | 2 - | 8,29E-05 - | 2 - | 8,41E-05 - | 2 - | 9,47E-05 - |
| 7500 | 1 | 3um_FIJJAFBF_625435 | Bacteria | Bacteroidetes | Flavobacteriia | Flavobacterial | Flavobacteriac | Arenitalea | Arenitalea lut | poorly charact | Function unkn | 09ARW | Zinc ribbon do | 1046627.BZAR | 1,2 | 0,4 | 6,34E-05 | 2,12E-05 | 1 - | 4,43E-05 - | 1 - | 5,50E-05 - | 2 - | 0,000104619 - | 1 - | 5,31E-05 - | 1 - | 5,98E-05 - |
| 7501 | 1 | 0.2um_NOKCPHJN_440248 | Bacteria | Bacteroidetes | unclassified | unclassified | unclassified | unclassified | Bacteroidetes | unclassified | unclassified | unclassified | unclassified | unclassified | 1 | 0 | 2,95E-05 | 2,80E-06 | 1 - | 2,47E-05 - | 1 - | 3,07E-05 - | 1 - | 2,92E-05 - | 1 - | 2,96E-05 - | 1 - | 3,34E-05 - |
| 7502 | 1 | 0.2um_NOKCPHJN_266025 | Bacteria | Proteobacteri | Alphaproteob | Rhodobactera | Rhodobactera | Sulfitobacter | Sulfitobacter s | unclassified | unclassified | unclassified | unclassified | unclassified | 9,8 | 1,833030278 | 0,000391182 | 4,12E-05 | 13 - | 0,000441449 - | 10 - | 0,000421653 - | 10 - | 0,000400835 - | 8 - | 0,000325404 - | 8 - | 0,000366567 - |
| 7503 | 1 | 0.2um_NOKCPHJN_1097287 | Eukaryota | Chromerida | unclassified | unclassified | unclassified | Vitrella | Vitrella brassi | unclassified | unclassified | unclassified | unclassified | unclassified | 0,4 | 0,489897949 | 2,83E-05 | 3,46E-05 | 0 - | 0 - | 0 - | 0 - | 1 - | 7,02E-05 - | 1 - | 7,12E-05 - | 0 - | 0 - |
| 7504 | 1 | 3um_FIJJAFBF_71026 | Bacteria | Bacteroidetes | Cytophagia | Cytophagales | Flammeovirga | unclassified | Flammeovirga | unclassified | unclassified | unclassified | unclassified | unclassified | 0,2 | 0,4 | 9,37E-06 | 1,87E-05 | 1 - | 4,68E-05 - | 0 - | 0 - | 0 - | 0 - | 0 - | 0 - | 0 - | 0 - |
| 7505 | 1 | 3um_FIJJAFBF_273615 | Bacteria | Bacteroidetes | Flavobacteriia | Flavobacterial | Flavobacteriac | Formosa | Formosa sp. H | unclassified | unclassified | unclassified | unclassified | unclassified | 1,2 | 0,748331477 | 7,67E-05 | 4,62E-05 | 2 - | 0,000112526 - | 1 - | 6,99E-05 - | 1 - | 6,64E-05 - | 2 - | 0,000134788 - | 0 - | 0 - |
| 7506 | 1 | 0.2um_NOKCPHJN_980145 | Eukaryota | unclassified | Bangiophycea | Bangiales | Bangiaceae | Porphyra | Porphyra umb | information st | Translation, ri | COG0088 | One of the pri | 45351.NEMVE | 0,6 | 0,489897949 | 2,02E-05 | 1,67E-05 | 1 - | 3,00E-05 - | 0 - | 0 - | 1 - | 3,54E-05 - | 1 - | 3,59E-05 - | 0 - | 0 - |
| 7507 | 1 | 0.2um_NOKCPHJN_258291 | Bacteria | unclassified | unclassified | unclassified | unclassified | unclassified | uncultured ba | unclassified | unclassified | unclassified | unclassified | unclassified | 0,6 | 0,489897949 | 1,17E-05 | 9,64E-06 | 1 - | 1,70E-05 - | 1 - | 2,11E-05 - | 0 - | 0 - | 1 - | 2,03E-05 - | 0 - | 0 - |
| 7508 | 4 | 0.2um_NOKCPHJN_288020;0.2um_NOKCPHJN_4 | Bacteria | Bacteroidetes | Flavobacteriia | Flavobacterial | Flavobacteriac | various | various | unclassified | unclassified | unclassified | unclassified | unclassified | 1 | 0 | 8,36E-06 | 7,94E-07 | 1 - | 7,00E-06 - | 1 - | 8,70E-06 - | 1 - | 8,27E-06 - | 1 - | 8,39E-06 - | 1 - | 9,45E-06 - |
| 7509 | 1 | 0.2um_NOKCPHJN_357798 | Bacteria | Proteobacteri | Alphaproteob | Pelagibacteral | Pelagibacterac | Candidatus Pe | Candidatus Pe | information st | Translation, ri | 17FIT | 50s ribosomal | 335992.SAR11 | 0,4 | 0,489897949 | 4,68E-05 | 5,76E-05 | 1 - | 0,000107268 - | 0 - | 0 - | 1 - | 0,000126619 - | 0 - | 0 - | 0 - | 0 - |
| 7510 | 3 | 0.2um_NOKCPHJN_327876;0.2um_NOKCPHJN_6 | Bacteria | Bacteroidetes | unclassified | unclassified | unclassified | unclassified | Bacteroidetes | unclassified | unclassified | unclassified | unclassified | unclassified | 0,8 | 0,748331477 | 3,25E-05 | 2,96E-05 | 0 - | 0 - | 1 - | 4,05E-05 - | 0 - | 0 - | 2 - | 7,81E-05 - | 1 - | 4,40E-05 - |
| 7511 | 1 | 3um_FIJJAFBF_168113 | Bacteria | Proteobacteri | Gammaprote | Oceanospirilla | Saccharospirill | Reinekea | Reinekea fors | cellular proces | Signal transdu | 0QTZG | Transcription | 314283.MED2 | 1,2 | 0,4 | 0,000124446 | 5,30E-05 | 1 - | 8,50E-05 - | 1 - | 0,00010557 - | 1 - | 0,000100357 - | 1 - | 0,000101839 - | 2 - | 0,000229444 - |
| 7512 | 1 | 3um_FIJJAFBF_732705 | Eukaryota | unclassified | Pelagophycea | Pelagomonad | unclassified | Aureococcus | Aureococcus a | unclassified | unclassified | unclassified | unclassified | unclassified | 0,6 | 0,489897949 | 6,74E-05 | 5,55E-05 | 1 - | 9,98E-05 - | 0 - | 0 - | 1 - | 0,000117811 - | 1 - | 0,000119551 - | 0 - | 0 - |
| 7513 | 1 | 3um_FIJJAFBF_502175 | Eukaryota | unclassified | unclassified | Prymnesiales | Chrysochromu | Chrysochromu | Chrysochromu | unclassified | unclassified | unclassified | unclassified | unclassified | 2 | 0 | 0,000185172 | 1,76E-05 | 2 - | 0,000155104 - | 2 - | 0,000192593 - | 2 - | 0,000183084 - | 2 - | 0,000185788 - | 2 - | 0,00020929 - |
| 7514 | 1 | 0.2um_NOKCPHJN_643785 | Bacteria | Proteobacteri | Alphaproteob | Pelagibacteral | Pelagibacterac | Candidatus Pe | Candidatus Pe | metabolism | Inorganic ion t | COG0841 | acriflavin resis | 335992.SAR11 | 1 | 0,632455532 | 1,88E-05 | 1,23E-05 | 1 - | 1,62E-05 - | 1 - | 2,01E-05 - | 1 - | 1,91E-05 - | 2 - | 3,87E-05 - | 0 - | 0 - |
| 7515 | 1 | 0.2um_NOKCPHJN_88830 | Bacteria | Bacteroidetes | Flavobacteriia | Flavobacterial | Flavobacteriac | Polaribacter | Polaribacter s | poorly charact | Function unkn | 0XSGQ | NA | 313598.MED1 | 0,4 | 0,8 | 9,15E-06 | 1,83E-05 | 2 - | 4,57E-05 - | 0 - | 0 - | 0 - | 0 - | 0 - | 0 - | 0 - | 0 - |
| 7516 | 1 | 3um_FIJJAFBF_583671 | Bacteria | Cyanobacteria | unclassified | Synechococca | Synechococca | Synechococcu | Synechococcu | poorly charact | Function unkn | 06QDP | C-phycoerythr | 32051.SynWH | 3 | 0,632455532 | 0,00024663 | 4,28E-05 | 4 - | 0,000278247 - | 3 - | 0,000259125 - | 2 - | 0,000164221 - | 3 - | 0,00024997 - | 3 - | 0,00028159 - |
| 7517 | 1 | 3um_FIJJAFBF_545136 | Bacteria | Proteobacteri | Gammaprote | Alteromonada | Colwelliaceae | Colwellia | Colwellia sp. P | information st | Translation, ri | 05KK9 | This protein bi | 167879.CPS_4 | 1 | 0 | 0,000133036 | 1,26E-05 | 1 - | 0,000111434 - | 1 - | 0,000138368 - | 1 - | 0,000131536 - | 1 - | 0,000133479 - | 1 - | 0,000150364 - |
| 7518 | 1 | 0.2um_NOKCPHJN_936837 | Bacteria | Bacteroidetes | Flavobacteriia | Flavobacterial | Flavobacteriac | Formosa | Formosa sp. H | information st | Transcription | COG0085 | DNA-depende | 391603.FBALC | 1 | 0 | 2,12E-05 | 2,01E-06 | 1 - | 1,78E-05 - | 1 - | 2,21E-05 - | 1 - | 2,10E-05 - | 1 - | 2,13E-05 - | 1 - | 2,40E-05 - |
| 7519 | 1 | 0.2um_NOKCPHJN_644973 | Bacteria | Proteobacteri | Gammaprote | Alteromonada | Colwelliaceae | Colwellia | Colwellia sp. P | poorly charact | Function unkn | 05C2A | tonB-depende | 283942.IL0582 | 0,2 | 0,4 | 2,21E-06 | 4,43E-06 | 1 - | 1,11E-05 - | 0 - | 0 - | 0 - | 0 - | 0 - | 0 - | 0 - | 0 - |
| 7520 | 1 | 3um_FIJJAFBF_600720 | Bacteria | Proteobacteri | Alphaproteob | Rhodospirillal | Rhodospirillac | unclassified | Rhodobactera | metabolism | Secondary me | 01Q86 | extracellular s | 367336.OM22 | 5,4 | 0,489897949 | 0,000202085 | 1,53E-05 | 6 - | 0,000189192 - | 5 - | 0,000195768 - | 5 - | 0,000186102 - | 6 - | 0,000226621 - | 5 - | 0,00021274 - |
| 7521 | 1 | 3um_FIJJAFBF_744840 | Bacteria | Proteobacteri | Gammaprote | Cellvibrionales | Cellvibrionace | Marinimicrobi | Marinimicrobi | information st | Translation, ri | 17FIH | One of the ear | 314287.GB220 | 0,8 | 0,4 | 0,000111766 | 5,74E-05 | 1 - | 0,000117119 - | 1 - | 0,000145427 - | 1 - | 0,000138247 - | 0 - | 0 - | 1 - | 0,000158035 - |
| 7522 | 1 | 3um_FIJJAFBF_08109 | Bacteria | Bacteroidetes | Flavobacteriia | Flavobacterial | Flavobacteriac | Tenacibaculum | Tenacibaculum | cellular proces | Posttranslatio | COG0330 | Band 7 protein | 313594.PI23P | 6,8 | 0,4 | 0,000336351 | 3,81E-05 | 7 - | 0,00029005 - | 7 - | 0,000360156 - | 7 - | 0,000342374 - | 6 - | 0,000297798 - | 7 - | 0,00039138 - |
| 7523 | 1 | 3um_FIJJAFBF_61063 | Eukaryota | Bacillariophyt | Coscinodiscop | Thalassiosirale | Thalassiosirac | Thalassiosira | Thalassiosira p | unclassified | unclassified | unclassified | unclassified | unclassified | 0,4 | 0,489897949 | 5,96E-06 | 7,30E-06 | 0 - | 0 - | 0 - | 0 - | 1 - | 1,48E-05 - | 1 - | 1,50E-05 - | 0 - | 0 - |
| 7524 | 1 | 0.2um_NOKCPHJN_106092 | Bacteria | Bacteroidetes | unclassified | unclassified | unclassified | unclassified | Bacteroidetes | metabolism | Inorganic ion t | 05C27 | TonB-depende | 886377.Murru | 1,8 | 0,4 | 2,39E-05 | 5,59E-06 | 2 - | 2,23E-05 - | 1 - | 1,39E-05 - | 2 - | 2,64E-05 - | 2 - | 2,67E-05 - | 2 - | 3,01E-05 - |
| 7525 | 1 | 0.2um_NOKCPHJN_636853 | Bacteria | Bacteroidetes | unclassified | unclassified | unclassified | unclassified | Bacteroidetes | poorly charact | Function unkn | 09DEA | Protein of unk | 926562.Oweh | 1 | 0 | 2,68E-05 | 2,55E-06 | 1 - | 2,25E-05 - | 1 - | 2,79E-05 - | 1 - | 2,65E-05 - | 1 - | 2,69E-05 - | 1 - | 3,03E-05 - |
| 7526 | 1 | 3um_FIJJAFBF_42576 | Bacteria | Proteobacteri | Gammaprote | Oceanospirilla | Saccharospirill | Reinekea | Reinekea blan | information st | Translation, ri | COG1185 | Involved in mF | 314283.MED2 | 10,4 | 1,356465997 | 0,000202982 | 3,62E-05 | 9 - | 0,000146524 - | 12 - | 0,000242585 - | 12 - | 0,000230608 - | 9 - | 0,000175511 - | 10 - | 0,00021968 - |
| 7527 | 1 | 0.2um_NOKCPHJN_231475 | Bacteria | Proteobacteri | Alphaproteob | Rhodobactera | Rhodobactera | unclassified | Rhodobactera | information st | Replication, re | COG0187 | DNA gyrase ne | 467661.RKLH1 | 1 | 0 | 2,07E-05 | 1,97E-06 | 1 - | 1,74E-05 - | 1 - | 2,16E-05 - | 1 - | 2,05E-05 - | 1 - | 2,08E-05 - | 1 - | 2,34E-05 - |
| 7528 | 1 | 3um_FIJJAFBF_06888 | Bacteria | Proteobacteri | Alphaproteob | Pelagibacteral | Pelagibacterac | Candidatus Pe | Candidatus Pe | unclassified | unclassified | unclassified | unclassified | unclassified | 5 | 0,632455532 | 0,000220882 | 3,40E-05 | 5 - | 0,000185124 - | 5 - | 0,000229869 - | 6 - | 0,000262224 - | 4 - | 0,000177398 - | 5 - | 0,000249798 - |
| 7529 | 1 | 3um_FIJJAFBF_806359 | Bacteria | Proteobacteri | Gammaprote | Oceanospirilla | Saccharospirill | Reinekea | Reinekea blan | metabolism | Carbohydrate | 0QKP9 | ABC-type suga | 314283.MED2 | 35,8 | 1,166190379 | 0,001181545 | 0,000116829 | 37 - | 0,001023311 - | 35 - | 0,001201966 - | 36 - | 0,001175268 - | 34 - | 0,001126369 - | 37 - | 0,001380809 - |
| 7530 | 1 | 0.2um_NOKCPHJN_173988 | Bacteria | Proteobacteri | Gammaprote | Oceanospirilla | Saccharospirill | Reinekea | Reinekea blan | information st | Translation, ri | 05C22 | threonyL-tRNA | 314283.MED2 | 0,4 | 0,489897949 | 8,39E-06 | 1,05E-05 | 1 - | 1,79E-05 - | 0 - | 0 - | 0 - | 0 - | 0 - | 0 - | 1 - | 2,41E-05 - |
| 7531 | 1 | 0.2um_NOKCPHJN_82401 | Bacteria | Bacteroidetes | Flavobacteriia | Flavobacterial | Flavobacteriac | Polaribacter | various | metabolism | Energy produc | 05C26 | Dehydrogenas | 156586.BBFL7 | 1 | 0,632455532 | 2,62E-05 | 1,77E-05 | 1 - | 2,24E-05 - | 2 - | 5,56E-05 - | 1 - | 2,64E-05 - | 1 - | 2,68E-05 - | 0 - | 0 - |
| 7532 | 2 | 0.2um_NOKCPHJN_1098985;3um_FIJJAFBF_8052 | Bacteria | Proteobacteri | Alphaproteob | Pelagibacteral | Pelagibacterac | Candidatus Pe | Candidatus Pe | poorly charact | Function unkn | 01QDW | Protein of unk | 335992.SAR11 | 1,4 | 0,489897949 | 9,24E-05 | 2,72E-05 | 2 - | 0,000113081 - | 1 - | 7,02E-05 - | 1 - | 6,67E-05 - | 2 - | 0,000135452 - | 1 - | 7,63E-05 - |
| 7533 | 1 | 0.2um_NOKCPHJN_295870 | Bacteria | Bacteroidetes | Flavobacteriia | Flavobacterial | Flavobacteriac | Flavobacterium | Flavobacterium | information st | Translation, ri | 05CRK | lysyL-tRNA syn | 50743.SCB49_ | 0,6 | 0,489897949 | 1,40E-05 | 1,16E-05 | 1 - | 2,04E-05 - | 1 - | 2,53E-05 - | 0 - | 0 - | 1 - | 2,44E-05 - | 0 - | 0 - |
| 7534 | 1 | 3um_FIJJAFBF_130195 | Bacteria | Proteobacteri | Betaproteoba | unclassified | unclassified | unclassified | Betaproteoba | metabolism | Energy produc | 16R80 | pyrophosphat | 887062.HGR_ | 0,2 | 0,4 | 1,75E-05 | 3,50E-05 | 0 - | 0 - | 0 - | 0 - | 0 - | 0 - | 0 - | 0 - | 1 - | 8,75E-05 - |
| 7535 | 1 | 0.2um_NOKCPHJN_10350 | Bacteria | Proteobacteri | Alphaproteob | Rhodobactera | Rhodobactera | Litorimicrobiu | Litorimicrobiu | unclassified | unclassified | unclassified | unclassified | unclassified | 6,8 | 1,166190379 | 0,000204347 | 2,86E-05 | 8 - | 0,000203595 - | 8 - | 0,000252805 - | 6 - | 0,000180242 - | 7 - | 0,000213389 - | 5 - | 0,000171701 - |
| 7536 | 1 | 3um_FIJJAFBF_175231 | Bacteria | Proteobacteri | Alphaproteob | Rhodobactera | Rhodobactera | Phaeobacter | Phaeobacter i | unclassified | unclassified | unclassified | unclassified | unclassified | 5 | 0,632455532 | 0,000273523 | 4,89E-05 | 4 - | 0,000182185 - | 5 - | 0,000282776 - | 6 - | 0,000322577 - | 5 - | 0,000272784 - | 5 - | 0,000307291 - |
| 7537 | 1 | 3um_FIJJAFBF_133795 | Bacteria | Proteobacteri | Alphaproteob | Pelagibacteral | Pelagibacterac | Candidatus Pe | Candidatus Pe | cellular proces | Cell wall/mem | 7257 | NA | 335992.SAR11 | 0,2 | 0,4 | 1,34E-05 | 2,68E-05 | 1 - | 6,71E-05 - | 0 - | 0 - | 0 - | 0 - | 0 - | 0 - | 0 - | 0 - |
| 7538 | 1 | 0.2um_NOKCPHJN_54732 | Eukaryota | Streptophyta | unclassified | Brassicales | Brassicaceae | Arabidopsis | various | metabolism | Energy produc | COG0055 | Produces ATP | 39947.LOC_O | 1 | 0 | 2,75E-05 | 2,61E-06 | 1 - | 2,30E-05 - | 1 - | 2,86E-05 - | 1 - | 2,72E-05 - | 1 - | 2,76E-05 - | 1 - | 3,11E-05 - |
| 7539 | 1 | 3um_FIJJAFBF_355591 | Bacteria | Proteobacteri | Alphaproteob | Pelagibacteral | Pelagibacterac | Candidatus Pe | Candidatus Pe | poorly charact | Function unkn | 17CZ8 | Lipoprotein | 335992.SAR11 | 3,6 | 1,959591794 | 0,000271997 | 0,000153204 | 3 - | 0,000189192 - | 1 - | 7,83E-05 - | 3 - | 0,000223322 - | 7 - | 0,000528782 - | 4 - | 0,000340383 - |
| 7540 | 1 | 3um_FIJJAFBF_97615 | Bacteria | Bacteroidetes | Flavobacteriia | Flavobacterial | Flavobacteriac | Polaribacter | various | information st | Translation, ri | 08UHY | One of the pri | 313598.MED1 | 8,4 | 1,019803903 | 0,00071588 | 2,96E-05 | 10 - | 0,000721866 - | 8 - | 0,000717076 - | 9 - | 0,000766881 - | 8 - | 0,00069174 - | 7 - | 0,000681837 - |
| 7541 | 1 | 0.2um_NOKCPHJN_183237 | unclassified | unclassified | unclassified | unclassified | unclassified | unclassified | unclassified | unclassified | unclassified | unclassified | unclassified | unclassified | 0,4 | 0,489897949 | 2,23E-05 | 2,75E-05 | 1 - | 5,12E-05 - | 0 - | 0 - | 1 - | 6,05E-05 - | 0 - | 0 - | 0 - | 0 - |
| 7542 | 1 | 3um_FIJJAFBF_514123 | Bacteria | Proteobacteri | Gammaprote | Oceanospirilla | Oceanospirilla | Nitrincola | Nitrincola laci | metabolism | Energy produc | 05CMV | Succinyl-CoA s | 207954.MED9 | 1,4 | 0,489897949 | 4,85E-05 | 1,52E-05 | 2 - | 5,90E-05 - | 2 - | 7,33E-05 - | 1 - | 3,48E-05 - | 1 - | 3,53E-05 - | 1 - | 3,98E-05 - |
| 7543 | 1 | 0.2um_NOKCPHJN_262676 | Bacteria | Bacteroidetes | Flavobacteriia | Flavobacterial | Flavobacteriac | Winogradskye | Winogradskye | information st | Translation, ri | 09FEM | Binds to the 2 | 391603.FBALC | 0,4 | 0,489897949 | 3,60E-05 | 4,48E-05 | 1 - | 7,65E-05 - | 0 - | 0 - | 0 - | 0 - | 0 - | 0 - | 1 - | 0,00010325 - |
| 7544 | 1 | 0.2um_NOKCPHJN_1060193 | Bacteria | Proteobacteri | Alphaproteob | unclassified | unclassified | unclassified | Alphaproteob | unclassified | unclassified | unclassified | unclassified | unclassified | 0,2 | 0,4 | 5,54E-06 | 1,11E-05 | 1 - | 2,77E-05 - | 0 - | 0 - | 0 - | 0 - | 0 - | 0 - | 0 - | 0 - |
| 7545 | 1 | 3um_FIJJAFBF_44827 | Bacteria | Bacteroidetes | Flavobacteriia | Flavobacterial | Flavobacteriac | Bizionia | Bizionia sp. AF | unclassified | unclassified | unclassified | unclassified | unclassified | 1,6 | 0,489897949 | 5,16E-05 | 1,35E-05 | 2 - | 5,52E-05 - | 1 - | 3,43E-05 - | 2 - | 6,51E-05 - | 2 - | 6,61E-05 - | 1 - | 3,72E-05 - |
| 7546 | 1 | 0.2um_NOKCPHJN_613245 | Bacteria | Proteobacteri | Gammaprote | unclassified | unclassified | unclassified | Gammaprote | information st | Translation, ri | 17C3B | The globular d | 207954.MED9 | 2 | 0,632455532 | 0,000248225 | 8,10E-05 | 2 - | 0,000208685 - | 1 - | 0,000129563 - | 2 - | 0,000246331 - | 3 - | 0,000374954 - | 2 - | 0,00028159 - |
| 7547 | 1 | 0.2um_NOKCPHJN_79535 | Bacteria | unclassified | unclassified | unclassified | unclassified | unclassified | uncultured ba | unclassified | unclassified | unclassified | unclassified | unclassified | 0,2 | 0,4 | 5,96E-06 | 1,19E-05 | 0 - | 0 - | 0 - | 0 - | 0 - | 0 - | 0 - | 0 - | 1 - | 2,98E-05 - |
| 7548 | 1 | 3um_FIJJAFBF_66745 | Bacteria | Proteobacteri | Gammaprote | Vibrionales | Vibrionaceae | Photobacteriu | various | unclassified | unclassified | unclassified | unclassified | unclassified | 8 | 1,095445115 | 0,00036055 | 4,60E-05 | 8 - | 0,000304044 - | 8 - | 0,000377533 - | 9 - | 0,000403755 - | 9 - | 0,000409718 - | 6 - | 0,000307698 - |
| 7549 | 1 | 3um_FIJJAFBF_424695 | Bacteria | Proteobacteri | Alphaproteob | Rhodobactera | Rhodobactera | Tropicibacter | Tropicibacter | information st | Replication, re | 05C68 | Can catalyze t | 252305.OB25 | 4,2 | 1,469693846 | 0,000158786 | 5,86E-05 | 3 - | 9,51E-05 - | 6 - | 0,000236219 - | 5 - | 0,00018713 - | 5 - | 0,000189894 - | 2 - | 8,56E-05 - |
| 7550 | 1 | 3um_FIJJAFBF_566351 | unclassified | unclassified | unclassified | unclassified | unclassified | unclassified | unclassified | unclassified | unclassified | unclassified | unclassified | unclassified | 0,2 | 0,4 | 1,70E-06 | 3,41E-06 | 0 - | 0 - | 0 - | 0 - | 1 - | 8,52E-06 - | 0 - | 0 - | 0 - | 0 - |
| 7551 | 1 | 3um_FIJJAFBF_37675 | Bacteria | Rhodotherma | unclassified | unclassified | unclassified | unclassified | Rhodotherma | cellular proces | Intracellular tr | 08SHB | Biopolymer tra | 643867.Ftrac_ | 1,4 | 1,019803903 | 0,000118531 | 7,67E-05 | 3 - | 0,000223591 - | 1 - | 9,25E-05 - | 2 - | 0,000175951 - | 0 - | 0 - | 1 - | 0,000100568 - |
| 7552 | 1 | 0.2um_NOKCPHJN_546028 | Bacteria | Proteobacteri | Gammaprote | unclassified | unclassified | unclassified | Gammaprote | unclassified | unclassified | unclassified | unclassified | unclassified | 25,2 | 2,481934729 | 0,000415405 | 6,63E-05 | 26 - | 0,000358247 - | 22 - | 0,0003764 - | 23 - | 0,000374081 - | 26 - | 0,00042912 - | 29 - | 0,000539179 - |
| 7553 | 1 | 3um_FIJJAFBF_337086 | Bacteria | Proteobacteri | Gammaprote | Oceanospirilla | Oceanospirilla | Motiliproteus | Motiliproteus | information st | Translation, ri | 05CEJ | Catalyzes the | 455436.GHTC | 4,2 | 1,720465053 | 8,00E-05 | 2,63E-05 | 7 - | 0,000115602 - | 5 - | 0,000102532 - | 4 - | 7,80E-05 - | 3 - | 5,93E-05 - | 2 - | 4,46E-05 - |
| 7554 | 1 | 0.2um_NOKCPHJN_256636 | Eukaryota | unclassified | Pelagophycea | Pelagomonad | unclassified | Aureococcus | Aureococcus a | unclassified | unclassified | unclassified | unclassified | unclassified | 3,8 | 0,748331477 | 6,00E-05 | 9,51E-06 | 5 - | 6,69E-05 - | 3 - | 4,98E-05 - | 4 - | 6,32E-05 - | 3 - | 4,81E-05 - | 4 - | 7,22E-05 - |
| 7555 | 1 | 3um_FIJJAFBF_40038 | Bacteria | Bacteroidetes | Flavobacteriia | Flavobacterial | Flavobacteriac | Maribacter | Maribacter pc | metabolism | Energy produc | 099Q7 | Catalyzes the | 983548.Krodi_ | 1 | 0 | 4,45E-05 | 4,22E-06 | 1 - | 3,73E-05 - | 1 - | 4,63E-05 - | 1 - | 4,40E-05 - | 1 - | 4,46E-05 - | 1 - | 5,03E-05 - |
| 7556 | 1 | 0.2um_NOKCPHJN_129777 | Bacteria | Bacteroidetes | Flavobacteriia | Flavobacterial | Flavobacteriac | Dokdonia | Dokdonia paci | unclassified | unclassified | unclassified | unclassified | unclassified | 1 | 0 | 2,05E-05 | 1,94E-06 | 1 - | 1,71E-05 - | 1 - | 2,13E-05 - | 1 - | 2,02E-05 - | 1 - | 2,05E-05 - | 1 - | 2,31E-05 - |
| 7557 | 1 | 3um_FIJJAFBF_246778 | Bacteria | Bacteroidetes | Flavobacteriia | Flavobacterial | Flavobacteriac | Flavobacterium | Flavobacterium | cellular proces | Cell wall/mem | 0PCMH | SusD family | 867900.Celly_ | 1 | 0 | 2,69E-05 | 2,55E-06 | 1 - | 2,25E-05 - | 1 - | 2,79E-05 - | 1 - | 2,66E-05 - | 1 - | 2,70E-05 - | 1 - | 3,04E-05 - |
| 7558 | 1 | 3um_FIJJAFBF_278068 | Bacteria | Proteobacteri | Alphaproteob | Rickettsiales | Candidatus Mi | Candidatus M | Candidatus M | metabolism | Energy produc | 16QAQ | C-type cytoch | 696127.midi_ | 0,2 | 0,4 | 9,79E-06 | 1,96E-05 | 0 - | 0 - | 0 - | 0 - | 0 - | 0 - | 1 - | 4,89E-05 - | 0 - | 0 - |
| 7559 | 1 | 3um_FIJJAFBF_607860 | Bacteria | Bacteroidetes | Flavobacteriia | Flavobacterial | Flavobacteriac | Formosa | Formosa sp. H | metabolism | Nucleotide tra | 0P85S | Plays an impo | 391603.FBALC | 1,8 | 0,748331477 | 5,74E-05 | 2,31E-05 | 2 - | 5,43E-05 - | 2 - | 6,74E-05 - | 3 - | 9,61E-05 - | 1 - | 3,25E-05 - | 1 - | 3,66E-05 - |
| 7560 | 1 | 3um_FIJJAFBF_699580 | Bacteria | Proteobacteri | Gammaprote | Cellvibrionales | Porticoccacea | unclassified | gamma prote | metabolism | Inorganic ion t | 16TQC | TONB-depend | 314287.GB220 | 0,2 | 0,4 | 5,22E-06 | 1,04E-05 | 0 - | 0 - | 0 - | 0 - | 0 - | 0 - | 1 - | 2,61E-05 - | 0 - | 0 - |
| 7561 | 1 | 0.2um_NOKCPHJN_347446 | Bacteria | Proteobacteri | Alphaproteob | Pelagibacteral | Pelagibacterac | unclassified | Pelagibacterac | cellular proces | Cell wall/mem | 7257 | NA | 439493.PB721 | 1,4 | 0,489897949 | 5,19E-05 | 2,11E-05 | 1 - | 3,05E-05 - | 1 - | 3,79E-05 - | 2 - | 7,21E-05 - | 1 - | 3,66E-05 - | 2 - | 8,24E-05 - |
| 7562 | 1 | 0.2um_NOKCPHJN_664804 | Bacteria | Proteobacteri | Alphaproteob | Pelagibacteral | Pelagibacterac | Candidatus Pe | Candidatus Pe | unclassified | unclassified | unclassified | unclassified | unclassified | 0,8 | 0,4 | 4,46E-05 | 2,24E-05 | 0 - | 0 - | 1 - | 5,57E-05 - | 1 - | 5,29E-05 - | 1 - | 5,37E-05 - | 1 - | 6,05E-05 - |
| 7563 | 1 | 3um_FIJJAFBF_306761 | Bacteria | Proteobacteri | Alphaproteob | Rhodobactera | Rhodobactera | Phaeobacter | Phaeobacter s | metabolism | Lipid transpor | COG2067 | lipid transport | 439496.RBY4I | 0,6 | 0,489897949 | 2,23E-05 | 1,85E-05 | 1 - | 3,14E-05 - | 0 - | 0 - | 0 - | 0 - | 1 - | 3,77E-05 - | 1 - | 4,24E-05 - |
| 7564 | 1 | 3um_FIJJAFBF_404360 | Bacteria | Bacteroidetes | Flavobacteriia | Flavobacterial | Flavobacteriac | Olleya | Olleya aquima | information st | Translation, ri | 08YY1 | This protein is | 313598.MED1 | 1,8 | 0,4 | 0,000211681 | 4,95E-05 | 2 - | 0,000197891 - | 1 - | 0,000122861 - | 2 - | 0,00023359 - | 2 - | 0,00023704 - | 2 - | 0,000267025 - |
| 7565 | 1 | 0.2um_NOKCPHJN_250021 | Bacteria | Proteobacteri | Alphaproteob | Pelagibacteral | Pelagibacterac | unclassified | alpha proteob | poorly charact | Function unkn | 0231W | FecR protein | 439493.PB721 | 0,4 | 0,489897949 | 5,15E-06 | 6,34E-06 | 1 - | 1,18E-05 - | 0 - | 0 - | 1 - | 1,39E-05 - | 0 - | 0 - | 0 - | 0 - |
| 7566 | 1 | 0.2um_NOKCPHJN_115346 | Bacteria | Bacteroidetes | Flavobacteriia | Flavobacterial | Flavobacteriac | Algibacter | Algibacter sp. | unclassified | unclassified | unclassified | unclassified | unclassified | 3,6 | 0,8 | 5,67E-05 | 1,47E-05 | 3 - | 3,94E-05 - | 3 - | 4,89E-05 - | 5 - | 7,74E-05 - | 3 - | 4,71E-05 - | 4 - | 7,08E-05 - |
| 7567 | 1 | 3um_FIJJAFBF_290327 | Bacteria | Proteobacteri | Gammaprote | Oceanospirilla | Saccharospirill | Saccharospiril | Saccharospiril | cellular proces | Signal transdu | 16Q0Y | phosphate reg | 999141.GME_ | 4,2 | 0,979795897 | 0,000241073 | 3,90E-05 | 6 - | 0,000293047 - | 4 - | 0,000242585 - | 4 - | 0,000230608 - | 3 - | 0,000175511 - | 4 - | 0,000263616 - |
| 7568 | 1 | 3um_FIJJAFBF_537673 | Eukaryota | unclassified | unclassified | Phaeocystales | Phaeocystace | Phaeocystis | Phaeocystis gl | metabolism | Energy produc | COG0055 | Produces ATP | 7739.JGI1257 | 5,2 | 1,166190379 | 0,000149162 | 3,27E-05 | 6 - | 0,000145287 - | 7 - | 0,000210471 - | 4 - | 0,000114331 - | 5 - | 0,000145025 - | 4 - | 0,000130696 - |
| 7569 | 2 | 3um_FIJJAFBF_29543;3um_FIJJAFBF_597393 | Bacteria | Bacteroidetes | Flavobacteriia | various | various | various | various | unclassified | unclassified | unclassified | unclassified | unclassified | 0,2 | 0,4 | 8,47E-06 | 1,69E-05 | 1 - | 4,24E-05 - | 0 - | 0 - | 0 - | 0 - | 0 - | 0 - | 0 - | 0 - |
| 7570 | 1 | 0.2um_NOKCPHJN_575659 | Bacteria | Bacteroidetes | Flavobacteriia | Flavobacterial | Cryomorphace | unclassified | Cryomorphace | unclassified | unclassified | unclassified | unclassified | unclassified | 0,2 | 0,4 | 1,36E-05 | 2,72E-05 | 1 - | 6,79E-05 - | 0 - | 0 - | 0 - | 0 - | 0 - | 0 - | 0 - | 0 - |
| 7571 | 1 | 0.2um_NOKCPHJN_491930 | Bacteria | Proteobacteri | Gammaprote | Alteromonada | Alteromonada | Paraglaciecola | Paraglaciecola | unclassified | unclassified | unclassified | unclassified | unclassified | 1,2 | 0,748331477 | 3,52E-05 | 2,39E-05 | 1 - | 2,39E-05 - | 2 - | 5,94E-05 - | 1 - | 2,82E-05 - | 0 - | 0 - | 2 - | 6,45E-05 - |
| 7572 | 1 | 3um_FIJJAFBF_44070 | unclassified | unclassified | unclassified | unclassified | unclassified | unclassified | unclassified | unclassified | unclassified | unclassified | unclassified | unclassified | 4,4 | 0,8 | 0,000399115 | 5,49E-05 | 5 - | 0,000385157 - | 4 - | 0,000382601 - | 5 - | 0,000454638 - | 5 - | 0,000461353 - | 3 - | 0,000311828 - |
| 7573 | 1 | 0.2um_NOKCPHJN_222058 | Bacteria | Bacteroidetes | Flavobacteriia | Flavobacterial | Flavobacteriac | unclassified | Flavobacteriac | metabolism | Inorganic ion t | 05C27 | TonB-depende | 742766.HMPR | 1 | 0 | 1,32E-05 | 1,25E-06 | 1 - | 1,10E-05 - | 1 - | 1,37E-05 - | 1 - | 1,30E-05 - | 1 - | 1,32E-05 - | 1 - | 1,49E-05 - |

| | | | | | | | | | | | | | | |
|---|---|---|---|---|---|---|---|---|---|---|---|---|---|---|
| 7574 | 1 | 3um_FIJJAFBF_94764 | unclassified | unclassified | unclassified | unclassified | unclassified | unclassified | unclassified | unclassified | unclassified | unclassified | unclassified | unclassified |
| 7575 | 1 | 3um_FIJJAFBF_407554 | Bacteria | Bacteroidetes | unclassified | unclassified | unclassified | unclassified | Bacteroidetes | unclassified | unclassified | unclassified | unclassified | unclassified |
| 7576 | 1 | 0.2um_NOKCPHJN_164119 | Bacteria | Bacteroidetes | Flavobacteriia | Flavobacterial | Flavobacteriac | Polaribacter | Polaribacter s | metabolism | Energy produc | 0P7S4 | Produces ATP | 313598.MED1 |
| 7577 | 1 | 0.2um_NOKCPHJN_167586 | Bacteria | Bacteroidetes | Saprospiria | Saprospirales | Saprospiracea | Aureispira | Aureispira sp. | unclassified | unclassified | unclassified | unclassified | unclassified |
| 7578 | 1 | 3um_FIJJAFBF_281980 | Bacteria | Bacteroidetes | Cytophagia | Cytophagales | Flammeovirga | unclassified | Flammeovirga | unclassified | unclassified | unclassified | unclassified | unclassified |
| 7579 | 1 | 3um_FIJJAFBF_79229 | Bacteria | Bacteroidetes | Flavobacteriia | Flavobacterial | Flavobacteriac | Polaribacter | Polaribacter s | unclassified | unclassified | unclassified | unclassified | unclassified |
| 7580 | 1 | 3um_FIJJAFBF_216066 | Bacteria | Proteobacteri | Gammaprotec | Alteromonada | various | various | various | information st | Translation, ril | 0QTKX | Seems to be tl | 203122.Sde_0 |
| 7581 | 3 | 0.2um_NOKCPHJN_833582;0.2um_NOKCPHJN_9 | Bacteria | unclassified | unclassified | unclassified | unclassified | unclassified | uncultured ba | various | various | various | various | various |
| 7582 | 1 | 0.2um_NOKCPHJN_263907 | Bacteria | Proteobacteri | Alphaproteob | Rickettsiales | unclassified | unclassified | Rickettsiales b | unclassified | unclassified | unclassified | unclassified | unclassified |
| 7583 | 1 | 3um_FIJJAFBF_20030 | Bacteria | unclassified | unclassified | unclassified | unclassified | unclassified | uncultured ba | metabolism | Amino acid tra | COG0436 | aminotransfer | 313598.MED1 |
| 7584 | 1 | 3um_FIJJAFBF_168181 | Bacteria | Bacteroidetes | Flavobacteriia | Flavobacterial | Flavobacteriac | Polaribacter | Polaribacter v | metabolism | Amino acid tra | 05C6M | Alpha-keto-be | 313594.PI23P |
| 7585 | 1 | 3um_FIJJAFBF_218819 | Bacteria | Proteobacteri | Gammaprotec | Oceanospirilla | Saccharospirill | Reinekea | Reinekea fors | poorly charact | Function unkn | 08UTA | Domain of unl | 325240.Sbal_( |
| 7586 | 1 | 0.2um_NOKCPHJN_49496 | Bacteria | Proteobacteri | Alphaproteob | Rhodospirillal | Rhodospirillac | unclassified | Rhodospirillac | metabolism | Secondary me | 01Q86 | extracellular s | 367336.OM22 |
| 7587 | 1 | 3um_FIJJAFBF_201862 | Bacteria | Bacteroidetes | Flavobacteriia | Flavobacterial | Flavobacteriac | unclassified | Flavobacteriac | unclassified | unclassified | unclassified | unclassified | unclassified |
| 7588 | 1 | 0.2um_NOKCPHJN_366538 | Bacteria | Bacteroidetes | unclassified | unclassified | unclassified | unclassified | Bacteroidetes | cellular proces | Intracellular tr | 05FGH | MotA TolQ ex | 391603.FBALC |
| 7589 | 1 | 0.2um_NOKCPHJN_67465 | Bacteria | Bacteroidetes | Flavobacteriia | Flavobacterial | Flavobacteriac | Gaetbulibacte | Gaetbulibacte | information st | Translation, ril | COG0088 | One of the pri | 1046627.BZAF |
| 7590 | 1 | 0.2um_NOKCPHJN_69345 | Bacteria | Bacteroidetes | Flavobacteriia | Flavobacterial | unclassified | unclassified | Flavobacterial | unclassified | unclassified | unclassified | unclassified | unclassified |
| 7591 | 1 | 3um_FIJJAFBF_438911 | Bacteria | Planctomycet | Phycisphaerae | Phycisphaeral | Phycisphaerac | unclassified | Phycisphaerac | unclassified | unclassified | unclassified | unclassified | unclassified |
| 7592 | 1 | 0.2um_NOKCPHJN_182727 | Bacteria | Bacteroidetes | unclassified | unclassified | unclassified | unclassified | Bacteroidetes | poorly charact | Function unkn | 11UE2 | S1 P1 Nucleas | 715451.ambt_ |
| 7593 | 2 | 3um_FIJJAFBF_352047;3um_FIJJAFBF_715031 | Bacteria | Proteobacteri | Alphaproteob | Rhodobactera | Rhodobactera | Cognatishimia | Cognatishimia | metabolism | Energy produc | 16RH6 | Produces ATP | 439496.RBY4I |
| 7594 | 1 | 3um_FIJJAFBF_196411 | Bacteria | Proteobacteri | Gammaprotec | Alteromonada | Alteromonada | Paraglaciecola | Paraglaciecola | unclassified | unclassified | unclassified | unclassified | unclassified |
| 7595 | 1 | 3um_FIJJAFBF_737463 | Bacteria | Proteobacteri | Gammaprotec | Oceanospirilla | Saccharospirill | Reinekea | Reinekea fors | information st | Translation, ril | 08UHH | Located on th | 314283.MED2 |
| 7596 | 1 | 3um_FIJJAFBF_53336 | Bacteria | Bacteroidetes | Flavobacteriia | Flavobacterial | Flavobacteriac | Polaribacter | Polaribacter s | unclassified | unclassified | unclassified | unclassified | unclassified |
| 7597 | 1 | 3um_FIJJAFBF_94123 | Bacteria | Proteobacteri | Gammaprotec | Arenicellales | Arenicellaceae | Arenicella | Arenicella xan | metabolism | Inorganic ion t | 05PWN | phosphate ab | 247634.GPB2 |
| 7598 | 1 | 3um_FIJJAFBF_188381 | Bacteria | Bacteroidetes | Flavobacteriia | Flavobacterial | Flavobacteriac | Formosa | Formosa sp. H | poorly charact | Function unkn | 0XPRT | NA | 391598.FBBAL |
| 7599 | 1 | 0.2um_NOKCPHJN_118513 | Bacteria | Proteobacteri | Gammaprotec | Oceanospirilla | Saccharospirill | Reinekea | Reinekea fors | unclassified | unclassified | unclassified | unclassified | unclassified |
| 7600 | 1 | 3um_FIJJAFBF_211172 | Bacteria | Proteobacteri | Betaproteoba | Burkholderial | Comamonada | Polaromonas | Polaromonas | unclassified | unclassified | unclassified | unclassified | unclassified |
| 7601 | 1 | 0.2um_NOKCPHJN_914416 | unclassified | unclassified | unclassified | unclassified | unclassified | unclassified | unclassified | unclassified | unclassified | unclassified | unclassified | unclassified |
| 7602 | 1 | 0.2um_NOKCPHJN_103660 | Eukaryota | unclassified | Oomycetes | Saprolegniales | Saprolegniace | Achlya | Achlya hypogy | information st | Translation, ril | COG5045 | Ribosomal prc | 67593.Physo1 |
| 7603 | 1 | 3um_FIJJAFBF_87433 | Bacteria | Proteobacteri | Alphaproteob | Rhodobactera | Rhodobactera | Sulfitobacter | Sulfitobacter g | unclassified | unclassified | unclassified | unclassified | unclassified |
| 7604 | 2 | 3um_FIJJAFBF_235100;3um_FIJJAFBF_750209 | Eukaryota | Bacillariophyt | various | various | various | various | various | cellular proces | Signal transdu | COG5040 | Tyrosine 3-mc | various |
| 7605 | 1 | 3um_FIJJAFBF_517423 | Bacteria | Bacteroidetes | Flavobacteriia | Flavobacterial | Flavobacteriac | Polaribacter | Polaribacter b | unclassified | unclassified | unclassified | unclassified | unclassified |
| 7606 | 1 | 3um_FIJJAFBF_245828 | Bacteria | Bacteroidetes | Flavobacteriia | Flavobacterial | Flavobacteriac | unclassified | Flavobacteriac | cellular proces | Posttranslatio | COG0330 | Band 7 protei | 313603.FB217 |
| 7607 | 1 | 3um_FIJJAFBF_639896 | Bacteria | Proteobacteri | Gammaprotec | Oceanospirilla | Saccharospirill | Reinekea | Reinekea fors | unclassified | unclassified | unclassified | unclassified | unclassified |
| 7608 | 1 | 3um_FIJJAFBF_81617 | Bacteria | Bacteroidetes | Flavobacteriia | Flavobacterial | Flavobacteriac | Winogradskye | Winogradskye | cellular proces | Cell wall/mem | 0P7KR | tyrosine-prote | 391603.FBALC |
| 7609 | 1 | 0.2um_NOKCPHJN_188899 | Bacteria | Proteobacteri | Alphaproteob | Rhodobactera | Rhodobactera | Sulfitobacter | Sulfitobacter s | information st | Translation, ril | 05G6W | One of the pri | 52598.EE36_0 |
| 7610 | 1 | 0.2um_NOKCPHJN_140287 | Bacteria | Proteobacteri | Alphaproteob | Rhodobactera | Rhodobactera | Actibacterium | Actibacterium | unclassified | unclassified | unclassified | unclassified | unclassified |
| 7611 | 1 | 3um_FIJJAFBF_191798 | Bacteria | Bacteroidetes | Flavobacteriia | Flavobacterial | Flavobacteriac | Cellulophaga | Cellulophaga t | poorly charact | Function unkn | 0PDYX | NA | 867900.Celly_ |
| 7612 | 1 | 3um_FIJJAFBF_527348 | Eukaryota | Bacillariophyt | Coscinodiscop | Coscinodiscale | Coscinodiscac | Coscinodiscus | Coscinodiscus | metabolism | Energy produc | COG0055 | Produces ATP | 7739.JGI1257 |
| 7613 | 1 | 3um_FIJJAFBF_76059 | Bacteria | Bacteroidetes | Saprospiria | Saprospirales | Saprospiracea | Aureispira | Aureispira sp. | unclassified | unclassified | unclassified | unclassified | unclassified |
| 7614 | 1 | 0.2um_NOKCPHJN_541382 | Bacteria | Proteobacteri | Alphaproteob | Rhodospirillal | Rhodospirillac | Azospirillum | Azospirillum s | unclassified | unclassified | unclassified | unclassified | unclassified |
| 7615 | 1 | 0.2um_NOKCPHJN_371584 | Bacteria | Proteobacteri | Gammaprotec | Alteromonada | Alteromonada | Paraglaciecola | Paraglaciecola | information st | Translation, ril | 05CEJ | Catalyzes the | 314282.PCNPT |
| 7616 | 1 | 0.2um_NOKCPHJN_779714 | Eukaryota | unclassified | Ichthyosporea | unclassified | unclassified | Capsaspora | Capsaspora ov | unclassified | unclassified | unclassified | unclassified | unclassified |
| 7617 | 1 | 0.2um_NOKCPHJN_256860 | Bacteria | Proteobacteri | Gammaprotec | Cellvibrionales | Porticoccacea | unclassified | SAR92 bacteri | information st | Replication, re | 05C68 | Can catalyze t | 498211.CJA_2 |
| 7618 | 1 | 0.2um_NOKCPHJN_1061371 | Bacteria | Proteobacteri | Alphaproteob | Pelagibacteral | Pelagibacterac | Candidatus Pe | Candidatus Pe | information st | Replication, re | 05C68 | Can catalyze t | 439493.PB721 |
| 7619 | 1 | 0.2um_NOKCPHJN_123345 | Bacteria | Proteobacteri | Gammaprotec | Oceanospirilla | Saccharospirill | Reinekea | Reinekea fors | unclassified | unclassified | unclassified | unclassified | unclassified |
| 7620 | 1 | 0.2um_NOKCPHJN_818504 | Bacteria | Proteobacteri | Alphaproteob | Rickettsiales | Anaplasmatac | Anaplasma | Anaplasma ph | unclassified | unclassified | unclassified | unclassified | unclassified |
| 7621 | 1 | 0.2um_NOKCPHJN_310343 | Eukaryota | Mucoromycot | Mucoromycet | Mucorales | Choanephorac | Choanephora | Choanephora | cellular proces | Intracellular tr | 0V837 | member RAS ( | 31033.ENSTRU |
| 7622 | 1 | 3um_FIJJAFBF_39637 | Bacteria | Bacteroidetes | Flavobacteriia | Flavobacterial | Flavobacteriac | unclassified | Flavobacteriac | metabolism | Carbohydrate | 099Q3 | phosphohexol | 391603.FBALC |
| 7623 | 1 | 0.2um_NOKCPHJN_67579 | Bacteria | Bacteroidetes | Flavobacteriia | Flavobacterial | Flavobacteriac | unclassified | Flavobacteriac | information st | Transcription | 09EME | Necessary for | 487796.Flav2A |
| 7624 | 1 | 0.2um_NOKCPHJN_27395 | Bacteria | unclassified | unclassified | unclassified | unclassified | unclassified | uncultured ba | unclassified | unclassified | unclassified | unclassified | unclassified |
| 7625 | 1 | 0.2um_NOKCPHJN_600197 | Bacteria | Proteobacteri | Gammaprotec | unclassified | unclassified | unclassified | SAR86 cluster | information st | Translation, ril | 05CEJ | Catalyzes the | 491952.Mar18 |
| 7626 | 1 | 0.2um_NOKCPHJN_379869 | Bacteria | Proteobacteri | Betaproteoba | Burkholderial | Comamonada | Hydrogenoph | Hydrogenoph | unclassified | unclassified | unclassified | unclassified | unclassified |
| 7627 | 1 | 3um_FIJJAFBF_582528 | unclassified | unclassified | unclassified | unclassified | unclassified | unclassified | unclassified | unclassified | unclassified | unclassified | unclassified | unclassified |
| 7628 | 1 | 0.2um_NOKCPHJN_355825 | Bacteria | Proteobacteri | Gammaprotec | unclassified | unclassified | unclassified | Gammaprotec | metabolism | Inorganic ion t | 173HE | CBS domain p | 443152.MDG8 |
| 7629 | 1 | 3um_FIJJAFBF_191136 | Bacteria | Proteobacteri | Alphaproteob | Rhodobactera | unclassified | unclassified | Rhodobactera | information st | Translation, ril | COG0088 | One of the pri | 367336.OM22 |
| 7630 | 1 | 0.2um_NOKCPHJN_668716 | Bacteria | Proteobacteri | Gammaprotec | Cellvibrionales | Porticoccacea | unclassified | SAR92 bacteri | poorly charact | Function unkn | 16R8I | Lipoprotein | 377629.TERTU |
| 7631 | 1 | 0.2um_NOKCPHJN_227309 | Bacteria | Bacteroidetes | Flavobacteriia | Flavobacterial | Flavobacteriac | Maribacter | Maribacter do | unclassified | unclassified | unclassified | unclassified | unclassified |
| 7632 | 1 | 3um_FIJJAFBF_444691 | Eukaryota | unclassified | unclassified | Isochrysidales | Noelaerhabda | Emiliania | Emiliania huxl | metabolism | Carbohydrate | COG1850 | RuBisCO catal | 45157.CMV01 |
| 7633 | 1 | 3um_FIJJAFBF_83506 | Bacteria | Proteobacteri | Betaproteoba | unclassified | unclassified | unclassified | Betaproteoba | unclassified | unclassified | unclassified | unclassified | unclassified |
| 7634 | 1 | 3um_FIJJAFBF_78180 | Bacteria | Bacteroidetes | Flavobacteriia | Flavobacterial | Flavobacteriac | Formosa | Formosa sp. H | metabolism | Amino acid tra | COG0436 | aminotransfer | 398720.MED2 |
| 7635 | 1 | 0.2um_NOKCPHJN_256291 | Bacteria | Proteobacteri | Alphaproteob | Rhodobactera | Rhodobactera | unclassified | Rhodobactera | metabolism | Carbohydrate | 0XPV5 | extracellular s | 744979.R2A13 |
| 7636 | 1 | 0.2um_NOKCPHJN_252950 | Bacteria | Cyanobacteria | unclassified | Oscillatoriales | Oscillatoriacea | Limnoraphis | Limnoraphis r | unclassified | unclassified | unclassified | unclassified | unclassified |
| 7637 | 1 | 0.2um_NOKCPHJN_178402 | Bacteria | Bacteroidetes | Flavobacteriia | Flavobacterial | Flavobacteriac | Polaribacter | various | poorly charact | Function unkn | 0PBMV | NA | 313603.FB217 |
| 7638 | 1 | 0.2um_NOKCPHJN_449387 | Bacteria | Proteobacteri | Alphaproteob | unclassified | unclassified | unclassified | Alphaproteob | metabolism | Carbohydrate | 16TFK | C4-dicarboxyl | 331869.BAL19 |
| 7639 | 1 | 0.2um_NOKCPHJN_224765 | Bacteria | Proteobacteri | Alphaproteob | Rhodospirillal | Rhodospirillac | unclassified | Rhodospirillac | metabolism | Amino acid tra | 05TIB | (ABC) transpo | 246200.SPO24 |
| 7640 | 1 | 0.2um_NOKCPHJN_559310 | unclassified | unclassified | unclassified | unclassified | unclassified | unclassified | unclassified | unclassified | unclassified | unclassified | unclassified | unclassified |
| 7641 | 1 | 0.2um_NOKCPHJN_1002588 | Bacteria | Proteobacteri | Alphaproteob | Pelagibacteral | Pelagibacterac | Candidatus Pe | Candidatus Pe | information st | Translation, ril | COG0050 | This protein p | 335992.SAR11 |
| 7642 | 1 | 0.2um_NOKCPHJN_483438 | Bacteria | Proteobacteri | Gammaprotec | unclassified | unclassified | unclassified | Gammaprotec | metabolism | Inorganic ion t | 0QJVD | TonB-depende | 327275.SOHN |
| 7643 | 1 | 0.2um_NOKCPHJN_552141 | Eukaryota | unclassified | Labyrinthulom | unclassified | Thraustochytr | Aurantiochytr | Aurantiochytr | unclassified | unclassified | unclassified | unclassified | unclassified |
| 7644 | 1 | 0.2um_NOKCPHJN_1061525 | Bacteria | Proteobacteri | Gammaprotec | Alteromonada | Pseudoalteror | Pseudoalteror | various | unclassified | unclassified | unclassified | unclassified | unclassified |
| 7645 | 1 | 3um_FIJJAFBF_395792 | Bacteria | Actinobacteri | Actinobacteri | Micrococcales | Demequinace | Lysinimicrobiu | Lysinimicrobiu | unclassified | unclassified | unclassified | unclassified | unclassified |
| 7646 | 1 | 0.2um_NOKCPHJN_467170 | Bacteria | Proteobacteri | Gammaprotec | unclassified | unclassified | unclassified | SAR86 cluster | unclassified | unclassified | unclassified | unclassified | unclassified |
| 7647 | 1 | 3um_FIJJAFBF_604254 | Bacteria | Proteobacteri | Gammaprotec | unclassified | unclassified | unclassified | Gammaprotec | information st | Translation, ril | COG0184 | Forms an inte | 247634.GPB2 |
| 7648 | 1 | 0.2um_NOKCPHJN_180480 | Bacteria | Proteobacteri | Alphaproteob | Rhodobactera | Rhodobactera | Planktotalea | Planktotalea f | cellular proces | Posttranslatio | COG0330 | Band 7 protei | 314270.RB208 |
| 7649 | 1 | 3um_FIJJAFBF_740444 | Bacteria | Proteobacteri | Alphaproteob | Pelagibacteral | Pelagibacterac | Candidatus Pe | Candidatus Pe | unclassified | unclassified | unclassified | unclassified | unclassified |
| 7650 | 1 | 3um_FIJJAFBF_345361 | Bacteria | Bacteroidetes | Flavobacteriia | Flavobacterial | Crocinitomica | unclassified | Crocinitomica | metabolism | Carbohydrate | 099YR | Catalyzes the | 755732.Fluta_ |
| 7651 | 1 | 3um_FIJJAFBF_561978 | Eukaryota | unclassified | Spirotrichea | Sporadotrichi | Oxytrichidae | Oxytricha | Oxytricha trifa | unclassified | unclassified | unclassified | unclassified | unclassified |
| 7652 | 1 | 3um_FIJJAFBF_181244 | Bacteria | Bacteroidetes | Flavobacteriia | Flavobacterial | Flavobacteriac | Leeuwenhoek | Leeuwenhoek | poorly charact | Function unkn | 11QQ6 | NA | 398720.MED2 |
| 7653 | 1 | 3um_FIJJAFBF_228001 | Eukaryota | Bacillariophyt | Bacillariophyc | Naviculales | Naviculaceae | Fistulifera | Fistulifera sola | unclassified | unclassified | unclassified | unclassified | unclassified |
| 7654 | 1 | 0.2um_NOKCPHJN_454362 | Bacteria | Proteobacteri | Alphaproteob | Rhodobactera | Rhodobactera | unclassified | Rhodobactera | unclassified | unclassified | unclassified | unclassified | unclassified |
| 7655 | 1 | 3um_FIJJAFBF_550979 | Bacteria | Proteobacteri | Alphaproteob | Rhodobactera | Rhodobactera | unclassified | Rhodobactera | unclassified | unclassified | unclassified | unclassified | unclassified |
| 7656 | 1 | 0.2um_NOKCPHJN_146467 | Bacteria | Bacteroidetes | Flavobacteriia | Flavobacterial | Flavobacteriac | Flavobacteriu | Flavobacteriu | metabolism | Energy produc | COG0055 | Produces ATP | 865938.Weev |
| 7657 | 1 | 0.2um_NOKCPHJN_138711 | Bacteria | Bacteroidetes | Flavobacteriia | Flavobacterial | Flavobacteriac | Aureicoccus | Aureicoccus m | unclassified | unclassified | unclassified | unclassified | unclassified |
| 7658 | 1 | 0.2um_NOKCPHJN_237508 | Bacteria | Proteobacteri | Gammaprotec | Cellvibrionales | unclassified | unclassified | Cellvibrionales | poorly charact | Function unkn | 0QNMQ | NA | 314287.GB22 |
| 7659 | 1 | 0.2um_NOKCPHJN_899405 | Bacteria | Proteobacteri | Alphaproteob | Pelagibacteral | Pelagibacterac | Candidatus Pe | Candidatus Pe | unclassified | unclassified | unclassified | unclassified | unclassified |
| 7660 | 1 | 3um_FIJJAFBF_579181 | Bacteria | Proteobacteri | Gammaprotec | Cellvibrionales | Porticoccacea | unclassified | gamma protec | unclassified | unclassified | unclassified | unclassified | unclassified |
| 7661 | 1 | 0.2um_NOKCPHJN_42568 | Bacteria | Proteobacteri | Alphaproteob | Rhodobactera | Rhodobactera | Litoreibacter | Litoreibacter p | unclassified | unclassified | unclassified | unclassified | unclassified |
| 7662 | 1 | 3um_FIJJAFBF_369630 | Bacteria | Proteobacteri | Alphaproteob | Rhodobactera | Rhodobactera | Thalassobium | Thalassobium | unclassified | unclassified | unclassified | unclassified | unclassified |
| 7663 | 1 | 0.2um_NOKCPHJN_457450 | Bacteria | Candidatus Po | unclassified | unclassified | unclassified | unclassified | Candidatus Po | poorly charact | Function unkn | 0ZJ76 | NA | 448385.sce87 |
| 7664 | 1 | 0.2um_NOKCPHJN_1061130 | Bacteria | Verrucomicrol | Opitutae | Opitutales | Opitutaceae | unclassified | Opitutaceae b | information st | Replication, re | 1DH1D | ATP-depender | 583355.Caka_ |
| 7665 | 1 | 3um_FIJJAFBF_711069 | Bacteria | Proteobacteri | Alphaproteob | Rhodobactera | Rhodobactera | Planktomarina | Planktomarina | poorly charact | Function unkn | 01UDD | YeeE YedE fan | 246200.SPO2 |
| 7666 | 1 | 0.2um_NOKCPHJN_568160 | Bacteria | Proteobacteri | Alphaproteob | Pelagibacteral | Pelagibacterac | Candidatus Pe | Candidatus Pe | metabolism | Amino acid tra | 01R4A | Polyamine AB | 335992.SAR11 |
| 7667 | 1 | 0.2um_NOKCPHJN_669445 | Bacteria | Proteobacteri | Gammaprotec | Cellvibrionales | Porticoccacea | unclassified | gamma protec | unclassified | unclassified | unclassified | unclassified | unclassified |
| 7668 | 1 | 3um_FIJJAFBF_459235 | Bacteria | Proteobacteri | Alphaproteob | various | various | various | various | information st | Translation, ril | 05VBI | Binds 16S rRN | 367336.OM22 |
| 7669 | 1 | 0.2um_NOKCPHJN_263427 | Bacteria | Bacteroidetes | Flavobacteriia | Flavobacterial | Flavobacteriac | Formosa | Formosa sp. H | unclassified | unclassified | unclassified | unclassified | unclassified |
| 7670 | 1 | 3um_FIJJAFBF_248985 | Bacteria | Actinobacteri | Actinobacteri | Micrococcales | Microbacteria | Candidatus Ac | Candidatus Ac | unclassified | unclassified | unclassified | unclassified | unclassified |
| 7671 | 2 | 3um_FIJJAFBF_18794;3um_FIJJAFBF_700800 | Bacteria | Proteobacteri | Alphaproteob | Rhodobactera | Rhodobactera | various | various | information st | Transcription | 178T0 | Transcriptiona | 314262.MED1 |
| 7672 | 1 | 0.2um_NOKCPHJN_405789 | Bacteria | Verrucomicrol | unclassified | unclassified | unclassified | unclassified | Verrucomicrol | unclassified | unclassified | unclassified | unclassified | unclassified |
| 7673 | 1 | 0.2um_NOKCPHJN_18843 | Bacteria | Proteobacteri | Gammaprotec | Oceanospirilla | Oceanospirilla | Neptunomona | Neptunomona | metabolism | Energy produc | COG0055 | Produces ATP | 207954.MED9 |
| 7674 | 1 | 0.2um_NOKCPHJN_260998 | Bacteria | Proteobacteri | Alphaproteob | Pelagibacteral | Pelagibacterac | unclassified | Pelagibacterac | unclassified | unclassified | unclassified | unclassified | unclassified |
| 7675 | 1 | 3um_FIJJAFBF_379396 | Viruses | unclassified | unclassified | unclassified | Phycodnavirid | unclassified | Pyramimonas | unclassified | unclassified | unclassified | unclassified | unclassified |
| 7676 | 1 | 3um_FIJJAFBF_486301 | Bacteria | Proteobacteri | Gammaprotec | Cellvibrionales | Halieaceae | Halioglobus | various | information st | Translation, ril | 0QQXI | This protein is | 247634.GPB2 |
| 7677 | 1 | 3um_FIJJAFBF_531098 | unclassified | unclassified | unclassified | unclassified | unclassified | unclassified | unclassified | unclassified | unclassified | unclassified | unclassified | unclassified |
| 7678 | 1 | 0.2um_NOKCPHJN_153284 | Bacteria | Proteobacteri | Gammaprotec | Alteromonada | Pseudoalteror | Pseudoalteror | Pseudoalteror | metabolism | Inorganic ion t | 0QN53 | TonB-depende | 983545.Glaag |
| 7679 | 1 | 0.2um_NOKCPHJN_27136 | Bacteria | Proteobacteri | Alphaproteob | Rhodobactera | Rhodobactera | Primorskyibac | Primorskyibac | metabolism | Carbohydrate | 16THB | transporter, d | 388399.SSE37 |
| 7680 | 2 | 0.2um_NOKCPHJN_206327;3um_FIJJAFBF_64841 | Eukaryota | Phaeophyceae | unclassified | Ectocarpales | Ectocarpaceae | Ectocarpus | Ectocarpus sili | unclassified | unclassified | unclassified | unclassified | unclassified |
| 7681 | 1 | 3um_FIJJAFBF_155472 | Bacteria | Proteobacteri | Gammaprotec | Oceanospirilla | Oceanospirilla | Balneatrix | Balneatrix alp | cellular proces | Signal transdu | COG1366 | stage II sporul | 83406.HDN1F |
| 7682 | 1 | 3um_FIJJAFBF_739990 | Eukaryota | unclassified | unclassified | Prymnesiales | Chrysochromu | Chrysochromu | various | unclassified | unclassified | unclassified | unclassified | unclassified |
| 7683 | 1 | 3um_FIJJAFBF_214611 | Bacteria | Proteobacteri | Alphaproteob | Rhodobactera | unclassified | unclassified | Rhodobactera | information st | Translation, ril | 177SZ | This protein bi | 391626.OA30 |
| 7684 | 1 | 3um_FIJJAFBF_140656 | Bacteria | Proteobacteri | Alphaproteob | Rhodobactera | unclassified | unclassified | Rhodobactera | cellular proces | Posttranslatio | COG0652 | peptidyl-proly | 367336.OM22 |
| 7685 | 1 | 3um_FIJJAFBF_264143 | Bacteria | Proteobacteri | Gammaprotec | Alteromonada | Alteromonada | Paraglaciecola | various | information st | Translation, ril | 0QVRF | 50S ribosomal | 342610.Patl_1 |
| 7686 | 1 | 0.2um_NOKCPHJN_09383 | Bacteria | Bacteroidetes | Flavobacteriia | Flavobacterial | Flavobacteriac | Polaribacter | Polaribacter d | cellular proces | Intracellular tr | 05FGH | MotA TolQ ex | 313598.MED1 |
| 7687 | 1 | 0.2um_NOKCPHJN_381565 | Bacteria | Proteobacteri | Alphaproteob | unclassified | unclassified | unclassified | Alphaproteob | information st | Translation, ril | COG0185 | Protein S19 fo | 488538.SAR11 |
| 7688 | 2 | 0.2um_NOKCPHJN_985727;3um_FIJJAFBF_45157 | unclassified | unclassified | unclassified | unclassified | unclassified | unclassified | unclassified | unclassified | unclassified | unclassified | unclassified | unclassified |
| 7689 | 1 | 3um_FIJJAFBF_157513 | unclassified | unclassified | unclassified | unclassified | unclassified | unclassified | unclassified | unclassified | unclassified | unclassified | unclassified | unclassified |
| 7690 | 1 | 3um_FIJJAFBF_254317 | unclassified | unclassified | unclassified | unclassified | unclassified | unclassified | unclassified | unclassified | unclassified | unclassified | unclassified | unclassified |
| 7691 | 1 | 3um_FIJJAFBF_411071 | Eukaryota | Phaeophyceae | unclassified | Ectocarpales | Ectocarpaceae | Ectocarpus | Ectocarpus sili | information st | Chromatin stri | KOG1744 | Core compone | 2850.Phatr54 |
| 7692 | 1 | 0.2um_NOKCPHJN_416915 | Bacteria | Proteobacteri | Alphaproteob | Pelagibacteral | Pelagibacterac | Candidatus Pe | Candidatus Pe | information st | Translation, ril | 01WSM | This is one of t | 335992.SAR11 |
| 7693 | 1 | 3um_FIJJAFBF_62817 | Bacteria | Bacteroidetes | Flavobacteriia | Flavobacterial | Flavobacteriac | Polaribacter | Polaribacter s | cellular proces | Cell wall/mem | 06B97 | phosphate-sel | 313594.PI23P |
| 7694 | 1 | 3um_FIJJAFBF_45932 | Eukaryota | Phaeophyceae | unclassified | Ectocarpales | Ectocarpaceae | Ectocarpus | Ectocarpus sili | unclassified | unclassified | unclassified | unclassified | unclassified |
| 7695 | 1 | 0.2um_NOKCPHJN_215618 | Eukaryota | unclassified | Pelagophycea | Pelagomonad | unclassified | Aureococcus | Aureococcus a | unclassified | unclassified | unclassified | unclassified | unclassified |
| 7696 | 1 | 0.2um_NOKCPHJN_607085 | Bacteria | Proteobacteri | Alphaproteob | unclassified | unclassified | Candidatus Pu | Candidatus Pu | unclassified | unclassified | unclassified | unclassified | unclassified |
| 7697 | 1 | 0.2um_NOKCPHJN_344166 | Bacteria | Proteobacteri | Alphaproteob | Rhodobactera | unclassified | unclassified | Rhodobactera | metabolism | Amino acid tra | COG0136 | Catalyzes the | 314232.SKA53 |
| 7698 | 1 | 3um_FIJJAFBF_557513 | Bacteria | Bacteroidetes | Flavobacteriia | Flavobacterial | Flavobacteriac | Tamlana | Tamlana agari | unclassified | unclassified | unclassified | unclassified | unclassified |
| 7699 | 1 | 3um_FIJJAFBF_26912 | Bacteria | Proteobacteri | Gammaprotec | Chromatiales | Granulosicocc | Granulosicocc | Granulosicocc | unclassified | unclassified | unclassified | unclassified | unclassified |
| 7700 | 1 | 3um_FIJJAFBF_201378 | Bacteria | Bacteroidetes | Flavobacteriia | Flavobacterial | Flavobacteriac | Polaribacter | Polaribacter fi | cellular proces | Cell wall/mem | 0PENK | NA | 391587.KAOT |
| 7701 | 1 | 0.2um_NOKCPHJN_21267 | Bacteria | Proteobacteri | unclassified | unclassified | unclassified | unclassified | uncultured ma | cellular proces | Posttranslatio | 01QDX | ATP binding to | 501479.CSE45 |
| 7702 | 1 | 0.2um_NOKCPHJN_785097 | Bacteria | Proteobacteri | Alphaproteob | Rhodobactera | Rhodobactera | Roseibium | Roseibium sp. | cellular proces | Posttranslatio | 16QU3 | Hemolysin-typ | 744980.TRICH |
| 7703 | 1 | 3um_FIJJAFBF_229664 | Bacteria | Bacteroidetes | Flavobacteriia | Flavobacterial | Flavobacteriac | Lutibacter | Lutibacter flav | metabolism | Inorganic ion t | 05C27 | TonB-depende | 313598.MED1 |
| 7704 | 1 | 3um_FIJJAFBF_346690 | Bacteria | Proteobacteri | Gammaprotec | Oceanospirilla | Oceanospirilla | Motiliproteus | Motiliproteus | metabolism | Energy produc | 16RH6 | Produces ATP | 207954.MED9 |
| 7705 | 2 | 0.2um_NOKCPHJN_436851;3um_FIJJAFBF_46386 | Bacteria | Proteobacteri | Gammaprotec | Cellvibrionales | Porticoccacea | unclassified | various | unclassified | unclassified | unclassified | unclassified | unclassified |
| 7706 | 1 | 3um_FIJJAFBF_152915 | Bacteria | Proteobacteri | Alphaproteob | Rhodobactera | Rhodobactera | Ruegeria | Ruegeria med | metabolism | Energy produc | 05C26 | Dehydrogenas | 314232.SKA53 |
| 7707 | 1 | 3um_FIJJAFBF_733378 | Viruses | unclassified | unclassified | unclassified | Mimiviridae | Cafeteriavirus | Cafeteria roen | unclassified | unclassified | unclassified | unclassified | unclassified |
| 7708 | 1 | 0.2um_NOKCPHJN_125762 | Bacteria | Bacteroidetes | Flavobacteriia | Flavobacterial | Flavobacteriac | Polaribacter | Polaribacter v | unclassified | unclassified | unclassified | unclassified | unclassified |
| 7709 | 1 | 0.2um_NOKCPHJN_91999 | Bacteria | Bacteroidetes | Flavobacteriia | Flavobacterial | Flavobacteriac | Ochrovirga | Ochrovirga pa | poorly charact | Function unkn | 0XY6Y | Pkd domain c | 391587.KAOT |
| 7710 | 1 | 3um_FIJJAFBF_53328 | Bacteria | Bacteroidetes | Flavobacteriia | Flavobacterial | Flavobacteriac | Polaribacter | Polaribacter s | metabolism | Nucleotide tra | 09B0C | Provides the p | 313594.PI23P |
| 7711 | 1 | 3um_FIJJAFBF_217605 | Bacteria | Bacteroidetes | Flavobacteriia | Flavobacterial | Flavobacteriac | Polaribacter | Polaribacter s | metabolism | Amino acid tra | 0P7Z9 | aminopeptida | 313598.MED1 |
| 7712 | 1 | 0.2um_NOKCPHJN_377561 | Eukaryota | Phaeophyceae | unclassified | Fucales | Fucaceae | Pelvetia | Pelvetia fastig | unclassified | unclassified | unclassified | unclassified | unclassified |
| 7713 | 1 | 3um_FIJJAFBF_774342 | Bacteria | Proteobacteri | Gammaprotec | Oceanospirilla | Saccharospirill | Reinekea | Reinekea fors | information st | Translation, ril | 0QSAC | Binds togethe | 314283.MED2 |
| 7714 | 1 | 0.2um_NOKCPHJN_686397 | Bacteria | Proteobacteri | Betaproteoba | Burkholderial | Comamonada | Limnohabitan | Limnohabitan | unclassified | unclassified | unclassified | unclassified | unclassified |
| 7715 | 1 | 0.2um_NOKCPHJN_472432 | Bacteria | Proteobacteri | Alphaproteob | Pelagibacteral | Pelagibacterac | Candidatus Pe | Candidatus Pe | metabolism | Amino acid tra | COG0683 | (ABC) transpo | 439493.PB721 |
| 7716 | 1 | 0.2um_NOKCPHJN_374504 | Bacteria | Proteobacteri | Gammaprotec | Cellvibrionales | Porticoccacea | unclassified | gamma protec | information st | Replication, re | 05XQP | Phage integra | 314287.GB22 |

| | | | | | | | | | | | | | | |
|---|---|---|---|---|---|---|---|---|---|---|---|---|---|---|
| 7574 | 0,4 | 0,489897949 | 3,78E-05 | 4,63E-05 | 0 - | 0 - | 1 - | 9,70E-05 - | 1 - | 9,22E-05 - | 0 - | 0 - | 0 - | 0 - |
| 7575 | 1,6 | 1,019803903 | 0,000221359 | 0,000144803 | 2 - | 0,000239118 - | 2 - | 0,000296914 - | 1 - | 0,000141127 - | 3 - | 0,000429635 - | 0 - | 0 - |
| 7576 | 0,8 | 0,4 | 3,68E-05 | 1,87E-05 | 1 - | 3,99E-05 - | 1 - | 4,95E-05 - | 1 - | 4,70E-05 - | 1 - | 4,77E-05 - | 0 - | 0 - |
| 7577 | 2,8 | 0,4 | 0,000223014 | 3,89E-05 | 3 - | 0,000200192 - | 3 - | 0,000248579 - | 3 - | 0,000236306 - | 2 - | 0,000159864 - | 3 - | 0,00027013 - |
| 7578 | 1 | 0 | 4,48E-05 | 4,25E-06 | 1 - | 3,75E-05 - | 1 - | 4,66E-05 - | 1 - | 4,43E-05 - | 1 - | 4,49E-05 - | 1 - | 5,06E-05 - |
| 7579 | 1 | 0 | 6,59E-05 | 6,26E-06 | 1 - | 5,52E-05 - | 1 - | 6,85E-05 - | 1 - | 6,51E-05 - | 1 - | 6,61E-05 - | 1 - | 7,45E-05 - |
| 7580 | 2,2 | 0,4 | 0,000240921 | 4,70E-05 | 2 - | 0,000183643 - | 2 - | 0,00022803 - | 3 - | 0,000325157 - | 2 - | 0,000219973 - | 2 - | 0,000247799 - |
| 7581 | 0,8 | 0,4 | 3,71E-05 | 1,88E-05 | 1 - | 4,01E-05 - | 1 - | 4,98E-05 - | 1 - | 4,74E-05 - | 1 - | 4,81E-05 - | 0 - | 0 - |
| 7582 | 0,2 | 0,4 | 5,71E-06 | 1,14E-05 | 1 - | 2,86E-05 - | 0 - | 0 - | 0 - | 0 - | 0 - | 0 - | 0 - | 0 - |
| 7583 | 6,6 | 1,019803903 | 0,000225745 | 1,75E-05 | 8 - | 0,000232459 - | 6 - | 0,000216484 - | 7 - | 0,000240095 - | 7 - | 0,000243641 - | 5 - | 0,000196044 - |
| 7584 | 3,4 | 0,8 | 9,43E-05 | 2,28E-05 | 3 - | 7,01E-05 - | 4 - | 0,000116105 - | 4 - | 0,000110372 - | 4 - | 0,000112003 - | 2 - | 6,31E-05 - |
| 7585 | 3,6 | 1,2 | 0,00028393 | 7,92E-05 | 5 - | 0,000337579 - | 2 - | 0,000167669 - | 5 - | 0,000398477 - | 3 - | 0,000242618 - | 3 - | 0,000273308 - |
| 7586 | 3,8 | 1,720465053 | 0,000139319 | 6,40E-05 | 3 - | 9,23E-05 - | 2 - | 7,64E-05 - | 4 - | 0,000145289 - | 7 - | 0,000258012 - | 3 - | 0,000124564 - |
| 7587 | 0,6 | 0,489897949 | 1,22E-05 | 1,00E-05 | 1 - | 1,81E-05 - | 0 - | 0 - | 1 - | 2,13E-05 - | 1 - | 2,17E-05 - | 0 - | 0 - |
| 7588 | 1,8 | 0,748331477 | 9,38E-05 | 3,33E-05 | 3 - | 0,000133981 - | 1 - | 5,55E-05 - | 2 - | 0,000105434 - | 1 - | 5,35E-05 - | 2 - | 0,000120525 - |
| 7589 | 1,4 | 0,489897949 | 8,97E-05 | 2,64E-05 | 2 - | 0,000109834 - | 1 - | 6,82E-05 - | 1 - | 6,48E-05 - | 2 - | 0,000131563 - | 1 - | 7,41E-05 - |
| 7590 | 1 | 0,894427191 | 2,26E-05 | 1,98E-05 | 0 - | 0 - | 2 - | 4,51E-05 - | 0 - | 0 - | 2 - | 4,35E-05 - | 1 - | 2,45E-05 - |
| 7591 | 0,4 | 0,489897949 | 6,37E-06 | 7,81E-06 | 0 - | 0 - | 1 - | 1,53E-05 - | 0 - | 0 - | 0 - | 0 - | 1 - | 1,66E-05 - |
| 7592 | 0,2 | 0,4 | 1,05E-05 | 2,11E-05 | 0 - | 0 - | 0 - | 0 - | 1 - | 5,27E-05 - | 0 - | 0 - | 0 - | 0 - |
| 7593 | 0,2 | 0,4 | 9,34E-06 | 1,87E-05 | 0 - | 0 - | 0 - | 0 - | 1 - | 4,67E-05 - | 0 - | 0 - | 0 - | 0 - |
| 7594 | 4 | 0,632455532 | 6,03E-05 | 1,08E-05 | 4 - | 5,05E-05 - | 4 - | 6,27E-05 - | 5 - | 7,45E-05 - | 3 - | 4,54E-05 - | 4 - | 6,82E-05 - |
| 7595 | 4,2 | 0,4 | 0,000445895 | 5,65E-05 | 4 - | 0,000355897 - | 4 - | 0,000441919 - | 5 - | 0,000525125 - | 4 - | 0,000426305 - | 4 - | 0,000480231 - |
| 7596 | 9,2 | 1,720465053 | 5,01E-05 | 5,64E-06 | 12 - | 5,56E-05 - | 10 - | 5,76E-05 - | 9 - | 4,93E-05 - | 8 - | 4,44E-05 - | 7 - | 4,38E-05 - |
| 7597 | 6,4 | 1,2 | 0,000280819 | 4,51E-05 | 7 - | 0,000260012 - | 5 - | 0,000230613 - | 7 - | 0,000306918 - | 8 - | 0,000355944 - | 5 - | 0,000250606 - |
| 7598 | 2 | 0,894427191 | 5,70E-05 | 2,37E-05 | 3 - | 7,34E-05 - | 3 - | 9,12E-05 - | 1 - | 2,89E-05 - | 2 - | 5,86E-05 - | 1 - | 3,30E-05 - |
| 7599 | 2,8 | 1,166190379 | 8,74E-05 | 3,39E-05 | 4 - | 0,000107519 - | 4 - | 0,000133507 - | 3 - | 9,52E-05 - | 2 - | 6,44E-05 - | 1 - | 3,63E-05 - |
| 7600 | 4 | 1,095445115 | 0,000302944 | 6,87E-05 | 5 - | 0,000324228 - | 4 - | 0,000322077 - | 5 - | 0,000382718 - | 4 - | 0,000310697 - | 2 - | 0,000174999 - |
| 7601 | 1,2 | 0,4 | 0,000225327 | 4,99E-05 | 2 - | 0,000323315 - | 1 - | 0,000200731 - | 1 - | 0,00019082 - | 1 - | 0,000193638 - | 1 - | 0,000218133 - |
| 7602 | 0,2 | 0,4 | 2,03E-05 | 4,06E-05 | 1 - | 0,000101572 - | 0 - | 0 - | 0 - | 0 - | 0 - | 0 - | 0 - | 0 - |
| 7603 | 0,6 | 0,489897949 | 3,46E-05 | 2,85E-05 | 1 - | 5,06E-05 - | 1 - | 6,28E-05 - | 1 - | 5,97E-05 - | 0 - | 0 - | 0 - | 0 - |
| 7604 | 0,4 | 0,489897949 | 2,28E-05 | 2,82E-05 | 1 - | 5,08E-05 - | 1 - | 6,31E-05 - | 0 - | 0 - | 0 - | 0 - | 0 - | 0 - |
| 7605 | 8,4 | 1,356465997 | 0,000672133 | 0,000122919 | 9 - | 0,000604088 - | 8 - | 0,000666755 - | 10 - | 0,000792294 - | 6 - | 0,000482398 - | 9 - | 0,000815129 - |
| 7606 | 0,2 | 0,4 | 8,55E-06 | 1,71E-05 | 0 - | 0 - | 0 - | 0 - | 1 - | 4,27E-05 - | 0 - | 0 - | 0 - | 0 - |
| 7607 | 5,8 | 0,748331477 | 0,000467607 | 8,76E-05 | 5 - | 0,000335605 - | 6 - | 0,000500066 - | 5 - | 0,000396147 - | 7 - | 0,000562797 - | 6 - | 0,000543419 - |
| 7608 | 0,2 | 0,4 | 3,97E-06 | 7,93E-06 | 0 - | 0 - | 0 - | 0 - | 0 - | 0 - | 0 - | 0 - | 1 - | 1,98E-05 - |
| 7609 | 1,4 | 0,8 | 8,78E-05 | 5,09E-05 | 2 - | 0,0001053 - | 1 - | 6,54E-05 - | 0 - | 0 - | 2 - | 0,000126131 - | 2 - | 0,000142087 - |
| 7610 | 0,4 | 0,489897949 | 1,10E-05 | 1,35E-05 | 1 - | 2,52E-05 - | 0 - | 0 - | 1 - | 2,97E-05 - | 0 - | 0 - | 0 - | 0 - |
| 7611 | 4 | 0,632455532 | 0,000148785 | 1,09E-05 | 5 - | 0,000158095 - | 4 - | 0,000157046 - | 4 - | 0,000149292 - | 4 - | 0,000151497 - | 3 - | 0,000127995 - |
| 7612 | 1 | 0,632455532 | 3,65E-05 | 2,41E-05 | 1 - | 3,04E-05 - | 2 - | 7,54E-05 - | 1 - | 3,58E-05 - | 0 - | 0 - | 1 - | 4,10E-05 - |
| 7613 | 1 | 0 | 2,26E-05 | 2,15E-06 | 1 - | 1,89E-05 - | 1 - | 2,35E-05 - | 1 - | 2,24E-05 - | 1 - | 2,27E-05 - | 1 - | 2,56E-05 - |
| 7614 | 2 | 1,095445115 | 7,20E-05 | 3,98E-05 | 1 - | 3,00E-05 - | 2 - | 7,46E-05 - | 2 - | 7,09E-05 - | 4 - | 0,000143962 - | 1 - | 4,05E-05 - |
| 7615 | 12,4 | 1,496662955 | 0,000241408 | 1,44E-05 | 15 - | 0,000247009 - | 12 - | 0,00024537 - | 11 - | 0,000213817 - | 13 - | 0,000256425 - | 11 - | 0,000244422 - |
| 7616 | 0,2 | 0,4 | 4,01E-06 | 8,01E-06 | 0 - | 0 - | 0 - | 0 - | 0 - | 0 - | 0 - | 0 - | 1 - | 2,00E-05 - |
| 7617 | 1 | 0 | 3,97E-05 | 3,77E-06 | 1 - | 3,33E-05 - | 1 - | 4,13E-05 - | 1 - | 3,93E-05 - | 1 - | 3,99E-05 - | 1 - | 4,49E-05 - |
| 7618 | 0,8 | 0,4 | 2,87E-05 | 1,47E-05 | 1 - | 3,00E-05 - | 1 - | 3,73E-05 - | 1 - | 3,55E-05 - | 0 - | 0 - | 1 - | 4,05E-05 - |
| 7619 | 2 | 0 | 0,000326255 | 3,10E-05 | 2 - | 0,000273278 - | 2 - | 0,000339331 - | 2 - | 0,000322577 - | 2 - | 0,000327341 - | 2 - | 0,000368749 - |
| 7620 | 0,2 | 0,4 | 1,12E-05 | 2,25E-05 | 0 - | 0 - | 0 - | 0 - | 1 - | 5,62E-05 - | 0 - | 0 - | 0 - | 0 - |
| 7621 | 1,6 | 0,489897949 | 9,98E-05 | 2,77E-05 | 2 - | 0,000106275 - | 2 - | 0,000131962 - | 2 - | 0,000125447 - | 1 - | 6,36E-05 - | 1 - | 7,17E-05 - |
| 7622 | 0,2 | 0,4 | 7,00E-06 | 1,40E-05 | 1 - | 3,50E-05 - | 0 - | 0 - | 0 - | 0 - | 0 - | 0 - | 0 - | 0 - |
| 7623 | 0,8 | 0,4 | 6,96E-05 | 3,57E-05 | 1 - | 7,26E-05 - | 1 - | 9,02E-05 - | 0 - | 0 - | 1 - | 8,70E-05 - | 1 - | 9,80E-05 - |
| 7624 | 11,4 | 1,356465997 | 0,000199177 | 2,82E-05 | 12 - | 0,000175903 - | 12 - | 0,00021842 - | 9 - | 0,000155727 - | 13 - | 0,000228261 - | 11 - | 0,000217576 - |
| 7625 | 1 | 0,632455532 | 2,08E-05 | 1,40E-05 | 1 - | 1,77E-05 - | 2 - | 4,41E-05 - | 1 - | 2,09E-05 - | 1 - | 2,12E-05 - | 0 - | 0 - |
| 7626 | 0,4 | 0,489897949 | 4,17E-05 | 5,10E-05 | 0 - | 0 - | 0 - | 0 - | 1 - | 0,000103422 - | 1 - | 0,000104949 - | 0 - | 0 - |
| 7627 | 0,2 | 0,4 | 5,17E-06 | 1,03E-05 | 1 - | 2,59E-05 - | 0 - | 0 - | 0 - | 0 - | 0 - | 0 - | 0 - | 0 - |
| 7628 | 0,8 | 0,4 | 2,60E-05 | 1,33E-05 | 1 - | 2,72E-05 - | 1 - | 3,38E-05 - | 1 - | 3,21E-05 - | 0 - | 0 - | 1 - | 3,67E-05 - |
| 7629 | 0,8 | 0,748331477 | 5,05E-05 | 4,39E-05 | 2 - | 0,000111434 - | 0 - | 0 - | 1 - | 6,58E-05 - | 0 - | 0 - | 1 - | 7,52E-05 - |
| 7630 | 4,8 | 0,748331477 | 9,64E-05 | 8,68E-06 | 6 - | 0,000102175 - | 5 - | 0,000105726 - | 5 - | 0,000100506 - | 4 - | 8,16E-05 - | 4 - | 9,19E-05 - |
| 7631 | 0,2 | 0,4 | 8,54E-07 | 1,71E-06 | 0 - | 0 - | 0 - | 0 - | 0 - | 0 - | 0 - | 0 - | 1 - | 4,27E-06 - |
| 7632 | 3 | 0 | 8,42E-05 | 8,00E-06 | 3 - | 7,06E-05 - | 3 - | 8,76E-05 - | 3 - | 8,33E-05 - | 3 - | 8,45E-05 - | 3 - | 9,52E-05 - |
| 7633 | 0,2 | 0,4 | 2,76E-05 | 5,53E-05 | 0 - | 0 - | 0 - | 0 - | 1 - | 0,000138247 - | 0 - | 0 - | 0 - | 0 - |
| 7634 | 0,2 | 0,4 | 5,98E-06 | 1,20E-05 | 1 - | 2,99E-05 - | 0 - | 0 - | 0 - | 0 - | 0 - | 0 - | 0 - | 0 - |
| 7635 | 0,8 | 0,4 | 2,75E-05 | 1,38E-05 | 0 - | 0 - | 1 - | 3,43E-05 - | 1 - | 3,26E-05 - | 1 - | 3,31E-05 - | 1 - | 3,73E-05 - |
| 7636 | 0,2 | 0,4 | 8,79E-07 | 1,76E-06 | 1 - | 4,39E-06 - | 0 - | 0 - | 0 - | 0 - | 0 - | 0 - | 0 - | 0 - |
| 7637 | 0,8 | 0,4 | 6,20E-05 | 3,15E-05 | 1 - | 6,71E-05 - | 1 - | 8,33E-05 - | 1 - | 7,92E-05 - | 1 - | 8,04E-05 - | 0 - | 0 - |
| 7638 | 1,4 | 0,489897949 | 5,72E-05 | 2,02E-05 | 1 - | 3,43E-05 - | 1 - | 4,25E-05 - | 2 - | 8,09E-05 - | 2 - | 8,21E-05 - | 1 - | 4,62E-05 - |
| 7639 | 0,2 | 0,4 | 7,81E-06 | 1,56E-05 | 0 - | 0 - | 0 - | 0 - | 1 - | 3,90E-05 - | 0 - | 0 - | 0 - | 0 - |
| 7640 | 0,8 | 0,4 | 3,21E-05 | 1,65E-05 | 1 - | 3,40E-05 - | 0 - | 0 - | 1 - | 4,01E-05 - | 1 - | 4,07E-05 - | 1 - | 4,58E-05 - |
| 7641 | 1 | 0 | 7,03E-05 | 6,67E-06 | 1 - | 5,89E-05 - | 1 - | 7,31E-05 - | 1 - | 6,95E-05 - | 1 - | 7,05E-05 - | 1 - | 7,94E-05 - |
| 7642 | 8,6 | 1,019803903 | 0,000139909 | 2,04E-05 | 9 - | 0,000122829 - | 7 - | 0,000118625 - | 8 - | 0,000128877 - | 10 - | 0,000163476 - | 9 - | 0,00016574 - |
| 7643 | 0,2 | 0,4 | 2,19E-06 | 4,39E-06 | 0 - | 0 - | 1 - | 1,10E-05 - | 0 - | 0 - | 0 - | 0 - | 0 - | 0 - |
| 7644 | 1 | 0 | 8,21E-05 | 7,79E-06 | 1 - | 6,87E-05 - | 1 - | 8,53E-05 - | 1 - | 8,11E-05 - | 1 - | 8,23E-05 - | 1 - | 9,27E-05 - |
| 7645 | 0,8 | 0,4 | 3,11E-05 | 1,60E-05 | 1 - | 3,25E-05 - | 1 - | 4,04E-05 - | 0 - | 0 - | 1 - | 3,89E-05 - | 1 - | 4,39E-05 - |
| 7646 | 3,6 | 0,8 | 9,09E-05 | 2,17E-05 | 3 - | 6,34E-05 - | 4 - | 0,000104986 - | 5 - | 0,000124753 - | 3 - | 7,60E-05 - | 3 - | 8,56E-05 - |
| 7647 | 0,4 | 0,489897949 | 5,78E-05 | 7,15E-05 | 1 - | 0,000128963 - | 1 - | 0,000160134 - | 0 - | 0 - | 0 - | 0 - | 0 - | 0 - |
| 7648 | 2,2 | 0,4 | 8,14E-05 | 1,59E-05 | 2 - | 6,20E-05 - | 2 - | 7,70E-05 - | 3 - | 0,00010985 - | 2 - | 7,43E-05 - | 2 - | 8,37E-05 - |
| 7649 | 1,8 | 0,4 | 0,00017004 | 4,17E-05 | 2 - | 0,000158313 - | 2 - | 0,000196578 - | 2 - | 0,000186872 - | 1 - | 9,48E-05 - | 2 - | 0,00021362 - |
| 7650 | 1 | 0 | 3,22E-05 | 3,06E-06 | 1 - | 2,70E-05 - | 1 - | 3,35E-05 - | 1 - | 3,19E-05 - | 1 - | 3,23E-05 - | 1 - | 3,64E-05 - |
| 7651 | 3,2 | 0,748331477 | 0,000319439 | 6,71E-05 | 4 - | 0,000337579 - | 3 - | 0,00031438 - | 2 - | 0,000199239 - | 4 - | 0,000404363 - | 3 - | 0,000341635 - |
| 7652 | 3,8 | 0,4 | 0,000165274 | 2,38E-05 | 4 - | 0,000145748 - | 4 - | 0,000180976 - | 4 - | 0,000172041 - | 3 - | 0,000130936 - | 4 - | 0,000196666 - |
| 7653 | 0,2 | 0,4 | 2,41E-06 | 4,82E-06 | 0 - | 0 - | 0 - | 0 - | 0 - | 0 - | 1 - | 1,20E-05 - | 0 - | 0 - |
| 7654 | 3,2 | 0,748331477 | 0,000154131 | 3,02E-05 | 4 - | 0,000163967 - | 4 - | 0,000203598 - | 3 - | 0,00014516 - | 3 - | 0,000147304 - | 2 - | 0,000110625 - |
| 7655 | 1,8 | 0,4 | 5,96E-05 | 1,65E-05 | 1 - | 2,73E-05 - | 2 - | 6,77E-05 - | 2 - | 6,44E-05 - | 2 - | 6,53E-05 - | 2 - | 7,36E-05 - |
| 7656 | 3,6 | 1,019803903 | 9,68E-05 | 2,24E-05 | 5 - | 0,000114548 - | 4 - | 0,000113788 - | 4 - | 0,000108169 - | 2 - | 5,49E-05 - | 3 - | 9,27E-05 - |
| 7657 | 2,2 | 0,4 | 6,08E-05 | 1,23E-05 | 2 - | 4,63E-05 - | 2 - | 5,75E-05 - | 2 - | 5,46E-05 - | 3 - | 8,32E-05 - | 2 - | 6,24E-05 - |
| 7658 | 0,8 | 0,4 | 9,51E-06 | 4,89E-06 | 1 - | 9,93E-06 - | 1 - | 1,23E-05 - | 0 - | 0 - | 1 - | 1,19E-05 - | 1 - | 1,34E-05 - |
| 7659 | 7,4 | 2,059126028 | 0,000232442 | 6,21E-05 | 8 - | 0,00021255 - | 5 - | 0,000164952 - | 7 - | 0,000219531 - | 11 - | 0,000350073 - | 6 - | 0,000215103 - |
| 7660 | 2,6 | 0,8 | 0,000247611 | 7,18E-05 | 3 - | 0,000244206 - | 3 - | 0,000303232 - | 3 - | 0,00028826 - | 3 - | 0,000292518 - | 1 - | 0,00010984 - |
| 7661 | 2,4 | 1,019803903 | 3,45E-05 | 1,47E-05 | 1 - | 1,19E-05 - | 2 - | 2,96E-05 - | 4 - | 5,62E-05 - | 3 - | 4,28E-05 - | 2 - | 3,21E-05 - |
| 7662 | 16,6 | 0,489897949 | 0,000358962 | 4,21E-05 | 16 - | 0,000289201 - | 17 - | 0,000381547 - | 16 - | 0,000341373 - | 17 - | 0,000368066 - | 17 - | 0,000414625 - |
| 7663 | 0,2 | 0,4 | 1,10E-05 | 2,19E-05 | 0 - | 0 - | 1 - | 5,48E-05 - | 0 - | 0 - | 0 - | 0 - | 0 - | 0 - |
| 7664 | 0,6 | 0,489897949 | 2,28E-05 | 1,87E-05 | 0 - | 0 - | 1 - | 3,76E-05 - | 1 - | 3,57E-05 - | 0 - | 0 - | 1 - | 4,09E-05 - |
| 7665 | 0,8 | 0,4 | 3,03E-05 | 1,54E-05 | 1 - | 3,28E-05 - | 1 - | 4,07E-05 - | 1 - | 3,87E-05 - | 1 - | 3,93E-05 - | 0 - | 0 - |
| 7666 | 0,4 | 0,489897949 | 1,45E-05 | 1,80E-05 | 1 - | 3,08E-05 - | 0 - | 0 - | 0 - | 0 - | 0 - | 0 - | 1 - | 4,15E-05 - |
| 7667 | 0,2 | 0,4 | 2,72E-06 | 5,45E-06 | 1 - | 1,36E-05 - | 0 - | 0 - | 0 - | 0 - | 0 - | 0 - | 0 - | 0 - |
| 7668 | 3,2 | 0,4 | 0,000429739 | 2,36E-05 | 4 - | 0,000454561 - | 3 - | 0,000423323 - | 3 - | 0,000402422 - | 3 - | 0,000408366 - | 3 - | 0,000460023 - |
| 7669 | 4,4 | 0,8 | 7,84E-05 | 1,48E-05 | 4 - | 5,99E-05 - | 5 - | 9,30E-05 - | 5 - | 8,84E-05 - | 5 - | 8,97E-05 - | 3 - | 6,07E-05 - |
| 7670 | 2,8 | 0,748331477 | 0,000387187 | 8,97E-05 | 4 - | 0,000468477 - | 3 - | 0,000436282 - | 2 - | 0,000276494 - | 2 - | 0,000280578 - | 3 - | 0,000474106 - |
| 7671 | 2 | 0 | 0,000158413 | 1,50E-05 | 2 - | 0,00013269 - | 2 - | 0,000164762 - | 2 - | 0,000156627 - | 2 - | 0,00015894 - | 2 - | 0,000179046 - |
| 7672 | 1,6 | 0,489897949 | 0,000234897 | 7,50E-05 | 2 - | 0,000246832 - | 1 - | 0,000153246 - | 1 - | 0,00014568 - | 2 - | 0,000295663 - | 2 - | 0,000333063 - |
| 7673 | 0,8 | 0,4 | 2,39E-05 | 1,23E-05 | 1 - | 2,51E-05 - | 1 - | 3,11E-05 - | 1 - | 2,96E-05 - | 0 - | 0 - | 1 - | 3,38E-05 - |
| 7674 | 0,4 | 0,489897949 | 1,28E-05 | 1,56E-05 | 0 - | 0 - | 0 - | 0 - | 1 - | 3,17E-05 - | 1 - | 3,21E-05 - | 0 - | 0 - |
| 7675 | 3 | 0,632455532 | 0,000107699 | 2,67E-05 | 3 - | 8,99E-05 - | 4 - | 0,000148845 - | 2 - | 7,07E-05 - | 3 - | 0,000107689 - | 3 - | 0,000121312 - |
| 7676 | 3 | 0,632455532 | 0,000343672 | 5,48E-05 | 4 - | 0,000389074 - | 2 - | 0,000241557 - | 3 - | 0,000344446 - | 3 - | 0,000349534 - | 3 - | 0,000393749 - |
| 7677 | 1 | 0 | 6,75E-05 | 6,41E-06 | 1 - | 5,65E-05 - | 1 - | 7,02E-05 - | 1 - | 6,67E-05 - | 1 - | 6,77E-05 - | 1 - | 7,63E-05 - |
| 7678 | 2 | 0,894427191 | 2,85E-05 | 1,18E-05 | 3 - | 3,67E-05 - | 3 - | 4,56E-05 - | 2 - | 2,89E-05 - | 1 - | 1,47E-05 - | 1 - | 1,65E-05 - |
| 7679 | 1 | 0,632455532 | 3,92E-05 | 2,57E-05 | 1 - | 3,37E-05 - | 1 - | 4,18E-05 - | 1 - | 3,97E-05 - | 2 - | 8,06E-05 - | 0 - | 0 - |
| 7680 | 0,2 | 0,4 | 1,09E-05 | 2,18E-05 | 1 - | 5,44E-05 - | 0 - | 0 - | 0 - | 0 - | 0 - | 0 - | 0 - | 0 - |
| 7681 | 0,2 | 0,4 | 2,27E-05 | 4,55E-05 | 1 - | 0,00011364 - | 0 - | 0 - | 0 - | 0 - | 0 - | 0 - | 0 - | 0 - |
| 7682 | 1 | 0 | 4,38E-05 | 4,16E-06 | 1 - | 3,67E-05 - | 1 - | 4,55E-05 - | 1 - | 4,33E-05 - | 1 - | 4,39E-05 - | 1 - | 4,95E-05 - |
| 7683 | 2 | 0 | 0,000194365 | 1,85E-05 | 2 - | 0,000162804 - | 2 - | 0,000202154 - | 2 - | 0,000192173 - | 2 - | 0,000195012 - | 2 - | 0,00021968 - |
| 7684 | 0,2 | 0,4 | 1,37E-05 | 2,73E-05 | 1 - | 6,83E-05 - | 0 - | 0 - | 0 - | 0 - | 0 - | 0 - | 0 - | 0 - |
| 7685 | 1 | 0 | 0,000228379 | 2,17E-05 | 1 - | 0,000191295 - | 1 - | 0,000237531 - | 1 - | 0,000225804 - | 1 - | 0,000229139 - | 1 - | 0,000258124 - |
| 7686 | 0,6 | 0,489897949 | 3,23E-05 | 2,66E-05 | 1 - | 4,78E-05 - | 0 - | 0 - | 1 - | 5,65E-05 - | 1 - | 5,73E-05 - | 0 - | 0 - |
| 7687 | 1 | 0 | 0,000150579 | 1,43E-05 | 1 - | 0,000126128 - | 1 - | 0,000156614 - | 1 - | 0,000148882 - | 1 - | 0,000151081 - | 1 - | 0,000170192 - |
| 7688 | 1,2 | 0,979795897 | 0,000168233 | 0,000139895 | 2 - | 0,000234238 - | 2 - | 0,000290855 - | 0 - | 0 - | 0 - | 0 - | 2 - | 0,00031607 - |
| 7689 | 0,8 | 0,4 | 0,000220942 | 0,000112218 | 1 - | 0,000239118 - | 1 - | 0,000296914 - | 1 - | 0,000282255 - | 1 - | 0,000286424 - | 0 - | 0 - |
| 7690 | 0,6 | 0,8 | 0,000188166 | 0,000260042 | 1 - | 0,000279943 - | 0 - | 0 - | 2 - | 0,000660889 - | 0 - | 0 - | 0 - | 0 - |
| 7691 | 3,8 | 0,748331477 | 0,000438113 | 5,44E-05 | 5 - | 0,000490499 - | 4 - | 0,000487244 - | 4 - | 0,000463187 - | 3 - | 0,000352521 - | 3 - | 0,000397114 - |
| 7692 | 2,4 | 0,489897949 | 0,000145669 | 3,88E-05 | 2 - | 0,000100681 - | 2 - | 0,000125017 - | 3 - | 0,000178266 - | 2 - | 0,000120599 - | 3 - | 0,000203782 - |
| 7693 | 0,2 | 0,4 | 6,74E-06 | 1,35E-05 | 0 - | 0 - | 0 - | 0 - | 1 - | 3,37E-05 - | 0 - | 0 - | 0 - | 0 - |
| 7694 | 0,4 | 0,489897949 | 7,94E-05 | 9,73E-05 | 0 - | 0 - | 1 - | 0,000203598 - | 1 - | 0,000193546 - | 0 - | 0 - | 0 - | 0 - |
| 7695 | 0,4 | 0,8 | 5,72E-06 | 1,14E-05 | 2 - | 2,86E-05 - | 0 - | 0 - | 0 - | 0 - | 0 - | 0 - | 0 - | 0 - |
| 7696 | 0,2 | 0,4 | 2,54E-05 | 5,08E-05 | 0 - | 0 - | 0 - | 0 - | 0 - | 0 - | 0 - | 0 - | 1 - | 0,000126946 - |
| 7697 | 0,8 | 0,748331477 | 3,09E-05 | 3,00E-05 | 1 - | 3,38E-05 - | 0 - | 0 - | 1 - | 3,98E-05 - | 2 - | 8,09E-05 - | 0 - | 0 - |
| 7698 | 1,6 | 0,489897949 | 6,94E-05 | 2,19E-05 | 2 - | 7,31E-05 - | 1 - | 4,54E-05 - | 2 - | 8,63E-05 - | 1 - | 4,38E-05 - | 2 - | 9,86E-05 - |
| 7699 | 0,6 | 0,8 | 1,63E-05 | 2,10E-05 | 2 - | 5,12E-05 - | 0 - | 0 - | 1 - | 3,02E-05 - | 0 - | 0 - | 0 - | 0 - |
| 7700 | 7,4 | 1,019803903 | 0,000172753 | 1,49E-05 | 9 - | 0,000177795 - | 6 - | 0,00014718 - | 8 - | 0,00018655 - | 7 - | 0,000165643 - | 7 - | 0,000186596 - |
| 7701 | 1 | 0,632455532 | 3,62E-05 | 2,38E-05 | 1 - | 3,00E-05 - | 2 - | 7,46E-05 - | 0 - | 0 - | 1 - | 3,60E-05 - | 1 - | 4,05E-05 - |
| 7702 | 1,4 | 0,489897949 | 2,77E-05 | 8,14E-06 | 2 - | 3,39E-05 - | 1 - | 2,10E-05 - | 1 - | 2,00E-05 - | 2 - | 4,06E-05 - | 1 - | 2,28E-05 - |
| 7703 | 0,4 | 0,489897949 | 6,56E-06 | 8,04E-06 | 0 - | 0 - | 1 - | 1,68E-05 - | 1 - | 1,60E-05 - | 0 - | 0 - | 0 - | 0 - |
| 7704 | 4,6 | 0,8 | 0,000217138 | 3,22E-05 | 5 - | 0,00019996 - | 5 - | 0,000248291 - | 5 - | 0,000236032 - | 5 - | 0,000239518 - | 3 - | 0,00016189 - |
| 7705 | 0,2 | 0,4 | 1,26E-05 | 2,52E-05 | 0 - | 0 - | 1 - | 6,31E-05 - | 0 - | 0 - | 0 - | 0 - | 0 - | 0 - |
| 7706 | 0,8 | 0,4 | 2,13E-05 | 1,08E-05 | 1 - | 2,30E-05 - | 1 - | 2,86E-05 - | 1 - | 2,72E-05 - | 1 - | 2,76E-05 - | 0 - | 0 - |
| 7707 | 3,4 | 0,8 | 7,96E-05 | 1,65E-05 | 4 - | 7,94E-05 - | 2 - | 4,93E-05 - | 4 - | 9,38E-05 - | 4 - | 9,51E-05 - | 3 - | 8,04E-05 - |
| 7708 | 2,2 | 1,166190379 | 3,69E-05 | 1,96E-05 | 3 - | 4,30E-05 - | 4 - | 7,12E-05 - | 2 - | 3,38E-05 - | 1 - | 1,72E-05 - | 1 - | 1,93E-05 - |
| 7709 | 3,4 | 0,8 | 9,74E-05 | 2,90E-05 | 2 - | 4,70E-05 - | 4 - | 0,000116819 - | 3 - | 8,33E-05 - | 4 - | 0,000112691 - | 4 - | 0,000126946 - |
| 7710 | 6,4 | 0,8 | 0,000146916 | 1,47E-05 | 7 - | 0,000135716 - | 7 - | 0,000168519 - | 7 - | 0,000160199 - | 6 - | 0,000139341 - | 5 - | 0,000130806 - |
| 7711 | 1,2 | 0,748331477 | 2,19E-05 | 1,38E-05 | 2 - | 3,07E-05 - | 1 - | 1,91E-05 - | 0 - | 0 - | 1 - | 1,84E-05 - | 2 - | 4,14E-05 - |
| 7712 | 2 | 0 | 6,05E-05 | 5,74E-06 | 2 - | 5,07E-05 - | 2 - | 6,29E-05 - | 2 - | 5,98E-05 - | 2 - | 6,07E-05 - | 2 - | 6,84E-05 - |
| 7713 | 5,6 | 0,8 | 0,000546128 | 0,000111677 | 6 - | 0,000488412 - | 5 - | 0,000505386 - | 5 - | 0,000480433 - | 5 - | 0,000487529 - | 7 - | 0,00076888 - |
| 7714 | 1 | 0 | 1,32E-05 | 1,26E-06 | 1 - | 1,11E-05 - | 1 - | 1,38E-05 - | 1 - | 1,31E-05 - | 1 - | 1,33E-05 - | 1 - | 1,50E-05 - |
| 7715 | 0,4 | 0,489897949 | 1,41E-05 | 1,72E-05 | 0 - | 0 - | 1 - | 3,61E-05 - | 1 - | 3,43E-05 - | 0 - | 0 - | 0 - | 0 - |
| 7716 | 0,2 | 0,4 | 1,40E-05 | 2,81E-05 | 0 - | 0 - | 0 - | 0 - | 1 - | 7,02E-05 - | 0 - | 0 - | 0 - | 0 - |

| | | | | | | | | | | | | | | | | | | | | | | | | | | | | |
|---|---|---|---|---|---|---|---|---|---|---|---|---|---|---|---|---|---|---|---|---|---|---|---|---|---|---|---|---|
| 7717 | 1 | 3um_FIJJAFBF_450004 | unclassified | unclassified | unclassified | unclassified | unclassified | unclassified | unclassified | unclassified | unclassified | unclassified | unclassified | unclassified | 0,4 | 0,489897949 | 9,21E-06 | 1,13E-05 | 0 - | 0 - | 0 - | 0 - | 1 - | 2,28E-05 - | 1 - | 2,32E-05 - | 0 - | 0 - |
| 7718 | 1 | 0.2um_NOKCPHJN_71525 | Bacteria | Proteobacteri | Alphaproteob | Rhodobactera | Rhodobactera | unclassified | Rhodobactera | unclassified | unclassified | unclassified | unclassified | unclassified | 2,4 | 0,489897949 | 6,13E-05 | 9,55E-06 | 3 - | 6,51E-05 - | 2 - | 5,39E-05 - | 2 - | 5,12E-05 - | 3 - | 7,80E-05 - | 2 - | 5,86E-05 - |
| 7719 | 1 | 0.2um_NOKCPHJN_273996 | unclassified | unclassified | unclassified | unclassified | unclassified | unclassified | unclassified | unclassified | unclassified | unclassified | unclassified | unclassified | 1 | 0 | 1,45E-05 | 1,37E-06 | 1 - | 1,21E-05 - | 1 - | 1,50E-05 - | 1 - | 1,43E-05 - | 1 - | 1,45E-05 - | 1 - | 1,63E-05 - |
| 7720 | 3 | 0.2um_NOKCPHJN_433132;0.2um_NOKCPHJN_8 | Bacteria | Proteobacteri | various | various | various | various | various | metabolism | Energy produc | 05C26 | Dehydrogenas | various | 0,8 | 0,4 | 1,86E-05 | 9,54E-06 | 1 - | 1,97E-05 - | 0 - | 0 - | 1 - | 2,32E-05 - | 1 - | 2,35E-05 - | 1 - | 2,65E-05 - |
| 7721 | 1 | 0.2um_NOKCPHJN_188895 | Bacteria | Proteobacteri | Alphaproteob | Rhodobactera | Rhodobactera | Planktomarina | Planktomarina | poorly charact | Function unkn | 0ZQA4 | motA TolQ Exl | 633131.TR2A6 | 0,2 | 0,4 | 7,05E-06 | 1,41E-05 | 0 - | 0 - | 0 - | 0 - | 0 - | 0 - | 1 - | 3,53E-05 - | 0 - | 0 - |
| 7722 | 1 | 3um_FIJJAFBF_539270 | Bacteria | Bacteroidetes | Flavobacteriia | Flavobacterial | Flavobacteriac | Ulvibacter | Ulvibacter sp. | unclassified | unclassified | unclassified | unclassified | unclassified | 0,2 | 0,4 | 2,91E-05 | 5,81E-05 | 1 - | 0,000145287 - | 0 - | 0 - | 0 - | 0 - | 0 - | 0 - | 0 - | 0 - |
| 7723 | 1 | 3um_FIJJAFBF_70347 | Bacteria | Proteobacteri | Betaproteoba | Burkholderiale | various | various | various | unclassified | unclassified | unclassified | unclassified | unclassified | 2 | 0 | 0,000251426 | 2,39E-05 | 2 - | 0,0002106 - | 2 - | 0,000261503 - | 2 - | 0,000248591 - | 2 - | 0,000252263 - | 2 - | 0,000284173 - |
| 7724 | 3 | 3um_FIJJAFBF_514002;3um_FIJJAFBF_55830;3ur | Bacteria | Bacteroidetes | Flavobacteriia | Flavobacterial | Flavobacteriac | Polaribacter | various | metabolism | Amino acid tra | 05CM2 | brancheD-cha | 313598.MED1 | 1,6 | 0,489897949 | 6,25E-05 | 2,07E-05 | 2 - | 6,54E-05 - | 2 - | 8,12E-05 - | 1 - | 3,86E-05 - | 1 - | 3,92E-05 - | 2 - | 8,82E-05 - |
| 7725 | 1 | 0.2um_NOKCPHJN_415238 | Bacteria | Proteobacteri | Alphaproteob | Rhodobactera | Rhodobactera | Planktomarina | Planktomarina | cellular proces | Cell wall/mem | 01XSD | Pfam:DUF583 | 52598.EE36_1 | 0,4 | 0,489897949 | 3,20E-05 | 3,95E-05 | 1 - | 7,13E-05 - | 1 - | 8,85E-05 - | 0 - | 0 - | 0 - | 0 - | 0 - | 0 - |
| 7726 | 1 | 3um_FIJJAFBF_672133 | Bacteria | Proteobacteri | Gammaprotec | Oceanospirilla | Saccharospirill | Reinekea | Reinekea blan | information st | Translation, ril | 0QHX6 | glycyl-tRNA sy | 314283.MED2 | 0,8 | 0,748331477 | 3,17E-05 | 2,79E-05 | 2 - | 7,15E-05 - | 1 - | 4,44E-05 - | 0 - | 0 - | 1 - | 4,28E-05 - | 0 - | 0 - |
| 7727 | 1 | 0.2um_NOKCPHJN_150777 | Bacteria | Bacteroidetes | Flavobacteriia | Flavobacterial | Crocinitomica | unclassified | Crocinitomica | poorly charact | Function unkn | 0P8NI | Inherit from b | 886377.Murru | 2,6 | 0,489897949 | 7,09E-05 | 1,03E-05 | 3 - | 6,94E-05 - | 2 - | 5,75E-05 - | 3 - | 8,19E-05 - | 3 - | 8,32E-05 - | 2 - | 6,24E-05 - |
| 7728 | 1 | 3um_FIJJAFBF_640408 | unclassified | unclassified | unclassified | unclassified | unclassified | unclassified | unclassified | unclassified | unclassified | unclassified | unclassified | unclassified | 0,4 | 0,489897949 | 2,42E-05 | 2,99E-05 | 1 - | 5,39E-05 - | 1 - | 6,69E-05 - | 0 - | 0 - | 0 - | 0 - | 0 - | 0 - |
| 7729 | 1 | 0.2um_NOKCPHJN_39806 | Bacteria | Bacteroidetes | Flavobacteriia | Flavobacterial | Flavobacteriac | Polaribacter | various | unclassified | unclassified | unclassified | unclassified | unclassified | 0,6 | 0,489897949 | 5,50E-05 | 4,50E-05 | 0 - | 0 - | 1 - | 9,44E-05 - | 1 - | 8,97E-05 - | 1 - | 9,10E-05 - | 0 - | 0 - |
| 7730 | 1 | 3um_FIJJAFBF_777841 | Bacteria | Bacteroidetes | Flavobacteriia | Flavobacterial | Flavobacteriac | Tenacibaculur | Tenacibaculur | unclassified | unclassified | unclassified | unclassified | unclassified | 1,6 | 0,489897949 | 7,40E-05 | 1,94E-05 | 2 - | 7,92E-05 - | 1 - | 4,91E-05 - | 2 - | 9,34E-05 - | 2 - | 9,48E-05 - | 1 - | 5,34E-05 - |
| 7731 | 1 | 0.2um_NOKCPHJN_61948 | Bacteria | Bacteroidetes | Flavobacteriia | Flavobacterial | Flavobacteriac | Kordia | Kordia algicida | unclassified | unclassified | unclassified | unclassified | unclassified | 0,4 | 0,8 | 3,97E-06 | 7,94E-06 | 2 - | 1,98E-05 - | 0 - | 0 - | 0 - | 0 - | 0 - | 0 - | 0 - | 0 - |
| 7732 | 4 | 0.2um_NOKCPHJN_389532;0.2um_NOKCPHJN_4 | Bacteria | Proteobacteri | Alphaproteob | Pelagibacteral | Pelagibacterac | Candidatus Pe | Candidatus Pe | unclassified | unclassified | unclassified | unclassified | unclassified | 2,8 | 0,4 | 0,000120669 | 1,44E-05 | 3 - | 0,000109311 - | 3 - | 0,000135732 - | 3 - | 0,000129031 - | 3 - | 0,000130936 - | 2 - | 9,83E-05 - |
| 7733 | 1 | 3um_FIJJAFBF_217630 | Bacteria | Bacteroidetes | Flavobacteriia | Flavobacterial | Flavobacteriac | Tenacibaculur | Tenacibaculur | cellular proces | Signal transdu | 09A4D | gtp-binding pr | 880070.Cycma | 1,2 | 0,748331477 | 2,70E-05 | 1,57E-05 | 2 - | 3,89E-05 - | 1 - | 2,42E-05 - | 2 - | 4,59E-05 - | 0 - | 0 - | 1 - | 2,62E-05 - |
| 7734 | 1 | 0.2um_NOKCPHJN_706271 | Bacteria | Proteobacteri | Gammaprotec | Alteromonada | Shewanellacea | Shewanella | Shewanella xia | poorly charact | Function unkn | 17A3F | NA | 311403.Arad_ | 0,4 | 0,489897949 | 1,16E-05 | 1,42E-05 | 0 - | 0 - | 0 - | 0 - | 1 - | 2,87E-05 - | 1 - | 2,91E-05 - | 0 - | 0 - |
| 7735 | 1 | 0.2um_NOKCPHJN_209633 | Bacteria | Actinobacteria | Actinobacteria | Corynebacteri | Mycobacteria | Mycobacteroi | Mycobacteroi | unclassified | unclassified | unclassified | unclassified | unclassified | 0,4 | 0,8 | 1,02E-06 | 2,03E-06 | 0 - | 0 - | 0 - | 0 - | 2 - | 5,08E-06 - | 0 - | 0 - | 0 - | 0 - |
| 7736 | 1 | 3um_FIJJAFBF_398158 | Bacteria | Bacteroidetes | Flavobacteriia | unclassified | unclassified | unclassified | Flavobacteria | unclassified | unclassified | unclassified | unclassified | unclassified | 0,6 | 0,489897949 | 5,27E-05 | 4,35E-05 | 1 - | 7,70E-05 - | 1 - | 9,57E-05 - | 1 - | 9,09E-05 - | 0 - | 0 - | 0 - | 0 - |
| 7737 | 1 | 3um_FIJJAFBF_103096 | Eukaryota | unclassified | unclassified | Isochrysidales | Noelaerhabda | Emiliania | Emiliania huxl | unclassified | unclassified | unclassified | unclassified | unclassified | 2,6 | 0,489897949 | 0,000156022 | 2,54E-05 | 3 - | 0,000152359 - | 3 - | 0,000189184 - | 3 - | 0,000179844 - | 2 - | 0,000121667 - | 2 - | 0,000137057 - |
| 7738 | 1 | 3um_FIJJAFBF_333553 | Bacteria | Proteobacteri | Alphaproteob | Rhodobactera | unclassified | unclassified | Rhodobactera | unclassified | unclassified | unclassified | unclassified | unclassified | 6,2 | 1,326649916 | 0,000160439 | 3,31E-05 | 6 - | 0,000131173 - | 6 - | 0,000162879 - | 7 - | 0,000180643 - | 8 - | 0,000209498 - | 4 - | 0,000118 - |
| 7739 | 1 | 3um_FIJJAFBF_397211 | Bacteria | Bacteroidetes | Cytophagia | Cytophagales | Flammeovirga | unclassified | Flammeovirga | unclassified | unclassified | unclassified | unclassified | unclassified | 0,2 | 0,4 | 1,11E-05 | 2,23E-05 | 1 - | 5,57E-05 - | 0 - | 0 - | 0 - | 0 - | 0 - | 0 - | 0 - | 0 - |
| 7740 | 1 | 0.2um_NOKCPHJN_648279 | Bacteria | Proteobacteri | Alphaproteob | Pelagibacteral | Pelagibacterac | Candidatus Pe | Candidatus Pe | information st | Translation, ril | 01QWB | One of the pri | 335992.SAR11 | 1 | 0 | 5,81E-05 | 5,51E-06 | 1 - | 4,86E-05 - | 1 - | 6,04E-05 - | 1 - | 5,74E-05 - | 1 - | 5,83E-05 - | 1 - | 6,56E-05 - |
| 7741 | 1 | 0.2um_NOKCPHJN_317015 | Bacteria | Proteobacteri | Gammaprotec | unclassified | unclassified | unclassified | Gammaprotec | unclassified | unclassified | unclassified | unclassified | unclassified | 0,4 | 0,489897949 | 3,23E-05 | 4,03E-05 | 1 - | 6,87E-05 - | 0 - | 0 - | 0 - | 0 - | 0 - | 0 - | 1 - | 9,27E-05 - |
| 7742 | 1 | 0.2um_NOKCPHJN_335007 | Eukaryota | Ascomycota | Dothideomyce | Pleosporales | Corynesporasc | Corynespora | Corynespora c | unclassified | unclassified | unclassified | unclassified | unclassified | 0,4 | 0,489897949 | 2,67E-06 | 3,33E-06 | 1 - | 5,68E-06 - | 0 - | 0 - | 0 - | 0 - | 0 - | 0 - | 1 - | 7,67E-06 - |
| 7743 | 1 | 0.2um_NOKCPHJN_132862 | Bacteria | Proteobacteri | Alphaproteob | Pelagibacteral | Pelagibacterac | Candidatus Pe | Candidatus Pe | metabolism | Energy produc | 172GV | Aconitate hyd | 439493.PB721 | 0,6 | 0,489897949 | 9,35E-06 | 7,64E-06 | 0 - | 0 - | 1 - | 1,60E-05 - | 1 - | 1,52E-05 - | 1 - | 1,55E-05 - | 0 - | 0 - |
| 7744 | 1 | 3um_FIJJAFBF_67857 | Bacteria | Bacteroidetes | unclassified | unclassified | unclassified | unclassified | Bacteroidetes | metabolism | Carbohydrate | 0P7K6 | Beta-glucosida | 376686.Fjoh_ | 0,6 | 0,489897949 | 1,11E-05 | 9,09E-06 | 0 - | 0 - | 0 - | 0 - | 1 - | 1,75E-05 - | 1 - | 1,78E-05 - | 1 - | 2,01E-05 - |
| 7745 | 1 | 3um_FIJJAFBF_654753 | Viruses | unclassified | unclassified | unclassified | unclassified | unclassified | uncultured ma | poorly charact | Function unkn | 0BBUF | Major capsid | 1097668.BYI2 | 2,2 | 0,748331477 | 7,39E-05 | 2,15E-05 | 3 - | 8,67E-05 - | 2 - | 7,18E-05 - | 2 - | 6,83E-05 - | 3 - | 0,000103892 - | 1 - | 3,90E-05 - |
| 7746 | 1 | 0.2um_NOKCPHJN_170444 | Bacteria | Proteobacteri | Alphaproteob | Rhodobactera | Rhodobactera | Sulfitobacter | Sulfitobacter s | metabolism | Amino acid tra | COG0059 | Alpha-keto-be | 391595.RLO14 | 1,6 | 0,8 | 6,62E-05 | 3,63E-05 | 1 - | 3,38E-05 - | 3 - | 0,000125752 - | 1 - | 3,98E-05 - | 1 - | 4,04E-05 - | 2 - | 9,11E-05 - |
| 7747 | 1 | 3um_FIJJAFBF_04044 | Bacteria | Bacteroidetes | Flavobacteriia | Flavobacterial | Flavobacteriac | Kordia | Kordia jejudor | unclassified | unclassified | unclassified | unclassified | unclassified | 0,2 | 0,4 | 1,30E-05 | 2,59E-05 | 1 - | 6,48E-05 - | 0 - | 0 - | 0 - | 0 - | 0 - | 0 - | 0 - | 0 - |
| 7748 | 1 | 3um_FIJJAFBF_452428 | Bacteria | Proteobacteri | Alphaproteob | Rhodospirillale | Rhodospirillac | unclassified | Rhodospirillac | metabolism | Energy produc | COG0056 | Produces ATP | 1082931.KKY_ | 2,2 | 0,748331477 | 5,70E-05 | 1,63E-05 | 3 - | 6,70E-05 - | 2 - | 5,55E-05 - | 3 - | 7,91E-05 - | 2 - | 5,35E-05 - | 1 - | 3,01E-05 - |
| 7749 | 1 | 0.2um_NOKCPHJN_368043 | Bacteria | Bacteroidetes | Flavobacteriia | Flavobacterial | Flavobacteriac | Muricauda | Muricauda sp. | metabolism | Energy produc | KOG0017 | Retrotranspos | 7668.SPU_013 | 0,2 | 0,4 | 7,81E-07 | 1,56E-06 | 1 - | 3,90E-06 - | 0 - | 0 - | 0 - | 0 - | 0 - | 0 - | 0 - | 0 - |
| 7750 | 1 | 0.2um_NOKCPHJN_285163 | Bacteria | Proteobacteri | Gammaprotec | unclassified | unclassified | unclassified | SAR86 cluster | information st | Transcription | 16PMT | DNA-depende | 205922.Pfl01_ | 1,8 | 0,4 | 7,32E-05 | 1,45E-05 | 2 - | 6,91E-05 - | 2 - | 8,59E-05 - | 2 - | 8,16E-05 - | 2 - | 8,28E-05 - | 1 - | 4,66E-05 - |
| 7751 | 1 | 0.2um_NOKCPHJN_410795 | Bacteria | Bacteroidetes | unclassified | unclassified | unclassified | unclassified | Bacteroidetes | poorly charact | Function unkn | 0XQTX | Chromosome | 926562.Oweh | 1,6 | 1,356465997 | 1,95E-05 | 1,50E-05 | 4 - | 4,39E-05 - | 2 - | 2,73E-05 - | 1 - | 1,30E-05 - | 1 - | 1,32E-05 - | 0 - | 0 - |
| 7752 | 1 | 0.2um_NOKCPHJN_208669 | Eukaryota | unclassified | Labyrinthulom | unclassified | Thraustochytr | Aurantiochytr | Aurantiochytr | cellular proces | Signal transdu | KOG0613 | fibronectin typ | 67593.Physo1 | 0,2 | 0,4 | 3,80E-07 | 7,60E-07 | 0 - | 0 - | 0 - | 0 - | 0 - | 0 - | 1 - | 1,90E-06 - | 0 - | 0 - |
| 7753 | 1 | 0.2um_NOKCPHJN_105399 | Bacteria | Proteobacteri | Gammaprotec | Oceanospirilla | Saccharospirill | Saccharospiril | Saccharospiril | information st | Translation, ril | 0QJEC | amino acids su | 314283.MED2 | 1,6 | 0,489897949 | 2,43E-05 | 8,86E-06 | 1 - | 1,24E-05 - | 2 - | 3,09E-05 - | 1 - | 1,47E-05 - | 2 - | 2,98E-05 - | 2 - | 3,36E-05 - |
| 7754 | 1 | 3um_FIJJAFBF_128395 | Bacteria | Bacteroidetes | Flavobacteriia | Flavobacterial | Flavobacteriac | Polaribacter | Polaribacter h | information st | Transcription | 0P7W9 | Transcription | 313594.PI23P | 0,6 | 0,489897949 | 1,88E-05 | 1,55E-05 | 1 - | 2,75E-05 - | 1 - | 3,41E-05 - | 1 - | 3,24E-05 - | 0 - | 0 - | 0 - | 0 - |
| 7755 | 1 | 3um_FIJJAFBF_804025 | Bacteria | Proteobacteri | Alphaproteob | Rhodobactera | Rhodobactera | Marivita | Marivita geoje | information st | Translation, ril | 16QAF | thus facilitatin | 246200.SPO0 | 0,2 | 0,4 | 5,53E-06 | 1,11E-05 | 0 - | 0 - | 0 - | 0 - | 0 - | 0 - | 0 - | 0 - | 1 - | 2,77E-05 - |
| 7756 | 1 | 0.2um_NOKCPHJN_40446 | Bacteria | Bacteroidetes | Flavobacteriia | Flavobacterial | Flavobacteriac | Algibacter | Algibacter sp. | unclassified | unclassified | unclassified | unclassified | unclassified | 2 | 0,894427191 | 3,33E-05 | 1,38E-05 | 3 - | 4,29E-05 - | 3 - | 5,32E-05 - | 1 - | 1,69E-05 - | 2 - | 3,42E-05 - | 1 - | 1,93E-05 - |
| 7757 | 1 | 3um_FIJJAFBF_598124 | Bacteria | Proteobacteri | Gammaprotec | unclassified | unclassified | unclassified | Gammaprotec | metabolism | Energy produc | 05CMV | Succinyl-CoA s | 351348.Maqu | 1,6 | 1,019803903 | 6,13E-05 | 3,62E-05 | 3 - | 0,000102175 - | 1 - | 4,23E-05 - | 2 - | 8,04E-05 - | 2 - | 8,16E-05 - | 0 - | 0 - |
| 7758 | 1 | 3um_FIJJAFBF_306723 | Bacteria | Proteobacteri | Alphaproteob | Rhodobactera | unclassified | unclassified | Rhodobactera | cellular proces | Posttranslatio | 01Y23 | cytochrome c | 439496.RBY4I | 0,2 | 0,4 | 1,46E-05 | 2,92E-05 | 1 - | 7,31E-05 - | 0 - | 0 - | 0 - | 0 - | 0 - | 0 - | 0 - | 0 - |
| 7759 | 1 | 3um_FIJJAFBF_347252 | Bacteria | Bacteroidetes | Flavobacteriia | Flavobacterial | Flavobacteriac | Polaribacter | Polaribacter ir | unclassified | unclassified | unclassified | unclassified | unclassified | 0,2 | 0,4 | 1,30E-05 | 2,59E-05 | 1 - | 6,48E-05 - | 0 - | 0 - | 0 - | 0 - | 0 - | 0 - | 0 - | 0 - |
| 7760 | 2 | 0.2um_NOKCPHJN_397544;3um_FIJJAFBF_60252 | Eukaryota | Platyhelminth | unclassified | Macrostomida | Macrostomida | Macrostomun | Macrostomun | unclassified | unclassified | unclassified | unclassified | unclassified | 0,2 | 0,4 | 9,38E-06 | 1,88E-05 | 0 - | 0 - | 0 - | 0 - | 0 - | 0 - | 1 - | 4,69E-05 - | 0 - | 0 - |
| 7761 | 1 | 3um_FIJJAFBF_187061 | Bacteria | Proteobacteri | Gammaprotec | Alteromonada | Alteromonada | Glaciecola | Glaciecola sp. | unclassified | unclassified | unclassified | unclassified | unclassified | 2 | 0,632455532 | 0,00014179 | 4,61E-05 | 2 - | 0,00011894 - | 2 - | 0,000147688 - | 3 - | 0,000210594 - | 1 - | 7,12E-05 - | 2 - | 0,000160492 - |
| 7762 | 1 | 3um_FIJJAFBF_760078 | Bacteria | Proteobacteri | Betaproteoba | unclassified | unclassified | unclassified | Betaproteoba | cellular proces | Posttranslatio | COG0330 | Band 7 protein | 296591.Bpro_ | 2,4 | 1,019803903 | 7,10E-05 | 3,20E-05 | 2 - | 4,93E-05 - | 4 - | 0,000122334 - | 3 - | 8,72E-05 - | 1 - | 2,95E-05 - | 2 - | 6,65E-05 - |
| 7763 | 7 | 0.2um_NOKCPHJN_122736;0.2um_NOKCPHJN_6 | Bacteria | Proteobacteri | Gammaprotec | Alteromonada | Colwelliaceae | Colwellia | various | unclassified | unclassified | unclassified | unclassified | unclassified | 0,2 | 0,4 | 1,04E-05 | 2,08E-05 | 0 - | 0 - | 1 - | 5,20E-05 - | 0 - | 0 - | 0 - | 0 - | 0 - | 0 - |
| 7764 | 1 | 3um_FIJJAFBF_149822 | Bacteria | Proteobacteri | Betaproteoba | unclassified | unclassified | unclassified | Betaproteoba | metabolism | Energy produc | COG0056 | Produces ATP | 543728.Vapar | 3,8 | 0,748331477 | 0,000101458 | 2,50E-05 | 3 - | 6,65E-05 - | 4 - | 0,000110053 - | 3 - | 7,85E-05 - | 5 - | 0,000132706 - | 4 - | 0,000119594 - |
| 7765 | 1 | 3um_FIJJAFBF_283891 | Bacteria | Proteobacteri | Alphaproteob | Rhodobactera | Rhodobactera | Planktomarina | Planktomarina | unclassified | unclassified | unclassified | unclassified | unclassified | 3,8 | 1,326649916 | 0,000292771 | 7,41E-05 | 6 - | 0,000400384 - | 3 - | 0,000248579 - | 4 - | 0,000315075 - | 4 - | 0,000319729 - | 2 - | 0,000180087 - |
| 7766 | 1 | 0.2um_NOKCPHJN_191029 | Bacteria | Proteobacteri | Alphaproteob | Rhodobactera | Rhodobactera | unclassified | Rhodobactera | metabolism | Amino acid tra | 01R4A | Polyamine AB | 388739.RSK20 | 0,4 | 0,489897949 | 1,24E-05 | 1,53E-05 | 1 - | 2,83E-05 - | 0 - | 0 - | 0 - | 0 - | 1 - | 3,39E-05 - | 0 - | 0 - |
| 7767 | 1 | 3um_FIJJAFBF_713065 | Bacteria | Proteobacteri | Gammaprotec | unclassified | unclassified | unclassified | OM182 bacter | unclassified | unclassified | unclassified | unclassified | unclassified | 1,8 | 0,4 | 0,000123694 | 3,42E-05 | 1 - | 5,65E-05 - | 2 - | 0,000140413 - | 2 - | 0,00013348 - | 2 - | 0,000135452 - | 2 - | 0,000152586 - |
| 7768 | 1 | 0.2um_NOKCPHJN_575859 | Bacteria | Proteobacteri | Alphaproteob | Pelagibacteral | Pelagibacterac | unclassified | Pelagibacterac | poorly charact | Function unkn | 0YZNV | NA | 93059.P9211_ | 1 | 0 | 1,63E-05 | 1,55E-06 | 1 - | 1,37E-05 - | 1 - | 1,70E-05 - | 1 - | 1,61E-05 - | 1 - | 1,64E-05 - | 1 - | 1,84E-05 - |
| 7769 | 1 | 3um_FIJJAFBF_612091 | Bacteria | Proteobacteri | Gammaprotec | Oceanospirilla | Saccharospirill | Reinekea | Reinekea fors | unclassified | unclassified | unclassified | unclassified | unclassified | 0,6 | 0,8 | 2,72E-05 | 3,52E-05 | 2 - | 8,57E-05 - | 0 - | 0 - | 1 - | 5,06E-05 - | 0 - | 0 - | 0 - | 0 - |
| 7770 | 1 | 3um_FIJJAFBF_777844 | Bacteria | Bacteroidetes | Flavobacteriia | Flavobacterial | Flavobacteriac | Polaribacter | Polaribacter s | unclassified | unclassified | unclassified | unclassified | unclassified | 14,2 | 2,785677655 | 0,000303759 | 4,78E-05 | 19 - | 0,000343968 - | 12 - | 0,000269752 - | 11 - | 0,000235064 - | 14 - | 0,000303591 - | 15 - | 0,000366422 - |
| 7771 | 1 | 0.2um_NOKCPHJN_767810 | Bacteria | Proteobacteri | Alphaproteob | Pelagibacteral | Pelagibacterac | Candidatus Pe | Candidatus Pe | information st | Transcription | COG0085 | DNA-depende | 335992.SAR11 | 2,4 | 0,8 | 3,81E-05 | 1,41E-05 | 2 - | 2,64E-05 - | 4 - | 6,56E-05 - | 2 - | 3,12E-05 - | 2 - | 3,16E-05 - | 2 - | 3,56E-05 - |
| 7772 | 1 | 0.2um_NOKCPHJN_58102 | Bacteria | Proteobacteri | Gammaprotec | Oceanospirilla | Saccharospirill | Reinekea | Reinekea blan | unclassified | unclassified | unclassified | unclassified | unclassified | 13,8 | 2,638181192 | 0,000116626 | 1,77E-05 | 18 - | 0,000128722 - | 15 - | 0,000133195 - | 10 - | 8,44E-05 - | 13 - | 0,000111357 - | 13 - | 0,000125444 - |
| 7773 | 1 | 3um_FIJJAFBF_416373 | Bacteria | Proteobacteri | Betaproteoba | unclassified | unclassified | unclassified | Betaproteoba | poorly charact | Function unkn | 0BH8N | BLUF domain | 232721.Ajs_0 | 0,2 | 0,4 | 1,63E-05 | 3,26E-05 | 1 - | 8,14E-05 - | 0 - | 0 - | 0 - | 0 - | 0 - | 0 - | 0 - | 0 - |
| 7774 | 1 | 3um_FIJJAFBF_532807 | Bacteria | Proteobacteri | Alphaproteob | Rhodobactera | unclassified | unclassified | Rhodobactera | cellular proces | Cell motility | 07W75 | Flagellin | 292414.TM10 | 0,2 | 0,4 | 9,64E-06 | 1,93E-05 | 0 - | 0 - | 0 - | 0 - | 1 - | 4,82E-05 - | 0 - | 0 - | 0 - | 0 - |
| 7775 | 1 | 3um_FIJJAFBF_262033 | Bacteria | Proteobacteri | Gammaprotec | unclassified | unclassified | unclassified | Gammaprotec | unclassified | unclassified | unclassified | unclassified | unclassified | 2,8 | 0,4 | 0,000279227 | 5,89E-05 | 2 - | 0,000165146 - | 3 - | 0,000307595 - | 3 - | 0,000292408 - | 3 - | 0,000296727 - | 3 - | 0,000334261 - |
| 7776 | 1 | 3um_FIJJAFBF_135425 | Bacteria | Proteobacteri | Alphaproteob | Rhodobactera | Rhodobactera | Planktomarina | Planktomarina | cellular proces | Intracellular tr | 05VIE | preprotein tra | 388739.RSK20 | 4,6 | 0,489897949 | 0,000727508 | 5,77E-05 | 5 - | 0,000667307 - | 4 - | 0,000662878 - | 5 - | 0,000787687 - | 5 - | 0,000799322 - | 4 - | 0,000720346 - |
| 7777 | 1 | 0.2um_NOKCPHJN_48124 | Bacteria | Proteobacteri | Gammaprotec | Cellvibrionales | Porticoccacea | unclassified | SAR92 bacteri | information st | Translation, ril | 179R6 | This protein is | 314287.GB220 | 0,6 | 0,489897949 | 5,85E-05 | 4,78E-05 | 0 - | 0 - | 1 - | 0,000100365 - | 1 - | 9,54E-05 - | 1 - | 9,68E-05 - | 0 - | 0 - |
| 7778 | 1 | 3um_FIJJAFBF_518688 | Bacteria | Bacteroidetes | Flavobacteriia | Flavobacterial | Flavobacteriac | Polaribacter | Polaribacter s | metabolism | Energy produc | 09A0E | phosphoenolp | 313598.MED1 | 6 | 1,414213562 | 9,47E-05 | 2,10E-05 | 7 - | 9,35E-05 - | 8 - | 0,00013273 - | 5 - | 7,89E-05 - | 6 - | 9,60E-05 - | 4 - | 7,21E-05 - |
| 7779 | 1 | 3um_FIJJAFBF_55181 | Bacteria | Proteobacteri | Gammaprotec | Oceanospirilla | Saccharospirill | Reinekea | Reinekea fors | poorly charact | Function unkn | 121Q1 | NA | 314283.MED2 | 3,6 | 0,489897949 | 0,000249301 | 2,48E-05 | 4 - | 0,000234238 - | 3 - | 0,000218141 - | 4 - | 0,000276494 - | 4 - | 0,000280578 - | 3 - | 0,000237053 - |
| 7780 | 1 | 0.2um_NOKCPHJN_296199 | Bacteria | Proteobacteri | Alphaproteob | Pelagibacteral | Pelagibacterac | Candidatus Pe | Candidatus Pe | metabolism | Energy produc | 16QPK | Succinate deh | 335992.SAR11 | 0,2 | 0,4 | 4,82E-06 | 9,65E-06 | 0 - | 0 - | 1 - | 2,41E-05 - | 0 - | 0 - | 0 - | 0 - | 0 - | 0 - |
| 7781 | 1 | 3um_FIJJAFBF_135181 | Bacteria | Bacteroidetes | Flavobacteriia | Flavobacterial | Cryomorphace | unclassified | Cryomorphace | unclassified | unclassified | unclassified | unclassified | unclassified | 0,8 | 0,4 | 4,33E-05 | 2,20E-05 | 1 - | 4,68E-05 - | 1 - | 5,82E-05 - | 1 - | 5,53E-05 - | 1 - | 5,61E-05 - | 0 - | 0 - |
| 7782 | 1 | 0.2um_NOKCPHJN_302189 | Eukaryota | unclassified | unclassified | unclassified | Apusomonadi | Thecamonas | Thecamonas t | unclassified | unclassified | unclassified | unclassified | unclassified | 0,8 | 0,4 | 2,99E-05 | 1,53E-05 | 1 - | 3,12E-05 - | 1 - | 3,87E-05 - | 0 - | 0 - | 1 - | 3,74E-05 - | 1 - | 4,21E-05 - |
| 7783 | 1 | 3um_FIJJAFBF_152852 | Bacteria | Bacteroidetes | Flavobacteriia | Flavobacterial | Flavobacteriac | Flaviramulus | Flaviramulus b | unclassified | unclassified | unclassified | unclassified | unclassified | 5,6 | 1,854723699 | 0,000117655 | 3,29E-05 | 8 - | 0,000144147 - | 7 - | 0,000156614 - | 4 - | 8,51E-05 - | 6 - | 0,000129498 - | 3 - | 7,29E-05 - |
| 7784 | 1 | 0.2um_NOKCPHJN_261541 | Bacteria | Bacteroidetes | Flavobacteriia | Flavobacterial | Flavobacteriac | Lutibacter | Lutibacter pro | metabolism | Inorganic ion t | 05C27 | TonB-depende | 487797.Flav30 | 25,8 | 2,227105745 | 0,000324688 | 1,44E-05 | 30 - | 0,000318529 - | 25 - | 0,0003296 - | 24 - | 0,000300793 - | 26 - | 0,000330672 - | 24 - | 0,000343847 - |
| 7785 | 1 | 3um_FIJJAFBF_534169 | Bacteria | Proteobacteri | Gammaprotec | Cellvibrionales | Halieaceae | unclassified | marine gamm | unclassified | unclassified | unclassified | unclassified | unclassified | 1,6 | 0,8 | 8,41E-05 | 4,18E-05 | 1 - | 4,41E-05 - | 1 - | 5,48E-05 - | 3 - | 0,000156326 - | 2 - | 0,000105756 - | 1 - | 5,96E-05 - |
| 7786 | 1 | 0.2um_NOKCPHJN_193463 | Bacteria | Proteobacteri | Alphaproteob | Rhodospirillale | Rhodospirillac | Candidatus En | Candidatus En | unclassified | unclassified | unclassified | unclassified | unclassified | 1,6 | 0,8 | 6,05E-05 | 2,42E-05 | 3 - | 9,89E-05 - | 1 - | 4,10E-05 - | 1 - | 3,89E-05 - | 2 - | 7,90E-05 - | 1 - | 4,45E-05 - |
| 7787 | 1 | 0.2um_NOKCPHJN_27669 | Bacteria | Actinobacteria | Actinobacteria | Micrococcales | Microbacteria | Candidatus Ac | Candidatus Ac | information st | Transcription | 00C46 | DNA-depende | 281090.Lxx20 | 3 | 1,673320053 | 3,47E-05 | 2,05E-05 | 2 - | 1,93E-05 - | 6 - | 7,20E-05 - | 3 - | 3,42E-05 - | 3 - | 3,47E-05 - | 1 - | 1,30E-05 - |
| 7788 | 1 | 0.2um_NOKCPHJN_668717 | Bacteria | Proteobacteri | Gammaprotec | Alteromonada | Alteromonada | Glaciecola | Glaciecola sp. | metabolism | Carbohydrate | 16QEQ | hydrolase, fan | 455436.GHTC | 0,8 | 0,4 | 4,00E-05 | 2,02E-05 | 0 - | 0 - | 1 - | 5,00E-05 - | 1 - | 4,75E-05 - | 1 - | 4,82E-05 - | 1 - | 5,43E-05 - |
| 7789 | 1 | 3um_FIJJAFBF_221204 | Bacteria | Proteobacteri | Gammaprotec | Oceanospirilla | Saccharospirill | Reinekea | Reinekea fors | information st | Translation, ril | 179R6 | This protein is | 491952.Mar18 | 5,2 | 0,4 | 0,000498656 | 2,47E-05 | 6 - | 0,000484972 - | 5 - | 0,000501827 - | 5 - | 0,00047705 - | 5 - | 0,000484096 - | 5 - | 0,000545333 - |
| 7790 | 1 | 3um_FIJJAFBF_632244 | Bacteria | Proteobacteri | Gammaprotec | Pseudomonad | Moraxellacea | Acinetobacter | various | unclassified | unclassified | unclassified | unclassified | unclassified | 0,4 | 0,489897949 | 4,47E-05 | 5,48E-05 | 0 - | 0 - | 0 - | 0 - | 1 - | 0,000111051 - | 1 - | 0,000112691 - | 0 - | 0 - |
| 7791 | 1 | 3um_FIJJAFBF_712629 | Bacteria | Proteobacteri | Gammaprotec | Oceanospirilla | Saccharospirill | Reinekea | Reinekea fors | poorly charact | Function unkn | 0QQPZ | UPF0325 prot | 314283.MED2 | 4,6 | 0,489897949 | 0,000496553 | 3,94E-05 | 5 - | 0,000455463 - | 4 - | 0,000452441 - | 5 - | 0,000537628 - | 5 - | 0,000545569 - | 4 - | 0,000491665 - |
| 7792 | 1 | 3um_FIJJAFBF_387026 | Bacteria | Bacteroidetes | Flavobacteriia | Flavobacterial | Flavobacteriac | Polaribacter | Polaribacter g | unclassified | unclassified | unclassified | unclassified | unclassified | 0,6 | 0,489897949 | 2,08E-05 | 1,71E-05 | 1 - | 3,08E-05 - | 0 - | 0 - | 1 - | 3,63E-05 - | 1 - | 3,69E-05 - | 0 - | 0 - |
| 7793 | 1 | 3um_FIJJAFBF_189729 | Bacteria | Bacteroidetes | Flavobacteriia | Flavobacterial | Flavobacteriac | Polaribacter | Polaribacter s | unclassified | unclassified | unclassified | unclassified | unclassified | 0,2 | 0,4 | 5,84E-06 | 1,17E-05 | 0 - | 0 - | 1 - | 2,92E-05 - | 0 - | 0 - | 0 - | 0 - | 0 - | 0 - |
| 7794 | 1 | 0.2um_NOKCPHJN_888294 | Eukaryota | unclassified | Pelagophycea | Pelagomonad | unclassified | Aureococcus | Aureococcus a | cellular proces | Signal transdu | KOG0613 | fibronectin typ | 65071.PYU1_T | 0,4 | 0,489897949 | 9,79E-06 | 1,20E-05 | 0 - | 0 - | 1 - | 2,51E-05 - | 1 - | 2,39E-05 - | 0 - | 0 - | 0 - | 0 - |
| 7795 | 1 | 3um_FIJJAFBF_786163 | Bacteria | Proteobacteri | Gammaprotec | Oceanospirilla | Saccharospirill | Reinekea | Reinekea fors | information st | Translation, ril | COG0051 | tRNA binding | 314283.MED2 | 3,2 | 0,748331477 | 0,000420029 | 8,22E-05 | 4 - | 0,000445735 - | 2 - | 0,000276736 - | 4 - | 0,000526145 - | 3 - | 0,000400437 - | 3 - | 0,000451091 - |
| 7796 | 1 | 3um_FIJJAFBF_332398 | Bacteria | Proteobacteri | Alphaproteob | Rhodobactera | Rhodobactera | Lentibacter | Lentibacter al | metabolism | Energy produc | COG0711 | F(1)F(0) ATP s | 388739.RSK20 | 1,8 | 0,979795897 | 0,000137384 | 7,67E-05 | 2 - | 0,000128963 - | 0 - | 0 - | 2 - | 0,000152227 - | 3 - | 0,000231713 - | 2 - | 0,000174016 - |
| 7797 | 1 | 0.2um_NOKCPHJN_359126 | Bacteria | Proteobacteri | Betaproteoba | Burkholderiale | unclassified | unclassified | various | cellular proces | Posttranslatio | 17FI0 | Binds to Cpn6 | 365046.Rta_3 | 1,2 | 0,4 | 0,000168403 | 3,73E-05 | 2 - | 0,000241635 - | 1 - | 0,00015002 - | 1 - | 0,000142613 - | 1 - | 0,000144719 - | 1 - | 0,000163026 - |
| 7798 | 2 | 0.2um_NOKCPHJN_455817;0.2um_NOKCPHJN_6 | Bacteria | Proteobacteri | Alphaproteob | Rhodobactera | Rhodobactera | Planktomarina | Planktomarina | information st | Translation, ril | 179RU | 30S ribosomal | 467661.RKLH1 | 1,6 | 0,489897949 | 0,000132454 | 3,47E-05 | 2 - | 0,0001417 - | 1 - | 8,80E-05 - | 2 - | 0,000167262 - | 2 - | 0,000169732 - | 1 - | 9,56E-05 - |
| 7799 | 1 | 0.2um_NOKCPHJN_174788 | Bacteria | Bacteroidetes | Flavobacteriia | Flavobacterial | Flavobacteriac | Polaribacter | Polaribacter s | unclassified | unclassified | unclassified | unclassified | unclassified | 0,4 | 0,8 | 1,30E-05 | 2,59E-05 | 2 - | 6,48E-05 - | 0 - | 0 - | 0 - | 0 - | 0 - | 0 - | 0 - | 0 - |
| 7800 | 1 | 3um_FIJJAFBF_556831 | Bacteria | Bacteroidetes | Flavobacteriia | Flavobacterial | Flavobacteriac | Dokdonia | various | information st | Translation, ril | COG0256 | This is one of t | 983548.Krodi_ | 0,2 | 0,4 | 2,34E-05 | 4,67E-05 | 0 - | 0 - | 0 - | 0 - | 1 - | 0,000116795 - | 0 - | 0 - | 0 - | 0 - |
| 7801 | 1 | 3um_FIJJAFBF_562472 | Bacteria | Proteobacteri | Alphaproteob | Rhodobactera | unclassified | unclassified | Rhodobactera | metabolism | Amino acid tra | 16RER | (ABC) transpo | 314262.MED1 | 1,6 | 0,8 | 8,20E-05 | 3,25E-05 | 3 - | 0,000134504 - | 1 - | 5,57E-05 - | 2 - | 0,000105845 - | 1 - | 5,37E-05 - | 1 - | 6,05E-05 - |
| 7802 | 1 | 0.2um_NOKCPHJN_224032 | Bacteria | Bacteroidetes | Flavobacteriia | Flavobacterial | Flavobacteriac | unclassified | Flavobacteriac | unclassified | unclassified | unclassified | unclassified | unclassified | 1,2 | 0,4 | 0,000143064 | 4,93E-05 | 1 - | 9,98E-05 - | 1 - | 0,000123929 - | 1 - | 0,000117811 - | 2 - | 0,000239101 - | 1 - | 0,000134673 - |
| 7803 | 1 | 3um_FIJJAFBF_806084 | Bacteria | Proteobacteri | Gammaprotec | Oceanospirilla | Saccharospirill | Reinekea | Reinekea fors | poorly charact | Function unkn | 17CZ8 | Lipoprotein | 314283.MED2 | 0,4 | 0,8 | 2,62E-05 | 5,25E-05 | 2 - | 0,000131173 - | 0 - | 0 - | 0 - | 0 - | 0 - | 0 - | 0 - | 0 - |
| 7804 | 3 | 0.2um_NOKCPHJN_112027;0.2um_NOKCPHJN_2 | Bacteria | Proteobacteri | Gammaprotec | Cellvibrionales | Porticoccacea | unclassified | gamma protec | cellular proces | Cell wall/mem | 0QTSX | Peptidoglycan | 314287.GB220 | 1 | 0 | 8,16E-05 | 7,74E-06 | 1 - | 6,83E-05 - | 1 - | 8,48E-05 - | 1 - | 8,06E-05 - | 1 - | 8,18E-05 - | 1 - | 9,22E-05 - |
| 7805 | 1 | 3um_FIJJAFBF_254299 | Bacteria | Proteobacteri | Gammaprotec | Cellvibrionales | Spongiibacter | Spongiibacter | Spongiibacter | information st | Translation, ril | 179ZI | One of the pri | 377629.TERTU | 3,6 | 1,496662955 | 0,000362643 | 0,000117834 | 6 - | 0,000525695 - | 2 - | 0,000217586 - | 4 - | 0,000413686 - | 4 - | 0,000419796 - | 2 - | 0,00023645 - |
| 7806 | 1 | 0.2um_NOKCPHJN_1113290 | Bacteria | Bacteroidetes | Bacteroidia | Bacteroidales | Prevotellaceae | Prevotella | various | unclassified | unclassified | unclassified | unclassified | unclassified | 1,2 | 0,4 | 3,80E-05 | 1,31E-05 | 1 - | 2,65E-05 - | 1 - | 3,29E-05 - | 1 - | 3,13E-05 - | 2 - | 6,35E-05 - | 1 - | 3,58E-05 - |
| 7807 | 1 | 0.2um_NOKCPHJN_817058 | Bacteria | Proteobacteri | Gammaprotec | unclassified | unclassified | unclassified | Gammaprotec | unclassified | unclassified | unclassified | unclassified | unclassified | 1,2 | 0,979795897 | 8,06E-05 | 6,64E-05 | 2 - | 0,00011772 - | 2 - | 0,000146173 - | 2 - | 0,000138956 - | 0 - | 0 - | 0 - | 0 - |
| 7808 | 1 | 0.2um_NOKCPHJN_326468 | Eukaryota | Phaeophyceae | unclassified | various | various | various | various | unclassified | unclassified | unclassified | unclassified | unclassified | 0,6 | 0,489897949 | 5,84E-05 | 4,78E-05 | 0 - | 0 - | 1 - | 9,57E-05 - | 0 - | 0 - | 1 - | 9,23E-05 - | 1 - | 0,000103943 - |
| 7809 | 1 | 3um_FIJJAFBF_175076 | Bacteria | Proteobacteri | Alphaproteob | Rhodobactera | Rhodobactera | various | various | metabolism | Amino acid tra | 01SDW | Spermidine pu | 439497.RR11_ | 8,2 | 1,6 | 0,000295686 | 3,91E-05 | 10 - | 0,00030689 - | 7 - | 0,000266747 - | 10 - | 0,000362252 - | 8 - | 0,000294082 - | 6 - | 0,000248462 - |
| 7810 | 1 | 0.2um_NOKCPHJN_805108 | unclassified | unclassified | unclassified | unclassified | unclassified | unclassified | unclassified | unclassified | unclassified | unclassified | unclassified | unclassified | 2,4 | 0,8 | 0,000139115 | 3,08E-05 | 4 - | 0,000199612 - | 2 - | 0,000123929 - | 2 - | 0,000117811 - | 2 - | 0,000119551 - | 2 - | 0,000134673 - |
| 7811 | 1 | 0.2um_NOKCPHJN_641068 | Eukaryota | Basidiomycota | Agaricomycet | Cantharellales | Ceratobasidia | Rhizoctonia | Rhizoctonia so | metabolism | Energy produc | KOG0017 | Retrotranspos | 67593.Physo1 | 0,2 | 0,4 | 2,38E-06 | 4,76E-06 | 0 - | 0 - | 0 - | 0 - | 0 - | 0 - | 1 - | 1,19E-05 - | 0 - | 0 - |
| 7812 | 1 | 0.2um_NOKCPHJN_05571 | Viruses | unclassified | unclassified | unclassified | Phycodnavirid | unclassified | Aureococcus a | unclassified | unclassified | unclassified | unclassified | unclassified | 3,6 | 0,8 | 9,71E-05 | 1,47E-05 | 5 - | 0,000114777 - | 4 - | 0,000114015 - | 3 - | 8,13E-05 - | 3 - | 8,25E-05 - | 3 - | 9,29E-05 - |
| 7813 | 1 | 3um_FIJJAFBF_176946 | Eukaryota | unclassified | Cryptophyta | Pyrenomonad | Geminigerace | Guillardia | Guillardia thet | unclassified | unclassified | unclassified | unclassified | unclassified | 1 | 0 | 4,31E-05 | 4,09E-06 | 1 - | 3,61E-05 - | 1 - | 4,48E-05 - | 1 - | 4,26E-05 - | 1 - | 4,32E-05 - | 1 - | 4,87E-05 - |
| 7814 | 1 | 3um_FIJJAFBF_401272 | Bacteria | Proteobacteri | Alphaproteob | Rhodobactera | Rhodobactera | Planktomarina | Planktomarina | cellular proces | Cell wall/mem | 08ZG0 | Ompa motb d | 89187.ISM_10 | 1,2 | 0,4 | 7,31E-05 | 1,62E-05 | 2 - | 0,000104819 - | 1 - | 6,51E-05 - | 1 - | 6,19E-05 - | 1 - | 6,28E-05 - | 1 - | 7,07E-05 - |
| 7815 | 1 | 0.2um_NOKCPHJN_384286 | Bacteria | Proteobacteri | Alphaproteob | Rhodobactera | Rhodobactera | unclassified | Rhodobactera | poorly charact | Function unkn | COG3181 | extra-cytoplas | 388401.RB215 | 0,4 | 0,489897949 | 1,60E-05 | 1,98E-05 | 1 - | 3,56E-05 - | 1 - | 4,43E-05 - | 0 - | 0 - | 0 - | 0 - | 0 - | 0 - |
| 7816 | 1 | 3um_FIJJAFBF_725633 | Bacteria | unclassified | unclassified | unclassified | unclassified | unclassified | bacterium TM | unclassified | unclassified | unclassified | unclassified | unclassified | 1 | 0 | 7,74E-05 | 7,35E-06 | 1 - | 6,48E-05 - | 1 - | 8,05E-05 - | 1 - | 7,65E-05 - | 1 - | 7,77E-05 - | 1 - | 8,75E-05 - |
| 7817 | 1 | 3um_FIJJAFBF_500138 | Bacteria | Bacteroidetes | Flavobacteriia | unclassified | unclassified | unclassified | Flavobacteria | unclassified | unclassified | unclassified | unclassified | unclassified | 2,4 | 0,489897949 | 0,00015288 | 2,25E-05 | 3 - | 0,00016242 - | 2 - | 0,000134452 - | 3 - | 0,00019172 - | 2 - | 0,000129701 - | 2 - | 0,000146108 - |
| 7818 | 1 | 3um_FIJJAFBF_118036 | Bacteria | Proteobacteri | Betaproteoba | Burkholderiale | Comamonada | Acidovorax | Acidovorax sp | unclassified | unclassified | unclassified | unclassified | unclassified | 4 | 1,095445115 | 0,000204023 | 3,43E-05 | 6 - | 0,000262848 - | 3 - | 0,00016319 - | 4 - | 0,000206843 - | 4 - | 0,000209898 - | 3 - | 0,000177337 - |
| 7819 | 1 | 3um_FIJJAFBF_710078 | Eukaryota | Eustigmatoph | unclassified | Eustigmatales | Monodopsida | Nannochlorop | Nannochlorop | metabolism | Energy produc | COG1290 | Component of | 35128.Thapsd | 1,4 | 0,8 | 0,000114336 | 5,06E-05 | 3 - | 0,000215206 - | 1 - | 8,91E-05 - | 1 - | 8,47E-05 - | 1 - | 8,59E-05 - | 1 - | 9,68E-05 - |
| 7820 | 1 | 0.2um_NOKCPHJN_165212 | Bacteria | Bacteroidetes | Flavobacteriia | Flavobacterial | Flavobacteriac | Formosa | Formosa sp. H | unclassified | unclassified | unclassified | unclassified | unclassified | 3,4 | 2,059126028 | 6,78E-05 | 4,14E-05 | 3 - | 5,07E-05 - | 1 - | 2,10E-05 - | 4 - | 7,98E-05 - | 7 - | 0,000141735 - | 2 - | 4,56E-05 - |
| 7821 | 2 | 0.2um_NOKCPHJN_57909;3um_FIJJAFBF_24784C | Bacteria | Proteobacteri | Alphaproteob | Rhodobactera | Rhodobactera | Planktomarina | Planktomarina | cellular proces | Cell wall/mem | COG2885 | Ompa motb d | various | 1,4 | 0,489897949 | 0,000110698 | 3,17E-05 | 2 - | 0,00013583 - | 1 - | 8,43E-05 - | 2 - | 0,000160334 - | 1 - | 8,14E-05 - | 1 - | 9,16E-05 - |
| 7822 | 1 | 3um_FIJJAFBF_571787 | Bacteria | Proteobacteri | Gammaprotec | Oceanospirilla | Saccharospirill | Reinekea | Reinekea fors | unclassified | unclassified | unclassified | unclassified | unclassified | 0,8 | 0,4 | 4,33E-05 | 2,22E-05 | 1 - | 4,52E-05 - | 1 - | 5,61E-05 - | 0 - | 0 - | 1 - | 5,41E-05 - | 1 - | 6,10E-05 - |
| 7823 | 1 | 3um_FIJJAFBF_20700 | Bacteria | Proteobacteri | Alphaproteob | Rhodobactera | unclassified | unclassified | Rhodobactera | cellular proces | Signal transdu | 13AP7 | NACHT domai | 7739.JGI9285 | 4 | 0 | 0,000326255 | 3,10E-05 | 4 - | 0,000273278 - | 4 - | 0,000339331 - | 4 - | 0,000322577 - | 4 - | 0,000327341 - | 4 - | 0,000368749 - |
| 7824 | 1 | 0.2um_NOKCPHJN_57093 | Bacteria | Proteobacteri | Betaproteoba | Burkholderiale | Alcaligenacea | Advenella | Advenella kas | unclassified | unclassified | unclassified | unclassified | unclassified | 4 | 0,632455532 | 0,000140083 | 3,43E-05 | 3 - | 8,67E-05 - | 4 - | 0,000143596 - | 4 - | 0,000136506 - | 4 - | 0,000138522 - | 5 - | 0,000195056 - |
| 7825 | 1 | 0.2um_NOKCPHJN_276317 | Bacteria | Bacteroidetes | unclassified | unclassified | unclassified | unclassified | Bacteroidetes | unclassified | unclassified | unclassified | unclassified | unclassified | 2 | 0,632455532 | 3,78E-05 | 8,76E-06 | 3 - | 4,90E-05 - | 2 - | 4,05E-05 - | 2 - | 3,85E-05 - | 2 - | 3,91E-05 - | 1 - | 2,20E-05 - |
| 7826 | 1 | 3um_FIJJAFBF_32971 | Bacteria | Bacteroidetes | Flavobacteriia | Flavobacterial | Flavobacteriac | Formosa | Formosa sp. H | cellular proces | Posttranslatio | 05C6P | ATP-depender | 1046627.BZAR | 1,4 | 1,356465997 | 2,35E-05 | 2,40E-05 | 1 - | 1,41E-05 - | 4 - | 6,99E-05 - | 1 - | 1,66E-05 - | 1 - | 1,69E-05 - | 0 - | 0 - |
| 7827 | 1 | 3um_FIJJAFBF_155851 | Bacteria | Proteobacteri | Gammaprotec | Oceanospirilla | Saccharospirill | Reinekea | Reinekea blan | metabolism | Amino acid tra | 0QIIB | Catalyzes the | 314283.MED2 | 0,8 | 0,4 | 3,73E-05 | 1,92E-05 | 1 - | 3,94E-05 - | 0 - | 0 - | 1 - | 4,66E-05 - | 1 - | 4,72E-05 - | 1 - | 5,32E-05 - |
| 7828 | 1 | 3um_FIJJAFBF_744830 | Bacteria | Proteobacteri | Gammaprotec | Pseudomonad | Pseudomonad | Pseudomonas | various | unclassified | unclassified | unclassified | unclassified | unclassified | 1,2 | 0,4 | 8,94E-05 | 1,98E-05 | 2 - | 0,000128242 - | 1 - | 7,96E-05 - | 1 - | 7,57E-05 - | 1 - | 7,68E-05 - | 1 - | 8,65E-05 - |
| 7829 | 1 | 3um_FIJJAFBF_100050 | Bacteria | Proteobacteri | Alphaproteob | Rhodospirillale | Rhodospirillac | Candidatus En | Candidatus En | unclassified | unclassified | unclassified | unclassified | unclassified | 1,4 | 1,019803903 | 4,03E-05 | 2,97E-05 | 2 - | 4,87E-05 - | 3 - | 9,08E-05 - | 0 - | 0 - | 1 - | 2,92E-05 - | 1 - | 3,29E-05 - |
| 7830 | 1 | 0.2um_NOKCPHJN_258100 | Bacteria | Bacteroidetes | Flavobacteriia | Flavobacterial | Flavobacteriac | Formosa | Formosa sp. H | unclassified | unclassified | unclassified | unclassified | unclassified | 1,6 | 1,019803903 | 0,000103797 | 6,14E-05 | 3 - | 0,00017303 - | 1 - | 7,16E-05 - | 2 - | 0,000136163 - | 2 - | 0,000138174 - | 0 - | 0 - |
| 7831 | 1 | 3um_FIJJAFBF_616663 | Bacteria | Proteobacteri | Gammaprotec | Chromatiales | unclassified | unclassified | Chromatiales | metabolism | Amino acid tra | COG0687 | polyamine bin | 391616.OA23 | 0,2 | 0,4 | 6,98E-06 | 1,40E-05 | 0 - | 0 - | 0 - | 0 - | 0 - | 0 - | 1 - | 3,49E-05 - | 0 - | 0 - |
| 7832 | 1 | 3um_FIJJAFBF_390144 | Eukaryota | Eustigmatoph | unclassified | Eustigmatales | Monodopsida | Nannochlorop | Nannochlorop | unclassified | unclassified | unclassified | unclassified | unclassified | 1,4 | 0,489897949 | 0,000269266 | 8,46E-05 | 2 - | 0,000327934 - | 2 - | 0,000407197 - | 1 - | 0,000193546 - | 1 - | 0,000196405 - | 1 - | 0,000221249 - |
| 7833 | 1 | 0.2um_NOKCPHJN_148412 | Bacteria | Proteobacteri | Alphaproteob | Pelagibacteral | Pelagibacterac | Candidatus Pe | Candidatus Pe | poorly charact | Function unkn | 05C2K | tail SHEATH pr | 644282.Deba_ | 1,6 | 1,019803903 | 1,91E-05 | 1,29E-05 | 1 - | 1,01E-05 - | 3 - | 3,76E-05 - | 2 - | 2,38E-05 - | 2 - | 2,42E-05 - | 0 - | 0 - |
| 7834 | 1 | 0.2um_NOKCPHJN_167906 | Bacteria | Proteobacteri | Gammaprotec | unclassified | unclassified | unclassified | various | metabolism | Carbohydrate | 07QQQ | Pfam:PQQ_N | 234267.Acid_ | 0,6 | 0,8 | 1,23E-05 | 1,65E-05 | 0 - | 0 - | 2 - | 4,16E-05 - | 1 - | 1,97E-05 - | 0 - | 0 - | 0 - | 0 - |
| 7835 | 1 | 3um_FIJJAFBF_715061 | Eukaryota | unclassified | Rhodellophyc | Rhodellales | Rhodellaceae | Corynoplastis | Corynoplastis | unclassified | unclassified | unclassified | unclassified | unclassified | 1 | 0,632455532 | 0,000174727 | 0,000115314 | 1 - | 0,000145287 - | 2 - | 0,000360807 - | 1 - | 0,000171496 - | 0 - | 0 - | 1 - | 0,000196044 - |
| 7836 | 1 | 0.2um_NOKCPHJN_143991 | Bacteria | Proteobacteri | Gammaprotec | Oceanospirilla | Saccharospirill | Reinekea | Reinekea blan | unclassified | unclassified | unclassified | unclassified | unclassified | 5 | 1,264911064 | 6,97E-05 | 1,16E-05 | 7 - | 8,37E-05 - | 5 - | 7,42E-05 - | 5 - | 7,06E-05 - | 5 - | 7,16E-05 - | 3 - | 4,84E-05 - |
| 7837 | 1 | 3um_FIJJAFBF_791191 | Eukaryota | Eustigmatoph | unclassified | Eustigmatales | Monodopsida | Nannochlorop | Nannochlorop | information st | Translation, ril | KOG3181 | ribosomal pro | 65071.PYU1_T | 0,4 | 0,489897949 | 3,55E-05 | 4,37E-05 | 1 - | 8,14E-05 - | 0 - | 0 - | 1 - | 9,61E-05 - | 0 - | 0 - | 0 - | 0 - |
| 7838 | 1 | 3um_FIJJAFBF_182040 | Bacteria | Proteobacteri | Alphaproteob | Rhodobactera | unclassified | unclassified | Rhodobactera | metabolism | Carbohydrate | 01RS7 | extracellular s | 367336.OM22 | 1,4 | 0,8 | 4,60E-05 | 2,76E-05 | 1 - | 2,79E-05 - | 2 - | 6,94E-05 - | 2 - | 6,59E-05 - | 2 - | 6,69E-05 - | 0 - | 0 - |
| 7839 | 1 | 3um_FIJJAFBF_291822 | Bacteria | Bacteroidetes | Flavobacteriia | Flavobacterial | Flavobacteriac | Polaribacter | Polaribacter d | poorly charact | Function unkn | 09A4Z | rhodopsin | 313598.MED1 | 1,2 | 0,4 | 6,84E-05 | 2,29E-05 | 1 - | 4,78E-05 - | 1 - | 5,94E-05 - | 2 - | 0,000112902 - | 1 - | 5,73E-05 - | 1 - | 6,45E-05 - |
| 7840 | 1 | 0.2um_NOKCPHJN_17420 | Bacteria | Proteobacteri | Gammaprotec | Alteromonada | Pseudoalteror | Pseudoalteror | Pseudoalteror | unclassified | unclassified | unclassified | unclassified | unclassified | 18 | 1,414213562 | 0,000264174 | 3,91E-05 | 17 - | 0,00020824 - | 18 - | 0,000273782 - | 16 - | 0,000231346 - | 20 - | 0,000293454 - | 19 - | 0,000314046 - |
| 7841 | 1 | 0.2um_NOKCPHJN_913365 | Eukaryota | unclassified | unclassified | Prymnesiales | Chrysochromu | Chrysochromu | Chrysochromu | unclassified | unclassified | unclassified | unclassified | unclassified | 0,8 | 0,4 | 6,01E-05 | 3,09E-05 | 1 - | 6,27E-05 - | 1 - | 7,79E-05 - | 0 - | 0 - | 1 - | 7,51E-05 - | 1 - | 8,46E-05 - |
| 7842 | 2 | 3um_FIJJAFBF_334866;3um_FIJJAFBF_458699 | Bacteria | Bacteroidetes | Flavobacteriia | Flavobacterial | Flavobacteriac | Formosa | Formosa sp. H | unclassified | unclassified | unclassified | unclassified | unclassified | 1 | 0 | 7,25E-05 | 6,88E-06 | 1 - | 6,07E-05 - | 1 - | 7,54E-05 - | 1 - | 7,17E-05 - | 1 - | 7,27E-05 - | 1 - | 8,19E-05 - |
| 7843 | 1 | 3um_FIJJAFBF_245504 | Bacteria | Bacteroidetes | Flavobacteriia | Flavobacterial | Flavobacteriac | Lutibacter | Lutibacter pro | unclassified | unclassified | unclassified | unclassified | unclassified | 6,8 | 0,4 | 0,000437603 | 5,10E-05 | 7 - | 0,000377201 - | 7 - | 0,000468372 - | 6 - | 0,00038164 - | 7 - | 0,000451823 - | 7 - | 0,000508977 - |
| 7844 | 1 | 3um_FIJJAFBF_111312 | Viruses | unclassified | unclassified | unclassified | various | various | various | information st | Transcription | 0WRII | DNA-depende | 339860.Msp_( | 1,2 | 0,4 | 4,52E-05 | 1,56E-05 | 1 - | 3,15E-05 - | 1 - | 3,92E-05 - | 1 - | 3,72E-05 - | 2 - | 7,55E-05 - | 1 - | 4,25E-05 - |
| 7845 | 1 | 3um_FIJJAFBF_793633 | Bacteria | Bacteroidetes | Flavobacteriia | Flavobacterial | Flavobacteriac | Tenacibaculur | Tenacibaculur | information st | Replication, re | 0PB0H | DNA-binding p | 313598.MED1 | 1 | 0 | 0,000152252 | 1,45E-05 | 1 - | 0,00012753 - | 1 - | 0,000158354 - | 1 - | 0,000150536 - | 1 - | 0,000152759 - | 1 - | 0,000172083 - |
| 7846 | 1 | 0.2um_NOKCPHJN_30295 | Bacteria | Proteobacteri | Gammaprotec | Oceanospirilla | Oceanospirilla | Neptunomona | Neptunomona | information st | Translation, ril | 179RU | 30S ribosomal | 207954.MED9 | 2,8 | 0,748331477 | 0,000291307 | 7,09E-05 | 3 - | 0,000264869 - | 2 - | 0,00021926 - | 4 - | 0,000416868 - | 3 - | 0,000317269 - | 2 - | 0,000238268 - |
| 7847 | 1 | 3um_FIJJAFBF_491685 | Bacteria | Proteobacteri | Gammaprotec | Cellvibrionales | Porticoccacea | unclassified | SAR92 bacteri | unclassified | unclassified | unclassified | unclassified | unclassified | 0,6 | 0,489897949 | 7,04E-05 | 5,75E-05 | 0 - | 0 - | 1 - | 0,000120779 - | 1 - | 0,000114815 - | 1 - | 0,000116511 - | 0 - | 0 - |
| 7848 | 1 | 0.2um_NOKCPHJN_506016 | Bacteria | Bacteroidetes | Flavobacteriia | Flavobacterial | Flavobacteriac | Gilvibacter | Gilvibacter sp. | poorly charact | Function unkn | 064XS | NA | 313595.P7007 | 1,8 | 0,4 | 8,94E-05 | 1,76E-05 | 2 - | 8,44E-05 - | 2 - | 0,000104793 - | 2 - | 9,96E-05 - | 2 - | 0,000101091 - | 1 - | 5,69E-05 - |
| 7849 | 1 | 0.2um_NOKCPHJN_235557 | Eukaryota | unclassified | Oomycetes | Peronosporale | Peronosporac | Phytophthora | Phytophthora | poorly charact | Function unkn | 0XNUF | Extracellular n | 340101.Vvad_ | 0,2 | 0,4 | 5,60E-07 | 1,12E-06 | 0 - | 0 - | 1 - | 2,80E-06 - | 0 - | 0 - | 0 - | 0 - | 0 - | 0 - |
| 7850 | 1 | 0.2um_NOKCPHJN_649562 | Bacteria | Proteobacteri | Alphaproteob | Pelagibacteral | Pelagibacterac | unclassified | Pelagibacterac | cellular proces | Cell wall/mem | 7257 | NA | 439493.PB721 | 1,4 | 0,489897949 | 8,88E-05 | 3,27E-05 | 1 - | 5,29E-05 - | 2 - | 0,000131354 - | 2 - | 0,000124868 - | 1 - | 6,34E-05 - | 1 - | 7,14E-05 - |
| 7851 | 1 | 3um_FIJJAFBF_653705 | Bacteria | Actinobacteria | Actinobacteria | Micrococcales | Microbacteria | Candidatus Ac | Candidatus Ac | cellular proces | Posttranslatio | 05DIW | FeS assembly | 979556.MTES | 0,4 | 0,489897949 | 1,06E-05 | 1,31E-05 | 1 - | 2,43E-05 - | 0 - | 0 - | 1 - | 2,87E-05 - | 0 - | 0 - | 0 - | 0 - |
| 7852 | 3 | 0.2um_NOKCPHJN_292421;0.2um_NOKCPHJN_8 | Bacteria | Bacteroidetes | Flavobacteriia | Flavobacterial | Crocinitomica | unclassified | various | unclassified | unclassified | unclassified | unclassified | unclassified | 0,6 | 0,489897949 | 0,00010564 | 8,67E-05 | 0 - | 0 - | 0 - | 0 - | 1 - | 0,000167262 - | 1 - | 0,000169732 - | 1 - | 0,000191203 - |
| 7853 | 1 | 3um_FIJJAFBF_09975 | Bacteria | Bacteroidetes | Flavobacteriia | Flavobacterial | Flavobacteriac | Polaribacter | Polaribacter s | unclassified | unclassified | unclassified | unclassified | unclassified | 1 | 0,632455532 | 8,48E-05 | 5,49E-05 | 1 - | 7,31E-05 - | 1 - | 9,08E-05 - | 2 - | 0,000172589 - | 1 - | 8,76E-05 - | 0 - | 0 - |
| 7854 | 1 | 0.2um_NOKCPHJN_72831 | Bacteria | Proteobacteri | Gammaprotec | Pseudomonad | Pseudomonad | Pseudomonas | Pseudomonas | information st | Replication, re | 05C68 | Can catalyze t | 390235.PputW | 2 | 0,632455532 | 7,23E-05 | 1,89E-05 | 3 - | 9,23E-05 - | 2 - | 7,64E-05 - | 2 - | 7,26E-05 - | 1 - | 3,69E-05 - | 2 - | 8,30E-05 - |
| 7855 | 1 | 0.2um_NOKCPHJN_409524 | Bacteria | Proteobacteri | Alphaproteob | Rhodospirillale | Rhodospirillac | unclassified | Rhodospirillac | unclassified | unclassified | unclassified | unclassified | unclassified | 2 | 0,894427191 | 8,37E-05 | 3,27E-05 | 3 - | 0,000108965 - | 1 - | 4,51E-05 - | 3 - | 0,000128622 - | 2 - | 8,70E-05 - | 1 - | 4,90E-05 - |
| 7856 | 1 | 0.2um_NOKCPHJN_451728 | Bacteria | Proteobacteri | Alphaproteob | Rhodobactera | unclassified | unclassified | Rhodobactera | unclassified | unclassified | unclassified | unclassified | unclassified | 0,8 | 0,4 | 5,83E-05 | 2,99E-05 | 1 - | 6,11E-05 - | 1 - | 7,58E-05 - | 1 - | 7,21E-05 - | 0 - | 0 - | 1 - | 8,24E-05 - |
| 7857 | 1 | 0.2um_NOKCPHJN_284053 | Bacteria | Bacteroidetes | Flavobacteriia | Flavobacterial | Flavobacteriac | Lutibacter | Lutibacter flav | poorly charact | Function unkn | 0P8NI | Inherit from b | 313594.PI23P | 3 | 0,632455532 | 6,39E-05 | 1,75E-05 | 3 - | 5,32E-05 - | 2 - | 4,41E-05 - | 3 - | 6,28E-05 - | 3 - | 6,37E-05 - | 4 - | 9,57E-05 - |
| 7858 | 1 | 3um_FIJJAFBF_491688 | Bacteria | Proteobacteri | Gammaprotec | Cellvibrionales | Porticoccacea | unclassified | SAR92 bacteri | information st | Transcription | 16PMT | DNA-depende | 314287.GB220 | 0,4 | 0,489897949 | 1,55E-05 | 1,92E-05 | 1 - | 3,47E-05 - | 1 - | 4,31E-05 - | 0 - | 0 - | 0 - | 0 - | 0 - | 0 - |
| 7859 | 1 | 3um_FIJJAFBF_263325 | Bacteria | Bacteroidetes | Flavobacteriia | Flavobacterial | Flavobacteriac | Polaribacter | Polaribacter s | metabolism | Energy produc | 09A0E | phosphoenolp | 313598.MED1 | 0,4 | 0,489897949 | 5,87E-06 | 7,24E-06 | 1 - | 1,34E-05 - | 0 - | 0 - | 0 - | 0 - | 1 - | 1,60E-05 - | 0 - | 0 - |

| | | | | | | | | | | | | | | | | | | | | | | | | | | | | |
|---|---|---|---|---|---|---|---|---|---|---|---|---|---|---|---|---|---|---|---|---|---|---|---|---|---|---|---|---|
| 7860 | 1 | 0.2um_NOKCPHJN_822824 | Eukaryota | unclassified | Spirotrichea | Sporadotrichi | Oxytrichidae | Stylonychia | Stylonychia le | metabolism | Energy produc | 0IFIU | Pyrophosphat | 412030.XP_00 | 0,8 | 0,4 | 4,84E-05 | 2,46E-05 | 1 - | 5,24E-05 - | 1 - | 6,51E-05 - | 1 - | 6,19E-05 - | 1 - | 6,28E-05 - | 0 - | 0 - |
| 7861 | 1 | 0.2um_NOKCPHJN_182121 | Bacteria | Bacteroidetes | Flavobacteriia | Flavobacterial | Flavobacteriac | unclassified | Flavobacteriac | metabolism | Inorganic ion t | COG0841 | acriflavin resis | 886377.Murru | 0,6 | 0,489897949 | 6,90E-06 | 5,69E-06 | 1 - | 1,00E-05 - | 1 - | 1,25E-05 - | 0 - | 0 - | 1 - | 1,20E-05 - | 0 - | 0 - |
| 7862 | 1 | 0.2um_NOKCPHJN_292104 | Bacteria | Proteobacteri | Gammaprote | unclassified | unclassified | unclassified | OM182 bacte | metabolism | Energy produc | 05CV4 | Subunits I and | 247633.GP214 | 0,8 | 0,4 | 2,32E-05 | 1,18E-05 | 1 - | 2,51E-05 - | 1 - | 3,12E-05 - | 1 - | 2,96E-05 - | 1 - | 3,01E-05 - | 0 - | 0 - |
| 7863 | 1 | 0.2um_NOKCPHJN_12460 | Bacteria | Bacteroidetes | Saprospiria | Saprospirales | Lewinellaceae | Lewinella | Lewinella pers | metabolism | Carbohydrate | 0NIRC | arabinogalact | 573061.Clocel | 7,8 | 1,469693846 | 0,00018591 | 3,02E-05 | 9 - | 0,000181545 - | 9 - | 0,000225425 - | 9 - | 0,000214295 - | 6 - | 0,000144974 - | 6 - | 0,000163312 - |
| 7864 | 1 | 3um_FIJJAFBF_676118 | Bacteria | Bacteroidetes | Cytophagia | Cytophagales | Flammeovirga | unclassified | Flammeovirga | unclassified | unclassified | unclassified | unclassified | unclassified | 2,2 | 0,4 | 0,00018033 | 3,52E-05 | 2 - | 0,000137457 - | 2 - | 0,000170681 - | 3 - | 0,000243381 - | 2 - | 0,000164651 - | 2 - | 0,000185478 - |
| 7865 | 1 | 3um_FIJJAFBF_726475 | Bacteria | Proteobacteri | Alphaproteob | Rhodobactera | unclassified | unclassified | Rhodobactera | metabolism | Energy produc | 01RNN | ferredoxin--na | 314271.RB265 | 0,2 | 0,4 | 8,00E-06 | 1,60E-05 | 1 - | 4,00E-05 - | 0 - | 0 - | 0 - | 0 - | 0 - | 0 - | 0 - | 0 - |
| 7866 | 1 | 3um_FIJJAFBF_645893 | Bacteria | Proteobacteri | Alphaproteob | Rhodobactera | Rhodobactera | Roseovarius | Roseovarius a | metabolism | Amino acid tra | COG0174 | glutamine syn | 89187.ISM_05 | 0,4 | 0,489897949 | 1,10E-05 | 1,36E-05 | 1 - | 2,45E-05 - | 1 - | 3,05E-05 - | 0 - | 0 - | 0 - | 0 - | 0 - | 0 - |
| 7867 | 1 | 3um_FIJJAFBF_645166 | Bacteria | Proteobacteri | Gammaprote | Alteromonada | various | various | various | information st | Translation, ri | 0QTKZ | 30S ribosomal | 342610.Patl_1 | 0,4 | 0,489897949 | 8,38E-05 | 0,000102748 | 0 - | 0 - | 1 - | 0,000200731 - | 0 - | 0 - | 0 - | 0 - | 1 - | 0,000218133 - |
| 7868 | 1 | 3um_FIJJAFBF_805938 | unclassified | unclassified | unclassified | unclassified | unclassified | unclassified | unclassified | unclassified | unclassified | unclassified | unclassified | unclassified | 1,6 | 0,489897949 | 0,000178168 | 4,94E-05 | 2 - | 0,000189714 - | 2 - | 0,000235568 - | 2 - | 0,000223938 - | 1 - | 0,000113623 - | 1 - | 0,000127995 - |
| 7869 | 1 | 0.2um_NOKCPHJN_845529 | Bacteria | Bacteroidetes | Flavobacteriia | Flavobacterial | Flavobacteriac | Tenacibaculur | Tenacibaculur | information st | Replication, re | COG0513 | purine NTP-de | 313598.MED1 | 1,4 | 0,8 | 4,30E-05 | 2,46E-05 | 2 - | 5,35E-05 - | 2 - | 6,64E-05 - | 2 - | 6,32E-05 - | 1 - | 3,20E-05 - | 0 - | 0 - |
| 7870 | 1 | 3um_FIJJAFBF_540523 | Bacteria | Proteobacteri | Alphaproteob | Rhodobactera | Rhodobactera | Planktomarina | Planktomarina | information st | Translation, ri | 177SZ | This protein bi | 644107.SL115 | 4,4 | 0,489897949 | 0,000405174 | 6,10E-05 | 4 - | 0,000308126 - | 5 - | 0,000478251 - | 5 - | 0,000454638 - | 4 - | 0,000369083 - | 4 - | 0,00041577 - |
| 7871 | 1 | 3um_FIJJAFBF_447102 | Bacteria | Proteobacteri | Betaproteoba | Burkholderial | Comamonada | Acidovorax | Acidovorax sp | metabolism | Energy produc | 0B8JG | Component of | 296591.Bpro_ | 0,4 | 0,489897949 | 5,57E-06 | 6,87E-06 | 1 - | 1,28E-05 - | 0 - | 0 - | 1 - | 1,51E-05 - | 0 - | 0 - | 0 - | 0 - |
| 7872 | 1 | 0.2um_NOKCPHJN_418692 | Bacteria | Proteobacteri | Alphaproteob | Pelagibacteral | Pelagibacterac | Candidatus Pe | Candidatus Pe | information st | Translation, ri | 08YY1 | This protein is | 335992.SAR11 | 0,4 | 0,489897949 | 3,22E-05 | 3,94E-05 | 0 - | 0 - | 1 - | 8,19E-05 - | 0 - | 0 - | 1 - | 7,90E-05 - | 0 - | 0 - |
| 7873 | 1 | 0.2um_NOKCPHJN_444454 | Bacteria | unclassified | unclassified | unclassified | unclassified | unclassified | uncultured ba | unclassified | unclassified | unclassified | unclassified | unclassified | 1,4 | 0,489897949 | 4,32E-05 | 1,77E-05 | 1 - | 2,53E-05 - | 1 - | 3,15E-05 - | 1 - | 2,99E-05 - | 2 - | 6,07E-05 - | 2 - | 6,84E-05 - |
| 7874 | 1 | 0.2um_NOKCPHJN_240819 | Bacteria | Bacteroidetes | Flavobacteriia | Flavobacterial | Flavobacteriac | Polaribacter | Polaribacter s | poorly charact | Function unkn | 09EUF | NA | 313598.MED1 | 2 | 0 | 0,000110953 | 1,05E-05 | 2 - | 9,29E-05 - | 2 - | 0,0001154 - | 2 - | 0,000109702 - | 2 - | 0,000111323 - | 2 - | 0,000125404 - |
| 7875 | 1 | 0.2um_NOKCPHJN_565444 | Bacteria | Proteobacteri | Alphaproteob | Rhodospirillal | Rhodospirillac | unclassified | Rhodospirillac | unclassified | unclassified | unclassified | unclassified | unclassified | 12,2 | 1,469693846 | 0,000534578 | 3,94E-05 | 15 - | 0,000555371 - | 11 - | 0,000505712 - | 12 - | 0,000524447 - | 11 - | 0,000487844 - | 12 - | 0,000599514 - |
| 7876 | 1 | 3um_FIJJAFBF_310874 | Bacteria | Bacteroidetes | Flavobacteriia | Flavobacterial | Crocinitomica | unclassified | Crocinitomica | unclassified | unclassified | unclassified | unclassified | unclassified | 1,6 | 0,489897949 | 0,000107255 | 2,97E-05 | 2 - | 0,000114206 - | 2 - | 0,00014181 - | 2 - | 0,000134808 - | 1 - | 6,84E-05 - | 1 - | 7,71E-05 - |
| 7877 | 1 | 0.2um_NOKCPHJN_571857 | Bacteria | Proteobacteri | Gammaprote | Cellvibrionale | Porticoccacea | unclassified | SAR92 bacteri | information st | Translation, ri | 0QTKX | Seems to be th | 314287.GB220 | 2 | 0 | 0,000217503 | 2,07E-05 | 2 - | 0,000182185 - | 2 - | 0,00022622 - | 2 - | 0,000215051 - | 2 - | 0,000218227 - | 2 - | 0,000245833 - |
| 7878 | 3 | 3um_FIJJAFBF_218562;3um_FIJJAFBF_301411;3u | Bacteria | Bacteroidetes | Flavobacteriia | Flavobacterial | Flavobacteriac | Formosa | various | poorly charact | Function unkn | 09AZX | Protein of unk | 391598.FBBAL | 1 | 0 | 4,22E-05 | 4,00E-06 | 1 - | 3,53E-05 - | 1 - | 4,39E-05 - | 1 - | 4,17E-05 - | 1 - | 4,23E-05 - | 1 - | 4,77E-05 - |
| 7879 | 1 | 3um_FIJJAFBF_497402 | Bacteria | Bacteroidetes | Flavobacteriia | Flavobacterial | Flavobacteriac | Arenibacter | Arenibacter c | metabolism | Carbohydrate | COG0057 | glyceraldehyd | 313603.FB217 | 3 | 0 | 0,000123078 | 1,17E-05 | 3 - | 0,000103093 - | 3 - | 0,000128011 - | 3 - | 0,000121691 - | 3 - | 0,000123488 - | 3 - | 0,000139109 - |
| 7880 | 1 | 3um_FIJJAFBF_527770 | Bacteria | Actinobacteri | Actinobacteri | Micrococcales | Microbacteria | Rhodoluna | Rhodoluna lac | unclassified | unclassified | unclassified | unclassified | unclassified | 19,4 | 0,489897949 | 0,000263302 | 2,93E-05 | 19 - | 0,000215703 - | 19 - | 0,00026784 - | 19 - | 0,000254615 - | 20 - | 0,000271975 - | 20 - | 0,000306379 - |
| 7881 | 1 | 3um_FIJJAFBF_225133 | Bacteria | Proteobacteri | Alphaproteob | Rhodobactera | Rhodobactera | Planktomarina | Planktomarina | information st | Translation, ri | 179ZI | One of the pri | 633131.TR2A6 | 6,6 | 0,489897949 | 0,000695069 | 8,19E-05 | 7 - | 0,000618029 - | 6 - | 0,000657779 - | 6 - | 0,000625303 - | 7 - | 0,000740295 - | 7 - | 0,000833939 - |
| 7882 | 1 | 0.2um_NOKCPHJN_539923 | Bacteria | Proteobacteri | Gammaprote | Cellvibrionale | Spongiibacter | Oceanicoccus | Oceanicoccus | cellular proces | Posttranslatio | 05C6P | ATP-depender | 247634.GPB21 | 0,2 | 0,4 | 3,40E-06 | 6,81E-06 | 0 - | 0 - | 0 - | 0 - | 0 - | 0 - | 1 - | 1,70E-05 - | 0 - | 0 - |
| 7883 | 1 | 3um_FIJJAFBF_350775 | Bacteria | Bacteroidetes | Flavobacteriia | Flavobacterial | Flavobacteriac | Arenitalea | Arenitalea lut | poorly charact | Function unkn | 09EUF | NA | 1046627.BZAR | 1,6 | 0,489897949 | 8,90E-05 | 2,80E-05 | 2 - | 9,37E-05 - | 1 - | 5,82E-05 - | 2 - | 0,000110598 - | 1 - | 5,61E-05 - | 2 - | 0,000126428 - |
| 7884 | 2 | 0.2um_NOKCPHJN_838813;3um_FIJJAFBF_34512 | Bacteria | Proteobacteri | Gammaprote | various | various | various | various | information st | Translation, ri | 0QTKW | Seems to be th | various | 0,6 | 0,8 | 7,04E-05 | 9,23E-05 | 0 - | 0 - | 2 - | 0,00022803 - | 0 - | 0 - | 0 - | 0 - | 1 - | 0,0001239 - |
| 7885 | 1 | 3um_FIJJAFBF_737462 | Bacteria | Proteobacteri | Gammaprote | Oceanospirilla | Saccharospirill | Reinekea | Reinekea blan | information st | Translation, ri | 05G6W | One of the pri | 314283.MED2 | 6,8 | 1,166190379 | 0,000445886 | 3,78E-05 | 9 - | 0,000501452 - | 6 - | 0,000415103 - | 6 - | 0,000394608 - | 7 - | 0,000467176 - | 6 - | 0,000451091 - |
| 7886 | 1 | 3um_FIJJAFBF_127758 | Bacteria | Bacteroidetes | Flavobacteriia | Flavobacterial | Flavobacteriac | Algibacter | Algibacter sp. | information st | Translation, ri | COG0254 | 50S ribosomal | 391603.FBALC | 1 | 0 | 0,000165093 | 1,57E-05 | 1 - | 0,000138285 - | 1 - | 0,000171709 - | 1 - | 0,000163232 - | 1 - | 0,000165643 - | 1 - | 0,000186596 - |
| 7887 | 1 | 3um_FIJJAFBF_30322 | Bacteria | Bacteroidetes | Flavobacteriia | Flavobacterial | Flavobacteriac | Polaribacter | Polaribacter b | information st | Transcription | 09EME | Necessary for | 313598.MED1 | 2,4 | 0,489897949 | 0,000209971 | 4,84E-05 | 2 - | 0,000146212 - | 3 - | 0,000272329 - | 3 - | 0,000258883 - | 2 - | 0,000175138 - | 2 - | 0,000197292 - |
| 7888 | 1 | 3um_FIJJAFBF_376034 | Bacteria | Bacteroidetes | Cytophagia | Cytophagales | Flammeovirga | Sediminitomi | Sediminitomi | unclassified | unclassified | unclassified | unclassified | unclassified | 0,4 | 0,489897949 | 9,55E-06 | 1,18E-05 | 1 - | 2,13E-05 - | 1 - | 2,64E-05 - | 0 - | 0 - | 0 - | 0 - | 0 - | 0 - |
| 7889 | 1 | 3um_FIJJAFBF_291320 | Bacteria | Proteobacteri | Alphaproteob | Rhodobactera | Rhodobactera | various | various | cellular proces | Posttranslatio | COG0330 | Band 7 protei | 391589.RGAI1 | 14,2 | 0,748331477 | 0,000493722 | 4,01E-05 | 15 - | 0,000438079 - | 13 - | 0,000471436 - | 14 - | 0,000482634 - | 15 - | 0,000524745 - | 14 - | 0,000551716 - |
| 7890 | 1 | 0.2um_NOKCPHJN_813798 | Bacteria | Bacteroidetes | Flavobacteriia | Flavobacterial | Flavobacteriac | Polaribacter | various | metabolism | Inorganic ion t | 0PA08 | Destroys radic | 391587.KAOT | 1 | 0 | 7,33E-05 | 6,96E-06 | 1 - | 6,14E-05 - | 1 - | 7,62E-05 - | 1 - | 7,25E-05 - | 1 - | 7,35E-05 - | 1 - | 8,28E-05 - |
| 7891 | 1 | 0.2um_NOKCPHJN_533955 | Eukaryota | unclassified | Oomycetes | Saprolegniale | Saprolegniace | Achlya | Achlya bisexu | metabolism | Carbohydrate | COG0057 | glyceraldehyd | 65071.PYU1_T | 1 | 0 | 3,98E-05 | 3,78E-06 | 1 - | 3,34E-05 - | 1 - | 4,14E-05 - | 1 - | 3,94E-05 - | 1 - | 4,00E-05 - | 1 - | 4,50E-05 - |
| 7892 | 1 | 3um_FIJJAFBF_551811 | unclassified | unclassified | unclassified | unclassified | unclassified | unclassified | unclassified | unclassified | unclassified | unclassified | unclassified | unclassified | 0,4 | 0,489897949 | 1,25E-05 | 1,54E-05 | 1 - | 2,87E-05 - | 0 - | 0 - | 1 - | 3,39E-05 - | 0 - | 0 - | 0 - | 0 - |
| 7893 | 1 | 0.2um_NOKCPHJN_515829 | Archaea | Euryarchaeota | unclassified | unclassified | unclassified | unclassified | Euryarchaeota | metabolism | Inorganic ion t | 09D2N | TonB depende | 484018.BACPI | 0,2 | 0,4 | 2,23E-06 | 4,45E-06 | 1 - | 1,11E-05 - | 0 - | 0 - | 0 - | 0 - | 0 - | 0 - | 0 - | 0 - |
| 7894 | 1 | 0.2um_NOKCPHJN_924353 | Bacteria | Bacteroidetes | Flavobacteriia | Flavobacterial | Flavobacteriac | Tenacibaculur | Tenacibaculur | poorly charact | Function unkn | 11P35 | NA | 983544.Lacal_ | 0,4 | 0,489897949 | 9,98E-06 | 1,22E-05 | 0 - | 0 - | 0 - | 0 - | 1 - | 2,48E-05 - | 1 - | 2,51E-05 - | 0 - | 0 - |
| 7895 | 1 | 3um_FIJJAFBF_532370 | Bacteria | Proteobacteri | Betaproteoba | unclassified | unclassified | unclassified | Betaproteoba | poorly charact | Function unkn | 17G76 | tol-pal system | 543728.Vapar | 0,4 | 0,489897949 | 1,97E-05 | 2,45E-05 | 1 - | 4,19E-05 - | 0 - | 0 - | 0 - | 0 - | 0 - | 0 - | 1 - | 5,65E-05 - |
| 7896 | 1 | 3um_FIJJAFBF_37762 | Eukaryota | Bacillariophyt | various | various | various | various | various | metabolism | Energy produc | 0J0AS | This b-type cy | 35128.Thapsd | 3 | 1,095445115 | 0,000485175 | 0,000172736 | 4 - | 0,000546556 - | 4 - | 0,000678661 - | 3 - | 0,000483865 - | 1 - | 0,000163671 - | 3 - | 0,000553123 - |
| 7897 | 1 | 3um_FIJJAFBF_811878 | Bacteria | Proteobacteri | Gammaprote | Oceanospirilla | Saccharospirill | Reinekea | Reinekea fors | poorly charact | Function unkn | 0QITB | UPF0070 prot | 314283.MED2 | 2,4 | 1,356465997 | 0,000149722 | 6,75E-05 | 5 - | 0,000273278 - | 1 - | 6,79E-05 - | 2 - | 0,000129031 - | 2 - | 0,000130936 - | 2 - | 0,0001475 - |
| 7898 | 1 | 0.2um_NOKCPHJN_442860 | Bacteria | Bacteroidetes | Flavobacteriia | Flavobacterial | unclassified | unclassified | Flavobacterial | unclassified | unclassified | unclassified | unclassified | unclassified | 0,4 | 0,489897949 | 1,08E-05 | 1,33E-05 | 0 - | 0 - | 1 - | 2,59E-05 - | 0 - | 0 - | 0 - | 0 - | 1 - | 2,82E-05 - |
| 7899 | 1 | 0.2um_NOKCPHJN_1032972 | Eukaryota | unclassified | Cryptophyta | Pyrenomonad | Geminigerace | Guillardia | Guillardia thet | metabolism | Lipid transpor | COG0439 | acetyl-CoA car | 5811.TGME49 | 0,8 | 0,4 | 5,25E-05 | 2,67E-05 | 1 - | 5,68E-05 - | 1 - | 7,06E-05 - | 1 - | 6,71E-05 - | 1 - | 6,81E-05 - | 0 - | 0 - |
| 7900 | 1 | 3um_FIJJAFBF_35352 | Bacteria | Bacteroidetes | Flavobacteriia | unclassified | unclassified | unclassified | Flavobacteria | cellular proces | Posttranslatio | COG0330 | Band 7 protei | 487796.Flav2A | 1,8 | 0,748331477 | 8,88E-05 | 3,18E-05 | 3 - | 0,000126592 - | 1 - | 5,24E-05 - | 1 - | 4,98E-05 - | 2 - | 0,000101091 - | 2 - | 0,000113878 - |
| 7901 | 1 | 0.2um_NOKCPHJN_924576 | Bacteria | Proteobacteri | Gammaprote | Pseudomonad | Pseudomonad | Pseudomonas | Pseudomonas | metabolism | Amino acid tra | 05CEU | Arginine dihyc | 746360.PFWH | 0,4 | 0,8 | 1,63E-05 | 3,25E-05 | 0 - | 0 - | 0 - | 0 - | 2 - | 8,14E-05 - | 0 - | 0 - | 0 - | 0 - |
| 7902 | 1 | 3um_FIJJAFBF_111302 | unclassified | unclassified | unclassified | unclassified | unclassified | unclassified | unclassified | unclassified | unclassified | unclassified | unclassified | unclassified | 1 | 0,632455532 | 3,03E-05 | 1,96E-05 | 1 - | 2,61E-05 - | 1 - | 3,25E-05 - | 2 - | 6,17E-05 - | 1 - | 3,13E-05 - | 0 - | 0 - |
| 7903 | 1 | 0.2um_NOKCPHJN_639776 | Bacteria | Bacteroidetes | Flavobacteriia | Flavobacterial | Flavobacteriac | Dokdonia | Dokdonia don | poorly charact | Function unkn | 05ERK | gliding motilit | 313590.MED1 | 12,4 | 1,743559577 | 0,000389472 | 4,82E-05 | 13 - | 0,000344595 - | 11 - | 0,000362057 - | 13 - | 0,00040676 - | 15 - | 0,00047627 - | 10 - | 0,000357678 - |
| 7904 | 1 | 3um_FIJJAFBF_241338 | Bacteria | Proteobacteri | Alphaproteob | Rhodobactera | unclassified | unclassified | Rhodobactera | information st | Translation, ri | 177SZ | This protein bi | 367336.OM22 | 1,2 | 0,748331477 | 9,69E-05 | 5,74E-05 | 2 - | 0,000139972 - | 0 - | 0 - | 1 - | 8,26E-05 - | 2 - | 0,000167663 - | 1 - | 9,44E-05 - |
| 7905 | 1 | 3um_FIJJAFBF_135024 | Bacteria | Bacteroidetes | Flavobacteriia | Flavobacterial | Flavobacteriac | Polaribacter | various | cellular proces | Posttranslatio | 07SFW | Trigger factor | 487797.Flav3C | 1,2 | 0,748331477 | 3,82E-05 | 2,60E-05 | 1 - | 2,60E-05 - | 2 - | 6,45E-05 - | 1 - | 3,07E-05 - | 0 - | 0 - | 2 - | 7,01E-05 - |
| 7906 | 1 | 3um_FIJJAFBF_569212 | Eukaryota | unclassified | Pelagophycea | Pelagomonad | unclassified | Aureococcus | Aureococcus a | metabolism | Carbohydrate | COG0057 | glyceraldehyd | 35128.Thaps3 | 2 | 1,095445115 | 0,000107607 | 5,89E-05 | 2 - | 9,07E-05 - | 1 - | 5,63E-05 - | 4 - | 0,000214201 - | 1 - | 5,43E-05 - | 2 - | 0,00012243 - |
| 7907 | 1 | 3um_FIJJAFBF_396484 | Bacteria | Proteobacteri | Gammaprote | Oceanospirilla | Saccharospirill | Reinekea | Reinekea fors | unclassified | unclassified | unclassified | unclassified | unclassified | 1 | 0 | 6,31E-05 | 6,00E-06 | 1 - | 5,29E-05 - | 1 - | 6,57E-05 - | 1 - | 6,24E-05 - | 1 - | 6,34E-05 - | 1 - | 7,14E-05 - |
| 7908 | 1 | 3um_FIJJAFBF_322026 | Bacteria | Proteobacteri | Gammaprote | Oceanospirilla | Saccharospirill | Reinekea | Reinekea fors | cellular proces | Posttranslatio | 0QW5X | glutaredoxin 3 | 314283.MED2 | 0,6 | 0,489897949 | 9,55E-05 | 7,80E-05 | 0 - | 0 - | 1 - | 0,000163815 - | 1 - | 0,000155727 - | 1 - | 0,000158027 - | 0 - | 0 - |
| 7909 | 1 | 0.2um_NOKCPHJN_726453 | Bacteria | Proteobacteri | Gammaprote | unclassified | unclassified | Candidatus Th | uncultured SU | information st | Translation, ri | 05CEJ | Catalyzes the | 413404.Rmag_ | 3 | 0 | 9,93E-05 | 9,43E-06 | 3 - | 8,32E-05 - | 3 - | 0,000103275 - | 3 - | 9,82E-05 - | 3 - | 9,96E-05 - | 3 - | 0,000112228 - |
| 7910 | 1 | 0.2um_NOKCPHJN_08438 | Bacteria | Proteobacteri | Gammaprote | Alteromonada | Alteromonada | Paraglaciecola | Paraglaciecola | cellular proces | Intracellular tr | 0QHXI | Signal peptida | 342610.Patl_3 | 0,2 | 0,4 | 7,63E-06 | 1,53E-05 | 1 - | 3,81E-05 - | 0 - | 0 - | 0 - | 0 - | 0 - | 0 - | 0 - | 0 - |
| 7911 | 1 | 0.2um_NOKCPHJN_554434 | Bacteria | Proteobacteri | Gammaprote | Pseudomonad | Pseudomonad | Pseudomonas | Pseudomonas | unclassified | unclassified | unclassified | unclassified | unclassified | 0,2 | 0,4 | 6,32E-06 | 1,26E-05 | 0 - | 0 - | 0 - | 0 - | 1 - | 3,16E-05 - | 0 - | 0 - | 0 - | 0 - |
| 7912 | 1 | 3um_FIJJAFBF_20032 | Bacteria | Bacteroidetes | Flavobacteriia | Flavobacterial | Flavobacteriac | Polaribacter | Polaribacter s | cellular proces | Cell wall/mem | 05D28 | Specifically me | 313594.PI23P | 1 | 1,095445115 | 5,96E-05 | 6,04E-05 | 3 - | 0,000164751 - | 1 - | 6,82E-05 - | 1 - | 6,48E-05 - | 0 - | 0 - | 0 - | 0 - |
| 7913 | 1 | 3um_FIJJAFBF_482001 | Bacteria | Proteobacteri | Alphaproteob | Rhodobactera | Rhodobactera | unclassified | Rhodobactera | unclassified | unclassified | unclassified | unclassified | unclassified | 4 | 1,414213562 | 9,39E-05 | 3,21E-05 | 5 - | 9,95E-05 - | 2 - | 4,94E-05 - | 3 - | 7,04E-05 - | 6 - | 0,000142964 - | 4 - | 0,000107365 - |
| 7914 | 1 | 0.2um_NOKCPHJN_57129 | Bacteria | Proteobacteri | Gammaprote | Oceanospirilla | Saccharospirill | Reinekea | Reinekea blan | information st | Replication, re | COG0187 | DNA gyrase ne | 314283.MED2 | 2,6 | 0,8 | 4,45E-05 | 1,51E-05 | 2 - | 2,85E-05 - | 4 - | 7,08E-05 - | 2 - | 3,37E-05 - | 3 - | 5,12E-05 - | 2 - | 3,85E-05 - |
| 7915 | 1 | 3um_FIJJAFBF_805172 | Bacteria | Proteobacteri | Alphaproteob | Rhodobactera | Rhodobactera | Jannaschia | Jannaschia sp | unclassified | unclassified | unclassified | unclassified | unclassified | 0,2 | 0,4 | 8,33E-06 | 1,67E-05 | 0 - | 0 - | 1 - | 4,17E-05 - | 0 - | 0 - | 0 - | 0 - | 0 - | 0 - |
| 7916 | 1 | 3um_FIJJAFBF_399524 | Bacteria | Proteobacteri | Gammaprote | Alteromonada | Alteromonada | Paraglaciecola | Paraglaciecola | metabolism | Lipid transpor | 0QIP1 | acyl-CoA dehy | 342610.Patl_0 | 1 | 0,632455532 | 3,39E-05 | 2,19E-05 | 1 - | 2,92E-05 - | 1 - | 3,63E-05 - | 2 - | 6,89E-05 - | 1 - | 3,50E-05 - | 0 - | 0 - |
| 7917 | 4 | 0.2um_NOKCPHJN_176161;0.2um_NOKCPHJN_3 | Bacteria | Bacteroidetes | Flavobacteriia | Flavobacterial | Flavobacteriac | Polaribacter | various | unclassified | unclassified | unclassified | unclassified | unclassified | 3 | 0 | 7,16E-05 | 6,80E-06 | 3 - | 6,00E-05 - | 3 - | 7,45E-05 - | 3 - | 7,08E-05 - | 3 - | 7,19E-05 - | 3 - | 8,09E-05 - |
| 7918 | 1 | 3um_FIJJAFBF_303920 | Bacteria | Bacteroidetes | Flavobacteriia | Flavobacterial | Crocinitomica | unclassified | Crocinitomica | cellular proces | Cell wall/mem | 09B44 | efflux transpo | 313596.RB250 | 1,8 | 0,748331477 | 7,17E-05 | 3,00E-05 | 1 - | 3,33E-05 - | 2 - | 8,26E-05 - | 3 - | 0,000117811 - | 2 - | 7,97E-05 - | 1 - | 4,49E-05 - |
| 7919 | 1 | 3um_FIJJAFBF_708513 | Bacteria | Cyanobacteria | unclassified | Synechococca | Synechococca | Synechococcu | Synechococcu | unclassified | unclassified | unclassified | unclassified | unclassified | 1,2 | 0,979795897 | 8,59E-05 | 6,07E-05 | 3 - | 0,000190238 - | 1 - | 7,87E-05 - | 1 - | 7,49E-05 - | 0 - | 0 - | 1 - | 8,56E-05 - |
| 7920 | 1 | 3um_FIJJAFBF_164710 | Bacteria | Actinobacteri | Actinobacteri | Micrococcales | Microbacteria | various | various | information st | Translation, ri | 0826X | This protein bi | 281090.Lxx08 | 2 | 0,632455532 | 0,000275181 | 0,000106218 | 2 - | 0,000227281 - | 2 - | 0,000282216 - | 1 - | 0,000134141 - | 2 - | 0,000272244 - | 3 - | 0,000460023 - |
| 7921 | 1 | 0.2um_NOKCPHJN_109745 | Bacteria | Bacteroidetes | Flavobacteriia | Flavobacterial | Flavobacteriac | Formosa | Formosa sp. H | unclassified | unclassified | unclassified | unclassified | unclassified | 1 | 0 | 0,000103808 | 9,86E-06 | 1 - | 8,70E-05 - | 1 - | 0,000107969 - | 1 - | 0,000102638 - | 1 - | 0,000104154 - | 1 - | 0,000117329 - |
| 7922 | 1 | 3um_FIJJAFBF_765131 | Bacteria | Proteobacteri | Betaproteoba | Burkholderial | unclassified | unclassified | various | unclassified | unclassified | unclassified | unclassified | unclassified | 1,2 | 1,166190379 | 0,000113748 | 0,000112546 | 2 - | 0,000168789 - | 0 - | 0 - | 3 - | 0,000298858 - | 1 - | 0,000101091 - | 0 - | 0 - |
| 7923 | 1 | 3um_FIJJAFBF_746721 | Eukaryota | Chlorophyta | Mamiellophyc | Mamiellales | Mamiellaceae | Micromonas | Micromonas c | unclassified | unclassified | unclassified | unclassified | unclassified | 1,8 | 0,4 | 0,000166493 | 3,29E-05 | 2 - | 0,000157228 - | 2 - | 0,000195231 - | 2 - | 0,000185592 - | 2 - | 0,000188333 - | 1 - | 0,000106078 - |
| 7924 | 1 | 3um_FIJJAFBF_163911 | Bacteria | Proteobacteri | Alphaproteob | Rhodobactera | Rhodobactera | unclassified | Rhodobactera | unclassified | unclassified | unclassified | unclassified | unclassified | 1,2 | 0,4 | 4,78E-05 | 1,65E-05 | 1 - | 3,34E-05 - | 1 - | 4,14E-05 - | 1 - | 3,94E-05 - | 2 - | 7,99E-05 - | 1 - | 4,50E-05 - |
| 7925 | 1 | 3um_FIJJAFBF_284884 | Bacteria | Proteobacteri | Gammaprote | Oceanospirilla | Saccharospirill | Reinekea | Reinekea fors | metabolism | Carbohydrate | COG1105 | 1-phosphofru | 177437.HRM2 | 0,4 | 0,489897949 | 1,64E-05 | 2,02E-05 | 1 - | 3,76E-05 - | 0 - | 0 - | 1 - | 4,44E-05 - | 0 - | 0 - | 0 - | 0 - |
| 7926 | 1 | 3um_FIJJAFBF_04972 | Bacteria | Proteobacteri | Alphaproteob | Rhodobactera | Rhodobactera | Planktomarina | Planktomarina | poorly charact | Function unkn | 01Y94 | Protein of unk | 633131.TR2A6 | 0,4 | 0,489897949 | 3,68E-05 | 4,55E-05 | 1 - | 8,20E-05 - | 1 - | 0,000101799 - | 0 - | 0 - | 0 - | 0 - | 0 - | 0 - |
| 7927 | 3 | 0.2um_NOKCPHJN_202118;0.2um_NOKCPHJN_4 | Viruses | unclassified | unclassified | unclassified | unclassified | unclassified | uncultured M | unclassified | unclassified | unclassified | unclassified | unclassified | 0,6 | 0,489897949 | 4,43E-05 | 3,65E-05 | 1 - | 6,56E-05 - | 0 - | 0 - | 1 - | 7,74E-05 - | 1 - | 7,86E-05 - | 0 - | 0 - |
| 7928 | 1 | 3um_FIJJAFBF_592385 | Bacteria | Proteobacteri | Alphaproteob | Rhodobactera | Rhodobactera | Natronohydro | Natronohydro | metabolism | Coenzyme tra | 01Q80 | Glutamate--cy | 391589.RGAI1 | 0,4 | 0,489897949 | 1,73E-05 | 2,13E-05 | 1 - | 3,93E-05 - | 0 - | 0 - | 0 - | 0 - | 1 - | 4,71E-05 - | 0 - | 0 - |
| 7929 | 1 | 0.2um_NOKCPHJN_663396 | Eukaryota | unclassified | Pelagophycea | Pelagomonad | unclassified | Aureococcus | Aureococcus a | unclassified | unclassified | unclassified | unclassified | unclassified | 0,4 | 0,489897949 | 1,21E-05 | 1,48E-05 | 0 - | 0 - | 0 - | 0 - | 1 - | 2,82E-05 - | 0 - | 0 - | 1 - | 3,22E-05 - |
| 7930 | 1 | 0.2um_NOKCPHJN_21852 | Bacteria | Bacteroidetes | Flavobacteriia | Flavobacterial | Flavobacteriac | Polaribacter | Polaribacter s | poorly charact | Function unkn | 0PBET | NA | 313594.PI23P | 3,4 | 1,019803903 | 0,000259427 | 5,96E-05 | 5 - | 0,00032607 - | 3 - | 0,00024293 - | 4 - | 0,000307914 - | 2 - | 0,000156231 - | 3 - | 0,000263991 - |
| 7931 | 1 | 3um_FIJJAFBF_21301 | Bacteria | Bacteroidetes | Flavobacteriia | Flavobacterial | Flavobacteriac | Polaribacter | Polaribacter s | metabolism | Carbohydrate | 07R4H | Transketolase | 313594.PI23P | 0,4 | 0,489897949 | 1,57E-05 | 1,93E-05 | 1 - | 3,60E-05 - | 0 - | 0 - | 1 - | 4,25E-05 - | 0 - | 0 - | 0 - | 0 - |
| 7932 | 1 | 0.2um_NOKCPHJN_20629 | Bacteria | unclassified | unclassified | unclassified | unclassified | unclassified | uncultured ma | metabolism | Lipid transpor | 05C9H | acyl-CoA dehy | 76869.PputGE | 0,8 | 0,4 | 2,63E-05 | 1,33E-05 | 1 - | 2,84E-05 - | 1 - | 3,53E-05 - | 1 - | 3,35E-05 - | 1 - | 3,40E-05 - | 0 - | 0 - |
| 7933 | 2 | 0.2um_NOKCPHJN_229752;3um_FIJJAFBF_77319 | Bacteria | Bacteroidetes | Cytophagia | Cytophagales | Cyclobacteriac | Cecembia | Cecembia lon | poorly charact | Function unkn | 0XP53 | Ragb susd dor | 388413.ALPR1 | 3 | 0 | 7,79E-05 | 7,39E-06 | 3 - | 6,52E-05 - | 3 - | 8,10E-05 - | 3 - | 7,70E-05 - | 3 - | 7,81E-05 - | 3 - | 8,80E-05 - |
| 7934 | 1 | 0.2um_NOKCPHJN_597103 | Bacteria | Proteobacteri | Alphaproteob | Pelagibacteral | Pelagibacterac | unclassified | Pelagibacterac | cellular proces | Cell wall/mem | 7257 | NA | 439493.PB721 | 1,8 | 0,4 | 0,000190349 | 4,45E-05 | 2 - | 0,000177948 - | 1 - | 0,00011048 - | 2 - | 0,00021005 - | 2 - | 0,000213152 - | 2 - | 0,000240115 - |
| 7935 | 1 | 0.2um_NOKCPHJN_446593 | Eukaryota | Phaeophyceae | unclassified | Ectocarpales | Ectocarpaceae | Ectocarpus | Ectocarpus sil | poorly charact | Function unkn | 0XT7F | HEAT repeat c | 35128.Thaps5 | 0,4 | 0,489897949 | 1,03E-05 | 1,27E-05 | 1 - | 2,37E-05 - | 0 - | 0 - | 1 - | 2,79E-05 - | 0 - | 0 - | 0 - | 0 - |
| 7936 | 1 | 3um_FIJJAFBF_599427 | Bacteria | Proteobacteri | Alphaproteob | Rhodospirillal | Rhodospirillac | unclassified | Rhodospirillac | unclassified | unclassified | unclassified | unclassified | unclassified | 1 | 0 | 2,47E-05 | 2,34E-06 | 1 - | 2,07E-05 - | 1 - | 2,57E-05 - | 1 - | 2,44E-05 - | 1 - | 2,48E-05 - | 1 - | 2,79E-05 - |
| 7937 | 1 | 3um_FIJJAFBF_163205 | Bacteria | Bacteroidetes | Flavobacteriia | Flavobacterial | unclassified | unclassified | Flavobacterial | unclassified | unclassified | unclassified | unclassified | unclassified | 0,6 | 0,489897949 | 3,13E-05 | 2,56E-05 | 0 - | 0 - | 1 - | 5,13E-05 - | 0 - | 0 - | 1 - | 4,95E-05 - | 1 - | 5,57E-05 - |
| 7938 | 1 | 0.2um_NOKCPHJN_467041 | Bacteria | Proteobacteri | Alphaproteob | Rhodobactera | unclassified | unclassified | Rhodobactera | unclassified | unclassified | unclassified | unclassified | unclassified | 4,2 | 0,748331477 | 0,00027394 | 6,15E-05 | 3 - | 0,00016242 - | 5 - | 0,000336129 - | 5 - | 0,000319534 - | 4 - | 0,000259402 - | 4 - | 0,000292216 - |
| 7939 | 1 | 0.2um_NOKCPHJN_12465 | Bacteria | Bacteroidetes | Flavobacteriia | Flavobacterial | Flavobacteriac | Formosa | Formosa sp. H | cellular proces | Signal transdu | 09DA6 | Histidine kinas | 483215.BACFI | 4,6 | 1,959591794 | 4,78E-05 | 1,59E-05 | 8 - | 7,24E-05 - | 5 - | 5,62E-05 - | 4 - | 4,27E-05 - | 4 - | 4,34E-05 - | 2 - | 2,44E-05 - |
| 7940 | 1 | 3um_FIJJAFBF_499745 | Bacteria | Cyanobacteria | unclassified | Synechococca | Synechococca | Synechococcu | Synechococcu | poorly charact | Function unkn | 13PBK | Conserved hy | 29850.GGTG_ | 0,4 | 0,489897949 | 1,01E-05 | 1,25E-05 | 1 - | 2,25E-05 - | 1 - | 2,79E-05 - | 0 - | 0 - | 0 - | 0 - | 0 - | 0 - |
| 7941 | 1 | 0.2um_NOKCPHJN_1013070 | Eukaryota | unclassified | Pelagophycea | Pelagomonad | unclassified | Aureococcus | Aureococcus a | information st | Translation, ri | 1520E | S25 ribosomal | 4787.PITG_13 | 0,4 | 0,489897949 | 5,15E-05 | 6,31E-05 | 0 - | 0 - | 0 - | 0 - | 1 - | 0,000127813 - | 1 - | 0,000129701 - | 0 - | 0 - |
| 7942 | 1 | 0.2um_NOKCPHJN_516222 | Bacteria | Bacteroidetes | Flavobacteriia | unclassified | unclassified | unclassified | Flavobacteria | unclassified | unclassified | unclassified | unclassified | unclassified | 11,4 | 0,8 | 0,000184932 | 2,13E-05 | 12 - | 0,00016319 - | 11 - | 0,000185747 - | 12 - | 0,000192629 - | 10 - | 0,000162895 - | 12 - | 0,000220201 - |
| 7943 | 1 | 3um_FIJJAFBF_741975 | Bacteria | Proteobacteri | Gammaprote | Alteromonada | Alteromonada | Paraglaciecola | Paraglaciecola | unclassified | unclassified | unclassified | unclassified | unclassified | 0,4 | 0,489897949 | 2,31E-05 | 2,83E-05 | 0 - | 0 - | 0 - | 0 - | 1 - | 5,74E-05 - | 1 - | 5,83E-05 - | 0 - | 0 - |
| 7944 | 1 | 0.2um_NOKCPHJN_01039 | Bacteria | Bacteroidetes | Flavobacteriia | Flavobacterial | Flavobacteriac | Formosa | Formosa sp. H | cellular proces | Posttranslatio | COG0330 | Band 7 protei | 983544.Lacal_ | 3,8 | 1,166190379 | 0,000190145 | 5,19E-05 | 5 - | 0,00021255 - | 4 - | 0,000211139 - | 2 - | 0,000100357 - | 5 - | 0,000254599 - | 3 - | 0,000172083 - |
| 7945 | 1 | 3um_FIJJAFBF_611290 | Bacteria | Proteobacteri | Gammaprote | Oceanospirilla | Saccharospirill | Reinekea | Reinekea blan | metabolism | Inorganic ion t | 16RE3 | may be the G1 | 314283.MED2 | 2,2 | 0,979795897 | 5,63E-05 | 2,69E-05 | 2 - | 4,25E-05 - | 4 - | 0,00010557 - | 1 - | 2,51E-05 - | 2 - | 5,09E-05 - | 2 - | 5,74E-05 - |
| 7946 | 1 | 0.2um_NOKCPHJN_76328 | Bacteria | Bacteroidetes | Flavobacteriia | Flavobacterial | Flavobacteriac | unclassified | Flavobacteriac | information st | Translation, ri | 09AI8 | Associates wit | 391603.FBALC | 2,6 | 1,2 | 0,00010768 | 4,38E-05 | 4 - | 0,000143024 - | 1 - | 4,44E-05 - | 4 - | 0,000168825 - | 2 - | 8,57E-05 - | 2 - | 9,65E-05 - |
| 7947 | 1 | 3um_FIJJAFBF_286875 | Bacteria | Proteobacteri | Gammaprote | Cellvibrionale | Porticoccacea | unclassified | SAR92 bacteri | unclassified | unclassified | unclassified | unclassified | unclassified | 1,2 | 0,748331477 | 3,80E-05 | 2,23E-05 | 2 - | 5,50E-05 - | 0 - | 0 - | 2 - | 6,50E-05 - | 1 - | 3,30E-05 - | 1 - | 3,71E-05 - |
| 7948 | 1 | 0.2um_NOKCPHJN_60612 | Bacteria | Bacteroidetes | Flavobacteriia | Flavobacterial | Flavobacteriac | Polaribacter | Polaribacter r | information st | Transcription | 0P7W9 | Transcription | 313598.MED1 | 0,2 | 0,4 | 6,58E-06 | 1,32E-05 | 0 - | 0 - | 0 - | 0 - | 0 - | 0 - | 1 - | 3,29E-05 - | 0 - | 0 - |
| 7949 | 1 | 3um_FIJJAFBF_211240 | Bacteria | Proteobacteri | Gammaprote | Arenicellales | Arenicellaceae | Arenicella | Arenicella xan | information st | Translation, ri | 0QQXI | This protein is | 722419.PH505 | 0,4 | 0,489897949 | 4,35E-05 | 5,36E-05 | 1 - | 9,98E-05 - | 0 - | 0 - | 1 - | 0,000117811 - | 0 - | 0 - | 0 - | 0 - |
| 7950 | 2 | 0.2um_NOKCPHJN_13852;0.2um_NOKCPHJN_64 | Bacteria | Bacteroidetes | Flavobacteriia | Flavobacterial | Crocinitomica | unclassified | Crocinitomica | poorly charact | Function unkn | 08M59 | Protein of unk | 269798.CHU_ | 0,4 | 0,489897949 | 1,53E-05 | 1,89E-05 | 1 - | 3,49E-05 - | 0 - | 0 - | 0 - | 0 - | 1 - | 4,18E-05 - | 0 - | 0 - |
| 7951 | 1 | 0.2um_NOKCPHJN_584203 | Bacteria | Proteobacteri | Gammaprote | unclassified | unclassified | unclassified | Gammaprote | poorly charact | Function unkn | 16R8I | Lipoprotein | 314287.GB220 | 6 | 0,632455532 | 0,000149322 | 1,57E-05 | 6 - | 0,000125668 - | 6 - | 0,000156043 - | 7 - | 0,000173061 - | 6 - | 0,000150529 - | 5 - | 0,000141309 - |
| 7952 | 1 | 0.2um_NOKCPHJN_12278 | Eukaryota | unclassified | Bangiophycea | Bangiales | Bangiaceae | Porphyra | Porphyra umb | unclassified | unclassified | unclassified | unclassified | unclassified | 0,4 | 0,489897949 | 9,80E-06 | 1,21E-05 | 1 - | 2,19E-05 - | 1 - | 2,71E-05 - | 0 - | 0 - | 0 - | 0 - | 0 - | 0 - |
| 7953 | 1 | 0.2um_NOKCPHJN_140603 | Bacteria | Bacteroidetes | Flavobacteriia | unclassified | unclassified | unclassified | Flavobacteria | cellular proces | Intracellular tr | 0BIEB | domain-conta | 1005048.CFU_ | 20 | 1,897366596 | 0,000280616 | 4,30E-05 | 21 - | 0,000246453 - | 18 - | 0,000262305 - | 20 - | 0,00027706 - | 18 - | 0,000253037 - | 23 - | 0,000364224 - |
| 7954 | 1 | 0.2um_NOKCPHJN_396629 | Bacteria | Proteobacteri | Alphaproteob | Pelagibacteral | Pelagibacterac | Candidatus Pe | Candidatus Pe | metabolism | Amino acid tra | COG0141 | Catalyzes the | 335992.SAR11 | 0,6 | 0,489897949 | 1,84E-05 | 1,51E-05 | 1 - | 2,68E-05 - | 1 - | 3,33E-05 - | 1 - | 3,17E-05 - | 0 - | 0 - | 0 - | 0 - |
| 7955 | 1 | 0.2um_NOKCPHJN_648226 | Bacteria | Bacteroidetes | Flavobacteriia | Flavobacterial | Flavobacteriac | Winogradskye | Winogradskye | poorly charact | Function unkn | 0XY6Y | Pkd domain co | 391587.KAOT | 2 | 0,894427191 | 2,70E-05 | 1,23E-05 | 3 - | 3,41E-05 - | 1 - | 1,41E-05 - | 2 - | 2,69E-05 - | 1 - | 1,36E-05 - | 3 - | 4,60E-05 - |
| 7956 | 1 | 0.2um_NOKCPHJN_144449 | Bacteria | Proteobacteri | Gammaprote | Oceanospirilla | Oceanospirilla | Marinomonas | Marinomonas | metabolism | Carbohydrate | 16SJ5 | Trap dicarbox | 225937.HP15_ | 9 | 2,097617696 | 0,000378465 | 6,21E-05 | 13 - | 0,000464828 - | 8 - | 0,000355187 - | 7 - | 0,000295444 - | 8 - | 0,000342638 - | 9 - | 0,000434228 - |
| 7957 | 1 | 3um_FIJJAFBF_08215 | unclassified | unclassified | unclassified | unclassified | unclassified | unclassified | unclassified | unclassified | unclassified | unclassified | unclassified | unclassified | 0,4 | 0,489897949 | 4,36E-06 | 5,38E-06 | 1 - | 9,93E-06 - | 0 - | 0 - | 0 - | 0 - | 1 - | 1,19E-05 - | 0 - | 0 - |
| 7958 | 1 | 0.2um_NOKCPHJN_474225 | unclassified | unclassified | unclassified | unclassified | unclassified | unclassified | unclassified | unclassified | unclassified | unclassified | unclassified | unclassified | 0,8 | 0,4 | 0,000224348 | 0,000115263 | 1 - | 0,000234238 - | 1 - | 0,000290855 - | 0 - | 0 - | 1 - | 0,000280578 - | 1 - | 0,00031607 - |
| 7959 | 2 | 0.2um_NOKCPHJN_363820;0.2um_NOKCPHJN_6 | Bacteria | unclassified | unclassified | unclassified | unclassified | unclassified | uncultured ba | unclassified | unclassified | unclassified | unclassified | unclassified | 1 | 0,632455532 | 4,99E-05 | 3,16E-05 | 1 - | 4,19E-05 - | 1 - | 5,20E-05 - | 2 - | 9,89E-05 - | 0 - | 0 - | 1 - | 5,65E-05 - |
| 7960 | 1 | 0.2um_NOKCPHJN_31742 | Bacteria | Proteobacteri | Betaproteoba | unclassified | unclassified | unclassified | Betaproteoba | information st | Transcription | 0B7RW | Facilitates tra | 399795.CtesD | 1 | 1,095445115 | 3,34E-05 | 3,72E-05 | 0 - | 0 - | 3 - | 0,000101799 - | 1 - | 3,23E-05 - | 1 - | 3,27E-05 - | 0 - | 0 - |
| 7961 | 1 | 0.2um_NOKCPHJN_431174 | Bacteria | Proteobacteri | Alphaproteob | Rhodobactera | Rhodobactera | Planktomarina | Planktomarina | unclassified | unclassified | unclassified | unclassified | unclassified | 0,8 | 0,4 | 2,37E-05 | 1,21E-05 | 1 - | 2,57E-05 - | 1 - | 3,19E-05 - | 1 - | 3,03E-05 - | 1 - | 3,08E-05 - | 0 - | 0 - |
| 7962 | 1 | 3um_FIJJAFBF_77421 | Viruses | unclassified | unclassified | unclassified | various | various | various | unclassified | unclassified | unclassified | unclassified | unclassified | 5,8 | 0,4 | 0,000149846 | 1,09E-05 | 6 - | 0,000130428 - | 6 - | 0,000161953 - | 6 - | 0,000153957 - | 6 - | 0,000156231 - | 5 - | 0,000146661 - |
| 7963 | 1 | 0.2um_NOKCPHJN_684260 | Bacteria | Proteobacteri | Alphaproteob | Pelagibacteral | Pelagibacterac | Candidatus Pe | Candidatus Pe | metabolism | Amino acid tra | 01R4A | Polyamine AB | 439493.PB721 | 1,8 | 0,4 | 8,41E-05 | 1,66E-05 | 2 - | 7,94E-05 - | 2 - | 9,86E-05 - | 2 - | 9,38E-05 - | 2 - | 9,51E-05 - | 1 - | 5,36E-05 - |
| 7964 | 1 | 0.2um_NOKCPHJN_564252 | Bacteria | Bacteroidetes | Flavobacteriia | Flavobacterial | Flavobacteriac | Polaribacter | Polaribacter fi | metabolism | Amino acid tra | 099QE | chorismate m | 983544.Lacal_ | 1,2 | 0,4 | 5,27E-05 | 1,82E-05 | 1 - | 3,68E-05 - | 1 - | 4,57E-05 - | 1 - | 4,34E-05 - | 2 - | 8,81E-05 - | 1 - | 4,96E-05 - |
| 7965 | 1 | 3um_FIJJAFBF_427347 | Bacteria | Proteobacteri | Gammaprote | Cellvibrionale | Porticoccacea | unclassified | gamma prote | cellular proces | Signal transdu | 16Q0Y | phosphate reg | 314287.GB220 | 0,6 | 0,489897949 | 3,39E-05 | 2,82E-05 | 1 - | 4,78E-05 - | 0 - | 0 - | 0 - | 0 - | 1 - | 5,73E-05 - | 1 - | 6,45E-05 - |
| 7966 | 1 | 0.2um_NOKCPHJN_884409 | Bacteria | Proteobacteri | Betaproteoba | Nitrosomonad | Nitrosomonad | Nitrosomonas | Nitrosomonas | unclassified | unclassified | unclassified | unclassified | unclassified | 3,8 | 0,748331477 | 7,54E-05 | 9,76E-06 | 5 - | 8,44E-05 - | 4 - | 8,38E-05 - | 3 - | 5,98E-05 - | 4 - | 8,09E-05 - | 3 - | 6,83E-05 - |
| 7967 | 1 | 0.2um_NOKCPHJN_756207 | Bacteria | Bacteroidetes | Flavobacteriia | Flavobacterial | Crocinitomica | unclassified | Crocinitomica | unclassified | unclassified | unclassified | unclassified | unclassified | 0,4 | 0,489897949 | 3,17E-05 | 3,90E-05 | 1 - | 7,26E-05 - | 0 - | 0 - | 1 - | 8,57E-05 - | 0 - | 0 - | 0 - | 0 - |
| 7968 | 1 | 0.2um_NOKCPHJN_237430 | Bacteria | Proteobacteri | Gammaprote | unclassified | unclassified | unclassified | SAR86 cluster | metabolism | Energy produc | COG0056 | Produces ATP | 247634.GPB21 | 2,4 | 0,8 | 6,22E-05 | 1,38E-05 | 4 - | 8,93E-05 - | 2 - | 5,55E-05 - | 2 - | 5,27E-05 - | 2 - | 5,35E-05 - | 2 - | 6,03E-05 - |
| 7969 | 1 | 3um_FIJJAFBF_169480 | Bacteria | Bacteroidetes | Flavobacteriia | Flavobacterial | Flavobacteriac | Aequorivita | Aequorivita su | unclassified | unclassified | unclassified | unclassified | unclassified | 0,6 | 0,489897949 | 2,53E-05 | 2,07E-05 | 0 - | 0 - | 1 - | 4,14E-05 - | 0 - | 0 - | 1 - | 4,00E-05 - | 1 - | 4,50E-05 - |
| 7970 | 1 | 3um_FIJJAFBF_208547 | Bacteria | Proteobacteri | Betaproteoba | Rhodocyclales | Zoogloeaceae | Thauera | Thauera terpe | cellular proces | Posttranslatio | 16QU3 | Hemolysin-typ | 744980.TRICH | 2,4 | 0,489897949 | 3,43E-05 | 7,40E-06 | 2 - | 2,39E-05 - | 2 - | 2,97E-05 - | 3 - | 4,24E-05 - | 3 - | 4,30E-05 - | 2 - | 3,23E-05 - |
| 7971 | 1 | 0.2um_NOKCPHJN_513502 | Bacteria | Proteobacteri | Alphaproteob | Rhodobactera | Rhodobactera | Planktomarina | Planktomarina | information st | Translation, ri | 08VZM | 50s ribosomal | 633131.TR2A6 | 4,2 | 0,4 | 0,00033592 | 1,59E-05 | 5 - | 0,000337579 - | 4 - | 0,000335339 - | 4 - | 0,000318782 - | 4 - | 0,00032349 - | 4 - | 0,000364411 - |
| 7972 | 1 | 0.2um_NOKCPHJN_102935 | Bacteria | Bacteroidetes | Flavobacteriia | Flavobacterial | Flavobacteriac | Algibacter | Algibacter sp. | metabolism | Carbohydrate | 0XNQK | Transporter | 391587.KAOT | 1,2 | 0,748331477 | 3,57E-05 | 2,26E-05 | 2 - | 5,00E-05 - | 1 - | 3,10E-05 - | 0 - | 0 - | 1 - | 3,00E-05 - | 2 - | 6,75E-05 - |
| 7973 | 1 | 0.2um_NOKCPHJN_173827 | Bacteria | Proteobacteri | Alphaproteob | Rhodobactera | Rhodobactera | Roseovarius | Roseovarius a | unclassified | unclassified | unclassified | unclassified | unclassified | 5,2 | 0,748331477 | 0,000196832 | 3,27E-05 | 6 - | 0,000190765 - | 4 - | 0,000157916 - | 5 - | 0,000187649 - | 5 - | 0,00019042 - | 6 - | 0,000257409 - |
| 7974 | 1 | 0.2um_NOKCPHJN_126580 | Bacteria | Verrucomicrol | Opitutae | unclassified | unclassified | unclassified | Opitutae bact | unclassified | unclassified | unclassified | unclassified | unclassified | 19 | 1,673320053 | 0,000541902 | 4,48E-05 | 22 - | 0,000528261 - | 19 - | 0,000566498 - | 18 - | 0,000510184 - | 17 - | 0,000488957 - | 19 - | 0,00061561 - |
| 7975 | 1 | 0.2um_NOKCPHJN_469918 | Bacteria | unclassified | unclassified | unclassified | unclassified | unclassified | bacterium B1 | unclassified | unclassified | unclassified | unclassified | unclassified | 1,4 | 0,489897949 | 3,36E-05 | 9,61E-06 | 2 - | 4,12E-05 - | 1 - | 2,56E-05 - | 2 - | 4,86E-05 - | 1 - | 2,47E-05 - | 1 - | 2,78E-05 - |
| 7976 | 1 | 0.2um_NOKCPHJN_112415 | Bacteria | Bacteroidetes | Flavobacteriia | Flavobacterial | Flavobacteriac | Marinirhabdu | Marinirhabdu | unclassified | unclassified | unclassified | unclassified | unclassified | 3 | 0,632455532 | 0,000323875 | 9,02E-05 | 3 - | 0,000269008 - | 3 - | 0,000334029 - | 3 - | 0,000317536 - | 2 - | 0,000214818 - | 4 - | 0,000483983 - |
| 7977 | 1 | 3um_FIJJAFBF_186974 | Bacteria | Proteobacteri | Gammaprote | Alteromonada | Alteromonada | Paraglaciecola | Paraglaciecola | unclassified | unclassified | unclassified | unclassified | unclassified | 0,2 | 0,4 | 1,34E-05 | 2,68E-05 | 1 - | 6,71E-05 - | 0 - | 0 - | 0 - | 0 - | 0 - | 0 - | 0 - | 0 - |
| 7978 | 1 | 3um_FIJJAFBF_315499 | Bacteria | Proteobacteri | Alphaproteob | unclassified | unclassified | unclassified | various | metabolism | Energy produc | 01RNF | Produces ATP | 269796.Rru_A | 1 | 0,632455532 | 4,58E-05 | 3,02E-05 | 1 - | 3,80E-05 - | 2 - | 9,44E-05 - | 0 - | 0 - | 1 - | 4,55E-05 - | 1 - | 5,13E-05 - |
| 7979 | 1 | 0.2um_NOKCPHJN_04178 | Bacteria | Bacteroidetes | Cytophagia | Cytophagales | Flammeovirga | Reichenbachie | Reichenbachie | unclassified | unclassified | unclassified | unclassified | unclassified | 0,4 | 0,8 | 6,09E-06 | 1,22E-05 | 2 - | 3,04E-05 - | 0 - | 0 - | 0 - | 0 - | 0 - | 0 - | 0 - | 0 - |
| 7980 | 1 | 0.2um_NOKCPHJN_91458 | Bacteria | Proteobacteri | Gammaprote | Oceanospirilla | Saccharospirill | Reinekea | Reinekea fors | metabolism | Carbohydrate | 0D2NF | Periplasmic bi | 765420.OSCT_ | 4,4 | 1,959591794 | 0,000185914 | 6,22E-05 | 8 - | 0,000294299 - | 3 - | 0,000137037 - | 5 - | 0,000217119 - | 3 - | 0,000132195 - | 3 - | 0,000148918 - |
| 7981 | 1 | 0.2um_NOKCPHJN_562408 | Bacteria | Proteobacteri | Betaproteoba | Nitrosomonad | unclassified | unclassified | Methylophilal | unclassified | unclassified | unclassified | unclassified | unclassified | 0,6 | 0,489897949 | 9,76E-05 | 8,12E-05 | 1 - | 0,000138285 - | 0 - | 0 - | 1 - | 0,000163232 - | 0 - | 0 - | 1 - | 0,000186596 - |
| 7982 | 1 | 0.2um_NOKCPHJN_62098 | Bacteria | Proteobacteri | Gammaprote | Cellvibrionale | Porticoccacea | unclassified | Gammaprote | poorly charact | Function unkn | 0QSE4 | Pfam:DUF400 | 314287.GB220 | 2 | 0 | 0,000159334 | 1,51E-05 | 2 - | 0,000133461 - | 2 - | 0,00016572 - | 2 - | 0,000157537 - | 2 - | 0,000159864 - | 2 - | 0,000180087 - |
| 7983 | 1 | 0.2um_NOKCPHJN_45933 | Bacteria | Bacteroidetes | Flavobacteriia | Flavobacterial | Flavobacteriac | Cellulophaga | Cellulophaga | poorly charact | Function unkn | 0XP53 | Ragb susd dor | 313598.MED1 | 5 | 1,095445115 | 0,000129781 | 3,94E-05 | 4 - | 8,57E-05 - | 4 - | 0,000106357 - | 5 - | 0,000126383 - | 5 - | 0,000128249 - | 7 - | 0,000202261 - |
| 7984 | 1 | 3um_FIJJAFBF_165509 | Bacteria | Proteobacteri | Gammaprote | Pseudomonad | Moraxellaceae | Psychrobacter | various | information st | Transcription | 05VEQ | Cold shock pr | 259536.Psyc_ | 1,2 | 0,4 | 0,000228546 | 5,06E-05 | 2 - | 0,000327934 - | 1 - | 0,000203598 - | 1 - | 0,000193546 - | 1 - | 0,000196405 - | 1 - | 0,000221249 - |
| 7985 | 1 | 3um_FIJJAFBF_48390 | Bacteria | Bacteroidetes | Cytophagia | Cytophagales | Flammeovirga | unclassified | Flammeovirga | unclassified | unclassified | unclassified | unclassified | unclassified | 1,4 | 0,489897949 | 6,74E-05 | 2,77E-05 | 1 - | 3,96E-05 - | 1 - | 4,91E-05 - | 1 - | 4,67E-05 - | 2 - | 9,48E-05 - | 2 - | 0,00010681 - |
| 7986 | 1 | 3um_FIJJAFBF_532856 | Bacteria | Proteobacteri | Alphaproteob | Rhodobactera | Rhodobactera | Leisingera | Leisingera sp. | cellular proces | Posttranslatio | COG0652 | peptidyl-proly | 439496.RBY4I | 1 | 0 | 8,16E-05 | 7,74E-06 | 1 - | 6,83E-05 - | 1 - | 8,48E-05 - | 1 - | 8,06E-05 - | 1 - | 8,18E-05 - | 1 - | 9,22E-05 - |
| 7987 | 1 | 0.2um_NOKCPHJN_383104 | Bacteria | Proteobacteri | Alphaproteob | unclassified | unclassified | unclassified | SAR116 cluste | unclassified | unclassified | unclassified | unclassified | unclassified | 0,6 | 0,8 | 1,46E-05 | 1,88E-05 | 2 - | 4,55E-05 - | 0 - | 0 - | 0 - | 0 - | 1 - | 2,73E-05 - | 0 - | 0 - |
| 7988 | 1 | 3um_FIJJAFBF_348506 | Bacteria | Bacteroidetes | Saprospiria | Saprospirales | Saprospiracea | Saprospira | Saprospira gra | poorly charact | Function unkn | 0XSVM | NA | 926562.Oweh | 1,8 | 1,166190379 | 4,77E-05 | 2,55E-05 | 4 - | 9,35E-05 - | 2 - | 5,81E-05 - | 1 - | 2,76E-05 - | 1 - | 2,80E-05 - | 1 - | 3,15E-05 - |
| 7989 | 2 | 0.2um_NOKCPHJN_131401;3um_FIJJAFBF_18782 | Bacteria | Bacteroidetes | Flavobacteriia | Flavobacterial | Flavobacteriac | Polaribacter | various | information st | Translation, ri | 09FE5 | One of the pri | 313594.PI23P | 2 | 0 | 0,000274054 | 2,60E-05 | 2 - | 0,000229554 - | 2 - | 0,000285038 - | 2 - | 0,000270964 - | 2 - | 0,000274967 - | 2 - | 0,000309749 - |
| 7990 | 1 | 3um_FIJJAFBF_196455 | Bacteria | Proteobacteri | Gammaprote | Cellvibrionale | Halieaceae | unclassified | marine gamm | unclassified | unclassified | unclassified | unclassified | unclassified | 11,2 | 0,748331477 | 0,000194691 | 1,06E-05 | 12 - | 0,000175679 - | 11 - | 0,000199963 - | 12 - | 0,000207371 - | 11 - | 0,000192897 - | 10 - | 0,000197544 - |
| 7991 | 1 | 3um_FIJJAFBF_405250 | Bacteria | Bacteroidetes | Flavobacteriia | Flavobacterial | Flavobacteriac | Polaribacter | Polaribacter ir | unclassified | unclassified | unclassified | unclassified | unclassified | 7,6 | 0,489897949 | 0,000314265 | 3,42E-05 | 8 - | 0,000277406 - | 7 - | 0,000301399 - | 8 - | 0,000327449 - | 7 - | 0,00029075 - | 8 - | 0,000374319 - |
| 7992 | 1 | 0.2um_NOKCPHJN_152943 | Bacteria | Proteobacteri | Alphaproteob | Rhodobactera | Rhodobactera | Litoreibacter | Litoreibacter a | cellular proces | Signal transdu | 01Q8R | gtp-binding pr | 391619.RGBS1 | 5,8 | 1,166190379 | 0,000131117 | 2,85E-05 | 5 - | 9,49E-05 - | 7 - | 0,000164898 - | 6 - | 0,000134363 - | 7 - | 0,000159072 - | 4 - | 0,000102396 - |
| 7993 | 1 | 0.2um_NOKCPHJN_785873 | Eukaryota | Bacillariophyt | Coscinodiscop | Thalassiosirale | Thalassiosirac | Thalassiosira | Thalassiosira c | information st | Chromatin str | KOG1744 | Core compone | 35128.Thaps6 | 4,2 | 0,748331477 | 0,000520354 | 6,22E-05 | 5 - | 0,000526499 - | 4 - | 0,000523005 - | 5 - | 0,000621478 - | 4 - | 0,000504526 - | 3 - | 0,00042626 - |
| 7994 | 1 | 0.2um_NOKCPHJN_175563 | Bacteria | Bacteroidetes | unclassified | unclassified | unclassified | unclassified | Bacteroidetes | unclassified | unclassified | unclassified | unclassified | unclassified | 0,6 | 0,489897949 | 1,48E-05 | 1,23E-05 | 1 - | 2,10E-05 - | 0 - | 0 - | 1 - | 2,48E-05 - | 0 - | 0 - | 1 - | 2,84E-05 - |
| 7995 | 1 | 3um_FIJJAFBF_744517 | Bacteria | Proteobacteri | Alphaproteob | Pelagibacteral | Pelagibacterac | Candidatus Pe | Candidatus Pe | unclassified | unclassified | unclassified | unclassified | unclassified | 3,4 | 0,489897949 | 0,000229977 | 4,82E-05 | 3 - | 0,000168789 - | 3 - | 0,000209587 - | 4 - | 0,000265651 - | 3 - | 0,000202181 - | 4 - | 0,000303675 - |
| 7996 | 1 | 0.2um_NOKCPHJN_28866 | Bacteria | Bacteroidetes | Flavobacteriia | Flavobacterial | Flavobacteriac | Polaribacter | Polaribacter b | metabolism | Carbohydrate | 05D2N | Aldolase | 391587.KAOT | 10 | 0,632455532 | 0,000385879 | 4,30E-05 | 10 - | 0,000323315 - | 10 - | 0,000401462 - | 11 - | 0,000419804 - | 9 - | 0,000348549 - | 10 - | 0,000436266 - |
| 7997 | 1 | 3um_FIJJAFBF_802884 | Bacteria | Proteobacteri | Gammaprote | Oceanospirilla | Saccharospirill | Reinekea | Reinekea fors | information st | Transcription | COG0217 | transcriptiona | 491952.Mar18 | 0,6 | 0,489897949 | 3,28E-05 | 2,70E-05 | 1 - | 4,76E-05 - | 1 - | 5,91E-05 - | 0 - | 0 - | 1 - | 5,70E-05 - | 0 - | 0 - |
| 7998 | 1 | 0.2um_NOKCPHJN_239174 | Bacteria | Bacteroidetes | Flavobacteriia | Flavobacterial | Flavobacteriac | unclassified | Flavobacteriac | unclassified | unclassified | unclassified | unclassified | unclassified | 3,2 | 0,4 | 5,73E-05 | 3,14E-06 | 4 - | 6,06E-05 - | 3 - | 5,65E-05 - | 3 - | 5,37E-05 - | 3 - | 5,45E-05 - | 3 - | 6,14E-05 - |
| 7999 | 1 | 0.2um_NOKCPHJN_479681 | Bacteria | Proteobacteri | Alphaproteob | Rhodobactera | Rhodobactera | Roseovarius | Roseovarius s | metabolism | Inorganic ion t | COG3221 | phosphonate | 375451.RD1_2 | 0,4 | 0,489897949 | 2,66E-05 | 3,26E-05 | 0 - | 0 - | 1 - | 6,82E-05 - | 1 - | 6,48E-05 - | 0 - | 0 - | 0 - | 0 - |
| 8000 | 1 | 0.2um_NOKCPHJN_50817 | Bacteria | Bacteroidetes | Flavobacteriia | Flavobacterial | Flavobacteriac | Polaribacter | various | unclassified | unclassified | unclassified | unclassified | unclassified | 0,8 | 0,4 | 8,29E-05 | 4,21E-05 | 1 - | 8,97E-05 - | 1 - | 0,000111343 - | 1 - | 0,000105845 - | 1 - | 0,000107409 - | 0 - | 0 - |
| 8001 | 1 | 3um_FIJJAFBF_404116 | Bacteria | Bacteroidetes | Flavobacteriia | Flavobacterial | Flavobacteriac | Formosa | Formosa sp. H | poorly charact | Function unkn | 11NCV | Protein of unk | 983544.Lacal_ | 0,4 | 0,489897949 | 3,30E-05 | 4,08E-05 | 1 - | 7,36E-05 - | 1 - | 9,14E-05 - | 0 - | 0 - | 0 - | 0 - | 0 - | 0 - |
| 8002 | 1 | 3um_FIJJAFBF_258136 | Bacteria | Proteobacteri | Gammaprote | unclassified | unclassified | unclassified | Gammaprote | cellular proces | Posttranslatio | COG0330 | Band 7 protei | 247634.GPB21 | 2,8 | 0,748331477 | 9,94E-05 | 1,87E-05 | 4 - | 0,000121457 - | 3 - | 0,00011311 - | 3 - | 0,000107526 - | 2 - | 7,27E-05 - | 2 - | 8,19E-05 - |

| | | | | | | | | | | | | | | | | | | | | | | | | | | | | |
|---|---|---|---|---|---|---|---|---|---|---|---|---|---|---|---|---|---|---|---|---|---|---|---|---|---|---|---|---|
| 8003 | 1 | 0.2um_NOKCPHJN_825917 | Bacteria | Proteobacteri | Gammaprotec | Oceanospirilla | Saccharospirill | Reinekea | Reinekea fors | metabolism | Secondary me | 0QQXG | DSBA Oxidore | 314283.MED2 | 0,8 | 0,4 | 4,96E-05 | 2,52E-05 | 1 - | 5,36E-05 - | 1 - | 6,66E-05 - | 1 - | 6,33E-05 - | 1 - | 6,42E-05 - | 0 - | 0 - |
| 8004 | 1 | 3um_FIJJAFBF_216195 | Bacteria | Bacteroidetes | Flavobacteriia | Flavobacterial | Flavobacteriac | Cellulophaga | Cellulophaga t | unclassified | unclassified | unclassified | unclassified | unclassified | 0,4 | 0,489897949 | 1,20E-05 | 1,47E-05 | 0 - | 0 - | 1 - | 3,08E-05 - | 1 - | 2,93E-05 - | 0 - | 0 - | 0 - | 0 - |
| 8005 | 1 | 3um_FIJJAFBF_609352 | Bacteria | Proteobacteri | Gammaprotec | Cellvibrionales | Halieaceae | unclassified | marine gamm | unclassified | unclassified | unclassified | unclassified | unclassified | 0,8 | 0,4 | 1,94E-05 | 9,77E-06 | 0 - | 0 - | 1 - | 2,42E-05 - | 1 - | 2,30E-05 - | 1 - | 2,34E-05 - | 1 - | 2,63E-05 - |
| 8006 | 1 | 0.2um_NOKCPHJN_426946 | Bacteria | Proteobacteri | Gammaprotec | unclassified | unclassified | unclassified | OM182 bacte | information st | Translation, ril | 08VZM | 50s ribosomal | 377629.TERTU | 0,6 | 0,489897949 | 4,46E-05 | 3,68E-05 | 1 - | 6,52E-05 - | 1 - | 8,10E-05 - | 1 - | 7,70E-05 - | 0 - | 0 - | 0 - | 0 - |
| 8007 | 1 | 3um_FIJJAFBF_409141 | Bacteria | Proteobacteri | Gammaprotec | unclassified | unclassified | unclassified | Gammaprotec | metabolism | Energy produc | COG1290 | Component of | 565045.NOR5 | 1 | 0 | 5,06E-05 | 4,80E-06 | 1 - | 4,24E-05 - | 1 - | 5,26E-05 - | 1 - | 5,00E-05 - | 1 - | 5,07E-05 - | 1 - | 5,71E-05 - |
| 8008 | 1 | 3um_FIJJAFBF_213100 | Bacteria | Bacteroidetes | Flavobacteriia | Flavobacterial | Flavobacteriac | Lutibacter | Lutibacter sp. | information st | Translation, ril | 05CA1 | Removes the | 313590.MED1 | 0,2 | 0,4 | 1,00E-05 | 2,01E-05 | 0 - | 0 - | 0 - | 0 - | 1 - | 5,02E-05 - | 0 - | 0 - | 0 - | 0 - |
| 8009 | 1 | 0.2um_NOKCPHJN_110814 | Bacteria | Bacteroidetes | Flavobacteriia | Flavobacterial | Flavobacteriac | Winogradskye | Winogradskye | unclassified | unclassified | unclassified | unclassified | unclassified | 0,4 | 0,489897949 | 2,36E-05 | 2,94E-05 | 1 - | 5,01E-05 - | 0 - | 0 - | 0 - | 0 - | 0 - | 0 - | 1 - | 6,76E-05 - |
| 8010 | 1 | 3um_FIJJAFBF_510918 | Bacteria | Actinobacteri | Actinobacteri | Micrococcales | Microbacteria | Candidatus Ac | Candidatus Ac | information st | Translation, ril | 08YY1 | This protein is | 288705.RSal3 | 2,4 | 0,489897949 | 0,000279747 | 4,36E-05 | 3 - | 0,000296836 - | 2 - | 0,000245722 - | 2 - | 0,00023359 - | 3 - | 0,00035556 - | 2 - | 0,000267025 - |
| 8011 | 1 | 3um_FIJJAFBF_60118 | Bacteria | unclassified | unclassified | unclassified | unclassified | unclassified | uncultured ba | information st | Translation, ril | COG0184 | Forms an inte | 313598.MED1 | 1,2 | 0,4 | 0,000184409 | 6,18E-05 | 1 - | 0,000128963 - | 1 - | 0,000160134 - | 2 - | 0,000304454 - | 1 - | 0,000154476 - | 1 - | 0,000174016 - |
| 8012 | 1 | 0.2um_NOKCPHJN_66479 | Bacteria | Bacteroidetes | unclassified | unclassified | unclassified | unclassified | Bacteroidetes | cellular proces | Cell wall/mem | 11TPU | outer membra | 926562.Oweh | 1,2 | 0,4 | 9,47E-05 | 2,10E-05 | 2 - | 0,00013583 - | 1 - | 8,43E-05 - | 1 - | 8,02E-05 - | 1 - | 8,14E-05 - | 1 - | 9,16E-05 - |
| 8013 | 1 | 0.2um_NOKCPHJN_36506 | Bacteria | Bacteroidetes | Cytophagia | Cytophagales | Flammeovirga | unclassified | Flammeovirga | unclassified | unclassified | unclassified | unclassified | unclassified | 0,2 | 0,4 | 8,03E-06 | 1,61E-05 | 1 - | 4,01E-05 - | 0 - | 0 - | 0 - | 0 - | 0 - | 0 - | 0 - | 0 - |
| 8014 | 1 | 3um_FIJJAFBF_457672 | Bacteria | Proteobacteri | Betaproteoba | unclassified | unclassified | unclassified | Betaproteoba | unclassified | unclassified | unclassified | unclassified | unclassified | 0,2 | 0,4 | 7,06E-06 | 1,41E-05 | 1 - | 3,53E-05 - | 0 - | 0 - | 0 - | 0 - | 0 - | 0 - | 0 - | 0 - |
| 8015 | 1 | 3um_FIJJAFBF_122379 | Bacteria | Bacteroidetes | Flavobacteriia | Flavobacterial | Flavobacteriac | Polaribacter | Polaribacter s | poorly charact | Function unkn | 09HDN | NA | 313598.MED1 | 1,4 | 0,8 | 0,000121151 | 5,36E-05 | 3 - | 0,000228033 - | 1 - | 9,44E-05 - | 1 - | 8,97E-05 - | 1 - | 9,10E-05 - | 1 - | 0,000102566 - |
| 8016 | 1 | 0.2um_NOKCPHJN_412145 | Bacteria | Proteobacteri | Alphaproteob | Rhodobactera | Rhodobactera | Acuticoccus | Acuticoccus y | unclassified | unclassified | unclassified | unclassified | unclassified | 0,2 | 0,4 | 4,09E-06 | 8,18E-06 | 1 - | 2,05E-05 - | 0 - | 0 - | 0 - | 0 - | 0 - | 0 - | 0 - | 0 - |
| 8017 | 1 | 3um_FIJJAFBF_689707 | Bacteria | Proteobacteri | Gammaprotec | Oceanospirilla | Oceanospirilla | Motiliproteus | Motiliproteus | unclassified | unclassified | unclassified | unclassified | unclassified | 0,4 | 0,489897949 | 2,29E-05 | 2,85E-05 | 1 - | 4,86E-05 - | 0 - | 0 - | 0 - | 0 - | 0 - | 0 - | 1 - | 6,56E-05 - |
| 8018 | 1 | 0.2um_NOKCPHJN_119310 | Bacteria | Proteobacteri | Gammaprotec | Cellvibrionales | Porticoccacea | unclassified | gamma prote | poorly charact | Function unkn | 16R8I | Lipoprotein | 314287.GB22 | 7,6 | 0,489897949 | 0,000189532 | 1,27E-05 | 8 - | 0,000167864 - | 7 - | 0,000182382 - | 8 - | 0,000198146 - | 8 - | 0,000201072 - | 7 - | 0,000198194 - |
| 8019 | 1 | 3um_FIJJAFBF_397801 | Bacteria | Proteobacteri | Betaproteoba | unclassified | unclassified | unclassified | Betaproteoba | metabolism | Amino acid tra | 0XPCZ | amino acid AB | 296591.Bpro_ | 6,6 | 1,019803903 | 0,000203668 | 3,91E-05 | 6 - | 0,000154755 - | 5 - | 0,000160134 - | 8 - | 0,000243564 - | 7 - | 0,000216266 - | 7 - | 0,000243623 - |
| 8020 | 1 | 0.2um_NOKCPHJN_332543 | Eukaryota | Bacillariophyt | Coscinodiscop | Thalassiosirale | Thalassiosirac | Thalassiosira | Thalassiosira | cellular proces | Posttranslatio | KOG0019 | heat shock pr | 35128.Thaps6 | 2 | 0 | 3,87E-05 | 3,68E-06 | 2 - | 3,24E-05 - | 2 - | 4,03E-05 - | 2 - | 3,83E-05 - | 2 - | 3,88E-05 - | 2 - | 4,37E-05 - |
| 8021 | 1 | 3um_FIJJAFBF_723900 | unclassified | unclassified | unclassified | unclassified | unclassified | unclassified | unclassified | unclassified | unclassified | unclassified | unclassified | unclassified | 0,4 | 0,489897949 | 2,38E-05 | 2,91E-05 | 0 - | 0 - | 1 - | 6,09E-05 - | 1 - | 5,79E-05 - | 0 - | 0 - | 0 - | 0 - |
| 8022 | 1 | 3um_FIJJAFBF_446710 | Bacteria | Bacteroidetes | Flavobacteriia | Flavobacterial | Flavobacteriac | Polaribacter | Polaribacter p | metabolism | Lipid transpor | 0P7W4 | reductase | 983544.Lacal_ | 1,2 | 0,4 | 6,45E-05 | 1,43E-05 | 2 - | 9,26E-05 - | 1 - | 5,75E-05 - | 1 - | 5,46E-05 - | 1 - | 5,54E-05 - | 1 - | 6,24E-05 - |
| 8023 | 1 | 0.2um_NOKCPHJN_216942 | Bacteria | Proteobacteri | Alphaproteob | Rhodobactera | unclassified | unclassified | Rhodobactera | unclassified | unclassified | unclassified | unclassified | unclassified | 0,4 | 0,489897949 | 2,31E-05 | 2,85E-05 | 1 - | 5,26E-05 - | 0 - | 0 - | 0 - | 0 - | 1 - | 6,31E-05 - | 0 - | 0 - |
| 8024 | 1 | 3um_FIJJAFBF_261510 | Bacteria | Bacteroidetes | Flavobacteriia | Flavobacterial | Flavobacteriac | Formosa | Formosa sp. H | poorly charact | Function unkn | 09A4Z | rhodopsin | 313594.PI23P | 0,8 | 0,4 | 4,42E-05 | 2,24E-05 | 1 - | 4,78E-05 - | 1 - | 5,94E-05 - | 1 - | 5,65E-05 - | 1 - | 5,73E-05 - | 0 - | 0 - |
| 8025 | 1 | 0.2um_NOKCPHJN_25800 | Bacteria | Bacteroidetes | Flavobacteriia | Flavobacterial | Flavobacteriac | Polaribacter | various | unclassified | unclassified | unclassified | unclassified | unclassified | 12,2 | 0,748331477 | 0,000144267 | 8,04E-06 | 13 - | 0,00012941 - | 12 - | 0,000148328 - | 13 - | 0,000152755 - | 12 - | 0,000143088 - | 11 - | 0,000147755 - |
| 8026 | 1 | 0.2um_NOKCPHJN_27666 | Bacteria | Actinobacteri | Actinobacteri | Micrococcales | Microbacteria | unclassified | Microbacteria | information st | Translation, ril | 08UHY | One of the pri | 281090.Lxx20 | 2,6 | 0,489897949 | 0,000228943 | 4,88E-05 | 2 - | 0,00014715 - | 3 - | 0,000274075 - | 3 - | 0,000260543 - | 3 - | 0,000264391 - | 2 - | 0,000198557 - |
| 8027 | 1 | 3um_FIJJAFBF_122435 | Bacteria | Bacteroidetes | Flavobacteriia | Flavobacterial | Flavobacteriac | Polaribacter | Polaribacter s | metabolism | Carbohydrate | COG1109 | Catalyzes the | 313598.MED1 | 1,2 | 0,748331477 | 2,92E-05 | 1,97E-05 | 1 - | 1,98E-05 - | 2 - | 4,91E-05 - | 0 - | 0 - | 1 - | 2,37E-05 - | 2 - | 5,33E-05 - |
| 8028 | 1 | 3um_FIJJAFBF_40402 | Bacteria | Proteobacteri | Gammaprotec | Oceanospirilla | Saccharospirill | Reinekea | Reinekea fors | cellular proces | Cell wall/mem | 0QTSX | Peptidoglycan | 314283.MED2 | 4,8 | 0,748331477 | 0,000349464 | 4,93E-05 | 6 - | 0,000368268 - | 5 - | 0,000381066 - | 4 - | 0,000289802 - | 4 - | 0,000294082 - | 5 - | 0,000414103 - |
| 8029 | 1 | 0.2um_NOKCPHJN_397687 | Eukaryota | Bacillariophyt | Bacillariophyc | Naviculales | Phaeodactylac | Phaeodactylu | Phaeodactylu | information st | Translation, ril | KOG3421 | ribosomal pro | 2850.Phatr25 | 3,8 | 0,4 | 0,000385921 | 3,44E-05 | 4 - | 0,000342617 - | 4 - | 0,000425429 - | 4 - | 0,000404425 - | 4 - | 0,000410398 - | 3 - | 0,000346734 - |
| 8030 | 1 | 0.2um_NOKCPHJN_1096186 | unclassified | unclassified | unclassified | unclassified | unclassified | unclassified | unclassified | unclassified | unclassified | unclassified | unclassified | unclassified | 0,2 | 0,4 | 7,49E-06 | 1,50E-05 | 0 - | 0 - | 0 - | 0 - | 0 - | 0 - | 1 - | 3,75E-05 - | 0 - | 0 - |
| 8031 | 1 | 0.2um_NOKCPHJN_959134 | Eukaryota | Phaeophyceae | unclassified | Ectocarpales | Ectocarpaceae | Ectocarpus | Ectocarpus sili | metabolism | Energy produc | COG0055 | Produces ATP | 35128.Thaps4 | 1 | 0 | 0,000106223 | 1,01E-05 | 1 - | 8,90E-05 - | 1 - | 0,00011048 - | 1 - | 0,000105025 - | 1 - | 0,000106576 - | 1 - | 0,000120058 - |
| 8032 | 1 | 0.2um_NOKCPHJN_984540 | Bacteria | Bacteroidetes | Flavobacteriia | Flavobacterial | unclassified | unclassified | Flavobacterial | unclassified | unclassified | unclassified | unclassified | unclassified | 0,6 | 0,489897949 | 1,99E-05 | 1,62E-05 | 0 - | 0 - | 1 - | 3,41E-05 - | 1 - | 3,24E-05 - | 1 - | 3,29E-05 - | 0 - | 0 - |
| 8033 | 1 | 3um_FIJJAFBF_202716 | Bacteria | Bacteroidetes | Flavobacteriia | unclassified | unclassified | unclassified | Flavobacteria | cellular proces | Posttranslatio | 0PA4Z | Glutathione p | 487796.Flav2 | 0,6 | 0,489897949 | 4,08E-05 | 3,36E-05 | 1 - | 6,04E-05 - | 0 - | 0 - | 1 - | 7,13E-05 - | 1 - | 7,24E-05 - | 0 - | 0 - |
| 8034 | 1 | 3um_FIJJAFBF_47863 | Bacteria | unclassified | unclassified | unclassified | unclassified | unclassified | uncultured ba | unclassified | unclassified | unclassified | unclassified | unclassified | 0,6 | 0,489897949 | 4,33E-05 | 3,60E-05 | 1 - | 6,11E-05 - | 0 - | 0 - | 0 - | 0 - | 1 - | 7,31E-05 - | 1 - | 8,24E-05 - |
| 8035 | 1 | 0.2um_NOKCPHJN_484153 | Eukaryota | unclassified | unclassified | Perkinsida | Perkinsidae | Perkinsus | Perkinsus mar | metabolism | Lipid transpor | KOG1180 | synthetase | 2850.Phatr20 | 0,4 | 0,489897949 | 3,87E-05 | 4,78E-05 | 1 - | 8,63E-05 - | 1 - | 0,000107157 - | 0 - | 0 - | 0 - | 0 - | 0 - | 0 - |
| 8036 | 1 | 0.2um_NOKCPHJN_851380 | Bacteria | Verrucomicrol | Opitutae | Opitutales | Opitutaceae | Opitutus | Opitutus terra | unclassified | unclassified | unclassified | unclassified | unclassified | 1,6 | 0,8 | 3,21E-05 | 1,65E-05 | 2 - | 3,36E-05 - | 2 - | 4,17E-05 - | 2 - | 3,97E-05 - | 0 - | 0 - | 2 - | 4,54E-05 - |
| 8037 | 1 | 3um_FIJJAFBF_469955 | Bacteria | Proteobacteri | Alphaproteob | Rhodobactera | Rhodobactera | unclassified | Rhodobactera | metabolism | Amino acid tra | 01R4A | Polyamine AB | 388739.RSK20 | 4,8 | 0,748331477 | 0,00017779 | 2,32E-05 | 6 - | 0,000187646 - | 4 - | 0,000155334 - | 4 - | 0,000147665 - | 5 - | 0,000187307 - | 5 - | 0,000211001 - |
| 8038 | 1 | 0.2um_NOKCPHJN_72789 | Bacteria | Proteobacteri | Gammaprotec | Oceanospirilla | Saccharospirill | Reinekea | Reinekea fors | cellular proces | Posttranslatio | 0QHT2 | Peroxiredoxin | 717774.Marm | 2 | 0,632455532 | 0,000155503 | 4,84E-05 | 2 - | 0,000131927 - | 2 - | 0,000163815 - | 2 - | 0,000155727 - | 3 - | 0,00023704 - | 1 - | 8,90E-05 - |
| 8039 | 1 | 0.2um_NOKCPHJN_129356 | Bacteria | Bacteroidetes | unclassified | unclassified | unclassified | unclassified | Bacteroidetes | unclassified | unclassified | unclassified | unclassified | unclassified | 0,2 | 0,4 | 6,48E-06 | 1,30E-05 | 1 - | 3,24E-05 - | 0 - | 0 - | 0 - | 0 - | 0 - | 0 - | 0 - | 0 - |
| 8040 | 1 | 3um_FIJJAFBF_44059 | Viruses | unclassified | unclassified | unclassified | Phycodnavirid | unclassified | Chrysochromu | unclassified | unclassified | unclassified | unclassified | unclassified | 4,2 | 0,4 | 0,00030925 | 3,92E-05 | 4 - | 0,000246832 - | 4 - | 0,000306492 - | 5 - | 0,0003642 - | 4 - | 0,000295663 - | 4 - | 0,000333063 - |
| 8041 | 1 | 3um_FIJJAFBF_481932 | Bacteria | Proteobacteri | Alphaproteob | Rhodobactera | Rhodobactera | Tateyamaria | Tateyamaria s | information st | Translation, ril | 05G6W | One of the pri | 52598.EE36_0 | 5,2 | 1,469693846 | 0,000338835 | 7,85E-05 | 7 - | 0,000390018 - | 6 - | 0,000415103 - | 4 - | 0,000263072 - | 6 - | 0,000400437 - | 3 - | 0,000225545 - |
| 8042 | 1 | 0.2um_NOKCPHJN_208316 | Eukaryota | Bacillariophyt | Bacillariophyc | Naviculales | Naviculaceae | Fistulifera | Fistulifera sola | information st | Translation, ril | 125NI | 60S acidic ribo | 65071.PYU1_ | 0,2 | 0,4 | 2,48E-05 | 4,96E-05 | 0 - | 0 - | 1 - | 0,000123929 - | 0 - | 0 - | 0 - | 0 - | 0 - | 0 - |
| 8043 | 1 | 0.2um_NOKCPHJN_751237 | Eukaryota | unclassified | unclassified | Prymnesiales | Chrysochromu | Chrysochromu | Chrysochromu | unclassified | unclassified | unclassified | unclassified | unclassified | 1 | 0 | 0,000161208 | 1,53E-05 | 1 - | 0,000135031 - | 1 - | 0,000167669 - | 1 - | 0,000159391 - | 1 - | 0,000161745 - | 1 - | 0,000182205 - |
| 8044 | 1 | 0.2um_NOKCPHJN_514099 | Bacteria | Proteobacteri | Betaproteoba | Nitrosomonad | Methylophilac | Candidatus M | Candidatus M | metabolism | Energy produc | COG0056 | Produces ATP | 383631.MB21 | 0,6 | 0,8 | 1,59E-05 | 2,12E-05 | 0 - | 0 - | 0 - | 0 - | 2 - | 5,28E-05 - | 1 - | 2,68E-05 - | 0 - | 0 - |
| 8045 | 1 | 3um_FIJJAFBF_409503 | Bacteria | Proteobacteri | Gammaprotec | Oceanospirilla | Oceanospirilla | Amphritea | Amphritea jap | cellular proces | Defense mech | 07SUK | Efflux transpo | 207954.MED9 | 1,8 | 1,469693846 | 5,78E-05 | 4,84E-05 | 1 - | 2,73E-05 - | 1 - | 3,39E-05 - | 3 - | 9,68E-05 - | 4 - | 0,000130936 - | 0 - | 0 - |
| 8046 | 1 | 3um_FIJJAFBF_770105 | Eukaryota | unclassified | Spirotrichea | Sporadotrichi | Oxytrichidae | Stylonychia | Stylonychia le | metabolism | Energy produc | 0IFIU | Pyrophosphat | 5911.EAS003 | 0,6 | 0,489897949 | 5,74E-05 | 4,73E-05 | 1 - | 8,50E-05 - | 0 - | 0 - | 1 - | 0,000100357 - | 1 - | 0,000101839 - | 0 - | 0 - |
| 8047 | 1 | 3um_FIJJAFBF_241622 | Viruses | unclassified | unclassified | unclassified | various | various | various | unclassified | unclassified | unclassified | unclassified | unclassified | 0,2 | 0,4 | 9,31E-06 | 1,86E-05 | 0 - | 0 - | 0 - | 0 - | 1 - | 4,66E-05 - | 0 - | 0 - | 0 - | 0 - |
| 8048 | 1 | 0.2um_NOKCPHJN_842153 | Bacteria | Proteobacteri | Gammaprotec | unclassified | unclassified | unclassified | Gammaprotec | unclassified | unclassified | unclassified | unclassified | unclassified | 4,8 | 1,166190379 | 8,59E-05 | 1,33E-05 | 7 - | 0,000106982 - | 5 - | 9,49E-05 - | 4 - | 7,22E-05 - | 4 - | 7,32E-05 - | 4 - | 8,25E-05 - |
| 8049 | 1 | 0.2um_NOKCPHJN_140231 | Bacteria | Bacteroidetes | unclassified | unclassified | unclassified | unclassified | Bacteroidetes | unclassified | unclassified | unclassified | unclassified | unclassified | 0,2 | 0,4 | 5,83E-06 | 1,17E-05 | 0 - | 0 - | 0 - | 0 - | 1 - | 2,91E-05 - | 0 - | 0 - | 0 - | 0 - |
| 8050 | 1 | 0.2um_NOKCPHJN_806252 | Bacteria | Proteobacteri | Alphaproteob | Pelagibacteral | Pelagibacterac | Candidatus Pe | Candidatus Pe | unclassified | unclassified | unclassified | unclassified | unclassified | 1,8 | 0,4 | 0,000116852 | 2,86E-05 | 2 - | 0,000108793 - | 2 - | 0,000135089 - | 2 - | 0,000128419 - | 1 - | 6,52E-05 - | 2 - | 0,0001468 - |
| 8051 | 1 | 0.2um_NOKCPHJN_893102 | Bacteria | Proteobacteri | Gammaprotec | unclassified | unclassified | unclassified | various | information st | Translation, ril | 05C64 | Binds directly | 207954.MED9 | 1,2 | 0,748331477 | 7,35E-05 | 4,48E-05 | 0 - | 0 - | 2 - | 0,000123393 - | 2 - | 0,000117301 - | 1 - | 5,95E-05 - | 1 - | 6,70E-05 - |
| 8052 | 1 | 0.2um_NOKCPHJN_46818 | Bacteria | Proteobacteri | Alphaproteob | Rhodobactera | Rhodobactera | Shimia | Shimia aestua | information st | Transcription | 05D27 | DNA-depende | 439497.RR11_ | 2 | 0,632455532 | 1,90E-05 | 4,84E-06 | 3 - | 2,44E-05 - | 1 - | 1,01E-05 - | 2 - | 1,92E-05 - | 2 - | 1,95E-05 - | 2 - | 2,19E-05 - |
| 8053 | 1 | 3um_FIJJAFBF_523711 | Bacteria | Proteobacteri | Gammaprotec | Oceanospirilla | Saccharospirill | Reinekea | Reinekea blan | metabolism | Nucleotide tra | 16Q4Q | Catalyzes the | 314283.MED2 | 1 | 0,632455532 | 2,87E-05 | 2,03E-05 | 1 - | 2,35E-05 - | 1 - | 2,91E-05 - | 1 - | 2,77E-05 - | 0 - | 0 - | 2 - | 6,33E-05 - |
| 8054 | 1 | 3um_FIJJAFBF_15367 | Bacteria | Bacteroidetes | Flavobacteriia | Flavobacterial | Flavobacteriac | Polaribacter | various | information st | Transcription | 0P7W9 | Transcription | 313590.MED1 | 8,2 | 2,039607805 | 0,000264581 | 4,74E-05 | 12 - | 0,000329503 - | 8 - | 0,000272763 - | 7 - | 0,000226884 - | 6 - | 0,000197344 - | 8 - | 0,00029641 - |
| 8055 | 1 | 0.2um_NOKCPHJN_1036776 | Bacteria | Proteobacteri | Alphaproteob | Pelagibacteral | Pelagibacterac | Candidatus Pe | Candidatus Pe | cellular proces | Cell wall/mem | 7257 | NA | 335992.SAR11 | 2,6 | 0,489897949 | 0,000100735 | 2,46E-05 | 2 - | 6,43E-05 - | 2 - | 7,98E-05 - | 3 - | 0,000113851 - | 3 - | 0,000115532 - | 3 - | 0,000130147 - |
| 8056 | 1 | 0.2um_NOKCPHJN_299589 | Bacteria | Proteobacteri | Alphaproteob | Pelagibacteral | Pelagibacterac | various | various | unclassified | unclassified | unclassified | unclassified | unclassified | 0,6 | 0,489897949 | 9,06E-05 | 7,53E-05 | 1 - | 0,000126128 - | 1 - | 0,000156614 - | 0 - | 0 - | 0 - | 0 - | 1 - | 0,000170192 - |
| 8057 | 1 | 3um_FIJJAFBF_488140 | Bacteria | Bacteroidetes | Flavobacteriia | Flavobacterial | Flavobacteriac | Algibacter | Algibacter sp. | poorly charact | Function unkn | 05ERK | gliding motilit | 1046627.BZAF | 1 | 0,894427191 | 2,54E-05 | 2,35E-05 | 1 - | 2,21E-05 - | 0 - | 0 - | 2 - | 5,21E-05 - | 2 - | 5,29E-05 - | 0 - | 0 - |
| 8058 | 1 | 3um_FIJJAFBF_267298 | Bacteria | Proteobacteri | Alphaproteob | Pelagibacteral | Pelagibacterac | Candidatus Pe | Candidatus Pe | metabolism | Amino acid tra | 08KMW | Glycine betain | 335992.SAR11 | 4,4 | 0,8 | 0,000193627 | 4,67E-05 | 4 - | 0,000146212 - | 5 - | 0,000226941 - | 3 - | 0,000129442 - | 5 - | 0,000218922 - | 5 - | 0,000246615 - |
| 8059 | 1 | 3um_FIJJAFBF_183990 | unclassified | unclassified | unclassified | unclassified | unclassified | unclassified | unclassified | unclassified | unclassified | unclassified | unclassified | unclassified | 1 | 0 | 5,98E-05 | 5,68E-06 | 1 - | 5,01E-05 - | 1 - | 6,22E-05 - | 1 - | 5,92E-05 - | 1 - | 6,00E-05 - | 1 - | 6,76E-05 - |
| 8060 | 1 | 0.2um_NOKCPHJN_656151 | Bacteria | Proteobacteri | Gammaprotec | Cellvibrionales | Halieaceae | Haliea | Haliea salexig | unclassified | unclassified | unclassified | unclassified | unclassified | 0,8 | 0,4 | 1,30E-05 | 6,53E-06 | 0 - | 0 - | 1 - | 1,62E-05 - | 1 - | 1,54E-05 - | 1 - | 1,56E-05 - | 1 - | 1,76E-05 - |
| 8061 | 1 | 0.2um_NOKCPHJN_37736 | Bacteria | Bacteroidetes | Flavobacteriia | Flavobacterial | unclassified | unclassified | Flavobacterial | metabolism | Inorganic ion t | 05C27 | TonB-depend | 313603.FB217 | 1,4 | 0,8 | 1,83E-05 | 1,03E-05 | 2 - | 2,29E-05 - | 1 - | 1,42E-05 - | 2 - | 2,71E-05 - | 2 - | 2,75E-05 - | 0 - | 0 - |
| 8062 | 1 | 3um_FIJJAFBF_132427 | Bacteria | Bacteroidetes | Flavobacteriia | Flavobacterial | Flavobacteriac | Aequorivita | Aequorivita sp | unclassified | unclassified | unclassified | unclassified | unclassified | 1,2 | 0,4 | 0,000260514 | 8,73E-05 | 1 - | 0,000182185 - | 1 - | 0,00022622 - | 2 - | 0,000430102 - | 1 - | 0,000218227 - | 1 - | 0,000245833 - |
| 8063 | 1 | 3um_FIJJAFBF_08681 | Bacteria | Bacteroidetes | Flavobacteriia | Flavobacterial | Flavobacteriac | Winogradskye | Winogradskye | poorly charact | Function unkn | 0904U | tail collar dom | 1046627.BZAF | 3,2 | 1,166190379 | 0,000232049 | 9,31E-05 | 2 - | 0,000119559 - | 5 - | 0,000371143 - | 2 - | 0,000141127 - | 4 - | 0,000286424 - | 3 - | 0,000241991 - |
| 8064 | 1 | 3um_FIJJAFBF_123201 | Viruses | unclassified | unclassified | unclassified | unclassified | unclassified | uncultured M | unclassified | unclassified | unclassified | unclassified | unclassified | 0,8 | 0,4 | 5,31E-05 | 2,73E-05 | 1 - | 5,54E-05 - | 1 - | 6,88E-05 - | 0 - | 0 - | 1 - | 6,64E-05 - | 1 - | 7,48E-05 - |
| 8065 | 1 | 3um_FIJJAFBF_01154 | Bacteria | Bacteroidetes | Flavobacteriia | Flavobacterial | Flavobacteriac | Polaribacter | Polaribacter a | cellular proces | Cell wall/mem | 09G63 | Channel that | 313598.MED1 | 1 | 0 | 0,000103808 | 9,86E-06 | 1 - | 8,70E-05 - | 1 - | 0,000107969 - | 1 - | 0,000102638 - | 1 - | 0,000104154 - | 1 - | 0,000117329 - |
| 8066 | 1 | 3um_FIJJAFBF_770533 | Bacteria | Cyanobacteria | unclassified | unclassified | unclassified | unclassified | Cyanobacteria | unclassified | unclassified | unclassified | unclassified | unclassified | 0,2 | 0,4 | 1,88E-05 | 3,76E-05 | 1 - | 9,41E-05 - | 0 - | 0 - | 0 - | 0 - | 0 - | 0 - | 0 - | 0 - |
| 8067 | 1 | 0.2um_NOKCPHJN_862589 | Bacteria | Proteobacteri | Gammaprotec | Cellvibrionales | Porticoccacea | unclassified | SAR92 bacteri | unclassified | unclassified | unclassified | unclassified | unclassified | 7 | 0,632455532 | 0,000132858 | 2,40E-05 | 6 - | 9,46E-05 - | 7 - | 0,000137037 - | 7 - | 0,000130271 - | 7 - | 0,000132195 - | 8 - | 0,000170192 - |
| 8068 | 1 | 0.2um_NOKCPHJN_317372 | Viruses | unclassified | unclassified | unclassified | unclassified | unclassified | uncultured vir | unclassified | unclassified | unclassified | unclassified | unclassified | 0,2 | 0,4 | 2,13E-05 | 4,25E-05 | 1 - | 0,000106275 - | 0 - | 0 - | 0 - | 0 - | 0 - | 0 - | 0 - | 0 - |
| 8069 | 1 | 0.2um_NOKCPHJN_290411 | Bacteria | Proteobacteri | Gammaprotec | Alteromonada | Colwelliaceae | Colwellia | various | information st | Translation, ril | COG0051 | tRNA binding | 167879.CPS_0 | 1 | 0 | 0,000133036 | 1,26E-05 | 1 - | 0,000111434 - | 1 - | 0,000138368 - | 1 - | 0,000131536 - | 1 - | 0,000133479 - | 1 - | 0,000150364 - |
| 8070 | 1 | 0.2um_NOKCPHJN_193556 | Bacteria | Verrucomicrol | Opitutae | unclassified | unclassified | unclassified | Opitutae bact | unclassified | unclassified | unclassified | unclassified | unclassified | 4,2 | 0,748331477 | 0,000118515 | 1,80E-05 | 5 - | 0,000119311 - | 3 - | 8,89E-05 - | 4 - | 0,000112667 - | 5 - | 0,000142914 - | 4 - | 0,000128794 - |
| 8071 | 1 | 0.2um_NOKCPHJN_495319 | Bacteria | Proteobacteri | Alphaproteob | Rhodobactera | Rhodobactera | Planktomarina | Planktomarina | metabolism | Amino acid tra | 01SEG | branched-chai | 314262.MED1 | 1,8 | 0,4 | 5,89E-05 | 1,63E-05 | 1 - | 2,69E-05 - | 2 - | 6,69E-05 - | 2 - | 6,36E-05 - | 2 - | 6,45E-05 - | 2 - | 7,27E-05 - |
| 8072 | 1 | 3um_FIJJAFBF_807137 | Viruses | unclassified | unclassified | unclassified | Mimiviridae | Cafeteriavirus | Cafeteria roen | unclassified | unclassified | unclassified | unclassified | unclassified | 0,6 | 0,489897949 | 1,60E-05 | 1,31E-05 | 0 - | 0 - | 1 - | 2,75E-05 - | 1 - | 2,62E-05 - | 1 - | 2,65E-05 - | 0 - | 0 - |
| 8073 | 1 | 0.2um_NOKCPHJN_05117 | Bacteria | Bacteroidetes | Flavobacteriia | Flavobacterial | Crocinitomica | unclassified | Crocinitomica | unclassified | unclassified | unclassified | unclassified | unclassified | 1 | 0 | 6,01E-05 | 5,71E-06 | 1 - | 5,03E-05 - | 1 - | 6,25E-05 - | 1 - | 5,94E-05 - | 1 - | 6,03E-05 - | 1 - | 6,79E-05 - |
| 8074 | 1 | 0.2um_NOKCPHJN_517649 | Viruses | unclassified | unclassified | unclassified | Mimiviridae | Indivirus | Indivirus ILV1 | information st | Replication, re | 0V9WU | polymerase (D | 7955.ENSDAR | 0,2 | 0,4 | 8,19E-06 | 1,64E-05 | 0 - | 0 - | 1 - | 4,10E-05 - | 0 - | 0 - | 0 - | 0 - | 0 - | 0 - |
| 8075 | 1 | 3um_FIJJAFBF_205191 | Eukaryota | Bacillariophyt | Coscinodiscop | Chaetocerotal | Acanthocerata | Acanthoceras | Acanthoceras | metabolism | Energy produc | 0IFBF | One of the tw | 45157.CMV08 | 12,4 | 0,489897949 | 0,000482843 | 3,64E-05 | 13 - | 0,000425099 - | 12 - | 0,000487244 - | 12 - | 0,000463187 - | 13 - | 0,000509197 - | 12 - | 0,000529485 - |
| 8076 | 1 | 0.2um_NOKCPHJN_1033107 | Bacteria | Proteobacteri | Gammaprotec | Oceanospirilla | Saccharospirill | Reinekea | Reinekea fors | unclassified | unclassified | unclassified | unclassified | unclassified | 1 | 0 | 3,99E-05 | 3,79E-06 | 1 - | 3,35E-05 - | 1 - | 4,16E-05 - | 1 - | 3,95E-05 - | 1 - | 4,01E-05 - | 1 - | 4,52E-05 - |
| 8077 | 1 | 3um_FIJJAFBF_31133 | Bacteria | Proteobacteri | Alphaproteob | Rhodobactera | Rhodobactera | Amylibacter | Amylibacter c | metabolism | Amino acid tra | COG0137 | Citrulline--asp | 351016.RAZW | 0,2 | 0,4 | 5,65E-06 | 1,13E-05 | 1 - | 2,83E-05 - | 0 - | 0 - | 0 - | 0 - | 0 - | 0 - | 0 - | 0 - |
| 8078 | 1 | 3um_FIJJAFBF_727986 | Bacteria | Bacteroidetes | Flavobacteriia | Flavobacterial | Flavobacteriac | Polaribacter | Polaribacter s | poorly charact | Function unkn | 05WQR | glycoside hydr | 313598.MED1 | 0,4 | 0,489897949 | 5,11E-05 | 6,29E-05 | 1 - | 0,000117119 - | 0 - | 0 - | 1 - | 0,000138247 - | 0 - | 0 - | 0 - | 0 - |
| 8079 | 1 | 3um_FIJJAFBF_215623 | Bacteria | Proteobacteri | Gammaprotec | Arenicellales | Arenicellaceae | Arenicella | Arenicella xan | unclassified | unclassified | unclassified | unclassified | unclassified | 1 | 0 | 9,65E-05 | 9,16E-06 | 1 - | 8,08E-05 - | 1 - | 0,000100365 - | 1 - | 9,54E-05 - | 1 - | 9,68E-05 - | 1 - | 0,000109067 - |
| 8080 | 1 | 3um_FIJJAFBF_738821 | Bacteria | Bacteroidetes | Flavobacteriia | Flavobacterial | Flavobacteriac | Arenitalea | Arenitalea lut | cellular proces | Cell wall/mem | 11TPU | outer membra | 1046627.BZAF | 2,2 | 0,748331477 | 0,000177433 | 5,43E-05 | 3 - | 0,000207428 - | 3 - | 0,000257564 - | 2 - | 0,000163232 - | 2 - | 0,000165643 - | 1 - | 9,33E-05 - |
| 8081 | 1 | 3um_FIJJAFBF_815478 | Eukaryota | Eustigmatoph | unclassified | Eustigmatales | Monodopsida | Nannochlorop | Nannochlorop | metabolism | Lipid transpor | COG0439 | acetyl-CoA ca | 35128.Thaps6 | 0,8 | 0,4 | 3,58E-05 | 1,84E-05 | 1 - | 3,74E-05 - | 1 - | 4,64E-05 - | 0 - | 0 - | 1 - | 4,48E-05 - | 1 - | 5,04E-05 - |
| 8082 | 1 | 3um_FIJJAFBF_175156 | Bacteria | Proteobacteri | Alphaproteob | Rhodobactera | Rhodobactera | unclassified | Rhodobactera | unclassified | unclassified | unclassified | unclassified | unclassified | 0,6 | 0,489897949 | 1,71E-05 | 1,41E-05 | 0 - | 0 - | 0 - | 0 - | 1 - | 2,72E-05 - | 1 - | 2,76E-05 - | 1 - | 3,10E-05 - |
| 8083 | 1 | 3um_FIJJAFBF_481948 | Bacteria | Proteobacteri | Alphaproteob | Rhodobactera | Rhodobactera | unclassified | Rhodobactera | metabolism | Energy produc | COG0712 | F(1)F(0) ATP s | 292414.TM10 | 2,4 | 0,489897949 | 0,000143328 | 3,30E-05 | 2 - | 9,98E-05 - | 3 - | 0,000185894 - | 3 - | 0,000176716 - | 2 - | 0,000119551 - | 2 - | 0,000134673 - |
| 8084 | 2 | 0.2um_NOKCPHJN_219941;0.2um_NOKCPHJN_8 | Bacteria | Proteobacteri | Gammaprotec | Cellvibrionales | unclassified | unclassified | Cellvibrionales | unclassified | unclassified | unclassified | unclassified | unclassified | 0,8 | 0,4 | 3,50E-05 | 1,78E-05 | 1 - | 3,79E-05 - | 1 - | 4,70E-05 - | 1 - | 4,47E-05 - | 1 - | 4,54E-05 - | 0 - | 0 - |
| 8085 | 1 | 0.2um_NOKCPHJN_447107 | Bacteria | Bacteroidetes | Flavobacteriia | Flavobacterial | Flavobacteriac | Formosa | Formosa sp. H | unclassified | unclassified | unclassified | unclassified | unclassified | 3,6 | 0,8 | 8,13E-05 | 1,61E-05 | 4 - | 7,68E-05 - | 4 - | 9,53E-05 - | 4 - | 9,06E-05 - | 4 - | 9,20E-05 - | 2 - | 5,18E-05 - |
| 8086 | 1 | 3um_FIJJAFBF_699203 | Bacteria | Proteobacteri | Gammaprotec | unclassified | unclassified | unclassified | Gammaprotec | unclassified | unclassified | unclassified | unclassified | unclassified | 1 | 0 | 2,91E-05 | 2,76E-06 | 1 - | 2,44E-05 - | 1 - | 3,03E-05 - | 1 - | 2,88E-05 - | 1 - | 2,92E-05 - | 1 - | 3,29E-05 - |
| 8087 | 1 | 3um_FIJJAFBF_402487 | Bacteria | Bacteroidetes | Flavobacteriia | various | various | various | various | information st | Translation, ril | COG0088 | One of the pri | 867900.Celly_ | 1,6 | 0,8 | 0,000102476 | 4,87E-05 | 2 - | 0,000109834 - | 1 - | 6,82E-05 - | 3 - | 0,000194472 - | 1 - | 6,58E-05 - | 1 - | 7,41E-05 - |
| 8088 | 1 | 0.2um_NOKCPHJN_911732 | Bacteria | Proteobacteri | Alphaproteob | Pelagibacteral | Pelagibacterac | Candidatus Pe | Candidatus Pe | unclassified | unclassified | unclassified | unclassified | unclassified | 2 | 0,632455532 | 0,000178011 | 5,54E-05 | 2 - | 0,000151022 - | 2 - | 0,000187525 - | 2 - | 0,000178266 - | 3 - | 0,000271349 - | 1 - | 0,000101891 - |
| 8089 | 1 | 3um_FIJJAFBF_336548 | Bacteria | Spirochaetes | Spirochaetia | unclassified | Leptospiracea | Leptospira | Leptospira sp. | unclassified | unclassified | unclassified | unclassified | unclassified | 4 | 1,095445115 | 7,96E-05 | 2,09E-05 | 5 - | 8,39E-05 - | 4 - | 8,33E-05 - | 2 - | 3,96E-05 - | 5 - | 0,000100499 - | 4 - | 9,06E-05 - |
| 8090 | 1 | 0.2um_NOKCPHJN_350722 | Bacteria | Proteobacteri | Alphaproteob | Rhodobactera | unclassified | unclassified | Rhodobactera | unclassified | unclassified | unclassified | unclassified | unclassified | 2 | 1,264911064 | 0,000169431 | 0,000108583 | 3 - | 0,00021387 - | 1 - | 8,85E-05 - | 0 - | 0 - | 3 - | 0,00025618 - | 3 - | 0,000288586 - |
| 8091 | 1 | 0.2um_NOKCPHJN_222811 | Bacteria | Bacteroidetes | unclassified | unclassified | unclassified | unclassified | Bacteroidetes | unclassified | unclassified | unclassified | unclassified | unclassified | 0,2 | 0,4 | 2,39E-06 | 4,77E-06 | 0 - | 0 - | 0 - | 0 - | 0 - | 0 - | 1 - | 1,19E-05 - | 0 - | 0 - |
| 8092 | 1 | 3um_FIJJAFBF_529379 | Bacteria | Proteobacteri | Alphaproteob | Pelagibacteral | Pelagibacterac | Candidatus Pe | Candidatus Pe | unclassified | unclassified | unclassified | unclassified | unclassified | 0,8 | 0,4 | 5,36E-05 | 2,70E-05 | 0 - | 0 - | 1 - | 6,69E-05 - | 1 - | 6,36E-05 - | 1 - | 6,45E-05 - | 1 - | 7,27E-05 - |
| 8093 | 1 | 3um_FIJJAFBF_10077 | Bacteria | Bacteroidetes | Flavobacteriia | Flavobacterial | Flavobacteriac | Formosa | Formosa sp. H | cellular proces | Posttranslatio | COG0330 | Band 7 protei | 983544.Lacal_ | 0,4 | 0,489897949 | 1,87E-05 | 2,30E-05 | 1 - | 4,25E-05 - | 0 - | 0 - | 0 - | 0 - | 1 - | 5,09E-05 - | 0 - | 0 - |
| 8094 | 1 | 0.2um_NOKCPHJN_813082 | Bacteria | Proteobacteri | Alphaproteob | Pelagibacteral | Pelagibacterac | Candidatus Pe | Candidatus Pe | metabolism | Inorganic ion t | 05PWN | phosphate ab | 335992.SAR11 | 4 | 0,894427191 | 0,000669552 | 0,000192798 | 3 - | 0,000414856 - | 4 - | 0,000686838 - | 5 - | 0,000816158 - | 3 - | 0,000496928 - | 5 - | 0,000932979 - |
| 8095 | 1 | 0.2um_NOKCPHJN_79998 | Bacteria | Proteobacteri | Gammaprotec | unclassified | unclassified | unclassified | Gammaprotec | metabolism | Energy produc | 05CV4 | Subunits I and | 314277.MED1 | 2,2 | 0,748331477 | 7,92E-05 | 2,66E-05 | 2 - | 6,11E-05 - | 2 - | 7,58E-05 - | 3 - | 0,000108098 - | 3 - | 0,000109694 - | 1 - | 4,12E-05 - |
| 8096 | 1 | 0.2um_NOKCPHJN_992072 | Eukaryota | Bacillariophyt | Coscinodiscop | Thalassiosirale | Thalassiosirac | Thalassiosira | Thalassiosira | metabolism | Energy produc | COG0377 | NDH-1 shuttle | 2850.Phatr41 | 1 | 0 | 6,62E-05 | 6,29E-06 | 1 - | 5,54E-05 - | 1 - | 6,88E-05 - | 1 - | 6,55E-05 - | 1 - | 6,64E-05 - | 1 - | 7,48E-05 - |
| 8097 | 1 | 3um_FIJJAFBF_124744 | Bacteria | Proteobacteri | Alphaproteob | Rhodobactera | Rhodobactera | Planktomarina | Planktomarina | unclassified | unclassified | unclassified | unclassified | unclassified | 4,4 | 0,8 | 0,000389062 | 8,28E-05 | 5 - | 0,000370248 - | 5 - | 0,000459738 - | 4 - | 0,000349632 - | 3 - | 0,000266097 - | 5 - | 0,000499595 - |
| 8098 | 1 | 3um_FIJJAFBF_607780 | Bacteria | Proteobacteri | Gammaprotec | Oceanospirilla | Saccharospirill | Reinekea | Reinekea fors | metabolism | Amino acid tra | 0QJVS | branched-chai | 999141.GME_ | 3,4 | 0,8 | 0,000115402 | 1,56E-05 | 5 - | 0,00014492 - | 3 - | 0,000107969 - | 3 - | 0,000102638 - | 3 - | 0,000104154 - | 3 - | 0,000117329 - |
| 8099 | 1 | 3um_FIJJAFBF_24165 | Bacteria | Bacteroidetes | Flavobacteriia | Flavobacterial | Flavobacteriac | Polaribacter | Polaribacter s | metabolism | Coenzyme tra | COG0171 | nh(3)-depend | 313598.MED1 | 2 | 0 | 0,000104601 | 9,93E-06 | 2 - | 8,76E-05 - | 2 - | 0,000108793 - | 2 - | 0,000103422 - | 2 - | 0,000104949 - | 2 - | 0,000118225 - |
| 8100 | 1 | 3um_FIJJAFBF_98072 | Bacteria | Proteobacteri | Gammaprotec | unclassified | unclassified | unclassified | OM182 bacte | cellular proces | Cell wall/mem | 0QTSX | Peptidoglycan | 351348.Maqu | 5,4 | 0,8 | 0,00041543 | 7,35E-05 | 6 - | 0,000386888 - | 5 - | 0,000400334 - | 4 - | 0,000304454 - | 6 - | 0,000463427 - | 6 - | 0,000522049 - |
| 8101 | 2 | 0.2um_NOKCPHJN_177948;0.2um_NOKCPHJN_8 | Eukaryota | unclassified | Oomycetes | Peronosporale | Peronosporac | Plasmopara | Plasmopara ha | cellular proces | Intracellular tr | 0YIKI | Small GTPase | various | 0,8 | 0,4 | 5,64E-05 | 2,87E-05 | 1 - | 6,11E-05 - | 1 - | 7,58E-05 - | 1 - | 7,21E-05 - | 1 - | 7,31E-05 - | 0 - | 0 - |
| 8102 | 1 | 0.2um_NOKCPHJN_944286 | Bacteria | Proteobacteri | Deltaproteoba | Desulfobacter | Desulfobacter | Desulfatibacill | Desulfatibacill | poorly charact | Function unkn | 0XNQ8 | alkaline phosp | 103690.alr529 | 0,2 | 0,4 | 1,84E-05 | 3,67E-05 | 1 - | 9,18E-05 - | 0 - | 0 - | 0 - | 0 - | 0 - | 0 - | 0 - | 0 - |
| 8103 | 1 | 3um_FIJJAFBF_350012 | Bacteria | Bacteroidetes | Cytophagia | Cytophagales | Flammeovirga | unclassified | Flammeovirga | unclassified | unclassified | unclassified | unclassified | unclassified | 5,6 | 0,8 | 0,001204037 | 8,80E-05 | 7 - | 0,001275297 - | 5 - | 0,001131102 - | 5 - | 0,001075256 - | 6 - | 0,001309365 - | 5 - | 0,001229163 - |
| 8104 | 1 | 3um_FIJJAFBF_292976 | Bacteria | Bacteroidetes | Flavobacteriia | Flavobacterial | Crocinitomica | unclassified | Crocinitomica | information st | Translation, ril | 09EM4 | Binds directly | 525257.HMPR | 1,4 | 0,489897949 | 0,000168971 | 6,23E-05 | 1 - | 0,000100681 - | 2 - | 0,000250033 - | 2 - | 0,000237688 - | 1 - | 0,000120599 - | 1 - | 0,000135855 - |
| 8105 | 1 | 3um_FIJJAFBF_451385 | Bacteria | Proteobacteri | Gammaprotec | Oceanospirilla | Saccharospirill | Saccharospiril | Saccharospiril | metabolism | Nucleotide tra | COG0015 | adenylosuccin | 247634.GPB2 | 2 | 0 | 5,98E-05 | 5,68E-06 | 2 - | 5,01E-05 - | 2 - | 6,22E-05 - | 2 - | 5,92E-05 - | 2 - | 6,00E-05 - | 2 - | 6,76E-05 - |
| 8106 | 1 | 3um_FIJJAFBF_80626 | Bacteria | Bacteroidetes | Flavobacteriia | Flavobacterial | Crocinitomica | unclassified | Crocinitomica | information st | Transcription | 0P7W9 | Transcription | 755732.Fluta_ | 0,2 | 0,4 | 7,55E-06 | 1,51E-05 | 0 - | 0 - | 0 - | 0 - | 0 - | 0 - | 0 - | 0 - | 1 - | 3,78E-05 - |
| 8107 | 1 | 0.2um_NOKCPHJN_608547 | Eukaryota | Phaeophyceae | unclassified | Ectocarpales | Ectocarpaceae | Ectocarpus | Ectocarpus sili | unclassified | unclassified | unclassified | unclassified | unclassified | 1 | 0,894427191 | 8,66E-05 | 8,05E-05 | 1 - | 7,40E-05 - | 2 - | 0,000183895 - | 2 - | 0,000174816 - | 0 - | 0 - | 0 - | 0 - |
| 8108 | 1 | 0.2um_NOKCPHJN_527312 | Bacteria | Bacteroidetes | Flavobacteriia | unclassified | unclassified | unclassified | Flavobacteria | cellular proces | Intracellular tr | 0BIEB | domain-conta | 1005048.CFU_ | 3,2 | 0,4 | 5,91E-05 | 1,25E-05 | 3 - | 4,60E-05 - | 3 - | 5,72E-05 - | 3 - | 5,43E-05 - | 3 - | 5,51E-05 - | 4 - | 8,28E-05 - |
| 8109 | 1 | 3um_FIJJAFBF_70541 | Bacteria | Bacteroidetes | unclassified | unclassified | unclassified | unclassified | Bacteroidetes | poorly charact | Function unkn | 11QQ6 | NA | 886377.Murru | 3 | 0 | 0,000130089 | 1,24E-05 | 3 - | 0,000108965 - | 3 - | 0,000135303 - | 3 - | 0,000128622 - | 3 - | 0,000130522 - | 3 - | 0,000147033 - |
| 8110 | 1 | 3um_FIJJAFBF_350298 | Bacteria | Proteobacteri | Gammaprotec | Cellvibrionales | Spongiibacter | Dasania | Dasania marin | information st | Translation, ril | COG0256 | This is one of | 1746360.PFWH | 4,4 | 0,489897949 | 0,000523512 | 7,56E-05 | 5 - | 0,000499029 - | 4 - | 0,000495718 - | 4 - | 0,000471243 - | 4 - | 0,000478203 - | 5 - | 0,000673367 - |
| 8111 | 1 | 3um_FIJJAFBF_14240 | Bacteria | Proteobacteri | Betaproteoba | unclassified | unclassified | unclassified | Betaproteoba | information st | Translation, ril | 17FIT | 50s ribosomal | 398578.Daci_ | 1 | 0 | 0,000177957 | 1,69E-05 | 1 - | 0,000149061 - | 1 - | 0,000185089 - | 1 - | 0,000175951 - | 1 - | 0,00017855 - | 1 - | 0,000201136 - |
| 8112 | 1 | 3um_FIJJAFBF_39296 | unclassified | unclassified | unclassified | unclassified | unclassified | unclassified | unclassified | unclassified | unclassified | unclassified | unclassified | unclassified | 1,4 | 0,489897949 | 4,02E-05 | 1,26E-05 | 2 - | 4,89E-05 - | 2 - | 6,08E-05 - | 1 - | 2,89E-05 - | 1 - | 2,93E-05 - | 1 - | 3,30E-05 - |
| 8113 | 1 | 3um_FIJJAFBF_184535 | Bacteria | Proteobacteri | Gammaprotec | Oceanospirilla | Saccharospirill | Reinekea | Reinekea fors | poorly charact | Function unkn | 05W3E | transcriptiona | 314283.MED2 | 0,6 | 0,489897949 | 9,26E-05 | 7,70E-05 | 1 - | 0,000128963 - | 1 - | 0,000160134 - | 0 - | 0 - | 0 - | 0 - | 1 - | 0,000174016 - |
| 8114 | 1 | 3um_FIJJAFBF_122459 | Bacteria | Bacteroidetes | Flavobacteriia | Flavobacterial | Flavobacteriac | Polaribacter | Polaribacter s | metabolism | Amino acid tra | 0P7S1 | Catalyzes the | 313598.MED1 | 0,8 | 0,4 | 3,78E-05 | 1,94E-05 | 1 - | 3,94E-05 - | 1 - | 4,90E-05 - | 0 - | 0 - | 1 - | 4,72E-05 - | 1 - | 5,32E-05 - |
| 8115 | 1 | 3um_FIJJAFBF_43620 | Bacteria | Proteobacteri | Alphaproteob | Rhodobactera | Rhodobactera | Primorskyibac | Primorskyibac | metabolism | Secondary me | 01Q86 | extracellular s | 391595.RLO14 | 1 | 1,095445115 | 3,51E-05 | 3,96E-05 | 1 - | 3,06E-05 - | 0 - | 0 - | 3 - | 0,000108386 - | 1 - | 3,67E-05 - | 0 - | 0 - |
| 8116 | 2 | 0.2um_NOKCPHJN_58705;0.2um_NOKCPHJN_96 | Bacteria | Proteobacteri | Alphaproteob | Pelagibacteral | Pelagibacterac | Candidatus Pe | Candidatus Pe | information st | Translation, ril | 082FS | 50S ribosomal | 335992.SAR11 | 1,2 | 0,4 | 0,000216479 | 7,46E-05 | 1 - | 0,000151022 - | 1 - | 0,000187525 - | 1 - | 0,000178266 - | 2 - | 0,000361798 - | 1 - | 0,000203782 - |
| 8117 | 1 | 0.2um_NOKCPHJN_224037 | Bacteria | unclassified | unclassified | unclassified | unclassified | unclassified | uncultured ba | unclassified | unclassified | unclassified | unclassified | unclassified | 3 | 1,095445115 | 5,15E-05 | 1,48E-05 | 5 - | 7,34E-05 - | 3 - | 5,47E-05 - | 2 - | 3,47E-05 - | 2 - | 3,52E-05 - | 3 - | 5,94E-05 - |
| 8118 | 1 | 3um_FIJJAFBF_30316 | Bacteria | unclassified | unclassified | unclassified | unclassified | unclassified | uncultured ba | unclassified | unclassified | unclassified | unclassified | unclassified | 1,8 | 0,748331477 | 5,12E-05 | 1,71E-05 | 3 - | 7,39E-05 - | 2 - | 6,12E-05 - | 2 - | 5,81E-05 - | 1 - | 2,95E-05 - | 1 - | 3,32E-05 - |
| 8119 | 1 | 3um_FIJJAFBF_506019 | Bacteria | Proteobacteri | Gammaprotec | Cellvibrionales | Porticoccacea | unclassified | gamma prote | cellular proces | Cell wall/mem | 0QTSX | Peptidoglycan | 314287.GB22 | 2,4 | 0,489897949 | 0,000199084 | 3,10E-05 | 3 - | 0,000211246 - | 2 - | 0,00017487 - | 2 - | 0,000166236 - | 3 - | 0,000253037 - | 2 - | 0,00019003 - |
| 8120 | 1 | 3um_FIJJAFBF_10622 | Bacteria | Bacteroidetes | Cytophagia | Cytophagales | Cytophagacea | Ekhidna | Ekhidna lutea | metabolism | Carbohydrate | 099YR | Catalyzes the | 388413.ALPR1 | 1,2 | 0,4 | 3,86E-05 | 1,33E-05 | 1 - | 2,69E-05 - | 1 - | 3,35E-05 - | 1 - | 3,18E-05 - | 2 - | 6,45E-05 - | 1 - | 3,64E-05 - |
| 8121 | 1 | 3um_FIJJAFBF_712024 | Bacteria | Proteobacteri | Gammaprotec | Oceanospirilla | Saccharospirill | Reinekea | Reinekea blan | cellular proces | Cell wall/mem | 16PEJ | Phospho-2-de | 314282.PCNP | 1,4 | 1,019803903 | 6,56E-05 | 4,29E-05 | 3 - | 0,000122537 - | 2 - | 0,000101437 - | 0 - | 0 - | 1 - | 4,89E-05 - | 1 - | 5,51E-05 - |
| 8122 | 1 | 0.2um_NOKCPHJN_179571 | Bacteria | Proteobacteri | Alphaproteob | Rhodobactera | Rhodobactera | Amylibacter | Amylibacter k | metabolism | Energy produc | 02EEZ | Catalyzes a 2- | 314262.MED1 | 2,4 | 0,489897949 | 2,89E-05 | 7,70E-06 | 2 - | 2,00E-05 - | 2 - | 2,48E-05 - | 3 - | 3,54E-05 - | 2 - | 2,40E-05 - | 3 - | 4,05E-05 - |
| 8123 | 1 | 0.2um_NOKCPHJN_209593 | Bacteria | Proteobacteri | Gammaprotec | Cellvibrionales | Porticoccacea | unclassified | gamma prote | metabolism | Energy produc | 0QJ3X | Aconitate hyd | 203122.Sde_1 | 0,4 | 0,489897949 | 5,89E-06 | 7,29E-06 | 1 - | 1,31E-05 - | 1 - | 1,63E-05 - | 0 - | 0 - | 0 - | 0 - | 0 - | 0 - |
| 8124 | 1 | 3um_FIJJAFBF_226844 | Bacteria | unclassified | unclassified | unclassified | unclassified | unclassified | uncultured ba | poorly charact | Function unkn | 05DPC | Ragb susd dor | 391598.FBBAL | 1,4 | 0,8 | 3,91E-05 | 2,30E-05 | 2 - | 4,68E-05 - | 2 - | 5,81E-05 - | 1 - | 2,76E-05 - | 0 - | 0 - | 2 - | 6,31E-05 - |
| 8125 | 1 | 3um_FIJJAFBF_627199 | Bacteria | Actinobacteri | unclassified | unclassified | unclassified | unclassified | Actinobacteri | unclassified | unclassified | unclassified | unclassified | unclassified | 0,6 | 0,8 | 2,75E-05 | 3,51E-05 | 2 - | 8,47E-05 - | 1 - | 5,26E-05 - | 0 - | 0 - | 0 - | 0 - | 0 - | 0 - |
| 8126 | 1 | 3um_FIJJAFBF_163905 | Bacteria | Proteobacteri | Alphaproteob | Rhodobactera | Rhodobactera | Tritonibacter | Tritonibacter | metabolism | Carbohydrate | 01QKD | xylose isomer | 292414.TM10 | 0,4 | 0,489897949 | 1,28E-05 | 1,56E-05 | 0 - | 0 - | 1 - | 3,27E-05 - | 1 - | 3,11E-05 - | 0 - | 0 - | 0 - | 0 - |
| 8127 | 1 | 3um_FIJJAFBF_242852 | Bacteria | various | various | various | various | various | various | metabolism | Amino acid tra | 099QE | chorismate m | 983544.Lacal_ | 11 | 1,788854382 | 0,000413441 | 4,70E-05 | 13 - | 0,000414472 - | 12 - | 0,000475063 - | 12 - | 0,000451607 - | 10 - | 0,000381898 - | 8 - | 0,000344166 - |
| 8128 | 1 | 0.2um_NOKCPHJN_137974 | Eukaryota | unclassified | unclassified | Kinetoplastida | Trypanosomat | Leishmania | Leishmania m | unclassified | unclassified | unclassified | unclassified | unclassified | 0,6 | 0,489897949 | 3,82E-06 | 3,12E-06 | 0 - | 0 - | 1 - | 6,54E-06 - | 1 - | 6,22E-06 - | 1 - | 6,31E-06 - | 0 - | 0 - |
| 8129 | 1 | 0.2um_NOKCPHJN_1092476 | Bacteria | Proteobacteri | Alphaproteob | Pelagibacteral | Pelagibacterac | Candidatus Pe | Candidatus Pe | unclassified | unclassified | unclassified | unclassified | unclassified | 1,2 | 0,748331477 | 3,37E-05 | 2,16E-05 | 0 - | 0 - | 1 - | 2,78E-05 - | 1 - | 2,65E-05 - | 2 - | 5,37E-05 - | 2 - | 6,05E-05 - |
| 8130 | 1 | 0.2um_NOKCPHJN_573253 | Eukaryota | Apicomplexa | Conoidasida | Eucoccidiorida | Eimeriidae | Eimeria | Eimeria mitis | poorly charact | Function unkn | 03Q02 | NA | 5022.CBX917 | 0,2 | 0,4 | 2,12E-06 | 4,24E-06 | 1 - | 1,06E-05 - | 0 - | 0 - | 0 - | 0 - | 0 - | 0 - | 0 - | 0 - |
| 8131 | 1 | 3um_FIJJAFBF_504068 | Bacteria | Proteobacteri | Gammaprotec | Cellvibrionales | Porticoccacea | unclassified | gamma prote | information st | Translation, ril | COG0050 | This protein p | 314287.GB22 | 0,2 | 0,4 | 3,57E-05 | 7,14E-05 | 0 - | 0 - | 0 - | 0 - | 0 - | 0 - | 1 - | 0,00017855 - | 0 - | 0 - |
| 8132 | 2 | 0.2um_NOKCPHJN_532236;0.2um_NOKCPHJN_6 | Eukaryota | unclassified | various | various | various | various | various | cellular proces | Posttranslatio | KOG0174 | The proteaso | various | 1 | 0 | 5,76E-05 | 5,47E-06 | 1 - | 4,82E-05 - | 1 - | 5,99E-05 - | 1 - | 5,69E-05 - | 1 - | 5,78E-05 - | 1 - | 6,51E-05 - |
| 8133 | 1 | 3um_FIJJAFBF_52009 | Bacteria | Proteobacteri | Alphaproteob | Rhodobactera | Rhodobactera | Cribrihabitans | Cribrihabitans | unclassified | unclassified | unclassified | unclassified | unclassified | 1 | 0 | 8,90E-05 | 8,45E-06 | 1 - | 7,45E-05 - | 1 - | 9,25E-05 - | 1 - | 8,80E-05 - | 1 - | 8,93E-05 - | 1 - | 0,000100568 - |
| 8134 | 1 | 3um_FIJJAFBF_100813 | Bacteria | Bacteroidetes | Flavobacteriia | Flavobacterial | Flavobacteriac | Polaribacter | Polaribacter s | metabolism | Carbohydrate | COG1501 | hydrolase, fan | 313594.PI23P | 4 | 1,264911064 | 6,82E-05 | 2,24E-05 | 3 - | 4,30E-05 - | 5 - | 8,90E-05 - | 5 - | 8,46E-05 - | 5 - | 8,58E-05 - | 2 - | 3,87E-05 - |
| 8135 | 1 | 3um_FIJJAFBF_391682 | Eukaryota | unclassified | Dinophyceae | Gymnodiniale | Gymnodiniace | Amphidinium | Amphidinium | metabolism | Coenzyme tra | 1DQD8 | adenosylhom | 3055.EDP033 | 0,4 | 0,489897949 | 2,20E-05 | 2,72E-05 | 1 - | 5,06E-05 - | 0 - | 0 - | 1 - | 5,97E-05 - | 0 - | 0 - | 0 - | 0 - |
| 8136 | 1 | 0.2um_NOKCPHJN_48598 | Bacteria | Firmicutes | Bacilli | Lactobacillales | Leuconostoca | Weissella | Weissella con | unclassified | unclassified | unclassified | unclassified | unclassified | 13,2 | 2,785677655 | 0,000103645 | 1,34E-05 | 18 - | 0,000120465 - | 14 - | 0,000116342 - | 11 - | 8,69E-05 - | 13 - | 0,000104215 - | 10 - | 9,03E-05 - |
| 8137 | 1 | 0.2um_NOKCPHJN_96410 | Bacteria | Bacteroidetes | Flavobacteriia | Flavobacterial | Flavobacteriac | Polaribacter | Polaribacter s | unclassified | unclassified | unclassified | unclassified | unclassified | 2 | 0 | 4,32E-05 | 4,10E-06 | 2 - | 3,62E-05 - | 2 - | 4,49E-05 - | 2 - | 4,27E-05 - | 2 - | 4,33E-05 - | 2 - | 4,88E-05 - |
| 8138 | 1 | 0.2um_NOKCPHJN_371590 | Bacteria | Bacteroidetes | Flavobacteriia | Flavobacterial | Flavobacteriac | Formosa | Formosa sp. H | metabolism | Carbohydrate | 09CFZ | Transporter | 391587.KAOT | 2 | 0,632455532 | 3,59E-05 | 1,29E-05 | 1 - | 1,48E-05 - | 3 - | 5,50E-05 - | 2 - | 3,48E-05 - | 2 - | 3,53E-05 - | 2 - | 3,98E-05 - |
| 8139 | 1 | 3um_FIJJAFBF_189720 | Bacteria | Bacteroidetes | Flavobacteriia | Flavobacterial | Flavobacteriac | Polaribacter | Polaribacter s | unclassified | unclassified | unclassified | unclassified | unclassified | 0,4 | 0,489897949 | 4,79E-05 | 5,89E-05 | 0 - | 0 - | 0 - | 0 - | 0 - | 0 - | 1 - | 0,000112691 - | 1 - | 0,000126946 - |
| 8140 | 1 | 0.2um_NOKCPHJN_83843 | Bacteria | Proteobacteri | Alphaproteob | Rhodobactera | Rhodobactera | Roseovarius | Roseovarius h | poorly charact | Function unkn | COG3181 | extra-cytoplas | 340100.Bpet0 | 1,2 | 0,748331477 | 4,88E-05 | 3,20E-05 | 1 - | 3,47E-05 - | 2 - | 8,61E-05 - | 2 - | 8,19E-05 - | 1 - | 4,15E-05 - | 0 - | 0 - |
| 8141 | 1 | 0.2um_NOKCPHJN_135912 | Bacteria | Proteobacteri | Alphaproteob | Rhodobactera | Rhodobactera | Actibacterium | Actibacterium | unclassified | unclassified | unclassified | unclassified | unclassified | 0,4 | 0,489897949 | 9,61E-06 | 1,18E-05 | 1 - | 2,20E-05 - | 0 - | 0 - | 1 - | 2,60E-05 - | 0 - | 0 - | 0 - | 0 - |
| 8142 | 1 | 3um_FIJJAFBF_03572 | Bacteria | Bacteroidetes | unclassified | unclassified | unclassified | unclassified | Bacteroidetes | unclassified | unclassified | unclassified | unclassified | unclassified | 1,4 | 0,8 | 0,0001492 | 8,59E-05 | 2 - | 0,000185124 - | 2 - | 0,000229869 - | 1 - | 0,00010926 - | 2 - | 0,000221747 - | 0 - | 0 - |
| 8143 | 1 | 3um_FIJJAFBF_520513 | Bacteria | Bacteroidetes | Flavobacteriia | Flavobacterial | Flavobacteriac | Polaribacter | Polaribacter s | metabolism | Amino acid tra | 09AVD | acetylornithin | 983544.Lacal_ | 1,2 | 0,4 | 4,35E-05 | 1,46E-05 | 1 - | 3,04E-05 - | 1 - | 3,78E-05 - | 2 - | 7,19E-05 - | 1 - | 3,65E-05 - | 1 - | 4,11E-05 - |
| 8144 | 1 | 0.2um_NOKCPHJN_584204 | Bacteria | Proteobacteri | Gammaprotec | unclassified | unclassified | unclassified | Gammaprotec | unclassified | unclassified | unclassified | unclassified | unclassified | 11,2 | 1,469693846 | 0,000215956 | 2,54E-05 | 13 - | 0,000211345 - | 13 - | 0,000262429 - | 10 - | 0,000191901 - | 10 - | 0,000194736 - | 10 - | 0,000219369 - |
| 8145 | 1 | 3um_FIJJAFBF_516962 | Bacteria | Proteobacteri | Gammaprotec | Oceanospirilla | Saccharospirill | Reinekea | Reinekea fors | metabolism | Coenzyme tra | 16QPW | GTP cyclohydr | 314283.MED2 | 1,2 | 0,748331477 | 7,35E-05 | 4,43E-05 | 2 - | 0,000107772 - | 1 - | 6,69E-05 - | 1 - | 6,36E-05 - | 2 - | 0,000129092 - | 0 - | 0 - |

| | | | | | | | | | | | | | | | | | | | | | | | | | | | | | | | | | | | | | | |
|---|---|---|---|---|---|---|---|---|---|---|---|---|---|---|---|---|---|---|---|---|---|---|---|---|---|---|---|---|---|---|---|---|---|---|---|---|---|---|
| 8146 | 1 | 3um_FIJJAFBF_413741 | Bacteria | Proteobacteri | Gammaprote | Oceanospirilla | Saccharospirill | Reinekea | Reinekea fors | unclassified | unclassified | unclassified | unclassified | unclassified | 2 | 0,632455532 | 5,18E-05 | 1,61E-05 | 2 | - | 4,40E-05 | - | 2 | - | 5,46E-05 | - | 2 | - | 5,19E-05 | - | 3 | - | 7,90E-05 | - | 1 | - | 2,97E-05 | - |
| 8147 | 1 | 3um_FIJJAFBF_556278 | Bacteria | Bacteroidetes | Flavobacteriia | Flavobacterial | Flavobacteriac | Tenacibaculur | Tenacibaculur | unclassified | unclassified | unclassified | unclassified | unclassified | 0,8 | 0,4 | 5,37E-05 | 2,76E-05 | 1 | - | 5,68E-05 | - | 0 | - | 0 | - | 1 | - | 6,71E-05 | - | 1 | - | 6,81E-05 | - | 1 | - | 7,67E-05 | - |
| 8148 | 1 | 3um_FIJJAFBF_562249 | Bacteria | Bacteroidetes | Flavobacteriia | Flavobacterial | Flavobacteriac | Algibacter | Algibacter pe | metabolism | Energy produc | 0P9HY | cytochrome C | 983544.Lacal_ | 1,2 | 0,4 | 9,18E-05 | 3,91E-05 | 1 | - | 6,27E-05 | - | 1 | - | 7,79E-05 | - | 1 | - | 7,40E-05 | - | 1 | - | 7,51E-05 | - | 2 | - | 0,000169262 | - |
| 8149 | 1 | 3um_FIJJAFBF_82222 | Bacteria | Bacteroidetes | Flavobacteriia | Flavobacterial | Flavobacteriac | Winogradskye | Winogradskye | metabolism | Carbohydrate | 0PF2Z | Transketolase | 867900.Celly_ | 0,4 | 0,489897949 | 1,98E-05 | 2,42E-05 | 0 | - | 0 | - | 1 | - | 5,07E-05 | - | 1 | - | 4,82E-05 | - | 0 | - | 0 | - | 0 | - | 0 | - |
| 8150 | 1 | 3um_FIJJAFBF_487028 | Bacteria | Proteobacteri | Betaproteoba | Burkholderial | Comamonada | Hydrogenoph | Hydrogenoph | metabolism | Inorganic ion t | 0BBBP | TRAP transpor | 296591.Bpro_ | 2,8 | 0,748331477 | 0,000114974 | 2,22E-05 | 4 | - | 0,0001404 | - | 3 | - | 0,000130751 | - | 2 | - | 8,29E-05 | - | 3 | - | 0,000126131 | - | 2 | - | 9,47E-05 | - |
| 8151 | 1 | 3um_FIJJAFBF_128454 | Bacteria | Bacteroidetes | Flavobacteriia | Flavobacterial | Flavobacteriac | Polaribacter | Polaribacter s | unclassified | unclassified | unclassified | unclassified | unclassified | 5,2 | 1,166190379 | 0,000345911 | 8,66E-05 | 6 | - | 0,000334301 | - | 6 | - | 0,000415103 | - | 5 | - | 0,00032884 | - | 3 | - | 0,000200218 | - | 6 | - | 0,000451091 | - |
| 8152 | 1 | 0.2um_NOKCPHJN_265067 | Eukaryota | Bacillariophyt | Coscinodiscop | Thalassiosiral | Thalassiosirac | Thalassiosira | Thalassiosira | information st | Translation, ri | KOG3181 | ribosomal pro | 35128.Thaps2 | 0,4 | 0,8 | 2,49E-05 | 4,97E-05 | 0 | - | 0 | - | 0 | - | 0 | - | 2 | - | 0,000124296 | - | 0 | - | 0 | - | 0 | - | 0 | - |
| 8153 | 1 | 3um_FIJJAFBF_158909 | Bacteria | Proteobacteri | Alphaproteob | Rhodobactera | Rhodobactera | Tateyamaria | Tateyamaria s | metabolism | Inorganic ion t | 05E2Z | phosphate AB | 252305.OB25 | 0,4 | 0,489897949 | 1,24E-05 | 1,51E-05 | 0 | - | 0 | - | 0 | - | 0 | - | 1 | - | 3,07E-05 | - | 1 | - | 3,11E-05 | - | 0 | - | 0 | - |
| 8154 | 1 | 3um_FIJJAFBF_95373 | Viruses | unclassified | unclassified | unclassified | Mimiviridae | Cafeteriavirus | Cafeteria roe | unclassified | unclassified | unclassified | unclassified | unclassified | 6 | 1,095445115 | 0,000156998 | 3,22E-05 | 6 | - | 0,000131675 | - | 8 | - | 0,000218002 | - | 5 | - | 0,000129524 | - | 6 | - | 0,000157725 | - | 5 | - | 0,000148064 | - |
| 8155 | 1 | 0.2um_NOKCPHJN_33189 | Bacteria | Bacteroidetes | Flavobacteriia | Flavobacterial | Flavobacteriac | Polaribacter | Polaribacter s | poorly charact | Function unkn | 0PE4G | Carbohydrate | 313598.MED1 | 0,8 | 0,4 | 7,23E-06 | 3,64E-06 | 0 | - | 0 | - | 1 | - | 9,03E-06 | - | 1 | - | 8,59E-06 | - | 1 | - | 8,71E-06 | - | 1 | - | 9,81E-06 | - |
| 8156 | 1 | 3um_FIJJAFBF_525962 | Bacteria | Bacteroidetes | Cytophagia | Cytophagales | Cytophagacea | Ekhidna | Ekhidna lutea | information st | Translation, ri | 09AI8 | Associates wit | 643867.Ftrac_ | 0,6 | 0,8 | 2,65E-05 | 3,41E-05 | 2 | - | 8,32E-05 | - | 0 | - | 0 | - | 1 | - | 4,91E-05 | - | 0 | - | 0 | - | 0 | - | 0 | - |
| 8157 | 1 | 3um_FIJJAFBF_420943 | Bacteria | Proteobacteri | Gammaprote | Cellvibrionale | Porticoccacea | unclassified | various | unclassified | unclassified | unclassified | unclassified | unclassified | 1,8 | 1,166190379 | 0,000130385 | 6,92E-05 | 4 | - | 0,000256484 | - | 1 | - | 7,96E-05 | - | 1 | - | 7,57E-05 | - | 2 | - | 0,000153613 | - | 1 | - | 8,65E-05 | - |
| 8158 | 1 | 0.2um_NOKCPHJN_192569 | Bacteria | Proteobacteri | Gammaprote | Alteromonada | Colwelliaceae | Colwellia | Colwellia sp. F | metabolism | Carbohydrate | 16QEQ | hydrolase, fan | 234831.PSM_ | 0,6 | 0,489897949 | 9,10E-06 | 7,57E-06 | 1 | - | 1,29E-05 | - | 0 | - | 0 | - | 1 | - | 1,52E-05 | - | 0 | - | 0 | - | 1 | - | 1,74E-05 | - |
| 8159 | 1 | 3um_FIJJAFBF_277423 | Bacteria | Proteobacteri | Gammaprote | Alteromonada | Alteromonada | Paraglaciecola | Paraglaciecola | metabolism | Carbohydrate | 05ECY | inositol mono | 342610.Patl_1 | 0,2 | 0,4 | 8,93E-06 | 1,79E-05 | 1 | - | 4,47E-05 | - | 0 | - | 0 | - | 0 | - | 0 | - | 0 | - | 0 | - | 0 | - | 0 | - |
| 8160 | 1 | 0.2um_NOKCPHJN_132873 | Bacteria | Proteobacteri | Alphaproteob | Pelagibacteral | Pelagibacterac | Candidatus Pe | Candidatus Pe | unclassified | unclassified | unclassified | unclassified | unclassified | 0,4 | 0,489897949 | 1,25E-05 | 1,53E-05 | 0 | - | 0 | - | 0 | - | 0 | - | 1 | - | 2,91E-05 | - | 0 | - | 0 | - | 1 | - | 3,32E-05 | - |
| 8161 | 1 | 3um_FIJJAFBF_43518 | Bacteria | Bacteroidetes | Flavobacteriia | Flavobacterial | Flavobacteriac | Polaribacter | various | poorly charact | Function unkn | 05VUE | NA | 867900.Celly_ | 1,6 | 0,8 | 0,000233816 | 9,35E-05 | 3 | - | 0,000382589 | - | 1 | - | 0,000158354 | - | 1 | - | 0,000150536 | - | 2 | - | 0,000305518 | - | 1 | - | 0,000172083 | - |
| 8162 | 2 | 0.2um_NOKCPHJN_1022817;3um_FIJJAFBF_4484 | Eukaryota | various | various | various | various | various | various | unclassified | unclassified | unclassified | unclassified | unclassified | 1 | 0 | 0,000147341 | 1,40E-05 | 1 | - | 0,000123416 | - | 1 | - | 0,000153246 | - | 1 | - | 0,00014568 | - | 1 | - | 0,000147832 | - | 1 | - | 0,000166532 | - |
| 8163 | 1 | 3um_FIJJAFBF_103147 | Bacteria | Bacteroidetes | Flavobacteriia | Flavobacterial | Flavobacteriac | Polaribacter | Polaribacter s | unclassified | unclassified | unclassified | unclassified | unclassified | 2,8 | 0,979795897 | 7,34E-05 | 2,59E-05 | 2 | - | 4,40E-05 | - | 2 | - | 5,46E-05 | - | 4 | - | 0,000103818 | - | 4 | - | 0,000105351 | - | 2 | - | 5,93E-05 | - |
| 8164 | 1 | 3um_FIJJAFBF_264531 | Bacteria | Proteobacteri | Alphaproteob | Rhodobactera | Rhodobactera | Planktomarina | Planktomarina | metabolism | Amino acid tra | 05CAI | ABC transport | 246200.SPO10 | 0,4 | 0,489897949 | 2,00E-05 | 2,47E-05 | 1 | - | 4,55E-05 | - | 0 | - | 0 | - | 0 | - | 0 | - | 1 | - | 5,46E-05 | - | 0 | - | 0 | - |
| 8165 | 1 | 3um_FIJJAFBF_306866 | Bacteria | Proteobacteri | Alphaproteob | Rhodobactera | unclassified | unclassified | Rhodobactera | metabolism | Amino acid tra | 16PK4 | phospho-2-de | 246200.SPO19 | 0,4 | 0,489897949 | 1,11E-05 | 1,36E-05 | 1 | - | 2,52E-05 | - | 0 | - | 0 | - | 0 | - | 0 | - | 1 | - | 3,01E-05 | - | 0 | - | 0 | - |
| 8166 | 1 | 0.2um_NOKCPHJN_396618 | Bacteria | Proteobacteri | Gammaprote | unclassified | unclassified | unclassified | Gammaprote | cellular proces | Posttranslatio | 0QJ3C | Acts as a chap | 717774.Marm | 2,2 | 0,748331477 | 4,65E-05 | 1,49E-05 | 3 | - | 5,37E-05 | - | 3 | - | 6,67E-05 | - | 2 | - | 4,23E-05 | - | 1 | - | 2,14E-05 | - | 2 | - | 4,83E-05 | - |
| 8167 | 1 | 0.2um_NOKCPHJN_72708 | Bacteria | Proteobacteri | Gammaprote | Cellvibrionale | Porticoccacea | unclassified | gamma prote | unclassified | unclassified | unclassified | unclassified | unclassified | 0,2 | 0,4 | 3,56E-06 | 7,13E-06 | 1 | - | 1,78E-05 | - | 0 | - | 0 | - | 0 | - | 0 | - | 0 | - | 0 | - | 0 | - | 0 | - |
| 8168 | 1 | 0.2um_NOKCPHJN_523501 | Bacteria | Bacteroidetes | Flavobacteriia | Flavobacterial | Flavobacteriac | Sediminicola | Sediminicola s | unclassified | unclassified | unclassified | unclassified | unclassified | 0,4 | 0,489897949 | 5,17E-06 | 6,39E-06 | 1 | - | 1,15E-05 | - | 1 | - | 1,43E-05 | - | 0 | - | 0 | - | 0 | - | 0 | - | 0 | - | 0 | - |
| 8169 | 1 | 0.2um_NOKCPHJN_169344 | Bacteria | Bacteroidetes | Flavobacteriia | Flavobacterial | Flavobacteriac | Polaribacter | Polaribacter s | metabolism | Inorganic ion t | 05C27 | TonB-depende | 313598.MED1 | 1,8 | 0,748331477 | 2,17E-05 | 7,05E-06 | 3 | - | 3,14E-05 | - | 1 | - | 1,30E-05 | - | 2 | - | 2,47E-05 | - | 2 | - | 2,51E-05 | - | 1 | - | 1,41E-05 | - |
| 8170 | 1 | 0.2um_NOKCPHJN_267722 | Bacteria | Proteobacteri | Betaproteoba | unclassified | unclassified | unclassified | Betaproteoba | cellular proces | Cell motility | 01YGF | Flagellin | 314270.RB208 | 0,8 | 0,4 | 3,32E-05 | 1,71E-05 | 1 | - | 3,48E-05 | - | 1 | - | 4,32E-05 | - | 1 | - | 4,11E-05 | - | 0 | - | 0 | - | 1 | - | 4,69E-05 | - |
| 8171 | 1 | 0.2um_NOKCPHJN_619316 | Eukaryota | Bacillariophyt | Bacillariophyc | Bacillariales | Bacillariaceae | Fragilariopsis | Fragilariopsis | unclassified | unclassified | unclassified | unclassified | unclassified | 4 | 0,632455532 | 0,00055934 | 6,55E-05 | 5 | - | 0,000591633 | - | 3 | - | 0,00044078 | - | 4 | - | 0,00055869 | - | 4 | - | 0,000566941 | - | 4 | - | 0,000638658 | - |
| 8172 | 1 | 0.2um_NOKCPHJN_255249 | Bacteria | Actinobacteria | Actinobacteria | Micrococcales | Cellulomonad | Cellulomonas | Cellulomonas | unclassified | unclassified | unclassified | unclassified | unclassified | 0,4 | 0,489897949 | 1,25E-05 | 1,53E-05 | 1 | - | 2,86E-05 | - | 0 | - | 0 | - | 1 | - | 3,37E-05 | - | 0 | - | 0 | - | 0 | - | 0 | - |
| 8173 | 1 | 0.2um_NOKCPHJN_15668 | Bacteria | Proteobacteri | Gammaprote | Alteromonada | Alteromonada | Glaciecola | Glaciecola pu | unclassified | unclassified | unclassified | unclassified | unclassified | 4,8 | 0,748331477 | 8,14E-05 | 7,80E-06 | 6 | - | 8,62E-05 | - | 5 | - | 8,92E-05 | - | 4 | - | 6,78E-05 | - | 5 | - | 8,60E-05 | - | 4 | - | 7,75E-05 | - |
| 8174 | 1 | 0.2um_NOKCPHJN_62798 | Bacteria | Bacteroidetes | Flavobacteriia | Flavobacterial | Crocinitomica | unclassified | Crocinitomica | poorly charact | Function unkn | 07EF7 | domain protei | 649349.Lbys_ | 1,4 | 0,489897949 | 2,35E-05 | 8,28E-06 | 1 | - | 1,40E-05 | - | 1 | - | 1,74E-05 | - | 2 | - | 3,32E-05 | - | 2 | - | 3,37E-05 | - | 1 | - | 1,90E-05 | - |
| 8175 | 1 | 0.2um_NOKCPHJN_767144 | Bacteria | Proteobacteri | Alphaproteob | Pelagibacteral | Pelagibacterac | Candidatus Pe | Candidatus Pe | unclassified | unclassified | unclassified | unclassified | unclassified | 12,4 | 0,489897949 | 0,000242356 | 2,39E-05 | 12 | - | 0,000196479 | - | 12 | - | 0,00024397 | - | 13 | - | 0,000251251 | - | 13 | - | 0,000254962 | - | 12 | - | 0,00026512 | - |
| 8176 | 1 | 3um_FIJJAFBF_780929 | Bacteria | Bacteroidetes | Flavobacteriia | Flavobacterial | Flavobacteriac | Polaribacter | Polaribacter s | metabolism | Energy produc | 05CMV | Succinyl-CoA s | 391587.KAOT | 0,4 | 0,489897949 | 1,37E-05 | 1,68E-05 | 0 | - | 0 | - | 0 | - | 0 | - | 1 | - | 3,40E-05 | - | 1 | - | 3,45E-05 | - | 0 | - | 0 | - |
| 8177 | 1 | 3um_FIJJAFBF_298536 | Bacteria | Proteobacteri | Alphaproteob | Rhodobactera | Rhodobactera | Rhodovulum | Rhodovulum s | information st | Translation, ri | 177SZ | This protein bi | 314271.RB265 | 0,8 | 0,748331477 | 7,20E-05 | 6,33E-05 | 2 | - | 0,000162804 | - | 1 | - | 0,000101077 | - | 1 | - | 9,61E-05 | - | 0 | - | 0 | - | 0 | - | 0 | - |
| 8178 | 1 | 0.2um_NOKCPHJN_162707 | Eukaryota | unclassified | Pelagophycea | Pelagomonad | unclassified | Aureococcus | Aureococcus a | unclassified | unclassified | unclassified | unclassified | unclassified | 0,8 | 0,4 | 1,67E-05 | 8,59E-06 | 1 | - | 1,75E-05 | - | 1 | - | 2,18E-05 | - | 1 | - | 2,07E-05 | - | 0 | - | 0 | - | 1 | - | 2,36E-05 | - |
| 8179 | 1 | 0.2um_NOKCPHJN_07554 | Bacteria | Bacteroidetes | Flavobacteriia | Flavobacterial | Flavobacteriac | unclassified | Flavobacteria | metabolism | Inorganic ion t | 05C27 | TonB-depende | 471870.BACIN | 5,2 | 1,326649916 | 6,63E-05 | 1,61E-05 | 6 | - | 6,50E-05 | - | 7 | - | 9,41E-05 | - | 5 | - | 6,39E-05 | - | 5 | - | 6,49E-05 | - | 3 | - | 4,38E-05 | - |
| 8180 | 1 | 3um_FIJJAFBF_353617 | Bacteria | Bacteroidetes | Cytophagia | Cytophagales | Flammeovirga | unclassified | Flammeovirga | unclassified | unclassified | unclassified | unclassified | unclassified | 0,8 | 0,748331477 | 0,000186759 | 0,000179653 | 1 | - | 0,000201363 | - | 1 | - | 0,000250033 | - | 0 | - | 0 | - | 2 | - | 0,000482398 | - | 0 | - | 0 | - |
| 8181 | 1 | 3um_FIJJAFBF_389123 | Bacteria | Bacteroidetes | Flavobacteriia | Flavobacterial | Flavobacteriac | Polaribacter | Polaribacter a | information st | Translation, ri | 05CAV | thus facilitatin | 391587.KAOT | 22,2 | 1,166190379 | 0,000493344 | 2,95E-05 | 24 | - | 0,000448639 | - | 22 | - | 0,000510654 | - | 23 | - | 0,000507507 | - | 21 | - | 0,00047022 | - | 21 | - | 0,000529701 | - |
| 8182 | 1 | 0.2um_NOKCPHJN_1092416 | Bacteria | Proteobacteri | Epsilonproteo | Campylobacte | Helicobactera | Sulfurimonas | Sulfurimonas | poorly charact | Function unkn | 0ZJXN | NA | 439483.CBGD | 0,4 | 0,489897949 | 1,38E-05 | 1,72E-05 | 1 | - | 2,93E-05 | - | 0 | - | 0 | - | 0 | - | 0 | - | 0 | - | 0 | - | 1 | - | 3,95E-05 | - |
| 8183 | 1 | 3um_FIJJAFBF_365448 | Bacteria | Bacteroidetes | Flavobacteriia | Flavobacterial | Flavobacteriac | Flavivirga | Flavivirga eck | poorly charact | Function unkn | 05YA2 | NA | 391603.FBALC | 2,4 | 0,489897949 | 4,55E-05 | 6,68E-06 | 3 | - | 4,83E-05 | - | 2 | - | 4,00E-05 | - | 3 | - | 5,70E-05 | - | 2 | - | 3,86E-05 | - | 2 | - | 4,34E-05 | - |
| 8184 | 1 | 0.2um_NOKCPHJN_445894 | Viruses | unclassified | unclassified | unclassified | Mimiviridae | Catovirus | Catovirus CTV | unclassified | unclassified | unclassified | unclassified | unclassified | 0,2 | 0,4 | 5,02E-06 | 1,00E-05 | 1 | - | 2,51E-05 | - | 0 | - | 0 | - | 0 | - | 0 | - | 0 | - | 0 | - | 0 | - | 0 | - |
| 8185 | 2 | 0.2um_NOKCPHJN_125622;3um_FIJJAFBF_49167 | Bacteria | Proteobacteri | Gammaprote | Cellvibrionale | Porticoccacea | unclassified | gamma prote | unclassified | unclassified | unclassified | unclassified | unclassified | 1,8 | 0,4 | 0,000179969 | 4,41E-05 | 2 | - | 0,000167557 | - | 2 | - | 0,000208057 | - | 2 | - | 0,000197784 | - | 1 | - | 0,000100353 | - | 2 | - | 0,000226094 | - |
| 8186 | 1 | 3um_FIJJAFBF_61247 | unclassified | unclassified | unclassified | unclassified | unclassified | unclassified | unclassified | unclassified | unclassified | unclassified | unclassified | unclassified | 3,2 | 1,166190379 | 8,00E-05 | 2,36E-05 | 5 | - | 0,000107469 | - | 4 | - | 0,000106756 | - | 3 | - | 7,61E-05 | - | 2 | - | 5,15E-05 | - | 2 | - | 5,80E-05 | - |
| 8187 | 1 | 3um_FIJJAFBF_155641 | Bacteria | Bacteroidetes | Flavobacteriia | Flavobacterial | Crocinitomica | unclassified | Crocinitomica | unclassified | unclassified | unclassified | unclassified | unclassified | 1,6 | 0,489897949 | 0,000146345 | 4,60E-05 | 2 | - | 0,000154063 | - | 1 | - | 9,57E-05 | - | 2 | - | 0,000181855 | - | 1 | - | 9,23E-05 | - | 2 | - | 0,000207885 | - |
| 8188 | 1 | 3um_FIJJAFBF_332580 | Bacteria | Bacteroidetes | Flavobacteriia | Flavobacterial | Flavobacteriac | Aequorivita | Aequorivita ar | unclassified | unclassified | unclassified | unclassified | unclassified | 1,8 | 0,4 | 0,00011797 | 2,89E-05 | 2 | - | 0,000109834 | - | 2 | - | 0,000136382 | - | 2 | - | 0,000129648 | - | 1 | - | 6,58E-05 | - | 2 | - | 0,000148205 | - |
| 8189 | 1 | 0.2um_NOKCPHJN_41285 | Bacteria | Bacteroidetes | Flavobacteriia | Flavobacterial | Flavobacteriac | Kordia | Kordia algicida | cellular proces | Cell wall/mem | 08ZG0 | Ompa motb d | 391587.KAOT | 0,6 | 0,8 | 1,20E-05 | 1,54E-05 | 2 | - | 3,71E-05 | - | 1 | - | 2,31E-05 | - | 0 | - | 0 | - | 0 | - | 0 | - | 0 | - | 0 | - |
| 8190 | 1 | 0.2um_NOKCPHJN_466971 | Bacteria | Bacteroidetes | Flavobacteriia | Flavobacterial | Flavobacteriac | Polaribacter | Polaribacter s | unclassified | unclassified | unclassified | unclassified | unclassified | 5,4 | 1,356465997 | 9,97E-05 | 1,98E-05 | 7 | - | 0,000110514 | - | 6 | - | 0,000117622 | - | 5 | - | 9,32E-05 | - | 6 | - | 0,000113466 | - | 3 | - | 6,39E-05 | - |
| 8191 | 7 | 0.2um_NOKCPHJN_156025;0.2um_NOKCPHJN_8 | Bacteria | Bacteroidetes | Flavobacteriia | Flavobacterial | Flavobacteriac | various | various | cellular proces | Intracellular tr | 0PBE9 | Preprotein tra | various | 0,4 | 0,489897949 | 5,31E-05 | 6,56E-05 | 1 | - | 0,000118327 | - | 1 | - | 0,000146927 | - | 0 | - | 0 | - | 0 | - | 0 | - | 0 | - | 0 | - |
| 8192 | 1 | 0.2um_NOKCPHJN_117727 | Eukaryota | Apicomplexa | Aconoidasida | Haemosporida | Plasmodiidae | Plasmodium | Plasmodium v | cellular proces | Signal transdu | KOG0613 | fibronectin typ | 135651.CBN1 | 0,2 | 0,4 | 4,08E-06 | 8,17E-06 | 1 | - | 2,04E-05 | - | 0 | - | 0 | - | 0 | - | 0 | - | 0 | - | 0 | - | 0 | - | 0 | - |
| 8193 | 1 | 3um_FIJJAFBF_786615 | Bacteria | Proteobacteri | Gammaprote | Oceanospirilla | Oceanospirilla | Motiliproteus | Motiliproteus | information st | Translation, ri | 05KK9 | This protein bi | 351348.Maqu | 5,6 | 0,489897949 | 0,000744235 | 9,39E-05 | 6 | - | 0,000668602 | - | 5 | - | 0,000691839 | - | 5 | - | 0,000657681 | - | 6 | - | 0,000800874 | - | 6 | - | 0,000902181 | - |
| 8194 | 1 | 3um_FIJJAFBF_554495 | Bacteria | Planctomycet | Planctomyceti | Planctomyceta | Planctomyceta | Rhodopirellula | Rhodopirellula | metabolism | Energy produc | 153I3 | Cytochrome c | 314230.DSM3 | 1 | 0 | 0,000121263 | 1,15E-05 | 1 | - | 0,000101572 | - | 1 | - | 0,000126123 | - | 1 | - | 0,000119896 | - | 1 | - | 0,000121667 | - | 1 | - | 0,000137057 | - |
| 8195 | 1 | 3um_FIJJAFBF_443625 | Bacteria | Proteobacteri | Alphaproteob | Rhodospirillal | Rhodospirillac | unclassified | Rhodobactera | information st | Transcription | 178T0 | Transcriptiona | 367336.OM22 | 1 | 0,632455532 | 7,73E-05 | 4,36E-05 | 2 | - | 0,000135031 | - | 0 | - | 0 | - | 1 | - | 7,97E-05 | - | 1 | - | 8,09E-05 | - | 1 | - | 9,11E-05 | - |
| 8196 | 1 | 3um_FIJJAFBF_505173 | Bacteria | Bacteroidetes | unclassified | unclassified | unclassified | unclassified | Bacteroidetes | unclassified | unclassified | unclassified | unclassified | unclassified | 5,8 | 1,720465053 | 0,000638434 | 0,000166597 | 7 | - | 0,000658555 | - | 6 | - | 0,000700912 | - | 8 | - | 0,000888408 | - | 5 | - | 0,000563456 | - | 3 | - | 0,000380839 | - |
| 8197 | 1 | 0.2um_NOKCPHJN_405981 | Bacteria | Proteobacteri | unclassified | unclassified | unclassified | unclassified | uncultured ma | unclassified | unclassified | unclassified | unclassified | unclassified | 0,4 | 0,8 | 7,23E-06 | 1,45E-05 | 2 | - | 3,62E-05 | - | 0 | - | 0 | - | 0 | - | 0 | - | 0 | - | 0 | - | 0 | - | 0 | - |
| 8198 | 1 | 0.2um_NOKCPHJN_841333 | Bacteria | Planctomycet | Phycisphaerae | Phycisphaeral | Phycisphaerac | Phycisphaera | Phycisphaera | cellular proces | Intracellular tr | COG0552 | Involved in tar | 634956.Geoth | 1 | 0 | 4,60E-05 | 4,37E-06 | 1 | - | 3,85E-05 | - | 1 | - | 4,78E-05 | - | 1 | - | 4,55E-05 | - | 1 | - | 4,61E-05 | - | 1 | - | 5,20E-05 | - |
| 8199 | 1 | 3um_FIJJAFBF_751786 | Eukaryota | unclassified | Pelagophycea | Pelagomonad | unclassified | Aureococcus | Aureococcus a | unclassified | unclassified | unclassified | unclassified | unclassified | 3,4 | 0,489897949 | 8,88E-05 | 9,43E-06 | 4 | - | 8,83E-05 | - | 3 | - | 8,22E-05 | - | 3 | - | 7,82E-05 | - | 4 | - | 0,000105756 | - | 3 | - | 8,94E-05 | - |
| 8200 | 1 | 0.2um_NOKCPHJN_460016 | Eukaryota | unclassified | Chrysophycea | Chromulinales | Chromulinace | Ochromonas | Ochromonas | unclassified | unclassified | unclassified | unclassified | unclassified | 0,8 | 0,4 | 3,24E-05 | 1,66E-05 | 1 | - | 3,43E-05 | - | 0 | - | 0 | - | 1 | - | 4,04E-05 | - | 1 | - | 4,10E-05 | - | 1 | - | 4,62E-05 | - |
| 8201 | 1 | 3um_FIJJAFBF_72483 | Bacteria | Bacteroidetes | Cytophagia | Cytophagales | Flammeovirga | unclassified | Flammeovirga | unclassified | unclassified | unclassified | unclassified | unclassified | 0,2 | 0,4 | 3,89E-05 | 7,78E-05 | 1 | - | 0,000194537 | - | 0 | - | 0 | - | 0 | - | 0 | - | 0 | - | 0 | - | 0 | - | 0 | - |
| 8202 | 1 | 0.2um_NOKCPHJN_229165 | Eukaryota | Cnidaria | Anthozoa | Scleractinia | Merulinidae | Orbicella | Orbicella fave | unclassified | unclassified | unclassified | unclassified | unclassified | 0,2 | 0,4 | 6,27E-07 | 1,25E-06 | 0 | - | 0 | - | 1 | - | 3,14E-06 | - | 0 | - | 0 | - | 0 | - | 0 | - | 0 | - | 0 | - |
| 8203 | 1 | 3um_FIJJAFBF_377212 | Bacteria | Bacteroidetes | Flavobacteriia | Flavobacterial | Flavobacteriac | Polaribacter | Polaribacter s | unclassified | unclassified | unclassified | unclassified | unclassified | 0,6 | 0,489897949 | 1,54E-05 | 1,27E-05 | 1 | - | 2,25E-05 | - | 1 | - | 2,80E-05 | - | 1 | - | 2,66E-05 | - | 0 | - | 0 | - | 0 | - | 0 | - |
| 8204 | 1 | 3um_FIJJAFBF_518388 | Bacteria | Bacteroidetes | Flavobacteriia | Flavobacterial | Flavobacteriac | Polaribacter | Polaribacter b | unclassified | unclassified | unclassified | unclassified | unclassified | 1 | 0,632455532 | 3,24E-05 | 2,19E-05 | 1 | - | 2,77E-05 | - | 2 | - | 6,87E-05 | - | 1 | - | 3,26E-05 | - | 1 | - | 3,31E-05 | - | 0 | - | 0 | - |
| 8205 | 1 | 3um_FIJJAFBF_554064 | Bacteria | Proteobacteri | Alphaproteob | Rhodobactera | Rhodobactera | Litoreibacter | Litoreibacter a | unclassified | unclassified | unclassified | unclassified | unclassified | 0,2 | 0,4 | 7,70E-06 | 1,54E-05 | 0 | - | 0 | - | 1 | - | 3,85E-05 | - | 0 | - | 0 | - | 0 | - | 0 | - | 0 | - | 0 | - |
| 8206 | 1 | 0.2um_NOKCPHJN_57910 | Bacteria | Proteobacteri | Alphaproteob | Rhodobactera | Rhodobactera | Pseudooceani | Pseudooceani | unclassified | unclassified | unclassified | unclassified | unclassified | 0,4 | 0,8 | 1,23E-05 | 2,46E-05 | 0 | - | 0 | - | 0 | - | 0 | - | 2 | - | 6,16E-05 | - | 0 | - | 0 | - | 0 | - | 0 | - |
| 8207 | 1 | 0.2um_NOKCPHJN_149519 | Bacteria | unclassified | unclassified | unclassified | unclassified | unclassified | uncultured ba | unclassified | unclassified | unclassified | unclassified | unclassified | 13 | 0 | 0,000191338 | 1,82E-05 | 13 | - | 0,000160268 | - | 13 | - | 0,000199006 | - | 13 | - | 0,00018918 | - | 13 | - | 0,000191975 | - | 13 | - | 0,000216259 | - |
| 8208 | 1 | 0.2um_NOKCPHJN_625942 | Bacteria | Proteobacteri | Gammaprote | Oceanospirilla | Saccharospirill | Reinekea | Reinekea fors | poorly charact | Function unkn | 061D9 | Protein with u | 247633.GP214 | 1 | 0 | 0,000123448 | 1,17E-05 | 1 | - | 0,000103402 | - | 1 | - | 0,000128395 | - | 1 | - | 0,000122056 | - | 1 | - | 0,000123859 | - | 1 | - | 0,000139527 | - |
| 8209 | 2 | 0.2um_NOKCPHJN_47587;3um_FIJJAFBF_442085 | Bacteria | various | various | various | various | various | various | metabolism | Energy produc | 0PEU8 | ATP synthase | 313603.FB217 | 1 | 0 | 0,000217503 | 2,07E-05 | 1 | - | 0,000182185 | - | 1 | - | 0,00022622 | - | 1 | - | 0,000215051 | - | 1 | - | 0,000218227 | - | 1 | - | 0,000245833 | - |
| 8210 | 1 | 3um_FIJJAFBF_276754 | Bacteria | Proteobacteri | Gammaprote | Alteromonada | Alteromonada | Paraglaciecola | Paraglaciecola | metabolism | Amino acid tra | 16STP | phosphoserine | 342610.Patl_1 | 0,2 | 0,4 | 6,20E-06 | 1,24E-05 | 1 | - | 3,10E-05 | - | 0 | - | 0 | - | 0 | - | 0 | - | 0 | - | 0 | - | 0 | - | 0 | - |
| 8211 | 1 | 3um_FIJJAFBF_414331 | Bacteria | Bacteroidetes | Cytophagia | Cytophagales | Flammeovirga | unclassified | Flammeovirga | unclassified | unclassified | unclassified | unclassified | unclassified | 0,6 | 0,489897949 | 3,20E-05 | 2,61E-05 | 0 | - | 0 | - | 1 | - | 5,48E-05 | - | 1 | - | 5,21E-05 | - | 1 | - | 5,29E-05 | - | 0 | - | 0 | - |
| 8212 | 1 | 0.2um_NOKCPHJN_42273 | Bacteria | Proteobacteri | Gammaprote | Cellvibrionale | Porticoccacea | unclassified | SAR92 bacteri | unclassified | unclassified | unclassified | unclassified | unclassified | 15,8 | 1,6 | 0,000255747 | 4,57E-05 | 14 | - | 0,0001886 | - | 14 | - | 0,000234186 | - | 16 | - | 0,000254427 | - | 17 | - | 0,000274321 | - | 18 | - | 0,0003272 | - |
| 8213 | 1 | 0.2um_NOKCPHJN_379860 | Bacteria | Proteobacteri | Betaproteoba | unclassified | unclassified | unclassified | Betaproteoba | unclassified | unclassified | unclassified | unclassified | unclassified | 1,2 | 0,4 | 9,77E-05 | 4,16E-05 | 1 | - | 6,67E-05 | - | 1 | - | 8,29E-05 | - | 1 | - | 7,88E-05 | - | 1 | - | 7,99E-05 | - | 2 | - | 0,000180087 | - |
| 8214 | 1 | 0.2um_NOKCPHJN_816703 | Bacteria | Proteobacteri | Betaproteoba | Nitrosomonad | unclassified | unclassified | various | poorly charact | Function unkn | 0XRND | NA | 314607.KB13_ | 3,2 | 0,4 | 0,000107216 | 1,84E-05 | 3 | - | 8,40E-05 | - | 4 | - | 0,000139043 | - | 3 | - | 9,91E-05 | - | 3 | - | 0,000100598 | - | 3 | - | 0,000113323 | - |
| 8215 | 1 | 0.2um_NOKCPHJN_812155 | Bacteria | Proteobacteri | Gammaprote | unclassified | unclassified | unclassified | Gammaprote | poorly charact | Function unkn | 0XY6Y | Pkd domain co | 314287.GB220 | 0,4 | 0,489897949 | 6,46E-06 | 7,95E-06 | 1 | - | 1,48E-05 | - | 0 | - | 0 | - | 1 | - | 1,75E-05 | - | 0 | - | 0 | - | 0 | - | 0 | - |
| 8216 | 1 | 0.2um_NOKCPHJN_35857 | Viruses | unclassified | unclassified | unclassified | Phycodnavirid | Prasinovirus | Micromonas s | unclassified | unclassified | unclassified | unclassified | unclassified | 1,2 | 0,979795897 | 3,85E-05 | 2,72E-05 | 3 | - | 8,52E-05 | - | 1 | - | 3,53E-05 | - | 1 | - | 3,35E-05 | - | 0 | - | 0 | - | 1 | - | 3,83E-05 | - |
| 8217 | 1 | 3um_FIJJAFBF_264894 | Bacteria | unclassified | unclassified | unclassified | unclassified | unclassified | uncultured ba | unclassified | unclassified | unclassified | unclassified | unclassified | 7,2 | 1,469693846 | 0,00010814 | 2,07E-05 | 9 | - | 0,000114143 | - | 5 | - | 7,87E-05 | - | 8 | - | 0,000119763 | - | 6 | - | 9,11E-05 | - | 8 | - | 0,000136906 | - |
| 8218 | 1 | 3um_FIJJAFBF_216288 | Bacteria | Proteobacteri | Alphaproteob | Rhodospirillal | Rhodospirillac | unclassified | Rhodobactera | metabolism | Carbohydrate | 01RS7 | extracellular s | 367336.OM22 | 0,6 | 0,489897949 | 1,87E-05 | 1,54E-05 | 1 | - | 2,77E-05 | - | 0 | - | 0 | - | 1 | - | 3,26E-05 | - | 1 | - | 3,31E-05 | - | 0 | - | 0 | - |
| 8219 | 1 | 0.2um_NOKCPHJN_164151 | Bacteria | Proteobacteri | Alphaproteob | Rhodobactera | Rhodobactera | Planktomarina | Planktomarina | information st | Translation, ri | 01ZNX | Binds as a het | 391595.RLO14 | 1 | 0 | 0,000182703 | 1,73E-05 | 1 | - | 0,000153036 | - | 1 | - | 0,000190025 | - | 1 | - | 0,000180643 | - | 1 | - | 0,000183311 | - | 1 | - | 0,000206499 | - |
| 8220 | 1 | 0.2um_NOKCPHJN_408470 | Bacteria | Bacteroidetes | Flavobacteriia | Flavobacterial | Flavobacteriac | Polaribacter | Polaribacter s | poorly charact | Function unkn | 05WTF | NA | 487797.Flav3 | 1,2 | 0,4 | 2,30E-05 | 7,69E-06 | 1 | - | 1,61E-05 | - | 1 | - | 1,99E-05 | - | 2 | - | 3,79E-05 | - | 1 | - | 1,92E-05 | - | 1 | - | 2,17E-05 | - |
| 8221 | 1 | 0.2um_NOKCPHJN_460846 | Eukaryota | Brachiopoda | Lingulata | Lingulida | Lingulidae | Lingula | Lingula anatin | unclassified | unclassified | unclassified | unclassified | unclassified | 0,2 | 0,4 | 3,28E-06 | 6,56E-06 | 0 | - | 0 | - | 0 | - | 0 | - | 0 | - | 0 | - | 0 | - | 0 | - | 1 | - | 1,64E-05 | - |
| 8222 | 1 | 0.2um_NOKCPHJN_870081 | Bacteria | Bacteroidetes | Flavobacteriia | Flavobacterial | Flavobacteriac | Aureitalea | Aureitalea ma | poorly charact | Function unkn | 11NCV | Protein of unk | 50743.SCB49_ | 8,6 | 0,8 | 0,00075267 | 7,78E-05 | 9 | - | 0,000662174 | - | 8 | - | 0,000730866 | - | 8 | - | 0,000694781 | - | 10 | - | 0,000881303 | - | 8 | - | 0,000794228 | - |
| 8223 | 1 | 3um_FIJJAFBF_38501 | Bacteria | Proteobacteri | Alphaproteob | Rhodobactera | various | various | various | information st | Translation, ri | COG0292 | Binds directly | 644076.SCH4 | 2,8 | 0,748331477 | 0,000315301 | 8,69E-05 | 3 | - | 0,00028457 | - | 4 | - | 0,000471137 | - | 2 | - | 0,000223938 | - | 3 | - | 0,000340868 | - | 2 | - | 0,000255991 | - |
| 8224 | 1 | 0.2um_NOKCPHJN_409669 | Bacteria | Proteobacteri | Gammaprote | unclassified | unclassified | unclassified | Gammaprote | unclassified | unclassified | unclassified | unclassified | unclassified | 15,2 | 2,785677655 | 0,000286469 | 7,20E-05 | 16 | - | 0,000250878 | - | 15 | - | 0,000292047 | - | 12 | - | 0,000222102 | - | 13 | - | 0,000244164 | - | 20 | - | 0,000423154 | - |
| 8225 | 1 | 0.2um_NOKCPHJN_73699 | Bacteria | Bacteroidetes | Flavobacteriia | Flavobacterial | Flavobacteriac | Winogradskye | Winogradskye | unclassified | unclassified | unclassified | unclassified | unclassified | 0,2 | 0,4 | 6,44E-06 | 1,29E-05 | 0 | - | 0 | - | 0 | - | 0 | - | 1 | - | 3,22E-05 | - | 0 | - | 0 | - | 0 | - | 0 | - |
| 8226 | 1 | 3um_FIJJAFBF_35653 | Bacteria | Proteobacteri | Gammaprote | Oceanospirilla | Saccharospirill | Reinekea | Reinekea fors | unclassified | unclassified | unclassified | unclassified | unclassified | 2 | 0 | 0,000149756 | 1,42E-05 | 2 | - | 0,000125439 | - | 2 | - | 0,000155758 | - | 2 | - | 0,000148068 | - | 2 | - | 0,000150255 | - | 2 | - | 0,000169262 | - |
| 8227 | 1 | 3um_FIJJAFBF_610542 | Bacteria | Bacteroidetes | Flavobacteriia | Flavobacterial | Flavobacteriac | Aequorivita | Aequorivita sp | unclassified | unclassified | unclassified | unclassified | unclassified | 6,4 | 1,019803903 | 0,000573126 | 8,92E-05 | 7 | - | 0,000528577 | - | 8 | - | 0,000750099 | - | 6 | - | 0,000534798 | - | 6 | - | 0,000542697 | - | 5 | - | 0,000509456 | - |
| 8228 | 1 | 0.2um_NOKCPHJN_09683 | Bacteria | Bacteroidetes | Flavobacteriia | Flavobacterial | Flavobacteriac | Formosa | Formosa sp. H | unclassified | unclassified | unclassified | unclassified | unclassified | 0,2 | 0,4 | 9,00E-06 | 1,80E-05 | 1 | - | 4,50E-05 | - | 0 | - | 0 | - | 0 | - | 0 | - | 0 | - | 0 | - | 0 | - | 0 | - |
| 8229 | 1 | 3um_FIJJAFBF_646408 | Bacteria | Bacteroidetes | Cytophagia | Cytophagales | Flammeovirga | unclassified | Flammeovirga | unclassified | unclassified | unclassified | unclassified | unclassified | 0,8 | 0,4 | 3,57E-05 | 1,81E-05 | 1 | - | 3,86E-05 | - | 1 | - | 4,80E-05 | - | 1 | - | 4,56E-05 | - | 1 | - | 4,63E-05 | - | 0 | - | 0 | - |
| 8230 | 1 | 0.2um_NOKCPHJN_585943 | Bacteria | Bacteroidetes | Flavobacteriia | Flavobacterial | Flavobacteriac | Polaribacter | Polaribacter p | unclassified | unclassified | unclassified | unclassified | unclassified | 4,4 | 0,489897949 | 6,69E-05 | 6,99E-06 | 5 | - | 6,40E-05 | - | 5 | - | 7,95E-05 | - | 4 | - | 6,05E-05 | - | 4 | - | 6,14E-05 | - | 4 | - | 6,91E-05 | - |
| 8231 | 1 | 0.2um_NOKCPHJN_877988 | Eukaryota | Mucoromycot | Mucoromycet | Mucorales | Choanephorac | Choanephora | Choanephora | unclassified | unclassified | unclassified | unclassified | unclassified | 0,4 | 0,489897949 | 2,86E-05 | 3,54E-05 | 1 | - | 6,38E-05 | - | 1 | - | 7,92E-05 | - | 0 | - | 0 | - | 0 | - | 0 | - | 0 | - | 0 | - |
| 8232 | 1 | 0.2um_NOKCPHJN_228000 | Bacteria | Proteobacteri | Gammaprote | unclassified | unclassified | unclassified | Gammaprote | information st | Translation, ri | COG0333 | 50s ribosomal | 314285.KT71_ | 1 | 0 | 0,000224635 | 2,13E-05 | 1 | - | 0,000188159 | - | 1 | - | 0,000233637 | - | 1 | - | 0,000222102 | - | 1 | - | 0,000225382 | - | 1 | - | 0,000253893 | - |
| 8233 | 1 | 0.2um_NOKCPHJN_217592 | Eukaryota | unclassified | unclassified | Kinetoplastida | Trypanosomat | Leishmania | Leishmania m | unclassified | unclassified | unclassified | unclassified | unclassified | 1 | 0 | 2,55E-06 | 2,42E-07 | 1 | - | 2,14E-06 | - | 1 | - | 2,65E-06 | - | 1 | - | 2,52E-06 | - | 1 | - | 2,56E-06 | - | 1 | - | 2,88E-06 | - |
| 8234 | 1 | 0.2um_NOKCPHJN_947667 | unclassified | unclassified | unclassified | unclassified | unclassified | unclassified | unclassified | unclassified | unclassified | unclassified | unclassified | unclassified | 1 | 0 | 0,000110506 | 1,05E-05 | 1 | - | 9,26E-05 | - | 1 | - | 0,000114935 | - | 1 | - | 0,00010926 | - | 1 | - | 0,000110874 | - | 1 | - | 0,000124899 | - |
| 8235 | 1 | 3um_FIJJAFBF_809550 | Bacteria | Bacteroidetes | Flavobacteriia | Flavobacterial | Flavobacteriac | various | various | information st | Translation, ri | COG0051 | tRNA binding | 313594.PI23P | 0,8 | 0,4 | 0,000107449 | 5,52E-05 | 1 | - | 0,00011364 | - | 0 | - | 0 | - | 1 | - | 0,000134141 | - | 1 | - | 0,000136122 | - | 1 | - | 0,000153341 | - |
| 8236 | 1 | 3um_FIJJAFBF_269247 | Bacteria | Bacteroidetes | Flavobacteriia | Flavobacterial | Flavobacteriac | Polaribacter | Polaribacter s | information st | Replication, re | COG0513 | purine NTP-de | 313594.PI23P | 0,4 | 0,489897949 | 8,49E-06 | 1,05E-05 | 1 | - | 1,89E-05 | - | 1 | - | 2,35E-05 | - | 0 | - | 0 | - | 0 | - | 0 | - | 0 | - | 0 | - |
| 8237 | 1 | 3um_FIJJAFBF_222227 | Bacteria | Proteobacteri | Alphaproteob | Rhodobactera | unclassified | unclassified | Rhodobactera | unclassified | unclassified | unclassified | unclassified | unclassified | 1 | 0 | 5,21E-05 | 4,95E-06 | 1 | - | 4,36E-05 | - | 1 | - | 5,42E-05 | - | 1 | - | 5,15E-05 | - | 1 | - | 5,23E-05 | - | 1 | - | 5,89E-05 | - |
| 8238 | 1 | 0.2um_NOKCPHJN_224040 | Bacteria | unclassified | unclassified | unclassified | unclassified | unclassified | uncultured ba | metabolism | Inorganic ion t | 05C27 | TonB-depende | 313596.RB250 | 15,6 | 2,939387691 | 0,000198448 | 2,56E-05 | 20 | - | 0,000215949 | - | 14 | - | 0,000187701 | - | 18 | - | 0,000229415 | - | 12 | - | 0,000155202 | - | 14 | - | 0,000203974 | - |
| 8239 | 1 | 3um_FIJJAFBF_662957 | Bacteria | Bacteroidetes | Flavobacteriia | Flavobacterial | Crocinitomica | unclassified | Crocinitomica | information st | Translation, ri | 05KK9 | This protein bi | 376686.Fjoh_ | 3,6 | 0,8 | 0,000369672 | 6,83E-05 | 5 | - | 0,00043476 | - | 3 | - | 0,000323907 | - | 3 | - | 0,000307914 | - | 3 | - | 0,000312462 | - | 4 | - | 0,000469317 | - |
| 8240 | 1 | 3um_FIJJAFBF_254301 | Bacteria | Proteobacteri | Gammaprote | Oceanospirilla | Endozoicomor | Endozoicomor | unclassified | information st | Translation, ri | COG0256 | This is one of t | 498211.CJA_0 | 0,4 | 0,489897949 | 4,79E-05 | 5,87E-05 | 0 | - | 0 | - | 1 | - | 0,000122861 | - | 1 | - | 0,000116795 | - | 0 | - | 0 | - | 0 | - | 0 | - |
| 8241 | 1 | 3um_FIJJAFBF_465005 | Bacteria | Proteobacteri | Gammaprote | Oceanospirilla | Saccharospirill | Reinekea | Reinekea blan | metabolism | Lipid transpor | 0QJ8V | Catalyzes the | 999141.GME_ | 1,2 | 0,748331477 | 3,98E-05 | 2,38E-05 | 2 | - | 5,65E-05 | - | 2 | - | 7,02E-05 | - | 0 | - | 0 | - | 1 | - | 3,39E-05 | - | 1 | - | 3,81E-05 | - |
| 8242 | 1 | 3um_FIJJAFBF_674281 | Bacteria | Bacteroidetes | Flavobacteriia | Flavobacterial | Crocinitomica | unclassified | Crocinitomica | information st | Translation, ri | COG0051 | tRNA binding | 398720.MED2 | 0,2 | 0,4 | 2,27E-05 | 4,55E-05 | 1 | - | 0,00011364 | - | 0 | - | 0 | - | 0 | - | 0 | - | 0 | - | 0 | - | 0 | - | 0 | - |
| 8243 | 1 | 0.2um_NOKCPHJN_487409 | Viruses | unclassified | unclassified | unclassified | unclassified | unclassified | uncultured M | unclassified | unclassified | unclassified | unclassified | unclassified | 0,2 | 0,4 | 1,21E-05 | 2,43E-05 | 1 | - | 6,07E-05 | - | 0 | - | 0 | - | 0 | - | 0 | - | 0 | - | 0 | - | 0 | - | 0 | - |
| 8244 | 1 | 3um_FIJJAFBF_52282 | Bacteria | Proteobacteri | Alphaproteob | Rhodobactera | unclassified | unclassified | Rhodobactera | poorly charact | Function unkn | 01Y94 | Protein of unk | 439497.RR11_ | 1,8 | 0,4 | 0,000173632 | 4,26E-05 | 2 | - | 0,000161657 | - | 2 | - | 0,000200731 | - | 2 | - | 0,00019082 | - | 1 | - | 9,68E-05 | - | 2 | - | 0,000218133 | - |
| 8245 | 1 | 3um_FIJJAFBF_600181 | Bacteria | Actinobacteria | Actinobacteria | Micrococcales | Microbacteria | unclassified | Microbacteria | metabolism | Inorganic ion t | 08UTW | Plays a role in | 312284.A20C1 | 3,2 | 0,748331477 | 0,000200291 | 3,92E-05 | 4 | - | 0,00021255 | - | 2 | - | 0,000131962 | - | 4 | - | 0,000250893 | - | 3 | - | 0,000190949 | - | 3 | - | 0,000215103 | - |
| 8246 | 1 | 3um_FIJJAFBF_295862 | Bacteria | Proteobacteri | Gammaprote | Oceanospirilla | Saccharospirill | Reinekea | Reinekea fors | metabolism | Secondary me | COG4181 | (ABC) transpo | 314283.MED2 | 0,2 | 0,4 | 1,00E-05 | 2,00E-05 | 1 | - | 5,01E-05 | - | 0 | - | 0 | - | 0 | - | 0 | - | 0 | - | 0 | - | 0 | - | 0 | - |
| 8247 | 1 | 3um_FIJJAFBF_81226 | Bacteria | Proteobacteri | Gammaprote | Cellvibrionale | Spongiibacter | Oceanicoccus | Oceanicoccus | information st | Translation, ri | 05CEJ | Catalyzes the | 247633.GP214 | 1,6 | 0,8 | 3,01E-05 | 1,23E-05 | 3 | - | 4,90E-05 | - | 2 | - | 4,06E-05 | - | 1 | - | 1,93E-05 | - | 1 | - | 1,96E-05 | - | 1 | - | 2,21E-05 | - |
| 8248 | 1 | 0.2um_NOKCPHJN_07544 | Bacteria | Bacteroidetes | Flavobacteriia | Flavobacterial | Flavobacteriac | Formosa | Formosa sp. H | poorly charact | Function unkn | 064XS | NA | 313598.MED1 | 2,6 | 0,489897949 | 0,00010665 | 1,80E-05 | 3 | - | 0,000104027 | - | 3 | - | 0,000129171 | - | 2 | - | 8,19E-05 | - | 3 | - | 0,000124607 | - | 2 | - | 9,36E-05 | - |
| 8249 | 1 | 0.2um_NOKCPHJN_262773 | Bacteria | Actinobacteria | Actinobacteria | Corynebacteri | Dietziaceae | Dietzia | Dietzia sp. WN | poorly charact | Function unkn | 0QKMK | Phage major c | 682634.SOD_ | 0,4 | 0,489897949 | 1,33E-05 | 1,62E-05 | 0 | - | 0 | - | 0 | - | 0 | - | 1 | - | 3,29E-05 | - | 1 | - | 3,34E-05 | - | 0 | - | 0 | - |
| 8250 | 1 | 0.2um_NOKCPHJN_43858 | Bacteria | Bacteroidetes | Flavobacteriia | Flavobacterial | Flavobacteriac | Algibacter | Algibacter aqu | metabolism | Energy produc | 0P7S4 | Produces ATP | 1046627.BZAF | 2 | 0,632455532 | 9,73E-05 | 3,29E-05 | 1 | - | 4,01E-05 | - | 2 | - | 9,97E-05 | - | 3 | - | 0,000142114 | - | 2 | - | 9,61E-05 | - | 2 | - | 0,000108304 | - |
| 8251 | 1 | 0.2um_NOKCPHJN_93313 | Bacteria | Proteobacteri | Betaproteoba | Burkholderial | Comamonada | Ramlibacter | Ramlibacter ta | metabolism | Lipid transpor | COG1024 | Enoyl-CoA hyd | 365046.Rta_0 | 0,2 | 0,4 | 9,04E-06 | 1,81E-05 | 1 | - | 4,52E-05 | - | 0 | - | 0 | - | 0 | - | 0 | - | 0 | - | 0 | - | 0 | - | 0 | - |
| 8252 | 1 | 3um_FIJJAFBF_21303 | Bacteria | Bacteroidetes | Flavobacteriia | Flavobacterial | Flavobacteriac | Polaribacter | Polaribacter s | metabolism | Nucleotide tra | COG0516 | Catalyzes the | 391587.KAOT | 1,8 | 0,4 | 5,03E-05 | 1,25E-05 | 2 | - | 4,68E-05 | - | 2 | - | 5,81E-05 | - | 1 | - | 2,76E-05 | - | 2 | - | 5,60E-05 | - | 2 | - | 6,31E-05 | - |
| 8253 | 1 | 3um_FIJJAFBF_540524 | Bacteria | Proteobacteri | Alphaproteob | Rhodobactera | Rhodobactera | Planktomarina | Planktomarina | information st | Translation, ri | 05C64 | Binds directly | 633131.TR2A | 2,4 | 0,489897949 | 0,000140307 | 2,48E-05 | 3 | - | 0,000148418 | - | 3 | - | 0,000184292 | - | 2 | - | 0,000116795 | - | 2 | - | 0,00011852 | - | 2 | - | 0,000133512 | - |
| 8254 | 1 | 0.2um_NOKCPHJN_22462 | Bacteria | Proteobacteri | Gammaprote | Alteromonada | Alteromonada | Paraglaciecola | Paraglaciecola | cellular proces | Intracellular tr | 0QITE | Outer membr | 1085623.GNIT | 3,8 | 1,469693846 | 0,000115343 | 4,54E-05 | 2 | - | 5,05E-05 | - | 3 | - | 9,40E-05 | - | 5 | - | 0,000148882 | - | 6 | - | 0,000181297 | - | 3 | - | 0,000102115 | - |
| 8255 | 1 | 3um_FIJJAFBF_47821 | Bacteria | Bacteroidetes | Flavobacteriia | Flavobacterial | Flavobacteriac | Polaribacter | Polaribacter s | unclassified | unclassified | unclassified | unclassified | unclassified | 0,4 | 0,489897949 | 2,12E-05 | 2,65E-05 | 1 | - | 4,52E-05 | - | 0 | - | 0 | - | 0 | - | 0 | - | 0 | - | 0 | - | 1 | - | 6,10E-05 | - |
| 8256 | 1 | 3um_FIJJAFBF_549856 | Bacteria | Bacteroidetes | Flavobacteriia | Flavobacterial | Crocinitomica | unclassified | Crocinitomica | unclassified | unclassified | unclassified | unclassified | unclassified | 1 | 0 | 0,000103808 | 9,86E-06 | 1 | - | 8,70E-05 | - | 1 | - | 0,000107969 | - | 1 | - | 0,000102638 | - | 1 | - | 0,000104154 | - | 1 | - | 0,000117329 | - |
| 8257 | 1 | 0.2um_NOKCPHJN_644066 | Bacteria | Bacteroidetes | Flavobacteriia | Flavobacterial | Flavobacteriac | Formosa | Formosa sp. H | poorly charact | Function unkn | 05C2A | tonB-depende | 688270.Celal_ | 0,8 | 0,4 | 1,22E-05 | 6,16E-06 | 0 | - | 0 | - | 1 | - | 1,53E-05 | - | 1 | - | 1,45E-05 | - | 1 | - | 1,48E-05 | - | 1 | - | 1,66E-05 | - |
| 8258 | 1 | 0.2um_NOKCPHJN_512717 | Eukaryota | Nematoda | Chromadorea | Diplogasterida | Neodiplogaste | Pristionchus | Pristionchus p | information st | Chromatin str | KOG3467 | Core compone | 7217.FBpp012 | 1,2 | 0,4 | 2,73E-05 | 6,04E-06 | 2 | - | 3,91E-05 | - | 1 | - | 2,43E-05 | - | 1 | - | 2,31E-05 | - | 1 | - | 2,34E-05 | - | 1 | - | 2,64E-05 | - |
| 8259 | 1 | 0.2um_NOKCPHJN_771178 | Eukaryota | Chordata | Aves | Bucerotiforme | Bucerotidae | Buceros | Buceros rhino | unclassified | unclassified | unclassified | unclassified | unclassified | 0,2 | 0,4 | 3,14E-06 | 6,27E-06 | 0 | - | 0 | - | 0 | - | 0 | - | 1 | - | 1,57E-05 | - | 0 | - | 0 | - | 0 | - | 0 | - |
| 8260 | 1 | 3um_FIJJAFBF_107569 | Bacteria | Proteobacteri | Alphaproteob | Rhodobactera | Rhodobactera | Planktotalea | Planktotalea f | unclassified | unclassified | unclassified | unclassified | unclassified | 0,6 | 0,489897949 | 4,79E-05 | 3,99E-05 | 1 | - | 6,67E-05 | - | 1 | - | 8,29E-05 | - | 0 | - | 0 | - | 0 | - | 0 | - | 1 | - | 9,00E-05 | - |
| 8261 | 1 | 0.2um_NOKCPHJN_438610 | Eukaryota | unclassified | Labyrinthulom | unclassified | Thraustochytr | Aurantiochytr | Aurantiochytr | unclassified | unclassified | unclassified | unclassified | unclassified | 14,8 | 0,748331477 | 0,000535298 | 4,49E-05 | 15 | - | 0,000455463 | - | 14 | - | 0,000527848 | - | 15 | - | 0,000537628 | - | 16 | - | 0,00058194 | - | 14 | - | 0,000573609 | - |
| 8262 | 1 | 0.2um_NOKCPHJN_476231 | Eukaryota | Phaeophyceae | unclassified | Ectocarpales | Ectocarpaceae | Ectocarpus | Ectocarpus sil | unclassified | unclassified | unclassified | unclassified | unclassified | 2,4 | 0,489897949 | 0,000244747 | 4,33E-05 | 3 | - | 0,000258895 | - | 3 | - | 0,000321471 | - | 2 | - | 0,000203733 | - | 2 | - | 0,000206742 | - | 2 | - | 0,000232894 | - |
| 8263 | 1 | 3um_FIJJAFBF_649754 | unclassified | unclassified | unclassified | unclassified | unclassified | unclassified | unclassified | unclassified | unclassified | unclassified | unclassified | unclassified | 0,2 | 0,4 | 4,53E-06 | 9,06E-06 | 0 | - | 0 | - | 0 | - | 0 | - | 0 | - | 0 | - | 1 | - | 2,26E-05 | - | 0 | - | 0 | - |
| 8264 | 1 | 0.2um_NOKCPHJN_16250 | Bacteria | Proteobacteri | Alphaproteob | Kiloniellales | Kiloniellaceae | Kiloniella | Kiloniella maj | poorly charact | Function unkn | COG1744 | basic membra | 911045.PSE_3 | 5,4 | 1,019803903 | 0,000207493 | 3,58E-05 | 7 | - | 0,00022696 | - | 5 | - | 0,000201298 | - | 5 | - | 0,000191359 | - | 4 | - | 0,000155348 | - | 6 | - | 0,000262499 | - |
| 8265 | 1 | 0.2um_NOKCPHJN_309859 | Bacteria | Proteobacteri | Gammaprote | Cellvibrionale | Halieaceae | unclassified | marine gamm | poorly charact | Function unkn | 16R8I | Lipoprotein | 247639.MGP2 | 0,8 | 0,748331477 | 1,44E-05 | 1,27E-05 | 2 | - | 3,26E-05 | - | 1 | - | 2,02E-05 | - | 1 | - | 1,92E-05 | - | 0 | - | 0 | - | 0 | - | 0 | - |
| 8266 | 1 | 3um_FIJJAFBF_387045 | Bacteria | Proteobacteri | Alphaproteob | Rhodobactera | Rhodobactera | unclassified | Rhodobactera | poorly charact | Function unkn | 0XR8Q | NA | 314265.R2601 | 0,2 | 0,4 | 4,50E-06 | 9,00E-06 | 1 | - | 2,25E-05 | - | 0 | - | 0 | - | 0 | - | 0 | - | 0 | - | 0 | - | 0 | - | 0 | - |
| 8267 | 1 | 3um_FIJJAFBF_193127 | Bacteria | Proteobacteri | Gammaprote | Oceanospirilla | Oceanospirilla | Motiliproteus | Motiliproteus | unclassified | unclassified | unclassified | unclassified | unclassified | 1 | 0 | 9,93E-05 | 9,43E-06 | 1 | - | 8,32E-05 | - | 1 | - | 0,000103275 | - | 1 | - | 9,82E-05 | - | 1 | - | 9,96E-05 | - | 1 | - | 0,000112228 | - |
| 8268 | 1 | 3um_FIJJAFBF_405912 | Bacteria | Proteobacteri | Gammaprote | Alteromonada | Alteromonada | Paraglaciecola | Paraglaciecola | information st | Transcription | COG0085 | DNA-depende | 342610.Patl_ | 8 | 0 | 8,17E-05 | 7,76E-06 | 8 | - | 6,84E-05 | - | 8 | - | 8,50E-05 | - | 8 | - | 8,08E-05 | - | 8 | - | 8,20E-05 | - | 8 | - | 9,23E-05 | - |
| 8269 | 1 | 0.2um_NOKCPHJN_96523 | Bacteria | Proteobacteri | Gammaprote | Oceanospirilla | Saccharospirill | Reinekea | Reinekea fors | poorly charact | Function unkn | 17EUC | FecR protein | 314283.MED2 | 0,4 | 0,8 | 6,81E-06 | 1,36E-05 | 2 | - | 3,41E-05 | - | 0 | - | 0 | - | 0 | - | 0 | - | 0 | - | 0 | - | 0 | - | 0 | - |
| 8270 | 1 | 3um_FIJJAFBF_16115 | Bacteria | Bacteroidetes | Cytophagia | Cytophagales | Flammeovirga | unclassified | Flammeovirga | unclassified | unclassified | unclassified | unclassified | unclassified | 0,8 | 0,748331477 | 5,91E-05 | 5,69E-05 | 1 | - | 6,48E-05 | - | 0 | - | 0 | - | 2 | - | 0,000153087 | - | 1 | - | 7,77E-05 | - | 0 | - | 0 | - |
| 8271 | 1 | 3um_FIJJAFBF_788764 | Bacteria | Proteobacteri | Alphaproteob | Rhodobactera | Rhodobactera | Yoonia | Yoonia marico | unclassified | unclassified | unclassified | unclassified | unclassified | 0,4 | 0,489897949 | 2,39E-05 | 2,95E-05 | 1 | - | 5,49E-05 | - | 0 | - | 0 | - | 1 | - | 6,48E-05 | - | 0 | - | 0 | - | 0 | - | 0 | - |
| 8272 | 1 | 3um_FIJJAFBF_137597 | Bacteria | Bacteroidetes | Flavobacteriia | Flavobacterial | Crocinitomica | unclassified | Crocinitomica | unclassified | unclassified | unclassified | unclassified | unclassified | 3 | 0,632455532 | 0,000307985 | 5,24E-05 | 4 | - | 0,000347808 | - | 3 | - | 0,000323907 | - | 3 | - | 0,000307914 | - | 2 | - | 0,000208308 | - | 3 | - | 0,000351987 | - |
| 8273 | 1 | 0.2um_NOKCPHJN_747889 | Bacteria | Proteobacteri | Gammaprote | Cellvibrionale | Spongiibacter | unclassified | marine gamm | metabolism | Inorganic ion t | 0QK8T | TonB-depende | 247633.GP214 | 17 | 1,264911064 | 0,00025785 | 1,36E-05 | 19 | - | 0,000242846 | - | 16 | - | 0,000253931 | - | 18 | - | 0,000271568 | - | 16 | - | 0,000244959 | - | 16 | - | 0,000275946 | - |
| 8274 | 1 | 3um_FIJJAFBF_705373 | Bacteria | Proteobacteri | Gammaprote | Alteromonada | Alteromonada | Paraglaciecola | Paraglaciecola | unclassified | unclassified | unclassified | unclassified | unclassified | 0,4 | 0,8 | 5,98E-06 | 1,20E-05 | 0 | - | 0 | - | 2 | - | 2,99E-05 | - | 0 | - | 0 | - | 0 | - | 0 | - | 0 | - | 0 | - |
| 8275 | 1 | 0.2um_NOKCPHJN_773140 | Bacteria | Bacteroidetes | Flavobacteriia | Flavobacterial | Flavobacteriac | Lacinutrix | Lacinutrix sp. | poorly charact | Function unkn | 1EB8T | NA | 436017.A4S9 | 0,6 | 0,489897949 | 9,94E-06 | 8,15E-06 | 0 | - | 0 | - | 0 | - | 0 | - | 1 | - | 1,57E-05 | - | 1 | - | 1,60E-05 | - | 1 | - | 1,80E-05 | - |
| 8276 | 1 | 3um_FIJJAFBF_581777 | Bacteria | Bacteroidetes | Flavobacteriia | Flavobacterial | Flavobacteriac | Polaribacter | Polaribacter s | cellular proces | Intracellular tr | 09S7I | Preprotein tra | 313594.PI23P | 0,6 | 0,489897949 | 7,11E-05 | 5,85E-05 | 1 | - | 0,0001053 | - | 0 | - | 0 | - | 1 | - | 0,000124296 | - | 1 | - | 0,000126131 | - | 0 | - | 0 | - |
| 8277 | 1 | 3um_FIJJAFBF_273643 | Bacteria | Bacteroidetes | Flavobacteriia | Flavobacterial | Flavobacteriac | Algibacter | Algibacter sp. | unclassified | unclassified | unclassified | unclassified | unclassified | 2 | 0,632455532 | 6,28E-05 | 2,01E-05 | 2 | - | 5,29E-05 | - | 1 | - | 3,28E-05 | - | 3 | - | 9,37E-05 | - | 2 | - | 6,34E-05 | - | 2 | - | 7,14E-05 | - |
| 8278 | 1 | 3um_FIJJAFBF_631848 | Bacteria | Bacteroidetes | Flavobacteriia | Flavobacterial | Flavobacteriac | Formosa | Formosa sp. H | cellular proces | Cell wall/mem | COG2885 | Ompa motb d | 983544.Lacal_ | 9,2 | 1,326649916 | 0,000265201 | 4,81E-05 | 9 | - | 0,000217015 | - | 11 | - | 0,00032935 | - | 10 | - | 0,000284627 | - | 7 | - | 0,000202181 | - | 9 | - | 0,00029283 | - |
| 8279 | 1 | 0.2um_NOKCPHJN_56921 | Bacteria | Bacteroidetes | Flavobacteriia | Flavobacterial | Flavobacteriac | Polaribacter | Polaribacter s | metabolism | Amino acid tra | 05C6M | Alpha-keto-be | 313598.MED1 | 1,6 | 0,489897949 | 4,38E-05 | 1,22E-05 | 2 | - | 4,67E-05 | - | 2 | - | 5,79E-05 | - | 2 | - | 5,51E-05 | - | 1 | - | 2,79E-05 | - | 1 | - | 3,15E-05 | - |
| 8280 | 1 | 0.2um_NOKCPHJN_701205 | Bacteria | Bacteroidetes | unclassified | unclassified | unclassified | unclassified | unidentified e | poorly charact | Function unkn | 05END | lipolytic prote | 50743.SCB49_ | 0,4 | 0,489897949 | 1,08E-05 | 1,32E-05 | 0 | - | 0 | - | 0 | - | 0 | - | 1 | - | 2,67E-05 | - | 1 | - | 2,71E-05 | - | 0 | - | 0 | - |
| 8281 | 1 | 3um_FIJJAFBF_539569 | Bacteria | Bacteroidetes | Flavobacteriia | Flavobacterial | Flavobacteriac | Nonlabens | Nonlabens do | poorly charact | Function unkn | 0PBEY | Pfam:DUF145 | 886377.Murru | 2,4 | 0,489897949 | 0,000484338 | 7,54E-05 | 3 | - | 0,000513926 | - | 2 | - | 0,000425429 | - | 2 | - | 0,000404425 | - | 3 | - | 0,000615597 | - | 2 | - | 0,000462312 | - |
| 8282 | 1 | 3um_FIJJAFBF_334134 | Bacteria | Proteobacteri | Alphaproteob | Rhizobiales | Cohaesibacter | Cohaesibacter | Cohaesibacter | unclassified | unclassified | unclassified | unclassified | unclassified | 2,2 | 0,748331477 | 9,79E-05 | 3,78E-05 | 2 | - | 7,40E-05 | - | 1 | - | 4,60E-05 | - | 3 | - | 0,000131112 | - | 2 | - | 8,87E-05 | - | 3 | - | 0,000149879 | - |
| 8283 | 1 | 3um_FIJJAFBF_10138 | Eukaryota | Bacillariophyt | Bacillariophyc | Naviculales | Phaeodactylac | Phaeodactylu | Phaeodactylu | information st | Translation, ri | KOG3421 | ribosomal pro | 2850.Phatr25 | 1,2 | 0,748331477 | 0,000121321 | 7,31E-05 | 2 | - | 0,000177948 | - | 1 | - | 0,00011048 | - | 1 | - | 0,000105025 | - | 2 | - | 0,000213152 | - | 0 | - | 0 | - |
| 8284 | 1 | 3um_FIJJAFBF_134065 | Bacteria | Bacteroidetes | Flavobacteriia | Flavobacterial | Crocinitomica | unclassified | Crocinitomica | unclassified | unclassified | unclassified | unclassified | unclassified | 1,2 | 0,748331477 | 0,00024088 | 0,000144083 | 2 | - | 0,000342617 | - | 2 | - | 0,000425429 | - | 0 | - | 0 | - | 1 | - | 0,000205199 | - | 1 | - | 0,000231156 | - |
| 8285 | 1 | 3um_FIJJAFBF_637811 | Bacteria | Proteobacteri | Gammaprote | Alteromonada | unclassified | unclassified | Alteromonada | unclassified | unclassified | unclassified | unclassified | unclassified | 1,4 | 1,2 | 2,42E-05 | 2,13E-05 | 2 | - | 2,87E-05 | - | 3 | - | 5,35E-05 | - | 0 | - | 0 | - | 0 | - | 0 | - | 2 | - | 3,88E-05 | - |
| 8286 | 1 | 0.2um_NOKCPHJN_361623 | Bacteria | Bacteroidetes | Flavobacteriia | Flavobacterial | Flavobacteriac | Polaribacter | various | unclassified | unclassified | unclassified | unclassified | unclassified | 1 | 0 | 0,000217503 | 2,07E-05 | 1 | - | 0,000182185 | - | 1 | - | 0,00022622 | - | 1 | - | 0,000215051 | - | 1 | - | 0,000218227 | - | 1 | - | 0,000245833 | - |
| 8287 | 2 | 3um_FIJJAFBF_453651;3um_FIJJAFBF_469687 | Bacteria | Bacteroidetes | Flavobacteriia | Flavobacterial | Flavobacteriac | Formosa | Formosa sp. H | information st | Transcription | 0P7W9 | Transcription | various | 2,2 | 0,748331477 | 7,47E-05 | 3,00E-05 | 2 | - | 5,60E-05 | - | 3 | - | 0,000104282 | - | 1 | - | 3,30E-05 | - | 2 | - | 6,71E-05 | - | 3 | - | 0,000113323 | - |
| 8288 | 1 | 3um_FIJJAFBF_564667 | Bacteria | Proteobacteri | Betaproteoba | unclassified | unclassified | unclassified | Betaproteoba | metabolism | Inorganic ion t | 07QMV | Part of the AB | 452638.Pnec_ | 10,4 | 0,489897949 | 0,001336215 | 0,000178395 | 10 | - | 0,00107268 | - | 11 | - | 0,001465147 | - | 10 | - | 0,001266189 | - | 10 | - | 0,001284891 | - | 11 | - | 0,001592168 | - |

| | | | | | | | | | | | | | | | | | | | | | | | | | | | | |
|---|---|---|---|---|---|---|---|---|---|---|---|---|---|---|---|---|---|---|---|---|---|---|---|---|---|---|---|---|
| 8289 | 1 | 3um_FIJJAFBF_106479 | Bacteria | Bacteroidetes | Flavobacteriia | Flavobacterial | Flavobacteriac | Flavivirga | Flavivirga eckl | information st | Translation, ri | 05C64 | Binds directly | 983544.Lacal_ | 1 | 0 | 5,96E-05 | 5,66E-06 | 1 - | 4,99E-05 - | 1 - | 6,20E-05 - | 1 - | 5,89E-05 - | 1 - | 5,98E-05 - | 1 - | 6,73E-05 - |
| 8290 | 1 | 0.2um_NOKCPHJN_191184 | Bacteria | Actinobacteria | Actinobacteria | unclassified | unclassified | unclassified | Actinobacteria | unclassified | unclassified | unclassified | unclassified | unclassified | 0,4 | 0,489897949 | 5,91E-05 | 7,25E-05 | 0 - | 0 - | 0 - | 0 - | 0 - | 0 - | 1 - | 0,000138872 - | 1 - | 0,000156439 - |
| 8291 | 1 | 3um_FIJJAFBF_337966 | Bacteria | Proteobacteri | Alphaproteob | Rhodobactera | unclassified | unclassified | Rhodobactera | unclassified | unclassified | unclassified | unclassified | unclassified | 1 | 0 | 9,86E-05 | 9,36E-06 | 1 - | 8,26E-05 - | 1 - | 0,000102532 - | 1 - | 9,75E-05 - | 1 - | 9,89E-05 - | 1 - | 0,00011142 - |
| 8292 | 1 | 0.2um_NOKCPHJN_645430 | Bacteria | Proteobacteri | Gammaprotec | Enterobactera | Enterobacteri | Candidatus Ri | Candidatus Ri | unclassified | unclassified | unclassified | unclassified | unclassified | 0,4 | 0,489897949 | 2,64E-05 | 3,23E-05 | 0 - | 0 - | 0 - | 0 - | 1 - | 6,55E-05 - | 1 - | 6,64E-05 - | 0 - | 0 - |
| 8293 | 1 | 3um_FIJJAFBF_206848 | Bacteria | Proteobacteri | Betaproteoba | Burkholderiale | Burkholderiac | Burkholderia | Burkholderia t | unclassified | unclassified | unclassified | unclassified | unclassified | 0,2 | 0,4 | 9,93E-06 | 1,99E-05 | 0 - | 0 - | 0 - | 0 - | 1 - | 4,96E-05 - | 0 - | 0 - | 0 - | 0 - |
| 8294 | 1 | 0.2um_NOKCPHJN_115030 | Viruses | unclassified | unclassified | Caudovirales | Myoviridae | unclassified | Synechococcu | unclassified | unclassified | unclassified | unclassified | unclassified | 0,2 | 0,4 | 5,86E-06 | 1,17E-05 | 0 - | 0 - | 0 - | 0 - | 0 - | 0 - | 1 - | 2,93E-05 - | 0 - | 0 - |
| 8295 | 1 | 3um_FIJJAFBF_214532 | Bacteria | Proteobacteri | Alphaproteob | Rhodobactera | unclassified | unclassified | Rhodobactera | unclassified | unclassified | unclassified | unclassified | unclassified | 1,4 | 1,019803903 | 6,73E-05 | 4,55E-05 | 3 - | 0,000128481 - | 2 - | 0,000106357 - | 1 - | 5,06E-05 - | 1 - | 5,13E-05 - | 0 - | 0 - |
| 8296 | 1 | 3um_FIJJAFBF_284366 | Bacteria | Bacteroidetes | Flavobacteriia | Flavobacterial | Flavobacteriac | Formosa | Formosa sp. H | information st | Translation, ri | 05KK9 | This protein bi | 313603.FB217 | 1,2 | 0,979795897 | 9,37E-05 | 7,79E-05 | 2 - | 0,000130428 - | 2 - | 0,000161953 - | 0 - | 0 - | 0 - | 0 - | 2 - | 0,000175994 - |
| 8297 | 1 | 3um_FIJJAFBF_348211 | Bacteria | Proteobacteri | Alphaproteob | unclassified | unclassified | Candidatus Pu | Candidatus Pu | metabolism | Energy produc | 01RNF | Produces ATP | 488538.SAR11 | 1,4 | 0,489897949 | 4,03E-05 | 1,49E-05 | 1 - | 2,40E-05 - | 2 - | 5,96E-05 - | 2 - | 5,67E-05 - | 1 - | 2,88E-05 - | 1 - | 3,24E-05 - |
| 8298 | 1 | 0.2um_NOKCPHJN_732605 | Bacteria | Bacteroidetes | unclassified | unclassified | unclassified | unclassified | Bacteroidetes | poorly charact | Function unkn | 16YV3 | Large repetitiv | 290398.Csal_ | 0,8 | 0,4 | 1,52E-05 | 7,83E-06 | 1 - | 1,60E-05 - | 1 - | 1,98E-05 - | 1 - | 1,88E-05 - | 0 - | 0 - | 1 - | 2,15E-05 - |
| 8299 | 1 | 0.2um_NOKCPHJN_517163 | Bacteria | Proteobacteri | Alphaproteob | Rhodobactera | Rhodobactera | Thalassococcu | Thalassococcu | metabolism | Carbohydrate | 08ERU | trap dicarboxy | 388739.RSK20 | 1,2 | 0,4 | 4,90E-05 | 1,64E-05 | 1 - | 3,43E-05 - | 1 - | 4,25E-05 - | 2 - | 8,09E-05 - | 1 - | 4,10E-05 - | 1 - | 4,62E-05 - |
| 8300 | 1 | 0.2um_NOKCPHJN_979126 | Eukaryota | unclassified | Cryptophyta | Pyrenomonad | Geminigerace | Guillardia | Guillardia thet | metabolism | Energy produc | 0IFIU | Pyrophosphat | 653948.CCA14 | 0,4 | 0,489897949 | 1,16E-05 | 1,42E-05 | 0 - | 0 - | 0 - | 0 - | 0 - | 0 - | 1 - | 2,72E-05 - | 1 - | 3,07E-05 - |
| 8301 | 1 | 3um_FIJJAFBF_754835 | Bacteria | Proteobacteri | Alphaproteob | Rhodobactera | Rhodobactera | unclassified | Rhodobactera | metabolism | Amino acid tra | 05CAS | nag kinase | 351016.RAZW | 0,2 | 0,4 | 9,44E-06 | 1,89E-05 | 0 - | 0 - | 0 - | 0 - | 1 - | 4,72E-05 - | 0 - | 0 - | 0 - | 0 - |
| 8302 | 1 | 3um_FIJJAFBF_75852 | Bacteria | Proteobacteri | Gammaprotec | unclassified | unclassified | unclassified | Gammaprotec | metabolism | Amino acid tra | COG0174 | glutamine syn | 504472.Slin_2 | 1 | 0 | 4,07E-05 | 3,86E-06 | 1 - | 3,41E-05 - | 1 - | 4,23E-05 - | 1 - | 4,02E-05 - | 1 - | 4,08E-05 - | 1 - | 4,60E-05 - |
| 8303 | 1 | 0.2um_NOKCPHJN_75525 | Bacteria | Proteobacteri | Gammaprotec | Oceanospirilla | Saccharospirill | Reinekea | Reinekea fors | information st | Replication, re | COG0513 | purine NTP-de | 247633.GP214 | 4,8 | 1,939071943 | 0,000141424 | 4,48E-05 | 8 - | 0,000204958 - | 5 - | 0,000159061 - | 5 - | 0,000151208 - | 4 - | 0,000122753 - | 2 - | 6,91E-05 - |
| 8304 | 1 | 0.2um_NOKCPHJN_198762 | Bacteria | Proteobacteri | Alphaproteob | Rhodobactera | Rhodobactera | Planktomarina | Planktomarina | unclassified | unclassified | unclassified | unclassified | unclassified | 0,6 | 0,8 | 2,41E-05 | 3,27E-05 | 0 - | 0 - | 1 - | 3,80E-05 - | 0 - | 0 - | 0 - | 0 - | 2 - | 8,26E-05 - |
| 8305 | 1 | 0.2um_NOKCPHJN_909610 | Eukaryota | Phaeophyceae | unclassified | Ectocarpales | Ectocarpaceae | Ectocarpus | Ectocarpus sili | unclassified | unclassified | unclassified | unclassified | unclassified | 8,4 | 0,489897949 | 0,000190644 | 1,59E-05 | 9 - | 0,000171593 - | 9 - | 0,000213068 - | 8 - | 0,000180043 - | 8 - | 0,000182702 - | 8 - | 0,000205813 - |
| 8306 | 1 | 0.2um_NOKCPHJN_952690 | Eukaryota | unclassified | Oomycetes | Pythiales | Pythiaceae | Pythium | Pythium insidi | unclassified | unclassified | unclassified | unclassified | unclassified | 1 | 0 | 7,41E-05 | 7,03E-06 | 1 - | 6,20E-05 - | 1 - | 7,70E-05 - | 1 - | 7,32E-05 - | 1 - | 7,43E-05 - | 1 - | 8,37E-05 - |
| 8307 | 1 | 0.2um_NOKCPHJN_759699 | Eukaryota | unclassified | Oomycetes | Peronosporale | Peronosporac | Phytophthora | various | information st | Chromatin stru | KOG1744 | Core compone | 164328.Phyra | 6,8 | 1,166190379 | 0,000794534 | 0,000106457 | 8 - | 0,000791564 - | 5 - | 0,000614305 - | 7 - | 0,000817565 - | 8 - | 0,000948161 - | 6 - | 0,000801075 - |
| 8308 | 1 | 0.2um_NOKCPHJN_80676 | Bacteria | Bacteroidetes | Flavobacteriia | Flavobacterial | Crocinitomica | unclassified | Crocinitomica | information st | Translation, ri | 05KK9 | This protein bi | 391603.FBALC | 0,4 | 0,489897949 | 3,82E-05 | 4,71E-05 | 1 - | 8,70E-05 - | 0 - | 0 - | 0 - | 0 - | 1 - | 0,000104154 - | 0 - | 0 - |
| 8309 | 1 | 3um_FIJJAFBF_520450 | Bacteria | Bacteroidetes | Flavobacteriia | Flavobacterial | Flavobacteriac | Polaribacter | Polaribacter s | cellular proces | Cell wall/mem | 05C2X | Arabinose 5-p | 313598.MED1 | 0,6 | 0,8 | 2,27E-05 | 2,93E-05 | 2 - | 7,13E-05 - | 0 - | 0 - | 1 - | 4,21E-05 - | 0 - | 0 - | 0 - | 0 - |
| 8310 | 1 | 0.2um_NOKCPHJN_166082 | Bacteria | Bacteroidetes | Flavobacteriia | Flavobacterial | Flavobacteriac | Formosa | Formosa sp. H | information st | Translation, ri | COG0018 | arginyL-tRNA s | 1046627.BZAR | 0,2 | 0,4 | 4,81E-06 | 9,63E-06 | 0 - | 0 - | 1 - | 2,41E-05 - | 0 - | 0 - | 0 - | 0 - | 0 - | 0 - |
| 8311 | 1 | 3um_FIJJAFBF_111340 | Viruses | unclassified | unclassified | unclassified | Phycodnavirid | unclassified | Chrysochromu | unclassified | unclassified | unclassified | unclassified | unclassified | 20,6 | 2,727636339 | 0,000527064 | 0,000100306 | 22 - | 0,000470221 - | 19 - | 0,000504257 - | 20 - | 0,000504589 - | 17 - | 0,000435236 - | 25 - | 0,000721017 - |
| 8312 | 1 | 3um_FIJJAFBF_269231 | Bacteria | Bacteroidetes | Flavobacteriia | Flavobacterial | Flavobacteriac | Polaribacter | Polaribacter ir | cellular proces | Posttranslatio | COG1225 | alkyl hydroper | 313594.PI23P | 0,6 | 0,489897949 | 5,24E-05 | 4,32E-05 | 1 - | 7,65E-05 - | 1 - | 9,50E-05 - | 1 - | 9,03E-05 - | 0 - | 0 - | 0 - | 0 - |
| 8313 | 1 | 3um_FIJJAFBF_504244 | Eukaryota | unclassified | Pelagophycea | Pelagomonad | unclassified | Aureoumbra | Aureoumbra l | metabolism | Energy produc | COG0055 | Produces ATP | 240292.Ava_2 | 1,6 | 0,8 | 4,46E-05 | 2,26E-05 | 2 - | 4,82E-05 - | 2 - | 5,99E-05 - | 2 - | 5,69E-05 - | 2 - | 5,78E-05 - | 0 - | 0 - |
| 8314 | 1 | 0.2um_NOKCPHJN_83701 | Bacteria | Bacteroidetes | Flavobacteriia | various | various | various | various | unclassified | unclassified | unclassified | unclassified | unclassified | 2 | 0,632455532 | 6,32E-05 | 1,46E-05 | 3 - | 8,18E-05 - | 2 - | 6,77E-05 - | 2 - | 6,44E-05 - | 2 - | 6,53E-05 - | 1 - | 3,68E-05 - |
| 8315 | 1 | 0.2um_NOKCPHJN_524232 | Bacteria | Proteobacteri | Betaproteoba | unclassified | unclassified | unclassified | Betaproteoba | unclassified | unclassified | unclassified | unclassified | unclassified | 1,8 | 0,748331477 | 0,000200071 | 8,51E-05 | 2 - | 0,000188159 - | 3 - | 0,000350456 - | 2 - | 0,000222102 - | 1 - | 0,000112691 - | 1 - | 0,000126946 - |
| 8316 | 1 | 3um_FIJJAFBF_21463 | Eukaryota | unclassified | Cryptophyta | Pyrenomonad | Pyrenomonad | Rhodomonas | Rhodomonas | metabolism | Energy produc | COG4451 | RuBisCO catal | 45157.CMV01 | 1 | 0 | 9,86E-05 | 9,36E-06 | 1 - | 8,26E-05 - | 1 - | 0,000102532 - | 1 - | 9,75E-05 - | 1 - | 9,89E-05 - | 1 - | 0,00011142 - |
| 8317 | 1 | 3um_FIJJAFBF_22495 | Bacteria | Proteobacteri | Gammaprotec | Alteromonada | Alteromonada | Paraglaciecola | Paraglaciecola | cellular proces | Cell wall/mem | 16QB0 | Part of the ou | 342610.Patl_1 | 0,2 | 0,4 | 2,78E-06 | 5,56E-06 | 1 - | 1,39E-05 - | 0 - | 0 - | 0 - | 0 - | 0 - | 0 - | 0 - | 0 - |
| 8318 | 1 | 0.2um_NOKCPHJN_313858 | Bacteria | Bacteroidetes | Flavobacteriia | Flavobacterial | Flavobacteriac | Formosa | Formosa sp. H | unclassified | unclassified | unclassified | unclassified | unclassified | 3,2 | 0,748331477 | 6,14E-05 | 1,56E-05 | 4 - | 6,44E-05 - | 3 - | 6,00E-05 - | 2 - | 3,80E-05 - | 3 - | 5,78E-05 - | 4 - | 8,69E-05 - |
| 8319 | 3 | 0.2um_NOKCPHJN_150702;0.2um_NOKCPHJN_2 | Bacteria | Proteobacteri | Alphaproteob | Rhodobactera | Rhodobactera | various | various | information st | Translation, ri | COG0012 | gtp-binding pr | various | 0,2 | 0,4 | 6,29E-06 | 1,26E-05 | 1 - | 3,14E-05 - | 0 - | 0 - | 0 - | 0 - | 0 - | 0 - | 0 - | 0 - |
| 8320 | 1 | 0.2um_NOKCPHJN_331120 | Bacteria | Proteobacteri | Gammaprotec | Pseudomonad | Moraxellaceae | Acinetobacter | Acinetobacter | unclassified | unclassified | unclassified | unclassified | unclassified | 0,4 | 0,489897949 | 2,40E-06 | 2,94E-06 | 0 - | 0 - | 1 - | 6,12E-06 - | 0 - | 0 - | 1 - | 5,90E-06 - | 0 - | 0 - |
| 8321 | 1 | 0.2um_NOKCPHJN_766226 | unclassified | unclassified | unclassified | unclassified | unclassified | unclassified | unclassified | unclassified | unclassified | unclassified | unclassified | unclassified | 0,8 | 0,4 | 2,16E-05 | 1,11E-05 | 1 - | 2,25E-05 - | 1 - | 2,79E-05 - | 0 - | 0 - | 1 - | 2,70E-05 - | 1 - | 3,04E-05 - |
| 8322 | 1 | 3um_FIJJAFBF_761859 | Bacteria | Planctomycete | Planctomyceti | Planctomyceta | Planctomyceta | unclassified | Planctomyceta | unclassified | unclassified | unclassified | unclassified | unclassified | 1 | 0 | 2,20E-05 | 2,09E-06 | 1 - | 1,84E-05 - | 1 - | 2,28E-05 - | 1 - | 2,17E-05 - | 1 - | 2,20E-05 - | 1 - | 2,48E-05 - |
| 8323 | 1 | 0.2um_NOKCPHJN_442695 | unclassified | unclassified | unclassified | unclassified | unclassified | unclassified | unclassified | unclassified | unclassified | unclassified | unclassified | unclassified | 1,8 | 1,166190379 | 5,19E-05 | 3,27E-05 | 3 - | 7,63E-05 - | 1 - | 3,16E-05 - | 2 - | 6,01E-05 - | 3 - | 9,15E-05 - | 0 - | 0 - |
| 8324 | 1 | 3um_FIJJAFBF_305074 | Bacteria | Proteobacteri | Alphaproteob | Rhodobactera | Rhodobactera | Octadecabact | Octadecabact | cellular proces | Posttranslatio | 16PM6 | feS assembly | 391616.OA23 | 1,4 | 0,489897949 | 7,77E-05 | 3,16E-05 | 1 - | 4,57E-05 - | 1 - | 5,68E-05 - | 2 - | 0,000107954 - | 1 - | 5,48E-05 - | 2 - | 0,000123406 - |
| 8325 | 1 | 3um_FIJJAFBF_789799 | Bacteria | Bacteroidetes | unclassified | unclassified | unclassified | unclassified | Bacteroidetes | cellular proces | Signal transdu | 09A4D | gtp-binding pr | 398720.MED2 | 0,4 | 0,489897949 | 9,76E-06 | 1,20E-05 | 0 - | 0 - | 0 - | 0 - | 0 - | 0 - | 1 - | 2,30E-05 - | 1 - | 2,59E-05 - |
| 8326 | 1 | 0.2um_NOKCPHJN_567099 | Bacteria | unclassified | unclassified | unclassified | unclassified | unclassified | uncultured ba | unclassified | unclassified | unclassified | unclassified | unclassified | 0,8 | 0,979795897 | 1,24E-05 | 1,52E-05 | 0 - | 0 - | 2 - | 3,18E-05 - | 2 - | 3,02E-05 - | 0 - | 0 - | 0 - | 0 - |
| 8327 | 1 | 3um_FIJJAFBF_158679 | Eukaryota | Arthropoda | Arachnida | Scorpiones | Buthidae | Centruroides | Centruroides s | information st | Translation, ri | KOG0886 | Ribosomal pro | 402676.XP_00 | 0,8 | 0,4 | 6,54E-05 | 3,36E-05 | 1 - | 6,91E-05 - | 0 - | 0 - | 1 - | 8,16E-05 - | 1 - | 8,28E-05 - | 1 - | 9,33E-05 - |
| 8328 | 1 | 3um_FIJJAFBF_452767 | Bacteria | Bacteroidetes | Flavobacteriia | Flavobacterial | Flavobacteriac | Polaribacter | Polaribacter t | unclassified | unclassified | unclassified | unclassified | unclassified | 2,2 | 0,748331477 | 0,000120898 | 3,72E-05 | 3 - | 0,000139972 - | 2 - | 0,000115869 - | 1 - | 5,51E-05 - | 3 - | 0,000167663 - | 2 - | 0,000125914 - |
| 8329 | 1 | 0.2um_NOKCPHJN_515768 | Bacteria | Proteobacteri | Gammaprotec | Alteromonada | Alteromonada | unclassified | Alteromonada | metabolism | Lipid transpor | 0QN9D | Carbamoyl-ph | 247633.GP214 | 0,2 | 0,4 | 2,10E-06 | 4,19E-06 | 1 - | 1,05E-05 - | 0 - | 0 - | 0 - | 0 - | 0 - | 0 - | 0 - | 0 - |
| 8330 | 1 | 0.2um_NOKCPHJN_1027022 | Eukaryota | Xanthophycea | unclassified | Botrydiales | Botrydiaceae | Botrydium | Botrydium gra | unclassified | unclassified | unclassified | unclassified | unclassified | 7 | 0 | 0,000507508 | 4,82E-05 | 7 - | 0,000425099 - | 7 - | 0,000527848 - | 7 - | 0,000501786 - | 7 - | 0,000509197 - | 7 - | 0,000573609 - |
| 8331 | 1 | 0.2um_NOKCPHJN_899665 | Bacteria | Proteobacteri | Gammaprotec | Oceanospirilla | Saccharospirill | Reinekea | Reinekea blan | metabolism | Carbohydrate | 05CZI | D-fructose-1,6 | 314283.MED2 | 0,2 | 0,4 | 6,83E-06 | 1,37E-05 | 1 - | 3,42E-05 - | 0 - | 0 - | 0 - | 0 - | 0 - | 0 - | 0 - | 0 - |
| 8332 | 1 | 3um_FIJJAFBF_205170 | Eukaryota | Bacillariophyta | Mediophycea | Cymatosirales | Cymatosiracea | Plagiogramma | Plagiogramma | metabolism | Energy produc | COG0055 | Produces ATP | 7739.JGI1257 | 0,2 | 0,4 | 5,80E-06 | 1,16E-05 | 0 - | 0 - | 0 - | 0 - | 0 - | 0 - | 1 - | 2,90E-05 - | 0 - | 0 - |
| 8333 | 1 | 3um_FIJJAFBF_450469 | Bacteria | Bacteroidetes | Cytophagia | Cytophagales | Flammeovirga | unclassified | Flammeovirga | cellular proces | Posttranslatio | 08V1T | peptidylprolyl | 523791.Kkor_ | 0,4 | 0,8 | 2,24E-05 | 4,48E-05 | 2 - | 0,000111977 - | 0 - | 0 - | 0 - | 0 - | 0 - | 0 - | 0 - | 0 - |
| 8334 | 1 | 0.2um_NOKCPHJN_755525 | Bacteria | Proteobacteri | Alphaproteob | Pelagibacteral | Pelagibacterac | Candidatus Pe | Candidatus Pe | unclassified | unclassified | unclassified | unclassified | unclassified | 1,6 | 0,489897949 | 9,54E-05 | 2,50E-05 | 2 - | 0,000102024 - | 1 - | 6,33E-05 - | 2 - | 0,000120429 - | 2 - | 0,000122207 - | 1 - | 6,88E-05 - |
| 8335 | 1 | 0.2um_NOKCPHJN_293110 | Bacteria | Proteobacteri | Gammaprotec | Cellvibrionales | Porticoccacea | unclassified | various | information st | Transcription | 05D27 | DNA-depende | 314287.GB220 | 0,2 | 0,4 | 2,47E-06 | 4,95E-06 | 0 - | 0 - | 0 - | 0 - | 0 - | 0 - | 1 - | 1,24E-05 - | 0 - | 0 - |
| 8336 | 1 | 3um_FIJJAFBF_191793 | Eukaryota | unclassified | Pelagophycea | Pelagomonad | unclassified | Pelagomonas | uncultured Pe | unclassified | unclassified | unclassified | unclassified | unclassified | 4 | 0 | 0,000227431 | 2,16E-05 | 4 - | 0,000190501 - | 4 - | 0,000236546 - | 4 - | 0,000224867 - | 4 - | 0,000228188 - | 4 - | 0,000257053 - |
| 8337 | 1 | 3um_FIJJAFBF_604546 | Bacteria | Proteobacteri | Gammaprotec | Alteromonada | Psychromonac | Psychromonas | Psychromonas | unclassified | unclassified | unclassified | unclassified | unclassified | 0,4 | 0,489897949 | 4,10E-05 | 5,05E-05 | 1 - | 9,41E-05 - | 0 - | 0 - | 1 - | 0,000111051 - | 0 - | 0 - | 0 - | 0 - |
| 8338 | 1 | 3um_FIJJAFBF_350306 | Bacteria | Proteobacteri | Gammaprotec | Oceanospirilla | Oceanospirilla | Motiliproteus | Motiliproteus | information st | Translation, ri | 05G6W | One of the pri | 207954.MED9 | 3 | 0,632455532 | 0,00019735 | 3,36E-05 | 4 - | 0,000222867 - | 3 - | 0,000207552 - | 3 - | 0,000197304 - | 2 - | 0,000133479 - | 3 - | 0,000225545 - |
| 8339 | 1 | 3um_FIJJAFBF_309215 | Bacteria | Bacteroidetes | Flavobacteriia | Flavobacterial | Flavobacteriac | Formosa | Formosa sp. H | unclassified | unclassified | unclassified | unclassified | unclassified | 2,4 | 0,8 | 0,000333297 | 7,38E-05 | 4 - | 0,000478236 - | 2 - | 0,000296914 - | 2 - | 0,000282255 - | 2 - | 0,000286424 - | 2 - | 0,000322655 - |
| 8340 | 1 | 0.2um_NOKCPHJN_61814 | Bacteria | Bacteroidetes | Flavobacteriia | Flavobacterial | Flavobacteriac | Polaribacter | Polaribacter fi | poorly charact | Function unkn | 0XRCJ | NA | 391587.KAOT | 25,4 | 2,727636339 | 0,00032709 | 1,87E-05 | 30 - | 0,000326379 - | 24 - | 0,000324214 - | 23 - | 0,000295364 - | 27 - | 0,000351853 - | 23 - | 0,000337641 - |
| 8341 | 1 | 3um_FIJJAFBF_359190 | Bacteria | Proteobacteri | Gammaprotec | Cellvibrionales | Porticoccacea | unclassified | Gammaprotec | information st | Translation, ri | 179RU | 30S ribosomal | 314287.GB220 | 3,2 | 1,469693846 | 0,000326315 | 0,000129293 | 5 - | 0,000441449 - | 2 - | 0,00021926 - | 5 - | 0,000521085 - | 2 - | 0,000211513 - | 2 - | 0,000238268 - |
| 8342 | 3 | 0.2um_NOKCPHJN_597293;3um_FIJJAFBF_2931 | Bacteria | Proteobacteri | Alphaproteob | Rhizobiales | Phyllobacteria | unclassified | Phyllobacteria | various | various | various | various | various | 2,6 | 0,8 | 0,000187415 | 6,11E-05 | 3 - | 0,000181226 - | 3 - | 0,00022503 - | 3 - | 0,000213919 - | 1 - | 7,24E-05 - | 3 - | 0,000244539 - |
| 8343 | 1 | 3um_FIJJAFBF_318784 | Eukaryota | Chlorophyta | Chlorophycea | Chlamydomor | Tetrabaenace | Tetrabaena | Tetrabaena so | unclassified | unclassified | unclassified | unclassified | unclassified | 4,4 | 0,489897949 | 0,000123332 | 9,42E-06 | 5 - | 0,000118327 - | 4 - | 0,000117541 - | 5 - | 0,000139672 - | 4 - | 0,000113388 - | 4 - | 0,000127732 - |
| 8344 | 1 | 0.2um_NOKCPHJN_143995 | Bacteria | Proteobacteri | Gammaprotec | Oceanospirilla | Saccharospirill | Reinekea | Reinekea fors | metabolism | Carbohydrate | 05CA9 | Pyruvate kinas | 675812.VHA_ | 3,8 | 1,166190379 | 0,000110846 | 3,43E-05 | 3 - | 7,34E-05 - | 3 - | 9,12E-05 - | 6 - | 0,000173325 - | 4 - | 0,000117257 - | 3 - | 9,91E-05 - |
| 8345 | 1 | 3um_FIJJAFBF_315281 | Bacteria | Proteobacteri | Alphaproteob | Rhodobactera | Rhodobactera | Ascidiaceihabi | Ascidiaceihabi | metabolism | Amino acid tra | 05C6M | Alpha-keto-be | 314270.RB208 | 1 | 0,632455532 | 3,92E-05 | 2,54E-05 | 1 - | 3,38E-05 - | 1 - | 4,19E-05 - | 2 - | 7,97E-05 - | 1 - | 4,04E-05 - | 0 - | 0 - |
| 8346 | 1 | 0.2um_NOKCPHJN_404169 | Eukaryota | unclassified | Oomycetes | Saprolegniales | Saprolegniace | Aphanomyces | Aphanomyces | unclassified | unclassified | unclassified | unclassified | unclassified | 0,6 | 0,489897949 | 5,92E-06 | 4,89E-06 | 1 - | 8,61E-06 - | 1 - | 1,07E-05 - | 0 - | 0 - | 1 - | 1,03E-05 - | 0 - | 0 - |
| 8347 | 1 | 3um_FIJJAFBF_687730 | Bacteria | Proteobacteri | Alphaproteob | Pelagibacteral | Pelagibacterac | Candidatus Pe | Candidatus Pe | unclassified | unclassified | unclassified | unclassified | unclassified | 2,8 | 0,748331477 | 0,000106456 | 2,91E-05 | 3 - | 9,59E-05 - | 2 - | 7,94E-05 - | 4 - | 0,000150955 - | 2 - | 7,66E-05 - | 3 - | 0,000129422 - |
| 8348 | 1 | 3um_FIJJAFBF_725808 | Eukaryota | Phaeophyceae | unclassified | Ectocarpales | Ectocarpaceae | Ectocarpus | Ectocarpus sili | metabolism | Energy produc | COG0056 | Produces ATP | 35128.Thaps2 | 3 | 0 | 0,000137027 | 1,30E-05 | 3 - | 0,000114777 - | 3 - | 0,000142519 - | 3 - | 0,000135482 - | 3 - | 0,000137483 - | 3 - | 0,000154874 - |
| 8349 | 1 | 3um_FIJJAFBF_112859 | Bacteria | Actinobacteria | Actinobacteria | Micrococcales | Microbacteria | Candidatus Ac | Candidatus Ac | unclassified | unclassified | unclassified | unclassified | unclassified | 2,6 | 0,489897949 | 0,000109308 | 2,67E-05 | 2 - | 6,98E-05 - | 2 - | 8,66E-05 - | 3 - | 0,00012354 - | 3 - | 0,000125365 - | 3 - | 0,000141223 - |
| 8350 | 1 | 3um_FIJJAFBF_540030 | Bacteria | Spirochaetes | Spirochaetia | unclassified | Leptospiracea | Leptospira | Leptospira ter | unclassified | unclassified | unclassified | unclassified | unclassified | 0,8 | 0,4 | 4,76E-05 | 2,44E-05 | 1 - | 4,97E-05 - | 1 - | 6,17E-05 - | 0 - | 0 - | 1 - | 5,95E-05 - | 1 - | 6,70E-05 - |
| 8351 | 1 | 0.2um_NOKCPHJN_33920 | Bacteria | Proteobacteri | Gammaprotec | Alteromonada | Alteromonada | Paraglaciecola | Paraglaciecola | metabolism | Inorganic ion t | 0QK8T | TonB-depende | 983545.Glaag | 9,8 | 1,166190379 | 0,000151588 | 2,13E-05 | 11 - | 0,000142822 - | 11 - | 0,000177342 - | 8 - | 0,000122608 - | 9 - | 0,000139972 - | 10 - | 0,000175197 - |
| 8352 | 1 | 3um_FIJJAFBF_593700 | Bacteria | Proteobacteri | Alphaproteob | Rhodobactera | Rhodobactera | Phaeobacter | Phaeobacter s | unclassified | unclassified | unclassified | unclassified | unclassified | 0,8 | 0,748331477 | 3,14E-05 | 2,73E-05 | 2 - | 6,94E-05 - | 0 - | 0 - | 1 - | 4,09E-05 - | 0 - | 0 - | 1 - | 4,68E-05 - |
| 8353 | 1 | 3um_FIJJAFBF_67777 | Eukaryota | unclassified | unclassified | Phaeocystales | Phaeocystace | Phaeocystis | Phaeocystis ar | unclassified | unclassified | unclassified | unclassified | unclassified | 0,2 | 0,4 | 1,73E-05 | 3,45E-05 | 0 - | 0 - | 0 - | 0 - | 1 - | 8,63E-05 - | 0 - | 0 - | 0 - | 0 - |
| 8354 | 1 | 0.2um_NOKCPHJN_143198 | Eukaryota | unclassified | Raphidophyce | Chattonellales | Chattonellace | Heterosigma | Heterosigma a | unclassified | unclassified | unclassified | unclassified | unclassified | 0,2 | 0,4 | 1,05E-05 | 2,10E-05 | 0 - | 0 - | 0 - | 0 - | 1 - | 5,25E-05 - | 0 - | 0 - | 0 - | 0 - |
| 8355 | 1 | 3um_FIJJAFBF_521179 | Bacteria | Proteobacteri | Alphaproteob | Rhodobactera | Rhodobactera | Planktotalea | Planktotalea f | unclassified | unclassified | unclassified | unclassified | unclassified | 0,2 | 0,4 | 2,35E-05 | 4,69E-05 | 0 - | 0 - | 0 - | 0 - | 0 - | 0 - | 0 - | 0 - | 1 - | 0,000117329 - |
| 8356 | 1 | 3um_FIJJAFBF_306499 | Bacteria | Proteobacteri | Betaproteoba | unclassified | unclassified | unclassified | Betaproteoba | unclassified | unclassified | unclassified | unclassified | unclassified | 8,8 | 1,326649916 | 0,000247311 | 3,34E-05 | 9 - | 0,000213428 - | 8 - | 0,000235568 - | 11 - | 0,000307914 - | 9 - | 0,000255651 - | 7 - | 0,000223992 - |
| 8357 | 1 | 3um_FIJJAFBF_198071 | Bacteria | Bacteroidetes | Cytophagia | Cytophagales | Flammeovirga | unclassified | Flammeovirga | metabolism | Inorganic ion t | 05PWN | phosphate ab | 391615.GP50 | 0,4 | 0,489897949 | 1,69E-05 | 2,08E-05 | 0 - | 0 - | 0 - | 0 - | 0 - | 0 - | 1 - | 3,97E-05 - | 1 - | 4,48E-05 - |
| 8358 | 1 | 0.2um_NOKCPHJN_05106 | Bacteria | Bacteroidetes | Flavobacteriia | Flavobacterial | Crocinitomica | unclassified | Crocinitomica | unclassified | unclassified | unclassified | unclassified | unclassified | 3,4 | 0,489897949 | 3,00E-05 | 2,91E-06 | 4 - | 2,98E-05 - | 3 - | 2,78E-05 - | 4 - | 3,52E-05 - | 3 - | 2,68E-05 - | 3 - | 3,02E-05 - |
| 8359 | 1 | 0.2um_NOKCPHJN_978660 | Bacteria | Proteobacteri | Gammaprotec | unclassified | unclassified | unclassified | Gammaprotec | unclassified | unclassified | unclassified | unclassified | unclassified | 0,4 | 0,489897949 | 8,78E-06 | 1,08E-05 | 1 - | 2,01E-05 - | 0 - | 0 - | 1 - | 2,38E-05 - | 0 - | 0 - | 0 - | 0 - |
| 8360 | 1 | 3um_FIJJAFBF_383207 | Bacteria | Proteobacteri | Deltaproteoba | Myxococcales | Nannocystace | Plesiocystis | Plesiocystis pa | unclassified | unclassified | unclassified | unclassified | unclassified | 2,8 | 0,748331477 | 0,000274427 | 5,30E-05 | 4 - | 0,000335115 - | 3 - | 0,000312085 - | 2 - | 0,000197784 - | 3 - | 0,000301058 - | 2 - | 0,000226094 - |
| 8361 | 1 | 0.2um_NOKCPHJN_242635 | Bacteria | Proteobacteri | Alphaproteob | unclassified | unclassified | unclassified | Alphaproteob | metabolism | Lipid transpor | COG2067 | lipid transport | 371731.Rsw2 | 0,4 | 0,489897949 | 1,42E-05 | 1,75E-05 | 1 - | 3,25E-05 - | 0 - | 0 - | 1 - | 3,84E-05 - | 0 - | 0 - | 0 - | 0 - |
| 8362 | 1 | 3um_FIJJAFBF_99502 | Bacteria | Bacteroidetes | Flavobacteriia | Flavobacterial | Flavobacteriac | Polaribacter | Polaribacter ir | cellular proces | Cell wall/mem | COG2885 | Ompa motb d | 313594.PI23P | 2,8 | 0,748331477 | 8,45E-05 | 2,31E-05 | 3 - | 7,63E-05 - | 4 - | 0,000126403 - | 3 - | 9,01E-05 - | 2 - | 6,10E-05 - | 2 - | 6,87E-05 - |
| 8363 | 1 | 0.2um_NOKCPHJN_170457 | Bacteria | Proteobacteri | Alphaproteob | Rhodobactera | Rhodobactera | Planktomarina | Planktomarina | unclassified | unclassified | unclassified | unclassified | unclassified | 0,2 | 0,4 | 1,41E-05 | 2,82E-05 | 0 - | 0 - | 0 - | 0 - | 0 - | 0 - | 0 - | 0 - | 1 - | 7,04E-05 - |
| 8364 | 1 | 3um_FIJJAFBF_53348 | Bacteria | Bacteroidetes | Flavobacteriia | Flavobacterial | Flavobacteriac | Polaribacter | Polaribacter s | unclassified | unclassified | unclassified | unclassified | unclassified | 3,6 | 0,489897949 | 0,000149736 | 1,77E-05 | 4 - | 0,0001404 - | 4 - | 0,000174335 - | 4 - | 0,000165727 - | 3 - | 0,000126131 - | 3 - | 0,000142087 - |
| 8365 | 1 | 3um_FIJJAFBF_280523 | unclassified | unclassified | unclassified | unclassified | unclassified | unclassified | unclassified | unclassified | unclassified | unclassified | unclassified | unclassified | 0,8 | 0,748331477 | 2,50E-05 | 2,46E-05 | 1 - | 2,68E-05 - | 2 - | 6,66E-05 - | 1 - | 3,17E-05 - | 0 - | 0 - | 0 - | 0 - |
| 8366 | 1 | 0.2um_NOKCPHJN_171971 | Eukaryota | unclassified | Oomycetes | Saprolegniales | Saprolegniace | Aphanomyces | Aphanomyces | information st | Translation, ri | KOG0052 | This protein p | 2850.Phatr28 | 1,6 | 0,489897949 | 5,18E-05 | 1,82E-05 | 1 - | 2,67E-05 - | 1 - | 3,31E-05 - | 2 - | 6,30E-05 - | 2 - | 6,39E-05 - | 2 - | 7,20E-05 - |
| 8367 | 1 | 3um_FIJJAFBF_778336 | Bacteria | Proteobacteri | Gammaprotec | Cellvibrionales | Porticoccacea | unclassified | SAR92 bacteri | unclassified | unclassified | unclassified | unclassified | unclassified | 2 | 0 | 0,000326255 | 3,10E-05 | 2 - | 0,000273278 - | 2 - | 0,000339331 - | 2 - | 0,000322577 - | 2 - | 0,000327341 - | 2 - | 0,000368749 - |
| 8368 | 1 | 3um_FIJJAFBF_15045 | Bacteria | Bacteroidetes | Flavobacteriia | Flavobacterial | Flavobacteriac | Polaribacter | Polaribacter s | unclassified | unclassified | unclassified | unclassified | unclassified | 1,2 | 0,979795897 | 1,81E-05 | 1,31E-05 | 3 - | 4,10E-05 - | 1 - | 1,70E-05 - | 1 - | 1,61E-05 - | 1 - | 1,64E-05 - | 0 - | 0 - |
| 8369 | 1 | 3um_FIJJAFBF_99520 | Viruses | unclassified | unclassified | Caudovirales | Siphoviridae | Cbastvirus | Cellulophaga v | unclassified | unclassified | unclassified | unclassified | unclassified | 2,4 | 0,489897949 | 0,000171485 | 2,52E-05 | 3 - | 0,000182185 - | 2 - | 0,000150814 - | 3 - | 0,000215051 - | 2 - | 0,000145485 - | 2 - | 0,000163888 - |
| 8370 | 1 | 0.2um_NOKCPHJN_221762 | Bacteria | Proteobacteri | Gammaprotec | unclassified | unclassified | unclassified | Gammaprotec | unclassified | unclassified | unclassified | unclassified | unclassified | 2,6 | 0,489897949 | 8,44E-05 | 1,37E-05 | 3 - | 8,24E-05 - | 3 - | 0,000102286 - | 3 - | 9,72E-05 - | 2 - | 6,58E-05 - | 2 - | 7,41E-05 - |
| 8371 | 1 | 0.2um_NOKCPHJN_379856 | Bacteria | Proteobacteri | Betaproteoba | unclassified | unclassified | unclassified | Betaproteoba | information st | Translation, ri | 05VBI | Binds 16S rRN | 595537.Varpa | 1,2 | 0,4 | 0,000158398 | 3,51E-05 | 2 - | 0,000227281 - | 1 - | 0,000141108 - | 1 - | 0,000134141 - | 1 - | 0,000136122 - | 1 - | 0,000153341 - |
| 8372 | 1 | 3um_FIJJAFBF_665394 | Bacteria | Bacteroidetes | Flavobacteriia | Flavobacterial | Flavobacteriac | Polaribacter | Polaribacter s | unclassified | unclassified | unclassified | unclassified | unclassified | 4,2 | 0,979795897 | 0,000137245 | 2,81E-05 | 5 - | 0,000138619 - | 5 - | 0,000172124 - | 5 - | 0,000163626 - | 3 - | 9,96E-05 - | 3 - | 0,000112228 - |
| 8373 | 1 | 0.2um_NOKCPHJN_23860 | Bacteria | Proteobacteri | Gammaprotec | Alteromonada | Alteromonada | Paraglaciecola | Paraglaciecola | unclassified | unclassified | unclassified | unclassified | unclassified | 0,8 | 0,748331477 | 0,00010634 | 9,91E-05 | 0 - | 0 - | 1 - | 0,000137037 - | 1 - | 0,000130271 - | 2 - | 0,000264391 - | 0 - | 0 - |
| 8374 | 1 | 3um_FIJJAFBF_203935 | Bacteria | Proteobacteri | Gammaprotec | Pseudomonad | Pseudomonad | Pseudomonas | Pseudomonas | unclassified | unclassified | unclassified | unclassified | unclassified | 5,8 | 1,166190379 | 0,000230726 | 3,37E-05 | 7 - | 0,000237002 - | 5 - | 0,000210205 - | 6 - | 0,000239792 - | 7 - | 0,000283889 - | 4 - | 0,000182743 - |
| 8375 | 1 | 0.2um_NOKCPHJN_58263 | Bacteria | Bacteroidetes | Flavobacteriia | Flavobacterial | Flavobacteriac | Cellulophaga | Cellulophaga s | information st | Transcription | 0P7SF | DNA-depende | 867900.Celly_ | 6 | 1,095445115 | 6,41E-05 | 8,67E-06 | 8 - | 7,24E-05 - | 6 - | 6,74E-05 - | 5 - | 5,34E-05 - | 5 - | 5,42E-05 - | 6 - | 7,32E-05 - |
| 8376 | 1 | 3um_FIJJAFBF_113088 | Bacteria | Proteobacteri | Gammaprotec | Cellvibrionales | Porticoccacea | unclassified | SAR92 bacteri | unclassified | unclassified | unclassified | unclassified | unclassified | 4 | 0 | 0,000394323 | 3,74E-05 | 4 - | 0,000330293 - | 4 - | 0,000410126 - | 4 - | 0,000389877 - | 4 - | 0,000395635 - | 4 - | 0,000445682 - |
| 8377 | 1 | 0.2um_NOKCPHJN_152692 | Bacteria | Proteobacteri | Gammaprotec | Cellvibrionales | Porticoccacea | unclassified | SAR92 bacteri | unclassified | unclassified | unclassified | unclassified | unclassified | 0,4 | 0,489897949 | 4,33E-06 | 5,34E-06 | 1 - | 9,86E-06 - | 0 - | 0 - | 0 - | 0 - | 1 - | 1,18E-05 - | 0 - | 0 - |
| 8378 | 1 | 3um_FIJJAFBF_634074 | Bacteria | Proteobacteri | Alphaproteob | Rhodobactera | unclassified | unclassified | Rhodobactera | unclassified | unclassified | unclassified | unclassified | unclassified | 1 | 0 | 7,49E-05 | 7,11E-06 | 1 - | 6,27E-05 - | 1 - | 7,79E-05 - | 1 - | 7,40E-05 - | 1 - | 7,51E-05 - | 1 - | 8,46E-05 - |
| 8379 | 1 | 3um_FIJJAFBF_180886 | Bacteria | Verrucomicrol | Verrucomicrol | Verrucomicrol | Rubritaleacea | Rubritalea | Rubritalea squ | unclassified | unclassified | unclassified | unclassified | unclassified | 0,4 | 0,489897949 | 1,11E-05 | 1,37E-05 | 1 - | 2,52E-05 - | 0 - | 0 - | 0 - | 0 - | 1 - | 3,02E-05 - | 0 - | 0 - |
| 8380 | 1 | 3um_FIJJAFBF_786154 | Bacteria | Proteobacteri | Gammaprotec | Oceanospirilla | Saccharospirill | Reinekea | Reinekea fors | information st | Translation, ri | COG0255 | 50s ribosomal | 314283.MED2 | 1,6 | 0,489897949 | 0,000346752 | 0,000110661 | 2 - | 0,000364371 - | 1 - | 0,00022622 - | 1 - | 0,000215051 - | 2 - | 0,000436455 - | 2 - | 0,000491665 - |
| 8381 | 1 | 0.2um_NOKCPHJN_464694 | Bacteria | Bacteroidetes | Flavobacteriia | Flavobacterial | Flavobacteriac | Polaribacter | Polaribacter s | unclassified | unclassified | unclassified | unclassified | unclassified | 1,8 | 0,4 | 3,50E-05 | 8,71E-06 | 2 - | 3,25E-05 - | 2 - | 4,04E-05 - | 1 - | 1,92E-05 - | 2 - | 3,89E-05 - | 2 - | 4,39E-05 - |
| 8382 | 1 | 3um_FIJJAFBF_30301 | Bacteria | Bacteroidetes | Flavobacteriia | Flavobacterial | Flavobacteriac | Polaribacter | Polaribacter s | unclassified | unclassified | unclassified | unclassified | unclassified | 6,4 | 0,8 | 0,000261821 | 4,28E-05 | 7 - | 0,000239832 - | 7 - | 0,000297801 - | 6 - | 0,000242655 - | 5 - | 0,000205199 - | 7 - | 0,000323618 - |
| 8383 | 1 | 0.2um_NOKCPHJN_307353 | Bacteria | Proteobacteri | Alphaproteob | Rhodobactera | Rhodobactera | Thalassobium | Thalassobium | metabolism | Carbohydrate | 01RW9 | glucose-6-pho | 391593.RCCS2 | 0,8 | 0,748331477 | 2,10E-05 | 1,85E-05 | 2 - | 4,75E-05 - | 1 - | 2,95E-05 - | 1 - | 2,81E-05 - | 0 - | 0 - | 0 - | 0 - |
| 8384 | 1 | 0.2um_NOKCPHJN_815085 | Viruses | unclassified | unclassified | unclassified | unclassified | unclassified | uncultured M | unclassified | unclassified | unclassified | unclassified | unclassified | 0,4 | 0,489897949 | 1,55E-05 | 1,91E-05 | 1 - | 3,55E-05 - | 0 - | 0 - | 1 - | 4,19E-05 - | 0 - | 0 - | 0 - | 0 - |
| 8385 | 1 | 0.2um_NOKCPHJN_07570 | Bacteria | unclassified | unclassified | unclassified | unclassified | unclassified | uncultured ba | metabolism | Inorganic ion t | 05C27 | TonB-depende | 313596.RB250 | 3,4 | 1,019803903 | 4,33E-05 | 9,10E-06 | 5 - | 5,49E-05 - | 3 - | 4,09E-05 - | 4 - | 5,18E-05 - | 3 - | 3,94E-05 - | 2 - | 2,96E-05 - |
| 8386 | 1 | 3um_FIJJAFBF_22756 | Bacteria | Proteobacteri | Gammaprotec | Cellvibrionales | Porticoccacea | unclassified | Gammaprotec | cellular proces | Posttranslatio | 17FMW | HeAt shock pr | 314287.GB220 | 0,2 | 0,4 | 1,21E-05 | 2,41E-05 | 0 - | 0 - | 0 - | 0 - | 0 - | 0 - | 1 - | 6,03E-05 - | 0 - | 0 - |
| 8387 | 1 | 3um_FIJJAFBF_726474 | Bacteria | Proteobacteri | Alphaproteob | Rhodobactera | unclassified | unclassified | Rhodobactera | poorly charact | Function unkn | 01YG5 | Bacterial prot | 314262.MED1 | 0,8 | 0,4 | 8,45E-05 | 4,25E-05 | 0 - | 0 - | 1 - | 0,00010557 - | 1 - | 0,000100357 - | 1 - | 0,000101839 - | 1 - | 0,000114722 - |
| 8388 | 1 | 0.2um_NOKCPHJN_33129 | Bacteria | Bacteroidetes | Flavobacteriia | Flavobacterial | Flavobacteriac | Polaribacter | Polaribacter s | unclassified | unclassified | unclassified | unclassified | unclassified | 0,6 | 0,489897949 | 5,86E-05 | 4,87E-05 | 1 - | 8,26E-05 - | 0 - | 0 - | 0 - | 0 - | 1 - | 9,89E-05 - | 1 - | 0,00011142 - |
| 8389 | 1 | 3um_FIJJAFBF_325787 | Bacteria | Proteobacteri | Gammaprotec | Cellvibrionales | unclassified | unclassified | Cellvibrionales | cellular proces | Posttranslatio | COG0330 | Band 7 protei | 314287.GB220 | 2 | 0,894427191 | 7,05E-05 | 2,83E-05 | 3 - | 9,11E-05 - | 2 - | 7,54E-05 - | 1 - | 3,58E-05 - | 3 - | 0,000109114 - | 1 - | 4,10E-05 - |
| 8390 | 1 | 3um_FIJJAFBF_542226 | Bacteria | Proteobacteri | Gammaprotec | Oceanospirilla | Saccharospirill | Reinekea | Reinekea fors | information st | Replication, re | COG0513 | purine NTP-de | 314283.MED2 | 9 | 1,095445115 | 0,000208071 | 3,11E-05 | 9 - | 0,000174492 - | 11 - | 0,000264815 - | 9 - | 0,00020597 - | 8 - | 0,000185788 - | 8 - | 0,00020929 - |
| 8391 | 1 | 0.2um_NOKCPHJN_963827 | Bacteria | Proteobacteri | Gammaprotec | Alteromonada | Alteromonada | Glaciecola | Glaciecola pal | unclassified | unclassified | unclassified | unclassified | unclassified | 0,8 | 0,4 | 7,77E-05 | 3,99E-05 | 1 - | 8,14E-05 - | 1 - | 0,000101077 - | 1 - | 9,61E-05 - | 0 - | 0 - | 1 - | 0,00010984 - |
| 8392 | 1 | 3um_FIJJAFBF_442207 | Bacteria | Bacteroidetes | Flavobacteriia | Flavobacterial | Flavobacteriac | Formosa | Formosa sp. H | cellular proces | Cell motility | COG1360 | Ompa motb d | 391603.FBALC | 7,4 | 1,019803903 | 0,000353039 | 2,72E-05 | 9 - | 0,000363729 - | 8 - | 0,000401462 - | 7 - | 0,000333935 - | 7 - | 0,000338867 - | 6 - | 0,0003272 - |
| 8393 | 1 | 0.2um_NOKCPHJN_637517 | Bacteria | Proteobacteri | Gammaprotec | Cellvibrionales | Porticoccacea | unclassified | Gammaprotec | unclassified | unclassified | unclassified | unclassified | unclassified | 0,2 | 0,4 | 7,26E-06 | 1,45E-05 | 0 - | 0 - | 0 - | 0 - | 1 - | 3,63E-05 - | 0 - | 0 - | 0 - | 0 - |
| 8394 | 1 | 0.2um_NOKCPHJN_383543 | Bacteria | Bacteroidetes | Flavobacteriia | unclassified | unclassified | unclassified | Flavobacteria | metabolism | Inorganic ion t | 05C27 | TonB-depende | 487796.Flav2 | 10,4 | 2,154065923 | 0,000129891 | 2,02E-05 | 13 - | 0,000138157 - | 12 - | 0,000158354 - | 11 - | 0,000137991 - | 9 - | 0,000114569 - | 7 - | 0,000100382 - |
| 8395 | 1 | 0.2um_NOKCPHJN_560383 | unclassified | unclassified | unclassified | unclassified | unclassified | unclassified | unclassified | unclassified | unclassified | unclassified | unclassified | unclassified | 0,2 | 0,4 | 6,20E-05 | 0,000123929 | 0 - | 0 - | 1 - | 0,000309824 - | 0 - | 0 - | 0 - | 0 - | 0 - | 0 - |
| 8396 | 1 | 0.2um_NOKCPHJN_140926 | Bacteria | Proteobacteri | Gammaprotec | Cellvibrionales | Spongiibacter | unclassified | marine gamm | unclassified | unclassified | unclassified | unclassified | unclassified | 1 | 0,894427191 | 1,63E-05 | 1,46E-05 | 2 - | 2,87E-05 - | 2 - | 3,56E-05 - | 1 - | 1,69E-05 - | 0 - | 0 - | 0 - | 0 - |
| 8397 | 1 | 3um_FIJJAFBF_29470 | Bacteria | Bacteroidetes | Flavobacteriia | Flavobacterial | Flavobacteriac | Formosa | Formosa sp. H | cellular proces | Posttranslatio | 099Y4 | peptidase, M1 | 391603.FBALC | 0,2 | 0,4 | 5,23E-06 | 1,05E-05 | 1 - | 2,61E-05 - | 0 - | 0 - | 0 - | 0 - | 0 - | 0 - | 0 - | 0 - |
| 8398 | 1 | 0.2um_NOKCPHJN_76179 | Eukaryota | Bacillariophyta | Coscinodiscop | Chaetocerotal | Attheyaceae | Attheya | Attheya longi | metabolism | Carbohydrate | COG1850 | RuBisCO catal | 35128.Thapsd | 2,6 | 0,489897949 | 7,20E-05 | 1,17E-05 | 3 - | 7,03E-05 - | 3 - | 8,73E-05 - | 3 - | 8,29E-05 - | 2 - | 5,61E-05 - | 2 - | 6,32E-05 - |
| 8399 | 1 | 3um_FIJJAFBF_474234 | Eukaryota | unclassified | Cryptophyta | Pyrenomonad | Geminigerace | Teleaulax | Teleaulax amp | unclassified | unclassified | unclassified | unclassified | unclassified | 4,8 | 0,748331477 | 0,000368413 | 4,67E-05 | 6 - | 0,000389074 - | 4 - | 0,000322077 - | 5 - | 0,000382718 - | 4 - | 0,000310697 - | 5 - | 0,000437499 - |
| 8400 | 1 | 0.2um_NOKCPHJN_307525 | Eukaryota | unclassified | Oomycetes | Saprolegniales | Saprolegniace | Thraustotheca | Thraustotheca | unclassified | unclassified | unclassified | unclassified | unclassified | 0,2 | 0,4 | 5,86E-06 | 1,17E-05 | 0 - | 0 - | 0 - | 0 - | 0 - | 0 - | 1 - | 2,93E-05 - | 0 - | 0 - |
| 8401 | 1 | 3um_FIJJAFBF_624552 | Bacteria | Bacteroidetes | Flavobacteriia | Flavobacterial | Flavobacteriac | Winogradskye | Winogradskye | cellular proces | Posttranslatio | COG0330 | Band 7 protei | 983544.Lacal_ | 1 | 0 | 4,46E-05 | 4,24E-06 | 1 - | 3,74E-05 - | 1 - | 4,64E-05 - | 1 - | 4,41E-05 - | 1 - | 4,48E-05 - | 1 - | 5,04E-05 - |
| 8402 | 1 | 3um_FIJJAFBF_768155 | Bacteria | Proteobacteri | Gammaprotec | Oceanospirilla | Oceanospirilla | Marinomonas | Marinomonas | information st | Translation, ri | 0QTKX | Seems to be t | 717774.Marm | 5,4 | 1,743559577 | 0,000586102 | 0,000185929 | 6 - | 0,000555371 - | 7 - | 0,000804542 - | 6 - | 0,000655559 - | 6 - | 0,000665242 - | 2 - | 0,000249798 - |
| 8403 | 1 | 0.2um_NOKCPHJN_210408 | Bacteria | various | various | various | various | various | various | metabolism | Energy produc | COG0055 | Produces ATP | 688270.Celal_ | 1,8 | 0,4 | 4,91E-05 | 1,22E-05 | 2 - | 4,56E-05 - | 2 - | 5,67E-05 - | 1 - | 2,69E-05 - | 2 - | 5,47E-05 - | 2 - | 6,16E-05 - |
| 8404 | 1 | 0.2um_NOKCPHJN_24039 | Viruses | unclassified | unclassified | unclassified | Phycodnavirid | unclassified | Chrysochromu | information st | Replication, re | 05C24 | DNA gyrase ne | 272633.MYPE | 1,2 | 0,4 | 7,34E-06 | 1,63E-06 | 2 - | 1,05E-05 - | 1 - | 6,54E-06 - | 1 - | 6,22E-06 - | 1 - | 6,31E-06 - | 1 - | 7,11E-06 - |
| 8405 | 1 | 3um_FIJJAFBF_161859 | Bacteria | Proteobacteri | Alphaproteob | Rhodobactera | Rhodobactera | Roseovarius | various | metabolism | Amino acid tra | COG0473 | 3-isopropylma | 314264.ROS2 | 2,2 | 0,4 | 8,19E-05 | 1,66E-05 | 2 - | 6,24E-05 - | 2 - | 7,75E-05 - | 2 - | 7,36E-05 - | 3 - | 0,000112079 - | 2 - | 8,42E-05 - |
| 8406 | 1 | 0.2um_NOKCPHJN_116287 | Bacteria | Proteobacteri | Alphaproteob | Rhodobactera | Rhodobactera | Thalassobium | Thalassobium | metabolism | Energy produc | 05C26 | Dehydrogenas | 765910.Marp | 6,2 | 0,4 | 0,000167838 | 1,83E-05 | 6 - | 0,000136099 - | 6 - | 0,000168995 - | 7 - | 0,000187426 - | 6 - | 0,000163024 - | 6 - | 0,000183646 - |
| 8407 | 1 | 3um_FIJJAFBF_485633 | Bacteria | Bacteroidetes | Flavobacteriia | Flavobacterial | Flavobacteriac | Tenacibaculum | Tenacibaculum | unclassified | unclassified | unclassified | unclassified | unclassified | 1,6 | 0,8 | 0,000141907 | 5,81E-05 | 3 - | 0,000231094 - | 2 - | 0,0001913 - | 1 - | 9,09E-05 - | 1 - | 9,23E-05 - | 1 - | 0,000103943 - |
| 8408 | 1 | 3um_FIJJAFBF_600652 | Bacteria | Actinobacteria | Actinobacteria | Micrococcales | Microbacteria | unclassified | Microbacteria | unclassified | unclassified | unclassified | unclassified | unclassified | 1,4 | 0,489897949 | 0,000120055 | 3,77E-05 | 2 - | 0,000146212 - | 2 - | 0,000181553 - | 1 - | 8,63E-05 - | 1 - | 8,76E-05 - | 1 - | 9,86E-05 - |
| 8409 | 1 | 0.2um_NOKCPHJN_892023 | Bacteria | Proteobacteri | Alphaproteob | Pelagibacteral | unclassified | unclassified | Pelagibacteral | cellular proces | Cell wall/mem | 7257 | NA | 439493.PB721 | 1 | 0 | 4,15E-05 | 3,94E-06 | 1 - | 3,48E-05 - | 1 - | 4,32E-05 - | 1 - | 4,11E-05 - | 1 - | 4,17E-05 - | 1 - | 4,69E-05 - |
| 8410 | 1 | 0.2um_NOKCPHJN_18194 | Eukaryota | Chordata | Mammalia | Rodentia | Muridae | Rattus | Rattus norveg | cellular proces | Signal transdu | 0DPD8 | gamma-aminc | 10116.ENSRN | 0,8 | 0,4 | 8,17E-06 | 4,20E-06 | 1 - | 8,53E-06 - | 1 - | 1,06E-05 - | 0 - | 0 - | 1 - | 1,02E-05 - | 1 - | 1,15E-05 - |
| 8411 | 1 | 0.2um_NOKCPHJN_949714 | Eukaryota | unclassified | Spirotrichea | Sporadotrichi | Oxytrichidae | Oxytricha | Oxytricha trifa | metabolism | Energy produc | COG0056 | Produces ATP | 412030.XP_00 | 1 | 0,632455532 | 2,88E-05 | 1,61E-05 | 2 - | 4,97E-05 - | 1 - | 3,08E-05 - | 0 - | 0 - | 1 - | 2,98E-05 - | 1 - | 3,35E-05 - |
| 8412 | 1 | 3um_FIJJAFBF_563699 | Bacteria | Proteobacteri | Gammaprotec | unclassified | unclassified | unclassified | Gammaprotec | unclassified | unclassified | unclassified | unclassified | unclassified | 2,4 | 0,8 | 0,000102188 | 3,89E-05 | 2 - | 7,04E-05 - | 3 - | 0,000131152 - | 3 - | 0,000124677 - | 1 - | 4,22E-05 - | 3 - | 0,000142523 - |
| 8413 | 1 | 3um_FIJJAFBF_180472 | Bacteria | Bacteroidetes | Flavobacteriia | Flavobacterial | Flavobacteriac | Formosa | Formosa sp. H | unclassified | unclassified | unclassified | unclassified | unclassified | 1,2 | 0,4 | 2,14E-05 | 7,18E-06 | 1 - | 1,50E-05 - | 1 - | 1,86E-05 - | 2 - | 3,54E-05 - | 1 - | 1,79E-05 - | 1 - | 2,02E-05 - |
| 8414 | 1 | 0.2um_NOKCPHJN_284549 | Bacteria | Proteobacteri | Gammaprotec | Cellvibrionales | Halieaceae | unclassified | marine gamm | information st | Transcription | 05D27 | DNA-depende | 876044.IMCC | 5,6 | 2,059126028 | 5,19E-05 | 1,50E-05 | 9 - | 7,17E-05 - | 6 - | 5,94E-05 - | 3 - | 2,82E-05 - | 6 - | 5,73E-05 - | 4 - | 4,30E-05 - |
| 8415 | 1 | 3um_FIJJAFBF_774033 | Bacteria | Proteobacteri | Betaproteoba | unclassified | unclassified | unclassified | Betaproteoba | metabolism | Carbohydrate | 08MVC | Trap dicarbox | 365046.Rta_1 | 2,6 | 0,489897949 | 0,000103823 | 2,21E-05 | 2 - | 6,67E-05 - | 3 - | 0,00012429 - | 3 - | 0,000118153 - | 3 - | 0,000119898 - | 2 - | 9,00E-05 - |
| 8416 | 1 | 0.2um_NOKCPHJN_1034378 | Eukaryota | Basidiomycota | Agaricomycet | Sebacinales | Serendipitace | Serendipita | Serendipita ve | unclassified | unclassified | unclassified | unclassified | unclassified | 1 | 0 | 8,01E-05 | 7,61E-06 | 1 - | 6,71E-05 - | 1 - | 8,33E-05 - | 1 - | 7,92E-05 - | 1 - | 8,04E-05 - | 1 - | 9,06E-05 - |
| 8417 | 1 | 3um_FIJJAFBF_573611 | Bacteria | Proteobacteri | Gammaprotec | Cellvibrionales | Porticoccacea | unclassified | gamma protec | unclassified | unclassified | unclassified | unclassified | unclassified | 9 | 1,414213562 | 0,000470056 | 6,24E-05 | 10 - | 0,000441449 - | 9 - | 0,000493335 - | 8 - | 0,000416868 - | 11 - | 0,00058166 - | 7 - | 0,00041697 - |
| 8418 | 1 | 3um_FIJJAFBF_675122 | Bacteria | Bacteroidetes | Flavobacteriia | Flavobacterial | Flavobacteriac | Formosa | Formosa sp. H | information st | Translation, ri | 0P97G | This is one of | 1046627.BZAR | 1 | 0 | 7,49E-05 | 7,11E-06 | 1 - | 6,27E-05 - | 1 - | 7,79E-05 - | 1 - | 7,40E-05 - | 1 - | 7,51E-05 - | 1 - | 8,46E-05 - |
| 8419 | 1 | 3um_FIJJAFBF_110764 | Bacteria | Proteobacteri | Gammaprotec | Cellvibrionales | Halieaceae | Halioglobus | Halioglobus sp | unclassified | unclassified | unclassified | unclassified | unclassified | 0,4 | 0,489897949 | 2,50E-05 | 3,07E-05 | 0 - | 0 - | 0 - | 0 - | 1 - | 6,21E-05 - | 1 - | 6,31E-05 - | 0 - | 0 - |
| 8420 | 1 | 3um_FIJJAFBF_271029 | unclassified | unclassified | unclassified | unclassified | unclassified | unclassified | unclassified | cellular proces | Intracellular tr | 116KZ | NA | 70448.Q019R | 0,6 | 0,489897949 | 7,38E-05 | 6,09E-05 | 1 - | 0,000107268 - | 1 - | 0,000133195 - | 0 - | 0 - | 1 - | 0,000128489 - | 0 - | 0 - |
| 8421 | 1 | 3um_FIJJAFBF_590239 | Bacteria | various | various | various | various | various | various | metabolism | Inorganic ion t | COG3221 | phosphonate | 675812.VHA_ | 34,2 | 2,315167381 | 0,001418671 | 0,000161923 | 37 - | 0,001286891 - | 32 - | 0,001382001 - | 33 - | 0,001354822 - | 32 - | 0,001333171 - | 37 - | 0,001736471 - |
| 8422 | 1 | 3um_FIJJAFBF_257403 | Bacteria | Bacteroidetes | Flavobacteriia | Flavobacterial | Flavobacteriac | Formosa | Formosa sp. H | cellular proces | Intracellular tr | 05FGH | MotA TolQ ex | 391603.FBALC | 5,6 | 1,019803903 | 0,000294244 | 4,18E-05 | 7 - | 0,00031141 - | 6 - | 0,000331439 - | 6 - | 0,000315075 - | 4 - | 0,000213152 - | 5 - | 0,000300144 - |
| 8423 | 1 | 3um_FIJJAFBF_214016 | Bacteria | Proteobacteri | Alphaproteob | Rhodobactera | Rhodobactera | Halocynthiiba | Halocynthiiba | metabolism | Amino acid tra | 16PK4 | phospho-2-de | 314271.RB265 | 1,4 | 1,019803903 | 4,12E-05 | 2,95E-05 | 2 - | 5,03E-05 - | 1 - | 3,13E-05 - | 0 - | 0 - | 3 - | 9,04E-05 - | 1 - | 3,40E-05 - |
| 8424 | 1 | 0.2um_NOKCPHJN_316719 | Eukaryota | Bacillariophyta | Coscinodiscop | Thalassiosirale | Thalassiosirac | Thalassiosira | Thalassiosira p | metabolism | Energy produc | COG0055 | Produces ATP | 35128.Thaps4 | 0,2 | 0,4 | 5,91E-06 | 1,18E-05 | 0 - | 0 - | 0 - | 0 - | 0 - | 0 - | 1 - | 2,96E-05 - | 0 - | 0 - |
| 8425 | 1 | 3um_FIJJAFBF_130521 | Bacteria | Proteobacteri | Gammaprotec | [illegible] | [illegible] | [illegible] | [illegible] | unclassified | unclassified | unclassified | unclassified | unclassified | 0,2 | 0,4 | 2,52E-05 | 5,05E-05 | 1 - | 0,000126128 - | 0 - | 0 - | 0 - | 0 - | 0 - | 0 - | 0 - | 0 - |
| 8426 | 1 | 0.2um_NOKCPHJN_57611 | Bacteria | Bacteroidetes | Flavobacteriia | Flavobacterial | Flavobacteriac | Polaribacter | various | unclassified | unclassified | unclassified | unclassified | unclassified | 0,2 | 0,4 | 1,95E-05 | 7,21E-06 | 1 - | 1,36E-05 - | 2 - | 3,37E-05 - | 1 - | 1,60E-05 - | 1 - | 1,62E-05 - | 1 - | 1,83E-05 - |
| 8427 | 1 | 3um_FIJJAFBF_542558 | Bacteria | Proteobacteri | Gammaprotec | Cellvibrionales | Porticoccacea | unclassified | gamma protec | cellular proces | Intracellular tr | 17SGA | One of the pr | 314287.GB220 | 1,4 | 0,489897949 | 0,000110127 | 3,89E-05 | 1 - | 6,60E-05 - | 1 - | 8,19E-05 - | 2 - | 0,000155727 - | 2 - | 0,000158027 - | 1 - | 8,90E-05 - |
| 8428 | 1 | 0.2um_NOKCPHJN_1056840 | Bacteria | Proteobacteri | Alphaproteob | Rhodobactera | Rhodobactera | Yoonia | Yoonia sedim | poorly charact | Function unkn | COG3970 | fumarylacetoa | 384765.SIAM | 1 | 0 | 4,46E-05 | 4,24E-06 | 1 - | 3,74E-05 - | 1 - | 4,64E-05 - | 1 - | 4,41E-05 - | 1 - | 4,48E-05 - | 1 - | 5,04E-05 - |
| 8429 | 1 | 0.2um_NOKCPHJN_72952 | Bacteria | Bacteroidetes | Flavobacteriia | unclassified | unclassified | unclassified | Flavobacteria | metabolism | Amino acid tra | COG0686 | alanine dehyd | 755732.Fluta | 0,4 | 0,489897949 | 1,27E-05 | 1,57E-05 | 1 - | 2,83E-05 - | 1 - | 3,51E-05 - | 0 - | 0 - | 0 - | 0 - | 0 - | 0 - |
| 8430 | 1 | 0.2um_NOKCPHJN_144331 | Bacteria | Proteobacteri | Alphaproteob | Rhodobactera | Rhodobactera | Roseovarius | Roseovarius a | unclassified | unclassified | unclassified | unclassified | unclassified | 0,8 | 0,748331477 | 1,50E-05 | 1,32E-05 | 2 - | 3,42E-05 - | 0 - | 0 - | 1 - | 2,02E-05 - | 1 - | 2,05E-05 - | 0 - | 0 - |
| 8431 | 2 | 3um_FIJJAFBF_181472;3um_FIJJAFBF_290030 | Bacteria | Bacteroidetes | Flavobacteriia | Flavobacterial | Flavobacteriac | Bizionia | Bizionia echin | cellular proces | Posttranslatio | COG0652 | peptidyl-proly | 983544.Lacal_ | 0,2 | 0,4 | 7,40E-06 | 1,48E-05 | 1 - | 3,70E-05 - | 0 - | 0 - | 0 - | 0 - | 0 - | 0 - | 0 - | 0 - |

| | | | | | | | | | | | | | | | | | | | | | | | | | | | | |
|---|---|---|---|---|---|---|---|---|---|---|---|---|---|---|---|---|---|---|---|---|---|---|---|---|---|---|---|---|
| 8432 | 1 | 3um_FIJJAFBF_366672 | Bacteria | Proteobacteri | Alphaproteob | Rhodobactera | Rhodobactera | Pseudorueger | Pseudorueger | metabolism | Amino acid tra | COG0436 | aminotransfer | 244592.SADFL | 0,6 | 0,489897949 | 2,13E-05 | 1,74E-05 | 0 - | 0 - | 1 - | 3,64E-05 - | 1 - | 3,47E-05 - | 1 - | 3,52E-05 - | 0 - | 0 - |
| 8433 | 1 | 3um_FIJJAFBF_564203 | Bacteria | Bacteroidetes | Cytophagia | Cytophagales | Flammeovirga | unclassified | Flammeovirga | information st | Translation, ri | 05KK9 | This protein bi | 388413.ALPR1 | 0,8 | 0,4 | 0,000107383 | 5,52E-05 | 1 - | 0,000112526 - | 1 - | 0,000139724 - | 1 - | 0,000132826 - | 0 - | 0 - | 1 - | 0,000151838 - |
| 8434 | 1 | 0.2um_NOKCPHJN_236556 | Bacteria | Proteobacteri | Gammaprotec | Alteromonada | Alteromonada | Paraglaciecola | Paraglaciecola | metabolism | Inorganic ion t | 16PIZ | Component of | 342610.Patl_4 | 0,2 | 0,4 | 4,73E-06 | 9,47E-06 | 0 - | 0 - | 1 - | 2,37E-05 - | 0 - | 0 - | 0 - | 0 - | 0 - | 0 - |
| 8435 | 1 | 0.2um_NOKCPHJN_447457 | unclassified | unclassified | unclassified | unclassified | unclassified | unclassified | unclassified | unclassified | unclassified | unclassified | unclassified | unclassified | 0,8 | 0,4 | 5,05E-05 | 2,54E-05 | 0 - | 0 - | 1 - | 6,31E-05 - | 1 - | 5,99E-05 - | 1 - | 6,08E-05 - | 1 - | 6,85E-05 - |
| 8436 | 1 | 3um_FIJJAFBF_227657 | Bacteria | Bacteroidetes | Flavobacteriia | unclassified | unclassified | unclassified | Flavobacteria | poorly charact | Function unkn | 09EUF | NA | 487796.Flav2A | 0,8 | 0,4 | 4,93E-05 | 2,51E-05 | 1 - | 5,34E-05 - | 1 - | 6,63E-05 - | 1 - | 6,30E-05 - | 1 - | 6,39E-05 - | 0 - | 0 - |
| 8437 | 1 | 3um_FIJJAFBF_231212 | Bacteria | Bacteroidetes | Flavobacteriia | Flavobacterial | Flavobacteriac | Polaribacter | Polaribacter s | unclassified | unclassified | unclassified | unclassified | unclassified | 0,6 | 0,489897949 | 2,30E-05 | 1,89E-05 | 1 - | 3,34E-05 - | 1 - | 4,14E-05 - | 0 - | 0 - | 1 - | 4,00E-05 - | 0 - | 0 - |
| 8438 | 1 | 0.2um_NOKCPHJN_840467 | Bacteria | Proteobacteri | Alphaproteob | Pelagibacteral | Pelagibacterac | Candidatus Pe | Candidatus Pe | information st | Transcription | COG0085 | DNA-depende | 335992.SAR11 | 3,6 | 1,019803903 | 7,89E-05 | 2,08E-05 | 5 - | 9,26E-05 - | 4 - | 9,19E-05 - | 3 - | 6,56E-05 - | 2 - | 4,43E-05 - | 4 - | 9,99E-05 - |
| 8439 | 1 | 0.2um_NOKCPHJN_296194 | Bacteria | Proteobacteri | Alphaproteob | Pelagibacteral | Pelagibacterac | Candidatus Pe | Candidatus Pe | unclassified | unclassified | unclassified | unclassified | unclassified | 9,6 | 0,8 | 0,000367396 | 4,72E-05 | 10 - | 0,000320605 - | 10 - | 0,000398097 - | 10 - | 0,000378442 - | 8 - | 0,000307225 - | 10 - | 0,00043261 - |
| 8440 | 1 | 3um_FIJJAFBF_357542 | Bacteria | Bacteroidetes | Flavobacteriia | Flavobacterial | Flavobacteriac | Polaribacter | Polaribacter h | unclassified | unclassified | unclassified | unclassified | unclassified | 1,4 | 0,489897949 | 0,000107094 | 3,37E-05 | 2 - | 0,000130428 - | 2 - | 0,000161953 - | 1 - | 7,70E-05 - | 1 - | 7,81E-05 - | 1 - | 8,80E-05 - |
| 8441 | 1 | 3um_FIJJAFBF_556665 | Bacteria | Proteobacteri | Gammaprotec | Oceanospirilla | Saccharospirill | Reinekea | Reinekea fors | unclassified | unclassified | unclassified | unclassified | unclassified | 0,2 | 0,4 | 2,31E-05 | 4,62E-05 | 0 - | 0 - | 0 - | 0 - | 0 - | 0 - | 1 - | 0,000115532 - | 0 - | 0 - |
| 8442 | 1 | 3um_FIJJAFBF_453245 | Bacteria | Proteobacteri | Betaproteoba | Burkholderial | Comamonada | Rhodoferax | Rhodoferax fe | metabolism | Inorganic ion t | COG3221 | phosphonate | 338969.Rfer_ | 1 | 0,894427191 | 4,11E-05 | 3,61E-05 | 2 - | 7,33E-05 - | 1 - | 4,55E-05 - | 2 - | 8,66E-05 - | 0 - | 0 - | 0 - | 0 - |
| 8443 | 1 | 0.2um_NOKCPHJN_450379 | Bacteria | Bacteroidetes | unclassified | unclassified | unclassified | unclassified | Bacteroidetes | cellular proces | Cell wall/mem | 09G29 | outer membra | 388413.ALPR1 | 0,4 | 0,489897949 | 2,54E-05 | 3,13E-05 | 1 - | 5,77E-05 - | 0 - | 0 - | 0 - | 0 - | 1 - | 6,91E-05 - | 0 - | 0 - |
| 8444 | 1 | 3um_FIJJAFBF_671251 | Bacteria | Proteobacteri | Betaproteoba | Burkholderial | Comamonada | Comamonas | Comamonas g | information st | Translation, ri | 0BCU4 | Binds directly | 688245.CtCNE | 1 | 0,632455532 | 0,000118629 | 7,23E-05 | 0 - | 0 - | 1 - | 0,000119764 - | 2 - | 0,000227701 - | 1 - | 0,000115532 - | 1 - | 0,000130147 - |
| 8445 | 1 | 3um_FIJJAFBF_60206 | Bacteria | Actinobacteria | unclassified | unclassified | unclassified | unclassified | Actinobacteria | cellular proces | Intracellular tr | 0QJ7H | motA TolQ Exl | 314285.KT71_ | 3 | 1,264911064 | 8,62E-05 | 3,45E-05 | 4 - | 9,89E-05 - | 4 - | 0,000122861 - | 4 - | 0,000116795 - | 2 - | 5,93E-05 - | 1 - | 3,34E-05 - |
| 8446 | 1 | 0.2um_NOKCPHJN_358473 | Bacteria | Proteobacteri | Alphaproteob | Rhodobactera | unclassified | unclassified | Rhodobactera | unclassified | unclassified | unclassified | unclassified | unclassified | 0,8 | 0,4 | 0,000116692 | 5,99E-05 | 1 - | 0,000123416 - | 0 - | 0 - | 1 - | 0,00014568 - | 1 - | 0,000147832 - | 1 - | 0,000166532 - |
| 8447 | 1 | 0.2um_NOKCPHJN_193775 | Eukaryota | unclassified | Oomycetes | Saprolegniales | Saprolegniace | Thraustotheca | Thraustotheca | unclassified | unclassified | unclassified | unclassified | unclassified | 0,2 | 0,4 | 5,29E-07 | 1,06E-06 | 1 - | 2,64E-06 - | 0 - | 0 - | 0 - | 0 - | 0 - | 0 - | 0 - | 0 - |
| 8448 | 1 | 3um_FIJJAFBF_251405 | Eukaryota | Bacillariophyta | Bacillariophyc | Bacillariales | Bacillariaceae | Fragilariopsis | Fragilariopsis | unclassified | unclassified | unclassified | unclassified | unclassified | 0,8 | 0,4 | 6,71E-05 | 3,41E-05 | 1 - | 7,26E-05 - | 1 - | 9,02E-05 - | 1 - | 8,57E-05 - | 1 - | 8,70E-05 - | 0 - | 0 - |
| 8449 | 1 | 0.2um_NOKCPHJN_1064535 | Bacteria | Proteobacteri | Gammaprotec | Cellvibrionales | Porticoccacea | unclassified | gamma prote | information st | Translation, ri | COG0050 | This protein p | 314287.GB22 | 0,6 | 0,489897949 | 2,28E-05 | 1,87E-05 | 0 - | 0 - | 1 - | 3,92E-05 - | 1 - | 3,72E-05 - | 1 - | 3,78E-05 - | 0 - | 0 - |
| 8450 | 1 | 0.2um_NOKCPHJN_824677 | Eukaryota | unclassified | Oomycetes | Saprolegniales | Saprolegniace | Aphanomyces | various | information st | Translation, ri | 0TJFM | 40S ribosomal | 4577.GRMZM | 4 | 0 | 0,00028253 | 2,68E-05 | 4 - | 0,000236653 - | 4 - | 0,000293853 - | 4 - | 0,000279345 - | 4 - | 0,000283471 - | 4 - | 0,000319329 - |
| 8451 | 1 | 3um_FIJJAFBF_319725 | Bacteria | Bacteroidetes | Flavobacteriia | Flavobacterial | Flavobacteriac | Polaribacter | Polaribacter g | unclassified | unclassified | unclassified | unclassified | unclassified | 4,2 | 0,748331477 | 0,000198317 | 2,37E-05 | 5 - | 0,000200659 - | 4 - | 0,000199327 - | 5 - | 0,000236857 - | 4 - | 0,000192284 - | 3 - | 0,000162456 - |
| 8452 | 2 | 0.2um_NOKCPHJN_63870;3um_FIJJAFBF_66657 | Bacteria | Bacteroidetes | Flavobacteriia | Flavobacterial | Flavobacteriac | various | various | unclassified | unclassified | unclassified | unclassified | unclassified | 1,4 | 0,489897949 | 5,52E-05 | 1,58E-05 | 2 - | 6,77E-05 - | 1 - | 4,20E-05 - | 2 - | 7,99E-05 - | 1 - | 4,06E-05 - | 1 - | 4,57E-05 - |
| 8453 | 1 | 0.2um_NOKCPHJN_1026858 | Bacteria | Proteobacteri | Gammaprotec | unclassified | unclassified | Candidatus Th | Candidatus Th | unclassified | unclassified | unclassified | unclassified | unclassified | 2 | 0 | 8,18E-05 | 7,77E-06 | 2 - | 6,85E-05 - | 2 - | 8,51E-05 - | 2 - | 8,09E-05 - | 2 - | 8,21E-05 - | 2 - | 9,25E-05 - |
| 8454 | 1 | 3um_FIJJAFBF_644938 | Bacteria | Proteobacteri | Alphaproteob | Rhodobactera | Rhodobactera | Sulfitobacter | Sulfitobacter g | metabolism | Nucleotide tra | 01Q8W | Plays an impo | 391589.RGAI1 | 1,2 | 0,4 | 3,72E-05 | 8,24E-06 | 2 - | 5,34E-05 - | 1 - | 3,31E-05 - | 1 - | 3,15E-05 - | 1 - | 3,20E-05 - | 1 - | 3,60E-05 - |
| 8455 | 1 | 3um_FIJJAFBF_239282 | Bacteria | Proteobacteri | Alphaproteob | Rhodobactera | various | various | various | unclassified | unclassified | unclassified | unclassified | unclassified | 1,6 | 0,489897949 | 0,000147983 | 3,88E-05 | 2 - | 0,000158313 - | 1 - | 9,83E-05 - | 2 - | 0,000186872 - | 2 - | 0,000189632 - | 1 - | 0,00010681 - |
| 8456 | 1 | 3um_FIJJAFBF_365804 | Bacteria | Bacteroidetes | unclassified | unclassified | unclassified | unclassified | Bacteroidetes | unclassified | unclassified | unclassified | unclassified | unclassified | 0,2 | 0,4 | 1,11E-05 | 2,23E-05 | 0 - | 0 - | 1 - | 5,57E-05 - | 0 - | 0 - | 0 - | 0 - | 0 - | 0 - |
| 8457 | 1 | 3um_FIJJAFBF_663759 | Bacteria | Proteobacteri | Alphaproteob | Rhodobactera | Rhodobactera | Phaeobacter | Phaeobacter g | metabolism | Energy produc | COG0377 | NDH-1 shuttle | 246200.SPO27 | 3,6 | 0,8 | 0,000266756 | 4,04E-05 | 5 - | 0,000315321 - | 4 - | 0,000313228 - | 3 - | 0,000223322 - | 3 - | 0,000226621 - | 3 - | 0,000255288 - |
| 8458 | 1 | 3um_FIJJAFBF_289836 | Bacteria | Proteobacteri | Gammaprotec | Oceanospirilla | Saccharospirill | Reinekea | Reinekea fors | metabolism | Amino acid tra | 0QIAJ | Glutamate syr | 314283.MED2 | 0,2 | 0,4 | 4,63E-06 | 9,26E-06 | 1 - | 2,31E-05 - | 0 - | 0 - | 0 - | 0 - | 0 - | 0 - | 0 - | 0 - |
| 8459 | 1 | 3um_FIJJAFBF_253702 | Bacteria | Bacteroidetes | Cytophagia | Cytophagales | Flammeovirga | unclassified | Flammeovirga | cellular proces | Posttranslatio | 08V1T | peptidylprolyl | 523791.Kkor_ | 0,6 | 1,2 | 3,36E-05 | 6,72E-05 | 3 - | 0,000167966 - | 0 - | 0 - | 0 - | 0 - | 0 - | 0 - | 0 - | 0 - |
| 8460 | 1 | 0.2um_NOKCPHJN_09520 | Bacteria | Bacteroidetes | Flavobacteriia | Flavobacterial | Flavobacteriac | Lutibacter | Lutibacter sp. | metabolism | Energy produc | 0P7SC | Delta-1-pyrrol | 391603.FBALC | 0,4 | 0,489897949 | 9,33E-06 | 1,15E-05 | 1 - | 2,12E-05 - | 0 - | 0 - | 0 - | 0 - | 1 - | 2,54E-05 - | 0 - | 0 - |
| 8461 | 1 | 3um_FIJJAFBF_323273 | Bacteria | Bacteroidetes | Flavobacteriia | Flavobacterial | Flavobacteriac | Polaribacter | Polaribacter b | poorly charact | Function unkn | 11NCV | Protein of unk | 313594.PI23P | 0,6 | 0,489897949 | 5,23E-05 | 4,35E-05 | 1 - | 7,40E-05 - | 0 - | 0 - | 1 - | 8,74E-05 - | 0 - | 0 - | 1 - | 9,99E-05 - |
| 8462 | 1 | 3um_FIJJAFBF_415536 | Bacteria | Proteobacteri | Gammaprotec | Alteromonada | Alteromonada | Paraglaciecola | Paraglaciecola | metabolism | Amino acid tra | 16PK4 | phospho-2-de | 715451.ambt_ | 0,2 | 0,4 | 6,15E-06 | 1,23E-05 | 0 - | 0 - | 0 - | 0 - | 0 - | 0 - | 1 - | 3,08E-05 - | 0 - | 0 - |
| 8463 | 1 | 3um_FIJJAFBF_487840 | various | various | various | various | various | various | various | unclassified | unclassified | unclassified | unclassified | unclassified | 1,8 | 1,166190379 | 0,000171869 | 0,000108792 | 3 - | 0,000251336 - | 2 - | 0,000208057 - | 1 - | 9,89E-05 - | 3 - | 0,000301058 - | 0 - | 0 - |
| 8464 | 1 | 0.2um_NOKCPHJN_677077 | Bacteria | Proteobacteri | Alphaproteob | Pelagibacteral | Pelagibacterac | Candidatus Pe | Candidatus Pe | information st | Translation, ri | COG1185 | Involved in mI | 439493.PB721 | 1 | 1,095445115 | 1,95E-05 | 1,93E-05 | 3 - | 5,23E-05 - | 1 - | 2,16E-05 - | 0 - | 0 - | 0 - | 0 - | 1 - | 2,35E-05 - |
| 8465 | 1 | 0.2um_NOKCPHJN_164344 | unclassified | unclassified | unclassified | unclassified | unclassified | unclassified | unclassified | unclassified | unclassified | unclassified | unclassified | unclassified | 0,4 | 0,489897949 | 4,93E-06 | 6,15E-06 | 1 - | 1,05E-05 - | 0 - | 0 - | 0 - | 0 - | 0 - | 0 - | 1 - | 1,42E-05 - |
| 8466 | 1 | 0.2um_NOKCPHJN_129924 | Bacteria | Proteobacteri | Gammaprotec | unclassified | unclassified | unclassified | Gammaprotec | cellular proces | Posttranslatio | 17FI0 | Binds to Cpn6 | 83406.HDN1F | 0,2 | 0,4 | 2,42E-05 | 4,83E-05 | 1 - | 0,000120818 - | 0 - | 0 - | 0 - | 0 - | 0 - | 0 - | 0 - | 0 - |
| 8467 | 1 | 3um_FIJJAFBF_244190 | Bacteria | Proteobacteri | Alphaproteob | Rhodospirillal | Rhodospirillac | unclassified | Rhodospirillac | unclassified | unclassified | unclassified | unclassified | unclassified | 1,4 | 0,489897949 | 5,99E-05 | 1,76E-05 | 2 - | 7,33E-05 - | 1 - | 4,55E-05 - | 1 - | 4,33E-05 - | 2 - | 8,78E-05 - | 1 - | 4,95E-05 - |
| 8468 | 1 | 3um_FIJJAFBF_173942 | Bacteria | Bacteroidetes | Flavobacteriia | Flavobacterial | Flavobacteriac | Tenacibaculur | Tenacibaculur | metabolism | Lipid transpor | 074D8 | Inherit from C | 573413.Spirs_ | 1 | 0,894427191 | 3,09E-05 | 2,87E-05 | 1 - | 2,69E-05 - | 0 - | 0 - | 2 - | 6,35E-05 - | 2 - | 6,44E-05 - | 0 - | 0 - |
| 8469 | 1 | 3um_FIJJAFBF_433328 | unclassified | unclassified | unclassified | unclassified | unclassified | unclassified | unclassified | unclassified | unclassified | unclassified | unclassified | unclassified | 0,6 | 0,489897949 | 4,68E-05 | 3,84E-05 | 0 - | 0 - | 0 - | 0 - | 1 - | 7,40E-05 - | 1 - | 7,51E-05 - | 1 - | 8,46E-05 - |
| 8470 | 1 | 3um_FIJJAFBF_158710 | Bacteria | Proteobacteri | Gammaprotec | Oceanospirilla | Oleiphilaceae | Oleiphilus | Oleiphilus sp. | information st | Replication, re | 0QUGS | DNA-binding | 349521.HCH_ | 0,4 | 0,489897949 | 6,50E-05 | 7,98E-05 | 0 - | 0 - | 0 - | 0 - | 0 - | 0 - | 1 - | 0,000152759 - | 1 - | 0,000172083 - |
| 8471 | 1 | 3um_FIJJAFBF_452433 | Bacteria | Proteobacteri | Gammaprotec | Cellvibrionales | Porticoccacea | unclassified | various | cellular proces | Cell wall/mem | COG2885 | Ompa motb d | 314287.GB22 | 1,2 | 0,4 | 5,16E-05 | 1,78E-05 | 1 - | 3,60E-05 - | 1 - | 4,47E-05 - | 1 - | 4,25E-05 - | 2 - | 8,62E-05 - | 1 - | 4,85E-05 - |
| 8472 | 1 | 3um_FIJJAFBF_127239 | Bacteria | Proteobacteri | Alphaproteob | Pelagibacteral | Pelagibacterac | Candidatus Pe | Candidatus Pe | unclassified | unclassified | unclassified | unclassified | unclassified | 0,2 | 0,4 | 4,85E-06 | 9,71E-06 | 1 - | 2,43E-05 - | 0 - | 0 - | 0 - | 0 - | 0 - | 0 - | 0 - | 0 - |
| 8473 | 1 | 3um_FIJJAFBF_783857 | Bacteria | Bacteroidetes | Flavobacteriia | Flavobacterial | Flavobacteriac | Tenacibaculur | Tenacibaculur | unclassified | unclassified | unclassified | unclassified | unclassified | 0,4 | 0,489897949 | 1,54E-05 | 1,89E-05 | 1 - | 3,52E-05 - | 0 - | 0 - | 1 - | 4,16E-05 - | 0 - | 0 - | 0 - | 0 - |
| 8474 | 1 | 3um_FIJJAFBF_297348 | Bacteria | Bacteroidetes | Flavobacteriia | Flavobacterial | Flavobacteriac | Formosa | Formosa sp. H | unclassified | unclassified | unclassified | unclassified | unclassified | 1,2 | 0,4 | 3,77E-05 | 8,36E-06 | 2 - | 5,41E-05 - | 1 - | 3,36E-05 - | 1 - | 3,20E-05 - | 1 - | 3,24E-05 - | 1 - | 3,65E-05 - |
| 8475 | 1 | 3um_FIJJAFBF_800083 | Bacteria | Proteobacteri | Gammaprotec | Cellvibrionales | Porticoccacea | unclassified | SAR92 bacteri | information st | Translation, ri | COG1185 | Involved in mI | 314287.GB22 | 1,2 | 0,748331477 | 2,21E-05 | 1,36E-05 | 2 - | 3,22E-05 - | 2 - | 4,00E-05 - | 1 - | 1,90E-05 - | 1 - | 1,93E-05 - | 0 - | 0 - |
| 8476 | 1 | 3um_FIJJAFBF_242310 | Bacteria | Proteobacteri | Gammaprotec | Oceanospirilla | Saccharospirill | Reinekea | Reinekea fors | unclassified | unclassified | unclassified | unclassified | unclassified | 5,2 | 0,748331477 | 0,000190436 | 1,82E-05 | 6 - | 0,000186124 - | 5 - | 0,000192593 - | 5 - | 0,000183084 - | 6 - | 0,000222946 - | 4 - | 0,000167432 - |
| 8477 | 1 | 0.2um_NOKCPHJN_922821 | Bacteria | Proteobacteri | Alphaproteob | Pelagibacteral | Pelagibacterac | Candidatus Pe | Candidatus Pe | unclassified | unclassified | unclassified | unclassified | unclassified | 1,8 | 0,748331477 | 9,37E-05 | 3,15E-05 | 3 - | 0,000135031 - | 2 - | 0,00011178 - | 1 - | 5,31E-05 - | 2 - | 0,00010783 - | 1 - | 6,07E-05 - |
| 8478 | 1 | 0.2um_NOKCPHJN_149063 | Bacteria | Bacteroidetes | Flavobacteriia | Flavobacterial | Flavobacteriac | Polaribacter | Polaribacter s | information st | Translation, ri | COG0088 | One of the pri | 313598.MED1 | 1 | 0 | 6,56E-05 | 6,23E-06 | 1 - | 5,49E-05 - | 1 - | 6,82E-05 - | 1 - | 6,48E-05 - | 1 - | 6,58E-05 - | 1 - | 7,41E-05 - |
| 8479 | 1 | 3um_FIJJAFBF_554178 | Eukaryota | unclassified | Oomycetes | Peronosporale | Peronosporac | Phytophthora | Phytophthora | unclassified | unclassified | unclassified | unclassified | unclassified | 1,8 | 0,4 | 6,34E-05 | 1,75E-05 | 1 - | 2,90E-05 - | 2 - | 7,20E-05 - | 2 - | 6,84E-05 - | 2 - | 6,94E-05 - | 2 - | 7,82E-05 - |
| 8480 | 1 | 3um_FIJJAFBF_57085 | Bacteria | Proteobacteri | Gammaprotec | Cellvibrionales | Porticoccacea | unclassified | Gammaprotec | poorly charact | Function unkn | 16R8I | Lipoprotein | 377629.TERTU | 13,4 | 1,854723699 | 0,000218735 | 2,47E-05 | 16 - | 0,00022046 - | 15 - | 0,000256637 - | 11 - | 0,000178908 - | 13 - | 0,00021456 - | 12 - | 0,000223108 - |
| 8481 | 1 | 3um_FIJJAFBF_353113 | Bacteria | Bacteroidetes | Cytophagia | Cytophagales | Flammeovirga | unclassified | Flammeovirga | unclassified | unclassified | unclassified | unclassified | unclassified | 0,6 | 0,8 | 3,54E-05 | 4,57E-05 | 2 - | 0,000111434 - | 0 - | 0 - | 1 - | 6,58E-05 - | 0 - | 0 - | 0 - | 0 - |
| 8482 | 1 | 0.2um_NOKCPHJN_222454 | Bacteria | Bacteroidetes | Flavobacteriia | Flavobacterial | Flavobacteriac | Marinirhabdu | Marinirhabdu | unclassified | unclassified | unclassified | unclassified | unclassified | 0,6 | 0,489897949 | 3,23E-06 | 2,66E-06 | 1 - | 4,72E-06 - | 1 - | 5,86E-06 - | 1 - | 5,57E-06 - | 0 - | 0 - | 0 - | 0 - |
| 8483 | 1 | 0.2um_NOKCPHJN_113685 | Bacteria | Bacteroidetes | Flavobacteriia | Flavobacterial | Flavobacteriac | Polaribacter | Polaribacter s | poorly charact | Function unkn | 0P8NI | Inherit from b | 867900.Celly_ | 0,8 | 0,4 | 1,44E-05 | 7,30E-06 | 1 - | 1,56E-05 - | 1 - | 1,93E-05 - | 1 - | 1,84E-05 - | 1 - | 1,86E-05 - | 0 - | 0 - |
| 8484 | 1 | 3um_FIJJAFBF_600549 | Bacteria | Bacteroidetes | Flavobacteriia | Flavobacterial | Flavobacteriac | Xanthomarina | Xanthomarina | cellular proces | Posttranslatio | 09ADN | alkyl hydroper | 867900.Celly_ | 1,4 | 0,489897949 | 8,84E-05 | 2,60E-05 | 2 - | 0,00010828 - | 1 - | 6,72E-05 - | 1 - | 6,39E-05 - | 2 - | 0,000129701 - | 1 - | 7,31E-05 - |
| 8485 | 1 | 0.2um_NOKCPHJN_170594 | Bacteria | Bacteroidetes | unclassified | unclassified | unclassified | unclassified | Bacteroidetes | unclassified | unclassified | unclassified | unclassified | unclassified | 4,2 | 0,748331477 | 0,000383964 | 9,59E-05 | 3 - | 0,000226533 - | 4 - | 0,00037505 - | 4 - | 0,000356532 - | 5 - | 0,000452248 - | 5 - | 0,000509456 - |
| 8486 | 2 | 0.2um_NOKCPHJN_656428;3um_FIJJAFBF_6032 | Bacteria | Bacteroidetes | Flavobacteriia | Flavobacterial | Flavobacteriac | Algibacter | Algibacter sp. | unclassified | unclassified | unclassified | unclassified | unclassified | 1,8 | 0,4 | 0,0001417 | 3,47E-05 | 2 - | 0,000131927 - | 2 - | 0,000163815 - | 2 - | 0,000155727 - | 1 - | 7,90E-05 - | 2 - | 0,000178017 - |
| 8487 | 1 | 3um_FIJJAFBF_294048 | Bacteria | Proteobacteri | Alphaproteob | Rhodobactera | Rhodobactera | Pelagimonas | Pelagimonas v | metabolism | Inorganic ion t | COG3221 | phosphonate | 391595.RLO14 | 0,2 | 0,4 | 9,73E-06 | 1,95E-05 | 1 - | 4,86E-05 - | 0 - | 0 - | 0 - | 0 - | 0 - | 0 - | 0 - | 0 - |
| 8488 | 1 | 0.2um_NOKCPHJN_191079 | Bacteria | Bacteroidetes | Flavobacteriia | Flavobacterial | Flavobacteriac | Formosa | Formosa sp. H | unclassified | unclassified | unclassified | unclassified | unclassified | 0,2 | 0,4 | 3,43E-05 | 6,87E-05 | 0 - | 0 - | 1 - | 0,000171709 - | 0 - | 0 - | 0 - | 0 - | 0 - | 0 - |
| 8489 | 1 | 3um_FIJJAFBF_447665 | Bacteria | Actinobacteria | Actinobacteria | Micrococcales | Microbacteria | Candidatus Ac | Candidatus Ac | unclassified | unclassified | unclassified | unclassified | unclassified | 4,2 | 0,748331477 | 5,31E-05 | 8,05E-06 | 5 - | 5,35E-05 - | 3 - | 3,98E-05 - | 4 - | 5,05E-05 - | 5 - | 6,41E-05 - | 4 - | 5,77E-05 - |
| 8490 | 1 | 0.2um_NOKCPHJN_674067 | Bacteria | Bacteroidetes | Cytophagia | Cytophagales | Flammeovirga | unclassified | Flammeovirga | unclassified | unclassified | unclassified | unclassified | unclassified | 1,4 | 0,489897949 | 2,76E-05 | 8,69E-06 | 2 - | 3,37E-05 - | 2 - | 4,18E-05 - | 1 - | 1,99E-05 - | 1 - | 2,02E-05 - | 1 - | 2,27E-05 - |
| 8491 | 1 | 0.2um_NOKCPHJN_1059776 | Viruses | unclassified | unclassified | unclassified | unclassified | unclassified | uncultured M | unclassified | unclassified | unclassified | unclassified | unclassified | 1 | 0 | 6,12E-05 | 5,81E-06 | 1 - | 5,12E-05 - | 1 - | 6,36E-05 - | 1 - | 6,05E-05 - | 1 - | 6,14E-05 - | 1 - | 6,91E-05 - |
| 8492 | 1 | 0.2um_NOKCPHJN_565448 | Bacteria | Proteobacteri | Alphaproteob | Pelagibacteral | Pelagibacterac | Candidatus Pe | Candidatus Pe | metabolism | Lipid transpor | 05VNB | Carrier of the | 335992.SAR11 | 0,6 | 0,8 | 9,87E-05 | 0,000136803 | 1 - | 0,000145287 - | 0 - | 0 - | 0 - | 0 - | 2 - | 0,000348059 - | 0 - | 0 - |
| 8493 | 2 | 0.2um_NOKCPHJN_970291;3um_FIJJAFBF_2033 | Bacteria | Bacteroidetes | Flavobacteriia | Flavobacterial | Flavobacteriac | various | various | unclassified | unclassified | unclassified | unclassified | unclassified | 1,2 | 0,4 | 0,000202509 | 4,49E-05 | 2 - | 0,000290574 - | 1 - | 0,000180404 - | 1 - | 0,000171496 - | 1 - | 0,00017403 - | 1 - | 0,000196044 - |
| 8494 | 1 | 3um_FIJJAFBF_256111 | Bacteria | Proteobacteri | Alphaproteob | Rhodospirillal | Rhodospirillac | Magnetospira | Magnetospira | poorly charact | Function unkn | 062M7 | phasin family | 414684.RC1_1 | 1 | 0,894427191 | 7,66E-05 | 6,81E-05 | 2 - | 0,000137457 - | 0 - | 0 - | 1 - | 8,11E-05 - | 2 - | 0,000164651 - | 0 - | 0 - |
| 8495 | 1 | 3um_FIJJAFBF_737106 | Bacteria | Proteobacteri | Alphaproteob | Pelagibacteral | Pelagibacterac | Candidatus Pe | Candidatus Pe | information st | Translation, ri | 08VZM | 50s ribosomal | 439493.PB721 | 0,4 | 0,489897949 | 2,92E-05 | 3,60E-05 | 1 - | 6,63E-05 - | 0 - | 0 - | 0 - | 0 - | 1 - | 7,95E-05 - | 0 - | 0 - |
| 8496 | 1 | 3um_FIJJAFBF_438665 | Bacteria | Proteobacteri | Gammaprotec | Oceanospirilla | Saccharospirill | Reinekea | Reinekea blan | metabolism | Amino acid tra | 16Q19 | homoserine d | 314283.MED2 | 0,6 | 0,489897949 | 1,92E-05 | 1,57E-05 | 0 - | 0 - | 1 - | 3,30E-05 - | 1 - | 3,14E-05 - | 1 - | 3,18E-05 - | 0 - | 0 - |
| 8497 | 1 | 0.2um_NOKCPHJN_420409 | Bacteria | Bacteroidetes | Cytophagia | Cytophagales | Flammeovirga | unclassified | Flammeovirga | poorly charact | Function unkn | 16R8I | Lipoprotein | 566466.NOR5 | 2 | 0,894427191 | 5,64E-05 | 2,79E-05 | 2 - | 4,68E-05 - | 1 - | 2,90E-05 - | 1 - | 2,76E-05 - | 3 - | 8,40E-05 - | 3 - | 9,46E-05 - |
| 8498 | 1 | 0.2um_NOKCPHJN_746240 | Bacteria | Proteobacteri | Gammaprotec | Cellvibrionales | Halieaceae | unclassified | marine gamm | unclassified | unclassified | unclassified | unclassified | unclassified | 0,6 | 0,489897949 | 9,18E-06 | 7,57E-06 | 1 - | 1,34E-05 - | 1 - | 1,66E-05 - | 1 - | 1,58E-05 - | 0 - | 0 - | 0 - | 0 - |
| 8499 | 1 | 0.2um_NOKCPHJN_726440 | unclassified | unclassified | unclassified | unclassified | unclassified | unclassified | unclassified | unclassified | unclassified | unclassified | unclassified | unclassified | 0,4 | 0,489897949 | 4,24E-05 | 5,22E-05 | 1 - | 9,73E-05 - | 0 - | 0 - | 1 - | 0,000114815 - | 0 - | 0 - | 0 - | 0 - |
| 8500 | 1 | 0.2um_NOKCPHJN_591136 | Eukaryota | Arthropoda | Insecta | Lepidoptera | Nymphalidae | Bicyclus | Bicyclus anyna | unclassified | unclassified | unclassified | unclassified | unclassified | 0,2 | 0,4 | 1,46E-05 | 2,93E-05 | 0 - | 0 - | 0 - | 0 - | 0 - | 0 - | 1 - | 7,31E-05 - | 0 - | 0 - |
| 8501 | 1 | 0.2um_NOKCPHJN_50272 | Viruses | unclassified | unclassified | unclassified | Phycodnavirid | unclassified | Aureococcus a | unclassified | unclassified | unclassified | unclassified | unclassified | 1,6 | 1,2 | 4,73E-05 | 3,51E-05 | 3 - | 7,79E-05 - | 3 - | 9,67E-05 - | 1 - | 3,07E-05 - | 1 - | 3,11E-05 - | 0 - | 0 - |
| 8502 | 1 | 0.2um_NOKCPHJN_52924 | Bacteria | Proteobacteri | Gammaprotec | Cellvibrionales | Porticoccacea | unclassified | gamma prote | unclassified | unclassified | unclassified | unclassified | unclassified | 1 | 0,894427191 | 3,60E-05 | 3,23E-05 | 0 - | 0 - | 2 - | 7,40E-05 - | 2 - | 7,04E-05 - | 1 - | 3,57E-05 - | 0 - | 0 - |
| 8503 | 1 | 3um_FIJJAFBF_233042 | Bacteria | Actinobacteria | Actinobacteria | Micrococcales | Microbacteria | unclassified | Microbacteria | information st | Translation, ri | 08UIK | This protein bi | 281090.Lxx02 | 1,6 | 0,489897949 | 0,000150053 | 3,93E-05 | 2 - | 0,000160527 - | 1 - | 9,97E-05 - | 2 - | 0,000189486 - | 2 - | 0,000192284 - | 1 - | 0,000108304 - |
| 8504 | 1 | 0.2um_NOKCPHJN_233281 | Bacteria | Bacteroidetes | Flavobacteriia | Flavobacterial | Flavobacteriac | Zhouia | Zhouia amylol | metabolism | Inorganic ion t | 05C27 | TonB-depende | 867900.Celly_ | 1 | 0 | 1,31E-05 | 1,25E-06 | 1 - | 1,10E-05 - | 1 - | 1,37E-05 - | 1 - | 1,30E-05 - | 1 - | 1,32E-05 - | 1 - | 1,48E-05 - |
| 8505 | 1 | 0.2um_NOKCPHJN_67766 | Bacteria | Proteobacteri | Alphaproteob | Rhodobactera | Rhodobactera | unclassified | Rhodobactera | information st | Translation, ri | 16PEA | histidyl-tRNA | 367336.OM22 | 0,6 | 0,489897949 | 1,59E-05 | 1,31E-05 | 1 - | 2,31E-05 - | 1 - | 2,87E-05 - | 0 - | 0 - | 1 - | 2,77E-05 - | 0 - | 0 - |
| 8506 | 1 | 3um_FIJJAFBF_73928 | Bacteria | Bacteroidetes | unclassified | unclassified | unclassified | unclassified | unidentified e | information st | Translation, ri | 09AI8 | Associates wit | 50743.SCB49_ | 3 | 0 | 0,000127665 | 1,21E-05 | 3 - | 0,000106935 - | 3 - | 0,000132782 - | 3 - | 0,000126226 - | 3 - | 0,00012809 - | 3 - | 0,000144293 - |
| 8507 | 1 | 0.2um_NOKCPHJN_13837 | Bacteria | Cyanobacteria | unclassified | Synechococca | Merismopedia | Aphanocapsa | Aphanocapsa | cellular proces | Posttranslatio | 08V1T | peptidylprolyl | 700598.Niako | 0,6 | 0,489897949 | 3,49E-05 | 2,90E-05 | 1 - | 4,86E-05 - | 1 - | 6,04E-05 - | 0 - | 0 - | 0 - | 0 - | 1 - | 6,56E-05 - |
| 8508 | 1 | 3um_FIJJAFBF_184544 | Bacteria | Proteobacteri | Alphaproteob | Rhodobactera | Rhodobactera | Roseobacter | uncultured Ro | metabolism | Amino acid tra | 01SEG | branched-cha | 314262.MED1 | 1,8 | 0,4 | 5,76E-05 | 1,35E-05 | 2 - | 5,39E-05 - | 1 - | 3,35E-05 - | 2 - | 6,36E-05 - | 2 - | 6,45E-05 - | 2 - | 7,27E-05 - |
| 8509 | 1 | 0.2um_NOKCPHJN_85189 | Bacteria | Proteobacteri | Alphaproteob | Rhodobactera | Rhodobactera | Amylibacter | Amylibacter k | metabolism | Carbohydrate | COG0057 | glyceraldehyd | 391619.RGBS1 | 4,4 | 1,356465997 | 0,000177953 | 5,01E-05 | 5 - | 0,000172857 - | 4 - | 0,000171709 - | 6 - | 0,000244847 - | 5 - | 0,000207053 - | 2 - | 9,33E-05 - |
| 8510 | 1 | 3um_FIJJAFBF_155128 | Bacteria | Proteobacteri | Gammaprotec | Cellvibrionales | Cellvibrionace | Marinimicrobi | Marinimicrobi | cellular proces | Posttranslatio | 17FI0 | Binds to Cpn6 | 203122.Sde_0 | 1,8 | 0,4 | 0,000256831 | 6,29E-05 | 2 - | 0,000239118 - | 2 - | 0,000296914 - | 2 - | 0,000282255 - | 1 - | 0,000143212 - | 2 - | 0,000322655 - |
| 8511 | 1 | 0.2um_NOKCPHJN_217909 | Bacteria | Bacteroidetes | Flavobacteriia | Flavobacterial | Flavobacteriac | Polaribacter | Polaribacter s | unclassified | unclassified | unclassified | unclassified | unclassified | 0,2 | 0,4 | 8,66E-06 | 1,73E-05 | 0 - | 0 - | 0 - | 0 - | 1 - | 4,33E-05 - | 0 - | 0 - | 0 - | 0 - |
| 8512 | 1 | 0.2um_NOKCPHJN_423481 | Bacteria | Bacteroidetes | Flavobacteriia | Flavobacterial | Flavobacteriac | Marixanthom | Marixanthom | unclassified | unclassified | unclassified | unclassified | unclassified | 1 | 0 | 8,36E-05 | 7,93E-06 | 1 - | 7,00E-05 - | 1 - | 8,69E-05 - | 1 - | 8,26E-05 - | 1 - | 8,38E-05 - | 1 - | 9,44E-05 - |
| 8513 | 1 | 0.2um_NOKCPHJN_15772 | Bacteria | Proteobacteri | Gammaprotec | Alteromonada | Alteromonada | Paraglaciecola | Paraglaciecola | metabolism | Carbohydrate | 0QHXA | transketolase | 675806.VII_00 | 0,2 | 0,4 | 3,45E-06 | 6,90E-06 | 1 - | 1,73E-05 - | 0 - | 0 - | 0 - | 0 - | 0 - | 0 - | 0 - | 0 - |
| 8514 | 1 | 0.2um_NOKCPHJN_662166 | Bacteria | Proteobacteri | Alphaproteob | Pelagibacteral | unclassified | unclassified | Pelagibacteral | cellular proces | Cell wall/mem | COG2885 | Ompa motb d | 335992.SAR11 | 1 | 0,894427191 | 9,24E-05 | 8,44E-05 | 0 - | 0 - | 2 - | 0,000180404 - | 1 - | 8,57E-05 - | 0 - | 0 - | 2 - | 0,000196044 - |
| 8515 | 1 | 3um_FIJJAFBF_475532 | Eukaryota | unclassified | Oomycetes | Peronosporale | Peronosporac | Plasmopara | Plasmopara h | information st | Translation, ri | COG0081 | Binds directly | 653948.CCA2 | 0,2 | 0,4 | 2,27E-05 | 4,55E-05 | 1 - | 0,00011364 - | 0 - | 0 - | 0 - | 0 - | 0 - | 0 - | 0 - | 0 - |
| 8516 | 1 | 3um_FIJJAFBF_687132 | Bacteria | Proteobacteri | Betaproteoba | unclassified | unclassified | unclassified | Betaproteoba | poorly charact | Function unkn | 05W4M | Transport-ass | 296591.Bpro_ | 4,2 | 0,4 | 0,000261639 | 3,47E-05 | 4 - | 0,000208685 - | 4 - | 0,000259125 - | 4 - | 0,000246331 - | 5 - | 0,000312462 - | 4 - | 0,00028159 - |
| 8517 | 1 | 0.2um_NOKCPHJN_36914 | Eukaryota | unclassified | Cryptophyta | Pyrenomonad | Pyrenomonad | Rhodomonas | Rhodomonas | metabolism | Carbohydrate | COG1850 | RuBisCO catal | 45157.CMV01 | 7,2 | 0,748331477 | 0,000200753 | 1,67E-05 | 8 - | 0,000188159 - | 8 - | 0,000233637 - | 7 - | 0,000194339 - | 7 - | 0,00019721 - | 6 - | 0,000190419 - |
| 8518 | 1 | 0.2um_NOKCPHJN_523562 | Eukaryota | Chlorophyta | Chlorophycea | Sphaeropleale | Selenastracea | Raphidocelis | Raphidocelis s | unclassified | unclassified | unclassified | unclassified | unclassified | 0,2 | 0,4 | 2,00E-06 | 4,01E-06 | 1 - | 1,00E-05 - | 0 - | 0 - | 0 - | 0 - | 0 - | 0 - | 0 - | 0 - |
| 8519 | 1 | 3um_FIJJAFBF_649064 | Bacteria | Proteobacteri | Gammaprotec | Oceanospirilla | Saccharospirill | Reinekea | Reinekea fors | unclassified | unclassified | unclassified | unclassified | unclassified | 0,4 | 0,489897949 | 2,43E-05 | 2,99E-05 | 1 - | 5,57E-05 - | 0 - | 0 - | 1 - | 6,58E-05 - | 0 - | 0 - | 0 - | 0 - |
| 8520 | 1 | 0.2um_NOKCPHJN_154986 | Bacteria | Proteobacteri | Gammaprotec | Oceanospirilla | Saccharospirill | Reinekea | Reinekea fors | cellular proces | Cell wall/mem | COG1686 | Carboxypeptic | 314283.MED2 | 3,2 | 0,748331477 | 0,000110931 | 2,17E-05 | 4 - | 0,00011772 - | 2 - | 7,31E-05 - | 4 - | 0,000138956 - | 3 - | 0,000105756 - | 3 - | 0,000119134 - |
| 8521 | 1 | 3um_FIJJAFBF_599962 | Bacteria | Proteobacteri | Gammaprotec | Oceanospirilla | Saccharospirill | Reinekea | Reinekea fors | metabolism | Lipid transpor | COG0511 | Acetyl-CoA ca | 314283.MED2 | 1,6 | 0,8 | 0,000137539 | 5,50E-05 | 3 - | 0,000225052 - | 1 - | 9,31E-05 - | 1 - | 8,86E-05 - | 2 - | 0,000179717 - | 1 - | 0,000101225 - |
| 8522 | 1 | 0.2um_NOKCPHJN_956058 | Bacteria | Proteobacteri | Gammaprotec | Cellvibrionales | Porticoccacea | unclassified | Gammaprotec | unclassified | unclassified | unclassified | unclassified | unclassified | 2 | 0 | 0,000119674 | 1,14E-05 | 2 - | 0,000100242 - | 2 - | 0,000124471 - | 2 - | 0,000118325 - | 2 - | 0,000120073 - | 2 - | 0,000135262 - |
| 8523 | 1 | 3um_FIJJAFBF_666765 | Eukaryota | Chlorophyta | Chlorophycea | Chlamydomor | Chlamydomor | Chlamydomor | Chlamydomor | unclassified | unclassified | unclassified | unclassified | unclassified | 0,2 | 0,4 | 4,10E-05 | 8,21E-05 | 0 - | 0 - | 0 - | 0 - | 0 - | 0 - | 1 - | 0,000205199 - | 0 - | 0 - |
| 8524 | 1 | 0.2um_NOKCPHJN_530269 | Bacteria | Proteobacteri | Alphaproteob | Pelagibacteral | Pelagibacterac | Candidatus Pe | Candidatus Pe | cellular proces | Cell wall/mem | 7257 | NA | 335992.SAR11 | 0,6 | 0,489897949 | 2,32E-05 | 1,92E-05 | 1 - | 3,28E-05 - | 0 - | 0 - | 1 - | 3,87E-05 - | 0 - | 0 - | 1 - | 4,42E-05 - |
| 8525 | 1 | 3um_FIJJAFBF_718088 | Bacteria | Bacteroidetes | Cytophagia | Cytophagales | Flammeovirga | unclassified | Flammeovirga | unclassified | unclassified | unclassified | unclassified | unclassified | 2,4 | 0,8 | 0,000258571 | 7,66E-05 | 3 - | 0,000277686 - | 2 - | 0,000229869 - | 3 - | 0,00032778 - | 3 - | 0,000332621 - | 1 - | 0,000124899 - |
| 8526 | 1 | 3um_FIJJAFBF_577836 | Eukaryota | Arthropoda | Insecta | Lepidoptera | Nymphalidae | Bicyclus | Bicyclus anyna | poorly charact | Function unkn | 140PW | NA | 400682.PAC_1 | 0,6 | 0,489897949 | 1,68E-05 | 1,38E-05 | 1 - | 2,49E-05 - | 0 - | 0 - | 1 - | 2,94E-05 - | 1 - | 2,98E-05 - | 0 - | 0 - |
| 8527 | 1 | 3um_FIJJAFBF_557880 | Bacteria | Bacteroidetes | Flavobacteriia | unclassified | unclassified | unclassified | Flavobacteria | unclassified | unclassified | unclassified | unclassified | unclassified | 1,2 | 0,4 | 6,13E-05 | 1,36E-05 | 2 - | 8,80E-05 - | 1 - | 5,46E-05 - | 1 - | 5,19E-05 - | 1 - | 5,27E-05 - | 1 - | 5,93E-05 - |
| 8528 | 1 | 3um_FIJJAFBF_397197 | Bacteria | Bacteroidetes | Cytophagia | Cytophagales | Flammeovirga | unclassified | Flammeovirga | unclassified | unclassified | unclassified | unclassified | unclassified | 4 | 1,264911064 | 0,000320653 | 0,000119316 | 3 - | 0,000197891 - | 5 - | 0,000409537 - | 2 - | 0,000155727 - | 5 - | 0,000395067 - | 5 - | 0,000445042 - |
| 8529 | 1 | 3um_FIJJAFBF_453403 | Bacteria | Proteobacteri | Alphaproteob | Rhodobactera | unclassified | unclassified | Rhodobactera | metabolism | Carbohydrate | 16SJ5 | Trap dicarbox | 314262.MED1 | 2 | 0 | 8,41E-05 | 7,98E-06 | 2 - | 7,04E-05 - | 2 - | 8,74E-05 - | 2 - | 8,31E-05 - | 2 - | 8,43E-05 - | 2 - | 9,50E-05 - |
| 8530 | 1 | 3um_FIJJAFBF_628527 | Bacteria | Proteobacteri | Alphaproteob | Rhodobactera | Rhodobactera | unclassified | Rhodobactera | unclassified | unclassified | unclassified | unclassified | unclassified | 0,2 | 0,4 | 4,11E-06 | 8,22E-06 | 0 - | 0 - | 0 - | 0 - | 0 - | 0 - | 0 - | 0 - | 1 - | 2,05E-05 - |
| 8531 | 2 | 0.2um_NOKCPHJN_207029;0.2um_NOKCPHJN_2 | Bacteria | Proteobacteri | Gammaprotec | Cellvibrionales | Porticoccacea | unclassified | various | information st | Translation, ri | COG0016 | phenylalanyl- | 314287.GB22 | 0,2 | 0,4 | 8,04E-06 | 1,61E-05 | 0 - | 0 - | 0 - | 0 - | 1 - | 4,02E-05 - | 0 - | 0 - | 0 - | 0 - |
| 8532 | 1 | 0.2um_NOKCPHJN_57904 | Bacteria | Proteobacteri | Alphaproteob | Rhodobactera | Rhodobactera | Tateyamaria | Tateyamaria s | unclassified | unclassified | unclassified | unclassified | unclassified | 5,6 | 1,2 | 0,000132802 | 2,60E-05 | 7 - | 0,000140216 - | 7 - | 0,000174107 - | 4 - | 9,46E-05 - | 5 - | 0,000119968 - | 5 - | 0,000135144 - |
| 8533 | 3 | 0.2um_NOKCPHJN_230057;3um_FIJJAFBF_1562 | Bacteria | unclassified | unclassified | unclassified | unclassified | unclassified | uncultured ba | various | various | various | various | various | 1 | 0 | 4,18E-05 | 3,97E-06 | 1 - | 3,50E-05 - | 1 - | 4,35E-05 - | 1 - | 4,13E-05 - | 1 - | 4,19E-05 - | 1 - | 4,72E-05 - |
| 8534 | 1 | 0.2um_NOKCPHJN_473382 | Bacteria | Proteobacteri | unclassified | unclassified | unclassified | unclassified | Proteobacteri | cellular proces | Cell wall/mem | 0HCHF | UDP-n-acetyln | 589865.DaAH | 0,2 | 0,4 | 4,48E-06 | 8,97E-06 | 1 - | 2,24E-05 - | 0 - | 0 - | 0 - | 0 - | 0 - | 0 - | 0 - | 0 - |
| 8535 | 1 | 0.2um_NOKCPHJN_540892 | Bacteria | Proteobacteri | Alphaproteob | Pelagibacteral | Pelagibacterac | Candidatus Pe | Candidatus Pe | metabolism | Energy produc | 16Q0J | alcohol dehyd | 335992.SAR11 | 2,6 | 1,019803903 | 9,93E-05 | 3,76E-05 | 3 - | 9,75E-05 - | 1 - | 4,04E-05 - | 3 - | 0,000115141 - | 4 - | 0,000155788 - | 2 - | 8,77E-05 - |
| 8536 | 1 | 3um_FIJJAFBF_700593 | unclassified | unclassified | unclassified | unclassified | unclassified | unclassified | unclassified | unclassified | unclassified | unclassified | unclassified | unclassified | 0,8 | 0,4 | 0,000115224 | 5,80E-05 | 0 - | 0 - | 1 - | 0,000143958 - | 1 - | 0,000136851 - | 1 - | 0,000138872 - | 1 - | 0,000156439 - |
| 8537 | 1 | 0.2um_NOKCPHJN_380276 | Viruses | unclassified | unclassified | Caudovirales | Siphoviridae | P12024virus | Nonlabens vir | unclassified | unclassified | unclassified | unclassified | unclassified | 0,2 | 0,4 | 3,92E-06 | 7,83E-06 | 0 - | 0 - | 0 - | 0 - | 1 - | 1,96E-05 - | 0 - | 0 - | 0 - | 0 - |
| 8538 | 1 | 3um_FIJJAFBF_504104 | Bacteria | Proteobacteri | Gammaprotec | Alteromonada | Psychromonac | Psychromonas | Psychromonas | poorly charact | Function unkn | 08VI5 | type VI secreti | 357804.Ping_ | 1,6 | 1,019803903 | 0,000120214 | 7,56E-05 | 2 - | 0,000128242 - | 2 - | 0,000159239 - | 3 - | 0,000227065 - | 0 - | 0 - | 1 - | 8,65E-05 - |
| 8539 | 1 | 0.2um_NOKCPHJN_774510 | Eukaryota | Bacillariophyta | Coscinodiscop | Thalassiosirale | Thalassiosirac | Thalassiosira | Thalassiosira | unclassified | unclassified | unclassified | unclassified | unclassified | 0,6 | 0,489897949 | 4,46E-05 | 3,68E-05 | 1 - | 6,52E-05 - | 1 - | 8,10E-05 - | 1 - | 7,70E-05 - | 0 - | 0 - | 0 - | 0 - |
| 8540 | 1 | 3um_FIJJAFBF_171078 | Bacteria | Proteobacteri | Gammaprotec | Oceanospirilla | Oceanospirilla | Amphritea | Amphritea jap | unclassified | unclassified | unclassified | unclassified | unclassified | 1 | 0 | 0,000101502 | 9,64E-06 | 1 - | 8,50E-05 - | 1 - | 0,00010557 - | 1 - | 0,000100357 - | 1 - | 0,000101839 - | 1 - | 0,000114722 - |
| 8541 | 1 | 3um_FIJJAFBF_715080 | Eukaryota | unclassified | Pelagophycea | Pelagomonad | unclassified | Pelagomonas | uncultured Pe | unclassified | unclassified | unclassified | unclassified | unclassified | 1 | 0 | 0,000119154 | 1,13E-05 | 1 - | 9,98E-05 - | 1 - | 0,000123929 - | 1 - | 0,000117811 - | 1 - | 0,000119551 - | 1 - | 0,000134673 - |
| 8542 | 1 | 3um_FIJJAFBF_410266 | Bacteria | Bacteroidetes | Flavobacteriia | Flavobacterial | Flavobacteriac | Polaribacter | various | metabolism | Carbohydrate | 05D2N | Aldolase | 391587.KAOT | 0,6 | 0,8 | 2,31E-05 | 3,09E-05 | 0 - | 0 - | 0 - | 0 - | 1 - | 3,82E-05 - | 2 - | 7,75E-05 - | 0 - | 0 - |
| 8543 | 1 | 0.2um_NOKCPHJN_158128 | Bacteria | Proteobacteri | Gammaprotec | Cellvibrionales | Porticoccacea | unclassified | various | unclassified | unclassified | unclassified | unclassified | unclassified | 0,4 | 0,489897949 | 4,36E-05 | 5,39E-05 | 1 - | 9,73E-05 - | 1 - | 0,000120779 - | 0 - | 0 - | 0 - | 0 - | 0 - | 0 - |
| 8544 | 2 | 3um_FIJJAFBF_224599;3um_FIJJAFBF_242857 | Bacteria | Bacteroidetes | Flavobacteriia | Flavobacterial | Flavobacteriac | various | various | unclassified | unclassified | unclassified | unclassified | unclassified | 0,4 | 0,8 | 7,68E-06 | 1,54E-05 | 2 - | 3,84E-05 - | 0 - | 0 - | 0 - | 0 - | 0 - | 0 - | 0 - | 0 - |
| 8545 | 1 | 0.2um_NOKCPHJN_880424 | Viruses | unclassified | unclassified | unclassified | unclassified | unclassified | uncultured vir | unclassified | unclassified | unclassified | unclassified | unclassified | 0,6 | 0,8 | 1,96E-05 | 2,51E-05 | 2 - | 6,04E-05 - | 1 - | 3,75E-05 - | 0 - | 0 - | 0 - | 0 - | 0 - | 0 - |
| 8546 | 1 | 3um_FIJJAFBF_201379 | Bacteria | Bacteroidetes | Flavobacteriia | Flavobacterial | Flavobacteriac | Polaribacter | Polaribacter s | unclassified | unclassified | unclassified | unclassified | unclassified | 3,2 | 0,4 | 0,000153887 | 2,39E-05 | 3 - | 0,000120818 - | 3 - | 0,00015002 - | 3 - | 0,000142613 - | 4 - | 0,000192959 - | 3 - | 0,000163026 - |
| 8547 | 1 | 0.2um_NOKCPHJN_118677 | Bacteria | Proteobacteri | Gammaprotec | Oceanospirilla | Saccharospirill | Reinekea | Reinekea fors | poorly charact | Function unkn | 0XNQ8 | alkaline phosp | 203122.Sde_3 | 18,8 | 2,039607805 | 0,000445154 | 5,51E-05 | 21 - | 0,000417732 - | 19 - | 0,0004693 - | 15 - | 0,000352207 - | 20 - | 0,000476545 - | 19 - | 0,000509985 - |
| 8548 | 1 | 3um_FIJJAFBF_328042 | Bacteria | Proteobacteri | Alphaproteob | Rhodobactera | Rhodobactera | unclassified | Rhodobactera | unclassified | unclassified | unclassified | unclassified | unclassified | 0,2 | 0,4 | 1,06E-05 | 2,12E-05 | 0 - | 0 - | 0 - | 0 - | 1 - | 5,29E-05 - | 0 - | 0 - | 0 - | 0 - |
| 8549 | 1 | 3um_FIJJAFBF_302304 | Bacteria | Proteobacteri | Betaproteoba | Burkholderial | Comamonada | Malikia | Malikia granos | metabolism | Amino acid tra | COG0174 | glutamine syn | 535289.Dtpsy | 3,6 | 1,2 | 0,0001019 | 2,26E-05 | 6 - | 0,000146212 - | 3 - | 9,08E-05 - | 3 - | 8,63E-05 - | 3 - | 8,76E-05 - | 3 - | 9,86E-05 - |
| 8550 | 1 | 0.2um_NOKCPHJN_906170 | unclassified | unclassified | unclassified | unclassified | unclassified | unclassified | unclassified | unclassified | unclassified | unclassified | unclassified | unclassified | 0,6 | 0,489897949 | 6,36E-05 | 5,29E-05 | 1 - | 8,97E-05 - | 0 - | 0 - | 0 - | 0 - | 1 - | 0,000107409 - | 1 - | 0,000120996 - |
| 8551 | 1 | 3um_FIJJAFBF_90781 | Eukaryota | unclassified | Spirotrichea | Sporadotrichi | Oxytrichidae | Stylonychia | Stylonychia le | unclassified | unclassified | unclassified | unclassified | unclassified | 1 | 0 | 0,000152252 | 1,45E-05 | 1 - | 0,00012753 - | 1 - | 0,000158354 - | 1 - | 0,000150536 - | 1 - | 0,000152759 - | 1 - | 0,000172083 - |
| 8552 | 1 | 3um_FIJJAFBF_66744 | Bacteria | Proteobacteri | Gammaprotec | Pseudomonad | Pseudomonad | Pseudomonas | Pseudomonas | metabolism | Inorganic ion t | 05ENV | phosphonate | 399739.Pmen | 11,2 | 1,6 | 0,000573566 | 2,76E-05 | 14 - | 0,000608665 - | 11 - | 0,000593829 - | 11 - | 0,000564509 - | 11 - | 0,000572847 - | 9 - | 0,000527981 - |
| 8553 | 1 | 0.2um_NOKCPHJN_228578 | Bacteria | Bacteroidetes | Flavobacteriia | Flavobacterial | Flavobacteriac | Formosa | Formosa sp. H | information st | Transcription | COG0085 | DNA-depende | 391603.FBALC | 0,2 | 0,4 | 2,66E-06 | 5,32E-06 | 1 - | 1,33E-05 - | 0 - | 0 - | 0 - | 0 - | 0 - | 0 - | 0 - | 0 - |
| 8554 | 1 | 3um_FIJJAFBF_242673 | Bacteria | Bacteroidetes | Saprospiria | Saprospirales | Haliscomenob | Haliscomenob | Haliscomenob | cellular proces | Posttranslatio | COG1404 | peptidase (S8 | 760192.Halhy | 1 | 0 | 3,77E-05 | 3,58E-06 | 1 - | 3,16E-05 - | 1 - | 3,93E-05 - | 1 - | 3,73E-05 - | 1 - | 3,79E-05 - | 1 - | 4,27E-05 - |
| 8555 | 1 | 3um_FIJJAFBF_496223 | Bacteria | Bacteroidetes | Flavobacteriia | Flavobacterial | unclassified | unclassified | Flavobacterial | unclassified | unclassified | unclassified | unclassified | unclassified | 2,6 | 0,489897949 | 0,000218704 | 4,82E-05 | 3 - | 0,000211246 - | 3 - | 0,000262305 - | 2 - | 0,000166236 - | 2 - | 0,000168691 - | 3 - | 0,000285045 - |
| 8556 | 1 | 3um_FIJJAFBF_653488 | Eukaryota | Bacillariophyta | Bacillariophyc | Bacillariales | Bacillariaceae | Fragilariopsis | Fragilariopsis | poorly charact | Function unkn | COG1100 | GTP-binding P | 2850.Phatr43 | 4,6 | 1,2 | 0,00042597 | 9,43E-05 | 6 - | 0,000471685 - | 4 - | 0,000390463 - | 3 - | 0,000278388 - | 6 - | 0,000565 - | 4 - | 0,000424314 - |
| 8557 | 1 | 0.2um_NOKCPHJN_106535 | Bacteria | Proteobacteri | Gammaprotec | Oceanospirilla | Saccharospirill | Reinekea | Reinekea blan | information st | Translation, ri | 05C6A | phenylalanyl-t | 314283.MED2 | 1 | 0,632455532 | 1,63E-05 | 9,30E-06 | 2 - | 2,90E-05 - | 1 - | 1,80E-05 - | 1 - | 1,71E-05 - | 1 - | 1,74E-05 - | 0 - | 0 - |
| 8558 | 1 | 0.2um_NOKCPHJN_285950 | Bacteria | Proteobacteri | Gammaprotec | Oceanospirilla | Saccharospirill | Reinekea | Reinekea fors | unclassified | unclassified | unclassified | unclassified | unclassified | 16,6 | 1,356465997 | 0,00023065 | 2,98E-05 | 18 - | 0,000209532 - | 17 - | 0,000245722 - | 15 - | 0,000206109 - | 15 - | 0,000209153 - | 18 - | 0,000282732 - |
| 8559 | 1 | 3um_FIJJAFBF_111286 | Viruses | unclassified | unclassified | unclassified | various | various | various | unclassified | unclassified | unclassified | unclassified | unclassified | 0,4 | 0,489897949 | 4,79E-05 | 5,87E-05 | 0 - | 0 - | 0 - | 0 - | 1 - | 0,000118844 - | 1 - | 0,000120599 - | 0 - | 0 - |
| 8560 | 1 | 3um_FIJJAFBF_314307 | Bacteria | Bacteroidetes | Flavobacteriia | Flavobacterial | Flavobacteriac | Polaribacter | Polaribacter s | cellular proces | Cell wall/mem | 11TPU | outer membra | 313598.MED1 | 3 | 0,894427191 | 0,000240156 | 6,04E-05 | 4 - | 0,000273278 - | 3 - | 0,000254498 - | 2 - | 0,000161288 - | 4 - | 0,000327341 - | 2 - | 0,000184374 - |
| 8561 | 1 | 3um_FIJJAFBF_277642 | Bacteria | Bacteroidetes | Flavobacteriia | Flavobacterial | Flavobacteriac | Polaribacter | various | poorly charact | Function unkn | 05FCH | NA | 487797.Flav3 | 0,6 | 0,8 | 3,13E-05 | 4,04E-05 | 2 - | 9,85E-05 - | 0 - | 0 - | 1 - | 5,81E-05 - | 0 - | 0 - | 0 - | 0 - |
| 8562 | 1 | 0.2um_NOKCPHJN_57126 | Bacteria | Proteobacteri | Gammaprotec | Oceanospirilla | Saccharospirill | Reinekea | Reinekea fors | cellular proces | Posttranslatio | 0QHT2 | Peroxiredoxin | 314283.MED2 | 1,6 | 0,489897949 | 0,000141774 | 5,18E-05 | 1 - | 7,26E-05 - | 2 - | 0,000180404 - | 1 - | 8,57E-05 - | 2 - | 0,00017403 - | 2 - | 0,000196044 - |
| 8563 | 1 | 0.2um_NOKCPHJN_123349 | Bacteria | Proteobacteri | Gammaprotec | Oceanospirilla | Saccharospirill | Reinekea | Reinekea fors | unclassified | unclassified | unclassified | unclassified | unclassified | 0,2 | 0,4 | 1,10E-05 | 2,20E-05 | 1 - | 5,49E-05 - | 0 - | 0 - | 0 - | 0 - | 0 - | 0 - | 0 - | 0 - |
| 8564 | 1 | 3um_FIJJAFBF_20171 | Bacteria | Proteobacteri | Gammaprotec | unclassified | unclassified | unclassified | various | unclassified | unclassified | unclassified | unclassified | unclassified | 0,6 | 0,489897949 | 6,92E-05 | 5,70E-05 | 1 - | 0,000102479 - | 0 - | 0 - | 1 - | 0,000120966 - | 1 - | 0,000122753 - | 0 - | 0 - |
| 8565 | 1 | 3um_FIJJAFBF_516961 | Bacteria | Proteobacteri | Gammaprotec | Oceanospirilla | Saccharospirill | Reinekea | Reinekea fors | metabolism | Coenzyme tra | 17FPS | D-erythro-7,8- | 314283.MED2 | 4,2 | 0,4 | 0,000487462 | 6,18E-05 | 4 - | 0,000389074 - | 4 - | 0,000483115 - | 5 - | 0,000574077 - | 4 - | 0,000466045 - | 4 - | 0,000524998 - |
| 8566 | 1 | 0.2um_NOKCPHJN_13537 | Bacteria | Bacteroidetes | Flavobacteriia | Flavobacterial | Flavobacteriac | unclassified | Flavobacteriac | metabolism | Inorganic ion t | 05C27 | TonB-depende | 313603.FB217 | 12 | 2,28035085 | 0,000152642 | 2,74E-05 | 12 - | 0,000128963 - | 13 - | 0,000173478 - | 15 - | 0,000190284 - | 12 - | 0,000154476 - | 8 - | 0,000116011 - |
| 8567 | 1 | 3um_FIJJAFBF_39851 | Bacteria | Proteobacteri | Gammaprotec | Cellvibrionales | Cellvibrionace | Saccharophag | Saccharophag | information st | Translation, ri | 0QTKX | Seems to be t | 203122.Sde_0 | 1,8 | 0,748331477 | 0,000192394 | 7,03E-05 | 3 - | 0,000273278 - | 2 - | 0,00022622 - | 1 - | 0,000107526 - | 1 - | 0,000109114 - | 2 - | 0,000245833 - |
| 8568 | 1 | 0.2um_NOKCPHJN_78753 | Bacteria | Proteobacteri | Alphaproteob | Rhodobactera | Rhodobactera | Confluentimic | Confluentimic | unclassified | unclassified | unclassified | unclassified | unclassified | 0,2 | 0,4 | 5,15E-06 | 1,03E-05 | 1 - | 2,57E-05 - | 0 - | 0 - | 0 - | 0 - | 0 - | 0 - | 0 - | 0 - |
| 8569 | 1 | 3um_FIJJAFBF_19843 | Bacteria | Bacteroidetes | Flavobacteriia | Flavobacterial | Flavobacteriac | Polaribacter | Polaribacter g | unclassified | unclassified | unclassified | unclassified | unclassified | 0,8 | 0,4 | 1,18E-05 | 5,98E-06 | 1 - | 1,28E-05 - | 1 - | 1,58E-05 - | 1 - | 1,51E-05 - | 1 - | 1,53E-05 - | 0 - | 0 - |
| 8570 | 1 | 3um_FIJJAFBF_443156 | Bacteria | Bacteroidetes | Flavobacteriia | Flavobacterial | Flavobacteriac | Muricauda | Muricauda an | unclassified | unclassified | unclassified | unclassified | unclassified | 7,8 | 1,720465053 | 0,000167094 | 3,45E-05 | 10 - | 0,000180751 - | 9 - | 0,000201995 - | 7 - | 0,00014935 - | 5 - | 0,000108255 - | 8 - | 0,000195117 - |
| 8571 | 1 | 3um_FIJJAFBF_484297 | Bacteria | Bacteroidetes | Flavobacteriia | Flavobacterial | Flavobacteriac | Gilvibacter | Gilvibacter sp. | unclassified | unclassified | unclassified | unclassified | unclassified | 0,8 | 0,4 | 1,43E-05 | 7,35E-06 | 1 - | 1,49E-05 - | 1 - | 1,86E-05 - | 0 - | 0 - | 1 - | 1,79E-05 - | 1 - | 2,02E-05 - |
| 8572 | 3 | 0.2um_NOKCPHJN_124809;0.2um_NOKCPHJN_4 | Bacteria | Bacteroidetes | Flavobacteriia | Flavobacterial | Flavobacteriac | Formosa | Formosa sp. H | information st | Translation, ri | COG0184 | Forms an inte | various | 1 | 0 | 0,000125713 | 1,19E-05 | 1 - | 0,0001053 - | 1 - | 0,000130751 - | 1 - | 0,000124296 - | 1 - | 0,000126131 - | 1 - | 0,000142087 - |
| 8573 | 1 | 3um_FIJJAFBF_489192 | Bacteria | Proteobacteri | Gammaprotec | Cellvibrionales | Porticoccacea | unclassified | SAR92 bacteri | unclassified | unclassified | unclassified | unclassified | unclassified | 1,2 | 0,4 | 0,000142375 | 6,06E-05 | 1 - | 9,73E-05 - | 1 - | 0,000120779 - | 1 - | 0,000114815 - | 1 - | 0,000116511 - | 2 - | 0,000262499 - |
| 8574 | 1 | 3um_FIJJAFBF_602616 | Bacteria | Proteobacteri | Gammaprotec | Oceanospirilla | Saccharospirill | Reinekea | Reinekea fors | cellular proces | Posttranslatio | 08V1T | peptidylprolyl | 314283.MED2 | 5,6 | 0,489897949 | 0,000321159 | 4,37E-05 | 6 - | 0,000288142 - | 6 - | 0,000357788 - | 5 - | 0,000283436 - | 5 - | 0,000287622 - | 6 - | 0,000388806 - |

| | | | | | | | | | | | | | | | | | | | | | | | | | | | | |
|---|---|---|---|---|---|---|---|---|---|---|---|---|---|---|---|---|---|---|---|---|---|---|---|---|---|---|---|---|
| 8575 | 1 | 0.2um_NOKCPHJN_191090 | Bacteria | Bacteroidetes | Flavobacteriia | Flavobacterial | Flavobacteriac | Formosa | Formosa sp. H | unclassified | unclassified | unclassified | unclassified | unclassified | 1 | 0,632455532 | 1,38E-05 | 9,14E-06 | 1 - | 1,15E-05 - | 2 - | 2,86E-05 - | 1 - | 1,36E-05 - | 0 - | 0 - | 1 - | 1,55E-05 - |
| 8576 | 1 | 3um_FIJJAFBF_316869 | Bacteria | Bacteroidetes | Flavobacteriia | Flavobacterial | Flavobacteriac | Polaribacter | Polaribacter b | information st | Translation, ril | 05C22 | threonyL-tRN | 313594.PI23P | 0,4 | 0,489897949 | 8,35E-06 | 1,04E-05 | 1 - | 1,78E-05 - | 0 - | 0 - | 0 - | 0 - | 0 - | 0 - | 1 - | 2,40E-05 - |
| 8577 | 1 | 3um_FIJJAFBF_788030 | Bacteria | Proteobacteri | Alphaproteob | Rhodobactera | Rhodobactera | unclassified | Rhodobactera | information st | Translation, ril | COG0256 | This is one of t | 644107.SL115 | 4,2 | 0,748331477 | 0,000490105 | 0,000121055 | 3 - | 0,000289353 - | 4 - | 0,000479055 - | 5 - | 0,000569253 - | 4 - | 0,000462129 - | 5 - | 0,000650733 - |
| 8578 | 1 | 0.2um_NOKCPHJN_1024190 | Bacteria | Proteobacteri | Alphaproteob | Rhodospirillal | Rhodospirillac | unclassified | Rhodospirillac | unclassified | unclassified | unclassified | unclassified | unclassified | 7,6 | 0,489897949 | 0,000473767 | 5,51E-05 | 7 - | 0,000365199 - | 8 - | 0,00051825 - | 8 - | 0,000492663 - | 8 - | 0,000499939 - | 7 - | 0,000492782 - |
| 8579 | 1 | 0.2um_NOKCPHJN_937431 | Bacteria | Verrucomicrol | unclassified | unclassified | unclassified | unclassified | Verrucomicrol | poorly charact | Function unkn | 1DIXS | Inherit from b | 382464.VDG1 | 2,8 | 0,4 | 0,000119621 | 2,14E-05 | 3 - | 0,000107268 - | 3 - | 0,000133195 - | 2 - | 8,44E-05 - | 3 - | 0,000128489 - | 3 - | 0,000144743 - |
| 8580 | 1 | 0.2um_NOKCPHJN_986771 | Bacteria | Proteobacteri | Gammaprote | Pseudomonad | Pseudomonad | Pseudomonas | Pseudomonas | metabolism | Carbohydrate | 05CR4 | methylisocitra | 1114970.PSF1 | 0,2 | 0,4 | 1,27E-05 | 2,53E-05 | 0 - | 0 - | 0 - | 0 - | 1 - | 6,33E-05 - | 0 - | 0 - | 0 - | 0 - |
| 8581 | 1 | 3um_FIJJAFBF_257121 | Eukaryota | unclassified | Cryptophyta | Pyrenomonad | Geminigerace | Guillardia | Guillardia thet | unclassified | unclassified | unclassified | unclassified | unclassified | 1,6 | 0,489897949 | 0,000178168 | 4,94E-05 | 2 - | 0,000189714 - | 2 - | 0,000235568 - | 2 - | 0,000223938 - | 1 - | 0,000113623 - | 1 - | 0,000127995 - |
| 8582 | 1 | 3um_FIJJAFBF_179844 | Bacteria | Bacteroidetes | Flavobacteriia | Flavobacterial | Flavobacteriac | Flavivirga | Flavivirga eckl | metabolism | Energy produc | COG0711 | F(1)F(0) ATP s | 391603.FBALC | 7,4 | 1,356465997 | 0,000600651 | 4,73E-05 | 10 - | 0,000691426 - | 7 - | 0,000600983 - | 7 - | 0,000571311 - | 7 - | 0,000579749 - | 6 - | 0,000559787 - |
| 8583 | 1 | 0.2um_NOKCPHJN_191603 | Eukaryota | Bacillariophyt | Bacillariophyc | Naviculales | Phaeodactylac | Phaeodactylu | Phaeodactylu | cellular proces | Cell cycle cont | 0IHFW | regulator of ch | 2850.Phatr31 | 1,8 | 0,4 | 3,92E-05 | 1,08E-05 | 1 - | 1,79E-05 - | 2 - | 4,45E-05 - | 2 - | 4,23E-05 - | 2 - | 4,29E-05 - | 2 - | 4,83E-05 - |
| 8584 | 1 | 0.2um_NOKCPHJN_277085 | Eukaryota | Phaeophyceae | unclassified | Ectocarpales | Ectocarpaceae | Ectocarpus | Ectocarpus sil | poorly charact | Function unkn | COG1100 | GTP-binding P | 2850.Phatr86 | 0,4 | 0,489897949 | 2,72E-05 | 3,35E-05 | 1 - | 6,24E-05 - | 0 - | 0 - | 1 - | 7,36E-05 - | 0 - | 0 - | 0 - | 0 - |
| 8585 | 1 | 0.2um_NOKCPHJN_515849 | Bacteria | Bacteroidetes | Flavobacteriia | Flavobacterial | Flavobacteriac | Formosa | Formosa sp. H | unclassified | unclassified | unclassified | unclassified | unclassified | 5,8 | 0,4 | 0,000399604 | 5,99E-05 | 5 - | 0,000286942 - | 6 - | 0,000427557 - | 6 - | 0,000406447 - | 6 - | 0,00041245 - | 6 - | 0,000464623 - |
| 8586 | 1 | 0.2um_NOKCPHJN_123233 | Bacteria | Bacteroidetes | unclassified | unclassified | unclassified | unclassified | Bacteroidetes | poorly charact | Function unkn | 0YECN | MAM domain | 926562.Oweh | 5 | 1,095445115 | 4,29E-05 | 1,01E-05 | 5 - | 3,60E-05 - | 4 - | 3,57E-05 - | 4 - | 3,40E-05 - | 7 - | 6,03E-05 - | 5 - | 4,85E-05 - |
| 8587 | 1 | 0.2um_NOKCPHJN_80879 | Bacteria | Proteobacteri | Gammaprote | Cellvibrionale | Porticoccacea | unclassified | Gammaprote | cellular proces | Cell wall/mem | 0QKII | polysaccharide | 377629.TERTU | 1,2 | 0,748331477 | 1,89E-05 | 1,14E-05 | 2 - | 2,70E-05 - | 2 - | 3,35E-05 - | 1 - | 1,59E-05 - | 0 - | 0 - | 1 - | 1,82E-05 - |
| 8588 | 1 | 0.2um_NOKCPHJN_306283 | Bacteria | Bacteroidetes | Flavobacteriia | Flavobacterial | Flavobacteriac | Winogradskye | Winogradskye | unclassified | unclassified | unclassified | unclassified | unclassified | 3,8 | 1,166190379 | 6,40E-05 | 1,84E-05 | 4 - | 5,73E-05 - | 3 - | 5,34E-05 - | 5 - | 8,46E-05 - | 5 - | 8,58E-05 - | 2 - | 3,87E-05 - |
| 8589 | 1 | 0.2um_NOKCPHJN_101862 | Bacteria | Proteobacteri | Gammaprote | Cellvibrionale | Porticoccacea | unclassified | gamma prote | unclassified | unclassified | unclassified | unclassified | unclassified | 2 | 0,632455532 | 3,01E-05 | 1,16E-05 | 2 - | 2,49E-05 - | 2 - | 3,09E-05 - | 1 - | 1,47E-05 - | 2 - | 2,98E-05 - | 3 - | 5,03E-05 - |
| 8590 | 1 | 3um_FIJJAFBF_143869 | Bacteria | Proteobacteri | Gammaprote | Oceanospirilla | Saccharospirill | Reinekea | Reinekea blan | unclassified | unclassified | unclassified | unclassified | unclassified | 4,6 | 1,2 | 0,000139487 | 2,97E-05 | 6 - | 0,000154755 - | 4 - | 0,000128107 - | 6 - | 0,000182673 - | 3 - | 9,27E-05 - | 4 - | 0,000139213 - |
| 8591 | 1 | 0.2um_NOKCPHJN_825697 | Bacteria | Bacteroidetes | Sphingobacter | Sphingobacter | Sphingobacter | Pedobacter | Pedobacter ar | unclassified | unclassified | unclassified | unclassified | unclassified | 1,4 | 0,489897949 | 4,87E-05 | 1,43E-05 | 2 - | 5,96E-05 - | 1 - | 3,70E-05 - | 1 - | 3,52E-05 - | 2 - | 7,14E-05 - | 1 - | 4,02E-05 - |
| 8592 | 1 | 3um_FIJJAFBF_788045 | Bacteria | Proteobacteri | Alphaproteob | Rhodobactera | unclassified | unclassified | Rhodobactera | information st | Translation, ril | 17C3B | The globular d | 391619.RGBS1 | 4,8 | 0,4 | 0,000522253 | 6,91E-05 | 5 - | 0,000455463 - | 5 - | 0,000565551 - | 4 - | 0,000430102 - | 5 - | 0,000545569 - | 5 - | 0,000614581 - |
| 8593 | 1 | 0.2um_NOKCPHJN_128516 | Bacteria | Bacteroidetes | Cytophagia | Cytophagales | Flammeovirga | unclassified | Flammeovirga | poorly charact | Function unkn | 11Q9F | Protein of unk | 388413.ALPR1 | 1,4 | 0,8 | 0,000103437 | 5,99E-05 | 2 - | 0,000124083 - | 1 - | 7,70E-05 - | 0 - | 0 - | 2 - | 0,000148631 - | 2 - | 0,000167432 - |
| 8594 | 1 | 0.2um_NOKCPHJN_505398 | unclassified | unclassified | unclassified | unclassified | unclassified | unclassified | unclassified | unclassified | unclassified | unclassified | unclassified | unclassified | 0,6 | 0,489897949 | 0,000147204 | 0,000122408 | 1 - | 0,000204958 - | 1 - | 0,000254498 - | 0 - | 0 - | 0 - | 0 - | 1 - | 0,000276562 - |
| 8595 | 1 | 0.2um_NOKCPHJN_1028808 | unclassified | unclassified | unclassified | unclassified | unclassified | unclassified | unclassified | unclassified | unclassified | unclassified | unclassified | unclassified | 0,6 | 0,489897949 | 3,92E-05 | 3,22E-05 | 1 - | 5,80E-05 - | 0 - | 0 - | 1 - | 6,84E-05 - | 1 - | 6,94E-05 - | 0 - | 0 - |
| 8596 | 1 | 0.2um_NOKCPHJN_815566 | Bacteria | Proteobacteri | Alphaproteob | Rhodobactera | Rhodobactera | Planktomarina | Planktomarina | information st | Translation, ril | 179R6 | This protein is | 391613.RTM1 | 7 | 0,632455532 | 0,000615122 | 8,24E-05 | 7 - | 0,000515024 - | 7 - | 0,000639508 - | 6 - | 0,000521085 - | 8 - | 0,000705043 - | 7 - | 0,00069495 - |
| 8597 | 1 | 3um_FIJJAFBF_739700 | Bacteria | Proteobacteri | Gammaprote | Cellvibrionale | Spongiibacter | Oceanicoccus | Oceanicoccus | metabolism | Energy produc | COG0055 | Produces ATP | 247634.GPB2 | 0,4 | 0,489897949 | 1,08E-05 | 1,34E-05 | 1 - | 2,47E-05 - | 0 - | 0 - | 0 - | 0 - | 1 - | 2,96E-05 - | 0 - | 0 - |
| 8598 | 1 | 3um_FIJJAFBF_276045 | Bacteria | Bacteroidetes | Flavobacteriia | Flavobacterial | Flavobacteriac | Polaribacter | various | metabolism | Amino acid tra | 09F6N | The glycine cle | 487797.Flav3 | 0,2 | 0,4 | 1,82E-05 | 3,64E-05 | 1 - | 9,11E-05 - | 0 - | 0 - | 0 - | 0 - | 0 - | 0 - | 0 - | 0 - |
| 8599 | 1 | 0.2um_NOKCPHJN_590601 | Bacteria | Proteobacteri | Alphaproteob | Pelagibacteral | Pelagibacterac | Candidatus Pe | Candidatus Pe | metabolism | Secondary me | 01Q86 | extracellular s | 439493.PB721 | 1,4 | 1,019803903 | 7,02E-05 | 5,27E-05 | 1 - | 4,27E-05 - | 2 - | 0,000105962 - | 3 - | 0,000151095 - | 1 - | 5,11E-05 - | 0 - | 0 - |
| 8600 | 1 | 3um_FIJJAFBF_777848 | Bacteria | Bacteroidetes | Flavobacteriia | Flavobacterial | Flavobacteriac | Polaribacter | Polaribacter s | information st | Transcription | 0P88Z | transcriptiona | 313598.MED1 | 4,6 | 0,489897949 | 0,000396568 | 5,97E-05 | 5 - | 0,000360933 - | 5 - | 0,000448173 - | 4 - | 0,000340836 - | 4 - | 0,00034587 - | 5 - | 0,000487027 - |
| 8601 | 2 | 0.2um_NOKCPHJN_241610;3um_FIJJAFBF_7846 | Bacteria | Proteobacteri | Alphaproteob | Pelagibacteral | Pelagibacterac | Candidatus Pe | Candidatus Pe | information st | Translation, ril | 01WSS | Interacts with various | | 2 | 0 | 0,000222808 | 2,12E-05 | 2 - | 0,000186629 - | 2 - | 0,000231738 - | 2 - | 0,000220296 - | 2 - | 0,00022355 - | 2 - | 0,000251828 - |
| 8602 | 1 | 0.2um_NOKCPHJN_57058 | Bacteria | Proteobacteri | Gammaprote | Oceanospirilla | Saccharospirill | Reinekea | Reinekea blan | poorly charact | Function unkn | 05DIH | basic membra | 314283.MED2 | 6,2 | 1,166190379 | 0,00025361 | 4,01E-05 | 8 - | 0,000276571 - | 7 - | 0,000300492 - | 5 - | 0,000204039 - | 5 - | 0,000207053 - | 6 - | 0,000279894 - |
| 8603 | 1 | 0.2um_NOKCPHJN_10097 | Bacteria | Bacteroidetes | Flavobacteriia | Flavobacterial | Flavobacteriac | Polaribacter | Polaribacter ir | unclassified | unclassified | unclassified | unclassified | unclassified | 3,4 | 0,489897949 | 7,36E-05 | 1,26E-05 | 4 - | 7,26E-05 - | 3 - | 6,77E-05 - | 3 - | 6,43E-05 - | 3 - | 6,53E-05 - | 4 - | 9,80E-05 - |
| 8604 | 1 | 3um_FIJJAFBF_198300 | Bacteria | Bacteroidetes | Flavobacteriia | Flavobacterial | Flavobacteriac | Polaribacter | Polaribacter s | metabolism | Inorganic ion t | 05C27 | TonB-depende | 487797.Flav3 | 14,4 | 1,019803903 | 0,000381336 | 4,20E-05 | 14 - | 0,000310807 - | 16 - | 0,000441064 - | 15 - | 0,000393082 - | 14 - | 0,000372295 - | 13 - | 0,000389433 - |
| 8605 | 1 | 3um_FIJJAFBF_560873 | Bacteria | Proteobacteri | Gammaprote | Cellvibrionale | Porticoccacea | unclassified | Gammaprote | metabolism | Inorganic ion t | 16TQC | TONB-depend | 314287.GB22 | 1,4 | 0,8 | 1,73E-05 | 9,90E-06 | 2 - | 2,15E-05 - | 2 - | 2,67E-05 - | 2 - | 2,54E-05 - | 1 - | 1,29E-05 - | 0 - | 0 - |
| 8606 | 1 | 0.2um_NOKCPHJN_418376 | Bacteria | Proteobacteri | Alphaproteob | Pelagibacteral | Pelagibacterac | Candidatus Pe | Candidatus Pe | information st | Replication, re | COG0178 | The UvrABC re | 439493.PB721 | 0,2 | 0,4 | 2,39E-06 | 4,79E-06 | 1 - | 1,20E-05 - | 0 - | 0 - | 0 - | 0 - | 0 - | 0 - | 0 - | 0 - |
| 8607 | 1 | 0.2um_NOKCPHJN_1025171 | Bacteria | Proteobacteri | Alphaproteob | Pelagibacteral | Pelagibacterac | Candidatus Pe | Candidatus Pe | metabolism | Energy produc | COG3474 | Electron carrie | 335992.SAR11 | 2,4 | 0,489897949 | 0,000171696 | 2,67E-05 | 3 - | 0,000182185 - | 2 - | 0,000150814 - | 2 - | 0,000143367 - | 3 - | 0,000218227 - | 2 - | 0,000163888 - |
| 8608 | 1 | 0.2um_NOKCPHJN_77522 | Bacteria | unclassified | unclassified | unclassified | unclassified | unclassified | bacterium BM | unclassified | unclassified | unclassified | unclassified | unclassified | 2 | 0 | 8,25E-05 | 7,84E-06 | 2 - | 6,91E-05 - | 2 - | 8,59E-05 - | 2 - | 8,16E-05 - | 2 - | 8,28E-05 - | 2 - | 9,33E-05 - |
| 8609 | 1 | 3um_FIJJAFBF_189609 | Bacteria | Bacteroidetes | Flavobacteriia | Flavobacterial | Flavobacteriac | Polaribacter | Polaribacter s | unclassified | unclassified | unclassified | unclassified | unclassified | 3,2 | 1,166190379 | 0,000214542 | 7,20E-05 | 5 - | 0,000284101 - | 3 - | 0,000211662 - | 2 - | 0,000134141 - | 2 - | 0,000136122 - | 4 - | 0,000306682 - |
| 8610 | 1 | 3um_FIJJAFBF_513826 | Viruses | unclassified | unclassified | unclassified | unclassified | unclassified | uncultured M | unclassified | unclassified | unclassified | unclassified | unclassified | 8,2 | 0,979795897 | 0,000361566 | 2,57E-05 | 10 - | 0,000372652 - | 7 - | 0,000323907 - | 8 - | 0,000351902 - | 8 - | 0,000357099 - | 8 - | 0,000402271 - |
| 8611 | 1 | 0.2um_NOKCPHJN_257141 | Bacteria | Bacteroidetes | Flavobacteriia | Flavobacterial | Flavobacteriac | Formosa | Formosa sp. H | unclassified | unclassified | unclassified | unclassified | unclassified | 17,6 | 1,019803903 | 0,0002847 | 3,51E-05 | 16 - | 0,000216305 - | 18 - | 0,00030216 - | 18 - | 0,000287241 - | 19 - | 0,000307678 - | 17 - | 0,000310114 - |
| 8612 | 1 | 3um_FIJJAFBF_02498 | Bacteria | Proteobacteri | Alphaproteob | Rhodobactera | Rhodobactera | Pseudooceani | Pseudooceani | information st | Translation, ril | 16Q16 | Binds the lowe | 644076.SCH4 | 1 | 0 | 5,83E-05 | 5,54E-06 | 1 - | 4,88E-05 - | 1 - | 6,06E-05 - | 1 - | 5,77E-05 - | 1 - | 5,85E-05 - | 1 - | 6,59E-05 - |
| 8613 | 1 | 0.2um_NOKCPHJN_60996 | Bacteria | Proteobacteri | Gammaprote | Cellvibrionale | Porticoccacea | unclassified | Gammaprote | unclassified | unclassified | unclassified | unclassified | unclassified | 0,8 | 0,4 | 5,87E-05 | 3,01E-05 | 1 - | 6,20E-05 - | 0 - | 0 - | 1 - | 7,32E-05 - | 1 - | 7,43E-05 - | 1 - | 8,37E-05 - |
| 8614 | 1 | 3um_FIJJAFBF_328085 | unclassified | unclassified | unclassified | unclassified | unclassified | unclassified | unclassified | unclassified | unclassified | unclassified | unclassified | unclassified | 0,2 | 0,4 | 7,51E-06 | 1,50E-05 | 0 - | 0 - | 0 - | 0 - | 0 - | 0 - | 1 - | 3,76E-05 - | 0 - | 0 - |
| 8615 | 1 | 0.2um_NOKCPHJN_81099 | Bacteria | Proteobacteri | Gammaprote | unclassified | unclassified | unclassified | Gammaprote | metabolism | Energy produc | COG0055 | Produces ATP | 498211.CJA_3 | 2,6 | 0,489897949 | 7,78E-05 | 1,72E-05 | 3 - | 7,52E-05 - | 3 - | 9,34E-05 - | 2 - | 5,92E-05 - | 2 - | 6,00E-05 - | 3 - | 0,000101446 - |
| 8616 | 1 | 3um_FIJJAFBF_457883 | Bacteria | Proteobacteri | Gammaprote | Cellvibrionale | Porticoccacea | unclassified | gamma prote | cellular proces | Cell wall/mem | 0QKXZ | Porin Gram-ne | 314287.GB22 | 0,8 | 0,748331477 | 3,52E-05 | 3,24E-05 | 0 - | 0 - | 0 - | 0 - | 1 - | 4,22E-05 - | 2 - | 8,57E-05 - | 1 - | 4,82E-05 - |
| 8617 | 1 | 3um_FIJJAFBF_122322 | unclassified | unclassified | unclassified | unclassified | unclassified | unclassified | unclassified | unclassified | unclassified | unclassified | unclassified | unclassified | 2,2 | 0,4 | 7,96E-05 | 7,30E-06 | 3 - | 9,23E-05 - | 2 - | 7,64E-05 - | 2 - | 7,26E-05 - | 2 - | 7,37E-05 - | 2 - | 8,30E-05 - |
| 8618 | 1 | 3um_FIJJAFBF_350085 | Bacteria | Proteobacteri | Alphaproteob | Rhodobactera | Rhodobactera | Ruegeria | Ruegeria aren | information st | Translation, ril | 01RJH | 30S ribosomal | 314270.RB208 | 2,8 | 1,166190379 | 0,000149321 | 6,34E-05 | 2 - | 9,00E-05 - | 3 - | 0,000167669 - | 4 - | 0,000212521 - | 4 - | 0,00021566 - | 1 - | 6,07E-05 - |
| 8619 | 1 | 3um_FIJJAFBF_237053 | Eukaryota | unclassified | Synurophycea | Parmales | Triparmaceae | Triparma | Triparma laevi | metabolism | Amino acid tra | 0FJ71 | Acetolactate s | 329726.AM1_ | 1 | 0,632455532 | 8,31E-05 | 5,86E-05 | 1 - | 6,79E-05 - | 1 - | 8,43E-05 - | 1 - | 8,02E-05 - | 0 - | 0 - | 2 - | 0,000183283 - |
| 8620 | 1 | 0.2um_NOKCPHJN_259107 | Eukaryota | Nematoda | Chromadorea | Diplogasterida | Neodiplogaste | Pristionchus | Pristionchus p | unclassified | unclassified | unclassified | unclassified | unclassified | 0,2 | 0,4 | 6,36E-07 | 1,27E-06 | 1 - | 3,18E-06 - | 0 - | 0 - | 0 - | 0 - | 0 - | 0 - | 0 - | 0 - |
| 8621 | 2 | 0.2um_NOKCPHJN_642868;3um_FIJJAFBF_66387 | unclassified | unclassified | unclassified | unclassified | unclassified | unclassified | unclassified | unclassified | unclassified | unclassified | unclassified | unclassified | 0,8 | 0,748331477 | 2,69E-05 | 2,64E-05 | 1 - | 2,87E-05 - | 2 - | 7,13E-05 - | 0 - | 0 - | 1 - | 3,44E-05 - | 0 - | 0 - |
| 8622 | 1 | 3um_FIJJAFBF_223667 | Bacteria | Proteobacteri | Alphaproteob | Rhodobactera | Rhodobactera | Planktomarina | Planktomarina | information st | Translation, ril | COG0251 | Endoribonucle | 314270.RB208 | 1,8 | 0,4 | 0,000160102 | 3,92E-05 | 2 - | 0,000149061 - | 2 - | 0,000185089 - | 2 - | 0,000175951 - | 1 - | 8,93E-05 - | 2 - | 0,000201136 - |
| 8623 | 1 | 3um_FIJJAFBF_303919 | Bacteria | Bacteroidetes | Flavobacteriia | Flavobacterial | Crocinitomica | unclassified | Crocinitomica | cellular proces | Cell wall/mem | 0P88B | Outer membra | 391598.FBBAL | 2 | 0,894427191 | 5,75E-05 | 2,47E-05 | 3 - | 7,31E-05 - | 3 - | 9,08E-05 - | 1 - | 2,88E-05 - | 1 - | 2,92E-05 - | 2 - | 6,58E-05 - |
| 8624 | 1 | 0.2um_NOKCPHJN_461489 | Bacteria | Bacteroidetes | Flavobacteriia | unclassified | unclassified | unclassified | Flavobacteria | poorly charact | Function unkn | 05DPC | Ragb susd dor | 487797.Flav3 | 13,4 | 2,059126028 | 0,000360326 | 5,02E-05 | 14 - | 0,000317564 - | 13 - | 0,000366155 - | 17 - | 0,000455177 - | 12 - | 0,000326047 - | 11 - | 0,000336684 - |
| 8625 | 3 | 3um_FIJJAFBF_156791;3um_FIJJAFBF_208153;3u | Bacteria | Bacteroidetes | Flavobacteriia | Flavobacterial | Flavobacteriac | Formosa | Formosa sp. H | unclassified | unclassified | unclassified | unclassified | unclassified | 1,2 | 0,4 | 0,000145439 | 3,22E-05 | 2 - | 0,000208685 - | 1 - | 0,000129563 - | 1 - | 0,000123166 - | 1 - | 0,000124985 - | 1 - | 0,000140795 - |
| 8626 | 1 | 3um_FIJJAFBF_420941 | Bacteria | Proteobacteri | Gammaprote | Cellvibrionale | Porticoccacea | unclassified | SAR92 bacteri | information st | Translation, ril | 179ZI | One of the pri | 314287.GB22 | 4 | 0,632455532 | 0,000418946 | 6,41E-05 | 4 - | 0,000353159 - | 4 - | 0,00043852 - | 4 - | 0,000416868 - | 5 - | 0,000528782 - | 3 - | 0,000357403 - |
| 8627 | 1 | 0.2um_NOKCPHJN_1116566 | unclassified | unclassified | unclassified | unclassified | unclassified | unclassified | unclassified | unclassified | unclassified | unclassified | unclassified | unclassified | 0,6 | 0,489897949 | 2,41E-05 | 2,00E-05 | 1 - | 3,40E-05 - | 0 - | 0 - | 0 - | 0 - | 1 - | 4,07E-05 - | 1 - | 4,58E-05 - |
| 8628 | 1 | 3um_FIJJAFBF_385223 | Bacteria | Proteobacteri | Alphaproteob | Rhodobactera | Rhodobactera | Thalassococcu | Thalassococcu | cellular proces | Cell wall/mem | 16QAK | protease | 388399.SSE37 | 3,6 | 1,019803903 | 0,000111548 | 3,72E-05 | 4 - | 0,00010317 - | 3 - | 9,61E-05 - | 2 - | 6,09E-05 - | 4 - | 0,00012358 - | 5 - | 0,000174016 - |
| 8629 | 1 | 3um_FIJJAFBF_293513 | Bacteria | Bacteroidetes | Cytophagia | Cytophagales | Flammeovirga | unclassified | Flammeovirga | poorly charact | Function unkn | 0900R | pur-alpha bet | 269798.CHU_ | 0,6 | 0,489897949 | 7,93E-05 | 6,59E-05 | 1 - | 0,000110362 - | 1 - | 0,000137037 - | 0 - | 0 - | 0 - | 0 - | 1 - | 0,000148918 - |
| 8630 | 1 | 3um_FIJJAFBF_788033 | Bacteria | Proteobacteri | Alphaproteob | Rhodobactera | unclassified | unclassified | Rhodobactera | information st | Translation, ril | 05VBI | Binds 16S rRN | 633131.TR2A | 1,4 | 1,019803903 | 0,000182124 | 0,000119612 | 3 - | 0,000340921 - | 2 - | 0,000282216 - | 1 - | 0,000134141 - | 0 - | 0 - | 1 - | 0,000153341 - |
| 8631 | 1 | 3um_FIJJAFBF_510697 | unclassified | unclassified | unclassified | unclassified | unclassified | unclassified | unclassified | unclassified | unclassified | unclassified | unclassified | unclassified | 0,4 | 0,489897949 | 4,33E-05 | 5,32E-05 | 0 - | 0 - | 0 - | 0 - | 0 - | 0 - | 1 - | 0,000101839 - | 1 - | 0,000114722 - |
| 8632 | 1 | 0.2um_NOKCPHJN_26721 | Bacteria | Proteobacteri | Betaproteoba | unclassified | unclassified | unclassified | Betaproteoba | metabolism | Energy produc | COG0372 | citrate syntha | 296591.Bpro_ | 1 | 0 | 3,14E-05 | 2,98E-06 | 1 - | 2,63E-05 - | 1 - | 3,27E-05 - | 1 - | 3,11E-05 - | 1 - | 3,15E-05 - | 1 - | 3,55E-05 - |
| 8633 | 1 | 3um_FIJJAFBF_712819 | Bacteria | Proteobacteri | Gammaprote | Oceanospirilla | Saccharospirill | Reinekea | Reinekea fors | cellular proces | Defense mech | 07SUK | Efflux transpo | 314283.MED2 | 0,4 | 0,489897949 | 1,12E-05 | 1,39E-05 | 1 - | 2,50E-05 - | 1 - | 3,10E-05 - | 0 - | 0 - | 0 - | 0 - | 0 - | 0 - |
| 8634 | 1 | 0.2um_NOKCPHJN_892527 | Bacteria | Bacteroidetes | Flavobacteriia | Flavobacterial | Flavobacteriac | Polaribacter | Polaribacter h | unclassified | unclassified | unclassified | unclassified | unclassified | 5,4 | 0,489897949 | 0,000156936 | 2,12E-05 | 5 - | 0,000121586 - | 6 - | 0,000181168 - | 6 - | 0,000172223 - | 5 - | 0,000145639 - | 5 - | 0,000164062 - |
| 8635 | 1 | 3um_FIJJAFBF_410806 | Bacteria | Proteobacteri | Gammaprote | Oceanospirilla | Oceanospirilla | Marinobacteri | Marinobacter | unclassified | unclassified | unclassified | unclassified | unclassified | 1 | 1,095445115 | 3,34E-05 | 3,56E-05 | 0 - | 0 - | 0 - | 0 - | 3 - | 9,72E-05 - | 1 - | 3,29E-05 - | 1 - | 3,71E-05 - |
| 8636 | 1 | 3um_FIJJAFBF_383274 | Eukaryota | Phaeophyceae | unclassified | Ectocarpales | Ectocarpaceae | Ectocarpus | Ectocarpus sil | cellular proces | Signal transdu | KOG0613 | fibronectin ty | 65071.PYU1_ | 0,8 | 0,748331477 | 1,66E-05 | 1,63E-05 | 1 - | 1,77E-05 - | 2 - | 4,41E-05 - | 0 - | 0 - | 1 - | 2,12E-05 - | 0 - | 0 - |
| 8637 | 1 | 3um_FIJJAFBF_103115 | Bacteria | Bacteroidetes | Flavobacteriia | Flavobacterial | Flavobacteriac | Polaribacter | Polaribacter s | unclassified | unclassified | unclassified | unclassified | unclassified | 2,4 | 1,019803903 | 2,92E-05 | 1,26E-05 | 3 - | 3,07E-05 - | 4 - | 5,09E-05 - | 1 - | 1,21E-05 - | 2 - | 2,45E-05 - | 2 - | 2,76E-05 - |
| 8638 | 1 | 0.2um_NOKCPHJN_224039 | Bacteria | unclassified | unclassified | unclassified | unclassified | unclassified | uncultured ba | unclassified | unclassified | unclassified | unclassified | unclassified | 6,6 | 1,959591794 | 0,00017436 | 3,94E-05 | 10 - | 0,000225495 - | 7 - | 0,000195998 - | 6 - | 0,000159704 - | 4 - | 0,000108042 - | 6 - | 0,000182563 - |
| 8639 | 1 | 3um_FIJJAFBF_642472 | Bacteria | Proteobacteri | Alphaproteob | Rhodobactera | Rhodobactera | Aestuariivita | Aestuariivita b | unclassified | unclassified | unclassified | unclassified | unclassified | 0,2 | 0,4 | 7,06E-06 | 1,41E-05 | 1 - | 3,53E-05 - | 0 - | 0 - | 0 - | 0 - | 0 - | 0 - | 0 - | 0 - |
| 8640 | 1 | 0.2um_NOKCPHJN_217489 | Bacteria | Proteobacteri | Gammaprote | Oceanospirilla | Saccharospirill | Reinekea | Reinekea fors | metabolism | Inorganic ion t | 0BF8H | Bacterial phos | 391597.LMED | 4,6 | 0,489897949 | 0,000281492 | 4,23E-05 | 5 - | 0,000256198 - | 5 - | 0,000318122 - | 4 - | 0,000241933 - | 4 - | 0,000245506 - | 5 - | 0,000345702 - |
| 8641 | 1 | 3um_FIJJAFBF_255679 | Bacteria | Bacteroidetes | Flavobacteriia | Flavobacterial | Flavobacteriac | Polaribacter | Polaribacter s | unclassified | unclassified | unclassified | unclassified | unclassified | 0,4 | 0,489897949 | 2,57E-05 | 3,16E-05 | 0 - | 0 - | 0 - | 0 - | 1 - | 5,99E-05 - | 0 - | 0 - | 1 - | 6,85E-05 - |
| 8642 | 1 | 3um_FIJJAFBF_494368 | Bacteria | Bacteroidetes | Flavobacteriia | Flavobacterial | Flavobacteriac | Polaribacter | Polaribacter s | unclassified | unclassified | unclassified | unclassified | unclassified | 0,6 | 0,489897949 | 1,51E-05 | 1,24E-05 | 1 - | 2,24E-05 - | 0 - | 0 - | 1 - | 2,64E-05 - | 1 - | 2,68E-05 - | 0 - | 0 - |
| 8643 | 1 | 0.2um_NOKCPHJN_600705 | Bacteria | Proteobacteri | Alphaproteob | Rhodobactera | Rhodobactera | Planktomarina | Planktomarina | cellular proces | Defense mech | 07SUK | Efflux transpo | 391593.RCCS2 | 1,6 | 0,489897949 | 9,67E-05 | 3,20E-05 | 2 - | 0,000101125 - | 2 - | 0,000125567 - | 1 - | 5,97E-05 - | 1 - | 6,06E-05 - | 2 - | 0,000136453 - |
| 8644 | 1 | 0.2um_NOKCPHJN_364494 | Bacteria | Proteobacteri | Gammaprote | Cellvibrionale | Halieaceae | unclassified | Halieaceae ba | cellular proces | Posttranslatio | 0QJ3C | Acts as a chap | 566466.NOR5 | 0,2 | 0,4 | 4,83E-06 | 9,66E-06 | 0 - | 0 - | 0 - | 0 - | 0 - | 0 - | 0 - | 0 - | 1 - | 2,42E-05 - |
| 8645 | 1 | 0.2um_NOKCPHJN_96771 | Bacteria | Proteobacteri | Betaproteoba | Burkholderial | unclassified | unclassified | various | unclassified | unclassified | unclassified | unclassified | unclassified | 1,2 | 0,748331477 | 6,08E-05 | 3,65E-05 | 2 - | 8,66E-05 - | 2 - | 0,000107561 - | 1 - | 5,11E-05 - | 0 - | 0 - | 1 - | 5,84E-05 - |
| 8646 | 1 | 3um_FIJJAFBF_177315 | Bacteria | Actinobacteri | Actinobacteri | Micrococcales | Microbacteria | unclassified | Microbacteria | information st | Translation, ril | 00IDP | 30S ribosomal | 930171.Asphe | 2,6 | 0,489897949 | 0,000219111 | 4,67E-05 | 2 - | 0,00014083 - | 3 - | 0,000262305 - | 3 - | 0,000249354 - | 3 - | 0,000253037 - | 2 - | 0,00019003 - |
| 8647 | 1 | 3um_FIJJAFBF_24379 | Bacteria | Bacteroidetes | Flavobacteriia | Flavobacterial | Flavobacteriac | Algibacter | Algibacter sp. | unclassified | unclassified | unclassified | unclassified | unclassified | 19 | 1,414213562 | 0,00050565 | 7,34E-05 | 17 - | 0,000377409 - | 21 - | 0,000578897 - | 20 - | 0,000524109 - | 18 - | 0,000478665 - | 19 - | 0,000569171 - |
| 8648 | 1 | 0.2um_NOKCPHJN_316057 | unclassified | unclassified | unclassified | unclassified | unclassified | unclassified | unclassified | unclassified | unclassified | unclassified | unclassified | unclassified | 1,2 | 0,4 | 4,22E-06 | 1,41E-06 | 1 - | 2,95E-06 - | 1 - | 3,67E-06 - | 2 - | 6,97E-06 - | 1 - | 3,54E-06 - | 1 - | 3,98E-06 - |
| 8649 | 1 | 0.2um_NOKCPHJN_245930 | Bacteria | Bacteroidetes | Flavobacteriia | Flavobacterial | Flavobacteriac | unclassified | Flavobacteria | unclassified | unclassified | unclassified | unclassified | unclassified | 1,8 | 0,4 | 3,85E-05 | 9,43E-06 | 2 - | 3,58E-05 - | 2 - | 4,45E-05 - | 2 - | 4,23E-05 - | 1 - | 2,14E-05 - | 2 - | 4,83E-05 - |
| 8650 | 1 | 0.2um_NOKCPHJN_687428 | Bacteria | Proteobacteri | Alphaproteob | Pelagibacteral | Pelagibacterac | Candidatus Pe | Candidatus Pe | metabolism | Energy produc | 01RN9 | Subunits I and | 335992.SAR11 | 0,2 | 0,4 | 9,37E-06 | 1,87E-05 | 1 - | 4,68E-05 - | 0 - | 0 - | 0 - | 0 - | 0 - | 0 - | 0 - | 0 - |
| 8651 | 1 | 3um_FIJJAFBF_465354 | Bacteria | Bacteroidetes | Flavobacteriia | Flavobacterial | Crocinitomica | unclassified | Crocinitomica | cellular proces | Posttranslatio | 099Y4 | peptidase, M1 | 643867.Ftrac_ | 1 | 0 | 3,13E-05 | 2,97E-06 | 1 - | 2,62E-05 - | 1 - | 3,25E-05 - | 1 - | 3,09E-05 - | 1 - | 3,14E-05 - | 1 - | 3,54E-05 - |
| 8652 | 1 | 0.2um_NOKCPHJN_530253 | Bacteria | Bacteroidetes | Flavobacteriia | Flavobacterial | Flavobacteriac | Formosa | Formosa sp. H | unclassified | unclassified | unclassified | unclassified | unclassified | 0,2 | 0,4 | 2,93E-06 | 5,85E-06 | 0 - | 0 - | 0 - | 0 - | 1 - | 1,46E-05 - | 0 - | 0 - | 0 - | 0 - |
| 8653 | 1 | 3um_FIJJAFBF_308824 | Bacteria | Proteobacteri | Alphaproteob | Pelagibacteral | Pelagibacterac | unclassified | Pelagibactera | unclassified | unclassified | unclassified | unclassified | unclassified | 11,8 | 1,166190379 | 0,000491051 | 6,73E-05 | 13 - | 0,000453525 - | 11 - | 0,000476507 - | 10 - | 0,0004118 - | 12 - | 0,000501459 - | 13 - | 0,000611966 - |
| 8654 | 1 | 3um_FIJJAFBF_579507 | Bacteria | Proteobacteri | Deltaproteoba | Desulfuromon | Geobacterace | Geobacter | Geobacter ura | cellular proces | Posttranslatio | 17FI0 | Binds to Cpn6 | 351605.Gura_ | 0,2 | 0,4 | 2,39E-05 | 4,78E-05 | 1 - | 0,000119559 - | 0 - | 0 - | 0 - | 0 - | 0 - | 0 - | 0 - | 0 - |
| 8655 | 2 | 0.2um_NOKCPHJN_329648;3um_FIJJAFBF_14239 | Bacteria | Proteobacteri | Alphaproteob | Rhodobactera | Rhodobactera | Actibacterium | Actibacterium | unclassified | unclassified | unclassified | unclassified | unclassified | 0,8 | 0,4 | 5,45E-05 | 2,80E-05 | 1 - | 5,77E-05 - | 0 - | 0 - | 1 - | 6,81E-05 - | 1 - | 6,91E-05 - | 1 - | 7,78E-05 - |
| 8656 | 1 | 3um_FIJJAFBF_44055 | Viruses | unclassified | unclassified | unclassified | various | various | various | unclassified | unclassified | unclassified | unclassified | unclassified | 8 | 1,095445115 | 0,000607712 | 0,000108477 | 7 - | 0,000443888 - | 8 - | 0,000629918 - | 10 - | 0,000748521 - | 7 - | 0,000531703 - | 8 - | 0,000684528 - |
| 8657 | 1 | 3um_FIJJAFBF_119765 | Bacteria | Bacteroidetes | Flavobacteriia | Flavobacterial | Flavobacteriac | Tamlana | Tamlana sp. U | unclassified | unclassified | unclassified | unclassified | unclassified | 1 | 0 | 0,000161208 | 1,53E-05 | 1 - | 0,000135031 - | 1 - | 0,000167669 - | 1 - | 0,000159391 - | 1 - | 0,000161745 - | 1 - | 0,000182205 - |
| 8658 | 1 | 0.2um_NOKCPHJN_416918 | Bacteria | Proteobacteri | Alphaproteob | Pelagibacteral | Pelagibacterac | Candidatus Pe | Candidatus Pe | unclassified | unclassified | unclassified | unclassified | unclassified | 1,8 | 0,4 | 9,39E-05 | 1,85E-05 | 2 - | 8,86E-05 - | 2 - | 0,000110053 - | 2 - | 0,000104619 - | 2 - | 0,000106165 - | 1 - | 5,98E-05 - |
| 8659 | 1 | 3um_FIJJAFBF_135696 | Bacteria | Bacteroidetes | Flavobacteriia | Flavobacterial | Flavobacteriac | Ulvibacter | Ulvibacter sp. | information st | Translation, ril | 05G6W | One of the pri | 983544.Lacal_ | 3 | 0 | 0,000204518 | 1,94E-05 | 3 - | 0,000171309 - | 3 - | 0,000212715 - | 3 - | 0,000202212 - | 3 - | 0,000205199 - | 3 - | 0,000231156 - |
| 8660 | 1 | 0.2um_NOKCPHJN_72816 | Bacteria | Proteobacteri | Gammaprote | Oceanospirilla | Saccharospirill | Reinekea | Reinekea blan | unclassified | unclassified | unclassified | unclassified | unclassified | 8,6 | 1,019803903 | 0,000271785 | 3,49E-05 | 10 - | 0,000265687 - | 9 - | 0,000296914 - | 8 - | 0,000250893 - | 7 - | 0,000222774 - | 9 - | 0,000322655 - |
| 8661 | 1 | 3um_FIJJAFBF_323224 | Viruses | unclassified | unclassified | unclassified | Phycodnavirid | Prasinovirus | Bathycoccus s | unclassified | unclassified | unclassified | unclassified | unclassified | 1,2 | 0,979795897 | 4,33E-05 | 3,52E-05 | 1 - | 3,04E-05 - | 1 - | 3,77E-05 - | 3 - | 0,000107526 - | 0 - | 0 - | 1 - | 4,10E-05 - |
| 8662 | 1 | 0.2um_NOKCPHJN_435179 | Bacteria | Proteobacteri | Gammaprote | unclassified | unclassified | unclassified | Gammaprote | unclassified | unclassified | unclassified | unclassified | unclassified | 2 | 0 | 0,000145002 | 1,38E-05 | 2 - | 0,000121457 - | 2 - | 0,000150814 - | 2 - | 0,000143367 - | 2 - | 0,000145485 - | 2 - | 0,000163888 - |
| 8663 | 1 | 3um_FIJJAFBF_737460 | Bacteria | Proteobacteri | Gammaprote | Oceanospirilla | Saccharospirill | Reinekea | Reinekea fors | unclassified | unclassified | unclassified | unclassified | unclassified | 3 | 0,894427191 | 0,000317373 | 9,88E-05 | 4 - | 0,000355897 - | 2 - | 0,000220959 - | 2 - | 0,00021005 - | 3 - | 0,000319729 - | 4 - | 0,000480231 - |
| 8664 | 1 | 3um_FIJJAFBF_282006 | Bacteria | Bacteroidetes | Flavobacteriia | Flavobacterial | Flavobacteriac | Psychroserper | Psychroserper | information st | Replication, re | COG0187 | DNA gyrase ne | 391603.FBALC | 3,8 | 0,748331477 | 8,04E-05 | 1,67E-05 | 4 - | 7,11E-05 - | 3 - | 6,62E-05 - | 5 - | 0,000104862 - | 3 - | 6,38E-05 - | 4 - | 9,59E-05 - |
| 8665 | 1 | 0.2um_NOKCPHJN_235595 | Bacteria | Proteobacteri | Gammaprote | Cellvibrionale | Porticoccacea | unclassified | gamma prote | information st | Transcription | COG0085 | DNA-depende | 314287.GB22 | 17,2 | 2,638181192 | 0,000169028 | 1,70E-05 | 22 - | 0,000182977 - | 18 - | 0,000185894 - | 15 - | 0,000147263 - | 15 - | 0,000149438 - | 16 - | 0,000179565 - |
| 8666 | 1 | 0.2um_NOKCPHJN_866594 | Bacteria | Proteobacteri | Alphaproteob | Pelagibacteral | Pelagibacterac | Candidatus Pe | Candidatus Pe | cellular proces | Cell wall/mem | 7257 | NA | 335992.SAR11 | 0,4 | 0,489897949 | 1,64E-05 | 2,03E-05 | 1 - | 3,66E-05 - | 1 - | 4,54E-05 - | 0 - | 0 - | 0 - | 0 - | 0 - | 0 - |
| 8667 | 1 | 3um_FIJJAFBF_135695 | Bacteria | Bacteroidetes | Flavobacteriia | Flavobacterial | Flavobacteriac | Ulvibacter | Ulvibacter sp. | unclassified | unclassified | unclassified | unclassified | unclassified | 3,8 | 1,326649916 | 0,000153955 | 4,15E-05 | 6 - | 0,000208685 - | 4 - | 0,00017275 - | 4 - | 0,000164221 - | 2 - | 8,33E-05 - | 3 - | 0,000140795 - |
| 8668 | 1 | 3um_FIJJAFBF_632631 | Bacteria | Proteobacteri | Alphaproteob | Rhodobactera | Rhodobactera | unclassified | Rhodobactera | metabolism | Inorganic ion t | 05ENH | Sulfatase | 391595.RLO14 | 1,4 | 1,019803903 | 3,59E-05 | 2,66E-05 | 2 - | 4,35E-05 - | 3 - | 8,10E-05 - | 1 - | 2,57E-05 - | 0 - | 0 - | 1 - | 2,93E-05 - |
| 8669 | 1 | 0.2um_NOKCPHJN_227645 | Eukaryota | Streptophyta | Bryopsida | Funariales | Funariaceae | Physcomitrell | Physcomitrell | information st | Replication, re | COG0258 | Structure-spec | 3218.PP1S10_ | 0,6 | 0,489897949 | 1,34E-05 | 1,10E-05 | 0 - | 0 - | 1 - | 2,19E-05 - | 0 - | 0 - | 1 - | 2,12E-05 - | 1 - | 2,38E-05 - |
| 8670 | 1 | 0.2um_NOKCPHJN_32624 | Bacteria | Verrucomicrol | Opitutae | unclassified | unclassified | unclassified | Opitutae bact | metabolism | Inorganic ion t | 177TK | phosphate-sel | 523791.Kkor_ | 4,8 | 0,748331477 | 0,000174533 | 2,42E-05 | 5 - | 0,000153445 - | 4 - | 0,000152427 - | 6 - | 0,000217351 - | 5 - | 0,000183801 - | 4 - | 0,000165641 - |
| 8671 | 1 | 3um_FIJJAFBF_56314 | Bacteria | Actinobacteri | Actinobacteri | Micrococcales | Microbacteria | Pontimonas | Pontimonas sa | metabolism | Amino acid tra | 00DZF | branched-chai | 312284.A20C1 | 2 | 0,632455532 | 6,53E-05 | 2,21E-05 | 1 - | 2,69E-05 - | 2 - | 6,69E-05 - | 3 - | 9,54E-05 - | 2 - | 6,45E-05 - | 2 - | 7,27E-05 - |
| 8672 | 1 | 3um_FIJJAFBF_208307 | Bacteria | Proteobacteri | Gammaprote | Oceanospirilla | Halomonadac | Salinicola | Salinicola peri | unclassified | unclassified | unclassified | unclassified | unclassified | 1,2 | 1,166190379 | 0,000117154 | 0,000113216 | 0 - | 0 - | 1 - | 0,000101077 - | 2 - | 0,000192173 - | 3 - | 0,000292518 - | 0 - | 0 - |
| 8673 | 1 | 3um_FIJJAFBF_66531 | Bacteria | Proteobacteri | Gammaprote | Oceanospirilla | Saccharospirill | Reinekea | Reinekea fors | metabolism | Energy produc | 16QM4 | alcohol dehyd | 314275.MADE | 1,6 | 0,8 | 6,60E-05 | 3,32E-05 | 1 - | 3,45E-05 - | 2 - | 8,56E-05 - | 3 - | 0,000122056 - | 1 - | 4,13E-05 - | 1 - | 4,65E-05 - |
| 8674 | 1 | 3um_FIJJAFBF_604554 | Bacteria | Proteobacteri | Gammaprote | Arenicellales | Arenicellaceae | Arenicella | Arenicella xan | information st | Translation, ril | 17FIH | One of the ear | 572477.Alvin_ | 0,4 | 0,489897949 | 5,05E-05 | 6,24E-05 | 1 - | 0,000112526 - | 1 - | 0,000139724 - | 0 - | 0 - | 0 - | 0 - | 0 - | 0 - |
| 8675 | 1 | 3um_FIJJAFBF_783318 | Eukaryota | unclassified | Pelagophycea | Pelagomonad | unclassified | Aureoumbra | Aureoumbra l | information st | Translation, ril | 05G6W | One of the pri | 696747.NIES3 | 0,6 | 0,489897949 | 4,02E-05 | 3,34E-05 | 1 - | 5,60E-05 - | 1 - | 6,95E-05 - | 0 - | 0 - | 0 - | 0 - | 1 - | 7,55E-05 - |
| 8676 | 1 | 3um_FIJJAFBF_226882 | Bacteria | Proteobacteri | Gammaprote | Alteromonada | Alteromonada | Paraglaciecola | various | information st | Transcription | 05VEQ | Cold shock pro | 983545.Glaag | 1,8 | 0,979795897 | 0,000345676 | 0,000177171 | 3 - | 0,000506368 - | 2 - | 0,000419173 - | 2 - | 0,000398477 - | 2 - | 0,000404363 - | 0 - | 0 - |
| 8677 | 1 | 0.2um_NOKCPHJN_397650 | Bacteria | Proteobacteri | Alphaproteob | Pelagibacteral | Pelagibacterac | unclassified | Pelagibactera | metabolism | Inorganic ion t | 16PII | TRAP transpor | 1002672.SAR1 | 0,4 | 0,489897949 | 1,57E-05 | 1,94E-05 | 1 - | 3,50E-05 - | 1 - | 4,35E-05 - | 0 - | 0 - | 0 - | 0 - | 0 - | 0 - |
| 8678 | 2 | 0.2um_NOKCPHJN_606024;3um_FIJJAFBF_26845 | Bacteria | Proteobacteri | Gammaprote | Cellvibrionale | Spongiibacter | Oceanicoccus | Oceanicoccus | cellular proces | Intracellular tr | COG2831 | Polypeptide-tr | 887062.HGR_ | 0,2 | 0,4 | 4,67E-06 | 9,34E-06 | 0 - | 0 - | 0 - | 0 - | 0 - | 0 - | 1 - | 2,33E-05 - | 0 - | 0 - |
| 8679 | 1 | 3um_FIJJAFBF_13867 | Bacteria | Proteobacteri | Alphaproteob | Rhodobactera | Rhodobactera | Planktomarina | Planktomarina | metabolism | Energy produc | 05FG6 | Dehydrogenas | 266779.Meso_ | 1 | 0,632455532 | 5,00E-05 | 2,82E-05 | 2 - | 8,73E-05 - | 0 - | 0 - | 1 - | 5,15E-05 - | 1 - | 5,23E-05 - | 1 - | 5,89E-05 - |
| 8680 | 1 | 3um_FIJJAFBF_163414 | Bacteria | Proteobacteri | Alphaproteob | Rhodospirillal | Rhodospirillac | unclassified | Rhodobactera | metabolism | Amino acid tra | 01Y25 | Nitrogen regu | 367336.OM22 | 0,8 | 0,4 | 8,72E-05 | 4,48E-05 | 1 - | 9,11E-05 - | 1 - | 0,00011311 - | 0 - | 0 - | 1 - | 0,000109114 - | 1 - | 0,000122916 - |
| 8681 | 1 | 0.2um_NOKCPHJN_149060 | Bacteria | Bacteroidetes | Flavobacteriia | Flavobacterial | Flavobacteriac | Polaribacter | various | information st | Translation, ril | 09GNN | Protein S19 fo | 313598.MED1 | 0,6 | 0,489897949 | 8,43E-05 | 6,94E-05 | 1 - | 0,000124757 - | 0 - | 0 - | 1 - | 0,000147263 - | 1 - | 0,000149438 - | 0 - | 0 - |
| 8682 | 1 | 0.2um_NOKCPHJN_91222 | Bacteria | Bacteroidetes | Flavobacteriia | Flavobacterial | Flavobacteriac | Formosa | Formosa sp. H | unclassified | unclassified | unclassified | unclassified | unclassified | 17 | 1,264911064 | 0,00025374 | 3,63E-05 | 17 - | 0,000212087 - | 15 - | 0,000232368 - | 17 - | 0,000250348 - | 17 - | 0,000254045 - | 19 - | 0,000319849 - |
| 8683 | 1 | 3um_FIJJAFBF_251384 | Bacteria | Bacteroidetes | Flavobacteriia | Flavobacterial | unclassified | unclassified | Flavobacterial | unclassified | unclassified | unclassified | unclassified | unclassified | 0,8 | 1,166190379 | 9,91E-06 | 1,44E-05 | 0 - | 0 - | 0 - | 0 - | 3 - | 3,70E-05 - | 1 - | 1,25E-05 - | 0 - | 0 - |
| 8684 | 1 | 0.2um_NOKCPHJN_547411 | Bacteria | Proteobacteri | Alphaproteob | Rhizobiales | unclassified | unclassified | Rhizobiales ba | unclassified | unclassified | unclassified | unclassified | unclassified | 0,4 | 0,489897949 | 4,17E-06 | 5,11E-06 | 0 - | 0 - | 0 - | 0 - | 1 - | 1,04E-05 - | 1 - | 1,05E-05 - | 0 - | 0 - |
| 8685 | 1 | 3um_FIJJAFBF_781290 | Bacteria | Proteobacteri | Gammaprote | Alteromonada | Pseudoalteror | Pseudoalteror | Pseudoalteror | unclassified | unclassified | unclassified | unclassified | unclassified | 0,2 | 0,4 | 1,09E-06 | 2,18E-06 | 1 - | 5,46E-06 - | 0 - | 0 - | 0 - | 0 - | 0 - | 0 - | 0 - | 0 - |
| 8686 | 1 | 3um_FIJJAFBF_764214 | Bacteria | Proteobacteri | Alphaproteob | Rhodobactera | Rhodobactera | Albimonas | Albimonas do | metabolism | Inorganic ion t | 16PAX | extracellular s | 644107.SL115 | 0,2 | 0,4 | 6,27E-06 | 1,25E-05 | 0 - | 0 - | 0 - | 0 - | 1 - | 3,14E-05 - | 0 - | 0 - | 0 - | 0 - |
| 8687 | 1 | 3um_FIJJAFBF_682764 | Bacteria | Bacteroidetes | Flavobacteriia | Flavobacterial | Flavobacteriac | Formosa | Formosa sp. H | metabolism | Inorganic ion t | 05C27 | TonB-depende | 983544.Lacal_ | 0,8 | 1,166190379 | 1,09E-05 | 1,59E-05 | 0 - | 0 - | 0 - | 0 - | 1 - | 1,34E-05 - | 3 - | 4,09E-05 - | 0 - | 0 - |
| 8688 | 1 | 0.2um_NOKCPHJN_624427 | Eukaryota | Ascomycota | Eurotiomycete | Eurotiales | Aspergillaceae | Aspergillus | Aspergillus te | cellular proces | Posttranslatio | 0JAJ8 | Dentin sialoph | 10020.ENSDO | 1 | 0 | 1,59E-05 | 1,51E-06 | 1 - | 1,33E-05 - | 1 - | 1,66E-05 - | 1 - | 1,58E-05 - | 1 - | 1,60E-05 - | 1 - | 1,80E-05 - |
| 8689 | 2 | 3um_FIJJAFBF_355827;3um_FIJJAFBF_51671 | Bacteria | Proteobacteri | Gammaprote | Cellvibrionale | unclassified | unclassified | Cellvibrionale | unclassified | unclassified | unclassified | unclassified | unclassified | 0,2 | 0,4 | 1,02E-05 | 2,03E-05 | 1 - | 5,08E-05 - | 0 - | 0 - | 0 - | 0 - | 0 - | 0 - | 0 - | 0 - |
| 8690 | 1 | 0.2um_NOKCPHJN_377232 | Bacteria | Bacteroidetes | Flavobacteriia | Flavobacterial | unclassified | unclassified | Flavobacterial | information st | Translation, ril | COG0012 | gtp-binding pr | 487796.Flav2 | 1 | 0 | 3,59E-05 | 3,41E-06 | 1 - | 3,00E-05 - | 1 - | 3,73E-05 - | 1 - | 3,55E-05 - | 1 - | 3,60E-05 - | 1 - | 4,05E-05 - |
| 8691 | 1 | 0.2um_NOKCPHJN_59638 | Bacteria | Bacteroidetes | Flavobacteriia | Flavobacterial | Crocinitomica | unclassified | Crocinitomica | unclassified | unclassified | unclassified | unclassified | unclassified | 1 | 0 | 2,09E-05 | 1,98E-06 | 1 - | 1,75E-05 - | 1 - | 2,17E-05 - | 1 - | 2,07E-05 - | 1 - | 2,10E-05 - | 1 - | 2,36E-05 - |
| 8692 | 1 | 3um_FIJJAFBF_189655 | Bacteria | Bacteroidetes | Flavobacteriia | Flavobacterial | Flavobacteriac | Polaribacter | Polaribacter s | cellular proces | Cell wall/mem | 09B44 | efflux transpo | 313598.MED1 | 1,8 | 0,4 | 6,32E-05 | 1,55E-05 | 2 - | 5,89E-05 - | 2 - | 7,31E-05 - | 2 - | 6,95E-05 - | 1 - | 3,53E-05 - | 2 - | 7,94E-05 - |
| 8693 | 1 | 0.2um_NOKCPHJN_684453 | Bacteria | Proteobacteri | Alphaproteob | Rhodobactera | Rhodobactera | Planktotalea | Planktotalea f | metabolism | Inorganic ion t | 05PWN | phosphate ab | 467661.RKLH1 | 3,2 | 0,4 | 0,000125808 | 6,90E-06 | 4 - | 0,000133075 - | 3 - | 0,000123929 - | 3 - | 0,000117811 - | 3 - | 0,000119551 - | 3 - | 0,000134673 - |
| 8694 | 1 | 3um_FIJJAFBF_729293 | Bacteria | Proteobacteri | Gammaprote | Cellvibrionale | unclassified | unclassified | Cellvibrionale | information st | Translation, ril | 08VZM | 50s ribosomal | 314287.GB22 | 1,8 | 0,4 | 0,00014009 | 3,43E-05 | 2 - | 0,000130428 - | 2 - | 0,000161953 - | 2 - | 0,000153957 - | 1 - | 7,81E-05 - | 2 - | 0,000175994 - |
| 8695 | 1 | 0.2um_NOKCPHJN_194183 | Bacteria | Proteobacteri | Gammaprote | unclassified | unclassified | unclassified | SAR86 cluster | metabolism | Energy produc | 0QJ3X | Aconitate hyd | 697282.Mettu | 0,8 | 0,4 | 1,32E-05 | 6,63E-06 | 0 - | 0 - | 1 - | 1,65E-05 - | 1 - | 1,56E-05 - | 1 - | 1,59E-05 - | 1 - | 1,79E-05 - |
| 8696 | 1 | 0.2um_NOKCPHJN_621198 | Bacteria | Bacteroidetes | Cytophagia | Cytophagales | Flammeovirga | unclassified | Flammeovirga | poorly charact | Function unkn | 09A4Z | rhodopsin | 760192.Halhy | 2,4 | 0,489897949 | 0,000136093 | 3,04E-05 | 3 - | 0,000142876 - | 2 - | 0,000118273 - | 2 - | 0,000112433 - | 2 - | 0,000114094 - | 3 - | 0,00019279 - |
| 8697 | 1 | 0.2um_NOKCPHJN_733220 | unclassified | unclassified | unclassified | unclassified | unclassified | unclassified | unclassified | unclassified | unclassified | unclassified | unclassified | unclassified | 1 | 0 | 2,04E-05 | 1,94E-06 | 1 - | 1,71E-05 - | 1 - | 2,12E-05 - | 1 - | 2,02E-05 - | 1 - | 2,05E-05 - | 1 - | 2,30E-05 - |
| 8698 | 1 | 0.2um_NOKCPHJN_607421 | Bacteria | Proteobacteri | Gammaprote | Oceanospirilla | Saccharospirill | Reinekea | Reinekea blan | metabolism | Nucleotide tra | 05CM0 | Catalyzes the | 314283.MED2 | 1,2 | 0,748331477 | 3,31E-05 | 1,98E-05 | 2 - | 4,86E-05 - | 1 - | 3,02E-05 - | 2 - | 5,74E-05 - | 1 - | 2,91E-05 - | 0 - | 0 - |
| 8699 | 1 | 0.2um_NOKCPHJN_01154 | Bacteria | Bacteroidetes | unclassified | unclassified | unclassified | unclassified | Bacteroidetes | poorly charact | Function unkn | 09PDM | NA | 860228.Ccan_ | 1,6 | 0,489897949 | 0,000115905 | 3,21E-05 | 2 - | 0,000123416 - | 2 - | 0,000153246 - | 2 - | 0,00014568 - | 1 - | 7,39E-05 - | 1 - | 8,33E-05 - |
| 8700 | 1 | 3um_FIJJAFBF_732143 | Bacteria | Proteobacteri | Alphaproteob | Rhizobiales | Hyphomicrobi | Devosia | Devosia enhy | unclassified | unclassified | unclassified | unclassified | unclassified | 5,6 | 0,8 | 0,000275482 | 3,80E-05 | 6 - | 0,000248614 - | 5 - | 0,000257254 - | 5 - | 0,000244553 - | 7 - | 0,000347431 - | 5 - | 0,000279557 - |
| 8701 | 1 | 3um_FIJJAFBF_630357 | Bacteria | Proteobacteri | Alphaproteob | Rhodobactera | Rhodobactera | unclassified | Rhodobactera | metabolism | Amino acid tra | 01R4A | Polyamine AB | 388739.RSK20 | 3,2 | 0,4 | 0,00011983 | 1,86E-05 | 3 - | 9,41E-05 - | 3 - | 0,000116819 - | 3 - | 0,000111051 - | 4 - | 0,000150255 - | 3 - | 0,000126946 - |
| 8702 | 1 | 0.2um_NOKCPHJN_300184 | Bacteria | Bacteroidetes | Flavobacteriia | Flavobacterial | unclassified | unclassified | Flavobacterial | metabolism | Carbohydrate | COG0057 | glyceraldehyd | 867900.Celly_ | 1 | 0 | 4,11E-05 | 3,91E-06 | 1 - | 3,45E-05 - | 1 - | 4,28E-05 - | 1 - | 4,07E-05 - | 1 - | 4,13E-05 - | 1 - | 4,65E-05 - |
| 8703 | 1 | 3um_FIJJAFBF_336547 | Bacteria | Proteobacteri | Deltaproteoba | Myxococcales | Anaeromyxob | Anaeromyxob | Anaeromyxob | unclassified | unclassified | unclassified | unclassified | unclassified | 0,6 | 0,489897949 | 1,74E-05 | 1,45E-05 | 1 - | 2,43E-05 - | 1 - | 3,01E-05 - | 0 - | 0 - | 0 - | 0 - | 1 - | 3,27E-05 - |
| 8704 | 1 | 3um_FIJJAFBF_349824 | Bacteria | Proteobacteri | Alphaproteob | Pelagibacteral | Pelagibacterac | unclassified | Pelagibactera | unclassified | unclassified | unclassified | unclassified | unclassified | 1 | 0 | 0,000141265 | 1,34E-05 | 1 - | 0,000118327 - | 1 - | 0,000146927 - | 1 - | 0,000139672 - | 1 - | 0,000141735 - | 1 - | 0,000159664 - |
| 8705 | 1 | 3um_FIJJAFBF_453300 | Bacteria | Proteobacteri | Alphaproteob | Rhodobactera | Rhodobactera | Ruegeria | Ruegeria halo | unclassified | unclassified | unclassified | unclassified | unclassified | 1,2 | 0,4 | 2,95E-05 | 1,02E-05 | 1 - | 2,06E-05 - | 1 - | 2,55E-05 - | 1 - | 2,43E-05 - | 2 - | 4,93E-05 - | 1 - | 2,78E-05 - |
| 8706 | 1 | 3um_FIJJAFBF_584721 | Eukaryota | Bacillariophyt | Coscinodiscop | Coscinodiscale | Coscinodiscac | Coscinodiscus | Coscinodiscus | information st | Translation, ril | 0FP0S | Seems to be t | 32049.SYNPC | 1 | 0,894427191 | 0,000105428 | 9,20E-05 | 2 - | 0,000180751 - | 2 - | 0,000224439 - | 0 - | 0 - | 0 - | 0 - | 1 - | 0,000121948 - |
| 8707 | 1 | 0.2um_NOKCPHJN_703721 | Viruses | unclassified | unclassified | unclassified | Phycodnavirid | Prasinovirus | Ostreococcus | unclassified | unclassified | unclassified | unclassified | unclassified | 0,8 | 0,4 | 2,71E-05 | 1,36E-05 | 0 - | 0 - | 1 - | 3,39E-05 - | 1 - | 3,22E-05 - | 1 - | 3,27E-05 - | 1 - | 3,68E-05 - |
| 8708 | 1 | 3um_FIJJAFBF_25410 | Bacteria | Bacteroidetes | Flavobacteriia | Flavobacterial | Flavobacteriac | Polaribacter | Polaribacter h | poorly charact | Function unkn | 09GJP | Protein of unk | 313598.MED1 | 0,6 | 0,489897949 | 5,79E-05 | 4,76E-05 | 1 - | 8,57E-05 - | 0 - | 0 - | 1 - | 0,000101106 - | 1 - | 0,000102599 - | 0 - | 0 - |
| 8709 | 1 | 0.2um_NOKCPHJN_45922 | Bacteria | Bacteroidetes | Flavobacteriia | Flavobacterial | Flavobacteriac | Arenitalea | Arenitalea lut | unclassified | unclassified | unclassified | unclassified | unclassified | 0,8 | 0,4 | 2,97E-05 | 1,51E-05 | 1 - | 3,22E-05 - | 1 - | 3,99E-05 - | 1 - | 3,80E-05 - | 1 - | 3,85E-05 - | 0 - | 0 - |
| 8710 | 1 | 3um_FIJJAFBF_72481 | Bacteria | Bacteroidetes | Cytophagia | Cytophagales | Flammeovirga | unclassified | Flammeovirga | information st | Translation, ril | COG0256 | This is one of t | 880070.Cycma | 1,2 | 0,4 | 0,000139115 | 3,08E-05 | 2 - | 0,000199612 - | 1 - | 0,000123929 - | 1 - | 0,000117811 - | 1 - | 0,000119551 - | 1 - | 0,000134673 - |
| 8711 | 1 | 0.2um_NOKCPHJN_271556 | Eukaryota | Apicomplexa | Conoidasida | Eucoccidiorida | Eimeriidae | Eimeria | Eimeria praec | unclassified | unclassified | unclassified | unclassified | unclassified | 11 | 2,28035085 | 0,000337433 | 5,93E-05 | 14 - | 0,000363546 - | 8 - | 0,000257953 - | 9 - | 0,000275869 - | 13 - | 0,000404363 - | 11 - | 0,000385434 - |
| 8712 | 1 | 0.2um_NOKCPHJN_217488 | Bacteria | Proteobacteri | Gammaprote | Oceanospirilla | Saccharospirill | Reinekea | Reinekea fors | metabolism | Inorganic ion t | 0GH39 | Phosphonate | 525897.Dbac_ | 5,8 | 0,748331477 | 0,000538799 | 3,83E-05 | 7 - | 0,0005503 - | 6 - | 0,000585694 - | 6 - | 0,000556776 - | 5 - | 0,000470833 - | 5 - | 0,000530392 - |
| 8713 | 1 | 3um_FIJJAFBF_193295 | Bacteria | Proteobacteri | Gammaprote | Oceanospirilla | Saccharospirill | Reinekea | Reinekea blan | metabolism | Amino acid tra | 16QEM | saccharopine | 314283.MED2 | 0,2 | 0,4 | 5,75E-06 | 1,15E-05 | 1 - | 2,88E-05 - | 0 - | 0 - | 0 - | 0 - | 0 - | 0 - | 0 - | 0 - |
| 8714 | 1 | 0.2um_NOKCPHJN_71917 | Bacteria | Proteobacteri | Gammaprote | Oceanospirilla | Saccharospirill | Reinekea | Reinekea blan | unclassified | unclassified | unclassified | unclassified | unclassified | 2,6 | 0,8 | 2,42E-05 | 8,16E-06 | 2 - | 1,55E-05 - | 4 - | 3,85E-05 - | 3 - | 2,74E-05 - | 2 - | 1,86E-05 - | 2 - | 2,09E-05 - |
| 8715 | 1 | 3um_FIJJAFBF_189639 | Bacteria | Bacteroidetes | Flavobacteriia | Flavobacterial | Flavobacteriac | Polaribacter | various | unclassified | unclassified | unclassified | unclassified | unclassified | 1,4 | 1,019803903 | 0,000105948 | 7,22E-05 | 3 - | 0,00019676 - | 0 - | 0 - | 1 - | 7,74E-05 - | 1 - | 7,86E-05 - | 2 - | 0,000176999 - |
| 8716 | 1 | 0.2um_NOKCPHJN_13536 | Bacteria | Proteobacteri | Gammaprote | Pseudomonad | Pseudomonad | Pseudomonas | Pseudomonas | unclassified | unclassified | unclassified | unclassified | unclassified | 0,2 | 0,4 | 5,11E-06 | 1,02E-05 | 0 - | 0 - | 0 - | 0 - | 0 - | 0 - | 1 - | 2,56E-05 - | 0 - | 0 - |
| 8717 | 1 | 0.2um_NOKCPHJN_960020 | Bacteria | Proteobacteri | Alphaproteob | Pelagibacteral | Pelagibacterac | Candidatus Pe | Candidatus Pe | poorly charact | Function unkn | 0ZNGQ | NA | 335992.SAR11 | 1,8 | 0,4 | 0,000285524 | 6,68E-05 | 2 - | 0,000266923 - | 1 - | 0,00016572 - | 2 - | 0,000315075 - | 2 - | 0,000319729 - | 2 - | 0,000360173 - |

| | | | | | | | | | | | | | | | | | | | | | | | | | | | | |
|---|---|---|---|---|---|---|---|---|---|---|---|---|---|---|---|---|---|---|---|---|---|---|---|---|---|---|---|---|
| 8718 | 1 | 0.2um_NOKCPHJN_902075 | Bacteria | Proteobacteri | Alphaproteob | Rhodobactera | Rhodobactera | Amylibacter | Amylibacter k | metabolism | Lipid transpor | 16QA8 | reductase | 252305.OB25 | 0,4 | 0,489897949 | 2,20E-05 | 2,75E-05 | 1 - | 4,68E-05 - | 0 - | 0 - | 0 - | 0 - | 0 - | 0 - | 1 - | 6,32E-05 - |
| 8719 | 1 | 3um_FIJJAFBF_00761 | Bacteria | Proteobacteri | Alphaproteob | Rhodobactera | Rhodobactera | Planktomarina | Planktomarina | unclassified | unclassified | unclassified | unclassified | unclassified | 0,8 | 0,4 | 3,66E-05 | 1,86E-05 | 1 - | 3,96E-05 - | 1 - | 4,91E-05 - | 1 - | 4,67E-05 - | 1 - | 4,74E-05 - | 0 - | 0 - |
| 8720 | 1 | 3um_FIJJAFBF_306777 | Bacteria | Proteobacteri | Alphaproteob | Rhodobactera | unclassified | unclassified | Rhodobactera | cellular proces | Signal transdu | 01RN6 | nitrogen | 439496.RBY4I | 0,2 | 0,4 | 5,96E-06 | 1,19E-05 | 0 - | 0 - | 0 - | 0 - | 1 - | 2,98E-05 - | 0 - | 0 - | 0 - | 0 - |
| 8721 | 1 | 3um_FIJJAFBF_551528 | Bacteria | Bacteroidetes | Flavobacteriia | Flavobacterial | Flavobacteriac | Winogradskye | Winogradskye | unclassified | unclassified | unclassified | unclassified | unclassified | 0,6 | 0,489897949 | 1,67E-05 | 1,38E-05 | 1 - | 2,35E-05 - | 0 - | 0 - | 0 - | 0 - | 1 - | 2,81E-05 - | 1 - | 3,17E-05 - |
| 8722 | 1 | 3um_FIJJAFBF_53342 | Bacteria | Bacteroidetes | Flavobacteriia | Flavobacterial | Flavobacteriac | Polaribacter | Polaribacter a | cellular proces | Cell wall/mem | 0P81E | outer membra | 313594.PI23P | 0,2 | 0,4 | 5,19E-06 | 1,04E-05 | 1 - | 2,60E-05 - | 0 - | 0 - | 0 - | 0 - | 0 - | 0 - | 0 - | 0 - |
| 8723 | 1 | 3um_FIJJAFBF_324100 | Bacteria | Proteobacteri | Gammaprotec | Alteromonada | Alteromonada | Aliiglaciecola | Aliiglaciecola l | information st | Transcription | 16PMT | DNA-depende | 314275.MADE | 5,2 | 1,166190379 | 0,000214902 | 4,71E-05 | 6 - | 0,000209319 - | 7 - | 0,000303232 - | 4 - | 0,00016472 - | 5 - | 0,000208941 - | 4 - | 0,000188297 - |
| 8724 | 1 | 3um_FIJJAFBF_745455 | Bacteria | Bacteroidetes | unclassified | unclassified | unclassified | unclassified | Bacteroidetes | information st | Replication, re | 17UKA | DNA ligase | 391625.PPSIR | 0,2 | 0,4 | 1,48E-05 | 2,96E-05 | 1 - | 7,40E-05 - | 0 - | 0 - | 0 - | 0 - | 0 - | 0 - | 0 - | 0 - |
| 8725 | 1 | 3um_FIJJAFBF_189605 | Bacteria | Bacteroidetes | Flavobacteriia | Flavobacterial | Flavobacteriac | Polaribacter | Polaribacter b | unclassified | unclassified | unclassified | unclassified | unclassified | 2,2 | 0,979795897 | 0,000248142 | 0,000121275 | 1 - | 9,18E-05 - | 3 - | 0,000342045 - | 3 - | 0,000325157 - | 1 - | 0,000109987 - | 3 - | 0,000371699 - |
| 8726 | 1 | 3um_FIJJAFBF_540522 | Bacteria | Proteobacteri | Alphaproteob | Rhodobactera | Rhodobactera | Planktomarina | Planktomarina | information st | Transcription | COG0250 | Participates in | 89187.ISM_01 | 4,4 | 0,8 | 0,000341906 | 7,73E-05 | 4 - | 0,000259383 - | 3 - | 0,000241557 - | 5 - | 0,000382718 - | 5 - | 0,000388371 - | 5 - | 0,000437499 - |
| 8727 | 1 | 3um_FIJJAFBF_718147 | Bacteria | Proteobacteri | Gammaprotec | Alteromonada | Alteromonada | Paraglaciecola | Paraglaciecola | poorly charact | Function unkn | 17DB1 | short-chain de | 983545.Glaag | 0,4 | 0,489897949 | 2,39E-05 | 2,93E-05 | 0 - | 0 - | 1 - | 6,09E-05 - | 0 - | 0 - | 1 - | 5,88E-05 - | 0 - | 0 - |
| 8728 | 1 | 3um_FIJJAFBF_741125 | Bacteria | Proteobacteri | Alphaproteob | Pelagibacteral | Pelagibacterac | unclassified | Pelagibacterac | poorly charact | Function unkn | 01SDU | solute sympor | 684719.HIMB | 6,6 | 0,489897949 | 0,000147242 | 1,29E-05 | 7 - | 0,000131281 - | 7 - | 0,000163012 - | 6 - | 0,000132826 - | 7 - | 0,000157252 - | 6 - | 0,000151838 - |
| 8729 | 1 | 0.2um_NOKCPHJN_290553 | Bacteria | Proteobacteri | Gammaprotec | Cellvibrionales | Halieaceae | Luminiphilus | Luminiphilus s | information st | Transcription | 05D27 | DNA-depende | 565045.NOR5 | 0,2 | 0,4 | 1,92E-06 | 3,84E-06 | 0 - | 0 - | 0 - | 0 - | 1 - | 9,61E-06 - | 0 - | 0 - | 0 - | 0 - |
| 8730 | 1 | 0.2um_NOKCPHJN_377061 | unclassified | unclassified | unclassified | unclassified | unclassified | unclassified | uncultured ma | unclassified | unclassified | unclassified | unclassified | unclassified | 0,4 | 0,489897949 | 1,09E-05 | 1,35E-05 | 1 - | 2,44E-05 - | 1 - | 3,03E-05 - | 0 - | 0 - | 0 - | 0 - | 0 - | 0 - |
| 8731 | 1 | 3um_FIJJAFBF_738497 | Bacteria | Proteobacteri | Gammaprotec | Alteromonada | Alteromonada | Paraglaciecola | Paraglaciecola | unclassified | unclassified | unclassified | unclassified | unclassified | 1,2 | 0,4 | 3,96E-05 | 1,37E-05 | 1 - | 2,77E-05 - | 1 - | 3,43E-05 - | 1 - | 3,26E-05 - | 2 - | 6,63E-05 - | 1 - | 3,73E-05 - |
| 8732 | 2 | 3um_FIJJAFBF_190674;3um_FIJJAFBF_378784 | Bacteria | Bacteroidetes | Flavobacteriia | Flavobacterial | Flavobacteriac | Ulvibacter | Ulvibacter lito | poorly charact | Function unkn | 09ARW | Zinc ribbon dc | 50743.SCB49_ | 0,4 | 0,489897949 | 1,94E-05 | 2,39E-05 | 1 - | 4,41E-05 - | 0 - | 0 - | 0 - | 0 - | 1 - | 5,29E-05 - | 0 - | 0 - |
| 8733 | 1 | 0.2um_NOKCPHJN_05633 | Bacteria | Bacteroidetes | Cytophagia | Cytophagales | Cytophagacea | Ekhidna | Ekhidna lutea | cellular proces | Cell wall/mem | COG2885 | Ompa motb d | 860228.Ccan_ | 5,6 | 0,8 | 9,82E-05 | 1,56E-05 | 6 - | 8,84E-05 - | 7 - | 0,000128066 - | 5 - | 8,70E-05 - | 5 - | 8,82E-05 - | 5 - | 9,94E-05 - |
| 8734 | 1 | 0.2um_NOKCPHJN_140577 | Bacteria | Proteobacteri | Gammaprotec | unclassified | unclassified | unclassified | OM182 bacter | unclassified | unclassified | unclassified | unclassified | unclassified | 0,6 | 0,489897949 | 2,13E-05 | 1,77E-05 | 1 - | 3,01E-05 - | 0 - | 0 - | 1 - | 3,56E-05 - | 0 - | 0 - | 1 - | 4,06E-05 - |
| 8735 | 1 | 3um_FIJJAFBF_257951 | Bacteria | Bacteroidetes | Flavobacteriia | Flavobacterial | Flavobacteriac | Olleya | Olleya sp. ITB | information st | Translation, ri | 05CEJ | Catalyzes the | 983548.Krodi_ | 0,6 | 0,489897949 | 1,29E-05 | 1,08E-05 | 1 - | 1,82E-05 - | 0 - | 0 - | 0 - | 0 - | 1 - | 2,19E-05 - | 1 - | 2,46E-05 - |
| 8736 | 1 | 3um_FIJJAFBF_328503 | Bacteria | Bacteroidetes | Flavobacteriia | Flavobacterial | Flavobacteriac | Arenitalea | Arenitalea lut | information st | Translation, ri | COG0088 | One of the pri | 983544.Lacal_ | 1,4 | 0,489897949 | 9,33E-05 | 3,79E-05 | 1 - | 5,49E-05 - | 1 - | 6,82E-05 - | 2 - | 0,000129648 - | 1 - | 6,58E-05 - | 2 - | 0,000148205 - |
| 8737 | 1 | 0.2um_NOKCPHJN_211484 | Bacteria | Proteobacteri | Gammaprotec | unclassified | unclassified | unclassified | various | poorly charact | Function unkn | 05C2A | tonB-depende | 566466.NOR5 | 33,6 | 2,8 | 0,000431083 | 5,24E-05 | 35 - | 0,000376494 - | 32 - | 0,000427423 - | 37 - | 0,000469807 - | 29 - | 0,000373666 - | 35 - | 0,000508023 - |
| 8738 | 1 | 3um_FIJJAFBF_218438 | Bacteria | Bacteroidetes | Flavobacteriia | Flavobacterial | Flavobacteriac | Formosa | Formosa sp. H | unclassified | unclassified | unclassified | unclassified | unclassified | 1 | 0 | 0,000217503 | 2,07E-05 | 1 - | 0,000182185 - | 1 - | 0,00022622 - | 1 - | 0,000215051 - | 1 - | 0,000218227 - | 1 - | 0,000245833 - |
| 8739 | 1 | 0.2um_NOKCPHJN_172059 | Bacteria | Bacteroidetes | unclassified | unclassified | unclassified | unclassified | Bacteroidetes | unclassified | unclassified | unclassified | unclassified | unclassified | 1 | 0 | 1,71E-05 | 1,62E-06 | 1 - | 1,43E-05 - | 1 - | 1,78E-05 - | 1 - | 1,69E-05 - | 1 - | 1,72E-05 - | 1 - | 1,93E-05 - |
| 8740 | 1 | 3um_FIJJAFBF_256444 | Bacteria | Proteobacteri | Alphaproteob | Rhodobactera | Rhodobactera | Roseobacter | Roseobacter d | poorly charact | Function unkn | COG3181 | extra-cytoplas | 367336.OM22 | 1,6 | 0,489897949 | 6,85E-05 | 2,50E-05 | 1 - | 3,51E-05 - | 2 - | 8,72E-05 - | 1 - | 4,14E-05 - | 2 - | 8,41E-05 - | 2 - | 9,47E-05 - |
| 8741 | 1 | 0.2um_NOKCPHJN_546626 | Bacteria | Proteobacteri | Alphaproteob | Pelagibacteral | Pelagibacterac | Candidatus Pe | Candidatus Pe | information st | Translation, ri | COG0251 | Endoribonucle | 335992.SAR11 | 1,8 | 0,4 | 0,000161149 | 3,95E-05 | 2 - | 0,000150035 - | 2 - | 0,000186299 - | 2 - | 0,000177101 - | 1 - | 8,99E-05 - | 2 - | 0,00020245 - |
| 8742 | 1 | 0.2um_NOKCPHJN_188517 | Bacteria | Bacteroidetes | Cytophagia | Cytophagales | Flammeovirga | Marivirga | Marivirga trac | metabolism | Inorganic ion t | 05C27 | TonB-depende | 643867.Ftrac_ | 1,4 | 1,019803903 | 1,73E-05 | 1,29E-05 | 1 - | 1,06E-05 - | 1 - | 1,31E-05 - | 3 - | 3,74E-05 - | 2 - | 2,53E-05 - | 0 - | 0 - |
| 8743 | 1 | 0.2um_NOKCPHJN_21219 | Bacteria | Proteobacteri | Alphaproteob | Rhodobactera | Rhodobactera | Planktotalea | Planktotalea a | metabolism | Energy produc | 05C6K | Malic enzyme | 314270.RB208 | 0,4 | 0,8 | 7,59E-06 | 1,52E-05 | 0 - | 0 - | 2 - | 3,80E-05 - | 0 - | 0 - | 0 - | 0 - | 0 - | 0 - |
| 8744 | 1 | 0.2um_NOKCPHJN_73077 | Bacteria | Bacteroidetes | unclassified | unclassified | unclassified | unclassified | Bacteroidetes | unclassified | unclassified | unclassified | unclassified | unclassified | 2 | 0,632455532 | 2,68E-05 | 8,15E-06 | 2 - | 2,28E-05 - | 2 - | 2,83E-05 - | 3 - | 4,03E-05 - | 2 - | 2,73E-05 - | 1 - | 1,54E-05 - |
| 8745 | 1 | 3um_FIJJAFBF_733805 | Bacteria | Proteobacteri | Gammaprotec | unclassified | unclassified | unclassified | Gammaprotec | information st | Translation, ri | 0QQXI | This protein is | 768066.HELO_ | 6 | 0 | 0,000702703 | 6,67E-05 | 6 - | 0,000588599 - | 6 - | 0,000730866 - | 6 - | 0,000694781 - | 6 - | 0,000705043 - | 6 - | 0,000794228 - |
| 8746 | 1 | 3um_FIJJAFBF_715090 | Eukaryota | unclassified | Pelagophycea | Pelagomonad | unclassified | Aureoumbra | Aureoumbra l | information st | Translation, ri | 0FJK5 | Binds the lowe | 197221.tlr008 | 1,8 | 0,979795897 | 0,00010933 | 5,60E-05 | 3 - | 0,000160154 - | 2 - | 0,000132576 - | 2 - | 0,00012603 - | 2 - | 0,000127891 - | 0 - | 0 - |
| 8747 | 1 | 3um_FIJJAFBF_454712 | Bacteria | Bacteroidetes | unclassified | unclassified | unclassified | unclassified | unidentified e | unclassified | unclassified | unclassified | unclassified | unclassified | 0,6 | 0,489897949 | 1,40E-05 | 1,15E-05 | 1 - | 2,04E-05 - | 1 - | 2,53E-05 - | 0 - | 0 - | 1 - | 2,44E-05 - | 0 - | 0 - |
| 8748 | 1 | 3um_FIJJAFBF_44069 | Viruses | unclassified | unclassified | unclassified | Phycodnavirid | unclassified | Chrysochromu | unclassified | unclassified | unclassified | unclassified | unclassified | 2,4 | 0,489897949 | 0,000174097 | 4,71E-05 | 2 - | 0,000120185 - | 2 - | 0,000149234 - | 2 - | 0,000141866 - | 3 - | 0,000215942 - | 3 - | 0,000243258 - |
| 8749 | 1 | 0.2um_NOKCPHJN_536531 | Bacteria | Proteobacteri | Gammaprotec | Xanthomonad | Rhodanobacte | Rhodanobacte | Rhodanobacte | unclassified | unclassified | unclassified | unclassified | unclassified | 0,2 | 0,4 | 1,58E-06 | 3,16E-06 | 1 - | 7,90E-06 - | 0 - | 0 - | 0 - | 0 - | 0 - | 0 - | 0 - | 0 - |
| 8750 | 1 | 3um_FIJJAFBF_637965 | Bacteria | Bacteroidetes | Flavobacteriia | Flavobacterial | Flavobacteriac | unclassified | Flavobacteriac | unclassified | unclassified | unclassified | unclassified | unclassified | 4,4 | 0,489897949 | 7,07E-05 | 7,39E-06 | 5 - | 6,77E-05 - | 5 - | 8,40E-05 - | 4 - | 6,39E-05 - | 4 - | 6,49E-05 - | 4 - | 7,31E-05 - |
| 8751 | 1 | 3um_FIJJAFBF_21306 | Bacteria | Bacteroidetes | Flavobacteriia | Flavobacterial | Flavobacteriac | Polaribacter | Polaribacter b | unclassified | unclassified | unclassified | unclassified | unclassified | 11,6 | 0,8 | 0,000377933 | 4,11E-05 | 12 - | 0,000327934 - | 10 - | 0,000339331 - | 12 - | 0,000387092 - | 12 - | 0,000392809 - | 12 - | 0,000442499 - |
| 8752 | 1 | 3um_FIJJAFBF_176642 | Bacteria | Bacteroidetes | unclassified | unclassified | unclassified | unclassified | Bacteroidetes | unclassified | unclassified | unclassified | unclassified | unclassified | 3 | 0,632455532 | 0,000310359 | 6,80E-05 | 3 - | 0,000260856 - | 2 - | 0,000215938 - | 4 - | 0,000410552 - | 3 - | 0,000312462 - | 3 - | 0,000351987 - |
| 8753 | 1 | 0.2um_NOKCPHJN_21215 | Bacteria | Proteobacteri | Alphaproteob | Rhodobactera | Rhodobactera | Ascidiaceihabi | Ascidiaceihabi | metabolism | Amino acid tra | COG0137 | Citrulline--asp | 398580.Dshi_ | 1,6 | 0,8 | 5,20E-05 | 2,08E-05 | 3 - | 8,50E-05 - | 1 - | 3,52E-05 - | 1 - | 3,35E-05 - | 2 - | 6,79E-05 - | 1 - | 3,82E-05 - |
| 8754 | 1 | 3um_FIJJAFBF_297882 | unclassified | unclassified | unclassified | unclassified | unclassified | unclassified | unclassified | unclassified | unclassified | unclassified | unclassified | unclassified | 0,6 | 0,489897949 | 1,62E-05 | 1,35E-05 | 1 - | 2,30E-05 - | 0 - | 0 - | 1 - | 2,72E-05 - | 0 - | 0 - | 1 - | 3,10E-05 - |
| 8755 | 1 | 3um_FIJJAFBF_347366 | Eukaryota | unclassified | Bangiophycea | Bangiales | Bangiaceae | Porphyra | Porphyra umb | unclassified | unclassified | unclassified | unclassified | unclassified | 11,2 | 0,4 | 0,000236122 | 2,37E-05 | 11 - | 0,000194238 - | 11 - | 0,000241186 - | 11 - | 0,000229278 - | 12 - | 0,000253815 - | 11 - | 0,000262095 - |
| 8756 | 1 | 0.2um_NOKCPHJN_39804 | Bacteria | Bacteroidetes | Flavobacteriia | Flavobacterial | Flavobacteriac | Polaribacter | Polaribacter ir | information st | Translation, ri | COG0052 | 30S ribosomal | 983548.Krodi_ | 1,8 | 1,166190379 | 7,96E-05 | 4,96E-05 | 3 - | 0,000114017 - | 3 - | 0,000141575 - | 0 - | 0 - | 2 - | 9,10E-05 - | 1 - | 5,13E-05 - |
| 8757 | 1 | 3um_FIJJAFBF_718692 | Eukaryota | unclassified | Cryptophyta | Pyrenomonad | Pyrenomonad | Rhodomonas | Rhodomonas | metabolism | Carbohydrate | 0IVA1 | Photosystem I | 3218.PP1S345 | 1,8 | 0,4 | 0,00013283 | 2,62E-05 | 2 - | 0,000125439 - | 2 - | 0,000155758 - | 2 - | 0,000148068 - | 2 - | 0,000150255 - | 1 - | 8,46E-05 - |
| 8758 | 1 | 3um_FIJJAFBF_715073 | Eukaryota | unclassified | Pelagophycea | Pelagomonad | unclassified | Aureoumbra | Aureoumbra l | information st | Translation, ri | 08UHY | One of the pri | 32049.SYNPC | 1,2 | 0,4 | 0,000105207 | 3,52E-05 | 1 - | 7,36E-05 - | 1 - | 9,14E-05 - | 2 - | 0,000173695 - | 1 - | 8,81E-05 - | 1 - | 9,93E-05 - |
| 8759 | 1 | 0.2um_NOKCPHJN_284928 | Bacteria | Bacteroidetes | Flavobacteriia | Flavobacterial | Flavobacteriac | Algibacter | Algibacter lect | metabolism | Coenzyme tra | 09A4R | May play a ke | 983544.Lacal_ | 2 | 0 | 6,26E-05 | 5,94E-06 | 2 - | 5,24E-05 - | 2 - | 6,51E-05 - | 2 - | 6,19E-05 - | 2 - | 6,28E-05 - | 2 - | 7,07E-05 - |
| 8760 | 1 | 3um_FIJJAFBF_338529 | Bacteria | Proteobacteri | Alphaproteob | unclassified | unclassified | unclassified | Alphaproteob | metabolism | Inorganic ion t | 16PII | TRAP transpor | 684719.HIMB | 0,2 | 0,4 | 8,56E-06 | 1,71E-05 | 0 - | 0 - | 1 - | 4,28E-05 - | 0 - | 0 - | 0 - | 0 - | 0 - | 0 - |
| 8761 | 1 | 3um_FIJJAFBF_520488 | Bacteria | Bacteroidetes | Flavobacteriia | Flavobacterial | Flavobacteriac | Polaribacter | Polaribacter s | cellular proces | Intracellular tr | COG0653 | Part of the Se | 313598.MED1 | 11,4 | 2,244994432 | 0,000138327 | 1,86E-05 | 15 - | 0,000154686 - | 13 - | 0,000166464 - | 10 - | 0,000121727 - | 10 - | 0,000123525 - | 9 - | 0,000125235 - |
| 8762 | 1 | 0.2um_NOKCPHJN_283743 | Bacteria | Proteobacteri | Alphaproteob | Rhodospirillal | Rhodospirillac | Candidatus En | Candidatus En | unclassified | unclassified | unclassified | unclassified | unclassified | 0,2 | 0,4 | 6,60E-06 | 1,32E-05 | 0 - | 0 - | 0 - | 0 - | 0 - | 0 - | 0 - | 0 - | 1 - | 3,30E-05 - |
| 8763 | 1 | 3um_FIJJAFBF_808729 | Bacteria | Proteobacteri | Gammaprotec | Alteromonada | Alteromonada | Paraglaciecola | Paraglaciecola | cellular proces | Intracellular tr | 175GA | One of the pro | 983545.Glaag | 0,8 | 0,4 | 6,35E-05 | 3,26E-05 | 1 - | 6,71E-05 - | 0 - | 0 - | 1 - | 7,92E-05 - | 1 - | 8,04E-05 - | 1 - | 9,06E-05 - |
| 8764 | 1 | 0.2um_NOKCPHJN_243019 | Eukaryota | Echinodermat | Echinoidea | Echinoida | Strongylocent | Strongylocent | Strongylocent | cellular proces | Cell wall/mem | KOG4177 | Ankyrin Repea | 5722.A2ERS8 | 1 | 0 | 2,08E-05 | 1,97E-06 | 1 - | 1,74E-05 - | 1 - | 2,16E-05 - | 1 - | 2,05E-05 - | 1 - | 2,08E-05 - | 1 - | 2,35E-05 - |
| 8765 | 1 | 0.2um_NOKCPHJN_123306 | Bacteria | Proteobacteri | Gammaprotec | Oceanospirilla | Saccharospirill | Reinekea | Reinekea blan | metabolism | Amino acid tra | COG0133 | The beta subu | 314283.MED2 | 1 | 0,894427191 | 3,06E-05 | 2,69E-05 | 2 - | 5,47E-05 - | 1 - | 3,39E-05 - | 2 - | 6,45E-05 - | 0 - | 0 - | 0 - | 0 - |
| 8766 | 1 | 0.2um_NOKCPHJN_227815 | Bacteria | Proteobacteri | Gammaprotec | Cellvibrionales | Porticoccacea | unclassified | SAR92 bacteri | unclassified | unclassified | unclassified | unclassified | unclassified | 0,2 | 0,4 | 3,29E-06 | 6,58E-06 | 0 - | 0 - | 1 - | 1,65E-05 - | 0 - | 0 - | 0 - | 0 - | 0 - | 0 - |
| 8767 | 1 | 3um_FIJJAFBF_445642 | unclassified | unclassified | unclassified | unclassified | unclassified | unclassified | unclassified | unclassified | unclassified | unclassified | unclassified | unclassified | 2 | 1,264911064 | 7,38E-05 | 4,52E-05 | 3 - | 9,65E-05 - | 0 - | 0 - | 3 - | 0,000113851 - | 3 - | 0,000115532 - | 1 - | 4,34E-05 - |
| 8768 | 1 | 3um_FIJJAFBF_29166 | Bacteria | Proteobacteri | Alphaproteob | Rhizobiales | Hyphomicrobi | Devosia | Devosia enhyc | unclassified | unclassified | unclassified | unclassified | unclassified | 1,6 | 0,489897949 | 0,000114054 | 3,68E-05 | 1 - | 5,95E-05 - | 2 - | 0,000147688 - | 2 - | 0,000140396 - | 2 - | 0,00014247 - | 1 - | 8,02E-05 - |
| 8769 | 1 | 0.2um_NOKCPHJN_96514 | Bacteria | Bacteroidetes | Flavobacteriia | unclassified | unclassified | unclassified | Flavobacteria | unclassified | unclassified | unclassified | unclassified | unclassified | 2,4 | 0,489897949 | 5,15E-05 | 8,02E-06 | 3 - | 5,47E-05 - | 2 - | 4,52E-05 - | 2 - | 4,30E-05 - | 3 - | 6,55E-05 - | 2 - | 4,92E-05 - |
| 8770 | 1 | 0.2um_NOKCPHJN_573220 | Bacteria | Proteobacteri | Alphaproteob | Pelagibacteral | Pelagibacterac | Candidatus Pe | Candidatus Pe | unclassified | unclassified | unclassified | unclassified | unclassified | 3,8 | 0,4 | 0,000568279 | 5,06E-05 | 4 - | 0,000504513 - | 4 - | 0,000626457 - | 4 - | 0,000595526 - | 4 - | 0,000604322 - | 3 - | 0,000510575 - |
| 8771 | 1 | 0.2um_NOKCPHJN_935668 | Eukaryota | Phaeophyceae | unclassified | Ectocarpales | Ectocarpaceae | Ectocarpus | Ectocarpus sili | unclassified | unclassified | unclassified | unclassified | unclassified | 0,2 | 0,4 | 4,41E-06 | 8,81E-06 | 0 - | 0 - | 1 - | 2,20E-05 - | 0 - | 0 - | 0 - | 0 - | 0 - | 0 - |
| 8772 | 1 | 0.2um_NOKCPHJN_924169 | Bacteria | Proteobacteri | Alphaproteob | Rhodobactera | Rhodobactera | Celeribacter | Celeribacter h | unclassified | unclassified | unclassified | unclassified | unclassified | 0,8 | 0,4 | 0,000126746 | 6,38E-05 | 0 - | 0 - | 1 - | 0,000158354 - | 1 - | 0,000150536 - | 1 - | 0,000152759 - | 1 - | 0,000172083 - |
| 8773 | 1 | 3um_FIJJAFBF_179795 | Bacteria | Proteobacteri | Alphaproteob | Rhodobactera | Rhodobactera | various | various | information st | Transcription | 05VEQ | Cold shock pro | 52598.EE36_0 | 1,2 | 0,4 | 0,000241947 | 8,34E-05 | 1 - | 0,000168789 - | 1 - | 0,000209587 - | 1 - | 0,000199239 - | 2 - | 0,000404363 - | 1 - | 0,000227757 - |
| 8774 | 1 | 3um_FIJJAFBF_359191 | Bacteria | Proteobacteri | Gammaprotec | Cellvibrionales | unclassified | unclassified | Cellvibrionales | information st | Translation, ri | 179R6 | This protein is | 314287.GB22 | 5,8 | 0,748331477 | 0,000560091 | 8,76E-05 | 5 - | 0,000404144 - | 6 - | 0,000602192 - | 7 - | 0,00066787 - | 6 - | 0,000580915 - | 5 - | 0,000545333 - |
| 8775 | 1 | 3um_FIJJAFBF_488139 | Bacteria | Bacteroidetes | Flavobacteriia | Flavobacterial | Flavobacteriac | Algibacter | Algibacter sp. | unclassified | unclassified | unclassified | unclassified | unclassified | 1,2 | 0,748331477 | 7,91E-05 | 5,35E-05 | 1 - | 5,44E-05 - | 0 - | 0 - | 1 - | 6,42E-05 - | 2 - | 0,000130316 - | 2 - | 0,0001468 - |
| 8776 | 1 | 0.2um_NOKCPHJN_606228 | Bacteria | Proteobacteri | Gammaprotec | Cellvibrionales | Porticoccacea | unclassified | gamma prote | metabolism | Energy produc | 16PVJ | Component of | 314287.GB22 | 0,8 | 0,748331477 | 1,27E-05 | 1,11E-05 | 2 - | 2,87E-05 - | 1 - | 1,78E-05 - | 1 - | 1,69E-05 - | 0 - | 0 - | 0 - | 0 - |
| 8777 | 1 | 0.2um_NOKCPHJN_107953 | Bacteria | Bacteroidetes | Flavobacteriia | Flavobacterial | Flavobacteriac | Polaribacter | Polaribacter s | metabolism | Inorganic ion t | 05C27 | TonB-depende | 313590.MED1 | 2 | 0,632455532 | 2,44E-05 | 7,42E-06 | 2 - | 2,07E-05 - | 2 - | 2,57E-05 - | 3 - | 3,67E-05 - | 2 - | 2,48E-05 - | 1 - | 1,40E-05 - |
| 8778 | 1 | 3um_FIJJAFBF_563594 | Bacteria | Proteobacteri | Gammaprotec | Oceanospirilla | Saccharospirill | Reinekea | Reinekea blan | metabolism | Energy produc | 16QX1 | ferredoxin-NA | 314283.MED2 | 3,2 | 0,979795897 | 0,000172532 | 6,03E-05 | 2 - | 8,90E-05 - | 5 - | 0,000276199 - | 3 - | 0,000157537 - | 3 - | 0,000159864 - | 3 - | 0,000180087 - |
| 8779 | 1 | 3um_FIJJAFBF_458071 | Bacteria | Bacteroidetes | Flavobacteriia | Flavobacterial | Flavobacteriac | unclassified | Flavobacteriac | unclassified | unclassified | unclassified | unclassified | unclassified | 0,2 | 0,4 | 2,33E-06 | 4,66E-06 | 1 - | 1,17E-05 - | 0 - | 0 - | 0 - | 0 - | 0 - | 0 - | 0 - | 0 - |
| 8780 | 1 | 3um_FIJJAFBF_756835 | unclassified | unclassified | unclassified | unclassified | unclassified | unclassified | unclassified | unclassified | unclassified | unclassified | unclassified | unclassified | 0,2 | 0,4 | 2,98E-05 | 5,96E-05 | 1 - | 0,000149061 - | 0 - | 0 - | 0 - | 0 - | 0 - | 0 - | 0 - | 0 - |
| 8781 | 1 | 3um_FIJJAFBF_744841 | Bacteria | Proteobacteri | Gammaprotec | Cellvibrionales | Spongiibacter | Zhongshania | Zhongshania s | information st | Translation, ri | COG0088 | One of the pri | 876044.IMCC | 0,4 | 0,489897949 | 2,51E-05 | 3,10E-05 | 1 - | 5,60E-05 - | 1 - | 6,95E-05 - | 0 - | 0 - | 0 - | 0 - | 0 - | 0 - |
| 8782 | 1 | 3um_FIJJAFBF_122292 | Bacteria | Proteobacteri | Gammaprotec | Oceanospirilla | Saccharospirill | Reinekea | Reinekea fors | information st | Translation, ri | COG0290 | IF-3 binds to t | 314283.MED2 | 1,2 | 0,979795897 | 0,000104802 | 7,60E-05 | 3 - | 0,000237469 - | 1 - | 9,83E-05 - | 1 - | 9,34E-05 - | 1 - | 9,48E-05 - | 0 - | 0 - |
| 8783 | 1 | 3um_FIJJAFBF_26304 | Bacteria | Proteobacteri | Alphaproteob | Rhodobactera | Rhodobactera | Amylibacter | Amylibacter k | metabolism | Nucleotide tra | COG0015 | adenylosuccin | 391595.RLO14 | 1,6 | 0,489897949 | 5,05E-05 | 1,63E-05 | 1 - | 2,63E-05 - | 2 - | 6,54E-05 - | 2 - | 6,21E-05 - | 2 - | 6,31E-05 - | 1 - | 3,55E-05 - |
| 8784 | 1 | 0.2um_NOKCPHJN_15492 | Bacteria | Bacteroidetes | Flavobacteriia | Flavobacterial | Flavobacteriac | Polaribacter | Polaribacter s | unclassified | unclassified | unclassified | unclassified | unclassified | 1 | 0 | 0,000107052 | 1,02E-05 | 1 - | 8,97E-05 - | 1 - | 0,000111343 - | 1 - | 0,000105845 - | 1 - | 0,000107409 - | 1 - | 0,000120996 - |
| 8785 | 1 | 0.2um_NOKCPHJN_366178 | Bacteria | Bacteroidetes | Flavobacteriia | Flavobacterial | Flavobacteriac | Winogradskye | Winogradskye | unclassified | unclassified | unclassified | unclassified | unclassified | 3,2 | 0,748331477 | 0,000118148 | 3,54E-05 | 2 - | 6,07E-05 - | 3 - | 0,00011311 - | 3 - | 0,000107526 - | 4 - | 0,000145485 - | 4 - | 0,000163888 - |
| 8786 | 1 | 3um_FIJJAFBF_678717 | Bacteria | Bacteroidetes | Bacteroidia | Bacteroidales | Bacteroidacea | Bacteroides | Bacteroides o | unclassified | unclassified | unclassified | unclassified | unclassified | 0,2 | 0,4 | 1,56E-06 | 3,13E-06 | 0 - | 0 - | 0 - | 0 - | 1 - | 7,82E-06 - | 0 - | 0 - | 0 - | 0 - |
| 8787 | 1 | 3um_FIJJAFBF_120533 | Bacteria | Proteobacteri | Gammaprotec | Oceanospirilla | Oceanospirilla | Oceanospirillu | Oceanospirillu | cellular proces | Intracellular tr | 05VIE | preprotein tra | 314276.OS145 | 0,6 | 0,8 | 7,05E-05 | 8,91E-05 | 2 - | 0,0002106 - | 0 - | 0 - | 0 - | 0 - | 0 - | 0 - | 1 - | 0,000142087 - |
| 8788 | 1 | 0.2um_NOKCPHJN_01987 | Bacteria | Proteobacteri | Alphaproteob | Rhodobactera | Rhodobactera | Sulfitobacter | Sulfitobacter r | information st | Translation, ri | COG0050 | This protein p | 391624.OIHEL | 22 | 2,607680962 | 0,000774707 | 0,000145439 | 22 - | 0,000645803 - | 21 - | 0,000765447 - | 18 - | 0,000623703 - | 23 - | 0,000808725 - | 26 - | 0,001029856 - |
| 8789 | 1 | 0.2um_NOKCPHJN_58200 | Bacteria | Bacteroidetes | Flavobacteriia | Flavobacterial | Flavobacteriac | Ulvibacter | Ulvibacter sp. | information st | Translation, ri | 0PA05 | 30S ribosomal | 886377.Murru | 0,4 | 0,489897949 | 2,16E-05 | 2,67E-05 | 1 - | 4,82E-05 - | 1 - | 5,99E-05 - | 0 - | 0 - | 0 - | 0 - | 0 - | 0 - |
| 8790 | 1 | 3um_FIJJAFBF_16993 | Bacteria | Proteobacteri | Gammaprotec | Cellvibrionales | unclassified | unclassified | Cellvibrionales | metabolism | Energy produc | 05CV4 | Subunits I and | 314287.GB22 | 4,8 | 1,326649916 | 0,000171417 | 4,90E-05 | 5 - | 0,000150626 - | 7 - | 0,000261846 - | 4 - | 0,000142239 - | 5 - | 0,000180424 - | 3 - | 0,000121948 - |
| 8791 | 1 | 0.2um_NOKCPHJN_264895 | Bacteria | Proteobacteri | Gammaprotec | Cellvibrionales | Porticoccacea | unclassified | various | information st | Transcription | 05D27 | DNA-depende | 314287.GB22 | 1 | 0 | 9,73E-06 | 9,24E-07 | 1 - | 8,15E-06 - | 1 - | 1,01E-05 - | 1 - | 9,62E-06 - | 1 - | 9,76E-06 - | 1 - | 1,10E-05 - |
| 8792 | 1 | 0.2um_NOKCPHJN_293114 | Bacteria | Proteobacteri | Gammaprotec | Cellvibrionales | Porticoccacea | unclassified | various | information st | Translation, ri | 05CEJ | Catalyzes the | 247634.GPB2 | 0,2 | 0,4 | 3,31E-06 | 6,62E-06 | 1 - | 1,65E-05 - | 0 - | 0 - | 0 - | 0 - | 0 - | 0 - | 0 - | 0 - |
| 8793 | 1 | 3um_FIJJAFBF_242580 | Bacteria | Bacteroidetes | Flavobacteriia | Flavobacterial | Flavobacteriac | Formosa | Formosa sp. H | unclassified | unclassified | unclassified | unclassified | unclassified | 0,2 | 0,4 | 2,94E-05 | 5,89E-05 | 1 - | 0,00014715 - | 0 - | 0 - | 0 - | 0 - | 0 - | 0 - | 0 - | 0 - |
| 8794 | 1 | 3um_FIJJAFBF_24302 | Bacteria | Proteobacteri | Gammaprotec | Oceanospirilla | Saccharospirill | Reinekea | Reinekea fors | metabolism | Inorganic ion t | COG0841 | acriflavin resis | 314283.MED2 | 3,2 | 0,748331477 | 4,16E-05 | 7,26E-06 | 4 - | 4,44E-05 - | 3 - | 4,14E-05 - | 4 - | 5,25E-05 - | 3 - | 3,99E-05 - | 2 - | 3,00E-05 - |
| 8795 | 1 | 3um_FIJJAFBF_306644 | Bacteria | Proteobacteri | Alphaproteob | Rhodobactera | Rhodobactera | unclassified | Rhodobactera | unclassified | unclassified | unclassified | unclassified | unclassified | 3,2 | 0,748331477 | 0,000215079 | 3,75E-05 | 4 - | 0,000229554 - | 3 - | 0,000213778 - | 4 - | 0,000270964 - | 3 - | 0,000206225 - | 2 - | 0,000154874 - |
| 8796 | 1 | 0.2um_NOKCPHJN_61813 | Bacteria | Bacteroidetes | Flavobacteriia | Flavobacterial | Flavobacteriac | Polaribacter | Polaribacter fi | poorly charact | Function unkn | 0XY6Y | Pkd domain cc | 391587.KAOT | 5,4 | 1,019803903 | 0,000163186 | 3,28E-05 | 6 - | 0,000152359 - | 7 - | 0,000220715 - | 5 - | 0,00014987 - | 4 - | 0,000121667 - | 5 - | 0,000171321 - |
| 8797 | 1 | 3um_FIJJAFBF_786151 | Bacteria | Proteobacteri | Gammaprotec | Oceanospirilla | Saccharospirill | Reinekea | Reinekea fors | unclassified | unclassified | unclassified | unclassified | unclassified | 3,4 | 1,019803903 | 0,000429859 | 0,000144274 | 3 - | 0,000315899 - | 2 - | 0,000261503 - | 3 - | 0,000372887 - | 5 - | 0,000630657 - | 4 - | 0,000568347 - |
| 8798 | 1 | 0.2um_NOKCPHJN_198405 | Bacteria | Bacteroidetes | Flavobacteriia | Flavobacterial | Flavobacteriac | Aureitalea | Aureitalea ma | metabolism | Inorganic ion t | 05C27 | TonB-depende | 398720.MED2 | 0,4 | 0,489897949 | 4,49E-06 | 5,54E-06 | 1 - | 1,02E-05 - | 0 - | 0 - | 0 - | 0 - | 1 - | 1,22E-05 - | 0 - | 0 - |
| 8799 | 1 | 0.2um_NOKCPHJN_40497 | Bacteria | Bacteroidetes | Flavobacteriia | Flavobacterial | Flavobacteriac | Algibacter | Algibacter sp. | poorly charact | Function unkn | COG3975 | peptidase m6 | 983544.Lacal_ | 1,8 | 0,979795897 | 3,94E-05 | 2,23E-05 | 2 - | 3,65E-05 - | 3 - | 6,80E-05 - | 2 - | 4,31E-05 - | 0 - | 0 - | 2 - | 4,92E-05 - |
| 8800 | 1 | 3um_FIJJAFBF_630544 | Eukaryota | unclassified | unclassified | Prymnesiales | Chrysochromu | Chrysochromu | Chrysochromu | poorly charact | Function unkn | 110WY | Chlorophyll A- | 2850.Phatr16 | 2,6 | 0,489897949 | 0,000271546 | 4,59E-05 | 3 - | 0,000264869 - | 3 - | 0,00032889 - | 2 - | 0,000208434 - | 3 - | 0,000317269 - | 2 - | 0,000238268 - |
| 8801 | 1 | 3um_FIJJAFBF_551806 | Bacteria | Proteobacteri | Alphaproteob | Rhodobactera | Rhodobactera | Hasllibacter | Hasllibacter ha | unclassified | unclassified | unclassified | unclassified | unclassified | 11,6 | 1,019803903 | 0,000381039 | 3,13E-05 | 13 - | 0,000359542 - | 12 - | 0,000412103 - | 12 - | 0,000391756 - | 10 - | 0,000331285 - | 11 - | 0,000410511 - |
| 8802 | 1 | 0.2um_NOKCPHJN_273609 | Eukaryota | unclassified | unclassified | Choanoflagelli | Salpingoecida | Monosiga | Monosiga bre | poorly charact | Function unkn | 0Y1WB | NA | 431895.XP_00 | 0,2 | 0,4 | 5,86E-06 | 1,17E-05 | 0 - | 0 - | 0 - | 0 - | 0 - | 0 - | 1 - | 2,93E-05 - | 0 - | 0 - |
| 8803 | 1 | 3um_FIJJAFBF_30211 | Bacteria | Proteobacteri | Alphaproteob | Rhodobactera | Rhodobactera | unclassified | Rhodobactera | unclassified | unclassified | unclassified | unclassified | unclassified | 0,2 | 0,4 | 1,49E-05 | 2,98E-05 | 0 - | 0 - | 0 - | 0 - | 1 - | 7,44E-05 - | 0 - | 0 - | 0 - | 0 - |
| 8804 | 1 | 3um_FIJJAFBF_222229 | Bacteria | Proteobacteri | Alphaproteob | Rhodobactera | Rhodobactera | various | various | unclassified | unclassified | unclassified | unclassified | unclassified | 2,2 | 0,4 | 0,000161909 | 3,16E-05 | 2 - | 0,000123416 - | 2 - | 0,000153246 - | 3 - | 0,00021852 - | 2 - | 0,000147832 - | 2 - | 0,000166532 - |
| 8805 | 1 | 0.2um_NOKCPHJN_39805 | Bacteria | Bacteroidetes | Flavobacteriia | Flavobacterial | Flavobacteriac | Polaribacter | various | unclassified | unclassified | unclassified | unclassified | unclassified | 3,2 | 0,748331477 | 0,000340191 | 7,58E-05 | 4 - | 0,000358677 - | 4 - | 0,000445371 - | 2 - | 0,000211691 - | 3 - | 0,000322226 - | 3 - | 0,000362987 - |
| 8806 | 2 | 0.2um_NOKCPHJN_545881;3um_FIJJAFBF_15582 | Bacteria | Proteobacteri | Alphaproteob | Rhodobactera | Rhodobactera | Planktomarina | Planktomarina | unclassified | unclassified | unclassified | unclassified | unclassified | 1,2 | 0,748331477 | 0,000105476 | 6,30E-05 | 2 - | 0,000155104 - | 1 - | 9,63E-05 - | 2 - | 0,000183084 - | 1 - | 9,29E-05 - | 0 - | 0 - |
| 8807 | 1 | 0.2um_NOKCPHJN_385093 | Bacteria | Proteobacteri | Alphaproteob | Rhodospirillal | Rhodospirillac | Tistlia | Tistlia consote | metabolism | Secondary me | 16Y20 | Inherit from C | 414684.RC1_3 | 0,2 | 0,4 | 8,26E-07 | 1,65E-06 | 1 - | 4,13E-06 - | 0 - | 0 - | 0 - | 0 - | 0 - | 0 - | 0 - | 0 - |
| 8808 | 1 | 3um_FIJJAFBF_356447 | Bacteria | Proteobacteri | Alphaproteob | Rhodobactera | Rhodobactera | Leisingera | Leisingera sp. | poorly charact | Function unkn | COG1744 | basic membra | 439497.RR11_ | 3,4 | 1,019803903 | 0,000128083 | 3,39E-05 | 4 - | 0,000128601 - | 3 - | 0,000119764 - | 5 - | 0,000189751 - | 3 - | 0,000115532 - | 2 - | 8,68E-05 - |
| 8809 | 1 | 3um_FIJJAFBF_259011 | Bacteria | Proteobacteri | Gammaprotec | Pseudomonad | Moraxellaceae | Psychrobacter | various | cellular proces | Cell wall/mem | 08ZG0 | Ompa motb d | 335284.Pcryo_ | 0,8 | 0,748331477 | 4,83E-05 | 4,18E-05 | 2 - | 0,0001053 - | 1 - | 6,54E-05 - | 0 - | 0 - | 0 - | 0 - | 1 - | 7,10E-05 - |
| 8810 | 1 | 0.2um_NOKCPHJN_106618 | Bacteria | Proteobacteri | Alphaproteob | Rhodobactera | Rhodobactera | Tateyamaria | Tateyamaria s | metabolism | Energy produc | 16QPK | Succinate deh | 391589.RGAI1 | 0,8 | 0,4 | 1,81E-05 | 9,27E-06 | 1 - | 1,91E-05 - | 0 - | 0 - | 1 - | 2,25E-05 - | 1 - | 2,29E-05 - | 1 - | 2,58E-05 - |
| 8811 | 1 | 3um_FIJJAFBF_520477 | Bacteria | Bacteroidetes | Flavobacteriia | Flavobacterial | Flavobacteriac | Polaribacter | Polaribacter a | cellular proces | Intracellular tr | 0PBE9 | Preprotein tra | 313598.MED1 | 3,8 | 0,4 | 0,000494862 | 6,53E-05 | 4 - | 0,000437245 - | 3 - | 0,000407197 - | 4 - | 0,000516123 - | 4 - | 0,000523746 - | 4 - | 0,000589998 - |
| 8812 | 1 | 0.2um_NOKCPHJN_803799 | Bacteria | Proteobacteri | Alphaproteob | Pelagibacteral | Pelagibacterac | Candidatus Pe | Candidatus Pe | cellular proces | Cell wall/mem | 7257 | NA | 335992.SAR11 | 0,8 | 0,748331477 | 3,36E-05 | 3,19E-05 | 0 - | 0 - | 2 - | 8,59E-05 - | 1 - | 4,08E-05 - | 1 - | 4,14E-05 - | 0 - | 0 - |
| 8813 | 1 | 0.2um_NOKCPHJN_123742 | unclassified | unclassified | unclassified | unclassified | unclassified | unclassified | unclassified | unclassified | unclassified | unclassified | unclassified | unclassified | 0,4 | 0,489897949 | 6,29E-05 | 7,71E-05 | 0 - | 0 - | 1 - | 0,000160134 - | 0 - | 0 - | 1 - | 0,000154476 - | 0 - | 0 - |
| 8814 | 1 | 3um_FIJJAFBF_786155 | Bacteria | Proteobacteri | Gammaprotec | Oceanospirilla | Saccharospirill | Reinekea | Reinekea fors | unclassified | unclassified | unclassified | unclassified | unclassified | 1,8 | 0,979795897 | 0,000179718 | 9,78E-05 | 1 - | 8,38E-05 - | 1 - | 0,000104028 - | 3 - | 0,000296676 - | 3 - | 0,000301058 - | 1 - | 0,000113047 - |
| 8815 | 1 | 3um_FIJJAFBF_278984 | Bacteria | Bacteroidetes | Flavobacteriia | Flavobacterial | Flavobacteriac | Polaribacter | Polaribacter a | metabolism | Lipid transpor | 07QM9 | Component of | 983544.Lacal_ | 2,6 | 1,019803903 | 0,000111886 | 4,64E-05 | 2 - | 7,24E-05 - | 4 - | 0,000179835 - | 3 - | 0,000128217 - | 3 - | 0,00013011 - | 1 - | 4,89E-05 - |
| 8816 | 1 | 3um_FIJJAFBF_367138 | Bacteria | Proteobacteri | Gammaprotec | Oceanospirilla | Saccharospirill | Reinekea | Reinekea blan | cellular proces | Cell cycle cont | 16Q4W | site-determini | 349521.HCH_ | 0,2 | 0,4 | 9,89E-06 | 1,98E-05 | 0 - | 0 - | 0 - | 0 - | 1 - | 4,94E-05 - | 0 - | 0 - | 0 - | 0 - |
| 8817 | 1 | 0.2um_NOKCPHJN_27907 | Bacteria | Bacteroidetes | Flavobacteriia | Flavobacterial | Flavobacteriac | Polaribacter | Polaribacter s | metabolism | Carbohydrate | 05CA9 | Pyruvate kina | 313594.PI23P | 0,2 | 0,4 | 6,52E-06 | 1,30E-05 | 0 - | 0 - | 0 - | 0 - | 0 - | 0 - | 0 - | 0 - | 1 - | 3,26E-05 - |
| 8818 | 1 | 3um_FIJJAFBF_386649 | unclassified | unclassified | unclassified | unclassified | unclassified | unclassified | unclassified | unclassified | unclassified | unclassified | unclassified | unclassified | 0,2 | 0,4 | 1,60E-05 | 3,20E-05 | 0 - | 0 - | 0 - | 0 - | 0 - | 0 - | 1 - | 7,99E-05 - | 0 - | 0 - |
| 8819 | 1 | 3um_FIJJAFBF_185502 | Bacteria | Proteobacteri | Alphaproteob | Pelagibacteral | Pelagibacterac | Candidatus Pe | Candidatus Pe | metabolism | Inorganic ion t | 05PWN | phosphate ab | 335992.SAR11 | 2 | 0,632455532 | 7,80E-05 | 1,81E-05 | 3 - | 0,000100977 - | 2 - | 8,36E-05 - | 2 - | 7,95E-05 - | 2 - | 8,06E-05 - | 1 - | 4,54E-05 - |
| 8820 | 1 | 0.2um_NOKCPHJN_115259 | Bacteria | Proteobacteri | Alphaproteob | Rhodobactera | Rhodobactera | Epibacterium | Epibacterium | metabolism | Inorganic ion t | 16PAX | extracellular s | 644076.SCH4 | 1,4 | 0,489897949 | 4,44E-05 | 1,57E-05 | 1 - | 2,66E-05 - | 1 - | 3,30E-05 - | 2 - | 6,27E-05 - | 2 - | 6,36E-05 - | 1 - | 3,59E-05 - |
| 8821 | 1 | 0.2um_NOKCPHJN_85830 | Bacteria | Proteobacteri | Alphaproteob | Rhodobactera | Hyphomonada | unclassified | Hyphomonada | unclassified | unclassified | unclassified | unclassified | unclassified | 1 | 0 | 2,16E-05 | 2,06E-06 | 1 - | 1,81E-05 - | 1 - | 2,25E-05 - | 1 - | 2,14E-05 - | 1 - | 2,17E-05 - | 1 - | 2,45E-05 - |
| 8822 | 1 | 0.2um_NOKCPHJN_291839 | Bacteria | Proteobacteri | Alphaproteob | Rhodobactera | unclassified | unclassified | Rhodobactera | unclassified | unclassified | unclassified | unclassified | unclassified | 3,6 | 1,019803903 | 0,000129815 | 3,76E-05 | 4 - | 0,000121457 - | 2 - | 7,54E-05 - | 5 - | 0,000179209 - | 3 - | 0,000109114 - | 4 - | 0,000163888 - |
| 8823 | 1 | 0.2um_NOKCPHJN_781170 | Eukaryota | unclassified | Raphidophyce | Chattonellales | Chattonellace | Heterosigma | Heterosigma a | unclassified | unclassified | unclassified | unclassified | unclassified | 1,8 | 0,4 | 0,000118575 | 2,34E-05 | 2 - | 0,000111977 - | 2 - | 0,000139043 - | 2 - | 0,000132178 - | 2 - | 0,00013413 - | 1 - | 7,55E-05 - |
| 8824 | 1 | 3um_FIJJAFBF_242061 | Bacteria | Proteobacteri | Alphaproteob | Rhodobactera | unclassified | unclassified | Rhodobactera | poorly charact | Function unkn | 0218I | Lipoprotein | 314262.MED1 | 0,4 | 0,489897949 | 2,92E-05 | 3,60E-05 | 1 - | 6,63E-05 - | 0 - | 0 - | 0 - | 0 - | 1 - | 7,95E-05 - | 0 - | 0 - |
| 8825 | 1 | 3um_FIJJAFBF_299529 | Bacteria | Proteobacteri | Alphaproteob | Rhodobactera | various | various | various | unclassified | unclassified | unclassified | unclassified | unclassified | 6,8 | 0,4 | 0,000644594 | 4,60E-05 | 7 - | 0,000557943 - | 7 - | 0,0006928 - | 7 - | 0,000658594 - | 7 - | 0,000668322 - | 6 - | 0,00064531 - |
| 8826 | 1 | 3um_FIJJAFBF_562790 | Bacteria | Proteobacteri | Gammaprotec | Oceanospirilla | Saccharospirill | Reinekea | Reinekea blan | metabolism | Nucleotide tra | 0QI1Y | Carbamoyl-ph | 314283.MED2 | 0,8 | 0,748331477 | 2,93E-05 | 2,71E-05 | 0 - | 0 - | 1 - | 3,79E-05 - | 2 - | 7,21E-05 - | 1 - | 3,66E-05 - | 0 - | 0 - |
| 8827 | 1 | 0.2um_NOKCPHJN_319419 | Bacteria | Bacteroidetes | Flavobacteriia | Flavobacterial | Flavobacteriac | Cellulophaga | Cellulophaga s | cellular proces | Posttranslatio | COG0612 | peptidase' | 487797.Flav3 | 0,8 | 0,748331477 | 1,46E-05 | 1,29E-05 | 2 - | 3,33E-05 - | 0 - | 0 - | 1 - | 1,97E-05 - | 1 - | 2,00E-05 - | 0 - | 0 - |
| 8828 | 1 | 3um_FIJJAFBF_06027 | Bacteria | Bacteroidetes | Flavobacteriia | Flavobacterial | Flavobacteriac | Siansivirga | Siansivirga zea | unclassified | unclassified | unclassified | unclassified | unclassified | 6,8 | 1,469693846 | 0,000145237 | 2,74E-05 | 8 - | 0,000144829 - | 8 - | 0,000179835 - | 8 - | 0,000170955 - | 5 - | 0,000108425 - | 5 - | 0,000122141 - |
| 8829 | 1 | 0.2um_NOKCPHJN_39142 | Eukaryota | Phaeophyceae | unclassified | Laminariales | Agaraceae | Costaria | Costaria costa | metabolism | Energy produc | COG4451 | RuBisCO catal | 288000.BBta_ | 14,4 | 1,356465997 | 0,001409356 | 8,75E-05 | 16 - | 0,001321171 - | 13 - | 0,00133291 - | 16 - | 0,001559508 - | 14 - | 0,001384724 - | 13 - | 0,001448466 - |
| 8830 | 1 | 3um_FIJJAFBF_266859 | Bacteria | Bacteroidetes | Flavobacteriia | Flavobacterial | Flavobacteriac | Spongiibacteri | Spongiibacteri | metabolism | Energy produc | COG0056 | Produces ATP | 688270.Celal_ | 3 | 1,095445115 | 7,79E-05 | 2,93E-05 | 4 - | 8,76E-05 - | 1 - | 2,72E-05 - | 3 - | 7,76E-05 - | 3 - | 7,87E-05 - | 4 - | 0,000118225 - |
| 8831 | 1 | 3um_FIJJAFBF_565140 | unclassified | unclassified | unclassified | unclassified | unclassified | unclassified | unclassified | unclassified | unclassified | unclassified | unclassified | unclassified | 1,2 | 0,748331477 | 0,000202566 | 0,000133554 | 1 - | 0,000143471 - | 2 - | 0,000356297 - | 1 - | 0,000169353 - | 2 - | 0,000343708 - | 0 - | 0 - |
| 8832 | 1 | 3um_FIJJAFBF_65172 | Bacteria | Bacteroidetes | Flavobacteriia | Flavobacterial | Flavobacteriac | Mesoflavibact | Mesoflavibact | poorly charact | Function unkn | 0PB02 | FN3 | 216432.CA25 | 5,8 | 1,166190379 | 0,000168337 | 2,52E-05 | 7 - | 0,000172782 - | 6 - | 0,000183895 - | 7 - | 0,000203952 - | 5 - | 0,000147832 - | 4 - | 0,000133225 - |
| 8833 | 1 | 0.2um_NOKCPHJN_154726 | Bacteria | Bacteroidetes | Flavobacteriia | Flavobacterial | Flavobacteriac | Formosa | Formosa sp. H | unclassified | unclassified | unclassified | unclassified | unclassified | 9,4 | 1,2 | 0,000128548 | 1,63E-05 | 10 - | 0,000115122 - | 10 - | 0,000142948 - | 8 - | 0,000108712 - | 11 - | 0,000151687 - | 8 - | 0,000124272 - |
| 8834 | 1 | 3um_FIJJAFBF_453277 | Bacteria | Proteobacteri | Alphaproteob | Rhodobactera | various | various | various | metabolism | Carbohydrate | COG0176 | Transaldolase | 246200.SPO3 | 0,8 | 0,4 | 5,05E-05 | 2,59E-05 | 1 - | 5,29E-05 - | 1 - | 6,57E-05 - | 1 - | 6,24E-05 - | 0 - | 0 - | 1 - | 7,14E-05 - |
| 8835 | 1 | 0.2um_NOKCPHJN_140944 | Bacteria | Bacteroidetes | unclassified | unclassified | unclassified | unclassified | Bacteroidetes | unclassified | unclassified | unclassified | unclassified | unclassified | 4,4 | 0,8 | 8,16E-05 | 1,60E-05 | 5 - | 7,80E-05 - | 3 - | 5,81E-05 - | 5 - | 9,20E-05 - | 4 - | 7,47E-05 - | 5 - | 0,000105214 - |
| 8836 | 1 | 3um_FIJJAFBF_270524 | Bacteria | Proteobacteri | Alphaproteob | Rhodobactera | Rhodobactera | unclassified | Rhodobactera | metabolism | Inorganic ion t | 16PAX | extracellular s | 388401.RB215 | 2,6 | 0,8 | 8,39E-05 | 2,66E-05 | 3 - | 8,16E-05 - | 1 - | 3,38E-05 - | 3 - | 9,63E-05 - | 3 - | 9,77E-05 - | 3 - | 0,0001101 - |
| 8837 | 1 | 0.2um_NOKCPHJN_675065 | Bacteria | Proteobacteri | Alphaproteob | Pelagibacteral | Pelagibacterac | Candidatus Pe | Candidatus Pe | metabolism | Secondary me | 0837G | toluene tolera | 335992.SAR11 | 0,4 | 0,489897949 | 2,54E-05 | 3,13E-05 | 1 - | 5,83E-05 - | 0 - | 0 - | 1 - | 6,88E-05 - | 0 - | 0 - | 0 - | 0 - |
| 8838 | 1 | 0.2um_NOKCPHJN_330360 | Eukaryota | Arthropoda | Insecta | Hymenoptera | Apidae | Eufriesea | Eufriesea mex | unclassified | unclassified | unclassified | unclassified | unclassified | 0,8 | 0,4 | 8,37E-06 | 4,30E-06 | 1 - | 8,85E-06 - | 0 - | 0 - | 1 - | 1,04E-05 - | 1 - | 1,06E-05 - | 1 - | 1,19E-05 - |
| 8839 | 1 | 0.2um_NOKCPHJN_356130 | Bacteria | Bacteroidetes | Flavobacteriia | unclassified | unclassified | unclassified | Flavobacteria | metabolism | Inorganic ion t | 05C27 | TonB-depende | 487796.Flav2 | 7,8 | 0,979795897 | 9,98E-05 | 8,44E-06 | 9 - | 9,74E-05 - | 7 - | 9,40E-05 - | 9 - | 0,000114924 - | 7 - | 9,07E-05 - | 7 - | 0,000102179 - |
| 8840 | 1 | 0.2um_NOKCPHJN_356379 | Bacteria | Proteobacteri | Gammaprotec | Cellvibrionales | Spongiibacter | unclassified | marine gamm | metabolism | Inorganic ion t | 16TQC | TONB-depend | 498211.CJA_0 | 1,2 | 0,4 | 1,56E-05 | 3,46E-06 | 2 - | 2,24E-05 - | 1 - | 1,39E-05 - | 1 - | 1,32E-05 - | 1 - | 1,34E-05 - | 1 - | 1,51E-05 - |
| 8841 | 1 | 3um_FIJJAFBF_227244 | Bacteria | Proteobacteri | Gammaprotec | unclassified | unclassified | unclassified | Gammaprotec | unclassified | unclassified | unclassified | unclassified | unclassified | 2,2 | 0,4 | 0,00014211 | 1,30E-05 | 3 - | 0,000164751 - | 2 - | 0,000136382 - | 2 - | 0,000129648 - | 2 - | 0,000131563 - | 2 - | 0,000148205 - |
| 8842 | 1 | 0.2um_NOKCPHJN_487262 | Bacteria | Bacteroidetes | Flavobacteriia | Flavobacterial | Flavobacteriac | Formosa | Formosa sp. H | unclassified | unclassified | unclassified | unclassified | unclassified | 1,8 | 0,748331477 | 0,000133261 | 4,44E-05 | 3 - | 0,000192363 - | 2 - | 0,000159239 - | 2 - | 0,000151377 - | 1 - | 7,68E-05 - | 1 - | 8,65E-05 - |
| 8843 | 1 | 0.2um_NOKCPHJN_386079 | Eukaryota | Bacillariophyt | Bacillariophyc | Naviculales | Phaeodactylac | Phaeodactylu | Phaeodactylu | cellular proces | Posttranslatio | COG1404 | peptidase (S8 | 2850.Phatr26 | 0,2 | 0,4 | 1,96E-06 | 3,92E-06 | 0 - | 0 - | 0 - | 0 - | 0 - | 0 - | 1 - | 9,81E-06 - | 0 - | 0 - |
| 8844 | 1 | 0.2um_NOKCPHJN_481955 | Bacteria | Bacteroidetes | Flavobacteriia | Flavobacterial | Flavobacteriac | Oceanihabitar | Oceanihabitar | unclassified | unclassified | unclassified | unclassified | unclassified | 0,2 | 0,4 | 5,28E-06 | 1,06E-05 | 0 - | 0 - | 0 - | 0 - | 0 - | 0 - | 1 - | 2,64E-05 - | 0 - | 0 - |
| 8845 | 1 | 3um_FIJJAFBF_333515 | Bacteria | Proteobacteri | Alphaproteob | Rhodobactera | unclassified | unclassified | Rhodobactera | metabolism | Lipid transpor | COG0511 | Acetyl-CoA ca | 439496.RBY4I | 0,6 | 0,8 | 5,52E-05 | 7,55E-05 | 0 - | 0 - | 0 - | 0 - | 0 - | 0 - | 1 - | 8,49E-05 - | 2 - | 0,000191203 - |
| 8846 | 1 | 0.2um_NOKCPHJN_176491 | Bacteria | unclassified | unclassified | unclassified | unclassified | unclassified | bacterium TM | metabolism | Energy produc | COG0056 | Produces ATP | 43989.cce_44 | 0,6 | 0,8 | 1,46E-05 | 1,88E-05 | 2 - | 4,59E-05 - | 0 - | 0 - | 1 - | 2,71E-05 - | 0 - | 0 - | 0 - | 0 - |
| 8847 | 1 | 3um_FIJJAFBF_19904 | Bacteria | Proteobacteri | Alphaproteob | Rhodobactera | Rhodobactera | Planktomarina | Planktomarina | unclassified | unclassified | unclassified | unclassified | unclassified | 0,4 | 0,489897949 | 5,36E-05 | 6,63E-05 | 1 - | 0,000119559 - | 1 - | 0,000148457 - | 0 - | 0 - | 0 - | 0 - | 0 - | 0 - |
| 8848 | 1 | 3um_FIJJAFBF_382727 | Bacteria | Cyanobacteria | unclassified | Oscillatoriales | Microcoleacea | Arthrospira | Arthrospira sp | information st | RNA processin | 0DKQ1 | Splicing regula | 13735.ENSPSI | 0,8 | 0,4 | 5,91E-05 | 3,04E-05 | 1 - | 6,17E-05 - | 1 - | 7,66E-05 - | 0 - | 0 - | 1 - | 7,39E-05 - | 1 - | 8,33E-05 - |
| 8849 | 1 | 0.2um_NOKCPHJN_597387 | Bacteria | Bacteroidetes | unclassified | unclassified | unclassified | unclassified | Bacteroidetes | metabolism | Amino acid tra | 09GJ6 | nitrogen regul | 688270.Celal_ | 1,8 | 0,748331477 | 0,000216443 | 7,90E-05 | 3 - | 0,000307438 - | 2 - | 0,000254498 - | 1 - | 0,000120966 - | 1 - | 0,000122753 - | 2 - | 0,000276562 - |
| 8850 | 1 | 3um_FIJJAFBF_293508 | Bacteria | Bacteroidetes | Cytophagia | Cytophagales | Flammeovirga | unclassified | Flammeovirga | metabolism | Amino acid tra | COG1748 | saccharopine | 742766.HMPR | 0,2 | 0,4 | 5,71E-06 | 1,14E-05 | 1 - | 2,86E-05 - | 0 - | 0 - | 0 - | 0 - | 0 - | 0 - | 0 - | 0 - |
| 8851 | 1 | 3um_FIJJAFBF_677249 | Bacteria | Proteobacteri | Gammaprotec | unclassified | unclassified | unclassified | Gammaprotec | metabolism | Energy produc | 0QIP8 | Electron trans | 207954.MED9 | 0,6 | 0,489897949 | 2,80E-05 | 2,30E-05 | 0 - | 0 - | 1 - | 4,61E-05 - | 1 - | 4,38E-05 - | 0 - | 0 - | 1 - | 5,01E-05 - |
| 8852 | 1 | 0.2um_NOKCPHJN_245726 | Eukaryota | unclassified | unclassified | unclassified | unclassified | Blastocystis | Blastocystis h | unclassified | unclassified | unclassified | unclassified | unclassified | 0,4 | 0,489897949 | 7,91E-06 | 9,75E-06 | 1 - | 1,80E-05 - | 0 - | 0 - | 0 - | 0 - | 1 - | 2,15E-05 - | 0 - | 0 - |
| 8853 | 1 | 0.2um_NOKCPHJN_556389 | Eukaryota | Chordata | Mammalia | Chiroptera | Phyllostomida | Desmodus | Desmodus rot | information st | Chromatin str | 13R2C | Histone H2A | 4932.YBL003C | 1 | 0 | 1,41E-05 | 1,34E-06 | 1 - | 1,18E-05 - | 1 - | 1,46E-05 - | 1 - | 1,39E-05 - | 1 - | 1,41E-05 - | 1 - | 1,59E-05 - |
| 8854 | 1 | 3um_FIJJAFBF_152376 | unclassified | unclassified | unclassified | unclassified | unclassified | unclassified | unclassified | unclassified | unclassified | unclassified | unclassified | unclassified | 6,4 | 0,489897949 | 0,000339571 | 4,09E-05 | 7 - | 0,00031141 - | 6 - | 0,000331439 - | 6 - | 0,000315075 - | 6 - | 0,000319729 - | 7 - | 0,000420202 - |
| 8855 | 1 | 3um_FIJJAFBF_440053 | Bacteria | Proteobacteri | Alphaproteob | Rhodobactera | Rhodobactera | Loktanella | Loktanella sp. | metabolism | Carbohydrate | COG0057 | glyceraldehyd | 391593.RCCS2 | 0,2 | 0,4 | 8,26E-06 | 1,65E-05 | 0 - | 0 - | 0 - | 0 - | 0 - | 0 - | 1 - | 4,13E-05 - | 0 - | 0 - |
| 8856 | 2 | 3um_FIJJAFBF_134824;3um_FIJJAFBF_732506 | Bacteria | Bacteroidetes | Flavobacteriia | Flavobacterial | Crocinitomica | unclassified | Crocinitomica | unclassified | unclassified | unclassified | unclassified | unclassified | 0,6 | 0,8 | 2,54E-05 | 3,37E-05 | 0 - | 0 - | 0 - | 0 - | 2 - | 8,42E-05 - | 1 - | 4,27E-05 - | 0 - | 0 - |
| 8857 | 1 | 3um_FIJJAFBF_40037 | Bacteria | Bacteroidetes | Flavobacteriia | Flavobacterial | Flavobacteriac | Cellulophaga | Cellulophaga s | information st | Replication, re | COG0187 | DNA gyrase ne | 487797.Flav3 | 1 | 0 | 2,12E-05 | 2,01E-06 | 1 - | 1,78E-05 - | 1 - | 2,21E-05 - | 1 - | 2,10E-05 - | 1 - | 2,13E-05 - | 1 - | 2,40E-05 - |
| 8858 | 1 | 3um_FIJJAFBF_149707 | Bacteria | Proteobacteri | Betaproteoba | unclassified | unclassified | unclassified | Betaproteoba | information st | Replication, re | COG0513 | purine NTP-de | 535289.Dtpsy | 0,2 | 0,4 | 4,50E-06 | 9,01E-06 | 0 - | 0 - | 1 - | 2,25E-05 - | 0 - | 0 - | 0 - | 0 - | 0 - | 0 - |
| 8859 | 1 | 3um_FIJJAFBF_306876 | Bacteria | Proteobacteri | Alphaproteob | Rhodobactera | unclassified | unclassified | Rhodobactera | unclassified | unclassified | unclassified | unclassified | unclassified | 1 | 0 | 5,15E-05 | 4,89E-06 | 1 - | 4,31E-05 - | 1 - | 5,36E-05 - | 1 - | 5,09E-05 - | 1 - | 5,17E-05 - | 1 - | 5,82E-05 - |
| 8860 | 1 | 3um_FIJJAFBF_394815 | Bacteria | Proteobacteri | Alphaproteob | various | various | various | various | unclassified | unclassified | unclassified | unclassified | unclassified | 0,6 | 0,489897949 | 0,000107706 | 8,86E-05 | 1 - | 0,000159412 - | 0 - | 0 - | 1 - | 0,00018817 - | 1 - | 0,000190949 - | 0 - | 0 - |

| | | | | | | | | | | | | | | | | | | | | | | | | | | | | |
|---|---|---|---|---|---|---|---|---|---|---|---|---|---|---|---|---|---|---|---|---|---|---|---|---|---|---|---|---|
| 8861 | 1 | 0.2um_NOKCPHJN_260410 | Bacteria | Proteobacteri | Gammaprotec | Oceanospirilla | Saccharospirill | Saccharospiril | Saccharospiril | unclassified | unclassified | unclassified | unclassified | unclassified | 2,4 | 1,019803903 | 7,76E-05 | 2,74E-05 | 4 - | 0,000112251 - | 3 - | 0,000104537 - | 2 - | 6,63E-05 - | 2 - | 6,72E-05 - | 1 - | 3,79E-05 - |
| 8862 | 1 | 3um_FIJJAFBF_102770 | unclassified | unclassified | unclassified | unclassified | unclassified | unclassified | unclassified | unclassified | unclassified | unclassified | unclassified | unclassified | 0,8 | 0,748331477 | 0,000155371 | 0,000148661 | 1 - | 0,000168789 - | 1 - | 0,000209587 - | 2 - | 0,000398477 - | 0 - | 0 - | 0 - | 0 - |
| 8863 | 1 | 3um_FIJJAFBF_557220 | Eukaryota | Chlorophyta | Ulvophyceae | Ulvales | Ulvaceae | Ulva | Ulva fasciata | metabolism | Energy produc | COG0055 | Produces ATP | 3702.ATCG00 | 1,6 | 0,489897949 | 4,52E-05 | 1,44E-05 | 2 - | 4,75E-05 - | 1 - | 2,95E-05 - | 1 - | 2,81E-05 - | 2 - | 5,69E-05 - | 2 - | 6,41E-05 - |
| 8864 | 1 | 3um_FIJJAFBF_368472 | Bacteria | Proteobacteri | Alphaproteob | Rhodospirillal | unclassified | unclassified | Rhodospirillal | metabolism | Inorganic ion t | 05PWN | phosphate ab | 156889.Mmc1 | 3,6 | 1,019803903 | 0,000143456 | 4,51E-05 | 4 - | 0,000133075 - | 5 - | 0,000206549 - | 2 - | 7,85E-05 - | 3 - | 0,000119551 - | 4 - | 0,000179565 - |
| 8865 | 1 | 3um_FIJJAFBF_629419 | Bacteria | Bacteroidetes | Flavobacteriia | Flavobacterial | Flavobacteriac | Polaribacter | Polaribacter t | metabolism | Inorganic ion t | 05C27 | TonB-depende | 873517.HMPR | 14,2 | 1,720465053 | 0,000192837 | 1,42E-05 | 17 - | 0,00019512 - | 14 - | 0,000199526 - | 15 - | 0,000203223 - | 12 - | 0,00016498 - | 13 - | 0,000201337 - |
| 8866 | 1 | 3um_FIJJAFBF_467049 | Bacteria | Bacteroidetes | Flavobacteriia | Flavobacterial | Flavobacteriac | Polaribacter | Polaribacter s | cellular proces | Intracellular tr | 0PBE9 | Preprotein tra | 313594.PI23P | 1 | 0 | 0,000129271 | 1,23E-05 | 1 - | 0,00010828 - | 1 - | 0,000134452 - | 1 - | 0,000127813 - | 1 - | 0,000129701 - | 1 - | 0,000146108 - |
| 8867 | 1 | 0.2um_NOKCPHJN_391505 | Bacteria | Proteobacteri | Alphaproteob | Rhodobactera | Rhodobactera | Aquimixticola | Aquimixticola | information st | Translation, ri | COG0173 | aspartyl-trna s | 314232.SKA53 | 0,8 | 0,4 | 1,86E-05 | 9,56E-06 | 1 - | 1,94E-05 - | 1 - | 2,41E-05 - | 0 - | 0 - | 1 - | 2,33E-05 - | 1 - | 2,62E-05 - |
| 8868 | 1 | 3um_FIJJAFBF_188372 | Bacteria | Bacteroidetes | unclassified | unclassified | unclassified | unclassified | Bacteroidetes | unclassified | unclassified | unclassified | unclassified | unclassified | 3 | 0 | 0,000153389 | 1,46E-05 | 3 - | 0,000128481 - | 3 - | 0,000159536 - | 3 - | 0,000151659 - | 3 - | 0,000153899 - | 3 - | 0,000173367 - |
| 8869 | 1 | 0.2um_NOKCPHJN_15784 | Bacteria | Proteobacteri | Gammaprotec | Alteromonada | Alteromonada | Paraglaciecola | Paraglaciecola | poorly charact | Function unkn | 16R8I | Lipoprotein | 1085623.GNIT | 14 | 1,095445115 | 0,000205459 | 2,59E-05 | 14 - | 0,000172042 - | 15 - | 0,000228885 - | 15 - | 0,000217584 - | 12 - | 0,000176638 - | 14 - | 0,000232146 - |
| 8870 | 1 | 3um_FIJJAFBF_183394 | Bacteria | Proteobacteri | Alphaproteob | Rhodobactera | Rhodobactera | Planktomarina | Planktomarina | information st | Transcription | 05VEQ | Cold shock pr | 412597.PaTRP | 2,6 | 0,489897949 | 0,000530337 | 0,000134641 | 2 - | 0,000337579 - | 3 - | 0,00062876 - | 3 - | 0,000597716 - | 2 - | 0,000404363 - | 3 - | 0,00068327 - |
| 8871 | 1 | 0.2um_NOKCPHJN_233299 | Bacteria | Proteobacteri | Gammaprotec | Alteromonada | Colwelliaceae | Colwellia | Colwellia sp. P | unclassified | unclassified | unclassified | unclassified | unclassified | 0,6 | 0,489897949 | 2,67E-05 | 2,19E-05 | 0 - | 0 - | 0 - | 0 - | 1 - | 4,23E-05 - | 1 - | 4,30E-05 - | 1 - | 4,84E-05 - |
| 8872 | 1 | 3um_FIJJAFBF_363430 | Bacteria | Bacteroidetes | Flavobacteriia | unclassified | unclassified | unclassified | Flavobacteria | unclassified | unclassified | unclassified | unclassified | unclassified | 5,8 | 0,748331477 | 0,000256422 | 3,00E-05 | 6 - | 0,000223591 - | 5 - | 0,000231362 - | 7 - | 0,000307914 - | 6 - | 0,000267825 - | 5 - | 0,00025142 - |
| 8873 | 3 | 3um_FIJJAFBF_255871;3um_FIJJAFBF_374667;3u | Bacteria | Proteobacteri | Alphaproteob | Rhodobactera | Rhodobactera | Planktomarina | Planktomarina | unclassified | unclassified | unclassified | unclassified | unclassified | 0,8 | 0,4 | 3,99E-05 | 2,02E-05 | 1 - | 4,31E-05 - | 1 - | 5,36E-05 - | 1 - | 5,09E-05 - | 1 - | 5,17E-05 - | 0 - | 0 - |
| 8874 | 1 | 3um_FIJJAFBF_606373 | Bacteria | Firmicutes | Clostridia | Clostridiales | unclassified | unclassified | Clostridiales b | unclassified | unclassified | unclassified | unclassified | unclassified | 0,6 | 0,8 | 6,54E-05 | 8,69E-05 | 0 - | 0 - | 0 - | 0 - | 2 - | 0,000216772 - | 1 - | 0,000109987 - | 0 - | 0 - |
| 8875 | 2 | 3um_FIJJAFBF_00830;3um_FIJJAFBF_777845 | Bacteria | Bacteroidetes | Flavobacteriia | Flavobacterial | Flavobacteriac | Polaribacter | various | metabolism | Lipid transpor | COG0332 | Catalyzes the | 313594.PI23P | 0,4 | 0,489897949 | 1,66E-05 | 2,03E-05 | 0 - | 0 - | 1 - | 4,25E-05 - | 1 - | 4,04E-05 - | 0 - | 0 - | 0 - | 0 - |
| 8876 | 1 | 0.2um_NOKCPHJN_53343 | Bacteria | Proteobacteri | Alphaproteob | Rhodospirillal | Rhodospirillac | Magnetospira | Magnetospira | cellular proces | Posttranslatio | 01QWG | Prevents misf | 342108.amb0 | 1,6 | 0,489897949 | 6,63E-05 | 2,19E-05 | 2 - | 6,94E-05 - | 2 - | 8,61E-05 - | 1 - | 4,09E-05 - | 1 - | 4,15E-05 - | 2 - | 9,36E-05 - |
| 8877 | 1 | 3um_FIJJAFBF_690505 | Bacteria | Proteobacteri | Alphaproteob | Rickettsiales | unclassified | unclassified | Rickettsiales b | unclassified | unclassified | unclassified | unclassified | unclassified | 1,2 | 0,979795897 | 7,89E-05 | 5,58E-05 | 3 - | 0,000174787 - | 1 - | 7,23E-05 - | 1 - | 6,88E-05 - | 0 - | 0 - | 1 - | 7,86E-05 - |
| 8878 | 2 | 0.2um_NOKCPHJN_441213;3um_FIJJAFBF_4217 | Bacteria | Proteobacteri | Gammaprotec | unclassified | unclassified | unclassified | various | cellular proces | Posttranslatio | COG1225 | alkyl hydroper | 391625.PPSIR | 1 | 0 | 4,13E-05 | 3,92E-06 | 1 - | 3,46E-05 - | 1 - | 4,29E-05 - | 1 - | 4,08E-05 - | 1 - | 4,14E-05 - | 1 - | 4,66E-05 - |
| 8879 | 1 | 3um_FIJJAFBF_104483 | Bacteria | Bacteroidetes | Flavobacteriia | Flavobacterial | Flavobacteriac | Algibacter | Algibacter sp. | cellular proces | Intracellular tr | 05FGH | MotA TolQ ex | 391603.FBALC | 0,6 | 0,489897949 | 3,18E-05 | 2,64E-05 | 1 - | 4,50E-05 - | 0 - | 0 - | 1 - | 5,31E-05 - | 0 - | 0 - | 1 - | 6,07E-05 - |
| 8880 | 1 | 3um_FIJJAFBF_48469 | Bacteria | Bacteroidetes | Flavobacteriia | Flavobacterial | Flavobacteriac | Algibacter | Algibacter sp. | cellular proces | Posttranslatio | COG0652 | peptidyl-proly | 391603.FBALC | 0,4 | 0,489897949 | 1,56E-05 | 1,93E-05 | 1 - | 3,55E-05 - | 0 - | 0 - | 0 - | 0 - | 1 - | 4,26E-05 - | 0 - | 0 - |
| 8881 | 1 | 3um_FIJJAFBF_547079 | Eukaryota | Ascomycota | Sordariomyce | Glomerellales | Glomerellacea | Colletotrichun | Colletotrichun | unclassified | unclassified | unclassified | unclassified | unclassified | 0,8 | 0,4 | 5,43E-05 | 2,79E-05 | 1 - | 5,74E-05 - | 0 - | 0 - | 1 - | 6,77E-05 - | 1 - | 6,87E-05 - | 1 - | 7,74E-05 - |
| 8882 | 1 | 0.2um_NOKCPHJN_07997 | Bacteria | Bacteroidetes | Flavobacteriia | Flavobacterial | Flavobacteriac | unclassified | Flavobacteriac | metabolism | Inorganic ion t | 05C27 | TonB-depende | 391603.FBALC | 5 | 2,449489743 | 6,05E-05 | 2,47E-05 | 9 - | 9,46E-05 - | 6 - | 7,83E-05 - | 5 - | 6,20E-05 - | 2 - | 2,52E-05 - | 3 - | 4,25E-05 - |
| 8883 | 1 | 3um_FIJJAFBF_46931 | Bacteria | Proteobacteri | Gammaprotec | Alteromonada | Alteromonada | Paraglaciecola | Paraglaciecola | unclassified | unclassified | unclassified | unclassified | unclassified | 0,8 | 0,4 | 5,17E-05 | 2,65E-05 | 1 - | 5,47E-05 - | 0 - | 0 - | 1 - | 6,45E-05 - | 1 - | 6,55E-05 - | 1 - | 7,37E-05 - |
| 8884 | 2 | 0.2um_NOKCPHJN_498504;0.2um_NOKCPHJN_6 | Bacteria | Proteobacteri | Alphaproteob | Pelagibacteral | Pelagibacterac | Candidatus Pe | Candidatus Pe | cellular proces | Posttranslatio | 16PM6 | feS assembly | 335992.SAR11 | 0,6 | 0,489897949 | 3,22E-05 | 2,66E-05 | 1 - | 4,70E-05 - | 1 - | 5,84E-05 - | 1 - | 5,55E-05 - | 0 - | 0 - | 0 - | 0 - |
| 8885 | 3 | 0.2um_NOKCPHJN_18896;0.2um_NOKCPHJN_53 | Bacteria | Proteobacteri | Gammaprotec | Cellvibrionales | Porticoccacea | unclassified | various | various | various | various | various | various | 0,2 | 0,4 | 9,18E-06 | 1,84E-05 | 1 - | 4,59E-05 - | 0 - | 0 - | 0 - | 0 - | 0 - | 0 - | 0 - | 0 - |
| 8886 | 1 | 0.2um_NOKCPHJN_734823 | Bacteria | Proteobacteri | Gammaprotec | Cellvibrionales | unclassified | unclassified | Cellvibrionales | information st | Translation, ri | 05C64 | Binds directly | 314287.GB22 | 1 | 0,632455532 | 5,91E-05 | 3,75E-05 | 1 - | 4,97E-05 - | 1 - | 6,17E-05 - | 2 - | 0,000117301 - | 0 - | 0 - | 1 - | 6,70E-05 - |
| 8887 | 1 | 0.2um_NOKCPHJN_206381 | Bacteria | Proteobacteri | Alphaproteob | Rhodobactera | Rhodobactera | Thalassobium | Thalassobium | information st | Transcription | 01R4Z | DNA-depende | 633131.TR2A | 28,4 | 3,2 | 0,000280932 | 2,92E-05 | 31 - | 0,000258394 - | 31 - | 0,000320849 - | 25 - | 0,000245974 - | 31 - | 0,000309512 - | 24 - | 0,000269934 - |
| 8888 | 1 | 3um_FIJJAFBF_400124 | Bacteria | Bacteroidetes | Flavobacteriia | Flavobacterial | Crocinitomica | unclassified | Crocinitomica | poorly charact | Function unkn | 11NCV | Protein of unk | 643867.Ftrac_ | 0,8 | 0,4 | 6,06E-05 | 3,08E-05 | 1 - | 6,56E-05 - | 1 - | 8,14E-05 - | 1 - | 7,74E-05 - | 1 - | 7,86E-05 - | 0 - | 0 - |
| 8889 | 1 | 0.2um_NOKCPHJN_311779 | Bacteria | Proteobacteri | Gammaprotec | Alteromonada | Shewanellace | Shewanella | Shewanella ar | poorly charact | Function unkn | 0QRAP | NA | 326297.Sama_ | 5 | 1,897366596 | 4,82E-05 | 1,98E-05 | 4 - | 3,19E-05 - | 7 - | 6,94E-05 - | 2 - | 1,89E-05 - | 7 - | 6,70E-05 - | 5 - | 5,39E-05 - |
| 8890 | 1 | 0.2um_NOKCPHJN_260411 | Bacteria | Proteobacteri | Gammaprotec | Oceanospirilla | Saccharospirill | Reinekea | Reinekea fors | cellular proces | Posttranslatio | COG0435 | glutathione St | 314283.MED2 | 0,4 | 0,8 | 1,66E-05 | 3,32E-05 | 0 - | 0 - | 0 - | 0 - | 0 - | 0 - | 2 - | 8,31E-05 - | 0 - | 0 - |
| 8891 | 1 | 0.2um_NOKCPHJN_795674 | Bacteria | Bacteroidetes | Flavobacteriia | unclassified | unclassified | unclassified | Flavobacteria | unclassified | unclassified | unclassified | unclassified | unclassified | 2,8 | 1,720465053 | 7,12E-05 | 3,57E-05 | 6 - | 0,000134242 - | 2 - | 5,56E-05 - | 3 - | 7,92E-05 - | 1 - | 2,68E-05 - | 2 - | 6,04E-05 - |
| 8892 | 1 | 0.2um_NOKCPHJN_221147 | Bacteria | Proteobacteri | Alphaproteob | Rhodobactera | Rhodobactera | Planktomarina | Planktomarina | cellular proces | Cell wall/mem | 16QB0 | Part of the ou | 633131.TR2A | 2,2 | 1,469693846 | 3,73E-05 | 2,38E-05 | 4 - | 5,88E-05 - | 4 - | 7,30E-05 - | 1 - | 1,73E-05 - | 1 - | 1,76E-05 - | 1 - | 1,98E-05 - |
| 8893 | 1 | 0.2um_NOKCPHJN_349052 | Bacteria | Bacteroidetes | Flavobacteriia | Flavobacterial | Flavobacteriac | Ulvibacter | Ulvibacter lito | metabolism | Amino acid tra | 09S3M | thermolysin | 50743.SCB49_ | 1,8 | 0,4 | 1,22E-05 | 2,86E-06 | 2 - | 1,14E-05 - | 1 - | 7,10E-06 - | 2 - | 1,35E-05 - | 2 - | 1,37E-05 - | 2 - | 1,54E-05 - |
| 8894 | 1 | 3um_FIJJAFBF_20890 | Bacteria | Bacteroidetes | Flavobacteriia | Flavobacterial | Crocinitomica | unclassified | Crocinitomica | cellular proces | Posttranslatio | 09GE9 | Thioredoxin | 525373.HMPR | 1 | 0 | 0,000129271 | 1,23E-05 | 1 - | 0,00010828 - | 1 - | 0,000134452 - | 1 - | 0,000127813 - | 1 - | 0,000129701 - | 1 - | 0,000146108 - |
| 8895 | 1 | 0.2um_NOKCPHJN_136874 | Eukaryota | unclassified | Oomycetes | Saprolegniales | Saprolegniace | Achlya | Achlya hypogy | unclassified | unclassified | unclassified | unclassified | unclassified | 0,2 | 0,4 | 2,56E-07 | 5,13E-07 | 0 - | 0 - | 0 - | 0 - | 1 - | 1,28E-06 - | 0 - | 0 - | 0 - | 0 - |
| 8896 | 1 | 3um_FIJJAFBF_38454 | Bacteria | Proteobacteri | Alphaproteob | Rhodobactera | Rhodobactera | unclassified | Rhodobactera | cellular proces | Cell wall/mem | 08ZG0 | Ompa motb d | 391619.RGBS | 2 | 0 | 0,000122894 | 1,17E-05 | 2 - | 0,000102939 - | 2 - | 0,00012782 - | 2 - | 0,000121509 - | 2 - | 0,000123303 - | 2 - | 0,000138901 - |
| 8897 | 1 | 3um_FIJJAFBF_463340 | Bacteria | Proteobacteri | Gammaprotec | Cellvibrionales | Porticoccacea | unclassified | Gammaprotec | unclassified | unclassified | unclassified | unclassified | unclassified | 2 | 0,632455532 | 0,000103659 | 2,71E-05 | 3 - | 0,000132435 - | 2 - | 0,00010963 - | 2 - | 0,000104217 - | 1 - | 5,29E-05 - | 2 - | 0,000119134 - |
| 8898 | 1 | 3um_FIJJAFBF_300638 | Bacteria | Bacteroidetes | Flavobacteriia | unclassified | unclassified | unclassified | Flavobacteria | unclassified | unclassified | unclassified | unclassified | unclassified | 11 | 2,607680962 | 0,000209854 | 3,95E-05 | 14 - | 0,000227603 - | 12 - | 0,000242242 - | 9 - | 0,000172711 - | 13 - | 0,000253156 - | 7 - | 0,000153558 - |
| 8899 | 1 | 0.2um_NOKCPHJN_1001141 | Bacteria | various | various | various | various | various | various | information st | Translation, ri | COG0051 | tRNA binding | 1114970.PSF1 | 1,8 | 0,748331477 | 0,00023801 | 9,95E-05 | 2 - | 0,000222867 - | 1 - | 0,000138368 - | 3 - | 0,000394608 - | 1 - | 0,000133479 - | 2 - | 0,000300727 - |
| 8900 | 1 | 0.2um_NOKCPHJN_803846 | Bacteria | Bacteroidetes | Flavobacteriia | Flavobacterial | Flavobacteriac | Polaribacter | Polaribacter s | unclassified | unclassified | unclassified | unclassified | unclassified | 0,4 | 0,489897949 | 1,04E-05 | 1,29E-05 | 1 - | 2,33E-05 - | 1 - | 2,89E-05 - | 0 - | 0 - | 0 - | 0 - | 0 - | 0 - |
| 8901 | 1 | 0.2um_NOKCPHJN_110052 | Bacteria | various | various | various | various | various | various | metabolism | Energy produc | COG0055 | Produces ATP | 865938.Weev | 1,6 | 1,854723699 | 3,95E-05 | 4,27E-05 | 5 - | 0,000113866 - | 2 - | 5,66E-05 - | 1 - | 2,69E-05 - | 0 - | 0 - | 0 - | 0 - |
| 8902 | 1 | 0.2um_NOKCPHJN_195815 | Bacteria | Proteobacteri | Gammaprotec | Oceanospirilla | Saccharospirill | Reinekea | Reinekea fors | metabolism | Energy produc | 07QSN | dihydrolipoam | 314283.MED2 | 4 | 1,095445115 | 8,49E-05 | 1,96E-05 | 5 - | 9,08E-05 - | 4 - | 9,02E-05 - | 4 - | 8,57E-05 - | 5 - | 0,000108768 - | 2 - | 4,90E-05 - |
| 8903 | 1 | 3um_FIJJAFBF_359189 | Bacteria | Proteobacteri | Gammaprotec | Cellvibrionales | Porticoccacea | unclassified | gamma protec | unclassified | unclassified | unclassified | unclassified | unclassified | 1 | 0 | 6,82E-05 | 6,47E-06 | 1 - | 5,71E-05 - | 1 - | 7,09E-05 - | 1 - | 6,74E-05 - | 1 - | 6,84E-05 - | 1 - | 7,71E-05 - |
| 8904 | 1 | 3um_FIJJAFBF_703920 | Bacteria | Proteobacteri | Gammaprotec | unclassified | unclassified | unclassified | Gammaprotec | metabolism | Inorganic ion t | 17AJ4 | TonB-depende | 247634.GPB2 | 1,8 | 0,748331477 | 2,55E-05 | 1,03E-05 | 2 - | 2,42E-05 - | 1 - | 1,50E-05 - | 2 - | 2,86E-05 - | 3 - | 4,35E-05 - | 1 - | 1,63E-05 - |
| 8905 | 1 | 0.2um_NOKCPHJN_590793 | Bacteria | Bacteroidetes | Flavobacteriia | Flavobacterial | Flavobacteriac | Polaribacter | Polaribacter t | poorly charact | Function unkn | 0XSVM | NA | 867900.Celly_ | 0,2 | 0,4 | 5,77E-06 | 1,15E-05 | 1 - | 2,88E-05 - | 0 - | 0 - | 0 - | 0 - | 0 - | 0 - | 0 - | 0 - |
| 8906 | 1 | 3um_FIJJAFBF_646507 | Bacteria | Proteobacteri | Alphaproteob | Rhizobiales | Bradyrhizobia | Bradyrhizobiu | Bradyrhizobiu | cellular proces | Intracellular tr | 0BIEB | domain-conta | 1005048.CFU_ | 6,8 | 0,748331477 | 0,000155954 | 2,68E-05 | 7 - | 0,00013413 - | 6 - | 0,000142757 - | 7 - | 0,000158326 - | 6 - | 0,000137713 - | 8 - | 0,000206844 - |
| 8907 | 1 | 0.2um_NOKCPHJN_911791 | Bacteria | Proteobacteri | Gammaprotec | Oceanospirilla | Halomonadac | Halomonas | Halomonas lic | unclassified | unclassified | unclassified | unclassified | unclassified | 3 | 0,632455532 | 6,77E-05 | 1,66E-05 | 3 - | 5,65E-05 - | 4 - | 9,36E-05 - | 3 - | 6,67E-05 - | 2 - | 4,52E-05 - | 3 - | 7,63E-05 - |
| 8908 | 4 | 3um_FIJJAFBF_122421;3um_FIJJAFBF_26095;3ur | Bacteria | Bacteroidetes | Flavobacteriia | Flavobacterial | Flavobacteriac | Polaribacter | various | unclassified | unclassified | unclassified | unclassified | unclassified | 1 | 0 | 0,000274054 | 2,60E-05 | 1 - | 0,000229554 - | 1 - | 0,000285038 - | 1 - | 0,000270964 - | 1 - | 0,000274967 - | 1 - | 0,000309749 - |
| 8909 | 1 | 3um_FIJJAFBF_393575 | Bacteria | Proteobacteri | Gammaprotec | Oceanospirilla | Saccharospirill | Reinekea | Reinekea fors | cellular proces | Cell wall/mem | 08ZG0 | Ompa motb d | 314283.MED2 | 0,4 | 0,489897949 | 2,25E-05 | 2,78E-05 | 1 - | 5,12E-05 - | 0 - | 0 - | 0 - | 0 - | 1 - | 6,14E-05 - | 0 - | 0 - |
| 8910 | 1 | 0.2um_NOKCPHJN_782497 | Eukaryota | Bacillariophyta | Coscinodiscop | Thalassiosirale | Thalassiosirac | Thalassiosira | Thalassiosira | cellular proces | Signal transdu | KOG0613 | fibronectin typ | 67593.Physo1 | 3,8 | 1,166190379 | 6,11E-05 | 1,84E-05 | 4 - | 5,43E-05 - | 3 - | 5,06E-05 - | 3 - | 4,81E-05 - | 6 - | 9,76E-05 - | 3 - | 5,50E-05 - |
| 8911 | 1 | 3um_FIJJAFBF_398122 | Bacteria | Bacteroidetes | Flavobacteriia | Flavobacterial | Flavobacteriac | Algibacter | Algibacter pe | unclassified | unclassified | unclassified | unclassified | unclassified | 0,2 | 0,4 | 9,71E-06 | 1,94E-05 | 0 - | 0 - | 0 - | 0 - | 0 - | 0 - | 0 - | 0 - | 1 - | 4,85E-05 - |
| 8912 | 1 | 3um_FIJJAFBF_733400 | Eukaryota | Phaeophyceae | unclassified | Ectocarpales | Ectocarpaceae | Ectocarpus | Ectocarpus sil | unclassified | unclassified | unclassified | unclassified | unclassified | 1,2 | 0,748331477 | 3,50E-05 | 2,31E-05 | 1 - | 2,48E-05 - | 2 - | 6,16E-05 - | 1 - | 2,93E-05 - | 2 - | 5,94E-05 - | 0 - | 0 - |
| 8913 | 2 | 0.2um_NOKCPHJN_359063;3um_FIJJAFBF_5349 | Bacteria | Proteobacteri | Alphaproteob | Rhodospirillal | Rhodospirillac | Candidatus En | Candidatus En | various | various | various | various | various | 0,4 | 0,489897949 | 1,09E-05 | 1,33E-05 | 0 - | 0 - | 1 - | 2,79E-05 - | 1 - | 2,65E-05 - | 0 - | 0 - | 0 - | 0 - |
| 8914 | 1 | 0.2um_NOKCPHJN_130205 | Eukaryota | unclassified | Synurophycea | Parmales | Triparmaceae | Triparma | Triparma laevi | cellular proces | Posttranslatio | 05C2Z | ATP-depender | 313612.L8106 | 0,2 | 0,4 | 3,31E-06 | 6,63E-06 | 0 - | 0 - | 0 - | 0 - | 0 - | 0 - | 1 - | 1,66E-05 - | 0 - | 0 - |
| 8915 | 1 | 3um_FIJJAFBF_214616 | Bacteria | Proteobacteri | Alphaproteob | Rhodobactera | unclassified | unclassified | Rhodobactera | unclassified | unclassified | unclassified | unclassified | unclassified | 4,2 | 0,4 | 0,00046428 | 2,20E-05 | 5 - | 0,000466572 - | 4 - | 0,000463476 - | 4 - | 0,000440593 - | 4 - | 0,0004471 - | 4 - | 0,000503657 - |
| 8916 | 1 | 3um_FIJJAFBF_570004 | Bacteria | Proteobacteri | Alphaproteob | Rhodobactera | unclassified | unclassified | Rhodobactera | metabolism | Inorganic ion t | COG3853 | Toxic anion re | 439496.RBY4 | 0,4 | 0,489897949 | 1,30E-05 | 1,61E-05 | 1 - | 2,90E-05 - | 1 - | 3,60E-05 - | 0 - | 0 - | 0 - | 0 - | 0 - | 0 - |
| 8917 | 1 | 0.2um_NOKCPHJN_495424 | Bacteria | Proteobacteri | Alphaproteob | Pelagibacteral | Pelagibacterac | Candidatus Pe | Candidatus Pe | unclassified | unclassified | unclassified | unclassified | unclassified | 3,4 | 0,489897949 | 0,00025784 | 2,74E-05 | 4 - | 0,000256484 - | 3 - | 0,000238858 - | 3 - | 0,000227065 - | 4 - | 0,000307225 - | 3 - | 0,000259566 - |
| 8918 | 1 | 3um_FIJJAFBF_513993 | Bacteria | Proteobacteri | Alphaproteob | Pelagibacteral | Pelagibacterac | Candidatus Pe | Candidatus Pe | cellular proces | Cell wall/mem | COG1686 | Carboxypeptic | 335992.SAR11 | 0,6 | 0,489897949 | 2,61E-05 | 2,15E-05 | 1 - | 3,79E-05 - | 1 - | 4,70E-05 - | 0 - | 0 - | 1 - | 4,54E-05 - | 0 - | 0 - |
| 8919 | 1 | 3um_FIJJAFBF_788016 | Bacteria | Proteobacteri | Alphaproteob | Rhodobactera | unclassified | unclassified | Rhodobactera | unclassified | unclassified | unclassified | unclassified | unclassified | 2 | 0,894427191 | 0,000194274 | 8,93E-05 | 2 - | 0,000163967 - | 3 - | 0,000305398 - | 1 - | 9,68E-05 - | 3 - | 0,000294607 - | 1 - | 0,000110625 - |
| 8920 | 2 | 3um_FIJJAFBF_224762;3um_FIJJAFBF_463528 | Bacteria | Bacteroidetes | Flavobacteriia | Flavobacterial | Flavobacteriac | Formosa | various | cellular proces | Posttranslatio | COG0652 | peptidyl-proly | 1046627.BZAF | 0,2 | 0,4 | 7,29E-06 | 1,46E-05 | 0 - | 0 - | 1 - | 3,64E-05 - | 0 - | 0 - | 0 - | 0 - | 0 - | 0 - |
| 8921 | 1 | 0.2um_NOKCPHJN_106480 | Bacteria | Bacteroidetes | Flavobacteriia | Flavobacterial | unclassified | unclassified | Flavobacterial | metabolism | Inorganic ion t | 05C27 | TonB-depende | 216432.CA25 | 6,2 | 0,748331477 | 8,28E-05 | 1,35E-05 | 6 - | 6,70E-05 - | 7 - | 9,70E-05 - | 5 - | 6,59E-05 - | 7 - | 9,36E-05 - | 6 - | 9,04E-05 - |
| 8922 | 1 | 3um_FIJJAFBF_300789 | Bacteria | Bacteroidetes | Flavobacteriia | Flavobacterial | Flavobacteriac | Formosa | Formosa sp. H | unclassified | unclassified | unclassified | unclassified | unclassified | 4,4 | 1,356465997 | 8,21E-05 | 2,03E-05 | 6 - | 9,63E-05 - | 4 - | 7,97E-05 - | 5 - | 9,47E-05 - | 5 - | 9,61E-05 - | 2 - | 4,33E-05 - |
| 8923 | 1 | 3um_FIJJAFBF_187828 | Bacteria | Bacteroidetes | Flavobacteriia | Flavobacterial | Flavobacteriac | Polaribacter | Polaribacter s | unclassified | unclassified | unclassified | unclassified | unclassified | 0,4 | 0,489897949 | 3,53E-05 | 4,35E-05 | 1 - | 8,03E-05 - | 0 - | 0 - | 0 - | 0 - | 1 - | 9,61E-05 - | 0 - | 0 - |
| 8924 | 1 | 0.2um_NOKCPHJN_205007 | Bacteria | Proteobacteri | Gammaprotec | Alteromonada | various | various | various | information st | Translation, ri | COG1185 | Involved in m | 342610.Patl_1 | 6,6 | 0,489897949 | 0,00012813 | 1,63E-05 | 7 - | 0,000113801 - | 7 - | 0,000141308 - | 6 - | 0,000115141 - | 6 - | 0,000116841 - | 7 - | 0,000153558 - |
| 8925 | 1 | 3um_FIJJAFBF_697218 | Bacteria | Bacteroidetes | Flavobacteriia | Flavobacterial | Flavobacteriac | Polaribacter | Polaribacter s | unclassified | unclassified | unclassified | unclassified | unclassified | 0,6 | 0,489897949 | 1,36E-05 | 1,12E-05 | 1 - | 1,99E-05 - | 1 - | 2,47E-05 - | 1 - | 2,35E-05 - | 0 - | 0 - | 0 - | 0 - |
| 8926 | 1 | 3um_FIJJAFBF_570134 | Bacteria | Proteobacteri | Alphaproteob | Rhodobactera | Rhodobactera | Epibacterium | Epibacterium | metabolism | Carbohydrate | 01QKM | inositol mono | 292414.TM10 | 0,2 | 0,4 | 1,04E-05 | 2,08E-05 | 0 - | 0 - | 0 - | 0 - | 1 - | 5,19E-05 - | 0 - | 0 - | 0 - | 0 - |
| 8927 | 1 | 0.2um_NOKCPHJN_968081 | Bacteria | Proteobacteri | Alphaproteob | Pelagibacteral | Pelagibacterac | Candidatus Pe | Candidatus Pe | metabolism | Secondary me | 01Q86 | extracellular s | 335992.SAR11 | 4,4 | 0,489897949 | 0,000644696 | 6,74E-05 | 5 - | 0,000617079 - | 5 - | 0,00076623 - | 4 - | 0,000582719 - | 4 - | 0,000591326 - | 4 - | 0,000666127 - |
| 8928 | 1 | 3um_FIJJAFBF_187037 | Bacteria | Proteobacteri | Gammaprotec | Alteromonada | Alteromonada | Paraglaciecola | Paraglaciecola | unclassified | unclassified | unclassified | unclassified | unclassified | 4,2 | 0,748331477 | 0,000207148 | 2,61E-05 | 5 - | 0,000209447 - | 4 - | 0,000208057 - | 4 - | 0,000197784 - | 5 - | 0,000250882 - | 3 - | 0,000169571 - |
| 8929 | 1 | 0.2um_NOKCPHJN_137415 | Bacteria | Bacteroidetes | Flavobacteriia | Flavobacterial | Flavobacteriac | Formosa | Formosa sp. H | metabolism | Nucleotide tra | COG0138 | bifunctional p | 983544.Lacal_ | 0,2 | 0,4 | 4,50E-06 | 9,00E-06 | 1 - | 2,25E-05 - | 0 - | 0 - | 0 - | 0 - | 0 - | 0 - | 0 - | 0 - |
| 8930 | 1 | 0.2um_NOKCPHJN_154964 | Bacteria | Bacteroidetes | unclassified | unclassified | unclassified | unclassified | Bacteroidetes | cellular proces | Cell wall/mem | 11TPU | outer membra | 487796.Flav2 | 3 | 0,632455532 | 0,000246527 | 6,53E-05 | 2 - | 0,00013583 - | 4 - | 0,000337323 - | 3 - | 0,000240501 - | 3 - | 0,000244053 - | 3 - | 0,000274925 - |
| 8931 | 1 | 3um_FIJJAFBF_746310 | Eukaryota | unclassified | Cryptophyta | Pyrenomonad | Geminigerace | Guillardia | Guillardia thet | metabolism | Energy produc | COG1290 | Component of | 35128.Thapsd | 1,2 | 0,4 | 7,44E-05 | 1,65E-05 | 2 - | 0,000106769 - | 1 - | 6,63E-05 - | 1 - | 6,30E-05 - | 1 - | 6,39E-05 - | 1 - | 7,20E-05 - |
| 8932 | 1 | 3um_FIJJAFBF_402420 | Bacteria | Bacteroidetes | Flavobacteriia | Flavobacterial | Flavobacteriac | Tenacibaculur | Tenacibaculun | unclassified | unclassified | unclassified | unclassified | unclassified | 1 | 0 | 0,000142737 | 1,36E-05 | 1 - | 0,000119559 - | 1 - | 0,000148457 - | 1 - | 0,000141127 - | 1 - | 0,000143212 - | 1 - | 0,000161328 - |
| 8933 | 1 | 3um_FIJJAFBF_189604 | Bacteria | Bacteroidetes | Flavobacteriia | Flavobacterial | Flavobacteriac | Polaribacter | Polaribacter s | unclassified | unclassified | unclassified | unclassified | unclassified | 4 | 1,673320053 | 4,71E-05 | 1,76E-05 | 6 - | 6,07E-05 - | 5 - | 6,28E-05 - | 2 - | 2,39E-05 - | 5 - | 6,06E-05 - | 2 - | 2,73E-05 - |
| 8934 | 1 | 0.2um_NOKCPHJN_915794 | Eukaryota | Ascomycota | Sordariomyce | Hypocreales | Nectriaceae | Neonectria | Neonectria dit | unclassified | unclassified | unclassified | unclassified | unclassified | 0,4 | 0,489897949 | 4,47E-05 | 5,48E-05 | 0 - | 0 - | 0 - | 0 - | 1 - | 0,000111051 - | 1 - | 0,000112691 - | 0 - | 0 - |
| 8935 | 1 | 0.2um_NOKCPHJN_83023 | Bacteria | Proteobacteri | Gammaprotec | unclassified | unclassified | unclassified | OM182 bacte | unclassified | unclassified | unclassified | unclassified | unclassified | 5,8 | 1,166190379 | 0,000111884 | 2,44E-05 | 5 - | 8,09E-05 - | 7 - | 0,00014071 - | 6 - | 0,000114654 - | 7 - | 0,000135738 - | 4 - | 8,74E-05 - |
| 8936 | 1 | 3um_FIJJAFBF_20047 | Bacteria | Bacteroidetes | Flavobacteriia | Flavobacterial | Flavobacteriac | Polaribacter | Polaribacter a | information st | Translation, ri | COG0173 | aspartyl-trna s | 313598.MED1 | 3,8 | 0,748331477 | 4,67E-05 | 9,53E-06 | 4 - | 4,14E-05 - | 5 - | 6,43E-05 - | 4 - | 4,89E-05 - | 3 - | 3,72E-05 - | 3 - | 4,19E-05 - |
| 8937 | 1 | 3um_FIJJAFBF_567865 | Eukaryota | unclassified | Florideophyce | Gigartinales | Gigartinaceae | Chondrus | Chondrus cris | unclassified | unclassified | unclassified | unclassified | unclassified | 0,8 | 0,4 | 5,97E-05 | 3,01E-05 | 0 - | 0 - | 1 - | 7,46E-05 - | 1 - | 7,09E-05 - | 1 - | 7,20E-05 - | 1 - | 8,11E-05 - |
| 8938 | 1 | 3um_FIJJAFBF_375446 | Eukaryota | Bacillariophyta | Bacillariophyc | Naviculales | Phaeodactylac | Phaeodactylu | Phaeodactylu | unclassified | unclassified | unclassified | unclassified | unclassified | 0,2 | 0,4 | 7,09E-06 | 1,42E-05 | 0 - | 0 - | 0 - | 0 - | 1 - | 3,55E-05 - | 0 - | 0 - | 0 - | 0 - |
| 8939 | 1 | 0.2um_NOKCPHJN_40258 | Bacteria | Proteobacteri | unclassified | unclassified | unclassified | unclassified | Proteobacteri | information st | Translation, ri | 05KK9 | This protein bi | 511062.GU3_ | 0,4 | 0,489897949 | 4,90E-05 | 6,04E-05 | 1 - | 0,000111434 - | 0 - | 0 - | 0 - | 0 - | 1 - | 0,000133479 - | 0 - | 0 - |
| 8940 | 1 | 0.2um_NOKCPHJN_335468 | Bacteria | Proteobacteri | Gammaprotec | Alteromonada | Colwelliaceae | Colwellia | Colwellia echi | metabolism | Amino acid tra | 16Q4U | Catalyzes the | 326442.PSHA | 0,6 | 0,489897949 | 1,99E-05 | 1,62E-05 | 0 - | 0 - | 1 - | 3,41E-05 - | 1 - | 3,24E-05 - | 1 - | 3,29E-05 - | 0 - | 0 - |
| 8941 | 1 | 3um_FIJJAFBF_342379 | Bacteria | Bacteroidetes | Flavobacteriia | Flavobacterial | Flavobacteriac | Polaribacter | Polaribacter s | information st | Translation, ri | 0PA05 | 30S ribosomal | 313594.PI23P | 0,8 | 0,4 | 5,55E-05 | 2,82E-05 | 1 - | 6,01E-05 - | 1 - | 7,46E-05 - | 1 - | 7,09E-05 - | 1 - | 7,20E-05 - | 0 - | 0 - |
| 8942 | 1 | 3um_FIJJAFBF_18413 | Bacteria | Bacteroidetes | Flavobacteriia | unclassified | unclassified | unclassified | Flavobacteria | metabolism | Secondary me | COG1228 | amidohydrola | 487797.Flav3 | 1 | 0 | 1,55E-05 | 1,47E-06 | 1 - | 1,30E-05 - | 1 - | 1,61E-05 - | 1 - | 1,53E-05 - | 1 - | 1,56E-05 - | 1 - | 1,75E-05 - |
| 8943 | 2 | 3um_FIJJAFBF_454368;3um_FIJJAFBF_808895 | Bacteria | Bacteroidetes | Bacteroidia | Marinilabiliale | unclassified | unclassified | Marinilabiliale | cellular proces | Cell wall/mem | 09BXT | Mechanosens | 313606.M231 | 0,2 | 0,4 | 1,08E-05 | 2,17E-05 | 0 - | 0 - | 0 - | 0 - | 0 - | 0 - | 0 - | 0 - | 1 - | 5,42E-05 - |
| 8944 | 1 | 0.2um_NOKCPHJN_338098 | Bacteria | Proteobacteri | Gammaprotec | Oceanospirilla | Saccharospirill | Reinekea | Reinekea fors | metabolism | Energy produc | 05C6V | Glycerol-3-ph | 314283.MED2 | 0,4 | 0,489897949 | 1,02E-05 | 1,27E-05 | 1 - | 2,17E-05 - | 0 - | 0 - | 0 - | 0 - | 0 - | 0 - | 1 - | 2,92E-05 - |
| 8945 | 1 | 0.2um_NOKCPHJN_295878 | Bacteria | Bacteroidetes | Flavobacteriia | Flavobacterial | Flavobacteriac | Ulvibacter | Ulvibacter lito | information st | Translation, ri | COG0013 | Catalyzes the | 983548.Krodi_ | 0,4 | 0,489897949 | 6,26E-06 | 7,67E-06 | 0 - | 0 - | 0 - | 0 - | 1 - | 1,55E-05 - | 1 - | 1,58E-05 - | 0 - | 0 - |
| 8946 | 1 | 3um_FIJJAFBF_478118 | Eukaryota | Bacillariophyta | Coscinodiscop | Thalassiosirale | Thalassiosirac | Thalassiosira | Thalassiosira | unclassified | unclassified | unclassified | unclassified | unclassified | 0,4 | 0,489897949 | 3,90E-05 | 4,78E-05 | 0 - | 0 - | 0 - | 0 - | 1 - | 9,68E-05 - | 1 - | 9,82E-05 - | 0 - | 0 - |
| 8947 | 1 | 0.2um_NOKCPHJN_109073 | Bacteria | Bacteroidetes | Flavobacteriia | Flavobacterial | Flavobacteriac | Polaribacter | Polaribacter s | information st | Translation, ri | 05CEJ | Catalyzes the | 983548.Krodi_ | 8,8 | 0,4 | 0,000171672 | 2,24E-05 | 8 - | 0,000130243 - | 9 - | 0,000181939 - | 9 - | 0,000172956 - | 9 - | 0,000175511 - | 9 - | 0,000197712 - |
| 8948 | 1 | 0.2um_NOKCPHJN_339415 | Eukaryota | unclassified | Oomycetes | various | various | various | various | unclassified | unclassified | unclassified | unclassified | unclassified | 4 | 1,414213562 | 0,000361824 | 0,000139557 | 4 - | 0,000302044 - | 2 - | 0,000187525 - | 3 - | 0,000267399 - | 6 - | 0,000542697 - | 5 - | 0,000509456 - |
| 8949 | 1 | 3um_FIJJAFBF_517421 | Bacteria | Bacteroidetes | Flavobacteriia | Flavobacterial | Flavobacteriac | Polaribacter | Polaribacter v | information st | Translation, ri | 08UIK | This protein bi | 313598.MED1 | 6,6 | 0,489897949 | 0,000626716 | 9,54E-05 | 6 - | 0,000474938 - | 7 - | 0,000688022 - | 7 - | 0,000654052 - | 6 - | 0,000568896 - | 7 - | 0,00074767 - |
| 8950 | 1 | 0.2um_NOKCPHJN_69844 | Bacteria | Proteobacteri | Betaproteoba | unclassified | unclassified | unclassified | Betaproteoba | cellular proces | Intracellular tr | COG0848 | Biopolymer tr | 338969.Rfer_ | 0,6 | 0,489897949 | 5,10E-05 | 4,20E-05 | 1 - | 7,55E-05 - | 0 - | 0 - | 1 - | 8,91E-05 - | 1 - | 9,04E-05 - | 0 - | 0 - |
| 8951 | 1 | 3um_FIJJAFBF_87434 | Bacteria | Proteobacteri | Alphaproteob | Rhodobactera | Rhodobactera | Loktanella | Loktanella sp. | metabolism | Inorganic ion t | 05ENV | phosphonate | 391595.RLO1 | 1,6 | 0,489897949 | 8,53E-05 | 2,72E-05 | 2 - | 8,97E-05 - | 1 - | 5,57E-05 - | 1 - | 5,29E-05 - | 2 - | 0,000107409 - | 2 - | 0,000120996 - |
| 8952 | 1 | 0.2um_NOKCPHJN_285186 | Bacteria | Proteobacteri | Gammaprotec | unclassified | unclassified | unclassified | various | information st | Translation, ri | COG0088 | One of the pri | 203122.Sde_0 | 1,2 | 0,979795897 | 7,50E-05 | 5,34E-05 | 3 - | 0,000167151 - | 0 - | 0 - | 1 - | 6,58E-05 - | 1 - | 6,67E-05 - | 1 - | 7,52E-05 - |
| 8953 | 1 | 3um_FIJJAFBF_390243 | Eukaryota | unclassified | Chrysophycea | Chromulinales | Chromulinace | Ochromonas | Ochromonas | unclassified | unclassified | unclassified | unclassified | unclassified | 2,4 | 1,019803903 | 5,09E-05 | 1,83E-05 | 4 - | 7,28E-05 - | 3 - | 6,78E-05 - | 2 - | 4,29E-05 - | 1 - | 2,18E-05 - | 2 - | 4,91E-05 - |
| 8954 | 1 | 3um_FIJJAFBF_277459 | Eukaryota | Bacillariophyta | Coscinodiscop | Rhizosoleniale | Rhizosoleniac | Proboscia | Proboscia sp. | unclassified | unclassified | unclassified | unclassified | unclassified | 2 | 0,632455532 | 0,000331661 | 0,000113968 | 1 - | 0,000136639 - | 2 - | 0,000339331 - | 2 - | 0,000322577 - | 3 - | 0,000491012 - | 2 - | 0,000368749 - |
| 8955 | 1 | 3um_FIJJAFBF_561464 | Bacteria | Proteobacteri | Gammaprotec | Oceanospirilla | Oceanospirilla | Neptuniibacte | Neptuniibacte | unclassified | unclassified | unclassified | unclassified | unclassified | 0,6 | 0,489897949 | 4,13E-05 | 3,38E-05 | 0 - | 0 - | 1 - | 7,09E-05 - | 1 - | 6,74E-05 - | 1 - | 6,84E-05 - | 0 - | 0 - |
| 8956 | 1 | 3um_FIJJAFBF_103124 | Bacteria | Bacteroidetes | Flavobacteriia | Flavobacterial | Flavobacteriac | Polaribacter | Polaribacter s | unclassified | unclassified | unclassified | unclassified | unclassified | 0,2 | 0,4 | 3,63E-06 | 7,26E-06 | 1 - | 1,82E-05 - | 0 - | 0 - | 0 - | 0 - | 0 - | 0 - | 0 - | 0 - |
| 8957 | 1 | 3um_FIJJAFBF_69920 | Bacteria | Bacteroidetes | Flavobacteriia | various | various | various | various | unclassified | unclassified | unclassified | unclassified | unclassified | 1 | 0 | 6,92E-05 | 6,57E-06 | 1 - | 5,80E-05 - | 1 - | 7,20E-05 - | 1 - | 6,84E-05 - | 1 - | 6,94E-05 - | 1 - | 7,82E-05 - |
| 8958 | 1 | 0.2um_NOKCPHJN_485592 | Bacteria | Proteobacteri | Alphaproteob | Rhodobactera | Rhodobactera | unclassified | Rhodobactera | unclassified | unclassified | unclassified | unclassified | unclassified | 0,8 | 0,4 | 3,20E-05 | 1,63E-05 | 1 - | 3,47E-05 - | 1 - | 4,31E-05 - | 1 - | 4,09E-05 - | 1 - | 4,15E-05 - | 0 - | 0 - |
| 8959 | 1 | 3um_FIJJAFBF_107304 | Bacteria | Bacteroidetes | Flavobacteriia | Flavobacterial | Flavobacteriac | Polaribacter | Polaribacter t | metabolism | Amino acid tra | 0P8GS | The glycine cle | 313594.PI23P | 0,2 | 0,4 | 2,42E-06 | 4,84E-06 | 1 - | 1,21E-05 - | 0 - | 0 - | 0 - | 0 - | 0 - | 0 - | 0 - | 0 - |
| 8960 | 1 | 0.2um_NOKCPHJN_57165 | Bacteria | Proteobacteri | Gammaprotec | Cellvibrionales | Porticoccacea | unclassified | gamma protec | poorly charact | Function unkn | 16T08 | saccharopine | 314287.GB22 | 0,2 | 0,4 | 7,27E-06 | 1,45E-05 | 0 - | 0 - | 1 - | 3,64E-05 - | 0 - | 0 - | 0 - | 0 - | 0 - | 0 - |
| 8961 | 1 | 0.2um_NOKCPHJN_180219 | Bacteria | Bacteroidetes | Cytophagia | Cytophagales | Flammeovirga | unclassified | Flammeovirga | cellular proces | Posttranslatio | 16QU3 | Hemolysin-typ | 744980.TRICH | 8,4 | 0,489897949 | 7,53E-05 | 8,82E-06 | 8 - | 6,00E-05 - | 9 - | 8,38E-05 - | 8 - | 7,08E-05 - | 9 - | 8,08E-05 - | 8 - | 8,09E-05 - |
| 8962 | 1 | 3um_FIJJAFBF_465283 | Eukaryota | Chlorophyta | Mamiellophyc | Mamiellales | Bathycoccace | Ostreococcus | Ostreococcus | unclassified | unclassified | unclassified | unclassified | unclassified | 2,4 | 1,019803903 | 0,000175411 | 6,18E-05 | 4 - | 0,00025365 - | 3 - | 0,000236219 - | 2 - | 0,000149704 - | 2 - | 0,000151915 - | 1 - | 8,56E-05 - |
| 8963 | 1 | 0.2um_NOKCPHJN_620638 | Bacteria | Proteobacteri | Gammaprotec | Oceanospirilla | Saccharospirill | Reinekea | Reinekea fors | metabolism | Inorganic ion t | 05PWN | phosphate ab | 314283.MED2 | 2,4 | 0,8 | 9,83E-05 | 3,77E-05 | 1 - | 3,34E-05 - | 3 - | 0,00012429 - | 2 - | 7,88E-05 - | 3 - | 0,000119898 - | 3 - | 0,000135065 - |
| 8964 | 1 | 3um_FIJJAFBF_707627 | unclassified | unclassified | unclassified | unclassified | unclassified | unclassified | unclassified | unclassified | unclassified | unclassified | unclassified | unclassified | 0,4 | 0,489897949 | 8,93E-06 | 1,10E-05 | 1 - | 1,99E-05 - | 1 - | 2,47E-05 - | 0 - | 0 - | 0 - | 0 - | 0 - | 0 - |
| 8965 | 1 | 3um_FIJJAFBF_584161 | Bacteria | Proteobacteri | Gammaprotec | Cellvibrionales | Microbulbifera | Microbulbifer | Microbulbifer | cellular proces | Signal transdu | 0QIPQ | gtp-binding pr | 247634.GPB2 | 0,4 | 0,489897949 | 8,09E-06 | 1,00E-05 | 1 - | 1,80E-05 - | 1 - | 2,24E-05 - | 0 - | 0 - | 0 - | 0 - | 0 - | 0 - |
| 8966 | 1 | 0.2um_NOKCPHJN_41540 | Bacteria | Bacteroidetes | Flavobacteriia | Flavobacterial | Crocinitomica | unclassified | Crocinitomica | metabolism | Amino acid tra | 0P7DH | glutamine syn | 755732.Fluta_ | 1,2 | 0,979795897 | 2,14E-05 | 1,51E-05 | 3 - | 4,72E-05 - | 1 - | 1,95E-05 - | 0 - | 0 - | 1 - | 1,89E-05 - | 1 - | 2,12E-05 - |
| 8967 | 1 | 0.2um_NOKCPHJN_62757 | Bacteria | Bacteroidetes | Flavobacteriia | Flavobacterial | Flavobacteriac | Polaribacter | various | cellular proces | Signal transdu | 09BM1 | Two compone | 886377.Murru | 0,4 | 0,8 | 1,95E-05 | 3,89E-05 | 2 - | 9,73E-05 - | 0 - | 0 - | 0 - | 0 - | 0 - | 0 - | 0 - | 0 - |
| 8968 | 1 | 0.2um_NOKCPHJN_06880 | Bacteria | Bacteroidetes | Cytophagia | Cytophagales | Flammeovirga | unclassified | Flammeovirga | information st | Translation, ri | 08UIK | This protein bi | 880070.Cycma | 1,6 | 0,489897949 | 0,000146345 | 4,60E-05 | 2 - | 0,000154063 - | 1 - | 9,57E-05 - | 2 - | 0,000181855 - | 1 - | 9,23E-05 - | 2 - | 0,000207885 - |
| 8969 | 1 | 3um_FIJJAFBF_773427 | Bacteria | Proteobacteri | Alphaproteob | Rhodospirillal | Rhodospirillac | Magnetospiril | Magnetospiril | unclassified | unclassified | unclassified | unclassified | unclassified | 0,8 | 0,4 | 1,57E-05 | 7,97E-06 | 1 - | 1,70E-05 - | 1 - | 2,11E-05 - | 1 - | 2,00E-05 - | 1 - | 2,03E-05 - | 0 - | 0 - |
| 8970 | 1 | 0.2um_NOKCPHJN_541968 | Bacteria | Proteobacteri | Gammaprotec | Cellvibrionales | Spongiibacter | unclassified | marine gamm | unclassified | unclassified | unclassified | unclassified | unclassified | 5 | 1,414213562 | 8,86E-05 | 2,10E-05 | 7 - | 0,000105577 - | 3 - | 5,62E-05 - | 4 - | 7,12E-05 - | 6 - | 0,000108397 - | 5 - | 0,000101757 - |
| 8971 | 1 | 0.2um_NOKCPHJN_156439 | Bacteria | Proteobacteri | Gammaprotec | Alteromonada | Pseudoalteror | Pseudoalteror | Pseudoalteror | unclassified | unclassified | unclassified | unclassified | unclassified | 0,2 | 0,4 | 1,54E-05 | 3,08E-05 | 1 - | 7,70E-05 - | 0 - | 0 - | 0 - | 0 - | 0 - | 0 - | 0 - | 0 - |
| 8972 | 1 | 3um_FIJJAFBF_617114 | Bacteria | Proteobacteri | Gammaprotec | Alteromonada | Pseudoalteror | Pseudoalteror | various | unclassified | unclassified | unclassified | unclassified | unclassified | 5,4 | 1,356465997 | 0,00081304 | 0,000219733 | 6 - | 0,00075677 - | 5 - | 0,000783071 - | 3 - | 0,000446645 - | 7 - | 0,001057564 - | 6 - | 0,00102115 - |
| 8973 | 1 | 3um_FIJJAFBF_755903 | Bacteria | Proteobacteri | Gammaprotec | Enterobactera | Yersiniaceae | unclassified | Yersiniaceae b | unclassified | unclassified | unclassified | unclassified | unclassified | 0,2 | 0,4 | 2,58E-05 | 5,16E-05 | 1 - | 0,000128963 - | 0 - | 0 - | 0 - | 0 - | 0 - | 0 - | 0 - | 0 - |
| 8974 | 1 | 3um_FIJJAFBF_649815 | Bacteria | Proteobacteri | Gammaprotec | Oceanospirilla | Saccharospirill | Reinekea | Reinekea fors | poorly charact | Function unkn | 0GF8Y | CBS domain co | 485915.Dret_ | 1 | 0 | 9,58E-05 | 9,10E-06 | 1 - | 8,03E-05 - | 1 - | 9,97E-05 - | 1 - | 9,47E-05 - | 1 - | 9,61E-05 - | 1 - | 0,000108304 - |
| 8975 | 1 | 3um_FIJJAFBF_118623 | Bacteria | Proteobacteri | Gammaprotec | Thiotrichales | Piscirickettsia | Cycloclasticus | Cycloclasticus | metabolism | Secondary me | 16QAT | Multicopper o | 314608.KT99_ | 0,6 | 0,489897949 | 3,15E-05 | 2,59E-05 | 1 - | 4,67E-05 - | 0 - | 0 - | 1 - | 5,51E-05 - | 1 - | 5,59E-05 - | 0 - | 0 - |
| 8976 | 1 | 3um_FIJJAFBF_28983 | Bacteria | Proteobacteri | Gammaprotec | Alteromonada | Shewanellace | Shewanella | Shewanella ha | unclassified | unclassified | unclassified | unclassified | unclassified | 0,2 | 0,4 | 1,92E-05 | 3,84E-05 | 0 - | 0 - | 0 - | 0 - | 1 - | 9,61E-05 - | 0 - | 0 - | 0 - | 0 - |
| 8977 | 1 | 3um_FIJJAFBF_73807 | Bacteria | Bacteroidetes | Flavobacteriia | Flavobacterial | Flavobacteriac | Polaribacter | Polaribacter b | cellular proces | Posttranslatio | COG1404 | peptidase (S8 | 313594.PI23P | 0,2 | 0,4 | 5,04E-06 | 1,01E-05 | 0 - | 0 - | 0 - | 0 - | 0 - | 0 - | 1 - | 2,52E-05 - | 0 - | 0 - |
| 8978 | 1 | 0.2um_NOKCPHJN_197130 | Viruses | unclassified | unclassified | unclassified | Phycodnavirid | Prasinovirus | Micromonas p | unclassified | unclassified | unclassified | unclassified | unclassified | 0,4 | 0,489897949 | 1,20E-05 | 1,48E-05 | 1 - | 2,68E-05 - | 1 - | 3,32E-05 - | 0 - | 0 - | 0 - | 0 - | 0 - | 0 - |
| 8979 | 1 | 3um_FIJJAFBF_403521 | Eukaryota | Bacillariophyta | Bacillariophyc | Bacillariales | Bacillariaceae | Fragilariopsis | Fragilariopsis | unclassified | unclassified | unclassified | unclassified | unclassified | 2 | 0,632455532 | 5,20E-05 | 1,36E-05 | 3 - | 6,65E-05 - | 2 - | 5,50E-05 - | 2 - | 5,23E-05 - | 1 - | 2,65E-05 - | 2 - | 5,98E-05 - |
| 8980 | 1 | 3um_FIJJAFBF_351766 | Bacteria | Actinobacteria | Actinobacteria | Bifidobacteria | Bifidobacteria | Bifidobacteriu | Bifidobacteriu | unclassified | unclassified | unclassified | unclassified | unclassified | 0,8 | 0,4 | 0,000127675 | 6,56E-05 | 1 - | 0,000135031 - | 0 - | 0 - | 1 - | 0,000159391 - | 1 - | 0,000161745 - | 1 - | 0,000182205 - |
| 8981 | 1 | 0.2um_NOKCPHJN_125613 | Bacteria | Proteobacteri | Gammaprotec | Cellvibrionales | Porticoccacea | unclassified | gamma protec | information st | Translation, ri | COG0256 | This is one of | 314287.GB22 | 0,4 | 0,489897949 | 4,83E-05 | 5,91E-05 | 0 - | 0 - | 1 - | 0,000122861 - | 0 - | 0 - | 1 - | 0,00011852 - | 0 - | 0 - |
| 8982 | 1 | 3um_FIJJAFBF_533443 | Bacteria | Bacteroidetes | Flavobacteriia | Flavobacterial | Crocinitomica | unclassified | Crocinitomica | metabolism | Coenzyme tra | 09A4R | May play a ke | 762903.Pedsa | 0,8 | 0,4 | 2,30E-05 | 1,17E-05 | 1 - | 2,49E-05 - | 1 - | 3,09E-05 - | 1 - | 2,94E-05 - | 1 - | 2,98E-05 - | 0 - | 0 - |
| 8983 | 1 | 3um_FIJJAFBF_323666 | Bacteria | Bacteroidetes | unclassified | Bacteroidetes | Rhodotherma | unclassified | Rhodotherma | unclassified | unclassified | unclassified | unclassified | unclassified | 0,2 | 0,4 | 9,37E-06 | 1,87E-05 | 1 - | 4,68E-05 - | 0 - | 0 - | 0 - | 0 - | 0 - | 0 - | 0 - | 0 - |
| 8984 | 1 | 0.2um_NOKCPHJN_611106 | Bacteria | Proteobacteri | Alphaproteob | Rhodospirillal | Rhodospirillac | unclassified | Rhodobactera | information st | Replication, re | 17B1W | resolvase | 388401.RB215 | 0,2 | 0,4 | 1,17E-05 | 2,33E-05 | 0 - | 0 - | 0 - | 0 - | 0 - | 0 - | 1 - | 5,83E-05 - | 0 - | 0 - |
| 8985 | 3 | 0.2um_NOKCPHJN_169037;0.2um_NOKCPHJN_6 | Bacteria | Bacteroidetes | various | various | various | various | various | unclassified | unclassified | unclassified | unclassified | unclassified | 0,2 | 0,4 | 1,25E-05 | 2,50E-05 | 0 - | 0 - | 0 - | 0 - | 0 - | 0 - | 0 - | 0 - | 1 - | 6,24E-05 - |
| 8986 | 2 | 0.2um_NOKCPHJN_787002;0.2um_NOKCPHJN_9 | Bacteria | Proteobacteri | Alphaproteob | Pelagibacteral | Pelagibacterac | unclassified | Pelagibacterac | unclassified | unclassified | unclassified | unclassified | unclassified | 1 | 0 | 8,90E-05 | 8,45E-06 | 1 - | 7,45E-05 - | 1 - | 9,25E-05 - | 1 - | 8,80E-05 - | 1 - | 8,93E-05 - | 1 - | 0,000100568 - |
| 8987 | 1 | 3um_FIJJAFBF_229418 | Bacteria | various | various | various | various | various | various | metabolism | Carbohydrate | 16QI2 | glycoside hydr | 314287.GB22 | 3,6 | 1,019803903 | 6,46E-05 | 2,12E-05 | 2 - | 2,95E-05 - | 4 - | 7,33E-05 - | 3 - | 5,22E-05 - | 5 - | 8,84E-05 - | 4 - | 7,96E-05 - |
| 8988 | 1 | 3um_FIJJAFBF_535056 | Bacteria | Bacteroidetes | Flavobacteriia | Flavobacterial | Flavobacteriac | Algibacter | Algibacter pe | poorly charact | Function unkn | 0PAPS | NA | 313603.FB217 | 2,2 | 0,4 | 0,000200767 | 3,91E-05 | 2 - | 0,000153036 - | 2 - | 0,000190025 - | 3 - | 0,000270964 - | 2 - | 0,000183311 - | 2 - | 0,000206499 - |
| 8989 | 1 | 3um_FIJJAFBF_292913 | Eukaryota | Bacillariophyta | Coscinodiscop | Coscinodiscale | Hemidiscacea | Actinocyclus | Actinocyclus s | metabolism | Energy produc | COG0056 | Produces ATP | 41431.PCC88 | 9,2 | 0,748331477 | 0,00025075 | 3,72E-05 | 8 - | 0,000181825 - | 10 - | 0,000282216 - | 9 - | 0,000241453 - | 10 - | 0,000272244 - | 9 - | 0,000276014 - |
| 8990 | 1 | 3um_FIJJAFBF_111224 | Bacteria | Bacteroidetes | Flavobacteriia | Flavobacterial | Flavobacteriac | Kordia | Kordia algicida | unclassified | unclassified | unclassified | unclassified | unclassified | 0,8 | 0,748331477 | 1,25E-05 | 1,16E-05 | 0 - | 0 - | 2 - | 3,09E-05 - | 0 - | 0 - | 1 - | 1,49E-05 - | 1 - | 1,68E-05 - |
| 8991 | 1 | 0.2um_NOKCPHJN_484615 | Eukaryota | unclassified | Pelagophycea | Pelagomonad | unclassified | Aureococcus | Aureococcus a | poorly charact | Function unkn | 01Q7A | thiJ PfpI | 366602.Caul_ | 0,6 | 0,489897949 | 4,82E-05 | 4,01E-05 | 1 - | 6,83E-05 - | 0 - | 0 - | 1 - | 8,06E-05 - | 0 - | 0 - | 1 - | 9,22E-05 - |
| 8992 | 1 | 3um_FIJJAFBF_08131 | Bacteria | Bacteroidetes | Flavobacteriia | Flavobacterial | Flavobacteriac | Polaribacter | Polaribacter s | unclassified | unclassified | unclassified | unclassified | unclassified | 1,2 | 0,4 | 4,34E-05 | 1,45E-05 | 1 - | 3,04E-05 - | 1 - | 3,77E-05 - | 2 - | 7,17E-05 - | 1 - | 3,64E-05 - | 1 - | 4,10E-05 - |
| 8993 | 1 | 3um_FIJJAFBF_195977 | Bacteria | Bacteroidetes | Flavobacteriia | Flavobacterial | Flavobacteriac | Polaribacter | Polaribacter b | metabolism | Nucleotide tra | COG0015 | adenylosuccin | 313594.PI23P | 4,4 | 1,019803903 | 0,000135462 | 3,61E-05 | 4 - | 0,000102709 - | 6 - | 0,0001913 - | 3 - | 9,09E-05 - | 5 - | 0,000153784 - | 4 - | 0,00013859 - |
| 8994 | 1 | 3um_FIJJAFBF_729200 | Bacteria | Proteobacteri | Gammaprotec | Oceanospirilla | Saccharospirill | Reinekea | Reinekea fors | unclassified | unclassified | unclassified | unclassified | unclassified | 0,4 | 0,489897949 | 2,86E-05 | 3,52E-05 | 1 - | 6,56E-05 - | 0 - | 0 - | 1 - | 7,74E-05 - | 0 - | 0 - | 0 - | 0 - |
| 8995 | 1 | 0.2um_NOKCPHJN_728723 | Bacteria | Proteobacteri | Alphaproteob | Rhodobactera | unclassified | unclassified | Rhodobactera | unclassified | unclassified | unclassified | unclassified | unclassified | 1,4 | 0,489897949 | 6,06E-05 | 2,23E-05 | 1 - | 3,61E-05 - | 2 - | 8,96E-05 - | 2 - | 8,52E-05 - | 1 - | 4,32E-05 - | 1 - | 4,87E-05 - |
| 8996 | 1 | 3um_FIJJAFBF_358211 | Eukaryota | Chlorophyta | Mamiellophyc | Mamiellales | Mamiellaceae | Micromonas | Micromonas c | unclassified | unclassified | unclassified | unclassified | unclassified | 0,6 | 0,489897949 | 1,17E-05 | 9,52E-06 | 0 - | 0 - | 1 - | 2,00E-05 - | 1 - | 1,90E-05 - | 1 - | 1,93E-05 - | 0 - | 0 - |
| 8997 | 1 | 0.2um_NOKCPHJN_404385 | Bacteria | Proteobacteri | Gammaprotec | unclassified | unclassified | unclassified | Gammaprotec | unclassified | unclassified | unclassified | unclassified | unclassified | 0,6 | 0,489897949 | 9,59E-05 | 7,97E-05 | 1 - | 0,000133461 - | 1 - | 0,00016572 - | 0 - | 0 - | 0 - | 0 - | 1 - | 0,000180087 - |
| 8998 | 1 | 0.2um_NOKCPHJN_108307 | Bacteria | Proteobacteri | Gammaprotec | Alteromonada | Alteromonada | Paraglaciecola | Paraglaciecola | unclassified | unclassified | unclassified | unclassified | unclassified | 1 | 0 | 9,65E-05 | 9,16E-06 | 1 - | 8,08E-05 - | 1 - | 0,000100365 - | 1 - | 9,54E-05 - | 1 - | 9,68E-05 - | 1 - | 0,000109067 - |
| 8999 | 1 | 0.2um_NOKCPHJN_674274 | Bacteria | Proteobacteri | Alphaproteob | Pelagibacteral | Pelagibacterac | Candidatus Pe | Candidatus Pe | metabolism | Carbohydrate | COG2721 | Dehydratase | 335992.SAR11 | 4,6 | 1,624807681 | 0,00015582 | 5,59E-05 | 5 - | 0,000143831 - | 7 - | 0,000250033 - | 5 - | 0,000169777 - | 4 - | 0,000137828 - | 2 - | 7,76E-05 - |
| 9000 | 1 | 3um_FIJJAFBF_445415 | Eukaryota | Echinodermat | Asteroidea | Valvatida | Acanthasterid | Acanthaster | Acanthaster p | unclassified | unclassified | unclassified | unclassified | unclassified | 0,2 | 0,4 | 3,52E-06 | 7,04E-06 | 0 - | 0 - | 0 - | 0 - | 0 - | 0 - | 1 - | 1,76E-05 - | 0 - | 0 - |
| 9001 | 1 | 3um_FIJJAFBF_573898 | Bacteria | Verrucomicrol | Opitutae | unclassified | unclassified | unclassified | Opitutae bact | unclassified | unclassified | unclassified | unclassified | unclassified | 2,8 | 0,979795897 | 8,10E-05 | 2,92E-05 | 3 - | 7,28E-05 - | 3 - | 9,04E-05 - | 4 - | 0,000114573 - | 1 - | 2,91E-05 - | 3 - | 9,82E-05 - |
| 9002 | 1 | 3um_FIJJAFBF_616971 | Bacteria | Bacteroidetes | unclassified | unclassified | unclassified | unclassified | Bacteroidetes | poorly charact | Function unkn | 0XR4Y | Ragb susd dor | 886377.Murru | 13,4 | 1,496662955 | 0,000322037 | 2,10E-05 | 15 - | 0,000304717 - | 13 - | 0,00032792 - | 15 - | 0,000359687 - | 13 - | 0,000316333 - | 11 - | 0,000301526 - |
| 9003 | 1 | 3um_FIJJAFBF_308252 | Bacteria | Bacteroidetes | Flavobacteriia | Flavobacterial | Flavobacteriac | Polaribacter | Polaribacter s | cellular proces | Posttranslatio | 09AIC | Glutamine cyc | 313594.PI23P | 0,6 | 0,489897949 | 2,51E-05 | 2,05E-05 | 0 - | 0 - | 1 - | 4,11E-05 - | 0 - | 0 - | 1 - | 3,96E-05 - | 1 - | 4,46E-05 - |

| | | | | | | | | | | | | | | | | | | | | | | | | | | | | |
|---|---|---|---|---|---|---|---|---|---|---|---|---|---|---|---|---|---|---|---|---|---|---|---|---|---|---|---|---|
| 9004 | 1 | 0.2um_NOKCPHJN_724163 | Bacteria | Proteobacteri | Gammaprotec | unclassified | unclassified | unclassified | Gammaprotec | metabolism | Inorganic ion t | 05F28 | tonB-depende | 565045.NOR5 | 12,6 | 1,2 | 0,000189541 | 1,10E-05 | 14 - | 0,000177752 - | 12 - | 0,000189184 - | 14 - | 0,000209818 - | 12 - | 0,0001825 - | 11 - | 0,000188453 - |
| 9005 | 1 | 3um_FIJJAFBF_131606 | Bacteria | Proteobacteri | Gammaprotec | Cellvibrionale | Porticoccacea | unclassified | SAR92 bacteri | unclassified | unclassified | unclassified | unclassified | unclassified | 21,4 | 2,332380758 | 0,000287731 | 2,80E-05 | 24 - | 0,000271929 - | 18 - | 0,000253242 - | 24 - | 0,000320985 - | 20 - | 0,000271438 - | 21 - | 0,000321063 - |
| 9006 | 1 | 3um_FIJJAFBF_793456 | Bacteria | Proteobacteri | Gammaprotec | unclassified | unclassified | unclassified | OM182 bacter | unclassified | unclassified | unclassified | unclassified | unclassified | 1,2 | 0,748331477 | 2,56E-05 | 1,69E-05 | 1 - | 1,82E-05 - | 2 - | 4,51E-05 - | 1 - | 2,14E-05 - | 2 - | 4,35E-05 - | 0 - | 0 - |
| 9007 | 1 | 0.2um_NOKCPHJN_430518 | Eukaryota | unclassified | Dinophyceae | Suessiales | Symbiodiniace | Symbiodinium | Symbiodinium | unclassified | unclassified | unclassified | unclassified | unclassified | 1,2 | 0,4 | 4,90E-05 | 1,64E-05 | 1 - | 3,43E-05 - | 1 - | 4,25E-05 - | 2 - | 8,09E-05 - | 1 - | 4,10E-05 - | 1 - | 4,62E-05 - |
| 9008 | 1 | 0.2um_NOKCPHJN_917246 | Bacteria | Proteobacteri | Alphaproteob | Rhodobactera | Rhodobactera | Planktomarina | Planktomarina | unclassified | unclassified | unclassified | unclassified | unclassified | 3,2 | 0,979795897 | 0,000101879 | 2,88E-05 | 4 - | 0,00010828 - | 4 - | 0,000134452 - | 2 - | 6,39E-05 - | 4 - | 0,000129701 - | 2 - | 7,31E-05 - |
| 9009 | 1 | 3um_FIJJAFBF_19845 | Bacteria | Bacteroidetes | Flavobacteriia | Flavobacterial | Flavobacteriac | Polaribacter | various | cellular proces | Posttranslatio | 09GE9 | Thioredoxin | 313594.PI23P | 4,6 | 0,489897949 | 0,000596823 | 5,82E-05 | 5 - | 0,000546556 - | 5 - | 0,000678661 - | 5 - | 0,000645153 - | 4 - | 0,000523746 - | 4 - | 0,000589998 - |
| 9010 | 1 | 3um_FIJJAFBF_660193 | Bacteria | Bacteroidetes | unclassified | unclassified | unclassified | unclassified | Bacteroidetes | unclassified | unclassified | unclassified | unclassified | unclassified | 1,2 | 0,4 | 0,000108956 | 3,76E-05 | 1 - | 7,60E-05 - | 1 - | 9,44E-05 - | 1 - | 8,97E-05 - | 2 - | 0,000182097 - | 1 - | 0,000102566 - |
| 9011 | 1 | 0.2um_NOKCPHJN_239108 | Bacteria | Bacteroidetes | Flavobacteriia | various | various | various | various | unclassified | unclassified | unclassified | unclassified | unclassified | 4 | 0,894427191 | 0,000368979 | 9,12E-05 | 3 - | 0,000231094 - | 5 - | 0,000478251 - | 5 - | 0,000454638 - | 4 - | 0,000369083 - | 3 - | 0,000311828 - |
| 9012 | 1 | 0.2um_NOKCPHJN_747877 | Bacteria | Proteobacteri | Alphaproteob | Rhodospirillal | Rhodospirillac | unclassified | Rhodospirillac | unclassified | unclassified | unclassified | unclassified | unclassified | 1 | 0 | 9,93E-05 | 9,43E-06 | 1 - | 8,32E-05 - | 1 - | 0,000103275 - | 1 - | 9,82E-05 - | 1 - | 9,96E-05 - | 1 - | 0,000112228 - |
| 9013 | 1 | 3um_FIJJAFBF_53337 | Bacteria | Bacteroidetes | Flavobacteriia | Flavobacterial | Flavobacteriac | Polaribacter | Polaribacter s | metabolism | Amino acid tra | 09F6N | The glycine cl | 313598.MED1 | 1,2 | 0,748331477 | 0,00012421 | 7,49E-05 | 2 - | 0,000182185 - | 1 - | 0,00011311 - | 1 - | 0,000107526 - | 2 - | 0,000218227 - | 0 - | 0 - |
| 9014 | 1 | 3um_FIJJAFBF_813731 | Bacteria | Proteobacteri | Gammaprotec | Cellvibrionale | Porticoccacea | unclassified | Gammaprotec | metabolism | Inorganic ion t | 16TQC | TONB-depend | 314287.GB22 | 5,8 | 0,4 | 0,000103694 | 7,53E-06 | 6 - | 9,03E-05 - | 6 - | 0,000112073 - | 6 - | 0,000106539 - | 6 - | 0,000108113 - | 5 - | 0,00010149 - |
| 9015 | 1 | 3um_FIJJAFBF_372780 | Bacteria | Proteobacteri | Alphaproteob | Rhodobactera | Rhodobactera | Tateyamaria | Tateyamaria s | metabolism | Energy produc | 01R8W | NDH-1 shuttle | 391595.RLO14 | 0,2 | 0,4 | 1,28E-05 | 2,56E-05 | 1 - | 6,41E-05 - | 0 - | 0 - | 0 - | 0 - | 0 - | 0 - | 0 - | 0 - |
| 9016 | 1 | 3um_FIJJAFBF_507285 | Bacteria | Proteobacteri | Alphaproteob | Rhodobactera | Rhodobactera | Planktomarina | Planktomarina | metabolism | Energy produc | COG0712 | F(1)F(0) ATP s | 391616.OA23 | 1 | 0,632455532 | 5,87E-05 | 3,31E-05 | 2 - | 0,000102479 - | 0 - | 0 - | 1 - | 6,05E-05 - | 1 - | 6,14E-05 - | 1 - | 6,91E-05 - |
| 9017 | 1 | 3um_FIJJAFBF_254297 | Bacteria | Proteobacteri | Gammaprotec | Cellvibrionale | Cellvibrionace | Simiduia | Simiduia agari | unclassified | unclassified | unclassified | unclassified | unclassified | 2 | 0 | 0,000152252 | 1,45E-05 | 2 - | 0,00012753 - | 2 - | 0,000158354 - | 2 - | 0,000150536 - | 2 - | 0,000152759 - | 2 - | 0,000172083 - |
| 9018 | 1 | 3um_FIJJAFBF_19836 | Bacteria | Bacteroidetes | Flavobacteriia | Flavobacterial | Flavobacteriac | Polaribacter | Polaribacter s | metabolism | Energy produc | 05CAY | ferredoxin--na | 391587.KAOT | 1,6 | 0,8 | 5,98E-05 | 2,37E-05 | 3 - | 9,81E-05 - | 1 - | 4,06E-05 - | 2 - | 7,72E-05 - | 1 - | 3,92E-05 - | 1 - | 4,41E-05 - |
| 9019 | 1 | 0.2um_NOKCPHJN_119890 | Bacteria | Proteobacteri | Alphaproteob | Rhodobactera | Rhodobactera | Labrenzia | various | metabolism | Amino acid tra | 01RBF | Branched-chai | 384765.SIAM | 0,6 | 0,489897949 | 1,86E-05 | 1,52E-05 | 0 - | 0 - | 1 - | 3,19E-05 - | 1 - | 3,03E-05 - | 1 - | 3,08E-05 - | 0 - | 0 - |
| 9020 | 1 | 3um_FIJJAFBF_149819 | Bacteria | Proteobacteri | various | various | various | various | various | unclassified | unclassified | unclassified | unclassified | unclassified | 0,6 | 0,489897949 | 0,000101338 | 8,28E-05 | 0 - | 0 - | 1 - | 0,000173803 - | 1 - | 0,000165222 - | 1 - | 0,000167663 - | 0 - | 0 - |
| 9021 | 1 | 0.2um_NOKCPHJN_374229 | Eukaryota | unclassified | Oomycetes | various | various | various | various | unclassified | unclassified | unclassified | unclassified | unclassified | 0,2 | 0,4 | 2,08E-05 | 4,16E-05 | 0 - | 0 - | 0 - | 0 - | 0 - | 0 - | 0 - | 0 - | 1 - | 0,000103943 - |
| 9022 | 1 | 3um_FIJJAFBF_285761 | Bacteria | unclassified | unclassified | unclassified | unclassified | unclassified | uncultured ba | information st | Translation, ri | 0PA05 | 30S ribosomal | 313594.PI23P | 5,8 | 0,748331477 | 0,000440108 | 6,32E-05 | 6 - | 0,000382589 - | 7 - | 0,00055424 - | 6 - | 0,000451607 - | 5 - | 0,000381898 - | 5 - | 0,000430207 - |
| 9023 | 1 | 3um_FIJJAFBF_486055 | Eukaryota | unclassified | Oomycetes | Peronosporale | Peronosporac | Phytophthora | Phytophthora | cellular proces | Signal transdu | 152IZ | EFh | 67593.Physo1 | 0,8 | 0,4 | 7,71E-05 | 3,88E-05 | 0 - | 0 - | 1 - | 9,63E-05 - | 1 - | 9,15E-05 - | 1 - | 9,29E-05 - | 1 - | 0,000104645 - |
| 9024 | 1 | 0.2um_NOKCPHJN_36238 | Eukaryota | unclassified | Cryptophyta | Cryptomonad | Cryptomonad | Cryptomonas | Cryptomonas | metabolism | Carbohydrate | COG1850 | RuBisCO catal | 45157.CMV01 | 1,4 | 0,489897949 | 3,93E-05 | 1,39E-05 | 1 - | 2,35E-05 - | 1 - | 2,92E-05 - | 2 - | 5,55E-05 - | 2 - | 5,63E-05 - | 1 - | 3,17E-05 - |
| 9025 | 1 | 0.2um_NOKCPHJN_58529 | Bacteria | Proteobacteri | Alphaproteob | Rhodobactera | Rhodobactera | unclassified | Rhodobactera | cellular proces | Cell motility | COG1749 | flagellar hook | 391626.OA30 | 1,6 | 0,8 | 1,26E-05 | 5,15E-06 | 3 - | 2,05E-05 - | 2 - | 1,69E-05 - | 1 - | 8,05E-06 - | 1 - | 8,17E-06 - | 1 - | 9,21E-06 - |
| 9026 | 1 | 3um_FIJJAFBF_497356 | Bacteria | Proteobacteri | Gammaprotec | Oceanospirilla | Saccharospirill | Reinekea | Reinekea fors | cellular proces | Cell motility | 174CZ | Pilus assembly | 314283.MED2 | 0,8 | 0,4 | 1,25E-05 | 6,34E-06 | 1 - | 1,35E-05 - | 1 - | 1,68E-05 - | 1 - | 1,59E-05 - | 1 - | 1,62E-05 - | 0 - | 0 - |
| 9027 | 1 | 3um_FIJJAFBF_786156 | Bacteria | Proteobacteri | Gammaprotec | Oceanospirilla | Saccharospirill | Saccharospiril | Saccharospirill | information st | Translation, ri | 16Q16 | Binds the lowe | 207954.MED9 | 7,6 | 1,356465997 | 0,000449201 | 3,53E-05 | 10 - | 0,000503407 - | 7 - | 0,000437558 - | 8 - | 0,000475376 - | 7 - | 0,000422098 - | 6 - | 0,000407564 - |
| 9028 | 1 | 3um_FIJJAFBF_196004 | Bacteria | Bacteroidetes | Flavobacteriia | Flavobacterial | Flavobacteriac | Polaribacter | Polaribacter s | cellular proces | Cell wall/mem | COG1368 | Sulfatase | 313594.PI23P | 0,8 | 0,4 | 1,65E-05 | 8,49E-06 | 1 - | 1,73E-05 - | 1 - | 2,14E-05 - | 0 - | 0 - | 1 - | 2,07E-05 - | 1 - | 2,33E-05 - |
| 9029 | 2 | 0.2um_NOKCPHJN_388854;3um_FIJJAFBF_8369 | Bacteria | Bacteroidetes | Flavobacteriia | Flavobacterial | Flavobacteriac | Polaribacter | various | poorly charact | Function unkn | 0PA0H | (Lipo)protein | 313594.PI23P | 1,2 | 0,4 | 9,07E-05 | 3,04E-05 | 1 - | 6,34E-05 - | 1 - | 7,87E-05 - | 2 - | 0,000149704 - | 1 - | 7,60E-05 - | 1 - | 8,56E-05 - |
| 9030 | 2 | 0.2um_NOKCPHJN_842037;3um_FIJJAFBF_55082 | Eukaryota | Chordata | Actinopteri | Carangiformes | Carangidae | Seriola | Seriola lalandi | unclassified | unclassified | unclassified | unclassified | unclassified | 3 | 0 | 0,000411081 | 3,90E-05 | 3 - | 0,00034433 - | 3 - | 0,000427557 - | 3 - | 0,000406447 - | 3 - | 0,00041245 - | 3 - | 0,000464623 - |
| 9031 | 1 | 3um_FIJJAFBF_08120 | Bacteria | Bacteroidetes | Flavobacteriia | Flavobacterial | Flavobacteriac | Polaribacter | Polaribacter s | metabolism | Amino acid tra | COG0019 | Specifically ca | 313598.MED1 | 1,8 | 0,4 | 6,09E-05 | 1,69E-05 | 1 - | 2,79E-05 - | 2 - | 6,92E-05 - | 2 - | 6,58E-05 - | 2 - | 6,67E-05 - | 2 - | 7,52E-05 - |
| 9032 | 1 | 3um_FIJJAFBF_297043 | unclassified | unclassified | unclassified | unclassified | unclassified | unclassified | unclassified | unclassified | unclassified | unclassified | unclassified | unclassified | 0,2 | 0,4 | 2,87E-05 | 5,74E-05 | 1 - | 0,000143471 - | 0 - | 0 - | 0 - | 0 - | 0 - | 0 - | 0 - | 0 - |
| 9033 | 1 | 0.2um_NOKCPHJN_10823 | Viruses | unclassified | unclassified | unclassified | Phycodnavirid | unclassified | Pyramimonas | unclassified | unclassified | unclassified | unclassified | unclassified | 0,4 | 0,489897949 | 1,58E-05 | 1,94E-05 | 0 - | 0 - | 1 - | 3,79E-05 - | 0 - | 0 - | 0 - | 0 - | 1 - | 4,12E-05 - |
| 9034 | 1 | 3um_FIJJAFBF_203490 | Bacteria | Proteobacteri | Gammaprotec | Alteromonada | Alteromonada | Paraglaciecola | Paraglaciecola | information st | Translation, ri | 16QAP | Associates wit | 455436.GHTC | 3 | 0 | 0,00013612 | 1,29E-05 | 3 - | 0,000114017 - | 3 - | 0,000141575 - | 3 - | 0,000134585 - | 3 - | 0,000136573 - | 3 - | 0,000153849 - |
| 9035 | 1 | 3um_FIJJAFBF_256256 | Bacteria | Proteobacteri | Alphaproteob | Rhodobactera | unclassified | unclassified | Rhodobactera | metabolism | Carbohydrate | 0XT7V | Trap dicarbox | 633131.TR2A | 0,4 | 0,489897949 | 1,32E-05 | 1,63E-05 | 1 - | 3,03E-05 - | 0 - | 0 - | 1 - | 3,57E-05 - | 0 - | 0 - | 0 - | 0 - |
| 9036 | 1 | 3um_FIJJAFBF_323242 | Bacteria | Bacteroidetes | Flavobacteriia | Flavobacterial | Flavobacteriac | Hyunsoonleel | Hyunsoonleel | cellular proces | Posttranslatio | 09GE9 | Thioredoxin | 391603.FBALC | 1,8 | 0,4 | 0,000235198 | 5,86E-05 | 2 - | 0,000218622 - | 2 - | 0,000271465 - | 1 - | 0,000129031 - | 2 - | 0,000261873 - | 2 - | 0,000294999 - |
| 9037 | 1 | 3um_FIJJAFBF_457156 | Bacteria | Actinobacteria | Actinobacteria | Streptomyceta | Streptomyceta | Streptomyces | Streptomyces | unclassified | unclassified | unclassified | unclassified | unclassified | 0,8 | 0,748331477 | 1,09E-05 | 1,07E-05 | 1 - | 1,17E-05 - | 2 - | 2,90E-05 - | 1 - | 1,38E-05 - | 0 - | 0 - | 0 - | 0 - |
| 9038 | 1 | 0.2um_NOKCPHJN_201839 | Bacteria | Bacteroidetes | unclassified | unclassified | unclassified | unclassified | Bacteroidetes | information st | Translation, ri | 05CAV | thus facilitatin | 755732.Fluta_ | 2,2 | 0,748331477 | 3,32E-05 | 1,01E-05 | 3 - | 3,88E-05 - | 3 - | 4,81E-05 - | 2 - | 3,05E-05 - | 2 - | 3,10E-05 - | 1 - | 1,74E-05 - |
| 9039 | 1 | 0.2um_NOKCPHJN_110551 | Bacteria | Proteobacteri | Gammaprotec | Cellvibrionale | Porticoccacea | unclassified | SAR92 bacteri | unclassified | unclassified | unclassified | unclassified | unclassified | 0,2 | 0,4 | 8,41E-06 | 1,68E-05 | 1 - | 4,20E-05 - | 0 - | 0 - | 0 - | 0 - | 0 - | 0 - | 0 - | 0 - |
| 9040 | 1 | 0.2um_NOKCPHJN_569876 | Bacteria | Bacteroidetes | unclassified | unclassified | unclassified | unclassified | Bacteroidetes | unclassified | unclassified | unclassified | unclassified | unclassified | 0,2 | 0,4 | 5,12E-06 | 1,02E-05 | 1 - | 2,56E-05 - | 0 - | 0 - | 0 - | 0 - | 0 - | 0 - | 0 - | 0 - |
| 9041 | 1 | 3um_FIJJAFBF_763633 | Bacteria | Proteobacteri | Betaproteoba | Burkholderiale | Comamonada | Limnohabitan | Limnohabitan | cellular proces | Cell cycle cont | 0B7CT | rod shape-det | 391597.LMED | 0,4 | 0,489897949 | 1,45E-05 | 1,79E-05 | 1 - | 3,31E-05 - | 0 - | 0 - | 0 - | 0 - | 1 - | 3,96E-05 - | 0 - | 0 - |
| 9042 | 1 | 3um_FIJJAFBF_220488 | Bacteria | Proteobacteri | Gammaprotec | Alteromonada | Alteromonada | Paraglaciecola | Paraglaciecola | information st | Translation, ri | COG0087 | One of the pri | 342610.Patl_ | 8,8 | 2,481934729 | 0,000554907 | 0,000117016 | 12 - | 0,00064968 - | 8 - | 0,000537807 - | 11 - | 0,000702974 - | 8 - | 0,000518805 - | 5 - | 0,00036527 - |
| 9043 | 1 | 3um_FIJJAFBF_673864 | Eukaryota | unclassified | Dinophyceae | Suessiales | Symbiodiniace | Symbiodinium | Symbiodinium | unclassified | unclassified | unclassified | unclassified | unclassified | 1 | 0 | 1,75E-05 | 1,66E-06 | 1 - | 1,46E-05 - | 1 - | 1,82E-05 - | 1 - | 1,73E-05 - | 1 - | 1,75E-05 - | 1 - | 1,98E-05 - |
| 9044 | 1 | 3um_FIJJAFBF_738679 | Bacteria | Bacteroidetes | Flavobacteriia | Flavobacterial | Flavobacteriac | Formosa | Formosa sp. H | unclassified | unclassified | unclassified | unclassified | unclassified | 1,8 | 0,748331477 | 2,63E-05 | 9,36E-06 | 3 - | 3,76E-05 - | 1 - | 1,56E-05 - | 2 - | 2,96E-05 - | 1 - | 1,50E-05 - | 2 - | 3,38E-05 - |
| 9045 | 1 | 3um_FIJJAFBF_336847 | Bacteria | Proteobacteri | Alphaproteob | Rhodobactera | Rhodobactera | Jannaschia | Jannaschia se | metabolism | Carbohydrate | 16THB | transporter, d | 744979.R2A13 | 3,8 | 0,748331477 | 0,000152504 | 2,48E-05 | 5 - | 0,000169788 - | 3 - | 0,000126496 - | 3 - | 0,00012025 - | 4 - | 0,000162702 - | 4 - | 0,000183283 - |
| 9046 | 1 | 3um_FIJJAFBF_327617 | Bacteria | Bacteroidetes | unclassified | unclassified | unclassified | unclassified | Bacteroidetes | unclassified | unclassified | unclassified | unclassified | unclassified | 1 | 0,632455532 | 8,66E-05 | 4,94E-05 | 2 - | 0,000154063 - | 1 - | 9,57E-05 - | 1 - | 9,09E-05 - | 1 - | 9,23E-05 - | 0 - | 0 - |
| 9047 | 1 | 0.2um_NOKCPHJN_1023200 | Bacteria | Bacteroidetes | Sphingobacter | Sphingobacter | Sphingobacter | Pedobacter | Pedobacter sp | unclassified | unclassified | unclassified | unclassified | unclassified | 1 | 0 | 4,22E-05 | 4,00E-06 | 1 - | 3,53E-05 - | 1 - | 4,39E-05 - | 1 - | 4,17E-05 - | 1 - | 4,23E-05 - | 1 - | 4,77E-05 - |
| 9048 | 1 | 3um_FIJJAFBF_338362 | Bacteria | Proteobacteri | Gammaprotec | Alteromonada | Alteromonada | Alteromonas | Alteromonas | information st | Translation, ri | COG0050 | This protein p | 675806.VII_0 | 10,8 | 0,748331477 | 0,000680028 | 6,35E-05 | 12 - | 0,00063471 - | 11 - | 0,000722446 - | 10 - | 0,000624342 - | 10 - | 0,000633564 - | 11 - | 0,000785078 - |
| 9049 | 1 | 0.2um_NOKCPHJN_1079231 | Bacteria | Proteobacteri | Alphaproteob | Pelagibacteral | Pelagibacterac | Candidatus Pe | Candidatus Pe | unclassified | unclassified | unclassified | unclassified | unclassified | 0,6 | 0,489897949 | 3,94E-05 | 3,24E-05 | 1 - | 5,83E-05 - | 0 - | 0 - | 1 - | 6,88E-05 - | 1 - | 6,98E-05 - | 0 - | 0 - |
| 9050 | 1 | 3um_FIJJAFBF_66597 | Eukaryota | Bacillariophyt | Bacillariophyc | Cymbellales | Gomphonema | Gomphoneis | Gomphoneis | metabolism | Energy produc | 1DYB8 | PsaA and PsaB | 3055.DAA014 | 0,2 | 0,4 | 3,05E-06 | 6,11E-06 | 1 - | 1,53E-05 - | 0 - | 0 - | 0 - | 0 - | 0 - | 0 - | 0 - | 0 - |
| 9051 | 1 | 3um_FIJJAFBF_716741 | Bacteria | Proteobacteri | Gammaprotec | Oceanospirilla | Saccharospirill | Reinekea | Reinekea fors | metabolism | Lipid transpor | COG1182 | Catalyzes the | 443152.MDG | 0,4 | 0,489897949 | 2,55E-05 | 3,14E-05 | 1 - | 5,80E-05 - | 0 - | 0 - | 0 - | 0 - | 1 - | 6,94E-05 - | 0 - | 0 - |
| 9052 | 2 | 0.2um_NOKCPHJN_23972;3um_FIJJAFBF_19055 | Eukaryota | Apicomplexa | Conoidasida | Eucoccidiorida | Eimeriidae | Eimeria | Eimeria necat | unclassified | unclassified | unclassified | unclassified | unclassified | 0,4 | 0,489897949 | 4,72E-06 | 5,83E-06 | 1 - | 1,05E-05 - | 1 - | 1,31E-05 - | 0 - | 0 - | 0 - | 0 - | 0 - | 0 - |
| 9053 | 1 | 0.2um_NOKCPHJN_103498 | Bacteria | Proteobacteri | Alphaproteob | Rhodobactera | unclassified | unclassified | Rhodobactera | unclassified | unclassified | unclassified | unclassified | unclassified | 0,4 | 0,489897949 | 4,31E-05 | 5,38E-05 | 1 - | 9,18E-05 - | 0 - | 0 - | 0 - | 0 - | 0 - | 0 - | 1 - | 0,0001239 - |
| 9054 | 1 | 0.2um_NOKCPHJN_372882 | Eukaryota | Arthropoda | Insecta | Diptera | Tephritidae | Zeugodacus | Zeugodacus c | unclassified | unclassified | unclassified | unclassified | unclassified | 0,2 | 0,4 | 1,12E-06 | 2,23E-06 | 0 - | 0 - | 0 - | 0 - | 0 - | 0 - | 1 - | 5,58E-06 - | 0 - | 0 - |
| 9055 | 1 | 0.2um_NOKCPHJN_231372 | Bacteria | Proteobacteri | Alphaproteob | Rhodobactera | Rhodobactera | Planktomarina | Planktomarina | unclassified | unclassified | unclassified | unclassified | unclassified | 0,4 | 0,489897949 | 1,93E-05 | 2,37E-05 | 0 - | 0 - | 0 - | 0 - | 0 - | 0 - | 1 - | 4,54E-05 - | 1 - | 5,11E-05 - |
| 9056 | 1 | 3um_FIJJAFBF_585924 | unclassified | unclassified | unclassified | unclassified | unclassified | unclassified | unclassified | unclassified | unclassified | unclassified | unclassified | unclassified | 0,2 | 0,4 | 1,54E-05 | 3,08E-05 | 0 - | 0 - | 1 - | 7,70E-05 - | 0 - | 0 - | 0 - | 0 - | 0 - | 0 - |
| 9057 | 1 | 3um_FIJJAFBF_01994 | Bacteria | Planctomycet | Planctomyceti | Planctomyceta | Planctomyceta | unclassified | Planctomyceta | unclassified | unclassified | unclassified | unclassified | unclassified | 3,4 | 1,2 | 0,000285509 | 9,91E-05 | 4 - | 0,00028516 - | 5 - | 0,000442605 - | 4 - | 0,000336602 - | 2 - | 0,000170787 - | 2 - | 0,000192391 - |
| 9058 | 1 | 0.2um_NOKCPHJN_144467 | Bacteria | Proteobacteri | Gammaprotec | Oceanospirilla | Saccharospirill | Reinekea | Reinekea blan | metabolism | Energy produc | COG0056 | Produces ATP | 314283.MED2 | 17,2 | 1,469693846 | 0,000456522 | 4,07E-05 | 18 - | 0,000401942 - | 15 - | 0,000415911 - | 18 - | 0,000474451 - | 19 - | 0,000508207 - | 16 - | 0,0004821 - |
| 9059 | 1 | 0.2um_NOKCPHJN_33601 | Bacteria | Bacteroidetes | Flavobacteriia | Flavobacterial | unclassified | unclassified | Flavobacterial | information st | Translation, ri | COG1185 | Involved in mF | 313603.FB217 | 2,6 | 1,356465997 | 4,58E-05 | 1,97E-05 | 5 - | 7,64E-05 - | 3 - | 5,69E-05 - | 2 - | 3,61E-05 - | 1 - | 1,83E-05 - | 2 - | 4,12E-05 - |
| 9060 | 1 | 0.2um_NOKCPHJN_139170 | Bacteria | Proteobacteri | Gammaprotec | Cellvibrionale | Porticoccacea | unclassified | Gammaprotec | unclassified | unclassified | unclassified | unclassified | unclassified | 0,2 | 0,4 | 3,72E-06 | 7,44E-06 | 1 - | 1,86E-05 - | 0 - | 0 - | 0 - | 0 - | 0 - | 0 - | 0 - | 0 - |
| 9061 | 1 | 0.2um_NOKCPHJN_1072705 | unclassified | unclassified | unclassified | unclassified | unclassified | unclassified | unclassified | unclassified | unclassified | unclassified | unclassified | unclassified | 0,6 | 0,489897949 | 7,18E-05 | 5,92E-05 | 1 - | 0,000104343 - | 1 - | 0,000129563 - | 0 - | 0 - | 1 - | 0,000124985 - | 0 - | 0 - |
| 9062 | 1 | 3um_FIJJAFBF_306833 | Bacteria | Proteobacteri | Alphaproteob | Rhodobactera | Rhodobactera | various | various | metabolism | Inorganic ion t | 16PII | TRAP transpor | 314262.MED1 | 3,2 | 0,979795897 | 0,000139565 | 5,10E-05 | 2 - | 7,15E-05 - | 4 - | 0,000177594 - | 2 - | 8,44E-05 - | 4 - | 0,000171319 - | 4 - | 0,00019299 - |
| 9063 | 1 | 3um_FIJJAFBF_13587 | Bacteria | Proteobacteri | Gammaprotec | Cellvibrionale | Porticoccacea | unclassified | various | information st | Translation, ri | 05KK9 | This protein bi | 314287.GB22 | 1,6 | 0,8 | 0,000210304 | 0,000104531 | 1 - | 0,000110362 - | 1 - | 0,000137037 - | 3 - | 0,000390814 - | 2 - | 0,000264391 - | 1 - | 0,000148918 - |
| 9064 | 1 | 0.2um_NOKCPHJN_689335 | Bacteria | Proteobacteri | Alphaproteob | Rhodobactera | Rhodobactera | Litoreibacter | Litoreibacter a | metabolism | Carbohydrate | 01RFA | ABC transport | 314232.SKA53 | 1,4 | 0,489897949 | 5,70E-05 | 2,32E-05 | 1 - | 3,36E-05 - | 1 - | 4,17E-05 - | 2 - | 7,92E-05 - | 1 - | 4,02E-05 - | 2 - | 9,06E-05 - |
| 9065 | 1 | 0.2um_NOKCPHJN_1029960 | Eukaryota | Phaeophyceae | unclassified | Ectocarpales | Ectocarpaceae | Ectocarpus | Ectocarpus sili | information st | RNA processin | 1DRWN | psbB mRNA m | 3694.POPTR_ | 0,4 | 0,489897949 | 9,96E-06 | 1,22E-05 | 0 - | 0 - | 1 - | 2,54E-05 - | 0 - | 0 - | 1 - | 2,45E-05 - | 0 - | 0 - |
| 9066 | 1 | 3um_FIJJAFBF_59808 | Viruses | unclassified | unclassified | unclassified | unclassified | unclassified | uncultured M | unclassified | unclassified | unclassified | unclassified | unclassified | 0,2 | 0,4 | 7,45E-06 | 1,49E-05 | 1 - | 3,73E-05 - | 0 - | 0 - | 0 - | 0 - | 0 - | 0 - | 0 - | 0 - |
| 9067 | 1 | 0.2um_NOKCPHJN_146211 | Bacteria | Bacteroidetes | Flavobacteriia | Flavobacterial | Flavobacteriac | Algibacter | Algibacter alg | cellular proces | Signal transdu | 09A4D | gtp-binding pr | 391587.KAOT | 1,2 | 1,166190379 | 2,56E-05 | 2,37E-05 | 3 - | 5,75E-05 - | 2 - | 4,76E-05 - | 0 - | 0 - | 1 - | 2,30E-05 - | 0 - | 0 - |
| 9068 | 1 | 3um_FIJJAFBF_646115 | Bacteria | Proteobacteri | Gammaprotec | unclassified | unclassified | unclassified | various | metabolism | Nucleotide tra | 08UTP | (2Fe-2S)-bindi | 644968.DFW1 | 1 | 0 | 7,06E-05 | 6,71E-06 | 1 - | 5,92E-05 - | 1 - | 7,35E-05 - | 1 - | 6,98E-05 - | 1 - | 7,09E-05 - | 1 - | 7,98E-05 - |
| 9069 | 1 | 3um_FIJJAFBF_274943 | Bacteria | Proteobacteri | Alphaproteob | Rhodobactera | Rhodobactera | unclassified | Rhodobactera | unclassified | unclassified | unclassified | unclassified | unclassified | 1 | 0,894427191 | 2,90E-05 | 2,53E-05 | 2 - | 4,97E-05 - | 2 - | 6,17E-05 - | 0 - | 0 - | 0 - | 0 - | 1 - | 3,35E-05 - |
| 9070 | 1 | 3um_FIJJAFBF_510233 | Bacteria | Bacteroidetes | Flavobacteriia | Flavobacterial | Flavobacteriac | Polaribacter | Polaribacter s | unclassified | unclassified | unclassified | unclassified | unclassified | 1,6 | 0,489897949 | 8,12E-05 | 2,86E-05 | 1 - | 4,19E-05 - | 1 - | 5,20E-05 - | 2 - | 9,89E-05 - | 2 - | 0,000100353 - | 2 - | 0,000113047 - |
| 9071 | 1 | 0.2um_NOKCPHJN_339287 | Bacteria | Bacteroidetes | Cytophagia | Cytophagales | unclassified | unclassified | Cytophagales | unclassified | unclassified | unclassified | unclassified | unclassified | 1 | 0 | 6,23E-05 | 5,91E-06 | 1 - | 5,22E-05 - | 1 - | 6,48E-05 - | 1 - | 6,16E-05 - | 1 - | 6,25E-05 - | 1 - | 7,04E-05 - |
| 9072 | 1 | 3um_FIJJAFBF_108911 | Bacteria | Proteobacteri | Gammaprotec | Alteromonada | Alteromonada | Paraglaciecola | various | cellular proces | Posttranslatio | 17FI0 | Binds to Cpn6 | 983545.Glaag | 2,4 | 0,489897949 | 0,000346381 | 9,37E-05 | 2 - | 0,000239118 - | 2 - | 0,000296914 - | 2 - | 0,000282255 - | 3 - | 0,000429635 - | 3 - | 0,000483983 - |
| 9073 | 2 | 3um_FIJJAFBF_30200;3um_FIJJAFBF_756301 | Bacteria | Proteobacteri | various | various | various | various | various | metabolism | Carbohydrate | 05CMT | Dihydroxyacet | various | 0,2 | 0,4 | 5,02E-06 | 1,00E-05 | 0 - | 0 - | 0 - | 0 - | 1 - | 2,51E-05 - | 0 - | 0 - | 0 - | 0 - |
| 9074 | 1 | 3um_FIJJAFBF_520481 | Bacteria | Bacteroidetes | Flavobacteriia | Flavobacterial | Flavobacteriac | Polaribacter | Polaribacter s | cellular proces | Signal transdu | 09BM1 | Two compone | 313598.MED1 | 0,2 | 0,4 | 9,85E-06 | 1,97E-05 | 1 - | 4,93E-05 - | 0 - | 0 - | 0 - | 0 - | 0 - | 0 - | 0 - | 0 - |
| 9075 | 1 | 0.2um_NOKCPHJN_574142 | Bacteria | Proteobacteri | Gammaprotec | Cellvibrionale | Porticoccacea | unclassified | gamma prote | poorly charact | Function unkn | 0QNMQ | NA | 314287.GB22 | 0,2 | 0,4 | 1,99E-06 | 3,98E-06 | 1 - | 9,95E-06 - | 0 - | 0 - | 0 - | 0 - | 0 - | 0 - | 0 - | 0 - |
| 9076 | 1 | 0.2um_NOKCPHJN_825274 | unclassified | unclassified | unclassified | unclassified | unclassified | unclassified | unclassified | unclassified | unclassified | unclassified | unclassified | unclassified | 1,2 | 0,4 | 0,000147859 | 4,95E-05 | 1 - | 0,000103402 - | 1 - | 0,000128395 - | 2 - | 0,000244112 - | 1 - | 0,000123859 - | 1 - | 0,000139527 - |
| 9077 | 1 | 3um_FIJJAFBF_604556 | Bacteria | Proteobacteri | Gammaprotec | Arenicellales | Arenicellaceae | Arenicella | Arenicella xan | information st | Translation, ri | COG0087 | One of the pri | 247634.GPB2 | 0,6 | 0,8 | 3,41E-05 | 4,40E-05 | 2 - | 0,000107268 - | 0 - | 0 - | 1 - | 6,33E-05 - | 0 - | 0 - | 0 - | 0 - |
| 9078 | 1 | 3um_FIJJAFBF_201380 | Bacteria | Bacteroidetes | Flavobacteriia | Flavobacterial | Flavobacteriac | Lacinutrix | Lacinutrix jang | poorly charact | Function unkn | 0XPRT | NA | 391598.FBBAL | 7 | 0 | 0,000192608 | 1,83E-05 | 7 - | 0,000161333 - | 7 - | 0,000200328 - | 7 - | 0,000190437 - | 7 - | 0,00019325 - | 7 - | 0,000217695 - |
| 9079 | 1 | 3um_FIJJAFBF_254415 | Bacteria | Bacteroidetes | unclassified | unclassified | unclassified | unclassified | Bacteroidetes | information st | Translation, ri | 05C64 | Binds directly | 575590.HMPR | 0,8 | 0,4 | 4,74E-05 | 2,44E-05 | 1 - | 4,97E-05 - | 1 - | 6,17E-05 - | 1 - | 5,87E-05 - | 0 - | 0 - | 1 - | 6,70E-05 - |
| 9080 | 1 | 3um_FIJJAFBF_345326 | Eukaryota | Apicomplexa | Conoidasida | Eucoccidiorida | Sarcocystidae | Hammondia | Hammondia h | unclassified | unclassified | unclassified | unclassified | unclassified | 0,2 | 0,4 | 1,89E-05 | 3,79E-05 | 0 - | 0 - | 0 - | 0 - | 1 - | 9,47E-05 - | 0 - | 0 - | 0 - | 0 - |
| 9081 | 1 | 3um_FIJJAFBF_494278 | Bacteria | Proteobacteri | Gammaprotec | Oceanospirilla | Saccharospirill | Reinekea | Reinekea fors | poorly charact | Function unkn | 05VQI | NA | 314292.VAS14 | 0,6 | 0,489897949 | 7,01E-05 | 5,78E-05 | 1 - | 0,000102479 - | 1 - | 0,000127249 - | 1 - | 0,000120966 - | 0 - | 0 - | 0 - | 0 - |
| 9082 | 1 | 3um_FIJJAFBF_103761 | Bacteria | Actinobacteria | Actinobacteria | Micrococcales | Microbacteria | Candidatus Ac | Candidatus Ac | metabolism | Inorganic ion t | 05C2J | ABC transport | 446462.Amir_ | 0,4 | 0,489897949 | 2,01E-05 | 2,48E-05 | 1 - | 4,61E-05 - | 0 - | 0 - | 1 - | 5,44E-05 - | 0 - | 0 - | 0 - | 0 - |
| 9083 | 1 | 0.2um_NOKCPHJN_91483 | Bacteria | Proteobacteri | Gammaprotec | Oceanospirilla | Saccharospirill | Reinekea | Reinekea fors | metabolism | Lipid transpor | COG1607 | thioesterase S | 314283.MED2 | 1 | 0 | 9,79E-05 | 9,29E-06 | 1 - | 8,20E-05 - | 1 - | 0,000101799 - | 1 - | 9,68E-05 - | 1 - | 9,82E-05 - | 1 - | 0,000110625 - |
| 9084 | 1 | 3um_FIJJAFBF_520456 | Bacteria | Bacteroidetes | Flavobacteriia | Flavobacterial | Flavobacteriac | Polaribacter | Polaribacter b | unclassified | unclassified | unclassified | unclassified | unclassified | 0,4 | 0,489897949 | 1,59E-05 | 1,94E-05 | 0 - | 0 - | 0 - | 0 - | 1 - | 3,94E-05 - | 1 - | 4,00E-05 - | 0 - | 0 - |
| 9085 | 1 | 3um_FIJJAFBF_203378 | Bacteria | Bacteroidetes | Flavobacteriia | Flavobacterial | Crocinitomica | unclassified | Crocinitomica | metabolism | Carbohydrate | COG0176 | Transaldolase | 313603.FB217 | 0,2 | 0,4 | 1,22E-05 | 2,44E-05 | 0 - | 0 - | 0 - | 0 - | 1 - | 6,10E-05 - | 0 - | 0 - | 0 - | 0 - |
| 9086 | 1 | 3um_FIJJAFBF_281086 | Bacteria | Proteobacteri | Gammaprotec | Alteromonada | Alteromonada | Paraglaciecola | Paraglaciecola | cellular proces | Posttranslatio | 08V1T | peptidylprolyl | 583345.Mmol | 9,2 | 0,979795897 | 0,000497827 | 7,65E-05 | 8 - | 0,000361502 - | 9 - | 0,000504988 - | 11 - | 0,000586734 - | 9 - | 0,000487146 - | 9 - | 0,000548768 - |
| 9087 | 1 | 0.2um_NOKCPHJN_26598 | Bacteria | Bacteroidetes | Flavobacteriia | Flavobacterial | Flavobacteriac | Cellulophaga | Cellulophaga | poorly charact | Function unkn | 0PDYX | NA | 867900.Celly_ | 0,8 | 0,748331477 | 2,94E-05 | 2,79E-05 | 1 - | 3,09E-05 - | 0 - | 0 - | 0 - | 0 - | 2 - | 7,41E-05 - | 1 - | 4,17E-05 - |
| 9088 | 1 | 3um_FIJJAFBF_104815 | Viruses | unclassified | unclassified | unclassified | unclassified | unclassified | uncultured M | unclassified | unclassified | unclassified | unclassified | unclassified | 1,8 | 0,4 | 4,71E-05 | 9,30E-06 | 2 - | 4,45E-05 - | 2 - | 5,52E-05 - | 2 - | 5,25E-05 - | 2 - | 5,33E-05 - | 1 - | 3,00E-05 - |
| 9089 | 1 | 0.2um_NOKCPHJN_212273 | Bacteria | Bacteroidetes | Flavobacteriia | Flavobacterial | Flavobacteriac | Polaribacter | Polaribacter s | unclassified | unclassified | unclassified | unclassified | unclassified | 1,4 | 0,8 | 4,69E-05 | 2,82E-05 | 1 - | 2,85E-05 - | 2 - | 7,07E-05 - | 2 - | 6,72E-05 - | 2 - | 6,82E-05 - | 0 - | 0 - |
| 9090 | 1 | 3um_FIJJAFBF_305232 | Bacteria | Proteobacteri | Gammaprotec | Oceanospirilla | Saccharospirill | Reinekea | Reinekea fors | information st | Transcription | 05VEQ | Cold shock pr | 945550.VISI12 | 4,8 | 0,4 | 0,000959682 | 0,000155061 | 4 - | 0,000665373 - | 5 - | 0,001032745 - | 5 - | 0,000981755 - | 5 - | 0,000996256 - | 5 - | 0,001122279 - |
| 9091 | 1 | 3um_FIJJAFBF_529752 | Bacteria | Proteobacteri | Gammaprotec | Oceanospirilla | Saccharospirill | Reinekea | Reinekea fors | unclassified | unclassified | unclassified | unclassified | unclassified | 0,6 | 0,489897949 | 4,12E-05 | 3,39E-05 | 1 - | 6,11E-05 - | 0 - | 0 - | 1 - | 7,21E-05 - | 1 - | 7,31E-05 - | 0 - | 0 - |
| 9092 | 1 | 3um_FIJJAFBF_111322 | Viruses | unclassified | unclassified | unclassified | various | various | various | unclassified | unclassified | unclassified | unclassified | unclassified | 0,8 | 0,4 | 7,14E-05 | 3,67E-05 | 1 - | 7,55E-05 - | 0 - | 0 - | 1 - | 8,91E-05 - | 1 - | 9,04E-05 - | 1 - | 0,000101891 - |
| 9093 | 1 | 0.2um_NOKCPHJN_101792 | Bacteria | Bacteroidetes | Flavobacteriia | Flavobacterial | Flavobacteriac | Polaribacter | Polaribacter s | unclassified | unclassified | unclassified | unclassified | unclassified | 4,2 | 0,748331477 | 0,0003501 | 5,41E-05 | 5 - | 0,000352076 - | 4 - | 0,00034974 - | 5 - | 0,00041559 - | 3 - | 0,000253037 - | 4 - | 0,00038006 - |
| 9094 | 1 | 3um_FIJJAFBF_188136 | Bacteria | Proteobacteri | Gammaprotec | Cellvibrionale | Porticoccacea | unclassified | SAR92 bacteri | information st | Translation, ri | 0QTKX | Seems to be t | 314287.GB22 | 1 | 0,632455532 | 0,000113255 | 6,98E-05 | 0 - | 0 - | 1 - | 0,000114015 - | 1 - | 0,000108386 - | 2 - | 0,000219973 - | 1 - | 0,0001239 - |
| 9095 | 1 | 0.2um_NOKCPHJN_207436 | Bacteria | Proteobacteri | Gammaprotec | Oceanospirilla | Saccharospirill | Reinekea | Reinekea fors | metabolism | Coenzyme tra | 0QPVY | molybdenum | 314283.MED2 | 2 | 0,894427191 | 0,000152621 | 6,87E-05 | 1 - | 6,38E-05 - | 2 - | 0,000158354 - | 3 - | 0,000225804 - | 3 - | 0,000229139 - | 1 - | 8,60E-05 - |
| 9096 | 1 | 0.2um_NOKCPHJN_122632 | Bacteria | Proteobacteri | Alphaproteob | Rhodobactera | Rhodobactera | Ruegeria | Ruegeria prof | information st | Replication, re | COG0187 | DNA gyrase ne | 467661.RKLH1 | 0,2 | 0,4 | 3,42E-06 | 6,83E-06 | 0 - | 0 - | 0 - | 0 - | 0 - | 0 - | 1 - | 1,71E-05 - | 0 - | 0 - |
| 9097 | 1 | 3um_FIJJAFBF_18799 | Bacteria | Proteobacteri | Alphaproteob | Rhodobactera | unclassified | unclassified | Rhodobactera | unclassified | unclassified | unclassified | unclassified | unclassified | 0,6 | 0,489897949 | 8,18E-05 | 6,75E-05 | 1 - | 0,000119559 - | 1 - | 0,000148457 - | 1 - | 0,000141127 - | 0 - | 0 - | 0 - | 0 - |
| 9098 | 6 | 0.2um_NOKCPHJN_87777;3um_FIJJAFBF_42772 | Bacteria | Bacteroidetes | Flavobacteriia | Flavobacterial | Flavobacteriac | Formosa | Formosa sp. H | unclassified | unclassified | unclassified | unclassified | unclassified | 1 | 0 | 0,000111404 | 1,06E-05 | 1 - | 9,33E-05 - | 1 - | 0,000115869 - | 1 - | 0,000110148 - | 1 - | 0,000111775 - | 1 - | 0,000125914 - |
| 9099 | 1 | 3um_FIJJAFBF_319726 | Bacteria | Bacteroidetes | Flavobacteriia | Flavobacterial | Flavobacteriac | Polaribacter | Polaribacter fi | poorly charact | Function unkn | 0PAEN | NA | 886377.Murru | 0,8 | 0,748331477 | 2,34E-05 | 2,22E-05 | 0 - | 0 - | 2 - | 5,98E-05 - | 1 - | 2,84E-05 - | 1 - | 2,88E-05 - | 0 - | 0 - |
| 9100 | 1 | 3um_FIJJAFBF_775048 | Eukaryota | unclassified | Cryptophyta | Pyrenomonad | Geminigerace | Teleaulax | Teleaulax amp | metabolism | Energy produc | 11HAV | photosystem i | 35128.Thapsd | 1 | 0 | 9,72E-05 | 9,23E-06 | 1 - | 8,14E-05 - | 1 - | 0,000101077 - | 1 - | 9,61E-05 - | 1 - | 9,75E-05 - | 1 - | 0,00010984 - |
| 9101 | 1 | 3um_FIJJAFBF_605906 | Bacteria | Proteobacteri | Betaproteoba | Nitrosomonad | Methylophilac | unclassified | Methylophilac | unclassified | unclassified | unclassified | unclassified | unclassified | 0,2 | 0,4 | 8,51E-06 | 1,70E-05 | 0 - | 0 - | 0 - | 0 - | 0 - | 0 - | 0 - | 0 - | 1 - | 4,25E-05 - |
| 9102 | 1 | 3um_FIJJAFBF_78895 | Bacteria | Proteobacteri | Gammaprotec | Cellvibrionale | Halieaceae | unclassified | gamma prote | unclassified | unclassified | unclassified | unclassified | unclassified | 9,6 | 1,019803903 | 0,000138636 | 1,46E-05 | 11 - | 0,000133744 - | 8 - | 0,000120779 - | 10 - | 0,000143519 - | 9 - | 0,000131075 - | 10 - | 0,000164062 - |
| 9103 | 2 | 0.2um_NOKCPHJN_106661;3um_FIJJAFBF_7861 | Bacteria | Proteobacteri | Gammaprotec | Cellvibrionale | unclassified | unclassified | Cellvibrionale | unclassified | unclassified | unclassified | unclassified | unclassified | 1 | 0 | 4,36E-05 | 4,14E-06 | 1 - | 3,66E-05 - | 1 - | 4,54E-05 - | 1 - | 4,31E-05 - | 1 - | 4,38E-05 - | 1 - | 4,93E-05 - |
| 9104 | 1 | 3um_FIJJAFBF_119800 | unclassified | unclassified | unclassified | unclassified | unclassified | unclassified | unclassified | unclassified | unclassified | unclassified | unclassified | unclassified | 1 | 0 | 9,86E-05 | 9,36E-06 | 1 - | 8,26E-05 - | 1 - | 0,000102532 - | 1 - | 9,75E-05 - | 1 - | 9,89E-05 - | 1 - | 0,00011142 - |
| 9105 | 1 | 0.2um_NOKCPHJN_381446 | Bacteria | Bacteroidetes | unclassified | unclassified | unclassified | unclassified | Bacteroidetes | poorly charact | Function unkn | 05ERK | gliding motilit | 487796.Flav2 | 2,6 | 1,356465997 | 6,56E-05 | 3,52E-05 | 3 - | 6,53E-05 - | 3 - | 8,11E-05 - | 3 - | 7,71E-05 - | 4 - | 0,000104352 - | 0 - | 0 - |
| 9106 | 1 | 3um_FIJJAFBF_46607 | Bacteria | Proteobacteri | Alphaproteob | unclassified | unclassified | unclassified | Alphaproteob | unclassified | unclassified | unclassified | unclassified | unclassified | 2 | 0 | 0,000115635 | 1,10E-05 | 2 - | 9,69E-05 - | 2 - | 0,000120269 - | 2 - | 0,000114331 - | 2 - | 0,00011602 - | 2 - | 0,000130696 - |
| 9107 | 1 | 3um_FIJJAFBF_56520 | Bacteria | Bacteroidetes | Flavobacteriia | Flavobacterial | Flavobacteriac | Polaribacter | Polaribacter s | metabolism | Carbohydrate | COG0057 | glyceraldehyd | 313598.MED1 | 1,6 | 0,489897949 | 6,55E-05 | 2,06E-05 | 2 - | 6,89E-05 - | 1 - | 4,28E-05 - | 2 - | 8,14E-05 - | 1 - | 4,13E-05 - | 2 - | 9,30E-05 - |
| 9108 | 1 | 3um_FIJJAFBF_165693 | Bacteria | Proteobacteri | Alphaproteob | Rhodobactera | unclassified | unclassified | Rhodobactera | information st | Translation, ri | 179RU | 30S ribosomal | 314262.MED1 | 1,8 | 0,748331477 | 0,00015123 | 6,21E-05 | 2 - | 0,00014258 - | 2 - | 0,000177042 - | 1 - | 8,42E-05 - | 3 - | 0,00025618 - | 1 - | 9,62E-05 - |
| 9109 | 1 | 3um_FIJJAFBF_703785 | Bacteria | Proteobacteri | Alphaproteob | Rhodobactera | Rhodobactera | Ascidiaceihabi | Ascidiaceihabi | metabolism | Amino acid tra | COG0131 | imidazoleglyc | 391589.RGAI1 | 0,2 | 0,4 | 1,71E-05 | 3,43E-05 | 1 - | 8,57E-05 - | 0 - | 0 - | 0 - | 0 - | 0 - | 0 - | 0 - | 0 - |
| 9110 | 1 | 3um_FIJJAFBF_145238 | Bacteria | Bacteroidetes | Flavobacteriia | Flavobacterial | Flavobacteriac | Flaviramulus | Flaviramulus b | metabolism | Energy produc | COG2838 | Isocitrate deh | 983544.Lacal_ | 0,2 | 0,4 | 3,12E-06 | 6,24E-06 | 1 - | 1,56E-05 - | 0 - | 0 - | 0 - | 0 - | 0 - | 0 - | 0 - | 0 - |
| 9111 | 1 | 0.2um_NOKCPHJN_728002 | Bacteria | Verrucomicrol | Verrucomicrol | Verrucomicrol | Rubritaleacea | Rubritalea | Rubritalea ma | unclassified | unclassified | unclassified | unclassified | unclassified | 0,4 | 0,489897949 | 1,16E-05 | 1,43E-05 | 0 - | 0 - | 1 - | 2,96E-05 - | 0 - | 0 - | 1 - | 2,86E-05 - | 0 - | 0 - |
| 9112 | 1 | 3um_FIJJAFBF_594168 | Eukaryota | unclassified | Pelagophycea | Pelagomonad | unclassified | Aureoumbra | Aureoumbra l | metabolism | Energy produc | COG0056 | Produces ATP | 35128.Thapsd | 18,8 | 0,748331477 | 0,000508208 | 4,25E-05 | 19 - | 0,00043098 - | 18 - | 0,000506984 - | 19 - | 0,000508728 - | 20 - | 0,000543412 - | 18 - | 0,000550937 - |
| 9113 | 1 | 0.2um_NOKCPHJN_127772 | Bacteria | Bacteroidetes | Flavobacteriia | Flavobacterial | Cryomorphace | unclassified | Cryomorphace | metabolism | Inorganic ion t | 05C27 | TonB-depende | 313596.RB250 | 1,2 | 0,4 | 1,55E-05 | 3,44E-06 | 2 - | 2,23E-05 - | 1 - | 1,38E-05 - | 1 - | 1,32E-05 - | 1 - | 1,33E-05 - | 1 - | 1,50E-05 - |
| 9114 | 1 | 3um_FIJJAFBF_45522 | Bacteria | Proteobacteri | Gammaprotec | unclassified | unclassified | unclassified | SAR86 cluster | metabolism | Amino acid tra | COG1104 | cysteine desul | 322710.Avin_ | 0,6 | 0,489897949 | 1,94E-05 | 1,60E-05 | 1 - | 2,84E-05 - | 1 - | 3,53E-05 - | 1 - | 3,35E-05 - | 0 - | 0 - | 0 - | 0 - |
| 9115 | 1 | 0.2um_NOKCPHJN_191831 | Bacteria | Proteobacteri | Gammaprotec | Oceanospirilla | Saccharospirill | Reinekea | Reinekea blan | cellular proces | Posttranslatio | COG0330 | Band 7 protei | 314283.MED2 | 2,2 | 0,748331477 | 9,41E-05 | 2,88E-05 | 3 - | 0,00011001 - | 3 - | 0,0001366 - | 2 - | 8,66E-05 - | 2 - | 8,78E-05 - | 1 - | 4,95E-05 - |
| 9116 | 2 | 0.2um_NOKCPHJN_498570;3um_FIJJAFBF_2122 | unclassified | unclassified | unclassified | unclassified | unclassified | unclassified | unclassified | unclassified | unclassified | unclassified | unclassified | unclassified | 0,6 | 0,489897949 | 2,66E-05 | 2,19E-05 | 1 - | 3,94E-05 - | 0 - | 0 - | 1 - | 4,66E-05 - | 1 - | 4,72E-05 - | 0 - | 0 - |
| 9117 | 2 | 0.2um_NOKCPHJN_915195;3um_FIJJAFBF_4719 | Eukaryota | Apicomplexa | Conoidasida | Eucoccidiorida | Eimeriidae | Eimeria | Eimeria necat | various | various | various | various | various | 0,2 | 0,4 | 1,10E-05 | 2,21E-05 | 1 - | 5,52E-05 - | 0 - | 0 - | 0 - | 0 - | 0 - | 0 - | 0 - | 0 - |
| 9118 | 1 | 3um_FIJJAFBF_49011 | Bacteria | Proteobacteri | Gammaprotec | Oceanospirilla | Saccharospirill | Reinekea | Reinekea blan | metabolism | Energy produc | COG0372 | citrate syntha | 629265.PMA4 | 1 | 0 | 3,23E-05 | 3,07E-06 | 1 - | 2,71E-05 - | 1 - | 3,36E-05 - | 1 - | 3,20E-05 - | 1 - | 3,24E-05 - | 1 - | 3,65E-05 - |
| 9119 | 1 | 0.2um_NOKCPHJN_01012 | Bacteria | Bacteroidetes | Flavobacteriia | Flavobacterial | Flavobacteriac | Formosa | Formosa sp. H | unclassified | unclassified | unclassified | unclassified | unclassified | 2 | 0,632455532 | 2,80E-05 | 7,34E-06 | 3 - | 3,58E-05 - | 2 - | 2,97E-05 - | 2 - | 2,82E-05 - | 1 - | 1,43E-05 - | 2 - | 3,22E-05 - |
| 9120 | 1 | 3um_FIJJAFBF_250838 | Bacteria | Proteobacteri | Betaproteoba | unclassified | unclassified | unclassified | Betaproteoba | information st | Replication, re | 0BE9U | DNA-binding p | 399795.CtesD | 1,2 | 0,748331477 | 0,000170113 | 0,000102544 | 2 - | 0,000249515 - | 1 - | 0,000154912 - | 1 - | 0,000147263 - | 2 - | 0,000298877 - | 0 - | 0 - |
| 9121 | 1 | 0.2um_NOKCPHJN_1077008 | Eukaryota | unclassified | Cryptophyta | Pyrenomonad | Geminigerace | Guillardia | Guillardia thet | unclassified | unclassified | unclassified | unclassified | unclassified | 1 | 0 | 3,90E-05 | 3,71E-06 | 1 - | 3,27E-05 - | 1 - | 4,06E-05 - | 1 - | 3,86E-05 - | 1 - | 3,92E-05 - | 1 - | 4,41E-05 - |
| 9122 | 1 | 0.2um_NOKCPHJN_63173 | Bacteria | Bacteroidetes | Flavobacteriia | Flavobacterial | Flavobacteriac | Flavobacteriu | Flavobacteriu | unclassified | unclassified | unclassified | unclassified | unclassified | 6,8 | 0,748331477 | 0,000177463 | 2,90E-05 | 6 - | 0,000130676 - | 6 - | 0,000162261 - | 7 - | 0,000179957 - | 8 - | 0,000208703 - | 7 - | 0,000205716 - |
| 9123 | 1 | 0.2um_NOKCPHJN_149058 | Bacteria | Bacteroidetes | Flavobacteriia | Flavobacterial | Flavobacteriac | Polaribacter | Polaribacter s | unclassified | unclassified | unclassified | unclassified | unclassified | 7,4 | 1,019803903 | 0,000415611 | 2,54E-05 | 9 - | 0,000428627 - | 7 - | 0,000413955 - | 7 - | 0,000393517 - | 8 - | 0,000456376 - | 6 - | 0,00038558 - |
| 9124 | 1 | 3um_FIJJAFBF_174679 | Bacteria | Bacteroidetes | Flavobacteriia | Flavobacterial | Flavobacteriac | Polaribacter | Polaribacter s | cellular proces | Posttranslatio | 08V1T | peptidylprolyl | 313594.PI23P | 11,2 | 0,979795897 | 0,000633523 | 8,00E-05 | 12 - | 0,000569141 - | 10 - | 0,000588921 - | 10 - | 0,000559844 - | 12 - | 0,000681735 - | 12 - | 0,000767973 - |
| 9125 | 1 | 3um_FIJJAFBF_674280 | Bacteria | Bacteroidetes | Flavobacteriia | Flavobacterial | Crocinitomica | unclassified | Crocinitomica | information st | Translation, ri | COG0087 | One of the pri | 755732.Fluta_ | 1,6 | 0,489897949 | 0,000104523 | 3,34E-05 | 2 - | 0,000109834 - | 1 - | 6,82E-05 - | 1 - | 6,48E-05 - | 2 - | 0,000131563 - | 2 - | 0,000148205 - |
| 9126 | 1 | 3um_FIJJAFBF_551529 | Bacteria | Bacteroidetes | Flavobacteriia | Flavobacterial | Flavobacteriac | Altibacter | Altibacter lent | unclassified | unclassified | unclassified | unclassified | unclassified | 0,4 | 0,489897949 | 5,53E-06 | 6,83E-06 | 1 - | 1,23E-05 - | 1 - | 1,53E-05 - | 0 - | 0 - | 0 - | 0 - | 0 - | 0 - |
| 9127 | 2 | 0.2um_NOKCPHJN_101061;0.2um_NOKCPHJN_8 | Bacteria | Proteobacteri | Gammaprotec | Alteromonada | Alteromonada | Glaciecola | Glaciecola sp. | metabolism | Carbohydrate | 16QEQ | hydrolase, fan | 455436.GHTC | 1,4 | 0,489897949 | 1,55E-05 | 4,57E-06 | 2 - | 1,90E-05 - | 1 - | 1,18E-05 - | 1 - | 1,12E-05 - | 2 - | 2,28E-05 - | 1 - | 1,28E-05 - |
| 9128 | 1 | 3um_FIJJAFBF_81319 | Bacteria | Proteobacteri | Gammaprotec | Alteromonada | Alteromonada | Paraglaciecola | Paraglaciecola | poorly charact | Function unkn | 16R01 | Extracellular li | 342610.Patl_2 | 0,4 | 0,489897949 | 5,83E-06 | 7,21E-06 | 1 - | 1,30E-05 - | 1 - | 1,62E-05 - | 0 - | 0 - | 0 - | 0 - | 0 - | 0 - |
| 9129 | 1 | 0.2um_NOKCPHJN_327210 | Bacteria | Proteobacteri | Gammaprotec | Cellvibrionale | Porticoccacea | unclassified | Gammaprotec | metabolism | Inorganic ion t | 16TQC | TONB-depend | 314287.GB22 | 1,2 | 0,979795897 | 1,58E-05 | 1,12E-05 | 3 - | 3,50E-05 - | 1 - | 1,45E-05 - | 0 - | 0 - | 1 - | 1,40E-05 - | 1 - | 1,57E-05 - |
| 9130 | 1 | 3um_FIJJAFBF_440992 | Bacteria | Proteobacteri | Betaproteoba | unclassified | unclassified | unclassified | Betaproteoba | metabolism | Amino acid tra | 16RER | (ABC) transpo | 338969.Rfer_ | 1,8 | 0,979795897 | 9,09E-05 | 4,40E-05 | 3 - | 0,000131927 - | 1 - | 5,46E-05 - | 3 - | 0,000155727 - | 1 - | 5,27E-05 - | 1 - | 5,93E-05 - |
| 9131 | 1 | 3um_FIJJAFBF_437597 | Bacteria | Bacteroidetes | Flavobacteriia | Flavobacterial | Flavobacteriac | Polaribacter | various | unclassified | unclassified | unclassified | unclassified | unclassified | 0,8 | 0,748331477 | 7,69E-05 | 6,76E-05 | 2 - | 0,000173904 - | 1 - | 0,000107969 - | 1 - | 0,000102638 - | 0 - | 0 - | 0 - | 0 - |
| 9132 | 1 | 0.2um_NOKCPHJN_661018 | Bacteria | Proteobacteri | Alphaproteob | Rhodospirillal | Rhodospirillac | Candidatus En | Candidatus En | unclassified | unclassified | unclassified | unclassified | unclassified | 0,2 | 0,4 | 5,03E-06 | 1,01E-05 | 0 - | 0 - | 1 - | 2,51E-05 - | 0 - | 0 - | 0 - | 0 - | 0 - | 0 - |
| 9133 | 1 | 0.2um_NOKCPHJN_306179 | Bacteria | Bacteroidetes | Flavobacteriia | Flavobacterial | Flavobacteriac | unclassified | Flavobacteriac | metabolism | Inorganic ion t | 05C27 | TonB-depende | 1034807.FBFL | 2,4 | 1,356465997 | 3,27E-05 | 1,92E-05 | 3 - | 3,52E-05 - | 4 - | 5,83E-05 - | 3 - | 4,16E-05 - | 2 - | 2,81E-05 - | 0 - | 0 - |
| 9134 | 1 | 0.2um_NOKCPHJN_957271 | Bacteria | Proteobacteri | Gammaprotec | Cellvibrionale | Porticoccacea | unclassified | SAR92 bacteri | metabolism | Inorganic ion t | 16TQC | TONB-depend | 314287.GB22 | 1,2 | 0,4 | 2,86E-05 | 1,06E-05 | 1 - | 1,98E-05 - | 2 - | 4,92E-05 - | 1 - | 2,34E-05 - | 1 - | 2,37E-05 - | 1 - | 2,67E-05 - |
| 9135 | 1 | 0.2um_NOKCPHJN_751138 | Bacteria | Proteobacteri | Alphaproteob | Pelagibacteral | Pelagibacterac | Candidatus Pe | Candidatus Pe | information st | Translation, ri | 16Q16 | Binds the lowe | 684719.HIMB | 2,8 | 1,166190379 | 0,000163724 | 4,96E-05 | 5 - | 0,000253931 - | 2 - | 0,000126123 - | 3 - | 0,000179844 - | 2 - | 0,000121667 - | 2 - | 0,000137057 - |
| 9136 | 1 | 3um_FIJJAFBF_58783 | Bacteria | Bacteroidetes | unclassified | unclassified | unclassified | unclassified | Bacteroidetes | unclassified | unclassified | unclassified | unclassified | unclassified | 0,4 | 0,489897949 | 6,75E-05 | 8,27E-05 | 0 - | 0 - | 1 - | 0,000171709 - | 0 - | 0 - | 1 - | 0,000165643 - | 0 - | 0 - |
| 9137 | 1 | 0.2um_NOKCPHJN_194408 | Bacteria | Proteobacteri | Gammaprotec | Oceanospirilla | Hahellaceae | Hahella | Hahella sp. CC | cellular proces | Cell motility | COG1749 | flagellar hook | 349521.HCH_ | 0,2 | 0,4 | 4,45E-06 | 8,89E-06 | 0 - | 0 - | 1 - | 2,22E-05 - | 0 - | 0 - | 0 - | 0 - | 0 - | 0 - |
| 9138 | 1 | 3um_FIJJAFBF_575459 | Bacteria | Bacteroidetes | Flavobacteriia | Flavobacterial | Flavobacteriac | Formosa | Formosa sp. H | metabolism | Carbohydrate | COG0057 | glyceraldehyd | 313598.MED1 | 1 | 0 | 4,11E-05 | 3,91E-06 | 1 - | 3,45E-05 - | 1 - | 4,28E-05 - | 1 - | 4,07E-05 - | 1 - | 4,13E-05 - | 1 - | 4,65E-05 - |
| 9139 | 1 | 3um_FIJJAFBF_332922 | Bacteria | Bacteroidetes | unclassified | unclassified | unclassified | unclassified | Bacteroidetes | unclassified | unclassified | unclassified | unclassified | unclassified | 0,8 | 0,979795897 | 9,17E-05 | 0,000113103 | 2 - | 0,000208685 - | 0 - | 0 - | 0 - | 0 - | 2 - | 0,00024997 - | 0 - | 0 - |
| 9140 | 1 | 0.2um_NOKCPHJN_100728 | Bacteria | Bacteroidetes | Flavobacteriia | Flavobacterial | Flavobacteriac | Winogradskye | Winogradskye | unclassified | unclassified | unclassified | unclassified | unclassified | 5,8 | 1,166190379 | 0,000164354 | 4,42E-05 | 5 - | 0,000117359 - | 7 - | 0,000204015 - | 6 - | 0,000166236 - | 4 - | 0,000112461 - | 7 - | 0,000221702 - |
| 9141 | 1 | 3um_FIJJAFBF_213858 | Bacteria | Proteobacteri | Gammaprotec | Oceanospirilla | Saccharospirill | Reinekea | Reinekea fors | cellular proces | Posttranslatio | COG0616 | Signal peptide | 314283.MED2 | 0,4 | 0,489897949 | 1,51E-05 | 1,86E-05 | 1 - | 3,43E-05 - | 0 - | 0 - | 0 - | 0 - | 1 - | 4,10E-05 - | 0 - | 0 - |
| 9142 | 1 | 3um_FIJJAFBF_526565 | Bacteria | Bacteroidetes | Flavobacteriia | Flavobacterial | Crocinitomica | unclassified | Crocinitomica | unclassified | unclassified | unclassified | unclassified | unclassified | 0,6 | 0,8 | 2,53E-05 | 3,24E-05 | 2 - | 7,81E-05 - | 1 - | 4,85E-05 - | 0 - | 0 - | 0 - | 0 - | 0 - | 0 - |
| 9143 | 1 | 3um_FIJJAFBF_420944 | Bacteria | Proteobacteri | Gammaprotec | Cellvibrionale | Porticoccacea | unclassified | various | unclassified | unclassified | unclassified | unclassified | unclassified | 2,4 | 0,8 | 0,000299652 | 8,88E-05 | 3 - | 0,000321804 - | 2 - | 0,00026639 - | 3 - | 0,000379857 - | 3 - | 0,000385467 - | 1 - | 0,000144743 - |
| 9144 | 1 | 3um_FIJJAFBF_792503 | Viruses | unclassified | unclassified | unclassified | unclassified | unclassified | uncultured M | unclassified | unclassified | unclassified | unclassified | unclassified | 0,2 | 0,4 | 5,60E-06 | 1,12E-05 | 0 - | 0 - | 1 - | 2,80E-05 - | 0 - | 0 - | 0 - | 0 - | 0 - | 0 - |
| 9145 | 1 | 0.2um_NOKCPHJN_356008 | Bacteria | Proteobacteri | Alphaproteob | Pelagibacteral | Pelagibacterac | Candidatus Pe | Candidatus Pe | metabolism | Energy produc | 17C3H | Produces ATP | 439493.PB72 | 0,8 | 0,4 | 8,43E-05 | 4,33E-05 | 1 - | 8,83E-05 - | 1 - | 0,00010963 - | 1 - | 0,000104217 - | 0 - | 0 - | 1 - | 0,000119134 - |
| 9146 | 1 | 0.2um_NOKCPHJN_654974 | Bacteria | Bacteroidetes | Flavobacteriia | Flavobacterial | Flavobacteriac | Polaribacter | Polaribacter s | unclassified | unclassified | unclassified | unclassified | unclassified | 0,4 | 0,489897949 | 3,91E-05 | 4,82E-05 | 1 - | 8,97E-05 - | 0 - | 0 - | 1 - | 0,000105845 - | 0 - | 0 - | 0 - | 0 - |

| | | | | | | | | | | | | | | | | | | | | | | | | | | | | |
|---|---|---|---|---|---|---|---|---|---|---|---|---|---|---|---|---|---|---|---|---|---|---|---|---|---|---|---|---|
| 9147 | 1 | 0.2um_NOKCPHJN_287506 | Bacteria | Proteobacteri | Gammaprote | unclassified | unclassified | unclassified | Gammaprote | metabolism | Energy produc | 0QIAU | Electron trans | 247633.GP214 | 0,2 | 0,4 | 1,14E-05 | 2,28E-05 | 0 - | 0 - | 1 - | 5,70E-05 - | 0 - | 0 - | 0 - | 0 - | 0 - | 0 - |
| 9148 | 8 | 0.2um_NOKCPHJN_655150;3um_FIJJAFBF_20521 | Eukaryota | various | various | various | various | various | various | metabolism | Energy produc | 123BF | photosystem I | 35128.Thapsd | 2 | 0 | 0,000315005 | 2,99E-05 | 2 - | 0,000263855 - | 2 - | 0,00032763 - | 2 - | 0,000311453 - | 2 - | 0,000316054 - | 2 - | 0,000356033 - |
| 9149 | 1 | 0.2um_NOKCPHJN_150691 | Bacteria | Proteobacteri | Alphaproteob | Rhodobactera | Rhodobactera | Planktomarina | Planktomarina | unclassified | unclassified | unclassified | unclassified | unclassified | 1,8 | 0,4 | 7,62E-05 | 1,50E-05 | 2 - | 7,20E-05 - | 2 - | 8,94E-05 - | 2 - | 8,49E-05 - | 2 - | 8,62E-05 - | 1 - | 4,85E-05 - |
| 9150 | 1 | 0.2um_NOKCPHJN_590541 | Bacteria | Proteobacteri | Alphaproteob | Rhodobactera | Rhodobactera | Planktomarina | Planktomarina | unclassified | unclassified | unclassified | unclassified | unclassified | 0,8 | 0,4 | 0,000141403 | 7,18E-05 | 1 - | 0,000153036 - | 1 - | 0,000190025 - | 1 - | 0,000180643 - | 1 - | 0,000183311 - | 0 - | 0 - |
| 9151 | 1 | 3um_FIJJAFBF_191139 | Bacteria | Proteobacteri | Alphaproteob | Rhodobactera | unclassified | unclassified | Rhodobactera | information st | Translation, ri | COG0185 | Protein S19 fo | 367336.OM22 | 1 | 0,632455532 | 0,000151081 | 0,000106529 | 1 - | 0,000123416 - | 1 - | 0,000153246 - | 1 - | 0,00014568 - | 0 - | 0 - | 2 - | 0,000333063 - |
| 9152 | 1 | 0.2um_NOKCPHJN_289090 | Eukaryota | unclassified | Oomycetes | Peronosporal | Peronosporac | Phytophthora | Phytophthora | unclassified | unclassified | unclassified | unclassified | unclassified | 0,2 | 0,4 | 1,39E-06 | 2,79E-06 | 0 - | 0 - | 0 - | 0 - | 0 - | 0 - | 0 - | 0 - | 1 - | 6,97E-06 - |
| 9153 | 1 | 3um_FIJJAFBF_122390 | Bacteria | Bacteroidetes | Flavobacteriia | Flavobacterial | Flavobacteriac | Polaribacter | Polaribacter a | information st | Replication, re | 05CZ8 | DNA polymera | 983544.Lacal_ | 7,8 | 1,469693846 | 0,000283357 | 3,73E-05 | 10 - | 0,00030854 - | 9 - | 0,000344804 - | 7 - | 0,00025494 - | 7 - | 0,000258705 - | 6 - | 0,000249798 - |
| 9154 | 1 | 0.2um_NOKCPHJN_131551 | Bacteria | Bacteroidetes | Flavobacteriia | Flavobacterial | Flavobacteriac | unclassified | Flavobacteriac | unclassified | unclassified | unclassified | unclassified | unclassified | 0,2 | 0,4 | 5,78E-06 | 1,16E-05 | 0 - | 0 - | 1 - | 2,89E-05 - | 0 - | 0 - | 0 - | 0 - | 0 - | 0 - |
| 9155 | 1 | 0.2um_NOKCPHJN_64986 | Bacteria | Bacteroidetes | Flavobacteriia | Flavobacterial | Flavobacteriac | Formosa | Formosa sp. H | information st | Translation, ri | COG0052 | 30S ribosomal | 867900.Celly_ | 0,6 | 0,489897949 | 2,68E-05 | 2,20E-05 | 0 - | 0 - | 0 - | 0 - | 1 - | 4,25E-05 - | 1 - | 4,31E-05 - | 1 - | 4,85E-05 - |
| 9156 | 1 | 0.2um_NOKCPHJN_235716 | Bacteria | Bacteroidetes | Flavobacteriia | Flavobacterial | Flavobacteriac | Formosa | Formosa sp. H | information st | Replication, re | COG0550 | Releases the s | 1046627.BZAF | 1,4 | 0,489897949 | 2,36E-05 | 9,61E-06 | 1 - | 1,39E-05 - | 1 - | 1,73E-05 - | 2 - | 3,28E-05 - | 1 - | 1,67E-05 - | 2 - | 3,75E-05 - |
| 9157 | 1 | 0.2um_NOKCPHJN_556776 | Bacteria | Bacteroidetes | Flavobacteriia | Flavobacterial | unclassified | unclassified | Flavobacterial | unclassified | unclassified | unclassified | unclassified | unclassified | 0,8 | 0,4 | 3,79E-05 | 1,91E-05 | 0 - | 0 - | 1 - | 4,73E-05 - | 1 - | 4,50E-05 - | 1 - | 4,57E-05 - | 1 - | 5,15E-05 - |
| 9158 | 1 | 0.2um_NOKCPHJN_172238 | Bacteria | Bacteroidetes | unclassified | unclassified | unclassified | unclassified | Bacteroidetes | unclassified | unclassified | unclassified | unclassified | unclassified | 1 | 0 | 4,58E-05 | 4,35E-06 | 1 - | 3,84E-05 - | 1 - | 4,77E-05 - | 1 - | 4,53E-05 - | 1 - | 4,60E-05 - | 1 - | 5,18E-05 - |
| 9159 | 1 | 3um_FIJJAFBF_177694 | Eukaryota | Bacillariophyt | Bacillariophyc | Bacillariales | Bacillariaceae | Fragilariopsis | Fragilariopsis | unclassified | unclassified | unclassified | unclassified | unclassified | 0,4 | 0,8 | 2,22E-05 | 4,44E-05 | 0 - | 0 - | 0 - | 0 - | 2 - | 0,000111051 - | 0 - | 0 - | 0 - | 0 - |
| 9160 | 1 | 3um_FIJJAFBF_172549 | Eukaryota | unclassified | unclassified | Phaeocystales | Phaeocystace | Phaeocystis | Phaeocystis gl | unclassified | unclassified | unclassified | unclassified | unclassified | 0,6 | 0,8 | 2,43E-05 | 3,36E-05 | 1 - | 3,62E-05 - | 0 - | 0 - | 2 - | 8,55E-05 - | 0 - | 0 - | 0 - | 0 - |
| 9161 | 1 | 0.2um_NOKCPHJN_709263 | unclassified | unclassified | unclassified | unclassified | unclassified | unclassified | unclassified | unclassified | unclassified | unclassified | unclassified | unclassified | 1 | 0 | 0,000148943 | 1,41E-05 | 1 - | 0,000124757 - | 1 - | 0,000154912 - | 1 - | 0,000147263 - | 1 - | 0,000149438 - | 1 - | 0,000168342 - |
| 9162 | 1 | 0.2um_NOKCPHJN_234570 | Bacteria | Bacteroidetes | Flavobacteriia | Flavobacterial | Flavobacteriac | Kordia | Kordia jejudor | information st | Translation, ri | 08UHY | One of the pri | 391587.KAOT | 1,4 | 0,489897949 | 0,000118404 | 3,39E-05 | 2 - | 0,000145287 - | 1 - | 9,02E-05 - | 2 - | 0,000171496 - | 1 - | 8,70E-05 - | 1 - | 9,80E-05 - |
| 9163 | 1 | 3um_FIJJAFBF_41555 | Eukaryota | Bacillariophyt | Coscinodiscop | Coscinodiscale | Coscinodiscac | Coscinodiscus | Coscinodiscus | metabolism | Carbohydrate | KOG1203 | Complex I inte | 35128.Thapsd | 0,4 | 0,8 | 1,44E-05 | 2,89E-05 | 2 - | 7,22E-05 - | 0 - | 0 - | 0 - | 0 - | 0 - | 0 - | 0 - | 0 - |
| 9164 | 1 | 0.2um_NOKCPHJN_609811 | Bacteria | Proteobacteri | Alphaproteob | Pelagibacteral | Pelagibacterac | Candidatus Pe | Candidatus Pe | metabolism | Energy produc | 07QM5 | aconitate hyd | 1002672.SAR1 | 0,2 | 0,4 | 2,75E-06 | 5,50E-06 | 1 - | 1,37E-05 - | 0 - | 0 - | 0 - | 0 - | 0 - | 0 - | 0 - | 0 - |
| 9165 | 13 | 0.2um_NOKCPHJN_156065;0.2um_NOKCPHJN_1 | Bacteria | various | various | various | various | various | various | unclassified | unclassified | unclassified | unclassified | unclassified | 0,6 | 0,489897949 | 5,04E-05 | 4,14E-05 | 1 - | 7,45E-05 - | 0 - | 0 - | 1 - | 8,80E-05 - | 1 - | 8,93E-05 - | 0 - | 0 - |
| 9166 | 1 | 3um_FIJJAFBF_350297 | Bacteria | Proteobacteri | Gammaprote | Oceanospirilla | Oceanospirilla | Nitrincola | Nitrincola sp. | information st | Translation, ri | 0QK14 | This protein bi | 717774.Marm | 3,8 | 0,4 | 0,000294357 | 4,37E-05 | 4 - | 0,000259383 - | 4 - | 0,000322077 - | 3 - | 0,000229631 - | 4 - | 0,000310697 - | 4 - | 0,000349999 - |
| 9167 | 1 | 3um_FIJJAFBF_14602 | Bacteria | Proteobacteri | Gammaprote | Oceanospirilla | Oceanospirilla | Oceanospirillu | Oceanospirillu | unclassified | unclassified | unclassified | unclassified | unclassified | 5,8 | 2,13541565 | 0,000226913 | 7,39E-05 | 9 - | 0,000299418 - | 7 - | 0,000289169 - | 4 - | 0,000157081 - | 3 - | 0,000119551 - | 6 - | 0,000269347 - |
| 9168 | 1 | 0.2um_NOKCPHJN_546978 | Bacteria | Proteobacteri | Gammaprote | Cellvibrionales | Porticoccacea | unclassified | SAR92 bacteri | metabolism | Inorganic ion t | 0QHT5 | TonB-depende | 314287.GB220 | 36,2 | 2,481934729 | 0,000571867 | 6,79E-05 | 39 - | 0,000516297 - | 36 - | 0,000591774 - | 34 - | 0,000531303 - | 33 - | 0,000523293 - | 39 - | 0,000696667 - |
| 9169 | 1 | 0.2um_NOKCPHJN_200776 | Bacteria | Proteobacteri | Gammaprote | Oceanospirilla | Saccharospirill | Reinekea | Reinekea blan | metabolism | Nucleotide tra | 0QKAC | Catalyzes the | 314283.MED2 | 0,2 | 0,4 | 5,24E-06 | 1,05E-05 | 1 - | 2,62E-05 - | 0 - | 0 - | 0 - | 0 - | 0 - | 0 - | 0 - | 0 - |
| 9170 | 1 | 3um_FIJJAFBF_399086 | Bacteria | Bacteroidetes | Flavobacteriia | Flavobacterial | Flavobacteriac | Polaribacter | Polaribacter s | unclassified | unclassified | unclassified | unclassified | unclassified | 6 | 0,632455532 | 0,000183884 | 3,24E-05 | 6 - | 0,000153377 - | 6 - | 0,000190448 - | 6 - | 0,000181045 - | 5 - | 0,000153099 - | 7 - | 0,000241452 - |
| 9171 | 1 | 0.2um_NOKCPHJN_50510 | Bacteria | Proteobacteri | Betaproteoba | Burkholderial | unclassified | unclassified | various | cellular proces | Cell wall/mem | COG2885 | Ompa motb d | 338969.Rfer_ | 3,4 | 0,489897949 | 0,000204041 | 1,98E-05 | 4 - | 0,000203145 - | 3 - | 0,000189184 - | 4 - | 0,000239792 - | 3 - | 0,0001825 - | 3 - | 0,000205586 - |
| 9172 | 1 | 0.2um_NOKCPHJN_172233 | Bacteria | Bacteroidetes | unclassified | unclassified | unclassified | unclassified | Bacteroidetes | unclassified | unclassified | unclassified | unclassified | unclassified | 2,2 | 0,4 | 0,000130978 | 2,92E-05 | 2 - | 9,94E-05 - | 3 - | 0,000185089 - | 2 - | 0,000117301 - | 2 - | 0,000119033 - | 2 - | 0,00013409 - |
| 9173 | 1 | 0.2um_NOKCPHJN_157072 | Eukaryota | Zoopagomyco | Basidiobolom | Basidiobolales | Basidiobolace | Basidiobolus | Basidiobolus r | unclassified | unclassified | unclassified | unclassified | unclassified | 1,8 | 0,748331477 | 0,000170384 | 5,68E-05 | 3 - | 0,00024595 - | 2 - | 0,000203598 - | 2 - | 0,000193546 - | 1 - | 9,82E-05 - | 1 - | 0,000110625 - |
| 9174 | 1 | 0.2um_NOKCPHJN_1088908 | Bacteria | Proteobacteri | Betaproteoba | Rhodocyclales | Zoogloeaceae | Thauera | Thauera terpe | unclassified | unclassified | unclassified | unclassified | unclassified | 1 | 0 | 2,64E-05 | 2,50E-06 | 1 - | 2,21E-05 - | 1 - | 2,74E-05 - | 1 - | 2,61E-05 - | 1 - | 2,64E-05 - | 1 - | 2,98E-05 - |
| 9175 | 1 | 0.2um_NOKCPHJN_414205 | Bacteria | Proteobacteri | Alphaproteob | Pelagibacteral | Pelagibacterac | Candidatus Pe | Candidatus Pe | cellular proces | Cell wall/mem | 7257 | NA | 335992.SAR11 | 1,2 | 0,4 | 4,78E-05 | 1,06E-05 | 2 - | 6,85E-05 - | 1 - | 4,25E-05 - | 1 - | 4,04E-05 - | 1 - | 4,10E-05 - | 1 - | 4,62E-05 - |
| 9176 | 1 | 3um_FIJJAFBF_389790 | Bacteria | Proteobacteri | Alphaproteob | unclassified | unclassified | unclassified | Alphaproteob | unclassified | unclassified | unclassified | unclassified | unclassified | 0,2 | 0,4 | 7,65E-06 | 1,53E-05 | 1 - | 3,83E-05 - | 0 - | 0 - | 0 - | 0 - | 0 - | 0 - | 0 - | 0 - |
| 9177 | 1 | 3um_FIJJAFBF_153101 | Bacteria | Bacteroidetes | Flavobacteriia | Flavobacterial | Flavobacteriac | Aquimarina | Aquimarina sp | unclassified | unclassified | unclassified | unclassified | unclassified | 0,6 | 0,8 | 2,98E-05 | 3,96E-05 | 0 - | 0 - | 0 - | 0 - | 2 - | 9,89E-05 - | 1 - | 5,02E-05 - | 0 - | 0 - |
| 9178 | 1 | 0.2um_NOKCPHJN_338389 | Bacteria | Verrucomicrol | Opitutae | Puniceicoccale | Puniceicoccac | Coraliomargar | Coraliomargar | unclassified | unclassified | unclassified | unclassified | unclassified | 1 | 0,632455532 | 3,44E-05 | 2,23E-05 | 1 - | 2,97E-05 - | 1 - | 3,68E-05 - | 2 - | 7,00E-05 - | 1 - | 3,55E-05 - | 0 - | 0 - |
| 9179 | 1 | 0.2um_NOKCPHJN_1018231 | Bacteria | Bacteroidetes | Flavobacteriia | Flavobacterial | Flavobacteriac | Polaribacter | Polaribacter s | metabolism | Inorganic ion t | 05C27 | TonB-depende | 313594.PI23P | 1,6 | 0,489897949 | 3,85E-05 | 1,24E-05 | 1 - | 2,01E-05 - | 2 - | 4,98E-05 - | 2 - | 4,74E-05 - | 2 - | 4,81E-05 - | 1 - | 2,71E-05 - |
| 9180 | 1 | 3um_FIJJAFBF_215936 | Bacteria | Bacteroidetes | Flavobacteriia | Flavobacterial | Crocinitomica | Crocinitomix | Crocinitomix s | information st | Translation, ri | COG0050 | This protein p | 391587.KAOT | 1,2 | 0,748331477 | 4,30E-05 | 2,62E-05 | 0 - | 0 - | 2 - | 7,22E-05 - | 2 - | 6,86E-05 - | 1 - | 3,48E-05 - | 1 - | 3,92E-05 - |
| 9181 | 1 | 0.2um_NOKCPHJN_527171 | Bacteria | Proteobacteri | Alphaproteob | Pelagibacteral | Pelagibacterac | Candidatus Pe | Candidatus Pe | information st | Replication, re | 05C2R | Single-strande | 335992.SAR11 | 0,2 | 0,4 | 4,88E-06 | 9,75E-06 | 0 - | 0 - | 0 - | 0 - | 0 - | 0 - | 1 - | 2,44E-05 - | 0 - | 0 - |
| 9182 | 1 | 0.2um_NOKCPHJN_529232 | Bacteria | Proteobacteri | Gammaprote | Cellvibrionales | Halieaceae | unclassified | marine gamm | metabolism | Inorganic ion t | 01UP7 | TonB-depende | 314254.OA26 | 0,4 | 0,8 | 5,66E-06 | 1,13E-05 | 2 - | 2,83E-05 - | 0 - | 0 - | 0 - | 0 - | 0 - | 0 - | 0 - | 0 - |
| 9183 | 1 | 0.2um_NOKCPHJN_282552 | Eukaryota | unclassified | Labyrinthulom | unclassified | Thraustochytr | Aurantiochytr | Aurantiochytr | unclassified | unclassified | unclassified | unclassified | unclassified | 12,2 | 0,979795897 | 0,000162709 | 1,94E-05 | 12 - | 0,000134111 - | 12 - | 0,000166526 - | 14 - | 0,000184689 - | 11 - | 0,000147256 - | 12 - | 0,000180963 - |
| 9184 | 1 | 3um_FIJJAFBF_132874 | Bacteria | Proteobacteri | Alphaproteob | Rhodobactera | Rhodobactera | Planktomarina | Planktomarina | unclassified | unclassified | unclassified | unclassified | unclassified | 2,6 | 1,624807681 | 0,000182768 | 0,000108417 | 4 - | 0,000245512 - | 1 - | 7,62E-05 - | 5 - | 0,000362252 - | 2 - | 0,000147041 - | 1 - | 8,28E-05 - |
| 9185 | 1 | 3um_FIJJAFBF_570109 | Bacteria | Proteobacteri | Alphaproteob | Rhodobactera | unclassified | unclassified | Rhodobactera | unclassified | unclassified | unclassified | unclassified | unclassified | 6,8 | 1,6 | 0,000626706 | 0,000139886 | 7 - | 0,000546556 - | 7 - | 0,000678661 - | 7 - | 0,000645153 - | 9 - | 0,000841735 - | 4 - | 0,000421427 - |
| 9186 | 1 | 3um_FIJJAFBF_467517 | Bacteria | Proteobacteri | Gammaprote | Alteromonada | Alteromonada | Paraglaciecola | Paraglaciecola | metabolism | Lipid transpor | 16QA8 | reductase | 342610.Patl_2 | 0,6 | 0,8 | 2,98E-05 | 3,85E-05 | 2 - | 9,37E-05 - | 0 - | 0 - | 1 - | 5,53E-05 - | 0 - | 0 - | 0 - | 0 - |
| 9187 | 1 | 3um_FIJJAFBF_324498 | Bacteria | Bacteroidetes | Cytophagia | Cytophagales | Flammeovirga | unclassified | Flammeovirga | poorly charact | Function unkn | 0P8NI | Inherit from b | 926562.Oweh | 2,2 | 0,979795897 | 7,17E-05 | 2,51E-05 | 4 - | 0,000112526 - | 2 - | 6,99E-05 - | 2 - | 6,64E-05 - | 1 - | 3,37E-05 - | 2 - | 7,59E-05 - |
| 9188 | 1 | 3um_FIJJAFBF_73797 | Bacteria | Bacteroidetes | Flavobacteriia | Flavobacterial | Flavobacteriac | Polaribacter | Polaribacter s | metabolism | Energy produc | 0P7WF | aldo keto redu | 313598.MED1 | 1 | 0 | 3,96E-05 | 3,76E-06 | 1 - | 3,32E-05 - | 1 - | 4,12E-05 - | 1 - | 3,92E-05 - | 1 - | 3,97E-05 - | 1 - | 4,48E-05 - |
| 9189 | 1 | 3um_FIJJAFBF_434751 | Bacteria | Verrucomicrol | Opitutae | Opitutales | unclassified | unclassified | Opitutales ba | unclassified | unclassified | unclassified | unclassified | unclassified | 0,4 | 0,489897949 | 3,79E-05 | 4,67E-05 | 1 - | 8,70E-05 - | 0 - | 0 - | 1 - | 0,000102638 - | 0 - | 0 - | 0 - | 0 - |
| 9190 | 1 | 0.2um_NOKCPHJN_393773 | Bacteria | Proteobacteri | Alphaproteob | Rhodospirillal | Rhodospirillac | unclassified | Rhodospirillac | unclassified | unclassified | unclassified | unclassified | unclassified | 4,8 | 0,748331477 | 0,000203327 | 1,47E-05 | 6 - | 0,000215881 - | 4 - | 0,000178707 - | 5 - | 0,000212355 - | 5 - | 0,000215491 - | 4 - | 0,0001942 - |
| 9191 | 2 | 0.2um_NOKCPHJN_592593;3um_FIJJAFBF_54681 | Bacteria | Bacteroidetes | Flavobacteriia | Flavobacterial | Flavobacteriac | various | various | metabolism | Amino acid tra | COG0473 | 3-isopropylma | 391603.FBALC | 0,4 | 0,489897949 | 1,35E-05 | 1,66E-05 | 1 - | 3,09E-05 - | 0 - | 0 - | 1 - | 3,64E-05 - | 0 - | 0 - | 0 - | 0 - |
| 9192 | 1 | 3um_FIJJAFBF_679391 | Bacteria | Bacteroidetes | Flavobacteriia | Flavobacterial | Flavobacteriac | Polaribacter | various | unclassified | unclassified | unclassified | unclassified | unclassified | 1 | 0,632455532 | 6,58E-05 | 4,67E-05 | 1 - | 5,41E-05 - | 0 - | 0 - | 1 - | 6,39E-05 - | 1 - | 6,49E-05 - | 2 - | 0,000146108 - |
| 9193 | 1 | 3um_FIJJAFBF_278990 | Bacteria | Bacteroidetes | Flavobacteriia | Flavobacterial | Flavobacteriac | Polaribacter | Polaribacter s | unclassified | unclassified | unclassified | unclassified | unclassified | 0,8 | 0,748331477 | 2,67E-05 | 2,49E-05 | 0 - | 0 - | 1 - | 3,44E-05 - | 1 - | 3,27E-05 - | 2 - | 6,64E-05 - | 0 - | 0 - |
| 9194 | 1 | 0.2um_NOKCPHJN_119206 | Bacteria | Actinobacteri | Actinobacteri | Actinomycetal | unclassified | unclassified | Actinomycetal | metabolism | Amino acid tra | 00DZF | branched-chai | 196162.Noca_ | 0,6 | 0,489897949 | 1,89E-05 | 1,54E-05 | 0 - | 0 - | 1 - | 3,24E-05 - | 1 - | 3,08E-05 - | 1 - | 3,12E-05 - | 0 - | 0 - |
| 9195 | 1 | 3um_FIJJAFBF_243622 | Bacteria | Bacteroidetes | Flavobacteriia | Flavobacterial | Flavobacteriac | Tenacibaculur | Tenacibaculur | information st | Translation, ri | 09EM4 | Binds directly | 313594.PI23P | 2 | 0 | 0,000240398 | 2,28E-05 | 2 - | 0,000201363 - | 2 - | 0,000250033 - | 2 - | 0,000237688 - | 2 - | 0,000241199 - | 2 - | 0,00027171 - |
| 9196 | 1 | 3um_FIJJAFBF_175473 | Bacteria | Proteobacteri | Gammaprote | Alteromonada | Alteromonada | Paraglaciecola | Paraglaciecola | unclassified | unclassified | unclassified | unclassified | unclassified | 1,4 | 0,489897949 | 4,30E-05 | 1,52E-05 | 1 - | 2,57E-05 - | 1 - | 3,20E-05 - | 2 - | 6,08E-05 - | 2 - | 6,17E-05 - | 1 - | 3,47E-05 - |
| 9197 | 1 | 0.2um_NOKCPHJN_163370 | Bacteria | Proteobacteri | Gammaprote | Cellvibrionales | Porticoccacea | unclassified | gamma prote | cellular proces | Intracellular tr | 0QJ7H | motA TolQ Exl | 314287.GB220 | 12 | 1,414213562 | 0,000362363 | 5,99E-05 | 13 - | 0,000328656 - | 10 - | 0,000313918 - | 11 - | 0,000328261 - | 12 - | 0,000363392 - | 14 - | 0,000477586 - |
| 9198 | 1 | 3um_FIJJAFBF_232377 | Eukaryota | Hemichordata | Enteropneust | unclassified | Harrimaniidae | Saccoglossus | Saccoglossus k | cellular proces | Posttranslatio | 1DGAQ | NHL repeat co | 9483.ENSCJAP | 0,4 | 0,489897949 | 1,47E-05 | 1,80E-05 | 0 - | 0 - | 1 - | 3,73E-05 - | 0 - | 0 - | 1 - | 3,60E-05 - | 0 - | 0 - |
| 9199 | 1 | 3um_FIJJAFBF_467264 | Bacteria | Proteobacteri | Gammaprote | unclassified | unclassified | unclassified | Gammaprote | unclassified | unclassified | unclassified | unclassified | unclassified | 1,2 | 0,748331477 | 3,56E-05 | 2,29E-05 | 1 - | 2,54E-05 - | 1 - | 3,16E-05 - | 2 - | 6,01E-05 - | 2 - | 6,10E-05 - | 0 - | 0 - |
| 9200 | 1 | 3um_FIJJAFBF_469914 | Bacteria | Proteobacteri | Alphaproteob | Rhodobactera | unclassified | unclassified | Rhodobactera | unclassified | unclassified | unclassified | unclassified | unclassified | 0,4 | 0,489897949 | 2,00E-05 | 2,47E-05 | 1 - | 4,59E-05 - | 0 - | 0 - | 1 - | 5,42E-05 - | 0 - | 0 - | 0 - | 0 - |
| 9201 | 1 | 3um_FIJJAFBF_155162 | Bacteria | Proteobacteri | Gammaprote | Cellvibrionales | Porticoccacea | unclassified | SAR92 bacteri | information st | Translation, ri | 179RU | 30S ribosomal | 314287.GB220 | 1,4 | 0,8 | 0,000142006 | 8,13E-05 | 2 - | 0,00017658 - | 2 - | 0,00021926 - | 2 - | 0,000208434 - | 1 - | 0,000105756 - | 0 - | 0 - |
| 9202 | 1 | 0.2um_NOKCPHJN_124517 | Bacteria | Proteobacteri | Alphaproteob | Rhodobactera | Rhodobactera | various | various | poorly charact | Function unkn | COG1744 | basic membra | 314262.MED1 | 3 | 0,894427191 | 0,000113931 | 3,53E-05 | 4 - | 0,000127885 - | 2 - | 7,94E-05 - | 3 - | 0,000113216 - | 2 - | 7,66E-05 - | 4 - | 0,000172562 - |
| 9203 | 1 | 3um_FIJJAFBF_748390 | Bacteria | Bacteroidetes | Flavobacteriia | Flavobacterial | Flavobacteriac | Tenacibaculur | various | cellular proces | Posttranslatio | 08V1J | Has antioxidar | 402612.FP159 | 1 | 0 | 8,30E-05 | 7,89E-06 | 1 - | 6,96E-05 - | 1 - | 8,64E-05 - | 1 - | 8,21E-05 - | 1 - | 8,33E-05 - | 1 - | 9,39E-05 - |
| 9204 | 1 | 0.2um_NOKCPHJN_642262 | Bacteria | Proteobacteri | Alphaproteob | Pelagibacteral | Pelagibacterac | Candidatus Pe | Candidatus Pe | unclassified | unclassified | unclassified | unclassified | unclassified | 2,6 | 0,489897949 | 0,000290581 | 4,21E-05 | 3 - | 0,00028457 - | 2 - | 0,000235568 - | 3 - | 0,000335906 - | 3 - | 0,000340868 - | 2 - | 0,000255991 - |
| 9205 | 1 | 3um_FIJJAFBF_328498 | Bacteria | various | various | various | various | various | various | unclassified | unclassified | unclassified | unclassified | unclassified | 1 | 0 | 0,000110506 | 1,05E-05 | 1 - | 9,26E-05 - | 1 - | 0,000114935 - | 1 - | 0,00010926 - | 1 - | 0,000110874 - | 1 - | 0,000124899 - |
| 9206 | 1 | 0.2um_NOKCPHJN_524593 | Eukaryota | Chlorophyta | Trebouxiophy | Chlorellales | Chlorellaceae | Micractinium | Micractinium | information st | Translation, ri | KOG3184 | Ribosomal pro | 436017.A4RU | 1,6 | 1,019803903 | 9,20E-05 | 5,96E-05 | 2 - | 9,98E-05 - | 2 - | 0,000123929 - | 3 - | 0,000176716 - | 1 - | 5,98E-05 - | 0 - | 0 - |
| 9207 | 1 | 0.2um_NOKCPHJN_435655 | Bacteria | Proteobacteri | Alphaproteob | Rhodobactera | Rhodobactera | Aestuariivita | Aestuariivita b | metabolism | Carbohydrate | COG0176 | Transaldolase | 388401.RB215 | 1,6 | 0,489897949 | 0,000103043 | 3,73E-05 | 1 - | 5,29E-05 - | 2 - | 0,000131354 - | 2 - | 0,000124868 - | 1 - | 6,34E-05 - | 2 - | 0,000142741 - |
| 9208 | 1 | 0.2um_NOKCPHJN_616511 | Eukaryota | unclassified | Pelagophycea | Pelagomonad | unclassified | Aureococcus | Aureococcus a | unclassified | unclassified | unclassified | unclassified | unclassified | 2,2 | 0,4 | 8,56E-05 | 7,85E-06 | 3 - | 9,92E-05 - | 2 - | 8,21E-05 - | 2 - | 7,81E-05 - | 2 - | 7,92E-05 - | 2 - | 8,93E-05 - |
| 9209 | 1 | 0.2um_NOKCPHJN_66824 | Bacteria | Proteobacteri | Gammaprote | Alteromonada | Pseudoalteror | Pseudoalteror | Pseudoalteror | metabolism | Carbohydrate | 16QEQ | hydrolase, fan | 247634.GPB2 | 1,6 | 1,019803903 | 2,12E-05 | 1,39E-05 | 2 - | 2,29E-05 - | 2 - | 2,85E-05 - | 1 - | 1,35E-05 - | 3 - | 4,12E-05 - | 0 - | 0 - |
| 9210 | 1 | 3um_FIJJAFBF_514039 | Bacteria | Bacteroidetes | Flavobacteriia | Flavobacterial | Flavobacteriac | Polaribacter | Polaribacter s | information st | Translation, ri | COG0256 | This is one of | 313598.MED1 | 2,2 | 0,979795897 | 0,00025126 | 8,73E-05 | 4 - | 0,000395782 - | 1 - | 0,000122861 - | 2 - | 0,00023359 - | 2 - | 0,00023704 - | 2 - | 0,000267025 - |
| 9211 | 1 | 0.2um_NOKCPHJN_435291 | Bacteria | unclassified | unclassified | unclassified | unclassified | unclassified | uncultured ba | unclassified | unclassified | unclassified | unclassified | unclassified | 10,4 | 1,496662955 | 0,000149496 | 1,67E-05 | 12 - | 0,000145903 - | 12 - | 0,000181168 - | 10 - | 0,000143519 - | 10 - | 0,000145639 - | 8 - | 0,00013125 - |
| 9212 | 1 | 0.2um_NOKCPHJN_285185 | Bacteria | Proteobacteri | Gammaprote | unclassified | unclassified | unclassified | various | information st | Translation, ri | 17FIH | One of the ea | 349521.HCH_ | 4,2 | 0,979795897 | 0,00057474 | 0,000121198 | 5 - | 0,000579681 - | 3 - | 0,000719792 - | 3 - | 0,000410552 - | 5 - | 0,00069436 - | 3 - | 0,000469317 - |
| 9213 | 2 | 0.2um_NOKCPHJN_261459;3um_FIJJAFBF_30372 | Bacteria | Bacteroidetes | unclassified | unclassified | unclassified | unclassified | Bacteroidetes | poorly charact | Function unkn | 0P884 | membrane | 926562.Oweh | 0,6 | 0,489897949 | 2,44E-05 | 2,01E-05 | 1 - | 3,61E-05 - | 0 - | 0 - | 1 - | 4,26E-05 - | 1 - | 4,32E-05 - | 0 - | 0 - |
| 9214 | 1 | 3um_FIJJAFBF_372266 | Bacteria | Proteobacteri | Alphaproteob | unclassified | unclassified | Candidatus Pu | Candidatus Pu | information st | Translation, ri | COG0335 | This protein is | 35128.Thapsd | 0,8 | 0,4 | 8,62E-05 | 4,38E-05 | 1 - | 9,33E-05 - | 1 - | 0,000115869 - | 1 - | 0,000110148 - | 1 - | 0,000111775 - | 0 - | 0 - |
| 9215 | 1 | 0.2um_NOKCPHJN_57273 | Bacteria | Proteobacteri | Deltaproteoba | Desulfuromon | Geobacterace | Geoalkalibact | Geoalkalibact | metabolism | Energy produc | COG0055 | Produces ATP | 378806.STAU | 0,6 | 0,8 | 1,78E-05 | 2,31E-05 | 0 - | 0 - | 0 - | 0 - | 2 - | 5,66E-05 - | 0 - | 0 - | 1 - | 3,23E-05 - |
| 9216 | 1 | 0.2um_NOKCPHJN_282230 | Bacteria | Bacteroidetes | Flavobacteriia | unclassified | unclassified | unclassified | Flavobacteria | unclassified | unclassified | unclassified | unclassified | unclassified | 3,6 | 0,8 | 0,000186347 | 3,69E-05 | 4 - | 0,000175232 - | 3 - | 0,00016319 - | 5 - | 0,000258554 - | 3 - | 0,000157424 - | 3 - | 0,000177337 - |
| 9217 | 1 | 3um_FIJJAFBF_12135 | Bacteria | Proteobacteri | Alphaproteob | Rhodobactera | Rhodobactera | Palleronia | Palleronia aby | metabolism | Inorganic ion t | 05EKV | Phosphonate | 375451.RD1_ | 3 | 1,414213562 | 0,000116624 | 6,26E-05 | 2 - | 6,34E-05 - | 3 - | 0,00011811 - | 4 - | 0,000149704 - | 1 - | 3,80E-05 - | 5 - | 0,000213915 - |
| 9218 | 1 | 3um_FIJJAFBF_89693 | Eukaryota | unclassified | Pelagophycea | Pelagomonad | unclassified | Aureococcus | Aureococcus a | cellular proces | Posttranslatio | KOG0730 | cell division cy | 35128.Thaps4 | 1 | 0 | 0,000150579 | 1,43E-05 | 1 - | 0,000126128 - | 1 - | 0,000156614 - | 1 - | 0,000148882 - | 1 - | 0,000151081 - | 1 - | 0,000170192 - |
| 9219 | 4 | 0.2um_NOKCPHJN_283787;3um_FIJJAFBF_54113 | Bacteria | Proteobacteri | Gammaprote | Cellvibrionales | Porticoccacea | unclassified | various | metabolism | Lipid transpor | 05VNB | Carrier of the | 314287.GB220 | 0,2 | 0,4 | 3,52E-05 | 7,04E-05 | 0 - | 0 - | 0 - | 0 - | 1 - | 0,000175951 - | 0 - | 0 - | 0 - | 0 - |
| 9220 | 1 | 3um_FIJJAFBF_144687 | Bacteria | Proteobacteri | Gammaprote | Oceanospirilla | Saccharospirill | Reinekea | Reinekea blan | metabolism | Amino acid tra | 0QJS2 | Catalyzes the | 314283.MED2 | 0,8 | 0,4 | 2,82E-05 | 1,43E-05 | 1 - | 3,05E-05 - | 1 - | 3,79E-05 - | 1 - | 3,60E-05 - | 1 - | 3,66E-05 - | 0 - | 0 - |
| 9221 | 1 | 3um_FIJJAFBF_788035 | Bacteria | Proteobacteri | Alphaproteob | Rhodobactera | various | various | various | unclassified | unclassified | unclassified | unclassified | unclassified | 1,8 | 0,4 | 0,000235999 | 4,66E-05 | 2 - | 0,000222867 - | 2 - | 0,000276736 - | 2 - | 0,000263072 - | 2 - | 0,000266958 - | 1 - | 0,000150364 - |
| 9222 | 1 | 3um_FIJJAFBF_161278 | Bacteria | Proteobacteri | Alphaproteob | Rhodobactera | Rhodobactera | Sulfitobacter | Sulfitobacter g | metabolism | Inorganic ion t | COG3221 | phosphonate | 375451.RD1_ | 4,8 | 1,166190379 | 0,000213845 | 3,82E-05 | 7 - | 0,000265161 - | 4 - | 0,000188144 - | 4 - | 0,000178854 - | 4 - | 0,000181496 - | 5 - | 0,000255568 - |
| 9223 | 1 | 3um_FIJJAFBF_466375 | Bacteria | Bacteroidetes | Flavobacteriia | Flavobacterial | Flavobacteriac | Formosa | Formosa sp. H | information st | Translation, ri | COG0254 | 50S ribosomal | 1046627.BZAF | 0,8 | 0,4 | 0,000132447 | 6,80E-05 | 1 - | 0,000138285 - | 1 - | 0,000171709 - | 0 - | 0 - | 1 - | 0,000165643 - | 1 - | 0,000186596 - |
| 9224 | 1 | 3um_FIJJAFBF_748937 | unclassified | unclassified | unclassified | unclassified | unclassified | unclassified | unclassified | unclassified | unclassified | unclassified | unclassified | unclassified | 0,2 | 0,4 | 1,32E-05 | 2,64E-05 | 1 - | 6,60E-05 - | 0 - | 0 - | 0 - | 0 - | 0 - | 0 - | 0 - | 0 - |
| 9225 | 1 | 0.2um_NOKCPHJN_62582 | Bacteria | Proteobacteri | Alphaproteob | Rhodobactera | Rhodobactera | Amylibacter | Amylibacter k | metabolism | Nucleotide tra | 16UID | reductase | 391613.RTM1 | 6,2 | 1,166190379 | 6,96E-05 | 1,36E-05 | 7 - | 6,61E-05 - | 7 - | 8,20E-05 - | 4 - | 4,46E-05 - | 7 - | 7,91E-05 - | 6 - | 7,64E-05 - |
| 9226 | 1 | 0.2um_NOKCPHJN_29661 | Bacteria | Proteobacteri | Gammaprote | Alteromonada | Alteromonada | Paraglaciecola | Paraglaciecola | unclassified | unclassified | unclassified | unclassified | unclassified | 6,4 | 1,356465997 | 9,84E-05 | 1,69E-05 | 8 - | 0,000104343 - | 4 - | 6,48E-05 - | 7 - | 0,00010777 - | 7 - | 0,000109362 - | 6 - | 0,000105596 - |
| 9227 | 1 | 0.2um_NOKCPHJN_785381 | Eukaryota | unclassified | unclassified | unclassified | Plasmodiopho | Plasmodiopho | Plasmodiopho | unclassified | unclassified | unclassified | unclassified | unclassified | 0,8 | 0,4 | 1,29E-05 | 6,58E-06 | 1 - | 1,40E-05 - | 1 - | 1,74E-05 - | 1 - | 1,65E-05 - | 1 - | 1,68E-05 - | 0 - | 0 - |
| 9228 | 1 | 0.2um_NOKCPHJN_272848 | Bacteria | Proteobacteri | Gammaprote | Oceanospirilla | Saccharospirill | Reinekea | Reinekea blan | unclassified | unclassified | unclassified | unclassified | unclassified | 1 | 0 | 5,86E-05 | 5,56E-06 | 1 - | 4,90E-05 - | 1 - | 6,09E-05 - | 1 - | 5,79E-05 - | 1 - | 5,88E-05 - | 1 - | 6,62E-05 - |
| 9229 | 1 | 0.2um_NOKCPHJN_787728 | Eukaryota | Basidiomycota | Exobasidiomy | Tilletiales | Tilletiaceae | Tilletia | Tilletia indica | poorly charact | Function unkn | 0XS1Q | Transmembra | 4558.Sb01g03 | 0,4 | 0,489897949 | 2,29E-05 | 2,81E-05 | 0 - | 0 - | 0 - | 0 - | 1 - | 5,33E-05 - | 0 - | 0 - | 1 - | 6,10E-05 - |
| 9230 | 2 | 3um_FIJJAFBF_416065;3um_FIJJAFBF_83281 | Bacteria | Proteobacteri | Alphaproteob | various | various | various | various | cellular proces | Cell wall/mem | COG2885 | Ompa motb d | 696127.midi_ | 0,6 | 0,489897949 | 5,00E-05 | 4,13E-05 | 1 - | 7,31E-05 - | 1 - | 9,08E-05 - | 1 - | 8,63E-05 - | 0 - | 0 - | 0 - | 0 - |
| 9231 | 1 | 3um_FIJJAFBF_01493 | Bacteria | Bacteroidetes | Flavobacteriia | Flavobacterial | Flavobacteriac | Polaribacter | Polaribacter s | cellular proces | Cell motility | COG1360 | Ompa motb d | 313598.MED1 | 4,2 | 0,4 | 0,000202505 | 9,58E-06 | 5 - | 0,000203505 - | 4 - | 0,000202154 - | 4 - | 0,000192173 - | 4 - | 0,000195012 - | 4 - | 0,00021968 - |
| 9232 | 1 | 3um_FIJJAFBF_50741 | Bacteria | Proteobacteri | Gammaprote | Cellvibrionales | Porticoccacea | unclassified | SAR92 bacteri | unclassified | unclassified | unclassified | unclassified | unclassified | 3 | 0,894427191 | 8,95E-05 | 2,55E-05 | 3 - | 7,58E-05 - | 2 - | 6,28E-05 - | 4 - | 0,000119368 - | 4 - | 0,000121131 - | 2 - | 6,82E-05 - |
| 9233 | 1 | 3um_FIJJAFBF_539523 | unclassified | unclassified | unclassified | unclassified | unclassified | unclassified | unclassified | unclassified | unclassified | unclassified | unclassified | unclassified | 0,6 | 0,489897949 | 4,65E-05 | 3,81E-05 | 0 - | 0 - | 0 - | 0 - | 1 - | 7,36E-05 - | 1 - | 7,47E-05 - | 1 - | 8,42E-05 - |
| 9234 | 1 | 3um_FIJJAFBF_694081 | Bacteria | Proteobacteri | Alphaproteob | Rhodobactera | Rhodobactera | Labrenzia | Labrenzia alex | unclassified | unclassified | unclassified | unclassified | unclassified | 3,2 | 0,748331477 | 0,000132073 | 2,39E-05 | 4 - | 0,00014083 - | 3 - | 0,000131152 - | 3 - | 0,000124677 - | 4 - | 0,000168691 - | 2 - | 9,50E-05 - |
| 9235 | 1 | 0.2um_NOKCPHJN_187255 | Bacteria | Bacteroidetes | Flavobacteriia | Flavobacterial | Flavobacteriac | Formosa | Formosa sp. H | cellular proces | Posttranslatio | 07RXU | peptidylprolyl | 391603.FBALC | 0,2 | 0,4 | 4,79E-06 | 9,58E-06 | 1 - | 2,40E-05 - | 0 - | 0 - | 0 - | 0 - | 0 - | 0 - | 0 - | 0 - |
| 9236 | 1 | 3um_FIJJAFBF_640522 | Eukaryota | unclassified | Spirotrichea | Euplotida | Euplotidae | Euplotes | various | information st | Chromatin str | KOG3467 | Core compone | 31033.ENSTRU | 1 | 0 | 0,000122346 | 1,16E-05 | 1 - | 0,000102479 - | 1 - | 0,000127249 - | 1 - | 0,000120966 - | 1 - | 0,000122753 - | 1 - | 0,000138281 - |
| 9237 | 1 | 3um_FIJJAFBF_345364 | Bacteria | Bacteroidetes | Flavobacteriia | Flavobacterial | Crocinitomica | unclassified | Crocinitomica | unclassified | unclassified | unclassified | unclassified | unclassified | 3 | 0,632455532 | 0,000123395 | 2,43E-05 | 3 - | 0,000104343 - | 3 - | 0,000129563 - | 4 - | 0,000164221 - | 3 - | 0,000124985 - | 2 - | 9,39E-05 - |
| 9238 | 1 | 3um_FIJJAFBF_119154 | Bacteria | Bacteroidetes | Flavobacteriia | Flavobacterial | Flavobacteriac | Polaribacter | Polaribacter b | poorly charact | Function unkn | 09ARW | Zinc ribbon do | 313594.PI23P | 3,6 | 1,496662955 | 0,000194108 | 9,32E-05 | 4 - | 0,000178641 - | 2 - | 0,00011091 - | 2 - | 0,000105434 - | 4 - | 0,000213982 - | 6 - | 0,000361575 - |
| 9239 | 2 | 3um_FIJJAFBF_491522;3um_FIJJAFBF_499654 | Bacteria | Bacteroidetes | Flavobacteriia | Flavobacterial | Flavobacteriac | Formosa | various | poorly charact | Function unkn | 11PIQ | YCE I like fami | 983544.Lacal_ | 1 | 0 | 7,29E-05 | 6,92E-06 | 1 - | 6,11E-05 - | 1 - | 7,58E-05 - | 1 - | 7,21E-05 - | 1 - | 7,31E-05 - | 1 - | 8,24E-05 - |
| 9240 | 1 | 3um_FIJJAFBF_286856 | Bacteria | Proteobacteri | Alphaproteob | Rhodobactera | Rhodobactera | Pseudopelagic | Pseudopelagic | metabolism | Carbohydrate | 16QKM | Fructose-1,6-b | 246200.SPO1 | 0,4 | 0,489897949 | 1,74E-05 | 2,14E-05 | 0 - | 0 - | 1 - | 4,44E-05 - | 0 - | 0 - | 1 - | 4,28E-05 - | 0 - | 0 - |
| 9241 | 1 | 3um_FIJJAFBF_491678 | Bacteria | Proteobacteri | Gammaprote | Cellvibrionales | Porticoccacea | unclassified | SAR92 bacteri | information st | Translation, ri | 179ZI | One of the pri | 314287.GB220 | 3,6 | 0,8 | 0,000375359 | 6,94E-05 | 5 - | 0,000441449 - | 3 - | 0,00032889 - | 3 - | 0,000312651 - | 3 - | 0,000317269 - | 4 - | 0,000476537 - |
| 9242 | 1 | 3um_FIJJAFBF_651104 | Bacteria | Proteobacteri | Betaproteoba | Nitrosomonad | Nitrosomonad | Nitrosomonas | Nitrosomonas | unclassified | unclassified | unclassified | unclassified | unclassified | 1 | 0 | 1,83E-05 | 1,74E-06 | 1 - | 1,53E-05 - | 1 - | 1,91E-05 - | 1 - | 1,81E-05 - | 1 - | 1,84E-05 - | 1 - | 2,07E-05 - |
| 9243 | 1 | 3um_FIJJAFBF_442084 | Bacteria | Bacteroidetes | unclassified | unclassified | unclassified | unclassified | Bacteroidetes | metabolism | Energy produc | COG0711 | F(1)F(0) ATP s | 313603.FB217 | 3,4 | 0,8 | 0,000276302 | 5,63E-05 | 4 - | 0,000276571 - | 4 - | 0,000343419 - | 4 - | 0,000326463 - | 3 - | 0,000248464 - | 2 - | 0,000186596 - |
| 9244 | 1 | 3um_FIJJAFBF_406347 | Bacteria | Bacteroidetes | Cytophagia | Cytophagales | Cytophagacea | Ekhidna | Ekhidna lutea | metabolism | Carbohydrate | 05D2N | Aldolase | 649349.Lbys_ | 0,8 | 0,4 | 3,07E-05 | 1,57E-05 | 1 - | 3,24E-05 - | 0 - | 0 - | 1 - | 3,83E-05 - | 1 - | 3,88E-05 - | 1 - | 4,37E-05 - |
| 9245 | 1 | 0.2um_NOKCPHJN_56842 | Bacteria | Proteobacteri | Gammaprote | Oceanospirilla | Saccharospirill | Reinekea | Reinekea blan | information st | Translation, ri | COG0172 | Catalyzes the | 314283.MED2 | 0,4 | 0,489897949 | 1,27E-05 | 1,55E-05 | 0 - | 0 - | 0 - | 0 - | 1 - | 3,14E-05 - | 1 - | 3,19E-05 - | 0 - | 0 - |
| 9246 | 1 | 3um_FIJJAFBF_191788 | Eukaryota | unclassified | Pelagophycea | Pelagomonad | unclassified | Aureoumbra | Aureoumbra l | metabolism | Energy produc | COG0056 | Produces ATP | 32049.SYNPC | 26,8 | 1,6 | 0,000728593 | 6,83E-05 | 28 - | 0,000638916 - | 28 - | 0,000793346 - | 28 - | 0,000754175 - | 24 - | 0,000655984 - | 26 - | 0,000800544 - |
| 9247 | 1 | 3um_FIJJAFBF_579414 | Bacteria | Bacteroidetes | Flavobacteriia | Flavobacterial | Flavobacteriac | Mesoflavibact | Mesoflavibact | metabolism | Lipid transpor | COG0439 | acetyl-CoA ca | 398720.MED2 | 0,2 | 0,4 | 5,11E-06 | 1,02E-05 | 1 - | 2,56E-05 - | 0 - | 0 - | 0 - | 0 - | 0 - | 0 - | 0 - | 0 - |
| 9248 | 1 | 3um_FIJJAFBF_107571 | Bacteria | Proteobacteri | Alphaproteob | Rhodobactera | Rhodobactera | Planktomarina | Planktomarina | metabolism | Lipid transpor | COG0511 | Acetyl-CoA ca | 439496.RBY4I | 1 | 0 | 8,36E-05 | 7,93E-06 | 1 - | 7,00E-05 - | 1 - | 8,69E-05 - | 1 - | 8,26E-05 - | 1 - | 8,38E-05 - | 1 - | 9,44E-05 - |
| 9249 | 1 | 3um_FIJJAFBF_430663 | Bacteria | Actinobacteri | Actinobacteri | Micrococcales | Microbacteria | unclassified | Microbacteria | unclassified | unclassified | unclassified | unclassified | unclassified | 4,2 | 0,4 | 0,000634515 | 3,00E-05 | 5 - | 0,000637649 - | 4 - | 0,000633417 - | 4 - | 0,000602143 - | 4 - | 0,000611037 - | 4 - | 0,000688331 - |
| 9250 | 1 | 3um_FIJJAFBF_332054 | unclassified | unclassified | unclassified | unclassified | unclassified | unclassified | unclassified | unclassified | unclassified | unclassified | unclassified | unclassified | 0,2 | 0,4 | 2,87E-05 | 5,74E-05 | 1 - | 0,000143471 - | 0 - | 0 - | 0 - | 0 - | 0 - | 0 - | 0 - | 0 - |
| 9251 | 1 | 3um_FIJJAFBF_756707 | Bacteria | Bacteroidetes | Flavobacteriia | Flavobacterial | Flavobacteriac | Lutibacter | Lutibacter sp. | poorly charact | Function unkn | COG0714 | ATPase activit | 376686.Fjoh_ | 0,6 | 0,8 | 2,94E-05 | 3,94E-05 | 0 - | 0 - | 0 - | 0 - | 1 - | 4,86E-05 - | 2 - | 9,86E-05 - | 0 - | 0 - |
| 9252 | 1 | 0.2um_NOKCPHJN_1052523 | Eukaryota | unclassified | Oomycetes | Peronosporal | Peronosporac | Plasmopara | Plasmopara ha | unclassified | unclassified | unclassified | unclassified | unclassified | 1 | 0 | 9,26E-05 | 8,79E-06 | 1 - | 7,76E-05 - | 1 - | 9,63E-05 - | 1 - | 9,15E-05 - | 1 - | 9,29E-05 - | 1 - | 0,000104645 - |
| 9253 | 1 | 3um_FIJJAFBF_646620 | Eukaryota | Bacillariophyt | Coscinodiscop | Thalassiosirale | Thalassiosirac | Thalassiosira | Thalassiosira c | information st | Transcription | COG1293 | Fibronectin-bi | 2850.Phatr42 | 0,2 | 0,4 | 2,88E-06 | 5,76E-06 | 0 - | 0 - | 1 - | 1,44E-05 - | 0 - | 0 - | 0 - | 0 - | 0 - | 0 - |
| 9254 | 1 | 0.2um_NOKCPHJN_62025 | Bacteria | Bacteroidetes | Cytophagia | Cytophagales | Flammeovirga | unclassified | Flammeovirga | cellular proces | Posttranslatio | 05E6M | Sulphatase-m | 643867.Ftrac_ | 2,6 | 0,489897949 | 9,53E-05 | 1,38E-05 | 3 - | 9,33E-05 - | 2 - | 7,72E-05 - | 3 - | 0,000110148 - | 3 - | 0,000111775 - | 2 - | 8,39E-05 - |
| 9255 | 1 | 3um_FIJJAFBF_01300 | Bacteria | Bacteroidetes | Flavobacteriia | Flavobacterial | Flavobacteriac | Polaribacter | Polaribacter s | unclassified | unclassified | unclassified | unclassified | unclassified | 19,4 | 1,496662955 | 0,000303062 | 2,78E-05 | 22 - | 0,000288912 - | 19 - | 0,000309824 - | 18 - | 0,000279025 - | 18 - | 0,000283146 - | 20 - | 0,000354404 - |
| 9256 | 1 | 0.2um_NOKCPHJN_938097 | Bacteria | Proteobacteri | Alphaproteob | Rhodobactera | Hyphomonad | Hyphomonas | various | metabolism | Energy produc | COG0055 | Produces ATP | 208348.PTTG_ | 1 | 0 | 0,000142737 | 1,36E-05 | 1 - | 0,000119559 - | 1 - | 0,000148457 - | 1 - | 0,000141127 - | 1 - | 0,000143212 - | 1 - | 0,000161328 - |
| 9257 | 1 | 3um_FIJJAFBF_161280 | Bacteria | Proteobacteri | Alphaproteob | Rhodobactera | Rhodobactera | Planktotalea | Planktotalea f | metabolism | Inorganic ion t | COG3639 | Phosphonate | 391595.RLO14 | 1,6 | 1,019803903 | 4,88E-05 | 3,07E-05 | 2 - | 5,25E-05 - | 0 - | 0 - | 3 - | 9,30E-05 - | 2 - | 6,29E-05 - | 1 - | 3,54E-05 - |
| 9258 | 1 | 0.2um_NOKCPHJN_795562 | Eukaryota | unclassified | Pelagophycea | Pelagomonad | unclassified | Aureococcus | Aureococcus a | cellular proces | Signal transdu | KOG0613 | fibronectin ty | 67593.Physo1 | 1 | 0 | 1,66E-05 | 1,58E-06 | 1 - | 1,39E-05 - | 1 - | 1,73E-05 - | 1 - | 1,64E-05 - | 1 - | 1,67E-05 - | 1 - | 1,88E-05 - |
| 9259 | 1 | 0.2um_NOKCPHJN_196592 | Bacteria | Proteobacteri | Betaproteoba | Nitrosomonad | Gallionellacea | Sulfuricella | Sulfuricella de | cellular proces | Signal transdu | KOG0613 | fibronectin ty | 7165.AGAP00 | 0,2 | 0,4 | 2,31E-06 | 4,61E-06 | 0 - | 0 - | 0 - | 0 - | 1 - | 1,15E-05 - | 0 - | 0 - | 0 - | 0 - |
| 9260 | 1 | 3um_FIJJAFBF_783325 | Eukaryota | Bacillariophyt | Mediophycea | Anaulales | Anaulaceae | Eunotogramm | Eunotogramm | information st | Translation, ri | 0FNCH | Binds directly | 65393.PCC742 | 4,4 | 0,489897949 | 0,000498467 | 3,81E-05 | 5 - | 0,000478236 - | 4 - | 0,000475063 - | 5 - | 0,000564509 - | 4 - | 0,000458278 - | 4 - | 0,000516248 - |
| 9261 | 1 | 3um_FIJJAFBF_473611 | Bacteria | Bacteroidetes | Flavobacteriia | Flavobacterial | Flavobacteriac | Formosa | Formosa sp. H | unclassified | unclassified | unclassified | unclassified | unclassified | 1,8 | 1,166190379 | 5,58E-05 | 2,98E-05 | 4 - | 0,000109311 - | 2 - | 6,79E-05 - | 1 - | 3,23E-05 - | 1 - | 3,27E-05 - | 1 - | 3,69E-05 - |
| 9262 | 1 | 0.2um_NOKCPHJN_648025 | Eukaryota | Cryptomycota | unclassified | unclassified | unclassified | Rozella | Rozella allomy | unclassified | unclassified | unclassified | unclassified | unclassified | 2,6 | 1,019803903 | 0,000206911 | 9,44E-05 | 2 - | 0,000131173 - | 1 - | 8,14E-05 - | 3 - | 0,000232255 - | 3 - | 0,000235686 - | 4 - | 0,000353999 - |
| 9263 | 1 | 3um_FIJJAFBF_175878 | Bacteria | Proteobacteri | Gammaprote | Oceanospirilla | Oceanospirilla | Marinobacteri | Marinobacter | information st | Translation, ri | 05C64 | Binds directly | 396588.Tgr7_ | 2 | 0,894427191 | 0,000116063 | 4,72E-05 | 3 - | 0,000149061 - | 1 - | 6,17E-05 - | 3 - | 0,000175951 - | 1 - | 5,95E-05 - | 2 - | 0,00013409 - |
| 9264 | 1 | 3um_FIJJAFBF_65170 | Bacteria | Bacteroidetes | Flavobacteriia | Flavobacterial | Flavobacteriac | Hyunsoonleell | Hyunsoonleell | unclassified | unclassified | unclassified | unclassified | unclassified | 4 | 0,894427191 | 8,02E-05 | 2,15E-05 | 4 - | 6,69E-05 - | 3 - | 6,23E-05 - | 5 - | 9,87E-05 - | 3 - | 6,01E-05 - | 5 - | 0,000112882 - |
| 9265 | 1 | 3um_FIJJAFBF_682054 | Bacteria | Bacteroidetes | Flavobacteriia | Flavobacterial | Flavobacteriac | Polaribacter | Polaribacter s | poorly charact | Function unkn | 0PAA9 | Inherit from b | 313598.MED1 | 1 | 0 | 6,72E-05 | 6,38E-06 | 1 - | 5,63E-05 - | 1 - | 6,99E-05 - | 1 - | 6,64E-05 - | 1 - | 6,74E-05 - | 1 - | 7,59E-05 - |
| 9266 | 1 | 0.2um_NOKCPHJN_594863 | Eukaryota | Phaeophyceae | unclassified | Ectocarpales | Ectocarpaceae | Ectocarpus | Ectocarpus sili | unclassified | unclassified | unclassified | unclassified | unclassified | 0,8 | 0,748331477 | 9,18E-06 | 9,30E-06 | 1 - | 9,37E-06 - | 0 - | 0 - | 0 - | 0 - | 1 - | 1,12E-05 - | 2 - | 2,53E-05 - |
| 9267 | 1 | 3um_FIJJAFBF_357546 | Bacteria | Bacteroidetes | Flavobacteriia | Flavobacterial | Flavobacteriac | Polaribacter | Polaribacter ir | unclassified | unclassified | unclassified | unclassified | unclassified | 0,6 | 0,489897949 | 2,01E-05 | 1,66E-05 | 1 - | 2,94E-05 - | 1 - | 3,64E-05 - | 1 - | 3,47E-05 - | 0 - | 0 - | 0 - | 0 - |
| 9268 | 1 | 3um_FIJJAFBF_688235 | Bacteria | Proteobacteri | Alphaproteob | Rhodobactera | Rhodobactera | Shimia | Shimia abyssi | unclassified | unclassified | unclassified | unclassified | unclassified | 0,6 | 0,489897949 | 2,34E-05 | 1,93E-05 | 1 - | 3,40E-05 - | 1 - | 4,22E-05 - | 0 - | 0 - | 1 - | 4,07E-05 - | 0 - | 0 - |
| 9269 | 1 | 0.2um_NOKCPHJN_59333 | Bacteria | Bacteroidetes | Flavobacteriia | Flavobacterial | Flavobacteriac | Lacinutrix | Lacinutrix ven | metabolism | Amino acid tra | 09GJ6 | nitrogen regul | 391603.FBALC | 3,2 | 0,4 | 0,00039123 | 5,79E-05 | 3 - | 0,000307438 - | 3 - | 0,000381747 - | 4 - | 0,000483865 - | 3 - | 0,000368259 - | 3 - | 0,000414842 - |
| 9270 | 1 | 3um_FIJJAFBF_147408 | Bacteria | Proteobacteri | Gammaprote | Alteromonada | unclassified | unclassified | gamma prote | information st | Transcription | 05VEQ | Cold shock pro | 983545.Glaag | 1,2 | 0,979795897 | 0,00022859 | 0,00016164 | 3 - | 0,000506368 - | 1 - | 0,000209587 - | 1 - | 0,000199239 - | 0 - | 0 - | 1 - | 0,000227757 - |
| 9271 | 1 | 0.2um_NOKCPHJN_479810 | Eukaryota | Chordata | Actinopteri | Gobiiformes | Gobiidae | Boleophthalm | Boleophthalm | unclassified | unclassified | unclassified | unclassified | unclassified | 0,4 | 0,489897949 | 3,96E-06 | 4,86E-06 | 0 - | 0 - | 1 - | 9,49E-06 - | 0 - | 0 - | 0 - | 0 - | 1 - | 1,03E-05 - |
| 9272 | 1 | 0.2um_NOKCPHJN_315798 | Bacteria | Bacteroidetes | unclassified | unclassified | unclassified | unclassified | Bacteroidetes | unclassified | unclassified | unclassified | unclassified | unclassified | 1 | 0 | 1,84E-05 | 1,75E-06 | 1 - | 1,54E-05 - | 1 - | 1,92E-05 - | 1 - | 1,82E-05 - | 1 - | 1,85E-05 - | 1 - | 2,08E-05 - |
| 9273 | 1 | 3um_FIJJAFBF_254310 | Bacteria | Proteobacteri | Gammaprote | Cellvibrionales | Spongiibacter | Oceanicoccus | Oceanicoccus | information st | Transcription | 16PMT | DNA-depende | 247633.GP214 | 6 | 1,264911064 | 0,000249486 | 6,72E-05 | 5 - | 0,000172337 - | 7 - | 0,000299589 - | 7 - | 0,000284797 - | 4 - | 0,000165145 - | 7 - | 0,000325562 - |
| 9274 | 1 | 0.2um_NOKCPHJN_16791 | Bacteria | Proteobacteri | Gammaprote | Alteromonada | Alteromonada | Paraglaciecola | various | information st | Translation, ri | 0QQXI | This protein is | 342610.Patl_1 | 4 | 1,095445115 | 0,00044629 | 7,89E-05 | 6 - | 0,000573884 - | 4 - | 0,000475063 - | 4 - | 0,000451607 - | 3 - | 0,000343708 - | 3 - | 0,000387186 - |
| 9275 | 1 | 3um_FIJJAFBF_183698 | Bacteria | Bacteroidetes | Flavobacteriia | Flavobacterial | Flavobacteriac | Algibacter | Algibacter sp. | unclassified | unclassified | unclassified | unclassified | unclassified | 2,2 | 0,748331477 | 5,36E-05 | 1,90E-05 | 3 - | 6,14E-05 - | 2 - | 5,08E-05 - | 2 - | 4,83E-05 - | 1 - | 2,45E-05 - | 3 - | 8,28E-05 - |
| 9276 | 2 | 0.2um_NOKCPHJN_370249;0.2um_NOKCPHJN_4 | Bacteria | Bacteroidetes | Flavobacteriia | Flavobacterial | Flavobacteriac | Polaribacter | Polaribacter s | cellular proces | Posttranslatio | 0XNQ9 | peptidase (S8 | 391598.FBBAL | 0,4 | 0,489897949 | 4,54E-06 | 5,59E-06 | 1 - | 1,03E-05 - | 0 - | 0 - | 0 - | 0 - | 1 - | 1,24E-05 - | 0 - | 0 - |
| 9277 | 2 | 0.2um_NOKCPHJN_18884;3um_FIJJAFBF_29160 | Bacteria | Proteobacteri | Gammaprote | Cellvibrionales | Porticoccacea | unclassified | SAR92 bacteri | cellular proces | Intracellular tr | 05VIE | preprotein tra | 314287.GB220 | 1 | 0 | 0,000126877 | 1,20E-05 | 1 - | 0,000106275 - | 1 - | 0,000131962 - | 1 - | 0,000125447 - | 1 - | 0,000127299 - | 1 - | 0,000143402 - |
| 9278 | 1 | 3um_FIJJAFBF_140603 | Bacteria | Proteobacteri | Gammaprote | Cellvibrionales | Halieaceae | unclassified | marine gamm | poorly charact | Function unkn | COG5430 | spore coat U c | 247634.GPB2 | 0,8 | 0,4 | 6,25E-05 | 3,21E-05 | 1 - | 6,52E-05 - | 1 - | 8,10E-05 - | 0 - | 0 - | 1 - | 7,81E-05 - | 1 - | 8,80E-05 - |
| 9279 | 1 | 0.2um_NOKCPHJN_87559 | Bacteria | Proteobacteri | Alphaproteob | Rhodospirillal | Rhodospirillac | Nisaea | Nisaea denitri | unclassified | unclassified | unclassified | unclassified | unclassified | 0,8 | 0,4 | 2,19E-05 | 1,12E-05 | 1 - | 2,29E-05 - | 1 - | 2,84E-05 - | 1 - | 2,70E-05 - | 0 - | 0 - | 1 - | 3,09E-05 - |
| 9280 | 1 | 3um_FIJJAFBF_471255 | Bacteria | Bacteroidetes | Flavobacteriia | Flavobacterial | Flavobacteriac | Formosa | Formosa sp. H | unclassified | unclassified | unclassified | unclassified | unclassified | 2,2 | 0,748331477 | 4,46E-05 | 1,58E-05 | 2 - | 3,43E-05 - | 3 - | 6,38E-05 - | 2 - | 4,04E-05 - | 3 - | 6,16E-05 - | 1 - | 2,31E-05 - |
| 9281 | 1 | 0.2um_NOKCPHJN_560588 | Bacteria | Bacteroidetes | Flavobacteriia | Flavobacterial | Flavobacteriac | Lutibacter | Lutibacter pro | poorly charact | Function unkn | 0P88U | NA | 313596.RB250 | 0,4 | 0,489897949 | 8,98E-06 | 1,11E-05 | 1 - | 2,00E-05 - | 1 - | 2,49E-05 - | 0 - | 0 - | 0 - | 0 - | 0 - | 0 - |
| 9282 | 1 | 3um_FIJJAFBF_377213 | Bacteria | Bacteroidetes | Flavobacteriia | Flavobacterial | Flavobacteriac | Polaribacter | Polaribacter t | unclassified | unclassified | unclassified | unclassified | unclassified | 2,8 | 0,748331477 | 0,000398935 | 0,000112358 | 3 - | 0,000358677 - | 2 - | 0,000296914 - | 2 - | 0,000282255 - | 4 - | 0,000572847 - | 3 - | 0,000483983 - |
| 9283 | 1 | 0.2um_NOKCPHJN_07978 | Bacteria | Bacteroidetes | Flavobacteriia | unclassified | unclassified | unclassified | Flavobacteria | unclassified | unclassified | unclassified | unclassified | unclassified | 0,4 | 0,489897949 | 2,81E-05 | 3,48E-05 | 1 - | 6,27E-05 - | 1 - | 7,79E-05 - | 0 - | 0 - | 0 - | 0 - | 0 - | 0 - |
| 9284 | 1 | 0.2um_NOKCPHJN_653095 | Bacteria | Proteobacteri | Gammaprote | Cellvibrionales | Halieaceae | unclassified | marine gamm | metabolism | Inorganic ion t | 16PDP | TonB-depende | 247634.GPB2 | 13,6 | 1,019803903 | 0,000203385 | 2,27E-05 | 14 - | 0,000175615 - | 12 - | 0,00018691 - | 15 - | 0,000222102 - | 13 - | 0,000195331 - | 14 - | 0,000236966 - |
| 9285 | 1 | 0.2um_NOKCPHJN_139037 | Bacteria | Bacteroidetes | Flavobacteriia | Flavobacterial | unclassified | unclassified | Flavobacterial | unclassified | unclassified | unclassified | unclassified | unclassified | 0,2 | 0,4 | 2,05E-06 | 4,10E-06 | 0 - | 0 - | 1 - | 1,02E-05 - | 0 - | 0 - | 0 - | 0 - | 0 - | 0 - |
| 9286 | 1 | 3um_FIJJAFBF_796563 | Eukaryota | unclassified | Bangiophycea | Bangiales | Bangiaceae | Bangia | Bangia fuscop | cellular proces | Posttranslatio | 0IEKR | One of the co | 35128.Thapsd | 2,4 | 0,489897949 | 7,13E-05 | 1,54E-05 | 2 - | 4,98E-05 - | 2 - | 6,18E-05 - | 3 - | 8,82E-05 - | 3 - | 8,95E-05 - | 2 - | 6,72E-05 - |
| 9287 | 1 | 0.2um_NOKCPHJN_161886 | Bacteria | unclassified | unclassified | unclassified | unclassified | unclassified | bacterium TM | unclassified | unclassified | unclassified | unclassified | unclassified | 1 | 0 | 5,93E-05 | 5,63E-06 | 1 - | 4,97E-05 - | 1 - | 6,17E-05 - | 1 - | 5,87E-05 - | 1 - | 5,95E-05 - | 1 - | 6,70E-05 - |
| 9288 | 1 | 3um_FIJJAFBF_397925 | Bacteria | Proteobacteri | Alphaproteob | Kordiimonada | unclassified | unclassified | Kordiimonada | information st | Translation, ri | 16Q16 | Binds the low | 536019.Meso | 0,2 | 0,4 | 1,15E-05 | 2,30E-05 | 0 - | 0 - | 0 - | 0 - | 1 - | 5,75E-05 - | 0 - | 0 - | 0 - | 0 - |
| 9289 | 1 | 3um_FIJJAFBF_386848 | Bacteria | Bacteroidetes | Flavobacteriia | unclassified | unclassified | unclassified | Flavobacteria | metabolism | Inorganic ion t | 0P78U | receptor | 487797.Flav3 | 2,6 | 0,489897949 | 4,81E-05 | 1,00E-05 | 3 - | 4,66E-05 - | 2 - | 3,86E-05 - | 2 - | 3,67E-05 - | 3 - | 5,58E-05 - | 3 - | 6,29E-05 - |

| | | | | | | | | | | | | | | | | | | | | | | | | | | | | |
|---|---|---|---|---|---|---|---|---|---|---|---|---|---|---|---|---|---|---|---|---|---|---|---|---|---|---|---|---|
| 9290 | 1 | 0.2um_NOKCPHJN_52330 | Bacteria | Bacteroidetes | Flavobacteriia | Flavobacterial | Flavobacteriac | Formosa | Formosa sp. H | unclassified | unclassified | unclassified | unclassified | unclassified | 1 | 0 | 1,61E-05 | 1,53E-06 | 1 - | 1,35E-05 - | 1 - | 1,68E-05 - | 1 - | 1,60E-05 - | 1 - | 1,62E-05 - | 1 - | 1,82E-05 - |
| 9291 | 1 | 3um_FIJJAFBF_689869 | Bacteria | Bacteroidetes | Flavobacteriia | Flavobacterial | Crocinitomica | unclassified | Crocinitomica | unclassified | unclassified | unclassified | unclassified | unclassified | 1 | 0,894427191 | 2,41E-05 | 2,09E-05 | 2 - | 4,20E-05 - | 0 - | 0 - | 0 - | 0 - | 2 - | 5,03E-05 - | 1 - | 2,83E-05 - |
| 9292 | 1 | 3um_FIJJAFBF_394404 | Bacteria | Proteobacteri | Alphaproteob | Rhodobactera | Rhodobactera | Planktomarina | Planktomarina | information st | Transcription | 01ZK0 | This protein is | 314270.RB208 | 1,4 | 0,489897949 | 0,000202673 | 6,37E-05 | 2 - | 0,000246832 - | 2 - | 0,000306492 - | 1 - | 0,00014568 - | 1 - | 0,000147832 - | 1 - | 0,000166532 - |
| 9293 | 1 | 0.2um_NOKCPHJN_695629 | Bacteria | Proteobacteri | Gammaprotec | Oceanospirilla | Saccharospirill | Reinekea | Reinekea fors | poorly charact | Function unkn | 0QXWX | NA | 391591.VSAK1 | 3 | 0,894427191 | 4,78E-05 | 1,16E-05 | 4 - | 5,45E-05 - | 3 - | 5,07E-05 - | 4 - | 6,43E-05 - | 2 - | 3,26E-05 - | 2 - | 3,67E-05 - |
| 9294 | 1 | 3um_FIJJAFBF_541150 | Bacteria | Bacteroidetes | Cytophagia | Cytophagales | Flammeovirga | unclassified | Flammeovirga | metabolism | Inorganic ion t | 05PWN | phosphate ab | 357804.Ping_ | 0,6 | 0,489897949 | 2,24E-05 | 1,84E-05 | 1 - | 3,32E-05 - | 0 - | 0 - | 1 - | 3,92E-05 - | 1 - | 3,97E-05 - | 0 - | 0 - |
| 9295 | 1 | 3um_FIJJAFBF_183932 | Eukaryota | unclassified | Pelagophycea | Pelagomonad | unclassified | Aureococcus | Aureococcus a | metabolism | Inorganic ion t | 1EKJE | calcium chann | 3218.PP1S45_ | 2,6 | 0,8 | 5,59E-05 | 1,90E-05 | 2 - | 3,58E-05 - | 4 - | 8,89E-05 - | 2 - | 4,23E-05 - | 3 - | 6,43E-05 - | 2 - | 4,83E-05 - |
| 9296 | 1 | 0.2um_NOKCPHJN_1079676 | Bacteria | Proteobacteri | Gammaprotec | Alteromonada | Pseudoalteror | Pseudoalteror | Pseudoalteror | unclassified | unclassified | unclassified | unclassified | unclassified | 0,8 | 0,4 | 2,02E-05 | 1,02E-05 | 1 - | 2,18E-05 - | 1 - | 2,71E-05 - | 1 - | 2,58E-05 - | 1 - | 2,61E-05 - | 0 - | 0 - |
| 9297 | 1 | 0.2um_NOKCPHJN_203950 | Eukaryota | unclassified | Oligohymenoр | Hymenostoma | Tetrahymenid | Tetrahymena | Tetrahymena | unclassified | unclassified | unclassified | unclassified | unclassified | 0,2 | 0,4 | 5,41E-06 | 1,08E-05 | 0 - | 0 - | 1 - | 2,70E-05 - | 0 - | 0 - | 0 - | 0 - | 0 - | 0 - |
| 9298 | 2 | 0.2um_NOKCPHJN_254901;0.2um_NOKCPHJN_4 | Bacteria | Bacteroidetes | various | various | various | various | various | metabolism | Secondary me | 08JEG | abc transporte | 487796.Flav2A | 0,2 | 0,4 | 1,06E-05 | 2,13E-05 | 0 - | 0 - | 0 - | 0 - | 1 - | 5,31E-05 - | 0 - | 0 - | 0 - | 0 - |
| 9299 | 1 | 3um_FIJJAFBF_350095 | Bacteria | Proteobacteri | Alphaproteob | Rhodobactera | Rhodobactera | Phaeobacter | Phaeobacter g | information st | Translation, ri | 05KK9 | This protein bi | 292414.TM10 | 4,8 | 0,748331477 | 0,000327582 | 2,36E-05 | 6 - | 0,000347808 - | 4 - | 0,000287917 - | 5 - | 0,000342127 - | 5 - | 0,00034718 - | 4 - | 0,000312878 - |
| 9300 | 1 | 3um_FIJJAFBF_278270 | Bacteria | Proteobacteri | various | various | various | various | various | cellular proces | Posttranslatio | 01ZS8 | Thioredoxin | 439496.RBY4I | 2,6 | 0,489897949 | 0,000333028 | 5,62E-05 | 3 - | 0,00032484 - | 3 - | 0,000403355 - | 2 - | 0,000255627 - | 3 - | 0,000389104 - | 2 - | 0,000292216 - |
| 9301 | 1 | 3um_FIJJAFBF_257113 | Eukaryota | unclassified | Florideophyce | Nemaliales | Liagoraceae | Yamadaella | Yamadaella ca | information st | Translation, ri | 0FNT6 | One of the ear | 391612.CY011 | 1,4 | 0,489897949 | 0,000173899 | 6,51E-05 | 1 - | 0,000103402 - | 2 - | 0,000256791 - | 1 - | 0,000122056 - | 2 - | 0,000247718 - | 1 - | 0,000139527 - |
| 9302 | 1 | 0.2um_NOKCPHJN_613724 | Bacteria | Bacteroidetes | unclassified | Bacteroidetes | Rhodotherma | Rhodothermu | Rhodothermu | unclassified | unclassified | unclassified | unclassified | unclassified | 0,6 | 0,489897949 | 1,09E-05 | 8,96E-06 | 1 - | 1,59E-05 - | 1 - | 1,97E-05 - | 1 - | 1,87E-05 - | 0 - | 0 - | 0 - | 0 - |
| 9303 | 1 | 0.2um_NOKCPHJN_94421 | Bacteria | Proteobacteri | Betaproteoba | unclassified | unclassified | unclassified | Betaproteoba | information st | Translation, ri | COG0088 | One of the pri | 296591.Bpro_ | 5,4 | 1,743559577 | 0,000349812 | 8,53E-05 | 8 - | 0,000445735 - | 3 - | 0,000207552 - | 6 - | 0,000394608 - | 6 - | 0,000400437 - | 4 - | 0,000300727 - |
| 9304 | 1 | 3um_FIJJAFBF_248948 | Bacteria | Bacteroidetes | Flavobacteriia | Flavobacterial | Flavobacteriac | Polaribacter | Polaribacter s | metabolism | Lipid transpor | 05VNB | Carrier of the | 313598.MED1 | 0,4 | 0,489897949 | 7,50E-05 | 9,21E-05 | 0 - | 0 - | 0 - | 0 - | 0 - | 0 - | 1 - | 0,000176261 - | 1 - | 0,000198557 - |
| 9305 | 1 | 0.2um_NOKCPHJN_312875 | Bacteria | Bacteroidetes | Flavobacteriia | Flavobacterial | Crocinitomica | unclassified | Crocinitomica | unclassified | unclassified | unclassified | unclassified | unclassified | 1 | 0 | 0,000108752 | 1,03E-05 | 1 - | 9,11E-05 - | 1 - | 0,00011311 - | 1 - | 0,000107526 - | 1 - | 0,000109114 - | 1 - | 0,000122916 - |
| 9306 | 1 | 0.2um_NOKCPHJN_325099 | Bacteria | Bacteroidetes | Flavobacteriia | Flavobacterial | Flavobacteriac | unclassified | Flavobacteriac | information st | Replication, re | 0PB0H | DNA-binding p | 487796.Flav2A | 1 | 0,632455532 | 0,000147651 | 8,26E-05 | 2 - | 0,000255059 - | 1 - | 0,000158354 - | 0 - | 0 - | 1 - | 0,000152759 - | 1 - | 0,000172083 - |
| 9307 | 1 | 3um_FIJJAFBF_332159 | Bacteria | Proteobacteri | Gammaprotec | Arenicellales | Arenicellaceae | Arenicella | Arenicella xan | information st | Translation, ri | 179RU | 30S ribosomal | 247634.GPB2 | 0,6 | 0,489897949 | 5,92E-05 | 4,87E-05 | 1 - | 8,76E-05 - | 0 - | 0 - | 1 - | 0,000103422 - | 1 - | 0,000104949 - | 0 - | 0 - |
| 9308 | 1 | 3um_FIJJAFBF_786143 | Bacteria | Proteobacteri | Gammaprotec | Oceanospirilla | Saccharospirill | Saccharospiril | Saccharospiril | unclassified | unclassified | unclassified | unclassified | unclassified | 4,8 | 1,166190379 | 0,00044801 | 8,42E-05 | 6 - | 0,000478236 - | 4 - | 0,000395886 - | 5 - | 0,000470424 - | 6 - | 0,000572847 - | 3 - | 0,000322655 - |
| 9309 | 1 | 3um_FIJJAFBF_151644 | Bacteria | Proteobacteri | Gammaprotec | Oceanospirilla | Saccharospirill | Reinekea | Reinekea blan | unclassified | unclassified | unclassified | unclassified | unclassified | 0,4 | 0,489897949 | 1,60E-05 | 1,97E-05 | 0 - | 0 - | 1 - | 4,08E-05 - | 0 - | 0 - | 1 - | 3,94E-05 - | 0 - | 0 - |
| 9310 | 1 | 3um_FIJJAFBF_366557 | unclassified | unclassified | unclassified | unclassified | unclassified | unclassified | unclassified | unclassified | unclassified | unclassified | unclassified | unclassified | 3,4 | 1,743559577 | 9,82E-05 | 4,31E-05 | 6 - | 0,000152359 - | 2 - | 6,31E-05 - | 4 - | 0,000119896 - | 4 - | 0,000121667 - | 1 - | 3,43E-05 - |
| 9311 | 1 | 3um_FIJJAFBF_195984 | Bacteria | Bacteroidetes | Flavobacteriia | Flavobacterial | Flavobacteriac | Polaribacter | various | unclassified | unclassified | unclassified | unclassified | unclassified | 2,8 | 0,748331477 | 0,000600693 | 0,000133497 | 4 - | 0,000728741 - | 2 - | 0,000452441 - | 2 - | 0,000430102 - | 3 - | 0,000654682 - | 3 - | 0,000737498 - |
| 9312 | 1 | 3um_FIJJAFBF_712785 | Eukaryota | unclassified | Pelagophycea | Pelagomonad | unclassified | Aureococcus | Aureococcus a | unclassified | unclassified | unclassified | unclassified | unclassified | 1,2 | 0,748331477 | 3,42E-05 | 2,25E-05 | 1 - | 2,42E-05 - | 2 - | 6,01E-05 - | 1 - | 2,86E-05 - | 2 - | 5,80E-05 - | 0 - | 0 - |
| 9313 | 1 | 0.2um_NOKCPHJN_372198 | Bacteria | Proteobacteri | Alphaproteob | Pelagibacteral | Pelagibacterac | unclassified | various | unclassified | unclassified | unclassified | unclassified | unclassified | 0,6 | 0,489897949 | 3,04E-05 | 2,50E-05 | 1 - | 4,50E-05 - | 0 - | 0 - | 1 - | 5,31E-05 - | 1 - | 5,39E-05 - | 0 - | 0 - |
| 9314 | 1 | 3um_FIJJAFBF_785348 | Bacteria | Proteobacteri | Betaproteoba | unclassified | unclassified | unclassified | Betaproteoba | unclassified | unclassified | unclassified | unclassified | unclassified | 8,4 | 0,8 | 0,000337404 | 4,57E-05 | 9 - | 0,00030293 - | 8 - | 0,000334355 - | 7 - | 0,000278116 - | 9 - | 0,000362859 - | 9 - | 0,00040876 - |
| 9315 | 1 | 0.2um_NOKCPHJN_509227 | Bacteria | Bacteroidetes | Flavobacteriia | unclassified | unclassified | unclassified | Flavobacteria | poorly charact | Function unkn | 0NVRZ | NA | 170187.SP_17 | 6,2 | 0,748331477 | 9,34E-05 | 1,43E-05 | 7 - | 8,84E-05 - | 6 - | 9,41E-05 - | 6 - | 8,94E-05 - | 5 - | 7,56E-05 - | 7 - | 0,000119265 - |
| 9316 | 1 | 3um_FIJJAFBF_769062 | Eukaryota | Phaeophyceae | unclassified | Ectocarpales | Ectocarpaceae | Ectocarpus | Ectocarpus sili | cellular proces | Signal transdu | KOG0613 | fibronectin typ | 67593.Physo1 | 2 | 0 | 5,76E-05 | 5,47E-06 | 2 - | 4,82E-05 - | 2 - | 5,99E-05 - | 2 - | 5,69E-05 - | 2 - | 5,78E-05 - | 2 - | 6,51E-05 - |
| 9317 | 1 | 0.2um_NOKCPHJN_110538 | Bacteria | Proteobacteri | Gammaprotec | Cellvibrionales | Porticoccacea | unclassified | SAR92 bacteri | information st | Translation, ri | COG0256 | This is one of t | 314287.GB22 | 2 | 0 | 0,000236254 | 2,24E-05 | 2 - | 0,000197891 - | 2 - | 0,000245722 - | 2 - | 0,00023359 - | 2 - | 0,00023704 - | 2 - | 0,000267025 - |
| 9318 | 1 | 0.2um_NOKCPHJN_43884 | Bacteria | Proteobacteri | Gammaprotec | Oceanospirilla | Oceanospirilla | Marinomonas | Marinomonas | information st | Translation, ri | 16QAP | Associates wit | 207954.MED9 | 1,4 | 0,8 | 6,42E-05 | 3,62E-05 | 2 - | 8,03E-05 - | 1 - | 4,98E-05 - | 2 - | 9,47E-05 - | 2 - | 9,61E-05 - | 0 - | 0 - |
| 9319 | 1 | 0.2um_NOKCPHJN_92442 | Bacteria | Proteobacteri | Gammaprotec | Oceanospirilla | Saccharospirill | Reinekea | Reinekea blan | metabolism | Amino acid tra | 16Q45 | Acetylornithin | 314283.MED2 | 0,8 | 0,4 | 2,84E-05 | 1,43E-05 | 0 - | 0 - | 1 - | 3,55E-05 - | 1 - | 3,37E-05 - | 1 - | 3,42E-05 - | 1 - | 3,85E-05 - |
| 9320 | 1 | 3um_FIJJAFBF_186279 | Bacteria | unclassified | unclassified | unclassified | unclassified | unclassified | uncultured ba | metabolism | Amino acid tra | COG2171 | Catalyzes the | 156586.BBFL7 | 0,4 | 0,8 | 1,52E-05 | 3,03E-05 | 2 - | 7,58E-05 - | 0 - | 0 - | 0 - | 0 - | 0 - | 0 - | 0 - | 0 - |
| 9321 | 1 | 3um_FIJJAFBF_228146 | Bacteria | Proteobacteri | Gammaprotec | unclassified | unclassified | unclassified | various | unclassified | unclassified | unclassified | unclassified | unclassified | 2 | 0 | 0,000145774 | 1,38E-05 | 2 - | 0,000122103 - | 2 - | 0,000151616 - | 2 - | 0,00014413 - | 2 - | 0,000146259 - | 2 - | 0,00016476 - |
| 9322 | 1 | 3um_FIJJAFBF_378522 | Archaea | Euryarchaeota | unclassified | unclassified | unclassified | unclassified | Euryarchaeota | unclassified | unclassified | unclassified | unclassified | unclassified | 0,2 | 0,4 | 2,02E-06 | 4,05E-06 | 0 - | 0 - | 1 - | 1,01E-05 - | 0 - | 0 - | 0 - | 0 - | 0 - | 0 - |
| 9323 | 1 | 3um_FIJJAFBF_155266 | Bacteria | Proteobacteri | Gammaprotec | Alteromonada | Pseudoalteror | Pseudoalteror | various | poorly charact | Function unkn | 129JZ | NA | 156578.ATW7 | 0,4 | 0,489897949 | 2,46E-05 | 3,03E-05 | 1 - | 5,60E-05 - | 0 - | 0 - | 0 - | 0 - | 1 - | 6,71E-05 - | 0 - | 0 - |
| 9324 | 1 | 3um_FIJJAFBF_601498 | Bacteria | Proteobacteri | Alphaproteob | unclassified | unclassified | unclassified | Alphaproteob | metabolism | Carbohydrate | 16W2E | Major Facilitat | 633131.TR2A6 | 0,2 | 0,4 | 5,80E-06 | 1,16E-05 | 0 - | 0 - | 0 - | 0 - | 1 - | 2,90E-05 - | 0 - | 0 - | 0 - | 0 - |
| 9325 | 1 | 3um_FIJJAFBF_470939 | Bacteria | Bacteroidetes | Cytophagia | Cytophagales | Flammeovirga | unclassified | Flammeovirga | unclassified | unclassified | unclassified | unclassified | unclassified | 1,4 | 0,489897949 | 0,000119763 | 4,23E-05 | 1 - | 7,17E-05 - | 1 - | 8,91E-05 - | 2 - | 0,000169353 - | 2 - | 0,000171854 - | 1 - | 9,68E-05 - |
| 9326 | 1 | 3um_FIJJAFBF_264567 | Bacteria | Proteobacteri | Alphaproteob | Pelagibacteral | Pelagibacterac | Candidatus Pe | Candidatus Pe | unclassified | unclassified | unclassified | unclassified | unclassified | 0,2 | 0,4 | 1,30E-05 | 2,59E-05 | 1 - | 6,48E-05 - | 0 - | 0 - | 0 - | 0 - | 0 - | 0 - | 0 - | 0 - |
| 9327 | 1 | 3um_FIJJAFBF_134489 | Bacteria | Bacteroidetes | Flavobacteriia | Flavobacterial | Flavobacteriac | Polaribacter | Polaribacter s | cellular proces | Intracellular tr | COG0653 | Part of the Sec | 313598.MED1 | 1 | 0,632455532 | 1,19E-05 | 6,67E-06 | 2 - | 2,06E-05 - | 1 - | 1,28E-05 - | 1 - | 1,22E-05 - | 0 - | 0 - | 1 - | 1,39E-05 - |
| 9328 | 2 | 0.2um_NOKCPHJN_394726;0.2um_NOKCPHJN_5 | Bacteria | unclassified | unclassified | unclassified | unclassified | unclassified | bacterium TM | cellular proces | Cell wall/mem | COG0438 | Glycosyl trans | 335992.SAR11 | 0,2 | 0,4 | 5,83E-06 | 1,17E-05 | 0 - | 0 - | 0 - | 0 - | 0 - | 0 - | 0 - | 0 - | 1 - | 2,92E-05 - |
| 9329 | 1 | 3um_FIJJAFBF_339842 | Bacteria | Bacteroidetes | Flavobacteriia | Flavobacterial | Flavobacteriac | Polaribacter | Polaribacter t | unclassified | unclassified | unclassified | unclassified | unclassified | 1 | 1,095445115 | 3,94E-05 | 4,45E-05 | 1 - | 3,44E-05 - | 0 - | 0 - | 3 - | 0,000121691 - | 1 - | 4,12E-05 - | 0 - | 0 - |
| 9330 | 1 | 0.2um_NOKCPHJN_1028962 | unclassified | unclassified | unclassified | unclassified | unclassified | unclassified | unclassified | unclassified | unclassified | unclassified | unclassified | unclassified | 0,8 | 0,4 | 4,87E-05 | 2,45E-05 | 0 - | 0 - | 1 - | 6,09E-05 - | 1 - | 5,79E-05 - | 1 - | 5,88E-05 - | 1 - | 6,62E-05 - |
| 9331 | 2 | 0.2um_NOKCPHJN_593576;0.2um_NOKCPHJN_7 | Bacteria | Bacteroidetes | Flavobacteriia | unclassified | unclassified | unclassified | Flavobacteria | unclassified | unclassified | unclassified | unclassified | unclassified | 2 | 0 | 0,000156602 | 1,49E-05 | 2 - | 0,000131173 - | 2 - | 0,000162879 - | 2 - | 0,000154837 - | 2 - | 0,000157124 - | 2 - | 0,000176999 - |
| 9332 | 1 | 0.2um_NOKCPHJN_591079 | Archaea | Euryarchaeota | unclassified | unclassified | unclassified | unclassified | Euryarchaeota | unclassified | unclassified | unclassified | unclassified | unclassified | 3,6 | 0,489897949 | 0,000148826 | 1,76E-05 | 4 - | 0,000139546 - | 4 - | 0,000173275 - | 4 - | 0,00016472 - | 3 - | 0,000125365 - | 3 - | 0,000141223 - |
| 9333 | 1 | 0.2um_NOKCPHJN_344710 | Bacteria | Bacteroidetes | Flavobacteriia | Flavobacterial | unclassified | unclassified | Flavobacterial | information st | Replication, re | 0PB0H | DNA-binding p | 755732.Fluta_ | 4 | 0 | 0,000609009 | 5,78E-05 | 4 - | 0,000510119 - | 4 - | 0,000633417 - | 4 - | 0,000602143 - | 4 - | 0,000611037 - | 4 - | 0,000688331 - |
| 9334 | 1 | 3um_FIJJAFBF_620000 | Bacteria | Bacteroidetes | Flavobacteriia | Flavobacterial | Flavobacteriac | Polaribacter | Polaribacter s | metabolism | Carbohydrate | 0PF2Z | Transketolase | 313594.PI23P | 0,4 | 0,489897949 | 1,98E-05 | 2,42E-05 | 0 - | 0 - | 1 - | 5,07E-05 - | 1 - | 4,82E-05 - | 0 - | 0 - | 0 - | 0 - |
| 9335 | 1 | 3um_FIJJAFBF_537763 | Bacteria | Bacteroidetes | Flavobacteriia | Flavobacterial | Crocinitomica | unclassified | Crocinitomica | information st | Translation, ri | COG0186 | One of the pri | 575590.HMPR | 1,4 | 0,8 | 0,00020038 | 0,000117636 | 2 - | 0,000239118 - | 2 - | 0,000296914 - | 0 - | 0 - | 1 - | 0,000143212 - | 2 - | 0,000322655 - |
| 9336 | 1 | 3um_FIJJAFBF_199361 | Bacteria | Proteobacteri | Gammaprotec | Oceanospirilla | Oceanospirilla | Nitrincola | Nitrincola sp. | unclassified | unclassified | unclassified | unclassified | unclassified | 2,2 | 0,748331477 | 0,000251177 | 0,000100936 | 2 - | 0,000188159 - | 3 - | 0,000350456 - | 1 - | 0,000111051 - | 2 - | 0,000225382 - | 3 - | 0,000380839 - |
| 9337 | 1 | 0.2um_NOKCPHJN_208347 | Bacteria | Proteobacteri | Gammaprotec | Cellvibrionales | Porticoccacea | unclassified | Gammaprotec | information st | Translation, ri | 179RU | 30S ribosomal | 314287.GB22 | 1 | 0 | 0,000105405 | 1,00E-05 | 1 - | 8,83E-05 - | 1 - | 0,00010963 - | 1 - | 0,000104217 - | 1 - | 0,000105756 - | 1 - | 0,000119134 - |
| 9338 | 1 | 3um_FIJJAFBF_649817 | Bacteria | Proteobacteri | Gammaprotec | Oceanospirilla | Saccharospirill | Reinekea | Reinekea blan | unclassified | unclassified | unclassified | unclassified | unclassified | 18,8 | 0,4 | 0,000484936 | 4,55E-05 | 19 - | 0,000410689 - | 18 - | 0,000483115 - | 19 - | 0,000484776 - | 19 - | 0,000491937 - | 19 - | 0,000554165 - |
| 9339 | 1 | 3um_FIJJAFBF_506115 | Bacteria | Proteobacteri | Gammaprotec | Oceanospirilla | Saccharospirill | Reinekea | Reinekea fors | poorly charact | Function unkn | 179VZ | UPF0234 prot | 314283.MED2 | 0,2 | 0,4 | 1,41E-05 | 2,82E-05 | 1 - | 7,04E-05 - | 0 - | 0 - | 0 - | 0 - | 0 - | 0 - | 0 - | 0 - |
| 9340 | 1 | 3um_FIJJAFBF_238323 | Bacteria | Bacteroidetes | Flavobacteriia | Flavobacterial | Flavobacteriac | Polaribacter | Polaribacter s | metabolism | Energy produc | 099Q7 | Catalyzes the | 313598.MED1 | 1 | 0 | 4,45E-05 | 4,22E-06 | 1 - | 3,73E-05 - | 1 - | 4,63E-05 - | 1 - | 4,40E-05 - | 1 - | 4,46E-05 - | 1 - | 5,03E-05 - |
| 9341 | 1 | 3um_FIJJAFBF_147679 | Bacteria | Proteobacteri | Alphaproteob | Rhodobactera | Rhodobactera | various | various | metabolism | Amino acid tra | COG0687 | polyamine bin | 314270.RB208 | 2,4 | 1,496662955 | 8,85E-05 | 4,64E-05 | 5 - | 0,000163035 - | 2 - | 8,10E-05 - | 3 - | 0,000115468 - | 1 - | 3,91E-05 - | 1 - | 4,40E-05 - |
| 9342 | 1 | 0.2um_NOKCPHJN_399203 | Eukaryota | Chordata | Actinopteri | Cypriniformes | Cyprinidae | Sinocyclocheil | Sinocyclocheil | cellular proces | Cell cycle cont | 0V9W7 | retinitis pigme | 31033.ENSTRU | 0,2 | 0,4 | 5,38E-06 | 1,08E-05 | 0 - | 0 - | 0 - | 0 - | 1 - | 2,69E-05 - | 0 - | 0 - | 0 - | 0 - |
| 9343 | 1 | 0.2um_NOKCPHJN_568703 | Bacteria | Planctomycet | Planctomyceti | Planctomycet | Gemmataceae | Gemmata | Gemmata sp. | poorly charact | Function unkn | 0SFVF | NA | 5507.FOXG_1 | 1 | 0 | 2,79E-05 | 2,65E-06 | 1 - | 2,34E-05 - | 1 - | 2,90E-05 - | 1 - | 2,76E-05 - | 1 - | 2,80E-05 - | 1 - | 3,15E-05 - |
| 9344 | 1 | 0.2um_NOKCPHJN_235590 | Bacteria | Proteobacteri | Gammaprotec | Cellvibrionales | Porticoccacea | unclassified | SAR92 bacteri | information st | Transcription | COG0250 | Participates in | 314285.KT71_ | 2,2 | 0,748331477 | 0,000171538 | 6,63E-05 | 2 - | 0,000129691 - | 1 - | 8,05E-05 - | 3 - | 0,000229631 - | 2 - | 0,000155348 - | 3 - | 0,000262499 - |
| 9345 | 1 | 3um_FIJJAFBF_177945 | Bacteria | Proteobacteri | Alphaproteob | Rhodobactera | Rhodobactera | Roseobacter | Roseobacter s | cellular proces | Cell cycle cont | 01Q8B | Required for c | 391593.RCCS2 | 0,8 | 0,4 | 9,99E-06 | 5,03E-06 | 0 - | 0 - | 1 - | 1,25E-05 - | 1 - | 1,19E-05 - | 1 - | 1,20E-05 - | 1 - | 1,36E-05 - |
| 9346 | 1 | 0.2um_NOKCPHJN_690213 | Bacteria | unclassified | unclassified | unclassified | unclassified | unclassified | uncultured ma | poorly charact | Function unkn | 11YJ6 | NA | 582744.Msip3 | 2 | 0,632455532 | 5,93E-05 | 1,95E-05 | 2 - | 5,01E-05 - | 3 - | 9,34E-05 - | 2 - | 5,92E-05 - | 2 - | 6,00E-05 - | 1 - | 3,38E-05 - |
| 9347 | 1 | 3um_FIJJAFBF_110703 | Bacteria | Bacteroidetes | Cytophagia | Cytophagales | Flammeovirga | unclassified | Flammeovirga | metabolism | Inorganic ion t | 0QJ3Y | Part of the AB | 572477.Alvin_ | 0,4 | 0,489897949 | 1,70E-05 | 2,09E-05 | 1 - | 3,89E-05 - | 0 - | 0 - | 1 - | 4,59E-05 - | 0 - | 0 - | 0 - | 0 - |
| 9348 | 1 | 3um_FIJJAFBF_508357 | Bacteria | Proteobacteri | various | various | various | various | various | unclassified | unclassified | unclassified | unclassified | unclassified | 1,2 | 0,748331477 | 0,000118862 | 7,79E-05 | 1 - | 8,44E-05 - | 2 - | 0,000209587 - | 2 - | 0,000199239 - | 1 - | 0,000101091 - | 0 - | 0 - |
| 9349 | 1 | 3um_FIJJAFBF_17217 | Bacteria | Proteobacteri | Alphaproteob | Rhodobactera | Rhodobactera | unclassified | Rhodobactera | unclassified | unclassified | unclassified | unclassified | unclassified | 3 | 1,095445115 | 9,15E-05 | 3,65E-05 | 3 - | 7,62E-05 - | 5 - | 0,000157654 - | 2 - | 5,99E-05 - | 2 - | 6,08E-05 - | 3 - | 0,000102793 - |
| 9350 | 1 | 3um_FIJJAFBF_467719 | Eukaryota | unclassified | Cryptophyta | Pyrenomonad | Geminigerace | Teleaulax | Teleaulax amp | metabolism | Energy produc | COG0711 | F(1)F(0) ATP s | 63737.Npun_ | 1,4 | 0,489897949 | 0,000104513 | 2,99E-05 | 2 - | 0,000128242 - | 1 - | 7,96E-05 - | 2 - | 0,000151377 - | 1 - | 7,68E-05 - | 1 - | 8,65E-05 - |
| 9351 | 1 | 0.2um_NOKCPHJN_865902 | Bacteria | Proteobacteri | Alphaproteob | Rhizobiales | unclassified | unclassified | Rhizobiales ba | metabolism | Secondary me | 01Q86 | extracellular s | 244592.SADFL | 2,8 | 0,4 | 0,000114544 | 1,85E-05 | 3 - | 0,000103093 - | 2 - | 8,53E-05 - | 3 - | 0,000121691 - | 3 - | 0,000123488 - | 3 - | 0,000139109 - |
| 9352 | 1 | 3um_FIJJAFBF_386738 | Bacteria | Verrucomicrol | Verrucomicrol | Verrucomicrol | Rubritaleacea | Rubritalea | Rubritalea squ | metabolism | Inorganic ion t | 1DH50 | phosphate bin | 240016.VspiD | 1 | 1,095445115 | 4,70E-05 | 4,68E-05 | 3 - | 0,000127059 - | 0 - | 0 - | 0 - | 0 - | 1 - | 5,07E-05 - | 1 - | 5,71E-05 - |
| 9353 | 1 | 0.2um_NOKCPHJN_733661 | Eukaryota | unclassified | Dinophyceae | Suessiales | Symbiodiniace | Symbiodinium | Symbiodinium | unclassified | unclassified | unclassified | unclassified | unclassified | 2 | 0 | 0,000232249 | 2,21E-05 | 2 - | 0,000194537 - | 2 - | 0,000241557 - | 2 - | 0,000229631 - | 2 - | 0,000233023 - | 2 - | 0,000262499 - |

**PROPHANE JOB INFORMATION**

**program versions**

prophane: 3.1.1
blast+: 2.7.1
diamond: 0.8.22
hmmer: 3.1b2
krona: 2.7

**tasks**

task: *taxonomic assignment*
task comment: tax_from_nr_20180808_qcover90
algorithm: diamond blastp
query: sequences without taxonomic classification (missing_taxa.faa)
database: NCBI NR database
database version: 2018-08-02
database comment: downloaded at 2018-08-02
database scope: tax
parameter:
- evalue 0.01
- query-cover 0.9
- max-target-seqs 1

task: *functional assignment*
task comment: fun_from_eggNog_4.5.1
algorithm: emapper
query: all sequences (all.faa)
database: eggmap database
database version: 4.5.1
database comment: includes euk, viruses, bakt, arch; downloaded at 2018-07-31
database scope: func
parameter:
- m diamond

| | | | taxonomic assignment | | | | | | | functional assignment | | | | | quantitative data mean | | | | quantitative data replicate 1 | | | | quantitative data replicate 2 | | | | quantitative data replicate 3 | | | | quantitative data replicate 4 | | | | quantitative data replicate 5 | | | |
|---|---|---|---|---|---|---|---|---|---|---|---|---|---|---|---|---|---|---|---|---|---|---|---|---|---|---|---|---|---|---|---|---|---|---|---|---|---|---|
| protein group | members_count | protein group: members_identifier | superkingdom | phylum | class | order | family | genus | species | main role | sub role | og | descr | prot | exclusive spetrum count | exclusive spetrum count, sd | NSAF | NSAF, sd | exclusive spetrum count | exclusive spetrum count, sd | NSAF | NSAF, sd | exclusive spetrum count | exclusive spetrum count, sd | NSAF | NSAF, sd | exclusive spetrum count | exclusive spetrum count, sd | NSAF | NSAF, sd | exclusive spetrum count | exclusive spetrum count, sd | NSAF | NSAF, sd | exclusive spetrum count | exclusive spetrum count, sd | NSAF | NSAF, sd |
| 0 | 1 | 0.2um_NOKCPHJN_245802 | Bacteria | Proteobacteri | Alphaproteob | Rhodobactera | Rhodobactera | Planktomarina | Planktomarina | information st | Transcription | COG0250 | Participates in | 351016.RAZW | 1 | 0 | 0,000491059 | 4,80E-05 | 1 | - | 0,000495445 | - | 1 | - | 0,000582881 | - | 1 | - | 0,000459756 | - | 1 | - | 0,000456866 | - | 1 | - | 0,000460347 | - |
| 1 | 1 | 3um_FIJJAFBF_445235 | unclassified | unclassified | unclassified | unclassified | unclassified | unclassified | unclassified | unclassified | unclassified | unclassified | unclassified | unclassified | 0,4 | 0,489897949 | 0,000579436 | 0,000709667 | 0 | - | 0 | - | 0 | - | 0 | - | 1 | - | 0,001453156 | - | 1 | - | 0,001444024 | - | 0 | - | 0 | - |
| 2 | 1 | 3um_FIJJAFBF_17424 | Eukaryota | Ascomycota | Eurotiomycete | Eurotiales | Aspergillaceae | Penicillium | Penicillium ste | unclassified | unclassified | unclassified | unclassified | unclassified | 0,6 | 0,489897949 | 8,46E-05 | 6,91E-05 | 0 | - | 0 | - | 0 | - | 0 | - | 1 | - | 0,000141279 | - | 1 | - | 0,000140391 | - | 1 | - | 0,000141461 | - |
| 3 | 2 | 3um_FIJJAFBF_135233;3um_FI | Eukaryota | Eustigmatoph | unclassified | Eustigmatales | Monodopsida | Nannochlorop | Nannochlorop | unclassified | unclassified | unclassified | unclassified | unclassified | 1,2 | 0,748331477 | 0,000248123 | 0,000151862 | 1 | - | 0,000220336 | - | 0 | - | 0 | - | 1 | - | 0,000204464 | - | 2 | - | 0,000406359 | - | 2 | - | 0,000409455 | - |
| 4 | 1 | 3um_FIJJAFBF_128453 | Bacteria | Bacteroidetes | Flavobacteriia | Flavobacterial | Flavobacteriac | Polaribacter | Polaribacter s | poorly charact | Function unkn | 11W7M | NA | 398720.MED2 | 0,8 | 0,4 | 0,000444735 | 0,000228939 | 0 | - | 0 | - | 1 | - | 0,000661345 | - | 1 | - | 0,000521646 | - | 1 | - | 0,000518368 | - | 1 | - | 0,000522317 | - |
| 5 | 1 | 0.2um_NOKCPHJN_551945 | Bacteria | Planctomycete | unclassified | unclassified | unclassified | unclassified | Planctomycete | poorly charact | Function unkn | 0ZPAX | Caulimovirus v | 4787.PITG_10 | 0,4 | 0,489897949 | 2,49E-05 | 3,04E-05 | 0 | - | 0 | - | 0 | - | 0 | - | 0 | - | 0 | - | 1 | - | 6,19E-05 | - | 1 | - | 6,24E-05 | - |
| 6 | 1 | 0.2um_NOKCPHJN_702692 | Bacteria | Bacteroidetes | Flavobacteriia | Flavobacterial | Flavobacteriac | Maribacter | Maribacter an | metabolism | Secondary me | 0P8JT | Phytoene des | 313595.P7007 | 0,8 | 0,4 | 0,000141589 | 7,29E-05 | 0 | - | 0 | - | 1 | - | 0,000210551 | - | 1 | - | 0,000166075 | - | 1 | - | 0,000165031 | - | 1 | - | 0,000166289 | - |
| 7 | 1 | 0.2um_NOKCPHJN_329715 | Eukaryota | unclassified | Oomycetes | Pythiales | Pythiaceae | Pythium | Pythium insidi | metabolism | Energy produc | COG0055 | Produces ATP | 65071.PYU1_ | 2 | 0 | 0,000354765 | 3,47E-05 | 2 | - | 0,000357934 | - | 2 | - | 0,000421102 | - | 2 | - | 0,00033215 | - | 2 | - | 0,000330063 | - | 2 | - | 0,000332578 | - |
| 8 | 1 | 0.2um_NOKCPHJN_809592 | unclassified | unclassified | unclassified | unclassified | unclassified | unclassified | unclassified | unclassified | unclassified | unclassified | unclassified | unclassified | 0,8 | 0,4 | 8,82E-05 | 4,52E-05 | 1 | - | 0,00010948 | - | 1 | - | 0,000128801 | - | 0 | - | 0 | - | 1 | - | 0,000100955 | - | 1 | - | 0,000101725 | - |
| 9 | 1 | 0.2um_NOKCPHJN_152857 | Bacteria | Bacteroidetes | Flavobacteriia | Flavobacterial | Crocinitomica | unclassified | Crocinitomica | unclassified | unclassified | unclassified | unclassified | unclassified | 0,2 | 0,4 | 0,00010197 | 0,000203939 | 1 | - | 0,000509848 | - | 0 | - | 0 | - | 0 | - | 0 | - | 0 | - | 0 | - | 0 | - | 0 | - |
| 10 | 1 | 0.2um_NOKCPHJN_566414 | unclassified | unclassified | unclassified | unclassified | unclassified | unclassified | unclassified | unclassified | unclassified | unclassified | unclassified | unclassified | 0,6 | 0,489897949 | 4,10E-05 | 3,34E-05 | 0 | - | 0 | - | 0 | - | 0 | - | 1 | - | 6,84E-05 | - | 1 | - | 6,80E-05 | - | 1 | - | 6,85E-05 | - |
| 11 | 1 | 3um_FIJJAFBF_444552 | Bacteria | Proteobacteri | Gammaprotec | Cellvibrionales | Porticoccacea | unclassified | SAR92 bacteri | poorly charact | Function unkn | 16R8I | Lipoprotein | 207949.RED6 | 2,6 | 1,019803903 | 0,000512863 | 0,000182519 | 3 | - | 0,000614677 | - | 1 | - | 0,000241051 | - | 2 | - | 0,000380265 | - | 3 | - | 0,000566813 | - | 4 | - | 0,000761509 | - |
| 12 | 1 | 3um_FIJJAFBF_594162 | Eukaryota | unclassified | Pelagophycea | Pelagomonad | unclassified | Pelagomonas | uncultured Pe | unclassified | unclassified | unclassified | unclassified | unclassified | 6,2 | 0,748331477 | 0,001660726 | 0,000199001 | 5 | - | 0,00135749 | - | 6 | - | 0,001916469 | - | 7 | - | 0,001763583 | - | 7 | - | 0,0017525 | - | 6 | - | 0,001513588 | - |
| 13 | 1 | 3um_FIJJAFBF_507105 | Eukaryota | Phaeophyceae | unclassified | Ectocarpales | Ectocarpaceae | Ectocarpus | Ectocarpus sil | unclassified | unclassified | unclassified | unclassified | unclassified | 1,6 | 0,489897949 | 0,000792069 | 0,00021792 | 2 | - | 0,001013802 | - | 1 | - | 0,000596358 | - | 2 | - | 0,000940772 | - | 1 | - | 0,00046743 | - | 2 | - | 0,000941983 | - |
| 14 | 1 | 3um_FIJJAFBF_161250 | Bacteria | Proteobacteri | Alphaproteob | Rhodospirillal | unclassified | unclassified | Rhodospirillal | unclassified | unclassified | unclassified | unclassified | unclassified | 1 | 0 | 0,000165873 | 1,62E-05 | 1 | - | 0,000167355 | - | 1 | - | 0,000196889 | - | 1 | - | 0,000155299 | - | 1 | - | 0,000154323 | - | 1 | - | 0,000155499 | - |
| 15 | 2 | 0.2um_NOKCPHJN_95248;3um | Eukaryota | unclassified | Chrysophycea | Chromulinales | Chromulinace | Ochromonas | Ochromonas s | unclassified | unclassified | unclassified | unclassified | unclassified | 0,2 | 0,4 | 6,35E-05 | 0,000127093 | 1 | - | 0,000317731 | - | 0 | - | 0 | - | 0 | - | 0 | - | 0 | - | 0 | - | 0 | - | 0 | - |
| 16 | 1 | 0.2um_NOKCPHJN_1085300 | Bacteria | Proteobacteri | Gammaprotec | unclassified | unclassified | unclassified | Gammaprotec | information st | Replication, re | COG0827 | DNA Methylas | 675815.VOA_ | 0,2 | 0,4 | 3,36E-05 | 6,72E-05 | 1 | - | 0,000167996 | - | 0 | - | 0 | - | 0 | - | 0 | - | 0 | - | 0 | - | 0 | - | 0 | - |
| 17 | 1 | 0.2um_NOKCPHJN_997351 | Eukaryota | unclassified | Oligohymenop | Philasterida | Philasteridae | Miamiensis | Miamiensis av | cellular proces | Signal transdu | KOG0699 | protein phosp | 3055.EDP008 | 0,6 | 0,489897949 | 0,000194226 | 0,000158833 | 1 | - | 0,000339899 | - | 0 | - | 0 | - | 1 | - | 0,000315414 | - | 0 | - | 0 | - | 1 | - | 0,00031582 | - |
| 18 | 1 | 3um_FIJJAFBF_715061 | Eukaryota | unclassified | Rhodellophyc | Rhodellales | Rhodellaceae | Corynoplastis | Corynoplastis | unclassified | unclassified | unclassified | unclassified | unclassified | 1,2 | 0,4 | 0,001322231 | 0,000461634 | 2 | - | 0,002220098 | - | 1 | - | 0,001305948 | - | 1 | - | 0,001030085 | - | 1 | - | 0,001023612 | - | 1 | - | 0,001031411 | - |
| 19 | 1 | 0.2um_NOKCPHJN_1013360 | Eukaryota | unclassified | Pelagophycea | Pelagomonad | unclassified | Aureococcus | Aureococcus a | unclassified | unclassified | unclassified | unclassified | unclassified | 1 | 0 | 0,000673779 | 6,59E-05 | 1 | - | 0,000679797 | - | 1 | - | 0,000799767 | - | 1 | - | 0,000630827 | - | 1 | - | 0,000626863 | - | 1 | - | 0,000631639 | - |
| 20 | 1 | 3um_FIJJAFBF_91013 | Eukaryota | unclassified | Florideophyce | Gigartinales | Gigartinaceae | Chondrus | Chondrus cris | information st | Translation, ri | KOG3280 | ribosomal pro | 35128.Thaps3 | 0,8 | 0,4 | 0,000619472 | 0,000310621 | 1 | - | 0,000819569 | - | 0 | - | 0 | - | 1 | - | 0,00076053 | - | 1 | - | 0,000755751 | - | 1 | - | 0,000761509 | - |
| 21 | 2 | 3um_FIJJAFBF_00233;3um_FIJ | Bacteria | Bacteroidetes | Flavobacteriia | Flavobacterial | Flavobacteriac | various | various | unclassified | unclassified | unclassified | unclassified | unclassified | 0,6 | 0,489897949 | 0,0005207 | 0,000425887 | 1 | - | 0,000913478 | - | 0 | - | 0 | - | 1 | - | 0,000847674 | - | 1 | - | 0,000842347 | - | 0 | - | 0 | - |
| 22 | 1 | 0.2um_NOKCPHJN_1058181 | Bacteria | Proteobacteri | Alphaproteob | Rhodospirillal | Rhodospirillac | Magnetovibrio | Magnetovibrio | metabolism | Carbohydrate | 05CV1 | Transketolase | 342108.amb1 | 0,4 | 0,489897949 | 6,15E-05 | 7,53E-05 | 0 | - | 0 | - | 0 | - | 0 | - | 1 | - | 0,000154123 | - | 1 | - | 0,000153154 | - | 0 | - | 0 | - |
| 23 | 1 | 3um_FIJJAFBF_391433 | Eukaryota | Apicomplexa | Conoidasida | Eucoccidiorida | Sarcocystidae | Besnoitia | Besnoitia besn | unclassified | unclassified | unclassified | unclassified | unclassified | 0,8 | 0,4 | 0,000257913 | 0,000132768 | 0 | - | 0 | - | 1 | - | 0,000383531 | - | 1 | - | 0,000302516 | - | 1 | - | 0,000300615 | - | 1 | - | 0,000302905 | - |
| 24 | 1 | 3um_FIJJAFBF_49815 | Eukaryota | Bacillariophyta | Bacillariophyc | Naviculales | Naviculaceae | Fistulifera | Fistulifera sola | unclassified | unclassified | unclassified | unclassified | unclassified | 3,6 | 0,8 | 0,00219574 | 0,000405983 | 3 | - | 0,001865827 | - | 3 | - | 0,002195104 | - | 5 | - | 0,0028857 | - | 3 | - | 0,001720539 | - | 4 | - | 0,002311532 | - |
| 25 | 1 | 3um_FIJJAFBF_640408 | unclassified | unclassified | unclassified | unclassified | unclassified | unclassified | unclassified | unclassified | unclassified | unclassified | unclassified | unclassified | 0,8 | 0,4 | 0,000325722 | 0,000167674 | 0 | - | 0 | - | 1 | - | 0,000484366 | - | 1 | - | 0,00038205 | - | 1 | - | 0,00037965 | - | 1 | - | 0,000382542 | - |
| 26 | 1 | 0.2um_NOKCPHJN_349772 | Bacteria | unclassified | unclassified | unclassified | unclassified | unclassified | uncultured ba | poorly charact | Function unkn | 08PIQ | CHU large pro | 50743.SCB49_ | 1,6 | 0,489897949 | 0,000105085 | 2,87E-05 | 2 | - | 0,000134603 | - | 1 | - | 7,92E-05 | - | 1 | - | 6,25E-05 | - | 2 | - | 0,000124122 | - | 2 | - | 0,000125068 | - |
| 27 | 1 | 0.2um_NOKCPHJN_757859 | Eukaryota | unclassified | Pelagophycea | Pelagomonad | unclassified | Aureococcus | Aureococcus a | unclassified | unclassified | unclassified | unclassified | unclassified | 0,2 | 0,4 | 2,44E-05 | 4,89E-05 | 0 | - | 0 | - | 0 | - | 0 | - | 1 | - | 0,000122187 | - | 0 | - | 0 | - | 0 | - | 0 | - |
| 28 | 1 | 3um_FIJJAFBF_118613 | unclassified | unclassified | unclassified | unclassified | unclassified | unclassified | unclassified | unclassified | unclassified | unclassified | unclassified | unclassified | 0,2 | 0,4 | 0,000123557 | 0,000247114 | 0 | - | 0 | - | 1 | - | 0,000617784 | - | 0 | - | 0 | - | 0 | - | 0 | - | 0 | - | 0 | - |
| 29 | 1 | 3um_FIJJAFBF_255966 | Bacteria | Candidatus W | unclassified | unclassified | unclassified | unclassified | Candidatus W | poorly charact | Function unkn | 1EB8T | NA | 436017.A4S9 | 0,6 | 0,489897949 | 3,55E-05 | 2,90E-05 | 0 | - | 0 | - | 0 | - | 0 | - | 1 | - | 5,92E-05 | - | 1 | - | 5,89E-05 | - | 1 | - | 5,93E-05 | - |
| 30 | 1 | 0.2um_NOKCPHJN_212384 | Eukaryota | unclassified | Oomycetes | Peronosporale | Peronosporac | Phytophthora | Phytophthora | poorly charact | Function unkn | COG5307 | and Sec7 dom | 65071.PYU1_ | 0,4 | 0,489897949 | 2,15E-05 | 2,65E-05 | 1 | - | 4,94E-05 | - | 1 | - | 5,81E-05 | - | 0 | - | 0 | - | 0 | - | 0 | - | 0 | - | 0 | - |
| 31 | 1 | 0.2um_NOKCPHJN_367359 | Bacteria | Firmicutes | Bacilli | Lactobacillales | Lactobacillace | Lactobacillus | Lactobacillus | poorly charact | Function unkn | 0NVRZ | NA | 170187.SP_17 | 0,6 | 0,489897949 | 6,02E-05 | 5,00E-05 | 0 | - | 0 | - | 1 | - | 0,000116708 | - | 1 | - | 9,21E-05 | - | 0 | - | 0 | - | 1 | - | 9,22E-05 | - |
| 32 | 1 | 3um_FIJJAFBF_257121 | Eukaryota | unclassified | Cryptophyta | Pyrenomonad | Geminigerace | Guillardia | Guillardia thet | unclassified | unclassified | unclassified | unclassified | unclassified | 0,6 | 0,489897949 | 0,000413117 | 0,000337894 | 1 | - | 0,000724743 | - | 0 | - | 0 | - | 1 | - | 0,000672535 | - | 1 | - | 0,000668309 | - | 0 | - | 0 | - |
| 33 | 1 | 3um_FIJJAFBF_62953 | Bacteria | Firmicutes | Clostridia | Clostridiales | unclassified | unclassified | Clostridiales b | metabolism | Inorganic ion t | COG1055 | arsenical pum | 431895.XP_00 | 0,2 | 0,4 | 0,000147386 | 0,000294771 | 0 | - | 0 | - | 1 | - | 0,000736928 | - | 0 | - | 0 | - | 0 | - | 0 | - | 0 | - | 0 | - |
| 34 | 1 | 3um_FIJJAFBF_203625 | Bacteria | Bacteroidetes | Flavobacteriia | Flavobacterial | Flavobacteriac | Winogradskye | Winogradskye | unclassified | unclassified | unclassified | unclassified | unclassified | 1 | 0 | 0,000595325 | 5,82E-05 | 1 | - | 0,000600643 | - | 1 | - | 0,000706643 | - | 1 | - | 0,000557375 | - | 1 | - | 0,000553872 | - | 1 | - | 0,000558092 | - |
| 35 | 1 | 3um_FIJJAFBF_80731 | Bacteria | Proteobacteri | Gammaprotec | Alteromonada | unclassified | unclassified | gamma protec | information st | Translation, ri | 05KK9 | This protein bi | 523791.Kkor_ | 1 | 0,632455532 | 0,000429461 | 0,000266726 | 1 | - | 0,000456739 | - | 0 | - | 0 | - | 1 | - | 0,000423837 | - | 2 | - | 0,000842347 | - | 1 | - | 0,000424383 | - |
| 36 | 1 | 3um_FIJJAFBF_08157 | Bacteria | Bacteroidetes | Flavobacteriia | Flavobacterial | Flavobacteriac | Polaribacter | various | metabolism | Inorganic ion t | 0P8Q4 | tonB-depende | 867900.Celly_ | 29,6 | 5,003998401 | 0,002600688 | 0,000277937 | 25 | - | 0,002246256 | - | 23 | - | 0,002431258 | - | 34 | - | 0,002834846 | - | 30 | - | 0,002485615 | - | 36 | - | 0,003005465 | - |
| 37 | 1 | 3um_FIJJAFBF_664428 | Bacteria | Proteobacteri | Alphaproteob | Rhodobactera | Rhodobactera | unclassified | Rhodobactera | unclassified | unclassified | unclassified | unclassified | unclassified | 0,8 | 0,4 | 7,29E-05 | 3,75E-05 | 0 | - | 0 | - | 1 | - | 0,000108372 | - | 1 | - | 8,55E-05 | - | 1 | - | 8,49E-05 | - | 1 | - | 8,56E-05 | - |
| 38 | 1 | 0.2um_NOKCPHJN_502677 | Eukaryota | Streptophyta | unclassified | Rosales | Rosaceae | Prunus | Prunus mume | unclassified | unclassified | unclassified | unclassified | unclassified | 0,2 | 0,4 | 4,09E-05 | 8,18E-05 | 0 | - | 0 | - | 0 | - | 0 | - | 1 | - | 0,000204464 | - | 0 | - | 0 | - | 0 | - | 0 | - |
| 39 | 1 | 3um_FIJJAFBF_566757 | Eukaryota | Bacillariophyta | Bacillariophyc | Naviculales | Naviculaceae | Fistulifera | Fistulifera sola | unclassified | unclassified | unclassified | unclassified | unclassified | 1 | 0 | 0,000279477 | 2,73E-05 | 1 | - | 0,000281974 | - | 1 | - | 0,000331736 | - | 1 | - | 0,000261662 | - | 1 | - | 0,000260017 | - | 1 | - | 0,000261998 | - |
| 40 | 1 | 0.2um_NOKCPHJN_532347 | Bacteria | Bacteroidetes | Flavobacteriia | Flavobacterial | Crocinitomica | unclassified | Crocinitomica | metabolism | Inorganic ion t | 0XP5Y | tonB-depende | 411477.PARM | 0,2 | 0,4 | 2,04E-05 | 4,08E-05 | 0 | - | 0 | - | 0 | - | 0 | - | 0 | - | 0 | - | 0 | - | 0 | - | 1 | - | 0,000101979 | - |
| 41 | 1 | 0.2um_NOKCPHJN_933811 | unclassified | unclassified | unclassified | unclassified | unclassified | unclassified | unclassified | unclassified | unclassified | unclassified | unclassified | unclassified | 0,4 | 0,489897949 | 0,000165054 | 0,000204609 | 0 | - | 0 | - | 1 | - | 0,000462645 | - | 0 | - | 0 | - | 1 | - | 0,000362625 | - | 0 | - | 0 | - |
| 42 | 1 | 3um_FIJJAFBF_135696 | Bacteria | Bacteroidetes | Flavobacteriia | Flavobacterial | Flavobacteriac | Ulvibacter | Ulvibacter sp. | information st | Translation, ri | 05G6W | One of the pri | 983544.Lacal_ | 0,8 | 0,4 | 0,000345168 | 0,000177684 | 0 | - | 0 | - | 1 | - | 0,000513283 | - | 1 | - | 0,000404859 | - | 1 | - | 0,000402315 | - | 1 | - | 0,000405381 | - |
| 43 | 1 | 3um_FIJJAFBF_235330 | Viruses | unclassified | unclassified | unclassified | unclassified | unclassified | uncultured M | unclassified | unclassified | unclassified | unclassified | unclassified | 0,4 | 0,489897949 | 0,000236609 | 0,000290111 | 1 | - | 0,000613244 | - | 0 | - | 0 | - | 0 | - | 0 | - | 0 | - | 0 | - | 1 | - | 0,000569801 | - |
| 44 | 1 | 3um_FIJJAFBF_222229 | Bacteria | Proteobacteri | Alphaproteob | Rhodobactera | Rhodobactera | various | various | unclassified | unclassified | unclassified | unclassified | unclassified | 0,8 | 0,4 | 0,000373004 | 0,000192013 | 0 | - | 0 | - | 1 | - | 0,000554677 | - | 1 | - | 0,000437509 | - | 1 | - | 0,00043476 | - | 1 | - | 0,000438073 | - |
| 45 | 1 | 0.2um_NOKCPHJN_718865 | Bacteria | Proteobacteri | Gammaprotec | Cellvibrionales | Porticoccacea | unclassified | gamma protec | poorly charact | Function unkn | 17NHM | NA | 203122.Sde_0 | 1 | 0 | 0,000371442 | 3,63E-05 | 1 | - | 0,00037476 | - | 1 | - | 0,000440897 | - | 1 | - | 0,000347764 | - | 1 | - | 0,000345578 | - | 1 | - | 0,000348212 | - |
| 46 | 1 | 3um_FIJJAFBF_526970 | unclassified | unclassified | unclassified | unclassified | unclassified | unclassified | unclassified | unclassified | unclassified | unclassified | unclassified | unclassified | 1 | 0 | 0,000127072 | 1,24E-05 | 1 | - | 0,000128207 | - | 1 | - | 0,000150833 | - | 1 | - | 0,000118972 | - | 1 | - | 0,000118224 | - | 1 | - | 0,000119125 | - |
| 47 | 1 | 0.2um_NOKCPHJN_409669 | Bacteria | Proteobacteri | Gammaprotec | unclassified | unclassified | unclassified | Gammaprotec | unclassified | unclassified | unclassified | unclassified | unclassified | 1,6 | 0,8 | 0,000185499 | 8,41E-05 | 1 | - | 0,0001198 | - | 1 | - | 0,000140942 | - | 2 | - | 0,000222341 | - | 1 | - | 0,000110472 | - | 3 | - | 0,000333941 | - |
| 48 | 1 | 3um_FIJJAFBF_28972 | Bacteria | Proteobacteri | Gammaprotec | unclassified | unclassified | unclassified | Gammaprotec | poorly charact | Function unkn | 070MQ | NA | 425104.Ssed_ | 3,2 | 0,748331477 | 0,000618641 | 0,000138624 | 3 | - | 0,000587236 | - | 3 | - | 0,00069087 | - | 4 | - | 0,000726578 | - | 2 | - | 0,000361006 | - | 4 | - | 0,000727513 | - |
| 49 | 1 | 3um_FIJJAFBF_566674 | Eukaryota | Nematoda | Chromadorea | Rhabditida | Diploscapteric | Diploscapter | Diploscapter p | poorly charact | Function unkn | 0IPY7 | Reverse trans | 7897.ENSLAC | 0,2 | 0,4 | 2,87E-05 | 5,74E-05 | 0 | - | 0 | - | 0 | - | 0 | - | 0 | - | 0 | - | 0 | - | 0 | - | 1 | - | 0,000143453 | - |
| 50 | 1 | 0.2um_NOKCPHJN_503156 | Viruses | unclassified | unclassified | unclassified | various | various | various | unclassified | unclassified | unclassified | unclassified | unclassified | 1 | 0 | 0,000172114 | 1,68E-05 | 1 | - | 0,000173651 | - | 1 | - | 0,000204297 | - | 1 | - | 0,000161142 | - | 1 | - | 0,000160129 | - | 1 | - | 0,000161349 | - |
| 51 | 1 | 0.2um_NOKCPHJN_822424 | Bacteria | Bacteroidetes | unclassified | unclassified | unclassified | unclassified | Bacteroidetes | metabolism | Inorganic ion t | COG0659 | sulfate transp | 926562.Oweh | 0,6 | 0,489897949 | 0,00015995 | 0,000132822 | 0 | - | 0 | - | 1 | - | 0,000310753 | - | 0 | - | 0 | - | 1 | - | 0,00024357 | - | 1 | - | 0,000245426 | - |
| 52 | 1 | 3um_FIJJAFBF_203258 | Bacteria | Proteobacteri | Gammaprotec | Arenicellales | Arenicellaceae | Arenicella | Arenicella xan | metabolism | Inorganic ion t | 16TQC | TONB-depend | 247639.MGP2 | 1,6 | 1,019803903 | 0,000141493 | 8,43E-05 | 2 | - | 0,000181373 | - | 1 | - | 0,000106691 | - | 0 | - | 0 | - | 3 | - | 0,000250875 | - | 2 | - | 0,000168524 | - |
| 53 | 1 | 3um_FIJJAFBF_191493 | Bacteria | Bacteroidetes | Flavobacteriia | Flavobacterial | Flavobacteriac | Polaribacter | Polaribacter s | unclassified | unclassified | unclassified | unclassified | unclassified | 0,6 | 0,489897949 | 5,97E-05 | 4,88E-05 | 1 | - | 0,000104647 | - | 0 | - | 0 | - | 0 | - | 0 | - | 1 | - | 9,65E-05 | - | 1 | - | 9,72E-05 | - |
| 54 | 1 | 3um_FIJJAFBF_226545 | Bacteria | Proteobacteri | Alphaproteob | Rhodospirillal | Rhodospirillac | Candidatus En | Candidatus En | unclassified | unclassified | unclassified | unclassified | unclassified | 0,4 | 0,8 | 6,73E-05 | 0,00013468 | 0 | - | 0 | - | 0 | - | 0 | - | 0 | - | 0 | - | 0 | - | 0 | - | 2 | - | 0,0003367 | - |
| 55 | 1 | 3um_FIJJAFBF_331695 | Bacteria | Bacteroidetes | Flavobacteriia | Flavobacterial | Crocinitomica | unclassified | Crocinitomica | poorly charact | Function unkn | 09AR5 | Inherit from b | 313606.M231 | 1 | 0 | 0,000167471 | 1,64E-05 | 1 | - | 0,000168967 | - | 1 | - | 0,000198786 | - | 1 | - | 0,000156795 | - | 1 | - | 0,00015581 | - | 1 | - | 0,000156997 | - |
| 56 | 1 | 3um_FIJJAFBF_358211 | Eukaryota | Chlorophyta | Mamiellophyc | Mamiellales | Mamiellaceae | Micromonas | Micromonas c | unclassified | unclassified | unclassified | unclassified | unclassified | 1,6 | 0,489897949 | 0,00019027 | 4,66E-05 | 1 | - | 0,000122993 | - | 1 | - | 0,000144698 | - | 2 | - | 0,000228266 | - | 2 | - | 0,000226831 | - | 2 | - | 0,00022856 | - |
| 57 | 1 | 3um_FIJJAFBF_390144 | Eukaryota | Eustigmatoph | unclassified | Eustigmatales | Monodopsida | Nannochlorop | Nannochlorop | unclassified | unclassified | unclassified | unclassified | unclassified | 1,2 | 0,748331477 | 0,001429966 | 0,000910939 | 2 | - | 0,002505539 | - | 0 | - | 0 | - | 2 | - | 0,00232505 | - | 1 | - | 0,001155219 | - | 1 | - | 0,001164021 | - |
| 58 | 1 | 0.2um_NOKCPHJN_888294 | Eukaryota | unclassified | Pelagophycea | Pelagomonad | unclassified | Aureococcus | Aureococcus a | cellular proces | Signal transdu | KOG0613 | fibronectin ty | 65071.PYU1_ | 0,8 | 0,4 | 0,000116696 | 5,85E-05 | 1 | - | 0,000154391 | - | 0 | - | 0 | - | 1 | - | 0,000143269 | - | 1 | - | 0,000142369 | - | 1 | - | 0,000143453 | - |
| 59 | 1 | 3um_FIJJAFBF_815479 | Eukaryota | unclassified | Cryptophyta | Pyrenomonad | Geminigerace | Guillardia | Guillardia thet | metabolism | Lipid transpor | COG0439 | acetyl-CoA ca | 2850.Phatr54 | 1 | 0,632455532 | 0,00063463 | 0,000387193 | 0 | - | 0 | - | 1 | - | 0,000764221 | - | 1 | - | 0,000602791 | - | 1 | - | 0,000599003 | - | 2 | - | 0,001207133 | - |
| 60 | 1 | 0.2um_NOKCPHJN_100732 | Bacteria | Bacteroidetes | Flavobacteriia | Flavobacterial | Flavobacteriac | Aureitalea | Aureitalea ma | unclassified | unclassified | unclassified | unclassified | unclassified | 7,8 | 0,748331477 | 0,000766604 | 0,00010196 | 7 | - | 0,000694408 | - | 8 | - | 0,000933664 | - | 8 | - | 0,000736441 | - | 9 | - | 0,00082329 | - | 7 | - | 0,000645215 | - |
| 61 | 1 | 0.2um_NOKCPHJN_206300 | Eukaryota | unclassified | Spirotrichea | Sporadotrichi | Oxytrichidae | Oxytricha | Oxytricha trifa | metabolism | Energy produc | COG0377 | NDH-1 shuttle | 35128.Thaps2 | 0,2 | 0,4 | 0,0001076 | 0,0002152 | 1 | - | 0,000537999 | - | 0 | - | 0 | - | 0 | - | 0 | - | 0 | - | 0 | - | 0 | - | 0 | - |
| 62 | 1 | 0.2um_NOKCPHJN_80485 | Bacteria | Bacteroidetes | Flavobacteriia | Flavobacterial | Flavobacteriac | Formosa | Formosa sp. H | cellular proces | Posttranslatio | 05E6M | Sulphatase-m | 391603.FBALC | 0,6 | 0,489897949 | 0,000108805 | 8,88E-05 | 0 | - | 0 | - | 0 | - | 0 | - | 1 | - | 0,000181645 | - | 1 | - | 0,000180503 | - | 1 | - | 0,000181878 | - |
| 63 | 1 | 3um_FIJJAFBF_311757 | Bacteria | Bacteroidetes | Flavobacteriia | Flavobacterial | Flavobacteriac | Polaribacter | Polaribacter a | information st | Translation, ri | COG0050 | This protein p | 313603.FB217 | 5,2 | 1,166190379 | 0,001158803 | 0,000336046 | 7 | - | 0,001554068 | - | 6 | - | 0,001567138 | - | 4 | - | 0,000824068 | - | 5 | - | 0,001023612 | - | 4 | - | 0,000825129 | - |
| 64 | 1 | 3um_FIJJAFBF_506614 | unclassified | unclassified | unclassified | unclassified | unclassified | unclassified | unclassified | unclassified | unclassified | unclassified | unclassified | unclassified | 0,6 | 0,489897949 | 0,00037677 | 0,000308112 | 1 | - | 0,000659352 | - | 0 | - | 0 | - | 1 | - | 0,000611855 | - | 0 | - | 0 | - | 1 | - | 0,000612643 | - |
| 65 | 6 | 0.2um_NOKCPHJN_1008923;0. | unclassified | unclassified | unclassified | unclassified | unclassified | unclassified | unclassified | unclassified | unclassified | unclassified | unclassified | unclassified | 1 | 0 | 0,00053986 | 5,28E-05 | 1 | - | 0,000544682 | - | 1 | - | 0,000640807 | - | 1 | - | 0,000505446 | - | 1 | - | 0,000502269 | - | 1 | - | 0,000506096 | - |
| 66 | 1 | 3um_FIJJAFBF_37832 | Bacteria | Proteobacteri | unclassified | unclassified | unclassified | unclassified | Proteobacteri | unclassified | unclassified | unclassified | unclassified | unclassified | 3,8 | 0,748331477 | 0,001068657 | 0,000307607 | 3 | - | 0,000840516 | - | 5 | - | 0,001648081 | - | 3 | - | 0,000779969 | - | 4 | - | 0,001033423 | - | 4 | - | 0,001041297 | - |
| 67 | 1 | 3um_FIJJAFBF_520274 | Bacteria | Firmicutes | Clostridia | Clostridiales | Clostridiaceae | Clostridium | Clostridium bo | cellular proces | Cell wall/mem | 125KQ | tolA protein | 322710.Avin_ | 0,6 | 0,489897949 | 0,000144215 | 0,000117753 | 0 | - | 0 | - | 0 | - | 0 | - | 1 | - | 0,00024076 | - | 1 | - | 0,000239247 | - | 1 | - | 0,00024107 | - |
| 68 | 1 | 0.2um_NOKCPHJN_932941 | Eukaryota | Arthropoda | Arachnida | Trombidiform | Tetranychidae | Tetranychus | Tetranychus u | unclassified | unclassified | unclassified | unclassified | unclassified | 1 | 0 | 0,000353323 | 3,46E-05 | 1 | - | 0,000356479 | - | 1 | - | 0,00041939 | - | 1 | - | 0,0003308 | - | 1 | - | 0,000328721 | - | 1 | - | 0,000331226 | - |
| 69 | 1 | 3um_FIJJAFBF_361450 | Eukaryota | unclassified | Pelagophycea | Pelagomonad | unclassified | Aureococcus | Aureococcus a | unclassified | unclassified | unclassified | unclassified | unclassified | 0,8 | 0,748331477 | 0,000450313 | 0,000433347 | 2 | - | 0,001169251 | - | 0 | - | 0 | - | 0 | - | 0 | - | 1 | - | 0,000539102 | - | 1 | - | 0,00054321 | - |
| 70 | 1 | 3um_FIJJAFBF_147450 | Bacteria | Bacteroidetes | Flavobacteriia | Flavobacterial | Flavobacteriac | Polaribacter | Polaribacter s | metabolism | Energy produc | 05D95 | Oxidoreducta | 382464.VDG1 | 0,2 | 0,4 | 3,56E-05 | 7,12E-05 | 0 | - | 0 | - | 1 | - | 0,000177879 | - | 0 | - | 0 | - | 0 | - | 0 | - | 0 | - | 0 | - |
| 71 | 1 | 3um_FIJJAFBF_489502 | Bacteria | Bacteroidetes | Flavobacteriia | Flavobacterial | Flavobacteriac | Polaribacter | various | unclassified | unclassified | unclassified | unclassified | unclassified | 3,8 | 0,748331477 | 0,000384953 | 5,38E-05 | 3 | - | 0,000310971 | - | 3 | - | 0,000365851 | - | 4 | - | 0,00038476 | - | 5 | - | 0,000477928 | - | 4 | - | 0,000385255 | - |
| 72 | 1 | 3um_FIJJAFBF_771871 | unclassified | unclassified | unclassified | unclassified | unclassified | unclassified | unclassified | unclassified | unclassified | unclassified | unclassified | unclassified | 0,4 | 0,489897949 | 0,000161539 | 0,000197847 | 0 | - | 0 | - | 0 | - | 0 | - | 0 | - | 0 | - | 1 | - | 0,000402315 | - | 1 | - | 0,000405381 | - |
| 73 | 1 | 0.2um_NOKCPHJN_1117233 | Eukaryota | unclassified | Oomycetes | Peronosporale | Peronosporac | Phytophthora | various | unclassified | unclassified | unclassified | unclassified | unclassified | 0,4 | 0,489897949 | 0,00042168 | 0,000516457 | 0 | - | 0 | - | 0 | - | 0 | - | 0 | - | 0 | - | 1 | - | 0,001050199 | - | 1 | - | 0,001058201 | - |
| 74 | 1 | 3um_FIJJAFBF_344729 | Eukaryota | Bacillariophyta | Coscinodiscop | Coscinodiscale | Coscinodiscac | Coscinodiscus | Coscinodiscus | unclassified | unclassified | unclassified | unclassified | unclassified | 0,4 | 0,489897949 | 0,000348815 | 0,000427689 | 1 | - | 0,00090406 | - | 0 | - | 0 | - | 0 | - | 0 | - | 0 | - | 0 | - | 1 | - | 0,000840015 | - |
| 75 | 1 | 3um_FIJJAFBF_272706 | Bacteria | Bacteroidetes | Flavobacteriia | Flavobacterial | unclassified | unclassified | Flavobacterial | poorly charact | Function unkn | 0XZ2H | Membrane as | 353153.XP_80 | 1,4 | 0,8 | 0,000335354 | 0,000196978 | 1 | - | 0,000237653 | - | 2 | - | 0,000559186 | - | 0 | - | 0 | - | 2 | - | 0,000438295 | - | 2 | - | 0,000441634 | - |
| 76 | 1 | 3um_FIJJAFBF_320826 | Bacteria | Proteobacteri | Gammaprotec | unclassified | unclassified | unclassified | Gammaprotec | information st | Replication, re | 06H78 | methyltransfe | 1026882.MAN | 1 | 0 | 0,000448028 | 4,38E-05 | 1 | - | 0,00045203 | - | 1 | - | 0,000531804 | - | 1 | - | 0,000419468 | - | 1 | - | 0,000416832 | - | 1 | - | 0,000420008 | - |
| 77 | 1 | 0.2um_NOKCPHJN_163581 | Eukaryota | unclassified | Oomycetes | Pythiales | Pythiaceae | Pythium | Pythium insidi | unclassified | unclassified | unclassified | unclassified | unclassified | 1 | 0 | 0,000147819 | 1,45E-05 | 1 | - | 0,000149139 | - | 1 | - | 0,000175459 | - | 1 | - | 0,000138396 | - | 1 | - | 0,000137526 | - | 1 | - | 0,000138574 | - |
| 78 | 1 | 0.2um_NOKCPHJN_959134 | Eukaryota | Phaeophyceae | unclassified | Ectocarpales | Ectocarpaceae | Ectocarpus | Ectocarpus sili | metabolism | Energy produc | COG0055 | Produces ATP | 35128.Thaps4 | 1 | 0 | 0,000673779 | 6,59E-05 | 1 | - | 0,000679797 | - | 1 | - | 0,000799767 | - | 1 | - | 0,000630827 | - | 1 | - | 0,000626863 | - | 1 | - | 0,000631639 | - |
| 79 | 1 | 0.2um_NOKCPHJN_1023367 | Bacteria | Bacteroidetes | Flavobacteriia | Flavobacterial | Flavobacteriac | Marinirhabdu | Marinirhabdu | metabolism | Inorganic ion t | 05C27 | TonB-depende | 983548.Krodi_ | 0,8 | 0,748331477 | 0,000164661 | 0,000154271 | 0 | - | 0 | - | 0 | - | 0 | - | 1 | - | 0,000206017 | - | 1 | - | 0,000204722 | - | 2 | - | 0,000412565 | - |
| 80 | 1 | 0.2um_NOKCPHJN_317337 | Eukaryota | unclassified | Pelagophycea | Pelagomonad | unclassified | Aureococcus | Aureococcus a | metabolism | Energy produc | 12NEE | ATPase, H tra | 6085.XP_0021 | 0,6 | 0,489897949 | 0,000120162 | 9,83E-05 | 1 | - | 0,000210803 | - | 0 | - | 0 | - | 1 | - | 0,000195617 | - | 1 | - | 0,000194388 | - | 0 | - | 0 | - |
| 81 | 1 | 3um_FIJJAFBF_310912 | Bacteria | Proteobacteri | Gammaprotec | Chromatiales | Chromatiacea | Candidatus Te | Candidatus Te | cellular proces | Cell wall/mem | 0GRGC | Protein of unk | 96561.Dole_2 | 1,2 | 0,4 | 0,000435413 | 0,000130155 | 1 | - | 0,000370016 | - | 1 | - | 0,000435316 | - | 2 | - | 0,000686724 | - | 1 | - | 0,000341204 | - | 1 | - | 0,000343804 | - |
| 82 | 1 | 0.2um_NOKCPHJN_138711 | Bacteria | Bacteroidetes | Flavobacteriia | Flavobacterial | Flavobacteriac | Aureicoccus | Aureicoccus m | unclassified | unclassified | unclassified | unclassified | unclassified | 1 | 0,894427191 | 0,000163738 | 0,000146381 | 0 | - | 0 | - | 0 | - | 0 | - | 1 | - | 0,000164066 | - | 2 | - | 0,00032607 | - | 2 | - | 0,000328554 | - |
| 83 | 1 | 3um_FIJJAFBF_205988 | Bacteria | Proteobacteri | Deltaproteoba | Myxococcales | Nannocystace | Plesiocystis | Plesiocystis pa | metabolism | Energy produc | 07T6V | glycerophosph | 391625.PPSIR | 1 | 0 | 0,000329233 | 3,22E-05 | 1 | - | 0,000332174 | - | 1 | - | 0,000390795 | - | 1 | - | 0,000308245 | - | 1 | - | 0,000306308 | - | 1 | - | 0,000308642 | - |
| 84 | 1 | 3um_FIJJAFBF_723333 | Bacteria | Proteobacteri | Alphaproteob | Rhodobactera | Rhodobactera | Marinosulfon | Marinosulfon | cellular proces | Cell wall/mem | 0RH7M | Glycosyl trans | 64091.VNG10 | 0,6 | 0,489897949 | 0,000141478 | 0,000116964 | 1 | - | 0,000227776 | - | 1 | - | 0,000267974 | - | 0 | - | 0 | - | 0 | - | 0 | - | 1 | - | 0,00021164 | - |
| 85 | 1 | 3um_FIJJAFBF_526565 | Bacteria | Bacteroidetes | Flavobacteriia | Flavobacterial | Crocinitomica | unclassified | Crocinitomica | unclassified | unclassified | unclassified | unclassified | unclassified | 0,6 | 0,8 | 0,000180204 | 0,000229517 | 0 | - | 0 | - | 1 | - | 0,000350918 | - | 0 | - | 0 | - | 2 | - | 0,000550104 | - | 0 | - | 0 | - |
| 86 | 1 | 3um_FIJJAFBF_644457 | Eukaryota | Chordata | Actinopteri | Cyprinodontif | Poeciliidae | Xiphophorus | Xiphophorus r | unclassified | unclassified | unclassified | unclassified | unclassified | 0,2 | 0,4 | 3,76E-05 | 7,52E-05 | 0 | - | 0 | - | 0 | - | 0 | - | 1 | - | 0,000187937 | - | 0 | - | 0 | - | 0 | - | 0 | - |
| 87 | 1 | 0.2um_NOKCPHJN_96088 | Bacteria | Proteobacteri | Gammaprotec | Alteromonada | Alteromonada | Paraglaciecola | Paraglaciecola | poorly charact | Function unkn | 05C2A | tonB-depende | 1085623.GNIT | 5,2 | 1,720465053 | 0,000409852 | 0,000141183 | 2 | - | 0,000159299 | - | 6 | - | 0,000562234 | - | 5 | - | 0,000369558 | - | 7 | - | 0,00051413 | - | 6 | - | 0,000444041 | - |
| 88 | 3 | 0.2um_NOKCPHJN_640868;3ur | Eukaryota | unclassified | unclassified | Prymnesiales | Chrysochromu | Chrysochromu | Chrysochromu | unclassified | unclassified | unclassified | unclassified | unclassified | 1 | 0 | 0,000398704 | 3,90E-05 | 1 | - | 0,000402265 | - | 1 | - | 0,000473256 | - | 1 | - | 0,000373288 | - | 1 | - | 0,000370942 | - | 1 | - | 0,000373768 | - |
| 89 | 1 | 3um_FIJJAFBF_570530 | unclassified | unclassified | unclassified | unclassified | unclassified | unclassified | unclassified | unclassified | unclassified | unclassified | unclassified | unclassified | 0,4 | 0,489897949 | 0,000151049 | 0,000185205 | 1 | - | 0,00039149 | - | 0 | - | 0 | - | 0 | - | 0 | - | 0 | - | 0 | - | 1 | - | 0,000363757 | - |
| 90 | 1 | 0.2um_NOKCPHJN_991012 | Eukaryota | unclassified | Cryptophyta | Pyrenomonad | Geminigerace | Guillardia | Guillardia thet | unclassified | unclassified | unclassified | unclassified | unclassified | 1,8 | 0,748331477 | 0,000569133 | 0,000207018 | 1 | - | 0,000328441 | - | 1 | - | 0,000386404 | - | 2 | - | 0,000609564 | - | 2 | - | 0,000605733 | - | 3 | - | 0,000915522 | - |
| 91 | 1 | 0.2um_NOKCPHJN_559288 | Eukaryota | unclassified | Labyrinthulom | unclassified | Thraustochytr | Aurantiochytr | Aurantiochytr | cellular proces | Posttranslatio | 0C0F6 | Domain of Un | 3711.Bra0278 | 0,2 | 0,4 | 1,43E-05 | 2,87E-05 | 0 | - | 0 | - | 0 | - | 0 | - | 0 | - | 0 | - | 0 | - | 0 | - | 1 | - | 7,17E-05 | - |
| 92 | 1 | 0.2um_NOKCPHJN_355727 | Bacteria | Bacteroidetes | Flavobacteriia | Flavobacterial | Flavobacteriac | Formosa | Formosa sp. H | metabolism | Inorganic ion t | 05C27 | TonB-depende | 983544.Lacal_ | 3 | 1,095445115 | 0,000252966 | 8,15E-05 | 3 | - | 0,000264138 | - | 1 | - | 0,000103584 | - | 3 | - | 0,000245111 | - | 4 | - | 0,00032476 | - | 4 | - | 0,000327235 | - |
| 93 | 1 | 3um_FIJJAFBF_477893 | Eukaryota | unclassified | unclassified | unclassified | unclassified | unclassified | uncultured pr | unclassified | unclassified | unclassified | unclassified | unclassified | 0,6 | 0,489897949 | 0,000104329 | 8,66E-05 | 0 | - | 0 | - | 1 | - | 0,000202691 | - | 0 | - | 0 | - | 1 | - | 0,000158871 | - | 1 | - | 0,000160082 | - |
| 94 | 1 | 0.2um_NOKCPHJN_358895 | Eukaryota | Eustigmatoph | unclassified | Eustigmatales | Monodopsida | Nannochlorop | Nannochlorop | unclassified | unclassified | unclassified | unclassified | unclassified | 0,4 | 0,489897949 | 2,83E-05 | 3,46E-05 | 0 | - | 0 | - | 0 | - | 0 | - | 1 | - | 7,09E-05 | - | 1 | - | 7,05E-05 | - | 0 | - | 0 | - |
| 95 | 1 | 0.2um_NOKCPHJN_899961 | Bacteria | unclassified | unclassified | unclassified | unclassified | unclassified | bacterium TM | information st | Translation, ri | KOG1936 | Histidyl-trna s | 436017.A4SAY | 0,2 | 0,4 | 5,46E-05 | 0,000109278 | 0 | - | 0 | - | 0 | - | 0 | - | 0 | - | 0 | - | 1 | - | 0,000273194 | - | 0 | - | 0 | - |
| 96 | 1 | 0.2um_NOKCPHJN_247996 | Bacteria | Proteobacteri | Gammaprotec | Alteromonada | Idiomarinacea | Idiomarina | various | metabolism | Inorganic ion t | 177TK | phosphate-sel | 455436.GHTC | 2,8 | 0,748331477 | 0,000787963 | 0,000163708 | 2 | - | 0,000578837 | - | 2 | - | 0,000680989 | - | 3 | - | 0,00080571 | - | 4 | - | 0,001067529 | - | 3 | - | 0,000806747 | - |
| 97 | 1 | 0.2um_NOKCPHJN_880073 | Bacteria | Cyanobacteria | unclassified | Synechococca | Synechococca | Synechococcu | Synechococcu | unclassified | unclassified | unclassified | unclassified | unclassified | 0,8 | 0,4 | 0,000144841 | 7,46E-05 | 0 | - | 0 | - | 1 | - | 0,000215386 | - | 1 | - | 0,000169889 | - | 1 | - | 0,000168821 | - | 1 | - | 0,000170107 | - |
| 98 | 6 | 0.2um_NOKCPHJN_1087161;0. | Eukaryota | various | various | various | various | various | various | various | various | various | various | various | 1 | 0 | 0,000211478 | 2,07E-05 | 1 | - | 0,000213367 | - | 1 | - | 0,000251022 | - | 1 | - | 0,000197997 | - | 1 | - | 0,000196753 | - | 1 | - | 0,000198252 | - |
| 99 | 1 | 3um_FIJJAFBF_544342 | Eukaryota | Chordata | Actinopteri | Cyprinodontif | Cyprinodontid | Cyprinodon | Cyprinodon va | unclassified | unclassified | unclassified | unclassified | unclassified | 0,2 | 0,4 | 1,76E-05 | 3,52E-05 | 0 | - | 0 | - | 0 | - | 0 | - | 0 | - | 0 | - | 0 | - | 0 | - | 1 | - | 8,80E-05 | - |
| 100 | 1 | 3um_FIJJAFBF_438578 | unclassified | unclassified | unclassified | unclassified | unclassified | unclassified | unclassified | unclassified | unclassified | unclassified | unclassified | unclassified | 2,4 | 1,356465997 | 0,000702117 | 0,000366914 | 1 | - | 0,000306622 | - | 1 | - | 0,000360734 | - | 4 | - | 0,001138136 | - | 2 | - | 0,000565492 | - | 4 | - | 0,001139601 | - |
| 101 | 3 | 0.2um_NOKCPHJN_537476;0.2 | various | various | various | various | various | various | various | unclassified | unclassified | unclassified | unclassified | unclassified | 1 | 0 | 0,000129341 | 1,27E-05 | 1 | - | 0,000130497 | - | 1 | - | 0,000153527 | - | 1 | - | 0,000121096 | - | 1 | - | 0,000120335 | - | 1 | - | 0,000121252 | - |
| 102 | 1 | 3um_FIJJAFBF_751778 | Eukaryota | Mollusca | Gastropoda | unclassified | Planorbidae | Biomphalaria | Biomphalaria | unclassified | unclassified | unclassified | unclassified | unclassified | 0,4 | 0,489897949 | 0,000224632 | 0,000275117 | 0 | - | 0 | - | 0 | - | 0 | - | 1 | - | 0,000561219 | - | 0 | - | 0 | - | 1 | - | 0,000561941 | - |
| 103 | 1 | 3um_FIJJAFBF_460908 | Bacteria | Bacteroidetes | Flavobacteriia | Flavobacterial | Flavobacteriac | Polaribacter | Polaribacter b | unclassified | unclassified | unclassified | unclassified | unclassified | 0,2 | 0,4 | 1,44E-05 | 2,89E-05 | 0 | - | 0 | - | 0 | - | 0 | - | 0 | - | 0 | - | 0 | - | 0 | - | 1 | - | 7,22E-05 | - |
| 104 | 1 | 3um_FIJJAFBF_81228 | Bacteria | Proteobacteri | Gammaprotec | Oceanospirilla | Endozoicomor | Endozoicomor | various | unclassified | unclassified | unclassified | unclassified | unclassified | 1,2 | 0,4 | 0,000839136 | 0,000251564 | 1 | - | 0,000712958 | - | 1 | - | 0,00083878 | - | 1 | - | 0,0006616 | - | 1 | - | 0,000657442 | - | 2 | - | 0,001324902 | - |
| 105 | 1 | 3um_FIJJAFBF_267386 | Eukaryota | Apicomplexa | Aconoidasida | Haemosporida | Plasmodiidae | Plasmodium | Plasmodium y | unclassified | unclassified | unclassified | unclassified | unclassified | 0,2 | 0,4 | 5,45E-05 | 0,00010891 | 0 | - | 0 | - | 0 | - | 0 | - | 0 | - | 0 | - | 1 | - | 0,000272274 | - | 0 | - | 0 | - |
| 106 | 1 | 3um_FIJJAFBF_243088 | Eukaryota | Bacillariophyta | Bacillariophyc | Naviculales | Phaeodactylac | Phaeodactylu | Phaeodactylu | unclassified | unclassified | unclassified | unclassified | unclassified | 1,2 | 0,4 | 0,000335042 | 0,000100152 | 1 | - | 0,00028472 | - | 1 | - | 0,000334967 | - | 2 | - | 0,00052842 | - | 1 | - | 0,00026255 | - | 1 | - | 0,00026455 | - |
| 107 | 1 | 3um_FIJJAFBF_470660 | Eukaryota | unclassified | Cryptophyta | Pyrenomonad | Geminigerace | Guillardia | Guillardia thet | metabolism | Energy produc | COG0711 | F(1)F(0) ATP s | 197221.tlr043 | 0,6 | 0,489897949 | 0,000271783 | 0,00022229 | 1 | - | 0,000476597 | - | 0 | - | 0 | - | 0 | - | 0 | - | 1 | - | 0,000439486 | - | 1 | - | 0,000442834 | - |
| 108 | 1 | 3um_FIJJAFBF_581779 | Bacteria | Bacteroidetes | Flavobacteriia | Flavobacterial | Flavobacteriac | Polaribacter | Polaribacter a | poorly charact | Function unkn | 11RX2 | NA | 313598.MED1 | 0,2 | 0,4 | 0,000104397 | 0,000208795 | 1 | - | 0,000521987 | - | 0 | - | 0 | - | 0 | - | 0 | - | 0 | - | 0 | - | 0 | - | 0 | - |
| 109 | 1 | 0.2um_NOKCPHJN_124941 | Bacteria | unclassified | unclassified | unclassified | unclassified | unclassified | bacterium TM | unclassified | unclassified | unclassified | unclassified | unclassified | 0,6 | 0,489897949 | 9,89E-05 | 8,07E-05 | 0 | - | 0 | - | 0 | - | 0 | - | 1 | - | 0,000165064 | - | 1 | - | 0,000164027 | - | 1 | - | 0,000165277 | - |
| 110 | 1 | 0.2um_NOKCPHJN_577911 | Viruses | unclassified | unclassified | unclassified | Phycodnavirid | unclassified | Aureococcus a | unclassified | unclassified | unclassified | unclassified | unclassified | 0,2 | 0,4 | 1,56E-05 | 3,12E-05 | 0 | - | 0 | - | 0 | - | 0 | - | 0 | - | 0 | - | 1 | - | 7,79E-05 | - | 0 | - | 0 | - |
| 111 | 1 | 0.2um_NOKCPHJN_216536 | Eukaryota | unclassified | Oomycetes | Peronosporale | Peronosporac | Phytophthora | Phytophthora | unclassified | unclassified | unclassified | unclassified | unclassified | 0,8 | 0,4 | 8,60E-05 | 4,43E-05 | 0 | - | 0 | - | 1 | - | 0,000127844 | - | 1 | - | 0,000100839 | - | 1 | - | 0,000100205 | - | 1 | - | 0,000100968 | - |
| 112 | 1 | 0.2um_NOKCPHJN_167586 | Bacteria | Bacteroidetes | Saprospiria | Saprospirales | Saprospiracea | Aureispira | Aureispira sp. | unclassified | unclassified | unclassified | unclassified | unclassified | 3 | 0,632455532 | 0,001516719 | 0,000336123 | 3 | - | 0,001529544 | - | 3 | - | 0,001799475 | - | 3 | - | 0,001419362 | - | 2 | - | 0,000940295 | - | 4 | - | 0,001894918 | - |
| 113 | 1 | 3um_FIJJAFBF_511166 | Eukaryota | Bacillariophyta | Coscinodiscop | Thalassiosirale | Thalassiosirac | Thalassiosira | Thalassiosira c | cellular proces | Signal transdu | 101WB | whole genom | 35128.Thaps4 | 0,8 | 0,4 | 7,47E-05 | 3,75E-05 | 1 | - | 9,89E-05 | - | 0 | - | 0 | - | 1 | - | 9,17E-05 | - | 1 | - | 9,12E-05 | - | 1 | - | 9,19E-05 | - |
| 114 | 1 | 0.2um_NOKCPHJN_322560 | Bacteria | Proteobacteri | Betaproteoba | Burkholderiale | Oxalobacterac | Collimonas | Collimonas pra | poorly charact | Function unkn | 04X2K | Mediates binc | 176279.SERP2 | 0,2 | 0,4 | 1,12E-05 | 2,24E-05 | 0 | - | 0 | - | 1 | - | 5,61E-05 | - | 0 | - | 0 | - | 0 | - | 0 | - | 0 | - | 0 | - |
| 115 | 2 | 0.2um_NOKCPHJN_164027;0.2 | Eukaryota | unclassified | Oomycetes | Peronosporale | Peronosporac | Phytophthora | various | unclassified | unclassified | unclassified | unclassified | unclassified | 1 | 0 | 0,000804791 | 7,87E-05 | 1 | - | 0,00081198 | - | 1 | - | 0,000955277 | - | 1 | - | 0,000753488 | - | 1 | - | 0,000748753 | - | 1 | - | 0,000754458 | - |
| 116 | 1 | 3um_FIJJAFBF_468768 | Bacteria | Proteobacteri | Gammaprotec | Oceanospirilla | Oceanospirilla | Marinomonas | Marinomonas | unclassified | unclassified | unclassified | unclassified | unclassified | 0,6 | 0,489897949 | 9,46E-05 | 7,82E-05 | 1 | - | 0,000152246 | - | 1 | - | 0,000179114 | - | 0 | - | 0 | - | 0 | - | 0 | - | 1 | - | 0,000141461 | - |
| 117 | 1 | 0.2um_NOKCPHJN_319780 | Eukaryota | unclassified | Oomycetes | Peronosporale | Peronosporac | Phytophthora | Phytophthora | unclassified | unclassified | unclassified | unclassified | unclassified | 1 | 0 | 0,000115736 | 1,13E-05 | 1 | - | 0,000116769 | - | 1 | - | 0,000137377 | - | 1 | - | 0,000108358 | - | 1 | - | 0,000107677 | - | 1 | - | 0,000108497 | - |
| 118 | 1 | 0.2um_NOKCPHJN_746047 | Bacteria | Bacteroidetes | Flavobacteriia | Flavobacterial | Flavobacteriac | Polaribacter | Polaribacter s | unclassified | unclassified | unclassified | unclassified | unclassified | 1 | 0,632455532 | 0,00015038 | 9,39E-05 | 1 | - | 0,000159734 | - | 0 | - | 0 | - | 2 | - | 0,000296454 | - | 1 | - | 0,000147296 | - | 1 | - | 0,000148418 | - |
| 119 | 1 | 3um_FIJJAFBF_442105 | Bacteria | Deinococcus-T | Deinococci | Deinococcales | Deinococcace | Deinococcus | Deinococcus a | unclassified | unclassified | unclassified | unclassified | unclassified | 0,4 | 0,489897949 | 8,47E-05 | 0,000103881 | 1 | - | 0,000220336 | - | 0 | - | 0 | - | 0 | - | 0 | - | 1 | - | 0,000203179 | - | 0 | - | 0 | - |
| 120 | 1 | 0.2um_NOKCPHJN_273533 | Eukaryota | Eustigmatoph | unclassified | Eustigmatales | Monodopsida | Nannochlorop | Nannochlorop | poorly charact | Function unkn | 14ZQ5 | ABC transport | 67593.Physo1 | 0,2 | 0,4 | 1,76E-05 | 3,51E-05 | 0 | - | 0 | - | 0 | - | 0 | - | 1 | - | 8,78E-05 | - | 0 | - | 0 | - | 0 | - | 0 | - |
| 121 | 1 | 0.2um_NOKCPHJN_727012 | Archaea | Euryarchaeota | unclassified | Candidatus Al | unclassified | unclassified | Candidatus Al | information st | Replication, re | 05CAR | Terminase, lar | 568816.Acin_ | 0,2 | 0,4 | 3,09E-05 | 6,17E-05 | 0 | - | 0 | - | 0 | - | 0 | - | 0 | - | 0 | - | 0 | - | 0 | - | 1 | - | 0,000154321 | - |
| 122 | 1 | 3um_FIJJAFBF_52009 | Bacteria | Proteobacteri | Alphaproteob | Rhodobactera | Rhodobactera | Cribrihabitans | Cribrihabitans | unclassified | unclassified | unclassified | unclassified | unclassified | 0,6 | 0,489897949 | 0,000316524 | 0,000258444 | 0 | - | 0 | - | 0 | - | 0 | - | 1 | - | 0,00052842 | - | 1 | - | 0,0005251 | - | 1 | - | 0,000529101 | - |
| 123 | 1 | 3um_FIJJAFBF_251768 | unclassified | unclassified | unclassified | unclassified | unclassified | unclassified | unclassified | unclassified | unclassified | unclassified | unclassified | unclassified | 0,2 | 0,4 | 0,00012802 | 0,00025604 | 1 | - | 0,000640101 | - | 0 | - | 0 | - | 0 | - | 0 | - | 0 | - | 0 | - | 0 | - | 0 | - |
| 124 | 1 | 3um_FIJJAFBF_306644 | Bacteria | Proteobacteri | Alphaproteob | Rhodobactera | Rhodobactera | unclassified | Rhodobactera | unclassified | unclassified | unclassified | unclassified | unclassified | 0,2 | 0,4 | 0,00010317 | 0,00020634 | 0 | - | 0 | - | 1 | - | 0,000515849 | - | 0 | - | 0 | - | 0 | - | 0 | - | 0 | - | 0 | - |
| 125 | 1 | 0.2um_NOKCPHJN_694005 | Eukaryota | unclassified | Bangiophycea | Bangiales | Bangiaceae | Wildemania | Wildemania s | unclassified | unclassified | unclassified | unclassified | unclassified | 1 | 0 | 0,000139964 | 1,37E-05 | 1 | - | 0,000141214 | - | 1 | - | 0,000166135 | - | 1 | - | 0,000131041 | - | 1 | - | 0,000130218 | - | 1 | - | 0,00013121 | - |
| 126 | 1 | 3um_FIJJAFBF_07962 | Viruses | unclassified | unclassified | unclassified | unclassified | unclassified | uncultured M | unclassified | unclassified | unclassified | unclassified | unclassified | 0,6 | 0,489897949 | 2,52E-05 | 2,06E-05 | 0 | - | 0 | - | 0 | - | 0 | - | 1 | - | 4,21E-05 | - | 1 | - | 4,19E-05 | - | 1 | - | 4,22E-05 | - |
| 127 | 1 | 0.2um_NOKCPHJN_503473 | Bacteria | Proteobacteri | Gammaprotec | Oceanospirilla | Oceanospirilla | Marinobacteri | Marinobacteri | metabolism | Energy produc | 05C26 | Dehydrogenas | 511062.GU3_ | 0,8 | 0,4 | 0,000148936 | 7,63E-05 | 1 | - | 0,000184619 | - | 1 | - | 0,0002172 | - | 1 | - | 0,000171319 | - | 0 | - | 0 | - | 1 | - | 0,00017154 | - |
| 128 | 1 | 3um_FIJJAFBF_44713 | Bacteria | Bacteroidetes | Flavobacteriia | Flavobacterial | Flavobacteriac | Polaribacter | Polaribacter s | cellular proces | Cell motility | COG1360 | Ompa motb d | 313598.MED1 | 0,6 | 0,489897949 | 0,00017726 | 0,000144983 | 1 | - | 0,000310971 | - | 0 | - | 0 | - | 1 | - | 0,00028857 | - | 1 | - | 0,000286757 | - | 0 | - | 0 | - |
| 129 | 1 | 3um_FIJJAFBF_287410 | Bacteria | Proteobacteri | Gammaprotec | Cellvibrionales | Spongiibacter | Oceanicoccus | Oceanicoccus | cellular proces | Intracellular tr | 0QITE | Outer membr | 247633.GP214 | 0,6 | 0,489897949 | 0,000105496 | 8,63E-05 | 1 | - | 0,000184619 | - | 0 | - | 0 | - | 1 | - | 0,000171319 | - | 0 | - | 0 | - | 1 | - | 0,00017154 | - |
| 130 | 1 | 3um_FIJJAFBF_329549 | Eukaryota | Bacillariophyta | Coscinodiscop | Thalassiosirale | Thalassiosirac | Thalassiosira | Thalassiosira p | unclassified | unclassified | unclassified | unclassified | unclassified | 0,4 | 0,489897949 | 0,000259205 | 0,000321322 | 0 | - | 0 | - | 1 | - | 0,000726549 | - | 0 | - | 0 | - | 1 | - | 0,000569474 | - | 0 | - | 0 | - |
| 131 | 1 | 0.2um_NOKCPHJN_13852 | Bacteria | Bacteroidetes | Flavobacteriia | Flavobacterial | Crocinitomica | unclassified | Crocinitomica | poorly charact | Function unkn | 08M59 | Protein of unk | 269798.CHU_ | 1 | 0,632455532 | 0,000273595 | 0,000201 | 0 | - | 0 | - | 2 | - | 0,000627173 | - | 1 | - | 0,000247346 | - | 1 | - | 0,000245791 | - | 1 | - | 0,000247664 | - |
| 132 | 1 | 3um_FIJJAFBF_304364 | Bacteria | Proteobacteri | Alphaproteob | Rhodobactera | Rhodobactera | Sulfitobacter | Sulfitobacter r | cellular proces | Cell wall/mem | 05EGS | Mechanosens | 391624.OIHEL | 0,6 | 0,489897949 | 9,75E-05 | 7,97E-05 | 1 | - | 0,000170943 | - | 0 | - | 0 | - | 0 | - | 0 | - | 1 | - | 0,000157632 | - | 1 | - | 0,000158833 | - |

| | | | | | | | | | | | | | | | | | | | | | | | | | | | | |
|---|---|---|---|---|---|---|---|---|---|---|---|---|---|---|---|---|---|---|---|---|---|---|---|---|---|---|---|---|
| 133 | 1 | 3um_FIJJAFBF_48889 | Eukaryota | unclassified | Oomycetes | Saprolegniale | Saprolegniace | Aphanomyces | various | unclassified | unclassified | unclassified | unclassified | unclassified | 1 | 0 | 0,000616436 | 6,03E-05 | 1 - | 0,000621942 - | 1 - | 0,000731701 - | 1 - | 0,00057714 - | 1 - | 0,000573513 - | 1 - | 0,000577883 - |
| 134 | 1 | 0.2um_NOKCPHJN_994281 | Eukaryota | Chlorophyta | unclassified | Pyramimonad | unclassified | Pyramimonas | Pyramimonas | unclassified | unclassified | unclassified | unclassified | unclassified | 0,6 | 0,489897949 | 0,000157241 | 0,000128389 | 0 - | 0 - | 0 - | 0 - | 1 - | 0,000262506 - | 1 - | 0,000260856 - | 1 - | 0,000262844 - |
| 135 | 1 | 0.2um_NOKCPHJN_797871 | Bacteria | Proteobacteri | Alphaproteob | Pelagibacteral | Pelagibacterac | unclassified | Pelagibacterac | information st | Replication, re | COG0470 | DNA polymera | 486041.XP_00 | 0,2 | 0,4 | 0,000111005 | 0,00022201 | 1 - | 0,000555024 - | 0 - | 0 - | 0 - | 0 - | 0 - | 0 - | 0 - | 0 - |
| 136 | 1 | 0.2um_NOKCPHJN_839592 | Viruses | unclassified | unclassified | unclassified | unclassified | unclassified | uncultured M | poorly charact | Function unkn | 0NGTE | Major head pr | 608534.GCWU | 1 | 0,632455532 | 0,000298046 | 0,000186035 | 1 - | 0,000316584 - | 0 - | 0 - | 2 - | 0,000587558 - | 1 - | 0,000291933 - | 1 - | 0,000294157 - |
| 137 | 1 | 0.2um_NOKCPHJN_417758 | Viruses | unclassified | unclassified | unclassified | unclassified | unclassified | Pleurochrysis | unclassified | unclassified | unclassified | unclassified | unclassified | 0,8 | 0,4 | 0,000134997 | 6,77E-05 | 1 - | 0,000178603 - | 0 - | 0 - | 1 - | 0,000165737 - | 1 - | 0,000164695 - | 1 - | 0,00016595 - |
| 138 | 1 | 0.2um_NOKCPHJN_25283 | Bacteria | unclassified | unclassified | unclassified | unclassified | unclassified | uncultured ba | unclassified | unclassified | unclassified | unclassified | unclassified | 0,8 | 0,4 | 7,30E-05 | 3,76E-05 | 0 - | 0 - | 1 - | 0,0001086 - | 1 - | 8,57E-05 - | 1 - | 8,51E-05 - | 1 - | 8,58E-05 - |
| 139 | 1 | 3um_FIJJAFBF_423716 | Eukaryota | Bacillariophyta | Coscinodiscop | Thalassiosirale | Thalassiosirac | Thalassiosira | Thalassiosira p | unclassified | unclassified | unclassified | unclassified | unclassified | 0,6 | 0,489897949 | 0,000281761 | 0,00023006 | 0 - | 0 - | 0 - | 0 - | 1 - | 0,000470386 - | 1 - | 0,00046743 - | 1 - | 0,000470991 - |
| 140 | 1 | 0.2um_NOKCPHJN_195162 | Eukaryota | unclassified | Labyrinthulon | unclassified | Thraustochytr | Aurantiochytr | Aurantiochytr | poorly charact | Function unkn | 0JBYD | C2 | 441375.XP_00 | 0,4 | 0,489897949 | 2,41E-05 | 2,98E-05 | 0 - | 0 - | 1 - | 6,72E-05 - | 0 - | 0 - | 0 - | 0 - | 1 - | 5,31E-05 - |
| 141 | 1 | 0.2um_NOKCPHJN_918711 | Eukaryota | unclassified | unclassified | Isochrysidales | Noelaerhabda | Emiliania | Emiliania huxl | cellular proces | Posttranslatio | KOG0731 | ATP-depender | 45157.CMQ29 | 1,2 | 0,4 | 0,000171451 | 5,14E-05 | 1 - | 0,000145671 - | 1 - | 0,000171379 - | 1 - | 0,000135177 - | 1 - | 0,000134328 - | 2 - | 0,000270703 - |
| 142 | 1 | 0.2um_NOKCPHJN_401250 | Eukaryota | Phaeophyceae | unclassified | Ectocarpales | Ectocarpaceae | Ectocarpus | Ectocarpus sili | metabolism | Carbohydrate | KOG2638 | UTP-glucose-1 | 164328.Phyra | 0,8 | 0,4 | 0,000102447 | 5,14E-05 | 1 - | 0,000135539 - | 0 - | 0 - | 1 - | 0,000125775 - | 1 - | 0,000124985 - | 1 - | 0,000125937 - |
| 143 | 1 | 0.2um_NOKCPHJN_564421 | Eukaryota | unclassified | Oomycetes | Peronosporale | Peronosporac | Phytophthora | Phytophthora | unclassified | unclassified | unclassified | unclassified | unclassified | 1,8 | 0,4 | 0,000598441 | 9,97E-05 | 2 - | 0,000685108 - | 1 - | 0,000403007 - | 2 - | 0,000635756 - | 2 - | 0,000631761 - | 2 - | 0,000636574 - |
| 144 | 1 | 0.2um_NOKCPHJN_576429 | Eukaryota | unclassified | unclassified | Kinetoplastida | Trypanosomat | Leishmania | Leishmania m | unclassified | unclassified | unclassified | unclassified | unclassified | 0,4 | 0,489897949 | 5,73E-05 | 7,03E-05 | 1 - | 0,000149139 - | 0 - | 0 - | 0 - | 0 - | 1 - | 0,000137526 - | 0 - | 0 - |
| 145 | 8 | 0.2um_NOKCPHJN_200646;0.2 | various | various | various | various | various | various | various | information st | Translation, ri | various | various | various | 0,2 | 0,4 | 0,000171319 | 0,000342639 | 0 - | 0 - | 0 - | 0 - | 1 - | 0,000856597 - | 0 - | 0 - | 0 - | 0 - |
| 146 | 1 | 3um_FIJJAFBF_39793 | Bacteria | Bacteroidetes | Flavobacteriia | Flavobacterial | Flavobacteriac | Polaribacter | Polaribacter s | unclassified | unclassified | unclassified | unclassified | unclassified | 0,8 | 0,4 | 0,000260959 | 0,000130852 | 1 - | 0,000345251 - | 0 - | 0 - | 1 - | 0,000320381 - | 1 - | 0,000318368 - | 1 - | 0,000320793 - |
| 147 | 1 | 0.2um_NOKCPHJN_1093443 | Bacteria | Proteobacteri | Alphaproteob | Pelagibacteral | Pelagibacterac | Candidatus Pe | Candidatus Pe | unclassified | unclassified | unclassified | unclassified | unclassified | 0,4 | 0,489897949 | 7,14E-05 | 8,84E-05 | 0 - | 0 - | 1 - | 0,000199555 - | 1 - | 0,000157402 - | 0 - | 0 - | 0 - | 0 - |
| 148 | 1 | 3um_FIJJAFBF_510235 | Bacteria | Bacteroidetes | Flavobacteriia | Flavobacterial | Flavobacteriac | Polaribacter | Polaribacter s | unclassified | unclassified | unclassified | unclassified | unclassified | 1 | 0 | 0,000634434 | 6,21E-05 | 1 - | 0,000640101 - | 1 - | 0,000753065 - | 1 - | 0,000593991 - | 1 - | 0,000590258 - | 1 - | 0,000594755 - |
| 149 | 2 | 0.2um_NOKCPHJN_393773;3ur | Bacteria | Proteobacteri | Alphaproteob | Rhodospirillal | Rhodospirillac | unclassified | Rhodospirillac | unclassified | unclassified | unclassified | unclassified | unclassified | 0,8 | 0,748331477 | 0,000203504 | 0,00018989 | 0 - | 0 - | 0 - | 0 - | 1 - | 0,0002551 - | 2 - | 0,000506993 - | 1 - | 0,000255428 - |
| 150 | 1 | 3um_FIJJAFBF_621085 | Bacteria | Actinobacteria | Actinobacteria | Micromonosp | Micromonosp | Actinoplanes | Actinoplanes f | unclassified | unclassified | unclassified | unclassified | unclassified | 0,2 | 0,4 | 7,90E-05 | 0,000158013 | 0 - | 0 - | 0 - | 0 - | 1 - | 0,000395033 - | 0 - | 0 - | 0 - | 0 - |
| 151 | 1 | 3um_FIJJAFBF_428565 | Bacteria | Proteobacteri | Betaproteoba | unclassified | unclassified | unclassified | Betaproteoba | unclassified | unclassified | unclassified | unclassified | unclassified | 0,2 | 0,4 | 1,16E-05 | 2,33E-05 | 1 - | 5,82E-05 - | 0 - | 0 - | 0 - | 0 - | 0 - | 0 - | 0 - | 0 - |
| 152 | 1 | 3um_FIJJAFBF_424915 | Eukaryota | Phaeophyceae | unclassified | Ectocarpales | Ectocarpaceae | Ectocarpus | Ectocarpus sili | unclassified | unclassified | unclassified | unclassified | unclassified | 1 | 0 | 0,001073055 | 0,000104992 | 1 - | 0,00108264 - | 1 - | 0,001273702 - | 1 - | 0,001004651 - | 1 - | 0,000998338 - | 1 - | 0,001005944 - |
| 153 | 1 | 3um_FIJJAFBF_253700 | Bacteria | Proteobacteri | Gammaprotec | Cellvibrionales | Halieaceae | Chromatocurv | Chromatocurv | unclassified | unclassified | unclassified | unclassified | unclassified | 3,6 | 0,489897949 | 0,00036212 | 6,86E-05 | 3 - | 0,000303439 - | 4 - | 0,000475986 - | 4 - | 0,000375441 - | 3 - | 0,000279811 - | 4 - | 0,000375924 - |
| 154 | 1 | 3um_FIJJAFBF_709097 | Eukaryota | unclassified | Oomycetes | Pythiales | Pythiaceae | Pythium | Pythium insidi | information st | Translation, ri | KOG0052 | This protein p | 65071.PYU1_T | 0,4 | 0,489897949 | 7,03E-05 | 8,62E-05 | 0 - | 0 - | 0 - | 0 - | 1 - | 0,00017576 - | 0 - | 0 - | 1 - | 0,000175986 - |
| 155 | 1 | 0.2um_NOKCPHJN_115354 | Bacteria | Bacteroidetes | Flavobacteriia | Flavobacterial | Flavobacteriac | Lutibacter | Lutibacter sp. | unclassified | unclassified | unclassified | unclassified | unclassified | 0,6 | 0,8 | 4,57E-05 | 6,11E-05 | 0 - | 0 - | 0 - | 0 - | 0 - | 0 - | 1 - | 7,59E-05 - | 2 - | 0,000152873 - |
| 156 | 1 | 0.2um_NOKCPHJN_997360 | Bacteria | Verrucomicrol | Verrucomicrol | Verrucomicrol | Rubritaleaceae | Rubritalea | Rubritalea squ | cellular proces | Cell wall/mem | 05C6Y | Udp-glucose 6 | 583355.Caka_ | 0,2 | 0,4 | 3,53E-05 | 7,06E-05 | 0 - | 0 - | 0 - | 0 - | 1 - | 0,000176522 - | 0 - | 0 - | 0 - | 0 - |
| 157 | 2 | 0.2um_NOKCPHJN_564820;3ur | Bacteria | Proteobacteri | Gammaprotec | various | various | various | various | information st | Translation, ri | 0QQXI | This protein is | various | 0,4 | 0,489897949 | 0,000223935 | 0,000274582 | 1 - | 0,000580754 - | 0 - | 0 - | 1 - | 0,000538919 - | 0 - | 0 - | 0 - | 0 - |
| 158 | 1 | 0.2um_NOKCPHJN_39209 | Eukaryota | unclassified | Pelagophycea | Pelagomonad | unclassified | Aureoumbra | Aureoumbra l | metabolism | Energy produc | COG0055 | Produces ATP | 489825.LYNGE | 32,6 | 3,382306905 | 0,005882205 | 0,000377402 | 31 - | 0,005687258 - | 28 - | 0,006043424 - | 36 - | 0,006128792 - | 31 - | 0,005244406 - | 37 - | 0,006307145 - |
| 159 | 1 | 0.2um_NOKCPHJN_595339 | Bacteria | Proteobacteri | Alphaproteob | Pelagibacteral | Pelagibacterac | Candidatus Pe | Candidatus Pe | unclassified | unclassified | unclassified | unclassified | unclassified | 0,4 | 0,489897949 | 0,000139317 | 0,00017156 | 1 - | 0,000320051 - | 1 - | 0,000376532 - | 0 - | 0 - | 0 - | 0 - | 0 - | 0 - |
| 160 | 1 | 0.2um_NOKCPHJN_1103368 | unclassified | unclassified | unclassified | unclassified | unclassified | unclassified | unclassified | unclassified | unclassified | unclassified | unclassified | unclassified | 0,2 | 0,4 | 0,000394465 | 0,00078893 | 0 - | 0 - | 0 - | 0 - | 0 - | 0 - | 1 - | 0,001972326 - | 0 - | 0 - |
| 161 | 1 | 0.2um_NOKCPHJN_1028497 | Viruses | unclassified | unclassified | unclassified | Phycodnavirid | unclassified | Chrysochromu | unclassified | unclassified | unclassified | unclassified | unclassified | 0,2 | 0,4 | 0,000110307 | 0,000220613 | 1 - | 0,000551534 - | 0 - | 0 - | 0 - | 0 - | 0 - | 0 - | 0 - | 0 - |
| 162 | 1 | 0.2um_NOKCPHJN_127444 | Bacteria | Proteobacteri | Alphaproteob | Rhizobiales | Aurantimonac | Martelella | Martelella me | unclassified | unclassified | unclassified | unclassified | unclassified | 0,4 | 0,489897949 | 1,59E-05 | 1,95E-05 | 1 - | 4,13E-05 - | 0 - | 0 - | 1 - | 3,83E-05 - | 0 - | 0 - | 0 - | 0 - |
| 163 | 1 | 0.2um_NOKCPHJN_27392 | Bacteria | unclassified | unclassified | unclassified | unclassified | unclassified | uncultured ba | metabolism | Inorganic ion t | 05C27 | TonB-depende | 313596.RB250 | 29,8 | 1,6 | 0,002476077 | 0,000254559 | 27 - | 0,002265774 - | 30 - | 0,002961815 - | 30 - | 0,002336174 - | 30 - | 0,002321493 - | 32 - | 0,002495127 - |
| 164 | 1 | 0.2um_NOKCPHJN_1057723 | Eukaryota | Bacillariophyta | Bacillariophyc | Naviculales | Phaeodactylac | Phaeodactylu | Phaeodactylu | unclassified | unclassified | unclassified | unclassified | unclassified | 0,2 | 0,4 | 3,82E-05 | 7,65E-05 | 0 - | 0 - | 0 - | 0 - | 0 - | 0 - | 1 - | 0,000191171 - | 0 - | 0 - |
| 165 | 1 | 0.2um_NOKCPHJN_949714 | Eukaryota | unclassified | Spirotrichea | Sporadotrichi | Oxytrichidae | Oxytricha | Oxytricha trifa | metabolism | Energy produc | COG0056 | Produces ATP | 412030.XP_00 | 0,4 | 0,489897949 | 7,30E-05 | 8,95E-05 | 1 - | 0,000189814 - | 0 - | 0 - | 0 - | 0 - | 1 - | 0,000175033 - | 0 - | 0 - |
| 166 | 1 | 3um_FIJJAFBF_673864 | Eukaryota | unclassified | Dinophyceae | Suessiales | Symbiodiniace | Symbiodinium | Symbiodinium | unclassified | unclassified | unclassified | unclassified | unclassified | 1 | 0 | 0,000110864 | 1,08E-05 | 1 - | 0,000111854 - | 1 - | 0,000131594 - | 1 - | 0,000103797 - | 1 - | 0,000103145 - | 1 - | 0,00010393 - |
| 167 | 1 | 3um_FIJJAFBF_102619 | unclassified | unclassified | unclassified | unclassified | unclassified | unclassified | unclassified | unclassified | unclassified | unclassified | unclassified | unclassified | 2,4 | 0,8 | 0,000689191 | 0,00020724 | 3 - | 0,000891802 - | 1 - | 0,000349728 - | 3 - | 0,00082756 - | 2 - | 0,00054824 - | 3 - | 0,000828625 - |
| 168 | 1 | 0.2um_NOKCPHJN_560573 | Bacteria | Proteobacteri | Alphaproteob | Rhodobactera | Rhodobactera | unclassified | Rhodobactera | unclassified | unclassified | unclassified | unclassified | unclassified | 1 | 0 | 0,000365199 | 3,57E-05 | 1 - | 0,000368462 - | 1 - | 0,000433487 - | 1 - | 0,000341919 - | 1 - | 0,00033977 - | 1 - | 0,000342359 - |
| 169 | 1 | 3um_FIJJAFBF_84361 | Eukaryota | Bacillariophyta | Mediophyceae | Cymatosirales | Cymatosiracea | Plagiogramma | Plagiogramma | metabolism | Energy produc | 0IHF0 | This is one of t | 45157.CMV04 | 11 | 1,095445115 | 0,002631655 | 9,44E-05 | 11 - | 0,002679534 - | 9 - | 0,002579247 - | 12 - | 0,002712558 - | 11 - | 0,002470886 - | 12 - | 0,00271605 - |
| 170 | 1 | 3um_FIJJAFBF_64866 | Bacteria | Bacteroidetes | Flavobacteriia | Flavobacterial | Flavobacteriac | Formosa | Formosa sp. H | poorly charact | Function unkn | 05C2A | tonB-depende | 688270.Celal_ | 0,4 | 0,489897949 | 3,05E-05 | 3,74E-05 | 0 - | 0 - | 0 - | 0 - | 1 - | 7,63E-05 - | 0 - | 0 - | 1 - | 7,64E-05 - |
| 171 | 1 | 3um_FIJJAFBF_437895 | Bacteria | Bacteroidetes | Saprospiria | Saprospirales | Lewinellaceae | unclassified | Lewinellaceae | metabolism | Inorganic ion t | 05C27 | TonB-depende | 391603.FBALC | 5,2 | 2,856571371 | 0,000409065 | 0,000230803 | 10 - | 0,000815757 - | 1 - | 9,60E-05 - | 5 - | 0,000378496 - | 5 - | 0,000376118 - | 5 - | 0,000378984 - |
| 172 | 1 | 3um_FIJJAFBF_510245 | Bacteria | Bacteroidetes | Flavobacteriia | Flavobacterial | Flavobacteriac | Polaribacter | various | unclassified | unclassified | unclassified | unclassified | unclassified | 3,2 | 0,4 | 0,00153054 | 0,000188931 | 3 - | 0,001453489 - | 3 - | 0,001709998 - | 4 - | 0,001798381 - | 3 - | 0,00134031 - | 3 - | 0,001350522 - |
| 173 | 1 | 3um_FIJJAFBF_564229 | Bacteria | Proteobacteri | Gammaprotec | Pseudomonad | Pseudomonad | Pseudomonas | Pseudomonas | cellular proces | Cell wall/mem | 0QX85 | 17 kDa surface | 205922.Pfl01_ | 1 | 0,894427191 | 0,001373792 | 0,001241482 | 2 - | 0,002875208 - | 0 - | 0 - | 2 - | 0,00266809 - | 1 - | 0,001325662 - | 0 - | 0 - |
| 174 | 1 | 3um_FIJJAFBF_435849 | Bacteria | Bacteroidetes | Saprospiria | Saprospirales | Saprospiracea | Aureispira | Aureispira sp. | unclassified | unclassified | unclassified | unclassified | unclassified | 0,4 | 0,489897949 | 0,000121987 | 0,000149404 | 0 - | 0 - | 0 - | 0 - | 1 - | 0,000305928 - | 1 - | 0,000304005 - | 0 - | 0 - |
| 175 | 1 | 3um_FIJJAFBF_103096 | Eukaryota | unclassified | unclassified | Isochrysidales | Noelaerhabda | Emiliania | Emiliania huxl | unclassified | unclassified | unclassified | unclassified | unclassified | 4,6 | 0,8 | 0,001740351 | 0,000193543 | 5 - | 0,001940129 - | 3 - | 0,001369512 - | 5 - | 0,00180037 - | 5 - | 0,001789056 - | 5 - | 0,001802688 - |
| 176 | 1 | 3um_FIJJAFBF_117751 | Eukaryota | Arthropoda | Arachnida | Araneae | Eresidae | Stegodyphus | Stegodyphus r | poorly charact | Function unkn | 0IRIW | Inherit from N | 7668.SPU_004 | 0,2 | 0,4 | 3,26E-05 | 6,52E-05 | 0 - | 0 - | 0 - | 0 - | 0 - | 0 - | 1 - | 0,000163035 - | 0 - | 0 - |
| 177 | 2 | 3um_FIJJAFBF_564495;3um_FI | Eukaryota | various | various | various | various | various | various | unclassified | unclassified | unclassified | unclassified | unclassified | 1,2 | 0,4 | 0,000982988 | 0,000294689 | 1 - | 0,00083518 - | 1 - | 0,00098257 - | 1 - | 0,000775017 - | 1 - | 0,000770146 - | 2 - | 0,001552028 - |
| 178 | 1 | 0.2um_NOKCPHJN_111289 | Bacteria | Proteobacteri | Alphaproteob | Rhodospirillal | Rhodospirillac | Candidatus En | Candidatus En | poorly charact | Function unkn | 9935 | NA | 214684.XP_56 | 0,2 | 0,4 | 1,61E-05 | 3,22E-05 | 0 - | 0 - | 1 - | 8,04E-05 - | 0 - | 0 - | 0 - | 0 - | 0 - | 0 - |
| 179 | 1 | 0.2um_NOKCPHJN_1027022 | Eukaryota | Xanthophycea | unclassified | Botrydiales | Botrydiaceae | Botrydium | Botrydium gra | unclassified | unclassified | unclassified | unclassified | unclassified | 6,2 | 1,166190379 | 0,002798844 | 0,000319873 | 6 - | 0,002783932 - | 4 - | 0,00218349 - | 7 - | 0,003013953 - | 7 - | 0,002995013 - | 7 - | 0,003017833 - |
| 180 | 1 | 0.2um_NOKCPHJN_350811 | Bacteria | Proteobacteri | Gammaprotec | Alteromonada | Alteromonada | Aliiglaciecola | Aliiglaciecola l | cellular proces | Signal transdu | 13AP7 | NACHT domai | 7739.JGI9285 | 0,2 | 0,4 | 3,65E-05 | 7,31E-05 | 1 - | 0,000182696 - | 0 - | 0 - | 0 - | 0 - | 0 - | 0 - | 0 - | 0 - |
| 181 | 1 | 3um_FIJJAFBF_35958 | Bacteria | Bacteroidetes | Flavobacteriia | Flavobacterial | Flavobacteriac | Arenitalea | Arenitalea lut | unclassified | unclassified | unclassified | unclassified | unclassified | 2,6 | 0,489897949 | 0,000571289 | 9,23E-05 | 3 - | 0,000671126 - | 2 - | 0,000526377 - | 3 - | 0,000622781 - | 2 - | 0,000412578 - | 3 - | 0,000623583 - |
| 182 | 1 | 0.2um_NOKCPHJN_563766 | Archaea | Euryarchaeota | Methanobact | Methanobact | Methanobact | Methanospha | Methanospha | cellular proces | Cell wall/mem | 0D8M5 | polymorphic c | 383372.Rcas_ | 1 | 0 | 8,97E-05 | 8,78E-06 | 1 - | 9,05E-05 - | 1 - | 0,00010647 - | 1 - | 8,40E-05 - | 1 - | 8,35E-05 - | 1 - | 8,41E-05 - |
| 183 | 1 | 3um_FIJJAFBF_713545 | Eukaryota | unclassified | unclassified | Prymnesiales | Chrysochromu | Chrysochromu | Chrysochromu | unclassified | unclassified | unclassified | unclassified | unclassified | 0,4 | 0,489897949 | 0,000232655 | 0,000284943 | 0 - | 0 - | 0 - | 0 - | 1 - | 0,000581262 - | 0 - | 0 - | 1 - | 0,000582011 - |
| 184 | 1 | 3um_FIJJAFBF_751786 | Eukaryota | unclassified | Pelagophycea | Pelagomonad | unclassified | Aureococcus | Aureococcus a | unclassified | unclassified | unclassified | unclassified | unclassified | 7 | 0,894427191 | 0,001167457 | 0,000166561 | 6 - | 0,001011852 - | 7 - | 0,001388825 - | 6 - | 0,000938962 - | 8 - | 0,001244082 - | 8 - | 0,001253561 - |
| 185 | 1 | 3um_FIJJAFBF_111340 | Viruses | unclassified | unclassified | unclassified | Phycodnavirid | unclassified | Chrysochromu | unclassified | unclassified | unclassified | unclassified | unclassified | 13,8 | 1,166190379 | 0,002222723 | 0,000178407 | 12 - | 0,001959639 - | 13 - | 0,002497595 - | 14 - | 0,002121554 - | 15 - | 0,002258809 - | 15 - | 0,002276019 - |
| 186 | 1 | 3um_FIJJAFBF_796563 | Eukaryota | unclassified | Bangiophycea | Bangiales | Bangiaceae | Bangia | Bangia fuscop | cellular proces | Posttranslatio | 0IEKR | One of the co | 35128.Thapsd | 2 | 0 | 0,000377082 | 3,69E-05 | 2 - | 0,000380451 - | 2 - | 0,000447592 - | 2 - | 0,000353044 - | 2 - | 0,000350826 - | 2 - | 0,000353499 - |
| 187 | 1 | 3um_FIJJAFBF_76725 | Bacteria | Proteobacteri | Alphaproteob | Rhodobactera | Rhodobactera | Loktanella | Loktanella sp. | unclassified | unclassified | unclassified | unclassified | unclassified | 0,6 | 0,8 | 0,000602948 | 0,000817912 | 2 - | 0,002063385 - | 0 - | 0 - | 0 - | 0 - | 1 - | 0,000951357 - | 0 - | 0 - |
| 188 | 1 | 3um_FIJJAFBF_296272 | Bacteria | Proteobacteri | Gammaprotec | Alteromonada | Alteromonada | Glaciecola | Glaciecola sp. | unclassified | unclassified | unclassified | unclassified | unclassified | 0,4 | 0,489897949 | 7,46E-05 | 9,14E-05 | 0 - | 0 - | 0 - | 0 - | 0 - | 0 - | 1 - | 0,000185897 - | 1 - | 0,000187314 - |
| 189 | 1 | 3um_FIJJAFBF_257124 | Eukaryota | unclassified | Florideophyce | Ceramiales | Delesseriacea | Membranopte | Membranopte | unclassified | unclassified | unclassified | unclassified | unclassified | 1 | 0 | 0,000634434 | 6,21E-05 | 1 - | 0,000640101 - | 1 - | 0,000753065 - | 1 - | 0,000593991 - | 1 - | 0,000590258 - | 1 - | 0,000594755 - |
| 190 | 1 | 3um_FIJJAFBF_140603 | Bacteria | Proteobacteri | Gammaprotec | Cellvibrionales | Halieaceae | unclassified | marine gamm | poorly charact | Function unkn | COG5430 | spore coat U c | 247634.GPB2 | 0,6 | 0,489897949 | 0,000284718 | 0,000232835 | 1 - | 0,000498261 - | 0 - | 0 - | 1 - | 0,000462368 - | 0 - | 0 - | 1 - | 0,000462963 - |
| 191 | 1 | 0.2um_NOKCPHJN_456112 | Bacteria | Proteobacteri | Betaproteoba | Burkholderiale | Oxalobacterac | Massilia | Massilia namu | unclassified | unclassified | unclassified | unclassified | unclassified | 0,8 | 0,4 | 6,09E-05 | 3,12E-05 | 1 - | 7,55E-05 - | 1 - | 8,88E-05 - | 1 - | 7,00E-05 - | 0 - | 0 - | 1 - | 7,01E-05 - |
| 192 | 1 | 3um_FIJJAFBF_156358 | Eukaryota | Bacillariophyta | Bacillariophyc | Naviculales | Phaeodactylac | Phaeodactylu | Phaeodactylu | unclassified | unclassified | unclassified | unclassified | unclassified | 0,2 | 0,4 | 4,93E-05 | 9,85E-05 | 1 - | 0,000246331 - | 0 - | 0 - | 0 - | 0 - | 0 - | 0 - | 0 - | 0 - |
| 193 | 1 | 0.2um_NOKCPHJN_423207 | Bacteria | Cyanobacteria | unclassified | Synechococca | Synechococca | Synechococcu | Synechococcu | unclassified | unclassified | unclassified | unclassified | unclassified | 0,2 | 0,4 | 6,90E-06 | 1,38E-05 | 0 - | 0 - | 0 - | 0 - | 1 - | 3,45E-05 - | 0 - | 0 - | 0 - | 0 - |
| 194 | 1 | 0.2um_NOKCPHJN_231475 | Bacteria | Proteobacteri | Alphaproteob | Rhodobactera | Rhodobactera | unclassified | Rhodobactera | information st | Replication, re | COG0187 | DNA gyrase ne | 467661.RKLH1 | 1 | 0 | 0,000131494 | 1,29E-05 | 1 - | 0,000132668 - | 1 - | 0,000156082 - | 1 - | 0,000123112 - | 1 - | 0,000122338 - | 1 - | 0,00012327 - |
| 195 | 1 | 0.2um_NOKCPHJN_428587 | Bacteria | Proteobacteri | Alphaproteob | unclassified | unclassified | unclassified | SAR116 cluste | metabolism | Lipid transpor | 01SHD | Short-chain de | 488538.SAR11 | 0,2 | 0,4 | 7,83E-05 | 0,000156695 | 0 - | 0 - | 0 - | 0 - | 0 - | 0 - | 0 - | 0 - | 1 - | 0,000391738 - |
| 196 | 1 | 0.2um_NOKCPHJN_56666 | Bacteria | Proteobacteri | Gammaprotec | Alteromonada | Alteromonada | Alteromonas | Alteromonas s | metabolism | Carbohydrate | 16QEQ | hydrolase, fan | 722419.PH505 | 1 | 0 | 8,11E-05 | 7,93E-06 | 1 - | 8,18E-05 - | 1 - | 9,62E-05 - | 1 - | 7,59E-05 - | 1 - | 7,54E-05 - | 1 - | 7,60E-05 - |
| 197 | 1 | 0.2um_NOKCPHJN_341284 | Bacteria | Bacteroidetes | Bacteroidia | Marinilabiliale | Marinilabiliac | Anaerophaga | Anaerophaga | unclassified | unclassified | unclassified | unclassified | unclassified | 0,8 | 0,4 | 0,000154871 | 7,94E-05 | 1 - | 0,000192311 - | 1 - | 0,00022625 - | 1 - | 0,000178458 - | 1 - | 0,000177336 - | 0 - | 0 - |
| 198 | 1 | 3um_FIJJAFBF_32817 | Bacteria | Proteobacteri | Alphaproteob | Rhodobactera | Rhodobactera | Pseudophaeol | Pseudophaeol | unclassified | unclassified | unclassified | unclassified | unclassified | 3,8 | 0,4 | 0,001004162 | 0,000163977 | 4 - | 0,001062956 - | 4 - | 0,001250544 - | 4 - | 0,000986385 - | 4 - | 0,000980186 - | 3 - | 0,000740741 - |
| 199 | 1 | 3um_FIJJAFBF_775040 | Eukaryota | unclassified | Cryptophyta | Pyrenomonad | Pyrenomonad | Rhodomonas | Rhodomonas | unclassified | unclassified | unclassified | unclassified | unclassified | 1,2 | 0,4 | 0,000325207 | 9,58E-05 | 1 - | 0,000276637 - | 1 - | 0,000325457 - | 1 - | 0,000256709 - | 2 - | 0,000510192 - | 1 - | 0,000257039 - |
| 200 | 1 | 3um_FIJJAFBF_743884 | Bacteria | Proteobacteri | Alphaproteob | Rhizobiales | unclassified | unclassified | Rhizobiales ba | poorly charact | Function unkn | 05C2A | tonB-depende | 1085623.GNIT | 3,6 | 0,489897949 | 0,000586301 | 6,74E-05 | 4 - | 0,00066184 - | 3 - | 0,000583981 - | 4 - | 0,000614164 - | 4 - | 0,000610305 - | 3 - | 0,000461216 - |
| 201 | 1 | 0.2um_NOKCPHJN_586048 | Bacteria | Bacteroidetes | unclassified | unclassified | unclassified | unclassified | Bacteroidetes | unclassified | unclassified | unclassified | unclassified | unclassified | 1,8 | 0,4 | 0,000120785 | 2,96E-05 | 1 - | 6,78E-05 - | 2 - | 0,000159459 - | 2 - | 0,000125775 - | 2 - | 0,000124985 - | 2 - | 0,000125937 - |
| 202 | 1 | 0.2um_NOKCPHJN_36914 | Eukaryota | unclassified | Cryptophyta | Pyrenomonad | Pyrenomonad | Rhodomonas | Rhodomonas | metabolism | Carbohydrate | COG1850 | RuBisCO catal | 45157.CMV01 | 1,2 | 1,166190379 | 0,000202405 | 0,000194011 | 1 - | 0,000179701 - | 0 - | 0 - | 0 - | 0 - | 2 - | 0,000331415 - | 3 - | 0,000500911 - |
| 203 | 1 | 0.2um_NOKCPHJN_616575 | unclassified | unclassified | unclassified | unclassified | unclassified | unclassified | unclassified | unclassified | unclassified | unclassified | unclassified | unclassified | 1 | 0 | 0,000155766 | 1,52E-05 | 1 - | 0,000157157 - | 1 - | 0,000184892 - | 1 - | 0,000145836 - | 1 - | 0,00014492 - | 1 - | 0,000146024 - |
| 204 | 1 | 3um_FIJJAFBF_715054 | Bacteria | Cyanobacteria | unclassified | Oscillatoriales | Cyanothecace | Cyanothece | various | unclassified | unclassified | unclassified | unclassified | unclassified | 1 | 0 | 0,000924654 | 9,05E-05 | 1 - | 0,000932913 - | 1 - | 0,001097552 - | 1 - | 0,00086571 - | 1 - | 0,00086027 - | 1 - | 0,000866824 - |
| 205 | 1 | 3um_FIJJAFBF_556069 | unclassified | unclassified | unclassified | unclassified | unclassified | unclassified | unclassified | unclassified | unclassified | unclassified | unclassified | unclassified | 1,4 | 0,489897949 | 0,001047069 | 0,000313292 | 1 - | 0,000769244 - | 1 - | 0,000904999 - | 2 - | 0,001427662 - | 2 - | 0,00141869 - | 1 - | 0,00071475 - |
| 206 | 1 | 3um_FIJJAFBF_510252 | Bacteria | Bacteroidetes | Flavobacteriia | Flavobacterial | Flavobacteriac | Flaviramulus | Flaviramulus i | unclassified | unclassified | unclassified | unclassified | unclassified | 0,6 | 0,489897949 | 0,000428253 | 0,000355619 | 0 - | 0 - | 1 - | 0,000832015 - | 0 - | 0 - | 1 - | 0,00065214 - | 1 - | 0,000657109 - |
| 207 | 1 | 0.2um_NOKCPHJN_1119963 | Eukaryota | unclassified | Oomycetes | Peronosporale | Peronosporac | Phytophthora | various | unclassified | unclassified | unclassified | unclassified | unclassified | 0,2 | 0,4 | 0,000224856 | 0,000449712 | 1 - | 0,00112428 - | 0 - | 0 - | 0 - | 0 - | 0 - | 0 - | 0 - | 0 - |
| 208 | 1 | 3um_FIJJAFBF_626507 | Bacteria | Chloroflexi | unclassified | unclassified | unclassified | unclassified | Chloroflexi ba | unclassified | unclassified | unclassified | unclassified | unclassified | 0,8 | 0,4 | 0,000107082 | 5,37E-05 | 1 - | 0,00014167 - | 0 - | 0 - | 1 - | 0,000131465 - | 1 - | 0,000130639 - | 1 - | 0,000131634 - |
| 209 | 1 | 0.2um_NOKCPHJN_438866 | Bacteria | Proteobacteri | Alphaproteob | Pelagibacteral | Pelagibacterac | unclassified | Pelagibacterac | poorly charact | Function unkn | COG2268 | Band 7 protei | 391600.BBAL3 | 0,2 | 0,4 | 2,47E-05 | 4,93E-05 | 0 - | 0 - | 0 - | 0 - | 1 - | 0,000123298 - | 0 - | 0 - | 0 - | 0 - |
| 210 | 1 | 0.2um_NOKCPHJN_995401 | Bacteria | Proteobacteri | Gammaprotec | Alteromonada | Colwelliaceae | Thalassotalea | Thalassotalea | information st | Transcription | COG0085 | DNA-depende | 167879.CPS_4 | 0,2 | 0,4 | 3,50E-05 | 6,99E-05 | 0 - | 0 - | 1 - | 0,000174864 - | 0 - | 0 - | 0 - | 0 - | 0 - | 0 - |
| 211 | 1 | 0.2um_NOKCPHJN_63759 | Bacteria | Bacteroidetes | Flavobacteriia | Flavobacterial | Crocinitomica | unclassified | Crocinitomica | poorly charact | Function unkn | 1AGHF | SusD family | 485918.Cpin_ | 0,4 | 0,489897949 | 6,52E-05 | 8,00E-05 | 1 - | 0,000169621 - | 0 - | 0 - | 0 - | 0 - | 1 - | 0,000156413 - | 0 - | 0 - |
| 212 | 1 | 3um_FIJJAFBF_606585 | Eukaryota | Basidiomycota | Microbotryom | Microbotryale | Microbotryace | Microbotryum | Microbotryum | information st | Chromatin str | KOG3467 | Core compone | 3218.PP1S133 | 1,4 | 0,489897949 | 0,000904287 | 0,000270571 | 1 - | 0,000664347 - | 1 - | 0,00078159 - | 2 - | 0,001232981 - | 2 - | 0,001225233 - | 1 - | 0,000617284 - |
| 213 | 1 | 3um_FIJJAFBF_70793 | Bacteria | Bacteroidetes | Flavobacteriia | Flavobacterial | Crocinitomica | unclassified | Crocinitomica | unclassified | unclassified | unclassified | unclassified | unclassified | 2,2 | 0,4 | 0,000661459 | 0,00019639 | 2 - | 0,000596557 - | 3 - | 0,001052754 - | 2 - | 0,000553583 - | 2 - | 0,000550104 - | 2 - | 0,000554296 - |
| 214 | 1 | 0.2um_NOKCPHJN_758975 | Bacteria | Proteobacteri | Alphaproteob | Pelagibacteral | Pelagibacterac | Candidatus Pe | Candidatus Pe | unclassified | unclassified | unclassified | unclassified | unclassified | 0,2 | 0,4 | 5,16E-05 | 0,000103141 | 0 - | 0 - | 0 - | 0 - | 0 - | 0 - | 0 - | 0 - | 1 - | 0,000257853 - |
| 215 | 1 | 3um_FIJJAFBF_518929 | Bacteria | Proteobacteri | Gammaprotec | Alteromonada | Alteromonada | Paraglaciecola | Paraglaciecola | information st | Transcription | 173HH | transcriptiona | 342610.Patl_2 | 1 | 0 | 0,000167471 | 1,64E-05 | 1 - | 0,000168967 - | 1 - | 0,000198786 - | 1 - | 0,000156795 - | 1 - | 0,00015581 - | 1 - | 0,000156997 - |
| 216 | 1 | 3um_FIJJAFBF_365122 | Bacteria | Bacteroidetes | Flavobacteriia | Flavobacterial | Flavobacteriac | Winogradskye | Winogradskye | unclassified | unclassified | unclassified | unclassified | unclassified | 0,8 | 0,4 | 0,000385437 | 0,000198414 | 0 - | 0 - | 1 - | 0,000573166 - | 1 - | 0,000452093 - | 1 - | 0,000449252 - | 1 - | 0,000452675 - |
| 217 | 1 | 3um_FIJJAFBF_187037 | Bacteria | Proteobacteri | Gammaprotec | Alteromonada | Alteromonada | Paraglaciecola | Paraglaciecola | unclassified | unclassified | unclassified | unclassified | unclassified | 1 | 0,632455532 | 0,000321752 | 0,000204545 | 2 - | 0,000640101 - | 1 - | 0,000376532 - | 1 - | 0,000296995 - | 1 - | 0,000295129 - | 0 - | 0 - |
| 218 | 1 | 0.2um_NOKCPHJN_296783 | Bacteria | Bacteroidetes | Flavobacteriia | Flavobacterial | Flavobacteriac | Flavobacteriu | Flavobacteriu | metabolism | Inorganic ion t | 05C27 | TonB-depende | 983548.Krodi_ | 0,8 | 0,4 | 9,07E-05 | 4,55E-05 | 1 - | 0,000119964 - | 0 - | 0 - | 1 - | 0,000111322 - | 1 - | 0,000110623 - | 1 - | 0,000111466 - |
| 219 | 1 | 3um_FIJJAFBF_348325 | Bacteria | Bacteroidetes | Flavobacteriia | Flavobacterial | Flavobacteriac | Algibacter | Algibacter lect | unclassified | unclassified | unclassified | unclassified | unclassified | 14 | 3,16227766 | 0,001307572 | 0,000206253 | 12 - | 0,001153866 - | 9 - | 0,001018124 - | 16 - | 0,001427662 - | 15 - | 0,001330022 - | 18 - | 0,001608187 - |
| 220 | 1 | 3um_FIJJAFBF_50289 | Bacteria | Bacteroidetes | Flavobacteriia | Flavobacterial | Flavobacteriac | Arenitalea | Arenitalea lut | metabolism | Carbohydrate | 05D2N | Aldolase | 313603.FB217 | 1 | 0,894427191 | 0,000228713 | 0,000204388 | 0 - | 0 - | 0 - | 0 - | 2 - | 0,000458461 - | 2 - | 0,000455579 - | 1 - | 0,000229525 - |
| 221 | 1 | 3um_FIJJAFBF_139789 | Bacteria | Acidobacteria | unclassified | unclassified | unclassified | unclassified | Acidobacteria | unclassified | unclassified | unclassified | unclassified | unclassified | 0,4 | 0,489897949 | 0,000156626 | 0,00019416 | 0 - | 0 - | 1 - | 0,000439021 - | 0 - | 0 - | 1 - | 0,000344108 - | 0 - | 0 - |
| 222 | 1 | 0.2um_NOKCPHJN_36238 | Eukaryota | unclassified | Cryptophyta | Pyrenomonad | Geminigerace | Teleaulax | Teleaulax amp | metabolism | Carbohydrate | COG1850 | RuBisCO catal | 45157.CMV01 | 0,6 | 0,489897949 | 9,99E-05 | 8,16E-05 | 0 - | 0 - | 0 - | 0 - | 1 - | 0,000166756 - | 1 - | 0,000165708 - | 1 - | 0,00016697 - |
| 223 | 1 | 0.2um_NOKCPHJN_152692 | Bacteria | Proteobacteri | Gammaprotec | Cellvibrionales | Porticoccacea | unclassified | SAR92 bacteri | unclassified | unclassified | unclassified | unclassified | unclassified | 0,8 | 0,4 | 5,96E-05 | 3,07E-05 | 0 - | 0 - | 1 - | 8,86E-05 - | 1 - | 6,99E-05 - | 1 - | 6,95E-05 - | 1 - | 7,00E-05 - |
| 224 | 1 | 3um_FIJJAFBF_12264 | Viruses | unclassified | unclassified | unclassified | unclassified | unclassified | uncultured M | poorly charact | Function unkn | 05BE3 | Phage portal p | 742766.HMPR | 0,6 | 0,489897949 | 8,65E-05 | 7,08E-05 | 1 - | 0,000151457 - | 0 - | 0 - | 1 - | 0,000140547 - | 0 - | 0 - | 1 - | 0,000140728 - |
| 225 | 1 | 3um_FIJJAFBF_804749 | Eukaryota | unclassified | unclassified | Isochrysidales | Noelaerhabda | Emiliania | Emiliania huxl | unclassified | unclassified | unclassified | unclassified | unclassified | 0,2 | 0,4 | 6,09E-05 | 0,000121797 | 1 - | 0,000304493 - | 0 - | 0 - | 0 - | 0 - | 0 - | 0 - | 0 - | 0 - |
| 226 | 1 | 0.2um_NOKCPHJN_219472 | Bacteria | Proteobacteri | Gammaprotec | Alteromonada | Pseudoalteror | Pseudoalteror | Pseudoalteror | metabolism | Inorganic ion t | 16PDP | TonB-depende | 983545.Glaag | 6,2 | 0,748331477 | 0,000980873 | 0,000119313 | 5 - | 0,000801589 - | 6 - | 0,001131662 - | 7 - | 0,001041384 - | 6 - | 0,000887006 - | 7 - | 0,001042725 - |
| 227 | 1 | 3um_FIJJAFBF_592540 | Bacteria | Proteobacteri | Gammaprotec | Alteromonada | Alteromonada | Glaciecola | Glaciecola sp. | unclassified | unclassified | unclassified | unclassified | unclassified | 3 | 0 | 0,000269651 | 2,64E-05 | 3 - | 0,00027206 - | 3 - | 0,000320072 - | 3 - | 0,000252461 - | 3 - | 0,000250875 - | 3 - | 0,000252786 - |
| 228 | 1 | 0.2um_NOKCPHJN_554855 | Bacteria | Proteobacteri | Gammaprotec | Enterobactera | Yersiniaceae | Serratia | Serratia marc | cellular proces | Signal transdu | KOG0613 | fibronectin typ | 7165.AGAP00 | 0,8 | 0,748331477 | 0,000108982 | 0,000110112 | 1 - | 0,000127462 - | 2 - | 0,000299912 - | 0 - | 0 - | 1 - | 0,000117537 - | 0 - | 0 - |
| 229 | 2 | 3um_FIJJAFBF_147391;3um_FI | Bacteria | Bacteroidetes | Flavobacteriia | Flavobacterial | Flavobacteriac | Polaribacter | various | unclassified | unclassified | unclassified | unclassified | unclassified | 1,4 | 1,019803903 | 0,000870492 | 0,000621537 | 1 - | 0,000664347 - | 0 - | 0 - | 2 - | 0,001232981 - | 3 - | 0,001837849 - | 1 - | 0,000617284 - |
| 230 | 1 | 0.2um_NOKCPHJN_407176 | Eukaryota | Mucoromycot | Mucoromycet | Mucorales | Syncephalastr | Syncephalastr | Syncephalastr | unclassified | unclassified | unclassified | unclassified | unclassified | 1 | 0 | 7,92E-05 | 7,75E-06 | 1 - | 7,99E-05 - | 1 - | 9,40E-05 - | 1 - | 7,41E-05 - | 1 - | 7,36E-05 - | 1 - | 7,42E-05 - |
| 231 | 1 | 0.2um_NOKCPHJN_966897 | Eukaryota | unclassified | unclassified | Isochrysidales | Noelaerhabda | Emiliania | Emiliania huxl | unclassified | unclassified | unclassified | unclassified | unclassified | 1 | 0 | 0,000308218 | 3,02E-05 | 1 - | 0,000310971 - | 1 - | 0,000365851 - | 1 - | 0,00028857 - | 1 - | 0,000286757 - | 1 - | 0,000288941 - |
| 232 | 1 | 0.2um_NOKCPHJN_94453 | Bacteria | Proteobacteri | Betaproteoba | Burkholderiale | Comamonada | Acidovorax | various | metabolism | Energy produc | COG0055 | Produces ATP | 420662.Mpe_ | 1,4 | 0,489897949 | 0,000257423 | 8,63E-05 | 2 - | 0,000373961 - | 1 - | 0,000219978 - | 2 - | 0,000347022 - | 1 - | 0,000172421 - | 1 - | 0,000173735 - |
| 233 | 1 | 0.2um_NOKCPHJN_137356 | Bacteria | Proteobacteri | Gammaprotec | Cellvibrionales | Halieaceae | Halioglobus | Halioglobus sp | unclassified | unclassified | unclassified | unclassified | unclassified | 0,4 | 0,489897949 | 3,52E-05 | 4,31E-05 | 0 - | 0 - | 0 - | 0 - | 0 - | 0 - | 1 - | 8,76E-05 - | 1 - | 8,83E-05 - |
| 234 | 1 | 0.2um_NOKCPHJN_26367 | Bacteria | Planctomycete | unclassified | unclassified | unclassified | unclassified | Planctomycete | information st | Translation, ri | COG0050 | This protein p | 487796.Flav2A | 1 | 0 | 0,000220044 | 2,15E-05 | 1 - | 0,00022201 - | 1 - | 0,00026119 - | 1 - | 0,000206017 - | 1 - | 0,000204722 - | 1 - | 0,000206282 - |
| 235 | 1 | 3um_FIJJAFBF_554090 | Bacteria | Verrucomicrol | Opitutae | unclassified | unclassified | unclassified | Opitutae bact | unclassified | unclassified | unclassified | unclassified | unclassified | 2,6 | 0,8 | 0,002447921 | 0,000759675 | 4 - | 0,003812776 - | 2 - | 0,002242824 - | 2 - | 0,00176906 - | 2 - | 0,001757942 - | 3 - | 0,002657005 - |
| 236 | 1 | 3um_FIJJAFBF_07267 | Bacteria | Planctomycete | Planctomyceti | Planctomyceta | Planctomyceta | Rhodopirellula | Rhodopirellula | unclassified | unclassified | unclassified | unclassified | unclassified | 0,8 | 0,4 | 0,000203324 | 0,000101952 | 1 - | 0,000269 - | 0 - | 0 - | 1 - | 0,000249622 - | 1 - | 0,000248053 - | 1 - | 0,000249943 - |
| 237 | 2 | 0.2um_NOKCPHJN_47587;3um | Bacteria | various | various | various | various | various | various | metabolism | Energy produc | 0PEU8 | ATP synthase | 313603.FB217 | 0,6 | 0,489897949 | 0,000862632 | 0,000713619 | 1 - | 0,001391966 - | 1 - | 0,001637617 - | 0 - | 0 - | 1 - | 0,001283577 - | 0 - | 0 - |
| 238 | 1 | 3um_FIJJAFBF_55089 | Bacteria | Bacteroidetes | Flavobacteriia | Flavobacterial | Flavobacteriac | Polaribacter | Polaribacter s | information st | Translation, ri | 0PA05 | 30S ribosomal | 313594.PI23P | 1 | 0 | 0,000480207 | 4,70E-05 | 1 - | 0,000484496 - | 1 - | 0,000569999 - | 1 - | 0,000449595 - | 1 - | 0,00044677 - | 1 - | 0,000450174 - |
| 239 | 1 | 0.2um_NOKCPHJN_536910 | Eukaryota | Streptophyta | unclassified | Sapindales | Rutaceae | Citrus | various | information st | RNA processin | 1BGEW | May be involv | 3218.PP1S162 | 0,2 | 0,4 | 4,12E-05 | 8,23E-05 | 0 - | 0 - | 0 - | 0 - | 0 - | 0 - | 1 - | 0,000205764 - | 0 - | 0 - |
| 240 | 1 | 3um_FIJJAFBF_527770 | Bacteria | Actinobacteria | Actinobacteria | Micrococcales | Microbacteria | Rhodoluna | Rhodoluna lac | unclassified | unclassified | unclassified | unclassified | unclassified | 1,6 | 2,727636339 | 0,000137555 | 0,000236877 | 7 - | 0,000607178 - | 0 - | 0 - | 0 - | 0 - | 0 - | 0 - | 1 - | 8,06E-05 - |
| 241 | 1 | 3um_FIJJAFBF_437894 | Bacteria | Bacteroidetes | unclassified | unclassified | unclassified | unclassified | Bacteroidetes | unclassified | unclassified | unclassified | unclassified | unclassified | 2 | 1,095445115 | 0,00039855 | 0,000231678 | 2 - | 0,000392366 - | 3 - | 0,000692415 - | 3 - | 0,000546153 - | 2 - | 0,000361814 - | 0 - | 0 - |
| 242 | 1 | 0.2um_NOKCPHJN_59638 | Bacteria | Bacteroidetes | Flavobacteriia | Flavobacterial | Crocinitomica | unclassified | Crocinitomica | unclassified | unclassified | unclassified | unclassified | unclassified | 1 | 0 | 0,000132496 | 1,30E-05 | 1 - | 0,00013368 - | 1 - | 0,000157271 - | 1 - | 0,00012405 - | 1 - | 0,00012327 - | 1 - | 0,00012421 - |
| 243 | 1 | 3um_FIJJAFBF_44700 | Bacteria | Bacteroidetes | Flavobacteriia | Flavobacterial | Flavobacteriac | Polaribacter | Polaribacter s | unclassified | unclassified | unclassified | unclassified | unclassified | 0,6 | 0,489897949 | 5,72E-05 | 4,67E-05 | 0 - | 0 - | 0 - | 0 - | 1 - | 9,55E-05 - | 1 - | 9,49E-05 - | 1 - | 9,56E-05 - |
| 244 | 1 | 3um_FIJJAFBF_744567 | Bacteria | unclassified | unclassified | unclassified | unclassified | unclassified | uncultured ba | unclassified | unclassified | unclassified | unclassified | unclassified | 9,8 | 3,059411708 | 0,00087575 | 0,000230461 | 10 - | 0,00092896 - | 4 - | 0,000437161 - | 11 - | 0,000948246 - | 11 - | 0,000942287 - | 13 - | 0,001122097 - |
| 245 | 1 | 0.2um_NOKCPHJN_946574 | Eukaryota | Chordata | Actinopteri | unclassified | Pomacentrida | Amphiprion | Amphiprion o | unclassified | unclassified | unclassified | unclassified | unclassified | 0,8 | 0,4 | 0,000217323 | 0,000108972 | 1 - | 0,000287521 - | 0 - | 0 - | 1 - | 0,000266809 - | 1 - | 0,000265132 - | 1 - | 0,000267152 - |
| 246 | 1 | 3um_FIJJAFBF_341378 | Bacteria | Bacteroidetes | Flavobacteriia | Flavobacterial | Flavobacteriac | Algibacter | Algibacter sp. | cellular proces | Cell wall/mem | COG2885 | Ompa motb d | 1046627.BZAF | 5,4 | 0,8 | 0,0010292 | 0,000206012 | 5 - | 0,000957356 - | 6 - | 0,001351571 - | 6 - | 0,001066071 - | 6 - | 0,001059371 - | 4 - | 0,000711629 - |
| 247 | 1 | 0.2um_NOKCPHJN_176004 | Eukaryota | unclassified | unclassified | Prymnesiales | Chrysochromu | Chrysochromu | various | unclassified | unclassified | unclassified | unclassified | unclassified | 0,2 | 0,4 | 2,17E-05 | 4,33E-05 | 0 - | 0 - | 0 - | 0 - | 0 - | 0 - | 0 - | 0 - | 1 - | 0,000108353 - |
| 248 | 1 | 3um_FIJJAFBF_744517 | Bacteria | Proteobacteri | Alphaproteob | Pelagibacteral | Pelagibacterac | Candidatus Pe | Candidatus Pe | unclassified | unclassified | unclassified | unclassified | unclassified | 0,4 | 0,489897949 | 0,000159665 | 0,000195549 | 0 - | 0 - | 0 - | 0 - | 1 - | 0,000398906 - | 0 - | 0 - | 1 - | 0,000399419 - |
| 249 | 1 | 3um_FIJJAFBF_209603 | Bacteria | Bacteroidetes | Flavobacteriia | Flavobacterial | Flavobacteriac | Flaviramulus | Flaviramulus i | unclassified | unclassified | unclassified | unclassified | unclassified | 1,6 | 0,8 | 0,000758237 | 0,000390322 | 0 - | 0 - | 2 - | 0,00112754 - | 2 - | 0,000889363 - | 2 - | 0,000883774 - | 2 - | 0,000890508 - |
| 250 | 1 | 0.2um_NOKCPHJN_244384 | Eukaryota | Streptophyta | Liliopsida | Poales | Poaceae | Aegilops | Aegilops taus | unclassified | unclassified | unclassified | unclassified | unclassified | 1 | 0 | 3,37E-05 | 3,30E-06 | 1 - | 3,40E-05 - | 1 - | 4,01E-05 - | 1 - | 3,16E-05 - | 1 - | 3,14E-05 - | 1 - | 3,16E-05 - |
| 251 | 1 | 3um_FIJJAFBF_461592 | Bacteria | Proteobacteri | Gammaprotec | Cellvibrionales | Porticoccacea | unclassified | Gammaprotec | metabolism | Inorganic ion t | 16TQC | TONB-depend | 314287.GB220 | 0,8 | 0,748331477 | 6,86E-05 | 6,60E-05 | 2 - | 0,000178059 - | 0 - | 0 - | 0 - | 0 - | 1 - | 8,21E-05 - | 1 - | 8,27E-05 - |
| 252 | 1 | 0.2um_NOKCPHJN_454728 | Eukaryota | unclassified | Labyrinthulon | unclassified | Thraustochytr | Aurantiochytr | Aurantiochytr | unclassified | unclassified | unclassified | unclassified | unclassified | 0,6 | 0,489897949 | 0,000190871 | 0,000156112 | 1 - | 0,000334709 - | 0 - | 0 - | 0 - | 0 - | 1 - | 0,000308646 - | 1 - | 0,000310998 - |
| 253 | 1 | 3um_FIJJAFBF_183700 | Bacteria | Bacteroidetes | Flavobacteriia | Flavobacterial | Flavobacteriac | Algibacter | Algibacter sp. | unclassified | unclassified | unclassified | unclassified | unclassified | 10,8 | 1,166190379 | 0,001009649 | 0,000138312 | 13 - | 0,001227148 - | 10 - | 0,001110548 - | 10 - | 0,000875961 - | 11 - | 0,000957501 - | 10 - | 0,000877088 - |
| 254 | 1 | 0.2um_NOKCPHJN_151232 | Bacteria | Bacteroidetes | Flavobacteriia | Flavobacterial | Flavobacteriac | Formosa | Formosa sp. H | unclassified | unclassified | unclassified | unclassified | unclassified | 2 | 0,894427191 | 0,000176054 | 6,96E-05 | 1 - | 9,18E-05 - | 1 - | 0,000108031 - | 2 - | 0,000170422 - | 3 - | 0,000254027 - | 3 - | 0,000255963 - |
| 255 | 1 | 0.2um_NOKCPHJN_617369 | Eukaryota | Chlorophyta | Ulvophyceae | Ulvales | Ulvaceae | Ulva | Ulva linza | unclassified | unclassified | unclassified | unclassified | unclassified | 1 | 0 | 0,000218936 | 2,14E-05 | 1 - | 0,000220891 - | 1 - | 0,000259874 - | 1 - | 0,000204979 - | 1 - | 0,000203691 - | 1 - | 0,000205243 - |
| 256 | 5 | 0.2um_NOKCPHJN_234612;0.2 | various | various | various | various | various | various | various | unclassified | unclassified | unclassified | unclassified | unclassified | 1 | 0 | 0,0002822 | 2,76E-05 | 1 - | 0,00028472 - | 1 - | 0,000334967 - | 1 - | 0,00026421 - | 1 - | 0,00026255 - | 1 - | 0,00026455 - |
| 257 | 2 | 0.2um_NOKCPHJN_299554;3ur | Bacteria | Proteobacteri | Gammaprotec | Pseudomonad | Pseudomonad | Pseudomonas | various | poorly charact | Function unkn | 061QR | Phosphatidylc | 216595.PFLU0 | 0,2 | 0,4 | 2,89E-05 | 5,79E-05 | 0 - | 0 - | 0 - | 0 - | 0 - | 0 - | 1 - | 0,000144661 - | 0 - | 0 - |
| 258 | 1 | 0.2um_NOKCPHJN_14938 | Bacteria | Bacteroidetes | Flavobacteriia | Flavobacterial | Flavobacteriac | Kordia | Kordia jejudor | unclassified | unclassified | unclassified | unclassified | unclassified | 1 | 0 | 0,000135809 | 1,33E-05 | 1 - | 0,000137022 - | 1 - | 0,000161203 - | 1 - | 0,000127151 - | 1 - | 0,000126352 - | 1 - | 0,000127315 - |
| 259 | 1 | 3um_FIJJAFBF_450895 | Bacteria | Bacteroidetes | Flavobacteriia | Flavobacterial | Flavobacteriac | Flaviramulus | Flaviramulus b | unclassified | unclassified | unclassified | unclassified | unclassified | 3,4 | 0,489897949 | 0,000864998 | 0,000187441 | 3 - | 0,000764772 - | 4 - | 0,00119965 - | 3 - | 0,000709681 - | 4 - | 0,000940295 - | 3 - | 0,000710594 - |
| 260 | 1 | 0.2um_NOKCPHJN_1033454 | unclassified | unclassified | unclassified | unclassified | unclassified | unclassified | unclassified | unclassified | unclassified | unclassified | unclassified | unclassified | 0,2 | 0,4 | 0,000103169 | 0,000206338 | 1 - | 0,000515846 - | 0 - | 0 - | 0 - | 0 - | 0 - | 0 - | 0 - | 0 - |
| 261 | 1 | 3um_FIJJAFBF_296937 | Bacteria | Proteobacteri | Alphaproteob | Rhodobactera | Rhodobactera | various | various | information st | Transcription | 05S3D | Transcriptiona | 439497.RR11_ | 0,6 | 0,489897949 | 0,000634309 | 0,00051872 | 1 - | 0,001110049 - | 0 - | 0 - | 1 - | 0,001030085 - | 0 - | 0 - | 1 - | 0,001031411 - |
| 262 | 1 | 3um_FIJJAFBF_546079 | Bacteria | Proteobacteri | unclassified | unclassified | unclassified | unclassified | Proteobacteri | unclassified | unclassified | unclassified | unclassified | unclassified | 0,4 | 0,489897949 | 0,000123458 | 0,000151206 | 0 - | 0 - | 0 - | 0 - | 0 - | 0 - | 1 - | 0,000307473 - | 1 - | 0,000309816 - |
| 263 | 1 | 3um_FIJJAFBF_183674 | Bacteria | Bacteroidetes | Flavobacteriia | Flavobacterial | Flavobacteriac | Jejuia | Jejuia pallidilu | poorly charact | Function unkn | 11P35 | NA | 398720.MED2 | 4,6 | 1,743559577 | 0,000734814 | 0,000336064 | 2 - | 0,000316584 - | 7 - | 0,001303591 - | 4 - | 0,000587558 - | 4 - | 0,000583865 - | 6 - | 0,000882471 - |
| 264 | 2 | 0.2um_NOKCPHJN_546137;3ur | various | various | various | various | various | various | various | various | various | various | various | various | 0,2 | 0,4 | 2,99E-05 | 5,97E-05 | 0 - | 0 - | 0 - | 0 - | 1 - | 0,000149315 - | 0 - | 0 - | 0 - | 0 - |
| 265 | 1 | 0.2um_NOKCPHJN_744384 | Bacteria | Proteobacteri | Alphaproteob | Pelagibacteral | Pelagibacterac | Candidatus Pe | Candidatus Pe | unclassified | unclassified | unclassified | unclassified | unclassified | 0,6 | 0,489897949 | 0,001020574 | 0,00083472 | 1 - | 0,00178967 - | 0 - | 0 - | 0 - | 0 - | 1 - | 0,001650313 - | 1 - | 0,001662888 - |
| 266 | 1 | 0.2um_NOKCPHJN_1041517 | Viruses | unclassified | unclassified | unclassified | various | various | various | unclassified | unclassified | unclassified | unclassified | unclassified | 0,2 | 0,4 | 7,70E-05 | 0,000153985 | 0 - | 0 - | 1 - | 0,000384962 - | 0 - | 0 - | 0 - | 0 - | 0 - | 0 - |
| 267 | 1 | 3um_FIJJAFBF_486409 | Bacteria | Proteobacteri | Gammaprotec | Cellvibrionales | Spongiibacter | Spongiibacter | Spongiibacter | poorly charact | Function unkn | 0ZX70 | NA | 525897.Dbac_ | 0,4 | 0,489897949 | 0,000328496 | 0,000402775 | 1 - | 0,000851397 - | 0 - | 0 - | 0 - | 0 - | 0 - | 0 - | 1 - | 0,000791082 - |
| 268 | 1 | 3um_FIJJAFBF_126100 | Eukaryota | Bacillariophyta | Coscinodiscop | Thalassiosirale | Thalassiosirac | Thalassiosira | Thalassiosira ( | unclassified | unclassified | unclassified | unclassified | unclassified | 1,8 | 0,4 | 0,000732738 | 0,000192106 | 2 - | 0,000815757 - | 2 - | 0,00095972 - | 2 - | 0,000756993 - | 2 - | 0,000752236 - | 1 - | 0,000378984 - |
| 269 | 1 | 0.2um_NOKCPHJN_539044 | Eukaryota | unclassified | unclassified | Kinetoplastida | Trypanosomat | Leishmania | Leishmania m | unclassified | unclassified | unclassified | unclassified | unclassified | 0,4 | 0,489897949 | 2,19E-05 | 2,68E-05 | 1 - | 5,69E-05 - | 0 - | 0 - | 0 - | 0 - | 1 - | 5,25E-05 - | 0 - | 0 - |

| | | | | | | | | | | | | | | | | | | | | | | | | | | | | |
|---|---|---|---|---|---|---|---|---|---|---|---|---|---|---|---|---|---|---|---|---|---|---|---|---|---|---|---|---|
| 270 | 1 | 0.2um_NOKCPHJN_573670 | unclassified | unclassified | unclassified | unclassified | unclassified | unclassified | unclassified | unclassified | unclassified | unclassified | unclassified | unclassified | 1 | 0 | 0,000154383 | 1,51E-05 | 1 - | 0,000155762 - | 1 - | 0,00018325 - | 1 - | 0,000144541 - | 1 - | 0,000143633 - | 1 - | 0,000144727 - |
| 271 | 1 | 0.2um_NOKCPHJN_512717 | Eukaryota | Nematoda | Chromadorea | Diplogasterida | Neodiplogaste | Pristionchus | Pristionchus p | information st | Chromatin str | KOG3467 | Core compone | 7217.FBpp012 | 1,2 | 0,979795897 | 0,000166081 | 0,000135606 | 0 - | 0 - | 0 - | 0 - | 2 - | 0,000277263 - | 2 - | 0,000275521 - | 2 - | 0,00027762 - |
| 272 | 1 | 3um_FIJJAFBF_755418 | Bacteria | Bacteroidetes | Flavobacteriia | Flavobacterial | Flavobacteriac | Flaviramulus | Flaviramulus b | cellular proces | Cell wall/mem | 06B97 | phosphate-sel | 391598.FBBAL | 0,4 | 0,489897949 | 8,48E-05 | 0,000105119 | 0 - | 0 - | 1 - | 0,000237172 - | 1 - | 0,000187073 - | 0 - | 0 - | 0 - | 0 - |
| 273 | 1 | 3um_FIJJAFBF_708514 | Bacteria | Cyanobacteria | unclassified | Synechococca | Synechococca | Synechococcu | Synechococcu | poorly charact | Function unkn | 06QDP | C-phycoerythr | 221359.RS991 | 1,4 | 0,489897949 | 0,000731087 | 0,000243421 | 2 - | 0,001062956 - | 1 - | 0,000625272 - | 1 - | 0,000493192 - | 2 - | 0,000980186 - | 1 - | 0,000493827 - |
| 274 | 1 | 3um_FIJJAFBF_521152 | Bacteria | Bacteroidetes | Flavobacteriia | Flavobacterial | Flavobacteriac | Gaetbulibacte | Gaetbulibacte | metabolism | Inorganic ion t | 0P8Q4 | tonB-depende | 867900.Celly_ | 1,6 | 0,8 | 0,000145896 | 6,60E-05 | 1 - | 9,43E-05 - | 1 - | 0,000110935 - | 1 - | 8,75E-05 - | 2 - | 0,000173904 - | 3 - | 0,000262844 - |
| 275 | 1 | 0.2um_NOKCPHJN_1007193 | Eukaryota | unclassified | Oomycetes | Peronosporale | Peronosporac | Plasmopara | Plasmopara ha | information st | Replication, re | COG0513 | purine NTP-de | 65071.PYU1_T | 2,8 | 0,748331477 | 0,000574201 | 0,000129356 | 3 - | 0,000629382 - | 2 - | 0,000493636 - | 2 - | 0,000389362 - | 4 - | 0,000773831 - | 3 - | 0,000584795 - |
| 276 | 1 | 3um_FIJJAFBF_128529 | unclassified | unclassified | unclassified | unclassified | unclassified | unclassified | unclassified | unclassified | unclassified | unclassified | unclassified | unclassified | 0,8 | 0,4 | 0,000338181 | 0,000169574 | 1 - | 0,000447418 - | 0 - | 0 - | 1 - | 0,000415187 - | 1 - | 0,000412578 - | 1 - | 0,000415722 - |
| 277 | 3 | 3um_FIJJAFBF_158178;3um_FI | Eukaryota | unclassified | Oomycetes | Pythiales | Pythiaceae | Pythium | Pythium insidi | information st | Translation, ri | COG0089 | One of the ear | various | 0,8 | 0,4 | 0,000460302 | 0,000230809 | 1 - | 0,000608985 - | 0 - | 0 - | 1 - | 0,000565116 - | 1 - | 0,000561565 - | 1 - | 0,000565844 - |
| 278 | 1 | 3um_FIJJAFBF_08158 | Bacteria | Bacteroidetes | Flavobacteriia | Flavobacterial | Flavobacteriac | Polaribacter | Polaribacter s | metabolism | Lipid transpor | COG4247 | Phytase | 688270.Celal_ | 2,2 | 0,4 | 0,000562518 | 9,34E-05 | 2 - | 0,000518899 - | 2 - | 0,000610473 - | 2 - | 0,000481519 - | 2 - | 0,000478493 - | 3 - | 0,000723209 - |
| 279 | 1 | 0.2um_NOKCPHJN_319552 | Eukaryota | Phaeophyceae | unclassified | Ectocarpales | Ectocarpaceae | Ectocarpus | Ectocarpus sili | metabolism | Energy produc | COG0055 | Produces ATP | 35128.Thaps4 | 4,4 | 0,489897949 | 0,000909669 | 8,69E-05 | 4 - | 0,000839176 - | 4 - | 0,000987272 - | 5 - | 0,000973406 - | 4 - | 0,000773831 - | 5 - | 0,000974659 - |
| 280 | 1 | 3um_FIJJAFBF_320548 | Bacteria | Proteobacteri | Alphaproteob | Rhodobactera | Rhodobactera | Sulfitobacter | Sulfitobacter s | unclassified | unclassified | unclassified | unclassified | unclassified | 0,2 | 0,4 | 6,79E-05 | 0,000135802 | 0 - | 0 - | 0 - | 0 - | 0 - | 0 - | 0 - | 0 - | 1 - | 0,000339506 - |
| 281 | 11 | 0.2um_NOKCPHJN_149835;0.2 | Bacteria | Proteobacteri | Gammaprotec | various | various | various | various | metabolism | Energy produc | 0QIP8 | Electron trans | various | 0,4 | 0,489897949 | 0,000104063 | 0,00012745 | 0 - | 0 - | 0 - | 0 - | 1 - | 0,00025999 - | 0 - | 0 - | 1 - | 0,000260324 - |
| 282 | 1 | 3um_FIJJAFBF_788009 | Bacteria | Proteobacteri | Alphaproteob | Rhodobactera | Rhodobactera | unclassified | Rhodobactera | unclassified | unclassified | unclassified | unclassified | unclassified | 5,8 | 0,748331477 | 0,001478128 | 0,000127106 | 5 - | 0,001297246 - | 5 - | 0,001526182 - | 7 - | 0,001685317 - | 6 - | 0,00143548 - | 6 - | 0,001446417 - |
| 283 | 1 | 0.2um_NOKCPHJN_579291 | Eukaryota | Basidiomycota | Agaricomycet | Gloeophyllale | Gloeophyllace | Neolentinus | Neolentinus le | unclassified | unclassified | unclassified | unclassified | unclassified | 0,2 | 0,4 | 2,54E-05 | 5,09E-05 | 0 - | 0 - | 0 - | 0 - | 0 - | 0 - | 1 - | 0,000127147 - | 0 - | 0 - |
| 284 | 1 | 3um_FIJJAFBF_672316 | Bacteria | Bacteroidetes | Cytophagia | Cytophagales | Flammeovirga | unclassified | Flammeovirga | unclassified | unclassified | unclassified | unclassified | unclassified | 0,8 | 0,748331477 | 0,000445542 | 0,000411347 | 1 - | 0,000588549 - | 0 - | 0 - | 2 - | 0,001092305 - | 0 - | 0 - | 1 - | 0,000546856 - |
| 285 | 1 | 0.2um_NOKCPHJN_91222 | Bacteria | Bacteroidetes | Flavobacteriia | Flavobacterial | Flavobacteriac | Formosa | Formosa sp. H | unclassified | unclassified | unclassified | unclassified | unclassified | 0,4 | 0,489897949 | 3,53E-05 | 4,32E-05 | 0 - | 0 - | 0 - | 0 - | 0 - | 0 - | 1 - | 8,79E-05 - | 1 - | 8,86E-05 - |
| 286 | 1 | 3um_FIJJAFBF_543287 | Bacteria | Bacteroidetes | Flavobacteriia | Flavobacterial | Flavobacteriac | Lacinutrix | Lacinutrix him | information st | Replication, re | 0PB0H | DNA-binding p | 983544.Lacal_ | 2,6 | 0,489897949 | 0,002486912 | 0,000394895 | 3 - | 0,002923128 - | 2 - | 0,002292664 - | 3 - | 0,002712558 - | 3 - | 0,002695512 - | 2 - | 0,0018107 - |
| 287 | 1 | 0.2um_NOKCPHJN_1094039 | Bacteria | Bacteroidetes | Flavobacteriia | Flavobacterial | Flavobacteriac | Formosa | Formosa sp. H | unclassified | unclassified | unclassified | unclassified | unclassified | 0,8 | 0,4 | 0,000184028 | 9,47E-05 | 0 - | 0 - | 1 - | 0,00027366 - | 1 - | 0,000215853 - | 1 - | 0,000214497 - | 1 - | 0,000216131 - |
| 288 | 1 | 3um_FIJJAFBF_254122 | unclassified | unclassified | unclassified | unclassified | unclassified | unclassified | unclassified | unclassified | unclassified | unclassified | unclassified | unclassified | 1 | 0 | 0,000924654 | 9,05E-05 | 1 - | 0,000932913 - | 1 - | 0,001097552 - | 1 - | 0,00086571 - | 1 - | 0,00086027 - | 1 - | 0,000866824 - |
| 289 | 1 | 3um_FIJJAFBF_98072 | Bacteria | Proteobacteri | Gammaprotec | unclassified | unclassified | unclassified | OM182 bacte | cellular proces | Cell wall/mem | 0QTSX | Peptidoglycan | 351348.Maqu | 1 | 0 | 0,0004883 | 4,78E-05 | 1 - | 0,000492662 - | 1 - | 0,000579606 - | 1 - | 0,000457173 - | 1 - | 0,0004543 - | 1 - | 0,000457761 - |
| 290 | 1 | 0.2um_NOKCPHJN_1116524 | unclassified | unclassified | unclassified | unclassified | unclassified | unclassified | unclassified | unclassified | unclassified | unclassified | unclassified | unclassified | 1 | 0 | 0,000955137 | 9,35E-05 | 1 - | 0,000963669 - | 1 - | 0,001133735 - | 1 - | 0,00089425 - | 1 - | 0,00088863 - | 1 - | 0,000895401 - |
| 291 | 1 | 3um_FIJJAFBF_576763 | Eukaryota | Bacillariophyt | Bacillariophyc | Bacillariales | Bacillariaceae | Fragilariopsis | Fragilariopsis | unclassified | unclassified | unclassified | unclassified | unclassified | 1 | 0 | 0,000413893 | 4,05E-05 | 1 - | 0,00041759 - | 1 - | 0,000491285 - | 1 - | 0,000387508 - | 1 - | 0,000385073 - | 1 - | 0,000388007 - |
| 292 | 1 | 3um_FIJJAFBF_504214 | Bacteria | Proteobacteri | Alphaproteob | unclassified | unclassified | Candidatus Pu | Candidatus Pu | unclassified | unclassified | unclassified | unclassified | unclassified | 0,2 | 0,4 | 2,60E-05 | 5,21E-05 | 0 - | 0 - | 0 - | 0 - | 0 - | 0 - | 1 - | 0,000130218 - | 0 - | 0 - |
| 293 | 3 | 0.2um_NOKCPHJN_09406;0.2u | Bacteria | Bacteroidetes | Flavobacteriia | Flavobacterial | Flavobacteriac | Polaribacter | Polaribacter b | metabolism | Carbohydrate | 05D2N | Aldolase | 391587.KAOT | 0,4 | 0,489897949 | 9,15E-05 | 0,00011202 | 0 - | 0 - | 0 - | 0 - | 0 - | 0 - | 1 - | 0,00022779 - | 1 - | 0,000229525 - |
| 294 | 1 | 0.2um_NOKCPHJN_785873 | Eukaryota | Bacillariophyt | Coscinodiscop | Thalassiosirale | Thalassiosirac | Thalassiosira | Thalassiosira ( | information st | Chromatin str | KOG1744 | Core compone | 35128.Thaps6 | 1 | 0 | 0,000797408 | 7,80E-05 | 1 - | 0,000804531 - | 1 - | 0,000946513 - | 1 - | 0,000746576 - | 1 - | 0,000741884 - | 1 - | 0,000747537 - |
| 295 | 1 | 3um_FIJJAFBF_249346 | Bacteria | Proteobacteri | Gammaprotec | unclassified | unclassified | unclassified | Gammaprotec | poorly charact | Function unkn | 17AGQ | NA | 765910.Marpu | 1,4 | 0,8 | 0,000260758 | 0,000139897 | 0 - | 0 - | 1 - | 0,000227748 - | 2 - | 0,000359279 - | 2 - | 0,000357021 - | 2 - | 0,000359742 - |
| 296 | 1 | 3um_FIJJAFBF_209012 | Eukaryota | Bacillariophyt | Coscinodiscop | Thalassiosirale | Thalassiosirac | Thalassiosira | Thalassiosira p | unclassified | unclassified | unclassified | unclassified | unclassified | 0,2 | 0,4 | 5,98E-05 | 0,000119672 | 0 - | 0 - | 0 - | 0 - | 1 - | 0,000299179 - | 0 - | 0 - | 0 - | 0 - |
| 297 | 1 | 3um_FIJJAFBF_399389 | Bacteria | Bacteroidetes | Flavobacteriia | Flavobacterial | Flavobacteriac | unclassified | Flavobacteriac | unclassified | unclassified | unclassified | unclassified | unclassified | 1 | 0 | 0,000208435 | 2,04E-05 | 1 - | 0,000210297 - | 1 - | 0,00024741 - | 1 - | 0,000195148 - | 1 - | 0,000193922 - | 1 - | 0,000195399 - |
| 298 | 1 | 0.2um_NOKCPHJN_207641 | Eukaryota | unclassified | Pelagophycea | Pelagomonad | unclassified | Aureococcus | Aureococcus a | unclassified | unclassified | unclassified | unclassified | unclassified | 1 | 0 | 0,000134547 | 1,32E-05 | 1 - | 0,000135749 - | 1 - | 0,000159706 - | 1 - | 0,00012597 - | 1 - | 0,000125179 - | 1 - | 0,000126132 - |
| 299 | 1 | 3um_FIJJAFBF_673146 | Eukaryota | Bacillariophyt | Coscinodiscop | Thalassiosirale | Thalassiosirac | Thalassiosira | Thalassiosira ( | unclassified | unclassified | unclassified | unclassified | unclassified | 0,8 | 0,4 | 0,000625316 | 0,000313552 | 1 - | 0,0008273 - | 0 - | 0 - | 1 - | 0,000767705 - | 1 - | 0,000762881 - | 1 - | 0,000768693 - |
| 300 | 1 | 3um_FIJJAFBF_565140 | unclassified | unclassified | unclassified | unclassified | unclassified | unclassified | unclassified | unclassified | unclassified | unclassified | unclassified | unclassified | 0,2 | 0,4 | 0,000202163 | 0,000404327 | 0 - | 0 - | 0 - | 0 - | 0 - | 0 - | 1 - | 0,001010817 - | 0 - | 0 - |
| 301 | 1 | 3um_FIJJAFBF_796580 | Bacteria | Bacteroidetes | unclassified | unclassified | unclassified | unclassified | Bacteroidetes | poorly charact | Function unkn | 11NCV | Protein of unk | 643867.Ftrac_ | 1 | 0,632455532 | 0,000469203 | 0,000293167 | 1 - | 0,000498261 - | 0 - | 0 - | 1 - | 0,000462368 - | 1 - | 0,000459462 - | 2 - | 0,000925926 - |
| 302 | 1 | 0.2um_NOKCPHJN_12180 | Bacteria | Proteobacteri | Gammaprotec | Cellvibrionales | Porticoccacea | unclassified | SAR92 bacteri | unclassified | unclassified | unclassified | unclassified | unclassified | 0,2 | 0,4 | 1,88E-05 | 3,75E-05 | 0 - | 0 - | 0 - | 0 - | 0 - | 0 - | 1 - | 9,38E-05 - | 0 - | 0 - |
| 303 | 3 | 0.2um_NOKCPHJN_530547;3ur | Eukaryota | various | various | various | various | various | various | information st | Chromatin str | KOG3467 | Core compone | various | 9,6 | 1,019803903 | 0,005631876 | 0,00038777 | 10 - | 0,005965568 - | 8 - | 0,005614688 - | 11 - | 0,006089416 - | 10 - | 0,005501044 - | 9 - | 0,004988663 - |
| 304 | 1 | 0.2um_NOKCPHJN_131710 | Bacteria | Bacteroidetes | unclassified | unclassified | unclassified | unclassified | Bacteroidetes | metabolism | Energy produc | 0NIET | alcohol dehyd | 656519.Halsa_ | 0,2 | 0,4 | 4,26E-05 | 8,51E-05 | 0 - | 0 - | 0 - | 0 - | 0 - | 0 - | 1 - | 0,000212804 - | 0 - | 0 - |
| 305 | 2 | 0.2um_NOKCPHJN_800807;0.2 | Bacteria | Proteobacteri | Gammaprotec | Methylococca | Methylococca | Methylomona | various | unclassified | unclassified | unclassified | unclassified | unclassified | 0,4 | 0,8 | 0,000165735 | 0,00033147 | 0 - | 0 - | 2 - | 0,000828674 - | 0 - | 0 - | 0 - | 0 - | 0 - | 0 - |
| 306 | 1 | 0.2um_NOKCPHJN_416858 | Bacteria | Proteobacteri | Alphaproteob | Rickettsiales | Anaplasmatac | Anaplasma | Anaplasma ph | unclassified | unclassified | unclassified | unclassified | unclassified | 1 | 0,632455532 | 4,59E-05 | 2,74E-05 | 1 - | 4,63E-05 - | 1 - | 5,45E-05 - | 1 - | 4,30E-05 - | 2 - | 8,55E-05 - | 0 - | 0 - |
| 307 | 1 | 0.2um_NOKCPHJN_971937 | unclassified | unclassified | unclassified | unclassified | unclassified | unclassified | unclassified | unclassified | unclassified | unclassified | unclassified | unclassified | 0,4 | 0,489897949 | 0,000113456 | 0,000138956 | 0 - | 0 - | 0 - | 0 - | 1 - | 0,000284534 - | 1 - | 0,000282746 - | 0 - | 0 - |
| 308 | 1 | 3um_FIJJAFBF_303919 | Bacteria | Bacteroidetes | Flavobacteriia | Flavobacterial | Crocinitomica | unclassified | Crocinitomica | cellular proces | Cell wall/mem | 0P88B | Outer membr | 391598.FBBAL | 2,2 | 0,748331477 | 0,000400776 | 0,000124521 | 1 - | 0,000186187 - | 2 - | 0,000438089 - | 2 - | 0,000345549 - | 3 - | 0,000515066 - | 3 - | 0,00051899 - |
| 309 | 1 | 0.2um_NOKCPHJN_953987 | Bacteria | Proteobacteri | Alphaproteob | Pelagibacteral | Pelagibactera | unclassified | Pelagibactera | metabolism | Energy produc | 05C26 | Dehydrogenas | 335992.SAR11 | 1 | 0 | 0,000176303 | 1,73E-05 | 1 - | 0,000177878 - | 1 - | 0,00020927 - | 1 - | 0,000165064 - | 1 - | 0,000164027 - | 1 - | 0,000165277 - |
| 310 | 1 | 0.2um_NOKCPHJN_157490 | Bacteria | Proteobacteri | Alphaproteob | Rhodobactera | Hyphomonad | Robiginitomac | Robiginitomac | unclassified | unclassified | unclassified | unclassified | unclassified | 1,2 | 0,4 | 0,000128382 | 3,78E-05 | 1 - | 0,000109208 - | 1 - | 0,000128481 - | 1 - | 0,000101341 - | 2 - | 0,000201408 - | 1 - | 0,000101471 - |
| 311 | 1 | 3um_FIJJAFBF_08771 | Eukaryota | unclassified | Raphidophyce | Chattonellales | Chattonellace | Heterosigma | Heterosigma a | metabolism | Energy produc | 0IFBF | One of the tw | 45157.CMV08 | 0,6 | 0,489897949 | 0,000138874 | 0,000113392 | 0 - | 0 - | 0 - | 0 - | 1 - | 0,000231843 - | 1 - | 0,000230386 - | 1 - | 0,000232141 - |
| 312 | 1 | 3um_FIJJAFBF_113804 | Bacteria | Proteobacteri | Alphaproteob | Rhizobiales | unclassified | unclassified | Rhizobiales ba | unclassified | unclassified | unclassified | unclassified | unclassified | 0,8 | 0,4 | 7,92E-05 | 3,97E-05 | 1 - | 0,000104772 - | 0 - | 0 - | 1 - | 9,72E-05 - | 1 - | 9,66E-05 - | 1 - | 9,73E-05 - |
| 313 | 1 | 3um_FIJJAFBF_477756 | Bacteria | Nitrospirae | unclassified | unclassified | unclassified | unclassified | Nitrospirae ba | unclassified | unclassified | unclassified | unclassified | unclassified | 0,6 | 0,489897949 | 8,85E-05 | 7,24E-05 | 1 - | 0,00015521 - | 0 - | 0 - | 1 - | 0,00014403 - | 1 - | 0,000143125 - | 0 - | 0 - |
| 314 | 1 | 3um_FIJJAFBF_607280 | Bacteria | Proteobacteri | Gammaprotec | unclassified | unclassified | unclassified | Gammaprotec | unclassified | unclassified | unclassified | unclassified | unclassified | 1 | 0 | 0,000223438 | 2,19E-05 | 1 - | 0,000225434 - | 1 - | 0,000265218 - | 1 - | 0,000209195 - | 1 - | 0,00020788 - | 1 - | 0,000209464 - |
| 315 | 1 | 0.2um_NOKCPHJN_17065 | Bacteria | Bacteroidetes | Flavobacteriia | Flavobacterial | Flavobacteriac | Polaribacter | Polaribacter t | poorly charact | Function unkn | 0ZVFQ | Pkd domain c | 269798.CHU_ | 1,2 | 0,979795897 | 1,39E-05 | 1,13E-05 | 0 - | 0 - | 0 - | 0 - | 2 - | 2,32E-05 - | 2 - | 2,30E-05 - | 2 - | 2,32E-05 - |
| 316 | 1 | 0.2um_NOKCPHJN_705309 | Eukaryota | Chordata | Aves | Passeriformes | Muscicapidae | Ficedula | Ficedula albic | poorly charact | Function unkn | 1EI7F | NA | 3702.AT4G362 | 0,4 | 0,489897949 | 3,27E-05 | 4,00E-05 | 0 - | 0 - | 0 - | 0 - | 1 - | 8,20E-05 - | 1 - | 8,14E-05 - | 0 - | 0 - |
| 317 | 1 | 3um_FIJJAFBF_211494 | Bacteria | Proteobacteri | Gammaprotec | Alteromonada | Alteromonada | Paraglaciecola | Paraglaciecola | metabolism | Inorganic ion t | 16TQC | TONB-depend | 87626.PTD2_2 | 5,4 | 1,496662955 | 0,000469684 | 0,000104604 | 4 - | 0,000357205 - | 4 - | 0,000420244 - | 5 - | 0,000414342 - | 8 - | 0,000658781 - | 6 - | 0,00049785 - |
| 318 | 1 | 3um_FIJJAFBF_760823 | Bacteria | Proteobacteri | Alphaproteob | Rhodobactera | Rhodobactera | Roseobacter | Roseobacter s | poorly charact | Function unkn | 090JY | secretion activ | 391593.RCCS2 | 0,2 | 0,4 | 8,02E-05 | 0,000160348 | 0 - | 0 - | 0 - | 0 - | 1 - | 0,000400871 - | 0 - | 0 - | 0 - | 0 - |
| 319 | 1 | 0.2um_NOKCPHJN_09839 | Bacteria | Proteobacteri | Gammaprotec | Alteromonada | Psychromonac | Psychromonas | Psychromonas | poorly charact | Function unkn | COG3517 | type VI secreti | 357804.Ping_ | 1,8 | 0,748331477 | 0,000316488 | 0,000118617 | 2 - | 0,000362371 - | 1 - | 0,000213161 - | 3 - | 0,000504401 - | 2 - | 0,000334154 - | 1 - | 0,00016835 - |
| 320 | 1 | 0.2um_NOKCPHJN_770232 | Eukaryota | unclassified | Pelagophycea | Pelagomonad | unclassified | Aureococcus | Aureococcus a | cellular proces | Posttranslatio | KOG0019 | heat shock pro | 272952.HpaP8 | 3 | 0,632455532 | 0,000331126 | 4,95E-05 | 3 - | 0,000339899 - | 2 - | 0,000266589 - | 3 - | 0,000315414 - | 4 - | 0,000417909 - | 3 - | 0,00031582 - |
| 321 | 1 | 3um_FIJJAFBF_453412 | Bacteria | Proteobacteri | Alphaproteob | Rhodobactera | unclassified | unclassified | Rhodobactera | poorly charact | Function unkn | COG3181 | extra-cytoplas | 388739.RSK20 | 0,6 | 0,489897949 | 0,000149066 | 0,000121714 | 0 - | 0 - | 0 - | 0 - | 1 - | 0,000248859 - | 1 - | 0,000247295 - | 1 - | 0,000249179 - |
| 322 | 1 | 0.2um_NOKCPHJN_849416 | Bacteria | Bacteroidetes | Flavobacteriia | Flavobacterial | Flavobacteriac | Muricauda | Muricauda sp. | unclassified | unclassified | unclassified | unclassified | unclassified | 0,2 | 0,4 | 2,62E-05 | 5,24E-05 | 1 - | 0,000131082 - | 0 - | 0 - | 0 - | 0 - | 0 - | 0 - | 0 - | 0 - |
| 323 | 1 | 3um_FIJJAFBF_665894 | unclassified | unclassified | unclassified | unclassified | unclassified | unclassified | unclassified | unclassified | unclassified | unclassified | unclassified | unclassified | 0,2 | 0,4 | 0,000188589 | 0,000377178 | 1 - | 0,000942945 - | 0 - | 0 - | 0 - | 0 - | 0 - | 0 - | 0 - | 0 - |
| 324 | 1 | 0.2um_NOKCPHJN_950149 | unclassified | unclassified | unclassified | unclassified | unclassified | unclassified | unclassified | unclassified | unclassified | unclassified | unclassified | unclassified | 0,2 | 0,4 | 2,56E-05 | 5,12E-05 | 0 - | 0 - | 0 - | 0 - | 0 - | 0 - | 1 - | 0,000127952 - | 0 - | 0 - |
| 325 | 1 | 3um_FIJJAFBF_500695 | Eukaryota | Bacillariophyt | Mediophyceae | Cymatosirales | Cymatosirace | Plagiogrammc | Plagiogrammc | metabolism | Energy produc | COG4451 | RuBisCO catal | 323097.Nham | 1,2 | 0,4 | 0,000741659 | 0,000218548 | 1 - | 0,000630891 - | 1 - | 0,000742229 - | 1 - | 0,000585444 - | 2 - | 0,00116353 - | 1 - | 0,000586198 - |
| 326 | 1 | 0.2um_NOKCPHJN_111274 | Bacteria | Proteobacteri | Gammaprotec | Cellvibrionales | Porticoccacea | unclassified | gamma prote | unclassified | unclassified | unclassified | unclassified | unclassified | 2,4 | 1,496662955 | 0,000229271 | 0,000133211 | 2 - | 0,000199078 - | 1 - | 0,000117105 - | 1 - | 9,24E-05 - | 3 - | 0,000275364 - | 5 - | 0,000462438 - |
| 327 | 1 | 0.2um_NOKCPHJN_306732 | Bacteria | Proteobacteri | Gammaprotec | Cellvibrionales | Porticoccacea | unclassified | Gammaprotec | unclassified | unclassified | unclassified | unclassified | unclassified | 0,2 | 0,4 | 4,47E-05 | 8,93E-05 | 0 - | 0 - | 1 - | 0,000223311 - | 0 - | 0 - | 0 - | 0 - | 0 - | 0 - |
| 328 | 6 | 3um_FIJJAFBF_124795;3um_FI | Eukaryota | unclassified | various | various | various | various | various | metabolism | Energy produc | 123BF | photosystem I | various | 0,6 | 0,489897949 | 0,000727533 | 0,000594036 | 0 - | 0 - | 0 - | 0 - | 1 - | 0,001214578 - | 1 - | 0,001206946 - | 1 - | 0,001216142 - |
| 329 | 1 | 0.2um_NOKCPHJN_627902 | unclassified | unclassified | unclassified | unclassified | unclassified | unclassified | unclassified | unclassified | unclassified | unclassified | unclassified | unclassified | 2,4 | 1,019803903 | 0,000601956 | 0,000265688 | 1 - | 0,000251994 - | 3 - | 0,000889396 - | 4 - | 0,000935365 - | 2 - | 0,000464743 - | 2 - | 0,000468284 - |
| 330 | 1 | 0.2um_NOKCPHJN_321141 | Eukaryota | unclassified | unclassified | Isochrysidales | Noelaerhabda | Emiliania | Emiliania huxl | unclassified | unclassified | unclassified | unclassified | unclassified | 0,2 | 0,4 | 2,17E-05 | 4,35E-05 | 0 - | 0 - | 0 - | 0 - | 1 - | 0,000108647 - | 0 - | 0 - | 0 - | 0 - |
| 331 | 1 | 3um_FIJJAFBF_135240 | Bacteria | Cyanobacteria | unclassified | Synechococca | Synechococca | Synechococcu | Synechococcu | unclassified | unclassified | unclassified | unclassified | unclassified | 1,2 | 0,4 | 0,000378072 | 0,000113342 | 1 - | 0,000321223 - | 1 - | 0,000377912 - | 1 - | 0,000298083 - | 1 - | 0,00029621 - | 2 - | 0,000596934 - |
| 332 | 1 | 0.2um_NOKCPHJN_15784 | Bacteria | Proteobacteri | Gammaprotec | Alteromonada | Alteromonada | Paraglaciecola | Paraglaciecola | poorly charact | Function unkn | 16R8I | Lipoprotein | 1085623.GNIT | 2,2 | 1,166190379 | 0,000206118 | 0,000109633 | 4 - | 0,000375563 - | 2 - | 0,000220921 - | 1 - | 8,71E-05 - | 3 - | 0,000259739 - | 1 - | 8,72E-05 - |
| 333 | 1 | 3um_FIJJAFBF_717710 | Eukaryota | Eustigmatoph | unclassified | Eustigmatales | Monodopsida | Nannochlorop | Nannochlorop | unclassified | unclassified | unclassified | unclassified | unclassified | 1,2 | 0,748331477 | 0,000599122 | 0,000367788 | 1 - | 0,000531478 - | 0 - | 0 - | 2 - | 0,000986385 - | 1 - | 0,000490093 - | 2 - | 0,000987654 - |
| 334 | 2 | 0.2um_NOKCPHJN_707303;3ur | Bacteria | Proteobacteri | Gammaprotec | Alteromonada | Colwelliaceae | Colwellia | Colwellia sp. P | unclassified | unclassified | unclassified | unclassified | unclassified | 0,2 | 0,4 | 4,68E-05 | 9,35E-05 | 0 - | 0 - | 0 - | 0 - | 1 - | 0,000233841 - | 0 - | 0 - | 0 - | 0 - |
| 335 | 1 | 0.2um_NOKCPHJN_598600 | Eukaryota | Chlorophyta | Chlorophycea | Sphaeropleale | Selenastracea | Raphidocelis | Raphidocelis s | information st | Chromatin str | KOG1862 | GRB10 interac | 7165.AGAP00 | 0,2 | 0,4 | 1,74E-05 | 3,48E-05 | 0 - | 0 - | 1 - | 8,70E-05 - | 0 - | 0 - | 0 - | 0 - | 0 - | 0 - |
| 336 | 1 | 3um_FIJJAFBF_262604 | unclassified | unclassified | unclassified | unclassified | unclassified | unclassified | unclassified | unclassified | unclassified | unclassified | unclassified | unclassified | 0,2 | 0,4 | 0,000121458 | 0,000242916 | 0 - | 0 - | 0 - | 0 - | 1 - | 0,000607289 - | 0 - | 0 - | 0 - | 0 - |
| 337 | 1 | 3um_FIJJAFBF_257116 | Eukaryota | unclassified | Cryptophyta | Pyrenomonad | Chroomonada | Chroomonas | various | unclassified | unclassified | unclassified | unclassified | unclassified | 1 | 0,632455532 | 0,000761062 | 0,000483016 | 2 - | 0,001511963 - | 1 - | 0,000889396 - | 1 - | 0,000701524 - | 0 - | 0 - | 1 - | 0,000702427 - |
| 338 | 1 | 0.2um_NOKCPHJN_565086 | Bacteria | Bacteroidetes | Cytophagia | Cytophagales | Flammeovirga | unclassified | Flammeovirga | unclassified | unclassified | unclassified | unclassified | unclassified | 0,4 | 0,489897949 | 5,65E-05 | 6,93E-05 | 1 - | 0,000146891 - | 0 - | 0 - | 0 - | 0 - | 1 - | 0,000135453 - | 0 - | 0 - |
| 339 | 1 | 0.2um_NOKCPHJN_1105265 | unclassified | unclassified | unclassified | unclassified | unclassified | unclassified | unclassified | unclassified | unclassified | unclassified | unclassified | unclassified | 0,4 | 0,489897949 | 8,18E-05 | 0,000100352 | 1 - | 0,000212849 - | 0 - | 0 - | 0 - | 0 - | 1 - | 0,000196275 - | 0 - | 0 - |
| 340 | 1 | 0.2um_NOKCPHJN_817880 | Eukaryota | unclassified | Pelagophycea | Pelagomonad | unclassified | Aureococcus | Aureococcus a | unclassified | unclassified | unclassified | unclassified | unclassified | 0,6 | 0,489897949 | 7,48E-05 | 6,10E-05 | 0 - | 0 - | 0 - | 0 - | 1 - | 0,000124811 - | 1 - | 0,000124027 - | 1 - | 0,000124972 - |
| 341 | 1 | 3um_FIJJAFBF_350095 | Bacteria | Proteobacteri | Alphaproteob | Rhodobactera | Rhodobactera | Phaeobacter | Phaeobacter g | information st | Translation, ri | 05KK9 | This protein b | 292414.TM10 | 0,8 | 0,4 | 0,000334765 | 0,000167861 | 1 - | 0,000442898 - | 0 - | 0 - | 1 - | 0,000410994 - | 1 - | 0,000408411 - | 1 - | 0,000411523 - |
| 342 | 1 | 3um_FIJJAFBF_67730 | Bacteria | Proteobacteri | Gammaprotec | Cellvibrionales | unclassified | unclassified | Cellvibrionales | unclassified | unclassified | unclassified | unclassified | unclassified | 0,4 | 0,489897949 | 0,000267459 | 0,000331354 | 0 - | 0 - | 1 - | 0,000747608 - | 1 - | 0,000589687 - | 0 - | 0 - | 0 - | 0 - |
| 343 | 1 | 0.2um_NOKCPHJN_635206 | Eukaryota | Arthropoda | Insecta | Hymenoptera | Apidae | Apis | Apis mellifera | poorly charact | Function unkn | 0KN5V | NA | 140110.Necha | 0,2 | 0,4 | 1,39E-05 | 2,77E-05 | 0 - | 0 - | 0 - | 0 - | 1 - | 6,93E-05 - | 0 - | 0 - | 0 - | 0 - |
| 344 | 3 | 0.2um_NOKCPHJN_202376;3ur | Bacteria | Bacteroidetes | various | various | various | various | various | poorly charact | Function unkn | COG2120 | ImbE family | various | 0,4 | 0,489897949 | 0,000135855 | 0,00016639 | 0 - | 0 - | 0 - | 0 - | 0 - | 0 - | 1 - | 0,000338349 - | 1 - | 0,000340927 - |
| 345 | 1 | 3um_FIJJAFBF_800231 | Bacteria | Bacteroidetes | Flavobacteriia | Flavobacterial | Flavobacteriac | Polaribacter | Polaribacter s | information st | Translation, ri | 09FEM | Binds to the 2 | 313594.PI23P | 1,8 | 0,4 | 0,001028194 | 0,000171231 | 2 - | 0,001177099 - | 1 - | 0,000692415 - | 2 - | 0,001092305 - | 2 - | 0,001085441 - | 2 - | 0,001093711 - |
| 346 | 1 | 0.2um_NOKCPHJN_170568 | unclassified | unclassified | unclassified | unclassified | unclassified | unclassified | unclassified | unclassified | unclassified | unclassified | unclassified | unclassified | 0,6 | 0,489897949 | 2,85E-05 | 2,36E-05 | 1 - | 4,59E-05 - | 1 - | 5,40E-05 - | 0 - | 0 - | 0 - | 0 - | 1 - | 4,26E-05 - |
| 347 | 1 | 0.2um_NOKCPHJN_76179 | Eukaryota | Bacillariophyt | Mediophyceae | Cymatosirales | Cymatosirace | Campylosira | Campylosira c | metabolism | Carbohydrate | COG1850 | RuBisCO catal | 35128.Thapsd | 1,4 | 0,489897949 | 0,000246391 | 8,26E-05 | 2 - | 0,000357934 - | 1 - | 0,000210551 - | 2 - | 0,00033215 - | 1 - | 0,000165031 - | 1 - | 0,000166289 - |
| 348 | 1 | 3um_FIJJAFBF_191786 | Eukaryota | unclassified | Florideophyce | Gracilariales | Gracilariaceae | Gracilaria | Gracilaria chile | unclassified | unclassified | unclassified | unclassified | unclassified | 1,8 | 0,4 | 0,00153201 | 0,000255134 | 2 - | 0,001753877 - | 1 - | 0,001031699 - | 2 - | 0,001627535 - | 2 - | 0,001617307 - | 2 - | 0,00162963 - |
| 349 | 1 | 3um_FIJJAFBF_294667 | Bacteria | Bacteroidetes | Flavobacteriia | Flavobacterial | Flavobacteriac | Olleya | Olleya sp. Bg1 | unclassified | unclassified | unclassified | unclassified | unclassified | 2,6 | 0,489897949 | 0,001840167 | 0,000464314 | 3 - | 0,002121625 - | 3 - | 0,002496046 - | 2 - | 0,001312528 - | 3 - | 0,00195642 - | 2 - | 0,001314218 - |
| 350 | 1 | 3um_FIJJAFBF_11563 | Bacteria | Bacteroidetes | Flavobacteriia | Flavobacterial | Flavobacteriac | Lutibacter | Lutibacter sp. | unclassified | unclassified | unclassified | unclassified | unclassified | 0,8 | 0,4 | 0,000168805 | 8,69E-05 | 0 - | 0 - | 1 - | 0,000251022 - | 1 - | 0,000197997 - | 1 - | 0,000196753 - | 1 - | 0,000198252 - |
| 351 | 1 | 0.2um_NOKCPHJN_785139 | Bacteria | unclassified | unclassified | unclassified | unclassified | unclassified | uncultured ba | metabolism | Inorganic ion t | 05C27 | TonB-depende | 313596.RB25C | 2,2 | 0,979795897 | 0,000235643 | 0,000105675 | 4 - | 0,000439018 - | 1 - | 0,000129124 - | 2 - | 0,000203696 - | 2 - | 0,000202416 - | 2 - | 0,000203959 - |
| 352 | 1 | 3um_FIJJAFBF_474234 | Eukaryota | unclassified | Cryptophyta | Pyrenomonad | Geminigerace | Teleaulax | Teleaulax amp | unclassified | unclassified | unclassified | unclassified | unclassified | 0,6 | 0,489897949 | 0,000275394 | 0,000224861 | 0 - | 0 - | 0 - | 0 - | 1 - | 0,000459756 - | 1 - | 0,000456866 - | 1 - | 0,000460347 - |
| 353 | 1 | 0.2um_NOKCPHJN_35610 | Eukaryota | unclassified | Oomycetes | Myzocytiopsic | Eurychasmata | Eurychasma | Eurychasma d | unclassified | unclassified | unclassified | unclassified | unclassified | 0,2 | 0,4 | 4,69E-07 | 9,39E-07 | 0 - | 0 - | 0 - | 0 - | 1 - | 2,35E-06 - | 0 - | 0 - | 0 - | 0 - |
| 354 | 1 | 0.2um_NOKCPHJN_671219 | unclassified | unclassified | unclassified | unclassified | unclassified | unclassified | unclassified | unclassified | unclassified | unclassified | unclassified | unclassified | 1 | 0 | 8,09E-05 | 7,92E-06 | 1 - | 8,17E-05 - | 1 - | 9,61E-05 - | 1 - | 7,58E-05 - | 1 - | 7,53E-05 - | 1 - | 7,59E-05 - |
| 355 | 1 | 3um_FIJJAFBF_06027 | Bacteria | Bacteroidetes | Flavobacteriia | Flavobacterial | Flavobacteriac | Algibacter | Algibacter algi | unclassified | unclassified | unclassified | unclassified | unclassified | 0,6 | 0,489897949 | 7,90E-05 | 6,46E-05 | 1 - | 0,000138318 - | 0 - | 0 - | 1 - | 0,000128354 - | 0 - | 0 - | 1 - | 0,00012852 - |
| 356 | 1 | 0.2um_NOKCPHJN_72831 | Bacteria | Proteobacteri | Gammaprotec | Pseudomonad | Pseudomonad | Pseudomonas | Pseudomonas | information st | Replication, re | 05C68 | Can catalyze t | 390235.PputW | 0,6 | 0,489897949 | 0,000134014 | 0,000109612 | 1 - | 0,000235104 - | 0 - | 0 - | 1 - | 0,000218168 - | 1 - | 0,000216797 - | 0 - | 0 - |
| 357 | 1 | 3um_FIJJAFBF_176995 | Eukaryota | unclassified | Bangiophycea | Porphyridiales | Porphyridiace | Porphyridium | Porphyridium | unclassified | unclassified | unclassified | unclassified | unclassified | 4,8 | 1,166190379 | 0,000811069 | 0,000168299 | 4 - | 0,000689146 - | 4 - | 0,000810765 - | 5 - | 0,000799379 - | 4 - | 0,000635484 - | 7 - | 0,001120571 - |
| 358 | 3 | 0.2um_NOKCPHJN_57009;0.2u | Bacteria | Bacteroidetes | Flavobacteriia | Flavobacterial | Flavobacteriac | various | various | information st | Translation, ri | 0PAI9 | Binds 16S rRN | various | 0,4 | 0,489897949 | 0,000379934 | 0,000465863 | 1 - | 0,000985324 - | 0 - | 0 - | 1 - | 0,000914345 - | 0 - | 0 - | 0 - | 0 - |
| 359 | 1 | 3um_FIJJAFBF_594483 | Bacteria | Bacteroidetes | unclassified | unclassified | unclassified | unclassified | various | unclassified | unclassified | unclassified | unclassified | unclassified | 0,2 | 0,4 | 3,54E-05 | 7,09E-05 | 0 - | 0 - | 0 - | 0 - | 0 - | 0 - | 0 - | 0 - | 1 - | 0,000177134 - |
| 360 | 1 | 0.2um_NOKCPHJN_437854 | Bacteria | Proteobacteri | Gammaprotec | Cellvibrionales | unclassified | unclassified | Cellvibrionales | cellular proces | Defense mech | 179PG | N-acetyl-anhy | 396588.Tgr7_ | 0,2 | 0,4 | 8,43E-05 | 0,000168656 | 0 - | 0 - | 0 - | 0 - | 1 - | 0,000421641 - | 0 - | 0 - | 0 - | 0 - |
| 361 | 1 | 3um_FIJJAFBF_24076 | Bacteria | unclassified | unclassified | unclassified | unclassified | unclassified | uncultured ba | unclassified | unclassified | unclassified | unclassified | unclassified | 0,2 | 0,4 | 7,67E-05 | 0,000153412 | 0 - | 0 - | 1 - | 0,000383531 - | 0 - | 0 - | 0 - | 0 - | 0 - | 0 - |
| 362 | 1 | 3um_FIJJAFBF_153653 | Eukaryota | Arthropoda | Insecta | Hemiptera | Delphacidae | Nilaparvata | Nilaparvata lu | unclassified | unclassified | unclassified | unclassified | unclassified | 0,2 | 0,4 | 2,78E-05 | 5,55E-05 | 1 - | 0,000138756 - | 0 - | 0 - | 0 - | 0 - | 0 - | 0 - | 0 - | 0 - |
| 363 | 1 | 0.2um_NOKCPHJN_868369 | Bacteria | Bacteroidetes | Flavobacteriia | Flavobacterial | Flavobacteriac | Polaribacter | Polaribacter s | poorly charact | Function unkn | 0P8NI | Inherit from b | 156586.BBFL7 | 1,8 | 0,4 | 0,000399734 | 9,79E-05 | 1 - | 0,000224281 - | 2 - | 0,000527723 - | 2 - | 0,000416249 - | 2 - | 0,000413634 - | 2 - | 0,000416785 - |
| 364 | 1 | 3um_FIJJAFBF_348211 | Bacteria | Proteobacteri | Alphaproteob | unclassified | unclassified | Candidatus Pu | Candidatus Pu | metabolism | Energy produc | 01RNF | Produces ATP | 488538.SAR11 | 9,4 | 1,019803903 | 0,001711182 | 0,000265987 | 8 - | 0,00146768 - | 10 - | 0,002158366 - | 9 - | 0,001532198 - | 9 - | 0,001522569 - | 11 - | 0,001875097 - |
| 365 | 1 | 3um_FIJJAFBF_382743 | Viruses | unclassified | unclassified | unclassified | Phycodnavirid | unclassified | Chrysochromu | unclassified | unclassified | unclassified | unclassified | unclassified | 0,8 | 0,4 | 0,000359475 | 0,000185049 | 0 - | 0 - | 1 - | 0,000534559 - | 1 - | 0,000421641 - | 1 - | 0,000418991 - | 1 - | 0,000422184 - |
| 366 | 1 | 0.2um_NOKCPHJN_390699 | Bacteria | Proteobacteri | Alphaproteob | Rhizobiales | Rhizobiaceae | Rhizobium | Rhizobium loe | poorly charact | Function unkn | 0ZKS3 | head-to-tail jo | 438753.AZC_3 | 0,4 | 0,489897949 | 5,98E-05 | 7,32E-05 | 0 - | 0 - | 0 - | 0 - | 1 - | 0,000149315 - | 0 - | 0 - | 1 - | 0,000149507 - |
| 367 | 1 | 0.2um_NOKCPHJN_477681 | Eukaryota | Nematoda | Enoplea | Trichinellida | Trichinellidae | Trichinella | Trichinella nel | poorly charact | Function unkn | 0MS4I | conserved hyp | 441960.XP_00 | 0,6 | 0,489897949 | 5,40E-05 | 4,41E-05 | 0 - | 0 - | 0 - | 0 - | 1 - | 9,01E-05 - | 1 - | 8,96E-05 - | 1 - | 9,02E-05 - |
| 368 | 1 | 0.2um_NOKCPHJN_65240 | Bacteria | Proteobacteri | Gammaprotec | Cellvibrionales | Porticoccacea | unclassified | SAR92 bacteri | unclassified | unclassified | unclassified | unclassified | unclassified | 4,2 | 0,979795897 | 0,000428589 | 7,83E-05 | 4 - | 0,000418087 - | 3 - | 0,000368903 - | 6 - | 0,000581955 - | 4 - | 0,000385532 - | 4 - | 0,00038847 - |
| 369 | 1 | 3um_FIJJAFBF_327766 | unclassified | unclassified | unclassified | unclassified | unclassified | unclassified | unclassified | unclassified | unclassified | unclassified | unclassified | unclassified | 0,2 | 0,4 | 0,000148634 | 0,000297267 | 1 - | 0,000743168 - | 0 - | 0 - | 0 - | 0 - | 0 - | 0 - | 0 - | 0 - |
| 370 | 1 | 3um_FIJJAFBF_201378 | Bacteria | Bacteroidetes | Flavobacteriia | Flavobacterial | Flavobacteriac | Polaribacter | Polaribacter fi | cellular proces | Cell wall/mem | 0PENK | NA | 391587.KAOT | 4 | 0,894427191 | 0,000588583 | 9,65E-05 | 3 - | 0,000452808 - | 3 - | 0,000532719 - | 4 - | 0,000560253 - | 5 - | 0,000695915 - | 5 - | 0,000701218 - |
| 371 | 1 | 3um_FIJJAFBF_557220 | Eukaryota | Chlorophyta | Ulvophyceae | Ulvales | Ulvaceae | Ulva | Ulva fasciata | metabolism | Energy produc | COG0055 | Produces ATP | 3702.ATCG00 | 1 | 0,894427191 | 0,000189233 | 0,000173355 | 1 - | 0,000181561 - | 2 - | 0,000427205 - | 0 - | 0 - | 0 - | 0 - | 2 - | 0,000337397 - |
| 372 | 1 | 3um_FIJJAFBF_728423 | Bacteria | Proteobacteri | Gammaprotec | Cellvibrionales | Spongiibacter | Oceanicoccus | Oceanicoccus | unclassified | unclassified | unclassified | unclassified | unclassified | 0,2 | 0,4 | 8,71E-05 | 0,000174126 | 0 - | 0 - | 1 - | 0,000435316 - | 0 - | 0 - | 0 - | 0 - | 0 - | 0 - |
| 373 | 1 | 0.2um_NOKCPHJN_586666 | Bacteria | Proteobacteri | Betaproteoba | unclassified | unclassified | unclassified | Betaproteoba | poorly charact | Function unkn | 0XRND | NA | 314607.KB13_ | 1 | 0 | 0,000395079 | 3,87E-05 | 1 - | 0,000398608 - | 1 - | 0,000468954 - | 1 - | 0,000369894 - | 1 - | 0,00036757 - | 1 - | 0,00037037 - |
| 374 | 1 | 3um_FIJJAFBF_26279 | Bacteria | Bacteroidetes | Flavobacteriia | Flavobacterial | Flavobacteriac | Flavobacteriu | Flavobacteriu | information st | Replication, re | 08IYB | Dead deah bo | 313628.LNTAR | 1 | 0 | 0,000102861 | 1,01E-05 | 1 - | 0,00010378 - | 1 - | 0,000122095 - | 1 - | 9,63E-05 - | 1 - | 9,57E-05 - | 1 - | 9,64E-05 - |
| 375 | 1 | 0.2um_NOKCPHJN_1059776 | Viruses | unclassified | unclassified | unclassified | unclassified | unclassified | uncultured M | unclassified | unclassified | unclassified | unclassified | unclassified | 1 | 0 | 0,000388024 | 3,80E-05 | 1 - | 0,00039149 - | 1 - | 0,00046058 - | 1 - | 0,000363289 - | 1 - | 0,000361006 - | 1 - | 0,000363757 - |
| 376 | 1 | 0.2um_NOKCPHJN_217488 | Bacteria | Proteobacteri | Gammaprotec | Oceanospirilla | Saccharospirill | Reinekea | Reinekea fors | metabolism | Inorganic ion t | 0GH39 | Phosphonate | 525897.Dbac_ | 1,6 | 0,489897949 | 0,000938547 | 0,00025822 | 2 - | 0,001201286 - | 1 - | 0,000706643 - | 2 - | 0,00111475 - | 1 - | 0,000553872 - | 2 - | 0,001116185 - |
| 377 | 1 | 0.2um_NOKCPHJN_369201 | Bacteria | Proteobacteri | Alphaproteob | Rhodobactera | Rhodobactera | Yoonia | Yoonia vestfol | information st | Transcription | 16QIX | transcriptiona | 391626.OA307 | 0,4 | 0,489897949 | 0,000110326 | 0,000135859 | 1 - | 0,00025345 - | 1 - | 0,000298179 - | 0 - | 0 - | 0 - | 0 - | 0 - | 0 - |
| 378 | 1 | 3um_FIJJAFBF_406358 | Bacteria | Bacteroidetes | Flavobacteriia | Flavobacterial | Flavobacteriac | Gaetbulibacte | Gaetbulibacte | poorly charact | Function unkn | 0PBET | NA | 655815.ZPR_3 | 0,8 | 0,4 | 0,000370299 | 0,000185679 | 1 - | 0,00048991 - | 0 - | 0 - | 1 - | 0,000454619 - | 1 - | 0,000451762 - | 1 - | 0,000455204 - |
| 379 | 1 | 0.2um_NOKCPHJN_219723 | Eukaryota | unclassified | Labyrinthulon | unclassified | Thraustochytr | Aurantiochytr | Aurantiochytr | metabolism | Coenzyme tra | KOG1426 | regulator of cl | 572307.XP_00 | 0,4 | 0,489897949 | 1,81E-05 | 2,22E-05 | 0 - | 0 - | 0 - | 0 - | 1 - | 4,53E-05 - | 0 - | 0 - | 1 - | 4,54E-05 - |
| 380 | 1 | 0.2um_NOKCPHJN_428374 | Viruses | unclassified | unclassified | unclassified | Phycodnavirid | unclassified | Aureococcus a | unclassified | unclassified | unclassified | unclassified | unclassified | 0,6 | 0,489897949 | 9,31E-05 | 7,61E-05 | 1 - | 0,000163303 - | 0 - | 0 - | 1 - | 0,00015154 - | 1 - | 0,000150587 - | 0 - | 0 - |
| 381 | 1 | 3um_FIJJAFBF_449836 | Eukaryota | Bacillariophyt | Bacillariophyc | Naviculales | Phaeodactylac | Phaeodactylu | Phaeodactylu | unclassified | unclassified | unclassified | unclassified | unclassified | 2,4 | 0,8 | 0,002470372 | 0,000728154 | 1 - | 0,001056552 - | 2 - | 0,002486021 - | 3 - | 0,002941328 - | 3 - | 0,002922844 - | 3 - | 0,002945114 - |
| 382 | 1 | 3um_FIJJAFBF_639191 | Bacteria | Planctomycet | unclassified | unclassified | unclassified | unclassified | Planctomycet | poorly charact | Function unkn | 0IRPE | NA | 43151.ADAR0 | 0,8 | 0,4 | 0,0001006 | 5,16E-05 | 1 - | 0,00012492 - | 1 - | 0,000146966 - | 1 - | 0,000115921 - | 1 - | 0,000115193 - | 0 - | 0 - |
| 383 | 1 | 0.2um_NOKCPHJN_194701 | Eukaryota | unclassified | Oomycetes | Saprolegniales | Saprolegniace | Aphanomyces | Aphanomyces | information st | Chromatin str | KOG0889 | Transformatio | 4787.PITG_03 | 0,2 | 0,4 | 3,85E-06 | 7,71E-06 | 0 - | 0 - | 0 - | 0 - | 0 - | 0 - | 0 - | 0 - | 1 - | 1,93E-05 - |
| 384 | 1 | 3um_FIJJAFBF_461043 | unclassified | unclassified | unclassified | unclassified | unclassified | unclassified | unclassified | unclassified | unclassified | unclassified | unclassified | unclassified | 0,6 | 0,489897949 | 9,24E-05 | 7,67E-05 | 0 - | 0 - | 1 - | 0,000179426 - | 0 - | 0 - | 1 - | 0,000140635 - | 1 - | 0,000141707 - |
| 385 | 1 | 3um_FIJJAFBF_616971 | Bacteria | Bacteroidetes | unclassified | unclassified | unclassified | unclassified | Bacteroidetes | poorly charact | Function unkn | 0XR4Y | Ragb susd dor | 886377.Murru | 1 | 0,632455532 | 0,000151637 | 9,25E-05 | 0 - | 0 - | 1 - | 0,000182602 - | 1 - | 0,00014403 - | 1 - | 0,000143125 - | 2 - | 0,00028843 - |
| 386 | 1 | 3um_FIJJAFBF_136477 | Bacteria | Proteobacteri | Gammaprotec | Alteromonada | Colwelliaceae | Colwellia | Colwellia sp. P | unclassified | unclassified | unclassified | unclassified | unclassified | 0,8 | 0,748331477 | 7,20E-05 | 6,72E-05 | 0 - | 0 - | 0 - | 0 - | 1 - | 9,02E-05 - | 2 - | 0,000179302 - | 1 - | 9,03E-05 - |
| 387 | 1 | 3um_FIJJAFBF_138590 | Bacteria | Bacteroidetes | Cytophagia | Cytophagales | Cytophagacea | Spirosoma | Spirosoma lut | unclassified | unclassified | unclassified | unclassified | unclassified | 0,4 | 0,489897949 | 0,000189879 | 0,000232557 | 0 - | 0 - | 0 - | 0 - | 0 - | 0 - | 1 - | 0,000472897 - | 1 - | 0,0004765 - |
| 388 | 1 | 3um_FIJJAFBF_722163 | Eukaryota | unclassified | Dinophyceae | Suessiales | Symbiodiniace | Symbiodinium | Symbiodinium | unclassified | unclassified | unclassified | unclassified | unclassified | 1,4 | 0,489897949 | 0,00058275 | 0,000194032 | 2 - | 0,000847284 - | 1 - | 0,000498405 - | 1 - | 0,000393124 - | 2 - | 0,000781308 - | 1 - | 0,00039363 - |
| 389 | 1 | 3um_FIJJAFBF_718642 | Bacteria | Verrucomicrol | Verrucomicrol | Verrucomicrol | Verrucomicrol | unclassified | Verrucomicro | unclassified | unclassified | unclassified | unclassified | unclassified | 1,8 | 1,469693846 | 0,000503368 | 0,000449869 | 3 - | 0,000790035 - | 4 - | 0,001239278 - | 0 - | 0 - | 1 - | 0,000242839 - | 1 - | 0,000244689 - |
| 390 | 1 | 3um_FIJJAFBF_698759 | Eukaryota | Bacillariophyt | Fragilariophyc | Licmophorales | Licmophorace | Licmophora | Licmophora sp | information st | Translation, ri | 0FNCH | Binds directly | 91464.S7335_ | 0,4 | 0,489897949 | 0,000294218 | 0,000360747 | 1 - | 0,000762555 - | 0 - | 0 - | 0 - | 0 - | 0 - | 0 - | 1 - | 0,000708535 - |
| 391 | 1 | 0.2um_NOKCPHJN_1010845 | Bacteria | Proteobacteri | Alphaproteob | Rhodobactera | Rhodobactera | unclassified | Rhodobactera | cellular proces | Cell wall/mem | 16RAR | cytidylyl-trans | 272942.RCAP_ | 0,2 | 0,4 | 7,63E-05 | 0,000152511 | 1 - | 0,000381278 - | 0 - | 0 - | 0 - | 0 - | 0 - | 0 - | 0 - | 0 - |
| 392 | 1 | 0.2um_NOKCPHJN_286046 | Eukaryota | unclassified | Pelagophycea | Pelagomonad | unclassified | Aureococcus | Aureococcus a | cellular proces | Cytoskeleton | 1514Z | Tubulin is the | 272952.HpaP8 | 0,8 | 0,4 | 0,000860825 | 0,000431643 | 1 - | 0,001138881 - | 0 - | 0 - | 1 - | 0,001056841 - | 1 - | 0,001050199 - | 1 - | 0,001058201 - |
| 393 | 1 | 0.2um_NOKCPHJN_137528 | Eukaryota | Chordata | Ascidiacea | Enterogona | Cionidae | Ciona | Ciona intestin | unclassified | unclassified | unclassified | unclassified | unclassified | 1 | 0 | 1,92E-05 | 1,88E-06 | 1 - | 1,94E-05 - | 1 - | 2,28E-05 - | 1 - | 1,80E-05 - | 1 - | 1,79E-05 - | 1 - | 1,80E-05 - |
| 394 | 1 | 3um_FIJJAFBF_235865 | Bacteria | Proteobacteri | Gammaprotec | Cellvibrionales | Spongiibacter | Oceanicoccus | Oceanicoccus | information st | Translation, ri | 0QSAC | Binds togethe | 247633.GP214 | 1,4 | 0,8 | 0,000831385 | 0,000480306 | 2 - | 0,001252769 - | 0 - | 0 - | 2 - | 0,001162525 - | 1 - | 0,00057761 - | 2 - | 0,001164021 - |
| 395 | 3 | 0.2um_NOKCPHJN_1019867;0. | Eukaryota | various | various | various | various | various | various | unclassified | unclassified | unclassified | unclassified | unclassified | 1 | 0 | 0,000896056 | 8,77E-05 | 1 - | 0,00090406 - | 1 - | 0,001063607 - | 1 - | 0,000838935 - | 1 - | 0,000833663 - | 1 - | 0,000840015 - |
| 396 | 1 | 3um_FIJJAFBF_786150 | Bacteria | Proteobacteri | Gammaprotec | Oceanospirilla | Saccharospirill | Reinekea | Reinekea fors | unclassified | unclassified | unclassified | unclassified | unclassified | 0,4 | 0,489897949 | 0,000181393 | 0,000222163 | 0 - | 0 - | 0 - | 0 - | 0 - | 0 - | 1 - | 0,000451762 - | 1 - | 0,000455204 - |
| 397 | 1 | 3um_FIJJAFBF_537784 | Eukaryota | Chlorophyta | Chlorophycea | Chlamydomor | Tetrabaenace | Tetrabaena | Tetrabaena so | unclassified | unclassified | unclassified | unclassified | unclassified | 1,2 | 0,748331477 | 0,000237963 | 0,000140633 | 0 - | 0 - | 1 - | 0,000241051 - | 1 - | 0,000190133 - | 2 - | 0,000377875 - | 2 - | 0,000380755 - |
| 398 | 1 | 3um_FIJJAFBF_586299 | Eukaryota | unclassified | Dinophyceae | Suessiales | Symbiodiniace | Symbiodinium | Symbiodinium | unclassified | unclassified | unclassified | unclassified | unclassified | 2,4 | 0,489897949 | 0,001109713 | 0,000183554 | 2 - | 0,000942945 - | 2 - | 0,001109354 - | 3 - | 0,001312528 - | 2 - | 0,00086952 - | 3 - | 0,001314218 - |
| 399 | 1 | 3um_FIJJAFBF_212346 | Bacteria | Bacteroidetes | Flavobacteriia | Flavobacterial | Crocinitomica | unclassified | Crocinitomica | unclassified | unclassified | unclassified | unclassified | unclassified | 0,6 | 0,489897949 | 6,15E-05 | 5,02E-05 | 0 - | 0 - | 0 - | 0 - | 1 - | 0,000102619 - | 1 - | 0,000101974 - | 1 - | 0,000102751 - |
| 400 | 1 | 3um_FIJJAFBF_556071 | Eukaryota | unclassified | Dinophyceae | Peridiniales | Heterocapsac | Heterocapsa | Heterocapsa t | unclassified | unclassified | unclassified | unclassified | unclassified | 1 | 0 | 0,001059969 | 0,000103712 | 1 - | 0,001069437 - | 1 - | 0,001258169 - | 1 - | 0,000992399 - | 1 - | 0,000986163 - | 1 - | 0,000993677 - |
| 401 | 1 | 0.2um_NOKCPHJN_1013197 | Eukaryota | Eustigmatoph | unclassified | Eustigmatales | Monodopsida | Nannochlorop | Nannochlorop | unclassified | unclassified | unclassified | unclassified | unclassified | 2,4 | 0,8 | 0,000375364 | 0,000104156 | 2 - | 0,000325999 - | 1 - | 0,000191766 - | 3 - | 0,000453774 - | 3 - | 0,000450922 - | 3 - | 0,000454358 - |
| 402 | 1 | 3um_FIJJAFBF_792638 | Eukaryota | unclassified | Synurophycea | Parmales | Triparmaceae | Triparma | Triparma laev | unclassified | unclassified | unclassified | unclassified | unclassified | 0,2 | 0,4 | 2,15E-05 | 4,30E-05 | 0 - | 0 - | 0 - | 0 - | 0 - | 0 - | 1 - | 0,000107534 - | 0 - | 0 - |
| 403 | 1 | 3um_FIJJAFBF_135487 | Bacteria | Proteobacteri | Deltaproteoba | Syntrophobac | Syntrophobac | Syntrophobac | Syntrophobac | cellular proces | Posttranslatio | 0HC06 | Prevents misf | 246197.MXAN | 0,4 | 0,489897949 | 6,53E-05 | 8,09E-05 | 0 - | 0 - | 1 - | 0,000182602 - | 1 - | 0,00014403 - | 0 - | 0 - | 0 - | 0 - |
| 404 | 1 | 3um_FIJJAFBF_58623 | unclassified | unclassified | unclassified | unclassified | unclassified | unclassified | unclassified | unclassified | unclassified | unclassified | unclassified | unclassified | 2 | 0,632455532 | 0,001078572 | 0,000326858 | 1 - | 0,000548087 - | 2 - | 0,001289624 - | 3 - | 0,001525814 - | 2 - | 0,001010817 - | 2 - | 0,001018519 - |
| 405 | 1 | 0.2um_NOKCPHJN_953225 | Bacteria | Bacteroidetes | Flavobacteriia | Flavobacterial | Flavobacteriac | Formosa | Formosa sp. H | metabolism | Inorganic ion t | 05C27 | TonB-depende | 886377.Murru | 0,2 | 0,4 | 2,98E-05 | 5,97E-05 | 0 - | 0 - | 0 - | 0 - | 0 - | 0 - | 0 - | 0 - | 1 - | 0,000149234 - |
| 406 | 1 | 0.2um_NOKCPHJN_261541 | Bacteria | Bacteroidetes | Flavobacteriia | Flavobacterial | Flavobacteriac | Lutibacter | Lutibacter pro | metabolism | Inorganic ion t | 05C27 | TonB-depende | 487797.Flav3C | 2,8 | 0,748331477 | 0,000223295 | 5,76E-05 | 4 - | 0,000324492 - | 2 - | 0,000190879 - | 2 - | 0,000150558 - | 3 - | 0,000224418 - | 3 - | 0,000226128 - |

| | | | | | | | | | | | | | | | | | | | | | | | | | | | | |
|---|---|---|---|---|---|---|---|---|---|---|---|---|---|---|---|---|---|---|---|---|---|---|---|---|---|---|---|---|
| 407 | 1 | 0.2um_NOKCPHJN_616670 | Eukaryota | Phaeophyceae | unclassified | Ectocarpales | Ectocarpaceae | Ectocarpus | Ectocarpus sil | cellular proces | Signal transdu | 14ZNA | Inherit from K | 164328.Phyra | 1 | 0 | 0,000459881 | 4,50E-05 | 1 - | 0,000463989 - | 1 - | 0,000545872 - | 1 - | 0,000430565 - | 1 - | 0,000427859 - | 1 - | 0,000431119 - |
| 408 | 1 | 3um_FIJJAFBF_697086 | Eukaryota | unclassified | Heterolobosea | Schizopyrenid | Vahlkampfiida | Naegleria | Naegleria grub | unclassified | unclassified | unclassified | unclassified | unclassified | 0,6 | 0,489897949 | 0,000130683 | 0,000106704 | 0 - | 0 - | 0 - | 0 - | 1 - | 0,000218168 - | 1 - | 0,000216797 - | 1 - | 0,000218449 - |
| 409 | 1 | 3um_FIJJAFBF_59958 | Eukaryota | Chlorophyta | unclassified | Pyramimonad | unclassified | Cymbomonas | Cymbomonas | metabolism | Energy produc | COG0711 | F(1)F(0) ATP s | 88036.ADH10 | 0,2 | 0,4 | 9,26E-05 | 0,000185185 | 0 - | 0 - | 0 - | 0 - | 0 - | 0 - | 0 - | 0 - | 1 - | 0,000462963 - |
| 410 | 1 | 0.2um_NOKCPHJN_310343 | Eukaryota | Mucoromycot | Mucoromycet | Mucorales | Choanephora | Choanephora | Choanephora | cellular proces | Intracellular tr | 0V837 | member RAS | 31033.ENSTRU | 0,6 | 0,489897949 | 0,000231519 | 0,000189358 | 1 - | 0,00040599 - | 0 - | 0 - | 0 - | 0 - | 1 - | 0,000374377 - | 1 - | 0,000377229 - |
| 411 | 1 | 3um_FIJJAFBF_299455 | Bacteria | Bacteroidetes | Saprospiria | Saprospirales | Saprospiracea | Aureispira | Aureispira sp. | poorly charact | Function unkn | 09PDM | NA | 755732.Fluta_ | 1 | 0,632455532 | 0,000320267 | 0,000231622 | 1 - | 0,000307698 - | 2 - | 0,000723999 - | 0 - | 0 - | 1 - | 0,000283738 - | 1 - | 0,0002859 - |
| 412 | 1 | 3um_FIJJAFBF_675681 | Viruses | unclassified | unclassified | unclassified | unclassified | unclassified | uncultured M | unclassified | unclassified | unclassified | unclassified | unclassified | 1,4 | 0,489897949 | 0,000459942 | 0,000190614 | 1 - | 0,000325999 - | 2 - | 0,000767062 - | 1 - | 0,000302516 - | 2 - | 0,000601229 - | 1 - | 0,000302905 - |
| 413 | 1 | 3um_FIJJAFBF_91115 | Eukaryota | unclassified | Raphidophyce | Chattonellales | Chattonellace | Heterosigma | Heterosigma a | unclassified | unclassified | unclassified | unclassified | unclassified | 15,2 | 1,6 | 0,004119807 | 0,000236158 | 14 - | 0,003860736 - | 13 - | 0,004217637 - | 17 - | 0,004350329 - | 15 - | 0,003814403 - | 17 - | 0,004355929 - |
| 414 | 1 | 0.2um_NOKCPHJN_247074 | Bacteria | Bacteroidetes | Flavobacteriia | Flavobacterial | Flavobacteriac | Polaribacter | Polaribacter fi | unclassified | unclassified | unclassified | unclassified | unclassified | 0,4 | 0,489897949 | 3,83E-05 | 4,69E-05 | 0 - | 0 - | 0 - | 0 - | 1 - | 9,56E-05 - | 0 - | 0 - | 1 - | 9,57E-05 - |
| 415 | 1 | 0.2um_NOKCPHJN_143029 | Bacteria | Bacteroidetes | Flavobacteriia | Flavobacterial | Flavobacteriac | Flavivirga | Flavivirga eckl | metabolism | Carbohydrate | COG0057 | glyceraldehyd | 313598.MED1 | 0,4 | 0,489897949 | 0,000100934 | 0,000123787 | 1 - | 0,000262556 - | 0 - | 0 - | 0 - | 0 - | 1 - | 0,000242112 - | 0 - | 0 - |
| 416 | 1 | 3um_FIJJAFBF_596874 | Bacteria | Bacteroidetes | Flavobacteriia | Flavobacterial | Flavobacteriac | Moheibacter | Moheibacter s | poorly charact | Function unkn | 06XI0 | Protein of unk | 313606.M231 | 2,2 | 0,4 | 0,001002417 | 0,000297622 | 2 - | 0,00090406 - | 3 - | 0,001595411 - | 2 - | 0,000838935 - | 2 - | 0,000833663 - | 2 - | 0,000840015 - |
| 417 | 1 | 3um_FIJJAFBF_280647 | Bacteria | Proteobacteri | Alphaproteob | Rhodobactera | unclassified | unclassified | Rhodobactera | information st | Replication, re | 05CVP | deoxyribo-dip | 367336.OM22 | 0,6 | 0,489897949 | 0,000107314 | 8,78E-05 | 1 - | 0,000188184 - | 0 - | 0 - | 0 - | 0 - | 1 - | 0,000173531 - | 1 - | 0,000174853 - |
| 418 | 1 | 3um_FIJJAFBF_67514 | Eukaryota | Bacillariophyta | Mediophyceae | Cymatosirales | Cymatosiracea | Plagiogrammo | Plagiogrammo | cellular proces | Posttranslatio | KOG0731 | ATP-depender | 35128.Thapsd | 0,8 | 0,748331477 | 0,000115176 | 0,00011637 | 1 - | 0,000134706 - | 2 - | 0,000316958 - | 0 - | 0 - | 1 - | 0,000124217 - | 0 - | 0 - |
| 419 | 1 | 3um_FIJJAFBF_504257 | Eukaryota | unclassified | Synurophycea | Parmales | Triparmaceae | Triparma | Triparma laev | metabolism | Energy produc | COG0712 | F(1)F(0) ATP s | 35128.Thapsd | 1 | 0 | 0,000474959 | 4,65E-05 | 1 - | 0,000479201 - | 1 - | 0,00056377 - | 1 - | 0,000444682 - | 1 - | 0,000441887 - | 1 - | 0,000445254 - |
| 420 | 1 | 3um_FIJJAFBF_183698 | Bacteria | Bacteroidetes | Flavobacteriia | Flavobacterial | Flavobacteriac | Algibacter | Algibacter sp. | unclassified | unclassified | unclassified | unclassified | unclassified | 0,2 | 0,4 | 2,88E-05 | 5,77E-05 | 0 - | 0 - | 0 - | 0 - | 0 - | 0 - | 1 - | 0,000144145 - | 0 - | 0 - |
| 421 | 1 | 3um_FIJJAFBF_552490 | Bacteria | Proteobacteri | Gammaprotec | Arenicellales | Arenicellaceae | Arenicella | Arenicella xan | unclassified | unclassified | unclassified | unclassified | unclassified | 0,2 | 0,4 | 2,14E-05 | 4,28E-05 | 1 - | 0,000107074 - | 0 - | 0 - | 0 - | 0 - | 0 - | 0 - | 0 - | 0 - |
| 422 | 1 | 0.2um_NOKCPHJN_111204 | Eukaryota | unclassified | Synurophycea | Parmales | Triparmaceae | Triparma | Triparma laev | metabolism | Energy produc | COG0055 | Produces ATP | 7739.JGI1257 | 1,6 | 0,8 | 0,000279677 | 0,000140238 | 2 - | 0,000370016 - | 0 - | 0 - | 2 - | 0,000343362 - | 2 - | 0,000341204 - | 2 - | 0,000343804 - |
| 423 | 1 | 3um_FIJJAFBF_758945 | Bacteria | Bacteroidetes | Flavobacteriia | Flavobacterial | Crocinitomica | unclassified | Crocinitomica | information st | Translation, ri | COG0087 | One of the pri | 755732.Fluta_ | 1 | 0 | 0,000415873 | 4,07E-05 | 1 - | 0,000419588 - | 1 - | 0,000493636 - | 1 - | 0,000389362 - | 1 - | 0,000386916 - | 1 - | 0,000389864 - |
| 424 | 1 | 3um_FIJJAFBF_737827 | unclassified | unclassified | unclassified | unclassified | unclassified | unclassified | unclassified | unclassified | unclassified | unclassified | unclassified | unclassified | 0,8 | 0,4 | 0,000302664 | 0,000151765 | 1 - | 0,000400429 - | 0 - | 0 - | 1 - | 0,000371583 - | 1 - | 0,000369248 - | 1 - | 0,000372062 - |
| 425 | 1 | 0.2um_NOKCPHJN_567099 | Bacteria | unclassified | unclassified | unclassified | unclassified | unclassified | uncultured ba | unclassified | unclassified | unclassified | unclassified | unclassified | 0,2 | 0,4 | 2,30E-05 | 4,60E-05 | 0 - | 0 - | 1 - | 0,000115017 - | 0 - | 0 - | 0 - | 0 - | 0 - | 0 - |
| 426 | 1 | 0.2um_NOKCPHJN_793458 | Bacteria | Cyanobacteria | unclassified | Nostocales | Nostocaceae | Trichormus | Trichormus sp | unclassified | unclassified | unclassified | unclassified | unclassified | 0,6 | 0,489897949 | 6,47E-05 | 5,37E-05 | 0 - | 0 - | 1 - | 0,000125664 - | 0 - | 0 - | 1 - | 9,85E-05 - | 1 - | 9,92E-05 - |
| 427 | 1 | 0.2um_NOKCPHJN_375927 | Eukaryota | unclassified | unclassified | unclassified | unclassified | Stygiella | Stygiella incar | information st | Translation, ri | KOG0052 | This protein p | 2850.Phatr18 | 1,8 | 0,4 | 0,000349656 | 8,56E-05 | 1 - | 0,000196183 - | 2 - | 0,00046161 - | 2 - | 0,000364102 - | 2 - | 0,000361814 - | 2 - | 0,00036457 - |
| 428 | 1 | 3um_FIJJAFBF_209610 | Bacteria | Bacteroidetes | Flavobacteriia | Flavobacterial | Flavobacteriac | Algibacter | Algibacter lect | unclassified | unclassified | unclassified | unclassified | unclassified | 0,8 | 0,4 | 0,000490989 | 0,000246196 | 1 - | 0,000649584 - | 0 - | 0 - | 1 - | 0,000602791 - | 1 - | 0,000599003 - | 1 - | 0,000603567 - |
| 429 | 1 | 3um_FIJJAFBF_254020 | Bacteria | Bacteroidetes | Flavobacteriia | Flavobacterial | Flavobacteriac | Arenitalea | Arenitalea lute | unclassified | unclassified | unclassified | unclassified | unclassified | 3,2 | 1,166190379 | 0,000703105 | 0,000276703 | 3 - | 0,000657704 - | 4 - | 0,001031699 - | 4 - | 0,000813767 - | 4 - | 0,000808654 - | 1 - | 0,000203704 - |
| 430 | 1 | 3um_FIJJAFBF_149840 | Eukaryota | Nematoda | Chromadorea | Rhabditida | Rhabditidae | Caenorhabditi | Caenorhabditi | unclassified | unclassified | unclassified | unclassified | unclassified | 0,2 | 0,4 | 5,75E-05 | 0,000115008 | 1 - | 0,000287521 - | 0 - | 0 - | 0 - | 0 - | 0 - | 0 - | 0 - | 0 - |
| 431 | 1 | 3um_FIJJAFBF_615123 | Bacteria | Proteobacteri | Gammaprotec | Cellvibrionales | Porticoccacea | unclassified | Gammaprotec | cellular proces | Cell wall/mem | 0QTSX | Peptidoglycan | 314287.GB22 | 1,6 | 0,8 | 0,0008702 | 0,000438329 | 1 - | 0,000544682 - | 2 - | 0,001281614 - | 3 - | 0,001516337 - | 1 - | 0,000502269 - | 1 - | 0,000506096 - |
| 432 | 1 | 3um_FIJJAFBF_533992 | Bacteria | Bacteroidetes | Flavobacteriia | Flavobacterial | Crocinitomica | unclassified | Crocinitomica | poorly charact | Function unkn | 0P884 | membrane | 926562.Oweh | 2,8 | 0,4 | 0,000688018 | 5,30E-05 | 3 - | 0,000753815 - | 2 - | 0,000591231 - | 3 - | 0,000699514 - | 3 - | 0,000695118 - | 3 - | 0,000700414 - |
| 433 | 1 | 3um_FIJJAFBF_125686 | Eukaryota | Bacillariophyta | Bacillariophyc | Naviculales | Phaeodactylac | Phaeodactylu | Phaeodactylu | unclassified | unclassified | unclassified | unclassified | unclassified | 1 | 0 | 0,000339521 | 3,32E-05 | 1 - | 0,000342554 - | 1 - | 0,000403007 - | 1 - | 0,000317878 - | 1 - | 0,00031588 - | 1 - | 0,000318287 - |
| 434 | 1 | 0.2um_NOKCPHJN_23322 | Eukaryota | unclassified | Pelagophyceae | Pelagomonad | unclassified | Pelagomonas | various | cellular proces | Posttranslatio | KOG0730 | cell division cy | 35128.Thapsd | 2,6 | 0,489897949 | 0,000767516 | 9,55E-05 | 2 - | 0,000604785 - | 2 - | 0,000711516 - | 3 - | 0,000841828 - | 3 - | 0,000836538 - | 3 - | 0,000842912 - |
| 435 | 1 | 0.2um_NOKCPHJN_1039177 | Bacteria | Proteobacteri | unclassified | unclassified | unclassified | unclassified | Proteobacteri | unclassified | unclassified | unclassified | unclassified | unclassified | 0,4 | 0,489897949 | 0,000341562 | 0,00041833 | 0 - | 0 - | 0 - | 0 - | 1 - | 0,000856597 - | 1 - | 0,000851214 - | 0 - | 0 - |
| 436 | 1 | 0.2um_NOKCPHJN_640148 | Eukaryota | unclassified | Pelagophyceae | Pelagomonad | unclassified | Aureococcus | Aureococcus a | cellular proces | Intracellular tr | KOG3342 | Catalytic comp | 2850.Phatr51 | 1,2 | 0,4 | 0,000446272 | 0,000205219 | 1 - | 0,000363875 - | 2 - | 0,000856182 - | 1 - | 0,000337663 - | 1 - | 0,000335541 - | 1 - | 0,000338097 - |
| 437 | 1 | 3um_FIJJAFBF_21583 | unclassified | unclassified | unclassified | unclassified | unclassified | unclassified | unclassified | unclassified | unclassified | unclassified | unclassified | unclassified | 0,4 | 0,489897949 | 0,000559232 | 0,000692831 | 0 - | 0 - | 1 - | 0,00156318 - | 1 - | 0,001232981 - | 0 - | 0 - | 0 - | 0 - |
| 438 | 1 | 3um_FIJJAFBF_319725 | Bacteria | Bacteroidetes | Flavobacteriia | Flavobacterial | Flavobacteriac | Polaribacter | Polaribacter g | unclassified | unclassified | unclassified | unclassified | unclassified | 0,6 | 0,489897949 | 0,000186034 | 0,000154411 | 0 - | 0 - | 1 - | 0,000360734 - | 1 - | 0,000284534 - | 0 - | 0 - | 1 - | 0,0002849 - |
| 439 | 2 | 0.2um_NOKCPHJN_109516;3ur | Bacteria | Bacteroidetes | Flavobacteriia | Flavobacterial | Crocinitomica | unclassified | Crocinitomica | unclassified | unclassified | unclassified | unclassified | unclassified | 0,4 | 0,489897949 | 2,68E-05 | 3,29E-05 | 0 - | 0 - | 0 - | 0 - | 0 - | 0 - | 1 - | 6,68E-05 - | 1 - | 6,73E-05 - |
| 440 | 1 | 3um_FIJJAFBF_248095 | Bacteria | Proteobacteri | Betaproteoba | Burkholderial | Comamonada | Hydrogenoph | various | unclassified | unclassified | unclassified | unclassified | unclassified | 0,2 | 0,4 | 3,81E-05 | 7,62E-05 | 0 - | 0 - | 0 - | 0 - | 1 - | 0,000190578 - | 0 - | 0 - | 0 - | 0 - |
| 441 | 1 | 3um_FIJJAFBF_130174 | unclassified | unclassified | unclassified | unclassified | unclassified | unclassified | unclassified | unclassified | unclassified | unclassified | unclassified | unclassified | 0,2 | 0,4 | 0,000254185 | 0,00050837 | 1 - | 0,001270925 - | 0 - | 0 - | 0 - | 0 - | 0 - | 0 - | 0 - | 0 - |
| 442 | 1 | 3um_FIJJAFBF_278417 | Eukaryota | unclassified | Rhodellophyc | Dixoniellales | Dixoniellaceae | Bulboplastis | Bulboplastis a | metabolism | Energy produc | 0IFBF | One of the tw | 45157.CMV08 | 0,8 | 0,4 | 0,00019766 | 0,000101751 | 0 - | 0 - | 1 - | 0,000293931 - | 1 - | 0,000231843 - | 1 - | 0,000230386 - | 1 - | 0,000232141 - |
| 443 | 1 | 3um_FIJJAFBF_715065 | Eukaryota | Bacillariophyta | Mediophyceae | Anaulales | Anaulaceae | Eunotogramm | Eunotogramm | unclassified | unclassified | unclassified | unclassified | unclassified | 2,2 | 0,748331477 | 0,002898052 | 0,0009795 | 3 - | 0,003986084 - | 2 - | 0,00312636 - | 1 - | 0,001232981 - | 3 - | 0,003675698 - | 2 - | 0,002469136 - |
| 444 | 1 | 0.2um_NOKCPHJN_725707 | Bacteria | Bacteroidetes | Flavobacteriia | Flavobacterial | Flavobacteriac | Formosa | Formosa sp. H | unclassified | unclassified | unclassified | unclassified | unclassified | 1,8 | 0,4 | 0,000259144 | 6,80E-05 | 2 - | 0,000288467 - | 2 - | 0,000339375 - | 1 - | 0,000133843 - | 2 - | 0,000266004 - | 2 - | 0,000268031 - |
| 445 | 1 | 0.2um_NOKCPHJN_157068 | Bacteria | unclassified | unclassified | unclassified | unclassified | unclassified | uncultured ba | poorly charact | Function unkn | 0P8NI | Inherit from b | 983544.Lacal_ | 2,6 | 0,8 | 0,000542488 | 0,000176797 | 1 - | 0,000210803 - | 3 - | 0,000744014 - | 3 - | 0,000586852 - | 3 - | 0,000583164 - | 3 - | 0,000587607 - |
| 446 | 1 | 3um_FIJJAFBF_163703 | Bacteria | Bacteroidetes | Flavobacteriia | Flavobacterial | Flavobacteriac | Lacinutrix | Lacinutrix sp. | unclassified | unclassified | unclassified | unclassified | unclassified | 1,6 | 0,489897949 | 0,000249329 | 8,60E-05 | 1 - | 0,000156039 - | 2 - | 0,000367153 - | 2 - | 0,000289597 - | 1 - | 0,000143889 - | 2 - | 0,00028997 - |
| 447 | 1 | 0.2um_NOKCPHJN_57013 | Bacteria | Bacteroidetes | Flavobacteriia | Flavobacterial | Flavobacteriac | Algibacter | Algibacter pec | unclassified | unclassified | unclassified | unclassified | unclassified | 3,4 | 0,8 | 0,001700172 | 0,000435269 | 5 - | 0,002519938 - | 3 - | 0,001778791 - | 3 - | 0,001403047 - | 3 - | 0,00139423 - | 3 - | 0,001404853 - |
| 448 | 1 | 3um_FIJJAFBF_349824 | Bacteria | Proteobacteri | Alphaproteob | Pelagibacteral | Pelagibacterac | unclassified | Pelagibacterac | unclassified | unclassified | unclassified | unclassified | unclassified | 1 | 0 | 0,000896056 | 8,77E-05 | 1 - | 0,00090406 - | 1 - | 0,001063607 - | 1 - | 0,000838935 - | 1 - | 0,000833663 - | 1 - | 0,000840015 - |
| 449 | 1 | 3um_FIJJAFBF_462596 | unclassified | unclassified | unclassified | unclassified | unclassified | unclassified | unclassified | information st | Replication, re | 0PFXX | DNA repair pr | 578458.XP_00 | 0,4 | 0,489897949 | 7,43E-05 | 9,15E-05 | 1 - | 0,000170611 - | 1 - | 0,00020072 - | 0 - | 0 - | 0 - | 0 - | 0 - | 0 - |
| 450 | 1 | 0.2um_NOKCPHJN_264390 | Viruses | unclassified | unclassified | unclassified | unclassified | unclassified | uncultured M | poorly charact | Function unkn | 0R2P9 | NA | 768066.HELO_ | 0,8 | 0,4 | 0,000106865 | 5,48E-05 | 1 - | 0,000132468 - | 1 - | 0,000155846 - | 1 - | 0,000122926 - | 0 - | 0 - | 1 - | 0,000123084 - |
| 451 | 1 | 3um_FIJJAFBF_84142 | Viruses | unclassified | unclassified | unclassified | Phycodnavirid | unclassified | Chrysochromu | cellular proces | Cell cycle cont | KOG0987 | DNA helicase | 44689.DDB02 | 0,8 | 0,4 | 0,00015253 | 7,82E-05 | 1 - | 0,000189404 - | 1 - | 0,000222829 - | 1 - | 0,00017576 - | 1 - | 0,000174655 - | 0 - | 0 - |
| 452 | 1 | 0.2um_NOKCPHJN_549424 | Eukaryota | Bacillariophyta | Coscinodiscop | Thalassiosirale | Thalassiosirac | Thalassiosira | Thalassiosira | metabolism | Inorganic ion | COG1226 | PotAssium vol | 2850.Phatr44 | 0,6 | 0,489897949 | 3,86E-05 | 3,15E-05 | 1 - | 6,77E-05 - | 0 - | 0 - | 1 - | 6,28E-05 - | 1 - | 6,24E-05 - | 0 - | 0 - |
| 453 | 1 | 3um_FIJJAFBF_162557 | Bacteria | Bacteroidetes | Flavobacteriia | Flavobacterial | Flavobacteriac | unclassified | Flavobacteriac | unclassified | unclassified | unclassified | unclassified | unclassified | 1,2 | 0,4 | 0,000338046 | 0,000118023 | 2 - | 0,000567598 - | 1 - | 0,000333883 - | 1 - | 0,000263355 - | 1 - | 0,0002617 - | 1 - | 0,000263694 - |
| 454 | 1 | 3um_FIJJAFBF_431586 | Viruses | unclassified | unclassified | unclassified | unclassified | unclassified | uncultured vir | unclassified | unclassified | unclassified | unclassified | unclassified | 0,2 | 0,4 | 1,12E-05 | 2,23E-05 | 0 - | 0 - | 0 - | 0 - | 0 - | 0 - | 0 - | 0 - | 1 - | 5,58E-05 - |
| 455 | 1 | 0.2um_NOKCPHJN_137368 | Bacteria | Bacteroidetes | Flavobacteriia | Flavobacterial | Crocinitomica | unclassified | Crocinitomica | unclassified | unclassified | unclassified | unclassified | unclassified | 1 | 0 | 5,74E-05 | 5,61E-06 | 1 - | 5,79E-05 - | 1 - | 6,81E-05 - | 1 - | 5,37E-05 - | 1 - | 5,34E-05 - | 1 - | 5,38E-05 - |
| 456 | 1 | 0.2um_NOKCPHJN_529682 | Bacteria | Proteobacteri | Alphaproteob | Rhodobactera | Rhodobactera | Tateyamaria | Tateyamaria s | unclassified | unclassified | unclassified | unclassified | unclassified | 5,4 | 0,8 | 0,001610821 | 0,000224608 | 5 - | 0,001511963 - | 5 - | 0,001778791 - | 7 - | 0,001964266 - | 5 - | 0,00139423 - | 5 - | 0,001404853 - |
| 457 | 1 | 3um_FIJJAFBF_520477 | Bacteria | Bacteroidetes | Flavobacteriia | Flavobacterial | Flavobacteriac | Polaribacter | Polaribacter a | cellular proces | Intracellular tr | 0PBE9 | Preprotein tra | 313598.MED1 | 0,2 | 0,4 | 0,000167036 | 0,000334072 | 1 - | 0,00083518 - | 0 - | 0 - | 0 - | 0 - | 0 - | 0 - | 0 - | 0 - |
| 458 | 1 | 0.2um_NOKCPHJN_469091 | unclassified | unclassified | unclassified | unclassified | unclassified | unclassified | unclassified | unclassified | unclassified | unclassified | unclassified | unclassified | 0,2 | 0,4 | 0,00015856 | 0,000317119 | 0 - | 0 - | 0 - | 0 - | 0 - | 0 - | 1 - | 0,000792798 - | 0 - | 0 - |
| 459 | 1 | 3um_FIJJAFBF_292927 | Eukaryota | Bacillariophyta | Coscinodiscop | Coscinodiscale | Coscinodiscac | Coscinodiscus | Coscinodiscus | cellular proces | Posttranslatio | KOG0731 | ATP-depender | 35128.Thapsd | 2,2 | 0,4 | 0,000298446 | 4,93E-05 | 2 - | 0,000275334 - | 2 - | 0,000323924 - | 3 - | 0,00038325 - | 2 - | 0,000253894 - | 2 - | 0,000255829 - |
| 460 | 1 | 3um_FIJJAFBF_20002 | Bacteria | Proteobacteri | Gammaprotec | Alteromonada | Colwelliaceae | Colwellia | Colwellia sp. C | metabolism | Inorganic ion | 16TQC | TONB-depend | 314287.GB22 | 5,6 | 1,019803903 | 0,000911071 | 0,000118482 | 6 - | 0,000998412 - | 4 - | 0,000783073 - | 5 - | 0,000772075 - | 7 - | 0,001074113 - | 6 - | 0,000927683 - |
| 461 | 1 | 3um_FIJJAFBF_748496 | Bacteria | Proteobacteri | Alphaproteob | Rhodobactera | Rhodobactera | Planktomarina | Planktomarina | unclassified | unclassified | unclassified | unclassified | unclassified | 1 | 0 | 0,00010372 | 1,01E-05 | 1 - | 0,000104647 - | 1 - | 0,000123114 - | 1 - | 9,71E-05 - | 1 - | 9,65E-05 - | 1 - | 9,72E-05 - |
| 462 | 1 | 0.2um_NOKCPHJN_938097 | Bacteria | Proteobacteri | Alphaproteob | Rhodobactera | Hyphomonada | Hyphomonas | various | metabolism | Energy produc | COG0055 | Produces ATP | 208348.PTTG_ | 0,8 | 0,4 | 0,000690453 | 0,000346213 | 1 - | 0,000913478 - | 0 - | 0 - | 1 - | 0,000847674 - | 1 - | 0,000842347 - | 1 - | 0,000848766 - |
| 463 | 1 | 3um_FIJJAFBF_808023 | Bacteria | Verrucomicrol | Verrucomicrol | Verrucomicrol | Akkermansiac | Akkermansia | Akkermansia | unclassified | unclassified | unclassified | unclassified | unclassified | 1 | 0 | 0,000222295 | 2,18E-05 | 1 - | 0,000224281 - | 1 - | 0,000263862 - | 1 - | 0,000208125 - | 1 - | 0,000206817 - | 1 - | 0,000208393 - |
| 464 | 1 | 0.2um_NOKCPHJN_721141 | Bacteria | Bacteroidetes | Flavobacteriia | Flavobacterial | Flavobacteriac | Altibacter | Altibacter lent | poorly charact | Function unkn | 0P8NI | Inherit from b | 313594.PI23P | 6 | 0,894427191 | 0,000808178 | 0,000150305 | 7 - | 0,000948774 - | 6 - | 0,000956753 - | 5 - | 0,000628877 - | 5 - | 0,000624925 - | 7 - | 0,000881562 - |
| 465 | 1 | 3um_FIJJAFBF_128443 | Bacteria | Bacteroidetes | Flavobacteriia | Flavobacterial | Flavobacteriac | Polaribacter | Polaribacter s | unclassified | unclassified | unclassified | unclassified | unclassified | 1 | 0 | 0,00011589 | 1,13E-05 | 1 - | 0,000116925 - | 1 - | 0,00013756 - | 1 - | 0,000108502 - | 1 - | 0,00010782 - | 1 - | 0,000108642 - |
| 466 | 1 | 3um_FIJJAFBF_118415 | Viruses | unclassified | unclassified | unclassified | Phycodnavirid | unclassified | Aureococcus a | unclassified | unclassified | unclassified | unclassified | unclassified | 1,4 | 0,8 | 0,000241771 | 0,000139396 | 2 - | 0,000364631 - | 0 - | 0 - | 1 - | 0,000169182 - | 2 - | 0,000336238 - | 2 - | 0,0003388 - |
| 467 | 1 | 3um_FIJJAFBF_651483 | Bacteria | Bacteroidetes | Flavobacteriia | Flavobacterial | Flavobacteriac | Formosa | Formosa sp. H | unclassified | unclassified | unclassified | unclassified | unclassified | 1,6 | 0,489897949 | 0,000498659 | 0,000171991 | 1 - | 0,000312078 - | 2 - | 0,000734305 - | 2 - | 0,000579194 - | 1 - | 0,000287777 - | 2 - | 0,000579939 - |
| 468 | 1 | 0.2um_NOKCPHJN_374229 | Eukaryota | unclassified | Oomycetes | various | various | various | various | unclassified | unclassified | unclassified | unclassified | unclassified | 0,2 | 0,4 | 0,000109231 | 0,000218461 | 0 - | 0 - | 0 - | 0 - | 1 - | 0,000546153 - | 0 - | 0 - | 0 - | 0 - |
| 469 | 1 | 3um_FIJJAFBF_322765 | Bacteria | Bacteroidetes | Flavobacteriia | Flavobacterial | Flavobacteriac | Psychroserpe | Psychroserpe | poorly charact | Function unkn | 0ZWUM | glyoxalase ble | 391587.KAOT | 0,4 | 0,489897949 | 0,000267554 | 0,000328134 | 1 - | 0,000695983 - | 0 - | 0 - | 0 - | 0 - | 1 - | 0,000641789 - | 0 - | 0 - |
| 470 | 1 | 0.2um_NOKCPHJN_227948 | Bacteria | Proteobacteri | Alphaproteob | Rhizobiales | unclassified | unclassified | Rhizobiales ba | cellular proces | Cell wall/mem | COG0438 | Glycosyl trans | 204773.HEAR | 0,2 | 0,4 | 4,07E-05 | 8,15E-05 | 0 - | 0 - | 0 - | 0 - | 0 - | 0 - | 1 - | 0,000203691 - | 0 - | 0 - |
| 471 | 1 | 0.2um_NOKCPHJN_926017 | Bacteria | Proteobacteri | Alphaproteob | Rhodobactera | Rhodobactera | unclassified | various | unclassified | unclassified | unclassified | unclassified | unclassified | 0,2 | 0,4 | 6,88E-05 | 0,000137559 | 1 - | 0,000343897 - | 0 - | 0 - | 0 - | 0 - | 0 - | 0 - | 0 - | 0 - |
| 472 | 1 | 0.2um_NOKCPHJN_100278 | Bacteria | Bacteroidetes | unclassified | unclassified | unclassified | unclassified | Bacteroidetes | metabolism | Inorganic ion | 05C27 | TonB-depende | 504472.Slin_4 | 0,6 | 1,2 | 4,70E-05 | 9,40E-05 | 3 - | 0,000235104 - | 0 - | 0 - | 0 - | 0 - | 0 - | 0 - | 0 - | 0 - |
| 473 | 1 | 0.2um_NOKCPHJN_904622 | Eukaryota | unclassified | Oomycetes | Peronosporale | Peronosporac | Phytophthora | Phytophthora | information st | Replication, re | COG0513 | purine NTP-de | 653948.CCA1 | 1 | 0 | 0,000211994 | 2,07E-05 | 1 - | 0,000213887 - | 1 - | 0,000251634 - | 1 - | 0,00019848 - | 1 - | 0,000197233 - | 1 - | 0,000198735 - |
| 474 | 1 | 3um_FIJJAFBF_115059 | Eukaryota | unclassified | unclassified | Phaeocystales | Phaeocystace | Phaeocystis | Phaeocystis gl | metabolism | Energy produc | COG0711 | F(1)F(0) ATP s | 91464.S7335_ | 1,2 | 0,4 | 0,000640836 | 0,000223737 | 2 - | 0,001075998 - | 1 - | 0,000632944 - | 1 - | 0,000499244 - | 1 - | 0,000496106 - | 1 - | 0,000499886 - |
| 475 | 1 | 3um_FIJJAFBF_300245 | Bacteria | Proteobacteri | Alphaproteob | Rhodobactera | Rhodobactera | Planktotalea | Planktotalea f | unclassified | unclassified | unclassified | unclassified | unclassified | 0,4 | 0,489897949 | 0,000107833 | 0,000132789 | 1 - | 0,000247723 - | 1 - | 0,00029144 - | 0 - | 0 - | 0 - | 0 - | 0 - | 0 - |
| 476 | 1 | 3um_FIJJAFBF_318784 | Eukaryota | Chlorophyta | Chlorophyceae | Chlamydomor | Tetrabaenace | Tetrabaena | Tetrabaena so | unclassified | unclassified | unclassified | unclassified | unclassified | 0,8 | 0,4 | 0,000143049 | 7,36E-05 | 0 - | 0 - | 1 - | 0,000212721 - | 1 - | 0,000167787 - | 1 - | 0,000166733 - | 1 - | 0,000168003 - |
| 477 | 1 | 3um_FIJJAFBF_328496 | Bacteria | Bacteroidetes | Flavobacteriia | Flavobacterial | Flavobacteriac | Arenitalea | Arenitalea lute | metabolism | Inorganic ion | 05C27 | TonB-depende | 886377.Murru | 25,8 | 2,712931993 | 0,002143342 | 0,000217399 | 30 - | 0,0025272 - | 22 - | 0,002180343 - | 24 - | 0,001876121 - | 26 - | 0,002019692 - | 27 - | 0,002113353 - |
| 478 | 1 | 3um_FIJJAFBF_775847 | unclassified | unclassified | unclassified | unclassified | unclassified | unclassified | unclassified | unclassified | unclassified | unclassified | unclassified | unclassified | 0,2 | 0,4 | 0,000140635 | 0,000281271 | 0 - | 0 - | 0 - | 0 - | 0 - | 0 - | 1 - | 0,000703177 - | 0 - | 0 - |
| 479 | 1 | 3um_FIJJAFBF_87014 | Bacteria | Proteobacteri | Alphaproteob | Rhodospirillale | Rhodospirillac | Azospirillum | Azospirillum b | unclassified | unclassified | unclassified | unclassified | unclassified | 1,2 | 0,4 | 6,79E-05 | 2,37E-05 | 2 - | 0,000113962 - | 1 - | 6,70E-05 - | 1 - | 5,29E-05 - | 1 - | 5,25E-05 - | 1 - | 5,29E-05 - |
| 480 | 1 | 3um_FIJJAFBF_24509 | Bacteria | Proteobacteri | Alphaproteob | Rhodobactera | unclassified | unclassified | Rhodobactera | unclassified | unclassified | unclassified | unclassified | unclassified | 1 | 0 | 0,000256394 | 2,51E-05 | 1 - | 0,000258684 - | 1 - | 0,000304336 - | 1 - | 0,000240049 - | 1 - | 0,000238541 - | 1 - | 0,000240358 - |
| 481 | 2 | 0.2um_NOKCPHJN_664786;3ur | Bacteria | Actinobacteria | Actinobacteria | Micrococcales | Microbacteria | Candidatus Ac | Candidatus Ac | various | various | various | various | various | 0,6 | 0,489897949 | 4,64E-05 | 3,79E-05 | 1 - | 8,11E-05 - | 0 - | 0 - | 1 - | 7,53E-05 - | 0 - | 0 - | 1 - | 7,54E-05 - |
| 482 | 1 | 3um_FIJJAFBF_194577 | Bacteria | Proteobacteri | Gammaprotec | Oceanospirilla | Saccharospiril | Reinekea | Reinekea fors | unclassified | unclassified | unclassified | unclassified | unclassified | 0,4 | 0,489897949 | 7,26E-05 | 8,99E-05 | 0 - | 0 - | 1 - | 0,000202691 - | 0 - | 0 - | 0 - | 0 - | 1 - | 0,000160082 - |
| 483 | 2 | 0.2um_NOKCPHJN_74285;3um | Bacteria | Bacteroidetes | Flavobacteriia | Flavobacterial | Flavobacteriac | Polaribacter | various | unclassified | unclassified | unclassified | unclassified | unclassified | 2,4 | 0,489897949 | 0,001674758 | 0,000333533 | 3 - | 0,002121625 - | 2 - | 0,00166403 - | 2 - | 0,001312528 - | 2 - | 0,00130428 - | 3 - | 0,001971326 - |
| 484 | 1 | 0.2um_NOKCPHJN_799555 | unclassified | unclassified | unclassified | unclassified | unclassified | unclassified | unclassified | unclassified | unclassified | unclassified | unclassified | unclassified | 0,2 | 0,4 | 5,09E-05 | 0,000101717 | 0 - | 0 - | 0 - | 0 - | 0 - | 0 - | 1 - | 0,000254294 - | 0 - | 0 - |
| 485 | 2 | 0.2um_NOKCPHJN_606024;3ur | Bacteria | Proteobacteri | Gammaprotec | Cellvibrionales | Spongiibacter | Oceanicoccus | Oceanicoccus | cellular proces | Intracellular tr | COG2831 | Polypeptide-tr | 887062.HGR_ | 0,2 | 0,4 | 2,75E-05 | 5,49E-05 | 0 - | 0 - | 0 - | 0 - | 0 - | 0 - | 1 - | 0,000137293 - | 0 - | 0 - |
| 486 | 1 | 0.2um_NOKCPHJN_555355 | Eukaryota | Chytridiomyc | Neocallimastig | Neocallimastig | Neocallimastig | Piromyces | Piromyces finn | unclassified | unclassified | unclassified | unclassified | unclassified | 0,2 | 0,4 | 3,75E-05 | 7,50E-05 | 1 - | 0,00018738 - | 0 - | 0 - | 0 - | 0 - | 0 - | 0 - | 0 - | 0 - |
| 487 | 1 | 3um_FIJJAFBF_442446 | Bacteria | Verrucomicrol | unclassified | unclassified | unclassified | unclassified | Verrucomicrol | unclassified | unclassified | unclassified | unclassified | unclassified | 0,2 | 0,4 | 4,05E-05 | 8,10E-05 | 0 - | 0 - | 0 - | 0 - | 1 - | 0,00020243 - | 0 - | 0 - | 0 - | 0 - |
| 488 | 1 | 0.2um_NOKCPHJN_458167 | Eukaryota | Arthropoda | Insecta | Hymenoptera | Halictidae | Dufourea | Dufourea nov | poorly charact | Function unkn | 0ANG0 | NA | 8364.ENSXETP | 0,6 | 0,489897949 | 9,69E-05 | 8,01E-05 | 1 - | 0,000156317 - | 1 - | 0,000183904 - | 0 - | 0 - | 1 - | 0,000144145 - | 0 - | 0 - |
| 489 | 1 | 0.2um_NOKCPHJN_45922 | Bacteria | Bacteroidetes | Flavobacteriia | Flavobacterial | Flavobacteriac | Arenitalea | Arenitalea lute | unclassified | unclassified | unclassified | unclassified | unclassified | 0,8 | 0,4 | 0,000185668 | 9,31E-05 | 1 - | 0,000245641 - | 0 - | 0 - | 1 - | 0,000227946 - | 1 - | 0,000226514 - | 1 - | 0,000228239 - |
| 490 | 1 | 3um_FIJJAFBF_167349 | Eukaryota | unclassified | Dinophyceae | Gymnodiniale | Gymnodiniace | Lepidodinium | Lepidodinium | metabolism | Carbohydrate | COG0057 | glyceraldehyd | 423536.XP_00 | 0,4 | 0,489897949 | 8,83E-05 | 0,000108318 | 1 - | 0,000228966 - | 0 - | 0 - | 0 - | 0 - | 0 - | 0 - | 1 - | 0,000212745 - |
| 491 | 1 | 0.2um_NOKCPHJN_810494 | Eukaryota | Chlorophyta | Mamiellophyc | Mamiellales | Bathycoccace | Ostreococcus | Ostreococcus | information st | Translation, ri | KOG0052 | This protein p | 436017.A4S6E | 2,2 | 1,166190379 | 0,000386812 | 0,0002068 | 3 - | 0,000557376 - | 0 - | 0 - | 2 - | 0,000344817 - | 3 - | 0,000513975 - | 3 - | 0,000517891 - |
| 492 | 1 | 0.2um_NOKCPHJN_478111 | Eukaryota | Streptophyta | unclassified | Fabales | Fabaceae | Glycine | Glycine max | cellular proces | Posttranslatio | KOG0019 | heat shock pr | 3847.GLYMA0 | 0,8 | 0,4 | 9,18E-05 | 4,60E-05 | 1 - | 0,00012146 - | 0 - | 0 - | 1 - | 0,00011271 - | 1 - | 0,000112002 - | 1 - | 0,000112855 - |
| 493 | 1 | 3um_FIJJAFBF_435972 | unclassified | unclassified | unclassified | unclassified | unclassified | unclassified | unclassified | unclassified | unclassified | unclassified | unclassified | unclassified | 0,6 | 0,489897949 | 0,000931638 | 0,000773628 | 0 - | 0 - | 1 - | 0,001809998 - | 0 - | 0 - | 1 - | 0,00141869 - | 1 - | 0,0014295 - |
| 494 | 1 | 0.2um_NOKCPHJN_243992 | Eukaryota | unclassified | Labyrinthulom | unclassified | Thraustochytr | Aurantiochytr | Aurantiochytr | unclassified | unclassified | unclassified | unclassified | unclassified | 0,2 | 0,4 | 2,14E-05 | 4,28E-05 | 0 - | 0 - | 1 - | 0,000106912 - | 0 - | 0 - | 0 - | 0 - | 0 - | 0 - |
| 495 | 1 | 3um_FIJJAFBF_201380 | Bacteria | Bacteroidetes | Flavobacteriia | Flavobacterial | Flavobacteriac | Lacinutrix | Lacinutrix jang | poorly charact | Function unkn | 0XPRT | NA | 391598.FBBAL | 3,8 | 0,979795897 | 0,000665245 | 0,000181328 | 4 - | 0,000704368 - | 4 - | 0,000828674 - | 2 - | 0,000326814 - | 5 - | 0,000811901 - | 4 - | 0,00065447 - |
| 496 | 1 | 0.2um_NOKCPHJN_130099 | Eukaryota | Arthropoda | Insecta | Hymenoptera | Apidae | Habropoda | Habropoda lab | poorly charact | Function unkn | 0F9Q7 | NA | 353151.XP_66 | 0,8 | 0,4 | 6,01E-06 | 3,01E-06 | 1 - | 7,95E-06 - | 0 - | 0 - | 1 - | 7,38E-06 - | 1 - | 7,33E-06 - | 1 - | 7,39E-06 - |
| 497 | 1 | 0.2um_NOKCPHJN_1070961 | Eukaryota | Streptophyta | unclassified | Cucurbitales | Cucurbitaceae | Cucurbita | various | information st | RNA processin | KOG1783 | Component of | 39946.BGIOS | 0,4 | 0,489897949 | 0,00035404 | 0,000433608 | 0 - | 0 - | 0 - | 0 - | 1 - | 0,00088453 - | 0 - | 0 - | 1 - | 0,000885668 - |
| 498 | 1 | 3um_FIJJAFBF_715069 | Eukaryota | unclassified | Compsopogon | Compsopogon | Compsopogon | Compsopogon | Compsopogon | unclassified | unclassified | unclassified | unclassified | unclassified | 0,8 | 0,4 | 0,001541476 | 0,000772942 | 1 - | 0,002039392 - | 0 - | 0 - | 1 - | 0,001892482 - | 1 - | 0,00188059 - | 1 - | 0,001894918 - |
| 499 | 1 | 0.2um_NOKCPHJN_956641 | Bacteria | Proteobacteri | Alphaproteob | Rhodobactera | unclassified | unclassified | Rhodobactera | unclassified | unclassified | unclassified | unclassified | unclassified | 1,2 | 0,4 | 0,000273721 | 8,18E-05 | 1 - | 0,00023261 - | 1 - | 0,00027366 - | 2 - | 0,000431707 - | 1 - | 0,000214497 - | 1 - | 0,000216131 - |
| 500 | 1 | 0.2um_NOKCPHJN_635290 | Eukaryota | unclassified | Pelagophyceae | Pelagomonad | unclassified | Aureococcus | Aureococcus a | unclassified | unclassified | unclassified | unclassified | unclassified | 1,6 | 0,489897949 | 0,000349645 | 8,56E-05 | 1 - | 0,000226015 - | 1 - | 0,000265902 - | 2 - | 0,000419468 - | 2 - | 0,000416832 - | 2 - | 0,000420008 - |
| 501 | 1 | 0.2um_NOKCPHJN_233889 | Bacteria | Proteobacteri | Alphaproteob | Rhizobiales | Bradyrhizobia | Bradyrhizobiu | Bradyrhizobiu | unclassified | unclassified | unclassified | unclassified | unclassified | 0,2 | 0,4 | 2,65E-05 | 5,29E-05 | 0 - | 0 - | 0 - | 0 - | 0 - | 0 - | 1 - | 0,000132349 - | 0 - | 0 - |
| 502 | 1 | 3um_FIJJAFBF_123736 | Bacteria | Proteobacteri | Alphaproteob | Rhizobiales | unclassified | unclassified | Rhizobiales ba | unclassified | unclassified | unclassified | unclassified | unclassified | 0,2 | 0,4 | 3,45E-05 | 6,91E-05 | 0 - | 0 - | 0 - | 0 - | 0 - | 0 - | 0 - | 0 - | 1 - | 0,00017263 - |
| 503 | 1 | 3um_FIJJAFBF_138049 | Viruses | unclassified | unclassified | unclassified | unclassified | unclassified | uncultured M | unclassified | unclassified | unclassified | unclassified | unclassified | 0,2 | 0,4 | 1,86E-05 | 3,72E-05 | 0 - | 0 - | 0 - | 0 - | 1 - | 9,30E-05 - | 0 - | 0 - | 0 - | 0 - |
| 504 | 1 | 0.2um_NOKCPHJN_372005 | Eukaryota | Phaeophyceae | unclassified | Ectocarpales | Ectocarpaceae | Ectocarpus | Ectocarpus sil | cellular proces | Signal transdu | KOG0613 | fibronectin typ | 653948.CCA1 | 0,2 | 0,4 | 1,14E-05 | 2,29E-05 | 0 - | 0 - | 0 - | 0 - | 1 - | 5,71E-05 - | 0 - | 0 - | 0 - | 0 - |
| 505 | 1 | 3um_FIJJAFBF_282298 | unclassified | unclassified | unclassified | unclassified | unclassified | unclassified | unclassified | unclassified | unclassified | unclassified | unclassified | unclassified | 1,8 | 1,166190379 | 0,001395325 | 0,000854751 | 0 - | 0 - | 1 - | 0,000955277 - | 3 - | 0,002260465 - | 2 - | 0,001497507 - | 3 - | 0,002263375 - |
| 506 | 2 | 0.2um_NOKCPHJN_80626;3um | Bacteria | Bacteroidetes | Flavobacteriia | Flavobacterial | Flavobacteriac | Polaribacter | Polaribacter s | unclassified | unclassified | unclassified | unclassified | unclassified | 0,6 | 0,489897949 | 0,000318523 | 0,000260518 | 1 - | 0,00055856 - | 0 - | 0 - | 0 - | 0 - | 1 - | 0,000515066 - | 1 - | 0,00051899 - |
| 507 | 1 | 3um_FIJJAFBF_37831 | Bacteria | Proteobacteri | unclassified | unclassified | unclassified | unclassified | Proteobacteri | unclassified | unclassified | unclassified | unclassified | unclassified | 6,6 | 1,019803903 | 0,001794427 | 0,00041526 | 5 - | 0,001361706 - | 8 - | 0,002563227 - | 7 - | 0,00176906 - | 6 - | 0,001506808 - | 7 - | 0,001771337 - |
| 508 | 1 | 0.2um_NOKCPHJN_97282 | Bacteria | Proteobacteri | Gammaprotec | Alteromonada | Alteromonada | Paraglaciecola | Paraglaciecola | metabolism | Energy produc | COG2838 | Isocitrate deh | 983545.Glaag | 1 | 0 | 0,000110442 | 1,08E-05 | 1 - | 0,000111428 - | 1 - | 0,000131093 - | 1 - | 0,000103401 - | 1 - | 0,000102751 - | 1 - | 0,000103534 - |
| 509 | 1 | 0.2um_NOKCPHJN_45933 | Bacteria | Bacteroidetes | Flavobacteriia | Flavobacterial | Flavobacteriac | Cellulophaga | Cellulophaga t | poorly charact | Function unkn | 0XP53 | Ragb susd dor | 313598.MED1 | 6,2 | 2,039607805 | 0,000990475 | 0,000284831 | 4 - | 0,000654432 - | 5 - | 0,000962406 - | 10 - | 0,001518223 - | 6 - | 0,000905209 - | 6 - | 0,000912106 - |
| 510 | 1 | 3um_FIJJAFBF_328495 | Bacteria | Bacteroidetes | Flavobacteriia | Flavobacterial | Flavobacteriac | Arenitalea | Arenitalea lute | poorly charact | Function unkn | 0PAEN | NA | 886377.Murru | 7,2 | 0,748331477 | 0,001315907 | 0,000248809 | 8 - | 0,001464615 - | 8 - | 0,001723088 - | 6 - | 0,001019333 - | 7 - | 0,001181748 - | 7 - | 0,001190752 - |
| 511 | 1 | 0.2um_NOKCPHJN_1113290 | Bacteria | Bacteroidetes | Bacteroidia | Bacteroidales | Prevotellaceae | Prevotella | various | unclassified | unclassified | unclassified | unclassified | unclassified | 1,4 | 0,489897949 | 0,00027859 | 9,28E-05 | 2 - | 0,000405052 - | 1 - | 0,000238268 - | 1 - | 0,000187937 - | 2 - | 0,000373512 - | 1 - | 0,000188179 - |
| 512 | 3 | 0.2um_NOKCPHJN_25800;0.2u | Bacteria | Bacteroidetes | Flavobacteriia | Flavobacterial | Flavobacteriac | Polaribacter | various | various | various | various | various | various | 0,2 | 0,4 | 1,41E-05 | 2,82E-05 | 0 - | 0 - | 0 - | 0 - | 1 - | 7,06E-05 - | 0 - | 0 - | 0 - | 0 - |
| 513 | 1 | 3um_FIJJAFBF_263708 | Eukaryota | Eustigmatoph | unclassified | Eustigmatales | Monodopsida | Nannochlorop | Nannochlorop | metabolism | Lipid transpor | COG0439 | acetyl-CoA ca | 35128.Thaps1 | 0,2 | 0,4 | 6,14E-05 | 0,000122821 | 0 - | 0 - | 1 - | 0,000307053 - | 0 - | 0 - | 0 - | 0 - | 0 - | 0 - |
| 514 | 1 | 3um_FIJJAFBF_165682 | Bacteria | Proteobacteri | Gammaprotec | Pseudomonad | Pseudomonad | Pseudomonas | Pseudomonas | unclassified | unclassified | unclassified | unclassified | unclassified | 1 | 0 | 0,000749288 | 7,33E-05 | 1 - | 0,000755981 - | 1 - | 0,000889396 - | 1 - | 0,000701524 - | 1 - | 0,000697115 - | 1 - | 0,000702427 - |
| 515 | 1 | 0.2um_NOKCPHJN_862435 | Eukaryota | unclassified | unclassified | Prymnesiales | Chrysochromu | Chrysochromu | Chrysochromu | unclassified | unclassified | unclassified | unclassified | unclassified | 0,8 | 0,748331477 | 0,000208133 | 0,000185329 | 1 - | 0,000257923 - | 1 - | 0,000303441 - | 0 - | 0 - | 0 - | 0 - | 2 - | 0,000479303 - |
| 516 | 1 | 3um_FIJJAFBF_37824 | Bacteria | Actinobacteria | unclassified | unclassified | unclassified | unclassified | Actinobacteria | unclassified | unclassified | unclassified | unclassified | unclassified | 5,8 | 1,166190379 | 0,000507969 | 6,70E-05 | 5 - | 0,000449712 - | 4 - | 0,000423261 - | 7 - | 0,000584243 - | 6 - | 0,000497633 - | 7 - | 0,000584995 - |
| 517 | 1 | 0.2um_NOKCPHJN_674215 | unclassified | unclassified | unclassified | unclassified | unclassified | unclassified | unclassified | unclassified | unclassified | unclassified | unclassified | unclassified | 0,2 | 0,4 | 0,00013468 | 0,00026936 | 0 - | 0 - | 0 - | 0 - | 0 - | 0 - | 0 - | 0 - | 1 - | 0,000673401 - |
| 518 | 1 | 0.2um_NOKCPHJN_291529 | Bacteria | Bacteroidetes | Flavobacteriia | Flavobacterial | Flavobacteriac | Formosa | Formosa sp. H | metabolism | Inorganic ion | 05C27 | TonB-depende | 742766.HMPR | 1 | 0,632455532 | 7,84E-05 | 4,90E-05 | 1 - | 8,33E-05 - | 0 - | 0 - | 1 - | 7,73E-05 - | 1 - | 7,68E-05 - | 2 - | 0,000154761 - |
| 519 | 1 | 3um_FIJJAFBF_755903 | Bacteria | Proteobacteri | Gammaprotec | Enterobactera | Yersiniaceae | unclassified | Yersiniaceae b | unclassified | unclassified | unclassified | unclassified | unclassified | 0,4 | 0,489897949 | 0,000378785 | 0,000464549 | 1 - | 0,000985324 - | 0 - | 0 - | 0 - | 0 - | 1 - | 0,000908599 - | 0 - | 0 - |
| 520 | 1 | 3um_FIJJAFBF_724142 | Eukaryota | Chordata | Actinopteri | Osteoglossifor | Osteoglossida | Scleropages | Scleropages fo | unclassified | unclassified | unclassified | unclassified | unclassified | 0,2 | 0,4 | 1,56E-05 | 3,12E-05 | 1 - | 7,80E-05 - | 0 - | 0 - | 0 - | 0 - | 0 - | 0 - | 0 - | 0 - |
| 521 | 1 | 3um_FIJJAFBF_106479 | Bacteria | Bacteroidetes | Flavobacteriia | Flavobacterial | Flavobacteriac | Flavivirga | Flavivirga eckl | information st | Translation, ri | 05C64 | Binds directly | 983544.Lacal_ | 0,4 | 0,489897949 | 0,000141616 | 0,000173443 | 0 - | 0 - | 0 - | 0 - | 1 - | 0,000353812 - | 0 - | 0 - | 1 - | 0,000354267 - |
| 522 | 1 | 0.2um_NOKCPHJN_122862 | Bacteria | Bacteroidetes | Flavobacteriia | Flavobacterial | Flavobacteriac | Cellulophaga | Cellulophaga t | unclassified | unclassified | unclassified | unclassified | unclassified | 0,4 | 0,489897949 | 4,66E-05 | 5,71E-05 | 0 - | 0 - | 0 - | 0 - | 1 - | 0,000116921 - | 1 - | 0,000116186 - | 0 - | 0 - |
| 523 | 1 | 0.2um_NOKCPHJN_1035812 | Bacteria | Proteobacteri | Gammaprotec | Oceanospirilla | Oceanospirilla | Neptuniibacte | Neptuniibacte | metabolism | Carbohydrate | 16PI5 | ribulose-phos | 207954.MED9 | 1 | 0 | 0,001059969 | 0,000103712 | 1 - | 0,001069437 - | 1 - | 0,001258169 - | 1 - | 0,000992399 - | 1 - | 0,000986163 - | 1 - | 0,000993677 - |
| 524 | 1 | 3um_FIJJAFBF_373756 | Bacteria | Bacteroidetes | Flavobacteriia | Flavobacterial | Flavobacteriac | Mesoflavibact | various | metabolism | Nucleotide tra | 0QPJ5 | Deoxycytidyla | 216895.VV2_ | 0,2 | 0,4 | 2,41E-05 | 4,81E-05 | 0 - | 0 - | 0 - | 0 - | 0 - | 0 - | 0 - | 0 - | 1 - | 0,000120357 - |
| 525 | 1 | 3um_FIJJAFBF_21475 | Eukaryota | unclassified | Cryptophyta | Pyrenomonad | Geminigerace | Guillardia | Guillardia thet | metabolism | Carbohydrate | 0IVA1 | Photosystem I | 3218.PP1S345 | 0,4 | 0,489897949 | 0,000184218 | 0,000225928 | 1 - | 0,000479201 - | 0 - | 0 - | 0 - | 0 - | 1 - | 0,000441887 - | 0 - | 0 - |
| 526 | 1 | 0.2um_NOKCPHJN_806559 | Eukaryota | Arthropoda | Collembola | Entomobryom | Isotomidae | Folsomia | Folsomia cand | poorly charact | Function unkn | 0WJDD | Inherit from N | 7227.FBpp029 | 0,4 | 0,8 | 6,50E-05 | 0,000129917 | 2 - | 0,000324792 - | 0 - | 0 - | 0 - | 0 - | 0 - | 0 - | 0 - | 0 - |
| 527 | 1 | 3um_FIJJAFBF_783325 | Eukaryota | Bacillariophyta | Mediophyceae | Anaulales | Anaulaceae | Eunotogramm | Eunotogramm | information st | Translation, ri | 0FNCH | Binds directly | 65393.PCC742 | 4,4 | 1,2 | 0,003105712 | 0,000700939 | 5 - | 0,00365391 - | 2 - | 0,001719498 - | 5 - | 0,003390698 - | 5 - | 0,00336939 - | 5 - | 0,003395062 - |
| 528 | 1 | 0.2um_NOKCPHJN_230839 | Eukaryota | Phaeophyceae | unclassified | Ectocarpales | Ectocarpaceae | Ectocarpus | Ectocarpus sil | unclassified | unclassified | unclassified | unclassified | unclassified | 0,2 | 0,4 | 3,40E-05 | 6,80E-05 | 0 - | 0 - | 0 - | 0 - | 0 - | 0 - | 0 - | 0 - | 1 - | 0,000170107 - |
| 529 | 1 | 0.2um_NOKCPHJN_888436 | Bacteria | Bacteroidetes | Flavobacteriia | unclassified | unclassified | unclassified | Flavobacteria | unclassified | unclassified | unclassified | unclassified | unclassified | 0,2 | 0,4 | 2,55E-05 | 5,10E-05 | 1 - | 0,000127462 - | 0 - | 0 - | 0 - | 0 - | 0 - | 0 - | 0 - | 0 - |
| 530 | 1 | 3um_FIJJAFBF_513056 | Bacteria | Proteobacteri | Alphaproteob | Rhizobiales | Hyphomicrobi | unclassified | Hyphomicrobi | unclassified | unclassified | unclassified | unclassified | unclassified | 0,2 | 0,4 | 7,50E-05 | 0,000149904 | 1 - | 0,00037476 - | 0 - | 0 - | 0 - | 0 - | 0 - | 0 - | 0 - | 0 - |
| 531 | 1 | 0.2um_NOKCPHJN_366105 | Viruses | unclassified | unclassified | unclassified | unclassified | unclassified | Tetraselmis vi | unclassified | unclassified | unclassified | unclassified | unclassified | 0,4 | 0,489897949 | 6,17E-05 | 7,57E-05 | 1 - | 0,000160025 - | 0 - | 0 - | 0 - | 0 - | 0 - | 0 - | 1 - | 0,000148689 - |
| 532 | 1 | 3um_FIJJAFBF_703717 | Bacteria | Bacteroidetes | Flavobacteriia | Flavobacterial | Flavobacteriac | Algibacter | Algibacter pec | cellular proces | Posttranslatio | 05E6M | Sulphatase-m | 867900.Celly_ | 0,8 | 0,4 | 0,000187005 | 9,63E-05 | 0 - | 0 - | 1 - | 0,000278086 - | 1 - | 0,000219344 - | 1 - | 0,000217966 - | 1 - | 0,000219627 - |
| 533 | 1 | 0.2um_NOKCPHJN_93295 | Bacteria | Proteobacteri | Alphaproteob | Pelagibacteral | Pelagibacterac | unclassified | various | cellular proces | Posttranslatio | 01QWG | Prevents misf | 439493.PB721 | 0,4 | 0,8 | 5,88E-05 | 0,000117512 | 0 - | 0 - | 0 - | 0 - | 2 - | 0,000293779 - | 0 - | 0 - | 0 - | 0 - |
| 534 | 1 | 3um_FIJJAFBF_121956 | Bacteria | Proteobacteri | Alphaproteob | Rhodobactera | Rhodobactera | Yoonia | Yoonia rosea | metabolism | Energy produc | 01QS0 | Aldehyde oxid | 467661.RKLH1 | 0,2 | 0,4 | 2,17E-05 | 4,34E-05 | 0 - | 0 - | 0 - | 0 - | 0 - | 0 - | 0 - | 0 - | 1 - | 0,000108497 - |
| 535 | 1 | 0.2um_NOKCPHJN_464767 | Eukaryota | unclassified | Cryptophyta | Pyrenomonad | Geminigerace | Guillardia | Guillardia thet | unclassified | unclassified | unclassified | unclassified | unclassified | 2 | 0,632455532 | 0,000810402 | 0,000212534 | 2 - | 0,000839176 - | 1 - | 0,000493636 - | 2 - | 0,000778725 - | 3 - | 0,001160747 - | 2 - | 0,000779727 - |
| 536 | 1 | 3um_FIJJAFBF_116593 | Bacteria | Cyanobacteria | unclassified | Synechococca | Synechococca | Synechococcu | Synechococcu | unclassified | unclassified | unclassified | unclassified | unclassified | 1 | 0 | 0,000335589 | 3,28E-05 | 1 - | 0,000338586 - | 1 - | 0,000398339 - | 1 - | 0,000314196 - | 1 - | 0,000312221 - | 1 - | 0,0003146 - |
| 537 | 1 | 0.2um_NOKCPHJN_557642 | Bacteria | Proteobacteri | Gammaprotec | Xanthomonad | Xanthomonad | Xanthomonas | Xanthomonas | poorly charact | Function unkn | 0NVRZ | NA | 170187.SP_17 | 2,4 | 0,489897949 | 0,000573334 | 0,000112605 | 3 - | 0,000726745 - | 2 - | 0,000569999 - | 2 - | 0,000449595 - | 3 - | 0,000670155 - | 2 - | 0,000450174 - |
| 538 | 1 | 0.2um_NOKCPHJN_215620 | Eukaryota | Chromerida | unclassified | unclassified | unclassified | Vitrella | Vitrella brassi | unclassified | unclassified | unclassified | unclassified | unclassified | 1 | 0 | 0,000164616 | 1,61E-05 | 1 - | 0,000166087 - | 1 - | 0,000195398 - | 1 - | 0,000154123 - | 1 - | 0,000153154 - | 1 - | 0,000154321 - |
| 539 | 10 | 0.2um_NOKCPHJN_1085110;0. | Bacteria | various | various | various | various | various | various | various | various | various | various | various | 1 | 0 | 6,85E-05 | 6,70E-06 | 1 - | 6,91E-05 - | 1 - | 8,13E-05 - | 1 - | 6,41E-05 - | 1 - | 6,37E-05 - | 1 - | 6,42E-05 - |
| 540 | 1 | 0.2um_NOKCPHJN_829615 | Bacteria | Proteobacteri | Gammaprotec | Pseudomonad | Pseudomonad | Pseudomonas | Pseudomonas | information st | Translation, ri | 16QAF | thus facilitatin | 205922.Pfl01_ | 0,8 | 0,4 | 0,000162062 | 8,13E-05 | 1 - | 0,00021441 - | 0 - | 0 - | 1 - | 0,000198965 - | 1 - | 0,000197715 - | 1 - | 0,000199221 - |
| 541 | 1 | 0.2um_NOKCPHJN_315471 | unclassified | unclassified | unclassified | unclassified | unclassified | unclassified | unclassified | unclassified | unclassified | unclassified | unclassified | unclassified | 2,8 | 0,4 | 0,000564612 | 0,000109585 | 3 - | 0,000607579 - | 3 - | 0,000714803 - | 2 - | 0,000375874 - | 3 - | 0,000560268 - | 3 - | 0,000564537 - |
| 542 | 1 | 3um_FIJJAFBF_510237 | Bacteria | Bacteroidetes | Flavobacteriia | Flavobacterial | Flavobacteriac | Polaribacter | Polaribacter s | unclassified | unclassified | unclassified | unclassified | unclassified | 0,4 | 0,489897949 | 0,000266942 | 0,000328723 | 1 - | 0,000613244 - | 1 - | 0,000721468 - | 0 - | 0 - | 0 - | 0 - | 0 - | 0 - |
| 543 | 2 | 0.2um_NOKCPHJN_326347;3ur | Eukaryota | unclassified | Oomycetes | Saprolegniales | Saprolegniace | Aphanomyces | Aphanomyces | cellular proces | Intracellular tr | various | various | various | 1 | 0 | 0,000436771 | 4,27E-05 | 1 - | 0,000440673 - | 1 - | 0,000518442 - | 1 - | 0,000408928 - | 1 - | 0,000406359 - | 1 - | 0,000409455 - |

| | | | | | | | | | | | | | | | | | | | | | | | | | | | | |
|---|---|---|---|---|---|---|---|---|---|---|---|---|---|---|---|---|---|---|---|---|---|---|---|---|---|---|---|---|
| 544 | 1 | 3um_FIJJAFBF_589704 | Eukaryota | unclassified | Cryptophyta | Pyrenomonad | Geminigerace | Teleaulax | Teleaulax amp | metabolism | Energy produc | COG4451 | RuBisCO catal | 45157.CMV01 | 1,6 | 0,8 | 0,000994751 | 0,0004977 | 3 - | 0,001892673 - | 1 - | 0,000742229 - | 2 - | 0,001170888 - | 1 - | 0,000581765 - | 1 - | 0,000586198 - |
| 545 | 1 | 0.2um_NOKCPHJN_92038 | Eukaryota | Basidiomycota | Dacrymycetes | Dacrymycetal | Dacrymycetac | Calocera | Calocera corn | cellular proces | Cell cycle cont | 19G3J | 14-3-3 protein | 644223.XP_00 | 0,4 | 0,489897949 | 0,000134718 | 0,000165187 | 1 - | 0,000349378 - | 0 - | 0 - | 1 - | 0,00032421 - | 0 - | 0 - | 0 - | 0 - |
| 546 | 1 | 0.2um_NOKCPHJN_458305 | Bacteria | unclassified | unclassified | unclassified | unclassified | unclassified | bacterium B1 | unclassified | unclassified | unclassified | unclassified | unclassified | 0,4 | 0,489897949 | 1,92E-05 | 2,35E-05 | 1 - | 4,99E-05 - | 0 - | 0 - | 0 - | 0 - | 1 - | 4,60E-05 - | 0 - | 0 - |
| 547 | 1 | 3um_FIJJAFBF_787562 | Eukaryota | unclassified | Pelagophycea | Pelagomonad | unclassified | Aureococcus | Aureococcus a | unclassified | unclassified | unclassified | unclassified | unclassified | 1 | 0 | 0,000776049 | 7,59E-05 | 1 - | 0,000782981 - | 1 - | 0,00092116 - | 1 - | 0,000726578 - | 1 - | 0,000722012 - | 1 - | 0,000727513 - |
| 548 | 1 | 3um_FIJJAFBF_412070 | Bacteria | Bacteroidetes | Flavobacteriia | Flavobacterial | Flavobacteriac | Arenitalea | Arenitalea lut | unclassified | unclassified | unclassified | unclassified | unclassified | 0,2 | 0,4 | 6,45E-05 | 0,000128962 | 1 - | 0,000322404 - | 0 - | 0 - | 0 - | 0 - | 0 - | 0 - | 0 - | 0 - |
| 549 | 1 | 0.2um_NOKCPHJN_891066 | Bacteria | Bacteroidetes | Flavobacteriia | unclassified | unclassified | unclassified | Flavobacteria | metabolism | Inorganic ion t | 05C27 | TonB-depende | 487797.Flav3 | 2,4 | 3,006659276 | 0,000387761 | 0,000498197 | 8 - | 0,001326183 - | 0 - | 0 - | 0 - | 0 - | 3 - | 0,000458594 - | 1 - | 0,000154029 - |
| 550 | 1 | 3um_FIJJAFBF_41553 | Eukaryota | Bacillariophyta | Coscinodiscop | Chaetocerotal | Chaetocerotac | Chaetoceros | Chaetoceros s | unclassified | unclassified | unclassified | unclassified | unclassified | 2 | 0 | 0,001151225 | 0,000112641 | 2 - | 0,001161508 - | 2 - | 0,001366489 - | 2 - | 0,001077838 - | 2 - | 0,001071064 - | 2 - | 0,001079225 - |
| 551 | 1 | 3um_FIJJAFBF_163900 | Bacteria | Proteobacteri | Alphaproteob | Rhodobactera | Rhodobactera | unclassified | Rhodobactera | metabolism | Carbohydrate | 01RFA | ABC transport | 644076.SCH4 | 2 | 1,095445115 | 0,000515977 | 0,000291343 | 4 - | 0,001025659 - | 2 - | 0,000603333 - | 1 - | 0,000237944 - | 1 - | 0,000236448 - | 2 - | 0,0004765 - |
| 552 | 1 | 0.2um_NOKCPHJN_330840 | Eukaryota | unclassified | Cryptophyta | Pyrenomonad | Geminigerace | Guillardia | Guillardia the | metabolism | Carbohydrate | COG1850 | RuBisCO catal | 35128.Thapsd | 1,2 | 0,4 | 0,000418488 | 0,000192443 | 1 - | 0,000341221 - | 2 - | 0,000802879 - | 1 - | 0,000316641 - | 1 - | 0,000314651 - | 1 - | 0,000317049 - |
| 553 | 2 | 3um_FIJJAFBF_275863;3um_FI | Bacteria | Bacteroidetes | Flavobacteriia | Flavobacterial | Flavobacteriac | Formosa | Formosa sp. H | unclassified | unclassified | unclassified | unclassified | unclassified | 0,6 | 0,489897949 | 0,000224152 | 0,000185314 | 1 - | 0,00036088 - | 1 - | 0,000424567 - | 0 - | 0 - | 0 - | 0 - | 1 - | 0,000335315 - |
| 554 | 1 | 3um_FIJJAFBF_21329 | Bacteria | Bacteroidetes | Flavobacteriia | Flavobacterial | Flavobacteriac | Polaribacter | Polaribacter s | cellular proces | Cell wall/mem | 09GJR | outer membra | 313598.MED1 | 0,4 | 0,489897949 | 0,000197743 | 0,000242467 | 1 - | 0,00051283 - | 0 - | 0 - | 1 - | 0,000475887 - | 0 - | 0 - | 0 - | 0 - |
| 555 | 1 | 3um_FIJJAFBF_475014 | Bacteria | Proteobacteri | Gammaprotec | Cellvibrionale | Cellvibrionace | Cellvibrio | Cellvibrio sp. | cellular proces | Cell wall/mem | 0QTSX | Peptidoglycan | 314287.GB22 | 4,8 | 0,4 | 0,002449428 | 7,43E-05 | 5 - | 0,002594493 - | 4 - | 0,002441891 - | 5 - | 0,002407596 - | 5 - | 0,002392466 - | 5 - | 0,002410695 - |
| 556 | 1 | 3um_FIJJAFBF_437252 | Bacteria | Bacteroidetes | Flavobacteriia | Flavobacterial | Flavobacteriac | Polaribacter | Polaribacter t | poorly charact | Function unkn | 0XSVM | NA | 867900.Celly_ | 1,2 | 0,4 | 0,00021522 | 6,34E-05 | 1 - | 0,000183077 - | 1 - | 0,000215386 - | 1 - | 0,000169889 - | 2 - | 0,000337642 - | 1 - | 0,000170107 - |
| 557 | 1 | 3um_FIJJAFBF_647835 | Viruses | unclassified | unclassified | unclassified | unclassified | unclassified | uncultured M | unclassified | unclassified | unclassified | unclassified | unclassified | 1 | 0 | 0,000323113 | 3,16E-05 | 1 - | 0,000325999 - | 1 - | 0,000383531 - | 1 - | 0,000302516 - | 1 - | 0,000300615 - | 1 - | 0,000302905 - |
| 558 | 1 | 0.2um_NOKCPHJN_24010 | Eukaryota | Arthropoda | Insecta | Lepidoptera | Noctuidae | Helicoverpa | Helicoverpa a | unclassified | unclassified | unclassified | unclassified | unclassified | 0,8 | 0,4 | 0,000276946 | 0,000142033 | 1 - | 0,000343897 - | 1 - | 0,000404588 - | 1 - | 0,000319124 - | 1 - | 0,000317119 - | 0 - | 0 - |
| 559 | 1 | 0.2um_NOKCPHJN_1041578 | unclassified | unclassified | unclassified | unclassified | unclassified | unclassified | unclassified | unclassified | unclassified | unclassified | unclassified | unclassified | 0,4 | 0,489897949 | 7,31E-05 | 9,06E-05 | 0 - | 0 - | 1 - | 0,000204297 - | 0 - | 0 - | 0 - | 0 - | 1 - | 0,000161349 - |
| 560 | 1 | 3um_FIJJAFBF_335656 | Bacteria | Planctomycete | Planctomyceti | Planctomyceta | Planctomyceta | Rubinisphaera | Rubinisphaera | unclassified | unclassified | unclassified | unclassified | unclassified | 1 | 0 | 0,000181078 | 1,77E-05 | 1 - | 0,000182696 - | 1 - | 0,000214937 - | 1 - | 0,000169535 - | 1 - | 0,000168469 - | 1 - | 0,000169753 - |
| 561 | 1 | 0.2um_NOKCPHJN_671436 | Bacteria | Verrucomicrol | Verrucomicrol | Verrucomicrol | Verrucomicrol | Pedosphaera | Pedosphaera | unclassified | unclassified | unclassified | unclassified | unclassified | 0,6 | 0,489897949 | 0,000837663 | 0,000692596 | 1 - | 0,001349136 - | 1 - | 0,001587229 - | 1 - | 0,00125195 - | 0 - | 0 - | 0 - | 0 - |
| 562 | 1 | 3um_FIJJAFBF_565482 | Eukaryota | unclassified | Pelagophycea | Pelagomonad | unclassified | Aureococcus | Aureococcus a | metabolism | Carbohydrate | KOG1203 | Complex I inte | 35128.Thapsd | 1,4 | 0,489897949 | 0,000374188 | 0,00011196 | 1 - | 0,000274902 - | 1 - | 0,000323417 - | 2 - | 0,000510199 - | 2 - | 0,000506993 - | 1 - | 0,000255428 - |
| 563 | 1 | 0.2um_NOKCPHJN_136446 | Eukaryota | Echinodermat | Echinoidea | Echinoida | Strongylocent | Strongylocent | Strongylocent | cellular proces | Cell wall/mem | 12RRB | ankyrin 3, nod | 7668.SPU_01 | 1 | 0 | 9,82E-05 | 9,61E-06 | 1 - | 9,91E-05 - | 1 - | 0,000116576 - | 1 - | 9,20E-05 - | 1 - | 9,14E-05 - | 1 - | 9,21E-05 - |
| 564 | 1 | 3um_FIJJAFBF_65359 | Bacteria | Bacteroidetes | Flavobacteriia | Flavobacterial | Flavobacteriac | Arenitalea | Arenitalea lut | poorly charact | Function unkn | 0PEU4 | NA | 1046627.BZAI | 0,2 | 0,4 | 5,31E-05 | 0,000106165 | 0 - | 0 - | 0 - | 0 - | 0 - | 0 - | 0 - | 0 - | 1 - | 0,000265412 - |
| 565 | 1 | 0.2um_NOKCPHJN_332245 | Eukaryota | unclassified | Labyrinthulom | unclassified | Thraustochytr | Aurantiochytr | Aurantiochytr | poorly charact | Function unkn | 0NVRZ | NA | 170187.SP_17 | 0,6 | 0,489897949 | 1,37E-05 | 1,11E-05 | 0 - | 0 - | 0 - | 0 - | 1 - | 2,28E-05 - | 1 - | 2,27E-05 - | 1 - | 2,28E-05 - |
| 566 | 1 | 3um_FIJJAFBF_346214 | Bacteria | Bacteroidetes | Flavobacteriia | Flavobacterial | Crocinitomica | unclassified | Crocinitomica | metabolism | Energy produc | COG0711 | F(1)F(0) ATP s | 926562.Oweh | 0,4 | 0,489897949 | 0,000197404 | 0,00024177 | 0 - | 0 - | 0 - | 0 - | 1 - | 0,000493192 - | 0 - | 0 - | 1 - | 0,000493827 - |
| 567 | 1 | 3um_FIJJAFBF_596860 | Bacteria | unclassified | unclassified | unclassified | unclassified | unclassified | uncultured ba | unclassified | unclassified | unclassified | unclassified | unclassified | 0,2 | 0,4 | 3,19E-05 | 6,38E-05 | 0 - | 0 - | 0 - | 0 - | 0 - | 0 - | 0 - | 0 - | 1 - | 0,000159455 - |
| 568 | 1 | 3um_FIJJAFBF_641954 | Bacteria | Proteobacteri | Gammaprotec | Cellvibrionale | Spongiibacter | Oceanicoccus | Oceanicoccus | metabolism | Inorganic ion t | 01UP7 | TonB-depende | 394221.Mmar | 8,6 | 1,356465997 | 0,000933451 | 9,19E-05 | 7 - | 0,000776052 - | 7 - | 0,000913008 - | 10 - | 0,001028783 - | 10 - | 0,001022318 - | 9 - | 0,000927097 - |
| 569 | 1 | 0.2um_NOKCPHJN_338400 | unclassified | unclassified | unclassified | unclassified | unclassified | unclassified | unclassified | unclassified | unclassified | unclassified | unclassified | unclassified | 0,8 | 0,4 | 0,000762403 | 0,000392467 | 0 - | 0 - | 1 - | 0,001133735 - | 1 - | 0,00089425 - | 1 - | 0,00088863 - | 1 - | 0,000895401 - |
| 570 | 1 | 3um_FIJJAFBF_557513 | Bacteria | Bacteroidetes | Flavobacteriia | Flavobacterial | Flavobacteriac | Tamlana | Tamlana agar | unclassified | unclassified | unclassified | unclassified | unclassified | 3 | 0 | 0,000830422 | 8,13E-05 | 3 - | 0,000837839 - | 3 - | 0,0009857 - | 3 - | 0,000777485 - | 3 - | 0,000772599 - | 3 - | 0,000778486 - |
| 571 | 1 | 3um_FIJJAFBF_255783 | Bacteria | Proteobacteri | Gammaprotec | Cellvibrionale | Porticoccacea | unclassified | Gammaprotec | unclassified | unclassified | unclassified | unclassified | unclassified | 1 | 0 | 0,000108106 | 1,06E-05 | 1 - | 0,000109072 - | 1 - | 0,000128321 - | 1 - | 0,000101215 - | 1 - | 0,000100579 - | 1 - | 0,000101345 - |
| 572 | 1 | 3um_FIJJAFBF_284683 | Eukaryota | Bacillariophyta | Coscinodiscop | Leptocylindral | Leptocylindrac | Leptocylindrus | Leptocylindrus | metabolism | Energy produc | 1DYB8 | PsaA and PsaB | 3055.DAA014 | 1,4 | 0,489897949 | 0,000158759 | 4,76E-05 | 1 - | 0,000116614 - | 1 - | 0,000137194 - | 1 - | 0,000108214 - | 2 - | 0,000215067 - | 2 - | 0,000216706 - |
| 573 | 2 | 0.2um_NOKCPHJN_180219;0.2 | Bacteria | Bacteroidetes | Cytophagia | Cytophagales | Flammeovirga | unclassified | Flammeovirga | cellular proces | Posttranslatio | 16QU3 | Hemolysin-typ | 744980.TRICH | 0,6 | 0,489897949 | 3,26E-05 | 2,66E-05 | 1 - | 5,71E-05 - | 0 - | 0 - | 0 - | 0 - | 1 - | 5,27E-05 - | 1 - | 5,31E-05 - |
| 574 | 1 | 0.2um_NOKCPHJN_358348 | Bacteria | Proteobacteri | Gammaprotec | unclassified | unclassified | unclassified | Gammaprotec | unclassified | unclassified | unclassified | unclassified | unclassified | 0,4 | 0,489897949 | 4,05E-05 | 4,96E-05 | 0 - | 0 - | 0 - | 0 - | 0 - | 0 - | 1 - | 0,000100955 - | 1 - | 0,000101725 - |
| 575 | 1 | 3um_FIJJAFBF_506077 | unclassified | unclassified | unclassified | unclassified | unclassified | unclassified | unclassified | unclassified | unclassified | unclassified | unclassified | unclassified | 1,8 | 0,748331477 | 0,000347712 | 0,000155352 | 3 - | 0,000575671 - | 2 - | 0,000451509 - | 2 - | 0,000356135 - | 1 - | 0,000176948 - | 1 - | 0,000178296 - |
| 576 | 1 | 3um_FIJJAFBF_372266 | Bacteria | Proteobacteri | Alphaproteob | unclassified | unclassified | Candidatus Pu | Candidatus Pu | information st | Translation, ri | COG0335 | This protein is | 35128.Thapsd | 0,2 | 0,4 | 0,000142592 | 0,000285183 | 1 - | 0,000712958 - | 0 - | 0 - | 0 - | 0 - | 0 - | 0 - | 0 - | 0 - |
| 577 | 1 | 3um_FIJJAFBF_673147 | unclassified | unclassified | unclassified | unclassified | unclassified | unclassified | unclassified | unclassified | unclassified | unclassified | unclassified | unclassified | 2 | 0 | 0,001524868 | 0,000149199 | 2 - | 0,001538489 - | 2 - | 0,001809998 - | 2 - | 0,001427662 - | 2 - | 0,00141869 - | 2 - | 0,0014295 - |
| 578 | 1 | 3um_FIJJAFBF_331698 | Bacteria | Proteobacteri | Alphaproteob | Rickettsiales | Candidatus M | Candidatus Jic | Candidatus Jic | metabolism | Energy produc | 01RNF | Produces ATP | 696127.midi_ | 0,4 | 0,489897949 | 7,20E-05 | 8,83E-05 | 1 - | 0,00018738 - | 0 - | 0 - | 0 - | 0 - | 1 - | 0,000172789 - | 0 - | 0 - |
| 579 | 1 | 3um_FIJJAFBF_100702 | Bacteria | Bacteroidetes | Flavobacteriia | Flavobacterial | Flavobacteriac | Formosa | Formosa sp. H | unclassified | unclassified | unclassified | unclassified | unclassified | 2,6 | 1,496662955 | 0,000255304 | 0,000135589 | 1 - | 0,000103291 - | 1 - | 0,000121519 - | 3 - | 0,00028755 - | 5 - | 0,000476239 - | 3 - | 0,00028792 - |
| 580 | 1 | 0.2um_NOKCPHJN_188311 | Bacteria | Proteobacteri | Gammaprotec | Cellvibrionale | Porticoccacea | unclassified | Gammaprotec | unclassified | unclassified | unclassified | unclassified | unclassified | 0,4 | 0,489897949 | 2,25E-05 | 2,75E-05 | 1 - | 5,83E-05 - | 0 - | 0 - | 1 - | 5,41E-05 - | 0 - | 0 - | 0 - | 0 - |
| 581 | 1 | 3um_FIJJAFBF_423475 | Eukaryota | Bacillariophyta | Coscinodiscop | Thalassiosirale | Thalassiosirac | Thalassiosira | Thalassiosira | unclassified | unclassified | unclassified | unclassified | unclassified | 2,6 | 0,489897949 | 0,000343487 | 4,27E-05 | 2 - | 0,00027066 - | 2 - | 0,000318426 - | 3 - | 0,000376744 - | 3 - | 0,000374377 - | 3 - | 0,000377229 - |
| 582 | 1 | 3um_FIJJAFBF_175076 | Bacteria | Proteobacteri | Alphaproteob | Rhodobactera | Rhodobactera | various | various | metabolism | Amino acid tra | 01SDW | Spermidine pu | 439497.RR11_ | 0,2 | 0,4 | 4,36E-05 | 8,71E-05 | 0 - | 0 - | 0 - | 0 - | 0 - | 0 - | 0 - | 0 - | 1 - | 0,000217865 - |
| 583 | 2 | 0.2um_NOKCPHJN_34939;3um | Bacteria | Bacteroidetes | Flavobacteriia | Flavobacterial | Flavobacteriac | Lutibacter | Lutibacter flav | poorly charact | Function unkn | 11P35 | NA | 867900.Celly_ | 0,4 | 0,489897949 | 9,15E-05 | 0,000112056 | 0 - | 0 - | 0 - | 0 - | 1 - | 0,000228586 - | 0 - | 0 - | 1 - | 0,000228881 - |
| 584 | 1 | 3um_FIJJAFBF_217104 | Bacteria | Bacteroidetes | Flavobacteriia | Flavobacterial | Crocinitomica | unclassified | Crocinitomica | unclassified | unclassified | unclassified | unclassified | unclassified | 4 | 0 | 0,001107229 | 0,000108336 | 4 - | 0,001117119 - | 4 - | 0,001314266 - | 4 - | 0,001036646 - | 4 - | 0,001030132 - | 4 - | 0,001037981 - |
| 585 | 1 | 3um_FIJJAFBF_112133 | Bacteria | Bacteroidetes | Flavobacteriia | Flavobacterial | Crocinitomica | unclassified | Crocinitomica | unclassified | unclassified | unclassified | unclassified | unclassified | 0,4 | 0,489897949 | 8,01E-05 | 9,83E-05 | 1 - | 0,000207805 - | 0 - | 0 - | 1 - | 0,000192836 - | 0 - | 0 - | 0 - | 0 - |
| 586 | 1 | 0.2um_NOKCPHJN_385913 | Bacteria | Proteobacteri | Alphaproteob | Rickettsiales | unclassified | unclassified | Rickettsiales b | unclassified | unclassified | unclassified | unclassified | unclassified | 5 | 0,894427191 | 0,000441671 | 5,36E-05 | 4 - | 0,000361252 - | 4 - | 0,000425005 - | 6 - | 0,000502843 - | 6 - | 0,000499683 - | 5 - | 0,000419575 - |
| 587 | 1 | 3um_FIJJAFBF_418831 | Eukaryota | Bacillariophyta | Bacillariophyc | Naviculales | Naviculaceae | Fistulifera | Fistulifera sola | unclassified | unclassified | unclassified | unclassified | unclassified | 0,6 | 0,8 | 8,19E-05 | 0,000109014 | 0 - | 0 - | 0 - | 0 - | 0 - | 0 - | 2 - | 0,000272274 - | 1 - | 0,000137174 - |
| 588 | 1 | 0.2um_NOKCPHJN_331161 | Viruses | unclassified | unclassified | unclassified | Phycodnavirid | unclassified | Chrysochromu | unclassified | unclassified | unclassified | unclassified | unclassified | 0,4 | 0,489897949 | 6,25E-05 | 7,67E-05 | 1 - | 0,000162096 - | 0 - | 0 - | 0 - | 0 - | 0 - | 0 - | 1 - | 0,000150613 - |
| 589 | 1 | 3um_FIJJAFBF_155642 | Bacteria | Bacteroidetes | Flavobacteriia | Flavobacterial | Crocinitomica | unclassified | Crocinitomica | unclassified | unclassified | unclassified | unclassified | unclassified | 1,6 | 0,8 | 0,001105381 | 0,00056647 | 2 - | 0,001370216 - | 2 - | 0,00161203 - | 2 - | 0,001271512 - | 0 - | 0 - | 2 - | 0,001273148 - |
| 590 | 4 | 0.2um_NOKCPHJN_176161;0.2 | Bacteria | Bacteroidetes | Flavobacteriia | Flavobacterial | Flavobacteriac | Polaribacter | various | unclassified | unclassified | unclassified | unclassified | unclassified | 0,2 | 0,4 | 3,06E-05 | 6,11E-05 | 1 - | 0,000152777 - | 0 - | 0 - | 0 - | 0 - | 0 - | 0 - | 0 - | 0 - |
| 591 | 1 | 3um_FIJJAFBF_574196 | unclassified | unclassified | unclassified | unclassified | unclassified | unclassified | unclassified | unclassified | unclassified | unclassified | unclassified | unclassified | 2 | 0 | 0,000823862 | 8,06E-05 | 2 - | 0,000831221 - | 2 - | 0,000977914 - | 2 - | 0,000771344 - | 2 - | 0,000766496 - | 2 - | 0,000772336 - |
| 592 | 1 | 3um_FIJJAFBF_216066 | Bacteria | Proteobacteri | Gammaprotec | Alteromonada | various | various | various | information st | Translation, ri | 0QTKX | Seems to be t | 203122.Sde_0 | 2 | 0 | 0,001390679 | 0,00013607 | 2 - | 0,001403102 - | 2 - | 0,001650718 - | 2 - | 0,001302028 - | 2 - | 0,001293846 - | 2 - | 0,001303704 - |
| 593 | 1 | 0.2um_NOKCPHJN_409097 | Eukaryota | Bacillariophyta | Coscinodiscop | Thalassiosirale | Thalassiosirac | Thalassiosira | Thalassiosira | cellular proces | Posttranslatio | KOG0019 | heat shock pr | 67593.Physo1 | 1 | 0 | 0,000122938 | 1,20E-05 | 1 - | 0,000124037 - | 1 - | 0,000145926 - | 1 - | 0,000115101 - | 1 - | 0,000114378 - | 1 - | 0,00011525 - |
| 594 | 1 | 3um_FIJJAFBF_263676 | Bacteria | Proteobacteri | Alphaproteob | unclassified | unclassified | Candidatus Pu | Candidatus Pu | metabolism | Carbohydrate | 0IVA1 | Photosystem I | 45157.CMV20 | 5,8 | 0,979795897 | 0,002729432 | 0,000459656 | 7 - | 0,003336179 - | 5 - | 0,00280353 - | 7 - | 0,003095854 - | 5 - | 0,002197428 - | 5 - | 0,002214171 - |
| 595 | 1 | 3um_FIJJAFBF_195324 | unclassified | unclassified | unclassified | unclassified | unclassified | unclassified | unclassified | unclassified | unclassified | unclassified | unclassified | unclassified | 7,8 | 1,326649916 | 0,001372372 | 0,000250383 | 6 - | 0,001067268 - | 8 - | 0,001674156 - | 7 - | 0,001155451 - | 8 - | 0,001312217 - | 10 - | 0,001652769 - |
| 596 | 1 | 0.2um_NOKCPHJN_954412 | Eukaryota | Arthropoda | Insecta | Hymenoptera | Formicidae | Harpegnathos | Harpegnathos | unclassified | unclassified | unclassified | unclassified | unclassified | 0,2 | 0,4 | 8,46E-05 | 0,000169131 | 0 - | 0 - | 1 - | 0,000422827 - | 0 - | 0 - | 0 - | 0 - | 0 - | 0 - |
| 597 | 1 | 3um_FIJJAFBF_450894 | Bacteria | Bacteroidetes | Flavobacteriia | Flavobacterial | Flavobacteriac | Hyunsoonleell | Hyunsoonleell | metabolism | Amino acid tra | 09GJ6 | nitrogen regul | 983548.Krodi_ | 1 | 0 | 0,000776049 | 7,59E-05 | 1 - | 0,000782981 - | 1 - | 0,00092116 - | 1 - | 0,000726578 - | 1 - | 0,000722012 - | 1 - | 0,000727513 - |
| 598 | 1 | 3um_FIJJAFBF_282599 | Eukaryota | unclassified | Pelagophycea | Pelagomonad | unclassified | Aureococcus | Aureococcus a | unclassified | unclassified | unclassified | unclassified | unclassified | 1 | 0 | 0,000469824 | 4,60E-05 | 1 - | 0,000474021 - | 1 - | 0,000557675 - | 1 - | 0,000439874 - | 1 - | 0,00043711 - | 1 - | 0,00044044 - |
| 599 | 7 | 0.2um_NOKCPHJN_861091;0.2 | Bacteria | various | various | various | various | various | various | unclassified | unclassified | unclassified | unclassified | unclassified | 0,4 | 0,489897949 | 0,000292931 | 0,000362911 | 0 - | 0 - | 1 - | 0,000818809 - | 1 - | 0,000645847 - | 0 - | 0 - | 0 - | 0 - |
| 600 | 1 | 0.2um_NOKCPHJN_574692 | Bacteria | Proteobacteri | Alphaproteob | Rhodospirillal | Rhodospirillac | Skermanella | Skermanella a | metabolism | Carbohydrate | 16PVY | Major Facilitat | 331869.BAL19 | 0,2 | 0,4 | 4,61E-05 | 9,22E-05 | 0 - | 0 - | 0 - | 0 - | 0 - | 0 - | 1 - | 0,000230386 - | 0 - | 0 - |
| 601 | 1 | 3um_FIJJAFBF_693487 | Bacteria | Bacteroidetes | unclassified | unclassified | unclassified | unclassified | Bacteroidetes | unclassified | unclassified | unclassified | unclassified | unclassified | 0,2 | 0,4 | 4,55E-05 | 9,11E-05 | 0 - | 0 - | 1 - | 0,000227748 - | 0 - | 0 - | 0 - | 0 - | 0 - | 0 - |
| 602 | 1 | 3um_FIJJAFBF_254304 | Bacteria | Proteobacteri | Gammaprotec | Cellvibrionale | Halieaceae | Pseudohaliea | Pseudohaliea | unclassified | unclassified | unclassified | unclassified | unclassified | 2,4 | 1,019803903 | 0,001411429 | 0,000603108 | 4 - | 0,002370104 - | 2 - | 0,001394188 - | 3 - | 0,001649529 - | 2 - | 0,001092775 - | 1 - | 0,000550551 - |
| 603 | 1 | 3um_FIJJAFBF_545173 | Eukaryota | Bacillariophyta | Bacillariophyc | Bacillariales | Bacillariaceae | Fragilariopsis | Fragilariopsis | unclassified | unclassified | unclassified | unclassified | unclassified | 0,8 | 0,4 | 0,000398728 | 0,000205256 | 0 - | 0 - | 1 - | 0,00059293 - | 1 - | 0,000467682 - | 1 - | 0,000464743 - | 1 - | 0,000468284 - |
| 604 | 1 | 3um_FIJJAFBF_498449 | Bacteria | Proteobacteri | Gammaprotec | Arenicellales | Arenicellaceae | Arenicella | Arenicella xan | unclassified | unclassified | unclassified | unclassified | unclassified | 0,2 | 0,4 | 1,96E-05 | 3,93E-05 | 0 - | 0 - | 0 - | 0 - | 0 - | 0 - | 0 - | 0 - | 1 - | 9,82E-05 - |
| 605 | 3 | 0.2um_NOKCPHJN_1061809;3u | Bacteria | Bacteroidetes | Flavobacteriia | Flavobacterial | Flavobacteriac | various | various | unclassified | unclassified | unclassified | unclassified | unclassified | 1 | 0 | 0,000459881 | 4,50E-05 | 1 - | 0,000463989 - | 1 - | 0,000545872 - | 1 - | 0,000430565 - | 1 - | 0,000427859 - | 1 - | 0,000431119 - |
| 606 | 1 | 3um_FIJJAFBF_532450 | Eukaryota | unclassified | Cryptophyta | Pyrenomonad | Geminigerace | Teleaulax | Teleaulax amp | metabolism | Energy produc | COG4451 | RuBisCO catal | 45157.CMV01 | 3 | 0,632455532 | 0,001876067 | 0,000414075 | 3 - | 0,001892673 - | 3 - | 0,002226688 - | 2 - | 0,001170888 - | 3 - | 0,001745295 - | 4 - | 0,002344791 - |
| 607 | 1 | 3um_FIJJAFBF_709591 | unclassified | unclassified | unclassified | unclassified | unclassified | unclassified | unclassified | unclassified | unclassified | unclassified | unclassified | unclassified | 0,8 | 0,4 | 0,000159334 | 8,17E-05 | 1 - | 0,000197509 - | 1 - | 0,000232365 - | 1 - | 0,000183281 - | 0 - | 0 - | 1 - | 0,000183517 - |
| 608 | 1 | 0.2um_NOKCPHJN_400172 | Eukaryota | unclassified | Oomycetes | Peronosporale | Peronosporac | Plasmopara | Plasmopara h | unclassified | unclassified | unclassified | unclassified | unclassified | 0,8 | 0,4 | 3,29E-05 | 1,65E-05 | 1 - | 4,35E-05 - | 0 - | 0 - | 1 - | 4,04E-05 - | 1 - | 4,01E-05 - | 1 - | 4,04E-05 - |
| 609 | 1 | 3um_FIJJAFBF_01111 | Bacteria | Bacteroidetes | Flavobacteriia | Flavobacterial | Flavobacteriac | various | various | unclassified | unclassified | unclassified | unclassified | unclassified | 0,8 | 0,748331477 | 0,001030441 | 0,000961507 | 0 - | 0 - | 0 - | 0 - | 1 - | 0,001291694 - | 2 - | 0,002567154 - | 1 - | 0,001293357 - |
| 610 | 1 | 0.2um_NOKCPHJN_135943 | Eukaryota | Apicomplexa | Aconoidasida | unclassified | Babesiidae | Babesia | Babesia ovata | unclassified | unclassified | unclassified | unclassified | unclassified | 0,2 | 0,4 | 1,66E-05 | 3,32E-05 | 1 - | 8,30E-05 - | 0 - | 0 - | 0 - | 0 - | 0 - | 0 - | 0 - | 0 - |
| 611 | 1 | 3um_FIJJAFBF_20059 | Bacteria | Bacteroidetes | Flavobacteriia | Flavobacterial | Flavobacteriac | Polaribacter | Polaribacter s | unclassified | unclassified | unclassified | unclassified | unclassified | 0,8 | 0,4 | 0,000468775 | 0,000241314 | 0 - | 0 - | 1 - | 0,000697094 - | 1 - | 0,000549843 - | 1 - | 0,000546388 - | 1 - | 0,000550551 - |
| 612 | 1 | 0.2um_NOKCPHJN_674145 | Eukaryota | unclassified | Pelagophycea | Pelagomonad | unclassified | Aureococcus | Aureococcus a | unclassified | unclassified | unclassified | unclassified | unclassified | 0,6 | 0,489897949 | 9,08E-05 | 7,41E-05 | 0 - | 0 - | 0 - | 0 - | 1 - | 0,00015154 - | 1 - | 0,000150587 - | 1 - | 0,000151735 - |
| 613 | 1 | 3um_FIJJAFBF_76059 | Bacteria | Bacteroidetes | Saprospiria | Saprospirales | Saprospiracea | Aureispira | Aureispira sp. | unclassified | unclassified | unclassified | unclassified | unclassified | 1 | 0 | 0,000143428 | 1,40E-05 | 1 - | 0,000144709 - | 1 - | 0,000170247 - | 1 - | 0,000134285 - | 1 - | 0,000133441 - | 1 - | 0,000134458 - |
| 614 | 1 | 3um_FIJJAFBF_329064 | Bacteria | Bacteroidetes | Flavobacteriia | Flavobacterial | Flavobacteriac | Psychroflexus | Psychroflexus | cellular proces | Posttranslatio | 05VJ6 | heat shock pr | 357804.Ping_ | 0,2 | 0,4 | 4,67E-05 | 9,34E-05 | 0 - | 0 - | 0 - | 0 - | 0 - | 0 - | 0 - | 0 - | 1 - | 0,000233471 - |
| 615 | 1 | 3um_FIJJAFBF_383942 | Archaea | Euryarchaeota | Methanobacte | Methanobacte | Methanobacte | Methanobrevi | Methanobrevi | unclassified | unclassified | unclassified | unclassified | unclassified | 0,4 | 0,489897949 | 9,97E-05 | 0,000123652 | 0 - | 0 - | 1 - | 0,000279593 - | 0 - | 0 - | 1 - | 0,000219147 - | 0 - | 0 - |
| 616 | 1 | 0.2um_NOKCPHJN_547411 | Bacteria | Proteobacteri | Alphaproteob | Rhizobiales | unclassified | unclassified | Rhizobiales ba | unclassified | unclassified | unclassified | unclassified | unclassified | 0,4 | 0,489897949 | 2,48E-05 | 3,04E-05 | 0 - | 0 - | 0 - | 0 - | 1 - | 6,22E-05 - | 1 - | 6,18E-05 - | 0 - | 0 - |
| 617 | 1 | 3um_FIJJAFBF_290796 | Bacteria | Proteobacteri | Alphaproteob | Rhodobactera | Rhodobactera | unclassified | Rhodobactera | unclassified | unclassified | unclassified | unclassified | unclassified | 0,4 | 0,8 | 7,16E-05 | 0,000143264 | 0 - | 0 - | 0 - | 0 - | 0 - | 0 - | 0 - | 0 - | 2 - | 0,00035816 - |
| 618 | 1 | 3um_FIJJAFBF_20710 | Eukaryota | various | various | various | various | various | various | unclassified | unclassified | unclassified | unclassified | unclassified | 3,8 | 0,748331477 | 0,004086408 | 0,000653553 | 4 - | 0,004384693 - | 3 - | 0,003868871 - | 4 - | 0,004068837 - | 5 - | 0,005054085 - | 3 - | 0,003055556 - |
| 619 | 1 | 3um_FIJJAFBF_176649 | Bacteria | Bacteroidetes | unclassified | unclassified | unclassified | unclassified | Bacteroidetes | unclassified | unclassified | unclassified | unclassified | unclassified | 0,4 | 0,489897949 | 0,000295275 | 0,000365815 | 0 - | 0 - | 1 - | 0,000825359 - | 1 - | 0,000651014 - | 0 - | 0 - | 0 - | 0 - |
| 620 | 1 | 3um_FIJJAFBF_549019 | Eukaryota | unclassified | Dinophyceae | Peridiniales | Kryptoperidini | Durinskia | Durinskia balt | unclassified | unclassified | unclassified | unclassified | unclassified | 1 | 0 | 0,000377902 | 3,70E-05 | 1 - | 0,000381278 - | 1 - | 0,000448565 - | 1 - | 0,000353812 - | 1 - | 0,000351588 - | 1 - | 0,000354267 - |
| 621 | 1 | 3um_FIJJAFBF_237055 | Eukaryota | Bacillariophyta | Bacillariophyc | Bacillariales | Bacillariaceae | Cylindrotheca | Cylindrotheca | metabolism | Energy produc | 0FJ7X | This is one of t | 103690.alr372 | 3,8 | 0,4 | 0,000917031 | 0,000132599 | 3 - | 0,000730782 - | 4 - | 0,001146332 - | 4 - | 0,000904186 - | 4 - | 0,000898504 - | 4 - | 0,00090535 - |
| 622 | 1 | 3um_FIJJAFBF_337088 | Bacteria | Proteobacteri | Gammaprotec | Oceanospirilla | Oceanospirilla | Motiliproteus | Motiliproteus | information st | Translation, ri | 177XJ | One of the pri | 205918.Psyr_ | 0,2 | 0,4 | 0,000104329 | 0,000208658 | 0 - | 0 - | 0 - | 0 - | 1 - | 0,000521646 - | 0 - | 0 - | 0 - | 0 - |
| 623 | 1 | 3um_FIJJAFBF_341696 | Bacteria | Bacteroidetes | Flavobacteriia | Flavobacterial | Crocinitomica | unclassified | Crocinitomica | poorly charact | Function unkn | 09N7H | NA | 50743.SCB49_ | 1 | 0 | 0,000290694 | 2,84E-05 | 1 - | 0,00029329 - | 1 - | 0,00034505 - | 1 - | 0,000272163 - | 1 - | 0,000270453 - | 1 - | 0,000272513 - |
| 624 | 1 | 0.2um_NOKCPHJN_678783 | Bacteria | unclassified | unclassified | unclassified | unclassified | unclassified | uncultured ba | metabolism | Inorganic ion t | 05C27 | TonB-depende | 313596.RB25 | 1,6 | 0,489897949 | 0,000139881 | 4,81E-05 | 1 - | 8,76E-05 - | 2 - | 0,000206134 - | 1 - | 8,13E-05 - | 2 - | 0,000161569 - | 2 - | 0,0001628 - |
| 625 | 1 | 3um_FIJJAFBF_815478 | Eukaryota | Eustigmatoph | unclassified | Eustigmatales | Monodopsida | Nannochlorop | Nannochlorop | metabolism | Lipid transpor | COG0439 | acetyl-CoA ca | 35128.Thaps6 | 1 | 0 | 0,000283119 | 2,77E-05 | 1 - | 0,000285648 - | 1 - | 0,000336058 - | 1 - | 0,000265071 - | 1 - | 0,000263405 - | 1 - | 0,000265412 - |
| 626 | 1 | 0.2um_NOKCPHJN_257142 | Bacteria | Bacteroidetes | Flavobacteriia | Flavobacterial | Flavobacteriac | Jejuia | Jejuia pallidilu | poorly charact | Function unkn | 11P35 | NA | 398720.MED2 | 0,8 | 0,748331477 | 0,000117191 | 0,000109797 | 0 - | 0 - | 0 - | 0 - | 1 - | 0,000146625 - | 1 - | 0,000145703 - | 2 - | 0,000293627 - |
| 627 | 1 | 3um_FIJJAFBF_516772 | Bacteria | Bacteroidetes | Flavobacteriia | Flavobacterial | unclassified | unclassified | Flavobacterial | poorly charact | Function unkn | 0XSVM | NA | 398720.MED2 | 0,2 | 0,4 | 3,71E-05 | 7,42E-05 | 1 - | 0,000185399 - | 0 - | 0 - | 0 - | 0 - | 0 - | 0 - | 0 - | 0 - |
| 628 | 1 | 3um_FIJJAFBF_229664 | Bacteria | Bacteroidetes | Flavobacteriia | Flavobacterial | Flavobacteriac | Lutibacter | Lutibacter flav | metabolism | Inorganic ion t | 05C27 | TonB-depende | 313598.MED1 | 0,2 | 0,4 | 2,07E-05 | 4,14E-05 | 1 - | 0,000103535 - | 0 - | 0 - | 0 - | 0 - | 0 - | 0 - | 0 - | 0 - |
| 629 | 1 | 0.2um_NOKCPHJN_221686 | unclassified | unclassified | unclassified | unclassified | unclassified | unclassified | unclassified | unclassified | unclassified | unclassified | unclassified | unclassified | 0,8 | 0,4 | 0,000328136 | 0,000164537 | 1 - | 0,000434128 - | 0 - | 0 - | 1 - | 0,000402855 - | 1 - | 0,000400324 - | 1 - | 0,000403374 - |
| 630 | 1 | 3um_FIJJAFBF_388951 | Bacteria | Bacteroidetes | Flavobacteriia | Flavobacterial | Flavobacteriac | Polaribacter | Polaribacter s | metabolism | Energy produc | COG0437 | 4fe-4S ferredo | 313598.MED1 | 0,8 | 0,748331477 | 6,40E-05 | 6,00E-05 | 0 - | 0 - | 0 - | 0 - | 1 - | 8,01E-05 - | 1 - | 7,96E-05 - | 2 - | 0,000160397 - |
| 631 | 1 | 0.2um_NOKCPHJN_268353 | Bacteria | Proteobacteri | Betaproteoba | Burkholderiale | Burkholderiac | Polynucleobac | Polynucleobac | unclassified | unclassified | unclassified | unclassified | unclassified | 0,2 | 0,4 | 2,88E-05 | 5,76E-05 | 1 - | 0,000143996 - | 0 - | 0 - | 0 - | 0 - | 0 - | 0 - | 0 - | 0 - |
| 632 | 1 | 3um_FIJJAFBF_336548 | Bacteria | Spirochaetes | Spirochaetia | unclassified | Leptospiracea | Leptospira | Leptospira sp. | unclassified | unclassified | unclassified | unclassified | unclassified | 3,4 | 0,489897949 | 0,000430683 | 6,41E-05 | 4 - | 0,00051283 - | 3 - | 0,0004525 - | 3 - | 0,000356916 - | 3 - | 0,000354673 - | 4 - | 0,0004765 - |
| 633 | 1 | 0.2um_NOKCPHJN_352204 | Eukaryota | Chordata | unclassified | unclassified | Branchiostom | Branchiostom | Branchiostom | unclassified | unclassified | unclassified | unclassified | unclassified | 0,8 | 0,4 | 3,39E-05 | 1,74E-05 | 1 - | 4,20E-05 - | 1 - | 4,95E-05 - | 1 - | 3,90E-05 - | 1 - | 3,88E-05 - | 0 - | 0 - |
| 634 | 6 | 0.2um_NOKCPHJN_302225;0.2 | Bacteria | Proteobacteri | Alphaproteob | various | various | various | various | unclassified | unclassified | unclassified | unclassified | unclassified | 0,6 | 0,489897949 | 0,000423867 | 0,000346091 | 0 - | 0 - | 0 - | 0 - | 1 - | 0,000707624 - | 1 - | 0,000703177 - | 1 - | 0,000708535 - |
| 635 | 1 | 0.2um_NOKCPHJN_522021 | unclassified | unclassified | unclassified | unclassified | unclassified | unclassified | unclassified | unclassified | unclassified | unclassified | unclassified | unclassified | 0,2 | 0,4 | 7,85E-05 | 0,000156912 | 0 - | 0 - | 1 - | 0,000392281 - | 0 - | 0 - | 0 - | 0 - | 0 - | 0 - |
| 636 | 1 | 3um_FIJJAFBF_142183 | Bacteria | Proteobacteri | Gammaprotec | Cellvibrionale | Halieaceae | Halioglobus | Halioglobus ja | metabolism | Energy produc | COG0056 | Produces ATP | 314285.KT71_ | 1 | 0,894427191 | 0,000158004 | 0,000141254 | 0 - | 0 - | 0 - | 0 - | 1 - | 0,000158321 - | 2 - | 0,000314651 - | 2 - | 0,000317049 - |
| 637 | 1 | 3um_FIJJAFBF_507403 | Eukaryota | Chromerida | unclassified | unclassified | unclassified | Vitrella | Vitrella brassi | poorly charact | Function unkn | 0VVDU | NA | 400682.PAC_ | 0,2 | 0,4 | 2,95E-05 | 5,90E-05 | 0 - | 0 - | 0 - | 0 - | 1 - | 0,000147422 - | 0 - | 0 - | 0 - | 0 - |
| 638 | 1 | 0.2um_NOKCPHJN_181378 | Eukaryota | unclassified | Bangiophycea | Cyanidiales | Cyanidiaceae | Cyanidioschyz | Cyanidioschyz | unclassified | unclassified | unclassified | unclassified | unclassified | 0,4 | 0,489897949 | 4,78E-06 | 5,85E-06 | 0 - | 0 - | 0 - | 0 - | 1 - | 1,20E-05 - | 1 - | 1,19E-05 - | 0 - | 0 - |
| 639 | 1 | 0.2um_NOKCPHJN_44818 | Eukaryota | unclassified | unclassified | Isochrysidales | Noelaerhabda | Emiliania | Emiliania huxl | metabolism | Energy produc | 0FJ7X | This is one of t | 32049.SYNPC | 1 | 0 | 0,000241437 | 2,36E-05 | 1 - | 0,000243594 - | 1 - | 0,000286583 - | 1 - | 0,000226047 - | 1 - | 0,000224626 - | 1 - | 0,000226337 - |
| 640 | 1 | 3um_FIJJAFBF_01139 | Bacteria | Bacteroidetes | Flavobacteriia | Flavobacterial | Flavobacteriac | Polaribacter | Polaribacter s | information st | Translation, ri | 05KK9 | This protein b | 313594.PI23P | 0,4 | 0,489897949 | 0,000202605 | 0,000248418 | 1 - | 0,000525113 - | 0 - | 0 - | 0 - | 0 - | 0 - | 0 - | 1 - | 0,000487913 - |
| 641 | 1 | 0.2um_NOKCPHJN_41914 | Bacteria | Bacteroidetes | unclassified | unclassified | unclassified | unclassified | Bacteroidetes | unclassified | unclassified | unclassified | unclassified | unclassified | 1 | 0 | 0,00010691 | 1,05E-05 | 1 - | 0,000107865 - | 1 - | 0,0001269 - | 1 - | 0,000100094 - | 1 - | 9,95E-05 - | 1 - | 0,000100223 - |
| 642 | 1 | 0.2um_NOKCPHJN_1098614 | Eukaryota | unclassified | Cryptophyta | Pyrenomonad | Geminigerace | Guillardia | Guillardia the | unclassified | unclassified | unclassified | unclassified | unclassified | 0,2 | 0,4 | 3,17E-05 | 6,35E-05 | 0 - | 0 - | 0 - | 0 - | 1 - | 0,000158629 - | 0 - | 0 - | 0 - | 0 - |
| 643 | 1 | 3um_FIJJAFBF_08681 | Bacteria | Bacteroidetes | Flavobacteriia | Flavobacterial | Flavobacteriac | Winogradskye | Winogradskye | poorly charact | Function unkn | 0904U | tail collar dom | 1046627.BZAI | 0,8 | 0,748331477 | 0,000345336 | 0,000319411 | 1 - | 0,000456739 - | 0 - | 0 - | 0 - | 0 - | 1 - | 0,000421174 - | 2 - | 0,000848766 - |
| 644 | 1 | 0.2um_NOKCPHJN_549879 | unclassified | unclassified | unclassified | unclassified | unclassified | unclassified | unclassified | unclassified | unclassified | unclassified | unclassified | unclassified | 0,2 | 0,4 | 0,000113023 | 0,000226047 | 0 - | 0 - | 0 - | 0 - | 1 - | 0,000565116 - | 0 - | 0 - | 0 - | 0 - |
| 645 | 2 | 0.2um_NOKCPHJN_112140;0.2 | unclassified | unclassified | unclassified | unclassified | unclassified | unclassified | unclassified | unclassified | unclassified | unclassified | unclassified | unclassified | 0,4 | 0,489897949 | 0,000131869 | 0,000161506 | 0 - | 0 - | 0 - | 0 - | 1 - | 0,00032946 - | 0 - | 0 - | 1 - | 0,000329885 - |
| 646 | 1 | 3um_FIJJAFBF_10380 | Eukaryota | Phaeophyceae | unclassified | Ectocarpales | Ectocarpaceae | Ectocarpus | Ectocarpus sil | unclassified | unclassified | unclassified | unclassified | unclassified | 1 | 0 | 0,0002324 | 2,27E-05 | 1 - | 0,000234476 - | 1 - | 0,000275855 - | 1 - | 0,000217585 - | 1 - | 0,000216218 - | 1 - | 0,000217865 - |
| 647 | 1 | 3um_FIJJAFBF_498939 | Viruses | unclassified | unclassified | Caudovirales | Myoviridae | unclassified | Pelagibacter p | unclassified | unclassified | unclassified | unclassified | unclassified | 0,2 | 0,4 | 0,000232505 | 0,00046501 | 0 - | 0 - | 0 - | 0 - | 1 - | 0,001162525 - | 0 - | 0 - | 0 - | 0 - |
| 648 | 1 | 3um_FIJJAFBF_625149 | Bacteria | Bacteroidetes | Bacteroidia | Marinilabiliale | unclassified | unclassified | Marinilabiliale | metabolism | Carbohydrate | 09G2S | Glycoside hyd | 388413.ALPR1 | 0,2 | 0,4 | 3,52E-05 | 7,03E-05 | 0 - | 0 - | 0 - | 0 - | 0 - | 0 - | 1 - | 0,000175794 - | 0 - | 0 - |
| 649 | 1 | 3um_FIJJAFBF_108764 | Bacteria | Proteobacteri | Alphaproteob | unclassified | unclassified | unclassified | Alphaproteob | poorly charact | Function unkn | 01UPA | von Willebran | 1082931.KKY_ | 0,4 | 0,489897949 | 0,000106997 | 0,000132639 | 0 - | 0 - | 1 - | 0,000299912 - | 0 - | 0 - | 1 - | 0,000235074 - | 0 - | 0 - |
| 650 | 1 | 3um_FIJJAFBF_355789 | Bacteria | Bacteroidetes | Flavobacteriia | Flavobacterial | Flavobacteriac | Polaribacter | Polaribacter s | poorly charact | Function unkn | 0P8NI | Inherit from b | 313594.PI23P | 1,4 | 0,489897949 | 0,000182426 | 5,52E-05 | 1 - | 0,000133884 - | 1 - | 0,000157511 - | 2 - | 0,000248479 - | 1 - | 0,000123459 - | 2 - | 0,000248798 - |
| 651 | 1 | 0.2um_NOKCPHJN_726929 | Bacteria | Verrucomicrol | Verrucomicrol | Verrucomicrol | unclassified | unclassified | Verrucomicro | unclassified | unclassified | unclassified | unclassified | unclassified | 0,6 | 0,489897949 | 0,000181789 | 0,00015097 | 0 - | 0 - | 1 - | 0,000353322 - | 1 - | 0,000278687 - | 1 - | 0,000276936 - | 0 - | 0 - |
| 652 | 1 | 0.2um_NOKCPHJN_356144 | Bacteria | Bacteroidetes | Flavobacteriia | Flavobacterial | Flavobacteriac | Gilvibacter | Gilvibacter sp. | poorly charact | Function unkn | 09C0E | Ragb susd dor | 411154.GFO_ | 1 | 0,632455532 | 0,000168159 | 0,000101022 | 1 - | 0,000169621 - | 1 - | 0,000199555 - | 0 - | 0 - | 1 - | 0,000156413 - | 2 - | 0,000315209 - |
| 653 | 1 | 0.2um_NOKCPHJN_636600 | Bacteria | Fusobacteria | Fusobacteriia | Fusobacteriale | Fusobacteriac | Fusobacterium | Fusobacterium | unclassified | unclassified | unclassified | unclassified | unclassified | 0,6 | 0,489897949 | 8,23E-05 | 6,73E-05 | 1 - | 0,000143996 - | 0 - | 0 - | 1 - | 0,000133624 - | 0 - | 0 - | 1 - | 0,000133796 - |
| 654 | 1 | 3um_FIJJAFBF_520476 | Bacteria | Bacteroidetes | Flavobacteriia | Flavobacterial | Flavobacteriac | Polaribacter | Polaribacter s | poorly charact | Function unkn | 11NCV | Protein of unk | 313594.PI23P | 1 | 0,632455532 | 0,000540789 | 0,000358406 | 2 - | 0,001131534 - | 0 - | 0 - | 1 - | 0,000525011 - | 1 - | 0,000521712 - | 1 - | 0,000525687 - |
| 655 | 1 | 0.2um_NOKCPHJN_483485 | Bacteria | Bacteroidetes | Flavobacteriia | Flavobacterial | Flavobacteriac | Arenitalea | Arenitalea lut | metabolism | Inorganic ion t | 05C27 | TonB-depende | 886377.Murru | 1,6 | 0,8 | 0,000139105 | 6,98E-05 | 2 - | 0,000184037 - | 0 - | 0 - | 2 - | 0,00017078 - | 2 - | 0,000169707 - | 2 - | 0,000171 - |
| 656 | 1 | 3um_FIJJAFBF_257123 | Eukaryota | Phaeophyceae | unclassified | Dictyotales | Dictyotaceae | Dictyopteris | Dictyopteris d | information st | Translation, ri | 0FKC1 | This is 1 of the | 551115.Aazo_ | 2,8 | 0,748331477 | 0,001334108 | 0,000304521 | 3 - | 0,001461564 - | 2 - | 0,001146332 - | 2 - | 0,000904186 - | 3 - | 0,001347756 - | 4 - | 0,0018107 - |
| 657 | 1 | 3um_FIJJAFBF_516119 | Eukaryota | Bacillariophyta | Coscinodiscop | Thalassiosirale | Thalassiosirac | Thalassiosira | Thalassiosira | cellular proces | Posttranslatio | KOG0730 | cell division cy | 35128.Thaps4 | 1,8 | 0,4 | 0,000481764 | 8,02E-05 | 2 - | 0,000551534 - | 1 - | 0,000324434 - | 2 - | 0,000511803 - | 2 - | 0,000508587 - | 2 - | 0,000512462 - |
| 658 | 1 | 0.2um_NOKCPHJN_23410 | Eukaryota | unclassified | Pelagophycea | Pelagomonad | unclassified | various | various | unclassified | unclassified | unclassified | unclassified | unclassified | 1,2 | 0,4 | 0,000964615 | 0,000289181 | 1 - | 0,000819569 - | 1 - | 0,000964205 - | 1 - | 0,00076053 - | 1 - | 0,000755751 - | 2 - | 0,001523019 - |
| 659 | 3 | 0.2um_NOKCPHJN_530304;3ur | Bacteria | Proteobacteri | Gammaprotec | Cellvibrionale | various | various | various | information st | Translation, ri | 0QTNF | One of the pri | various | 0,2 | 0,4 | 0,000197065 | 0,00039413 | 1 - | 0,000985324 - | 0 - | 0 - | 0 - | 0 - | 0 - | 0 - | 0 - | 0 - |
| 660 | 1 | 3um_FIJJAFBF_112130 | Bacteria | Bacteroidetes | Flavobacteriia | Flavobacterial | Crocinitomica | unclassified | Crocinitomica | unclassified | unclassified | unclassified | unclassified | unclassified | 0,6 | 0,489897949 | 8,32E-05 | 6,79E-05 | 0 - | 0 - | 0 - | 0 - | 1 - | 0,000138868 - | 1 - | 0,000137995 - | 1 - | 0,000139047 - |
| 661 | 1 | 0.2um_NOKCPHJN_45917 | Bacteria | Bacteroidetes | Flavobacteriia | Flavobacterial | Flavobacteriac | Algibacter | Algibacter sp. | metabolism | Energy produc | COG0437 | 4fe-4S ferredo | 983544.Lacal_ | 2 | 0,632455532 | 0,00016454 | 4,37E-05 | 2 - | 0,000170279 - | 1 - | 0,000100165 - | 3 - | 0,00023702 - | 2 - | 0,00015702 - | 2 - | 0,000158216 - |
| 662 | 1 | 3um_FIJJAFBF_560542 | Eukaryota | Chlorophyta | Chlorophycea | Chlamydomor | Volvocaceae | Volvox | Volvox carteri | unclassified | unclassified | unclassified | unclassified | unclassified | 0,2 | 0,4 | 7,69E-05 | 0,000153739 | 0 - | 0 - | 0 - | 0 - | 0 - | 0 - | 0 - | 0 - | 1 - | 0,000384347 - |
| 663 | 1 | 3um_FIJJAFBF_306761 | Bacteria | Proteobacteri | Alphaproteob | Rhodobactera | Rhodobactera | Phaeobacter | Phaeobacter s | metabolism | Lipid transpor | COG2067 | lipid transport | 439496.RBY4I | 1 | 0 | 0,00023813 | 2,33E-05 | 1 - | 0,000240257 - | 1 - | 0,000282657 - | 1 - | 0,00022295 - | 1 - | 0,000221549 - | 1 - | 0,000223237 - |
| 664 | 1 | 3um_FIJJAFBF_75485 | unclassified | unclassified | unclassified | unclassified | unclassified | unclassified | unclassified | unclassified | unclassified | unclassified | unclassified | unclassified | 0,2 | 0,4 | 5,99E-05 | 0,000119801 | 0 - | 0 - | 0 - | 0 - | 0 - | 0 - | 1 - | 0,000299501 - | 0 - | 0 - |
| 665 | 1 | 3um_FIJJAFBF_76654 | Bacteria | Proteobacteri | Alphaproteob | Rhodobactera | Rhodobactera | Tateyamaria | Tateyamaria s | unclassified | unclassified | unclassified | unclassified | unclassified | 1 | 0,894427191 | 0,000128057 | 0,000114697 | 0 - | 0 - | 0 - | 0 - | 2 - | 0,000256305 - | 1 - | 0,000127347 - | 2 - | 0,000256635 - |
| 666 | 1 | 3um_FIJJAFBF_305834 | Eukaryota | Chlorophyta | unclassified | Pyramimonad | unclassified | Cymbomonas | Cymbomonas | metabolism | Energy produc | COG1290 | Component of | 70448.Q0P3P | 0,6 | 0,489897949 | 0,000233072 | 0,0001906 | 1 - | 0,000407878 - | 0 - | 0 - | 1 - | 0,000378496 - | 0 - | 0 - | 1 - | 0,000378984 - |
| 667 | 1 | 3um_FIJJAFBF_537683 | Eukaryota | Bacillariophyta | Bacillariophyc | Naviculales | Naviculaceae | Fistulifera | Fistulifera sola | metabolism | Energy produc | 1DYB8 | PsaA and PsaB | 3055.DAA014 | 7,2 | 1,326649916 | 0,000822347 | 0,000126439 | 9 - | 0,001049527 - | 5 - | 0,00068597 - | 7 - | 0,000757496 - | 8 - | 0,00086027 - | 7 - | 0,000758471 - |
| 668 | 1 | 3um_FIJJAFBF_186059 | Viruses | unclassified | unclassified | unclassified | Phycodnavirid | unclassified | Aureococcus a | unclassified | unclassified | unclassified | unclassified | unclassified | 0,4 | 0,489897949 | 4,34E-05 | 5,32E-05 | 0 - | 0 - | 0 - | 0 - | 1 - | 0,000108502 - | 0 - | 0 - | 1 - | 0,000108642 - |
| 669 | 2 | 0.2um_NOKCPHJN_228484;3ur | Bacteria | Proteobacteri | Alphaproteob | Rhodobactera | Rhodobactera | various | various | metabolism | Carbohydrate | 05WYW | fructose-bisph | various | 0,2 | 0,4 | 5,48E-05 | 0,000109598 | 0 - | 0 - | 0 - | 0 - | 1 - | 0,000273996 - | 0 - | 0 - | 0 - | 0 - |
| 670 | 1 | 3um_FIJJAFBF_564667 | Bacteria | Proteobacteri | Betaproteoba | unclassified | unclassified | unclassified | Betaproteoba | metabolism | Inorganic ion t | 07QMV | Part of the AB | 452638.Pnec_ | 1,2 | 0,4 | 0,000963463 | 0,000283908 | 1 - | 0,000819569 - | 1 - | 0,000964205 - | 1 - | 0,00076053 - | 2 - | 0,001511502 - | 1 - | 0,000761509 - |
| 671 | 1 | 3um_FIJJAFBF_238099 | Bacteria | Bacteroidetes | Flavobacteriia | Flavobacterial | Flavobacteriac | Polaribacter | Polaribacter in | metabolism | Inorganic ion t | 05C27 | TonB-depende | 313594.PI23P | 0,4 | 0,8 | 3,32E-05 | 6,64E-05 | 2 - | 0,000166087 - | 0 - | 0 - | 0 - | 0 - | 0 - | 0 - | 0 - | 0 - |
| 672 | 1 | 0.2um_NOKCPHJN_400612 | Eukaryota | Ascomycota | Leotiomycetes | Helotiales | Sclerotiniacea | Botrytis | Botrytis ciner | metabolism | Energy produc | KOG0017 | Retrotranspos | 10090.ENSMU | 0,6 | 0,489897949 | 8,46E-05 | 6,91E-05 | 0 - | 0 - | 0 - | 0 - | 1 - | 0,000141279 - | 1 - | 0,000140391 - | 1 - | 0,000141461 - |
| 673 | 1 | 0.2um_NOKCPHJN_556144 | Eukaryota | unclassified | Oomycetes | Peronosporale | Peronosporac | Phytophthora | various | information st | Translation, ri | COG0052 | 30S ribosomal | 486041.XP_00 | 1 | 0 | 0,000294635 | 2,88E-05 | 1 - | 0,000297267 - | 1 - | 0,000349728 - | 1 - | 0,000275853 - | 1 - | 0,00027412 - | 1 - | 0,000276208 - |
| 674 | 1 | 0.2um_NOKCPHJN_165212 | Bacteria | Bacteroidetes | Flavobacteriia | Flavobacterial | Flavobacteriac | Formosa | Formosa sp. H | unclassified | unclassified | unclassified | unclassified | unclassified | 5,4 | 1,624807681 | 0,000673041 | 0,000159657 | 4 - | 0,000516606 - | 3 - | 0,000455832 - | 6 - | 0,000719088 - | 7 - | 0,000833663 - | 7 - | 0,000840015 - |
| 675 | 2 | 3um_FIJJAFBF_232873;3um_FI | Eukaryota | various | various | various | various | various | various | poorly charact | Function unkn | COG1100 | GTP-binding P | various | 2 | 0 | 0,000819976 | 8,02E-05 | 2 - | 0,0008273 - | 2 - | 0,000973301 - | 2 - | 0,000767705 - | 2 - | 0,000762881 - | 2 - | 0,000768693 - |
| 676 | 1 | 3um_FIJJAFBF_521113 | Bacteria | Proteobacteri | Gammaprotec | unclassified | unclassified | unclassified | various | unclassified | unclassified | unclassified | unclassified | unclassified | 0,8 | 0,4 | 0,000271051 | 0,000138905 | 1 - | 0,000335992 - | 1 - | 0,000395287 - | 1 - | 0,000311788 - | 0 - | 0 - | 1 - | 0,00031219 - |
| 677 | 1 | 3um_FIJJAFBF_288420 | unclassified | unclassified | unclassified | unclassified | unclassified | unclassified | unclassified | unclassified | unclassified | unclassified | unclassified | unclassified | 1 | 0 | 0,000955137 | 9,35E-05 | 1 - | 0,000963669 - | 1 - | 0,001133735 - | 1 - | 0,00089425 - | 1 - | 0,00088863 - | 1 - | 0,000895401 - |
| 678 | 1 | 3um_FIJJAFBF_172360 | Bacteria | Proteobacteri | Gammaprotec | Oceanospirilla | Saccharospiril | Reinekea | Reinekea fors | information st | Translation, ri | 08VZM | 50s ribosomal | 314283.MED2 | 0,2 | 0,4 | 0,000106944 | 0,000213887 | 1 - | 0,000534719 - | 0 - | 0 - | 0 - | 0 - | 0 - | 0 - | 0 - | 0 - |
| 679 | 1 | 3um_FIJJAFBF_212746 | Bacteria | Bacteroidetes | Flavobacteriia | Flavobacterial | Flavobacteriac | Polaribacter | Polaribacter s | poorly charact | Function unkn | 05ERK | gliding motilit | 313594.PI23P | 0,2 | 0,4 | 3,14E-05 | 6,27E-05 | 0 - | 0 - | 0 - | 0 - | 1 - | 0,000156795 - | 0 - | 0 - | 0 - | 0 - |
| 680 | 1 | 3um_FIJJAFBF_336984 | Bacteria | Proteobacteri | Gammaprotec | Cellvibrionale | Cellvibrionace | Cellvibrio | Cellvibrio sp. | poorly charact | Function unkn | 129NV | NA | 62977.ACIAD3 | 0,8 | 0,4 | 0,000243434 | 0,000125314 | 0 - | 0 - | 1 - | 0,000362 - | 1 - | 0,000285532 - | 1 - | 0,000283738 - | 1 - | 0,0002859 - |

| | | | | | | | | | | | | | | | | | | | | | | | | | | | | |
|---|---|---|---|---|---|---|---|---|---|---|---|---|---|---|---|---|---|---|---|---|---|---|---|---|---|---|---|---|
| 681 | 1 | 0.2um_NOKCPHJN_604581 | Bacteria | Proteobacteri | Alphaproteob | Rickettsiales | unclassified | unclassified | Rickettsiales b | unclassified | unclassified | unclassified | unclassified | unclassified | 1 | 0 | 0,000176662 | 1,73E-05 | 1 - | 0,00017824 - | 1 - | 0,000209695 - | 1 - | 0,0001654 - | 1 - | 0,00016436 - | 1 - | 0,000165613 - |
| 682 | 1 | 0.2um_NOKCPHJN_90233 | unclassified | unclassified | unclassified | unclassified | unclassified | unclassified | unclassified | unclassified | unclassified | unclassified | unclassified | unclassified | 0,2 | 0,4 | 2,36E-05 | 4,72E-05 | 0 - | 0 - | 0 - | 0 - | 0 - | 0 - | 0 - | 0 - | 1 - | 0,000117918 - |
| 683 | 1 | 0.2um_NOKCPHJN_635778 | Eukaryota | Ascomycota | Saccharomyce | Saccharomyce | Phaffomyceta | Komagataella | Komagataella | poorly charact | Function unkn | 19A1Y | Cell wall prote | 436907.XP_00 | 0,8 | 0,4 | 5,94E-05 | 2,98E-05 | 1 - | 7,86E-05 - | 0 - | 0 - | 1 - | 7,29E-05 - | 1 - | 7,25E-05 - | 1 - | 7,30E-05 - |
| 684 | 1 | 3um_FIJJAFBF_663734 | Eukaryota | Bacillariophyta | Bacillariophyc | Naviculales | Naviculaceae | Fistulifera | Fistulifera sola | poorly charact | Function unkn | 110WY | Chlorophyll A- | 2850.Phatr18 | 1 | 0 | 0,000827785 | 8,10E-05 | 1 - | 0,00083518 - | 1 - | 0,00098257 - | 1 - | 0,000775017 - | 1 - | 0,000770146 - | 1 - | 0,000776014 - |
| 685 | 1 | 0.2um_NOKCPHJN_479684 | Bacteria | Bacteroidetes | Saprospiria | Saprospirales | Haliscomenob | Haliscomenob | Haliscomenob | unclassified | unclassified | unclassified | unclassified | unclassified | 0,6 | 0,489897949 | 6,41E-05 | 5,24E-05 | 0 - | 0 - | 0 - | 0 - | 1 - | 0,000107075 - | 1 - | 0,000106402 - | 1 - | 0,000107212 - |
| 686 | 1 | 3um_FIJJAFBF_91113 | Eukaryota | unclassified | Florideophyce | Ceramiales | Rhodomelace | Polysiphonia | Polysiphonia s | unclassified | unclassified | unclassified | unclassified | unclassified | 3,8 | 0,748331477 | 0,002468132 | 0,000353136 | 3 - | 0,001993042 - | 3 - | 0,00234477 - | 4 - | 0,002465962 - | 4 - | 0,002450465 - | 5 - | 0,00308642 - |
| 687 | 1 | 3um_FIJJAFBF_154390 | Bacteria | unclassified | unclassified | unclassified | unclassified | unclassified | bacterium B1 | unclassified | unclassified | unclassified | unclassified | unclassified | 1,6 | 1,2 | 0,000243377 | 0,000179379 | 1 - | 0,000162697 - | 0 - | 0 - | 3 - | 0,000452932 - | 3 - | 0,000450085 - | 1 - | 0,000151172 - |
| 688 | 2 | 0.2um_NOKCPHJN_56987;3um | Bacteria | Bacteroidetes | Flavobacteriia | Flavobacterial | Flavobacteria | Mangrovimon | Mangrovimon | poorly charact | Function unkn | 1015T | domain protei | various | 2 | 0,632455532 | 0,000397881 | 0,0001058 | 2 - | 0,000411708 - | 1 - | 0,000242183 - | 2 - | 0,00038205 - | 2 - | 0,00037965 - | 3 - | 0,000573813 - |
| 689 | 1 | 0.2um_NOKCPHJN_728308 | Archaea | Euryarchaeota | unclassified | unclassified | unclassified | unclassified | Euryarchaeota | unclassified | unclassified | unclassified | unclassified | unclassified | 0,4 | 0,489897949 | 0,000855978 | 0,001061109 | 0 - | 0 - | 1 - | 0,0023993 - | 0 - | 0 - | 1 - | 0,00188059 - | 0 - | 0 - |
| 690 | 1 | 3um_FIJJAFBF_319817 | unclassified | unclassified | unclassified | unclassified | unclassified | unclassified | unclassified | unclassified | unclassified | unclassified | unclassified | unclassified | 0,4 | 0,489897949 | 0,000162076 | 0,000198773 | 1 - | 0,000421605 - | 0 - | 0 - | 0 - | 0 - | 1 - | 0,000388776 - | 0 - | 0 - |
| 691 | 1 | 0.2um_NOKCPHJN_374320 | Bacteria | Proteobacteri | Alphaproteob | Rhodospirillal | unclassified | unclassified | Rhodospirillal | metabolism | Amino acid tra | COG0683 | (ABC) transpo | 504832.OCA5 | 0,8 | 0,4 | 0,000161975 | 8,31E-05 | 1 - | 0,000201133 - | 1 - | 0,000236628 - | 1 - | 0,000186644 - | 1 - | 0,000185471 - | 0 - | 0 - |
| 692 | 1 | 0.2um_NOKCPHJN_977312 | Eukaryota | Phaeophyceae | unclassified | Ectocarpales | Ectocarpaceae | Ectocarpus | Ectocarpus sil | information st | Translation, ri | 0TJFM | 40S ribosomal | 4641.GSMUA_ | 1,2 | 0,4 | 0,000806358 | 0,000241737 | 1 - | 0,000685108 - | 1 - | 0,000806015 - | 1 - | 0,000635756 - | 1 - | 0,000631761 - | 2 - | 0,001273148 - |
| 693 | 1 | 3um_FIJJAFBF_110764 | Bacteria | Proteobacteri | Gammaprotec | Cellvibrionales | Halieaceae | Halioglobus | Halioglobus sp | unclassified | unclassified | unclassified | unclassified | unclassified | 0,2 | 0,4 | 7,48E-05 | 0,000149507 | 0 - | 0 - | 0 - | 0 - | 0 - | 0 - | 0 - | 0 - | 1 - | 0,000373768 - |
| 694 | 1 | 0.2um_NOKCPHJN_431475 | Bacteria | Proteobacteri | Alphaproteob | Rhodospirillal | Rhodospirillac | unclassified | Rhodospirillac | unclassified | unclassified | unclassified | unclassified | unclassified | 0,8 | 0,4 | 3,10E-05 | 1,59E-05 | 1 - | 3,84E-05 - | 1 - | 4,52E-05 - | 1 - | 3,56E-05 - | 0 - | 0 - | 1 - | 3,57E-05 - |
| 695 | 1 | 3um_FIJJAFBF_617422 | Eukaryota | unclassified | Dictyochophy | Florenciellales | unclassified | Pseudochatto | Pseudochatto | cellular proces | Cytoskeleton | 1514Z | Tubulin is the | 4787.PITG_00 | 1,6 | 0,489897949 | 0,001085297 | 0,000265705 | 1 - | 0,000701551 - | 1 - | 0,000825359 - | 2 - | 0,001302028 - | 2 - | 0,001293846 - | 2 - | 0,001303704 - |
| 696 | 1 | 3um_FIJJAFBF_481949 | Bacteria | Proteobacteri | Alphaproteob | Rhodobactera | Rhodobactera | Pseudooceani | Pseudooceani | metabolism | Energy produc | COG0056 | Produces ATP | 398580.Dshi_ | 1,6 | 0,489897949 | 0,000267392 | 7,29E-05 | 2 - | 0,000342554 - | 1 - | 0,000201504 - | 2 - | 0,000317878 - | 2 - | 0,00031588 - | 1 - | 0,000159144 - |
| 697 | 1 | 0.2um_NOKCPHJN_1094406 | Bacteria | Proteobacteri | Alphaproteob | Rhodobactera | Rhodobactera | Shimia | Shimia abyssi | unclassified | unclassified | unclassified | unclassified | unclassified | 1 | 0 | 0,000493849 | 4,83E-05 | 1 - | 0,000498261 - | 1 - | 0,000586193 - | 1 - | 0,000462368 - | 1 - | 0,000459462 - | 1 - | 0,000462963 - |
| 698 | 1 | 0.2um_NOKCPHJN_49737 | Bacteria | Proteobacteri | Alphaproteob | Rhodospirillal | Rhodospirillac | unclassified | Rhodospirillac | information st | RNA processin | 0PGMY | DEAD DEAH b | 5141.EFNCRP | 0,2 | 0,4 | 2,67E-05 | 5,33E-05 | 0 - | 0 - | 0 - | 0 - | 0 - | 0 - | 0 - | 0 - | 1 - | 0,000133358 - |
| 699 | 1 | 3um_FIJJAFBF_788016 | Bacteria | Proteobacteri | Alphaproteob | Rhodobactera | unclassified | unclassified | Rhodobactera | unclassified | unclassified | unclassified | unclassified | unclassified | 0,4 | 0,489897949 | 0,000240799 | 0,00029532 | 1 - | 0,000626385 - | 0 - | 0 - | 0 - | 0 - | 1 - | 0,00057761 - | 0 - | 0 - |
| 700 | 1 | 3um_FIJJAFBF_602360 | Bacteria | Proteobacteri | Alphaproteob | Rhodospirillal | Rhodospirillac | Candidatus En | Candidatus En | unclassified | unclassified | unclassified | unclassified | unclassified | 0,8 | 0,4 | 0,000803914 | 0,000411978 | 1 - | 0,000996521 - | 1 - | 0,001172385 - | 1 - | 0,000924736 - | 0 - | 0 - | 1 - | 0,000925926 - |
| 701 | 1 | 0.2um_NOKCPHJN_222208 | Eukaryota | Phaeophyceae | unclassified | Ectocarpales | Ectocarpaceae | Ectocarpus | Ectocarpus sil | unclassified | unclassified | unclassified | unclassified | unclassified | 0,8 | 0,4 | 1,56E-05 | 8,04E-06 | 0 - | 0 - | 1 - | 2,32E-05 - | 1 - | 1,83E-05 - | 1 - | 1,82E-05 - | 1 - | 1,83E-05 - |
| 702 | 1 | 0.2um_NOKCPHJN_193556 | Bacteria | Verrucomicrol | Opitutae | unclassified | unclassified | unclassified | Opitutae bact | unclassified | unclassified | unclassified | unclassified | unclassified | 5,2 | 0,979795897 | 0,000927858 | 0,000126801 | 5 - | 0,000911578 - | 4 - | 0,000857962 - | 5 - | 0,000845912 - | 7 - | 0,001176835 - | 5 - | 0,000847001 - |
| 703 | 1 | 3um_FIJJAFBF_524441 | Bacteria | Bacteroidetes | Flavobacteriia | Flavobacterial | Flavobacteria | Formosa | Formosa sp. H | unclassified | unclassified | unclassified | unclassified | unclassified | 0,8 | 0,4 | 0,00040355 | 0,000206962 | 1 - | 0,000501108 - | 1 - | 0,000589542 - | 1 - | 0,00046501 - | 1 - | 0,000462088 - | 0 - | 0 - |
| 704 | 1 | 3um_FIJJAFBF_67793 | Eukaryota | unclassified | unclassified | Phaeocystales | Phaeocystace | Phaeocystis | Phaeocystis an | metabolism | Energy produc | 11HAV | photosystem i | 35128.Thapsd | 0,2 | 0,4 | 0,000124388 | 0,000248777 | 1 - | 0,000621942 - | 0 - | 0 - | 0 - | 0 - | 0 - | 0 - | 0 - | 0 - |
| 705 | 1 | 0.2um_NOKCPHJN_33511 | Bacteria | unclassified | unclassified | unclassified | unclassified | unclassified | bacterium B1 | unclassified | unclassified | unclassified | unclassified | unclassified | 0,6 | 0,489897949 | 8,93E-05 | 7,29E-05 | 0 - | 0 - | 0 - | 0 - | 1 - | 0,000149042 - | 1 - | 0,000148105 - | 1 - | 0,000149234 - |
| 706 | 1 | 3um_FIJJAFBF_761547 | Viruses | unclassified | unclassified | Caudovirales | Myoviridae | unclassified | Shewanella sp | unclassified | unclassified | unclassified | unclassified | unclassified | 1,2 | 0,979795897 | 0,000625079 | 0,000580855 | 1 - | 0,00048991 - | 3 - | 0,001729104 - | 1 - | 0,000454619 - | 1 - | 0,000451762 - | 0 - | 0 - |
| 707 | 1 | 0.2um_NOKCPHJN_881016 | Bacteria | Proteobacteri | Gammaprotec | Cellvibrionales | Porticoccacea | unclassified | gamma protec | metabolism | Inorganic ion t | 0QHT5 | TonB-depende | 314287.GB22 | 0,2 | 0,4 | 2,89E-05 | 5,79E-05 | 1 - | 0,000144709 - | 0 - | 0 - | 0 - | 0 - | 0 - | 0 - | 0 - | 0 - |
| 708 | 1 | 3um_FIJJAFBF_191785 | Eukaryota | unclassified | Raphidophyce | Chattonellales | Chattonellace | Heterosigma | Heterosigma a | unclassified | unclassified | unclassified | unclassified | unclassified | 1,2 | 0,748331477 | 0,000500593 | 0,000319065 | 2 - | 0,000876939 - | 0 - | 0 - | 1 - | 0,000406884 - | 1 - | 0,000404327 - | 2 - | 0,000814815 - |
| 709 | 2 | 0.2um_NOKCPHJN_443521;3ur | various | various | various | various | various | various | various | various | various | various | various | various | 0,4 | 0,489897949 | 5,50E-05 | 6,74E-05 | 0 - | 0 - | 0 - | 0 - | 1 - | 0,000137461 - | 0 - | 0 - | 1 - | 0,000137638 - |
| 710 | 1 | 3um_FIJJAFBF_128630 | Eukaryota | unclassified | Raphidophyce | Chattonellales | Chattonellace | Heterosigma | Heterosigma a | unclassified | unclassified | unclassified | unclassified | unclassified | 1,4 | 1,019803903 | 0,000516464 | 0,000420037 | 1 - | 0,000353604 - | 3 - | 0,001248023 - | 0 - | 0 - | 2 - | 0,00065214 - | 1 - | 0,000328554 - |
| 711 | 1 | 3um_FIJJAFBF_684627 | Bacteria | Bacteroidetes | Flavobacteriia | Flavobacterial | Flavobacteria | Polaribacter | Polaribacter s | poorly charact | Function unkn | 0P8NI | Inherit from b | 487797.Flav3 | 0,4 | 0,489897949 | 9,98E-05 | 0,000122281 | 0 - | 0 - | 0 - | 0 - | 1 - | 0,00025039 - | 1 - | 0,000248816 - | 0 - | 0 - |
| 712 | 1 | 3um_FIJJAFBF_52276 | Bacteria | Proteobacteri | Alphaproteob | Rhodobactera | unclassified | unclassified | Rhodobactera | metabolism | Secondary me | 01Q86 | extracellular s | 644107.SL115 | 0,8 | 1,166190379 | 0,000192945 | 0,000266783 | 0 - | 0 - | 1 - | 0,000286583 - | 3 - | 0,00067814 - | 0 - | 0 - | 0 - | 0 - |
| 713 | 1 | 3um_FIJJAFBF_189700 | Bacteria | Bacteroidetes | Flavobacteriia | Flavobacterial | Flavobacteria | Polaribacter | Polaribacter s | unclassified | unclassified | unclassified | unclassified | unclassified | 1 | 0,632455532 | 0,000137061 | 8,23E-05 | 1 - | 0,000138318 - | 1 - | 0,000162729 - | 2 - | 0,000256709 - | 1 - | 0,000127548 - | 0 - | 0 - |
| 714 | 1 | 3um_FIJJAFBF_279887 | unclassified | unclassified | unclassified | unclassified | unclassified | unclassified | unclassified | poorly charact | Function unkn | 006NJ | Inherit from b | 204669.Acid3 | 1,2 | 0,748331477 | 0,000181196 | 0,00012004 | 1 - | 0,000147882 - | 2 - | 0,000347959 - | 0 - | 0 - | 2 - | 0,000272733 - | 1 - | 0,000137406 - |
| 715 | 1 | 3um_FIJJAFBF_426302 | Bacteria | Proteobacteri | Gammaprotec | Cellvibrionales | Halieaceae | Luminiphilus | Luminiphilus s | information st | Translation, ri | COG0050 | This protein p | 565045.NOR5 | 0,2 | 0,4 | 4,00E-05 | 8,00E-05 | 0 - | 0 - | 0 - | 0 - | 1 - | 0,000199943 - | 0 - | 0 - | 0 - | 0 - |
| 716 | 1 | 3um_FIJJAFBF_715096 | Eukaryota | unclassified | Pelagophycea | Pelagomonad | unclassified | Pelagomonas | uncultured Pe | metabolism | Energy produc | 0IE88 | oxidoreductas | 35128.Thapsd | 2,6 | 0,489897949 | 0,000304963 | 4,86E-05 | 3 - | 0,000358422 - | 2 - | 0,000281117 - | 2 - | 0,000221735 - | 3 - | 0,000330512 - | 3 - | 0,000333031 - |
| 717 | 1 | 0.2um_NOKCPHJN_681701 | Eukaryota | Arthropoda | Insecta | Hymenoptera | Agaonidae | Ceratosolen | Ceratosolen s | unclassified | unclassified | unclassified | unclassified | unclassified | 0,2 | 0,4 | 2,84E-05 | 5,67E-05 | 0 - | 0 - | 0 - | 0 - | 0 - | 0 - | 1 - | 0,000141869 - | 0 - | 0 - |
| 718 | 1 | 3um_FIJJAFBF_12816 | Bacteria | Bacteroidetes | Flavobacteriia | Flavobacterial | Flavobacteria | Flavivirga | Flavivirga eckl | metabolism | Energy produc | 099Q7 | Catalyzes the | 391603.FBALC | 1,2 | 0,748331477 | 0,000334642 | 0,000194063 | 1 - | 0,00028472 - | 1 - | 0,000334967 - | 2 - | 0,00052842 - | 2 - | 0,0005251 - | 0 - | 0 - |
| 719 | 1 | 3um_FIJJAFBF_708117 | Eukaryota | unclassified | Cryptophyta | Pyrenomonad | Geminigerace | Guillardia | Guillardia thet | unclassified | unclassified | unclassified | unclassified | unclassified | 1 | 0 | 0,000526773 | 5,15E-05 | 1 - | 0,000531478 - | 1 - | 0,000625272 - | 1 - | 0,000493192 - | 1 - | 0,000490093 - | 1 - | 0,000493827 - |
| 720 | 1 | 3um_FIJJAFBF_104815 | Viruses | unclassified | unclassified | unclassified | unclassified | unclassified | uncultured M | unclassified | unclassified | unclassified | unclassified | unclassified | 0,6 | 0,489897949 | 0,000103112 | 8,56E-05 | 0 - | 0 - | 1 - | 0,000199942 - | 1 - | 0,000157707 - | 0 - | 0 - | 1 - | 0,00015791 - |
| 721 | 1 | 3um_FIJJAFBF_715091 | Eukaryota | unclassified | Pelagophycea | Pelagomonad | unclassified | Aureococcus | Aureococcus a | unclassified | unclassified | unclassified | unclassified | unclassified | 1 | 0 | 0,000653515 | 6,39E-05 | 1 - | 0,000659352 - | 1 - | 0,000775713 - | 1 - | 0,000611855 - | 1 - | 0,00060801 - | 1 - | 0,000612643 - |
| 722 | 1 | 0.2um_NOKCPHJN_709263 | unclassified | unclassified | unclassified | unclassified | unclassified | unclassified | unclassified | unclassified | unclassified | unclassified | unclassified | unclassified | 1 | 0 | 0,000944755 | 9,24E-05 | 1 - | 0,000953194 - | 1 - | 0,001121412 - | 1 - | 0,00088453 - | 1 - | 0,000878971 - | 1 - | 0,000885668 - |
| 723 | 1 | 0.2um_NOKCPHJN_575859 | Bacteria | Proteobacteri | Alphaproteob | Pelagibacteral | Pelagibactera | unclassified | Pelagibactera | poorly charact | Function unkn | 0YZNV | NA | 93059.P9211_ | 1 | 0 | 0,000103473 | 1,01E-05 | 1 - | 0,000104397 - | 1 - | 0,000122821 - | 1 - | 9,69E-05 - | 1 - | 9,63E-05 - | 1 - | 9,70E-05 - |
| 724 | 1 | 3um_FIJJAFBF_189702 | Bacteria | Bacteroidetes | Flavobacteriia | Flavobacterial | Flavobacteria | Polaribacter | Polaribacter fi | poorly charact | Function unkn | 09A4Z | rhodopsin | 313598.MED1 | 0,6 | 0,489897949 | 0,000203103 | 0,000165835 | 0 - | 0 - | 0 - | 0 - | 1 - | 0,00033907 - | 1 - | 0,000336939 - | 1 - | 0,000339506 - |
| 725 | 1 | 0.2um_NOKCPHJN_835021 | Eukaryota | Streptophyta | unclassified | Solanales | Convolvulacea | Cuscuta | Cuscuta austr | cellular proces | Intracellular tr | KOG1241 | importin | 3218.PP1S332 | 0,2 | 0,4 | 4,57E-05 | 9,14E-05 | 0 - | 0 - | 0 - | 0 - | 1 - | 0,000228586 - | 0 - | 0 - | 0 - | 0 - |
| 726 | 1 | 3um_FIJJAFBF_405250 | Bacteria | Bacteroidetes | Flavobacteriia | Flavobacterial | Flavobacteria | Polaribacter | Polaribacter ir | unclassified | unclassified | unclassified | unclassified | unclassified | 6,2 | 0,979795897 | 0,001615117 | 0,000214539 | 7 - | 0,001854553 - | 5 - | 0,001558458 - | 5 - | 0,001229256 - | 7 - | 0,001710143 - | 7 - | 0,001723174 - |
| 727 | 1 | 3um_FIJJAFBF_27824 | Bacteria | Proteobacteri | Gammaprotec | Alteromonada | Colwelliaceae | Colwellia | various | information st | Transcription | 05VEQ | Cold shock pro | 156578.ATW7 | 0,6 | 0,8 | 0,000793801 | 0,001012061 | 0 - | 0 - | 1 - | 0,001539849 - | 2 - | 0,002429157 - | 0 - | 0 - | 0 - | 0 - |
| 728 | 1 | 0.2um_NOKCPHJN_34249 | Eukaryota | unclassified | Labyrinthulom | unclassified | Thraustochytr | Aurantiochytr | Aurantiochytr | unclassified | unclassified | unclassified | unclassified | unclassified | 1 | 0 | 8,51E-05 | 8,33E-06 | 1 - | 8,59E-05 - | 1 - | 0,000101048 - | 1 - | 7,97E-05 - | 1 - | 7,92E-05 - | 1 - | 7,98E-05 - |
| 729 | 1 | 0.2um_NOKCPHJN_39631 | Bacteria | Bacteroidetes | Flavobacteriia | Flavobacterial | Flavobacteria | Polaribacter | Polaribacter s | unclassified | unclassified | unclassified | unclassified | unclassified | 15 | 2,28035085 | 0,001435045 | 0,000153058 | 17 - | 0,001660129 - | 11 - | 0,001263774 - | 14 - | 0,00126868 - | 17 - | 0,001530859 - | 16 - | 0,001451786 - |
| 730 | 1 | 3um_FIJJAFBF_49868 | Bacteria | Proteobacteri | Alphaproteob | Rhodobactera | Rhodobactera | Sulfitobacter | various | metabolism | Energy produc | 01RJJ | NADH flavin o | 314264.ROS2 | 0,8 | 0,4 | 0,000619472 | 0,000310621 | 1 - | 0,000819569 - | 0 - | 0 - | 1 - | 0,00076053 - | 1 - | 0,000755751 - | 1 - | 0,000761509 - |
| 731 | 1 | 3um_FIJJAFBF_669336 | Bacteria | Proteobacteri | Gammaprotec | unclassified | unclassified | unclassified | Gammaprotec | unclassified | unclassified | unclassified | unclassified | unclassified | 1,2 | 0,4 | 0,000115549 | 4,03E-05 | 2 - | 0,000194013 - | 1 - | 0,000114126 - | 1 - | 9,00E-05 - | 1 - | 8,95E-05 - | 1 - | 9,01E-05 - |
| 732 | 1 | 0.2um_NOKCPHJN_138597 | Bacteria | Bacteroidetes | Flavobacteriia | Flavobacterial | Crocinitomica | Lishizhenia | Lishizhenia tia | unclassified | unclassified | unclassified | unclassified | unclassified | 0,2 | 0,4 | 3,48E-05 | 6,96E-05 | 0 - | 0 - | 0 - | 0 - | 0 - | 0 - | 1 - | 0,000173904 - | 0 - | 0 - |
| 733 | 1 | 3um_FIJJAFBF_119125 | Bacteria | Bacteroidetes | Flavobacteriia | Flavobacterial | Flavobacteria | Polaribacter | various | information st | Translation, ri | 08YY1 | This protein is | 313594.PI23P | 1 | 0 | 0,000673779 | 6,59E-05 | 1 - | 0,000679797 - | 1 - | 0,000799767 - | 1 - | 0,000630827 - | 1 - | 0,000626863 - | 1 - | 0,000631639 - |
| 734 | 1 | 0.2um_NOKCPHJN_08633 | Bacteria | Proteobacteri | Gammaprotec | Alteromonada | Alteromonada | Paraglaciecola | Paraglaciecola | unclassified | unclassified | unclassified | unclassified | unclassified | 1 | 0,894427191 | 0,000360474 | 0,000335009 | 2 - | 0,000682442 - | 2 - | 0,000802879 - | 0 - | 0 - | 0 - | 0 - | 1 - | 0,000317049 - |
| 735 | 1 | 3um_FIJJAFBF_346691 | Bacteria | Proteobacteri | Gammaprotec | Oceanospirilla | Oceanospirilla | Amphritea | Amphritea atl | metabolism | Energy produc | COG0056 | Produces ATP | 207954.MED9 | 0,8 | 0,748331477 | 0,000131414 | 0,000126463 | 2 - | 0,000341221 - | 0 - | 0 - | 0 - | 0 - | 1 - | 0,000157326 - | 1 - | 0,000158524 - |
| 736 | 1 | 3um_FIJJAFBF_17219 | Bacteria | Proteobacteri | Alphaproteob | Rhodobactera | Rhodobactera | Phaeobacter | Phaeobacter g | cellular proces | Posttranslatio | COG0330 | Band 7 protei | 246200.SPO1 | 1,2 | 0,748331477 | 0,000367141 | 0,000243751 | 1 - | 0,000299296 - | 2 - | 0,000704231 - | 0 - | 0 - | 1 - | 0,000275991 - | 2 - | 0,000556188 - |
| 737 | 1 | 0.2um_NOKCPHJN_231997 | Bacteria | Proteobacteri | Betaproteoba | Nitrosomonad | unclassified | unclassified | beta proteoba | unclassified | unclassified | unclassified | unclassified | unclassified | 0,6 | 1,2 | 0,000261102 | 0,000522204 | 0 - | 0 - | 0 - | 0 - | 3 - | 0,001305509 - | 0 - | 0 - | 0 - | 0 - |
| 738 | 1 | 3um_FIJJAFBF_481353 | Bacteria | Proteobacteri | Gammaprotec | Alteromonada | Colwelliaceae | Colwellia | various | unclassified | unclassified | unclassified | unclassified | unclassified | 0,2 | 0,4 | 5,23E-05 | 0,000104665 | 0 - | 0 - | 0 - | 0 - | 1 - | 0,000261662 - | 0 - | 0 - | 0 - | 0 - |
| 739 | 1 | 3um_FIJJAFBF_653259 | Bacteria | Bacteroidetes | Flavobacteriia | Flavobacterial | Flavobacteria | Polaribacter | Polaribacter t | metabolism | Inorganic ion t | 05C27 | TonB-depende | 867900.Celly_ | 0,2 | 0,4 | 1,52E-05 | 3,03E-05 | 0 - | 0 - | 0 - | 0 - | 0 - | 0 - | 0 - | 0 - | 1 - | 7,59E-05 - |
| 740 | 1 | 0.2um_NOKCPHJN_21340 | Bacteria | Proteobacteri | Alphaproteob | Rhodospirillal | unclassified | unclassified | Rhodospirillal | information st | Translation, ri | COG1185 | Involved in m | 1064539.AZO | 0,2 | 0,4 | 2,19E-05 | 4,38E-05 | 0 - | 0 - | 0 - | 0 - | 0 - | 0 - | 1 - | 0,000109574 - | 0 - | 0 - |
| 741 | 1 | 0.2um_NOKCPHJN_183442 | Eukaryota | Eustigmatoph | unclassified | Eustigmatales | Monodopsida | Nannochlorop | Nannochlorop | cellular proces | Posttranslatio | KOG0738 | katanin p60 | 400682.PAC_ | 0,2 | 0,4 | 2,60E-05 | 5,20E-05 | 0 - | 0 - | 0 - | 0 - | 0 - | 0 - | 1 - | 0,000130009 - | 0 - | 0 - |
| 742 | 1 | 0.2um_NOKCPHJN_144610 | Bacteria | Proteobacteri | Gammaprotec | Cellvibrionales | Porticoccacea | unclassified | SAR92 bacteri | unclassified | unclassified | unclassified | unclassified | unclassified | 0,8 | 0,4 | 0,000299046 | 0,000153942 | 0 - | 0 - | 1 - | 0,000444698 - | 1 - | 0,000350762 - | 1 - | 0,000348558 - | 1 - | 0,000351213 - |
| 743 | 1 | 3um_FIJJAFBF_640407 | unclassified | unclassified | unclassified | unclassified | unclassified | unclassified | unclassified | unclassified | unclassified | unclassified | unclassified | unclassified | 1 | 0 | 0,000363671 | 3,56E-05 | 1 - | 0,00036692 - | 1 - | 0,000431673 - | 1 - | 0,000340488 - | 1 - | 0,000338349 - | 1 - | 0,000340927 - |
| 744 | 1 | 0.2um_NOKCPHJN_902013 | Eukaryota | unclassified | unclassified | Prymnesiales | Chrysochromu | Chrysochromu | Chrysochromu | unclassified | unclassified | unclassified | unclassified | unclassified | 1,2 | 0,4 | 0,000433675 | 0,000199427 | 1 - | 0,000353604 - | 2 - | 0,000832015 - | 1 - | 0,000328132 - | 1 - | 0,00032607 - | 1 - | 0,000328554 - |
| 745 | 1 | 0.2um_NOKCPHJN_413021 | Bacteria | Proteobacteri | Betaproteoba | Burkholderiale | Oxalobacterac | Duganella | Duganella sac | poorly charact | Function unkn | 08QSJ | NA | 225937.HP15_ | 0,6 | 0,489897949 | 0,000147711 | 0,000120607 | 0 - | 0 - | 0 - | 0 - | 1 - | 0,000246596 - | 1 - | 0,000245047 - | 1 - | 0,000246914 - |
| 746 | 1 | 3um_FIJJAFBF_203229 | Bacteria | Proteobacteri | Gammaprotec | Arenicellales | Arenicellaceae | Arenicella | Arenicella xan | cellular proces | Posttranslatio | 17FI0 | Binds to Cpn6 | 572477.Alvin_ | 0,2 | 0,4 | 0,000171319 | 0,000342639 | 0 - | 0 - | 0 - | 0 - | 1 - | 0,000856597 - | 0 - | 0 - | 0 - | 0 - |
| 747 | 1 | 3um_FIJJAFBF_346921 | unclassified | unclassified | unclassified | unclassified | unclassified | unclassified | unclassified | unclassified | unclassified | unclassified | unclassified | unclassified | 0,4 | 0,489897949 | 0,000217915 | 0,000266894 | 0 - | 0 - | 0 - | 0 - | 0 - | 0 - | 1 - | 0,00054272 - | 1 - | 0,000546856 - |
| 748 | 1 | 0.2um_NOKCPHJN_841344 | Eukaryota | Chlorophyta | Trebouxiophy | Chlorellales | Chlorellaceae | Chlorella | Chlorella soro | cellular proces | Posttranslatio | KOG3085 | Haloacid deha | 9986.ENSOCU | 1 | 0 | 0,000116045 | 1,14E-05 | 1 - | 0,000117081 - | 1 - | 0,000137744 - | 1 - | 0,000108647 - | 1 - | 0,000107964 - | 1 - | 0,000108787 - |
| 749 | 1 | 0.2um_NOKCPHJN_92345 | Bacteria | Bacteroidetes | Flavobacteriia | Flavobacterial | Flavobacteria | Polaribacter | Polaribacter b | unclassified | unclassified | unclassified | unclassified | unclassified | 1,2 | 0,4 | 0,000863275 | 0,000301397 | 2 - | 0,001449485 - | 1 - | 0,000852644 - | 1 - | 0,000672535 - | 1 - | 0,000668309 - | 1 - | 0,000673401 - |
| 750 | 1 | 3um_FIJJAFBF_639894 | Bacteria | Proteobacteri | Gammaprotec | Oceanospirilla | Saccharospiril | Reinekea | Reinekea fors | poorly charact | Function unkn | 08VI5 | type VI secreti | 83406.HDN1F | 0,6 | 0,489897949 | 0,000331896 | 0,000275605 | 0 - | 0 - | 1 - | 0,000644812 - | 0 - | 0 - | 1 - | 0,000505408 - | 1 - | 0,000509259 - |
| 751 | 1 | 0.2um_NOKCPHJN_310436 | unclassified | unclassified | unclassified | unclassified | unclassified | unclassified | unclassified | unclassified | unclassified | unclassified | unclassified | unclassified | 0,4 | 0,489897949 | 9,40E-05 | 0,000115782 | 1 - | 0,000215995 - | 1 - | 0,000254113 - | 0 - | 0 - | 0 - | 0 - | 0 - | 0 - |
| 752 | 1 | 0.2um_NOKCPHJN_657137 | Eukaryota | unclassified | Cryptophyta | Pyrenomonad | Geminigerace | Guillardia | Guillardia thet | unclassified | unclassified | unclassified | unclassified | unclassified | 6,2 | 0,4 | 0,004556592 | 0,000421637 | 6 - | 0,004459009 - | 6 - | 0,005245927 - | 6 - | 0,0041378 - | 7 - | 0,004797097 - | 6 - | 0,004143127 - |
| 753 | 1 | 0.2um_NOKCPHJN_481465 | Eukaryota | Ascomycota | Saccharomyce | Saccharomyce | Metschnikowi | Clavispora | [Candida] inte | unclassified | unclassified | unclassified | unclassified | unclassified | 1 | 0,632455532 | 0,000258895 | 0,00016431 | 2 - | 0,000514333 - | 1 - | 0,000302551 - | 1 - | 0,000238641 - | 0 - | 0 - | 1 - | 0,000238949 - |
| 754 | 2 | 0.2um_NOKCPHJN_21880;0.2u | Bacteria | Bacteroidetes | Flavobacteriia | Flavobacterial | Flavobacteria | Polaribacter | various | unclassified | unclassified | unclassified | unclassified | unclassified | 1,6 | 0,489897949 | 0,000898425 | 0,000219954 | 1 - | 0,000580754 - | 1 - | 0,000683244 - | 2 - | 0,001077838 - | 2 - | 0,001071064 - | 2 - | 0,001079225 - |
| 755 | 1 | 3um_FIJJAFBF_783318 | Eukaryota | unclassified | Pelagophycea | Pelagomonad | unclassified | Aureoumbra | Aureoumbra l | information st | Translation, ri | 05G6W | One of the pri | 696747.NIES3 | 0,8 | 0,748331477 | 0,000317273 | 0,000297254 | 0 - | 0 - | 0 - | 0 - | 1 - | 0,00039696 - | 1 - | 0,000394465 - | 2 - | 0,000794941 - |
| 756 | 1 | 0.2um_NOKCPHJN_05106 | Bacteria | Bacteroidetes | Flavobacteriia | Flavobacterial | Crocinitomica | unclassified | Crocinitomica | unclassified | unclassified | unclassified | unclassified | unclassified | 2,4 | 0,489897949 | 0,000134117 | 2,22E-05 | 2 - | 0,000113962 - | 2 - | 0,000134074 - | 3 - | 0,000158629 - | 2 - | 0,000105088 - | 3 - | 0,000158833 - |
| 757 | 1 | 0.2um_NOKCPHJN_376481 | Bacteria | Bacteroidetes | Cytophagia | Cytophagales | Flammeovirga | unclassified | Flammeovirga | poorly charact | Function unkn | 0XR4Y | Ragb susd dor | 886377.Murru | 0,2 | 0,4 | 3,13E-05 | 6,26E-05 | 1 - | 0,000156596 - | 0 - | 0 - | 0 - | 0 - | 0 - | 0 - | 0 - | 0 - |
| 758 | 1 | 0.2um_NOKCPHJN_217813 | Bacteria | unclassified | unclassified | unclassified | unclassified | unclassified | bacterium B1 | unclassified | unclassified | unclassified | unclassified | unclassified | 0,6 | 0,489897949 | 8,72E-05 | 7,12E-05 | 0 - | 0 - | 0 - | 0 - | 1 - | 0,000145576 - | 1 - | 0,000144661 - | 1 - | 0,000145763 - |
| 759 | 1 | 3um_FIJJAFBF_518449 | Bacteria | Bacteroidetes | Flavobacteriia | Flavobacterial | Flavobacteria | Ochrovirga | Ochrovirga pa | information st | Translation, ri | COG0254 | 50S ribosomal | 313598.MED1 | 0,6 | 0,489897949 | 0,000639799 | 0,000531287 | 0 - | 0 - | 1 - | 0,001243011 - | 0 - | 0 - | 1 - | 0,000974281 - | 1 - | 0,000981705 - |
| 760 | 1 | 0.2um_NOKCPHJN_106480 | Bacteria | Bacteroidetes | Flavobacteriia | Flavobacterial | unclassified | unclassified | Flavobacterial | metabolism | Inorganic ion t | 05C27 | TonB-depende | 216432.CA25 | 0,4 | 0,489897949 | 3,59E-05 | 4,45E-05 | 0 - | 0 - | 1 - | 0,00010036 - | 0 - | 0 - | 0 - | 0 - | 1 - | 7,93E-05 - |
| 761 | 1 | 0.2um_NOKCPHJN_975273 | unclassified | unclassified | unclassified | unclassified | unclassified | unclassified | unclassified | unclassified | unclassified | unclassified | unclassified | unclassified | 0,4 | 0,489897949 | 0,000179171 | 0,000221975 | 0 - | 0 - | 1 - | 0,000500825 - | 1 - | 0,000395033 - | 0 - | 0 - | 0 - | 0 - |
| 762 | 1 | 0.2um_NOKCPHJN_583156 | Bacteria | Proteobacteri | Alphaproteob | Pelagibacteral | unclassified | unclassified | Pelagibacteral | cellular proces | Cell wall/mem | 7257 | NA | 335992.SAR11 | 1 | 0 | 0,0002618 | 2,56E-05 | 1 - | 0,000264138 - | 1 - | 0,000310753 - | 1 - | 0,000245111 - | 1 - | 0,00024357 - | 1 - | 0,000245426 - |
| 763 | 1 | 0.2um_NOKCPHJN_348376 | Eukaryota | Streptophyta | unclassified | Asterales | Asteraceae | Cynara | Cynara cardur | unclassified | unclassified | unclassified | unclassified | unclassified | 0,2 | 0,4 | 1,17E-05 | 2,34E-05 | 0 - | 0 - | 0 - | 0 - | 0 - | 0 - | 1 - | 5,85E-05 - | 0 - | 0 - |
| 764 | 1 | 3um_FIJJAFBF_636516 | Bacteria | Bacteroidetes | Flavobacteriia | Flavobacterial | Flavobacteria | Polaribacter | Polaribacter s | metabolism | Inorganic ion t | 0P8Q4 | tonB-depende | 867900.Celly_ | 3,6 | 1,356465997 | 0,000427277 | 0,000136796 | 2 - | 0,000246678 - | 2 - | 0,000290211 - | 4 - | 0,000457816 - | 5 - | 0,000568673 - | 5 - | 0,000573006 - |
| 765 | 1 | 3um_FIJJAFBF_343895 | Eukaryota | unclassified | unclassified | Phaeocystales | Phaeocystace | Phaeocystis | Phaeocystis gl | metabolism | Energy produc | COG0711 | F(1)F(0) ATP s | 32049.SYNPC | 2 | 0 | 0,001073055 | 0,000104992 | 2 - | 0,00108264 - | 2 - | 0,001273702 - | 2 - | 0,001004651 - | 2 - | 0,000998338 - | 2 - | 0,001005944 - |
| 766 | 1 | 3um_FIJJAFBF_60207 | Bacteria | Actinobacteria | unclassified | unclassified | unclassified | unclassified | Actinobacteria | unclassified | unclassified | unclassified | unclassified | unclassified | 2 | 0,894427191 | 0,000632826 | 0,000270478 | 3 - | 0,000974376 - | 1 - | 0,000382111 - | 2 - | 0,000602791 - | 1 - | 0,000299501 - | 3 - | 0,00090535 - |
| 767 | 1 | 3um_FIJJAFBF_292915 | Eukaryota | Bacillariophyta | Coscinodiscop | Coscinodiscale | Coscinodiscac | Coscinodiscus | Coscinodiscus | metabolism | Energy produc | COG0711 | F(1)F(0) ATP s | 35128.Thapsd | 5,6 | 0,8 | 0,002682887 | 0,000206583 | 6 - | 0,002939459 - | 4 - | 0,002305472 - | 6 - | 0,002727712 - | 6 - | 0,00271057 - | 6 - | 0,002731223 - |
| 768 | 1 | 0.2um_NOKCPHJN_290418 | Bacteria | Proteobacteri | Gammaprotec | Alteromonada | Colwelliaceae | Colwellia | Colwellia sp. P | information st | Translation, ri | 16Q16 | Binds the low | 167879.CPS_ | 0,4 | 0,489897949 | 0,000140469 | 0,000172041 | 0 - | 0 - | 0 - | 0 - | 1 - | 0,00035228 - | 1 - | 0,000350066 - | 0 - | 0 - |
| 769 | 1 | 0.2um_NOKCPHJN_494189 | Bacteria | Proteobacteri | Alphaproteob | Pelagibacteral | Pelagibactera | Candidatus Pe | Candidatus Pe | metabolism | Energy produc | 01RNF | Produces ATP | 331869.BAL1 | 1,8 | 0,4 | 0,000332323 | 5,53E-05 | 2 - | 0,000380451 - | 1 - | 0,000223796 - | 2 - | 0,000353044 - | 2 - | 0,000350826 - | 2 - | 0,000353499 - |
| 770 | 1 | 0.2um_NOKCPHJN_329566 | Eukaryota | Chytridiomyc | Neocallimasti | Neocallimasti | Neocallimasti | Anaeromyces | Anaeromyces | unclassified | unclassified | unclassified | unclassified | unclassified | 0,2 | 0,4 | 0,000101089 | 0,000202178 | 0 - | 0 - | 0 - | 0 - | 1 - | 0,000505446 - | 0 - | 0 - | 0 - | 0 - |
| 771 | 1 | 3um_FIJJAFBF_558999 | Bacteria | Proteobacteri | Gammaprotec | Cellvibrionales | Cellvibrionace | Cellvibrio | Cellvibrio sp. | unclassified | unclassified | unclassified | unclassified | unclassified | 0,2 | 0,4 | 5,82E-05 | 0,000116402 | 0 - | 0 - | 0 - | 0 - | 0 - | 0 - | 0 - | 0 - | 1 - | 0,000291005 - |
| 772 | 1 | 3um_FIJJAFBF_220488 | Bacteria | Proteobacteri | Gammaprotec | Alteromonada | Alteromonada | Paraglaciecola | Paraglaciecola | information st | Translation, ri | COG0087 | One of the pri | 342610.Patl_ | 0,8 | 0,748331477 | 0,000313239 | 0,000289394 | 1 - | 0,00041365 - | 0 - | 0 - | 1 - | 0,000383853 - | 0 - | 0 - | 2 - | 0,000768693 - |
| 773 | 1 | 3um_FIJJAFBF_234946 | Eukaryota | Bacillariophyta | Bacillariophyc | Bacillariales | Bacillariaceae | Fragilariopsis | Fragilariopsis | unclassified | unclassified | unclassified | unclassified | unclassified | 0,4 | 0,489897949 | 0,000254241 | 0,000311743 | 1 - | 0,000659352 - | 0 - | 0 - | 1 - | 0,000611855 - | 0 - | 0 - | 0 - | 0 - |
| 774 | 1 | 3um_FIJJAFBF_594159 | Eukaryota | unclassified | Pelagophycea | Pelagomonad | unclassified | Aureoumbra | Aureoumbra l | metabolism | Energy produc | COG0055 | Produces ATP | 7739.JGI1257 | 5,8 | 0,748331477 | 0,001796781 | 0,00014971 | 5 - | 0,001577228 - | 5 - | 0,001855574 - | 6 - | 0,001756333 - | 7 - | 0,002036178 - | 6 - | 0,001758593 - |
| 775 | 1 | 0.2um_NOKCPHJN_929635 | Bacteria | Proteobacteri | Gammaprotec | Cellvibrionales | Halieaceae | Luminiphilus | Luminiphilus s | unclassified | unclassified | unclassified | unclassified | unclassified | 0,2 | 0,4 | 5,26E-05 | 0,000105137 | 0 - | 0 - | 0 - | 0 - | 0 - | 0 - | 0 - | 0 - | 1 - | 0,000262844 - |
| 776 | 1 | 0.2um_NOKCPHJN_602244 | Bacteria | Bacteroidetes | Flavobacteriia | Flavobacterial | Flavobacteria | unclassified | Flavobacteria | unclassified | unclassified | unclassified | unclassified | unclassified | 1 | 0 | 0,000104218 | 1,02E-05 | 1 - | 0,000105149 - | 1 - | 0,000123705 - | 1 - | 9,76E-05 - | 1 - | 9,70E-05 - | 1 - | 9,77E-05 - |
| 777 | 1 | 0.2um_NOKCPHJN_917897 | Bacteria | Proteobacteri | Alphaproteob | Rhodospirillal | Rhodospirillac | unclassified | Rhodospirillac | cellular proces | Cell motility | 16UR5 | flagellar hook- | 269796.Rru_A | 0,2 | 0,4 | 0,000131427 | 0,000262853 | 0 - | 0 - | 1 - | 0,000657133 - | 0 - | 0 - | 0 - | 0 - | 0 - | 0 - |
| 778 | 1 | 0.2um_NOKCPHJN_302339 | Bacteria | Proteobacteri | Betaproteoba | Burkholderiale | Comamonada | Polaromonas | Polaromonas | metabolism | Energy produc | 05CMV | Succinyl-CoA s | 365046.Rta_3 | 0,2 | 0,4 | 3,86E-05 | 7,71E-05 | 0 - | 0 - | 0 - | 0 - | 1 - | 0,000192836 - | 0 - | 0 - | 0 - | 0 - |
| 779 | 1 | 3um_FIJJAFBF_156011 | Eukaryota | Chlorophyta | Chlorophycea | Chlamydomor | Volvocaceae | Gonium | Gonium pecto | unclassified | unclassified | unclassified | unclassified | unclassified | 1,8 | 0,979795897 | 0,000329315 | 0,000162872 | 1 - | 0,000190225 - | 1 - | 0,000223796 - | 3 - | 0,000529567 - | 3 - | 0,000526239 - | 1 - | 0,000176749 - |
| 780 | 1 | 0.2um_NOKCPHJN_420409 | Bacteria | Bacteroidetes | Cytophagia | Cytophagales | Flammeovirga | unclassified | Flammeovirga | poorly charact | Function unkn | 16R8I | Lipoprotein | 566466.NOR5 | 3,6 | 1,019803903 | 0,000621506 | 0,000137349 | 3 - | 0,000535808 - | 2 - | 0,000420244 - | 4 - | 0,000662947 - | 4 - | 0,000658781 - | 5 - | 0,00082975 - |
| 781 | 1 | 3um_FIJJAFBF_46932 | Bacteria | Proteobacteri | Gammaprotec | Alteromonada | Alteromonada | Paraglaciecola | various | information st | Translation, ri | 179RU | 30S ribosomal | 342610.Patl_ | 0,6 | 0,489897949 | 0,000384517 | 0,000314501 | 1 - | 0,000674568 - | 0 - | 0 - | 1 - | 0,000625975 - | 1 - | 0,000622041 - | 0 - | 0 - |
| 782 | 1 | 3um_FIJJAFBF_503363 | unclassified | unclassified | unclassified | unclassified | unclassified | unclassified | unclassified | unclassified | unclassified | unclassified | unclassified | unclassified | 0,4 | 0,489897949 | 0,000444786 | 0,000544756 | 0 - | 0 - | 0 - | 0 - | 0 - | 0 - | 1 - | 0,001107745 - | 1 - | 0,001116185 - |
| 783 | 1 | 3um_FIJJAFBF_143821 | Bacteria | Proteobacteri | Gammaprotec | Cellvibrionales | Halieaceae | Halioglobus | Halioglobus ja | unclassified | unclassified | unclassified | unclassified | unclassified | 1 | 0 | 0,000161257 | 1,58E-05 | 1 - | 0,000162697 - | 1 - | 0,00019141 - | 1 - | 0,000150977 - | 1 - | 0,000150028 - | 1 - | 0,000151172 - |
| 784 | 1 | 3um_FIJJAFBF_49461 | Bacteria | Actinobacteria | Acidimicrobiia | Acidimicrobial | Acidimicrobia | unclassified | Acidimicrobia | cellular proces | Posttranslatio | COG1225 | alkyl hydroper | 269798.CHU_ | 0,8 | 0,4 | 0,000424894 | 0,000213054 | 1 - | 0,00056214 - | 0 - | 0 - | 1 - | 0,000521646 - | 1 - | 0,000518368 - | 1 - | 0,000522317 - |
| 785 | 1 | 0.2um_NOKCPHJN_1099431 | unclassified | unclassified | unclassified | unclassified | unclassified | unclassified | unclassified | unclassified | unclassified | unclassified | unclassified | unclassified | 0,4 | 0,489897949 | 0,0001358 | 0,000166513 | 1 - | 0,000352184 - | 0 - | 0 - | 1 - | 0,000326814 - | 0 - | 0 - | 0 - | 0 - |
| 786 | 1 | 0.2um_NOKCPHJN_296782 | Bacteria | Bacteroidetes | Flavobacteriia | Flavobacterial | Flavobacteria | Marinirhabdu | Marinirhabdu | unclassified | unclassified | unclassified | unclassified | unclassified | 0,6 | 0,8 | 9,30E-05 | 0,000122171 | 1 - | 0,000162697 - | 0 - | 0 - | 0 - | 0 - | 0 - | 0 - | 2 - | 0,000302343 - |
| 787 | 1 | 3um_FIJJAFBF_37426 | Bacteria | Proteobacteri | Alphaproteob | Rhodobactera | Rhodobactera | Loktanella | Loktanella sp. | metabolism | Energy produc | 0259B | 2-oxoacid deh | 991905.SL003 | 0,8 | 0,4 | 0,000216808 | 0,000111608 | 0 - | 0 - | 1 - | 0,000322406 - | 1 - | 0,000254302 - | 1 - | 0,000252704 - | 1 - | 0,00025463 - |
| 788 | 1 | 0.2um_NOKCPHJN_191187 | Eukaryota | Platyhelminth | unclassified | Macrostomida | Macrostomida | Macrostomum | Macrostomum | poorly charact | Function unkn | 0NVRZ | NA | 170187.SP_17 | 1 | 0 | 3,77E-05 | 3,68E-06 | 1 - | 3,80E-05 - | 1 - | 4,47E-05 - | 1 - | 3,53E-05 - | 1 - | 3,50E-05 - | 1 - | 3,53E-05 - |
| 789 | 1 | 3um_FIJJAFBF_192887 | Bacteria | Proteobacteri | Alphaproteob | Rhizobiales | unclassified | unclassified | Rhizobiales ba | poorly charact | Function unkn | 05C2A | tonB-depende | 1085623.GNIT | 1,6 | 0,489897949 | 0,000394425 | 0,000135523 | 1 - | 0,000247025 - | 2 - | 0,000581239 - | 1 - | 0,00022923 - | 2 - | 0,000455579 - | 2 - | 0,000459051 - |
| 790 | 1 | 0.2um_NOKCPHJN_605600 | Eukaryota | unclassified | Oomycetes | Saprolegniales | Saprolegniace | Thraustotheca | Thraustotheca | unclassified | unclassified | unclassified | unclassified | unclassified | 0,2 | 0,4 | 3,05E-05 | 6,10E-05 | 0 - | 0 - | 0 - | 0 - | 0 - | 0 - | 1 - | 0,000152576 - | 0 - | 0 - |
| 791 | 1 | 0.2um_NOKCPHJN_487588 | Bacteria | Verrucomicrol | unclassified | unclassified | unclassified | unclassified | Verrucomicro | unclassified | unclassified | unclassified | unclassified | unclassified | 1 | 0 | 0,000755804 | 7,40E-05 | 1 - | 0,000762555 - | 1 - | 0,000897129 - | 1 - | 0,000707624 - | 1 - | 0,000703177 - | 1 - | 0,000708535 - |
| 792 | 1 | 0.2um_NOKCPHJN_994909 | Bacteria | Proteobacteri | Alphaproteob | Rhodobactera | Rhodobactera | Planktomarina | Planktomarina | unclassified | unclassified | unclassified | unclassified | unclassified | 0,8 | 0,4 | 0,000237782 | 0,000121948 | 1 - | 0,000295265 - | 1 - | 0,000347373 - | 1 - | 0,000273996 - | 1 - | 0,000272274 - | 0 - | 0 - |
| 793 | 1 | 0.2um_NOKCPHJN_725744 | Bacteria | Proteobacteri | Betaproteoba | unclassified | unclassified | unclassified | Betaproteoba | unclassified | unclassified | unclassified | unclassified | unclassified | 0,2 | 0,4 | 8,23E-05 | 0,000164683 | 1 - | 0,000411708 - | 0 - | 0 - | 0 - | 0 - | 0 - | 0 - | 0 - | 0 - |
| 794 | 1 | 3um_FIJJAFBF_257126 | Eukaryota | unclassified | Florideophyce | Nemaliales | Liagoraceae | Dermonema | Dermonema v | unclassified | unclassified | unclassified | unclassified | unclassified | 1,4 | 0,489897949 | 0,000656534 | 0,000198577 | 1 - | 0,000481834 - | 1 - | 0,000566868 - | 2 - | 0,00089425 - | 1 - | 0,000444315 - | 2 - | 0,000895401 - |
| 795 | 1 | 3um_FIJJAFBF_242749 | Bacteria | Bacteroidetes | Flavobacteriia | unclassified | unclassified | unclassified | Flavobacteria | unclassified | unclassified | unclassified | unclassified | unclassified | 3,6 | 2,244994432 | 0,000397859 | 0,000252793 | 6 - | 0,000697829 - | 0 - | 0 - | 3 - | 0,00032378 - | 6 - | 0,000643491 - | 3 - | 0,000324197 - |
| 796 | 1 | 3um_FIJJAFBF_282387 | Eukaryota | Bacillariophyta | Coscinodiscop | Thalassiosirale | Thalassiosirac | Thalassiosira | Thalassiosira p | unclassified | unclassified | unclassified | unclassified | unclassified | 0,6 | 0,489897949 | 0,000113141 | 9,25E-05 | 1 - | 0,000198402 - | 0 - | 0 - | 0 - | 0 - | 1 - | 0,000182953 - | 1 - | 0,000184347 - |
| 797 | 1 | 3um_FIJJAFBF_65825 | Eukaryota | unclassified | Dinophyceae | Suessiales | Symbiodiniace | Symbiodinium | Symbiodinium | information st | Replication, re | 08Z6R | Alkylated DNA | 523791.Kkor_ | 1 | 0 | 0,000168445 | 1,65E-05 | 1 - | 0,000169949 - | 1 - | 0,000199942 - | 1 - | 0,000157707 - | 1 - | 0,000156716 - | 1 - | 0,00015791 - |
| 798 | 1 | 3um_FIJJAFBF_510230 | Bacteria | Bacteroidetes | Flavobacteriia | Flavobacterial | Flavobacteria | Polaribacter | Polaribacter s | information st | Translation, ri | COG0087 | One of the pri | 313594.PI23P | 1 | 0,894427191 | 0,000396064 | 0,00035394 | 0 - | 0 - | 0 - | 0 - | 2 - | 0,000793919 - | 2 - | 0,00078893 - | 1 - | 0,000397471 - |
| 799 | 1 | 0.2um_NOKCPHJN_440998 | Eukaryota | Bacillariophyta | Coscinodiscop | Thalassiosirale | Thalassiosirac | Thalassiosira | Thalassiosira p | information st | RNA processin | KOG1596 | rRNA 2'-O-me | 35128.Thaps9 | 0,6 | 0,489897949 | 0,000214908 | 0,000178475 | 0 - | 0 - | 1 - | 0,000417692 - | 1 - | 0,00032946 - | 1 - | 0,00032739 - | 0 - | 0 - |
| 800 | 1 | 3um_FIJJAFBF_103105 | Bacteria | Bacteroidetes | Flavobacteriia | Flavobacterial | Flavobacteria | Polaribacter | Polaribacter s | cellular proces | Cell wall/mem | COG2885 | Ompa motb d | 313594.PI23P | 5,8 | 1,326649916 | 0,0010698 | 0,000220181 | 6 - | 0,00112428 - | 5 - | 0,001102242 - | 4 - | 0,000695528 - | 8 - | 0,001382314 - | 6 - | 0,001044635 - |
| 801 | 1 | 3um_FIJJAFBF_205216 | Eukaryota | Bacillariophyta | Mediophyceae | Cymatosirales | Cymatosirace | Plagiogramm | Plagiogramm | metabolism | Energy produc | COG0056 | Produces ATP | 41431.PCC88 | 0,8 | 0,4 | 0,000140442 | 7,20E-05 | 1 - | 0,000174342 - | 1 - | 0,000205109 - | 0 - | 0 - | 1 - | 0,000160766 - | 1 - | 0,000161991 - |
| 802 | 1 | 3um_FIJJAFBF_25828 | unclassified | unclassified | unclassified | unclassified | unclassified | unclassified | unclassified | unclassified | unclassified | unclassified | unclassified | unclassified | 0,6 | 0,489897949 | 0,000288196 | 0,000238261 | 1 - | 0,000463989 - | 1 - | 0,000545872 - | 0 - | 0 - | 0 - | 0 - | 1 - | 0,000431119 - |
| 803 | 2 | 0.2um_NOKCPHJN_727059;3ur | unclassified | unclassified | unclassified | unclassified | unclassified | unclassified | unclassified | unclassified | unclassified | unclassified | unclassified | unclassified | 0,4 | 0,489897949 | 8,60E-05 | 0,000106607 | 0 - | 0 - | 1 - | 0,000241051 - | 0 - | 0 - | 1 - | 0,000188938 - | 0 - | 0 - |
| 804 | 1 | 3um_FIJJAFBF_63332 | Bacteria | Proteobacteri | Gammaprotec | Alteromonada | Alteromonada | Glaciecola | Glaciecola sp. | metabolism | Inorganic ion t | 0QJVD | TonB-depende | 455436.GHTC | 7 | 1,264911064 | 0,00068829 | 8,18E-05 | 7 - | 0,000704773 - | 5 - | 0,00059225 - | 7 - | 0,000654004 - | 9 - | 0,000835578 - | 7 - | 0,000654846 - |
| 805 | 1 | 3um_FIJJAFBF_12818 | Bacteria | Bacteroidetes | Flavobacteriia | Flavobacterial | Flavobacteria | Arenitalea | Arenitalea lut | unclassified | unclassified | unclassified | unclassified | unclassified | 3,6 | 2,653299832 | 0,00029529 | 0,000215557 | 1 - | 8,81E-05 - | 0 - | 0 - | 7 - | 0,0005725 - | 5 - | 0,000406359 - | 5 - | 0,000409455 - |
| 806 | 1 | 3um_FIJJAFBF_175283 | Eukaryota | Apicomplexa | Conoidasida | Eucoccidiorida | Sarcocystidae | Hammondia | Hammondia h | unclassified | unclassified | unclassified | unclassified | unclassified | 0,2 | 0,4 | 2,36E-05 | 4,72E-05 | 0 - | 0 - | 0 - | 0 - | 0 - | 0 - | 1 - | 0,00011788 - | 0 - | 0 - |
| 807 | 1 | 3um_FIJJAFBF_313317 | Bacteria | Proteobacteri | Gammaprotec | Oceanospirilla | Saccharospiril | Reinekea | Reinekea fors | cellular proces | Posttranslatio | COG0330 | Band 7 protei | 314283.MED2 | 0,2 | 0,4 | 4,45E-05 | 8,90E-05 | 1 - | 0,000222573 - | 0 - | 0 - | 0 - | 0 - | 0 - | 0 - | 0 - | 0 - |
| 808 | 1 | 3um_FIJJAFBF_524119 | Bacteria | Fibrobacteres | Fibrobacteria | Fibrobacterale | Fibrobacterac | Fibrobacter | Fibrobacter sp | unclassified | unclassified | unclassified | unclassified | unclassified | 1 | 0 | 0,000220603 | 2,16E-05 | 1 - | 0,000222573 - | 1 - | 0,000261853 - | 1 - | 0,00020654 - | 1 - | 0,000205242 - | 1 - | 0,000206806 - |
| 809 | 1 | 0.2um_NOKCPHJN_170714 | Bacteria | Bacteroidetes | unclassified | unclassified | unclassified | unclassified | Bacteroidetes | poorly charact | Function unkn | 05WQ7 | NA | 313595.P7007 | 0,2 | 0,4 | 3,89E-05 | 7,78E-05 | 0 - | 0 - | 0 - | 0 - | 0 - | 0 - | 1 - | 0,000194388 - | 0 - | 0 - |
| 810 | 1 | 3um_FIJJAFBF_95716 | Eukaryota | unclassified | Raphidophyce | Chattonellales | Chattonellace | Heterosigma | Heterosigma a | cellular proces | Posttranslatio | 0IEKR | One of the co | 35128.Thapsd | 11,6 | 1,019803903 | 0,002184496 | 8,28E-05 | 11 - | 0,002106184 - | 10 - | 0,002252618 - | 12 - | 0,002132142 - | 12 - | 0,002118743 - | 13 - | 0,002312793 - |
| 811 | 1 | 3um_FIJJAFBF_351766 | Bacteria | Actinobacteria | Actinobacteria | Bifidobacteria | Bifidobacteria | Bifidobacteriu | Bifidobacteriu | unclassified | unclassified | unclassified | unclassified | unclassified | 0,2 | 0,4 | 0,000190271 | 0,000380543 | 0 - | 0 - | 0 - | 0 - | 0 - | 0 - | 1 - | 0,000951357 - | 0 - | 0 - |
| 812 | 1 | 3um_FIJJAFBF_403052 | Eukaryota | Chlorophyta | Trebouxiophy | Chlorellales | Chlorellaceae | Chlorella | Chlorella soro | information st | Chromatin str | KOG3467 | Core compone | 3218.PP1S133 | 1 | 0 | 0,000843859 | 8,26E-05 | 1 - | 0,000851397 - | 1 - | 0,001001649 - | 1 - | 0,000790065 - | 1 - | 0,000785101 - | 1 - | 0,000791082 - |
| 813 | 1 | 3um_FIJJAFBF_274555 | unclassified | unclassified | unclassified | unclassified | unclassified | unclassified | unclassified | unclassified | unclassified | unclassified | unclassified | unclassified | 0,6 | 0,489897949 | 5,58E-05 | 4,56E-05 | 0 - | 0 - | 0 - | 0 - | 1 - | 9,32E-05 - | 1 - | 9,26E-05 - | 1 - | 9,33E-05 - |
| 814 | 1 | 3um_FIJJAFBF_06900 | Bacteria | Bacteroidetes | unclassified | unclassified | unclassified | unclassified | Bacteroidetes | unclassified | unclassified | unclassified | unclassified | unclassified | 0,2 | 0,4 | 3,64E-05 | 7,28E-05 | 0 - | 0 - | 0 - | 0 - | 0 - | 0 - | 0 - | 0 - | 1 - | 0,000181878 - |
| 815 | 1 | 0.2um_NOKCPHJN_266842 | Bacteria | Bacteroidetes | Flavobacteriia | unclassified | unclassified | unclassified | Flavobacteria | poorly charact | Function unkn | 0P8NI | Inherit from b | 487796.Flav2 | 0,6 | 0,489897949 | 0,000111289 | 9,09E-05 | 0 - | 0 - | 0 - | 0 - | 1 - | 0,000185792 - | 1 - | 0,000184624 - | 1 - | 0,000186031 - |
| 816 | 1 | 3um_FIJJAFBF_139372 | Bacteria | Bacteroidetes | unclassified | unclassified | unclassified | unclassified | Bacteroidetes | metabolism | Inorganic ion t | 05C27 | TonB-depende | 391603.FBALC | 0,2 | 0,4 | 1,49E-05 | 2,98E-05 | 0 - | 0 - | 0 - | 0 - | 0 - | 0 - | 0 - | 0 - | 1 - | 7,46E-05 - |
| 817 | 1 | 0.2um_NOKCPHJN_485725 | Bacteria | Proteobacteri | Gammaprotec | Cellvibrionales | Spongiibacter | Oceanicoccus | Oceanicoccus | cellular proces | Cell wall/mem | 0QTSX | Peptidoglycan | 247634.GPB2 | 1 | 0 | 0,000455065 | 4,45E-05 | 1 - | 0,00045913 - | 1 - | 0,000540156 - | 1 - | 0,000426056 - | 1 - | 0,000423379 - | 1 - | 0,000426605 - |

| | | | | | | | | | | | | | | | | | | | | | | | | | | | | |
|---|---|---|---|---|---|---|---|---|---|---|---|---|---|---|---|---|---|---|---|---|---|---|---|---|---|---|---|---|
| 818 | 1 | 3um_FIJJAFBF_237981 | Eukaryota | unclassified | Compsopogor | Erythropeltida | Erythrotrichia | Erythrotrichia | Erythrotrichia | information st | Translation, ril | COG0333 | 50s ribosomal | 696747.NIES3 | 0,6 | 0,489897949 | 0,000812412 | 0,000663341 | 0 - | 0 - | 0 - | 0 - | 1 - | 0,001356279 - | 1 - | 0,001347756 - | 1 - | 0,001358025 - |
| 819 | 1 | 3um_FIJJAFBF_711375 | Bacteria | Bacteroidetes | Flavobacteriia | Flavobacterial | Flavobacteria | Aquimarina | Aquimarina sp | poorly charact | Function unkn | 0PAPS | NA | 873517.HMPR | 1 | 0 | 0,00069534 | 6,80E-05 | 1 - | 0,000701551 - | 1 - | 0,000825359 - | 1 - | 0,000651014 - | 1 - | 0,000646923 - | 1 - | 0,000651852 - |
| 820 | 1 | 3um_FIJJAFBF_551806 | Bacteria | Proteobacteri | Alphaproteob | Rhodobactera | Rhodobactera | Hasllibacter | Hasllibacter h | unclassified | unclassified | unclassified | unclassified | unclassified | 3 | 0,632455532 | 0,000628269 | 0,00013829 | 3 - | 0,000633931 - | 3 - | 0,000745806 - | 4 - | 0,000784354 - | 3 - | 0,000584569 - | 2 - | 0,000392682 - |
| 821 | 1 | 0.2um_NOKCPHJN_134680 | Bacteria | Bacteroidetes | Flavobacteriia | Flavobacterial | Flavobacteria | Flaviramulus | Flaviramulus i | unclassified | unclassified | unclassified | unclassified | unclassified | 1 | 0,632455532 | 0,000408811 | 0,000255432 | 1 - | 0,000434128 - | 0 - | 0 - | 1 - | 0,000402855 - | 1 - | 0,000400324 - | 2 - | 0,000806747 - |
| 822 | 1 | 0.2um_NOKCPHJN_418551 | Eukaryota | Chordata | unclassified | unclassified | Branchiostom | Branchiostom | Branchiostom | metabolism | Secondary me | KOG0054 | ATP-binding c | 7739.JGI2562 | 1 | 0 | 6,50E-05 | 6,36E-06 | 1 - | 6,56E-05 - | 1 - | 7,72E-05 - | 1 - | 6,09E-05 - | 1 - | 6,05E-05 - | 1 - | 6,09E-05 - |
| 823 | 1 | 0.2um_NOKCPHJN_76427 | Bacteria | Bacteroidetes | Flavobacteriia | Flavobacterial | Flavobacteria | unclassified | Flavobacteria | metabolism | Inorganic ion t | 0P7D0 | tonB-depende | 487796.Flav2A | 0,2 | 0,4 | 2,14E-05 | 4,29E-05 | 1 - | 0,000107205 - | 0 - | 0 - | 0 - | 0 - | 0 - | 0 - | 0 - | 0 - |
| 824 | 1 | 0.2um_NOKCPHJN_479664 | Eukaryota | Streptophyta | Bryopsida | Funariales | Funariaceae | Physcomitrell | Physcomitrell | poorly charact | Function unkn | 0NVRZ | NA | 170187.SP_17 | 1 | 0 | 4,59E-05 | 4,49E-06 | 1 - | 4,63E-05 - | 1 - | 5,44E-05 - | 1 - | 4,29E-05 - | 1 - | 4,27E-05 - | 1 - | 4,30E-05 - |
| 825 | 1 | 0.2um_NOKCPHJN_66855 | Bacteria | Proteobacteri | Gammaprotec | Arenicellales | Arenicellaceae | Arenicella | Arenicella xan | unclassified | unclassified | unclassified | unclassified | unclassified | 0,6 | 0,489897949 | 0,000203103 | 0,000165835 | 0 - | 0 - | 0 - | 0 - | 1 - | 0,00033907 - | 1 - | 0,000336939 - | 1 - | 0,000339506 - |
| 826 | 1 | 0.2um_NOKCPHJN_1045255 | Bacteria | Proteobacteri | Alphaproteob | Rhodobactera | Hyphomonad | Hyphomonas | Hyphomonas | poorly charact | Function unkn | 11H20 | NA | 483219.LILAB | 0,4 | 0,489897949 | 0,000123031 | 0,000152423 | 0 - | 0 - | 1 - | 0,0003439 - | 1 - | 0,000271256 - | 0 - | 0 - | 0 - | 0 - |
| 827 | 2 | 3um_FIJJAFBF_01992;3um_FIJJ | unclassified | unclassified | unclassified | unclassified | unclassified | unclassified | unclassified | unclassified | unclassified | unclassified | unclassified | unclassified | 0,2 | 0,4 | 0,000167036 | 0,000334072 | 1 - | 0,00083518 - | 0 - | 0 - | 0 - | 0 - | 0 - | 0 - | 0 - | 0 - |
| 828 | 1 | 3um_FIJJAFBF_569212 | Eukaryota | unclassified | Pelagophycea | Pelagomonad | unclassified | Aureococcus | Aureococcus a | metabolism | Carbohydrate | COG0057 | glyceraldehyd | 35128.Thaps3 | 0,4 | 0,8 | 0,00012785 | 0,000255701 | 0 - | 0 - | 0 - | 0 - | 0 - | 0 - | 2 - | 0,000639252 - | 0 - | 0 - |
| 829 | 1 | 0.2um_NOKCPHJN_36762 | Viruses | unclassified | unclassified | unclassified | various | various | various | unclassified | unclassified | unclassified | unclassified | unclassified | 0,8 | 0,4 | 0,00012946 | 6,49E-05 | 1 - | 0,000171277 - | 0 - | 0 - | 1 - | 0,000158939 - | 1 - | 0,00015794 - | 1 - | 0,000159144 - |
| 830 | 1 | 0.2um_NOKCPHJN_296182 | Bacteria | Proteobacteri | Alphaproteob | Rhodobactera | Rhodobactera | Monaibacteriu | Monaibacteriu | metabolism | Lipid transpor | 08IQF | Catalyzes the | 381666.H16_A | 0,4 | 0,489897949 | 5,87E-05 | 7,19E-05 | 0 - | 0 - | 0 - | 0 - | 0 - | 0 - | 1 - | 0,00014623 - | 1 - | 0,000147344 - |
| 831 | 1 | 0.2um_NOKCPHJN_184643 | Bacteria | Bacteroidetes | Flavobacteriia | Flavobacterial | Cryomorphace | Owenweeksia | Owenweeksia | poorly charact | Function unkn | 082FF | Colicin v prod | 177437.HRM2 | 0,6 | 0,489897949 | 0,000314517 | 0,000257241 | 1 - | 0,000551534 - | 0 - | 0 - | 0 - | 0 - | 1 - | 0,000508587 - | 1 - | 0,000512462 - |
| 832 | 1 | 3um_FIJJAFBF_95704 | Eukaryota | unclassified | Raphidophyce | Chattonellales | Chattonellace | Heterosigma | Heterosigma a | unclassified | unclassified | unclassified | unclassified | unclassified | 10,8 | 1,6 | 0,001749331 | 0,000118768 | 10 - | 0,001657729 - | 8 - | 0,001560225 - | 12 - | 0,001845975 - | 12 - | 0,001834375 - | 12 - | 0,001848351 - |
| 833 | 1 | 0.2um_NOKCPHJN_1015854 | unclassified | unclassified | unclassified | unclassified | unclassified | unclassified | unclassified | unclassified | unclassified | unclassified | unclassified | unclassified | 0,8 | 0,4 | 0,00031414 | 0,000157519 | 1 - | 0,000415611 - | 0 - | 0 - | 1 - | 0,000385672 - | 1 - | 0,000383248 - | 1 - | 0,000386168 - |
| 834 | 1 | 0.2um_NOKCPHJN_507733 | Eukaryota | Phaeophyceae | unclassified | Ectocarpales | Ectocarpaceae | Ectocarpus | Ectocarpus sili | poorly charact | Function unkn | 102D8 | PAP_fibrillin | 2850.Phatr46 | 2,8 | 0,979795897 | 0,001180347 | 0,000410547 | 1 - | 0,000427775 - | 3 - | 0,001509803 - | 3 - | 0,001190879 - | 3 - | 0,001183395 - | 4 - | 0,001589883 - |
| 835 | 1 | 3um_FIJJAFBF_114132 | unclassified | unclassified | unclassified | unclassified | unclassified | unclassified | unclassified | unclassified | unclassified | unclassified | unclassified | unclassified | 2 | 0 | 0,00117456 | 0,000114924 | 2 - | 0,001185052 - | 2 - | 0,001394188 - | 2 - | 0,001099686 - | 2 - | 0,001092775 - | 2 - | 0,001101101 - |
| 836 | 2 | 0.2um_NOKCPHJN_1043665;3u | Bacteria | Proteobacteri | Gammaprotec | Cellvibrionales | various | various | various | unclassified | unclassified | unclassified | unclassified | unclassified | 0,2 | 0,4 | 7,66E-05 | 0,000153177 | 1 - | 0,000382943 - | 0 - | 0 - | 0 - | 0 - | 0 - | 0 - | 0 - | 0 - |
| 837 | 1 | 3um_FIJJAFBF_39791 | Bacteria | Bacteroidetes | Flavobacteriia | Flavobacterial | Cryomorphace | unclassified | Cryomorphace | unclassified | unclassified | unclassified | unclassified | unclassified | 1,8 | 0,979795897 | 0,000579604 | 0,00028695 | 1 - | 0,000334709 - | 1 - | 0,000393778 - | 1 - | 0,000310598 - | 3 - | 0,000925939 - | 3 - | 0,000932994 - |
| 838 | 1 | 0.2um_NOKCPHJN_07200 | Viruses | unclassified | unclassified | unclassified | unclassified | unclassified | Pacmanvirus A | unclassified | unclassified | unclassified | unclassified | unclassified | 0,2 | 0,4 | 2,51E-05 | 5,02E-05 | 0 - | 0 - | 0 - | 0 - | 0 - | 0 - | 0 - | 0 - | 1 - | 0,000125549 - |
| 839 | 1 | 0.2um_NOKCPHJN_112415 | Bacteria | Bacteroidetes | Flavobacteriia | Flavobacterial | Flavobacteria | Marinirhabdu | Marinirhabdu | unclassified | unclassified | unclassified | unclassified | unclassified | 0,2 | 0,4 | 0,000137022 | 0,000274043 | 1 - | 0,000685108 - | 0 - | 0 - | 0 - | 0 - | 0 - | 0 - | 0 - | 0 - |
| 840 | 1 | 3um_FIJJAFBF_269222 | Bacteria | Bacteroidetes | Flavobacteriia | Flavobacterial | Flavobacteria | Polaribacter | Polaribacter v | unclassified | unclassified | unclassified | unclassified | unclassified | 0,4 | 0,489897949 | 0,000187791 | 0,000232794 | 0 - | 0 - | 1 - | 0,000526377 - | 0 - | 0 - | 1 - | 0,000412578 - | 0 - | 0 - |
| 841 | 1 | 3um_FIJJAFBF_202445 | Bacteria | Proteobacteri | Alphaproteob | Rhodobactera | Rhodobactera | Oceanicola | Oceanicola sp | metabolism | Amino acid tra | COG3931 | N-formylgluta | 272943.RSP_1 | 0,2 | 0,4 | 6,32E-05 | 0,000126352 | 0 - | 0 - | 0 - | 0 - | 0 - | 0 - | 1 - | 0,00031588 - | 0 - | 0 - |
| 842 | 1 | 0.2um_NOKCPHJN_1096773 | Viruses | unclassified | unclassified | unclassified | unclassified | unclassified | uncultured M | information st | Replication, re | 12UUK | crossover junc | 7029.ACYPI00 | 0,2 | 0,4 | 6,66E-05 | 0,000133112 | 0 - | 0 - | 0 - | 0 - | 0 - | 0 - | 1 - | 0,000332779 - | 0 - | 0 - |
| 843 | 1 | 3um_FIJJAFBF_738679 | Bacteria | Bacteroidetes | Flavobacteriia | Flavobacterial | Flavobacteria | Formosa | Formosa sp. H | unclassified | unclassified | unclassified | unclassified | unclassified | 0,2 | 0,4 | 1,78E-05 | 3,55E-05 | 0 - | 0 - | 0 - | 0 - | 1 - | 8,88E-05 - | 0 - | 0 - | 0 - | 0 - |
| 844 | 1 | 3um_FIJJAFBF_501887 | Bacteria | Proteobacteri | Alphaproteob | Rhodobactera | Rhodobactera | Yoonia | Yoonia litorea | poorly charact | Function unkn | 11PVH | #NAME? | 391593.RCCS2 | 0,2 | 0,4 | 0,000106512 | 0,000213024 | 0 - | 0 - | 0 - | 0 - | 0 - | 0 - | 0 - | 0 - | 1 - | 0,000532559 - |
| 845 | 1 | 3um_FIJJAFBF_294668 | Bacteria | Bacteroidetes | Flavobacteriia | Flavobacterial | Flavobacteria | Algibacter | Algibacter sp. | unclassified | unclassified | unclassified | unclassified | unclassified | 1,2 | 0,4 | 0,00059649 | 0,000178305 | 1 - | 0,000506901 - | 1 - | 0,000596358 - | 2 - | 0,000940772 - | 1 - | 0,00046743 - | 1 - | 0,000470991 - |
| 846 | 1 | 3um_FIJJAFBF_720178 | Bacteria | Proteobacteri | Alphaproteob | Sphingomona | Erythrobacter | Porphyrobact | Porphyrobact | information st | Translation, ril | COG0050 | This protein p | 237727.NAP1 | 0,6 | 0,489897949 | 0,000134046 | 0,000111322 | 0 - | 0 - | 1 - | 0,00026053 - | 1 - | 0,000205497 - | 1 - | 0,000204205 - | 0 - | 0 - |
| 847 | 1 | 3um_FIJJAFBF_456402 | Eukaryota | Ascomycota | Leotiomycetes | unclassified | Pseudeurotiac | Pseudogymno | Pseudogymno | unclassified | unclassified | unclassified | unclassified | unclassified | 0,8 | 0,4 | 0,000212076 | 0,000108764 | 1 - | 0,000263345 - | 1 - | 0,000309819 - | 1 - | 0,000244375 - | 1 - | 0,000242839 - | 0 - | 0 - |
| 848 | 1 | 0.2um_NOKCPHJN_310259 | Bacteria | Proteobacteri | Gammaprotec | unclassified | unclassified | unclassified | Gammaprotec | unclassified | unclassified | unclassified | unclassified | unclassified | 0,2 | 0,4 | 7,21E-05 | 0,000144294 | 0 - | 0 - | 1 - | 0,000360734 - | 0 - | 0 - | 0 - | 0 - | 0 - | 0 - |
| 849 | 1 | 3um_FIJJAFBF_94430 | Bacteria | Proteobacteri | Alphaproteob | Pelagibacteral | Pelagibactera | Candidatus Pe | Candidatus Pe | unclassified | unclassified | unclassified | unclassified | unclassified | 0,2 | 0,4 | 2,71E-05 | 5,41E-05 | 1 - | 0,00013533 - | 0 - | 0 - | 0 - | 0 - | 0 - | 0 - | 0 - | 0 - |
| 850 | 1 | 0.2um_NOKCPHJN_116666 | Bacteria | Proteobacteri | Gammaprotec | Cellvibrionales | Porticoccacea | unclassified | Gammaprotec | metabolism | Inorganic ion t | 16TQC | TONB-depend | 314287.GB22 | 6,4 | 1,624807681 | 0,000566112 | 0,00013395 | 9 - | 0,000813654 - | 4 - | 0,000425443 - | 7 - | 0,000587255 - | 6 - | 0,000500198 - | 6 - | 0,000504009 - |
| 851 | 1 | 0.2um_NOKCPHJN_114165 | Bacteria | Bacteroidetes | Flavobacteriia | Flavobacterial | Flavobacteria | Formosa | Formosa sp. H | poorly charact | Function unkn | 06E9D | NA | 391598.FBBAL | 0,8 | 0,4 | 0,000116287 | 5,83E-05 | 1 - | 0,000153849 - | 0 - | 0 - | 1 - | 0,000142766 - | 1 - | 0,000141869 - | 1 - | 0,00014295 - |
| 852 | 1 | 0.2um_NOKCPHJN_249844 | Eukaryota | unclassified | Pelagophycea | Pelagomonad | unclassified | Aureococcus | Aureococcus a | poorly charact | Function unkn | 0XT7J | HYDIN, axone | 67593.Physo1 | 0,4 | 0,489897949 | 7,10E-06 | 8,69E-06 | 0 - | 0 - | 0 - | 0 - | 1 - | 1,77E-05 - | 0 - | 0 - | 1 - | 1,78E-05 - |
| 853 | 3 | 0.2um_NOKCPHJN_58032;3um | Eukaryota | unclassified | Chrysophycea | Chromulinales | Chromulinace | Ochromonas | Ochromonas s | unclassified | unclassified | unclassified | unclassified | unclassified | 0 | 0 | 0 | 0 | 0 - | 0 - | 0 - | 0 - | 0 - | 0 - | 0 - | 0 - | 0 - | 0 - |
| 854 | 1 | 3um_FIJJAFBF_92868 | Eukaryota | unclassified | Cryptophyta | Pyrenomonad | Pyrenomonad | Rhodomonas | Rhodomonas | unclassified | unclassified | unclassified | unclassified | unclassified | 1,2 | 0,748331477 | 0,000315831 | 0,000201303 | 2 - | 0,000553274 - | 0 - | 0 - | 1 - | 0,000256709 - | 1 - | 0,000255096 - | 2 - | 0,000514079 - |
| 855 | 2 | 0.2um_NOKCPHJN_152866;3un | Bacteria | Bacteroidetes | various | various | various | various | various | poorly charact | Function unkn | 09A4Z | rhodopsin | various | 0,4 | 0,489897949 | 0,000130925 | 0,000160351 | 0 - | 0 - | 0 - | 0 - | 0 - | 0 - | 1 - | 0,00032607 - | 1 - | 0,000328554 - |
| 856 | 1 | 0.2um_NOKCPHJN_975818 | Bacteria | Proteobacteri | Gammaprotec | Pseudomonad | Pseudomonad | Pseudomonas | various | unclassified | unclassified | unclassified | unclassified | unclassified | 0,6 | 0,489897949 | 0,000432294 | 0,000357391 | 1 - | 0,000695983 - | 1 - | 0,000818809 - | 0 - | 0 - | 0 - | 0 - | 1 - | 0,000646679 - |
| 857 | 1 | 3um_FIJJAFBF_235861 | Bacteria | Proteobacteri | Gammaprotec | Cellvibrionales | Microbulbifer | Microbulbifer | Microbulbifer | unclassified | unclassified | unclassified | unclassified | unclassified | 2,2 | 0,4 | 0,001275908 | 0,000210971 | 2 - | 0,001177099 - | 2 - | 0,001384831 - | 3 - | 0,001638458 - | 2 - | 0,001085441 - | 2 - | 0,001093711 - |
| 858 | 1 | 3um_FIJJAFBF_110706 | Bacteria | Bacteroidetes | Cytophagia | Cytophagales | Flammeovirga | unclassified | Flammeovirga | metabolism | Inorganic ion t | 05PWN | phosphate ab | 391615.GP501 | 1,4 | 0,489897949 | 0,000343994 | 0,000102926 | 1 - | 0,00025272 - | 1 - | 0,00029732 - | 2 - | 0,00046903 - | 2 - | 0,000466083 - | 1 - | 0,000234817 - |
| 859 | 1 | 3um_FIJJAFBF_194075 | Bacteria | Bacteroidetes | Flavobacteriia | Flavobacterial | Flavobacteria | Polaribacter | Polaribacter s | cellular proces | Posttranslatio | 07SFW | Trigger factor | 313598.MED1 | 0,2 | 0,4 | 3,68E-05 | 7,35E-05 | 0 - | 0 - | 0 - | 0 - | 0 - | 0 - | 1 - | 0,000183785 - | 0 - | 0 - |
| 860 | 1 | 0.2um_NOKCPHJN_546028 | Bacteria | Proteobacteri | Gammaprotec | unclassified | unclassified | unclassified | Gammaprotec | unclassified | unclassified | unclassified | unclassified | unclassified | 13 | 1,264911064 | 0,00134599 | 7,43E-05 | 13 - | 0,001368572 - | 11 - | 0,001362387 - | 15 - | 0,001465368 - | 13 - | 0,001262004 - | 13 - | 0,00127162 - |
| 861 | 1 | 0.2um_NOKCPHJN_546259 | Bacteria | Proteobacteri | Gammaprotec | unclassified | unclassified | unclassified | various | unclassified | unclassified | unclassified | unclassified | unclassified | 0,8 | 0,4 | 0,000265493 | 0,000136159 | 1 - | 0,000329676 - | 1 - | 0,000387857 - | 1 - | 0,000305928 - | 1 - | 0,000304005 - | 0 - | 0 - |
| 862 | 1 | 3um_FIJJAFBF_238480 | Bacteria | Proteobacteri | Gammaprotec | Oceanospirilla | Oceanospirilla | Pseudospirillu | Pseudospirillu | information st | Translation, ril | COG0050 | This protein p | 207954.MED9 | 2 | 0 | 0,000437871 | 4,28E-05 | 2 - | 0,000441783 - | 2 - | 0,000519748 - | 2 - | 0,000409958 - | 2 - | 0,000407382 - | 2 - | 0,000410486 - |
| 863 | 2 | 0.2um_NOKCPHJN_411418;0.2 | Bacteria | Proteobacteri | Alphaproteob | Rhodobactera | Rhodobactera | various | various | unclassified | unclassified | unclassified | unclassified | unclassified | 0,2 | 0,4 | 4,59E-05 | 9,19E-05 | 0 - | 0 - | 0 - | 0 - | 0 - | 0 - | 1 - | 0,000229731 - | 0 - | 0 - |
| 864 | 1 | 3um_FIJJAFBF_595134 | Eukaryota | Basidiomycota | Ustilaginomyc | Violaceomyce | Violaceomyce | Violaceomyce | Violaceomyce | unclassified | unclassified | unclassified | unclassified | unclassified | 0,2 | 0,4 | 0,00017154 | 0,00034308 | 0 - | 0 - | 0 - | 0 - | 0 - | 0 - | 0 - | 0 - | 1 - | 0,0008577 - |
| 865 | 3 | 0.2um_NOKCPHJN_1064534;3u | Bacteria | Proteobacteri | Gammaprotec | Cellvibrionales | various | various | various | information st | Translation, ril | COG0051 | tRNA binding | various | 0,6 | 0,489897949 | 0,000485516 | 0,0003971 | 1 - | 0,000851397 - | 0 - | 0 - | 0 - | 0 - | 1 - | 0,000785101 - | 1 - | 0,000791082 - |
| 866 | 1 | 3um_FIJJAFBF_552359 | Bacteria | Proteobacteri | Alphaproteob | Rhodobactera | Rhodobactera | unclassified | Rhodobactera | unclassified | unclassified | unclassified | unclassified | unclassified | 1,8 | 0,979795897 | 0,000814687 | 0,000493301 | 2 - | 0,000881345 - | 3 - | 0,001555325 - | 2 - | 0,000817857 - | 0 - | 0 - | 2 - | 0,000818909 - |
| 867 | 1 | 3um_FIJJAFBF_783366 | Eukaryota | Ascomycota | Sordariomyce | Glomerellales | Plectosphaere | Verticillium | Verticillium lo | cellular proces | Posttranslatio | COG1225 | alkyl hydroper | 391625.PPSIR | 0,2 | 0,4 | 5,02E-05 | 0,000100409 | 0 - | 0 - | 1 - | 0,000251022 - | 0 - | 0 - | 0 - | 0 - | 0 - | 0 - |
| 868 | 1 | 0.2um_NOKCPHJN_996696 | Bacteria | Proteobacteri | Alphaproteob | Rhodobactera | Rhodobactera | Ruegeria | Ruegeria sp. A | metabolism | Amino acid tra | COG0436 | aminotransfer | 467661.RKLH1 | 0,2 | 0,4 | 5,42E-05 | 0,000108315 | 0 - | 0 - | 1 - | 0,000270787 - | 0 - | 0 - | 0 - | 0 - | 0 - | 0 - |
| 869 | 1 | 3um_FIJJAFBF_302191 | Bacteria | Proteobacteri | Gammaprotec | Oceanospirilla | Saccharospiril | Reinekea | Reinekea fors | cellular proces | Signal transdu | 0R28X | 7TMR-DISM e | 314283.MED2 | 0,2 | 0,4 | 2,63E-05 | 5,26E-05 | 0 - | 0 - | 1 - | 0,000131427 - | 0 - | 0 - | 0 - | 0 - | 0 - | 0 - |
| 870 | 1 | 3um_FIJJAFBF_387035 | Bacteria | Proteobacteri | Alphaproteob | unclassified | unclassified | unclassified | Alphaproteob | unclassified | unclassified | unclassified | unclassified | unclassified | 11,6 | 0,8 | 0,003391488 | 0,000466166 | 12 - | 0,003531296 - | 12 - | 0,004154492 - | 12 - | 0,003276916 - | 10 - | 0,002713602 - | 12 - | 0,003281134 - |
| 871 | 1 | 0.2um_NOKCPHJN_260227 | Eukaryota | Phaeophyceae | unclassified | Ectocarpales | Ectocarpaceae | Ectocarpus | Ectocarpus sili | information st | Translation, ril | 09V1S | Sep (O-phosph | 8128.ENSONI | 0,4 | 0,489897949 | 9,76E-05 | 0,00012097 | 0 - | 0 - | 1 - | 0,000272936 - | 1 - | 0,000215282 - | 0 - | 0 - | 0 - | 0 - |
| 872 | 1 | 3um_FIJJAFBF_592170 | Bacteria | Bacteroidetes | Flavobacteriia | Flavobacterial | Crocinitomica | unclassified | Crocinitomica | unclassified | unclassified | unclassified | unclassified | unclassified | 1 | 0 | 0,000591275 | 5,79E-05 | 1 - | 0,000596557 - | 1 - | 0,000701836 - | 1 - | 0,000553583 - | 1 - | 0,000550104 - | 1 - | 0,000554296 - |
| 873 | 1 | 3um_FIJJAFBF_442115 | unclassified | unclassified | unclassified | unclassified | unclassified | unclassified | unclassified | unclassified | unclassified | unclassified | unclassified | unclassified | 1,8 | 0,4 | 0,000997213 | 0,000261688 | 2 - | 0,001110049 - | 2 - | 0,001305948 - | 1 - | 0,000515043 - | 2 - | 0,001023612 - | 2 - | 0,001031411 - |
| 874 | 1 | 3um_FIJJAFBF_348329 | Bacteria | Bacteroidetes | Flavobacteriia | Flavobacterial | Flavobacteria | Algibacter | Algibacter sp. | unclassified | unclassified | unclassified | unclassified | unclassified | 0,6 | 0,489897949 | 6,90E-05 | 5,73E-05 | 0 - | 0 - | 1 - | 0,000134161 - | 1 - | 0,000105822 - | 1 - | 0,000105157 - | 0 - | 0 - |
| 875 | 1 | 3um_FIJJAFBF_250215 | Bacteria | Bacteroidetes | Saprospiria | Saprospirales | Saprospiracea | Aureispira | Aureispira sp. | unclassified | unclassified | unclassified | unclassified | unclassified | 1,4 | 0,489897949 | 0,000498367 | 0,000217486 | 2 - | 0,000698756 - | 2 - | 0,000822071 - | 1 - | 0,00032421 - | 1 - | 0,000322173 - | 1 - | 0,000324627 - |
| 876 | 1 | 0.2um_NOKCPHJN_589037 | Eukaryota | Streptophyta | Liliopsida | Poales | Poaceae | Hordeum | Hordeum vulg | unclassified | unclassified | unclassified | unclassified | unclassified | 2 | 0 | 0,000569951 | 5,58E-05 | 2 - | 0,000575042 - | 2 - | 0,000676524 - | 2 - | 0,000533618 - | 2 - | 0,000530265 - | 2 - | 0,000534305 - |
| 877 | 1 | 3um_FIJJAFBF_315499 | Bacteria | Proteobacteri | Alphaproteob | unclassified | unclassified | unclassified | various | metabolism | Energy produc | 01RNF | Produces ATP | 269796.Rru_A | 1,6 | 0,489897949 | 0,000449213 | 0,000109977 | 1 - | 0,000290377 - | 1 - | 0,000341622 - | 2 - | 0,000538919 - | 2 - | 0,000535532 - | 2 - | 0,000539613 - |
| 878 | 1 | 3um_FIJJAFBF_574558 | Bacteria | Proteobacteri | Gammaprotec | Alteromonada | Pseudoalteror | Pseudoalteror | unclassified | cellular proces | Defense mech | COG2367 | Beta-lactamas | 351348.Maqu | 0,2 | 0,4 | 0,000130428 | 0,000260856 | 0 - | 0 - | 0 - | 0 - | 0 - | 0 - | 1 - | 0,00065214 - | 0 - | 0 - |
| 879 | 1 | 0.2um_NOKCPHJN_499401 | Eukaryota | unclassified | Labyrinthulom | unclassified | Thraustochytr | Aurantiochytr | Aurantiochytr | unclassified | unclassified | unclassified | unclassified | unclassified | 0,2 | 0,4 | 7,50E-05 | 0,000150065 | 0 - | 0 - | 1 - | 0,000375163 - | 0 - | 0 - | 0 - | 0 - | 0 - | 0 - |
| 880 | 1 | 0.2um_NOKCPHJN_552431 | Bacteria | Proteobacteri | Betaproteoba | Nitrosomonad | Sterolibacteri | Methyloversa | Methyloversa | unclassified | unclassified | unclassified | unclassified | unclassified | 3 | 2,28035085 | 0,000516364 | 0,000397151 | 7 - | 0,001232644 - | 1 - | 0,000207168 - | 4 - | 0,000653628 - | 2 - | 0,00032476 - | 1 - | 0,000163617 - |
| 881 | 1 | 3um_FIJJAFBF_182249 | Bacteria | Proteobacteri | Gammaprotec | Pseudomonad | Moraxellaceae | Psychrobacter | Psychrobacter | metabolism | Carbohydrate | COG0176 | Transaldolase | 335284.Pcryo | 0,2 | 0,4 | 5,18E-05 | 0,000103665 | 0 - | 0 - | 0 - | 0 - | 1 - | 0,000259162 - | 0 - | 0 - | 0 - | 0 - |
| 882 | 1 | 3um_FIJJAFBF_32479 | Eukaryota | Bacillariophyta | Bacillariophyc | unclassified | Entomoneidac | Entomoneis | Entomoneis sp | unclassified | unclassified | unclassified | unclassified | unclassified | 0,2 | 0,4 | 3,20E-05 | 6,40E-05 | 0 - | 0 - | 0 - | 0 - | 0 - | 0 - | 0 - | 0 - | 1 - | 0,000160082 - |
| 883 | 1 | 3um_FIJJAFBF_10655 | Bacteria | Bacteroidetes | Flavobacteriia | Flavobacterial | Flavobacteria | Flavobacteriu | Flavobacteriu | unclassified | unclassified | unclassified | unclassified | unclassified | 1,6 | 0,489897949 | 0,00025974 | 8,92E-05 | 1 - | 0,000162697 - | 2 - | 0,00038282 - | 2 - | 0,000301955 - | 2 - | 0,000300057 - | 1 - | 0,000151172 - |
| 884 | 1 | 3um_FIJJAFBF_715080 | Eukaryota | unclassified | Pelagophycea | Pelagomonad | unclassified | Pelagomonas | uncultured Pe | unclassified | unclassified | unclassified | unclassified | unclassified | 1 | 0 | 0,000755804 | 7,40E-05 | 1 - | 0,000762555 - | 1 - | 0,000897129 - | 1 - | 0,000707624 - | 1 - | 0,000703177 - | 1 - | 0,000708535 - |
| 885 | 1 | 0.2um_NOKCPHJN_48324 | Bacteria | Bacteroidetes | Cytophagia | Cytophagales | Flammeovirga | Reichenbachie | Reichenbachie | unclassified | unclassified | unclassified | unclassified | unclassified | 0,4 | 0,489897949 | 0,000208003 | 0,000254752 | 0 - | 0 - | 0 - | 0 - | 1 - | 0,000521646 - | 1 - | 0,000518368 - | 0 - | 0 - |
| 886 | 1 | 0.2um_NOKCPHJN_115345 | Bacteria | Bacteroidetes | Flavobacteriia | Flavobacterial | Flavobacteria | Olleya | Olleya namha | unclassified | unclassified | unclassified | unclassified | unclassified | 12,4 | 3,006659276 | 0,001150524 | 0,000212795 | 10 - | 0,000952159 - | 9 - | 0,001008175 - | 11 - | 0,000971926 - | 16 - | 0,001404827 - | 16 - | 0,001415531 - |
| 887 | 1 | 0.2um_NOKCPHJN_137064 | Bacteria | Bacteroidetes | Flavobacteriia | Flavobacterial | Crocinitomica | unclassified | Crocinitomica | unclassified | unclassified | unclassified | unclassified | unclassified | 0,2 | 0,4 | 9,91E-05 | 0,000198178 | 1 - | 0,000495445 - | 0 - | 0 - | 0 - | 0 - | 0 - | 0 - | 0 - | 0 - |
| 888 | 4 | 0.2um_NOKCPHJN_865894;3ur | Eukaryota | various | various | various | various | various | various | information st | various | various | various | various | 1 | 0 | 0,000234279 | 2,29E-05 | 1 - | 0,000236372 - | 1 - | 0,000278086 - | 1 - | 0,000219344 - | 1 - | 0,000217966 - | 1 - | 0,000219627 - |
| 889 | 1 | 0.2um_NOKCPHJN_379868 | Bacteria | Proteobacteri | Betaproteoba | unclassified | unclassified | unclassified | Betaproteoba | information st | Transcription | 16PMT | DNA-depende | 596154.Alide2 | 0,2 | 0,4 | 4,93E-05 | 9,86E-05 | 0 - | 0 - | 0 - | 0 - | 1 - | 0,000246596 - | 0 - | 0 - | 0 - | 0 - |
| 890 | 1 | 3um_FIJJAFBF_469955 | Bacteria | Proteobacteri | Alphaproteob | Rhodobactera | Rhodobactera | unclassified | Rhodobactera | metabolism | Amino acid tra | 01R4A | Polyamine AB | 388739.RSK20 | 1,8 | 0,4 | 0,000425875 | 0,000104302 | 1 - | 0,000238948 - | 2 - | 0,000562234 - | 2 - | 0,00044347 - | 2 - | 0,000440683 - | 2 - | 0,000444041 - |
| 891 | 2 | 3um_FIJJAFBF_583281;3um_FI | Eukaryota | unclassified | unclassified | Phaeocystales | Phaeocystace | Phaeocystis | various | information st | Translation, ril | 0FKC1 | This is 1 of the | various | 0,2 | 0,4 | 9,69E-05 | 0,000193799 | 1 - | 0,000484496 - | 0 - | 0 - | 0 - | 0 - | 0 - | 0 - | 0 - | 0 - |
| 892 | 1 | 3um_FIJJAFBF_767728 | Eukaryota | Chordata | Actinopteri | Cyprinodontif | Cyprinodontid | Cyprinodon | Cyprinodon va | unclassified | unclassified | unclassified | unclassified | unclassified | 0,6 | 0,489897949 | 9,96E-05 | 8,27E-05 | 0 - | 0 - | 1 - | 0,000193565 - | 1 - | 0,000152677 - | 1 - | 0,000151717 - | 0 - | 0 - |
| 893 | 1 | 0.2um_NOKCPHJN_127050 | Bacteria | Bacteroidetes | Flavobacteriia | Flavobacterial | Flavobacteria | Formosa | Formosa sp. H | metabolism | Secondary me | COG1228 | amidohydrola | 313603.FB217 | 0,4 | 0,489897949 | 0,000103736 | 0,000127052 | 0 - | 0 - | 0 - | 0 - | 0 - | 0 - | 1 - | 0,000258356 - | 1 - | 0,000260324 - |
| 894 | 1 | 3um_FIJJAFBF_262967 | Bacteria | unclassified | unclassified | unclassified | unclassified | unclassified | uncultured ba | metabolism | Carbohydrate | 16QEQ | hydrolase, fan | 983545.Glaag | 1,6 | 0,489897949 | 0,000333459 | 0,000120742 | 2 - | 0,00041365 - | 2 - | 0,00048665 - | 1 - | 0,000191926 - | 1 - | 0,00019072 - | 2 - | 0,000384347 - |
| 895 | 1 | 0.2um_NOKCPHJN_400179 | Bacteria | Proteobacteri | Gammaprotec | Oceanospirilla | Saccharospiril | Reinekea | Reinekea blan | unclassified | unclassified | unclassified | unclassified | unclassified | 0,8 | 0,4 | 0,000124359 | 6,24E-05 | 1 - | 0,000164529 - | 0 - | 0 - | 1 - | 0,000152677 - | 1 - | 0,000151717 - | 1 - | 0,000152873 - |
| 896 | 1 | 3um_FIJJAFBF_269831 | Eukaryota | unclassified | Oomycetes | Saprolegniales | Saprolegniace | various | various | unclassified | unclassified | unclassified | unclassified | unclassified | 5,6 | 1,496662955 | 0,001262061 | 0,000270579 | 6 - | 0,001395658 - | 3 - | 0,000820981 - | 5 - | 0,001079267 - | 7 - | 0,001501479 - | 7 - | 0,001512919 - |
| 897 | 1 | 0.2um_NOKCPHJN_108842 | Bacteria | Proteobacteri | Alphaproteob | Rhodobactera | Rhodobactera | Ruegeria | Ruegeria sp. 6 | metabolism | Energy produc | 16Q0H | fad dependen | 391619.RGBS1 | 0,2 | 0,4 | 3,75E-05 | 7,51E-05 | 0 - | 0 - | 0 - | 0 - | 0 - | 0 - | 0 - | 0 - | 1 - | 0,000187745 - |
| 898 | 2 | 0.2um_NOKCPHJN_968216;3ur | Eukaryota | various | various | various | various | various | various | information st | Chromatin str | various | various | various | 0,6 | 0,489897949 | 0,000353559 | 0,000292329 | 1 - | 0,000569441 - | 1 - | 0,000669934 - | 1 - | 0,00052842 - | 0 - | 0 - | 0 - | 0 - |
| 899 | 1 | 3um_FIJJAFBF_48832 | Eukaryota | unclassified | Chrysophycea | Chromulinales | Chromulinace | Ochromonas | Ochromonas s | information st | Transcription | 05D27 | DNA-depende | 118168.MC74 | 1 | 0 | 0,000126887 | 1,24E-05 | 1 - | 0,00012802 - | 1 - | 0,000150613 - | 1 - | 0,000118798 - | 1 - | 0,000118052 - | 1 - | 0,000118951 - |
| 900 | 1 | 3um_FIJJAFBF_691249 | Eukaryota | unclassified | Chrysophycea | Chromulinales | Chromulinace | Ochromonas | Ochromonas s | information st | Translation, ril | COG0088 | One of the pri | 756067.Micva | 0,2 | 0,4 | 8,73E-05 | 0,000174515 | 1 - | 0,000436288 - | 0 - | 0 - | 0 - | 0 - | 0 - | 0 - | 0 - | 0 - |
| 901 | 1 | 3um_FIJJAFBF_545217 | Bacteria | Bacteroidetes | Flavobacteriia | Flavobacterial | Flavobacteria | Arenitalea | Arenitalea lut | information st | Translation, ril | COG0256 | This is one of | 983544.Lacal | 1 | 0 | 0,000736589 | 7,21E-05 | 1 - | 0,000743168 - | 1 - | 0,000874321 - | 1 - | 0,000689633 - | 1 - | 0,0006853 - | 1 - | 0,000690521 - |
| 902 | 1 | 3um_FIJJAFBF_788032 | Bacteria | Proteobacteri | Alphaproteob | Rhodobactera | Rhodobactera | unclassified | Rhodobactera | information st | Translation, ril | 179ZI | One of the pri | 52598.EE36_0 | 1,4 | 0,489897949 | 0,000928704 | 0,000311352 | 2 - | 0,001349136 - | 1 - | 0,000793615 - | 2 - | 0,00125195 - | 1 - | 0,000622041 - | 1 - | 0,000626781 - |
| 903 | 3 | 0.2um_NOKCPHJN_202118;0.2 | Viruses | unclassified | unclassified | unclassified | unclassified | unclassified | uncultured M | unclassified | unclassified | unclassified | unclassified | unclassified | 1 | 0 | 0,000496671 | 4,86E-05 | 1 - | 0,000501108 - | 1 - | 0,000589542 - | 1 - | 0,00046501 - | 1 - | 0,000462088 - | 1 - | 0,000465609 - |
| 904 | 1 | 3um_FIJJAFBF_303505 | unclassified | unclassified | unclassified | unclassified | unclassified | unclassified | unclassified | unclassified | unclassified | unclassified | unclassified | unclassified | 0,8 | 0,4 | 0,000279677 | 0,000140238 | 1 - | 0,000370016 - | 0 - | 0 - | 1 - | 0,000343362 - | 1 - | 0,000341204 - | 1 - | 0,000343804 - |
| 905 | 1 | 3um_FIJJAFBF_348506 | Bacteria | Bacteroidetes | Saprospiria | Saprospirales | Saprospiracea | Saprospira | Saprospira gra | poorly charact | Function unkn | 0XSVM | NA | 926562.Oweh | 0,8 | 0,4 | 0,000143874 | 7,38E-05 | 1 - | 0,000178603 - | 1 - | 0,000210122 - | 0 - | 0 - | 1 - | 0,000164695 - | 1 - | 0,00016595 - |
| 906 | 1 | 3um_FIJJAFBF_78500 | Bacteria | Proteobacteri | Alphaproteob | Rhodobactera | Rhodobactera | Jannaschia | Jannaschia fae | cellular proces | Posttranslatio | 17DF7 | Glutathione S | 203122.Sde_3 | 0,2 | 0,4 | 7,02E-05 | 0,000140305 | 0 - | 0 - | 0 - | 0 - | 1 - | 0,000350762 - | 0 - | 0 - | 0 - | 0 - |
| 907 | 1 | 3um_FIJJAFBF_69734 | Bacteria | Proteobacteri | Gammaprotec | Pseudomonad | Pseudomonad | Pseudomonas | various | unclassified | unclassified | unclassified | unclassified | unclassified | 12,8 | 1,6 | 0,00227694 | 0,000485718 | 13 - | 0,002312414 - | 15 - | 0,003139043 - | 14 - | 0,002310901 - | 11 - | 0,001804298 - | 11 - | 0,001818045 - |
| 908 | 2 | 0.2um_NOKCPHJN_160356;3ur | Eukaryota | unclassified | Pelagophycea | Pelagomonad | unclassified | Pelagomonas | uncultured Pe | unclassified | unclassified | unclassified | unclassified | unclassified | 3,4 | 0,8 | 0,001198602 | 0,000198028 | 3 - | 0,001091625 - | 2 - | 0,000856182 - | 4 - | 0,001350651 - | 4 - | 0,001342164 - | 4 - | 0,00135239 - |
| 909 | 1 | 0.2um_NOKCPHJN_278109 | Eukaryota | unclassified | Bangiophycea | Bangiales | Bangiaceae | Porphyra | Porphyra umb | unclassified | unclassified | unclassified | unclassified | unclassified | 0,4 | 0,489897949 | 1,17E-05 | 1,43E-05 | 1 - | 3,04E-05 - | 0 - | 0 - | 0 - | 0 - | 1 - | 2,80E-05 - | 0 - | 0 - |
| 910 | 1 | 3um_FIJJAFBF_662957 | Bacteria | Bacteroidetes | Flavobacteriia | Flavobacterial | Crocinitomica | unclassified | Crocinitomica | information st | Translation, ril | 05KK9 | This protein b | 376686.Fjoh | 0,8 | 0,4 | 0,000535168 | 0,000274429 | 1 - | 0,000664347 - | 1 - | 0,00078159 - | 0 - | 0 - | 1 - | 0,000612616 - | 1 - | 0,000617284 - |
| 911 | 1 | 3um_FIJJAFBF_348326 | Bacteria | Bacteroidetes | Flavobacteriia | Flavobacterial | Flavobacteria | Algibacter | Algibacter lect | poorly charact | Function unkn | 0P8NI | Inherit from b | 313594.PI23P | 2,2 | 0,748331477 | 0,000280901 | 8,73E-05 | 1 - | 0,000130497 - | 2 - | 0,000307053 - | 2 - | 0,000242193 - | 3 - | 0,000361006 - | 3 - | 0,000363757 - |
| 912 | 1 | 3um_FIJJAFBF_328499 | Bacteria | Bacteroidetes | Flavobacteriia | Flavobacterial | Flavobacteria | Arenitalea | Arenitalea lut | information st | Translation, ril | 08UHY | One of the pri | 1046627.BZAR | 2,4 | 0,8 | 0,001286781 | 0,000386935 | 3 - | 0,001665073 - | 1 - | 0,000652974 - | 3 - | 0,001545128 - | 2 - | 0,001023612 - | 3 - | 0,001547117 - |
| 913 | 1 | 3um_FIJJAFBF_390243 | Eukaryota | unclassified | Chrysophycea | Chromulinales | Chromulinace | Ochromonas | Ochromonas c | unclassified | unclassified | unclassified | unclassified | unclassified | 1,8 | 0,4 | 0,000242791 | 4,04E-05 | 2 - | 0,000277952 - | 1 - | 0,000163502 - | 2 - | 0,000257929 - | 2 - | 0,000256309 - | 2 - | 0,000258261 - |
| 914 | 1 | 3um_FIJJAFBF_254285 | Bacteria | Proteobacteri | Gammaprotec | Cellvibrionales | Spongiibacter | Oceanicoccus | Oceanicoccus | information st | Translation, ril | COG0087 | One of the pri | 247633.GP214 | 1,2 | 1,166190379 | 0,000531233 | 0,000551833 | 2 - | 0,000823416 - | 3 - | 0,001453097 - | 0 - | 0 - | 1 - | 0,00037965 - | 0 - | 0 - |
| 915 | 1 | 3um_FIJJAFBF_492315 | Eukaryota | unclassified | Cryptophyta | Pyrenomonad | Geminigerace | Teleaulax | Teleaulax amp | unclassified | unclassified | unclassified | unclassified | unclassified | 2,4 | 0,489897949 | 0,001165443 | 0,000188775 | 2 - | 0,000990891 - | 2 - | 0,001165761 - | 3 - | 0,001379267 - | 3 - | 0,001370599 - | 2 - | 0,000920695 - |
| 916 | 1 | 3um_FIJJAFBF_785179 | Eukaryota | Euglenida | unclassified | Euglenales | Euglenaceae | Trachelomona | Trachelomona | cellular proces | Posttranslatio | 0IEKR | One of the co | 3055.DAA009 | 1 | 0 | 0,000452695 | 4,43E-05 | 1 - | 0,000456739 - | 1 - | 0,000537343 - | 1 - | 0,000423837 - | 1 - | 0,000421174 - | 1 - | 0,000424383 - |
| 917 | 1 | 0.2um_NOKCPHJN_97273 | Bacteria | Proteobacteri | Gammaprotec | Alteromonada | Alteromonada | Paraglaciecola | Paraglaciecola | metabolism | Carbohydrate | 0QJ3A | Catalyzes the | 342610.Patl_1 | 0,2 | 0,4 | 2,06E-05 | 4,12E-05 | 0 - | 0 - | 0 - | 0 - | 1 - | 0,000103009 - | 0 - | 0 - | 0 - | 0 - |
| 918 | 1 | 3um_FIJJAFBF_348971 | Bacteria | Bacteroidetes | Flavobacteriia | Flavobacterial | Flavobacteria | Kordia | Kordia algicida | unclassified | unclassified | unclassified | unclassified | unclassified | 0,8 | 0,4 | 6,50E-05 | 3,26E-05 | 1 - | 8,60E-05 - | 0 - | 0 - | 1 - | 7,98E-05 - | 1 - | 7,93E-05 - | 1 - | 7,99E-05 - |
| 919 | 1 | 3um_FIJJAFBF_369614 | Bacteria | Bacteroidetes | Flavobacteriia | Flavobacterial | Flavobacteria | Winogradskye | Winogradskye | unclassified | unclassified | unclassified | unclassified | unclassified | 1,6 | 0,489897949 | 0,000671595 | 0,000164421 | 1 - | 0,000434128 - | 1 - | 0,000510742 - | 2 - | 0,00080571 - | 2 - | 0,000800647 - | 2 - | 0,000806747 - |
| 920 | 1 | 0.2um_NOKCPHJN_102880 | Bacteria | Proteobacteri | Alphaproteob | unclassified | unclassified | unclassified | SAR116 cluste | information st | Transcription | 05D27 | DNA-depende | 488538.SAR11 | 0,4 | 0,489897949 | 2,32E-05 | 2,84E-05 | 0 - | 0 - | 0 - | 0 - | 1 - | 5,82E-05 - | 1 - | 5,78E-05 - | 0 - | 0 - |
| 921 | 1 | 0.2um_NOKCPHJN_588085 | unclassified | unclassified | unclassified | unclassified | unclassified | unclassified | unclassified | unclassified | unclassified | unclassified | unclassified | unclassified | 0,4 | 0,489897949 | 6,55E-05 | 8,11E-05 | 0 - | 0 - | 1 - | 0,000182925 - | 0 - | 0 - | 0 - | 0 - | 1 - | 0,000144471 - |
| 922 | 1 | 3um_FIJJAFBF_300355 | Eukaryota | Apicomplexa | Conoidasida | Eucoccidiorida | Eimeriidae | Eimeria | Eimeria acerv | unclassified | unclassified | unclassified | unclassified | unclassified | 5,2 | 0,4 | 0,00160899 | 0,000200514 | 6 - | 0,001872467 - | 5 - | 0,001835763 - | 5 - | 0,001447985 - | 5 - | 0,001438885 - | 5 - | 0,001449849 - |
| 923 | 1 | 0.2um_NOKCPHJN_485486 | Bacteria | Proteobacteri | Gammaprotec | Cellvibrionales | Porticoccacea | unclassified | Gammaprotec | cellular proces | Posttranslatio | COG2844 | Modifies, by u | 314287.GB22 | 0,8 | 0,4 | 7,81E-05 | 4,01E-05 | 1 - | 9,70E-05 - | 1 - | 0,000114126 - | 1 - | 9,00E-05 - | 1 - | 8,95E-05 - | 0 - | 0 - |
| 924 | 1 | 0.2um_NOKCPHJN_747889 | Bacteria | Proteobacteri | Gammaprotec | Cellvibrionales | Spongiibacter | unclassified | marine gamm | metabolism | Inorganic ion t | 0QK8T | TonB-depende | 247633.GP214 | 4,6 | 1,019803903 | 0,000437858 | 7,43E-05 | 5 - | 0,000488273 - | 3 - | 0,000344666 - | 5 - | 0,0004531 - | 6 - | 0,000540303 - | 4 - | 0,000362947 - |
| 925 | 1 | 0.2um_NOKCPHJN_39143 | Eukaryota | unclassified | Pelagophycea | Pelagomonad | unclassified | Aureococcus | Aureococcus a | metabolism | Carbohydrate | COG1850 | RuBisCO catal | 45157.CMV01 | 15,8 | 0,979795897 | 0,002805134 | 0,000223658 | 15 - | 0,002695508 - | 15 - | 0,003171206 - | 17 - | 0,002834846 - | 15 - | 0,002485615 - | 17 - | 0,002838495 - |
| 926 | 1 | 3um_FIJJAFBF_531533 | Bacteria | unclassified | unclassified | unclassified | unclassified | unclassified | bacterium BM | unclassified | unclassified | unclassified | unclassified | unclassified | 0,4 | 0,489897949 | 6,96E-05 | 8,53E-05 | 0 - | 0 - | 0 - | 0 - | 1 - | 0,000174628 - | 1 - | 0,000173531 - | 0 - | 0 - |
| 927 | 1 | 3um_FIJJAFBF_33368 | Bacteria | Proteobacteri | Alphaproteob | Rhodobactera | Rhodobactera | Roseobacter | Roseobacter s | information st | Replication, re | 01QWN | helicase RecD | 351016.RAZW | 1 | 0 | 0,000118094 | 1,16E-05 | 1 - | 0,000119149 - | 1 - | 0,000140176 - | 1 - | 0,000110566 - | 1 - | 0,000109871 - | 1 - | 0,000110709 - |
| 928 | 1 | 3um_FIJJAFBF_195099 | Eukaryota | unclassified | unclassified | Prymnesiales | Chrysochromu | Chrysochromu | Chrysochromu | unclassified | unclassified | unclassified | unclassified | unclassified | 0,8 | 0,4 | 0,000340092 | 0,000175071 | 0 - | 0 - | 1 - | 0,000505735 - | 1 - | 0,000398906 - | 1 - | 0,000396399 - | 1 - | 0,000399419 - |
| 929 | 1 | 3um_FIJJAFBF_673918 | Eukaryota | unclassified | Oomycetes | Peronosporale | Peronosporac | Phytophthora | various | unclassified | unclassified | unclassified | unclassified | unclassified | 2,8 | 0,748331477 | 0,000659327 | 0,000148369 | 3 - | 0,000722752 - | 2 - | 0,000566868 - | 3 - | 0,000670687 - | 4 - | 0,00088863 - | 2 - | 0,000447701 - |
| 930 | 1 | 0.2um_NOKCPHJN_103568 | Bacteria | Proteobacteri | Gammaprotec | Alteromonada | Alteromonada | Paraglaciecola | Paraglaciecola | unclassified | unclassified | unclassified | unclassified | unclassified | 0,8 | 0,4 | 0,000232574 | 0,000116619 | 1 - | 0,000307698 - | 0 - | 0 - | 1 - | 0,000285532 - | 1 - | 0,000283738 - | 1 - | 0,0002859 - |
| 931 | 1 | 0.2um_NOKCPHJN_383112 | Bacteria | Bacteroidetes | unclassified | unclassified | unclassified | unclassified | Bacteroidetes | information st | Translation, ril | 09AI8 | Associates wit | 487796.Flav2A | 0,2 | 0,4 | 6,91E-05 | 0,000138231 | 0 - | 0 - | 0 - | 0 - | 0 - | 0 - | 1 - | 0,000345578 - | 0 - | 0 - |
| 932 | 1 | 3um_FIJJAFBF_19624 | Bacteria | Proteobacteri | Gammaprotec | Cellvibrionales | Porticoccacea | unclassified | gamma protec | metabolism | Inorganic ion t | 16TQC | TONB-depend | 314287.GB22 | 1,8 | 0,979795897 | 0,00015067 | 7,95E-05 | 3 - | 0,000256415 - | 1 - | 0,000100555 - | 1 - | 7,93E-05 - | 1 - | 7,88E-05 - | 3 - | 0,00023825 - |
| 933 | 1 | 3um_FIJJAFBF_305344 | Eukaryota | unclassified | Dinophyceae | Gonyaulacales | unclassified | Lingulodinium | Lingulodinium | poorly charact | Function unkn | 0XRVT | NA | 35128.Thaps3 | 0,8 | 0,4 | 0,000191126 | 9,84E-05 | 0 - | 0 - | 1 - | 0,000284215 - | 1 - | 0,000224178 - | 1 - | 0,00022277 - | 1 - | 0,000224467 - |
| 934 | 6 | 0.2um_NOKCPHJN_859809;3ur | Eukaryota | various | various | various | various | various | various | unclassified | unclassified | unclassified | unclassified | unclassified | 1 | 0 | 0,002414374 | 0,000236232 | 1 - | 0,00243594 - | 1 - | 0,00286583 - | 1 - | 0,002260465 - | 1 - | 0,00224626 - | 1 - | 0,002263375 - |
| 935 | 1 | 0.2um_NOKCPHJN_78103 | Viruses | unclassified | unclassified | unclassified | Lavidaviridae | Sputnikvirus | Mimivirus-dep | unclassified | unclassified | unclassified | unclassified | unclassified | 0,8 | 0,4 | 0,000161605 | 8,29E-05 | 1 - | 0,000200672 - | 1 - | 0,000236087 - | 1 - | 0,000186217 - | 1 - | 0,000185047 - | 0 - | 0 - |
| 936 | 1 | 0.2um_NOKCPHJN_1079676 | Bacteria | Proteobacteri | Gammaprotec | Alteromonada | Pseudoalteror | Pseudoalteror | Pseudoalteror | unclassified | unclassified | unclassified | unclassified | unclassified | 1,2 | 0,4 | 0,000198586 | 6,93E-05 | 2 - | 0,000333437 - | 1 - | 0,00019614 - | 1 - | 0,000154709 - | 1 - | 0,000153736 - | 1 - | 0,000154908 - |
| 937 | 1 | 0.2um_NOKCPHJN_128221 | Bacteria | Bacteroidetes | Cytophagia | Cytophagales | Amoebophilac | Candidatus Ar | Candidatus Ar | unclassified | unclassified | unclassified | unclassified | unclassified | 0,6 | 0,489897949 | 4,16E-05 | 3,40E-05 | 0 - | 0 - | 0 - | 0 - | 1 - | 6,94E-05 - | 1 - | 6,90E-05 - | 1 - | 6,95E-05 - |
| 938 | 1 | 0.2um_NOKCPHJN_341818 | Bacteria | Verrucomicrol | Verrucomicrol | Verrucomicrol | Verrucomicrol | Haloferula | Haloferula sp. | unclassified | unclassified | unclassified | unclassified | unclassified | 0,4 | 0,8 | 7,98E-05 | 0,000159562 | 0 - | 0 - | 0 - | 0 - | 2 - | 0,000398906 - | 0 - | 0 - | 0 - | 0 - |
| 939 | 1 | 0.2um_NOKCPHJN_139936 | Bacteria | Bacteroidetes | unclassified | unclassified | unclassified | unclassified | Bacteroidetes | metabolism | Inorganic ion t | 05C27 | TonB-depende | 983544.Lacal | 0,6 | 0,489897949 | 4,77E-05 | 3,89E-05 | 0 - | 0 - | 0 - | 0 - | 1 - | 7,96E-05 - | 1 - | 7,91E-05 - | 1 - | 7,97E-05 - |
| 940 | 1 | 3um_FIJJAFBF_92098 | Bacteria | Proteobacteri | Gammaprotec | unclassified | unclassified | unclassified | Gammaprotec | unclassified | unclassified | unclassified | unclassified | unclassified | 1 | 0 | 5,66E-05 | 5,54E-06 | 1 - | 5,71E-05 - | 1 - | 6,72E-05 - | 1 - | 5,30E-05 - | 1 - | 5,27E-05 - | 1 - | 5,31E-05 - |
| 941 | 1 | 0.2um_NOKCPHJN_40987 | Bacteria | Actinobacteria | Actinobacteria | Micrococcales | Microbacteria | Candidatus Ac | Candidatus Ac | unclassified | unclassified | unclassified | unclassified | unclassified | 1 | 0 | 0,00024211 | 2,37E-05 | 1 - | 0,000244273 - | 1 - | 0,000287381 - | 1 - | 0,000226676 - | 1 - | 0,000225252 - | 1 - | 0,000226968 - |
| 942 | 1 | 3um_FIJJAFBF_539671 | Bacteria | unclassified | unclassified | unclassified | unclassified | unclassified | bacterium (Ca | cellular proces | Posttranslatio | 066HJ | ATP-depender | 445970.ALIPU | 0,8 | 0,4 | 0,000164795 | 8,48E-05 | 0 - | 0 - | 1 - | 0,000245059 - | 1 - | 0,000193294 - | 1 - | 0,000192079 - | 1 - | 0,000193543 - |
| 943 | 1 | 0.2um_NOKCPHJN_21852 | Bacteria | Bacteroidetes | Flavobacteriia | Flavobacterial | Flavobacteria | Polaribacter | Polaribacter s | poorly charact | Function unkn | 0PBET | NA | 313594.PI23P | 0,2 | 0,4 | 9,97E-05 | 0,000199304 | 1 - | 0,000498261 - | 0 - | 0 - | 0 - | 0 - | 0 - | 0 - | 0 - | 0 - |
| 944 | 11 | 0.2um_NOKCPHJN_1011604;0. | Viruses | unclassified | unclassified | unclassified | unclassified | unclassified | uncultured M | unclassified | unclassified | unclassified | unclassified | unclassified | 1 | 0 | 0,000474959 | 4,65E-05 | 1 - | 0,000479201 - | 1 - | 0,00056377 - | 1 - | 0,000444682 - | 1 - | 0,000441887 - | 1 - | 0,000445254 - |
| 945 | 1 | 3um_FIJJAFBF_788036 | Bacteria | Proteobacteri | Alphaproteob | Rhodobactera | Rhodobactera | Pseudophaeol | Pseudophaeol | unclassified | unclassified | unclassified | unclassified | unclassified | 1 | 0 | 0,000712438 | 6,97E-05 | 1 - | 0,000718802 - | 1 - | 0,000845655 - | 1 - | 0,000667022 - | 1 - | 0,000662831 - | 1 - | 0,000667881 - |
| 946 | 1 | 3um_FIJJAFBF_251385 | Bacteria | Bacteroidetes | Flavobacteriia | Flavobacterial | Flavobacteria | Polaribacter | Polaribacter s | unclassified | unclassified | unclassified | unclassified | unclassified | 3 | 0,632455532 | 0,000541842 | 8,10E-05 | 3 - | 0,000556198 - | 2 - | 0,000436236 - | 3 - | 0,000516132 - | 4 - | 0,000683851 - | 3 - | 0,000516796 - |
| 947 | 1 | 3um_FIJJAFBF_411697 | unclassified | unclassified | unclassified | unclassified | unclassified | unclassified | unclassified | unclassified | unclassified | unclassified | unclassified | unclassified | 0,8 | 0,4 | 0,000630733 | 0,000323435 | 1 - | 0,000782981 - | 1 - | 0,00092116 - | 0 - | 0 - | 1 - | 0,000722012 - | 1 - | 0,000727513 - |
| 948 | 1 | 3um_FIJJAFBF_579113 | Bacteria | Bacteroidetes | Flavobacteriia | Flavobacterial | unclassified | unclassified | Flavobacterial | metabolism | Energy produc | 0PBIY | subunit C | 926562.Oweh | 1 | 0 | 0,001259673 | 0,000123252 | 1 - | 0,001270925 - | 1 - | 0,001495216 - | 1 - | 0,001179373 - | 1 - | 0,001171962 - | 1 - | 0,001180891 - |
| 949 | 1 | 3um_FIJJAFBF_545215 | Bacteria | Bacteroidetes | Flavobacteriia | Flavobacterial | Flavobacteria | Flavivirga | Flavivirga eckl | unclassified | unclassified | unclassified | unclassified | unclassified | 2 | 0 | 0,001316931 | 0,000128854 | 2 - | 0,001328695 - | 2 - | 0,00156318 - | 2 - | 0,001232981 - | 2 - | 0,001225233 - | 2 - | 0,001234568 - |
| 950 | 1 | 0.2um_NOKCPHJN_66824 | Bacteria | Proteobacteri | Gammaprotec | Alteromonada | Pseudoalteror | Pseudoalteror | Pseudoalteror | metabolism | Carbohydrate | 16QEQ | hydrolase, fan | 247634.GPB2 | 0,6 | 0,489897949 | 5,00E-05 | 4,09E-05 | 1 - | 8,76E-05 - | 0 - | 0 - | 0 - | 0 - | 1 - | 8,08E-05 - | 1 - | 8,14E-05 - |
| 951 | 1 | 0.2um_NOKCPHJN_01012 | Bacteria | Bacteroidetes | Flavobacteriia | Flavobacterial | Flavobacteria | Formosa | Formosa sp. H | unclassified | unclassified | unclassified | unclassified | unclassified | 0,6 | 0,489897949 | 5,20E-05 | 4,25E-05 | 1 - | 9,13E-05 - | 0 - | 0 - | 1 - | 8,47E-05 - | 1 - | 8,41E-05 - | 0 - | 0 - |
| 952 | 1 | 3um_FIJJAFBF_135237 | Bacteria | Cyanobacteria | unclassified | Synechococca | Synechococca | Synechococcu | Synechococcu | metabolism | Energy produc | COG1290 | Component of | 70448.Q0P3F | 0,8 | 0,4 | 0,000173517 | 8,70E-05 | 1 - | 0,000229565 - | 0 - | 0 - | 1 - | 0,000213028 - | 1 - | 0,000211689 - | 1 - | 0,000213302 - |
| 953 | 1 | 3um_FIJJAFBF_412012 | Bacteria | Proteobacteri | Gammaprotec | Alteromonada | Alteromonada | unclassified | Alteromonada | information st | Translation, ril | COG1185 | Involved in m | 377629.TERTU | 0,8 | 0,748331477 | 9,49E-05 | 8,77E-05 | 1 - | 0,000125456 - | 0 - | 0 - | 0 - | 0 - | 1 - | 0,000115687 - | 2 - | 0,000233137 - |
| 954 | 1 | 3um_FIJJAFBF_537191 | Eukaryota | unclassified | unclassified | Isochrysidales | Noelaerhabda | Emiliania | Emiliania huxl | poorly charact | Function unkn | 0XSKP | von Willebran | 65071.PYU1_ | 0,2 | 0,4 | 1,52E-05 | 3,04E-05 | 0 - | 0 - | 0 - | 0 - | 1 - | 7,59E-05 - | 0 - | 0 - | 0 - | 0 - |

| | | | | | | | | | | | | | | | | | | | | | | | | | | | | |
|---|---|---|---|---|---|---|---|---|---|---|---|---|---|---|---|---|---|---|---|---|---|---|---|---|---|---|---|---|
| 955 | 1 | 0.2um_NOKCPHJN_698224 | Eukaryota | Bacillariophyta | Coscinodiscop | Thalassiosirale | Thalassiosirac | Thalassiosira | Thalassiosira c | metabolism | Amino acid tra | COG0010 | hydrolase acti | 2850.Phatr385 | 0,6 | 0,489897949 | 0,0001417 | 0,000115699 | 0 - | 0 - | 0 - | 0 - | 1 - | 0,00023656 - | 1 - | 0,000235074 - | 1 - | 0,000236865 - |
| 956 | 1 | 0.2um_NOKCPHJN_1061905 | unclassified | unclassified | unclassified | unclassified | unclassified | unclassified | unclassified | unclassified | unclassified | unclassified | unclassified | unclassified | 1 | 0 | 0,000436771 | 4,27E-05 | 1 - | 0,000440673 - | 1 - | 0,000518442 - | 1 - | 0,000408928 - | 1 - | 0,000406359 - | 1 - | 0,000409455 - |
| 957 | 1 | 0.2um_NOKCPHJN_683378 | Eukaryota | Phaeophyceae | unclassified | Ectocarpales | Ectocarpaceae | Ectocarpus | Ectocarpus sili | unclassified | unclassified | unclassified | unclassified | unclassified | 0,8 | 0,4 | 8,25E-05 | 4,14E-05 | 1 - | 0,000109208 - | 0 - | 0 - | 1 - | 0,000101341 - | 1 - | 0,000100704 - | 1 - | 0,000101471 - |
| 958 | 1 | 3um_FIJJAFBF_261098 | Bacteria | Bacteroidetes | Flavobacteriia | Flavobacterial | Flavobacteriac | Lutibacter | Lutibacter aga | metabolism | Amino acid tra | 0P7S1 | Catalyzes the | 376686.Fjoh_ | 1 | 0 | 0,000296647 | 2,90E-05 | 1 - | 0,000299296 - | 1 - | 0,000352116 - | 1 - | 0,000277736 - | 1 - | 0,000275991 - | 1 - | 0,000278094 - |
| 959 | 3 | 0.2um_NOKCPHJN_175137;0.2 | Bacteria | Bacteroidetes | unclassified | unclassified | unclassified | unclassified | Bacteroidetes | unclassified | unclassified | unclassified | unclassified | unclassified | 0,2 | 0,4 | 1,80E-05 | 3,60E-05 | 1 - | 8,99E-05 - | 0 - | 0 - | 0 - | 0 - | 0 - | 0 - | 0 - | 0 - |
| 960 | 2 | 0.2um_NOKCPHJN_546865;3ur | Bacteria | Bacteroidetes | Cytophagia | Cytophagales | Flammeovirga | unclassified | Flammeovirga | cellular proces | Cell wall/mem | 0QKXZ | Porin Gram-ne | 247633.GP214 | 3,4 | 0,8 | 0,000921037 | 0,000175076 | 4 - | 0,001113573 - | 2 - | 0,000655047 - | 3 - | 0,000775017 - | 4 - | 0,001026862 - | 4 - | 0,001034686 - |
| 961 | 1 | 3um_FIJJAFBF_715082 | Eukaryota | unclassified | Pelagophyceae | Pelagomonada | unclassified | Aureoumbra | Aureoumbra l | unclassified | unclassified | unclassified | unclassified | unclassified | 1,2 | 0,748331477 | 0,000606779 | 0,000386746 | 2 - | 0,001062956 - | 0 - | 0 - | 1 - | 0,000493192 - | 1 - | 0,000490093 - | 2 - | 0,000987654 - |
| 962 | 1 | 3um_FIJJAFBF_305779 | Eukaryota | Bacillariophyta | Bacillariophyc | Bacillariales | Bacillariaceae | Fragilariopsis | Fragilariopsis | poorly charact | Function unkn | 1DV37 | plastid-lipid-a | 436017.A4RTN | 0,2 | 0,4 | 0,000158723 | 0,000317446 | 0 - | 0 - | 1 - | 0,000793615 - | 0 - | 0 - | 0 - | 0 - | 0 - | 0 - |
| 963 | 1 | 0.2um_NOKCPHJN_155316 | Eukaryota | unclassified | unclassified | Trichomonadi | Trichomonadi | Trichomonas | Trichomonas v | cellular proces | Signal transdu | KOG0613 | fibronectin typ | 7070.TC00296 | 0,6 | 0,489897949 | 3,45E-05 | 2,82E-05 | 1 - | 6,03E-05 - | 0 - | 0 - | 1 - | 5,60E-05 - | 0 - | 0 - | 1 - | 5,60E-05 - |
| 964 | 1 | 3um_FIJJAFBF_246767 | Bacteria | Bacteroidetes | Flavobacteriia | Flavobacterial | Flavobacteriac | Bergeyella | Bergeyella zoo | poorly charact | Function unkn | 0FXH3 | Inherit from b | 643867.Ftrac_ | 1 | 0,632455532 | 0,00020411 | 0,000149953 | 0 - | 0 - | 2 - | 0,000467891 - | 1 - | 0,000184528 - | 1 - | 0,000183368 - | 1 - | 0,000184765 - |
| 965 | 1 | 0.2um_NOKCPHJN_898151 | Viruses | unclassified | unclassified | unclassified | unclassified | unclassified | uncultured M | unclassified | unclassified | unclassified | unclassified | unclassified | 0,6 | 0,489897949 | 0,000154281 | 0,000126189 | 1 - | 0,00027066 - | 0 - | 0 - | 1 - | 0,000251163 - | 1 - | 0,000249584 - | 0 - | 0 - |
| 966 | 1 | 0.2um_NOKCPHJN_276939 | Bacteria | Proteobacteria | Betaproteobac | Burkholderiale | Comamonada | Acidovorax | Acidovorax sp | metabolism | Inorganic ion t | 05EKV | Phosphonate | 543728.Vapar | 0,4 | 0,489897949 | 0,000101523 | 0,000125019 | 1 - | 0,000233228 - | 1 - | 0,000274388 - | 0 - | 0 - | 0 - | 0 - | 0 - | 0 - |
| 967 | 1 | 0.2um_NOKCPHJN_893102 | Bacteria | Proteobacteria | Gammaprotec | unclassified | unclassified | unclassified | various | information st | Translation, ri | 05C64 | Binds directly | 207954.MED9 | 0,6 | 0,489897949 | 0,000211016 | 0,000172296 | 0 - | 0 - | 0 - | 0 - | 1 - | 0,00035228 - | 1 - | 0,000350066 - | 1 - | 0,000352734 - |
| 968 | 1 | 0.2um_NOKCPHJN_705037 | Bacteria | Proteobacteria | Betaproteobac | Nitrosomonad | Methylophilac | Methylotener | Methylotener | unclassified | unclassified | unclassified | unclassified | unclassified | 0,2 | 0,4 | 3,82E-05 | 7,64E-05 | 0 - | 0 - | 0 - | 0 - | 1 - | 0,000191025 - | 0 - | 0 - | 0 - | 0 - |
| 969 | 1 | 0.2um_NOKCPHJN_729548 | Bacteria | Proteobacteria | Gammaprotec | Alteromonada | Alteromonada | Paraglaciecola | Paraglaciecola | metabolism | Inorganic ion t | 0QJE8 | receptor | 342610.Patl_2 | 0,4 | 0,489897949 | 4,59E-05 | 5,69E-05 | 0 - | 0 - | 1 - | 0,000128321 - | 1 - | 0,000101215 - | 0 - | 0 - | 0 - | 0 - |
| 970 | 1 | 0.2um_NOKCPHJN_1119393 | Eukaryota | Arthropoda | Insecta | Lepidoptera | Pyralidae | Amyelois | Amyelois tran | unclassified | unclassified | unclassified | unclassified | unclassified | 1 | 0,632455532 | 0,000178699 | 0,00011154 | 1 - | 0,000189814 - | 0 - | 0 - | 2 - | 0,00035228 - | 1 - | 0,000175033 - | 1 - | 0,000176367 - |
| 971 | 1 | 3um_FIJJAFBF_513826 | Viruses | unclassified | unclassified | unclassified | unclassified | unclassified | uncultured M | unclassified | unclassified | unclassified | unclassified | unclassified | 6 | 0,894427191 | 0,001702847 | 0,000365121 | 5 - | 0,001423601 - | 7 - | 0,00234477 - | 6 - | 0,001585261 - | 7 - | 0,001837849 - | 5 - | 0,001322752 - |
| 972 | 1 | 3um_FIJJAFBF_270535 | unclassified | unclassified | unclassified | unclassified | unclassified | unclassified | unclassified | unclassified | unclassified | unclassified | unclassified | unclassified | 1 | 0 | 0,000144142 | 1,41E-05 | 1 - | 0,000145429 - | 1 - | 0,000171094 - | 1 - | 0,000134953 - | 1 - | 0,000134105 - | 1 - | 0,000135127 - |
| 973 | 1 | 3um_FIJJAFBF_594150 | Eukaryota | unclassified | Pelagophyceae | Pelagomonada | unclassified | Aureoumbra | Aureoumbra l | metabolism | Energy produc | COG4451 | RuBisCO catal | 543728.Vapar | 5 | 0,632455532 | 0,003125792 | 0,00047522 | 5 - | 0,003154455 - | 5 - | 0,003711147 - | 4 - | 0,002341777 - | 6 - | 0,003490591 - | 5 - | 0,002930989 - |
| 974 | 1 | 3um_FIJJAFBF_493300 | Bacteria | Proteobacteria | Gammaprotec | unclassified | unclassified | unclassified | OM182 bacter | unclassified | unclassified | unclassified | unclassified | unclassified | 2 | 0,894427191 | 0,000636883 | 0,000319832 | 1 - | 0,000315446 - | 3 - | 0,001113344 - | 2 - | 0,000585444 - | 1 - | 0,000290883 - | 3 - | 0,000879297 - |
| 975 | 1 | 3um_FIJJAFBF_313476 | unclassified | unclassified | unclassified | unclassified | unclassified | unclassified | unclassified | unclassified | unclassified | unclassified | unclassified | unclassified | 0,8 | 0,4 | 0,00021451 | 0,000107561 | 1 - | 0,000283799 - | 0 - | 0 - | 1 - | 0,000263355 - | 1 - | 0,0002617 - | 1 - | 0,000263694 - |
| 976 | 1 | 3um_FIJJAFBF_13125 | Eukaryota | Bacillariophyta | Coscinodiscop | Chaetocerotal | Acanthocerata | Acanthoceras | Acanthoceras | metabolism | Energy produc | 0IFBF | One of the tw | 45157.CMV08 | 0,8 | 0,748331477 | 0,000204799 | 0,000189441 | 2 - | 0,00049968 - | 1 - | 0,000293931 - | 0 - | 0 - | 1 - | 0,000230386 - | 0 - | 0 - |
| 977 | 1 | 3um_FIJJAFBF_157067 | Bacteria | Bacteroidetes | Flavobacteriia | unclassified | unclassified | unclassified | Flavobacteria | cellular proces | Intracellular tr | 18ATJ | Pfam:YadA | 1005048.CFU_ | 0,8 | 0,4 | 6,92E-05 | 3,47E-05 | 1 - | 9,15E-05 - | 0 - | 0 - | 1 - | 8,49E-05 - | 1 - | 8,44E-05 - | 1 - | 8,51E-05 - |
| 978 | 1 | 3um_FIJJAFBF_209608 | Bacteria | Bacteroidetes | Flavobacteriia | Flavobacterial | Flavobacteriac | Tamlana | various | unclassified | unclassified | unclassified | unclassified | unclassified | 1 | 0 | 0,000625306 | 6,12E-05 | 1 - | 0,000630891 - | 1 - | 0,000742229 - | 1 - | 0,000585444 - | 1 - | 0,000581765 - | 1 - | 0,000586198 - |
| 979 | 1 | 3um_FIJJAFBF_74383 | Eukaryota | unclassified | Cryptophyta | Pyrenomonad | Geminigerace | Teleaulax | Teleaulax amp | metabolism | Energy produc | COG1290 | Component of | 3055.DAA009 | 1,2 | 0,4 | 0,000500239 | 0,000230036 | 1 - | 0,000407878 - | 2 - | 0,00095972 - | 1 - | 0,000378496 - | 1 - | 0,000376118 - | 1 - | 0,000378984 - |
| 980 | 1 | 0.2um_NOKCPHJN_448283 | Eukaryota | Chordata | Ascidiacea | Enterogona | Cionidae | Ciona | Ciona intestin | poorly charact | Function unkn | 0Q0SF | SRP40, C-term | 29850.GGTG_ | 0,4 | 0,489897949 | 3,37E-05 | 4,17E-05 | 0 - | 0 - | 1 - | 9,44E-05 - | 0 - | 0 - | 1 - | 7,40E-05 - | 0 - | 0 - |
| 981 | 1 | 0.2um_NOKCPHJN_698927 | Bacteria | Proteobacteria | Alphaproteob | Rhodobactera | Rhodobactera | Planktotalea | Planktotalea f | metabolism | Energy produc | 0HMD2 | Inherit from p | 596151.Desfr | 1 | 0 | 0,000266618 | 2,61E-05 | 1 - | 0,000269 - | 1 - | 0,000316472 - | 1 - | 0,000249622 - | 1 - | 0,000248053 - | 1 - | 0,000249943 - |
| 982 | 1 | 0.2um_NOKCPHJN_149872 | Bacteria | Bacteroidetes | Flavobacteriia | Flavobacterial | Crocinitomica | unclassified | Crocinitomica | unclassified | unclassified | unclassified | unclassified | unclassified | 0,4 | 0,489897949 | 3,34E-05 | 4,10E-05 | 0 - | 0 - | 0 - | 0 - | 0 - | 0 - | 1 - | 8,33E-05 - | 1 - | 8,39E-05 - |
| 983 | 1 | 3um_FIJJAFBF_510246 | Bacteria | Bacteroidetes | Flavobacteriia | Flavobacterial | Flavobacteriac | Polaribacter | Polaribacter s | information st | Translation, ri | COG0256 | This is one of | 313598.MED1 | 1 | 0 | 0,000749288 | 7,33E-05 | 1 - | 0,000755981 - | 1 - | 0,000889396 - | 1 - | 0,000701524 - | 1 - | 0,000697115 - | 1 - | 0,000702427 - |
| 984 | 1 | 3um_FIJJAFBF_214623 | Bacteria | Proteobacteria | Alphaproteob | Rhodobactera | unclassified | unclassified | Rhodobactera | information st | Translation, ri | 177XJ | One of the pri | 89187.ISM_01 | 0,2 | 0,4 | 0,000104329 | 0,000208658 | 0 - | 0 - | 0 - | 0 - | 1 - | 0,000521646 - | 0 - | 0 - | 0 - | 0 - |
| 985 | 1 | 3um_FIJJAFBF_789445 | Bacteria | Proteobacteria | Gammaprotec | Alteromonada | Alteromonada | Paraglaciecola | Paraglaciecola | cellular proces | Cell wall/mem | 0QTSX | Peptidoglycan | 983545.Glaag | 0,4 | 0,489897949 | 0,000195287 | 0,000241941 | 0 - | 0 - | 1 - | 0,000545872 - | 1 - | 0,000430565 - | 0 - | 0 - | 0 - | 0 - |
| 986 | 1 | 3um_FIJJAFBF_594169 | Eukaryota | unclassified | Synurophycea | Parmales | Triparmaceae | Triparma | Triparma laev | unclassified | unclassified | unclassified | unclassified | unclassified | 0,6 | 0,489897949 | 8,17E-05 | 6,68E-05 | 1 - | 0,000143291 - | 0 - | 0 - | 0 - | 0 - | 1 - | 0,000132133 - | 1 - | 0,00013314 - |
| 987 | 1 | 3um_FIJJAFBF_60140 | Viruses | unclassified | unclassified | unclassified | unclassified | unclassified | uncultured M | unclassified | unclassified | unclassified | unclassified | unclassified | 1 | 0 | 0,000147068 | 1,44E-05 | 1 - | 0,000148382 - | 1 - | 0,000174568 - | 1 - | 0,000137693 - | 1 - | 0,000136828 - | 1 - | 0,000137871 - |
| 988 | 1 | 3um_FIJJAFBF_586277 | Eukaryota | unclassified | Pelagophyceae | Pelagomonada | unclassified | Aureococcus | Aureococcus a | unclassified | unclassified | unclassified | unclassified | unclassified | 0,2 | 0,4 | 0,00016546 | 0,00033092 | 1 - | 0,0008273 - | 0 - | 0 - | 0 - | 0 - | 0 - | 0 - | 0 - | 0 - |
| 989 | 2 | 0.2um_NOKCPHJN_228707;3ur | Bacteria | Bacteroidetes | Flavobacteriia | Flavobacterial | Crocinitomica | unclassified | Crocinitomica | unclassified | unclassified | unclassified | unclassified | unclassified | 0,6 | 0,489897949 | 0,00035962 | 0,000294138 | 1 - | 0,000630891 - | 0 - | 0 - | 1 - | 0,000585444 - | 1 - | 0,000581765 - | 0 - | 0 - |
| 990 | 1 | 0.2um_NOKCPHJN_881844 | Bacteria | Proteobacteria | Alphaproteob | Pelagibacteral | Pelagibacterac | Candidatus Pe | Candidatus Pe | metabolism | Amino acid tra | 01R4A | Polyamine AB | 335992.SAR11 | 1 | 0 | 0,000233023 | 2,28E-05 | 1 - | 0,000235104 - | 1 - | 0,000276595 - | 1 - | 0,000218168 - | 1 - | 0,000216797 - | 1 - | 0,000218449 - |
| 991 | 2 | 0.2um_NOKCPHJN_933735;0.2 | various | various | various | various | various | various | various | unclassified | unclassified | unclassified | unclassified | unclassified | 0,4 | 0,489897949 | 5,43E-05 | 6,66E-05 | 1 - | 0,000141214 - | 0 - | 0 - | 0 - | 0 - | 1 - | 0,000130218 - | 0 - | 0 - |
| 992 | 1 | 0.2um_NOKCPHJN_428163 | Eukaryota | unclassified | Oomycetes | Peronosporale | Peronosporac | Phytophthora | Phytophthora | metabolism | Amino acid tra | KOG0540 | carboxylase, b | 164328.Phyra | 0,8 | 0,4 | 0,000126143 | 6,49E-05 | 0 - | 0 - | 1 - | 0,000187582 - | 1 - | 0,000147958 - | 1 - | 0,000147028 - | 1 - | 0,000148148 - |
| 993 | 1 | 3um_FIJJAFBF_296064 | Bacteria | Proteobacteria | Gammaprotec | unclassified | unclassified | unclassified | Gammaprotec | metabolism | Energy produc | COG0056 | Produces ATP | 349521.HCH_ | 0,8 | 0,4 | 0,000137436 | 7,05E-05 | 1 - | 0,000170611 - | 1 - | 0,00020072 - | 0 - | 0 - | 1 - | 0,000157326 - | 1 - | 0,000158524 - |
| 994 | 1 | 3um_FIJJAFBF_102824 | Bacteria | Bacteroidetes | Flavobacteriia | Flavobacterial | Flavobacteriac | Arenitalea | Arenitalea lut | unclassified | unclassified | unclassified | unclassified | unclassified | 2,6 | 1,356465997 | 0,000434296 | 0,000206001 | 2 - | 0,000349378 - | 1 - | 0,000205518 - | 2 - | 0,00032421 - | 5 - | 0,000805432 - | 3 - | 0,000486941 - |
| 995 | 1 | 0.2um_NOKCPHJN_372789 | Eukaryota | unclassified | Oomycetes | Peronosporale | Peronosporac | Phytophthora | Phytophthora | unclassified | unclassified | unclassified | unclassified | unclassified | 0,6 | 0,489897949 | 8,51E-05 | 6,95E-05 | 0 - | 0 - | 0 - | 0 - | 1 - | 0,000142019 - | 1 - | 0,000141126 - | 1 - | 0,000142202 - |
| 996 | 2 | 3um_FIJJAFBF_56398;3um_FIJ | Eukaryota | unclassified | Dinophyceae | Gonyaulacales | various | various | various | metabolism | Energy produc | 0FJ7X | This is one of | various | 1 | 0 | 0,000247628 | 2,42E-05 | 1 - | 0,00024984 - | 1 - | 0,000293931 - | 1 - | 0,000231843 - | 1 - | 0,000230386 - | 1 - | 0,000232141 - |
| 997 | 1 | 3um_FIJJAFBF_725808 | Eukaryota | Phaeophyceae | unclassified | Ectocarpales | Ectocarpaceae | Ectocarpus | Ectocarpus sili | metabolism | Energy produc | COG0056 | Produces ATP | 35128.Thaps2 | 2,8 | 0,4 | 0,000815264 | 0,000159321 | 3 - | 0,000876939 - | 3 - | 0,001031699 - | 3 - | 0,000813767 - | 2 - | 0,000539102 - | 3 - | 0,000814815 - |
| 998 | 1 | 0.2um_NOKCPHJN_728863 | Eukaryota | unclassified | Labyrinthulom | unclassified | Thraustochytr | Aurantiochytr | Aurantiochytr | poorly charact | Function unkn | 0A4HT | Coiled-coil do | 7668.SPU_011 | 0,6 | 0,489897949 | 8,79E-05 | 7,26E-05 | 1 - | 0,000141442 - | 1 - | 0,000166403 - | 0 - | 0 - | 0 - | 0 - | 1 - | 0,000131422 - |
| 999 | 1 | 0.2um_NOKCPHJN_252641 | Bacteria | Proteobacteria | Alphaproteob | Rhodobactera | Rhodobactera | unclassified | Rhodobactera | unclassified | unclassified | unclassified | unclassified | unclassified | 5 | 1,264911064 | 0,000500619 | 9,22E-05 | 4 - | 0,00041365 - | 3 - | 0,000364988 - | 6 - | 0,000575779 - | 6 - | 0,000572161 - | 6 - | 0,00057652 - |
| 1000 | 1 | 0.2um_NOKCPHJN_792463 | Bacteria | Proteobacteria | Gammaprotec | Enterobactera | Enterobacteri | Citrobacter | Citrobacter w | cellular proces | Intracellular tr | 0224F | Hep_Hag repe | 237727.NAP1_ | 0,6 | 0,489897949 | 7,19E-05 | 5,88E-05 | 1 - | 0,000126178 - | 0 - | 0 - | 1 - | 0,000117089 - | 1 - | 0,000116353 - | 0 - | 0 - |
| 1001 | 1 | 0.2um_NOKCPHJN_1092416 | Bacteria | Proteobacteria | Epsilonproteo | Campylobacte | Helicobactera | Sulfurimonas | Sulfurimonas | poorly charact | Function unkn | 0ZJXN | NA | 439483.CBGD | 0,4 | 0,489897949 | 9,74E-05 | 0,000119917 | 1 - | 0,000223709 - | 1 - | 0,000263188 - | 0 - | 0 - | 0 - | 0 - | 0 - | 0 - |
| 1002 | 1 | 0.2um_NOKCPHJN_27620 | Bacteria | Actinobacteria | Actinobacteria | Micrococcales | Microbacteria | Candidatus Ac | Candidatus Ac | metabolism | Amino acid tra | 00DZF | branched-chai | 312284.A20C1 | 1,6 | 0,489897949 | 0,00033126 | 0,000114254 | 1 - | 0,000207314 - | 2 - | 0,000487801 - | 2 - | 0,00038476 - | 1 - | 0,000191171 - | 2 - | 0,000385255 - |
| 1003 | 1 | 0.2um_NOKCPHJN_151069 | Eukaryota | unclassified | Labyrinthulom | unclassified | Thraustochytr | Aurantiochytr | Aurantiochytr | unclassified | unclassified | unclassified | unclassified | unclassified | 0,2 | 0,4 | 1,37E-05 | 2,74E-05 | 1 - | 6,86E-05 - | 0 - | 0 - | 0 - | 0 - | 0 - | 0 - | 0 - | 0 - |
| 1004 | 1 | 3um_FIJJAFBF_95715 | Eukaryota | unclassified | Raphidophyce | Chattonellales | Chattonellace | Heterosigma | Heterosigma a | metabolism | Energy produc | 0IFBF | One of the tw | 45157.CMV08 | 2,8 | 0,4 | 0,000652047 | 0,000113733 | 2 - | 0,000470208 - | 3 - | 0,000829785 - | 3 - | 0,000654505 - | 3 - | 0,000650392 - | 3 - | 0,000655347 - |
| 1005 | 1 | 3um_FIJJAFBF_365652 | Bacteria | Bacteroidetes | Flavobacteriia | Flavobacterial | Flavobacteriac | Aureitalea | Aureitalea ma | unclassified | unclassified | unclassified | unclassified | unclassified | 3,6 | 1,624807681 | 0,000343931 | 0,000137015 | 2 - | 0,000198627 - | 2 - | 0,00023368 - | 3 - | 0,000276478 - | 6 - | 0,000549481 - | 5 - | 0,00046139 - |
| 1006 | 1 | 0.2um_NOKCPHJN_568137 | Bacteria | Proteobacteria | Alphaproteob | Pelagibacteral | Pelagibacterac | Candidatus Pe | Candidatus Pe | cellular proces | Cell wall/mem | 16RAV | Exopolysaccha | 439493.PB721 | 0,2 | 0,4 | 2,28E-05 | 4,56E-05 | 0 - | 0 - | 0 - | 0 - | 1 - | 0,000113973 - | 0 - | 0 - | 0 - | 0 - |
| 1007 | 2 | 0.2um_NOKCPHJN_482977;3ur | various | various | various | various | various | various | various | unclassified | unclassified | unclassified | unclassified | unclassified | 0,2 | 0,4 | 1,64E-05 | 3,28E-05 | 0 - | 0 - | 0 - | 0 - | 1 - | 8,21E-05 - | 0 - | 0 - | 0 - | 0 - |
| 1008 | 1 | 3um_FIJJAFBF_405471 | Bacteria | Proteobacteria | Gammaprotec | Alteromonada | Alteromonada | Paraglaciecola | Paraglaciecola | poorly charact | Function unkn | 0NMWQ | Collagen triple | 324057.Pjdr2_ | 0,4 | 0,489897949 | 5,09E-05 | 6,23E-05 | 0 - | 0 - | 0 - | 0 - | 0 - | 0 - | 1 - | 0,000126748 - | 1 - | 0,000127714 - |
| 1009 | 1 | 3um_FIJJAFBF_17218 | Bacteria | Proteobacteria | Alphaproteob | Rhodobactera | unclassified | unclassified | Rhodobactera | cellular proces | Posttranslatio | COG0330 | Band 7 protein | 439496.RBY4I | 0,6 | 0,8 | 0,000127326 | 0,000170037 | 0 - | 0 - | 0 - | 0 - | 0 - | 0 - | 1 - | 0,000211137 - | 2 - | 0,000425491 - |
| 1010 | 1 | 0.2um_NOKCPHJN_12460 | Bacteria | Bacteroidetes | Saprospiria | Saprospirales | Lewinellaceae | Lewinella | Lewinella pers | metabolism | Carbohydrate | 0NIRC | arabinogalacta | 573061.Clocel | 1,4 | 0,489897949 | 0,000212219 | 7,12E-05 | 2 - | 0,000308238 - | 1 - | 0,000181318 - | 1 - | 0,000143017 - | 1 - | 0,000142118 - | 2 - | 0,000286402 - |
| 1011 | 1 | 3um_FIJJAFBF_20039 | Bacteria | Bacteroidetes | Flavobacteriia | Flavobacterial | Flavobacteriac | Polaribacter | Polaribacter s | unclassified | unclassified | unclassified | unclassified | unclassified | 2,6 | 1,356465997 | 0,000217195 | 0,000112495 | 3 - | 0,000265739 - | 0 - | 0 - | 3 - | 0,000246596 - | 4 - | 0,000326729 - | 3 - | 0,000246914 - |
| 1012 | 1 | 3um_FIJJAFBF_499026 | Bacteria | Proteobacteria | Betaproteobac | unclassified | unclassified | unclassified | Betaproteoba | poorly charact | Function unkn | COG4147 | solute sympor | 887062.HGR_ | 0,2 | 0,4 | 2,53E-05 | 5,05E-05 | 1 - | 0,00012636 - | 0 - | 0 - | 0 - | 0 - | 0 - | 0 - | 0 - | 0 - |
| 1013 | 1 | 3um_FIJJAFBF_143869 | Bacteria | Proteobacteria | Gammaprotec | Oceanospirilla | Saccharospirill | Reinekea | Reinekea blan | unclassified | unclassified | unclassified | unclassified | unclassified | 1 | 0 | 0,00019532 | 1,91E-05 | 1 - | 0,000197065 - | 1 - | 0,000231842 - | 1 - | 0,000182869 - | 1 - | 0,00018172 - | 1 - | 0,000183104 - |
| 1014 | 1 | 3um_FIJJAFBF_313117 | Bacteria | Cyanobacteria | unclassified | Synechococca | Synechococca | Synechococcu | Synechococcu | unclassified | unclassified | unclassified | unclassified | unclassified | 2,2 | 0,979795897 | 0,000849801 | 0,000335554 | 1 - | 0,000404119 - | 1 - | 0,000475437 - | 3 - | 0,001125024 - | 3 - | 0,001117954 - | 3 - | 0,001126472 - |
| 1015 | 1 | 0.2um_NOKCPHJN_505891 | Bacteria | Proteobacteria | Gammaprotec | Cellvibrionales | Spongiibacter | unclassified | marine gamm | cellular proces | Cell wall/mem | 0QKXZ | Porin Gram-ne | 247633.GP214 | 0,8 | 0,748331477 | 0,000217832 | 0,000201496 | 2 - | 0,000531478 - | 1 - | 0,000312636 - | 0 - | 0 - | 1 - | 0,000245047 - | 0 - | 0 - |
| 1016 | 1 | 3um_FIJJAFBF_87923 | Bacteria | Bacteroidetes | Flavobacteriia | Flavobacterial | Crocinitomica | unclassified | Crocinitomica | unclassified | unclassified | unclassified | unclassified | unclassified | 0,8 | 0,4 | 0,000101283 | 5,21E-05 | 0 - | 0 - | 1 - | 0,000150613 - | 1 - | 0,000118798 - | 1 - | 0,000118052 - | 1 - | 0,000118951 - |
| 1017 | 1 | 3um_FIJJAFBF_414690 | Bacteria | Bacteroidetes | Flavobacteriia | Flavobacterial | Flavobacteriac | Jejuia | Jejuia pallidilu | unclassified | unclassified | unclassified | unclassified | unclassified | 0,2 | 0,4 | 3,99E-05 | 7,99E-05 | 0 - | 0 - | 0 - | 0 - | 0 - | 0 - | 0 - | 0 - | 1 - | 0,00019971 - |
| 1018 | 1 | 3um_FIJJAFBF_715072 | Eukaryota | Bacillariophyta | Mediophyceae | Triceratiales | Triceratiaceae | Triceratium | Triceratium d | information st | Translation, ri | COG0050 | This protein p | 32049.SYNPC | 5 | 0,894427191 | 0,001054743 | 0,00016944 | 6 - | 0,001286462 - | 4 - | 0,001008996 - | 4 - | 0,000795861 - | 6 - | 0,001186289 - | 5 - | 0,000996106 - |
| 1019 | 2 | 0.2um_NOKCPHJN_05117;0.2u | Bacteria | Bacteroidetes | Flavobacteriia | Flavobacterial | Crocinitomica | unclassified | Crocinitomica | unclassified | unclassified | unclassified | unclassified | unclassified | 0,6 | 0,489897949 | 0,000238899 | 0,000197506 | 1 - | 0,000384622 - | 1 - | 0,0004525 - | 0 - | 0 - | 0 - | 0 - | 1 - | 0,000357375 - |
| 1020 | 1 | 3um_FIJJAFBF_318783 | Eukaryota | Chlorophyta | Chlorophyceae | Chlamydomor | Chlamydomor | Chlamydomor | Chlamydomor | unclassified | unclassified | unclassified | unclassified | unclassified | 0,4 | 0,489897949 | 7,23E-05 | 8,90E-05 | 1 - | 0,000166087 - | 1 - | 0,000195398 - | 0 - | 0 - | 0 - | 0 - | 0 - | 0 - |
| 1021 | 1 | 0.2um_NOKCPHJN_925580 | Eukaryota | unclassified | Synurophycea | Synurales | Mallomonada | Mallomonas | Mallomonas r | metabolism | Carbohydrate | COG0149 | triosephospha | 35128.Thaps2 | 0,6 | 0,489897949 | 0,000158778 | 0,000129643 | 0 - | 0 - | 0 - | 0 - | 1 - | 0,000265071 - | 1 - | 0,000263405 - | 1 - | 0,000265412 - |
| 1022 | 1 | 0.2um_NOKCPHJN_492269 | Eukaryota | Nematoda | Enoplea | Trichinellida | Trichinellidae | Trichinella | Trichinella pse | cellular proces | Intracellular tr | 0V6YH | RAN member | 6334.EFV5555 | 1 | 0 | 0,000421929 | 4,13E-05 | 1 - | 0,000425698 - | 1 - | 0,000500825 - | 1 - | 0,000395033 - | 1 - | 0,00039255 - | 1 - | 0,000395541 - |
| 1023 | 1 | 3um_FIJJAFBF_145781 | Eukaryota | Bacillariophyta | Coscinodiscop | Chaetocerotal | Chaetocerotac | Chaetoceros | Chaetoceros g | unclassified | unclassified | unclassified | unclassified | unclassified | 0,2 | 0,4 | 0,000296296 | 0,000592593 | 0 - | 0 - | 0 - | 0 - | 0 - | 0 - | 0 - | 0 - | 1 - | 0,001481482 - |
| 1024 | 1 | 3um_FIJJAFBF_486301 | Bacteria | Proteobacteria | Gammaprotec | Cellvibrionales | Halieaceae | Halioglobus | various | information st | Translation, ri | 0QQXI | This protein is | 247634.GPB2 | 1,4 | 1,019803903 | 0,000986211 | 0,000717347 | 2 - | 0,001486336 - | 0 - | 0 - | 3 - | 0,0020689 - | 1 - | 0,0006853 - | 1 - | 0,000690521 - |
| 1025 | 1 | 3um_FIJJAFBF_196001 | Bacteria | Bacteroidetes | Flavobacteriia | Flavobacterial | Flavobacteriac | Polaribacter | Polaribacter s | cellular proces | Posttranslatio | 08V1T | peptidylprolyl | 313594.PI23P | 1,8 | 0,748331477 | 0,000659818 | 0,000245513 | 2 - | 0,000755981 - | 1 - | 0,000444698 - | 2 - | 0,000701524 - | 3 - | 0,001045673 - | 1 - | 0,000351213 - |
| 1026 | 1 | 0.2um_NOKCPHJN_723505 | Bacteria | Actinobacteria | Actinobacteria | Corynebacteri | Mycobacteria | Mycobacteriu | Mycobacteriu | unclassified | unclassified | unclassified | unclassified | unclassified | 0,4 | 0,489897949 | 6,65E-05 | 8,14E-05 | 0 - | 0 - | 0 - | 0 - | 1 - | 0,000166756 - | 1 - | 0,000165708 - | 0 - | 0 - |
| 1027 | 1 | 0.2um_NOKCPHJN_968394 | Eukaryota | unclassified | Cryptophyta | Pyrenomonad | various | various | various | unclassified | unclassified | unclassified | unclassified | unclassified | 4,4 | 1,496662955 | 0,004564264 | 0,0014221 | 3 - | 0,003169657 - | 4 - | 0,004972043 - | 7 - | 0,006863099 - | 5 - | 0,004871407 - | 3 - | 0,002945114 - |
| 1028 | 1 | 0.2um_NOKCPHJN_698497 | Bacteria | Proteobacteria | Alphaproteob | Rhizobiales | unclassified | unclassified | Rhizobiales ba | unclassified | unclassified | unclassified | unclassified | unclassified | 0,2 | 0,4 | 1,83E-05 | 3,65E-05 | 0 - | 0 - | 0 - | 0 - | 1 - | 9,13E-05 - | 0 - | 0 - | 0 - | 0 - |
| 1029 | 1 | 0.2um_NOKCPHJN_937652 | unclassified | unclassified | unclassified | unclassified | unclassified | unclassified | unclassified | unclassified | unclassified | unclassified | unclassified | unclassified | 0,2 | 0,4 | 0,000246914 | 0,000493827 | 0 - | 0 - | 0 - | 0 - | 0 - | 0 - | 0 - | 0 - | 1 - | 0,001234568 - |
| 1030 | 3 | 0.2um_NOKCPHJN_1045591;3u | Eukaryota | various | various | various | various | various | various | metabolism | Energy produc | COG1145 | 4Fe-4S ferredo | various | 1,8 | 0,4 | 0,001683527 | 0,000280367 | 2 - | 0,001927337 - | 1 - | 0,001133735 - | 2 - | 0,0017885 - | 2 - | 0,001777261 - | 2 - | 0,001790802 - |
| 1031 | 1 | 0.2um_NOKCPHJN_61811 | Bacteria | Bacteroidetes | Flavobacteriia | Flavobacterial | Flavobacteriac | Algibacter | Algibacter sp. | metabolism | Inorganic ion t | 05C27 | TonB-depende | 391587.KAOT | 19 | 1,549193338 | 0,001582545 | 6,43E-05 | 20 - | 0,001692931 - | 16 - | 0,001593357 - | 20 - | 0,00157098 - | 20 - | 0,001561107 - | 19 - | 0,001494352 - |
| 1032 | 1 | 0.2um_NOKCPHJN_119224 | Bacteria | Actinobacteria | Actinobacteria | Actinomycetal | unclassified | unclassified | Actinomyceta | metabolism | Carbohydrate | 00BJE | Glyceraldehyd | 469371.Tbis_1 | 1,4 | 0,489897949 | 0,000359907 | 0,000108858 | 1 - | 0,000264138 - | 1 - | 0,000310753 - | 2 - | 0,000490221 - | 1 - | 0,00024357 - | 2 - | 0,000490852 - |
| 1033 | 1 | 3um_FIJJAFBF_316866 | Bacteria | Bacteroidetes | Flavobacteriia | Flavobacterial | Flavobacteriac | Algibacter | Algibacter alg | information st | Translation, ri | 09EM4 | Binds directly | 1046627.BZAF | 0,8 | 0,4 | 0,000581434 | 0,000291548 | 1 - | 0,000769244 - | 0 - | 0 - | 1 - | 0,000713831 - | 1 - | 0,000709345 - | 1 - | 0,00071475 - |
| 1034 | 2 | 3um_FIJJAFBF_549291;3um_FI | Bacteria | various | various | various | various | various | various | various | various | various | various | various | 0,2 | 0,4 | 8,44E-06 | 1,69E-05 | 0 - | 0 - | 0 - | 0 - | 0 - | 0 - | 1 - | 4,22E-05 - | 0 - | 0 - |
| 1035 | 1 | 0.2um_NOKCPHJN_285950 | Bacteria | Proteobacteria | Gammaprotec | Oceanospirilla | Saccharospirill | Reinekea | Reinekea fors | unclassified | unclassified | unclassified | unclassified | unclassified | 0,2 | 0,4 | 1,78E-05 | 3,56E-05 | 1 - | 8,89E-05 - | 0 - | 0 - | 0 - | 0 - | 0 - | 0 - | 0 - | 0 - |
| 1036 | 1 | 0.2um_NOKCPHJN_941524 | Eukaryota | unclassified | Spirotrichea | Sporadotrichi | Oxytrichidae | Oxytricha | Oxytricha trifa | unclassified | unclassified | unclassified | unclassified | unclassified | 4,6 | 1,959591794 | 0,000858487 | 0,000341821 | 5 - | 0,000978726 - | 1 - | 0,00023029 - | 7 - | 0,001271512 - | 5 - | 0,000902515 - | 5 - | 0,000909392 - |
| 1037 | 1 | 3um_FIJJAFBF_24644 | Bacteria | Proteobacteria | Alphaproteob | Rhodobactera | Rhodobactera | unclassified | Rhodobactera | unclassified | unclassified | unclassified | unclassified | unclassified | 0,2 | 0,4 | 3,75E-05 | 7,50E-05 | 1 - | 0,00018738 - | 0 - | 0 - | 0 - | 0 - | 0 - | 0 - | 0 - | 0 - |
| 1038 | 1 | 3um_FIJJAFBF_189664 | Bacteria | Bacteroidetes | Flavobacteriia | Flavobacterial | Flavobacteriac | Polaribacter | Polaribacter s | information st | Translation, ri | 0PBIW | 50S ribosomal | 313594.PI23P | 0,4 | 0,489897949 | 0,000507334 | 0,000621362 | 0 - | 0 - | 0 - | 0 - | 0 - | 0 - | 1 - | 0,001263521 - | 1 - | 0,001273148 - |
| 1039 | 1 | 0.2um_NOKCPHJN_283743 | Bacteria | Proteobacteria | Alphaproteob | Rhodospirillal | Rhodospirillac | Candidatus En | Candidatus En | unclassified | unclassified | unclassified | unclassified | unclassified | 0,4 | 0,489897949 | 7,21E-05 | 8,84E-05 | 1 - | 0,00018698 - | 0 - | 0 - | 1 - | 0,000173511 - | 0 - | 0 - | 0 - | 0 - |
| 1040 | 1 | 3um_FIJJAFBF_254310 | Bacteria | Proteobacteria | Gammaprotec | Cellvibrionales | Spongiibacter | Oceanicoccus | Oceanicoccus | information st | Transcription | 16PMT | DNA-depende | 247633.GP214 | 2,2 | 0,748331477 | 0,000566863 | 0,000176125 | 1 - | 0,000263345 - | 2 - | 0,000619639 - | 2 - | 0,000488749 - | 3 - | 0,000728517 - | 3 - | 0,000734067 - |
| 1041 | 1 | 0.2um_NOKCPHJN_359261 | Bacteria | Bacteroidetes | Flavobacteriia | Flavobacterial | Flavobacteriac | Formosa | Formosa sp. H | metabolism | Inorganic ion t | 05C27 | TonB-depende | 742766.HMPR | 0,2 | 0,4 | 1,55E-05 | 3,10E-05 | 0 - | 0 - | 0 - | 0 - | 0 - | 0 - | 0 - | 0 - | 1 - | 7,74E-05 - |
| 1042 | 1 | 3um_FIJJAFBF_189626 | Bacteria | Bacteroidetes | Flavobacteriia | Flavobacterial | Flavobacteriac | Polaribacter | Polaribacter b | unclassified | unclassified | unclassified | unclassified | unclassified | 6,2 | 0,748331477 | 0,00141962 | 0,000262447 | 5 - | 0,001150838 - | 7 - | 0,00189551 - | 7 - | 0,001495111 - | 6 - | 0,00127347 - | 6 - | 0,001283173 - |
| 1043 | 2 | 0.2um_NOKCPHJN_786247;0.2 | Bacteria | Bacteroidetes | Flavobacteriia | Flavobacterial | Flavobacteriac | Formosa | Formosa sp. H | unclassified | unclassified | unclassified | unclassified | unclassified | 1,2 | 0,4 | 0,000596893 | 0,000208395 | 2 - | 0,001002215 - | 1 - | 0,000589542 - | 1 - | 0,00046501 - | 1 - | 0,000462088 - | 1 - | 0,000465609 - |
| 1044 | 1 | 3um_FIJJAFBF_539569 | Bacteria | Bacteroidetes | Flavobacteriia | Flavobacterial | Flavobacteriac | Nonlabens | Nonlabens do | poorly charact | Function unkn | 0PBEY | Pfam:DUF145 | 886377.Murru | 1,6 | 0,489897949 | 0,002024809 | 0,000495718 | 1 - | 0,001308863 - | 1 - | 0,001539849 - | 2 - | 0,002429157 - | 2 - | 0,002413891 - | 2 - | 0,002432283 - |
| 1045 | 2 | 3um_FIJJAFBF_331179;3um_FI | Viruses | unclassified | unclassified | unclassified | unclassified | unclassified | uncultured M | unclassified | unclassified | unclassified | unclassified | unclassified | 0,2 | 0,4 | 4,95E-05 | 9,89E-05 | 0 - | 0 - | 0 - | 0 - | 0 - | 0 - | 1 - | 0,000247295 - | 0 - | 0 - |
| 1046 | 1 | 0.2um_NOKCPHJN_291421 | Bacteria | Bacteroidetes | unclassified | unclassified | unclassified | unclassified | Bacteroidetes | poorly charact | Function unkn | 05EV9 | Alpha Beta Hy | 391603.FBALC | 0,2 | 0,4 | 6,37E-05 | 0,000127347 | 0 - | 0 - | 0 - | 0 - | 0 - | 0 - | 1 - | 0,000318368 - | 0 - | 0 - |
| 1047 | 1 | 0.2um_NOKCPHJN_115346 | Bacteria | Bacteroidetes | Flavobacteriia | Flavobacterial | Flavobacteriac | Algibacter | Algibacter sp. | unclassified | unclassified | unclassified | unclassified | unclassified | 5,4 | 2,244994432 | 0,000519905 | 0,000195218 | 6 - | 0,000601329 - | 2 - | 0,000235817 - | 5 - | 0,00046501 - | 9 - | 0,000831758 - | 5 - | 0,000465609 - |
| 1048 | 2 | 0.2um_NOKCPHJN_91999;3um | Bacteria | Bacteroidetes | Flavobacteriia | Flavobacterial | various | various | various | poorly charact | Function unkn | various | various | various | 0,2 | 0,4 | 2,80E-05 | 5,61E-05 | 0 - | 0 - | 1 - | 0,000140176 - | 0 - | 0 - | 0 - | 0 - | 0 - | 0 - |
| 1049 | 9 | 0.2um_NOKCPHJN_445402;3ur | various | various | various | various | various | various | various | various | various | various | various | various | 0,4 | 0,489897949 | 0,000182823 | 0,000226471 | 0 - | 0 - | 1 - | 0,000510742 - | 0 - | 0 - | 0 - | 0 - | 1 - | 0,000403374 - |
| 1050 | 1 | 3um_FIJJAFBF_81743 | Eukaryota | unclassified | unclassified | Kinetoplastida | Trypanosomat | Trypanosoma | Trypanosoma | unclassified | unclassified | unclassified | unclassified | unclassified | 0,2 | 0,4 | 3,88E-05 | 7,76E-05 | 1 - | 0,000194013 - | 0 - | 0 - | 0 - | 0 - | 0 - | 0 - | 0 - | 0 - |
| 1051 | 1 | 3um_FIJJAFBF_77475 | Bacteria | Nitrospirae | Nitrospira | Nitrospirales | Nitrospiraceae | Nitrospira | Nitrospira sp. | unclassified | unclassified | unclassified | unclassified | unclassified | 0,2 | 0,4 | 9,13E-05 | 0,000182602 | 0 - | 0 - | 1 - | 0,000456504 - | 0 - | 0 - | 0 - | 0 - | 0 - | 0 - |
| 1052 | 1 | 3um_FIJJAFBF_609352 | Bacteria | Proteobacteria | Gammaprotec | Cellvibrionales | Halieaceae | unclassified | marine gamm | unclassified | unclassified | unclassified | unclassified | unclassified | 0,8 | 0,4 | 0,000117991 | 6,07E-05 | 0 - | 0 - | 1 - | 0,000175459 - | 1 - | 0,000138396 - | 1 - | 0,000137526 - | 1 - | 0,000138574 - |
| 1053 | 1 | 0.2um_NOKCPHJN_897706 | Bacteria | Proteobacteria | Gammaprotec | unclassified | unclassified | unclassified | SAR86 cluster | information st | Translation, ri | 16Q16 | Binds the low | 498211.CJA_0 | 1 | 0 | 0,000505334 | 4,94E-05 | 1 - | 0,000509848 - | 1 - | 0,000599825 - | 1 - | 0,000473121 - | 1 - | 0,000470147 - | 1 - | 0,00047373 - |
| 1054 | 1 | 3um_FIJJAFBF_01994 | Bacteria | Planctomycet | Planctomyceti | Planctomycet | Planctomycet | unclassified | Planctomycet | unclassified | unclassified | unclassified | unclassified | unclassified | 3 | 0,632455532 | 0,001638805 | 0,000490589 | 2 - | 0,001089365 - | 4 - | 0,002563227 - | 3 - | 0,001516337 - | 3 - | 0,001506808 - | 3 - | 0,001518289 - |
| 1055 | 1 | 0.2um_NOKCPHJN_356926 | Bacteria | Bacteroidetes | unclassified | unclassified | unclassified | unclassified | Bacteroidetes | poorly charact | Function unkn | 08QKU | NA | 755732.Fluta_ | 0,2 | 0,4 | 1,15E-05 | 2,30E-05 | 0 - | 0 - | 0 - | 0 - | 0 - | 0 - | 1 - | 5,75E-05 - | 0 - | 0 - |
| 1056 | 1 | 0.2um_NOKCPHJN_469875 | Bacteria | Proteobacteria | Gammaprotec | Alteromonada | Pseudoalteror | Pseudoalteror | Pseudoalteror | poorly charact | Function unkn | 05NCR | NA | 269799.Gmet_ | 1,6 | 0,489897949 | 0,000360626 | 0,000130491 | 2 - | 0,000447418 - | 2 - | 0,000526377 - | 2 - | 0,000415187 - | 1 - | 0,000206289 - | 1 - | 0,000207861 - |
| 1057 | 1 | 0.2um_NOKCPHJN_584203 | Bacteria | Proteobacteria | Gammaprotec | unclassified | unclassified | unclassified | Gammaprotec | poorly charact | Function unkn | 16R8I | Lipoprotein | 314287.GB220 | 1,4 | 0,8 | 0,000212398 | 0,000122706 | 2 - | 0,000320051 - | 0 - | 0 - | 2 - | 0,000296995 - | 1 - | 0,000147565 - | 2 - | 0,000297378 - |
| 1058 | 1 | 0.2um_NOKCPHJN_291839 | Bacteria | Proteobacteria | Alphaproteob | Rhodobactera | unclassified | unclassified | Rhodobactera | unclassified | unclassified | unclassified | unclassified | unclassified | 1 | 0 | 0,00022994 | 2,25E-05 | 1 - | 0,000231994 - | 1 - | 0,000272936 - | 1 - | 0,000215282 - | 1 - | 0,00021393 - | 1 - | 0,00021556 - |
| 1059 | 1 | 3um_FIJJAFBF_66744 | Bacteria | Proteobacteria | Gammaprotec | Pseudomonad | Pseudomonad | Pseudomonas | Pseudomonas | metabolism | Inorganic ion t | 05ENV | phosphonate | 399739.Pmen | 0,4 | 0,489897949 | 0,000144594 | 0,000178058 | 1 - | 0,000332174 - | 1 - | 0,000390795 - | 0 - | 0 - | 0 - | 0 - | 0 - | 0 - |
| 1060 | 2 | 3um_FIJJAFBF_165523;3um_FI | Bacteria | Proteobacteria | Gammaprotec | various | various | various | various | poorly charact | Function unkn | 0QRAP | NA | 393595.ABO_ | 0,8 | 0,979795897 | 7,17E-05 | 8,78E-05 | 0 - | 0 - | 0 - | 0 - | 0 - | 0 - | 2 - | 0,000178511 - | 2 - | 0,000179871 - |
| 1061 | 1 | 3um_FIJJAFBF_390935 | Viruses | unclassified | unclassified | unclassified | Phycodnavirid | unclassified | Organic Lake p | unclassified | unclassified | unclassified | unclassified | unclassified | 2 | 0 | 0,002045117 | 0,000200103 | 2 - | 0,002063385 - | 2 - | 0,002427527 - | 2 - | 0,001914747 - | 2 - | 0,001902714 - | 2 - | 0,001917212 - |
| 1062 | 1 | 3um_FIJJAFBF_243622 | Bacteria | Bacteroidetes | Flavobacteriia | Flavobacterial | Flavobacteriac | Tenacibaculum | Tenacibaculum | information st | Translation, ri | 09EM4 | Binds directly | 313594.PI23P | 2,4 | 0,489897949 | 0,001821667 | 0,000362791 | 3 - | 0,002307733 - | 2 - | 0,001809998 - | 2 - | 0,001427662 - | 2 - | 0,00141869 - | 3 - | 0,00214425 - |
| 1063 | 1 | 3um_FIJJAFBF_788047 | Bacteria | Proteobacteria | Alphaproteob | unclassified | unclassified | unclassified | Alphaproteob | unclassified | unclassified | unclassified | unclassified | unclassified | 0,6 | 0,489897949 | 0,000174088 | 0,000142144 | 0 - | 0 - | 0 - | 0 - | 1 - | 0,000290631 - | 1 - | 0,000288805 - | 1 - | 0,000291005 - |
| 1064 | 1 | 3um_FIJJAFBF_465472 | Eukaryota | Cnidaria | Anthozoa | Scleractinia | Acroporidae | Acropora | Acropora digit | poorly charact | Function unkn | 13CT1 | Reverse trans | 8128.ENSONI | 0,2 | 0,4 | 3,20E-05 | 6,41E-05 | 0 - | 0 - | 0 - | 0 - | 1 - | 0,00016019 - | 0 - | 0 - | 0 - | 0 - |
| 1065 | 1 | 3um_FIJJAFBF_658713 | Eukaryota | unclassified | unclassified | Isochrysidales | Noelaerhabda | Emiliania | Emiliania hux | unclassified | unclassified | unclassified | unclassified | unclassified | 1,4 | 0,489897949 | 0,000511681 | 0,000212865 | 1 - | 0,000362371 - | 2 - | 0,000852644 - | 2 - | 0,000672535 - | 1 - | 0,000334154 - | 1 - | 0,0003367 - |
| 1066 | 1 | 3um_FIJJAFBF_115452 | Bacteria | Proteobacteria | Gammaprotec | unclassified | unclassified | unclassified | SAR86 cluster | information st | Translation, ri | 179RU | 30S ribosomal | 566466.NOR5 | 0,4 | 0,489897949 | 0,000249764 | 0,000305901 | 0 - | 0 - | 0 - | 0 - | 0 - | 0 - | 1 - | 0,000622041 - | 1 - | 0,000626781 - |
| 1067 | 1 | 3um_FIJJAFBF_65172 | Bacteria | Bacteroidetes | Flavobacteriia | Flavobacterial | Flavobacteriac | Mesoflavibact | Mesoflavibact | poorly charact | Function unkn | 0PB02 | FN3 | 216432.CA25 | 1,8 | 0,748331477 | 0,000341775 | 0,000152734 | 3 - | 0,000565767 - | 2 - | 0,000443741 - | 1 - | 0,000175004 - | 1 - | 0,000173904 - | 2 - | 0,000350458 - |
| 1068 | 1 | 3um_FIJJAFBF_115558 | Eukaryota | unclassified | unclassified | Kinetoplastida | Trypanosomat | Leishmania | Leishmania in | unclassified | unclassified | unclassified | unclassified | unclassified | 0,4 | 0,489897949 | 2,67E-05 | 3,30E-05 | 0 - | 0 - | 1 - | 7,45E-05 - | 1 - | 5,88E-05 - | 0 - | 0 - | 0 - | 0 - |
| 1069 | 1 | 0.2um_NOKCPHJN_892782 | unclassified | unclassified | unclassified | unclassified | unclassified | unclassified | unclassified | unclassified | unclassified | unclassified | unclassified | unclassified | 3 | 0,894427191 | 0,001002078 | 0,000273951 | 4 - | 0,001364885 - | 2 - | 0,000802879 - | 3 - | 0,000949923 - | 4 - | 0,001258605 - | 2 - | 0,000634097 - |
| 1070 | 1 | 3um_FIJJAFBF_128441 | Bacteria | Bacteroidetes | Flavobacteriia | Flavobacterial | Flavobacteriac | Polaribacter | Polaribacter s | unclassified | unclassified | unclassified | unclassified | unclassified | 24,2 | 3,6 | 0,002212214 | 0,000203658 | 20 - | 0,001865827 - | 20 - | 0,002195104 - | 25 - | 0,002164275 - | 28 - | 0,002408755 - | 28 - | 0,002427108 - |
| 1071 | 1 | 3um_FIJJAFBF_254313 | Bacteria | Proteobacteria | Gammaprotec | Cellvibrionales | Spongiibacter | Spongiibacter | Spongiibacter | information st | Replication, re | 16W6D | single-strande | 207954.MED9 | 0,4 | 0,489897949 | 0,000207577 | 0,000254515 | 1 - | 0,000537999 - | 0 - | 0 - | 0 - | 0 - | 0 - | 0 - | 1 - | 0,000499886 - |
| 1072 | 1 | 3um_FIJJAFBF_321054 | unclassified | unclassified | unclassified | unclassified | unclassified | unclassified | unclassified | unclassified | unclassified | unclassified | unclassified | unclassified | 0,4 | 0,489897949 | 0,000339288 | 0,000415541 | 0 - | 0 - | 0 - | 0 - | 1 - | 0,000847674 - | 0 - | 0 - | 1 - | 0,000848766 - |
| 1073 | 3 | 0.2um_NOKCPHJN_439880;0.2 | various | various | various | various | various | various | various | information st | Replication, re | COG0513 | purine NTP-de | various | 1 | 0 | 0,00023178 | 2,27E-05 | 1 - | 0,00023385 - | 1 - | 0,00027512 - | 1 - | 0,000217005 - | 1 - | 0,000215641 - | 1 - | 0,000217284 - |
| 1074 | 1 | 3um_FIJJAFBF_434438 | Eukaryota | Bacillariophyta | Bacillariophyc | Naviculales | Naviculaceae | Fistulifera | Fistulifera sola | information st | Transcription | COG5108 | DNA-depende | 3218.PP1S256 | 0,2 | 0,4 | 3,72E-05 | 7,43E-05 | 0 - | 0 - | 0 - | 0 - | 1 - | 0,000185792 - | 0 - | 0 - | 0 - | 0 - |
| 1075 | 1 | 3um_FIJJAFBF_196455 | Bacteria | Proteobacteria | Gammaprotec | Cellvibrionales | Halieaceae | unclassified | marine gamm | unclassified | unclassified | unclassified | unclassified | unclassified | 1,8 | 0,4 | 0,000199357 | 4,88E-05 | 1 - | 0,000111854 - | 2 - | 0,000263188 - | 2 - | 0,000207594 - | 2 - | 0,000206289 - | 2 - | 0,000207861 - |
| 1076 | 1 | 0.2um_NOKCPHJN_548160 | Eukaryota | Phaeophyceae | unclassified | Ectocarpales | Ectocarpaceae | Ectocarpus | Ectocarpus sili | cellular proces | Signal transdu | 151NR | Pfam:efhand | 67593.Physo1 | 1,6 | 0,489897949 | 0,000832283 | 0,000203761 | 1 - | 0,000537999 - | 1 - | 0,000632944 - | 2 - | 0,000998488 - | 2 - | 0,000992213 - | 2 - | 0,000999773 - |
| 1077 | 1 | 3um_FIJJAFBF_254286 | Bacteria | Proteobacteria | Gammaprotec | Cellvibrionales | Spongiibacter | Spongiibacter | Spongiibacter | information st | Translation, ri | COG0088 | One of the pri | 876044.IMCC3 | 0,8 | 0,748331477 | 0,000351258 | 0,000324443 | 2 - | 0,00085555 - | 1 - | 0,000503268 - | 0 - | 0 - | 0 - | 0 - | 1 - | 0,000397471 - |
| 1078 | 1 | 0.2um_NOKCPHJN_699240 | Eukaryota | Streptophyta | unclassified | Solanales | Convolvulacea | Ipomoea | Ipomoea nil | unclassified | unclassified | unclassified | unclassified | unclassified | 0,2 | 0,4 | 2,59E-05 | 5,18E-05 | 0 - | 0 - | 1 - | 0,00012961 - | 0 - | 0 - | 0 - | 0 - | 0 - | 0 - |
| 1079 | 1 | 0.2um_NOKCPHJN_315798 | Bacteria | Bacteroidetes | unclassified | unclassified | unclassified | unclassified | Bacteroidetes | unclassified | unclassified | unclassified | unclassified | unclassified | 1 | 0 | 0,000116825 | 1,14E-05 | 1 - | 0,000117868 - | 1 - | 0,000138669 - | 1 - | 0,000109377 - | 1 - | 0,00010869 - | 1 - | 0,000109518 - |
| 1080 | 1 | 3um_FIJJAFBF_263679 | Bacteria | Proteobacteria | Alphaproteob | unclassified | unclassified | Candidatus Pu | Candidatus Pu | information st | Translation, ri | 0FN1J | This protein is | 533247.CRD_ | 1,2 | 0,748331477 | 0,00084222 | 0,000508757 | 2 - | 0,001414417 - | 1 - | 0,000832015 - | 2 - | 0,001312528 - | 1 - | 0,00065214 - | 0 - | 0 - |
| 1081 | 1 | 0.2um_NOKCPHJN_26421 | Bacteria | Bacteroidetes | Flavobacteriia | unclassified | unclassified | unclassified | Flavobacteria | metabolism | Inorganic ion t | 05C27 | TonB-depende | 487796.Flav2 | 0,2 | 0,4 | 1,66E-05 | 3,32E-05 | 1 - | 8,29E-05 - | 0 - | 0 - | 0 - | 0 - | 0 - | 0 - | 0 - | 0 - |
| 1082 | 1 | 0.2um_NOKCPHJN_257407 | Eukaryota | Apicomplexa | Aconoidasida | Haemosporida | Plasmodiidae | Plasmodium | Plasmodium fa | cellular proces | Signal transdu | KOG0613 | fibronectin typ | 7460.GB1563 | 0,8 | 0,4 | 0,000125537 | 6,29E-05 | 1 - | 0,000166087 - | 0 - | 0 - | 1 - | 0,000154123 - | 1 - | 0,000153154 - | 1 - | 0,000154321 - |
| 1083 | 1 | 3um_FIJJAFBF_632635 | Bacteria | Proteobacteria | Alphaproteob | Rhodobactera | Rhodobactera | Epibacterium | Epibacterium | unclassified | unclassified | unclassified | unclassified | unclassified | 0,2 | 0,4 | 3,47E-05 | 6,94E-05 | 0 - | 0 - | 0 - | 0 - | 1 - | 0,000173511 - | 0 - | 0 - | 0 - | 0 - |
| 1084 | 1 | 0.2um_NOKCPHJN_996157 | Eukaryota | unclassified | Oomycetes | Peronosporale | Peronosporac | Phytophthora | Phytophthora | unclassified | unclassified | unclassified | unclassified | unclassified | 0,2 | 0,4 | 3,12E-05 | 6,23E-05 | 0 - | 0 - | 0 - | 0 - | 0 - | 0 - | 0 - | 0 - | 1 - | 0,000155796 - |
| 1085 | 1 | 0.2um_NOKCPHJN_312745 | Eukaryota | unclassified | Labyrinthulom | unclassified | Thraustochytr | Aurantiochytr | Aurantiochytr | cellular proces | Signal transdu | KOG0613 | fibronectin typ | 7165.AGAP00 | 0,8 | 0,4 | 2,17E-05 | 1,09E-05 | 1 - | 2,87E-05 - | 0 - | 0 - | 1 - | 2,66E-05 - | 1 - | 2,64E-05 - | 1 - | 2,66E-05 - |
| 1086 | 2 | 0.2um_NOKCPHJN_656428;3ur | Bacteria | Bacteroidetes | Flavobacteriia | Flavobacterial | Flavobacteriac | Algibacter | Algibacter sp. | unclassified | unclassified | unclassified | unclassified | unclassified | 0,4 | 0,489897949 | 0,000194454 | 0,000238425 | 1 - | 0,000503988 - | 0 - | 0 - | 0 - | 0 - | 0 - | 0 - | 1 - | 0,000468284 - |
| 1087 | 1 | 0.2um_NOKCPHJN_556776 | Bacteria | Bacteroidetes | Flavobacteriia | Flavobacterial | unclassified | unclassified | Flavobacterial | unclassified | unclassified | unclassified | unclassified | unclassified | 1,4 | 0,489897949 | 0,000415582 | 0,000181359 | 2 - | 0,000582683 - | 2 - | 0,000685514 - | 1 - | 0,000270355 - | 1 - | 0,000268656 - | 1 - | 0,000270703 - |
| 1088 | 1 | 3um_FIJJAFBF_734730 | Eukaryota | Chromerida | unclassified | unclassified | unclassified | Vitrella | Vitrella brassi | poorly charact | Function unkn | 0XPG5 | KH domain co | 67593.Physo1 | 2,8 | 1,166190379 | 0,000427726 | 0,000157055 | 2 - | 0,000320051 - | 1 - | 0,000188266 - | 4 - | 0,000593991 - | 4 - | 0,000590258 - | 3 - | 0,000446067 - |
| 1089 | 1 | 3um_FIJJAFBF_106477 | Bacteria | Bacteroidetes | Flavobacteriia | Flavobacterial | Flavobacteriac | Mangrovimon | Mangrovimon | unclassified | unclassified | unclassified | unclassified | unclassified | 1 | 0 | 0,00069534 | 6,80E-05 | 1 - | 0,000701551 - | 1 - | 0,000825359 - | 1 - | 0,000651014 - | 1 - | 0,000646923 - | 1 - | 0,000651852 - |
| 1090 | 1 | 3um_FIJJAFBF_711074 | Bacteria | Bacteroidetes | Flavobacteriia | Flavobacterial | Flavobacteriac | Flaviramulus | Flaviramulus i | unclassified | unclassified | unclassified | unclassified | unclassified | 14,2 | 3,059411708 | 0,001272218 | 0,000185865 | 13 - | 0,001198759 - | 9 - | 0,000976371 - | 18 - | 0,001540254 - | 16 - | 0,001360511 - | 15 - | 0,001285197 - |
| 1091 | 1 | 3um_FIJJAFBF_269224 | Bacteria | Bacteroidetes | Flavobacteriia | Flavobacterial | Flavobacteriac | Riemerella | Riemerella co | unclassified | unclassified | unclassified | unclassified | unclassified | 3,4 | 0,489897949 | 0,001679355 | 0,000395436 | 4 - | 0,001970648 - | 4 - | 0,002318424 - | 3 - | 0,001371518 - | 3 - | 0,001362899 - | 3 - | 0,001373284 - |

| | | | | | | | | | | | | | | | | | | | | | | | | | | | | |
|---|---|---|---|---|---|---|---|---|---|---|---|---|---|---|---|---|---|---|---|---|---|---|---|---|---|---|---|---|
| 1092 | 1 | 0.2um_NOKCPHJN_469346 | Eukaryota | unclassified | Ichthyosporea | unclassified | unclassified | Capsaspora | Capsaspora o | cellular proces | Defense mech | COG0842 | (ABC) transpo | 5786.XP_0032 | 0,2 | 0,4 | 4,96E-05 | 9,92E-05 | 0 - | 0 - | 1 - | 0,000248005 - | 0 - | 0 - | 0 - | 0 - | 0 - | 0 - |
| 1093 | 1 | 0.2um_NOKCPHJN_912086 | Viruses | unclassified | unclassified | Caudovirales | Podoviridae | unclassified | Cellulophaga | unclassified | unclassified | unclassified | unclassified | unclassified | 1 | 0 | 0,000175946 | 1,72E-05 | 1 - | 0,000177518 - | 1 - | 0,000208846 - | 1 - | 0,00016473 - | 1 - | 0,000163695 - | 1 - | 0,000164942 - |
| 1094 | 2 | 0.2um_NOKCPHJN_188037;3ur | Eukaryota | various | various | various | various | various | various | poorly charact | Function unkn | 0VCF1 | chromosome | 45351.NEMVE | 1 | 0 | 0,000317217 | 3,10E-05 | 1 - | 0,000320051 - | 1 - | 0,000376532 - | 1 - | 0,000296995 - | 1 - | 0,000295129 - | 1 - | 0,000297378 - |
| 1095 | 1 | 0.2um_NOKCPHJN_1065485 | unclassified | unclassified | unclassified | unclassified | unclassified | unclassified | unclassified | unclassified | unclassified | unclassified | unclassified | unclassified | 1 | 0 | 0,000679043 | 6,64E-05 | 1 - | 0,000685108 - | 1 - | 0,000806015 - | 1 - | 0,000635756 - | 1 - | 0,000631761 - | 1 - | 0,000636574 - |
| 1096 | 1 | 3um_FIJJAFBF_253585 | Viruses | unclassified | unclassified | unclassified | unclassified | unclassified | uncultured M | unclassified | unclassified | unclassified | unclassified | unclassified | 3 | 0 | 0,000841137 | 8,23E-05 | 3 - | 0,00084865 - | 3 - | 0,000998418 - | 3 - | 0,000787517 - | 3 - | 0,000782568 - | 3 - | 0,000788531 - |
| 1097 | 1 | 3um_FIJJAFBF_51134 | unclassified | unclassified | unclassified | unclassified | unclassified | unclassified | unclassified | unclassified | unclassified | unclassified | unclassified | unclassified | 1 | 0 | 0,001849308 | 0,000180944 | 1 - | 0,001865827 - | 1 - | 0,002195104 - | 1 - | 0,00173142 - | 1 - | 0,001720539 - | 1 - | 0,001733649 - |
| 1098 | 1 | 3um_FIJJAFBF_47498 | Bacteria | Bacteroidetes | unclassified | unclassified | unclassified | unclassified | unidentified e | information st | Replication, re | 09GNB | DNA-binding | 50743.SCB49_ | 1,8 | 0,748331477 | 0,001717407 | 0,000702491 | 3 - | 0,002923128 - | 1 - | 0,001146332 - | 2 - | 0,001808372 - | 1 - | 0,000898504 - | 2 - | 0,0018107 - |
| 1099 | 1 | 3um_FIJJAFBF_642735 | Eukaryota | Bacillariophyta | Bacillariophyc | Naviculales | Phaeodactylac | Phaeodactylu | Phaeodactylu | unclassified | unclassified | unclassified | unclassified | unclassified | 0,6 | 0,489897949 | 0,000257909 | 0,000210584 | 0 - | 0 - | 0 - | 0 - | 1 - | 0,000430565 - | 1 - | 0,000427859 - | 1 - | 0,000431119 - |
| 1100 | 4 | 0.2um_NOKCPHJN_138709;0.2 | Bacteria | unclassified | unclassified | unclassified | unclassified | unclassified | uncultured ba | unclassified | unclassified | unclassified | unclassified | unclassified | 0,4 | 0,489897949 | 5,95E-05 | 7,29E-05 | 0 - | 0 - | 0 - | 0 - | 1 - | 0,000148769 - | 0 - | 0 - | 1 - | 0,000148961 - |
| 1101 | 1 | 0.2um_NOKCPHJN_381260 | Eukaryota | Chlorophyta | Chlorophyceae | Chlamydomor | Volvocaceae | Gonium | Gonium pecto | information st | Replication, re | 0XNUT | helicase activi | 3055.EDP0692 | 0,4 | 0,489897949 | 2,00E-05 | 2,48E-05 | 0 - | 0 - | 1 - | 5,59E-05 - | 0 - | 0 - | 0 - | 0 - | 1 - | 4,42E-05 - |
| 1102 | 2 | 0.2um_NOKCPHJN_93629;3um | Eukaryota | Xanthophycea | unclassified | various | various | various | various | metabolism | Carbohydrate | COG1850 | RuBisCO catal | various | 2,2 | 0,4 | 0,00038719 | 6,40E-05 | 2 - | 0,000357205 - | 2 - | 0,000420244 - | 3 - | 0,00049721 - | 2 - | 0,00032939 - | 2 - | 0,0003319 - |
| 1103 | 2 | 3um_FIJJAFBF_257128;3um_FI | various | various | various | various | various | various | various | unclassified | unclassified | unclassified | unclassified | unclassified | 1 | 0 | 0,000712438 | 6,97E-05 | 1 - | 0,000718802 - | 1 - | 0,000845655 - | 1 - | 0,000667022 - | 1 - | 0,000662831 - | 1 - | 0,000667881 - |
| 1104 | 1 | 3um_FIJJAFBF_164074 | Bacteria | Bacteroidetes | Flavobacteriia | Flavobacterial | Flavobacteriac | Formosa | Formosa sp. H | unclassified | unclassified | unclassified | unclassified | unclassified | 0,8 | 0,4 | 0,000297763 | 0,000153281 | 0 - | 0 - | 1 - | 0,000442789 - | 1 - | 0,000349256 - | 1 - | 0,000347062 - | 1 - | 0,000349706 - |
| 1105 | 1 | 0.2um_NOKCPHJN_538433 | Bacteria | Proteobacteri | Gammaprotec | unclassified | unclassified | unclassified | Gammaprotec | unclassified | unclassified | unclassified | unclassified | unclassified | 0,8 | 0,4 | 0,000161283 | 8,27E-05 | 1 - | 0,000200214 - | 1 - | 0,000235548 - | 0 - | 0 - | 1 - | 0,000184624 - | 1 - | 0,000186031 - |
| 1106 | 1 | 3um_FIJJAFBF_375705 | unclassified | unclassified | unclassified | unclassified | unclassified | unclassified | unclassified | unclassified | unclassified | unclassified | unclassified | unclassified | 0,2 | 0,4 | 0,00021951 | 0,000439021 | 0 - | 0 - | 1 - | 0,001097552 - | 0 - | 0 - | 0 - | 0 - | 0 - | 0 - |
| 1107 | 1 | 3um_FIJJAFBF_483811 | Bacteria | Actinobacteria | Actinobacteria | Micrococcales | Microbacteria | Candidatus Ac | Candidatus Ac | unclassified | unclassified | unclassified | unclassified | unclassified | 0,4 | 0,8 | 3,09E-05 | 6,18E-05 | 2 - | 0,000154527 - | 0 - | 0 - | 0 - | 0 - | 0 - | 0 - | 0 - | 0 - |
| 1108 | 1 | 3um_FIJJAFBF_128394 | Bacteria | Bacteroidetes | Flavobacteriia | Flavobacterial | Flavobacteriac | Polaribacter | Polaribacter s | unclassified | unclassified | unclassified | unclassified | unclassified | 0,2 | 0,4 | 2,17E-05 | 4,35E-05 | 0 - | 0 - | 1 - | 0,000108714 - | 0 - | 0 - | 0 - | 0 - | 0 - | 0 - |
| 1109 | 1 | 0.2um_NOKCPHJN_551609 | Bacteria | Bacteroidetes | Flavobacteriia | Flavobacterial | Flavobacteriac | Formosa | Formosa algae | poorly charact | Function unkn | 09SDX | Inherit from N | 391598.FBBAL | 1 | 0 | 6,62E-05 | 6,48E-06 | 1 - | 6,68E-05 - | 1 - | 7,86E-05 - | 1 - | 6,20E-05 - | 1 - | 6,16E-05 - | 1 - | 6,21E-05 - |
| 1110 | 1 | 3um_FIJJAFBF_40656 | Eukaryota | unclassified | Pelagophycea | Pelagomonad | unclassified | Aureococcus | Aureococcus a | unclassified | unclassified | unclassified | unclassified | unclassified | 0,4 | 0,489897949 | 0,000676445 | 0,000828483 | 0 - | 0 - | 0 - | 0 - | 0 - | 0 - | 1 - | 0,001684695 - | 1 - | 0,001697531 - |
| 1111 | 1 | 0.2um_NOKCPHJN_116081 | Bacteria | Proteobacteri | Alphaproteob | Rhodobactera | Rhodobactera | unclassified | Rhodobactera | unclassified | unclassified | unclassified | unclassified | unclassified | 0,4 | 0,489897949 | 9,12E-05 | 0,000111706 | 0 - | 0 - | 0 - | 0 - | 0 - | 0 - | 1 - | 0,00022715 - | 1 - | 0,000228881 - |
| 1112 | 1 | 3um_FIJJAFBF_734855 | unclassified | unclassified | unclassified | unclassified | unclassified | unclassified | unclassified | unclassified | unclassified | unclassified | unclassified | unclassified | 0,2 | 0,4 | 1,30E-05 | 2,60E-05 | 0 - | 0 - | 0 - | 0 - | 1 - | 6,50E-05 - | 0 - | 0 - | 0 - | 0 - |
| 1113 | 1 | 0.2um_NOKCPHJN_710508 | Eukaryota | Eustigmatoph | unclassified | Eustigmatales | Monodopsida | Nannochlorop | Nannochlorop | metabolism | Energy produc | 1BFZ8 | Cytochrome b | 3218.PP1S35_ | 3,2 | 0,4 | 0,002269881 | 0,000273076 | 3 - | 0,002156406 - | 3 - | 0,002536964 - | 3 - | 0,002001067 - | 4 - | 0,002651323 - | 3 - | 0,002003643 - |
| 1114 | 1 | 0.2um_NOKCPHJN_488159 | Bacteria | Proteobacteri | Alphaproteob | Sphingomona | various | various | various | unclassified | unclassified | unclassified | unclassified | unclassified | 0,4 | 0,8 | 0,000220042 | 0,000440084 | 0 - | 0 - | 0 - | 0 - | 0 - | 0 - | 2 - | 0,001100209 - | 0 - | 0 - |
| 1115 | 1 | 3um_FIJJAFBF_380757 | Bacteria | Proteobacteri | Gammaprotec | Thiotrichales | Piscirickettsia | Methylophaga | Methylophaga | information st | Translation, ri | 05CRK | lysyL-tRNA syr | 637616.MDM | 0,8 | 0,4 | 0,000142137 | 7,29E-05 | 1 - | 0,000176446 - | 1 - | 0,000207585 - | 0 - | 0 - | 1 - | 0,000162707 - | 1 - | 0,000163947 - |
| 1116 | 1 | 3um_FIJJAFBF_452428 | Bacteria | Proteobacteri | Alphaproteob | Rhodospirilla | Rhodospirillac | unclassified | Rhodospirilla | metabolism | Energy produc | COG0056 | Produces ATP | 1082931.KKY_ | 0,6 | 0,8 | 9,99E-05 | 0,000135361 | 2 - | 0,000341221 - | 0 - | 0 - | 0 - | 0 - | 0 - | 0 - | 1 - | 0,000158524 - |
| 1117 | 1 | 0.2um_NOKCPHJN_484873 | Bacteria | Proteobacteri | Gammaprotec | Chromatiales | Chromatiacea | Candidatus Te | Candidatus Te | unclassified | unclassified | unclassified | unclassified | unclassified | 1 | 0 | 0,000196646 | 1,92E-05 | 1 - | 0,000198402 - | 1 - | 0,000233416 - | 1 - | 0,00018411 - | 1 - | 0,000182953 - | 1 - | 0,000184347 - |
| 1118 | 1 | 0.2um_NOKCPHJN_559310 | unclassified | unclassified | unclassified | unclassified | unclassified | unclassified | unclassified | unclassified | unclassified | unclassified | unclassified | unclassified | 1 | 0 | 0,000257152 | 2,52E-05 | 1 - | 0,000259449 - | 1 - | 0,000305236 - | 1 - | 0,00024076 - | 1 - | 0,000239247 - | 1 - | 0,00024107 - |
| 1119 | 1 | 0.2um_NOKCPHJN_443530 | Bacteria | Bacteroidetes | Flavobacteriia | Flavobacterial | Flavobacteriac | unclassified | Flavobacteria | cellular proces | Cell wall/mem | 0P81E | outer membra | 313598.MED1 | 0,2 | 0,4 | 3,72E-05 | 7,44E-05 | 0 - | 0 - | 0 - | 0 - | 0 - | 0 - | 1 - | 0,000185897 - | 0 - | 0 - |
| 1120 | 1 | 3um_FIJJAFBF_543375 | Eukaryota | Bacillariophyta | Bacillariophyc | Naviculales | Naviculaceae | Fistulifera | Fistulifera sola | information st | Replication, re | COG0513 | purine NTP-de | 2850.Phatr257 | 1,2 | 0,4 | 0,001214033 | 0,000362902 | 1 - | 0,001031692 - | 1 - | 0,001213763 - | 2 - | 0,001914747 - | 1 - | 0,000951357 - | 1 - | 0,000958606 - |
| 1121 | 1 | 3um_FIJJAFBF_309030 | Eukaryota | unclassified | Dinophyceae | Suessiales | Symbiodiniace | Symbiodinium | Symbiodinium | unclassified | unclassified | unclassified | unclassified | unclassified | 0,2 | 0,4 | 4,28E-05 | 8,57E-05 | 0 - | 0 - | 0 - | 0 - | 1 - | 0,000214149 - | 0 - | 0 - | 0 - | 0 - |
| 1122 | 1 | 3um_FIJJAFBF_407604 | Eukaryota | Bacillariophyta | Bacillariophyc | Naviculales | Phaeodactylac | Phaeodactylu | Phaeodactylu | cellular proces | Posttranslatio | KOG0543 | Peptidyl-proly | 2850.Phatr196 | 0,6 | 0,489897949 | 0,000245035 | 0,000200418 | 1 - | 0,000429872 - | 0 - | 0 - | 1 - | 0,000398906 - | 1 - | 0,000396399 - | 0 - | 0 - |
| 1123 | 1 | 0.2um_NOKCPHJN_732467 | Eukaryota | unclassified | Oomycetes | Saprolegniales | Saprolegniace | Thraustotheca | Thraustotheca | information st | Translation, ri | KOG3184 | Ribosomal prc | 10228.TriadP3 | 0,2 | 0,4 | 8,39E-05 | 0,000167756 | 0 - | 0 - | 1 - | 0,00041939 - | 0 - | 0 - | 0 - | 0 - | 0 - | 0 - |
| 1124 | 1 | 0.2um_NOKCPHJN_1001141 | Bacteria | various | various | various | various | various | various | information st | Translation, ri | COG0051 | tRNA binding | 205922.Pfl01_ | 1,4 | 0,489897949 | 0,001214468 | 0,00052999 | 2 - | 0,001702793 - | 2 - | 0,002003299 - | 1 - | 0,000790065 - | 1 - | 0,000785101 - | 1 - | 0,000791082 - |
| 1125 | 1 | 0.2um_NOKCPHJN_54710 | Eukaryota | Streptophyta | unclassified | Ericales | Primulaceae | Androsace | Androsace lax | cellular proces | Posttranslatio | 0IEKR | One of the co | 59689.fgenesh | 1,4 | 0,489897949 | 0,000259195 | 7,84E-05 | 1 - | 0,000190225 - | 1 - | 0,000223796 - | 2 - | 0,000353044 - | 1 - | 0,000175413 - | 2 - | 0,000353499 - |
| 1126 | 1 | 0.2um_NOKCPHJN_100353 | Bacteria | Proteobacteri | Alphaproteob | Rhodobactera | Rhodobactera | Thalassococcu | Thalassococcu | information st | Replication, re | 01QHS | DNA polymera | 314270.RB208 | 0,8 | 0,4 | 5,67E-05 | 2,84E-05 | 1 - | 7,50E-05 - | 0 - | 0 - | 1 - | 6,96E-05 - | 1 - | 6,91E-05 - | 1 - | 6,96E-05 - |
| 1127 | 1 | 3um_FIJJAFBF_163782 | Bacteria | Bacteroidetes | Flavobacteriia | Flavobacterial | unclassified | unclassified | Flavobacterial | unclassified | unclassified | unclassified | unclassified | unclassified | 1 | 0 | 0,000303907 | 2,97E-05 | 1 - | 0,000306622 - | 1 - | 0,000360734 - | 1 - | 0,000284534 - | 1 - | 0,000282746 - | 1 - | 0,0002849 - |
| 1128 | 1 | 0.2um_NOKCPHJN_523564 | Bacteria | Proteobacteri | unclassified | unclassified | unclassified | unclassified | Proteobacteri | unclassified | unclassified | unclassified | unclassified | unclassified | 1 | 0 | 0,000129921 | 1,27E-05 | 1 - | 0,000131082 - | 1 - | 0,000154215 - | 1 - | 0,000121639 - | 1 - | 0,000120875 - | 1 - | 0,000121796 - |
| 1129 | 1 | 0.2um_NOKCPHJN_221102 | Bacteria | Proteobacteri | Gammaprotec | Cellvibrionales | Halieaceae | Congregibacte | Congregibacte | poorly charact | Function unkn | 0XNQ8 | alkaline phos | 314285.KT71_ | 1,4 | 0,8 | 0,000184636 | 0,000103367 | 1 - | 0,000140761 - | 0 - | 0 - | 2 - | 0,000261242 - | 2 - | 0,0002596 - | 2 - | 0,000261578 - |
| 1130 | 1 | 3um_FIJJAFBF_279094 | unclassified | unclassified | unclassified | unclassified | unclassified | unclassified | unclassified | unclassified | unclassified | unclassified | unclassified | unclassified | 1 | 0 | 0,000607814 | 5,95E-05 | 1 - | 0,000613244 - | 1 - | 0,000721468 - | 1 - | 0,000569068 - | 1 - | 0,000565492 - | 1 - | 0,000569801 - |
| 1131 | 1 | 0.2um_NOKCPHJN_30655 | Bacteria | Bacteroidetes | Flavobacteriia | Flavobacterial | unclassified | unclassified | Flavobacterial | poorly charact | Function unkn | 11P35 | NA | 391603.FBALC | 1 | 0 | 0,000157459 | 1,54E-05 | 1 - | 0,000158866 - | 1 - | 0,000186902 - | 1 - | 0,000147422 - | 1 - | 0,000146495 - | 1 - | 0,000147611 - |
| 1132 | 1 | 3um_FIJJAFBF_361016 | Bacteria | Cyanobacteria | unclassified | Synechococca | Synechococca | Synechococcu | Synechococcu | unclassified | unclassified | unclassified | unclassified | unclassified | 0,8 | 0,4 | 0,000130057 | 6,67E-05 | 1 - | 0,000161499 - | 1 - | 0,00019 - | 1 - | 0,000149865 - | 1 - | 0,000148923 - | 0 - | 0 - |
| 1133 | 1 | 3um_FIJJAFBF_507104 | Eukaryota | unclassified | Cryptophyta | Pyrenomonad | Geminigerace | Guillardia | Guillardia thet | unclassified | unclassified | unclassified | unclassified | unclassified | 6,4 | 0,8 | 0,004617478 | 0,000572926 | 6 - | 0,004384693 - | 6 - | 0,005158494 - | 6 - | 0,004068837 - | 6 - | 0,004043268 - | 8 - | 0,005432099 - |
| 1134 | 1 | 3um_FIJJAFBF_453383 | Bacteria | Proteobacteri | Alphaproteob | Rhodobactera | Rhodobactera | unclassified | Rhodobactera | poorly charact | Function unkn | 177FT | NA | 411684.HPDF | 0,2 | 0,4 | 4,52E-05 | 9,05E-05 | 0 - | 0 - | 1 - | 0,00022625 - | 0 - | 0 - | 0 - | 0 - | 0 - | 0 - |
| 1135 | 1 | 0.2um_NOKCPHJN_933150 | Eukaryota | unclassified | Pelagophycea | Pelagomonad | unclassified | Aureococcus | Aureococcus a | unclassified | unclassified | unclassified | unclassified | unclassified | 1,2 | 0,4 | 0,000374395 | 0,000130713 | 2 - | 0,00062863 - | 1 - | 0,000369785 - | 1 - | 0,000291673 - | 1 - | 0,00028984 - | 1 - | 0,000292048 - |
| 1136 | 1 | 3um_FIJJAFBF_87379 | Viruses | unclassified | unclassified | unclassified | unclassified | unclassified | uncultured M | poorly charact | Function unkn | 17BR7 | Erf family | 749222.Nitsa_ | 0,2 | 0,4 | 7,76E-05 | 0,000155203 | 0 - | 0 - | 0 - | 0 - | 0 - | 0 - | 0 - | 0 - | 1 - | 0,000388007 - |
| 1137 | 1 | 3um_FIJJAFBF_281830 | Bacteria | Bacteroidetes | unclassified | unclassified | unclassified | unclassified | Bacteroidetes | information st | Replication, re | 0PB0H | DNA-binding | 487796.Flav2A | 0,6 | 0,8 | 0,000577234 | 0,000781747 | 2 - | 0,001970648 - | 0 - | 0 - | 0 - | 0 - | 0 - | 0 - | 1 - | 0,000915522 - |
| 1138 | 1 | 3um_FIJJAFBF_99520 | Viruses | unclassified | unclassified | Caudovirales | Siphoviridae | Cbastvirus | Cellulophaga | unclassified | unclassified | unclassified | unclassified | unclassified | 2,6 | 1,019803903 | 0,001168519 | 0,000444715 | 4 - | 0,001855955 - | 1 - | 0,000545872 - | 3 - | 0,001291694 - | 2 - | 0,000855718 - | 3 - | 0,001293357 - |
| 1139 | 1 | 3um_FIJJAFBF_355093 | Eukaryota | unclassified | Cryptophyta | Pyrenomonad | Geminigerace | Guillardia | Guillardia thet | unclassified | unclassified | unclassified | unclassified | unclassified | 1,6 | 0,489897949 | 0,002524385 | 0,00091344 | 2 - | 0,003131923 - | 2 - | 0,003684639 - | 2 - | 0,002906312 - | 1 - | 0,001444024 - | 1 - | 0,001455027 - |
| 1140 | 1 | 3um_FIJJAFBF_490907 | Bacteria | Bacteroidetes | Flavobacteriia | unclassified | unclassified | unclassified | Flavobacteria | metabolism | Inorganic ion t | 05C27 | TonB-depend | 487797.Flav3 | 0,8 | 0,4 | 6,32E-05 | 3,24E-05 | 1 - | 7,85E-05 - | 1 - | 9,24E-05 - | 0 - | 0 - | 1 - | 7,24E-05 - | 1 - | 7,29E-05 - |
| 1141 | 1 | 3um_FIJJAFBF_119151 | Bacteria | Bacteroidetes | Flavobacteriia | Flavobacterial | Flavobacteriac | Tenacibaculur | Tenacibaculur | unclassified | unclassified | unclassified | unclassified | unclassified | 1 | 0,894427191 | 0,000345592 | 0,000308957 | 0 - | 0 - | 0 - | 0 - | 1 - | 0,000346284 - | 2 - | 0,000688216 - | 2 - | 0,00069346 - |
| 1142 | 1 | 0.2um_NOKCPHJN_405347 | Eukaryota | unclassified | Oomycetes | Peronosporale | Peronosporac | Phytophthora | Phytophthora | unclassified | unclassified | unclassified | unclassified | unclassified | 0,2 | 0,4 | 2,77E-05 | 5,54E-05 | 0 - | 0 - | 0 - | 0 - | 0 - | 0 - | 0 - | 0 - | 1 - | 0,000138574 - |
| 1143 | 2 | 0.2um_NOKCPHJN_290410;0.2 | Bacteria | Proteobacteri | Gammaprotec | Alteromonada | Colwelliaceae | Colwellia | various | information st | Translation, ri | 05CEJ | Catalyzes the | various | 0,2 | 0,4 | 2,50E-05 | 5,00E-05 | 1 - | 0,000125098 - | 0 - | 0 - | 0 - | 0 - | 0 - | 0 - | 0 - | 0 - |
| 1144 | 1 | 3um_FIJJAFBF_87924 | Bacteria | Bacteroidetes | Flavobacteriia | Flavobacterial | Crocinitomica | unclassified | Crocinitomica | unclassified | unclassified | unclassified | unclassified | unclassified | 0,2 | 0,4 | 2,54E-05 | 5,09E-05 | 0 - | 0 - | 0 - | 0 - | 1 - | 0,000127151 - | 0 - | 0 - | 0 - | 0 - |
| 1145 | 1 | 3um_FIJJAFBF_594167 | Eukaryota | unclassified | Pelagophycea | Pelagomonad | unclassified | Aureococcus | Aureococcus a | metabolism | Energy produc | 0FN1Q | F(1)F(0) ATP s | 497965.Cyan7 | 7,6 | 0,489897949 | 0,003544873 | 0,000513399 | 8 - | 0,003751609 - | 8 - | 0,004413685 - | 7 - | 0,003046188 - | 7 - | 0,003027045 - | 8 - | 0,003485839 - |
| 1146 | 1 | 3um_FIJJAFBF_653488 | Eukaryota | Bacillariophyta | Bacillariophyc | Bacillariales | Bacillariaceae | Fragilariopsis | Fragilariopsis | poorly charact | Function unkn | COG1100 | GTP-binding P | 2850.Phatr43 | 2,8 | 0,4 | 0,001644647 | 0,000126638 | 3 - | 0,001801928 - | 2 - | 0,001413286 - | 3 - | 0,001672125 - | 3 - | 0,001661617 - | 3 - | 0,001674277 - |
| 1147 | 1 | 0.2um_NOKCPHJN_569319 | Bacteria | Proteobacteri | Alphaproteob | Pelagibacteral | Pelagibacterac | Candidatus Pe | Candidatus Pe | unclassified | unclassified | unclassified | unclassified | unclassified | 0,6 | 0,489897949 | 9,09E-05 | 7,52E-05 | 1 - | 0,0001464 - | 1 - | 0,000172237 - | 1 - | 0,000135854 - | 0 - | 0 - | 0 - | 0 - |
| 1148 | 1 | 0.2um_NOKCPHJN_820029 | Bacteria | Proteobacteri | Gammaprotec | Alteromonada | Colwelliaceae | Thalassomona | Thalassomona | unclassified | unclassified | unclassified | unclassified | unclassified | 0,6 | 0,489897949 | 0,000435273 | 0,000361449 | 0 - | 0 - | 1 - | 0,000845655 - | 0 - | 0 - | 1 - | 0,000662831 - | 1 - | 0,000667881 - |
| 1149 | 1 | 0.2um_NOKCPHJN_27395 | Bacteria | unclassified | unclassified | unclassified | unclassified | unclassified | uncultured ba | unclassified | unclassified | unclassified | unclassified | unclassified | 1,8 | 0,979795897 | 0,000193942 | 9,60E-05 | 1 - | 0,000111997 - | 1 - | 0,000131762 - | 1 - | 0,000103929 - | 3 - | 0,000309829 - | 3 - | 0,00031219 - |
| 1150 | 1 | 0.2um_NOKCPHJN_140603 | Bacteria | Bacteroidetes | Flavobacteriia | unclassified | unclassified | unclassified | Flavobacteria | cellular proces | Intracellular tr | 0BIEB | domain-conta | 1005048.CFU_ | 5 | 3,577708764 | 0,000444576 | 0,000321816 | 12 - | 0,001075998 - | 3 - | 0,000316472 - | 2 - | 0,000166415 - | 4 - | 0,000330738 - | 4 - | 0,000333258 - |
| 1151 | 1 | 3um_FIJJAFBF_799495 | unclassified | unclassified | unclassified | unclassified | unclassified | unclassified | unclassified | unclassified | unclassified | unclassified | unclassified | unclassified | 0,8 | 0,4 | 0,000323334 | 0,000162129 | 1 - | 0,000427775 - | 0 - | 0 - | 1 - | 0,00039696 - | 1 - | 0,000394465 - | 1 - | 0,000397471 - |
| 1152 | 1 | 0.2um_NOKCPHJN_330360 | Eukaryota | Arthropoda | Insecta | Hymenoptera | Apidae | Eufriesea | Eufriesea mex | unclassified | unclassified | unclassified | unclassified | unclassified | 0,2 | 0,4 | 1,25E-05 | 2,51E-05 | 0 - | 0 - | 0 - | 0 - | 1 - | 6,27E-05 - | 0 - | 0 - | 0 - | 0 - |
| 1153 | 1 | 3um_FIJJAFBF_745490 | Eukaryota | Bacillariophyta | Coscinodiscop | Thalassiosirale | Thalassiosirac | Thalassiosira | Thalassiosira | cellular proces | Posttranslatio | KOG0731 | ATP-depender | 2850.Phatr17 | 1,6 | 0,489897949 | 0,000997516 | 0,000244214 | 1 - | 0,000644808 - | 1 - | 0,000758602 - | 2 - | 0,001196717 - | 2 - | 0,001189196 - | 2 - | 0,001198257 - |
| 1154 | 1 | 3um_FIJJAFBF_742841 | Eukaryota | unclassified | Labyrinthulon | unclassified | Thraustochytr | Aurantiochytr | Aurantiochytr | unclassified | unclassified | unclassified | unclassified | unclassified | 0,2 | 0,4 | 2,66E-05 | 5,32E-05 | 0 - | 0 - | 0 - | 0 - | 1 - | 0,000132969 - | 0 - | 0 - | 0 - | 0 - |
| 1155 | 2 | 0.2um_NOKCPHJN_887992;3ur | Bacteria | Bacteroidetes | unclassified | unclassified | unclassified | unclassified | Bacteroidetes | unclassified | unclassified | unclassified | unclassified | unclassified | 0,2 | 0,4 | 3,16E-05 | 6,33E-05 | 0 - | 0 - | 0 - | 0 - | 0 - | 0 - | 1 - | 0,000158249 - | 0 - | 0 - |
| 1156 | 1 | 3um_FIJJAFBF_537682 | Eukaryota | Phaeophyceae | unclassified | Laminariales | Agaraceae | Thalassiophyll | Thalassiophyll | metabolism | Energy produc | 0IE88 | oxidoreductas | 35128.Thapsd | 8,6 | 1,019803903 | 0,001011243 | 0,000100404 | 10 - | 0,001194739 - | 7 - | 0,000983909 - | 8 - | 0,00088694 - | 9 - | 0,000991537 - | 9 - | 0,000999092 - |
| 1157 | 1 | 0.2um_NOKCPHJN_869431 | Eukaryota | unclassified | Pelagophycea | Pelagomonad | unclassified | Aureococcus | Aureococcus a | unclassified | unclassified | unclassified | unclassified | unclassified | 1 | 0 | 0,00031042 | 3,04E-05 | 1 - | 0,000313192 - | 1 - | 0,000368464 - | 1 - | 0,000290631 - | 1 - | 0,000288805 - | 1 - | 0,000291005 - |
| 1158 | 1 | 3um_FIJJAFBF_246857 | Eukaryota | Bacillariophyta | Mediophyceae | Biddulphiales | Biddulphiacea | Biddulphia | Biddulphia bio | information st | Translation, ri | 082FX | This protein b | 197221.tll016 | 1,2 | 0,4 | 0,000991255 | 0,000292098 | 1 - | 0,00084321 - | 1 - | 0,000992018 - | 1 - | 0,000782469 - | 2 - | 0,001555103 - | 1 - | 0,000783476 - |
| 1159 | 1 | 0.2um_NOKCPHJN_239679 | Bacteria | Proteobacteri | Alphaproteob | Rhodobactera | Rhodobactera | Marivivens | Marivivens niv | cellular proces | Signal transdu | 172JY | phosphoenolp | 391589.RGAI1 | 0,6 | 0,489897949 | 6,51E-05 | 5,31E-05 | 0 - | 0 - | 0 - | 0 - | 1 - | 0,000108647 - | 1 - | 0,000107964 - | 1 - | 0,000108787 - |
| 1160 | 1 | 3um_FIJJAFBF_582148 | Bacteria | Bacteroidetes | Flavobacteriia | Flavobacterial | Crocinitomica | unclassified | Crocinitomica | poorly charact | Function unkn | 09AR5 | Inherit from b | 313606.M231 | 1 | 0 | 0,000170426 | 1,67E-05 | 1 - | 0,000171949 - | 1 - | 0,000202294 - | 1 - | 0,000159562 - | 1 - | 0,00015856 - | 1 - | 0,000159768 - |
| 1161 | 2 | 3um_FIJJAFBF_327843;3um_FI | Bacteria | Bacteroidetes | Flavobacteriia | Flavobacterial | Flavobacteriac | various | various | information st | Translation, ri | COG0051 | tRNA binding | various | 1,4 | 0,489897949 | 0,001182048 | 0,000354342 | 1 - | 0,000868256 - | 1 - | 0,001021484 - | 1 - | 0,00080571 - | 2 - | 0,001601294 - | 2 - | 0,001613495 - |
| 1162 | 3 | 0.2um_NOKCPHJN_99117;3um | Bacteria | Bacteroidetes | Flavobacteriia | Flavobacterial | Flavobacteriac | Polaribacter | various | metabolism | Lipid transpor | 0P7W4 | reductase | various | 0,2 | 0,4 | 7,07E-05 | 0,000141442 | 1 - | 0,000353604 - | 0 - | 0 - | 0 - | 0 - | 0 - | 0 - | 0 - | 0 - |
| 1163 | 1 | 3um_FIJJAFBF_375489 | Bacteria | Bacteroidetes | Flavobacteriia | Flavobacterial | Flavobacteriac | Tamlana | Tamlana sp. U | metabolism | Carbohydrate | 099YR | Catalyzes the | 391603.FBALC | 2 | 0,632455532 | 0,000394131 | 0,000104559 | 2 - | 0,000407878 - | 1 - | 0,00023993 - | 3 - | 0,000567745 - | 2 - | 0,000376118 - | 2 - | 0,000378984 - |
| 1164 | 1 | 3um_FIJJAFBF_56137 | Bacteria | Bacteroidetes | Flavobacteriia | Flavobacterial | Flavobacteriac | Polaribacter | Polaribacter s | metabolism | Energy produc | COG0711 | F(1)F(0) ATP s | 313598.MED1 | 0,4 | 0,489897949 | 0,000221729 | 0,000274866 | 0 - | 0 - | 1 - | 0,000621505 - | 0 - | 0 - | 1 - | 0,000487141 - | 0 - | 0 - |
| 1165 | 1 | 3um_FIJJAFBF_273643 | Bacteria | Bacteroidetes | Flavobacteriia | Flavobacterial | Flavobacteriac | Algibacter | Algibacter sp. | unclassified | unclassified | unclassified | unclassified | unclassified | 1,6 | 0,8 | 0,000312537 | 0,000141051 | 1 - | 0,00020206 - | 1 - | 0,000237719 - | 3 - | 0,000562512 - | 2 - | 0,000372651 - | 1 - | 0,000187745 - |
| 1166 | 1 | 3um_FIJJAFBF_254302 | Bacteria | Proteobacteri | Gammaprotec | unclassified | unclassified | unclassified | SAR86 cluster | unclassified | unclassified | unclassified | unclassified | unclassified | 0,4 | 0,489897949 | 0,000194866 | 0,000238987 | 1 - | 0,000506901 - | 0 - | 0 - | 0 - | 0 - | 1 - | 0,00046743 - | 0 - | 0 - |
| 1167 | 1 | 0.2um_NOKCPHJN_915793 | unclassified | unclassified | unclassified | unclassified | unclassified | unclassified | unclassified | unclassified | unclassified | unclassified | unclassified | unclassified | 1 | 0 | 0,000860569 | 8,42E-05 | 1 - | 0,000868256 - | 1 - | 0,001021484 - | 1 - | 0,00080571 - | 1 - | 0,000800647 - | 1 - | 0,000806747 - |
| 1168 | 1 | 3um_FIJJAFBF_577836 | Eukaryota | Arthropoda | Insecta | Lepidoptera | Nymphalidae | Bicyclus | Bicyclus anyna | poorly charact | Function unkn | 140PW | NA | 400682.PAC_1 | 1 | 0 | 0,000188541 | 1,84E-05 | 1 - | 0,000190225 - | 1 - | 0,000223796 - | 1 - | 0,000176522 - | 1 - | 0,000175413 - | 1 - | 0,000176749 - |
| 1169 | 1 | 3um_FIJJAFBF_350453 | Eukaryota | Bacillariophyta | Bacillariophyc | Naviculales | Phaeodactylac | Phaeodactylu | Phaeodactylu | unclassified | unclassified | unclassified | unclassified | unclassified | 1 | 0 | 0,000852132 | 8,34E-05 | 1 - | 0,000859744 - | 1 - | 0,001011469 - | 1 - | 0,000797811 - | 1 - | 0,000792798 - | 1 - | 0,000798838 - |
| 1170 | 1 | 0.2um_NOKCPHJN_55584 | Bacteria | Bacteroidetes | Flavobacteriia | Flavobacterial | Flavobacteriac | Polaribacter | Polaribacter g | poorly charact | Function unkn | 06E9D | NA | 983544.Lacal_ | 1 | 0,894427191 | 0,000309188 | 0,000276931 | 0 - | 0 - | 0 - | 0 - | 2 - | 0,000618835 - | 1 - | 0,000307473 - | 2 - | 0,000619631 - |
| 1171 | 1 | 3um_FIJJAFBF_359334 | Bacteria | Bacteroidetes | Sphingobacter | unclassified | unclassified | unclassified | Sphingobacte | cellular proces | Cytoskeleton | 0F97C | NA | 441375.XP_00 | 0,4 | 0,489897949 | 8,79E-05 | 0,000107755 | 1 - | 0,000227776 - | 0 - | 0 - | 0 - | 0 - | 0 - | 0 - | 1 - | 0,00021164 - |
| 1172 | 1 | 3um_FIJJAFBF_315982 | Bacteria | Bacteroidetes | Flavobacteriia | Flavobacterial | Flavobacteriac | Flaviramulus | Flaviramulus b | poorly charact | Function unkn | 0P8G1 | peptidylprolyl | 391603.FBALC | 0,6 | 0,489897949 | 0,000154745 | 0,000126351 | 0 - | 0 - | 0 - | 0 - | 1 - | 0,000258339 - | 1 - | 0,000256715 - | 1 - | 0,000258671 - |
| 1173 | 1 | 3um_FIJJAFBF_292964 | Bacteria | Proteobacteri | Alphaproteob | Rhodobactera | Rhodobactera | Tateyamaria | Tateyamaria s | unclassified | unclassified | unclassified | unclassified | unclassified | 5,8 | 0,4 | 0,001658513 | 5,57E-05 | 6 - | 0,001742262 - | 5 - | 0,001708111 - | 6 - | 0,001616756 - | 6 - | 0,001606596 - | 6 - | 0,001618838 - |
| 1174 | 1 | 3um_FIJJAFBF_639236 | Bacteria | Bacteroidetes | Flavobacteriia | Flavobacterial | Flavobacteriac | Polaribacter | Polaribacter s | unclassified | unclassified | unclassified | unclassified | unclassified | 1 | 0 | 0,000201665 | 1,97E-05 | 1 - | 0,000203466 - | 1 - | 0,000239373 - | 1 - | 0,000188809 - | 1 - | 0,000187623 - | 1 - | 0,000189052 - |
| 1175 | 1 | 3um_FIJJAFBF_607840 | Eukaryota | unclassified | Dinophyceae | Suessiales | Symbiodiniace | Symbiodinium | Symbiodinium | unclassified | unclassified | unclassified | unclassified | unclassified | 5,2 | 0,4 | 0,001165273 | 0,000145218 | 6 - | 0,00135609 - | 5 - | 0,001329509 - | 5 - | 0,001048669 - | 5 - | 0,001042079 - | 5 - | 0,001050019 - |
| 1176 | 1 | 0.2um_NOKCPHJN_557692 | Eukaryota | Eustigmatoph | unclassified | Eustigmatales | Monodopsida | Nannochlorop | Nannochlorop | unclassified | unclassified | unclassified | unclassified | unclassified | 0,2 | 0,4 | 2,43E-05 | 4,86E-05 | 0 - | 0 - | 0 - | 0 - | 1 - | 0,000121458 - | 0 - | 0 - | 0 - | 0 - |
| 1177 | 1 | 0.2um_NOKCPHJN_1022212 | Eukaryota | unclassified | Spirotrichea | Sporadotrichi | Oxytrichidae | Oxytricha | Oxytricha trifa | unclassified | unclassified | unclassified | unclassified | unclassified | 0,2 | 0,4 | 0,000216528 | 0,000433056 | 1 - | 0,00108264 - | 0 - | 0 - | 0 - | 0 - | 0 - | 0 - | 0 - | 0 - |
| 1178 | 1 | 3um_FIJJAFBF_654925 | Bacteria | Bacteroidetes | Flavobacteriia | various | various | various | various | information st | Replication, re | 0PB0H | DNA-binding | 487797.Flav3 | 0,8 | 0,4 | 0,000736483 | 0,000369294 | 1 - | 0,000974376 - | 0 - | 0 - | 1 - | 0,000904186 - | 1 - | 0,000898504 - | 1 - | 0,00090535 - |
| 1179 | 1 | 0.2um_NOKCPHJN_667152 | Eukaryota | unclassified | Oomycetes | Peronosporale | Peronosporac | Phytophthora | various | unclassified | unclassified | unclassified | unclassified | unclassified | 0,4 | 0,489897949 | 0,000596449 | 0,000734489 | 1 - | 0,001370216 - | 1 - | 0,00161203 - | 0 - | 0 - | 0 - | 0 - | 0 - | 0 - |
| 1180 | 1 | 3um_FIJJAFBF_715085 | Eukaryota | unclassified | Pelagophycea | Pelagomonad | unclassified | Aureoumbra | Aureoumbra l | information st | Translation, ri | 0FKC1 | This is 1 of the | 449447.MAE_ | 2,2 | 0,4 | 0,001062186 | 0,000176419 | 2 - | 0,00097982 - | 2 - | 0,001152736 - | 2 - | 0,000909237 - | 2 - | 0,000903523 - | 3 - | 0,001365612 - |
| 1181 | 1 | 3um_FIJJAFBF_21306 | Bacteria | Bacteroidetes | Flavobacteriia | Flavobacterial | Flavobacteriac | Polaribacter | Polaribacter b | unclassified | unclassified | unclassified | unclassified | unclassified | 3,8 | 0,979795897 | 0,000782775 | 0,000202855 | 2 - | 0,00041759 - | 4 - | 0,00098257 - | 4 - | 0,000775017 - | 5 - | 0,000962683 - | 4 - | 0,000776014 - |
| 1182 | 1 | 0.2um_NOKCPHJN_23471 | Bacteria | Proteobacteri | Gammaprotec | Arenicellales | Arenicellaceae | Arenicella | Arenicella xan | metabolism | Energy produc | 0QJ3X | Aconitate hyd | 878320.AcavA | 1,6 | 0,489897949 | 0,000158597 | 4,36E-05 | 2 - | 0,000202995 - | 1 - | 0,00011941 - | 2 - | 0,000188372 - | 1 - | 9,36E-05 - | 2 - | 0,000188615 - |
| 1183 | 1 | 0.2um_NOKCPHJN_54434 | Bacteria | Proteobacteri | Gammaprotec | Cellvibrionales | Spongiibacter | Oceanicoccus | Oceanicoccus | cellular proces | Cell wall/mem | COG2885 | Ompa motb d | 247633.GP214 | 2 | 1,264911064 | 0,000434411 | 0,000269953 | 1 - | 0,00023261 - | 0 - | 0 - | 3 - | 0,00064756 - | 3 - | 0,000643491 - | 3 - | 0,000648394 - |
| 1184 | 1 | 0.2um_NOKCPHJN_163795 | Bacteria | Proteobacteri | Gammaprotec | Vibrionales | Vibrionaceae | Vibrio | Vibrio vulnific | metabolism | Inorganic ion t | 16SSZ | Involved in co | 216895.VV1_ | 0,8 | 0,4 | 8,76E-05 | 4,49E-05 | 1 - | 0,000108801 - | 1 - | 0,000128002 - | 0 - | 0 - | 1 - | 0,000100329 - | 1 - | 0,000101094 - |
| 1185 | 1 | 0.2um_NOKCPHJN_241771 | Bacteria | Bacteroidetes | Flavobacteriia | Flavobacterial | Flavobacteriac | unclassified | Flavobacteriac | metabolism | Amino acid tra | COG0473 | 3-isopropylma | 886377.Murru | 0,8 | 0,4 | 0,000228564 | 0,000114609 | 1 - | 0,000302393 - | 0 - | 0 - | 1 - | 0,000280609 - | 1 - | 0,000278846 - | 1 - | 0,000280971 - |
| 1186 | 2 | 0.2um_NOKCPHJN_237440;0.2 | Bacteria | Proteobacteri | Gammaprotec | unclassified | unclassified | unclassified | various | unclassified | unclassified | unclassified | unclassified | unclassified | 0,2 | 0,4 | 4,05E-05 | 8,09E-05 | 0 - | 0 - | 1 - | 0,000202294 - | 0 - | 0 - | 0 - | 0 - | 0 - | 0 - |
| 1187 | 1 | 3um_FIJJAFBF_117655 | Eukaryota | various | various | various | various | various | various | metabolism | Energy produc | COG1145 | 4Fe-4S ferred | 35128.Thapsd | 0,8 | 0,4 | 0,000856527 | 0,00044092 | 0 - | 0 - | 1 - | 0,001273702 - | 1 - | 0,001004651 - | 1 - | 0,000998338 - | 1 - | 0,001005944 - |
| 1188 | 1 | 0.2um_NOKCPHJN_210688 | Bacteria | Bacteroidetes | Flavobacteriia | Flavobacterial | Flavobacteriac | Flavobacteriu | Flavobacteriu | metabolism | Inorganic ion t | 05C27 | TonB-depend | 983548.Krodi_ | 4 | 1,414213562 | 0,000313375 | 9,93E-05 | 5 - | 0,000404119 - | 2 - | 0,000190175 - | 3 - | 0,000225005 - | 6 - | 0,000447182 - | 4 - | 0,000300393 - |
| 1189 | 1 | 3um_FIJJAFBF_263677 | Eukaryota | Bacillariophyta | Coscinodiscop | Rhizosoleniale | Rhizosoleniac | Rhizosolenia | Rhizosolenia s | metabolism | Energy produc | 11IIK | Low-potential | 35128.Thapsd | 5,4 | 0,489897949 | 0,002900973 | 0,000242543 | 5 - | 0,002723412 - | 5 - | 0,003204034 - | 5 - | 0,002527228 - | 6 - | 0,003013616 - | 6 - | 0,003036577 - |
| 1190 | 1 | 0.2um_NOKCPHJN_589718 | Eukaryota | unclassified | Oomycetes | Peronosporale | Peronosporac | Phytophthora | Phytophthora | poorly charact | Function unkn | 1AYEZ | NLR family, C | 9601.ENSPPY | 0,4 | 0,489897949 | 5,38E-05 | 6,62E-05 | 1 - | 0,000123512 - | 1 - | 0,00014531 - | 0 - | 0 - | 0 - | 0 - | 0 - | 0 - |
| 1191 | 1 | 3um_FIJJAFBF_67526 | unclassified | unclassified | unclassified | unclassified | unclassified | unclassified | unclassified | unclassified | unclassified | unclassified | unclassified | unclassified | 0,6 | 0,489897949 | 0,000576983 | 0,000479167 | 0 - | 0 - | 1 - | 0,001121412 - | 1 - | 0,00088453 - | 1 - | 0,000878971 - | 0 - | 0 - |
| 1192 | 1 | 0.2um_NOKCPHJN_738405 | Bacteria | Actinobacteria | Actinobacteria | Streptomycet | Streptomycet | Streptomyces | Streptomyces | unclassified | unclassified | unclassified | unclassified | unclassified | 0,2 | 0,4 | 3,49E-05 | 6,97E-05 | 0 - | 0 - | 1 - | 0,000174273 - | 0 - | 0 - | 0 - | 0 - | 0 - | 0 - |
| 1193 | 1 | 0.2um_NOKCPHJN_909610 | Eukaryota | Phaeophyceae | unclassified | Ectocarpales | Ectocarpaceae | Ectocarpus | Ectocarpus sil | unclassified | unclassified | unclassified | unclassified | unclassified | 6,4 | 0,489897949 | 0,000920188 | 7,29E-05 | 6 - | 0,000874025 - | 6 - | 0,001028271 - | 7 - | 0,000946241 - | 7 - | 0,000940295 - | 6 - | 0,000812108 - |
| 1194 | 2 | 0.2um_NOKCPHJN_654066;0.2 | Bacteria | Proteobacteri | Alphaproteob | Rhodobactera | unclassified | unclassified | Rhodobactera | unclassified | unclassified | unclassified | unclassified | unclassified | 1 | 0 | 0,000274188 | 2,68E-05 | 1 - | 0,000276637 - | 1 - | 0,000325457 - | 1 - | 0,000256709 - | 1 - | 0,000255096 - | 1 - | 0,000257039 - |
| 1195 | 1 | 3um_FIJJAFBF_806703 | Eukaryota | unclassified | Oomycetes | Saprolegniales | Saprolegniace | Thraustotheca | Thraustotheca | unclassified | unclassified | unclassified | unclassified | unclassified | 0,2 | 0,4 | 3,90E-05 | 7,81E-05 | 0 - | 0 - | 0 - | 0 - | 1 - | 0,000195148 - | 0 - | 0 - | 0 - | 0 - |
| 1196 | 1 | 3um_FIJJAFBF_545826 | Bacteria | Bacteroidetes | Flavobacteriia | Flavobacterial | Flavobacteriac | Polaribacter | Polaribacter s | cellular proces | Intracellular tr | 0P8JN | The central su | 313598.MED1 | 1 | 0 | 0,000197092 | 1,93E-05 | 1 - | 0,000198852 - | 1 - | 0,000233945 - | 1 - | 0,000184528 - | 1 - | 0,000183368 - | 1 - | 0,000184765 - |
| 1197 | 1 | 0.2um_NOKCPHJN_394615 | Bacteria | Bacteroidetes | Flavobacteriia | Flavobacterial | Crocinitomica | Fluviicola | various | metabolism | Energy produc | COG0056 | Produces ATP | 755732.Fluta_ | 0,4 | 0,489897949 | 8,26E-05 | 0,000101747 | 1 - | 0,000189814 - | 1 - | 0,000223311 - | 0 - | 0 - | 0 - | 0 - | 0 - | 0 - |
| 1198 | 1 | 3um_FIJJAFBF_444690 | Eukaryota | unclassified | unclassified | Phaeocystales | Phaeocystace | Phaeocystis | Phaeocystis ar | metabolism | Energy produc | COG4451 | RuBisCO catal | 977880.RALTA | 0,4 | 0,489897949 | 0,000243418 | 0,000298459 | 1 - | 0,000630891 - | 0 - | 0 - | 0 - | 0 - | 0 - | 0 - | 1 - | 0,000586198 - |
| 1199 | 1 | 3um_FIJJAFBF_693942 | unclassified | unclassified | unclassified | unclassified | unclassified | unclassified | unclassified | unclassified | unclassified | unclassified | unclassified | unclassified | 1 | 0 | 0,000152487 | 1,49E-05 | 1 - | 0,000153849 - | 1 - | 0,000181 - | 1 - | 0,000142766 - | 1 - | 0,000141869 - | 1 - | 0,00014295 - |
| 1200 | 1 | 3um_FIJJAFBF_264641 | Bacteria | Bacteroidetes | Flavobacteriia | Flavobacterial | Flavobacteriac | Polaribacter | Polaribacter t | poorly charact | Function unkn | 05END | lipolytic prote | 313598.MED1 | 9 | 0,894427191 | 0,001548616 | 0,000276272 | 9 - | 0,001553631 - | 10 - | 0,002030903 - | 10 - | 0,001601904 - | 8 - | 0,00127347 - | 8 - | 0,001283173 - |
| 1201 | 1 | 3um_FIJJAFBF_739161 | Bacteria | Proteobacteri | Gammaprotec | Cellvibrionales | Porticoccacea | unclassified | gamma prote | metabolism | Inorganic ion t | 16TQC | TONB-depend | 314287.GB22 | 0,4 | 0,489897949 | 3,28E-05 | 4,02E-05 | 1 - | 8,53E-05 - | 0 - | 0 - | 0 - | 0 - | 1 - | 7,87E-05 - | 0 - | 0 - |
| 1202 | 1 | 3um_FIJJAFBF_254287 | Bacteria | Proteobacteri | Gammaprotec | Cellvibrionales | Halieaceae | Haliea | Haliea salexig | information st | Translation, ri | 17FIH | One of the ea | 566466.NOR5 | 1 | 0 | 0,000886913 | 8,68E-05 | 1 - | 0,000894835 - | 1 - | 0,001052754 - | 1 - | 0,000830375 - | 1 - | 0,000825157 - | 1 - | 0,000831444 - |
| 1203 | 1 | 3um_FIJJAFBF_292379 | Bacteria | Bacteroidetes | Cytophagia | Cytophagales | Flammeovirga | unclassified | Flammeovirga | poorly charact | Function unkn | 16R8I | Lipoprotein | 314260.PB25 | 14,4 | 0,8 | 0,002562732 | 0,000211136 | 13 - | 0,002340904 - | 14 - | 0,002965869 - | 15 - | 0,002506471 - | 15 - | 0,002490719 - | 15 - | 0,002509697 - |
| 1204 | 1 | 0.2um_NOKCPHJN_382781 | unclassified | unclassified | unclassified | unclassified | unclassified | unclassified | unclassified | unclassified | unclassified | unclassified | unclassified | unclassified | 0,6 | 0,489897949 | 0,000374799 | 0,000310055 | 1 - | 0,000604785 - | 1 - | 0,000711516 - | 0 - | 0 - | 1 - | 0,000557692 - | 0 - | 0 - |
| 1205 | 2 | 3um_FIJJAFBF_176961;3um_FI | Eukaryota | unclassified | Rhodellophyc | Rhodellales | Rhodellaceae | Corynoplastis | Corynoplastis | unclassified | unclassified | unclassified | unclassified | unclassified | 2,4 | 1,019803903 | 0,000268709 | 0,000101674 | 2 - | 0,000233228 - | 1 - | 0,000137194 - | 3 - | 0,000324641 - | 2 - | 0,000215067 - | 4 - | 0,000433412 - |
| 1206 | 1 | 3um_FIJJAFBF_12251 | Bacteria | Proteobacteri | Gammaprotec | Cellvibrionales | Porticoccacea | unclassified | Gammaprotec | unclassified | unclassified | unclassified | unclassified | unclassified | 2,8 | 0,4 | 0,000237271 | 1,83E-05 | 3 - | 0,000259962 - | 2 - | 0,000203893 - | 3 - | 0,000241235 - | 3 - | 0,000239719 - | 3 - | 0,000241546 - |
| 1207 | 1 | 0.2um_NOKCPHJN_846361 | Bacteria | Proteobacteri | Alphaproteob | unclassified | unclassified | unclassified | Alphaproteob | metabolism | Energy produc | 01QH6 | CoA-binding d | 331869.BAL19 | 0,2 | 0,4 | 2,66E-05 | 5,32E-05 | 0 - | 0 - | 0 - | 0 - | 1 - | 0,000132969 - | 0 - | 0 - | 0 - | 0 - |
| 1208 | 1 | 3um_FIJJAFBF_501851 | Bacteria | Proteobacteri | Alphaproteob | Rhodobactera | Rhodobactera | Leisingera | Leisingera sp. | information st | Replication, re | 16RMN | DNA helicase | 252305.OB25 | 0,2 | 0,4 | 2,36E-05 | 4,72E-05 | 0 - | 0 - | 0 - | 0 - | 1 - | 0,000117937 - | 0 - | 0 - | 0 - | 0 - |
| 1209 | 1 | 3um_FIJJAFBF_177760 | Viruses | unclassified | unclassified | unclassified | Phycodnavirid | unclassified | Phaeocystis p | unclassified | unclassified | unclassified | unclassified | unclassified | 0,8 | 0,4 | 0,000124593 | 6,25E-05 | 1 - | 0,000164838 - | 0 - | 0 - | 1 - | 0,000152964 - | 1 - | 0,000152003 - | 1 - | 0,000153161 - |
| 1210 | 3 | 0.2um_NOKCPHJN_230057;3ur | Bacteria | unclassified | unclassified | unclassified | unclassified | unclassified | uncultured ba | various | various | various | various | various | 0,8 | 0,4 | 0,000202084 | 0,000101331 | 1 - | 0,000267359 - | 0 - | 0 - | 1 - | 0,0002481 - | 1 - | 0,000246541 - | 1 - | 0,000248419 - |
| 1211 | 1 | 3um_FIJJAFBF_692267 | Eukaryota | Mollusca | Gastropoda | unclassified | Lottiidae | Lottia | Lottia gigante | unclassified | unclassified | unclassified | unclassified | unclassified | 0,8 | 0,4 | 0,000113573 | 5,82E-05 | 1 - | 0,000140987 - | 1 - | 0,000165868 - | 0 - | 0 - | 1 - | 0,000130009 - | 1 - | 0,000130999 - |
| 1212 | 1 | 3um_FIJJAFBF_305831 | Eukaryota | Chlorophyta | Nephroselmid | unclassified | unclassified | Nephroselmis | Nephroselmis | unclassified | unclassified | unclassified | unclassified | unclassified | 1,8 | 0,4 | 0,00030767 | 7,54E-05 | 1 - | 0,000172626 - | 2 - | 0,000406181 - | 2 - | 0,000320381 - | 2 - | 0,000318368 - | 2 - | 0,000320793 - |
| 1213 | 1 | 0.2um_NOKCPHJN_753709 | Eukaryota | unclassified | Cryptophyta | Pyrenomonad | Geminigerace | Guillardia | Guillardia thet | cellular proces | Posttranslatio | KOG0171 | Mitochondrial | 35128.Thaps3 | 1 | 0 | 0,0003863 | 3,78E-05 | 1 - | 0,00038975 - | 1 - | 0,000458533 - | 1 - | 0,000361674 - | 1 - | 0,000359402 - | 1 - | 0,00036214 - |
| 1214 | 1 | 3um_FIJJAFBF_467719 | Eukaryota | unclassified | Cryptophyta | Pyrenomonad | Geminigerace | Teleaulax | Teleaulax amp | metabolism | Energy produc | COG0711 | F(1)F(0) ATP s | 63737.Npun_ | 0,8 | 0,4 | 0,000370299 | 0,000185679 | 1 - | 0,00048991 - | 0 - | 0 - | 1 - | 0,000454619 - | 1 - | 0,000451762 - | 1 - | 0,000455204 - |
| 1215 | 1 | 0.2um_NOKCPHJN_124256 | Eukaryota | unclassified | Labyrinthulon | unclassified | Thraustochytr | Aurantiochytr | Aurantiochytr | unclassified | unclassified | unclassified | unclassified | unclassified | 0,4 | 0,489897949 | 7,25E-06 | 8,89E-06 | 1 - | 1,88E-05 - | 0 - | 0 - | 1 - | 1,75E-05 - | 0 - | 0 - | 0 - | 0 - |
| 1216 | 1 | 3um_FIJJAFBF_497373 | Bacteria | Planctomycet | Phycisphaerae | Phycisphaeral | Phycisphaerac | Phycisphaera | Phycisphaera | unclassified | unclassified | unclassified | unclassified | unclassified | 0,6 | 0,489897949 | 6,28E-05 | 5,13E-05 | 0 - | 0 - | 0 - | 0 - | 1 - | 0,000104867 - | 1 - | 0,000104208 - | 1 - | 0,000105002 - |
| 1217 | 1 | 0.2um_NOKCPHJN_115240 | Bacteria | Actinobacteria | Actinobacteria | Actinomycetal | unclassified | unclassified | Actinomyceta | information st | Transcription | 05D27 | DNA-depende | 266940.Krad_ | 0,2 | 0,4 | 1,25E-05 | 2,50E-05 | 0 - | 0 - | 0 - | 0 - | 0 - | 0 - | 1 - | 6,26E-05 - | 0 - | 0 - |
| 1218 | 1 | 0.2um_NOKCPHJN_277085 | Eukaryota | Phaeophyceae | unclassified | Ectocarpales | Ectocarpaceae | Ectocarpus | Ectocarpus sili | poorly charact | Function unkn | COG1100 | GTP-binding P | 2850.Phatr86 | 0,4 | 0,489897949 | 0,000183886 | 0,000225467 | 1 - | 0,000476597 - | 0 - | 0 - | 0 - | 0 - | 0 - | 0 - | 1 - | 0,000442834 - |
| 1219 | 3 | 0.2um_NOKCPHJN_122736;3ur | Bacteria | Proteobacteri | Gammaprotec | Alteromonada | Colwelliaceae | Colwellia | various | unclassified | unclassified | unclassified | unclassified | unclassified | 0,4 | 0,489897949 | 0,000139827 | 0,000172188 | 1 - | 0,000321223 - | 1 - | 0,000377912 - | 0 - | 0 - | 0 - | 0 - | 0 - | 0 - |
| 1220 | 1 | 0.2um_NOKCPHJN_277638 | Bacteria | Bacteroidetes | Flavobacteriia | Flavobacterial | Cryomorphace | Owenweeksia | Owenweeksia | information st | Translation, ri | 0FWTQ | This protein is | 504472.Slin_4 | 0,4 | 0,489897949 | 0,000259679 | 0,000319778 | 1 - | 0,000596557 - | 1 - | 0,000701836 - | 0 - | 0 - | 0 - | 0 - | 0 - | 0 - |
| 1221 | 1 | 3um_FIJJAFBF_92871 | Eukaryota | unclassified | Cryptophyta | Pyrenomonad | Geminigerace | Guillardia | Guillardia thet | metabolism | Energy produc | COG0055 | Produces ATP | 4081.Solyc01g | 1,2 | 0,4 | 0,000217292 | 6,51E-05 | 1 - | 0,000184619 - | 1 - | 0,0002172 - | 1 - | 0,000171319 - | 1 - | 0,000170243 - | 2 - | 0,00034308 - |
| 1222 | 1 | 3um_FIJJAFBF_347377 | Bacteria | Proteobacteri | Gammaprotec | Enterobactera | Enterobacteri | Escherichia | Escherichia co | unclassified | unclassified | unclassified | unclassified | unclassified | 0,6 | 0,489897949 | 0,000122677 | 0,000101485 | 1 - | 0,000197955 - | 1 - | 0,000232889 - | 0 - | 0 - | 1 - | 0,00018254 - | 0 - | 0 - |
| 1223 | 1 | 3um_FIJJAFBF_105513 | Bacteria | Bacteroidetes | Flavobacteriia | Flavobacterial | Flavobacteriac | Polaribacter | Polaribacter b | unclassified | unclassified | unclassified | unclassified | unclassified | 0,6 | 0,489897949 | 0,000312373 | 0,000259393 | 0 - | 0 - | 1 - | 0,000606882 - | 0 - | 0 - | 1 - | 0,000475679 - | 1 - | 0,000479303 - |
| 1224 | 1 | 0.2um_NOKCPHJN_445514 | Eukaryota | unclassified | unclassified | Isochrysidales | Noelaerhabda | Emiliania | Emiliania huxl | poorly charact | Function unkn | COG2340 | Scp-like extra | 6085.XP_0021 | 1 | 0 | 0,000142956 | 1,40E-05 | 1 - | 0,000144233 - | 1 - | 0,000169687 - | 1 - | 0,000133843 - | 1 - | 0,000133002 - | 1 - | 0,000134016 - |
| 1225 | 1 | 3um_FIJJAFBF_131607 | Bacteria | Proteobacteri | Gammaprotec | Cellvibrionales | Porticoccacea | unclassified | SAR92 bacteri | poorly charact | Function unkn | 16R8I | Lipoprotein | 377629.TERTU | 4 | 1,095445115 | 0,000373586 | 0,000102797 | 6 - | 0,000568211 - | 3 - | 0,000334244 - | 4 - | 0,000351519 - | 3 - | 0,000261983 - | 4 - | 0,000351972 - |
| 1226 | 1 | 0.2um_NOKCPHJN_396986 | Eukaryota | Eustigmatoph | unclassified | Eustigmatales | Monodopsida | Nannochlorop | Nannochlorop | unclassified | unclassified | unclassified | unclassified | unclassified | 2,4 | 0,8 | 0,000344422 | 0,000103044 | 3 - | 0,000445901 - | 1 - | 0,000174864 - | 2 - | 0,000275853 - | 3 - | 0,00041118 - | 3 - | 0,000414313 - |
| 1227 | 1 | 0.2um_NOKCPHJN_41537 | Bacteria | Bacteroidetes | Cytophagia | Cytophagales | Persicobacter | Persicobacter | Persicobacter | poorly charact | Function unkn | 11TE0 | Domain-Conta | 7668.SPU_002 | 0,6 | 0,489897949 | 1,48E-05 | 1,23E-05 | 0 - | 0 - | 1 - | 2,87E-05 - | 1 - | 2,26E-05 - | 1 - | 2,25E-05 - | 0 - | 0 - |
| 1228 | 1 | 3um_FIJJAFBF_121184 | Bacteria | Proteobacteri | Gammaprotec | Alteromonada | Alteromonada | Paraglaciecola | Paraglaciecola | unclassified | unclassified | unclassified | unclassified | unclassified | 1,2 | 0,748331477 | 0,000110958 | 6,79E-05 | 1 - | 9,85E-05 - | 0 - | 0 - | 1 - | 9,14E-05 - | 2 - | 0,00018172 - | 2 - | 0,000183104 - |

| | | | | | | | | | | | | | | | | | | | | | | | | | | | | |
|---|---|---|---|---|---|---|---|---|---|---|---|---|---|---|---|---|---|---|---|---|---|---|---|---|---|---|---|---|
| 1229 | 1 | 3um_FIJJAFBF_510254 | Bacteria | Bacteroidetes | Flavobacteriia | Flavobacterial | Flavobacteriac | Flaviramulus | Flaviramulus i | information st | Translation, ri | 05G6W | One of the pri | 983544.Lacal_ | 0,2 | 0,4 | 8,11E-05 | 0,000162152 | 0 - | 0 - | 0 - | 0 - | 0 - | 0 - | 0 - | 0 - | 1 - | 0,000405381 - |
| 1230 | 1 | 3um_FIJJAFBF_311149 | Eukaryota | unclassified | Chrysophycea | Chromulinales | Chromulinace | Ochromonas | Ochromonas s | metabolism | Energy produc | COG1290 | Component of | 35128.Thapsd | 0,2 | 0,4 | 7,52E-05 | 0,000150447 | 0 - | 0 - | 0 - | 0 - | 0 - | 0 - | 1 - | 0,000376118 - | 0 - | 0 - |
| 1231 | 1 | 3um_FIJJAFBF_559087 | Bacteria | Proteobacteri | Gammaprotec | Alteromonada | Alteromonada | Paraglaciecola | Paraglaciecola | unclassified | unclassified | unclassified | unclassified | unclassified | 5,2 | 1,326649916 | 0,000535063 | 0,00010328 | 5 - | 0,000530834 - | 3 - | 0,000374709 - | 5 - | 0,000492595 - | 7 - | 0,0006853 - | 6 - | 0,000591875 - |
| 1232 | 1 | 3um_FIJJAFBF_08772 | Eukaryota | unclassified | Pelagophycea | Pelagomonad | unclassified | Aureococcus | Aureococcus a | cellular proces | Posttranslatio | 0IEKR | One of the co | 35128.Thapsd | 1 | 0 | 0,000189776 | 1,86E-05 | 1 - | 0,000191471 - | 1 - | 0,000225262 - | 1 - | 0,000177678 - | 1 - | 0,000176562 - | 1 - | 0,000177907 - |
| 1233 | 1 | 3um_FIJJAFBF_292921 | Eukaryota | Bacillariophyta | Coscinodiscop | Coscinodiscale | Coscinodiscac | Stellarima | Stellarima mic | metabolism | Carbohydrate | COG1850 | RuBisCO catal | 261292.Nit79 | 3,2 | 0,748331477 | 0,000562618 | 0,00011902 | 2 - | 0,000357934 - | 3 - | 0,000631652 - | 3 - | 0,000498225 - | 4 - | 0,000660125 - | 4 - | 0,000665155 - |
| 1234 | 1 | 3um_FIJJAFBF_594171 | Eukaryota | unclassified | Dinophyceae | Peridiniales | Kryptoperidini | Kryptoperidini | Kryptoperidin | metabolism | Energy produc | 11HAV | photosystem i | 35128.Thapsd | 5,4 | 0,8 | 0,003199734 | 0,000320951 | 6 - | 0,003628711 - | 4 - | 0,002846066 - | 5 - | 0,002806095 - | 6 - | 0,003346153 - | 6 - | 0,003371648 - |
| 1235 | 1 | 3um_FIJJAFBF_517421 | Bacteria | Bacteroidetes | Flavobacteriia | Flavobacterial | Flavobacteriac | Polaribacter | Polaribacter v | information st | Translation, ri | 08UIK | This protein b | 313598.MED1 | 1,2 | 0,4 | 0,000711675 | 0,000212736 | 1 - | 0,000604785 - | 1 - | 0,000711516 - | 2 - | 0,001122438 - | 1 - | 0,000557692 - | 1 - | 0,000561941 - |
| 1236 | 1 | 0.2um_NOKCPHJN_82975 | Bacteria | Proteobacteri | Alphaproteob | Pelagibacteral | Pelagibacterac | Candidatus Pe | Candidatus Pe | unclassified | unclassified | unclassified | unclassified | unclassified | 0,4 | 0,489897949 | 5,76E-05 | 7,06E-05 | 0 - | 0 - | 0 - | 0 - | 1 - | 0,00014403 - | 0 - | 0 - | 1 - | 0,000144215 - |
| 1237 | 1 | 3um_FIJJAFBF_191791 | Eukaryota | Bacillariophyta | Coscinodiscop | Rhizosoleniale | Rhizosoleniace | Rhizosolenia | Rhizosolenia i | unclassified | unclassified | unclassified | unclassified | unclassified | 8,6 | 1,019803903 | 0,00459457 | 0,000239439 | 8 - | 0,004357458 - | 7 - | 0,004485647 - | 10 - | 0,005054456 - | 9 - | 0,004520423 - | 9 - | 0,004554866 - |
| 1238 | 1 | 0.2um_NOKCPHJN_105761 | Bacteria | Proteobacteri | Alphaproteob | unclassified | unclassified | unclassified | SAR116 cluste | unclassified | unclassified | unclassified | unclassified | unclassified | 0,2 | 0,4 | 2,20E-05 | 4,41E-05 | 1 - | 0,000110168 - | 0 - | 0 - | 0 - | 0 - | 0 - | 0 - | 0 - | 0 - |
| 1239 | 1 | 0.2um_NOKCPHJN_740335 | Bacteria | Bacteroidetes | Flavobacteriia | Flavobacterial | Flavobacteriac | Polaribacter | Polaribacter s | unclassified | unclassified | unclassified | unclassified | unclassified | 8,4 | 1,356465997 | 0,000910716 | 9,59E-05 | 9 - | 0,000997781 - | 6 - | 0,000782578 - | 8 - | 0,000823026 - | 9 - | 0,000920086 - | 10 - | 0,001030107 - |
| 1240 | 2 | 0.2um_NOKCPHJN_63870;3um | Bacteria | Bacteroidetes | Flavobacteriia | Flavobacterial | Flavobacteriac | various | various | unclassified | unclassified | unclassified | unclassified | unclassified | 1 | 0 | 0,000256394 | 2,51E-05 | 1 - | 0,000258684 - | 1 - | 0,000304336 - | 1 - | 0,000240049 - | 1 - | 0,000238541 - | 1 - | 0,000240358 - |
| 1241 | 1 | 0.2um_NOKCPHJN_86824 | Bacteria | Bacteroidetes | unclassified | unclassified | unclassified | unclassified | Bacteroidetes | unclassified | unclassified | unclassified | unclassified | unclassified | 0,8 | 0,4 | 8,56E-05 | 4,29E-05 | 1 - | 0,0001133 - | 0 - | 0 - | 1 - | 0,000105138 - | 1 - | 0,000104477 - | 1 - | 0,000105273 - |
| 1242 | 1 | 0.2um_NOKCPHJN_906556 | Bacteria | Bacteroidetes | Cytophagia | Cytophagales | Flammeovirga | Sediminitomix | Sediminitomix | information st | Translation, ri | COG0180 | tryptophanylt | 926562.Oweh | 0,8 | 0,4 | 0,000205212 | 0,000102899 | 1 - | 0,000271498 - | 0 - | 0 - | 1 - | 0,00025194 - | 1 - | 0,000250357 - | 1 - | 0,000252265 - |
| 1243 | 1 | 3um_FIJJAFBF_250545 | Bacteria | Proteobacteri | Alphaproteob | Rhodobactera | unclassified | unclassified | Rhodobactera | poorly charact | Function unkn | 01VGS | ParB-like nucl | 314262.MED1 | 0,4 | 0,489897949 | 9,47E-05 | 0,000116137 | 1 - | 0,000246331 - | 0 - | 0 - | 0 - | 0 - | 1 - | 0,00022715 - | 0 - | 0 - |
| 1244 | 1 | 0.2um_NOKCPHJN_234273 | Bacteria | Proteobacteri | Gammaprotec | Cellvibrionales | Halieaceae | unclassified | marine gamm | metabolism | Inorganic ion t | 16PDP | TonB-depende | 247639.MGP2 | 0,2 | 0,4 | 1,69E-05 | 3,38E-05 | 0 - | 0 - | 0 - | 0 - | 0 - | 0 - | 1 - | 8,46E-05 - | 0 - | 0 - |
| 1245 | 1 | 3um_FIJJAFBF_642657 | Bacteria | Proteobacteri | Alphaproteob | Rhodobactera | Rhodobactera | unclassified | Rhodobactera | metabolism | Coenzyme tra | 05CVM | Molybdenum | 314256.OG25 | 0,8 | 0,4 | 0,000132402 | 6,82E-05 | 0 - | 0 - | 1 - | 0,000196889 - | 1 - | 0,000155299 - | 1 - | 0,000154323 - | 1 - | 0,000155499 - |
| 1246 | 1 | 3um_FIJJAFBF_222361 | Bacteria | Proteobacteri | Alphaproteob | Rhodobactera | unclassified | unclassified | Rhodobactera | cellular proces | Intracellular tr | 175GA | One of the pro | 439497.RR11_ | 0,4 | 0,489897949 | 0,000226438 | 0,000280533 | 0 - | 0 - | 1 - | 0,000632944 - | 1 - | 0,000499244 - | 0 - | 0 - | 0 - | 0 - |
| 1247 | 1 | 0.2um_NOKCPHJN_1082065 | Bacteria | Verrucomicrol | Verrucomicrol | Verrucomicrol | Rubritaleacea | Rubritalea | Rubritalea squ | poorly charact | Function unkn | 1DMX1 | Permease Ygj | 349741.Amuc | 0,8 | 0,748331477 | 0,000195871 | 0,000181182 | 2 - | 0,000477896 - | 1 - | 0,000281117 - | 0 - | 0 - | 1 - | 0,000220342 - | 0 - | 0 - |
| 1248 | 1 | 3um_FIJJAFBF_813295 | Eukaryota | Bacillariophyta | Bacillariophyc | Bacillariales | Bacillariaceae | Fragilariopsis | Fragilariopsis | cellular proces | Posttranslatio | KOG0840 | ATP-depender | 2850.Phatr40 | 0,2 | 0,4 | 0,000153849 | 0,000307698 | 1 - | 0,000769244 - | 0 - | 0 - | 0 - | 0 - | 0 - | 0 - | 0 - | 0 - |
| 1249 | 1 | 3um_FIJJAFBF_510233 | Bacteria | Bacteroidetes | Flavobacteriia | Flavobacterial | Flavobacteriac | Polaribacter | Polaribacter s | unclassified | unclassified | unclassified | unclassified | unclassified | 0,4 | 0,489897949 | 0,000118425 | 0,000145041 | 0 - | 0 - | 0 - | 0 - | 1 - | 0,000296995 - | 1 - | 0,000295129 - | 0 - | 0 - |
| 1250 | 1 | 0.2um_NOKCPHJN_12324 | Eukaryota | unclassified | Pelagophycea | Pelagomonad | unclassified | Aureococcus | Aureococcus a | unclassified | unclassified | unclassified | unclassified | unclassified | 0,2 | 0,4 | 2,44E-05 | 4,88E-05 | 0 - | 0 - | 0 - | 0 - | 0 - | 0 - | 1 - | 0,000121969 - | 0 - | 0 - |
| 1251 | 1 | 3um_FIJJAFBF_175545 | Bacteria | Proteobacteri | Gammaprotec | Oceanospirilla | Oceanospirilla | Bacterioplane | Bacterioplane | unclassified | unclassified | unclassified | unclassified | unclassified | 0,2 | 0,4 | 0,00018107 | 0,00036214 | 0 - | 0 - | 0 - | 0 - | 0 - | 0 - | 0 - | 0 - | 1 - | 0,00090535 - |
| 1252 | 1 | 3um_FIJJAFBF_786162 | Bacteria | Proteobacteri | Gammaprotec | Oceanospirilla | Saccharospiril | Reinekea | Reinekea fors | information st | Translation, ri | COG0087 | One of the pri | 314283.MED2 | 0,6 | 0,8 | 0,000273529 | 0,000384613 | 0 - | 0 - | 2 - | 0,00098257 - | 0 - | 0 - | 1 - | 0,000385073 - | 0 - | 0 - |
| 1253 | 1 | 3um_FIJJAFBF_742492 | Eukaryota | Bacillariophyta | Bacillariophyc | Naviculales | Naviculaceae | Fistulifera | Fistulifera sola | unclassified | unclassified | unclassified | unclassified | unclassified | 0,4 | 0,489897949 | 0,000159665 | 0,000195549 | 0 - | 0 - | 0 - | 0 - | 1 - | 0,000398906 - | 0 - | 0 - | 1 - | 0,000399419 - |
| 1254 | 1 | 0.2um_NOKCPHJN_319100 | Bacteria | Proteobacteri | Alphaproteob | Rhodobactera | Rhodobactera | Litorimicrobiu | Litorimicrobiu | metabolism | Energy produc | 01RNF | Produces ATP | 388401.RB215 | 0,6 | 0,489897949 | 0,000112032 | 9,30E-05 | 0 - | 0 - | 1 - | 0,000217658 - | 0 - | 0 - | 1 - | 0,000170602 - | 1 - | 0,000171902 - |
| 1255 | 1 | 3um_FIJJAFBF_39170 | Eukaryota | Chordata | Actinopteri | Osteoglossifor | Mormyridae | Paramormyro | Paramormyro | unclassified | unclassified | unclassified | unclassified | unclassified | 2,2 | 0,748331477 | 0,000523852 | 0,000164748 | 3 - | 0,000736923 - | 1 - | 0,000288991 - | 3 - | 0,000683838 - | 2 - | 0,000453027 - | 2 - | 0,000456479 - |
| 1256 | 1 | 3um_FIJJAFBF_610024 | Eukaryota | Bacillariophyta | Coscinodiscop | Thalassiosirale | Thalassiosirac | Thalassiosira | Thalassiosira p | unclassified | unclassified | unclassified | unclassified | unclassified | 0,4 | 0,489897949 | 7,82E-05 | 9,69E-05 | 0 - | 0 - | 1 - | 0,00021858 - | 1 - | 0,000172408 - | 0 - | 0 - | 0 - | 0 - |
| 1257 | 1 | 3um_FIJJAFBF_155127 | Bacteria | Bacteroidetes | Cytophagia | Cytophagales | Flammeovirga | unclassified | Flammeovirga | unclassified | unclassified | unclassified | unclassified | unclassified | 6,2 | 1,469693846 | 0,000957746 | 0,000161698 | 5 - | 0,00079577 - | 4 - | 0,000748965 - | 8 - | 0,001181514 - | 7 - | 0,001027328 - | 7 - | 0,001035155 - |
| 1258 | 1 | 3um_FIJJAFBF_170869 | Bacteria | Bacteroidetes | Flavobacteriia | Flavobacterial | Crocinitomica | unclassified | Crocinitomica | unclassified | unclassified | unclassified | unclassified | unclassified | 2,6 | 0,489897949 | 0,001544494 | 0,000364178 | 2 - | 0,001193114 - | 3 - | 0,002105508 - | 3 - | 0,00166075 - | 2 - | 0,001100209 - | 3 - | 0,001662888 - |
| 1259 | 1 | 0.2um_NOKCPHJN_564677 | unclassified | unclassified | unclassified | unclassified | unclassified | unclassified | unclassified | unclassified | unclassified | unclassified | unclassified | unclassified | 1,2 | 0,4 | 0,00023699 | 6,98E-05 | 1 - | 0,000201595 - | 1 - | 0,000237172 - | 1 - | 0,000187073 - | 2 - | 0,000371795 - | 1 - | 0,000187314 - |
| 1260 | 1 | 3um_FIJJAFBF_44702 | Bacteria | Bacteroidetes | Flavobacteriia | Flavobacterial | Flavobacteriac | Polaribacter | Polaribacter s | cellular proces | Cell wall/mem | 11TPU | outer membra | 313598.MED1 | 2,4 | 1,019803903 | 0,001272854 | 0,00066107 | 1 - | 0,000521987 - | 4 - | 0,002456426 - | 3 - | 0,001453156 - | 2 - | 0,000962683 - | 2 - | 0,000970018 - |
| 1261 | 1 | 3um_FIJJAFBF_305842 | Eukaryota | Chlorophyta | unclassified | Pyramimonad | unclassified | Pyramimonas | Pyramimonas | metabolism | Energy produc | 0DCW7 | Produces ATP | 3055.DAA009 | 3 | 1,095445115 | 0,000529991 | 0,000164317 | 2 - | 0,000364631 - | 2 - | 0,000428981 - | 3 - | 0,000507547 - | 5 - | 0,000840596 - | 3 - | 0,000508201 - |
| 1262 | 1 | 0.2um_NOKCPHJN_162707 | Eukaryota | unclassified | Pelagophycea | Pelagomonad | unclassified | Aureococcus | Aureococcus a | unclassified | unclassified | unclassified | unclassified | unclassified | 0,2 | 0,4 | 2,48E-05 | 4,97E-05 | 0 - | 0 - | 0 - | 0 - | 1 - | 0,000124239 - | 0 - | 0 - | 0 - | 0 - |
| 1263 | 1 | 0.2um_NOKCPHJN_216802 | Eukaryota | unclassified | Oomycetes | Saprolegniales | Saprolegniace | Saprolegnia | Saprolegnia di | unclassified | unclassified | unclassified | unclassified | unclassified | 0,2 | 0,4 | 5,55E-06 | 1,11E-05 | 1 - | 2,78E-05 - | 0 - | 0 - | 0 - | 0 - | 0 - | 0 - | 0 - | 0 - |
| 1264 | 1 | 3um_FIJJAFBF_191792 | Eukaryota | unclassified | Raphidophyce | Chattonellales | Chattonellace | Heterosigma | Heterosigma a | unclassified | unclassified | unclassified | unclassified | unclassified | 2 | 0,894427191 | 0,002050129 | 0,000809626 | 1 - | 0,001069437 - | 1 - | 0,001258169 - | 3 - | 0,002977198 - | 3 - | 0,002958489 - | 2 - | 0,001987353 - |
| 1265 | 1 | 0.2um_NOKCPHJN_1095134 | Eukaryota | unclassified | unclassified | Euglyphida | Paulinellidae | Paulinella | Paulinella mic | unclassified | unclassified | unclassified | unclassified | unclassified | 0,6 | 0,489897949 | 0,000184314 | 0,000153067 | 0 - | 0 - | 1 - | 0,000358229 - | 1 - | 0,000282558 - | 1 - | 0,000280782 - | 0 - | 0 - |
| 1266 | 1 | 3um_FIJJAFBF_229359 | unclassified | unclassified | unclassified | unclassified | unclassified | unclassified | unclassified | cellular proces | Posttranslatio | 0BTMU | Cytosine speci | 59689.fgenes | 0,2 | 0,4 | 4,53E-05 | 9,06E-05 | 1 - | 0,000226599 - | 0 - | 0 - | 0 - | 0 - | 0 - | 0 - | 0 - | 0 - |
| 1267 | 1 | 0.2um_NOKCPHJN_513056 | Bacteria | Bacteroidetes | Flavobacteriia | Flavobacterial | Flavobacteriac | Gramella | Gramella flava | metabolism | Energy produc | 099Q7 | Catalyzes the | 655815.ZPR_3 | 0,4 | 0,489897949 | 9,84E-05 | 0,000120598 | 1 - | 0,000254924 - | 0 - | 0 - | 0 - | 0 - | 0 - | 0 - | 1 - | 0,000236865 - |
| 1268 | 1 | 0.2um_NOKCPHJN_968745 | Eukaryota | unclassified | Cryptophyta | Pyrenomonad | Geminigerace | Guillardia | Guillardia thet | unclassified | unclassified | unclassified | unclassified | unclassified | 0,8 | 0,4 | 0,000219703 | 0,00011259 | 1 - | 0,000272341 - | 1 - | 0,000320403 - | 1 - | 0,000252723 - | 0 - | 0 - | 1 - | 0,000253048 - |
| 1269 | 1 | 3um_FIJJAFBF_584729 | Eukaryota | Bacillariophyta | Fragilariophyc | Fragilariales | Fragilariaceae | Asterionella | Asterionella fo | metabolism | Energy produc | 11IIK | Low-potential | 35128.Thapsd | 1,8 | 0,4 | 0,000939883 | 0,000156524 | 2 - | 0,001075998 - | 1 - | 0,000632944 - | 2 - | 0,000998488 - | 2 - | 0,000992213 - | 2 - | 0,000999773 - |
| 1270 | 1 | 3um_FIJJAFBF_344296 | Bacteria | Proteobacteri | Gammaprotec | unclassified | unclassified | unclassified | Gammaprotec | metabolism | Inorganic ion t | 0ZZVV | tonB-depende | 314275.MADE | 10,8 | 1,469693846 | 0,001201029 | 0,000222436 | 8 - | 0,000895978 - | 12 - | 0,001581148 - | 11 - | 0,001143224 - | 11 - | 0,001136039 - | 12 - | 0,001248759 - |
| 1271 | 1 | 3um_FIJJAFBF_20041 | Bacteria | Bacteroidetes | Flavobacteriia | Flavobacterial | Flavobacteriac | Polaribacter | Polaribacter h | unclassified | unclassified | unclassified | unclassified | unclassified | 3,4 | 0,489897949 | 0,001785252 | 0,000264914 | 4 - | 0,002125912 - | 3 - | 0,001875816 - | 4 - | 0,00197277 - | 3 - | 0,001470279 - | 3 - | 0,001481482 - |
| 1272 | 2 | 3um_FIJJAFBF_115057;3um_FI | Eukaryota | unclassified | unclassified | Phaeocystales | Phaeocystace | Phaeocystis | various | metabolism | Energy produc | 0FN1Q | F(1)F(0) ATP s | various | 1 | 0 | 0,000474959 | 4,65E-05 | 1 - | 0,000479201 - | 1 - | 0,00056377 - | 1 - | 0,000444682 - | 1 - | 0,000441887 - | 1 - | 0,000445254 - |
| 1273 | 1 | 3um_FIJJAFBF_30331 | Bacteria | Bacteroidetes | Flavobacteriia | Flavobacterial | Flavobacteriac | Polaribacter | Polaribacter s | metabolism | Inorganic ion t | 09B4C | tonB-depende | 313594.PI23P | 0,6 | 0,489897949 | 6,23E-05 | 5,09E-05 | 0 - | 0 - | 0 - | 0 - | 1 - | 0,000104062 - | 1 - | 0,000103408 - | 1 - | 0,000104196 - |
| 1274 | 1 | 0.2um_NOKCPHJN_326123 | Eukaryota | unclassified | Oomycetes | Saprolegniales | Saprolegniace | Aphanomyces | Aphanomyces | unclassified | unclassified | unclassified | unclassified | unclassified | 0,2 | 0,4 | 1,47E-05 | 2,95E-05 | 0 - | 0 - | 0 - | 0 - | 0 - | 0 - | 1 - | 7,37E-05 - | 0 - | 0 - |
| 1275 | 1 | 3um_FIJJAFBF_531098 | unclassified | unclassified | unclassified | unclassified | unclassified | unclassified | unclassified | unclassified | unclassified | unclassified | unclassified | unclassified | 1 | 0 | 0,000428165 | 4,19E-05 | 1 - | 0,000431989 - | 1 - | 0,000508226 - | 1 - | 0,000400871 - | 1 - | 0,000398351 - | 1 - | 0,000401387 - |
| 1276 | 2 | 0.2um_NOKCPHJN_1098470;0. | Bacteria | Proteobacteri | various | various | various | various | various | unclassified | unclassified | unclassified | unclassified | unclassified | 0,6 | 0,489897949 | 0,000186049 | 0,00015191 | 0 - | 0 - | 0 - | 0 - | 1 - | 0,000310598 - | 1 - | 0,000308646 - | 1 - | 0,000310998 - |
| 1277 | 1 | 0.2um_NOKCPHJN_108160 | Bacteria | Proteobacteri | Gammaprotec | Alteromonada | Alteromonada | Alishewanella | Alishewanella | cellular proces | Cell wall/mem | 0QITS | Nad-depende | 553480.PAU_ | 1 | 0 | 0,000276807 | 2,71E-05 | 1 - | 0,00027928 - | 1 - | 0,000328567 - | 1 - | 0,000259162 - | 1 - | 0,000257533 - | 1 - | 0,000259495 - |
| 1278 | 1 | 0.2um_NOKCPHJN_559480 | Eukaryota | Eustigmatoph | unclassified | Eustigmatales | Monodopsida | Nannochlorop | Nannochlorop | poorly charact | Function unkn | 0ZI4C | NUF2, NDC80 | 65071.PYU1_ | 0,2 | 0,4 | 3,45E-05 | 6,91E-05 | 0 - | 0 - | 0 - | 0 - | 0 - | 0 - | 0 - | 0 - | 1 - | 0,00017263 - |
| 1279 | 1 | 3um_FIJJAFBF_323298 | Bacteria | Bacteroidetes | Saprospiria | Saprospirales | Saprospiracea | Aureispira | Aureispira sp. | unclassified | unclassified | unclassified | unclassified | unclassified | 0,4 | 0,489897949 | 0,000549973 | 0,000673582 | 0 - | 0 - | 0 - | 0 - | 1 - | 0,001379267 - | 1 - | 0,001370599 - | 0 - | 0 - |
| 1280 | 1 | 3um_FIJJAFBF_401626 | unclassified | unclassified | unclassified | unclassified | unclassified | unclassified | unclassified | unclassified | unclassified | unclassified | unclassified | unclassified | 0,2 | 0,4 | 6,32E-05 | 0,000126328 | 0 - | 0 - | 0 - | 0 - | 0 - | 0 - | 0 - | 0 - | 1 - | 0,00031582 - |
| 1281 | 1 | 3um_FIJJAFBF_207309 | Bacteria | Proteobacteri | Alphaproteob | Sphingomona | Erythrobacter | Croceicoccus | Croceicoccus | metabolism | Energy produc | COG2224 | Isocitrate lyas | 279238.Saro_ | 1,2 | 0,748331477 | 0,000303962 | 0,000179527 | 0 - | 0 - | 1 - | 0,00030797 - | 2 - | 0,000485831 - | 2 - | 0,000482778 - | 1 - | 0,000243228 - |
| 1282 | 1 | 3um_FIJJAFBF_379396 | Viruses | unclassified | unclassified | unclassified | Phycodnavirid | unclassified | Pyramimonas | unclassified | unclassified | unclassified | unclassified | unclassified | 0,2 | 0,4 | 4,25E-05 | 8,51E-05 | 0 - | 0 - | 0 - | 0 - | 0 - | 0 - | 0 - | 0 - | 1 - | 0,000212745 - |
| 1283 | 1 | 3um_FIJJAFBF_347891 | Eukaryota | unclassified | Cryptophyta | Pyrenomonad | Geminigerace | Guillardia | Guillardia thet | unclassified | unclassified | unclassified | unclassified | unclassified | 1,4 | 0,489897949 | 0,000502055 | 0,000151853 | 1 - | 0,000368462 - | 1 - | 0,000433487 - | 2 - | 0,000683838 - | 1 - | 0,00033977 - | 2 - | 0,000684718 - |
| 1284 | 2 | 0.2um_NOKCPHJN_84487;0.2u | various | various | various | various | various | various | various | unclassified | unclassified | unclassified | unclassified | unclassified | 0,4 | 0,489897949 | 0,000206678 | 0,00025313 | 0 - | 0 - | 0 - | 0 - | 1 - | 0,000518323 - | 1 - | 0,000515066 - | 0 - | 0 - |
| 1285 | 2 | 3um_FIJJAFBF_139825;3um_FI | Bacteria | Bacteroidetes | Flavobacteriia | Flavobacterial | Flavobacteriac | various | various | unclassified | unclassified | unclassified | unclassified | unclassified | 0,2 | 0,4 | 7,83E-05 | 0,000156596 | 1 - | 0,00039149 - | 0 - | 0 - | 0 - | 0 - | 0 - | 0 - | 0 - | 0 - |
| 1286 | 1 | 3um_FIJJAFBF_05397 | Bacteria | Bacteroidetes | Flavobacteriia | Flavobacterial | Crocinitomica | unclassified | Crocinitomica | poorly charact | Function unkn | 055DA | Tetratricopep | 709991.Odosp | 1 | 0 | 0,000153294 | 1,50E-05 | 1 - | 0,000154663 - | 1 - | 0,000181957 - | 1 - | 0,000143522 - | 1 - | 0,00014262 - | 1 - | 0,000143706 - |
| 1287 | 1 | 3um_FIJJAFBF_788557 | Bacteria | Bacteroidetes | Flavobacteriia | Flavobacterial | Flavobacteriac | Flavobacteriu | Flavobacteriu | unclassified | unclassified | unclassified | unclassified | unclassified | 0,4 | 0,489897949 | 0,000205052 | 0,000254038 | 0 - | 0 - | 1 - | 0,000573166 - | 1 - | 0,000452093 - | 0 - | 0 - | 0 - | 0 - |
| 1288 | 1 | 0.2um_NOKCPHJN_198182 | Bacteria | Proteobacteri | Gammaprotec | Alteromonada | Alteromonada | unclassified | Alteromonada | cellular proces | Cell wall/mem | COG2885 | Ompa motb d | 247633.GP214 | 1,6 | 1,356465997 | 0,000372222 | 0,000306448 | 0 - | 0 - | 2 - | 0,00055319 - | 4 - | 0,000872673 - | 1 - | 0,000216797 - | 1 - | 0,000218449 - |
| 1289 | 2 | 0.2um_NOKCPHJN_284237;0.2 | Bacteria | Proteobacteri | Gammaprotec | unclassified | unclassified | unclassified | various | information st | Translation, ri | COG0256 | This is one of | 999141.GME_ | 0,2 | 0,4 | 0,000140305 | 0,000280609 | 0 - | 0 - | 0 - | 0 - | 1 - | 0,000701524 - | 0 - | 0 - | 0 - | 0 - |
| 1290 | 2 | 0.2um_NOKCPHJN_177948;0.2 | Eukaryota | unclassified | Oomycetes | Peronosporale | Peronosporac | Plasmopara | Plasmopara ha | cellular proces | Intracellular tr | 0YIKI | Small GTPase | various | 1 | 0 | 0,000462327 | 4,52E-05 | 1 - | 0,000466457 - | 1 - | 0,000548776 - | 1 - | 0,000432855 - | 1 - | 0,000430135 - | 1 - | 0,000433412 - |
| 1291 | 1 | 3um_FIJJAFBF_175086 | Bacteria | Proteobacteri | Alphaproteob | Rhodobactera | Rhodobactera | Phaeobacter | Phaeobacter g | metabolism | Inorganic ion t | 16PAX | extracellular s | 439496.RBY4I | 1,8 | 0,979795897 | 0,000354814 | 0,000190816 | 0 - | 0 - | 2 - | 0,000472173 - | 2 - | 0,000372434 - | 2 - | 0,000370093 - | 3 - | 0,00055937 - |
| 1292 | 1 | 3um_FIJJAFBF_348593 | Bacteria | Acidobacteria | unclassified | unclassified | unclassified | unclassified | Acidobacteria | metabolism | Energy produc | 07EQ1 | H -transportin | 682795.AciX8 | 0,8 | 0,4 | 0,000631271 | 0,000316538 | 1 - | 0,00083518 - | 0 - | 0 - | 1 - | 0,000775017 - | 1 - | 0,000770146 - | 1 - | 0,000776014 - |
| 1293 | 4 | 0.2um_NOKCPHJN_487028;0.2 | Eukaryota | unclassified | various | various | various | various | various | unclassified | unclassified | unclassified | unclassified | unclassified | 0,6 | 0,489897949 | 0,000202685 | 0,000168308 | 0 - | 0 - | 1 - | 0,000393778 - | 0 - | 0 - | 1 - | 0,000308646 - | 1 - | 0,000310998 - |
| 1294 | 1 | 3um_FIJJAFBF_545216 | Bacteria | Bacteroidetes | Flavobacteriia | Flavobacterial | Flavobacteriac | Flavivirga | Flavivirga eckl | unclassified | unclassified | unclassified | unclassified | unclassified | 1,2 | 0,748331477 | 0,000556214 | 0,000354517 | 2 - | 0,000974376 - | 0 - | 0 - | 1 - | 0,000452093 - | 1 - | 0,000449252 - | 2 - | 0,00090535 - |
| 1295 | 1 | 3um_FIJJAFBF_306499 | Bacteria | Proteobacteri | Betaproteoba | unclassified | unclassified | unclassified | Betaproteoba | unclassified | unclassified | unclassified | unclassified | unclassified | 6,8 | 2,039607805 | 0,001197266 | 0,000295271 | 4 - | 0,000724743 - | 5 - | 0,001065805 - | 7 - | 0,001176936 - | 9 - | 0,001503695 - | 9 - | 0,001515152 - |
| 1296 | 1 | 3um_FIJJAFBF_292916 | Eukaryota | Bacillariophyta | Coscinodiscop | Coscinodiscale | Coscinodiscac | Coscinodiscus | Coscinodiscus | unclassified | unclassified | unclassified | unclassified | unclassified | 3,4 | 0,489897949 | 0,001888381 | 0,000281043 | 4 - | 0,00224856 - | 3 - | 0,001984036 - | 3 - | 0,001564937 - | 3 - | 0,001555103 - | 4 - | 0,002089269 - |
| 1297 | 1 | 0.2um_NOKCPHJN_349565 | Eukaryota | Ascomycota | Eurotiomycete | Chaetothyriale | Herpotrichiell | Exophiala | Exophiala aqu | information st | Replication, re | 0XESV | DNA helicase | 6239.K02F3.1 | 0,2 | 0,4 | 5,72E-06 | 1,14E-05 | 0 - | 0 - | 0 - | 0 - | 0 - | 0 - | 1 - | 2,86E-05 - | 0 - | 0 - |
| 1298 | 5 | 0.2um_NOKCPHJN_223179;0.2 | Bacteria | Bacteroidetes | Flavobacteriia | Flavobacterial | various | various | various | unclassified | unclassified | unclassified | unclassified | unclassified | 1 | 0,632455532 | 0,000250242 | 0,000156356 | 1 - | 0,000265739 - | 0 - | 0 - | 1 - | 0,000246596 - | 1 - | 0,000245047 - | 2 - | 0,000493827 - |
| 1299 | 1 | 0.2um_NOKCPHJN_335418 | Bacteria | Proteobacteri | Alphaproteob | unclassified | unclassified | unclassified | alpha proteob | cellular proces | Posttranslatio | 16QU3 | Hemolysin-typ | 744980.TRICH | 0,4 | 0,489897949 | 2,98E-05 | 3,65E-05 | 0 - | 0 - | 0 - | 0 - | 1 - | 7,45E-05 - | 0 - | 0 - | 1 - | 7,45E-05 - |
| 1300 | 3 | 0.2um_NOKCPHJN_130385;0.2 | Bacteria | Proteobacteri | Gammaprotec | various | various | various | various | unclassified | unclassified | unclassified | unclassified | unclassified | 0,6 | 0,489897949 | 0,000211934 | 0,000173045 | 0 - | 0 - | 0 - | 0 - | 1 - | 0,000353812 - | 1 - | 0,000351588 - | 1 - | 0,000354267 - |
| 1301 | 1 | 0.2um_NOKCPHJN_62391 | Bacteria | Bacteroidetes | Flavobacteriia | Flavobacterial | Cryomorphace | unclassified | Cryomorphace | metabolism | Carbohydrate | 16QEQ | hydrolase, fan | 455436.GHTC | 0,4 | 0,489897949 | 8,69E-05 | 0,000106379 | 0 - | 0 - | 0 - | 0 - | 1 - | 0,000217005 - | 0 - | 0 - | 1 - | 0,000217284 - |
| 1302 | 1 | 3um_FIJJAFBF_715060 | Eukaryota | Bacillariophyta | Coscinodiscop | Coscinodiscale | Coscinodiscac | Coscinodiscus | Coscinodiscus | metabolism | Energy produc | 11HAV | photosystem i | 35128.Thapsd | 1 | 0 | 0,000629837 | 6,16E-05 | 1 - | 0,000635463 - | 1 - | 0,000747608 - | 1 - | 0,000589687 - | 1 - | 0,000585981 - | 1 - | 0,000590446 - |
| 1303 | 1 | 0.2um_NOKCPHJN_724920 | Bacteria | Bacteroidetes | Flavobacteriia | Flavobacterial | Flavobacteriac | Polaribacter | Polaribacter s | metabolism | Inorganic ion t | 0P8Q4 | tonB-depende | 688270.Celal_ | 14,4 | 4,45421149 | 0,001427712 | 0,000358919 | 11 - | 0,001125592 - | 9 - | 0,001083464 - | 13 - | 0,00123442 - | 18 - | 0,001698455 - | 21 - | 0,001996629 - |
| 1304 | 1 | 3um_FIJJAFBF_350308 | Bacteria | Proteobacteri | Gammaprotec | Oceanospirilla | Oceanospirilla | Bermanella | Bermanella m | unclassified | unclassified | unclassified | unclassified | unclassified | 1 | 0 | 0,000700947 | 6,86E-05 | 1 - | 0,000707208 - | 1 - | 0,000832015 - | 1 - | 0,000656264 - | 1 - | 0,00065214 - | 1 - | 0,000657109 - |
| 1305 | 1 | 3um_FIJJAFBF_306873 | Bacteria | Proteobacteri | Alphaproteob | Rhodobactera | unclassified | unclassified | Rhodobactera | metabolism | Inorganic ion t | 05PWN | phosphate ab | 439497.RR11_ | 1,2 | 0,4 | 0,000293997 | 8,79E-05 | 1 - | 0,00024984 - | 1 - | 0,000293931 - | 2 - | 0,000463685 - | 1 - | 0,000230386 - | 1 - | 0,000232141 - |
| 1306 | 1 | 0.2um_NOKCPHJN_363833 | Bacteria | Bacteroidetes | Flavobacteriia | Flavobacterial | Flavobacteriac | Formosa | Formosa sp. H | information st | Translation, ri | COG0256 | This is one of | 983544.Lacal_ | 0,6 | 0,489897949 | 0,000461602 | 0,000381621 | 1 - | 0,000743168 - | 1 - | 0,000874321 - | 0 - | 0 - | 0 - | 0 - | 1 - | 0,000690521 - |
| 1307 | 1 | 0.2um_NOKCPHJN_1110152 | unclassified | unclassified | unclassified | unclassified | unclassified | unclassified | unclassified | unclassified | unclassified | unclassified | unclassified | unclassified | 1 | 0 | 0,000171774 | 1,68E-05 | 1 - | 0,000173308 - | 1 - | 0,000203893 - | 1 - | 0,000160824 - | 1 - | 0,000159813 - | 1 - | 0,000161031 - |
| 1308 | 1 | 3um_FIJJAFBF_328503 | Bacteria | Bacteroidetes | Flavobacteriia | Flavobacterial | Flavobacteriac | Arenitalea | Arenitalea lut | information st | Translation, ri | COG0088 | One of the pri | 983544.Lacal_ | 0,8 | 0,4 | 0,000317146 | 0,000159026 | 1 - | 0,000419588 - | 0 - | 0 - | 1 - | 0,000389362 - | 1 - | 0,000386916 - | 1 - | 0,000389864 - |
| 1309 | 1 | 3um_FIJJAFBF_676357 | Eukaryota | Chordata | Actinopteri | Cyprinodontif | Nothobranchi | Nothobranchi | Nothobranchi | information st | Chromatin str | KOG3467 | Core compone | 6085.XP_0021 | 1,8 | 0,4 | 0,001187604 | 0,000197778 | 2 - | 0,001359595 - | 1 - | 0,000799767 - | 2 - | 0,001261655 - | 2 - | 0,001253726 - | 2 - | 0,001263279 - |
| 1310 | 1 | 0.2um_NOKCPHJN_27322 | Bacteria | Bacteroidetes | Flavobacteriia | Flavobacterial | Flavobacteriac | Formosa | Formosa sp. H | metabolism | Inorganic ion t | 05C27 | TonB-depende | 742766.HMPR | 18,6 | 3,006659276 | 0,001516685 | 0,000164972 | 19 - | 0,00158232 - | 14 - | 0,001371679 - | 17 - | 0,001313775 - | 20 - | 0,001535904 - | 23 - | 0,001779748 - |
| 1311 | 1 | 3um_FIJJAFBF_341105 | Viruses | unclassified | unclassified | unclassified | unclassified | unclassified | uncultured vir | unclassified | unclassified | unclassified | unclassified | unclassified | 0,8 | 0,4 | 0,000155943 | 8,00E-05 | 1 - | 0,000193585 - | 1 - | 0,000227748 - | 0 - | 0 - | 1 - | 0,000178511 - | 1 - | 0,000179871 - |
| 1312 | 1 | 0.2um_NOKCPHJN_816703 | Bacteria | Proteobacteri | Betaproteoba | Nitrosomonad | unclassified | unclassified | various | poorly charact | Function unkn | 0XRND | NA | 314607.KB13_ | 3 | 0 | 0,000635981 | 6,22E-05 | 3 - | 0,000641662 - | 3 - | 0,000754902 - | 3 - | 0,00059544 - | 3 - | 0,000591698 - | 3 - | 0,000596206 - |
| 1313 | 1 | 3um_FIJJAFBF_231203 | Bacteria | Bacteroidetes | Flavobacteriia | Flavobacterial | Flavobacteriac | Polaribacter | Polaribacter h | unclassified | unclassified | unclassified | unclassified | unclassified | 0,4 | 0,489897949 | 0,000142858 | 0,000174965 | 0 - | 0 - | 0 - | 0 - | 1 - | 0,000356916 - | 0 - | 0 - | 1 - | 0,000357375 - |
| 1314 | 1 | 3um_FIJJAFBF_807669 | Eukaryota | Bacillariophyta | Coscinodiscop | Thalassiosirale | Thalassiosirac | Thalassiosira | Thalassiosira d | unclassified | unclassified | unclassified | unclassified | unclassified | 0,2 | 0,4 | 4,89E-05 | 9,79E-05 | 0 - | 0 - | 0 - | 0 - | 0 - | 0 - | 0 - | 0 - | 1 - | 0,000244689 - |
| 1315 | 1 | 0.2um_NOKCPHJN_07570 | Bacteria | unclassified | unclassified | unclassified | unclassified | unclassified | uncultured ba | metabolism | Inorganic ion t | 05C27 | TonB-depende | 313596.RB250 | 0,8 | 0,4 | 6,34E-05 | 3,18E-05 | 1 - | 8,38E-05 - | 0 - | 0 - | 1 - | 7,78E-05 - | 1 - | 7,73E-05 - | 1 - | 7,79E-05 - |
| 1316 | 1 | 0.2um_NOKCPHJN_622978 | Eukaryota | unclassified | Labyrinthulom | unclassified | Thraustochytr | Aurantiochytr | Aurantiochytr | cellular proces | Signal transdu | KOG0614 | cgmp-depend | 653948.CCA2 | 0,8 | 0,4 | 9,91E-05 | 5,08E-05 | 1 - | 0,000122993 - | 1 - | 0,000144698 - | 0 - | 0 - | 1 - | 0,000113416 - | 1 - | 0,00011428 - |
| 1317 | 1 | 3um_FIJJAFBF_669258 | Eukaryota | Bacillariophyta | various | various | various | various | various | metabolism | Energy produc | 0J0AS | This b-type cy | 35128.Thapsd | 5,6 | 0,8 | 0,00580681 | 0,001061863 | 5 - | 0,005219872 - | 6 - | 0,007369278 - | 7 - | 0,006781395 - | 5 - | 0,004813414 - | 5 - | 0,004850089 - |
| 1318 | 1 | 3um_FIJJAFBF_746721 | Eukaryota | Chlorophyta | Mamiellophyc | Mamiellales | Mamiellaceae | Micromonas | Micromonas c | unclassified | unclassified | unclassified | unclassified | unclassified | 1 | 0 | 0,000595325 | 5,82E-05 | 1 - | 0,000600643 - | 1 - | 0,000706643 - | 1 - | 0,000557375 - | 1 - | 0,000553872 - | 1 - | 0,000558092 - |
| 1319 | 1 | 0.2um_NOKCPHJN_724163 | Bacteria | Proteobacteri | Gammaprotec | unclassified | unclassified | unclassified | Gammaprotec | metabolism | Inorganic ion t | 05F28 | tonB-depende | 565045.NOR5 | 1,6 | 0,489897949 | 0,000150069 | 3,67E-05 | 1 - | 9,70E-05 - | 1 - | 0,000114126 - | 2 - | 0,000180037 - | 2 - | 0,000178906 - | 2 - | 0,000180269 - |
| 1320 | 2 | 0.2um_NOKCPHJN_34045;0.2u | Bacteria | Bacteroidetes | Flavobacteriia | Flavobacterial | Flavobacteriac | unclassified | various | poorly charact | Function unkn | 0P7WW | NA | 487796.Flav2 | 0,6 | 0,489897949 | 9,17E-05 | 7,50E-05 | 1 - | 0,000160906 - | 0 - | 0 - | 1 - | 0,000149315 - | 1 - | 0,000148377 - | 0 - | 0 - |
| 1321 | 1 | 3um_FIJJAFBF_118448 | Bacteria | Proteobacteri | Alphaproteob | Rhodobactera | Rhodobactera | unclassified | Rhodobactera | unclassified | unclassified | unclassified | unclassified | unclassified | 0,6 | 0,489897949 | 0,000108081 | 8,82E-05 | 0 - | 0 - | 0 - | 0 - | 1 - | 0,000180436 - | 1 - | 0,000179302 - | 1 - | 0,000180668 - |
| 1322 | 1 | 3um_FIJJAFBF_22823 | Eukaryota | Bacillariophyta | Bacillariophyc | Naviculales | Phaeodactylac | Phaeodactylu | Phaeodactylu | unclassified | unclassified | unclassified | unclassified | unclassified | 1 | 0 | 0,000216752 | 2,12E-05 | 1 - | 0,000218688 - | 1 - | 0,000257282 - | 1 - | 0,000202935 - | 1 - | 0,000201659 - | 1 - | 0,000203196 - |
| 1323 | 1 | 3um_FIJJAFBF_08764 | Eukaryota | Bacillariophyta | various | various | various | various | various | metabolism | Energy produc | 0J0AS | This b-type cy | 35128.Thapsd | 2,8 | 0,748331477 | 0,002895151 | 0,000793401 | 2 - | 0,002087949 - | 3 - | 0,003684639 - | 4 - | 0,003875083 - | 3 - | 0,002888048 - | 2 - | 0,001940036 - |
| 1324 | 1 | 3um_FIJJAFBF_788052 | Bacteria | Proteobacteri | Alphaproteob | Rhodobactera | Rhodobactera | unclassified | Rhodobactera | information st | Translation, ri | 01QWB | One of the pri | 391619.RGBS | 1,8 | 0,748331477 | 0,0006362 | 0,000239404 | 2 - | 0,00072775 - | 1 - | 0,000428091 - | 2 - | 0,000675326 - | 1 - | 0,000335541 - | 3 - | 0,001014292 - |
| 1325 | 1 | 0.2um_NOKCPHJN_689502 | Bacteria | Actinobacteria | Actinobacteria | Streptomyceta | Streptomyceta | Streptomyces | Streptomyces | unclassified | unclassified | unclassified | unclassified | unclassified | 1 | 0 | 0,000543234 | 5,32E-05 | 1 - | 0,000548087 - | 1 - | 0,000644812 - | 1 - | 0,000508605 - | 1 - | 0,000505408 - | 1 - | 0,000509259 - |
| 1326 | 1 | 3um_FIJJAFBF_488345 | Eukaryota | Bacillariophyta | Coscinodiscop | Thalassiosirale | Thalassiosirac | Thalassiosira | Thalassiosira c | information st | Translation, ri | KOG0886 | Ribosomal pro | 35128.Thaps3 | 0,8 | 0,4 | 0,000430745 | 0,000220882 | 1 - | 0,000534719 - | 1 - | 0,000629085 - | 0 - | 0 - | 1 - | 0,000493081 - | 1 - | 0,000496838 - |
| 1327 | 1 | 3um_FIJJAFBF_214150 | Bacteria | Bacteroidetes | Flavobacteriia | Flavobacterial | Flavobacteriac | Olleya | various | information st | Translation, ri | COG0050 | This protein p | 1046627.BZAI | 3,4 | 0,8 | 0,000734757 | 0,000140915 | 4 - | 0,000888039 - | 2 - | 0,000522379 - | 4 - | 0,000824068 - | 3 - | 0,000614167 - | 4 - | 0,000825129 - |
| 1328 | 1 | 3um_FIJJAFBF_254297 | Bacteria | Proteobacteri | Gammaprotec | Cellvibrionales | Spongiibacter | Oceanicoccus | Oceanicoccus | unclassified | unclassified | unclassified | unclassified | unclassified | 0,8 | 0,4 | 0,000368242 | 0,000184647 | 1 - | 0,000487188 - | 0 - | 0 - | 1 - | 0,000452093 - | 1 - | 0,000449252 - | 1 - | 0,000452675 - |
| 1329 | 1 | 0.2um_NOKCPHJN_681420 | Bacteria | Proteobacteri | Gammaprotec | Alteromonada | Colwelliaceae | Colwellia | Colwellia psyc | unclassified | unclassified | unclassified | unclassified | unclassified | 0,2 | 0,4 | 2,14E-05 | 4,27E-05 | 1 - | 0,000106813 - | 0 - | 0 - | 0 - | 0 - | 0 - | 0 - | 0 - | 0 - |
| 1330 | 1 | 0.2um_NOKCPHJN_782497 | Eukaryota | Bacillariophyta | Coscinodiscop | Thalassiosirale | Thalassiosirac | Thalassiosira | Thalassiosira c | cellular proces | Signal transdu | KOG0613 | fibronectin ty | 67593.Physo1 | 0,8 | 1,166190379 | 7,70E-05 | 0,000112429 | 0 - | 0 - | 0 - | 0 - | 0 - | 0 - | 1 - | 9,57E-05 - | 3 - | 0,000289283 - |
| 1331 | 2 | 0.2um_NOKCPHJN_119310;0.2 | Bacteria | Proteobacteri | Gammaprotec | Cellvibrionales | Porticoccacea | unclassified | gamma protec | poorly charact | Function unkn | 16R8I | Lipoprotein | 314287.GB22 | 1 | 0,632455532 | 0,000156402 | 9,49E-05 | 0 - | 0 - | 1 - | 0,00018861 - | 1 - | 0,000148769 - | 2 - | 0,000295669 - | 1 - | 0,000148961 - |
| 1332 | 1 | 0.2um_NOKCPHJN_770496 | Eukaryota | Bacillariophyta | Coscinodiscop | Thalassiosirale | Thalassiosirac | Thalassiosira | Thalassiosira c | unclassified | unclassified | unclassified | unclassified | unclassified | 0,4 | 0,489897949 | 0,000151726 | 0,000185828 | 0 - | 0 - | 0 - | 0 - | 0 - | 0 - | 1 - | 0,000377875 - | 1 - | 0,000380755 - |
| 1333 | 1 | 0.2um_NOKCPHJN_741193 | Eukaryota | Mucoromycot | Mortierellomy | Mortierellales | Mortierellacea | Lobosporangiu | Lobosporangiu | unclassified | unclassified | unclassified | unclassified | unclassified | 0,2 | 0,4 | 2,08E-05 | 4,16E-05 | 0 - | 0 - | 0 - | 0 - | 1 - | 0,000104062 - | 0 - | 0 - | 0 - | 0 - |
| 1334 | 1 | 3um_FIJJAFBF_212745 | Bacteria | Bacteroidetes | Flavobacteriia | Flavobacterial | Flavobacteriac | Polaribacter | Polaribacter t | unclassified | unclassified | unclassified | unclassified | unclassified | 1,2 | 0,748331477 | 0,00046194 | 0,000283575 | 1 - | 0,000409784 - | 0 - | 0 - | 2 - | 0,00076053 - | 1 - | 0,000377875 - | 2 - | 0,000761509 - |
| 1335 | 1 | 3um_FIJJAFBF_283757 | unclassified | unclassified | unclassified | unclassified | unclassified | unclassified | unclassified | unclassified | unclassified | unclassified | unclassified | unclassified | 1 | 0 | 0,001010668 | 9,89E-05 | 1 - | 0,001019696 - | 1 - | 0,00119965 - | 1 - | 0,000946241 - | 1 - | 0,000940295 - | 1 - | 0,000947459 - |
| 1336 | 4 | 0.2um_NOKCPHJN_469736;3u | Bacteria | Proteobacteri | Gammaprotec | Cellvibrionales | various | various | various | cellular proces | Signal transdu | various | various | various | 0,8 | 0,748331477 | 0,000307438 | 0,000284035 | 1 - | 0,00040599 - | 0 - | 0 - | 1 - | 0,000376744 - | 0 - | 0 - | 2 - | 0,000754458 - |
| 1337 | 1 | 0.2um_NOKCPHJN_192568 | Bacteria | Proteobacteri | Gammaprotec | Alteromonada | Colwelliaceae | Colwellia | Colwellia sp. E | metabolism | Inorganic ion t | 16TQC | TONB-depend | 87626.PTD2_2 | 7 | 2 | 0,000604629 | 0,000157744 | 7 - | 0,000614471 - | 6 - | 0,000619639 - | 4 - | 0,000325833 - | 8 - | 0,00064757 - | 10 - | 0,000815631 - |
| 1338 | 1 | 3um_FIJJAFBF_689038 | Eukaryota | Mucoromycot | Mucoromycet | Mucorales | Choanephorac | Choanephora | Choanephora | unclassified | unclassified | unclassified | unclassified | unclassified | 1 | 0 | 0,000730399 | 7,15E-05 | 1 - | 0,000736923 - | 1 - | 0,000866974 - | 1 - | 0,000683838 - | 1 - | 0,000679541 - | 1 - | 0,000684718 - |
| 1339 | 1 | 3um_FIJJAFBF_505134 | unclassified | unclassified | unclassified | unclassified | unclassified | unclassified | unclassified | unclassified | unclassified | unclassified | unclassified | unclassified | 0,4 | 0,489897949 | 0,000131455 | 0,000161001 | 0 - | 0 - | 0 - | 0 - | 0 - | 0 - | 1 - | 0,00032739 - | 1 - | 0,000329885 - |
| 1340 | 1 | 3um_FIJJAFBF_482441 | Eukaryota | Bacillariophyta | Coscinodiscop | Thalassiosirale | Thalassiosirac | Thalassiosira | Thalassiosira c | unclassified | unclassified | unclassified | unclassified | unclassified | 0,2 | 0,4 | 1,51E-05 | 3,01E-05 | 0 - | 0 - | 0 - | 0 - | 1 - | 7,53E-05 - | 0 - | 0 - | 0 - | 0 - |
| 1341 | 1 | 0.2um_NOKCPHJN_126580 | Bacteria | Verrucomicrol | Opitutae | unclassified | unclassified | unclassified | Opitutae bact | unclassified | unclassified | unclassified | unclassified | unclassified | 18,4 | 1,854723699 | 0,003335168 | 0,000375663 | 16 - | 0,002935359 - | 18 - | 0,003885059 - | 20 - | 0,003404885 - | 21 - | 0,003552662 - | 17 - | 0,002897877 - |
| 1342 | 1 | 3um_FIJJAFBF_91116 | Eukaryota | unclassified | Pelagophycea | Pelagomonad | unclassified | Aureococcus | Aureococcus a | metabolism | Carbohydrate | KOG1203 | Complex I inte | 35128.Thapsd | 1,6 | 0,8 | 0,000421376 | 0,000190516 | 1 - | 0,000272341 - | 1 - | 0,000320403 - | 1 - | 0,000252723 - | 2 - | 0,000502269 - | 3 - | 0,000759144 - |
| 1343 | 1 | 3um_FIJJAFBF_733449 | unclassified | unclassified | unclassified | unclassified | unclassified | unclassified | unclassified | unclassified | unclassified | unclassified | unclassified | unclassified | 0,4 | 0,489897949 | 0,000269695 | 0,000330759 | 1 - | 0,000701551 - | 0 - | 0 - | 0 - | 0 - | 1 - | 0,000646923 - | 0 - | 0 - |
| 1344 | 1 | 0.2um_NOKCPHJN_845630 | Bacteria | Actinobacteria | Actinobacteria | Corynebacteri | Mycobacteriac | Mycobacteriu | Mycobacteriu | poorly charact | Function unkn | 0IQEP | NA | 5057.CADACL | 0,4 | 0,489897949 | 6,69E-05 | 8,20E-05 | 1 - | 0,000173996 - | 0 - | 0 - | 0 - | 0 - | 1 - | 0,000160447 - | 0 - | 0 - |
| 1345 | 3 | 0.2um_NOKCPHJN_158130;0.2 | Bacteria | Proteobacteri | Gammaprotec | Cellvibrionales | Porticoccacea | unclassified | various | information st | Translation, ri | 05G6W | One of the pri | various | 1 | 0 | 0,000421929 | 4,13E-05 | 1 - | 0,000425698 - | 1 - | 0,000500825 - | 1 - | 0,000395033 - | 1 - | 0,00039255 - | 1 - | 0,000395541 - |
| 1346 | 1 | 0.2um_NOKCPHJN_667434 | Bacteria | Proteobacteri | Betaproteoba | Nitrosomonad | unclassified | unclassified | Methylophilal | metabolism | Inorganic ion t | COG0841 | acriflavin resis | 383631.MB21 | 0,2 | 0,4 | 1,61E-05 | 3,23E-05 | 0 - | 0 - | 0 - | 0 - | 0 - | 0 - | 0 - | 0 - | 1 - | 8,07E-05 - |
| 1347 | 1 | 3um_FIJJAFBF_721624 | Eukaryota | unclassified | Oomycetes | Pythiales | Pythiaceae | Pythium | Pythium insidi | unclassified | unclassified | unclassified | unclassified | unclassified | 0,6 | 0,489897949 | 0,000527478 | 0,000431357 | 1 - | 0,000923093 - | 0 - | 0 - | 1 - | 0,000856597 - | 0 - | 0 - | 1 - | 0,0008577 - |
| 1348 | 1 | 0.2um_NOKCPHJN_84217 | Eukaryota | unclassified | Oomycetes | Saprolegniales | Saprolegniace | Achlya | Achlya hypogy | information st | Translation, ri | COG0052 | 30S ribosomal | 653948.CCA1 | 0,8 | 0,4 | 0,000199364 | 0,000102628 | 0 - | 0 - | 1 - | 0,000296465 - | 1 - | 0,000233841 - | 1 - | 0,000232372 - | 1 - | 0,000234142 - |
| 1349 | 1 | 3um_FIJJAFBF_807524 | Bacteria | Proteobacteri | Alphaproteob | Rhodobactera | Rhodobactera | Planktomarina | Planktomarina | cellular proces | Cell wall/mem | 16PAV | penicillin-bind | 644107.SL115 | 1 | 0 | 0,000122247 | 1,20E-05 | 1 - | 0,000123339 - | 1 - | 0,000145105 - | 1 - | 0,000114454 - | 1 - | 0,000113735 - | 1 - | 0,000114601 - |
| 1350 | 1 | 3um_FIJJAFBF_99818 | Bacteria | Planctomycete | Phycisphaerae | Phycisphaeral | Phycisphaerac | Phycisphaera | Phycisphaera | unclassified | unclassified | unclassified | unclassified | unclassified | 0,2 | 0,4 | 6,05E-05 | 0,00012102 | 0 - | 0 - | 1 - | 0,000302551 - | 0 - | 0 - | 0 - | 0 - | 0 - | 0 - |
| 1351 | 1 | 0.2um_NOKCPHJN_1077861 | Eukaryota | unclassified | Cryptophyta | Pyrenomonad | Geminigerace | Guillardia | Guillardia thet | unclassified | unclassified | unclassified | unclassified | unclassified | 1,2 | 0,4 | 0,001120549 | 0,000330198 | 1 - | 0,000953194 - | 1 - | 0,001121412 - | 1 - | 0,00088453 - | 2 - | 0,001757942 - | 1 - | 0,000885668 - |
| 1352 | 1 | 3um_FIJJAFBF_66815 | Eukaryota | Chlorophyta | Mamiellophyc | Mamiellales | Bathycoccacea | Bathycoccus | Bathycoccus p | unclassified | unclassified | unclassified | unclassified | unclassified | 0,4 | 0,489897949 | 0,000337118 | 0,000413448 | 1 - | 0,000876939 - | 0 - | 0 - | 0 - | 0 - | 1 - | 0,000808654 - | 0 - | 0 - |
| 1353 | 1 | 0.2um_NOKCPHJN_27323 | Bacteria | Bacteroidetes | unclassified | unclassified | unclassified | unclassified | Bacteroidetes | poorly charact | Function unkn | 0Y63T | Ragb susd dor | 471854.Dfer_ | 8 | 3,16227766 | 0,001335037 | 0,000455986 | 4 - | 0,000691865 - | 5 - | 0,001017455 - | 8 - | 0,001284051 - | 12 - | 0,001913973 - | 11 - | 0,001767843 - |
| 1354 | 1 | 0.2um_NOKCPHJN_12461 | Bacteria | Bacteroidetes | Flavobacteriia | Flavobacterial | Flavobacteriac | Formosa | Formosa sp. H | unclassified | unclassified | unclassified | unclassified | unclassified | 1 | 0 | 0,000790159 | 7,73E-05 | 1 - | 0,000797217 - | 1 - | 0,000937908 - | 1 - | 0,000739789 - | 1 - | 0,00073514 - | 1 - | 0,000740741 - |
| 1355 | 1 | 0.2um_NOKCPHJN_506016 | Bacteria | Bacteroidetes | Flavobacteriia | Flavobacterial | Flavobacteriac | Gilvibacter | Gilvibacter sp. | poorly charact | Function unkn | 064XS | NA | 313595.P7007 | 1,2 | 0,4 | 0,00039541 | 0,00018183 | 1 - | 0,000322404 - | 2 - | 0,000758602 - | 1 - | 0,000299179 - | 1 - | 0,000297299 - | 1 - | 0,000299564 - |
| 1356 | 1 | 0.2um_NOKCPHJN_344034 | Bacteria | Proteobacteri | Gammaprotec | Cellvibrionales | Porticoccacea | unclassified | gamma protec | unclassified | unclassified | unclassified | unclassified | unclassified | 1 | 0 | 0,000248336 | 2,43E-05 | 1 - | 0,000250554 - | 1 - | 0,000294771 - | 1 - | 0,000232505 - | 1 - | 0,000231044 - | 1 - | 0,000232804 - |
| 1357 | 1 | 3um_FIJJAFBF_231373 | unclassified | unclassified | unclassified | unclassified | unclassified | unclassified | unclassified | unclassified | unclassified | unclassified | unclassified | unclassified | 0,8 | 0,4 | 0,000492047 | 0,000253294 | 0 - | 0 - | 1 - | 0,000731701 - | 1 - | 0,00057714 - | 1 - | 0,000573513 - | 1 - | 0,000577883 - |
| 1358 | 3 | 0.2um_NOKCPHJN_833582;0.2 | Bacteria | unclassified | unclassified | unclassified | unclassified | unclassified | uncultured ba | various | various | various | various | various | 1 | 0 | 0,000303907 | 2,97E-05 | 1 - | 0,000306622 - | 1 - | 0,000360734 - | 1 - | 0,000284534 - | 1 - | 0,000282746 - | 1 - | 0,0002849 - |
| 1359 | 1 | 3um_FIJJAFBF_566364 | Bacteria | Proteobacteri | Gammaprotec | Pseudomonad | Moraxellaceae | Psychrobacter | various | cellular proces | Cell wall/mem | 07Q6S | Gram-negativ | 335284.Pcryo_ | 0,4 | 0,489897949 | 8,80E-05 | 0,00010777 | 0 - | 0 - | 0 - | 0 - | 0 - | 0 - | 1 - | 0,000219147 - | 1 - | 0,000220817 - |
| 1360 | 1 | 3um_FIJJAFBF_507106 | Eukaryota | Bacillariophyta | Coscinodiscop | Thalassiosirale | Thalassiosirac | Thalassiosira | Thalassiosira c | unclassified | unclassified | unclassified | unclassified | unclassified | 3 | 0 | 0,002929802 | 0,000286664 | 3 - | 0,002955973 - | 3 - | 0,003477637 - | 3 - | 0,002743036 - | 3 - | 0,002725798 - | 3 - | 0,002746567 - |
| 1361 | 1 | 3um_FIJJAFBF_730452 | Bacteria | Bacteroidetes | Flavobacteriia | Flavobacterial | unclassified | unclassified | Flavobacterial | poorly charact | Function unkn | 09AR5 | Inherit from b | 388413.ALPR1 | 0,8 | 0,4 | 0,000170883 | 8,80E-05 | 0 - | 0 - | 1 - | 0,000254113 - | 1 - | 0,000200435 - | 1 - | 0,000199176 - | 1 - | 0,000200693 - |
| 1362 | 7 | 0.2um_NOKCPHJN_454363;0.2 | Bacteria | Proteobacteri | Alphaproteob | Rhodobactera | various | various | various | information st | Translation, ri | COG0185 | Protein S19 fo | various | 0,8 | 0,4 | 0,000746007 | 0,000384027 | 0 - | 0 - | 1 - | 0,001109354 - | 1 - | 0,000875019 - | 1 - | 0,00086952 - | 1 - | 0,000876145 - |
| 1363 | 1 | 3um_FIJJAFBF_17186 | Bacteria | Proteobacteri | Alphaproteob | Rhodobactera | unclassified | unclassified | Rhodobactera | unclassified | unclassified | unclassified | unclassified | unclassified | 1,2 | 0,748331477 | 0,000227819 | 0,000151243 | 1 - | 0,000185792 - | 2 - | 0,000437161 - | 2 - | 0,000344817 - | 1 - | 0,000171325 - | 0 - | 0 - |
| 1364 | 1 | 3um_FIJJAFBF_792634 | Eukaryota | Bacillariophyta | Bacillariophyc | Naviculales | Phaeodactylac | Phaeodactylu | Phaeodactylu | unclassified | unclassified | unclassified | unclassified | unclassified | 1,8 | 0,4 | 0,000300984 | 5,01E-05 | 2 - | 0,000344573 - | 1 - | 0,000202691 - | 2 - | 0,000319751 - | 2 - | 0,000317742 - | 2 - | 0,000320163 - |
| 1365 | 1 | 3um_FIJJAFBF_594168 | Eukaryota | unclassified | Pelagophycea | Pelagomonad | unclassified | Aureoumbra | Aureoumbra l | metabolism | Energy produc | COG0056 | Produces ATP | 35128.Thapsd | 16,4 | 2,332380758 | 0,002784333 | 0,000223352 | 15 - | 0,00259962 - | 13 - | 0,00265061 - | 17 - | 0,002734001 - | 17 - | 0,00271682 - | 20 - | 0,003220612 - |

| | | | | | | | | | | | | | | | | | | | | | | | | | | | | |
|---|---|---|---|---|---|---|---|---|---|---|---|---|---|---|---|---|---|---|---|---|---|---|---|---|---|---|---|---|
| 1366 | 1 | 0.2um_NOKCPHJN_892804 | Eukaryota | Bacillariophyta | Coscinodiscop | Thalassiosirale | Thalassiosirac | Conticribra | Conticribra we | unclassified | unclassified | unclassified | unclassified | unclassified | 7,6 | 1,019803903 | 0,001241083 | 0,000109578 | 8 - | 0,001331216 - | 6 - | 0,00117461 - | 9 - | 0,001389736 - | 8 - | 0,001227558 - | 7 - | 0,001082297 - |
| 1367 | 1 | 3um_FIJJAFBF_205215 | Eukaryota | unclassified | Raphidophyce | Chattonellales | Chattonellace | Heterosigma | Heterosigma a | unclassified | unclassified | unclassified | unclassified | unclassified | 1,6 | 0,489897949 | 0,00026921 | 7,41E-05 | 2 - | 0,000344573 - | 1 - | 0,000202691 - | 2 - | 0,000319751 - | 1 - | 0,000158871 - | 2 - | 0,000320163 - |
| 1368 | 1 | 3um_FIJJAFBF_734480 | Eukaryota | Platyhelminth | Trematoda | Opisthorchiida | Opisthorchiida | Opisthorchis | Opisthorchis v | unclassified | unclassified | unclassified | unclassified | unclassified | 0,4 | 0,8 | 0,000127093 | 0,000254185 | 2 - | 0,000635463 - | 0 - | 0 - | 0 - | 0 - | 0 - | 0 - | 0 - | 0 - |
| 1369 | 1 | 3um_FIJJAFBF_89693 | Eukaryota | unclassified | Pelagophycea | Pelagomonad | unclassified | Aureococcus | Aureococcus a | cellular proces | Posttranslatio | KOG0730 | cell division cy | 35128.Thaps4 | 1,8 | 0,748331477 | 0,001669873 | 0,000607404 | 1 - | 0,000963669 - | 1 - | 0,001133735 - | 2 - | 0,0017885 - | 2 - | 0,001777261 - | 3 - | 0,002686203 - |
| 1370 | 1 | 0.2um_NOKCPHJN_503409 | Eukaryota | unclassified | Oomycetes | Saprolegniales | Saprolegniace | Saprolegnia | Saprolegnia p | unclassified | unclassified | unclassified | unclassified | unclassified | 0,8 | 0,4 | 6,39E-05 | 3,20E-05 | 1 - | 8,45E-05 - | 0 - | 0 - | 1 - | 7,84E-05 - | 1 - | 7,79E-05 - | 1 - | 7,85E-05 - |
| 1371 | 1 | 0.2um_NOKCPHJN_632309 | unclassified | unclassified | unclassified | unclassified | unclassified | unclassified | unclassified | unclassified | unclassified | unclassified | unclassified | unclassified | 1,4 | 0,489897949 | 0,000239088 | 9,95E-05 | 1 - | 0,000169293 - | 2 - | 0,000398339 - | 1 - | 0,000157098 - | 1 - | 0,000156111 - | 2 - | 0,0003146 - |
| 1372 | 1 | 0.2um_NOKCPHJN_1072291 | unclassified | unclassified | unclassified | unclassified | unclassified | unclassified | unclassified | unclassified | unclassified | unclassified | unclassified | unclassified | 0,2 | 0,4 | 0,000152844 | 0,000305689 | 0 - | 0 - | 1 - | 0,000764221 - | 0 - | 0 - | 0 - | 0 - | 0 - | 0 - |
| 1373 | 1 | 3um_FIJJAFBF_119808 | unclassified | unclassified | unclassified | unclassified | unclassified | unclassified | unclassified | unclassified | unclassified | unclassified | unclassified | unclassified | 2 | 0 | 0,001485769 | 0,000145374 | 2 - | 0,00149904 - | 2 - | 0,001763588 - | 2 - | 0,001391055 - | 2 - | 0,001382314 - | 2 - | 0,001392846 - |
| 1374 | 1 | 3um_FIJJAFBF_633909 | Bacteria | unclassified | unclassified | unclassified | unclassified | unclassified | uncultured ba | information st | Translation, ri | 09AI8 | Associates wit | 391603.FBALC | 0,4 | 0,489897949 | 0,000105021 | 0,0001288 | 1 - | 0,00027319 - | 0 - | 0 - | 0 - | 0 - | 1 - | 0,000251917 - | 0 - | 0 - |
| 1375 | 2 | 0.2um_NOKCPHJN_409524;3ur | Bacteria | Proteobacteri | Alphaproteob | Rhodospirillal | Rhodospirillac | unclassified | Rhodospirillac | unclassified | unclassified | unclassified | unclassified | unclassified | 0,6 | 0,489897949 | 0,000167518 | 0,000139107 | 0 - | 0 - | 1 - | 0,000325457 - | 0 - | 0 - | 1 - | 0,000255096 - | 1 - | 0,000257039 - |
| 1376 | 1 | 3um_FIJJAFBF_259311 | Bacteria | Planctomycet | Phycisphaerae | Phycisphaeral | Phycisphaerac | Phycisphaera | Phycisphaera | cellular proces | Cell wall/mem | 06B97 | phosphate-sel | 443144.GM21 | 3,4 | 1,496662955 | 0,0006275 | 0,00024234 | 2 - | 0,000382943 - | 2 - | 0,000450524 - | 3 - | 0,000533035 - | 6 - | 0,001059371 - | 4 - | 0,000711629 - |
| 1377 | 1 | 3um_FIJJAFBF_670752 | Bacteria | Bacteroidetes | Flavobacteriia | Flavobacterial | Flavobacteriac | Arenitalea | Arenitalea lut | metabolism | Inorganic ion t | 05C27 | TonB-depende | 886377.Murru | 4,4 | 1,2 | 0,00053772 | 0,000125369 | 5 - | 0,000625491 - | 3 - | 0,000441526 - | 3 - | 0,00034826 - | 6 - | 0,000692143 - | 5 - | 0,00058118 - |
| 1378 | 1 | 3um_FIJJAFBF_465283 | Eukaryota | Chlorophyta | Mamiellophyc | Mamiellales | Bathycoccace | Ostreococcus | Ostreococcus | unclassified | unclassified | unclassified | unclassified | unclassified | 1,2 | 0,4 | 0,000577106 | 0,000201486 | 2 - | 0,000968993 - | 1 - | 0,000569999 - | 1 - | 0,000449595 - | 1 - | 0,00044677 - | 1 - | 0,000450174 - |
| 1379 | 1 | 0.2um_NOKCPHJN_1017248 | Eukaryota | unclassified | Spirotrichea | Sporadotrichi | Oxytrichidae | Stylonychia | Stylonychia le | cellular proces | Signal transdu | COG5040 | Tyrosine 3-mo | 412030.XP_00 | 0,6 | 0,489897949 | 0,000203953 | 0,000166529 | 0 - | 0 - | 0 - | 0 - | 1 - | 0,000340488 - | 1 - | 0,000338349 - | 1 - | 0,000340927 - |
| 1380 | 1 | 0.2um_NOKCPHJN_604778 | Bacteria | Proteobacteri | Alphaproteob | Rhodospirillal | Rhodospirillac | Fodinicurvata | Fodinicurvata | unclassified | unclassified | unclassified | unclassified | unclassified | 0,2 | 0,4 | 3,82E-05 | 7,64E-05 | 0 - | 0 - | 1 - | 0,000191055 - | 0 - | 0 - | 0 - | 0 - | 0 - | 0 - |
| 1381 | 1 | 3um_FIJJAFBF_56092 | Eukaryota | Bacillariophyta | Bacillariophyc | Naviculales | Naviculaceae | Fistulifera | Fistulifera sola | metabolism | Energy produc | COG0055 | Produces ATP | 2850.Phatr54 | 2 | 0 | 0,000687095 | 6,72E-05 | 2 - | 0,000693232 - | 2 - | 0,000815572 - | 2 - | 0,000643294 - | 2 - | 0,000639252 - | 2 - | 0,000644122 - |
| 1382 | 1 | 0.2um_NOKCPHJN_148310 | Bacteria | Bacteroidetes | Flavobacteriia | Flavobacterial | Flavobacteriac | Flavobacteriu | Flavobacteriu | poorly charact | Function unkn | 0XZ2H | Membrane as | 353153.XP_80 | 2,4 | 0,489897949 | 0,000364033 | 6,02E-05 | 2 - | 0,000309326 - | 2 - | 0,000363915 - | 3 - | 0,000430565 - | 2 - | 0,000285239 - | 3 - | 0,000431119 - |
| 1383 | 1 | 3um_FIJJAFBF_627897 | Eukaryota | Phaeophyceae | unclassified | Ectocarpales | Ectocarpaceae | Ectocarpus | Ectocarpus sili | unclassified | unclassified | unclassified | unclassified | unclassified | 0,8 | 0,748331477 | 0,000676415 | 0,00068257 | 1 - | 0,000790035 - | 2 - | 0,001858917 - | 1 - | 0,000733124 - | 0 - | 0 - | 0 - | 0 - |
| 1384 | 1 | 0.2um_NOKCPHJN_144817 | Bacteria | Bacteroidetes | Flavobacteriia | Flavobacterial | Flavobacteriac | Polaribacter | Polaribacter g | poorly charact | Function unkn | 06HJQ | Hyalin repeat | 696747.NIES3 | 1,8 | 1,166190379 | 0,000308786 | 0,000196855 | 1 - | 0,00018346 - | 0 - | 0 - | 3 - | 0,000510733 - | 2 - | 0,000338349 - | 3 - | 0,00051139 - |
| 1385 | 1 | 3um_FIJJAFBF_292922 | Eukaryota | Bacillariophyta | Coscinodiscop | Coscinodiscale | Coscinodiscac | Coscinodiscus | Coscinodiscus | metabolism | Energy produc | COG4451 | RuBisCO catal | 977880.RALTA | 5,4 | 0,8 | 0,003351031 | 0,00042133 | 4 - | 0,002523564 - | 5 - | 0,003711147 - | 6 - | 0,003512665 - | 6 - | 0,003490591 - | 6 - | 0,003517187 - |
| 1386 | 1 | 3um_FIJJAFBF_226619 | Bacteria | Bacteroidetes | Flavobacteriia | Flavobacterial | Flavobacteriac | Formosa | Formosa sp. H | metabolism | Inorganic ion t | COG3119 | Sulfatase | 655815.ZPR_3 | 0,6 | 0,489897949 | 9,28E-05 | 7,58E-05 | 0 - | 0 - | 0 - | 0 - | 1 - | 0,000155003 - | 1 - | 0,000154029 - | 1 - | 0,000155203 - |
| 1387 | 1 | 0.2um_NOKCPHJN_538357 | Bacteria | Bacteroidetes | Flavobacteriia | Flavobacterial | Cryomorphace | unclassified | Cryomorphace | unclassified | unclassified | unclassified | unclassified | unclassified | 0,8 | 0,4 | 0,000102936 | 5,30E-05 | 0 - | 0 - | 1 - | 0,000153071 - | 1 - | 0,000120737 - | 1 - | 0,000119978 - | 1 - | 0,000120892 - |
| 1388 | 1 | 3um_FIJJAFBF_216064 | Bacteria | Proteobacteri | Gammaprotec | Cellvibrionales | Microbulbifer | Microbulbifer | Microbulbifer | information st | Translation, ri | 05C64 | Binds directly | 314277.MED1 | 1 | 0,632455532 | 0,000381735 | 0,000242609 | 2 - | 0,000759254 - | 1 - | 0,000446623 - | 0 - | 0 - | 1 - | 0,000350066 - | 1 - | 0,000352734 - |
| 1389 | 1 | 3um_FIJJAFBF_201379 | Bacteria | Bacteroidetes | Flavobacteriia | Flavobacterial | Flavobacteriac | Polaribacter | Polaribacter s | unclassified | unclassified | unclassified | unclassified | unclassified | 1,6 | 0,489897949 | 0,000480441 | 0,000131265 | 2 - | 0,000615395 - | 1 - | 0,000362 - | 1 - | 0,000285532 - | 2 - | 0,000567476 - | 2 - | 0,0005718 - |
| 1390 | 1 | 3um_FIJJAFBF_20314 | Bacteria | Bacteroidetes | Flavobacteriia | Flavobacterial | Flavobacteriac | Polaribacter | Polaribacter h | metabolism | Carbohydrate | COG0057 | glyceraldehyd | 762903.Pedsa | 1,6 | 0,489897949 | 0,000424214 | 0,000152998 | 2 - | 0,00052669 - | 2 - | 0,000619639 - | 1 - | 0,000244375 - | 2 - | 0,000485678 - | 1 - | 0,000244689 - |
| 1391 | 1 | 0.2um_NOKCPHJN_591136 | Eukaryota | Arthropoda | Insecta | Lepidoptera | Nymphalidae | Bicyclus | Bicyclus anyna | unclassified | unclassified | unclassified | unclassified | unclassified | 0,4 | 0,489897949 | 0,000173253 | 0,000212191 | 0 - | 0 - | 0 - | 0 - | 1 - | 0,000432855 - | 0 - | 0 - | 1 - | 0,000433412 - |
| 1392 | 1 | 3um_FIJJAFBF_817538 | unclassified | unclassified | unclassified | unclassified | unclassified | unclassified | unclassified | unclassified | unclassified | unclassified | unclassified | unclassified | 0,4 | 0,489897949 | 0,000213476 | 0,000261456 | 0 - | 0 - | 0 - | 0 - | 1 - | 0,000535373 - | 1 - | 0,000532009 - | 0 - | 0 - |
| 1393 | 1 | 3um_FIJJAFBF_326228 | Bacteria | Proteobacteri | Alphaproteob | Rhodobactera | Rhodobactera | Ruegeria | Ruegeria med | metabolism | Amino acid tra | 01SEG | branched-chai | 439496.RBY4I | 0,4 | 0,489897949 | 7,60E-05 | 9,31E-05 | 0 - | 0 - | 0 - | 0 - | 0 - | 0 - | 1 - | 0,00018938 - | 1 - | 0,000190823 - |
| 1394 | 1 | 0.2um_NOKCPHJN_380154 | Eukaryota | unclassified | Oomycetes | Pythiales | Pythiaceae | Pythium | Pythium insidi | information st | Translation, ri | KOG3184 | Ribosomal prc | 65071.PYU1_T | 0,4 | 0,489897949 | 0,000147383 | 0,000180507 | 0 - | 0 - | 0 - | 0 - | 1 - | 0,000368221 - | 0 - | 0 - | 1 - | 0,000368695 - |
| 1395 | 2 | 3um_FIJJAFBF_532815;3um_FI | Bacteria | Proteobacteri | Alphaproteob | Rhodobactera | various | various | various | unclassified | unclassified | unclassified | unclassified | unclassified | 0,8 | 0,4 | 0,000552362 | 0,000276971 | 1 - | 0,000730782 - | 0 - | 0 - | 1 - | 0,00067814 - | 1 - | 0,000673878 - | 1 - | 0,000679012 - |
| 1396 | 1 | 0.2um_NOKCPHJN_392180 | Eukaryota | unclassified | Oomycetes | Saprolegniales | Saprolegniace | Thraustotheca | Thraustotheca | cellular proces | Defense mech | KOG1823 | UTP20, small s | 67593.Physo1 | 0,6 | 0,489897949 | 1,95E-05 | 1,59E-05 | 0 - | 0 - | 0 - | 0 - | 1 - | 3,26E-05 - | 1 - | 3,24E-05 - | 1 - | 3,26E-05 - |
| 1397 | 1 | 0.2um_NOKCPHJN_1111840 | Eukaryota | unclassified | Cryptophyta | Pyrenomonad | Geminigerace | Guillardia | Guillardia thet | cellular proces | Cell wall/mem | 17CVN | Cell envelope | 502347.ESCAE | 0,4 | 0,489897949 | 0,000242309 | 0,00029677 | 0 - | 0 - | 0 - | 0 - | 0 - | 0 - | 1 - | 0,000603473 - | 1 - | 0,000608071 - |
| 1398 | 1 | 0.2um_NOKCPHJN_594863 | Eukaryota | Phaeophyceae | unclassified | Ectocarpales | Ectocarpaceae | Ectocarpus | Ectocarpus sili | unclassified | unclassified | unclassified | unclassified | unclassified | 0,8 | 0,4 | 5,78E-05 | 2,96E-05 | 1 - | 7,16E-05 - | 1 - | 8,42E-05 - | 1 - | 6,64E-05 - | 0 - | 0 - | 1 - | 6,65E-05 - |
| 1399 | 1 | 0.2um_NOKCPHJN_855581 | Eukaryota | unclassified | Pelagophycea | Pelagomonad | unclassified | Aureococcus | Aureococcus a | unclassified | unclassified | unclassified | unclassified | unclassified | 1 | 0 | 0,000174533 | 1,71E-05 | 1 - | 0,000176092 - | 1 - | 0,000207168 - | 1 - | 0,000163407 - | 1 - | 0,00016238 - | 1 - | 0,000163617 - |
| 1400 | 1 | 0.2um_NOKCPHJN_766226 | unclassified | unclassified | unclassified | unclassified | unclassified | unclassified | unclassified | unclassified | unclassified | unclassified | unclassified | unclassified | 1 | 0 | 0,000170426 | 1,67E-05 | 1 - | 0,000171949 - | 1 - | 0,000202294 - | 1 - | 0,000159562 - | 1 - | 0,00015856 - | 1 - | 0,000159768 - |
| 1401 | 1 | 3um_FIJJAFBF_335138 | Bacteria | Proteobacteri | Alphaproteob | Rhizobiales | unclassified | unclassified | Rhizobiales ba | unclassified | unclassified | unclassified | unclassified | unclassified | 0,8 | 0,4 | 0,000236727 | 0,000118702 | 1 - | 0,000313192 - | 0 - | 0 - | 1 - | 0,000290631 - | 1 - | 0,000288805 - | 1 - | 0,000291005 - |
| 1402 | 1 | 0.2um_NOKCPHJN_815085 | Viruses | unclassified | unclassified | unclassified | unclassified | unclassified | uncultured M | unclassified | unclassified | unclassified | unclassified | unclassified | 1,4 | 0,489897949 | 0,000369554 | 0,000110574 | 1 - | 0,000271498 - | 1 - | 0,000319411 - | 2 - | 0,000503881 - | 2 - | 0,000500714 - | 1 - | 0,000252265 - |
| 1403 | 1 | 0.2um_NOKCPHJN_487497 | Eukaryota | Arthropoda | Insecta | Lepidoptera | Nymphalidae | Bicyclus | Bicyclus anyna | unclassified | unclassified | unclassified | unclassified | unclassified | 0,2 | 0,4 | 3,08E-05 | 6,16E-05 | 1 - | 0,000154119 - | 0 - | 0 - | 0 - | 0 - | 0 - | 0 - | 0 - | 0 - |
| 1404 | 1 | 3um_FIJJAFBF_513507 | Bacteria | Actinobacteria | Actinobacteria | Streptomycet | Streptomycet | Streptomyces | Streptomyces | unclassified | unclassified | unclassified | unclassified | unclassified | 0,2 | 0,4 | 4,37E-05 | 8,74E-05 | 0 - | 0 - | 0 - | 0 - | 0 - | 0 - | 1 - | 0,000218555 - | 0 - | 0 - |
| 1405 | 1 | 0.2um_NOKCPHJN_828264 | Eukaryota | Chlorophyta | unclassified | Pyramimonad | unclassified | Cymbomonas | Cymbomonas | metabolism | Energy produc | 0IHF0 | This is one of t | 70448.Q0P3J1 | 0,8 | 0,4 | 0,000187772 | 9,42E-05 | 1 - | 0,000248425 - | 0 - | 0 - | 1 - | 0,000230529 - | 1 - | 0,00022908 - | 1 - | 0,000230826 - |
| 1406 | 1 | 0.2um_NOKCPHJN_1021094 | Bacteria | Proteobacteri | Betaproteoba | Nitrosomonad | unclassified | unclassified | Methylophilal | cellular proces | Signal transdu | 0B8J2 | Signal transdu | 383631.MB21 | 0,4 | 0,489897949 | 0,000100841 | 0,000125007 | 0 - | 0 - | 1 - | 0,000282657 - | 0 - | 0 - | 1 - | 0,000221549 - | 0 - | 0 - |
| 1407 | 1 | 0.2um_NOKCPHJN_227146 | Viruses | unclassified | unclassified | unclassified | Phycodnavirid | unclassified | Organic Lake p | information st | Transcription | COG0086 | DNA-depende | 88036.EFJ103 | 1 | 0 | 5,68E-05 | 5,56E-06 | 1 - | 5,74E-05 - | 1 - | 6,75E-05 - | 1 - | 5,32E-05 - | 1 - | 5,29E-05 - | 1 - | 5,33E-05 - |
| 1408 | 1 | 0.2um_NOKCPHJN_146801 | Eukaryota | Apicomplexa | Aconoidasida | Haemosporida | Plasmodiidae | Plasmodium | Plasmodium c | unclassified | unclassified | unclassified | unclassified | unclassified | 0,2 | 0,4 | 9,89E-06 | 1,98E-05 | 0 - | 0 - | 1 - | 4,95E-05 - | 0 - | 0 - | 0 - | 0 - | 0 - | 0 - |
| 1409 | 1 | 0.2um_NOKCPHJN_435831 | Bacteria | Proteobacteri | Gammaprotec | Alteromonada | Alteromonada | Bowmanella | Bowmanella d | metabolism | Inorganic ion t | 0QJVD | TonB-depende | 223926.VP016 | 0,8 | 0,4 | 8,03E-05 | 4,12E-05 | 1 - | 9,98E-05 - | 1 - | 0,000117372 - | 1 - | 9,26E-05 - | 1 - | 9,20E-05 - | 0 - | 0 - |
| 1410 | 1 | 3um_FIJJAFBF_238677 | Bacteria | Proteobacteri | Alphaproteob | Rhodobactera | Rhodobactera | unclassified | Rhodobactera | unclassified | unclassified | unclassified | unclassified | unclassified | 0,4 | 0,489897949 | 0,000212071 | 0,000261152 | 1 - | 0,000487188 - | 1 - | 0,000573166 - | 0 - | 0 - | 0 - | 0 - | 0 - | 0 - |
| 1411 | 1 | 3um_FIJJAFBF_231278 | Bacteria | Proteobacteri | Gammaprotec | Oceanospirilla | Oceanospirilla | Oceanobacter | Oceanobacter | metabolism | Carbohydrate | 0QKP9 | ABC-type suga | 400668.Mmw | 0,2 | 0,4 | 4,23E-05 | 8,45E-05 | 1 - | 0,00021131 - | 0 - | 0 - | 0 - | 0 - | 0 - | 0 - | 0 - | 0 - |
| 1412 | 1 | 0.2um_NOKCPHJN_175435 | Eukaryota | unclassified | Oomycetes | Peronosporale | Peronosporac | Phytophthora | Phytophthora | cellular proces | Cytoskeleton | 0DGVY | Dynein, axone | 7897.ENSLACF | 0,2 | 0,4 | 2,40E-06 | 4,80E-06 | 0 - | 0 - | 0 - | 0 - | 0 - | 0 - | 1 - | 1,20E-05 - | 0 - | 0 - |
| 1413 | 1 | 0.2um_NOKCPHJN_1052523 | Eukaryota | unclassified | Oomycetes | Peronosporale | Peronosporac | Plasmopara | Plasmopara ha | unclassified | unclassified | unclassified | unclassified | unclassified | 1 | 0 | 0,00058728 | 5,75E-05 | 1 - | 0,000592526 - | 1 - | 0,000697094 - | 1 - | 0,000549843 - | 1 - | 0,000546388 - | 1 - | 0,000550551 - |
| 1414 | 1 | 0.2um_NOKCPHJN_1062283 | Eukaryota | unclassified | Oomycetes | Peronosporale | Peronosporac | Phytophthora | Phytophthora | unclassified | unclassified | unclassified | unclassified | unclassified | 1,6 | 0,489897949 | 0,000279969 | 7,64E-05 | 2 - | 0,000358666 - | 1 - | 0,000210981 - | 2 - | 0,000332829 - | 2 - | 0,000330738 - | 1 - | 0,000166629 - |
| 1415 | 1 | 3um_FIJJAFBF_355317 | Bacteria | Proteobacteri | Gammaprotec | Cellvibrionales | Halieaceae | Halioglobus | Halioglobus lu | metabolism | Lipid transpor | 0RADM | Putative lysop | 247634.GPB2 | 0,4 | 0,489897949 | 0,000132086 | 0,000162655 | 1 - | 0,000303439 - | 1 - | 0,000356989 - | 0 - | 0 - | 0 - | 0 - | 0 - | 0 - |
| 1416 | 1 | 3um_FIJJAFBF_461253 | Bacteria | Proteobacteri | Alphaproteob | Rhodobactera | Rhodobactera | unclassified | Rhodobactera | unclassified | unclassified | unclassified | unclassified | unclassified | 0,6 | 0,489897949 | 0,000149066 | 0,000121714 | 0 - | 0 - | 0 - | 0 - | 1 - | 0,000248859 - | 1 - | 0,000247295 - | 1 - | 0,000249179 - |
| 1417 | 1 | 3um_FIJJAFBF_39296 | unclassified | unclassified | unclassified | unclassified | unclassified | unclassified | unclassified | unclassified | unclassified | unclassified | unclassified | unclassified | 0,2 | 0,4 | 3,74E-05 | 7,48E-05 | 1 - | 0,00018698 - | 0 - | 0 - | 0 - | 0 - | 0 - | 0 - | 0 - | 0 - |
| 1418 | 1 | 3um_FIJJAFBF_161769 | unclassified | unclassified | unclassified | unclassified | unclassified | unclassified | unclassified | unclassified | unclassified | unclassified | unclassified | unclassified | 0,2 | 0,4 | 5,45E-05 | 0,00010891 | 0 - | 0 - | 0 - | 0 - | 0 - | 0 - | 1 - | 0,000272274 - | 0 - | 0 - |
| 1419 | 1 | 0.2um_NOKCPHJN_122191 | Bacteria | Proteobacteri | Gammaprotec | unclassified | unclassified | unclassified | Gammaprotec | metabolism | Lipid transpor | COG1960 | acyl-CoA dehy | 632772.ROP_ | 1 | 0 | 0,000224014 | 2,19E-05 | 1 - | 0,000226015 - | 1 - | 0,000265902 - | 1 - | 0,000209734 - | 1 - | 0,000208416 - | 1 - | 0,000210004 - |
| 1420 | 1 | 0.2um_NOKCPHJN_820822 | Bacteria | Proteobacteri | Alphaproteob | Rickettsiales | unclassified | unclassified | Rickettsiales b | unclassified | unclassified | unclassified | unclassified | unclassified | 0,8 | 0,4 | 0,000169523 | 8,50E-05 | 1 - | 0,000224281 - | 0 - | 0 - | 1 - | 0,000208125 - | 1 - | 0,000206817 - | 1 - | 0,000208393 - |
| 1421 | 1 | 3um_FIJJAFBF_99324 | Bacteria | Bacteroidetes | Flavobacteriia | Flavobacterial | Flavobacteriac | unclassified | various | metabolism | Inorganic ion t | 05C27 | TonB-depende | 313598.MED1 | 23,8 | 1,939071943 | 0,002075819 | 0,000192755 | 21 - | 0,001854553 - | 23 - | 0,002389635 - | 24 - | 0,001966809 - | 27 - | 0,002198756 - | 24 - | 0,001969341 - |
| 1422 | 1 | 3um_FIJJAFBF_314913 | Bacteria | Proteobacteri | Gammaprotec | Alteromonada | Alteromonada | Paraglaciecola | Paraglaciecola | unclassified | unclassified | unclassified | unclassified | unclassified | 0,8 | 0,4 | 9,58E-05 | 4,80E-05 | 1 - | 0,000126725 - | 0 - | 0 - | 1 - | 0,000117596 - | 1 - | 0,000116857 - | 1 - | 0,000117748 - |
| 1423 | 1 | 0.2um_NOKCPHJN_859297 | Bacteria | Proteobacteri | Gammaprotec | Cellvibrionales | Porticoccacea | unclassified | gamma protec | poorly charact | Function unkn | 0XY6Y | Pkd domain cc | 314287.GB22 | 0,8 | 0,748331477 | 9,89E-05 | 9,52E-05 | 2 - | 0,00025679 - | 0 - | 0 - | 0 - | 0 - | 1 - | 0,000118397 - | 1 - | 0,000119299 - |
| 1424 | 1 | 3um_FIJJAFBF_254284 | Bacteria | Proteobacteri | Gammaprotec | Oceanospirilla | Oceanospirilla | Marinobacteri | various | information st | Translation, ri | COG0051 | tRNA binding | 876044.IMCC3 | 0,8 | 0,4 | 0,000667103 | 0,000343409 | 0 - | 0 - | 1 - | 0,000992018 - | 1 - | 0,000782469 - | 1 - | 0,000777551 - | 1 - | 0,000783476 - |
| 1425 | 1 | 3um_FIJJAFBF_244013 | Eukaryota | Chlorophyta | unclassified | Pyramimonad | unclassified | Cymbomonas | Cymbomonas | metabolism | Energy produc | COG1290 | Component o | 4538.ORGLA0 | 0,6 | 0,489897949 | 0,000212859 | 0,000173801 | 0 - | 0 - | 0 - | 0 - | 1 - | 0,000355357 - | 1 - | 0,000353124 - | 1 - | 0,000355814 - |
| 1426 | 1 | 3um_FIJJAFBF_664332 | Bacteria | Proteobacteri | Gammaprotec | Pseudomonad | Moraxellaceae | Moraxella | Moraxella boe | unclassified | unclassified | unclassified | unclassified | unclassified | 1,4 | 0,489897949 | 0,001205716 | 0,000360761 | 1 - | 0,000885796 - | 1 - | 0,00104212 - | 2 - | 0,001643975 - | 2 - | 0,001633643 - | 1 - | 0,000823045 - |
| 1427 | 1 | 0.2um_NOKCPHJN_983105 | Eukaryota | Chlorophyta | Ulvophyceae | Ulvales | Ulvaceae | Ulva | Ulva linza | unclassified | unclassified | unclassified | unclassified | unclassified | 0,6 | 0,489897949 | 0,000260311 | 0,000216161 | 0 - | 0 - | 1 - | 0,000505735 - | 0 - | 0 - | 1 - | 0,000396399 - | 1 - | 0,000399419 - |
| 1428 | 8 | 0.2um_NOKCPHJN_655150;3ur | Eukaryota | various | various | various | various | various | various | metabolism | Energy produc | 123BF | photosystem I | 35128.Thapsd | 2 | 0 | 0,001998103 | 0,000195503 | 2 - | 0,002015951 - | 2 - | 0,002371722 - | 2 - | 0,00187073 - | 2 - | 0,001858974 - | 2 - | 0,001873138 - |
| 1429 | 2 | 0.2um_NOKCPHJN_842037;3ur | Eukaryota | Chordata | Actinopteri | Carangiformes | Carangidae | Seriola | Seriola lalandi | unclassified | unclassified | unclassified | unclassified | unclassified | 0,8 | 0,4 | 0,000662835 | 0,000332365 | 1 - | 0,000876939 - | 0 - | 0 - | 1 - | 0,000813767 - | 1 - | 0,000808654 - | 1 - | 0,000814815 - |
| 1430 | 2 | 0.2um_NOKCPHJN_947652;3ur | Eukaryota | various | various | various | various | various | various | various | various | various | various | various | 0,2 | 0,4 | 0,000104397 | 0,000208795 | 1 - | 0,000521987 - | 0 - | 0 - | 0 - | 0 - | 0 - | 0 - | 0 - | 0 - |
| 1431 | 1 | 0.2um_NOKCPHJN_372100 | Bacteria | Proteobacteri | Gammaprotec | unclassified | unclassified | unclassified | Gammaprotec | unclassified | unclassified | unclassified | unclassified | unclassified | 1,6 | 0,489897949 | 0,000465525 | 0,000160563 | 1 - | 0,000291342 - | 2 - | 0,000685514 - | 2 - | 0,000540709 - | 1 - | 0,000268656 - | 2 - | 0,000541405 - |
| 1432 | 1 | 0.2um_NOKCPHJN_278290 | Eukaryota | Phaeophyceae | unclassified | Ectocarpales | Ectocarpaceae | Ectocarpus | Ectocarpus sili | metabolism | Energy produc | COG0055 | Produces ATP | 35128.Thaps4 | 4,8 | 1,469693846 | 0,00079775 | 0,000205623 | 5 - | 0,000863128 - | 2 - | 0,000406181 - | 6 - | 0,000961143 - | 5 - | 0,000795919 - | 6 - | 0,00096238 - |
| 1433 | 1 | 3um_FIJJAFBF_470068 | Bacteria | Bacteroidetes | Flavobacteriia | Flavobacterial | Flavobacteriac | Algibacter | Algibacter sp. | unclassified | unclassified | unclassified | unclassified | unclassified | 1,6 | 1,019803903 | 0,000186833 | 0,000118104 | 2 - | 0,000247373 - | 0 - | 0 - | 1 - | 0,000114777 - | 3 - | 0,000342167 - | 2 - | 0,000229849 - |
| 1434 | 1 | 0.2um_NOKCPHJN_279514 | unclassified | unclassified | unclassified | unclassified | unclassified | unclassified | unclassified | unclassified | unclassified | unclassified | unclassified | unclassified | 0,6 | 0,489897949 | 0,000291278 | 0,000240809 | 1 - | 0,000468951 - | 1 - | 0,000551711 - | 0 - | 0 - | 0 - | 0 - | 1 - | 0,00043573 - |
| 1435 | 1 | 3um_FIJJAFBF_58538 | Eukaryota | Phaeophyceae | unclassified | Ectocarpales | Ectocarpaceae | Ectocarpus | Ectocarpus sili | information st | Translation, ri | 1508W | Ribosomal prc | 272952.HpaP8 | 0,8 | 0,4 | 0,000660749 | 0,000340138 | 0 - | 0 - | 1 - | 0,00098257 - | 1 - | 0,000775017 - | 1 - | 0,000770146 - | 1 - | 0,000776014 - |
| 1436 | 1 | 0.2um_NOKCPHJN_411814 | Eukaryota | Chlorophyta | Trebouxiophy | Chlorellales | Chlorellaceae | Micractinium | Micractinium | cellular proces | Intracellular tr | KOG1109 | vacuole meml | 65071.PYU1_T | 0,4 | 0,489897949 | 7,17E-05 | 8,89E-05 | 0 - | 0 - | 1 - | 0,000201111 - | 0 - | 0 - | 1 - | 0,000157632 - | 0 - | 0 - |
| 1437 | 1 | 3um_FIJJAFBF_21463 | Eukaryota | unclassified | Cryptophyta | Pyrenomonad | Pyrenomonad | Rhodomonas | Rhodomonas | metabolism | Energy produc | COG4451 | RuBisCO catal | 45157.CMV01 | 1,2 | 0,4 | 0,000773751 | 0,000355811 | 1 - | 0,000630891 - | 2 - | 0,001484459 - | 1 - | 0,000585444 - | 1 - | 0,000581765 - | 1 - | 0,000586198 - |
| 1438 | 1 | 0.2um_NOKCPHJN_575170 | Bacteria | Proteobacteri | Gammaprotec | Pseudomonad | Moraxellaceae | Psychrobacter | Psychrobacter | unclassified | unclassified | unclassified | unclassified | unclassified | 0,2 | 0,4 | 7,32E-05 | 0,00014634 | 0 - | 0 - | 1 - | 0,000365851 - | 0 - | 0 - | 0 - | 0 - | 0 - | 0 - |
| 1439 | 1 | 3um_FIJJAFBF_358760 | Eukaryota | unclassified | Pelagophycea | Pelagomonad | unclassified | Aureococcus | Aureococcus a | unclassified | unclassified | unclassified | unclassified | unclassified | 1 | 0 | 0,00117456 | 0,000114924 | 1 - | 0,001185052 - | 1 - | 0,001394188 - | 1 - | 0,001099686 - | 1 - | 0,001092775 - | 1 - | 0,001101101 - |
| 1440 | 1 | 3um_FIJJAFBF_666764 | Bacteria | Proteobacteri | Epsilonproteo | Campylobacte | Campylobacte | Arcobacter | Arcobacter sp | metabolism | Carbohydrate | 05D2N | Aldolase | 469613.HMPR | 1 | 0 | 0,000526773 | 5,15E-05 | 1 - | 0,000531478 - | 1 - | 0,000625272 - | 1 - | 0,000493192 - | 1 - | 0,000490093 - | 1 - | 0,000493827 - |
| 1441 | 1 | 3um_FIJJAFBF_806359 | Bacteria | Proteobacteri | Gammaprotec | Oceanospirilla | Saccharospiril | Reinekea | Reinekea blan | metabolism | Carbohydrate | 0QKP9 | ABC-type suga | 314283.MED2 | 0,2 | 0,4 | 4,23E-05 | 8,45E-05 | 1 - | 0,00021131 - | 0 - | 0 - | 0 - | 0 - | 0 - | 0 - | 0 - | 0 - |
| 1442 | 1 | 0.2um_NOKCPHJN_245658 | Viruses | unclassified | unclassified | unclassified | unclassified | unclassified | Vibrio phage 1 | poorly charact | Function unkn | COG1783 | phage termina | 572265.HDEF_ | 0,8 | 0,4 | 0,000116902 | 5,86E-05 | 1 - | 0,000154663 - | 0 - | 0 - | 1 - | 0,000143522 - | 1 - | 0,00014262 - | 1 - | 0,000143706 - |
| 1443 | 1 | 3um_FIJJAFBF_452346 | unclassified | unclassified | unclassified | unclassified | unclassified | unclassified | unclassified | unclassified | unclassified | unclassified | unclassified | unclassified | 1 | 0,632455532 | 0,000457496 | 0,000277527 | 0 - | 0 - | 1 - | 0,000551711 - | 1 - | 0,00043517 - | 2 - | 0,00086487 - | 1 - | 0,00043573 - |
| 1444 | 1 | 0.2um_NOKCPHJN_688594 | Bacteria | Proteobacteri | Alphaproteob | Rhodospirillal | Rhodospirillac | Azospirillum | Azospirillum s | metabolism | Lipid transpor | 16PEQ | Amp-depende | 1064539.AZO | 1 | 0 | 0,000295638 | 2,89E-05 | 1 - | 0,000298278 - | 1 - | 0,000350918 - | 1 - | 0,000276792 - | 1 - | 0,000275052 - | 1 - | 0,000277148 - |
| 1445 | 1 | 0.2um_NOKCPHJN_359127 | Bacteria | Proteobacteri | Betaproteoba | various | various | various | various | unclassified | unclassified | unclassified | unclassified | unclassified | 2,6 | 1,743559577 | 0,000396193 | 0,000249158 | 2 - | 0,000318308 - | 1 - | 0,000187241 - | 6 - | 0,000886135 - | 2 - | 0,000293522 - | 2 - | 0,000295759 - |
| 1446 | 1 | 3um_FIJJAFBF_498429 | Bacteria | Proteobacteri | Gammaprotec | Arenicellales | Arenicellaceae | Arenicella | Arenicella xan | metabolism | Inorganic ion t | 177TK | phosphate-sel | 455436.GHTC | 2,6 | 0,489897949 | 0,000660606 | 0,000106684 | 3 - | 0,000776052 - | 2 - | 0,000608672 - | 3 - | 0,000720148 - | 2 - | 0,000477082 - | 3 - | 0,000721075 - |
| 1447 | 1 | 3um_FIJJAFBF_545834 | Bacteria | Bacteroidetes | Flavobacteriia | Flavobacterial | Flavobacteriac | Polaribacter | Polaribacter s | metabolism | Carbohydrate | 099YR | Catalyzes the | 313598.MED1 | 1,2 | 0,4 | 0,000240304 | 7,08E-05 | 1 - | 0,000204415 - | 1 - | 0,000240489 - | 1 - | 0,000189689 - | 2 - | 0,000376995 - | 1 - | 0,000189934 - |
| 1448 | 1 | 0.2um_NOKCPHJN_567304 | Eukaryota | unclassified | Pelagophycea | Pelagomonad | unclassified | Aureococcus | Aureococcus a | unclassified | unclassified | unclassified | unclassified | unclassified | 9,4 | 0,489897949 | 0,003258906 | 0,000255602 | 9 - | 0,003156979 - | 9 - | 0,003714116 - | 9 - | 0,002929563 - | 10 - | 0,003234614 - | 10 - | 0,00325926 - |
| 1449 | 1 | 3um_FIJJAFBF_625435 | Bacteria | Bacteroidetes | Flavobacteriia | Flavobacterial | Flavobacteriac | Arenitalea | Arenitalea lut | poorly charact | Function unkn | 09ARW | Zinc ribbon dc | 1046627.BZAF | 1,2 | 0,4 | 0,000398509 | 0,000119469 | 1 - | 0,000338586 - | 1 - | 0,000398339 - | 1 - | 0,000314196 - | 1 - | 0,000312221 - | 2 - | 0,000629201 - |
| 1450 | 1 | 0.2um_NOKCPHJN_1024190 | Bacteria | Proteobacteri | Alphaproteob | Rhodospirillal | Rhodospirillac | unclassified | Rhodospirillac | unclassified | unclassified | unclassified | unclassified | unclassified | 1 | 1,095445115 | 0,000389801 | 0,000407127 | 0 - | 0 - | 1 - | 0,000468954 - | 3 - | 0,001109683 - | 0 - | 0 - | 1 - | 0,00037037 - |
| 1451 | 5 | 0.2um_NOKCPHJN_310378;3ur | Bacteria | Bacteroidetes | Flavobacteriia | various | various | various | various | unclassified | unclassified | unclassified | unclassified | unclassified | 0,8 | 0,4 | 0,001159744 | 0,000594329 | 1 - | 0,001437604 - | 1 - | 0,00169131 - | 1 - | 0,001334045 - | 0 - | 0 - | 1 - | 0,001335762 - |
| 1452 | 2 | 0.2um_NOKCPHJN_145187;0.2 | Bacteria | Bacteroidetes | unclassified | unclassified | unclassified | unclassified | Bacteroidetes | unclassified | unclassified | unclassified | unclassified | unclassified | 0,6 | 0,489897949 | 0,000135814 | 0,00011278 | 0 - | 0 - | 1 - | 0,000263862 - | 0 - | 0 - | 1 - | 0,000206817 - | 1 - | 0,000208393 - |
| 1453 | 1 | 0.2um_NOKCPHJN_239171 | Bacteria | Bacteroidetes | unclassified | unclassified | unclassified | unclassified | Bacteroidetes | poorly charact | Function unkn | 0P8NI | Inherit from b | 313590.MED1 | 0,4 | 0,489897949 | 8,59E-05 | 0,000106389 | 0 - | 0 - | 1 - | 0,00023993 - | 0 - | 0 - | 0 - | 0 - | 1 - | 0,000189492 - |
| 1454 | 2 | 0.2um_NOKCPHJN_576092;3ur | Bacteria | Proteobacteri | Gammaprotec | unclassified | unclassified | unclassified | Gammaprotec | metabolism | Energy produc | COG0372 | citrate syntha | 351348.Maqu | 0,2 | 0,4 | 4,18E-05 | 8,37E-05 | 0 - | 0 - | 0 - | 0 - | 1 - | 0,000209195 - | 0 - | 0 - | 0 - | 0 - |
| 1455 | 1 | 3um_FIJJAFBF_737884 | Bacteria | Proteobacteri | Betaproteoba | unclassified | unclassified | unclassified | Betaproteoba | unclassified | unclassified | unclassified | unclassified | unclassified | 2 | 0 | 0,001287666 | 0,000125991 | 2 - | 0,001299168 - | 2 - | 0,001528443 - | 2 - | 0,001205581 - | 2 - | 0,001198005 - | 2 - | 0,001207133 - |
| 1456 | 1 | 0.2um_NOKCPHJN_728627 | Eukaryota | Bacillariophyta | Coscinodiscop | Thalassiosirale | Thalassiosirac | Thalassiosira | Thalassiosira p | unclassified | unclassified | unclassified | unclassified | unclassified | 0,6 | 0,489897949 | 0,000354668 | 0,00029008 | 1 - | 0,000621942 - | 0 - | 0 - | 0 - | 0 - | 1 - | 0,000573513 - | 1 - | 0,000577883 - |
| 1457 | 1 | 3um_FIJJAFBF_497706 | Bacteria | Proteobacteri | Alphaproteob | unclassified | unclassified | unclassified | Alphaproteob | unclassified | unclassified | unclassified | unclassified | unclassified | 1 | 0 | 0,000194446 | 1,90E-05 | 1 - | 0,000196183 - | 1 - | 0,000230805 - | 1 - | 0,000182051 - | 1 - | 0,000180907 - | 1 - | 0,000182285 - |
| 1458 | 1 | 3um_FIJJAFBF_526759 | Bacteria | Proteobacteri | Gammaprotec | Cellvibrionales | Halieaceae | unclassified | gamma protec | metabolism | Inorganic ion t | 16TQC | TONB-depend | 326297.Sama_ | 0,2 | 0,4 | 1,66E-05 | 3,32E-05 | 1 - | 8,30E-05 - | 0 - | 0 - | 0 - | 0 - | 0 - | 0 - | 0 - | 0 - |
| 1459 | 1 | 3um_FIJJAFBF_578962 | Eukaryota | Bacillariophyta | Coscinodiscop | Leptocylindral | Leptocylindrac | Leptocylindrus | Leptocylindrus | information st | Translation, ri | 0FP0S | Seems to be tl | 1140.Synpcc7 | 0,4 | 0,489897949 | 0,000266253 | 0,000326471 | 1 - | 0,000690503 - | 0 - | 0 - | 1 - | 0,000640762 - | 0 - | 0 - | 0 - | 0 - |
| 1460 | 1 | 3um_FIJJAFBF_335857 | Bacteria | Firmicutes | Bacilli | Bacillales | Bacillaceae | Bacillus | various | information st | RNA processin | 09V5W | DEAH (Asp-Glu | 8083.ENSXMA | 1,2 | 0,748331477 | 0,001195538 | 0,000812262 | 2 - | 0,001927337 - | 2 - | 0,00226747 - | 1 - | 0,00089425 - | 1 - | 0,00088863 - | 0 - | 0 - |
| 1461 | 2 | 0.2um_NOKCPHJN_312154;0.2 | Bacteria | Bacteroidetes | Flavobacteriia | Flavobacterial | Crocinitomica | unclassified | Crocinitomica | unclassified | unclassified | unclassified | unclassified | unclassified | 0,2 | 0,4 | 5,07E-05 | 0,00010138 | 1 - | 0,00025345 - | 0 - | 0 - | 0 - | 0 - | 0 - | 0 - | 0 - | 0 - |
| 1462 | 1 | 3um_FIJJAFBF_470658 | Eukaryota | unclassified | Cryptophyta | Pyrenomonad | Chroomonada | Chroomonas | Chroomonas r | metabolism | Energy produc | COG0056 | Produces ATP | 43989.cce_44 | 0,8 | 0,748331477 | 0,000134759 | 0,000129529 | 2 - | 0,000349378 - | 0 - | 0 - | 1 - | 0,000162105 - | 0 - | 0 - | 1 - | 0,000162314 - |
| 1463 | 1 | 3um_FIJJAFBF_765335 | Eukaryota | Bacillariophyta | Bacillariophyc | Naviculales | Phaeodactylac | Phaeodactylu | Phaeodactylu | unclassified | unclassified | unclassified | unclassified | unclassified | 0,4 | 0,489897949 | 5,62E-05 | 6,88E-05 | 0 - | 0 - | 0 - | 0 - | 0 - | 0 - | 1 - | 0,000139905 - | 1 - | 0,000140971 - |
| 1464 | 1 | 0.2um_NOKCPHJN_817058 | Bacteria | Proteobacteri | Gammaprotec | unclassified | unclassified | unclassified | Gammaprotec | unclassified | unclassified | unclassified | unclassified | unclassified | 0,2 | 0,4 | 8,99E-05 | 0,000179885 | 1 - | 0,000449712 - | 0 - | 0 - | 0 - | 0 - | 0 - | 0 - | 0 - | 0 - |
| 1465 | 1 | 3um_FIJJAFBF_122300 | Bacteria | Bacteroidetes | Flavobacteriia | Flavobacterial | Flavobacteriac | Formosa | Formosa algae | metabolism | Inorganic ion t | 05C27 | TonB-depende | 867900.Celly_ | 0,2 | 0,4 | 1,57E-05 | 3,14E-05 | 0 - | 0 - | 0 - | 0 - | 0 - | 0 - | 0 - | 0 - | 1 - | 7,86E-05 - |
| 1466 | 1 | 0.2um_NOKCPHJN_254831 | Eukaryota | Arthropoda | Hexanauplia | Calanoida | Temoridae | Eurytemora | Eurytemora af | unclassified | unclassified | unclassified | unclassified | unclassified | 0,4 | 0,489897949 | 9,39E-05 | 0,000115172 | 1 - | 0,000243594 - | 0 - | 0 - | 1 - | 0,000226047 - | 0 - | 0 - | 0 - | 0 - |
| 1467 | 5 | 0.2um_NOKCPHJN_70575;3um | Bacteria | Bacteroidetes | Flavobacteriia | Flavobacterial | Flavobacteriac | Polaribacter | various | metabolism | Amino acid tra | 08IQK | o-acetylhomo | various | 0,4 | 0,489897949 | 7,94E-05 | 9,74E-05 | 1 - | 0,000205854 - | 0 - | 0 - | 0 - | 0 - | 0 - | 0 - | 1 - | 0,000191271 - |
| 1468 | 1 | 3um_FIJJAFBF_521304 | Bacteria | Bacteroidetes | Flavobacteriia | Flavobacterial | Flavobacteriac | Salegentibacte | Salegentibacte | poorly charact | Function unkn | 080GT | oxidoreductas | 411154.GFO_ | 0,2 | 0,4 | 5,73E-05 | 0,000114632 | 1 - | 0,000286581 - | 0 - | 0 - | 0 - | 0 - | 0 - | 0 - | 0 - | 0 - |
| 1469 | 1 | 0.2um_NOKCPHJN_728002 | Bacteria | Verrucomicrol | Verrucomicrol | Verrucomicrol | Rubritaleacea | Rubritalea | Rubritalea ma | unclassified | unclassified | unclassified | unclassified | unclassified | 1,4 | 0,489897949 | 0,000250789 | 8,35E-05 | 2 - | 0,000364631 - | 1 - | 0,00021449 - | 1 - | 0,000169182 - | 2 - | 0,000336238 - | 1 - | 0,0001694 - |
| 1470 | 1 | 3um_FIJJAFBF_657588 | unclassified | unclassified | unclassified | unclassified | unclassified | unclassified | unclassified | unclassified | unclassified | unclassified | unclassified | unclassified | 1 | 0 | 0,000508289 | 4,97E-05 | 1 - | 0,00051283 - | 1 - | 0,000603333 - | 1 - | 0,000475887 - | 1 - | 0,000472897 - | 1 - | 0,0004765 - |
| 1471 | 1 | 0.2um_NOKCPHJN_21801 | Bacteria | Bacteroidetes | Flavobacteriia | Flavobacterial | Flavobacteriac | Polaribacter | Polaribacter s | poorly charact | Function unkn | 0P9SU | NA | 313598.MED1 | 0,4 | 0,489897949 | 9,44E-05 | 0,000115629 | 0 - | 0 - | 0 - | 0 - | 1 - | 0,000235875 - | 0 - | 0 - | 1 - | 0,000236178 - |
| 1472 | 1 | 3um_FIJJAFBF_135258 | Eukaryota | unclassified | Synurophycea | Parmales | Triparmaceae | Triparma | Triparma laevi | metabolism | Energy produc | COG0711 | F(1)F(0) ATP s | 314260.PB250 | 2,2 | 0,4 | 0,001439455 | 0,000232768 | 2 - | 0,001328695 - | 2 - | 0,00156318 - | 2 - | 0,001232981 - | 3 - | 0,001837849 - | 2 - | 0,001234568 - |
| 1473 | 1 | 3um_FIJJAFBF_502558 | Bacteria | Proteobacteri | Alphaproteob | unclassified | unclassified | unclassified | Alphaproteob | poorly charact | Function unkn | 11T16 | Pilus assembly | 985867.COdyt | 1 | 0 | 0,000351893 | 3,44E-05 | 1 - | 0,000355036 - | 1 - | 0,000417692 - | 1 - | 0,00032946 - | 1 - | 0,00032739 - | 1 - | 0,000329885 - |
| 1474 | 1 | 0.2um_NOKCPHJN_1115200 | Eukaryota | Chlorophyta | Chlorophycea | Chlamydomor | Chlamydomor | Chlamydomor | Chlamydomor | information st | Translation, ri | KOG3181 | ribosomal pro | 4558.Sb02g03 | 2,4 | 0,489897949 | 0,000948838 | 0,000259589 | 2 - | 0,000790035 - | 3 - | 0,001394188 - | 2 - | 0,000733124 - | 3 - | 0,001092775 - | 2 - | 0,000734067 - |
| 1475 | 1 | 3um_FIJJAFBF_734139 | Eukaryota | Streptophyta | Liliopsida | Poales | Poaceae | Dichanthelium | Dichanthelium | unclassified | unclassified | unclassified | unclassified | unclassified | 0,8 | 0,748331477 | 0,000329664 | 0,000304781 | 1 - | 0,000436288 - | 0 - | 0 - | 2 - | 0,000809719 - | 1 - | 0,000402315 - | 0 - | 0 - |
| 1476 | 1 | 3um_FIJJAFBF_578937 | Bacteria | Proteobacteri | Alphaproteob | Rhodobactera | Rhodobactera | Roseobacter | Roseobacter s | metabolism | Lipid transpor | 01R4W | acetyl-coa ace | 391593.RCCS2 | 0,2 | 0,4 | 4,57E-05 | 9,13E-05 | 1 - | 0,000228369 - | 0 - | 0 - | 0 - | 0 - | 0 - | 0 - | 0 - | 0 - |
| 1477 | 1 | 0.2um_NOKCPHJN_334766 | Bacteria | Proteobacteri | Alphaproteob | Rhodospirillal | Rhodospirillac | Candidatus En | Candidatus En | unclassified | unclassified | unclassified | unclassified | unclassified | 0,4 | 0,489897949 | 7,15E-05 | 8,77E-05 | 1 - | 0,000185399 - | 0 - | 0 - | 1 - | 0,000172044 - | 0 - | 0 - | 0 - | 0 - |
| 1478 | 2 | 3um_FIJJAFBF_08375;3um_FIJJ | Bacteria | Bacteroidetes | Flavobacteriia | Flavobacterial | Crocinitomica | unclassified | Crocinitomica | unclassified | unclassified | unclassified | unclassified | unclassified | 0,8 | 0,4 | 0,000504587 | 0,000258748 | 1 - | 0,000626385 - | 1 - | 0,000736928 - | 0 - | 0 - | 1 - | 0,00057761 - | 1 - | 0,000582011 - |
| 1479 | 1 | 3um_FIJJAFBF_07325 | Viruses | unclassified | unclassified | Caudovirales | Siphoviridae | unclassified | Tenacibaculur | unclassified | unclassified | unclassified | unclassified | unclassified | 2,8 | 0,4 | 0,000420785 | 7,34E-05 | 2 - | 0,000303439 - | 3 - | 0,000535484 - | 3 - | 0,000422371 - | 3 - | 0,000419716 - | 3 - | 0,000422914 - |
| 1480 | 1 | 0.2um_NOKCPHJN_270820 | unclassified | unclassified | unclassified | unclassified | unclassified | unclassified | unclassified | unclassified | unclassified | unclassified | unclassified | unclassified | 1 | 0 | 7,55E-05 | 7,39E-06 | 1 - | 7,62E-05 - | 1 - | 8,96E-05 - | 1 - | 7,07E-05 - | 1 - | 7,03E-05 - | 1 - | 7,08E-05 - |
| 1481 | 1 | 0.2um_NOKCPHJN_576524 | Bacteria | Bacteroidetes | Flavobacteriia | Flavobacterial | Flavobacteriac | Tenacibaculur | Tenacibaculur | unclassified | unclassified | unclassified | unclassified | unclassified | 0,2 | 0,4 | 3,13E-05 | 6,26E-05 | 1 - | 0,000156596 - | 0 - | 0 - | 0 - | 0 - | 0 - | 0 - | 0 - | 0 - |
| 1482 | 2 | 3um_FIJJAFBF_151896;3um_FI | various | various | various | various | various | various | various | unclassified | unclassified | unclassified | unclassified | unclassified | 0,8 | 0,4 | 0,000266199 | 0,00013348 | 1 - | 0,000352184 - | 0 - | 0 - | 1 - | 0,000326814 - | 1 - | 0,00032476 - | 1 - | 0,000327235 - |
| 1483 | 1 | 3um_FIJJAFBF_156949 | unclassified | unclassified | unclassified | unclassified | unclassified | unclassified | unclassified | unclassified | unclassified | unclassified | unclassified | unclassified | 0,2 | 0,4 | 1,35E-05 | 2,69E-05 | 0 - | 0 - | 0 - | 0 - | 1 - | 6,73E-05 - | 0 - | 0 - | 0 - | 0 - |
| 1484 | 1 | 3um_FIJJAFBF_85816 | Viruses | unclassified | unclassified | unclassified | various | various | various | unclassified | unclassified | unclassified | unclassified | unclassified | 0,4 | 0,489897949 | 0,000204132 | 0,000251376 | 1 - | 0,000468951 - | 1 - | 0,000551711 - | 0 - | 0 - | 0 - | 0 - | 0 - | 0 - |
| 1485 | 1 | 3um_FIJJAFBF_744568 | Bacteria | unclassified | unclassified | unclassified | unclassified | unclassified | uncultured ba | poorly charact | Function unkn | 0P8NI | Inherit from b | 983544.Lacal_ | 5,8 | 0,748331477 | 0,001218913 | 0,000133519 | 6 - | 0,001280202 - | 5 - | 0,001255108 - | 7 - | 0,001385979 - | 5 - | 0,000983763 - | 6 - | 0,001189511 - |
| 1486 | 1 | 3um_FIJJAFBF_39858 | Bacteria | Proteobacteri | Gammaprotec | Cellvibrionales | Spongiibacter | Oceanicoccus | Oceanicoccus | information st | Translation, ri | COG0050 | This protein p | 247633.GP214 | 0,4 | 0,489897949 | 0,000114695 | 0,00014063 | 1 - | 0,000297267 - | 0 - | 0 - | 0 - | 0 - | 0 - | 0 - | 1 - | 0,000276208 - |
| 1487 | 1 | 3um_FIJJAFBF_560942 | Eukaryota | unclassified | Florideophyce | Ceramiales | Rhodomelace | Polysiphonia | Polysiphonia t | metabolism | Energy produc | 0FJ7X | This is one of t | 240292.Ava_3 | 1,8 | 0,4 | 0,000434156 | 0,00010633 | 1 - | 0,000243594 - | 2 - | 0,000573166 - | 2 - | 0,000452093 - | 2 - | 0,000449252 - | 2 - | 0,000452675 - |
| 1488 | 1 | 0.2um_NOKCPHJN_726022 | Bacteria | Proteobacteri | Gammaprotec | unclassified | unclassified | Candidatus Th | Candidatus Th | unclassified | unclassified | unclassified | unclassified | unclassified | 1 | 0 | 0,000268264 | 2,62E-05 | 1 - | 0,00027066 - | 1 - | 0,000318426 - | 1 - | 0,000251163 - | 1 - | 0,000249584 - | 1 - | 0,000251486 - |
| 1489 | 2 | 3um_FIJJAFBF_32275;3um_FIJJ | Bacteria | Proteobacteri | Alphaproteob | Rhodobactera | Rhodobactera | various | various | poorly charact | Function unkn | 177T7 | NA | 391593.RCCS2 | 0,2 | 0,4 | 5,75E-05 | 0,000115008 | 1 - | 0,000287521 - | 0 - | 0 - | 0 - | 0 - | 0 - | 0 - | 0 - | 0 - |
| 1490 | 1 | 0.2um_NOKCPHJN_13684 | Bacteria | Bacteroidetes | Flavobacteriia | unclassified | unclassified | unclassified | Flavobacteria | unclassified | unclassified | unclassified | unclassified | unclassified | 0,4 | 0,489897949 | 4,17E-05 | 5,10E-05 | 0 - | 0 - | 0 - | 0 - | 0 - | 0 - | 1 - | 0,000103807 - | 1 - | 0,000104598 - |
| 1491 | 1 | 3um_FIJJAFBF_629419 | Bacteria | Bacteroidetes | Flavobacteriia | Flavobacterial | Flavobacteriac | Polaribacter | Polaribacter ta | metabolism | Inorganic ion t | 05C27 | TonB-depende | 873517.HMPR | 0,4 | 0,489897949 | 3,38E-05 | 4,15E-05 | 1 - | 8,77E-05 - | 0 - | 0 - | 1 - | 8,14E-05 - | 0 - | 0 - | 0 - | 0 - |
| 1492 | 1 | 0.2um_NOKCPHJN_23025 | Bacteria | unclassified | unclassified | unclassified | unclassified | unclassified | uncultured ba | unclassified | unclassified | unclassified | unclassified | unclassified | 0,8 | 0,748331477 | 6,98E-05 | 6,54E-05 | 0 - | 0 - | 0 - | 0 - | 1 - | 8,73E-05 - | 1 - | 8,68E-05 - | 2 - | 0,000174853 - |
| 1493 | 1 | 3um_FIJJAFBF_739631 | Eukaryota | unclassified | Chrysophycea | Chromulinales | Chromulinace | Ochromonas | Ochromonas s | unclassified | unclassified | unclassified | unclassified | unclassified | 0,4 | 0,489897949 | 6,66E-05 | 8,20E-05 | 1 - | 0,000153043 - | 1 - | 0,000180052 - | 0 - | 0 - | 0 - | 0 - | 0 - | 0 - |
| 1494 | 1 | 0.2um_NOKCPHJN_157840 | unclassified | unclassified | unclassified | unclassified | unclassified | unclassified | unclassified | unclassified | unclassified | unclassified | unclassified | unclassified | 1 | 0 | 0,0001162 | 1,14E-05 | 1 - | 0,000117238 - | 1 - | 0,000137928 - | 1 - | 0,000108792 - | 1 - | 0,000108109 - | 1 - | 0,000108932 - |
| 1495 | 1 | 0.2um_NOKCPHJN_115733 | Bacteria | Proteobacteri | Alphaproteob | unclassified | unclassified | unclassified | Alphaproteob | metabolism | Energy produc | COG0056 | Produces ATP | 488538.SAR11 | 0,2 | 0,4 | 3,18E-05 | 6,37E-05 | 0 - | 0 - | 0 - | 0 - | 1 - | 0,00015925 - | 0 - | 0 - | 0 - | 0 - |
| 1496 | 1 | 0.2um_NOKCPHJN_372004 | Eukaryota | Phaeophyceae | unclassified | Ectocarpales | Ectocarpaceae | Ectocarpus | Ectocarpus sili | unclassified | unclassified | unclassified | unclassified | unclassified | 0,6 | 0,489897949 | 8,40E-05 | 6,86E-05 | 0 - | 0 - | 0 - | 0 - | 1 - | 0,000140305 - | 1 - | 0,000139423 - | 1 - | 0,000140485 - |
| 1497 | 1 | 3um_FIJJAFBF_807627 | Bacteria | Bacteroidetes | Flavobacteriia | Flavobacterial | Flavobacteriac | Chryseobacte | Chryseobacte | unclassified | unclassified | unclassified | unclassified | unclassified | 0,8 | 0,4 | 0,000277515 | 0,000142858 | 0 - | 0 - | 1 - | 0,00041268 - | 1 - | 0,000325507 - | 1 - | 0,000323461 - | 1 - | 0,000325926 - |
| 1498 | 1 | 0.2um_NOKCPHJN_316399 | Bacteria | Proteobacteri | Gammaprotec | unclassified | unclassified | unclassified | SAR86 cluster | metabolism | Inorganic ion t | 0QK8T | TonB-depende | 247633.GP214 | 0,8 | 0,4 | 5,91E-05 | 2,96E-05 | 1 - | 7,82E-05 - | 0 - | 0 - | 1 - | 7,26E-05 - | 1 - | 7,21E-05 - | 1 - | 7,27E-05 - |
| 1499 | 1 | 3um_FIJJAFBF_25653 | Bacteria | Proteobacteri | Gammaprotec | Alteromonada | Alteromonada | Paraglaciecola | Paraglaciecola | unclassified | unclassified | unclassified | unclassified | unclassified | 3,2 | 0,748331477 | 0,00029279 | 4,97E-05 | 3 - | 0,00028258 - | 2 - | 0,000221632 - | 3 - | 0,000262224 - | 4 - | 0,000347434 - | 4 - | 0,000350082 - |
| 1500 | 1 | 0.2um_NOKCPHJN_483486 | Bacteria | Bacteroidetes | Flavobacteriia | Flavobacterial | Flavobacteriac | Arenitalea | Arenitalea lut | poorly charact | Function unkn | 0PAEN | NA | 886377.Murru | 2 | 1,414213562 | 0,000347429 | 0,000232847 | 0 - | 0 - | 1 - | 0,000214937 - | 2 - | 0,00033907 - | 4 - | 0,000673878 - | 3 - | 0,000509259 - |
| 1501 | 1 | 0.2um_NOKCPHJN_891697 | Eukaryota | unclassified | Pelagophycea | Pelagomonad | unclassified | Aureococcus | Aureococcus a | unclassified | unclassified | unclassified | unclassified | unclassified | 18,6 | 1,019803903 | 0,009081334 | 0,001020455 | 17 - | 0,008375255 - | 19 - | 0,011012516 - | 19 - | 0,008686282 - | 18 - | 0,008177395 - | 20 - | 0,009155224 - |
| 1502 | 3 | 0.2um_NOKCPHJN_741458;3ur | Bacteria | Actinobacteria | Actinobacteria | Micrococcales | Microbacteria | various | various | various | various | various | various | various | 0,2 | 0,4 | 5,52E-05 | 0,000110342 | 0 - | 0 - | 1 - | 0,000275855 - | 0 - | 0 - | 0 - | 0 - | 0 - | 0 - |

| | | | | | | | | | | | | | | | | | | | | | | | | | | | | |
|---|---|---|---|---|---|---|---|---|---|---|---|---|---|---|---|---|---|---|---|---|---|---|---|---|---|---|---|---|
| 1503 | 1 | 3um_FIJJAFBF_496263 | Bacteria | Bacteroidetes | Flavobacteriia | Flavobacterial | Flavobacteriac | Winogradskye | Winogradskye | metabolism | Energy produc | COG0055 | Produces ATP | 391603.FBALC | 5,6 | 1,854723699 | 0,000955952 | 0,000277668 | 6 - | 0,001048134 - | 4 - | 0,000822071 - | 5 - | 0,000810525 - | 9 - | 0,001449777 - | 4 - | 0,000649255 - |
| 1504 | 1 | 3um_FIJJAFBF_251917 | Eukaryota | Bacillariophyta | Coscinodiscop | Thalassiosirale | Thalassiosirac | Thalassiosira | Thalassiosira c | unclassified | unclassified | unclassified | unclassified | unclassified | 0,2 | 0,4 | 0,000332806 | 0,000665612 | 0 - | 0 - | 1 - | 0,00166403 - | 0 - | 0 - | 0 - | 0 - | 0 - | 0 - |
| 1505 | 2 | 0.2um_NOKCPHJN_101567;3ur | Bacteria | Bacteroidetes | Flavobacteriia | Flavobacterial | Flavobacteriac | Polaribacter | various | information st | Replication, re | 0PB0H | DNA-binding p | 313598.MED1 | 0,2 | 0,4 | 0,00018107 | 0,00036214 | 0 - | 0 - | 0 - | 0 - | 0 - | 0 - | 0 - | 0 - | 1 - | 0,00090535 - |
| 1506 | 3 | 0.2um_NOKCPHJN_237470;0.2 | Bacteria | Bacteroidetes | Flavobacteriia | Flavobacterial | Flavobacteriac | Tenacibaculur | various | metabolism | Energy produc | COG2838 | Isocitrate deh | various | 0,2 | 0,4 | 2,21E-05 | 4,42E-05 | 0 - | 0 - | 0 - | 0 - | 0 - | 0 - | 0 - | 0 - | 1 - | 0,000110409 - |
| 1507 | 1 | 3um_FIJJAFBF_156303 | Viruses | unclassified | unclassified | unclassified | various | various | various | information st | Replication, re | 1BHM5 | DNA polymera | 88036.EFJ152 | 0,8 | 0,4 | 4,55E-05 | 2,28E-05 | 1 - | 6,02E-05 - | 0 - | 0 - | 1 - | 5,59E-05 - | 1 - | 5,55E-05 - | 1 - | 5,59E-05 - |
| 1508 | 1 | 3um_FIJJAFBF_737020 | Bacteria | Proteobacteri | Alphaproteob | Rhodospirillale | unclassified | unclassified | Rhodospirillale | cellular proces | Cell wall/mem | COG2885 | Ompa motb d | 331869.BAL19 | 0,4 | 0,489897949 | 0,000195457 | 0,000239664 | 1 - | 0,000506901 - | 0 - | 0 - | 1 - | 0,000470386 - | 0 - | 0 - | 0 - | 0 - |
| 1509 | 1 | 3um_FIJJAFBF_788045 | Bacteria | Proteobacteri | Alphaproteob | Rhodobactera | unclassified | unclassified | Rhodobactera | information st | Translation, ri | 17C3B | The globular d | 391619.RGBS1 | 0,4 | 0,489897949 | 0,000257527 | 0,000315408 | 0 - | 0 - | 0 - | 0 - | 1 - | 0,000645847 - | 1 - | 0,000641789 - | 0 - | 0 - |
| 1510 | 2 | 3um_FIJJAFBF_326268;3um_FI | Bacteria | Proteobacteri | Alphaproteob | various | various | various | various | information st | Translation, ri | COG0184 | Forms an inte | various | 0,2 | 0,4 | 0,000197065 | 0,00039413 | 1 - | 0,000985324 - | 0 - | 0 - | 0 - | 0 - | 0 - | 0 - | 0 - | 0 - |
| 1511 | 1 | 3um_FIJJAFBF_63092 | unclassified | unclassified | unclassified | unclassified | unclassified | unclassified | unclassified | unclassified | unclassified | unclassified | unclassified | unclassified | 0,2 | 0,4 | 3,18E-05 | 6,37E-05 | 0 - | 0 - | 0 - | 0 - | 0 - | 0 - | 0 - | 0 - | 1 - | 0,000159144 - |
| 1512 | 1 | 0.2um_NOKCPHJN_326468 | Eukaryota | Phaeophyceae | unclassified | various | various | various | various | unclassified | unclassified | unclassified | unclassified | unclassified | 1,8 | 0,4 | 0,001048968 | 0,000256904 | 1 - | 0,000588549 - | 2 - | 0,001384831 - | 2 - | 0,001092305 - | 2 - | 0,001085441 - | 2 - | 0,001093711 - |
| 1513 | 1 | 0.2um_NOKCPHJN_59333 | Bacteria | Bacteroidetes | Flavobacteriia | Flavobacterial | Flavobacteriac | Lacinutrix | Lacinutrix ven | metabolism | Amino acid tra | 09GJ6 | nitrogen regul | 391603.FBALC | 2,6 | 0,489897949 | 0,001998599 | 0,00031829 | 3 - | 0,002348942 - | 2 - | 0,001842319 - | 2 - | 0,001453156 - | 3 - | 0,002166036 - | 3 - | 0,00218254 - |
| 1514 | 1 | 3um_FIJJAFBF_761774 | Bacteria | Proteobacteri | Gammaprotec | Aeromonadale | Aeromonadac | Aeromonas | Aeromonas ve | unclassified | unclassified | unclassified | unclassified | unclassified | 0,2 | 0,4 | 2,53E-05 | 5,06E-05 | 1 - | 0,000126542 - | 0 - | 0 - | 0 - | 0 - | 0 - | 0 - | 0 - | 0 - |
| 1515 | 1 | 0.2um_NOKCPHJN_857856 | Eukaryota | unclassified | Oomycetes | Saprolegniales | Saprolegniace | Thraustotheca | Thraustotheca | unclassified | unclassified | unclassified | unclassified | unclassified | 1 | 0,894427191 | 0,000221896 | 0,000198374 | 0 - | 0 - | 0 - | 0 - | 1 - | 0,000222341 - | 2 - | 0,000441887 - | 2 - | 0,000445254 - |
| 1516 | 1 | 0.2um_NOKCPHJN_85563 | Bacteria | Firmicutes | Clostridia | Clostridiales | Clostridiaceae | Clostridium | Clostridium sp | unclassified | unclassified | unclassified | unclassified | unclassified | 0,4 | 0,489897949 | 0,000471885 | 0,000584971 | 0 - | 0 - | 1 - | 0,001322691 - | 0 - | 0 - | 1 - | 0,001036735 - | 0 - | 0 - |
| 1517 | 1 | 3um_FIJJAFBF_344571 | Bacteria | various | various | various | various | various | various | metabolism | Amino acid tra | 0FNCB | carbon dioxide | 84588.SYNW1 | 1,8 | 0,748331477 | 0,001488381 | 0,000559263 | 2 - | 0,001702793 - | 1 - | 0,001001649 - | 3 - | 0,002370196 - | 1 - | 0,000785101 - | 2 - | 0,001582165 - |
| 1518 | 1 | 3um_FIJJAFBF_582313 | Eukaryota | Eustigmatophy | unclassified | Eustigmatales | Monodopsida | Nannochlorop | Nannochlorop | cellular proces | Signal transdu | COG5040 | Tyrosine 3-mc | 35128.Thaps2 | 2,6 | 0,489897949 | 0,001598157 | 0,00037459 | 2 - | 0,001235125 - | 3 - | 0,002179646 - | 2 - | 0,001146151 - | 3 - | 0,001708423 - | 3 - | 0,00172144 - |
| 1519 | 1 | 3um_FIJJAFBF_331637 | Eukaryota | unclassified | Pelagophycea | Pelagomonad | unclassified | Aureococcus | Aureococcus a | cellular proces | Posttranslatio | 1DQ9N | cysteine prote | 4113.PGSC000 | 0,8 | 0,4 | 0,000246024 | 0,000126647 | 0 - | 0 - | 1 - | 0,000365851 - | 1 - | 0,00028857 - | 1 - | 0,000286757 - | 1 - | 0,000288941 - |
| 1520 | 1 | 0.2um_NOKCPHJN_167800 | Bacteria | Proteobacteri | Gammaprotec | Cellvibrionales | Porticoccacea | unclassified | SAR92 bacteri | unclassified | unclassified | unclassified | unclassified | unclassified | 0,6 | 0,489897949 | 0,00035962 | 0,000294138 | 1 - | 0,000630891 - | 0 - | 0 - | 1 - | 0,000585444 - | 1 - | 0,000581765 - | 0 - | 0 - |
| 1521 | 1 | 0.2um_NOKCPHJN_71343 | Bacteria | Proteobacteri | Gammaprotec | Cellvibrionales | unclassified | unclassified | Cellvibrionales | unclassified | unclassified | unclassified | unclassified | unclassified | 0,4 | 0,489897949 | 0,000144859 | 0,000177417 | 0 - | 0 - | 0 - | 0 - | 1 - | 0,000363289 - | 1 - | 0,000361006 - | 0 - | 0 - |
| 1522 | 1 | 3um_FIJJAFBF_594166 | Eukaryota | unclassified | Pelagophycea | Pelagomonad | unclassified | Aureococcus | Aureococcus a | metabolism | Energy produc | COG0711 | F(1)F(0) ATP s | 35128.Thapsd | 5,4 | 1,019803903 | 0,002655013 | 0,000474972 | 5 - | 0,002491303 - | 5 - | 0,002930963 - | 6 - | 0,002774207 - | 4 - | 0,001837849 - | 7 - | 0,003240741 - |
| 1523 | 1 | 3um_FIJJAFBF_116542 | Eukaryota | Chordata | Actinopteri | Carangiformes | Carangidae | Seriola | Seriola lalandi | unclassified | unclassified | unclassified | unclassified | unclassified | 1,2 | 0,748331477 | 0,00147052 | 0,000934344 | 2 - | 0,002579231 - | 0 - | 0 - | 1 - | 0,001196717 - | 2 - | 0,002378393 - | 1 - | 0,001198257 - |
| 1524 | 1 | 3um_FIJJAFBF_715066 | Eukaryota | unclassified | Florideophyce | Gigartinales | Peyssonneliac | Riquetophycu | Riquetophycu | unclassified | unclassified | unclassified | unclassified | unclassified | 6,6 | 0,8 | 0,000877076 | 0,000143479 | 6 - | 0,000803302 - | 7 - | 0,001102579 - | 6 - | 0,000745436 - | 8 - | 0,000987668 - | 6 - | 0,000746395 - |
| 1525 | 1 | 3um_FIJJAFBF_306072 | Eukaryota | unclassified | Bangiophycea | Bangiales | Bangiaceae | Pyropia | Pyropia endivi | cellular proces | Posttranslatio | 0IEKR | One of the cor | 35128.Thapsd | 0,4 | 0,489897949 | 7,07E-05 | 8,66E-05 | 0 - | 0 - | 0 - | 0 - | 0 - | 0 - | 1 - | 0,000176177 - | 1 - | 0,00017752 - |
| 1526 | 1 | 3um_FIJJAFBF_437595 | Bacteria | Bacteroidetes | Flavobacteriia | Flavobacterial | Flavobacteriac | Polaribacter | Polaribacter s | unclassified | unclassified | unclassified | unclassified | unclassified | 2 | 0 | 0,000949918 | 9,29E-05 | 2 - | 0,000958403 - | 2 - | 0,00112754 - | 2 - | 0,000889363 - | 2 - | 0,000883774 - | 2 - | 0,000890508 - |
| 1527 | 1 | 0.2um_NOKCPHJN_187364 | Bacteria | Proteobacteri | Alphaproteob | Rhodobactera | Hyphomonad | unclassified | Hyphomonad | information st | Translation, ri | COG0617 | Catalyzes the | 755732.Fluta_ | 0,2 | 0,4 | 3,26E-05 | 6,51E-05 | 0 - | 0 - | 0 - | 0 - | 1 - | 0,000162753 - | 0 - | 0 - | 0 - | 0 - |
| 1528 | 1 | 0.2um_NOKCPHJN_981236 | Bacteria | Proteobacteri | Alphaproteob | Pelagibacteral | Pelagibacterac | Candidatus Pe | Candidatus Pe | metabolism | Amino acid tra | 05CM2 | brancheD-cha | 439493.PB721 | 0,4 | 0,489897949 | 0,000185579 | 0,000229885 | 0 - | 0 - | 1 - | 0,000518442 - | 0 - | 0 - | 0 - | 0 - | 1 - | 0,000409455 - |
| 1529 | 1 | 3um_FIJJAFBF_280809 | Bacteria | Bacteroidetes | Flavobacteriia | Flavobacterial | Flavobacteriac | Arenitalea | Arenitalea lut | cellular proces | Cell motility | COG1360 | Ompa motb d | 391587.KAOT1 | 7,2 | 1,939071943 | 0,002192051 | 0,000464235 | 7 - | 0,002200204 - | 4 - | 0,001479138 - | 7 - | 0,00204171 - | 8 - | 0,00231872 - | 10 - | 0,002920484 - |
| 1530 | 1 | 0.2um_NOKCPHJN_321072 | unclassified | unclassified | unclassified | unclassified | unclassified | unclassified | unclassified | unclassified | unclassified | unclassified | unclassified | unclassified | 0,2 | 0,4 | 3,15E-05 | 6,30E-05 | 1 - | 0,00015744 - | 0 - | 0 - | 0 - | 0 - | 0 - | 0 - | 0 - | 0 - |
| 1531 | 1 | 3um_FIJJAFBF_556902 | Bacteria | unclassified | unclassified | unclassified | unclassified | unclassified | uncultured ba | information st | Replication, re | COG0178 | The UvrABC re | 50743.SCB49_ | 0,2 | 0,4 | 1,86E-05 | 3,72E-05 | 1 - | 9,30E-05 - | 0 - | 0 - | 0 - | 0 - | 0 - | 0 - | 0 - | 0 - |
| 1532 | 1 | 0.2um_NOKCPHJN_123118 | Bacteria | Proteobacteri | Alphaproteob | Rhodobactera | Rhodobactera | Planktomarina | Planktomarina | metabolism | Energy produc | 01RNF | Produces ATP | 388401.RB215 | 0,2 | 0,4 | 3,70E-05 | 7,40E-05 | 1 - | 0,000185008 - | 0 - | 0 - | 0 - | 0 - | 0 - | 0 - | 0 - | 0 - |
| 1533 | 2 | 0.2um_NOKCPHJN_117309;3ur | Bacteria | Bacteroidetes | Flavobacteriia | Flavobacterial | Flavobacteriac | Polaribacter | various | unclassified | unclassified | unclassified | unclassified | unclassified | 0,2 | 0,4 | 3,95E-05 | 7,91E-05 | 0 - | 0 - | 1 - | 0,000197643 - | 0 - | 0 - | 0 - | 0 - | 0 - | 0 - |
| 1534 | 1 | 3um_FIJJAFBF_151367 | Eukaryota | Annelida | Polychaeta | Golfingiida | Sipunculidae | Sipunculus | Sipunculus nu | information st | Chromatin str | KOG1744 | Core compone | 400682.PAC_1 | 2,6 | 0,8 | 0,001830795 | 0,000567769 | 4 - | 0,002851833 - | 2 - | 0,001677559 - | 3 - | 0,001984799 - | 2 - | 0,001314884 - | 2 - | 0,001324902 - |
| 1535 | 1 | 0.2um_NOKCPHJN_273568 | Eukaryota | unclassified | Oomycetes | Saprolegniales | Saprolegniace | Aphanomyces | Aphanomyces | unclassified | unclassified | unclassified | unclassified | unclassified | 0,2 | 0,4 | 3,49E-05 | 6,97E-05 | 1 - | 0,000174342 - | 0 - | 0 - | 0 - | 0 - | 0 - | 0 - | 0 - | 0 - |
| 1536 | 1 | 0.2um_NOKCPHJN_256507 | Bacteria | Bacteroidetes | Flavobacteriia | Flavobacterial | Flavobacteriac | Polaribacter | Polaribacter s | metabolism | Inorganic ion t | 05C27 | TonB-depende | 471870.BACIN | 0,8 | 0,4 | 6,15E-05 | 3,09E-05 | 1 - | 8,14E-05 - | 0 - | 0 - | 1 - | 7,56E-05 - | 1 - | 7,51E-05 - | 1 - | 7,57E-05 - |
| 1537 | 1 | 3um_FIJJAFBF_68872 | Eukaryota | unclassified | unclassified | unclassified | unclassified | Ancoracysta | Ancoracysta t | unclassified | unclassified | unclassified | unclassified | unclassified | 0,8 | 0,4 | 0,000247358 | 0,000126763 | 1 - | 0,000306622 - | 1 - | 0,000360734 - | 1 - | 0,000284534 - | 0 - | 0 - | 1 - | 0,0002849 - |
| 1538 | 1 | 3um_FIJJAFBF_800979 | Eukaryota | Bacillariophyta | Mediophyceae | Triceratiales | Triceratiaceae | Trieres | Trieres chinen | unclassified | unclassified | unclassified | unclassified | unclassified | 2,2 | 0,4 | 0,000953279 | 0,000283033 | 2 - | 0,000859744 - | 3 - | 0,001517204 - | 2 - | 0,000797811 - | 2 - | 0,000792798 - | 2 - | 0,000798838 - |
| 1539 | 1 | 3um_FIJJAFBF_40378 | Eukaryota | Bacillariophyta | Coscinodiscop | Thalassiosirale | Thalassiosirac | Thalassiosira | Thalassiosira c | metabolism | Energy produc | 11HAV | photosystem i | 35128.Thapsd | 0,2 | 0,4 | 0,000126178 | 0,000252356 | 1 - | 0,000630891 - | 0 - | 0 - | 0 - | 0 - | 0 - | 0 - | 0 - | 0 - |
| 1540 | 1 | 3um_FIJJAFBF_517515 | Bacteria | Proteobacteri | Gammaprotec | Cellvibrionales | Spongiibacter | Dasania | Dasania marin | metabolism | Amino acid tra | 05C6M | Alpha-keto-be | 314285.KT71_ | 0,4 | 0,489897949 | 6,65E-05 | 8,14E-05 | 0 - | 0 - | 0 - | 0 - | 1 - | 0,000166075 - | 0 - | 0 - | 1 - | 0,000166289 - |
| 1541 | 1 | 0.2um_NOKCPHJN_89225 | Eukaryota | unclassified | Oomycetes | Saprolegniales | Saprolegniace | Saprolegnia | Saprolegnia di | unclassified | unclassified | unclassified | unclassified | unclassified | 1 | 0 | 5,47E-05 | 5,35E-06 | 1 - | 5,52E-05 - | 1 - | 6,49E-05 - | 1 - | 5,12E-05 - | 1 - | 5,09E-05 - | 1 - | 5,12E-05 - |
| 1542 | 1 | 3um_FIJJAFBF_135101 | unclassified | unclassified | unclassified | unclassified | unclassified | unclassified | unclassified | unclassified | unclassified | unclassified | unclassified | unclassified | 0,6 | 0,489897949 | 0,000363578 | 0,000301941 | 0 - | 0 - | 1 - | 0,000706643 - | 1 - | 0,000557375 - | 1 - | 0,000553872 - | 0 - | 0 - |
| 1543 | 1 | 0.2um_NOKCPHJN_21267 | Bacteria | Proteobacteri | unclassified | unclassified | unclassified | unclassified | uncultured ma | cellular proces | Posttranslatio | 01QDX | ATP binding to | 501479.CSE45 | 0,4 | 0,489897949 | 8,85E-05 | 0,000108539 | 1 - | 0,000229565 - | 0 - | 0 - | 1 - | 0,000213028 - | 0 - | 0 - | 0 - | 0 - |
| 1544 | 1 | 0.2um_NOKCPHJN_56842 | Bacteria | Proteobacteri | Gammaprotec | Oceanospirilla | Saccharospirill | Reinekea | Reinekea blan | information st | Translation, ri | COG0172 | Catalyzes the | 314283.MED2 | 0,2 | 0,4 | 3,78E-05 | 7,55E-05 | 0 - | 0 - | 0 - | 0 - | 1 - | 0,000188809 - | 0 - | 0 - | 0 - | 0 - |
| 1545 | 1 | 3um_FIJJAFBF_57085 | Bacteria | Proteobacteri | Gammaprotec | Cellvibrionales | Porticoccacea | unclassified | Gammaprotec | poorly charact | Function unkn | 16R8I | Lipoprotein | 377629.TERTU | 2,8 | 0,4 | 0,000293465 | 5,69E-05 | 3 - | 0,000315824 - | 3 - | 0,00037156 - | 3 - | 0,000293074 - | 3 - | 0,000291232 - | 2 - | 0,000195634 - |
| 1546 | 1 | 0.2um_NOKCPHJN_624427 | Eukaryota | Ascomycota | Eurotiomycete | Eurotiales | Aspergillaceae | Aspergillus | Aspergillus te | cellular proces | Posttranslatio | 0JAJ8 | Dentin sialoph | 10020.ENSDO | 1 | 0 | 0,000101067 | 9,89E-06 | 1 - | 0,00010197 - | 1 - | 0,000119965 - | 1 - | 9,46E-05 - | 1 - | 9,40E-05 - | 1 - | 9,47E-05 - |
| 1547 | 1 | 3um_FIJJAFBF_386845 | unclassified | unclassified | unclassified | unclassified | unclassified | unclassified | unclassified | unclassified | unclassified | unclassified | unclassified | unclassified | 0,2 | 0,4 | 0,000139419 | 0,000278838 | 0 - | 0 - | 1 - | 0,000697094 - | 0 - | 0 - | 0 - | 0 - | 0 - | 0 - |
| 1548 | 1 | 3um_FIJJAFBF_753878 | Bacteria | Proteobacteri | Alphaproteob | Rhodospirillale | Rhodospirillac | unclassified | Rhodospirillac | metabolism | Inorganic ion t | COG0841 | acriflavin resis | 391593.RCCS2 | 0,6 | 0,489897949 | 6,07E-05 | 5,01E-05 | 1 - | 9,77E-05 - | 1 - | 0,000114889 - | 0 - | 0 - | 0 - | 0 - | 1 - | 9,07E-05 - |
| 1549 | 1 | 3um_FIJJAFBF_66140 | Bacteria | Bacteroidetes | Flavobacteriia | Flavobacterial | Flavobacteriac | Polaribacter | various | unclassified | unclassified | unclassified | unclassified | unclassified | 0,2 | 0,4 | 0,00015521 | 0,000310421 | 1 - | 0,000776052 - | 0 - | 0 - | 0 - | 0 - | 0 - | 0 - | 0 - | 0 - |
| 1550 | 1 | 3um_FIJJAFBF_129163 | Bacteria | Proteobacteri | Gammaprotec | unclassified | unclassified | unclassified | OM182 bacter | unclassified | unclassified | unclassified | unclassified | unclassified | 0,6 | 0,8 | 5,19E-05 | 7,04E-05 | 2 - | 0,000177518 - | 0 - | 0 - | 0 - | 0 - | 1 - | 8,18E-05 - | 0 - | 0 - |
| 1551 | 1 | 3um_FIJJAFBF_660323 | Bacteria | Proteobacteri | Alphaproteob | Rhodobactera | Rhodobactera | Pseudodonghi | Pseudodonghi | unclassified | unclassified | unclassified | unclassified | unclassified | 0,4 | 0,489897949 | 0,000251084 | 0,000311067 | 0 - | 0 - | 1 - | 0,000701836 - | 1 - | 0,000553583 - | 0 - | 0 - | 0 - | 0 - |
| 1552 | 1 | 3um_FIJJAFBF_157376 | Bacteria | Bacteroidetes | Flavobacteriia | Flavobacterial | Flavobacteriac | Algibacter | Algibacter aqu | information st | Translation, ri | COG0052 | 30S ribosomal | 487797.Flav3C | 1,6 | 1,019803903 | 0,000488656 | 0,000310223 | 0 - | 0 - | 2 - | 0,000726549 - | 1 - | 0,000286538 - | 2 - | 0,000569474 - | 3 - | 0,00086072 - |
| 1553 | 1 | 3um_FIJJAFBF_419336 | Bacteria | Bacteroidetes | unclassified | unclassified | unclassified | unclassified | Bacteroidetes | poorly charact | Function unkn | 05DPE | Conserved Pro | 329726.AM1_ | 0,6 | 0,489897949 | 9,14E-05 | 7,59E-05 | 0 - | 0 - | 1 - | 0,000177268 - | 1 - | 0,000139823 - | 0 - | 0 - | 1 - | 0,000140003 - |
| 1554 | 1 | 3um_FIJJAFBF_580261 | Bacteria | Proteobacteri | Gammaprotec | Pseudomonad | Pseudomonad | Pseudomonas | Pseudomonas | poorly charact | Function unkn | 17B58 | Inherit from b | 471874.PROST | 2,8 | 0,4 | 0,000726011 | 0,000126634 | 2 - | 0,000523545 - | 3 - | 0,000923909 - | 3 - | 0,000728747 - | 3 - | 0,000724167 - | 3 - | 0,000729685 - |
| 1555 | 1 | 3um_FIJJAFBF_730482 | unclassified | unclassified | unclassified | unclassified | unclassified | unclassified | unclassified | unclassified | unclassified | unclassified | unclassified | unclassified | 0,8 | 0,4 | 0,000444735 | 0,000228939 | 0 - | 0 - | 1 - | 0,000661345 - | 1 - | 0,000521646 - | 1 - | 0,000518368 - | 1 - | 0,000522317 - |
| 1556 | 1 | 0.2um_NOKCPHJN_832213 | Bacteria | Verrucomicrob | Opitutae | unclassified | unclassified | unclassified | Opitutae bact | unclassified | unclassified | unclassified | unclassified | unclassified | 0,2 | 0,4 | 0,000170243 | 0,000340486 | 0 - | 0 - | 0 - | 0 - | 0 - | 0 - | 1 - | 0,000851214 - | 0 - | 0 - |
| 1557 | 1 | 3um_FIJJAFBF_574434 | Bacteria | Proteobacteri | Alphaproteob | Rhodobactera | Rhodobactera | Pseudophaeol | Pseudophaeol | information st | Transcription | 069B1 | acetyltransfer | 314262.MED1 | 1 | 0 | 0,000485572 | 4,75E-05 | 1 - | 0,00048991 - | 1 - | 0,000576368 - | 1 - | 0,000454619 - | 1 - | 0,000451762 - | 1 - | 0,000455204 - |
| 1558 | 1 | 0.2um_NOKCPHJN_1036455 | unclassified | unclassified | unclassified | unclassified | unclassified | unclassified | unclassified | unclassified | unclassified | unclassified | unclassified | unclassified | 0,2 | 0,4 | 3,70E-05 | 7,40E-05 | 0 - | 0 - | 1 - | 0,000184892 - | 0 - | 0 - | 0 - | 0 - | 0 - | 0 - |
| 1559 | 1 | 3um_FIJJAFBF_586240 | Bacteria | Bacteroidetes | Flavobacteriia | Flavobacterial | Flavobacteriac | Lacinutrix | Lacinutrix algi | metabolism | Amino acid tra | COG0174 | glutamine syn | 313594.PI23P | 0,8 | 0,4 | 0,000205262 | 0,000105664 | 0 - | 0 - | 1 - | 0,000305236 - | 1 - | 0,00024076 - | 1 - | 0,000239247 - | 1 - | 0,00024107 - |
| 1560 | 1 | 3um_FIJJAFBF_257125 | Bacteria | Cyanobacteria | unclassified | Oscillatoriales | Microcoleacea | Arthrospira | various | unclassified | unclassified | unclassified | unclassified | unclassified | 3,6 | 1,2 | 0,001732342 | 0,000511977 | 2 - | 0,000985324 - | 3 - | 0,001738818 - | 5 - | 0,002285864 - | 3 - | 0,001362899 - | 5 - | 0,002288806 - |
| 1561 | 1 | 0.2um_NOKCPHJN_43232 | Bacteria | Actinobacteria | Actinobacteria | Micrococcales | Microbacteria | Candidatus Ac | Candidatus Ac | information st | Replication, re | COG0470 | DNA polymera | 743718.Isova_ | 0,4 | 0,489897949 | 8,78E-05 | 0,000107693 | 1 - | 0,000227776 - | 0 - | 0 - | 1 - | 0,000211368 - | 0 - | 0 - | 0 - | 0 - |
| 1562 | 1 | 3um_FIJJAFBF_313534 | Bacteria | Actinobacteria | Actinobacteria | Streptomyceta | Streptomyceta | Streptomyces | Streptomyces | unclassified | unclassified | unclassified | unclassified | unclassified | 1 | 0 | 0,000480207 | 4,70E-05 | 1 - | 0,000484496 - | 1 - | 0,000569999 - | 1 - | 0,000449595 - | 1 - | 0,00044677 - | 1 - | 0,000450174 - |
| 1563 | 1 | 0.2um_NOKCPHJN_216476 | unclassified | unclassified | unclassified | unclassified | unclassified | unclassified | unclassified | information st | Transcription | KOG2252 | homeobox | 7460.GB13918 | 0,4 | 0,489897949 | 7,04E-06 | 8,62E-06 | 0 - | 0 - | 0 - | 0 - | 1 - | 1,76E-05 - | 1 - | 1,75E-05 - | 0 - | 0 - |
| 1564 | 1 | 3um_FIJJAFBF_187829 | Bacteria | Bacteroidetes | Flavobacteriia | Flavobacterial | Flavobacteriac | Polaribacter | Polaribacter s | unclassified | unclassified | unclassified | unclassified | unclassified | 0,4 | 0,489897949 | 0,000136485 | 0,000167388 | 1 - | 0,000355036 - | 0 - | 0 - | 0 - | 0 - | 1 - | 0,00032739 - | 0 - | 0 - |
| 1565 | 1 | 3um_FIJJAFBF_48303 | Bacteria | Proteobacteri | Gammaprotec | Cellvibrionales | Halieaceae | Halioglobus | Halioglobus pa | poorly charact | Function unkn | 05ZFT | NA | 314285.KT71_ | 1 | 0,632455532 | 0,000172769 | 0,000126927 | 0 - | 0 - | 2 - | 0,000396046 - | 1 - | 0,000156193 - | 1 - | 0,000155212 - | 1 - | 0,000156394 - |
| 1566 | 1 | 3um_FIJJAFBF_346215 | Bacteria | Bacteroidetes | Flavobacteriia | Flavobacterial | Crocinitomica | unclassified | Crocinitomica | unclassified | unclassified | unclassified | unclassified | unclassified | 2 | 0 | 0,002287302 | 0,000223799 | 2 - | 0,002307733 - | 2 - | 0,002714997 - | 2 - | 0,002141493 - | 2 - | 0,002128036 - | 2 - | 0,00214425 - |
| 1567 | 1 | 0.2um_NOKCPHJN_944186 | Eukaryota | unclassified | Heterotrichea | Heterotrichida | Stentoridae | Stentor | Stentor coeru | information st | Chromatin str | KOG2130 | jumonji doma | 572307.XP_00 | 0,4 | 0,489897949 | 8,54E-05 | 0,000105849 | 0 - | 0 - | 1 - | 0,000238819 - | 1 - | 0,000188372 - | 0 - | 0 - | 0 - | 0 - |
| 1568 | 1 | 3um_FIJJAFBF_36386 | Eukaryota | unclassified | unclassified | Prymnesiales | Chrysochromu | Chrysochromu | Chrysochromu | unclassified | unclassified | unclassified | unclassified | unclassified | 0,2 | 0,4 | 3,09E-05 | 6,17E-05 | 0 - | 0 - | 0 - | 0 - | 0 - | 0 - | 0 - | 0 - | 1 - | 0,000154321 - |
| 1569 | 1 | 3um_FIJJAFBF_440532 | Eukaryota | Chordata | Mammalia | Primates | Cercopithecid | Cercocebus | Cercocebus at | cellular proces | Cell cycle cont | 0V9W7 | retinitis pigme | 9606.ENSP000 | 0,4 | 0,489897949 | 2,95E-05 | 3,66E-05 | 0 - | 0 - | 1 - | 8,25E-05 - | 0 - | 0 - | 0 - | 0 - | 1 - | 6,52E-05 - |
| 1570 | 1 | 0.2um_NOKCPHJN_84326 | Bacteria | Bacteroidetes | Flavobacteriia | Flavobacterial | Flavobacteriac | Algibacter | Algibacter sp. | poorly charact | Function unkn | 11NCV | Protein of unk | 1046627.BZAR | 3 | 0,632455532 | 0,001656813 | 0,000508997 | 3 - | 0,00164426 - | 4 - | 0,002579247 - | 3 - | 0,001525814 - | 3 - | 0,001516225 - | 2 - | 0,001018519 - |
| 1571 | 1 | 3um_FIJJAFBF_00853 | Bacteria | Bacteroidetes | Flavobacteriia | Flavobacterial | Flavobacteriac | Polaribacter | Polaribacter s | unclassified | unclassified | unclassified | unclassified | unclassified | 1 | 0 | 0,000679043 | 6,64E-05 | 1 - | 0,000685108 - | 1 - | 0,000806015 - | 1 - | 0,000635756 - | 1 - | 0,000631761 - | 1 - | 0,000636574 - |
| 1572 | 1 | 3um_FIJJAFBF_225879 | Bacteria | Bacteroidetes | Flavobacteriia | Flavobacterial | Flavobacteriac | Flaviramulus | Flaviramulus b | cellular proces | Cell wall/mem | 06B97 | phosphate-sel | 391598.FBBAL | 1,8 | 0,4 | 0,000380528 | 9,98E-05 | 2 - | 0,000423642 - | 2 - | 0,000498405 - | 2 - | 0,000393124 - | 2 - | 0,000390654 - | 1 - | 0,000196815 - |
| 1573 | 2 | 0.2um_NOKCPHJN_220959;0.2 | Bacteria | unclassified | unclassified | unclassified | unclassified | unclassified | uncultured ba | metabolism | Inorganic ion t | COG4206 | tonB-depende | 1046627.BZAR | 0,6 | 0,489897949 | 8,03E-05 | 6,56E-05 | 0 - | 0 - | 0 - | 0 - | 1 - | 0,000134064 - | 1 - | 0,000133221 - | 1 - | 0,000134236 - |
| 1574 | 1 | 3um_FIJJAFBF_712785 | Eukaryota | unclassified | Pelagophycea | Pelagomonad | unclassified | Aureococcus | Aureococcus a | unclassified | unclassified | unclassified | unclassified | unclassified | 0,4 | 0,489897949 | 7,14E-05 | 8,75E-05 | 1 - | 0,000185008 - | 0 - | 0 - | 0 - | 0 - | 0 - | 0 - | 1 - | 0,000171902 - |
| 1575 | 1 | 3um_FIJJAFBF_91117 | Eukaryota | Bacillariophyta | Coscinodiscop | Leptocylindral | Leptocylindrac | Leptocylindrus | Leptocylindrus | unclassified | unclassified | unclassified | unclassified | unclassified | 1 | 0 | 0,000852132 | 8,34E-05 | 1 - | 0,000859744 - | 1 - | 0,001011469 - | 1 - | 0,000797811 - | 1 - | 0,000792798 - | 1 - | 0,000798838 - |
| 1576 | 1 | 3um_FIJJAFBF_287676 | Bacteria | Bacteroidetes | Flavobacteriia | Flavobacterial | Flavobacteriac | various | various | information st | Translation, ri | 0PBIW | 50S ribosomal | 391603.FBALC | 0,6 | 0,489897949 | 0,000727533 | 0,000594036 | 0 - | 0 - | 0 - | 0 - | 1 - | 0,001214578 - | 1 - | 0,001206946 - | 1 - | 0,001216142 - |
| 1577 | 1 | 0.2um_NOKCPHJN_712059 | Eukaryota | unclassified | Ichthyosporea | Ichthyophonic | unclassified | Sphaeroforma | Sphaeroforma | unclassified | unclassified | unclassified | unclassified | unclassified | 0,4 | 0,489897949 | 0,000252501 | 0,000309596 | 1 - | 0,000654432 - | 0 - | 0 - | 0 - | 0 - | 0 - | 0 - | 1 - | 0,000608071 - |
| 1578 | 1 | 3um_FIJJAFBF_503090 | Bacteria | Verrucomicrob | Opitutae | unclassified | unclassified | unclassified | Opitutae bact | unclassified | unclassified | unclassified | unclassified | unclassified | 3 | 0,632455532 | 0,000631111 | 0,000129342 | 2 - | 0,000426734 - | 3 - | 0,000753065 - | 3 - | 0,000593991 - | 4 - | 0,000787011 - | 3 - | 0,000594755 - |
| 1579 | 1 | 0.2um_NOKCPHJN_987129 | unclassified | unclassified | unclassified | unclassified | unclassified | unclassified | unclassified | unclassified | unclassified | unclassified | unclassified | unclassified | 0,6 | 0,489897949 | 0,000594448 | 0,000485371 | 0 - | 0 - | 0 - | 0 - | 1 - | 0,000992399 - | 1 - | 0,000986163 - | 1 - | 0,000993677 - |
| 1580 | 1 | 3um_FIJJAFBF_242895 | Bacteria | Bacteroidetes | Flavobacteriia | Flavobacterial | Flavobacteriac | Polaribacter | Polaribacter d | poorly charact | Function unkn | 08ME3 | Alpha amylase | 983548.Krodi_ | 0,4 | 0,489897949 | 9,44E-05 | 0,000115629 | 0 - | 0 - | 0 - | 0 - | 1 - | 0,000235875 - | 0 - | 0 - | 1 - | 0,000236178 - |
| 1581 | 1 | 0.2um_NOKCPHJN_155794 | Eukaryota | unclassified | Raphidophyce | Chattonellales | Chattonellace | Heterosigma | Heterosigma a | unclassified | unclassified | unclassified | unclassified | unclassified | 0,8 | 0,4 | 0,000138987 | 7,12E-05 | 1 - | 0,000172287 - | 1 - | 0,000202691 - | 1 - | 0,000159876 - | 0 - | 0 - | 1 - | 0,000160082 - |
| 1582 | 1 | 0.2um_NOKCPHJN_311381 | Eukaryota | unclassified | Oomycetes | Saprolegniales | Saprolegniace | Aphanomyces | Aphanomyces | unclassified | unclassified | unclassified | unclassified | unclassified | 0,4 | 0,489897949 | 2,12E-05 | 2,61E-05 | 1 - | 4,86E-05 - | 1 - | 5,72E-05 - | 0 - | 0 - | 0 - | 0 - | 0 - | 0 - |
| 1583 | 1 | 0.2um_NOKCPHJN_469918 | Bacteria | unclassified | unclassified | unclassified | unclassified | unclassified | bacterium B1 | unclassified | unclassified | unclassified | unclassified | unclassified | 0,6 | 0,489897949 | 8,75E-05 | 7,15E-05 | 0 - | 0 - | 0 - | 0 - | 1 - | 0,000146098 - | 1 - | 0,00014518 - | 1 - | 0,000146286 - |
| 1584 | 1 | 0.2um_NOKCPHJN_504894 | Bacteria | Proteobacteri | Alphaproteob | Rhodobactera | Rhodobactera | Oceaniovalibu | Oceaniovalibu | unclassified | unclassified | unclassified | unclassified | unclassified | 4,8 | 0,979795897 | 0,000962233 | 0,000233211 | 6 - | 0,001206796 - | 5 - | 0,001183141 - | 5 - | 0,00093322 - | 5 - | 0,000927355 - | 3 - | 0,000560652 - |
| 1585 | 1 | 0.2um_NOKCPHJN_668827 | Bacteria | Proteobacteri | Alphaproteob | Pelagibacteral | Pelagibacterac | unclassified | Pelagibacterac | cellular proces | Cell wall/mem | 7257 | NA | 439493.PB721 | 0,2 | 0,4 | 4,89E-05 | 9,77E-05 | 0 - | 0 - | 0 - | 0 - | 0 - | 0 - | 1 - | 0,000244306 - | 0 - | 0 - |
| 1586 | 1 | 3um_FIJJAFBF_198300 | Bacteria | Bacteroidetes | Flavobacteriia | Flavobacterial | Flavobacteriac | Polaribacter | Polaribacter s | metabolism | Inorganic ion t | 05C27 | TonB-depende | 487797.Flav3C | 3,2 | 1,6 | 0,000526307 | 0,000267321 | 6 - | 0,001017724 - | 1 - | 0,000199555 - | 3 - | 0,000472205 - | 3 - | 0,000469238 - | 3 - | 0,000472813 - |
| 1587 | 1 | 3um_FIJJAFBF_414092 | Bacteria | Bacteroidetes | Flavobacteriia | Flavobacterial | Flavobacteriac | Polaribacter | various | metabolism | Carbohydrate | COG0149 | triosephospha | 313598.MED1 | 0,8 | 0,4 | 0,000275312 | 0,000141724 | 0 - | 0 - | 1 - | 0,000409404 - | 1 - | 0,000322924 - | 1 - | 0,000320894 - | 1 - | 0,000323339 - |
| 1588 | 1 | 3um_FIJJAFBF_313972 | Bacteria | Proteobacteri | Gammaprotec | Chromatiales | Granulosicocc | Granulosicocc | Granulosicocc | cellular proces | Cell wall/mem | 177PY | Molecular cha | 187272.Mlg_1 | 0,2 | 0,4 | 8,34E-05 | 0,000166733 | 0 - | 0 - | 0 - | 0 - | 0 - | 0 - | 1 - | 0,000416832 - | 0 - | 0 - |
| 1589 | 1 | 0.2um_NOKCPHJN_343102 | Eukaryota | Cnidaria | Anthozoa | Scleractinia | Merulinidae | Orbicella | Orbicella fave | unclassified | unclassified | unclassified | unclassified | unclassified | 0,8 | 0,4 | 5,24E-05 | 2,69E-05 | 1 - | 6,50E-05 - | 1 - | 7,65E-05 - | 0 - | 0 - | 1 - | 5,99E-05 - | 1 - | 6,04E-05 - |
| 1590 | 2 | 0.2um_NOKCPHJN_1022817;3u | Eukaryota | various | various | various | various | various | various | unclassified | unclassified | unclassified | unclassified | unclassified | 1 | 0 | 0,000934596 | 9,14E-05 | 1 - | 0,000942945 - | 1 - | 0,001109354 - | 1 - | 0,000875019 - | 1 - | 0,00086952 - | 1 - | 0,000876145 - |
| 1591 | 1 | 3um_FIJJAFBF_41558 | Eukaryota | Bacillariophyta | Mediophyceae | Lithodesmiale | Lithodesmiace | Lithodesmium | Lithodesmium | metabolism | Carbohydrate | 0IVA1 | Photosystem I | 35128.Thapsd | 3 | 0,632455532 | 0,001386026 | 0,000212476 | 3 - | 0,001422062 - | 2 - | 0,00111535 - | 3 - | 0,001319623 - | 3 - | 0,00131133 - | 4 - | 0,001761762 - |
| 1592 | 1 | 0.2um_NOKCPHJN_421216 | Bacteria | Verrucomicrob | unclassified | unclassified | unclassified | unclassified | Verrucomicro | unclassified | unclassified | unclassified | unclassified | unclassified | 0,4 | 0,489897949 | 4,31E-05 | 5,28E-05 | 1 - | 0,000111712 - | 0 - | 0 - | 1 - | 0,000103665 - | 0 - | 0 - | 0 - | 0 - |
| 1593 | 1 | 0.2um_NOKCPHJN_338143 | Eukaryota | unclassified | Bangiophycea | Bangiales | Bangiaceae | Porphyra | Porphyra umb | cellular proces | Posttranslatio | KOG0734 | YME1-like 1 (S | 45157.CMI157 | 1 | 0 | 0,000198442 | 1,94E-05 | 1 - | 0,000200214 - | 1 - | 0,000235548 - | 1 - | 0,000185792 - | 1 - | 0,000184624 - | 1 - | 0,000186031 - |
| 1594 | 1 | 3um_FIJJAFBF_532805 | Bacteria | Proteobacteri | Alphaproteob | Rhodobactera | Rhodobactera | unclassified | Rhodobactera | unclassified | unclassified | unclassified | unclassified | unclassified | 8,6 | 0,489897949 | 0,002017642 | 0,000256906 | 8 - | 0,001890973 - | 9 - | 0,002502774 - | 9 - | 0,001974099 - | 8 - | 0,001743727 - | 9 - | 0,00197664 - |
| 1595 | 1 | 0.2um_NOKCPHJN_161249 | Archaea | Euryarchaeota | unclassified | unclassified | unclassified | unclassified | Euryarchaeota | unclassified | unclassified | unclassified | unclassified | unclassified | 1 | 0 | 0,000331746 | 3,25E-05 | 1 - | 0,000334709 - | 1 - | 0,000393778 - | 1 - | 0,000310598 - | 1 - | 0,000308646 - | 1 - | 0,000310998 - |
| 1596 | 2 | 0.2um_NOKCPHJN_551016;3ur | Bacteria | Bacteroidetes | Cytophagia | Cytophagales | Flammeovirga | Fabibacter | Fabibacter sp. | unclassified | unclassified | unclassified | unclassified | unclassified | 0,2 | 0,4 | 3,65E-05 | 7,31E-05 | 1 - | 0,000182696 - | 0 - | 0 - | 0 - | 0 - | 0 - | 0 - | 0 - | 0 - |
| 1597 | 1 | 3um_FIJJAFBF_535169 | Bacteria | Proteobacteri | Alphaproteob | Rhodobactera | unclassified | unclassified | Rhodobactera | unclassified | unclassified | unclassified | unclassified | unclassified | 1 | 0 | 0,000523599 | 5,12E-05 | 1 - | 0,000528276 - | 1 - | 0,000621505 - | 1 - | 0,000490221 - | 1 - | 0,000487141 - | 1 - | 0,000490852 - |
| 1598 | 1 | 0.2um_NOKCPHJN_663999 | Bacteria | Proteobacteri | Gammaprotec | Chromatiales | Ectothiorhodc | Thioalkalivibri | Thioalkalivibri | unclassified | unclassified | unclassified | unclassified | unclassified | 0,4 | 0,489897949 | 0,000106041 | 0,000129874 | 0 - | 0 - | 0 - | 0 - | 1 - | 0,000265937 - | 1 - | 0,000264266 - | 0 - | 0 - |
| 1599 | 1 | 3um_FIJJAFBF_417061 | Bacteria | Proteobacteri | Gammaprotec | Alteromonada | Alteromonada | Paraglaciecola | various | information st | Translation, ri | 179ZI | One of the pri | 342610.Patl_1 | 0,4 | 0,489897949 | 0,000249764 | 0,000305901 | 0 - | 0 - | 0 - | 0 - | 0 - | 0 - | 1 - | 0,000622041 - | 1 - | 0,000626781 - |
| 1600 | 1 | 3um_FIJJAFBF_220489 | Bacteria | Proteobacteri | Gammaprotec | Alteromonada | Alteromonada | Paraglaciecola | Paraglaciecola | information st | Translation, ri | COG0088 | One of the pri | 983545.Glaag | 0,4 | 0,489897949 | 0,000161539 | 0,000197847 | 0 - | 0 - | 0 - | 0 - | 0 - | 0 - | 1 - | 0,000402315 - | 1 - | 0,000405381 - |
| 1601 | 1 | 3um_FIJJAFBF_690863 | Viruses | unclassified | unclassified | unclassified | unclassified | unclassified | uncultured M | unclassified | unclassified | unclassified | unclassified | unclassified | 0,4 | 0,489897949 | 0,000208594 | 0,000256871 | 1 - | 0,000479201 - | 1 - | 0,00056377 - | 0 - | 0 - | 0 - | 0 - | 0 - | 0 - |
| 1602 | 1 | 3um_FIJJAFBF_426980 | Bacteria | Bacteroidetes | Flavobacteriia | Flavobacterial | Flavobacteriac | Mesoflavibact | Mesoflavibact | unclassified | unclassified | unclassified | unclassified | unclassified | 1 | 0 | 0,000186119 | 1,82E-05 | 1 - | 0,000187781 - | 1 - | 0,000220921 - | 1 - | 0,000174254 - | 1 - | 0,000173159 - | 1 - | 0,000174479 - |
| 1603 | 1 | 3um_FIJJAFBF_147734 | Bacteria | Bacteroidetes | Flavobacteriia | Flavobacterial | Crocinitomica | unclassified | Crocinitomica | unclassified | unclassified | unclassified | unclassified | unclassified | 0,4 | 0,489897949 | 5,10E-05 | 6,24E-05 | 0 - | 0 - | 0 - | 0 - | 0 - | 0 - | 1 - | 0,000126947 - | 1 - | 0,000127914 - |
| 1604 | 1 | 3um_FIJJAFBF_577814 | Eukaryota | Chlorophyta | Chlorophycea | Sphaeropleale | Selenastracea | Raphidocelis | Raphidocelis s | information st | Replication, re | 0XNUT | helicase activi | 3055.EDP0692 | 0,2 | 0,4 | 3,70E-05 | 7,40E-05 | 1 - | 0,000185008 - | 0 - | 0 - | 0 - | 0 - | 0 - | 0 - | 0 - | 0 - |
| 1605 | 1 | 0.2um_NOKCPHJN_237508 | Bacteria | Proteobacteri | Gammaprotec | Cellvibrionales | unclassified | unclassified | Cellvibrionales | poorly charact | Function unkn | 0QNMQ | NA | 314287.GB22C | 0,8 | 0,4 | 5,73E-05 | 2,88E-05 | 1 - | 7,59E-05 - | 0 - | 0 - | 1 - | 7,04E-05 - | 1 - | 7,00E-05 - | 1 - | 7,05E-05 - |
| 1606 | 1 | 3um_FIJJAFBF_03426 | Bacteria | Bacteroidetes | Flavobacteriia | Flavobacterial | Flavobacteriac | Algibacter | Algibacter aqu | information st | Transcription | 05D27 | DNA-depende | 983544.Lacal_ | 1,8 | 0,979795897 | 0,000103796 | 5,59E-05 | 2 - | 0,000122392 - | 0 - | 0 - | 2 - | 0,000113575 - | 3 - | 0,000169292 - | 2 - | 0,000113722 - |
| 1607 | 1 | 0.2um_NOKCPHJN_356814 | Bacteria | Proteobacteri | Gammaprotec | unclassified | unclassified | unclassified | Gammaprotec | metabolism | Energy produc | 0QIP8 | Electron trans | 220664.PFL_1 | 0,8 | 0,4 | 0,00021451 | 0,000107561 | 1 - | 0,000283799 - | 0 - | 0 - | 1 - | 0,000263355 - | 1 - | 0,0002617 - | 1 - | 0,000263694 - |
| 1608 | 1 | 3um_FIJJAFBF_367967 | Bacteria | Proteobacteri | Gammaprotec | Enterobactera | Enterobacteri | Kosakonia | Kosakonia ory | unclassified | unclassified | unclassified | unclassified | unclassified | 0,4 | 0,489897949 | 0,000161246 | 0,000197485 | 0 - | 0 - | 0 - | 0 - | 1 - | 0,000402855 - | 0 - | 0 - | 1 - | 0,000403374 - |
| 1609 | 1 | 0.2um_NOKCPHJN_54732 | Eukaryota | Streptophyta | unclassified | Brassicales | Brassicaceae | Arabidopsis | various | metabolism | Energy produc | COG0055 | Produces ATP | 39947.LOC_Os | 0,2 | 0,4 | 3,52E-05 | 7,04E-05 | 1 - | 0,000176092 - | 0 - | 0 - | 0 - | 0 - | 0 - | 0 - | 0 - | 0 - |
| 1610 | 1 | 0.2um_NOKCPHJN_604534 | Bacteria | Proteobacteri | Alphaproteob | Rhodobactera | unclassified | unclassified | Rhodobactera | poorly charact | Function unkn | 175CM | beta-lactamas | 991905.SL003 | 1 | 0 | 0,000242786 | 2,38E-05 | 1 - | 0,000244955 - | 1 - | 0,000288184 - | 1 - | 0,000227309 - | 1 - | 0,000225881 - | 1 - | 0,000227602 - |
| 1611 | 1 | 0.2um_NOKCPHJN_1002686 | unclassified | unclassified | unclassified | unclassified | unclassified | unclassified | unclassified | unclassified | unclassified | unclassified | unclassified | unclassified | 1 | 0 | 0,001010668 | 9,89E-05 | 1 - | 0,001019696 - | 1 - | 0,00119965 - | 1 - | 0,000946241 - | 1 - | 0,000940295 - | 1 - | 0,000947459 - |
| 1612 | 1 | 0.2um_NOKCPHJN_870446 | Bacteria | Proteobacteri | Gammaprotec | unclassified | unclassified | unclassified | Gammaprotec | poorly charact | Function unkn | 16R8I | Lipoprotein | 314287.GB22C | 0,2 | 0,4 | 3,37E-05 | 6,75E-05 | 0 - | 0 - | 0 - | 0 - | 0 - | 0 - | 0 - | 0 - | 1 - | 0,000168699 - |
| 1613 | 1 | 0.2um_NOKCPHJN_356143 | Bacteria | Bacteroidetes | Flavobacteriia | Flavobacterial | Cryomorphac | unclassified | Cryomorphac | metabolism | Inorganic ion t | 05C27 | TonB-depende | 313596.RB250 | 0,4 | 0,489897949 | 3,28E-05 | 4,02E-05 | 1 - | 8,51E-05 - | 0 - | 0 - | 0 - | 0 - | 0 - | 0 - | 1 - | 7,90E-05 - |
| 1614 | 1 | 0.2um_NOKCPHJN_11291 | Bacteria | Proteobacteri | Alphaproteob | unclassified | unclassified | unclassified | SAR116 cluste | cellular proces | Defense mech | 01QSH | ABC transport | 488538.SAR11 | 0,2 | 0,4 | 2,74E-05 | 5,48E-05 | 0 - | 0 - | 0 - | 0 - | 0 - | 0 - | 1 - | 0,00013706 - | 0 - | 0 - |
| 1615 | 1 | 3um_FIJJAFBF_407842 | Bacteria | Proteobacteri | Alphaproteob | Rhodobactera | Rhodobactera | Roseivivax | Roseivivax len | unclassified | unclassified | unclassified | unclassified | unclassified | 0,2 | 0,4 | 4,16E-05 | 8,31E-05 | 1 - | 0,000207805 - | 0 - | 0 - | 0 - | 0 - | 0 - | 0 - | 0 - | 0 - |
| 1616 | 1 | 0.2um_NOKCPHJN_789774 | unclassified | unclassified | unclassified | unclassified | unclassified | unclassified | unclassified | unclassified | unclassified | unclassified | unclassified | unclassified | 1 | 0 | 0,000188133 | 1,84E-05 | 1 - | 0,000189814 - | 1 - | 0,000223311 - | 1 - | 0,00017614 - | 1 - | 0,000175033 - | 1 - | 0,000176367 - |
| 1617 | 1 | 3um_FIJJAFBF_180780 | Bacteria | Verrucomicrob | Opitutae | Opitutales | unclassified | unclassified | Opitutales ba | unclassified | unclassified | unclassified | unclassified | unclassified | 1 | 0 | 0,000658466 | 6,44E-05 | 1 - | 0,000664347 - | 1 - | 0,00078159 - | 1 - | 0,00061649 - | 1 - | 0,000612616 - | 1 - | 0,000617284 - |
| 1618 | 1 | 0.2um_NOKCPHJN_233772 | Bacteria | Proteobacteri | Alphaproteob | Rhodobactera | Rhodobactera | Amylibacter | Amylibacter c | unclassified | unclassified | unclassified | unclassified | unclassified | 0,6 | 0,489897949 | 0,000168667 | 0,000137718 | 0 - | 0 - | 0 - | 0 - | 1 - | 0,00028158 - | 1 - | 0,000279811 - | 1 - | 0,000281943 - |
| 1619 | 1 | 3um_FIJJAFBF_84528 | Bacteria | Bacteroidetes | Flavobacteriia | Flavobacterial | Flavobacteriac | Polaribacter | Polaribacter r | cellular proces | Posttranslatio | COG0330 | Band 7 protei | 391603.FBALC | 1,6 | 0,489897949 | 0,000447804 | 0,000123203 | 2 - | 0,000573162 - | 1 - | 0,000337156 - | 2 - | 0,000531874 - | 1 - | 0,000264266 - | 2 - | 0,000532559 - |
| 1620 | 1 | 0.2um_NOKCPHJN_33920 | Bacteria | Proteobacteri | Gammaprotec | Alteromonada | Alteromonada | Paraglaciecola | Paraglaciecola | metabolism | Inorganic ion t | 0QK8T | TonB-depende | 983545.Glaag | 0,8 | 0,4 | 7,50E-05 | 3,76E-05 | 1 - | 9,92E-05 - | 0 - | 0 - | 1 - | 9,21E-05 - | 1 - | 9,15E-05 - | 1 - | 9,22E-05 - |
| 1621 | 1 | 0.2um_NOKCPHJN_420122 | Eukaryota | Apicomplexa | Conoidasida | Eucoccidiorida | Cryptosporidii | Cryptosporidii | Cryptosporidii | unclassified | unclassified | unclassified | unclassified | unclassified | 0,6 | 0,489897949 | 0,000133272 | 0,000108986 | 1 - | 0,000233228 - | 0 - | 0 - | 1 - | 0,000216428 - | 0 - | 0 - | 1 - | 0,000216706 - |
| 1622 | 1 | 0.2um_NOKCPHJN_106487 | Bacteria | Bacteroidetes | Flavobacteriia | Flavobacterial | Flavobacteriac | Aquimarina | Aquimarina m | information st | Translation, ri | 05CEJ | Catalyzes the | 50743.SCB49_ | 1,6 | 1,019803903 | 0,000187067 | 0,000118231 | 2 - | 0,000247723 - | 0 - | 0 - | 2 - | 0,000229878 - | 3 - | 0,00034265 - | 1 - | 0,000115087 - |
| 1623 | 1 | 3um_FIJJAFBF_698996 | Bacteria | Proteobacteri | Alphaproteob | Rhodobactera | Rhodobactera | Tateyamaria | Tateyamaria s | metabolism | Amino acid tra | COG1932 | Catalyzes the | 391589.RGAI1 | 0,8 | 0,4 | 0,00017489 | 8,77E-05 | 1 - | 0,000231382 - | 0 - | 0 - | 1 - | 0,000214714 - | 1 - | 0,000213365 - | 1 - | 0,000214991 - |
| 1624 | 1 | 0.2um_NOKCPHJN_242718 | Bacteria | Chloroflexi | Chloroflexia | Herpetosiphoi | Herpetosiphoi | Herpetosiphoi | Herpetosipho | cellular proces | Cytoskeleton | COG5245 | heavy chain | 7668.SPU_020 | 0,2 | 0,4 | 1,91E-05 | 3,82E-05 | 0 - | 0 - | 0 - | 0 - | 0 - | 0 - | 0 - | 0 - | 1 - | 9,54E-05 - |
| 1625 | 1 | 3um_FIJJAFBF_362249 | Viruses | unclassified | unclassified | unclassified | Phycodnavirid | unclassified | Chrysochromu | unclassified | unclassified | unclassified | unclassified | unclassified | 1,2 | 0,4 | 0,000338337 | 0,000101137 | 1 - | 0,000287521 - | 1 - | 0,000338262 - | 2 - | 0,000533618 - | 1 - | 0,000265132 - | 1 - | 0,000267152 - |
| 1626 | 1 | 3um_FIJJAFBF_735065 | Bacteria | Actinobacteria | Actinobacteria | Micrococcales | Microbacteria | unclassified | Microbacteria | unclassified | unclassified | unclassified | unclassified | unclassified | 0,8 | 0,4 | 0,000188018 | 9,68E-05 | 0 - | 0 - | 1 - | 0,000279593 - | 1 - | 0,000220533 - | 1 - | 0,000219147 - | 1 - | 0,000220817 - |
| 1627 | 1 | 0.2um_NOKCPHJN_192042 | Bacteria | Bacteroidetes | Flavobacteriia | Flavobacterial | Flavobacteriac | Lutibacter | Lutibacter sp. | metabolism | Carbohydrate | 05D2N | Aldolase | 391587.KAOT1 | 0,4 | 0,489897949 | 0,000104029 | 0,000128865 | 0 - | 0 - | 1 - | 0,000290619 - | 0 - | 0 - | 0 - | 0 - | 1 - | 0,000229525 - |
| 1628 | 2 | 0.2um_NOKCPHJN_985727;3ur | unclassified | unclassified | unclassified | unclassified | unclassified | unclassified | unclassified | unclassified | unclassified | unclassified | unclassified | unclassified | 0,6 | 0,8 | 0,000511545 | 0,000671943 | 1 - | 0,000894835 - | 0 - | 0 - | 0 - | 0 - | 0 - | 0 - | 2 - | 0,001662888 - |
| 1629 | 1 | 0.2um_NOKCPHJN_198494 | Bacteria | Bacteroidetes | Flavobacteriia | Flavobacterial | Flavobacteriac | Algibacter | Algibacter sp. | cellular proces | Intracellular tr | 05FGH | MotA TolQ exl | 391603.FBALC | 2,4 | 0,489897949 | 0,000809438 | 0,000133887 | 2 - | 0,000687795 - | 2 - | 0,000809176 - | 3 - | 0,000957373 - | 2 - | 0,000634238 - | 3 - | 0,000958606 - |
| 1630 | 1 | 3um_FIJJAFBF_667394 | Bacteria | Proteobacteri | Alphaproteob | Rhodobactera | Rhodobactera | Sulfitobacter | Sulfitobacter g | unclassified | unclassified | unclassified | unclassified | unclassified | 1,8 | 0,4 | 0,000545116 | 0,000142916 | 2 - | 0,000606878 - | 2 - | 0,000713978 - | 2 - | 0,000563161 - | 2 - | 0,000559622 - | 1 - | 0,000281943 - |
| 1631 | 1 | 0.2um_NOKCPHJN_172999 | Bacteria | Bacteroidetes | unclassified | unclassified | unclassified | unclassified | Bacteroidetes | poorly charact | Function unkn | 05DES | (LipO)protein | 860228.Ccan_ | 0,2 | 0,4 | 2,96E-05 | 5,92E-05 | 0 - | 0 - | 0 - | 0 - | 0 - | 0 - | 0 - | 0 - | 1 - | 0,000147879 - |
| 1632 | 1 | 0.2um_NOKCPHJN_489599 | Eukaryota | unclassified | Pelagophycea | Pelagomonad | unclassified | Aureococcus | Aureococcus a | unclassified | unclassified | unclassified | unclassified | unclassified | 2 | 0 | 0,001151225 | 0,000112641 | 2 - | 0,001161508 - | 2 - | 0,001366489 - | 2 - | 0,001077838 - | 2 - | 0,001071064 - | 2 - | 0,001079225 - |
| 1633 | 1 | 3um_FIJJAFBF_750093 | Eukaryota | Bacillariophyta | Bacillariophyc | Naviculales | Phaeodactylac | Phaeodactylu | Phaeodactylu | unclassified | unclassified | unclassified | unclassified | unclassified | 1 | 0 | 0,000514305 | 5,03E-05 | 1 - | 0,000518899 - | 1 - | 0,000610473 - | 1 - | 0,000481519 - | 1 - | 0,000478493 - | 1 - | 0,000482139 - |
| 1634 | 1 | 3um_FIJJAFBF_755414 | Bacteria | Bacteroidetes | Flavobacteriia | Flavobacterial | Flavobacteriac | Psychroflexus | Psychroflexus | poorly charact | Function unkn | 05X9M | NA | 313595.P7007 | 0,2 | 0,4 | 2,86E-05 | 5,72E-05 | 0 - | 0 - | 0 - | 0 - | 0 - | 0 - | 0 - | 0 - | 1 - | 0,00014295 - |
| 1635 | 1 | 3um_FIJJAFBF_111336 | Viruses | unclassified | unclassified | unclassified | Phycodnavirid | unclassified | Organic Lake p | unclassified | unclassified | unclassified | unclassified | unclassified | 1,2 | 0,4 | 0,001272723 | 0,000375039 | 1 - | 0,00108264 - | 1 - | 0,001273702 - | 1 - | 0,001004651 - | 2 - | 0,001996675 - | 1 - | 0,001005944 - |
| 1636 | 1 | 3um_FIJJAFBF_454585 | Viruses | unclassified | unclassified | Caudovirales | Siphoviridae | unclassified | Tenacibaculur | unclassified | unclassified | unclassified | unclassified | unclassified | 0,2 | 0,4 | 3,55E-05 | 7,09E-05 | 0 - | 0 - | 1 - | 0,000177268 - | 0 - | 0 - | 0 - | 0 - | 0 - | 0 - |
| 1637 | 1 | 3um_FIJJAFBF_405813 | unclassified | unclassified | unclassified | unclassified | unclassified | unclassified | unclassified | unclassified | unclassified | unclassified | unclassified | unclassified | 1,6 | 0,8 | 0,001077496 | 0,000487563 | 1 - | 0,000695983 - | 1 - | 0,000818809 - | 3 - | 0,001937541 - | 1 - | 0,000641789 - | 2 - | 0,001293357 - |
| 1638 | 1 | 3um_FIJJAFBF_140885 | Bacteria | Proteobacteri | Gammaprotec | unclassified | Competibacte | Plasticicumula | Plasticicumula | unclassified | unclassified | unclassified | unclassified | unclassified | 2,8 | 0,748331477 | 0,000821508 | 0,000302593 | 3 - | 0,000868256 - | 4 - | 0,001361979 - | 3 - | 0,00080571 - | 2 - | 0,000533765 - | 2 - | 0,000537832 - |
| 1639 | 1 | 3um_FIJJAFBF_161768 | Bacteria | Bacteroidetes | Flavobacteriia | Flavobacterial | unclassified | unclassified | Flavobacterial | unclassified | unclassified | unclassified | unclassified | unclassified | 0,2 | 0,4 | 2,56E-05 | 5,13E-05 | 0 - | 0 - | 0 - | 0 - | 0 - | 0 - | 1 - | 0,000128154 - | 0 - | 0 - |

| | | | | | | | | | | | | | | | | | | | | | | | | | | | | | | | | | |
|---|---|---|---|---|---|---|---|---|---|---|---|---|---|---|---|---|---|---|---|---|---|---|---|---|---|---|---|---|---|---|---|---|---|
| 1640 | 1 | 0.2um_NOKCPHJN_88818 | Bacteria | Bacteroidetes | Flavobacteriia | Flavobacterial | Flavobacteriac | Ulvibacter | Ulvibacter lito | information st | Replication, re | COG0178 | The UvrABC re | 50743.SCB49_ | 0,2 | 0,4 | 1,86E-05 | 3,71E-05 | 1 | 9,28E-05 | - | 0 | 0 | - | 0 | 0 | - | 0 | 0 | - | 0 | 0 | - |
| 1641 | 1 | 0.2um_NOKCPHJN_947787 | Eukaryota | unclassified | Oomycetes | Peronosporale | Peronosporac | Phytophthora | Phytophthora | unclassified | unclassified | unclassified | unclassified | unclassified | 0,6 | 0,489897949 | 0,000118669 | 9,81E-05 | 1 | 0,000191054 | - | 1 | 0,000224771 | - | 0 | 0 | - | 0 | 0 | - | 1 | 0,00017752 | - |
| 1642 | 1 | 0.2um_NOKCPHJN_490937 | Bacteria | Proteobacteri | Gammaprotec | unclassified | unclassified | Candidatus Th | Candidatus Th | metabolism | Energy produc | COG0055 | Produces ATP | 412965.COSY_ | 0,4 | 0,489897949 | 8,05E-05 | 9,97E-05 | 0 | 0 | - | 1 | 0,000224771 | - | 0 | 0 | - | 0 | 0 | - | 1 | 0,00017752 | - |
| 1643 | 2 | 3um_FIJJAFBF_582844;3um_FI | Bacteria | Bacteroidetes | Flavobacteriia | Flavobacterial | Flavobacteriac | various | various | unclassified | unclassified | unclassified | unclassified | unclassified | 2,8 | 0,748331477 | 0,000822013 | 0,000226168 | 2 | 0,000592526 | - | 3 | 0,001045641 | - | 4 | 0,001099686 | - | 2 | 0,000546388 | - | 3 | 0,000825826 | - |
| 1644 | 1 | 0.2um_NOKCPHJN_1115842 | Bacteria | Bacteroidetes | Flavobacteriia | Flavobacterial | Flavobacteriac | Flavobacteriu | Flavobacteriu | cellular proces | Cell wall/mem | 09ARE | ompa motb d | 376686.Fjoh_ | 1 | 0,632455532 | 0,000218087 | 0,00015751 | 1 | 0,000209293 | - | 2 | 0,000492458 | - | 1 | 0,000194217 | - | 0 | 0 | - | 1 | 0,000194467 | - |
| 1645 | 1 | 0.2um_NOKCPHJN_39231 | Eukaryota | Bacillariophyt | Coscinodiscop | Rhizosoleniale | Rhizosoleniac | Rhizosolenia | Rhizosolenia s | information st | Translation, ri | COG0050 | This protein p | 91464.S7335_ | 5 | 0,894427191 | 0,001048616 | 0,000127603 | 4 | 0,000857642 | - | 4 | 0,001008996 | - | 5 | 0,000994826 | - | 6 | 0,001186289 | - | 6 | 0,001195328 | - |
| 1646 | 1 | 3um_FIJJAFBF_579438 | Eukaryota | Bacillariophyt | Bacillariophyc | Bacillariales | Bacillariaceae | Fragilariopsis | Fragilariopsis | cellular proces | Cell cycle cont | 0GTVD | BRCA1 C Term | 7260.FBpp024 | 0,8 | 0,4 | 8,85E-05 | 4,56E-05 | 0 | 0 | - | 1 | 0,000131594 | - | 1 | 0,000103797 | - | 1 | 0,000103145 | - | 1 | 0,00010393 | - |
| 1647 | 1 | 3um_FIJJAFBF_87388 | Bacteria | Bacteroidetes | Flavobacteriia | Flavobacterial | Flavobacteriac | unclassified | Flavobacteriac | poorly charact | Function unkn | 0M0RX | Carboxymuco | 309800.HVO_ | 0,8 | 0,4 | 0,00047686 | 0,000239111 | 1 | 0,000630891 | - | 0 | 0 | - | 1 | 0,000585444 | - | 1 | 0,000581765 | - | 1 | 0,000586198 | - |
| 1648 | 1 | 0.2um_NOKCPHJN_547183 | Eukaryota | Chlorophyta | Chlorophycea | Sphaeropleale | Selenastracea | Monoraphidiu | Monoraphidiu | unclassified | unclassified | unclassified | unclassified | unclassified | 0,4 | 0,489897949 | 0,000225336 | 0,000275982 | 0 | 0 | - | 0 | 0 | - | 1 | 0,000565116 | - | 1 | 0,000561565 | - | 0 | 0 | - |
| 1649 | 1 | 0.2um_NOKCPHJN_212074 | Eukaryota | unclassified | Spirotrichea | Sporadotrichi | Oxytrichidae | Stylonychia | Stylonychia le | poorly charact | Function unkn | 0JPIP | Inherit from N | 5722.A2EVM7 | 0,2 | 0,4 | 3,46E-05 | 6,91E-05 | 0 | 0 | - | 0 | 0 | - | 1 | 0,000172774 | - | 0 | 0 | - | 0 | 0 | - |
| 1650 | 1 | 0.2um_NOKCPHJN_751237 | Eukaryota | unclassified | unclassified | Prymnesiales | Chrysochromu | Chrysochromu | Chrysochromu | unclassified | unclassified | unclassified | unclassified | unclassified | 0,4 | 0,489897949 | 0,00039661 | 0,00048641 | 1 | 0,001031692 | - | 0 | 0 | - | 0 | 0 | - | 1 | 0,000951357 | - | 0 | 0 | - |
| 1651 | 1 | 3um_FIJJAFBF_176964 | Eukaryota | unclassified | Cryptophyta | Pyrenomonad | Pyrenomonad | Storeatula | Storeatula sp. | unclassified | unclassified | unclassified | unclassified | unclassified | 0,2 | 0,4 | 2,57E-05 | 5,14E-05 | 0 | 0 | - | 0 | 0 | - | 0 | 0 | - | 1 | 0,000128562 | - | 0 | 0 | - |
| 1652 | 2 | 0.2um_NOKCPHJN_339415;0.2 | Eukaryota | various | various | various | various | various | various | unclassified | unclassified | unclassified | unclassified | unclassified | 0,4 | 0,489897949 | 0,000202308 | 0,000247776 | 0 | 0 | - | 0 | 0 | - | 1 | 0,000505446 | - | 0 | 0 | - | 1 | 0,000506096 | - |
| 1653 | 1 | 3um_FIJJAFBF_08025 | Bacteria | Bacteroidetes | Flavobacteriia | Flavobacterial | Flavobacteriac | Psychroserper | Psychroserper | cellular proces | Cell wall/mem | COG0438 | Glycosyl trans | 31964.CMS_0 | 0,4 | 0,489897949 | 9,51E-05 | 0,000117853 | 0 | 0 | - | 1 | 0,000265902 | - | 1 | 0,000209734 | - | 0 | 0 | - | 0 | 0 | - |
| 1654 | 1 | 0.2um_NOKCPHJN_07567 | Bacteria | unclassified | unclassified | unclassified | unclassified | unclassified | uncultured ba | unclassified | unclassified | unclassified | unclassified | unclassified | 0,6 | 0,489897949 | 6,23E-05 | 5,08E-05 | 0 | 0 | - | 0 | 0 | - | 1 | 0,000103929 | - | 1 | 0,000103276 | - | 1 | 0,000104063 | - |
| 1655 | 1 | 3um_FIJJAFBF_774536 | Eukaryota | unclassified | Cryptophyta | Pyrenomonad | Geminigerace | Guillardia | Guillardia thet | unclassified | unclassified | unclassified | unclassified | unclassified | 3,4 | 1,356465997 | 0,001607488 | 0,000574191 | 3 | 0,001486336 | - | 1 | 0,000582881 | - | 4 | 0,001839022 | - | 4 | 0,001827466 | - | 5 | 0,002301737 | - |
| 1656 | 1 | 0.2um_NOKCPHJN_284613 | Eukaryota | Phaeophyceae | unclassified | Ectocarpales | Ectocarpaceae | Ectocarpus | Ectocarpus sili | metabolism | Energy produc | arCOG00097 | formylmethan | 634498.mru_ | 0,4 | 0,489897949 | 1,55E-05 | 1,89E-05 | 0 | 0 | - | 0 | 0 | - | 1 | 3,86E-05 | - | 0 | 0 | - | 1 | 3,87E-05 | - |
| 1657 | 1 | 3um_FIJJAFBF_493815 | Eukaryota | Bacillariophyt | Coscinodiscop | Thalassiosirale | Thalassiosirac | Thalassiosira | Thalassiosira p | poorly charact | Function unkn | 0XS5J | prolyl 4-hydro | 35128.Thaps2 | 0,2 | 0,4 | 3,49E-05 | 6,99E-05 | 1 | 0,000174689 | - | 0 | 0 | - | 0 | 0 | - | 0 | 0 | - | 0 | 0 | - |
| 1658 | 1 | 3um_FIJJAFBF_507802 | Bacteria | Proteobacteri | Gammaprotec | Chromatiales | Ectothiorhodc | Ectothiorhodc | Ectothiorhodo | unclassified | unclassified | unclassified | unclassified | unclassified | 1 | 0,632455532 | 0,000763855 | 0,00046337 | 0 | 0 | - | 1 | 0,00092116 | - | 1 | 0,000726578 | - | 2 | 0,001444024 | - | 1 | 0,000727513 | - |
| 1659 | 1 | 3um_FIJJAFBF_715063 | Eukaryota | unclassified | Raphidophyce | Chattonellales | Chattonellace | Heterosigma | Heterosigma a | cellular proces | Posttranslatio | 0IEKR | One of the co | 35128.Thapsd | 5 | 0,894427191 | 0,000936382 | 0,000113541 | 4 | 0,000765885 | - | 4 | 0,000901047 | - | 6 | 0,001066071 | - | 6 | 0,001059371 | - | 5 | 0,000889536 | - |
| 1660 | 1 | 0.2um_NOKCPHJN_673538 | Bacteria | Bacteroidetes | Flavobacteriia | Flavobacterial | Flavobacteriac | Muricauda | Muricauda sp. | unclassified | unclassified | unclassified | unclassified | unclassified | 1 | 0 | 8,15E-05 | 7,97E-06 | 1 | 8,22E-05 | - | 1 | 9,67E-05 | - | 1 | 7,63E-05 | - | 1 | 7,58E-05 | - | 1 | 7,64E-05 | - |
| 1661 | 1 | 0.2um_NOKCPHJN_84840 | Bacteria | Verrucomicrol | unclassified | unclassified | unclassified | unclassified | Verrucomicrol | cellular proces | Posttranslatio | 1DME7 | Molecular cha | 382464.VDG1 | 0,2 | 0,4 | 2,63E-05 | 5,27E-05 | 0 | 0 | - | 0 | 0 | - | 1 | 0,000131678 | - | 0 | 0 | - | 0 | 0 | - |
| 1662 | 1 | 0.2um_NOKCPHJN_269537 | Bacteria | Bacteroidetes | Flavobacteriia | Flavobacterial | Flavobacteriac | unclassified | Flavobacteriac | information st | Replication, re | 0PDAJ | Inherit from b | 1041826.FCOL | 0,2 | 0,4 | 8,05E-05 | 0,000160906 | 1 | 0,000402265 | - | 0 | 0 | - | 0 | 0 | - | 0 | 0 | - | 0 | 0 | - |
| 1663 | 1 | 3um_FIJJAFBF_477845 | Eukaryota | unclassified | Bangiophycea | Porphyridiales | Porphyridiace | Flintiella | Flintiella sang | information st | Translation, ri | COG0051 | tRNA binding | 321332.CYB_0 | 0,2 | 0,4 | 0,000162753 | 0,000325507 | 0 | 0 | - | 0 | 0 | - | 1 | 0,000813767 | - | 0 | 0 | - | 0 | 0 | - |
| 1664 | 1 | 3um_FIJJAFBF_247264 | unclassified | unclassified | unclassified | unclassified | unclassified | unclassified | unclassified | unclassified | unclassified | unclassified | unclassified | unclassified | 0,4 | 0,489897949 | 0,000408967 | 0,000506974 | 0 | 0 | - | 1 | 0,001146332 | - | 0 | 0 | - | 1 | 0,000898504 | - | 0 | 0 | - |
| 1665 | 1 | 0.2um_NOKCPHJN_221762 | Bacteria | Proteobacteri | Gammaprotec | unclassified | unclassified | unclassified | Gammaprotec | unclassified | unclassified | unclassified | unclassified | unclassified | 1,6 | 0,489897949 | 0,00033795 | 0,000121886 | 2 | 0,000419588 | - | 2 | 0,000493636 | - | 1 | 0,000194681 | - | 2 | 0,000386916 | - | 1 | 0,000194932 | - |
| 1666 | 1 | 0.2um_NOKCPHJN_263318 | Eukaryota | Phaeophyceae | unclassified | Ectocarpales | Ectocarpaceae | Ectocarpus | Ectocarpus sili | unclassified | unclassified | unclassified | unclassified | unclassified | 2 | 0,632455532 | 0,000261871 | 7,91E-05 | 3 | 0,000403499 | - | 1 | 0,000158236 | - | 2 | 0,000249622 | - | 2 | 0,000248053 | - | 2 | 0,000249943 | - |
| 1667 | 1 | 3um_FIJJAFBF_573898 | Bacteria | Verrucomicrol | Opitutae | unclassified | unclassified | unclassified | Opitutae bact | unclassified | unclassified | unclassified | unclassified | unclassified | 0,4 | 0,489897949 | 7,13E-05 | 8,74E-05 | 1 | 0,000185399 | - | 0 | 0 | - | 0 | 0 | - | 1 | 0,000170963 | - | 0 | 0 | - |
| 1668 | 1 | 0.2um_NOKCPHJN_329634 | Eukaryota | Chordata | Actinopteri | Cyprinodontifi | Poeciliidae | Poecilia | Poecilia latipir | unclassified | unclassified | unclassified | unclassified | unclassified | 0,8 | 0,4 | 0,000219482 | 0,000110055 | 1 | 0,000290377 | - | 0 | 0 | - | 1 | 0,000269459 | - | 1 | 0,000267766 | - | 1 | 0,000269806 | - |
| 1669 | 1 | 0.2um_NOKCPHJN_1064122 | Eukaryota | Chordata | Mammalia | Rodentia | Muridae | Rattus | Rattus norveg | unclassified | unclassified | unclassified | unclassified | unclassified | 0,6 | 0,489897949 | 0,000181169 | 0,000150455 | 0 | 0 | - | 1 | 0,000352116 | - | 1 | 0,000277736 | - | 1 | 0,000275991 | - | 0 | 0 | - |
| 1670 | 1 | 0.2um_NOKCPHJN_537008 | Bacteria | Bacteroidetes | Flavobacteriia | Flavobacterial | Flavobacteriac | Formosa | Formosa sp. H | unclassified | unclassified | unclassified | unclassified | unclassified | 0,6 | 0,489897949 | 0,000168049 | 0,000139547 | 0 | 0 | - | 1 | 0,000326487 | - | 0 | 0 | - | 1 | 0,000255903 | - | 1 | 0,000257853 | - |
| 1671 | 1 | 3um_FIJJAFBF_24379 | Bacteria | Bacteroidetes | Flavobacteriia | Flavobacterial | Flavobacteriac | Algibacter | Algibacter sp. | unclassified | unclassified | unclassified | unclassified | unclassified | 6,6 | 1,624807681 | 0,001088859 | 0,000212404 | 7 | 0,001187344 | - | 4 | 0,00079822 | - | 7 | 0,001101813 | - | 6 | 0,000938476 | - | 9 | 0,00141844 | - |
| 1672 | 1 | 3um_FIJJAFBF_128464 | Bacteria | Bacteroidetes | Flavobacteriia | Flavobacterial | Flavobacteriac | Polaribacter | Polaribacter s | information st | Translation, ri | COG0052 | 30S ribosomal | 487797.Flav3 | 0,2 | 0,4 | 5,24E-05 | 0,000104709 | 1 | 0,000261773 | - | 0 | 0 | - | 0 | 0 | - | 0 | 0 | - | 0 | 0 | - |
| 1673 | 1 | 0.2um_NOKCPHJN_373383 | Bacteria | Proteobacteri | Alphaproteob | Pelagibacteral | Pelagibacterac | Candidatus Pe | Candidatus Pe | unclassified | unclassified | unclassified | unclassified | unclassified | 0,2 | 0,4 | 1,99E-05 | 3,98E-05 | 0 | 0 | - | 1 | 9,94E-05 | - | 0 | 0 | - | 0 | 0 | - | 0 | 0 | - |
| 1674 | 1 | 3um_FIJJAFBF_45671 | Bacteria | Proteobacteri | Alphaproteob | Sphingomona | Sphingomona | Sphingobium | Sphingobium | information st | Translation, ri | 05C6A | phenylalanyl-t | 627192.SLG_3 | 0,2 | 0,4 | 3,12E-05 | 6,24E-05 | 1 | 0,000156039 | - | 0 | 0 | - | 0 | 0 | - | 0 | 0 | - | 0 | 0 | - |
| 1675 | 1 | 0.2um_NOKCPHJN_133072 | Bacteria | Bacteroidetes | Flavobacteriia | Flavobacterial | Crocinitomica | unclassified | Crocinitomica | unclassified | unclassified | unclassified | unclassified | unclassified | 1 | 0 | 0,000276807 | 2,71E-05 | 1 | 0,00027928 | - | 1 | 0,000328567 | - | 1 | 0,000259162 | - | 1 | 0,000257533 | - | 1 | 0,000259495 | - |
| 1676 | 1 | 3um_FIJJAFBF_65170 | Bacteria | Bacteroidetes | Flavobacteriia | Flavobacterial | Flavobacteriac | Hyunsoonleell | Hyunsoonleell | unclassified | unclassified | unclassified | unclassified | unclassified | 0,6 | 0,489897949 | 7,29E-05 | 5,96E-05 | 1 | 0,000127834 | - | 0 | 0 | - | 0 | 0 | - | 1 | 0,00011788 | - | 1 | 0,000118778 | - |
| 1677 | 1 | 3um_FIJJAFBF_53313 | Bacteria | Bacteroidetes | Flavobacteriia | Flavobacterial | Flavobacteriac | Lutibacter | Lutibacter sp. | unclassified | unclassified | unclassified | unclassified | unclassified | 4,4 | 1,496662955 | 0,000374181 | 0,000115708 | 3 | 0,000259962 | - | 4 | 0,000407786 | - | 3 | 0,000241235 | - | 7 | 0,000559345 | - | 5 | 0,000402577 | - |
| 1678 | 1 | 3um_FIJJAFBF_282388 | Eukaryota | Bacillariophyt | Bacillariophyc | Naviculales | Phaeodactylac | Phaeodactylu | Phaeodactylu | unclassified | unclassified | unclassified | unclassified | unclassified | 2 | 0,894427191 | 0,001411625 | 0,000567153 | 2 | 0,001461564 | - | 1 | 0,000859749 | - | 1 | 0,00067814 | - | 3 | 0,002021634 | - | 3 | 0,002037037 | - |
| 1679 | 1 | 0.2um_NOKCPHJN_1072722 | unclassified | unclassified | unclassified | unclassified | unclassified | unclassified | unclassified | unclassified | unclassified | unclassified | unclassified | unclassified | 1,6 | 0,489897949 | 0,001818452 | 0,000624812 | 1 | 0,001138881 | - | 2 | 0,002679737 | - | 1 | 0,001056841 | - | 2 | 0,002100399 | - | 2 | 0,002116402 | - |
| 1680 | 1 | 0.2um_NOKCPHJN_923670 | Bacteria | Proteobacteri | Alphaproteob | unclassified | unclassified | unclassified | Alphaproteob | poorly charact | Function unkn | 0HE85 | Hydrolase Coc | 690850.Desaf | 0,2 | 0,4 | 2,66E-05 | 5,31E-05 | 0 | 0 | - | 0 | 0 | - | 0 | 0 | - | 1 | 0,000132784 | - | 0 | 0 | - |
| 1681 | 1 | 0.2um_NOKCPHJN_494079 | Bacteria | Cyanobacteria | unclassified | Oscillatoriales | Cyanothecace | Cyanothece | various | cellular proces | Intracellular tr | 0DNW5 | translocated p | 13616.ENSMO | 0,2 | 0,4 | 4,90E-05 | 9,80E-05 | 0 | 0 | - | 1 | 0,000245059 | - | 0 | 0 | - | 0 | 0 | - | 0 | 0 | - |
| 1682 | 1 | 0.2um_NOKCPHJN_136728 | Bacteria | Bacteroidetes | Flavobacteriia | unclassified | unclassified | unclassified | Flavobacteria | metabolism | Inorganic ion t | 05C27 | TonB-depende | 487796.Flav2A | 0,6 | 1,2 | 4,92E-05 | 9,83E-05 | 3 | 0,000245871 | - | 0 | 0 | - | 0 | 0 | - | 0 | 0 | - | 0 | 0 | - |
| 1683 | 1 | 3um_FIJJAFBF_457406 | Eukaryota | Bacillariophyt | Coscinodiscop | Thalassiosirale | Thalassiosirac | Thalassiosira | Thalassiosira c | unclassified | unclassified | unclassified | unclassified | unclassified | 0,6 | 0,489897949 | 0,000276172 | 0,000225885 | 1 | 0,000484496 | - | 0 | 0 | - | 1 | 0,000449595 | - | 1 | 0,00044677 | - | 0 | 0 | - |
| 1684 | 1 | 3um_FIJJAFBF_37634 | Bacteria | Proteobacteri | Gammaprotec | Oceanospirilla | Oceanospirilla | Motiliproteus | Motiliproteus | information st | Replication, re | 0QKMF | The RuvA-Ruv | 223926.VP105 | 0,4 | 0,489897949 | 0,000188043 | 0,000231563 | 1 | 0,000431989 | - | 1 | 0,000508226 | - | 0 | 0 | - | 0 | 0 | - | 0 | 0 | - |
| 1685 | 1 | 3um_FIJJAFBF_552358 | Viruses | unclassified | unclassified | unclassified | unclassified | unclassified | uncultured vir | unclassified | unclassified | unclassified | unclassified | unclassified | 7 | 0,632455532 | 0,001976713 | 0,000366886 | 7 | 0,001980184 | - | 8 | 0,002662449 | - | 6 | 0,001575034 | - | 7 | 0,001825992 | - | 7 | 0,001839905 | - |
| 1686 | 1 | 0.2um_NOKCPHJN_332543 | Eukaryota | Bacillariophyt | Coscinodiscop | Thalassiosirale | Thalassiosirac | Thalassiosira | Thalassiosira c | cellular proces | Posttranslatio | KOG0019 | heat shock pro | 35128.Thaps6 | 2 | 0 | 0,00024553 | 2,40E-05 | 2 | 0,000247723 | - | 2 | 0,00029144 | - | 2 | 0,000229878 | - | 2 | 0,000228433 | - | 2 | 0,000230174 | - |
| 1687 | 3 | 3um_FIJJAFBF_237980;3um_FI | Eukaryota | various | various | various | various | various | various | unclassified | unclassified | unclassified | unclassified | unclassified | 1 | 0 | 0,002172937 | 0,000212609 | 1 | 0,002192346 | - | 1 | 0,002579247 | - | 1 | 0,002034419 | - | 1 | 0,002021634 | - | 1 | 0,002037037 | - |
| 1688 | 1 | 0.2um_NOKCPHJN_31647 | Bacteria | Proteobacteri | Betaproteoba | unclassified | unclassified | unclassified | Betaproteoba | cellular proces | Cell wall/mem | COG2885 | Ompa motb d | 365044.Pnap_ | 2,4 | 0,489897949 | 0,001103373 | 0,000303527 | 2 | 0,00091826 | - | 3 | 0,001620469 | - | 3 | 0,001278169 | - | 2 | 0,000846758 | - | 2 | 0,000853209 | - |
| 1689 | 1 | 0.2um_NOKCPHJN_42273 | Bacteria | Proteobacteri | Gammaprotec | Cellvibrionales | Porticoccacea | unclassified | SAR92 bacteri | unclassified | unclassified | unclassified | unclassified | unclassified | 1 | 0,632455532 | 9,84E-05 | 6,52E-05 | 2 | 0,000205854 | - | 0 | 0 | - | 1 | 9,55E-05 | - | 1 | 9,49E-05 | - | 1 | 9,56E-05 | - |
| 1690 | 1 | 0.2um_NOKCPHJN_470890 | unclassified | unclassified | unclassified | unclassified | unclassified | unclassified | unclassified | unclassified | unclassified | unclassified | unclassified | unclassified | 0,6 | 0,489897949 | 0,000331041 | 0,000270763 | 1 | 0,000580754 | - | 0 | 0 | - | 1 | 0,000538919 | - | 1 | 0,000535532 | - | 0 | 0 | - |
| 1691 | 1 | 3um_FIJJAFBF_191789 | Eukaryota | unclassified | Synurophycea | Parmales | Triparmaceae | Triparma | Triparma laev | metabolism | Energy produc | COG0712 | F(1)F(0) ATP s | 35128.Thapsd | 13,8 | 1,166190379 | 0,0063829 | 0,000512324 | 12 | 0,005627413 | - | 13 | 0,007172238 | - | 14 | 0,006092376 | - | 15 | 0,006486526 | - | 15 | 0,006535949 | - |
| 1692 | 1 | 0.2um_NOKCPHJN_191603 | Eukaryota | Bacillariophyt | Bacillariophyc | Naviculales | Phaeodactylac | Phaeodactylu | Phaeodactylu | cellular proces | Cell cycle cont | 0IHFW | regulator of c | 2850.Phatr31 | 1,6 | 0,489897949 | 0,000211641 | 5,18E-05 | 1 | 0,000136808 | - | 1 | 0,000160951 | - | 2 | 0,000253906 | - | 2 | 0,00025231 | - | 2 | 0,000254232 | - |
| 1693 | 1 | 0.2um_NOKCPHJN_316872 | Bacteria | Proteobacteri | Alphaproteob | unclassified | unclassified | unclassified | various | metabolism | Energy produc | 01RNF | Produces ATP | 862719.AZOLI | 0,8 | 0,4 | 0,000147929 | 7,62E-05 | 0 | 0 | - | 1 | 0,000219978 | - | 1 | 0,000173511 | - | 1 | 0,000172421 | - | 1 | 0,000173735 | - |
| 1694 | 1 | 0.2um_NOKCPHJN_653095 | Bacteria | Proteobacteri | Gammaprotec | Cellvibrionales | Halieaceae | unclassified | marine gamm | metabolism | Inorganic ion t | 16PDP | TonB-depende | 247634.GPB2 | 1,8 | 0,4 | 0,000172173 | 4,51E-05 | 2 | 0,000191681 | - | 2 | 0,000225508 | - | 2 | 0,000177873 | - | 2 | 0,000176755 | - | 1 | 8,91E-05 | - |
| 1695 | 1 | 3um_FIJJAFBF_652364 | Bacteria | Bacteroidetes | Flavobacteriia | Flavobacterial | Flavobacteriac | Lutibacter | Lutibacter aga | poorly charact | Function unkn | 0XRCJ | NA | 391587.KAOT | 0,8 | 0,4 | 6,27E-05 | 3,14E-05 | 1 | 8,30E-05 | - | 0 | 0 | - | 1 | 7,70E-05 | - | 1 | 7,65E-05 | - | 1 | 7,71E-05 | - |
| 1696 | 1 | 3um_FIJJAFBF_190036 | Bacteria | Proteobacteri | Alphaproteob | Rhodobactera | Rhodobactera | Planktomarina | Planktomarina | unclassified | unclassified | unclassified | unclassified | unclassified | 1 | 0 | 0,000214082 | 2,09E-05 | 1 | 0,000215995 | - | 1 | 0,000254113 | - | 1 | 0,000200435 | - | 1 | 0,000199176 | - | 1 | 0,000200693 | - |
| 1697 | 1 | 3um_FIJJAFBF_151368 | Eukaryota | Annelida | Polychaeta | Golfingiida | Sipunculidae | Sipunculus | Sipunculus nu | unclassified | unclassified | unclassified | unclassified | unclassified | 1,4 | 0,489897949 | 0,000962797 | 0,000288617 | 1 | 0,000707208 | - | 1 | 0,000832015 | - | 1 | 0,000656264 | - | 2 | 0,00130428 | - | 2 | 0,001314218 | - |
| 1698 | 1 | 0.2um_NOKCPHJN_530253 | Bacteria | Bacteroidetes | Flavobacteriia | Flavobacterial | Flavobacteriac | Formosa | Formosa sp. H | unclassified | unclassified | unclassified | unclassified | unclassified | 2 | 1,673320053 | 0,000176617 | 0,000145269 | 1 | 9,47E-05 | - | 0 | 0 | - | 2 | 0,00017576 | - | 5 | 0,000436638 | - | 2 | 0,000175986 | - |
| 1699 | 1 | 3um_FIJJAFBF_448400 | Eukaryota | Bacillariophyt | Bacillariophyc | Naviculales | Phaeodactylac | Phaeodactylu | Phaeodactylu | unclassified | unclassified | unclassified | unclassified | unclassified | 1,6 | 0,489897949 | 0,003802907 | 0,001037507 | 2 | 0,004871881 | - | 1 | 0,00286583 | - | 2 | 0,00452093 | - | 2 | 0,00449252 | - | 1 | 0,002263375 | - |
| 1700 | 1 | 3um_FIJJAFBF_337762 | Bacteria | Verrucomicrol | Verrucomicrol | Verrucomicrol | Verrucomicrol | unclassified | Verrucomicrol | information st | Translation, ri | COG0185 | Protein S19 fo | 240016.VspiD | 0,8 | 0,4 | 0,000794881 | 0,000407349 | 1 | 0,000985324 | - | 1 | 0,001159212 | - | 1 | 0,000914345 | - | 0 | 0 | - | 1 | 0,000915522 | - |
| 1701 | 1 | 0.2um_NOKCPHJN_220621 | Bacteria | Bacteroidetes | unclassified | unclassified | unclassified | unclassified | Bacteroidetes | metabolism | Inorganic ion t | 05C27 | TonB-depende | 886377.Murru | 3 | 1,264911064 | 0,000250307 | 0,000105074 | 5 | 0,00043114 | - | 1 | 0,000101445 | - | 3 | 0,000240049 | - | 3 | 0,000238541 | - | 3 | 0,000240358 | - |
| 1702 | 1 | 3um_FIJJAFBF_45623 | Bacteria | Planctomycete | Phycisphaerae | Phycisphaeral | Phycisphaerac | Phycisphaera | Phycisphaera | unclassified | unclassified | unclassified | unclassified | unclassified | 1,6 | 0,489897949 | 0,000447804 | 0,000123203 | 2 | 0,000573162 | - | 1 | 0,000337156 | - | 2 | 0,000531874 | - | 1 | 0,000264266 | - | 2 | 0,000532559 | - |
| 1703 | 1 | 3um_FIJJAFBF_111353 | unclassified | unclassified | unclassified | unclassified | unclassified | unclassified | unclassified | unclassified | unclassified | unclassified | unclassified | unclassified | 0,2 | 0,4 | 5,14E-05 | 0,000102816 | 0 | 0 | - | 0 | 0 | - | 0 | 0 | - | 0 | 0 | - | 1 | 0,000257039 | - |
| 1704 | 9 | 0.2um_NOKCPHJN_196703;0.2 | Bacteria | Bacteroidetes | various | various | various | various | various | information st | Translation, ri | 05CAV | thus facilitatin | various | 0,2 | 0,4 | 2,61E-05 | 5,22E-05 | 0 | 0 | - | 0 | 0 | - | 0 | 0 | - | 0 | 0 | - | 1 | 0,000130579 | - |
| 1705 | 1 | 0.2um_NOKCPHJN_63174 | Bacteria | Bacteroidetes | Flavobacteriia | Flavobacterial | Flavobacteriac | Flavobacteriu | Flavobacteriu | metabolism | Inorganic ion t | 05C27 | TonB-depende | 983548.Krodi_ | 0,2 | 0,4 | 1,62E-05 | 3,23E-05 | 1 | 8,08E-05 | - | 0 | 0 | - | 0 | 0 | - | 0 | 0 | - | 0 | 0 | - |
| 1706 | 1 | 3um_FIJJAFBF_437253 | Bacteria | Bacteroidetes | Flavobacteriia | Flavobacterial | Flavobacteriac | Polaribacter | Polaribacter s | metabolism | Inorganic ion t | 05C27 | TonB-depende | 867900.Celly_ | 7,2 | 1,720465053 | 0,000567055 | 0,000128182 | 10 | 0,000802323 | - | 5 | 0,000471957 | - | 6 | 0,000446716 | - | 7 | 0,000517893 | - | 8 | 0,000596388 | - |
| 1707 | 1 | 0.2um_NOKCPHJN_102627 | Viruses | unclassified | unclassified | unclassified | unclassified | unclassified | uncultured vir | unclassified | unclassified | unclassified | unclassified | unclassified | 1,4 | 0,489897949 | 0,000243074 | 0,000100737 | 1 | 0,000172287 | - | 2 | 0,000405383 | - | 1 | 0,000159876 | - | 2 | 0,000317742 | - | 1 | 0,000160082 | - |
| 1708 | 1 | 3um_FIJJAFBF_344737 | Eukaryota | Bacillariophyt | Mediophyceae | Toxariales | Toxariaceae | Toxarium | Toxarium und | information st | Translation, ri | COG0050 | This protein p | 321332.CYB_0 | 0,4 | 0,489897949 | 8,27E-05 | 0,000101432 | 1 | 0,00021441 | - | 0 | 0 | - | 0 | 0 | - | 0 | 0 | - | 1 | 0,000199221 | - |
| 1709 | 1 | 0.2um_NOKCPHJN_132742 | Eukaryota | unclassified | Oomycetes | Albuginales | Albuginaceae | Albugo | Albugo candid | unclassified | unclassified | unclassified | unclassified | unclassified | 4,6 | 0,489897949 | 0,000886834 | 5,14E-05 | 4 | 0,000784732 | - | 4 | 0,00092322 | - | 5 | 0,000910254 | - | 5 | 0,000904534 | - | 5 | 0,000911426 | - |
| 1710 | 1 | 3um_FIJJAFBF_174375 | Viruses | unclassified | unclassified | unclassified | Phycodnavirid | unclassified | Chrysochromu | unclassified | unclassified | unclassified | unclassified | unclassified | 1 | 0 | 0,000148323 | 1,45E-05 | 1 | 0,000149648 | - | 1 | 0,000176058 | - | 1 | 0,000138868 | - | 1 | 0,000137995 | - | 1 | 0,000139047 | - |
| 1711 | 1 | 3um_FIJJAFBF_594160 | Eukaryota | unclassified | Bangiophycea | Bangiales | Bangiaceae | Wildemania | Wildemania s | unclassified | unclassified | unclassified | unclassified | unclassified | 3 | 1,264911064 | 0,001706878 | 0,000673689 | 4 | 0,002323016 | - | 2 | 0,001366489 | - | 5 | 0,002694594 | - | 2 | 0,001071064 | - | 2 | 0,001079225 | - |
| 1712 | 1 | 3um_FIJJAFBF_263212 | unclassified | unclassified | unclassified | unclassified | unclassified | unclassified | unclassified | unclassified | unclassified | unclassified | unclassified | unclassified | 0,2 | 0,4 | 8,26E-05 | 0,000165232 | 0 | 0 | - | 0 | 0 | - | 1 | 0,00041308 | - | 0 | 0 | - | 0 | 0 | - |
| 1713 | 2 | 0.2um_NOKCPHJN_885492;3ur | Bacteria | Bacteroidetes | Flavobacteriia | Flavobacterial | Flavobacteriac | Tenacibaculur | Tenacibaculur | poorly charact | Function unkn | 0P8NI | Inherit from b | various | 1 | 0 | 0,000233649 | 2,29E-05 | 1 | 0,000235736 | - | 1 | 0,000277338 | - | 1 | 0,000218755 | - | 1 | 0,00021738 | - | 1 | 0,000219036 | - |
| 1714 | 1 | 0.2um_NOKCPHJN_648025 | Eukaryota | Cryptomycota | unclassified | unclassified | unclassified | Rozella | Rozella allomy | unclassified | unclassified | unclassified | unclassified | unclassified | 1,6 | 0,489897949 | 0,000775212 | 0,000189789 | 1 | 0,000501108 | - | 1 | 0,000589542 | - | 2 | 0,00093002 | - | 2 | 0,000924175 | - | 2 | 0,000931217 | - |
| 1715 | 1 | 3um_FIJJAFBF_185356 | Bacteria | Bacteroidetes | Flavobacteriia | Flavobacterial | Flavobacteriac | Formosa | Formosa sp. H | unclassified | unclassified | unclassified | unclassified | unclassified | 1,2 | 0,4 | 0,000881569 | 0,000405392 | 1 | 0,000718802 | - | 2 | 0,00169131 | - | 1 | 0,000667022 | - | 1 | 0,000662831 | - | 1 | 0,000667881 | - |
| 1716 | 1 | 0.2um_NOKCPHJN_240819 | Bacteria | Bacteroidetes | Flavobacteriia | Flavobacterial | Flavobacteriac | Polaribacter | Polaribacter s | poorly charact | Function unkn | 09EUF | NA | 313598.MED1 | 1 | 0 | 0,000351893 | 3,44E-05 | 1 | 0,000355036 | - | 1 | 0,000417692 | - | 1 | 0,00032946 | - | 1 | 0,00032739 | - | 1 | 0,000329885 | - |
| 1717 | 1 | 3um_FIJJAFBF_115058 | Eukaryota | unclassified | unclassified | Phaeocystales | Phaeocystace | Phaeocystis | Phaeocystis gl | metabolism | Energy produc | COG0711 | F(1)F(0) ATP s | 63737.Npun_ | 2,8 | 0,4 | 0,001389792 | 0,000242413 | 2 | 0,001002215 | - | 3 | 0,001768627 | - | 3 | 0,00139503 | - | 3 | 0,001386263 | - | 3 | 0,001396826 | - |
| 1718 | 1 | 3um_FIJJAFBF_505169 | Bacteria | Proteobacteri | Alphaproteob | Rhodobactera | Hyphomonad | Robiginitomac | Robiginitomac | unclassified | unclassified | unclassified | unclassified | unclassified | 1 | 0 | 0,000210965 | 2,06E-05 | 1 | 0,000212849 | - | 1 | 0,000250412 | - | 1 | 0,000197516 | - | 1 | 0,000196275 | - | 1 | 0,000197771 | - |
| 1719 | 1 | 0.2um_NOKCPHJN_513503 | Bacteria | Proteobacteri | Alphaproteob | Rhodobactera | Rhodobactera | Planktomarina | Planktomarina | unclassified | unclassified | unclassified | unclassified | unclassified | 1,4 | 0,8 | 0,001000554 | 0,000600301 | 2 | 0,001403102 | - | 2 | 0,001650718 | - | 2 | 0,001302028 | - | 1 | 0,000646923 | - | 0 | 0 | - |
| 1720 | 1 | 3um_FIJJAFBF_56139 | Bacteria | Bacteroidetes | Flavobacteriia | Flavobacterial | Flavobacteriac | Polaribacter | Polaribacter s | metabolism | Energy produc | COG0056 | Produces ATP | 313598.MED1 | 2,4 | 0,8 | 0,000387065 | 0,000115802 | 3 | 0,000501108 | - | 1 | 0,000196514 | - | 2 | 0,000310007 | - | 3 | 0,000462088 | - | 3 | 0,000465609 | - |
| 1721 | 1 | 3um_FIJJAFBF_442870 | Eukaryota | unclassified | Cryptophyta | Cryptomonad | Hemiselmidac | Hemiselmis | Hemiselmis an | unclassified | unclassified | unclassified | unclassified | unclassified | 1 | 0 | 0,000648638 | 6,35E-05 | 1 | 0,000654432 | - | 1 | 0,000769925 | - | 1 | 0,000607289 | - | 1 | 0,000603473 | - | 1 | 0,000608071 | - |
| 1722 | 1 | 0.2um_NOKCPHJN_449563 | unclassified | unclassified | unclassified | unclassified | unclassified | unclassified | unclassified | unclassified | unclassified | unclassified | unclassified | unclassified | 0,6 | 0,489897949 | 8,79E-05 | 7,19E-05 | 1 | 0,000153849 | - | 0 | 0 | - | 1 | 0,000142766 | - | 0 | 0 | - | 1 | 0,00014295 | - |
| 1723 | 1 | 3um_FIJJAFBF_681094 | Eukaryota | Mollusca | Bivalvia | Ostreoida | Ostreidae | Crassostrea | Crassostrea vi | unclassified | unclassified | unclassified | unclassified | unclassified | 0,8 | 0,4 | 9,47E-05 | 4,75E-05 | 1 | 0,000125277 | - | 0 | 0 | - | 1 | 0,000116252 | - | 1 | 0,000115522 | - | 1 | 0,000116402 | - |
| 1724 | 1 | 0.2um_NOKCPHJN_556777 | Bacteria | Bacteroidetes | Flavobacteriia | Flavobacterial | unclassified | unclassified | Flavobacterial | unclassified | unclassified | unclassified | unclassified | unclassified | 0,6 | 0,489897949 | 0,000241844 | 0,000200735 | 0 | 0 | - | 1 | 0,000468954 | - | 1 | 0,000369894 | - | 0 | 0 | - | 1 | 0,00037037 | - |
| 1725 | 1 | 0.2um_NOKCPHJN_530493 | Eukaryota | unclassified | Labyrinthulom | unclassified | Thraustochytr | Aurantiochytr | Aurantiochytr | unclassified | unclassified | unclassified | unclassified | unclassified | 0,4 | 0,8 | 2,07E-05 | 4,15E-05 | 0 | 0 | - | 0 | 0 | - | 0 | 0 | - | 0 | 0 | - | 2 | 0,000103732 | - |
| 1726 | 1 | 0.2um_NOKCPHJN_328778 | Bacteria | Proteobacteri | Gammaprotec | Nevskiales | Algiphilaceae | Algiphilus | Algiphilus aro | poorly charact | Function unkn | 0XSV1 | NA | 204773.HEAR | 1 | 0 | 2,86E-05 | 2,80E-06 | 1 | 2,88E-05 | - | 1 | 3,39E-05 | - | 1 | 2,68E-05 | - | 1 | 2,66E-05 | - | 1 | 2,68E-05 | - |
| 1727 | 1 | 3um_FIJJAFBF_321055 | unclassified | unclassified | unclassified | unclassified | unclassified | unclassified | unclassified | unclassified | unclassified | unclassified | unclassified | unclassified | 0,2 | 0,4 | 0,000385073 | 0,000770146 | 0 | 0 | - | 0 | 0 | - | 0 | 0 | - | 1 | 0,001925366 | - | 0 | 0 | - |
| 1728 | 1 | 0.2um_NOKCPHJN_109945 | Eukaryota | unclassified | unclassified | Acytosteliales | Acytosteliacea | Acytostelium | Acytostelium | unclassified | unclassified | unclassified | unclassified | unclassified | 0,2 | 0,4 | 1,08E-05 | 2,15E-05 | 0 | 0 | - | 0 | 0 | - | 0 | 0 | - | 0 | 0 | - | 1 | 5,38E-05 | - |
| 1729 | 1 | 0.2um_NOKCPHJN_252102 | Eukaryota | Apicomplexa | Conoidasida | Eucoccidiorida | Eimeriidae | Eimeria | Eimeria acerv | unclassified | unclassified | unclassified | unclassified | unclassified | 1 | 0 | 9,42E-05 | 9,21E-06 | 1 | 9,50E-05 | - | 1 | 0,000111777 | - | 1 | 8,82E-05 | - | 1 | 8,76E-05 | - | 1 | 8,83E-05 | - |
| 1730 | 1 | 3um_FIJJAFBF_594165 | Eukaryota | unclassified | Pelagophycea | Pelagomonad | unclassified | Aureococcus | Aureococcus a | unclassified | unclassified | unclassified | unclassified | unclassified | 4,6 | 1,019803903 | 0,002549497 | 0,000471967 | 3 | 0,0016973 | - | 4 | 0,002662449 | - | 5 | 0,002625056 | - | 5 | 0,00260856 | - | 6 | 0,003154122 | - |
| 1731 | 2 | 0.2um_NOKCPHJN_567918;3ur | Eukaryota | unclassified | Pelagophycea | Pelagomonad | unclassified | Aureococcus | Aureococcus a | cellular proces | Signal transdu | KOG0613 | fibronectin typ | various | 0,6 | 0,489897949 | 4,45E-05 | 3,63E-05 | 0 | 0 | - | 0 | 0 | - | 1 | 7,43E-05 | - | 1 | 7,38E-05 | - | 1 | 7,44E-05 | - |
| 1732 | 1 | 0.2um_NOKCPHJN_814531 | unclassified | unclassified | unclassified | unclassified | unclassified | unclassified | unclassified | unclassified | unclassified | unclassified | unclassified | unclassified | 0,6 | 0,489897949 | 0,000273729 | 0,000227303 | 0 | 0 | - | 1 | 0,000531804 | - | 0 | 0 | - | 1 | 0,000416832 | - | 1 | 0,000420008 | - |
| 1733 | 1 | 3um_FIJJAFBF_11305 | Bacteria | Bacteroidetes | unclassified | unclassified | unclassified | unclassified | Bacteroidetes | cellular proces | Cell cycle cont | 05CR9 | regulator of c | 391587.KAOT | 1,6 | 0,489897949 | 0,000307624 | 7,53E-05 | 1 | 0,000198852 | - | 1 | 0,000233945 | - | 2 | 0,000369056 | - | 2 | 0,000366736 | - | 2 | 0,000369531 | - |
| 1734 | 1 | 3um_FIJJAFBF_15942 | Eukaryota | unclassified | Pelagophycea | Pelagomonad | unclassified | Aureococcus | Aureococcus a | unclassified | unclassified | unclassified | unclassified | unclassified | 1 | 0 | 0,000274188 | 2,68E-05 | 1 | 0,000276637 | - | 1 | 0,000325457 | - | 1 | 0,000256709 | - | 1 | 0,000255096 | - | 1 | 0,000257039 | - |
| 1735 | 1 | 0.2um_NOKCPHJN_582029 | Bacteria | Proteobacteri | Betaproteoba | Burkholderiale | Comamonada | Rhodoferax | Rhodoferax sp | unclassified | unclassified | unclassified | unclassified | unclassified | 0,6 | 0,489897949 | 0,000218751 | 0,000180849 | 1 | 0,000352184 | - | 1 | 0,000414337 | - | 0 | 0 | - | 0 | 0 | - | 1 | 0,000327235 | - |
| 1736 | 1 | 0.2um_NOKCPHJN_227614 | Bacteria | Bacteroidetes | Flavobacteriia | Flavobacterial | Flavobacteriac | Maribacter | Maribacter sp | metabolism | Inorganic ion t | 05C27 | TonB-depende | 714943.Mucp | 0,6 | 0,489897949 | 4,34E-05 | 3,55E-05 | 1 | 7,61E-05 | - | 0 | 0 | - | 0 | 0 | - | 1 | 7,01E-05 | - | 1 | 7,07E-05 | - |
| 1737 | 1 | 0.2um_NOKCPHJN_29615 | Bacteria | Proteobacteri | Gammaprotec | Alteromonada | Colwelliaceae | Thalassotalea | Thalassotalea | metabolism | Carbohydrate | 16QEQ | hydrolase, fan | 392500.Swoo_ | 0,4 | 0,489897949 | 3,56E-05 | 4,37E-05 | 1 | 9,26E-05 | - | 0 | 0 | - | 0 | 0 | - | 1 | 8,54E-05 | - | 0 | 0 | - |
| 1738 | 1 | 3um_FIJJAFBF_142181 | Bacteria | Proteobacteri | Gammaprotec | Cellvibrionales | Spongiibacter | Oceanicoccus | Oceanicoccus | metabolism | Energy produc | COG0055 | Produces ATP | 247639.MGP2 | 3,4 | 1,496662955 | 0,000611663 | 0,000244129 | 3 | 0,000565767 | - | 1 | 0,000221871 | - | 3 | 0,000525011 | - | 5 | 0,00086952 | - | 5 | 0,000876145 | - |
| 1739 | 1 | 3um_FIJJAFBF_182957 | Bacteria | Proteobacteri | Gammaprotec | Alteromonada | Alteromonada | Paraglaciecola | Paraglaciecola | metabolism | Inorganic ion t | 16TQC | TONB-depend | 314287.GB22 | 10,2 | 2,315167381 | 0,000842027 | 0,000142722 | 11 | 0,000934721 | - | 6 | 0,000599825 | - | 11 | 0,000867388 | - | 13 | 0,001018653 | - | 10 | 0,000789549 | - |
| 1740 | 1 | 3um_FIJJAFBF_62637 | Bacteria | Proteobacteri | Alphaproteob | Pelagibacteral | Pelagibacterac | Candidatus Pe | Candidatus Pe | metabolism | Inorganic ion t | 05PWN | phosphate ab | 335992.SAR1 | 0,4 | 0,489897949 | 0,00015801 | 0,000193747 | 1 | 0,000409784 | - | 0 | 0 | - | 1 | 0,000380265 | - | 0 | 0 | - | 0 | 0 | - |
| 1741 | 1 | 0.2um_NOKCPHJN_477555 | unclassified | unclassified | unclassified | unclassified | unclassified | unclassified | unclassified | unclassified | unclassified | unclassified | unclassified | unclassified | 0,2 | 0,4 | 6,00E-05 | 0,000119965 | 0 | 0 | - | 1 | 0,000299912 | - | 0 | 0 | - | 0 | 0 | - | 0 | 0 | - |
| 1742 | 1 | 0.2um_NOKCPHJN_494513 | Bacteria | Bacteroidetes | Flavobacteriia | Flavobacterial | Flavobacteriac | Cellulophaga | Cellulophaga t | poorly charact | Function unkn | 0P8NI | Inherit from b | 867900.Celly_ | 0,2 | 0,4 | 2,18E-05 | 4,35E-05 | 0 | 0 | - | 0 | 0 | - | 0 | 0 | - | 0 | 0 | - | 1 | 0,000108787 | - |
| 1743 | 1 | 0.2um_NOKCPHJN_782226 | Bacteria | Proteobacteri | Gammaprotec | Alteromonada | Pseudoalteror | Pseudoalteror | Pseudoalteror | metabolism | Inorganic ion t | 16PDP | TonB-depende | 637905.SVI_1 | 4,4 | 1,624807681 | 0,000827913 | 0,000258929 | 3 | 0,000588549 | - | 2 | 0,00046161 | - | 5 | 0,000910254 | - | 6 | 0,001085441 | - | 6 | 0,001093711 | - |
| 1744 | 1 | 0.2um_NOKCPHJN_166763 | unclassified | unclassified | unclassified | unclassified | unclassified | unclassified | synthetic cons | unclassified | unclassified | unclassified | unclassified | unclassified | 0,4 | 0,489897949 | 0,000170186 | 0,000210817 | 0 | 0 | - | 1 | 0,000475437 | - | 0 | 0 | - | 0 | 0 | - | 1 | 0,000375491 | - |
| 1745 | 1 | 3um_FIJJAFBF_513825 | Viruses | unclassified | unclassified | unclassified | unclassified | unclassified | uncultured M | unclassified | unclassified | unclassified | unclassified | unclassified | 1 | 0 | 0,000496671 | 4,86E-05 | 1 | 0,000501108 | - | 1 | 0,000589542 | - | 1 | 0,00046501 | - | 1 | 0,000462088 | - | 1 | 0,000465609 | - |
| 1746 | 1 | 3um_FIJJAFBF_404360 | Bacteria | Bacteroidetes | Flavobacteriia | Flavobacterial | Flavobacteriac | Olleya | Olleya aquima | information st | Translation, ri | 08YY1 | This protein is | 313598.MED1 | 0,8 | 0,4 | 0,000571409 | 0,000286521 | 1 | 0,000755981 | - | 0 | 0 | - | 1 | 0,000701524 | - | 1 | 0,000697115 | - | 1 | 0,000702427 | - |
| 1747 | 1 | 3um_FIJJAFBF_777597 | Bacteria | Verrucomicrol | Opitutae | Puniceicoccale | Puniceicoccac | Coraliomargar | Coraliomargar | poorly charact | Function unkn | 1DIXS | Inherit from b | 382464.VDG1 | 1,8 | 0,4 | 0,000441515 | 0,000108132 | 1 | 0,000247723 | - | 2 | 0,000582881 | - | 2 | 0,000459756 | - | 2 | 0,000456866 | - | 2 | 0,000460347 | - |
| 1748 | 1 | 3um_FIJJAFBF_254413 | Bacteria | Bacteroidetes | Flavobacteriia | Flavobacterial | Crocinitomica | unclassified | Crocinitomica | unclassified | unclassified | unclassified | unclassified | unclassified | 0,2 | 0,4 | 0,000129385 | 0,000258769 | 0 | 0 | - | 0 | 0 | - | 0 | 0 | - | 1 | 0,000646923 | - | 0 | 0 | - |
| 1749 | 1 | 3um_FIJJAFBF_26690 | Bacteria | Bacteroidetes | Flavobacteriia | Flavobacterial | Flavobacteriac | Polaribacter | Polaribacter s | unclassified | unclassified | unclassified | unclassified | unclassified | 0,8 | 0,4 | 0,000212167 | 0,000109219 | 0 | 0 | - | 1 | 0,000315504 | - | 1 | 0,000248859 | - | 1 | 0,000247295 | - | 1 | 0,000249179 | - |
| 1750 | 1 | 3um_FIJJAFBF_328494 | Bacteria | Bacteroidetes | Flavobacteriia | Flavobacterial | Flavobacteriac | Arenitalea | Arenitalea lute | unclassified | unclassified | unclassified | unclassified | unclassified | 1 | 0 | 0,000284975 | 2,79E-05 | 1 | 0,000287521 | - | 1 | 0,000338262 | - | 1 | 0,000266809 | - | 1 | 0,000265132 | - | 1 | 0,000267152 | - |
| 1751 | 1 | 0.2um_NOKCPHJN_669120 | Bacteria | Proteobacteri | Alphaproteob | Pelagibacteral | Pelagibacterac | Candidatus Pe | Candidatus Pe | poorly charact | Function unkn | 01QDW | Protein of unk | 1002672.SAR1 | 1 | 0 | 0,000415873 | 4,07E-05 | 1 | 0,000419588 | - | 1 | 0,000493636 | - | 1 | 0,000389362 | - | 1 | 0,000386916 | - | 1 | 0,000389864 | - |
| 1752 | 1 | 3um_FIJJAFBF_799075 | Bacteria | Proteobacteri | unclassified | unclassified | unclassified | unclassified | Proteobacteri | metabolism | Lipid transpor | 08IQF | Catalyzes the | 991905.SL003 | 0,8 | 0,4 | 0,000129857 | 6,66E-05 | 1 | 0,000161202 | - | 1 | 0,000189651 | - | 0 | 0 | - | 1 | 0,00014865 | - | 1 | 0,000149782 | - |
| 1753 | 1 | 3um_FIJJAFBF_714940 | Bacteria | Proteobacteri | Alphaproteob | unclassified | unclassified | Candidatus Pu | Candidatus Pu | metabolism | Carbohydrate | 0IVA1 | Photosystem I | 35128.Thapsd | 0,6 | 0,8 | 0,00027703 | 0,000375798 | 2 | 0,000948042 | - | 0 | 0 | - | 0 | 0 | - | 1 | 0,00043711 | - | 0 | 0 | - |
| 1754 | 1 | 0.2um_NOKCPHJN_926215 | Eukaryota | unclassified | unclassified | Isochrysidales | Isochrysidacea | Chrysotila | Chrysotila cart | unclassified | unclassified | unclassified | unclassified | unclassified | 3 | 0 | 0,005014469 | 0,000490637 | 3 | 0,005059261 | - | 3 | 0,005952109 | - | 3 | 0,004694812 | - | 3 | 0,004665309 | - | 3 | 0,004700855 | - |
| 1755 | 1 | 3um_FIJJAFBF_347183 | Bacteria | Proteobacteri | Alphaproteob | Rhodospirillale | Rhodospirillac | unclassified | Rhodospirillac | cellular proces | Cell wall/mem | COG2982 | asmA family | 744980.TRICH | 0,2 | 0,4 | 2,25E-05 | 4,50E-05 | 0 | 0 | - | 1 | 0,000112385 | - | 0 | 0 | - | 0 | 0 | - | 0 | 0 | - |
| 1756 | 2 | 0.2um_NOKCPHJN_993350;3ur | Bacteria | Bacteroidetes | Cytophagia | Cytophagales | Flammeovirga | unclassified | Flammeovirga | unclassified | unclassified | unclassified | unclassified | unclassified | 0,2 | 0,4 | 2,85E-05 | 5,70E-05 | 0 | 0 | - | 0 | 0 | - | 0 | 0 | - | 0 | 0 | - | 1 | 0,00014245 | - |
| 1757 | 1 | 3um_FIJJAFBF_145237 | Bacteria | Bacteroidetes | Flavobacteriia | Flavobacterial | Flavobacteriac | Hyunsoonleell | Hyunsoonleell | information st | Translation, ri | 08YY1 | This protein is | 391603.FBALC | 2,6 | 0,8 | 0,00138152 | 0,000428439 | 4 | 0,002151996 | - | 2 | 0,001265888 | - | 3 | 0,001497731 | - | 2 | 0,000992213 | - | 2 | 0,000999773 | - |
| 1758 | 1 | 0.2um_NOKCPHJN_347401 | Bacteria | unclassified | unclassified | unclassified | unclassified | Thermobaculu | Thermobaculu | unclassified | unclassified | unclassified | unclassified | unclassified | 0,2 | 0,4 | 3,12E-05 | 6,25E-05 | 0 | 0 | - | 0 | 0 | - | 1 | 0,000156193 | - | 0 | 0 | - | 0 | 0 | - |
| 1759 | 1 | 3um_FIJJAFBF_283091 | Bacteria | Bacteroidetes | Flavobacteriia | Flavobacterial | Flavobacteriac | Algibacter | Algibacter sp. | poorly charact | Function unkn | 08KQQ | NA | 391603.FBALC | 0,6 | 0,8 | 0,000183635 | 0,000244782 | 0 | 0 | - | 0 | 0 | - | 2 | 0,000611855 | - | 0 | 0 | - | 1 | 0,000306321 | - |
| 1760 | 1 | 0.2um_NOKCPHJN_349749 | Bacteria | Firmicutes | Bacilli | Lactobacillales | Lactobacillace | Lactobacillus | Lactobacillus r | unclassified | unclassified | unclassified | unclassified | unclassified | 0,2 | 0,4 | 5,44E-06 | 1,09E-05 | 1 | 2,72E-05 | - | 0 | 0 | - | 0 | 0 | - | 0 | 0 | - | 0 | 0 | - |
| 1761 | 1 | 0.2um_NOKCPHJN_35230 | Bacteria | Proteobacteri | Alphaproteob | unclassified | unclassified | unclassified | SAR116 cluste | metabolism | Energy produc | 01RNF | Produces ATP | 1064539.AZO | 1,2 | 0,4 | 0,000217706 | 6,51E-05 | 1 | 0,000185008 | - | 1 | 0,000217658 | - | 2 | 0,000343362 | - | 1 | 0,000170602 | - | 1 | 0,000171902 | - |
| 1762 | 1 | 3um_FIJJAFBF_216063 | Bacteria | Proteobacteri | Gammaprotec | Cellvibrionales | unclassified | unclassified | Cellvibrionales | information st | Translation, ri | 177SZ | This protein b | 247633.GP214 | 1,6 | 0,489897949 | 0,000964117 | 0,00026303 | 2 | 0,001235125 | - | 1 | 0,000726549 | - | 2 | 0,001146151 | - | 2 | 0,001138949 | - | 1 | 0,000573813 | - |
| 1763 | 1 | 0.2um_NOKCPHJN_259382 | Bacteria | Proteobacteri | Alphaproteob | Rhodobactera | Rhodobactera | Sulfitobacter | Sulfitobacter g | unclassified | unclassified | unclassified | unclassified | unclassified | 0,4 | 0,489897949 | 4,14E-05 | 5,13E-05 | 0 | 0 | - | 1 | 0,000115791 | - | 1 | 9,13E-05 | - | 0 | 0 | - | 0 | 0 | - |
| 1764 | 1 | 3um_FIJJAFBF_268268 | Eukaryota | unclassified | unclassified | Choanoflagelli | Salpingoecida | Salpingoeca | Salpingoeca r | unclassified | unclassified | unclassified | unclassified | unclassified | 0,4 | 0,489897949 | 6,25E-05 | 7,67E-05 | 1 | 0,000162096 | - | 0 | 0 | - | 0 | 0 | - | 0 | 0 | - | 1 | 0,000150613 | - |
| 1765 | 1 | 0.2um_NOKCPHJN_783159 | Bacteria | Proteobacteri | Gammaprotec | unclassified | unclassified | unclassified | OM182 bacter | metabolism | Energy produc | 17C3H | Produces ATP | 379731.PST_4 | 0,4 | 0,489897949 | 0,000269564 | 0,000333921 | 0 | 0 | - | 1 | 0,000753065 | - | 0 | 0 | - | 0 | 0 | - | 1 | 0,000594755 | - |
| 1766 | 1 | 0.2um_NOKCPHJN_52330 | Bacteria | Bacteroidetes | Flavobacteriia | Flavobacterial | Flavobacteriac | Formosa | Formosa sp. H | unclassified | unclassified | unclassified | unclassified | unclassified | 1 | 0 | 0,000102376 | 1,00E-05 | 1 | 0,000103291 | - | 1 | 0,000121519 | - | 1 | 9,59E-05 | - | 1 | 9,52E-05 | - | 1 | 9,60E-05 | - |
| 1767 | 1 | 0.2um_NOKCPHJN_359987 | Bacteria | Proteobacteri | Gammaprotec | Oceanospirilla | unclassified | unclassified | Oceanospirilla | cellular proces | Posttranslatio | 0IEKR | One of the co | 35128.Thapsd | 0,4 | 0,489897949 | 7,04E-05 | 8,62E-05 | 0 | 0 | - | 0 | 0 | - | 1 | 0,000176522 | - | 1 | 0,000175413 | - | 0 | 0 | - |
| 1768 | 1 | 3um_FIJJAFBF_60206 | Bacteria | Actinobacteria | unclassified | unclassified | unclassified | unclassified | Actinobacteria | cellular proces | Intracellular tr | 0QJ7H | motA TolQ Exl | 314285.KT71_ | 0,8 | 0,4 | 0,000142852 | 7,16E-05 | 1 | 0,000188995 | - | 0 | 0 | - | 1 | 0,000175381 | - | 1 | 0,000174279 | - | 1 | 0,000175607 | - |
| 1769 | 1 | 3um_FIJJAFBF_348505 | Bacteria | Bacteroidetes | Cytophagia | Cytophagales | Flammeovirga | Marivirga | Marivirga seri | metabolism | Inorganic ion t | 05C27 | TonB-depende | 643867.Ftrac_ | 4,8 | 2,315167381 | 0,000380429 | 0,000171996 | 5 | 0,0004164 | - | 1 | 9,80E-05 | - | 4 | 0,000309123 | - | 8 | 0,000614362 | - | 6 | 0,000464282 | - |
| 1770 | 1 | 3um_FIJJAFBF_191788 | Eukaryota | unclassified | Pelagophycea | Pelagomonad | unclassified | Aureoumbra | Aureoumbra l | metabolism | Energy produc | COG0056 | Produces ATP | 32049.SYNPC | 18,8 | 0,979795897 | 0,003233274 | 0,000155224 | 19 | 0,003312491 | - | 17 | 0,003486855 | - | 19 | 0,003073873 | - | 20 | 0,003215322 | - | 19 | 0,00307783 | - |
| 1771 | 1 | 3um_FIJJAFBF_147847 | Eukaryota | unclassified | Oomycetes | Peronosporale | Peronosporac | Phytophthora | Phytophthora | unclassified | unclassified | unclassified | unclassified | unclassified | 1,2 | 0,748331477 | 0,000151618 | 9,31E-05 | 1 | 0,0001345 | - | 0 | 0 | - | 2 | 0,000249622 | - | 1 | 0,000124027 | - | 2 | 0,000249943 | - |
| 1772 | 1 | 3um_FIJJAFBF_40363 | Eukaryota | unclassified | Raphidophyce | Chattonellales | Chattonellace | Heterosigma | Heterosigma a | unclassified | unclassified | unclassified | unclassified | unclassified | 1,8 | 0,748331477 | 0,000298502 | 0,000108403 | 1 | 0,000172287 | - | 1 | 0,000202691 | - | 3 | 0,000479627 | - | 2 | 0,000317742 | - | 2 | 0,000320163 | - |
| 1773 | 1 | 0.2um_NOKCPHJN_321483 | Bacteria | Proteobacteri | Gammaprotec | Cellvibrionales | Halieaceae | unclassified | marine gamm | metabolism | Inorganic ion t | 0QJVD | TonB-depende | 247639.MGP2 | 0,4 | 0,489897949 | 3,73E-05 | 4,57E-05 | 1 | 9,69E-05 | - | 0 | 0 | - | 0 | 0 | - | 1 | 8,94E-05 | - | 0 | 0 | - |
| 1774 | 1 | 0.2um_NOKCPHJN_818235 | unclassified | unclassified | unclassified | unclassified | unclassified | unclassified | unclassified | unclassified | unclassified | unclassified | unclassified | unclassified | 0,4 | 0,489897949 | 0,000164135 | 0,000203321 | 0 | 0 | - | 1 | 0,000458533 | - | 0 | 0 | - | 0 | 0 | - | 1 | 0,00036214 | - |
| 1775 | 1 | 3um_FIJJAFBF_92885 | Eukaryota | unclassified | Cryptophyta | Pyrenomonad | Pyrenomonad | Storeatula | Storeatula sp. | metabolism | Energy produc | COG0711 | F(1)F(0) ATP s | 91464.S7335_ | 0,8 | 0,4 | 0,000368242 | 0,000184647 | 1 | 0,000487188 | - | 0 | 0 | - | 1 | 0,000452093 | - | 1 | 0,000449252 | - | 1 | 0,000452675 | - |
| 1776 | 1 | 0.2um_NOKCPHJN_1027277 | Eukaryota | unclassified | Florideophyce | Gigartinales | Gigartinaceae | Chondrus | Chondrus cris | unclassified | unclassified | unclassified | unclassified | unclassified | 0,8 | 0,4 | 0,000396907 | 0,000199021 | 1 | 0,000525113 | - | 0 | 0 | - | 1 | 0,000487286 | - | 1 | 0,000484224 | - | 1 | 0,000487913 | - |

| | | | | | | | | | | | | | | | | | | | | | | | | | | | | |
|---|---|---|---|---|---|---|---|---|---|---|---|---|---|---|---|---|---|---|---|---|---|---|---|---|---|---|---|---|
| 1777 | 1 | 3um_FIJJAFBF_729268 | Eukaryota | unclassified | unclassified | Phaeocystales | Phaeocystace | Phaeocystis | Phaeocystis a | metabolism | Energy produc | COG1290 | Component ol | 70448.Q0P3F3 | 0,2 | 0,4 | 4,24E-05 | 8,48E-05 | 0 - | 0 - | 0 - | 0 - | 1 - | 0,000211919 - | 0 - | 0 - | 0 - | 0 - |
| 1778 | 1 | 0.2um_NOKCPHJN_1008157 | Eukaryota | unclassified | Cryptophyta | Pyrenomonad | Geminigerace | Guillardia | Guillardia the | cellular proces | Cytoskeleton | 0DKI0 | asp (abnorma | 7739.JGI8877 | 0,2 | 0,4 | 3,56E-05 | 7,12E-05 | 0 - | 0 - | 1 - | 0,000177879 - | 0 - | 0 - | 0 - | 0 - | 0 - | 0 - |
| 1779 | 2 | 3um_FIJJAFBF_333770;3um_FI | Bacteria | Bacteroidetes | Flavobacteriia | Flavobacterial | Flavobacteria | Polaribacter | various | poorly charact | Function unkn | 09EUF | NA | 313598.MED1 | 0,4 | 0,489897949 | 0,000124404 | 0,000152365 | 0 - | 0 - | 0 - | 0 - | 0 - | 0 - | 1 - | 0,000309829 - | 1 - | 0,00031219 - |
| 1780 | 1 | 0.2um_NOKCPHJN_978568 | Eukaryota | unclassified | Oligohymenop | Hymenostoma | Tetrahymenid | Tetrahymena | Tetrahymena | unclassified | unclassified | unclassified | unclassified | unclassified | 2,4 | 0,489897949 | 0,000461442 | 9,17E-05 | 3 - | 0,000584626 - | 2 - | 0,000458533 - | 3 - | 0,000542512 - | 2 - | 0,000359402 - | 2 - | 0,00036214 - |
| 1781 | 1 | 0.2um_NOKCPHJN_571842 | Bacteria | Proteobacteri | Alphaproteob | Pelagibacteral | Pelagibactera | unclassified | alpha proteob | cellular proces | Intracellular tr | 16Q1A | type II secreti | 684719.HIMB | 0,8 | 0,4 | 0,000114085 | 5,72E-05 | 1 - | 0,000150936 - | 0 - | 0 - | 1 - | 0,000140063 - | 1 - | 0,000139183 - | 1 - | 0,000140244 - |
| 1782 | 1 | 0.2um_NOKCPHJN_604311 | Bacteria | Proteobacteri | Alphaproteob | Pelagibacteral | Pelagibactera | Candidatus Pe | Candidatus Pe | metabolism | Energy produc | COG0056 | Produces ATP | 684719.HIMB | 0,2 | 0,4 | 3,63E-05 | 7,26E-05 | 1 - | 0,000181561 - | 0 - | 0 - | 0 - | 0 - | 0 - | 0 - | 0 - | 0 - |
| 1783 | 1 | 0.2um_NOKCPHJN_301289 | Bacteria | Firmicutes | Bacilli | Lactobacillales | Lactobacillace | Lactobacillus | Lactobacillus | unclassified | unclassified | unclassified | unclassified | unclassified | 0,4 | 0,489897949 | 1,00E-05 | 1,23E-05 | 1 - | 2,61E-05 - | 0 - | 0 - | 0 - | 0 - | 1 - | 2,40E-05 - | 0 - | 0 - |
| 1784 | 1 | 3um_FIJJAFBF_92950 | Bacteria | Proteobacteri | Gammaprotec | Cellvibrionales | Porticoccacea | unclassified | various | cellular proces | Cell wall/mem | COG2885 | Ompa motb d | 314287.GB22 | 1 | 0 | 0,000276807 | 2,71E-05 | 1 - | 0,00027928 - | 1 - | 0,000328567 - | 1 - | 0,000259162 - | 1 - | 0,000257533 - | 1 - | 0,000259495 - |
| 1785 | 1 | 3um_FIJJAFBF_01300 | Bacteria | Bacteroidetes | Flavobacteriia | Flavobacterial | Flavobacteria | Polaribacter | Polaribacter s | unclassified | unclassified | unclassified | unclassified | unclassified | 1 | 0 | 9,94E-05 | 9,73E-06 | 1 - | 0,000100336 - | 1 - | 0,000118043 - | 1 - | 9,31E-05 - | 1 - | 9,25E-05 - | 1 - | 9,32E-05 - |
| 1786 | 1 | 3um_FIJJAFBF_774175 | Eukaryota | unclassified | Pelagophycea | Pelagomonad | unclassified | Aureococcus | Aureococcus a | unclassified | unclassified | unclassified | unclassified | unclassified | 1,4 | 0,489897949 | 0,000241674 | 7,24E-05 | 1 - | 0,000177518 - | 1 - | 0,000208846 - | 1 - | 0,00016473 - | 2 - | 0,00032739 - | 2 - | 0,000329885 - |
| 1787 | 1 | 0.2um_NOKCPHJN_990892 | Eukaryota | unclassified | unclassified | Prymnesiales | Chrysochromu | Chrysochromu | Chrysochromu | unclassified | unclassified | unclassified | unclassified | unclassified | 0,6 | 0,489897949 | 0,000248692 | 0,000203409 | 1 - | 0,000436288 - | 0 - | 0 - | 1 - | 0,000404859 - | 1 - | 0,000402315 - | 0 - | 0 - |
| 1788 | 1 | 0.2um_NOKCPHJN_523420 | Bacteria | Bacteroidetes | Flavobacteriia | Flavobacterial | Crocinitomica | Fluviicola | Fluviicola taff | unclassified | unclassified | unclassified | unclassified | unclassified | 0,8 | 0,4 | 0,000286942 | 0,000143881 | 1 - | 0,000379627 - | 0 - | 0 - | 1 - | 0,00035228 - | 1 - | 0,000350066 - | 1 - | 0,000352734 - |
| 1789 | 1 | 3um_FIJJAFBF_17872 | Bacteria | Proteobacteri | Gammaprotec | Vibrionales | Vibrionaceae | Photobacteriu | Photobacteriu | unclassified | unclassified | unclassified | unclassified | unclassified | 0,4 | 0,489897949 | 0,000112741 | 0,00013808 | 0 - | 0 - | 0 - | 0 - | 0 - | 0 - | 1 - | 0,000280782 - | 1 - | 0,000282922 - |
| 1790 | 1 | 3um_FIJJAFBF_704767 | unclassified | unclassified | unclassified | unclassified | unclassified | unclassified | unclassified | unclassified | unclassified | unclassified | unclassified | unclassified | 0,6 | 0,489897949 | 0,000100712 | 8,22E-05 | 0 - | 0 - | 0 - | 0 - | 1 - | 0,000168134 - | 1 - | 0,000167077 - | 1 - | 0,00016835 - |
| 1791 | 1 | 3um_FIJJAFBF_410461 | Bacteria | Candidatus At | unclassified | unclassified | unclassified | unclassified | Candidatus At | information st | Translation, ri | COG0256 | This is one of | 684719.HIMB | 1 | 0 | 0,000730399 | 7,15E-05 | 1 - | 0,000736923 - | 1 - | 0,000866974 - | 1 - | 0,000683838 - | 1 - | 0,000679541 - | 1 - | 0,000684718 - |
| 1792 | 1 | 0.2um_NOKCPHJN_371014 | Bacteria | Proteobacteri | Betaproteoba | unclassified | unclassified | unclassified | Betaproteoba | metabolism | Inorganic ion | 07QMV | Part of the AB | 261292.Nit79 | 0,8 | 0,4 | 0,000878211 | 0,000452082 | 0 - | 0 - | 1 - | 0,001305948 - | 1 - | 0,001030085 - | 1 - | 0,001023612 - | 1 - | 0,001031411 - |
| 1793 | 1 | 0.2um_NOKCPHJN_351457 | Bacteria | Proteobacteri | Alphaproteob | Rhodospirillal | Rhodospirillac | unclassified | Rhodospirillac | unclassified | unclassified | unclassified | unclassified | unclassified | 0,8 | 0,4 | 0,000224527 | 0,000115581 | 0 - | 0 - | 1 - | 0,000333883 - | 1 - | 0,000263355 - | 1 - | 0,0002617 - | 1 - | 0,000263694 - |
| 1794 | 1 | 3um_FIJJAFBF_405912 | Bacteria | Proteobacteri | Gammaprotec | Alteromonada | Alteromonada | Paraglaciecola | Paraglaciecola | information st | Transcription | COG0085 | DNA-depende | 342610.Patl_ | 1,8 | 1,939071943 | 0,000123616 | 0,00013208 | 5 - | 0,000326728 - | 3 - | 0,000230633 - | 0 - | 0 - | 0 - | 0 - | 1 - | 6,07E-05 - |
| 1795 | 1 | 3um_FIJJAFBF_634956 | Bacteria | Proteobacteri | Gammaprotec | Oceanospirilla | Oceanospirilla | Motiliproteus | Motiliproteus | information st | Translation, ri | COG1185 | Involved in m | 207954.MED9 | 0,6 | 0,489897949 | 7,20E-05 | 5,89E-05 | 1 - | 0,000125997 - | 0 - | 0 - | 1 - | 0,000116921 - | 0 - | 0 - | 1 - | 0,000117071 - |
| 1796 | 1 | 0.2um_NOKCPHJN_422582 | Bacteria | Proteobacteri | Alphaproteob | Rhizobiales | unclassified | unclassified | Rhizobiales ba | unclassified | unclassified | unclassified | unclassified | unclassified | 0,2 | 0,4 | 4,17E-05 | 8,34E-05 | 0 - | 0 - | 0 - | 0 - | 0 - | 0 - | 1 - | 0,000208416 - | 0 - | 0 - |
| 1797 | 1 | 3um_FIJJAFBF_565197 | unclassified | unclassified | unclassified | unclassified | unclassified | unclassified | unclassified | unclassified | unclassified | unclassified | unclassified | unclassified | 0,8 | 0,4 | 0,001142819 | 0,000573043 | 1 - | 0,001511963 - | 0 - | 0 - | 1 - | 0,001403047 - | 1 - | 0,00139423 - | 1 - | 0,001404853 - |
| 1798 | 1 | 3um_FIJJAFBF_31340 | Bacteria | Proteobacteri | Alphaproteob | Rhodobactera | Rhodobactera | Roseobacter | Roseobacter s | metabolism | Inorganic ion | COG0841 | acriflavin resis | 391593.RCCS2 | 1 | 0 | 0,000175946 | 1,72E-05 | 1 - | 0,000177518 - | 1 - | 0,000208846 - | 1 - | 0,00016473 - | 1 - | 0,000163695 - | 1 - | 0,000164942 - |
| 1799 | 1 | 3um_FIJJAFBF_316515 | Bacteria | Proteobacteri | Gammaprotec | Alteromonada | Alteromonada | Paraglaciecola | Paraglaciecola | unclassified | unclassified | unclassified | unclassified | unclassified | 1,2 | 0,748331477 | 0,000125555 | 7,44E-05 | 0 - | 0 - | 1 - | 0,000127057 - | 2 - | 0,000200435 - | 1 - | 9,96E-05 - | 2 - | 0,000200693 - |
| 1800 | 1 | 0.2um_NOKCPHJN_367726 | Eukaryota | unclassified | unclassified | Trichomonadi | Trichomonadi | Trichomonas | Trichomonas | cellular proces | Cytoskeleton | 0F97C | NA | 353152.XP_62 | 1 | 0,894427191 | 3,57E-05 | 3,05E-05 | 0 - | 0 - | 1 - | 4,31E-05 - | 2 - | 6,80E-05 - | 2 - | 6,76E-05 - | 0 - | 0 - |
| 1801 | 1 | 3um_FIJJAFBF_630544 | Eukaryota | unclassified | unclassified | Prymnesiales | Chrysochromu | Chrysochromu | Chrysochromu | poorly charact | Function unkn | 110WY | Chlorophyll A- | 2850.Phatr16 | 1,2 | 0,4 | 0,000793791 | 0,000237282 | 1 - | 0,000674568 - | 1 - | 0,000793615 - | 2 - | 0,00125195 - | 1 - | 0,000622041 - | 1 - | 0,000626781 - |
| 1802 | 1 | 0.2um_NOKCPHJN_476838 | Eukaryota | unclassified | Oomycetes | Saprolegniales | Saprolegniace | Aphanomyces | Aphanomyces | unclassified | unclassified | unclassified | unclassified | unclassified | 1 | 0 | 9,31E-05 | 9,11E-06 | 1 - | 9,39E-05 - | 1 - | 0,00011046 - | 1 - | 8,71E-05 - | 1 - | 8,66E-05 - | 1 - | 8,72E-05 - |
| 1803 | 1 | 0.2um_NOKCPHJN_65231 | Bacteria | Proteobacteri | Gammaprotec | Cellvibrionales | Porticoccacea | unclassified | gamma prote | unclassified | unclassified | unclassified | unclassified | unclassified | 0,4 | 0,489897949 | 5,19E-05 | 6,37E-05 | 1 - | 0,000134706 - | 0 - | 0 - | 1 - | 0,000125003 - | 0 - | 0 - | 0 - | 0 - |
| 1804 | 1 | 3um_FIJJAFBF_674421 | Viruses | unclassified | unclassified | unclassified | unclassified | unclassified | uncultured M | unclassified | unclassified | unclassified | unclassified | unclassified | 2 | 0 | 0,00085633 | 8,38E-05 | 2 - | 0,000863979 - | 2 - | 0,001016452 - | 2 - | 0,000801741 - | 2 - | 0,000796703 - | 2 - | 0,000802773 - |
| 1805 | 1 | 3um_FIJJAFBF_337584 | unclassified | unclassified | unclassified | unclassified | unclassified | unclassified | unclassified | unclassified | unclassified | unclassified | unclassified | unclassified | 0,2 | 0,4 | 0,00033092 | 0,00066184 | 1 - | 0,001654601 - | 0 - | 0 - | 0 - | 0 - | 0 - | 0 - | 0 - | 0 - |
| 1806 | 1 | 3um_FIJJAFBF_176595 | Eukaryota | Phaeophyceae | unclassified | Ectocarpales | Ectocarpaceae | Ectocarpus | Ectocarpus sil | cellular proces | Posttranslatio | 0XRR7 | Calreticulin | 67593.Physo1 | 0,2 | 0,4 | 0,000121602 | 0,000243204 | 0 - | 0 - | 0 - | 0 - | 0 - | 0 - | 1 - | 0,00060801 - | 0 - | 0 - |
| 1807 | 1 | 3um_FIJJAFBF_510231 | Bacteria | Bacteroidetes | Flavobacteriia | Flavobacterial | Flavobacteria | Polaribacter | Polaribacter r | information st | Translation, ri | COG0088 | One of the pri | 313598.MED1 | 1,4 | 0,8 | 0,000571818 | 0,000301363 | 1 - | 0,000419588 - | 1 - | 0,000493636 - | 1 - | 0,000389362 - | 1 - | 0,000386916 - | 3 - | 0,001169591 - |
| 1808 | 1 | 3um_FIJJAFBF_79252 | Bacteria | Bacteroidetes | Flavobacteriia | Flavobacterial | Flavobacteria | Polaribacter | Polaribacter s | information st | Replication, re | 0PB0H | DNA-binding | 313598.MED1 | 2,8 | 0,748331477 | 0,00266775 | 0,000606927 | 3 - | 0,002923128 - | 2 - | 0,002292664 - | 4 - | 0,003616744 - | 3 - | 0,002695512 - | 2 - | 0,0018107 - |
| 1809 | 1 | 0.2um_NOKCPHJN_684761 | Bacteria | Bacteroidetes | Flavobacteriia | Flavobacterial | Flavobacteria | Formosa | Formosa sp. H | poorly charact | Function unkn | 06VZ7 | NA | 688270.Celal_ | 0,4 | 0,489897949 | 0,00010535 | 0,000129203 | 1 - | 0,000274043 - | 0 - | 0 - | 0 - | 0 - | 1 - | 0,000252704 - | 0 - | 0 - |
| 1810 | 1 | 3um_FIJJAFBF_715094 | Eukaryota | Bacillariophyt | Bacillariophyc | Eunotiales | Eunotiaceae | Eunotia | Eunotia naege | information st | Translation, ri | 0FNT6 | One of the ea | 497965.Cyan7 | 1,2 | 0,4 | 0,001132863 | 0,000333826 | 1 - | 0,000963669 - | 1 - | 0,001133735 - | 1 - | 0,00089425 - | 2 - | 0,001777261 - | 1 - | 0,000895401 - |
| 1811 | 1 | 3um_FIJJAFBF_684040 | unclassified | unclassified | unclassified | unclassified | unclassified | unclassified | unclassified | unclassified | unclassified | unclassified | unclassified | unclassified | 0,6 | 0,489897949 | 7,44E-05 | 6,09E-05 | 1 - | 0,000130497 - | 0 - | 0 - | 0 - | 0 - | 1 - | 0,000120335 - | 1 - | 0,000121252 - |
| 1812 | 1 | 0.2um_NOKCPHJN_77102 | Eukaryota | Arthropoda | Arachnida | Trombidiform | Tetranychidae | Tetranychus | Tetranychus c | cellular proces | Posttranslatio | 0FI38 | heat shock pr | 441375.XP_0 | 0,4 | 0,489897949 | 4,79E-05 | 5,87E-05 | 1 - | 0,000124037 - | 0 - | 0 - | 0 - | 0 - | 0 - | 0 - | 1 - | 0,00011525 - |
| 1813 | 1 | 3um_FIJJAFBF_186254 | Bacteria | Bacteroidetes | Flavobacteriia | Flavobacterial | Flavobacteria | Formosa | Formosa sp. H | unclassified | unclassified | unclassified | unclassified | unclassified | 0,4 | 0,489897949 | 0,000111507 | 0,000136568 | 0 - | 0 - | 0 - | 0 - | 1 - | 0,000279645 - | 1 - | 0,000277888 - | 0 - | 0 - |
| 1814 | 1 | 0.2um_NOKCPHJN_129918 | Bacteria | Proteobacteri | Gammaprotec | unclassified | unclassified | unclassified | Gammaprotec | metabolism | Energy produc | 16PVJ | Component ol | 349521.HCH_ | 0,4 | 0,489897949 | 3,65E-05 | 4,47E-05 | 0 - | 0 - | 0 - | 0 - | 0 - | 0 - | 1 - | 9,09E-05 - | 1 - | 9,16E-05 - |
| 1815 | 1 | 0.2um_NOKCPHJN_440914 | Bacteria | Proteobacteri | Gammaprotec | Cellvibrionales | Porticoccacea | unclassified | various | unclassified | unclassified | unclassified | unclassified | unclassified | 1,4 | 1,2 | 0,000133245 | 0,000113144 | 2 - | 0,000201133 - | 0 - | 0 - | 0 - | 0 - | 3 - | 0,000278206 - | 2 - | 0,000186884 - |
| 1816 | 1 | 0.2um_NOKCPHJN_402249 | Bacteria | Proteobacteri | Alphaproteob | Pelagibacteral | Pelagibactera | unclassified | Pelagibactera | cellular proces | Cell wall/mem | 7257 | NA | 439493.PB721 | 0,2 | 0,4 | 4,76E-05 | 9,51E-05 | 0 - | 0 - | 0 - | 0 - | 0 - | 0 - | 1 - | 0,000237839 - | 0 - | 0 - |
| 1817 | 3 | 0.2um_NOKCPHJN_116238;0.2 | Bacteria | Bacteroidetes | Flavobacteriia | Flavobacterial | Crocinitomica | unclassified | various | unclassified | unclassified | unclassified | unclassified | unclassified | 1,4 | 0,489897949 | 0,000982919 | 0,000409224 | 1 - | 0,000695983 - | 2 - | 0,001637617 - | 1 - | 0,000645847 - | 1 - | 0,000641789 - | 2 - | 0,001293357 - |
| 1818 | 1 | 3um_FIJJAFBF_41560 | Eukaryota | Bacillariophyt | Coscinodiscop | Chaetocerotal | Chaetocerota | Chaetoceros | Chaetoceros s | metabolism | Energy produc | 0IE88 | oxidoreductas | 45157.CMV13 | 1,8 | 0,748331477 | 0,00020731 | 7,54E-05 | 1 - | 0,000119637 - | 1 - | 0,00014075 - | 2 - | 0,000222037 - | 2 - | 0,000220642 - | 3 - | 0,000333485 - |
| 1819 | 1 | 3um_FIJJAFBF_259312 | Bacteria | Planctomycet | Phycisphaerae | Phycisphaeral | Phycisphaerac | Phycisphaera | Phycisphaera | cellular proces | Cell wall/mem | 06B97 | phosphate-sel | 316067.Geob_ | 1 | 0,632455532 | 0,000173186 | 0,000114779 | 2 - | 0,000362371 - | 0 - | 0 - | 1 - | 0,000168134 - | 1 - | 0,000167077 - | 1 - | 0,00016835 - |
| 1820 | 1 | 3um_FIJJAFBF_414105 | Bacteria | Proteobacteri | Gammaprotec | Alteromonada | Alteromonada | unclassified | Alteromonada | unclassified | unclassified | unclassified | unclassified | unclassified | 0,4 | 0,489897949 | 0,000121991 | 0,000149408 | 0 - | 0 - | 0 - | 0 - | 1 - | 0,000304782 - | 0 - | 0 - | 1 - | 0,000305174 - |
| 1821 | 1 | 0.2um_NOKCPHJN_27340 | Bacteria | Bacteroidetes | Flavobacteriia | Flavobacterial | Flavobacteria | Flavobacteriu | Flavobacteriu | unclassified | unclassified | unclassified | unclassified | unclassified | 2 | 0,632455532 | 0,000198952 | 6,78E-05 | 3 - | 0,000298956 - | 2 - | 0,000234477 - | 2 - | 0,000184947 - | 2 - | 0,000183785 - | 1 - | 9,26E-05 - |
| 1822 | 1 | 0.2um_NOKCPHJN_658935 | Bacteria | Proteobacteri | Deltaproteoba | Myxococcales | Myxococcacea | Myxococcus | Myxococcus s | cellular proces | Posttranslatio | 05JRK | feS assembly | 290397.Adeh_ | 0,4 | 0,489897949 | 0,000137312 | 0,000169091 | 1 - | 0,000315446 - | 1 - | 0,000371115 - | 0 - | 0 - | 0 - | 0 - | 0 - | 0 - |
| 1823 | 1 | 3um_FIJJAFBF_180252 | unclassified | unclassified | unclassified | unclassified | unclassified | unclassified | unclassified | unclassified | unclassified | unclassified | unclassified | unclassified | 0,2 | 0,4 | 2,13E-05 | 4,27E-05 | 0 - | 0 - | 0 - | 0 - | 0 - | 0 - | 1 - | 0,000106683 - | 0 - | 0 - |
| 1824 | 1 | 3um_FIJJAFBF_271409 | Bacteria | Bacteroidetes | Flavobacteriia | Flavobacterial | Flavobacteria | Flavobacteriu | Flavobacteriu | unclassified | unclassified | unclassified | unclassified | unclassified | 0,2 | 0,4 | 2,39E-05 | 4,78E-05 | 0 - | 0 - | 0 - | 0 - | 1 - | 0,000119496 - | 0 - | 0 - | 0 - | 0 - |
| 1825 | 1 | 3um_FIJJAFBF_513536 | Bacteria | Bacteroidetes | Saprospiria | Saprospirales | Saprospiracea | various | various | poorly charact | Function unkn | 1AINX | Phage tail she | 760192.Halhy | 0,2 | 0,4 | 3,53E-05 | 7,06E-05 | 1 - | 0,000176446 - | 0 - | 0 - | 0 - | 0 - | 0 - | 0 - | 0 - | 0 - |
| 1826 | 1 | 3um_FIJJAFBF_618324 | Bacteria | Bacteroidetes | Flavobacteriia | Flavobacterial | Flavobacteria | Algibacter | Algibacter sp. | metabolism | Carbohydrate | 0XNQK | Transporter | 391587.KAOT | 0,2 | 0,4 | 3,55E-05 | 7,10E-05 | 0 - | 0 - | 0 - | 0 - | 0 - | 0 - | 0 - | 0 - | 1 - | 0,00017752 - |
| 1827 | 2 | 0.2um_NOKCPHJN_936620;3ur | Eukaryota | unclassified | Oomycetes | Peronosporale | Peronosporac | Phytophthora | various | cellular proces | Signal transdu | 152IZ | EFh | 67593.Physo1 | 2,6 | 0,8 | 0,001332074 | 0,000345924 | 2 - | 0,001050226 - | 2 - | 0,001235568 - | 3 - | 0,001461858 - | 4 - | 0,001936895 - | 2 - | 0,000975826 - |
| 1828 | 1 | 0.2um_NOKCPHJN_39203 | Eukaryota | Bacillariophyt | Fragilariophyc | Licmophorales | Licmophorace | Licmophora | Licmophora sp | metabolism | Energy produc | COG1290 | Component ol | 35128.Thapsd | 5,2 | 0,748331477 | 0,002117882 | 0,000455439 | 5 - | 0,002039392 - | 6 - | 0,00287916 - | 5 - | 0,001892482 - | 4 - | 0,001504472 - | 6 - | 0,002273902 - |
| 1829 | 1 | 0.2um_NOKCPHJN_316030 | Viruses | unclassified | unclassified | unclassified | unclassified | unclassified | uncultured M | unclassified | unclassified | unclassified | unclassified | unclassified | 2,2 | 0,748331477 | 0,000606299 | 0,000250896 | 3 - | 0,000814494 - | 3 - | 0,000958234 - | 2 - | 0,000503881 - | 1 - | 0,000250357 - | 2 - | 0,000504529 - |
| 1830 | 1 | 0.2um_NOKCPHJN_835463 | Eukaryota | Apicomplexa | Conoidasida | Eucoccidiorida | Eimeriidae | Eimeria | Eimeria mitis | poorly charact | Function unkn | 0NF4H | copper amine | 479436.Vpar_ | 1 | 0 | 0,000149087 | 1,46E-05 | 1 - | 0,000150418 - | 1 - | 0,000176964 - | 1 - | 0,000139583 - | 1 - | 0,000138706 - | 1 - | 0,000139762 - |
| 1831 | 1 | 3um_FIJJAFBF_31984 | Bacteria | Proteobacteri | Gammaprotec | Cellvibrionales | Halieaceae | Halioglobus | Halioglobus lu | unclassified | unclassified | unclassified | unclassified | unclassified | 0,2 | 0,4 | 3,02E-05 | 6,04E-05 | 1 - | 0,000150936 - | 0 - | 0 - | 0 - | 0 - | 0 - | 0 - | 0 - | 0 - |
| 1832 | 1 | 3um_FIJJAFBF_102176 | Bacteria | Proteobacteri | Alphaproteob | Rhodobactera | Rhodobactera | Ruegeria | Ruegeria med | metabolism | Inorganic ion | COG3221 | phosphonate | 388739.RSK20 | 3,6 | 0,489897949 | 0,001029008 | 0,000118778 | 4 - | 0,001161508 - | 3 - | 0,001024866 - | 3 - | 0,000808378 - | 4 - | 0,001071064 - | 4 - | 0,001079225 - |
| 1833 | 1 | 0.2um_NOKCPHJN_430990 | Bacteria | Verrucomicrol | Opitutae | Puniceicoccale | Puniceicoccac | Coraliomargar | Coraliomargar | unclassified | unclassified | unclassified | unclassified | unclassified | 1 | 0 | 0,000284975 | 2,79E-05 | 1 - | 0,000287521 - | 1 - | 0,000338262 - | 1 - | 0,000266809 - | 1 - | 0,000265132 - | 1 - | 0,000267152 - |
| 1834 | 1 | 3um_FIJJAFBF_116533 | Eukaryota | Bacillariophyt | Coscinodiscop | Thalassiosirale | Stephanodisca | Cyclotella | Cyclotella pse | metabolism | Energy produc | 11IIK | Low-potential | 35128.Thapsd | 3,6 | 0,8 | 0,001933718 | 0,000350109 | 3 - | 0,00164426 - | 3 - | 0,001934435 - | 3 - | 0,001525814 - | 5 - | 0,002527042 - | 4 - | 0,002037037 - |
| 1835 | 2 | 0.2um_NOKCPHJN_27393;3um | Bacteria | unclassified | unclassified | unclassified | unclassified | unclassified | uncultured ba | unclassified | unclassified | unclassified | unclassified | unclassified | 3,2 | 0,4 | 0,0005443 | 6,75E-05 | 3 - | 0,00051686 - | 3 - | 0,000608074 - | 3 - | 0,000479627 - | 3 - | 0,000476613 - | 4 - | 0,000640326 - |
| 1836 | 1 | 3um_FIJJAFBF_85545 | Bacteria | Proteobacteri | Gammaprotec | Alteromonada | Alteromonada | Alteromonas | Alteromonas | poorly charact | Function unkn | 17TP6 | PEP-CTERM m | 335283.Neut_ | 1 | 0 | 0,0002822 | 2,76E-05 | 1 - | 0,00028472 - | 1 - | 0,000334967 - | 1 - | 0,00026421 - | 1 - | 0,00026255 - | 1 - | 0,00026455 - |
| 1837 | 1 | 0.2um_NOKCPHJN_1001140 | Bacteria | Proteobacteri | Gammaprotec | Pseudomonad | Pseudomonad | Pseudomonas | various | information st | Translation, ri | COG0087 | One of the pri | 629265.PMA4 | 2 | 0,632455532 | 0,000803305 | 0,000213606 | 2 - | 0,000831221 - | 1 - | 0,000488957 - | 2 - | 0,000771344 - | 2 - | 0,000766496 - | 3 - | 0,001158505 - |
| 1838 | 1 | 0.2um_NOKCPHJN_469467 | Bacteria | Bacteroidetes | Flavobacteriia | Flavobacterial | Flavobacteria | unclassified | Flavobacteria | poorly charact | Function unkn | 178AZ | NADPH dehyd | 335992.SAR11 | 0,2 | 0,4 | 9,13E-05 | 0,000182696 | 1 - | 0,000456739 - | 0 - | 0 - | 0 - | 0 - | 0 - | 0 - | 0 - | 0 - |
| 1839 | 1 | 0.2um_NOKCPHJN_919228 | Eukaryota | unclassified | Pelagophycea | Pelagomonad | unclassified | Aureococcus | Aureococcus a | metabolism | Energy produc | COG0055 | Produces ATP | 2850.Phatr54 | 1 | 0,632455532 | 0,000917553 | 0,000573304 | 1 - | 0,000974376 - | 0 - | 0 - | 1 - | 0,000904186 - | 1 - | 0,000898504 - | 2 - | 0,0018107 - |
| 1840 | 1 | 3um_FIJJAFBF_746307 | Eukaryota | unclassified | Bangiophycea | Bangiales | Bangiaceae | various | various | metabolism | Energy produc | 0J0AS | This b-type cy | 35128.Thapsd | 0,2 | 0,4 | 0,000192537 | 0,000385073 | 0 - | 0 - | 0 - | 0 - | 0 - | 0 - | 1 - | 0,000962683 - | 0 - | 0 - |
| 1841 | 1 | 3um_FIJJAFBF_77421 | Viruses | unclassified | unclassified | unclassified | various | various | various | unclassified | unclassified | unclassified | unclassified | unclassified | 12,8 | 1,720465053 | 0,002093795 | 0,000250189 | 10 - | 0,001660868 - | 12 - | 0,00234477 - | 15 - | 0,002311839 - | 13 - | 0,001991003 - | 14 - | 0,002160494 - |
| 1842 | 1 | 3um_FIJJAFBF_176110 | Bacteria | Proteobacteri | Gammaprotec | Alteromonada | Ferrimonadac | Paraferrimona | Paraferrimona | cellular proces | Intracellular tr | 175GA | One of the pr | 323850.Shew_ | 0,2 | 0,4 | 0,000105655 | 0,00021131 | 1 - | 0,000528276 - | 0 - | 0 - | 0 - | 0 - | 0 - | 0 - | 0 - | 0 - |
| 1843 | 1 | 0.2um_NOKCPHJN_54716 | Eukaryota | Streptophyta | unclassified | Ericales | Primulaceae | Androsace | Androsace lax | metabolism | Energy produc | 1DQDB | PsaA and PsaE | 59689.fgenesh | 0,8 | 0,4 | 9,03E-05 | 4,53E-05 | 1 - | 0,000119474 - | 0 - | 0 - | 1 - | 0,000110867 - | 1 - | 0,000110171 - | 1 - | 0,00011101 - |
| 1844 | 1 | 0.2um_NOKCPHJN_805964 | Bacteria | Proteobacteri | Alphaproteob | Rhodobactera | Rhodobactera | unclassified | Rhodobactera | poorly charact | Function unkn | 17P8K | NA | 856793.MICA_ | 1 | 0,632455532 | 0,000309296 | 0,000188704 | 0 - | 0 - | 1 - | 0,000372454 - | 1 - | 0,000293779 - | 1 - | 0,000291933 - | 2 - | 0,000588314 - |
| 1845 | 1 | 0.2um_NOKCPHJN_241055 | Eukaryota | Ascomycota | Sordariomyce | Hypocreales | Nectriaceae | Fusarium | Fusarium gran | cellular proces | Posttranslatio | 0V7QW | ankyrin repea | 7668.SPU_024 | 1 | 0 | 6,31E-05 | 6,18E-06 | 1 - | 6,37E-05 - | 1 - | 7,49E-05 - | 1 - | 5,91E-05 - | 1 - | 5,87E-05 - | 1 - | 5,92E-05 - |
| 1846 | 1 | 3um_FIJJAFBF_13890 | unclassified | unclassified | unclassified | unclassified | unclassified | unclassified | unclassified | unclassified | unclassified | unclassified | unclassified | unclassified | 0,8 | 0,4 | 0,000793732 | 0,000407019 | 1 - | 0,000985324 - | 1 - | 0,001159212 - | 0 - | 0 - | 1 - | 0,000908599 - | 1 - | 0,000915522 - |
| 1847 | 1 | 3um_FIJJAFBF_71019 | Bacteria | Cyanobacteria | unclassified | Oscillatoriales | Microcoleacea | Arthrospira | various | information st | Translation, ri | 0FP0S | Seems to be t | 513049.Amax | 0,6 | 0,8 | 0,000377704 | 0,000504268 | 0 - | 0 - | 0 - | 0 - | 2 - | 0,001261655 - | 1 - | 0,000626863 - | 0 - | 0 - |
| 1848 | 1 | 3um_FIJJAFBF_71348 | Bacteria | Bacteroidetes | Cytophagia | Cytophagales | Flammeovirga | Marivirga | Marivirga trac | poorly charact | Function unkn | 09AR5 | Inherit from b | 643867.Ftrac_ | 2 | 0 | 0,000397792 | 3,89E-05 | 2 - | 0,000401345 - | 2 - | 0,000472173 - | 2 - | 0,000372434 - | 2 - | 0,000370093 - | 2 - | 0,000372913 - |
| 1849 | 1 | 0.2um_NOKCPHJN_793302 | Bacteria | Proteobacteri | Gammaprotec | Alteromonada | Pseudoalteror | Pseudoalteror | Pseudoalteror | poorly charact | Function unkn | COG4681 | Yaeq family | 357804.Ping_ | 0,4 | 0,489897949 | 0,00023858 | 0,000293796 | 1 - | 0,000548087 - | 1 - | 0,000644812 - | 0 - | 0 - | 0 - | 0 - | 0 - | 0 - |
| 1850 | 1 | 3um_FIJJAFBF_338127 | Bacteria | Bacteroidetes | Saprospiria | Saprospirales | Haliscomenob | Phaeodactylib | Phaeodactylib | information st | Translation, ri | 1AFYF | This protein p | 485917.Phep_ | 0,6 | 0,489897949 | 0,000141932 | 0,00011787 | 0 - | 0 - | 1 - | 0,000275855 - | 1 - | 0,000217585 - | 1 - | 0,000216218 - | 0 - | 0 - |
| 1851 | 1 | 3um_FIJJAFBF_722063 | unclassified | unclassified | unclassified | unclassified | unclassified | unclassified | unclassified | unclassified | unclassified | unclassified | unclassified | unclassified | 3,2 | 0,748331477 | 0,001058221 | 0,000258559 | 4 - | 0,001333747 - | 3 - | 0,001176843 - | 2 - | 0,000618835 - | 3 - | 0,000922418 - | 4 - | 0,001239262 - |
| 1852 | 1 | 3um_FIJJAFBF_74634 | Viruses | unclassified | unclassified | unclassified | Phycodnavirid | Coccolithoviru | Emiliania hux | poorly charact | Function unkn | 0JUYB | Mycoplasma | 70448.Q00TR | 0,2 | 0,4 | 1,93E-05 | 3,87E-05 | 1 - | 9,67E-05 - | 0 - | 0 - | 0 - | 0 - | 0 - | 0 - | 0 - | 0 - |
| 1853 | 1 | 3um_FIJJAFBF_371763 | Bacteria | Bacteroidetes | Flavobacteriia | Flavobacterial | Flavobacteria | Flaviramulus | Flaviramulus i | metabolism | Inorganic ion | 05C27 | TonB-depend | 688270.Celal_ | 4,8 | 1,720465053 | 0,000402516 | 0,000131203 | 6 - | 0,000520954 - | 2 - | 0,000204297 - | 4 - | 0,000322284 - | 5 - | 0,000400324 - | 7 - | 0,000564723 - |
| 1854 | 1 | 3um_FIJJAFBF_542441 | Bacteria | unclassified | unclassified | unclassified | unclassified | unclassified | bacterium B1 | unclassified | unclassified | unclassified | unclassified | unclassified | 1,2 | 0,979795897 | 0,000202279 | 0,000190661 | 0 - | 0 - | 3 - | 0,000565831 - | 1 - | 0,000148769 - | 1 - | 0,000147834 - | 1 - | 0,000148961 - |
| 1855 | 1 | 0.2um_NOKCPHJN_72300 | Bacteria | Proteobacteri | Gammaprotec | Oceanospirilla | Saccharospiril | Reinekea | Reinekea fors | unclassified | unclassified | unclassified | unclassified | unclassified | 1 | 0 | 0,00025489 | 2,49E-05 | 1 - | 0,000257167 - | 1 - | 0,000302551 - | 1 - | 0,000238641 - | 1 - | 0,000237142 - | 1 - | 0,000238949 - |
| 1856 | 1 | 3um_FIJJAFBF_257406 | Bacteria | Bacteroidetes | Flavobacteriia | Flavobacterial | Flavobacteria | Algibacter | Algibacter sp. | cellular proces | Intracellular tr | 08SHB | Biopolymer tr | 411154.GFO_ | 0,6 | 0,489897949 | 0,000317155 | 0,00025936 | 1 - | 0,000555024 - | 0 - | 0 - | 1 - | 0,000515043 - | 0 - | 0 - | 1 - | 0,000515706 - |
| 1857 | 1 | 3um_FIJJAFBF_33587 | Bacteria | Proteobacteri | Alphaproteob | Sphingomona | Sphingomona | Sphingomona | Sphingomona | unclassified | unclassified | unclassified | unclassified | unclassified | 2,6 | 0,489897949 | 0,000297953 | 7,03E-05 | 2 - | 0,000230168 - | 3 - | 0,000406181 - | 3 - | 0,000320381 - | 2 - | 0,000212245 - | 3 - | 0,000320793 - |
| 1858 | 1 | 0.2um_NOKCPHJN_237004 | Eukaryota | unclassified | unclassified | Prymnesiales | Chrysochromu | Chrysochromu | Chrysochromu | poorly charact | Function unkn | COG4102 | (twin-arginine | 35128.Thaps2 | 0,8 | 0,4 | 3,00E-05 | 1,54E-05 | 1 - | 3,72E-05 - | 1 - | 4,38E-05 - | 1 - | 3,46E-05 - | 1 - | 3,43E-05 - | 0 - | 0 - |
| 1859 | 1 | 0.2um_NOKCPHJN_635596 | unclassified | unclassified | unclassified | unclassified | unclassified | unclassified | unclassified | unclassified | unclassified | unclassified | unclassified | unclassified | 1 | 0 | 0,000157174 | 1,54E-05 | 1 - | 0,000158578 - | 1 - | 0,000186564 - | 1 - | 0,000147155 - | 1 - | 0,00014623 - | 1 - | 0,000147344 - |
| 1860 | 1 | 0.2um_NOKCPHJN_450502 | Eukaryota | unclassified | Labyrinthulom | unclassified | Thraustochytr | Aurantiochytr | Aurantiochytr | unclassified | unclassified | unclassified | unclassified | unclassified | 1 | 0,632455532 | 0,000227199 | 0,000138481 | 0 - | 0 - | 1 - | 0,00027366 - | 2 - | 0,000431707 - | 1 - | 0,000214497 - | 1 - | 0,000216131 - |
| 1861 | 2 | 0.2um_NOKCPHJN_100728;3ur | Bacteria | Bacteroidetes | Flavobacteriia | Flavobacterial | Flavobacteria | Winogradskye | Winogradskye | unclassified | unclassified | unclassified | unclassified | unclassified | 2,2 | 0,748331477 | 0,000385981 | 0,000119778 | 1 - | 0,000179333 - | 2 - | 0,000421963 - | 3 - | 0,000499244 - | 3 - | 0,000496106 - | 2 - | 0,000333258 - |
| 1862 | 1 | 3um_FIJJAFBF_395548 | Eukaryota | unclassified | unclassified | Isochrysidales | Noelaerhabda | Emiliania | Emiliania hux | poorly charact | Function unkn | 17F80 | NA | 391597.LMED | 0,2 | 0,4 | 0,000173142 | 0,000346284 | 0 - | 0 - | 0 - | 0 - | 1 - | 0,00086571 - | 0 - | 0 - | 0 - | 0 - |
| 1863 | 1 | 3um_FIJJAFBF_385889 | Bacteria | Proteobacteri | Alphaproteob | Rhodospirillal | Rhodospirillac | Oceanibaculu | Oceanibaculu | metabolism | Energy produc | COG0056 | Produces ATP | 414684.RC1_ | 1 | 0 | 0,000171435 | 1,68E-05 | 1 - | 0,000172966 - | 1 - | 0,000203491 - | 1 - | 0,000160506 - | 1 - | 0,000159498 - | 1 - | 0,000160713 - |
| 1864 | 1 | 0.2um_NOKCPHJN_279402 | Eukaryota | unclassified | Oomycetes | Peronosporale | Peronosporac | Phytophthora | Phytophthora | unclassified | unclassified | unclassified | unclassified | unclassified | 0,2 | 0,4 | 1,48E-05 | 2,96E-05 | 0 - | 0 - | 1 - | 7,40E-05 - | 0 - | 0 - | 0 - | 0 - | 0 - | 0 - |
| 1865 | 1 | 3um_FIJJAFBF_208051 | Bacteria | Proteobacteri | Gammaprotec | Oceanospirilla | Saccharospiril | Reinekea | Reinekea fors | metabolism | Inorganic ion | 0QI15 | sulfurtransfer | 314283.MED2 | 0,2 | 0,4 | 5,69E-05 | 0,000113814 | 0 - | 0 - | 0 - | 0 - | 1 - | 0,000284534 - | 0 - | 0 - | 0 - | 0 - |
| 1866 | 1 | 3um_FIJJAFBF_393524 | Bacteria | Bacteroidetes | Flavobacteriia | Flavobacterial | Flavobacteria | Confluentibac | Confluentibac | metabolism | Secondary me | 099V5 | phenylacetate | 1046627.BZAF | 1 | 0 | 0,000201198 | 1,97E-05 | 1 - | 0,000202995 - | 1 - | 0,000238819 - | 1 - | 0,000188372 - | 1 - | 0,000187188 - | 1 - | 0,000188615 - |
| 1867 | 1 | 3um_FIJJAFBF_495044 | Bacteria | Bacteroidetes | Flavobacteriia | Flavobacterial | Flavobacteria | Cellulophaga | Cellulophaga | poorly charact | Function unkn | 0XP53 | Ragb susd dor | 313598.MED1 | 1,2 | 0,748331477 | 0,000714252 | 0,000453824 | 2 - | 0,001252769 - | 0 - | 0 - | 1 - | 0,000581262 - | 2 - | 0,001155219 - | 1 - | 0,000582011 - |
| 1868 | 1 | 0.2um_NOKCPHJN_154905 | Eukaryota | unclassified | Oomycetes | Peronosporale | Peronosporac | Phytophthora | Phytophthora | unclassified | unclassified | unclassified | unclassified | unclassified | 2 | 0,894427191 | 0,000371377 | 0,000147526 | 1 - | 0,000193585 - | 1 - | 0,000227748 - | 3 - | 0,000538919 - | 2 - | 0,000357021 - | 3 - | 0,000539613 - |
| 1869 | 1 | 3um_FIJJAFBF_276754 | Bacteria | Proteobacteri | Gammaprotec | Alteromonada | Alteromonada | Paraglaciecola | Paraglaciecola | metabolism | Amino acid tra | 16STP | phosphoserin | 342610.Patl_1 | 0,4 | 0,489897949 | 9,98E-05 | 0,000123641 | 0 - | 0 - | 1 - | 0,000278838 - | 0 - | 0 - | 0 - | 0 - | 1 - | 0,00022022 - |
| 1870 | 1 | 3um_FIJJAFBF_444691 | Eukaryota | unclassified | unclassified | Isochrysidales | Noelaerhabda | Emiliania | Emiliania hux | metabolism | Carbohydrate | COG1850 | RuBisCO catal | 45157.CMV01 | 1,2 | 0,748331477 | 0,000208662 | 0,000123241 | 0 - | 0 - | 1 - | 0,000211414 - | 2 - | 0,000333511 - | 2 - | 0,000331415 - | 1 - | 0,00016697 - |
| 1871 | 2 | 0.2um_NOKCPHJN_1056495;3u | unclassified | unclassified | unclassified | unclassified | unclassified | unclassified | unclassified | unclassified | unclassified | unclassified | unclassified | unclassified | 22 | 1,264911064 | 0,014879744 | 0,00097804 | 23 - | 0,015757489 - | 20 - | 0,016120295 - | 23 - | 0,014622384 - | 23 - | 0,014530493 - | 21 - | 0,013368057 - |
| 1872 | 1 | 0.2um_NOKCPHJN_546978 | Bacteria | Proteobacteri | Gammaprotec | Cellvibrionales | Porticoccacea | unclassified | SAR92 bacteri | metabolism | Inorganic ion | 0QHT5 | TonB-depend | 314287.GB22 | 9,2 | 2,13541565 | 0,000916662 | 0,000210001 | 13 - | 0,001314902 - | 7 - | 0,000832975 - | 8 - | 0,000750881 - | 10 - | 0,000932703 - | 8 - | 0,000751848 - |
| 1873 | 1 | 3um_FIJJAFBF_392122 | Bacteria | Proteobacteri | Alphaproteob | unclassified | unclassified | unclassified | Alphaproteob | information st | Replication, re | COG0513 | purine NTP-de | 414684.RC1_ | 0,2 | 0,4 | 1,64E-05 | 3,28E-05 | 0 - | 0 - | 0 - | 0 - | 0 - | 0 - | 1 - | 8,19E-05 - | 0 - | 0 - |
| 1874 | 1 | 0.2um_NOKCPHJN_271368 | Bacteria | Proteobacteri | Deltaproteoba | unclassified | unclassified | unclassified | Deltaproteoba | information st | Translation, ri | 0NJE3 | Located at the | 760568.Desku | 0,2 | 0,4 | 0,000134507 | 0,000269014 | 0 - | 0 - | 0 - | 0 - | 1 - | 0,000672535 - | 0 - | 0 - | 0 - | 0 - |
| 1875 | 1 | 3um_FIJJAFBF_344754 | Eukaryota | Bacillariophyt | Coscinodiscop | Coscinodiscale | Coscinodiscac | Coscinodiscus | Coscinodiscus | unclassified | unclassified | unclassified | unclassified | unclassified | 0,2 | 0,4 | 0,000144949 | 0,000289897 | 1 - | 0,000724743 - | 0 - | 0 - | 0 - | 0 - | 0 - | 0 - | 0 - | 0 - |
| 1876 | 1 | 0.2um_NOKCPHJN_243019 | Eukaryota | Echinodermat | Echinoidea | Echinoida | Strongylocent | Strongylocent | Strongylocent | cellular proces | Cell wall/mem | KOG4177 | Ankyrin Repea | 5722.A2ERS8 | 1 | 0 | 0,000131693 | 1,29E-05 | 1 - | 0,000132869 - | 1 - | 0,000156318 - | 1 - | 0,000123298 - | 1 - | 0,000122523 - | 1 - | 0,000123457 - |
| 1877 | 1 | 0.2um_NOKCPHJN_78018 | Bacteria | Bacteroidetes | Flavobacteriia | Flavobacterial | Crocinitomica | unclassified | Crocinitomica | unclassified | unclassified | unclassified | unclassified | unclassified | 1 | 0 | 0,000124881 | 1,22E-05 | 1 - | 0,000125997 - | 1 - | 0,000148233 - | 1 - | 0,000116921 - | 1 - | 0,000116186 - | 1 - | 0,000117071 - |
| 1878 | 1 | 3um_FIJJAFBF_203100 | Bacteria | Bacteroidetes | Flavobacteriia | Flavobacterial | Crocinitomica | unclassified | Crocinitomica | cellular proces | Intracellular tr | 05FGH | MotA TolQ ex | 655815.ZPR_4 | 0,8 | 0,748331477 | 0,000217845 | 0,000203272 | 0 - | 0 - | 0 - | 0 - | 1 - | 0,000273076 - | 2 - | 0,00054272 - | 1 - | 0,000273428 - |
| 1879 | 1 | 3um_FIJJAFBF_254288 | Bacteria | Proteobacteri | Gammaprotec | Cellvibrionales | Spongiibacter | Zhongshania | Zhongshania s | unclassified | unclassified | unclassified | unclassified | unclassified | 0,4 | 0,8 | 0,000127555 | 0,000255109 | 2 - | 0,000637773 - | 0 - | 0 - | 0 - | 0 - | 0 - | 0 - | 0 - | 0 - |
| 1880 | 1 | 0.2um_NOKCPHJN_287456 | Bacteria | Proteobacteri | Gammaprotec | Alteromonada | Colwelliaceae | Colwellia | Colwellia sp. 3 | unclassified | unclassified | unclassified | unclassified | unclassified | 4,6 | 1,356465997 | 0,000453749 | 0,00011979 | 6 - | 0,000611818 - | 2 - | 0,00023993 - | 5 - | 0,000473121 - | 5 - | 0,000470147 - | 5 - | 0,00047373 - |
| 1881 | 1 | 0.2um_NOKCPHJN_900409 | Bacteria | Proteobacteri | Gammaprotec | Pseudomonad | Pseudomonad | Pseudomonas | various | information st | Translation, ri | 179ZI | One of the pri | 220664.PFL_5 | 1 | 0,632455532 | 0,000667809 | 0,000398902 | 1 - | 0,000674568 - | 1 - | 0,000793615 - | 0 - | 0 - | 2 - | 0,001244082 - | 1 - | 0,000626781 - |
| 1882 | 1 | 3um_FIJJAFBF_254301 | Bacteria | Proteobacteri | Gammaprotec | Oceanospirilla | Endozoicomor | Endozoicomor | unclassified | information st | Translation, ri | COG0256 | This is one of | 498211.CJA_0 | 0,8 | 0,748331477 | 0,000609685 | 0,000542678 | 1 - | 0,000755981 - | 1 - | 0,000889396 - | 2 - | 0,001403047 - | 0 - | 0 - | 0 - | 0 - |
| 1883 | 1 | 0.2um_NOKCPHJN_198779 | Bacteria | Proteobacteri | Gammaprotec | Vibrionales | Vibrionaceae | Vibrio | Vibrio breoga | unclassified | unclassified | unclassified | unclassified | unclassified | 0,2 | 0,4 | 3,94E-05 | 7,88E-05 | 0 - | 0 - | 0 - | 0 - | 1 - | 0,000197038 - | 0 - | 0 - | 0 - | 0 - |
| 1884 | 1 | 0.2um_NOKCPHJN_975731 | Eukaryota | Arthropoda | Insecta | Hymenoptera | Formicidae | Harpegnathos | Harpegnathos | cellular proces | Signal transdu | 0N554 | cadherin, EGF | 31033.ENSTRU | 0,2 | 0,4 | 7,50E-05 | 0,000149904 | 1 - | 0,00037476 - | 0 - | 0 - | 0 - | 0 - | 0 - | 0 - | 0 - | 0 - |
| 1885 | 1 | 3um_FIJJAFBF_788038 | Bacteria | Proteobacteri | Alphaproteob | Rhodobactera | unclassified | unclassified | Rhodobactera | information st | Translation, ri | COG0255 | 50s ribosomal | 644076.SCH4 | 0,4 | 0,489897949 | 0,000578375 | 0,000712232 | 1 - | 0,001328695 - | 1 - | 0,00156318 - | 0 - | 0 - | 0 - | 0 - | 0 - | 0 - |
| 1886 | 1 | 3um_FIJJAFBF_630935 | Viruses | unclassified | unclassified | unclassified | unclassified | unclassified | uncultured M | unclassified | unclassified | unclassified | unclassified | unclassified | 0,2 | 0,4 | 3,43E-05 | 6,87E-05 | 0 - | 0 - | 1 - | 0,000171664 - | 0 - | 0 - | 0 - | 0 - | 0 - | 0 - |
| 1887 | 1 | 3um_FIJJAFBF_486460 | Eukaryota | Streptophyta | Liliopsida | Arecales | Arecaceae | various | various | information st | Translation, ri | KOG0887 | Ribosomal pro | 3847.GLYMA0 | 0,6 | 0,489897949 | 0,000355393 | 0,00029063 | 1 - | 0,000621942 - | 0 - | 0 - | 1 - | 0,00057714 - | 0 - | 0 - | 1 - | 0,000577883 - |
| 1888 | 1 | 3um_FIJJAFBF_226993 | Bacteria | unclassified | unclassified | unclassified | unclassified | unclassified | uncultured ba | metabolism | Inorganic ion | 09C0C | TonB depende | 742767.HMPR | 0,8 | 0,4 | 0,000149846 | 7,71E-05 | 0 - | 0 - | 1 - | 0,000222829 - | 1 - | 0,00017576 - | 1 - | 0,000174655 - | 1 - | 0,000175986 - |
| 1889 | 1 | 3um_FIJJAFBF_09654 | Bacteria | Proteobacteri | Gammaprotec | Cellvibrionales | Spongiibacter | Spongiibacter | Spongiibacter | cellular proces | Intracellular tr | 05E4G | Type IV pilus | 247633.GP214 | 0,4 | 0,8 | 4,69E-05 | 9,38E-05 | 2 - | 0,000234476 - | 0 - | 0 - | 0 - | 0 - | 0 - | 0 - | 0 - | 0 - |
| 1890 | 1 | 3um_FIJJAFBF_389027 | Eukaryota | Bacillariophyt | Coscinodiscop | Thalassiosirale | Thalassiosirac | Thalassiosira | Thalassiosira | unclassified | unclassified | unclassified | unclassified | unclassified | 0,2 | 0,4 | 5,07E-05 | 0,000101404 | 0 - | 0 - | 0 - | 0 - | 1 - | 0,00025351 - | 0 - | 0 - | 0 - | 0 - |
| 1891 | 1 | 3um_FIJJAFBF_654287 | Bacteria | Proteobacteri | Gammaprotec | Cellvibrionales | Spongiibacter | various | various | poorly charact | Function unkn | 129NV | NA | 575588.HMPR | 0,8 | 0,4 | 0,00022469 | 0,000112666 | 1 - | 0,000297267 - | 0 - | 0 - | 1 - | 0,000275853 - | 1 - | 0,00027412 - | 1 - | 0,000276208 - |
| 1892 | 1 | 0.2um_NOKCPHJN_851899 | Eukaryota | Bacillariophyt | Bacillariophyc | Bacillariales | Bacillariaceae | Fragilariopsis | Fragilariopsis | unclassified | unclassified | unclassified | unclassified | unclassified | 4,6 | 0,8 | 0,001589071 | 0,00043545 | 4 - | 0,00137559 - | 6 - | 0,002427527 - | 5 - | 0,001595622 - | 4 - | 0,001268476 - | 4 - | 0,001278141 - |
| 1893 | 1 | 3um_FIJJAFBF_191787 | Eukaryota | unclassified | Dictyochophy | Pedinellales | unclassified | Apedinella | Apedinella rad | cellular proces | Posttranslatio | KOG0730 | cell division cy | 45157.CMV01 | 5,6 | 0,8 | 0,001600029 | 0,000185562 | 5 - | 0,001451885 - | 5 - | 0,001708111 - | 6 - | 0,001616756 - | 7 - | 0,001874362 - | 5 - | 0,001349031 - |
| 1894 | 1 | 0.2um_NOKCPHJN_312733 | Eukaryota | Phaeophyceae | unclassified | Ectocarpales | Ectocarpaceae | Ectocarpus | Ectocarpus sil | cellular proces | Signal transdu | KOG0613 | fibronectin typ | 67593.Physo1 | 0,4 | 0,489897949 | 8,54E-06 | 1,05E-05 | 1 - | 2,21E-05 - | 0 - | 0 - | 0 - | 0 - | 0 - | 0 - | 1 - | 2,06E-05 - |
| 1895 | 1 | 0.2um_NOKCPHJN_515544 | Bacteria | Proteobacteri | Deltaproteoba | unclassified | unclassified | unclassified | Deltaproteoba | cellular proces | Signal transdu | 16QED | Histidine kina | 526222.Desal_ | 0,2 | 0,4 | 2,92E-05 | 5,84E-05 | 0 - | 0 - | 1 - | 0,000145926 - | 0 - | 0 - | 0 - | 0 - | 0 - | 0 - |
| 1896 | 1 | 0.2um_NOKCPHJN_866315 | unclassified | unclassified | unclassified | unclassified | unclassified | unclassified | unclassified | unclassified | unclassified | unclassified | unclassified | unclassified | 0,8 | 0,4 | 0,000221684 | 0,000111159 | 1 - | 0,00029329 - | 0 - | 0 - | 1 - | 0,000272163 - | 1 - | 0,000270453 - | 1 - | 0,000272513 - |
| 1897 | 1 | 3um_FIJJAFBF_772645 | unclassified | unclassified | unclassified | unclassified | unclassified | unclassified | unclassified | unclassified | unclassified | unclassified | unclassified | unclassified | 0,2 | 0,4 | 3,27E-05 | 6,55E-05 | 0 - | 0 - | 0 - | 0 - | 0 - | 0 - | 1 - | 0,000163695 - | 0 - | 0 - |
| 1898 | 1 | 0.2um_NOKCPHJN_344904 | Bacteria | Bacteroidetes | Flavobacteriia | Flavobacterial | Cryomorphace | unclassified | Cryomorphace | metabolism | Nucleotide tra | COG0138 | bifunctional p | 655815.ZPR_0 | 0,2 | 0,4 | 6,32E-05 | 0,000126328 | 0 - | 0 - | 0 - | 0 - | 0 - | 0 - | 0 - | 0 - | 1 - | 0,00031582 - |
| 1899 | 1 | 3um_FIJJAFBF_537760 | Bacteria | Bacteroidetes | Flavobacteriia | Flavobacterial | Crocinitomica | unclassified | Crocinitomica | unclassified | unclassified | unclassified | unclassified | unclassified | 0,4 | 0,489897949 | 0,000150233 | 0,000186236 | 0 - | 0 - | 1 - | 0,000421102 - | 0 - | 0 - | 1 - | 0,000330063 - | 0 - | 0 - |
| 1900 | 2 | 0.2um_NOKCPHJN_05620;3um | Bacteria | Bacteroidetes | Cytophagia | Cytophagales | Flammeovirga | unclassified | Flammeovirga | cellular proces | Cell cycle cont | 05CR9 | regulator of c | 391587.KAOT | 0,2 | 0,4 | 5,82E-05 | 0,000116402 | 0 - | 0 - | 0 - | 0 - | 0 - | 0 - | 0 - | 0 - | 1 - | 0,000291005 - |
| 1901 | 5 | 3um_FIJJAFBF_124352;3um_FI | Bacteria | Proteobacteri | Alphaproteob | Rhodobactera | various | various | various | information st | Transcription | 05VEQ | Cold shock pro | various | 0,4 | 0,489897949 | 0,000497575 | 0,000610086 | 1 - | 0,001289615 - | 0 - | 0 - | 0 - | 0 - | 0 - | 0 - | 1 - | 0,001198257 - |
| 1902 | 1 | 3um_FIJJAFBF_424816 | Bacteria | Proteobacteri | Alphaproteob | Rhodobactera | Rhodobactera | unclassified | Rhodobactera | unclassified | unclassified | unclassified | unclassified | unclassified | 0,4 | 0,489897949 | 5,09E-05 | 6,23E-05 | 0 - | 0 - | 0 - | 0 - | 1 - | 0,000127151 - | 0 - | 0 - | 1 - | 0,000127315 - |
| 1903 | 1 | 3um_FIJJAFBF_343898 | Eukaryota | unclassified | unclassified | Isochrysidales | Noelaerhabda | Emiliania | Emiliania hux | metabolism | Energy produc | COG0056 | Produces ATP | 41431.PCC88 | 1,4 | 0,489897949 | 0,000241023 | 8,09E-05 | 2 - | 0,000350075 - | 1 - | 0,000205928 - | 1 - | 0,000162429 - | 1 - | 0,000161408 - | 2 - | 0,000325275 - |
| 1904 | 1 | 0.2um_NOKCPHJN_1108412 | Eukaryota | unclassified | unclassified | Kinetoplastida | Trypanosomat | Leishmania | Leishmania m | unclassified | unclassified | unclassified | unclassified | unclassified | 0,8 | 0,4 | 0,000130737 | 6,56E-05 | 1 - | 0,000172966 - | 0 - | 0 - | 1 - | 0,000160506 - | 1 - | 0,000159498 - | 1 - | 0,000160713 - |
| 1905 | 3 | 0.2um_NOKCPHJN_292421;0.2 | Bacteria | Bacteroidetes | Flavobacteriia | Flavobacterial | Crocinitomica | unclassified | various | unclassified | unclassified | unclassified | unclassified | unclassified | 0,8 | 0,4 | 0,000818315 | 0,000410327 | 1 - | 0,00108264 - | 0 - | 0 - | 1 - | 0,001004651 - | 1 - | 0,000998338 - | 1 - | 0,001005944 - |
| 1906 | 1 | 0.2um_NOKCPHJN_62798 | Bacteria | Bacteroidetes | Flavobacteriia | Flavobacterial | Crocinitomica | unclassified | Crocinitomica | poorly charact | Function unkn | 07EF7 | domain protei | 649349.Lbys_ | 0,6 | 0,489897949 | 6,12E-05 | 5,00E-05 | 1 - | 0,000107336 - | 0 - | 0 - | 1 - | 9,96E-05 - | 1 - | 9,90E-05 - | 0 - | 0 - |
| 1907 | 1 | 0.2um_NOKCPHJN_570486 | Eukaryota | Phaeophyceae | unclassified | Ectocarpales | Ectocarpaceae | Ectocarpus | Ectocarpus sil | cellular proces | Posttranslatio | 14ZJ0 | STI1 | 67593.Physo1 | 0,6 | 0,489897949 | 0,000240192 | 0,000199472 | 0 - | 0 - | 1 - | 0,000466832 - | 1 - | 0,000368221 - | 1 - | 0,000365907 - | 0 - | 0 - |
| 1908 | 1 | 3um_FIJJAFBF_535056 | Bacteria | Bacteroidetes | Flavobacteriia | Flavobacterial | Flavobacteria | Algibacter | Algibacter pe | poorly charact | Function unkn | 0PAPS | NA | 313603.FB217 | 1,6 | 1,019803903 | 0,000942297 | 0,000593178 | 2 - | 0,001169251 - | 2 - | 0,001375599 - | 3 - | 0,001627535 - | 1 - | 0,000539102 - | 0 - | 0 - |
| 1909 | 1 | 0.2um_NOKCPHJN_947667 | unclassified | unclassified | unclassified | unclassified | unclassified | unclassified | unclassified | unclassified | unclassified | unclassified | unclassified | unclassified | 1 | 0 | 0,000700947 | 6,86E-05 | 1 - | 0,000707208 - | 1 - | 0,000832015 - | 1 - | 0,000656264 - | 1 - | 0,00065214 - | 1 - | 0,000657109 - |
| 1910 | 1 | 0.2um_NOKCPHJN_262403 | Eukaryota | unclassified | Oomycetes | Peronosporale | Peronosporac | Phytophthora | Phytophthora | unclassified | unclassified | unclassified | unclassified | unclassified | 0,2 | 0,4 | 2,46E-05 | 4,92E-05 | 0 - | 0 - | 0 - | 0 - | 1 - | 0,000123112 - | 0 - | 0 - | 0 - | 0 - |
| 1911 | 1 | 3um_FIJJAFBF_323987 | Bacteria | Proteobacteri | Gammaprotec | Alteromonada | Aestuariibacte | Aestuariibacte | bacterium TM | unclassified | unclassified | 0YV5D | NA | 203122.Sde_3 | 0,4 | 0,489897949 | 0,000206821 | 0,000253649 | 1 - | 0,000537999 - | 0 - | 0 - | 0 - | 0 - | 1 - | 0,000496106 - | 0 - | 0 - |
| 1912 | 1 | 3um_FIJJAFBF_725633 | unclassified | unclassified | unclassified | unclassified | unclassified | unclassified | unclassified | unclassified | unclassified | unclassified | unclassified | unclassified | 0,8 | 0,4 | 0,000374483 | 0,000187777 | 1 - | 0,000495445 - | 0 - | 0 - | 1 - | 0,000459756 - | 1 - | 0,000456866 - | 1 - | 0,000460347 - |
| 1913 | 1 | 0.2um_NOKCPHJN_790587 | unclassified | unclassified | unclassified | unclassified | unclassified | unclassified | unclassified | unclassified | unclassified | unclassified | unclassified | unclassified | 0,6 | 0,489897949 | 0,000168907 | 0,000140196 | 0 - | 0 - | 1 - | 0,000327523 - | 1 - | 0,000258339 - | 0 - | 0 - | 1 - | 0,000258671 - |

| | | | | | | | | | | | | | | | | | | | | | | | | | | | | |
|---|---|---|---|---|---|---|---|---|---|---|---|---|---|---|---|---|---|---|---|---|---|---|---|---|---|---|---|---|
| 1914 | 1 | 0.2um_NOKCPHJN_75313 | Bacteria | Proteobacteri | Gammaprotec | Alteromonada | Alteromonada | Glaciecola | Glaciecola sp. | metabolism | Carbohydrate | 16QEQ | hydrolase, fan | 455436.GHTC | 0,8 | 0,748331477 | 7,64E-05 | 7,35E-05 | 2 - | 0,000198402 - | 0 - | 0 - | 0 - | 0 - | 1 - | 9,15E-05 - | 1 - | 9,22E-05 - |
| 1915 | 1 | 3um_FIJJAFBF_564131 | Bacteria | Bacteroidetes | Flavobacteriia | Flavobacterial | Flavobacteria | Mesoflavibact | various | metabolism | Carbohydrate | COG0057 | glyceraldehyd | 391587.KAOT | 0,4 | 0,489897949 | 9,80E-05 | 0,000120065 | 0 - | 0 - | 0 - | 0 - | 1 - | 0,000245851 - | 1 - | 0,000244306 - | 0 - | 0 - |
| 1916 | 1 | 0.2um_NOKCPHJN_460006 | Bacteria | Planctomycet | Phycisphaerae | Phycisphaeral | Phycisphaerac | Phycisphaera | Phycisphaera | metabolism | Energy produc | 05C26 | Dehydrogenas | 502025.Hoch_ | 0,6 | 0,489897949 | 0,000105942 | 8,66E-05 | 1 - | 0,000185399 - | 0 - | 0 - | 1 - | 0,000172044 - | 0 - | 0 - | 1 - | 0,000172265 - |
| 1917 | 1 | 3um_FIJJAFBF_683762 | Bacteria | Proteobacteri | Gammaprotec | unclassified | unclassified | unclassified | Gammaprotec | cellular proces | Intracellular tr | 0QITE | Outer membr | 207949.RED6 | 0,2 | 0,4 | 3,42E-05 | 6,83E-05 | 0 - | 0 - | 0 - | 0 - | 0 - | 0 - | 0 - | 0 - | 1 - | 0,000170821 - |
| 1918 | 1 | 3um_FIJJAFBF_753362 | Bacteria | Cyanobacteria | unclassified | Oscillatoriales | Oscillatoriace | Moorea | Moorea produ | cellular proces | Signal transdu | 1AK23 | HisKA | 760192.Halhy | 0,4 | 0,489897949 | 7,06E-05 | 8,65E-05 | 0 - | 0 - | 0 - | 0 - | 0 - | 0 - | 1 - | 0,000175794 - | 1 - | 0,000177134 - |
| 1919 | 1 | 3um_FIJJAFBF_269588 | Bacteria | Proteobacteri | Alphaproteob | Rhodobactera | Rhodobactera | unclassified | Rhodobactera | cellular proces | Defense mech | COG0610 | Type I site-spe | 388401.RB21 | 1 | 0 | 7,97E-05 | 7,80E-06 | 1 - | 8,04E-05 - | 1 - | 9,46E-05 - | 1 - | 7,46E-05 - | 1 - | 7,41E-05 - | 1 - | 7,47E-05 - |
| 1920 | 1 | 0.2um_NOKCPHJN_1028962 | unclassified | unclassified | unclassified | unclassified | unclassified | unclassified | unclassified | unclassified | unclassified | unclassified | unclassified | unclassified | 1 | 0 | 0,000371442 | 3,63E-05 | 1 - | 0,00037476 - | 1 - | 0,000440897 - | 1 - | 0,000347764 - | 1 - | 0,000345578 - | 1 - | 0,000348212 - |
| 1921 | 1 | 3um_FIJJAFBF_177926 | Eukaryota | Bacillariophyt | Coscinodiscop | Coscinodiscale | Coscinodiscac | Coscinodiscus | Coscinodiscus | unclassified | unclassified | unclassified | unclassified | unclassified | 3,8 | 0,748331477 | 0,001041908 | 0,000171281 | 4 - | 0,001117119 - | 3 - | 0,0009857 - | 4 - | 0,001036646 - | 3 - | 0,000772599 - | 5 - | 0,001297476 - |
| 1922 | 1 | 3um_FIJJAFBF_438692 | Eukaryota | Bacillariophyt | Coscinodiscop | Thalassiosirale | Thalassiosirac | Thalassiosira | Thalassiosira | unclassified | unclassified | unclassified | unclassified | unclassified | 2,2 | 0,748331477 | 0,001054945 | 0,000302069 | 2 - | 0,000996521 - | 1 - | 0,000586193 - | 2 - | 0,000924736 - | 3 - | 0,001378387 - | 3 - | 0,001388889 - |
| 1923 | 1 | 0.2um_NOKCPHJN_63758 | Bacteria | Bacteroidetes | Flavobacteriia | Flavobacterial | Crocinitomica | unclassified | Crocinitomica | metabolism | Inorganic ion | 05C27 | TonB-depend | 470145.BACC | 0,4 | 0,489897949 | 3,19E-05 | 3,91E-05 | 0 - | 0 - | 0 - | 0 - | 0 - | 0 - | 1 - | 7,95E-05 - | 1 - | 8,01E-05 - |
| 1924 | 1 | 0.2um_NOKCPHJN_596448 | Bacteria | Bacteroidetes | Bacteroidia | Marinilabiliale | Prolixibacter | Prolixibacter | Prolixibacter | unclassified | unclassified | unclassified | unclassified | unclassified | 1 | 0 | 0,000265803 | 2,60E-05 | 1 - | 0,000268177 - | 1 - | 0,000315504 - | 1 - | 0,000248859 - | 1 - | 0,000247295 - | 1 - | 0,000249179 - |
| 1925 | 1 | 3um_FIJJAFBF_104645 | Bacteria | Proteobacteri | Gammaprotec | Alteromonada | Pseudoalteror | Pseudoalteror | Pseudoalteror | unclassified | unclassified | unclassified | unclassified | unclassified | 0,2 | 0,4 | 8,81E-05 | 0,000176269 | 1 - | 0,000440673 - | 0 - | 0 - | 0 - | 0 - | 0 - | 0 - | 0 - | 0 - |
| 1926 | 1 | 0.2um_NOKCPHJN_938882 | Viruses | unclassified | unclassified | unclassified | Phycodnavirid | Prasinovirus | Bathycoccus s | unclassified | unclassified | unclassified | unclassified | unclassified | 0,6 | 0,489897949 | 0,000279375 | 0,000228499 | 1 - | 0,00048991 - | 0 - | 0 - | 0 - | 0 - | 1 - | 0,000451762 - | 1 - | 0,000455204 - |
| 1927 | 1 | 3um_FIJJAFBF_257135 | Eukaryota | Bacillariophyt | Mediophyceae | Biddulphiales | Biddulphiacea | Biddulphia | Biddulphia tri | information st | Translation, ri | 08UHY | One of the pri | 489825.LYNG | 2,6 | 1,019803903 | 0,001398853 | 0,000478529 | 2 - | 0,00112428 - | 1 - | 0,000661345 - | 4 - | 0,002086583 - | 3 - | 0,001555103 - | 3 - | 0,001566952 - |
| 1928 | 1 | 3um_FIJJAFBF_715097 | Eukaryota | unclassified | Pelagophycea | Pelagomonad | unclassified | Aureococcus | Aureococcus | metabolism | Energy produc | 1DYB8 | PsaA and PsaE | 3055.DAA014 | 10 | 2,19089023 | 0,001136561 | 0,000180411 | 9 - | 0,001049527 - | 7 - | 0,000960358 - | 9 - | 0,000973924 - | 12 - | 0,001290405 - | 13 - | 0,00140859 - |
| 1929 | 1 | 3um_FIJJAFBF_708523 | Bacteria | Cyanobacteria | unclassified | Synechococca | Synechococca | Synechococcu | Synechococcu | poorly charact | Function unkn | 11FY4 | C-phycoerythr | 64471.sync_0 | 0,6 | 0,489897949 | 0,000305551 | 0,000249871 | 1 - | 0,000534719 - | 0 - | 0 - | 1 - | 0,0004962 - | 0 - | 0 - | 1 - | 0,000496838 - |
| 1930 | 1 | 3um_FIJJAFBF_514244 | unclassified | unclassified | unclassified | unclassified | unclassified | unclassified | unclassified | unclassified | unclassified | unclassified | unclassified | unclassified | 0,2 | 0,4 | 0,000103688 | 0,000207377 | 0 - | 0 - | 1 - | 0,000518442 - | 0 - | 0 - | 0 - | 0 - | 0 - | 0 - |
| 1931 | 1 | 3um_FIJJAFBF_99502 | Bacteria | Bacteroidetes | Flavobacteriia | Flavobacterial | Flavobacteria | Polaribacter | Polaribacter i | cellular proces | Cell wall/mem | COG2885 | Ompa motb d | 313594.PI23P | 0,2 | 0,4 | 3,89E-05 | 7,78E-05 | 1 - | 0,000194443 - | 0 - | 0 - | 0 - | 0 - | 0 - | 0 - | 0 - | 0 - |
| 1932 | 1 | 0.2um_NOKCPHJN_60577 | Bacteria | Bacteroidetes | Flavobacteriia | Flavobacterial | Crocinitomica | unclassified | Crocinitomica | poorly charact | Function unkn | 05CZH | peptidase | 926562.Oweh | 0,2 | 0,4 | 2,20E-05 | 4,39E-05 | 0 - | 0 - | 0 - | 0 - | 1 - | 0,00010982 - | 0 - | 0 - | 0 - | 0 - |
| 1933 | 1 | 3um_FIJJAFBF_576716 | Bacteria | Proteobacteri | Alphaproteob | Sphingomona | Sphingomona | Sphingomona | various | unclassified | unclassified | unclassified | unclassified | unclassified | 0,2 | 0,4 | 1,56E-05 | 3,13E-05 | 0 - | 0 - | 0 - | 0 - | 0 - | 0 - | 1 - | 7,81E-05 - | 0 - | 0 - |
| 1934 | 1 | 3um_FIJJAFBF_550914 | Eukaryota | Bacillariophyt | Coscinodiscop | Thalassiosirale | Thalassiosirac | Thalassiosira | Thalassiosira | unclassified | unclassified | unclassified | unclassified | unclassified | 2,2 | 1,166190379 | 0,002277891 | 0,001103966 | 1 - | 0,00108264 - | 1 - | 0,001273702 - | 4 - | 0,004018605 - | 2 - | 0,001996675 - | 3 - | 0,003017833 - |
| 1935 | 1 | 0.2um_NOKCPHJN_300636 | Bacteria | Proteobacteri | Gammaprotec | Oceanospirilla | unclassified | unclassified | Oceanospirilla | information st | Translation, ri | COG0050 | This protein p | 553480.PAU_ | 1,2 | 0,4 | 0,000265117 | 9,26E-05 | 2 - | 0,000445146 - | 1 - | 0,000261853 - | 1 - | 0,00020654 - | 1 - | 0,000205242 - | 1 - | 0,000206806 - |
| 1936 | 2 | 0.2um_NOKCPHJN_779798;3u | Bacteria | Bacteroidetes | Flavobacteriia | Flavobacterial | Flavobacteria | various | various | various | various | various | various | various | 0,8 | 0,979795897 | 8,57E-05 | 0,000104927 | 0 - | 0 - | 0 - | 0 - | 0 - | 0 - | 2 - | 0,000213365 - | 2 - | 0,000214991 - |
| 1937 | 1 | 3um_FIJJAFBF_38454 | Bacteria | Proteobacteri | Alphaproteob | Rhodobactera | Rhodobactera | unclassified | Rhodobactera | cellular proces | Cell wall/mem | 08ZG0 | Ompa motb d | 391619.RGBS | 0,6 | 0,489897949 | 0,000224158 | 0,000183342 | 1 - | 0,000393246 - | 0 - | 0 - | 1 - | 0,000364918 - | 1 - | 0,000362625 - | 0 - | 0 - |
| 1938 | 2 | 0.2um_NOKCPHJN_854801;0.2 | Bacteria | Bacteroidetes | Flavobacteriia | Flavobacterial | Flavobacteria | Polaribacter | Polaribacter s | metabolism | Coenzyme tra | 099YY | Nicotinate-nu | 313598.MED1 | 0,2 | 0,4 | 7,21E-05 | 0,000144294 | 0 - | 0 - | 1 - | 0,000360734 - | 0 - | 0 - | 0 - | 0 - | 0 - | 0 - |
| 1939 | 1 | 3um_FIJJAFBF_204503 | Bacteria | Actinobacteria | Acidimicrobiia | Acidimicrobial | Acidimicrobia | Acidimicrobiu | Acidimicrobiu | unclassified | unclassified | unclassified | unclassified | unclassified | 0,8 | 0,4 | 0,000241031 | 0,00012086 | 1 - | 0,000318887 - | 0 - | 0 - | 1 - | 0,000295915 - | 1 - | 0,000294056 - | 1 - | 0,000296296 - |
| 1940 | 1 | 3um_FIJJAFBF_16490 | Bacteria | Proteobacteri | Gammaprotec | Alteromonada | Alteromonada | Paraglaciecola | Paraglaciecola | information st | Translation, ri | 0QTKX | Seems to be t | 1085623.GNIT | 0,6 | 0,489897949 | 0,000442838 | 0,000366108 | 1 - | 0,000712958 - | 1 - | 0,00083878 - | 0 - | 0 - | 0 - | 0 - | 1 - | 0,000662451 - |
| 1941 | 1 | 3um_FIJJAFBF_715058 | Eukaryota | Bacillariophyt | Mediophyceae | Cymatosirales | Cymatosiracea | Plagiogrammo | Plagiogrammo | information st | Translation, ri | COG0052 | 30S ribosomal | 35128.Thapsd | 0,2 | 0,4 | 7,03E-05 | 0,000140635 | 0 - | 0 - | 0 - | 0 - | 0 - | 0 - | 1 - | 0,000351588 - | 0 - | 0 - |
| 1942 | 1 | 0.2um_NOKCPHJN_153284 | Bacteria | Proteobacteri | Gammaprotec | Alteromonada | Pseudoalteror | Pseudoalteror | Pseudoalteror | metabolism | Inorganic ion | 0QN53 | TonB-depend | 983545.Glaag | 4,6 | 0,489897949 | 0,000424525 | 4,15E-05 | 5 - | 0,00046795 - | 4 - | 0,000440426 - | 5 - | 0,000434241 - | 4 - | 0,00034521 - | 5 - | 0,0004348 - |
| 1943 | 1 | 3um_FIJJAFBF_163897 | Bacteria | Proteobacteri | Alphaproteob | Rhodobactera | unclassified | unclassified | Rhodobactera | unclassified | unclassified | unclassified | unclassified | unclassified | 0,4 | 0,489897949 | 0,000104565 | 0,000129624 | 0 - | 0 - | 1 - | 0,000293096 - | 0 - | 0 - | 1 - | 0,000229731 - | 0 - | 0 - |
| 1944 | 1 | 0.2um_NOKCPHJN_1100554 | unclassified | unclassified | unclassified | unclassified | unclassified | unclassified | unclassified | unclassified | unclassified | unclassified | unclassified | unclassified | 0,2 | 0,4 | 5,53E-05 | 0,000110638 | 0 - | 0 - | 1 - | 0,000276595 - | 0 - | 0 - | 0 - | 0 - | 0 - | 0 - |
| 1945 | 1 | 3um_FIJJAFBF_322515 | unclassified | unclassified | unclassified | unclassified | unclassified | unclassified | unclassified | unclassified | unclassified | unclassified | unclassified | unclassified | 0,2 | 0,4 | 3,29E-05 | 6,58E-05 | 1 - | 0,000164529 - | 0 - | 0 - | 0 - | 0 - | 0 - | 0 - | 0 - | 0 - |
| 1946 | 1 | 3um_FIJJAFBF_100050 | Bacteria | Proteobacteri | Alphaproteob | Rhodospirillal | Rhodospirillac | Candidatus En | Candidatus En | unclassified | unclassified | unclassified | unclassified | unclassified | 3,6 | 0,489897949 | 0,00066926 | 0,000138266 | 4 - | 0,000744746 - | 4 - | 0,000876177 - | 3 - | 0,000518323 - | 3 - | 0,000515066 - | 4 - | 0,000691987 - |
| 1947 | 1 | 3um_FIJJAFBF_545830 | Bacteria | Bacteroidetes | Flavobacteriia | Flavobacterial | Flavobacteria | Polaribacter | Polaribacter h | information st | Translation, ri | 05G6W | One of the pri | 313594.PI23P | 1,8 | 0,748331477 | 0,000756012 | 0,000274994 | 1 - | 0,000436288 - | 1 - | 0,000513283 - | 2 - | 0,000809719 - | 2 - | 0,00080463 - | 3 - | 0,001216142 - |
| 1948 | 1 | 3um_FIJJAFBF_177155 | unclassified | unclassified | unclassified | unclassified | unclassified | unclassified | unclassified | unclassified | unclassified | unclassified | unclassified | unclassified | 1 | 0 | 0,000294635 | 2,88E-05 | 1 - | 0,000297267 - | 1 - | 0,000349728 - | 1 - | 0,000275853 - | 1 - | 0,00027412 - | 1 - | 0,000276208 - |
| 1949 | 1 | 0.2um_NOKCPHJN_619316 | Eukaryota | Bacillariophyt | Bacillariophyc | Bacillariales | Bacillariaceae | Fragilariopsis | Fragilariopsis | unclassified | unclassified | unclassified | unclassified | unclassified | 2,2 | 0,4 | 0,001972925 | 0,000409033 | 3 - | 0,002712181 - | 2 - | 0,002127214 - | 2 - | 0,001677871 - | 2 - | 0,001667327 - | 2 - | 0,001680031 - |
| 1950 | 1 | 3um_FIJJAFBF_469510 | Bacteria | Bacteroidetes | Flavobacteriia | Flavobacterial | Flavobacteria | Mesoflavibact | Mesoflavibact | unclassified | unclassified | unclassified | unclassified | unclassified | 0,2 | 0,4 | 8,64E-05 | 0,000172796 | 1 - | 0,000431989 - | 0 - | 0 - | 0 - | 0 - | 0 - | 0 - | 0 - | 0 - |
| 1951 | 1 | 3um_FIJJAFBF_594156 | Eukaryota | unclassified | Raphidophyce | Chattonellales | Chattonellace | Heterosigma | Heterosigma | metabolism | Energy produc | COG0055 | Produces ATP | 35128.Thapsd | 5 | 0 | 0,002556396 | 0,000250129 | 5 - | 0,002579231 - | 5 - | 0,003034408 - | 5 - | 0,002393434 - | 5 - | 0,002378393 - | 5 - | 0,002396514 - |
| 1952 | 1 | 0.2um_NOKCPHJN_938763 | Eukaryota | unclassified | Oomycetes | Peronosporale | Peronosporac | Phytophthora | various | unclassified | unclassified | unclassified | unclassified | unclassified | 1 | 0 | 0,000406156 | 3,97E-05 | 1 - | 0,000409784 - | 1 - | 0,000482102 - | 1 - | 0,000380265 - | 1 - | 0,000377875 - | 1 - | 0,000380755 - |
| 1953 | 1 | 3um_FIJJAFBF_769062 | Eukaryota | Phaeophyceae | unclassified | Ectocarpales | Ectocarpaceae | Ectocarpus | Ectocarpus sili | cellular proces | Signal transdu | KOG0613 | fibronectin typ | 67593.Physo1 | 0,4 | 0,489897949 | 7,11E-05 | 8,72E-05 | 1 - | 0,000184231 - | 0 - | 0 - | 0 - | 0 - | 0 - | 0 - | 1 - | 0,00017118 - |
| 1954 | 1 | 0.2um_NOKCPHJN_112963 | Bacteria | Proteobacteri | Alphaproteob | Rhodobactera | Rhodobactera | Thalassobius | Thalassobius s | cellular proces | Posttranslatio | 01QWG | Prevents misf | 391619.RGBS | 1,8 | 0,748331477 | 0,000280297 | 0,000114429 | 3 - | 0,000477462 - | 1 - | 0,000187241 - | 2 - | 0,000295378 - | 2 - | 0,000293522 - | 1 - | 0,000147879 - |
| 1955 | 1 | 0.2um_NOKCPHJN_341236 | Eukaryota | unclassified | Pelagophycea | Pelagomonad | unclassified | Aureococcus | Aureococcus | unclassified | unclassified | unclassified | unclassified | unclassified | 0,8 | 0,4 | 0,000107327 | 5,50E-05 | 1 - | 0,000133273 - | 1 - | 0,000156793 - | 1 - | 0,000123673 - | 1 - | 0,000122896 - | 0 - | 0 - |
| 1956 | 2 | 0.2um_NOKCPHJN_954744;3u | Eukaryota | Eustigmatoph | unclassified | Eustigmatales | Monodopsida | Nannochlorop | Nannochlorop | unclassified | unclassified | unclassified | unclassified | unclassified | 1 | 0 | 0,000234912 | 2,30E-05 | 1 - | 0,00023701 - | 1 - | 0,000278838 - | 1 - | 0,000219937 - | 1 - | 0,000218555 - | 1 - | 0,00022022 - |
| 1957 | 1 | 0.2um_NOKCPHJN_512123 | Bacteria | Bacteroidetes | Flavobacteriia | Flavobacterial | Flavobacteria | Polaribacter | Polaribacter s | poorly charact | Function unkn | 05DPC | Ragb susd dor | 487797.Flav3 | 0,6 | 0,489897949 | 9,88E-05 | 8,08E-05 | 1 - | 0,000173308 - | 0 - | 0 - | 0 - | 0 - | 1 - | 0,000159813 - | 1 - | 0,000161031 - |
| 1958 | 2 | 0.2um_NOKCPHJN_355825;0.2 | Bacteria | Proteobacteri | Gammaprotec | unclassified | unclassified | unclassified | Gammaprotec | metabolism | Inorganic ion | 173HE | CBS domain p | various | 0,6 | 0,489897949 | 0,00011739 | 9,60E-05 | 1 - | 0,000205854 - | 0 - | 0 - | 0 - | 0 - | 1 - | 0,000189825 - | 1 - | 0,000191271 - |
| 1959 | 1 | 0.2um_NOKCPHJN_192650 | Bacteria | Proteobacteri | Gammaprotec | Arenicellales | Arenicellaceae | Arenicella | Arenicella xan | information st | Translation, ri | 05CEJ | Catalyzes the | 153948.NAL2 | 0,2 | 0,4 | 2,34E-05 | 4,68E-05 | 0 - | 0 - | 0 - | 0 - | 1 - | 0,000116921 - | 0 - | 0 - | 0 - | 0 - |
| 1960 | 1 | 3um_FIJJAFBF_97615 | Bacteria | Bacteroidetes | Flavobacteriia | Flavobacterial | Flavobacteria | Polaribacter | various | information st | Translation, ri | 08UHY | One of the pri | 313598.MED1 | 0,6 | 0,489897949 | 0,000306571 | 0,000250317 | 0 - | 0 - | 0 - | 0 - | 1 - | 0,000511803 - | 1 - | 0,000508587 - | 1 - | 0,000512462 - |
| 1961 | 1 | 3um_FIJJAFBF_583671 | Bacteria | Cyanobacteria | unclassified | Synechococca | Synechococca | Synechococcu | Synechococcu | poorly charact | Function unkn | 06QDP | C-phycoerythr | 32051.SynWH | 0,4 | 0,489897949 | 0,000223693 | 0,000277132 | 0 - | 0 - | 1 - | 0,000625272 - | 1 - | 0,000493192 - | 0 - | 0 - | 0 - | 0 - |
| 1962 | 1 | 3um_FIJJAFBF_646004 | unclassified | unclassified | unclassified | unclassified | unclassified | unclassified | unclassified | unclassified | unclassified | unclassified | unclassified | unclassified | 0,8 | 0,4 | 0,000289448 | 0,000145138 | 1 - | 0,000382943 - | 0 - | 0 - | 1 - | 0,000355357 - | 1 - | 0,000353124 - | 1 - | 0,000355814 - |
| 1963 | 1 | 3um_FIJJAFBF_350452 | Eukaryota | Bacillariophyt | Coscinodiscop | Thalassiosirale | Thalassiosirac | Thalassiosira | Thalassiosira | unclassified | unclassified | unclassified | unclassified | unclassified | 0,2 | 0,4 | 4,28E-05 | 8,55E-05 | 0 - | 0 - | 0 - | 0 - | 0 - | 0 - | 0 - | 0 - | 1 - | 0,000213862 - |
| 1964 | 1 | 3um_FIJJAFBF_254291 | Bacteria | Proteobacteri | Gammaprotec | Cellvibrionales | Spongiibacter | Oceanicoccus | Oceanicoccus | information st | Translation, ri | 16Q16 | Binds the low | 247634.GPB2 | 0,8 | 0,4 | 0,000298574 | 0,000149714 | 1 - | 0,000395017 - | 0 - | 0 - | 1 - | 0,000366562 - | 1 - | 0,000364258 - | 1 - | 0,000367034 - |
| 1965 | 1 | 3um_FIJJAFBF_254290 | Bacteria | Proteobacteri | Gammaprotec | Cellvibrionales | Spongiibacter | Oceanicoccus | Oceanicoccus | information st | Translation, ri | 17C3B | The globular d | 247633.GP21 | 1,4 | 1,019803903 | 0,001079425 | 0,000817636 | 3 - | 0,002413592 - | 0 - | 0 - | 1 - | 0,000746576 - | 1 - | 0,000741884 - | 2 - | 0,001495073 - |
| 1966 | 1 | 3um_FIJJAFBF_295198 | Bacteria | unclassified | unclassified | unclassified | unclassified | unclassified | candidate divi | unclassified | unclassified | unclassified | unclassified | unclassified | 0,4 | 0,489897949 | 5,09E-05 | 6,23E-05 | 0 - | 0 - | 0 - | 0 - | 1 - | 0,00012755 - | 1 - | 0,000126748 - | 0 - | 0 - |
| 1967 | 1 | 3um_FIJJAFBF_74738 | Bacteria | Proteobacteri | Alphaproteob | Rhizobiales | unclassified | Bauldia | Bauldia litoral | poorly charact | Function unkn | 0268K | NA | 349163.Acry_ | 0,4 | 0,489897949 | 0,000113954 | 0,000141262 | 0 - | 0 - | 1 - | 0,000319411 - | 0 - | 0 - | 1 - | 0,000250357 - | 0 - | 0 - |
| 1968 | 1 | 3um_FIJJAFBF_454939 | Bacteria | Bacteroidetes | Flavobacteriia | Flavobacterial | Flavobacteria | Marinirhabdu | Marinirhabdu | unclassified | unclassified | unclassified | unclassified | unclassified | 1 | 0 | 0,000153836 | 1,51E-05 | 1 - | 0,00015521 - | 1 - | 0,000182602 - | 1 - | 0,00014403 - | 1 - | 0,000143125 - | 1 - | 0,000144215 - |
| 1969 | 1 | 0.2um_NOKCPHJN_804742 | Eukaryota | unclassified | Oomycetes | Peronosporale | Peronosporac | Phytophthora | Phytophthora | information st | Translation, ri | COG0052 | 30S ribosomal | 67593.Physo1 | 1,8 | 0,4 | 0,000573786 | 9,56E-05 | 2 - | 0,000656883 - | 1 - | 0,000386404 - | 2 - | 0,000609564 - | 2 - | 0,000605733 - | 2 - | 0,000610348 - |
| 1970 | 1 | 0.2um_NOKCPHJN_215618 | Eukaryota | unclassified | Pelagophycea | Pelagomonad | unclassified | Aureococcus | Aureococcus | unclassified | unclassified | unclassified | unclassified | unclassified | 1,4 | 0,489897949 | 0,000148676 | 4,46E-05 | 1 - | 0,000109208 - | 1 - | 0,000128481 - | 1 - | 0,000101341 - | 2 - | 0,000201408 - | 2 - | 0,000202943 - |
| 1971 | 1 | 0.2um_NOKCPHJN_1013357 | Eukaryota | Nematoda | Chromadorea | Rhabditida | Haemonchida | Haemonchus | Haemonchus | cellular proces | Cell cycle cont | 0V9W7 | retinitis pigme | 8128.ENSONI | 1 | 0 | 0,000294635 | 2,88E-05 | 1 - | 0,000297267 - | 1 - | 0,000349728 - | 1 - | 0,000275853 - | 1 - | 0,00027412 - | 1 - | 0,000276208 - |
| 1972 | 1 | 0.2um_NOKCPHJN_445696 | Bacteria | Proteobacteri | Alphaproteob | Pelagibacteral | Pelagibactera | unclassified | Pelagibactera | cellular proces | Cell wall/mem | 0PD4C | Cytidylyltrans | 313598.MED1 | 0,2 | 0,4 | 0,000101089 | 0,000202178 | 0 - | 0 - | 0 - | 0 - | 1 - | 0,000505446 - | 0 - | 0 - | 0 - | 0 - |
| 1973 | 1 | 0.2um_NOKCPHJN_1105933 | Eukaryota | unclassified | Cryptophyta | Pyrenomonad | Geminigerace | Guillardia | Guillardia thet | poorly charact | Function unkn | arCOG02812 | bacteriorhodo | 272569.rrnAC | 0,8 | 0,4 | 0,000654516 | 0,000336929 | 0 - | 0 - | 1 - | 0,000973301 - | 1 - | 0,000767705 - | 1 - | 0,000762881 - | 1 - | 0,000768693 - |
| 1974 | 1 | 3um_FIJJAFBF_228661 | Bacteria | Bacteroidetes | Flavobacteriia | Flavobacterial | Flavobacteria | Formosa | Formosa sp. H | poorly charact | Function unkn | 0P8NI | Inherit from b | 983544.Lacal_ | 4,2 | 0,748331477 | 0,000885592 | 0,000218111 | 4 - | 0,00084321 - | 5 - | 0,001240023 - | 5 - | 0,000978086 - | 3 - | 0,000583164 - | 4 - | 0,000783476 - |
| 1975 | 1 | 3um_FIJJAFBF_594175 | Eukaryota | Phaeophyceae | unclassified | Ectocarpales | various | various | various | metabolism | Energy produc | COG1290 | Component of | 35128.Thapsd | 4,4 | 0,489897949 | 0,001768089 | 0,000164238 | 4 - | 0,001631514 - | 4 - | 0,00191944 - | 4 - | 0,001513986 - | 5 - | 0,00188059 - | 5 - | 0,001894918 - |
| 1976 | 1 | 0.2um_NOKCPHJN_227678 | Bacteria | Bacteroidetes | Flavobacteriia | Flavobacterial | Flavobacteria | Formosa | Formosa sp. H | metabolism | Inorganic ion | 05C27 | TonB-depend | 867900.Celly_ | 5,2 | 1,939071943 | 0,000435737 | 0,000141403 | 5 - | 0,000436722 - | 2 - | 0,000205518 - | 6 - | 0,000486315 - | 8 - | 0,000644345 - | 5 - | 0,000405784 - |
| 1977 | 1 | 3um_FIJJAFBF_801470 | Eukaryota | Arthropoda | Insecta | Hemiptera | Aphididae | Melanaphis | Melanaphis sa | poorly charact | Function unkn | 0SVN9 | NA | 7029.ACYPI00 | 0,2 | 0,4 | 5,13E-05 | 0,000102566 | 1 - | 0,000256415 - | 0 - | 0 - | 0 - | 0 - | 0 - | 0 - | 0 - | 0 - |
| 1978 | 1 | 3um_FIJJAFBF_213931 | Bacteria | Bacteroidetes | Flavobacteriia | Flavobacterial | Flavobacteria | Formosa | Formosa sp. H | poorly charact | Function unkn | 0XPRT | NA | 391598.FBBA | 0,8 | 0,4 | 0,000136106 | 6,82E-05 | 1 - | 0,00018007 - | 0 - | 0 - | 1 - | 0,000167098 - | 1 - | 0,000166048 - | 1 - | 0,000167313 - |
| 1979 | 1 | 0.2um_NOKCPHJN_861422 | unclassified | unclassified | unclassified | unclassified | unclassified | unclassified | unclassified | unclassified | unclassified | unclassified | unclassified | unclassified | 0,4 | 0,489897949 | 0,00019437 | 0,000240775 | 0 - | 0 - | 1 - | 0,000542999 - | 0 - | 0 - | 0 - | 0 - | 1 - | 0,00042885 - |
| 1980 | 1 | 0.2um_NOKCPHJN_608547 | Eukaryota | Phaeophyceae | unclassified | Ectocarpales | Ectocarpaceae | Ectocarpus | Ectocarpus sili | unclassified | unclassified | unclassified | unclassified | unclassified | 1,6 | 0,489897949 | 0,00090336 | 0,00031039 | 1 - | 0,000565767 - | 2 - | 0,001331224 - | 1 - | 0,000525011 - | 2 - | 0,001043424 - | 2 - | 0,001051374 - |
| 1981 | 3 | 0.2um_NOKCPHJN_150646;0.2 | Bacteria | Proteobacteri | Gammaprotec | Cellvibrionales | Porticoccacea | unclassified | Gammaprotec | cellular proces | Intracellular tr | 16Q1A | type II secreti | various | 0,8 | 0,4 | 0,000145013 | 7,44E-05 | 1 - | 0,00018007 - | 1 - | 0,000211848 - | 1 - | 0,000167098 - | 1 - | 0,000166048 - | 0 - | 0 - |
| 1982 | 1 | 3um_FIJJAFBF_292913 | Eukaryota | Bacillariophyt | Coscinodiscop | Coscinodiscale | Hemidiscacea | Actinocyclus | Actinocyclus s | metabolism | Energy produc | COG0056 | Produces ATP | 41431.PCC88 | 6,6 | 2,497999199 | 0,001099972 | 0,000384928 | 8 - | 0,00138921 - | 2 - | 0,000408594 - | 8 - | 0,001289137 - | 6 - | 0,000960776 - | 9 - | 0,001452145 - |
| 1983 | 1 | 0.2um_NOKCPHJN_399402 | Bacteria | Proteobacteri | Alphaproteob | Rhodobactera | Rhodobactera | Puniceibacteri | Puniceibacteri | poorly charact | Function unkn | 0NVRZ | NA | 170187.SP_17 | 0,2 | 0,4 | 1,85E-05 | 3,69E-05 | 0 - | 0 - | 0 - | 0 - | 0 - | 0 - | 0 - | 0 - | 1 - | 9,23E-05 - |
| 1984 | 1 | 3um_FIJJAFBF_411071 | Eukaryota | Phaeophyceae | unclassified | Ectocarpales | Ectocarpaceae | Ectocarpus | Ectocarpus sili | information st | Chromatin str | KOG1744 | Core compone | 2850.Phatr54 | 6,4 | 1,2 | 0,004685896 | 0,000589492 | 5 - | 0,0037476 - | 5 - | 0,00440897 - | 7 - | 0,004868694 - | 8 - | 0,005529255 - | 7 - | 0,004874961 - |
| 1985 | 1 | 3um_FIJJAFBF_348266 | Bacteria | Proteobacteri | Alphaproteob | Rhodobactera | Rhodobactera | Sulfitobacter | Sulfitobacter g | cellular proces | Signal transdu | COG3279 | Regulator | 314262.MED1 | 0,6 | 0,489897949 | 0,000196551 | 0,000160486 | 0 - | 0 - | 0 - | 0 - | 1 - | 0,000328132 - | 1 - | 0,00032607 - | 1 - | 0,000328554 - |
| 1986 | 1 | 3um_FIJJAFBF_172525 | Bacteria | Proteobacteri | Betaproteoba | Burkholderial | Comamonada | Hydrogenoph | various | unclassified | unclassified | unclassified | unclassified | unclassified | 0,4 | 0,489897949 | 0,000187856 | 0,000230343 | 1 - | 0,000487188 - | 0 - | 0 - | 1 - | 0,000452093 - | 0 - | 0 - | 0 - | 0 - |
| 1987 | 1 | 3um_FIJJAFBF_615878 | Bacteria | Bacteroidetes | Flavobacteriia | Flavobacterial | Flavobacteria | Polaribacter | Polaribacter s | poorly charact | Function unkn | 0P8NI | Inherit from b | 313594.PI23P | 3,2 | 0,4 | 0,000417242 | 5,18E-05 | 3 - | 0,000396207 - | 3 - | 0,000466129 - | 3 - | 0,000367666 - | 3 - | 0,000365356 - | 4 - | 0,000490852 - |
| 1988 | 1 | 0.2um_NOKCPHJN_710509 | Eukaryota | Streptophyta | unclassified | Asterales | Asteraceae | Lactuca | Lactuca sativa | metabolism | Energy produc | 1BFZ8 | Cytochrome b | 3847.GLYMA1 | 1,4 | 0,489897949 | 0,000947348 | 0,000283455 | 1 - | 0,000695983 - | 1 - | 0,000818809 - | 2 - | 0,001291694 - | 2 - | 0,001283577 - | 1 - | 0,000646679 - |
| 1989 | 1 | 3um_FIJJAFBF_546283 | Eukaryota | unclassified | Oomycetes | Saprolegniales | Saprolegniace | Thraustotheca | Thraustotheca | information st | Translation, ri | 11UVJ | Ribosomal pro | 29760.VIT_05 | 2,2 | 1,166190379 | 0,001574805 | 0,000843474 | 3 - | 0,002267944 - | 0 - | 0 - | 3 - | 0,002104571 - | 2 - | 0,00139423 - | 3 - | 0,00210728 - |
| 1990 | 1 | 3um_FIJJAFBF_515657 | Bacteria | Proteobacteri | Alphaproteob | Rhodobactera | Rhodobactera | Ruegeria | Ruegeria sp. T | unclassified | unclassified | unclassified | unclassified | unclassified | 6,2 | 0,4 | 0,001712676 | 0,000161798 | 6 - | 0,001675679 - | 6 - | 0,001971399 - | 7 - | 0,001814131 - | 6 - | 0,001545198 - | 6 - | 0,001556971 - |
| 1991 | 1 | 3um_FIJJAFBF_92861 | Eukaryota | unclassified | Cryptophyta | Pyrenomonad | Pyrenomonad | Rhodomonas | Rhodomonas | metabolism | Energy produc | 11HAV | photosystem i | 35128.Thapsd | 0,8 | 0,4 | 0,000470096 | 0,00023572 | 1 - | 0,000621942 - | 0 - | 0 - | 1 - | 0,00057714 - | 1 - | 0,000573513 - | 1 - | 0,000577883 - |
| 1992 | 1 | 3um_FIJJAFBF_628820 | Bacteria | Actinobacteria | Actinobacteria | Micrococcales | Intrasporangia | Janibacter | Janibacter hoy | unclassified | unclassified | unclassified | unclassified | unclassified | 0,4 | 0,489897949 | 0,000192732 | 0,000236047 | 0 - | 0 - | 0 - | 0 - | 1 - | 0,000481519 - | 0 - | 0 - | 1 - | 0,000482139 - |
| 1993 | 1 | 0.2um_NOKCPHJN_165213 | Bacteria | Bacteroidetes | Flavobacteriia | Flavobacterial | Flavobacteria | Flaviramulus | Flaviramulus i | unclassified | unclassified | unclassified | unclassified | unclassified | 3,6 | 1,019803903 | 0,000616484 | 0,000136239 | 3 - | 0,000531478 - | 2 - | 0,000416848 - | 4 - | 0,00065759 - | 4 - | 0,000653457 - | 5 - | 0,000823045 - |
| 1994 | 1 | 0.2um_NOKCPHJN_316057 | unclassified | unclassified | unclassified | unclassified | unclassified | unclassified | unclassified | unclassified | unclassified | unclassified | unclassified | unclassified | 1 | 0 | 2,24E-05 | 2,19E-06 | 1 - | 2,26E-05 - | 1 - | 2,65E-05 - | 1 - | 2,09E-05 - | 1 - | 2,08E-05 - | 1 - | 2,10E-05 - |
| 1995 | 1 | 3um_FIJJAFBF_459065 | unclassified | unclassified | unclassified | unclassified | unclassified | unclassified | unclassified | unclassified | unclassified | unclassified | unclassified | unclassified | 0,8 | 0,4 | 0,000148285 | 7,44E-05 | 1 - | 0,000196183 - | 0 - | 0 - | 1 - | 0,000182051 - | 1 - | 0,000180907 - | 1 - | 0,000182285 - |
| 1996 | 1 | 0.2um_NOKCPHJN_194408 | Bacteria | Proteobacteri | Gammaprotec | Oceanospirilla | Hahellaceae | Hahella | Hahella sp. CC | cellular proces | Cell motility | COG1749 | flagellar hook | 349521.HCH_ | 1,8 | 0,748331477 | 0,000250631 | 0,000128215 | 1 - | 0,000136808 - | 3 - | 0,000482854 - | 1 - | 0,000126953 - | 2 - | 0,00025231 - | 2 - | 0,000254232 - |
| 1997 | 1 | 3um_FIJJAFBF_257132 | Eukaryota | unclassified | Florideophyce | Plocamiales | Plocamiaceae | Plocamium | Plocamium ca | unclassified | unclassified | unclassified | unclassified | unclassified | 0,6 | 0,489897949 | 0,000366502 | 0,000299251 | 0 - | 0 - | 0 - | 0 - | 1 - | 0,000611855 - | 1 - | 0,00060801 - | 1 - | 0,000612643 - |
| 1998 | 1 | 0.2um_NOKCPHJN_61813 | Bacteria | Bacteroidetes | Flavobacteriia | Flavobacterial | Flavobacteria | Polaribacter | Polaribacter fi | poorly charact | Function unkn | 0XY6Y | Pkd domain c | 391587.KAOT | 3,2 | 0,4 | 0,000612893 | 7,57E-05 | 3 - | 0,000582039 - | 3 - | 0,000684756 - | 4 - | 0,000720148 - | 3 - | 0,000536717 - | 3 - | 0,000540806 - |
| 1999 | 1 | 0.2um_NOKCPHJN_274455 | Eukaryota | Arthropoda | Insecta | Hymenoptera | Apidae | Apis | Apis cerana | unclassified | unclassified | unclassified | unclassified | unclassified | 0,8 | 0,4 | 9,74E-05 | 5,02E-05 | 0 - | 0 - | 1 - | 0,000144902 - | 1 - | 0,000114293 - | 1 - | 0,000113575 - | 1 - | 0,00011444 - |
| 2000 | 1 | 0.2um_NOKCPHJN_263907 | Bacteria | Proteobacteri | Alphaproteob | Rickettsiales | unclassified | unclassified | Rickettsiales b | unclassified | unclassified | unclassified | unclassified | unclassified | 1,6 | 0,489897949 | 0,000337468 | 8,26E-05 | 1 - | 0,000218144 - | 1 - | 0,000256642 - | 2 - | 0,000404859 - | 2 - | 0,000402315 - | 2 - | 0,000405381 - |
| 2001 | 1 | 0.2um_NOKCPHJN_907916 | Viruses | unclassified | unclassified | unclassified | Circoviridae | unclassified | Avon-Heathco | unclassified | unclassified | unclassified | unclassified | unclassified | 0,6 | 0,489897949 | 0,000188631 | 0,000154284 | 1 - | 0,00033092 - | 0 - | 0 - | 1 - | 0,000307082 - | 1 - | 0,000305152 - | 0 - | 0 - |
| 2002 | 1 | 0.2um_NOKCPHJN_18132 | Viruses | unclassified | unclassified | unclassified | Phycodnavirid | unclassified | Chrysochrom | information st | Transcription | 1BKE5 | SWIB complex | 3711.Bra0223 | 0,2 | 0,4 | 7,43E-05 | 0,000148633 | 0 - | 0 - | 0 - | 0 - | 1 - | 0,000371583 - | 0 - | 0 - | 0 - | 0 - |
| 2003 | 1 | 3um_FIJJAFBF_633970 | unclassified | unclassified | unclassified | unclassified | unclassified | unclassified | unclassified | unclassified | unclassified | unclassified | unclassified | unclassified | 0,6 | 0,489897949 | 7,58E-05 | 6,19E-05 | 0 - | 0 - | 0 - | 0 - | 1 - | 0,000126558 - | 1 - | 0,000125763 - | 1 - | 0,000126721 - |
| 2004 | 1 | 0.2um_NOKCPHJN_535342 | Bacteria | Actinobacteria | Actinobacteria | Micrococcales | Microbacteria | Candidatus Ac | Candidatus Ad | unclassified | unclassified | unclassified | unclassified | unclassified | 0,2 | 0,4 | 1,73E-05 | 3,45E-05 | 1 - | 8,63E-05 - | 0 - | 0 - | 0 - | 0 - | 0 - | 0 - | 0 - | 0 - |
| 2005 | 1 | 3um_FIJJAFBF_338021 | Bacteria | Acidobacteria | unclassified | unclassified | unclassified | unclassified | Acidobacteria | cellular proces | Posttranslatio | COG0652 | peptidyl-proly | 2850.Phatr14 | 0,6 | 0,489897949 | 0,001085829 | 0,000901257 | 0 - | 0 - | 1 - | 0,002105508 - | 1 - | 0,00166075 - | 0 - | 0 - | 1 - | 0,001662888 - |
| 2006 | 1 | 0.2um_NOKCPHJN_409279 | Eukaryota | Phaeophyceae | unclassified | Ectocarpales | Ectocarpaceae | Ectocarpus | Ectocarpus sili | unclassified | unclassified | unclassified | unclassified | unclassified | 0,6 | 0,489897949 | 4,84E-05 | 4,00E-05 | 1 - | 7,80E-05 - | 1 - | 9,17E-05 - | 1 - | 7,23E-05 - | 0 - | 0 - | 0 - | 0 - |
| 2007 | 1 | 0.2um_NOKCPHJN_614920 | Eukaryota | unclassified | unclassified | Choanoflagelli | Salpingoecida | Salpingoeca | Salpingoeca r | poorly charact | Function unkn | 1EJ2N | NA | 436017.A4S4 | 0,2 | 0,4 | 2,71E-05 | 5,43E-05 | 1 - | 0,000135749 - | 0 - | 0 - | 0 - | 0 - | 0 - | 0 - | 0 - | 0 - |
| 2008 | 1 | 3um_FIJJAFBF_715070 | Eukaryota | unclassified | Raphidophyce | Chattonellales | Chattonellace | Heterosigma | Heterosigma | unclassified | unclassified | unclassified | unclassified | unclassified | 25,6 | 1,743559577 | 0,004359471 | 0,000399818 | 23 - | 0,003962591 - | 25 - | 0,005067283 - | 25 - | 0,003996893 - | 28 - | 0,004448389 - | 27 - | 0,004322201 - |
| 2009 | 1 | 0.2um_NOKCPHJN_730596 | Eukaryota | unclassified | Oomycetes | Saprolegniales | Saprolegniace | Achlya | Achlya hypogy | poorly charact | Function unkn | COG1100 | GTP-binding P | 272952.HpaP | 0,4 | 0,489897949 | 0,000149532 | 0,00018314 | 0 - | 0 - | 0 - | 0 - | 1 - | 0,000375008 - | 1 - | 0,000372651 - | 0 - | 0 - |
| 2010 | 1 | 0.2um_NOKCPHJN_127271 | Bacteria | Bacteroidetes | Flavobacteriia | Flavobacterial | Flavobacteria | Arenitalea | Arenitalea lut | cellular proces | Posttranslatio | 08V1T | peptidylprolyl | 313594.PI23P | 1 | 0 | 0,000371442 | 3,63E-05 | 1 - | 0,00037476 - | 1 - | 0,000440897 - | 1 - | 0,000347764 - | 1 - | 0,000345578 - | 1 - | 0,000348212 - |
| 2011 | 1 | 0.2um_NOKCPHJN_1100803 | Bacteria | Verrucomicrol | Opitutae | Opitutales | Opitutaceae | unclassified | Opitutaceae b | unclassified | unclassified | unclassified | unclassified | unclassified | 0,2 | 0,4 | 3,26E-05 | 6,52E-05 | 0 - | 0 - | 0 - | 0 - | 0 - | 0 - | 0 - | 0 - | 1 - | 0,000162963 - |
| 2012 | 1 | 3um_FIJJAFBF_102730 | Bacteria | Bacteroidetes | Flavobacteriia | Flavobacterial | Flavobacteria | Aquimarina | Aquimarina at | metabolism | Amino acid tra | 07SUW | oxidoreductas | 313590.MED1 | 1 | 0 | 0,000123113 | 1,20E-05 | 1 - | 0,000124212 - | 1 - | 0,000146133 - | 1 - | 0,000115265 - | 1 - | 0,00011454 - | 1 - | 0,000115413 - |
| 2013 | 1 | 3um_FIJJAFBF_26346 | Bacteria | Bacteroidetes | Flavobacteriia | Flavobacterial | Flavobacteria | Seonamhaeic | Seonamhaeic | poorly charact | Function unkn | 09AVF | Protein of unk | 391587.KAOT | 1,8 | 0,748331477 | 0,000500216 | 0,000204609 | 3 - | 0,000851397 - | 1 - | 0,000333883 - | 2 - | 0,00052671 - | 1 - | 0,0002617 - | 2 - | 0,000527388 - |
| 2014 | 1 | 3um_FIJJAFBF_100818 | Bacteria | Bacteroidetes | Flavobacteriia | Flavobacterial | Flavobacteria | Polaribacter | various | information st | Translation, ri | 09EM4 | Binds directly | 313598.MED1 | 1 | 0 | 0,000762434 | 7,46E-05 | 1 - | 0,000769244 - | 1 - | 0,000904999 - | 1 - | 0,000713831 - | 1 - | 0,000709345 - | 1 - | 0,00071475 - |
| 2015 | 1 | 3um_FIJJAFBF_166626 | Bacteria | Firmicutes | Tissierellia | Tissierellales | Peptoniphila | Anaerococcus | Anaerococcus | unclassified | unclassified | unclassified | unclassified | unclassified | 1 | 0 | 0,000160069 | 1,57E-05 | 1 - | 0,000161499 - | 1 - | 0,00019 - | 1 - | 0,000149865 - | 1 - | 0,000148923 - | 1 - | 0,000150058 - |
| 2016 | 1 | 3um_FIJJAFBF_501038 | Bacteria | Bacteroidetes | Flavobacteriia | Flavobacterial | Flavobacteria | Flavobacteriu | Flavobacteriu | cellular proces | Cell wall/mem | 06B97 | phosphate-sel | 886377.Murr | 0,6 | 0,489897949 | 0,000150884 | 0,00012474 | 1 - | 0,000242919 - | 1 - | 0,000285789 - | 0 - | 0 - | 0 - | 0 - | 1 - | 0,000225711 - |
| 2017 | 1 | 0.2um_NOKCPHJN_887041 | Bacteria | Proteobacteri | Gammaprotec | unclassified | unclassified | unclassified | Gammaprotec | unclassified | unclassified | unclassified | unclassified | unclassified | 0,4 | 0,489897949 | 0,000111291 | 0,000137048 | 1 - | 0,000255667 - | 1 - | 0,000300787 - | 0 - | 0 - | 0 - | 0 - | 0 - | 0 - |
| 2018 | 1 | 0.2um_NOKCPHJN_211484 | Bacteria | Proteobacteri | Gammaprotec | unclassified | unclassified | unclassified | various | poorly charact | Function unkn | 05C2A | tonB-depend | 566466.NOR5 | 4,6 | 1,496662955 | 0,000363257 | 9,76E-05 | 4 - | 0,000328749 - | 2 - | 0,000193383 - | 6 - | 0,000457601 - | 6 - | 0,000454726 - | 5 - | 0,000381825 - |
| 2019 | 2 | 0.2um_NOKCPHJN_392967;3u | Eukaryota | Chordata | unclassified | unclassified | Branchiostom | Branchiostom | Branchiostom | unclassified | unclassified | unclassified | unclassified | unclassified | 0,6 | 0,489897949 | 6,17E-05 | 5,04E-05 | 0 - | 0 - | 0 - | 0 - | 1 - | 0,000103009 - | 1 - | 0,000102361 - | 1 - | 0,000103141 - |
| 2020 | 1 | 3um_FIJJAFBF_352965 | Bacteria | Proteobacteri | Gammaprotec | Alteromonada | Alteromonada | Glaciecola | Glaciecola sp. | metabolism | Inorganic ion | 0QJVD | TonB-depend | 455436.GHTC | 12 | 1,414213562 | 0,001189599 | 0,000147008 | 13 - | 0,001304371 - | 11 - | 0,001298477 - | 10 - | 0,000931084 - | 14 - | 0,001295326 - | 12 - | 0,001118739 - |
| 2021 | 1 | 3um_FIJJAFBF_56880 | Bacteria | Proteobacteri | Alphaproteob | Rhodobactera | Rhodobactera | Sulfitobacter | Sulfitobacter | metabolism | Lipid transpor | 01SS3 | Dehydrogenas | 52598.EE36_1 | 0,6 | 0,489897949 | 0,000487447 | 0,000398004 | 0 - | 0 - | 0 - | 0 - | 1 - | 0,000813767 - | 1 - | 0,000808654 - | 1 - | 0,000814815 - |
| 2022 | 1 | 3um_FIJJAFBF_735995 | Bacteria | Actinobacteria | Actinobacteria | Streptomyceta | Streptomyceta | Streptomyces | Streptomyces | unclassified | unclassified | unclassified | unclassified | unclassified | 0,6 | 0,489897949 | 0,000754803 | 0,000624416 | 1 - | 0,00121797 - | 1 - | 0,001432915 - | 0 - | 0 - | 1 - | 0,00112313 - | 0 - | 0 - |
| 2023 | 2 | 3um_FIJJAFBF_15541;3um_FIJ | Bacteria | Bacteroidetes | various | various | various | various | various | poorly charact | Function unkn | 11RX2 | NA | various | 1 | 0 | 0,000514305 | 5,03E-05 | 1 - | 0,000518899 - | 1 - | 0,000610473 - | 1 - | 0,000481519 - | 1 - | 0,000478493 - | 1 - | 0,000482139 - |
| 2024 | 1 | 3um_FIJJAFBF_46520 | Bacteria | Proteobacteri | Gammaprotec | Oceanospirilla | Oceanospirilla | Amphritea | Amphritea jap | information st | Translation, ri | COG0087 | One of the pri | 377629.TERT | 0,2 | 0,4 | 7,69E-05 | 0,000153739 | 0 - | 0 - | 0 - | 0 - | 0 - | 0 - | 0 - | 0 - | 1 - | 0,000384347 - |
| 2025 | 1 | 3um_FIJJAFBF_128452 | Bacteria | Bacteroidetes | Flavobacteriia | Flavobacterial | Flavobacteria | Polaribacter | Polaribacter s | cellular proces | Intracellular tr | 05FGH | MotA TolQ ex | 313594.PI23P | 4 | 0,894427191 | 0,00142486 | 0,00023362 | 3 - | 0,001096173 - | 3 - | 0,001289624 - | 4 - | 0,001356279 - | 5 - | 0,001684695 - | 5 - | 0,001697531 - |
| 2026 | 1 | 0.2um_NOKCPHJN_795562 | Eukaryota | unclassified | Pelagophycea | Pelagomonad | unclassified | Aureococcus | Aureococcus | cellular proces | Signal transdu | KOG0613 | fibronectin typ | 67593.Physo1 | 0,8 | 0,4 | 8,04E-05 | 4,03E-05 | 1 - | 0,000106425 - | 0 - | 0 - | 1 - | 9,88E-05 - | 1 - | 9,81E-05 - | 1 - | 9,89E-05 - |
| 2027 | 1 | 3um_FIJJAFBF_360602 | Bacteria | Tenericutes | Mollicutes | Mycoplasmata | Mycoplasmata | Mycoplasma | Mycoplasma | poorly charact | Function unkn | 03BSV | Inherit from e | 7070.TC0147 | 1 | 0 | 7,37E-05 | 7,21E-06 | 1 - | 7,44E-05 - | 1 - | 8,75E-05 - | 1 - | 6,90E-05 - | 1 - | 6,86E-05 - | 1 - | 6,91E-05 - |
| 2028 | 1 | 3um_FIJJAFBF_148493 | Eukaryota | Bacillariophyt | Coscinodiscop | Thalassiosirale | Thalassiosirac | Thalassiosira | Thalassiosira | poorly charact | Function unkn | 0ISAJ | BP28CT (NUC | 2850.Phatr46 | 0,8 | 0,4 | 0,00013533 | 6,94E-05 | 1 - | 0,000167996 - | 1 - | 0,000197643 - | 0 - | 0 - | 1 - | 0,000154914 - | 1 - | 0,000156095 - |
| 2029 | 1 | 0.2um_NOKCPHJN_1031253 | Eukaryota | Chordata | unclassified | unclassified | Branchiostom | Branchiostom | Branchiostom | unclassified | unclassified | unclassified | unclassified | unclassified | 0,4 | 0,489897949 | 0,000237749 | 0,000291182 | 0 - | 0 - | 0 - | 0 - | 1 - | 0,000593991 - | 0 - | 0 - | 1 - | 0,000594755 - |
| 2030 | 1 | 3um_FIJJAFBF_197554 | Bacteria | Verrucomicrol | Verrucomicrol | Verrucomicrol | Rubritaleacea | Rubritalea | Rubritalea squ | metabolism | Nucleotide tra | 07XAE | Catalyzes the | 349741.Amuc | 0,2 | 0,4 | 9,28E-05 | 0,000185595 | 1 - | 0,000463989 - | 0 - | 0 - | 0 - | 0 - | 0 - | 0 - | 0 - | 0 - |
| 2031 | 1 | 3um_FIJJAFBF_112857 | unclassified | unclassified | unclassified | unclassified | unclassified | unclassified | unclassified | unclassified | unclassified | unclassified | unclassified | unclassified | 1 | 0 | 0,001524868 | 0,000149199 | 1 - | 0,001538489 - | 1 - | 0,001809998 - | 1 - | 0,001427662 - | 1 - | 0,00141869 - | 1 - | 0,0014295 - |
| 2032 | 1 | 0.2um_NOKCPHJN_876272 | Eukaryota | unclassified | Pelagophycea | Pelagomonad | unclassified | Aureococcus | Aureococcus | cellular proces | Posttranslatio | 08V1T | peptidylprolyl | 513049.Amax | 1 | 0,632455532 | 0,000489571 | 0,000298692 | 0 - | 0 - | 1 - | 0,000589542 - | 1 - | 0,00046501 - | 1 - | 0,000462088 - | 2 - | 0,000931217 - |
| 2033 | 2 | 0.2um_NOKCPHJN_205147;0.2 | Bacteria | Bacteroidetes | Flavobacteriia | Flavobacterial | Crocinitomica | unclassified | various | metabolism | Inorganic ion | 05C27 | TonB-depend | various | 0,2 | 0,4 | 1,58E-05 | 3,17E-05 | 0 - | 0 - | 0 - | 0 - | 0 - | 0 - | 0 - | 0 - | 1 - | 7,92E-05 - |
| 2034 | 1 | 0.2um_NOKCPHJN_282157 | Eukaryota | Bacillariophyt | Fragilariophyc | Fragilariales | Fragilariaceae | Ardissonea | Ardissonea cr | unclassified | unclassified | unclassified | unclassified | unclassified | 3,8 | 0,4 | 0,000440684 | 7,21E-05 | 4 - | 0,000466457 - | 4 - | 0,000548776 - | 3 - | 0,000324641 - | 4 - | 0,000430135 - | 4 - | 0,000433412 - |
| 2035 | 1 | 0.2um_NOKCPHJN_754061 | Bacteria | Bacteroidetes | unclassified | unclassified | unclassified | unclassified | Bacteroidetes | poorly charact | Function unkn | 0QNU1 | Uncharacteriz | 187272.Mlg_1 | 1 | 0 | 0,000141329 | 1,38E-05 | 1 - | 0,000142592 - | 1 - | 0,000167756 - | 1 - | 0,00013232 - | 1 - | 0,000131488 - | 1 - | 0,00013249 - |
| 2036 | 1 | 0.2um_NOKCPHJN_383104 | Bacteria | Proteobacteri | Alphaproteob | unclassified | unclassified | unclassified | SAR116 cluste | unclassified | unclassified | unclassified | unclassified | unclassified | 0,4 | 0,489897949 | 6,71E-05 | 8,23E-05 | 1 - | 0,000173996 - | 0 - | 0 - | 1 - | 0,000161462 - | 0 - | 0 - | 0 - | 0 - |
| 2037 | 1 | 0.2um_NOKCPHJN_446856 | Bacteria | Proteobacteri | Alphaproteob | Pelagibacteral | Pelagibactera | Candidatus Pe | Candidatus Pe | unclassified | unclassified | unclassified | unclassified | unclassified | 0,6 | 0,489897949 | 0,000276853 | 0,000226403 | 1 - | 0,000484496 - | 0 - | 0 - | 1 - | 0,000449595 - | 0 - | 0 - | 1 - | 0,000450174 - |
| 2038 | 1 | 3um_FIJJAFBF_807864 | Eukaryota | Chlorophyta | Chlorophycea | Chlamydomor | Tetrabaenace | Tetrabaena | Tetrabaena so | information st | Replication, re | COG0592 | DNA polymera | 3847.GLYMA0 | 0,8 | 0,4 | 0,0002717 | 0,000139326 | 1 - | 0,000337284 - | 1 - | 0,000396807 - | 0 - | 0 - | 1 - | 0,000311021 - | 1 - | 0,00031339 - |
| 2039 | 1 | 3um_FIJJAFBF_753922 | Bacteria | Proteobacteri | Alphaproteob | Rhizobiales | unclassified | unclassified | Rhizobiales ba | metabolism | Inorganic ion | 05F28 | tonB-depend | 247633.GP21 | 1,2 | 1,166190379 | 0,000120652 | 0,000117007 | 0 - | 0 - | 0 - | 0 - | 2 - | 0,000201677 - | 3 - | 0,000300615 - | 1 - | 0,000100968 - |
| 2040 | 2 | 3um_FIJJAFBF_169112;3um_FI | Eukaryota | various | various | various | various | various | various | information st | Translation, ri | COG0087 | One of the pri | various | 1 | 0 | 0,000415873 | 4,07E-05 | 1 - | 0,000419588 - | 1 - | 0,000493636 - | 1 - | 0,000389362 - | 1 - | 0,000386916 - | 1 - | 0,000389864 - |
| 2041 | 1 | 3um_FIJJAFBF_383125 | Eukaryota | Bacillariophyt | Coscinodiscop | Thalassiosirale | Thalassiosirac | Thalassiosira | Thalassiosira | cellular proces | Signal transdu | KOG0613 | fibronectin typ | 67593.Physo1 | 0,6 | 0,489897949 | 8,13E-05 | 6,65E-05 | 1 - | 0,000142592 - | 0 - | 0 - | 0 - | 0 - | 1 - | 0,000131488 - | 1 - | 0,00013249 - |
| 2042 | 1 | 3um_FIJJAFBF_257113 | Eukaryota | unclassified | Florideophyce | Nemaliales | Liagoraceae | Yamadaella | Yamadaella ca | information st | Translation, ri | 0FNT6 | One of the ea | 391612.CY011 | 0,4 | 0,489897949 | 0,000292328 | 0,00035803 | 0 - | 0 - | 0 - | 0 - | 1 - | 0,000733124 - | 1 - | 0,000728517 - | 0 - | 0 - |
| 2043 | 2 | 3um_FIJJAFBF_283034;3um_FI | Bacteria | Bacteroidetes | Flavobacteriia | Flavobacterial | Flavobacteria | various | various | poorly charact | Function unkn | 08ZIU | NA | 398720.MED2 | 0,8 | 0,4 | 0,000433259 | 0,000222199 | 1 - | 0,000537999 - | 1 - | 0,000632944 - | 1 - | 0,000499244 - | 1 - | 0,000496106 - | 0 - | 0 - |
| 2044 | 1 | 0.2um_NOKCPHJN_786900 | unclassified | unclassified | unclassified | unclassified | unclassified | unclassified | unclassified | unclassified | unclassified | unclassified | unclassified | unclassified | 1 | 0 | 0,000103844 | 1,02E-05 | 1 - | 0,000104772 - | 1 - | 0,000123262 - | 1 - | 9,72E-05 - | 1 - | 9,66E-05 - | 1 - | 9,73E-05 - |
| 2045 | 1 | 3um_FIJJAFBF_142185 | Bacteria | Proteobacteri | Gammaprotec | Oceanospirilla | Oceanospirilla | Marinomonas | Marinomonas | metabolism | Energy produc | 17CGT | F(1)F(0) ATP s | 717774.Marm | 0,8 | 0,4 | 0,000444735 | 0,000228939 | 0 - | 0 - | 1 - | 0,000661345 - | 1 - | 0,000521646 - | 1 - | 0,000518368 - | 1 - | 0,000522317 - |
| 2046 | 1 | 0.2um_NOKCPHJN_565444 | Bacteria | Proteobacteri | Alphaproteob | Rhodospirillal | Rhodospirillac | unclassified | Rhodospirillac | unclassified | unclassified | unclassified | unclassified | unclassified | 7,2 | 0,4 | 0,002029214 | 0,000321642 | 7 - | 0,001980184 - | 8 - | 0,002662449 - | 7 - | 0,001837539 - | 7 - | 0,001825992 - | 7 - | 0,001839905 - |
| 2047 | 1 | 3um_FIJJAFBF_458234 | Bacteria | Proteobacteri | Alphaproteob | Rhodobactera | Rhodobactera | Marinosulfon | Marinosulfon | information st | Translation, ri | COG0051 | tRNA binding | 367336.OM22 | 1 | 0 | 0,000852132 | 8,34E-05 | 1 - | 0,000859744 - | 1 - | 0,001011469 - | 1 - | 0,000797811 - | 1 - | 0,000792798 - | 1 - | 0,000798838 - |
| 2048 | 1 | 3um_FIJJAFBF_176621 | Bacteria | Proteobacteri | Gammaprotec | Alteromonada | Colwelliaceae | Colwellia | Colwellia psyc | poorly charact | Function unkn | 0XNQ8 | alkaline phos | 167879.CPS_1 | 0,2 | 0,4 | 4,28E-05 | 8,55E-05 | 0 - | 0 - | 0 - | 0 - | 0 - | 0 - | 0 - | 0 - | 1 - | 0,000213862 - |
| 2049 | 1 | 3um_FIJJAFBF_20171 | Bacteria | Proteobacteri | Gammaprotec | unclassified | unclassified | unclassified | various | unclassified | unclassified | unclassified | unclassified | unclassified | 0,2 | 0,4 | 0,000156596 | 0,000313192 | 1 - | 0,000782981 - | 0 - | 0 - | 0 - | 0 - | 0 - | 0 - | 0 - | 0 - |
| 2050 | 1 | 3um_FIJJAFBF_501912 | Bacteria | Proteobacteri | Alphaproteob | Rhodospirillal | Rhodospirillac | unclassified | Rhodospirillac | unclassified | unclassified | unclassified | unclassified | unclassified | 0,2 | 0,4 | 5,23E-05 | 0,00010468 | 0 - | 0 - | 0 - | 0 - | 0 - | 0 - | 1 - | 0,0002617 - | 0 - | 0 - |

| | | | | | | | | | | | | | | | | | | | | | | | | | | | | |
|---|---|---|---|---|---|---|---|---|---|---|---|---|---|---|---|---|---|---|---|---|---|---|---|---|---|---|---|---|
| 2051 | 1 | 3um_FIJJAFBF_263680 | Bacteria | Proteobacteri | Alphaproteob | unclassified | unclassified | Candidatus Pu | Candidatus Pu | cellular proces | Posttranslatio | 05C2Z | ATP-depender | 313624.N941 | 3,8 | 0,979795897 | 0,00037523 | 7,63E-05 | 4 - | 0,000407878 - | 2 - | 0,00023993 - | 4 - | 0,000378496 - | 4 - | 0,000376118 - | 5 - | 0,00047373 - |
| 2052 | 1 | 0.2um_NOKCPHJN_224040 | Bacteria | unclassified | unclassified | unclassified | unclassified | unclassified | uncultured ba | metabolism | Inorganic ion t | 05C27 | TonB-depende | 313596.RB250 | 4 | 0 | 0,000327065 | 3,20E-05 | 4 - | 0,000329986 - | 4 - | 0,000388222 - | 4 - | 0,000306215 - | 4 - | 0,000304291 - | 4 - | 0,00030661 - |
| 2053 | 1 | 3um_FIJJAFBF_631848 | Bacteria | Bacteroidetes | Flavobacteriia | Flavobacterial | Flavobacteriac | Formosa | Formosa sp. H | cellular proces | Cell wall/mem | COG2885 | Ompa motb d | 983544.Lacal_ | 1 | 0,632455532 | 0,000179731 | 0,000109028 | 0 - | 0 - | 1 - | 0,000216743 - | 1 - | 0,00017096 - | 2 - | 0,00033977 - | 1 - | 0,00017118 - |
| 2054 | 1 | 0.2um_NOKCPHJN_232960 | Eukaryota | unclassified | Labyrinthulom | unclassified | Thraustochytr | Aurantiochytr | Aurantiochytr | unclassified | unclassified | unclassified | unclassified | unclassified | 0,4 | 0,489897949 | 1,99E-05 | 2,44E-05 | 1 - | 5,17E-05 - | 0 - | 0 - | 1 - | 4,80E-05 - | 0 - | 0 - | 0 - | 0 - |
| 2055 | 1 | 0.2um_NOKCPHJN_547497 | Bacteria | unclassified | unclassified | unclassified | unclassified | unclassified | uncultured ba | metabolism | Inorganic ion t | 05C27 | TonB-depende | 700598.Niako | 0,6 | 0,489897949 | 5,46E-05 | 4,46E-05 | 0 - | 0 - | 0 - | 0 - | 1 - | 9,12E-05 - | 1 - | 9,07E-05 - | 1 - | 9,13E-05 - |
| 2056 | 1 | 3um_FIJJAFBF_639895 | Bacteria | Proteobacteri | Gammaprotec | Oceanospirilla | various | various | various | poorly charact | Function unkn | COG3517 | type VI secreti | 83406.HDN1F | 1 | 0 | 0,000181078 | 1,77E-05 | 1 - | 0,000182696 - | 1 - | 0,000214937 - | 1 - | 0,000169535 - | 1 - | 0,000168469 - | 1 - | 0,000169753 - |
| 2057 | 1 | 3um_FIJJAFBF_465281 | Eukaryota | Bacillariophyt | Bacillariophyc | Bacillariales | Bacillariaceae | Fragilariopsis | Fragilariopsis | unclassified | unclassified | unclassified | unclassified | unclassified | 0,4 | 0,489897949 | 0,000294218 | 0,000360747 | 1 - | 0,000762555 - | 0 - | 0 - | 0 - | 0 - | 0 - | 0 - | 1 - | 0,000708535 - |
| 2058 | 1 | 3um_FIJJAFBF_185959 | Bacteria | Bacteroidetes | Flavobacteriia | Flavobacterial | Flavobacteriac | Formosa | Formosa sp. H | poorly charact | Function unkn | 08PAV | RagB SusD dor | 983544.Lacal_ | 1,4 | 0,489897949 | 0,00025275 | 0,000105229 | 1 - | 0,000178967 - | 2 - | 0,000421102 - | 1 - | 0,000166075 - | 1 - | 0,000165031 - | 2 - | 0,000332578 - |
| 2059 | 1 | 3um_FIJJAFBF_69380 | Eukaryota | Chlorophyta | Mamiellophyc | Mamiellales | Bathycoccace | Ostreococcus | Ostreococcus | metabolism | Energy produc | COG0056 | Produces ATP | 70448.Q0P3H | 1 | 0 | 0,000198896 | 1,95E-05 | 1 - | 0,000200672 - | 1 - | 0,000236087 - | 1 - | 0,000186217 - | 1 - | 0,000185047 - | 1 - | 0,000186457 - |
| 2060 | 1 | 0.2um_NOKCPHJN_955130 | Eukaryota | Bacillariophyt | Bacillariophyc | Naviculales | Naviculaceae | Fistulifera | Fistulifera sola | metabolism | Energy produc | COG0056 | Produces ATP | 13037.EHJ650 | 3,6 | 0,489897949 | 0,000785526 | 6,09E-05 | 3 - | 0,00066772 - | 3 - | 0,000785558 - | 4 - | 0,00082616 - | 4 - | 0,000820968 - | 4 - | 0,000827223 - |
| 2061 | 1 | 3um_FIJJAFBF_317844 | Bacteria | Bacteroidetes | Flavobacteriia | Flavobacterial | Crocinitomica | unclassified | Crocinitomica | unclassified | unclassified | unclassified | unclassified | unclassified | 4 | 1,414213562 | 0,000452024 | 0,000139021 | 2 - | 0,00023261 - | 3 - | 0,00041049 - | 5 - | 0,000539634 - | 4 - | 0,000428994 - | 6 - | 0,000648394 - |
| 2062 | 1 | 0.2um_NOKCPHJN_316226 | Bacteria | Bacteroidetes | Flavobacteriia | Flavobacterial | Flavobacteriac | Muricauda | Muricauda sp. | unclassified | unclassified | unclassified | unclassified | unclassified | 1 | 0 | 3,71E-05 | 3,63E-06 | 1 - | 3,75E-05 - | 1 - | 4,41E-05 - | 1 - | 3,48E-05 - | 1 - | 3,46E-05 - | 1 - | 3,48E-05 - |
| 2063 | 1 | 0.2um_NOKCPHJN_483426 | Eukaryota | Chordata | Aves | Bucerotiforme | Bucerotidae | Buceros | Buceros rhino | unclassified | unclassified | unclassified | unclassified | unclassified | 1,6 | 0,489897949 | 0,000233836 | 6,43E-05 | 2 - | 0,000299296 - | 1 - | 0,000176058 - | 2 - | 0,000277736 - | 1 - | 0,000137995 - | 2 - | 0,000278094 - |
| 2064 | 3 | 0.2um_NOKCPHJN_573220;0.2 | Bacteria | Proteobacteri | Alphaproteob | Pelagibacteral | Pelagibacterac | various | various | unclassified | unclassified | unclassified | unclassified | unclassified | 0,6 | 0,489897949 | 0,00054931 | 0,000449288 | 1 - | 0,000963669 - | 0 - | 0 - | 1 - | 0,00089425 - | 1 - | 0,00088863 - | 0 - | 0 - |
| 2065 | 1 | 0.2um_NOKCPHJN_728405 | Bacteria | Proteobacteri | Alphaproteob | Rhizobiales | unclassified | unclassified | Rhizobiales ba | metabolism | Inorganic ion t | 05F28 | tonB-depende | 247634.GPB2 | 0,6 | 0,489897949 | 6,29E-05 | 5,14E-05 | 0 - | 0 - | 0 - | 0 - | 1 - | 0,000105002 - | 1 - | 0,000104342 - | 1 - | 0,000105137 - |
| 2066 | 1 | 0.2um_NOKCPHJN_861812 | Eukaryota | Bacillariophyt | Bacillariophyc | Naviculales | Naviculaceae | Fistulifera | Fistulifera sola | poorly charact | Function unkn | 110WY | Chlorophyll A- | 2850.Phatr18 | 0,6 | 0,489897949 | 0,000541608 | 0,000442227 | 0 - | 0 - | 0 - | 0 - | 1 - | 0,000904186 - | 1 - | 0,000898504 - | 1 - | 0,00090535 - |
| 2067 | 2 | 3um_FIJJAFBF_500683;3um_FI | Eukaryota | Bacillariophyt | Mediophyceae | various | various | various | various | metabolism | Energy produc | 11HAV | photosystem i | 35128.Thapsd | 6,2 | 0,748331477 | 0,002309275 | 0,000141795 | 6 - | 0,002277762 - | 5 - | 0,002233114 - | 7 - | 0,002465962 - | 6 - | 0,002100399 - | 7 - | 0,002469136 - |
| 2068 | 2 | 3um_FIJJAFBF_260030;3um_FI | Bacteria | Verrucomicrol | Verrucomicrol | unclassified | unclassified | unclassified | Verrucomicrol | poorly charact | Function unkn | 0ZMMU | Prokaryotic N- | 240016.VspiD | 1,2 | 0,4 | 0,000407877 | 0,000121924 | 1 - | 0,000346616 - | 1 - | 0,000407786 - | 2 - | 0,000643294 - | 1 - | 0,000319626 - | 1 - | 0,000322061 - |
| 2069 | 2 | 0.2um_NOKCPHJN_1085971;0. | Viruses | unclassified | unclassified | unclassified | unclassified | unclassified | uncultured M | unclassified | unclassified | unclassified | unclassified | unclassified | 0,4 | 0,489897949 | 0,000377307 | 0,000462109 | 0 - | 0 - | 0 - | 0 - | 1 - | 0,000946241 - | 1 - | 0,000940295 - | 0 - | 0 - |
| 2070 | 1 | 3um_FIJJAFBF_561978 | Eukaryota | unclassified | Spirotrichea | Sporadotrichi | Oxytrichidae | Oxytricha | Oxytricha trifa | unclassified | unclassified | unclassified | unclassified | unclassified | 1,6 | 0,489897949 | 0,000997516 | 0,000244214 | 1 - | 0,000644808 - | 1 - | 0,000758602 - | 2 - | 0,001196717 - | 2 - | 0,001189196 - | 2 - | 0,001198257 - |
| 2071 | 1 | 0.2um_NOKCPHJN_1024349 | Bacteria | Proteobacteri | Alphaproteob | Pelagibacteral | Pelagibacterac | Candidatus Pe | Candidatus Pe | metabolism | Secondary me | 01Q86 | extracellular s | 439493.PB72 | 0,6 | 0,489897949 | 0,000138196 | 0,000114705 | 0 - | 0 - | 1 - | 0,000267974 - | 1 - | 0,000211368 - | 0 - | 0 - | 1 - | 0,00021164 - |
| 2072 | 1 | 0.2um_NOKCPHJN_603468 | unclassified | unclassified | unclassified | unclassified | unclassified | unclassified | unclassified | unclassified | unclassified | unclassified | unclassified | unclassified | 0,4 | 0,489897949 | 0,000646996 | 0,000796734 | 1 - | 0,001486336 - | 1 - | 0,001748642 - | 0 - | 0 - | 0 - | 0 - | 0 - | 0 - |
| 2073 | 1 | 3um_FIJJAFBF_689592 | unclassified | unclassified | unclassified | unclassified | unclassified | unclassified | unclassified | unclassified | unclassified | unclassified | unclassified | unclassified | 0,2 | 0,4 | 0,000224282 | 0,000448565 | 0 - | 0 - | 1 - | 0,001121412 - | 0 - | 0 - | 0 - | 0 - | 0 - | 0 - |
| 2074 | 1 | 0.2um_NOKCPHJN_39166 | Eukaryota | unclassified | Synurophycea | Parmales | Triparmaceae | Triparma | Triparma laev | unclassified | unclassified | unclassified | unclassified | unclassified | 4 | 0,632455532 | 0,002400063 | 0,000232781 | 4 - | 0,002452975 - | 3 - | 0,002164403 - | 4 - | 0,002276273 - | 5 - | 0,00282746 - | 4 - | 0,002279203 - |
| 2075 | 1 | 3um_FIJJAFBF_191790 | Eukaryota | Bacillariophyt | Fragilariophyc | Fragilariales | Fragilariaceae | Asterionellops | Asterionellops | metabolism | Energy produc | COG0711 | F(1)F(0) ATP s | 35128.Thapsd | 7,6 | 0,8 | 0,003490985 | 0,000458999 | 9 - | 0,004175898 - | 7 - | 0,003821107 - | 7 - | 0,003013953 - | 7 - | 0,002995013 - | 8 - | 0,003448952 - |
| 2076 | 1 | 0.2um_NOKCPHJN_434902 | Eukaryota | Streptophyta | Chlorokyboph | Chlorokybales | Chlorokybace | Chlorokybus | Chlorokybus a | unclassified | unclassified | unclassified | unclassified | unclassified | 0,6 | 0,489897949 | 0,000115629 | 9,57E-05 | 1 - | 0,000186583 - | 1 - | 0,00021951 - | 0 - | 0 - | 1 - | 0,000172054 - | 0 - | 0 - |
| 2077 | 1 | 3um_FIJJAFBF_582068 | Bacteria | Verrucomicrol | Opitutae | unclassified | unclassified | unclassified | Opitutae bact | unclassified | unclassified | unclassified | unclassified | unclassified | 0,8 | 0,4 | 0,000144462 | 7,41E-05 | 1 - | 0,000179333 - | 1 - | 0,000210981 - | 0 - | 0 - | 1 - | 0,000165369 - | 1 - | 0,000166629 - |
| 2078 | 1 | 0.2um_NOKCPHJN_235200 | Eukaryota | unclassified | Labyrinthulom | unclassified | Thraustochytr | Aurantiochytr | Aurantiochytr | cellular proces | Cell wall/mem | 12RRB | ankyrin 3, nod | 400682.PAC_1 | 1 | 0 | 0,000103105 | 1,01E-05 | 1 - | 0,000104026 - | 1 - | 0,000122384 - | 1 - | 9,65E-05 - | 1 - | 9,59E-05 - | 1 - | 9,67E-05 - |
| 2079 | 1 | 3um_FIJJAFBF_732705 | Eukaryota | unclassified | Pelagophycea | Pelagomonad | unclassified | Aureococcus | Aureococcus a | unclassified | unclassified | unclassified | unclassified | unclassified | 0,6 | 0,489897949 | 0,000435743 | 0,000356338 | 1 - | 0,000762555 - | 0 - | 0 - | 1 - | 0,000707624 - | 0 - | 0 - | 1 - | 0,000708535 - |
| 2080 | 1 | 3um_FIJJAFBF_744828 | Bacteria | Proteobacteri | Gammaprotec | Pseudomonad | Pseudomonad | Azotobacter | Azotobacter v | information st | Translation, ri | 179ZI | One of the pri | 223283.PSPTC | 0,2 | 0,4 | 0,000133884 | 0,000267767 | 1 - | 0,000669419 - | 0 - | 0 - | 0 - | 0 - | 0 - | 0 - | 0 - | 0 - |
| 2081 | 1 | 3um_FIJJAFBF_116531 | Bacteria | Proteobacteri | Alphaproteob | unclassified | unclassified | Candidatus Pu | Candidatus Pu | metabolism | Carbohydrate | 0IVA1 | Photosystem I | 35128.Thapsd | 6,4 | 0,489897949 | 0,002908516 | 0,000455771 | 6 - | 0,002740433 - | 7 - | 0,003761402 - | 6 - | 0,002543023 - | 6 - | 0,002527042 - | 7 - | 0,002970679 - |
| 2082 | 1 | 0.2um_NOKCPHJN_01987 | Bacteria | Proteobacteri | Alphaproteob | Rhodobactera | Rhodobactera | Sulfitobacter | Sulfitobacter r | information st | Translation, ri | COG0050 | This protein p | 391624.OIHEL | 1,2 | 0,748331477 | 0,000252826 | 0,000155205 | 1 - | 0,000224281 - | 0 - | 0 - | 2 - | 0,000416249 - | 1 - | 0,000206817 - | 2 - | 0,000416785 - |
| 2083 | 1 | 0.2um_NOKCPHJN_957389 | Eukaryota | unclassified | Oomycetes | Saprolegniales | Saprolegniace | Aphanomyces | Aphanomyces | information st | Chromatin str | KOG1744 | Core compone | 4787.PITG_03 | 2 | 0 | 0,001485769 | 0,000145374 | 2 - | 0,00149904 - | 2 - | 0,001763588 - | 2 - | 0,001391055 - | 2 - | 0,001382314 - | 2 - | 0,001392846 - |
| 2084 | 1 | 3um_FIJJAFBF_122322 | unclassified | unclassified | unclassified | unclassified | unclassified | unclassified | unclassified | unclassified | unclassified | unclassified | unclassified | unclassified | 4,6 | 0,8 | 0,001086362 | 0,000297694 | 4 - | 0,000940417 - | 6 - | 0,001659569 - | 5 - | 0,001090841 - | 4 - | 0,000867189 - | 4 - | 0,000873796 - |
| 2085 | 1 | 0.2um_NOKCPHJN_529099 | unclassified | unclassified | unclassified | unclassified | unclassified | unclassified | unclassified | unclassified | unclassified | unclassified | unclassified | unclassified | 0,2 | 0,4 | 3,74E-05 | 7,49E-05 | 0 - | 0 - | 0 - | 0 - | 0 - | 0 - | 1 - | 0,000187188 - | 0 - | 0 - |
| 2086 | 1 | 3um_FIJJAFBF_523829 | Bacteria | Proteobacteri | Gammaprotec | Arenicellales | Arenicellaceae | Arenicella | Arenicella xan | unclassified | unclassified | unclassified | unclassified | unclassified | 4,2 | 0,979795897 | 0,000574449 | 0,000123087 | 3 - | 0,000416928 - | 4 - | 0,000654009 - | 4 - | 0,000515859 - | 6 - | 0,000768926 - | 4 - | 0,000516523 - |
| 2087 | 1 | 3um_FIJJAFBF_577614 | unclassified | unclassified | unclassified | unclassified | unclassified | unclassified | unclassified | unclassified | unclassified | unclassified | unclassified | unclassified | 0,4 | 0,489897949 | 0,000507334 | 0,000621362 | 0 - | 0 - | 0 - | 0 - | 0 - | 0 - | 1 - | 0,001263521 - | 1 - | 0,001273148 - |
| 2088 | 1 | 0.2um_NOKCPHJN_310213 | unclassified | unclassified | unclassified | unclassified | unclassified | unclassified | unclassified | unclassified | unclassified | unclassified | unclassified | unclassified | 0,4 | 0,489897949 | 0,000354861 | 0,000435209 | 1 - | 0,000923093 - | 0 - | 0 - | 0 - | 0 - | 1 - | 0,000851214 - | 0 - | 0 - |
| 2089 | 1 | 0.2um_NOKCPHJN_226763 | Bacteria | Proteobacteri | Alphaproteob | Rhodobactera | Rhodobactera | Thalassobium | Thalassobium | metabolism | Lipid transpor | COG1960 | acyl-CoA dehy | 643867.Ftrac_ | 0,4 | 0,489897949 | 7,77E-05 | 9,63E-05 | 0 - | 0 - | 1 - | 0,0002172 - | 1 - | 0,000171319 - | 0 - | 0 - | 0 - | 0 - |
| 2090 | 1 | 3um_FIJJAFBF_150012 | Bacteria | Proteobacteri | Betaproteoba | Burkholderiale | Comamonada | Limnohabitan | Limnohabitan | information st | Translation, ri | COG0050 | This protein p | 339670.Bamb | 0,8 | 1,6 | 0,0001665 | 0,000332999 | 0 - | 0 - | 0 - | 0 - | 4 - | 0,000832499 - | 0 - | 0 - | 0 - | 0 - |
| 2091 | 1 | 0.2um_NOKCPHJN_372676 | Eukaryota | Chordata | Mammalia | unclassified | Bovidae | Bubalus | Bubalus bubal | metabolism | Amino acid tra | 0UPXV | aminopeptida | 9986.ENSOCU | 0,2 | 0,4 | 1,95E-05 | 3,89E-05 | 0 - | 0 - | 1 - | 9,73E-05 - | 0 - | 0 - | 0 - | 0 - | 0 - | 0 - |
| 2092 | 1 | 3um_FIJJAFBF_386500 | Viruses | unclassified | unclassified | unclassified | unclassified | unclassified | uncultured vir | unclassified | unclassified | unclassified | unclassified | unclassified | 0,4 | 0,489897949 | 7,09E-05 | 8,69E-05 | 1 - | 0,000183845 - | 0 - | 0 - | 1 - | 0,000170601 - | 0 - | 0 - | 0 - | 0 - |
| 2093 | 1 | 0.2um_NOKCPHJN_282552 | Eukaryota | unclassified | Labyrinthulom | unclassified | Thraustochytr | Aurantiochytr | Aurantiochytr | unclassified | unclassified | unclassified | unclassified | unclassified | 1,4 | 1,2 | 0,000110774 | 9,50E-05 | 0 - | 0 - | 0 - | 0 - | 3 - | 0,000237712 - | 2 - | 0,000157479 - | 2 - | 0,000158679 - |
| 2094 | 1 | 3um_FIJJAFBF_128440 | Bacteria | Bacteroidetes | Flavobacteriia | Flavobacterial | Flavobacteriac | Polaribacter | Polaribacter s | poorly charact | Function unkn | 0P8NI | Inherit from b | 313594.PI23P | 13,6 | 1,2 | 0,001776329 | 0,000216289 | 15 - | 0,001978057 - | 13 - | 0,002016855 - | 15 - | 0,001835566 - | 13 - | 0,001580826 - | 12 - | 0,001470343 - |
| 2095 | 1 | 0.2um_NOKCPHJN_299100 | Bacteria | Candidatus Pa | unclassified | unclassified | unclassified | unclassified | Candidatus Pa | unclassified | unclassified | unclassified | unclassified | unclassified | 1,4 | 0,489897949 | 0,000346636 | 0,000115415 | 2 - | 0,000503988 - | 1 - | 0,000296465 - | 1 - | 0,000233841 - | 2 - | 0,000464743 - | 1 - | 0,000234142 - |
| 2096 | 1 | 3um_FIJJAFBF_273735 | unclassified | unclassified | unclassified | unclassified | unclassified | unclassified | unclassified | unclassified | unclassified | unclassified | unclassified | unclassified | 0,2 | 0,4 | 0,00023701 | 0,000474021 | 1 - | 0,001185052 - | 0 - | 0 - | 0 - | 0 - | 0 - | 0 - | 0 - | 0 - |
| 2097 | 1 | 3um_FIJJAFBF_209605 | Bacteria | Bacteroidetes | Flavobacteriia | Flavobacterial | Flavobacteriac | various | various | unclassified | unclassified | unclassified | unclassified | unclassified | 0,8 | 0,4 | 0,000543307 | 0,00027243 | 1 - | 0,000718802 - | 0 - | 0 - | 1 - | 0,000667022 - | 1 - | 0,000662831 - | 1 - | 0,000667881 - |
| 2098 | 1 | 0.2um_NOKCPHJN_562145 | Bacteria | Bacteroidetes | Flavobacteriia | Flavobacterial | Crocinitomica | unclassified | Crocinitomica | poorly charact | Function unkn | 16R8I | Lipoprotein | 715451.ambt_ | 3 | 0,894427191 | 0,000199702 | 5,75E-05 | 2 - | 0,000134914 - | 3 - | 0,000238084 - | 4 - | 0,00025039 - | 2 - | 0,000124408 - | 4 - | 0,000250712 - |
| 2099 | 1 | 3um_FIJJAFBF_599427 | Bacteria | Proteobacteri | Alphaproteob | Rhodospirillal | Rhodospirillac | unclassified | Rhodospirillac | unclassified | unclassified | unclassified | unclassified | unclassified | 0,6 | 0,489897949 | 9,01E-05 | 7,37E-05 | 1 - | 0,000158007 - | 0 - | 0 - | 0 - | 0 - | 1 - | 0,000145703 - | 1 - | 0,000146813 - |
| 2100 | 1 | 0.2um_NOKCPHJN_473189 | Bacteria | Proteobacteri | Gammaprotec | unclassified | unclassified | unclassified | Gammaprotec | poorly charact | Function unkn | 070MQ | NA | 425104.Ssed_ | 0,6 | 0,489897949 | 0,00016559 | 0,000135435 | 1 - | 0,000290377 - | 0 - | 0 - | 0 - | 0 - | 1 - | 0,000267766 - | 1 - | 0,000269806 - |
| 2101 | 1 | 3um_FIJJAFBF_652365 | Bacteria | Bacteroidetes | Flavobacteriia | Flavobacterial | Flavobacteriac | Gaetbulibacte | Gaetbulibacte | poorly charact | Function unkn | 0XPRT | NA | 391598.FBBAL | 1,4 | 0,489897949 | 0,000434341 | 0,000189545 | 2 - | 0,000608985 - | 2 - | 0,000716458 - | 1 - | 0,000282558 - | 1 - | 0,000280782 - | 1 - | 0,000282922 - |
| 2102 | 1 | 3um_FIJJAFBF_259313 | Bacteria | Planctomycete | Phycisphaerae | Phycisphaeral | Phycisphaerac | Phycisphaera | Phycisphaera | unclassified | unclassified | unclassified | unclassified | unclassified | 4,2 | 0,4 | 0,001368294 | 0,000146123 | 4 - | 0,001318705 - | 4 - | 0,001551427 - | 4 - | 0,00122371 - | 4 - | 0,00121602 - | 5 - | 0,001531607 - |
| 2103 | 1 | 3um_FIJJAFBF_663182 | Bacteria | Proteobacteri | Gammaprotec | unclassified | unclassified | unclassified | SAR86 cluster | information st | Translation, ri | COG1186 | Peptide chain | 247633.GP21 | 0,8 | 0,4 | 0,000212167 | 0,000109219 | 0 - | 0 - | 1 - | 0,000315504 - | 1 - | 0,000248859 - | 1 - | 0,000247295 - | 1 - | 0,000249179 - |
| 2104 | 1 | 0.2um_NOKCPHJN_36508 | Bacteria | Bacteroidetes | Cytophagia | Cytophagales | Flammeovirga | unclassified | Flammeovirga | poorly charact | Function unkn | 1AINX | Phage tail she | 760192.Halhy | 3,4 | 0,489897949 | 0,00059862 | 6,98E-05 | 3 - | 0,000536901 - | 3 - | 0,000631652 - | 4 - | 0,0006643 - | 3 - | 0,000495094 - | 4 - | 0,000665155 - |
| 2105 | 1 | 3um_FIJJAFBF_383207 | Bacteria | Proteobacteri | Deltaproteoba | Myxococcales | Nannocystace | Plesiocystis | Plesiocystis pa | unclassified | unclassified | unclassified | unclassified | unclassified | 1 | 0,894427191 | 0,000592804 | 0,000529962 | 0 - | 0 - | 0 - | 0 - | 1 - | 0,000593991 - | 2 - | 0,001180516 - | 2 - | 0,001189511 - |
| 2106 | 1 | 3um_FIJJAFBF_798846 | Bacteria | Proteobacteri | Gammaprotec | Oceanospirilla | Oceanospirilla | Motiliproteus | Motiliproteus | unclassified | unclassified | unclassified | unclassified | unclassified | 1 | 0 | 0,000269094 | 2,63E-05 | 1 - | 0,000271498 - | 1 - | 0,000319411 - | 1 - | 0,00025194 - | 1 - | 0,000250357 - | 1 - | 0,000252265 - |
| 2107 | 1 | 0.2um_NOKCPHJN_700347 | Eukaryota | Eustigmatoph | unclassified | Eustigmatales | Monodopsida | Nannochlorop | Nannochlorop | metabolism | Carbohydrate | KOG2638 | UTP-glucose-1 | 653948.CCA1 | 4,8 | 1,469693846 | 0,000443739 | 0,000113778 | 5 - | 0,000480251 - | 2 - | 0,000226002 - | 6 - | 0,000534787 - | 6 - | 0,000531426 - | 5 - | 0,000446229 - |
| 2108 | 1 | 3um_FIJJAFBF_715087 | Eukaryota | Bacillariophyt | Bacillariophyc | Naviculales | Phaeodactylac | Phaeodactylu | Phaeodactylu | unclassified | unclassified | unclassified | unclassified | unclassified | 1,4 | 0,489897949 | 0,001023362 | 0,000425729 | 1 - | 0,000724743 - | 2 - | 0,001705287 - | 2 - | 0,00134507 - | 1 - | 0,000668309 - | 1 - | 0,000673401 - |
| 2109 | 1 | 0.2um_NOKCPHJN_145730 | Bacteria | Proteobacteri | Betaproteoba | Burkholderiale | Comamonada | Hydrogenoph | various | information st | Transcription | COG0086 | DNA-depende | 596154.Alide2 | 0,2 | 0,4 | 1,15E-05 | 2,30E-05 | 0 - | 0 - | 0 - | 0 - | 1 - | 5,76E-05 - | 0 - | 0 - | 0 - | 0 - |
| 2110 | 1 | 0.2um_NOKCPHJN_479544 | Eukaryota | unclassified | Labyrinthulom | unclassified | Thraustochytr | Aurantiochytr | Aurantiochytr | poorly charact | Function unkn | 0XNR8 | NIPA-like dom | 67593.Physo1 | 0,2 | 0,4 | 5,84E-05 | 0,000116819 | 0 - | 0 - | 0 - | 0 - | 0 - | 0 - | 0 - | 0 - | 1 - | 0,000292048 - |
| 2111 | 1 | 0.2um_NOKCPHJN_405135 | Bacteria | Proteobacteri | various | various | various | various | various | cellular proces | Cell wall/mem | 0B7GS | outer membra | 583345.Mmol | 1 | 0 | 0,000204994 | 2,01E-05 | 1 - | 0,000206825 - | 1 - | 0,000243325 - | 1 - | 0,000191926 - | 1 - | 0,00019072 - | 1 - | 0,000192173 - |
| 2112 | 1 | 3um_FIJJAFBF_277459 | Eukaryota | Bacillariophyt | Mediophyceae | Triceratiales | Plagiogramma | Plagiogramma | Plagiogramma | unclassified | unclassified | unclassified | unclassified | unclassified | 1 | 0 | 0,001034732 | 0,000101242 | 1 - | 0,001043974 - | 1 - | 0,001228213 - | 1 - | 0,000968771 - | 1 - | 0,000962683 - | 1 - | 0,000970018 - |
| 2113 | 1 | 0.2um_NOKCPHJN_759699 | Eukaryota | unclassified | Oomycetes | Peronosporale | Peronosporac | Phytophthora | various | information st | Chromatin str | KOG1744 | Core compone | 164328.Phyra | 1,2 | 0,979795897 | 0,000840426 | 0,000686215 | 0 - | 0 - | 0 - | 0 - | 2 - | 0,001403047 - | 2 - | 0,00139423 - | 2 - | 0,001404853 - |
| 2114 | 1 | 0.2um_NOKCPHJN_951164 | unclassified | unclassified | unclassified | unclassified | unclassified | unclassified | unclassified | unclassified | unclassified | unclassified | unclassified | unclassified | 1 | 0 | 0,000835745 | 8,18E-05 | 1 - | 0,00084321 - | 1 - | 0,000992018 - | 1 - | 0,000782469 - | 1 - | 0,000777551 - | 1 - | 0,000783476 - |
| 2115 | 1 | 3um_FIJJAFBF_114517 | unclassified | unclassified | unclassified | unclassified | unclassified | unclassified | unclassified | unclassified | unclassified | unclassified | unclassified | unclassified | 0,2 | 0,4 | 3,99E-05 | 7,99E-05 | 0 - | 0 - | 0 - | 0 - | 0 - | 0 - | 0 - | 0 - | 1 - | 0,00019971 - |
| 2116 | 1 | 3um_FIJJAFBF_107495 | Bacteria | Proteobacteri | Alphaproteob | Rhodobactera | Rhodobactera | Planktomarina | Planktomarina | metabolism | Inorganic ion t | 16SYE | High-affinity z | 391616.OA23 | 0,2 | 0,4 | 0,000129773 | 0,000259547 | 0 - | 0 - | 1 - | 0,000648867 - | 0 - | 0 - | 0 - | 0 - | 0 - | 0 - |
| 2117 | 1 | 3um_FIJJAFBF_44059 | Viruses | unclassified | unclassified | unclassified | Phycodnavirid | unclassified | Chrysochromu | unclassified | unclassified | unclassified | unclassified | unclassified | 3,2 | 0,748331477 | 0,001482717 | 0,000316291 | 2 - | 0,000942945 - | 3 - | 0,00166403 - | 4 - | 0,001750037 - | 3 - | 0,00130428 - | 4 - | 0,00175229 - |
| 2118 | 1 | 0.2um_NOKCPHJN_864503 | Bacteria | Bacteroidetes | Flavobacteriia | Flavobacterial | Flavobacteriac | Formosa | Formosa sp. H | metabolism | Inorganic ion t | 153M9 | iduronate-2-s | 756272.Plabr_ | 1 | 0,632455532 | 0,00020516 | 0,000124454 | 0 - | 0 - | 1 - | 0,00024741 - | 1 - | 0,000195148 - | 2 - | 0,000387843 - | 1 - | 0,000195399 - |
| 2119 | 1 | 3um_FIJJAFBF_105537 | Eukaryota | Ascomycota | Saccharomyce | Saccharomyce | Saccharomyce | Saccharomyce | Saccharomyce | poorly charact | Function unkn | 0NVRZ | NA | 170187.SP_17 | 7,6 | 1,019803903 | 0,001803055 | 0,000112448 | 7 - | 0,001695737 - | 6 - | 0,001709998 - | 8 - | 0,001798381 - | 9 - | 0,002010465 - | 8 - | 0,001800696 - |
| 2120 | 1 | 0.2um_NOKCPHJN_695286 | Bacteria | Proteobacteri | Alphaproteob | Rhodospirillal | Rhodospirillac | Candidatus En | Candidatus En | unclassified | unclassified | unclassified | unclassified | unclassified | 0,8 | 0,4 | 0,000289936 | 0,000148582 | 1 - | 0,000359401 - | 1 - | 0,000422827 - | 1 - | 0,000333511 - | 0 - | 0 - | 1 - | 0,000333941 - |
| 2121 | 1 | 0.2um_NOKCPHJN_75279 | Bacteria | Bacteroidetes | unclassified | unclassified | unclassified | unclassified | Bacteroidetes | unclassified | unclassified | unclassified | unclassified | unclassified | 1,6 | 1,2 | 0,000102767 | 7,33E-05 | 0 - | 0 - | 1 - | 7,88E-05 - | 3 - | 0,000186501 - | 1 - | 6,18E-05 - | 3 - | 0,000186741 - |
| 2122 | 1 | 3um_FIJJAFBF_99521 | Viruses | unclassified | unclassified | Caudovirales | Siphoviridae | Cbastvirus | Cellulophaga v | unclassified | unclassified | unclassified | unclassified | unclassified | 1,6 | 0,489897949 | 0,000426829 | 0,00014654 | 1 - | 0,000267359 - | 2 - | 0,000629085 - | 2 - | 0,0004962 - | 2 - | 0,000493081 - | 1 - | 0,000248419 - |
| 2123 | 1 | 3um_FIJJAFBF_619427 | Bacteria | Proteobacteri | Gammaprotec | Alteromonada | Colwelliaceae | Thalassomona | Thalassomona | cellular proces | Intracellular tr | 08ZUW | Major pilin sul | 259536.Psyc_ | 0,4 | 0,489897949 | 0,000211504 | 0,000259039 | 0 - | 0 - | 0 - | 0 - | 1 - | 0,00052842 - | 0 - | 0 - | 1 - | 0,000529101 - |
| 2124 | 1 | 3um_FIJJAFBF_807137 | Viruses | unclassified | unclassified | unclassified | Mimiviridae | Cafeteriavirus | Cafeteria roer | unclassified | unclassified | unclassified | unclassified | unclassified | 0,6 | 0,489897949 | 0,000104915 | 8,68E-05 | 1 - | 0,000169293 - | 1 - | 0,00019917 - | 0 - | 0 - | 1 - | 0,000156111 - | 0 - | 0 - |
| 2125 | 1 | 0.2um_NOKCPHJN_48598 | Bacteria | Firmicutes | Bacilli | Lactobacillales | Leuconostoca | Weissella | Weissella con | unclassified | unclassified | unclassified | unclassified | unclassified | 0,2 | 0,4 | 1,20E-05 | 2,41E-05 | 0 - | 0 - | 1 - | 6,02E-05 - | 0 - | 0 - | 0 - | 0 - | 0 - | 0 - |
| 2126 | 1 | 3um_FIJJAFBF_73737 | Bacteria | Proteobacteri | Gammaprotec | Alteromonada | Alteromonada | Alteromonas | Alteromonas s | unclassified | unclassified | unclassified | unclassified | unclassified | 1 | 0 | 0,000819976 | 8,02E-05 | 1 - | 0,0008273 - | 1 - | 0,000973301 - | 1 - | 0,000767705 - | 1 - | 0,000762881 - | 1 - | 0,000768693 - |
| 2127 | 1 | 0.2um_NOKCPHJN_243077 | Bacteria | Bacteroidetes | Flavobacteriia | Flavobacterial | Flavobacteriac | Muricauda | Muricauda sp. | cellular proces | Intracellular tr | 18ATJ | Pfam:YadA | 626418.bglu_ | 0,8 | 0,4 | 6,10E-05 | 3,13E-05 | 1 - | 7,58E-05 - | 1 - | 8,92E-05 - | 1 - | 7,03E-05 - | 1 - | 6,99E-05 - | 0 - | 0 - |
| 2128 | 1 | 3um_FIJJAFBF_815477 | Eukaryota | Bacillariophyt | Coscinodiscop | Thalassiosirale | Stephanodisca | Cyclotella | Cyclotella cry | metabolism | Lipid transpor | COG0439 | acetyl-CoA ca | 2850.Phatr54 | 2,4 | 1,019803903 | 0,001358703 | 0,000528051 | 1 - | 0,000580754 - | 2 - | 0,001366489 - | 3 - | 0,001616756 - | 2 - | 0,001071064 - | 4 - | 0,00215845 - |
| 2129 | 1 | 0.2um_NOKCPHJN_266660 | Bacteria | Acidobacteria | unclassified | unclassified | unclassified | unclassified | Acidobacteria | information st | Replication, re | COG0210 | helicase | 498848.TaqD | 0,8 | 0,4 | 0,000131008 | 6,71E-05 | 1 - | 0,000162396 - | 1 - | 0,000191055 - | 1 - | 0,000150698 - | 0 - | 0 - | 1 - | 0,000150892 - |
| 2130 | 1 | 3um_FIJJAFBF_152376 | unclassified | unclassified | unclassified | unclassified | unclassified | unclassified | unclassified | unclassified | unclassified | unclassified | unclassified | unclassified | 10,2 | 0,748331477 | 0,003443577 | 0,000495047 | 9 - | 0,003059088 - | 11 - | 0,004398716 - | 10 - | 0,003154137 - | 11 - | 0,003447748 - | 10 - | 0,003158197 - |
| 2131 | 1 | 0.2um_NOKCPHJN_447107 | Bacteria | Bacteroidetes | Flavobacteriia | Flavobacterial | Flavobacteriac | Formosa | Formosa sp. H | unclassified | unclassified | unclassified | unclassified | unclassified | 0,4 | 0,489897949 | 5,66E-05 | 6,94E-05 | 1 - | 0,000146645 - | 0 - | 0 - | 0 - | 0 - | 0 - | 0 - | 1 - | 0,000136257 - |
| 2132 | 1 | 0.2um_NOKCPHJN_56999 | Bacteria | Bacteroidetes | Flavobacteriia | Flavobacterial | Flavobacteriac | Algibacter | Algibacter lect | unclassified | unclassified | unclassified | unclassified | unclassified | 0,6 | 0,489897949 | 0,000193027 | 0,000160303 | 0 - | 0 - | 1 - | 0,000375163 - | 1 - | 0,000295915 - | 1 - | 0,000294056 - | 0 - | 0 - |
| 2133 | 1 | 3um_FIJJAFBF_459209 | Eukaryota | unclassified | unclassified | Isochrysidales | Noelaerhabda | Emiliania | Emiliania hux | cellular proces | Posttranslatio | KOG0730 | cell division cy | 35128.Thapsd | 0,4 | 0,489897949 | 0,00013163 | 0,000162094 | 1 - | 0,000302393 - | 1 - | 0,000355758 - | 0 - | 0 - | 0 - | 0 - | 0 - | 0 - |
| 2134 | 1 | 3um_FIJJAFBF_155390 | Eukaryota | Chytridiomyc | Monoblephari | Monoblephari | Gonapodyace | Gonapodya | Gonapodya pr | unclassified | unclassified | unclassified | unclassified | unclassified | 1 | 0 | 0,000648638 | 6,35E-05 | 1 - | 0,000654432 - | 1 - | 0,000769925 - | 1 - | 0,000607289 - | 1 - | 0,000603473 - | 1 - | 0,000608071 - |
| 2135 | 1 | 3um_FIJJAFBF_635344 | Eukaryota | unclassified | unclassified | unclassified | Apusomonadi | Thecamonas | Thecamonas t | poorly charact | Function unkn | 0T9G1 | kinetoplast-as | 353153.XP_8 | 0,2 | 0,4 | 0,000105706 | 0,000211413 | 0 - | 0 - | 0 - | 0 - | 0 - | 0 - | 1 - | 0,000528532 - | 0 - | 0 - |
| 2136 | 1 | 0.2um_NOKCPHJN_285267 | unclassified | unclassified | unclassified | unclassified | unclassified | unclassified | unclassified | unclassified | unclassified | unclassified | unclassified | unclassified | 0,6 | 0,489897949 | 3,02E-05 | 2,47E-05 | 0 - | 0 - | 0 - | 0 - | 1 - | 5,05E-05 - | 1 - | 5,02E-05 - | 1 - | 5,05E-05 - |
| 2137 | 1 | 3um_FIJJAFBF_205192 | Eukaryota | Bacillariophyt | Mediophyceae | Toxariales | Toxariaceae | Toxarium | Toxarium und | cellular proces | Posttranslatio | 0IEKR | One of the cor | 45157.CMV08 | 1,2 | 0,4 | 0,000224598 | 6,62E-05 | 1 - | 0,000191054 - | 1 - | 0,000224771 - | 1 - | 0,000177291 - | 2 - | 0,000352354 - | 1 - | 0,00017752 - |
| 2138 | 1 | 3um_FIJJAFBF_158698 | Bacteria | Proteobacteri | Gammaprotec | Oceanospirilla | Alcanivoracac | Ketobacter | Ketobacter alk | unclassified | unclassified | unclassified | unclassified | unclassified | 0,2 | 0,4 | 6,82E-05 | 0,000136488 | 1 - | 0,000341221 - | 0 - | 0 - | 0 - | 0 - | 0 - | 0 - | 0 - | 0 - |
| 2139 | 1 | 0.2um_NOKCPHJN_843197 | Viruses | unclassified | unclassified | unclassified | various | various | various | unclassified | unclassified | unclassified | unclassified | unclassified | 0,6 | 0,489897949 | 0,000164852 | 0,000136905 | 0 - | 0 - | 1 - | 0,000320403 - | 1 - | 0,000252723 - | 1 - | 0,000251135 - | 0 - | 0 - |
| 2140 | 1 | 3um_FIJJAFBF_430380 | Bacteria | Proteobacteri | Alphaproteob | Rhodobactera | unclassified | unclassified | Rhodobactera | cellular proces | Cell wall/mem | COG2885 | Ompa motb d | 388739.RSK20 | 0,6 | 0,489897949 | 0,000266545 | 0,000217973 | 1 - | 0,000466457 - | 0 - | 0 - | 1 - | 0,000432855 - | 0 - | 0 - | 1 - | 0,000433412 - |
| 2141 | 2 | 0.2um_NOKCPHJN_489442;3ur | Eukaryota | various | various | various | various | various | various | poorly charact | Function unkn | COG1100 | GTP-binding P | various | 1 | 0 | 0,000634434 | 6,21E-05 | 1 - | 0,000640101 - | 1 - | 0,000753065 - | 1 - | 0,000593991 - | 1 - | 0,000590258 - | 1 - | 0,000594755 - |
| 2142 | 1 | 0.2um_NOKCPHJN_530548 | Eukaryota | Bacillariophyt | Coscinodiscop | Thalassiosirale | Thalassiosirac | Thalassiosira | Thalassiosira | information st | Chromatin str | KOG1744 | Core compone | 2850.Phatr54 | 1,4 | 0,489897949 | 0,001067472 | 0,00044408 | 1 - | 0,000755981 - | 2 - | 0,001778791 - | 2 - | 0,001403047 - | 1 - | 0,000697115 - | 1 - | 0,000702427 - |
| 2143 | 1 | 0.2um_NOKCPHJN_281234 | Eukaryota | unclassified | unclassified | Kinetoplastida | Trypanosomat | Trypanosoma | Trypanosoma | unclassified | unclassified | unclassified | unclassified | unclassified | 0,6 | 0,489897949 | 0,00010332 | 8,45E-05 | 1 - | 0,000180812 - | 0 - | 0 - | 1 - | 0,000167787 - | 0 - | 0 - | 1 - | 0,000168003 - |
| 2144 | 1 | 0.2um_NOKCPHJN_312807 | Eukaryota | Ascomycota | Saccharomyce | Saccharomyce | Saccharomyce | Nakaseomyce | [Candida] glab | unclassified | unclassified | unclassified | unclassified | unclassified | 0,2 | 0,4 | 1,04E-05 | 2,09E-05 | 1 - | 5,22E-05 - | 0 - | 0 - | 0 - | 0 - | 0 - | 0 - | 0 - | 0 - |
| 2145 | 1 | 3um_FIJJAFBF_241394 | Eukaryota | Bacillariophyt | Coscinodiscop | Thalassiosirale | Thalassiosirac | Thalassiosira | Thalassiosira | unclassified | unclassified | unclassified | unclassified | unclassified | 1,4 | 0,489897949 | 0,000622328 | 0,000208638 | 2 - | 0,00090406 - | 1 - | 0,000531804 - | 2 - | 0,000838935 - | 1 - | 0,000416832 - | 1 - | 0,000420008 - |
| 2146 | 1 | 0.2um_NOKCPHJN_175750 | Eukaryota | unclassified | Pelagophycea | Pelagomonad | unclassified | Aureococcus | Aureococcus a | unclassified | unclassified | unclassified | unclassified | unclassified | 0,6 | 0,489897949 | 0,000244948 | 0,000200002 | 0 - | 0 - | 0 - | 0 - | 1 - | 0,000408928 - | 1 - | 0,000406359 - | 1 - | 0,000409455 - |
| 2147 | 1 | 0.2um_NOKCPHJN_373963 | Eukaryota | unclassified | Pelagophycea | Pelagomonad | unclassified | Aureococcus | Aureococcus a | poorly charact | Function unkn | 0J2G0 | C-Myc binding | 67593.Physo1 | 0,2 | 0,4 | 0,000229266 | 0,000458533 | 0 - | 0 - | 1 - | 0,001146332 - | 0 - | 0 - | 0 - | 0 - | 0 - | 0 - |
| 2148 | 1 | 0.2um_NOKCPHJN_1101372 | unclassified | unclassified | unclassified | unclassified | unclassified | unclassified | unclassified | unclassified | unclassified | unclassified | unclassified | unclassified | 1 | 0 | 0,000234279 | 2,29E-05 | 1 - | 0,000236372 - | 1 - | 0,000278086 - | 1 - | 0,000219344 - | 1 - | 0,000217966 - | 1 - | 0,000219627 - |
| 2149 | 1 | 3um_FIJJAFBF_70346 | Bacteria | Proteobacteri | Betaproteoba | unclassified | unclassified | unclassified | Betaproteoba | information st | Translation, ri | 16Q16 | Binds the low | 887062.HGR_ | 0,2 | 0,4 | 5,67E-05 | 0,000113417 | 0 - | 0 - | 0 - | 0 - | 1 - | 0,000283543 - | 0 - | 0 - | 0 - | 0 - |
| 2150 | 1 | 3um_FIJJAFBF_123244 | Viruses | unclassified | unclassified | unclassified | various | various | various | unclassified | unclassified | unclassified | unclassified | unclassified | 3,8 | 0,748331477 | 0,000619379 | 8,86E-05 | 3 - | 0,000500155 - | 3 - | 0,000588421 - | 4 - | 0,000618835 - | 4 - | 0,000614946 - | 5 - | 0,000774539 - |
| 2151 | 1 | 3um_FIJJAFBF_636517 | Bacteria | Bacteroidetes | Flavobacteriia | Flavobacterial | Flavobacteriac | Flavivirga | Flavivirga eck | metabolism | Lipid transpor | COG4247 | Phytase | 867900.Celly_ | 1,6 | 0,489897949 | 0,000407394 | 9,97E-05 | 1 - | 0,000263345 - | 1 - | 0,000309819 - | 2 - | 0,000488749 - | 2 - | 0,000485678 - | 2 - | 0,000489378 - |
| 2152 | 1 | 3um_FIJJAFBF_38797 | Viruses | unclassified | unclassified | unclassified | Phycodnavirid | unclassified | Chrysochromu | cellular proces | Posttranslatio | 1BMP7 | DnaJ homolog | 3694.POPTR_ | 0,2 | 0,4 | 4,56E-05 | 9,11E-05 | 0 - | 0 - | 0 - | 0 - | 0 - | 0 - | 1 - | 0,00022779 - | 0 - | 0 - |
| 2153 | 1 | 3um_FIJJAFBF_594149 | Eukaryota | unclassified | Pelagophycea | Pelagomonad | unclassified | Aureoumbra | Aureoumbra l | metabolism | Carbohydrate | COG1850 | RuBisCO catal | 45157.CMV01 | 19,2 | 1,833030278 | 0,003397686 | 0,000130689 | 19 - | 0,003421321 - | 16 - | 0,003389565 - | 21 - | 0,003509059 - | 19 - | 0,003154911 - | 21 - | 0,003513576 - |
| 2154 | 1 | 3um_FIJJAFBF_452605 | Bacteria | Proteobacteri | Alphaproteob | Sphingomona | Erythrobacter | Altererythrob | Altererythrob | cellular proces | Signal transdu | 01WWM | Histidine kina | 314225.ELI_1 | 0,2 | 0,4 | 4,08E-05 | 8,16E-05 | 0 - | 0 - | 1 - | 0,000203893 - | 0 - | 0 - | 0 - | 0 - | 0 - | 0 - |
| 2155 | 1 | 3um_FIJJAFBF_197145 | Viruses | unclassified | unclassified | unclassified | various | various | various | unclassified | unclassified | unclassified | unclassified | unclassified | 0,6 | 0,489897949 | 7,33E-05 | 5,99E-05 | 0 - | 0 - | 0 - | 0 - | 1 - | 0,000122371 - | 1 - | 0,000121602 - | 1 - | 0,000122529 - |
| 2156 | 1 | 0.2um_NOKCPHJN_39142 | Eukaryota | Phaeophyceae | unclassified | Laminariales | Agaraceae | Costaria | Costaria costa | metabolism | Energy produc | COG4451 | RuBisCO catal | 288000.BBta_ | 9,8 | 1,166190379 | 0,006118674 | 0,000876208 | 8 - | 0,005047128 - | 10 - | 0,007422294 - | 11 - | 0,006439886 - | 9 - | 0,005235886 - | 11 - | 0,006448176 - |
| 2157 | 1 | 0.2um_NOKCPHJN_433505 | Bacteria | Bacteroidetes | Flavobacteriia | Flavobacterial | unclassified | unclassified | Flavobacterial | poorly charact | Function unkn | 0NVRZ | NA | 170187.SP_17 | 0,6 | 0,489897949 | 4,38E-05 | 3,62E-05 | 1 - | 7,06E-05 - | 1 - | 8,30E-05 - | 1 - | 6,55E-05 - | 0 - | 0 - | 0 - | 0 - |
| 2158 | 1 | 3um_FIJJAFBF_183675 | Bacteria | Bacteroidetes | Flavobacteriia | Flavobacterial | Flavobacteriac | Arenitalea | Arenitalea lut | unclassified | unclassified | unclassified | unclassified | unclassified | 17,6 | 1,356465997 | 0,001805959 | 9,87E-05 | 16 - | 0,001666391 - | 16 - | 0,001960473 - | 19 - | 0,001836292 - | 19 - | 0,001824753 - | 18 - | 0,001741885 - |
| 2159 | 1 | 0.2um_NOKCPHJN_86147 | Eukaryota | unclassified | Oomycetes | Peronosporale | Peronosporac | Phytophthora | Phytophthora | cellular proces | Cell cycle cont | 0WI8M | Echinoderm m | 8128.ENSONI | 0,2 | 0,4 | 1,34E-05 | 2,67E-05 | 0 - | 0 - | 0 - | 0 - | 1 - | 6,69E-05 - | 0 - | 0 - | 0 - | 0 - |
| 2160 | 1 | 3um_FIJJAFBF_741972 | unclassified | unclassified | unclassified | unclassified | unclassified | unclassified | unclassified | unclassified | unclassified | unclassified | unclassified | unclassified | 0,2 | 0,4 | 3,09E-05 | 6,17E-05 | 0 - | 0 - | 0 - | 0 - | 0 - | 0 - | 1 - | 0,000154323 - | 0 - | 0 - |
| 2161 | 2 | 0.2um_NOKCPHJN_13837;0.2u | Bacteria | various | various | various | various | various | various | cellular proces | Posttranslatio | various | various | various | 1,2 | 0,4 | 0,000437346 | 0,000131112 | 1 - | 0,000371584 - | 1 - | 0,000437161 - | 1 - | 0,000344817 - | 1 - | 0,00034265 - | 2 - | 0,000690521 - |
| 2162 | 1 | 3um_FIJJAFBF_183439 | Bacteria | Bacteroidetes | Cytophagia | Cytophagales | Flammeovirga | unclassified | Flammeovirga | metabolism | Inorganic ion t | 05C27 | TonB-depende | 688270.Celal_ | 2,2 | 2,13541565 | 0,000179394 | 0,000180257 | 6 - | 0,000505926 - | 0 - | 0 - | 1 - | 7,82E-05 - | 1 - | 7,78E-05 - | 3 - | 0,000235043 - |
| 2163 | 1 | 3um_FIJJAFBF_115900 | Bacteria | Proteobacteri | Gammaprotec | Chromatiales | Chromatiacea | Thiorhodovibr | Thiorhodovibr | information st | Translation, ri | COG0050 | This protein p | 572477.Alvin_ | 0,2 | 0,4 | 6,33E-05 | 0,000126656 | 0 - | 0 - | 0 - | 0 - | 1 - | 0,000316641 - | 0 - | 0 - | 0 - | 0 - |
| 2164 | 1 | 3um_FIJJAFBF_353321 | Bacteria | Proteobacteri | Alphaproteob | Rhodobactera | Rhodobactera | unclassified | Rhodobactera | unclassified | unclassified | unclassified | unclassified | unclassified | 1 | 0 | 0,00019576 | 1,92E-05 | 1 - | 0,000197509 - | 1 - | 0,000232365 - | 1 - | 0,000183281 - | 1 - | 0,000182129 - | 1 - | 0,000183517 - |
| 2165 | 1 | 0.2um_NOKCPHJN_343386 | Bacteria | Proteobacteri | Gammaprotec | unclassified | unclassified | unclassified | Gammaprotec | metabolism | Inorganic ion t | 16TQC | TONB-depend | 314287.GB22 | 1 | 0,632455532 | 0,000105443 | 6,55E-05 | 1 - | 0,00011214 - | 0 - | 0 - | 1 - | 0,000104062 - | 2 - | 0,000206817 - | 1 - | 0,000104196 - |
| 2166 | 1 | 3um_FIJJAFBF_187036 | Bacteria | Proteobacteri | Gammaprotec | Alteromonada | Alteromonada | Paraglaciecola | Paraglaciecola | unclassified | unclassified | unclassified | unclassified | unclassified | 0,4 | 0,489897949 | 0,000118442 | 0,000145062 | 0 - | 0 - | 0 - | 0 - | 1 - | 0,000295915 - | 0 - | 0 - | 1 - | 0,000296296 - |
| 2167 | 1 | 0.2um_NOKCPHJN_309398 | Bacteria | Proteobacteri | Alphaproteob | Rhodobactera | Rhodobactera | unclassified | Rhodobactera | metabolism | Energy produc | 01RNF | Produces ATP | 314264.ROS2 | 1 | 0 | 0,00018337 | 1,79E-05 | 1 - | 0,000185008 - | 1 - | 0,000217658 - | 1 - | 0,000171681 - | 1 - | 0,000170602 - | 1 - | 0,000171902 - |
| 2168 | 2 | 0.2um_NOKCPHJN_784123;3ur | various | various | various | various | various | various | various | unclassified | unclassified | unclassified | unclassified | unclassified | 0,2 | 0,4 | 4,73E-05 | 9,47E-05 | 0 - | 0 - | 1 - | 0,000236628 - | 0 - | 0 - | 0 - | 0 - | 0 - | 0 - |
| 2169 | 1 | 0.2um_NOKCPHJN_87559 | Bacteria | Proteobacteri | Alphaproteob | Rhodospirillal | Rhodospirillac | Nisaea | Nisaea denitri | unclassified | unclassified | unclassified | unclassified | unclassified | 0,8 | 0,4 | 0,000141002 | 7,23E-05 | 1 - | 0,000175038 - | 1 - | 0,000205928 - | 0 - | 0 - | 1 - | 0,000161408 - | 1 - | 0,000162638 - |
| 2170 | 1 | 3um_FIJJAFBF_356540 | Bacteria | Proteobacteri | Alphaproteob | Rhodobactera | various | various | various | information st | Transcription | 05VEQ | Cold shock pro | 391619.RGBS | 0,8 | 0,4 | 0,000974757 | 0,000488772 | 1 - | 0,001289615 - | 0 - | 0 - | 1 - | 0,001196717 - | 1 - | 0,001189196 - | 1 - | 0,001198257 - |
| 2171 | 2 | 3um_FIJJAFBF_145240;3um_FI | Bacteria | Bacteroidetes | Flavobacteriia | Flavobacterial | Flavobacteriac | Algibacter | Algibacter sp. | unclassified | unclassified | unclassified | unclassified | unclassified | 0,6 | 0,489897949 | 0,000195156 | 0,000162071 | 0 - | 0 - | 1 - | 0,000379301 - | 1 - | 0,000299179 - | 1 - | 0,000297299 - | 0 - | 0 - |
| 2172 | 1 | 3um_FIJJAFBF_467720 | Eukaryota | unclassified | Cryptophyta | Pyrenomonad | Geminigerace | Teleaulax | Teleaulax amp | metabolism | Energy produc | COG0711 | F(1)F(0) ATP s | 118168.MC74 | 0,8 | 0,4 | 0,000423041 | 0,000217771 | 0 - | 0 - | 1 - | 0,000629085 - | 1 - | 0,0004962 - | 1 - | 0,000493081 - | 1 - | 0,000496838 - |
| 2173 | 1 | 0.2um_NOKCPHJN_787782 | unclassified | unclassified | unclassified | unclassified | unclassified | unclassified | unclassified | unclassified | unclassified | unclassified | unclassified | unclassified | 0,6 | 0,489897949 | 0,002717291 | 0,002247899 | 1 - | 0,004384693 - | 1 - | 0,005158494 - | 0 - | 0 - | 1 - | 0,004043268 - | 0 - | 0 - |
| 2174 | 1 | 0.2um_NOKCPHJN_1011565 | Viruses | unclassified | unclassified | unclassified | unclassified | unclassified | uncultured vir | unclassified | unclassified | unclassified | unclassified | unclassified | 0,6 | 0,489897949 | 0,000210644 | 0,000174934 | 0 - | 0 - | 1 - | 0,000409404 - | 1 - | 0,000322924 - | 1 - | 0,000320894 - | 0 - | 0 - |
| 2175 | 1 | 0.2um_NOKCPHJN_912697 | Eukaryota | Basidiomycota | Agaricomycet | Auriculariales | Auriculariacea | Auricularia | Auricularia sul | unclassified | unclassified | unclassified | unclassified | unclassified | 1 | 0,632455532 | 0,000181848 | 0,000113506 | 1 - | 0,000193158 - | 0 - | 0 - | 2 - | 0,000358488 - | 1 - | 0,000178118 - | 1 - | 0,000179475 - |
| 2176 | 1 | 3um_FIJJAFBF_783854 | Bacteria | Bacteroidetes | Flavobacteriia | Flavobacterial | Flavobacteriac | Tenacibaculur | Tenacibaculur | poorly charact | Function unkn | 11P35 | NA | 983544.Lacal_ | 2,6 | 0,489897949 | 0,000405427 | 5,04E-05 | 2 - | 0,000319468 - | 2 - | 0,000375847 - | 3 - | 0,000444682 - | 3 - | 0,000441887 - | 3 - | 0,000445254 - |
| 2177 | 1 | 0.2um_NOKCPHJN_908003 | Bacteria | Proteobacteri | Gammaprotec | Aeromonadale | Aeromonadac | Aeromonas | Aeromonas sp | unclassified | unclassified | unclassified | unclassified | unclassified | 1 | 0 | 0,000259455 | 2,54E-05 | 1 - | 0,000261773 - | 1 - | 0,00030797 - | 1 - | 0,000242916 - | 1 - | 0,000241389 - | 1 - | 0,000243228 - |
| 2178 | 2 | 0.2um_NOKCPHJN_130260;3ur | Eukaryota | various | various | various | various | various | various | metabolism | Energy produc | COG1290 | Component of | 35128.Thapsd | 2 | 0 | 0,000808535 | 7,91E-05 | 2 - | 0,000815757 - | 2 - | 0,00095972 - | 2 - | 0,000756993 - | 2 - | 0,000752236 - | 2 - | 0,000757967 - |
| 2179 | 1 | 3um_FIJJAFBF_485317 | Eukaryota | Phaeophyceae | unclassified | Ectocarpales | Ectocarpaceae | Ectocarpus | Ectocarpus sili | information st | Translation, ri | 151B4 | Ribosomal fan | 67593.Physo1 | 0,6 | 0,489897949 | 0,000206216 | 0,000168637 | 1 - | 0,00036088 - | 0 - | 0 - | 1 - | 0,000334884 - | 0 - | 0 - | 1 - | 0,000335315 - |
| 2180 | 1 | 3um_FIJJAFBF_118447 | Bacteria | Proteobacteri | Alphaproteob | Rhodobactera | Rhodobactera | unclassified | Rhodobactera | unclassified | unclassified | unclassified | unclassified | unclassified | 2 | 0,632455532 | 0,000372251 | 9,76E-05 | 2 - | 0,000385467 - | 1 - | 0,000226747 - | 2 - | 0,0003577 - | 3 - | 0,000533178 - | 2 - | 0,00035816 - |
| 2181 | 1 | 0.2um_NOKCPHJN_168610 | Bacteria | Spirochaetes | Spirochaetia | unclassified | Leptospiracea | Leptospira | Leptospira sp. | unclassified | unclassified | unclassified | unclassified | unclassified | 0,2 | 0,4 | 5,35E-05 | 0,000106912 | 0 - | 0 - | 1 - | 0,00026728 - | 0 - | 0 - | 0 - | 0 - | 0 - | 0 - |
| 2182 | 2 | 0.2um_NOKCPHJN_18165;3um | Bacteria | Proteobacteri | Epsilonproteo | Nautiliales | Nautiliaceae | Nautilia | Nautilia profu | various | various | various | various | various | 0,4 | 0,489897949 | 0,000242309 | 0,00029677 | 0 - | 0 - | 0 - | 0 - | 0 - | 0 - | 1 - | 0,000603473 - | 1 - | 0,000608071 - |
| 2183 | 1 | 0.2um_NOKCPHJN_430131 | Bacteria | Proteobacteri | Alphaproteob | Rhodobactera | Rhodobactera | Paracoccus | Paracoccus pa | unclassified | unclassified | unclassified | unclassified | unclassified | 1 | 0 | 0,000445731 | 4,36E-05 | 1 - | 0,000449712 - | 1 - | 0,000529076 - | 1 - | 0,000417317 - | 1 - | 0,000414694 - | 1 - | 0,000417854 - |
| 2184 | 1 | 3um_FIJJAFBF_783853 | Bacteria | Bacteroidetes | Flavobacteriia | Flavobacterial | Flavobacteriac | Polaribacter | Polaribacter s | unclassified | unclassified | unclassified | unclassified | unclassified | 2 | 0,632455532 | 0,000203985 | 6,18E-05 | 1 - | 0,000103657 - | 2 - | 0,0002439 - | 3 - | 0,00028857 - | 2 - | 0,000191171 - | 2 - | 0,000192628 - |
| 2185 | 1 | 3um_FIJJAFBF_169756 | Bacteria | Bacteroidetes | Saprospiria | Saprospirales | Lewinellaceae | Lewinella | Lewinella pers | metabolism | Inorganic ion t | 05C27 | TonB-depende | 391603.FBALC | 1 | 0 | 8,09E-05 | 7,91E-06 | 1 - | 8,16E-05 - | 1 - | 9,60E-05 - | 1 - | 7,57E-05 - | 1 - | 7,52E-05 - | 1 - | 7,58E-05 - |
| 2186 | 1 | 0.2um_NOKCPHJN_51590 | Bacteria | Proteobacteri | Gammaprotec | Cellvibrionales | Porticoccacea | unclassified | Gammaprotec | metabolism | Inorganic ion t | 16TQC | TONB-depend | 314287.GB22 | 0,6 | 0,489897949 | 5,17E-05 | 4,22E-05 | 1 - | 9,04E-05 - | 0 - | 0 - | 1 - | 8,39E-05 - | 0 - | 0 - | 1 - | 8,40E-05 - |
| 2187 | 1 | 3um_FIJJAFBF_242673 | Bacteria | Bacteroidetes | Saprospiria | Saprospirales | Haliscomenob | Haliscomenob | Haliscomenob | cellular proces | Posttranslatio | COG1404 | peptidase (S8 | 760192.Halhy | 0,8 | 0,4 | 0,000182599 | 9,16E-05 | 1 - | 0,000241581 - | 0 - | 0 - | 1 - | 0,000224178 - | 1 - | 0,00022277 - | 1 - | 0,000224467 - |

| | | | | | | | | | | | | | | | | | | | | | | | | | | | | |
|---|---|---|---|---|---|---|---|---|---|---|---|---|---|---|---|---|---|---|---|---|---|---|---|---|---|---|---|---|
| 2188 | 1 | 3um_FIJJAFBF_145240 | Bacteria | Bacteroidetes | Flavobacteriia | Flavobacterial | Flavobacteriac | Algibacter | Algibacter sp. | unclassified | unclassified | unclassified | unclassified | unclassified | 0,2 | 0,4 | 6,01E-05 | 0,000120268 | 0 - | 0 - | 0 - | 0 - | 0 - | 0 - | 0 - | 0 - | 1 - | 0,00030067 - |
| 2189 | 1 | 3um_FIJJAFBF_779814 | Eukaryota | unclassified | unclassified | Kinetoplastida | Trypanosomat | Trypanosoma | Trypanosoma | poorly charact | Function unkn | 0T9G1 | kinetoplast-as | 353153.XP_80 | 0,6 | 0,489897949 | 0,000357201 | 0,000292152 | 1 - | 0,000626385 - | 0 - | 0 - | 0 - | 0 - | 1 - | 0,00057761 - | 1 - | 0,000582011 - |
| 2190 | 1 | 0.2um_NOKCPHJN_17420 | Bacteria | Proteobacteri | Gammaprotec | Alteromonada | Pseudoalteror | Pseudoalteror | Pseudoalteror | unclassified | unclassified | unclassified | unclassified | unclassified | 1 | 0 | 9,28E-05 | 9,08E-06 | 1 - | 9,36E-05 - | 1 - | 0,000110107 - | 1 - | 8,68E-05 - | 1 - | 8,63E-05 - | 1 - | 8,70E-05 - |
| 2191 | 1 | 0.2um_NOKCPHJN_408548 | Bacteria | Proteobacteri | Gammaprotec | Alteromonada | Alteromonada | Bowmanella | Bowmanella d | metabolism | Inorganic ion t | 0QJVD | TonB-depende | 743721.Psesu | 1 | 0 | 9,89E-05 | 9,68E-06 | 1 - | 9,98E-05 - | 1 - | 0,000117372 - | 1 - | 9,26E-05 - | 1 - | 9,20E-05 - | 1 - | 9,27E-05 - |
| 2192 | 1 | 3um_FIJJAFBF_560873 | Bacteria | Proteobacteri | Gammaprotec | Cellvibrionales | Porticoccacea | unclassified | Gammaprotec | metabolism | Inorganic ion t | 16TQC | TONB-depend | 314287.GB22( | 2,8 | 0,979795897 | 0,000220768 | 6,83E-05 | 3 - | 0,000246331 - | 1 - | 9,66E-05 - | 3 - | 0,000228586 - | 3 - | 0,00022715 - | 4 - | 0,000305174 - |
| 2193 | 1 | 0.2um_NOKCPHJN_411656 | Eukaryota | Arthropoda | Insecta | Hymenoptera | Braconidae | Diachasma | Diachasma all | unclassified | unclassified | unclassified | unclassified | unclassified | 0,6 | 0,489897949 | 5,64E-05 | 4,61E-05 | 1 - | 9,89E-05 - | 0 - | 0 - | 1 - | 9,17E-05 - | 1 - | 9,12E-05 - | 0 - | 0 - |
| 2194 | 1 | 0.2um_NOKCPHJN_435179 | Bacteria | Proteobacteri | Gammaprotec | unclassified | unclassified | unclassified | Gammaprotec | unclassified | unclassified | unclassified | unclassified | unclassified | 0,4 | 0,489897949 | 0,00017837 | 0,000218756 | 1 - | 0,000463989 - | 0 - | 0 - | 0 - | 0 - | 1 - | 0,000427859 - | 0 - | 0 - |
| 2195 | 1 | 3um_FIJJAFBF_405742 | unclassified | unclassified | unclassified | unclassified | unclassified | unclassified | unclassified | unclassified | unclassified | unclassified | unclassified | unclassified | 0,2 | 0,4 | 0,000187582 | 0,000375163 | 0 - | 0 - | 1 - | 0,000937908 - | 0 - | 0 - | 0 - | 0 - | 0 - | 0 - |
| 2196 | 1 | 3um_FIJJAFBF_592424 | Bacteria | Actinobacteria | Actinobacteria | unclassified | unclassified | unclassified | actinobacteriu | cellular proces | Posttranslatio | 00CWM | ATP-depender | 312284.A20C1 | 0,4 | 0,489897949 | 7,66E-05 | 9,38E-05 | 0 - | 0 - | 0 - | 0 - | 0 - | 0 - | 1 - | 0,00019072 - | 1 - | 0,000192173 - |
| 2197 | 1 | 3um_FIJJAFBF_257117 | Eukaryota | Bacillariophyta | Fragilariophyc | Fragilariales | Fragilariaceae | Asterionellops | Asterionellops | information st | Translation, ri | 0FJK5 | Binds the low | 197221.tlr008 | 1 | 0 | 0,000395079 | 3,87E-05 | 1 - | 0,000398608 - | 1 - | 0,000468954 - | 1 - | 0,000369894 - | 1 - | 0,00036757 - | 1 - | 0,00037037 - |
| 2198 | 1 | 0.2um_NOKCPHJN_326393 | Bacteria | Bacteroidetes | unclassified | unclassified | unclassified | unclassified | Bacteroidetes | metabolism | Inorganic ion t | 05C27 | TonB-depende | 391603.FBALC | 0,4 | 0,489897949 | 3,12E-05 | 3,82E-05 | 1 - | 8,10E-05 - | 0 - | 0 - | 0 - | 0 - | 1 - | 7,47E-05 - | 0 - | 0 - |
| 2199 | 1 | 0.2um_NOKCPHJN_1062322 | Bacteria | Proteobacteri | Alphaproteob | Pelagibacteral | Pelagibactera | Candidatus Pe | Candidatus Pe | unclassified | unclassified | unclassified | unclassified | unclassified | 0,6 | 0,489897949 | 0,000209895 | 0,000174296 | 0 - | 0 - | 1 - | 0,000407786 - | 0 - | 0 - | 1 - | 0,000319626 - | 1 - | 0,000322061 - |
| 2200 | 1 | 3um_FIJJAFBF_280514 | Eukaryota | unclassified | Oomycetes | Saprolegniales | Saprolegniace | Saprolegnia | Saprolegnia pa | unclassified | unclassified | unclassified | unclassified | unclassified | 0,2 | 0,4 | 5,10E-05 | 0,00010197 | 1 - | 0,000254924 - | 0 - | 0 - | 0 - | 0 - | 0 - | 0 - | 0 - | 0 - |
| 2201 | 1 | 3um_FIJJAFBF_256676 | Bacteria | Bacteroidetes | Flavobacteriia | Flavobacterial | Flavobacteriac | Algibacter | Algibacter sp. | cellular proces | Cell wall/mem | 0P9R9 | TonB family | 1046627.BZAR | 0,4 | 0,489897949 | 0,000141569 | 0,000173581 | 1 - | 0,00036692 - | 0 - | 0 - | 0 - | 0 - | 0 - | 0 - | 1 - | 0,000340927 - |
| 2202 | 1 | 3um_FIJJAFBF_235815 | Bacteria | Bacteroidetes | Flavobacteriia | unclassified | unclassified | unclassified | Flavobacteria | metabolism | Inorganic ion t | 05C27 | TonB-depende | 487797.Flav3( | 2 | 2,607680962 | 0,000171414 | 0,000229217 | 7 - | 0,000613857 - | 0 - | 0 - | 0 - | 0 - | 2 - | 0,000161731 - | 1 - | 8,15E-05 - |
| 2203 | 1 | 3um_FIJJAFBF_142849 | Bacteria | Proteobacteri | Alphaproteob | Rhodobactera | unclassified | unclassified | Rhodobactera | cellular proces | Cell wall/mem | 16QB0 | Part of the ou | 439497.RR11_ | 1,8 | 1,166190379 | 0,000189961 | 0,000121103 | 1 - | 0,000112862 - | 0 - | 0 - | 3 - | 0,000314196 - | 2 - | 0,000208148 - | 3 - | 0,0003146 - |
| 2204 | 1 | 0.2um_NOKCPHJN_259848 | Bacteria | Proteobacteri | Gammaprotec | unclassified | unclassified | unclassified | Gammaprotec | cellular proces | Defense mech | COG1680 | Beta-lactamas | 247633.GP214 | 1 | 0 | 0,000212512 | 2,08E-05 | 1 - | 0,00021441 - | 1 - | 0,000252249 - | 1 - | 0,000198965 - | 1 - | 0,000197715 - | 1 - | 0,000199221 - |
| 2205 | 1 | 0.2um_NOKCPHJN_364077 | Archaea | Thaumarchae | unclassified | unclassified | unclassified | unclassified | Thaumarchae | information st | Transcription | COG0086 | DNA-depende | 436308.Nmar_ | 1 | 0 | 7,00E-05 | 6,85E-06 | 1 - | 7,07E-05 - | 1 - | 8,31E-05 - | 1 - | 6,56E-05 - | 1 - | 6,52E-05 - | 1 - | 6,57E-05 - |
| 2206 | 1 | 0.2um_NOKCPHJN_352637 | Bacteria | Proteobacteri | Gammaprotec | Cellvibrionales | Cellvibrionace | Simiduia | Simiduia agari | metabolism | Inorganic ion t | 16TQC | TONB-depend | 190485.XCC39 | 3,8 | 1,720465053 | 0,000345194 | 0,000180748 | 1 - | 9,00E-05 - | 6 - | 0,000635543 - | 3 - | 0,000250647 - | 5 - | 0,00041512 - | 4 - | 0,000334626 - |
| 2207 | 1 | 0.2um_NOKCPHJN_401629 | Bacteria | Firmicutes | Clostridia | Clostridiales | unclassified | unclassified | Clostridiales b | poorly charact | Function unkn | 07F5I | NA | 760192.Halhy | 1,4 | 0,8 | 0,000104016 | 6,01E-05 | 2 - | 0,000156736 - | 0 - | 0 - | 2 - | 0,000145445 - | 1 - | 7,23E-05 - | 2 - | 0,000145633 - |
| 2208 | 1 | 0.2um_NOKCPHJN_310980 | Eukaryota | unclassified | unclassified | unclassified | unclassified | Gymnochlora | Gymnochlora | information st | Chromatin stru | KOG3467 | Core compone | 2850.Phatr349 | 2,2 | 0,4 | 0,002043097 | 0,00033038 | 2 - | 0,001885889 - | 2 - | 0,002218707 - | 2 - | 0,001750037 - | 3 - | 0,00260856 - | 2 - | 0,00175229 - |
| 2209 | 1 | 3um_FIJJAFBF_502175 | Eukaryota | unclassified | unclassified | Prymnesiales | Chrysochromu | Chrysochromu | Chrysochromu | unclassified | unclassified | unclassified | unclassified | unclassified | 1,8 | 0,4 | 0,001035142 | 0,000172388 | 2 - | 0,001185052 - | 1 - | 0,000697094 - | 2 - | 0,001099686 - | 2 - | 0,001092775 - | 2 - | 0,001101101 - |
| 2210 | 1 | 3um_FIJJAFBF_362977 | unclassified | unclassified | unclassified | unclassified | unclassified | unclassified | unclassified | unclassified | unclassified | unclassified | unclassified | unclassified | 0,2 | 0,4 | 3,07E-05 | 6,14E-05 | 0 - | 0 - | 0 - | 0 - | 1 - | 0,000153541 - | 0 - | 0 - | 0 - | 0 - |
| 2211 | 1 | 0.2um_NOKCPHJN_61814 | Bacteria | Bacteroidetes | Flavobacteriia | Flavobacterial | Flavobacteriac | Polaribacter | Polaribacter fi | poorly charact | Function unkn | 0XRCJ | NA | 391587.KAOT | 8 | 2,898275349 | 0,00064276 | 0,000203104 | 8 - | 0,000664977 - | 4 - | 0,000391165 - | 8 - | 0,000617075 - | 7 - | 0,000536547 - | 13 - | 0,001004037 - |
| 2212 | 1 | 0.2um_NOKCPHJN_756783 | Bacteria | Proteobacteri | Gammaprotec | unclassified | unclassified | unclassified | OM182 bacte | unclassified | unclassified | unclassified | unclassified | unclassified | 6,6 | 0,489897949 | 0,000641451 | 3,31E-05 | 6 - | 0,00059186 - | 6 - | 0,00069631 - | 7 - | 0,000640762 - | 7 - | 0,000636735 - | 7 - | 0,000641587 - |
| 2213 | 1 | 0.2um_NOKCPHJN_833975 | Eukaryota | Phaeophyceae | unclassified | Ectocarpales | Ectocarpaceae | Ectocarpus | Ectocarpus sili | unclassified | unclassified | unclassified | unclassified | unclassified | 0,4 | 0,489897949 | 4,93E-05 | 6,04E-05 | 1 - | 0,000127648 - | 0 - | 0 - | 0 - | 0 - | 0 - | 0 - | 1 - | 0,000118605 - |
| 2214 | 1 | 3um_FIJJAFBF_37522 | Bacteria | Bacteroidetes | Flavobacteriia | Flavobacterial | Crocinitomica | unclassified | Crocinitomica | information st | Translation, ri | 08UHY | One of the pri | 755732.Fluta_ | 0,2 | 0,4 | 0,000105002 | 0,000210004 | 0 - | 0 - | 0 - | 0 - | 1 - | 0,000525011 - | 0 - | 0 - | 0 - | 0 - |
| 2215 | 2 | 0.2um_NOKCPHJN_168916;0.2 | Eukaryota | Bacillariophyta | Coscinodiscop | Thalassiosirale | Thalassiosirac | Thalassiosira | various | metabolism | Energy produc | COG0377 | NDH-1 shuttle | various | 1,2 | 1,166190379 | 0,000471559 | 0,000459013 | 0 - | 0 - | 0 - | 0 - | 2 - | 0,000786249 - | 1 - | 0,000390654 - | 3 - | 0,001180891 - |
| 2216 | 1 | 0.2um_NOKCPHJN_1004726 | Bacteria | Bacteroidetes | Flavobacteriia | Flavobacterial | Flavobacteriac | Polaribacter | Polaribacter s | poorly charact | Function unkn | 0P8NI | Inherit from b | 983548.Krodi_ | 1 | 0 | 0,000187322 | 1,83E-05 | 1 - | 0,000188995 - | 1 - | 0,000222349 - | 1 - | 0,000175381 - | 1 - | 0,000174279 - | 1 - | 0,000175607 - |
| 2217 | 12 | 0.2um_NOKCPHJN_1085758;0. | Bacteria | Bacteroidetes | various | various | various | various | various | cellular proces | Posttranslatio | 09GE9 | Thioredoxin | various | 1 | 0 | 0,000769181 | 7,53E-05 | 1 - | 0,000776052 - | 1 - | 0,000913008 - | 1 - | 0,000720148 - | 1 - | 0,000715623 - | 1 - | 0,000721075 - |
| 2218 | 1 | 3um_FIJJAFBF_128631 | Eukaryota | Chlorophyta | Mamiellophyc | Mamiellales | Mamiellaceae | Micromonas | Micromonas c | unclassified | unclassified | unclassified | unclassified | unclassified | 2 | 0 | 0,001524868 | 0,000149199 | 2 - | 0,001538489 - | 2 - | 0,001809998 - | 2 - | 0,001427662 - | 2 - | 0,00141869 - | 2 - | 0,0014295 - |
| 2219 | 1 | 0.2um_NOKCPHJN_818796 | Bacteria | Proteobacteri | Alphaproteob | Pelagibacteral | Pelagibactera | unclassified | alpha proteob | metabolism | Energy produc | 01RNF | Produces ATP | 331869.BAL19 | 0,2 | 0,4 | 4,35E-05 | 8,71E-05 | 0 - | 0 - | 1 - | 0,000217658 - | 0 - | 0 - | 0 - | 0 - | 0 - | 0 - |
| 2220 | 1 | 3um_FIJJAFBF_237053 | Eukaryota | unclassified | Synurophycea | Parmales | Triparmaceae | Triparma | Triparma laevi | metabolism | Amino acid tra | 0FJ71 | Acetolactate s | 329726.AM1_ | 1 | 1,095445115 | 0,000529737 | 0,00057004 | 3 - | 0,001556696 - | 1 - | 0,000610473 - | 1 - | 0,000481519 - | 0 - | 0 - | 0 - | 0 - |
| 2221 | 1 | 0.2um_NOKCPHJN_283114 | Bacteria | Proteobacteri | Gammaprotec | Pasteurellales | Pasteurellacea | Haemophilus | Haemophilus ( | unclassified | unclassified | unclassified | unclassified | unclassified | 0,2 | 0,4 | 1,28E-05 | 2,56E-05 | 0 - | 0 - | 1 - | 6,40E-05 - | 0 - | 0 - | 0 - | 0 - | 0 - | 0 - |
| 2222 | 1 | 0.2um_NOKCPHJN_509418 | unclassified | unclassified | unclassified | unclassified | unclassified | unclassified | unclassified | unclassified | unclassified | unclassified | unclassified | unclassified | 1 | 0 | 0,000139739 | 1,37E-05 | 1 - | 0,000140987 - | 1 - | 0,000165868 - | 1 - | 0,000130831 - | 1 - | 0,000130009 - | 1 - | 0,000130999 - |
| 2223 | 1 | 0.2um_NOKCPHJN_292026 | unclassified | unclassified | unclassified | unclassified | unclassified | unclassified | unclassified | unclassified | unclassified | unclassified | unclassified | unclassified | 0,6 | 0,489897949 | 7,20E-05 | 5,89E-05 | 1 - | 0,00012636 - | 0 - | 0 - | 1 - | 0,000117258 - | 1 - | 0,000116521 - | 0 - | 0 - |
| 2224 | 1 | 3um_FIJJAFBF_633503 | Viruses | unclassified | unclassified | unclassified | unclassified | unclassified | uncultured M | unclassified | unclassified | unclassified | unclassified | unclassified | 0,4 | 0,489897949 | 0,000101401 | 0,000124192 | 0 - | 0 - | 0 - | 0 - | 1 - | 0,000254302 - | 1 - | 0,000252704 - | 0 - | 0 - |
| 2225 | 1 | 3um_FIJJAFBF_284143 | Bacteria | Actinobacteria | Actinobacteria | Actinomycetal | unclassified | unclassified | Actinomyceta | cellular proces | Cell cycle cont | COG3599 | Divisome com | 351607.Acel_ | 0,6 | 0,489897949 | 0,000287991 | 0,000235511 | 1 - | 0,000503988 - | 0 - | 0 - | 1 - | 0,000467682 - | 0 - | 0 - | 1 - | 0,000468284 - |
| 2226 | 1 | 3um_FIJJAFBF_161153 | Viruses | unclassified | unclassified | unclassified | unclassified | unclassified | Vibrio phage 1 | unclassified | unclassified | unclassified | unclassified | unclassified | 4,2 | 0,979795897 | 0,001112552 | 0,000188074 | 3 - | 0,000817023 - | 3 - | 0,00096121 - | 5 - | 0,001263614 - | 5 - | 0,001255673 - | 5 - | 0,001265241 - |
| 2227 | 1 | 3um_FIJJAFBF_379962 | Eukaryota | unclassified | unclassified | Isochrysidales | Noelaerhabda | Emiliania | Emiliania huxl | information st | Replication, re | 1E8C4 | Inherit from N | 436017.A4S7F | 0,6 | 0,489897949 | 0,000169842 | 0,000138677 | 0 - | 0 - | 0 - | 0 - | 1 - | 0,000283543 - | 1 - | 0,000281761 - | 1 - | 0,000283908 - |
| 2228 | 1 | 0.2um_NOKCPHJN_480383 | Bacteria | Bacteroidetes | Flavobacteriia | Flavobacterial | Cryomorphace | Owenweeksia | Owenweeksia | metabolism | Amino acid tra | COG2171 | Catalyzes the | 926562.Oweh | 1 | 0 | 0,000462327 | 4,52E-05 | 1 - | 0,000466457 - | 1 - | 0,000548776 - | 1 - | 0,000432855 - | 1 - | 0,000430135 - | 1 - | 0,000433412 - |
| 2229 | 2 | 0.2um_NOKCPHJN_276340;0.2 | Bacteria | Bacteroidetes | Flavobacteriia | Flavobacterial | Flavobacteriac | Altibacter | Altibacter lent | metabolism | Lipid transpor | COG2067 | lipid transport | 50743.SCB49_ | 1,4 | 0,8 | 0,000290484 | 0,000174219 | 0 - | 0 - | 2 - | 0,000488957 - | 2 - | 0,000385672 - | 1 - | 0,000191624 - | 2 - | 0,000386168 - |
| 2230 | 1 | 0.2um_NOKCPHJN_129777 | Bacteria | Bacteroidetes | Flavobacteriia | Flavobacterial | Flavobacteriac | Dokdonia | Dokdonia paci | unclassified | unclassified | unclassified | unclassified | unclassified | 1 | 0 | 0,000129728 | 1,27E-05 | 1 - | 0,000130886 - | 1 - | 0,000153985 - | 1 - | 0,000121458 - | 1 - | 0,000120695 - | 1 - | 0,000121614 - |
| 2231 | 1 | 3um_FIJJAFBF_176097 | Bacteria | Proteobacteri | Gammaprotec | Cellvibrionales | Spongiibacter | Oceanicoccus | Oceanicoccus | cellular proces | Intracellular tr | 124XR | MSHA pilin pr | 903510.vfu_A | 2 | 0 | 0,001066472 | 0,000104348 | 2 - | 0,001075998 - | 2 - | 0,001265888 - | 2 - | 0,000998488 - | 2 - | 0,000992213 - | 2 - | 0,000999773 - |
| 2232 | 1 | 3um_FIJJAFBF_182748 | Viruses | unclassified | unclassified | unclassified | various | various | various | unclassified | unclassified | unclassified | unclassified | unclassified | 0,6 | 0,489897949 | 0,000285502 | 0,00023708 | 0 - | 0 - | 1 - | 0,000554677 - | 0 - | 0 - | 1 - | 0,00043476 - | 1 - | 0,000438073 - |
| 2233 | 1 | 3um_FIJJAFBF_277464 | Eukaryota | unclassified | Pelagophycea | Pelagomonad | unclassified | Pelagomonas | uncultured Pe | metabolism | Energy produc | COG1290 | Component of | 35128.Thapsd | 0,2 | 0,4 | 0,000101721 | 0,000203442 | 0 - | 0 - | 0 - | 0 - | 1 - | 0,000508605 - | 0 - | 0 - | 0 - | 0 - |
| 2234 | 1 | 3um_FIJJAFBF_00462 | Bacteria | Bacteroidetes | unclassified | unclassified | unclassified | unclassified | Bacteroidetes | metabolism | Energy produc | 0PEUJ | ATP synthase | 391603.FBALC | 1 | 0 | 0,00117456 | 0,000114924 | 1 - | 0,001185052 - | 1 - | 0,001394188 - | 1 - | 0,001099686 - | 1 - | 0,001092775 - | 1 - | 0,001101101 - |
| 2235 | 1 | 0.2um_NOKCPHJN_675062 | Bacteria | Bacteroidetes | Flavobacteriia | Flavobacterial | Flavobacteriac | Tenacibaculur | Tenacibaculur | unclassified | unclassified | unclassified | unclassified | unclassified | 0,6 | 0,489897949 | 5,21E-05 | 4,26E-05 | 1 - | 9,14E-05 - | 0 - | 0 - | 0 - | 0 - | 1 - | 8,43E-05 - | 1 - | 8,50E-05 - |
| 2236 | 1 | 0.2um_NOKCPHJN_592028 | Eukaryota | unclassified | Pelagophycea | Pelagomonad | unclassified | Aureococcus | Aureococcus a | cellular proces | Posttranslatio | KOG0019 | heat shock pr | 67593.Physo1 | 0,4 | 0,489897949 | 4,75E-05 | 5,82E-05 | 1 - | 0,000123166 - | 0 - | 0 - | 1 - | 0,000114293 - | 0 - | 0 - | 0 - | 0 - |
| 2237 | 1 | 3um_FIJJAFBF_437251 | Bacteria | Bacteroidetes | Flavobacteriia | Flavobacterial | Flavobacteriac | Flaviramulus | Flaviramulus i | unclassified | unclassified | unclassified | unclassified | unclassified | 0,4 | 0,8 | 3,51E-05 | 7,03E-05 | 2 - | 0,000175739 - | 0 - | 0 - | 0 - | 0 - | 0 - | 0 - | 0 - | 0 - |
| 2238 | 1 | 3um_FIJJAFBF_205191 | Eukaryota | Bacillariophyta | Coscinodiscop | Chaetocerotal | Acanthocerat | Acanthoceras | Acanthoceras | metabolism | Energy produc | 0IFBF | One of the tw | 45157.CMV08 | 12 | 0,894427191 | 0,00295558 | 0,000181015 | 11 - | 0,00274824 - | 11 - | 0,003233244 - | 13 - | 0,003013953 - | 12 - | 0,002764627 - | 13 - | 0,003017833 - |
| 2239 | 1 | 3um_FIJJAFBF_711075 | Bacteria | Bacteroidetes | Flavobacteriia | Flavobacterial | Flavobacteriac | Flaviramulus | Flaviramulus i | poorly charact | Function unkn | 0P8NI | Inherit from b | 391603.FBALC | 2,4 | 0,489897949 | 0,000512175 | 8,47E-05 | 2 - | 0,000435205 - | 2 - | 0,000512009 - | 3 - | 0,000605782 - | 2 - | 0,000401317 - | 3 - | 0,000606562 - |
| 2240 | 1 | 0.2um_NOKCPHJN_267389 | Bacteria | Proteobacteri | Gammaprotec | Cellvibrionales | Porticoccacea | unclassified | SAR92 bacteri | poorly charact | Function unkn | 16R8I | Lipoprotein | 377629.TERTU | 0,2 | 0,4 | 1,74E-05 | 3,48E-05 | 0 - | 0 - | 0 - | 0 - | 0 - | 0 - | 0 - | 0 - | 1 - | 8,70E-05 - |
| 2241 | 1 | 0.2um_NOKCPHJN_435291 | Bacteria | unclassified | unclassified | unclassified | unclassified | unclassified | uncultured ba | unclassified | unclassified | unclassified | unclassified | unclassified | 2 | 0,894427191 | 0,000178105 | 7,04E-05 | 1 - | 9,29E-05 - | 1 - | 0,00010929 - | 2 - | 0,000172408 - | 3 - | 0,000256987 - | 3 - | 0,000258945 - |
| 2242 | 1 | 0.2um_NOKCPHJN_206968 | Eukaryota | unclassified | Oomycetes | Peronosporale | Peronosporac | Phytophthora | Phytophthora | unclassified | unclassified | unclassified | unclassified | unclassified | 0,2 | 0,4 | 7,60E-06 | 1,52E-05 | 0 - | 0 - | 0 - | 0 - | 0 - | 0 - | 0 - | 0 - | 1 - | 3,80E-05 - |
| 2243 | 1 | 3um_FIJJAFBF_768267 | Bacteria | Cyanobacteria | unclassified | Synechococca | Synechococca | Synechococcu | Synechococcu | poorly charact | Function unkn | 129GX | NA | 56780.SYN_0C | 0,4 | 0,489897949 | 0,000139827 | 0,000172188 | 1 - | 0,000321223 - | 1 - | 0,000377912 - | 0 - | 0 - | 0 - | 0 - | 0 - | 0 - |
| 2244 | 1 | 0.2um_NOKCPHJN_889433 | Bacteria | Proteobacteri | Alphaproteob | Rhodobactera | Rhodobactera | Pseudooctade | Pseudooctade | poorly charact | Function unkn | 0J6G5 | NA | 306901.XP_0C | 0,6 | 0,489897949 | 0,000266365 | 0,000217489 | 0 - | 0 - | 0 - | 0 - | 1 - | 0,000444682 - | 1 - | 0,000441887 - | 1 - | 0,000445254 - |
| 2245 | 1 | 3um_FIJJAFBF_81542 | Bacteria | Proteobacteri | Alphaproteob | Rhizobiales | Methylobacte | Microvirga | Microvirga gu | metabolism | Energy produc | COG0056 | Produces ATP | 460265.Mnod | 1 | 0 | 0,000169761 | 1,66E-05 | 1 - | 0,000171277 - | 1 - | 0,000201504 - | 1 - | 0,000158939 - | 1 - | 0,00015794 - | 1 - | 0,000159144 - |
| 2246 | 1 | 3um_FIJJAFBF_257111 | Eukaryota | Bacillariophyta | Coscinodiscop | Leptocylindral | Leptocylindrac | Leptocylindrus | Leptocylindrus | information st | Translation, ri | COG0087 | One of the pri | 489825.LYNGE | 2,6 | 0,489897949 | 0,001022019 | 0,000162286 | 3 - | 0,001201286 - | 2 - | 0,000942191 - | 3 - | 0,00111475 - | 3 - | 0,001107745 - | 2 - | 0,000744123 - |
| 2247 | 1 | 3um_FIJJAFBF_96040 | Bacteria | Proteobacteri | Gammaprotec | Enterobactera | Enterobacteri | Citrobacter | Citrobacter w | unclassified | unclassified | unclassified | unclassified | unclassified | 1 | 0 | 0,000119228 | 1,17E-05 | 1 - | 0,000120293 - | 1 - | 0,000141522 - | 1 - | 0,000111628 - | 1 - | 0,000110926 - | 1 - | 0,000111772 - |
| 2248 | 1 | 3um_FIJJAFBF_405248 | Bacteria | Bacteroidetes | Flavobacteriia | Flavobacterial | Flavobacteriac | Polaribacter | Polaribacter s | metabolism | Amino acid tra | 09GJ6 | nitrogen regul | 313598.MED1 | 2,6 | 0,489897949 | 0,002026048 | 0,000474292 | 2 - | 0,001565962 - | 3 - | 0,002763479 - | 3 - | 0,002179734 - | 3 - | 0,002166036 - | 2 - | 0,001455027 - |
| 2249 | 1 | 0.2um_NOKCPHJN_669311 | unclassified | unclassified | unclassified | unclassified | unclassified | unclassified | unclassified | unclassified | unclassified | unclassified | unclassified | unclassified | 0,4 | 0,489897949 | 0,000153922 | 0,000189546 | 1 - | 0,000353604 - | 1 - | 0,000416008 - | 0 - | 0 - | 0 - | 0 - | 0 - | 0 - |
| 2250 | 1 | 0.2um_NOKCPHJN_1094357 | unclassified | unclassified | unclassified | unclassified | unclassified | unclassified | unclassified | unclassified | unclassified | unclassified | unclassified | unclassified | 0,4 | 0,489897949 | 0,000136829 | 0,00016962 | 0 - | 0 - | 1 - | 0,000383531 - | 0 - | 0 - | 1 - | 0,000300615 - | 0 - | 0 - |
| 2251 | 1 | 0.2um_NOKCPHJN_492829 | Bacteria | Bacteroidetes | Flavobacteriia | Flavobacterial | Flavobacteriac | Formosa | Formosa sp. H | poorly charact | Function unkn | 0PAEN | NA | 886377.Murru | 1,4 | 0,489897949 | 0,000257759 | 0,000106823 | 1 - | 0,000182696 - | 2 - | 0,000429875 - | 1 - | 0,000169535 - | 2 - | 0,000336939 - | 1 - | 0,000169753 - |
| 2252 | 1 | 3um_FIJJAFBF_135693 | Bacteria | Bacteroidetes | Flavobacteriia | Flavobacterial | Flavobacteriac | Algibacter | Algibacter aqu | poorly charact | Function unkn | 0ZWIA | NA | 457570.Nther | 0,2 | 0,4 | 5,05E-05 | 0,000101088 | 1 - | 0,00025272 - | 0 - | 0 - | 0 - | 0 - | 0 - | 0 - | 0 - | 0 - |
| 2253 | 1 | 0.2um_NOKCPHJN_700962 | Bacteria | Proteobacteri | Alphaproteob | Rhodospirillale | Rhodospirillac | unclassified | Rhodobactera | metabolism | Inorganic ion t | 05PWN | phosphate ab | 765910.Marpu | 0,8 | 0,979795897 | 0,000194866 | 0,000238987 | 2 - | 0,000506901 - | 0 - | 0 - | 0 - | 0 - | 2 - | 0,00046743 - | 0 - | 0 - |
| 2254 | 1 | 3um_FIJJAFBF_566201 | unclassified | unclassified | unclassified | unclassified | unclassified | unclassified | unclassified | unclassified | unclassified | unclassified | unclassified | unclassified | 0,8 | 0,4 | 0,000526059 | 0,000263782 | 1 - | 0,000695983 - | 0 - | 0 - | 1 - | 0,000645847 - | 1 - | 0,000641789 - | 1 - | 0,000646679 - |
| 2255 | 1 | 3um_FIJJAFBF_545136 | Bacteria | Proteobacteri | Gammaprotec | Alteromonada | Colwelliaceae | Colwellia | Colwellia sp. P | information st | Translation, ri | 05KK9 | This protein bi | 167879.CPS_4 | 0,2 | 0,4 | 0,00020033 | 0,00040066 | 0 - | 0 - | 1 - | 0,001001649 - | 0 - | 0 - | 0 - | 0 - | 0 - | 0 - |
| 2256 | 1 | 3um_FIJJAFBF_555528 | Bacteria | Proteobacteri | Alphaproteob | Rhodobactera | Hyphomonada | Hyphomonas | Hyphomonas | cellular proces | Cell wall/mem | COG3637 | Membrane | 411684.HPDFI | 1,4 | 0,489897949 | 0,000436084 | 0,000181557 | 1 - | 0,000308781 - | 2 - | 0,000726549 - | 1 - | 0,000286538 - | 1 - | 0,000284737 - | 2 - | 0,000573813 - |
| 2257 | 1 | 3um_FIJJAFBF_581298 | Bacteria | Bacteroidetes | Flavobacteriia | Flavobacterial | Crocinitomica | unclassified | Crocinitomica | information st | Replication, re | 0PB0H | DNA-binding p | 755732.Fluta_ | 2,8 | 0,4 | 0,002231318 | 0,000389195 | 2 - | 0,001609061 - | 3 - | 0,002839538 - | 3 - | 0,002239727 - | 3 - | 0,002225652 - | 3 - | 0,00224261 - |
| 2258 | 1 | 0.2um_NOKCPHJN_812308 | unclassified | unclassified | unclassified | unclassified | unclassified | unclassified | unclassified | unclassified | unclassified | unclassified | unclassified | unclassified | 1 | 0 | 0,000742884 | 7,27E-05 | 1 - | 0,00074952 - | 1 - | 0,000881794 - | 1 - | 0,000695528 - | 1 - | 0,000691157 - | 1 - | 0,000696423 - |
| 2259 | 1 | 3um_FIJJAFBF_306833 | Bacteria | Proteobacteri | Alphaproteob | Rhodobactera | Rhodobactera | various | various | metabolism | Inorganic ion t | 16PII | TRAP transpor | 314262.MED1 | 0,8 | 0,748331477 | 0,000210745 | 0,000202565 | 2 - | 0,000546379 - | 0 - | 0 - | 1 - | 0,00025351 - | 0 - | 0 - | 1 - | 0,000253836 - |
| 2260 | 3 | 0.2um_NOKCPHJN_597293;3ur | Bacteria | Proteobacteri | Alphaproteob | Rhizobiales | Phyllobacteria | unclassified | Phyllobacteria | various | various | various | various | various | 0,8 | 0,4 | 0,00037169 | 0,000190623 | 1 - | 0,000461547 - | 1 - | 0,000542999 - | 1 - | 0,000428299 - | 1 - | 0,000425607 - | 0 - | 0 - |
| 2261 | 1 | 3um_FIJJAFBF_352546 | Eukaryota | unclassified | Pelagophycea | Pelagomonad | unclassified | Aureococcus | Aureococcus a | unclassified | unclassified | unclassified | unclassified | unclassified | 1,4 | 0,489897949 | 0,00036392 | 0,000108888 | 1 - | 0,000267359 - | 1 - | 0,000314542 - | 2 - | 0,0004962 - | 2 - | 0,000493081 - | 1 - | 0,000248419 - |
| 2262 | 1 | 3um_FIJJAFBF_624005 | Bacteria | Bacteroidetes | Flavobacteriia | Flavobacterial | Flavobacteriac | Marixanthom | Marixanthom | poorly charact | Function unkn | 0P8NI | Inherit from b | 313594.PI23P | 0,6 | 0,489897949 | 7,52E-05 | 6,14E-05 | 0 - | 0 - | 0 - | 0 - | 1 - | 0,000125581 - | 1 - | 0,000124792 - | 1 - | 0,000125743 - |

**PROPHANE JOB INFORMATION**

**program versions**

prophane: 3.1.1
blast+: 2.7.1
diamond: 0.8.22
hmmer: 3.1b2
krona: 2.7

**tasks**

task: *taxonomic assignment*
task comment: tax_from_nr_20180808_qcover90
algorithm: diamond blastp
query: sequences without taxonomic classification (missing_taxa.faa)
database: NCBI NR database
database version: 2018-08-02
database comment: downloaded at 2018-08-02
database scope: tax
parameter:
- evalue 0.01
- query-cover 0.9
- max-target-seqs 1

task: *functional assignment*
task comment: fun_from_eggNog_4.5.1
algorithm: emapper
query: all sequences (all.faa)
database: eggmap database
database version: 4.5.1
database comment: includes euk, viruses, bakt, arch; downloaded at 2018-07-31
database scope: func
parameter:
- m diamond

| protein group | members_count | protein group: members_identifier | taxonomic assignment<br>superkingdom | phylum | class | order | family | genus | species | functional assignment<br>main role | sub role | og | descr | prot | quantitative data mean<br>exclusive spetrum count | exclusive spetrum count, sd | NSAF | NSAF, sd | quantitative data replicate 1<br>exclusive spetrum count | exclusive spetrum count, sd | NSAF | NSAF, sd | quantitative data replicate 2<br>exclusive spetrum count | exclusive spetrum count, sd | NSAF | NSAF, sd | quantitative data replicate 3<br>exclusive spetrum count | exclusive spetrum count, sd | NSAF | NSAF, sd |
|---|---|---|---|---|---|---|---|---|---|---|---|---|---|---|---|---|---|---|---|---|---|---|---|---|---|---|---|---|---|---|
| 0 | 1 | 3um_FIJJAFBF_594168 | Eukaryota | unclassified | Pelagophycea | Pelagomonad | unclassified | Aureoumbra | Aureoumbra l | metabolism | Energy produc | COG0056 | Produces ATP | 35128.Thapsd | 18,4 | 2,332380758 | 0,002262507 | 0,000145723 | 14 | - | 0,002070898 | - | 20 | - | 0,002473317 | - | 18 | - | 0,002240856 | - |
| 1 | 1 | 3um_FIJJAFBF_669279 | Eukaryota | Bacillariophyt | Fragilariophyc | Fragilariales | Fragilariaceae | Asterionella | Asterionella fo | unclassified | unclassified | unclassified | unclassified | unclassified | 1,6 | 0,489897949 | 0,000277343 | 8,24E-05 | 1 | - | 0,000207336 | - | 2 | - | 0,000346675 | - | 2 | - | 0,000348991 | - |
| 2 | 1 | 0.2um_NOKCPHJN_464263 | Eukaryota | Arthropoda | Insecta | Diptera | Culicidae | Aedes | Aedes aegypti | information st | Chromatin str | KOG1245 | bromodomain | 7668.SPU_013 | 0,2 | 0,4 | 6,54E-05 | 0,000130739 | 1 | - | 0,000326848 | - | 0 | - | 0 | - | 0 | - | 0 | - |
| 3 | 1 | 0.2um_NOKCPHJN_717763 | Viruses | unclassified | unclassified | Ortervirales | Retroviridae | Lentivirus | Caprine arthri | unclassified | unclassified | unclassified | unclassified | unclassified | 0,4 | 0,8 | 4,02E-05 | 8,05E-05 | 0 | - | 0 | - | 2 | - | 0,000201206 | - | 0 | - | 0 | - |
| 4 | 1 | 3um_FIJJAFBF_292913 | Eukaryota | Bacillariophyt | Coscinodiscop | Coscinodiscale | Hemidiscacea | Actinocyclus | Actinocyclus s | metabolism | Energy produc | COG0056 | Produces ATP | 41431.PCC880 | 9,8 | 1,6 | 0,001201585 | 8,85E-05 | 7 | - | 0,0010375 | - | 10 | - | 0,001239107 | - | 10 | - | 0,001247385 | - |
| 5 | 1 | 0.2um_NOKCPHJN_227146 | Viruses | unclassified | unclassified | unclassified | Phycodnavirid | unclassified | Organic Lake p | information st | Transcription | COG0086 | DNA-depende | 88036.EFJ103 | 1 | 0 | 4,12E-05 | 4,36E-06 | 1 | - | 4,90E-05 | - | 1 | - | 4,09E-05 | - | 1 | - | 4,12E-05 | - |
| 6 | 1 | 0.2um_NOKCPHJN_150079 | Bacteria | Proteobacteri | Betaproteoba | unclassified | unclassified | unclassified | Betaproteoba | unclassified | unclassified | unclassified | unclassified | unclassified | 0,2 | 0,4 | 2,92E-05 | 5,85E-05 | 0 | - | 0 | - | 0 | - | 0 | - | 0 | - | 0 | - |
| 7 | 1 | 0.2um_NOKCPHJN_31505 | Bacteria | Bacteroidetes | unclassified | unclassified | unclassified | unclassified | Bacteroidetes | poorly charact | Function unkn | 11RX2 | NA | 762903.Pedsa | 0,6 | 0,489897949 | 0,000240019 | 0,000197744 | 1 | - | 0,000448193 | - | 1 | - | 0,0003747 | - | 1 | - | 0,000377203 | - |
| 8 | 1 | 3um_FIJJAFBF_596874 | Bacteria | Bacteroidetes | Flavobacteriia | Flavobacterial | Flavobacteria | Moheibacter | Moheibacter s | poorly charact | Function unkn | 06XI0 | Protein of unk | 313606.M231 | 1 | 0 | 0,000324702 | 3,43E-05 | 1 | - | 0,000385815 | - | 1 | - | 0,000322551 | - | 1 | - | 0,000324706 | - |
| 9 | 1 | 0.2um_NOKCPHJN_411988 | Bacteria | Proteobacteri | Gammaprotec | Cellvibrionales | Cellvibrionace | Cellvibrio | Cellvibrio sp. F | poorly charact | Function unkn | 178PC | Domain of Un | 498211.CJA_2 | 0,6 | 0,489897949 | 4,03E-05 | 3,36E-05 | 1 | - | 7,95E-05 | - | 0 | - | 0 | - | 0 | - | 0 | - |
| 10 | 1 | 3um_FIJJAFBF_557220 | Eukaryota | Chlorophyta | Ulvophyceae | Ulvales | Ulvaceae | Ulva | Ulva fasciata | metabolism | Energy produc | COG0055 | Produces ATP | 3702.ATCG00 | 2,2 | 0,4 | 0,000286748 | 5,79E-05 | 2 | - | 0,00030993 | - | 3 | - | 0,000388664 | - | 2 | - | 0,00026084 | - |
| 11 | 1 | 0.2um_NOKCPHJN_315471 | unclassified | unclassified | unclassified | unclassified | unclassified | unclassified | unclassified | unclassified | unclassified | unclassified | unclassified | unclassified | 2,8 | 0,748331477 | 0,000400709 | 9,07E-05 | 2 | - | 0,000345719 | - | 2 | - | 0,00028903 | - | 3 | - | 0,000436441 | - |
| 12 | 1 | 3um_FIJJAFBF_510235 | Bacteria | Bacteroidetes | Flavobacteriia | Flavobacterial | Flavobacteria | Polaribacter | Polaribacter s | unclassified | unclassified | unclassified | unclassified | unclassified | 1 | 0 | 0,000459797 | 4,86E-05 | 1 | - | 0,000546337 | - | 1 | - | 0,000456751 | - | 1 | - | 0,000459802 | - |
| 13 | 1 | 0.2um_NOKCPHJN_656677 | unclassified | unclassified | unclassified | unclassified | unclassified | unclassified | unclassified | unclassified | unclassified | unclassified | unclassified | unclassified | 0,4 | 0,489897949 | 2,33E-05 | 2,91E-05 | 1 | - | 6,73E-05 | - | 0 | - | 0 | - | 0 | - | 0 | - |
| 14 | 1 | 0.2um_NOKCPHJN_442140 | Bacteria | Actinobacteri | Actinobacteri | unclassified | unclassified | unclassified | Actinobacteri | unclassified | unclassified | unclassified | unclassified | unclassified | 0,2 | 0,4 | 2,32E-05 | 4,64E-05 | 0 | - | 0 | - | 1 | - | 0,000115879 | - | 0 | - | 0 | - |
| 15 | 1 | 0.2um_NOKCPHJN_288060 | Viruses | unclassified | unclassified | unclassified | Phycodnavirid | Prasinovirus | Micromonas p | unclassified | unclassified | unclassified | unclassified | unclassified | 0,2 | 0,4 | 2,30E-05 | 4,61E-05 | 0 | - | 0 | - | 0 | - | 0 | - | 0 | - | 0 | - |
| 16 | 1 | 3um_FIJJAFBF_277463 | Bacteria | Cyanobacteria | unclassified | Synechococca | Leptolyngbyac | Nodosilinea | Nodosilinea n | information st | Translation, ri | 0FP0S | Seems to be tl | 513049.Amax | 1 | 0 | 0,000488311 | 5,16E-05 | 1 | - | 0,000580218 | - | 1 | - | 0,000485077 | - | 1 | - | 0,000488317 | - |
| 17 | 1 | 0.2um_NOKCPHJN_483159 | Eukaryota | Bacillariophyt | Bacillariophyc | Naviculales | Phaeodactylac | Phaeodactylu | Phaeodactylu | unclassified | unclassified | unclassified | unclassified | unclassified | 1 | 0 | 0,000181012 | 1,91E-05 | 1 | - | 0,000215081 | - | 1 | - | 0,000179813 | - | 1 | - | 0,000181014 | - |
| 18 | 1 | 3um_FIJJAFBF_348505 | Bacteria | Bacteroidetes | Cytophagia | Cytophagales | Flammeovirga | Marivirga | Marivirga seri | metabolism | Inorganic ion t | 05C27 | TonB-depende | 643867.Ftrac_ | 3,4 | 1,356465997 | 0,000195098 | 6,75E-05 | 1 | - | 7,11E-05 | - | 3 | - | 0,000178276 | - | 4 | - | 0,000239289 | - |
| 19 | 2 | 0.2um_NOKCPHJN_132117;3um_FIJJAF | various | various | various | various | various | various | various | various | various | various | various | various | 0,2 | 0,4 | 2,61E-05 | 5,21E-05 | 0 | - | 0 | - | 0 | - | 0 | - | 0 | - | 0 | - |
| 20 | 1 | 0.2um_NOKCPHJN_608547 | Eukaryota | Phaeophyceae | unclassified | Ectocarpales | Ectocarpaceae | Ectocarpus | Ectocarpus sili | unclassified | unclassified | unclassified | unclassified | unclassified | 3 | 0 | 0,001219203 | 0,000128934 | 3 | - | 0,001448675 | - | 3 | - | 0,001211127 | - | 3 | - | 0,001219218 | - |
| 21 | 1 | 3um_FIJJAFBF_520840 | Eukaryota | Bacillariophyt | Bacillariophyc | Naviculales | Phaeodactylac | Phaeodactylu | Phaeodactylu | metabolism | Amino acid tra | COG0473 | 3-isopropylma | 2850.Phatr20 | 0,4 | 0,489897949 | 0,000104295 | 0,000127795 | 0 | - | 0 | - | 0 | - | 0 | - | 1 | - | 0,000266919 | - |
| 22 | 1 | 3um_FIJJAFBF_692267 | Eukaryota | Mollusca | Gastropoda | unclassified | Lottiidae | Lottia | Lottia gigante | unclassified | unclassified | unclassified | unclassified | unclassified | 1,2 | 0,748331477 | 0,000114038 | 6,83E-05 | 0 | - | 0 | - | 1 | - | 0,000100603 | - | 1 | - | 0,000101275 | - |
| 23 | 1 | 0.2um_NOKCPHJN_26367 | Bacteria | Planctomycet | unclassified | unclassified | unclassified | unclassified | Planctomycet | information st | Translation, ri | COG0050 | This protein p | 487796.Flav2A | 1,2 | 0,4 | 0,000191369 | 6,60E-05 | 1 | - | 0,000189489 | - | 1 | - | 0,000158418 | - | 2 | - | 0,000318952 | - |
| 24 | 3 | 0.2um_NOKCPHJN_116238;0.2um_NO | Bacteria | Bacteroidetes | Flavobacteriia | Flavobacterial | Crocinitomica | unclassified | various | unclassified | unclassified | unclassified | unclassified | unclassified | 0,8 | 0,4 | 0,000381131 | 0,000192086 | 0 | - | 0 | - | 1 | - | 0,000496626 | - | 1 | - | 0,000499944 | - |
| 25 | 1 | 0.2um_NOKCPHJN_592342 | Eukaryota | Bacillariophyt | Bacillariophyc | Naviculales | Phaeodactylac | Phaeodactylu | Phaeodactylu | cellular proces | Posttranslatio | 0XRV1 | UnCoupling Pr | 2850.Phatr49 | 0,6 | 0,489897949 | 0,000151546 | 0,000123803 | 0 | - | 0 | - | 1 | - | 0,000255408 | - | 1 | - | 0,000257114 | - |
| 26 | 1 | 3um_FIJJAFBF_584729 | Eukaryota | Bacillariophyt | Fragilariophyc | Fragilariales | Fragilariaceae | Asterionella | Asterionella fo | metabolism | Energy produc | 11IIK | Low-potential | 35128.Thapsd | 1 | 0 | 0,000386455 | 4,09E-05 | 1 | - | 0,000459191 | - | 1 | - | 0,000383895 | - | 1 | - | 0,00038646 | - |
| 27 | 1 | 3um_FIJJAFBF_332003 | Bacteria | Proteobacteri | Gammaprotec | Chromatiales | Granulosicocc | Granulosicocc | Granulosicocc | unclassified | unclassified | unclassified | unclassified | unclassified | 0,2 | 0,4 | 2,55E-05 | 5,09E-05 | 0 | - | 0 | - | 0 | - | 0 | - | 0 | - | 0 | - |
| 28 | 1 | 0.2um_NOKCPHJN_93629 | Eukaryota | Xanthophycea | unclassified | Mischococcale | Pleurochlorida | Pleurochloris | Pleurochloris | metabolism | Carbohydrate | COG1850 | RuBisCO catal | 45157.CMV01 | 3,2 | 0,4 | 0,000409571 | 3,65E-05 | 3 | - | 0,000460132 | - | 3 | - | 0,000384682 | - | 3 | - | 0,000387252 | - |
| 29 | 1 | 0.2um_NOKCPHJN_21880 | Bacteria | Bacteroidetes | Flavobacteriia | Flavobacterial | Flavobacteria | Polaribacter | various | unclassified | unclassified | unclassified | unclassified | unclassified | 3,6 | 0,8 | 0,00147536 | 0,000212301 | 3 | - | 0,00148705 | - | 3 | - | 0,00124321 | - | 3 | - | 0,001251515 | - |
| 30 | 1 | 3um_FIJJAFBF_746159 | Eukaryota | Chordata | Actinopteri | Cypriniformes | Cyprinidae | Sinocyclocheil | Sinocyclocheil | poorly charact | Function unkn | 0EBNZ | NA | 135651.CBN2 | 0,4 | 0,489897949 | 7,67E-05 | 9,48E-05 | 1 | - | 0,000212637 | - | 0 | - | 0 | - | 0 | - | 0 | - |
| 31 | 1 | 3um_FIJJAFBF_744567 | Bacteria | unclassified | unclassified | unclassified | unclassified | unclassified | uncultured ba | unclassified | unclassified | unclassified | unclassified | unclassified | 7,8 | 1,469693846 | 0,000516609 | 9,10E-05 | 7 | - | 0,000555018 | - | 10 | - | 0,00066287 | - | 6 | - | 0,000400379 | - |
| 32 | 1 | 0.2um_NOKCPHJN_254094 | Bacteria | Proteobacteri | Gammaprotec | Cellvibrionales | Porticoccacea | unclassified | gamma protec | cellular proces | Cell wall/mem | 16QA6 | penicillin-bind | 314287.GB22 | 0,6 | 0,489897949 | 6,35E-05 | 5,25E-05 | 1 | - | 0,000120335 | - | 1 | - | 0,000100603 | - | 0 | - | 0 | - |
| 33 | 1 | 3um_FIJJAFBF_282598 | Eukaryota | unclassified | Pelagophycea | Pelagomonad | unclassified | Aureococcus | Aureococcus a | unclassified | unclassified | unclassified | unclassified | unclassified | 1 | 0 | 0,000250965 | 2,65E-05 | 1 | - | 0,0002982 | - | 1 | - | 0,000249302 | - | 1 | - | 0,000250968 | - |
| 34 | 1 | 3um_FIJJAFBF_564667 | Bacteria | Proteobacteri | Betaproteoba | unclassified | unclassified | unclassified | Betaproteoba | metabolism | Inorganic ion t | 07QMV | Part of the AB | 452638.Pnec_ | 1,2 | 0,979795897 | 0,000663693 | 0,000544317 | 0 | - | 0 | - | 0 | - | 0 | - | 2 | - | 0,001177438 | - |
| 35 | 1 | 3um_FIJJAFBF_221402 | Viruses | unclassified | unclassified | unclassified | unclassified | unclassified | uncultured M | unclassified | unclassified | unclassified | unclassified | unclassified | 0,2 | 0,4 | 3,65E-05 | 7,30E-05 | 1 | - | 0,000182557 | - | 0 | - | 0 | - | 0 | - | 0 | - |
| 36 | 1 | 3um_FIJJAFBF_641954 | Bacteria | Proteobacteri | Gammaprotec | Cellvibrionales | Spongiibacter | Oceanicoccus | Oceanicoccus | metabolism | Inorganic ion t | 01UP7 | TonB-depende | 394221.Mmar | 5 | 0,632455532 | 0,000393028 | 1,29E-05 | 4 | - | 0,000378499 | - | 5 | - | 0,000395543 | - | 5 | - | 0,000398185 | - |
| 37 | 1 | 3um_FIJJAFBF_589604 | Bacteria | Actinobacteri | Actinobacteri | unclassified | unclassified | unclassified | Actinobacteri | unclassified | unclassified | unclassified | unclassified | unclassified | 0,2 | 0,4 | 2,24E-05 | 4,47E-05 | 0 | - | 0 | - | 0 | - | 0 | - | 0 | - | 0 | - |
| 38 | 1 | 3um_FIJJAFBF_760215 | Bacteria | Proteobacteri | Alphaproteob | Rhodobactera | Rhodobactera | Sulfitobacter | Sulfitobacter g | metabolism | Carbohydrate | 05WYW | fructose-bispł | 391589.RGAI1 | 0,2 | 0,4 | 4,24E-05 | 8,48E-05 | 0 | - | 0 | - | 0 | - | 0 | - | 1 | - | 0,000212097 | - |
| 39 | 1 | 3um_FIJJAFBF_439855 | Bacteria | Bacteroidetes | Flavobacteriia | Flavobacterial | Flavobacteria | Flaviramulus | Flaviramulus b | unclassified | unclassified | unclassified | unclassified | unclassified | 0,6 | 0,489897949 | 3,68E-05 | 3,08E-05 | 1 | - | 7,28E-05 | - | 0 | - | 0 | - | 0 | - | 0 | - |
| 40 | 1 | 3um_FIJJAFBF_116589 | Eukaryota | Eustigmatoph | unclassified | Eustigmatales | Monodopsida | Nannochlorop | various | unclassified | unclassified | unclassified | unclassified | unclassified | 3 | 0,632455532 | 0,000468858 | 8,86E-05 | 2 | - | 0,000376122 | - | 3 | - | 0,00047167 | - | 4 | - | 0,000633095 | - |
| 41 | 1 | 0.2um_NOKCPHJN_1084374 | Eukaryota | unclassified | Oomycetes | Pythiales | Pythiaceae | Pythium | Pythium insidi | information st | Replication, re | COG0513 | purine NTP-de | 65071.PYU1_ | 1 | 0 | 0,000396177 | 4,19E-05 | 1 | - | 0,000470743 | - | 1 | - | 0,000393553 | - | 1 | - | 0,000396182 | - |
| 42 | 1 | 0.2um_NOKCPHJN_702692 | Bacteria | Bacteroidetes | Flavobacteriia | Flavobacterial | Flavobacteria | Maribacter | Maribacter an | metabolism | Secondary me | 0P8JT | Phytoene des | 313595.P7007 | 1 | 0 | 0,000128555 | 1,36E-05 | 1 | - | 0,000152751 | - | 1 | - | 0,000127704 | - | 1 | - | 0,000128557 | - |
| 43 | 1 | 0.2um_NOKCPHJN_488159 | Bacteria | Proteobacteri | Alphaproteob | Sphingomona | various | various | various | unclassified | unclassified | unclassified | unclassified | unclassified | 0,2 | 0,4 | 8,17E-05 | 0,000163471 | 0 | - | 0 | - | 0 | - | 0 | - | 0 | - | 0 | - |
| 44 | 1 | 0.2um_NOKCPHJN_07567 | Bacteria | unclassified | unclassified | unclassified | unclassified | unclassified | uncultured ba | unclassified | unclassified | unclassified | unclassified | unclassified | 0,4 | 0,489897949 | 2,93E-05 | 3,59E-05 | 0 | - | 0 | - | 0 | - | 0 | - | 0 | - | 0 | - |
| 45 | 1 | 3um_FIJJAFBF_510245 | Bacteria | Bacteroidetes | Flavobacteriia | Flavobacterial | Flavobacteria | Polaribacter | various | unclassified | unclassified | unclassified | unclassified | unclassified | 3,2 | 0,4 | 0,001104256 | 9,84E-05 | 3 | - | 0,001240578 | - | 3 | - | 0,001037153 | - | 3 | - | 0,001044082 | - |
| 46 | 1 | 0.2um_NOKCPHJN_825274 | unclassified | unclassified | unclassified | unclassified | unclassified | unclassified | unclassified | unclassified | unclassified | unclassified | unclassified | unclassified | 0,6 | 0,8 | 0,00030453 | 0,000398104 | 0 | - | 0 | - | 0 | - | 0 | - | 0 | - | 0 | - |
| 47 | 1 | 0.2um_NOKCPHJN_115345 | Bacteria | Bacteroidetes | Flavobacteriia | Flavobacterial | Flavobacteria | Olleya | Olleya namha | unclassified | unclassified | unclassified | unclassified | unclassified | 5,8 | 1,469693846 | 0,000388567 | 7,16E-05 | 4 | - | 0,000325074 | - | 5 | - | 0,000339712 | - | 7 | - | 0,000478774 | - |
| 48 | 1 | 3um_FIJJAFBF_150253 | Bacteria | Bacteroidetes | Saprospiria | Saprospirales | Saprospiracea | Aureispira | Aureispira sp. | cellular proces | Signal transdu | arCOG02393 | Response regu | 868132.Metb | 0 | 0 | 0 | 0 | 0 | - | 0 | - | 0 | - | 0 | - | 0 | - | 0 | - |
| 49 | 3 | 3um_FIJJAFBF_537646;3um_FIJJAFBF_ | Eukaryota | Bacillariophyt | various | various | various | various | various | information st | Translation, ri | various | various | various | 0,4 | 0,489897949 | 0,000223738 | 0,000275221 | 0 | - | 0 | - | 0 | - | 0 | - | 1 | - | 0,000599933 | - |
| 50 | 1 | 0.2um_NOKCPHJN_1006223 | Eukaryota | Phaeophyceae | unclassified | Ectocarpales | Ectocarpaceae | Ectocarpus | Ectocarpus sili | metabolism | Amino acid tra | COG0436 | aminotransfer | 35128.Thaps3 | 1 | 0 | 0,000170711 | 1,81E-05 | 1 | - | 0,000202841 | - | 1 | - | 0,00016958 | - | 1 | - | 0,000170713 | - |
| 51 | 1 | 0.2um_NOKCPHJN_780172 | Eukaryota | unclassified | Cryptophyta | Pyrenomonad | Geminigerace | Guillardia | Guillardia thet | metabolism | Amino acid tra | COG0473 | 3-isopropylma | 2850.Phatr20 | 2 | 0 | 0,000386455 | 4,09E-05 | 2 | - | 0,000459191 | - | 2 | - | 0,000383895 | - | 2 | - | 0,00038646 | - |
| 52 | 1 | 3um_FIJJAFBF_353183 | Eukaryota | Streptophyta | Liliopsida | Poales | Poaceae | Oryza | Oryza sativa | unclassified | unclassified | unclassified | unclassified | unclassified | 1,2 | 0,4 | 0,000719912 | 0,000248222 | 1 | - | 0,00071284 | - | 1 | - | 0,000595952 | - | 2 | - | 0,001199865 | - |
| 53 | 1 | 0.2um_NOKCPHJN_657137 | Eukaryota | unclassified | Cryptophyta | Pyrenomonad | Geminigerace | Guillardia | Guillardia thet | unclassified | unclassified | unclassified | unclassified | unclassified | 6 | 0,632455532 | 0,003168452 | 6,04E-05 | 5 | - | 0,003171533 | - | 6 | - | 0,003181776 | - | 6 | - | 0,003203031 | - |
| 54 | 1 | 3um_FIJJAFBF_280809 | Bacteria | Bacteroidetes | Flavobacteriia | Flavobacterial | Flavobacteria | Arenitalea | Arenitalea lut | cellular proces | Cell motility | COG1360 | Ompa motb d | 391587.KAOT | 3,8 | 0,748331477 | 0,000855569 | 0,000174288 | 3 | - | 0,000804819 | - | 4 | - | 0,000897131 | - | 5 | - | 0,001128906 | - |
| 55 | 1 | 3um_FIJJAFBF_556071 | Eukaryota | unclassified | Dinophyceae | Peridiniales | Heterocapsac | Heterocapsa | Heterocapsa t | unclassified | unclassified | unclassified | unclassified | unclassified | 2,6 | 0,489897949 | 0,001969413 | 0,000276486 | 2 | - | 0,001825565 | - | 2 | - | 0,001526218 | - | 3 | - | 0,00230462 | - |
| 56 | 1 | 0.2um_NOKCPHJN_249844 | Eukaryota | unclassified | Pelagophycea | Pelagomonad | unclassified | Aureococcus | Aureococcus a | poorly charact | Function unkn | 0XT7J | HYDIN, axonei | 67593.Physo1 | 0,4 | 0,489897949 | 6,01E-06 | 7,40E-06 | 1 | - | 1,63E-05 | - | 0 | - | 0 | - | 1 | - | 1,37E-05 | - |
| 57 | 1 | 0.2um_NOKCPHJN_469021 | Bacteria | Bacteroidetes | Flavobacteriia | Flavobacterial | Flavobacteria | Lacinutrix | Lacinutrix sp. | unclassified | unclassified | unclassified | unclassified | unclassified | 0,4 | 0,489897949 | 2,70E-05 | 3,31E-05 | 0 | - | 0 | - | 0 | - | 0 | - | 0 | - | 0 | - |
| 58 | 1 | 0.2um_NOKCPHJN_12988 | Bacteria | Bacteroidetes | Flavobacteriia | Flavobacterial | Flavobacteria | Formosa | Formosa sp. H | information st | Translation, ri | 0PA05 | 30S ribosomal | 983544.Lacal_ | 0,6 | 0,489897949 | 0,000173658 | 0,0001448 | 1 | - | 0,00033868 | - | 1 | - | 0,000283144 | - | 0 | - | 0 | - |
| 59 | 1 | 0.2um_NOKCPHJN_597321 | Eukaryota | unclassified | unclassified | Isochrysidales | Noelaerhabda | Emiliania | Emiliania huxl | unclassified | unclassified | unclassified | unclassified | unclassified | 1 | 0 | 0,000228233 | 2,41E-05 | 1 | - | 0,000271189 | - | 1 | - | 0,000226721 | - | 1 | - | 0,000228235 | - |
| 60 | 1 | 3um_FIJJAFBF_612753 | Bacteria | Proteobacteri | Alphaproteob | Rhodobactera | Rhodobactera | Labrenzia | Labrenzia agg | unclassified | unclassified | unclassified | unclassified | unclassified | 0,2 | 0,4 | 2,88E-05 | 5,76E-05 | 0 | - | 0 | - | 0 | - | 0 | - | 0 | - | 0 | - |
| 61 | 1 | 3um_FIJJAFBF_669282 | Eukaryota | unclassified | Pelagophycea | Pelagomonad | unclassified | various | various | metabolism | Energy produc | 0J0AS | This b-type cy | 35128.Thapsd | 1,8 | 0,748331477 | 0,001363178 | 0,000581479 | 2 | - | 0,0017821 | - | 2 | - | 0,001489879 | - | 1 | - | 0,000749916 | - |
| 62 | 1 | 3um_FIJJAFBF_179604 | Eukaryota | Bacillariophyt | Bacillariophyc | Naviculales | Phaeodactylac | Phaeodactylu | Phaeodactylu | unclassified | unclassified | unclassified | unclassified | unclassified | 0,8 | 0,979795897 | 8,71E-05 | 0,00010676 | 0 | - | 0 | - | 0 | - | 0 | - | 2 | - | 0,000222984 | - |
| 63 | 3 | 0.2um_NOKCPHJN_276340;0.2um_NO | Bacteria | Bacteroidetes | Flavobacteriia | Flavobacterial | Flavobacteria | various | various | metabolism | Lipid transpor | COG2067 | lipid transport | 50743.SCB49_ | 0,6 | 0,489897949 | 8,95E-05 | 7,48E-05 | 1 | - | 0,000176946 | - | 0 | - | 0 | - | 0 | - | 0 | - |
| 64 | 1 | 3um_FIJJAFBF_140885 | Bacteria | Proteobacteri | Gammaprotec | unclassified | Competibacte | Plasticicumula | Plasticicumula | unclassified | unclassified | unclassified | unclassified | unclassified | 0,8 | 0,748331477 | 0,00015314 | 0,000137504 | 0 | - | 0 | - | 0 | - | 0 | - | 1 | - | 0,000207897 | - |
| 65 | 1 | 0.2um_NOKCPHJN_121297 | Bacteria | Proteobacteri | Alphaproteob | Rhodobactera | Rhodobactera | Planktomarina | Planktomarina | unclassified | unclassified | unclassified | unclassified | unclassified | 0,4 | 0,489897949 | 2,53E-05 | 3,11E-05 | 0 | - | 0 | - | 0 | - | 0 | - | 1 | - | 6,78E-05 | - |
| 66 | 1 | 0.2um_NOKCPHJN_201800 | Eukaryota | Mollusca | Cephalopoda | Octopoda | Octopodidae | Octopus | Octopus bima | cellular proces | Cell cycle cont | 12QMH | septin 2 | 7955.ENSDAR | 0,2 | 0,4 | 1,86E-05 | 3,71E-05 | 0 | - | 0 | - | 0 | - | 0 | - | 1 | - | 9,28E-05 | - |
| 67 | 1 | 3um_FIJJAFBF_689436 | Bacteria | Proteobacteri | Gammaprotec | Alteromonada | Colwelliaceae | Colwellia | Colwellia pola | unclassified | unclassified | unclassified | unclassified | unclassified | 0,4 | 0,489897949 | 0,000134008 | 0,000167488 | 1 | - | 0,000387814 | - | 0 | - | 0 | - | 0 | - | 0 | - |
| 68 | 1 | 0.2um_NOKCPHJN_35230 | Bacteria | Proteobacteri | Alphaproteob | unclassified | unclassified | unclassified | SAR116 cluste | metabolism | Energy produc | 01RNF | Produces ATP | 1064539.AZO | 1 | 0 | 0,000132895 | 1,41E-05 | 1 | - | 0,000157908 | - | 1 | - | 0,000132015 | - | 1 | - | 0,000132896 | - |
| 69 | 1 | 3um_FIJJAFBF_507802 | Bacteria | Proteobacteri | Gammaprotec | Chromatiales | Ectothiorhodc | Ectothiorhodc | Ectothiorhodc | unclassified | unclassified | unclassified | unclassified | unclassified | 0,8 | 0,4 | 0,000455153 | 0,00023486 | 1 | - | 0,000668287 | - | 1 | - | 0,000558705 | - | 1 | - | 0,000562437 | - |
| 70 | 1 | 3um_FIJJAFBF_377522 | Bacteria | Bacteroidetes | Flavobacteriia | Flavobacterial | Flavobacteria | Kordia | Kordia sp. SM | information st | Replication, re | COG0550 | Releases the s | 655815.ZPR_4 | 0,2 | 0,4 | 1,51E-05 | 3,01E-05 | 0 | - | 0 | - | 0 | - | 0 | - | 1 | - | 7,53E-05 | - |
| 71 | 1 | 3um_FIJJAFBF_599427 | Bacteria | Proteobacteri | Alphaproteob | Rhodospirillal | Rhodospirillac | unclassified | Rhodospirillac | unclassified | unclassified | unclassified | unclassified | unclassified | 0,8 | 0,4 | 8,65E-05 | 4,36E-05 | 0 | - | 0 | - | 1 | - | 0,000112748 | - | 1 | - | 0,000113501 | - |
| 72 | 1 | 0.2um_NOKCPHJN_652146 | Eukaryota | unclassified | Oomycetes | Albuginales | Albuginaceae | Albugo | Albugo candid | unclassified | unclassified | unclassified | unclassified | unclassified | 8,4 | 0,489897949 | 0,001188864 | 0,000183724 | 9 | - | 0,00150701 | - | 9 | - | 0,001259898 | - | 8 | - | 0,00112739 | - |
| 73 | 1 | 0.2um_NOKCPHJN_34584 | Eukaryota | unclassified | Bangiophycea | Cyanidiales | Cyanidiaceae | Galdieria | Galdieria sulpl | cellular proces | Posttranslatio | KOG0731 | ATP-depender | 59689.scaffold | 0,6 | 0,489897949 | 5,69E-05 | 4,74E-05 | 1 | - | 0,000110886 | - | 1 | - | 9,27E-05 | - | 0 | - | 0 | - |
| 74 | 3 | 0.2um_NOKCPHJN_833582;0.2um_NO | Bacteria | unclassified | unclassified | unclassified | unclassified | unclassified | uncultured ba | various | various | various | various | various | 0,6 | 0,489897949 | 0,000134191 | 0,000111891 | 1 | - | 0,000261707 | - | 1 | - | 0,000218793 | - | 0 | - | 0 | - |
| 75 | 1 | 3um_FIJJAFBF_325931 | Eukaryota | Bacillariophyt | Coscinodiscop | Chaetocerotal | Chaetocerota | Chaetoceros | Chaetoceros s | cellular proces | Posttranslatio | KOG0731 | ATP-depender | 35128.Thapsd | 1,2 | 0,4 | 0,000280696 | 9,55E-05 | 1 | - | 0,000278246 | - | 2 | - | 0,000465241 | - | 1 | - | 0,000234174 | - |
| 76 | 1 | 3um_FIJJAFBF_111262 | Bacteria | Proteobacteri | Alphaproteob | Rhodospirillal | Rhodospirillac | Candidatus En | Candidatus En | unclassified | unclassified | unclassified | unclassified | unclassified | 2 | 0,632455532 | 0,001261244 | 0,000358584 | 2 | - | 0,001512085 | - | 2 | - | 0,00126414 | - | 2 | - | 0,001272585 | - |
| 77 | 1 | 3um_FIJJAFBF_391433 | Eukaryota | Apicomplexa | Conoidasida | Eucoccidiorida | Sarcocystidae | Besnoitia | Besnoitia besr | unclassified | unclassified | unclassified | unclassified | unclassified | 1 | 0 | 0,000234172 | 2,48E-05 | 1 | - | 0,000278246 | - | 1 | - | 0,000232621 | - | 1 | - | 0,000234174 | - |
| 78 | 1 | 0.2um_NOKCPHJN_862562 | Bacteria | Bacteroidetes | Flavobacteriia | Flavobacterial | Flavobacteria | Aureitalea | Aureitalea sp. | metabolism | Energy produc | COG2838 | Isocitrate deh | 983544.Lacal_ | 0,6 | 0,489897949 | 5,26E-05 | 4,32E-05 | 0 | - | 0 | - | 1 | - | 9,15E-05 | - | 1 | - | 9,21E-05 | - |
| 79 | 1 | 0.2um_NOKCPHJN_257407 | Eukaryota | Apicomplexa | Aconoidasida | Haemosporida | Plasmodiidae | Plasmodium | Plasmodium fa | cellular proces | Signal transdu | KOG0613 | fibronectin typ | 7460.GB1563 | 1 | 0 | 0,000119303 | 1,26E-05 | 1 | - | 0,000141758 | - | 1 | - | 0,000118513 | - | 1 | - | 0,000119305 | - |
| 80 | 2 | 3um_FIJJAFBF_01992;3um_FIJJAFBF_1 | unclassified | unclassified | unclassified | unclassified | unclassified | unclassified | unclassified | unclassified | unclassified | unclassified | unclassified | unclassified | 0,8 | 0,4 | 0,000457358 | 0,000230503 | 0 | - | 0 | - | 1 | - | 0,000595952 | - | 1 | - | 0,000599933 | - |
| 81 | 1 | 0.2um_NOKCPHJN_1104412 | Eukaryota | Arthropoda | Insecta | Coleoptera | Tenebrionidae | Tribolium | Tribolium cast | metabolism | Lipid transpor | COG0439 | acetyl-CoA car | 7070.TC01561 | 0,2 | 0,4 | 6,88E-05 | 0,000137527 | 0 | - | 0 | - | 1 | - | 0,000343818 | - | 0 | - | 0 | - |
| 82 | 1 | 0.2um_NOKCPHJN_315798 | Bacteria | Bacteroidetes | unclassified | unclassified | unclassified | unclassified | Bacteroidetes | unclassified | unclassified | unclassified | unclassified | unclassified | 0,6 | 0,489897949 | 4,76E-05 | 3,90E-05 | 0 | - | 0 | - | 1 | - | 8,41E-05 | - | 0 | - | 0 | - |
| 83 | 1 | 3um_FIJJAFBF_355789 | Bacteria | Bacteroidetes | Flavobacteriia | Flavobacterial | Flavobacteria | Polaribacter | Polaribacter s | poorly charact | Function unkn | 0P8NI | Inherit from b | 313594.PI23P | 0,4 | 0,489897949 | 3,50E-05 | 4,29E-05 | 0 | - | 0 | - | 0 | - | 0 | - | 0 | - | 0 | - |
| 84 | 1 | 3um_FIJJAFBF_715073 | Eukaryota | unclassified | Pelagophycea | Pelagomonad | unclassified | Aureoumbra | Aureoumbra l | information st | Translation, ri | 08UHY | One of the pri | 32049.SYNPC | 2,2 | 0,4 | 0,000888352 | 0,000182714 | 2 | - | 0,000959592 | - | 2 | - | 0,000802243 | - | 3 | - | 0,001211403 | - |
| 85 | 1 | 3um_FIJJAFBF_257125 | Bacteria | Cyanobacteria | unclassified | Oscillatoriales | Microcoleacea | Arthrospira | various | unclassified | unclassified | unclassified | unclassified | unclassified | 4,8 | 0,4 | 0,001685344 | 8,54E-05 | 4 | - | 0,001681982 | - | 5 | - | 0,001757722 | - | 5 | - | 0,001769465 | - |
| 86 | 2 | 0.2um_NOKCPHJN_635586;3um_FIJJAF | unclassified | unclassified | unclassified | unclassified | unclassified | unclassified | unclassified | unclassified | unclassified | unclassified | unclassified | unclassified | 0,6 | 0,489897949 | 0,000157994 | 0,000129072 | 0 | - | 0 | - | 1 | - | 0,000266276 | - | 1 | - | 0,000268055 | - |
| 87 | 1 | 3um_FIJJAFBF_437250 | Bacteria | Bacteroidetes | Cytophagia | Cytophagales | Flammeovirga | Fabibacter | Fabibacter sp. | unclassified | unclassified | unclassified | unclassified | unclassified | 0,6 | 0,8 | 7,04E-05 | 9,21E-05 | 0 | - | 0 | - | 0 | - | 0 | - | 0 | - | 0 | - |
| 88 | 1 | 0.2um_NOKCPHJN_539670 | Eukaryota | Chordata | Actinopteri | Pleuronectifor | Paralichthyida | Paralichthys | Paralichthys o | unclassified | unclassified | unclassified | unclassified | unclassified | 0,2 | 0,4 | 0,000122702 | 0,000245404 | 1 | - | 0,00061351 | - | 0 | - | 0 | - | 0 | - | 0 | - |
| 89 | 1 | 3um_FIJJAFBF_211494 | Bacteria | Proteobacteri | Gammaprotec | Alteromonada | Alteromonada | Paraglaciecola | Paraglaciecola | metabolism | Inorganic ion t | 16TQC | TONB-depend | 87626.PTD2_ | 3,6 | 1,2 | 0,000226354 | 6,30E-05 | 3 | - | 0,00022866 | - | 2 | - | 0,000127444 | - | 3 | - | 0,000192443 | - |
| 90 | 1 | 3um_FIJJAFBF_292927 | Eukaryota | Bacillariophyt | Coscinodiscop | Coscinodiscale | Coscinodiscac | Coscinodiscus | Coscinodiscus | cellular proces | Posttranslatio | KOG0731 | ATP-depender | 35128.Thapsd | 2,4 | 0,489897949 | 0,000237202 | 5,21E-05 | 2 | - | 0,000235002 | - | 3 | - | 0,000294701 | - | 3 | - | 0,00029667 | - |
| 91 | 1 | 3um_FIJJAFBF_49741 | Eukaryota | unclassified | Cryptophyta | Pyrenomonad | Geminigerace | Guillardia | Guillardia thet | cellular proces | Posttranslatio | 09YID | Heat shock 70 | 135651.CBN1 | 1 | 0 | 9,69E-05 | 1,02E-05 | 1 | - | 0,000115151 | - | 1 | - | 9,63E-05 | - | 1 | - | 9,69E-05 | - |
| 92 | 1 | 3um_FIJJAFBF_103164 | Bacteria | Bacteroidetes | Flavobacteriia | Flavobacterial | Flavobacteria | Polaribacter | Polaribacter b | information st | Translation, ri | 0P97G | This is one of t | 313598.MED1 | 0,4 | 0,489897949 | 0,000125551 | 0,000156918 | 1 | - | 0,000363341 | - | 0 | - | 0 | - | 0 | - | 0 | - |
| 93 | 1 | 0.2um_NOKCPHJN_206649 | unclassified | unclassified | unclassified | unclassified | unclassified | unclassified | unclassified | information st | RNA processir | 1C5YF | RNA recogniti | 3711.Bra0187 | 0,2 | 0,4 | 9,11E-06 | 1,82E-05 | 0 | - | 0 | - | 0 | - | 0 | - | 0 | - | 0 | - |
| 94 | 1 | 0.2um_NOKCPHJN_969995 | Eukaryota | unclassified | Ichthyosporea | Ichthyophonic | unclassified | Sphaeroforma | Sphaeroforma | unclassified | unclassified | unclassified | unclassified | unclassified | 0,6 | 0,489897949 | 0,000131998 | 0,000108256 | 0 | - | 0 | - | 0 | - | 0 | - | 1 | - | 0,000234174 | - |
| 95 | 1 | 3um_FIJJAFBF_22824 | Eukaryota | Eustigmatoph | unclassified | Eustigmatales | Monodopsida | Nannochlorop | Nannochlorop | unclassified | unclassified | unclassified | unclassified | unclassified | 0,6 | 0,489897949 | 0,000152393 | 0,000124983 | 0 | - | 0 | - | 0 | - | 0 | - | 1 | - | 0,000270356 | - |
| 96 | 1 | 0.2um_NOKCPHJN_417984 | Eukaryota | unclassified | unclassified | Protosteliales | Cavosteliacea | Planoprotoste | Planoprotoste | unclassified | unclassified | unclassified | unclassified | unclassified | 0,4 | 0,489897949 | 2,31E-05 | 2,88E-05 | 1 | - | 6,67E-05 | - | 0 | - | 0 | - | 0 | - | 0 | - |

| | | | | | | | | | | | | | | | | | | | | | | | | |
|---|---|---|---|---|---|---|---|---|---|---|---|---|---|---|---|---|---|---|---|---|---|---|---|---|
| 97 | 1 | 3um_FIJJAFBF_182957 | Bacteria | Proteobacteri | Gammaprotec | Alteromonada | Alteromonada | Paraglaciecola | Paraglaciecola | metabolism | Inorganic ion t | 16TQC | TONB-depend | 314287.GB22( | 6,2 | 1,166190379 | 0,000372356 | 4,16E-05 | 5 - | 0,000362637 - | 5 - | 0,000303173 - | 6 - | 0,000366238 - |
| 98 | 1 | 3um_FIJJAFBF_142849 | Bacteria | Proteobacteri | Alphaproteob | Rhodobactera | unclassified | unclassified | Rhodobactera | cellular proces | Cell wall/mem | 16QB0 | Part of the ou | 439497.RR11_ | 0,6 | 0,489897949 | 4,87E-05 | 4,07E-05 | 1 - | 9,63E-05 - | 0 - | 0 - | 0 - | 0 - |
| 99 | 1 | 0.2um_NOKCPHJN_381565 | Bacteria | Proteobacteri | Alphaproteob | unclassified | unclassified | unclassified | Alphaproteob | information st | Translation, ri | COG0185 | Protein S19 fo | 488538.SAR11 | 1 | 0 | 0,000692222 | 7,32E-05 | 1 - | 0,000822508 - | 1 - | 0,000687636 - | 1 - | 0,00069223 - |
| 100 | 1 | 3um_FIJJAFBF_440713 | Bacteria | Bacteroidetes | Flavobacteriia | Flavobacterial | Flavobacteria | Flaviramulus | Flaviramulus b | metabolism | Energy produc | 05D26 | NQR complex | 983544.Lacal_ | 1,6 | 0,489897949 | 0,000236955 | 8,84E-05 | 2 - | 0,00034413 - | 2 - | 0,000287701 - | 2 - | 0,000289623 - |
| 101 | 1 | 0.2um_NOKCPHJN_1100803 | Bacteria | Verrucomicrol | Opitutae | Opitutales | Opitutaceae | unclassified | Opitutaceae b | unclassified | unclassified | unclassified | unclassified | unclassified | 0,4 | 0,489897949 | 5,50E-05 | 6,78E-05 | 1 - | 0,000149696 - | 1 - | 0,00012515 - | 0 - | 0 - |
| 102 | 1 | 0.2um_NOKCPHJN_710509 | Eukaryota | Streptophyta | unclassified | Asterales | Asteraceae | Lactuca | Lactuca sativa | metabolism | Energy produc | 1BFZ8 | Cytochrome b | 3847.GLYMA1 | 2 | 0 | 0,000999876 | 0,00010574 | 2 - | 0,001188066 - | 2 - | 0,000993253 - | 2 - | 0,000999888 - |
| 103 | 1 | 3um_FIJJAFBF_632919 | Bacteria | Planctomycet | Phycisphaerae | Phycisphaeral | Phycisphaerac | Phycisphaera | Phycisphaera | unclassified | unclassified | unclassified | unclassified | unclassified | 0,2 | 0,4 | 3,33E-05 | 6,66E-05 | 0 - | 0 - | 0 - | 0 - | 0 - | 0 - |
| 104 | 1 | 0.2um_NOKCPHJN_184643 | Bacteria | Bacteroidetes | Flavobacteriia | Flavobacterial | Cryomorphac | Owenweeksia | Owenweeksia | poorly charact | Function unkn | 082FF | Colicin v prod | 177437.HRM2 | 0,2 | 0,4 | 7,56E-05 | 0,000151133 | 0 - | 0 - | 0 - | 0 - | 0 - | 0 - |
| 105 | 1 | 0.2um_NOKCPHJN_1032972 | Eukaryota | unclassified | Cryptophyta | Pyrenomonad | Geminigerace | Guillardia | Guillardia thet | metabolism | Lipid transpor | COG0439 | acetyl-CoA car | 5811.TGME49 | 3,8 | 1,166190379 | 0,001159687 | 0,000262516 | 3 - | 0,001111607 - | 3 - | 0,00092933 - | 4 - | 0,001247385 - |
| 106 | 1 | 0.2um_NOKCPHJN_154141 | Eukaryota | Streptophyta | Liliopsida | Poales | Poaceae | Hordeum | Hordeum vulg | unclassified | unclassified | unclassified | unclassified | unclassified | 3,4 | 0,489897949 | 0,000701608 | 6,81E-05 | 3 - | 0,000743525 - | 3 - | 0,000621605 - | 3 - | 0,000625758 - |
| 107 | 1 | 0.2um_NOKCPHJN_15951 | Bacteria | Bacteroidetes | Flavobacteriia | Flavobacterial | Flavobacteria | Formosa | Formosa sp. H | information st | Translation, ri | COG1185 | Involved in mI | 688270.Celal_ | 0,8 | 0,4 | 6,57E-05 | 3,31E-05 | 0 - | 0 - | 1 - | 8,56E-05 - | 1 - | 8,62E-05 - |
| 108 | 1 | 0.2um_NOKCPHJN_165212 | Bacteria | Bacteroidetes | Flavobacteriia | Flavobacterial | Flavobacteria | Formosa | Formosa sp. H | unclassified | unclassified | unclassified | unclassified | unclassified | 1,6 | 0,489897949 | 0,000151068 | 5,51E-05 | 2 - | 0,000220466 - | 1 - | 9,22E-05 - | 2 - | 0,000185546 - |
| 109 | 1 | 0.2um_NOKCPHJN_528916 | Bacteria | Bacteroidetes | Cytophagia | Cytophagales | Cytophagacea | Flectobacillus | Flectobacillus | cellular proces | Cell wall/mem | 1AJ2I | UDPG_MGDP_ | 485918.Cpin_ | 0,2 | 0,4 | 2,85E-05 | 5,71E-05 | 0 - | 0 - | 0 - | 0 - | 0 - | 0 - |
| 110 | 1 | 3um_FIJJAFBF_326272 | Bacteria | Proteobacteri | Alphaproteob | Rhodobactera | Rhodobactera | Phaeobacter | Phaeobacter s | information st | Translation, ri | COG1185 | Involved in mI | 391595.RLO14 | 0,4 | 0,489897949 | 3,31E-05 | 4,07E-05 | 0 - | 0 - | 1 - | 8,85E-05 - | 0 - | 0 - |
| 111 | 1 | 3um_FIJJAFBF_669262 | Eukaryota | unclassified | Synurophycea | Parmales | Triparmaceae | Triparma | Triparma laev | unclassified | unclassified | unclassified | unclassified | unclassified | 0,4 | 0,489897949 | 4,19E-05 | 5,14E-05 | 0 - | 0 - | 0 - | 0 - | 1 - | 0,000107313 - |
| 112 | 1 | 0.2um_NOKCPHJN_1004726 | Bacteria | Bacteroidetes | Flavobacteriia | Flavobacterial | Flavobacteria | Polaribacter | Polaribacter s | poorly charact | Function unkn | 0P8NI | Inherit from b | 983548.Krodi_ | 1 | 0 | 0,000135759 | 1,44E-05 | 1 - | 0,000161311 - | 1 - | 0,00013486 - | 1 - | 0,000135761 - |
| 113 | 1 | 0.2um_NOKCPHJN_786900 | unclassified | unclassified | unclassified | unclassified | unclassified | unclassified | unclassified | unclassified | unclassified | unclassified | unclassified | unclassified | 1 | 0 | 7,53E-05 | 7,96E-06 | 1 - | 8,94E-05 - | 1 - | 7,48E-05 - | 1 - | 7,53E-05 - |
| 114 | 1 | 3um_FIJJAFBF_431946 | unclassified | unclassified | unclassified | unclassified | unclassified | unclassified | unclassified | unclassified | unclassified | unclassified | unclassified | unclassified | 0,8 | 0,748331477 | 0,001026508 | 0,000895915 | 1 - | 0,001559337 - | 1 - | 0,001303644 - | 0 - | 0 - |
| 115 | 1 | 0.2um_NOKCPHJN_257141 | Bacteria | Bacteroidetes | Flavobacteriia | Flavobacterial | Flavobacteria | Formosa | Formosa sp. H | unclassified | unclassified | unclassified | unclassified | unclassified | 0,4 | 0,489897949 | 2,76E-05 | 3,39E-05 | 0 - | 0 - | 1 - | 7,37E-05 - | 0 - | 0 - |
| 116 | 1 | 0.2um_NOKCPHJN_632604 | Viruses | unclassified | unclassified | unclassified | unclassified | unclassified | uncultured M | unclassified | unclassified | unclassified | unclassified | unclassified | 0,4 | 0,489897949 | 0,000140977 | 0,000172721 | 0 - | 0 - | 1 - | 0,000359626 - | 0 - | 0 - |
| 117 | 1 | 0.2um_NOKCPHJN_987129 | unclassified | unclassified | unclassified | unclassified | unclassified | unclassified | unclassified | unclassified | unclassified | unclassified | unclassified | unclassified | 1 | 0 | 0,000768197 | 8,12E-05 | 1 - | 0,000912783 - | 1 - | 0,000763109 - | 1 - | 0,000768207 - |
| 118 | 1 | 0.2um_NOKCPHJN_216573 | Bacteria | Bacteroidetes | Flavobacteriia | Flavobacterial | Flavobacteria | Lutibacter | Lutibacter sp. | unclassified | unclassified | unclassified | unclassified | unclassified | 1 | 0 | 6,34E-05 | 6,71E-06 | 1 - | 7,54E-05 - | 1 - | 6,30E-05 - | 1 - | 6,34E-05 - |
| 119 | 2 | 0.2um_NOKCPHJN_56987;3um_FIJJAFE | Bacteria | Bacteroidetes | Flavobacteriia | Flavobacterial | Flavobacteria | Mangrovimon | Mangrovimon | poorly charact | Function unkn | 1015T | domain protei | various | 1,2 | 0,4 | 0,000173441 | 4,29E-05 | 1 - | 0,0001757 - | 1 - | 0,000146889 - | 1 - | 0,000147871 - |
| 120 | 1 | 3um_FIJJAFBF_257114 | Eukaryota | Bacillariophyt | Mediophycea | Biddulphiales | Biddulphiacea | Biddulphia | Biddulphia tri | unclassified | unclassified | unclassified | unclassified | unclassified | 1 | 0 | 0,000229062 | 2,42E-05 | 1 - | 0,000272175 - | 1 - | 0,000227545 - | 1 - | 0,000229065 - |
| 121 | 1 | 0.2um_NOKCPHJN_431584 | Bacteria | Proteobacteri | Gammaprotec | Enterobactera | Enterobacteri | Salmonella | Salmonella en | cellular proces | Cell cycle cont | 0V9W7 | retinitis pigme | 9606.ENSP000 | 0,2 | 0,4 | 1,51E-05 | 3,01E-05 | 0 - | 0 - | 0 - | 0 - | 1 - | 7,53E-05 - |
| 122 | 1 | 0.2um_NOKCPHJN_363833 | Bacteria | Bacteroidetes | Flavobacteriia | Flavobacterial | Flavobacteria | Formosa | Formosa sp. H | information st | Translation, ri | COG0256 | This is one of t | 983544.Lacal_ | 1,4 | 0,489897949 | 0,000746659 | 0,000265297 | 1 - | 0,000634307 - | 2 - | 0,001060592 - | 2 - | 0,001067677 - |
| 123 | 1 | 3um_FIJJAFBF_573648 | Eukaryota | unclassified | unclassified | Isochrysidales | Noelaerhabda | Emiliania | Emiliania huxl | unclassified | unclassified | unclassified | unclassified | unclassified | 0,4 | 0,8 | 0,000163818 | 0,000327635 | 0 - | 0 - | 0 - | 0 - | 0 - | 0 - |
| 124 | 1 | 3um_FIJJAFBF_713652 | Eukaryota | Bacillariophyt | Bacillariophyc | Bacillariales | Bacillariaceae | Fragilariopsis | Fragilariopsis | unclassified | unclassified | unclassified | unclassified | unclassified | 1 | 0 | 0,000212095 | 2,24E-05 | 1 - | 0,000252014 - | 1 - | 0,00021069 - | 1 - | 0,000212097 - |
| 125 | 1 | 0.2um_NOKCPHJN_912086 | Viruses | unclassified | unclassified | Caudovirales | Podoviridae | unclassified | Cellulophaga | unclassified | unclassified | unclassified | unclassified | unclassified | 1 | 0 | 0,000127515 | 1,35E-05 | 1 - | 0,000151515 - | 1 - | 0,00012667 - | 1 - | 0,000127516 - |
| 126 | 1 | 0.2um_NOKCPHJN_117608 | Bacteria | Proteobacteri | Betaproteoba | Burkholderial | Comamonada | Hydrogenoph | Hydrogenoph | unclassified | unclassified | unclassified | unclassified | unclassified | 0,6 | 0,489897949 | 3,89E-05 | 3,25E-05 | 1 - | 7,68E-05 - | 0 - | 0 - | 0 - | 0 - |
| 127 | 1 | 0.2um_NOKCPHJN_484213 | Bacteria | Proteobacteri | Gammaprotec | Thiotrichales | Piscirickettsia | Thiomicrospir | Thiomicrospir | unclassified | unclassified | unclassified | unclassified | unclassified | 0,4 | 0,489897949 | 9,05E-05 | 0,000111121 | 0 - | 0 - | 0 - | 0 - | 0 - | 0 - |
| 128 | 1 | 0.2um_NOKCPHJN_312799 | Eukaryota | Ascomycota | Eurotiomycet | Chaetothyriale | Herpotrichiell | Fonsecaea | Fonsecaea ere | information st | Transcription | 1DA2B | Homeobox do | 10090.ENSMU | 0,2 | 0,4 | 5,02E-05 | 0,000100467 | 1 - | 0,000251168 - | 0 - | 0 - | 0 - | 0 - |
| 129 | 1 | 3um_FIJJAFBF_58538 | Eukaryota | Phaeophyceae | unclassified | Ectocarpales | Ectocarpaceae | Ectocarpus | Ectocarpus sili | information st | Translation, ri | 1508W | Ribosomal prc | 272952.HpaP8 | 1 | 0 | 0,000599925 | 6,34E-05 | 1 - | 0,00071284 - | 1 - | 0,000595952 - | 1 - | 0,000599933 - |
| 130 | 1 | 0.2um_NOKCPHJN_487848 | Bacteria | Proteobacteri | Alphaproteob | Rhodobactera | Rhodobactera | Roseobacter | Roseobacter c | information st | Translation, ri | COG0050 | This protein p | 375451.RD1_1 | 0,4 | 0,489897949 | 6,27E-05 | 7,69E-05 | 0 - | 0 - | 1 - | 0,000160038 - | 0 - | 0 - |
| 131 | 1 | 3um_FIJJAFBF_671982 | Bacteria | Proteobacteri | Alphaproteob | Rhodospirillal | Rhodospirillac | Thalassospira | Thalassospira | metabolism | Energy produc | 01QSC | Dihydrolipoyl | 331869.BAL19 | 1 | 0 | 0,000174494 | 1,85E-05 | 1 - | 0,000207336 - | 1 - | 0,000173338 - | 1 - | 0,000174496 - |
| 132 | 1 | 3um_FIJJAFBF_614416 | Bacteria | Bacteroidetes | Flavobacteriia | Flavobacterial | Flavobacteria | Zobellia | various | poorly charact | Function unkn | 09ARW | Zinc ribbon dc | 313603.FB217 | 0,2 | 0,4 | 4,64E-05 | 9,28E-05 | 0 - | 0 - | 0 - | 0 - | 0 - | 0 - |
| 133 | 1 | 3um_FIJJAFBF_393923 | Eukaryota | Bacillariophyt | Bacillariophyc | Naviculales | Phaeodactylac | Phaeodactylu | Phaeodactylu | unclassified | unclassified | unclassified | unclassified | unclassified | 0,2 | 0,4 | 3,16E-05 | 6,32E-05 | 0 - | 0 - | 0 - | 0 - | 0 - | 0 - |
| 134 | 2 | 3um_FIJJAFBF_243622;3um_FIJJAFBF_ | Bacteria | Bacteroidetes | Flavobacteriia | Flavobacterial | Flavobacteria | various | various | information st | Translation, ri | 09EM4 | Binds directly | 313594.PI23P | 2,4 | 0,489897949 | 0,00132542 | 0,00029129 | 2 - | 0,001313126 - | 3 - | 0,001646708 - | 3 - | 0,001657709 - |
| 135 | 1 | 3um_FIJJAFBF_28972 | Bacteria | Proteobacteri | Gammaprotec | unclassified | unclassified | unclassified | Gammaprotec | poorly charact | Function unkn | 070MQ | NA | 425104.Ssed_ | 1,6 | 0,489897949 | 0,000226274 | 7,82E-05 | 2 - | 0,000334144 - | 2 - | 0,000279352 - | 1 - | 0,000140609 - |
| 136 | 1 | 3um_FIJJAFBF_352965 | Bacteria | Proteobacteri | Gammaprotec | Alteromonada | Alteromonada | Glaciecola | Glaciecola sp. | metabolism | Inorganic ion t | 0QJVD | TonB-depende | 455436.GHTC | 5,4 | 1,2 | 0,000383103 | 6,71E-05 | 4 - | 0,000342555 - | 6 - | 0,000429576 - | 4 - | 0,000288297 - |
| 137 | 1 | 0.2um_NOKCPHJN_163581 | Eukaryota | unclassified | Oomycetes | Pythiales | Pythiaceae | Pythium | Pythium insidi | unclassified | unclassified | unclassified | unclassified | unclassified | 1 | 0 | 0,00010713 | 1,13E-05 | 1 - | 0,000127293 - | 1 - | 0,00010642 - | 1 - | 0,000107131 - |
| 138 | 1 | 3um_FIJJAFBF_142185 | Bacteria | Proteobacteri | Gammaprotec | Oceanospirilla | Oceanospirilla | Marinomonas | Marinomonas | metabolism | Energy produc | 17CGT | F(1)F(0) ATP s | 717774.Marm | 0,4 | 0,489897949 | 0,00015778 | 0,000193331 | 0 - | 0 - | 0 - | 0 - | 1 - | 0,000403801 - |
| 139 | 1 | 3um_FIJJAFBF_721624 | Eukaryota | unclassified | Oomycetes | Pythiales | Pythiaceae | Pythium | Pythium insidi | unclassified | unclassified | unclassified | unclassified | unclassified | 1 | 0 | 0,000663076 | 7,01E-05 | 1 - | 0,000787876 - | 1 - | 0,000658683 - | 1 - | 0,000663084 - |
| 140 | 1 | 0.2um_NOKCPHJN_504794 | Eukaryota | unclassified | Oomycetes | Saprolegniales | Saprolegniace | Thraustotheca | Thraustotheca | unclassified | unclassified | unclassified | unclassified | unclassified | 0,6 | 0,489897949 | 3,99E-05 | 3,29E-05 | 1 - | 7,45E-05 - | 1 - | 6,23E-05 - | 1 - | 6,27E-05 - |
| 141 | 1 | 3um_FIJJAFBF_434038 | Eukaryota | Bacillariophyt | Coscinodiscop | Thalassiosirale | Stephanodisca | Cyclotella | Cyclotella sp. | unclassified | unclassified | unclassified | unclassified | unclassified | 1 | 0 | 0,000874891 | 9,25E-05 | 1 - | 0,001039558 - | 1 - | 0,000869096 - | 1 - | 0,000874902 - |
| 142 | 1 | 0.2um_NOKCPHJN_388322 | unclassified | unclassified | unclassified | unclassified | unclassified | unclassified | unclassified | unclassified | unclassified | unclassified | unclassified | unclassified | 0,4 | 0,489897949 | 7,01E-05 | 8,60E-05 | 0 - | 0 - | 0 - | 0 - | 0 - | 0 - |
| 143 | 1 | 0.2um_NOKCPHJN_1026862 | Bacteria | Proteobacteri | Gammaprotec | Alteromonada | Alteromonada | Glaciecola | Glaciecola sp. | information st | Translation, ri | COG0050 | This protein p | 345073.VC395 | 1 | 0 | 0,000159879 | 1,69E-05 | 1 - | 0,00018997 - | 1 - | 0,00015882 - | 1 - | 0,000159881 - |
| 144 | 1 | 3um_FIJJAFBF_44713 | Bacteria | Bacteroidetes | Flavobacteriia | Flavobacterial | Flavobacteria | Polaribacter | Polaribacter s | cellular proces | Cell motility | COG1360 | Ompa motb d | 313598.MED1 | 2,2 | 0,4 | 0,000485384 | 5,94E-05 | 2 - | 0,000530838 - | 2 - | 0,000443794 - | 2 - | 0,000446758 - |
| 145 | 1 | 0.2um_NOKCPHJN_869431 | Eukaryota | unclassified | Pelagophycea | Pelagomonad | unclassified | Aureococcus | Aureococcus a | unclassified | unclassified | unclassified | unclassified | unclassified | 1 | 0 | 0,000224972 | 2,38E-05 | 1 - | 0,000267315 - | 1 - | 0,000223482 - | 1 - | 0,000224975 - |
| 146 | 1 | 3um_FIJJAFBF_103744 | Viruses | unclassified | unclassified | unclassified | Phycodnavirid | unclassified | Aureococcus a | unclassified | unclassified | unclassified | unclassified | unclassified | 0,8 | 0,748331477 | 0,000633802 | 0,000565399 | 0 - | 0 - | 1 - | 0,000845607 - | 1 - | 0,000851256 - |
| 147 | 1 | 3um_FIJJAFBF_344296 | Bacteria | Proteobacteri | Gammaprotec | unclassified | unclassified | unclassified | Gammaprotec | metabolism | Inorganic ion t | 0ZZVV | tonB-depende | 314275.MADE | 7,4 | 1,854723699 | 0,000583486 | 0,000108829 | 5 - | 0,000477958 - | 7 - | 0,000559418 - | 9 - | 0,000724057 - |
| 148 | 1 | 3um_FIJJAFBF_393738 | Bacteria | Bacteroidetes | Flavobacteriia | Flavobacterial | Flavobacteria | Lutibacter | Lutibacter sp. | unclassified | unclassified | unclassified | unclassified | unclassified | 1 | 0 | 9,33E-05 | 9,87E-06 | 1 - | 0,000110886 - | 1 - | 9,27E-05 - | 1 - | 9,33E-05 - |
| 149 | 1 | 3um_FIJJAFBF_684040 | unclassified | unclassified | unclassified | unclassified | unclassified | unclassified | unclassified | unclassified | unclassified | unclassified | unclassified | unclassified | 0,8 | 0,4 | 7,75E-05 | 3,95E-05 | 1 - | 0,000111381 - | 1 - | 9,31E-05 - | 1 - | 9,37E-05 - |
| 150 | 1 | 0.2um_NOKCPHJN_60178 | Bacteria | Bacteroidetes | Flavobacteriia | Flavobacterial | Flavobacteria | unclassified | Flavobacteria | metabolism | Energy produc | COG1053 | Succinate DeH | 487796.Flav2A | 1,2 | 1,166190379 | 0,000116635 | 0,000109593 | 1 - | 0,000112216 - | 2 - | 0,000187631 - | 3 - | 0,000283327 - |
| 151 | 1 | 3um_FIJJAFBF_69734 | Bacteria | Proteobacteri | Gammaprotec | Pseudomonad | Pseudomonac | Pseudomonas | various | unclassified | unclassified | unclassified | unclassified | unclassified | 5,6 | 1,496662955 | 0,000696459 | 0,000138605 | 3 - | 0,000455466 - | 6 - | 0,000761561 - | 5 - | 0,000638874 - |
| 152 | 1 | 3um_FIJJAFBF_784152 | Eukaryota | unclassified | unclassified | Prymnesiales | Chrysochrom | Chrysochrom | Chrysochrom | unclassified | unclassified | unclassified | unclassified | unclassified | 0,6 | 0,8 | 0,000280655 | 0,000385542 | 0 - | 0 - | 2 - | 0,000977733 - | 0 - | 0 - |
| 153 | 1 | 3um_FIJJAFBF_402284 | unclassified | unclassified | unclassified | unclassified | unclassified | unclassified | unclassified | unclassified | unclassified | unclassified | unclassified | unclassified | 0,8 | 0,4 | 0,000302259 | 0,0001563 | 1 - | 0,000448193 - | 0 - | 0 - | 1 - | 0,000377203 - |
| 154 | 1 | 3um_FIJJAFBF_416695 | unclassified | unclassified | unclassified | unclassified | unclassified | unclassified | unclassified | unclassified | unclassified | unclassified | unclassified | unclassified | 0,8 | 0,4 | 0,000657843 | 0,000331546 | 0 - | 0 - | 1 - | 0,000857191 - | 1 - | 0,000862917 - |
| 155 | 2 | 0.2um_NOKCPHJN_207569;3um_FIJJAF | Bacteria | Bacteroidetes | Cytophagia | Cytophagales | Flammeovirga | unclassified | Flammeovirga | cellular proces | Intracellular tr | 05VIE | preprotein tra | various | 0,6 | 0,489897949 | 0,00032734 | 0,000268696 | 0 - | 0 - | 1 - | 0,000568863 - | 1 - | 0,000572663 - |
| 156 | 1 | 0.2um_NOKCPHJN_39631 | Bacteria | Bacteroidetes | Flavobacteriia | Flavobacterial | Flavobacteria | Polaribacter | Polaribacter s | unclassified | unclassified | unclassified | unclassified | unclassified | 14,6 | 1,854723699 | 0,001013637 | 8,12E-05 | 12 - | 0,001000198 - | 16 - | 0,001114921 - | 15 - | 0,001052221 - |
| 157 | 1 | 0.2um_NOKCPHJN_332245 | Eukaryota | unclassified | Labyrinthulom | unclassified | Thraustochytr | Aurantiochytr | Aurantiochytr | poorly charact | Function unkn | 0NVRZ | NA | 170187.SP_17 | 0,8 | 0,4 | 1,41E-05 | 7,31E-06 | 1 - | 2,10E-05 - | 0 - | 0 - | 1 - | 1,76E-05 - |
| 158 | 1 | 0.2um_NOKCPHJN_299100 | Bacteria | Candidatus Pa | unclassified | unclassified | unclassified | unclassified | Candidatus Pa | unclassified | unclassified | unclassified | unclassified | unclassified | 1 | 0 | 0,000181012 | 1,91E-05 | 1 - | 0,000215081 - | 1 - | 0,000179813 - | 1 - | 0,000181014 - |
| 159 | 1 | 3um_FIJJAFBF_663907 | Eukaryota | Bacillariophyt | Coscinodiscop | Thalassiosirale | Thalassiosirac | Thalassiosira | Thalassiosira c | unclassified | unclassified | unclassified | unclassified | unclassified | 2,2 | 0,4 | 0,000677663 | 0,000203052 | 3 - | 0,001079541 - | 2 - | 0,000601682 - | 2 - | 0,000605701 - |
| 160 | 1 | 0.2um_NOKCPHJN_1059394 | Eukaryota | unclassified | Oomycetes | Saprolegniales | Saprolegniace | Saprolegnia | Saprolegnia di | unclassified | unclassified | unclassified | unclassified | unclassified | 1 | 0 | 0,000133176 | 1,41E-05 | 1 - | 0,000158241 - | 1 - | 0,000132294 - | 1 - | 0,000133177 - |
| 161 | 1 | 3um_FIJJAFBF_586299 | Eukaryota | unclassified | Dinophyceae | Suessiales | Symbiodiniace | Symbiodinium | Symbiodinium | unclassified | unclassified | unclassified | unclassified | unclassified | 3,6 | 0,489897949 | 0,001206454 | 0,000113309 | 3 - | 0,001207229 - | 4 - | 0,001345697 - | 3 - | 0,001016015 - |
| 162 | 1 | 3um_FIJJAFBF_99779 | Bacteria | Bacteroidetes | Flavobacteriia | Flavobacterial | Flavobacteria | Formosa | Formosa sp. H | unclassified | unclassified | unclassified | unclassified | unclassified | 0,6 | 0,489897949 | 8,03E-05 | 6,63E-05 | 1 - | 0,000151822 - | 0 - | 0 - | 1 - | 0,000127775 - |
| 163 | 1 | 3um_FIJJAFBF_569212 | Eukaryota | unclassified | Pelagophycea | Pelagomonad | unclassified | Aureococcus | Aureococcus a | metabolism | Carbohydrate | COG0057 | glyceraldehyd | 35128.Thaps3 | 7 | 1,673320053 | 0,00170424 | 0,000246472 | 5 - | 0,001479213 - | 6 - | 0,00148399 - | 7 - | 0,001742888 - |
| 164 | 1 | 0.2um_NOKCPHJN_709263 | unclassified | unclassified | unclassified | unclassified | unclassified | unclassified | unclassified | unclassified | unclassified | unclassified | unclassified | unclassified | 1 | 0 | 0,000684698 | 7,24E-05 | 1 - | 0,000813567 - | 1 - | 0,000680162 - | 1 - | 0,000684706 - |
| 165 | 1 | 3um_FIJJAFBF_128452 | Bacteria | Bacteroidetes | Flavobacteriia | Flavobacterial | Flavobacteria | Polaribacter | Polaribacter s | cellular proces | Intracellular tr | 05FGH | MotA TolQ exl | 313594.PI23P | 4,4 | 0,489897949 | 0,001152078 | 0,000149717 | 4 - | 0,00124747 - | 5 - | 0,001303644 - | 4 - | 0,001049882 - |
| 166 | 1 | 3um_FIJJAFBF_687052 | Eukaryota | Streptophyta | Liliopsida | Poales | Poaceae | Setaria | Setaria italica | cellular proces | Posttranslatio | 033V7 | ubiquitin carb | 7070.TC00279 | 0,2 | 0,4 | 2,26E-05 | 4,51E-05 | 0 - | 0 - | 0 - | 0 - | 0 - | 0 - |
| 167 | 2 | 0.2um_NOKCPHJN_59639;3um_FIJJAFE | Bacteria | Bacteroidetes | Flavobacteriia | Flavobacterial | Crocinitomica | unclassified | Crocinitomica | unclassified | unclassified | unclassified | unclassified | unclassified | 0,6 | 0,489897949 | 5,26E-05 | 4,31E-05 | 0 - | 0 - | 1 - | 9,14E-05 - | 1 - | 9,20E-05 - |
| 168 | 1 | 0.2um_NOKCPHJN_1043521 | unclassified | unclassified | unclassified | unclassified | unclassified | unclassified | unclassified | unclassified | unclassified | unclassified | unclassified | unclassified | 0,8 | 0,4 | 0,000717987 | 0,000370483 | 1 - | 0,0010542 - | 1 - | 0,000881337 - | 1 - | 0,000887224 - |
| 169 | 1 | 3um_FIJJAFBF_254308 | Bacteria | Proteobacteri | Gammaprotec | Cellvibrionales | Halieaceae | Chromatocurv | Chromatocurv | information st | Translation, ri | 08UHH | Located on th | 566466.NOR5 | 0,6 | 0,8 | 0,000277623 | 0,000381937 | 0 - | 0 - | 0 - | 0 - | 2 - | 0,000969122 - |
| 170 | 1 | 3um_FIJJAFBF_254301 | Bacteria | Proteobacteri | Gammaprotec | Oceanospirilla | Endozoicomoı | Endozoicomoı | unclassified | information st | Translation, ri | COG0256 | This is one of t | 498211.CJA_0 | 0,6 | 0,489897949 | 0,000305379 | 0,000250366 | 0 - | 0 - | 1 - | 0,000539439 - | 0 - | 0 - |
| 171 | 3 | 0.2um_NOKCPHJN_27919;3um_FIJJAFE | Bacteria | Bacteroidetes | Flavobacteriia | Flavobacterial | Flavobacteria | Formosa | various | unclassified | unclassified | unclassified | unclassified | unclassified | 1 | 0 | 0,000364117 | 3,85E-05 | 1 - | 0,000432648 - | 1 - | 0,000361705 - | 1 - | 0,000364121 - |
| 172 | 1 | 0.2um_NOKCPHJN_868369 | Bacteria | Bacteroidetes | Flavobacteriia | Flavobacterial | Flavobacteria | Polaribacter | Polaribacter s | poorly charact | Function unkn | 0P8NI | Inherit from b | 156586.BBFL7 | 1,4 | 0,489897949 | 0,000225334 | 8,01E-05 | 1 - | 0,000191428 - | 2 - | 0,000320076 - | 2 - | 0,000322215 - |
| 173 | 1 | 3um_FIJJAFBF_280545 | unclassified | unclassified | unclassified | unclassified | unclassified | unclassified | unclassified | unclassified | unclassified | unclassified | unclassified | unclassified | 0,8 | 0,4 | 0,000560197 | 0,000285458 | 1 - | 0,000804819 - | 1 - | 0,000672849 - | 1 - | 0,000677343 - |
| 174 | 1 | 0.2um_NOKCPHJN_811890 | Eukaryota | unclassified | Cryptophyta | Pyrenomonad | Geminigerace | Guillardia | Guillardia thet | metabolism | Energy produc | KOG2799 | beta' subunit | 423536.XP_00 | 0,4 | 0,489897949 | 7,54E-05 | 9,25E-05 | 0 - | 0 - | 0 - | 0 - | 0 - | 0 - |
| 175 | 1 | 0.2um_NOKCPHJN_523102 | unclassified | unclassified | unclassified | unclassified | unclassified | unclassified | unclassified | unclassified | unclassified | unclassified | unclassified | unclassified | 1,4 | 0,489897949 | 0,001022633 | 0,00028645 | 1 - | 0,00089105 - | 1 - | 0,00074494 - | 1 - | 0,000749916 - |
| 176 | 1 | 3um_FIJJAFBF_403052 | Eukaryota | Chlorophyta | Trebouxiophy | Chlorellales | Chlorellaceae | Chlorella | Chlorella soro | information st | Chromatin str | KOG3467 | Core compone | 3218.PP1S133 | 1 | 0 | 0,000611575 | 6,47E-05 | 1 - | 0,000726681 - | 1 - | 0,000607523 - | 1 - | 0,000611582 - |
| 177 | 1 | 3um_FIJJAFBF_152852 | Bacteria | Bacteroidetes | Flavobacteriia | Flavobacterial | Flavobacteria | Flaviramulus | Flaviramulus b | unclassified | unclassified | unclassified | unclassified | unclassified | 1 | 0 | 9,89E-05 | 1,05E-05 | 1 - | 0,000117501 - | 1 - | 9,82E-05 - | 1 - | 9,89E-05 - |
| 178 | 1 | 0.2um_NOKCPHJN_443356 | Eukaryota | Phaeophyceae | unclassified | Ectocarpales | Ectocarpaceae | Ectocarpus | Ectocarpus sili | metabolism | Energy produc | 0IFIU | Pyrophosphat | 2850.Phatr21 | 0,2 | 0,4 | 1,95E-05 | 3,90E-05 | 0 - | 0 - | 1 - | 9,75E-05 - | 0 - | 0 - |
| 179 | 1 | 0.2um_NOKCPHJN_220576 | Eukaryota | unclassified | Oomycetes | Saprolegniales | Saprolegniace | Thraustotheca | Thraustotheca | unclassified | unclassified | unclassified | unclassified | unclassified | 1 | 0 | 0,000237706 | 2,51E-05 | 1 - | 0,000282446 - | 1 - | 0,000236132 - | 1 - | 0,000237709 - |
| 180 | 1 | 0.2um_NOKCPHJN_11364 | Bacteria | Proteobacteri | Alphaproteob | unclassified | unclassified | unclassified | SAR116 cluste | unclassified | unclassified | unclassified | unclassified | unclassified | 0,6 | 0,489897949 | 9,30E-05 | 7,62E-05 | 0 - | 0 - | 0 - | 0 - | 1 - | 0,000164903 - |
| 181 | 1 | 3um_FIJJAFBF_209608 | Bacteria | Bacteroidetes | Flavobacteriia | Flavobacterial | Flavobacteria | Tamlana | various | unclassified | unclassified | unclassified | unclassified | unclassified | 0,8 | 0,4 | 0,000366742 | 0,00018924 | 1 - | 0,000538476 - | 1 - | 0,000450179 - | 1 - | 0,000453187 - |
| 182 | 1 | 0.2um_NOKCPHJN_413021 | Bacteria | Proteobacteri | Betaproteoba | Burkholderiale | Oxalobacterac | Duganella | Duganella sac | poorly charact | Function unkn | 08QSJ | NA | 225937.HP15_ | 0,8 | 0,4 | 0,000157874 | 8,04E-05 | 1 - | 0,000226813 - | 1 - | 0,000189621 - | 1 - | 0,000190888 - |
| 183 | 1 | 0.2um_NOKCPHJN_364780 | Eukaryota | Arthropoda | Insecta | Diptera | Culicidae | Aedes | Aedes albopic | poorly charact | Function unkn | 0IRPM | Retrotranspos | 4081.Solyc11g | 0,2 | 0,4 | 7,42E-06 | 1,48E-05 | 0 - | 0 - | 0 - | 0 - | 1 - | 3,71E-05 - |
| 184 | 1 | 3um_FIJJAFBF_128631 | Eukaryota | Chlorophyta | Mamiellophyc | Mamiellales | Mamiellaceae | Micromonas | Micromonas c | unclassified | unclassified | unclassified | unclassified | unclassified | 2,8 | 0,4 | 0,001552293 | 0,000302232 | 3 - | 0,001969689 - | 3 - | 0,001646708 - | 3 - | 0,001657709 - |
| 185 | 1 | 3um_FIJJAFBF_468776 | Bacteria | Bacteroidetes | Flavobacteriia | Flavobacterial | Flavobacteria | Polaribacter | Polaribacter s | metabolism | Amino acid tra | 0P85E | aminopeptida | 313598.MED1 | 0,2 | 0,4 | 1,41E-05 | 2,82E-05 | 0 - | 0 - | 0 - | 0 - | 0 - | 0 - |
| 186 | 1 | 3um_FIJJAFBF_235253 | Eukaryota | unclassified | Dinophyceae | Gymnodiniale | Kareniaceae | Karlodinium | Karlodinium v | unclassified | unclassified | unclassified | unclassified | unclassified | 0,6 | 0,8 | 0,000359605 | 0,000470101 | 0 - | 0 - | 0 - | 0 - | 0 - | 0 - |
| 187 | 1 | 3um_FIJJAFBF_447157 | unclassified | unclassified | unclassified | unclassified | unclassified | unclassified | unclassified | unclassified | unclassified | unclassified | unclassified | unclassified | 0,4 | 0,489897949 | 0,000391541 | 0,000481638 | 0 - | 0 - | 0 - | 0 - | 1 - | 0,001049882 - |
| 188 | 1 | 3um_FIJJAFBF_594169 | Eukaryota | unclassified | Synurophycea | Parmales | Triparmaceae | Triparma | Triparma laev | unclassified | unclassified | unclassified | unclassified | unclassified | 6 | 2,366431913 | 0,000607507 | 0,000218771 | 5 - | 0,000611505 - | 3 - | 0,00030674 - | 5 - | 0,000514648 - |
| 189 | 1 | 0.2um_NOKCPHJN_340184 | Eukaryota | unclassified | Oomycetes | Albuginales | Albuginaceae | Albugo | Albugo laibacl | unclassified | unclassified | unclassified | unclassified | unclassified | 1 | 0,632455532 | 0,000105796 | 6,98E-05 | 0 - | 0 - | 1 - | 0,000109206 - | 2 - | 0,000219871 - |
| 190 | 1 | 3um_FIJJAFBF_744832 | Bacteria | Proteobacteri | Gammaprotec | Cellvibrionales | Spongiibacter | various | various | unclassified | unclassified | unclassified | unclassified | unclassified | 0,8 | 0,4 | 0,000413062 | 0,000213627 | 1 - | 0,00061351 - | 1 - | 0,000512909 - | 0 - | 0 - |
| 191 | 1 | 0.2um_NOKCPHJN_583758 | Eukaryota | unclassified | Oomycetes | Peronosporale | Peronosporac | Phytophthora | Phytophthora | unclassified | unclassified | unclassified | unclassified | unclassified | 0,2 | 0,4 | 4,42E-05 | 8,83E-05 | 0 - | 0 - | 0 - | 0 - | 0 - | 0 - |
| 192 | 1 | 0.2um_NOKCPHJN_374735 | Viruses | unclassified | unclassified | unclassified | unclassified | unclassified | uncultured M | poorly charact | Function unkn | 09QDG | peptidase M1 | 926562.Oweh | 0,8 | 0,4 | 0,000302029 | 0,000152219 | 0 - | 0 - | 1 - | 0,000393553 - | 1 - | 0,000396182 - |
| 193 | 1 | 3um_FIJJAFBF_155390 | Eukaryota | Chytridiomyco | Monoblephari | Monoblephari | Gonapodyace | Gonapodya | Gonapodya pr | unclassified | unclassified | unclassified | unclassified | unclassified | 1 | 0 | 0,000470091 | 4,97E-05 | 1 - | 0,000558569 - | 1 - | 0,000466977 - | 1 - | 0,000470097 - |
| 194 | 1 | 3um_FIJJAFBF_786591 | unclassified | unclassified | unclassified | unclassified | unclassified | unclassified | unclassified | unclassified | unclassified | unclassified | unclassified | unclassified | 0,2 | 0,4 | 0,00017821 | 0,00035642 | 1 - | 0,00089105 - | 0 - | 0 - | 0 - | 0 - |
| 195 | 1 | 3um_FIJJAFBF_261098 | Bacteria | Bacteroidetes | Flavobacteriia | Flavobacterial | Flavobacteria | Lutibacter | Lutibacter aga | metabolism | Amino acid tra | 0P7S1 | Catalyzes the | 376686.Fjoh_ | 1 | 0 | 0,000214999 | 2,27E-05 | 1 - | 0,000255455 - | 1 - | 0,000213566 - | 1 - | 0,000214993 - |
| 196 | 1 | 0.2um_NOKCPHJN_45864 | Bacteria | Bacteroidetes | Flavobacteriia | Flavobacterial | Crocinitomica | unclassified | Crocinitomica | information st | Replication, re | COG0513 | purine NTP-de | 926562.Oweh | 0,6 | 0,489897949 | 6,35E-05 | 5,18E-05 | 0 - | 0 - | 1 - | 0,000106966 - | 1 - | 0,00010768 - |
| 197 | 1 | 0.2um_NOKCPHJN_622978 | Eukaryota | unclassified | Labyrinthulom | unclassified | Thraustochytr | Aurantiochytr | Aurantiochytr | cellular proces | Signal transdu | KOG0614 | cgmp-depend | 653948.CCA21 | 1 | 0 | 8,83E-05 | 9,34E-06 | 1 - | 0,000104976 - | 1 - | 8,78E-05 - | 1 - | 8,83E-05 - |

| | | | | | | | | | | | | | | | | | | | | | | | | |
|---|---|---|---|---|---|---|---|---|---|---|---|---|---|---|---|---|---|---|---|---|---|---|---|---|
| 198 | 1 | 0.2um_NOKCPHJN_392072 | Eukaryota | Arthropoda | Insecta | Hemiptera | Liviidae | Diaphorina | Diaphorina cit | cellular proces | Posttranslatio | 007GN | Inherit from a | 5833.PFL0585 | 1,8 | 0,4 | 0,000446401 | 0,000112213 | 2 - | 0,000589356 - | 2 - | 0,000492716 - | 1 - | 0,000248004 - |
| 199 | 1 | 3um_FIJJAFBF_372257 | Eukaryota | Phaeophyceae | unclassified | Dictyotales | Dictyotaceae | Dictyopteris | Dictyopteris d | unclassified | unclassified | unclassified | unclassified | unclassified | 0,4 | 0,489897949 | 0,000113975 | 0,000139869 | 0 - | 0 - | 0 - | 0 - | 0 - | 0 - |
| 200 | 1 | 0.2um_NOKCPHJN_267523 | Eukaryota | Cnidaria | Anthozoa | Scleractinia | Acroporidae | Acropora | Acropora digit | unclassified | unclassified | unclassified | unclassified | unclassified | 1 | 0 | 0,000331538 | 3,51E-05 | 1 - | 0,000393938 - | 1 - | 0,000329342 - | 1 - | 0,000331542 - |
| 201 | 1 | 0.2um_NOKCPHJN_1034378 | Eukaryota | Basidiomycota | Agaricomycete | Sebacinales | Serendipitace | Serendipita | Serendipita ve | unclassified | unclassified | unclassified | unclassified | unclassified | 1 | 0 | 0,000368375 | 3,90E-05 | 1 - | 0,000437709 - | 1 - | 0,000365935 - | 1 - | 0,00036838 - |
| 202 | 1 | 3um_FIJJAFBF_40823 | Bacteria | Proteobacteri | Alphaproteob | Rhodobactera | Rhodobactera | Pseudooceani | Pseudooceani | metabolism | Energy produc | COG0056 | Produces ATP | 375451.RD1_3 | 1 | 0 | 0,000124245 | 1,31E-05 | 1 - | 0,00014763 - | 1 - | 0,000123422 - | 1 - | 0,000124246 - |
| 203 | 1 | 3um_FIJJAFBF_609410 | Bacteria | unclassified | unclassified | unclassified | unclassified | unclassified | uncultured ba | unclassified | unclassified | unclassified | unclassified | unclassified | 1 | 0 | 0,000215727 | 2,28E-05 | 1 - | 0,000256329 - | 1 - | 0,000214298 - | 1 - | 0,000215729 - |
| 204 | 1 | 0.2um_NOKCPHJN_625007 | Bacteria | Bacteroidetes | Cytophagia | Cytophagales | Persicobacter | Persicobacter | Persicobacter | metabolism | Inorganic ion t | COG0659 | sulfate transp | 926562.Oweh | 0,6 | 0,489897949 | 6,86E-05 | 5,73E-05 | 1 - | 0,000135595 - | 0 - | 0 - | 0 - | 0 - |
| 205 | 1 | 3um_FIJJAFBF_671855 | Eukaryota | Bacillariophyta | Bacillariophyc | Naviculales | Phaeodactylac | Phaeodactylu | Phaeodactylu | information st | Replication, re | COG0513 | purine NTP-de | 35128.Thaps9 | 0,4 | 0,489897949 | 8,08E-05 | 9,96E-05 | 1 - | 0,000219496 - | 0 - | 0 - | 1 - | 0,00018473 - |
| 206 | 1 | 3um_FIJJAFBF_692602 | Eukaryota | Chlorophyta | unclassified | Pyramimonad | unclassified | Pyramimonas | Pyramimonas | metabolism | Carbohydrate | COG1850 | RuBisCO catal | 3055.DAA009 | 0,8 | 0,4 | 0,000124334 | 6,42E-05 | 1 - | 0,000182557 - | 1 - | 0,000152622 - | 1 - | 0,000153641 - |
| 207 | 1 | 3um_FIJJAFBF_257130 | Eukaryota | Bacillariophyta | Coscinodiscop | Chaetocerotal | Attheyaceae | Attheya | Attheya longi | unclassified | unclassified | unclassified | unclassified | unclassified | 0,2 | 0,4 | 3,47E-05 | 6,94E-05 | 0 - | 0 - | 0 - | 0 - | 0 - | 0 - |
| 208 | 1 | 0.2um_NOKCPHJN_468774 | Bacteria | Firmicutes | Bacilli | Bacillales | Bacillaceae | Bacillus | Bacillus soli | unclassified | unclassified | unclassified | unclassified | unclassified | 0,2 | 0,4 | 5,52E-06 | 1,10E-05 | 0 - | 0 - | 0 - | 0 - | 0 - | 0 - |
| 209 | 1 | 0.2um_NOKCPHJN_647559 | Bacteria | Proteobacteri | Gammaprotec | unclassified | unclassified | unclassified | OM182 bacte | unclassified | unclassified | unclassified | unclassified | unclassified | 0,2 | 0,4 | 1,32E-05 | 2,64E-05 | 0 - | 0 - | 0 - | 0 - | 0 - | 0 - |
| 210 | 1 | 3um_FIJJAFBF_737827 | unclassified | unclassified | unclassified | unclassified | unclassified | unclassified | unclassified | unclassified | unclassified | unclassified | unclassified | unclassified | 1 | 0 | 0,000287636 | 3,04E-05 | 1 - | 0,000341773 - | 1 - | 0,00028573 - | 1 - | 0,000287639 - |
| 211 | 1 | 0.2um_NOKCPHJN_424897 | unclassified | unclassified | unclassified | unclassified | unclassified | unclassified | unclassified | unclassified | unclassified | unclassified | unclassified | unclassified | 1,4 | 0,489897949 | 0,000644161 | 0,000218607 | 1 - | 0,000550354 - | 1 - | 0,00046011 - | 2 - | 0,000926367 - |
| 212 | 1 | 0.2um_NOKCPHJN_1008844 | unclassified | unclassified | unclassified | unclassified | unclassified | unclassified | unclassified | unclassified | unclassified | unclassified | unclassified | unclassified | 0,8 | 0,4 | 0,000100189 | 5,11E-05 | 1 - | 0,000143939 - | 1 - | 0,000120336 - | 1 - | 0,00012114 - |
| 213 | 1 | 0.2um_NOKCPHJN_61191 | Viruses | unclassified | unclassified | unclassified | Phycodnavirid | Prasinovirus | Ostreococcus | unclassified | unclassified | unclassified | unclassified | unclassified | 1 | 0 | 0,000263566 | 2,79E-05 | 1 - | 0,000313172 - | 1 - | 0,00026182 - | 1 - | 0,000263569 - |
| 214 | 1 | 0.2um_NOKCPHJN_161227 | Bacteria | Bacteroidetes | Flavobacteriia | unclassified | unclassified | unclassified | Flavobacteria | unclassified | unclassified | unclassified | unclassified | unclassified | 0,2 | 0,4 | 1,19E-05 | 2,38E-05 | 0 - | 0 - | 0 - | 0 - | 0 - | 0 - |
| 215 | 1 | 3um_FIJJAFBF_594162 | Eukaryota | unclassified | Pelagophycea | Pelagomonad | unclassified | Pelagomonas | uncultured Pe | unclassified | unclassified | unclassified | unclassified | unclassified | 7,8 | 0,979795897 | 0,001501214 | 6,10E-05 | 6 - | 0,001390369 - | 8 - | 0,001549843 - | 8 - | 0,001560197 - |
| 216 | 1 | 0.2um_NOKCPHJN_415195 | Eukaryota | unclassified | unclassified | Prymnesiales | Chrysochrom | Chrysochrom | Chrysochrom | poorly charact | Function unkn | 017A6 | Pfam:DUF395 | 486041.XP_00 | 0,2 | 0,4 | 2,01E-05 | 4,01E-05 | 0 - | 0 - | 0 - | 0 - | 0 - | 0 - |
| 217 | 1 | 3um_FIJJAFBF_191794 | Eukaryota | Bacillariophyta | Coscinodiscop | Rhizosoleniale | Rhizosoleniac | Rhizosolenia | Rhizosolenia f | unclassified | unclassified | unclassified | unclassified | unclassified | 0,8 | 1,166190379 | 0,000128439 | 0,000183199 | 0 - | 0 - | 0 - | 0 - | 0 - | 0 - |
| 218 | 1 | 0.2um_NOKCPHJN_286911 | unclassified | unclassified | unclassified | unclassified | unclassified | unclassified | unclassified | unclassified | unclassified | unclassified | unclassified | unclassified | 0,4 | 0,489897949 | 1,25E-05 | 1,55E-05 | 1 - | 3,48E-05 - | 0 - | 0 - | 0 - | 0 - |
| 219 | 1 | 0.2um_NOKCPHJN_1023367 | Bacteria | Bacteroidetes | Flavobacteriia | Flavobacterial | Flavobacteriac | Marinirhabdu | Marinirhabdu | metabolism | Inorganic ion t | 05C27 | TonB-depende | 983548.Krodi_ | 1 | 0,632455532 | 0,000158208 | 9,77E-05 | 1 - | 0,000189489 - | 0 - | 0 - | 1 - | 0,000159476 - |
| 220 | 1 | 3um_FIJJAFBF_31679 | unclassified | unclassified | unclassified | unclassified | unclassified | unclassified | unclassified | unclassified | unclassified | unclassified | unclassified | unclassified | 1 | 0 | 0,000732467 | 7,75E-05 | 1 - | 0,000870328 - | 1 - | 0,000727615 - | 1 - | 0,000732476 - |
| 221 | 1 | 3um_FIJJAFBF_292914 | Eukaryota | Bacillariophyta | Coscinodiscop | Coscinodiscale | Coscinodiscac | Coscinodiscus | Coscinodiscus | metabolism | Energy produc | COG0712 | F(1)F(0) ATP s | 35128.Thapsd | 0,4 | 0,489897949 | 0,000125181 | 0,000153926 | 0 - | 0 - | 1 - | 0,000334625 - | 0 - | 0 - |
| 222 | 1 | 3um_FIJJAFBF_586240 | Bacteria | Bacteroidetes | Flavobacteriia | Flavobacterial | Flavobacteriac | Lacinutrix | Lacinutrix algi | metabolism | Amino acid tra | COG0174 | glutamine syn | 313594.PI23P_ | 1,8 | 0,4 | 0,000340504 | 9,47E-05 | 2 - | 0,000442889 - | 2 - | 0,000370266 - | 2 - | 0,000372739 - |
| 223 | 1 | 3um_FIJJAFBF_673918 | Eukaryota | unclassified | Oomycetes | Peronosporale | Peronosporac | Phytophthora | various | unclassified | unclassified | unclassified | unclassified | unclassified | 1,4 | 0,489897949 | 0,000248563 | 0,000107707 | 2 - | 0,000411254 - | 2 - | 0,000343818 - | 1 - | 0,000173058 - |
| 224 | 1 | 3um_FIJJAFBF_79252 | Bacteria | Bacteroidetes | Flavobacteriia | Flavobacterial | Flavobacteriac | Polaribacter | Polaribacter s | information st | Replication, re | 0PB0H | DNA-binding p | 313598.MED1 | 3,4 | 1,019803903 | 0,002365812 | 0,000688353 | 3 - | 0,002494939 - | 4 - | 0,002781107 - | 2 - | 0,001399843 - |
| 225 | 1 | 3um_FIJJAFBF_716599 | Eukaryota | Bacillariophyta | Bacillariophyc | Bacillariales | Bacillariaceae | Fragilariopsis | Fragilariopsis | poorly charact | Function unkn | COG1100 | GTP-binding P | 63737.Npun_ | 0,2 | 0,4 | 4,13E-05 | 8,26E-05 | 0 - | 0 - | 0 - | 0 - | 1 - | 0,000206534 - |
| 226 | 1 | 3um_FIJJAFBF_143869 | Bacteria | Proteobacteri | Gammaprotec | Oceanospirilla | Saccharospiril | Reinekea | Reinekea blan | unclassified | unclassified | unclassified | unclassified | unclassified | 0,8 | 0,4 | 0,000107916 | 5,44E-05 | 0 - | 0 - | 1 - | 0,000140618 - | 1 - | 0,000141557 - |
| 227 | 1 | 3um_FIJJAFBF_115452 | Bacteria | Proteobacteri | Gammaprotec | unclassified | unclassified | unclassified | SAR86 cluster | information st | Translation, ri | 179RU | 30S ribosomal | 566466.NOR5 | 1,2 | 0,4 | 0,000580824 | 0,000197684 | 1 - | 0,000575755 - | 2 - | 0,000962691 - | 1 - | 0,000484561 - |
| 228 | 1 | 0.2um_NOKCPHJN_601842 | Bacteria | Proteobacteri | Alphaproteob | Pelagibacteral | Pelagibacterac | Candidatus Pe | Candidatus Pe | information st | Replication, re | 05CVP | deoxyribo-dip | 335992.SAR11 | 0,4 | 0,489897949 | 4,96E-05 | 6,10E-05 | 0 - | 0 - | 1 - | 0,000132574 - | 0 - | 0 - |
| 229 | 1 | 0.2um_NOKCPHJN_1057723 | Eukaryota | Bacillariophyta | Bacillariophyc | Naviculales | Phaeodactylac | Phaeodactylu | Phaeodactylu | unclassified | unclassified | unclassified | unclassified | unclassified | 1,6 | 1,2 | 0,00022745 | 0,000158298 | 1 - | 0,000176946 - | 1 - | 0,000147931 - | 0 - | 0 - |
| 230 | 1 | 3um_FIJJAFBF_627437 | Eukaryota | unclassified | Dinophyceae | Suessiales | Symbiodiniace | Symbiodinium | Symbiodinium | metabolism | Energy produc | KOG1647 | subunit D | 423536.XP_00 | 0,2 | 0,4 | 5,03E-05 | 0,000100545 | 0 - | 0 - | 0 - | 0 - | 0 - | 0 - |
| 231 | 2 | 0.2um_NOKCPHJN_60647;3um_FIJJAF | Bacteria | Bacteroidetes | Flavobacteriia | Flavobacterial | Flavobacteriac | Polaribacter | various | information st | Translation, ri | 09AI8 | Associates wit | 313598.MED1 | 0,8 | 0,4 | 0,000148677 | 7,49E-05 | 0 - | 0 - | 1 - | 0,00019373 - | 1 - | 0,000195025 - |
| 232 | 1 | 3um_FIJJAFBF_254290 | Bacteria | Proteobacteri | Gammaprotec | Cellvibrionales | Spongiibacter | Oceanicoccus | Oceanicoccus | information st | Translation, ri | 17C3B | The globular d | 247633.GP214 | 1,8 | 0,4 | 0,00104559 | 0,000274485 | 2 - | 0,001373361 - | 2 - | 0,001148164 - | 2 - | 0,001155834 - |
| 233 | 1 | 0.2um_NOKCPHJN_39143 | Eukaryota | unclassified | Pelagophycea | Pelagomonad | unclassified | Aureococcus | Aureococcus a | metabolism | Carbohydrate | COG1850 | RuBisCO catal | 45157.CMV01 | 34,6 | 0,8 | 0,004465054 | 0,000464714 | 34 - | 0,005214832 - | 34 - | 0,004359728 - | 36 - | 0,00464702 - |
| 234 | 1 | 3um_FIJJAFBF_783325 | Eukaryota | Bacillariophyta | Mediophyceae | Anaulales | Anaulaceae | Eunotogramm | Eunotogramm | information st | Translation, ri | 0FNCH | Binds directly | 65393.PCC742 | 5,2 | 0,4 | 0,002728965 | 0,000341941 | 5 - | 0,003118674 - | 6 - | 0,003128746 - | 5 - | 0,002624706 - |
| 235 | 1 | 3um_FIJJAFBF_357686 | Bacteria | Proteobacteri | Deltaproteoba | Myxococcales | Anaeromyxob | Anaeromyxob | Anaeromyxob | unclassified | unclassified | unclassified | unclassified | unclassified | 0,6 | 0,489897949 | 2,21E-05 | 1,81E-05 | 0 - | 0 - | 0 - | 0 - | 1 - | 3,92E-05 - |
| 236 | 1 | 3um_FIJJAFBF_561978 | Eukaryota | unclassified | Spirotrichea | Sporadotrichic | Oxytrichidae | Oxytricha | Oxytricha trifa | unclassified | unclassified | unclassified | unclassified | unclassified | 1,4 | 0,489897949 | 0,000647836 | 0,000230184 | 1 - | 0,000550354 - | 2 - | 0,000920219 - | 2 - | 0,000926367 - |
| 237 | 1 | 3um_FIJJAFBF_282298 | unclassified | unclassified | unclassified | unclassified | unclassified | unclassified | unclassified | unclassified | unclassified | unclassified | unclassified | unclassified | 6,8 | 0,979795897 | 0,00392149 | 0,000423628 | 5 - | 0,003465194 - | 8 - | 0,004635179 - | 7 - | 0,004082876 - |
| 238 | 1 | 3um_FIJJAFBF_161153 | Viruses | unclassified | unclassified | unclassified | unclassified | unclassified | Vibrio phage 1 | unclassified | unclassified | unclassified | unclassified | unclassified | 1,8 | 0,748331477 | 0,000339732 | 0,000113632 | 1 - | 0,000232448 - | 1 - | 0,000194332 - | 2 - | 0,00039126 - |
| 239 | 1 | 3um_FIJJAFBF_344400 | Bacteria | Bacteroidetes | Flavobacteriia | Flavobacterial | Crocinitomica | unclassified | Crocinitomica | unclassified | unclassified | unclassified | unclassified | unclassified | 1 | 0 | 9,39E-05 | 9,93E-06 | 1 - | 0,000111547 - | 1 - | 9,33E-05 - | 1 - | 9,39E-05 - |
| 240 | 1 | 3um_FIJJAFBF_438578 | unclassified | unclassified | unclassified | unclassified | unclassified | unclassified | unclassified | unclassified | unclassified | unclassified | unclassified | unclassified | 3 | 0,632455532 | 0,000654797 | 0,000122445 | 3 - | 0,000785121 - | 3 - | 0,00065638 - | 2 - | 0,00044051 - |
| 241 | 2 | 0.2um_NOKCPHJN_332638;3um_FIJJAF | Viruses | unclassified | unclassified | unclassified | various | various | various | unclassified | unclassified | unclassified | unclassified | unclassified | 1 | 0,632455532 | 0,000306314 | 0,000194471 | 0 - | 0 - | 1 - | 0,00031926 - | 1 - | 0,000321393 - |
| 242 | 1 | 3um_FIJJAFBF_640407 | unclassified | unclassified | unclassified | unclassified | unclassified | unclassified | unclassified | unclassified | unclassified | unclassified | unclassified | unclassified | 1 | 0 | 0,000263566 | 2,79E-05 | 1 - | 0,000313172 - | 1 - | 0,00026182 - | 1 - | 0,000263569 - |
| 243 | 1 | 3um_FIJJAFBF_535056 | Bacteria | Bacteroidetes | Flavobacteriia | Flavobacterial | Flavobacteriac | Algibacter | Algibacter pec | poorly charact | Function unkn | 0PAPS | NA | 313603.FB217 | 1,2 | 0,4 | 0,000519745 | 0,000239972 | 2 - | 0,000997976 - | 1 - | 0,000417166 - | 1 - | 0,000419953 - |
| 244 | 1 | 3um_FIJJAFBF_487611 | Bacteria | Proteobacteri | Alphaproteob | Rhodobactera | Rhodobactera | Planktotalea | Planktotalea f | unclassified | unclassified | unclassified | unclassified | unclassified | 0,2 | 0,4 | 2,43E-05 | 4,85E-05 | 0 - | 0 - | 1 - | 0,000121269 - | 0 - | 0 - |
| 245 | 1 | 3um_FIJJAFBF_711128 | Eukaryota | Bacillariophyta | Coscinodiscop | Chaetocerotal | Chaetocerotac | Chaetoceros | Chaetoceros s | information st | Translation, ri | 08UIK | This protein bi | 1140.Synpcc7 | 0,6 | 0,489897949 | 0,000255372 | 0,000209621 | 0 - | 0 - | 1 - | 0,000443794 - | 1 - | 0,000446758 - |
| 246 | 1 | 3um_FIJJAFBF_577814 | Eukaryota | Chlorophyta | Chlorophyceae | Sphaeropleale | Selenastracea | Raphidocelis | Raphidocelis s | information st | Replication, re | 0XNUT | helicase activi | 3055.EDP0692 | 0,4 | 0,489897949 | 5,46E-05 | 6,82E-05 | 1 - | 0,000157908 - | 0 - | 0 - | 0 - | 0 - |
| 247 | 1 | 3um_FIJJAFBF_676357 | Eukaryota | Chordata | Actinopteri | Cyprinodontif | Nothobranchi | Nothobranchi | Nothobranchi | information st | Chromatin str | KOG3467 | Core compone | 6085.XP_0021 | 2,2 | 0,4 | 0,001073638 | 0,000216932 | 2 - | 0,001160437 - | 3 - | 0,001455231 - | 2 - | 0,000976635 - |
| 248 | 1 | 3um_FIJJAFBF_715083 | Eukaryota | unclassified | Pelagophycea | Pelagomonad | unclassified | Aureoumbra | Aureoumbra l | unclassified | unclassified | unclassified | unclassified | unclassified | 1,2 | 0,4 | 0,000419035 | 0,000131224 | 1 - | 0,000418146 - | 1 - | 0,000349581 - | 1 - | 0,000351916 - |
| 249 | 1 | 3um_FIJJAFBF_145237 | Bacteria | Bacteroidetes | Flavobacteriia | Flavobacterial | Flavobacteriac | Hyunsoonleell | Hyunsoonleell | information st | Translation, ri | 08YY1 | This protein is | 391603.FBALC | 2,4 | 0,489897949 | 0,000916523 | 0,00015199 | 2 - | 0,000918383 - | 3 - | 0,001151686 - | 2 - | 0,000772919 - |
| 250 | 1 | 0.2um_NOKCPHJN_877855 | Bacteria | Proteobacteri | Gammaprotec | unclassified | unclassified | Candidatus Th | Candidatus Th | metabolism | Energy produc | 16QEJ | Key enzyme in | 411684.HPDFI | 0,2 | 0,4 | 0,00010587 | 0,000211741 | 0 - | 0 - | 0 - | 0 - | 1 - | 0,000529352 - |
| 251 | 2 | 0.2um_NOKCPHJN_362980;3um_FIJJAF | Bacteria | Proteobacteri | Gammaprotec | unclassified | unclassified | unclassified | Gammaprotec | various | various | various | various | various | 0,6 | 0,489897949 | 0,000145755 | 0,000120431 | 1 - | 0,000276193 - | 1 - | 0,000230904 - | 0 - | 0 - |
| 252 | 1 | 3um_FIJJAFBF_44458 | Bacteria | Bacteroidetes | Flavobacteriia | Flavobacterial | Flavobacteriac | Flavobacteriu | Flavobacteriu | metabolism | Coenzyme tra | 09A4R | May play a ke | 376686.Fjoh_1 | 0,2 | 0,4 | 3,82E-05 | 7,63E-05 | 0 - | 0 - | 1 - | 0,000190777 - | 0 - | 0 - |
| 253 | 1 | 0.2um_NOKCPHJN_153362 | Eukaryota | Ascomycota | Dothideomyce | unclassified | Gloniaceae | Cenococcum | Cenococcum g | cellular proces | Posttranslatio | COG0652 | peptidyl-proly | 30611.ENSOG | 0,2 | 0,4 | 1,91E-05 | 3,81E-05 | 0 - | 0 - | 0 - | 0 - | 0 - | 0 - |
| 254 | 1 | 3um_FIJJAFBF_305779 | Eukaryota | Bacillariophyta | Bacillariophyc | Bacillariales | Bacillariaceae | Fragilariopsis | Fragilariopsis | poorly charact | Function unkn | 1DV37 | plastid-lipid-a | 436017.A4RTN | 1,2 | 0,4 | 0,000576979 | 0,000180685 | 1 - | 0,000575755 - | 1 - | 0,000481346 - | 1 - | 0,000484561 - |
| 255 | 1 | 0.2um_NOKCPHJN_83083 | Viruses | unclassified | unclassified | unclassified | various | various | various | unclassified | unclassified | unclassified | unclassified | unclassified | 1 | 0 | 3,32E-05 | 3,52E-06 | 1 - | 3,95E-05 - | 1 - | 3,30E-05 - | 1 - | 3,32E-05 - |
| 256 | 1 | 3um_FIJJAFBF_669267 | Eukaryota | unclassified | Pelagophycea | Pelagomonad | unclassified | Aureococcus | Aureococcus a | unclassified | unclassified | unclassified | unclassified | unclassified | 1 | 0,632455532 | 0,000123546 | 8,10E-05 | 0 - | 0 - | 2 - | 0,000255408 - | 1 - | 0,000128557 - |
| 257 | 1 | 0.2um_NOKCPHJN_331909 | Eukaryota | unclassified | Labyrinthulom | unclassified | Thraustochytr | Aurantiochytr | Aurantiochytr | metabolism | Energy produc | COG1053 | Succinate DeH | 431895.XP_00 | 1 | 0 | 9,90E-05 | 1,05E-05 | 1 - | 0,000117686 - | 1 - | 9,84E-05 - | 1 - | 9,90E-05 - |
| 258 | 1 | 3um_FIJJAFBF_774028 | Bacteria | Proteobacteri | Alphaproteob | Sphingomona | Erythrobacter | Altererythrob | Altererythrob | unclassified | unclassified | unclassified | unclassified | unclassified | 0,4 | 0,489897949 | 0,000738958 | 0,000923574 | 1 - | 0,00213852 - | 0 - | 0 - | 0 - | 0 - |
| 259 | 1 | 3um_FIJJAFBF_440532 | Eukaryota | Chordata | Mammalia | Primates | Cercopithecid | Cercocebus | Cercocebus at | cellular proces | Cell cycle cont | 0V9W7 | retinitis pigme | 9606.ENSP000 | 0,2 | 0,4 | 1,01E-05 | 2,02E-05 | 0 - | 0 - | 0 - | 0 - | 1 - | 5,04E-05 - |
| 260 | 1 | 3um_FIJJAFBF_594171 | Eukaryota | unclassified | Dinophyceae | Peridiniales | Kryptoperidini | Kryptoperidini | Kryptoperidini | metabolism | Energy produc | 11HAV | photosystem i | 35128.Thapsd | 8,6 | 0,8 | 0,003732067 | 0,000477372 | 8 - | 0,004129555 - | 8 - | 0,003452409 - | 10 - | 0,004344341 - |
| 261 | 1 | 3um_FIJJAFBF_43159 | unclassified | unclassified | unclassified | unclassified | unclassified | unclassified | unclassified | unclassified | unclassified | unclassified | unclassified | unclassified | 1 | 0 | 0,000663076 | 7,01E-05 | 1 - | 0,000787876 - | 1 - | 0,000658683 - | 1 - | 0,000663084 - |
| 262 | 1 | 0.2um_NOKCPHJN_925569 | unclassified | unclassified | unclassified | unclassified | unclassified | unclassified | unclassified | unclassified | unclassified | unclassified | unclassified | unclassified | 0,6 | 0,489897949 | 0,000203432 | 0,000166986 | 0 - | 0 - | 1 - | 0,000353531 - | 1 - | 0,000355892 - |
| 263 | 1 | 3um_FIJJAFBF_235910 | Eukaryota | Bacillariophyta | Bacillariophyc | Naviculales | Naviculaceae | Fistulifera | Fistulifera sola | metabolism | Carbohydrate | 09WWU | ER degradatio | 31033.ENSTRU | 0,8 | 0,748331477 | 0,000349919 | 0,000338418 | 0 - | 0 - | 2 - | 0,000913502 - | 0 - | 0 - |
| 264 | 1 | 3um_FIJJAFBF_557699 | Bacteria | Proteobacteri | Gammaprotec | Chromatiales | Ectothiorhodc | Thioalkalivibri | Thioalkalivibri | cellular proces | Cell wall/mem | 12RRB | ankyrin 3, nod | 7668.SPU_017 | 0,6 | 0,489897949 | 7,38E-05 | 6,17E-05 | 1 - | 0,000145903 - | 0 - | 0 - | 0 - | 0 - |
| 265 | 1 | 0.2um_NOKCPHJN_561246 | unclassified | unclassified | unclassified | unclassified | unclassified | unclassified | unclassified | unclassified | unclassified | unclassified | unclassified | unclassified | 0,2 | 0,4 | 3,70E-05 | 7,39E-05 | 0 - | 0 - | 0 - | 0 - | 0 - | 0 - |
| 266 | 1 | 3um_FIJJAFBF_11153 | Bacteria | Bacteroidetes | Flavobacteriia | Flavobacterial | Flavobacteriac | Polaribacter | Polaribacter p | cellular proces | Posttranslatio | 07SFW | Trigger factor | 313598.MED1 | 0,6 | 0,489897949 | 8,05E-05 | 6,60E-05 | 0 - | 0 - | 1 - | 0,000142216 - | 0 - | 0 - |
| 267 | 1 | 3um_FIJJAFBF_216063 | Bacteria | Proteobacteri | Gammaprotec | Cellvibrionales | unclassified | unclassified | Cellvibrionales | information st | Translation, ri | 177SZ | This protein bi | 247633.GP214 | 1,2 | 0,748331477 | 0,000503627 | 0,000310064 | 0 - | 0 - | 1 - | 0,000440668 - | 2 - | 0,000887224 - |
| 268 | 1 | 3um_FIJJAFBF_582352 | Bacteria | Cyanobacteria | unclassified | Synechococca | Synechococca | Synechococcu | Synechococcu | poorly charact | Function unkn | 11FY4 | C-phycoerythr | 91464.S7335_ | 0,2 | 0,4 | 9,13E-05 | 0,000182557 | 1 - | 0,000456391 - | 0 - | 0 - | 0 - | 0 - |
| 269 | 1 | 0.2um_NOKCPHJN_576041 | unclassified | unclassified | unclassified | unclassified | unclassified | unclassified | unclassified | unclassified | unclassified | unclassified | unclassified | unclassified | 0,2 | 0,4 | 2,16E-05 | 4,33E-05 | 0 - | 0 - | 0 - | 0 - | 1 - | 0,000108235 - |
| 270 | 1 | 3um_FIJJAFBF_323298 | Bacteria | Bacteroidetes | Saprospiria | Saprospirales | Saprospiracea | Aureispira | Aureispira sp. | unclassified | unclassified | unclassified | unclassified | unclassified | 0,2 | 0,4 | 0,000184642 | 0,000369284 | 0 - | 0 - | 0 - | 0 - | 0 - | 0 - |
| 271 | 1 | 3um_FIJJAFBF_181271 | Eukaryota | Nematoda | Chromadorea | Rhabditida | Ancylostomati | Ancylostoma | Ancylostoma c | unclassified | unclassified | unclassified | unclassified | unclassified | 0,8 | 0,4 | 0,000104768 | 5,42E-05 | 1 - | 0,00015561 - | 1 - | 0,000130093 - | 0 - | 0 - |
| 272 | 1 | 0.2um_NOKCPHJN_547562 | unclassified | unclassified | unclassified | unclassified | unclassified | unclassified | unclassified | unclassified | unclassified | unclassified | unclassified | unclassified | 0,6 | 0,489897949 | 0,000738048 | 0,000615399 | 1 - | 0,001439388 - | 1 - | 0,001203364 - | 0 - | 0 - |
| 273 | 1 | 0.2um_NOKCPHJN_63946 | Bacteria | Bacteroidetes | unclassified | unclassified | unclassified | unclassified | Bacteroidetes | unclassified | unclassified | unclassified | unclassified | unclassified | 1 | 0 | 7,70E-05 | 8,14E-06 | 1 - | 9,15E-05 - | 1 - | 7,65E-05 - | 1 - | 7,70E-05 - |
| 274 | 1 | 3um_FIJJAFBF_411071 | Eukaryota | Phaeophyceae | unclassified | Ectocarpales | Ectocarpaceae | Ectocarpus | Ectocarpus sil | information st | Chromatin str | KOG1744 | Core compone | 2850.Phatr543 | 10,2 | 0,979795897 | 0,005452519 | 0,000417223 | 9 - | 0,005757553 - | 9 - | 0,004813455 - | 11 - | 0,005922413 - |
| 275 | 3 | 0.2um_NOKCPHJN_164868;0.2um_NO | Bacteria | Bacteroidetes | Flavobacteriia | Flavobacterial | Flavobacteriac | Polaribacter | various | poorly charact | Function unkn | 0900R | pur-alpha beta | various | 0,6 | 0,489897949 | 0,000283523 | 0,000232729 | 0 - | 0 - | 1 - | 0,000492716 - | 1 - | 0,000496007 - |
| 276 | 1 | 3um_FIJJAFBF_579113 | Bacteria | Bacteroidetes | Flavobacteriia | Flavobacterial | unclassified | unclassified | Flavobacterial | metabolism | Energy produc | 0PBIY | subunit C | 926562.Oweh | 1 | 0 | 0,00091293 | 9,65E-05 | 1 - | 0,001084756 - | 1 - | 0,000906883 - | 1 - | 0,000912941 - |
| 277 | 1 | 3um_FIJJAFBF_135240 | Bacteria | Cyanobacteria | unclassified | Synechococca | Synechococca | Synechococcu | Synechococcu | unclassified | unclassified | unclassified | unclassified | unclassified | 1,6 | 0,489897949 | 0,000360499 | 9,13E-05 | 1 - | 0,000274169 - | 2 - | 0,000458424 - | 1 - | 0,000230743 - |
| 278 | 1 | 0.2um_NOKCPHJN_525206 | Bacteria | Proteobacteri | Alphaproteob | unclassified | unclassified | unclassified | Alphaproteob | unclassified | unclassified | unclassified | unclassified | unclassified | 0 | 0 | 0 | 0 | 0 - | 0 - | 0 - | 0 - | 0 - | 0 - |
| 279 | 1 | 3um_FIJJAFBF_265695 | unclassified | unclassified | unclassified | unclassified | unclassified | unclassified | unclassified | unclassified | unclassified | unclassified | unclassified | unclassified | 24,2 | 1,939071943 | 0,011886672 | 0,001329799 | 23 - | 0,013449283 - | 23 - | 0,011243931 - | 24 - | 0,011811176 - |
| 280 | 1 | 3um_FIJJAFBF_207965 | Eukaryota | unclassified | unclassified | Prymnesiales | Chrysochrom | Chrysochrom | Chrysochrom | unclassified | unclassified | unclassified | unclassified | unclassified | 0,2 | 0,4 | 4,14E-05 | 8,29E-05 | 0 - | 0 - | 0 - | 0 - | 0 - | 0 - |
| 281 | 1 | 0.2um_NOKCPHJN_673538 | Bacteria | Bacteroidetes | Flavobacteriia | Flavobacterial | Flavobacteriac | Muricauda | Muricauda sp. | unclassified | unclassified | unclassified | unclassified | unclassified | 1 | 0 | 5,90E-05 | 6,24E-06 | 1 - | 7,01E-05 - | 1 - | 5,86E-05 - | 1 - | 5,90E-05 - |
| 282 | 1 | 3um_FIJJAFBF_414291 | unclassified | unclassified | unclassified | unclassified | unclassified | unclassified | unclassified | unclassified | unclassified | unclassified | unclassified | unclassified | 1 | 0 | 0,00104987 | 0,000111027 | 1 - | 0,00124747 - | 1 - | 0,001042915 - | 1 - | 0,001049882 - |
| 283 | 1 | 0.2um_NOKCPHJN_300833 | Bacteria | Proteobacteri | Alphaproteob | Rhodospirillale | Rhodospirillac | Candidatus En | Candidatus En | unclassified | unclassified | unclassified | unclassified | unclassified | 0,6 | 0,489897949 | 5,36E-05 | 4,41E-05 | 1 - | 0,000100064 - | 1 - | 8,37E-05 - | 1 - | 8,42E-05 - |
| 284 | 1 | 3um_FIJJAFBF_560797 | Eukaryota | Bacillariophyta | Coscinodiscop | Thalassiosirale | Thalassiosirac | Thalassiosira | Thalassiosira p | poorly charact | Function unkn | 0IHC7 | Thylakoid mer | 35128.Thaps4 | 0,8 | 0,4 | 0,000228404 | 0,000118109 | 1 - | 0,00033868 - | 0 - | 0 - | 1 - | 0,000285036 - |
| 285 | 1 | 3um_FIJJAFBF_116591 | Eukaryota | unclassified | Synurophycea | Parmales | Triparmaceae | Triparma | Triparma laevi | metabolism | Energy produc | 01TRE | NADH dehydr | 360095.BARBA | 0,4 | 0,489897949 | 0,000116807 | 0,00014306 | 0 - | 0 - | 1 - | 0,000291046 - | 1 - | 0,00029299 - |
| 286 | 1 | 0.2um_NOKCPHJN_987138 | unclassified | unclassified | unclassified | unclassified | unclassified | unclassified | unclassified | unclassified | unclassified | unclassified | unclassified | unclassified | 0,6 | 0,489897949 | 0,000276436 | 0,000227746 | 1 - | 0,000516194 - | 1 - | 0,000431551 - | 1 - | 0,000434434 - |
| 287 | 1 | 3um_FIJJAFBF_437253 | Bacteria | Bacteroidetes | Flavobacteriia | Flavobacterial | Flavobacteriac | Polaribacter | Polaribacter s | metabolism | Inorganic ion t | 05C27 | TonB-depende | 867900.Celly_ | 7,8 | 1,166190379 | 0,0004431 | 3,32E-05 | 6 - | 0,000410877 - | 8 - | 0,000458005 - | 7 - | 0,000403431 - |
| 288 | 1 | 0.2um_NOKCPHJN_926215 | Eukaryota | unclassified | unclassified | Isochrysidales | Isochrysidacea | Chrysotila | Chrysotila cart | unclassified | unclassified | unclassified | unclassified | unclassified | 1,4 | 0,489897949 | 0,001739938 | 0,00075395 | 2 - | 0,002878776 - | 2 - | 0,002406728 - | 1 - | 0,001211403 - |
| 289 | 1 | 0.2um_NOKCPHJN_457075 | Bacteria | Bacteroidetes | Flavobacteriia | Flavobacterial | Flavobacteriac | Aequorivita | Aequorivita ca | unclassified | unclassified | unclassified | unclassified | unclassified | 0,4 | 0,489897949 | 0,000155843 | 0,000191249 | 0 - | 0 - | 0 - | 0 - | 0 - | 0 - |
| 290 | 1 | 0.2um_NOKCPHJN_454728 | Eukaryota | unclassified | Labyrinthulom | unclassified | Thraustochytr | Aurantiochytr | Aurantiochytr | unclassified | unclassified | unclassified | unclassified | unclassified | 1,4 | 0,489897949 | 0,000327867 | 9,18E-05 | 1 - | 0,00028568 - | 1 - | 0,000238836 - | 1 - | 0,000240431 - |
| 291 | 1 | 3um_FIJJAFBF_715066 | Eukaryota | unclassified | Florideophyce | Gigartinales | Peyssonneliac | Riquetophycu | Riquetophycu | unclassified | unclassified | unclassified | unclassified | unclassified | 16,4 | 2,244994432 | 0,00155854 | 0,000111308 | 14 - | 0,001599808 - | 17 - | 0,001624082 - | 14 - | 0,001346414 - |
| 292 | 1 | 3um_FIJJAFBF_178015 | Bacteria | Proteobacteri | Gammaprotec | Cellvibrionales | Halieaceae | unclassified | marine gamm | poorly charact | Function unkn | 05WYC | Short-chain de | 316056.RPC_3 | 1 | 0 | 0,000247028 | 2,61E-05 | 1 - | 0,000293522 - | 1 - | 0,000245392 - | 1 - | 0,000247031 - |
| 293 | 1 | 3um_FIJJAFBF_405248 | Bacteria | Bacteroidetes | Flavobacteriia | Flavobacterial | Flavobacteriac | Polaribacter | Polaribacter s | metabolism | Amino acid tra | 09GJ6 | nitrogen regul | 313598.MED1 | 2,6 | 0,489897949 | 0,001446355 | 0,000229182 | 2 - | 0,001336575 - | 3 - | 0,001676114 - | 3 - | 0,001687311 - |
| 294 | 1 | 0.2um_NOKCPHJN_802229 | unclassified | unclassified | unclassified | unclassified | unclassified | unclassified | unclassified | unclassified | unclassified | unclassified | unclassified | unclassified | 0,2 | 0,4 | 2,63E-05 | 5,25E-05 | 0 - | 0 - | 0 - | 0 - | 0 - | 0 - |
| 295 | 1 | 3um_FIJJAFBF_80731 | Bacteria | Proteobacteri | Gammaprotec | Alteromonada | unclassified | unclassified | gamma protec | information st | Translation, ri | 05KK9 | This protein bi | 523791.Kkor_ | 2,4 | 0,8 | 0,000763137 | 0,00021872 | 1 - | 0,000389834 - | 2 - | 0,000651822 - | 3 - | 0,000984265 - |
| 296 | 1 | 3um_FIJJAFBF_91113 | Eukaryota | unclassified | Florideophyce | Ceramiales | Rhodomelace | Polysiphonia | Polysiphonia s | unclassified | unclassified | unclassified | unclassified | unclassified | 3,2 | 0,4 | 0,001526451 | 0,000238914 | 3 - | 0,001701095 - | 4 - | 0,00189621 - | 3 - | 0,001431658 - |
| 297 | 1 | 3um_FIJJAFBF_56092 | Eukaryota | Bacillariophyta | Bacillariophyc | Naviculales | Naviculaceae | Fistulifera | Fistulifera sola | metabolism | Energy produc | COG0055 | Produces ATP | 2850.Phatr540 | 2 | 0 | 0,000497962 | 5,27E-05 | 2 - | 0,000591685 - | 2 - | 0,000494663 - | 2 - | 0,000497968 - |
| 298 | 1 | 0.2um_NOKCPHJN_338143 | Eukaryota | unclassified | Bangiophycea | Bangiales | Bangiaceae | Porphyra | Porphyra umb | cellular proces | Posttranslatio | KOG0734 | YME1-like 1 (S | 45157.CMI157 | 0,6 | 0,489897949 | 9,04E-05 | 7,46E-05 | 1 - | 0,000170886 - | 0 - | 0 - | 1 - | 0,000143819 - |

| | | | | | | | | | | | | | | | | | | | | | | | | |
|---|---|---|---|---|---|---|---|---|---|---|---|---|---|---|---|---|---|---|---|---|---|---|---|---|
| 299 | 1 | 3um_FIJJAFBF_269222 | Bacteria | Bacteroidetes | Flavobacteriia | Flavobacterial | Flavobacteriac | Polaribacter | Polaribacter v | unclassified | unclassified | unclassified | unclassified | unclassified | 0,4 | 0,489897949 | 0,00011986 | 0,00014744 | 0 - | 0 - | 0 - | 0 - | 1 - | 0,000321393 - |
| 300 | 1 | 3um_FIJJAFBF_400754 | Bacteria | Proteobacteri | Alphaproteob | Rhizobiales | unclassified | Methyloceanil | Methyloceanil | unclassified | unclassified | unclassified | unclassified | unclassified | 0,8 | 0,4 | 0,000315938 | 0,000159229 | 0 - | 0 - | 1 - | 0,000411677 - | 1 - | 0,000414427 - |
| 301 | 1 | 3um_FIJJAFBF_99324 | Bacteria | Bacteroidetes | Flavobacteriia | Flavobacterial | Flavobacteriac | unclassified | various | metabolism | Inorganic ion t | 05C27 | TonB-depende | 313598.MED1 | 21,4 | 2,576819745 | 0,001342582 | 8,78E-05 | 17 - | 0,001281389 - | 24 - | 0,001512385 - | 21 - | 0,001332177 - |
| 302 | 1 | 3um_FIJJAFBF_491869 | Bacteria | Bacteroidetes | Flavobacteriia | Flavobacterial | Flavobacteriac | Flavobacteriu | Flavobacteriu | unclassified | unclassified | unclassified | unclassified | unclassified | 0,2 | 0,4 | 4,76E-05 | 9,51E-05 | 0 - | 0 - | 0 - | 0 - | 0 - | 0 - |
| 303 | 1 | 3um_FIJJAFBF_715069 | Eukaryota | unclassified | Compsopogon | Compsopogon | Compsopogon | Compsopogon | Compsopogon | unclassified | unclassified | unclassified | unclassified | unclassified | 1 | 0 | 0,001464934 | 0,000154921 | 1 - | 0,001740655 - | 1 - | 0,001455231 - | 1 - | 0,001464952 - |
| 304 | 1 | 0.2um_NOKCPHJN_857856 | Eukaryota | unclassified | Oomycetes | Saprolegniales | Saprolegniace | Thraustotheca | Thraustotheca | unclassified | unclassified | unclassified | unclassified | unclassified | 1 | 0 | 0,00017211 | 1,82E-05 | 1 - | 0,000204503 - | 1 - | 0,00017097 - | 1 - | 0,000172112 - |
| 305 | 1 | 3um_FIJJAFBF_437895 | Bacteria | Bacteroidetes | Saprospiria | Saprospirales | Lewinellaceae | unclassified | Lewinellaceae | metabolism | Inorganic ion t | 05C27 | TonB-depende | 391603.FBALC | 0,4 | 0,489897949 | 2,34E-05 | 2,86E-05 | 0 - | 0 - | 1 - | 5,82E-05 - | 1 - | 5,86E-05 - |
| 306 | 1 | 0.2um_NOKCPHJN_211484 | Bacteria | Proteobacteri | Gammaprotec | unclassified | unclassified | unclassified | various | poorly charact | Function unkn | 05C2A | tonB-depende | 566466.NOR5 | 2,4 | 0,8 | 0,000143285 | 5,48E-05 | 3 - | 0,000210445 - | 3 - | 0,000175937 - | 1 - | 5,90E-05 - |
| 307 | 1 | 3um_FIJJAFBF_675681 | Viruses | unclassified | unclassified | unclassified | unclassified | unclassified | uncultured M | unclassified | unclassified | unclassified | unclassified | unclassified | 2,8 | 0,4 | 0,000655991 | 0,000120873 | 3 - | 0,000834738 - | 2 - | 0,000465241 - | 3 - | 0,000702523 - |
| 308 | 1 | 3um_FIJJAFBF_91579 | Bacteria | Bacteroidetes | Flavobacteriia | Flavobacterial | Flavobacteriac | Lutibacter | Lutibacter sp. | unclassified | unclassified | unclassified | unclassified | unclassified | 0,4 | 0,489897949 | 2,57E-05 | 3,15E-05 | 0 - | 0 - | 0 - | 0 - | 1 - | 6,58E-05 - |
| 309 | 1 | 3um_FIJJAFBF_41558 | Eukaryota | Bacillariophyt | Mediophyceae | Lithodesmiale | Lithodesmiace | Lithodesmium | Lithodesmium | metabolism | Carbohydrate | 0IVA1 | Photosystem I | 35128.Thapsd | 2,4 | 0,489897949 | 0,000814043 | 0,00016634 | 2 - | 0,00080917 - | 2 - | 0,000676486 - | 3 - | 0,001021507 - |
| 310 | 1 | 0.2um_NOKCPHJN_390598 | Bacteria | Proteobacteri | Alphaproteob | Rhodobactera | Rhodobactera | Sulfitobacter | Sulfitobacter s | metabolism | Energy produc | 01RNF | Produces ATP | 391595.RLO14 | 0,8 | 0,4 | 0,000106092 | 5,49E-05 | 1 - | 0,000157575 - | 1 - | 0,000131737 - | 0 - | 0 - |
| 311 | 1 | 3um_FIJJAFBF_801206 | Eukaryota | Bacillariophyt | Mediophyceae | Triceratiales | Plagiogramma | Plagiogramma | Plagiogramma | cellular proces | Posttranslatio | 16R8D | CbbX protein | 472759.Nhal_ | 2,6 | 0,489897949 | 0,000618423 | 8,47E-05 | 2 - | 0,000573549 - | 3 - | 0,000719252 - | 2 - | 0,000482705 - |
| 312 | 1 | 3um_FIJJAFBF_253585 | Viruses | unclassified | unclassified | unclassified | unclassified | unclassified | uncultured M | unclassified | unclassified | unclassified | unclassified | unclassified | 1,6 | 1,2 | 0,000313585 | 0,000223043 | 1 - | 0,000241446 - | 3 - | 0,000605564 - | 0 - | 0 - |
| 313 | 1 | 0.2um_NOKCPHJN_503156 | Viruses | unclassified | unclassified | unclassified | various | various | various | unclassified | unclassified | unclassified | unclassified | unclassified | 1 | 0 | 0,000124737 | 1,32E-05 | 1 - | 0,000148214 - | 1 - | 0,000123911 - | 1 - | 0,000124738 - |
| 314 | 1 | 0.2um_NOKCPHJN_785823 | Bacteria | Proteobacteri | Gammaprotec | Cellvibrionales | unclassified | unclassified | Cellvibrionales | metabolism | Energy produc | 0QIIJ | Succinyl-CoA l | 377629.TERTU | 1,4 | 0,8 | 0,000295923 | 0,000159856 | 1 - | 0,000258097 - | 2 - | 0,000431551 - | 0 - | 0 - |
| 315 | 1 | 0.2um_NOKCPHJN_118112 | Bacteria | Bacteroidetes | Flavobacteriia | Flavobacterial | Flavobacteriac | Maribacter | Maribacter sp | poorly charact | Function unkn | 0P8GC | NA | 487797.Flav3C | 0,2 | 0,4 | 1,55E-05 | 3,09E-05 | 0 - | 0 - | 1 - | 7,73E-05 - | 0 - | 0 - |
| 316 | 1 | 3um_FIJJAFBF_390144 | Eukaryota | Eustigmatoph | unclassified | Eustigmatales | Monodopsida | Nannochlorop | Nannochlorop | unclassified | unclassified | unclassified | unclassified | unclassified | 1,4 | 0,489897949 | 0,001235495 | 0,00037629 | 1 - | 0,00106926 - | 1 - | 0,000893927 - | 2 - | 0,001799798 - |
| 317 | 1 | 0.2um_NOKCPHJN_130099 | Eukaryota | Arthropoda | Insecta | Hymenoptera | Apidae | Habropoda | Habropoda la | poorly charact | Function unkn | 0F9Q7 | NA | 353151.XP_66 | 0,8 | 0,4 | 4,62E-06 | 2,38E-06 | 1 - | 6,78E-06 - | 1 - | 5,67E-06 - | 1 - | 5,71E-06 - |
| 318 | 1 | 0.2um_NOKCPHJN_62798 | Bacteria | Bacteroidetes | Flavobacteriia | Flavobacterial | Crocinitomica | unclassified | Crocinitomica | poorly charact | Function unkn | 07EF7 | domain protei | 649349.Lbys_( | 0,2 | 0,4 | 1,53E-05 | 3,06E-05 | 0 - | 0 - | 1 - | 7,66E-05 - | 0 - | 0 - |
| 319 | 1 | 3um_FIJJAFBF_405912 | Bacteria | Proteobacteri | Gammaprotec | Alteromonada | Alteromonada | Paraglaciecola | Paraglaciecola | information st | Transcription | COG0085 | DNA-depende | 342610.Patl_( | 0,4 | 0,489897949 | 1,83E-05 | 2,25E-05 | 0 - | 0 - | 0 - | 0 - | 1 - | 4,69E-05 - |
| 320 | 1 | 3um_FIJJAFBF_383207 | Bacteria | Proteobacteri | Deltaproteoba | Myxococcales | Nannocystace | Plesiocystis | Plesiocystis pa | unclassified | unclassified | unclassified | unclassified | unclassified | 2,2 | 0,4 | 0,001011554 | 0,000208054 | 2 - | 0,001092674 - | 2 - | 0,000913502 - | 3 - | 0,001379407 - |
| 321 | 1 | 0.2um_NOKCPHJN_390699 | Bacteria | Proteobacteri | Alphaproteob | Rhizobiales | Rhizobiaceae | Rhizobium | Rhizobium loe | poorly charact | Function unkn | 0ZKS3 | head-to-tail jo | 438753.AZC_3 | 0,4 | 0,489897949 | 4,20E-05 | 5,16E-05 | 0 - | 0 - | 0 - | 0 - | 0 - | 0 - |
| 322 | 1 | 0.2um_NOKCPHJN_977312 | Eukaryota | Phaeophyceae | unclassified | Ectocarpales | Ectocarpaceae | Ectocarpus | Ectocarpus sili | information st | Translation, ri | 0TJFM | 40S ribosomal | 4641.GSMUA_ | 1 | 0 | 0,000492126 | 5,20E-05 | 1 - | 0,000584751 - | 1 - | 0,000488867 - | 1 - | 0,000492132 - |
| 323 | 1 | 0.2um_NOKCPHJN_592028 | Eukaryota | unclassified | Pelagophyceae | Pelagomonada | unclassified | Aureococcus | Aureococcus a | cellular proces | Posttranslatio | KOG0019 | heat shock pro | 67593.Physo1 | 1 | 0 | 8,85E-05 | 9,36E-06 | 1 - | 0,000105124 - | 1 - | 8,79E-05 - | 1 - | 8,85E-05 - |
| 324 | 1 | 0.2um_NOKCPHJN_559310 | unclassified | unclassified | unclassified | unclassified | unclassified | unclassified | unclassified | unclassified | unclassified | unclassified | unclassified | unclassified | 1 | 0 | 0,000186367 | 1,97E-05 | 1 - | 0,000221444 - | 1 - | 0,000185133 - | 1 - | 0,00018637 - |
| 325 | 1 | 3um_FIJJAFBF_281830 | Bacteria | Bacteroidetes | unclassified | unclassified | unclassified | unclassified | Bacteroidetes | information st | Replication, re | 0PB0H | DNA-binding p | 487796.Flav2A | 1,2 | 0,748331477 | 0,0008026 | 0,000492209 | 0 - | 0 - | 2 - | 0,001406178 - | 1 - | 0,000707786 - |
| 326 | 1 | 0.2um_NOKCPHJN_709264 | unclassified | unclassified | unclassified | unclassified | unclassified | unclassified | unclassified | unclassified | unclassified | unclassified | unclassified | unclassified | 1,8 | 0,4 | 0,002134898 | 0,000366873 | 1 - | 0,001439388 - | 2 - | 0,002406728 - | 2 - | 0,002422805 - |
| 327 | 1 | 0.2um_NOKCPHJN_220377 | Bacteria | Proteobacteri | Gammaprotec | Cellvibrionales | Halieaceae | Halioglobus | Halioglobus pa | unclassified | unclassified | unclassified | unclassified | unclassified | 0,2 | 0,4 | 4,46E-05 | 8,93E-05 | 0 - | 0 - | 0 - | 0 - | 0 - | 0 - |
| 328 | 1 | 3um_FIJJAFBF_365812 | unclassified | unclassified | unclassified | unclassified | unclassified | unclassified | unclassified | unclassified | unclassified | unclassified | unclassified | unclassified | 0,2 | 0,4 | 6,07E-05 | 0,000121365 | 0 - | 0 - | 0 - | 0 - | 0 - | 0 - |
| 329 | 1 | 0.2um_NOKCPHJN_52148 | Bacteria | Proteobacteri | Gammaprotec | Cellvibrionales | Halieaceae | Chromatocurv | Chromatocurv | unclassified | unclassified | unclassified | unclassified | unclassified | 0,6 | 0,489897949 | 0,000112523 | 9,24E-05 | 0 - | 0 - | 1 - | 0,000195547 - | 1 - | 0,000196853 - |
| 330 | 1 | 0.2um_NOKCPHJN_619986 | Bacteria | Proteobacteri | Gammaprotec | Cellvibrionales | Cellvibrionace | Teredinibacte | Teredinibacte | cellular proces | Nuclear struct | 19I79 | SRP40, C-term | 1071381.XP_( | 0,2 | 0,4 | 2,38E-05 | 4,76E-05 | 0 - | 0 - | 0 - | 0 - | 0 - | 0 - |
| 331 | 1 | 3um_FIJJAFBF_785067 | unclassified | unclassified | unclassified | unclassified | unclassified | unclassified | unclassified | unclassified | unclassified | unclassified | unclassified | unclassified | 0,4 | 0,489897949 | 0,000175619 | 0,000215091 | 0 - | 0 - | 1 - | 0,000437587 - | 1 - | 0,00044051 - |
| 332 | 1 | 0.2um_NOKCPHJN_448361 | Bacteria | Proteobacteri | Betaproteoba | Neisseriales | Neisseriaceae | Neisseria | Neisseria lacta | cellular proces | Cell wall/mem | COG3637 | Membrane | 489653.NLA_4 | 0,2 | 0,4 | 7,77E-06 | 1,55E-05 | 0 - | 0 - | 0 - | 0 - | 0 - | 0 - |
| 333 | 1 | 0.2um_NOKCPHJN_1000769 | unclassified | unclassified | unclassified | unclassified | unclassified | unclassified | unclassified | unclassified | unclassified | unclassified | unclassified | unclassified | 0,6 | 0,8 | 0,000228307 | 0,000287523 | 1 - | 0,000464896 - | 0 - | 0 - | 0 - | 0 - |
| 334 | 1 | 3um_FIJJAFBF_205760 | Bacteria | Proteobacteri | Gammaprotec | unclassified | unclassified | unclassified | Gammaprotec | unclassified | unclassified | unclassified | unclassified | unclassified | 0,4 | 0,489897949 | 3,68E-05 | 4,53E-05 | 0 - | 0 - | 1 - | 9,84E-05 - | 0 - | 0 - |
| 335 | 1 | 0.2um_NOKCPHJN_88358 | Bacteria | Proteobacteri | Gammaprotec | Thiotrichales | Thiotrichaceae | Thiothrix | Thiothrix nive | unclassified | unclassified | unclassified | unclassified | unclassified | 1 | 0 | 0,000104812 | 1,11E-05 | 1 - | 0,000124539 - | 1 - | 0,000104118 - | 1 - | 0,000104814 - |
| 336 | 1 | 3um_FIJJAFBF_130932 | unclassified | unclassified | unclassified | unclassified | unclassified | unclassified | unclassified | unclassified | unclassified | unclassified | unclassified | unclassified | 0,2 | 0,4 | 1,65E-05 | 3,31E-05 | 0 - | 0 - | 1 - | 8,27E-05 - | 0 - | 0 - |
| 337 | 1 | 0.2um_NOKCPHJN_176471 | Eukaryota | unclassified | unclassified | Isochrysidales | Noelaerhabda | Emiliania | Emiliania huxl | metabolism | Carbohydrate | COG1850 | RuBisCO catal | 45157.CMV01 | 1 | 0 | 0,000129082 | 1,37E-05 | 1 - | 0,000153377 - | 1 - | 0,000128227 - | 1 - | 0,000129084 - |
| 338 | 1 | 0.2um_NOKCPHJN_11384 | Bacteria | Proteobacteri | Alphaproteob | unclassified | unclassified | unclassified | SAR116 cluste | metabolism | Amino acid tra | 07QMX | Aminotransfer | 488538.SAR11 | 0,6 | 0,489897949 | 0,000103132 | 8,52E-05 | 1 - | 0,000195426 - | 1 - | 0,000163381 - | 0 - | 0 - |
| 339 | 1 | 3um_FIJJAFBF_41553 | Eukaryota | Bacillariophyt | Coscinodiscop | Chaetocerotal | Chaetocerota | Chaetoceros | Chaetoceros s | unclassified | unclassified | unclassified | unclassified | unclassified | 2 | 0 | 0,000834333 | 8,82E-05 | 2 - | 0,000991367 - | 2 - | 0,000828807 - | 2 - | 0,000834344 - |
| 340 | 1 | 3um_FIJJAFBF_350452 | Eukaryota | Bacillariophyt | Coscinodiscop | Thalassiosirale | Thalassiosirac | Thalassiosira | Thalassiosira p | unclassified | unclassified | unclassified | unclassified | unclassified | 4,6 | 1,356465997 | 0,000740333 | 0,00018851 | 2 - | 0,000392904 - | 5 - | 0,000821193 - | 5 - | 0,000826679 - |
| 341 | 1 | 0.2um_NOKCPHJN_237508 | Bacteria | Proteobacteri | Gammaprotec | Cellvibrionales | unclassified | unclassified | Cellvibrionales | poorly charact | Function unkn | 0QNMQ | NA | 314287.GB22( | 1 | 0 | 5,45E-05 | 5,76E-06 | 1 - | 6,47E-05 - | 1 - | 5,41E-05 - | 1 - | 5,45E-05 - |
| 342 | 1 | 3um_FIJJAFBF_628367 | Eukaryota | Bacillariophyt | Coscinodiscop | Chaetocerotal | Attheyaceae | Attheya | Attheya longi | metabolism | Energy produc | COG0056 | Produces ATP | 41431.PCC88( | 1,8 | 1,469693846 | 0,00021343 | 0,000176264 | 0 - | 0 - | 4 - | 0,000497614 - | 1 - | 0,000125234 - |
| 343 | 1 | 3um_FIJJAFBF_810755 | Eukaryota | Bacillariophyt | Coscinodiscop | Thalassiosirale | Thalassiosirac | Thalassiosira | Thalassiosira p | metabolism | Nucleotide tra | COG0299 | phosphoribos | 35128.Thaps2 | 0,2 | 0,4 | 2,04E-05 | 4,08E-05 | 0 - | 0 - | 1 - | 0,00010208 - | 0 - | 0 - |
| 344 | 1 | 3um_FIJJAFBF_20710 | Eukaryota | various | various | various | various | various | various | unclassified | unclassified | unclassified | unclassified | unclassified | 5 | 0 | 0,003937011 | 0,00041635 | 5 - | 0,004678011 - | 5 - | 0,003910932 - | 5 - | 0,003937059 - |
| 345 | 1 | 3um_FIJJAFBF_372266 | Bacteria | Proteobacteri | Alphaproteob | unclassified | unclassified | Candidatus Pu | Candidatus Pu | information st | Translation, ri | COG0335 | This protein is | 35128.Thapsd | 1 | 0 | 0,000512132 | 5,42E-05 | 1 - | 0,000608522 - | 1 - | 0,000508739 - | 1 - | 0,000512138 - |
| 346 | 1 | 0.2um_NOKCPHJN_616575 | unclassified | unclassified | unclassified | unclassified | unclassified | unclassified | unclassified | unclassified | unclassified | unclassified | unclassified | unclassified | 0,4 | 0,489897949 | 4,20E-05 | 5,16E-05 | 0 - | 0 - | 1 - | 0,000112141 - | 0 - | 0 - |
| 347 | 1 | 0.2um_NOKCPHJN_159500 | Bacteria | Bacteroidetes | Flavobacteriia | Flavobacterial | Cryomorphace | unclassified | Cryomorphace | metabolism | Amino acid tra | COG0141 | Catalyzes the | 761193.Runsl_ | 0,2 | 0,4 | 2,81E-05 | 5,61E-05 | 0 - | 0 - | 0 - | 0 - | 0 - | 0 - |
| 348 | 1 | 0.2um_NOKCPHJN_235200 | Eukaryota | unclassified | Labyrinthulom | unclassified | Thraustochytr | Aurantiochytr | Aurantiochytr | cellular proces | Cell wall/mem | 12RRB | ankyrin 3, nod | 400682.PAC_1 | 1 | 0 | 7,47E-05 | 7,90E-06 | 1 - | 8,88E-05 - | 1 - | 7,42E-05 - | 1 - | 7,47E-05 - |
| 349 | 1 | 0.2um_NOKCPHJN_1102021 | Bacteria | Verrucomicrol | Verrucomicrol | Verrucomicrol | Rubritaleacea | Rubritalea | Rubritalea squ | cellular proces | Posttranslatio | 1DME7 | Molecular cha | 240016.VspiD | 0,4 | 0,489897949 | 4,94E-05 | 6,06E-05 | 0 - | 0 - | 0 - | 0 - | 1 - | 0,000126492 - |
| 350 | 1 | 0.2um_NOKCPHJN_153284 | Bacteria | Proteobacteri | Gammaprotec | Alteromonada | Pseudoalteror | Pseudoalteror | Pseudoalteror | metabolism | Inorganic ion t | 0QN53 | TonB-depende | 983545.Glaag | 0,8 | 0,4 | 5,38E-05 | 2,78E-05 | 1 - | 7,99E-05 - | 1 - | 6,68E-05 - | 0 - | 0 - |
| 351 | 1 | 3um_FIJJAFBF_505119 | Eukaryota | Chlorophyta | Mamiellophyc | Mamiellales | Mamiellaceae | Micromonas | Micromonas p | unclassified | unclassified | unclassified | unclassified | unclassified | 0,2 | 0,4 | 4,52E-05 | 9,03E-05 | 0 - | 0 - | 0 - | 0 - | 0 - | 0 - |
| 352 | 1 | 3um_FIJJAFBF_434829 | Eukaryota | Streptophyta | unclassified | Asterales | Asteraceae | Lactuca | Lactuca sativa | poorly charact | Function unkn | 0IRPM | Retrotranspos | 400682.PAC_1 | 0,2 | 0,4 | 2,12E-05 | 4,25E-05 | 0 - | 0 - | 1 - | 0,000106239 - | 0 - | 0 - |
| 353 | 1 | 3um_FIJJAFBF_321054 | unclassified | unclassified | unclassified | unclassified | unclassified | unclassified | unclassified | unclassified | unclassified | unclassified | unclassified | unclassified | 0,8 | 0,748331477 | 0,000508047 | 0,000446466 | 1 - | 0,000779669 - | 0 - | 0 - | 0 - | 0 - |
| 354 | 1 | 3um_FIJJAFBF_734730 | Eukaryota | Chromerida | unclassified | unclassified | unclassified | Vitrella | Vitrella brassi | poorly charact | Function unkn | 0XPG5 | KH domain co | 67593.Physo1 | 1,2 | 0,748331477 | 0,000132548 | 8,49E-05 | 0 - | 0 - | 1 - | 0,000114188 - | 2 - | 0,000229901 - |
| 355 | 1 | 3um_FIJJAFBF_448400 | Eukaryota | Bacillariophyt | Bacillariophyc | Naviculales | Phaeodactylac | Phaeodactylu | Phaeodactylu | unclassified | unclassified | unclassified | unclassified | unclassified | 2,4 | 1,019803903 | 0,004131302 | 0,001758493 | 1 - | 0,002079116 - | 3 - | 0,005214577 - | 4 - | 0,006999215 - |
| 356 | 1 | 3um_FIJJAFBF_331637 | Eukaryota | unclassified | Pelagophyceae | Pelagomonada | unclassified | Aureococcus | Aureococcus a | cellular proces | Posttranslatio | 1DQ9N | cysteine prote | 4113.PGSC000 | 2 | 0,632455532 | 0,000436276 | 0,000120825 | 1 - | 0,000265419 - | 2 - | 0,000443794 - | 2 - | 0,000446758 - |
| 357 | 1 | 3um_FIJJAFBF_189664 | Bacteria | Bacteroidetes | Flavobacteriia | Flavobacterial | Flavobacteriac | Polaribacter | Polaribacter s | information st | Translation, ri | 0PBIW | 50S ribosomal | 313598.MED1 | 0,8 | 0,4 | 0,000750352 | 0,000378169 | 0 - | 0 - | 1 - | 0,000977733 - | 1 - | 0,000984265 - |
| 358 | 2 | 0.2um_NOKCPHJN_403347;0.2um_NO | Bacteria | Proteobacteri | Gammaprotec | Cellvibrionales | Halieaceae | unclassified | marine gamm | metabolism | Inorganic ion t | 16PDP | TonB-depende | 247634.GPB2 | 0,4 | 0,489897949 | 2,48E-05 | 3,05E-05 | 0 - | 0 - | 1 - | 6,64E-05 - | 0 - | 0 - |
| 359 | 1 | 3um_FIJJAFBF_03427 | Bacteria | Bacteroidetes | Flavobacteriia | Flavobacterial | Flavobacteriac | Flavivirga | Flavivirga eckl | information st | Transcription | COG0085 | DNA-depende | 391603.FBALC | 0,2 | 0,4 | 9,82E-06 | 1,96E-05 | 0 - | 0 - | 1 - | 4,91E-05 - | 0 - | 0 - |
| 360 | 1 | 3um_FIJJAFBF_99349 | Bacteria | Actinobacteria | Actinobacteria | Micrococcales | Microbacteria | unclassified | Microbacteria | unclassified | unclassified | unclassified | unclassified | unclassified | 0,2 | 0,4 | 3,28E-05 | 6,55E-05 | 1 - | 0,000163782 - | 0 - | 0 - | 0 - | 0 - |
| 361 | 1 | 0.2um_NOKCPHJN_551231 | Bacteria | Bacteroidetes | Flavobacteriia | Flavobacterial | Flavobacteriac | Flavobacteriu | Flavobacteriu | information st | Translation, ri | 09FEM | Binds to the 2 | 376686.Fjoh_( | 1,6 | 0,489897949 | 0,000675804 | 0,000233547 | 2 - | 0,000997976 - | 2 - | 0,000834332 - | 1 - | 0,000419953 - |
| 362 | 1 | 3um_FIJJAFBF_85816 | Viruses | unclassified | unclassified | unclassified | various | various | various | unclassified | unclassified | unclassified | unclassified | unclassified | 2,6 | 0,489897949 | 0,000879393 | 0,000210141 | 3 - | 0,001200773 - | 2 - | 0,00066925 - | 3 - | 0,001010582 - |
| 363 | 1 | 0.2um_NOKCPHJN_461918 | unclassified | unclassified | unclassified | unclassified | unclassified | unclassified | unclassified | unclassified | unclassified | unclassified | unclassified | unclassified | 0,2 | 0,4 | 0,000133138 | 0,000266276 | 0 - | 0 - | 1 - | 0,000665691 - | 0 - | 0 - |
| 364 | 1 | 3um_FIJJAFBF_128529 | unclassified | unclassified | unclassified | unclassified | unclassified | unclassified | unclassified | unclassified | unclassified | unclassified | unclassified | unclassified | 1 | 0 | 0,000321389 | 3,40E-05 | 1 - | 0,000381878 - | 1 - | 0,00031926 - | 1 - | 0,000321393 - |
| 365 | 1 | 3um_FIJJAFBF_353321 | Bacteria | Proteobacteri | Alphaproteob | Rhodobactera | Rhodobactera | unclassified | Rhodobactera | unclassified | unclassified | unclassified | unclassified | unclassified | 0,2 | 0,4 | 2,45E-05 | 4,91E-05 | 0 - | 0 - | 0 - | 0 - | 0 - | 0 - |
| 366 | 1 | 3um_FIJJAFBF_449301 | Bacteria | Proteobacteri | Alphaproteob | Rhodobactera | Rhodobactera | Defluviimonas | various | information st | Replication, re | 176NF | Crispr-associa | 488538.SAR11 | 0,4 | 0,489897949 | 2,18E-05 | 2,68E-05 | 0 - | 0 - | 0 - | 0 - | 1 - | 5,84E-05 - |
| 367 | 1 | 0.2um_NOKCPHJN_774080 | Eukaryota | Streptophyta | unclassified | Apiales | Apiaceae | Daucus | Daucus carota | information st | RNA processir | 04DVS | nucleolin prot | 306902.XP_00 | 0,6 | 0,489897949 | 9,50E-05 | 7,92E-05 | 1 - | 0,000185268 - | 1 - | 0,000154888 - | 0 - | 0 - |
| 368 | 1 | 3um_FIJJAFBF_593424 | Eukaryota | unclassified | Oomycetes | Peronosporale | Peronosporac | Phytophthora | Phytophthora | unclassified | unclassified | unclassified | unclassified | unclassified | 0,4 | 0,489897949 | 0,000105305 | 0,000129826 | 1 - | 0,000286775 - | 1 - | 0,000239751 - | 0 - | 0 - |
| 369 | 3 | 0.2um_NOKCPHJN_505529;3um_FIJJAF | various | various | various | various | various | various | various | various | various | various | various | various | 0,2 | 0,4 | 2,89E-05 | 5,78E-05 | 0 - | 0 - | 0 - | 0 - | 1 - | 0,000144479 - |
| 370 | 1 | 3um_FIJJAFBF_304886 | unclassified | unclassified | unclassified | unclassified | unclassified | unclassified | unclassified | unclassified | unclassified | unclassified | unclassified | unclassified | 0,8 | 0,4 | 0,000614182 | 0,000316919 | 1 - | 0,000901785 - | 1 - | 0,000753915 - | 1 - | 0,000758951 - |
| 371 | 1 | 3um_FIJJAFBF_715090 | Eukaryota | unclassified | Pelagophyceae | Pelagomonada | unclassified | Aureoumbra | Aureoumbra l | information st | Translation, ri | 0FJK5 | Binds the lowe | 197221.tlr008 | 4,8 | 0,4 | 0,001395308 | 7,07E-05 | 4 - | 0,001392524 - | 5 - | 0,001455231 - | 5 - | 0,001464952 - |
| 372 | 1 | 0.2um_NOKCPHJN_171762 | Bacteria | Bacteroidetes | Flavobacteriia | Flavobacterial | unclassified | unclassified | Flavobacterial | information st | Translation, ri | 0P7DE | amino acids su | 755732.Fluta_ | 0,4 | 0,489897949 | 3,09E-05 | 3,82E-05 | 1 - | 8,57E-05 - | 0 - | 0 - | 0 - | 0 - |
| 373 | 1 | 3um_FIJJAFBF_718023 | Bacteria | Cyanobacteria | unclassified | Synechococca | Leptolyngbya | Phormidesmis | Phormidesmis | unclassified | unclassified | unclassified | unclassified | unclassified | 0,4 | 0,489897949 | 4,33E-05 | 5,34E-05 | 1 - | 0,000117871 - | 1 - | 9,85E-05 - | 0 - | 0 - |
| 374 | 2 | 0.2um_NOKCPHJN_86190;3um_FIJJAFE | Bacteria | Bacteroidetes | Flavobacteriia | Flavobacterial | Crocinitomica | unclassified | Crocinitomica | poorly charact | Function unkn | 08WMU | Domain of unl | 382464.VDG1 | 0,2 | 0,4 | 8,38E-05 | 0,000167598 | 0 - | 0 - | 0 - | 0 - | 0 - | 0 - |
| 375 | 1 | 3um_FIJJAFBF_53313 | Bacteria | Bacteroidetes | Flavobacteriia | Flavobacterial | Flavobacteriac | Lutibacter | Lutibacter sp. | unclassified | unclassified | unclassified | unclassified | unclassified | 4 | 0,894427191 | 0,000251818 | 7,51E-05 | 5 - | 0,000369803 - | 5 - | 0,000309165 - | 3 - | 0,000186738 - |
| 376 | 1 | 0.2um_NOKCPHJN_733887 | Eukaryota | unclassified | unclassified | Prymnesiales | Chrysochromu | Chrysochromu | Chrysochromu | information st | Chromatin str | KOG1347 | Mate efflux fa | 272952.HpaP8 | 0,4 | 0,489897949 | 4,36E-05 | 5,37E-05 | 1 - | 0,000118244 - | 0 - | 0 - | 1 - | 9,95E-05 - |
| 377 | 1 | 0.2um_NOKCPHJN_712059 | Eukaryota | unclassified | Ichthyosporea | Ichthyophonic | unclassified | Sphaeroforma | Sphaeroforma | unclassified | unclassified | unclassified | unclassified | unclassified | 0,2 | 0,4 | 0,000111714 | 0,000223427 | 1 - | 0,000558569 - | 0 - | 0 - | 0 - | 0 - |
| 378 | 1 | 0.2um_NOKCPHJN_912697 | Eukaryota | Basidiomycota | Agaricomycete | Auriculariales | Auriculariacea | Auricularia | Auricularia sul | unclassified | unclassified | unclassified | unclassified | unclassified | 0,4 | 0,489897949 | 6,07E-05 | 7,48E-05 | 1 - | 0,000164864 - | 0 - | 0 - | 1 - | 0,000138751 - |
| 379 | 1 | 0.2um_NOKCPHJN_133072 | Bacteria | Bacteroidetes | Flavobacteriia | Flavobacterial | Crocinitomica | unclassified | Crocinitomica | unclassified | unclassified | unclassified | unclassified | unclassified | 1,2 | 0,748331477 | 0,000225897 | 0,00013535 | 0 - | 0 - | 1 - | 0,000199283 - | 1 - | 0,000200614 - |
| 380 | 1 | 3um_FIJJAFBF_666765 | Eukaryota | Chlorophyta | Chlorophyceae | Chlamydomor | Chlamydomor | Chlamydomor | Chlamydomor | unclassified | unclassified | unclassified | unclassified | unclassified | 1,2 | 0,4 | 0,001163609 | 0,000537251 | 2 - | 0,002234274 - | 1 - | 0,000933954 - | 1 - | 0,000940193 - |
| 381 | 1 | 0.2um_NOKCPHJN_648025 | Eukaryota | Cryptomycota | unclassified | unclassified | unclassified | Rozella | Rozella allomy | unclassified | unclassified | unclassified | unclassified | unclassified | 2,2 | 0,4 | 0,000782161 | 9,58E-05 | 2 - | 0,000855408 - | 2 - | 0,000715142 - | 2 - | 0,000719919 - |
| 382 | 1 | 0.2um_NOKCPHJN_202839 | Eukaryota | unclassified | unclassified | Acytosteliales | Acytosteliacea | Acytostelium | Acytostelium s | unclassified | unclassified | unclassified | unclassified | unclassified | 0,2 | 0,4 | 1,09E-05 | 2,19E-05 | 0 - | 0 - | 0 - | 0 - | 1 - | 5,47E-05 - |
| 383 | 1 | 0.2um_NOKCPHJN_134680 | Bacteria | Bacteroidetes | Flavobacteriia | Flavobacterial | Flavobacteriac | Flaviramulus | Flaviramulus i | unclassified | unclassified | unclassified | unclassified | unclassified | 0,4 | 0,489897949 | 0,000116299 | 0,000143061 | 0 - | 0 - | 0 - | 0 - | 1 - | 0,000311846 - |
| 384 | 2 | 3um_FIJJAFBF_195084;3um_FIJJAFBF_ | Bacteria | Proteobacteri | Alphaproteob | Rhodobactera | various | various | various | unclassified | unclassified | unclassified | unclassified | unclassified | 0,2 | 0,4 | 7,72E-05 | 0,000154326 | 1 - | 0,000385815 - | 0 - | 0 - | 0 - | 0 - |
| 385 | 1 | 3um_FIJJAFBF_56139 | Bacteria | Bacteroidetes | Flavobacteriia | Flavobacterial | Flavobacteriac | Polaribacter | Polaribacter s | metabolism | Energy produc | COG0056 | Produces ATP | 313598.MED1 | 2,6 | 0,8 | 0,000306492 | 8,05E-05 | 2 - | 0,000285136 - | 2 - | 0,000238381 - | 2 - | 0,000239973 - |
| 386 | 1 | 0.2um_NOKCPHJN_884992 | Eukaryota | unclassified | Cryptophyta | Pyrenomonad | Geminigerace | Guillardia | Guillardia thet | unclassified | unclassified | unclassified | unclassified | unclassified | 0,2 | 0,4 | 2,53E-05 | 5,06E-05 | 0 - | 0 - | 0 - | 0 - | 1 - | 0,000126492 - |
| 387 | 1 | 0.2um_NOKCPHJN_131719 | Bacteria | unclassified | unclassified | unclassified | unclassified | unclassified | uncultured ba | unclassified | unclassified | unclassified | unclassified | unclassified | 0,4 | 0,489897949 | 8,29E-05 | 0,000101532 | 0 - | 0 - | 1 - | 0,000211402 - | 0 - | 0 - |
| 388 | 1 | 0.2um_NOKCPHJN_774510 | Eukaryota | Bacillariophyt | Coscinodiscop | Thalassiosirale | Thalassiosirac | Thalassiosira | Thalassiosira d | unclassified | unclassified | unclassified | unclassified | unclassified | 1 | 0 | 0,00035791 | 3,78E-05 | 1 - | 0,000425274 - | 1 - | 0,000355539 - | 1 - | 0,000357914 - |
| 389 | 1 | 0.2um_NOKCPHJN_798730 | Eukaryota | unclassified | Cryptophyta | Pyrenomonad | Geminigerace | Guillardia | Guillardia thet | metabolism | Energy produc | 0SQ0U | V-type proton | 43151.ADAR0 | 1,2 | 0,4 | 0,000165293 | 4,08E-05 | 1 - | 0,000167446 - | 1 - | 0,000139989 - | 1 - | 0,000140924 - |
| 390 | 1 | 3um_FIJJAFBF_282599 | Eukaryota | unclassified | Pelagophyceae | Pelagomonada | unclassified | Aureococcus | Aureococcus a | unclassified | unclassified | unclassified | unclassified | unclassified | 1,4 | 0,489897949 | 0,000467485 | 0,00014238 | 1 - | 0,000404585 - | 1 - | 0,000338243 - | 2 - | 0,000681005 - |
| 391 | 1 | 3um_FIJJAFBF_254020 | Bacteria | Bacteroidetes | Flavobacteriia | Flavobacterial | Flavobacteriac | Arenitalea | Arenitalea lut | unclassified | unclassified | unclassified | unclassified | unclassified | 0,2 | 0,4 | 2,72E-05 | 5,45E-05 | 0 - | 0 - | 0 - | 0 - | 0 - | 0 - |
| 392 | 1 | 3um_FIJJAFBF_779814 | Eukaryota | unclassified | unclassified | Kinetoplastida | Trypanosomat | Trypanosoma | Trypanosoma | poorly charact | Function unkn | 0T9G1 | kinetoplast-as | 353153.XP_80 | 0,8 | 0,4 | 0,000372131 | 0,000189626 | 1 - | 0,00053463 - | 1 - | 0,000446964 - | 1 - | 0,00044995 - |
| 393 | 1 | 3um_FIJJAFBF_03426 | Bacteria | Bacteroidetes | Flavobacteriia | Flavobacterial | Flavobacteriac | Algibacter | Algibacter aqu | information st | Transcription | 05D27 | DNA-depende | 983544.Lacal_ | 1,4 | 0,489897949 | 6,15E-05 | 2,18E-05 | 1 - | 5,22E-05 - | 2 - | 8,73E-05 - | 2 - | 8,79E-05 - |
| 394 | 1 | 3um_FIJJAFBF_91115 | Eukaryota | unclassified | Raphidophyce | Chattonellales | Chattonellace | Heterosigma | Heterosigma a | unclassified | unclassified | unclassified | unclassified | unclassified | 17,6 | 0,8 | 0,003486358 | 0,000409011 | 18 - | 0,00423669 - | 18 - | 0,003541977 - | 16 - | 0,003169456 - |
| 395 | 1 | 3um_FIJJAFBF_166626 | Bacteria | Firmicutes | Tissierellia | Tissierellales | Peptoniphilac | Anaerococcus | Anaerococcus | unclassified | unclassified | unclassified | unclassified | unclassified | 0,6 | 0,489897949 | 6,54E-05 | 5,36E-05 | 0 - | 0 - | 0 - | 0 - | 1 - | 0,000116009 - |
| 396 | 1 | 0.2um_NOKCPHJN_206084 | Viruses | unclassified | unclassified | unclassified | unclassified | Pandoravirus | Pandoravirus | cellular proces | Signal transdu | 15M6X | PAS fold | 15368.BRADI4 | 0,8 | 0,4 | 8,21E-05 | 4,24E-05 | 1 - | 0,000121704 - | 0 - | 0 - | 1 - | 0,000102428 - |
| 397 | 1 | 3um_FIJJAFBF_565482 | Eukaryota | unclassified | Pelagophyceae | Pelagomonada | unclassified | Aureococcus | Aureococcus a | metabolism | Carbohydrate | KOG1203 | Complex I inte | 35128.Thapsd | 5 | 0,894427191 | 0,000976129 | 0,000137015 | 4 - | 0,000938535 - | 6 - | 0,001176958 - | 5 - | 0,00098735 - |
| 398 | 1 | 3um_FIJJAFBF_405471 | Bacteria | Proteobacteri | Gammaprotec | Alteromonada | Alteromonada | Paraglaciecola | Paraglaciecola | poorly charact | Function unkn | 0NMWQ | Collagen triple | 324057.Pjdr2_ | 0,2 | 0,4 | 1,88E-05 | 3,77E-05 | 0 - | 0 - | 0 - | 0 - | 0 - | 0 - |
| 399 | 1 | 0.2um_NOKCPHJN_42710 | Bacteria | Proteobacteri | Alphaproteob | Rhodobactera | Rhodobactera | Thalassococcu | Thalassococcu | information st | Replication, re | 01QHS | DNA polymera | 314270.RB208 | 0,2 | 0,4 | 1,07E-05 | 2,14E-05 | 0 - | 0 - | 1 - | 5,35E-05 - | 0 - | 0 - |

| | | | | | | | | | | | | | | | | | | | | | | | | | | | | | | |
|---|---|---|---|---|---|---|---|---|---|---|---|---|---|---|---|---|---|---|---|---|---|---|---|---|---|---|---|---|---|---|
| 400 | 1 | 0.2um_NOKCPHJN_83239 | Eukaryota | unclassified | Spirotrichea | Sporadotrichi | Oxytrichidae | Stylonychia | Stylonychia le | unclassified | unclassified | unclassified | unclassified | unclassified | 0,2 | 0,4 | 1,75E-05 | 3,50E-05 | 0 | - | 0 | - | 0 | - | 0 | - | 0 | - | 0 | - |
| 401 | 1 | 3um_FIJJAFBF_651104 | Bacteria | Proteobacteri | Betaproteoba | Nitrosomonad | Nitrosomonac | Nitrosomonas | Nitrosomonas | unclassified | unclassified | unclassified | unclassified | unclassified | 0,4 | 0,489897949 | 3,06E-05 | 3,76E-05 | 0 | - | 0 | - | 0 | - | 0 | - | 0 | - | 0 | - |
| 402 | 1 | 3um_FIJJAFBF_636517 | Bacteria | Bacteroidetes | Flavobacteriia | Flavobacterial | Flavobacteria | Flavivirga | Flavivirga eckl | metabolism | Lipid transpor | COG4247 | Phytase | 867900.Celly_ | 0,6 | 0,489897949 | 0,00012037 | 9,92E-05 | 1 | - | 0,000224769 | - | 1 | - | 0,000187913 | - | 1 | - | 0,000189168 | - |
| 403 | 1 | 0.2um_NOKCPHJN_441897 | Eukaryota | Arthropoda | Insecta | Coleoptera | Tenebrionidae | Tribolium | Tribolium cast | unclassified | unclassified | unclassified | unclassified | unclassified | 0,2 | 0,4 | 2,02E-05 | 4,05E-05 | 0 | - | 0 | - | 0 | - | 0 | - | 0 | - | 0 | - |
| 404 | 1 | 3um_FIJJAFBF_513826 | Viruses | unclassified | unclassified | unclassified | unclassified | unclassified | uncultured M | unclassified | unclassified | unclassified | unclassified | unclassified | 4,8 | 0,748331477 | 0,000978055 | 0,000160768 | 5 | - | 0,001215068 | - | 5 | - | 0,001015827 | - | 4 | - | 0,00081809 | - |
| 405 | 1 | 3um_FIJJAFBF_763981 | Eukaryota | unclassified | Oomycetes | Pythiales | Pythiaceae | Pythium | Pythium insidi | information st | Translation, ri | COG0089 | One of the ea | 653948.CCA24 | 1 | 0 | 0,000453181 | 4,79E-05 | 1 | - | 0,000538476 | - | 1 | - | 0,000450179 | - | 1 | - | 0,000453187 | - |
| 406 | 1 | 3um_FIJJAFBF_456402 | Eukaryota | Ascomycota | Leotiomycetes | unclassified | Pseudeurotiac | Pseudogymno | Pseudogymno | unclassified | unclassified | unclassified | unclassified | unclassified | 0,8 | 0,4 | 0,000151583 | 7,84E-05 | 1 | - | 0,000224769 | - | 0 | - | 0 | - | 1 | - | 0,000189168 | - |
| 407 | 1 | 3um_FIJJAFBF_400977 | unclassified | unclassified | unclassified | unclassified | unclassified | unclassified | unclassified | poorly charact | Function unkn | 0IPYB | Reverse trans | 44689.DDB02 | 0,2 | 0,4 | 1,76E-05 | 3,52E-05 | 0 | - | 0 | - | 0 | - | 0 | - | 0 | - | 0 | - |
| 408 | 1 | 0.2um_NOKCPHJN_409026 | Eukaryota | Basidiomycota | Agaricomycete | Agaricales | Mycenaceae | Mycena | Mycena chlor | unclassified | unclassified | unclassified | unclassified | unclassified | 0,6 | 0,489897949 | 5,41E-05 | 4,44E-05 | 0 | - | 0 | - | 1 | - | 9,40E-05 | - | 1 | - | 9,46E-05 | - |
| 409 | 1 | 3um_FIJJAFBF_184996 | Eukaryota | unclassified | Oomycetes | Albuginales | Albuginaceae | Albugo | Albugo laibacl | information st | Translation, ri | KOG1749 | Ribosomal pro | 653948.CCA18 | 0,2 | 0,4 | 8,63E-05 | 0,00017262 | 0 | - | 0 | - | 1 | - | 0,000431551 | - | 0 | - | 0 | - |
| 410 | 1 | 0.2um_NOKCPHJN_366105 | Viruses | unclassified | unclassified | unclassified | unclassified | unclassified | Tetraselmis vi | unclassified | unclassified | unclassified | unclassified | unclassified | 0,6 | 0,489897949 | 6,78E-05 | 5,54E-05 | 0 | - | 0 | - | 1 | - | 0,000114188 | - | 1 | - | 0,000114951 | - |
| 411 | 1 | 0.2um_NOKCPHJN_112574 | Viruses | unclassified | unclassified | unclassified | Phycodnavirid | unclassified | Aureococcus a | information st | Translation, ri | KOG0052 | This protein p | 65071.PYU1_T | 1,6 | 0,489897949 | 0,000223476 | 8,15E-05 | 2 | - | 0,000326136 | - | 1 | - | 0,000136329 | - | 2 | - | 0,000274479 | - |
| 412 | 1 | 0.2um_NOKCPHJN_509146 | unclassified | unclassified | unclassified | unclassified | unclassified | unclassified | unclassified | unclassified | unclassified | unclassified | unclassified | unclassified | 0,4 | 0,489897949 | 0,000122508 | 0,00015034 | 0 | - | 0 | - | 0 | - | 0 | - | 0 | - | 0 | - |
| 413 | 2 | 0.2um_NOKCPHJN_18962;3um_FIJJAF | Bacteria | Proteobacteri | Gammaprotec | various | various | various | various | information st | Transcription | 0ZZMP | Helix-turn-hel | 717773.Thicy_ | 0,2 | 0,4 | 0,000136173 | 0,000272347 | 0 | - | 0 | - | 0 | - | 0 | - | 0 | - | 0 | - |
| 414 | 1 | 0.2um_NOKCPHJN_575550 | Eukaryota | Phaeophyceae | unclassified | Ectocarpales | Ectocarpaceae | Ectocarpus | Ectocarpus sil | information st | Translation, ri | COG5045 | Ribosomal pro | 67593.Physo1 | 0,2 | 0,4 | 4,54E-05 | 9,08E-05 | 0 | - | 0 | - | 0 | - | 0 | - | 0 | - | 0 | - |
| 415 | 1 | 3um_FIJJAFBF_407843 | Bacteria | Proteobacteri | Alphaproteob | Rhodobactera | Rhodobactera | Rhodosalinus | Rhodosalinus | unclassified | unclassified | unclassified | unclassified | unclassified | 0,2 | 0,4 | 4,60E-05 | 9,20E-05 | 0 | - | 0 | - | 1 | - | 0,000230055 | - | 0 | - | 0 | - |
| 416 | 1 | 0.2um_NOKCPHJN_915793 | unclassified | unclassified | unclassified | unclassified | unclassified | unclassified | unclassified | unclassified | unclassified | unclassified | unclassified | unclassified | 1 | 0 | 0,000623685 | 6,60E-05 | 1 | - | 0,000741071 | - | 1 | - | 0,000619554 | - | 1 | - | 0,000623692 | - |
| 417 | 1 | 3um_FIJJAFBF_570821 | Bacteria | Proteobacteri | Deltaproteoba | Myxococcales | unclassified | Minicystis | Minicystis ros | poorly charact | Function unkn | 12323 | NA | 423536.XP_00 | 0,8 | 0,4 | 0,000102426 | 5,30E-05 | 1 | - | 0,00015213 | - | 1 | - | 0,000127185 | - | 0 | - | 0 | - |
| 418 | 1 | 3um_FIJJAFBF_135258 | Eukaryota | unclassified | Synurophycea | Parmales | Triparmaceae | Triparma | Triparma laev | metabolism | Energy produ | COG0711 | F(1)F(0) ATP s | 314260.PB250 | 2,6 | 0,489897949 | 0,001227211 | 0,000194457 | 2 | - | 0,001134063 | - | 3 | - | 0,001422157 | - | 3 | - | 0,001431658 | - |
| 419 | 1 | 3um_FIJJAFBF_624005 | Bacteria | Bacteroidetes | Flavobacteriia | Flavobacterial | Flavobacteria | Marixanthom | Marixanthom | poorly charact | Function unkn | 0P8NI | Inherit from b | 313594.PI23P | 0,4 | 0,489897949 | 3,54E-05 | 4,34E-05 | 0 | - | 0 | - | 0 | - | 0 | - | 0 | - | 0 | - |
| 420 | 1 | 0.2um_NOKCPHJN_1006750 | unclassified | unclassified | unclassified | unclassified | unclassified | unclassified | unclassified | unclassified | unclassified | unclassified | unclassified | unclassified | 0,4 | 0,489897949 | 4,22E-05 | 5,17E-05 | 0 | - | 0 | - | 1 | - | 0,000107702 | - | 0 | - | 0 | - |
| 421 | 1 | 0.2um_NOKCPHJN_537861 | Bacteria | Proteobacteri | Betaproteoba | Burkholderiale | Comamonada | Limnohabitan | Limnohabitan | poorly charact | Function unkn | 0KN5V | NA | 140110.Necha | 0,6 | 0,489897949 | 2,59E-05 | 2,13E-05 | 1 | - | 4,84E-05 | - | 1 | - | 4,04E-05 | - | 1 | - | 4,07E-05 | - |
| 422 | 1 | 0.2um_NOKCPHJN_86330 | Bacteria | Bacteroidetes | Flavobacteriia | Flavobacterial | Crocinitomica | unclassified | Crocinitomica | unclassified | unclassified | unclassified | unclassified | unclassified | 1,2 | 0,4 | 9,83E-05 | 2,43E-05 | 1 | - | 9,95E-05 | - | 1 | - | 8,32E-05 | - | 1 | - | 8,38E-05 | - |
| 423 | 1 | 0.2um_NOKCPHJN_696789 | Bacteria | Proteobacteri | Betaproteoba | Neisseriales | Neisseriaceae | Snodgrassella | Snodgrassella | unclassified | unclassified | unclassified | unclassified | unclassified | 0,2 | 0,4 | 1,22E-05 | 2,44E-05 | 0 | - | 0 | - | 0 | - | 0 | - | 0 | - | 0 | - |
| 424 | 1 | 3um_FIJJAFBF_476362 | Eukaryota | unclassified | unclassified | Prymnesiales | Chrysochrom | Chrysochrom | various | unclassified | unclassified | unclassified | unclassified | unclassified | 2,4 | 0,489897949 | 0,000296852 | 6,52E-05 | 2 | - | 0,000294099 | - | 3 | - | 0,000368811 | - | 3 | - | 0,000371275 | - |
| 425 | 1 | 0.2um_NOKCPHJN_915378 | Eukaryota | unclassified | Cryptophyta | Pyrenomonad | Geminigerace | Guillardia | Guillardia the | unclassified | unclassified | unclassified | unclassified | unclassified | 0,2 | 0,4 | 2,13E-05 | 4,26E-05 | 0 | - | 0 | - | 0 | - | 0 | - | 0 | - | 0 | - |
| 426 | 1 | 3um_FIJJAFBF_361450 | Eukaryota | unclassified | Pelagophycea | Pelagomonad | unclassified | Aureococcus | Aureococcus a | unclassified | unclassified | unclassified | unclassified | unclassified | 2,4 | 0,489897949 | 0,001023126 | 0,000299237 | 3 | - | 0,001496964 | - | 3 | - | 0,001251498 | - | 2 | - | 0,000839906 | - |
| 427 | 1 | 3um_FIJJAFBF_472206 | Bacteria | Bacteroidetes | Flavobacteriia | Flavobacterial | Flavobacteria | Lutibacter | Lutibacter aga | metabolism | Inorganic ion t | COG3119 | Sulfatase | 485917.Phep_ | 0,2 | 0,4 | 1,27E-05 | 2,55E-05 | 0 | - | 0 | - | 0 | - | 0 | - | 0 | - | 0 | - |
| 428 | 1 | 0.2um_NOKCPHJN_312154 | Bacteria | Bacteroidetes | Flavobacteriia | Flavobacterial | Crocinitomica | unclassified | Crocinitomica | unclassified | unclassified | unclassified | unclassified | unclassified | 0,2 | 0,4 | 3,47E-05 | 6,95E-05 | 0 | - | 0 | - | 0 | - | 0 | - | 0 | - | 0 | - |
| 429 | 1 | 0.2um_NOKCPHJN_624427 | Eukaryota | Ascomycota | Eurotiomycete | Eurotiales | Aspergillaceae | Aspergillus | Aspergillus te | cellular proces | Posttranslatio | 0JAJ8 | Dentin sialoph | 10020.ENSDO | 1 | 0 | 7,32E-05 | 7,75E-06 | 1 | - | 8,70E-05 | - | 1 | - | 7,28E-05 | - | 1 | - | 7,32E-05 | - |
| 430 | 1 | 3um_FIJJAFBF_67828 | Eukaryota | unclassified | Spirotrichea | Sporadotrichi | Oxytrichidae | Oxytricha | Oxytricha trifa | unclassified | unclassified | unclassified | unclassified | unclassified | 1,4 | 0,8 | 0,000249344 | 0,000141315 | 0 | - | 0 | - | 1 | - | 0,000186791 | - | 2 | - | 0,000376077 | - |
| 431 | 1 | 3um_FIJJAFBF_697086 | Eukaryota | unclassified | Heterolobosea | Schizopyrenid | Vahlkampfiida | Naegleria | Naegleria grul | unclassified | unclassified | unclassified | unclassified | unclassified | 1,4 | 0,489897949 | 0,000230298 | 6,45E-05 | 1 | - | 0,000200665 | - | 1 | - | 0,000167761 | - | 1 | - | 0,000168882 | - |
| 432 | 1 | 3um_FIJJAFBF_715087 | Eukaryota | Bacillariophyt | Bacillariophyc | Naviculales | Phaeodactyla | Phaeodactylu | Phaeodactylu | unclassified | unclassified | unclassified | unclassified | unclassified | 2 | 0,632455532 | 0,001021598 | 0,000306574 | 1 | - | 0,00061858 | - | 2 | - | 0,001034296 | - | 3 | - | 0,001561808 | - |
| 433 | 1 | 0.2um_NOKCPHJN_31647 | Bacteria | Proteobacteri | Betaproteoba | unclassified | unclassified | unclassified | Betaproteoba | cellular proces | Cell wall/mem | COG2885 | Ompa motb d | 365044.Pnap_ | 1 | 0 | 0,000329802 | 3,49E-05 | 1 | - | 0,000391875 | - | 1 | - | 0,000327617 | - | 1 | - | 0,000329806 | - |
| 434 | 1 | 3um_FIJJAFBF_600643 | unclassified | unclassified | unclassified | unclassified | unclassified | unclassified | unclassified | unclassified | unclassified | unclassified | unclassified | unclassified | 0,2 | 0,4 | 3,74E-05 | 7,47E-05 | 0 | - | 0 | - | 1 | - | 0,000186791 | - | 0 | - | 0 | - |
| 435 | 1 | 3um_FIJJAFBF_344729 | Eukaryota | Bacillariophyt | Coscinodiscop | Coscinodiscale | Coscinodiscac | Coscinodiscus | Coscinodiscus | unclassified | unclassified | unclassified | unclassified | unclassified | 0,2 | 0,4 | 0,000123867 | 0,000247734 | 0 | - | 0 | - | 0 | - | 0 | - | 0 | - | 0 | - |
| 436 | 1 | 0.2um_NOKCPHJN_176491 | Bacteria | unclassified | unclassified | unclassified | unclassified | unclassified | bacterium TM | metabolism | Energy produ | COG0056 | Produces ATP | 43989.cce_44 | 0,6 | 0,489897949 | 7,68E-05 | 6,40E-05 | 1 | - | 0,000149696 | - | 1 | - | 0,00012515 | - | 0 | - | 0 | - |
| 437 | 1 | 3um_FIJJAFBF_737020 | Bacteria | Proteobacteri | Alphaproteob | Rhodospirillal | unclassified | unclassified | Rhodospirillal | cellular proces | Cell wall/mem | COG2885 | Ompa motb d | 331869.BAL19 | 0,4 | 0,489897949 | 0,0001495 | 0,00018685 | 1 | - | 0,000432648 | - | 0 | - | 0 | - | 0 | - | 0 | - |
| 438 | 1 | 0.2um_NOKCPHJN_857717 | unclassified | unclassified | unclassified | unclassified | unclassified | unclassified | unclassified | unclassified | unclassified | unclassified | unclassified | unclassified | 0,2 | 0,4 | 1,48E-05 | 2,96E-05 | 0 | - | 0 | - | 0 | - | 0 | - | 0 | - | 0 | - |
| 439 | 1 | 3um_FIJJAFBF_539523 | unclassified | unclassified | unclassified | unclassified | unclassified | unclassified | unclassified | unclassified | unclassified | unclassified | unclassified | unclassified | 0,6 | 0,489897949 | 0,000192976 | 0,000158266 | 0 | - | 0 | - | 0 | - | 0 | - | 1 | - | 0,000342353 | - |
| 440 | 1 | 0.2um_NOKCPHJN_770496 | Eukaryota | Bacillariophyt | Coscinodiscop | Thalassiosirale | Thalassiosirac | Thalassiosira | Thalassiosira | unclassified | unclassified | unclassified | unclassified | unclassified | 1,2 | 0,4 | 0,000352837 | 0,000120088 | 1 | - | 0,000349758 | - | 2 | - | 0,000584812 | - | 1 | - | 0,00029436 | - |
| 441 | 1 | 0.2um_NOKCPHJN_117065 | Eukaryota | Bacillariophyt | Bacillariophyc | Naviculales | Naviculaceae | Fistulifera | Fistulifera sola | metabolism | Energy produ | 0IFIU | Pyrophosphat | 2850.Phatr15 | 0,6 | 0,489897949 | 4,51E-05 | 3,70E-05 | 0 | - | 0 | - | 1 | - | 7,97E-05 | - | 0 | - | 0 | - |
| 442 | 1 | 0.2um_NOKCPHJN_653211 | Bacteria | Proteobacteri | Alphaproteob | Pelagibacteral | Pelagibactera | unclassified | Pelagibactera | unclassified | unclassified | unclassified | unclassified | unclassified | 1 | 0 | 0,00061757 | 6,53E-05 | 1 | - | 0,000733806 | - | 1 | - | 0,00061348 | - | 1 | - | 0,000617578 | - |
| 443 | 1 | 3um_FIJJAFBF_815478 | Eukaryota | Eustigmatoph | unclassified | Eustigmatales | Monodopsida | Nannochlorop | Nannochlorop | metabolism | Lipid transpor | COG0439 | acetyl-CoA ca | 35128.Thaps6 | 3,2 | 0,4 | 0,000656324 | 0,000102725 | 3 | - | 0,000731415 | - | 4 | - | 0,000815308 | - | 3 | - | 0,000615566 | - |
| 444 | 1 | 3um_FIJJAFBF_216059 | Bacteria | Proteobacteri | Gammaprotec | Xanthomonad | Rhodanobacte | Oleiagrimonas | Oleiagrimonas | information st | Translation, ri | COG0050 | This protein p | 523791.Kkor_ | 0,8 | 0,4 | 0,000878915 | 0,000453522 | 1 | - | 0,001290486 | - | 1 | - | 0,001078878 | - | 1 | - | 0,001086085 | - |
| 445 | 1 | 0.2um_NOKCPHJN_192042 | Bacteria | Bacteroidetes | Flavobacteriia | Flavobacterial | Flavobacteria | Lutibacter | Lutibacter sp. | metabolism | Carbohydrate | 05D2N | Aldolase | 391587.KAOT | 0,8 | 0,4 | 0,000135275 | 6,82E-05 | 0 | - | 0 | - | 1 | - | 0,000176267 | - | 1 | - | 0,000177445 | - |
| 446 | 1 | 3um_FIJJAFBF_469420 | Bacteria | Bacteroidetes | Cytophagia | Cytophagales | Flammeovirga | unclassified | Flammeovirga | metabolism | Lipid transpor | COG0657 | alpha beta hy | 504472.Slin_1 | 0,4 | 0,489897949 | 7,44E-05 | 9,13E-05 | 0 | - | 0 | - | 0 | - | 0 | - | 0 | - | 0 | - |
| 447 | 1 | 0.2um_NOKCPHJN_1116522 | unclassified | unclassified | unclassified | unclassified | unclassified | unclassified | unclassified | unclassified | unclassified | unclassified | unclassified | unclassified | 1 | 0 | 0,000887214 | 9,38E-05 | 1 | - | 0,0010542 | - | 1 | - | 0,000881337 | - | 1 | - | 0,000887224 | - |
| 448 | 1 | 3um_FIJJAFBF_146773 | Bacteria | various | various | various | various | various | various | unclassified | unclassified | unclassified | unclassified | unclassified | 0,8 | 0,748331477 | 0,000203036 | 0,000181123 | 0 | - | 0 | - | 1 | - | 0,000270887 | - | 1 | - | 0,000272697 | - |
| 449 | 1 | 0.2um_NOKCPHJN_730596 | Eukaryota | unclassified | Oomycetes | Saprolegniales | Saprolegniace | Achlya | Achlya hypogy | poorly charact | Function unkn | COG1100 | GTP-binding P | 272952.HpaP8 | 1,6 | 1,019803903 | 0,000451612 | 0,000274989 | 1 | - | 0,000344923 | - | 0 | - | 0 | - | 2 | - | 0,00058058 | - |
| 450 | 1 | 3um_FIJJAFBF_788032 | Bacteria | Proteobacteri | Alphaproteob | Rhodobactera | Rhodobactera | unclassified | Rhodobactera | information st | Translation, ri | 179ZI | One of the pri | 52598.EE36_0 | 1 | 0 | 0,000484555 | 5,12E-05 | 1 | - | 0,000575755 | - | 1 | - | 0,000481346 | - | 1 | - | 0,000484561 | - |
| 451 | 1 | 3um_FIJJAFBF_121610 | unclassified | unclassified | unclassified | unclassified | unclassified | unclassified | unclassified | unclassified | unclassified | unclassified | unclassified | unclassified | 0,2 | 0,4 | 6,95E-05 | 0,000138903 | 0 | - | 0 | - | 0 | - | 0 | - | 0 | - | 0 | - |
| 452 | 1 | 0.2um_NOKCPHJN_86824 | Bacteria | Bacteroidetes | unclassified | unclassified | unclassified | unclassified | Bacteroidetes | unclassified | unclassified | unclassified | unclassified | unclassified | 0,6 | 0,489897949 | 5,11E-05 | 4,22E-05 | 1 | - | 9,67E-05 | - | 0 | - | 0 | - | 1 | - | 8,14E-05 | - |
| 453 | 1 | 3um_FIJJAFBF_306873 | Bacteria | Proteobacteri | Alphaproteob | Rhodobactera | unclassified | unclassified | Rhodobactera | metabolism | Inorganic ion t | 05PWN | phosphate ab | 439497.RR11_ | 1,2 | 0,4 | 0,00021512 | 7,32E-05 | 1 | - | 0,000213243 | - | 2 | - | 0,000356552 | - | 1 | - | 0,000179467 | - |
| 454 | 1 | 0.2um_NOKCPHJN_858740 | unclassified | unclassified | unclassified | unclassified | unclassified | unclassified | unclassified | unclassified | unclassified | unclassified | unclassified | unclassified | 0,2 | 0,4 | 0,000391093 | 0,000782186 | 0 | - | 0 | - | 1 | - | 0,001955466 | - | 0 | - | 0 | - |
| 455 | 1 | 0.2um_NOKCPHJN_1032283 | Eukaryota | unclassified | Oomycetes | Saprolegniales | Saprolegniace | Aphanomyces | Aphanomyces | information st | Replication, re | COG0513 | purine NTP-de | 4787.PITG_03 | 0,8 | 0,4 | 0,000125713 | 6,34E-05 | 0 | - | 0 | - | 1 | - | 0,000163809 | - | 1 | - | 0,000164903 | - |
| 456 | 1 | 3um_FIJJAFBF_08158 | Bacteria | Bacteroidetes | Flavobacteriia | Flavobacterial | Flavobacteria | Polaribacter | Polaribacter s | metabolism | Lipid transpor | COG4247 | Phytase | 688270.Celal_ | 3,6 | 0,8 | 0,000665386 | 0,000139242 | 3 | - | 0,000664333 | - | 5 | - | 0,000925664 | - | 3 | - | 0,000559109 | - |
| 457 | 1 | 0.2um_NOKCPHJN_416546 | Eukaryota | unclassified | unclassified | unclassified | unclassified | Stygiella | Stygiella incar | information st | Translation, ri | KOG0052 | This protein p | 5664.LmjF.17. | 2 | 0 | 0,000266352 | 2,82E-05 | 2 | - | 0,000316483 | - | 2 | - | 0,000264587 | - | 2 | - | 0,000266355 | - |
| 458 | 2 | 0.2um_NOKCPHJN_480841;3um_FIJJAF | Bacteria | Bacteroidetes | Flavobacteriia | Flavobacterial | Flavobacteria | various | various | various | various | various | various | various | 0,2 | 0,4 | 2,17E-05 | 4,35E-05 | 0 | - | 0 | - | 0 | - | 0 | - | 0 | - | 0 | - |
| 459 | 1 | 3um_FIJJAFBF_119125 | Bacteria | Bacteroidetes | Flavobacteriia | Flavobacterial | Flavobacteria | Polaribacter | various | information st | Translation, ri | 08YY1 | This protein is | 313594.PI23P | 2,8 | 0,4 | 0,001348891 | 0,000117786 | 2 | - | 0,001160437 | - | 3 | - | 0,001455231 | - | 3 | - | 0,001464952 | - |
| 460 | 1 | 0.2um_NOKCPHJN_913570 | Bacteria | Proteobacteri | Alphaproteob | Pelagibacteral | Pelagibactera | Candidatus Pe | Candidatus Pe | unclassified | unclassified | unclassified | unclassified | unclassified | 1 | 0 | 9,40E-05 | 9,94E-06 | 1 | - | 0,000111714 | - | 1 | - | 9,34E-05 | - | 1 | - | 9,40E-05 | - |
| 461 | 1 | 3um_FIJJAFBF_176097 | Bacteria | Proteobacteri | Gammaprotec | Cellvibrionales | Spongiibacter | Oceanicoccus | Oceanicoccus | cellular proces | Intracellular tr | 124XR | MSHA pilin pr | 903510.vfu_A | 1,2 | 0,4 | 0,000463747 | 0,000159897 | 1 | - | 0,000459191 | - | 1 | - | 0,000383895 | - | 2 | - | 0,000772919 | - |
| 462 | 1 | 0.2um_NOKCPHJN_1014683 | Eukaryota | Basidiomycota | Exobasidiomy | Microstromat | unclassified | Jaminaea | Jaminaea rose | poorly charact | Function unkn | 121QZ | Glycine-rich R | 3711.Bra0140 | 0,4 | 0,489897949 | 7,44E-05 | 9,11E-05 | 0 | - | 0 | - | 0 | - | 0 | - | 1 | - | 0,000190311 | - |
| 463 | 1 | 3um_FIJJAFBF_72763 | Bacteria | Actinobacteria | Actinobacteria | Micrococcales | Microbacteria | Candidatus Ac | Candidatus Ac | unclassified | unclassified | unclassified | unclassified | unclassified | 0,2 | 0,4 | 2,70E-05 | 5,41E-05 | 0 | - | 0 | - | 0 | - | 0 | - | 0 | - | 0 | - |
| 464 | 1 | 0.2um_NOKCPHJN_241055 | Eukaryota | Ascomycota | Sordariomyce | Hypocreales | Nectriaceae | Fusarium | Fusarium gran | cellular proces | Posttranslatio | 0V7QW | ankyrin repea | 7668.SPU_024 | 1 | 0 | 4,57E-05 | 4,84E-06 | 1 | - | 5,44E-05 | - | 1 | - | 4,54E-05 | - | 1 | - | 4,57E-05 | - |
| 465 | 1 | 3um_FIJJAFBF_24024 | Viruses | unclassified | unclassified | unclassified | Phycodnavirid | unclassified | Chrysochrom | unclassified | unclassified | unclassified | unclassified | unclassified | 1 | 0 | 8,29E-05 | 8,77E-06 | 1 | - | 9,85E-05 | - | 1 | - | 8,23E-05 | - | 1 | - | 8,29E-05 | - |
| 466 | 1 | 3um_FIJJAFBF_487685 | Eukaryota | Bacillariophyt | Bacillariophyc | Bacillariales | Bacillariaceae | Nitzschia | Nitzschia sp. I | unclassified | unclassified | unclassified | unclassified | unclassified | 0,2 | 0,4 | 3,15E-05 | 6,31E-05 | 0 | - | 0 | - | 0 | - | 0 | - | 0 | - | 0 | - |
| 467 | 1 | 3um_FIJJAFBF_217104 | Bacteria | Bacteroidetes | Flavobacteriia | Flavobacterial | Crocinitomica | unclassified | Crocinitomica | unclassified | unclassified | unclassified | unclassified | unclassified | 3,4 | 1,2 | 0,000659426 | 0,000214058 | 1 | - | 0,00023837 | - | 4 | - | 0,000797133 | - | 4 | - | 0,000802458 | - |
| 468 | 1 | 3um_FIJJAFBF_715079 | Eukaryota | unclassified | Pelagophycea | Pelagomonad | unclassified | Pelagomonas | uncultured Pe | information st | Translation, ri | 08UHH | Located on th | 580331.Thit_1 | 0,4 | 0,489897949 | 0,000180711 | 0,000222294 | 0 | - | 0 | - | 0 | - | 0 | - | 1 | - | 0,000484561 | - |
| 469 | 1 | 3um_FIJJAFBF_465667 | Eukaryota | unclassified | Oomycetes | Albuginales | Albuginaceae | Albugo | Albugo candid | cellular proces | Intracellular tr | KOG0092 | member RAS | 653948.CCA18 | 1,4 | 0,489897949 | 0,000434596 | 0,000132363 | 1 | - | 0,000376122 | - | 1 | - | 0,000314447 | - | 2 | - | 0,000633095 | - |
| 470 | 1 | 3um_FIJJAFBF_495044 | Bacteria | Bacteroidetes | Flavobacteriia | Flavobacterial | Flavobacteria | Cellulophaga | Cellulophaga t | poorly charact | Function unkn | 0XP53 | Ragb susd dor | 313598.MED1 | 0,6 | 0,489897949 | 0,000257196 | 0,000211119 | 0 | - | 0 | - | 1 | - | 0,000446964 | - | 1 | - | 0,00044995 | - |
| 471 | 1 | 0.2um_NOKCPHJN_374320 | Bacteria | Proteobacteri | Alphaproteob | Rhodospirillal | unclassified | unclassified | Rhodospirillal | metabolism | Amino acid tra | COG0683 | (ABC) transpo | 504832.OCA5 | 0,8 | 0,4 | 0,000110143 | 5,55E-05 | 0 | - | 0 | - | 1 | - | 0,00014352 | - | 1 | - | 0,000144479 | - |
| 472 | 1 | 3um_FIJJAFBF_175076 | Bacteria | Proteobacteri | Alphaproteob | Rhodobactera | Rhodobactera | various | various | metabolism | Amino acid tra | 01SDW | Spermidine pu | 439497.RR11_ | 0,4 | 0,489897949 | 6,13E-05 | 7,52E-05 | 0 | - | 0 | - | 0 | - | 0 | - | 0 | - | 0 | - |
| 473 | 1 | 0.2um_NOKCPHJN_284549 | Bacteria | Proteobacteri | Gammaprotec | Cellvibrionales | Halieaceae | unclassified | marine gamm | information st | Transcription | 05D27 | DNA-depende | 876044.IMCC3 | 4,6 | 1,743559577 | 0,000206859 | 9,68E-05 | 7 | - | 0,000363845 | - | 6 | - | 0,000260729 | - | 4 | - | 0,00017498 | - |
| 474 | 1 | 0.2um_NOKCPHJN_76785 | Bacteria | Proteobacteri | Deltaproteoba | Myxococcales | Polyangiaceae | Chondromyce | Chondromyce | unclassified | unclassified | unclassified | unclassified | unclassified | 1,4 | 0,8 | 0,000409858 | 0,000245646 | 2 | - | 0,000689845 | - | 0 | - | 0 | - | 2 | - | 0,00058058 | - |
| 475 | 1 | 3um_FIJJAFBF_181303 | Eukaryota | unclassified | Pelagophycea | Pelagomonad | unclassified | Aureococcus | Aureococcus a | unclassified | unclassified | unclassified | unclassified | unclassified | 1 | 0 | 0,000173532 | 1,84E-05 | 1 | - | 0,000206193 | - | 1 | - | 0,000172383 | - | 1 | - | 0,000173534 | - |
| 476 | 1 | 0.2um_NOKCPHJN_291436 | Bacteria | Proteobacteri | Alphaproteob | Rhodobactera | Rhodobactera | Planktomarina | Planktomarina | unclassified | unclassified | unclassified | unclassified | unclassified | 0,8 | 0,4 | 0,000377608 | 0,000194847 | 1 | - | 0,000554431 | - | 1 | - | 0,000463518 | - | 1 | - | 0,000466614 | - |
| 477 | 1 | 3um_FIJJAFBF_546136 | unclassified | unclassified | unclassified | unclassified | unclassified | unclassified | unclassified | unclassified | unclassified | unclassified | unclassified | unclassified | 1 | 0,632455532 | 9,38E-05 | 5,80E-05 | 1 | - | 0,000112554 | - | 1 | - | 9,41E-05 | - | 0 | - | 0 | - |
| 478 | 1 | 0.2um_NOKCPHJN_160361 | Eukaryota | unclassified | Pelagophycea | Pelagomonad | unclassified | Aureoumbra | Aureoumbra l | metabolism | Energy produ | COG0056 | Produces ATP | 41431.PCC880 | 2 | 0 | 0,000250466 | 2,65E-05 | 2 | - | 0,000297607 | - | 2 | - | 0,000248807 | - | 2 | - | 0,000250469 | - |
| 479 | 1 | 0.2um_NOKCPHJN_915792 | unclassified | unclassified | unclassified | unclassified | unclassified | unclassified | unclassified | unclassified | unclassified | unclassified | unclassified | unclassified | 0,4 | 0,8 | 0,00015026 | 0,000300521 | 0 | - | 0 | - | 0 | - | 0 | - | 0 | - | 0 | - |
| 480 | 1 | 0.2um_NOKCPHJN_243019 | Eukaryota | Echinodermat | Echinoidea | Echinoida | Strongylocent | Strongylocent | Strongylocent | cellular proces | Cell wall/mem | KOG4177 | Ankyrin Repea | 5722.A2ERS8 | 1 | 0 | 9,54E-05 | 1,01E-05 | 1 | - | 0,000113406 | - | 1 | - | 9,48E-05 | - | 1 | - | 9,54E-05 | - |
| 481 | 1 | 3um_FIJJAFBF_537530 | Eukaryota | Bacillariophyt | Coscinodiscop | Chaetocerotal | Chaetocerota | Chaetoceros | Chaetoceros s | metabolism | Carbohydrate | 0IVA1 | Photosystem I | 35128.Thapsd | 2 | 0 | 0,000940182 | 9,94E-05 | 2 | - | 0,001117137 | - | 2 | - | 0,000933954 | - | 2 | - | 0,000940193 | - |
| 482 | 1 | 3um_FIJJAFBF_653488 | Eukaryota | Bacillariophyt | Bacillariophyc | Bacillariales | Bacillariaceae | Fragilariopsis | Fragilariopsis | poorly charact | Function unkn | COG1100 | GTP-binding P | 2850.Phatr43 | 6 | 0,632455532 | 0,002568483 | 0,000203276 | 5 | - | 0,002563294 | - | 6 | - | 0,002571572 | - | 6 | - | 0,002588751 | - |
| 483 | 1 | 3um_FIJJAFBF_362915 | Bacteria | Proteobacteri | Candidatus La | unclassified | unclassified | unclassified | Candidatus La | cellular proces | Cell wall/mem | 17CVN | Cell envelope | 634499.EpC_2 | 0,2 | 0,4 | 7,15E-05 | 0,000143037 | 0 | - | 0 | - | 0 | - | 0 | - | 0 | - | 0 | - |
| 484 | 1 | 0.2um_NOKCPHJN_1028962 | unclassified | unclassified | unclassified | unclassified | unclassified | unclassified | unclassified | unclassified | unclassified | unclassified | unclassified | unclassified | 0,8 | 0,4 | 0,000215357 | 0,000111378 | 1 | - | 0,000319864 | - | 1 | - | 0,000267414 | - | 0 | - | 0 | - |
| 485 | 1 | 3um_FIJJAFBF_450894 | Bacteria | Bacteroidetes | Flavobacteriia | Flavobacterial | Flavobacteria | Hyunsoonleell | Hyunsoonleell | metabolism | Amino acid tra | 09GJ6 | nitrogen regul | 983548.Krodi_ | 1 | 0 | 0,00056243 | 5,95E-05 | 1 | - | 0,000668287 | - | 1 | - | 0,000558705 | - | 1 | - | 0,000562437 | - |
| 486 | 1 | 3um_FIJJAFBF_71348 | Bacteria | Bacteroidetes | Cytophagia | Cytophagales | Flammeovirga | Marivirga | Marivirga trac | poorly charact | Function unkn | 09AR5 | Inherit from b | 643867.Ftrac_ | 1,6 | 0,489897949 | 0,0002254 | 5,77E-05 | 1 | - | 0,000171277 | - | 1 | - | 0,000143192 | - | 2 | - | 0,000288297 | - |
| 487 | 1 | 3um_FIJJAFBF_00853 | Bacteria | Bacteroidetes | Flavobacteriia | Flavobacterial | Flavobacteria | Polaribacter | Polaribacter s | unclassified | unclassified | unclassified | unclassified | unclassified | 3,8 | 1,166190379 | 0,001836937 | 0,000530201 | 2 | - | 0,001169503 | - | 5 | - | 0,002444333 | - | 5 | - | 0,002460662 | - |
| 488 | 1 | 0.2um_NOKCPHJN_483238 | Eukaryota | Arthropoda | Insecta | Lepidoptera | Pyralidae | Amyelois | Amyelois tran | unclassified | unclassified | unclassified | unclassified | unclassified | 0,6 | 0,489897949 | 2,03E-05 | 1,69E-05 | 1 | - | 4,01E-05 | - | 0 | - | 0 | - | 0 | - | 0 | - |
| 489 | 1 | 3um_FIJJAFBF_518929 | Bacteria | Proteobacteri | Gammaprotec | Alteromonada | Alteromonada | Paraglaciecola | Paraglaciecola | information st | Transcription | 173HH | transcriptiona | 342610.Patl_2 | 1 | 0 | 0,000121372 | 1,28E-05 | 1 | - | 0,000144216 | - | 1 | - | 0,000120568 | - | 1 | - | 0,000121374 | - |
| 490 | 1 | 3um_FIJJAFBF_271734 | Eukaryota | Cryptomycota | unclassified | unclassified | unclassified | Rozella | Rozella allomy | cellular proces | Posttranslatio | 033RP | heat shock | 6945.ISCW017 | 0,2 | 0,4 | 1,68E-05 | 3,35E-05 | 0 | - | 0 | - | 0 | - | 0 | - | 0 | - | 0 | - |
| 491 | 1 | 0.2um_NOKCPHJN_457094 | Bacteria | Bacteroidetes | Flavobacteriia | Flavobacterial | Flavobacteria | unclassified | Flavobacteria | unclassified | unclassified | unclassified | unclassified | unclassified | 0,6 | 0,489897949 | 0,000180555 | 0,000148753 | 1 | - | 0,000337154 | - | 1 | - | 0,000281869 | - | 1 | - | 0,000283752 | - |
| 492 | 1 | 3um_FIJJAFBF_287410 | Bacteria | Proteobacteri | Gammaprotec | Cellvibrionales | Spongiibacter | Oceanicoccus | Oceanicoccus | cellular proces | Intracellular tr | 0QITE | Outer membr | 247633.GP214 | 0,4 | 0,8 | 5,06E-05 | 0,00010118 | 0 | - | 0 | - | 0 | - | 0 | - | 0 | - | 0 | - |
| 493 | 1 | 0.2um_NOKCPHJN_181378 | Eukaryota | unclassified | Bangiophycea | Cyanidiales | Cyanidiaceae | Cyanidioschyz | Cyanidioschyz | unclassified | unclassified | unclassified | unclassified | unclassified | 0,4 | 0,489897949 | 3,45E-06 | 4,24E-06 | 0 | - | 0 | - | 1 | - | 9,22E-06 | - | 0 | - | 0 | - |
| 494 | 1 | 3um_FIJJAFBF_285698 | Eukaryota | Bacillariophyt | Bacillariophyc | Naviculales | Naviculaceae | Fistulifera | Fistulifera sola | unclassified | unclassified | unclassified | unclassified | unclassified | 0,6 | 0,489897949 | 0,000513391 | 0,000420905 | 0 | - | 0 | - | 1 | - | 0,000906883 | - | 0 | - | 0 | - |
| 495 | 1 | 0.2um_NOKCPHJN_315401 | Bacteria | Bacteroidetes | Flavobacteriia | Flavobacterial | Flavobacteria | Lutibacter | Lutibacter pro | poorly charact | Function unkn | 0P88U | NA | 313596.RB250 | 0,6 | 0,489897949 | 6,37E-05 | 5,23E-05 | 0 | - | 0 | - | 1 | - | 0,000110752 | - | 1 | - | 0,000111492 | - |
| 496 | 1 | 3um_FIJJAFBF_702557 | Eukaryota | Bacillariophyt | Coscinodiscop | Chaetocerotal | Chaetocerota | Chaetoceros | Chaetoceros s | information st | Translation, ri | 0FNXB | Protein S19 fo | 65393.PCC742 | 1 | 0 | 0,000684698 | 7,24E-05 | 1 | - | 0,000813567 | - | 1 | - | 0,000680162 | - | 1 | - | 0,000684706 | - |
| 497 | 1 | 3um_FIJJAFBF_135278 | Bacteria | Cyanobacteria | unclassified | Synechococca | Synechococca | Synechococcu | Synechococcu | metabolism | Energy produ | 05CMZ | NDH-1 shuttle | 331869.BAL19 | 1,2 | 0,4 | 0,000231164 | 7,97E-05 | 1 | - | 0,000228894 | - | 1 | - | 0,000191361 | - | 2 | - | 0,000385278 | - |
| 498 | 1 | 0.2um_NOKCPHJN_251837 | Eukaryota | unclassified | Oomycetes | Peronosporale | Peronosporac | Phytophthora | Phytophthora | information st | Translation, ri | 14XM1 | EFG_IV | 4787.PITG_10 | 0,6 | 0,489897949 | 4,06E-05 | 3,31E-05 | 0 | - | 0 | - | 1 | - | 6,84E-05 | - | 1 | - | 6,88E-05 | - |
| 499 | 1 | 3um_FIJJAFBF_375705 | unclassified | unclassified | unclassified | unclassified | unclassified | unclassified | unclassified | unclassified | unclassified | unclassified | unclassified | unclassified | 0,6 | 0,489897949 | 0,000421099 | 0,000347833 | 1 | - | 0,000796257 | - | 0 | - | 0 | - | 1 | - | 0,000670138 | - |
| 500 | 1 | 0.2um_NOKCPHJN_233299 | Bacteria | Proteobacteri | Gammaprotec | Alteromonada | Colwelliaceae | Colwellia | Colwellia sp. P | unclassified | unclassified | unclassified | unclassified | unclassified | 0,2 | 0,4 | 3,91E-05 | 7,82E-05 | 0 | - | 0 | - | 1 | - | 0,000195547 | - | 0 | - | 0 | - |

| | | | | | | | | | | | | | | | | | | | | | | | | |
|---|---|---|---|---|---|---|---|---|---|---|---|---|---|---|---|---|---|---|---|---|---|---|---|---|
| 501 | 1 | 3um_FIJJAFBF_350095 | Bacteria | Proteobacteri | Alphaproteob | Rhodobactera | Rhodobactera | Phaeobacter | Phaeobacter g | information st | Translation, ri | 05KK9 | This protein bi | 292414.TM10 | 1 | 0 | 0,000318142 | 3,36E-05 | 1 - | 0,000378021 - | 1 - | 0,000316035 - | 1 - | 0,000318146 - |
| 502 | 1 | 3um_FIJJAFBF_707424 | Eukaryota | Bacillariophyt | Coscinodiscop | Thalassiosirale | Thalassiosirac | Thalassiosira | Thalassiosira p | cellular proces | Intracellular tr | COG5101 | exportin | 35128.Thaps2 | 0,4 | 0,489897949 | 3,34E-05 | 4,10E-05 | 0 - | 0 - | 0 - | 0 - | 0 - | 0 - |
| 503 | 1 | 0.2um_NOKCPHJN_754361 | Eukaryota | unclassified | Oomycetes | Saprolegniales | Saprolegniace | Achlya | Achlya hypogy | cellular proces | Intracellular tr | 9190 | (GTP)-binding | 578458.XP_00 | 1,8 | 0,979795897 | 0,000576478 | 0,000339431 | 3 - | 0,001128365 - | 1 - | 0,000314447 - | 1 - | 0,000316547 - |
| 504 | 1 | 3um_FIJJAFBF_658500 | unclassified | unclassified | unclassified | unclassified | unclassified | unclassified | unclassified | unclassified | unclassified | unclassified | unclassified | unclassified | 0,8 | 0,4 | 0,000111429 | 5,76E-05 | 1 - | 0,000165228 - | 0 - | 0 - | 1 - | 0,000139057 - |
| 505 | 1 | 3um_FIJJAFBF_76059 | Bacteria | Bacteroidetes | Saprospiria | Saprospirales | Saprospiracea | Aureispira | Aureispira sp. | unclassified | unclassified | unclassified | unclassified | unclassified | 0,6 | 0,489897949 | 6,25E-05 | 5,22E-05 | 1 - | 0,000123512 - | 0 - | 0 - | 0 - | 0 - |
| 506 | 1 | 0.2um_NOKCPHJN_753649 | unclassified | unclassified | unclassified | unclassified | unclassified | unclassified | unclassified | unclassified | unclassified | unclassified | unclassified | unclassified | 0,4 | 0,489897949 | 8,45E-05 | 0,00010374 | 0 - | 0 - | 0 - | 0 - | 0 - | 0 - |
| 507 | 1 | 3um_FIJJAFBF_138631 | unclassified | unclassified | unclassified | unclassified | unclassified | unclassified | unclassified | unclassified | unclassified | unclassified | unclassified | unclassified | 0,4 | 0,489897949 | 8,39E-05 | 0,000103169 | 0 - | 0 - | 1 - | 0,000224283 - | 0 - | 0 - |
| 508 | 1 | 3um_FIJJAFBF_255966 | Bacteria | Candidatus W | unclassified | unclassified | unclassified | unclassified | Candidatus W | poorly charact | Function unkn | 1EB8T | NA | 436017.A4S90 | 1 | 0 | 4,58E-05 | 4,85E-06 | 1 - | 5,45E-05 - | 1 - | 4,55E-05 - | 1 - | 4,58E-05 - |
| 509 | 1 | 3um_FIJJAFBF_254284 | Bacteria | Proteobacteri | Gammaprotec | Oceanospirilla | Oceanospirilla | Marinobacteri | various | information st | Translation, ri | COG0051 | tRNA binding | 876044.IMCC3 | 0,8 | 0,748331477 | 0,000460951 | 0,0004458 | 0 - | 0 - | 2 - | 0,001203364 - | 0 - | 0 - |
| 510 | 2 | 0.2um_NOKCPHJN_385173;3um_FIJJAF | various | various | various | various | various | various | various | unclassified | unclassified | unclassified | unclassified | unclassified | 0,2 | 0,4 | 9,58E-06 | 1,92E-05 | 0 - | 0 - | 1 - | 4,79E-05 - | 0 - | 0 - |
| 511 | 1 | 3um_FIJJAFBF_594156 | Eukaryota | unclassified | Raphidophyce | Chattonellales | Chattonellace | Heterosigma | Heterosigma a | metabolism | Energy produc | COG0055 | Produces ATP | 35128.Thapsd | 7 | 0,632455532 | 0,002594287 | 0,000363777 | 7 - | 0,003081984 - | 6 - | 0,002208527 - | 8 - | 0,002964374 - |
| 512 | 1 | 0.2um_NOKCPHJN_17065 | Bacteria | Bacteroidetes | Flavobacteriia | Flavobacterial | Flavobacteria | Polaribacter | Polaribacter t | poorly charact | Function unkn | 0ZVFQ | Pkd domain cc | 269798.CHU_ | 0,6 | 0,489897949 | 5,63E-06 | 4,65E-06 | 1 - | 1,07E-05 - | 1 - | 8,92E-06 - | 0 - | 0 - |
| 513 | 1 | 0.2um_NOKCPHJN_477124 | Eukaryota | Chordata | Mammalia | Dasyuromorph | Dasyuridae | Sarcophilus | Sarcophilus ha | cellular proces | Cytoskeleton | 0DGS2 | Tubulin is the | 9305.ENSSHA | 0,8 | 0,4 | 0,000105544 | 5,32E-05 | 0 - | 0 - | 1 - | 0,000137527 - | 1 - | 0,000138446 - |
| 514 | 1 | 0.2um_NOKCPHJN_252641 | Bacteria | Proteobacteri | Alphaproteob | Rhodobactera | Rhodobactera | unclassified | Rhodobactera | unclassified | unclassified | unclassified | unclassified | unclassified | 1 | 0,632455532 | 7,08E-05 | 4,49E-05 | 0 - | 0 - | 1 - | 7,38E-05 - | 1 - | 7,43E-05 - |
| 515 | 1 | 3um_FIJJAFBF_149822 | Bacteria | Proteobacteri | Betaproteoba | unclassified | unclassified | unclassified | Betaproteoba | metabolism | Energy produc | COG0056 | Produces ATP | 543728.Vapar | 1 | 0 | 0,000121607 | 1,29E-05 | 1 - | 0,000144495 - | 1 - | 0,000120801 - | 1 - | 0,000121608 - |
| 516 | 1 | 0.2um_NOKCPHJN_539429 | Viruses | unclassified | unclassified | unclassified | unclassified | unclassified | uncultured M | unclassified | unclassified | unclassified | unclassified | unclassified | 0,2 | 0,4 | 5,59E-05 | 0,000111714 | 1 - | 0,000279284 - | 0 - | 0 - | 0 - | 0 - |
| 517 | 1 | 0.2um_NOKCPHJN_285498 | Bacteria | Bacteroidetes | unclassified | unclassified | unclassified | unclassified | Bacteroidetes | poorly charact | Function unkn | 05ERK | gliding motilit | 755732.Fluta_ | 0,2 | 0,4 | 1,88E-05 | 3,77E-05 | 0 - | 0 - | 0 - | 0 - | 1 - | 9,42E-05 - |
| 518 | 1 | 0.2um_NOKCPHJN_351970 | Eukaryota | Ascomycota | Eurotiomycete | Eurotiales | Trichocomace | Talaromyces | Talaromyces a | unclassified | unclassified | unclassified | unclassified | unclassified | 0,4 | 0,489897949 | 1,18E-05 | 1,44E-05 | 0 - | 0 - | 1 - | 3,01E-05 - | 0 - | 0 - |
| 519 | 1 | 3um_FIJJAFBF_284366 | Bacteria | Bacteroidetes | Flavobacteriia | Flavobacterial | Flavobacteria | Formosa | Formosa sp. H | information st | Translation, ri | 05KK9 | This protein bi | 313603.FB217 | 1,2 | 0,748331477 | 0,000423122 | 0,000248048 | 1 - | 0,000425274 - | 1 - | 0,000355539 - | 2 - | 0,000715829 - |
| 520 | 1 | 0.2um_NOKCPHJN_885933 | Eukaryota | Bacillariophyt | Coscinodiscop | Thalassiosirale | Thalassiosirac | Thalassiosira | Thalassiosira c | information st | Translation, ri | COG0251 | Endoribonucle | 35128.Thaps2 | 0,2 | 0,4 | 8,13E-05 | 0,000162595 | 0 - | 0 - | 0 - | 0 - | 0 - | 0 - |
| 521 | 1 | 0.2um_NOKCPHJN_24897 | Bacteria | Proteobacteri | Gammaprotec | Cellvibrionales | Porticoccacea | unclassified | gamma protec | unclassified | unclassified | unclassified | unclassified | unclassified | 0,2 | 0,4 | 5,02E-05 | 0,000100387 | 0 - | 0 - | 0 - | 0 - | 1 - | 0,000250968 - |
| 522 | 1 | 3um_FIJJAFBF_116253 | Eukaryota | unclassified | Oomycetes | Saprolegniales | Saprolegniace | Thraustotheca | Thraustotheca | cellular proces | Intracellular tr | COG5541 | vesicle-mediat | 4787.PITG_00 | 0,4 | 0,489897949 | 0,00036164 | 0,000445851 | 1 - | 0,000984845 - | 1 - | 0,000823354 - | 0 - | 0 - |
| 523 | 1 | 0.2um_NOKCPHJN_505695 | Eukaryota | unclassified | Oomycetes | Albuginales | Albuginaceae | Albugo | Albugo candid | unclassified | unclassified | unclassified | unclassified | unclassified | 0,2 | 0,4 | 1,12E-05 | 2,23E-05 | 0 - | 0 - | 1 - | 5,58E-05 - | 0 - | 0 - |
| 524 | 1 | 3um_FIJJAFBF_652364 | Bacteria | Bacteroidetes | Flavobacteriia | Flavobacterial | Flavobacteria | Lutibacter | Lutibacter aga | poorly charact | Function unkn | 0XRCJ | NA | 391587.KAOT | 0,2 | 0,4 | 1,03E-05 | 2,06E-05 | 0 - | 0 - | 0 - | 0 - | 0 - | 0 - |
| 525 | 1 | 0.2um_NOKCPHJN_1039154 | Bacteria | Actinobacteria | Actinobacteria | Micromonosp | Micromonosp | Salinispora | Salinispora pa | unclassified | unclassified | unclassified | unclassified | unclassified | 0,2 | 0,4 | 2,26E-05 | 4,53E-05 | 0 - | 0 - | 0 - | 0 - | 0 - | 0 - |
| 526 | 1 | 0.2um_NOKCPHJN_667152 | Eukaryota | unclassified | Oomycetes | Peronosporale | Peronosporac | Phytophthora | various | unclassified | unclassified | unclassified | unclassified | unclassified | 0,6 | 0,489897949 | 0,000599664 | 0,000500012 | 1 - | 0,001169503 - | 1 - | 0,000977733 - | 0 - | 0 - |
| 527 | 1 | 0.2um_NOKCPHJN_352637 | Bacteria | Proteobacteri | Gammaprotec | Cellvibrionales | Cellvibrionace | Simiduia | Simiduia agari | metabolism | Inorganic ion t | 16TQC | TONB-depend | 190485.XCC39 | 3,6 | 1,854723699 | 0,000222489 | 0,000116833 | 0 - | 0 - | 4 - | 0,000256981 - | 5 - | 0,000323372 - |
| 528 | 1 | 3um_FIJJAFBF_08764 | Eukaryota | Bacillariophyt | various | various | various | various | various | metabolism | Energy produc | 0J0AS | This b-type cy | 35128.Thapsd | 1,2 | 0,4 | 0,000892944 | 0,000279632 | 1 - | 0,00089105 - | 1 - | 0,00074494 - | 1 - | 0,000749916 - |
| 529 | 1 | 0.2um_NOKCPHJN_681331 | Bacteria | Bacteroidetes | Flavobacteriia | Flavobacterial | Flavobacteria | Salinimicrobiu | Salinimicrobiu | cellular proces | Defense mech | 16WMC | Type III restric | 1002672.SAR1 | 0,4 | 0,489897949 | 2,89E-05 | 3,56E-05 | 1 - | 7,87E-05 - | 1 - | 6,58E-05 - | 0 - | 0 - |
| 530 | 1 | 3um_FIJJAFBF_134225 | Eukaryota | unclassified | Rhodellophyce | Rhodellales | Rhodellaceae | Corynoplastis | Corynoplastis | metabolism | Energy produc | 1DYB8 | PsaA and PsaE | 3055.DAA014 | 0,8 | 0,4 | 6,70E-05 | 3,47E-05 | 1 - | 9,95E-05 - | 1 - | 8,32E-05 - | 0 - | 0 - |
| 531 | 1 | 0.2um_NOKCPHJN_300745 | Bacteria | Firmicutes | Bacilli | Bacillales | Paenibacillace | Paenibacillus | Paenibacillus | poorly charact | Function unkn | COG3236 | Swarming mo | 471881.PROP | 0,2 | 0,4 | 9,80E-06 | 1,96E-05 | 0 - | 0 - | 0 - | 0 - | 1 - | 4,90E-05 - |
| 532 | 1 | 3um_FIJJAFBF_205175 | Eukaryota | Bacillariophyt | Mediophyceae | Cymatosirales | Cymatosiraces | Plagiogrammo | Plagiogrammo | information st | Translation, ri | COG0335 | This protein is | 35128.Thapsd | 0,8 | 0,4 | 0,000424809 | 0,000219202 | 1 - | 0,000623735 - | 1 - | 0,000521458 - | 1 - | 0,000524941 - |
| 533 | 1 | 3um_FIJJAFBF_112857 | unclassified | unclassified | unclassified | unclassified | unclassified | unclassified | unclassified | unclassified | unclassified | unclassified | unclassified | unclassified | 1 | 0 | 0,001105126 | 0,00011687 | 1 - | 0,001313126 - | 1 - | 0,001097806 - | 1 - | 0,001105139 - |
| 534 | 1 | 0.2um_NOKCPHJN_93296 | Bacteria | Proteobacteri | Alphaproteob | Pelagibacteral | Pelagibactera | Candidatus Pe | Candidatus Pe | information st | Replication, re | COG0513 | purine NTP-de | 439493.PB721 | 0,2 | 0,4 | 2,47E-05 | 4,93E-05 | 0 - | 0 - | 0 - | 0 - | 1 - | 0,000123274 - |
| 535 | 1 | 3um_FIJJAFBF_755990 | Eukaryota | Bacillariophyt | Coscinodiscop | Thalassiosirale | Thalassiosirac | Thalassiosira | Thalassiosira p | unclassified | unclassified | unclassified | unclassified | unclassified | 1 | 0 | 0,000288955 | 3,06E-05 | 1 - | 0,00034334 - | 1 - | 0,000287041 - | 1 - | 0,000288958 - |
| 536 | 2 | 0.2um_NOKCPHJN_123927;3um_FIJJAF | Bacteria | Bacteroidetes | Flavobacteriia | Flavobacterial | Flavobacteria | various | various | unclassified | unclassified | unclassified | unclassified | unclassified | 0,2 | 0,4 | 2,99E-05 | 5,98E-05 | 0 - | 0 - | 0 - | 0 - | 0 - | 0 - |
| 537 | 1 | 0.2um_NOKCPHJN_315969 | Eukaryota | unclassified | Oligohymeno | Philasterida | Pseudocohnile | Pseudocohnile | Pseudocohnile | metabolism | Energy produc | 0D9WN | Mitochondrial | 436017.A4S9X | 1 | 0 | 0,000181534 | 1,92E-05 | 1 - | 0,000215701 - | 1 - | 0,000180331 - | 1 - | 0,000181536 - |
| 538 | 1 | 0.2um_NOKCPHJN_78018 | Bacteria | Bacteroidetes | Flavobacteriia | Flavobacterial | Crocinitomica | unclassified | Crocinitomica | unclassified | unclassified | unclassified | unclassified | unclassified | 1 | 0 | 9,05E-05 | 9,57E-06 | 1 - | 0,00010754 - | 1 - | 8,99E-05 - | 1 - | 9,05E-05 - |
| 539 | 1 | 3um_FIJJAFBF_369614 | Bacteria | Bacteroidetes | Flavobacteriia | Flavobacterial | Flavobacteria | Winogradskye | Winogradskye | unclassified | unclassified | unclassified | unclassified | unclassified | 1,4 | 0,8 | 0,000413515 | 0,000234358 | 0 - | 0 - | 1 - | 0,000309777 - | 2 - | 0,000623692 - |
| 540 | 1 | 0.2um_NOKCPHJN_100732 | Bacteria | Bacteroidetes | Flavobacteriia | Flavobacterial | Flavobacteria | Aureitalea | Aureitalea ma | unclassified | unclassified | unclassified | unclassified | unclassified | 2 | 1,095445115 | 0,000135129 | 7,31E-05 | 0 - | 0 - | 3 - | 0,000212358 - | 2 - | 0,000142518 - |
| 541 | 1 | 0.2um_NOKCPHJN_263327 | unclassified | unclassified | unclassified | unclassified | unclassified | unclassified | unclassified | unclassified | unclassified | unclassified | unclassified | unclassified | 0,2 | 0,4 | 3,90E-06 | 7,80E-06 | 0 - | 0 - | 0 - | 0 - | 0 - | 0 - |
| 542 | 1 | 3um_FIJJAFBF_750093 | Eukaryota | Bacillariophyt | Bacillariophyc | Naviculales | Phaeodactyla | Phaeodactylu | Phaeodactylu | unclassified | unclassified | unclassified | unclassified | unclassified | 1,8 | 0,979795897 | 0,000643847 | 0,00030714 | 1 - | 0,000442889 - | 1 - | 0,000370266 - | 1 - | 0,000372739 - |
| 543 | 1 | 3um_FIJJAFBF_366219 | Eukaryota | unclassified | unclassified | Prymnesiales | Chrysochromu | Chrysochromu | Chrysochromu | unclassified | unclassified | unclassified | unclassified | unclassified | 1 | 0 | 0,000828844 | 8,77E-05 | 1 - | 0,000984845 - | 1 - | 0,000823354 - | 1 - | 0,000828854 - |
| 544 | 1 | 3um_FIJJAFBF_744836 | Bacteria | Proteobacteri | Gammaprotec | Cellvibrionales | Spongiibacter | unclassified | marine gamm | information st | Translation, ri | 16Q16 | Binds the lowe | 247639.MGP2 | 0,2 | 0,4 | 5,68E-05 | 0,000113501 | 0 - | 0 - | 0 - | 0 - | 1 - | 0,000283752 - |
| 545 | 1 | 3um_FIJJAFBF_487840 | various | various | various | various | various | various | various | unclassified | unclassified | unclassified | unclassified | unclassified | 0,2 | 0,4 | 7,95E-05 | 0,000159035 | 0 - | 0 - | 0 - | 0 - | 0 - | 0 - |
| 546 | 1 | 3um_FIJJAFBF_269588 | Bacteria | Proteobacteri | Alphaproteob | Rhodobactera | Rhodobactera | unclassified | Rhodobactera | cellular proces | Defense mech | COG0610 | Type I site-spe | 388401.RB215 | 1,2 | 0,4 | 6,77E-05 | 1,67E-05 | 1 - | 6,86E-05 - | 1 - | 5,74E-05 - | 1 - | 5,77E-05 - |
| 547 | 1 | 3um_FIJJAFBF_123244 | Viruses | unclassified | unclassified | unclassified | various | various | various | unclassified | unclassified | unclassified | unclassified | unclassified | 2,6 | 0,489897949 | 0,000307019 | 4,31E-05 | 2 - | 0,000284594 - | 2 - | 0,000237927 - | 3 - | 0,000359275 - |
| 548 | 1 | 0.2um_NOKCPHJN_416399 | Bacteria | Proteobacteri | Betaproteoba | Nitrosomonad | unclassified | unclassified | Methylophilal | metabolism | Energy produc | 05CMV | Succinyl-CoA s | 383631.MB21 | 0,2 | 0,4 | 3,26E-05 | 6,51E-05 | 0 - | 0 - | 0 - | 0 - | 1 - | 0,000162772 - |
| 549 | 1 | 3um_FIJJAFBF_497706 | Bacteria | Proteobacteri | Alphaproteob | unclassified | unclassified | unclassified | Alphaproteob | unclassified | unclassified | unclassified | unclassified | unclassified | 1 | 0 | 0,000140922 | 1,49E-05 | 1 - | 0,000167446 - | 1 - | 0,000139989 - | 1 - | 0,000140924 - |
| 550 | 1 | 0.2um_NOKCPHJN_130645 | Eukaryota | unclassified | Dinophyceae | Suessiales | Symbiodiniace | Symbiodinium | Symbiodinium | poorly charact | Function unkn | 0E9KN | von Willebran | 10090.ENSMU | 0,6 | 0,489897949 | 1,88E-05 | 1,56E-05 | 1 - | 3,66E-05 - | 1 - | 3,06E-05 - | 0 - | 0 - |
| 551 | 1 | 3um_FIJJAFBF_594483 | Bacteria | Bacteroidetes | unclassified | unclassified | unclassified | unclassified | various | unclassified | unclassified | unclassified | unclassified | unclassified | 0,4 | 0,489897949 | 5,46E-05 | 6,69E-05 | 0 - | 0 - | 1 - | 0,000136032 - | 1 - | 0,000136941 - |
| 552 | 1 | 3um_FIJJAFBF_669743 | Bacteria | Bacteroidetes | Flavobacteriia | Flavobacterial | Flavobacteria | Maribacter | Maribacter ar | metabolism | Carbohydrate | 05DEI | Mannitol deh | 487797.Flav3 | 1 | 0 | 0,000127515 | 1,35E-05 | 1 - | 0,000151515 - | 1 - | 0,00012667 - | 1 - | 0,000127516 - |
| 553 | 1 | 3um_FIJJAFBF_372336 | unclassified | unclassified | unclassified | unclassified | unclassified | unclassified | unclassified | unclassified | unclassified | unclassified | unclassified | unclassified | 0,2 | 0,4 | 8,29E-05 | 0,000165761 | 0 - | 0 - | 1 - | 0,000414403 - | 0 - | 0 - |
| 554 | 2 | 0.2um_NOKCPHJN_852052;3um_FIJJAF | Eukaryota | unclassified | unclassified | Isochrysidales | Noelaerhabda | Emiliania | Emiliania huxl | unclassified | unclassified | unclassified | unclassified | unclassified | 1,2 | 0,4 | 0,000297564 | 0,000137389 | 2 - | 0,00057136 - | 1 - | 0,000238836 - | 1 - | 0,000240431 - |
| 555 | 1 | 3um_FIJJAFBF_662957 | Bacteria | Bacteroidetes | Flavobacteriia | Flavobacterial | Crocinitomica | unclassified | Crocinitomica | information st | Translation, ri | 05KK9 | This protein bi | 376686.Fjoh_ | 0,6 | 0,489897949 | 0,000268997 | 0,000220613 | 0 - | 0 - | 0 - | 0 - | 1 - | 0,000477219 - |
| 556 | 1 | 0.2um_NOKCPHJN_82975 | Bacteria | Proteobacteri | Alphaproteob | Pelagibacteral | Pelagibactera | Candidatus Pe | Candidatus Pe | unclassified | unclassified | unclassified | unclassified | unclassified | 0,6 | 0,489897949 | 7,09E-05 | 5,84E-05 | 1 - | 0,000132475 - | 1 - | 0,000110752 - | 1 - | 0,000111492 - |
| 557 | 1 | 3um_FIJJAFBF_110706 | Bacteria | Bacteroidetes | Cytophagia | Cytophagales | Flammeovirga | unclassified | Flammeovirga | metabolism | Inorganic ion t | 05PWN | phosphate ab | 391615.GP501 | 0,6 | 0,8 | 0,000104009 | 0,000143089 | 0 - | 0 - | 0 - | 0 - | 2 - | 0,000363072 - |
| 558 | 1 | 0.2um_NOKCPHJN_850367 | Eukaryota | unclassified | Oomycetes | Peronosporale | Peronosporac | Phytophthora | Phytophthora | cellular proces | Cytoskeleton | KOG0443 | capping prote | 67593.Physo1 | 0,2 | 0,4 | 1,68E-05 | 3,37E-05 | 0 - | 0 - | 1 - | 8,42E-05 - | 0 - | 0 - |
| 559 | 1 | 0.2um_NOKCPHJN_319552 | Eukaryota | Phaeophyceae | unclassified | Ectocarpales | Ectocarpaceae | Ectocarpus | Ectocarpus sili | metabolism | Energy produc | COG0055 | Produces ATP | 35128.Thaps4 | 3,6 | 0,489897949 | 0,000542716 | 9,86E-05 | 4 - | 0,000716251 - | 3 - | 0,000449102 - | 3 - | 0,000452102 - |
| 560 | 1 | 3um_FIJJAFBF_651156 | Eukaryota | Bacillariophyt | Bacillariophyc | Cymbellales | Gomphonema | Didymosphen | Didymosphen | information st | Translation, ri | COG0050 | This protein p | 91464.S7335_ | 0,8 | 0,4 | 0,00026386 | 0,000132983 | 0 - | 0 - | 1 - | 0,000343818 - | 1 - | 0,000346115 - |
| 561 | 1 | 3um_FIJJAFBF_358728 | Bacteria | Proteobacteri | Gammaprotec | unclassified | unclassified | unclassified | gamma protec | unclassified | unclassified | unclassified | unclassified | unclassified | 0,8 | 0,4 | 0,000564971 | 0,000284739 | 0 - | 0 - | 1 - | 0,000736176 - | 1 - | 0,000741093 - |
| 562 | 1 | 3um_FIJJAFBF_271573 | Bacteria | Bacteroidetes | Cytophagia | Cytophagales | Catalimonada | Catalinimonas | Catalinimonas | poorly charact | Function unkn | 087ST | AAA domain ( | 391587.KAOT | 0,2 | 0,4 | 2,80E-05 | 5,60E-05 | 0 - | 0 - | 1 - | 0,000139989 - | 0 - | 0 - |
| 563 | 1 | 0.2um_NOKCPHJN_919228 | Eukaryota | unclassified | Pelagophycea | Pelagomonad | unclassified | Aureococcus | Aureococcus a | metabolism | Energy produc | COG0055 | Produces ATP | 2850.Phatr54 | 1,2 | 0,4 | 0,000833414 | 0,00026099 | 1 - | 0,000831646 - | 1 - | 0,000695277 - | 1 - | 0,000699922 - |
| 564 | 1 | 3um_FIJJAFBF_121956 | Bacteria | Proteobacteri | Alphaproteob | Rhodobactera | Rhodobactera | Yoonia | Yoonia rosea | metabolism | Energy produc | 01QS0 | Aldehyde oxid | 467661.RKLH1 | 0,2 | 0,4 | 1,67E-05 | 3,33E-05 | 0 - | 0 - | 1 - | 8,33E-05 - | 0 - | 0 - |
| 565 | 1 | 3um_FIJJAFBF_296064 | Bacteria | Proteobacteri | Gammaprotec | unclassified | unclassified | unclassified | Gammaprotec | metabolism | Energy produc | COG0056 | Produces ATP | 349521.HCH_ | 0,2 | 0,4 | 2,12E-05 | 4,24E-05 | 0 - | 0 - | 0 - | 0 - | 0 - | 0 - |
| 566 | 1 | 0.2um_NOKCPHJN_718865 | Bacteria | Proteobacteri | Gammaprotec | Cellvibrionales | Porticoccacea | unclassified | gamma protec | poorly charact | Function unkn | 17NHM | NA | 203122.Sde_0 | 1 | 0 | 0,000269197 | 2,85E-05 | 1 - | 0,000319864 - | 1 - | 0,000267414 - | 1 - | 0,000269201 - |
| 567 | 1 | 3um_FIJJAFBF_111979 | Bacteria | Proteobacteri | Gammaprotec | Cellvibrionales | Halieaceae | unclassified | marine gamm | metabolism | Energy produc | COG1290 | Component of | 565045.NOR5 | 0,2 | 0,4 | 1,61E-05 | 3,22E-05 | 0 - | 0 - | 0 - | 0 - | 0 - | 0 - |
| 568 | 1 | 3um_FIJJAFBF_135261 | Bacteria | Cyanobacteria | unclassified | Synechococca | Synechococca | Synechococcu | Synechococcu | unclassified | unclassified | unclassified | unclassified | unclassified | 1 | 0 | 0,000127 | 1,34E-05 | 1 - | 0,000150904 - | 1 - | 0,000126159 - | 1 - | 0,000127002 - |
| 569 | 1 | 3um_FIJJAFBF_190036 | Bacteria | Proteobacteri | Alphaproteob | Rhodobactera | Rhodobactera | Planktomarina | Planktomarina | unclassified | unclassified | unclassified | unclassified | unclassified | 1 | 0 | 0,000155153 | 1,64E-05 | 1 - | 0,000184355 - | 1 - | 0,000154125 - | 1 - | 0,000155155 - |
| 570 | 1 | 0.2um_NOKCPHJN_700301 | Bacteria | Actinobacteria | Thermoleophi | Solirubrobacte | Patulibacterac | Patulibacter | Patulibacter m | poorly charact | Function unkn | 0ER2W | Short-chain de | 643648.Slip_0 | 0,2 | 0,4 | 4,41E-05 | 8,82E-05 | 0 - | 0 - | 0 - | 0 - | 0 - | 0 - |
| 571 | 1 | 3um_FIJJAFBF_507105 | Eukaryota | Phaeophyceae | unclassified | Ectocarpales | Ectocarpaceae | Ectocarpus | Ectocarpus sili | unclassified | unclassified | unclassified | unclassified | unclassified | 1,6 | 0,489897949 | 0,000585958 | 0,000202498 | 2 - | 0,000865297 - | 2 - | 0,000723409 - | 1 - | 0,000364121 - |
| 572 | 1 | 0.2um_NOKCPHJN_438431 | unclassified | unclassified | unclassified | unclassified | unclassified | unclassified | unclassified | unclassified | unclassified | unclassified | unclassified | unclassified | 0,4 | 0,489897949 | 0,00020885 | 0,00025736 | 1 - | 0,000567032 - | 0 - | 0 - | 1 - | 0,000477219 - |
| 573 | 1 | 0.2um_NOKCPHJN_1023097 | Bacteria | Proteobacteri | Alphaproteob | Sphingomona | unclassified | unclassified | Sphingomona | cellular proces | Defense mech | 16Q0U | Type III restric | 391613.RTM1 | 0,2 | 0,4 | 3,00E-05 | 5,99E-05 | 0 - | 0 - | 0 - | 0 - | 0 - | 0 - |
| 574 | 1 | 0.2um_NOKCPHJN_740335 | Bacteria | Bacteroidetes | Flavobacteriia | Flavobacterial | Flavobacteria | Polaribacter | Polaribacter s | unclassified | unclassified | unclassified | unclassified | unclassified | 5,2 | 1,326649916 | 0,000406638 | 8,92E-05 | 3 - | 0,000283874 - | 5 - | 0,000395543 - | 6 - | 0,000477823 - |
| 575 | 1 | 3um_FIJJAFBF_243088 | Eukaryota | Bacillariophyt | Bacillariophyc | Naviculales | Phaeodactyla | Phaeodactylu | Phaeodactylu | unclassified | unclassified | unclassified | unclassified | unclassified | 3 | 0 | 0,00061356 | 6,49E-05 | 3 - | 0,000729041 - | 3 - | 0,000609496 - | 3 - | 0,000613568 - |
| 576 | 1 | 3um_FIJJAFBF_594167 | Eukaryota | unclassified | Pelagophycea | Pelagomonad | unclassified | Aureococcus | Aureococcus a | metabolism | Energy produc | 0FN1Q | F(1)F(0) ATP s | 497965.Cyan7 | 10 | 1,095445115 | 0,003340774 | 0,000242943 | 9 - | 0,003602319 - | 10 - | 0,003346252 - | 10 - | 0,003368606 - |
| 577 | 1 | 3um_FIJJAFBF_520476 | Bacteria | Bacteroidetes | Flavobacteriia | Flavobacterial | Flavobacteria | Polaribacter | Polaribacter s | poorly charact | Function unkn | 11NCV | Protein of unk | 313594.PI23P | 1 | 0 | 0,000406401 | 4,30E-05 | 1 - | 0,000482892 - | 1 - | 0,000403709 - | 1 - | 0,000406406 - |
| 578 | 1 | 3um_FIJJAFBF_128464 | Bacteria | Bacteroidetes | Flavobacteriia | Flavobacterial | Flavobacteria | Polaribacter | Polaribacter s | information st | Translation, ri | COG0052 | 30S ribosomal | 487797.Flav3 | 2 | 1,414213562 | 0,000359676 | 0,000261518 | 0 - | 0 - | 4 - | 0,000747163 - | 1 - | 0,000188039 - |
| 579 | 1 | 3um_FIJJAFBF_754316 | Eukaryota | Phaeophyceae | unclassified | Ectocarpales | Ectocarpaceae | Ectocarpus | Ectocarpus sili | unclassified | unclassified | unclassified | unclassified | unclassified | 1 | 0 | 0,000466609 | 4,93E-05 | 1 - | 0,000554431 - | 1 - | 0,000463518 - | 1 - | 0,000466614 - |
| 580 | 1 | 0.2um_NOKCPHJN_452177 | Bacteria | Proteobacteri | Gammaprotec | unclassified | unclassified | unclassified | various | metabolism | Inorganic ion t | 0QK8T | TonB-depende | 566466.NOR5 | 0,4 | 0,489897949 | 2,57E-05 | 3,15E-05 | 0 - | 0 - | 0 - | 0 - | 0 - | 0 - |
| 581 | 1 | 0.2um_NOKCPHJN_428374 | Viruses | unclassified | unclassified | unclassified | Phycodnavirid | unclassified | Aureococcus a | unclassified | unclassified | unclassified | unclassified | unclassified | 0,6 | 0,8 | 6,84E-05 | 8,62E-05 | 1 - | 0,000139382 - | 0 - | 0 - | 0 - | 0 - |
| 582 | 1 | 0.2um_NOKCPHJN_15784 | Bacteria | Proteobacteri | Gammaprotec | Alteromonada | Alteromonada | Paraglaciecola | Paraglaciecola | poorly charact | Function unkn | 16R8I | Lipoprotein | 1085623.GNIT | 2,4 | 0,489897949 | 0,000161775 | 3,56E-05 | 2 - | 0,000160274 - | 3 - | 0,00020099 - | 3 - | 0,000202333 - |
| 583 | 1 | 3um_FIJJAFBF_531098 | unclassified | unclassified | unclassified | unclassified | unclassified | unclassified | unclassified | unclassified | unclassified | unclassified | unclassified | unclassified | 1 | 0 | 0,000310306 | 3,28E-05 | 1 - | 0,00036871 - | 1 - | 0,000308251 - | 1 - | 0,00031031 - |
| 584 | 1 | 3um_FIJJAFBF_628210 | Bacteria | Bacteroidetes | Cytophagia | Cytophagales | Flammeovirga | Roseivirga | Roseivirga sec | metabolism | Coenzyme tra | 09A4R | May play a ke | 525373.HMPR | 0,8 | 0,4 | 0,000110651 | 5,58E-05 | 0 - | 0 - | 1 - | 0,000144182 - | 1 - | 0,000145145 - |
| 585 | 1 | 0.2um_NOKCPHJN_1081893 | unclassified | unclassified | unclassified | unclassified | unclassified | unclassified | unclassified | unclassified | unclassified | unclassified | unclassified | unclassified | 1 | 0 | 0,000340498 | 3,60E-05 | 1 - | 0,000404585 - | 1 - | 0,000338243 - | 1 - | 0,000340502 - |
| 586 | 1 | 0.2um_NOKCPHJN_27395 | Bacteria | unclassified | unclassified | unclassified | unclassified | unclassified | uncultured ba | unclassified | unclassified | unclassified | unclassified | unclassified | 3,6 | 0,8 | 0,000283562 | 4,87E-05 | 2 - | 0,000191183 - | 4 - | 0,000319668 - | 4 - | 0,000321803 - |
| 587 | 1 | 3um_FIJJAFBF_461082 | Eukaryota | Arthropoda | Arachnida | Trombidiform | Tetranychidae | Tetranychus | Tetranychus u | metabolism | Inorganic ion t | 0DGSY | ATPase Na K t | 9305.ENSSHA | 1,2 | 0,4 | 0,000287417 | 9,91E-05 | 1 - | 0,000284594 - | 1 - | 0,000237927 - | 2 - | 0,000479034 - |
| 588 | 1 | 3um_FIJJAFBF_423716 | Eukaryota | Bacillariophyt | Coscinodiscop | Thalassiosirale | Thalassiosirac | Thalassiosira | Thalassiosira p | unclassified | unclassified | unclassified | unclassified | unclassified | 1 | 0 | 0,000364117 | 3,85E-05 | 1 - | 0,000432648 - | 1 - | 0,000361705 - | 1 - | 0,000364121 - |
| 589 | 2 | 0.2um_NOKCPHJN_602920;3um_FIJJAF | Bacteria | Proteobacteri | Alphaproteob | Rhodobactera | Rhodobactera | various | various | unclassified | unclassified | unclassified | unclassified | unclassified | 0,4 | 0,489897949 | 6,77E-05 | 8,33E-05 | 0 - | 0 - | 0 - | 0 - | 1 - | 0,000181536 - |
| 590 | 1 | 3um_FIJJAFBF_191493 | Bacteria | Bacteroidetes | Flavobacteriia | Flavobacterial | Flavobacteria | Polaribacter | Polaribacter s | unclassified | unclassified | unclassified | unclassified | unclassified | 0,8 | 0,4 | 5,73E-05 | 2,89E-05 | 0 - | 0 - | 1 - | 7,47E-05 - | 1 - | 7,52E-05 - |
| 591 | 1 | 3um_FIJJAFBF_424915 | Eukaryota | Phaeophyceae | unclassified | Ectocarpales | Ectocarpaceae | Ectocarpus | Ectocarpus sili | unclassified | unclassified | unclassified | unclassified | unclassified | 1 | 0 | 0,000777681 | 8,22E-05 | 1 - | 0,000924052 - | 1 - | 0,00077253 - | 1 - | 0,000777691 - |
| 592 | 1 | 3um_FIJJAFBF_85808 | Bacteria | Proteobacteri | Betaproteoba | Burkholderiale | Comamonada | Ramlibacter | Ramlibacter ta | poorly charact | Function unkn | 0XNMM | collagen, type | 45351.NEMVE | 4,6 | 0,489897949 | 0,001127358 | 9,07E-05 | 4 - | 0,001174089 - | 4 - | 0,000981567 - | 5 - | 0,001235156 - |
| 593 | 1 | 3um_FIJJAFBF_43848 | Bacteria | Bacteroidetes | unclassified | unclassified | unclassified | unclassified | Bacteroidetes | unclassified | unclassified | unclassified | unclassified | unclassified | 0,8 | 0,4 | 0,000205553 | 0,000106066 | 1 - | 0,000301807 - | 1 - | 0,000252318 - | 1 - | 0,000254004 - |
| 594 | 1 | 3um_FIJJAFBF_755903 | Bacteria | Proteobacteri | Gammaprotec | Enterobactera | Yersiniaceae | unclassified | Yersiniaceae b | unclassified | unclassified | unclassified | unclassified | unclassified | 0,8 | 0,748331477 | 0,000533962 | 0,000504474 | 0 - | 0 - | 0 - | 0 - | 1 - | 0,000707786 - |
| 595 | 1 | 0.2um_NOKCPHJN_1090264 | Eukaryota | Phaeophyceae | unclassified | Ectocarpales | Ectocarpaceae | Ectocarpus | Ectocarpus sili | cellular proces | Cytoskeleton | 0DFD2 | Kinesin family | 31033.ENSTRU | 1,2 | 0,4 | 0,000180839 | 6,24E-05 | 1 - | 0,000179063 - | 1 - | 0,000149701 - | 2 - | 0,000301402 - |
| 596 | 2 | 0.2um_NOKCPHJN_196290;3um_FIJJAF | Viruses | unclassified | unclassified | unclassified | Phycodnavirid | unclassified | Chrysochromu | information st | Transcription | 1BKE5 | SWIB complex | 3847.GLYMA0 | 2,6 | 0,489897949 | 0,000782569 | 0,000124002 | 2 - | 0,000723171 - | 3 - | 0,000906883 - | 3 - | 0,000912941 - |
| 597 | 1 | 0.2um_NOKCPHJN_36287 | Bacteria | Bacteroidetes | unclassified | unclassified | unclassified | unclassified | Bacteroidetes | metabolism | Energy produc | COG0056 | Produces ATP | 926562.Oweh | 0,2 | 0,4 | 2,40E-05 | 4,81E-05 | 0 - | 0 - | 0 - | 0 - | 1 - | 0,000120216 - |
| 598 | 1 | 0.2um_NOKCPHJN_1062283 | Eukaryota | unclassified | Oomycetes | Peronosporale | Peronosporac | Phytophthora | Phytophthora | unclassified | unclassified | unclassified | unclassified | unclassified | 1,2 | 1,166190379 | 0,000141568 | 0,000132822 | 0 - | 0 - | 1 - | 0,000127965 - | 0 - | 0 - |
| 599 | 1 | 0.2um_NOKCPHJN_612534 | Eukaryota | Chlorophyta | Mamiellophyc | Mamiellales | Bathycoccace | Ostreococcus | Ostreococcus | information st | Translation, ri | KOG0886 | Ribosomal prc | 436017.A4RV | 1 | 0 | 0,000366234 | 3,87E-05 | 1 - | 0,000435164 - | 1 - | 0,000363808 - | 1 - | 0,000366238 - |
| 600 | 1 | 0.2um_NOKCPHJN_530548 | Eukaryota | Bacillariophyt | Coscinodiscop | Thalassiosirale | Thalassiosirac | Thalassiosira | Thalassiosira c | information st | Chromatin str | KOG1744 | Core compone | 2850.Phatr54 | 1,6 | 0,489897949 | 0,000884271 | 0,000322362 | 2 - | 0,001290486 - | 1 - | 0,000539439 - | 2 - | 0,001086085 - |
| 601 | 1 | 3um_FIJJAFBF_118447 | Bacteria | Proteobacteri | Alphaproteob | Rhodobactera | Rhodobactera | unclassified | Rhodobactera | unclassified | unclassified | unclassified | unclassified | unclassified | 1,6 | 0,8 | 0,000215201 | 9,78E-05 | 1 - | 0,000164502 - | 1 - | 0,000137527 - | 1 - | 0,000138446 - |

| | | | | | | | | | | | | | | | | | | | | | | | | |
|---|---|---|---|---|---|---|---|---|---|---|---|---|---|---|---|---|---|---|---|---|---|---|---|---|
| 602 | 1 | 3um_FIJJAFBF_191790 | Eukaryota | Bacillariophyta | Fragilariophyc | Fragilariales | Fragilariaceae | Asterionellops | Asterionellops | metabolism | Energy produc | COG0711 | F(1)F(0) ATP s | 35128.Thapsd | 9,8 | 1,469693846 | 0,003219605 | 0,000287109 | 7 - | 0,002772155 - | 10 - | 0,003310842 - | 11 - | 0,003666256 - |
| 603 | 1 | 3um_FIJJAFBF_277458 | Eukaryota | Bacillariophyta | Coscinodiscop | Thalassiosirale | Thalassiosirac | Thalassiosira | Thalassiosira c | information st | Translation, ri | 082FX | This protein bi | 197221.tll016 | 1 | 0 | 0,000583261 | 6,17E-05 | 1 - | 0,000693039 - | 1 - | 0,000579397 - | 1 - | 0,000583268 - |
| 604 | 1 | 0.2um_NOKCPHJN_676453 | Bacteria | Proteobacteria | Gammaprotec | Pseudomonad | Pseudomonac | Pseudomonas | Pseudomonas | cellular proces | Signal transdu | 07H8Q | Histidine kinas | 216595.PFLU3 | 0,8 | 0,4 | 0,000133929 | 6,82E-05 | 1 - | 0,000192412 - | 1 - | 0,000160861 - | 1 - | 0,000161936 - |
| 605 | 1 | 0.2um_NOKCPHJN_548160 | Eukaryota | Phaeophyceae | unclassified | Ectocarpales | Ectocarpaceae | Ectocarpus | Ectocarpus sili | cellular proces | Signal transdu | 151NR | Pfam:efhand | 67593.Physo1 | 1 | 0 | 0,000386455 | 4,09E-05 | 1 - | 0,000459191 - | 1 - | 0,000383895 - | 1 - | 0,00038646 - |
| 606 | 1 | 3um_FIJJAFBF_248053 | Bacteria | Proteobacteria | Gammaprotec | Pseudomonad | Pseudomonac | Pseudomonas | Pseudomonas | unclassified | unclassified | unclassified | unclassified | unclassified | 0,4 | 0,489897949 | 0,000143103 | 0,000175346 | 0 - | 0 - | 0 - | 0 - | 1 - | 0,000366238 - |
| 607 | 1 | 3um_FIJJAFBF_41560 | Eukaryota | Bacillariophyta | Coscinodiscop | Chaetocerotal | Chaetocerotac | Chaetoceros | Chaetoceros s | metabolism | Energy produc | 0IE88 | oxidoreductas | 45157.CMV13 | 8,6 | 0,8 | 0,000737357 | 8,60E-05 | 8 - | 0,000816897 - | 9 - | 0,000768314 - | 8 - | 0,000687508 - |
| 608 | 2 | 0.2um_NOKCPHJN_160385;3um_FIJJAF | Eukaryota | Phaeophyceae | unclassified | Laminariales | Agaraceae | Thalassiophyll | Thalassiophyll | metabolism | Energy produc | 0IE88 | oxidoreductas | 35128.Thapsd | 7 | 0,632455532 | 0,000598534 | 6,56E-05 | 7 - | 0,000713811 - | 6 - | 0,000511512 - | 7 - | 0,00060075 - |
| 609 | 1 | 3um_FIJJAFBF_27824 | Bacteria | Proteobacteria | Gammaprotec | Alteromonada | Colwelliaceae | Colwellia | various | information st | Transcription | 05VEQ | Cold shock pro | 156578.ATW7 | 0,8 | 0,748331477 | 0,000796241 | 0,000819168 | 2 - | 0,002234274 - | 1 - | 0,000933954 - | 0 - | 0 - |
| 610 | 1 | 0.2um_NOKCPHJN_23471 | Bacteria | Proteobacteria | Gammaprotec | Arenicellales | Arenicellaceae | Arenicella | Arenicella xan | metabolism | Energy produc | 0QJ3X | Aconitate hyd | 878320.AcavA | 2 | 0 | 0,000145815 | 1,54E-05 | 2 - | 0,00017326 - | 2 - | 0,000144849 - | 2 - | 0,000145817 - |
| 611 | 1 | 0.2um_NOKCPHJN_992072 | Eukaryota | Bacillariophyta | Coscinodiscop | Thalassiosirale | Thalassiosirac | Thalassiosira | Thalassiosira c | metabolism | Energy produc | COG0377 | NDH-1 shuttle | 2850.Phatr41 | 2,8 | 0,4 | 0,000852471 | 0,000157077 | 3 - | 0,001084756 - | 2 - | 0,000604589 - | 3 - | 0,000912941 - |
| 612 | 1 | 3um_FIJJAFBF_647442 | Eukaryota | unclassified | Dinophyceae | Gymnodiniales | Gymnodiniace | Cochlodinium | Cochlodinium | unclassified | unclassified | unclassified | unclassified | unclassified | 1 | 0 | 0,000707777 | 7,48E-05 | 1 - | 0,000840991 - | 1 - | 0,000703089 - | 1 - | 0,000707786 - |
| 613 | 1 | 3um_FIJJAFBF_296272 | Bacteria | Proteobacteria | Gammaprotec | Alteromonada | Alteromonada | Glaciecola | Glaciecola sp. | unclassified | unclassified | unclassified | unclassified | unclassified | 0,2 | 0,4 | 2,50E-05 | 5,01E-05 | 0 - | 0 - | 0 - | 0 - | 0 - | 0 - |
| 614 | 1 | 3um_FIJJAFBF_95719 | Bacteria | Cyanobacteria | unclassified | Chroococcales | Cyanobacteria | Cyanobacteriu | Cyanobacteriu | information st | Translation, ri | 05G6W | One of the pri | 197221.tlr014 | 1,4 | 0,489897949 | 0,000440529 | 0,000156525 | 1 - | 0,000374241 - | 2 - | 0,000625749 - | 2 - | 0,000629929 - |
| 615 | 1 | 0.2um_NOKCPHJN_300628 | Bacteria | Proteobacteria | Gammaprotec | unclassified | unclassified | unclassified | Gammaprotec | information st | Translation, ri | 16Q16 | Binds the lowe | 208964.PA425 | 0,8 | 0,4 | 0,000234917 | 0,000121218 | 1 - | 0,000344923 - | 1 - | 0,000288364 - | 1 - | 0,00029029 - |
| 616 | 1 | 3um_FIJJAFBF_808656 | Eukaryota | Bacillariophyta | Bacillariophyc | Naviculales | Phaeodactylac | Phaeodactylum | Phaeodactylum | unclassified | unclassified | unclassified | unclassified | unclassified | 0,2 | 0,4 | 7,11E-05 | 0,00014219 | 0 - | 0 - | 0 - | 0 - | 0 - | 0 - |
| 617 | 1 | 0.2um_NOKCPHJN_787620 | Eukaryota | unclassified | unclassified | Isochrysidales | Noelaerhabda | Emiliania | Emiliania huxl | unclassified | unclassified | unclassified | unclassified | unclassified | 0,4 | 0,489897949 | 0,000143613 | 0,00017659 | 0 - | 0 - | 1 - | 0,000383895 - | 0 - | 0 - |
| 618 | 1 | 0.2um_NOKCPHJN_479664 | Eukaryota | Streptophyta | Bryopsida | Funariales | Funariaceae | Physcomitrella | Physcomitrella | poorly charact | Function unkn | 0NVRZ | NA | 170187.SP_17 | 1 | 0 | 3,32E-05 | 3,52E-06 | 1 - | 3,95E-05 - | 1 - | 3,30E-05 - | 1 - | 3,32E-05 - |
| 619 | 1 | 3um_FIJJAFBF_58647 | Eukaryota | unclassified | Spirotrichea | Sporadotrichic | Oxytrichidae | Stylonychia | Stylonychia le | unclassified | unclassified | unclassified | unclassified | unclassified | 0,2 | 0,4 | 5,99E-05 | 0,000119713 | 0 - | 0 - | 0 - | 0 - | 0 - | 0 - |
| 620 | 1 | 0.2um_NOKCPHJN_224040 | Bacteria | unclassified | unclassified | unclassified | unclassified | unclassified | uncultured ba | metabolism | Inorganic ion t | 05C27 | TonB-depende | 313596.RB250 | 1,2 | 0,4 | 7,10E-05 | 2,42E-05 | 1 - | 7,04E-05 - | 2 - | 0,000117733 - | 1 - | 5,93E-05 - |
| 621 | 1 | 0.2um_NOKCPHJN_410795 | Bacteria | Bacteroidetes | unclassified | unclassified | unclassified | unclassified | Bacteroidetes | poorly charact | Function unkn | 0XQTX | Chromosome | 926562.Oweh | 0,2 | 0,4 | 1,04E-05 | 2,08E-05 | 0 - | 0 - | 0 - | 0 - | 0 - | 0 - |
| 622 | 1 | 3um_FIJJAFBF_707273 | Eukaryota | unclassified | Oomycetes | Peronosporale | Peronosporac | Phytophthora | Phytophthora | unclassified | unclassified | unclassified | unclassified | unclassified | 5,8 | 0,748331477 | 0,000523679 | 4,46E-05 | 5 - | 0,000541593 - | 6 - | 0,000543342 - | 6 - | 0,000546972 - |
| 623 | 1 | 3um_FIJJAFBF_715084 | Eukaryota | Xanthophycea | unclassified | Vaucheriales | Vaucheriaceae | Vaucheria | Vaucheria lito | unclassified | unclassified | unclassified | unclassified | unclassified | 2,6 | 0,489897949 | 0,001222858 | 0,000276897 | 3 - | 0,001675706 - | 2 - | 0,000933954 - | 2 - | 0,000940193 - |
| 624 | 1 | 3um_FIJJAFBF_750048 | Bacteria | Proteobacteria | Alphaproteob | Rhizobiales | Rhizobiaceae | Sinorhizobium | Sinorhizobium | unclassified | unclassified | unclassified | unclassified | unclassified | 0,6 | 0,489897949 | 5,76E-05 | 4,81E-05 | 1 - | 0,000113751 - | 0 - | 0 - | 0 - | 0 - |
| 625 | 1 | 3um_FIJJAFBF_594150 | Eukaryota | unclassified | Pelagophycea | Pelagomonada | unclassified | Aureoumbra | Aureoumbra l | metabolism | Energy produc | COG4451 | RuBisCO catal | 543728.Vapar | 7,4 | 1,743559577 | 0,003277835 | 0,000490429 | 5 - | 0,002692381 - | 8 - | 0,003601434 - | 6 - | 0,00271912 - |
| 626 | 1 | 0.2um_NOKCPHJN_907240 | Eukaryota | Phaeophyceae | unclassified | Ectocarpales | Ectocarpaceae | Ectocarpus | Ectocarpus sili | information st | Replication, re | COG0513 | purine NTP-de | 2850.Phatrdra | 1,2 | 0,748331477 | 0,000138801 | 9,17E-05 | 2 - | 0,000272175 - | 0 - | 0 - | 1 - | 0,000114533 - |
| 627 | 1 | 3um_FIJJAFBF_175545 | Bacteria | Proteobacteria | Gammaprotec | Oceanospirilla | Oceanospirilla | Bacterioplane | Bacterioplane | unclassified | unclassified | unclassified | unclassified | unclassified | 0,2 | 0,4 | 0,000121043 | 0,000242086 | 0 - | 0 - | 0 - | 0 - | 0 - | 0 - |
| 628 | 1 | 0.2um_NOKCPHJN_400172 | Eukaryota | unclassified | Oomycetes | Peronosporale | Peronosporac | Plasmopara | Plasmopara ha | unclassified | unclassified | unclassified | unclassified | unclassified | 0,8 | 0,4 | 2,38E-05 | 1,20E-05 | 0 - | 0 - | 1 - | 3,11E-05 - | 1 - | 3,13E-05 - |
| 629 | 1 | 0.2um_NOKCPHJN_418551 | Eukaryota | Chordata | unclassified | unclassified | Branchiostom | Branchiostom | Branchiostom | metabolism | Secondary me | KOG0054 | ATP-binding ca | 7739.JGI25620 | 1 | 0 | 4,71E-05 | 4,98E-06 | 1 - | 5,60E-05 - | 1 - | 4,68E-05 - | 1 - | 4,71E-05 - |
| 630 | 1 | 0.2um_NOKCPHJN_774908 | Viruses | unclassified | unclassified | unclassified | unclassified | unclassified | uncultured M | unclassified | unclassified | unclassified | unclassified | unclassified | 0,2 | 0,4 | 2,04E-05 | 4,07E-05 | 0 - | 0 - | 0 - | 0 - | 0 - | 0 - |
| 631 | 1 | 0.2um_NOKCPHJN_487588 | Bacteria | Verrucomicrob | unclassified | unclassified | unclassified | unclassified | Verrucomicro | unclassified | unclassified | unclassified | unclassified | unclassified | 0,8 | 0,4 | 0,000438932 | 0,000226975 | 1 - | 0,000650854 - | 0 - | 0 - | 1 - | 0,000547765 - |
| 632 | 1 | 3um_FIJJAFBF_220961 | Bacteria | Proteobacteria | Gammaprotec | Oceanospirilla | Oceanospirilla | Motiliproteus | Motiliproteus | unclassified | unclassified | unclassified | unclassified | unclassified | 0,6 | 0,489897949 | 0,000270412 | 0,000221698 | 0 - | 0 - | 1 - | 0,000477671 - | 0 - | 0 - |
| 633 | 2 | 0.2um_NOKCPHJN_1100070;0.2um_NO | various | various | various | various | various | various | various | unclassified | unclassified | unclassified | unclassified | unclassified | 0,4 | 0,489897949 | 5,07E-05 | 6,23E-05 | 0 - | 0 - | 1 - | 0,000135444 - | 0 - | 0 - |
| 634 | 1 | 0.2um_NOKCPHJN_125506 | Bacteria | Proteobacteria | Alphaproteob | unclassified | unclassified | unclassified | Alphaproteob | metabolism | Secondary me | 0837G | toluene tolera | 137722.AZL_0 | 0,2 | 0,4 | 5,80E-05 | 0,000116088 | 0 - | 0 - | 0 - | 0 - | 0 - | 0 - |
| 635 | 1 | 3um_FIJJAFBF_191789 | Eukaryota | unclassified | Synurophycea | Parmales | Triparmaceae | Triparma | Triparma laevi | metabolism | Energy produc | COG0712 | F(1)F(0) ATP s | 35128.Thapsd | 15 | 0,894427191 | 0,005048838 | 0,000550034 | 14 - | 0,005603607 - | 16 - | 0,005354004 - | 16 - | 0,00538977 - |
| 636 | 1 | 3um_FIJJAFBF_491393 | Eukaryota | unclassified | Oomycetes | Peronosporale | Peronosporac | Phytophthora | Phytophthora | unclassified | unclassified | unclassified | unclassified | unclassified | 0,4 | 0,489897949 | 1,42E-05 | 1,74E-05 | 0 - | 0 - | 0 - | 0 - | 1 - | 3,79E-05 - |
| 637 | 1 | 3um_FIJJAFBF_449836 | Eukaryota | Bacillariophyta | Bacillariophyc | Naviculales | Phaeodactylac | Phaeodactylum | Phaeodactylum | unclassified | unclassified | unclassified | unclassified | unclassified | 5,4 | 0,489897949 | 0,004097282 | 0,000539263 | 5 - | 0,004508927 - | 6 - | 0,004523488 - | 6 - | 0,004553706 - |
| 638 | 1 | 3um_FIJJAFBF_67727 | Bacteria | Proteobacteria | Gammaprotec | Cellvibrionales | unclassified | unclassified | Cellvibrionales | unclassified | unclassified | unclassified | unclassified | unclassified | 0,2 | 0,4 | 7,36E-05 | 0,000147235 | 0 - | 0 - | 1 - | 0,000368088 - | 0 - | 0 - |
| 639 | 1 | 0.2um_NOKCPHJN_899834 | Eukaryota | Basidiomycota | Agaricomycete | Polyporales | Phanerochaet | Phlebiopsis | Phlebiopsis gi | information st | Translation, ri | 14Y9H | Ribosomal Pro | 4787.PITG_06 | 0,8 | 0,4 | 0,000315747 | 0,000160895 | 1 - | 0,000453625 - | 1 - | 0,000379242 - | 1 - | 0,000381775 - |
| 640 | 1 | 0.2um_NOKCPHJN_263318 | Eukaryota | Phaeophyceae | unclassified | Ectocarpales | Ectocarpaceae | Ectocarpus | Ectocarpus sili | unclassified | unclassified | unclassified | unclassified | unclassified | 2 | 0,632455532 | 0,000186976 | 4,32E-05 | 1 - | 0,000114798 - | 2 - | 0,000191948 - | 2 - | 0,00019323 - |
| 641 | 1 | 0.2um_NOKCPHJN_140603 | Bacteria | Bacteroidetes | Flavobacteriia | unclassified | unclassified | unclassified | Flavobacteria | cellular proces | Intracellular tr | 0BIEB | domain-conta | 1005048.CFU_ | 1,4 | 0,489897949 | 8,95E-05 | 3,01E-05 | 1 - | 7,65E-05 - | 2 - | 0,000127965 - | 1 - | 6,44E-05 - |
| 642 | 1 | 3um_FIJJAFBF_388951 | Bacteria | Bacteroidetes | Flavobacteriia | Flavobacterial | Flavobacteriac | Polaribacter | Polaribacter s | metabolism | Energy produc | COG0437 | 4fe-4S ferredo | 313598.MED1 | 1,6 | 0,8 | 9,90E-05 | 4,95E-05 | 1 - | 7,37E-05 - | 3 - | 0,000184768 - | 2 - | 0,000124002 - |
| 643 | 1 | 3um_FIJJAFBF_08084 | Bacteria | Bacteroidetes | Flavobacteriia | Flavobacterial | Flavobacteriac | Polaribacter | Polaribacter s | metabolism | Amino acid tra | 05C65 | Catalyzes the | 313598.MED1 | 0,2 | 0,4 | 2,95E-05 | 5,90E-05 | 0 - | 0 - | 1 - | 0,000147582 - | 0 - | 0 - |
| 644 | 1 | 0.2um_NOKCPHJN_187364 | Bacteria | Proteobacteria | Alphaproteob | Rhodobactera | Hyphomonada | unclassified | Hyphomonada | information st | Translation, ri | COG0617 | Catalyzes the | 755732.Fluta_ | 0,2 | 0,4 | 2,50E-05 | 5,01E-05 | 0 - | 0 - | 1 - | 0,00012515 - | 0 - | 0 - |
| 645 | 1 | 0.2um_NOKCPHJN_766226 | unclassified | unclassified | unclassified | unclassified | unclassified | unclassified | unclassified | unclassified | unclassified | unclassified | unclassified | unclassified | 1 | 0 | 0,000123514 | 1,31E-05 | 1 - | 0,000146761 - | 1 - | 0,000122696 - | 1 - | 0,000123516 - |
| 646 | 1 | 3um_FIJJAFBF_715096 | Eukaryota | unclassified | Pelagophycea | Pelagomonada | unclassified | Pelagomonas | uncultured Pe | metabolism | Energy produc | 0IE88 | oxidoreductas | 35128.Thapsd | 4,6 | 0,8 | 0,000395001 | 8,31E-05 | 5 - | 0,000509865 - | 3 - | 0,000255756 - | 5 - | 0,000429107 - |
| 647 | 1 | 3um_FIJJAFBF_05987 | Eukaryota | Streptophyta | unclassified | Fabales | Fabaceae | Phaseolus | Phaseolus vul | information st | Translation, ri | 0TRRE | ribosomal pro | 4641.GSMUA_ | 0,4 | 0,489897949 | 0,000305453 | 0,000374848 | 0 - | 0 - | 0 - | 0 - | 0 - | 0 - |
| 648 | 1 | 0.2um_NOKCPHJN_918711 | Eukaryota | unclassified | unclassified | Isochrysidales | Noelaerhabda | Emiliania | Emiliania huxl | cellular proces | Posttranslatio | KOG0731 | ATP-depender | 45157.CMQ29 | 3,2 | 0,4 | 0,000332011 | 2,96E-05 | 3 - | 0,000372998 - | 3 - | 0,000311835 - | 3 - | 0,000313918 - |
| 649 | 1 | 0.2um_NOKCPHJN_594863 | Eukaryota | Phaeophyceae | unclassified | Ectocarpales | Ectocarpaceae | Ectocarpus | Ectocarpus sili | unclassified | unclassified | unclassified | unclassified | unclassified | 0,6 | 0,489897949 | 3,22E-05 | 2,66E-05 | 1 - | 6,11E-05 - | 1 - | 5,11E-05 - | 0 - | 0 - |
| 650 | 1 | 3um_FIJJAFBF_63332 | Bacteria | Proteobacteria | Gammaprotec | Alteromonada | Alteromonada | Glaciecola | Glaciecola sp. | metabolism | Inorganic ion t | 0QJVD | TonB-depende | 455436.GHTC | 7 | 1,673320053 | 0,000496415 | 8,66E-05 | 5 - | 0,000429668 - | 5 - | 0,000359213 - | 8 - | 0,00057858 - |
| 651 | 1 | 0.2um_NOKCPHJN_694005 | Eukaryota | unclassified | Bangiophycea | Bangiales | Bangiaceae | Wildemania | Wildemania s | unclassified | unclassified | unclassified | unclassified | unclassified | 1 | 0 | 0,000101437 | 1,07E-05 | 1 - | 0,000120528 - | 1 - | 0,000100765 - | 1 - | 0,000101438 - |
| 652 | 1 | 3um_FIJJAFBF_802634 | Eukaryota | Bacillariophyta | Coscinodiscop | Thalassiosirale | Thalassiosirac | Thalassiosira | Thalassiosira p | cellular proces | Cell wall/mem | COG0438 | Glycosyl trans | 35128.Thaps6 | 0,6 | 0,489897949 | 0,000108456 | 8,90E-05 | 0 - | 0 - | 1 - | 0,000188479 - | 1 - | 0,000189738 - |
| 653 | 1 | 0.2um_NOKCPHJN_764447 | Eukaryota | Chordata | Actinopteri | Siluriformes | Ictaluridae | Ictalurus | Ictalurus punc | unclassified | unclassified | unclassified | unclassified | unclassified | 0,6 | 0,489897949 | 0,000117288 | 9,63E-05 | 0 - | 0 - | 1 - | 0,000203827 - | 1 - | 0,000205189 - |
| 654 | 1 | 0.2um_NOKCPHJN_387207 | Eukaryota | unclassified | unclassified | unclassified | unclassified | Blastocystis | Blastocystis h | poorly charact | Function unkn | 0Y9AP | NA | 65071.PYU1_T | 0,6 | 0,489897949 | 3,89E-05 | 3,18E-05 | 0 - | 0 - | 1 - | 6,55E-05 - | 1 - | 6,60E-05 - |
| 655 | 1 | 3um_FIJJAFBF_95703 | Eukaryota | unclassified | Pelagophycea | Pelagomonada | unclassified | Pelagomonas | uncultured Pe | information st | Translation, ri | COG0051 | tRNA binding | 123214.PERM | 0,8 | 0,4 | 0,000432635 | 0,000218043 | 0 - | 0 - | 1 - | 0,000563738 - | 1 - | 0,000567504 - |
| 656 | 1 | 0.2um_NOKCPHJN_525304 | Eukaryota | Annelida | Polychaeta | Capitellida | Capitellidae | Capitella | Capitella telet | cellular proces | Signal transdu | KOG0613 | fibronectin typ | 7070.TC00296 | 0,2 | 0,4 | 8,20E-06 | 1,64E-05 | 0 - | 0 - | 0 - | 0 - | 0 - | 0 - |
| 657 | 1 | 3um_FIJJAFBF_344571 | Bacteria | various | various | various | various | various | various | metabolism | Amino acid tra | 0FNCB | carbon dioxide | 84588.SYNW1 | 1,2 | 0,748331477 | 0,000693509 | 0,000425307 | 0 - | 0 - | 2 - | 0,001215047 - | 1 - | 0,000611582 - |
| 658 | 1 | 3um_FIJJAFBF_346921 | unclassified | unclassified | unclassified | unclassified | unclassified | unclassified | unclassified | unclassified | unclassified | unclassified | unclassified | unclassified | 1 | 0 | 0,000422766 | 4,47E-05 | 1 - | 0,000502337 - | 1 - | 0,000419966 - | 1 - | 0,000422771 - |
| 659 | 1 | 0.2um_NOKCPHJN_413933 | Eukaryota | Streptophyta | unclassified | Fagales | Fagaceae | Quercus | Quercus sube | unclassified | unclassified | unclassified | unclassified | unclassified | 0,8 | 0,4 | 2,54E-05 | 1,31E-05 | 1 - | 3,72E-05 - | 1 - | 3,11E-05 - | 1 - | 3,13E-05 - |
| 660 | 1 | 0.2um_NOKCPHJN_909475 | Bacteria | Verrucomicrob | Verrucomicrob | Verrucomicrob | Rubritaleacea | Rubritalea | Rubritalea ma | poorly charact | Function unkn | 129P6 | NA | 644107.SL115 | 0,2 | 0,4 | 4,30E-05 | 8,60E-05 | 1 - | 0,000215081 - | 0 - | 0 - | 0 - | 0 - |
| 661 | 1 | 3um_FIJJAFBF_360602 | Bacteria | Tenericutes | Mollicutes | Mycoplasmata | Mycoplasmata | Mycoplasma | Mycoplasma h | poorly charact | Function unkn | 03BSV | Inherit from e | 7070.TC01476 | 0,4 | 0,489897949 | 2,09E-05 | 2,56E-05 | 0 - | 0 - | 0 - | 0 - | 1 - | 5,34E-05 - |
| 662 | 1 | 0.2um_NOKCPHJN_1115200 | Eukaryota | Chlorophyta | Chlorophyceae | Chlamydomor | Chlamydomor | Chlamydomor | Chlamydomor | information st | Translation, ri | KOG3181 | ribosomal pro | 4558.Sb02g03 | 2,6 | 0,489897949 | 0,000740373 | 0,000175572 | 3 - | 0,001011462 - | 3 - | 0,000845607 - | 2 - | 0,000567504 - |
| 663 | 1 | 0.2um_NOKCPHJN_996159 | Eukaryota | Chlorophyta | Trebouxiophy | Chlorellales | Chlorellaceae | Helicosporidiu | Helicosporidiu | cellular proces | Posttranslatio | KOG3083 | prohibitin | 3218.PP1S24_ | 0,8 | 0,4 | 0,000221302 | 0,000111534 | 0 - | 0 - | 1 - | 0,000288364 - | 1 - | 0,00029029 - |
| 664 | 1 | 3um_FIJJAFBF_111294 | Viruses | unclassified | unclassified | unclassified | Mimiviridae | unclassified | Yellowstone la | unclassified | unclassified | unclassified | unclassified | unclassified | 0,2 | 0,4 | 6,81E-05 | 0,000136173 | 0 - | 0 - | 0 - | 0 - | 0 - | 0 - |
| 665 | 1 | 0.2um_NOKCPHJN_14938 | Bacteria | Bacteroidetes | Flavobacteriia | Flavobacterial | Flavobacteriac | Kordia | Kordia jejudor | unclassified | unclassified | unclassified | unclassified | unclassified | 1 | 0 | 9,84E-05 | 1,04E-05 | 1 - | 0,00011695 - | 1 - | 9,78E-05 - | 1 - | 9,84E-05 - |
| 666 | 1 | 0.2um_NOKCPHJN_440251 | Bacteria | Bacteroidetes | Flavobacteriia | Flavobacterial | Flavobacteriac | Flaviramulus | Flaviramulus b | metabolism | Amino acid tra | COG0473 | 3-isopropylma | 714943.Mucp | 1 | 0 | 0,000178955 | 1,89E-05 | 1 - | 0,000212637 - | 1 - | 0,00017777 - | 1 - | 0,000178957 - |
| 667 | 1 | 0.2um_NOKCPHJN_742903 | Bacteria | Bacteroidetes | Flavobacteriia | Flavobacterial | Flavobacteriac | Polaribacter | Polaribacter s | metabolism | Carbohydrate | 0PEN0 | Carbohydrate | 313594.PI23P | 0,4 | 0,489897949 | 5,86E-05 | 7,19E-05 | 0 - | 0 - | 0 - | 0 - | 0 - | 0 - |
| 668 | 1 | 3um_FIJJAFBF_390511 | Bacteria | Bacteroidetes | Flavobacteriia | Flavobacterial | Flavobacteriac | Leeuwenhoek | Leeuwenhoek | unclassified | unclassified | unclassified | unclassified | unclassified | 0,2 | 0,4 | 2,72E-05 | 5,43E-05 | 0 - | 0 - | 0 - | 0 - | 0 - | 0 - |
| 669 | 1 | 3um_FIJJAFBF_187826 | Bacteria | Bacteroidetes | Flavobacteriia | Flavobacterial | Flavobacteriac | Polaribacter | various | information st | Translation, ri | 0PAE2 | One of the pri | 313594.PI23P | 0,4 | 0,489897949 | 0,000269517 | 0,000330748 | 0 - | 0 - | 0 - | 0 - | 0 - | 0 - |
| 670 | 1 | 0.2um_NOKCPHJN_642351 | Eukaryota | unclassified | Dinophyceae | Suessiales | Symbiodiniace | Symbiodinium | Symbiodinium | metabolism | Amino acid tra | 05C6M | Alpha-keto-be | 500637.PROV | 2 | 0 | 0,000591476 | 6,26E-05 | 2 - | 0,0007028 - | 2 - | 0,000587558 - | 2 - | 0,000591483 - |
| 671 | 1 | 0.2um_NOKCPHJN_955130 | Eukaryota | Bacillariophyta | Bacillariophyc | Naviculales | Naviculaceae | Fistulifera | Fistulifera sola | metabolism | Energy produc | COG0056 | Produces ATP | 13037.EHJ650 | 3,2 | 0,4 | 0,0005114 | 8,00E-05 | 3 - | 0,00056991 - | 4 - | 0,000635278 - | 3 - | 0,000479642 - |
| 672 | 1 | 0.2um_NOKCPHJN_64273 | Bacteria | Verrucomicrob | unclassified | unclassified | unclassified | unclassified | Verrucomicro | metabolism | Coenzyme tra | 05DPM | folylpolyglutar | 583355.Caka_ | 0,6 | 0,489897949 | 9,01E-05 | 7,39E-05 | 0 - | 0 - | 0 - | 0 - | 1 - | 0,000159881 - |
| 673 | 1 | 3um_FIJJAFBF_286087 | Eukaryota | Apicomplexa | Aconoidasida | Haemosporida | Plasmodiidae | Plasmodium | Plasmodium r | unclassified | unclassified | unclassified | unclassified | unclassified | 0,4 | 0,489897949 | 6,90E-05 | 8,47E-05 | 0 - | 0 - | 0 - | 0 - | 0 - | 0 - |
| 674 | 1 | 0.2um_NOKCPHJN_369494 | Eukaryota | Echinodermat | Echinoidea | Echinoida | Strongylocent | Strongylocent | Strongylocent | cellular proces | Posttranslatio | KOG0019 | heat shock pro | 281687.CJA14 | 1 | 0 | 7,54E-05 | 7,98E-06 | 1 - | 8,96E-05 - | 1 - | 7,49E-05 - | 1 - | 7,54E-05 - |
| 675 | 1 | 3um_FIJJAFBF_145432 | unclassified | unclassified | unclassified | unclassified | unclassified | unclassified | unclassified | unclassified | unclassified | unclassified | unclassified | unclassified | 0,2 | 0,4 | 0,000115892 | 0,000231785 | 0 - | 0 - | 0 - | 0 - | 0 - | 0 - |
| 676 | 1 | 3um_FIJJAFBF_686924 | Eukaryota | Eustigmatoph | unclassified | Eustigmatales | Monodopsida | Nannochlorop | Nannochlorop | unclassified | unclassified | unclassified | unclassified | unclassified | 0,8 | 0,4 | 0,000372268 | 0,000187619 | 0 - | 0 - | 1 - | 0,000485077 - | 1 - | 0,000488317 - |
| 677 | 1 | 0.2um_NOKCPHJN_747889 | Bacteria | Proteobacteria | Gammaprotec | Cellvibrionales | Spongiibacter | unclassified | marine gamm | metabolism | Inorganic ion t | 0QK8T | TonB-depende | 247633.GP214 | 0,4 | 0,489897949 | 2,74E-05 | 3,36E-05 | 0 - | 0 - | 0 - | 0 - | 1 - | 7,01E-05 - |
| 678 | 1 | 0.2um_NOKCPHJN_505398 | unclassified | unclassified | unclassified | unclassified | unclassified | unclassified | unclassified | unclassified | unclassified | unclassified | unclassified | unclassified | 0,6 | 0,489897949 | 0,000715772 | 0,0005897 | 1 - | 0,001336575 - | 1 - | 0,001117409 - | 1 - | 0,001124874 - |
| 679 | 1 | 0.2um_NOKCPHJN_532535 | Eukaryota | Chordata | Chondrichthye | Orectolobiforr | Rhincodontida | Rhincodon | Rhincodon typ | unclassified | unclassified | unclassified | unclassified | unclassified | 0,4 | 0,489897949 | 0,000128433 | 0,00015834 | 1 - | 0,000349758 - | 1 - | 0,000292406 - | 0 - | 0 - |
| 680 | 1 | 3um_FIJJAFBF_280837 | Bacteria | Bacteroidetes | Flavobacteriia | Flavobacterial | Crocinitomica | unclassified | Crocinitomica | information st | Translation, ri | COG0184 | Forms an inter | 398720.MED2 | 0,4 | 0,489897949 | 0,000257404 | 0,000315883 | 0 - | 0 - | 0 - | 0 - | 0 - | 0 - |
| 681 | 1 | 3um_FIJJAFBF_47498 | Bacteria | Bacteroidetes | unclassified | unclassified | unclassified | unclassified | unidentified e | information st | Replication, re | 09GNB | DNA-binding p | 50743.SCB49_ | 3,8 | 0,4 | 0,002659668 | 0,000407469 | 4 - | 0,003326586 - | 4 - | 0,002781107 - | 3 - | 0,002099765 - |
| 682 | 1 | 0.2um_NOKCPHJN_636600 | Bacteria | Fusobacteria | Fusobacteriia | Fusobacteriale | Fusobacteriac | Fusobacterium | Fusobacterium | unclassified | unclassified | unclassified | unclassified | unclassified | 0,4 | 0,489897949 | 4,25E-05 | 5,31E-05 | 1 - | 0,000122903 - | 0 - | 0 - | 0 - | 0 - |
| 683 | 1 | 0.2um_NOKCPHJN_684944 | Eukaryota | unclassified | Cryptophyta | Pyrenomonad | Geminigerace | Guillardia | Guillardia thet | unclassified | unclassified | unclassified | unclassified | unclassified | 0,4 | 0,489897949 | 0,000114749 | 0,000141099 | 0 - | 0 - | 1 - | 0,00030674 - | 0 - | 0 - |
| 684 | 1 | 3um_FIJJAFBF_13125 | Eukaryota | Bacillariophyta | Coscinodiscop | Chaetocerotal | Acanthocerata | Acanthoceras | Acanthoceras | metabolism | Energy produc | 0IFBF | One of the tw | 45157.CMV08 | 1,6 | 0,489897949 | 0,000292 | 0,000105988 | 2 - | 0,000426485 - | 2 - | 0,000356552 - | 1 - | 0,000179467 - |
| 685 | 1 | 0.2um_NOKCPHJN_222208 | Eukaryota | Phaeophyceae | unclassified | Ectocarpales | Ectocarpaceae | Ectocarpus | Ectocarpus sili | unclassified | unclassified | unclassified | unclassified | unclassified | 0,8 | 0,4 | 1,14E-05 | 5,88E-06 | 1 - | 1,69E-05 - | 0 - | 0 - | 1 - | 1,42E-05 - |
| 686 | 1 | 0.2um_NOKCPHJN_221762 | Bacteria | Proteobacteria | Gammaprotec | unclassified | unclassified | unclassified | Gammaprotec | unclassified | unclassified | unclassified | unclassified | unclassified | 1,6 | 0,489897949 | 0,000235645 | 6,04E-05 | 1 - | 0,000179063 - | 1 - | 0,000149701 - | 2 - | 0,000301402 - |
| 687 | 1 | 3um_FIJJAFBF_738739 | Eukaryota | Chordata | unclassified | unclassified | Branchiostom | Branchiostom | Branchiostom | cellular proces | Signal transdu | 1DF4H | Low density li | 9031.ENSGAL | 1 | 0,632455532 | 0,000121287 | 6,97E-05 | 1 - | 0,000147921 - | 0 - | 0 - | 1 - | 0,000124492 - |
| 688 | 1 | 0.2um_NOKCPHJN_812308 | unclassified | unclassified | unclassified | unclassified | unclassified | unclassified | unclassified | unclassified | unclassified | unclassified | unclassified | unclassified | 1 | 0 | 0,000538395 | 5,69E-05 | 1 - | 0,000639728 - | 1 - | 0,000534828 - | 1 - | 0,000538401 - |
| 689 | 1 | 0.2um_NOKCPHJN_650697 | Eukaryota | unclassified | Oomycetes | Saprolegniales | Saprolegniace | Saprolegnia | Saprolegnia di | metabolism | Coenzyme tra | COG0214 | Involved in the | 65071.PYU1_T | 0,4 | 0,489897949 | 8,18E-05 | 0,000100178 | 0 - | 0 - | 1 - | 0,000208583 - | 0 - | 0 - |
| 690 | 1 | 3um_FIJJAFBF_65171 | Bacteria | Bacteroidetes | Flavobacteriia | Flavobacterial | Flavobacteriac | unclassified | Flavobacteriac | unclassified | unclassified | unclassified | unclassified | unclassified | 0,6 | 0,489897949 | 0,00031865 | 0,000261563 | 0 - | 0 - | 1 - | 0,00055376 - | 1 - | 0,00055746 - |
| 691 | 1 | 0.2um_NOKCPHJN_724888 | Eukaryota | Bacillariophyta | Mediophyceae | Lithodesmiale | Lithodesmiace | Ditylum | Ditylum bright | metabolism | Carbohydrate | COG0057 | glyceraldehyd | 2850.Phatr22 | 0,6 | 0,489897949 | 0,000101551 | 8,48E-05 | 1 - | 0,000200665 - | 0 - | 0 - | 0 - | 0 - |
| 692 | 1 | 3um_FIJJAFBF_423475 | Eukaryota | Bacillariophyta | Coscinodiscop | Thalassiosirale | Thalassiosirac | Thalassiosira | Thalassiosira c | unclassified | unclassified | unclassified | unclassified | unclassified | 2 | 0 | 0,00019442 | 2,06E-05 | 2 - | 0,000231013 - | 2 - | 0,000193132 - | 2 - | 0,000194423 - |
| 693 | 1 | 0.2um_NOKCPHJN_575859 | Bacteria | Proteobacteria | Alphaproteob | Pelagibacteral | Pelagibactera | unclassified | Pelagibactera | poorly charact | Function unkn | 0YZNV | NA | 93059.P9211_ | 1 | 0 | 7,50E-05 | 7,93E-06 | 1 - | 8,91E-05 - | 1 - | 7,45E-05 - | 1 - | 7,50E-05 - |
| 694 | 1 | 3um_FIJJAFBF_788036 | Bacteria | Proteobacteria | Alphaproteob | Rhodobactera | Rhodobactera | Pseudophaeol | Pseudophaeol | unclassified | unclassified | unclassified | unclassified | unclassified | 0,8 | 0,979795897 | 0,000375557 | 0,000460879 | 0 - | 0 - | 0 - | 0 - | 0 - | 0 - |
| 695 | 1 | 3um_FIJJAFBF_387035 | Bacteria | Proteobacteria | Alphaproteob | unclassified | unclassified | unclassified | Alphaproteob | unclassified | unclassified | unclassified | unclassified | unclassified | 5,4 | 0,489897949 | 0,001149146 | 0,000211707 | 6 - | 0,00150701 - | 6 - | 0,001259898 - | 5 - | 0,001056928 - |
| 696 | 1 | 3um_FIJJAFBF_52276 | Bacteria | Proteobacteria | Alphaproteob | Rhodobactera | unclassified | unclassified | Rhodobactera | metabolism | Secondary me | 01Q86 | extracellular s | 644107.SL115 | 1 | 0,894427191 | 0,000168854 | 0,000143966 | 1 - | 0,000207912 - | 0 - | 0 - | 0 - | 0 - |
| 697 | 1 | 3um_FIJJAFBF_190021 | Bacteria | Proteobacteria | Gammaprotec | Cellvibrionales | Spongiibacter | various | various | metabolism | Amino acid tra | 0RATG | O-acetylhomo | 400668.Mmw | 0,4 | 0,489897949 | 5,55E-05 | 6,83E-05 | 0 - | 0 - | 0 - | 0 - | 1 - | 0,000148919 - |
| 698 | 1 | 0.2um_NOKCPHJN_428163 | Eukaryota | unclassified | Oomycetes | Peronosporale | Peronosporac | Phytophthora | Phytophthora | metabolism | Amino acid tra | KOG0540 | carboxylase, b | 164328.Phyra | 0,2 | 0,4 | 2,29E-05 | 4,58E-05 | 0 - | 0 - | 0 - | 0 - | 1 - | 0,000114533 - |
| 699 | 1 | 0.2um_NOKCPHJN_18132 | Viruses | unclassified | unclassified | unclassified | Phycodnavirid | unclassified | Chrysochromu | information st | Transcription | 1BKE5 | SWIB complex | 3711.Bra0223 | 2 | 0,632455532 | 0,00055666 | 0,00012862 | 1 - | 0,000341773 - | 2 - | 0,00057146 - | 2 - | 0,000575278 - |
| 700 | 1 | 0.2um_NOKCPHJN_91817 | Bacteria | Bacteroidetes | Flavobacteriia | Flavobacterial | Flavobacteriac | Kordia | Kordia jejudor | unclassified | unclassified | unclassified | unclassified | unclassified | 0,4 | 0,489897949 | 5,39E-05 | 6,60E-05 | 0 - | 0 - | 0 - | 0 - | 1 - | 0,00013784 - |
| 701 | 4 | 0.2um_NOKCPHJN_168847;0.2um_NO | Eukaryota | various | various | various | various | various | various | unclassified | unclassified | unclassified | unclassified | unclassified | 0,4 | 0,489897949 | 9,47E-05 | 0,000116172 | 0 - | 0 - | 0 - | 0 - | 0 - | 0 - |
| 702 | 1 | 3um_FIJJAFBF_475014 | Bacteria | Proteobacteria | Gammaprotec | Cellvibrionales | Cellvibrionace | Cellvibrio | Cellvibrio sp. F | cellular proces | Cell wall/mem | 0QTSX | Peptidoglycan | 314287.GB22 | 0,8 | 0,748331477 | 0,000277523 | 0,000247571 | 0 - | 0 - | 1 - | 0,000370266 - | 1 - | 0,000372739 - |

| | | | | | | | | | | | | | | | | | | | | | | | | | | | | | | |
|---|---|---|---|---|---|---|---|---|---|---|---|---|---|---|---|---|---|---|---|---|---|---|---|---|---|---|---|---|---|---|
| 703 | 1 | 0.2um_NOKCPHJN_59638 | Bacteria | Bacteroidetes | Flavobacteriia | Flavobacterial | Crocinitomica | unclassified | Crocinitomica | unclassified | unclassified | unclassified | unclassified | unclassified | 1 | 0 | 9,60E-05 | 1,02E-05 | 1 | - | 0,000114098 | - | 1 | - | 9,54E-05 | - | 1 | - | 9,60E-05 | - |
| 704 | 1 | 0.2um_NOKCPHJN_131710 | Bacteria | Bacteroidetes | unclassified | unclassified | unclassified | unclassified | Bacteroidetes | metabolism | Energy produc | 0NIET | alcohol dehyd | 656519.Halsa_ | 0,2 | 0,4 | 3,16E-05 | 6,32E-05 | 0 | - | 0 | - | 0 | - | 0 | - | 0 | - | 0 | - |
| 705 | 1 | 0.2um_NOKCPHJN_316030 | Viruses | unclassified | unclassified | unclassified | unclassified | unclassified | uncultured M | unclassified | unclassified | unclassified | unclassified | unclassified | 1 | 0,632455532 | 0,000190003 | 0,000109153 | 1 | - | 0,000231728 | - | 0 | - | 0 | - | 1 | - | 0,000195025 | - |
| 706 | 1 | 0.2um_NOKCPHJN_368946 | Bacteria | Candidatus M | unclassified | unclassified | unclassified | unclassified | Candidatus M | information st | Replication, re | COG0210 | helicase | 523850.TON_ | 1 | 0 | 0,000128818 | 1,36E-05 | 1 | - | 0,000153064 | - | 1 | - | 0,000127965 | - | 1 | - | 0,00012882 | - |
| 707 | 1 | 3um_FIJJAFBF_350453 | Eukaryota | Bacillariophyta | Bacillariophyc | Naviculales | Phaeodactylac | Phaeodactylu | Phaeodactylu | unclassified | unclassified | unclassified | unclassified | unclassified | 1 | 0 | 0,00061757 | 6,53E-05 | 1 | - | 0,000733806 | - | 1 | - | 0,00061348 | - | 1 | - | 0,000617578 | - |
| 708 | 1 | 3um_FIJJAFBF_711074 | Bacteria | Bacteroidetes | Flavobacteriia | Flavobacterial | Flavobacteriac | Flaviramulus | Flaviramulus i | unclassified | unclassified | unclassified | unclassified | unclassified | 9,2 | 2,039607805 | 0,000596005 | 8,27E-05 | 6 | - | 0,000472228 | - | 8 | - | 0,000526393 | - | 10 | - | 0,000662386 | - |
| 709 | 1 | 3um_FIJJAFBF_285696 | Eukaryota | Bacillariophyta | Coscinodiscop | Thalassiosirale | Thalassiosirac | Thalassiosira | Thalassiosira c | unclassified | unclassified | unclassified | unclassified | unclassified | 1 | 0 | 0,000543036 | 5,74E-05 | 1 | - | 0,000645243 | - | 1 | - | 0,000539439 | - | 1 | - | 0,000543043 | - |
| 710 | 1 | 0.2um_NOKCPHJN_775606 | Bacteria | Proteobacteria | Alphaproteoba | unclassified | unclassified | Candidatus Pu | Candidatus Pu | unclassified | unclassified | unclassified | unclassified | unclassified | 0,4 | 0,489897949 | 3,57E-05 | 4,37E-05 | 0 | - | 0 | - | 1 | - | 8,89E-05 | - | 1 | - | 8,95E-05 | - |
| 711 | 1 | 3um_FIJJAFBF_507106 | Eukaryota | Bacillariophyta | Coscinodiscop | Thalassiosirale | Thalassiosirac | Thalassiosira | Thalassiosira c | unclassified | unclassified | unclassified | unclassified | unclassified | 3,2 | 0,4 | 0,002245735 | 0,000200057 | 3 | - | 0,002522972 | - | 3 | - | 0,002109267 | - | 3 | - | 0,002123357 | - |
| 712 | 1 | 0.2um_NOKCPHJN_1003860 | unclassified | unclassified | unclassified | unclassified | unclassified | unclassified | unclassified | unclassified | unclassified | unclassified | unclassified | unclassified | 1 | 0 | 0,000109361 | 1,16E-05 | 1 | - | 0,000129945 | - | 1 | - | 0,000108637 | - | 1 | - | 0,000109363 | - |
| 713 | 1 | 0.2um_NOKCPHJN_1082065 | Bacteria | Verrucomicrol | Verrucomicrol | Verrucomicrol | Rubritaleacea | Rubritalea | Rubritalea squ | poorly charact | Function unkn | 1DMX1 | Permease YjgF | 349741.Amuc_ | 0,8 | 0,748331477 | 0,000144001 | 0,000149344 | 2 | - | 0,000407892 | - | 0 | - | 0 | - | 0 | - | 0 | - |
| 714 | 1 | 3um_FIJJAFBF_207484 | unclassified | unclassified | unclassified | unclassified | unclassified | unclassified | unclassified | unclassified | unclassified | unclassified | unclassified | unclassified | 2,8 | 0,4 | 0,000471563 | 4,12E-05 | 2 | - | 0,000405681 | - | 3 | - | 0,000508739 | - | 3 | - | 0,000512138 | - |
| 715 | 1 | 3um_FIJJAFBF_182748 | Viruses | unclassified | unclassified | unclassified | various | various | various | unclassified | unclassified | unclassified | unclassified | unclassified | 1,2 | 0,4 | 0,000397237 | 9,82E-05 | 1 | - | 0,00040241 | - | 1 | - | 0,000336424 | - | 1 | - | 0,000338672 | - |
| 716 | 1 | 3um_FIJJAFBF_412012 | Bacteria | Proteobacteria | Gammaprotec | Alteromonada | Alteromonada | unclassified | Alteromonada | information st | Translation, ri | COG1185 | Involved in mF | 377629.TERTU | 3 | 0,894427191 | 0,000268445 | 7,66E-05 | 2 | - | 0,000214158 | - | 4 | - | 0,000358083 | - | 3 | - | 0,000270356 | - |
| 717 | 1 | 0.2um_NOKCPHJN_236546 | Eukaryota | Phaeophyceae | unclassified | Laminariales | Laminariaceae | Saccharina | Saccharina jap | metabolism | Energy produc | KOG0369 | Catalyzes a 2-s | 65071.PYU1_T | 0,6 | 0,489897949 | 3,02E-05 | 2,48E-05 | 0 | - | 0 | - | 1 | - | 5,24E-05 | - | 1 | - | 5,28E-05 | - |
| 718 | 1 | 0.2um_NOKCPHJN_263907 | Bacteria | Proteobacteria | Alphaproteoba | Rickettsiales | unclassified | unclassified | Rickettsiales b | unclassified | unclassified | unclassified | unclassified | unclassified | 0,6 | 0,489897949 | 8,96E-05 | 7,35E-05 | 0 | - | 0 | - | 1 | - | 0,000155659 | - | 1 | - | 0,000156699 | - |
| 719 | 1 | 0.2um_NOKCPHJN_907916 | Viruses | unclassified | unclassified | unclassified | Circoviridae | unclassified | Avon-Heathco | unclassified | unclassified | unclassified | unclassified | unclassified | 1,8 | 0,748331477 | 0,000412806 | 0,000138074 | 1 | - | 0,000282446 | - | 1 | - | 0,000236132 | - | 2 | - | 0,000475418 | - |
| 720 | 1 | 0.2um_NOKCPHJN_873188 | Eukaryota | Phaeophyceae | unclassified | Ectocarpales | Ectocarpaceae | Ectocarpus | Ectocarpus sili | information st | Translation, ri | COG0052 | 30S ribosomal | 2850.Phatr136 | 1 | 0 | 0,000207895 | 2,20E-05 | 1 | - | 0,000247024 | - | 1 | - | 0,000206518 | - | 1 | - | 0,000207897 | - |
| 721 | 1 | 0.2um_NOKCPHJN_403354 | Eukaryota | Arthropoda | Hexanauplia | Calanoida | Temoridae | Eurytemora | Eurytemora af | unclassified | unclassified | unclassified | unclassified | unclassified | 0,4 | 0,8 | 4,46E-05 | 8,91E-05 | 0 | - | 0 | - | 0 | - | 0 | - | 0 | - | 0 | - |
| 722 | 1 | 0.2um_NOKCPHJN_333033 | Eukaryota | unclassified | Oomycetes | Peronosporale | Peronosporac | Phytophthora | Phytophthora | unclassified | unclassified | unclassified | unclassified | unclassified | 0,4 | 0,489897949 | 7,32E-05 | 8,98E-05 | 0 | - | 0 | - | 0 | - | 0 | - | 0 | - | 0 | - |
| 723 | 1 | 0.2um_NOKCPHJN_341641 | Bacteria | Proteobacteria | Gammaprotec | Cellvibrionales | Porticoccacea | unclassified | SAR92 bacteri | information st | Translation, ri | 05G6W | One of the pri | 314287.GB220 | 1 | 0 | 0,000305787 | 3,23E-05 | 1 | - | 0,000363341 | - | 1 | - | 0,000303762 | - | 1 | - | 0,000305791 | - |
| 724 | 1 | 3um_FIJJAFBF_751786 | Eukaryota | unclassified | Pelagophycea | Pelagomonada | unclassified | Aureococcus | Aureococcus a | unclassified | unclassified | unclassified | unclassified | unclassified | 4,2 | 0,4 | 0,000505505 | 4,04E-05 | 4 | - | 0,000575755 | - | 4 | - | 0,000481346 | - | 4 | - | 0,000484561 | - |
| 725 | 1 | 3um_FIJJAFBF_711075 | Bacteria | Bacteroidetes | Flavobacteriia | Flavobacterial | Flavobacteriac | Flaviramulus | Flaviramulus i | poorly charact | Function unkn | 0P8NI | Inherit from b | 391603.FBALC | 1 | 0 | 0,000156308 | 1,65E-05 | 1 | - | 0,000185728 | - | 1 | - | 0,000155273 | - | 1 | - | 0,00015631 | - |
| 726 | 1 | 3um_FIJJAFBF_549760 | Eukaryota | unclassified | Oomycetes | Saprolegniales | Saprolegniace | Aphanomyces | Aphanomyces | unclassified | unclassified | unclassified | unclassified | unclassified | 0,2 | 0,4 | 3,40E-05 | 6,80E-05 | 0 | - | 0 | - | 1 | - | 0,000170041 | - | 0 | - | 0 | - |
| 727 | 1 | 0.2um_NOKCPHJN_902013 | Eukaryota | unclassified | unclassified | Prymnesiales | Chrysochromu | Chrysochromu | Chrysochromu | unclassified | unclassified | unclassified | unclassified | unclassified | 0,6 | 0,489897949 | 0,000143176 | 0,000117423 | 0 | - | 0 | - | 0 | - | 0 | - | 1 | - | 0,000254004 | - |
| 728 | 1 | 0.2um_NOKCPHJN_280545 | Bacteria | Proteobacteria | Alphaproteoba | Pelagibacteral | Pelagibacterac | unclassified | Pelagibactera | unclassified | unclassified | unclassified | unclassified | unclassified | 0,4 | 0,489897949 | 3,95E-05 | 4,84E-05 | 0 | - | 0 | - | 1 | - | 9,84E-05 | - | 1 | - | 9,90E-05 | - |
| 729 | 1 | 0.2um_NOKCPHJN_1017248 | Eukaryota | unclassified | Spirotrichea | Sporadotrichic | Oxytrichidae | Stylonychia | Stylonychia le | cellular proces | Signal transdu | COG5040 | Tyrosine 3-mc | 412030.XP_00 | 0,8 | 0,4 | 0,000200931 | 0,000101267 | 0 | - | 0 | - | 1 | - | 0,00026182 | - | 1 | - | 0,000263569 | - |
| 730 | 1 | 3um_FIJJAFBF_237053 | Eukaryota | unclassified | Synurophycea | Parmales | Triparmaceae | Triparma | Triparma laevi | metabolism | Amino acid tra | 0FJ71 | Acetolactate s | 329726.AM1_ | 2 | 0,632455532 | 0,000759994 | 0,000313434 | 3 | - | 0,001328666 | - | 1 | - | 0,000370266 | - | 2 | - | 0,000745479 | - |
| 731 | 1 | 3um_FIJJAFBF_93294 | Eukaryota | unclassified | Dinophyceae | Suessiales | Symbiodiniace | Symbiodinium | Symbiodinium | unclassified | unclassified | unclassified | unclassified | unclassified | 0,2 | 0,4 | 1,69E-05 | 3,38E-05 | 0 | - | 0 | - | 0 | - | 0 | - | 0 | - | 0 | - |
| 732 | 1 | 3um_FIJJAFBF_585224 | Bacteria | unclassified | unclassified | unclassified | unclassified | unclassified | uncultured ma | unclassified | unclassified | unclassified | unclassified | unclassified | 0,2 | 0,4 | 3,19E-05 | 6,39E-05 | 0 | - | 0 | - | 0 | - | 0 | - | 0 | - | 0 | - |
| 733 | 1 | 0.2um_NOKCPHJN_189632 | Bacteria | Bacteroidetes | unclassified | unclassified | unclassified | unclassified | Bacteroidetes | unclassified | unclassified | unclassified | unclassified | unclassified | 0,4 | 0,489897949 | 7,35E-05 | 9,00E-05 | 0 | - | 0 | - | 0 | - | 0 | - | 1 | - | 0,000188039 | - |
| 734 | 1 | 0.2um_NOKCPHJN_836267 | unclassified | unclassified | unclassified | unclassified | unclassified | unclassified | unclassified | unclassified | unclassified | unclassified | unclassified | unclassified | 0,6 | 0,8 | 0,000136957 | 0,000186468 | 0 | - | 0 | - | 0 | - | 0 | - | 0 | - | 0 | - |
| 735 | 1 | 3um_FIJJAFBF_738132 | Eukaryota | Phaeophyceae | unclassified | Ectocarpales | Ectocarpaceae | Ectocarpus | Ectocarpus sili | information st | Translation, ri | KOG1242 | GCN1 general | 35128.Thaps2 | 1,6 | 0,489897949 | 0,000360805 | 9,24E-05 | 1 | - | 0,000274169 | - | 1 | - | 0,000229212 | - | 2 | - | 0,000461487 | - |
| 736 | 1 | 3um_FIJJAFBF_344748 | Eukaryota | Bacillariophyta | Coscinodiscop | Coscinodiscale | Coscinodiscac | Coscinodiscus | Coscinodiscus | unclassified | unclassified | unclassified | unclassified | unclassified | 1 | 0 | 0,000353889 | 3,74E-05 | 1 | - | 0,000420495 | - | 1 | - | 0,000351544 | - | 1 | - | 0,000353893 | - |
| 737 | 1 | 3um_FIJJAFBF_18637 | Bacteria | Proteobacteria | Gammaprotec | Alteromonada | Alteromonada | Paraglaciecola | Paraglaciecola | information st | Replication, re | 08IYS | Site-specific ty | 342610.Patl_4 | 0,6 | 0,489897949 | 0,00012083 | 9,92E-05 | 0 | - | 0 | - | 1 | - | 0,000209983 | - | 1 | - | 0,000211386 | - |
| 738 | 1 | 0.2um_NOKCPHJN_669311 | unclassified | unclassified | unclassified | unclassified | unclassified | unclassified | unclassified | unclassified | unclassified | unclassified | unclassified | unclassified | 1 | 0 | 0,000254001 | 2,69E-05 | 1 | - | 0,000301807 | - | 1 | - | 0,000252318 | - | 1 | - | 0,000254004 | - |
| 739 | 1 | 3um_FIJJAFBF_394608 | Eukaryota | Bacillariophyta | Bacillariophyc | Naviculales | Phaeodactylac | Phaeodactylu | Phaeodactylu | cellular proces | Cell wall/mem | KOG0916 | synthase | 65071.PYU1_T | 0,4 | 0,489897949 | 1,48E-05 | 1,81E-05 | 0 | - | 0 | - | 1 | - | 3,69E-05 | - | 1 | - | 3,71E-05 | - |
| 740 | 1 | 3um_FIJJAFBF_20059 | Bacteria | Bacteroidetes | Flavobacteriia | Flavobacterial | Flavobacteriac | Polaribacter | Polaribacter s | unclassified | unclassified | unclassified | unclassified | unclassified | 0,6 | 0,489897949 | 0,000239916 | 0,000196763 | 0 | - | 0 | - | 0 | - | 0 | - | 1 | - | 0,000425628 | - |
| 741 | 1 | 3um_FIJJAFBF_385206 | Bacteria | Proteobacteria | Gammaprotec | Alteromonada | Pseudoalteror | Pseudoalteror | Pseudoalteror | unclassified | unclassified | unclassified | unclassified | unclassified | 0,4 | 0,8 | 0,000149231 | 0,000298462 | 0 | - | 0 | - | 0 | - | 0 | - | 0 | - | 0 | - |
| 742 | 1 | 3um_FIJJAFBF_02965 | Bacteria | Proteobacteria | Gammaprotec | unclassified | unclassified | Thiolapillus | Thiolapillus br | unclassified | unclassified | unclassified | unclassified | unclassified | 0,6 | 0,489897949 | 6,12E-05 | 5,11E-05 | 1 | - | 0,000120918 | - | 0 | - | 0 | - | 0 | - | 0 | - |
| 743 | 1 | 3um_FIJJAFBF_414731 | Bacteria | Bacteroidetes | Flavobacteriia | Flavobacterial | Flavobacteriac | Flavobacterium | Flavobacteriu | unclassified | unclassified | unclassified | unclassified | unclassified | 0,8 | 0,4 | 0,000111986 | 5,79E-05 | 1 | - | 0,000166329 | - | 1 | - | 0,000139055 | - | 0 | - | 0 | - |
| 744 | 1 | 3um_FIJJAFBF_328499 | Bacteria | Bacteroidetes | Flavobacteriia | Flavobacterial | Flavobacteriac | Arenitalea | Arenitalea lute | information st | Translation, ri | 08UHY | One of the pri | 1046627.BZAF | 1,6 | 0,489897949 | 0,000622887 | 0,000157799 | 1 | - | 0,000473723 | - | 2 | - | 0,000792088 | - | 1 | - | 0,000398689 | - |
| 745 | 1 | 0.2um_NOKCPHJN_796646 | Eukaryota | Chlorophyta | Mamiellophyc | Mamiellales | Bathycoccace | Ostreococcus | Ostreococcus | unclassified | unclassified | unclassified | unclassified | unclassified | 0,4 | 0,489897949 | 0,000269457 | 0,000332202 | 1 | - | 0,000733806 | - | 1 | - | 0,00061348 | - | 0 | - | 0 | - |
| 746 | 1 | 3um_FIJJAFBF_549856 | Bacteria | Bacteroidetes | Flavobacteriia | Flavobacterial | Crocinitomica | unclassified | Crocinitomica | unclassified | unclassified | unclassified | unclassified | unclassified | 0,8 | 0,4 | 0,000363807 | 0,000183355 | 0 | - | 0 | - | 1 | - | 0,000474052 | - | 1 | - | 0,000477219 | - |
| 747 | 1 | 3um_FIJJAFBF_545826 | Bacteria | Bacteroidetes | Flavobacteriia | Flavobacterial | Flavobacteriac | Polaribacter | Polaribacter s | cellular proces | Intracellular tr | 0P8JN | The central su | 313598.MED1 | 1,2 | 0,4 | 0,000171408 | 5,91E-05 | 1 | - | 0,000169724 | - | 1 | - | 0,000141893 | - | 2 | - | 0,000285682 | - |
| 748 | 1 | 3um_FIJJAFBF_557513 | Bacteria | Bacteroidetes | Flavobacteriia | Flavobacterial | Flavobacteriac | Tamlana | Tamlana agari | unclassified | unclassified | unclassified | unclassified | unclassified | 2,4 | 0,8 | 0,000470612 | 0,000116299 | 2 | - | 0,00047674 | - | 2 | - | 0,000398566 | - | 2 | - | 0,000401229 | - |
| 749 | 1 | 0.2um_NOKCPHJN_119224 | Bacteria | Actinobacteria | Actinobacteria | Actinomycetal | unclassified | unclassified | Actinomyceta | metabolism | Carbohydrate | 00BJE | Glyceraldehyd | 469371.Tbis_1 | 2 | 0 | 0,000379471 | 4,01E-05 | 2 | - | 0,000450893 | - | 2 | - | 0,000376957 | - | 2 | - | 0,000379476 | - |
| 750 | 1 | 0.2um_NOKCPHJN_1002686 | unclassified | unclassified | unclassified | unclassified | unclassified | unclassified | unclassified | unclassified | unclassified | unclassified | unclassified | unclassified | 1 | 0 | 0,000732467 | 7,75E-05 | 1 | - | 0,000870328 | - | 1 | - | 0,000727615 | - | 1 | - | 0,000732476 | - |
| 751 | 1 | 0.2um_NOKCPHJN_283743 | Bacteria | Proteobacteria | Alphaproteoba | Rhodospirillale | Rhodospirillac | Candidatus En | Candidatus En | unclassified | unclassified | unclassified | unclassified | unclassified | 0,4 | 0,489897949 | 4,88E-05 | 5,99E-05 | 0 | - | 0 | - | 0 | - | 0 | - | 0 | - | 0 | - |
| 752 | 1 | 0.2um_NOKCPHJN_919867 | Bacteria | Proteobacteria | Epsilonproteo | Campylobacte | Helicobactera | Helicobacter | Helicobacter s | unclassified | unclassified | unclassified | unclassified | unclassified | 0,4 | 0,489897949 | 6,31E-05 | 7,74E-05 | 0 | - | 0 | - | 0 | - | 0 | - | 0 | - | 0 | - |
| 753 | 1 | 3um_FIJJAFBF_91471 | Eukaryota | Bacillariophyta | Coscinodiscop | Thalassiosirale | Thalassiosirac | Thalassiosira | Thalassiosira p | metabolism | Inorganic ion t | COG0175 | Reduction of a | 35128.Thaps2 | 0,2 | 0,4 | 2,84E-05 | 5,69E-05 | 0 | - | 0 | - | 1 | - | 0,000142216 | - | 0 | - | 0 | - |
| 754 | 1 | 0.2um_NOKCPHJN_976115 | unclassified | unclassified | unclassified | unclassified | unclassified | unclassified | unclassified | unclassified | unclassified | unclassified | unclassified | unclassified | 0,2 | 0,4 | 7,12E-05 | 0,000142404 | 0 | - | 0 | - | 0 | - | 0 | - | 0 | - | 0 | - |
| 755 | 1 | 3um_FIJJAFBF_810150 | Eukaryota | unclassified | Oomycetes | Pythiales | Pythiaceae | Pythium | Pythium insidi | unclassified | unclassified | unclassified | unclassified | unclassified | 2,4 | 0,8 | 0,000295408 | 8,47E-05 | 1 | - | 0,000150904 | - | 2 | - | 0,000252318 | - | 3 | - | 0,000381006 | - |
| 756 | 1 | 3um_FIJJAFBF_545832 | Bacteria | Bacteroidetes | Flavobacteriia | Flavobacterial | Flavobacteriac | Polaribacter | Polaribacter s | unclassified | unclassified | unclassified | unclassified | unclassified | 0,2 | 0,4 | 7,97E-05 | 0,000159427 | 0 | - | 0 | - | 1 | - | 0,000398566 | - | 0 | - | 0 | - |
| 757 | 2 | 3um_FIJJAFBF_235100;3um_FIJJAFBF_ | Eukaryota | Bacillariophyta | various | various | various | various | various | cellular proces | Signal transdu | COG5040 | Tyrosine 3-mc | various | 2,2 | 0,748331477 | 0,000620372 | 0,000221061 | 2 | - | 0,000662373 | - | 3 | - | 0,000830641 | - | 3 | - | 0,000836189 | - |
| 758 | 1 | 3um_FIJJAFBF_111082 | Bacteria | Proteobacteria | Deltaproteoba | unclassified | unclassified | unclassified | Deltaproteoba | cellular proces | Signal transdu | 0HFDJ | Histidine kinas | 391625.PPSIR | 0,2 | 0,4 | 1,09E-05 | 2,18E-05 | 0 | - | 0 | - | 1 | - | 5,46E-05 | - | 0 | - | 0 | - |
| 759 | 1 | 3um_FIJJAFBF_466108 | unclassified | unclassified | unclassified | unclassified | unclassified | unclassified | unclassified | unclassified | unclassified | unclassified | unclassified | unclassified | 1 | 0 | 0,000129614 | 1,37E-05 | 1 | - | 0,000154009 | - | 1 | - | 0,000128755 | - | 1 | - | 0,000129615 | - |
| 760 | 1 | 0.2um_NOKCPHJN_669120 | Bacteria | Proteobacteria | Alphaproteoba | Pelagibacteral | Pelagibacterac | Candidatus Pe | Candidatus Pe | poorly charact | Function unkn | 01QDW | Protein of unk | 1002672.SAR1 | 1 | 0 | 0,000301398 | 3,19E-05 | 1 | - | 0,000358125 | - | 1 | - | 0,000299402 | - | 1 | - | 0,000301402 | - |
| 761 | 2 | 3um_FIJJAFBF_263771;3um_FIJJAFBF_ | Bacteria | Actinobacteria | unclassified | unclassified | unclassified | unclassified | Actinobacteria | unclassified | unclassified | unclassified | unclassified | unclassified | 0,8 | 0,4 | 0,000390427 | 0,000196771 | 0 | - | 0 | - | 1 | - | 0,000508739 | - | 1 | - | 0,000512138 | - |
| 762 | 1 | 3um_FIJJAFBF_588338 | Bacteria | unclassified | unclassified | unclassified | unclassified | unclassified | uncultured ba | unclassified | unclassified | unclassified | unclassified | unclassified | 1 | 0 | 8,04E-05 | 8,51E-06 | 1 | - | 9,56E-05 | - | 1 | - | 7,99E-05 | - | 1 | - | 8,05E-05 | - |
| 763 | 1 | 3um_FIJJAFBF_191787 | Eukaryota | unclassified | Dictyochophy | Pedinellales | unclassified | Apedinella | Apedinella rad | cellular proces | Posttranslatio | KOG0730 | cell division cy | 45157.CMV01 | 13 | 2,097617696 | 0,002675362 | 0,000287829 | 10 | - | 0,002478417 | - | 11 | - | 0,002279219 | - | 15 | - | 0,003128788 | - |
| 764 | 1 | 3um_FIJJAFBF_257113 | Eukaryota | unclassified | Florideophyce | Nemaliales | Liagoraceae | Yamadaella | Yamadaella ca | information st | Translation, ri | 0FNT6 | One of the ear | 391612.CY011 | 1,6 | 0,8 | 0,000891135 | 0,000429528 | 1 | - | 0,000674308 | - | 3 | - | 0,001691214 | - | 1 | - | 0,000567504 | - |
| 765 | 1 | 3um_FIJJAFBF_201377 | Bacteria | Bacteroidetes | Flavobacteriia | Flavobacterial | Flavobacteriac | Polaribacter | Polaribacter s | unclassified | unclassified | unclassified | unclassified | unclassified | 0,2 | 0,4 | 1,04E-05 | 2,09E-05 | 0 | - | 0 | - | 0 | - | 0 | - | 0 | - | 0 | - |
| 766 | 1 | 3um_FIJJAFBF_405813 | unclassified | unclassified | unclassified | unclassified | unclassified | unclassified | unclassified | unclassified | unclassified | unclassified | unclassified | unclassified | 1 | 0 | 0,000499938 | 5,29E-05 | 1 | - | 0,000594033 | - | 1 | - | 0,000496626 | - | 1 | - | 0,000499944 | - |
| 767 | 1 | 3um_FIJJAFBF_37823 | Bacteria | Actinobacteria | unclassified | unclassified | unclassified | unclassified | Actinobacteria | unclassified | unclassified | unclassified | unclassified | unclassified | 0,4 | 0,489897949 | 4,07E-05 | 4,99E-05 | 0 | - | 0 | - | 0 | - | 0 | - | 0 | - | 0 | - |
| 768 | 1 | 0.2um_NOKCPHJN_477681 | Eukaryota | Nematoda | Enoplea | Trichinellida | Trichinellidae | Trichinella | Trichinella nel | poorly charact | Function unkn | 0MS4I | conserved hyp | 441960.XP_00 | 0,4 | 0,489897949 | 2,99E-05 | 3,70E-05 | 1 | - | 8,29E-05 | - | 0 | - | 0 | - | 0 | - | 0 | - |
| 769 | 1 | 0.2um_NOKCPHJN_369201 | Bacteria | Proteobacteria | Alphaproteoba | Rhodobactera | Rhodobactera | Yoonia | Yoonia vestfol | information st | Transcription | 16QIX | transcriptiona | 391626.OA307 | 0,8 | 0,4 | 0,000138793 | 7,00E-05 | 0 | - | 0 | - | 1 | - | 0,000180852 | - | 1 | - | 0,000182061 | - |
| 770 | 2 | 3um_FIJJAFBF_167713;3um_FIJJAFBF_ | unclassified | unclassified | unclassified | unclassified | unclassified | unclassified | unclassified | unclassified | unclassified | unclassified | unclassified | unclassified | 0,2 | 0,4 | 5,61E-05 | 0,000112242 | 0 | - | 0 | - | 1 | - | 0,000280605 | - | 0 | - | 0 | - |
| 771 | 1 | 0.2um_NOKCPHJN_216066 | Bacteria | Proteobacteria | Deltaproteoba | Desulfobacter | Desulfobacter | Desulfatibacill | Desulfatibacill | poorly charact | Function unkn | 071EU | NA | 439235.Dalk_ | 0,4 | 0,489897949 | 3,91E-05 | 4,84E-05 | 1 | - | 0,000108476 | - | 0 | - | 0 | - | 0 | - | 0 | - |
| 772 | 1 | 0.2um_NOKCPHJN_451306 | Bacteria | Proteobacteria | Alphaproteoba | Pelagibacteral | Pelagibacterac | Candidatus Pe | Candidatus Pe | poorly charact | Function unkn | 179ZR | VWA-like dom | 550540.Fbal_ | 0,2 | 0,4 | 2,85E-05 | 5,71E-05 | 0 | - | 0 | - | 0 | - | 0 | - | 0 | - | 0 | - |
| 773 | 1 | 3um_FIJJAFBF_10545 | Eukaryota | unclassified | Pelagophycea | Pelagomonada | unclassified | Aureococcus | Aureococcus a | information st | Transcription | 12N88 | dna-directed r | 6238.CBG1819 | 0,2 | 0,4 | 9,49E-06 | 1,90E-05 | 0 | - | 0 | - | 0 | - | 0 | - | 0 | - | 0 | - |
| 774 | 1 | 3um_FIJJAFBF_228001 | Eukaryota | Bacillariophyta | Bacillariophyc | Naviculales | Naviculaceae | Fistulifera | Fistulifera sola | unclassified | unclassified | unclassified | unclassified | unclassified | 0,6 | 0,489897949 | 3,46E-05 | 2,86E-05 | 1 | - | 6,56E-05 | - | 1 | - | 5,48E-05 | - | 0 | - | 0 | - |
| 775 | 1 | 0.2um_NOKCPHJN_835463 | Eukaryota | Apicomplexa | Conoidasida | Eucoccidiorida | Eimeriidae | Eimeria | Eimeria mitis | poorly charact | Function unkn | 0NF4H | copper amine | 479436.Vpar_ | 1 | 0 | 0,000108048 | 1,14E-05 | 1 | - | 0,000128385 | - | 1 | - | 0,000107333 | - | 1 | - | 0,00010805 | - |
| 776 | 1 | 0.2um_NOKCPHJN_1094406 | Bacteria | Proteobacteria | Alphaproteoba | Rhodobactera | Rhodobactera | Shimia | Shimia abyssi | unclassified | unclassified | unclassified | unclassified | unclassified | 1 | 0 | 0,00035791 | 3,78E-05 | 1 | - | 0,000425274 | - | 1 | - | 0,000355539 | - | 1 | - | 0,000357914 | - |
| 777 | 1 | 0.2um_NOKCPHJN_38349 | Bacteria | Proteobacteria | Gammaprotec | Pseudomonad | Pseudomonac | Pseudomonas | Pseudomonas | metabolism | Nucleotide tra | 08Z6B | deaminase | 1042876.PPS_ | 0,6 | 0,489897949 | 0,000267098 | 0,000222669 | 1 | - | 0,000519779 | - | 0 | - | 0 | - | 1 | - | 0,000437451 | - |
| 778 | 1 | 0.2um_NOKCPHJN_322238 | Bacteria | Proteobacteria | Betaproteobac | Burkholderiale | Alcaligenacea | Bordetella | Bordetella ger | unclassified | unclassified | unclassified | unclassified | unclassified | 0,6 | 0,489897949 | 0,000364217 | 0,000304195 | 1 | - | 0,000719694 | - | 0 | - | 0 | - | 0 | - | 0 | - |
| 779 | 1 | 0.2um_NOKCPHJN_215618 | Eukaryota | unclassified | Pelagophycea | Pelagomonada | unclassified | Aureococcus | Aureococcus a | unclassified | unclassified | unclassified | unclassified | unclassified | 4,4 | 0,489897949 | 0,000344332 | 4,47E-05 | 4 | - | 0,000372843 | - | 5 | - | 0,000389632 | - | 4 | - | 0,000313788 | - |
| 780 | 1 | 3um_FIJJAFBF_337088 | Bacteria | Proteobacteria | Gammaprotec | Oceanospirilla | Oceanospirilla | Motiliproteus | Motiliproteus | information st | Translation, ri | 177XJ | One of the pri | 205918.Psyr_4 | 0,6 | 0,489897949 | 0,000246552 | 0,000205541 | 1 | - | 0,000479796 | - | 0 | - | 0 | - | 1 | - | 0,000403801 | - |
| 781 | 2 | 0.2um_NOKCPHJN_138709;0.2um_NO | Bacteria | unclassified | unclassified | unclassified | unclassified | unclassified | uncultured ba | unclassified | unclassified | unclassified | unclassified | unclassified | 0,2 | 0,4 | 2,00E-05 | 4,01E-05 | 0 | - | 0 | - | 0 | - | 0 | - | 0 | - | 0 | - |
| 782 | 1 | 0.2um_NOKCPHJN_283237 | Eukaryota | Arthropoda | Insecta | Lepidoptera | Papilionidae | Papilio | Papilio macha | unclassified | unclassified | unclassified | unclassified | unclassified | 0,2 | 0,4 | 9,67E-06 | 1,93E-05 | 0 | - | 0 | - | 0 | - | 0 | - | 1 | - | 4,83E-05 | - |
| 783 | 1 | 0.2um_NOKCPHJN_690840 | Bacteria | Proteobacteria | Alphaproteoba | Rhodobactera | unclassified | unclassified | Rhodobactera | unclassified | unclassified | unclassified | unclassified | unclassified | 0,6 | 0,489897949 | 0,000153223 | 0,000125773 | 0 | - | 0 | - | 1 | - | 0,000266276 | - | 1 | - | 0,000268055 | - |
| 784 | 1 | 0.2um_NOKCPHJN_65240 | Bacteria | Proteobacteria | Gammaprotec | Cellvibrionales | Porticoccacea | unclassified | SAR92 bacteri | unclassified | unclassified | unclassified | unclassified | unclassified | 2,6 | 0,8 | 0,000189556 | 5,13E-05 | 1 | - | 8,92E-05 | - | 3 | - | 0,000223748 | - | 3 | - | 0,000225243 | - |
| 785 | 1 | 0.2um_NOKCPHJN_157068 | Bacteria | unclassified | unclassified | unclassified | unclassified | unclassified | uncultured ba | poorly charact | Function unkn | 0P8NI | Inherit from b | 983544.Lacal_ | 0,8 | 0,748331477 | 0,000114237 | 0,000107928 | 0 | - | 0 | - | 0 | - | 0 | - | 1 | - | 0,000151425 | - |
| 786 | 1 | 3um_FIJJAFBF_126100 | Eukaryota | Bacillariophyta | Coscinodiscop | Thalassiosirale | Thalassiosirac | Thalassiosira | Thalassiosira c | unclassified | unclassified | unclassified | unclassified | unclassified | 6 | 0,894427191 | 0,001779203 | 0,000426215 | 7 | - | 0,002436918 | - | 7 | - | 0,002037323 | - | 6 | - | 0,001757942 | - |
| 787 | 1 | 3um_FIJJAFBF_735995 | Bacteria | Actinobacteria | Actinobacteria | Streptomyceta | Streptomyceta | Streptomyces | Streptomyces | unclassified | unclassified | unclassified | unclassified | unclassified | 0,6 | 0,489897949 | 0,00049316 | 0,000404458 | 0 | - | 0 | - | 0 | - | 0 | - | 1 | - | 0,000874902 | - |
| 788 | 1 | 3um_FIJJAFBF_713545 | Eukaryota | unclassified | unclassified | Prymnesiales | Chrysochromu | Chrysochromu | Chrysochromu | unclassified | unclassified | unclassified | unclassified | unclassified | 0,2 | 0,4 | 9,00E-05 | 0,00017998 | 0 | - | 0 | - | 0 | - | 0 | - | 1 | - | 0,00044995 | - |
| 789 | 1 | 3um_FIJJAFBF_715071 | Eukaryota | unclassified | Florideophyce | Ceramiales | Rhodomelacea | Rhodomela | Rhodomela co | information st | Translation, ri | COG0051 | tRNA binding | 41431.PCC880 | 2,2 | 0,4 | 0,001332528 | 0,000274071 | 2 | - | 0,001439388 | - | 2 | - | 0,001203364 | - | 3 | - | 0,001817104 | - |
| 790 | 1 | 0.2um_NOKCPHJN_86872 | Bacteria | Proteobacteria | Gammaprotec | unclassified | unclassified | unclassified | Gammaprotec | information st | Transcription | 05D27 | DNA-depende | 208964.PA426 | 1,4 | 0,489897949 | 6,45E-05 | 2,79E-05 | 2 | - | 0,000106697 | - | 2 | - | 8,92E-05 | - | 1 | - | 4,49E-05 | - |
| 791 | 1 | 0.2um_NOKCPHJN_563536 | Eukaryota | Phaeophyceae | unclassified | Ectocarpales | Ectocarpaceae | Ectocarpus | Ectocarpus sili | metabolism | Carbohydrate | KOG2638 | UTP-glucose-1 | 653948.CCA18 | 0,8 | 0,4 | 4,76E-05 | 2,46E-05 | 1 | - | 7,07E-05 | - | 1 | - | 5,91E-05 | - | 0 | - | 0 | - |
| 792 | 1 | 0.2um_NOKCPHJN_39203 | Eukaryota | Bacillariophyta | Fragilariophyc | Licmophorales | Licmophorace | Licmophora | Licmophora sp | metabolism | Energy produc | COG1290 | Component of | 35128.Thapsd | 5,8 | 0,4 | 0,001699323 | 0,000219757 | 6 | - | 0,002088787 | - | 6 | - | 0,001746277 | - | 5 | - | 0,001464952 | - |
| 793 | 1 | 3um_FIJJAFBF_813295 | Eukaryota | Bacillariophyta | Bacillariophyc | Bacillariales | Bacillariaceae | Fragilariopsis | Fragilariopsis c | cellular proces | Posttranslatio | KOG0840 | ATP-depender | 2850.Phatr409 | 0,2 | 0,4 | 9,56E-05 | 0,000191121 | 0 | - | 0 | - | 0 | - | 0 | - | 0 | - | 0 | - |
| 794 | 1 | 3um_FIJJAFBF_437251 | Bacteria | Bacteroidetes | Flavobacteriia | Flavobacterial | Flavobacteriac | Flaviramulus | Flaviramulus i | unclassified | unclassified | unclassified | unclassified | unclassified | 0,2 | 0,4 | 1,26E-05 | 2,52E-05 | 0 | - | 0 | - | 0 | - | 0 | - | 1 | - | 6,31E-05 | - |
| 795 | 1 | 3um_FIJJAFBF_502175 | Eukaryota | unclassified | unclassified | Prymnesiales | Chrysochromu | Chrysochromu | Chrysochromu | unclassified | unclassified | unclassified | unclassified | unclassified | 2,8 | 0,4 | 0,001195685 | 0,0002328 | 3 | - | 0,001517193 | - | 3 | - | 0,001268411 | - | 3 | - | 0,001276884 | - |
| 796 | 1 | 0.2um_NOKCPHJN_87772 | Eukaryota | unclassified | Labyrinthulom | unclassified | Thraustochytr | Aurantiochytr | Aurantiochytr | cellular proces | Cytoskeleton | COG5245 | heavy chain | 67593.Physo1 | 0,4 | 0,489897949 | 5,51E-06 | 6,78E-06 | 0 | - | 0 | - | 0 | - | 0 | - | 1 | - | 1,48E-05 | - |
| 797 | 1 | 3um_FIJJAFBF_652999 | Eukaryota | unclassified | unclassified | Isochrysidales | Noelaerhabda | Emiliania | Emiliania huxl | unclassified | unclassified | unclassified | unclassified | unclassified | 0,2 | 0,4 | 7,32E-05 | 0,000146495 | 0 | - | 0 | - | 0 | - | 0 | - | 1 | - | 0,000366238 | - |
| 798 | 1 | 0.2um_NOKCPHJN_304690 | Bacteria | Bacteroidetes | Flavobacteriia | Flavobacterial | Flavobacteriac | Lutibacter | Lutibacter flav | unclassified | unclassified | unclassified | unclassified | unclassified | 0,8 | 0,4 | 0,0001376 | 6,93E-05 | 0 | - | 0 | - | 1 | - | 0,000179298 | - | 1 | - | 0,000180496 | - |
| 799 | 1 | 3um_FIJJAFBF_403521 | Eukaryota | Bacillariophyta | Bacillariophyc | Bacillariales | Bacillariaceae | Fragilariopsis | Fragilariopsis c | unclassified | unclassified | unclassified | unclassified | unclassified | 1 | 0 | 0,000121607 | 1,29E-05 | 1 | - | 0,000144495 | - | 1 | - | 0,000120801 | - | 1 | - | 0,000121608 | - |
| 800 | 1 | 0.2um_NOKCPHJN_474497 | Eukaryota | unclassified | Oomycetes | Saprolegniales | Saprolegniace | Aphanomyces | Aphanomyces | unclassified | unclassified | unclassified | unclassified | unclassified | 0,2 | 0,4 | 8,81E-06 | 1,76E-05 | 0 | - | 0 | - | 1 | - | 4,40E-05 | - | 0 | - | 0 | - |
| 801 | 2 | 0.2um_NOKCPHJN_710197;3um_FIJJAF | Bacteria | various | various | various | various | various | various | unclassified | unclassified | unclassified | unclassified | unclassified | 0,2 | 0,4 | 3,53E-05 | 7,05E-05 | 0 | - | 0 | - | 0 | - | 0 | - | 0 | - | 0 | - |
| 802 | 1 | 3um_FIJJAFBF_126699 | Eukaryota | unclassified | Oomycetes | Peronosporale | Peronosporac | Phytophthora | various | unclassified | unclassified | unclassified | unclassified | unclassified | 1 | 0 | 0,000807592 | 8,54E-05 | 1 | - | 0,000959592 | - | 1 | - | 0,000802243 | - | 1 | - | 0,000807602 | - |
| 803 | 1 | 3um_FIJJAFBF_385889 | Bacteria | Proteobacteria | Alphaproteoba | Rhodospirillale | Rhodospirillac | Oceanibaculu | Oceanibaculu | metabolism | Energy produc | COG0056 | Produces ATP | 414684.RC1_2 | 1 | 0 | 0,000124245 | 1,31E-05 | 1 | - | 0,00014763 | - | 1 | - | 0,000123422 | - | 1 | - | 0,000124246 | - |

| | | | | | | | | | | | | | | | | | | | | | | | | |
|---|---|---|---|---|---|---|---|---|---|---|---|---|---|---|---|---|---|---|---|---|---|---|---|---|
| 804 | 1 | 3um_FIJJAFBF_546283 | Eukaryota | unclassified | Oomycetes | Saprolegniales | Saprolegniace | Thraustotheca | Thraustotheca | information st | Translation, ri | 11UVJ | Ribosomal prc | 29760.VIT_05 | 3 | 0 | 0,001629108 | 0,000172283 | 3 - | 0,001935729 - | 3 - | 0,001618317 - | 3 - | 0,001629128 - |
| 805 | 1 | 3um_FIJJAFBF_462696 | Bacteria | Bacteroidetes | Sphingobacter | Sphingobacter | Sphingobacter | Pedobacter | Pedobacter nu | unclassified | unclassified | unclassified | unclassified | unclassified | 0,4 | 0,489897949 | 4,59E-05 | 5,62E-05 | 0 - | 0 - | 1 - | 0,000114397 - | 1 - | 0,000115161 - |
| 806 | 1 | 3um_FIJJAFBF_814035 | Bacteria | Bacteroidetes | Flavobacteriia | Flavobacterial | Flavobacteriac | Algibacter | Algibacter lect | unclassified | unclassified | unclassified | unclassified | unclassified | 1 | 0 | 0,000111887 | 1,18E-05 | 1 - | 0,000132945 - | 1 - | 0,000111146 - | 1 - | 0,000111888 - |
| 807 | 1 | 3um_FIJJAFBF_40378 | Eukaryota | Bacillariophyta | Coscinodiscop | Thalassiosirale | Thalassiosirac | Thalassiosira | Thalassiosira | metabolism | Energy produc | 11HAV | photosystem i | 35128.Thapsd | 1 | 0 | 0,000453181 | 4,79E-05 | 1 - | 0,000538476 - | 1 - | 0,000450179 - | 1 - | 0,000453187 - |
| 808 | 1 | 0.2um_NOKCPHJN_340028 | Bacteria | Proteobacteri | Alphaproteob | Rhodobactera | Rhodobactera | Roseobacter | Roseobacter s | metabolism | Energy produc | 05C6V | Glycerol-3-ph | 391589.RGAI1 | 0,2 | 0,4 | 2,06E-05 | 4,12E-05 | 0 - | 0 - | 0 - | 0 - | 0 - | 0 - |
| 809 | 1 | 3um_FIJJAFBF_15726 | Viruses | unclassified | unclassified | unclassified | Mimiviridae | unclassified | Yellowstone la | unclassified | unclassified | unclassified | unclassified | unclassified | 0,2 | 0,4 | 1,36E-05 | 2,73E-05 | 0 - | 0 - | 1 - | 6,82E-05 - | 0 - | 0 - |
| 810 | 2 | 0.2um_NOKCPHJN_96608;3um_FIJJAFB | Bacteria | Bacteroidetes | Flavobacteriia | Flavobacterial | Flavobacteriac | various | various | unclassified | unclassified | unclassified | unclassified | unclassified | 1 | 0 | 0,000508001 | 5,37E-05 | 1 - | 0,000603614 - | 1 - | 0,000504636 - | 1 - | 0,000508008 - |
| 811 | 5 | 0.2um_NOKCPHJN_70575;3um_FIJJAFB | Bacteria | Bacteroidetes | Flavobacteriia | Flavobacterial | Flavobacteriac | Polaribacter | various | metabolism | Amino acid tra | 08IQK | o-acetylhomo | various | 0,8 | 0,748331477 | 0,000112925 | 0,000109488 | 0 - | 0 - | 0 - | 0 - | 2 - | 0,000295741 - |
| 812 | 1 | 0.2um_NOKCPHJN_899987 | Bacteria | Proteobacteri | Gammaprotec | Cellvibrionales | Porticoccacea | unclassified | various | unclassified | unclassified | unclassified | unclassified | unclassified | 0,8 | 0,4 | 0,000388793 | 0,000198116 | 1 - | 0,000558569 - | 1 - | 0,000466977 - | 1 - | 0,000470097 - |
| 813 | 1 | 3um_FIJJAFBF_250490 | Eukaryota | Chytridiomyco | Chytridiomyce | Rhizophydiale | unclassified | Batrachochytr | Batrachochytr | metabolism | Amino acid tra | COG0137 | Citrulline--asp | 2850.Phatr21 | 0,2 | 0,4 | 6,81E-05 | 0,000136201 | 0 - | 0 - | 0 - | 0 - | 1 - | 0,000340502 - |
| 814 | 1 | 0.2um_NOKCPHJN_195670 | unclassified | unclassified | unclassified | unclassified | unclassified | unclassified | unclassified | unclassified | unclassified | unclassified | unclassified | unclassified | 0,2 | 0,4 | 4,05E-05 | 8,10E-05 | 0 - | 0 - | 0 - | 0 - | 0 - | 0 - |
| 815 | 1 | 3um_FIJJAFBF_32817 | Bacteria | Proteobacteri | Alphaproteob | Rhodobactera | Rhodobactera | Pseudophaeol | Pseudophaeol | unclassified | unclassified | unclassified | unclassified | unclassified | 2,8 | 0,748331477 | 0,000522381 | 9,73E-05 | 2 - | 0,000453625 - | 2 - | 0,000379242 - | 3 - | 0,000572663 - |
| 816 | 1 | 3um_FIJJAFBF_628214 | Eukaryota | Eustigmatoph | unclassified | Eustigmatales | Monodopsida | Nannochlorop | Nannochlorop | unclassified | unclassified | unclassified | unclassified | unclassified | 0,4 | 0,489897949 | 6,06E-05 | 7,47E-05 | 1 - | 0,000164502 - | 0 - | 0 - | 1 - | 0,000138446 - |
| 817 | 1 | 0.2um_NOKCPHJN_43232 | Bacteria | Actinobacteria | Actinobacteria | Micrococcales | Microbacteria | Candidatus Ac | Candidatus Ac | information st | Replication, re | COG0470 | DNA polymera | 743718.Isova_ | 0,8 | 0,4 | 0,000132408 | 6,83E-05 | 1 - | 0,000194411 - | 1 - | 0,000162532 - | 1 - | 0,000163618 - |
| 818 | 1 | 3um_FIJJAFBF_789081 | Eukaryota | Bacillariophyta | Coscinodiscop | Thalassiosirale | Thalassiosirac | Thalassiosira | Thalassiosira p | poorly charact | Function unkn | 0XP1S | solute carrier | 35128.Thaps7 | 2 | 0,632455532 | 0,00031704 | 8,80E-05 | 2 - | 0,000381878 - | 2 - | 0,00031926 - | 1 - | 0,000160696 - |
| 819 | 1 | 0.2um_NOKCPHJN_191792 | Eukaryota | unclassified | Pelagophycea | Pelagomonad | unclassified | Aureococcus | Aureococcus a | cellular proces | Signal transdu | 0XNQP | NIMA-related | 65071.PYU1_T | 0,8 | 0,4 | 4,86E-05 | 2,51E-05 | 1 - | 7,14E-05 - | 1 - | 5,97E-05 - | 1 - | 6,01E-05 - |
| 820 | 1 | 3um_FIJJAFBF_103676 | unclassified | unclassified | unclassified | unclassified | unclassified | unclassified | unclassified | unclassified | unclassified | unclassified | unclassified | unclassified | 1,2 | 0,4 | 0,000490244 | 0,000153523 | 1 - | 0,000489204 - | 1 - | 0,000408986 - | 1 - | 0,000411719 - |
| 821 | 1 | 0.2um_NOKCPHJN_274455 | Eukaryota | Arthropoda | Insecta | Hymenoptera | Apidae | Apis | Apis cerana | unclassified | unclassified | unclassified | unclassified | unclassified | 0,8 | 0,4 | 7,16E-05 | 3,69E-05 | 1 - | 0,000105124 - | 1 - | 8,79E-05 - | 1 - | 8,85E-05 - |
| 822 | 7 | 0.2um_NOKCPHJN_716770;0.2um_NO | various | various | various | various | various | various | various | unclassified | unclassified | unclassified | unclassified | unclassified | 0,2 | 0,4 | 0,000139984 | 0,000279969 | 0 - | 0 - | 0 - | 0 - | 1 - | 0,000699922 - |
| 823 | 1 | 0.2um_NOKCPHJN_56666 | Bacteria | Proteobacteri | Gammaprotec | Alteromonada | Alteromonada | Alteromonas | Alteromonas s | metabolism | Carbohydrate | 16QEQ | hydrolase, fan | 722419.PH505 | 0,6 | 0,489897949 | 3,53E-05 | 2,95E-05 | 1 - | 6,98E-05 - | 0 - | 0 - | 0 - | 0 - |
| 824 | 1 | 3um_FIJJAFBF_81226 | Bacteria | Proteobacteri | Gammaprotec | Cellvibrionales | Spongiibacter | Oceanicoccus | Oceanicoccus | information st | Translation, ri | 05CEJ | Catalyzes the | 247633.GP214 | 1,2 | 0,4 | 0,000111057 | 5,13E-05 | 2 - | 0,000213243 - | 1 - | 8,91E-05 - | 1 - | 8,97E-05 - |
| 825 | 1 | 0.2um_NOKCPHJN_179233 | Eukaryota | Eustigmatoph | unclassified | Eustigmatales | Monodopsida | Nannochlorop | Nannochlorop | metabolism | Amino acid tra | 15H0A | Interconversic | 15368.BRADI1 | 0,8 | 0,4 | 0,000101805 | 5,27E-05 | 1 - | 0,000151208 - | 1 - | 0,000126414 - | 0 - | 0 - |
| 826 | 1 | 0.2um_NOKCPHJN_72952 | Bacteria | Bacteroidetes | Flavobacteriia | unclassified | unclassified | unclassified | Flavobacteriia | metabolism | Amino acid tra | COG0686 | alanine dehyd | 755732.Fluta_ | 0,2 | 0,4 | 3,10E-05 | 6,21E-05 | 0 - | 0 - | 0 - | 0 - | 1 - | 0,000155155 - |
| 827 | 1 | 3um_FIJJAFBF_269230 | Bacteria | Bacteroidetes | Flavobacteriia | Flavobacterial | Flavobacteriac | Polaribacter | Polaribacter s | unclassified | unclassified | unclassified | unclassified | unclassified | 0,2 | 0,4 | 1,47E-05 | 2,94E-05 | 0 - | 0 - | 0 - | 0 - | 0 - | 0 - |
| 828 | 1 | 0.2um_NOKCPHJN_328356 | Bacteria | Proteobacteri | Alphaproteob | Rhodospirillale | Rhodospirillac | unclassified | Rhodobactera | unclassified | unclassified | unclassified | unclassified | unclassified | 1 | 0 | 0,00026579 | 2,81E-05 | 1 - | 0,000315815 - | 1 - | 0,000264029 - | 1 - | 0,000265793 - |
| 829 | 1 | 0.2um_NOKCPHJN_219635 | Eukaryota | Porifera | Demospongiae | Haplosclerida | Niphatidae | Amphimedon | Amphimedon | metabolism | Lipid transpor | 0V6XM | High density li | 6669.DappuP2 | 0,2 | 0,4 | 9,82E-06 | 1,96E-05 | 0 - | 0 - | 0 - | 0 - | 0 - | 0 - |
| 830 | 1 | 3um_FIJJAFBF_95715 | Eukaryota | unclassified | Raphidophyce | Chattonellales | Chattonellace | Heterosigma | Heterosigma a | metabolism | Energy produc | 0IFBF | One of the tw | 45157.CMV08 | 11,2 | 0,979795897 | 0,001891901 | 0,00027373 | 12 - | 0,002407984 - | 10 - | 0,001677612 - | 10 - | 0,001688819 - |
| 831 | 1 | 3um_FIJJAFBF_407604 | Eukaryota | Bacillariophyta | Bacillariophyc | Naviculales | Phaeodactylac | Phaeodactylu | Phaeodactylu | cellular proces | Posttranslatio | KOG0543 | Peptidyl-proly | 2850.Phatr19 | 0,4 | 0,489897949 | 0,000126782 | 0,000158456 | 1 - | 0,000366903 - | 0 - | 0 - | 0 - | 0 - |
| 832 | 1 | 0.2um_NOKCPHJN_307745 | Eukaryota | Phaeophyceae | unclassified | Ectocarpales | Ectocarpaceae | Ectocarpus | Ectocarpus sili | unclassified | unclassified | unclassified | unclassified | unclassified | 0,8 | 0,4 | 0,000221025 | 0,000114309 | 1 - | 0,000328282 - | 1 - | 0,000274451 - | 0 - | 0 - |
| 833 | 1 | 0.2um_NOKCPHJN_971461 | Eukaryota | unclassified | Oomycetes | Peronosporale | Peronosporac | Phytophthora | Phytophthora | unclassified | unclassified | unclassified | unclassified | unclassified | 0,4 | 0,489897949 | 5,28E-05 | 6,60E-05 | 1 - | 0,000152751 - | 0 - | 0 - | 0 - | 0 - |
| 834 | 1 | 3um_FIJJAFBF_248998 | Bacteria | Bacteroidetes | Flavobacteriia | Flavobacterial | Flavobacteriac | Polaribacter | various | information st | Translation, ri | COG0254 | 50S ribosomal | 313598.MED1 | 0,8 | 0,748331477 | 0,000572692 | 0,000555263 | 0 - | 0 - | 0 - | 0 - | 2 - | 0,001499832 - |
| 835 | 3 | 0.2um_NOKCPHJN_464803;0.2um_NO | various | various | various | various | various | various | various | unclassified | unclassified | unclassified | unclassified | unclassified | 0,2 | 0,4 | 1,39E-05 | 2,78E-05 | 0 - | 0 - | 0 - | 0 - | 0 - | 0 - |
| 836 | 1 | 3um_FIJJAFBF_411697 | unclassified | unclassified | unclassified | unclassified | unclassified | unclassified | unclassified | unclassified | unclassified | unclassified | unclassified | unclassified | 1 | 0 | 0,00056243 | 5,95E-05 | 1 - | 0,000668287 - | 1 - | 0,000558705 - | 1 - | 0,000562437 - |
| 837 | 1 | 3um_FIJJAFBF_700428 | Eukaryota | Cnidaria | Hydrozoa | Anthoathecata | Hydridae | Hydra | Hydra vulgaris | information st | Replication, re | KOG0442 | Excision repai | 6085.XP_0021 | 0,2 | 0,4 | 2,06E-05 | 4,12E-05 | 0 - | 0 - | 1 - | 0,000102919 - | 0 - | 0 - |
| 838 | 1 | 0.2um_NOKCPHJN_310540 | Eukaryota | Chordata | Aves | Galliformes | Phasianidae | Gallus | Gallus gallus | poorly charact | Function unkn | 0PXWZ | ChtBD1 | 318829.MGG_ | 0,6 | 0,489897949 | 9,32E-06 | 7,61E-06 | 0 - | 0 - | 1 - | 1,57E-05 - | 1 - | 1,58E-05 - |
| 839 | 1 | 3um_FIJJAFBF_566201 | unclassified | unclassified | unclassified | unclassified | unclassified | unclassified | unclassified | unclassified | unclassified | unclassified | unclassified | unclassified | 1,6 | 0,489897949 | 0,00078108 | 0,000197874 | 1 - | 0,000594033 - | 2 - | 0,000993253 - | 1 - | 0,000499944 - |
| 840 | 1 | 3um_FIJJAFBF_322481 | Eukaryota | Bacillariophyta | Bacillariophyc | Naviculales | Phaeodactylac | Phaeodactylu | Phaeodactylu | unclassified | unclassified | unclassified | unclassified | unclassified | 0,4 | 0,489897949 | 9,37E-05 | 0,000114769 | 0 - | 0 - | 1 - | 0,000233489 - | 1 - | 0,000235048 - |
| 841 | 1 | 0.2um_NOKCPHJN_324069 | Bacteria | Proteobacteri | Gammaprotec | unclassified | unclassified | unclassified | SAR86 cluster | metabolism | Energy produc | COG0055 | Produces ATP | 314287.GB22 | 0,4 | 0,489897949 | 5,29E-05 | 6,49E-05 | 0 - | 0 - | 0 - | 0 - | 1 - | 0,000135469 - |
| 842 | 1 | 3um_FIJJAFBF_674374 | unclassified | unclassified | unclassified | unclassified | unclassified | unclassified | unclassified | unclassified | unclassified | unclassified | unclassified | unclassified | 0,2 | 0,4 | 0,000196969 | 0,000393937 | 0 - | 0 - | 0 - | 0 - | 0 - | 0 - |
| 843 | 1 | 0.2um_NOKCPHJN_698927 | Bacteria | Proteobacteri | Alphaproteob | Rhodobactera | Rhodobactera | Planktotalea | Planktotalea f | metabolism | Energy produc | 0HMD2 | Inherit from p | 596151.Desfr | 0,6 | 0,489897949 | 0,000121421 | 0,000100295 | 1 - | 0,000229596 - | 0 - | 0 - | 1 - | 0,00019323 - |
| 844 | 1 | 3um_FIJJAFBF_191788 | Eukaryota | unclassified | Pelagophycea | Pelagomonad | unclassified | Aureoumbra | Aureoumbra l | metabolism | Energy produc | COG0056 | Produces ATP | 32049.SYNPC | 20,6 | 3,322649545 | 0,00254689 | 0,000248918 | 18 - | 0,002678464 - | 18 - | 0,002239261 - | 18 - | 0,00225422 - |
| 845 | 1 | 3um_FIJJAFBF_377020 | Eukaryota | Bacillariophyta | Coscinodiscop | Thalassiosirale | Thalassiosirac | Thalassiosira | Thalassiosira p | unclassified | unclassified | unclassified | unclassified | unclassified | 1,2 | 0,748331477 | 0,000595956 | 0,000344075 | 1 - | 0,000608522 - | 1 - | 0,000508739 - | 0 - | 0 - |
| 846 | 1 | 3um_FIJJAFBF_492315 | Eukaryota | unclassified | Cryptophyta | Pyrenomonad | Geminigerace | Teleaulax | Teleaulax amp | unclassified | unclassified | unclassified | unclassified | unclassified | 3,8 | 0,748331477 | 0,001338505 | 0,000234237 | 3 - | 0,001268613 - | 5 - | 0,001767653 - | 3 - | 0,001067677 - |
| 847 | 1 | 3um_FIJJAFBF_509785 | Eukaryota | Bacillariophyta | Coscinodiscop | Thalassiosirale | Thalassiosirac | Thalassiosira | Thalassiosira p | metabolism | Carbohydrate | KOG2670 | enolase (EC 4. | 35128.Thaps4 | 1,2 | 0,4 | 0,000346883 | 8,57E-05 | 1 - | 0,0003514 - | 1 - | 0,000293779 - | 1 - | 0,000295741 - |
| 848 | 1 | 3um_FIJJAFBF_01300 | Bacteria | Bacteroidetes | Flavobacteriia | Flavobacterial | Flavobacteriac | Polaribacter | Polaribacter s | unclassified | unclassified | unclassified | unclassified | unclassified | 1 | 0 | 7,21E-05 | 7,62E-06 | 1 - | 8,56E-05 - | 1 - | 7,16E-05 - | 1 - | 7,21E-05 - |
| 849 | 2 | 3um_FIJJAFBF_147391;3um_FIJJAFBF_ | Bacteria | Bacteroidetes | Flavobacteriia | Flavobacterial | Flavobacteriac | Polaribacter | various | unclassified | unclassified | unclassified | unclassified | unclassified | 1,8 | 0,4 | 0,000841021 | 0,000144526 | 1 - | 0,000567032 - | 2 - | 0,000948105 - | 2 - | 0,000954438 - |
| 850 | 1 | 0.2um_NOKCPHJN_364077 | Archaea | Thaumarchaeo | unclassified | unclassified | unclassified | unclassified | Thaumarchaeo | information st | Transcription | COG0086 | DNA-depende | 436308.Nmar_ | 1 | 0 | 5,08E-05 | 5,37E-06 | 1 - | 6,03E-05 - | 1 - | 5,04E-05 - | 1 - | 5,08E-05 - |
| 851 | 1 | 3um_FIJJAFBF_715070 | Eukaryota | unclassified | Raphidophyce | Chattonellales | Chattonellace | Heterosigma | Heterosigma a | unclassified | unclassified | unclassified | unclassified | unclassified | 14,8 | 0,748331477 | 0,001825959 | 0,000154093 | 14 - | 0,002058693 - | 14 - | 0,001721118 - | 15 - | 0,001856373 - |
| 852 | 1 | 0.2um_NOKCPHJN_881171 | Bacteria | Bacteroidetes | Sphingobacter | Sphingobacter | Sphingobacter | Mucilaginibac | Mucilaginibac | cellular proces | Signal transdu | KOG0613 | fibronectin ty | 54126.PPA28 | 0,8 | 0,4 | 0,000104942 | 5,43E-05 | 1 - | 0,00015561 - | 0 - | 0 - | 1 - | 0,000130962 - |
| 853 | 1 | 3um_FIJJAFBF_08157 | Bacteria | Bacteroidetes | Flavobacteriia | Flavobacterial | Flavobacteriac | Polaribacter | various | metabolism | Inorganic ion t | 0P8Q4 | tonB-depende | 867900.Celly_ | 32 | 5,138093031 | 0,002032119 | 0,00014962 | 24 - | 0,001840529 - | 29 - | 0,001859296 - | 33 - | 0,002129884 - |
| 854 | 1 | 3um_FIJJAFBF_07325 | Viruses | unclassified | unclassified | Caudovirales | Siphoviridae | unclassified | Tenacibaculur | unclassified | unclassified | unclassified | unclassified | unclassified | 0,2 | 0,4 | 1,88E-05 | 3,77E-05 | 0 - | 0 - | 0 - | 0 - | 0 - | 0 - |
| 855 | 1 | 3um_FIJJAFBF_314913 | Bacteria | Proteobacteri | Gammaprotec | Alteromonada | Alteromonada | Paraglaciecola | Paraglaciecola | unclassified | unclassified | unclassified | unclassified | unclassified | 1 | 0 | 9,10E-05 | 9,63E-06 | 1 - | 0,000108162 - | 1 - | 9,04E-05 - | 1 - | 9,10E-05 - |
| 856 | 1 | 0.2um_NOKCPHJN_329715 | Eukaryota | unclassified | Oomycetes | Pythiales | Pythiaceae | Pythium | Pythium insidi | metabolism | Energy produc | COG0055 | Produces ATP | 65071.PYU1_T | 2,2 | 0,4 | 0,000287661 | 8,62E-05 | 3 - | 0,000458254 - | 2 - | 0,000255408 - | 2 - | 0,000257114 - |
| 857 | 1 | 3um_FIJJAFBF_251385 | Bacteria | Bacteroidetes | Flavobacteriia | Flavobacterial | Flavobacteriac | Polaribacter | Polaribacter s | unclassified | unclassified | unclassified | unclassified | unclassified | 2,2 | 1,469693846 | 0,000278476 | 0,000169006 | 1 - | 0,000158241 - | 1 - | 0,000132294 - | 1 - | 0,000133177 - |
| 858 | 2 | 0.2um_NOKCPHJN_482392;3um_FIJJAF | various | various | various | various | various | various | various | unclassified | unclassified | unclassified | unclassified | unclassified | 0,2 | 0,4 | 1,63E-05 | 3,25E-05 | 0 - | 0 - | 1 - | 8,13E-05 - | 0 - | 0 - |
| 859 | 1 | 0.2um_NOKCPHJN_400612 | Eukaryota | Ascomycota | Leotiomycetes | Helotiales | Sclerotiniacea | Botrytis | Botrytis cinere | metabolism | Energy produc | KOG0017 | Retrotranspos | 10090.ENSMU | 0,6 | 0,489897949 | 6,96E-05 | 5,73E-05 | 1 - | 0,000129945 - | 1 - | 0,000108637 - | 1 - | 0,000109363 - |
| 860 | 1 | 3um_FIJJAFBF_431402 | Eukaryota | Arthropoda | Insecta | Diptera | Drosophilidae | Drosophila | Drosophila me | cellular proces | Signal transdu | 033RB | inositol 1,4,5-t | 7245.FBpp027 | 0,4 | 0,489897949 | 3,29E-05 | 4,07E-05 | 1 - | 9,12E-05 - | 0 - | 0 - | 0 - | 0 - |
| 861 | 1 | 0.2um_NOKCPHJN_254296 | Eukaryota | unclassified | Labyrinthulom | unclassified | Thraustochytr | Aurantiochytr | Aurantiochytr | unclassified | unclassified | unclassified | unclassified | unclassified | 0,6 | 0,489897949 | 1,20E-05 | 9,81E-06 | 0 - | 0 - | 0 - | 0 - | 1 - | 2,12E-05 - |
| 862 | 1 | 3um_FIJJAFBF_479496 | Eukaryota | Bacillariophyta | Bacillariophyc | Naviculales | Naviculaceae | Fistulifera | Fistulifera sola | information st | Replication, re | KOG0981 | topoisomeras | 35128.Thaps9 | 0,8 | 0,4 | 7,23E-05 | 3,65E-05 | 0 - | 0 - | 1 - | 9,42E-05 - | 1 - | 9,49E-05 - |
| 863 | 1 | 3um_FIJJAFBF_45139 | Eukaryota | unclassified | Oligohymenop | Peniculida | Parameciidae | Paramecium | Paramecium t | unclassified | unclassified | unclassified | unclassified | unclassified | 0,2 | 0,4 | 2,35E-05 | 4,70E-05 | 0 - | 0 - | 0 - | 0 - | 0 - | 0 - |
| 864 | 1 | 0.2um_NOKCPHJN_124256 | Eukaryota | unclassified | Labyrinthulom | unclassified | Thraustochytr | Aurantiochytr | Aurantiochytr | unclassified | unclassified | unclassified | unclassified | unclassified | 0,2 | 0,4 | 2,58E-06 | 5,15E-06 | 0 - | 0 - | 0 - | 0 - | 0 - | 0 - |
| 865 | 1 | 0.2um_NOKCPHJN_1119963 | Eukaryota | unclassified | Oomycetes | Peronosporale | Peronosporac | Phytophthora | various | unclassified | unclassified | unclassified | unclassified | unclassified | 1,6 | 0,8 | 0,001255336 | 0,000570397 | 1 - | 0,000959592 - | 1 - | 0,000802243 - | 1 - | 0,000807602 - |
| 866 | 1 | 3um_FIJJAFBF_551806 | Bacteria | Proteobacteri | Alphaproteob | Rhodobactera | Rhodobactera | Hasllibacter | Hasllibacter h | unclassified | unclassified | unclassified | unclassified | unclassified | 2 | 0 | 0,000303577 | 3,21E-05 | 2 - | 0,000360714 - | 2 - | 0,000301566 - | 2 - | 0,00030358 - |
| 867 | 2 | 0.2um_NOKCPHJN_607027;3um_FIJJAF | Eukaryota | Apicomplexa | Aconoidasida | Haemosporida | Plasmodiidae | Plasmodium | various | various | various | various | various | various | 0,8 | 0,4 | 4,34E-05 | 2,21E-05 | 1 - | 6,23E-05 - | 1 - | 5,21E-05 - | 1 - | 5,25E-05 - |
| 868 | 1 | 3um_FIJJAFBF_168836 | Bacteria | Proteobacteri | Alphaproteob | Rhodobactera | Hyphomonada | Hyphobacteri | Hyphobacteri | cellular proces | Cell wall/mem | 01TJH | Udp-sulfoquin | 394221.Mmar | 1 | 0 | 0,000213533 | 2,26E-05 | 1 - | 0,000253723 - | 1 - | 0,000212118 - | 1 - | 0,000213535 - |
| 869 | 1 | 3um_FIJJAFBF_332470 | Bacteria | Bacteroidetes | Flavobacteriia | Flavobacterial | Flavobacteriac | Polaribacter | Polaribacter ir | metabolism | Carbohydrate | 08IUS | Transketolase | 487796.Flav2A | 0,6 | 0,489897949 | 0,000117125 | 9,57E-05 | 0 - | 0 - | 1 - | 0,000197397 - | 1 - | 0,000198716 - |
| 870 | 1 | 3um_FIJJAFBF_208005 | Eukaryota | Phaeophyceae | unclassified | Ectocarpales | Ectocarpaceae | Ectocarpus | Ectocarpus sili | information st | Translation, ri | COG0050 | This protein p | 321327.CYA_1 | 0,2 | 0,4 | 5,75E-05 | 0,000115056 | 0 - | 0 - | 0 - | 0 - | 1 - | 0,000287639 - |
| 871 | 2 | 0.2um_NOKCPHJN_1012642;3um_FIJJA | Eukaryota | unclassified | unclassified | Prymnesiales | Chrysochrom | Chrysochrom | Chrysochrom | poorly charact | Function unkn | 110WY | Chlorophyll A- | 2850.Phatr16 | 1,8 | 0,4 | 0,000872841 | 0,000220931 | 2 - | 0,001151511 - | 1 - | 0,000481346 - | 2 - | 0,000969122 - |
| 872 | 1 | 3um_FIJJAFBF_277464 | Eukaryota | unclassified | Pelagophycea | Pelagomonad | unclassified | Pelagomonas | uncultured Pe | metabolism | Energy produc | COG1290 | Component of | 35128.Thapsd | 1,6 | 0,489897949 | 0,000615623 | 0,000157699 | 1 - | 0,000467801 - | 1 - | 0,000391093 - | 2 - | 0,000787412 - |
| 873 | 1 | 0.2um_NOKCPHJN_409279 | Eukaryota | Phaeophyceae | unclassified | Ectocarpales | Ectocarpaceae | Ectocarpus | Ectocarpus sili | unclassified | unclassified | unclassified | unclassified | unclassified | 0,6 | 0,489897949 | 3,15E-05 | 2,58E-05 | 0 - | 0 - | 1 - | 5,56E-05 - | 0 - | 0 - |
| 874 | 1 | 3um_FIJJAFBF_517420 | Bacteria | Bacteroidetes | Flavobacteriia | Flavobacterial | Flavobacteriac | Polaribacter | various | information st | Transcription | COG0250 | Participates in | 313598.MED1 | 0,6 | 0,489897949 | 0,000206987 | 0,000172876 | 1 - | 0,000409006 - | 0 - | 0 - | 0 - | 0 - |
| 875 | 1 | 3um_FIJJAFBF_183932 | Eukaryota | unclassified | Pelagophycea | Pelagomonad | unclassified | Aureococcus | Aureococcus a | metabolism | Inorganic ion t | 1EKJE | calcium chann | 3218.PP1S45_ | 2 | 0 | 0,000196543 | 2,08E-05 | 2 - | 0,000233536 - | 2 - | 0,000195242 - | 2 - | 0,000196546 - |
| 876 | 1 | 3um_FIJJAFBF_239813 | Bacteria | Bacteroidetes | Flavobacteriia | Flavobacterial | Flavobacteriac | Formosa | Formosa sp. H | unclassified | unclassified | unclassified | unclassified | unclassified | 1 | 0 | 0,00034422 | 3,64E-05 | 1 - | 0,000409006 - | 1 - | 0,000341939 - | 1 - | 0,000344224 - |
| 877 | 1 | 3um_FIJJAFBF_313117 | Bacteria | Cyanobacteria | unclassified | Synechococca | Synechococca | Synechococcu | Synechococcu | unclassified | unclassified | unclassified | unclassified | unclassified | 1 | 0,894427191 | 0,000271119 | 0,000242387 | 0 - | 0 - | 2 - | 0,000576727 - | 0 - | 0 - |
| 878 | 1 | 0.2um_NOKCPHJN_555603 | unclassified | unclassified | unclassified | unclassified | unclassified | unclassified | unclassified | unclassified | unclassified | unclassified | unclassified | unclassified | 3 | 0,632455532 | 0,00107089 | 0,000235993 | 3 - | 0,001275821 - | 2 - | 0,000711079 - | 3 - | 0,001073743 - |
| 879 | 1 | 3um_FIJJAFBF_783854 | Bacteria | Bacteroidetes | Flavobacteriia | Flavobacterial | Flavobacteriac | Tenacibaculur | Tenacibaculur | poorly charact | Function unkn | 11P35 | NA | 983544.Lacal_ | 2,8 | 0,748331477 | 0,00031589 | 7,10E-05 | 2 - | 0,000272671 - | 3 - | 0,000341939 - | 2 - | 0,000229482 - |
| 880 | 1 | 3um_FIJJAFBF_05630 | Eukaryota | Phaeophyceae | unclassified | Ectocarpales | Ectocarpaceae | Ectocarpus | Ectocarpus sili | unclassified | unclassified | unclassified | unclassified | unclassified | 1 | 0 | 0,000508001 | 5,37E-05 | 1 - | 0,000603614 - | 1 - | 0,000504636 - | 1 - | 0,000508008 - |
| 881 | 1 | 3um_FIJJAFBF_545173 | Eukaryota | Bacillariophyta | Bacillariophyc | Bacillariales | Bacillariaceae | Fragilariopsis | Fragilariopsis | unclassified | unclassified | unclassified | unclassified | unclassified | 0,6 | 0,489897949 | 0,000213383 | 0,000174321 | 0 - | 0 - | 1 - | 0,000359626 - | 1 - | 0,000362028 - |
| 882 | 1 | 0.2um_NOKCPHJN_956102 | Bacteria | Proteobacteri | Gammaprotec | Chromatiales | Chromatiacea | Nitrosococcus | Nitrosococcus | unclassified | unclassified | unclassified | unclassified | unclassified | 0,2 | 0,4 | 0,000184044 | 0,000368088 | 0 - | 0 - | 1 - | 0,000920219 - | 0 - | 0 - |
| 883 | 1 | 0.2um_NOKCPHJN_142020 | Bacteria | Nitrospinae | unclassified | unclassified | unclassified | unclassified | Nitrospinae ba | information st | Translation, ri | 16T1C | amidotransfer | 331869.BAL19 | 0,2 | 0,4 | 2,70E-05 | 5,40E-05 | 0 - | 0 - | 0 - | 0 - | 1 - | 0,000134889 - |
| 884 | 1 | 0.2um_NOKCPHJN_902159 | Eukaryota | Bacillariophyta | Bacillariophyc | Bacillariales | Bacillariaceae | Fragilariopsis | Fragilariopsis | poorly charact | Function unkn | 0XRVT | NA | 35128.Thaps3 | 0,8 | 0,748331477 | 0,000371211 | 0,00033331 | 0 - | 0 - | 0 - | 0 - | 1 - | 0,000503944 - |
| 885 | 1 | 3um_FIJJAFBF_163897 | Bacteria | Proteobacteri | Alphaproteob | Rhodobactera | unclassified | unclassified | Rhodobactera | unclassified | unclassified | unclassified | unclassified | unclassified | 1 | 0 | 0,000178955 | 1,89E-05 | 1 - | 0,000212637 - | 1 - | 0,00017777 - | 1 - | 0,000178957 - |
| 886 | 3 | 3um_FIJJAFBF_237980;3um_FIJJAFBF_ | Eukaryota | various | various | various | various | various | various | unclassified | unclassified | unclassified | unclassified | unclassified | 1 | 0 | 0,001574804 | 0,00016654 | 1 - | 0,001871205 - | 1 - | 0,001564373 - | 1 - | 0,001574823 - |
| 887 | 2 | 0.2um_NOKCPHJN_78071;3um_FIJJAFB | Eukaryota | unclassified | Oomycetes | Peronosporale | Peronosporac | Phytophthora | various | information st | Translation, ri | KOG1145 | One of the ess | various | 0,2 | 0,4 | 8,66E-06 | 1,73E-05 | 0 - | 0 - | 0 - | 0 - | 1 - | 4,33E-05 - |
| 888 | 1 | 3um_FIJJAFBF_594332 | Bacteria | Proteobacteri | Alphaproteob | Rhodobactera | Rhodobactera | Roseobacter | Roseobacter s | unclassified | unclassified | unclassified | unclassified | unclassified | 1 | 0 | 0,000839896 | 8,88E-05 | 1 - | 0,000997976 - | 1 - | 0,000834332 - | 1 - | 0,000839906 - |
| 889 | 1 | 3um_FIJJAFBF_106302 | Bacteria | Proteobacteri | Alphaproteob | Rhodobactera | Rhodobactera | unclassified | Rhodobactera | unclassified | unclassified | unclassified | unclassified | unclassified | 0,4 | 0,489897949 | 9,15E-05 | 0,00011214 | 0 - | 0 - | 1 - | 0,000233489 - | 0 - | 0 - |
| 890 | 1 | 0.2um_NOKCPHJN_869722 | Bacteria | Proteobacteri | Alphaproteob | Pelagibacteral | Pelagibacterac | Candidatus Pe | Candidatus Pe | unclassified | unclassified | unclassified | unclassified | unclassified | 0,6 | 0,489897949 | 0,000103042 | 8,49E-05 | 1 - | 0,000192412 - | 1 - | 0,000160861 - | 1 - | 0,000161936 - |
| 891 | 1 | 3um_FIJJAFBF_366991 | Eukaryota | Bacillariophyta | Bacillariophyc | Bacillariales | Bacillariaceae | Fragilariopsis | Fragilariopsis | unclassified | unclassified | unclassified | unclassified | unclassified | 1,6 | 0,489897949 | 0,000456017 | 0,000116814 | 1 - | 0,000346519 - | 1 - | 0,000289699 - | 2 - | 0,000583268 - |
| 892 | 1 | 0.2um_NOKCPHJN_1097045 | Bacteria | Proteobacteri | Alphaproteob | Rhodobactera | Rhodobactera | Maritimibacte | Maritimibacte | unclassified | unclassified | unclassified | unclassified | unclassified | 0,2 | 0,4 | 3,35E-05 | 6,69E-05 | 0 - | 0 - | 1 - | 0,000167313 - | 0 - | 0 - |
| 893 | 1 | 3um_FIJJAFBF_116580 | Bacteria | Cyanobacteria | unclassified | Synechococca | Synechococca | Synechococcu | Synechococcu | unclassified | unclassified | unclassified | unclassified | unclassified | 2 | 0 | 0,000684698 | 7,24E-05 | 2 - | 0,000813567 - | 2 - | 0,000680162 - | 2 - | 0,000684706 - |
| 894 | 1 | 0.2um_NOKCPHJN_501796 | Bacteria | Proteobacteri | Alphaproteob | Pelagibacteral | Pelagibacterac | Candidatus Pe | Candidatus Pe | unclassified | unclassified | unclassified | unclassified | unclassified | 0,2 | 0,4 | 8,39E-06 | 1,68E-05 | 0 - | 0 - | 0 - | 0 - | 1 - | 4,20E-05 - |
| 895 | 1 | 0.2um_NOKCPHJN_372415 | Eukaryota | Chlorophyta | Chlorophycea | Chlamydomor | Volvocaceae | Volvox | Volvox carteri | unclassified | unclassified | unclassified | unclassified | unclassified | 0,8 | 0,4 | 5,93E-05 | 2,99E-05 | 0 - | 0 - | 1 - | 7,73E-05 - | 1 - | 7,78E-05 - |
| 896 | 1 | 3um_FIJJAFBF_60207 | Bacteria | Actinobacteria | unclassified | unclassified | unclassified | unclassified | Actinobacteria | unclassified | unclassified | unclassified | unclassified | unclassified | 3 | 0,632455532 | 0,000700223 | 0,000165946 | 3 - | 0,000831646 - | 2 - | 0,000463518 - | 4 - | 0,000933229 - |
| 897 | 1 | 0.2um_NOKCPHJN_281234 | Eukaryota | unclassified | unclassified | Kinetoplastida | Trypanosomat | Trypanosoma | Trypanosoma | unclassified | unclassified | unclassified | unclassified | unclassified | 1 | 0 | 0,000129881 | 1,37E-05 | 1 - | 0,000154326 - | 1 - | 0,00012902 - | 1 - | 0,000129882 - |
| 898 | 1 | 0.2um_NOKCPHJN_269537 | Bacteria | Bacteroidetes | Flavobacteriia | Flavobacterial | Flavobacteriac | unclassified | Flavobacteriac | information st | Replication, re | 0PDAJ | Inherit from b | 1041826.FCOL | 0,4 | 0,489897949 | 0,00010738 | 0,000132038 | 0 - | 0 - | 1 - | 0,000287041 - | 0 - | 0 - |
| 899 | 1 | 3um_FIJJAFBF_110539 | Bacteria | Bacteroidetes | Flavobacteriia | Flavobacterial | Flavobacteriac | unclassified | Flavobacteriac | unclassified | unclassified | unclassified | unclassified | unclassified | 0,2 | 0,4 | 2,42E-05 | 4,84E-05 | 0 - | 0 - | 0 - | 0 - | 0 - | 0 - |
| 900 | 1 | 3um_FIJJAFBF_239272 | unclassified | unclassified | unclassified | unclassified | unclassified | unclassified | unclassified | unclassified | unclassified | unclassified | unclassified | unclassified | 1 | 0 | 0,000543036 | 5,74E-05 | 1 - | 0,000645243 - | 1 - | 0,000539439 - | 1 - | 0,000543043 - |
| 901 | 1 | 0.2um_NOKCPHJN_76179 | Eukaryota | Bacillariophyta | Mediophyceae | Cymatosirales | Cymatosiracea | Campylosira | Campylosira c | metabolism | Carbohydrate | COG1850 | RuBisCO catal | 35128.Thapsd | 5,2 | 1,469693846 | 0,000654481 | 0,000140713 | 4 - | 0,000611006 - | 4 - | 0,000510816 - | 4 - | 0,000514228 - |
| 902 | 1 | 3um_FIJJAFBF_220319 | Eukaryota | Bacillariophyta | Coscinodiscop | Thalassiosirale | Thalassiosirac | Thalassiosira | Thalassiosira a | metabolism | Carbohydrate | COG1850 | RuBisCO catal | 35128.Thapsd | 1,6 | 0,489897949 | 0,000443688 | 0,000161046 | 2 - | 0,000648036 - | 2 - | 0,000541774 - | 1 - | 0,000272697 - |
| 903 | 1 | 3um_FIJJAFBF_647835 | Viruses | unclassified | unclassified | unclassified | unclassified | unclassified | uncultured M | unclassified | unclassified | unclassified | unclassified | unclassified | 1 | 0 | 0,000234172 | 2,48E-05 | 1 - | 0,000278246 - | 1 - | 0,000232621 - | 1 - | 0,000234174 - |
| 904 | 1 | 0.2um_NOKCPHJN_36762 | Viruses | unclassified | unclassified | unclassified | various | various | various | unclassified | unclassified | unclassified | unclassified | unclassified | 0,8 | 0,4 | 9,38E-05 | 4,73E-05 | 0 - | 0 - | 1 - | 0,000122217 - | 1 - | 0,000123033 - |

| | | | | | | | | | | | | | | | | | | | | | | | | |
|---|---|---|---|---|---|---|---|---|---|---|---|---|---|---|---|---|---|---|---|---|---|---|---|---|
| 905 | 1 | 0.2um_NOKCPHJN_655398 | Eukaryota | Eustigmatoph | unclassified | Eustigmatales | Monodopsida | Nannochlorop | Nannochlorop | information st | Translation, ri | COG0017 | asparaginyl-tR | 2850.Phatr42 | 0,4 | 0,489897949 | 4,99E-05 | 6,11E-05 | 0 - | 0 - | 1 - | 0,000124403 - | 1 - | 0,000125234 - |
| 906 | 1 | 3um_FIJJAFBF_581444 | Bacteria | Proteobacteri | Gammaprotec | Arenicellales | Arenicellaceae | Arenicella | Arenicella xan | unclassified | unclassified | unclassified | unclassified | unclassified | 0,6 | 0,489897949 | 6,28E-05 | 5,19E-05 | 1 - | 0,000118996 - | 1 - | 9,95E-05 - | 0 - | 0 - |
| 907 | 1 | 3um_FIJJAFBF_551491 | Eukaryota | unclassified | Pelagophycea | Pelagomonad | unclassified | Aureococcus | Aureococcus a | unclassified | unclassified | unclassified | unclassified | unclassified | 0,2 | 0,4 | 6,09E-05 | 0,000121719 | 0 - | 0 - | 0 - | 0 - | 0 - | 0 - |
| 908 | 1 | 3um_FIJJAFBF_304770 | Bacteria | Bacteroidetes | Flavobacteriia | Flavobacterial | Flavobacteria | Kordia | Kordia zhangz | unclassified | unclassified | unclassified | unclassified | unclassified | 0,2 | 0,4 | 9,24E-06 | 1,85E-05 | 0 - | 0 - | 0 - | 0 - | 0 - | 0 - |
| 909 | 1 | 0.2um_NOKCPHJN_611919 | Viruses | unclassified | unclassified | unclassified | unclassified | unclassified | uncultured M | unclassified | unclassified | unclassified | unclassified | unclassified | 0,4 | 0,489897949 | 3,22E-05 | 3,96E-05 | 0 - | 0 - | 1 - | 8,62E-05 - | 0 - | 0 - |
| 910 | 1 | 0.2um_NOKCPHJN_678783 | Bacteria | unclassified | unclassified | unclassified | unclassified | unclassified | uncultured ba | metabolism | Inorganic ion t | 05C27 | TonB-depende | 313596.RB250 | 1,2 | 0,4 | 7,79E-05 | 3,60E-05 | 2 - | 0,000149547 - | 1 - | 6,25E-05 - | 1 - | 6,29E-05 - |
| 911 | 1 | 0.2um_NOKCPHJN_889433 | Bacteria | Proteobacteri | Alphaproteob | Rhodobactera | Rhodobactera | Pseudooctade | Pseudooctade | poorly charact | Function unkn | 0J6G5 | NA | 306901.XP_00 | 1,2 | 0,4 | 0,000409876 | 0,000128356 | 1 - | 0,000409006 - | 1 - | 0,000341939 - | 1 - | 0,000344224 - |
| 912 | 1 | 0.2um_NOKCPHJN_499496 | Eukaryota | unclassified | unclassified | unclassified | Plasmodiopho | Plasmodiopho | Plasmodiopho | cellular proces | Cytoskeleton | 0V4MW | Myosin VIIA | 7070.TC01374 | 0,4 | 0,489897949 | 1,65E-05 | 2,03E-05 | 0 - | 0 - | 0 - | 0 - | 0 - | 0 - |
| 913 | 2 | 0.2um_NOKCPHJN_50907;3um_FIJJAFE | Bacteria | Bacteroidetes | Flavobacteriia | Flavobacterial | Crocinitomica | unclassified | Crocinitomica | unclassified | unclassified | unclassified | unclassified | unclassified | 0,2 | 0,4 | 1,88E-05 | 3,75E-05 | 1 - | 9,38E-05 - | 0 - | 0 - | 0 - | 0 - |
| 914 | 1 | 3um_FIJJAFBF_552358 | Viruses | unclassified | unclassified | unclassified | unclassified | unclassified | uncultured vir | unclassified | unclassified | unclassified | unclassified | unclassified | 4,4 | 0,489897949 | 0,000888315 | 8,86E-05 | 4 - | 0,000965783 - | 5 - | 0,001009273 - | 4 - | 0,000812812 - |
| 915 | 1 | 3um_FIJJAFBF_418657 | Bacteria | Proteobacteri | Gammaprotec | Alteromonada | Colwelliaceae | Colwellia | Colwellia sp. P | metabolism | Coenzyme tra | 08UTT | Catalyzes the | 167879.CPS_1 | 0,6 | 0,489897949 | 0,000260281 | 0,000214436 | 1 - | 0,000486027 - | 1 - | 0,000406331 - | 1 - | 0,000409045 - |
| 916 | 1 | 0.2um_NOKCPHJN_469181 | Bacteria | Bacteroidetes | unclassified | unclassified | unclassified | unclassified | Bacteroidetes | unclassified | unclassified | unclassified | unclassified | unclassified | 0,2 | 0,4 | 0,000189621 | 0,000379242 | 0 - | 0 - | 1 - | 0,000948105 - | 0 - | 0 - |
| 917 | 1 | 3um_FIJJAFBF_537078 | Viruses | unclassified | unclassified | unclassified | Phycodnavirid | unclassified | Chrysochromu | information st | Translation, ri | 1BUH7 | glycine-rich pr | 3218.PP1S103 | 0,4 | 0,489897949 | 0,000150717 | 0,000184958 | 0 - | 0 - | 0 - | 0 - | 0 - | 0 - |
| 918 | 1 | 3um_FIJJAFBF_269224 | Bacteria | Bacteroidetes | Flavobacteriia | Flavobacterial | Flavobacteria | Riemerella | Riemerella col | unclassified | unclassified | unclassified | unclassified | unclassified | 2,6 | 0,8 | 0,000946284 | 0,000398798 | 4 - | 0,001681982 - | 3 - | 0,001054633 - | 2 - | 0,000707786 - |
| 919 | 1 | 3um_FIJJAFBF_96040 | Bacteria | Proteobacteri | Gammaprotec | Enterobactera | Enterobacteri | Citrobacter | Citrobacter w | unclassified | unclassified | unclassified | unclassified | unclassified | 1 | 0 | 8,64E-05 | 9,14E-06 | 1 - | 0,000102672 - | 1 - | 8,58E-05 - | 1 - | 8,64E-05 - |
| 920 | 1 | 3um_FIJJAFBF_241394 | Eukaryota | Bacillariophyt | Coscinodiscop | Thalassiosirale | Thalassiosirac | Thalassiosira | Thalassiosira c | unclassified | unclassified | unclassified | unclassified | unclassified | 1,6 | 0,489897949 | 0,000507731 | 0,000130061 | 1 - | 0,000385815 - | 1 - | 0,000322551 - | 2 - | 0,000649412 - |
| 921 | 1 | 0.2um_NOKCPHJN_495718 | Eukaryota | Eustigmatoph | unclassified | Eustigmatales | Monodopsida | Nannochlorop | Nannochlorop | metabolism | Inorganic ion t | COG3119 | Sulfatase | 400682.PAC_1 | 0,4 | 0,489897949 | 3,07E-05 | 3,84E-05 | 1 - | 8,89E-05 - | 0 - | 0 - | 0 - | 0 - |
| 922 | 1 | 3um_FIJJAFBF_236963 | Bacteria | Bacteroidetes | Flavobacteriia | Flavobacterial | Flavobacteria | Formosa | Formosa sp. H | information st | Replication, re | COG0513 | purine NTP-de | 1046627.BZAF | 0,2 | 0,4 | 2,04E-05 | 4,09E-05 | 0 - | 0 - | 1 - | 0,000102247 - | 0 - | 0 - |
| 923 | 1 | 0.2um_NOKCPHJN_1058181 | Bacteria | Proteobacteri | Alphaproteob | Rhodospirillal | Rhodospirillac | Magnetovibrio | Magnetovibrio | metabolism | Carbohydrate | 05CV1 | Transketolase | 342108.amb1 | 1 | 0 | 0,000119303 | 1,26E-05 | 1 - | 0,000141758 - | 1 - | 0,000118513 - | 1 - | 0,000119305 - |
| 924 | 1 | 0.2um_NOKCPHJN_86863 | Bacteria | Proteobacteri | Gammaprotec | unclassified | unclassified | unclassified | uncultured ga | information st | Translation, ri | COG0050 | This protein p | 314285.KT71_ | 2 | 0 | 0,000310306 | 3,28E-05 | 2 - | 0,00036871 - | 2 - | 0,000308251 - | 2 - | 0,00031031 - |
| 925 | 1 | 0.2um_NOKCPHJN_586666 | Bacteria | Proteobacteri | Betaproteoba | unclassified | unclassified | unclassified | Betaproteoba | poorly charact | Function unkn | 0XRND | NA | 314607.KB13_ | 1 | 0 | 0,000286328 | 3,03E-05 | 1 - | 0,000340219 - | 1 - | 0,000284431 - | 1 - | 0,000286332 - |
| 926 | 1 | 0.2um_NOKCPHJN_450502 | Eukaryota | unclassified | Labyrinthulom | unclassified | Thraustochytr | Aurantiochytr | Aurantiochytr | unclassified | unclassified | unclassified | unclassified | unclassified | 3,8 | 1,166190379 | 0,000631738 | 0,000192887 | 4 - | 0,000794145 - | 5 - | 0,000829906 - | 2 - | 0,00033418 - |
| 927 | 1 | 0.2um_NOKCPHJN_305592 | Bacteria | Proteobacteri | Alphaproteob | Rhodobactera | Hyphomonada | unclassified | Hyphomonada | metabolism | Nucleotide tra | 0FW5M | adenylosuccin | 761193.Runsl_ | 1,8 | 0,4 | 0,000261211 | 4,49E-05 | 1 - | 0,000176113 - | 2 - | 0,00029447 - | 2 - | 0,000296437 - |
| 928 | 1 | 3um_FIJJAFBF_257135 | Eukaryota | Bacillariophyt | Mediophycea | Biddulphiales | Biddulphiacea | Biddulphia | Biddulphia tri | information st | Translation, ri | 08UHY | One of the pri | 489825.LYNGI | 4,2 | 0,4 | 0,001695944 | 0,000235092 | 4 - | 0,001919184 - | 4 - | 0,001604485 - | 5 - | 0,002019004 - |
| 929 | 1 | 0.2um_NOKCPHJN_430840 | Viruses | unclassified | unclassified | unclassified | Phycodnavirid | unclassified | Organic Lake p | unclassified | unclassified | unclassified | unclassified | unclassified | 0,2 | 0,4 | 2,22E-05 | 4,43E-05 | 0 - | 0 - | 0 - | 0 - | 0 - | 0 - |
| 930 | 2 | 3um_FIJJAFBF_310091;3um_FIJJAFBF_ | Bacteria | Proteobacteri | Gammaprotec | Cellvibrionales | Halieaceae | unclassified | marine gamm | metabolism | Nucleotide tra | 0QIPM | dihydroorotas | 247634.GPB2 | 0,2 | 0,4 | 3,61E-05 | 7,21E-05 | 0 - | 0 - | 1 - | 0,000180331 - | 0 - | 0 - |
| 931 | 1 | 0.2um_NOKCPHJN_866315 | unclassified | unclassified | unclassified | unclassified | unclassified | unclassified | unclassified | unclassified | unclassified | unclassified | unclassified | unclassified | 1,2 | 0,4 | 0,000247111 | 6,11E-05 | 1 - | 0,000250328 - | 1 - | 0,000209281 - | 1 - | 0,000210679 - |
| 932 | 1 | 3um_FIJJAFBF_457406 | Eukaryota | Bacillariophyt | Bacillariophyc | Naviculales | Naviculaceae | Fistulifera | Fistulifera sola | unclassified | unclassified | unclassified | unclassified | unclassified | 1,2 | 0,4 | 0,000414405 | 0,000129774 | 1 - | 0,000413526 - | 1 - | 0,000345718 - | 1 - | 0,000348027 - |
| 933 | 1 | 3um_FIJJAFBF_30117 | Eukaryota | Chromerida | unclassified | unclassified | unclassified | Vitrella | Vitrella brassi | unclassified | unclassified | unclassified | unclassified | unclassified | 0,2 | 0,4 | 8,23E-05 | 0,000164671 | 0 - | 0 - | 1 - | 0,000411677 - | 0 - | 0 - |
| 934 | 1 | 0.2um_NOKCPHJN_1035812 | Bacteria | Proteobacteri | Gammaprotec | Oceanospirilla | Oceanospirilla | Neptuniibacte | Neptuniibacte | metabolism | Carbohydrate | 16PI5 | ribulose-phos | 207954.MED9 | 3,8 | 0,4 | 0,002890232 | 0,000166564 | 3 - | 0,002738348 - | 4 - | 0,003052435 - | 4 - | 0,003072826 - |
| 935 | 1 | 3um_FIJJAFBF_26346 | Bacteria | Bacteroidetes | Flavobacteriia | Flavobacterial | Flavobacteria | Seonamhaeic | Seonamhaeic | poorly charact | Function unkn | 09AVF | Protein of unk | 391587.KAOT | 0,4 | 0,8 | 7,05E-05 | 0,000141021 | 0 - | 0 - | 0 - | 0 - | 0 - | 0 - |
| 936 | 1 | 3um_FIJJAFBF_740349 | unclassified | unclassified | unclassified | unclassified | unclassified | unclassified | unclassified | unclassified | unclassified | unclassified | unclassified | unclassified | 0,8 | 0,4 | 9,23E-05 | 4,77E-05 | 1 - | 0,000135595 - | 1 - | 0,00011336 - | 1 - | 0,000114118 - |
| 937 | 1 | 0.2um_NOKCPHJN_53803 | Bacteria | Fibrobacteres | Fibrobacteria | Fibrobacterale | Fibrobacterac | Fibrobacter | Fibrobacter sp | unclassified | unclassified | unclassified | unclassified | unclassified | 0,8 | 0,4 | 8,03E-05 | 4,05E-05 | 0 - | 0 - | 1 - | 0,00010464 - | 1 - | 0,000105339 - |
| 938 | 1 | 0.2um_NOKCPHJN_45030 | Bacteria | Proteobacteri | Gammaprotec | unclassified | unclassified | unclassified | Gammaprotec | unclassified | unclassified | unclassified | unclassified | unclassified | 0,4 | 0,489897949 | 7,28E-05 | 8,92E-05 | 0 - | 0 - | 1 - | 0,000181377 - | 1 - | 0,000182588 - |
| 939 | 1 | 3um_FIJJAFBF_111336 | Viruses | unclassified | unclassified | unclassified | Phycodnavirid | unclassified | Organic Lake p | unclassified | unclassified | unclassified | unclassified | unclassified | 3 | 1,095445115 | 0,00226168 | 0,000614126 | 2 - | 0,001848103 - | 3 - | 0,00231759 - | 2 - | 0,001555381 - |
| 940 | 1 | 3um_FIJJAFBF_732705 | Eukaryota | unclassified | Pelagophycea | Pelagomonad | unclassified | Aureococcus | Aureococcus a | unclassified | unclassified | unclassified | unclassified | unclassified | 2,4 | 0,489897949 | 0,001299071 | 0,000215429 | 2 - | 0,001301708 - | 3 - | 0,001632389 - | 2 - | 0,001095529 - |
| 941 | 1 | 3um_FIJJAFBF_189626 | Bacteria | Bacteroidetes | Flavobacteriia | Flavobacterial | Flavobacteria | Polaribacter | Polaribacter b | unclassified | unclassified | unclassified | unclassified | unclassified | 4 | 0 | 0,000661335 | 6,99E-05 | 4 - | 0,000785808 - | 4 - | 0,000656955 - | 4 - | 0,000661343 - |
| 942 | 1 | 3um_FIJJAFBF_656239 | Eukaryota | Apicomplexa | Aconoidasida | Haemosporida | Plasmodiidae | Plasmodium | Plasmodium f | unclassified | unclassified | unclassified | unclassified | unclassified | 0,2 | 0,4 | 4,65E-05 | 9,30E-05 | 0 - | 0 - | 0 - | 0 - | 1 - | 0,000232446 - |
| 943 | 1 | 0.2um_NOKCPHJN_208494 | Eukaryota | Arthropoda | Insecta | Lepidoptera | Papilionidae | Papilio | Papilio polyte | information st | Translation, ri | COG0081 | Binds directly | 65071.PYU1_ | 0,4 | 0,489897949 | 0,000205109 | 0,000252871 | 1 - | 0,000558569 - | 1 - | 0,000466977 - | 0 - | 0 - |
| 944 | 1 | 0.2um_NOKCPHJN_308920 | Eukaryota | unclassified | Pelagophycea | Pelagomonad | unclassified | Aureococcus | Aureococcus a | unclassified | unclassified | unclassified | unclassified | unclassified | 2 | 0,632455532 | 5,14E-05 | 1,19E-05 | 1 - | 3,15E-05 - | 2 - | 5,27E-05 - | 2 - | 5,31E-05 - |
| 945 | 1 | 3um_FIJJAFBF_570974 | Bacteria | Cyanobacteria | unclassified | Synechococca | Prochloraceae | Prochlorococc | Prochlorococc | information st | Translation, ri | COG0185 | Protein S19 fo | 452637.Oter_ | 1 | 0 | 0,000663076 | 7,01E-05 | 1 - | 0,000787876 - | 1 - | 0,000658683 - | 1 - | 0,000663084 - |
| 946 | 1 | 0.2um_NOKCPHJN_464767 | Eukaryota | unclassified | Cryptophyta | Pyrenomonad | Geminigerace | Guillardia | Guillardia thet | unclassified | unclassified | unclassified | unclassified | unclassified | 1,2 | 0,4 | 0,000361278 | 0,000122961 | 1 - | 0,000358125 - | 2 - | 0,000598803 - | 1 - | 0,000301402 - |
| 947 | 1 | 0.2um_NOKCPHJN_759699 | Eukaryota | unclassified | Oomycetes | Peronosporale | Peronosporac | Phytophthora | various | information st | Chromatin str | KOG1744 | Core compone | 164328.Phyra | 2,6 | 0,489897949 | 0,001406147 | 0,000263813 | 2 - | 0,001290486 - | 3 - | 0,001618317 - | 3 - | 0,001629128 - |
| 948 | 1 | 0.2um_NOKCPHJN_99167 | Eukaryota | Bacillariophyt | Bacillariophyc | Naviculales | Naviculaceae | Fistulifera | Fistulifera sola | information st | Transcription | KOG1474 | bromodomain | 35128.Thaps2 | 1 | 0 | 4,33E-05 | 4,58E-06 | 1 - | 5,14E-05 - | 1 - | 4,30E-05 - | 1 - | 4,33E-05 - |
| 949 | 1 | 3um_FIJJAFBF_673864 | Eukaryota | unclassified | Dinophyceae | Suessiales | Symbiodiniace | Symbiodinium | Symbiodinium | unclassified | unclassified | unclassified | unclassified | unclassified | 1 | 0 | 8,03E-05 | 8,50E-06 | 1 - | 9,55E-05 - | 1 - | 7,98E-05 - | 1 - | 8,03E-05 - |
| 950 | 2 | 3um_FIJJAFBF_02505;3um_FIJJAFBF_2 | various | various | various | various | various | various | various | various | various | various | various | various | 1 | 0 | 0,000104465 | 1,10E-05 | 1 - | 0,000124126 - | 1 - | 0,000103773 - | 1 - | 0,000104466 - |
| 951 | 1 | 3um_FIJJAFBF_158698 | Bacteria | Proteobacteri | Gammaprotec | Oceanospirilla | Alcanivoraca | Ketobacter | Ketobacter alk | unclassified | unclassified | unclassified | unclassified | unclassified | 0,4 | 0,489897949 | 8,91E-05 | 0,000109391 | 0 - | 0 - | 0 - | 0 - | 0 - | 0 - |
| 952 | 1 | 0.2um_NOKCPHJN_938763 | Eukaryota | unclassified | Oomycetes | Peronosporale | Peronosporac | Phytophthora | various | unclassified | unclassified | unclassified | unclassified | unclassified | 1 | 0 | 0,000294356 | 3,11E-05 | 1 - | 0,000349758 - | 1 - | 0,000292406 - | 1 - | 0,00029436 - |
| 953 | 1 | 3um_FIJJAFBF_254297 | Bacteria | Proteobacteri | Gammaprotec | Cellvibrionales | Spongiibacter | Oceanicoccus | Oceanicoccus | unclassified | unclassified | unclassified | unclassified | unclassified | 0,8 | 0,4 | 0,000283206 | 0,000146135 | 1 - | 0,000415823 - | 1 - | 0,000347638 - | 1 - | 0,000349961 - |
| 954 | 1 | 0.2um_NOKCPHJN_277177 | unclassified | unclassified | unclassified | unclassified | unclassified | unclassified | unclassified | unclassified | unclassified | unclassified | unclassified | unclassified | 0,2 | 0,4 | 3,05E-06 | 6,10E-06 | 0 - | 0 - | 0 - | 0 - | 0 - | 0 - |
| 955 | 1 | 3um_FIJJAFBF_416696 | unclassified | unclassified | unclassified | unclassified | unclassified | unclassified | unclassified | unclassified | unclassified | unclassified | unclassified | unclassified | 0,4 | 0,489897949 | 0,000151564 | 0,00018644 | 0 - | 0 - | 0 - | 0 - | 1 - | 0,000406406 - |
| 956 | 1 | 0.2um_NOKCPHJN_409404 | Bacteria | Proteobacteri | Alphaproteob | Pelagibacteral | Pelagibactera | Candidatus Pe | Candidatus Pe | unclassified | unclassified | unclassified | unclassified | unclassified | 1 | 0 | 7,29E-05 | 7,71E-06 | 1 - | 8,66E-05 - | 1 - | 7,24E-05 - | 1 - | 7,29E-05 - |
| 957 | 1 | 0.2um_NOKCPHJN_1070961 | Eukaryota | Streptophyta | unclassified | Cucurbitales | Cucurbitaceae | Cucurbita | various | information st | RNA processir | KOG1783 | Component of | 39946.BGIOS | 1 | 0 | 0,000684698 | 7,24E-05 | 1 - | 0,000813567 - | 1 - | 0,000680162 - | 1 - | 0,000684706 - |
| 958 | 1 | 0.2um_NOKCPHJN_372100 | Bacteria | Proteobacteri | Gammaprotec | unclassified | unclassified | unclassified | Gammaprotec | unclassified | unclassified | unclassified | unclassified | unclassified | 0,6 | 0,489897949 | 0,000125843 | 0,000105104 | 1 - | 0,000248665 - | 0 - | 0 - | 0 - | 0 - |
| 959 | 1 | 3um_FIJJAFBF_771871 | unclassified | unclassified | unclassified | unclassified | unclassified | unclassified | unclassified | unclassified | unclassified | unclassified | unclassified | unclassified | 0,4 | 0,489897949 | 0,000116878 | 0,000143772 | 0 - | 0 - | 0 - | 0 - | 1 - | 0,000313398 - |
| 960 | 1 | 3um_FIJJAFBF_576166 | Bacteria | Proteobacteri | Alphaproteob | Rhodobactera | Rhodobactera | Flavimaricola | Flavimaricola | poorly charact | Function unkn | 05D2Z | Putative cycla | 290400.Jann_ | 0,6 | 0,489897949 | 0,000229347 | 0,000188259 | 0 - | 0 - | 1 - | 0,000398566 - | 1 - | 0,000401229 - |
| 961 | 1 | 3um_FIJJAFBF_91013 | Eukaryota | unclassified | Florideophyce | Gigartinales | Gigartinaceae | Chondrus | Chondrus cris | information st | Translation, ri | KOG3280 | ribosomal pro | 35128.Thaps3 | 1 | 0 | 0,000588712 | 6,23E-05 | 1 - | 0,000699516 - | 1 - | 0,000584812 - | 1 - | 0,000588719 - |
| 962 | 1 | 3um_FIJJAFBF_259312 | Bacteria | Planctomycete | Phycisphaerae | Phycisphaeral | Phycisphaerac | Phycisphaera | Phycisphaera | cellular proces | Cell wall/mem | 06B97 | phosphate-sel | 316067.Geob_ | 0,4 | 0,489897949 | 5,07E-05 | 6,21E-05 | 0 - | 0 - | 1 - | 0,000129287 - | 0 - | 0 - |
| 963 | 2 | 0.2um_NOKCPHJN_1003586;3um_FIJJA | Bacteria | Proteobacteri | Gammaprotec | Thiotrichales | Piscirickettsia | Methylophaga | Methylophaga | unclassified | unclassified | unclassified | unclassified | unclassified | 1 | 0 | 0,000144477 | 1,53E-05 | 1 - | 0,00017167 - | 1 - | 0,00014352 - | 1 - | 0,000144479 - |
| 964 | 1 | 0.2um_NOKCPHJN_974541 | Bacteria | Proteobacteri | Betaproteoba | Ferrovales | Ferrovaceae | Ferrovum | Ferrovum sp. | unclassified | unclassified | unclassified | unclassified | unclassified | 1 | 0 | 0,000104638 | 1,11E-05 | 1 - | 0,000124333 - | 1 - | 0,000103945 - | 1 - | 0,000104639 - |
| 965 | 1 | 0.2um_NOKCPHJN_1116524 | unclassified | unclassified | unclassified | unclassified | unclassified | unclassified | unclassified | unclassified | unclassified | unclassified | unclassified | unclassified | 1 | 0 | 0,000692222 | 7,32E-05 | 1 - | 0,000822508 - | 1 - | 0,000687636 - | 1 - | 0,00069223 - |
| 966 | 1 | 3um_FIJJAFBF_765335 | Eukaryota | Bacillariophyt | Bacillariophyc | Naviculales | Phaeodactylac | Phaeodactylu | Phaeodactylu | unclassified | unclassified | unclassified | unclassified | unclassified | 7,2 | 0,748331477 | 0,000781584 | 9,24E-05 | 7 - | 0,000906466 - | 8 - | 0,000866089 - | 6 - | 0,000653906 - |
| 967 | 1 | 3um_FIJJAFBF_24379 | Bacteria | Bacteroidetes | Flavobacteriia | Flavobacterial | Flavobacteria | Algibacter | Algibacter sp. | unclassified | unclassified | unclassified | unclassified | unclassified | 4,2 | 0,4 | 0,000508438 | 4,07E-05 | 4 - | 0,000579096 - | 4 - | 0,000484139 - | 4 - | 0,000487373 - |
| 968 | 1 | 3um_FIJJAFBF_372262 | Eukaryota | unclassified | Pelagophycea | Pelagomonad | unclassified | Aureococcus | Aureococcus a | information st | Translation, ri | 08UIK | This protein bi | 497965.Cyan7 | 0,2 | 0,4 | 7,51E-05 | 0,00015026 | 0 - | 0 - | 0 - | 0 - | 0 - | 0 - |
| 969 | 3 | 0.2um_NOKCPHJN_151309;3um_FIJJAI | Bacteria | Bacteroidetes | Flavobacteriia | Flavobacterial | unclassified | unclassified | Flavobacterial | unclassified | unclassified | unclassified | unclassified | unclassified | 0,2 | 0,4 | 3,51E-05 | 7,03E-05 | 0 - | 0 - | 0 - | 0 - | 0 - | 0 - |
| 970 | 1 | 3um_FIJJAFBF_418333 | Bacteria | Verrucomicrol | Opitutae | Puniceicoccale | Puniceicoccac | Coraliomargar | Coraliomargar | poorly charact | Function unkn | 1DIXS | Inherit from b | 382464.VDG1 | 1 | 0 | 0,000174012 | 1,84E-05 | 1 - | 0,000206763 - | 1 - | 0,000172859 - | 1 - | 0,000174014 - |
| 971 | 1 | 0.2um_NOKCPHJN_292273 | Eukaryota | Echinodermat | Holothuroidea | Aspidochirotic | Stichopodidae | Apostichopus | Apostichopus | unclassified | unclassified | unclassified | unclassified | unclassified | 0,2 | 0,4 | 1,61E-05 | 3,23E-05 | 1 - | 8,07E-05 - | 0 - | 0 - | 0 - | 0 - |
| 972 | 1 | 0.2um_NOKCPHJN_769317 | Eukaryota | Chordata | Mammalia | Proboscidea | Elephantidae | Loxodonta | Loxodonta afr | cellular proces | Cytoskeleton | 09W8V | centromere p | 9785.ENSLAFF | 0,4 | 0,489897949 | 2,84E-05 | 3,48E-05 | 0 - | 0 - | 0 - | 0 - | 1 - | 7,27E-05 - |
| 973 | 1 | 0.2um_NOKCPHJN_07570 | Bacteria | unclassified | unclassified | unclassified | unclassified | unclassified | uncultured ba | metabolism | Inorganic ion t | 05C27 | TonB-depende | 313596.RB250 | 0,6 | 0,8 | 3,28E-05 | 4,25E-05 | 0 - | 0 - | 1 - | 5,98E-05 - | 0 - | 0 - |
| 974 | 1 | 0.2um_NOKCPHJN_98378 | Eukaryota | unclassified | Oomycetes | Peronosporale | Peronosporac | Phytophthora | Phytophthora | unclassified | unclassified | unclassified | unclassified | unclassified | 0,6 | 0,489897949 | 7,52E-06 | 6,17E-06 | 0 - | 0 - | 1 - | 1,33E-05 - | 0 - | 0 - |
| 975 | 1 | 0.2um_NOKCPHJN_165213 | Bacteria | Bacteroidetes | Flavobacteriia | Flavobacterial | Flavobacteria | Flaviramulus | Flaviramulus i | unclassified | unclassified | unclassified | unclassified | unclassified | 0,2 | 0,4 | 2,20E-05 | 4,40E-05 | 0 - | 0 - | 0 - | 0 - | 0 - | 0 - |
| 976 | 1 | 0.2um_NOKCPHJN_49275 | Viruses | unclassified | unclassified | unclassified | Phycodnavirid | unclassified | Organic Lake p | unclassified | unclassified | unclassified | unclassified | unclassified | 0,2 | 0,4 | 2,02E-05 | 4,05E-05 | 0 - | 0 - | 0 - | 0 - | 0 - | 0 - |
| 977 | 1 | 3um_FIJJAFBF_594159 | Eukaryota | unclassified | Pelagophycea | Pelagomonad | unclassified | Aureoumbra | Aureoumbra l | metabolism | Energy produc | COG0055 | Produces ATP | 7739.JGI1257 | 8,2 | 0,4 | 0,001851911 | 0,000153849 | 8 - | 0,002153905 - | 8 - | 0,001800717 - | 8 - | 0,001812746 - |
| 978 | 1 | 0.2um_NOKCPHJN_231004 | Eukaryota | Phaeophyceae | unclassified | Ectocarpales | Ectocarpaceae | Ectocarpus | Ectocarpus sili | unclassified | unclassified | unclassified | unclassified | unclassified | 2 | 0 | 0,000656168 | 6,94E-05 | 2 - | 0,000779669 - | 2 - | 0,000651822 - | 2 - | 0,000656176 - |
| 979 | 1 | 0.2um_NOKCPHJN_616670 | Eukaryota | Phaeophyceae | unclassified | Ectocarpales | Ectocarpaceae | Ectocarpus | Ectocarpus sili | cellular proces | Signal transdu | 14ZNA | Inherit from K | 164328.Phyra | 1 | 0 | 0,000333292 | 3,52E-05 | 1 - | 0,000396022 - | 1 - | 0,000331084 - | 1 - | 0,000333296 - |
| 980 | 1 | 0.2um_NOKCPHJN_968394 | Eukaryota | unclassified | Cryptophyta | Pyrenomonad | various | various | various | unclassified | unclassified | unclassified | unclassified | unclassified | 3 | 0,632455532 | 0,002305392 | 0,000699695 | 4 - | 0,003607141 - | 3 - | 0,002261744 - | 2 - | 0,001517902 - |
| 981 | 1 | 3um_FIJJAFBF_33943 | Eukaryota | unclassified | unclassified | Isochrysidales | Noelaerhabda | Emiliania | Emiliania huxl | information st | Replication, re | 0WNHR | Domain of unl | 990316.MCON | 0,6 | 0,489897949 | 4,76E-05 | 3,98E-05 | 1 - | 9,41E-05 - | 0 - | 0 - | 0 - | 0 - |
| 982 | 2 | 3um_FIJJAFBF_226580;3um_FIJJAFBF_ | Bacteria | Proteobacteri | Alphaproteob | various | various | various | various | unclassified | unclassified | unclassified | unclassified | unclassified | 0,8 | 0,4 | 0,000505809 | 0,000257744 | 1 - | 0,000726681 - | 1 - | 0,000607523 - | 1 - | 0,000611582 - |
| 983 | 1 | 3um_FIJJAFBF_698996 | Bacteria | Proteobacteri | Alphaproteob | Rhodobactera | Rhodobactera | Tateyamaria | Tateyamaria s | metabolism | Amino acid tra | COG1932 | Catalyzes the | 391589.RGAI1 | 0,4 | 0,489897949 | 7,25E-05 | 8,94E-05 | 1 - | 0,000197489 - | 1 - | 0,000165105 - | 0 - | 0 - |
| 984 | 1 | 0.2um_NOKCPHJN_271001 | Eukaryota | unclassified | Labyrinthulom | unclassified | Thraustochytr | Aurantiochytr | Aurantiochytr | metabolism | Coenzyme tra | KOG1426 | regulator of cl | 65071.PYU1_ | 0,8 | 0,4 | 2,87E-05 | 1,48E-05 | 1 - | 4,26E-05 - | 1 - | 3,56E-05 - | 0 - | 0 - |
| 985 | 1 | 0.2um_NOKCPHJN_348376 | Eukaryota | Streptophyta | unclassified | Asterales | Asteraceae | Cynara | Cynara cardur | unclassified | unclassified | unclassified | unclassified | unclassified | 0,2 | 0,4 | 8,69E-06 | 1,74E-05 | 0 - | 0 - | 0 - | 0 - | 0 - | 0 - |
| 986 | 1 | 3um_FIJJAFBF_251780 | Bacteria | Proteobacteri | Alphaproteob | Rhodobactera | Rhodobactera | Rhodobacter | Candidatus Rh | unclassified | unclassified | unclassified | unclassified | unclassified | 0,8 | 0,4 | 0,000130376 | 6,73E-05 | 1 - | 0,000191428 - | 1 - | 0,000160038 - | 1 - | 0,000161107 - |
| 987 | 1 | 0.2um_NOKCPHJN_699789 | Eukaryota | Mucoromycot | Mortierellomy | Mortierellales | Mortierellacea | Mortierella | Mortierella ve | unclassified | unclassified | unclassified | unclassified | unclassified | 0,2 | 0,4 | 1,97E-05 | 3,94E-05 | 0 - | 0 - | 0 - | 0 - | 0 - | 0 - |
| 988 | 1 | 3um_FIJJAFBF_449534 | Eukaryota | Bacillariophyt | Coscinodiscop | Chaetocerotal | Chaetocerota | Chaetoceros | Chaetoceros s | unclassified | unclassified | unclassified | unclassified | unclassified | 0,4 | 0,489897949 | 0,00025116 | 0,000307751 | 0 - | 0 - | 0 - | 0 - | 1 - | 0,000642785 - |
| 989 | 1 | 3um_FIJJAFBF_100818 | Bacteria | Bacteroidetes | Flavobacteriia | Flavobacterial | Flavobacteria | Polaribacter | various | information st | Translation, ri | 09EM4 | Binds directly | 313598.MED1 | 1 | 0 | 0,000552563 | 5,84E-05 | 1 - | 0,000656563 - | 1 - | 0,000548903 - | 1 - | 0,00055257 - |
| 990 | 1 | 3um_FIJJAFBF_201913 | Bacteria | Bacteroidetes | Flavobacteriia | Flavobacterial | Flavobacteria | Hyunsoonleell | Hyunsoonleell | poorly charact | Function unkn | COG2322 | membrAne | 391603.FBALC | 0,4 | 0,489897949 | 0,000136584 | 0,000168013 | 0 - | 0 - | 0 - | 0 - | 1 - | 0,000366238 - |
| 991 | 2 | 0.2um_NOKCPHJN_18114;3um_FIJJAFE | Eukaryota | Mucoromycot | Glomeromyce | Glomerales | Glomeraceae | Rhizophagus | Rhizophagus c | unclassified | unclassified | unclassified | unclassified | unclassified | 0,4 | 0,489897949 | 9,92E-05 | 0,000121967 | 0 - | 0 - | 1 - | 0,000265148 - | 0 - | 0 - |
| 992 | 1 | 3um_FIJJAFBF_445608 | Eukaryota | Eustigmatoph | unclassified | Eustigmatales | Monodopsida | Nannochlorop | Nannochlorop | metabolism | Lipid transpor | COG0439 | acetyl-CoA car | 35128.Thaps6 | 4,2 | 1,166190379 | 0,000601623 | 0,000173279 | 5 - | 0,000852485 - | 4 - | 0,000570159 - | 3 - | 0,000430476 - |
| 993 | 1 | 3um_FIJJAFBF_485317 | Eukaryota | Phaeophyceae | unclassified | Ectocarpales | Ectocarpaceae | Ectocarpus | Ectocarpus sili | information st | Translation, ri | 151B4 | Ribosomal fan | 67593.Physo1 | 1,8 | 0,4 | 0,000456851 | 7,85E-05 | 1 - | 0,000308017 - | 2 - | 0,00051502 - | 2 - | 0,00051846 - |
| 994 | 1 | 0.2um_NOKCPHJN_82662 | Eukaryota | Bacillariophyt | Bacillariophyc | Naviculales | Phaeodactylac | Phaeodactylu | Phaeodactylu | information st | Replication, re | COG0513 | purine NTP-de | 281687.CJA01 | 0,2 | 0,4 | 2,09E-05 | 4,18E-05 | 0 - | 0 - | 0 - | 0 - | 1 - | 0,000104466 - |
| 995 | 1 | 0.2um_NOKCPHJN_479085 | Bacteria | Proteobacteri | Alphaproteob | Pelagibacteral | Pelagibactera | unclassified | Pelagibactera | unclassified | unclassified | unclassified | unclassified | unclassified | 0,2 | 0,4 | 0,000116651 | 0,000233303 | 0 - | 0 - | 0 - | 0 - | 0 - | 0 - |
| 996 | 1 | 3um_FIJJAFBF_250215 | Bacteria | Bacteroidetes | Saprospiria | Saprospirales | Saprospiracea | Aureispira | Aureispira sp. | unclassified | unclassified | unclassified | unclassified | unclassified | 0,4 | 0,489897949 | 9,13E-05 | 0,000112006 | 0 - | 0 - | 0 - | 0 - | 0 - | 0 - |
| 997 | 1 | 0.2um_NOKCPHJN_644123 | unclassified | unclassified | unclassified | unclassified | unclassified | unclassified | unclassified | unclassified | unclassified | unclassified | unclassified | unclassified | 0,6 | 0,489897949 | 4,25E-05 | 3,55E-05 | 1 - | 8,30E-05 - | 1 - | 6,94E-05 - | 0 - | 0 - |
| 998 | 1 | 3um_FIJJAFBF_805938 | unclassified | unclassified | unclassified | unclassified | unclassified | unclassified | unclassified | unclassified | unclassified | unclassified | unclassified | unclassified | 1,6 | 0,489897949 | 0,000851863 | 0,000317881 | 2 - | 0,00123716 - | 2 - | 0,001034296 - | 2 - | 0,001041206 - |
| 999 | 1 | 3um_FIJJAFBF_277459 | Eukaryota | Bacillariophyt | Coscinodiscop | Rhizosoleniale | Rhizosoleniac | Proboscia | Proboscia sp. | unclassified | unclassified | unclassified | unclassified | unclassified | 1,4 | 0,489897949 | 0,001042927 | 0,000353935 | 1 - | 0,00089105 - | 1 - | 0,00074494 - | 2 - | 0,001499832 - |
| 1000 | 1 | 0.2um_NOKCPHJN_573978 | Eukaryota | Eustigmatoph | unclassified | Eustigmatales | Monodopsida | Nannochlorop | Nannochlorop | unclassified | unclassified | unclassified | unclassified | unclassified | 1 | 0,632455532 | 9,51E-05 | 5,91E-05 | 1 - | 0,000110071 - | 1 - | 9,20E-05 - | 2 - | 0,000185273 - |
| 1001 | 1 | 3um_FIJJAFBF_526970 | unclassified | unclassified | unclassified | unclassified | unclassified | unclassified | unclassified | unclassified | unclassified | unclassified | unclassified | unclassified | 1,4 | 0,489897949 | 0,000125586 | 3,52E-05 | 1 - | 0,000109427 - | 1 - | 9,15E-05 - | 1 - | 9,21E-05 - |
| 1002 | 1 | 3um_FIJJAFBF_453302 | Bacteria | Proteobacteri | Alphaproteob | Rhodobactera | unclassified | unclassified | Rhodobactera | unclassified | unclassified | unclassified | unclassified | unclassified | 0,6 | 0,489897949 | 0,000128356 | 0,000107026 | 1 - | 0,000250328 - | 1 - | 0,000209281 - | 0 - | 0 - |
| 1003 | 1 | 3um_FIJJAFBF_274555 | unclassified | unclassified | unclassified | unclassified | unclassified | unclassified | unclassified | unclassified | unclassified | unclassified | unclassified | unclassified | 1 | 0 | 7,22E-05 | 7,63E-06 | 1 - | 8,57E-05 - | 1 - | 7,17E-05 - | 1 - | 7,22E-05 - |
| 1004 | 1 | 3um_FIJJAFBF_118448 | Bacteria | Proteobacteri | Alphaproteob | Rhodobactera | Rhodobactera | unclassified | Rhodobactera | unclassified | unclassified | unclassified | unclassified | unclassified | 0,6 | 0,489897949 | 7,87E-05 | 6,46E-05 | 0 - | 0 - | 0 - | 0 - | 1 - | 0,000139674 - |
| 1005 | 2 | 3um_FIJJAFBF_756834;3um_FIJJAFBF_ | unclassified | unclassified | unclassified | unclassified | unclassified | unclassified | unclassified | unclassified | unclassified | unclassified | unclassified | unclassified | 1 | 0 | 0,00057791 | 6,11E-05 | 1 - | 0,000686681 - | 1 - | 0,000574082 - | 1 - | 0,000577917 - |

| | | | | | | | | | | | | | | | | | | | | | | | | |
|---|---|---|---|---|---|---|---|---|---|---|---|---|---|---|---|---|---|---|---|---|---|---|---|---|
| 1006 | 1 | 3um_FIJJAFBF_10571 | unclassified | unclassified | unclassified | unclassified | unclassified | unclassified | unclassified | unclassified | unclassified | unclassified | unclassified | unclassified | 0,6 | 0,489897949 | 0,000407172 | 0,000333821 | 0 - | 0 - | 1 - | 0,000719252 - | 0 - | 0 - |
| 1007 | 1 | 3um_FIJJAFBF_33587 | Bacteria | Proteobacteri | Alphaproteob | Sphingomona | Sphingomona | Sphingomona | Sphingomona | unclassified | unclassified | unclassified | unclassified | unclassified | 1,8 | 0,748331477 | 0,000147816 | 6,07E-05 | 1 - | 9,82E-05 - | 3 - | 0,000246358 - | 2 - | 0,000165336 - |
| 1008 | 1 | 0.2um_NOKCPHJN_23322 | Eukaryota | unclassified | Pelagophycea | Pelagomonad | unclassified | Pelagomonas | various | cellular proces | Posttranslatio | KOG0730 | cell division cy | 35128.Thapsd | 6,2 | 0,4 | 0,001340851 | 0,000106736 | 6 - | 0,001548583 - | 6 - | 0,001294653 - | 6 - | 0,001303302 - |
| 1009 | 1 | 3um_FIJJAFBF_122637 | Bacteria | Actinobacteri | Acidimicrobiia | Acidimicrobial | unclassified | unclassified | Acidimicrobial | unclassified | unclassified | unclassified | unclassified | unclassified | 0,2 | 0,4 | 2,61E-05 | 5,23E-05 | 0 - | 0 - | 0 - | 0 - | 1 - | 0,000130691 - |
| 1010 | 1 | 3um_FIJJAFBF_119800 | unclassified | unclassified | unclassified | unclassified | unclassified | unclassified | unclassified | unclassified | unclassified | unclassified | unclassified | unclassified | 0,8 | 0,4 | 0,000362544 | 0,0001875 | 1 - | 0,000538476 - | 1 - | 0,000450179 - | 0 - | 0 - |
| 1011 | 1 | 0.2um_NOKCPHJN_548395 | Bacteria | Bacteroidetes | Flavobacteriia | Flavobacterial | Flavobacteria | Polaribacter | Polaribacter a | poorly charact | Function unkn | 0ZVFE | GH3 auxin-res | 313598.MED1 | 0,2 | 0,4 | 2,52E-05 | 5,05E-05 | 0 - | 0 - | 0 - | 0 - | 1 - | 0,000126238 - |
| 1012 | 1 | 3um_FIJJAFBF_205191 | Eukaryota | Bacillariophyt | Coscinodiscop | Chaetocerotal | Acanthocerat | Acanthoceras | Acanthoceras | metabolism | Energy produc | 0IFBF | One of the tw | 45157.CMV08 | 4,8 | 0,4 | 0,000854676 | 4,33E-05 | 4 - | 0,000852971 - | 5 - | 0,000891381 - | 5 - | 0,000897335 - |
| 1013 | 1 | 3um_FIJJAFBF_263677 | Eukaryota | Bacillariophyt | Coscinodiscop | Rhizosoleniale | Rhizosoleniac | Rhizosolenia | Rhizosolenia s | metabolism | Energy produc | 11IIK | Low-potential | 35128.Thapsd | 6 | 0,632455532 | 0,002332288 | 0,000224179 | 5 - | 0,002324478 - | 7 - | 0,002720649 - | 6 - | 0,002347563 - |
| 1014 | 1 | 3um_FIJJAFBF_400448 | Eukaryota | Bacillariophyt | Bacillariophyc | Naviculales | Naviculaceae | Fistulifera | Fistulifera sola | cellular proces | Posttranslatio | KOG1641 | 10 kda heat sh | 45157.CMS18 | 0,8 | 0,748331477 | 0,000345737 | 0,000310256 | 1 - | 0,000502337 - | 1 - | 0,000419966 - | 0 - | 0 - |
| 1015 | 1 | 0.2um_NOKCPHJN_352204 | Eukaryota | Chordata | unclassified | unclassified | Branchiostom | Branchiostom | Branchiostom | unclassified | unclassified | unclassified | unclassified | unclassified | 0,4 | 0,489897949 | 1,13E-05 | 1,39E-05 | 0 - | 0 - | 0 - | 0 - | 1 - | 3,02E-05 - |
| 1016 | 1 | 0.2um_NOKCPHJN_458305 | Bacteria | unclassified | unclassified | unclassified | unclassified | unclassified | bacterium B17 | unclassified | unclassified | unclassified | unclassified | unclassified | 0,4 | 0,489897949 | 1,33E-05 | 1,64E-05 | 0 - | 0 - | 1 - | 3,56E-05 - | 0 - | 0 - |
| 1017 | 1 | 0.2um_NOKCPHJN_547411 | Bacteria | Proteobacteri | Alphaproteob | Rhizobiales | unclassified | unclassified | Rhizobiales ba | unclassified | unclassified | unclassified | unclassified | unclassified | 0,4 | 0,489897949 | 1,88E-05 | 2,31E-05 | 0 - | 0 - | 0 - | 0 - | 1 - | 4,82E-05 - |
| 1018 | 1 | 3um_FIJJAFBF_285697 | Eukaryota | Phaeophyceae | unclassified | Ectocarpales | Ectocarpaceae | Ectocarpus | Ectocarpus sili | unclassified | unclassified | unclassified | unclassified | unclassified | 1 | 0 | 0,000732467 | 7,75E-05 | 1 - | 0,000870328 - | 1 - | 0,000727615 - | 1 - | 0,000732476 - |
| 1019 | 1 | 3um_FIJJAFBF_399389 | Bacteria | Bacteroidetes | Flavobacteriia | Flavobacterial | Flavobacteria | unclassified | Flavobacteria | unclassified | unclassified | unclassified | unclassified | unclassified | 1 | 0 | 0,00015106 | 1,60E-05 | 1 - | 0,000179492 - | 1 - | 0,00015006 - | 1 - | 0,000151062 - |
| 1020 | 1 | 3um_FIJJAFBF_715061 | Eukaryota | unclassified | Rhodellophyce | Rhodellales | Rhodellaceae | Corynoplastis | Corynoplastis | unclassified | unclassified | unclassified | unclassified | unclassified | 2,2 | 0,748331477 | 0,001791614 | 0,000740439 | 3 - | 0,002842336 - | 2 - | 0,001584175 - | 3 - | 0,002392137 - |
| 1021 | 1 | 0.2um_NOKCPHJN_275213 | unclassified | unclassified | unclassified | unclassified | unclassified | unclassified | unclassified | unclassified | unclassified | unclassified | unclassified | unclassified | 0,2 | 0,4 | 2,82E-05 | 5,64E-05 | 0 - | 0 - | 0 - | 0 - | 0 - | 0 - |
| 1022 | 1 | 0.2um_NOKCPHJN_627902 | unclassified | unclassified | unclassified | unclassified | unclassified | unclassified | unclassified | unclassified | unclassified | unclassified | unclassified | unclassified | 1,2 | 0,4 | 0,000212316 | 5,25E-05 | 1 - | 0,000215081 - | 1 - | 0,000179813 - | 1 - | 0,000181014 - |
| 1023 | 1 | 0.2um_NOKCPHJN_316872 | Bacteria | Proteobacteri | Alphaproteob | unclassified | unclassified | unclassified | various | metabolism | Energy produc | 01RNF | Produces ATP | 862719.AZOLI | 1 | 0 | 0,000134312 | 1,42E-05 | 1 - | 0,000159591 - | 1 - | 0,000133422 - | 1 - | 0,000134313 - |
| 1024 | 1 | 3um_FIJJAFBF_57085 | Bacteria | Proteobacteri | Gammaprotec | Cellvibrionales | Porticoccacea | unclassified | Gammaprotec | poorly charact | Function unkn | 16R8I | Lipoprotein | 377629.TERTU | 2,2 | 0,4 | 0,00016432 | 2,01E-05 | 2 - | 0,000179708 - | 2 - | 0,00015024 - | 2 - | 0,000151244 - |
| 1025 | 1 | 0.2um_NOKCPHJN_665331 | Bacteria | unclassified | unclassified | unclassified | unclassified | unclassified | uncultured ma | metabolism | Amino acid tra | 01SDM | gamma-glutar | 367336.OM22 | 0,2 | 0,4 | 2,39E-05 | 4,78E-05 | 0 - | 0 - | 1 - | 0,000119418 - | 0 - | 0 - |
| 1026 | 1 | 0.2um_NOKCPHJN_796234 | Eukaryota | Chromerida | unclassified | unclassified | unclassified | Vitrella | Vitrella brassic | unclassified | unclassified | unclassified | unclassified | unclassified | 0,4 | 0,489897949 | 0,00047727 | 0,0005857 | 0 - | 0 - | 0 - | 0 - | 0 - | 0 - |
| 1027 | 1 | 3um_FIJJAFBF_788045 | Bacteria | Proteobacteri | Alphaproteob | Rhodobactera | unclassified | unclassified | Rhodobactera | information st | Translation, ri | 17C3B | The globular d | 439497.RR11_ | 0,2 | 0,4 | 1,00E-04 | 0,000199978 | 0 - | 0 - | 0 - | 0 - | 1 - | 0,000499944 - |
| 1028 | 1 | 3um_FIJJAFBF_100811 | unclassified | unclassified | unclassified | unclassified | unclassified | unclassified | unclassified | unclassified | unclassified | unclassified | unclassified | unclassified | 0,4 | 0,489897949 | 3,07E-05 | 3,76E-05 | 0 - | 0 - | 0 - | 0 - | 0 - | 0 - |
| 1029 | 1 | 0.2um_NOKCPHJN_987128 | unclassified | unclassified | unclassified | unclassified | unclassified | unclassified | unclassified | unclassified | unclassified | unclassified | unclassified | unclassified | 1,2 | 0,4 | 0,00045892 | 0,00011341 | 1 - | 0,000464896 - | 1 - | 0,000388664 - | 1 - | 0,00039126 - |
| 1030 | 1 | 0.2um_NOKCPHJN_864502 | Bacteria | unclassified | unclassified | unclassified | unclassified | unclassified | uncultured ba | unclassified | unclassified | unclassified | unclassified | unclassified | 1 | 0 | 0,00020452 | 2,16E-05 | 1 - | 0,000243014 - | 1 - | 0,000203165 - | 1 - | 0,000204523 - |
| 1031 | 1 | 3um_FIJJAFBF_756153 | Eukaryota | Bacillariophyt | Coscinodiscop | Thalassiosirale | Thalassiosirac | Thalassiosira | Thalassiosira p | cellular proces | Signal transdu | COG5040 | Tyrosine 3-mo | 35128.Thaps2 | 0,2 | 0,4 | 0,000107888 | 0,000215776 | 0 - | 0 - | 1 - | 0,000539439 - | 0 - | 0 - |
| 1032 | 1 | 0.2um_NOKCPHJN_613397 | Bacteria | Proteobacteri | Betaproteoba | Nitrosomonad | Thiobacillacea | Thiobacillus | Thiobacillus d | unclassified | unclassified | unclassified | unclassified | unclassified | 0,2 | 0,4 | 2,14E-05 | 4,29E-05 | 0 - | 0 - | 0 - | 0 - | 0 - | 0 - |
| 1033 | 1 | 3um_FIJJAFBF_513825 | Viruses | unclassified | unclassified | unclassified | unclassified | unclassified | uncultured M | unclassified | unclassified | unclassified | unclassified | unclassified | 1 | 0 | 0,000359955 | 3,81E-05 | 1 - | 0,000427704 - | 1 - | 0,000357571 - | 1 - | 0,00035996 - |
| 1034 | 1 | 3um_FIJJAFBF_128440 | Bacteria | Bacteroidetes | Flavobacteriia | Flavobacterial | Flavobacteria | Polaribacter | Polaribacter s | poorly charact | Function unkn | 0P8NI | Inherit from b | 313594.PI23P | 10,8 | 1,469693846 | 0,001008893 | 6,30E-05 | 8 - | 0,000900429 - | 11 - | 0,001035074 - | 11 - | 0,001041988 - |
| 1035 | 1 | 3um_FIJJAFBF_545834 | Bacteria | Bacteroidetes | Flavobacteriia | Flavobacterial | Flavobacteria | Polaribacter | Polaribacter s | metabolism | Carbohydrate | 099YR | Catalyzes the | 313598.MED1 | 2,8 | 0,4 | 0,000411332 | 7,58E-05 | 3 - | 0,000523414 - | 2 - | 0,000291725 - | 3 - | 0,00044051 - |
| 1036 | 1 | 3um_FIJJAFBF_555595 | Bacteria | Cyanobacteria | unclassified | Oscillatoriales | Microcoleace | Planktothricoi | Planktothricoi | poorly charact | Function unkn | 12527 | NA | 513049.Amax | 1 | 0 | 0,000101764 | 1,08E-05 | 1 - | 0,000120918 - | 1 - | 0,00010109 - | 1 - | 0,000101766 - |
| 1037 | 1 | 3um_FIJJAFBF_156011 | Eukaryota | Chlorophyta | Chlorophycea | Chlamydomor | Volvocaceae | Gonium | Gonium pecto | unclassified | unclassified | unclassified | unclassified | unclassified | 0,4 | 0,489897949 | 5,61E-05 | 7,01E-05 | 1 - | 0,00016236 - | 0 - | 0 - | 0 - | 0 - |
| 1038 | 1 | 3um_FIJJAFBF_629419 | Bacteria | Bacteroidetes | Flavobacteriia | Flavobacterial | Flavobacteria | Polaribacter | Polaribacter ta | metabolism | Inorganic ion t | 05C27 | TonB-depende | 873517.HMPR | 0,2 | 0,4 | 1,09E-05 | 2,18E-05 | 0 - | 0 - | 0 - | 0 - | 0 - | 0 - |
| 1039 | 1 | 0.2um_NOKCPHJN_446749 | Eukaryota | unclassified | Oligohymenop | Hymenostoma | Tetrahymenid | Tetrahymena | Tetrahymena | cellular proces | Signal transdu | KOG0613 | fibronectin typ | 7165.AGAP00 | 0,4 | 0,489897949 | 1,36E-05 | 1,67E-05 | 0 - | 0 - | 0 - | 0 - | 0 - | 0 - |
| 1040 | 2 | 0.2um_NOKCPHJN_968216;3um_FIJJAF | Eukaryota | various | various | various | various | various | various | information st | Chromatin str | various | various | various | 1 | 0 | 0,00040904 | 4,33E-05 | 1 - | 0,000486027 - | 1 - | 0,000406331 - | 1 - | 0,000409045 - |
| 1041 | 1 | 3um_FIJJAFBF_706909 | unclassified | unclassified | unclassified | unclassified | unclassified | unclassified | unclassified | unclassified | unclassified | unclassified | unclassified | unclassified | 0,8 | 0,4 | 0,000458124 | 0,000236932 | 1 - | 0,000680438 - | 1 - | 0,000568863 - | 0 - | 0 - |
| 1042 | 1 | 3um_FIJJAFBF_333454 | Bacteria | Proteobacteri | Gammaprotec | Cellvibrionales | unclassified | unclassified | Cellvibrionales | unclassified | unclassified | unclassified | unclassified | unclassified | 0,2 | 0,4 | 3,07E-05 | 6,15E-05 | 0 - | 0 - | 0 - | 0 - | 1 - | 0,000153641 - |
| 1043 | 1 | 3um_FIJJAFBF_277461 | Bacteria | Cyanobacteria | unclassified | Oscillatoriales | unclassified | unclassified | filamentous cy | information st | Translation, ri | 08UIK | This protein bi | 1140.Synpcc7 | 1,6 | 1,019803903 | 0,000669674 | 0,000405524 | 0 - | 0 - | 1 - | 0,000443794 - | 2 - | 0,000893517 - |
| 1044 | 1 | 0.2um_NOKCPHJN_785873 | Eukaryota | Bacillariophyt | Coscinodiscop | Thalassiosirale | Thalassiosirac | Thalassiosira | Thalassiosira c | information st | Chromatin str | KOG1744 | Core compone | 35128.Thaps6 | 1 | 0 | 0,00057791 | 6,11E-05 | 1 - | 0,000686681 - | 1 - | 0,000574082 - | 1 - | 0,000577917 - |
| 1045 | 1 | 0.2um_NOKCPHJN_700348 | Eukaryota | Bacillariophyt | Bacillariophyc | Naviculales | Naviculaceae | Fistulifera | Fistulifera sola | metabolism | Carbohydrate | KOG2638 | UTP-glucose-1 | 35128.Thaps2 | 1,2 | 0,4 | 0,000912174 | 0,00022542 | 1 - | 0,000924052 - | 1 - | 0,00077253 - | 1 - | 0,000777691 - |
| 1046 | 1 | 3um_FIJJAFBF_85802 | Viruses | unclassified | unclassified | unclassified | Phycodnavirid | unclassified | Chrysochromu | information st | Replication, re | COG0513 | purine NTP-de | 7159.AAEL014 | 0,8 | 0,4 | 0,000126509 | 6,54E-05 | 1 - | 0,000187589 - | 0 - | 0 - | 1 - | 0,000157877 - |
| 1047 | 1 | 3um_FIJJAFBF_715054 | Bacteria | Cyanobacteria | unclassified | Oscillatoriales | Cyanothecace | Cyanothece | various | unclassified | unclassified | unclassified | unclassified | unclassified | 1 | 0 | 0,00067013 | 7,09E-05 | 1 - | 0,000796257 - | 1 - | 0,000665691 - | 1 - | 0,000670138 - |
| 1048 | 1 | 3um_FIJJAFBF_205192 | Eukaryota | Bacillariophyt | Mediophyceae | Toxariales | Toxariaceae | Toxarium | Toxarium und | cellular proces | Posttranslatio | 0IEKR | One of the cor | 45157.CMV08 | 2,8 | 0,748331477 | 0,000376657 | 7,43E-05 | 2 - | 0,000326136 - | 3 - | 0,000408986 - | 3 - | 0,000411719 - |
| 1049 | 1 | 3um_FIJJAFBF_329053 | Bacteria | Bacteroidetes | Flavobacteriia | Flavobacterial | Flavobacteria | Formosa | Formosa sp. H | information st | Translation, ri | COG0184 | Forms an inte | 983544.Lacal_ | 0,6 | 0,489897949 | 0,000352096 | 0,000293585 | 1 - | 0,000686681 - | 1 - | 0,000574082 - | 0 - | 0 - |
| 1050 | 1 | 0.2um_NOKCPHJN_123174 | Eukaryota | Mollusca | Bivalvia | Ostreoida | Ostreidae | Crassostrea | Crassostrea gi | unclassified | unclassified | unclassified | unclassified | unclassified | 0,2 | 0,4 | 1,52E-05 | 3,04E-05 | 0 - | 0 - | 1 - | 7,60E-05 - | 0 - | 0 - |
| 1051 | 1 | 3um_FIJJAFBF_715086 | Eukaryota | unclassified | Pelagophycea | Pelagomonad | unclassified | Aureoumbra | Aureoumbra l | unclassified | unclassified | unclassified | unclassified | unclassified | 0,4 | 0,489897949 | 0,0002437 | 0,00029861 | 0 - | 0 - | 0 - | 0 - | 1 - | 0,000623692 - |
| 1052 | 1 | 0.2um_NOKCPHJN_1065485 | unclassified | unclassified | unclassified | unclassified | unclassified | unclassified | unclassified | unclassified | unclassified | unclassified | unclassified | unclassified | 0,4 | 0,489897949 | 0,000178976 | 0,000219637 | 0 - | 0 - | 0 - | 0 - | 0 - | 0 - |
| 1053 | 1 | 0.2um_NOKCPHJN_509418 | unclassified | unclassified | unclassified | unclassified | unclassified | unclassified | unclassified | unclassified | unclassified | unclassified | unclassified | unclassified | 1 | 0 | 0,000101274 | 1,07E-05 | 1 - | 0,000120335 - | 1 - | 0,000100603 - | 1 - | 0,000101275 - |
| 1054 | 3 | 0.2um_NOKCPHJN_504918;3um_FIJJAF | Eukaryota | various | various | various | various | various | various | unclassified | unclassified | unclassified | unclassified | unclassified | 0,8 | 0,4 | 0,000392131 | 0,000202341 | 1 - | 0,000575755 - | 1 - | 0,000481346 - | 1 - | 0,000484561 - |
| 1055 | 1 | 3um_FIJJAFBF_393524 | Bacteria | Bacteroidetes | Flavobacteriia | Flavobacterial | Flavobacteria | Confluentibac | Confluentibac | metabolism | Secondary me | 099V5 | phenylacetate | 1046627.BZAR | 1 | 0 | 0,000145815 | 1,54E-05 | 1 - | 0,00017326 - | 1 - | 0,000144849 - | 1 - | 0,000145817 - |
| 1056 | 1 | 0.2um_NOKCPHJN_862435 | Eukaryota | unclassified | unclassified | Prymnesiales | Chrysochromu | Chrysochromu | Chrysochromu | unclassified | unclassified | unclassified | unclassified | unclassified | 3 | 0,894427191 | 0,000549329 | 0,000148534 | 3 - | 0,000660425 - | 2 - | 0,000368088 - | 2 - | 0,000370547 - |
| 1057 | 1 | 3um_FIJJAFBF_565197 | unclassified | unclassified | unclassified | unclassified | unclassified | unclassified | unclassified | unclassified | unclassified | unclassified | unclassified | unclassified | 1 | 0,632455532 | 0,001076012 | 0,000665384 | 1 - | 0,001290486 - | 1 - | 0,001078878 - | 0 - | 0 - |
| 1058 | 1 | 0.2um_NOKCPHJN_995236 | unclassified | unclassified | unclassified | unclassified | unclassified | unclassified | unclassified | poorly charact | Function unkn | 0A7Q8 | Inherit from N | 7668.SPU_004 | 0,6 | 0,489897949 | 6,70E-05 | 5,53E-05 | 1 - | 0,000126647 - | 0 - | 0 - | 1 - | 0,000106587 - |
| 1059 | 1 | 0.2um_NOKCPHJN_865894 | Eukaryota | unclassified | Oomycetes | Saprolegniales | Saprolegniace | Aphanomyces | Aphanomyces | information st | Replication, re | COG0513 | purine NTP-de | 4787.PITG_03 | 1 | 0 | 0,00016979 | 1,80E-05 | 1 - | 0,000201747 - | 1 - | 0,000168666 - | 1 - | 0,000169792 - |
| 1060 | 2 | 0.2um_NOKCPHJN_1117093;3um_FIJJA | Eukaryota | Bacillariophyt | various | various | various | various | various | unclassified | unclassified | unclassified | unclassified | unclassified | 0,6 | 0,489897949 | 6,44E-05 | 5,32E-05 | 1 - | 0,000122101 - | 1 - | 0,00010208 - | 0 - | 0 - |
| 1061 | 1 | 0.2um_NOKCPHJN_794309 | Bacteria | Proteobacteri | Deltaproteoba | unclassified | unclassified | Candidatus Ba | Candidatus Ba | cellular proces | Cell wall/mem | KOG4177 | Ankyrin Repea | 5722.A2EK94 | 0,8 | 0,4 | 0,000148007 | 7,54E-05 | 1 - | 0,000212637 - | 1 - | 0,00017777 - | 1 - | 0,000178957 - |
| 1062 | 1 | 3um_FIJJAFBF_178652 | unclassified | unclassified | unclassified | unclassified | unclassified | unclassified | unclassified | unclassified | unclassified | unclassified | unclassified | unclassified | 0,2 | 0,4 | 0,000166866 | 0,000333733 | 0 - | 0 - | 1 - | 0,000834332 - | 0 - | 0 - |
| 1063 | 1 | 0.2um_NOKCPHJN_1001141 | Bacteria | various | various | various | various | various | various | information st | Translation, ri | COG0051 | tRNA binding | 205922.Pfl01_ | 1 | 0 | 0,000611575 | 6,47E-05 | 1 - | 0,000726681 - | 1 - | 0,000607523 - | 1 - | 0,000611582 - |
| 1064 | 1 | 3um_FIJJAFBF_62816 | Bacteria | Bacteroidetes | Flavobacteriia | Flavobacterial | Flavobacteria | Polaribacter | Polaribacter s | unclassified | unclassified | unclassified | unclassified | unclassified | 0,6 | 0,489897949 | 5,28E-05 | 4,36E-05 | 1 - | 9,98E-05 - | 0 - | 0 - | 1 - | 8,40E-05 - |
| 1065 | 5 | 0.2um_NOKCPHJN_45711;0.2um_NOK | Bacteria | various | various | various | various | various | various | information st | Translation, ri | 0PAE2 | One of the pri | various | 0,6 | 0,489897949 | 0,000423617 | 0,000347725 | 0 - | 0 - | 1 - | 0,000736176 - | 1 - | 0,000741093 - |
| 1066 | 1 | 3um_FIJJAFBF_468363 | Bacteria | Proteobacteri | Gammaprotec | unclassified | unclassified | unclassified | Gammaprotec | unclassified | unclassified | unclassified | unclassified | unclassified | 0,4 | 0,489897949 | 8,15E-05 | 9,98E-05 | 0 - | 0 - | 1 - | 0,00020789 - | 0 - | 0 - |
| 1067 | 1 | 0.2um_NOKCPHJN_836194 | Eukaryota | unclassified | Pelagophycea | Pelagomonad | unclassified | Aureococcus | Aureococcus a | poorly charact | Function unkn | 0XQGZ | guanine nucle | 164328.Phyra | 1 | 0,894427191 | 0,000297082 | 0,000263123 | 0 - | 0 - | 2 - | 0,000625749 - | 1 - | 0,000314965 - |
| 1068 | 1 | 3um_FIJJAFBF_639225 | Bacteria | Planctomycete | Phycisphaerae | Phycisphaeral | Phycisphaerac | Phycisphaera | Phycisphaera | unclassified | unclassified | unclassified | unclassified | unclassified | 0,2 | 0,4 | 3,18E-05 | 6,36E-05 | 0 - | 0 - | 0 - | 0 - | 0 - | 0 - |
| 1069 | 1 | 0.2um_NOKCPHJN_292026 | unclassified | unclassified | unclassified | unclassified | unclassified | unclassified | unclassified | unclassified | unclassified | unclassified | unclassified | unclassified | 0,8 | 0,4 | 7,26E-05 | 3,76E-05 | 1 - | 0,00010785 - | 1 - | 9,02E-05 - | 0 - | 0 - |
| 1070 | 1 | 0.2um_NOKCPHJN_720672 | unclassified | unclassified | unclassified | unclassified | unclassified | unclassified | unclassified | unclassified | unclassified | unclassified | unclassified | unclassified | 0,8 | 0,4 | 5,01E-05 | 2,52E-05 | 0 - | 0 - | 1 - | 6,53E-05 - | 1 - | 6,57E-05 - |
| 1071 | 1 | 0.2um_NOKCPHJN_107953 | Bacteria | Bacteroidetes | Flavobacteriia | Flavobacterial | Flavobacteria | Polaribacter | Polaribacter s | metabolism | Inorganic ion t | 05C27 | TonB-depende | 313590.MED1 | 0,2 | 0,4 | 1,08E-05 | 2,17E-05 | 0 - | 0 - | 0 - | 0 - | 0 - | 0 - |
| 1072 | 1 | 0.2um_NOKCPHJN_597809 | Bacteria | Proteobacteri | Epsilonproteo | Campylobacte | Campylobacte | Sulfurospirillu | Sulfurospirillu | unclassified | unclassified | unclassified | unclassified | unclassified | 1 | 0 | 0,000250965 | 2,65E-05 | 1 - | 0,0002982 - | 1 - | 0,000249302 - | 1 - | 0,000250968 - |
| 1073 | 1 | 0.2um_NOKCPHJN_967210 | Bacteria | Firmicutes | Bacilli | Lactobacillales | Lactobacillace | Lactobacillus | Lactobacillus k | unclassified | unclassified | unclassified | unclassified | unclassified | 1 | 0 | 0,000143164 | 1,51E-05 | 1 - | 0,00017011 - | 1 - | 0,000142216 - | 1 - | 0,000143166 - |
| 1074 | 1 | 3um_FIJJAFBF_44059 | Viruses | unclassified | unclassified | unclassified | Phycodnavirid | unclassified | Chrysochromu | unclassified | unclassified | unclassified | unclassified | unclassified | 8,6 | 0,489897949 | 0,002902929 | 0,000248988 | 8 - | 0,003219277 - | 9 - | 0,003027819 - | 9 - | 0,003048045 - |
| 1075 | 1 | 0.2um_NOKCPHJN_596098 | Eukaryota | unclassified | Oomycetes | Peronosporale | Peronosporac | Phytophthora | Phytophthora | information st | Translation, ri | COG0052 | 30S ribosomal | 65071.PYU1_T | 1 | 0 | 0,00020452 | 2,16E-05 | 1 - | 0,000243014 - | 1 - | 0,000203165 - | 1 - | 0,000204523 - |
| 1076 | 1 | 3um_FIJJAFBF_215318 | Bacteria | Bacteroidetes | Flavobacteriia | Flavobacterial | Flavobacteria | Kordia | Kordia algicida | metabolism | Amino acid tra | 0P8J9 | DegT DnrJ Ery | 391587.KAOT | 1 | 0 | 0,000167088 | 1,77E-05 | 1 - | 0,000198536 - | 1 - | 0,000165981 - | 1 - | 0,00016709 - |
| 1077 | 1 | 0.2um_NOKCPHJN_123309 | Bacteria | Proteobacteri | Gammaprotec | Oceanospirilla | Saccharospiril | Reinekea | Reinekea fors | poorly charact | Function unkn | 08W7N | glutathione-de | 203122.Sde_0 | 0,2 | 0,4 | 8,71E-05 | 0,000174132 | 0 - | 0 - | 0 - | 0 - | 0 - | 0 - |
| 1078 | 1 | 0.2um_NOKCPHJN_30971 | Bacteria | Proteobacteri | Alphaproteob | Rhodospirillale | Rhodospirillac | unclassified | Rhodobactera | metabolism | Inorganic ion t | 05CAU | Arsenical resis | 367336.OM22 | 0,2 | 0,4 | 5,13E-05 | 0,000102693 | 0 - | 0 - | 0 - | 0 - | 0 - | 0 - |
| 1079 | 2 | 0.2um_NOKCPHJN_708367;0.2um_NO | unclassified | unclassified | unclassified | unclassified | unclassified | unclassified | unclassified | unclassified | unclassified | unclassified | unclassified | unclassified | 1,2 | 0,748331477 | 0,000430813 | 0,000265235 | 0 - | 0 - | 1 - | 0,000376957 - | 2 - | 0,000758951 - |
| 1080 | 1 | 3um_FIJJAFBF_156949 | unclassified | unclassified | unclassified | unclassified | unclassified | unclassified | unclassified | unclassified | unclassified | unclassified | unclassified | unclassified | 0,2 | 0,4 | 1,04E-05 | 2,07E-05 | 0 - | 0 - | 1 - | 5,18E-05 - | 0 - | 0 - |
| 1081 | 1 | 0.2um_NOKCPHJN_851921 | Bacteria | Proteobacteri | Gammaprotec | unclassified | unclassified | unclassified | OM182 bacter | metabolism | Amino acid tra | 07QZM | n-acetylglutan | 575586.HMPR | 0,2 | 0,4 | 2,60E-05 | 5,19E-05 | 0 - | 0 - | 0 - | 0 - | 0 - | 0 - |
| 1082 | 1 | 3um_FIJJAFBF_357687 | unclassified | unclassified | unclassified | unclassified | unclassified | unclassified | unclassified | unclassified | unclassified | unclassified | unclassified | unclassified | 0,8 | 0,4 | 9,65E-05 | 4,99E-05 | 1 - | 0,000143113 - | 0 - | 0 - | 1 - | 0,000120445 - |
| 1083 | 1 | 3um_FIJJAFBF_342671 | Bacteria | Bacteroidetes | Flavobacteriia | Flavobacterial | Flavobacteria | Formosa | Formosa sp. H | metabolism | Amino acid tra | 05CRQ | synthase (Con | 1046627.BZAR | 0,4 | 0,489897949 | 5,34E-05 | 6,55E-05 | 0 - | 0 - | 0 - | 0 - | 0 - | 0 - |
| 1084 | 1 | 3um_FIJJAFBF_171859 | unclassified | unclassified | unclassified | unclassified | unclassified | unclassified | unclassified | unclassified | unclassified | unclassified | unclassified | unclassified | 0,2 | 0,4 | 6,71E-05 | 0,000134247 | 0 - | 0 - | 0 - | 0 - | 0 - | 0 - |
| 1085 | 1 | 3um_FIJJAFBF_719178 | Bacteria | unclassified | unclassified | unclassified | unclassified | unclassified | bacterium TM | poorly charact | Function unkn | 0HJ78 | Uncharacteris | 56780.SYN_00 | 0,2 | 0,4 | 2,68E-05 | 5,36E-05 | 0 - | 0 - | 0 - | 0 - | 1 - | 0,000134028 - |
| 1086 | 1 | 0.2um_NOKCPHJN_103568 | Bacteria | Proteobacteri | Gammaprotec | Alteromonada | Alteromonada | Paraglaciecola | Paraglaciecola | unclassified | unclassified | unclassified | unclassified | unclassified | 1 | 0 | 0,000221025 | 2,34E-05 | 1 - | 0,000262625 - | 1 - | 0,000219561 - | 1 - | 0,000221028 - |
| 1087 | 1 | 3um_FIJJAFBF_183675 | Bacteria | Bacteroidetes | Flavobacteriia | Flavobacterial | Flavobacteria | Arenitalea | Arenitalea lut | unclassified | unclassified | unclassified | unclassified | unclassified | 12,6 | 3,136877428 | 0,000928018 | 0,000181283 | 11 - | 0,000977827 - | 8 - | 0,000594536 - | 12 - | 0,000897762 - |
| 1088 | 1 | 3um_FIJJAFBF_163900 | Bacteria | Proteobacteri | Alphaproteob | Rhodobactera | Rhodobactera | unclassified | Rhodobactera | metabolism | Carbohydrate | 01RFA | ABC transport | 644076.SCH4E | 0,2 | 0,4 | 3,51E-05 | 7,03E-05 | 0 - | 0 - | 0 - | 0 - | 0 - | 0 - |
| 1089 | 1 | 0.2um_NOKCPHJN_601798 | Eukaryota | Chordata | Aves | Passeriformes | Estrildidae | Lonchura | Lonchura stria | cellular proces | Cell cycle cont | 0V9W7 | retinitis pigme | 31033.ENSTRU | 0,8 | 0,4 | 9,69E-05 | 5,01E-05 | 1 - | 0,000143939 - | 1 - | 0,000120336 - | 0 - | 0 - |
| 1090 | 1 | 3um_FIJJAFBF_538012 | Eukaryota | Bacillariophyt | Bacillariophyc | Naviculales | Naviculaceae | Fistulifera | Fistulifera sola | unclassified | unclassified | unclassified | unclassified | unclassified | 2 | 0 | 0,000367301 | 3,88E-05 | 2 - | 0,000436433 - | 2 - | 0,000364868 - | 2 - | 0,000367306 - |
| 1091 | 1 | 0.2um_NOKCPHJN_160386 | Eukaryota | Bacillariophyt | Mediophyceae | Triceratiales | Triceratiaceae | Triceratium | Triceratium du | metabolism | Energy produc | 1DYB8 | PsaA and PsaE | 3055.DAA014 | 1,4 | 0,489897949 | 0,000120315 | 5,21E-05 | 2 - | 0,000199064 - | 2 - | 0,000166423 - | 1 - | 8,38E-05 - |
| 1092 | 1 | 0.2um_NOKCPHJN_370026 | Eukaryota | unclassified | Cryptophyta | Pyrenomonad | Geminigerace | Guillardia | Guillardia thet | cellular proces | Cytoskeleton | 0DKI0 | asp (abnorma | 99883.ENSTNI | 0,2 | 0,4 | 5,13E-06 | 1,03E-05 | 0 - | 0 - | 0 - | 0 - | 1 - | 2,56E-05 - |
| 1093 | 1 | 3um_FIJJAFBF_103105 | Bacteria | Bacteroidetes | Flavobacteriia | Flavobacterial | Flavobacteria | Polaribacter | Polaribacter s | cellular proces | Cell wall/mem | COG2885 | Ompa motb d | 313594.PI23P | 4,8 | 2,13541565 | 0,000639841 | 0,000269375 | 4 - | 0,000639728 - | 4 - | 0,000534828 - | 3 - | 0,000403801 - |
| 1094 | 1 | 3um_FIJJAFBF_395027 | unclassified | unclassified | unclassified | unclassified | unclassified | unclassified | unclassified | unclassified | unclassified | unclassified | unclassified | unclassified | 0,2 | 0,4 | 3,18E-05 | 6,37E-05 | 0 - | 0 - | 1 - | 0,000159224 - | 0 - | 0 - |
| 1095 | 1 | 0.2um_NOKCPHJN_785684 | Eukaryota | Ascomycota | Dothideomyce | unclassified | unclassified | Coniosporium | Coniosporium | unclassified | unclassified | unclassified | unclassified | unclassified | 0,8 | 0,748331477 | 5,52E-05 | 4,96E-05 | 0 - | 0 - | 1 - | 7,46E-05 - | 0 - | 0 - |
| 1096 | 1 | 3um_FIJJAFBF_574025 | Bacteria | Bacteroidetes | Flavobacteriia | Flavobacterial | Flavobacteria | Flavivirga | Flavivirga eckl | metabolism | Amino acid tra | COG0436 | aminotransfer | 398720.MED2 | 0,4 | 0,489897949 | 6,36E-05 | 7,79E-05 | 0 - | 0 - | 1 - | 0,000158418 - | 1 - | 0,000159476 - |
| 1097 | 1 | 0.2um_NOKCPHJN_565444 | Bacteria | Proteobacteri | Alphaproteob | Rhodospirillale | Rhodospirillac | unclassified | Rhodospirillac | unclassified | unclassified | unclassified | unclassified | unclassified | 4,4 | 0,8 | 0,000878784 | 0,00010331 | 3 - | 0,000724337 - | 5 - | 0,001009273 - | 4 - | 0,000812812 - |
| 1098 | 2 | 3um_FIJJAFBF_147114;3um_FIJJAFBF_ | Bacteria | Proteobacteri | Alphaproteob | Rhodobactera | Rhodobactera | various | various | metabolism | Energy produc | COG0056 | Produces ATP | 391593.RCCS2 | 1 | 0 | 0,000120214 | 1,27E-05 | 1 - | 0,00014284 - | 1 - | 0,000119418 - | 1 - | 0,000120216 - |
| 1099 | 1 | 0.2um_NOKCPHJN_266447 | Eukaryota | unclassified | Labyrinthulom | unclassified | Thraustochytr | Aurantiochytr | Aurantiochytr | unclassified | unclassified | unclassified | unclassified | unclassified | 0,2 | 0,4 | 4,83E-06 | 9,65E-06 | 0 - | 0 - | 0 - | 0 - | 0 - | 0 - |
| 1100 | 1 | 3um_FIJJAFBF_410461 | Bacteria | Candidatus At | unclassified | unclassified | unclassified | unclassified | Candidatus At | information st | Translation, ri | COG0256 | This is one of t | 684719.HIMB | 1 | 0 | 0,000529346 | 5,60E-05 | 1 - | 0,000628976 - | 1 - | 0,00052584 - | 1 - | 0,000529352 - |
| 1101 | 1 | 0.2um_NOKCPHJN_892804 | Eukaryota | Bacillariophyt | Coscinodiscop | Thalassiosirale | Thalassiosirac | Conticribra | Conticribra we | unclassified | unclassified | unclassified | unclassified | unclassified | 6,4 | 0,489897949 | 0,000766255 | 0,000114767 | 7 - | 0,000994188 - | 6 - | 0,000712428 - | 6 - | 0,000717187 - |
| 1102 | 1 | 3um_FIJJAFBF_415869 | Bacteria | Bacteroidetes | Flavobacteriia | Flavobacterial | Flavobacteria | Polaribacter | Polaribacter s | cellular proces | Posttranslatio | 09GE9 | Thioredoxin | 688270.Celal_ | 1 | 0 | 0,000599925 | 6,34E-05 | 1 - | 0,00071284 - | 1 - | 0,000595952 - | 1 - | 0,000599933 - |
| 1103 | 1 | 3um_FIJJAFBF_290586 | Eukaryota | Bacillariophyt | Coscinodiscop | Chaetocerotal | Chaetocerota | Chaetoceros | Chaetoceros s | unclassified | unclassified | unclassified | unclassified | unclassified | 0,8 | 0,4 | 0,000307837 | 0,000155146 | 0 - | 0 - | 1 - | 0,000401121 - | 1 - | 0,000403801 - |
| 1104 | 1 | 3um_FIJJAFBF_158710 | Bacteria | Proteobacteri | Gammaprotec | Oceanospirilla | Oleiphilaceae | Oleiphilus | Oleiphilus sp. | information st | Replication, re | 0QUGS | DNA-binding p | 349521.HCH_ | 0,2 | 0,4 | 0,000121043 | 0,000242086 | 0 - | 0 - | 0 - | 0 - | 0 - | 0 - |
| 1105 | 1 | 3um_FIJJAFBF_156303 | Viruses | unclassified | unclassified | unclassified | various | various | various | information st | Replication, re | 1BHM5 | DNA polymera | 88036.EFJ152 | 0,8 | 0,4 | 3,30E-05 | 1,66E-05 | 0 - | 0 - | 1 - | 4,29E-05 - | 1 - | 4,32E-05 - |
| 1106 | 1 | 3um_FIJJAFBF_651483 | Bacteria | Bacteroidetes | Flavobacteriia | Flavobacterial | Flavobacteria | Formosa | Formosa sp. H | unclassified | unclassified | unclassified | unclassified | unclassified | 1 | 0,894427191 | 0,000213955 | 0,000196232 | 0 - | 0 - | 0 - | 0 - | 2 - | 0,000448348 - |

| | | | | | | | | | | | | | | | | | | | | | | | | |
|---|---|---|---|---|---|---|---|---|---|---|---|---|---|---|---|---|---|---|---|---|---|---|---|---|
| 1107 | 1 | 3um_FIJJAFBF_539671 | Bacteria | unclassified | unclassified | unclassified | unclassified | unclassified | bacterium (Ca | cellular proces | Posttranslatio | 066HJ | ATP-depender | 445970.ALIPU | 1 | 0 | 0,000149625 | 1,58E-05 | 1 - | 0,000177787 - | 1 - | 0,000148634 - | 1 - | 0,000149627 - |
| 1108 | 2 | 0.2um_NOKCPHJN_12281;3um_FIJJAFB | various | various | various | various | various | various | various | unclassified | unclassified | unclassified | unclassified | unclassified | 0,8 | 0,4 | 0,000200094 | 0,000100845 | 0 - | 0 - | 1 - | 0,000260729 - | 1 - | 0,000262471 - |
| 1109 | 1 | 0.2um_NOKCPHJN_205463 | Bacteria | Bacteroidetes | Flavobacteriia | Flavobacterial | Flavobacteria | Tenacibaculur | Tenacibaculur | unclassified | unclassified | unclassified | unclassified | unclassified | 0,2 | 0,4 | 1,56E-05 | 3,11E-05 | 0 - | 0 - | 1 - | 7,78E-05 - | 0 - | 0 - |
| 1110 | 1 | 0.2um_NOKCPHJN_1068544 | Eukaryota | Nematoda | Enoplea | Trichinellida | Trichinellidae | Trichinella | Trichinella mu | unclassified | unclassified | unclassified | unclassified | unclassified | 1 | 0 | 0,000207895 | 2,20E-05 | 1 - | 0,000247024 - | 1 - | 0,000206518 - | 1 - | 0,000207897 - |
| 1111 | 1 | 0.2um_NOKCPHJN_1073503 | Bacteria | Bacteroidetes | Flavobacteriia | Flavobacterial | Cryomorphac | Owenweeksia | Owenweeksia | information st | Translation, ri | 0P78W | Pseudouridine | 926562.Oweh | 0,2 | 0,4 | 4,87E-05 | 9,74E-05 | 0 - | 0 - | 1 - | 0,000243482 - | 0 - | 0 - |
| 1112 | 1 | 3um_FIJJAFBF_390935 | Viruses | unclassified | unclassified | unclassified | Phycodnavirid | unclassified | Organic Lake p | unclassified | unclassified | unclassified | unclassified | unclassified | 1,6 | 0,489897949 | 0,001192596 | 0,000412142 | 2 - | 0,001761134 - | 2 - | 0,001472351 - | 1 - | 0,000741093 - |
| 1113 | 1 | 3um_FIJJAFBF_352289 | Eukaryota | unclassified | Cryptophyta | Cryptomonad | Hemiselmidac | Hemiselmis | Hemiselmis ar | information st | Replication, re | COG0513 | purine NTP-de | 45157.CMK02 | 1,4 | 0,489897949 | 0,000214395 | 6,46E-05 | 1 - | 0,000185728 - | 2 - | 0,000310546 - | 1 - | 0,00015631 - |
| 1114 | 3 | 0.2um_NOKCPHJN_1064534;3um_FIJJA | Bacteria | Proteobacteri | Gammaprotec | Cellvibrionales | various | various | various | information st | Translation, ri | COG0051 | tRNA binding | various | 0,2 | 0,4 | 0,000105766 | 0,000211532 | 0 - | 0 - | 0 - | 0 - | 0 - | 0 - |
| 1115 | 1 | 3um_FIJJAFBF_743482 | Bacteria | Proteobacteri | Gammaprotec | Cellvibrionales | Porticoccacea | unclassified | various | information st | Translation, ri | COG0292 | Binds directly | 314287.GB22( | 0,2 | 0,4 | 0,000106768 | 0,000213535 | 0 - | 0 - | 0 - | 0 - | 1 - | 0,000533838 - |
| 1116 | 1 | 3um_FIJJAFBF_607840 | Eukaryota | unclassified | Dinophyceae | Suessiales | Symbiodiniace | Symbiodinium | Symbiodinium | unclassified | unclassified | unclassified | unclassified | unclassified | 5,2 | 0,4 | 0,000839832 | 6,60E-05 | 5 - | 0,000964538 - | 5 - | 0,000806378 - | 5 - | 0,000811765 - |
| 1117 | 1 | 3um_FIJJAFBF_255783 | Bacteria | Proteobacteri | Gammaprotec | Cellvibrionales | Porticoccacea | unclassified | Gammaprotec | unclassified | unclassified | unclassified | unclassified | unclassified | 1 | 0 | 7,83E-05 | 8,29E-06 | 1 - | 9,31E-05 - | 1 - | 7,78E-05 - | 1 - | 7,83E-05 - |
| 1118 | 1 | 3um_FIJJAFBF_698759 | Eukaryota | Bacillariophyta | Fragilariophyc | Licmophorales | Licmophorace | Licmophora | Licmophora sp | information st | Translation, ri | 0FNCH | Binds directly | 91464.S7335_ | 1 | 0 | 0,000547758 | 5,79E-05 | 1 - | 0,000650854 - | 1 - | 0,00054413 - | 1 - | 0,000547765 - |
| 1119 | 1 | 0.2um_NOKCPHJN_309398 | Bacteria | Proteobacteri | Alphaproteob | Rhodobactera | Rhodobactera | unclassified | Rhodobactera | metabolism | Energy produc | 01RNF | Produces ATP | 314264.ROS21 | 1,4 | 0,489897949 | 0,000190879 | 8,27E-05 | 2 - | 0,000315815 - | 2 - | 0,000264029 - | 1 - | 0,000132896 - |
| 1120 | 2 | 0.2um_NOKCPHJN_954744;3um_FIJJAF | Eukaryota | Eustigmatoph | unclassified | Eustigmatales | Monodopsida | Nannochlorop | Nannochlorop | unclassified | unclassified | unclassified | unclassified | unclassified | 1 | 0 | 0,000170249 | 1,80E-05 | 1 - | 0,000202292 - | 1 - | 0,000169121 - | 1 - | 0,000170251 - |
| 1121 | 1 | 0.2um_NOKCPHJN_479681 | Bacteria | Proteobacteri | Alphaproteob | Rhodobactera | Rhodobactera | Roseovarius | Roseovarius s | metabolism | Inorganic ion t | COG3221 | phosphonate . | 375451.RD1_2 | 0,2 | 0,4 | 5,99E-05 | 0,000119761 | 0 - | 0 - | 1 - | 0,000299402 - | 0 - | 0 - |
| 1122 | 1 | 3um_FIJJAFBF_654925 | Bacteria | Bacteroidetes | Flavobacteriia | various | various | various | various | information st | Replication, re | 0PB0H | DNA-binding p | 487797.Flav3( | 0,8 | 0,4 | 0,000533584 | 0,00026892 | 0 - | 0 - | 1 - | 0,000695277 - | 1 - | 0,000699922 - |
| 1123 | 1 | 3um_FIJJAFBF_414690 | Bacteria | Bacteroidetes | Flavobacteriia | Flavobacterial | Flavobacteria | Jejuia | Jejuia pallidilu | unclassified | unclassified | unclassified | unclassified | unclassified | 1 | 0 | 0,000154393 | 1,63E-05 | 1 - | 0,000183451 - | 1 - | 0,00015337 - | 1 - | 0,000154394 - |
| 1124 | 1 | 0.2um_NOKCPHJN_608964 | Bacteria | Proteobacteri | Gammaprotec | Thiotrichales | Piscirickettsia | Methylophaga | Methylophaga | poorly charact | Function unkn | 05VQX | NA | 565045.NOR5 | 0,2 | 0,4 | 2,67E-05 | 5,34E-05 | 0 - | 0 - | 0 - | 0 - | 0 - | 0 - |
| 1125 | 1 | 0.2um_NOKCPHJN_778194 | Eukaryota | unclassified | Spirotrichea | Sporadotrichic | Oxytrichidae | Stylonychia | Stylonychia le | unclassified | unclassified | unclassified | unclassified | unclassified | 1 | 0 | 0,00012649 | 1,34E-05 | 1 - | 0,000150298 - | 1 - | 0,000125652 - | 1 - | 0,000126492 - |
| 1126 | 1 | 0.2um_NOKCPHJN_52330 | Bacteria | Bacteroidetes | Flavobacteriia | Flavobacterial | Flavobacteria | Formosa | Formosa sp. H | unclassified | unclassified | unclassified | unclassified | unclassified | 0,4 | 0,489897949 | 2,76E-05 | 3,39E-05 | 0 - | 0 - | 1 - | 7,37E-05 - | 0 - | 0 - |
| 1127 | 2 | 3um_FIJJAFBF_232873;3um_FIJJAFBF_ | Eukaryota | various | various | various | various | various | various | poorly charact | Function unkn | COG1100 | GTP-binding P | various | 2 | 0 | 0,000594266 | 6,28E-05 | 2 - | 0,000706115 - | 2 - | 0,000590329 - | 2 - | 0,000594273 - |
| 1128 | 1 | 0.2um_NOKCPHJN_670407 | Bacteria | Firmicutes | Negativicutes | Veillonellales | Veillonellacea | Megasphaera | Megasphaera | unclassified | unclassified | unclassified | unclassified | unclassified | 0,6 | 0,489897949 | 7,23E-05 | 5,93E-05 | 0 - | 0 - | 0 - | 0 - | 1 - | 0,000128295 - |
| 1129 | 1 | 0.2um_NOKCPHJN_324343 | Eukaryota | unclassified | Oomycetes | Peronosporale | Peronosporac | Phytophthora | Phytophthora | poorly charact | Function unkn | 0IG8C | kiaa1009 | 7897.ENSLACF | 1,2 | 0,4 | 5,48E-05 | 1,86E-05 | 1 - | 5,43E-05 - | 2 - | 9,08E-05 - | 1 - | 4,57E-05 - |
| 1130 | 1 | 3um_FIJJAFBF_162557 | Bacteria | Bacteroidetes | Flavobacteriia | Flavobacterial | Flavobacteria | unclassified | Flavobacteria | unclassified | unclassified | unclassified | unclassified | unclassified | 1,4 | 0,489897949 | 0,000279886 | 8,52E-05 | 1 - | 0,000242227 - | 1 - | 0,000202508 - | 2 - | 0,000407721 - |
| 1131 | 1 | 3um_FIJJAFBF_517421 | Bacteria | Bacteroidetes | Flavobacteriia | Flavobacterial | Flavobacteria | Polaribacter | Polaribacter v | information st | Translation, ri | 08UIK | This protein bi | 313598.MED1 | 2 | 0,632455532 | 0,000872305 | 0,000292717 | 2 - | 0,001032389 - | 3 - | 0,001294653 - | 2 - | 0,000868868 - |
| 1132 | 1 | 0.2um_NOKCPHJN_640148 | Eukaryota | unclassified | Pelagophycea | Pelagomonad | unclassified | Aureococcus | Aureococcus a | cellular proces | Intracellular tr | KOG3342 | Catalytic comp | 2850.Phatr512 | 3,4 | 0,489897949 | 0,000885919 | 0,000135158 | 3 - | 0,00093172 - | 4 - | 0,001038588 - | 3 - | 0,000784144 - |
| 1133 | 1 | 0.2um_NOKCPHJN_344034 | Bacteria | Proteobacteri | Gammaprotec | Cellvibrionales | Porticoccacea | unclassified | gamma protec | unclassified | unclassified | unclassified | unclassified | unclassified | 1 | 0 | 0,000179978 | 1,90E-05 | 1 - | 0,000213852 - | 1 - | 0,000178785 - | 1 - | 0,00017998 - |
| 1134 | 1 | 3um_FIJJAFBF_791054 | Eukaryota | unclassified | Pelagophycea | Pelagomonad | unclassified | Aureococcus | Aureococcus a | unclassified | unclassified | unclassified | unclassified | unclassified | 2 | 0 | 0,001105126 | 0,00011687 | 2 - | 0,001313126 - | 2 - | 0,001097806 - | 2 - | 0,001105139 - |
| 1135 | 1 | 3um_FIJJAFBF_688570 | Bacteria | Bacteroidetes | Flavobacteriia | Flavobacterial | Flavobacteria | Flavobacteriu | Flavobacteriu | metabolism | Energy produc | COG2838 | Isocitrate deh | 391598.FBBAL | 2 | 0 | 0,000196543 | 2,08E-05 | 2 - | 0,000233536 - | 2 - | 0,000195242 - | 2 - | 0,000196546 - |
| 1136 | 1 | 0.2um_NOKCPHJN_378970 | Eukaryota | unclassified | Labyrinthulom | unclassified | Thraustochytr | Aurantiochytr | Aurantiochytr | metabolism | Nucleotide tra | 0IUMV | carbamoylphc | 65071.PYU1_T | 0,4 | 0,489897949 | 2,50E-05 | 3,06E-05 | 0 - | 0 - | 1 - | 6,22E-05 - | 1 - | 6,26E-05 - |
| 1137 | 1 | 0.2um_NOKCPHJN_39166 | Eukaryota | unclassified | Synurophycea | Parmales | Triparmaceae | Triparma | Triparma laevi | unclassified | unclassified | unclassified | unclassified | unclassified | 5,4 | 1,019803903 | 0,002334224 | 0,000219847 | 4 - | 0,002093655 - | 5 - | 0,002187934 - | 5 - | 0,00220255 - |
| 1138 | 1 | 0.2um_NOKCPHJN_12874 | Bacteria | Bacteroidetes | Flavobacteriia | Flavobacterial | Flavobacteria | Hyunsoonleell | Hyunsoonleell | information st | Translation, ri | COG0050 | This protein p | 688270.Celal_ | 2,6 | 0,489897949 | 0,000420424 | 0,000111998 | 3 - | 0,000568467 - | 3 - | 0,000475253 - | 3 - | 0,000478427 - |
| 1139 | 1 | 0.2um_NOKCPHJN_128221 | Bacteria | Bacteroidetes | Cytophagia | Cytophagales | Amoebophila | Candidatus Ar | Candidatus Ar | unclassified | unclassified | unclassified | unclassified | unclassified | 0,6 | 0,489897949 | 3,03E-05 | 2,48E-05 | 0 - | 0 - | 0 - | 0 - | 1 - | 5,37E-05 - |
| 1140 | 1 | 3um_FIJJAFBF_567865 | Eukaryota | unclassified | Florideophyce | Gigartinales | Gigartinaceae | Chondrus | Chondrus cris | unclassified | unclassified | unclassified | unclassified | unclassified | 1 | 0 | 0,000329802 | 3,49E-05 | 1 - | 0,000391875 - | 1 - | 0,000327617 - | 1 - | 0,000329806 - |
| 1141 | 1 | 3um_FIJJAFBF_462350 | unclassified | unclassified | unclassified | unclassified | unclassified | unclassified | unclassified | unclassified | unclassified | unclassified | unclassified | unclassified | 0,4 | 0,489897949 | 0,000261758 | 0,000322711 | 1 - | 0,00071284 - | 1 - | 0,000595952 - | 0 - | 0 - |
| 1142 | 1 | 0.2um_NOKCPHJN_1108412 | Eukaryota | unclassified | unclassified | Kinetoplastida | Trypanosomat | Leishmania | Leishmania m | unclassified | unclassified | unclassified | unclassified | unclassified | 0,6 | 0,489897949 | 6,99E-05 | 5,73E-05 | 0 - | 0 - | 1 - | 0,000123422 - | 0 - | 0 - |
| 1143 | 1 | 3um_FIJJAFBF_39793 | Bacteria | Bacteroidetes | Flavobacteriia | Flavobacterial | Flavobacteria | Polaribacter | Polaribacter s | unclassified | unclassified | unclassified | unclassified | unclassified | 1 | 0 | 0,000248001 | 2,62E-05 | 1 - | 0,000294678 - | 1 - | 0,000246358 - | 1 - | 0,000248004 - |
| 1144 | 1 | 3um_FIJJAFBF_437459 | Eukaryota | unclassified | Cryptophyta | Pyrenomonad | Chroomonada | Chroomonas | Chroomonas r | unclassified | unclassified | unclassified | unclassified | unclassified | 0,8 | 0,4 | 0,000141957 | 7,23E-05 | 1 - | 0,000203946 - | 1 - | 0,000170504 - | 1 - | 0,000171643 - |
| 1145 | 1 | 0.2um_NOKCPHJN_216536 | Eukaryota | unclassified | Oomycetes | Peronosporale | Peronosporac | Phytophthora | Phytophthora | unclassified | unclassified | unclassified | unclassified | unclassified | 2 | 0 | 0,000156114 | 1,65E-05 | 2 - | 0,000185497 - | 2 - | 0,00015508 - | 2 - | 0,000156116 - |
| 1146 | 1 | 3um_FIJJAFBF_484231 | Eukaryota | Bacillariophyta | Coscinodiscop | Thalassiosirale | Thalassiosirac | Thalassiosira | Thalassiosira c | unclassified | unclassified | unclassified | unclassified | unclassified | 0,2 | 0,4 | 5,16E-05 | 0,000103134 | 0 - | 0 - | 0 - | 0 - | 0 - | 0 - |
| 1147 | 1 | 0.2um_NOKCPHJN_1116704 | Viruses | unclassified | unclassified | unclassified | unclassified | unclassified | uncultured M | unclassified | unclassified | unclassified | unclassified | unclassified | 0,2 | 0,4 | 2,39E-05 | 4,79E-05 | 0 - | 0 - | 0 - | 0 - | 0 - | 0 - |
| 1148 | 1 | 3um_FIJJAFBF_191791 | Eukaryota | Bacillariophyta | Coscinodiscop | Rhizosoleniale | Rhizosoleniac | Rhizosolenia | Rhizosolenia i | unclassified | unclassified | unclassified | unclassified | unclassified | 8,6 | 0,489897949 | 0,003368422 | 0,000445076 | 9 - | 0,00418406 - | 9 - | 0,003497977 - | 8 - | 0,003130084 - |
| 1149 | 1 | 3um_FIJJAFBF_254313 | Bacteria | Proteobacteri | Gammaprotec | Cellvibrionales | Spongiibacter | Spongiibacter | Spongiibacter | information st | Replication, re | 16W6D | single-strande | 207954.MED9 | 0,8 | 0,4 | 0,000294617 | 0,000148484 | 0 - | 0 - | 1 - | 0,000383895 - | 1 - | 0,00038646 - |
| 1150 | 1 | 3um_FIJJAFBF_135972 | Bacteria | Proteobacteri | Gammaprotec | Thiotrichales | Piscirickettsia | Methylophaga | various | metabolism | Carbohydrate | COG0176 | Transaldolase | 1026882.MAN | 0,8 | 0,4 | 0,000146858 | 7,40E-05 | 0 - | 0 - | 1 - | 0,000191361 - | 1 - | 0,000192639 - |
| 1151 | 1 | 0.2um_NOKCPHJN_220621 | Bacteria | Bacteroidetes | unclassified | unclassified | unclassified | unclassified | Bacteroidetes | metabolism | Inorganic ion t | 05C27 | TonB-depende | 886377.Murru | 0,4 | 0,489897949 | 2,25E-05 | 2,76E-05 | 0 - | 0 - | 0 - | 0 - | 0 - | 0 - |
| 1152 | 1 | 3um_FIJJAFBF_06549 | unclassified | unclassified | unclassified | unclassified | unclassified | unclassified | unclassified | unclassified | unclassified | unclassified | unclassified | unclassified | 0,8 | 0,4 | 0,000183292 | 9,24E-05 | 0 - | 0 - | 1 - | 0,000238836 - | 1 - | 0,000240431 - |
| 1153 | 1 | 3um_FIJJAFBF_244104 | Eukaryota | unclassified | unclassified | Trichomonadi | Trichomonadi | Trichomonas | Trichomonas v | metabolism | Lipid transpor | COG2267 | alpha beta hy | 44689.DDB02 | 0,2 | 0,4 | 2,11E-05 | 4,21E-05 | 0 - | 0 - | 0 - | 0 - | 0 - | 0 - |
| 1154 | 1 | 3um_FIJJAFBF_254286 | Bacteria | Proteobacteri | Gammaprotec | Cellvibrionales | Spongiibacter | Spongiibacter | Spongiibacter | information st | Translation, ri | COG0088 | One of the pri | 876044.IMCC3 | 2 | 0,632455532 | 0,000617404 | 0,000208836 | 2 - | 0,000730226 - | 2 - | 0,000610487 - | 3 - | 0,000921848 - |
| 1155 | 1 | 0.2um_NOKCPHJN_267970 | unclassified | unclassified | unclassified | unclassified | unclassified | unclassified | unclassified | unclassified | unclassified | unclassified | unclassified | unclassified | 0,2 | 0,4 | 0,00027307 | 0,00054614 | 0 - | 0 - | 0 - | 0 - | 0 - | 0 - |
| 1156 | 1 | 3um_FIJJAFBF_40656 | Eukaryota | unclassified | Pelagophycea | Pelagomonad | unclassified | Aureococcus | Aureococcus a | unclassified | unclassified | unclassified | unclassified | unclassified | 1 | 0 | 0,001312337 | 0,000138783 | 1 - | 0,001559337 - | 1 - | 0,001303644 - | 1 - | 0,001312353 - |
| 1157 | 6 | 3um_FIJJAFBF_124795;3um_FIJJAFBF_ | Eukaryota | unclassified | various | various | various | various | various | metabolism | Energy produc | 123BF | photosystem I | various | 1,2 | 0,4 | 0,00112822 | 0,000389004 | 1 - | 0,001117137 - | 1 - | 0,000933954 - | 2 - | 0,001880386 - |
| 1158 | 1 | 0.2um_NOKCPHJN_1011565 | Viruses | unclassified | unclassified | unclassified | unclassified | unclassified | uncultured vir | unclassified | unclassified | unclassified | unclassified | unclassified | 0,2 | 0,4 | 4,32E-05 | 8,65E-05 | 0 - | 0 - | 0 - | 0 - | 0 - | 0 - |
| 1159 | 1 | 3um_FIJJAFBF_560873 | Bacteria | Proteobacteri | Gammaprotec | Cellvibrionales | Porticoccacea | unclassified | Gammaprotec | metabolism | Inorganic ion t | 16TQC | TONB-depend | 314287.GB22( | 0,6 | 0,489897949 | 3,32E-05 | 2,72E-05 | 0 - | 0 - | 1 - | 5,86E-05 - | 0 - | 0 - |
| 1160 | 1 | 3um_FIJJAFBF_545864 | Eukaryota | Bacillariophyta | Coscinodiscop | Thalassiosirale | Thalassiosirac | Thalassiosira | Thalassiosira c | poorly charact | Function unkn | COG1100 | GTP-binding P | 35128.Thaps3 | 1 | 0 | 0,000459797 | 4,86E-05 | 1 - | 0,000546337 - | 1 - | 0,000456751 - | 1 - | 0,000459802 - |
| 1161 | 1 | 3um_FIJJAFBF_290585 | Eukaryota | Bacillariophyta | Coscinodiscop | Chaetocerotal | Chaetocerota | Chaetoceros | Chaetoceros s | metabolism | Energy produc | COG0711 | F(1)F(0) ATP s | 35128.Thapsd | 2,4 | 1,2 | 0,000804847 | 0,000405634 | 0 - | 0 - | 3 - | 0,001048742 - | 3 - | 0,001055748 - |
| 1162 | 1 | 0.2um_NOKCPHJN_546028 | Bacteria | Proteobacteri | Gammaprotec | unclassified | unclassified | unclassified | Gammaprotec | unclassified | unclassified | unclassified | unclassified | unclassified | 9,4 | 2,416609195 | 0,000693156 | 0,000136653 | 5 - | 0,000449269 - | 10 - | 0,0007512 - | 9 - | 0,000680596 - |
| 1163 | 1 | 0.2um_NOKCPHJN_161604 | unclassified | unclassified | unclassified | unclassified | unclassified | unclassified | unclassified | unclassified | unclassified | unclassified | unclassified | unclassified | 1 | 0 | 0,000154393 | 1,63E-05 | 1 - | 0,000183451 - | 1 - | 0,00015337 - | 1 - | 0,000154394 - |
| 1164 | 1 | 0.2um_NOKCPHJN_979126 | Eukaryota | unclassified | Cryptophyta | Pyrenomonad | Geminigerace | Guillardia | Guillardia thet | metabolism | Energy produc | 0IFIU | Pyrophosphat | 653948.CCA14 | 0,8 | 0,4 | 0,000103165 | 5,26E-05 | 1 - | 0,000148214 - | 1 - | 0,000123911 - | 1 - | 0,000124738 - |
| 1165 | 1 | 3um_FIJJAFBF_663734 | Eukaryota | Bacillariophyta | Bacillariophyc | Naviculales | Naviculaceae | Fistulifera | Fistulifera sola | poorly charact | Function unkn | 110WY | Chlorophyll A- | 2850.Phatr18 | 1,8 | 0,4 | 0,001057283 | 0,000181689 | 1 - | 0,00071284 - | 2 - | 0,001191903 - | 2 - | 0,001199865 - |
| 1166 | 1 | 3um_FIJJAFBF_175283 | Eukaryota | Apicomplexa | Conoidasida | Eucoccidiorida | Sarcocystidae | Hammondia | Hammondia h | unclassified | unclassified | unclassified | unclassified | unclassified | 0,4 | 0,489897949 | 3,93E-05 | 4,87E-05 | 1 - | 0,000109108 - | 0 - | 0 - | 0 - | 0 - |
| 1167 | 1 | 3um_FIJJAFBF_128441 | Bacteria | Bacteroidetes | Flavobacteriia | Flavobacterial | Flavobacteria | Polaribacter | Polaribacter s | unclassified | unclassified | unclassified | unclassified | unclassified | 31,4 | 3,666060556 | 0,002085978 | 0,000178104 | 27 - | 0,002149895 - | 27 - | 0,001797365 - | 35 - | 0,002345482 - |
| 1168 | 1 | 0.2um_NOKCPHJN_807201 | Bacteria | Proteobacteri | Gammaprotec | Alteromonada | Colwelliaceae | Colwellia | Colwellia pola | cellular proces | Defense mech | COG0842 | (ABC) transpo | 87626.PTD2_1 | 1 | 0 | 0,000384099 | 4,06E-05 | 1 - | 0,000456391 - | 1 - | 0,000381554 - | 1 - | 0,000384103 - |
| 1169 | 1 | 0.2um_NOKCPHJN_152692 | Bacteria | Proteobacteri | Gammaprotec | Cellvibrionales | Porticoccacea | unclassified | SAR92 bacteri | unclassified | unclassified | unclassified | unclassified | unclassified | 0,6 | 0,489897949 | 3,30E-05 | 2,75E-05 | 1 - | 6,43E-05 - | 0 - | 0 - | 1 - | 5,41E-05 - |
| 1170 | 1 | 0.2um_NOKCPHJN_744827 | Eukaryota | unclassified | Cryptophyta | Pyrenomonad | Geminigerace | Guillardia | Guillardia thet | unclassified | unclassified | unclassified | unclassified | unclassified | 1 | 0 | 0,000492126 | 5,20E-05 | 1 - | 0,000584751 - | 1 - | 0,000488867 - | 1 - | 0,000492132 - |
| 1171 | 1 | 3um_FIJJAFBF_145781 | Eukaryota | Bacillariophyta | Coscinodiscop | Chaetocerotal | Chaetocerota | Chaetoceros | Chaetoceros g | unclassified | unclassified | unclassified | unclassified | unclassified | 0,8 | 0,4 | 0,000873137 | 0,000440051 | 0 - | 0 - | 1 - | 0,001137726 - | 1 - | 0,001145326 - |
| 1172 | 1 | 0.2um_NOKCPHJN_717491 | Eukaryota | unclassified | Cryptophyta | Pyrenomonad | Geminigerace | Guillardia | Guillardia thet | cellular proces | Posttranslatio | 0TJU0 | Hsp90 protein | 4555.Si00587 | 1 | 0 | 9,18E-05 | 9,71E-06 | 1 - | 0,000109108 - | 1 - | 9,12E-05 - | 1 - | 9,18E-05 - |
| 1173 | 1 | 3um_FIJJAFBF_486301 | Bacteria | Proteobacteri | Gammaprotec | Cellvibrionales | Halieaceae | Halioglobus | various | information st | Translation, ri | 0QQXI | This protein is | 247634.GPB2 | 2,4 | 0,489897949 | 0,001261808 | 0,00018103 | 2 - | 0,001268613 - | 2 - | 0,001060592 - | 2 - | 0,001067677 - |
| 1174 | 1 | 3um_FIJJAFBF_73823 | Bacteria | Bacteroidetes | Flavobacteriia | Flavobacterial | Flavobacteria | Polaribacter | Polaribacter s | unclassified | unclassified | unclassified | unclassified | unclassified | 0,6 | 0,8 | 8,02E-05 | 0,000103847 | 0 - | 0 - | 0 - | 0 - | 1 - | 0,000146837 - |
| 1175 | 1 | 3um_FIJJAFBF_378479 | Bacteria | Candidatus Ha | unclassified | unclassified | unclassified | unclassified | Candidatus Ha | unclassified | unclassified | unclassified | unclassified | unclassified | 0,6 | 0,489897949 | 0,000174828 | 0,000145747 | 1 - | 0,000340219 - | 0 - | 0 - | 1 - | 0,000286332 - |
| 1176 | 1 | 0.2um_NOKCPHJN_709000 | Eukaryota | Chlorophyta | Chlorophycea | Chlamydomor | Chlamydomor | Chlamydomor | Chlamydomor | metabolism | Amino acid tra | 1BDGU | amino acid tra | 4533.OB04G1 | 0,2 | 0,4 | 3,74E-05 | 7,48E-05 | 1 - | 0,00018712 - | 0 - | 0 - | 0 - | 0 - |
| 1177 | 1 | 3um_FIJJAFBF_745913 | Bacteria | Proteobacteri | Gammaprotec | Cellvibrionales | Halieaceae | Halioglobus | Halioglobus p | unclassified | unclassified | unclassified | unclassified | unclassified | 0,8 | 0,4 | 0,000461755 | 0,000232719 | 0 - | 0 - | 1 - | 0,000601682 - | 1 - | 0,000605701 - |
| 1178 | 1 | 3um_FIJJAFBF_636077 | Eukaryota | Bacillariophyta | Coscinodiscop | Thalassiosirale | Thalassiosirac | Thalassiosira | Thalassiosira p | unclassified | unclassified | unclassified | unclassified | unclassified | 0,6 | 0,489897949 | 0,000163629 | 0,000134198 | 0 - | 0 - | 0 - | 0 - | 1 - | 0,00029029 - |
| 1179 | 1 | 3um_FIJJAFBF_447354 | Bacteria | Planctomycet | Phycisphaerae | Phycisphaeral | Phycisphaerac | Phycisphaera | Phycisphaera | information st | Translation, ri | COG0050 | This protein p | 314230.DSM3 | 0,2 | 0,4 | 2,70E-05 | 5,41E-05 | 0 - | 0 - | 0 - | 0 - | 0 - | 0 - |
| 1180 | 1 | 3um_FIJJAFBF_640408 | unclassified | unclassified | unclassified | unclassified | unclassified | unclassified | unclassified | unclassified | unclassified | unclassified | unclassified | unclassified | 2 | 0,632455532 | 0,000603 | 0,000248687 | 3 - | 0,0010542 - | 1 - | 0,000293779 - | 2 - | 0,000591483 - |
| 1181 | 2 | 0.2um_NOKCPHJN_485750;0.2um_NO | Eukaryota | Bacillariophyta | Coscinodiscop | Thalassiosirale | Thalassiosirac | Thalassiosira | Thalassiosira c | unclassified | unclassified | unclassified | unclassified | unclassified | 0,6 | 0,489897949 | 6,99E-05 | 5,74E-05 | 0 - | 0 - | 1 - | 0,000121505 - | 1 - | 0,000122316 - |
| 1182 | 1 | 3um_FIJJAFBF_95709 | Eukaryota | unclassified | Florideophyce | Gelidiales | Gelidiaceae | Gelidium | Gelidium eleg | unclassified | unclassified | unclassified | unclassified | unclassified | 0,8 | 0,4 | 0,000490026 | 0,000246968 | 0 - | 0 - | 1 - | 0,00063852 - | 1 - | 0,000642785 - |
| 1183 | 1 | 3um_FIJJAFBF_74442 | Bacteria | Bacteroidetes | unclassified | unclassified | unclassified | unclassified | Bacteroidetes | information st | Translation, ri | 05CAV | thus facilitatin | 313594.PI23P | 0,4 | 0,489897949 | 3,80E-05 | 4,67E-05 | 0 - | 0 - | 1 - | 0,000101583 - | 0 - | 0 - |
| 1184 | 1 | 0.2um_NOKCPHJN_556304 | Bacteria | Bacteroidetes | Flavobacteriia | Flavobacterial | Flavobacteria | Flavobacteriu | Flavobacteriu | metabolism | Amino acid tra | 0NIVA | O-Acetyltrans | 562983.HMPR | 0,2 | 0,4 | 5,56E-05 | 0,000111162 | 0 - | 0 - | 0 - | 0 - | 0 - | 0 - |
| 1185 | 1 | 3um_FIJJAFBF_478118 | Eukaryota | Bacillariophyta | Coscinodiscop | Thalassiosirale | Thalassiosirac | Thalassiosira | Thalassiosira p | unclassified | unclassified | unclassified | unclassified | unclassified | 1 | 0 | 0,000449944 | 4,76E-05 | 1 - | 0,00053463 - | 1 - | 0,000446964 - | 1 - | 0,00044995 - |
| 1186 | 2 | 3um_FIJJAFBF_278688;3um_FIJJAFBF_ | Bacteria | Bacteroidetes | Flavobacteriia | Flavobacterial | Flavobacteria | Polaribacter | various | metabolism | Carbohydrate | COG0149 | triosephospha | various | 1 | 0 | 0,000249969 | 2,64E-05 | 1 - | 0,000297017 - | 1 - | 0,000248313 - | 1 - | 0,000249972 - |
| 1187 | 1 | 3um_FIJJAFBF_32479 | Eukaryota | Bacillariophyta | Bacillariophyc | unclassified | Entomoneida | Entomoneis | Entomoneis sp | unclassified | unclassified | unclassified | unclassified | unclassified | 0,8 | 0,4 | 9,92E-05 | 5,13E-05 | 1 - | 0,000147049 - | 0 - | 0 - | 1 - | 0,000123758 - |
| 1188 | 1 | 3um_FIJJAFBF_776242 | Eukaryota | unclassified | Cryptophyta | Pyrenomonad | Geminigerace | Guillardia | Guillardia thet | unclassified | unclassified | unclassified | unclassified | unclassified | 0,2 | 0,4 | 6,29E-05 | 0,000125812 | 0 - | 0 - | 0 - | 0 - | 0 - | 0 - |
| 1189 | 1 | 3um_FIJJAFBF_722063 | unclassified | unclassified | unclassified | unclassified | unclassified | unclassified | unclassified | unclassified | unclassified | unclassified | unclassified | unclassified | 3 | 0,632455532 | 0,000716641 | 0,000157927 | 3 - | 0,000853782 - | 2 - | 0,000475855 - | 3 - | 0,000718551 - |
| 1190 | 1 | 0.2um_NOKCPHJN_997360 | Bacteria | Verrucomicrol | Verrucomicrol | Verrucomicrol | Rubritaleacea | Rubritalea | Rubritalea squ | cellular proces | Cell wall/mem | 05C6Y | Udp-glucose 6 | 583355.Caka_ | 1 | 0 | 0,000136642 | 1,45E-05 | 1 - | 0,00016236 - | 1 - | 0,000135737 - | 1 - | 0,000136644 - |
| 1191 | 1 | 0.2um_NOKCPHJN_1016645 | Eukaryota | unclassified | Actinophryida | unclassified | unclassified | Actinophrys | Actinophrys s | unclassified | unclassified | unclassified | unclassified | unclassified | 0,2 | 0,4 | 8,93E-05 | 0,000178588 | 0 - | 0 - | 0 - | 0 - | 0 - | 0 - |
| 1192 | 1 | 3um_FIJJAFBF_554298 | Eukaryota | unclassified | Oomycetes | Saprolegniales | Saprolegniace | Saprolegnia | Saprolegnia p | poorly charact | Function unkn | 0VXSK | NA | 7739.JGI1044 | 0,8 | 0,4 | 0,000151491 | 7,63E-05 | 0 - | 0 - | 1 - | 0,000197397 - | 1 - | 0,000198716 - |
| 1193 | 1 | 0.2um_NOKCPHJN_1094039 | Bacteria | Bacteroidetes | Flavobacteriia | Flavobacterial | Flavobacteria | Formosa | Formosa sp. H | unclassified | unclassified | unclassified | unclassified | unclassified | 0,2 | 0,4 | 3,97E-05 | 7,94E-05 | 1 - | 0,000198536 - | 0 - | 0 - | 0 - | 0 - |
| 1194 | 1 | 3um_FIJJAFBF_237055 | Eukaryota | Bacillariophyta | Bacillariophyc | Bacillariales | Bacillariaceae | Cylindrotheca | Cylindrotheca | metabolism | Energy produc | 0FJ7X | This is one of t | 103690.alr372 | 11,8 | 0,748331477 | 0,00206024 | 0,00021969 | 12 - | 0,002494939 - | 11 - | 0,001912011 - | 11 - | 0,001924784 - |
| 1195 | 1 | 3um_FIJJAFBF_16760 | Bacteria | Bacteroidetes | Flavobacteriia | Flavobacterial | Flavobacteria | Algibacter | Algibacter alg | metabolism | Amino acid tra | 05C6M | Alpha-keto-be | 867900.Celly_ | 0,4 | 0,489897949 | 5,50E-05 | 6,80E-05 | 1 - | 0,00015244 - | 0 - | 0 - | 0 - | 0 - |
| 1196 | 1 | 3um_FIJJAFBF_66276 | Bacteria | Bacteroidetes | Flavobacteriia | Flavobacterial | Crocinitomica | unclassified | Crocinitomica | information st | Translation, ri | 05C22 | threonyL-tRN | 755732.Fluta_ | 0,4 | 0,489897949 | 3,62E-05 | 4,45E-05 | 0 - | 0 - | 1 - | 9,67E-05 - | 0 - | 0 - |
| 1197 | 1 | 3um_FIJJAFBF_20041 | Bacteria | Bacteroidetes | Flavobacteriia | Flavobacterial | Flavobacteria | Polaribacter | Polaribacter h | unclassified | unclassified | unclassified | unclassified | unclassified | 2,6 | 0,489897949 | 0,000978232 | 0,000133954 | 2 - | 0,000907251 - | 3 - | 0,001137726 - | 2 - | 0,000763551 - |
| 1198 | 1 | 3um_FIJJAFBF_566757 | Eukaryota | Bacillariophyta | Bacillariophyc | Naviculales | Naviculaceae | Fistulifera | Fistulifera sola | unclassified | unclassified | unclassified | unclassified | unclassified | 4 | 0,632455532 | 0,000802296 | 0,000108849 | 3 - | 0,000722008 - | 5 - | 0,001006028 - | 4 - | 0,000810199 - |
| 1199 | 1 | 3um_FIJJAFBF_630544 | Eukaryota | unclassified | unclassified | Prymnesiales | Chrysochromu | Chrysochromu | Chrysochromu | poorly charact | Function unkn | 110WY | Chlorophyll A- | 2850.Phatr16 | 0,8 | 0,4 | 0,000369404 | 0,000186176 | 0 - | 0 - | 1 - | 0,000481346 - | 1 - | 0,000484561 - |
| 1200 | 3 | 0.2um_NOKCPHJN_1045591;3um_FIJJA | Eukaryota | various | various | various | various | various | various | metabolism | Energy produc | COG1145 | 4Fe-4S ferred | various | 1 | 0 | 0,000692222 | 7,32E-05 | 1 - | 0,000822508 - | 1 - | 0,000687636 - | 1 - | 0,00069223 - |
| 1201 | 1 | 3um_FIJJAFBF_86326 | Eukaryota | Chlorophyta | Chlorophycea | Chlamydomor | Chlamydomor | Chlamydomor | Chlamydomor | poorly charact | Function unkn | 0ZP14 | NA | 9685.ENSFCAF | 0,2 | 0,4 | 9,39E-05 | 0,000187825 | 0 - | 0 - | 0 - | 0 - | 0 - | 0 - |
| 1202 | 1 | 0.2um_NOKCPHJN_405885 | unclassified | unclassified | unclassified | unclassified | unclassified | unclassified | unclassified | unclassified | unclassified | unclassified | unclassified | unclassified | 0,8 | 0,4 | 3,91E-05 | 1,99E-05 | 1 - | 5,61E-05 - | 1 - | 4,69E-05 - | 1 - | 4,72E-05 - |
| 1203 | 1 | 3um_FIJJAFBF_257116 | Eukaryota | unclassified | Cryptophyta | Pyrenomonad | Chroomonada | Chroomonas | various | unclassified | unclassified | unclassified | unclassified | unclassified | 3 | 0,894427191 | 0,001593251 | 0,000387387 | 2 - | 0,001290486 - | 4 - | 0,002157756 - | 2 - | 0,001086085 - |
| 1204 | 1 | 3um_FIJJAFBF_56982 | unclassified | unclassified | unclassified | unclassified | unclassified | unclassified | unclassified | unclassified | unclassified | unclassified | unclassified | unclassified | 0,4 | 0,489897949 | 0,000449195 | 0,000551247 | 0 - | 0 - | 0 - | 0 - | 0 - | 0 - |
| 1205 | 1 | 0.2um_NOKCPHJN_790587 | unclassified | unclassified | unclassified | unclassified | unclassified | unclassified | unclassified | unclassified | unclassified | unclassified | unclassified | unclassified | 0,2 | 0,4 | 3,97E-05 | 7,95E-05 | 0 - | 0 - | 1 - | 0,000198651 - | 0 - | 0 - |
| 1206 | 1 | 3um_FIJJAFBF_452346 | unclassified | unclassified | unclassified | unclassified | unclassified | unclassified | unclassified | unclassified | unclassified | unclassified | unclassified | unclassified | 0,2 | 0,4 | 6,74E-05 | 0,000134744 | 0 - | 0 - | 0 - | 0 - | 1 - | 0,000336861 - |
| 1207 | 1 | 0.2um_NOKCPHJN_40987 | Bacteria | Actinobacteri | Actinobacteri | Micrococcales | Microbacteria | Candidatus Ac | Candidatus Ac | unclassified | unclassified | unclassified | unclassified | unclassified | 1 | 0 | 0,000175466 | 1,86E-05 | 1 - | 0,000208491 - | 1 - | 0,000174303 - | 1 - | 0,000175468 - |

| | | | | | | | | | | | | | | | | | | | | | | | | |
|---|---|---|---|---|---|---|---|---|---|---|---|---|---|---|---|---|---|---|---|---|---|---|---|---|
| 1208 | 1 | 3um_FIJJAFBF_558999 | Bacteria | Proteobacteri | Gammaprotec | Cellvibrionales | Cellvibrionace | Cellvibrio | Cellvibrio sp. 7 | unclassified | unclassified | unclassified | unclassified | unclassified | 0,6 | 0,489897949 | 0,000135281 | 0,000112987 | 1 - | 0,000267315 - | 0 - | 0 - | 0 - | 0 - |
| 1209 | 1 | 3um_FIJJAFBF_235863 | Bacteria | Proteobacteri | Gammaprotec | Alteromonada | unclassified | unclassified | Alteromonada | unclassified | unclassified | unclassified | unclassified | unclassified | 0,6 | 0,8 | 0,000458485 | 0,000594005 | 0 - | 0 - | 0 - | 0 - | 1 - | 0,000839906 - |
| 1210 | 1 | 0.2um_NOKCPHJN_765960 | Eukaryota | unclassified | unclassified | Longamoebia | Acanthamoeb | Acanthamoeb | Acanthamoeb | unclassified | unclassified | unclassified | unclassified | unclassified | 0,2 | 0,4 | 1,90E-05 | 3,80E-05 | 1 - | 9,50E-05 - | 0 - | 0 - | 0 - | 0 - |
| 1211 | 1 | 0.2um_NOKCPHJN_1065032 | Eukaryota | unclassified | Spirotrichea | Sporadotrichi | Oxytrichidae | Stylonychia | Stylonychia le | metabolism | Carbohydrate | COG0057 | glyceraldehyd | 423536.XP_00 | 1,4 | 0,489897949 | 0,000249713 | 6,99E-05 | 1 - | 0,000217582 - | 1 - | 0,000181904 - | 1 - | 0,000183119 - |
| 1212 | 1 | 0.2um_NOKCPHJN_312733 | Eukaryota | Phaeophyceae | unclassified | Ectocarpales | Ectocarpaceae | Ectocarpus | Ectocarpus sili | cellular proces | Signal transdu | KOG0613 | fibronectin typ | 67593.Physo1 | 0,2 | 0,4 | 2,75E-06 | 5,50E-06 | 0 - | 0 - | 0 - | 0 - | 0 - | 0 - |
| 1213 | 1 | 3um_FIJJAFBF_85799 | Viruses | unclassified | unclassified | unclassified | unclassified | unclassified | uncultured M | unclassified | unclassified | unclassified | unclassified | unclassified | 0,2 | 0,4 | 7,89E-05 | 0,000157882 | 0 - | 0 - | 0 - | 0 - | 0 - | 0 - |
| 1214 | 1 | 3um_FIJJAFBF_515763 | Eukaryota | Bacillariophyta | Coscinodiscop | Thalassiosirale | Thalassiosirac | Thalassiosira | Thalassiosira c | metabolism | Amino acid tra | COG0436 | aminotransfer | 35128.Thaps3 | 0,2 | 0,4 | 7,19E-05 | 0,000143893 | 0 - | 0 - | 0 - | 0 - | 0 - | 0 - |
| 1215 | 1 | 0.2um_NOKCPHJN_740862 | Eukaryota | Arthropoda | Insecta | Hymenoptera | Apidae | Bombus | Bombus impat | information st | Replication, re | 0V9WU | polymerase (D | 12957.ACEP_0 | 0,6 | 0,489897949 | 0,000139794 | 0,000114649 | 0 - | 0 - | 0 - | 0 - | 1 - | 0,000248004 - |
| 1216 | 1 | 0.2um_NOKCPHJN_882893 | Eukaryota | unclassified | Cryptophyta | Pyrenomonad | Geminigerace | Guillardia | Guillardia thet | unclassified | unclassified | unclassified | unclassified | unclassified | 0,2 | 0,4 | 3,80E-05 | 7,59E-05 | 0 - | 0 - | 0 - | 0 - | 0 - | 0 - |
| 1217 | 2 | 3um_FIJJAFBF_11563;3um_FIJJAFBF_4 | Bacteria | Bacteroidetes | Flavobacteriia | Flavobacterial | Flavobacteriac | Lutibacter | Lutibacter sp. | unclassified | unclassified | unclassified | unclassified | unclassified | 0,8 | 0,4 | 0,000122517 | 6,34E-05 | 1 - | 0,00018167 - | 0 - | 0 - | 1 - | 0,000152895 - |
| 1218 | 1 | 3um_FIJJAFBF_437595 | Bacteria | Bacteroidetes | Flavobacteriia | Flavobacterial | Flavobacteriac | Polaribacter | Polaribacter s | unclassified | unclassified | unclassified | unclassified | unclassified | 1,8 | 0,4 | 0,000606638 | 0,000104248 | 1 - | 0,000409006 - | 2 - | 0,000683879 - | 2 - | 0,000688447 - |
| 1219 | 1 | 3um_FIJJAFBF_500695 | Eukaryota | Bacillariophyta | Mediophyceae | Cymatosirales | Cymatosiracea | Plagiogrammc | Plagiogrammc | metabolism | Energy produc | COG4451 | RuBisCO catal | 323097.Nham | 3,2 | 1,166190379 | 0,001409196 | 0,000411699 | 2 - | 0,001076952 - | 2 - | 0,000900359 - | 4 - | 0,001812746 - |
| 1220 | 1 | 0.2um_NOKCPHJN_1077861 | Eukaryota | unclassified | Cryptophyta | Pyrenomonad | Geminigerace | Guillardia | Guillardia thet | unclassified | unclassified | unclassified | unclassified | unclassified | 1 | 0 | 0,000684698 | 7,24E-05 | 1 - | 0,000813567 - | 1 - | 0,000680162 - | 1 - | 0,000684706 - |
| 1221 | 1 | 0.2um_NOKCPHJN_06883 | Bacteria | Bacteroidetes | Cytophagia | Cytophagales | Cytophagacea | Ekhidna | Ekhidna lutea | information st | Translation, ri | 0FVR5 | This protein p | 643867.Ftrac_ | 0,4 | 0,8 | 6,38E-05 | 0,000127581 | 0 - | 0 - | 0 - | 0 - | 2 - | 0,000318952 - |
| 1222 | 1 | 0.2um_NOKCPHJN_1049390 | Bacteria | Bacteroidetes | Flavobacteriia | Flavobacterial | Cryomorphace | Owenweeksia | Owenweeksia | metabolism | Amino acid tra | 0P7SN | Catalyzes the | 888059.HMPR | 0,2 | 0,4 | 4,74E-05 | 9,47E-05 | 0 - | 0 - | 0 - | 0 - | 0 - | 0 - |
| 1223 | 1 | 0.2um_NOKCPHJN_485725 | Bacteria | Proteobacteri | Gammaprotec | Cellvibrionales | Spongiibacter | Oceanicoccus | Oceanicoccus | cellular proces | Cell wall/mem | 0QTSX | Peptidoglycan | 247634.GPB2 | 4,2 | 0,4 | 0,001384731 | 0,000188391 | 4 - | 0,001567501 - | 5 - | 0,001638087 - | 4 - | 0,001319224 - |
| 1224 | 1 | 3um_FIJJAFBF_151368 | Eukaryota | Annelida | Polychaeta | Golfingiida | Sipunculidae | Sipunculus | Sipunculus nu | unclassified | unclassified | unclassified | unclassified | unclassified | 2,2 | 0,748331477 | 0,001093777 | 0,000348768 | 1 - | 0,000603614 - | 2 - | 0,001009273 - | 3 - | 0,001524023 - |
| 1225 | 1 | 0.2um_NOKCPHJN_855491 | Eukaryota | Phaeophyceae | unclassified | Ectocarpales | Ectocarpaceae | Ectocarpus | Ectocarpus sili | unclassified | unclassified | unclassified | unclassified | unclassified | 1 | 0 | 9,83E-05 | 1,04E-05 | 1 - | 0,000116768 - | 1 - | 9,76E-05 - | 1 - | 9,83E-05 - |
| 1226 | 1 | 3um_FIJJAFBF_67730 | Bacteria | Proteobacteri | Gammaprotec | Cellvibrionales | unclassified | unclassified | Cellvibrionales | unclassified | unclassified | unclassified | unclassified | unclassified | 0,8 | 0,4 | 0,000365777 | 0,000189146 | 1 - | 0,000542378 - | 0 - | 0 - | 1 - | 0,000456471 - |
| 1227 | 1 | 0.2um_NOKCPHJN_1033454 | unclassified | unclassified | unclassified | unclassified | unclassified | unclassified | unclassified | unclassified | unclassified | unclassified | unclassified | unclassified | 0,2 | 0,4 | 6,41E-05 | 0,000128163 | 0 - | 0 - | 0 - | 0 - | 0 - | 0 - |
| 1228 | 6 | 0.2um_NOKCPHJN_454363;0.2um_NO | Bacteria | Proteobacteri | Alphaproteob | Rhodobactera | Rhodobactera | various | various | information st | Translation, ri | COG0185 | Protein S19 fo | various | 1 | 0 | 0,000677335 | 7,16E-05 | 1 - | 0,000804819 - | 1 - | 0,000672849 - | 1 - | 0,000677343 - |
| 1229 | 1 | 0.2um_NOKCPHJN_809592 | unclassified | unclassified | unclassified | unclassified | unclassified | unclassified | unclassified | unclassified | unclassified | unclassified | unclassified | unclassified | 1 | 0 | 7,86E-05 | 8,32E-06 | 1 - | 9,34E-05 - | 1 - | 7,81E-05 - | 1 - | 7,86E-05 - |
| 1230 | 1 | 0.2um_NOKCPHJN_512717 | Eukaryota | Nematoda | Chromadorea | Diplogasterida | Neodiplogaste | Pristionchus | Pristionchus p | information st | Chromatin str | KOG3467 | Core compone | 7217.FBpp012 | 1,8 | 0,4 | 0,000196066 | 5,45E-05 | 2 - | 0,000255019 - | 2 - | 0,000213202 - | 2 - | 0,000214627 - |
| 1231 | 1 | 3um_FIJJAFBF_715095 | Eukaryota | Bacillariophyta | Coscinodiscop | Rhizosoleniale | Rhizosoleniac | Rhizosolenia | Rhizosolenia i | information st | Translation, ri | COG0087 | One of the pri | 118168.MC74 | 1,6 | 0,8 | 0,000476655 | 0,000226626 | 1 - | 0,000358125 - | 1 - | 0,000299402 - | 2 - | 0,000602803 - |
| 1232 | 1 | 0.2um_NOKCPHJN_468509 | Eukaryota | Chordata | unclassified | Squamata | Gekkonidae | Gekko | Gekko japonic | unclassified | unclassified | unclassified | unclassified | unclassified | 1 | 0 | 9,95E-05 | 1,05E-05 | 1 - | 0,000118244 - | 1 - | 9,89E-05 - | 1 - | 9,95E-05 - |
| 1233 | 2 | 0.2um_NOKCPHJN_486326;0.2um_NO | Eukaryota | various | various | various | various | various | various | unclassified | unclassified | unclassified | unclassified | unclassified | 0,8 | 0,4 | 0,000390427 | 0,000196771 | 0 - | 0 - | 1 - | 0,000508739 - | 1 - | 0,000512138 - |
| 1234 | 1 | 0.2um_NOKCPHJN_750086 | Bacteria | Proteobacteri | Alphaproteob | Pelagibacteral | Pelagibacterac | Candidatus Pe | Candidatus Pe | metabolism | Energy produc | 01RNF | Produces ATP | 439493.PB721 | 1,2 | 0,4 | 0,000158914 | 4,98E-05 | 1 - | 0,000158577 - | 1 - | 0,000132574 - | 1 - | 0,00013346 - |
| 1235 | 1 | 0.2um_NOKCPHJN_84217 | Eukaryota | unclassified | Oomycetes | Saprolegniales | Saprolegniace | Achlya | Achlya hypogy | information st | Translation, ri | COG0052 | 30S ribosomal | 653948.CCA13 | 0,2 | 0,4 | 3,13E-05 | 6,26E-05 | 0 - | 0 - | 0 - | 0 - | 0 - | 0 - |
| 1236 | 1 | 0.2um_NOKCPHJN_904616 | Eukaryota | unclassified | Pelagophyceae | Pelagomonada | unclassified | Aureococcus | Aureococcus a | cellular proces | Posttranslatio | KOG3083 | prohibitin | 294747.XP_00 | 1,6 | 0,489897949 | 0,000895452 | 0,00022938 | 1 - | 0,000680438 - | 1 - | 0,000568863 - | 2 - | 0,001145326 - |
| 1237 | 1 | 3um_FIJJAFBF_504209 | Bacteria | Proteobacteri | Alphaproteob | unclassified | unclassified | Candidatus Pu | Candidatus Pu | cellular proces | Posttranslatio | 0FJCG | Prevents misfo | 321327.CYA_1 | 1,6 | 0,489897949 | 0,00019501 | 7,11E-05 | 2 - | 0,000284594 - | 1 - | 0,000118964 - | 2 - | 0,000239517 - |
| 1238 | 1 | 3um_FIJJAFBF_390383 | Eukaryota | Bacillariophyta | Bacillariophyc | Naviculales | Naviculaceae | Fistulifera | Fistulifera sola | unclassified | unclassified | unclassified | unclassified | unclassified | 0,6 | 0,489897949 | 6,49E-05 | 5,30E-05 | 0 - | 0 - | 1 - | 0,000109397 - | 1 - | 0,000110128 - |
| 1239 | 1 | 0.2um_NOKCPHJN_234273 | Bacteria | Proteobacteri | Gammaprotec | Cellvibrionales | Halieaceae | unclassified | marine gamm | metabolism | Inorganic ion t | 16PDP | TonB-depende | 247639.MGP2 | 0,4 | 0,489897949 | 2,40E-05 | 2,94E-05 | 0 - | 0 - | 0 - | 0 - | 0 - | 0 - |
| 1240 | 4 | 0.2um_NOKCPHJN_108598;0.2um_NO | Bacteria | Bacteroidetes | Flavobacteriia | Flavobacterial | Crocinitomica | unclassified | Crocinitomica | metabolism | Nucleotide tra | 0PF2Y | Catalyzes two | various | 0 | 0 | 0 | 0 | 0 - | 0 - | 0 - | 0 - | 0 - | 0 - |
| 1241 | 1 | 3um_FIJJAFBF_593283 | unclassified | unclassified | unclassified | unclassified | unclassified | unclassified | unclassified | unclassified | unclassified | unclassified | unclassified | unclassified | 1 | 0 | 0,000319757 | 3,38E-05 | 1 - | 0,00037994 - | 1 - | 0,000317639 - | 1 - | 0,000319761 - |
| 1242 | 1 | 0.2um_NOKCPHJN_792829 | Bacteria | Proteobacteri | Alphaproteob | Rhodobactera | unclassified | unclassified | Rhodobactera | unclassified | unclassified | unclassified | unclassified | unclassified | 0,8 | 0,4 | 7,40E-05 | 3,77E-05 | 1 - | 0,000106318 - | 1 - | 8,89E-05 - | 1 - | 8,95E-05 - |
| 1243 | 1 | 0.2um_NOKCPHJN_114353 | Bacteria | Proteobacteri | Alphaproteob | Pelagibacteral | Pelagibacterac | Candidatus Pe | Candidatus Pe | unclassified | unclassified | unclassified | unclassified | unclassified | 0,2 | 0,4 | 3,13E-05 | 6,26E-05 | 0 - | 0 - | 1 - | 0,000156437 - | 0 - | 0 - |
| 1244 | 1 | 3um_FIJJAFBF_328495 | Bacteria | Bacteroidetes | Flavobacteriia | Flavobacterial | Flavobacteriac | Arenitalea | Arenitalea lut | poorly charact | Function unkn | 0PAEN | NA | 886377.Murru | 5 | 1,095445115 | 0,000644079 | 0,000103518 | 3 - | 0,000468778 - | 5 - | 0,000653183 - | 6 - | 0,000789056 - |
| 1245 | 1 | 3um_FIJJAFBF_240437 | Archaea | unclassified | unclassified | unclassified | unclassified | unclassified | archaeon GW | unclassified | unclassified | unclassified | unclassified | unclassified | 0,4 | 0,489897949 | 8,16E-05 | 0,000100548 | 1 - | 0,000222101 - | 1 - | 0,000185682 - | 0 - | 0 - |
| 1246 | 1 | 3um_FIJJAFBF_799495 | unclassified | unclassified | unclassified | unclassified | unclassified | unclassified | unclassified | unclassified | unclassified | unclassified | unclassified | unclassified | 0,4 | 0,489897949 | 0,000120067 | 0,00014712 | 0 - | 0 - | 0 - | 0 - | 1 - | 0,000307283 - |
| 1247 | 1 | 0.2um_NOKCPHJN_87559 | Bacteria | Proteobacteri | Alphaproteob | Rhodospirillale | Rhodospirillac | Nisaea | Nisaea denitri | unclassified | unclassified | unclassified | unclassified | unclassified | 0,8 | 0,4 | 9,59E-05 | 4,83E-05 | 0 - | 0 - | 1 - | 0,0001249 - | 1 - | 0,000125734 - |
| 1248 | 1 | 0.2um_NOKCPHJN_132742 | Eukaryota | unclassified | Oomycetes | Albuginales | Albuginaceae | Albugo | Albugo candid | unclassified | unclassified | unclassified | unclassified | unclassified | 5 | 0 | 0,00070461 | 7,45E-05 | 5 - | 0,000837228 - | 5 - | 0,000699943 - | 5 - | 0,000704619 - |
| 1249 | 1 | 3um_FIJJAFBF_658713 | Eukaryota | unclassified | unclassified | Isochrysidales | Noelaerhabda | Emiliania | Emiliania huxl | unclassified | unclassified | unclassified | unclassified | unclassified | 2,6 | 0,489897949 | 0,000679531 | 0,000162382 | 3 - | 0,00092787 - | 2 - | 0,000517148 - | 3 - | 0,000780904 - |
| 1250 | 1 | 3um_FIJJAFBF_111260 | unclassified | unclassified | unclassified | unclassified | unclassified | unclassified | unclassified | unclassified | unclassified | unclassified | unclassified | unclassified | 1,2 | 0,748331477 | 0,000514496 | 0,000316756 | 0 - | 0 - | 1 - | 0,000450179 - | 2 - | 0,000906373 - |
| 1251 | 1 | 3um_FIJJAFBF_01138 | Bacteria | Bacteroidetes | Flavobacteriia | Flavobacterial | Flavobacteriac | Polaribacter | various | information st | Translation, ri | 09G6S | 50S ribosomal | 313594.PI23P | 0,6 | 0,489897949 | 0,000446262 | 0,000372102 | 1 - | 0,000870328 - | 1 - | 0,000727615 - | 0 - | 0 - |
| 1252 | 2 | 0.2um_NOKCPHJN_970291;3um_FIJJAF | Bacteria | Bacteroidetes | Flavobacteriia | Flavobacterial | Flavobacteriac | various | various | unclassified | unclassified | unclassified | unclassified | unclassified | 0,2 | 0,4 | 0,000137897 | 0,000275794 | 0 - | 0 - | 0 - | 0 - | 0 - | 0 - |
| 1253 | 1 | 0.2um_NOKCPHJN_951164 | unclassified | unclassified | unclassified | unclassified | unclassified | unclassified | unclassified | unclassified | unclassified | unclassified | unclassified | unclassified | 1 | 0 | 0,000605694 | 6,41E-05 | 1 - | 0,000719694 - | 1 - | 0,000601682 - | 1 - | 0,000605701 - |
| 1254 | 1 | 3um_FIJJAFBF_111300 | unclassified | unclassified | unclassified | unclassified | unclassified | unclassified | unclassified | unclassified | unclassified | unclassified | unclassified | unclassified | 0,2 | 0,4 | 1,38E-05 | 2,76E-05 | 0 - | 0 - | 0 - | 0 - | 0 - | 0 - |
| 1255 | 1 | 3um_FIJJAFBF_444417 | unclassified | unclassified | unclassified | unclassified | unclassified | unclassified | unclassified | unclassified | unclassified | unclassified | unclassified | unclassified | 0,2 | 0,4 | 2,93E-05 | 5,86E-05 | 1 - | 0,000146474 - | 0 - | 0 - | 0 - | 0 - |
| 1256 | 1 | 3um_FIJJAFBF_122480 | Bacteria | Bacteroidetes | Flavobacteriia | Flavobacterial | Flavobacteriac | Polaribacter | Polaribacter v | information st | Transcription | 05D27 | DNA-depende | 313594.PI23P | 7 | 2 | 0,000303369 | 7,45E-05 | 4 - | 0,000210248 - | 8 - | 0,000351544 - | 6 - | 0,00026542 - |
| 1257 | 1 | 3um_FIJJAFBF_527148 | Eukaryota | Bacillariophyta | Coscinodiscop | Thalassiosirale | Thalassiosirac | Thalassiosira | Thalassiosira c | poorly charact | Function unkn | COG1100 | GTP-binding P | 35128.Thaps2 | 1 | 0 | 0,000290287 | 3,07E-05 | 1 - | 0,000344923 - | 1 - | 0,000288364 - | 1 - | 0,00029029 - |
| 1258 | 1 | 3um_FIJJAFBF_257126 | Eukaryota | unclassified | Florideophyce | Nemaliales | Liagoraceae | Dermonema | Dermonema v | unclassified | unclassified | unclassified | unclassified | unclassified | 3,6 | 0,489897949 | 0,001255823 | 0,000264828 | 4 - | 0,001645015 - | 3 - | 0,001031455 - | 4 - | 0,00138446 - |
| 1259 | 1 | 0.2um_NOKCPHJN_959134 | Eukaryota | Phaeophyceae | unclassified | Ectocarpales | Ectocarpaceae | Ectocarpus | Ectocarpus sili | metabolism | Energy produc | COG0055 | Produces ATP | 35128.Thaps4 | 1 | 0 | 0,000488311 | 5,16E-05 | 1 - | 0,000580218 - | 1 - | 0,000485077 - | 1 - | 0,000488317 - |
| 1260 | 1 | 0.2um_NOKCPHJN_28866 | Bacteria | Bacteroidetes | Flavobacteriia | Flavobacterial | Flavobacteriac | Polaribacter | Polaribacter b | metabolism | Carbohydrate | 05D2N | Aldolase | 391587.KAOT | 1,8 | 0,748331477 | 0,000314126 | 0,000126067 | 1 - | 0,00021084 - | 3 - | 0,000528802 - | 2 - | 0,00035489 - |
| 1261 | 1 | 0.2um_NOKCPHJN_627366 | Eukaryota | Mollusca | Gastropoda | Architaenioglc | Ampullariidae | Pomacea | Pomacea cana | unclassified | unclassified | unclassified | unclassified | unclassified | 0,2 | 0,4 | 9,62E-06 | 1,92E-05 | 0 - | 0 - | 0 - | 0 - | 0 - | 0 - |
| 1262 | 1 | 3um_FIJJAFBF_14623 | Bacteria | Proteobacteri | Alphaproteob | Rhodobactera | Rhodobactera | Actibacterium | Actibacterium | cellular proces | Cell wall/mem | 01VG7 | Salt-stress ind | 89187.ISM_10 | 0,6 | 0,489897949 | 0,000126361 | 0,000103633 | 0 - | 0 - | 0 - | 0 - | 1 - | 0,000224174 - |
| 1263 | 1 | 0.2um_NOKCPHJN_59333 | Bacteria | Bacteroidetes | Flavobacteriia | Flavobacterial | Flavobacteriac | Lacinutrix | Lacinutrix ven | metabolism | Amino acid tra | 09GJ6 | nitrogen regul | 391603.FBALC | 2 | 0,894427191 | 0,001099227 | 0,000468992 | 1 - | 0,000668287 - | 1 - | 0,000558705 - | 3 - | 0,001687311 - |
| 1264 | 1 | 0.2um_NOKCPHJN_1029958 | Bacteria | Bacteroidetes | Flavobacteriia | Flavobacterial | Flavobacteriac | Flavobacteriu | Flavobacteriu | poorly charact | Function unkn | 0Z0C5 | Family of unkr | 762903.Pedsa | 0,4 | 0,489897949 | 9,14E-05 | 0,000114223 | 1 - | 0,000264481 - | 0 - | 0 - | 0 - | 0 - |
| 1265 | 1 | 3um_FIJJAFBF_503090 | Bacteria | Verrucomicrol | Opitutae | unclassified | unclassified | unclassified | Opitutae bact | unclassified | unclassified | unclassified | unclassified | unclassified | 1 | 0 | 0,000153266 | 1,62E-05 | 1 - | 0,000182112 - | 1 - | 0,00015225 - | 1 - | 0,000153267 - |
| 1266 | 1 | 3um_FIJJAFBF_73819 | Bacteria | Bacteroidetes | Flavobacteriia | Flavobacterial | Flavobacteriac | Polaribacter | Polaribacter s | unclassified | unclassified | unclassified | unclassified | unclassified | 0,2 | 0,4 | 1,80E-05 | 3,60E-05 | 0 - | 0 - | 1 - | 8,99E-05 - | 0 - | 0 - |
| 1267 | 1 | 0.2um_NOKCPHJN_219472 | Bacteria | Proteobacteri | Gammaprotec | Alteromonada | Pseudoalteror | Pseudoalteror | Pseudoalteror | metabolism | Inorganic ion t | 16PDP | TonB-depende | 983545.Glaag | 4,8 | 0,748331477 | 0,000547516 | 6,90E-05 | 4 - | 0,000547336 - | 4 - | 0,000457586 - | 5 - | 0,000575804 - |
| 1268 | 1 | 0.2um_NOKCPHJN_487408 | Bacteria | Proteobacteri | Gammaprotec | Alteromonada | Pseudoalteror | Pseudoalteror | Pseudoalteror | unclassified | unclassified | unclassified | unclassified | unclassified | 0,4 | 0,489897949 | 6,92E-05 | 8,49E-05 | 0 - | 0 - | 0 - | 0 - | 0 - | 0 - |
| 1269 | 1 | 3um_FIJJAFBF_448401 | Eukaryota | Bacillariophyta | Bacillariophyc | Bacillariales | Bacillariaceae | Fragilariopsis | Fragilariopsis | unclassified | unclassified | unclassified | unclassified | unclassified | 1,4 | 0,489897949 | 0,002202593 | 0,000616969 | 1 - | 0,001919184 - | 1 - | 0,001604485 - | 1 - | 0,001615204 - |
| 1270 | 1 | 0.2um_NOKCPHJN_395741 | Eukaryota | unclassified | Oomycetes | Saprolegniales | Saprolegniace | Thraustotheca | Thraustotheca | unclassified | unclassified | unclassified | unclassified | unclassified | 2,4 | 0,489897949 | 0,000306997 | 4,40E-05 | 2 - | 0,000308652 - | 2 - | 0,000258041 - | 2 - | 0,000259765 - |
| 1271 | 3 | 0.2um_NOKCPHJN_249529;0.2um_NO | Bacteria | Proteobacteri | Alphaproteob | Rhodobactera | various | various | various | unclassified | unclassified | unclassified | unclassified | unclassified | 0,2 | 0,4 | 5,78E-05 | 0,000115596 | 1 - | 0,000288989 - | 0 - | 0 - | 0 - | 0 - |
| 1272 | 1 | 3um_FIJJAFBF_284683 | Eukaryota | Bacillariophyta | Coscinodiscop | Leptocylindral | Leptocylindrac | Leptocylindrus | Leptocylindrus | metabolism | Energy produc | 1DYB8 | PsaA and PsaE | 3055.DAA014 | 1,6 | 0,8 | 0,00012772 | 6,44E-05 | 0 - | 0 - | 2 - | 0,000166423 - | 2 - | 0,000167534 - |
| 1273 | 1 | 0.2um_NOKCPHJN_1004298 | Bacteria | Proteobacteri | Gammaprotec | Cellvibrionales | unclassified | unclassified | Cellvibrionales | unclassified | unclassified | unclassified | unclassified | unclassified | 0,4 | 0,489897949 | 0,000100921 | 0,000123848 | 0 - | 0 - | 0 - | 0 - | 0 - | 0 - |
| 1274 | 1 | 3um_FIJJAFBF_225879 | Bacteria | Bacteroidetes | Flavobacteriia | Flavobacterial | Flavobacteriac | Flaviramulus | Flaviramulus b | cellular proces | Cell wall/mem | 06B97 | phosphate-sel | 391598.FBBAL | 0,6 | 0,489897949 | 8,70E-05 | 7,14E-05 | 0 - | 0 - | 1 - | 0,000151147 - | 1 - | 0,000152157 - |
| 1275 | 1 | 3um_FIJJAFBF_114132 | unclassified | unclassified | unclassified | unclassified | unclassified | unclassified | unclassified | unclassified | unclassified | unclassified | unclassified | unclassified | 2 | 0 | 0,000851246 | 9,00E-05 | 2 - | 0,001011462 - | 2 - | 0,000845607 - | 2 - | 0,000851256 - |
| 1276 | 1 | 3um_FIJJAFBF_504214 | Bacteria | Proteobacteri | Alphaproteob | unclassified | unclassified | Candidatus Pu | Candidatus Pu | unclassified | unclassified | unclassified | unclassified | unclassified | 3,4 | 0,8 | 0,000344616 | 8,59E-05 | 3 - | 0,000361585 - | 5 - | 0,000503824 - | 3 - | 0,000304314 - |
| 1277 | 1 | 0.2um_NOKCPHJN_898151 | Viruses | unclassified | unclassified | unclassified | unclassified | unclassified | uncultured M | unclassified | unclassified | unclassified | unclassified | unclassified | 0,8 | 0,748331477 | 0,000144757 | 0,000129134 | 0 - | 0 - | 1 - | 0,000193132 - | 1 - | 0,000194423 - |
| 1278 | 1 | 0.2um_NOKCPHJN_546398 | Eukaryota | Arthropoda | Insecta | Hymenoptera | Formicidae | Lasius | Lasius niger | unclassified | unclassified | unclassified | unclassified | unclassified | 0,6 | 0,489897949 | 2,83E-05 | 2,32E-05 | 0 - | 0 - | 1 - | 5,01E-05 - | 0 - | 0 - |
| 1279 | 1 | 0.2um_NOKCPHJN_891630 | Bacteria | Proteobacteri | Alphaproteob | Rhizobiales | Methylobacte | Methylorubru | Methylorubru | unclassified | unclassified | unclassified | unclassified | unclassified | 1 | 0 | 0,000777681 | 8,22E-05 | 1 - | 0,000924052 - | 1 - | 0,00077253 - | 1 - | 0,000777691 - |
| 1280 | 1 | 0.2um_NOKCPHJN_577036 | Eukaryota | Bacillariophyta | Coscinodiscop | Thalassiosirale | Thalassiosirac | Thalassiosira | Thalassiosira c | unclassified | unclassified | unclassified | unclassified | unclassified | 1 | 0 | 0,000470091 | 4,97E-05 | 1 - | 0,000558569 - | 1 - | 0,000466977 - | 1 - | 0,000470097 - |
| 1281 | 1 | 3um_FIJJAFBF_344737 | Eukaryota | Bacillariophyta | Mediophyceae | Toxariales | Toxariaceae | Toxarium | Toxarium und | information st | Translation, ri | COG0050 | This protein p | 321332.CYB_0 | 3 | 0,632455532 | 0,000454821 | 7,57E-05 | 2 - | 0,000366006 - | 3 - | 0,000458985 - | 3 - | 0,000462051 - |
| 1282 | 1 | 0.2um_NOKCPHJN_553159 | unclassified | unclassified | unclassified | unclassified | unclassified | unclassified | unclassified | unclassified | unclassified | unclassified | unclassified | unclassified | 0,2 | 0,4 | 3,11E-05 | 6,23E-05 | 0 - | 0 - | 0 - | 0 - | 0 - | 0 - |
| 1283 | 1 | 0.2um_NOKCPHJN_290418 | Bacteria | Proteobacteri | Gammaprotec | Alteromonada | Colwelliaceae | Colwellia | Colwellia sp. P | information st | Translation, ri | 16Q16 | Binds the lowe | 167879.CPS_0 | 0,2 | 0,4 | 5,45E-05 | 0,000109079 | 0 - | 0 - | 0 - | 0 - | 1 - | 0,000272697 - |
| 1284 | 1 | 3um_FIJJAFBF_715082 | Eukaryota | unclassified | Pelagophyceae | Pelagomonada | unclassified | Aureoumbra | Aureoumbra l | unclassified | unclassified | unclassified | unclassified | unclassified | 4,8 | 0,979795897 | 0,001810827 | 0,000300295 | 4 - | 0,001814501 - | 6 - | 0,002275452 - | 4 - | 0,001527102 - |
| 1285 | 1 | 3um_FIJJAFBF_37474 | Eukaryota | Bacillariophyta | Coscinodiscop | Thalassiosirale | Thalassiosirac | Porosira | Porosira glacia | metabolism | Carbohydrate | COG1850 | RuBisCO catal | 35128.Thapsd | 0,8 | 0,4 | 0,000241598 | 0,000124665 | 1 - | 0,000354731 - | 1 - | 0,000296564 - | 1 - | 0,000298545 - |
| 1286 | 1 | 3um_FIJJAFBF_768334 | Eukaryota | unclassified | Oomycetes | Saprolegniales | Saprolegniace | Aphanomyces | Aphanomyces | unclassified | unclassified | unclassified | unclassified | unclassified | 1 | 0 | 0,000206532 | 2,18E-05 | 1 - | 0,000245404 - | 1 - | 0,000205164 - | 1 - | 0,000206534 - |
| 1287 | 1 | 3um_FIJJAFBF_91116 | Eukaryota | unclassified | Pelagophyceae | Pelagomonada | unclassified | Aureococcus | Aureococcus a | metabolism | Carbohydrate | KOG1203 | Complex I inte | 35128.Thapsd | 7 | 2,19089023 | 0,001333492 | 0,000294984 | 5 - | 0,001162239 - | 7 - | 0,001360324 - | 5 - | 0,000978151 - |
| 1288 | 1 | 3um_FIJJAFBF_364989 | Bacteria | Proteobacteri | Betaproteoba | Burkholderiale | unclassified | unclassified | various | poorly charact | Function unkn | COG0714 | ATPase activit | 187272.Mlg_1 | 0,2 | 0,4 | 4,69E-05 | 9,39E-05 | 1 - | 0,000234634 - | 0 - | 0 - | 0 - | 0 - |
| 1289 | 1 | 3um_FIJJAFBF_802740 | Bacteria | Proteobacteri | Alphaproteob | Rhodobactera | Rhodobactera | Marinovum | Marinovum al | cellular proces | Cell wall/mem | 16RAV | Exopolysaccha | 314256.OG25 | 0,8 | 0,4 | 7,05E-05 | 3,64E-05 | 1 - | 0,000103524 - | 1 - | 8,65E-05 - | 1 - | 8,71E-05 - |
| 1290 | 2 | 0.2um_NOKCPHJN_45852;3um_FIJJAFE | Bacteria | Bacteroidetes | Flavobacteriia | Flavobacterial | Crocinitomica | unclassified | Crocinitomica | metabolism | Nucleotide tra | 0PF2Y | Catalyzes two | 313590.MED1 | 0,2 | 0,4 | 3,73E-05 | 7,47E-05 | 1 - | 0,000186654 - | 0 - | 0 - | 0 - | 0 - |
| 1291 | 1 | 0.2um_NOKCPHJN_390653 | Eukaryota | Phaeophyceae | unclassified | Ectocarpales | Ectocarpaceae | Ectocarpus | Ectocarpus sili | unclassified | unclassified | unclassified | unclassified | unclassified | 1 | 0 | 0,000206532 | 2,18E-05 | 1 - | 0,000245404 - | 1 - | 0,000205164 - | 1 - | 0,000206534 - |
| 1292 | 1 | 3um_FIJJAFBF_196455 | Bacteria | Proteobacteri | Gammaprotec | Cellvibrionales | Halieaceae | unclassified | marine gamm | unclassified | unclassified | unclassified | unclassified | unclassified | 2,2 | 0,748331477 | 0,000173845 | 5,21E-05 | 2 - | 0,000190939 - | 2 - | 0,00015963 - | 1 - | 8,03E-05 - |
| 1293 | 1 | 3um_FIJJAFBF_592540 | Bacteria | Proteobacteri | Gammaprotec | Alteromonada | Alteromonada | Glaciecola | Glaciecola sp. | unclassified | unclassified | unclassified | unclassified | unclassified | 1,6 | 1,019803903 | 9,88E-05 | 6,29E-05 | 0 - | 0 - | 1 - | 6,47E-05 - | 2 - | 0,000130285 - |
| 1294 | 1 | 0.2um_NOKCPHJN_562145 | Bacteria | Bacteroidetes | Flavobacteriia | Flavobacterial | Crocinitomica | unclassified | Crocinitomica | poorly charact | Function unkn | 16R8I | Lipoprotein | 715451.ambt_ | 1,6 | 0,8 | 7,48E-05 | 3,12E-05 | 1 - | 5,76E-05 - | 2 - | 9,63E-05 - | 1 - | 4,85E-05 - |
| 1295 | 1 | 0.2um_NOKCPHJN_61813 | Bacteria | Bacteroidetes | Flavobacteriia | Flavobacterial | Flavobacteriac | Polaribacter | Polaribacter fi | poorly charact | Function unkn | 0XY6Y | Pkd domain cc | 391587.KAOT | 2,8 | 0,748331477 | 0,000385156 | 9,49E-05 | 2 - | 0,000331187 - | 2 - | 0,00027688 - | 4 - | 0,00055746 - |
| 1296 | 1 | 3um_FIJJAFBF_129173 | Bacteria | Bacteroidetes | Flavobacteriia | Flavobacterial | Flavobacteriac | Polaribacter | Polaribacter s | unclassified | unclassified | unclassified | unclassified | unclassified | 0,2 | 0,4 | 1,26E-05 | 2,53E-05 | 0 - | 0 - | 0 - | 0 - | 0 - | 0 - |
| 1297 | 1 | 3um_FIJJAFBF_532202 | Bacteria | Proteobacteri | Gammaprotec | Cellvibrionales | Halieaceae | Haliea | Haliea salexige | metabolism | Amino acid tra | 05C6M | Alpha-keto-be | 247633.GP214 | 0,2 | 0,4 | 2,55E-05 | 5,11E-05 | 0 - | 0 - | 1 - | 0,000127704 - | 0 - | 0 - |
| 1298 | 1 | 3um_FIJJAFBF_490002 | Bacteria | Bacteroidetes | unclassified | unclassified | unclassified | unclassified | Bacteroidetes | metabolism | Inorganic ion t | COG0659 | sulfate transp | 391603.FBALC | 0,2 | 0,4 | 2,33E-05 | 4,67E-05 | 0 - | 0 - | 0 - | 0 - | 1 - | 0,000116654 - |
| 1299 | 1 | 0.2um_NOKCPHJN_301303 | Eukaryota | unclassified | unclassified | unclassified | Apusomonadi | Thecamonas | Thecamonas t | unclassified | unclassified | unclassified | unclassified | unclassified | 0,2 | 0,4 | 3,34E-05 | 6,68E-05 | 0 - | 0 - | 0 - | 0 - | 0 - | 0 - |
| 1300 | 1 | 0.2um_NOKCPHJN_745046 | Bacteria | Bacteroidetes | Flavobacteriia | Flavobacterial | Flavobacteriac | unclassified | Flavobacteriac | unclassified | unclassified | unclassified | unclassified | unclassified | 0,4 | 0,489897949 | 3,87E-05 | 4,74E-05 | 0 - | 0 - | 0 - | 0 - | 1 - | 9,90E-05 - |
| 1301 | 1 | 0.2um_NOKCPHJN_604260 | unclassified | unclassified | unclassified | unclassified | unclassified | unclassified | unclassified | unclassified | unclassified | unclassified | unclassified | unclassified | 0,8 | 0,4 | 3,96E-05 | 2,05E-05 | 1 - | 5,88E-05 - | 1 - | 4,91E-05 - | 0 - | 0 - |
| 1302 | 1 | 3um_FIJJAFBF_804749 | Eukaryota | unclassified | unclassified | Isochrysidales | Noelaerhabda | Emiliania | Emiliania huxl | unclassified | unclassified | unclassified | unclassified | unclassified | 0,8 | 1,166190379 | 0,00016819 | 0,000252667 | 0 - | 0 - | 3 - | 0,000651822 - | 0 - | 0 - |
| 1303 | 3 | 3um_FIJJAFBF_260030;3um_FIJJAFBF_ | Bacteria | Verrucomicrol | Verrucomicrol | unclassified | unclassified | unclassified | various | poorly charact | Function unkn | 0ZMMU | Prokaryotic N- | 240016.VspiD | 1 | 0 | 0,000248981 | 2,63E-05 | 1 - | 0,000295843 - | 1 - | 0,000247332 - | 1 - | 0,000248984 - |
| 1304 | 1 | 0.2um_NOKCPHJN_959461 | Eukaryota | Bacillariophyta | Coscinodiscop | Thalassiosirale | Thalassiosirac | Thalassiosira | Thalassiosira p | metabolism | Carbohydrate | 0IP3Z | Solute carrier | 35128.Thaps2 | 0,2 | 0,4 | 2,67E-05 | 5,34E-05 | 0 - | 0 - | 0 - | 0 - | 0 - | 0 - |
| 1305 | 1 | 3um_FIJJAFBF_711586 | Bacteria | Cyanobacteria | unclassified | Synechococca | Synechococca | Synechococcu | Candidatus Sy | unclassified | unclassified | unclassified | unclassified | unclassified | 0,6 | 0,489897949 | 0,000157189 | 0,000129503 | 1 - | 0,000293522 - | 1 - | 0,000245392 - | 1 - | 0,000247031 - |
| 1306 | 1 | 0.2um_NOKCPHJN_1079676 | Bacteria | Proteobacteri | Gammaprotec | Alteromonada | Pseudoalteror | Pseudoalteror | Pseudoalteror | unclassified | unclassified | unclassified | unclassified | unclassified | 0,6 | 0,489897949 | 6,75E-05 | 5,54E-05 | 0 - | 0 - | 0 - | 0 - | 1 - | 0,000119758 - |
| 1307 | 1 | 3um_FIJJAFBF_462347 | unclassified | unclassified | unclassified | unclassified | unclassified | unclassified | unclassified | unclassified | unclassified | unclassified | unclassified | unclassified | 1,6 | 0,489897949 | 0,000435426 | 0,00015124 | 2 - | 0,000642474 - | 1 - | 0,000268562 - | 2 - | 0,000540712 - |
| 1308 | 1 | 0.2um_NOKCPHJN_03604 | Viruses | unclassified | unclassified | unclassified | Phycodnavirid | unclassified | Chrysochrom | unclassified | unclassified | unclassified | unclassified | unclassified | 0,4 | 0,489897949 | 2,24E-05 | 2,74E-05 | 0 - | 0 - | 1 - | 5,58E-05 - | 1 - | 5,62E-05 - |

| | | | | | | | | | | | | | | | | | | | | | | | | |
|---|---|---|---|---|---|---|---|---|---|---|---|---|---|---|---|---|---|---|---|---|---|---|---|---|
| 1309 | 1 | 0.2um_NOKCPHJN_420409 | Bacteria | Bacteroidetes | Cytophagia | Cytophagales | Flammeovirga | unclassified | Flammeovirga | poorly charact | Function unkn | 16R8I | Lipoprotein | 566466.NOR5 | 3,8 | 0,748331477 | 0,000489799 | 0,000114297 | 4 - | 0,000609761 - | 4 - | 0,000509775 - | 3 - | 0,000384886 - |
| 1310 | 1 | 0.2um_NOKCPHJN_278478 | Bacteria | Proteobacteri | Gammaprotec | Alteromonada | Colwelliaceae | Thalassomona | various | unclassified | unclassified | unclassified | unclassified | unclassified | 0,8 | 0,748331477 | 0,000539096 | 0,000480914 | 0 - | 0 - | 1 - | 0,000719252 - | 1 - | 0,000724057 - |
| 1311 | 1 | 0.2um_NOKCPHJN_529682 | Bacteria | Proteobacteri | Alphaproteob | Rhodobactera | Rhodobactera | Tateyamaria | Tateyamaria s | unclassified | unclassified | unclassified | unclassified | unclassified | 2,8 | 0,748331477 | 0,000594145 | 0,000109414 | 2 - | 0,000516194 - | 3 - | 0,000647327 - | 2 - | 0,000434434 - |
| 1312 | 1 | 3um_FIJJAFBF_510252 | Bacteria | Bacteroidetes | Flavobacteriia | Flavobacterial | Flavobacteria | Flaviramulus | Flaviramulus i | unclassified | unclassified | unclassified | unclassified | unclassified | 1 | 0,894427191 | 0,000484849 | 0,000444687 | 0 - | 0 - | 0 - | 0 - | 2 - | 0,001016015 - |
| 1313 | 1 | 3um_FIJJAFBF_693166 | Bacteria | Proteobacteri | Gammaprotec | unclassified | unclassified | unclassified | Gammaprotec | information st | Translation, ri | COG0184 | Forms an inter | 203122.Sde_2 | 0,2 | 0,4 | 0,000135001 | 0,000270002 | 0 - | 0 - | 0 - | 0 - | 0 - | 0 - |
| 1314 | 1 | 3um_FIJJAFBF_24076 | Bacteria | unclassified | unclassified | unclassified | unclassified | unclassified | uncultured ba | unclassified | unclassified | unclassified | unclassified | unclassified | 0,4 | 0,489897949 | 0,000102173 | 0,000125965 | 1 - | 0,000278246 - | 1 - | 0,000232621 - | 0 - | 0 - |
| 1315 | 1 | 3um_FIJJAFBF_625309 | Eukaryota | Bacillariophyta | Coscinodiscop | Chaetocerotal | Acanthocerata | Acanthoceras | Acanthoceras | metabolism | Lipid transpor | 05VNB | Carrier of the | 699218.HMPR | 0,6 | 0,489897949 | 0,000455791 | 0,000374134 | 0 - | 0 - | 1 - | 0,000792088 - | 1 - | 0,000797379 - |
| 1316 | 1 | 0.2um_NOKCPHJN_347404 | Bacteria | Proteobacteri | Alphaproteob | Pelagibacteral | Pelagibactera | Candidatus Pe | Candidatus Pe | unclassified | unclassified | unclassified | unclassified | unclassified | 0,2 | 0,4 | 9,51E-06 | 1,90E-05 | 0 - | 0 - | 0 - | 0 - | 0 - | 0 - |
| 1317 | 1 | 0.2um_NOKCPHJN_897328 | Eukaryota | Chordata | Amphibia | Anura | Pipidae | Xenopus | Xenopus laevi | unclassified | unclassified | unclassified | unclassified | unclassified | 1,2 | 0,4 | 0,000365889 | 0,000114581 | 1 - | 0,000365113 - | 1 - | 0,000305244 - | 1 - | 0,000307283 - |
| 1318 | 1 | 3um_FIJJAFBF_205173 | Eukaryota | Bacillariophyta | Mediophyceae | Cymatosirales | Cymatosirace | Plagiogrammc | Plagiogrammc | unclassified | unclassified | unclassified | unclassified | unclassified | 0,2 | 0,4 | 4,01E-05 | 8,02E-05 | 0 - | 0 - | 0 - | 0 - | 1 - | 0,000200614 - |
| 1319 | 1 | 0.2um_NOKCPHJN_957388 | Eukaryota | unclassified | Labyrinthulom | unclassified | Thraustochytr | Aurantiochytr | Aurantiochytr | unclassified | unclassified | unclassified | unclassified | unclassified | 0,6 | 0,8 | 0,000224716 | 0,00029672 | 0 - | 0 - | 0 - | 0 - | 1 - | 0,00038646 - |
| 1320 | 1 | 3um_FIJJAFBF_343083 | Bacteria | Proteobacteri | Alphaproteob | unclassified | unclassified | unclassified | Alphaproteob | unclassified | unclassified | unclassified | unclassified | unclassified | 0,2 | 0,4 | 7,20E-05 | 0,000143939 | 1 - | 0,000359847 - | 0 - | 0 - | 0 - | 0 - |
| 1321 | 1 | 0.2um_NOKCPHJN_736695 | Eukaryota | unclassified | Pelagophycea | Pelagomonad | unclassified | Aureococcus | Aureococcus a | unclassified | unclassified | unclassified | unclassified | unclassified | 0,8 | 0,748331477 | 0,000213221 | 0,000196381 | 1 - | 0,000319864 - | 0 - | 0 - | 0 - | 0 - |
| 1322 | 1 | 3um_FIJJAFBF_110488 | Eukaryota | Chlorophyta | Trebouxiophy | Chlorellales | Chlorellaceae | Micractinium | Micractinium | unclassified | unclassified | unclassified | unclassified | unclassified | 1 | 0 | 0,000221803 | 2,35E-05 | 1 - | 0,00026355 - | 1 - | 0,000220334 - | 1 - | 0,000221806 - |
| 1323 | 1 | 3um_FIJJAFBF_369771 | unclassified | unclassified | unclassified | unclassified | unclassified | unclassified | unclassified | unclassified | unclassified | unclassified | unclassified | unclassified | 0,2 | 0,4 | 0,000114672 | 0,000229345 | 0 - | 0 - | 0 - | 0 - | 0 - | 0 - |
| 1324 | 1 | 0.2um_NOKCPHJN_206300 | Eukaryota | unclassified | Spirotrichea | Sporadotrichi | Oxytrichidae | Oxytricha | Oxytricha trifa | metabolism | Energy produc | COG0377 | NDH-1 shuttle | 35128.Thaps2 | 0,2 | 0,4 | 7,37E-05 | 0,000147424 | 0 - | 0 - | 0 - | 0 - | 0 - | 0 - |
| 1325 | 1 | 3um_FIJJAFBF_22826 | Eukaryota | Bacillariophyta | Coscinodiscop | Thalassiosirale | Thalassiosirac | Thalassiosira | Thalassiosira c | unclassified | unclassified | unclassified | unclassified | unclassified | 0,2 | 0,4 | 0,00015179 | 0,00030358 | 0 - | 0 - | 0 - | 0 - | 1 - | 0,000758951 - |
| 1326 | 1 | 0.2um_NOKCPHJN_1013197 | Eukaryota | Eustigmatoph | unclassified | Eustigmatales | Monodopsida | Nannochlorop | Nannochlorop | unclassified | unclassified | unclassified | unclassified | unclassified | 5,4 | 0,8 | 0,000625927 | 6,31E-05 | 5 - | 0,000695615 - | 5 - | 0,000581551 - | 5 - | 0,000585436 - |
| 1327 | 1 | 3um_FIJJAFBF_25653 | Bacteria | Proteobacteri | Gammaprotec | Alteromonada | Alteromonada | Paraglaciecola | Paraglaciecola | unclassified | unclassified | unclassified | unclassified | unclassified | 2 | 0,894427191 | 0,000130317 | 5,00E-05 | 1 - | 8,04E-05 - | 2 - | 0,000134425 - | 1 - | 6,77E-05 - |
| 1328 | 1 | 0.2um_NOKCPHJN_491012 | Eukaryota | unclassified | Pelagophycea | Pelagomonad | unclassified | Aureococcus | Aureococcus a | unclassified | unclassified | unclassified | unclassified | unclassified | 1 | 0 | 0,000103776 | 1,10E-05 | 1 - | 0,000123308 - | 1 - | 0,000103089 - | 1 - | 0,000103777 - |
| 1329 | 4 | 0.2um_NOKCPHJN_411418;0.2um_NO | Bacteria | Proteobacteri | Alphaproteob | Rhodobactera | Rhodobactera | various | various | unclassified | unclassified | unclassified | unclassified | unclassified | 0,6 | 0,489897949 | 0,000112453 | 9,29E-05 | 1 - | 0,000212637 - | 0 - | 0 - | 1 - | 0,000178957 - |
| 1330 | 1 | 3um_FIJJAFBF_373756 | Bacteria | Bacteroidetes | Flavobacteriia | Flavobacterial | Flavobacteria | Mesoflavibact | various | metabolism | Nucleotide tra | 0QPJ5 | Deoxycytidyla | 216895.VV2_( | 0,2 | 0,4 | 1,61E-05 | 3,22E-05 | 0 - | 0 - | 0 - | 0 - | 0 - | 0 - |
| 1331 | 1 | 0.2um_NOKCPHJN_845694 | Eukaryota | unclassified | Pelagophycea | Pelagomonad | unclassified | Aureococcus | Aureococcus a | cellular proces | Posttranslatio | KOG0730 | cell division cy | 272952.HpaP8 | 0,8 | 0,748331477 | 0,000123514 | 0,000111016 | 0 - | 0 - | 1 - | 0,000166866 - | 0 - | 0 - |
| 1332 | 1 | 0.2um_NOKCPHJN_111274 | Bacteria | Proteobacteri | Gammaprotec | Cellvibrionales | Porticoccacea | unclassified | gamma protec | unclassified | unclassified | unclassified | unclassified | unclassified | 1,4 | 1,019803903 | 9,56E-05 | 6,26E-05 | 1 - | 8,50E-05 - | 1 - | 7,10E-05 - | 0 - | 0 - |
| 1333 | 1 | 3um_FIJJAFBF_720342 | unclassified | unclassified | unclassified | unclassified | unclassified | unclassified | unclassified | unclassified | unclassified | unclassified | unclassified | unclassified | 1 | 0 | 0,000272693 | 2,88E-05 | 1 - | 0,000324018 - | 1 - | 0,000270887 - | 1 - | 0,000272697 - |
| 1334 | 1 | 0.2um_NOKCPHJN_193556 | Bacteria | Verrucomicrol | Opitutae | unclassified | unclassified | unclassified | Opitutae bact | unclassified | unclassified | unclassified | unclassified | unclassified | 2,6 | 1,019803903 | 0,000329599 | 0,0001167 | 1 - | 0,00015561 - | 2 - | 0,000260187 - | 3 - | 0,000392887 - |
| 1335 | 1 | 3um_FIJJAFBF_101509 | Bacteria | Proteobacteri | Gammaprotec | Thiotrichales | Piscirickettsia | Cycloclasticus | Cycloclasticus | unclassified | unclassified | unclassified | unclassified | unclassified | 0,2 | 0,4 | 1,94E-05 | 3,88E-05 | 0 - | 0 - | 0 - | 0 - | 0 - | 0 - |
| 1336 | 1 | 3um_FIJJAFBF_249852 | Bacteria | Proteobacteri | Gammaprotec | unclassified | unclassified | unclassified | gamma protec | unclassified | unclassified | unclassified | unclassified | unclassified | 0,2 | 0,4 | 2,49E-05 | 4,99E-05 | 0 - | 0 - | 0 - | 0 - | 1 - | 0,000124738 - |
| 1337 | 1 | 0.2um_NOKCPHJN_193078 | Eukaryota | Eustigmatoph | unclassified | Eustigmatales | Monodopsida | Nannochlorop | Nannochlorop | cellular proces | Posttranslatio | KOG0175 | The proteasor | 2850.Phatr20( | 0,4 | 0,8 | 9,12E-05 | 0,000182324 | 0 - | 0 - | 0 - | 0 - | 0 - | 0 - |
| 1338 | 1 | 0.2um_NOKCPHJN_295621 | Bacteria | Firmicutes | Bacilli | Lactobacillales | Lactobacillace | Lactobacillus | Lactobacillus z | unclassified | unclassified | unclassified | unclassified | unclassified | 0,6 | 0,489897949 | 3,09E-05 | 2,53E-05 | 0 - | 0 - | 1 - | 5,46E-05 - | 0 - | 0 - |
| 1339 | 1 | 3um_FIJJAFBF_594175 | Eukaryota | Phaeophyceae | unclassified | Ectocarpales | various | various | various | metabolism | Energy produc | COG1290 | Component of | 35128.Thapsd | 6,2 | 0,4 | 0,001816519 | 0,000219768 | 6 - | 0,002088787 - | 6 - | 0,001746277 - | 7 - | 0,002050933 - |
| 1340 | 1 | 0.2um_NOKCPHJN_644547 | unclassified | unclassified | unclassified | unclassified | unclassified | unclassified | unclassified | unclassified | unclassified | unclassified | unclassified | unclassified | 0,2 | 0,4 | 1,69E-05 | 3,38E-05 | 0 - | 0 - | 0 - | 0 - | 0 - | 0 - |
| 1341 | 1 | 0.2um_NOKCPHJN_113026 | unclassified | unclassified | unclassified | unclassified | unclassified | unclassified | unclassified | unclassified | unclassified | unclassified | unclassified | unclassified | 0,4 | 0,489897949 | 0,000145917 | 0,000179067 | 0 - | 0 - | 0 - | 0 - | 0 - | 0 - |
| 1342 | 1 | 0.2um_NOKCPHJN_492829 | Bacteria | Bacteroidetes | Flavobacteriia | Flavobacterial | Flavobacteria | Formosa | Formosa sp. H | poorly charact | Function unkn | 0PAEN | NA | 886377.Murru | 1 | 0 | 0,000131234 | 1,39E-05 | 1 - | 0,000155934 - | 1 - | 0,000130364 - | 1 - | 0,000131235 - |
| 1343 | 1 | 3um_FIJJAFBF_165682 | Bacteria | Proteobacteri | Gammaprotec | Pseudomonad | Pseudomonac | Pseudomonas | Pseudomonas | unclassified | unclassified | unclassified | unclassified | unclassified | 1,2 | 0,4 | 0,000646614 | 0,000202492 | 1 - | 0,000645243 - | 1 - | 0,000539439 - | 1 - | 0,000543043 - |
| 1344 | 1 | 3um_FIJJAFBF_715085 | Eukaryota | unclassified | Pelagophycea | Pelagomonad | unclassified | Aureoumbra | Aureoumbra l | information st | Translation, ri | 0FKC1 | This is 1 of the | 449447.MAE_ | 4 | 0,632455532 | 0,001391141 | 0,000162239 | 3 - | 0,001254439 - | 4 - | 0,001398322 - | 4 - | 0,001407663 - |
| 1345 | 1 | 0.2um_NOKCPHJN_754061 | Bacteria | Bacteroidetes | unclassified | unclassified | unclassified | unclassified | Bacteroidetes | poorly charact | Function unkn | 0QNU1 | Uncharacteriz | 187272.Mlg_1 | 0,8 | 0,4 | 8,47E-05 | 4,32E-05 | 1 - | 0,000121704 - | 1 - | 0,000101748 - | 1 - | 0,000102428 - |
| 1346 | 1 | 0.2um_NOKCPHJN_351629 | Eukaryota | Chordata | Actinopteri | Characiformes | Characidae | Astyanax | Astyanax mex | cellular proces | Intracellular tr | COG0552 | Involved in tar | 367336.OM22 | 0,2 | 0,4 | 4,65E-05 | 9,30E-05 | 1 - | 0,000232448 - | 0 - | 0 - | 0 - | 0 - |
| 1347 | 2 | 3um_FIJJAFBF_425116;3um_FIJJAFBF_ | Bacteria | Proteobacteri | Gammaprotec | Cellvibrionales | Halieaceae | unclassified | various | cellular proces | Signal transdu | various | various | various | 1,2 | 0,4 | 0,000348871 | 0,000109252 | 1 - | 0,000348131 - | 1 - | 0,000291046 - | 1 - | 0,00029299 - |
| 1348 | 1 | 0.2um_NOKCPHJN_819165 | Eukaryota | Apicomplexa | Conoidasida | Eucoccidiorida | Eimeriidae | Eimeria | Eimeria mitis | unclassified | unclassified | unclassified | unclassified | unclassified | 0,2 | 0,4 | 2,01E-05 | 4,02E-05 | 0 - | 0 - | 0 - | 0 - | 0 - | 0 - |
| 1349 | 1 | 3um_FIJJAFBF_773582 | Eukaryota | Bacillariophyta | Bacillariophyc | Naviculales | Naviculaceae | Fistulifera | Fistulifera sola | unclassified | unclassified | unclassified | unclassified | unclassified | 0,8 | 0,4 | 0,000355723 | 0,00017928 | 0 - | 0 - | 1 - | 0,000463518 - | 1 - | 0,000466614 - |
| 1350 | 1 | 3um_FIJJAFBF_426183 | unclassified | unclassified | unclassified | unclassified | unclassified | unclassified | unclassified | unclassified | unclassified | unclassified | unclassified | unclassified | 0,2 | 0,4 | 5,03E-05 | 0,000100522 | 0 - | 0 - | 1 - | 0,000251305 - | 0 - | 0 - |
| 1351 | 1 | 3um_FIJJAFBF_343937 | Eukaryota | Hemichordata | Enteropneusta | unclassified | Harrimaniidae | Saccoglossus | Saccoglossus k | poorly charact | Function unkn | 0XTGE | Polycystic kidr | 65071.PYU1_T | 1,4 | 1,019803903 | 0,000107257 | 7,03E-05 | 1 - | 9,53E-05 - | 1 - | 7,97E-05 - | 0 - | 0 - |
| 1352 | 1 | 0.2um_NOKCPHJN_523420 | Bacteria | Bacteroidetes | Flavobacteriia | Flavobacterial | Crocinitomica | Fluviicola | Fluviicola taffe | unclassified | unclassified | unclassified | unclassified | unclassified | 0,8 | 0,4 | 0,00022068 | 0,000113871 | 1 - | 0,000324018 - | 1 - | 0,000270887 - | 1 - | 0,000272697 - |
| 1353 | 1 | 3um_FIJJAFBF_102271 | Bacteria | Bacteroidetes | Flavobacteriia | Flavobacterial | Flavobacteria | Gaetbulibacte | Gaetbulibacte | metabolism | Amino acid tra | COG1363 | Peptidase m4 | 313594.PI23P | 0,2 | 0,4 | 3,32E-05 | 6,64E-05 | 0 - | 0 - | 0 - | 0 - | 0 - | 0 - |
| 1354 | 1 | 3um_FIJJAFBF_205310 | Eukaryota | Bacillariophyta | Coscinodiscop | Thalassiosirale | Thalassiosirac | Thalassiosira | Thalassiosira c | cellular proces | Posttranslatio | COG0330 | Band 7 protei | 489825.LYNGE | 0,6 | 0,8 | 0,000265802 | 0,000357967 | 0 - | 0 - | 0 - | 0 - | 2 - | 0,000899899 - |
| 1355 | 2 | 0.2um_NOKCPHJN_101567;3um_FIJJAF | Bacteria | Bacteroidetes | Flavobacteriia | Flavobacterial | Flavobacteria | Polaribacter | various | information st | Replication, re | 0PB0H | DNA-binding p | 313598.MED1 | 1,4 | 0,8 | 0,000946124 | 0,000559639 | 0 - | 0 - | 2 - | 0,001390554 - | 2 - | 0,001399843 - |
| 1356 | 1 | 3um_FIJJAFBF_466392 | Bacteria | Proteobacteri | Alphaproteob | Rhodospirillal | Rhodospirillac | Candidatus En | Candidatus En | unclassified | unclassified | unclassified | unclassified | unclassified | 0,6 | 0,489897949 | 0,000109779 | 9,01E-05 | 0 - | 0 - | 1 - | 0,000190777 - | 1 - | 0,000192052 - |
| 1357 | 1 | 3um_FIJJAFBF_163782 | Bacteria | Bacteroidetes | Flavobacteriia | Flavobacterial | unclassified | unclassified | Flavobacterial | unclassified | unclassified | unclassified | unclassified | unclassified | 0,6 | 0,489897949 | 0,0001259 | 0,000103345 | 0 - | 0 - | 1 - | 0,000218793 - | 1 - | 0,000220255 - |
| 1358 | 2 | 0.2um_NOKCPHJN_296782;3um_FIJJAF | Bacteria | Bacteroidetes | Flavobacteriia | Flavobacterial | Flavobacteria | various | various | unclassified | unclassified | unclassified | unclassified | unclassified | 0,2 | 0,4 | 2,32E-05 | 4,64E-05 | 0 - | 0 - | 1 - | 0,000116094 - | 0 - | 0 - |
| 1359 | 1 | 3um_FIJJAFBF_292922 | Eukaryota | Bacillariophyta | Coscinodiscop | Coscinodiscale | Coscinodiscac | Coscinodiscus | Coscinodiscus | metabolism | Energy produc | COG4451 | RuBisCO catal | 977880.RALTA | 4 | 0 | 0,001812724 | 0,000191701 | 4 - | 0,002153905 - | 4 - | 0,001800717 - | 4 - | 0,001812746 - |
| 1360 | 1 | 3um_FIJJAFBF_715063 | Eukaryota | unclassified | Raphidophyce | Chattonellales | Chattonellace | Heterosigma | Heterosigma a | cellular proces | Posttranslatio | 0IEKR | One of the cor | 35128.Thapsd | 5,4 | 0,489897949 | 0,000737707 | 5,44E-05 | 5 - | 0,00081712 - | 5 - | 0,000683132 - | 5 - | 0,000687696 - |
| 1361 | 1 | 0.2um_NOKCPHJN_526672 | Bacteria | Bacteroidetes | Bacteroidia | Marinilabiliale | Marinilabiliac | Saccharicrinis | Saccharicrinis | unclassified | unclassified | unclassified | unclassified | unclassified | 0,8 | 0,4 | 9,74E-05 | 4,96E-05 | 1 - | 0,000139903 - | 1 - | 0,000116962 - | 1 - | 0,000117744 - |
| 1362 | 1 | 3um_FIJJAFBF_451551 | Bacteria | Firmicutes | Bacilli | Bacillales | Bacillaceae | Bacillus | Bacillus marin | information st | Translation, ri | 05KK9 | This protein bi | 479434.Sthe_ | 0,8 | 0,4 | 0,000494055 | 0,000255515 | 1 - | 0,000733806 - | 1 - | 0,00061348 - | 0 - | 0 - |
| 1363 | 1 | 0.2um_NOKCPHJN_767494 | unclassified | unclassified | unclassified | unclassified | unclassified | unclassified | unclassified | unclassified | unclassified | unclassified | unclassified | unclassified | 0,4 | 0,489897949 | 2,69E-05 | 3,30E-05 | 0 - | 0 - | 1 - | 7,18E-05 - | 0 - | 0 - |
| 1364 | 1 | 3um_FIJJAFBF_606153 | unclassified | unclassified | unclassified | unclassified | unclassified | unclassified | unclassified | unclassified | unclassified | unclassified | unclassified | unclassified | 0,8 | 0,4 | 0,000354008 | 0,000182669 | 1 - | 0,000519779 - | 1 - | 0,000434548 - | 1 - | 0,000437451 - |
| 1365 | 1 | 0.2um_NOKCPHJN_24031 | Viruses | unclassified | unclassified | unclassified | Phycodnavirid | unclassified | Chrysochromu | unclassified | unclassified | unclassified | unclassified | unclassified | 4,4 | 0,8 | 0,000946204 | 0,000113823 | 3 - | 0,000779669 - | 4 - | 0,000869096 - | 5 - | 0,001093627 - |
| 1366 | 1 | 0.2um_NOKCPHJN_828264 | Eukaryota | Chlorophyta | unclassified | Pyramimonad | unclassified | Cymbomonas | Cymbomonas | metabolism | Energy produc | 0IHF0 | This is one of | 70448.Q0P3J1 | 1 | 0 | 0,000178448 | 1,89E-05 | 1 - | 0,000212035 - | 1 - | 0,000177266 - | 1 - | 0,00017845 - |
| 1367 | 1 | 0.2um_NOKCPHJN_1002571 | Eukaryota | Eustigmatoph | unclassified | Eustigmatales | Monodopsida | Nannochlorop | Nannochlorop | unclassified | unclassified | unclassified | unclassified | unclassified | 0,2 | 0,4 | 0,000222324 | 0,000444648 | 0 - | 0 - | 0 - | 0 - | 0 - | 0 - |
| 1368 | 1 | 0.2um_NOKCPHJN_398174 | Eukaryota | Phaeophyceae | unclassified | various | various | various | various | unclassified | unclassified | unclassified | unclassified | unclassified | 4 | 0,894427191 | 0,001536603 | 0,000241092 | 3 - | 0,001394687 - | 3 - | 0,001165992 - | 4 - | 0,001565042 - |
| 1369 | 1 | 3um_FIJJAFBF_257128 | Bacteria | Firmicutes | Clostridia | Clostridiales | Clostridiaceae | Clostridium | various | unclassified | unclassified | unclassified | unclassified | unclassified | 1 | 0 | 0,000516329 | 5,46E-05 | 1 - | 0,00061351 - | 1 - | 0,000512909 - | 1 - | 0,000516336 - |
| 1370 | 1 | 3um_FIJJAFBF_216067 | Bacteria | Proteobacteri | Gammaprotec | Cellvibrionales | Microbulbifer | Microbulbifer | Microbulbifer | information st | Transcription | COG0085 | DNA-depende | 247633.GP214 | 2,8 | 1,166190379 | 0,000125011 | 4,67E-05 | 1 - | 5,50E-05 - | 2 - | 9,20E-05 - | 4 - | 0,000185273 - |
| 1371 | 1 | 3um_FIJJAFBF_746721 | Eukaryota | Chlorophyta | Mamiellophyc | Mamiellales | Mamiellaceae | Micromonas | Micromonas c | unclassified | unclassified | unclassified | unclassified | unclassified | 1,6 | 0,489897949 | 0,000674083 | 0,000170768 | 1 - | 0,000512659 - | 2 - | 0,000857191 - | 1 - | 0,000431458 - |
| 1372 | 1 | 3um_FIJJAFBF_405250 | Bacteria | Bacteroidetes | Flavobacteriia | Flavobacterial | Flavobacteria | Polaribacter | Polaribacter ir | unclassified | unclassified | unclassified | unclassified | unclassified | 4,8 | 0,4 | 0,000906318 | 4,59E-05 | 4 - | 0,00090451 - | 5 - | 0,00094524 - | 5 - | 0,000951555 - |
| 1373 | 1 | 3um_FIJJAFBF_292915 | Eukaryota | Bacillariophyta | Coscinodiscop | Coscinodiscale | Coscinodiscac | Coscinodiscus | Coscinodiscus | metabolism | Energy produc | COG0711 | F(1)F(0) ATP s | 35128.Thapsd | 2 | 1,095445115 | 0,000671997 | 0,000305498 | 1 - | 0,000418146 - | 1 - | 0,000349581 - | 2 - | 0,000703832 - |
| 1374 | 1 | 3um_FIJJAFBF_630539 | Bacteria | Bacteroidetes | Flavobacteriia | Flavobacterial | Flavobacteria | Flavobacteriu | Flavobacteriu | cellular proces | Signal transdu | 07UWE | serine threoni | 376686.Fjoh_( | 0,4 | 0,489897949 | 9,71E-05 | 0,000119437 | 0 - | 0 - | 1 - | 0,000259647 - | 0 - | 0 - |
| 1375 | 1 | 0.2um_NOKCPHJN_259529 | Bacteria | Proteobacteri | Betaproteoba | Burkholderiale | Burkholderiac | various | various | unclassified | unclassified | unclassified | unclassified | unclassified | 0,2 | 0,4 | 2,29E-05 | 4,58E-05 | 0 - | 0 - | 0 - | 0 - | 0 - | 0 - |
| 1376 | 1 | 3um_FIJJAFBF_552359 | Bacteria | Proteobacteri | Alphaproteob | Rhodobactera | Rhodobactera | unclassified | Rhodobactera | unclassified | unclassified | unclassified | unclassified | unclassified | 1,2 | 0,4 | 0,000371287 | 9,18E-05 | 1 - | 0,000376122 - | 1 - | 0,000314447 - | 1 - | 0,000316547 - |
| 1377 | 1 | 3um_FIJJAFBF_49815 | Eukaryota | Bacillariophyta | Bacillariophyc | Naviculales | Naviculaceae | Fistulifera | Fistulifera sola | unclassified | unclassified | unclassified | unclassified | unclassified | 4,6 | 0,489897949 | 0,0020592 | 0,000345221 | 5 - | 0,002654191 - | 5 - | 0,002218969 - | 4 - | 0,001787034 - |
| 1378 | 1 | 3um_FIJJAFBF_142183 | Bacteria | Proteobacteri | Gammaprotec | Cellvibrionales | Halieaceae | Halioglobus | Halioglobus ja | metabolism | Energy produc | COG0056 | Produces ATP | 314285.KT71_ | 1 | 0,632455532 | 0,000117777 | 7,72E-05 | 0 - | 0 - | 2 - | 0,000243482 - | 1 - | 0,000122554 - |
| 1379 | 1 | 3um_FIJJAFBF_39170 | Eukaryota | Chordata | Actinopteri | Osteoglossifor | Mormyridae | Paramormyro | Paramormyro | unclassified | unclassified | unclassified | unclassified | unclassified | 3 | 0 | 0,000529346 | 5,60E-05 | 3 - | 0,000628976 - | 3 - | 0,00052584 - | 3 - | 0,000529352 - |
| 1380 | 1 | 0.2um_NOKCPHJN_782930 | Eukaryota | unclassified | Cryptophyta | Pyrenomonad | Geminigerace | Guillardia | Guillardia thet | cellular proces | Posttranslatio | 0V8PF | Proteasome (F | 6669.DappuP5 | 0,4 | 0,489897949 | 6,43E-05 | 7,91E-05 | 0 - | 0 - | 1 - | 0,000171909 - | 0 - | 0 - |
| 1381 | 1 | 0.2um_NOKCPHJN_196523 | Eukaryota | unclassified | Pelagophycea | Pelagomonad | unclassified | Aureococcus | Aureococcus a | unclassified | unclassified | unclassified | unclassified | unclassified | 1 | 0,632455532 | 0,000130911 | 8,27E-05 | 1 - | 0,000154326 - | 2 - | 0,000258041 - | 1 - | 0,000129882 - |
| 1382 | 1 | 0.2um_NOKCPHJN_476231 | Eukaryota | Phaeophyceae | unclassified | Ectocarpales | Ectocarpaceae | Ectocarpus | Ectocarpus sili | unclassified | unclassified | unclassified | unclassified | unclassified | 1,6 | 0,8 | 0,000722143 | 0,000363952 | 0 - | 0 - | 2 - | 0,000940976 - | 2 - | 0,000947262 - |
| 1383 | 1 | 3um_FIJJAFBF_111353 | unclassified | unclassified | unclassified | unclassified | unclassified | unclassified | unclassified | unclassified | unclassified | unclassified | unclassified | unclassified | 3,8 | 1,166190379 | 0,000736351 | 0,000189336 | 2 - | 0,000472228 - | 5 - | 0,000986986 - | 4 - | 0,000794864 - |
| 1384 | 1 | 3um_FIJJAFBF_510237 | Bacteria | Bacteroidetes | Flavobacteriia | Flavobacterial | Flavobacteria | Polaribacter | Polaribacter s | unclassified | unclassified | unclassified | unclassified | unclassified | 1,2 | 0,748331477 | 0,000511441 | 0,00033342 | 0 - | 0 - | 2 - | 0,000875174 - | 2 - | 0,00088102 - |
| 1385 | 1 | 3um_FIJJAFBF_220667 | Eukaryota | Bacillariophyta | Coscinodiscop | Thalassiosirale | Thalassiosirac | Thalassiosira | Thalassiosira c | metabolism | Secondary me | KOG0055 | (ABC) transpo | 35128.Thaps2 | 0,8 | 0,4 | 3,68E-05 | 1,90E-05 | 1 - | 5,46E-05 - | 1 - | 4,57E-05 - | 0 - | 0 - |
| 1386 | 1 | 0.2um_NOKCPHJN_277085 | Eukaryota | Phaeophyceae | unclassified | Ectocarpales | Ectocarpaceae | Ectocarpus | Ectocarpus sili | poorly charact | Function unkn | COG1100 | GTP-binding P | 2850.Phatr86 | 0,6 | 0,489897949 | 0,000192521 | 0,000157839 | 0 - | 0 - | 1 - | 0,000340081 - | 0 - | 0 - |
| 1387 | 1 | 3um_FIJJAFBF_254287 | Bacteria | Proteobacteri | Gammaprotec | Cellvibrionales | Halieaceae | Haliea | Haliea salexige | information st | Translation, ri | 17FIH | One of the ear | 566466.NOR5 | 1 | 0 | 0,000642777 | 6,80E-05 | 1 - | 0,000763757 - | 1 - | 0,00063852 - | 1 - | 0,000642785 - |
| 1388 | 1 | 3um_FIJJAFBF_348329 | Bacteria | Bacteroidetes | Flavobacteriia | Flavobacterial | Flavobacteria | Algibacter | Algibacter sp. | unclassified | unclassified | unclassified | unclassified | unclassified | 1,2 | 0,748331477 | 9,43E-05 | 6,02E-05 | 0 - | 0 - | 2 - | 0,000162744 - | 1 - | 8,19E-05 - |
| 1389 | 1 | 3um_FIJJAFBF_65170 | Bacteria | Bacteroidetes | Flavobacteriia | Flavobacterial | Flavobacteria | Hyunsoonleell | Hyunsoonleell | unclassified | unclassified | unclassified | unclassified | unclassified | 0,2 | 0,4 | 1,82E-05 | 3,65E-05 | 0 - | 0 - | 1 - | 9,12E-05 - | 0 - | 0 - |
| 1390 | 1 | 3um_FIJJAFBF_276224 | unclassified | unclassified | unclassified | unclassified | unclassified | unclassified | unclassified | unclassified | unclassified | unclassified | unclassified | unclassified | 0,2 | 0,4 | 4,70E-05 | 9,40E-05 | 0 - | 0 - | 0 - | 0 - | 1 - | 0,000235048 - |
| 1391 | 1 | 3um_FIJJAFBF_41613 | Eukaryota | unclassified | unclassified | Choanoflagelli | Stephanoecid | Diaphanoeca | Diaphanoeca g | information st | Translation, ri | KOG0052 | This protein p | 3055.EDP0854 | 0,8 | 0,4 | 0,000738798 | 0,000381222 | 1 - | 0,001084756 - | 1 - | 0,000906883 - | 1 - | 0,000912941 - |
| 1392 | 1 | 3um_FIJJAFBF_494604 | Eukaryota | unclassified | Pelagophycea | Pelagomonad | unclassified | Aureococcus | Aureococcus a | poorly charact | Function unkn | 110WY | Chlorophyll A- | 35128.Thaps3 | 0,6 | 0,489897949 | 0,000336653 | 0,000280708 | 1 - | 0,000656563 - | 1 - | 0,000548903 - | 0 - | 0 - |
| 1393 | 1 | 3um_FIJJAFBF_567532 | Bacteria | Proteobacteri | Alphaproteob | Rhodobactera | Rhodobactera | Albidovulum | Albidovulum i | poorly charact | Function unkn | 0XZ2H | Membrane as | 353153.XP_80 | 0,4 | 0,489897949 | 3,19E-05 | 3,94E-05 | 1 - | 8,67E-05 - | 0 - | 0 - | 1 - | 7,30E-05 - |
| 1394 | 1 | 3um_FIJJAFBF_167349 | Eukaryota | unclassified | Dinophyceae | Gymnodiniale | Gymnodiniace | Lepidodinium | Lepidodinium | metabolism | Carbohydrate | COG0057 | glyceraldehyd | 423536.XP_00 | 3,8 | 0,748331477 | 0,000623029 | 0,000125488 | 3 - | 0,000586278 - | 5 - | 0,000816905 - | 4 - | 0,00065789 - |
| 1395 | 1 | 3um_FIJJAFBF_725633 | Bacteria | unclassified | unclassified | unclassified | unclassified | unclassified | bacterium TM | unclassified | unclassified | unclassified | unclassified | unclassified | 1 | 0 | 0,000355888 | 3,76E-05 | 1 - | 0,000422871 - | 1 - | 0,000353531 - | 1 - | 0,000355892 - |
| 1396 | 1 | 0.2um_NOKCPHJN_01987 | Bacteria | Proteobacteri | Alphaproteob | Rhodobactera | Rhodobactera | Sulfitobacter | Sulfitobacter r | information st | Translation, ri | COG0050 | This protein p | 391624.OIHEL | 2,4 | 1,019803903 | 0,000379426 | 0,000139227 | 2 - | 0,000382855 - | 2 - | 0,000320076 - | 3 - | 0,000483322 - |
| 1397 | 1 | 3um_FIJJAFBF_20002 | Bacteria | Proteobacteri | Gammaprotec | Alteromonada | Colwelliaceae | Colwellia | Colwellia sp. C | metabolism | Inorganic ion t | 16TQC | TONB-depend | 314287.GB22( | 3 | 0,894427191 | 0,000349748 | 7,99E-05 | 2 - | 0,000284054 - | 3 - | 0,000356214 - | 2 - | 0,000239062 - |
| 1398 | 1 | 0.2um_NOKCPHJN_726453 | Bacteria | Proteobacteri | Gammaprotec | unclassified | unclassified | Candidatus Th | uncultured SU | information st | Translation, ri | 05CEJ | Catalyzes the | 413404.Rmag_ | 1 | 0 | 0,000152155 | 1,61E-05 | 1 - | 0,000180793 - | 1 - | 0,000151147 - | 1 - | 0,000152157 - |
| 1399 | 1 | 3um_FIJJAFBF_220310 | Eukaryota | Bacillariophyta | Coscinodiscop | Thalassiosirale | Thalassiosirac | Thalassiosira | Thalassiosira c | unclassified | unclassified | unclassified | unclassified | unclassified | 1 | 0 | 0,000425623 | 4,50E-05 | 1 - | 0,000505731 - | 1 - | 0,000422804 - | 1 - | 0,000425628 - |
| 1400 | 1 | 0.2um_NOKCPHJN_508302 | unclassified | unclassified | unclassified | unclassified | unclassified | unclassified | unclassified | unclassified | unclassified | unclassified | unclassified | unclassified | 0,6 | 0,489897949 | 0,000455491 | 0,000375264 | 1 - | 0,000850548 - | 1 - | 0,000711079 - | 1 - | 0,000715829 - |
| 1401 | 1 | 3um_FIJJAFBF_738104 | Eukaryota | unclassified | unclassified | Prymnesiales | Chrysochromu | Chrysochromu | Chrysochromu | cellular proces | Posttranslatio | COG0652 | peptidyl-proly | 423536.XP_00 | 1,4 | 0,489897949 | 0,000471233 | 0,000204195 | 2 - | 0,000779669 - | 2 - | 0,000651822 - | 1 - | 0,000328088 - |
| 1402 | 1 | 3um_FIJJAFBF_410760 | Bacteria | Proteobacteri | Alphaproteob | Pelagibacteral | Pelagibactera | Candidatus Pe | Candidatus Pe | unclassified | unclassified | unclassified | unclassified | unclassified | 0,4 | 0,489897949 | 8,42E-05 | 0,000103359 | 0 - | 0 - | 0 - | 0 - | 0 - | 0 - |
| 1403 | 1 | 3um_FIJJAFBF_279094 | unclassified | unclassified | unclassified | unclassified | unclassified | unclassified | unclassified | unclassified | unclassified | unclassified | unclassified | unclassified | 1 | 0 | 0,000440505 | 4,66E-05 | 1 - | 0,000523414 - | 1 - | 0,000437587 - | 1 - | 0,00044051 - |
| 1404 | 1 | 0.2um_NOKCPHJN_1039968 | Eukaryota | Ascomycota | Eurotiomycete | Eurotiales | Aspergillaceae | Aspergillus | Aspergillus tu | cellular proces | Posttranslatio | KOG0880 | PPIases accele | 515849.XP_00 | 0,2 | 0,4 | 0,000110039 | 0,000220078 | 0 - | 0 - | 0 - | 0 - | 0 - | 0 - |
| 1405 | 1 | 3um_FIJJAFBF_348022 | unclassified | unclassified | unclassified | unclassified | unclassified | unclassified | unclassified | unclassified | unclassified | unclassified | unclassified | unclassified | 1,6 | 0,489897949 | 0,001539058 | 0,000394247 | 1 - | 0,001169503 - | 1 - | 0,000977733 - | 2 - | 0,001968529 - |
| 1406 | 1 | 3um_FIJJAFBF_97614 | Bacteria | unclassified | unclassified | unclassified | unclassified | unclassified | uncultured ba | information st | Translation, ri | 05CEJ | Catalyzes the | 391587.KAOT1 | 5 | 1,414213562 | 0,000560783 | 0,000119701 | 3 - | 0,000409753 - | 6 - | 0,000685127 - | 5 - | 0,000574753 - |
| 1407 | 1 | 3um_FIJJAFBF_142181 | Bacteria | Proteobacteri | Gammaprotec | Cellvibrionales | Spongiibacter | Oceanicoccus | Oceanicoccus | metabolism | Energy produc | COG0055 | Produces ATP | 247639.MGP2 | 2,6 | 0,489897949 | 0,000347294 | 4,88E-05 | 2 - | 0,000321928 - | 2 - | 0,000269139 - | 3 - | 0,000406406 - |
| 1408 | 1 | 0.2um_NOKCPHJN_114836 | Bacteria | Bacteroidetes | Flavobacteriia | unclassified | unclassified | unclassified | Flavobacteria | metabolism | Energy produc | 0P7SJ | Fe-S oxidored | 487796.Flav2A | 0,2 | 0,4 | 4,17E-05 | 8,35E-05 | 0 - | 0 - | 0 - | 0 - | 0 - | 0 - |
| 1409 | 1 | 0.2um_NOKCPHJN_698224 | Eukaryota | Bacillariophyta | Coscinodiscop | Thalassiosirale | Thalassiosirac | Thalassiosira | Thalassiosira c | metabolism | Amino acid tra | COG0010 | hydrolase acti | 2850.Phatr38 | 0,4 | 0,489897949 | 7,30E-05 | 8,94E-05 | 0 - | 0 - | 1 - | 0,000181904 - | 1 - | 0,000183119 - |

| | | | | | | | | | | | | | | | | | | | | | | | | |
|---|---|---|---|---|---|---|---|---|---|---|---|---|---|---|---|---|---|---|---|---|---|---|---|---|
| 1410 | 1 | 0.2um_NOKCPHJN_417822 | Bacteria | Proteobacteri | Alphaproteob | Pelagibacteral | Pelagibacterac | unclassified | Pelagibacterac | information st | Replication, re | 07SNH | Integrase | 945681.APO_( | 0,2 | 0,4 | 2,42E-05 | 4,84E-05 | 0 - | 0 - | 0 - | 0 - | 0 - | 0 - |
| 1411 | 1 | 0.2um_NOKCPHJN_674145 | Eukaryota | unclassified | Pelagophycea | Pelagomonad | unclassified | Aureococcus | Aureococcus a | unclassified | unclassified | unclassified | unclassified | unclassified | 1 | 0 | 0,000117304 | 1,24E-05 | 1 - | 0,000139382 - | 1 - | 0,000116527 - | 1 - | 0,000117305 - |
| 1412 | 1 | 3um_FIJJAFBF_39296 | unclassified | unclassified | unclassified | unclassified | unclassified | unclassified | unclassified | unclassified | unclassified | unclassified | unclassified | unclassified | 3 | 1,095445115 | 0,000391854 | 0,000109126 | 2 - | 0,000319182 - | 3 - | 0,000400266 - | 3 - | 0,00040294 - |
| 1413 | 1 | 0.2um_NOKCPHJN_350811 | Bacteria | Proteobacteri | Gammaprotec | Alteromonada | Alteromonada | Aliiglaciecola | Aliiglaciecola l | cellular proces | Signal transdu | 13AP7 | NACHT domai | 7739.JGI92858 | 0,4 | 0,489897949 | 4,89E-05 | 6,02E-05 | 0 - | 0 - | 0 - | 0 - | 1 - | 0,000131235 - |
| 1414 | 1 | 0.2um_NOKCPHJN_692237 | Bacteria | Bacteroidetes | Flavobacteriia | Flavobacterial | Flavobacteriac | Muricauda | Muricauda sp. | unclassified | unclassified | unclassified | unclassified | unclassified | 0,4 | 0,489897949 | 2,29E-05 | 2,82E-05 | 0 - | 0 - | 1 - | 6,12E-05 - | 0 - | 0 - |
| 1415 | 1 | 0.2um_NOKCPHJN_191603 | Eukaryota | Bacillariophyta | Bacillariophyc | Naviculales | Phaeodactylac | Phaeodactylu | Phaeodactylu | cellular proces | Cell cycle cont | 0IHFW | regulator of ch | 2850.Phatr318 | 2,6 | 0,489897949 | 0,000254466 | 4,77E-05 | 2 - | 0,000233536 - | 3 - | 0,000292862 - | 3 - | 0,000294819 - |
| 1416 | 1 | 3um_FIJJAFBF_295360 | Bacteria | Bacteroidetes | Flavobacteriia | Flavobacterial | Flavobacteriac | Flagellimonas | Flagellimonas | metabolism | Energy produc | 0PEUJ | ATP synthase | 391603.FBALC | 0,8 | 0,4 | 0,000788706 | 0,000407845 | 1 - | 0,001169503 - | 0 - | 0 - | 1 - | 0,000984265 - |
| 1417 | 1 | 0.2um_NOKCPHJN_371590 | Bacteria | Bacteroidetes | Flavobacteriia | Flavobacterial | Flavobacteriac | Formosa | Formosa sp. H | metabolism | Carbohydrate | 09CFZ | Transporter | 391587.KAOT1 | 0,6 | 0,489897949 | 4,87E-05 | 4,07E-05 | 1 - | 9,62E-05 - | 0 - | 0 - | 0 - | 0 - |
| 1418 | 1 | 0.2um_NOKCPHJN_1037148 | unclassified | unclassified | unclassified | unclassified | unclassified | unclassified | unclassified | unclassified | unclassified | unclassified | unclassified | unclassified | 1 | 0 | 0,000294356 | 3,11E-05 | 1 - | 0,000349758 - | 1 - | 0,000292406 - | 1 - | 0,00029436 - |
| 1419 | 1 | 3um_FIJJAFBF_560542 | Eukaryota | Chlorophyta | Chlorophycea | Chlamydomor | Volvocaceae | Volvox | Volvox carteri | unclassified | unclassified | unclassified | unclassified | unclassified | 1 | 0 | 0,000297133 | 3,14E-05 | 1 - | 0,000353057 - | 1 - | 0,000295165 - | 1 - | 0,000297137 - |
| 1420 | 1 | 3um_FIJJAFBF_226619 | Bacteria | Bacteroidetes | Flavobacteriia | Flavobacterial | Flavobacteriac | Formosa | Formosa sp. H | metabolism | Inorganic ion t | COG3119 | Sulfatase | 655815.ZPR_3 | 0,6 | 0,8 | 6,65E-05 | 9,06E-05 | 0 - | 0 - | 0 - | 0 - | 0 - | 0 - |
| 1421 | 1 | 3um_FIJJAFBF_329475 | unclassified | unclassified | unclassified | unclassified | unclassified | unclassified | unclassified | unclassified | unclassified | unclassified | unclassified | unclassified | 0,8 | 0,4 | 0,000585944 | 0,000302348 | 1 - | 0,000860324 - | 1 - | 0,000719252 - | 1 - | 0,000724057 - |
| 1422 | 1 | 0.2um_NOKCPHJN_246253 | Bacteria | Bacteroidetes | Flavobacteriia | Flavobacterial | Flavobacteriac | Capnocytopha | Capnocytopha | poorly charact | Function unkn | 09IR6 | Peptidase M1 | 313606.M231 | 0,8 | 0,4 | 0,000360551 | 0,000186444 | 1 - | 0,00053463 - | 0 - | 0 - | 1 - | 0,00044995 - |
| 1423 | 1 | 3um_FIJJAFBF_520477 | Bacteria | Bacteroidetes | Flavobacteriia | Flavobacterial | Flavobacteriac | Polaribacter | Polaribacter a | cellular proces | Intracellular tr | 0PBE9 | Preprotein tra | 313598.MED1 | 0,8 | 0,748331477 | 0,000441918 | 0,000396798 | 0 - | 0 - | 0 - | 0 - | 1 - | 0,000599933 - |
| 1424 | 1 | 0.2um_NOKCPHJN_798656 | Eukaryota | Arthropoda | Branchiopoda | Diplostraca | Daphniidae | Daphnia | Daphnia pulex | metabolism | Nucleotide tra | KOG0888 | nucleoside dip | 6669.DappuP3 | 0,2 | 0,4 | 7,07E-05 | 0,000141354 | 0 - | 0 - | 0 - | 0 - | 0 - | 0 - |
| 1425 | 1 | 3um_FIJJAFBF_292379 | Bacteria | Bacteroidetes | Cytophagia | Cytophagales | Flammeovirga | unclassified | Flammeovirga | poorly charact | Function unkn | 16R8I | Lipoprotein | 314260.PB250 | 13,6 | 1,496662955 | 0,001749557 | 0,000192991 | 13 - | 0,001998001 - | 14 - | 0,001798868 - | 11 - | 0,001422838 - |
| 1426 | 1 | 3um_FIJJAFBF_538586 | unclassified | unclassified | unclassified | unclassified | unclassified | unclassified | unclassified | unclassified | unclassified | unclassified | unclassified | unclassified | 0,4 | 0,489897949 | 0,000111162 | 0,000136982 | 1 - | 0,000301807 - | 0 - | 0 - | 1 - | 0,000254004 - |
| 1427 | 1 | 3um_FIJJAFBF_818329 | Bacteria | Bacteroidetes | Flavobacteriia | Flavobacterial | Flavobacteriac | Formosa | Formosa sp. H | cellular proces | Posttranslatio | 07S5S | Peptidyl-proly | 391603.FBALC | 0,2 | 0,4 | 1,95E-05 | 3,89E-05 | 0 - | 0 - | 0 - | 0 - | 1 - | 9,74E-05 - |
| 1428 | 1 | 3um_FIJJAFBF_205170 | Eukaryota | Bacillariophyta | Mediophyceae | Cymatosirales | Cymatosiracea | Plagiogrammc | Plagiogrammc | metabolism | Energy produc | COG0055 | Produces ATP | 7739.JGI12576 | 3,8 | 0,4 | 0,000506231 | 8,35E-05 | 4 - | 0,00063163 - | 4 - | 0,000528058 - | 4 - | 0,000531586 - |
| 1429 | 3 | 0.2um_NOKCPHJN_596126;3um_FIJJAF | Bacteria | various | various | various | various | various | various | unclassified | unclassified | unclassified | unclassified | unclassified | 0,6 | 0,489897949 | 0,000118057 | 9,69E-05 | 0 - | 0 - | 1 - | 0,000205164 - | 1 - | 0,000206534 - |
| 1430 | 1 | 3um_FIJJAFBF_257131 | Eukaryota | unclassified | Raphidophyce | Chattonellales | Chattonellace | Heterosigma | Heterosigma a | unclassified | unclassified | unclassified | unclassified | unclassified | 0,2 | 0,4 | 7,67E-05 | 0,000153435 | 0 - | 0 - | 0 - | 0 - | 0 - | 0 - |
| 1431 | 1 | 3um_FIJJAFBF_372168 | Bacteria | Proteobacteri | Alphaproteob | Rhodobactera | Rhodobactera | Sulfitobacter | Sulfitobacter s | unclassified | unclassified | unclassified | unclassified | unclassified | 0,2 | 0,4 | 4,86E-05 | 9,73E-05 | 0 - | 0 - | 0 - | 0 - | 1 - | 0,000243216 - |
| 1432 | 1 | 0.2um_NOKCPHJN_600010 | Eukaryota | Phaeophyceae | unclassified | Ectocarpales | Ectocarpaceae | Ectocarpus | Ectocarpus sili | unclassified | unclassified | unclassified | unclassified | unclassified | 1 | 0 | 0,001399826 | 0,000148035 | 1 - | 0,001663293 - | 1 - | 0,001390554 - | 1 - | 0,001399843 - |
| 1433 | 1 | 0.2um_NOKCPHJN_430131 | Bacteria | Proteobacteri | Alphaproteob | Rhodobactera | Rhodobactera | Paracoccus | Paracoccus pa | unclassified | unclassified | unclassified | unclassified | unclassified | 1 | 0 | 0,000323037 | 3,42E-05 | 1 - | 0,000383837 - | 1 - | 0,000320897 - | 1 - | 0,000323041 - |
| 1434 | 3 | 0.2um_NOKCPHJN_1061809;3um_FIJJA | Bacteria | Bacteroidetes | Flavobacteriia | Flavobacterial | Flavobacteriac | various | various | unclassified | unclassified | unclassified | unclassified | unclassified | 1 | 0 | 0,000333292 | 3,52E-05 | 1 - | 0,000396022 - | 1 - | 0,000331084 - | 1 - | 0,000333296 - |
| 1435 | 2 | 0.2um_NOKCPHJN_654066;0.2um_NO | Bacteria | Proteobacteri | Alphaproteob | Rhodobactera | unclassified | unclassified | Rhodobactera | unclassified | unclassified | unclassified | unclassified | unclassified | 1 | 0 | 0,000198713 | 2,10E-05 | 1 - | 0,000236114 - | 1 - | 0,000197397 - | 1 - | 0,000198716 - |
| 1436 | 1 | 0.2um_NOKCPHJN_117986 | Eukaryota | unclassified | Oomycetes | Peronosporale | Peronosporac | Phytophthora | Phytophthora | unclassified | unclassified | unclassified | unclassified | unclassified | 0,8 | 0,4 | 9,06E-05 | 4,69E-05 | 1 - | 0,000134619 - | 1 - | 0,000112545 - | 0 - | 0 - |
| 1437 | 1 | 3um_FIJJAFBF_119808 | unclassified | unclassified | unclassified | unclassified | unclassified | unclassified | unclassified | unclassified | unclassified | unclassified | unclassified | unclassified | 2,6 | 0,489897949 | 0,001384545 | 0,000219388 | 2 - | 0,001279456 - | 3 - | 0,001604485 - | 3 - | 0,001615204 - |
| 1438 | 1 | 3um_FIJJAFBF_445086 | Bacteria | Bacteroidetes | Flavobacteriia | Flavobacterial | Flavobacteriac | Algibacter | Algibacter sp. | metabolism | Lipid transpor | 0PF5H | long-chain-fat | 391603.FBALC | 0,4 | 0,489897949 | 4,26E-05 | 5,21E-05 | 0 - | 0 - | 1 - | 0,000106059 - | 1 - | 0,000106768 - |
| 1439 | 1 | 0.2um_NOKCPHJN_726022 | Bacteria | Proteobacteri | Gammaprotec | unclassified | unclassified | Candidatus Th | Candidatus Th | unclassified | unclassified | unclassified | unclassified | unclassified | 1 | 0 | 0,00019442 | 2,06E-05 | 1 - | 0,000231013 - | 1 - | 0,000193132 - | 1 - | 0,000194423 - |
| 1440 | 1 | 0.2um_NOKCPHJN_185702 | Bacteria | Proteobacteri | Oligoflexia | Oligoflexales | Pseudobacteri | Pseudobacteri | Pseudobacter | metabolism | Energy produc | COG2838 | Isocitrate deh | 331678.Cphar | 0,2 | 0,4 | 1,68E-05 | 3,35E-05 | 0 - | 0 - | 1 - | 8,38E-05 - | 0 - | 0 - |
| 1441 | 1 | 0.2um_NOKCPHJN_168665 | unclassified | unclassified | unclassified | unclassified | unclassified | unclassified | unclassified | unclassified | unclassified | unclassified | unclassified | unclassified | 0,8 | 0,4 | 0,000148027 | 7,65E-05 | 1 - | 0,000219496 - | 0 - | 0 - | 1 - | 0,00018473 - |
| 1442 | 2 | 0.2um_NOKCPHJN_862145;3um_FIJJAF | unclassified | unclassified | unclassified | unclassified | unclassified | unclassified | unclassified | unclassified | unclassified | unclassified | unclassified | unclassified | 1 | 0 | 0,000299963 | 3,17E-05 | 1 - | 0,00035642 - | 1 - | 0,000297976 - | 1 - | 0,000299966 - |
| 1443 | 1 | 3um_FIJJAFBF_220318 | Eukaryota | Bacillariophyta | Coscinodiscop | Thalassiosirale | Thalassiosirac | Thalassiosira | Thalassiosira c | metabolism | Energy produc | COG4451 | RuBisCO catal | 323097.Nham | 4,4 | 0,8 | 0,001969471 | 0,000241122 | 4 - | 0,002153905 - | 4 - | 0,001800717 - | 4 - | 0,001812746 - |
| 1444 | 1 | 0.2um_NOKCPHJN_864503 | Bacteria | Bacteroidetes | Flavobacteriia | Flavobacterial | Flavobacteriac | Formosa | Formosa sp. H | metabolism | Inorganic ion t | 153M9 | iduronate-2-su | 756272.Plabr_ | 1,8 | 0,4 | 0,000272109 | 6,89E-05 | 2 - | 0,000358984 - | 1 - | 0,00015006 - | 2 - | 0,000302124 - |
| 1445 | 1 | 3um_FIJJAFBF_316866 | Bacteria | Bacteroidetes | Flavobacteriia | Flavobacterial | Flavobacteriac | Algibacter | Algibacter alg | information st | Translation, ri | 09EM4 | Binds directly | 1046627.BZAF | 0,2 | 0,4 | 0,000109781 | 0,000219561 | 0 - | 0 - | 1 - | 0,000548903 - | 0 - | 0 - |
| 1446 | 1 | 0.2um_NOKCPHJN_389419 | Bacteria | Bacteroidetes | Saprospiria | Saprospirales | Saprospiracea | Aureispira | Aureispira sp. | unclassified | unclassified | unclassified | unclassified | unclassified | 0,6 | 0,489897949 | 0,000344203 | 0,000284315 | 1 - | 0,000650854 - | 0 - | 0 - | 1 - | 0,000547765 - |
| 1447 | 1 | 0.2um_NOKCPHJN_874072 | Eukaryota | Chlorophyta | Mamiellophyc | Mamiellales | Bathycoccacea | Ostreococcus | Ostreococcus | information st | Transcription | 1EBZF | SNF2 family N | 70448.Q00XM | 0,4 | 0,489897949 | 3,48E-05 | 4,27E-05 | 0 - | 0 - | 1 - | 8,89E-05 - | 0 - | 0 - |
| 1448 | 3 | 0.2um_NOKCPHJN_230057;3um_FIJJAF | Bacteria | unclassified | unclassified | unclassified | unclassified | unclassified | uncultured ba | various | various | various | various | various | 1 | 0 | 0,000192049 | 2,03E-05 | 1 - | 0,000228196 - | 1 - | 0,000190777 - | 1 - | 0,000192052 - |
| 1449 | 1 | 3um_FIJJAFBF_758338 | Eukaryota | Bacillariophyta | Coscinodiscop | Thalassiosirale | Thalassiosirac | Thalassiosira | Thalassiosira p | unclassified | unclassified | unclassified | unclassified | unclassified | 2 | 0,632455532 | 0,00150646 | 0,000447122 | 1 - | 0,000912783 - | 3 - | 0,002289326 - | 2 - | 0,001536413 - |
| 1450 | 1 | 0.2um_NOKCPHJN_678021 | Bacteria | Proteobacteri | Gammaprotec | Pseudomonad | Pseudomonac | Pseudomonas | various | metabolism | Energy produc | 0QIP8 | Electron trans | 746360.PFWH | 0,6 | 0,8 | 0,000113023 | 0,000153881 | 0 - | 0 - | 0 - | 0 - | 0 - | 0 - |
| 1451 | 1 | 3um_FIJJAFBF_586153 | Eukaryota | Bacillariophyta | Bacillariophyc | Naviculales | Phaeodactylac | Phaeodactylu | Phaeodactylu | cellular proces | Posttranslatio | KOG0176 | The proteasor | 2850.Phatr282 | 0,6 | 0,489897949 | 0,000330849 | 0,000275869 | 1 - | 0,000645243 - | 1 - | 0,000539439 - | 0 - | 0 - |
| 1452 | 1 | 3um_FIJJAFBF_426980 | Bacteria | Bacteroidetes | Flavobacteriia | Flavobacterial | Flavobacteriac | Mesoflavibact | Mesoflavibact | unclassified | unclassified | unclassified | unclassified | unclassified | 0,8 | 0,4 | 0,000102832 | 5,18E-05 | 0 - | 0 - | 1 - | 0,000133993 - | 1 - | 0,000134889 - |
| 1453 | 1 | 0.2um_NOKCPHJN_833833 | Eukaryota | unclassified | Oligohymenop | Philasterida | Pseudocohnile | Pseudocohnile | Pseudocohnile | unclassified | unclassified | unclassified | unclassified | unclassified | 0,4 | 0,489897949 | 0,000378672 | 0,000463994 | 0 - | 0 - | 0 - | 0 - | 1 - | 0,000969122 - |
| 1454 | 1 | 3um_FIJJAFBF_348325 | Bacteria | Bacteroidetes | Flavobacteriia | Flavobacterial | Flavobacteriac | Algibacter | Algibacter lect | unclassified | unclassified | unclassified | unclassified | unclassified | 9,2 | 2,925747768 | 0,000615794 | 0,000142392 | 5 - | 0,000410352 - | 8 - | 0,000548903 - | 10 - | 0,000690712 - |
| 1455 | 1 | 0.2um_NOKCPHJN_530154 | Eukaryota | Chlorophyta | Chlorophycea | Chlamydomor | Chlamydomor | Chlamydomor | Chlamydomor | unclassified | unclassified | unclassified | unclassified | unclassified | 0,2 | 0,4 | 1,28E-05 | 2,55E-05 | 0 - | 0 - | 0 - | 0 - | 0 - | 0 - |
| 1456 | 1 | 3um_FIJJAFBF_90637 | Bacteria | Proteobacteri | Gammaprotec | Alteromonada | Alteromonada | Paraglaciecola | Paraglaciecola | cellular proces | Intracellular tr | 05VIE | preprotein tra | 983545.Glaag_ | 0,6 | 0,489897949 | 0,000324391 | 0,000266276 | 0 - | 0 - | 1 - | 0,000563738 - | 1 - | 0,000567504 - |
| 1457 | 3 | 0.2um_NOKCPHJN_253384;0.2um_NO | Bacteria | Proteobacteri | Alphaproteob | various | various | various | various | various | various | various | various | various | 0,2 | 0,4 | 2,79E-05 | 5,57E-05 | 0 - | 0 - | 0 - | 0 - | 1 - | 0,000139365 - |
| 1458 | 1 | 3um_FIJJAFBF_670752 | Bacteria | Bacteroidetes | Flavobacteriia | Flavobacterial | Flavobacteriac | Arenitalea | Arenitalea lute | metabolism | Inorganic ion t | 05C27 | TonB-depende | 886377.Murru | 3 | 0,632455532 | 0,000263767 | 3,11E-05 | 2 - | 0,000213547 - | 3 - | 0,000267796 - | 3 - | 0,000269585 - |
| 1459 | 1 | 3um_FIJJAFBF_469955 | Bacteria | Proteobacteri | Alphaproteob | Rhodobactera | Rhodobactera | unclassified | Rhodobactera | metabolism | Amino acid tra | 01R4A | Polyamine AB | 388739.RSK20 | 1,6 | 1,019803903 | 0,00026703 | 0,000162595 | 1 - | 0,000203946 - | 0 - | 0 - | 2 - | 0,000343286 - |
| 1460 | 1 | 3um_FIJJAFBF_475533 | Eukaryota | unclassified | Oomycetes | Albuginales | Albuginaceae | Albugo | Albugo laibach | information st | Translation, ri | COG0081 | Binds directly | 65071.PYU1_T | 1,4 | 0,489897949 | 0,000692751 | 0,000194047 | 1 - | 0,000603614 - | 1 - | 0,000504636 - | 1 - | 0,000508008 - |
| 1461 | 1 | 0.2um_NOKCPHJN_1072722 | unclassified | unclassified | unclassified | unclassified | unclassified | unclassified | unclassified | unclassified | unclassified | unclassified | unclassified | unclassified | 1,8 | 0,4 | 0,00144175 | 0,000247758 | 1 - | 0,000972054 - | 2 - | 0,001625323 - | 2 - | 0,00163618 - |
| 1462 | 2 | 0.2um_NOKCPHJN_307535;3um_FIJJAF | Bacteria | Bacteroidetes | Flavobacteriia | Flavobacterial | Flavobacteriac | Polaribacter | various | cellular proces | Posttranslatio | 08V1T | peptidylprolyl | 313594.PI23P_ | 0,6 | 0,489897949 | 0,000153878 | 0,00012631 | 0 - | 0 - | 1 - | 0,000267414 - | 1 - | 0,000269201 - |
| 1463 | 1 | 3um_FIJJAFBF_117655 | Eukaryota | various | various | various | various | various | various | metabolism | Energy produc | COG1145 | 4Fe-4S ferred | 35128.Thapsd | 1 | 0 | 0,000777681 | 8,22E-05 | 1 - | 0,000924052 - | 1 - | 0,00077253 - | 1 - | 0,000777691 - |
| 1464 | 2 | 0.2um_NOKCPHJN_47587;3um_FIJJAFE | Bacteria | various | various | various | various | various | various | metabolism | Energy produc | 0PEU8 | ATP synthase | 313603.FB217 | 0,6 | 0,489897949 | 0,000562285 | 0,000460991 | 0 - | 0 - | 1 - | 0,000993253 - | 0 - | 0 - |
| 1465 | 1 | 0.2um_NOKCPHJN_409097 | Eukaryota | Bacillariophyta | Coscinodiscop | Thalassiosirale | Thalassiosirac | Thalassiosira | Thalassiosira c | cellular proces | Posttranslatio | KOG0019 | heat shock pro | 67593.Physo1 | 1 | 0 | 8,91E-05 | 9,42E-06 | 1 - | 0,000105867 - | 1 - | 8,85E-05 - | 1 - | 8,91E-05 - |
| 1466 | 2 | 0.2um_NOKCPHJN_766095;3um_FIJJAF | Eukaryota | Chlorophyta | Mamiellophyc | Mamiellales | Bathycoccacea | various | various | metabolism | Energy produc | COG0056 | Produces ATP | 70448.Q0P3H | 1 | 0 | 0,000144147 | 1,52E-05 | 1 - | 0,000171277 - | 1 - | 0,000143192 - | 1 - | 0,000144149 - |
| 1467 | 1 | 0.2um_NOKCPHJN_780235 | Bacteria | Proteobacteri | Gammaprotec | unclassified | unclassified | unclassified | various | metabolism | Energy produc | 0QJ3X | Aconitate hyd | 697282.Mettu | 0,2 | 0,4 | 1,38E-05 | 2,76E-05 | 0 - | 0 - | 0 - | 0 - | 0 - | 0 - |
| 1468 | 1 | 3um_FIJJAFBF_586277 | Eukaryota | unclassified | Pelagophycea | Pelagomonad | unclassified | Aureococcus | Aureococcus a | unclassified | unclassified | unclassified | unclassified | unclassified | 1,2 | 1,166190379 | 0,000653083 | 0,000612733 | 0 - | 0 - | 1 - | 0,000590329 - | 0 - | 0 - |
| 1469 | 1 | 3um_FIJJAFBF_651155 | Eukaryota | Bacillariophyta | Coscinodiscop | Chaetocerotal | Chaetocerota | Chaetoceros | Chaetoceros s | information st | Translation, ri | 08UHY | One of the pri | 32049.SYNPC0 | 2,8 | 0,748331477 | 0,001118633 | 0,000281574 | 2 - | 0,000959592 - | 4 - | 0,001604485 - | 3 - | 0,001211403 - |
| 1470 | 1 | 0.2um_NOKCPHJN_210688 | Bacteria | Bacteroidetes | Flavobacteriia | Flavobacterial | Flavobacteriac | Flavobacteriu | Flavobacteriu | metabolism | Inorganic ion t | 05C27 | TonB-depende | 983548.Krodi_ | 3 | 1,095445115 | 0,000168844 | 4,58E-05 | 2 - | 0,000137969 - | 3 - | 0,000173018 - | 2 - | 0,000116116 - |
| 1471 | 1 | 3um_FIJJAFBF_175086 | Bacteria | Proteobacteri | Alphaproteob | Rhodobactera | Rhodobactera | Phaeobacter | Phaeobacter g | metabolism | Inorganic ion t | 16PAX | extracellular s | 439496.RBY4I_ | 0,8 | 0,4 | 0,000109891 | 5,54E-05 | 0 - | 0 - | 1 - | 0,000143192 - | 1 - | 0,000144149 - |
| 1472 | 1 | 0.2um_NOKCPHJN_890902 | Bacteria | Proteobacteri | Gammaprotec | Alteromonada | Pseudoalteror | Algicola | Algicola sagan | metabolism | Energy produc | 0QJAP | Catalyzes the | 506534.Rhein_ | 0,2 | 0,4 | 4,02E-05 | 8,05E-05 | 0 - | 0 - | 1 - | 0,000201206 - | 0 - | 0 - |
| 1473 | 1 | 3um_FIJJAFBF_715060 | Eukaryota | Bacillariophyta | Coscinodiscop | Coscinodiscale | Coscinodiscac | Coscinodiscus | Coscinodiscus | metabolism | Energy produc | 11HAV | photosystem i | 35128.Thapsd | 1 | 0 | 0,000456465 | 4,83E-05 | 1 - | 0,000542378 - | 1 - | 0,000453441 - | 1 - | 0,000456471 - |
| 1474 | 1 | 3um_FIJJAFBF_292916 | Eukaryota | Bacillariophyta | Coscinodiscop | Coscinodiscale | Coscinodiscac | Coscinodiscus | Coscinodiscus | unclassified | unclassified | unclassified | unclassified | unclassified | 3,4 | 0,8 | 0,001358776 | 0,000288584 | 3 - | 0,001439388 - | 2 - | 0,000802243 - | 4 - | 0,001615204 - |
| 1475 | 1 | 3um_FIJJAFBF_662596 | Bacteria | Bacteroidetes | Flavobacteriia | Flavobacterial | Flavobacteriac | Formosa | Formosa sp. H | unclassified | unclassified | unclassified | unclassified | unclassified | 0,6 | 0,489897949 | 0,00013501 | 0,000110726 | 0 - | 0 - | 0 - | 0 - | 1 - | 0,000239517 - |
| 1476 | 1 | 0.2um_NOKCPHJN_971779 | Eukaryota | unclassified | Labyrinthulom | unclassified | Thraustochytr | Aurantiochytr | Aurantiochytr | metabolism | Energy produc | COG0372 | citrate syntha | 272952.HpaP8 | 1,2 | 0,4 | 0,000199448 | 6,88E-05 | 1 - | 0,000197489 - | 1 - | 0,000165105 - | 2 - | 0,000332417 - |
| 1477 | 1 | 0.2um_NOKCPHJN_728863 | Eukaryota | unclassified | Labyrinthulom | unclassified | Thraustochytr | Aurantiochytr | Aurantiochytr | poorly charact | Function unkn | 0A4HT | Coiled-coil do | 7668.SPU_011 | 0,4 | 0,489897949 | 3,96E-05 | 4,85E-05 | 0 - | 0 - | 1 - | 0,000100927 - | 0 - | 0 - |
| 1478 | 1 | 0.2um_NOKCPHJN_155316 | Eukaryota | unclassified | unclassified | Trichomonadi | Trichomonadi | Trichomonas | Trichomonas v | cellular proces | Signal transdu | KOG0613 | fibronectin typ | 7070.TC00296 | 1 | 0 | 4,33E-05 | 4,58E-06 | 1 - | 5,15E-05 - | 1 - | 4,30E-05 - | 1 - | 4,33E-05 - |
| 1479 | 1 | 0.2um_NOKCPHJN_912152 | Eukaryota | unclassified | Spirotrichea | Sporadotrichi | Oxytrichidae | Stylonychia | Stylonychia le | cellular proces | Posttranslatio | 0FI38 | heat shock pro | 441375.XP_00 | 0,8 | 0,748331477 | 7,48E-05 | 6,52E-05 | 1 - | 0,000113406 - | 0 - | 0 - | 1 - | 9,54E-05 - |
| 1480 | 1 | 0.2um_NOKCPHJN_846217 | Eukaryota | unclassified | unclassified | Prymnesiales | Chrysochromu | Chrysochromu | Chrysochromu | unclassified | unclassified | unclassified | unclassified | unclassified | 0,2 | 0,4 | 0,000139665 | 0,00027933 | 0 - | 0 - | 0 - | 0 - | 0 - | 0 - |
| 1481 | 1 | 0.2um_NOKCPHJN_949714 | Eukaryota | unclassified | Spirotrichea | Sporadotrichi | Oxytrichidae | Oxytricha | Oxytricha trifa | metabolism | Energy produc | COG0056 | Produces ATP | 412030.XP_00 | 1 | 0,632455532 | 0,000127525 | 7,50E-05 | 0 - | 0 - | 1 - | 0,000135444 - | 1 - | 0,000136348 - |
| 1482 | 1 | 0.2um_NOKCPHJN_200139 | Eukaryota | Chordata | Actinopteri | Cyprinodontif | Fundulidae | Fundulus | Fundulus hete | metabolism | Carbohydrate | 09XUV | Family with se | 10020.ENSDO | 0,2 | 0,4 | 2,56E-05 | 5,13E-05 | 1 - | 0,000128165 - | 0 - | 0 - | 0 - | 0 - |
| 1483 | 1 | 0.2um_NOKCPHJN_982466 | unclassified | unclassified | unclassified | unclassified | unclassified | unclassified | unclassified | unclassified | unclassified | unclassified | unclassified | unclassified | 1 | 0 | 0,000699913 | 7,40E-05 | 1 - | 0,000831646 - | 1 - | 0,000695277 - | 1 - | 0,000699922 - |
| 1484 | 1 | 3um_FIJJAFBF_176595 | Eukaryota | Phaeophyceae | unclassified | Ectocarpales | Ectocarpaceae | Ectocarpus | Ectocarpus sili | cellular proces | Posttranslatio | 0XRR7 | Calreticulin | 67593.Physo1 | 1 | 0 | 0,000473625 | 5,01E-05 | 1 - | 0,000562768 - | 1 - | 0,000470488 - | 1 - | 0,000473631 - |
| 1485 | 1 | 3um_FIJJAFBF_556069 | unclassified | unclassified | unclassified | unclassified | unclassified | unclassified | unclassified | unclassified | unclassified | unclassified | unclassified | unclassified | 2,2 | 0,748331477 | 0,00120142 | 0,000380005 | 2 - | 0,001313126 - | 1 - | 0,000548903 - | 3 - | 0,001657709 - |
| 1486 | 1 | 0.2um_NOKCPHJN_1041578 | unclassified | unclassified | unclassified | unclassified | unclassified | unclassified | unclassified | unclassified | unclassified | unclassified | unclassified | unclassified | 0,2 | 0,4 | 2,38E-05 | 4,76E-05 | 0 - | 0 - | 0 - | 0 - | 0 - | 0 - |
| 1487 | 1 | 0.2um_NOKCPHJN_1055434 | unclassified | unclassified | unclassified | unclassified | unclassified | unclassified | unclassified | unclassified | unclassified | unclassified | unclassified | unclassified | 0,6 | 0,489897949 | 6,63E-05 | 5,44E-05 | 0 - | 0 - | 1 - | 0,000115239 - | 1 - | 0,000116009 - |
| 1488 | 1 | 3um_FIJJAFBF_347377 | Bacteria | Proteobacteri | Gammaprotec | Enterobactera | Enterobacteri | Escherichia | Escherichia co | unclassified | unclassified | unclassified | unclassified | unclassified | 8 | 1,095445115 | 0,001122815 | 8,47E-05 | 6 - | 0,001013745 - | 9 - | 0,001271274 - | 8 - | 0,00113757 - |
| 1489 | 1 | 3um_FIJJAFBF_37634 | Bacteria | Proteobacteri | Gammaprotec | Oceanospirilla | Oceanospirilla | Motiliproteus | Motiliproteus | information st | Replication, re | 0QKMF | The RuvA-Ruv | 223926.VP105 | 0,6 | 0,489897949 | 0,00019458 | 0,000160772 | 1 - | 0,00036871 - | 1 - | 0,000308251 - | 0 - | 0 - |
| 1490 | 1 | 3um_FIJJAFBF_237981 | Eukaryota | unclassified | Compsopogor | Erythropeltida | Erythrotrichia | Erythrotrichia | Erythrotrichia | information st | Translation, ri | COG0333 | 50s ribosomal | 696747.NIES3 | 1 | 0 | 0,00104987 | 0,000111027 | 1 - | 0,00124747 - | 1 - | 0,001042915 - | 1 - | 0,001049882 - |
| 1491 | 1 | 3um_FIJJAFBF_438911 | Bacteria | Planctomycete | Phycisphaerae | Phycisphaeral | Phycisphaerac | unclassified | Phycisphaerac | unclassified | unclassified | unclassified | unclassified | unclassified | 0,8 | 0,4 | 5,14E-05 | 2,59E-05 | 0 - | 0 - | 1 - | 6,70E-05 - | 1 - | 6,74E-05 - |
| 1492 | 1 | 3um_FIJJAFBF_177183 | Eukaryota | unclassified | unclassified | Prymnesiales | Chrysochromu | Chrysochromu | Chrysochromu | unclassified | unclassified | unclassified | unclassified | unclassified | 1 | 0 | 0,000125733 | 1,33E-05 | 1 - | 0,000149398 - | 1 - | 0,0001249 - | 1 - | 0,000125734 - |
| 1493 | 1 | 0.2um_NOKCPHJN_643038 | Viruses | unclassified | unclassified | unclassified | unclassified | unclassified | Pleurochrysis | unclassified | unclassified | unclassified | unclassified | unclassified | 0,8 | 0,748331477 | 0,000103993 | 9,73E-05 | 1 - | 0,00015244 - | 0 - | 0 - | 2 - | 0,00025659 - |
| 1494 | 1 | 0.2um_NOKCPHJN_459990 | Bacteria | Proteobacteri | Gammaprotec | Oceanospirilla | unclassified | unclassified | Oceanospirilla | information st | Replication, re | 0Z2SC | Phage integra | 390235.PputW | 0,2 | 0,4 | 2,56E-05 | 5,11E-05 | 0 - | 0 - | 0 - | 0 - | 1 - | 0,000127775 - |
| 1495 | 1 | 0.2um_NOKCPHJN_666805 | unclassified | unclassified | unclassified | unclassified | unclassified | unclassified | unclassified | unclassified | unclassified | unclassified | unclassified | unclassified | 0,4 | 0,489897949 | 4,83E-05 | 5,93E-05 | 0 - | 0 - | 1 - | 0,00012902 - | 0 - | 0 - |
| 1496 | 1 | 0.2um_NOKCPHJN_644182 | Eukaryota | unclassified | Oomycetes | Pythiales | Pythiaceae | Pythium | Pythium insidi | cellular proces | Posttranslatio | KOG0019 | heat shock pro | 4787.PITG_05 | 0,4 | 0,489897949 | 7,43E-05 | 9,10E-05 | 0 - | 0 - | 1 - | 0,000185133 - | 1 - | 0,00018637 - |
| 1497 | 2 | 0.2um_NOKCPHJN_596724;3um_FIJJAF | Eukaryota | Basidiomycota | various | various | various | various | various | metabolism | Energy produc | COG1053 | Succinate DeH | various | 0,8 | 0,4 | 8,41E-05 | 4,24E-05 | 0 - | 0 - | 1 - | 0,000109588 - | 1 - | 0,00011032 - |
| 1498 | 1 | 3um_FIJJAFBF_215382 | Viruses | unclassified | unclassified | unclassified | unclassified | unclassified | uncultured vir | unclassified | unclassified | unclassified | unclassified | unclassified | 0,6 | 0,489897949 | 0,000144588 | 0,000118541 | 0 - | 0 - | 1 - | 0,000255408 - | 0 - | 0 - |
| 1499 | 1 | 0.2um_NOKCPHJN_300331 | Eukaryota | unclassified | Pelagophycea | Pelagomonad | unclassified | Aureococcus | Aureococcus a | unclassified | unclassified | unclassified | unclassified | unclassified | 0,8 | 0,4 | 2,15E-05 | 1,11E-05 | 1 - | 3,20E-05 - | 1 - | 2,67E-05 - | 0 - | 0 - |
| 1500 | 1 | 0.2um_NOKCPHJN_291839 | Bacteria | Proteobacteri | Alphaproteob | Rhodobactera | unclassified | unclassified | Rhodobactera | unclassified | unclassified | unclassified | unclassified | unclassified | 0,4 | 0,489897949 | 7,27E-05 | 8,96E-05 | 1 - | 0,000198011 - | 1 - | 0,000165542 - | 0 - | 0 - |
| 1501 | 1 | 0.2um_NOKCPHJN_917897 | Bacteria | Proteobacteri | Alphaproteob | Rhodospirillal | Rhodospirillac | unclassified | Rhodospirillac | cellular proces | Cell motility | 16UR5 | flagellar hook- | 269796.Rru_A | 0,4 | 0,489897949 | 0,000149101 | 0,000183339 | 0 - | 0 - | 1 - | 0,000398566 - | 0 - | 0 - |
| 1502 | 4 | 0.2um_NOKCPHJN_1097847;0.2um_N( | Eukaryota | various | various | various | various | various | various | various | various | various | various | various | 0,2 | 0,4 | 6,26E-05 | 0,000125157 | 0 - | 0 - | 0 - | 0 - | 0 - | 0 - |
| 1503 | 1 | 3um_FIJJAFBF_79208 | Bacteria | Proteobacteri | Alphaproteob | Rhodobactera | Rhodobactera | Shimia | Shimia aestua | information st | Translation, ri | 16Q16 | Binds the lowe | 367336.OM22 | 1 | 0 | 0,000263566 | 2,79E-05 | 1 - | 0,000313172 - | 1 - | 0,00026182 - | 1 - | 0,000263569 - |
| 1504 | 1 | 3um_FIJJAFBF_529327 | Bacteria | Bacteroidetes | Flavobacteriia | Flavobacterial | Flavobacteriac | Algibacter | Algibacter sp. | poorly charact | Function unkn | 09BCJ | secretion prot | 313598.MED1 | 0,6 | 0,489897949 | 9,11E-05 | 7,60E-05 | 1 - | 0,000179924 - | 0 - | 0 - | 0 - | 0 - |
| 1505 | 1 | 3um_FIJJAFBF_564229 | Bacteria | Proteobacteri | Gammaprotec | Pseudomonad | Pseudomonac | Pseudomonas | Pseudomonas | cellular proces | Cell wall/mem | 0QX85 | 17 kDa surface | 205922.Pfl01_ | 1,2 | 0,748331477 | 0,001190758 | 0,00076229 | 0 - | 0 - | 1 - | 0,001025818 - | 2 - | 0,002065342 - |
| 1506 | 1 | 3um_FIJJAFBF_660512 | Eukaryota | Bacillariophyta | Coscinodiscop | Thalassiosirale | Thalassiosirac | Thalassiosira | Thalassiosira p | unclassified | unclassified | unclassified | unclassified | unclassified | 1 | 0 | 0,000153266 | 1,62E-05 | 1 - | 0,000182112 - | 1 - | 0,00015225 - | 1 - | 0,000153267 - |
| 1507 | 1 | 3um_FIJJAFBF_100050 | Bacteria | Proteobacteri | Alphaproteob | Rhodospirillal | Rhodospirillac | Candidatus En | Candidatus En | unclassified | unclassified | unclassified | unclassified | unclassified | 0,4 | 0,489897949 | 4,86E-05 | 5,97E-05 | 0 - | 0 - | 0 - | 0 - | 0 - | 0 - |
| 1508 | 1 | 0.2um_NOKCPHJN_938596 | Eukaryota | Eustigmatoph | unclassified | Eustigmatales | Monodopsida | Nannochlorop | Nannochlorop | metabolism | Lipid transpor | KOG1175 | synthetase | 65071.PYU1_T | 0,8 | 0,4 | 0,000100151 | 5,17E-05 | 1 - | 0,000147049 - | 1 - | 0,000122937 - | 1 - | 0,000123758 - |
| 1509 | 1 | 3um_FIJJAFBF_507104 | Eukaryota | unclassified | Cryptophyta | Pyrenomonad | Geminigerace | Guillardia | Guillardia thet | unclassified | unclassified | unclassified | unclassified | unclassified | 4,6 | 0,489897949 | 0,002433766 | 0,000469833 | 5 - | 0,003118674 - | 5 - | 0,002607288 - | 5 - | 0,002624706 - |
| 1510 | 1 | 3um_FIJJAFBF_758228 | Eukaryota | Chordata | Mammalia | Primates | Hominidae | Pongo | Pongo abelii | information st | Chromatin str | KOG3467 | Core compone | 7217.FBpp012 | 0,6 | 0,489897949 | 0,000368999 | 0,000302525 | 0 - | 0 - | 1 - | 0,000651822 - | 0 - | 0 - |

| | | | | | | | | | | | | | | | | | | | | | | | | |
|---|---|---|---|---|---|---|---|---|---|---|---|---|---|---|---|---|---|---|---|---|---|---|---|---|
| 1511 | 1 | 3um_FIJJAFBF_204899 | unclassified | unclassified | unclassified | unclassified | unclassified | unclassified | unclassified | unclassified | unclassified | unclassified | unclassified | unclassified | 0,2 | 0,4 | 8,01E-05 | 0,000160201 | 0 - | 0 - | 0 - | 0 - | 0 - | 0 - |
| 1512 | 2 | 3um_FIJJAFBF_355968;3um_FIJJAFBF_ | Eukaryota | unclassified | Cryptophyta | Pyrenomonad | Geminigerace | Teleaulax | Teleaulax amp | metabolism | Energy produc | COG0711 | F(1)F(0) ATP s | various | 0,2 | 0,4 | 6,09E-05 | 0,000121719 | 0 - | 0 - | 0 - | 0 - | 0 - | 0 - |
| 1513 | 1 | 0.2um_NOKCPHJN_721141 | Bacteria | Bacteroidetes | Flavobacteriia | Flavobacterial | Flavobacteria | Altibacter | Altibacter lent | poorly charact | Function unkn | 0P8NI | Inherit from b | 313594.PI23P_ | 3,8 | 1,166190379 | 0,000361867 | 9,89E-05 | 2 - | 0,00023137 - | 3 - | 0,000290146 - | 5 - | 0,000486808 - |
| 1514 | 1 | 0.2um_NOKCPHJN_671436 | Bacteria | Verrucomicrol | Verrucomicrol | Verrucomicrol | Verrucomicrol | Pedosphaera | Pedosphaera | unclassified | unclassified | unclassified | unclassified | unclassified | 0,6 | 0,489897949 | 0,000544984 | 0,000446807 | 0 - | 0 - | 1 - | 0,000962691 - | 0 - | 0 - |
| 1515 | 1 | 3um_FIJJAFBF_465283 | Eukaryota | Chlorophyta | Mamiellophyc | Mamiellales | Bathycoccace | Ostreococcus | Ostreococcus | unclassified | unclassified | unclassified | unclassified | unclassified | 2,2 | 0,748331477 | 0,00075301 | 0,00022567 | 2 - | 0,000827052 - | 2 - | 0,000691436 - | 1 - | 0,000348027 - |
| 1516 | 2 | 0.2um_NOKCPHJN_144039;3um_FIJJAF | Eukaryota | Bacillariophyta | Coscinodiscop | Thalassiosirale | Thalassiosirac | Thalassiosira | Thalassiosira c | cellular proces | Intracellular tr | KOG0096 | GTP-binding n | 35128.Thaps3 | 1,2 | 0,4 | 0,000351585 | 0,000121224 | 1 - | 0,000348131 - | 1 - | 0,000291046 - | 2 - | 0,000585981 - |
| 1517 | 1 | 3um_FIJJAFBF_68872 | Eukaryota | unclassified | unclassified | unclassified | unclassified | Ancoracysta | Ancoracysta tw | unclassified | unclassified | unclassified | unclassified | unclassified | 1 | 0 | 0,000220252 | 2,33E-05 | 1 - | 0,000261707 - | 1 - | 0,000218793 - | 1 - | 0,000220255 - |
| 1518 | 1 | 0.2um_NOKCPHJN_610453 | unclassified | unclassified | unclassified | unclassified | unclassified | unclassified | unclassified | unclassified | unclassified | unclassified | unclassified | unclassified | 0,2 | 0,4 | 3,36E-05 | 6,72E-05 | 0 - | 0 - | 0 - | 0 - | 0 - | 0 - |
| 1519 | 1 | 0.2um_NOKCPHJN_892782 | unclassified | unclassified | unclassified | unclassified | unclassified | unclassified | unclassified | unclassified | unclassified | unclassified | unclassified | unclassified | 6 | 1,788854382 | 0,001434228 | 0,000310958 | 4 - | 0,001164952 - | 7 - | 0,001704375 - | 5 - | 0,001225544 - |
| 1520 | 1 | 0.2um_NOKCPHJN_770232 | Eukaryota | unclassified | Pelagophycea | Pelagomonad | unclassified | Aureococcus | Aureococcus a | cellular proces | Posttranslatio | KOG0019 | heat shock pro | 272952.HpaP8 | 8 | 1,673320053 | 0,000638182 | 9,30E-05 | 5 - | 0,000483515 - | 8 - | 0,000646769 - | 8 - | 0,00065109 - |
| 1521 | 1 | 3um_FIJJAFBF_688127 | Eukaryota | Microsporidia | unclassified | unclassified | unclassified | Pseudoloma | Pseudoloma n | cellular proces | Intracellular tr | 19BK2 | Gtr1/RagA G p | 644223.XP_00 | 0,6 | 0,489897949 | 0,000181889 | 0,000151662 | 1 - | 0,000354731 - | 1 - | 0,000296564 - | 0 - | 0 - |
| 1522 | 2 | 0.2um_NOKCPHJN_309389;3um_FIJJAF | Bacteria | Bacteroidetes | Flavobacteriia | Flavobacterial | Flavobacteria | Polaribacter | Polaribacter s | metabolism | Energy produc | 09A0E | phosphoenolp | 313598.MED1 | 0,2 | 0,4 | 1,46E-05 | 2,91E-05 | 0 - | 0 - | 1 - | 7,28E-05 - | 0 - | 0 - |
| 1523 | 1 | 3um_FIJJAFBF_812317 | unclassified | unclassified | unclassified | unclassified | unclassified | unclassified | unclassified | unclassified | unclassified | unclassified | unclassified | unclassified | 0,6 | 0,489897949 | 0,000172101 | 0,000142158 | 1 - | 0,000325427 - | 0 - | 0 - | 1 - | 0,000273882 - |
| 1524 | 1 | 0.2um_NOKCPHJN_356712 | Eukaryota | unclassified | Oomycetes | Peronosporale | Peronosporac | Phytophthora | Phytophthora | unclassified | unclassified | unclassified | unclassified | unclassified | 0,2 | 0,4 | 1,66E-05 | 3,32E-05 | 0 - | 0 - | 0 - | 0 - | 0 - | 0 - |
| 1525 | 1 | 0.2um_NOKCPHJN_538279 | Bacteria | Bacteroidetes | Flavobacteriia | Flavobacterial | Flavobacteria | Mangrovimon | Mangrovimon | metabolism | Energy produc | 099Q7 | Catalyzes the | 391603.FBALC | 1,2 | 0,4 | 0,000245425 | 8,46E-05 | 1 - | 0,000243014 - | 1 - | 0,000203165 - | 2 - | 0,000409045 - |
| 1526 | 1 | 3um_FIJJAFBF_594165 | Eukaryota | unclassified | Pelagophycea | Pelagomonad | unclassified | Aureococcus | Aureococcus a | unclassified | unclassified | unclassified | unclassified | unclassified | 12 | 1,264911064 | 0,004831996 | 0,000306223 | 10 - | 0,004828915 - | 11 - | 0,004440801 - | 13 - | 0,005283279 - |
| 1527 | 1 | 0.2um_NOKCPHJN_310980 | Eukaryota | unclassified | unclassified | unclassified | unclassified | Gymnochlora | Gymnochlora | information st | Chromatin str | KOG3467 | Core compone | 2850.Phatr349 | 2,4 | 0,489897949 | 0,001644829 | 0,000463905 | 3 - | 0,002414458 - | 2 - | 0,001345697 - | 2 - | 0,001354687 - |
| 1528 | 1 | 3um_FIJJAFBF_542763 | Bacteria | Bacteroidetes | Sphingobacter | Sphingobacter | Sphingobacter | Parapedobact | Parapedobact | information st | Replication, re | 0P9S1 | Inherit from b | 531844.FIC_01 | 0,2 | 0,4 | 2,23E-05 | 4,45E-05 | 0 - | 0 - | 1 - | 0,000111343 - | 0 - | 0 - |
| 1529 | 1 | 0.2um_NOKCPHJN_574936 | unclassified | unclassified | unclassified | unclassified | unclassified | unclassified | unclassified | unclassified | unclassified | unclassified | unclassified | unclassified | 0,2 | 0,4 | 0,000108939 | 0,000217878 | 0 - | 0 - | 0 - | 0 - | 0 - | 0 - |
| 1530 | 1 | 3um_FIJJAFBF_669273 | Eukaryota | unclassified | Synurophycea | Parmales | Triparmaceae | Triparma | Triparma laev | cellular proces | Posttranslatio | 0FJCR | cysteine desul | 197221.tll049 | 0,6 | 0,489897949 | 7,38E-05 | 6,05E-05 | 0 - | 0 - | 0 - | 0 - | 1 - | 0,000130962 - |
| 1531 | 1 | 3um_FIJJAFBF_497373 | Bacteria | Planctomycet | Phycisphaerae | Phycisphaeral | Phycisphaerac | Phycisphaera | Phycisphaera | unclassified | unclassified | unclassified | unclassified | unclassified | 0,6 | 0,489897949 | 4,64E-05 | 3,81E-05 | 0 - | 0 - | 1 - | 8,06E-05 - | 1 - | 8,12E-05 - |
| 1532 | 1 | 3um_FIJJAFBF_259311 | Bacteria | Planctomycet | Phycisphaerae | Phycisphaeral | Phycisphaerac | Phycisphaera | Phycisphaera | cellular proces | Cell wall/mem | 06B97 | phosphate-sel | 443144.GM21 | 0,8 | 0,4 | 0,000104853 | 5,28E-05 | 0 - | 0 - | 1 - | 0,000136626 - | 1 - | 0,000137539 - |
| 1533 | 1 | 3um_FIJJAFBF_180780 | Bacteria | Verrucomicrol | Opitutae | Opitutales | unclassified | unclassified | Opitutales bac | unclassified | unclassified | unclassified | unclassified | unclassified | 1 | 0 | 0,000477213 | 5,05E-05 | 1 - | 0,000567032 - | 1 - | 0,000474052 - | 1 - | 0,000477219 - |
| 1534 | 1 | 0.2um_NOKCPHJN_385377 | Eukaryota | unclassified | unclassified | Isochrysidales | Noelaerhabda | Emiliania | Emiliania huxl | unclassified | unclassified | unclassified | unclassified | unclassified | 0,2 | 0,4 | 6,40E-06 | 1,28E-05 | 0 - | 0 - | 0 - | 0 - | 1 - | 3,20E-05 - |
| 1535 | 1 | 0.2um_NOKCPHJN_247996 | Bacteria | Proteobacteri | Gammaprotec | Alteromonada | Idiomarinacea | Idiomarina | various | metabolism | Inorganic ion t | 177TK | phosphate-sel | 455436.GHTC | 2,4 | 0,489897949 | 0,000498673 | 0,000109594 | 2 - | 0,000494047 - | 3 - | 0,000619554 - | 3 - | 0,000623692 - |
| 1536 | 1 | 3um_FIJJAFBF_151367 | Eukaryota | Annelida | Polychaeta | Golfingiida | Sipunculidae | Sipunculus | Sipunculus nu | information st | Chromatin str | KOG1744 | Core compone | 400682.PAC_1 | 3,6 | 0,489897949 | 0,001825074 | 0,000178146 | 3 - | 0,001825565 - | 3 - | 0,001526218 - | 4 - | 0,002048551 - |
| 1537 | 1 | 3um_FIJJAFBF_197145 | Viruses | unclassified | unclassified | unclassified | various | various | various | unclassified | unclassified | unclassified | unclassified | unclassified | 1 | 0 | 9,47E-05 | 1,00E-05 | 1 - | 0,000112554 - | 1 - | 9,41E-05 - | 1 - | 9,47E-05 - |
| 1538 | 1 | 3um_FIJJAFBF_170869 | Bacteria | Bacteroidetes | Flavobacteriia | Flavobacterial | Crocinitomica | unclassified | Crocinitomica | unclassified | unclassified | unclassified | unclassified | unclassified | 1,4 | 0,489897949 | 0,000587762 | 0,000176972 | 1 - | 0,000509171 - | 2 - | 0,000851359 - | 1 - | 0,000428523 - |
| 1539 | 1 | 0.2um_NOKCPHJN_479810 | Eukaryota | Chordata | Actinopteri | Gobiiformes | Gobiidae | Boleophthalm | Boleophthalm | unclassified | unclassified | unclassified | unclassified | unclassified | 1 | 0 | 4,20E-05 | 4,44E-06 | 1 - | 4,99E-05 - | 1 - | 4,17E-05 - | 1 - | 4,20E-05 - |
| 1540 | 1 | 0.2um_NOKCPHJN_215620 | Eukaryota | Chromerida | unclassified | unclassified | unclassified | Vitrella | Vitrella brassi | unclassified | unclassified | unclassified | unclassified | unclassified | 1 | 0 | 0,000119303 | 1,26E-05 | 1 - | 0,000141758 - | 1 - | 0,000118513 - | 1 - | 0,000119305 - |
| 1541 | 1 | 0.2um_NOKCPHJN_147128 | Eukaryota | Mucoromycot | Mortierellomy | Mortierellales | Mortierellacea | Mortierella | Mortierella ve | unclassified | unclassified | unclassified | unclassified | unclassified | 0,4 | 0,489897949 | 5,53E-05 | 6,84E-05 | 1 - | 0,000153377 - | 0 - | 0 - | 0 - | 0 - |
| 1542 | 1 | 3um_FIJJAFBF_640522 | Eukaryota | unclassified | Spirotrichea | Euplotida | Euplotidae | Euplotes | various | information st | Chromatin str | KOG3467 | Core compone | 31033.ENSTRU | 1,4 | 0,8 | 0,000750268 | 0,000432026 | 0 - | 0 - | 2 - | 0,001117409 - | 2 - | 0,001124874 - |
| 1543 | 1 | 3um_FIJJAFBF_669258 | Eukaryota | Bacillariophyta | various | various | various | various | various | metabolism | Energy produc | 0J0AS | This b-type cy | 35128.Thapsd | 6,4 | 1,019803903 | 0,004784519 | 0,000806038 | 6 - | 0,005346299 - | 7 - | 0,005214577 - | 5 - | 0,00374958 - |
| 1544 | 1 | 0.2um_NOKCPHJN_347401 | Bacteria | unclassified | unclassified | unclassified | unclassified | Thermobacul | Thermobaculu | unclassified | unclassified | unclassified | unclassified | unclassified | 0,4 | 0,489897949 | 4,40E-05 | 5,40E-05 | 0 - | 0 - | 0 - | 0 - | 0 - | 0 - |
| 1545 | 1 | 0.2um_NOKCPHJN_724437 | Bacteria | Bacteroidetes | Flavobacteriia | Flavobacterial | Flavobacteria | Chryseobacter | Chryseobacter | unclassified | unclassified | unclassified | unclassified | unclassified | 0,4 | 0,489897949 | 7,68E-05 | 9,44E-05 | 0 - | 0 - | 1 - | 0,000205164 - | 0 - | 0 - |
| 1546 | 1 | 3um_FIJJAFBF_399086 | Bacteria | Bacteroidetes | Flavobacteriia | Flavobacterial | Flavobacteria | Polaribacter | Polaribacter s | unclassified | unclassified | unclassified | unclassified | unclassified | 0,2 | 0,4 | 3,33E-05 | 6,67E-05 | 1 - | 0,0001667 - | 0 - | 0 - | 0 - | 0 - |
| 1547 | 1 | 0.2um_NOKCPHJN_469361 | Bacteria | Bacteroidetes | unclassified | unclassified | unclassified | unclassified | unidentified e | unclassified | unclassified | unclassified | unclassified | unclassified | 1 | 0 | 4,93E-05 | 5,21E-06 | 1 - | 5,85E-05 - | 1 - | 4,89E-05 - | 1 - | 4,93E-05 - |
| 1548 | 1 | 3um_FIJJAFBF_799075 | Bacteria | Proteobacteri | unclassified | unclassified | unclassified | unclassified | Proteobacteri | metabolism | Lipid transpor | 08IQF | Catalyzes the | 991905.SL003 | 1 | 0 | 0,000115794 | 1,22E-05 | 1 - | 0,000137589 - | 1 - | 0,000115027 - | 1 - | 0,000115796 - |
| 1549 | 1 | 3um_FIJJAFBF_282387 | Eukaryota | Bacillariophyta | Coscinodiscop | Thalassiosirale | Thalassiosirac | Thalassiosira | Thalassiosira p | unclassified | unclassified | unclassified | unclassified | unclassified | 5,8 | 0,4 | 0,000821229 | 4,32E-05 | 5 - | 0,000846699 - | 6 - | 0,000849433 - | 6 - | 0,000855108 - |
| 1550 | 1 | 0.2um_NOKCPHJN_114165 | Bacteria | Bacteroidetes | Flavobacteriia | Flavobacterial | Flavobacteria | Formosa | Formosa sp. H | poorly charact | Function unkn | 06E9D | NA | 391598.FBBAL | 0,2 | 0,4 | 2,21E-05 | 4,42E-05 | 0 - | 0 - | 0 - | 0 - | 1 - | 0,000110514 - |
| 1551 | 1 | 3um_FIJJAFBF_579438 | Eukaryota | Bacillariophyta | Bacillariophyc | Bacillariales | Bacillariaceae | Fragilariopsis | Fragilariopsis | cellular proces | Cell cycle cont | 0GTVD | BRCA1 C Term | 7260.FBpp024 | 0,2 | 0,4 | 1,53E-05 | 3,07E-05 | 0 - | 0 - | 0 - | 0 - | 0 - | 0 - |
| 1552 | 1 | 3um_FIJJAFBF_137701 | Eukaryota | Eustigmatoph | unclassified | Eustigmatales | Monodopsida | Nannochlorop | Nannochlorop | unclassified | unclassified | unclassified | unclassified | unclassified | 1,4 | 0,489897949 | 0,000391581 | 0,000139134 | 1 - | 0,000332659 - | 2 - | 0,000556221 - | 2 - | 0,000559937 - |
| 1553 | 1 | 3um_FIJJAFBF_628820 | Bacteria | Actinobacteria | Actinobacteria | Micrococcales | Intrasporangia | Janibacter | Janibacter hoy | unclassified | unclassified | unclassified | unclassified | unclassified | 2,8 | 0,4 | 0,001029627 | 8,99E-05 | 2 - | 0,000885777 - | 3 - | 0,001110797 - | 3 - | 0,001118218 - |
| 1554 | 1 | 3um_FIJJAFBF_389027 | Eukaryota | Bacillariophyta | Coscinodiscop | Thalassiosirale | Thalassiosirac | Thalassiosira | Thalassiosira c | unclassified | unclassified | unclassified | unclassified | unclassified | 1 | 0 | 0,000196237 | 2,08E-05 | 1 - | 0,000233172 - | 1 - | 0,000194937 - | 1 - | 0,00019624 - |
| 1555 | 1 | 0.2um_NOKCPHJN_981456 | unclassified | unclassified | unclassified | unclassified | unclassified | unclassified | unclassified | unclassified | unclassified | unclassified | unclassified | unclassified | 0,4 | 0,489897949 | 0,000295454 | 0,000361859 | 0 - | 0 - | 1 - | 0,000736176 - | 1 - | 0,000741093 - |
| 1556 | 1 | 0.2um_NOKCPHJN_384647 | Bacteria | Proteobacteri | Alphaproteob | Rickettsiales | unclassified | unclassified | Rickettsiales b | unclassified | unclassified | unclassified | unclassified | unclassified | 0,8 | 0,4 | 0,000106954 | 5,39E-05 | 0 - | 0 - | 1 - | 0,000139365 - | 1 - | 0,000140296 - |
| 1557 | 1 | 0.2um_NOKCPHJN_183170 | Bacteria | Proteobacteri | Gammaprotec | Cellvibrionales | Porticoccacea | unclassified | Porticoccacea | unclassified | unclassified | unclassified | unclassified | unclassified | 0,4 | 0,489897949 | 2,83E-05 | 3,50E-05 | 1 - | 7,85E-05 - | 0 - | 0 - | 0 - | 0 - |
| 1558 | 1 | 3um_FIJJAFBF_464071 | Eukaryota | Brachiopoda | Lingulata | Lingulida | Lingulidae | Lingula | Lingula anatin | unclassified | unclassified | unclassified | unclassified | unclassified | 0,2 | 0,4 | 5,86E-05 | 0,00011722 | 0 - | 0 - | 0 - | 0 - | 0 - | 0 - |
| 1559 | 1 | 0.2um_NOKCPHJN_782226 | Bacteria | Proteobacteri | Gammaprotec | Alteromonada | Pseudoalteror | Pseudoalteror | Pseudoalteror | metabolism | Inorganic ion t | 16PDP | TonB-depende | 637905.SVI_1 | 1,6 | 0,8 | 0,000219052 | 9,95E-05 | 1 - | 0,000167446 - | 1 - | 0,000139989 - | 1 - | 0,000140924 - |
| 1560 | 1 | 3um_FIJJAFBF_348211 | Bacteria | Proteobacteri | Alphaproteob | unclassified | unclassified | Candidatus Pu | Candidatus Pu | metabolism | Energy produc | 01RNF | Produces ATP | 488538.SAR11 | 8,2 | 0,748331477 | 0,001088971 | 0,000201788 | 9 - | 0,001409275 - | 9 - | 0,001178189 - | 8 - | 0,001054275 - |
| 1561 | 1 | 0.2um_NOKCPHJN_851899 | Eukaryota | Bacillariophyta | Bacillariophyc | Bacillariales | Bacillariaceae | Fragilariopsis | Fragilariopsis | unclassified | unclassified | unclassified | unclassified | unclassified | 4,4 | 0,8 | 0,001068653 | 0,000128553 | 3 - | 0,000880567 - | 4 - | 0,000981567 - | 5 - | 0,001235156 - |
| 1562 | 1 | 3um_FIJJAFBF_594166 | Eukaryota | unclassified | Pelagophycea | Pelagomonad | unclassified | Aureococcus | Aureococcus a | metabolism | Energy produc | COG0711 | F(1)F(0) ATP s | 35128.Thapsd | 10,4 | 1,019803903 | 0,003707845 | 0,000410215 | 9 - | 0,003827464 - | 12 - | 0,004266472 - | 11 - | 0,003937059 - |
| 1563 | 1 | 3um_FIJJAFBF_177945 | Bacteria | Proteobacteri | Alphaproteob | Rhodobactera | Rhodobactera | Roseobacter | Roseobacter s | cellular proces | Cell cycle cont | 01Q8B | Required for c | 391593.RCCS2 | 0,6 | 0,489897949 | 3,32E-05 | 2,77E-05 | 1 - | 6,55E-05 - | 0 - | 0 - | 0 - | 0 - |
| 1564 | 1 | 3um_FIJJAFBF_05988 | Eukaryota | unclassified | Ichthyosporea | unclassified | unclassified | Capsaspora | Capsaspora ov | unclassified | unclassified | unclassified | unclassified | unclassified | 1 | 0 | 0,000588712 | 6,23E-05 | 1 - | 0,000699516 - | 1 - | 0,000584812 - | 1 - | 0,000588719 - |
| 1565 | 1 | 3um_FIJJAFBF_212746 | Bacteria | Bacteroidetes | Flavobacteriia | Flavobacterial | Flavobacteria | Polaribacter | Polaribacter s | poorly charact | Function unkn | 05ERK | gliding motilit | 313594.PI23P_ | 0,6 | 0,489897949 | 7,15E-05 | 5,84E-05 | 0 - | 0 - | 1 - | 0,000120568 - | 1 - | 0,000121374 - |
| 1566 | 1 | 3um_FIJJAFBF_615878 | Bacteria | Bacteroidetes | Flavobacteriia | Flavobacterial | Flavobacteria | Polaribacter | Polaribacter s | poorly charact | Function unkn | 0P8NI | Inherit from b | 313594.PI23P_ | 1,6 | 0,8 | 0,000145776 | 5,94E-05 | 1 - | 0,000112723 - | 1 - | 9,42E-05 - | 1 - | 9,49E-05 - |
| 1567 | 1 | 0.2um_NOKCPHJN_978568 | Eukaryota | unclassified | Oligohymenop | Hymenostoma | Tetrahymenid | Tetrahymena | Tetrahymena | unclassified | unclassified | unclassified | unclassified | unclassified | 7,2 | 1,720465053 | 0,000995585 | 0,000197125 | 7 - | 0,001164305 - | 5 - | 0,000695277 - | 6 - | 0,000839906 - |
| 1568 | 1 | 0.2um_NOKCPHJN_847711 | Eukaryota | Arthropoda | Insecta | Diptera | Tephritidae | Ceratitis | Ceratitis capit | unclassified | unclassified | unclassified | unclassified | unclassified | 0,2 | 0,4 | 2,29E-05 | 4,58E-05 | 0 - | 0 - | 0 - | 0 - | 0 - | 0 - |
| 1569 | 1 | 0.2um_NOKCPHJN_44818 | Eukaryota | unclassified | unclassified | Isochrysidales | Noelaerhabda | Emiliania | Emiliania huxl | metabolism | Energy produc | 0FJ7X | This is one of t | 32049.SYNPC | 1 | 0 | 0,000174978 | 1,85E-05 | 1 - | 0,000207912 - | 1 - | 0,000173819 - | 1 - | 0,00017498 - |
| 1570 | 1 | 0.2um_NOKCPHJN_227842 | Bacteria | Proteobacteri | Alphaproteob | Rhodospirillal | Rhodospirillac | unclassified | Rhodobactera | information st | Translation, ri | COG2265 | catalyzes the t | 367336.OM22 | 1 | 0 | 0,000154772 | 1,64E-05 | 1 - | 0,000183902 - | 1 - | 0,000153747 - | 1 - | 0,000154774 - |
| 1571 | 2 | 0.2um_NOKCPHJN_188037;3um_FIJJAF | Eukaryota | various | various | various | various | various | various | poorly charact | Function unkn | 0VCF1 | chromosome | 45351.NEMVE | 1 | 0 | 0,000229898 | 2,43E-05 | 1 - | 0,000273169 - | 1 - | 0,000228376 - | 1 - | 0,000229901 - |
| 1572 | 1 | 3um_FIJJAFBF_328039 | Bacteria | Bacteroidetes | Flavobacteriia | Flavobacterial | Flavobacteria | unclassified | Flavobacteria | poorly charact | Function unkn | 08W7B | integral memb | 391603.FBALC | 0,2 | 0,4 | 4,27E-05 | 8,54E-05 | 0 - | 0 - | 0 - | 0 - | 1 - | 0,000213535 - |
| 1573 | 1 | 3um_FIJJAFBF_576763 | Eukaryota | Bacillariophyta | Bacillariophyc | Bacillariales | Bacillariaceae | Fragilariopsis | Fragilariopsis | unclassified | unclassified | unclassified | unclassified | unclassified | 1 | 0 | 0,000299963 | 3,17E-05 | 1 - | 0,00035642 - | 1 - | 0,000297976 - | 1 - | 0,000299966 - |
| 1574 | 1 | 3um_FIJJAFBF_356540 | Bacteria | Proteobacteri | Alphaproteob | Rhodobactera | various | various | various | information st | Transcription | 05VEQ | Cold shock pro | 391619.RGBS1 | 0,4 | 0,489897949 | 0,000344248 | 0,000423298 | 0 - | 0 - | 1 - | 0,000920219 - | 0 - | 0 - |
| 1575 | 1 | 0.2um_NOKCPHJN_228601 | Bacteria | Bacteroidetes | Cytophagia | Cytophagales | Amoebophila | Candidatus Ar | Candidatus Ar | cellular proces | Posttranslatio | 0V7QW | ankyrin repeat | 7668.SPU_010 | 0,2 | 0,4 | 1,06E-05 | 2,11E-05 | 0 - | 0 - | 0 - | 0 - | 0 - | 0 - |
| 1576 | 1 | 3um_FIJJAFBF_754547 | Bacteria | Proteobacteri | Alphaproteob | Rhodobactera | Rhodobactera | Sulfitobacter | Sulfitobacter s | unclassified | unclassified | unclassified | unclassified | unclassified | 1 | 0 | 0,000134026 | 1,42E-05 | 1 - | 0,000159251 - | 1 - | 0,000133138 - | 1 - | 0,000134028 - |
| 1577 | 1 | 0.2um_NOKCPHJN_635596 | unclassified | unclassified | unclassified | unclassified | unclassified | unclassified | unclassified | unclassified | unclassified | unclassified | unclassified | unclassified | 1 | 0 | 0,00011391 | 1,20E-05 | 1 - | 0,000135349 - | 1 - | 0,000113155 - | 1 - | 0,000113911 - |
| 1578 | 1 | 0.2um_NOKCPHJN_485499 | Eukaryota | Phaeophyceae | unclassified | Ectocarpales | Ectocarpaceae | Ectocarpus | Ectocarpus sil | metabolism | Energy produc | COG1143 | NDH-1 shuttle | 88036.EFJ152 | 0,2 | 0,4 | 4,97E-05 | 9,95E-05 | 0 - | 0 - | 0 - | 0 - | 0 - | 0 - |
| 1579 | 1 | 3um_FIJJAFBF_708117 | Eukaryota | unclassified | Cryptophyta | Pyrenomonad | Geminigerace | Guillardia | Guillardia thet | unclassified | unclassified | unclassified | unclassified | unclassified | 1 | 0 | 0,000381771 | 4,04E-05 | 1 - | 0,000453625 - | 1 - | 0,000379242 - | 1 - | 0,000381775 - |
| 1580 | 1 | 3um_FIJJAFBF_783318 | Eukaryota | unclassified | Pelagophycea | Pelagomonad | unclassified | Aureoumbra | Aureoumbra l | information st | Translation, ri | 05G6W | One of the pri | 696747.NIES3 | 0,6 | 0,489897949 | 0,000187212 | 0,000156101 | 1 - | 0,000365113 - | 1 - | 0,000305244 - | 0 - | 0 - |
| 1581 | 1 | 3um_FIJJAFBF_49461 | Bacteria | Actinobacteria | Acidimicrobiia | Acidimicrobial | Acidimicrobia | unclassified | Acidimicrobia | cellular proces | Posttranslatio | COG1225 | alkyl hydroper | 269798.CHU_ | 0,8 | 0,4 | 0,000307837 | 0,000155146 | 0 - | 0 - | 1 - | 0,000401121 - | 1 - | 0,000403801 - |
| 1582 | 1 | 3um_FIJJAFBF_77421 | Viruses | unclassified | unclassified | unclassified | various | various | various | unclassified | unclassified | unclassified | unclassified | unclassified | 10,4 | 1,019803903 | 0,001233193 | 0,000111439 | 10 - | 0,001417579 - | 10 - | 0,001185131 - | 9 - | 0,001073743 - |
| 1583 | 2 | 0.2um_NOKCPHJN_164027;0.2um_NO | Eukaryota | unclassified | Oomycetes | Peronosporale | Peronosporac | Phytophthora | various | unclassified | unclassified | unclassified | unclassified | unclassified | 0,8 | 0,4 | 0,000467381 | 0,000241686 | 1 - | 0,000693039 - | 0 - | 0 - | 1 - | 0,000583268 - |
| 1584 | 1 | 0.2um_NOKCPHJN_393294 | Eukaryota | unclassified | Oomycetes | Saprolegniales | Saprolegniace | Aphanomyces | Aphanomyces | information st | Translation, ri | 1524W | Ribosomal S1 | 164328.Phyra | 0,8 | 0,4 | 0,00031803 | 0,000160284 | 0 - | 0 - | 1 - | 0,000414403 - | 1 - | 0,000417172 - |
| 1585 | 1 | 0.2um_NOKCPHJN_108597 | Bacteria | Bacteroidetes | Flavobacteriia | Flavobacterial | Crocinitomica | unclassified | Crocinitomica | unclassified | unclassified | unclassified | unclassified | unclassified | 0,2 | 0,4 | 2,58E-05 | 5,16E-05 | 0 - | 0 - | 0 - | 0 - | 0 - | 0 - |
| 1586 | 1 | 3um_FIJJAFBF_73880 | Eukaryota | Bacillariophyta | Coscinodiscop | Thalassiosirale | Thalassiosirac | Thalassiosira | Thalassiosira p | unclassified | unclassified | unclassified | unclassified | unclassified | 1 | 0 | 0,000599925 | 6,34E-05 | 1 - | 0,00071284 - | 1 - | 0,000595952 - | 1 - | 0,000599933 - |
| 1587 | 1 | 0.2um_NOKCPHJN_1055588 | Eukaryota | unclassified | Bangiophycea | Bangiales | Bangiaceae | Porphyra | Porphyra umb | cellular proces | Cell wall/mem | 00D84 | UDP-sulfoquir | 479433.Caci_ | 0,8 | 0,4 | 0,000410223 | 0,000209036 | 1 - | 0,000589356 - | 1 - | 0,000492716 - | 1 - | 0,000496007 - |
| 1588 | 1 | 3um_FIJJAFBF_183700 | Bacteria | Bacteroidetes | Flavobacteriia | Flavobacterial | Flavobacteria | Algibacter | Algibacter sp. | unclassified | unclassified | unclassified | unclassified | unclassified | 5 | 1,788854382 | 0,000328732 | 0,000105662 | 2 - | 0,000161137 - | 7 - | 0,000471501 - | 4 - | 0,000271229 - |
| 1589 | 1 | 3um_FIJJAFBF_216064 | Bacteria | Proteobacteri | Gammaprotec | Cellvibrionales | Microbulbifer | Microbulbifer | Microbulbifer | information st | Translation, ri | 05C64 | Binds directly | 314277.MED1 | 0,8 | 0,4 | 0,000225534 | 0,000114925 | 1 - | 0,000324018 - | 1 - | 0,000270887 - | 1 - | 0,000272697 - |
| 1590 | 1 | 0.2um_NOKCPHJN_761973 | unclassified | unclassified | unclassified | unclassified | unclassified | unclassified | unclassified | unclassified | unclassified | unclassified | unclassified | unclassified | 0,2 | 0,4 | 1,45E-05 | 2,90E-05 | 0 - | 0 - | 0 - | 0 - | 0 - | 0 - |
| 1591 | 1 | 0.2um_NOKCPHJN_957389 | Eukaryota | unclassified | Oomycetes | Saprolegniales | Saprolegniace | Aphanomyces | Aphanomyces | information st | Chromatin str | KOG1744 | Core compone | 4787.PITG_03 | 2 | 0 | 0,001076789 | 0,000113873 | 2 - | 0,001279456 - | 2 - | 0,001069657 - | 2 - | 0,001076802 - |
| 1592 | 1 | 3um_FIJJAFBF_631848 | Bacteria | Bacteroidetes | Flavobacteriia | Flavobacterial | Flavobacteria | Formosa | Formosa sp. H | cellular proces | Cell wall/mem | COG2885 | Ompa motb d | 983544.Lacal_ | 0,8 | 0,748331477 | 9,98E-05 | 9,43E-05 | 0 - | 0 - | 0 - | 0 - | 1 - | 0,000132338 - |
| 1593 | 1 | 3um_FIJJAFBF_470068 | Bacteria | Bacteroidetes | Flavobacteriia | Flavobacterial | Flavobacteria | Algibacter | Algibacter sp. | unclassified | unclassified | unclassified | unclassified | unclassified | 0,4 | 0,489897949 | 3,23E-05 | 3,97E-05 | 0 - | 0 - | 0 - | 0 - | 0 - | 0 - |
| 1594 | 1 | 3um_FIJJAFBF_503027 | Bacteria | Proteobacteri | Alphaproteob | unclassified | unclassified | unclassified | Alphaproteob | unclassified | unclassified | unclassified | unclassified | unclassified | 0,2 | 0,4 | 0,000112554 | 0,000225107 | 1 - | 0,000562768 - | 0 - | 0 - | 0 - | 0 - |
| 1595 | 1 | 0.2um_NOKCPHJN_238398 | Eukaryota | Phaeophyceae | unclassified | Ectocarpales | Ectocarpaceae | Ectocarpus | Ectocarpus sil | unclassified | unclassified | unclassified | unclassified | unclassified | 0,2 | 0,4 | 2,52E-05 | 5,04E-05 | 0 - | 0 - | 1 - | 0,000125905 - | 0 - | 0 - |
| 1596 | 1 | 3um_FIJJAFBF_715094 | Eukaryota | Bacillariophyta | Bacillariophyc | Eunotiales | Eunotiaceae | Eunotia | Eunotia naege | information st | Translation, ri | 0FNT6 | One of the ear | 497965.Cyan7 | 1,8 | 0,4 | 0,00126473 | 0,000351579 | 2 - | 0,001645015 - | 2 - | 0,001375273 - | 2 - | 0,00138446 - |
| 1597 | 1 | 0.2um_NOKCPHJN_348474 | Bacteria | Proteobacteri | Alphaproteob | Rhodobactera | Rhodobactera | unclassified | Rhodobactera | unclassified | unclassified | unclassified | unclassified | unclassified | 0,2 | 0,4 | 1,48E-05 | 2,95E-05 | 0 - | 0 - | 0 - | 0 - | 0 - | 0 - |
| 1598 | 1 | 0.2um_NOKCPHJN_805138 | Eukaryota | unclassified | unclassified | Kinetoplastida | Trypanosomat | Trypanosoma | Trypanosoma | unclassified | unclassified | unclassified | unclassified | unclassified | 0,6 | 0,489897949 | 0,000217693 | 0,000181818 | 1 - | 0,000430162 - | 0 - | 0 - | 0 - | 0 - |
| 1599 | 1 | 3um_FIJJAFBF_539569 | Bacteria | Bacteroidetes | Flavobacteriia | Flavobacterial | Flavobacteria | Nonlabens | Nonlabens do | poorly charact | Function unkn | 0PBEY | Pfam:DUF145 | 886377.Murru | 2 | 0 | 0,001880363 | 0,000198854 | 2 - | 0,002234274 - | 2 - | 0,001867908 - | 2 - | 0,001880386 - |
| 1600 | 3 | 3um_FIJJAFBF_178126;3um_FIJJAFBF_ | various | various | various | various | various | various | various | various | various | various | various | various | 0,2 | 0,4 | 3,74E-05 | 7,48E-05 | 0 - | 0 - | 0 - | 0 - | 1 - | 0,000186923 - |
| 1601 | 1 | 0.2um_NOKCPHJN_192568 | Bacteria | Proteobacteri | Gammaprotec | Alteromonada | Colwelliaceae | Colwellia | Colwellia sp. E | metabolism | Inorganic ion t | 16TQC | TONB-depend | 87626.PTD2_2 | 2 | 1,414213562 | 0,000117367 | 7,86E-05 | 0 - | 0 - | 2 - | 0,000125275 - | 1 - | 6,31E-05 - |
| 1602 | 4 | 0.2um_NOKCPHJN_1042306;0.2um_NO | Bacteria | Bacteroidetes | unclassified | unclassified | unclassified | unclassified | Bacteroidetes | unclassified | unclassified | unclassified | unclassified | unclassified | 1 | 0,894427191 | 5,14E-05 | 4,73E-05 | 2 - | 0,000113838 - | 1 - | 4,76E-05 - | 2 - | 9,58E-05 - |
| 1603 | 1 | 0.2um_NOKCPHJN_419460 | Eukaryota | Mucoromycot | Mortierellomy | Mortierellales | Mortierellacea | Mortierella | Mortierella ve | unclassified | unclassified | unclassified | unclassified | unclassified | 0,8 | 0,4 | 4,39E-05 | 2,27E-05 | 1 - | 6,51E-05 - | 1 - | 5,45E-05 - | 0 - | 0 - |
| 1604 | 1 | 0.2um_NOKCPHJN_241962 | Bacteria | Bacteroidetes | Flavobacteriia | Flavobacterial | Flavobacteria | Flaviramulus | Flaviramulus b | information st | Translation, ri | 05CAV | thus facilitatin | 983544.Lacal_ | 1,4 | 0,489897949 | 0,000139582 | 4,20E-05 | 1 - | 0,000120918 - | 2 - | 0,000202181 - | 1 - | 0,000101766 - |
| 1605 | 1 | 3um_FIJJAFBF_593596 | Bacteria | Proteobacteri | Alphaproteob | Rhodobactera | Rhodobactera | Puniceibacteri | Puniceibacteri | unclassified | unclassified | unclassified | unclassified | unclassified | 0,4 | 0,489897949 | 8,10E-05 | 9,92E-05 | 0 - | 0 - | 0 - | 0 - | 1 - | 0,000207214 - |
| 1606 | 1 | 3um_FIJJAFBF_521152 | Bacteria | Bacteroidetes | Flavobacteriia | Flavobacterial | Flavobacteria | Gaetbulibacte | Gaetbulibacte | metabolism | Inorganic ion t | 0P8Q4 | tonB-depende | 867900.Celly_ | 1,2 | 0,4 | 7,94E-05 | 1,96E-05 | 1 - | 8,05E-05 - | 1 - | 6,73E-05 - | 1 - | 6,77E-05 - |
| 1607 | 1 | 0.2um_NOKCPHJN_36981 | Eukaryota | unclassified | Rhodellophyc | Rhodellales | Rhodellaceae | Corynoplastis | Corynoplastis | unclassified | unclassified | unclassified | unclassified | unclassified | 2,6 | 0,489897949 | 0,000218568 | 5,18E-05 | 3 - | 0,000298596 - | 3 - | 0,000249634 - | 2 - | 0,000167534 - |
| 1608 | 1 | 3um_FIJJAFBF_92098 | Bacteria | Proteobacteri | Gammaprotec | unclassified | unclassified | unclassified | Gammaprotec | unclassified | unclassified | unclassified | unclassified | unclassified | 1 | 0 | 4,10E-05 | 4,34E-06 | 1 - | 4,88E-05 - | 1 - | 4,08E-05 - | 1 - | 4,10E-05 - |
| 1609 | 2 | 0.2um_NOKCPHJN_426473;0.2um_NO | Bacteria | various | various | various | various | various | various | various | various | various | various | various | 0,4 | 0,489897949 | 4,76E-05 | 5,87E-05 | 1 - | 0,00012972 - | 1 - | 0,000108449 - | 0 - | 0 - |
| 1610 | 1 | 0.2um_NOKCPHJN_11896 | Bacteria | Actinobacteria | Actinobacteria | Micrococcales | Microbacteria | unclassified | Microbacteria | unclassified | unclassified | unclassified | unclassified | unclassified | 1 | 0 | 0,000160285 | 1,70E-05 | 1 - | 0,000190453 - | 1 - | 0,000159224 - | 1 - | 0,000160287 - |
| 1611 | 1 | 3um_FIJJAFBF_282388 | Eukaryota | Bacillariophyta | Bacillariophyc | Naviculales | Phaeodactyla | Phaeodactylu | Phaeodactylu | unclassified | unclassified | unclassified | unclassified | unclassified | 4,4 | 0,489897949 | 0,002290647 | 0,000182055 | 4 - | 0,002494939 - | 4 - | 0,002085831 - | 4 - | 0,002099765 - |

| | | | | | | | | | | | | | | | | | | | | | | | | | | | | | | |
|---|---|---|---|---|---|---|---|---|---|---|---|---|---|---|---|---|---|---|---|---|---|---|---|---|---|---|---|---|---|---|
| 1612 | 2 | 0.2um_NOKCPHJN_160356;3um_FIJJAI | Eukaryota | unclassified | Pelagophycea | Pelagomonada | unclassified | Pelagomonas | uncultured Pe | unclassified | unclassified | unclassified | unclassified | unclassified | 2,6 | 0,8 | 0,000663017 | 0,000146542 | 2 | - | 0,000621147 | - | 2 | - | 0,000519294 | - | 2 | - | 0,000522763 | - |
| 1613 | 1 | 0.2um_NOKCPHJN_505060 | Eukaryota | Phaeophyceae | unclassified | Ectocarpales | Ectocarpaceae | Ectocarpus | Ectocarpus sili | information st | Translation, ri | 14XM1 | EFG_IV | 4787.PITG_10 | 0,6 | 0,489897949 | 4,10E-05 | 3,37E-05 | 0 | - | 0 | - | 0 | - | 0 | - | 1 | - | 7,28E-05 | - |
| 1614 | 4 | 0.2um_NOKCPHJN_105152;0.2um_NO | Bacteria | Bacteroidetes | Flavobacteriia | Flavobacterial | Flavobacteria | Polaribacter | various | unclassified | unclassified | unclassified | unclassified | unclassified | 2 | 0,632455532 | 0,001031927 | 0,000293387 | 2 | - | 0,00123716 | - | 2 | - | 0,001034296 | - | 2 | - | 0,001041206 | - |
| 1615 | 1 | 3um_FIJJAFBF_692160 | Eukaryota | unclassified | Oomycetes | Saprolegniales | Saprolegniace | Achlya | Achlya hypogy | unclassified | unclassified | unclassified | unclassified | unclassified | 1 | 0,632455532 | 0,000572441 | 0,000342782 | 1 | - | 0,000668287 | - | 1 | - | 0,000558705 | - | 1 | - | 0,000562437 | - |
| 1616 | 1 | 3um_FIJJAFBF_116533 | Eukaryota | Bacillariophyta | Coscinodiscop | Thalassiosirale | Stephanodisca | Cyclotella | Cyclotella pse | metabolism | Energy produc | 11IIK | Low-potential | 35128.Thapsd | 6,2 | 0,979795897 | 0,002423286 | 0,000330106 | 6 | - | 0,002806807 | - | 6 | - | 0,002346559 | - | 6 | - | 0,002362235 | - |
| 1617 | 1 | 0.2um_NOKCPHJN_375427 | Eukaryota | unclassified | unclassified | Kinetoplastida | Trypanosomat | Leishmania | Leishmania m | unclassified | unclassified | unclassified | unclassified | unclassified | 1 | 0 | 6,40E-05 | 6,77E-06 | 1 | - | 7,61E-05 | - | 1 | - | 6,36E-05 | - | 1 | - | 6,40E-05 | - |
| 1618 | 1 | 3um_FIJJAFBF_305831 | Eukaryota | Chlorophyta | Nephroselmid | unclassified | unclassified | Nephroselmis | Nephroselmis | unclassified | unclassified | unclassified | unclassified | unclassified | 0,8 | 0,4 | 9,45E-05 | 4,76E-05 | 0 | - | 0 | - | 1 | - | 0,000123179 | - | 1 | - | 0,000124002 | - |
| 1619 | 2 | 3um_FIJJAFBF_167273;3um_FIJJAFBF_ | Bacteria | Bacteroidetes | Flavobacteriia | Flavobacterial | Flavobacteria | Dokdonia | Dokdonia paci | unclassified | unclassified | unclassified | unclassified | unclassified | 0,8 | 0,4 | 0,000154911 | 7,81E-05 | 0 | - | 0 | - | 1 | - | 0,000201855 | - | 1 | - | 0,000203203 | - |
| 1620 | 1 | 0.2um_NOKCPHJN_816703 | Bacteria | Proteobacteri | Betaproteoba | Nitrosomonad | unclassified | unclassified | various | poorly charact | Function unkn | 0XRND | NA | 314607.KB13_ | 1,8 | 0,979795897 | 0,000265391 | 0,000126602 | 1 | - | 0,000182557 | - | 1 | - | 0,000152622 | - | 1 | - | 0,000153641 | - |
| 1621 | 1 | 3um_FIJJAFBF_262967 | Bacteria | unclassified | unclassified | unclassified | unclassified | unclassified | uncultured ba | metabolism | Carbohydrate | 16QEQ | hydrolase, fan | 983545.Glaag_ | 1,6 | 0,489897949 | 0,000237903 | 7,98E-05 | 2 | - | 0,000353057 | - | 1 | - | 0,000147582 | - | 1 | - | 0,000148568 | - |
| 1622 | 1 | 3um_FIJJAFBF_259313 | Bacteria | Planctomycete | Phycisphaerae | Phycisphaeral | Phycisphaerac | Phycisphaera | Phycisphaera | unclassified | unclassified | unclassified | unclassified | unclassified | 1 | 0,632455532 | 0,00022149 | 0,000130281 | 0 | - | 0 | - | 1 | - | 0,000235244 | - | 1 | - | 0,000236816 | - |
| 1623 | 1 | 3um_FIJJAFBF_15942 | Eukaryota | unclassified | Pelagophycea | Pelagomonada | unclassified | Aureococcus | Aureococcus a | unclassified | unclassified | unclassified | unclassified | unclassified | 3,2 | 0,4 | 0,000635884 | 0,000101455 | 3 | - | 0,000708342 | - | 3 | - | 0,000592192 | - | 4 | - | 0,000794864 | - |
| 1624 | 1 | 0.2um_NOKCPHJN_05106 | Bacteria | Bacteroidetes | Flavobacteriia | Flavobacterial | Crocinitomica | unclassified | Crocinitomica | unclassified | unclassified | unclassified | unclassified | unclassified | 2 | 0 | 8,19E-05 | 8,66E-06 | 2 | - | 9,73E-05 | - | 2 | - | 8,13E-05 | - | 2 | - | 8,19E-05 | - |
| 1625 | 1 | 3um_FIJJAFBF_20039 | Bacteria | Bacteroidetes | Flavobacteriia | Flavobacterial | Flavobacteria | Polaribacter | Polaribacter s | unclassified | unclassified | unclassified | unclassified | unclassified | 2 | 0,894427191 | 0,000124188 | 5,26E-05 | 1 | - | 7,56E-05 | - | 3 | - | 0,000189621 | - | 1 | - | 6,36E-05 | - |
| 1626 | 1 | 3um_FIJJAFBF_346215 | Bacteria | Bacteroidetes | Flavobacteriia | Flavobacterial | Crocinitomica | unclassified | Crocinitomica | unclassified | unclassified | unclassified | unclassified | unclassified | 1,8 | 0,4 | 0,00146072 | 0,000251018 | 1 | - | 0,000984845 | - | 2 | - | 0,001646708 | - | 2 | - | 0,001657709 | - |
| 1627 | 1 | 3um_FIJJAFBF_723333 | Bacteria | Proteobacteri | Alphaproteob | Rhodobactera | Rhodobactera | Marinosulfon | Marinosulfon | cellular proces | Cell wall/mem | 0RH7M | Glycosyl trans | 64091.VNG10 | 0,6 | 0,489897949 | 0,000102814 | 8,49E-05 | 1 | - | 0,000194411 | - | 0 | - | 0 | - | 1 | - | 0,000163618 | - |
| 1628 | 1 | 3um_FIJJAFBF_161768 | Bacteria | Bacteroidetes | Flavobacteriia | Flavobacterial | unclassified | unclassified | Flavobacterial | unclassified | unclassified | unclassified | unclassified | unclassified | 0,2 | 0,4 | 1,73E-05 | 3,45E-05 | 0 | - | 0 | - | 0 | - | 0 | - | 0 | - | 0 | - |
| 1629 | 1 | 3um_FIJJAFBF_39791 | Bacteria | Bacteroidetes | Flavobacteriia | Flavobacterial | Cryomorphace | unclassified | Cryomorphace | unclassified | unclassified | unclassified | unclassified | unclassified | 0,8 | 0,748331477 | 0,000177105 | 0,000159022 | 0 | - | 0 | - | 0 | - | 0 | - | 1 | - | 0,000240431 | - |
| 1630 | 1 | 0.2um_NOKCPHJN_776268 | Eukaryota | Streptophyta | unclassified | Brassicales | Brassicaceae | Brassica | various | unclassified | unclassified | unclassified | unclassified | unclassified | 1,8 | 0,4 | 0,000536303 | 9,22E-05 | 1 | - | 0,000361585 | - | 2 | - | 0,000604589 | - | 2 | - | 0,000608627 | - |
| 1631 | 1 | 0.2um_NOKCPHJN_227402 | Eukaryota | Phaeophyceae | unclassified | Ectocarpales | Ectocarpaceae | Ectocarpus | Ectocarpus sili | cellular proces | Cytoskeleton | COG5245 | heavy chain | 65071.PYU1_T | 0,2 | 0,4 | 2,87E-06 | 5,74E-06 | 0 | - | 0 | - | 0 | - | 0 | - | 0 | - | 0 | - |
| 1632 | 1 | 3um_FIJJAFBF_121184 | Bacteria | Proteobacteri | Gammaproteo | Alteromonada | Alteromonada | Paraglaciecola | Paraglaciecola | unclassified | unclassified | unclassified | unclassified | unclassified | 0,2 | 0,4 | 1,42E-05 | 2,83E-05 | 0 | - | 0 | - | 0 | - | 0 | - | 1 | - | 7,08E-05 | - |
| 1633 | 1 | 3um_FIJJAFBF_20315 | Bacteria | Bacteroidetes | Flavobacteriia | Flavobacterial | Flavobacteria | Polaribacter | Polaribacter s | metabolism | Carbohydrate | 099Q3 | phosphohexol | 313594.PI23P_ | 0,2 | 0,4 | 3,33E-05 | 6,66E-05 | 0 | - | 0 | - | 0 | - | 0 | - | 0 | - | 0 | - |
| 1634 | 1 | 3um_FIJJAFBF_276754 | Bacteria | Proteobacteri | Gammaproteo | Alteromonada | Alteromonada | Paraglaciecola | Paraglaciecola | metabolism | Amino acid tra | 16STP | phosphoserine | 342610.Patl_1 | 0,2 | 0,4 | 3,25E-05 | 6,49E-05 | 0 | - | 0 | - | 0 | - | 0 | - | 0 | - | 0 | - |
| 1635 | 1 | 3um_FIJJAFBF_737884 | Bacteria | Proteobacteri | Betaproteoba | unclassified | unclassified | unclassified | Betaproteoba | unclassified | unclassified | unclassified | unclassified | unclassified | 1,4 | 0,489897949 | 0,000666496 | 0,000281417 | 2 | - | 0,001108862 | - | 1 | - | 0,000463518 | - | 1 | - | 0,000466614 | - |
| 1636 | 1 | 3um_FIJJAFBF_519979 | Bacteria | Bacteroidetes | Flavobacteriia | Flavobacterial | Flavobacteria | Flavobacteriu | Flavobacteriu | cellular proces | Signal transdu | COG3275 | Histidine kinas | 376686.Fjoh_4 | 0,6 | 0,489897949 | 0,000115514 | 9,52E-05 | 1 | - | 0,000215701 | - | 1 | - | 0,000180331 | - | 1 | - | 0,000181536 | - |
| 1637 | 1 | 3um_FIJJAFBF_257121 | Eukaryota | unclassified | Cryptophyta | Pyrenomonad | Geminigerace | Guillardia | Guillardia thet | unclassified | unclassified | unclassified | unclassified | unclassified | 1,6 | 0,489897949 | 0,00084704 | 0,000307452 | 2 | - | 0,00123716 | - | 2 | - | 0,001034296 | - | 1 | - | 0,000520603 | - |
| 1638 | 1 | 0.2um_NOKCPHJN_620209 | Eukaryota | Bacillariophyta | Bacillariophyc | Bacillariales | Bacillariaceae | Fragilariopsis | Fragilariopsis | unclassified | unclassified | unclassified | unclassified | unclassified | 1,6 | 0,489897949 | 0,001778433 | 0,000614598 | 2 | - | 0,002626252 | - | 2 | - | 0,002195611 | - | 1 | - | 0,001105139 | - |
| 1639 | 1 | 0.2um_NOKCPHJN_521598 | Bacteria | Proteobacteri | Gammaproteo | Cellvibrionales | unclassified | unclassified | Cellvibrionales | poorly charact | Function unkn | 16SJR | amidohydrola | 314285.KT71_ | 0,4 | 0,489897949 | 4,61E-05 | 5,65E-05 | 0 | - | 0 | - | 1 | - | 0,000117622 | - | 0 | - | 0 | - |
| 1640 | 1 | 0.2um_NOKCPHJN_573914 | Bacteria | Cyanobacteria | unclassified | Oscillatoriales | Oscillatoriace | Lyngbya | Lyngbya sp. P | poorly charact | Function unkn | 0XRC0 | fg-gap repeat | 313612.L8106 | 0,2 | 0,4 | 9,09E-06 | 1,82E-05 | 0 | - | 0 | - | 0 | - | 0 | - | 1 | - | 4,54E-05 | - |
| 1641 | 1 | 0.2um_NOKCPHJN_997351 | Eukaryota | unclassified | Oligohymenop | Philasterida | Philasteridae | Miamiensis | Miamiensis av | cellular proces | Signal transdu | KOG0699 | protein phosp | 3055.EDP0089 | 0,4 | 0,489897949 | 9,07E-05 | 0,000111567 | 0 | - | 0 | - | 1 | - | 0,000242538 | - | 0 | - | 0 | - |
| 1642 | 1 | 0.2um_NOKCPHJN_805291 | Eukaryota | unclassified | Oomycetes | Pythiales | Pythiaceae | Pythium | Pythium insidi | unclassified | unclassified | unclassified | unclassified | unclassified | 1,2 | 0,4 | 0,000401555 | 9,92E-05 | 1 | - | 0,000406784 | - | 1 | - | 0,000340081 | - | 1 | - | 0,000342353 | - |
| 1643 | 2 | 0.2um_NOKCPHJN_1042131;3um_FIJJA | Eukaryota | unclassified | Oomycetes | Peronosporale | Peronosporac | Plasmopara | Plasmopara ha | information st | Replication, re | COG0513 | purine NTP-de | 65071.PYU1_T | 5,4 | 0,489897949 | 0,000854187 | 0,000158475 | 6 | - | 0,001119923 | - | 5 | - | 0,000780236 | - | 6 | - | 0,000942538 | - |
| 1644 | 1 | 0.2um_NOKCPHJN_938882 | Viruses | unclassified | unclassified | unclassified | Phycodnavirid | Prasinovirus | Bathycoccus s | unclassified | unclassified | unclassified | unclassified | unclassified | 1 | 0 | 0,000351912 | 3,72E-05 | 1 | - | 0,000418146 | - | 1 | - | 0,000349581 | - | 1 | - | 0,000351916 | - |
| 1645 | 1 | 0.2um_NOKCPHJN_853407 | unclassified | unclassified | unclassified | unclassified | unclassified | unclassified | unclassified | unclassified | unclassified | unclassified | unclassified | unclassified | 0,2 | 0,4 | 5,42E-05 | 0,000108355 | 0 | - | 0 | - | 1 | - | 0,000270887 | - | 0 | - | 0 | - |
| 1646 | 1 | 3um_FIJJAFBF_673146 | Eukaryota | Bacillariophyta | Coscinodiscop | Thalassiosirale | Thalassiosirac | Thalassiosira | Thalassiosira | unclassified | unclassified | unclassified | unclassified | unclassified | 2,8 | 0,748331477 | 0,001626281 | 0,000302839 | 2 | - | 0,00141223 | - | 2 | - | 0,001180659 | - | 3 | - | 0,001782819 | - |
| 1647 | 1 | 3um_FIJJAFBF_66597 | Eukaryota | Bacillariophyta | Bacillariophyc | Cymbellales | Gomphonema | Gomphoneis | Gomphoneis r | metabolism | Energy produc | 1DYB8 | PsaA and PsaE | 3055.DAA014 | 3,6 | 0,8 | 0,000300561 | 6,79E-05 | 3 | - | 0,000298596 | - | 5 | - | 0,000416057 | - | 3 | - | 0,000251302 | - |
| 1648 | 1 | 3um_FIJJAFBF_111312 | Viruses | unclassified | unclassified | unclassified | various | various | various | information st | Transcription | 0WRII | DNA-depende | 339860.Msp_ | 0,2 | 0,4 | 2,99E-05 | 5,99E-05 | 0 | - | 0 | - | 0 | - | 0 | - | 0 | - | 0 | - |
| 1649 | 1 | 3um_FIJJAFBF_807864 | Eukaryota | Chlorophyta | Chlorophycea | Chlamydomor | Tetrabaenace | Tetrabaena | Tetrabaena sc | information st | Replication, re | COG0592 | DNA polymera | 3847.GLYMA0 | 1 | 0 | 0,000242278 | 2,56E-05 | 1 | - | 0,000287878 | - | 1 | - | 0,000240673 | - | 1 | - | 0,000242281 | - |
| 1650 | 1 | 3um_FIJJAFBF_99521 | Viruses | unclassified | unclassified | Caudovirales | Siphoviridae | Cbastvirus | Cellulophaga v | unclassified | unclassified | unclassified | unclassified | unclassified | 0,8 | 0,4 | 0,00014641 | 7,38E-05 | 0 | - | 0 | - | 1 | - | 0,000190777 | - | 1 | - | 0,000192052 | - |
| 1651 | 1 | 0.2um_NOKCPHJN_39142 | Eukaryota | Phaeophyceae | unclassified | Laminariales | Agaraceae | Costaria | Costaria costa | metabolism | Energy produc | COG4451 | RuBisCO catal | 288000.BBta_ | 13 | 0 | 0,005891355 | 0,000623027 | 13 | - | 0,00700019 | - | 13 | - | 0,005852331 | - | 13 | - | 0,005891426 | - |
| 1652 | 1 | 3um_FIJJAFBF_204879 | unclassified | unclassified | unclassified | unclassified | unclassified | unclassified | unclassified | unclassified | unclassified | unclassified | unclassified | unclassified | 0,4 | 0,8 | 0,000131252 | 0,000262503 | 0 | - | 0 | - | 0 | - | 0 | - | 0 | - | 0 | - |
| 1653 | 2 | 3um_FIJJAFBF_126542;3um_FIJJAFBF_ | Eukaryota | various | various | various | various | various | various | information st | Translation, ri | KOG3181 | ribosomal pro | various | 1 | 0 | 0,000282476 | 2,99E-05 | 1 | - | 0,000335642 | - | 1 | - | 0,000280605 | - | 1 | - | 0,00028248 | - |
| 1654 | 1 | 3um_FIJJAFBF_598844 | Bacteria | Bacteroidetes | Flavobacteriia | Flavobacterial | Flavobacteria | Polaribacter | Polaribacter b | metabolism | Nucleotide tra | COG0138 | bifunctional p | 313598.MED1 | 0,8 | 0,4 | 0,000100349 | 5,18E-05 | 1 | - | 0,000147339 | - | 1 | - | 0,000123179 | - | 1 | - | 0,000124002 | - |
| 1655 | 1 | 0.2um_NOKCPHJN_144610 | Bacteria | Proteobacteri | Gammaproteo | Cellvibrionales | Porticoccacea | unclassified | SAR92 bacteri | unclassified | unclassified | unclassified | unclassified | unclassified | 0,4 | 0,489897949 | 9,87E-05 | 0,000121179 | 0 | - | 0 | - | 0 | - | 0 | - | 0 | - | 0 | - |
| 1656 | 1 | 0.2um_NOKCPHJN_819662 | Bacteria | Proteobacteri | Gammaproteo | Vibrionales | Vibrionaceae | Vibrio | Vibrio sp. G98 | unclassified | unclassified | unclassified | unclassified | unclassified | 0,4 | 0,8 | 6,26E-05 | 0,000125157 | 0 | - | 0 | - | 0 | - | 0 | - | 0 | - | 0 | - |
| 1657 | 1 | 3um_FIJJAFBF_732530 | Bacteria | Bacteroidetes | Cytophagia | Cytophagales | Flammeovirga | unclassified | Flammeovirga | cellular proces | Cell wall/mem | 0QKXZ | Porin Gram-ne | 247633.GP214 | 4,6 | 1,854723699 | 0,000905216 | 0,000338304 | 3 | - | 0,00071284 | - | 3 | - | 0,000595952 | - | 5 | - | 0,000999888 | - |
| 1658 | 1 | 3um_FIJJAFBF_122322 | unclassified | unclassified | unclassified | unclassified | unclassified | unclassified | unclassified | unclassified | unclassified | unclassified | unclassified | unclassified | 3,2 | 0,4 | 0,000535846 | 4,77E-05 | 3 | - | 0,000601996 | - | 3 | - | 0,000503284 | - | 3 | - | 0,000506646 | - |
| 1659 | 1 | 0.2um_NOKCPHJN_751237 | Eukaryota | unclassified | unclassified | Prymnesiales | Chrysochromu | Chrysochromu | Chrysochromu | unclassified | unclassified | unclassified | unclassified | unclassified | 1 | 0 | 0,000741084 | 7,84E-05 | 1 | - | 0,000880567 | - | 1 | - | 0,000736176 | - | 1 | - | 0,000741093 | - |
| 1660 | 1 | 0.2um_NOKCPHJN_728658 | Bacteria | Proteobacteri | Gammaproteo | Pseudomonad | Pseudomonac | Pseudomonas | various | unclassified | unclassified | unclassified | unclassified | unclassified | 0,6 | 0,489897949 | 0,000234992 | 0,000191974 | 0 | - | 0 | - | 1 | - | 0,000396044 | - | 1 | - | 0,000398689 | - |
| 1661 | 1 | 0.2um_NOKCPHJN_96088 | Bacteria | Proteobacteri | Gammaproteo | Alteromonada | Alteromonada | Paraglaciecola | Paraglaciecola | poorly charact | Function unkn | 05C2A | tonB-depende | 1085623.GNIT | 4 | 1,095445115 | 0,000226777 | 6,03E-05 | 3 | - | 0,000203946 | - | 3 | - | 0,000170504 | - | 6 | - | 0,000343286 | - |
| 1662 | 1 | 3um_FIJJAFBF_753922 | Bacteria | Proteobacteri | Alphaproteob | Rhizobiales | unclassified | unclassified | Rhizobiales ba | metabolism | Inorganic ion t | 05F28 | tonB-depende | 247633.GP214 | 1 | 0,894427191 | 7,43E-05 | 6,81E-05 | 0 | - | 0 | - | 2 | - | 0,00015508 | - | 0 | - | 0 | - |
| 1663 | 1 | 0.2um_NOKCPHJN_663091 | Eukaryota | Bacillariophyta | Bacillariophyc | Bacillariales | Bacillariaceae | Fragilariopsis | Fragilariopsis | metabolism | Inorganic ion t | 0INF7 | Two pore segr | 2850.Phatr15 | 0,8 | 0,4 | 0,000100466 | 5,06E-05 | 0 | - | 0 | - | 1 | - | 0,00013091 | - | 1 | - | 0,000131784 | - |
| 1664 | 1 | 3um_FIJJAFBF_335656 | Bacteria | Planctomycete | Planctomyceti | Planctomyceta | Planctomyceta | Rubinisphaera | Rubinisphaera | unclassified | unclassified | unclassified | unclassified | unclassified | 0,8 | 0,4 | 0,000105161 | 5,44E-05 | 1 | - | 0,000155934 | - | 0 | - | 0 | - | 1 | - | 0,000131235 | - |
| 1665 | 1 | 0.2um_NOKCPHJN_439969 | Bacteria | Proteobacteri | Alphaproteob | Pelagibacteral | Pelagibactera | Candidatus Pe | Candidatus Pe | unclassified | unclassified | unclassified | unclassified | unclassified | 0,2 | 0,4 | 1,54E-05 | 3,08E-05 | 0 | - | 0 | - | 0 | - | 0 | - | 1 | - | 7,71E-05 | - |
| 1666 | 1 | 3um_FIJJAFBF_97615 | Bacteria | Bacteroidetes | Flavobacteriia | Flavobacterial | Flavobacteria | Polaribacter | various | information st | Translation, ri | 08UHY | One of the pri | 313598.MED1 | 3,2 | 1,166190379 | 0,001245804 | 0,000422603 | 2 | - | 0,000941487 | - | 3 | - | 0,001180659 | - | 5 | - | 0,00198091 | - |
| 1667 | 1 | 3um_FIJJAFBF_668498 | Eukaryota | Bacillariophyta | Bacillariophyc | Bacillariales | Bacillariaceae | Fragilariopsis | Fragilariopsis | unclassified | unclassified | unclassified | unclassified | unclassified | 1,2 | 0,748331477 | 0,000580845 | 0,000360563 | 1 | - | 0,00057136 | - | 2 | - | 0,000955342 | - | 2 | - | 0,000961724 | - |
| 1668 | 1 | 0.2um_NOKCPHJN_328778 | Bacteria | Proteobacteri | Gammaproteo | Nevskiales | Algiphilaceae | Algiphilus | Algiphilus aro | poorly charact | Function unkn | 0XSV1 | NA | 204773.HEAR | 1 | 0 | 2,07E-05 | 2,19E-06 | 1 | - | 2,46E-05 | - | 1 | - | 2,06E-05 | - | 1 | - | 2,07E-05 | - |
| 1669 | 1 | 3um_FIJJAFBF_257112 | Eukaryota | unclassified | Dinophyceae | Peridiniales | Kryptoperidini | Durinskia | Durinskia balt | information st | Translation, ri | COG0088 | One of the pri | 329726.AM1_ | 1,6 | 0,8 | 0,000448809 | 0,000226195 | 0 | - | 0 | - | 2 | - | 0,000584812 | - | 2 | - | 0,000588719 | - |
| 1670 | 1 | 3um_FIJJAFBF_545707 | Bacteria | Bacteroidetes | Saprospiria | Saprospirales | Saprospiracea | Aureispira | Aureispira sp. | metabolism | Energy produc | COG0055 | Produces ATP | 391596.PBAL3 | 2,2 | 0,4 | 0,000274966 | 5,66E-05 | 2 | - | 0,000297017 | - | 2 | - | 0,000248313 | - | 3 | - | 0,000374958 | - |
| 1671 | 1 | 3um_FIJJAFBF_442115 | unclassified | unclassified | unclassified | unclassified | unclassified | unclassified | unclassified | unclassified | unclassified | unclassified | unclassified | unclassified | 2,4 | 0,489897949 | 0,000952623 | 0,000191935 | 2 | - | 0,000947445 | - | 3 | - | 0,001188131 | - | 2 | - | 0,000797379 | - |
| 1672 | 1 | 0.2um_NOKCPHJN_226948 | unclassified | unclassified | unclassified | unclassified | unclassified | unclassified | unclassified | unclassified | unclassified | unclassified | unclassified | unclassified | 0,2 | 0,4 | 2,61E-05 | 5,21E-05 | 0 | - | 0 | - | 1 | - | 0,000130364 | - | 0 | - | 0 | - |
| 1673 | 1 | 0.2um_NOKCPHJN_416858 | Bacteria | Proteobacteri | Alphaproteob | Rickettsiales | Anaplasmatac | Anaplasma | Anaplasma ph | unclassified | unclassified | unclassified | unclassified | unclassified | 0,4 | 0,489897949 | 1,37E-05 | 1,71E-05 | 1 | - | 3,96E-05 | - | 0 | - | 0 | - | 0 | - | 0 | - |
| 1674 | 1 | 3um_FIJJAFBF_582313 | Eukaryota | Eustigmatoph | unclassified | Eustigmatales | Monodopsida | Nannochlorop | Nannochlorop | cellular proces | Signal transdu | COG5040 | Tyrosine 3-mo | 35128.Thaps2 | 4,4 | 0,8 | 0,00197209 | 0,000497949 | 5 | - | 0,002635499 | - | 5 | - | 0,002203342 | - | 5 | - | 0,002218061 | - |
| 1675 | 1 | 0.2um_NOKCPHJN_1046856 | Viruses | unclassified | unclassified | unclassified | Phycodnavirid | unclassified | Chrysochromu | unclassified | unclassified | unclassified | unclassified | unclassified | 1 | 0 | 0,000238607 | 2,52E-05 | 1 | - | 0,000283516 | - | 1 | - | 0,000237026 | - | 1 | - | 0,00023861 | - |
| 1676 | 3 | 0.2um_NOKCPHJN_738060;0.2um_NO | Bacteria | Bacteroidetes | Flavobacteriia | Flavobacterial | Flavobacteria | Formosa | Formosa sp. H | unclassified | unclassified | unclassified | unclassified | unclassified | 0,8 | 0,4 | 0,000291298 | 0,00015031 | 1 | - | 0,000427704 | - | 1 | - | 0,000357571 | - | 1 | - | 0,00035996 | - |
| 1677 | 1 | 0.2um_NOKCPHJN_725707 | Bacteria | Bacteroidetes | Flavobacteriia | Flavobacterial | Flavobacteria | Formosa | Formosa sp. H | unclassified | unclassified | unclassified | unclassified | unclassified | 1,4 | 0,489897949 | 0,000143951 | 4,84E-05 | 1 | - | 0,000123106 | - | 2 | - | 0,000205839 | - | 1 | - | 0,000103607 | - |
| 1678 | 1 | 3um_FIJJAFBF_57641 | Bacteria | Proteobacteri | Alphaproteob | Rhodospirillal | Rhodospirillac | unclassified | Rhodospirillac | poorly charact | Function unkn | 0YPRH | FBG | 6085.XP_0021 | 0,2 | 0,4 | 2,10E-05 | 4,21E-05 | 0 | - | 0 | - | 0 | - | 0 | - | 0 | - | 0 | - |
| 1679 | 1 | 0.2um_NOKCPHJN_90233 | unclassified | unclassified | unclassified | unclassified | unclassified | unclassified | unclassified | unclassified | unclassified | unclassified | unclassified | unclassified | 0,6 | 0,489897949 | 5,55E-05 | 4,63E-05 | 1 | - | 0,000108319 | - | 1 | - | 9,06E-05 | - | 0 | - | 0 | - |
| 1680 | 1 | 0.2um_NOKCPHJN_157509 | Eukaryota | Phaeophyceae | unclassified | Ectocarpales | Ectocarpaceae | Ectocarpus | Ectocarpus sili | unclassified | unclassified | unclassified | unclassified | unclassified | 0,8 | 0,4 | 7,74E-05 | 4,00E-05 | 1 | - | 0,000114974 | - | 1 | - | 9,61E-05 | - | 0 | - | 0 | - |
| 1681 | 1 | 3um_FIJJAFBF_168406 | Bacteria | Bacteroidetes | Flavobacteriia | Flavobacterial | Flavobacteria | unclassified | Flavobacteria | poorly charact | Function unkn | 0ZMES | Caspase doma | 272559.BF343 | 0,4 | 0,489897949 | 2,58E-05 | 3,16E-05 | 0 | - | 0 | - | 1 | - | 6,58E-05 | - | 0 | - | 0 | - |
| 1682 | 1 | 3um_FIJJAFBF_459209 | Eukaryota | unclassified | unclassified | Isochrysidales | Noelaerhabda | Emiliania | Emiliania huxl | cellular proces | Posttranslatio | KOG0730 | cell division cy | 35128.Thapsd | 0,4 | 0,489897949 | 8,46E-05 | 0,000103633 | 0 | - | 0 | - | 1 | - | 0,000215776 | - | 0 | - | 0 | - |
| 1683 | 1 | 0.2um_NOKCPHJN_1015854 | unclassified | unclassified | unclassified | unclassified | unclassified | unclassified | unclassified | unclassified | unclassified | unclassified | unclassified | unclassified | 1 | 0 | 0,000298541 | 3,16E-05 | 1 | - | 0,000354731 | - | 1 | - | 0,000296564 | - | 1 | - | 0,000298545 | - |
| 1684 | 1 | 0.2um_NOKCPHJN_1011380 | unclassified | unclassified | unclassified | unclassified | unclassified | unclassified | unclassified | unclassified | unclassified | unclassified | unclassified | unclassified | 0,2 | 0,4 | 2,08E-05 | 4,16E-05 | 0 | - | 0 | - | 0 | - | 0 | - | 0 | - | 0 | - |
| 1685 | 1 | 3um_FIJJAFBF_627897 | Eukaryota | Phaeophyceae | unclassified | Ectocarpales | Ectocarpaceae | Ectocarpus | Ectocarpus sili | unclassified | unclassified | unclassified | unclassified | unclassified | 0,8 | 0,4 | 0,000453996 | 0,000234798 | 1 | - | 0,000674308 | - | 1 | - | 0,000563738 | - | 0 | - | 0 | - |
| 1686 | 1 | 0.2um_NOKCPHJN_270820 | unclassified | unclassified | unclassified | unclassified | unclassified | unclassified | unclassified | unclassified | unclassified | unclassified | unclassified | unclassified | 1 | 0 | 5,47E-05 | 5,79E-06 | 1 | - | 6,50E-05 | - | 1 | - | 5,44E-05 | - | 1 | - | 5,47E-05 | - |
| 1687 | 1 | 0.2um_NOKCPHJN_724920 | Bacteria | Bacteroidetes | Flavobacteriia | Flavobacterial | Flavobacteria | Polaribacter | Polaribacter s | metabolism | Inorganic ion t | 0P8Q4 | tonB-depende | 688270.Celal_ | 9,6 | 2,576819745 | 0,000688887 | 0,00012189 | 7 | - | 0,000611362 | - | 8 | - | 0,00058413 | - | 8 | - | 0,000588032 | - |
| 1688 | 1 | 3um_FIJJAFBF_714542 | Bacteria | Proteobacteri | Alphaproteob | Rhodobactera | Rhodobactera | Sulfitobacter | Sulfitobacter s | metabolism | Energy produc | 01R4M | Acetoacetyl-C | 375451.RD1_3 | 0,4 | 0,489897949 | 9,79E-05 | 0,000120409 | 0 | - | 0 | - | 0 | - | 0 | - | 1 | - | 0,000262471 | - |
| 1689 | 3 | 0.2um_NOKCPHJN_292421;0.2um_NO | Bacteria | Bacteroidetes | Flavobacteriia | Flavobacterial | Crocinitomica | unclassified | various | unclassified | unclassified | unclassified | unclassified | unclassified | 0,6 | 0,489897949 | 0,000474841 | 0,000395856 | 1 | - | 0,000924052 | - | 0 | - | 0 | - | 1 | - | 0,000777691 | - |
| 1690 | 1 | 3um_FIJJAFBF_711587 | Eukaryota | unclassified | Pelagophycea | Pelagomonada | unclassified | Aureococcus | Aureococcus a | unclassified | unclassified | unclassified | unclassified | unclassified | 0,6 | 0,489897949 | 0,000511873 | 0,000427518 | 1 | - | 0,001011462 | - | 0 | - | 0 | - | 0 | - | 0 | - |
| 1691 | 1 | 3um_FIJJAFBF_370013 | Bacteria | Bacteroidetes | unclassified | unclassified | unclassified | unclassified | Bacteroidetes | poorly charact | Function unkn | 09DAT | Pfam:NPD | 983548.Krodi_ | 0,6 | 0,489897949 | 0,000131068 | 0,000109468 | 1 | - | 0,00025899 | - | 0 | - | 0 | - | 0 | - | 0 | - |
| 1692 | 1 | 0.2um_NOKCPHJN_1054080 | unclassified | unclassified | unclassified | unclassified | unclassified | unclassified | unclassified | unclassified | unclassified | unclassified | unclassified | unclassified | 0,6 | 0,489897949 | 6,51E-05 | 5,34E-05 | 0 | - | 0 | - | 1 | - | 0,000115027 | - | 0 | - | 0 | - |
| 1693 | 1 | 0.2um_NOKCPHJN_701777 | Eukaryota | Apicomplexa | Conoidasida | Eucoccidiorida | Eimeriidae | Eimeria | Eimeria praec | unclassified | unclassified | unclassified | unclassified | unclassified | 0,2 | 0,4 | 4,01E-05 | 8,01E-05 | 0 | - | 0 | - | 0 | - | 0 | - | 0 | - | 0 | - |
| 1694 | 1 | 0.2um_NOKCPHJN_617369 | Eukaryota | Chlorophyta | Ulvophyceae | Ulvales | Ulvaceae | Ulva | Ulva linza | unclassified | unclassified | unclassified | unclassified | unclassified | 1,4 | 0,489897949 | 0,000220459 | 7,42E-05 | 1 | - | 0,000188534 | - | 2 | - | 0,000315239 | - | 1 | - | 0,000158672 | - |
| 1695 | 1 | 3um_FIJJAFBF_516119 | Eukaryota | Bacillariophyta | Coscinodiscop | Thalassiosirale | Thalassiosirac | Thalassiosira | Thalassiosira o | cellular proces | Posttranslatio | KOG0730 | cell division cy | 35128.Thaps4 | 2 | 0 | 0,000396177 | 4,19E-05 | 2 | - | 0,000470743 | - | 2 | - | 0,000393553 | - | 2 | - | 0,000396182 | - |
| 1696 | 1 | 0.2um_NOKCPHJN_268023 | Bacteria | Proteobacteri | Alphaproteob | unclassified | unclassified | Candidatus Pu | Candidatus Pu | unclassified | unclassified | unclassified | unclassified | unclassified | 0,8 | 0,4 | 8,30E-05 | 4,28E-05 | 1 | - | 0,000121903 | - | 1 | - | 0,000101914 | - | 1 | - | 0,000102594 | - |
| 1697 | 1 | 3um_FIJJAFBF_31880 | Bacteria | unclassified | unclassified | unclassified | unclassified | unclassified | uncultured ba | metabolism | Coenzyme tra | 09A4R | May play a ke | 391587.KAOT | 0,6 | 0,489897949 | 8,76E-05 | 7,31E-05 | 1 | - | 0,000170886 | - | 1 | - | 0,000142865 | - | 0 | - | 0 | - |
| 1698 | 1 | 0.2um_NOKCPHJN_728627 | Eukaryota | Bacillariophyta | Coscinodiscop | Thalassiosirale | Thalassiosirac | Thalassiosira | Thalassiosira p | unclassified | unclassified | unclassified | unclassified | unclassified | 2 | 0 | 0,000893506 | 9,45E-05 | 2 | - | 0,001061676 | - | 2 | - | 0,000887587 | - | 2 | - | 0,000893517 | - |
| 1699 | 2 | 0.2um_NOKCPHJN_74285;3um_FIJJAFE | Bacteria | Bacteroidetes | Flavobacteriia | Flavobacterial | Flavobacteria | Polaribacter | various | unclassified | unclassified | unclassified | unclassified | unclassified | 1 | 0 | 0,000508001 | 5,37E-05 | 1 | - | 0,000603614 | - | 1 | - | 0,000504636 | - | 1 | - | 0,000508008 | - |
| 1700 | 1 | 3um_FIJJAFBF_467310 | Eukaryota | unclassified | Pelagophycea | Pelagomonada | unclassified | Aureococcus | Aureococcus a | metabolism | Amino acid tra | 05C6M | Alpha-keto-be | 867900.Celly_ | 0,8 | 0,4 | 0,000275832 | 0,000142634 | 1 | - | 0,000409006 | - | 0 | - | 0 | - | 1 | - | 0,000344224 | - |
| 1701 | 1 | 3um_FIJJAFBF_205215 | Eukaryota | unclassified | Raphidophyce | Chattonellales | Chattonellace | Heterosigma | Heterosigma a | unclassified | unclassified | unclassified | unclassified | unclassified | 2 | 0 | 0,000247513 | 2,62E-05 | 2 | - | 0,000294099 | - | 2 | - | 0,000245874 | - | 2 | - | 0,000247516 | - |
| 1702 | 1 | 0.2um_NOKCPHJN_223179 | Bacteria | Bacteroidetes | Flavobacteriia | Flavobacterial | Flavobacteria | Tenacibaculum | Tenacibaculum | unclassified | unclassified | unclassified | unclassified | unclassified | 2,6 | 1,019803903 | 0,000478533 | 0,000159057 | 1 | - | 0,000226813 | - | 3 | - | 0,000568863 | - | 3 | - | 0,000572663 | - |
| 1703 | 1 | 3um_FIJJAFBF_254288 | Bacteria | Proteobacteri | Gammaproteo | Cellvibrionales | Spongiibacter | Zhongshania | Zhongshania s | unclassified | unclassified | unclassified | unclassified | unclassified | 0,8 | 0,4 | 0,000183249 | 9,48E-05 | 1 | - | 0,000272175 | - | 1 | - | 0,000227545 | - | 0 | - | 0 | - |
| 1704 | 1 | 3um_FIJJAFBF_85545 | Bacteria | Proteobacteri | Gammaproteo | Alteromonada | Alteromonada | Alteromonas | Alteromonas s | poorly charact | Function unkn | 17TP6 | PEP-CTERM m | 335283.Neut_ | 0,2 | 0,4 | 3,90E-05 | 7,80E-05 | 0 | - | 0 | - | 0 | - | 0 | - | 0 | - | 0 | - |
| 1705 | 1 | 0.2um_NOKCPHJN_421076 | Bacteria | Bacteroidetes | Flavobacteriia | Flavobacterial | Flavobacteria | Cellulophaga | Cellulophaga a | metabolism | Inorganic ion t | 09HYP | Sodium/calciu | 688270.Celal_ | 0,6 | 0,489897949 | 9,58E-05 | 7,86E-05 | 0 | - | 0 | - | 1 | - | 0,000166423 | - | 1 | - | 0,000167534 | - |
| 1706 | 1 | 0.2um_NOKCPHJN_914725 | unclassified | unclassified | unclassified | unclassified | unclassified | unclassified | unclassified | unclassified | unclassified | unclassified | unclassified | unclassified | 0,2 | 0,4 | 8,13E-05 | 0,000162595 | 0 | - | 0 | - | 0 | - | 0 | - | 0 | - | 0 | - |
| 1707 | 1 | 0.2um_NOKCPHJN_560343 | Eukaryota | Bacillariophyta | Coscinodiscop | Thalassiosirale | Thalassiosirac | Thalassiosira | Thalassiosira p | unclassified | unclassified | unclassified | unclassified | unclassified | 0,2 | 0,4 | 3,26E-05 | 6,52E-05 | 0 | - | 0 | - | 0 | - | 0 | - | 0 | - | 0 | - |
| 1708 | 1 | 0.2um_NOKCPHJN_127271 | Bacteria | Bacteroidetes | Flavobacteriia | Flavobacterial | Flavobacteria | Arenitalea | Arenitalea lut | cellular proces | Posttranslatio | 08V1T | peptidylprolyl | 313594.PI23P_ | 0,8 | 0,4 | 0,000205225 | 0,000103431 | 0 | - | 0 | - | 1 | - | 0,000267414 | - | 1 | - | 0,000269201 | - |
| 1709 | 1 | 0.2um_NOKCPHJN_671818 | unclassified | unclassified | unclassified | unclassified | unclassified | unclassified | unclassified | unclassified | unclassified | unclassified | unclassified | unclassified | 1 | 0 | 0,000212095 | 2,24E-05 | 1 | - | 0,000252014 | - | 1 | - | 0,00021069 | - | 1 | - | 0,000212097 | - |
| 1710 | 1 | 3um_FIJJAFBF_768481 | unclassified | unclassified | unclassified | unclassified | unclassified | unclassified | unclassified | unclassified | unclassified | unclassified | unclassified | unclassified | 0,4 | 0,489897949 | 3,66E-05 | 4,57E-05 | 1 | - | 0,000105867 | - | 0 | - | 0 | - | 0 | - | 0 | - |
| 1711 | 1 | 0.2um_NOKCPHJN_405347 | Eukaryota | unclassified | Oomycetes | Peronosporale | Peronosporac | Phytophthora | Phytophthora | unclassified | unclassified | unclassified | unclassified | unclassified | 0,6 | 0,489897949 | 6,44E-05 | 5,38E-05 | 1 | - | 0,000127293 | - | 0 | - | 0 | - | 0 | - | 0 | - |
| 1712 | 1 | 0.2um_NOKCPHJN_316226 | Bacteria | Bacteroidetes | Flavobacteriia | Flavobacterial | Flavobacteria | Muricauda | Muricauda sp. | unclassified | unclassified | unclassified | unclassified | unclassified | 1 | 0 | 2,69E-05 | 2,85E-06 | 1 | - | 3,20E-05 | - | 1 | - | 2,67E-05 | - | 1 | - | 2,69E-05 | - |

| | | | | | | | | | | | | | | | | | | | | | | | | |
|---|---|---|---|---|---|---|---|---|---|---|---|---|---|---|---|---|---|---|---|---|---|---|---|---|
| 1713 | 1 | 0.2um_NOKCPHJN_12462 | Bacteria | Bacteroidetes | Flavobacteriia | Flavobacterial | Flavobacteriac | Formosa | Formosa sp. H | metabolism | Inorganic ion t | COG3119 | Sulfatase | 313628.LNTAF | 1 | 0 | 0,000131508 | 1,39E-05 | 1 - | 0,000156259 - | 1 - | 0,000130637 - | 1 - | 0,000131509 - |
| 1714 | 1 | 0.2um_NOKCPHJN_359572 | Bacteria | Proteobacteri | Alphaproteob | Rhodospirillal | unclassified | unclassified | various | poorly charact | Function unkn | 01QWE | Integrase | 398580.Dshi_ | 0,2 | 0,4 | 3,18E-05 | 6,37E-05 | 0 - | 0 - | 1 - | 0,000159224 - | 0 - | 0 - |
| 1715 | 1 | 3um_FIJJAFBF_205988 | Bacteria | Proteobacteri | Deltaproteoba | Myxococcales | Nannocystace | Plesiocystis | Plesiocystis pa | metabolism | Energy produc | 07T6V | glycerophosph | 391625.PPSIR | 1 | 0 | 0,000238607 | 2,52E-05 | 1 - | 0,000283516 - | 1 - | 0,000237026 - | 1 - | 0,00023861 - |
| 1716 | 1 | 0.2um_NOKCPHJN_188581 | Bacteria | Bacteroidetes | unclassified | unclassified | unclassified | unclassified | unidentified e | information st | Translation, ri | COG0018 | arginyL-tRNA s | 50743.SCB49_ | 0,4 | 0,8 | 4,26E-05 | 8,53E-05 | 0 - | 0 - | 0 - | 0 - | 2 - | 0,000213174 - |
| 1717 | 1 | 3um_FIJJAFBF_559551 | unclassified | unclassified | unclassified | unclassified | unclassified | unclassified | unclassified | unclassified | unclassified | unclassified | unclassified | unclassified | 0,6 | 0,489897949 | 0,00068533 | 0,000571442 | 1 - | 0,001336575 - | 1 - | 0,001117409 - | 0 - | 0 - |
| 1718 | 1 | 3um_FIJJAFBF_310912 | Bacteria | Proteobacteri | Gammaprotec | Chromatiales | Chromatiacea | Candidatus Te | Candidatus Te | cellular proces | Cell wall/mem | 0GRGC | Protein of unk | 96561.Dole_2 | 1,2 | 0,4 | 0,000316486 | 9,91E-05 | 1 - | 0,000315815 - | 1 - | 0,000264029 - | 1 - | 0,000265793 - |
| 1719 | 1 | 3um_FIJJAFBF_235865 | Bacteria | Proteobacteri | Gammaprotec | Cellvibrionales | Spongiibacter | Oceanicoccus | Oceanicoccus | information st | Translation, ri | 0QSAC | Binds togethe | 247633.GP214 | 1 | 0 | 0,000449944 | 4,76E-05 | 1 - | 0,00053463 - | 1 - | 0,000446964 - | 1 - | 0,00044995 - |
| 1720 | 2 | 3um_FIJJAFBF_447548;3um_FIJJAFBF_ | various | various | various | various | various | various | various | various | various | various | various | various | 0,8 | 0,4 | 0,000145965 | 7,36E-05 | 0 - | 0 - | 1 - | 0,000190197 - | 1 - | 0,000191468 - |
| 1721 | 1 | 3um_FIJJAFBF_01139 | Bacteria | Bacteroidetes | Flavobacteriia | Flavobacterial | Flavobacteriac | Polaribacter | Polaribacter s | information st | Translation, ri | 05KK9 | This protein bi | 313594.PI23P | 0,4 | 0,489897949 | 0,000140173 | 0,000172361 | 0 - | 0 - | 1 - | 0,0003747 - | 0 - | 0 - |
| 1722 | 1 | 3um_FIJJAFBF_390314 | Eukaryota | Nematoda | Chromadorea | Rhabditida | Ancylostomat | Ancylostoma | Ancylostoma c | cellular proces | Signal transdu | 0A7BN | Inherit from m | 54126.PPA002 | 0,4 | 0,489897949 | 6,91E-05 | 8,50E-05 | 0 - | 0 - | 0 - | 0 - | 1 - | 0,000185273 - |
| 1723 | 1 | 3um_FIJJAFBF_594160 | Eukaryota | unclassified | Bangiophycea | Bangiales | Bangiaceae | Wildemania | Wildemania s | unclassified | unclassified | unclassified | unclassified | unclassified | 5,4 | 1,019803903 | 0,002245528 | 0,000436451 | 5 - | 0,002478417 - | 4 - | 0,001657614 - | 6 - | 0,002503031 - |
| 1724 | 1 | 3um_FIJJAFBF_111396 | Bacteria | Chlorobi | Chlorobia | Chlorobiales | Chlorobiaceae | Prosthecochlo | Prosthecochlo | cellular proces | Cell wall/mem | 05D2T | synthase | 517417.Cpar_ | 2,8 | 0,4 | 0,000639731 | 5,59E-05 | 2 - | 0,000550354 - | 3 - | 0,000690165 - | 3 - | 0,000694775 - |
| 1725 | 1 | 0.2um_NOKCPHJN_1023200 | Bacteria | Bacteroidetes | Sphingobacter | Sphingobacter | Sphingobacter | Pedobacter | Pedobacter sp | unclassified | unclassified | unclassified | unclassified | unclassified | 1 | 0 | 0,000193822 | 2,05E-05 | 1 - | 0,000230302 - | 1 - | 0,000192538 - | 1 - | 0,000193824 - |
| 1726 | 1 | 3um_FIJJAFBF_545215 | Bacteria | Bacteroidetes | Flavobacteriia | Flavobacterial | Flavobacteriac | Flavivirga | Flavivirga eckl | unclassified | unclassified | unclassified | unclassified | unclassified | 2,8 | 0,4 | 0,00133683 | 0,000246325 | 3 - | 0,001701095 - | 2 - | 0,000948105 - | 3 - | 0,001431658 - |
| 1727 | 2 | 3um_FIJJAFBF_49184;3um_FIJJAFBF_5 | Bacteria | Bacteroidetes | Flavobacteriia | Flavobacterial | Flavobacteriac | Polaribacter | various | unclassified | unclassified | unclassified | unclassified | unclassified | 0,2 | 0,4 | 1,38E-05 | 2,77E-05 | 0 - | 0 - | 0 - | 0 - | 1 - | 6,91E-05 - |
| 1728 | 1 | 3um_FIJJAFBF_292921 | Eukaryota | Bacillariophyta | Coscinodiscop | Coscinodiscale | Coscinodiscac | Stellarima | Stellarima mic | metabolism | Carbohydrate | COG1850 | RuBisCO catal | 261292.Nit79 | 8,4 | 1,854723699 | 0,001061112 | 0,000147308 | 7 - | 0,00106926 - | 8 - | 0,001021631 - | 7 - | 0,000899899 - |
| 1729 | 2 | 0.2um_NOKCPHJN_836546;0.2um_NO | various | various | various | various | various | various | various | information st | Replication, re | various | various | various | 0,2 | 0,4 | 7,13E-05 | 0,000142568 | 1 - | 0,00035642 - | 0 - | 0 - | 0 - | 0 - |
| 1730 | 1 | 3um_FIJJAFBF_269831 | Eukaryota | unclassified | Oomycetes | Saprolegniales | Saprolegniace | various | various | unclassified | unclassified | unclassified | unclassified | unclassified | 17,4 | 2,059126028 | 0,002881408 | 0,000225765 | 16 - | 0,003176581 - | 17 - | 0,002821681 - | 15 - | 0,00250635 - |
| 1731 | 1 | 3um_FIJJAFBF_195099 | Eukaryota | unclassified | unclassified | Prymnesiales | Chrysochromu | Chrysochromu | Chrysochromu | unclassified | unclassified | unclassified | unclassified | unclassified | 1 | 0 | 0,000308785 | 3,27E-05 | 1 - | 0,000366903 - | 1 - | 0,00030674 - | 1 - | 0,000308789 - |
| 1732 | 1 | 0.2um_NOKCPHJN_1001222 | unclassified | unclassified | unclassified | unclassified | unclassified | unclassified | unclassified | unclassified | unclassified | unclassified | unclassified | unclassified | 0,2 | 0,4 | 0,000375472 | 0,000750943 | 0 - | 0 - | 0 - | 0 - | 0 - | 0 - |
| 1733 | 1 | 3um_FIJJAFBF_319475 | Bacteria | Bacteroidetes | Flavobacteriia | Flavobacterial | Flavobacteriac | Flavobacteriu | Flavobacteriu | unclassified | unclassified | unclassified | unclassified | unclassified | 0,2 | 0,4 | 5,94E-05 | 0,000118855 | 0 - | 0 - | 0 - | 0 - | 1 - | 0,000297137 - |
| 1734 | 1 | 0.2um_NOKCPHJN_206707 | Bacteria | Proteobacteri | Alphaproteob | Pelagibacteral | Pelagibacterac | Candidatus Pe | Candidatus Pe | cellular proces | Cell wall/mem | COG0438 | Glycosyl trans | 656519.Halsa_ | 0,4 | 0,489897949 | 6,45E-05 | 7,94E-05 | 0 - | 0 - | 0 - | 0 - | 1 - | 0,000173058 - |
| 1735 | 1 | 3um_FIJJAFBF_310979 | unclassified | unclassified | unclassified | unclassified | unclassified | unclassified | unclassified | unclassified | unclassified | unclassified | unclassified | unclassified | 0,2 | 0,4 | 0,000129195 | 0,000258389 | 0 - | 0 - | 0 - | 0 - | 0 - | 0 - |
| 1736 | 1 | 0.2um_NOKCPHJN_417758 | Viruses | unclassified | unclassified | unclassified | unclassified | unclassified | Pleurochrysis | unclassified | unclassified | unclassified | unclassified | unclassified | 1 | 0 | 0,000128294 | 1,36E-05 | 1 - | 0,00015244 - | 1 - | 0,000127444 - | 1 - | 0,000128295 - |
| 1737 | 1 | 0.2um_NOKCPHJN_602110 | Eukaryota | unclassified | Oomycetes | Saprolegniales | Saprolegniace | Thraustotheca | Thraustotheca | unclassified | unclassified | unclassified | unclassified | unclassified | 0,6 | 0,489897949 | 5,86E-05 | 4,90E-05 | 1 - | 0,000115864 - | 0 - | 0 - | 0 - | 0 - |
| 1738 | 1 | 0.2um_NOKCPHJN_1117233 | Eukaryota | unclassified | Oomycetes | Peronosporale | Peronosporac | Phytophthora | various | unclassified | unclassified | unclassified | unclassified | unclassified | 0,6 | 1,2 | 0,00046812 | 0,000936241 | 0 - | 0 - | 0 - | 0 - | 0 - | 0 - |
| 1739 | 1 | 3um_FIJJAFBF_89693 | Eukaryota | unclassified | Pelagophycea | Pelagomonad | unclassified | Aureococcus | Aureococcus a | cellular proces | Posttranslatio | KOG0730 | cell division cy | 35128.Thaps4 | 2,8 | 0,4 | 0,001956952 | 0,000415724 | 3 - | 0,002467523 - | 3 - | 0,002062909 - | 3 - | 0,00207669 - |
| 1740 | 1 | 3um_FIJJAFBF_332981 | unclassified | unclassified | unclassified | unclassified | unclassified | unclassified | unclassified | unclassified | unclassified | unclassified | unclassified | unclassified | 0,2 | 0,4 | 2,81E-05 | 5,61E-05 | 0 - | 0 - | 0 - | 0 - | 0 - | 0 - |
| 1741 | 1 | 0.2um_NOKCPHJN_538830 | Eukaryota | unclassified | unclassified | Kinetoplastida | Trypanosomat | Leishmania | Leishmania m | unclassified | unclassified | unclassified | unclassified | unclassified | 0,2 | 0,4 | 3,44E-05 | 6,87E-05 | 0 - | 0 - | 0 - | 0 - | 0 - | 0 - |
| 1742 | 1 | 3um_FIJJAFBF_610542 | Bacteria | Bacteroidetes | Flavobacteriia | Flavobacterial | Flavobacteriac | Aequorivita | Aequorivita sp | unclassified | unclassified | unclassified | unclassified | unclassified | 0,4 | 0,489897949 | 0,000170155 | 0,000212665 | 1 - | 0,000492422 - | 0 - | 0 - | 0 - | 0 - |
| 1743 | 2 | 0.2um_NOKCPHJN_404857;3um_FIJJAF | Bacteria | Firmicutes | Bacilli | Bacillales | Paenibacillace | Paenibacillus | various | unclassified | unclassified | unclassified | unclassified | unclassified | 0,4 | 0,489897949 | 2,58E-05 | 3,18E-05 | 0 - | 0 - | 0 - | 0 - | 1 - | 6,93E-05 - |
| 1744 | 1 | 3um_FIJJAFBF_168628 | Bacteria | Proteobacteri | Gammaprotec | Cellvibrionales | Halieaceae | Halioglobus | Halioglobus lu | information st | Replication, re | 05C24 | DNA gyrase ne | 247634.GPB2 | 0,2 | 0,4 | 1,27E-05 | 2,54E-05 | 0 - | 0 - | 0 - | 0 - | 0 - | 0 - |
| 1745 | 1 | 3um_FIJJAFBF_800979 | Eukaryota | Bacillariophyta | Mediophyceae | Triceratiales | Triceratiaceae | Trieres | Trieres chinen | unclassified | unclassified | unclassified | unclassified | unclassified | 2 | 0 | 0,00061757 | 6,53E-05 | 2 - | 0,000733806 - | 2 - | 0,00061348 - | 2 - | 0,000617578 - |
| 1746 | 1 | 0.2um_NOKCPHJN_541868 | Eukaryota | Apicomplexa | Conoidasida | Eucoccidiorida | Eimeriidae | Eimeria | Eimeria brune | poorly charact | Function unkn | 0KN5V | NA | 140110.Necha | 0,2 | 0,4 | 1,77E-05 | 3,55E-05 | 1 - | 8,87E-05 - | 0 - | 0 - | 0 - | 0 - |
| 1747 | 8 | 0.2um_NOKCPHJN_11819;3um_FIJJAFB | Bacteria | Bacteroidetes | Flavobacteriia | Flavobacterial | Flavobacteriac | various | various | poorly charact | Function unkn | various | various | various | 0,6 | 0,489897949 | 0,000254162 | 0,000211925 | 1 - | 0,000495683 - | 1 - | 0,000414403 - | 0 - | 0 - |
| 1748 | 1 | 3um_FIJJAFBF_456031 | Bacteria | Bacteroidetes | Cytophagia | Cytophagales | Cytophagacea | Leadbetterella | Leadbetterella | unclassified | unclassified | unclassified | unclassified | unclassified | 0,8 | 0,4 | 0,000189812 | 9,57E-05 | 0 - | 0 - | 1 - | 0,000247332 - | 1 - | 0,000248984 - |
| 1749 | 1 | 3um_FIJJAFBF_715055 | Eukaryota | unclassified | Cryptophyta | Pyrenomonad | Pyrenomonad | Rhodomonas | Rhodomonas | information st | Transcription | COG0085 | DNA-depende | 395961.Cyan7 | 0,2 | 0,4 | 1,15E-05 | 2,29E-05 | 0 - | 0 - | 1 - | 5,73E-05 - | 0 - | 0 - |
| 1750 | 1 | 3um_FIJJAFBF_214150 | Bacteria | Bacteroidetes | Flavobacteriia | Flavobacterial | Flavobacteriac | Olleya | various | information st | Translation, ri | COG0050 | This protein p | 1046627.BZAF | 1,8 | 0,979795897 | 0,00027357 | 0,00014977 | 0 - | 0 - | 2 - | 0,000316835 - | 2 - | 0,000318952 - |
| 1751 | 1 | 0.2um_NOKCPHJN_719164 | unclassified | unclassified | unclassified | unclassified | unclassified | unclassified | unclassified | unclassified | unclassified | unclassified | unclassified | unclassified | 0,2 | 0,4 | 0,000138446 | 0,000276892 | 0 - | 0 - | 0 - | 0 - | 1 - | 0,00069223 - |
| 1752 | 1 | 0.2um_NOKCPHJN_316893 | Bacteria | Bacteroidetes | unclassified | unclassified | unclassified | unclassified | Bacteroidetes | metabolism | Carbohydrate | 07R46 | aggregation fa | 344747.PM87 | 0,2 | 0,4 | 1,59E-05 | 3,18E-05 | 0 - | 0 - | 0 - | 0 - | 1 - | 7,94E-05 - |
| 1753 | 1 | 3um_FIJJAFBF_532815 | Bacteria | Proteobacteri | Alphaproteob | Rhodobactera | unclassified | unclassified | Rhodobactera | unclassified | unclassified | unclassified | unclassified | unclassified | 0,8 | 0,4 | 0,000410449 | 0,000206862 | 0 - | 0 - | 1 - | 0,000534828 - | 1 - | 0,000538401 - |
| 1754 | 1 | 3um_FIJJAFBF_629059 | Eukaryota | Chytridiomyco | Chytridiomyce | Rhizophydiale | unclassified | Batrachochytr | Batrachochytr | information st | Translation, ri | KOG1242 | GCN1 general | 436017.A4S74 | 0,4 | 0,489897949 | 6,50E-05 | 8,04E-05 | 1 - | 0,000180357 - | 0 - | 0 - | 0 - | 0 - |
| 1755 | 1 | 0.2um_NOKCPHJN_774804 | Archaea | Euryarchaeota | unclassified | unclassified | unclassified | unclassified | Marine Group | unclassified | unclassified | unclassified | unclassified | unclassified | 0,8 | 0,4 | 8,98E-05 | 4,58E-05 | 1 - | 0,000129049 - | 1 - | 0,000107888 - | 1 - | 0,000108609 - |
| 1756 | 1 | 0.2um_NOKCPHJN_136913 | Eukaryota | unclassified | Oomycetes | Saprolegniales | Saprolegniace | Saprolegnia | Saprolegnia di | cellular proces | Cytoskeleton | COG5245 | heavy chain | 4787.PITG_08 | 0,8 | 0,4 | 1,19E-05 | 5,99E-06 | 0 - | 0 - | 1 - | 1,55E-05 - | 1 - | 1,56E-05 - |
| 1757 | 1 | 3um_FIJJAFBF_242673 | Bacteria | Bacteroidetes | Saprospiria | Saprospirales | Haliscomenob | Haliscomenob | Haliscomenob | cellular proces | Posttranslatio | COG1404 | peptidase (S8 | 760192.Halhy | 1 | 0 | 0,000173532 | 1,84E-05 | 1 - | 0,000206193 - | 1 - | 0,000172383 - | 1 - | 0,000173534 - |
| 1758 | 1 | 0.2um_NOKCPHJN_137528 | Eukaryota | Chordata | Ascidiacea | Enterogona | Cionidae | Ciona | Ciona intestin | unclassified | unclassified | unclassified | unclassified | unclassified | 1 | 0 | 1,39E-05 | 1,47E-06 | 1 - | 1,65E-05 - | 1 - | 1,38E-05 - | 1 - | 1,39E-05 - |
| 1759 | 1 | 0.2um_NOKCPHJN_915794 | Eukaryota | Ascomycota | Sordariomycet | Hypocreales | Nectriaceae | Neonectria | Neonectria dit | unclassified | unclassified | unclassified | unclassified | unclassified | 1 | 0 | 0,000516329 | 5,46E-05 | 1 - | 0,00061351 - | 1 - | 0,000512909 - | 1 - | 0,000516336 - |
| 1760 | 1 | 0.2um_NOKCPHJN_1027022 | Eukaryota | Xanthophycea | unclassified | Botrydiales | Botrydiaceae | Botrydium | Botrydium gra | unclassified | unclassified | unclassified | unclassified | unclassified | 6,8 | 0,4 | 0,002253839 | 0,000128093 | 6 - | 0,002376133 - | 7 - | 0,00231759 - | 7 - | 0,002333072 - |
| 1761 | 1 | 3um_FIJJAFBF_213038 | Bacteria | Bacteroidetes | Flavobacteriia | Flavobacterial | Flavobacteriac | Muricauda | Muricauda sp. | unclassified | unclassified | unclassified | unclassified | unclassified | 0,4 | 0,489897949 | 0,000231771 | 0,000284992 | 0 - | 0 - | 1 - | 0,000619554 - | 0 - | 0 - |
| 1762 | 1 | 0.2um_NOKCPHJN_89203 | Bacteria | Bacteroidetes | Flavobacteriia | Flavobacterial | Flavobacteriac | Kordia | Kordia zhangz | unclassified | unclassified | unclassified | unclassified | unclassified | 0,2 | 0,4 | 2,96E-05 | 5,92E-05 | 0 - | 0 - | 1 - | 0,000147931 - | 0 - | 0 - |
| 1763 | 2 | 0.2um_NOKCPHJN_1000825;0.2um_N | Bacteria | various | various | various | various | various | various | unclassified | unclassified | unclassified | unclassified | unclassified | 0,4 | 0,489897949 | 4,88E-05 | 6,00E-05 | 0 - | 0 - | 1 - | 0,000130364 - | 0 - | 0 - |
| 1764 | 1 | 0.2um_NOKCPHJN_675926 | unclassified | unclassified | unclassified | unclassified | unclassified | unclassified | unclassified | unclassified | unclassified | unclassified | unclassified | unclassified | 0,8 | 0,4 | 0,000121374 | 6,26E-05 | 1 - | 0,00017821 - | 1 - | 0,000148988 - | 1 - | 0,000149983 - |
| 1765 | 1 | 3um_FIJJAFBF_102176 | Bacteria | Proteobacteri | Alphaproteob | Rhodobactera | Rhodobactera | Ruegeria | Ruegeria med | metabolism | Inorganic ion t | COG3221 | phosphonate | 388739.RSK20 | 2,8 | 0,979795897 | 0,000562686 | 0,000163061 | 1 - | 0,000247842 - | 3 - | 0,000621605 - | 3 - | 0,000625758 - |
| 1766 | 1 | 3um_FIJJAFBF_314150 | Bacteria | Proteobacteri | Gammaprotec | Oceanospirilla | unclassified | unclassified | Oceanospirilla | unclassified | unclassified | unclassified | unclassified | unclassified | 0,2 | 0,4 | 2,48E-05 | 4,96E-05 | 0 - | 0 - | 0 - | 0 - | 0 - | 0 - |
| 1767 | 1 | 3um_FIJJAFBF_407704 | Bacteria | Proteobacteri | Gammaprotec | Oceanospirilla | Halomonadac | Halomonas | Halomonas ve | unclassified | unclassified | unclassified | unclassified | unclassified | 0,2 | 0,4 | 0,000137527 | 0,000275055 | 0 - | 0 - | 1 - | 0,000687636 - | 0 - | 0 - |
| 1768 | 1 | 3um_FIJJAFBF_666764 | Bacteria | Proteobacteri | Epsilonproteo | Campylobacte | Campylobacte | Arcobacter | Arcobacter sp | metabolism | Carbohydrate | 05D2N | Aldolase | 469613.HMPR | 1,4 | 0,489897949 | 0,000520613 | 0,000145829 | 1 - | 0,000453625 - | 1 - | 0,000379242 - | 1 - | 0,000381775 - |
| 1769 | 1 | 0.2um_NOKCPHJN_1027277 | Eukaryota | unclassified | Florideophyce | Gigartinales | Gigartinaceae | Chondrus | Chondrus cris | unclassified | unclassified | unclassified | unclassified | unclassified | 1 | 0 | 0,000377199 | 3,99E-05 | 1 - | 0,000448193 - | 1 - | 0,0003747 - | 1 - | 0,000377203 - |
| 1770 | 1 | 3um_FIJJAFBF_321055 | unclassified | unclassified | unclassified | unclassified | unclassified | unclassified | unclassified | unclassified | unclassified | unclassified | unclassified | unclassified | 0,4 | 0,489897949 | 0,000545452 | 0,000669371 | 0 - | 0 - | 0 - | 0 - | 0 - | 0 - |
| 1771 | 1 | 0.2um_NOKCPHJN_106200 | Bacteria | Bacteroidetes | Cytophagia | Cytophagales | Cytophagacea | Dyadobacter | Dyadobacter f | poorly charact | Function unkn | 08NCR | NA | 471854.Dfer_ | 0,2 | 0,4 | 1,35E-05 | 2,69E-05 | 0 - | 0 - | 1 - | 6,73E-05 - | 0 - | 0 - |
| 1772 | 1 | 3um_FIJJAFBF_257111 | Eukaryota | Bacillariophyta | Coscinodiscop | Leptocylindral | Leptocylindrac | Leptocylindrus | Leptocylindrus | information st | Translation, ri | COG0087 | One of the pri | 489825.LYNG | 6 | 0,632455532 | 0,00172353 | 0,000239209 | 6 - | 0,002050635 - | 5 - | 0,001428651 - | 6 - | 0,001725834 - |
| 1773 | 1 | 3um_FIJJAFBF_609352 | Bacteria | Proteobacteri | Gammaprotec | Cellvibrionales | Halieaceae | unclassified | marine gamm | unclassified | unclassified | unclassified | unclassified | unclassified | 0,4 | 0,489897949 | 4,19E-05 | 5,13E-05 | 0 - | 0 - | 0 - | 0 - | 1 - | 0,000107131 - |
| 1774 | 1 | 3um_FIJJAFBF_163703 | Bacteria | Bacteroidetes | Flavobacteriia | Flavobacterial | Flavobacteriac | Lacinutrix | Lacinutrix sp. | unclassified | unclassified | unclassified | unclassified | unclassified | 0,8 | 0,4 | 8,54E-05 | 4,31E-05 | 0 - | 0 - | 1 - | 0,000111343 - | 1 - | 0,000112087 - |
| 1775 | 1 | 3um_FIJJAFBF_177760 | Viruses | unclassified | unclassified | unclassified | Phycodnavirid | unclassified | Phaeocystis p | unclassified | unclassified | unclassified | unclassified | unclassified | 0,8 | 1,166190379 | 8,40E-05 | 0,000119837 | 0 - | 0 - | 0 - | 0 - | 0 - | 0 - |
| 1776 | 1 | 3um_FIJJAFBF_391682 | Eukaryota | unclassified | Dinophyceae | Gymnodiniale | Gymnodiniace | Amphidinium | Amphidinium | metabolism | Coenzyme tra | 1DQD8 | adenosylhomc | 3055.EDP0336 | 1 | 0,632455532 | 0,000275296 | 0,000170013 | 1 - | 0,000329728 - | 0 - | 0 - | 1 - | 0,000277502 - |
| 1777 | 1 | 3um_FIJJAFBF_26690 | Bacteria | Bacteroidetes | Flavobacteriia | Flavobacterial | Flavobacteriac | Polaribacter | Polaribacter s | unclassified | unclassified | unclassified | unclassified | unclassified | 0,6 | 0,489897949 | 0,000108586 | 8,91E-05 | 0 - | 0 - | 0 - | 0 - | 1 - | 0,000192639 - |
| 1778 | 1 | 3um_FIJJAFBF_787562 | Eukaryota | unclassified | Pelagophycea | Pelagomonad | unclassified | Aureococcus | Aureococcus a | unclassified | unclassified | unclassified | unclassified | unclassified | 1 | 0 | 0,00056243 | 5,95E-05 | 1 - | 0,000668287 - | 1 - | 0,000558705 - | 1 - | 0,000562437 - |
| 1779 | 1 | 0.2um_NOKCPHJN_281073 | Bacteria | Bacteroidetes | Flavobacteriia | Flavobacterial | unclassified | unclassified | Flavobacterial | unclassified | unclassified | unclassified | unclassified | unclassified | 0,4 | 0,489897949 | 1,27E-05 | 1,56E-05 | 0 - | 0 - | 1 - | 3,38E-05 - | 0 - | 0 - |
| 1780 | 1 | 3um_FIJJAFBF_758945 | Bacteria | Bacteroidetes | Flavobacteriia | Flavobacterial | Crocinitomica | unclassified | Crocinitomica | information st | Translation, ri | COG0087 | One of the pri | 755732.Fluta_ | 1 | 0 | 0,000301398 | 3,19E-05 | 1 - | 0,000358125 - | 1 - | 0,000299402 - | 1 - | 0,000301402 - |
| 1781 | 1 | 0.2um_NOKCPHJN_1088455 | Eukaryota | Chlorophyta | Mamiellophyc | Mamiellales | Bathycoccace | Bathycoccus | Bathycoccus p | metabolism | Energy produc | COG0055 | Produces ATP | 436017.A4RSS | 1,2 | 0,4 | 0,000697038 | 0,000172255 | 1 - | 0,000706115 - | 1 - | 0,000590329 - | 1 - | 0,000594273 - |
| 1782 | 1 | 0.2um_NOKCPHJN_678825 | Bacteria | Proteobacteri | Gammaprotec | unclassified | unclassified | unclassified | SAR86 cluster | unclassified | unclassified | unclassified | unclassified | unclassified | 0,6 | 0,489897949 | 0,000223667 | 0,000182722 | 0 - | 0 - | 1 - | 0,000376957 - | 1 - | 0,000379476 - |
| 1783 | 1 | 0.2um_NOKCPHJN_715413 | Bacteria | Proteobacteri | Alphaproteob | Pelagibacteral | unclassified | unclassified | Pelagibacteral | metabolism | Carbohydrate | 5,00E+29 | HAD-superfan | 439493.PB721 | 0,2 | 0,4 | 5,18E-05 | 0,000103596 | 1 - | 0,00025899 - | 0 - | 0 - | 0 - | 0 - |
| 1784 | 1 | 0.2um_NOKCPHJN_1096728 | Bacteria | unclassified | unclassified | unclassified | unclassified | unclassified | bacterium TM | metabolism | Lipid transpor | COG0439 | acetyl-CoA car | 355276.LBL_1 | 0,6 | 0,489897949 | 0,000118475 | 9,71E-05 | 0 - | 0 - | 1 - | 0,000209281 - | 0 - | 0 - |
| 1785 | 1 | 3um_FIJJAFBF_577836 | Eukaryota | Arthropoda | Insecta | Lepidoptera | Nymphalidae | Bicyclus | Bicyclus anyna | poorly charact | Function unkn | 140PW | NA | 400682.PAC_1 | 1 | 0 | 0,000136642 | 1,45E-05 | 1 - | 0,00016236 - | 1 - | 0,000135737 - | 1 - | 0,000136644 - |
| 1786 | 1 | 0.2um_NOKCPHJN_1033377 | Eukaryota | Apicomplexa | Aconoidasida | Haemosporida | Plasmodiidae | Plasmodium | Plasmodium v | unclassified | unclassified | unclassified | unclassified | unclassified | 0,2 | 0,4 | 2,23E-05 | 4,47E-05 | 0 - | 0 - | 1 - | 0,000111741 - | 0 - | 0 - |
| 1787 | 1 | 0.2um_NOKCPHJN_363820 | Bacteria | unclassified | unclassified | unclassified | unclassified | unclassified | uncultured ba | unclassified | unclassified | unclassified | unclassified | unclassified | 0,4 | 0,489897949 | 8,98E-05 | 0,000110071 | 0 - | 0 - | 0 - | 0 - | 1 - | 0,000229901 - |
| 1788 | 1 | 0.2um_NOKCPHJN_154905 | Eukaryota | unclassified | Oomycetes | Peronosporale | Peronosporac | Phytophthora | Phytophthora | unclassified | unclassified | unclassified | unclassified | unclassified | 4,4 | 0,489897949 | 0,000610373 | 7,93E-05 | 4 - | 0,000660911 - | 5 - | 0,000690672 - | 4 - | 0,000556229 - |
| 1789 | 10 | 0.2um_NOKCPHJN_1011604;0.2um_N | Viruses | unclassified | unclassified | unclassified | unclassified | unclassified | uncultured M | unclassified | unclassified | unclassified | unclassified | unclassified | 1 | 0 | 0,00034422 | 3,64E-05 | 1 - | 0,000409006 - | 1 - | 0,000341939 - | 1 - | 0,000344224 - |
| 1790 | 1 | 3um_FIJJAFBF_13890 | unclassified | unclassified | unclassified | unclassified | unclassified | unclassified | unclassified | unclassified | unclassified | unclassified | unclassified | unclassified | 1,4 | 0,489897949 | 0,000970798 | 0,000292302 | 1 - | 0,000840991 - | 2 - | 0,001406178 - | 1 - | 0,000707786 - |
| 1791 | 1 | 0.2um_NOKCPHJN_243505 | Eukaryota | Streptophyta | Klebsormidiop | Klebsormidial | Klebsormidiac | Klebsormidiur | Klebsormidiur | cellular proces | Signal transdu | 1BD7E | serine threoni | 39946.BGIOS | 0,2 | 0,4 | 6,54E-06 | 1,31E-05 | 0 - | 0 - | 0 - | 0 - | 0 - | 0 - |
| 1792 | 1 | 3um_FIJJAFBF_458234 | Bacteria | Proteobacteri | Alphaproteob | Rhodobactera | Rhodobactera | Marinosulfonc | Marinosulfonc | information st | Translation, ri | COG0051 | tRNA binding | 314265.R2601 | 1 | 0 | 0,00061757 | 6,53E-05 | 1 - | 0,000733806 - | 1 - | 0,00061348 - | 1 - | 0,000617578 - |
| 1793 | 1 | 3um_FIJJAFBF_254311 | Bacteria | Proteobacteri | Gammaprotec | Cellvibrionales | Halieaceae | Luminiphilus | Luminiphilus s | unclassified | unclassified | unclassified | unclassified | unclassified | 1,2 | 0,748331477 | 0,000537427 | 0,000342897 | 0 - | 0 - | 2 - | 0,000927036 - | 1 - | 0,000466614 - |
| 1794 | 1 | 3um_FIJJAFBF_715091 | Eukaryota | unclassified | Pelagophycea | Pelagomonad | unclassified | Aureococcus | Aureococcus a | unclassified | unclassified | unclassified | unclassified | unclassified | 1 | 0 | 0,000473625 | 5,01E-05 | 1 - | 0,000562768 - | 1 - | 0,000470488 - | 1 - | 0,000473631 - |
| 1795 | 1 | 0.2um_NOKCPHJN_278290 | Eukaryota | Phaeophyceae | unclassified | Ectocarpales | Ectocarpaceae | Ectocarpus | Ectocarpus sili | metabolism | Energy produc | COG0055 | Produces ATP | 35128.Thaps4 | 5 | 1,673320053 | 0,000598139 | 0,00016033 | 2 - | 0,000294678 - | 5 - | 0,000615895 - | 5 - | 0,000620009 - |
| 1796 | 1 | 0.2um_NOKCPHJN_308846 | Eukaryota | Phaeophyceae | unclassified | Ectocarpales | Ectocarpaceae | Ectocarpus | Ectocarpus sili | unclassified | unclassified | unclassified | unclassified | unclassified | 0,2 | 0,4 | 2,72E-05 | 5,43E-05 | 1 - | 0,000135841 - | 0 - | 0 - | 0 - | 0 - |
| 1797 | 1 | 3um_FIJJAFBF_263676 | Bacteria | Proteobacteri | Alphaproteob | unclassified | unclassified | Candidatus Pu | Candidatus Pu | metabolism | Carbohydrate | 0IVA1 | Photosystem I | 45157.CMV20 | 7 | 0,894427191 | 0,002371119 | 0,000180799 | 6 - | 0,002440702 - | 7 - | 0,002380568 - | 6 - | 0,002054118 - |
| 1798 | 1 | 0.2um_NOKCPHJN_876272 | Eukaryota | unclassified | Pelagophycea | Pelagomonad | unclassified | Aureococcus | Aureococcus a | cellular proces | Posttranslatio | 08V1T | peptidylprolyl | 513049.Amax | 2 | 0 | 0,000719911 | 7,61E-05 | 2 - | 0,000855408 - | 2 - | 0,000715142 - | 2 - | 0,000719919 - |
| 1799 | 1 | 0.2um_NOKCPHJN_128597 | Eukaryota | unclassified | Labyrinthulom | unclassified | Thraustochytr | Aurantiochytr | Aurantiochytr | cellular proces | Posttranslatio | KOG0417 | ubiquitin-conj | 9601.ENSPPYF | 0,8 | 0,4 | 0,00035441 | 0,000180596 | 1 - | 0,000509171 - | 1 - | 0,00042568 - | 1 - | 0,000428523 - |
| 1800 | 1 | 0.2um_NOKCPHJN_17420 | Bacteria | Proteobacteri | Gammaprotec | Alteromonada | Pseudoalteror | Pseudoalteror | Pseudoalteror | unclassified | unclassified | unclassified | unclassified | unclassified | 3 | 0,632455532 | 0,000199863 | 3,74E-05 | 3 - | 0,000239642 - | 3 - | 0,000200347 - | 2 - | 0,000134457 - |
| 1801 | 1 | 3um_FIJJAFBF_183674 | Bacteria | Bacteroidetes | Flavobacteriia | Flavobacterial | Flavobacteriac | Jejuia | Jejuia pallidilu | poorly charact | Function unkn | 11P35 | NA | 398720.MED2 | 2 | 0 | 0,000227409 | 2,40E-05 | 2 - | 0,00027021 - | 2 - | 0,000225902 - | 2 - | 0,000227411 - |
| 1802 | 1 | 3um_FIJJAFBF_535169 | Bacteria | Proteobacteri | Alphaproteob | Rhodobactera | unclassified | unclassified | Rhodobactera | unclassified | unclassified | unclassified | unclassified | unclassified | 1 | 0 | 0,000379471 | 4,01E-05 | 1 - | 0,000450893 - | 1 - | 0,000376957 - | 1 - | 0,000379476 - |
| 1803 | 1 | 3um_FIJJAFBF_07737 | Eukaryota | Bacillariophyta | Coscinodiscop | Thalassiosirale | Thalassiosirac | Thalassiosira | Thalassiosira p | unclassified | unclassified | unclassified | unclassified | unclassified | 0,2 | 0,4 | 2,20E-05 | 4,39E-05 | 0 - | 0 - | 0 - | 0 - | 0 - | 0 - |
| 1804 | 1 | 3um_FIJJAFBF_105537 | Eukaryota | Ascomycota | Saccharomyce | Saccharomyce | Saccharomyce | Saccharomyce | Saccharomyce | poorly charact | Function unkn | 0NVRZ | NA | 170187.SP_17 | 5,6 | 0,8 | 0,000965048 | 9,72E-05 | 5 - | 0,001033815 - | 5 - | 0,000864294 - | 6 - | 0,001044082 - |
| 1805 | 1 | 3um_FIJJAFBF_498449 | Bacteria | Proteobacteri | Gammaprotec | Arenicellales | Arenicellaceae | Arenicella | Arenicella xan | unclassified | unclassified | unclassified | unclassified | unclassified | 0,8 | 0,4 | 5,79E-05 | 2,92E-05 | 0 - | 0 - | 1 - | 7,54E-05 - | 1 - | 7,59E-05 - |
| 1806 | 1 | 3um_FIJJAFBF_111367 | Bacteria | Bacteroidetes | Cytophagia | Cytophagales | Cyclobacteriac | Echinicola | Echinicola pac | metabolism | Nucleotide tra | 09B0C | Provides the p | 504472.Slin_6 | 0,2 | 0,4 | 1,63E-05 | 3,26E-05 | 0 - | 0 - | 0 - | 0 - | 1 - | 8,15E-05 - |
| 1807 | 1 | 0.2um_NOKCPHJN_198182 | Bacteria | Proteobacteri | Gammaprotec | Alteromonada | Alteromonada | unclassified | Alteromonada | cellular proces | Cell wall/mem | COG2885 | Ompa motb d | 247633.GP214 | 0,6 | 0,8 | 9,06E-05 | 0,00011847 | 0 - | 0 - | 0 - | 0 - | 0 - | 0 - |
| 1808 | 1 | 0.2um_NOKCPHJN_304300 | Bacteria | Bacteroidetes | Flavobacteriia | unclassified | unclassified | unclassified | Flavobacteria | unclassified | unclassified | unclassified | unclassified | unclassified | 0,2 | 0,4 | 1,92E-05 | 3,84E-05 | 0 - | 0 - | 1 - | 9,61E-05 - | 0 - | 0 - |
| 1809 | 1 | 3um_FIJJAFBF_510254 | Bacteria | Bacteroidetes | Flavobacteriia | Flavobacterial | Flavobacteriac | Flaviramulus | Flaviramulus i | information st | Translation, ri | 05G6W | One of the pri | 983544.Lacal_ | 1,2 | 0,748331477 | 0,00037856 | 0,000234994 | 1 - | 0,000372379 - | 2 - | 0,000622636 - | 2 - | 0,000626795 - |
| 1810 | 1 | 3um_FIJJAFBF_793426 | Eukaryota | Bacillariophyta | Coscinodiscop | Thalassiosirale | Thalassiosirac | Thalassiosira | Thalassiosira p | unclassified | unclassified | unclassified | unclassified | unclassified | 0,2 | 0,4 | 1,80E-05 | 3,60E-05 | 0 - | 0 - | 0 - | 0 - | 0 - | 0 - |
| 1811 | 1 | 3um_FIJJAFBF_533445 | Bacteria | Bacteroidetes | Flavobacteriia | Flavobacterial | Flavobacteriac | Flavobacteriu | Flavobacteriu | information st | Replication, re | COG3513 | Crispr-associa | 1034807.FBFL | 0,4 | 0,489897949 | 1,63E-05 | 2,01E-05 | 0 - | 0 - | 0 - | 0 - | 1 - | 4,37E-05 - |
| 1812 | 1 | 0.2um_NOKCPHJN_760455 | unclassified | unclassified | unclassified | unclassified | unclassified | unclassified | unclassified | unclassified | unclassified | unclassified | unclassified | unclassified | 0,4 | 0,489897949 | 6,67E-05 | 8,20E-05 | 0 - | 0 - | 1 - | 0,000178276 - | 0 - | 0 - |
| 1813 | 1 | 0.2um_NOKCPHJN_515980 | unclassified | unclassified | unclassified | unclassified | unclassified | unclassified | unclassified | unclassified | unclassified | unclassified | unclassified | unclassified | 0,2 | 0,4 | 7,73E-05 | 0,000154523 | 0 - | 0 - | 0 - | 0 - | 0 - | 0 - |

| | | | | | | | | | | | | | | | | | | | | | | | | |
|---|---|---|---|---|---|---|---|---|---|---|---|---|---|---|---|---|---|---|---|---|---|---|---|---|
| 1814 | 1 | 3um_FIJJAFBF_379962 | Eukaryota | unclassified | unclassified | Isochrysidales | Noelaerhabda | Emiliania | Emiliania huxl | information st | Replication, re | 1E8C4 | Inherit from N | 436017.A4S7F | 1,2 | 0,4 | 0,000263091 | 8,95E-05 | 1 - | 0,000260795 - | 2 - | 0,000436062 - | 1 - | 0,000219488 - |
| 1815 | 1 | 3um_FIJJAFBF_556534 | Eukaryota | unclassified | unclassified | Choanoflagelli | Salpingoecida | Monosiga | Monosiga bre | unclassified | unclassified | unclassified | unclassified | unclassified | 1 | 0 | 5,21E-05 | 5,51E-06 | 1 - | 6,19E-05 - | 1 - | 5,18E-05 - | 1 - | 5,21E-05 - |
| 1816 | 1 | 0.2um_NOKCPHJN_167586 | Bacteria | Bacteroidetes | Saprospiria | Saprospirales | Saprospiracea | Aureispira | Aureispira sp. | unclassified | unclassified | unclassified | unclassified | unclassified | 2,6 | 0,489897949 | 0,000938906 | 0,000131813 | 2 - | 0,000870328 - | 2 - | 0,000727615 - | 3 - | 0,001098714 - |
| 1817 | 1 | 0.2um_NOKCPHJN_1110152 | unclassified | unclassified | unclassified | unclassified | unclassified | unclassified | unclassified | unclassified | unclassified | unclassified | unclassified | unclassified | 0,4 | 0,489897949 | 5,33E-05 | 6,60E-05 | 1 - | 0,000147921 - | 0 - | 0 - | 0 - | 0 - |
| 1818 | 1 | 3um_FIJJAFBF_20040 | Bacteria | Bacteroidetes | Flavobacteriia | Flavobacterial | Flavobacteriac | Polaribacter | Polaribacter s | metabolism | Energy produc | 099Q7 | Catalyzes the | 313594.PI23P | 0,4 | 0,489897949 | 7,44E-05 | 9,13E-05 | 0 - | 0 - | 0 - | 0 - | 0 - | 0 - |
| 1819 | 1 | 3um_FIJJAFBF_807137 | Viruses | unclassified | unclassified | unclassified | Mimiviridae | Cafeteriavirus | Cafeteria roen | unclassified | unclassified | unclassified | unclassified | unclassified | 0,8 | 0,4 | 9,27E-05 | 4,67E-05 | 0 - | 0 - | 1 - | 0,000120801 - | 1 - | 0,000121608 - |
| 1820 | 5 | 0.2um_NOKCPHJN_329666;0.2um_NO | Bacteria | Bacteroidetes | Flavobacteriia | various | various | various | various | unclassified | unclassified | unclassified | unclassified | unclassified | 0,8 | 0,4 | 0,000710946 | 0,000367635 | 1 - | 0,0010542 - | 0 - | 0 - | 1 - | 0,000887224 - |
| 1821 | 1 | 0.2um_NOKCPHJN_237169 | Bacteria | Proteobacteri | Alphaproteob | Rhodobactera | Rhodobactera | Sulfitobacter | Sulfitobacter d | unclassified | unclassified | unclassified | unclassified | unclassified | 0,8 | 0,4 | 0,000204852 | 0,000105945 | 1 - | 0,000304261 - | 1 - | 0,00025437 - | 0 - | 0 - |
| 1822 | 1 | 0.2um_NOKCPHJN_27392 | Bacteria | unclassified | unclassified | unclassified | unclassified | unclassified | uncultured ba | metabolism | Inorganic ion t | 05C27 | TonB-depende | 313596.RB250 | 26,6 | 3,826225294 | 0,001586002 | 0,000152081 | 23 - | 0,001647376 - | 22 - | 0,001317367 - | 26 - | 0,001567288 - |
| 1823 | 1 | 3um_FIJJAFBF_257122 | Bacteria | Candidatus Ke | unclassified | unclassified | unclassified | unclassified | various | information st | Translation, ri | 0EU2Y | One of the pro | 1042156.CXIV | 0,6 | 0,489897949 | 0,000364006 | 0,000297371 | 0 - | 0 - | 1 - | 0,00061348 - | 1 - | 0,000617578 - |
| 1824 | 1 | 3um_FIJJAFBF_468844 | Eukaryota | Bacillariophyta | Bacillariophyc | Naviculales | Phaeodactylac | Phaeodactylu | Phaeodactylu | metabolism | Carbohydrate | COG0057 | glyceraldehyd | 2850.Phatr327 | 0,8 | 0,4 | 0,000217297 | 0,000109515 | 0 - | 0 - | 1 - | 0,000283144 - | 1 - | 0,000285036 - |
| 1825 | 1 | 0.2um_NOKCPHJN_431905 | unclassified | unclassified | unclassified | unclassified | unclassified | unclassified | unclassified | unclassified | unclassified | unclassified | unclassified | unclassified | 0,2 | 0,4 | 1,75E-05 | 3,50E-05 | 0 - | 0 - | 1 - | 8,75E-05 - | 0 - | 0 - |
| 1826 | 1 | 0.2um_NOKCPHJN_111852 | Bacteria | Proteobacteri | Gammaprotec | unclassified | unclassified | unclassified | Gammaprotec | metabolism | Energy produc | COG0554 | Key enzyme in | 83406.HDN1F | 0,2 | 0,4 | 2,56E-05 | 5,11E-05 | 0 - | 0 - | 0 - | 0 - | 1 - | 0,000127775 - |
| 1827 | 1 | 0.2um_NOKCPHJN_832919 | unclassified | unclassified | unclassified | unclassified | unclassified | unclassified | unclassified | unclassified | unclassified | unclassified | unclassified | unclassified | 0,6 | 0,489897949 | 9,61E-05 | 7,94E-05 | 1 - | 0,00018167 - | 0 - | 0 - | 1 - | 0,000152895 - |
| 1828 | 1 | 3um_FIJJAFBF_300355 | Eukaryota | Apicomplexa | Conoidasida | Eucoccidiorida | Eimeriidae | Eimeria | Eimeria acerv | unclassified | unclassified | unclassified | unclassified | unclassified | 5 | 0 | 0,001120857 | 0,000118534 | 5 - | 0,001331818 - | 5 - | 0,001113433 - | 5 - | 0,001120871 - |
| 1829 | 1 | 3um_FIJJAFBF_186254 | Bacteria | Bacteroidetes | Flavobacteriia | Flavobacterial | Flavobacteriac | Formosa | Formosa sp. H | unclassified | unclassified | unclassified | unclassified | unclassified | 1 | 0 | 0,000216468 | 2,29E-05 | 1 - | 0,00025721 - | 1 - | 0,000215034 - | 1 - | 0,000216471 - |
| 1830 | 1 | 3um_FIJJAFBF_371763 | Bacteria | Bacteroidetes | Flavobacteriia | Flavobacterial | Flavobacteriac | Flaviramulus | Flaviramulus i | metabolism | Inorganic ion t | 05C27 | TonB-depende | 688270.Celal_ | 0,6 | 0,489897949 | 3,57E-05 | 2,93E-05 | 0 - | 0 - | 1 - | 6,20E-05 - | 1 - | 6,24E-05 - |
| 1831 | 1 | 0.2um_NOKCPHJN_586723 | Viruses | unclassified | unclassified | Caudovirales | Podoviridae | unclassified | Synechococcu | unclassified | unclassified | unclassified | unclassified | unclassified | 0,2 | 0,4 | 1,37E-05 | 2,75E-05 | 0 - | 0 - | 0 - | 0 - | 0 - | 0 - |
| 1832 | 1 | 3um_FIJJAFBF_618344 | unclassified | unclassified | unclassified | unclassified | unclassified | unclassified | unclassified | unclassified | unclassified | unclassified | unclassified | unclassified | 0,2 | 0,4 | 9,93E-05 | 0,000198597 | 0 - | 0 - | 0 - | 0 - | 0 - | 0 - |
| 1833 | 1 | 3um_FIJJAFBF_372531 | Bacteria | Bacteroidetes | Flavobacteriia | Flavobacterial | Flavobacteriac | Flavobacteriu | Flavobacteriu | metabolism | Inorganic ion t | 1AFXQ | Inherit from b | 760192.Halhy_ | 0,2 | 0,4 | 1,69E-05 | 3,37E-05 | 0 - | 0 - | 1 - | 8,43E-05 - | 0 - | 0 - |
| 1834 | 1 | 0.2um_NOKCPHJN_807312 | unclassified | unclassified | unclassified | unclassified | unclassified | unclassified | unclassified | unclassified | unclassified | unclassified | unclassified | unclassified | 0,8 | 0,4 | 0,000787255 | 0,000396768 | 0 - | 0 - | 1 - | 0,001025818 - | 1 - | 0,001032671 - |
| 1835 | 1 | 3um_FIJJAFBF_735694 | Bacteria | Cyanobacteria | unclassified | Synechococca | Synechococca | Synechococcu | Synechococcu | unclassified | unclassified | unclassified | unclassified | unclassified | 0,4 | 0,489897949 | 5,76E-05 | 7,06E-05 | 0 - | 0 - | 0 - | 0 - | 1 - | 0,000147524 - |
| 1836 | 1 | 0.2um_NOKCPHJN_220829 | Eukaryota | unclassified | unclassified | unclassified | Apusomonadi | Thecamonas | Thecamonas t | unclassified | unclassified | unclassified | unclassified | unclassified | 0,2 | 0,4 | 5,53E-06 | 1,11E-05 | 0 - | 0 - | 0 - | 0 - | 0 - | 0 - |
| 1837 | 1 | 0.2um_NOKCPHJN_1093171 | Bacteria | Actinobacteria | Actinobacteria | Corynebacteri | Mycobacteria | Mycobacteriu | Mycobacteriu | unclassified | unclassified | unclassified | unclassified | unclassified | 0,2 | 0,4 | 0,000143037 | 0,000286074 | 0 - | 0 - | 0 - | 0 - | 0 - | 0 - |
| 1838 | 1 | 0.2um_NOKCPHJN_12460 | Bacteria | Bacteroidetes | Saprospiria | Saprospirales | Lewinellaceae | Lewinella | Lewinella pers | metabolism | Carbohydrate | 0NIRC | arabinogalact | 573061.Clocel | 2,6 | 0,8 | 0,000287545 | 9,04E-05 | 2 - | 0,000263087 - | 4 - | 0,000439894 - | 3 - | 0,000332124 - |
| 1839 | 1 | 3um_FIJJAFBF_321849 | Viruses | unclassified | unclassified | unclassified | unclassified | unclassified | uncultured vir | unclassified | unclassified | unclassified | unclassified | unclassified | 0,8 | 0,4 | 9,92E-05 | 5,13E-05 | 1 - | 0,000147049 - | 0 - | 0 - | 1 - | 0,000123758 - |
| 1840 | 1 | 3um_FIJJAFBF_715093 | Eukaryota | unclassified | Pelagophycea | Pelagomonad | unclassified | Pelagomonas | uncultured Pe | unclassified | unclassified | unclassified | unclassified | unclassified | 4 | 0 | 0,00091625 | 9,69E-05 | 4 - | 0,001088701 - | 4 - | 0,000910181 - | 4 - | 0,000916261 - |
| 1841 | 1 | 3um_FIJJAFBF_04289 | Bacteria | Bacteroidetes | Flavobacteriia | Flavobacterial | Flavobacteriac | Kordia | Kordia jejudor | metabolism | Energy produc | 0P7W5 | TLC ATP/ADP t | 391587.KAOT1 | 0,4 | 0,489897949 | 2,65E-05 | 3,25E-05 | 0 - | 0 - | 1 - | 6,61E-05 - | 1 - | 6,66E-05 - |
| 1842 | 1 | 3um_FIJJAFBF_329124 | Eukaryota | Phaeophyceae | unclassified | Fucales | Sargassaceae | Sargassum | Sargassum thu | unclassified | unclassified | unclassified | unclassified | unclassified | 1,6 | 0,489897949 | 0,001091229 | 0,000397809 | 2 - | 0,001592515 - | 1 - | 0,000665691 - | 2 - | 0,001340275 - |
| 1843 | 1 | 0.2um_NOKCPHJN_109160 | Eukaryota | Chlorophyta | Chlorophycea | Sphaeropleale | Selenastracea | Monoraphidiu | Monoraphidiu | unclassified | unclassified | unclassified | unclassified | unclassified | 0,8 | 0,4 | 0,000349955 | 0,00018099 | 1 - | 0,000519779 - | 1 - | 0,000434548 - | 0 - | 0 - |
| 1844 | 1 | 0.2um_NOKCPHJN_385913 | Bacteria | Proteobacteri | Alphaproteob | Rickettsiales | unclassified | unclassified | Rickettsiales b | unclassified | unclassified | unclassified | unclassified | unclassified | 2,4 | 0,8 | 0,000150812 | 4,29E-05 | 1 - | 7,71E-05 - | 3 - | 0,000193331 - | 2 - | 0,000129749 - |
| 1845 | 1 | 0.2um_NOKCPHJN_853396 | Eukaryota | Bacillariophyta | Coscinodiscop | Thalassiosirale | Thalassiosirac | Thalassiosira | Thalassiosira p | unclassified | unclassified | unclassified | unclassified | unclassified | 0,6 | 0,489897949 | 0,000205233 | 0,000171127 | 1 - | 0,000400258 - | 1 - | 0,000334625 - | 0 - | 0 - |
| 1846 | 1 | 3um_FIJJAFBF_39858 | Bacteria | Proteobacteri | Gammaprotec | Cellvibrionales | Spongiibacter | Oceanicoccus | Oceanicoccus | information st | Translation, ri | COG0050 | This protein p | 247633.GP214 | 2 | 0 | 0,000427066 | 4,52E-05 | 2 - | 0,000507445 - | 2 - | 0,000424237 - | 2 - | 0,000427071 - |
| 1847 | 1 | 0.2um_NOKCPHJN_496300 | Bacteria | Proteobacteri | Alphaproteob | Pelagibacteral | Pelagibactera | Candidatus Pe | Candidatus Pe | metabolism | Energy produc | 01R8W | NDH-1 shuttle | 439493.PB721 | 1 | 0 | 0,000313394 | 3,31E-05 | 1 - | 0,000372379 - | 1 - | 0,000311318 - | 1 - | 0,000313398 - |
| 1848 | 1 | 0.2um_NOKCPHJN_655476 | Eukaryota | Bacillariophyta | Coscinodiscop | Thalassiosirale | Thalassiosirac | Thalassiosira | Thalassiosira c | cellular proces | Posttranslatio | KOG0179 | The proteasor | 35128.Thaps2 | 0,4 | 0,489897949 | 0,000341857 | 0,000418883 | 0 - | 0 - | 0 - | 0 - | 1 - | 0,000874902 - |
| 1849 | 1 | 0.2um_NOKCPHJN_195247 | Bacteria | Firmicutes | Bacilli | Lactobacillales | Lactobacillace | Lactobacillus | Lactobacillus k | unclassified | unclassified | unclassified | unclassified | unclassified | 0,2 | 0,4 | 1,72E-05 | 3,44E-05 | 0 - | 0 - | 0 - | 0 - | 1 - | 8,59E-05 - |
| 1850 | 1 | 3um_FIJJAFBF_337197 | unclassified | unclassified | unclassified | unclassified | unclassified | unclassified | unclassified | unclassified | unclassified | unclassified | unclassified | unclassified | 0,2 | 0,4 | 0,000393706 | 0,000787412 | 0 - | 0 - | 0 - | 0 - | 1 - | 0,001968529 - |
| 1851 | 1 | 0.2um_NOKCPHJN_261541 | Bacteria | Bacteroidetes | Flavobacteriia | Flavobacterial | Flavobacteriac | Lutibacter | Lutibacter pro | metabolism | Inorganic ion t | 05C27 | TonB-depende | 487797.Flav30 | 0,4 | 0,8 | 2,32E-05 | 4,63E-05 | 0 - | 0 - | 2 - | 0,000115772 - | 0 - | 0 - |
| 1852 | 1 | 3um_FIJJAFBF_722163 | Eukaryota | unclassified | Dinophyceae | Suessiales | Symbiodiniace | Symbiodinium | Symbiodinium | unclassified | unclassified | unclassified | unclassified | unclassified | 5,6 | 1,019803903 | 0,001684445 | 0,000235917 | 5 - | 0,001807927 - | 6 - | 0,001813766 - | 4 - | 0,001217255 - |
| 1853 | 1 | 0.2um_NOKCPHJN_727290 | Bacteria | unclassified | unclassified | unclassified | unclassified | unclassified | uncultured ba | unclassified | unclassified | unclassified | unclassified | unclassified | 0,8 | 0,4 | 6,13E-05 | 3,09E-05 | 0 - | 0 - | 1 - | 7,99E-05 - | 1 - | 8,05E-05 - |
| 1854 | 1 | 3um_FIJJAFBF_436969 | Eukaryota | unclassified | unclassified | unclassified | Apusomonadi | Thecamonas | Thecamonas t | unclassified | unclassified | unclassified | unclassified | unclassified | 0,2 | 0,4 | 4,74E-05 | 9,47E-05 | 0 - | 0 - | 0 - | 0 - | 0 - | 0 - |
| 1855 | 1 | 3um_FIJJAFBF_594155 | Eukaryota | unclassified | Florideophyce | Ceramiales | Delesseriacea | Acrosorium | Acrosorium ci | unclassified | unclassified | unclassified | unclassified | unclassified | 0,4 | 0,489897949 | 0,000263321 | 0,000323145 | 0 - | 0 - | 0 - | 0 - | 0 - | 0 - |
| 1856 | 1 | 0.2um_NOKCPHJN_1052523 | Eukaryota | unclassified | Oomycetes | Peronosporale | Peronosporac | Plasmopara | Plasmopara ha | unclassified | unclassified | unclassified | unclassified | unclassified | 1 | 0 | 0,000425623 | 4,50E-05 | 1 - | 0,000505731 - | 1 - | 0,000422804 - | 1 - | 0,000425628 - |
| 1857 | 1 | 3um_FIJJAFBF_102619 | unclassified | unclassified | unclassified | unclassified | unclassified | unclassified | unclassified | unclassified | unclassified | unclassified | unclassified | unclassified | 1 | 0,894427191 | 0,000199433 | 0,000178298 | 0 - | 0 - | 2 - | 0,000424237 - | 0 - | 0 - |
| 1858 | 1 | 0.2um_NOKCPHJN_704660 | Eukaryota | unclassified | unclassified | Prymnesiales | Chrysochromu | Chrysochromu | Chrysochromu | unclassified | unclassified | unclassified | unclassified | unclassified | 1 | 0 | 0,00071582 | 7,57E-05 | 1 - | 0,000850548 - | 1 - | 0,000711079 - | 1 - | 0,000715829 - |
| 1859 | 1 | 3um_FIJJAFBF_131607 | Bacteria | Proteobacteri | Gammaprotec | Cellvibrionales | Porticoccacea | unclassified | SAR92 bacteri | poorly charact | Function unkn | 16R8I | Lipoprotein | 377629.TERTU | 1,4 | 0,489897949 | 9,46E-05 | 3,21E-05 | 1 - | 8,08E-05 - | 1 - | 6,76E-05 - | 2 - | 0,000136054 - |
| 1860 | 1 | 0.2um_NOKCPHJN_27620 | Bacteria | Actinobacteria | Actinobacteria | Micrococcales | Microbacteria | Candidatus Ac | Candidatus Ac | metabolism | Amino acid tra | 00DZF | branched-chai | 312284.A20C1 | 1 | 0 | 0,000148918 | 1,57E-05 | 1 - | 0,000176946 - | 1 - | 0,000147931 - | 1 - | 0,000148919 - |
| 1861 | 1 | 3um_FIJJAFBF_683762 | Bacteria | Proteobacteri | Gammaprotec | unclassified | unclassified | unclassified | Gammaprotec | cellular proces | Intracellular tr | 0QITE | Outer membra | 207949.RED65 | 1,6 | 0,489897949 | 0,000212517 | 7,34E-05 | 2 - | 0,000313829 - | 2 - | 0,000262369 - | 1 - | 0,000132061 - |
| 1862 | 1 | 3um_FIJJAFBF_469902 | Bacteria | Proteobacteri | Alphaproteob | Rhodobactera | unclassified | unclassified | Rhodobactera | information st | Translation, ri | 17FUG | Binds directly | 439496.RBY4I | 0,2 | 0,4 | 0,000125217 | 0,000250434 | 0 - | 0 - | 0 - | 0 - | 0 - | 0 - |
| 1863 | 1 | 3um_FIJJAFBF_176419 | Bacteria | Bacteroidetes | Flavobacteriia | Flavobacterial | Crocinitomica | unclassified | Crocinitomica | information st | Translation, ri | COG1185 | Involved in mF | 743722.Sph21 | 0,2 | 0,4 | 1,66E-05 | 3,33E-05 | 0 - | 0 - | 0 - | 0 - | 0 - | 0 - |
| 1864 | 1 | 3um_FIJJAFBF_712785 | Eukaryota | unclassified | Pelagophycea | Pelagomonad | unclassified | Aureococcus | Aureococcus a | unclassified | unclassified | unclassified | unclassified | unclassified | 1,6 | 0,489897949 | 0,00020886 | 5,67E-05 | 1 - | 0,000157908 - | 2 - | 0,000264029 - | 2 - | 0,000265793 - |
| 1865 | 1 | 0.2um_NOKCPHJN_148310 | Bacteria | Bacteroidetes | Flavobacteriia | Flavobacterial | Flavobacteriac | Flavobacteriu | Flavobacteriu | poorly charact | Function unkn | 0XZ2H | Membrane as | 353153.XP_80 | 0,6 | 0,489897949 | 6,26E-05 | 5,14E-05 | 0 - | 0 - | 0 - | 0 - | 1 - | 0,000111099 - |
| 1866 | 4 | 0.2um_NOKCPHJN_727195;0.2um_NO | Eukaryota | unclassified | various | various | various | various | various | various | various | various | various | various | 0,8 | 0,4 | 0,00025275 | 0,000127383 | 0 - | 0 - | 1 - | 0,000329342 - | 1 - | 0,000331542 - |
| 1867 | 1 | 0.2um_NOKCPHJN_579370 | Bacteria | Bacteroidetes | Flavobacteriia | Flavobacterial | Flavobacteriac | Polaribacter | Polaribacter fi | poorly charact | Function unkn | 0PDH4 | NA | 313598.MED1 | 0,2 | 0,4 | 3,45E-05 | 6,90E-05 | 0 - | 0 - | 0 - | 0 - | 1 - | 0,000172583 - |
| 1868 | 1 | 0.2um_NOKCPHJN_216476 | unclassified | unclassified | unclassified | unclassified | unclassified | unclassified | unclassified | information st | Transcription | KOG2252 | homeobox | 7460.GB13918 | 0,6 | 0,489897949 | 7,70E-06 | 6,31E-06 | 0 - | 0 - | 0 - | 0 - | 1 - | 1,37E-05 - |
| 1869 | 1 | 0.2um_NOKCPHJN_359403 | Eukaryota | unclassified | unclassified | Prymnesiales | Chrysochromu | Chrysochromu | Chrysochromu | metabolism | Lipid transpor | COG0657 | alpha beta hyc | 706436.HMPR | 0,4 | 0,489897949 | 5,90E-05 | 7,38E-05 | 1 - | 0,000170886 - | 0 - | 0 - | 0 - | 0 - |
| 1870 | 1 | 3um_FIJJAFBF_420046 | unclassified | unclassified | unclassified | unclassified | unclassified | unclassified | unclassified | unclassified | unclassified | unclassified | unclassified | unclassified | 1 | 0 | 0,000777681 | 8,22E-05 | 1 - | 0,000924052 - | 1 - | 0,00077253 - | 1 - | 0,000777691 - |
| 1871 | 1 | 0.2um_NOKCPHJN_445514 | Eukaryota | unclassified | unclassified | Isochrysidales | Noelaerhabda | Emiliania | Emiliania huxl | poorly charact | Function unkn | COG2340 | Scp-like extrac | 6085.XP_0021 | 1 | 0 | 0,000103606 | 1,10E-05 | 1 - | 0,000123106 - | 1 - | 0,000102919 - | 1 - | 0,000103607 - |
| 1872 | 1 | 3um_FIJJAFBF_316185 | unclassified | unclassified | unclassified | unclassified | unclassified | unclassified | unclassified | unclassified | unclassified | unclassified | unclassified | unclassified | 0,4 | 0,8 | 5,01E-05 | 0,00010012 | 0 - | 0 - | 2 - | 0,0002503 - | 0 - | 0 - |
| 1873 | 1 | 3um_FIJJAFBF_212745 | Bacteria | Bacteroidetes | Flavobacteriia | Flavobacterial | Flavobacteriac | Polaribacter | Polaribacter ta | unclassified | unclassified | unclassified | unclassified | unclassified | 1 | 0 | 0,000294356 | 3,11E-05 | 1 - | 0,000349758 - | 1 - | 0,000292406 - | 1 - | 0,00029436 - |
| 1874 | 1 | 3um_FIJJAFBF_98072 | Bacteria | Proteobacteri | Gammaprotec | unclassified | unclassified | unclassified | OM182 bacte | cellular proces | Cell wall/mem | 0QTSX | Peptidoglycan | 351348.Maqu | 0,6 | 0,489897949 | 0,000202289 | 0,000166048 | 0 - | 0 - | 1 - | 0,000351544 - | 1 - | 0,000353893 - |
| 1875 | 1 | 3um_FIJJAFBF_254291 | Bacteria | Proteobacteri | Gammaprotec | Cellvibrionales | Spongiibacter | Oceanicoccus | Oceanicoccus | information st | Translation, ri | 16Q16 | Binds the lowe | 247634.GPB21 | 1 | 0 | 0,000283749 | 3,00E-05 | 1 - | 0,000337154 - | 1 - | 0,000281869 - | 1 - | 0,000283752 - |
| 1876 | 1 | 0.2um_NOKCPHJN_564928 | Eukaryota | Phaeophyceae | unclassified | Ectocarpales | Ectocarpaceae | Ectocarpus | Ectocarpus sili | information st | Chromatin str | KOG1744 | Core compone | 2850.Phatr543 | 1 | 0 | 0,000529346 | 5,60E-05 | 1 - | 0,000628976 - | 1 - | 0,00052584 - | 1 - | 0,000529352 - |
| 1877 | 1 | 0.2um_NOKCPHJN_891697 | Eukaryota | unclassified | Pelagophycea | Pelagomonad | unclassified | Aureococcus | Aureococcus a | unclassified | unclassified | unclassified | unclassified | unclassified | 17,6 | 1,019803903 | 0,006201327 | 0,000455311 | 16 - | 0,006727927 - | 19 - | 0,006679345 - | 17 - | 0,006016179 - |
| 1878 | 1 | 3um_FIJJAFBF_663803 | Bacteria | Bacteroidetes | Flavobacteriia | Flavobacterial | Flavobacteriac | Polaribacter | unclassified | information st | Replication, re | 0PB0H | DNA-binding p | 313598.MED1 | 0,4 | 0,489897949 | 0,000273485 | 0,000335107 | 0 - | 0 - | 0 - | 0 - | 1 - | 0,000699922 - |
| 1879 | 1 | 0.2um_NOKCPHJN_265381 | Bacteria | Proteobacteri | Betaproteoba | Burkholderiale | Burkholderiac | Caballeronia | Caballeronia s | cellular proces | Cell wall/mem | 16PZG | Glycosyltransf | 945681.APO_ | 0,6 | 0,489897949 | 2,35E-05 | 1,96E-05 | 1 - | 4,58E-05 - | 0 - | 0 - | 1 - | 3,86E-05 - |
| 1880 | 1 | 3um_FIJJAFBF_470660 | Eukaryota | unclassified | Cryptophyta | Pyrenomonad | Geminigerace | Guillardia | Guillardia thet | metabolism | Energy produc | COG0711 | F(1)F(0) ATP s | 197221.tlr043 | 0,2 | 0,4 | 6,80E-05 | 0,000136032 | 0 - | 0 - | 1 - | 0,000340081 - | 0 - | 0 - |
| 1881 | 2 | 0.2um_NOKCPHJN_1009653;3um_FIJJA | Eukaryota | unclassified | various | various | various | various | various | various | various | various | various | various | 0,2 | 0,4 | 3,43E-05 | 6,87E-05 | 0 - | 0 - | 0 - | 0 - | 0 - | 0 - |
| 1882 | 1 | 3um_FIJJAFBF_214089 | Bacteria | Proteobacteri | Gammaprotec | Thiotrichales | Francisellacea | unclassified | Francisella en | metabolism | Nucleotide tra | 16Q4Q | Catalyzes the | 573569.F7308 | 0,2 | 0,4 | 6,92E-05 | 0,000138446 | 0 - | 0 - | 0 - | 0 - | 1 - | 0,000346115 - |
| 1883 | 1 | 0.2um_NOKCPHJN_36238 | Eukaryota | unclassified | Cryptophyta | Pyrenomonad | Geminigerace | Teleaulax | Teleaulax amp | metabolism | Carbohydrate | COG1850 | RuBisCO catal | 45157.CMV01 | 2 | 0 | 0,000258165 | 2,73E-05 | 2 - | 0,000306755 - | 2 - | 0,000256455 - | 2 - | 0,000258168 - |
| 1884 | 1 | 0.2um_NOKCPHJN_27322 | Bacteria | Bacteroidetes | Flavobacteriia | Flavobacterial | Flavobacteriac | Formosa | Formosa sp. H | metabolism | Inorganic ion t | 05C27 | TonB-depende | 742766.HMPR | 13,8 | 3,370459909 | 0,000804524 | 0,000130768 | 8 - | 0,000568647 - | 13 - | 0,00077253 - | 14 - | 0,000837513 - |
| 1885 | 1 | 3um_FIJJAFBF_37832 | Bacteria | Proteobacteri | unclassified | unclassified | unclassified | unclassified | Proteobacteri | unclassified | unclassified | unclassified | unclassified | unclassified | 2,8 | 0,4 | 0,000555933 | 4,85E-05 | 2 - | 0,000478263 - | 3 - | 0,00059976 - | 3 - | 0,000603766 - |
| 1886 | 1 | 0.2um_NOKCPHJN_340558 | Eukaryota | Phaeophyceae | unclassified | Ectocarpales | Ectocarpaceae | Ectocarpus | Ectocarpus sili | cellular proces | Cytoskeleton | COG5245 | heavy chain | 65071.PYU1_T | 0,2 | 0,4 | 5,32E-06 | 1,06E-05 | 0 - | 0 - | 0 - | 0 - | 0 - | 0 - |
| 1887 | 1 | 0.2um_NOKCPHJN_613335 | Eukaryota | Chlorophyta | Chlorophycea | Chlamydomor | Tetrabaenace | Tetrabaena | Tetrabaena so | unclassified | unclassified | unclassified | unclassified | unclassified | 0,2 | 0,4 | 2,48E-05 | 4,96E-05 | 1 - | 0,000123921 - | 0 - | 0 - | 0 - | 0 - |
| 1888 | 1 | 3um_FIJJAFBF_264641 | Bacteria | Bacteroidetes | Flavobacteriia | Flavobacterial | Flavobacteriac | Polaribacter | Polaribacter ta | poorly charact | Function unkn | 05END | lipolytic prote | 313598.MED1 | 5,4 | 0,489897949 | 0,000674105 | 0,00012419 | 6 - | 0,000884034 - | 6 - | 0,000739074 - | 5 - | 0,000620009 - |
| 1889 | 1 | 3um_FIJJAFBF_270756 | Bacteria | Bacteroidetes | Flavobacteriia | Flavobacterial | Flavobacteriac | Lacinutrix | Lacinutrix him | information st | Translation, ri | 08YY1 | This protein is | 391603.FBALC | 0,6 | 0,489897949 | 0,000320075 | 0,000261481 | 0 - | 0 - | 1 - | 0,000539439 - | 1 - | 0,000543043 - |
| 1890 | 1 | 0.2um_NOKCPHJN_161983 | Eukaryota | unclassified | Cryptophyta | Pyrenomonad | Geminigerace | Guillardia | Guillardia thet | metabolism | Energy produc | 0IFIU | Pyrophosphat | 65071.PYU1_T | 1,2 | 0,4 | 8,19E-05 | 2,82E-05 | 1 - | 8,11E-05 - | 1 - | 6,78E-05 - | 2 - | 0,000136496 - |
| 1891 | 1 | 3um_FIJJAFBF_128453 | Bacteria | Bacteroidetes | Flavobacteriia | Flavobacterial | Flavobacteriac | Polaribacter | Polaribacter s | poorly charact | Function unkn | 11W7M | NA | 398720.MED2 | 0,8 | 0,4 | 0,000307837 | 0,000155146 | 0 - | 0 - | 1 - | 0,000401121 - | 1 - | 0,000403801 - |
| 1892 | 1 | 3um_FIJJAFBF_753878 | Bacteria | Proteobacteri | Alphaproteob | Rhodospirillale | Rhodospirillac | unclassified | Rhodospirillac | metabolism | Inorganic ion t | COG0841 | acriflavin resis | 391593.RCCS2 | 0,6 | 0,489897949 | 3,95E-05 | 3,24E-05 | 0 - | 0 - | 0 - | 0 - | 1 - | 7,01E-05 - |
| 1893 | 1 | 3um_FIJJAFBF_01994 | Bacteria | Planctomycete | Planctomyceti | Planctomyceta | Planctomyceta | unclassified | Planctomyceta | unclassified | unclassified | unclassified | unclassified | unclassified | 3,4 | 1,2 | 0,001286086 | 0,000417479 | 1 - | 0,000464896 - | 4 - | 0,001554656 - | 4 - | 0,001565042 - |
| 1894 | 1 | 3um_FIJJAFBF_540030 | Bacteria | Spirochaetes | Spirochaetia | unclassified | Leptospiracea | Leptospira | Leptospira ter | unclassified | unclassified | unclassified | unclassified | unclassified | 0,6 | 0,489897949 | 0,000163977 | 0,000136954 | 1 - | 0,000324018 - | 0 - | 0 - | 0 - | 0 - |
| 1895 | 1 | 3um_FIJJAFBF_161250 | Bacteria | Proteobacteri | Alphaproteob | Rhodospirillale | unclassified | unclassified | Rhodospirillale | unclassified | unclassified | unclassified | unclassified | unclassified | 0,8 | 0,4 | 9,94E-05 | 5,07E-05 | 1 - | 0,00014284 - | 1 - | 0,000119418 - | 1 - | 0,000120216 - |
| 1896 | 1 | 3um_FIJJAFBF_476392 | Bacteria | Proteobacteri | Gammaprotec | Oceanospirilla | Halomonadac | Halomonas | Halomonas sp | metabolism | Coenzyme tra | 0CZE3 | Together with | 517418.Ctha_ | 1 | 0 | 0,000391256 | 4,14E-05 | 1 - | 0,000464896 - | 1 - | 0,000388664 - | 1 - | 0,00039126 - |
| 1897 | 1 | 0.2um_NOKCPHJN_34980 | Archaea | Euryarchaeota | unclassified | unclassified | unclassified | unclassified | Euryarchaeota | unclassified | unclassified | unclassified | unclassified | unclassified | 0,2 | 0,4 | 5,65E-06 | 1,13E-05 | 0 - | 0 - | 0 - | 0 - | 1 - | 2,82E-05 - |
| 1898 | 1 | 3um_FIJJAFBF_323161 | Bacteria | Bacteroidetes | Cytophagia | Cytophagales | Flammeovirga | unclassified | Flammeovirga | information st | Translation, ri | 0FX08 | Interacts with | 643867.Ftrac_ | 0,2 | 0,4 | 9,93E-05 | 0,000198651 | 0 - | 0 - | 1 - | 0,000496626 - | 0 - | 0 - |
| 1899 | 1 | 0.2um_NOKCPHJN_318710 | Eukaryota | unclassified | Labyrinthulom | unclassified | Thraustochytr | Aurantiochytr | Aurantiochytr | unclassified | unclassified | unclassified | unclassified | unclassified | 0,2 | 0,4 | 4,96E-06 | 9,93E-06 | 0 - | 0 - | 1 - | 2,48E-05 - | 0 - | 0 - |
| 1900 | 1 | 3um_FIJJAFBF_532958 | unclassified | unclassified | unclassified | unclassified | unclassified | unclassified | unclassified | unclassified | unclassified | unclassified | unclassified | unclassified | 0,2 | 0,4 | 5,57E-05 | 0,000111492 | 0 - | 0 - | 0 - | 0 - | 1 - | 0,00027873 - |
| 1901 | 2 | 0.2um_NOKCPHJN_214009;0.2um_NO | Eukaryota | Chlorophyta | Chlorophycea | Chlamydomor | Chlamydomor | Chlamydomor | Chlamydomor | cellular proces | Posttranslatio | KOG0543 | Peptidyl-proly | 35128.Thaps1 | 1 | 0 | 0,000247028 | 2,61E-05 | 1 - | 0,000293522 - | 1 - | 0,000245392 - | 1 - | 0,000247031 - |
| 1902 | 1 | 3um_FIJJAFBF_731710 | Eukaryota | Phaeophyceae | unclassified | Ectocarpales | Ectocarpaceae | Ectocarpus | Ectocarpus sili | unclassified | unclassified | unclassified | unclassified | unclassified | 1 | 0 | 0,000707777 | 7,48E-05 | 1 - | 0,000840991 - | 1 - | 0,000703089 - | 1 - | 0,000707786 - |
| 1903 | 1 | 3um_FIJJAFBF_64397 | Eukaryota | Bacillariophyta | Coscinodiscop | Thalassiosirale | Thalassiosirac | Thalassiosira | Thalassiosira p | unclassified | unclassified | unclassified | unclassified | unclassified | 0,6 | 0,489897949 | 0,00024263 | 0,000198921 | 0 - | 0 - | 1 - | 0,000428595 - | 0 - | 0 - |
| 1904 | 1 | 3um_FIJJAFBF_115134 | Eukaryota | unclassified | Labyrinthulom | unclassified | Thraustochytr | Aurantiochytr | Aurantiochytr | unclassified | unclassified | unclassified | unclassified | unclassified | 0,4 | 0,489897949 | 0,000298971 | 0,000366167 | 0 - | 0 - | 1 - | 0,00074494 - | 1 - | 0,000749916 - |
| 1905 | 1 | 0.2um_NOKCPHJN_619316 | Eukaryota | Bacillariophyta | Bacillariophyc | Bacillariales | Bacillariaceae | Fragilariopsis | Fragilariopsis c | unclassified | unclassified | unclassified | unclassified | unclassified | 3 | 0,632455532 | 0,00199023 | 0,000628661 | 4 - | 0,003086523 - | 3 - | 0,001935307 - | 3 - | 0,001948235 - |
| 1906 | 1 | 0.2um_NOKCPHJN_557642 | Bacteria | Proteobacteri | Gammaprotec | Xanthomonad | Xanthomonad | Xanthomonas | Xanthomonas | poorly charact | Function unkn | 0NVRZ | NA | 170187.SP_17 | 1,4 | 0,8 | 0,000230515 | 0,000130262 | 0 - | 0 - | 2 - | 0,000345718 - | 1 - | 0,000174014 - |
| 1907 | 1 | 3um_FIJJAFBF_737098 | Bacteria | Bacteroidetes | Flavobacteriia | Flavobacterial | Flavobacteriac | Polaribacter | Polaribacter g | cellular proces | Signal transdu | COG3275 | Histidine kinas | 50743.SCB49_ | 0,2 | 0,4 | 2,42E-05 | 4,84E-05 | 1 - | 0,000121114 - | 0 - | 0 - | 0 - | 0 - |
| 1908 | 1 | 3um_FIJJAFBF_157376 | Bacteria | Bacteroidetes | Flavobacteriia | Flavobacterial | Flavobacteriac | Algibacter | Algibacter aqu | information st | Translation, ri | COG0052 | 30S ribosomal | 487797.Flav30 | 0,2 | 0,4 | 4,23E-05 | 8,46E-05 | 0 - | 0 - | 0 - | 0 - | 0 - | 0 - |
| 1909 | 1 | 3um_FIJJAFBF_201379 | Bacteria | Bacteroidetes | Flavobacteriia | Flavobacterial | Flavobacteriac | Polaribacter | Polaribacter s | unclassified | unclassified | unclassified | unclassified | unclassified | 1,6 | 1,2 | 0,000329265 | 0,000238169 | 0 - | 0 - | 1 - | 0,000219561 - | 1 - | 0,000221028 - |
| 1910 | 1 | 0.2um_NOKCPHJN_194408 | Bacteria | Proteobacteri | Gammaprotec | Oceanospirilla | Hahellaceae | Hahella | Hahella sp. CC | cellular proces | Cell motility | COG1749 | flagellar hook | 349521.HCH_ | 0,2 | 0,4 | 1,70E-05 | 3,40E-05 | 0 - | 0 - | 0 - | 0 - | 0 - | 0 - |
| 1911 | 1 | 0.2um_NOKCPHJN_704656 | Eukaryota | unclassified | unclassified | Prymnesiales | Chrysochromu | Chrysochromu | Chrysochromu | unclassified | unclassified | unclassified | unclassified | unclassified | 1 | 0 | 0,000529346 | 5,60E-05 | 1 - | 0,000628976 - | 1 - | 0,00052584 - | 1 - | 0,000529352 - |
| 1912 | 1 | 3um_FIJJAFBF_111335 | Viruses | unclassified | unclassified | unclassified | Phycodnavirid | unclassified | Chrysochromu | unclassified | unclassified | unclassified | unclassified | unclassified | 1,4 | 0,8 | 0,000300167 | 0,000169622 | 0 - | 0 - | 2 - | 0,000450179 - | 1 - | 0,000226593 - |
| 1913 | 1 | 0.2um_NOKCPHJN_470890 | unclassified | unclassified | unclassified | unclassified | unclassified | unclassified | unclassified | unclassified | unclassified | unclassified | unclassified | unclassified | 1,6 | 0,489897949 | 0,000652316 | 0,000167098 | 1 - | 0,000495683 - | 1 - | 0,000414403 - | 2 - | 0,000834344 - |
| 1914 | 1 | 3um_FIJJAFBF_289952 | Bacteria | Proteobacteri | Gammaprotec | Alteromonada | Alteromonada | Alteromonas | Alteromonas r | cellular proces | Intracellular tr | 08ZUW | Major pilin sul | 697282.Mettu | 1,2 | 0,748331477 | 0,000518225 | 0,000319051 | 0 - | 0 - | 1 - | 0,000453441 - | 2 - | 0,000912941 - |

| | | | | | | | | | | | | | | | | | | | | | | | | |
|---|---|---|---|---|---|---|---|---|---|---|---|---|---|---|---|---|---|---|---|---|---|---|---|---|
| 1915 | 1 | 3um_FIJJAFBF_382743 | Viruses | unclassified | unclassified | unclassified | Phycodnavirid | unclassified | Chrysochromu | unclassified | unclassified | unclassified | unclassified | unclassified | 1,8 | 0,748331477 | 0,000566373 | 0,000188646 | 1 - | 0,000387814 - | 2 - | 0,000648445 - | 1 - | 0,000326388 - |
| 1916 | 1 | 3um_FIJJAFBF_783853 | Bacteria | Bacteroidetes | Flavobacteriia | Flavobacterial | Flavobacteria | Polaribacter | Polaribacter s | unclassified | unclassified | unclassified | unclassified | unclassified | 3,4 | 0,489897949 | 0,000250456 | 2,43E-05 | 3 - | 0,000265419 - | 3 - | 0,000221897 - | 3 - | 0,000223379 - |
| 1917 | 1 | 0.2um_NOKCPHJN_312903 | Bacteria | Proteobacteri | Alphaproteob | Rhodobactera | Rhodobactera | Cribrihabitans | Cribrihabitans | metabolism | Inorganic ion t | 05C2T | Binding-protei | 246200.SPO15 | 0,4 | 0,489897949 | 8,76E-05 | 0,00010837 | 1 - | 0,000243014 - | 0 - | 0 - | 0 - | 0 - |
| 1918 | 1 | 3um_FIJJAFBF_532806 | Bacteria | Proteobacteri | Alphaproteob | Rhodobactera | Rhodobactera | unclassified | Rhodobactera | unclassified | unclassified | unclassified | unclassified | unclassified | 0,2 | 0,4 | 2,83E-05 | 5,65E-05 | 0 - | 0 - | 1 - | 0,000141253 - | 0 - | 0 - |
| 1919 | 1 | 3um_FIJJAFBF_652362 | Bacteria | Bacteroidetes | Flavobacteriia | Flavobacterial | Flavobacteria | Confluentibac | Confluentibac | unclassified | unclassified | unclassified | unclassified | unclassified | 0,6 | 0,489897949 | 0,000129657 | 0,000108111 | 1 - | 0,000252865 - | 1 - | 0,000211402 - | 0 - | 0 - |
| 1920 | 1 | 3um_FIJJAFBF_26279 | Bacteria | Bacteroidetes | Flavobacteriia | Flavobacterial | Flavobacteria | Flavobacteriu | Flavobacteriu | information st | Replication, re | 08IYB | Dead deah bo | 313628.LNTAF | 1 | 0 | 7,45E-05 | 7,88E-06 | 1 - | 8,86E-05 - | 1 - | 7,41E-05 - | 1 - | 7,45E-05 - |
| 1921 | 1 | 0.2um_NOKCPHJN_909610 | Eukaryota | Phaeophyceae | unclassified | Ectocarpales | Ectocarpaceae | Ectocarpus | Ectocarpus sili | unclassified | unclassified | unclassified | unclassified | unclassified | 3 | 0,632455532 | 0,000316607 | 7,96E-05 | 3 - | 0,000372998 - | 4 - | 0,00041578 - | 3 - | 0,000313918 - |
| 1922 | 4 | 0.2um_NOKCPHJN_1034679;0.2um_NO | Bacteria | Proteobacteri | Alphaproteob | various | various | various | various | unclassified | unclassified | unclassified | unclassified | unclassified | 0,2 | 0,4 | 8,18E-05 | 0,000163618 | 0 - | 0 - | 0 - | 0 - | 1 - | 0,000409045 - |
| 1923 | 1 | 3um_FIJJAFBF_65172 | Bacteria | Bacteroidetes | Flavobacteriia | Flavobacterial | Flavobacteria | Mesoflavibact | Mesoflavibact | poorly charact | Function unkn | 0PB02 | FN3 | 216432.CA255 | 1,4 | 0,489897949 | 0,000191088 | 7,63E-05 | 2 - | 0,000321928 - | 1 - | 0,00013457 - | 1 - | 0,000135469 - |
| 1924 | 1 | 3um_FIJJAFBF_609869 | Eukaryota | Bacillariophyta | Coscinodiscop | Thalassiosirale | Thalassiosirac | Thalassiosira | Thalassiosira | unclassified | unclassified | unclassified | unclassified | unclassified | 1,6 | 0,489897949 | 0,00039301 | 0,000143272 | 2 - | 0,000573549 - | 1 - | 0,000239751 - | 2 - | 0,000482705 - |
| 1925 | 2 | 0.2um_NOKCPHJN_593480;3um_FIJJAF | various | various | various | various | various | various | various | unclassified | unclassified | unclassified | unclassified | unclassified | 0,2 | 0,4 | 1,40E-05 | 2,79E-05 | 0 - | 0 - | 0 - | 0 - | 1 - | 6,98E-05 - |
| 1926 | 1 | 3um_FIJJAFBF_506812 | Bacteria | Bacteroidetes | Flavobacteriia | Flavobacterial | Flavobacteria | Formosa | Formosa sp. H | metabolism | Carbohydrate | 0P8NX | Glyceraldehyd | 1046627.BZAF | 0,2 | 0,4 | 3,82E-05 | 7,64E-05 | 0 - | 0 - | 0 - | 0 - | 1 - | 0,000190888 - |
| 1927 | 1 | 0.2um_NOKCPHJN_442404 | Eukaryota | unclassified | Pelagophycea | Pelagomonada | unclassified | Aureococcus | Aureococcus a | unclassified | unclassified | unclassified | unclassified | unclassified | 0,2 | 0,4 | 1,09E-05 | 2,18E-05 | 1 - | 5,46E-05 - | 0 - | 0 - | 0 - | 0 - |
| 1928 | 1 | 0.2um_NOKCPHJN_136446 | Eukaryota | Echinodermat | Echinoidea | Echinoida | Strongylocent | Strongylocent | Strongylocent | cellular proces | Cell wall/mem | 12RRB | ankyrin 3, nod | 7668.SPU_015 | 1 | 0 | 7,12E-05 | 7,53E-06 | 1 - | 8,46E-05 - | 1 - | 7,07E-05 - | 1 - | 7,12E-05 - |
| 1929 | 1 | 3um_FIJJAFBF_247264 | unclassified | unclassified | unclassified | unclassified | unclassified | unclassified | unclassified | unclassified | unclassified | unclassified | unclassified | unclassified | 2 | 0 | 0,001399826 | 0,000148035 | 2 - | 0,001663293 - | 2 - | 0,001390554 - | 2 - | 0,001399843 - |
| 1930 | 1 | 0.2um_NOKCPHJN_1031253 | Eukaryota | Chordata | unclassified | unclassified | Branchiostom | Branchiostom | Branchiostom | unclassified | unclassified | unclassified | unclassified | unclassified | 0,8 | 0,4 | 0,000350529 | 0,000176663 | 0 - | 0 - | 1 - | 0,000456751 - | 1 - | 0,000459802 - |
| 1931 | 1 | 0.2um_NOKCPHJN_1084528 | Eukaryota | unclassified | Oomycetes | Saprolegniales | Saprolegniace | Saprolegnia | Saprolegnia pa | unclassified | unclassified | unclassified | unclassified | unclassified | 1 | 0,632455532 | 0,000135777 | 7,80E-05 | 1 - | 0,000165593 - | 0 - | 0 - | 1 - | 0,000139365 - |
| 1932 | 1 | 0.2um_NOKCPHJN_327556 | Eukaryota | Ascomycota | Sordariomycet | Hypocreales | Hypocreaceae | Trichoderma | Trichoderma v | cellular proces | Signal transdu | 09UWE | dual-specificit | 12957.ACEP_( | 0,2 | 0,4 | 6,84E-06 | 1,37E-05 | 0 - | 0 - | 1 - | 3,42E-05 - | 0 - | 0 - |
| 1933 | 1 | 0.2um_NOKCPHJN_832554 | Eukaryota | Phaeophyceae | unclassified | Ectocarpales | Ectocarpaceae | Ectocarpus | Ectocarpus sili | unclassified | unclassified | unclassified | unclassified | unclassified | 0,2 | 0,4 | 1,64E-05 | 3,28E-05 | 0 - | 0 - | 0 - | 0 - | 1 - | 8,19E-05 - |
| 1934 | 1 | 3um_FIJJAFBF_348326 | Bacteria | Bacteroidetes | Flavobacteriia | Flavobacterial | Flavobacteria | Algibacter | Algibacter lect | poorly charact | Function unkn | 0P8NI | Inherit from b | 313594.PI23P_ | 1,4 | 0,8 | 0,000125045 | 7,20E-05 | 0 - | 0 - | 2 - | 0,000186235 - | 2 - | 0,000187479 - |
| 1935 | 1 | 0.2um_NOKCPHJN_440998 | Eukaryota | Bacillariophyta | Coscinodiscop | Thalassiosirale | Thalassiosirac | Thalassiosira | Thalassiosira p | information st | RNA processin | KOG1596 | rRNA 2'-O-me | 35128.Thaps9 | 1 | 0 | 0,000255029 | 2,70E-05 | 1 - | 0,000303029 - | 1 - | 0,00025334 - | 1 - | 0,000255032 - |
| 1936 | 1 | 0.2um_NOKCPHJN_39231 | Eukaryota | Bacillariophyta | Coscinodiscop | Rhizosoleniale | Rhizosoleniac | Rhizosolenia | Rhizosolenia s | information st | Translation, ri | COG0050 | This protein p | 91464.S7335_ | 10 | 1,673320053 | 0,001518183 | 0,000180537 | 7 - | 0,00128102 - | 12 - | 0,001835939 - | 10 - | 0,00154017 - |
| 1937 | 1 | 0.2um_NOKCPHJN_793117 | unclassified | unclassified | unclassified | unclassified | unclassified | unclassified | unclassified | unclassified | unclassified | unclassified | unclassified | unclassified | 0,2 | 0,4 | 3,01E-05 | 6,02E-05 | 0 - | 0 - | 0 - | 0 - | 0 - | 0 - |
| 1938 | 1 | 0.2um_NOKCPHJN_346572 | Bacteria | Proteobacteri | Alphaproteob | Pelagibacteral | Pelagibactera | Candidatus Pe | Candidatus Pe | metabolism | Coenzyme tra | COG0611 | Catalyzes the | 439493.PB721 | 0,2 | 0,4 | 3,42E-05 | 6,83E-05 | 0 - | 0 - | 0 - | 0 - | 0 - | 0 - |
| 1939 | 1 | 3um_FIJJAFBF_136575 | Bacteria | Proteobacteri | Gammaprotec | Cellvibrionales | Spongiibacter | unclassified | marine gamm | poorly charact | Function unkn | 05N42 | UPF0307 prot | 247633.GP214 | 0,4 | 0,489897949 | 0,000111869 | 0,000137611 | 0 - | 0 - | 0 - | 0 - | 1 - | 0,000299966 - |
| 1940 | 1 | 3um_FIJJAFBF_257123 | Eukaryota | Phaeophyceae | unclassified | Dictyotales | Dictyotaceae | Dictyopteris | Dictyopteris d | information st | Translation, ri | 0FKC1 | This is 1 of the | 551115.Aazo_ | 3 | 0 | 0,00104987 | 0,000111027 | 3 - | 0,00124747 - | 3 - | 0,001042915 - | 3 - | 0,001049882 - |
| 1941 | 1 | 3um_FIJJAFBF_717710 | Eukaryota | Eustigmatoph | unclassified | Eustigmatales | Monodopsida | Nannochlorop | Nannochlorop | unclassified | unclassified | unclassified | unclassified | unclassified | 1,2 | 0,4 | 0,000458126 | 0,000157959 | 1 - | 0,000453625 - | 1 - | 0,000379242 - | 2 - | 0,000763551 - |
| 1942 | 1 | 3um_FIJJAFBF_10655 | Bacteria | Bacteroidetes | Flavobacteriia | Flavobacterial | Flavobacteria | Flavobacteriu | Flavobacteriu | unclassified | unclassified | unclassified | unclassified | unclassified | 1,2 | 0,4 | 0,00013708 | 3,39E-05 | 1 - | 0,000138865 - | 1 - | 0,000116094 - | 1 - | 0,00011687 - |
| 1943 | 1 | 0.2um_NOKCPHJN_330360 | Eukaryota | Arthropoda | Insecta | Hymenoptera | Apidae | Eufriesea | Eufriesea mex | unclassified | unclassified | unclassified | unclassified | unclassified | 0,6 | 0,489897949 | 2,96E-05 | 2,47E-05 | 1 - | 5,77E-05 - | 1 - | 4,82E-05 - | 0 - | 0 - |
| 1944 | 1 | 0.2um_NOKCPHJN_401476 | Bacteria | Bacteroidetes | Flavobacteriia | Flavobacterial | unclassified | unclassified | Flavobacterial | unclassified | unclassified | unclassified | unclassified | unclassified | 1 | 0 | 0,000116652 | 1,23E-05 | 1 - | 0,000138608 - | 1 - | 0,000115879 - | 1 - | 0,000116654 - |
| 1945 | 1 | 0.2um_NOKCPHJN_789774 | unclassified | unclassified | unclassified | unclassified | unclassified | unclassified | unclassified | unclassified | unclassified | unclassified | unclassified | unclassified | 1 | 0 | 0,000136347 | 1,44E-05 | 1 - | 0,000162009 - | 1 - | 0,000135444 - | 1 - | 0,000136348 - |
| 1946 | 2 | 0.2um_NOKCPHJN_606024;3um_FIJJAF | Bacteria | Proteobacteri | Gammaprotec | Cellvibrionales | Spongiibacter | Oceanicoccus | Oceanicoccus | cellular proces | Intracellular tr | COG2831 | Polypeptide-tr | 887062.HGR_( | 0,2 | 0,4 | 2,04E-05 | 4,08E-05 | 0 - | 0 - | 0 - | 0 - | 0 - | 0 - |
| 1947 | 1 | 0.2um_NOKCPHJN_956764 | unclassified | unclassified | unclassified | unclassified | unclassified | unclassified | unclassified | unclassified | unclassified | unclassified | unclassified | unclassified | 0,2 | 0,4 | 0,000160201 | 0,000320402 | 0 - | 0 - | 0 - | 0 - | 0 - | 0 - |
| 1948 | 1 | 3um_FIJJAFBF_436031 | Eukaryota | Mollusca | Gastropoda | unclassified | various | various | various | poorly charact | Function unkn | 11NK2 | dynein, light c | 272952.HpaP8 | 0,6 | 0,489897949 | 0,000297508 | 0,000248068 | 1 - | 0,000580218 - | 1 - | 0,000485077 - | 0 - | 0 - |
| 1949 | 1 | 0.2um_NOKCPHJN_423660 | Eukaryota | Chordata | Mammalia | Scandentia | Tupaiidae | Tupaia | Tupaia chinen | unclassified | unclassified | unclassified | unclassified | unclassified | 0,6 | 0,489897949 | 2,55E-05 | 2,13E-05 | 1 - | 5,04E-05 - | 0 - | 0 - | 0 - | 0 - |
| 1950 | 1 | 0.2um_NOKCPHJN_671219 | unclassified | unclassified | unclassified | unclassified | unclassified | unclassified | unclassified | unclassified | unclassified | unclassified | unclassified | unclassified | 1 | 0 | 5,87E-05 | 6,20E-06 | 1 - | 6,97E-05 - | 1 - | 5,83E-05 - | 1 - | 5,87E-05 - |
| 1951 | 1 | 3um_FIJJAFBF_458152 | Bacteria | Verrucomicrol | unclassified | unclassified | unclassified | unclassified | Verrucomicrol | poorly charact | Function unkn | 153IU | Inherit from N | 344747.PM87 | 0,4 | 0,489897949 | 2,37E-05 | 2,94E-05 | 1 - | 6,58E-05 - | 0 - | 0 - | 0 - | 0 - |
| 1952 | 1 | 3um_FIJJAFBF_475532 | Eukaryota | unclassified | Oomycetes | Peronosporale | Peronosporac | Plasmopara | Plasmopara ha | information st | Translation, ri | COG0081 | Binds directly | 653948.CCA20 | 1 | 0 | 0,000623685 | 6,60E-05 | 1 - | 0,000741071 - | 1 - | 0,000619554 - | 1 - | 0,000623692 - |
| 1953 | 1 | 0.2um_NOKCPHJN_222203 | Bacteria | Firmicutes | Bacilli | Bacillales | Staphylococca | Staphylococcu | Staphylococcu | poorly charact | Function unkn | 0ZPU4 | NA | 5786.XP_0032 | 0,8 | 0,748331477 | 4,91E-05 | 4,64E-05 | 0 - | 0 - | 0 - | 0 - | 1 - | 6,51E-05 - |
| 1954 | 1 | 3um_FIJJAFBF_450895 | Bacteria | Bacteroidetes | Flavobacteriia | Flavobacterial | Flavobacteria | Flaviramulus | Flaviramulus b | unclassified | unclassified | unclassified | unclassified | unclassified | 2,8 | 0,4 | 0,000514423 | 0,000100158 | 3 - | 0,000652746 - | 3 - | 0,000545711 - | 3 - | 0,000549357 - |
| 1955 | 1 | 3um_FIJJAFBF_257132 | Eukaryota | unclassified | Florideophyce | Plocamiales | Plocamiaceae | Plocamium | Plocamium ca | unclassified | unclassified | unclassified | unclassified | unclassified | 2,2 | 0,4 | 0,00102916 | 0,000126 | 2 - | 0,001125537 - | 2 - | 0,000940976 - | 2 - | 0,000947262 - |
| 1956 | 1 | 0.2um_NOKCPHJN_115346 | Bacteria | Bacteroidetes | Flavobacteriia | Flavobacterial | Flavobacteria | Algibacter | Algibacter sp. | unclassified | unclassified | unclassified | unclassified | unclassified | 1,4 | 1,356465997 | 9,57E-05 | 8,82E-05 | 1 - | 8,55E-05 - | 0 - | 0 - | 0 - | 0 - |
| 1957 | 1 | 3um_FIJJAFBF_379396 | Viruses | unclassified | unclassified | unclassified | Phycodnavirid | unclassified | Pyramimonas | unclassified | unclassified | unclassified | unclassified | unclassified | 0,8 | 0,4 | 0,000131576 | 6,80E-05 | 1 - | 0,000195426 - | 1 - | 0,000163381 - | 0 - | 0 - |
| 1958 | 1 | 3um_FIJJAFBF_257124 | Eukaryota | unclassified | Florideophyce | Ceramiales | Delesseriacea | Membranopte | Membranopte | unclassified | unclassified | unclassified | unclassified | unclassified | 1,2 | 0,4 | 0,000539314 | 0,000133277 | 1 - | 0,000546337 - | 1 - | 0,000456751 - | 1 - | 0,000459802 - |
| 1959 | 1 | 0.2um_NOKCPHJN_355727 | Bacteria | Bacteroidetes | Flavobacteriia | Flavobacterial | Flavobacteria | Formosa | Formosa sp. H | metabolism | Inorganic ion t | 05C27 | TonB-depende | 983544.Lacal_ | 2,8 | 0,748331477 | 0,000175208 | 4,41E-05 | 2 - | 0,000150298 - | 4 - | 0,000251305 - | 3 - | 0,000189738 - |
| 1960 | 1 | 3um_FIJJAFBF_118044 | Bacteria | Bacteroidetes | unclassified | unclassified | unclassified | unclassified | Bacteroidetes | unclassified | unclassified | unclassified | unclassified | unclassified | 0,4 | 0,489897949 | 7,59E-05 | 9,31E-05 | 0 - | 0 - | 0 - | 0 - | 0 - | 0 - |
| 1961 | 2 | 3um_FIJJAFBF_423989;3um_FIJJAFBF_( | Bacteria | Proteobacteri | Alphaproteob | Rhodobactera | Rhodobactera | Roseovarius | Roseovarius a | metabolism | Energy produc | 16RMM | alcohol dehyd | 89187.ISM_11 | 1 | 0 | 0,000225778 | 2,39E-05 | 1 - | 0,000268273 - | 1 - | 0,000224283 - | 1 - | 0,000225781 - |
| 1962 | 1 | 3um_FIJJAFBF_44700 | Bacteria | Bacteroidetes | Flavobacteriia | Flavobacterial | Flavobacteria | Polaribacter | Polaribacter s | unclassified | unclassified | unclassified | unclassified | unclassified | 0,8 | 0,4 | 5,64E-05 | 2,84E-05 | 0 - | 0 - | 1 - | 7,34E-05 - | 1 - | 7,39E-05 - |
| 1963 | 1 | 0.2um_NOKCPHJN_403622 | Bacteria | Proteobacteri | Alphaproteob | Rhodobactera | Rhodobactera | unclassified | Rhodobactera | unclassified | unclassified | unclassified | unclassified | unclassified | 0,8 | 0,4 | 0,000110755 | 5,73E-05 | 1 - | 0,000164502 - | 1 - | 0,000137527 - | 0 - | 0 - |
| 1964 | 1 | 3um_FIJJAFBF_205161 | Eukaryota | Bacillariophyta | Coscinodiscop | Rhizosoleniale | Rhizosoleniac | Guinardia | Guinardia stria | metabolism | Energy produc | 0IE88 | oxidoreductas | 35128.Thapsd | 1,2 | 0,4 | 0,0001008 | 2,49E-05 | 1 - | 0,000102112 - | 1 - | 8,54E-05 - | 1 - | 8,59E-05 - |
| 1965 | 1 | 3um_FIJJAFBF_257127 | Eukaryota | unclassified | Synurophycea | Parmales | Triparmaceae | Triparma | Triparma laevi | unclassified | unclassified | unclassified | unclassified | unclassified | 0,2 | 0,4 | 3,14E-05 | 6,27E-05 | 0 - | 0 - | 1 - | 0,000156829 - | 0 - | 0 - |
| 1966 | 1 | 3um_FIJJAFBF_231373 | unclassified | unclassified | unclassified | unclassified | unclassified | unclassified | unclassified | unclassified | unclassified | unclassified | unclassified | unclassified | 1 | 0 | 0,000446753 | 4,72E-05 | 1 - | 0,000530838 - | 1 - | 0,000443794 - | 1 - | 0,000446758 - |
| 1967 | 1 | 3um_FIJJAFBF_288420 | unclassified | unclassified | unclassified | unclassified | unclassified | unclassified | unclassified | unclassified | unclassified | unclassified | unclassified | unclassified | 2 | 0,894427191 | 0,001371204 | 0,000583129 | 2 - | 0,001645015 - | 3 - | 0,002062909 - | 1 - | 0,00069223 - |
| 1968 | 1 | 3um_FIJJAFBF_369772 | unclassified | unclassified | unclassified | unclassified | unclassified | unclassified | unclassified | unclassified | unclassified | unclassified | unclassified | unclassified | 1 | 0,632455532 | 0,000688876 | 0,000454245 | 0 - | 0 - | 1 - | 0,000711079 - | 2 - | 0,001431658 - |
| 1969 | 2 | 0.2um_NOKCPHJN_1000581;3um_FIJJA | Bacteria | Proteobacteri | various | various | various | various | various | various | various | various | various | various | 0,4 | 0,489897949 | 7,60E-05 | 9,35E-05 | 0 - | 0 - | 1 - | 0,000203165 - | 0 - | 0 - |
| 1970 | 1 | 3um_FIJJAFBF_506019 | Bacteria | Proteobacteri | Gammaprotec | Cellvibrionales | Porticoccacea | unclassified | gamma protec | cellular proces | Cell wall/mem | 0QTSX | Peptidoglycan | 314287.GB22( | 1 | 0 | 0,000386455 | 4,09E-05 | 1 - | 0,000459191 - | 1 - | 0,000383895 - | 1 - | 0,00038646 - |
| 1971 | 1 | 0.2um_NOKCPHJN_251593 | Eukaryota | Chlorophyta | Trebouxiophy | Chlorellales | Chlorellaceae | Chlorella | Chlorella varia | metabolism | Energy produc | COG3474 | Electron carrie | 45351.NEMVE | 0,8 | 0,4 | 0,000500945 | 0,000255265 | 1 - | 0,000719694 - | 1 - | 0,000601682 - | 1 - | 0,000605701 - |
| 1972 | 1 | 3um_FIJJAFBF_523829 | Bacteria | Proteobacteri | Gammaprotec | Arenicellales | Arenicellaceae | Arenicella | Arenicella xan | unclassified | unclassified | unclassified | unclassified | unclassified | 3,6 | 1,2 | 0,000353514 | 0,000110132 | 2 - | 0,000237237 - | 5 - | 0,000495839 - | 3 - | 0,000299491 - |
| 1973 | 1 | 0.2um_NOKCPHJN_265132 | Bacteria | Bacteroidetes | unclassified | unclassified | unclassified | unclassified | Bacteroidetes | metabolism | Energy produc | COG0056 | Produces ATP | 525373.HMPR | 0,8 | 0,4 | 0,000106323 | 5,42E-05 | 1 - | 0,000152751 - | 1 - | 0,000127704 - | 1 - | 0,000128557 - |
| 1974 | 1 | 3um_FIJJAFBF_784243 | Eukaryota | Bacillariophyta | Bacillariophyc | Bacillariales | Bacillariaceae | Fragilariopsis | Fragilariopsis | metabolism | Carbohydrate | 1255K | major facilitat | 2850.Phatr444 | 1 | 0 | 0,000396177 | 4,19E-05 | 1 - | 0,000470743 - | 1 - | 0,000393553 - | 1 - | 0,000396182 - |
| 1975 | 1 | 3um_FIJJAFBF_536896 | Eukaryota | unclassified | Dinophyceae | Gymnodiniales | Gymnodiniace | Amphidinium | Amphidinium | poorly charact | Function unkn | COG1100 | GTP-binding P | 423536.XP_00 | 2,6 | 0,489897949 | 0,001106109 | 0,000155287 | 2 - | 0,001025318 - | 2 - | 0,000857191 - | 3 - | 0,001294375 - |
| 1976 | 1 | 3um_FIJJAFBF_154705 | Bacteria | Bacteroidetes | Flavobacteriia | Flavobacterial | Flavobacteria | Arenitalea | Arenitalea lute | unclassified | unclassified | unclassified | unclassified | unclassified | 1 | 0 | 0,000403796 | 4,27E-05 | 1 - | 0,000479796 - | 1 - | 0,000401121 - | 1 - | 0,000403801 - |
| 1977 | 1 | 0.2um_NOKCPHJN_54732 | Eukaryota | Streptophyta | unclassified | Brassicales | Brassicaceae | Arabidopsis | various | metabolism | Energy produc | COG0055 | Produces ATP | 39947.LOC_Os | 1 | 0 | 0,00012649 | 1,34E-05 | 1 - | 0,000150298 - | 1 - | 0,000125652 - | 1 - | 0,000126492 - |
| 1978 | 1 | 3um_FIJJAFBF_213931 | Bacteria | Bacteroidetes | Flavobacteriia | Flavobacterial | Flavobacteria | Formosa | Formosa sp. H | poorly charact | Function unkn | 0XPRT | NA | 391598.FBBAL | 0,2 | 0,4 | 2,47E-05 | 4,93E-05 | 0 - | 0 - | 0 - | 0 - | 0 - | 0 - |
| 1979 | 1 | 0.2um_NOKCPHJN_718641 | Eukaryota | Arthropoda | Insecta | Hemiptera | Liviidae | Diaphorina | Diaphorina cit | cellular proces | Posttranslatio | 007GN | Inherit from a | 5833.PFL0585 | 3,8 | 1,166190379 | 0,003243127 | 0,001033382 | 4 - | 0,004045848 - | 3 - | 0,002536821 - | 3 - | 0,002553768 - |
| 1980 | 1 | 3um_FIJJAFBF_590488 | Eukaryota | Streptophyta | unclassified | Ranunculales | Ranunculacea | Aquilegia | Aquilegia coer | unclassified | unclassified | unclassified | unclassified | unclassified | 1 | 0 | 0,000349957 | 3,70E-05 | 1 - | 0,000415823 - | 1 - | 0,000347638 - | 1 - | 0,000349961 - |
| 1981 | 1 | 0.2um_NOKCPHJN_536213 | Eukaryota | unclassified | Oomycetes | Peronosporale | Peronosporac | Phytophthora | Phytophthora | unclassified | unclassified | unclassified | unclassified | unclassified | 0,2 | 0,4 | 2,42E-05 | 4,84E-05 | 0 - | 0 - | 0 - | 0 - | 0 - | 0 - |
| 1982 | 1 | 3um_FIJJAFBF_302476 | Eukaryota | unclassified | Cryptophyta | Pyrenomonad | Geminigerace | Guillardia | Guillardia thet | unclassified | unclassified | unclassified | unclassified | unclassified | 0,8 | 0,4 | 0,0004869 | 0,000248108 | 1 - | 0,000699516 - | 1 - | 0,000584812 - | 1 - | 0,000588719 - |
| 1983 | 1 | 3um_FIJJAFBF_808023 | Bacteria | Verrucomicrol | Verrucomicrol | Verrucomicrol | Akkermansiac | Akkermansia | Akkermansia r | unclassified | unclassified | unclassified | unclassified | unclassified | 1 | 0 | 0,000161105 | 1,70E-05 | 1 - | 0,000191428 - | 1 - | 0,000160038 - | 1 - | 0,000161107 - |
| 1984 | 1 | 0.2um_NOKCPHJN_489599 | Eukaryota | unclassified | Pelagophycea | Pelagomonada | unclassified | Aureococcus | Aureococcus a | unclassified | unclassified | unclassified | unclassified | unclassified | 2 | 0 | 0,000834333 | 8,82E-05 | 2 - | 0,000991367 - | 2 - | 0,000828807 - | 2 - | 0,000834344 - |
| 1985 | 1 | 0.2um_NOKCPHJN_949177 | Eukaryota | Phaeophyceae | unclassified | Ectocarpales | Ectocarpaceae | Ectocarpus | Ectocarpus sili | metabolism | Carbohydrate | 0V5IG | enolase 2, gan | 400682.PAC_1 | 0,8 | 0,4 | 0,000119757 | 6,04E-05 | 0 - | 0 - | 1 - | 0,000156047 - | 1 - | 0,00015709 - |
| 1986 | 1 | 0.2um_NOKCPHJN_783728 | Eukaryota | Mucoromycot | Mucoromycet | Mucorales | Choanephora | Choanephora | Choanephora | poorly charact | Function unkn | 0YY9G | WD repeat do | 4787.PITG_12 | 0,2 | 0,4 | 2,57E-05 | 5,14E-05 | 0 - | 0 - | 0 - | 0 - | 1 - | 0,000128557 - |
| 1987 | 1 | 3um_FIJJAFBF_669269 | Eukaryota | Bacillariophyta | Coscinodiscop | Coscinodiscale | Coscinodiscac | Coscinodiscus | Coscinodiscus | cellular proces | Intracellular tr | COG0653 | Part of the Se | 197221.tll185 | 0,4 | 0,489897949 | 2,56E-05 | 3,15E-05 | 0 - | 0 - | 1 - | 6,85E-05 - | 0 - | 0 - |
| 1988 | 1 | 3um_FIJJAFBF_220309 | Eukaryota | Bacillariophyta | Coscinodiscop | Thalassiosirale | Thalassiosirac | Thalassiosira | Thalassiosira | unclassified | unclassified | unclassified | unclassified | unclassified | 0,4 | 0,489897949 | 0,000152311 | 0,000187688 | 1 - | 0,000413526 - | 0 - | 0 - | 1 - | 0,000348027 - |
| 1989 | 1 | 0.2um_NOKCPHJN_405135 | Bacteria | Proteobacteri | various | various | various | various | various | cellular proces | Cell wall/mem | 0B7GS | outer membra | 583345.Mmol | 1 | 0 | 0,000148566 | 1,57E-05 | 1 - | 0,000176529 - | 1 - | 0,000147582 - | 1 - | 0,000148568 - |
| 1990 | 1 | 3um_FIJJAFBF_30301 | Bacteria | Bacteroidetes | Flavobacteriia | Flavobacterial | Flavobacteria | Polaribacter | Polaribacter s | unclassified | unclassified | unclassified | unclassified | unclassified | 1,2 | 1,166190379 | 0,000209994 | 0,000204736 | 0 - | 0 - | 1 - | 0,000186791 - | 0 - | 0 - |
| 1991 | 1 | 3um_FIJJAFBF_580261 | Bacteria | Proteobacteri | Gammaprotec | Pseudomonad | Pseudomonac | Pseudomonas | Pseudomonas | poorly charact | Function unkn | 17B58 | Inherit from b | 471874.PROST | 1 | 0,632455532 | 0,00017587 | 0,000103447 | 0 - | 0 - | 1 - | 0,000186791 - | 1 - | 0,000188039 - |
| 1992 | 1 | 3um_FIJJAFBF_216066 | Bacteria | Proteobacteri | Gammaprotec | Alteromonada | various | various | various | information st | Translation, ri | 0QTKX | Seems to be tl | 203122.Sde_0 | 2 | 0 | 0,001007875 | 0,000106586 | 2 - | 0,001197571 - | 2 - | 0,001001199 - | 2 - | 0,001007887 - |
| 1993 | 1 | 3um_FIJJAFBF_112505 | Bacteria | Proteobacteri | Deltaproteoba | unclassified | unclassified | unclassified | various | poorly charact | Function unkn | 0HFRS | MotA TolQ Exl | 391625.PPSIR | 0,8 | 0,4 | 0,000219281 | 0,000110515 | 0 - | 0 - | 1 - | 0,00028573 - | 1 - | 0,000287639 - |
| 1994 | 1 | 3um_FIJJAFBF_21329 | Bacteria | Bacteroidetes | Flavobacteriia | Flavobacterial | Flavobacteria | Polaribacter | Polaribacter s | cellular proces | Cell wall/mem | 09GJR | outer membra | 313598.MED1 | 0,4 | 0,489897949 | 0,000137383 | 0,000168996 | 0 - | 0 - | 0 - | 0 - | 1 - | 0,00036838 - |
| 1995 | 1 | 0.2um_NOKCPHJN_551609 | Bacteria | Bacteroidetes | Flavobacteriia | Flavobacterial | Flavobacteria | Formosa | Formosa algae | poorly charact | Function unkn | 09SDX | Inherit from N | 391598.FBBAL | 1 | 0 | 4,80E-05 | 5,08E-06 | 1 - | 5,70E-05 - | 1 - | 4,77E-05 - | 1 - | 4,80E-05 - |
| 1996 | 1 | 3um_FIJJAFBF_21302 | Bacteria | Bacteroidetes | Flavobacteriia | Flavobacterial | Flavobacteria | Polaribacter | Polaribacter s | unclassified | unclassified | unclassified | unclassified | unclassified | 0,8 | 0,748331477 | 0,000253201 | 0,000236315 | 1 - | 0,000372379 - | 2 - | 0,000622636 - | 0 - | 0 - |
| 1997 | 1 | 3um_FIJJAFBF_341378 | Bacteria | Bacteroidetes | Flavobacteriia | Flavobacterial | Flavobacteria | Algibacter | Algibacter sp. | cellular proces | Cell wall/mem | COG2885 | Ompa motb d | 1046627.BZAF | 2 | 1,264911064 | 0,00026381 | 0,000154203 | 1 - | 0,000163424 - | 1 - | 0,000136626 - | 1 - | 0,000137539 - |
| 1998 | 3 | 0.2um_NOKCPHJN_597293;3um_FIJJAF | Bacteria | Proteobacteri | Alphaproteob | Rhizobiales | Phyllobacteria | unclassified | Phyllobacteria | various | various | various | various | various | 0,2 | 0,4 | 5,73E-05 | 0,000114672 | 0 - | 0 - | 0 - | 0 - | 0 - | 0 - |
| 1999 | 1 | 3um_FIJJAFBF_55089 | Bacteria | Bacteroidetes | Flavobacteriia | Flavobacterial | Flavobacteria | Polaribacter | Polaribacter s | information st | Translation, ri | 0PA05 | 30S ribosomal | 313594.PI23P_ | 2,8 | 0,748331477 | 0,000955169 | 0,000188529 | 2 - | 0,000827052 - | 3 - | 0,001037153 - | 3 - | 0,001044082 - |
| 2000 | 1 | 0.2um_NOKCPHJN_559361 | Bacteria | Spirochaetes | Spirochaetia | Brachyspirales | Brachyspirace | Brachyspira | Brachyspira in | unclassified | unclassified | unclassified | unclassified | unclassified | 0,4 | 0,489897949 | 1,87E-05 | 2,30E-05 | 0 - | 0 - | 1 - | 5,01E-05 - | 0 - | 0 - |
| 2001 | 1 | 3um_FIJJAFBF_196001 | Bacteria | Bacteroidetes | Flavobacteriia | Flavobacterial | Flavobacteria | Polaribacter | Polaribacter s | cellular proces | Posttranslatio | 08V1T | peptidylprolyl | 313594.PI23P_ | 3 | 0 | 0,000814554 | 8,61E-05 | 3 - | 0,000967864 - | 3 - | 0,000809158 - | 3 - | 0,000814564 - |
| 2002 | 1 | 3um_FIJJAFBF_670830 | Bacteria | Proteobacteri | Betaproteoba | unclassified | unclassified | unclassified | Betaproteoba | metabolism | Coenzyme tra | 16Q82 | coproporphyri | 399795.CtesD | 0,4 | 0,489897949 | 5,44E-05 | 6,68E-05 | 0 - | 0 - | 0 - | 0 - | 0 - | 0 - |
| 2003 | 1 | 3um_FIJJAFBF_40363 | Eukaryota | unclassified | Raphidophyce | Chattonellales | Chattonellace | Heterosigma | Heterosigma a | unclassified | unclassified | unclassified | unclassified | unclassified | 5,8 | 0,4 | 0,00071313 | 3,75E-05 | 5 - | 0,000735247 - | 6 - | 0,000737622 - | 6 - | 0,000742549 - |
| 2004 | 1 | 3um_FIJJAFBF_155642 | Bacteria | Bacteroidetes | Flavobacteriia | Flavobacterial | Crocinitomica | unclassified | Crocinitomica | unclassified | unclassified | unclassified | unclassified | unclassified | 0,6 | 0,489897949 | 0,000281308 | 0,000230911 | 0 - | 0 - | 1 - | 0,000488867 - | 1 - | 0,000492132 - |
| 2005 | 1 | 3um_FIJJAFBF_280872 | Archaea | Euryarchaeota | unclassified | unclassified | unclassified | unclassified | Marine Group | unclassified | unclassified | unclassified | unclassified | unclassified | 1 | 0 | 0,000132615 | 1,40E-05 | 1 - | 0,000157575 - | 1 - | 0,000131737 - | 1 - | 0,000132617 - |
| 2006 | 1 | 3um_FIJJAFBF_514974 | Eukaryota | unclassified | Pelagophycea | Pelagomonada | unclassified | Aureococcus | Aureococcus a | unclassified | unclassified | unclassified | unclassified | unclassified | 2 | 0 | 0,000332413 | 3,52E-05 | 2 - | 0,000394977 - | 2 - | 0,000330211 - | 2 - | 0,000332417 - |
| 2007 | 1 | 3um_FIJJAFBF_592170 | Bacteria | Bacteroidetes | Flavobacteriia | Flavobacterial | Crocinitomica | unclassified | Crocinitomica | unclassified | unclassified | unclassified | unclassified | unclassified | 1 | 0 | 0,000428518 | 4,53E-05 | 1 - | 0,000509171 - | 1 - | 0,00042568 - | 1 - | 0,000428523 - |
| 2008 | 1 | 0.2um_NOKCPHJN_240819 | Bacteria | Bacteroidetes | Flavobacteriia | Flavobacterial | Flavobacteria | Polaribacter | Polaribacter s | poorly charact | Function unkn | 09EUF | NA | 313598.MED1 | 1 | 0 | 0,000255029 | 2,70E-05 | 1 - | 0,000303029 - | 1 - | 0,00025334 - | 1 - | 0,000255032 - |
| 2009 | 1 | 0.2um_NOKCPHJN_804742 | Eukaryota | unclassified | Oomycetes | Peronosporale | Peronosporac | Phytophthora | Phytophthora | information st | Translation, ri | COG0052 | 30S ribosomal | 67593.Physo1 | 2 | 0 | 0,000471851 | 4,99E-05 | 2 - | 0,000560661 - | 2 - | 0,000468726 - | 2 - | 0,000471857 - |
| 2010 | 2 | 0.2um_NOKCPHJN_783672;0.2um_NO | various | various | various | various | various | various | various | unclassified | unclassified | unclassified | unclassified | unclassified | 1 | 0 | 0,000177443 | 1,88E-05 | 1 - | 0,00021084 - | 1 - | 0,000176267 - | 1 - | 0,000177445 - |
| 2011 | 1 | 3um_FIJJAFBF_24509 | Bacteria | Proteobacteri | Alphaproteob | Rhodobactera | unclassified | unclassified | Rhodobactera | unclassified | unclassified | unclassified | unclassified | unclassified | 0,2 | 0,4 | 3,54E-05 | 7,09E-05 | 0 - | 0 - | 0 - | 0 - | 0 - | 0 - |
| 2012 | 2 | 0.2um_NOKCPHJN_985727;3um_FIJJAF | unclassified | unclassified | unclassified | unclassified | unclassified | unclassified | unclassified | unclassified | unclassified | unclassified | unclassified | unclassified | 1,4 | 0,8 | 0,000851495 | 0,000481172 | 0 - | 0 - | 2 - | 0,001277039 - | 1 - | 0,000642785 - |
| 2013 | 1 | 0.2um_NOKCPHJN_530425 | Eukaryota | unclassified | unclassified | Dictyosteliales | Raperosteliac | Tieghemosteli | Tieghemosteli | information st | Transcription | KOG3282 | Peptidyl-trna l | 164328.Phyra | 0,8 | 0,4 | 8,75E-05 | 4,52E-05 | 1 - | 0,000129945 - | 1 - | 0,000108637 - | 0 - | 0 - |
| 2014 | 1 | 3um_FIJJAFBF_290584 | Eukaryota | Bacillariophyta | Coscinodiscop | Chaetocerotal | Chaetocerota | Chaetoceros | Chaetoceros s | metabolism | Energy produc | COG0712 | F(1)F(0) ATP s | 35128.Thapsd | 1,8 | 0,4 | 0,000606788 | 0,000153588 | 2 - | 0,000800515 - | 1 - | 0,000334625 - | 2 - | 0,000673721 - |
| 2015 | 1 | 0.2um_NOKCPHJN_1111441 | unclassified | unclassified | unclassified | unclassified | unclassified | unclassified | unclassified | unclassified | unclassified | unclassified | unclassified | unclassified | 0,2 | 0,4 | 4,40E-05 | 8,81E-05 | 1 - | 0,000220142 - | 0 - | 0 - | 0 - | 0 - |

| | | | | | | | | | | | | | | | | | | | | | | | | |
|---|---|---|---|---|---|---|---|---|---|---|---|---|---|---|---|---|---|---|---|---|---|---|---|---|
| 2016 | 1 | 3um_FIJJAFBF_08771 | Eukaryota | unclassified | Raphidophyce | Chattonellales | Chattonellace | Heterosigma | Heterosigma a | metabolism | Energy produc | 0IFBF | One of the tw | 45157.CMV08 | 1,6 | 0,489897949 | 0,000288805 | 9,98E-05 | 2 - | 0,000426485 - | 2 - | 0,000356552 - | 1 - | 0,000179467 - |
| 2017 | 1 | 0.2um_NOKCPHJN_279899 | Bacteria | Cyanobacteria | unclassified | Oscillatoriales | Oscillatoriace | Phormidium | Phormidium s | metabolism | Carbohydrate | 0FJZR | phosphoglyce | 1148.SYNGTS_ | 0,6 | 0,489897949 | 7,71E-05 | 6,33E-05 | 0 - | 0 - | 1 - | 0,000133993 - | 1 - | 0,000134889 - |
| 2018 | 1 | 3um_FIJJAFBF_37831 | Bacteria | Proteobacteri | unclassified | unclassified | unclassified | unclassified | Proteobacteri | unclassified | unclassified | unclassified | unclassified | unclassified | 4,6 | 0,489897949 | 0,000897818 | 0,000112317 | 4 - | 0,000929791 - | 5 - | 0,00097166 - | 5 - | 0,000978151 - |
| 2019 | 1 | 3um_FIJJAFBF_715097 | Eukaryota | unclassified | Pelagophycea | Pelagomonad | unclassified | Aureococcus | Aureococcus a | metabolism | Energy produc | 1DYB8 | PsaA and PsaE | 3055.DAA014 | 14,6 | 1,356465997 | 0,001220165 | 0,00015357 | 15 - | 0,001492982 - | 13 - | 0,001081747 - | 13 - | 0,001088974 - |
| 2020 | 1 | 3um_FIJJAFBF_108764 | Bacteria | Proteobacteri | Alphaproteob | unclassified | unclassified | unclassified | Alphaproteob | poorly charact | Function unkn | 01UPA | von Willebran | 1082931.KKY_ | 1 | 0 | 0,000183117 | 1,94E-05 | 1 - | 0,000217582 - | 1 - | 0,000181904 - | 1 - | 0,000183119 - |
| 2021 | 1 | 3um_FIJJAFBF_792634 | Eukaryota | Bacillariophyt | Bacillariophyc | Naviculales | Phaeodactylac | Phaeodactylu | Phaeodactylu | unclassified | unclassified | unclassified | unclassified | unclassified | 1,2 | 0,748331477 | 0,000144013 | 8,31E-05 | 1 - | 0,000147049 - | 1 - | 0,000122937 - | 0 - | 0 - |
| 2022 | 1 | 0.2um_NOKCPHJN_273533 | Eukaryota | Eustigmatoph | unclassified | Eustigmatales | Monodopsida | Nannochlorop | Nannochlorop | poorly charact | Function unkn | 14ZQ5 | ABC transport | 67593.Physo1 | 0,4 | 0,489897949 | 2,71E-05 | 3,32E-05 | 0 - | 0 - | 1 - | 6,75E-05 - | 1 - | 6,80E-05 - |
| 2023 | 1 | 0.2um_NOKCPHJN_402166 | Bacteria | Bacteroidetes | Flavobacteriia | Flavobacterial | Crocinitomica | unclassified | Crocinitomica | unclassified | unclassified | unclassified | unclassified | unclassified | 0,2 | 0,4 | 1,98E-05 | 3,96E-05 | 0 - | 0 - | 0 - | 0 - | 1 - | 9,90E-05 - |
| 2024 | 1 | 3um_FIJJAFBF_81228 | Bacteria | Proteobacteri | Gammaprotec | Oceanospirilla | Endozoicomo | Endozoicomo | various | unclassified | unclassified | unclassified | unclassified | unclassified | 1 | 0,894427191 | 0,00050197 | 0,000447726 | 0 - | 0 - | 1 - | 0,000508739 - | 2 - | 0,001024275 - |
| 2025 | 1 | 3um_FIJJAFBF_695952 | Eukaryota | unclassified | Oomycetes | Peronosporale | Peronosporac | Phytophthora | various | unclassified | unclassified | unclassified | unclassified | unclassified | 2 | 0 | 0,000363067 | 3,84E-05 | 2 - | 0,000431402 - | 2 - | 0,000360662 - | 2 - | 0,000363072 - |
| 2026 | 1 | 3um_FIJJAFBF_328503 | Bacteria | Bacteroidetes | Flavobacteriia | Flavobacterial | Flavobacteria | Arenitalea | Arenitalea lut | information st | Translation, ri | COG0088 | One of the pri | 983544.Lacal_ | 0,4 | 0,489897949 | 0,000109612 | 0,000134515 | 0 - | 0 - | 0 - | 0 - | 0 - | 0 - |
| 2027 | 1 | 3um_FIJJAFBF_666897 | unclassified | unclassified | unclassified | unclassified | unclassified | unclassified | unclassified | unclassified | unclassified | unclassified | unclassified | unclassified | 1 | 0 | 0,000807592 | 8,54E-05 | 1 - | 0,000959592 - | 1 - | 0,000802243 - | 1 - | 0,000807602 - |
| 2028 | 1 | 0.2um_NOKCPHJN_421642 | Bacteria | Proteobacteri | Alphaproteob | Rickettsiales | unclassified | unclassified | Rickettsiales b | unclassified | unclassified | unclassified | unclassified | unclassified | 0,6 | 0,489897949 | 6,63E-05 | 5,53E-05 | 1 - | 0,000129271 - | 1 - | 0,000108074 - | 0 - | 0 - |
| 2029 | 1 | 0.2um_NOKCPHJN_734407 | Bacteria | Proteobacteri | Gammaprotec | Cellvibrionales | Spongiibacter | Oceanicoccus | Oceanicoccus | poorly charact | Function unkn | 16PAT | VWA containi | 247633.GP214 | 0,4 | 0,489897949 | 0,00012002 | 0,000147968 | 1 - | 0,000326848 - | 1 - | 0,000273253 - | 0 - | 0 - |
| 2030 | 1 | 0.2um_NOKCPHJN_12324 | Eukaryota | unclassified | Pelagophycea | Pelagomonad | unclassified | Aureococcus | Aureococcus a | unclassified | unclassified | unclassified | unclassified | unclassified | 3,4 | 0,489897949 | 0,000322911 | 5,47E-05 | 3 - | 0,00033868 - | 4 - | 0,000377526 - | 4 - | 0,000380048 - |
| 2031 | 1 | 0.2um_NOKCPHJN_452099 | Bacteria | unclassified | unclassified | unclassified | unclassified | unclassified | uncultured ba | metabolism | Inorganic ion t | COG3119 | Sulfatase | 655815.ZPR_3 | 0,6 | 0,489897949 | 6,57E-05 | 5,39E-05 | 0 - | 0 - | 1 - | 0,000116094 - | 0 - | 0 - |
| 2032 | 1 | 3um_FIJJAFBF_115451 | Bacteria | Proteobacteri | Gammaprotec | Cellvibrionales | Halieaceae | Congregibacte | Congregibacte | information st | Translation, ri | 179R6 | This protein is | 314285.KT71_ | 0,4 | 0,8 | 0,000153435 | 0,00030687 | 0 - | 0 - | 0 - | 0 - | 0 - | 0 - |
| 2033 | 1 | 3um_FIJJAFBF_232387 | Eukaryota | unclassified | Dinophyceae | Suessiales | Symbiodiniace | Symbiodinium | Symbiodinium | metabolism | Amino acid tra | COG0591 | symporter | 35128.Thaps2 | 2,2 | 0,748331477 | 0,00029241 | 7,96E-05 | 1 - | 0,000163424 - | 2 - | 0,000273253 - | 2 - | 0,000275078 - |
| 2034 | 1 | 3um_FIJJAFBF_248454 | Bacteria | Proteobacteri | Gammaprotec | Oceanospirilla | Saccharospiril | Reinekea | Reinekea fors | metabolism | Nucleotide tra | 173P7 | UPF0207 prot | 349521.HCH_ | 0,4 | 0,489897949 | 0,000111751 | 0,000137139 | 0 - | 0 - | 0 - | 0 - | 0 - | 0 - |
| 2035 | 1 | 3um_FIJJAFBF_715089 | Eukaryota | Phaeophyceae | unclassified | Dictyotales | Dictyotaceae | Dictyopteris | Dictyopteris d | unclassified | unclassified | unclassified | unclassified | unclassified | 1 | 0 | 0,000456465 | 4,83E-05 | 1 - | 0,000542378 - | 1 - | 0,000453441 - | 1 - | 0,000456471 - |
| 2036 | 1 | 3um_FIJJAFBF_338021 | Bacteria | Acidobacteria | unclassified | unclassified | unclassified | unclassified | Acidobacteria | cellular proce | Posttranslatio | COG0652 | peptidyl-proly | 2850.Phatr14 | 0,6 | 0,489897949 | 0,000724644 | 0,000594305 | 0 - | 0 - | 0 - | 0 - | 1 - | 0,00128557 - |
| 2037 | 1 | 0.2um_NOKCPHJN_115733 | Bacteria | Proteobacteri | Alphaproteob | unclassified | unclassified | unclassified | Alphaproteob | metabolism | Energy produc | COG0056 | Produces ATP | 488538.SAR11 | 0,6 | 0,489897949 | 7,27E-05 | 5,94E-05 | 0 - | 0 - | 1 - | 0,000122456 - | 1 - | 0,000123274 - |
| 2038 | 1 | 3um_FIJJAFBF_724267 | Bacteria | Proteobacteri | Gammaprotec | unclassified | Competibacte | Plasticicumula | Plasticicumula | information st | Replication, re | 05C68 | Can catalyze t | 1002339.HMP | 1 | 0 | 0,000252981 | 2,68E-05 | 1 - | 0,000300595 - | 1 - | 0,000251305 - | 1 - | 0,000252984 - |
| 2039 | 1 | 3um_FIJJAFBF_254317 | unclassified | unclassified | unclassified | unclassified | unclassified | unclassified | unclassified | unclassified | unclassified | unclassified | unclassified | unclassified | 0,6 | 0,489897949 | 0,000977639 | 0,000805444 | 1 - | 0,001825565 - | 1 - | 0,001526218 - | 1 - | 0,001536413 - |
| 2040 | 1 | 0.2um_NOKCPHJN_428843 | Bacteria | Bacteroidetes | Flavobacteriia | Flavobacterial | Cryomorphace | Owenweeksia | Owenweeksia | metabolism | Energy produc | COG0056 | Produces ATP | 926562.Oweh | 0,8 | 0,4 | 9,69E-05 | 5,00E-05 | 1 - | 0,000142297 - | 1 - | 0,000118964 - | 1 - | 0,000119758 - |
| 2041 | 1 | 3um_FIJJAFBF_369510 | Bacteria | Bacteroidetes | Flavobacteriia | Flavobacterial | Flavobacteria | Polaribacter | Polaribacter s | metabolism | Carbohydrate | COG0057 | glyceraldehyd | 992406.RIA_1 | 1 | 0 | 0,000189166 | 2,00E-05 | 1 - | 0,000224769 - | 1 - | 0,000187913 - | 1 - | 0,000189168 - |
| 2042 | 1 | 3um_FIJJAFBF_203258 | Bacteria | Proteobacteri | Gammaprotec | Arenicellales | Arenicellaceae | Arenicella | Arenicella xan | metabolism | Inorganic ion t | 16TQC | TONB-depend | 247639.MGP2 | 0,6 | 0,489897949 | 3,97E-05 | 3,31E-05 | 1 - | 7,74E-05 - | 1 - | 6,47E-05 - | 0 - | 0 - |
| 2043 | 1 | 3um_FIJJAFBF_335138 | Bacteria | Proteobacteri | Alphaproteob | Rhizobiales | unclassified | unclassified | Rhizobiales ba | unclassified | unclassified | unclassified | unclassified | unclassified | 1 | 0 | 0,000224972 | 2,38E-05 | 1 - | 0,000267315 - | 1 - | 0,000223482 - | 1 - | 0,000224975 - |
| 2044 | 1 | 0.2um_NOKCPHJN_302555 | Bacteria | Actinobacteria | Actinobacteria | Pseudonocard | Pseudonocard | Actinophytoc | Actinophytoc | unclassified | unclassified | unclassified | unclassified | unclassified | 0,2 | 0,4 | 3,00E-05 | 6,00E-05 | 0 - | 0 - | 0 - | 0 - | 1 - | 0,000149983 - |
| 2045 | 1 | 0.2um_NOKCPHJN_981236 | Bacteria | Proteobacteri | Alphaproteob | Pelagibacteral | Pelagibactera | Candidatus Pe | Candidatus Pe | metabolism | Amino acid tra | 05CM2 | brancheD-cha | 439493.PB721 | 0,6 | 0,489897949 | 0,00017801 | 0,000145942 | 0 - | 0 - | 1 - | 0,000314447 - | 0 - | 0 - |
| 2046 | 1 | 3um_FIJJAFBF_570530 | unclassified | unclassified | unclassified | unclassified | unclassified | unclassified | unclassified | unclassified | unclassified | unclassified | unclassified | unclassified | 0,4 | 0,489897949 | 0,000123072 | 0,000151659 | 1 - | 0,000334144 - | 0 - | 0 - | 1 - | 0,000281218 - |
| 2047 | 1 | 3um_FIJJAFBF_279280 | Eukaryota | Cnidaria | Anthozoa | Scleractinia | Pocilloporidae | Stylophora | Stylophora pis | unclassified | unclassified | unclassified | unclassified | unclassified | 0,8 | 0,4 | 0,000134141 | 6,76E-05 | 0 - | 0 - | 1 - | 0,00017479 - | 1 - | 0,000175958 - |
| 2048 | 1 | 0.2um_NOKCPHJN_239593 | Bacteria | Proteobacteri | Alphaproteob | Pelagibacteral | Pelagibactera | Candidatus Pe | Candidatus Pe | unclassified | unclassified | unclassified | unclassified | unclassified | 0,2 | 0,4 | 4,43E-05 | 8,86E-05 | 0 - | 0 - | 0 - | 0 - | 0 - | 0 - |
| 2049 | 1 | 0.2um_NOKCPHJN_70512 | Bacteria | Bacteroidetes | Flavobacteriia | Flavobacterial | Flavobacteria | Polaribacter | Polaribacter ir | unclassified | unclassified | unclassified | unclassified | unclassified | 3 | 0,632455532 | 0,000233736 | 5,68E-05 | 3 - | 0,000276874 - | 3 - | 0,000231473 - | 4 - | 0,000310693 - |
| 2050 | 1 | 0.2um_NOKCPHJN_863379 | Bacteria | Bacteroidetes | Flavobacteriia | Flavobacterial | Flavobacteria | Gilvibacter | Gilvibacter sp. | unclassified | unclassified | unclassified | unclassified | unclassified | 0,2 | 0,4 | 2,66E-05 | 5,33E-05 | 0 - | 0 - | 1 - | 0,000133138 - | 0 - | 0 - |
| 2051 | 1 | 3um_FIJJAFBF_254365 | Viruses | unclassified | unclassified | unclassified | Phycodnavirid | unclassified | Pyramimonas | unclassified | unclassified | unclassified | unclassified | unclassified | 1,4 | 1,019803903 | 0,000227636 | 0,000149228 | 1 - | 0,000199595 - | 1 - | 0,000166866 - | 2 - | 0,000335962 - |
| 2052 | 1 | 3um_FIJJAFBF_263680 | Bacteria | Proteobacteri | Alphaproteob | unclassified | unclassified | Candidatus Pu | Candidatus Pu | cellular proce | Posttranslatio | 05C2Z | ATP-depender | 313624.N9414 | 9,4 | 1,854723699 | 0,000674927 | 8,52E-05 | 6 - | 0,000522197 - | 9 - | 0,000654854 - | 10 - | 0,000732476 - |
| 2053 | 1 | 3um_FIJJAFBF_401626 | unclassified | unclassified | unclassified | unclassified | unclassified | unclassified | unclassified | unclassified | unclassified | unclassified | unclassified | unclassified | 0,4 | 0,489897949 | 8,88E-05 | 0,000108967 | 0 - | 0 - | 0 - | 0 - | 0 - | 0 - |
| 2054 | 1 | 0.2um_NOKCPHJN_440914 | Bacteria | Proteobacteri | Gammaprotec | Cellvibrionales | Porticoccacea | unclassified | various | unclassified | unclassified | unclassified | unclassified | unclassified | 0,2 | 0,4 | 1,38E-05 | 2,76E-05 | 0 - | 0 - | 0 - | 0 - | 0 - | 0 - |
| 2055 | 1 | 0.2um_NOKCPHJN_217592 | Eukaryota | unclassified | unclassified | Kinetoplastida | Trypanosomat | Leishmania | Leishmania m | unclassified | unclassified | unclassified | unclassified | unclassified | 0,2 | 0,4 | 2,35E-06 | 4,69E-06 | 0 - | 0 - | 0 - | 0 - | 1 - | 1,17E-05 - |
| 2056 | 1 | 3um_FIJJAFBF_19624 | Bacteria | Proteobacteri | Gammaprotec | Cellvibrionales | Porticoccacea | unclassified | gamma prote | metabolism | Inorganic ion t | 16TQC | TONB-depend | 314287.GB22 | 0,4 | 0,489897949 | 2,45E-05 | 3,00E-05 | 0 - | 0 - | 1 - | 6,10E-05 - | 1 - | 6,14E-05 - |
| 2057 | 1 | 3um_FIJJAFBF_805908 | Bacteria | unclassified | unclassified | unclassified | unclassified | unclassified | candidate divi | unclassified | unclassified | unclassified | unclassified | unclassified | 0,6 | 0,489897949 | 0,000341419 | 0,000280009 | 0 - | 0 - | 0 - | 0 - | 1 - | 0,000605701 - |
| 2058 | 1 | 3um_FIJJAFBF_352546 | Eukaryota | unclassified | Pelagophycea | Pelagomonad | unclassified | Aureococcus | Aureococcus a | unclassified | unclassified | unclassified | unclassified | unclassified | 1 | 0 | 0,000192049 | 2,03E-05 | 1 - | 0,000228196 - | 1 - | 0,000190777 - | 1 - | 0,000192052 - |
| 2059 | 1 | 3um_FIJJAFBF_226618 | Bacteria | Bacteroidetes | Flavobacteriia | Flavobacterial | Flavobacteria | Formosa | Formosa sp. H | unclassified | unclassified | unclassified | unclassified | unclassified | 1,2 | 0,4 | 0,000168178 | 5,27E-05 | 1 - | 0,000167821 - | 1 - | 0,000140303 - | 1 - | 0,00014124 - |
| 2060 | 2 | 0.2um_NOKCPHJN_96622;3um_FIJJAFE | Bacteria | Bacteroidetes | Flavobacteriia | Flavobacterial | Flavobacteria | Polaribacter | various | unclassified | unclassified | unclassified | unclassified | unclassified | 0,8 | 0,4 | 0,000417845 | 0,000215609 | 1 - | 0,00061351 - | 1 - | 0,000512909 - | 1 - | 0,000516336 - |
| 2061 | 3 | 0.2um_NOKCPHJN_1053285;0.2um_N | Bacteria | Proteobacteri | Alphaproteob | Rhodobactera | unclassified | unclassified | Rhodobactera | unclassified | unclassified | unclassified | unclassified | unclassified | 1 | 0 | 0,000167088 | 1,77E-05 | 1 - | 0,000198536 - | 1 - | 0,000165981 - | 1 - | 0,00016709 - |
| 2062 | 1 | 0.2um_NOKCPHJN_111222 | Eukaryota | unclassified | Synurophycea | Parmales | Triparmaceae | Triparma | Triparma laev | metabolism | Carbohydrate | COG1850 | RuBisCO catal | 35128.Thapsd | 1,4 | 0,489897949 | 0,000176499 | 5,38E-05 | 1 - | 0,000152751 - | 1 - | 0,000127704 - | 2 - | 0,000257114 - |
| 2063 | 1 | 3um_FIJJAFBF_438692 | Eukaryota | Bacillariophyt | Coscinodiscop | Thalassiosirale | Thalassiosirac | Thalassiosira | Thalassiosira c | unclassified | unclassified | unclassified | unclassified | unclassified | 5 | 0 | 0,00178955 | 0,00018925 | 5 - | 0,002126369 - | 5 - | 0,001777697 - | 5 - | 0,001789572 - |
| 2064 | 1 | 3um_FIJJAFBF_545830 | Bacteria | Bacteroidetes | Flavobacteriia | Flavobacterial | Flavobacteria | Polaribacter | Polaribacter h | information st | Translation, ri | 05G6W | One of the pri | 313594.PI23P | 1 | 0 | 0,000313394 | 3,31E-05 | 1 - | 0,000372379 - | 1 - | 0,000311318 - | 1 - | 0,000313398 - |
| 2065 | 1 | 3um_FIJJAFBF_815756 | Bacteria | Planctomycet | Phycisphaerae | Phycisphaeral | Phycisphaerac | Phycisphaera | Phycisphaera | information st | Transcription | 153I6 | DNA-depende | 575540.Isop_ | 1 | 0 | 4,29E-05 | 4,54E-06 | 1 - | 5,10E-05 - | 1 - | 4,27E-05 - | 1 - | 4,29E-05 - |
| 2066 | 1 | 3um_FIJJAFBF_21306 | Bacteria | Bacteroidetes | Flavobacteriia | Flavobacterial | Flavobacteria | Polaribacter | Polaribacter b | unclassified | unclassified | unclassified | unclassified | unclassified | 0,8 | 0,4 | 0,000121374 | 6,26E-05 | 1 - | 0,00017821 - | 1 - | 0,000148988 - | 1 - | 0,000149983 - |
| 2067 | 1 | 3um_FIJJAFBF_328494 | Bacteria | Bacteroidetes | Flavobacteriia | Flavobacterial | Flavobacteria | Arenitalea | Arenitalea lut | unclassified | unclassified | unclassified | unclassified | unclassified | 1 | 0 | 0,000206532 | 2,18E-05 | 1 - | 0,000245404 - | 1 - | 0,000205164 - | 1 - | 0,000206534 - |
| 2068 | 1 | 3um_FIJJAFBF_308981 | Bacteria | Proteobacteri | Alphaproteob | Pelagibacteral | unclassified | unclassified | Pelagibacteral | unclassified | unclassified | unclassified | unclassified | unclassified | 0,4 | 0,489897949 | 8,67E-05 | 0,000106829 | 1 - | 0,000235372 - | 0 - | 0 - | 1 - | 0,000198091 - |
| 2069 | 1 | 3um_FIJJAFBF_597540 | Eukaryota | Bacillariophyt | Bacillariophyc | Naviculales | Naviculaceae | Fistulifera | Fistulifera sola | unclassified | unclassified | unclassified | unclassified | unclassified | 2,6 | 0,489897949 | 0,000299567 | 6,78E-05 | 3 - | 0,000410502 - | 2 - | 0,000228793 - | 2 - | 0,000230322 - |
| 2070 | 1 | 3um_FIJJAFBF_116592 | Bacteria | Cyanobacteria | unclassified | Synechococca | Synechococca | Synechococcu | Synechococcu | metabolism | Energy produc | 1BCA9 | NADH dehydr | 3694.POPTR_ | 1 | 0,894427191 | 8,39E-05 | 7,50E-05 | 0 - | 0 - | 2 - | 0,00017853 - | 0 - | 0 - |
| 2071 | 1 | 3um_FIJJAFBF_788009 | Bacteria | Proteobacteri | Alphaproteob | Rhodobactera | Rhodobactera | unclassified | Rhodobactera | unclassified | unclassified | unclassified | unclassified | unclassified | 4,6 | 0,8 | 0,000863812 | 0,000202815 | 5 - | 0,001107222 - | 6 - | 0,001110797 - | 4 - | 0,000745479 - |
| 2072 | 4 | 0.2um_NOKCPHJN_108877;0.2um_NO | Bacteria | various | various | various | various | various | various | information st | Translation, ri | 08YY1 | This protein is | various | 0,2 | 0,4 | 0,000100967 | 0,000201934 | 0 - | 0 - | 0 - | 0 - | 0 - | 0 - |
| 2073 | 2 | 0.2um_NOKCPHJN_177948;0.2um_NO | Eukaryota | unclassified | Oomycetes | Peronosporale | Peronosporac | Plasmopara | Plasmopara ha | cellular proce | Intracellular tr | 0YIKI | Small GTPase | various | 1 | 0 | 0,000335065 | 3,54E-05 | 1 - | 0,000398129 - | 1 - | 0,000332845 - | 1 - | 0,000335069 - |
| 2074 | 1 | 0.2um_NOKCPHJN_251566 | Eukaryota | Phaeophyceae | unclassified | Ectocarpales | Ectocarpaceae | Ectocarpus | Ectocarpus sili | unclassified | unclassified | unclassified | unclassified | unclassified | 1 | 0 | 0,000342349 | 3,62E-05 | 1 - | 0,000406784 - | 1 - | 0,000340081 - | 1 - | 0,000342353 - |
| 2075 | 1 | 3um_FIJJAFBF_778664 | unclassified | unclassified | unclassified | unclassified | unclassified | unclassified | unclassified | unclassified | unclassified | unclassified | unclassified | unclassified | 1 | 0 | 0,002333044 | 0,000246726 | 1 - | 0,002772155 - | 1 - | 0,00231759 - | 1 - | 0,002333072 - |
| 2076 | 1 | 0.2um_NOKCPHJN_08458 | Bacteria | Proteobacteri | Gammaprotec | Alteromonada | Alteromonada | Paraglaciecola | Paraglaciecola | cellular proce | Posttranslatio | 05C6P | ATP-depender | 314275.MADE | 0,4 | 0,489897949 | 3,49E-05 | 4,30E-05 | 1 - | 9,50E-05 - | 1 - | 7,94E-05 - | 0 - | 0 - |
| 2077 | 1 | 3um_FIJJAFBF_84361 | Eukaryota | Bacillariophyt | Mediophyceae | Cymatosirales | Cymatosiracea | Plagiogrammo | Plagiogrammo | metabolism | Energy produc | 0IHF0 | This is one of | 45157.CMV04 | 4 | 0 | 0,000699913 | 7,40E-05 | 4 - | 0,000831646 - | 4 - | 0,000695277 - | 4 - | 0,000699922 - |
| 2078 | 1 | 3um_FIJJAFBF_817776 | Bacteria | Proteobacteri | Alphaproteob | Rhodobactera | Rhodobactera | Tateyamaria | Tateyamaria s | metabolism | Energy produc | 01RNF | Produces ATP | 391613.RTM1 | 0,4 | 0,489897949 | 4,94E-05 | 6,07E-05 | 0 - | 0 - | 1 - | 0,000132015 - | 0 - | 0 - |
| 2079 | 1 | 0.2um_NOKCPHJN_1105444 | Eukaryota | Bacillariophyt | Coscinodiscop | Thalassiosirale | Thalassiosirac | Thalassiosira | Thalassiosira p | unclassified | unclassified | unclassified | unclassified | unclassified | 0,8 | 0,4 | 0,000102354 | 5,22E-05 | 1 - | 0,000147049 - | 1 - | 0,000122937 - | 1 - | 0,000123758 - |
| 2080 | 1 | 0.2um_NOKCPHJN_163825 | Bacteria | Bacteroidetes | Cytophagia | Cytophagales | Flammeovirga | unclassified | Flammeovirga | unclassified | unclassified | unclassified | unclassified | unclassified | 0,2 | 0,4 | 2,64E-05 | 5,28E-05 | 0 - | 0 - | 0 - | 0 - | 0 - | 0 - |
| 2081 | 1 | 0.2um_NOKCPHJN_800496 | Eukaryota | unclassified | Oomycetes | Saprolegniales | Saprolegniace | Saprolegnia | Saprolegnia pa | cellular proce | Posttranslatio | KOG0019 | heat shock pro | 67593.Physo1 | 2,2 | 0,748331477 | 0,000263815 | 8,76E-05 | 1 - | 0,000145055 - | 3 - | 0,000363808 - | 3 - | 0,000366238 - |
| 2082 | 1 | 3um_FIJJAFBF_113994 | Viruses | unclassified | unclassified | unclassified | unclassified | unclassified | uncultured vir | cellular proce | Cell wall/mem | 01Y9Y | glycosyl transf | 488538.SAR11 | 0,8 | 0,4 | 0,000105869 | 5,48E-05 | 1 - | 0,000157244 - | 1 - | 0,00013146 - | 0 - | 0 - |
| 2083 | 1 | 0.2um_NOKCPHJN_1102032 | Eukaryota | Ascomycota | Pneumocystid | Pneumocystid | Pneumocystid | Pneumocystis | Pneumocystis | cellular proce | Posttranslatio | 09YID | Heat shock 70 | 7425.NV1371 | 1 | 0 | 0,000623685 | 6,60E-05 | 1 - | 0,000741071 - | 1 - | 0,000619554 - | 1 - | 0,000623692 - |
| 2084 | 1 | 3um_FIJJAFBF_247267 | Eukaryota | Apicomplexa | Conoidasida | Eucoccidiorida | Sarcocystidae | Toxoplasma | Toxoplasma g | unclassified | unclassified | unclassified | unclassified | unclassified | 0,4 | 0,489897949 | 0,000186068 | 0,000232555 | 1 - | 0,000538476 - | 0 - | 0 - | 0 - | 0 - |
| 2085 | 1 | 3um_FIJJAFBF_387408 | Bacteria | Bacteroidetes | Flavobacteriia | Flavobacterial | Flavobacteria | Polaribacter | Polaribacter s | unclassified | unclassified | unclassified | unclassified | unclassified | 0,8 | 0,4 | 7,88E-05 | 4,07E-05 | 1 - | 0,000115685 - | 1 - | 9,67E-05 - | 1 - | 9,74E-05 - |
| 2086 | 3 | 0.2um_NOKCPHJN_359057;0.2um_NO | Bacteria | Proteobacteri | Alphaproteob | Rhodospirillale | Rhodospirillac | Candidatus En | Candidatus En | unclassified | unclassified | unclassified | unclassified | unclassified | 0,2 | 0,4 | 0,000118066 | 0,000236132 | 0 - | 0 - | 1 - | 0,000590329 - | 0 - | 0 - |
| 2087 | 1 | 0.2um_NOKCPHJN_570486 | Eukaryota | Phaeophyceae | unclassified | Ectocarpales | Ectocarpaceae | Ectocarpus | Ectocarpus sili | cellular proce | Posttranslatio | 14ZJ0 | STI1 | 67593.Physo1 | 0,4 | 0,489897949 | 0,000113636 | 0,000139176 | 0 - | 0 - | 1 - | 0,000283144 - | 1 - | 0,000285036 - |
| 2088 | 2 | 0.2um_NOKCPHJN_157072;0.2um_NO | Eukaryota | various | various | various | various | various | various | unclassified | unclassified | unclassified | unclassified | unclassified | 1,6 | 0,489897949 | 0,000674656 | 0,000172821 | 1 - | 0,000512659 - | 1 - | 0,000428595 - | 2 - | 0,000862917 - |
| 2089 | 1 | 0.2um_NOKCPHJN_888611 | Eukaryota | unclassified | Pelagophycea | Pelagomonad | unclassified | Aureococcus | Aureococcus a | unclassified | unclassified | unclassified | unclassified | unclassified | 0,2 | 0,4 | 1,56E-05 | 3,13E-05 | 0 - | 0 - | 0 - | 0 - | 0 - | 0 - |
| 2090 | 1 | 3um_FIJJAFBF_263351 | unclassified | unclassified | unclassified | unclassified | unclassified | unclassified | unclassified | unclassified | unclassified | unclassified | unclassified | unclassified | 1 | 0 | 0,000232443 | 2,46E-05 | 1 - | 0,000276193 - | 1 - | 0,000230904 - | 1 - | 0,000232446 - |
| 2091 | 1 | 0.2um_NOKCPHJN_420410 | Bacteria | Bacteroidetes | Cytophagia | Cytophagales | Flammeovirga | unclassified | Flammeovirga | unclassified | unclassified | unclassified | unclassified | unclassified | 0,6 | 0,489897949 | 3,76E-05 | 3,09E-05 | 0 - | 0 - | 1 - | 6,53E-05 - | 1 - | 6,58E-05 - |
| 2092 | 1 | 3um_FIJJAFBF_10572 | Eukaryota | unclassified | Oomycetes | Albuginales | Albuginaceae | Albugo | Albugo candid | unclassified | unclassified | unclassified | unclassified | unclassified | 2 | 1,264911064 | 0,000422041 | 0,000237369 | 1 - | 0,000261707 - | 3 - | 0,00065638 - | 1 - | 0,000220255 - |
| 2093 | 1 | 3um_FIJJAFBF_641634 | unclassified | unclassified | unclassified | unclassified | unclassified | unclassified | unclassified | unclassified | unclassified | unclassified | unclassified | unclassified | 0,4 | 0,489897949 | 4,73E-05 | 5,81E-05 | 0 - | 0 - | 1 - | 0,000126414 - | 0 - | 0 - |
| 2094 | 1 | 3um_FIJJAFBF_122379 | Bacteria | Bacteroidetes | Flavobacteriia | Flavobacterial | Flavobacteria | Polaribacter | Polaribacter s | poorly charact | Function unkn | 09HDN | NA | 313598.MED1 | 1 | 0 | 0,000417167 | 4,41E-05 | 1 - | 0,000495683 - | 1 - | 0,000414403 - | 1 - | 0,000417172 - |
| 2095 | 8 | 0.2um_NOKCPHJN_655150;3um_FIJJA | Eukaryota | various | various | various | various | various | various | metabolism | Energy produc | 123BF | photosystem I | 35128.Thapsd | 2 | 0 | 0,001448096 | 0,00015314 | 2 - | 0,001720648 - | 2 - | 0,001438504 - | 2 - | 0,001448114 - |
| 2096 | 1 | 3um_FIJJAFBF_565140 | unclassified | unclassified | unclassified | unclassified | unclassified | unclassified | unclassified | unclassified | unclassified | unclassified | unclassified | unclassified | 0,6 | 0,489897949 | 0,000464108 | 0,000379148 | 0 - | 0 - | 1 - | 0,000782186 - | 1 - | 0,000787412 - |
| 2097 | 1 | 3um_FIJJAFBF_362249 | Viruses | unclassified | unclassified | unclassified | Phycodnavirid | unclassified | Chrysochromu | unclassified | unclassified | unclassified | unclassified | unclassified | 2,4 | 0,489897949 | 0,000501538 | 0,000141453 | 3 - | 0,000736212 - | 2 - | 0,000410327 - | 2 - | 0,000413068 - |
| 2098 | 1 | 3um_FIJJAFBF_263679 | Bacteria | Proteobacteri | Alphaproteob | unclassified | unclassified | Candidatus Pu | Candidatus Pu | information st | Translation, ri | 0FN1J | This protein is | 533247.CRD_ | 2 | 0 | 0,001016003 | 0,000107445 | 2 - | 0,001207229 - | 2 - | 0,001009273 - | 2 - | 0,001016015 - |
| 2099 | 1 | 0.2um_NOKCPHJN_545392 | Archaea | unclassified | unclassified | unclassified | unclassified | unclassified | archaeon GW | poorly charact | Function unkn | 07GSP | RhoGEF Guani | 203122.Sde_0 | 0,2 | 0,4 | 1,98E-05 | 3,97E-05 | 0 - | 0 - | 1 - | 9,92E-05 - | 0 - | 0 - |
| 2100 | 2 | 0.2um_NOKCPHJN_18872;0.2um_NOK | Bacteria | Proteobacteri | Gammaprotec | Cellvibrionales | Porticoccacea | unclassified | various | metabolism | Lipid transpor | 16QA8 | reductase | 314287.GB22 | 0,2 | 0,4 | 5,07E-05 | 0,000101336 | 0 - | 0 - | 1 - | 0,00025334 - | 0 - | 0 - |
| 2101 | 1 | 3um_FIJJAFBF_191786 | Eukaryota | unclassified | Florideophyce | Gracilariales | Gracilariaceae | Gracilaria | Gracilaria chil | unclassified | unclassified | unclassified | unclassified | unclassified | 2 | 0 | 0,001259844 | 0,000133232 | 2 - | 0,001496964 - | 2 - | 0,001251498 - | 2 - | 0,001259859 - |
| 2102 | 1 | 0.2um_NOKCPHJN_94453 | Bacteria | Proteobacteri | Betaproteoba | Burkholderiale | Comamonada | Acidovorax | various | metabolism | Energy produc | COG0055 | Produces ATP | 420662.Mpe_ | 1 | 0 | 0,000134312 | 1,42E-05 | 1 - | 0,000159591 - | 1 - | 0,000133422 - | 1 - | 0,000134313 - |
| 2103 | 1 | 0.2um_NOKCPHJN_1007957 | Eukaryota | unclassified | Labyrinthulom | unclassified | Thraustochytr | Aurantiochytr | Aurantiochytr | metabolism | Inorganic ion t | COG0475 | Sodium hydro | 35128.Thaps2 | 0,6 | 0,489897949 | 7,96E-05 | 6,53E-05 | 0 - | 0 - | 1 - | 0,000140618 - | 0 - | 0 - |
| 2104 | 1 | 3um_FIJJAFBF_715058 | Eukaryota | Bacillariophyt | Mediophyceae | Cymatosirales | Cymatosiracea | Plagiogrammo | Plagiogrammo | information st | Translation, ri | COG0052 | 30S ribosomal | 35128.Thapsd | 3 | 0,632455532 | 0,000803916 | 9,47E-05 | 2 - | 0,000650854 - | 3 - | 0,000816195 - | 3 - | 0,000821647 - |
| 2105 | 1 | 3um_FIJJAFBF_22823 | Eukaryota | Bacillariophyt | Bacillariophyc | Naviculales | Phaeodactylac | Phaeodactylu | Phaeodactylu | unclassified | unclassified | unclassified | unclassified | unclassified | 1,2 | 0,4 | 0,000188297 | 6,41E-05 | 1 - | 0,000186654 - | 2 - | 0,000312094 - | 1 - | 0,00015709 - |
| 2106 | 1 | 3um_FIJJAFBF_111340 | Viruses | unclassified | unclassified | unclassified | Phycodnavirid | unclassified | Chrysochromu | unclassified | unclassified | unclassified | unclassified | unclassified | 17,4 | 2,870540019 | 0,002023439 | 0,000271096 | 17 - | 0,002369496 - | 15 - | 0,001747903 - | 16 - | 0,001876885 - |
| 2107 | 1 | 0.2um_NOKCPHJN_709035 | Eukaryota | unclassified | Labyrinthulom | unclassified | Thraustochytr | Aurantiochytr | Aurantiochytr | unclassified | unclassified | unclassified | unclassified | unclassified | 0,2 | 0,4 | 0,00010012 | 0,00020024 | 0 - | 0 - | 1 - | 0,000500599 - | 0 - | 0 - |
| 2108 | 1 | 0.2um_NOKCPHJN_479684 | Bacteria | Bacteroidetes | Saprospiria | Saprospirales | Haliscomenob | Haliscomenob | Haliscomenob | unclassified | unclassified | unclassified | unclassified | unclassified | 1 | 0 | 8,29E-05 | 8,77E-06 | 1 - | 9,85E-05 - | 1 - | 8,23E-05 - | 1 - | 8,29E-05 - |
| 2109 | 3 | 0.2um_NOKCPHJN_106262;0.2um_NO | Bacteria | Bacteroidetes | Flavobacteriia | Flavobacterial | Flavobacteria | Polaribacter | Polaribacter s | poorly charact | Function unkn | 0XQTX | Chromosome | 313594.PI23P | 0,4 | 0,489897949 | 2,18E-05 | 2,68E-05 | 0 - | 0 - | 0 - | 0 - | 0 - | 0 - |
| 2110 | 1 | 0.2um_NOKCPHJN_30588 | Bacteria | unclassified | unclassified | unclassified | unclassified | unclassified | uncultured ba | unclassified | unclassified | unclassified | unclassified | unclassified | 1,4 | 0,489897949 | 9,03E-05 | 2,75E-05 | 1 - | 7,81E-05 - | 1 - | 6,53E-05 - | 2 - | 0,000131509 - |
| 2111 | 1 | 3um_FIJJAFBF_715065 | Eukaryota | Bacillariophyt | Mediophyceae | Anaulales | Anaulaceae | Eunotogramm | Eunotogramm | unclassified | unclassified | unclassified | unclassified | unclassified | 1,6 | 0,8 | 0,001466591 | 0,000597595 | 1 - | 0,001134063 - | 1 - | 0,000948105 - | 1 - | 0,000954438 - |
| 2112 | 1 | 0.2um_NOKCPHJN_1092416 | Bacteria | Proteobacteri | Epsilonproteo | Campylobacte | Helicobactera | Sulfurimonas | Sulfurimonas | poorly charact | Function unkn | 0ZJXN | NA | 439483.CBGD | 0,2 | 0,4 | 3,07E-05 | 6,13E-05 | 0 - | 0 - | 0 - | 0 - | 0 - | 0 - |
| 2113 | 1 | 0.2um_NOKCPHJN_350434 | Bacteria | Bacteroidetes | Flavobacteriia | Flavobacterial | unclassified | unclassified | Flavobacterial | information st | Replication, re | 0EQ45 | snf2 family | 720554.Clocl_ | 1 | 0 | 0,000572656 | 6,06E-05 | 1 - | 0,000680438 - | 1 - | 0,000568863 - | 1 - | 0,000572663 - |
| 2114 | 1 | 0.2um_NOKCPHJN_513690 | Eukaryota | Apicomplexa | Conoidasida | Eucoccidiorida | Eimeriidae | Eimeria | Eimeria maxin | unclassified | unclassified | unclassified | unclassified | unclassified | 0,6 | 0,489897949 | 2,96E-05 | 2,43E-05 | 0 - | 0 - | 1 - | 5,23E-05 - | 0 - | 0 - |
| 2115 | 1 | 3um_FIJJAFBF_198300 | Bacteria | Bacteroidetes | Flavobacteriia | Flavobacterial | Flavobacteria | Polaribacter | Polaribacter s | metabolism | Inorganic ion t | 05C27 | TonB-depende | 487797.Flav3 | 0,6 | 0,489897949 | 7,44E-05 | 6,20E-05 | 1 - | 0,000144774 - | 0 - | 0 - | 1 - | 0,000121843 - |
| 2116 | 1 | 3um_FIJJAFBF_46437 | unclassified | unclassified | unclassified | unclassified | unclassified | unclassified | unclassified | unclassified | unclassified | unclassified | unclassified | unclassified | 0,4 | 0,489897949 | 0,000523695 | 0,000641693 | 0 - | 0 - | 0 - | 0 - | 1 - | 0,001340275 - |

| | | | | | | | | | | | | | | | | | | | | | | | | |
|---|---|---|---|---|---|---|---|---|---|---|---|---|---|---|---|---|---|---|---|---|---|---|---|---|
| 2117 | 1 | 0.2um_NOKCPHJN_126580 | Bacteria | Verrucomicrol | Opitutae | unclassified | unclassified | unclassified | Opitutae bact | unclassified | unclassified | unclassified | unclassified | unclassified | 10,6 | 0,8 | 0,001389766 | 0,000108882 | 10 - | 0,001565862 - | 10 - | 0,001309099 - | 11 - | 0,001449628 - |
| 2118 | 1 | 0.2um_NOKCPHJN_289944 | Bacteria | unclassified | unclassified | unclassified | unclassified | unclassified | bacterium BM | cellular proce: | Signal transdu | KOG0613 | fibronectin typ | 7070.TC00296 | 0,2 | 0,4 | 8,46E-06 | 1,69E-05 | 0 - | 0 - | 1 - | 4,23E-05 - | 0 - | 0 - |
| 2119 | 1 | 0.2um_NOKCPHJN_953987 | Bacteria | Proteobacteri | Alphaproteob | Pelagibacteral | Pelagibacterac | unclassified | Pelagibacterac | metabolism | Energy produc | 05C26 | Dehydrogenas | 335992.SAR11 | 0,8 | 0,4 | 9,74E-05 | 4,91E-05 | 0 - | 0 - | 1 - | 0,000126927 - | 1 - | 0,000127775 - |
| 2120 | 1 | 0.2um_NOKCPHJN_288576 | Eukaryota | unclassified | Labyrinthulom | unclassified | Thraustochytr | Aurantiochytr | Aurantiochytr | poorly charact | Function unkn | 0Z05D | TUDOR | 67593.Physo1 | 0,2 | 0,4 | 2,42E-06 | 4,83E-06 | 0 - | 0 - | 0 - | 0 - | 0 - | 0 - |
| 2121 | 1 | 0.2um_NOKCPHJN_1034651 | Eukaryota | unclassified | Dinophyceae | Suessiales | Symbiodiniace | Symbiodinium | Symbiodinium | unclassified | unclassified | unclassified | unclassified | unclassified | 0,6 | 0,489897949 | 0,0001107 | 9,08E-05 | 0 - | 0 - | 1 - | 0,000195547 - | 0 - | 0 - |
| 2122 | 1 | 0.2um_NOKCPHJN_933735 | Bacteria | Actinobacteri | Actinobacteri | Streptomycet | Streptomycet | Streptomyces | Streptomyces | unclassified | unclassified | unclassified | unclassified | unclassified | 1 | 0 | 0,000101437 | 1,07E-05 | 1 - | 0,000120528 - | 1 - | 0,000100765 - | 1 - | 0,000101438 - |
| 2123 | 1 | 0.2um_NOKCPHJN_632309 | unclassified | unclassified | unclassified | unclassified | unclassified | unclassified | unclassified | unclassified | unclassified | unclassified | unclassified | unclassified | 0,6 | 0,489897949 | 6,84E-05 | 5,61E-05 | 0 - | 0 - | 1 - | 0,000120801 - | 0 - | 0 - |
| 2124 | 1 | 3um_FIJJAFBF_510231 | Bacteria | Bacteroidetes | Flavobacteriia | Flavobacterial | Flavobacteria | Polaribacter | Polaribacter r | information st | Translation, ri | COG0088 | One of the pri | 313598.MED1 | 1,4 | 0,489897949 | 0,00041101 | 0,000115128 | 1 - | 0,000358125 - | 1 - | 0,000299402 - | 1 - | 0,000301402 - |
| 2125 | 1 | 3um_FIJJAFBF_457728 | Eukaryota | Mollusca | Gastropoda | Architaeniogl | Ampullariidae | Pomacea | Pomacea cana | unclassified | unclassified | unclassified | unclassified | unclassified | 1,8 | 1,166190379 | 7,46E-05 | 4,93E-05 | 0 - | 0 - | 1 - | 4,32E-05 - | 3 - | 0,00013033 - |
| 2126 | 1 | 3um_FIJJAFBF_522166 | Eukaryota | unclassified | unclassified | Isochrysidales | Noelaerhabda | Emiliania | Emiliania huxl | unclassified | unclassified | unclassified | unclassified | unclassified | 1 | 0 | 0,000533832 | 5,65E-05 | 1 - | 0,000634307 - | 1 - | 0,000530296 - | 1 - | 0,000533838 - |
| 2127 | 1 | 3um_FIJJAFBF_152376 | unclassified | unclassified | unclassified | unclassified | unclassified | unclassified | unclassified | unclassified | unclassified | unclassified | unclassified | unclassified | 11,6 | 2,244994432 | 0,002801433 | 0,000413585 | 11 - | 0,003191202 - | 11 - | 0,002667923 - | 10 - | 0,002441587 - |
| 2128 | 1 | 3um_FIJJAFBF_82638 | unclassified | unclassified | unclassified | unclassified | unclassified | unclassified | unclassified | unclassified | unclassified | unclassified | unclassified | unclassified | 1 | 0 | 0,000169334 | 1,79E-05 | 1 - | 0,000201205 - | 1 - | 0,000168212 - | 1 - | 0,000169336 - |
| 2129 | 1 | 3um_FIJJAFBF_49868 | Bacteria | Proteobacteri | Alphaproteob | Rhodobactera | Rhodobactera | Sulfitobacter | various | metabolism | Energy produc | 01RJJ | NADH flavin o | 314264.ROS2 | 1 | 0 | 0,000588712 | 6,23E-05 | 1 - | 0,000699516 - | 1 - | 0,000584812 - | 1 - | 0,000588719 - |
| 2130 | 1 | 0.2um_NOKCPHJN_682989 | Eukaryota | Phaeophyceae | unclassified | Ectocarpales | Ectocarpaceae | Ectocarpus | Ectocarpus sili | unclassified | unclassified | unclassified | unclassified | unclassified | 0,8 | 0,4 | 0,000118675 | 6,05E-05 | 1 - | 0,000170497 - | 1 - | 0,00014254 - | 1 - | 0,000143492 - |
| 2131 | 1 | 3um_FIJJAFBF_657472 | Bacteria | Proteobacteri | Alphaproteob | Rhodobactera | Rhodobactera | Yoonia | Yoonia marico | metabolism | Amino acid tra | 16S09 | Synthesizes se | 391593.RCCS2 | 0,4 | 0,489897949 | 9,63E-05 | 0,000118454 | 0 - | 0 - | 1 - | 0,00025751 - | 0 - | 0 - |
| 2132 | 1 | 0.2um_NOKCPHJN_21852 | Bacteria | Bacteroidetes | Flavobacteriia | Flavobacterial | Flavobacteria | Polaribacter | Polaribacter s | poorly charact | Function unkn | 0PBET | NA | 313594.PI23P | 0,2 | 0,4 | 7,11E-05 | 0,000142216 | 0 - | 0 - | 1 - | 0,000355539 - | 0 - | 0 - |
| 2133 | 1 | 0.2um_NOKCPHJN_596448 | Bacteria | Bacteroidetes | Bacteroidia | Marinilabiliale | Prolixibactera | Prolixibacter | Prolixibacter c | unclassified | unclassified | unclassified | unclassified | unclassified | 0,8 | 0,4 | 0,000155893 | 8,04E-05 | 1 - | 0,000228894 - | 1 - | 0,000191361 - | 1 - | 0,000192639 - |
| 2134 | 1 | 0.2um_NOKCPHJN_504894 | Bacteria | Proteobacteri | Alphaproteob | Rhodobactera | Rhodobactera | Oceaniovalibu | Oceaniovalibu | unclassified | unclassified | unclassified | unclassified | unclassified | 1,6 | 1,356465997 | 0,000223432 | 0,00018272 | 1 - | 0,00017167 - | 0 - | 0 - | 1 - | 0,000144479 - |
| 2135 | 1 | 3um_FIJJAFBF_44066 | unclassified | unclassified | unclassified | unclassified | unclassified | unclassified | unclassified | unclassified | unclassified | unclassified | unclassified | unclassified | 1 | 0 | 0,001105126 | 0,00011687 | 1 - | 0,001313126 - | 1 - | 0,001097806 - | 1 - | 0,001105139 - |
| 2136 | 1 | 3um_FIJJAFBF_503363 | unclassified | unclassified | unclassified | unclassified | unclassified | unclassified | unclassified | unclassified | unclassified | unclassified | unclassified | unclassified | 1 | 0 | 0,000862907 | 9,13E-05 | 1 - | 0,001025318 - | 1 - | 0,000857191 - | 1 - | 0,000862917 - |
| 2137 | 1 | 3um_FIJJAFBF_154096 | Eukaryota | Bacillariophyt | Bacillariophyc | Naviculales | Naviculaceae | Fistulifera | Fistulifera sola | unclassified | unclassified | unclassified | unclassified | unclassified | 0,8 | 0,4 | 0,000440574 | 0,000222044 | 0 - | 0 - | 1 - | 0,000574082 - | 1 - | 0,000577917 - |
| 2138 | 1 | 0.2um_NOKCPHJN_106716 | Bacteria | Proteobacteri | Gammaprotec | Cellvibrionales | Spongiibacter | Oceanicoccus | Oceanicoccus | poorly charact | Function unkn | 0R9NA | TPR | 857087.Metm | 0,6 | 0,8 | 4,34E-05 | 5,63E-05 | 0 - | 0 - | 1 - | 7,92E-05 - | 0 - | 0 - |
| 2139 | 1 | 3um_FIJJAFBF_51134 | unclassified | unclassified | unclassified | unclassified | unclassified | unclassified | unclassified | unclassified | unclassified | unclassified | unclassified | unclassified | 2 | 1,264911064 | 0,002546893 | 0,001404672 | 1 - | 0,001592515 - | 1 - | 0,001331381 - | 1 - | 0,001340275 - |
| 2140 | 1 | 3um_FIJJAFBF_177926 | Eukaryota | Bacillariophyt | Coscinodiscop | Coscinodiscale | Coscinodiscac | Coscinodiscus | Coscinodiscus | unclassified | unclassified | unclassified | unclassified | unclassified | 2 | 1,095445115 | 0,000384673 | 0,000175991 | 1 - | 0,00023837 - | 2 - | 0,000398566 - | 2 - | 0,000401229 - |
| 2141 | 1 | 0.2um_NOKCPHJN_688594 | Bacteria | Proteobacteri | Alphaproteob | Rhodospirillal | Rhodospirillac | Azospirillum | Azospirillum s | metabolism | Lipid transpor | 16PEQ | Amp-depende | 1064539.AZOI | 1 | 0 | 0,000214259 | 2,27E-05 | 1 - | 0,000254586 - | 1 - | 0,00021284 - | 1 - | 0,000214262 - |
| 2142 | 1 | 3um_FIJJAFBF_684334 | Eukaryota | unclassified | Cryptophyta | Pyrenomonad | Geminigerace | Guillardia | Guillardia thet | cellular proce: | Cytoskeleton | 0XPVJ | GTPase activit | 45157.CMD10 | 1,8 | 0,4 | 0,00047048 | 0,000118266 | 2 - | 0,000621147 - | 2 - | 0,000519294 - | 1 - | 0,000261381 - |
| 2143 | 1 | 3um_FIJJAFBF_260595 | Bacteria | Bacteroidetes | Saprospiria | Saprospirales | Saprospiracea | Aureispira | Aureispira sp. | information st | Translation, ri | 09EM4 | Binds directly | 926562.Oweh | 0,2 | 0,4 | 0,000109781 | 0,000219561 | 0 - | 0 - | 1 - | 0,000548903 - | 0 - | 0 - |
| 2144 | 1 | 0.2um_NOKCPHJN_794140 | unclassified | unclassified | unclassified | unclassified | unclassified | unclassified | unclassified | unclassified | unclassified | unclassified | unclassified | unclassified | 0,4 | 0,489897949 | 0,000121446 | 0,000149654 | 1 - | 0,000329728 - | 0 - | 0 - | 1 - | 0,000277502 - |
| 2145 | 1 | 0.2um_NOKCPHJN_243308 | Eukaryota | unclassified | unclassified | Longamoebia | Acanthamoeb | Acanthamoeb | Acanthamoeb | cellular proce: | Defense mech | COG1680 | Beta-lactamas | 44689.DDB01 | 0,2 | 0,4 | 2,39E-05 | 4,79E-05 | 0 - | 0 - | 0 - | 0 - | 0 - | 0 - |
| 2146 | 1 | 3um_FIJJAFBF_400812 | Bacteria | Firmicutes | Clostridia | Clostridiales | Clostridiaceae | Clostridium | Clostridium sp | metabolism | Carbohydrate | COG0057 | glyceraldehyd | 281687.CJA04 | 1 | 0,632455532 | 0,000181036 | 0,000104001 | 1 - | 0,000220791 - | 0 - | 0 - | 1 - | 0,00018582 - |
| 2147 | 1 | 0.2um_NOKCPHJN_341236 | Eukaryota | unclassified | Pelagophycea | Pelagomonad | unclassified | Aureococcus | Aureococcus a | unclassified | unclassified | unclassified | unclassified | unclassified | 3 | 0 | 0,000287198 | 3,04E-05 | 3 - | 0,000341253 - | 3 - | 0,000285296 - | 3 - | 0,000287202 - |
| 2148 | 1 | 0.2um_NOKCPHJN_20493 | Bacteria | Proteobacteri | Gammaprotec | Cellvibrionales | Spongiibacter | unclassified | marine gamm | metabolism | Amino acid tra | COG0174 | glutamine syn | 247633.GP21 | 2,8 | 0,4 | 0,000376876 | 6,87E-05 | 3 - | 0,000479796 - | 3 - | 0,000401121 - | 2 - | 0,000269201 - |
| 2149 | 1 | 3um_FIJJAFBF_45623 | Bacteria | Planctomycet | Phycisphaerae | Phycisphaeral | Phycisphaerac | Phycisphaera | Phycisphaera | unclassified | unclassified | unclassified | unclassified | unclassified | 0,8 | 0,748331477 | 0,00015503 | 0,000146633 | 0 - | 0 - | 1 - | 0,000204493 - | 0 - | 0 - |
| 2150 | 1 | 0.2um_NOKCPHJN_880073 | Bacteria | Cyanobacteria | unclassified | Synechococca | Synechococca | Synechococcu | Synechococcu | unclassified | unclassified | unclassified | unclassified | unclassified | 0,8 | 0,748331477 | 0,000101821 | 8,95E-05 | 1 - | 0,000156259 - | 0 - | 0 - | 0 - | 0 - |
| 2151 | 1 | 0.2um_NOKCPHJN_523104 | Eukaryota | Phaeophyceae | unclassified | Ectocarpales | Ectocarpaceae | Ectocarpus | Ectocarpus sili | cellular proce: | Intracellular tr | KOG2514 | Anoctamin | 65071.PYU1_T | 1 | 0 | 0,000176944 | 1,87E-05 | 1 - | 0,000210248 - | 1 - | 0,000175772 - | 1 - | 0,000176946 - |
| 2152 | 1 | 3um_FIJJAFBF_442784 | Eukaryota | Bacillariophyt | Bacillariophyc | Naviculales | Naviculaceae | Fistulifera | Fistulifera sola | unclassified | unclassified | unclassified | unclassified | unclassified | 1,2 | 0,4 | 0,000272642 | 6,74E-05 | 1 - | 0,000276193 - | 1 - | 0,000230904 - | 1 - | 0,000232446 - |
| 2153 | 1 | 0.2um_NOKCPHJN_934182 | Bacteria | Proteobacteri | Gammaprotec | Pseudomonad | Pseudomonac | Pseudomonas | Pseudomonas | unclassified | unclassified | unclassified | unclassified | unclassified | 0,2 | 0,4 | 3,75E-05 | 7,50E-05 | 1 - | 0,000187589 - | 0 - | 0 - | 0 - | 0 - |
| 2154 | 1 | 3um_FIJJAFBF_119151 | Bacteria | Bacteroidetes | Flavobacteriia | Flavobacterial | Flavobacteria | Tenacibaculur | Tenacibaculur | unclassified | unclassified | unclassified | unclassified | unclassified | 1 | 0 | 0,000268052 | 2,83E-05 | 1 - | 0,000318503 - | 1 - | 0,000266276 - | 1 - | 0,000268055 - |
| 2155 | 1 | 3um_FIJJAFBF_267386 | Eukaryota | Apicomplexa | Aconoidasida | Haemosporida | Plasmodiidae | Plasmodium | Plasmodium y | unclassified | unclassified | unclassified | unclassified | unclassified | 4,4 | 0,489897949 | 0,000927479 | 9,54E-05 | 4 - | 0,001008056 - | 4 - | 0,00084276 - | 5 - | 0,001060487 - |
| 2156 | 1 | 0.2um_NOKCPHJN_496935 | unclassified | unclassified | unclassified | unclassified | unclassified | unclassified | unclassified | unclassified | unclassified | unclassified | unclassified | unclassified | 0,8 | 0,4 | 0,000266633 | 0,000137897 | 1 - | 0,000396022 - | 1 - | 0,000331084 - | 0 - | 0 - |
| 2157 | 1 | 0.2um_NOKCPHJN_741825 | Eukaryota | unclassified | Spirotrichea | Sporadotrichi | Oxytrichidae | Stylonychia | Stylonychia le | cellular proce: | Posttranslatio | KOG0019 | heat shock pro | 423536.XP_00 | 0,4 | 0,489897949 | 4,75E-05 | 5,93E-05 | 1 - | 0,000137336 - | 0 - | 0 - | 0 - | 0 - |
| 2158 | 1 | 3um_FIJJAFBF_253700 | Bacteria | Proteobacteri | Gammaprotec | Cellvibrionales | Halieaceae | Chromatocurv | Chromatocurv | unclassified | unclassified | unclassified | unclassified | unclassified | 2 | 0,894427191 | 0,000143869 | 6,24E-05 | 1 - | 8,63E-05 - | 2 - | 0,000144348 - | 3 - | 0,000217969 - |
| 2159 | 1 | 3um_FIJJAFBF_734480 | Eukaryota | Platyhelminth | Trematoda | Opisthorchiida | Opisthorchiida | Opisthorchis | Opisthorchis v | unclassified | unclassified | unclassified | unclassified | unclassified | 3 | 0 | 0,000684698 | 7,24E-05 | 3 - | 0,000813567 - | 3 - | 0,000680162 - | 3 - | 0,000684706 - |
| 2160 | 1 | 3um_FIJJAFBF_10380 | Eukaryota | Phaeophyceae | unclassified | Ectocarpales | Ectocarpaceae | Ectocarpus | Ectocarpus sili | unclassified | unclassified | unclassified | unclassified | unclassified | 1 | 0 | 0,000168428 | 1,78E-05 | 1 - | 0,000200129 - | 1 - | 0,000167313 - | 1 - | 0,00016843 - |
| 2161 | 1 | 3um_FIJJAFBF_321781 | Eukaryota | unclassified | Oomycetes | Peronosporale | Peronosporac | Phytophthora | Phytophthora | information st | Translation, ri | 151Y2 | Ribosomal pro | 272952.HpaP8 | 0,4 | 0,489897949 | 0,00018498 | 0,000227545 | 0 - | 0 - | 0 - | 0 - | 1 - | 0,000496007 - |
| 2162 | 1 | 3um_FIJJAFBF_690863 | Viruses | unclassified | unclassified | unclassified | unclassified | unclassified | uncultured M | unclassified | unclassified | unclassified | unclassified | unclassified | 0,2 | 0,4 | 5,95E-05 | 0,000119059 | 0 - | 0 - | 0 - | 0 - | 0 - | 0 - |
| 2163 | 1 | 0.2um_NOKCPHJN_385093 | Bacteria | Proteobacteri | Alphaproteob | Rhodospirillal | Rhodospirillac | Tistlia | Tistlia consote | metabolism | Secondary me | 16Y20 | Inherit from C | 414684.RC1_3 | 1 | 0 | 2,27E-05 | 2,40E-06 | 1 - | 2,69E-05 - | 1 - | 2,25E-05 - | 1 - | 2,27E-05 - |
| 2164 | 1 | 0.2um_NOKCPHJN_710508 | Eukaryota | Eustigmatoph | unclassified | Eustigmatales | Monodopsida | Nannochlorop | Nannochlorop | metabolism | Energy produc | 1BFZ8 | Cytochrome b | 3218.PP1S35_ | 3,6 | 0,489897949 | 0,001840033 | 0,000179606 | 3 - | 0,001840529 - | 3 - | 0,001538728 - | 4 - | 0,002065342 - |
| 2165 | 1 | 3um_FIJJAFBF_816773 | Eukaryota | unclassified | Spirotrichea | Sporadotrichi | Oxytrichidae | Laurentiella | Laurentiella st | unclassified | unclassified | unclassified | unclassified | unclassified | 0,2 | 0,4 | 2,86E-05 | 5,72E-05 | 0 - | 0 - | 0 - | 0 - | 0 - | 0 - |
| 2166 | 1 | 3um_FIJJAFBF_300655 | Eukaryota | Bacillariophyt | Coscinodiscop | Thalassiosirale | Thalassiosirac | Thalassiosira | Thalassiosira c | unclassified | unclassified | unclassified | unclassified | unclassified | 0,8 | 0,4 | 0,000167698 | 8,67E-05 | 1 - | 0,000248665 - | 0 - | 0 - | 1 - | 0,000209279 - |
| 2167 | 1 | 3um_FIJJAFBF_518684 | Bacteria | Bacteroidetes | Flavobacteriia | Flavobacterial | Flavobacteria | Polaribacter | Polaribacter s | metabolism | Carbohydrate | 05CA9 | Pyruvate kina: | 313594.PI23P | 0,4 | 0,489897949 | 5,29E-05 | 6,48E-05 | 0 - | 0 - | 1 - | 0,000131737 - | 1 - | 0,000132617 - |
| 2168 | 1 | 3um_FIJJAFBF_590394 | Eukaryota | unclassified | Pelagophycea | Pelagomonad | unclassified | Aureococcus | Aureococcus a | unclassified | unclassified | unclassified | unclassified | unclassified | 1 | 0 | 0,000177944 | 1,88E-05 | 1 - | 0,000211436 - | 1 - | 0,000176765 - | 1 - | 0,000177946 - |
| 2169 | 1 | 3um_FIJJAFBF_581298 | Bacteria | Bacteroidetes | Flavobacteriia | Flavobacterial | Crocinitomica | unclassified | Crocinitomica | information st | Replication, re | 0PB0H | DNA-binding p | 755732.Fluta_ | 3,2 | 0,4 | 0,001833673 | 0,000163349 | 3 - | 0,002060042 - | 3 - | 0,001722245 - | 3 - | 0,001733751 - |
| 2170 | 1 | 3um_FIJJAFBF_564131 | Bacteria | Bacteroidetes | Flavobacteriia | Flavobacterial | Flavobacteria | Mesoflavibact | various | metabolism | Carbohydrate | COG0057 | glyceraldehyd | 391587.KAOT | 0,2 | 0,4 | 3,81E-05 | 7,61E-05 | 0 - | 0 - | 0 - | 0 - | 1 - | 0,000190311 - |
| 2171 | 1 | 3um_FIJJAFBF_328496 | Bacteria | Bacteroidetes | Flavobacteriia | Flavobacterial | Flavobacteria | Arenitalea | Arenitalea lut | metabolism | Inorganic ion t | 05C27 | TonB-depende | 886377.Murru | 19 | 3,405877273 | 0,001132692 | 0,00013121 | 15 - | 0,001078504 - | 15 - | 0,000901656 - | 20 - | 0,001210239 - |
| 2172 | 1 | 0.2um_NOKCPHJN_36508 | Bacteria | Bacteroidetes | Cytophagia | Cytophagales | Flammeovirga | unclassified | Flammeovirga | poorly charact | Function unkn | 1AINX | Phage tail she | 760192.Halhy | 1,6 | 0,8 | 0,000210529 | 0,000113278 | 2 - | 0,000305503 - | 1 - | 0,000127704 - | 3 - | 0,000385671 - |
| 2173 | 1 | 3um_FIJJAFBF_125686 | Eukaryota | Bacillariophyt | Bacillariophyc | Naviculales | Phaeodactyla | Phaeodactylu | Phaeodactylu | unclassified | unclassified | unclassified | unclassified | unclassified | 2 | 0 | 0,000492126 | 5,20E-05 | 2 - | 0,000584751 - | 2 - | 0,000488867 - | 2 - | 0,000492132 - |
| 2174 | 1 | 3um_FIJJAFBF_01154 | Bacteria | Bacteroidetes | Flavobacteriia | Flavobacterial | Flavobacteria | Polaribacter | Polaribacter a | cellular proce: | Cell wall/mem | 09G63 | Channel that c | 313598.MED1 | 0,2 | 0,4 | 8,25E-05 | 0,000165059 | 0 - | 0 - | 0 - | 0 - | 0 - | 0 - |
| 2175 | 1 | 3um_FIJJAFBF_774536 | Eukaryota | unclassified | Cryptophyta | Pyrenomonad | Geminigerace | Guillardia | Guillardia thet | unclassified | unclassified | unclassified | unclassified | unclassified | 2,8 | 0,4 | 0,00098309 | 8,58E-05 | 2 - | 0,000845742 - | 3 - | 0,001060592 - | 3 - | 0,001067677 - |
| 2176 | 1 | 3um_FIJJAFBF_269266 | Eukaryota | Arthropoda | Insecta | Lepidoptera | Papilionidae | Papilio | Papilio macha | poorly charact | Function unkn | 0IRPM | Retrotranspos | 6085.XP_0021 | 0,2 | 0,4 | 2,26E-05 | 4,53E-05 | 0 - | 0 - | 0 - | 0 - | 0 - | 0 - |
| 2177 | 1 | 0.2um_NOKCPHJN_664508 | Eukaryota | Eustigmatoph | unclassified | Eustigmatales | Monodopsida | Nannochlorop | Nannochlorop | cellular proce: | Posttranslatio | KOG0179 | The proteasor | 35128.Thaps2 | 0,4 | 0,489897949 | 9,85E-05 | 0,000120638 | 0 - | 0 - | 0 - | 0 - | 1 - | 0,000251972 - |
| 2178 | 1 | 0.2um_NOKCPHJN_899537 | Bacteria | Proteobacteri | Gammaprotec | Thiotrichales | Piscirickettsia | Thiomicrospir | Thiomicrospir | cellular proce: | Cell motility | COG1360 | Ompa motb d | 394.NGR_c05 | 0,6 | 0,489897949 | 0,000157289 | 0,000131151 | 1 - | 0,000306755 - | 1 - | 0,000256455 - | 0 - | 0 - |
| 2179 | 1 | 3um_FIJJAFBF_273735 | unclassified | unclassified | unclassified | unclassified | unclassified | unclassified | unclassified | unclassified | unclassified | unclassified | unclassified | unclassified | 1 | 0 | 0,000851246 | 9,00E-05 | 1 - | 0,001011462 - | 1 - | 0,000845607 - | 1 - | 0,000851256 - |
| 2180 | 1 | 3um_FIJJAFBF_545217 | Bacteria | Bacteroidetes | Flavobacteriia | Flavobacterial | Flavobacteria | Arenitalea | Arenitalea lut | information st | Translation, ri | COG0256 | This is one of | 983544.Lacal_ | 0,6 | 0,489897949 | 0,000305148 | 0,00025048 | 0 - | 0 - | 1 - | 0,000530296 - | 1 - | 0,000533838 - |
| 2181 | 1 | 3um_FIJJAFBF_302477 | Eukaryota | unclassified | Pelagophycea | Pelagomonad | unclassified | Aureococcus | Aureococcus a | unclassified | unclassified | unclassified | unclassified | unclassified | 1 | 0 | 0,000642777 | 6,80E-05 | 1 - | 0,000763757 - | 1 - | 0,00063852 - | 1 - | 0,000642785 - |
| 2182 | 1 | 0.2um_NOKCPHJN_904618 | Eukaryota | unclassified | Pelagophycea | Pelagomonad | unclassified | Aureococcus | Aureococcus a | cellular proce: | Posttranslatio | KOG3083 | prohibitin | 653948.CCA24 | 1 | 0,632455532 | 0,000467843 | 0,000269106 | 1 - | 0,00057136 - | 1 - | 0,000477671 - | 0 - | 0 - |
| 2183 | 1 | 0.2um_NOKCPHJN_1113290 | Bacteria | Bacteroidetes | Bacteroidia | Bacteroidales | Prevotellacea | Prevotella | various | unclassified | unclassified | unclassified | unclassified | unclassified | 2 | 0 | 0,000290957 | 3,08E-05 | 2 - | 0,000345719 - | 2 - | 0,00028903 - | 2 - | 0,00029096 - |
| 2184 | 1 | 0.2um_NOKCPHJN_1021094 | Bacteria | Proteobacteri | Betaproteoba | Nitrosomonad | unclassified | unclassified | Methylophilal | cellular proce: | Signal transdu | 0B8J2 | Signal transdu | 383631.MB21 | 0,2 | 0,4 | 3,45E-05 | 6,90E-05 | 0 - | 0 - | 0 - | 0 - | 1 - | 0,000172583 - |
| 2185 | 1 | 0.2um_NOKCPHJN_78044 | Eukaryota | Bacillariophyt | Bacillariophyc | Bacillariales | Bacillariaceae | Nitzschia | Nitzschia sp. I | cellular proce: | Posttranslatio | KOG0730 | cell division cy | 2850.Phatr54 | 1 | 0 | 7,86E-05 | 8,32E-06 | 1 - | 9,34E-05 - | 1 - | 7,81E-05 - | 1 - | 7,86E-05 - |
| 2186 | 1 | 3um_FIJJAFBF_798417 | Eukaryota | unclassified | unclassified | Kinetoplastida | Trypanosomat | Leishmania | Leishmania m | unclassified | unclassified | unclassified | unclassified | unclassified | 0,2 | 0,4 | 1,18E-05 | 2,36E-05 | 0 - | 0 - | 0 - | 0 - | 0 - | 0 - |
| 2187 | 1 | 0.2um_NOKCPHJN_170568 | unclassified | unclassified | unclassified | unclassified | unclassified | unclassified | unclassified | unclassified | unclassified | unclassified | unclassified | unclassified | 0,6 | 0,8 | 1,83E-05 | 2,49E-05 | 0 - | 0 - | 0 - | 0 - | 0 - | 0 - |
| 2188 | 1 | 0.2um_NOKCPHJN_1003141 | Eukaryota | Nemertea | Pilidiophora | Heteronemert | Cerebratulida | Cerebratulus | Cerebratulus l | unclassified | unclassified | unclassified | unclassified | unclassified | 1,8 | 0,748331477 | 0,000990032 | 0,000338661 | 1 - | 0,000674308 - | 2 - | 0,001127476 - | 2 - | 0,001135008 - |
| 2189 | 1 | 3um_FIJJAFBF_60206 | Bacteria | Actinobacteri | unclassified | unclassified | unclassified | unclassified | Actinobacteri | cellular proce: | Intracellular tr | 0QJ7H | motA TolQ Exl | 314285.KT71_ | 2,4 | 0,8 | 0,000315601 | 8,98E-05 | 1 - | 0,000161311 - | 3 - | 0,000404579 - | 2 - | 0,000271521 - |
| 2190 | 1 | 0.2um_NOKCPHJN_398524 | Eukaryota | unclassified | Oomycetes | Saprolegniales | Saprolegniace | Saprolegnia | Saprolegnia di | unclassified | unclassified | unclassified | unclassified | unclassified | 0,2 | 0,4 | 3,97E-05 | 7,93E-05 | 0 - | 0 - | 0 - | 0 - | 0 - | 0 - |
| 2191 | 1 | 3um_FIJJAFBF_189700 | Bacteria | Bacteroidetes | Flavobacteriia | Flavobacterial | Flavobacteria | Polaribacter | Polaribacter s | unclassified | unclassified | unclassified | unclassified | unclassified | 1 | 0 | 9,94E-05 | 1,05E-05 | 1 - | 0,000118057 - | 1 - | 9,87E-05 - | 1 - | 9,94E-05 - |
| 2192 | 1 | 0.2um_NOKCPHJN_1060146 | Eukaryota | unclassified | unclassified | Choanoflagelli | Salpingoecida | Salpingoeca | Salpingoeca r | unclassified | unclassified | unclassified | unclassified | unclassified | 0,4 | 0,489897949 | 4,33E-05 | 5,33E-05 | 0 - | 0 - | 1 - | 0,000115879 - | 0 - | 0 - |
| 2193 | 1 | 3um_FIJJAFBF_235861 | Bacteria | Proteobacteri | Gammaprotec | Cellvibrionales | Microbulbifer | Microbulbifer | Microbulbifer | unclassified | unclassified | unclassified | unclassified | unclassified | 1,2 | 0,4 | 0,00049588 | 0,000122544 | 1 - | 0,000502337 - | 1 - | 0,000419966 - | 1 - | 0,000422771 - |
| 2194 | 1 | 3um_FIJJAFBF_500683 | Eukaryota | Bacillariophyt | Mediophycea | Cymatosirales | Cymatosiracea | Plagiogrammc | Plagiogrammc | metabolism | Energy produc | 11HAV | photosystem i | 35128.Thapsd | 8,2 | 1,469693846 | 0,003682294 | 0,00055554 | 7 - | 0,003769333 - | 9 - | 0,004051613 - | 9 - | 0,004078679 - |
| 2195 | 1 | 3um_FIJJAFBF_70793 | Bacteria | Bacteroidetes | Flavobacteriia | Flavobacterial | Crocinitomica | unclassified | Crocinitomica | unclassified | unclassified | unclassified | unclassified | unclassified | 0,8 | 0,979795897 | 0,000155843 | 0,000191249 | 0 - | 0 - | 0 - | 0 - | 0 - | 0 - |
| 2196 | 1 | 3um_FIJJAFBF_100702 | Bacteria | Bacteroidetes | Flavobacteriia | Flavobacterial | Flavobacteria | Formosa | Formosa sp. H | unclassified | unclassified | unclassified | unclassified | unclassified | 1,2 | 0,4 | 8,70E-05 | 2,15E-05 | 1 - | 8,82E-05 - | 1 - | 7,37E-05 - | 1 - | 7,42E-05 - |
| 2197 | 2 | 3um_FIJJAFBF_164232;3um_FIJJAFBF_ | Bacteria | Bacteroidetes | Flavobacteriia | Flavobacterial | Crocinitomica | unclassified | Crocinitomica | poorly charact | Function unkn | 0P884 | membrane | 926562.Oweh | 1 | 0,632455532 | 0,000175609 | 0,000101011 | 1 - | 0,000214465 - | 1 - | 0,000179298 - | 0 - | 0 - |
| 2198 | 1 | 3um_FIJJAFBF_135245 | Bacteria | Cyanobacteria | unclassified | Synechococca | Synechococca | Synechococcu | Synechococcu | unclassified | unclassified | unclassified | unclassified | unclassified | 1,4 | 0,489897949 | 0,000129343 | 3,89E-05 | 1 - | 0,000112048 - | 2 - | 0,00018735 - | 1 - | 9,43E-05 - |
| 2199 | 1 | 3um_FIJJAFBF_530726 | unclassified | unclassified | unclassified | unclassified | unclassified | unclassified | unclassified | unclassified | unclassified | unclassified | unclassified | unclassified | 0,2 | 0,4 | 2,02E-05 | 4,03E-05 | 0 - | 0 - | 0 - | 0 - | 0 - | 0 - |
| 2200 | 1 | 3um_FIJJAFBF_360713 | Bacteria | Bacteroidetes | Flavobacteriia | Flavobacterial | Flavobacteria | Polaribacter | Polaribacter s | unclassified | unclassified | unclassified | unclassified | unclassified | 0,8 | 0,4 | 5,31E-05 | 2,74E-05 | 1 - | 7,87E-05 - | 0 - | 0 - | 1 - | 6,62E-05 - |
| 2201 | 1 | 3um_FIJJAFBF_412096 | Bacteria | Proteobacteri | Alphaproteob | Rhodobactera | Rhodobactera | Aquimixticola | Aquimixticola | metabolism | Amino acid tra | 05CZ6 | Can catalyze t | 999141.GME_ | 0,6 | 0,489897949 | 6,42E-05 | 5,26E-05 | 0 - | 0 - | 1 - | 0,00011336 - | 0 - | 0 - |
| 2202 | 1 | 3um_FIJJAFBF_494165 | Bacteria | Bacteroidetes | Flavobacteriia | Flavobacterial | Flavobacteria | Polaribacter | various | unclassified | unclassified | unclassified | unclassified | unclassified | 0,2 | 0,4 | 6,02E-05 | 0,000120374 | 0 - | 0 - | 0 - | 0 - | 0 - | 0 - |
| 2203 | 1 | 3um_FIJJAFBF_201380 | Bacteria | Bacteroidetes | Flavobacteriia | Flavobacterial | Flavobacteria | Lacinutrix | Lacinutrix jang | poorly charact | Function unkn | 0XPRT | NA | 391598.FBBAL | 2,2 | 0,4 | 0,000278111 | 5,62E-05 | 2 - | 0,000300595 - | 3 - | 0,000376957 - | 2 - | 0,000252984 - |
| 2204 | 1 | 3um_FIJJAFBF_85780 | unclassified | unclassified | unclassified | unclassified | unclassified | unclassified | unclassified | unclassified | unclassified | unclassified | unclassified | unclassified | 4 | 1,095445115 | 0,000651513 | 0,000120974 | 3 - | 0,000594033 - | 3 - | 0,000496626 - | 4 - | 0,000666592 - |
| 2205 | 1 | 0.2um_NOKCPHJN_09839 | Bacteria | Proteobacteri | Gammaprotec | Alteromonada | Psychromona | Psychromonas | Psychromonas | poorly charact | Function unkn | COG3517 | type VI secreti | 357804.Ping_ | 1 | 0,632455532 | 0,000121728 | 7,16E-05 | 0 - | 0 - | 1 - | 0,000129287 - | 1 - | 0,000130151 - |
| 2206 | 1 | 0.2um_NOKCPHJN_228743 | Eukaryota | Blastocladiom | Blastocladiom | Blastocladiale | Blastocladiace | Allomyces | Allomyces ma | unclassified | unclassified | unclassified | unclassified | unclassified | 2,2 | 0,4 | 0,000678918 | 0,000137178 | 2 - | 0,000733806 - | 3 - | 0,000920219 - | 2 - | 0,000617578 - |
| 2207 | 1 | 0.2um_NOKCPHJN_469204 | Bacteria | Proteobacteri | Alphaproteob | Pelagibacteral | Pelagibacterac | Candidatus Pe | Candidatus Pe | poorly charact | Function unkn | 16XWJ | Domain of unl | 335992.SAR11 | 1 | 0 | 0,000353889 | 3,74E-05 | 1 - | 0,000420495 - | 1 - | 0,000351544 - | 1 - | 0,000353893 - |
| 2208 | 11 | 0.2um_NOKCPHJN_234612;0.2um_NO | various | various | various | various | various | various | various | unclassified | unclassified | unclassified | unclassified | unclassified | 0,4 | 0,489897949 | 7,44E-05 | 9,13E-05 | 0 - | 0 - | 0 - | 0 - | 0 - | 0 - |
| 2209 | 3 | 3um_FIJJAFBF_195466;3um_FIJJAFBF_ | Bacteria | Proteobacteri | Alphaproteob | Rhodobactera | Rhodobactera | various | various | metabolism | Nucleotide tra | 05CAT | Cytidine mon | various | 0,2 | 0,4 | 5,39E-05 | 0,000107759 | 0 - | 0 - | 0 - | 0 - | 0 - | 0 - |
| 2210 | 1 | 0.2um_NOKCPHJN_128184 | Bacteria | Proteobacteri | Alphaproteob | Rhodobactera | Rhodobactera | unclassified | Rhodobactera | unclassified | unclassified | unclassified | unclassified | unclassified | 0,6 | 0,489897949 | 0,000179817 | 0,000147423 | 0 - | 0 - | 1 - | 0,000317639 - | 0 - | 0 - |
| 2211 | 2 | 3um_FIJJAFBF_250160;3um_FIJJAFBF_ | Eukaryota | Bacillariophyt | Coscinodiscop | Coscinodiscale | Hemidiscacea | Actinocyclus | Actinocyclus s | unclassified | unclassified | unclassified | unclassified | unclassified | 1 | 0 | 0,000496001 | 5,25E-05 | 1 - | 0,000589356 - | 1 - | 0,000492716 - | 1 - | 0,000496007 - |
| 2212 | 1 | 3um_FIJJAFBF_292964 | Bacteria | Proteobacteri | Alphaproteob | Rhodobactera | Rhodobactera | Tateyamaria | Tateyamaria s | unclassified | unclassified | unclassified | unclassified | unclassified | 4,8 | 0,979795897 | 0,000999419 | 0,000214336 | 5 - | 0,001239208 - | 5 - | 0,001036009 - | 5 - | 0,001042929 - |
| 2213 | 1 | 3um_FIJJAFBF_221603 | Eukaryota | unclassified | Pelagophycea | Pelagomonad | unclassified | Aureococcus | Aureococcus a | information st | Translation, ri | 1520E | S25 ribosomal | 4787.PITG_13 | 1 | 0,894427191 | 0,000568499 | 0,000522274 | 2 - | 0,001257953 - | 1 - | 0,00052584 - | 2 - | 0,001058705 - |
| 2214 | 1 | 0.2um_NOKCPHJN_1095133 | Eukaryota | Platyhelminth | Trematoda | Plagiorchiida | Fasciolidae | Fasciola | Fasciola hepat | cellular proce: | Cytoskeleton | 0VQU4 | ACTIN | 7897.ENSLACP | 1 | 0 | 0,000414422 | 4,38E-05 | 1 - | 0,000492422 - | 1 - | 0,000411677 - | 1 - | 0,000414427 - |
| 2215 | 1 | 3um_FIJJAFBF_693899 | Bacteria | Bacteroidetes | Flavobacteriia | Flavobacterial | Flavobacteria | Formosa | Formosa sp. H | unclassified | unclassified | unclassified | unclassified | unclassified | 0,2 | 0,4 | 2,51E-05 | 5,03E-05 | 0 - | 0 - | 0 - | 0 - | 1 - | 0,000125734 - |
| 2216 | 1 | 0.2um_NOKCPHJN_524511 | Eukaryota | unclassified | Cryptophyta | Pyrenomonad | Geminigerace | Guillardia | Guillardia thet | unclassified | unclassified | unclassified | unclassified | unclassified | 0,4 | 0,489897949 | 4,35E-05 | 5,43E-05 | 1 - | 0,000125795 - | 0 - | 0 - | 0 - | 0 - |
| 2217 | 2 | 0.2um_NOKCPHJN_885492;3um_FIJJAI | Bacteria | Bacteroidetes | Flavobacteriia | Flavobacterial | Flavobacteria | Tenacibaculur | Tenacibaculur | poorly charact | Function unkn | 0P8NI | Inherit from b | various | 0,6 | 0,489897949 | 0,000103168 | 8,60E-05 | 1 - | 0,000201205 - | 1 - | 0,000168212 - | 0 - | 0 - |

| | | | | | | | | | | | | | | | | | | | | | | | | |
|---|---|---|---|---|---|---|---|---|---|---|---|---|---|---|---|---|---|---|---|---|---|---|---|---|
| 2218 | 3 | 0.2um_NOKCPHJN_58213;3um_FIJJAFE | Bacteria | Bacteroidetes | Flavobacteriia | Flavobacterial | Flavobacteriac | Marinirhabdu | Marinirhabdu | unclassified | unclassified | unclassified | unclassified | unclassified | 0,6 | 0,489897949 | 6,28E-05 | 5,15E-05 | 0 - | 0 - | 0 - | 0 - | 1 - | 0,000111492 - |
| 2219 | 1 | 3um_FIJJAFBF_715318 | Eukaryota | Microsporidia | unclassified | unclassified | unclassified | Mitosporidium | Mitosporidium | unclassified | unclassified | unclassified | unclassified | unclassified | 0,4 | 0,489897949 | 0,000216749 | 0,000266521 | 0 - | 0 - | 1 - | 0,000579397 - | 0 - | 0 - |
| 2220 | 1 | 3um_FIJJAFBF_715076 | Eukaryota | unclassified | Pelagophyceae | Pelagomonad | unclassified | Aureoumbra | Aureoumbra l | unclassified | unclassified | unclassified | unclassified | unclassified | 0,4 | 0,489897949 | 0,000164283 | 0,000202086 | 0 - | 0 - | 0 - | 0 - | 1 - | 0,00044051 - |
| 2221 | 1 | 3um_FIJJAFBF_726401 | Bacteria | Proteobacteri | Betaproteoba | Burkholderiale | Burkholderiac | Cupriavidus | Cupriavidus m | unclassified | unclassified | unclassified | unclassified | unclassified | 0,6 | 0,489897949 | 0,000339688 | 0,000279858 | 1 - | 0,000634307 - | 1 - | 0,000530296 - | 1 - | 0,000533838 - |
| 2222 | 1 | 3um_FIJJAFBF_444690 | Eukaryota | unclassified | unclassified | Phaeocystales | Phaeocystace | Phaeocystis | Phaeocystis ar | metabolism | Energy produc | COG4451 | RuBisCO catal | 977880.RALTA | 0,4 | 0,489897949 | 0,000168409 | 0,000207081 | 0 - | 0 - | 1 - | 0,000450179 - | 0 - | 0 - |
| 2223 | 1 | 3um_FIJJAFBF_391108 | Bacteria | Proteobacteri | Alphaproteob | Pelagibacteral | Pelagibacterac | Candidatus Pe | Candidatus Pe | unclassified | unclassified | unclassified | unclassified | unclassified | 0,2 | 0,4 | 7,28E-05 | 0,000145523 | 0 - | 0 - | 1 - | 0,000363808 - | 0 - | 0 - |
| 2224 | 1 | 0.2um_NOKCPHJN_503483 | Eukaryota | Chytridiomyco | Chytridiomyce | Chytridiales | Chytriomyceta | Rhizoclosmati | Rhizoclosmati | cellular proces | Posttranslatio | 0JAJ8 | Dentin sialoph | 10116.ENSRN | 0,4 | 0,489897949 | 2,17E-05 | 2,66E-05 | 0 - | 0 - | 1 - | 5,53E-05 - | 0 - | 0 - |
| 2225 | 1 | 0.2um_NOKCPHJN_930429 | Bacteria | Proteobacteri | Alphaproteob | unclassified | unclassified | Emcibacter | Emcibacter co | cellular proces | Cell wall/mem | COG4948 | mandelate rac | 313606.M231 | 0,4 | 0,489897949 | 7,07E-05 | 8,68E-05 | 0 - | 0 - | 0 - | 0 - | 0 - | 0 - |
| 2226 | 1 | 3um_FIJJAFBF_344329 | unclassified | unclassified | unclassified | unclassified | unclassified | unclassified | unclassified | unclassified | unclassified | unclassified | unclassified | unclassified | 0,2 | 0,4 | 3,12E-05 | 6,24E-05 | 0 - | 0 - | 0 - | 0 - | 0 - | 0 - |
| 2227 | 1 | 0.2um_NOKCPHJN_175750 | Eukaryota | unclassified | Pelagophyceae | Pelagomonad | unclassified | Aureococcus | Aureococcus a | unclassified | unclassified | unclassified | unclassified | unclassified | 1 | 0 | 0,000316544 | 3,35E-05 | 1 - | 0,000376122 - | 1 - | 0,000314447 - | 1 - | 0,000316547 - |
| 2228 | 1 | 3um_FIJJAFBF_242749 | Bacteria | Bacteroidetes | Flavobacteriia | unclassified | unclassified | unclassified | Flavobacteria | unclassified | unclassified | unclassified | unclassified | unclassified | 0,8 | 0,748331477 | 6,30E-05 | 5,95E-05 | 0 - | 0 - | 0 - | 0 - | 1 - | 8,35E-05 - |
| 2229 | 1 | 3um_FIJJAFBF_155127 | Bacteria | Bacteroidetes | Cytophagia | Cytophagales | Flammeovirga | unclassified | Flammeovirga | unclassified | unclassified | unclassified | unclassified | unclassified | 3 | 0 | 0,00034297 | 3,63E-05 | 3 - | 0,000407522 - | 3 - | 0,000340698 - | 3 - | 0,000342974 - |
| 2230 | 1 | 0.2um_NOKCPHJN_74284 | Bacteria | Bacteroidetes | Flavobacteriia | Flavobacterial | Flavobacteriac | Polaribacter | Polaribacter ta | information st | Transcription | 0P7SF | DNA-depende | 313598.MED1 | 6,6 | 1,854723699 | 0,000329976 | 9,32E-05 | 7 - | 0,000418146 - | 7 - | 0,000349581 - | 3 - | 0,000150821 - |
| 2231 | 1 | 3um_FIJJAFBF_84528 | Bacteria | Bacteroidetes | Flavobacteriia | Flavobacterial | Flavobacteriac | Polaribacter | Polaribacter r | cellular proces | Posttranslatio | COG0330 | Band 7 protei | 391603.FBALC | 1,2 | 0,4 | 0,000247029 | 8,52E-05 | 1 - | 0,000244602 - | 1 - | 0,000204493 - | 2 - | 0,000411719 - |
| 2232 | 1 | 0.2um_NOKCPHJN_159101 | Bacteria | Bacteroidetes | unclassified | unclassified | unclassified | unclassified | Bacteroidetes | cellular proces | Intracellular tr | COG0653 | Part of the Sec | 926562.Oweh | 0,8 | 0,748331477 | 4,43E-05 | 4,08E-05 | 0 - | 0 - | 1 - | 5,64E-05 - | 1 - | 5,68E-05 - |
| 2233 | 1 | 3um_FIJJAFBF_48824 | Eukaryota | unclassified | Chrysophycea | Chromulinales | Chromulinace | Ochromonas | Ochromonas s | metabolism | Energy produc | COG0056 | Produces ATP | 41431.PCC880 | 1 | 0 | 0,000125482 | 1,33E-05 | 1 - | 0,0001491 - | 1 - | 0,000124651 - | 1 - | 0,000125484 - |
| 2234 | 1 | 0.2um_NOKCPHJN_760115 | Eukaryota | unclassified | Cryptophyta | Pyrenomonad | Geminigerace | Guillardia | Guillardia thet | information st | RNA processir | 0VB7C | Mago-nashi h | 45351.NEMVE | 1 | 0 | 0,000529346 | 5,60E-05 | 1 - | 0,000628976 - | 1 - | 0,00052584 - | 1 - | 0,000529352 - |
| 2235 | 2 | 0.2um_NOKCPHJN_530547;3um_FIJJAI | Eukaryota | various | various | various | various | various | various | information st | Chromatin str | KOG3467 | Core compone | various | 12 | 1,095445115 | 0,005146756 | 0,000746321 | 12 - | 0,006110056 - | 11 - | 0,004682477 - | 14 - | 0,005999327 - |
| 2236 | 7 | 0.2um_NOKCPHJN_861091;0.2um_NO | Bacteria | various | various | various | various | various | various | unclassified | unclassified | unclassified | unclassified | unclassified | 0,2 | 0,4 | 1,00E-04 | 0,000199978 | 0 - | 0 - | 0 - | 0 - | 1 - | 0,000499944 - |
| 2237 | 1 | 3um_FIJJAFBF_574196 | unclassified | unclassified | unclassified | unclassified | unclassified | unclassified | unclassified | unclassified | unclassified | unclassified | unclassified | unclassified | 2 | 0 | 0,000597082 | 6,31E-05 | 2 - | 0,000709461 - | 2 - | 0,000593127 - | 2 - | 0,000597089 - |
| 2238 | 1 | 3um_FIJJAFBF_366559 | Viruses | unclassified | unclassified | unclassified | Phycodnavirid | unclassified | Chrysochromu | information st | Transcription | 12NXR | DNA-depende | 4950.XP_0036 | 0,4 | 0,489897949 | 0,000118335 | 0,000144998 | 0 - | 0 - | 0 - | 0 - | 1 - | 0,000302851 - |
| 2239 | 1 | 0.2um_NOKCPHJN_72555 | Eukaryota | unclassified | Labyrinthulom | unclassified | Thraustochytr | Aurantiochytr | Aurantiochytr | cellular proces | Signal transdu | KOG0613 | fibronectin typ | 67593.Physo1 | 0,2 | 0,4 | 7,82E-07 | 1,56E-06 | 0 - | 0 - | 0 - | 0 - | 1 - | 3,91E-06 - |
| 2240 | 1 | 0.2um_NOKCPHJN_506016 | Bacteria | Bacteroidetes | Flavobacteriia | Flavobacterial | Flavobacteriac | Gilvibacter | Gilvibacter sp. | poorly charact | Function unkn | 064XS | NA | 313595.P7007 | 0,2 | 0,4 | 4,42E-05 | 8,83E-05 | 0 - | 0 - | 0 - | 0 - | 0 - | 0 - |
| 2241 | 1 | 3um_FIJJAFBF_543025 | unclassified | unclassified | unclassified | unclassified | unclassified | unclassified | unclassified | unclassified | unclassified | unclassified | unclassified | unclassified | 0,6 | 0,489897949 | 0,000196813 | 0,000164106 | 1 - | 0,000383837 - | 1 - | 0,000320897 - | 0 - | 0 - |
| 2242 | 1 | 3um_FIJJAFBF_257117 | Eukaryota | Bacillariophyta | Fragilariophyc | Fragilariales | Fragilariaceae | Asterionellops | Asterionellops | information st | Translation, ri | 0FJK5 | Binds the lowe | 197221.tlr008 | 2,2 | 0,4 | 0,00062727 | 0,000112703 | 2 - | 0,000680438 - | 2 - | 0,000568863 - | 2 - | 0,000572663 - |
| 2243 | 1 | 0.2um_NOKCPHJN_814372 | Eukaryota | Chlorophyta | Trebouxiophy | Chlorellales | Chlorellaceae | Micractinium | Micractinium | poorly charact | Function unkn | 08WXZ | Peptidoglycan | 649638.Trad_ | 0,8 | 0,4 | 0,000105325 | 5,43E-05 | 1 - | 0,000154645 - | 1 - | 0,000129287 - | 1 - | 0,000130151 - |
| 2244 | 1 | 3um_FIJJAFBF_37824 | Bacteria | Actinobacteria | unclassified | unclassified | unclassified | unclassified | Actinobacteria | unclassified | unclassified | unclassified | unclassified | unclassified | 9,6 | 0,489897949 | 0,000617798 | 4,63E-05 | 9 - | 0,000690906 - | 10 - | 0,000641794 - | 9 - | 0,000581473 - |
| 2245 | 1 | 3um_FIJJAFBF_67514 | Eukaryota | Bacillariophyta | Mediophyceae | Cymatosirales | Cymatosiracea | Plagiogrammc | Plagiogrammc | cellular proces | Posttranslatio | KOG0731 | ATP-depender | 35128.Thapsd | 2,6 | 0,489897949 | 0,000255096 | 6,80E-05 | 3 - | 0,000344923 - | 3 - | 0,000288364 - | 3 - | 0,00029029 - |
| 2246 | 1 | 0.2um_NOKCPHJN_349277 | unclassified | unclassified | unclassified | unclassified | unclassified | unclassified | unclassified | unclassified | unclassified | unclassified | unclassified | unclassified | 0,6 | 0,489897949 | 4,00E-05 | 3,30E-05 | 1 - | 7,56E-05 - | 0 - | 0 - | 1 - | 6,36E-05 - |
| 2247 | 1 | 3um_FIJJAFBF_259506 | Eukaryota | Bacillariophyta | Coscinodiscop | Thalassiosirale | Stephanodisca | Cyclotella | Cyclotella pse | cellular proces | Posttranslatio | KOG0731 | ATP-depender | 35128.Thapsd | 0,4 | 0,489897949 | 3,91E-05 | 4,79E-05 | 0 - | 0 - | 1 - | 9,75E-05 - | 1 - | 9,81E-05 - |
| 2248 | 1 | 3um_FIJJAFBF_744568 | Bacteria | unclassified | unclassified | unclassified | unclassified | unclassified | uncultured ba | poorly charact | Function unkn | 0P8NI | Inherit from b | 983544.Lacal_ | 4,8 | 0,4 | 0,000735878 | 0,000102887 | 5 - | 0,000910562 - | 4 - | 0,000609002 - | 5 - | 0,000766337 - |
| 2249 | 1 | 3um_FIJJAFBF_470344 | Eukaryota | Bacillariophyta | Bacillariophyc | Bacillariales | Bacillariaceae | Fragilariopsis | Fragilariopsis | unclassified | unclassified | unclassified | unclassified | unclassified | 0,4 | 0,489897949 | 0,000238968 | 0,000292812 | 0 - | 0 - | 0 - | 0 - | 1 - | 0,000611582 - |
| 2250 | 1 | 3um_FIJJAFBF_715057 | Eukaryota | Xanthophycea | unclassified | Vaucheriales | Vaucheriaceae | Vaucheria | Vaucheria lito | information st | Transcription | 05D27 | DNA-depende | 329726.AM1_ | 0,6 | 0,489897949 | 3,02E-05 | 2,47E-05 | 0 - | 0 - | 1 - | 5,09E-05 - | 1 - | 5,12E-05 - |
| 2251 | 1 | 3um_FIJJAFBF_720178 | Bacteria | Proteobacteri | Alphaproteob | Sphingomona | Erythrobacter | Porphyrobact | Porphyrobact | information st | Translation, ri | COG0050 | This protein p | 237727.NAP1_ | 0,4 | 0,8 | 6,36E-05 | 0,000127258 | 0 - | 0 - | 0 - | 0 - | 2 - | 0,000318146 - |
| 2252 | 1 | 3um_FIJJAFBF_552490 | Bacteria | Proteobacteri | Gammaprotec | Arenicellales | Arenicellaceae | Arenicella | Arenicella xan | unclassified | unclassified | unclassified | unclassified | unclassified | 1 | 0,632455532 | 7,48E-05 | 4,30E-05 | 1 - | 9,14E-05 - | 1 - | 7,64E-05 - | 0 - | 0 - |
| 2253 | 1 | 0.2um_NOKCPHJN_991243 | Eukaryota | unclassified | unclassified | Acytosteliales | Acytosteliacea | Heterostelium | Heterostelium | poorly charact | Function unkn | 0JMN7 | NA | 5786.XP_0032 | 0,4 | 0,489897949 | 0,000220279 | 0,000270323 | 0 - | 0 - | 0 - | 0 - | 0 - | 0 - |
| 2254 | 1 | 0.2um_NOKCPHJN_666584 | Eukaryota | Bacillariophyta | Bacillariophyc | Bacillariales | Bacillariaceae | Nitzschia | Nitzschia sp. I | cellular proces | Posttranslatio | KOG0730 | cell division cy | 35128.Thaps2 | 1 | 0 | 8,11E-05 | 8,57E-06 | 1 - | 9,63E-05 - | 1 - | 8,05E-05 - | 1 - | 8,11E-05 - |
| 2255 | 1 | 0.2um_NOKCPHJN_1097287 | Eukaryota | Chromerida | unclassified | unclassified | unclassified | Vitrella | Vitrella brassic | unclassified | unclassified | unclassified | unclassified | unclassified | 0,6 | 0,489897949 | 0,000196262 | 0,000163919 | 1 - | 0,000387814 - | 0 - | 0 - | 0 - | 0 - |
| 2256 | 1 | 3um_FIJJAFBF_594149 | Eukaryota | unclassified | Pelagophyceae | Pelagomonad | unclassified | Aureoumbra | Aureoumbra l | metabolism | Carbohydrate | COG1850 | RuBisCO catal | 45157.CMV01 | 22,2 | 0,748331477 | 0,002866985 | 0,000276036 | 22 - | 0,003381232 - | 21 - | 0,002698302 - | 22 - | 0,002845677 - |
| 2257 | 1 | 3um_FIJJAFBF_525903 | Bacteria | Bacteroidetes | Flavobacteriia | Flavobacterial | Flavobacteriac | Polaribacter | Polaribacter v | unclassified | unclassified | unclassified | unclassified | unclassified | 1 | 0 | 9,17E-05 | 9,70E-06 | 1 - | 0,000108949 - | 1 - | 9,11E-05 - | 1 - | 9,17E-05 - |
| 2258 | 1 | 3um_FIJJAFBF_715080 | Eukaryota | unclassified | Pelagophyceae | Pelagomonad | unclassified | Pelagomonas | uncultured Pe | unclassified | unclassified | unclassified | unclassified | unclassified | 1 | 0 | 0,000547758 | 5,79E-05 | 1 - | 0,000650854 - | 1 - | 0,00054413 - | 1 - | 0,000547765 - |
| 2259 | 1 | 3um_FIJJAFBF_207078 | Bacteria | Proteobacteri | Alphaproteob | Rhodospirillal | Rhodospirillac | Candidatus En | Candidatus En | unclassified | unclassified | unclassified | unclassified | unclassified | 0,6 | 0,489897949 | 0,000225022 | 0,000183829 | 0 - | 0 - | 1 - | 0,000379242 - | 1 - | 0,000381775 - |
| 2260 | 1 | 3um_FIJJAFBF_228661 | Bacteria | Bacteroidetes | Flavobacteriia | Flavobacterial | Flavobacteriac | Formosa | Formosa sp. H | poorly charact | Function unkn | 0P8NI | Inherit from b | 983544.Lacal_ | 2,4 | 0,489897949 | 0,000357917 | 5,13E-05 | 2 - | 0,000359847 - | 2 - | 0,000300841 - | 2 - | 0,000302851 - |
| 2261 | 2 | 0.2um_NOKCPHJN_130064;0.2um_NO | Bacteria | Proteobacteri | Alphaproteob | Rhodobactera | various | various | various | metabolism | Carbohydrate | various | various | various | 0,4 | 0,489897949 | 9,15E-05 | 0,000113146 | 1 - | 0,000253723 - | 0 - | 0 - | 0 - | 0 - |
| 2262 | 1 | 3um_FIJJAFBF_486409 | Bacteria | Proteobacteri | Gammaprotec | Cellvibrionale | Spongiibacter | Spongiibacter | Spongiibacter | poorly charact | Function unkn | 0ZX70 | NA | 525897.Dbac_ | 1 | 0 | 0,000611575 | 6,47E-05 | 1 - | 0,000726681 - | 1 - | 0,000607523 - | 1 - | 0,000611582 - |
| 2263 | 1 | 0.2um_NOKCPHJN_278109 | Eukaryota | unclassified | Bangiophycea | Bangiales | Bangiaceae | Porphyra | Porphyra umb | unclassified | unclassified | unclassified | unclassified | unclassified | 0,6 | 0,489897949 | 1,23E-05 | 1,01E-05 | 0 - | 0 - | 0 - | 0 - | 1 - | 2,18E-05 - |
| 2264 | 1 | 0.2um_NOKCPHJN_546978 | Bacteria | Proteobacteri | Gammaprotec | Cellvibrionale | Porticoccacea | unclassified | SAR92 bacteri | metabolism | Inorganic ion t | 0QHT5 | TonB-depende | 314287.GB220 | 4,8 | 2,227105745 | 0,000331908 | 0,000135262 | 1 - | 8,63E-05 - | 5 - | 0,00036087 - | 5 - | 0,000363281 - |
| 2265 | 1 | 0.2um_NOKCPHJN_227678 | Bacteria | Bacteroidetes | Flavobacteriia | Flavobacterial | Flavobacteriac | Formosa | Formosa sp. H | metabolism | Inorganic ion t | 05C27 | TonB-depende | 867900.Celly_ | 3,4 | 1,019803903 | 0,000209343 | 4,99E-05 | 3 - | 0,00022365 - | 3 - | 0,000186977 - | 2 - | 0,000125484 - |
| 2266 | 1 | 0.2um_NOKCPHJN_958977 | Eukaryota | unclassified | Spirotrichea | Sporadotrichi | Oxytrichidae | Sterkiella | Sterkiella nova | information st | Translation, ri | KOG0052 | This protein p | 67593.Physo1 | 2,8 | 0,4 | 0,00039547 | 3,45E-05 | 2 - | 0,000340219 - | 3 - | 0,000426647 - | 3 - | 0,000429497 - |
| 2267 | 1 | 0.2um_NOKCPHJN_175435 | Eukaryota | unclassified | Oomycetes | Peronosporale | Peronosporac | Phytophthora | Phytophthora | cellular proces | Cytoskeleton | 0DGVY | Dynein, axone | 7897.ENSLACI | 0,4 | 0,489897949 | 4,09E-06 | 5,04E-06 | 1 - | 1,11E-05 - | 0 - | 0 - | 1 - | 9,34E-06 - |
| 2268 | 1 | 0.2um_NOKCPHJN_438308 | Eukaryota | unclassified | Oomycetes | Saprolegniales | Saprolegniace | Aphanomyces | Aphanomyces | unclassified | unclassified | unclassified | unclassified | unclassified | 0,8 | 0,4 | 0,00027407 | 0,000141421 | 1 - | 0,00040241 - | 1 - | 0,000336424 - | 1 - | 0,000338672 - |
| 2269 | 1 | 3um_FIJJAFBF_559087 | Bacteria | Proteobacteri | Gammaprotec | Alteromonada | Alteromonada | Paraglaciecola | Paraglaciecola | unclassified | unclassified | unclassified | unclassified | unclassified | 4,4 | 1,019803903 | 0,000330718 | 5,91E-05 | 4 - | 0,000362461 - | 4 - | 0,000303026 - | 3 - | 0,000228788 - |
| 2270 | 2 | 3um_FIJJAFBF_13108;3um_FIJJAFBF_7 | Eukaryota | unclassified | Cryptophyta | Pyrenomonad | Geminigerace | various | various | metabolism | Energy produc | COG1290 | Component of | various | 0,6 | 0,489897949 | 0,000164763 | 0,000135081 | 0 - | 0 - | 1 - | 0,000291046 - | 0 - | 0 - |
| 2271 | 1 | 3um_FIJJAFBF_391683 | Eukaryota | unclassified | Dinophyceae | Suessiales | Symbiodiniace | Symbiodinium | Symbiodinium | unclassified | unclassified | unclassified | unclassified | unclassified | 3,2 | 0,748331477 | 0,000812215 | 0,00022855 | 4 - | 0,00120238 - | 3 - | 0,000753915 - | 2 - | 0,000505967 - |
| 2272 | 1 | 0.2um_NOKCPHJN_663396 | Eukaryota | unclassified | Pelagophyceae | Pelagomonad | unclassified | Aureococcus | Aureococcus a | unclassified | unclassified | unclassified | unclassified | unclassified | 0,4 | 0,489897949 | 5,61E-05 | 6,94E-05 | 1 - | 0,00015561 - | 0 - | 0 - | 0 - | 0 - |
| 2273 | 1 | 3um_FIJJAFBF_814041 | Bacteria | Bacteroidetes | Flavobacteriia | Flavobacterial | Flavobacteriac | Winogradskye | Winogradskye | metabolism | Inorganic ion t | 0P7D0 | tonB-depende | 391587.KAOT | 0,4 | 0,489897949 | 2,74E-05 | 3,37E-05 | 0 - | 0 - | 0 - | 0 - | 0 - | 0 - |
| 2274 | 1 | 3um_FIJJAFBF_510233 | Bacteria | Bacteroidetes | Flavobacteriia | Flavobacterial | Flavobacteriac | Polaribacter | Polaribacter s | unclassified | unclassified | unclassified | unclassified | unclassified | 1,6 | 0,8 | 0,000357359 | 0,000162376 | 1 - | 0,000273169 - | 1 - | 0,000228376 - | 1 - | 0,000229901 - |
| 2275 | 1 | 3um_FIJJAFBF_748937 | unclassified | unclassified | unclassified | unclassified | unclassified | unclassified | unclassified | unclassified | unclassified | unclassified | unclassified | unclassified | 1 | 0 | 0,000362024 | 3,83E-05 | 1 - | 0,000430162 - | 1 - | 0,000359626 - | 1 - | 0,000362028 - |
| 2276 | 1 | 3um_FIJJAFBF_442105 | Bacteria | Deinococcus-T | Deinococci | Deinococcales | Deinococcace | Deinococcus | Deinococcus a | unclassified | unclassified | unclassified | unclassified | unclassified | 1,2 | 0,4 | 0,000185643 | 4,59E-05 | 1 - | 0,000188061 - | 1 - | 0,000157223 - | 1 - | 0,000158274 - |
| 2277 | 1 | 0.2um_NOKCPHJN_823660 | Eukaryota | Chlorophyta | Mamiellophyc | Mamiellales | Mamiellaceae | Micromonas | Micromonas p | cellular proces | Posttranslatio | KOG0019 | heat shock pro | 4558.Sb02g02 | 0,4 | 0,489897949 | 4,60E-05 | 5,66E-05 | 0 - | 0 - | 1 - | 0,000122937 - | 0 - | 0 - |
| 2278 | 1 | 3um_FIJJAFBF_313476 | unclassified | unclassified | unclassified | unclassified | unclassified | unclassified | unclassified | unclassified | unclassified | unclassified | unclassified | unclassified | 1 | 0 | 0,000203858 | 2,16E-05 | 1 - | 0,000242227 - | 1 - | 0,000202508 - | 1 - | 0,000203861 - |
| 2279 | 1 | 0.2um_NOKCPHJN_1095134 | Eukaryota | unclassified | unclassified | Euglyphida | Paulinellidae | Paulinella | Paulinella mic | unclassified | unclassified | unclassified | unclassified | unclassified | 1,6 | 0,8 | 0,000339987 | 0,000154483 | 1 - | 0,00025989 - | 1 - | 0,000217274 - | 1 - | 0,000218725 - |
| 2280 | 1 | 0.2um_NOKCPHJN_991012 | Eukaryota | unclassified | Cryptophyta | Pyrenomonad | Geminigerace | Guillardia | Guillardia thet | unclassified | unclassified | unclassified | unclassified | unclassified | 5 | 0 | 0,001179629 | 0,000124749 | 5 - | 0,001401651 - | 5 - | 0,001171815 - | 5 - | 0,001179643 - |
| 2281 | 1 | 3um_FIJJAFBF_175150 | Bacteria | Proteobacteri | Alphaproteob | Rhodobactera | Rhodobactera | unclassified | Rhodobactera | unclassified | unclassified | unclassified | unclassified | unclassified | 0,8 | 0,4 | 0,000406971 | 0,000205109 | 0 - | 0 - | 1 - | 0,000530296 - | 1 - | 0,000533838 - |
| 2282 | 1 | 3um_FIJJAFBF_320586 | Bacteria | Bacteroidetes | Flavobacteriia | Flavobacterial | Flavobacteriac | Polaribacter | Polaribacter s | metabolism | Energy produc | COG0055 | Produces ATP | 411154.GFO_ | 4,2 | 2,039607805 | 0,000510787 | 0,000216648 | 3 - | 0,0004473 - | 5 - | 0,000623256 - | 1 - | 0,000125484 - |
| 2283 | 1 | 0.2um_NOKCPHJN_279514 | unclassified | unclassified | unclassified | unclassified | unclassified | unclassified | unclassified | unclassified | unclassified | unclassified | unclassified | unclassified | 0,4 | 0,489897949 | 0,000131177 | 0,000160714 | 0 - | 0 - | 1 - | 0,000334625 - | 0 - | 0 - |
| 2284 | 1 | 3um_FIJJAFBF_165677 | Bacteria | Actinobacteria | Acidimicrobiia | Acidimicrobial | Microthrixace | Candidatus M | Candidatus M | metabolism | Carbohydrate | 00BJE | Glyceraldehyd | 675635.Psed_ | 0,2 | 0,4 | 3,25E-05 | 6,50E-05 | 0 - | 0 - | 0 - | 0 - | 0 - | 0 - |
| 2285 | 1 | 0.2um_NOKCPHJN_1059776 | Viruses | unclassified | unclassified | unclassified | unclassified | unclassified | uncultured M | unclassified | unclassified | unclassified | unclassified | unclassified | 1 | 0 | 0,000281215 | 2,97E-05 | 1 - | 0,000334144 - | 1 - | 0,000279352 - | 1 - | 0,000281218 - |
| 2286 | 1 | 0.2um_NOKCPHJN_600276 | Bacteria | Lentisphaerae | Lentisphaeria | Lentisphaerale | Lentisphaerac | Lentisphaera | Lentisphaera a | unclassified | unclassified | unclassified | unclassified | unclassified | 0,2 | 0,4 | 4,60E-05 | 9,20E-05 | 0 - | 0 - | 0 - | 0 - | 1 - | 0,000229901 - |
| 2287 | 1 | 0.2um_NOKCPHJN_925580 | Eukaryota | unclassified | Synurophycea | Synurales | Mallomonada | Mallomonas | Mallomonas r | metabolism | Carbohydrate | COG0149 | triosephospha | 35128.Thaps2 | 1 | 0 | 0,000205186 | 2,17E-05 | 1 - | 0,000243805 - | 1 - | 0,000203827 - | 1 - | 0,000205189 - |
| 2288 | 1 | 3um_FIJJAFBF_191792 | Eukaryota | unclassified | Raphidophyce | Chattonellales | Chattonellace | Heterosigma | Heterosigma a | unclassified | unclassified | unclassified | unclassified | unclassified | 2,8 | 0,748331477 | 0,002135709 | 0,000535084 | 2 - | 0,001825565 - | 3 - | 0,002289326 - | 3 - | 0,00230462 - |
| 2289 | 1 | 0.2um_NOKCPHJN_396965 | Eukaryota | unclassified | Oligohymenop | Hymenostoma | Tetrahymenid | Tetrahymena | Tetrahymena | poorly charact | Function unkn | 0XZ2H | Membrane as | 929439.XP_00 | 0,6 | 0,489897949 | 2,93E-05 | 2,40E-05 | 0 - | 0 - | 1 - | 5,18E-05 - | 0 - | 0 - |
| 2290 | 1 | 0.2um_NOKCPHJN_899961 | Bacteria | unclassified | unclassified | unclassified | unclassified | unclassified | bacterium TM | information st | Translation, ri | KOG1936 | Histidyl-trna s | 436017.A4SAY | 1 | 0 | 0,000212811 | 2,25E-05 | 1 - | 0,000252865 - | 1 - | 0,000211402 - | 1 - | 0,000212814 - |
| 2291 | 1 | 0.2um_NOKCPHJN_861812 | Eukaryota | Bacillariophyta | Bacillariophyc | Naviculales | Naviculaceae | Fistulifera | Fistulifera sola | poorly charact | Function unkn | 110WY | Chlorophyll A- | 2850.Phatr18 | 1 | 0 | 0,000699913 | 7,40E-05 | 1 - | 0,000831646 - | 1 - | 0,000695277 - | 1 - | 0,000699922 - |
| 2292 | 1 | 3um_FIJJAFBF_351766 | Bacteria | Actinobacteria | Actinobacteria | Bifidobacteria | Bifidobacteria | Bifidobacteriu | Bifidobacteriu | unclassified | unclassified | unclassified | unclassified | unclassified | 0,4 | 0,489897949 | 0,000275398 | 0,000338638 | 0 - | 0 - | 1 - | 0,000736176 - | 0 - | 0 - |
| 2293 | 1 | 0.2um_NOKCPHJN_770497 | Eukaryota | Eustigmatoph | unclassified | Eustigmatales | Monodopsida | Nannochlorop | Nannochlorop | unclassified | unclassified | unclassified | unclassified | unclassified | 0,2 | 0,4 | 8,87E-05 | 0,000177445 | 0 - | 0 - | 0 - | 0 - | 1 - | 0,000443612 - |
| 2294 | 1 | 0.2um_NOKCPHJN_241147 | Eukaryota | Streptophyta | unclassified | Brassicales | Brassicaceae | Arabis | Arabis alpina | cellular proces | Posttranslatio | KOG0010 | Ubiquilin | 44689.DDB01 | 0,2 | 0,4 | 2,05E-05 | 4,10E-05 | 0 - | 0 - | 1 - | 0,000102414 - | 0 - | 0 - |
| 2295 | 1 | 0.2um_NOKCPHJN_248636 | Eukaryota | unclassified | Oomycetes | Saprolegniales | Saprolegniace | Achlya | Achlya hypogy | unclassified | unclassified | unclassified | unclassified | unclassified | 0,2 | 0,4 | 3,37E-06 | 6,75E-06 | 0 - | 0 - | 0 - | 0 - | 1 - | 1,69E-05 - |
| 2296 | 1 | 0.2um_NOKCPHJN_73952 | Bacteria | Proteobacteri | Gammaprotec | Cellvibrionale | Porticoccacea | unclassified | Gammaprotec | cellular proces | Cell motility | 16VRE | flagellar moto | 314287.GB220 | 0,2 | 0,4 | 4,56E-05 | 9,13E-05 | 1 - | 0,000228196 - | 0 - | 0 - | 0 - | 0 - |
| 2297 | 1 | 0.2um_NOKCPHJN_363822 | Bacteria | unclassified | unclassified | unclassified | unclassified | unclassified | uncultured ba | unclassified | unclassified | unclassified | unclassified | unclassified | 0,2 | 0,4 | 8,83E-05 | 0,000176692 | 0 - | 0 - | 0 - | 0 - | 0 - | 0 - |
| 2298 | 1 | 3um_FIJJAFBF_758079 | unclassified | unclassified | unclassified | unclassified | unclassified | unclassified | unclassified | unclassified | unclassified | unclassified | unclassified | unclassified | 0,2 | 0,4 | 7,63E-05 | 0,000152622 | 0 - | 0 - | 1 - | 0,000381554 - | 0 - | 0 - |
| 2299 | 1 | 3um_FIJJAFBF_506077 | unclassified | unclassified | unclassified | unclassified | unclassified | unclassified | unclassified | unclassified | unclassified | unclassified | unclassified | unclassified | 2,4 | 1,496662955 | 0,000313157 | 0,000168023 | 1 - | 0,000163782 - | 2 - | 0,000273851 - | 1 - | 0,00013784 - |
| 2300 | 1 | 3um_FIJJAFBF_175693 | Bacteria | Proteobacteri | Alphaproteob | Rhodobactera | Rhodobactera | Planktotalea | Planktotalea f | unclassified | unclassified | unclassified | unclassified | unclassified | 0,2 | 0,4 | 4,77E-05 | 9,54E-05 | 0 - | 0 - | 0 - | 0 - | 1 - | 0,00023861 - |
| 2301 | 1 | 3um_FIJJAFBF_135241 | Eukaryota | unclassified | unclassified | unclassified | Histionidae | Reclinomonas | Reclinomonas | metabolism | Energy produc | 12T8S | NADH dehydr | 9305.ENSSHA | 0,8 | 0,4 | 7,60E-05 | 3,92E-05 | 1 - | 0,000111547 - | 1 - | 9,33E-05 - | 1 - | 9,39E-05 - |
| 2302 | 1 | 3um_FIJJAFBF_462349 | Eukaryota | unclassified | Oomycetes | Saprolegniales | Saprolegniace | Aphanomyces | Aphanomyces | cellular proces | Intracellular tr | KOG2514 | Anoctamin | 65071.PYU1_ | 1 | 0 | 0,000203858 | 2,16E-05 | 1 - | 0,000242227 - | 1 - | 0,000202508 - | 1 - | 0,000203861 - |
| 2303 | 1 | 0.2um_NOKCPHJN_451227 | unclassified | unclassified | unclassified | unclassified | unclassified | unclassified | unclassified | unclassified | unclassified | unclassified | unclassified | unclassified | 0,2 | 0,4 | 6,23E-05 | 0,000124509 | 0 - | 0 - | 0 - | 0 - | 0 - | 0 - |
| 2304 | 1 | 0.2um_NOKCPHJN_855581 | Eukaryota | unclassified | Pelagophyceae | Pelagomonad | unclassified | Aureococcus | Aureococcus a | unclassified | unclassified | unclassified | unclassified | unclassified | 0,8 | 0,4 | 0,000101192 | 5,23E-05 | 1 - | 0,000150298 - | 1 - | 0,000125652 - | 0 - | 0 - |
| 2305 | 1 | 0.2um_NOKCPHJN_469875 | Bacteria | Proteobacteri | Gammaprotec | Alteromonada | Pseudoalteror | Pseudoalteror | Pseudoalteror | poorly charact | Function unkn | 05NCR | NA | 269799.Gmet_ | 1,8 | 0,4 | 0,000290738 | 7,63E-05 | 2 - | 0,000381878 - | 2 - | 0,00031926 - | 2 - | 0,000321393 - |
| 2306 | 1 | 0.2um_NOKCPHJN_25462 | Viruses | unclassified | unclassified | unclassified | Phycodnavirid | unclassified | Chrysochromu | poorly charact | Function unkn | 007GH | NA | 5833.PFI1000 | 0,2 | 0,4 | 8,18E-06 | 1,64E-05 | 0 - | 0 - | 0 - | 0 - | 1 - | 4,09E-05 - |
| 2307 | 1 | 0.2um_NOKCPHJN_66824 | Bacteria | Proteobacteri | Gammaprotec | Alteromonada | Pseudoalteror | Pseudoalteror | Pseudoalteror | metabolism | Carbohydrate | 16QEQ | hydrolase, fan | 247634.GPB2 | 0,4 | 0,489897949 | 2,35E-05 | 2,89E-05 | 0 - | 0 - | 0 - | 0 - | 1 - | 6,29E-05 - |
| 2308 | 1 | 0.2um_NOKCPHJN_797213 | Eukaryota | unclassified | Oomycetes | Saprolegniales | Saprolegniace | Achlya | Achlya hypogy | poorly charact | Function unkn | 0J2G0 | C-Myc binding | 67593.Physo1 | 0,2 | 0,4 | 0,000196969 | 0,000393937 | 0 - | 0 - | 0 - | 0 - | 0 - | 0 - |
| 2309 | 1 | 3um_FIJJAFBF_638938 | Bacteria | Proteobacteri | Alphaproteob | Rhodobactera | Rhodobactera | unclassified | Rhodobactera | unclassified | unclassified | unclassified | unclassified | unclassified | 0,4 | 0,489897949 | 4,05E-05 | 4,97E-05 | 0 - | 0 - | 0 - | 0 - | 0 - | 0 - |
| 2310 | 1 | 3um_FIJJAFBF_339167 | Bacteria | Bacteroidetes | Flavobacteriia | Flavobacterial | Flavobacteriac | Polaribacter | Polaribacter s | cellular proces | Cell wall/mem | 11TPU | outer membra | 313594.PI23P | 0,8 | 0,4 | 0,000285182 | 0,00014747 | 1 - | 0,000422871 - | 0 - | 0 - | 1 - | 0,000355892 - |
| 2311 | 1 | 0.2um_NOKCPHJN_709837 | Eukaryota | unclassified | Spirotrichea | Sporadotrichi | Oxytrichidae | Oxytricha | Oxytricha trifa | unclassified | unclassified | unclassified | unclassified | unclassified | 0,8 | 0,4 | 5,74E-05 | 2,89E-05 | 0 - | 0 - | 1 - | 7,48E-05 - | 1 - | 7,53E-05 - |
| 2312 | 1 | 0.2um_NOKCPHJN_820335 | Bacteria | Proteobacteri | Alphaproteob | Rhodobactera | Rhodobactera | Roseovarius | Roseovarius a | unclassified | unclassified | unclassified | unclassified | unclassified | 0,4 | 0,489897949 | 3,41E-05 | 4,19E-05 | 0 - | 0 - | 0 - | 0 - | 1 - | 9,14E-05 - |
| 2313 | 1 | 3um_FIJJAFBF_50289 | Bacteria | Bacteroidetes | Flavobacteriia | Flavobacterial | Flavobacteriac | Arenitalea | Arenitalea lut | metabolism | Carbohydrate | 05D2N | Aldolase | 313603.FB217 | 0,6 | 0,489897949 | 0,000101429 | 8,33E-05 | 0 - | 0 - | 1 - | 0,000176267 - | 1 - | 0,000177445 - |
| 2314 | 1 | 3um_FIJJAFBF_402154 | Bacteria | Proteobacteri | Alphaproteob | Sphingomona | Sphingomona | Sphingomonas | Sphingomona | unclassified | unclassified | unclassified | unclassified | unclassified | 0,2 | 0,4 | 3,05E-05 | 6,10E-05 | 0 - | 0 - | 1 - | 0,000152622 - | 0 - | 0 - |
| 2315 | 1 | 3um_FIJJAFBF_254304 | Bacteria | Proteobacteri | Gammaprotec | Cellvibrionale | Halieaceae | Pseudohaliea | Pseudohaliea | unclassified | unclassified | unclassified | unclassified | unclassified | 2,4 | 0,8 | 0,000990015 | 0,000283745 | 1 - | 0,000505731 - | 2 - | 0,000845607 - | 3 - | 0,001276884 - |
| 2316 | 1 | 0.2um_NOKCPHJN_723505 | Bacteria | Actinobacteria | Actinobacteria | Corynebacteri | Mycobacteria | Mycobacteriu | Mycobacteriu | unclassified | unclassified | unclassified | unclassified | unclassified | 1 | 0 | 0,000129082 | 1,37E-05 | 1 - | 0,000153377 - | 1 - | 0,000128227 - | 1 - | 0,000129084 - |
| 2317 | 1 | 3um_FIJJAFBF_461081 | Eukaryota | unclassified | Bangiophycea | Bangiales | Bangiaceae | Porphyra | Porphyra umb | metabolism | Inorganic ion t | 1E1DU | Cation transp | 70448.Q01GZ | 2,4 | 0,8 | 0,000682597 | 0,000228283 | 2 - | 0,000671284 - | 3 - | 0,000841815 - | 3 - | 0,000847439 - |
| 2318 | 1 | 3um_FIJJAFBF_742492 | Eukaryota | Bacillariophyta | Bacillariophyc | Naviculales | Naviculaceae | Fistulifera | Fistulifera sola | unclassified | unclassified | unclassified | unclassified | unclassified | 0,8 | 0,4 | 0,000235405 | 0,000118641 | 0 - | 0 - | 1 - | 0,00030674 - | 1 - | 0,000308789 - |

| | | | | | | | | | | | | | | | | | | | | | | | | |
|---|---|---|---|---|---|---|---|---|---|---|---|---|---|---|---|---|---|---|---|---|---|---|---|---|
| 2319 | 1 | 3um_FIJJAFBF_301944 | Eukaryota | unclassified | Oomycetes | Peronosporale | Peronosporac | Phytophthora | various | metabolism | Inorganic ion t | 0IG0G | Calcium-bindi | 2850.Phatr313 | 2,2 | 0,748331477 | 0,00039086 | 0,000124632 | 1 - | 0,000215701 - | 2 - | 0,000360662 - | 3 - | 0,000544608 - |
| 2320 | 1 | 3um_FIJJAFBF_09654 | Bacteria | Proteobacteri | Gammaprotec | Cellvibrionale | Spongiibacter | Spongiibacter | Spongiibacter | cellular proce | Intracellular tr | 05E4G | Type IV pilus | 247633.GP214 | 0,4 | 0,489897949 | 3,69E-05 | 4,54E-05 | 1 - | 0,000100064 - | 0 - | 0 - | 1 - | 8,42E-05 - |
| 2321 | 1 | 3um_FIJJAFBF_191785 | Eukaryota | unclassified | Raphidophyce | Chattonellales | Chattonellace | Heterosigma | Heterosigma a | unclassified | unclassified | unclassified | unclassified | unclassified | 2,8 | 0,4 | 0,000882308 | 0,000162575 | 3 - | 0,001122723 - | 2 - | 0,000625749 - | 3 - | 0,000944894 - |
| 2322 | 1 | 3um_FIJJAFBF_20314 | Bacteria | Bacteroidetes | Flavobacteriia | Flavobacterial | Flavobacteria | Polaribacter | Polaribacter h | metabolism | Carbohydrate | COG0057 | glyceraldehyd | 762903.Pedsa | 2,6 | 0,8 | 0,000481594 | 0,000113142 | 2 - | 0,000449539 - | 2 - | 0,000375825 - | 3 - | 0,000567504 - |
| 2323 | 1 | 3um_FIJJAFBF_814534 | unclassified | unclassified | unclassified | unclassified | unclassified | unclassified | unclassified | unclassified | unclassified | unclassified | unclassified | unclassified | 0,2 | 0,4 | 0,000100967 | 0,000201934 | 0 - | 0 - | 0 - | 0 - | 0 - | 0 - |
| 2324 | 1 | 3um_FIJJAFBF_577614 | unclassified | unclassified | unclassified | unclassified | unclassified | unclassified | unclassified | unclassified | unclassified | unclassified | unclassified | unclassified | 0,2 | 0,4 | 0,000187736 | 0,000375472 | 0 - | 0 - | 0 - | 0 - | 0 - | 0 - |
| 2325 | 1 | 0.2um_NOKCPHJN_282157 | Eukaryota | Bacillariophyta | Fragilariophyc | Fragilariales | Fragilariaceae | Ardissonea | Ardissonea cry | unclassified | unclassified | unclassified | unclassified | unclassified | 2,8 | 0,4 | 0,000236812 | 5,03E-05 | 3 - | 0,000298596 - | 3 - | 0,000249634 - | 3 - | 0,000251302 - |
| 2326 | 1 | 3um_FIJJAFBF_510795 | Viruses | unclassified | unclassified | unclassified | Phycodnavirid | unclassified | Phaeocystis p | unclassified | unclassified | unclassified | unclassified | unclassified | 0,2 | 0,4 | 2,28E-05 | 4,57E-05 | 0 - | 0 - | 0 - | 0 - | 0 - | 0 - |
| 2327 | 1 | 3um_FIJJAFBF_596860 | Bacteria | unclassified | unclassified | unclassified | unclassified | unclassified | uncultured ba | unclassified | unclassified | unclassified | unclassified | unclassified | 0,8 | 0,4 | 9,86E-05 | 5,10E-05 | 1 - | 0,000146474 - | 1 - | 0,000122456 - | 0 - | 0 - |
| 2328 | 1 | 3um_FIJJAFBF_357541 | Bacteria | Bacteroidetes | Flavobacteriia | Flavobacterial | Flavobacteria | Polaribacter | Polaribacter s | metabolism | Amino acid tra | 05C6M | Alpha-keto-be | 313594.PI23P | 2,2 | 0,748331477 | 0,000276229 | 8,81E-05 | 1 - | 0,00015244 - | 2 - | 0,000254888 - | 3 - | 0,000384886 - |
| 2329 | 1 | 3um_FIJJAFBF_807610 | Eukaryota | Bacillariophyta | Bacillariophyc | Naviculales | Phaeodactyla | Phaeodactylu | Phaeodactylu | information st | Translation, ri | KOG2671 | tRNA methyltr | 35128.Thaps2 | 0,4 | 0,489897949 | 3,36E-05 | 4,13E-05 | 0 - | 0 - | 0 - | 0 - | 0 - | 0 - |
| 2330 | 1 | 3um_FIJJAFBF_673147 | unclassified | unclassified | unclassified | unclassified | unclassified | unclassified | unclassified | unclassified | unclassified | unclassified | unclassified | unclassified | 1,6 | 0,489897949 | 0,000864033 | 0,000221332 | 1 - | 0,000656563 - | 1 - | 0,000548903 - | 2 - | 0,001105139 - |
| 2331 | 2 | 0.2um_NOKCPHJN_702776;3um_FIJJAF | Bacteria | Proteobacteri | Gammaprotec | Alteromonada | various | various | various | unclassified | unclassified | unclassified | unclassified | unclassified | 0,2 | 0,4 | 4,05E-05 | 8,10E-05 | 0 - | 0 - | 0 - | 0 - | 1 - | 0,00020255 - |
| 2332 | 1 | 0.2um_NOKCPHJN_1101737 | unclassified | unclassified | unclassified | unclassified | unclassified | unclassified | unclassified | unclassified | unclassified | unclassified | unclassified | unclassified | 0,2 | 0,4 | 0,000162532 | 0,000325065 | 0 - | 0 - | 1 - | 0,000812661 - | 0 - | 0 - |
| 2333 | 1 | 3um_FIJJAFBF_99520 | Viruses | unclassified | unclassified | Caudovirales | Siphoviridae | Cbastvirus | Cellulophaga v | unclassified | unclassified | unclassified | unclassified | unclassified | 1 | 0,632455532 | 0,000324272 | 0,000186523 | 1 - | 0,000396022 - | 1 - | 0,000331084 - | 0 - | 0 - |
| 2334 | 1 | 3um_FIJJAFBF_214727 | Eukaryota | Bacillariophyta | Coscinodiscop | Thalassiosirale | Thalassiosirac | Thalassiosira | Thalassiosira p | metabolism | Carbohydrate | KOG2670 | enolase (EC 4. | 35128.Thaps4 | 0,4 | 0,489897949 | 8,43E-05 | 0,000103286 | 0 - | 0 - | 0 - | 0 - | 1 - | 0,000215729 - |
| 2335 | 1 | 3um_FIJJAFBF_271029 | unclassified | unclassified | unclassified | unclassified | unclassified | unclassified | unclassified | cellular proces | Intracellular tr | 116KZ | NA | 70448.Q019R5 | 0,4 | 0,489897949 | 0,000214102 | 0,000262744 | 0 - | 0 - | 0 - | 0 - | 0 - | 0 - |
| 2336 | 1 | 3um_FIJJAFBF_125762 | Eukaryota | unclassified | Pelagophycea | Pelagomonada | unclassified | Aureococcus | Aureococcus a | metabolism | Energy produc | 0IFBF | One of the tw | 45157.CMV08 | 1 | 0 | 0,000186367 | 1,97E-05 | 1 - | 0,000221444 - | 1 - | 0,000185133 - | 1 - | 0,00018637 - |
| 2337 | 1 | 0.2um_NOKCPHJN_115137 | Bacteria | Bacteroidetes | Flavobacteriia | Flavobacterial | Crocinitomica | unclassified | Crocinitomica | unclassified | unclassified | unclassified | unclassified | unclassified | 1 | 0,632455532 | 0,000127152 | 7,31E-05 | 1 - | 0,000155287 - | 1 - | 0,000129823 - | 0 - | 0 - |
| 2338 | 1 | 3um_FIJJAFBF_411145 | unclassified | unclassified | unclassified | unclassified | unclassified | unclassified | unclassified | unclassified | unclassified | unclassified | unclassified | unclassified | 1 | 0 | 0,000503937 | 5,33E-05 | 1 - | 0,000598785 - | 1 - | 0,000500599 - | 1 - | 0,000503944 - |
| 2339 | 1 | 0.2um_NOKCPHJN_983105 | Eukaryota | Chlorophyta | Ulvophyceae | Ulvales | Ulvaceae | Ulva | Ulva linza | unclassified | unclassified | unclassified | unclassified | unclassified | 1,2 | 0,4 | 0,000362187 | 8,95E-05 | 1 - | 0,000366903 - | 1 - | 0,00030674 - | 1 - | 0,000308789 - |
| 2340 | 1 | 0.2um_NOKCPHJN_535135 | Bacteria | Proteobacteri | Alphaproteob | Rickettsiales | Candidatus M | Candidatus Jic | Candidatus Jic | information st | Transcription | 05D27 | DNA-depende | 862719.AZOLI | 1 | 0 | 6,29E-05 | 6,65E-06 | 1 - | 7,47E-05 - | 1 - | 6,25E-05 - | 1 - | 6,29E-05 - |
| 2341 | 1 | 0.2um_NOKCPHJN_333506 | Eukaryota | unclassified | Labyrinthulom | unclassified | Thraustochytr | Aurantiochytr | Aurantiochytr | poorly charact | Function unkn | 0NVRZ | NA | 170187.SP_17 | 0,6 | 0,489897949 | 1,00E-05 | 8,22E-06 | 0 - | 0 - | 0 - | 0 - | 1 - | 1,78E-05 - |
| 2342 | 1 | 3um_FIJJAFBF_348593 | Bacteria | Acidobacteria | unclassified | unclassified | unclassified | unclassified | Acidobacteria | metabolism | Energy produc | 07EQ1 | H -transportin | 682795.AciX8 | 0,6 | 0,489897949 | 0,000381745 | 0,000314507 | 1 - | 0,00071284 - | 1 - | 0,000595952 - | 1 - | 0,000599933 - |
| 2343 | 1 | 3um_FIJJAFBF_37031 | Eukaryota | unclassified | Oomycetes | Saprolegniales | Saprolegniace | Aphanomyces | Aphanomyces | cellular proces | Signal transdu | KOG0613 | fibronectin typ | 135651.CBN15 | 0,8 | 0,748331477 | 7,53E-05 | 6,77E-05 | 0 - | 0 - | 1 - | 0,000101748 - | 0 - | 0 - |
| 2344 | 1 | 3um_FIJJAFBF_777843 | Bacteria | Bacteroidetes | Flavobacteriia | Flavobacterial | Flavobacteria | Polaribacter | Polaribacter s | metabolism | Lipid transpor | 0P7W4 | reductase | 1046627.BZAF | 0,4 | 0,489897949 | 9,24E-05 | 0,000113361 | 0 - | 0 - | 0 - | 0 - | 0 - | 0 - |
| 2345 | 3 | 0.2um_NOKCPHJN_1019867;0.2um_NO | Eukaryota | various | various | various | various | various | various | unclassified | unclassified | unclassified | unclassified | unclassified | 1 | 0 | 0,000649404 | 6,87E-05 | 1 - | 0,000771631 - | 1 - | 0,000645102 - | 1 - | 0,000649412 - |
| 2346 | 1 | 0.2um_NOKCPHJN_739197 | Bacteria | Proteobacteri | Gammaprotec | Cellvibrionale | Porticoccacea | unclassified | Gammaprotec | metabolism | Energy produc | COG2224 | Isocitrate lyas | 314287.GB220 | 1,2 | 0,748331477 | 0,000186842 | 0,000110395 | 1 - | 0,000189489 - | 2 - | 0,000316835 - | 0 - | 0 - |
| 2347 | 1 | 3um_FIJJAFBF_743884 | Bacteria | Proteobacteri | Alphaproteob | Rhizobiales | unclassified | unclassified | Rhizobiales ba | poorly charact | Function unkn | 05C2A | tonB-depende | 1085623.GNIT | 0,2 | 0,4 | 2,27E-05 | 4,53E-05 | 0 - | 0 - | 0 - | 0 - | 0 - | 0 - |
| 2348 | 1 | 3um_FIJJAFBF_798846 | Bacteria | Proteobacteri | Gammaprotec | Oceanospirilla | Oceanospirilla | Motiliproteus | Motiliproteus | unclassified | unclassified | unclassified | unclassified | unclassified | 1,6 | 0,489897949 | 0,000312293 | 0,000104734 | 2 - | 0,000463456 - | 1 - | 0,00019373 - | 1 - | 0,000195025 - |
| 2349 | 1 | 3um_FIJJAFBF_645987 | Bacteria | Actinobacteria | Actinobacteria | unclassified | unclassified | unclassified | Actinobacteria | unclassified | unclassified | unclassified | unclassified | unclassified | 0,4 | 0,489897949 | 9,12E-05 | 0,000111702 | 0 - | 0 - | 0 - | 0 - | 1 - | 0,000233307 - |
| 2350 | 1 | 3um_FIJJAFBF_44702 | Bacteria | Bacteroidetes | Flavobacteriia | Flavobacterial | Flavobacteria | Polaribacter | Polaribacter s | cellular proces | Cell wall/mem | 11TPU | outer membra | 313598.MED1 | 3,2 | 0,4 | 0,001199852 | 0,000191436 | 3 - | 0,001336575 - | 3 - | 0,001117409 - | 4 - | 0,001499832 - |
| 2351 | 1 | 0.2um_NOKCPHJN_116666 | Bacteria | Proteobacteri | Gammaprotec | Cellvibrionale | Porticoccacea | unclassified | Gammaprotec | metabolism | Inorganic ion t | 16TQC | TONB-depend | 314287.GB220 | 0,8 | 1,166190379 | 4,61E-05 | 6,57E-05 | 0 - | 0 - | 0 - | 0 - | 0 - | 0 - |
| 2352 | 1 | 3um_FIJJAFBF_116593 | Bacteria | Cyanobacteria | unclassified | Synechococca | Synechococca | Synechococcu | Synechococcu | unclassified | unclassified | unclassified | unclassified | unclassified | 1 | 0 | 0,000243213 | 2,57E-05 | 1 - | 0,000288989 - | 1 - | 0,000241602 - | 1 - | 0,000243216 - |
| 2353 | 1 | 3um_FIJJAFBF_242895 | Bacteria | Bacteroidetes | Flavobacteriia | Flavobacterial | Flavobacteria | Polaribacter | Polaribacter d | poorly charact | Function unkn | 08ME3 | Alpha amylase | 983548.Krodi_ | 0,4 | 0,489897949 | 7,50E-05 | 9,37E-05 | 1 - | 0,000216951 - | 0 - | 0 - | 0 - | 0 - |
| 2354 | 1 | 0.2um_NOKCPHJN_1016379 | Eukaryota | Arthropoda | Insecta | Hymenoptera | Apidae | Melipona | Melipona qua | information st | RNA processir | 1DWIU | Pre-mRNA-spl | 3055.EDO970 | 0,2 | 0,4 | 5,56E-05 | 0,000111162 | 0 - | 0 - | 0 - | 0 - | 0 - | 0 - |
| 2355 | 1 | 3um_FIJJAFBF_323687 | Eukaryota | unclassified | Oomycetes | Peronosporale | Peronosporac | Phytophthora | Phytophthora | unclassified | unclassified | unclassified | unclassified | unclassified | 1 | 0 | 9,40E-05 | 9,94E-06 | 1 - | 0,000111714 - | 1 - | 9,34E-05 - | 1 - | 9,40E-05 - |
| 2356 | 1 | 0.2um_NOKCPHJN_689502 | Bacteria | Actinobacteria | Actinobacteria | Streptomyceta | Streptomyceta | Streptomyces | Streptomyces | unclassified | unclassified | unclassified | unclassified | unclassified | 1 | 0 | 0,000393701 | 4,16E-05 | 1 - | 0,000467801 - | 1 - | 0,000391093 - | 1 - | 0,000393706 - |
| 2357 | 1 | 0.2um_NOKCPHJN_418168 | Bacteria | Verrucomicrol | Opitutae | unclassified | unclassified | unclassified | Opitutae bact | unclassified | unclassified | unclassified | unclassified | unclassified | 0,2 | 0,4 | 2,49E-05 | 4,98E-05 | 0 - | 0 - | 0 - | 0 - | 0 - | 0 - |
| 2358 | 1 | 0.2um_NOKCPHJN_225033 | Eukaryota | unclassified | Labyrinthulom | unclassified | Thraustochytr | Aurantiochytr | Aurantiochytr | poorly charact | Function unkn | 0XR4D | armadillo repe | 7719.ENSCINP | 0,2 | 0,4 | 6,56E-06 | 1,31E-05 | 0 - | 0 - | 0 - | 0 - | 0 - | 0 - |
| 2359 | 1 | 0.2um_NOKCPHJN_538650 | unclassified | unclassified | unclassified | unclassified | unclassified | unclassified | unclassified | unclassified | unclassified | unclassified | unclassified | unclassified | 0,2 | 0,4 | 6,12E-05 | 0,000122316 | 0 - | 0 - | 0 - | 0 - | 1 - | 0,000305791 - |
| 2360 | 1 | 3um_FIJJAFBF_135101 | unclassified | unclassified | unclassified | unclassified | unclassified | unclassified | unclassified | unclassified | unclassified | unclassified | unclassified | unclassified | 0,2 | 0,4 | 7,46E-05 | 0,000149231 | 0 - | 0 - | 0 - | 0 - | 0 - | 0 - |
| 2361 | 1 | 0.2um_NOKCPHJN_904622 | Eukaryota | unclassified | Oomycetes | Peronosporale | Peronosporac | Phytophthora | Phytophthora | information st | Replication, re | COG0513 | purine NTP-de | 653948.CCA19 | 2,8 | 0,748331477 | 0,000427455 | 0,000108341 | 3 - | 0,00054767 - | 3 - | 0,000457865 - | 2 - | 0,000307283 - |
| 2362 | 2 | 3um_FIJJAFBF_551628;3um_FIJJAFBF_ | Bacteria | Proteobacteri | Gammaprotec | Pseudomonad | Moraxellaceae | various | various | information st | Translation, ri | 0QQXI | This protein is | various | 0,2 | 0,4 | 8,81E-05 | 0,000176204 | 0 - | 0 - | 0 - | 0 - | 1 - | 0,00044051 - |
| 2363 | 1 | 0.2um_NOKCPHJN_484873 | Bacteria | Proteobacteri | Gammaprotec | Chromatiales | Chromatiacea | Candidatus Te | Candidatus Te | unclassified | unclassified | unclassified | unclassified | unclassified | 1 | 0 | 0,000142516 | 1,51E-05 | 1 - | 0,00016934 - | 1 - | 0,000141572 - | 1 - | 0,000142518 - |
| 2364 | 1 | 3um_FIJJAFBF_245869 | Eukaryota | unclassified | Dinophyceae | Suessiales | Symbiodiniace | Symbiodinium | Symbiodinium | unclassified | unclassified | unclassified | unclassified | unclassified | 0,4 | 0,489897949 | 1,36E-05 | 1,66E-05 | 0 - | 0 - | 0 - | 0 - | 0 - | 0 - |
| 2365 | 1 | 3um_FIJJAFBF_143488 | Bacteria | Actinobacteria | Actinobacteria | Micrococcales | Microbacteria | unclassified | Microbacteria | poorly charact | Function unkn | 00QCW | NA | 312284.A20C1 | 0,6 | 0,489897949 | 0,000232076 | 0,000190333 | 0 - | 0 - | 0 - | 0 - | 1 - | 0,000411719 - |
| 2366 | 1 | 0.2um_NOKCPHJN_525992 | Eukaryota | unclassified | Cryptophyta | Pyrenomonad | Geminigerace | Guillardia | Guillardia thet | unclassified | unclassified | unclassified | unclassified | unclassified | 1 | 0 | 0,000128818 | 1,36E-05 | 1 - | 0,000153064 - | 1 - | 0,000127965 - | 1 - | 0,00012882 - |
| 2367 | 1 | 3um_FIJJAFBF_135237 | Bacteria | Cyanobacteria | unclassified | Synechococca | Synechococca | Synechococcu | Synechococcu | metabolism | Energy produc | COG1290 | Component of | 70448.Q0P3F3 | 1,4 | 0,489897949 | 0,000226181 | 6,81E-05 | 1 - | 0,000195938 - | 2 - | 0,000327617 - | 1 - | 0,000164903 - |
| 2368 | 1 | 3um_FIJJAFBF_657588 | unclassified | unclassified | unclassified | unclassified | unclassified | unclassified | unclassified | unclassified | unclassified | unclassified | unclassified | unclassified | 1 | 0 | 0,000368375 | 3,90E-05 | 1 - | 0,000437709 - | 1 - | 0,000365935 - | 1 - | 0,00036838 - |
| 2369 | 1 | 0.2um_NOKCPHJN_597353 | Eukaryota | Bacillariophyta | Bacillariophyc | Naviculales | Naviculaceae | Fistulifera | Fistulifera sola | metabolism | Energy produc | KOG0017 | Retrotranspos | 39947.LOC_O | 0,4 | 0,489897949 | 2,55E-05 | 3,14E-05 | 0 - | 0 - | 1 - | 6,82E-05 - | 0 - | 0 - |
| 2370 | 1 | 3um_FIJJAFBF_607280 | Bacteria | Proteobacteri | Gammaprotec | unclassified | unclassified | unclassified | Gammaprotec | unclassified | unclassified | unclassified | unclassified | unclassified | 0,8 | 0,4 | 0,000133929 | 6,82E-05 | 1 - | 0,000192412 - | 1 - | 0,000160861 - | 1 - | 0,000161936 - |
| 2371 | 1 | 0.2um_NOKCPHJN_430403 | Eukaryota | Nematoda | Enoplea | Trichinellida | Trichuridae | Trichuris | Trichuris trich | cellular proces | Signal transdu | 0AFXJ | Pfam:efhand | 6334.EFV5491 | 0,2 | 0,4 | 1,85E-05 | 3,71E-05 | 0 - | 0 - | 0 - | 0 - | 1 - | 9,26E-05 - |
| 2372 | 1 | 0.2um_NOKCPHJN_452764 | Bacteria | Proteobacteri | Gammaprotec | Pseudomonad | Moraxellaceae | Acinetobacter | Acinetobacter | unclassified | unclassified | unclassified | unclassified | unclassified | 0,8 | 0,4 | 0,000176494 | 9,13E-05 | 1 - | 0,000261707 - | 0 - | 0 - | 1 - | 0,000220255 - |
| 2373 | 1 | 3um_FIJJAFBF_489502 | Bacteria | Bacteroidetes | Flavobacteriia | Flavobacterial | Flavobacteria | Polaribacter | various | unclassified | unclassified | unclassified | unclassified | unclassified | 4,2 | 0,979795897 | 0,000305895 | 4,18E-05 | 3 - | 0,000265419 - | 4 - | 0,000295862 - | 4 - | 0,000297839 - |
| 2374 | 1 | 3um_FIJJAFBF_128630 | Eukaryota | unclassified | Raphidophyce | Chattonellales | Chattonellace | Heterosigma | Heterosigma a | unclassified | unclassified | unclassified | unclassified | unclassified | 6,6 | 1,019803903 | 0,001650745 | 0,000116903 | 5 - | 0,001509036 - | 6 - | 0,001513909 - | 7 - | 0,001778026 - |
| 2375 | 1 | 3um_FIJJAFBF_517423 | Bacteria | Bacteroidetes | Flavobacteriia | Flavobacterial | Flavobacteria | Polaribacter | Polaribacter b | unclassified | unclassified | unclassified | unclassified | unclassified | 0,6 | 0,489897949 | 0,000207647 | 0,000170298 | 0 - | 0 - | 0 - | 0 - | 1 - | 0,00036838 - |
| 2376 | 1 | 3um_FIJJAFBF_778055 | Bacteria | Bacteroidetes | Flavobacteriia | Flavobacterial | Flavobacteria | Polaribacter | Polaribacter s | metabolism | Inorganic ion t | 09B4C | tonB-depende | 313594.PI23P | 0,2 | 0,4 | 1,52E-05 | 3,04E-05 | 0 - | 0 - | 0 - | 0 - | 0 - | 0 - |
| 2377 | 1 | 0.2um_NOKCPHJN_642959 | Eukaryota | Phaeophyceae | unclassified | Ectocarpales | Ectocarpaceae | Ectocarpus | Ectocarpus sili | unclassified | unclassified | unclassified | unclassified | unclassified | 1 | 0 | 8,13E-05 | 8,60E-06 | 1 - | 9,66E-05 - | 1 - | 8,07E-05 - | 1 - | 8,13E-05 - |
| 2378 | 1 | 3um_FIJJAFBF_774175 | Eukaryota | unclassified | Pelagophycea | Pelagomonada | unclassified | Aureococcus | Aureococcus a | unclassified | unclassified | unclassified | unclassified | unclassified | 3,4 | 0,489897949 | 0,000428918 | 4,16E-05 | 3 - | 0,000454544 - | 3 - | 0,00038001 - | 3 - | 0,000382548 - |
| 2379 | 2 | 0.2um_NOKCPHJN_1045583;3um_FIJJA | Eukaryota | various | various | various | various | various | various | cellular proces | Intracellular tr | KOG0096 | GTP-binding n | various | 1 | 0 | 0,000294356 | 3,11E-05 | 1 - | 0,000349758 - | 1 - | 0,000292406 - | 1 - | 0,00029436 - |
| 2380 | 1 | 3um_FIJJAFBF_116542 | Eukaryota | Chordata | Actinopteri | Carangiformes | Carangidae | Seriola | Seriola lalandi | unclassified | unclassified | unclassified | unclassified | unclassified | 2,8 | 0,748331477 | 0,002557695 | 0,000619346 | 2 - | 0,002201417 - | 4 - | 0,003680878 - | 2 - | 0,001852733 - |
| 2381 | 1 | 3um_FIJJAFBF_336548 | Bacteria | Spirochaetes | Spirochaetia | unclassified | Leptospiracea | Leptospira | Leptospira sp. | unclassified | unclassified | unclassified | unclassified | unclassified | 4 | 0,632455532 | 0,000367522 | 6,45E-05 | 4 - | 0,000437709 - | 4 - | 0,000365935 - | 3 - | 0,000276285 - |
| 2382 | 1 | 3um_FIJJAFBF_525182 | Eukaryota | Apicomplexa | Aconoidasida | Haemosporida | Plasmodiidae | Plasmodium | Plasmodium r | unclassified | unclassified | unclassified | unclassified | unclassified | 0,2 | 0,4 | 1,73E-05 | 3,47E-05 | 0 - | 0 - | 0 - | 0 - | 1 - | 8,66E-05 - |
| 2383 | 1 | 0.2um_NOKCPHJN_349413 | Bacteria | Verrucomicrol | unclassified | unclassified | unclassified | unclassified | Verrucomicrol | unclassified | unclassified | unclassified | unclassified | unclassified | 0,2 | 0,4 | 1,70E-05 | 3,39E-05 | 1 - | 8,49E-05 - | 0 - | 0 - | 0 - | 0 - |
| 2384 | 1 | 0.2um_NOKCPHJN_503363 | Bacteria | unclassified | unclassified | unclassified | unclassified | unclassified | uncultured ba | cellular proces | Posttranslatio | 05E6M | Sulphatase-m | 391603.FBALC | 1 | 0 | 0,000131234 | 1,39E-05 | 1 - | 0,000155934 - | 1 - | 0,000130364 - | 1 - | 0,000131235 - |
| 2385 | 1 | 3um_FIJJAFBF_657951 | Bacteria | unclassified | unclassified | unclassified | unclassified | unclassified | bacterium B17 | poorly charact | Function unkn | 0Y3P1 | NA | 44689.DDB01 | 1 | 0 | 9,28E-05 | 9,81E-06 | 1 - | 0,000110233 - | 1 - | 9,22E-05 - | 1 - | 9,28E-05 - |
| 2386 | 1 | 3um_FIJJAFBF_327843 | Bacteria | Bacteroidetes | Flavobacteriia | Flavobacterial | Flavobacteria | Polaribacter | various | information st | Translation, ri | COG0051 | tRNA binding | 313598.MED1 | 1,8 | 0,4 | 0,001099156 | 0,000188885 | 1 - | 0,000741071 - | 2 - | 0,001239107 - | 2 - | 0,001247385 - |
| 2387 | 3 | 3um_FIJJAFBF_447188;3um_FIJJAFBF_ | Bacteria | Proteobacteri | Gammaprotec | Alteromonada | Alteromonada | Paraglaciecola | Paraglaciecola | unclassified | unclassified | unclassified | unclassified | unclassified | 0,4 | 0,489897949 | 3,22E-05 | 3,95E-05 | 0 - | 0 - | 0 - | 0 - | 1 - | 8,25E-05 - |
| 2388 | 1 | 3um_FIJJAFBF_417001 | unclassified | unclassified | unclassified | unclassified | unclassified | unclassified | unclassified | unclassified | unclassified | unclassified | unclassified | unclassified | 0,4 | 0,489897949 | 0,000293705 | 0,000360431 | 0 - | 0 - | 0 - | 0 - | 0 - | 0 - |
| 2389 | 1 | 3um_FIJJAFBF_264684 | Bacteria | Proteobacteri | Gammaprotec | Oceanospirilla | Kangiellaceae | Kangiella | Kangiella spor | information st | Translation, ri | 17FIT | 50s ribosomal | 247639.MGP2 | 0,6 | 0,489897949 | 0,000461634 | 0,000378931 | 0 - | 0 - | 1 - | 0,000802243 - | 1 - | 0,000807602 - |
| 2390 | 1 | 3um_FIJJAFBF_747678 | unclassified | unclassified | unclassified | unclassified | unclassified | unclassified | unclassified | unclassified | unclassified | unclassified | unclassified | unclassified | 0,4 | 0,489897949 | 0,000168448 | 0,000206718 | 0 - | 0 - | 0 - | 0 - | 0 - | 0 - |
| 2391 | 1 | 0.2um_NOKCPHJN_169297 | Eukaryota | Chordata | Actinopteri | Clupeiformes | Clupeidae | Clupea | Clupea hareng | unclassified | unclassified | unclassified | unclassified | unclassified | 1 | 0 | 2,07E-05 | 2,19E-06 | 1 - | 2,46E-05 - | 1 - | 2,06E-05 - | 1 - | 2,07E-05 - |
| 2392 | 1 | 0.2um_NOKCPHJN_936620 | Eukaryota | unclassified | Oomycetes | Peronosporale | Peronosporac | Phytophthora | Phytophthora | cellular proces | Signal transdu | 152IZ | EFh | 67593.Physo1 | 3,6 | 0,8 | 0,001334008 | 0,000191961 | 3 - | 0,001344578 - | 3 - | 0,0011241 - | 3 - | 0,00113161 - |
| 2393 | 1 | 0.2um_NOKCPHJN_867614 | Eukaryota | Ascomycota | Eurotiomycete | Eurotiales | Aspergillaceae | Penicillium | Penicillium na | unclassified | unclassified | unclassified | unclassified | unclassified | 0,8 | 0,4 | 6,99E-05 | 3,62E-05 | 1 - | 0,000103668 - | 0 - | 0 - | 1 - | 8,72E-05 - |
| 2394 | 1 | 3um_FIJJAFBF_510247 | Bacteria | Bacteroidetes | Flavobacteriia | Flavobacterial | Flavobacteria | Polaribacter | Polaribacter s | unclassified | unclassified | unclassified | unclassified | unclassified | 2,2 | 0,748331477 | 0,000777015 | 0,00022395 | 1 - | 0,000432648 - | 3 - | 0,001085114 - | 2 - | 0,000728242 - |
| 2395 | 1 | 3um_FIJJAFBF_669336 | Bacteria | Proteobacteri | Gammaprotec | unclassified | unclassified | unclassified | Gammaprotec | unclassified | unclassified | unclassified | unclassified | unclassified | 0,8 | 0,4 | 5,58E-05 | 2,89E-05 | 1 - | 8,28E-05 - | 0 - | 0 - | 1 - | 6,97E-05 - |
| 2396 | 1 | 0.2um_NOKCPHJN_700347 | Eukaryota | Eustigmatoph | unclassified | Eustigmatales | Monodopsida | Nannochlorop | Nannochlorop | metabolism | Carbohydrate | KOG2638 | UTP-glucose-1 | 653948.CCA18 | 10,4 | 1,356465997 | 0,000709845 | 6,41E-05 | 8 - | 0,000655844 - | 10 - | 0,000685377 - | 12 - | 0,000827947 - |
| 2397 | 1 | 0.2um_NOKCPHJN_27323 | Bacteria | Bacteroidetes | unclassified | unclassified | unclassified | unclassified | Bacteroidetes | poorly charact | Function unkn | 0Y63T | Ragb susd dor | 471854.Dfer_ | 6 | 1,414213562 | 0,000728241 | 0,000107735 | 4 - | 0,000590518 - | 6 - | 0,000740532 - | 5 - | 0,000621232 - |
| 2398 | 1 | 3um_FIJJAFBF_451103 | unclassified | unclassified | unclassified | unclassified | unclassified | unclassified | unclassified | unclassified | unclassified | unclassified | unclassified | unclassified | 0,4 | 0,489897949 | 9,55E-05 | 0,000117473 | 0 - | 0 - | 0 - | 0 - | 1 - | 0,000256069 - |
| 2399 | 1 | 3um_FIJJAFBF_156358 | Eukaryota | Bacillariophyta | Bacillariophyc | Naviculales | Phaeodactyla | Phaeodactylu | Phaeodactylu | unclassified | unclassified | unclassified | unclassified | unclassified | 0,2 | 0,4 | 3,52E-05 | 7,03E-05 | 0 - | 0 - | 1 - | 0,000175772 - | 0 - | 0 - |
| 2400 | 1 | 3um_FIJJAFBF_555528 | Bacteria | Proteobacteri | Alphaproteob | Rhodobactera | Hyphomonad | Hyphomonas | Hyphomonas | cellular proces | Cell wall/mem | COG3637 | Membrane | 411684.HPDFI | 3,2 | 0,4 | 0,000709477 | 0,000111044 | 3 - | 0,00079065 - | 4 - | 0,000881337 - | 3 - | 0,000665418 - |
| 2401 | 1 | 0.2um_NOKCPHJN_1064791 | Eukaryota | Bacillariophyta | Bacillariophyc | Naviculales | Phaeodactyla | Phaeodactylu | Phaeodactylu | cellular proces | Intracellular tr | 11QUX | nuclear transp | 35128.Thaps2 | 0,8 | 0,4 | 0,000326684 | 0,000164645 | 0 - | 0 - | 1 - | 0,00042568 - | 1 - | 0,000428523 - |
| 2402 | 1 | 3um_FIJJAFBF_358760 | Eukaryota | unclassified | Pelagophycea | Pelagomonada | unclassified | Aureococcus | Aureococcus a | unclassified | unclassified | unclassified | unclassified | unclassified | 1,8 | 0,4 | 0,001500199 | 0,000257803 | 1 - | 0,001011462 - | 2 - | 0,001691214 - | 2 - | 0,001702512 - |
| 2403 | 1 | 3um_FIJJAFBF_155128 | Bacteria | Proteobacteri | Gammaprotec | Cellvibrionale | Cellvibrionace | Marinimicrobi | Marinimicrobi | cellular proces | Posttranslatio | 17FI0 | Binds to Cpn6 | 203122.Sde_0 | 0,2 | 0,4 | 0,000130364 | 0,000260729 | 0 - | 0 - | 1 - | 0,000651822 - | 0 - | 0 - |
| 2404 | 1 | 3um_FIJJAFBF_205216 | Eukaryota | Bacillariophyta | Mediophyceae | Cymatosirales | Cymatosiracea | Plagiogrammo | Plagiogrammo | metabolism | Energy produc | COG0056 | Produces ATP | 41431.PCC880 | 1,2 | 0,4 | 0,000146891 | 3,63E-05 | 1 - | 0,000148804 - | 1 - | 0,000124403 - | 1 - | 0,000125234 - |
| 2405 | 1 | 3um_FIJJAFBF_12251 | Bacteria | Proteobacteri | Gammaprotec | Cellvibrionale | Porticoccacea | unclassified | Gammaprotec | unclassified | unclassified | unclassified | unclassified | unclassified | 0,6 | 0,489897949 | 3,51E-05 | 2,88E-05 | 0 - | 0 - | 0 - | 0 - | 1 - | 6,22E-05 - |
| 2406 | 3 | 0.2um_NOKCPHJN_192220;0.2um_NO | Eukaryota | various | various | various | various | various | various | information st | Translation, ri | KOG0886 | Ribosomal pro | various | 1 | 0 | 0,000381771 | 4,04E-05 | 1 - | 0,000453625 - | 1 - | 0,000379242 - | 1 - | 0,000381775 - |
| 2407 | 1 | 3um_FIJJAFBF_201378 | Bacteria | Bacteroidetes | Flavobacteriia | Flavobacterial | Flavobacteria | Polaribacter | Polaribacter fi | cellular proces | Cell wall/mem | 0PENK | NA | 391587.KAOT | 3,4 | 0,489897949 | 0,000364691 | 3,54E-05 | 3 - | 0,000386479 - | 3 - | 0,000323106 - | 3 - | 0,000325265 - |
| 2408 | 1 | 0.2um_NOKCPHJN_168317 | Eukaryota | Phaeophyceae | unclassified | Ectocarpales | Ectocarpaceae | Ectocarpus | Ectocarpus sili | unclassified | unclassified | unclassified | unclassified | unclassified | 0,4 | 0,489897949 | 2,02E-05 | 2,48E-05 | 0 - | 0 - | 0 - | 0 - | 0 - | 0 - |
| 2409 | 1 | 3um_FIJJAFBF_452605 | Bacteria | Proteobacteri | Alphaproteob | Sphingomona | Erythrobacter | Altererythrob | Altererythrob | cellular proces | Signal transdu | 01WWM | Histidine kina | 314225.ELI_11 | 0,2 | 0,4 | 2,37E-05 | 4,75E-05 | 0 - | 0 - | 0 - | 0 - | 0 - | 0 - |
| 2410 | 5 | 0.2um_NOKCPHJN_1097847;0.2um_NO | Eukaryota | various | various | various | various | various | various | various | various | various | various | various | 0,2 | 0,4 | 6,56E-05 | 0,000131235 | 0 - | 0 - | 0 - | 0 - | 1 - | 0,000328088 - |
| 2411 | 1 | 3um_FIJJAFBF_742304 | Eukaryota | Chordata | Amphibia | Anura | Ranidae | Rana | Rana catesbei | metabolism | Inorganic ion t | KOG2332 | Stores iron in | 3711.Bra0056 | 1 | 0 | 0,000377199 | 3,99E-05 | 1 - | 0,000448193 - | 1 - | 0,0003747 - | 1 - | 0,000377203 - |
| 2412 | 1 | 3um_FIJJAFBF_574558 | Bacteria | Proteobacteri | Gammaprotec | Alteromonada | Pseudoalteror | Pseudoalteror | unclassified | cellular proces | Defense mech | COG2367 | Beta-lactamas | 351348.Maqu | 0,2 | 0,4 | 0,000101602 | 0,000203203 | 0 - | 0 - | 0 - | 0 - | 1 - | 0,000508008 - |
| 2413 | 1 | 0.2um_NOKCPHJN_635290 | Eukaryota | unclassified | Pelagophycea | Pelagomonada | unclassified | Aureococcus | Aureococcus a | unclassified | unclassified | unclassified | unclassified | unclassified | 1,4 | 0,489897949 | 0,000227077 | 8,07E-05 | 1 - | 0,000192908 - | 2 - | 0,000322551 - | 2 - | 0,000324706 - |
| 2414 | 1 | 0.2um_NOKCPHJN_110537 | Bacteria | Proteobacteri | Gammaprotec | Cellvibrionale | Porticoccacea | unclassified | various | unclassified | unclassified | unclassified | unclassified | unclassified | 0,8 | 0,4 | 0,000298187 | 0,000154216 | 1 - | 0,000442889 - | 1 - | 0,000370266 - | 0 - | 0 - |
| 2415 | 1 | 0.2um_NOKCPHJN_09370 | Bacteria | Bacteroidetes | Flavobacteriia | Flavobacterial | Flavobacteria | Polaribacter | Polaribacter s | cellular proces | Cell wall/mem | COG2885 | Ompa motb d | 313598.MED1 | 0,4 | 0,8 | 4,82E-05 | 9,64E-05 | 0 - | 0 - | 0 - | 0 - | 0 - | 0 - |
| 2416 | 1 | 3um_FIJJAFBF_406308 | Bacteria | various | various | various | various | various | various | information st | Translation, ri | 05KK9 | This protein bi | 411154.GFO_ | 0,4 | 0,489897949 | 0,000137699 | 0,000169319 | 0 - | 0 - | 1 - | 0,000368088 - | 0 - | 0 - |
| 2417 | 1 | 3um_FIJJAFBF_744840 | Bacteria | Proteobacteri | Gammaprotec | Cellvibrionale | Cellvibrionace | Marinimicrobi | Marinimicrobi | information st | Translation, ri | 17FIH | One of the ear | 314287.GB220 | 0,2 | 0,4 | 0,000111162 | 0,000222324 | 0 - | 0 - | 0 - | 0 - | 0 - | 0 - |
| 2418 | 1 | 0.2um_NOKCPHJN_403322 | Eukaryota | Chlorophyta | Chlorophycea | Chlamydomor | Chlamydomor | Chlamydomor | Chlamydomor | unclassified | unclassified | unclassified | unclassified | unclassified | 0,2 | 0,4 | 1,94E-05 | 3,87E-05 | 0 - | 0 - | 0 - | 0 - | 1 - | 9,68E-05 - |
| 2419 | 1 | 0.2um_NOKCPHJN_505891 | Bacteria | Proteobacteri | Gammaprotec | Cellvibrionale | Spongiibacter | unclassified | marine gamm | cellular proces | Cell wall/mem | 0QKXZ | Porin Gram-ne | 247633.GP214 | 1,4 | 0,8 | 0,00026056 | 0,000140912 | 1 - | 0,000226813 - | 0 - | 0 - | 2 - | 0,000381775 - |

| | | | | | | | | | | | | | | | | | | | | | | | | |
|---|---|---|---|---|---|---|---|---|---|---|---|---|---|---|---|---|---|---|---|---|---|---|---|---|
| 2420 | 1 | 0.2um_NOKCPHJN_1083545 | Eukaryota | unclassified | unclassified | Prymnesiales | Chrysochromu | Chrysochromu | Chrysochromu | cellular proces | Posttranslatio | 11VAU | glutathione St | 70448.Q01GE | 0,4 | 0,489897949 | 9,97E-05 | 0,000122056 | 0 - | 0 - | 1 - | 0,000248313 - | 1 - | 0,000249972 - |
| 2421 | 1 | 0.2um_NOKCPHJN_24010 | Eukaryota | Arthropoda | Insecta | Lepidoptera | Noctuidae | Helicoverpa | Helicoverpa a | unclassified | unclassified | unclassified | unclassified | unclassified | 2,4 | 0,489897949 | 0,000585856 | 9,72E-05 | 2 - | 0,000587045 - | 3 - | 0,000736176 - | 2 - | 0,000494062 - |
| 2422 | 1 | 0.2um_NOKCPHJN_1119393 | Eukaryota | Arthropoda | Insecta | Lepidoptera | Pyralidae | Amyelois | Amyelois tran | unclassified | unclassified | unclassified | unclassified | unclassified | 0,8 | 0,4 | 0,00011034 | 5,69E-05 | 1 - | 0,000162009 - | 1 - | 0,000135444 - | 1 - | 0,000136348 - |
| 2423 | 1 | 0.2um_NOKCPHJN_1043614 | Eukaryota | Euglenida | unclassified | Eutreptiales | unclassified | Eutreptiella | Eutreptiella b | unclassified | unclassified | unclassified | unclassified | unclassified | 1 | 0 | 0,000174978 | 1,85E-05 | 1 - | 0,000207912 - | 1 - | 0,000173819 - | 1 - | 0,00017498 - |
| 2424 | 1 | 0.2um_NOKCPHJN_354376 | Bacteria | Proteobacteri | Betaproteoba | Burkholderial | Comamonada | Candidatus Sy | Candidatus Sy | metabolism | Amino acid tr | 07SUW | oxidoreductas | 314256.OG25 | 0,4 | 0,489897949 | 3,26E-05 | 4,00E-05 | 0 - | 0 - | 0 - | 0 - | 0 - | 0 - |
| 2425 | 1 | 3um_FIJJAFBF_402705 | unclassified | unclassified | unclassified | unclassified | unclassified | unclassified | unclassified | unclassified | unclassified | unclassified | unclassified | unclassified | 0,2 | 0,4 | 0,000155627 | 0,000311254 | 0 - | 0 - | 0 - | 0 - | 0 - | 0 - |
| 2426 | 1 | 3um_FIJJAFBF_258847 | Bacteria | Proteobacteri | Gammaprotec | Alteromonada | Colwelliaceae | Colwellia | Colwellia sp. F | unclassified | unclassified | unclassified | unclassified | unclassified | 0,6 | 0,489897949 | 0,000227092 | 0,000189354 | 1 - | 0,000442889 - | 1 - | 0,000370266 - | 0 - | 0 - |
| 2427 | 1 | 3um_FIJJAFBF_776741 | Bacteria | Bacteroidetes | Flavobacteriia | Flavobacterial | Flavobacteria | Cellulophaga | Cellulophaga f | unclassified | unclassified | unclassified | unclassified | unclassified | 0,4 | 0,489897949 | 9,51E-05 | 0,000116654 | 0 - | 0 - | 0 - | 0 - | 0 - | 0 - |
| 2428 | 1 | 0.2um_NOKCPHJN_36914 | Eukaryota | unclassified | Cryptophyta | Pyrenomonad | Pyrenomonad | Rhodomonas | Rhodomonas | metabolism | Carbohydrate | COG1850 | RuBisCO catal | 45157.CMV01 | 2 | 0 | 0,000258165 | 2,73E-05 | 2 - | 0,000306755 - | 2 - | 0,000256455 - | 2 - | 0,000258168 - |
| 2429 | 1 | 3um_FIJJAFBF_617422 | Eukaryota | unclassified | Dictyochophy | Florenciellales | unclassified | Pseudochatto | Pseudochatto | cellular proces | Cytoskeleton | 1514Z | Tubulin is the | 4787.PITG_00 | 2,8 | 0,4 | 0,001411692 | 0,00026012 | 3 - | 0,001796356 - | 2 - | 0,001001199 - | 3 - | 0,001511831 - |
| 2430 | 1 | 3um_FIJJAFBF_215516 | Eukaryota | Bacillariophyt | Bacillariophyc | Bacillariales | Bacillariaceae | Fragilariopsis | Fragilariopsis | metabolism | Inorganic ion t | COG0659 | sulfate transp | 4787.PITG_20 | 0,4 | 0,489897949 | 3,44E-05 | 4,23E-05 | 0 - | 0 - | 1 - | 9,20E-05 - | 0 - | 0 - |
| 2431 | 2 | 3um_FIJJAFBF_137599;3um_FIJJAFBF_ | Bacteria | Bacteroidetes | Flavobacteriia | Flavobacterial | Crocinitomica | unclassified | Crocinitomica | information st | Translation, ri | COG0256 | This is one of t | 888832.HMPR | 0,2 | 0,4 | 9,15E-05 | 0,00018309 | 0 - | 0 - | 0 - | 0 - | 0 - | 0 - |
| 2432 | 1 | 0.2um_NOKCPHJN_941524 | Eukaryota | unclassified | Spirotrichea | Sporadotrichi | Oxytrichidae | Oxytricha | Oxytricha trifa | unclassified | unclassified | unclassified | unclassified | unclassified | 3,2 | 0,748331477 | 0,000439544 | 6,84E-05 | 2 - | 0,000334144 - | 3 - | 0,000419028 - | 3 - | 0,000421828 - |
| 2433 | 1 | 3um_FIJJAFBF_715072 | Eukaryota | Bacillariophyt | Mediophyceae | Triceratiales | Triceratiaceae | Triceratium | Triceratium d | information st | Translation, ri | COG0050 | This protein p | 32049.SYNPC | 8,2 | 2,315167381 | 0,001233326 | 0,000275697 | 5 - | 0,000915014 - | 6 - | 0,000917969 - | 9 - | 0,001386153 - |
| 2434 | 1 | 0.2um_NOKCPHJN_443911 | unclassified | unclassified | unclassified | unclassified | unclassified | unclassified | unclassified | unclassified | unclassified | unclassified | unclassified | unclassified | 0,2 | 0,4 | 0,000115892 | 0,000231785 | 0 - | 0 - | 0 - | 0 - | 0 - | 0 - |
| 2435 | 1 | 0.2um_NOKCPHJN_111204 | Eukaryota | unclassified | Synurophycea | Parmales | Triparmaceae | Triparma | Triparma laev | metabolism | Energy produc | COG0055 | Produces ATP | 7739.JGI1257 | 2,6 | 0,8 | 0,000341578 | 9,89E-05 | 2 - | 0,000315815 - | 4 - | 0,000528058 - | 2 - | 0,000265793 - |
| 2436 | 2 | 3um_FIJJAFBF_135233;3um_FIJJAFBF_ | Eukaryota | Eustigmatoph | unclassified | Eustigmatales | Monodopsida | Nannochlorop | Nannochlorop | unclassified | unclassified | unclassified | unclassified | unclassified | 3,6 | 0,489897949 | 0,000565286 | 6,57E-05 | 3 - | 0,000564182 - | 4 - | 0,000628894 - | 4 - | 0,000633095 - |
| 2437 | 1 | 3um_FIJJAFBF_436970 | Bacteria | Proteobacteri | Gammaprotec | Cellvibrionale | Halieaceae | unclassified | marine gamm | unclassified | unclassified | unclassified | unclassified | unclassified | 0,2 | 0,4 | 2,69E-05 | 5,38E-05 | 1 - | 0,000134619 - | 0 - | 0 - | 0 - | 0 - |
| 2438 | 1 | 0.2um_NOKCPHJN_129689 | Bacteria | Proteobacteri | Betaproteoba | unclassified | unclassified | unclassified | Betaproteoba | information st | Replication, re | 0B7G8 | ATP-depender | 397945.Aave_ | 0,2 | 0,4 | 9,01E-06 | 1,80E-05 | 0 - | 0 - | 0 - | 0 - | 0 - | 0 - |
| 2439 | 2 | 0.2um_NOKCPHJN_123861;3um_FIJJA | Eukaryota | unclassified | Florideophyce | Ceramiales | Rhodomelace | Polysiphonia | Polysiphonia t | metabolism | Energy produc | 0FJ7X | This is one of t | various | 1,6 | 0,489897949 | 0,000284932 | 0,000103872 | 2 - | 0,000415823 - | 1 - | 0,000173819 - | 2 - | 0,000349961 - |
| 2440 | 1 | 0.2um_NOKCPHJN_151232 | Bacteria | Bacteroidetes | Flavobacteriia | Flavobacterial | Flavobacteria | Formosa | Formosa sp. H | unclassified | unclassified | unclassified | unclassified | unclassified | 0,8 | 0,4 | 5,03E-05 | 2,53E-05 | 0 - | 0 - | 1 - | 6,55E-05 - | 1 - | 6,60E-05 - |
| 2441 | 1 | 0.2um_NOKCPHJN_355365 | Eukaryota | unclassified | Oomycetes | Peronosporale | Peronosporac | Phytophthora | Phytophthora | unclassified | unclassified | unclassified | unclassified | unclassified | 1 | 0 | 5,93E-05 | 6,27E-06 | 1 - | 7,04E-05 - | 1 - | 5,89E-05 - | 1 - | 5,93E-05 - |
| 2442 | 3 | 0.2um_NOKCPHJN_566389;3um_FIJJA | Eukaryota | various | various | various | various | various | various | unclassified | unclassified | unclassified | unclassified | unclassified | 0,2 | 0,4 | 3,10E-05 | 6,21E-05 | 0 - | 0 - | 0 - | 0 - | 1 - | 0,000155155 - |
| 2443 | 1 | 0.2um_NOKCPHJN_829615 | Bacteria | Proteobacteri | Gammaprotec | Pseudomonad | Pseudomonac | Pseudomonas | Pseudomonas | information st | Translation, ri | 16QAF | thus facilitatin | 205922.Pfl01_ | 0,8 | 0,4 | 0,00012738 | 6,49E-05 | 1 - | 0,000183003 - | 1 - | 0,000152995 - | 1 - | 0,000154017 - |
| 2444 | 1 | 0.2um_NOKCPHJN_198494 | Bacteria | Bacteroidetes | Flavobacteriia | Flavobacterial | Flavobacteria | Algibacter | Algibacter sp. | cellular proces | Intracellular tr | 05FGH | MotA TolQ ex | 391603.FBALC | 2,6 | 0,489897949 | 0,000632974 | 8,67E-05 | 2 - | 0,000587045 - | 3 - | 0,000736176 - | 2 - | 0,000494062 - |
| 2445 | 2 | 0.2um_NOKCPHJN_1000666;3um_FIJJA | Bacteria | Bacteroidetes | Flavobacteriia | Flavobacterial | Flavobacteria | Polaribacter | Polaribacter s | poorly charact | Function unkn | 11P35 | NA | 867900.Celly_ | 0,4 | 0,489897949 | 6,59E-05 | 8,11E-05 | 0 - | 0 - | 1 - | 0,000176267 - | 0 - | 0 - |
| 2446 | 1 | 0.2um_NOKCPHJN_810494 | Eukaryota | Chlorophyta | Mamiellophyc | Mamiellales | Bathycoccace | Ostreococcus | Ostreococcus | information st | Translation, ri | KOG0052 | This protein p | 436017.A4S6E | 6 | 1,095445115 | 0,000788501 | 8,19E-05 | 5 - | 0,000792883 - | 6 - | 0,000795444 - | 5 - | 0,000667298 - |
| 2447 | 1 | 0.2um_NOKCPHJN_947787 | Eukaryota | unclassified | Oomycetes | Peronosporale | Peronosporac | Phytophthora | Phytophthora | unclassified | unclassified | unclassified | unclassified | unclassified | 1 | 0 | 0,000137238 | 1,45E-05 | 1 - | 0,000163068 - | 1 - | 0,000136329 - | 1 - | 0,00013724 - |
| 2448 | 1 | 3um_FIJJAFBF_115057 | Eukaryota | unclassified | unclassified | Phaeocystales | Phaeocystace | Phaeocystis | Phaeocystis gl | metabolism | Energy produc | 0FN1Q | F(1)F(0) ATP s | 65393.PCC742 | 0,6 | 0,489897949 | 0,000219034 | 0,000180455 | 1 - | 0,000409006 - | 1 - | 0,000341939 - | 1 - | 0,000344224 - |
| 2449 | 1 | 3um_FIJJAFBF_745490 | Eukaryota | Bacillariophyt | Coscinodiscop | Thalassiosirale | Thalassiosirac | Thalassiosira | Thalassiosira | cellular proces | Posttranslatio | KOG0731 | ATP-depender | 2850.Phatr17 | 2,8 | 0,748331477 | 0,001266928 | 0,000233309 | 2 - | 0,001100709 - | 3 - | 0,001380329 - | 2 - | 0,000926367 - |
| 2450 | 1 | 0.2um_NOKCPHJN_45933 | Bacteria | Bacteroidetes | Flavobacteriia | Flavobacterial | Flavobacteria | Cellulophaga | Cellulophaga t | poorly charact | Function unkn | 0XP53 | Ragb susd dor | 313598.MED1 | 3,6 | 0,489897949 | 0,000421838 | 6,13E-05 | 3 - | 0,000418926 - | 4 - | 0,000466977 - | 4 - | 0,000470097 - |
| 2451 | 1 | 0.2um_NOKCPHJN_366226 | Eukaryota | unclassified | Oomycetes | Peronosporale | Peronosporac | Phytophthora | Phytophthora | unclassified | unclassified | unclassified | unclassified | unclassified | 0,4 | 0,489897949 | 8,69E-06 | 1,06E-05 | 0 - | 0 - | 1 - | 2,22E-05 - | 0 - | 0 - |
| 2452 | 1 | 3um_FIJJAFBF_645627 | Eukaryota | unclassified | Pelagophycea | Pelagomonad | unclassified | Aureococcus | Aureococcus a | information st | Replication, re | COG0513 | purine NTP-de | 4787.PITG_19 | 0,4 | 0,489897949 | 0,000199208 | 0,000244466 | 0 - | 0 - | 0 - | 0 - | 0 - | 0 - |
| 2453 | 1 | 0.2um_NOKCPHJN_437854 | Bacteria | Proteobacteri | Gammaprotec | Cellvibrionale | unclassified | unclassified | Cellvibrionale | cellular proces | Defense mech | 179PG | N-acetyl-anhy | 396588.Tgr7_ | 0,2 | 0,4 | 7,76E-05 | 0,000155126 | 1 - | 0,000387814 - | 0 - | 0 - | 0 - | 0 - |
| 2454 | 1 | 0.2um_NOKCPHJN_648272 | Eukaryota | unclassified | Oomycetes | Saprolegniales | Saprolegniace | Achlya | Achlya hypog | metabolism | Inorganic ion t | COG0053 | cation diffusio | 4787.PITG_13 | 0,8 | 0,4 | 0,000108141 | 5,59E-05 | 1 - | 0,000160618 - | 1 - | 0,000134281 - | 0 - | 0 - |
| 2455 | 1 | 3um_FIJJAFBF_633503 | Viruses | unclassified | unclassified | unclassified | unclassified | unclassified | uncultured M | unclassified | unclassified | unclassified | unclassified | unclassified | 0,2 | 0,4 | 3,94E-05 | 7,87E-05 | 0 - | 0 - | 0 - | 0 - | 1 - | 0,000196853 - |
| 2456 | 1 | 0.2um_NOKCPHJN_723310 | unclassified | unclassified | unclassified | unclassified | unclassified | unclassified | unclassified | unclassified | unclassified | unclassified | unclassified | unclassified | 0,4 | 0,489897949 | 0,000344248 | 0,000423298 | 0 - | 0 - | 1 - | 0,000920219 - | 0 - | 0 - |
| 2457 | 3 | 0.2um_NOKCPHJN_1038489;0.2um_N | various | various | various | various | various | various | various | various | various | various | various | various | 0,4 | 0,489897949 | 3,49E-05 | 4,36E-05 | 1 - | 0,00010101 - | 0 - | 0 - | 0 - | 0 - |
| 2458 | 1 | 3um_FIJJAFBF_543375 | Eukaryota | Bacillariophyt | Bacillariophyc | Naviculales | Naviculaceae | Fistulifera | Fistulifera sol | information st | Replication, re | COG0513 | purine NTP-de | 2850.Phatr25 | 1,4 | 0,489897949 | 0,001017466 | 0,000309886 | 1 - | 0,000880567 - | 1 - | 0,000736176 - | 2 - | 0,001482187 - |
| 2459 | 1 | 3um_FIJJAFBF_252264 | Bacteria | Proteobacteri | Gammaprotec | Cellvibrionale | Cellvibrionace | Gilvimarinus | Gilvimarinus c | information st | Translation, ri | 177XJ | One of the pri | 377629.TERTU | 0,8 | 0,748331477 | 0,000315849 | 0,000275666 | 1 - | 0,000479796 - | 1 - | 0,000401121 - | 0 - | 0 - |
| 2460 | 1 | 0.2um_NOKCPHJN_966220 | Eukaryota | Eustigmatoph | unclassified | Eustigmatales | Monodopsida | Nannochlorop | Nannochlorop | unclassified | unclassified | unclassified | unclassified | unclassified | 0,2 | 0,4 | 7,00E-05 | 0,000139984 | 0 - | 0 - | 0 - | 0 - | 1 - | 0,000349961 - |
| 2461 | 1 | 0.2um_NOKCPHJN_756783 | Bacteria | Proteobacteri | Gammaprotec | unclassified | unclassified | unclassified | OM182 bacte | unclassified | unclassified | unclassified | unclassified | unclassified | 4,6 | 0,8 | 0,000324776 | 5,94E-05 | 5 - | 0,000420968 - | 4 - | 0,000281552 - | 4 - | 0,000283433 - |
| 2462 | 1 | 3um_FIJJAFBF_606585 | Eukaryota | Basidiomycota | Microbotryom | Microbotryale | Microbotryace | Microbotryum | Microbotryum | information st | Chromatin str | KOG3467 | Core compone | 3218.PP1S133 | 1 | 0 | 0,000477213 | 5,05E-05 | 1 - | 0,000567032 - | 1 - | 0,000474052 - | 1 - | 0,000477219 - |
| 2463 | 1 | 3um_FIJJAFBF_245504 | Bacteria | Bacteroidetes | Flavobacteriia | Flavobacterial | Flavobacteria | Lutibacter | Lutibacter pro | unclassified | unclassified | unclassified | unclassified | unclassified | 2,6 | 0,489897949 | 0,000757785 | 0,000103767 | 2 - | 0,0007028 - | 3 - | 0,000881337 - | 2 - | 0,000591483 - |
| 2464 | 1 | 0.2um_NOKCPHJN_681420 | Bacteria | Proteobacteri | Gammaprotec | Alteromonada | Colwelliaceae | Colwellia | Colwellia psyc | unclassified | unclassified | unclassified | unclassified | unclassified | 0,8 | 0,4 | 6,14E-05 | 3,17E-05 | 1 - | 9,12E-05 - | 1 - | 7,62E-05 - | 0 - | 0 - |
| 2465 | 1 | 3um_FIJJAFBF_725808 | Eukaryota | Phaeophyceae | unclassified | Ectocarpales | Ectocarpaceae | Ectocarpus | Ectocarpus sil | metabolism | Energy produc | COG0056 | Produces ATP | 35128.Thaps2 | 2,8 | 0,4 | 0,000593609 | 0,000126103 | 3 - | 0,000748482 - | 3 - | 0,000625749 - | 3 - | 0,000629929 - |
| 2466 | 1 | 3um_FIJJAFBF_730482 | unclassified | unclassified | unclassified | unclassified | unclassified | unclassified | unclassified | unclassified | unclassified | unclassified | unclassified | unclassified | 1 | 0 | 0,000403796 | 4,27E-05 | 1 - | 0,000479796 - | 1 - | 0,000401121 - | 1 - | 0,000403801 - |
| 2467 | 1 | 3um_FIJJAFBF_540416 | Bacteria | Proteobacteri | Deltaproteoba | unclassified | unclassified | unclassified | Deltaproteoba | metabolism | Lipid transpor | 0WKF6 | mevalonate ki | 192952.MM_ | 0,8 | 0,4 | 0,000180131 | 9,29E-05 | 1 - | 0,000264481 - | 1 - | 0,000221113 - | 1 - | 0,00022259 - |
| 2468 | 1 | 3um_FIJJAFBF_513507 | Bacteria | Actinobacteri | Actinobacteri | Streptomycet | Streptomycet | Streptomyces | Streptomyces | unclassified | unclassified | unclassified | unclassified | unclassified | 0,6 | 0,489897949 | 9,57E-05 | 7,85E-05 | 0 - | 0 - | 1 - | 0,000169121 - | 0 - | 0 - |
| 2469 | 1 | 0.2um_NOKCPHJN_129265 | Eukaryota | Chordata | Mammalia | Rodentia | Muridae | Mus | Mus musculus | poorly charact | Function unkn | 0T9G1 | kinetoplast-as | 353153.XP_80 | 0,8 | 0,4 | 6,61E-05 | 3,41E-05 | 1 - | 9,71E-05 - | 1 - | 8,12E-05 - | 1 - | 8,17E-05 - |
| 2470 | 1 | 0.2um_NOKCPHJN_619645 | Eukaryota | unclassified | unclassified | Dictyosteliales | Dictyosteliace | Dictyostelium | Dictyostelium | cellular proces | Signal transdu | COG0515 | Serine Threon | 44689.DDB02 | 0,2 | 0,4 | 2,26E-05 | 4,52E-05 | 1 - | 0,000113064 - | 0 - | 0 - | 0 - | 0 - |
| 2471 | 1 | 0.2um_NOKCPHJN_112415 | Bacteria | Bacteroidetes | Flavobacteriia | Flavobacterial | Flavobacteria | Marinirhabdu | Marinirhabdu | unclassified | unclassified | unclassified | unclassified | unclassified | 0,2 | 0,4 | 8,51E-05 | 0,000170217 | 0 - | 0 - | 0 - | 0 - | 0 - | 0 - |
| 2472 | 1 | 3um_FIJJAFBF_177694 | Eukaryota | Bacillariophyt | Bacillariophyc | Bacillariales | Bacillariaceae | Fragilariopsis | Fragilariopsis | unclassified | unclassified | unclassified | unclassified | unclassified | 1,4 | 0,489897949 | 0,000361089 | 0,000128299 | 1 - | 0,000306755 - | 2 - | 0,000512909 - | 2 - | 0,000516336 - |
| 2473 | 1 | 0.2um_NOKCPHJN_521157 | Bacteria | Proteobacteri | Alphaproteob | unclassified | unclassified | unclassified | Alphaproteob | unclassified | unclassified | unclassified | unclassified | unclassified | 0,4 | 0,489897949 | 2,23E-05 | 2,74E-05 | 0 - | 0 - | 0 - | 0 - | 0 - | 0 - |
| 2474 | 2 | 0.2um_NOKCPHJN_913579;3um_FIJJA | Eukaryota | Bacillariophyt | Coscinodiscop | Thalassiosirale | Thalassiosirac | Thalassiosira | Thalassiosira | cellular proces | Signal transdu | KOG0613 | fibronectin ty | 67593.Physo1 | 1 | 0 | 9,40E-05 | 9,94E-06 | 1 - | 0,000111714 - | 1 - | 9,34E-05 - | 1 - | 9,40E-05 - |
| 2475 | 2 | 0.2um_NOKCPHJN_07665;3um_FIJJAF | Bacteria | Bacteroidetes | Flavobacteriia | Flavobacterial | Flavobacteria | various | various | metabolism | Energy produc | 099YH | Pyruvate dehy | various | 0,4 | 0,489897949 | 7,05E-05 | 8,65E-05 | 0 - | 0 - | 0 - | 0 - | 0 - | 0 - |
| 2476 | 1 | 3um_FIJJAFBF_451308 | unclassified | unclassified | unclassified | unclassified | unclassified | unclassified | unclassified | unclassified | unclassified | unclassified | unclassified | unclassified | 1 | 0 | 0,000153639 | 1,62E-05 | 1 - | 0,000182557 - | 1 - | 0,000152622 - | 1 - | 0,000153641 - |
| 2477 | 1 | 0.2um_NOKCPHJN_823005 | Eukaryota | Chromerida | unclassified | unclassified | unclassified | Vitrella | Vitrella brassi | unclassified | unclassified | unclassified | unclassified | unclassified | 0,8 | 0,4 | 0,000327231 | 0,000169237 | 1 - | 0,000486027 - | 1 - | 0,000406331 - | 0 - | 0 - |
| 2478 | 1 | 0.2um_NOKCPHJN_191187 | Eukaryota | Platyhelminth | unclassified | Macrostomida | Macrostomida | Macrostomun | Macrostomun | poorly charact | Function unkn | 0NVRZ | NA | 170187.SP_17 | 0,8 | 0,4 | 2,26E-05 | 1,15E-05 | 1 - | 3,24E-05 - | 1 - | 2,71E-05 - | 1 - | 2,73E-05 - |
| 2479 | 1 | 3um_FIJJAFBF_815477 | Eukaryota | Bacillariophyt | Coscinodiscop | Thalassiosirale | Stephanodisca | Cyclotella | Cyclotella cry | metabolism | Lipid transpor | COG0439 | acetyl-CoA ca | 2850.Phatr54 | 5 | 0,632455532 | 0,002058842 | 6,73E-05 | 4 - | 0,001982733 - | 5 - | 0,002072017 - | 5 - | 0,002085859 - |
| 2480 | 1 | 3um_FIJJAFBF_313534 | Bacteria | Actinobacteri | Actinobacteri | Streptomycet | Streptomycet | Streptomyces | Streptomyces | unclassified | unclassified | unclassified | unclassified | unclassified | 0,4 | 0,489897949 | 0,000135987 | 0,000166628 | 0 - | 0 - | 0 - | 0 - | 1 - | 0,000348027 - |
| 2481 | 1 | 0.2um_NOKCPHJN_817880 | Eukaryota | unclassified | Pelagophycea | Pelagomonad | unclassified | Aureococcus | Aureococcus a | unclassified | unclassified | unclassified | unclassified | unclassified | 1 | 0 | 9,66E-05 | 1,02E-05 | 1 - | 0,000114798 - | 1 - | 9,60E-05 - | 1 - | 9,66E-05 - |
| 2482 | 1 | 0.2um_NOKCPHJN_538589 | Bacteria | Bacteroidetes | Flavobacteriia | Flavobacterial | Flavobacteria | Formosa | Formosa sp. H | unclassified | unclassified | unclassified | unclassified | unclassified | 1,4 | 0,489897949 | 9,28E-05 | 2,60E-05 | 1 - | 8,08E-05 - | 1 - | 6,76E-05 - | 1 - | 6,80E-05 - |
| 2483 | 1 | 0.2um_NOKCPHJN_523103 | Eukaryota | Phaeophyceae | unclassified | Ectocarpales | Ectocarpaceae | Ectocarpus | Ectocarpus sil | cellular proces | Intracellular tr | KOG2514 | Anoctamin | 4787.PITG_09 | 2 | 0 | 0,000510058 | 5,39E-05 | 2 - | 0,000606058 - | 2 - | 0,00050668 - | 2 - | 0,000510064 - |
| 2484 | 1 | 0.2um_NOKCPHJN_106480 | Bacteria | Bacteroidetes | Flavobacteriia | Flavobacterial | unclassified | unclassified | Flavobacterial | metabolism | Inorganic ion t | 05C27 | TonB-depend | 216432.CA25 | 0,2 | 0,4 | 1,06E-05 | 2,12E-05 | 0 - | 0 - | 0 - | 0 - | 0 - | 0 - |
| 2485 | 2 | 0.2um_NOKCPHJN_1001159;3um_FIJJA | Eukaryota | various | various | various | various | various | various | unclassified | unclassified | unclassified | unclassified | unclassified | 1 | 0 | 0,000437446 | 4,63E-05 | 1 - | 0,000519779 - | 1 - | 0,000434548 - | 1 - | 0,000437451 - |
| 2486 | 1 | 0.2um_NOKCPHJN_483485 | Bacteria | Bacteroidetes | Flavobacteriia | Flavobacterial | Flavobacteria | Arenitalea | Arenitalea lut | metabolism | Inorganic ion t | 05C27 | TonB-depend | 886377.Murru | 0,4 | 0,489897949 | 2,58E-05 | 3,16E-05 | 0 - | 0 - | 0 - | 0 - | 1 - | 6,61E-05 - |
| 2487 | 1 | 0.2um_NOKCPHJN_349772 | Bacteria | unclassified | unclassified | unclassified | unclassified | unclassified | uncultured ba | poorly charact | Function unkn | 08PIQ | CHU large pro | 50743.SCB49_ | 1,2 | 0,979795897 | 5,54E-05 | 4,03E-05 | 1 - | 5,74E-05 - | 1 - | 4,80E-05 - | 0 - | 0 - |
| 2488 | 2 | 0.2um_NOKCPHJN_228707;3um_FIJJA | Bacteria | Bacteroidetes | Flavobacteriia | Flavobacterial | Crocinitomica | unclassified | Crocinitomica | unclassified | unclassified | unclassified | unclassified | unclassified | 0,4 | 0,489897949 | 0,000169011 | 0,000207901 | 0 - | 0 - | 0 - | 0 - | 1 - | 0,000453187 - |
| 2489 | 1 | 0.2um_NOKCPHJN_63870 | Bacteria | Bacteroidetes | Flavobacteriia | Flavobacterial | Flavobacteria | unclassified | various | unclassified | unclassified | unclassified | unclassified | unclassified | 1 | 0 | 0,000185818 | 1,97E-05 | 1 - | 0,000220791 - | 1 - | 0,000184587 - | 1 - | 0,00018582 - |
| 2490 | 1 | 0.2um_NOKCPHJN_371014 | Bacteria | Proteobacteri | Betaproteoba | unclassified | unclassified | unclassified | Betaproteoba | metabolism | Inorganic ion t | 07QMV | Part of the AB | 261292.Nit79 | 1 | 0 | 0,000797369 | 8,43E-05 | 1 - | 0,000947445 - | 1 - | 0,000792088 - | 1 - | 0,000797379 - |
| 2491 | 2 | 0.2um_NOKCPHJN_24000;3um_FIJJAF | various | various | various | various | various | various | various | various | various | various | various | various | 5,2 | 0,4 | 0,000972701 | 7,64E-05 | 5 - | 0,001117137 - | 5 - | 0,000933954 - | 5 - | 0,000940193 - |
| 2492 | 1 | 3um_FIJJAFBF_56601 | Bacteria | unclassified | unclassified | unclassified | unclassified | unclassified | magneto-ovoi | unclassified | unclassified | unclassified | unclassified | unclassified | 0,8 | 0,4 | 0,000243851 | 0,000126097 | 1 - | 0,000361585 - | 0 - | 0 - | 1 - | 0,000304314 - |
| 2493 | 1 | 0.2um_NOKCPHJN_420122 | Eukaryota | Apicomplexa | Conoidasida | Eucoccidiorida | Cryptosporidii | Cryptosporidiu | Cryptosporidiu | unclassified | unclassified | unclassified | unclassified | unclassified | 0,2 | 0,4 | 3,98E-05 | 7,96E-05 | 1 - | 0,000199064 - | 0 - | 0 - | 0 - | 0 - |
| 2494 | 1 | 0.2um_NOKCPHJN_66855 | Bacteria | Proteobacteri | Gammaprotec | Arenicellales | Arenicellaceae | Arenicella | Arenicella xan | unclassified | unclassified | unclassified | unclassified | unclassified | 0,6 | 0,489897949 | 0,000147948 | 0,000121337 | 0 - | 0 - | 0 - | 0 - | 1 - | 0,000262471 - |
| 2495 | 1 | 3um_FIJJAFBF_174375 | Viruses | unclassified | unclassified | unclassified | Phycodnavirid | unclassified | Chrysochromu | unclassified | unclassified | unclassified | unclassified | unclassified | 1 | 0 | 0,000107495 | 1,14E-05 | 1 - | 0,000127727 - | 1 - | 0,000106783 - | 1 - | 0,000107496 - |
| 2496 | 1 | 0.2um_NOKCPHJN_07024 | Bacteria | Bacteroidetes | Cytophagia | Cytophagales | Flammeovirga | unclassified | Flammeovirga | unclassified | unclassified | unclassified | unclassified | unclassified | 1 | 0,632455532 | 0,000178083 | 0,000113076 | 1 - | 0,000209659 - | 1 - | 0,00017528 - | 2 - | 0,000352902 - |
| 2497 | 1 | 0.2um_NOKCPHJN_286046 | Eukaryota | unclassified | Pelagophycea | Pelagomonad | unclassified | Aureococcus | Aureococcus a | cellular proces | Cytoskeleton | 1514Z | Tubulin is the | 272952.HpaP | 1,2 | 0,4 | 0,000959559 | 0,00023713 | 1 - | 0,000972054 - | 1 - | 0,000812661 - | 1 - | 0,00081809 - |
| 2498 | 1 | 3um_FIJJAFBF_226993 | Bacteria | unclassified | unclassified | unclassified | unclassified | unclassified | uncultured ba | metabolism | Inorganic ion t | 09C0C | TonB depende | 742767.HMPR | 1,2 | 0,4 | 0,000159581 | 3,94E-05 | 1 - | 0,000161659 - | 1 - | 0,000135151 - | 1 - | 0,000136054 - |
| 2499 | 1 | 3um_FIJJAFBF_807524 | Bacteria | Proteobacteri | Alphaproteob | Rhodobactera | Rhodobactera | Planktomarina | Planktomarina | cellular proces | Cell wall/mem | 16PAV | penicillin-bind | 644107.SL115 | 1 | 0 | 8,86E-05 | 9,37E-06 | 1 - | 0,000105272 - | 1 - | 8,80E-05 - | 1 - | 8,86E-05 - |
| 2500 | 1 | 0.2um_NOKCPHJN_45922 | Bacteria | Bacteroidetes | Flavobacteriia | Flavobacterial | Flavobacteria | Arenitalea | Arenitalea lut | unclassified | unclassified | unclassified | unclassified | unclassified | 0,6 | 0,489897949 | 0,000110878 | 9,16E-05 | 1 - | 0,000209659 - | 0 - | 0 - | 1 - | 0,000176451 - |
| 2501 | 1 | 3um_FIJJAFBF_07267 | Bacteria | Planctomycet | Planctomyceti | Planctomycet | Planctomycet | Rhodopirellul | Rhodopirellul | unclassified | unclassified | unclassified | unclassified | unclassified | 1 | 0 | 0,000193228 | 2,04E-05 | 1 - | 0,000229596 - | 1 - | 0,000191948 - | 1 - | 0,00019323 - |
| 2502 | 1 | 0.2um_NOKCPHJN_257951 | Eukaryota | Chordata | Mammalia | Proboscidea | Elephantidae | Loxodonta | Loxodonta afr | unclassified | unclassified | unclassified | unclassified | unclassified | 0,2 | 0,4 | 1,73E-05 | 3,46E-05 | 0 - | 0 - | 0 - | 0 - | 0 - | 0 - |
| 2503 | 2 | 0.2um_NOKCPHJN_842037;3um_FIJJA | Eukaryota | Chordata | Actinopteri | Carangiformes | Carangidae | Seriola | Seriola laland | unclassified | unclassified | unclassified | unclassified | unclassified | 1,6 | 0,8 | 0,000979162 | 0,00044491 | 1 - | 0,000748482 - | 1 - | 0,000625749 - | 1 - | 0,000629929 - |
| 2504 | 1 | 0.2um_NOKCPHJN_517197 | Archaea | Euryarchaeota | Methanomicr | Methanosarci | Candidatus M | Candidatus M | Candidatus M | poorly charact | Function unkn | COG0714 | ATPase activit | 439481.Aboo_ | 0,2 | 0,4 | 1,17E-05 | 2,34E-05 | 0 - | 0 - | 0 - | 0 - | 1 - | 5,85E-05 - |
| 2505 | 1 | 0.2um_NOKCPHJN_947667 | unclassified | unclassified | unclassified | unclassified | unclassified | unclassified | unclassified | unclassified | unclassified | unclassified | unclassified | unclassified | 1 | 0 | 0,000508001 | 5,37E-05 | 1 - | 0,000603614 - | 1 - | 0,000504636 - | 1 - | 0,000508008 - |
| 2506 | 1 | 0.2um_NOKCPHJN_748716 | Eukaryota | Basidiomycota | Agaricomycet | Sebacinales | Serendipitace | Serendipita | Serendipita v | information st | Replication, re | COG0513 | purine NTP-de | 272952.HpaP | 1 | 0 | 0,000594266 | 6,28E-05 | 1 - | 0,000706115 - | 1 - | 0,000590329 - | 1 - | 0,000594273 - |
| 2507 | 1 | 3um_FIJJAFBF_642666 | Eukaryota | Bacillariophyt | Bacillariophyc | Bacillariales | Bacillariaceae | Fragilariopsis | Fragilariopsis | unclassified | unclassified | unclassified | unclassified | unclassified | 1,2 | 0,748331477 | 0,000778044 | 0,000507224 | 0 - | 0 - | 2 - | 0,001331381 - | 2 - | 0,001340275 - |
| 2508 | 1 | 0.2um_NOKCPHJN_09296 | Eukaryota | Chlorophyta | Chlorophycea | Chlamydomor | Volvocaceae | Volvox | Volvox carteri | information st | Replication, re | 0XNUT | helicase activi | 3055.EDP069 | 0,2 | 0,4 | 9,31E-06 | 1,86E-05 | 0 - | 0 - | 0 - | 0 - | 0 - | 0 - |
| 2509 | 1 | 0.2um_NOKCPHJN_476838 | Eukaryota | unclassified | Oomycetes | Saprolegniales | Saprolegniace | Aphanomyces | Aphanomyces | unclassified | unclassified | unclassified | unclassified | unclassified | 1 | 0 | 6,74E-05 | 7,13E-06 | 1 - | 8,01E-05 - | 1 - | 6,70E-05 - | 1 - | 6,74E-05 - |
| 2510 | 1 | 0.2um_NOKCPHJN_112963 | Bacteria | Proteobacteri | Alphaproteob | Rhodobactera | Rhodobactera | Thalassobius | Thalassobius s | cellular proces | Posttranslatio | 01QWG | Prevents misf | 391619.RGBS | 1 | 0 | 0,000114323 | 1,21E-05 | 1 - | 0,000135841 - | 1 - | 0,000113566 - | 1 - | 0,000114325 - |
| 2511 | 1 | 0.2um_NOKCPHJN_84326 | Bacteria | Bacteroidetes | Flavobacteriia | Flavobacterial | Flavobacteria | Algibacter | Algibacter sp. | poorly charact | Function unkn | 11NCV | Protein of unk | 1046627.BZA | 1,8 | 0,979795897 | 0,000675376 | 0,000369746 | 0 - | 0 - | 2 - | 0,000782186 - | 2 - | 0,000787412 - |
| 2512 | 1 | 3um_FIJJAFBF_463195 | Eukaryota | unclassified | unclassified | Kinetoplastida | Trypanosomat | Leishmania | Leishmania m | poorly charact | Function unkn | 0J56E | membrane as | 5664.LmjF.29. | 0,4 | 0,489897949 | 2,79E-05 | 3,42E-05 | 0 - | 0 - | 0 - | 0 - | 0 - | 0 - |
| 2513 | 1 | 3um_FIJJAFBF_254310 | Bacteria | Proteobacteri | Gammaprotec | Cellvibrionale | Spongiibacter | Oceanicoccus | Oceanicoccus | information st | Transcription | 16PMT | DNA-depende | 247633.GP21 | 1 | 0,632455532 | 0,000182045 | 0,000120041 | 0 - | 0 - | 1 - | 0,000187913 - | 2 - | 0,000378336 - |
| 2514 | 1 | 0.2um_NOKCPHJN_507733 | Eukaryota | Phaeophyceae | unclassified | Ectocarpales | Ectocarpaceae | Ectocarpus | Ectocarpus sil | poorly charact | Function unkn | 102D8 | PAP_fibrillin | 2850.Phatr46 | 3,8 | 0,4 | 0,001156093 | 6,66E-05 | 3 - | 0,001095339 - | 4 - | 0,001220974 - | 4 - | 0,001229131 - |
| 2515 | 1 | 3um_FIJJAFBF_155266 | Bacteria | Proteobacteri | Gammaprotec | Alteromonada | Pseudoalteror | Pseudoalteror | various | poorly charact | Function unkn | 129JZ | NA | 156578.ATW7 | 0,6 | 0,489897949 | 0,000173208 | 0,000142053 | 0 - | 0 - | 0 - | 0 - | 1 - | 0,000307283 - |
| 2516 | 1 | 3um_FIJJAFBF_316515 | Bacteria | Proteobacteri | Gammaprotec | Alteromonada | Alteromonada | Paraglaciecola | Paraglaciecola | unclassified | unclassified | unclassified | unclassified | unclassified | 0,6 | 0,489897949 | 4,36E-05 | 3,58E-05 | 0 - | 0 - | 1 - | 7,71E-05 - | 0 - | 0 - |
| 2517 | 1 | 0.2um_NOKCPHJN_54434 | Bacteria | Proteobacteri | Gammaprotec | Cellvibrionale | Spongiibacter | Oceanicoccus | Oceanicoccus | cellular proces | Cell wall/mem | COG2885 | Ompa motb d | 247633.GP21 | 0,8 | 0,748331477 | 0,000144703 | 0,000145797 | 2 - | 0,000397073 - | 0 - | 0 - | 1 - | 0,00016709 - |
| 2518 | 1 | 3um_FIJJAFBF_372258 | Eukaryota | unclassified | Pelagophycea | Pelagomonad | unclassified | Aureococcus | Aureococcus a | cellular proces | Posttranslatio | 0FJCG | Prevents misf | 313624.N941 | 0,6 | 0,489897949 | 6,62E-05 | 5,43E-05 | 0 - | 0 - | 1 - | 0,000116962 - | 0 - | 0 - |
| 2519 | 1 | 0.2um_NOKCPHJN_262361 | Bacteria | Proteobacteri | Gammaprotec | unclassified | unclassified | unclassified | Gammaprotec | unclassified | unclassified | unclassified | unclassified | unclassified | 0,2 | 0,4 | 1,67E-05 | 3,34E-05 | 0 - | 0 - | 0 - | 0 - | 1 - | 8,35E-05 - |
| 2520 | 1 | 0.2um_NOKCPHJN_310213 | unclassified | unclassified | unclassified | unclassified | unclassified | unclassified | unclassified | unclassified | unclassified | unclassified | unclassified | unclassified | 0,6 | 0,489897949 | 0,000404864 | 0,000337519 | 1 - | 0,000787876 - | 0 - | 0 - | 1 - | 0,000663084 - |

| | | | | | | | | | | | | | | | | | | | | | | | | |
|---|---|---|---|---|---|---|---|---|---|---|---|---|---|---|---|---|---|---|---|---|---|---|---|---|
| 2521 | 1 | 0.2um_NOKCPHJN_419974 | Bacteria | Proteobacteri | Gammaprotec | Oceanospirilla | unclassified | unclassified | Oceanospirilla | cellular proces | Posttranslatio | 0IEKR | One of the cor | 35128.Thapsd | 0,4 | 0,489897949 | 5,34E-05 | 6,54E-05 | 0 - | 0 - | 0 - | 0 - | 1 - | 0,000136644 - |
| 2522 | 1 | 3um_FIJJAFBF_54806 | Bacteria | Bacteroidetes | Flavobacteriia | Flavobacterial | Flavobacteriac | Polaribacter | Polaribacter b | cellular proces | Posttranslatio | 09ADN | alkyl hydroper | 391587.KAOT | 0,6 | 0,489897949 | 0,000166702 | 0,000136718 | 0 - | 0 - | 0 - | 0 - | 1 - | 0,000295741 - |
| 2523 | 1 | 3um_FIJJAFBF_715077 | Bacteria | Cyanobacteria | unclassified | Synechococca | Leptolyngbyac | Leptolyngbya | Leptolyngbya | unclassified | unclassified | unclassified | unclassified | unclassified | 0,8 | 0,4 | 0,00033349 | 0,000168075 | 0 - | 0 - | 1 - | 0,000434548 - | 1 - | 0,000437451 - |
| 2524 | 1 | 3um_FIJJAFBF_571878 | Bacteria | Bacteroidetes | Flavobacteriia | Flavobacterial | Flavobacteriac | Flavobacteriu | Flavobacteriu | poorly charact | Function unkn | 09IJP | Heavy-metal-a | 398720.MED2 | 0,8 | 0,4 | 0,000717987 | 0,000370483 | 1 - | 0,0010542 - | 1 - | 0,000881337 - | 1 - | 0,000887224 - |
| 2525 | 1 | 0.2um_NOKCPHJN_287456 | Bacteria | Proteobacteri | Gammaprotec | Alteromonada | Colwelliaceae | Colwellia | Colwellia sp. 3 | unclassified | unclassified | unclassified | unclassified | unclassified | 3,8 | 0,748331477 | 0,000274902 | 4,36E-05 | 3 - | 0,000261098 - | 4 - | 0,000291046 - | 3 - | 0,000219743 - |
| 2526 | 1 | 0.2um_NOKCPHJN_364766 | Bacteria | Proteobacteri | Betaproteoba | Nitrosomonad | unclassified | unclassified | Methylophilal | cellular proces | Posttranslatio | 16TCX | Peptidase, M1 | 383631.MB21 | 0,2 | 0,4 | 2,92E-05 | 5,85E-05 | 0 - | 0 - | 0 - | 0 - | 1 - | 0,000146155 - |
| 2527 | 1 | 3um_FIJJAFBF_674421 | Viruses | unclassified | unclassified | unclassified | unclassified | unclassified | uncultured M | unclassified | unclassified | unclassified | unclassified | unclassified | 1 | 0,894427191 | 0,000304371 | 0,000261074 | 1 - | 0,00036871 - | 2 - | 0,000616502 - | 0 - | 0 - |
| 2528 | 2 | 0.2um_NOKCPHJN_24089;3um_FIJJAFE | Viruses | unclassified | unclassified | unclassified | Phycodnavirid | unclassified | Chrysochromu | unclassified | unclassified | unclassified | unclassified | unclassified | 0,8 | 0,4 | 0,000240458 | 0,000124077 | 1 - | 0,000353057 - | 1 - | 0,000295165 - | 1 - | 0,000297137 - |
| 2529 | 1 | 3um_FIJJAFBF_305842 | Eukaryota | Chlorophyta | unclassified | Pyramimonad | unclassified | Pyramimonas | Pyramimonas | metabolism | Energy produc | 0DCW7 | Produces ATP | 3055.DAA009 | 4 | 0 | 0,000523843 | 5,54E-05 | 4 - | 0,000622438 - | 4 - | 0,000520374 - | 4 - | 0,00052385 - |
| 2530 | 1 | 3um_FIJJAFBF_81227 | Bacteria | Proteobacteri | Gammaprotec | Cellvibrionales | Cellvibrionace | Marinagarivor | Marinagarivor | information st | Translation, ri | 177XJ | One of the pri | 377629.TERTU | 0,8 | 0,4 | 0,000307837 | 0,000155146 | 0 - | 0 - | 1 - | 0,000401121 - | 1 - | 0,000403801 - |
| 2531 | 1 | 0.2um_NOKCPHJN_883238 | Bacteria | Firmicutes | Bacilli | Bacillales | Bacillaceae | Bacillus | Bacillus sp. CC | metabolism | Amino acid tra | 0XSVU | dehydrogenas | 358681.BBR47 | 0,2 | 0,4 | 2,71E-05 | 5,43E-05 | 0 - | 0 - | 1 - | 0,000135737 - | 0 - | 0 - |
| 2532 | 1 | 3um_FIJJAFBF_414471 | Eukaryota | Bacillariophyta | Coscinodiscop | Thalassiosirale | Thalassiosirac | Thalassiosira | Thalassiosira d | metabolism | Energy produc | COG0654 | oxidoreductas | 35128.Thaps2 | 0,2 | 0,4 | 4,39E-05 | 8,77E-05 | 0 - | 0 - | 0 - | 0 - | 0 - | 0 - |
| 2533 | 1 | 3um_FIJJAFBF_355093 | Eukaryota | unclassified | Cryptophyta | Pyrenomonad | Geminigerace | Guillardia | Guillardia thet | unclassified | unclassified | unclassified | unclassified | unclassified | 1,8 | 0,748331477 | 0,002044767 | 0,000872219 | 2 - | 0,002673149 - | 2 - | 0,002234819 - | 1 - | 0,001124874 - |
| 2534 | 1 | 0.2um_NOKCPHJN_911240 | unclassified | unclassified | unclassified | unclassified | unclassified | unclassified | unclassified | unclassified | unclassified | unclassified | unclassified | unclassified | 0,4 | 0,489897949 | 0,000242872 | 0,000297559 | 0 - | 0 - | 1 - | 0,000619554 - | 0 - | 0 - |
| 2535 | 1 | 3um_FIJJAFBF_305344 | Eukaryota | unclassified | Dinophyceae | Gonyaulacales | unclassified | Lingulodinium | Lingulodinium | poorly charact | Function unkn | 0XRVT | NA | 35128.Thaps3 | 1 | 0 | 0,000173532 | 1,84E-05 | 1 - | 0,000206193 - | 1 - | 0,000172383 - | 1 - | 0,000173534 - |
| 2536 | 2 | 0.2um_NOKCPHJN_406222;3um_FIJJAF | Eukaryota | various | various | various | various | various | various | unclassified | unclassified | unclassified | unclassified | unclassified | 0,2 | 0,4 | 2,10E-05 | 4,19E-05 | 0 - | 0 - | 0 - | 0 - | 0 - | 0 - |
| 2537 | 1 | 0.2um_NOKCPHJN_531699 | Bacteria | Proteobacteri | Gammaprotec | Cellvibrionales | Halieaceae | unclassified | Halieaceae ba | unclassified | unclassified | unclassified | unclassified | unclassified | 0,2 | 0,4 | 1,09E-05 | 2,19E-05 | 0 - | 0 - | 1 - | 5,47E-05 - | 0 - | 0 - |
| 2538 | 1 | 0.2um_NOKCPHJN_225231 | Eukaryota | unclassified | Oomycetes | Albuginales | Albuginaceae | Albugo | Albugo candid | cellular proces | Posttranslatio | KOG0734 | YME1-like 1 (S | 65071.PYU1_T | 1 | 0 | 9,75E-05 | 1,03E-05 | 1 - | 0,000115864 - | 1 - | 9,69E-05 - | 1 - | 9,75E-05 - |
| 2539 | 1 | 0.2um_NOKCPHJN_467912 | Eukaryota | unclassified | Oomycetes | Saprolegniales | Saprolegniace | various | various | unclassified | unclassified | unclassified | unclassified | unclassified | 1,2 | 0,748331477 | 0,000935063 | 0,000637036 | 2 - | 0,0017821 - | 1 - | 0,00074494 - | 2 - | 0,001499832 - |
| 2540 | 1 | 0.2um_NOKCPHJN_446826 | Bacteria | Bacteroidetes | Flavobacteriia | Flavobacterial | Flavobacteriac | unclassified | Flavobacteriac | unclassified | unclassified | unclassified | unclassified | unclassified | 0,4 | 0,489897949 | 0,000532767 | 0,000653804 | 0 - | 0 - | 0 - | 0 - | 0 - | 0 - |
| 2541 | 1 | 0.2um_NOKCPHJN_124651 | Bacteria | Proteobacteri | Gammaprotec | Cellvibrionales | Porticoccacea | unclassified | gamma protec | unclassified | unclassified | unclassified | unclassified | unclassified | 0,2 | 0,4 | 2,59E-05 | 5,17E-05 | 0 - | 0 - | 0 - | 0 - | 1 - | 0,000129349 - |
| 2542 | 1 | 3um_FIJJAFBF_194067 | Bacteria | Bacteroidetes | Flavobacteriia | Flavobacterial | Flavobacteriac | Polaribacter | Polaribacter h | information st | Translation, ri | COG1185 | Involved in mF | 688270.Celal_ | 0,2 | 0,4 | 1,70E-05 | 3,39E-05 | 0 - | 0 - | 0 - | 0 - | 1 - | 8,48E-05 - |
| 2543 | 1 | 0.2um_NOKCPHJN_326468 | Eukaryota | Phaeophyceae | unclassified | various | various | various | various | unclassified | unclassified | unclassified | unclassified | unclassified | 0,6 | 0,489897949 | 0,000237745 | 0,000194916 | 0 - | 0 - | 1 - | 0,000419966 - | 0 - | 0 - |
| 2544 | 1 | 3um_FIJJAFBF_344568 | Bacteria | Cyanobacteria | unclassified | Synechococca | Synechococca | Synechococcu | Synechococcu | unclassified | unclassified | unclassified | unclassified | unclassified | 0,2 | 0,4 | 3,30E-05 | 6,60E-05 | 0 - | 0 - | 0 - | 0 - | 0 - | 0 - |
| 2545 | 1 | 3um_FIJJAFBF_669718 | Eukaryota | Bacillariophyta | Coscinodiscop | Thalassiosirale | Thalassiosirac | Thalassiosira | Thalassiosira c | unclassified | unclassified | unclassified | unclassified | unclassified | 2,2 | 0,4 | 0,00068783 | 8,42E-05 | 2 - | 0,000752243 - | 2 - | 0,000628894 - | 2 - | 0,000633095 - |
| 2546 | 1 | 0.2um_NOKCPHJN_108160 | Bacteria | Proteobacteri | Gammaprotec | Alteromonada | Alteromonada | Alishewanella | Alishewanella | cellular proces | Cell wall/mem | 0QITS | Nad-depender | 553480.PAU_ | 0,8 | 0,4 | 0,000160489 | 8,30E-05 | 1 - | 0,00023837 - | 1 - | 0,000199283 - | 0 - | 0 - |
| 2547 | 1 | 3um_FIJJAFBF_669270 | unclassified | unclassified | unclassified | unclassified | unclassified | unclassified | unclassified | unclassified | unclassified | unclassified | unclassified | unclassified | 0,2 | 0,4 | 4,01E-05 | 8,02E-05 | 0 - | 0 - | 1 - | 0,000200561 - | 0 - | 0 - |
| 2548 | 1 | 0.2um_NOKCPHJN_23410 | Eukaryota | unclassified | Pelagophycea | Pelagomonada | unclassified | various | various | unclassified | unclassified | unclassified | unclassified | unclassified | 1,2 | 0,4 | 0,000728615 | 0,000336409 | 2 - | 0,001399031 - | 1 - | 0,000584812 - | 1 - | 0,000588719 - |
| 2549 | 1 | 0.2um_NOKCPHJN_264279 | Eukaryota | Phaeophyceae | unclassified | Ectocarpales | Ectocarpaceae | Ectocarpus | Ectocarpus sili | metabolism | Coenzyme tra | KOG2794 | delta-aminole | 2850.Phatr41 | 1 | 0 | 0,000247028 | 2,61E-05 | 1 - | 0,000293522 - | 1 - | 0,000245392 - | 1 - | 0,000247031 - |
| 2550 | 1 | 0.2um_NOKCPHJN_744345 | Eukaryota | Streptophyta | unclassified | Vitales | Vitaceae | Vitis | Vitis vinifera | unclassified | unclassified | unclassified | unclassified | unclassified | 0,4 | 0,489897949 | 4,16E-05 | 5,10E-05 | 0 - | 0 - | 0 - | 0 - | 0 - | 0 - |
| 2551 | 1 | 0.2um_NOKCPHJN_560573 | Bacteria | Proteobacteri | Alphaproteob | Rhodobactera | Rhodobactera | unclassified | Rhodobactera | unclassified | unclassified | unclassified | unclassified | unclassified | 1 | 0 | 0,000264673 | 2,80E-05 | 1 - | 0,000314488 - | 1 - | 0,00026292 - | 1 - | 0,000264676 - |
| 2552 | 1 | 3um_FIJJAFBF_388958 | Bacteria | Bacteroidetes | Flavobacteriia | Flavobacterial | Flavobacteriac | Polaribacter | Polaribacter s | cellular proces | Cell wall/mem | COG0750 | Membrane-as | 313594.PI23P | 0,2 | 0,4 | 2,79E-05 | 5,59E-05 | 0 - | 0 - | 1 - | 0,000139676 - | 0 - | 0 - |
| 2553 | 1 | 0.2um_NOKCPHJN_504510 | Bacteria | Bacteroidetes | unclassified | unclassified | unclassified | unclassified | Bacteroidetes | cellular proces | Cell wall/mem | 05Q51 | Glycosyl trans | 398720.MED2 | 1 | 0 | 0,000147523 | 1,56E-05 | 1 - | 0,000175288 - | 1 - | 0,000146545 - | 1 - | 0,000147524 - |
| 2554 | 1 | 0.2um_NOKCPHJN_89225 | Eukaryota | unclassified | Oomycetes | Saprolegniales | Saprolegniace | Saprolegnia | Saprolegnia di | unclassified | unclassified | unclassified | unclassified | unclassified | 0,8 | 0,4 | 3,02E-05 | 1,52E-05 | 0 - | 0 - | 1 - | 3,94E-05 - | 1 - | 3,96E-05 - |
| 2555 | 1 | 3um_FIJJAFBF_577920 | unclassified | unclassified | unclassified | unclassified | unclassified | unclassified | unclassified | unclassified | unclassified | unclassified | unclassified | unclassified | 0,2 | 0,4 | 2,04E-05 | 4,07E-05 | 0 - | 0 - | 0 - | 0 - | 0 - | 0 - |
| 2556 | 1 | 3um_FIJJAFBF_521113 | Bacteria | Proteobacteri | Gammaprotec | unclassified | unclassified | unclassified | various | unclassified | unclassified | unclassified | unclassified | unclassified | 0,8 | 0,4 | 0,000193399 | 0,000100008 | 1 - | 0,000286775 - | 0 - | 0 - | 1 - | 0,000241352 - |
| 2557 | 1 | 0.2um_NOKCPHJN_798900 | Bacteria | Proteobacteri | Alphaproteob | Pelagibacteral | Pelagibacterac | Candidatus Pe | Candidatus Pe | cellular proces | Posttranslatio | 0251V | PPIC-type PPI | 335992.SAR11 | 0,8 | 0,4 | 0,000171942 | 8,76E-05 | 1 - | 0,000247024 - | 1 - | 0,000206518 - | 1 - | 0,000207897 - |
| 2558 | 1 | 0.2um_NOKCPHJN_916986 | Eukaryota | Annelida | Polychaeta | Capitellida | Capitellidae | Capitella | Capitella telet | unclassified | unclassified | unclassified | unclassified | unclassified | 0,2 | 0,4 | 2,38E-05 | 4,76E-05 | 0 - | 0 - | 0 - | 0 - | 0 - | 0 - |
| 2559 | 2 | 0.2um_NOKCPHJN_633339;3um_FIJJAF | Eukaryota | unclassified | unclassified | Prymnesiales | Chrysochromu | Chrysochromu | Chrysochromu | poorly charact | Function unkn | 11UER | mitochondrial | 2850.Phatr14 | 0,6 | 0,489897949 | 0,000113124 | 9,43E-05 | 1 - | 0,000220142 - | 0 - | 0 - | 1 - | 0,000185273 - |
| 2560 | 1 | 0.2um_NOKCPHJN_933150 | Eukaryota | unclassified | Pelagophycea | Pelagomonada | unclassified | Aureococcus | Aureococcus a | unclassified | unclassified | unclassified | unclassified | unclassified | 1,4 | 0,489897949 | 0,000307889 | 8,62E-05 | 1 - | 0,000268273 - | 1 - | 0,000224283 - | 1 - | 0,000225781 - |
| 2561 | 2 | 0.2um_NOKCPHJN_27393;3um_FIJJAFE | Bacteria | unclassified | unclassified | unclassified | unclassified | unclassified | uncultured ba | unclassified | unclassified | unclassified | unclassified | unclassified | 3,2 | 0,4 | 0,000394875 | 5,44E-05 | 3 - | 0,000441148 - | 3 - | 0,000368811 - | 3 - | 0,000371275 - |
| 2562 | 1 | 3um_FIJJAFBF_115137 | Eukaryota | Streptophyta | Liliopsida | Poales | Poaceae | Hordeum | Hordeum vulg | unclassified | unclassified | unclassified | unclassified | unclassified | 2,8 | 0,4 | 0,003393491 | 0,000625288 | 3 - | 0,004318164 - | 2 - | 0,002406728 - | 3 - | 0,003634208 - |
| 2563 | 1 | 0.2um_NOKCPHJN_466965 | Bacteria | various | various | various | various | various | various | information st | Translation, ri | 17FIT | 50s ribosomal | 413404.Rmag | 0,2 | 0,4 | 0,000160449 | 0,000320897 | 0 - | 0 - | 1 - | 0,000802243 - | 0 - | 0 - |
| 2564 | 1 | 3um_FIJJAFBF_256129 | Bacteria | Planctomycete | Phycisphaerae | Phycisphaeral | Phycisphaerac | Phycisphaera | Phycisphaera | poorly charact | Function unkn | 154D9 | NA | 314230.DSM3 | 0,4 | 0,489897949 | 9,29E-05 | 0,000114475 | 1 - | 0,000252865 - | 1 - | 0,000211402 - | 0 - | 0 - |
| 2565 | 1 | 3um_FIJJAFBF_558283 | unclassified | unclassified | unclassified | unclassified | unclassified | unclassified | unclassified | unclassified | unclassified | unclassified | unclassified | unclassified | 0,8 | 0,4 | 0,000463428 | 0,00023913 | 1 - | 0,000680438 - | 1 - | 0,000568863 - | 1 - | 0,000572663 - |
| 2566 | 1 | 3um_FIJJAFBF_128443 | Bacteria | Bacteroidetes | Flavobacteriia | Flavobacterial | Flavobacteriac | Polaribacter | Polaribacter s | unclassified | unclassified | unclassified | unclassified | unclassified | 2,6 | 0,489897949 | 0,000215323 | 3,02E-05 | 2 - | 0,000199595 - | 2 - | 0,000166866 - | 3 - | 0,000251972 - |
| 2567 | 1 | 0.2um_NOKCPHJN_878846 | Eukaryota | unclassified | Oomycetes | Peronosporale | Peronosporac | Plasmopara | Plasmopara ha | information st | Replication, re | KOG2093 | REV1, polyme | 4787.PITG_08 | 0,2 | 0,4 | 1,54E-05 | 3,07E-05 | 0 - | 0 - | 0 - | 0 - | 0 - | 0 - |
| 2568 | 1 | 3um_FIJJAFBF_345733 | Bacteria | Verrucomicrol | Verrucomicrol | Verrucomicrol | Rubritaleacea | Rubritalea | Rubritalea ma | metabolism | Amino acid tra | 05CAS | nag kinase | 349741.Amuc | 0,6 | 0,489897949 | 0,000127153 | 0,000103876 | 0 - | 0 - | 1 - | 0,000214298 - | 1 - | 0,000215729 - |
| 2569 | 1 | 0.2um_NOKCPHJN_301289 | Bacteria | Firmicutes | Bacilli | Lactobacillales | Lactobacillace | Lactobacillus | Lactobacillus r | unclassified | unclassified | unclassified | unclassified | unclassified | 0,4 | 0,489897949 | 7,31E-06 | 8,96E-06 | 0 - | 0 - | 0 - | 0 - | 1 - | 1,87E-05 - |
| 2570 | 1 | 0.2um_NOKCPHJN_125068 | Eukaryota | Streptophyta | Klebsormidiop | Klebsormidial | Klebsormidiac | Klebsormidiur | Klebsormidiur | cellular proces | Signal transdu | 1BCNW | calcium-deper | 3218.PP1S316 | 0,2 | 0,4 | 9,74E-06 | 1,95E-05 | 0 - | 0 - | 0 - | 0 - | 0 - | 0 - |
| 2571 | 1 | 0.2um_NOKCPHJN_244384 | Eukaryota | Streptophyta | Liliopsida | Poales | Poaceae | Aegilops | Aegilops tausc | unclassified | unclassified | unclassified | unclassified | unclassified | 1 | 0 | 2,45E-05 | 2,59E-06 | 1 - | 2,91E-05 - | 1 - | 2,43E-05 - | 1 - | 2,45E-05 - |
| 2572 | 1 | 3um_FIJJAFBF_510236 | Bacteria | Bacteroidetes | Flavobacteriia | Flavobacterial | Flavobacteriac | Polaribacter | various | unclassified | unclassified | unclassified | unclassified | unclassified | 2,2 | 1,166190379 | 0,000545515 | 0,000286735 | 0 - | 0 - | 3 - | 0,000778941 - | 2 - | 0,000522763 - |
| 2573 | 1 | 3um_FIJJAFBF_151895 | unclassified | unclassified | unclassified | unclassified | unclassified | unclassified | unclassified | unclassified | unclassified | unclassified | unclassified | unclassified | 0,8 | 0,4 | 0,000245822 | 0,000127134 | 1 - | 0,000365113 - | 1 - | 0,000305244 - | 0 - | 0 - |
| 2574 | 1 | 0.2um_NOKCPHJN_277851 | Eukaryota | unclassified | Dinophyceae | Gymnodiniale | Kareniaceae | Karlodinium | Karlodinium v | unclassified | unclassified | unclassified | unclassified | unclassified | 1 | 0 | 8,69E-05 | 9,19E-06 | 1 - | 0,000103239 - | 1 - | 8,63E-05 - | 1 - | 8,69E-05 - |
| 2575 | 1 | 3um_FIJJAFBF_17424 | Eukaryota | Ascomycota | Eurotiomycete | Eurotiales | Aspergillaceae | Penicillium | Penicillium ste | unclassified | unclassified | unclassified | unclassified | unclassified | 1 | 0 | 0,000109361 | 1,16E-05 | 1 - | 0,000129945 - | 1 - | 0,000108637 - | 1 - | 0,000109363 - |
| 2576 | 1 | 3um_FIJJAFBF_681880 | Eukaryota | Chlorophyta | unclassified | Pyramimonad | unclassified | Cymbomonas | Cymbomonas | metabolism | Energy produc | COG0055 | Produces ATP | 39947.LOC_O | 0,2 | 0,4 | 2,50E-05 | 5,00E-05 | 0 - | 0 - | 0 - | 0 - | 0 - | 0 - |
| 2577 | 1 | 3um_FIJJAFBF_451925 | unclassified | unclassified | unclassified | unclassified | unclassified | unclassified | unclassified | unclassified | unclassified | unclassified | unclassified | unclassified | 1 | 0 | 0,000473625 | 5,01E-05 | 1 - | 0,000562768 - | 1 - | 0,000470488 - | 1 - | 0,000473631 - |
| 2578 | 1 | 3um_FIJJAFBF_490907 | Bacteria | Bacteroidetes | Flavobacteriia | unclassified | unclassified | unclassified | Flavobacteria | metabolism | Inorganic ion t | 05C27 | TonB-depende | 487797.Flav3 | 0,6 | 0,489897949 | 3,22E-05 | 2,65E-05 | 0 - | 0 - | 1 - | 5,60E-05 - | 1 - | 5,64E-05 - |
| 2579 | 1 | 3um_FIJJAFBF_11305 | Bacteria | Bacteroidetes | unclassified | unclassified | unclassified | unclassified | Bacteroidetes | cellular proces | Cell cycle cont | 05CR9 | regulator of ch | 391587.KAOT | 0,6 | 0,489897949 | 9,09E-05 | 7,49E-05 | 1 - | 0,000169724 - | 1 - | 0,000141893 - | 1 - | 0,000142841 - |
| 2580 | 1 | 3um_FIJJAFBF_347891 | Eukaryota | unclassified | Cryptophyta | Pyrenomonad | Geminigerace | Guillardia | Guillardia thet | unclassified | unclassified | unclassified | unclassified | unclassified | 1,8 | 0,4 | 0,000478862 | 0,000125709 | 2 - | 0,000628976 - | 2 - | 0,00052584 - | 2 - | 0,000529352 - |
| 2581 | 2 | 0.2um_NOKCPHJN_381924;3um_FIJJAF | various | various | various | various | various | various | various | unclassified | unclassified | unclassified | unclassified | unclassified | 0,2 | 0,4 | 0,000100789 | 0,000201577 | 0 - | 0 - | 0 - | 0 - | 1 - | 0,000503944 - |
| 2582 | 1 | 0.2um_NOKCPHJN_779575 | Eukaryota | Bacillariophyta | Coscinodiscop | Thalassiosirale | Thalassiosirac | Thalassiosira | Thalassiosira p | information st | Translation, ri | KOG0052 | This protein p | 35128.Thaps4 | 1,8 | 0,4 | 0,000357424 | 9,94E-05 | 2 - | 0,000464896 - | 2 - | 0,000388664 - | 2 - | 0,00039126 - |
| 2583 | 1 | 3um_FIJJAFBF_277462 | Eukaryota | Bacillariophyta | Coscinodiscop | Thalassiosirale | Stephanodisca | Cyclotella | Cyclotella sp. | information st | Translation, ri | KOG1569 | ribosomal pro | 35128.Thapsd | 0,4 | 0,489897949 | 0,000103037 | 0,000126747 | 0 - | 0 - | 0 - | 0 - | 1 - | 0,000276285 - |
| 2584 | 1 | 0.2um_NOKCPHJN_54710 | Eukaryota | Streptophyta | unclassified | Ericales | Primulaceae | Androsace | Androsace lax | cellular proces | Posttranslatio | 0IEKR | One of the cor | 59689.fgenesh | 1 | 0 | 0,000136642 | 1,45E-05 | 1 - | 0,00016236 - | 1 - | 0,000135737 - | 1 - | 0,000136644 - |
| 2585 | 1 | 3um_FIJJAFBF_144667 | unclassified | unclassified | unclassified | unclassified | unclassified | unclassified | unclassified | unclassified | unclassified | unclassified | unclassified | unclassified | 0,2 | 0,4 | 4,74E-05 | 9,47E-05 | 0 - | 0 - | 0 - | 0 - | 1 - | 0,000236816 - |
| 2586 | 1 | 3um_FIJJAFBF_335857 | Bacteria | Firmicutes | Bacilli | Bacillales | Bacillaceae | Bacillus | various | information st | RNA processir | 09V5W | DEAH (Asp-Glu | 8083.ENSXMA | 0,4 | 0,489897949 | 0,000302948 | 0,000373314 | 1 - | 0,000822508 - | 0 - | 0 - | 1 - | 0,00069223 - |
| 2587 | 1 | 3um_FIJJAFBF_789459 | Eukaryota | Bacillariophyta | Coscinodiscop | Thalassiosirale | Thalassiosirac | Thalassiosira | Thalassiosira p | unclassified | unclassified | unclassified | unclassified | unclassified | 0,8 | 0,4 | 0,000717987 | 0,000370483 | 1 - | 0,0010542 - | 1 - | 0,000881337 - | 1 - | 0,000887224 - |
| 2588 | 1 | 3um_FIJJAFBF_186025 | Bacteria | Bacteroidetes | Saprospiria | Saprospirales | Lewinellaceae | Lewinella | Lewinella agar | poorly charact | Function unkn | 05MT5 | tonB-depende | 50743.SCB49_ | 0,6 | 0,489897949 | 5,47E-05 | 4,57E-05 | 1 - | 0,000108006 - | 0 - | 0 - | 0 - | 0 - |
| 2589 | 1 | 0.2um_NOKCPHJN_999000 | Bacteria | Spirochaetes | Spirochaetia | unclassified | Leptospiracea | Leptospira | Leptospira sp. | unclassified | unclassified | unclassified | unclassified | unclassified | 0,6 | 0,489897949 | 7,45E-05 | 6,14E-05 | 1 - | 0,000139123 - | 1 - | 0,00011631 - | 1 - | 0,000117087 - |
| 2590 | 2 | 0.2um_NOKCPHJN_640868;3um_FIJJAF | Eukaryota | unclassified | unclassified | Prymnesiales | Chrysochromu | Chrysochromu | Chrysochromu | unclassified | unclassified | unclassified | unclassified | unclassified | 3 | 0,894427191 | 0,000848552 | 0,000209381 | 2 - | 0,000686681 - | 2 - | 0,000574082 - | 4 - | 0,001155834 - |
| 2591 | 1 | 3um_FIJJAFBF_573534 | Eukaryota | Eustigmatoph | unclassified | Eustigmatales | Monodopsida | Nannochlorop | Nannochlorop | unclassified | unclassified | unclassified | unclassified | unclassified | 0,8 | 0,748331477 | 6,75E-05 | 6,53E-05 | 0 - | 0 - | 2 - | 0,000176267 - | 0 - | 0 - |
| 2592 | 1 | 0.2um_NOKCPHJN_262403 | Eukaryota | unclassified | Oomycetes | Peronosporale | Peronosporac | Phytophthora | Phytophthora | unclassified | unclassified | unclassified | unclassified | unclassified | 0,8 | 0,4 | 7,62E-05 | 3,94E-05 | 1 - | 0,000113235 - | 1 - | 9,47E-05 - | 0 - | 0 - |
| 2593 | 1 | 0.2um_NOKCPHJN_990892 | Eukaryota | unclassified | unclassified | Prymnesiales | Chrysochromu | Chrysochromu | Chrysochromu | unclassified | unclassified | unclassified | unclassified | unclassified | 1 | 0 | 0,000313394 | 3,31E-05 | 1 - | 0,000372379 - | 1 - | 0,000311318 - | 1 - | 0,000313398 - |
| 2594 | 1 | 0.2um_NOKCPHJN_68677 | Bacteria | Proteobacteri | Alphaproteob | Rhodobactera | Rhodobactera | Planktomarina | Planktomarina | cellular proces | Intracellular tr | COG0552 | Involved in tar | 314270.RB208 | 0,8 | 0,4 | 0,000122506 | 6,17E-05 | 0 - | 0 - | 1 - | 0,00015963 - | 1 - | 0,000160696 - |
| 2595 | 1 | 3um_FIJJAFBF_510230 | Bacteria | Bacteroidetes | Flavobacteriia | Flavobacterial | Flavobacteriac | Polaribacter | Polaribacter s | information st | Translation, ri | COG0087 | One of the pri | 313594.PI23P | 1,4 | 0,489897949 | 0,000427346 | 0,000145027 | 1 - | 0,000365113 - | 1 - | 0,000305244 - | 2 - | 0,000614565 - |
| 2596 | 1 | 3um_FIJJAFBF_592424 | Bacteria | Actinobacteria | Actinobacteria | unclassified | unclassified | unclassified | actinobacteriu | cellular proces | Posttranslatio | 00CWM | ATP-depender | 312284.A20C1 | 1 | 0 | 0,000148566 | 1,57E-05 | 1 - | 0,000176529 - | 1 - | 0,000147582 - | 1 - | 0,000148568 - |
| 2597 | 1 | 0.2um_NOKCPHJN_731847 | unclassified | unclassified | unclassified | unclassified | unclassified | unclassified | unclassified | unclassified | unclassified | unclassified | unclassified | unclassified | 0,2 | 0,4 | 5,85E-05 | 0,000116962 | 0 - | 0 - | 1 - | 0,000292406 - | 0 - | 0 - |
| 2598 | 1 | 0.2um_NOKCPHJN_563766 | Archaea | Euryarchaeota | Methanobacte | Methanobacte | Methanobacte | Methanospha | Methanospha | cellular proces | Cell wall/mem | 0D8M5 | polymorphic c | 383372.Rcas_ | 0,6 | 0,489897949 | 3,96E-05 | 3,30E-05 | 1 - | 7,72E-05 - | 1 - | 6,46E-05 - | 0 - | 0 - |
| 2599 | 1 | 0.2um_NOKCPHJN_678579 | Eukaryota | Eustigmatoph | unclassified | Eustigmatales | Monodopsida | Nannochlorop | Nannochlorop | poorly charact | Function unkn | 0XYGB | EH-domain co | 35128.Thaps2 | 1 | 0 | 0,000425623 | 4,50E-05 | 1 - | 0,000505731 - | 1 - | 0,000422804 - | 1 - | 0,000425628 - |
| 2600 | 1 | 3um_FIJJAFBF_252631 | Eukaryota | Chordata | Amphibia | Anura | Pipidae | Xenopus | Xenopus tropi | unclassified | unclassified | unclassified | unclassified | unclassified | 0,4 | 0,489897949 | 9,59E-05 | 0,00011763 | 0 - | 0 - | 0 - | 0 - | 0 - | 0 - |
| 2601 | 1 | 0.2um_NOKCPHJN_285110 | Eukaryota | Phaeophyceae | unclassified | Ectocarpales | Ectocarpaceae | Ectocarpus | Ectocarpus sili | cellular proces | Posttranslatio | KOG0361 | t-complex pro | 272952.HpaP8 | 0,4 | 0,489897949 | 4,70E-05 | 5,79E-05 | 1 - | 0,000127946 - | 1 - | 0,000106966 - | 0 - | 0 - |
| 2602 | 1 | 0.2um_NOKCPHJN_30432 | Bacteria | Proteobacteri | Alphaproteob | Rhodospirillale | unclassified | unclassified | Rhodospirillale | information st | Translation, ri | COG0013 | Catalyzes the | 269796.Rru_A | 0,4 | 0,489897949 | 2,93E-05 | 3,66E-05 | 1 - | 8,49E-05 - | 0 - | 0 - | 0 - | 0 - |
| 2603 | 1 | 3um_FIJJAFBF_748496 | Bacteria | Proteobacteri | Alphaproteob | Rhodobactera | Rhodobactera | Planktomarina | Planktomarina | unclassified | unclassified | unclassified | unclassified | unclassified | 1 | 0 | 7,52E-05 | 7,95E-06 | 1 - | 8,93E-05 - | 1 - | 7,47E-05 - | 1 - | 7,52E-05 - |
| 2604 | 2 | 0.2um_NOKCPHJN_211756;3um_FIJJAF | Bacteria | Bacteroidetes | Flavobacteriia | Flavobacterial | Flavobacteriac | unclassified | Flavobacteriac | unclassified | unclassified | unclassified | unclassified | unclassified | 0,6 | 0,489897949 | 0,00011824 | 9,74E-05 | 1 - | 0,000220791 - | 1 - | 0,000184587 - | 1 - | 0,00018582 - |
| 2605 | 1 | 3um_FIJJAFBF_532450 | Eukaryota | unclassified | Cryptophyta | Pyrenomonad | Geminigerace | Teleaulax | Teleaulax amp | metabolism | Energy produc | COG4451 | RuBisCO catal | 45157.CMV01 | 1,2 | 0,4 | 0,000560876 | 0,000258963 | 2 - | 0,001076952 - | 1 - | 0,000450179 - | 1 - | 0,000453187 - |
| 2606 | 1 | 3um_FIJJAFBF_457156 | Bacteria | Actinobacteria | Actinobacteria | Streptomyceta | Streptomyceta | Streptomyces | Streptomyces | unclassified | unclassified | unclassified | unclassified | unclassified | 0,2 | 0,4 | 1,52E-05 | 3,04E-05 | 1 - | 7,61E-05 - | 0 - | 0 - | 0 - | 0 - |
| 2607 | 1 | 0.2um_NOKCPHJN_538357 | Bacteria | Bacteroidetes | Flavobacteriia | Flavobacterial | Cryomorphace | unclassified | Cryomorphace | unclassified | unclassified | unclassified | unclassified | unclassified | 0,8 | 0,4 | 7,13E-05 | 3,59E-05 | 0 - | 0 - | 1 - | 9,28E-05 - | 1 - | 9,35E-05 - |
| 2608 | 1 | 3um_FIJJAFBF_269861 | Bacteria | Proteobacteri | Alphaproteob | Pelagibacteral | Pelagibacterac | unclassified | Pelagibacterac | unclassified | unclassified | unclassified | unclassified | unclassified | 0,6 | 0,489897949 | 0,000175804 | 0,000144839 | 1 - | 0,000328282 - | 1 - | 0,000274451 - | 1 - | 0,000276285 - |
| 2609 | 1 | 3um_FIJJAFBF_530420 | Bacteria | Bacteroidetes | Saprospiria | Saprospirales | Saprospiracea | Saprospira | Saprospira gra | poorly charact | Function unkn | 1AK27 | OmpA MotB d | 700598.Niako | 0,6 | 0,489897949 | 6,64E-05 | 5,44E-05 | 0 - | 0 - | 0 - | 0 - | 1 - | 0,000117744 - |
| 2610 | 1 | 3um_FIJJAFBF_636516 | Bacteria | Bacteroidetes | Flavobacteriia | Flavobacterial | Flavobacteriac | Polaribacter | Polaribacter s | metabolism | Inorganic ion t | 0P8Q4 | tonB-depende | 867900.Celly_ | 2,2 | 0,979795897 | 0,000186783 | 6,56E-05 | 1 - | 0,000105272 - | 2 - | 0,000176019 - | 2 - | 0,000177195 - |
| 2611 | 1 | 3um_FIJJAFBF_545076 | Bacteria | Verrucomicrol | Opitutae | unclassified | unclassified | unclassified | Opitutae bact | unclassified | unclassified | unclassified | unclassified | unclassified | 0,2 | 0,4 | 2,88E-05 | 5,77E-05 | 1 - | 0,000144216 - | 0 - | 0 - | 0 - | 0 - |
| 2612 | 1 | 0.2um_NOKCPHJN_119122 | Bacteria | Bacteroidetes | Flavobacteriia | Flavobacterial | Flavobacteriac | Polaribacter | Polaribacter s | unclassified | unclassified | unclassified | unclassified | unclassified | 0,4 | 0,489897949 | 2,51E-05 | 3,09E-05 | 0 - | 0 - | 1 - | 6,72E-05 - | 0 - | 0 - |
| 2613 | 1 | 0.2um_NOKCPHJN_1060599 | Bacteria | Proteobacteri | Alphaproteob | Pelagibacteral | Pelagibacterac | Candidatus Pe | Candidatus Pe | poorly charact | Function unkn | 0QJ8F | Phospholipid | 207954.MED9 | 1 | 0 | 0,000243213 | 2,57E-05 | 1 - | 0,000288989 - | 1 - | 0,000241602 - | 1 - | 0,000243216 - |
| 2614 | 1 | 3um_FIJJAFBF_474234 | Eukaryota | unclassified | Cryptophyta | Pyrenomonad | Geminigerace | Teleaulax | Teleaulax amp | unclassified | unclassified | unclassified | unclassified | unclassified | 1,2 | 0,4 | 0,000427066 | 0,00014725 | 1 - | 0,000422871 - | 1 - | 0,000353531 - | 2 - | 0,000711785 - |
| 2615 | 1 | 0.2um_NOKCPHJN_785139 | Bacteria | unclassified | unclassified | unclassified | unclassified | unclassified | uncultured ba | metabolism | Inorganic ion t | 05C27 | TonB-depende | 313596.RB250 | 1,8 | 0,4 | 0,000138942 | 2,39E-05 | 1 - | 9,37E-05 - | 2 - | 0,000156633 - | 2 - | 0,000157679 - |
| 2616 | 1 | 3um_FIJJAFBF_730097 | unclassified | unclassified | unclassified | unclassified | unclassified | unclassified | unclassified | unclassified | unclassified | unclassified | unclassified | unclassified | 0,6 | 0,489897949 | 0,000610758 | 0,000500731 | 0 - | 0 - | 1 - | 0,001078878 - | 0 - | 0 - |
| 2617 | 1 | 0.2um_NOKCPHJN_48861 | Bacteria | Bacteroidetes | Flavobacteriia | Flavobacterial | Flavobacteriac | Polaribacter | Polaribacter s | cellular proces | Posttranslatio | 05DIW | FeS assembly | 313598.MED1 | 0,4 | 0,489897949 | 5,60E-05 | 6,92E-05 | 1 - | 0,000155287 - | 0 - | 0 - | 0 - | 0 - |
| 2618 | 1 | 3um_FIJJAFBF_453081 | Eukaryota | Basidiomycota | Exobasidiomy | Tilletiales | Tilletiaceae | Tilletia | Tilletia indica | poorly charact | Function unkn | 01I70 | Reverse trans | 5306.JGI1000 | 0,2 | 0,4 | 8,14E-06 | 1,63E-05 | 0 - | 0 - | 0 - | 0 - | 0 - | 0 - |
| 2619 | 1 | 0.2um_NOKCPHJN_456112 | Bacteria | Proteobacteri | Betaproteoba | Burkholderiale | Oxalobacterac | Massilia | Massilia namu | unclassified | unclassified | unclassified | unclassified | unclassified | 0,8 | 0,4 | 4,48E-05 | 2,28E-05 | 1 - | 6,44E-05 - | 1 - | 5,39E-05 - | 1 - | 5,42E-05 - |
| 2620 | 1 | 0.2um_NOKCPHJN_248889 | Eukaryota | Chordata | Mammalia | unclassified | Chrysochlorid | Chrysochloris | Chrysochloris | unclassified | unclassified | unclassified | unclassified | unclassified | 1 | 0 | 0,000126237 | 1,33E-05 | 1 - | 0,000149996 - | 1 - | 0,000125401 - | 1 - | 0,000126238 - |
| 2621 | 1 | 0.2um_NOKCPHJN_205757 | unclassified | unclassified | unclassified | unclassified | unclassified | unclassified | unclassified | unclassified | unclassified | unclassified | unclassified | unclassified | 0,8 | 0,4 | 0,000132659 | 6,69E-05 | 0 - | 0 - | 1 - | 0,000172859 - | 1 - | 0,000174014 - |

| | | | | | | | | | | | | | | | | | | | | | | | | |
|---|---|---|---|---|---|---|---|---|---|---|---|---|---|---|---|---|---|---|---|---|---|---|---|---|
| 2622 | 1 | 3um_FIJJAFBF_721778 | Eukaryota | unclassified | unclassified | Isochrysidales | Noelaerhabda | Emiliania | Emiliania huxl | unclassified | unclassified | unclassified | unclassified | unclassified | 0,4 | 0,489897949 | 0,000222244 | 0,000272195 | 0 - | 0 - | 1 - | 0,00055376 - | 1 - | 0,00055746 - |
| 2623 | 1 | 3um_FIJJAFBF_510246 | Bacteria | Bacteroidetes | Flavobacteriia | Flavobacterial | Flavobacteriac | Polaribacter | Polaribacter s | information st | Translation, ri | COG0256 | This is one of t | 313598.MED1 | 1 | 0 | 0,000543036 | 5,74E-05 | 1 - | 0,000645243 - | 1 - | 0,000539439 - | 1 - | 0,000543043 - |
| 2624 | 1 | 0.2um_NOKCPHJN_416465 | Bacteria | Bacteroidetes | various | various | various | various | various | metabolism | Nucleotide tra | COG0756 | This enzyme is | 867900.Celly_ | 0,8 | 0,4 | 0,000348119 | 0,000180015 | 1 - | 0,000516194 - | 0 - | 0 - | 1 - | 0,000434434 - |
| 2625 | 1 | 3um_FIJJAFBF_397384 | Bacteria | unclassified | unclassified | unclassified | unclassified | unclassified | bacterium TM | unclassified | unclassified | unclassified | unclassified | unclassified | 0,2 | 0,4 | 2,85E-05 | 5,70E-05 | 0 - | 0 - | 1 - | 0,00014254 - | 0 - | 0 - |
| 2626 | 1 | 3um_FIJJAFBF_370622 | Bacteria | Proteobacteri | Alphaproteob | unclassified | unclassified | unclassified | Alphaproteob | unclassified | unclassified | unclassified | unclassified | unclassified | 1 | 0,632455532 | 0,000172028 | 0,000109216 | 0 - | 0 - | 1 - | 0,000179298 - | 1 - | 0,000180496 - |
| 2627 | 1 | 3um_FIJJAFBF_95708 | Eukaryota | unclassified | Pelagophycea | Pelagomonad | unclassified | Aureococcus | Aureococcus a | information st | Translation, ri | COG0052 | 30S ribosomal | 32049.SYNPC( | 3,8 | 0,4 | 0,000839982 | 0,000152973 | 4 - | 0,00104318 - | 4 - | 0,000872124 - | 4 - | 0,00087795 - |
| 2628 | 1 | 0.2um_NOKCPHJN_491503 | Bacteria | Proteobacteri | Alphaproteob | Pelagibacteral | unclassified | unclassified | Pelagibacteral | metabolism | Energy produc | COG0056 | Produces ATP | 335992.SAR11 | 2,8 | 0,748331477 | 0,000337847 | 6,22E-05 | 2 - | 0,000293522 - | 3 - | 0,000368088 - | 2 - | 0,000247031 - |
| 2629 | 1 | 3um_FIJJAFBF_589764 | Eukaryota | unclassified | unclassified | unclassified | unclassified | Gymnochlora | Gymnochlora | cellular proces | Posttranslatio | KOG0417 | ubiquitin-conj | 423536.XP_00 | 1 | 0 | 0,000422766 | 4,47E-05 | 1 - | 0,000502337 - | 1 - | 0,000419966 - | 1 - | 0,000422771 - |
| 2630 | 1 | 3um_FIJJAFBF_713603 | Bacteria | Proteobacteri | Alphaproteob | Rhodobactera | Rhodobactera | various | various | information st | Replication, re | 01QWU | Transposase | 314265.R2601 | 0,2 | 0,4 | 2,31E-05 | 4,61E-05 | 0 - | 0 - | 0 - | 0 - | 1 - | 0,000115372 - |
| 2631 | 1 | 0.2um_NOKCPHJN_1001140 | Bacteria | Proteobacteri | Gammaprotec | Pseudomonad | Pseudomonac | Pseudomonas | various | information st | Translation, ri | COG0087 | One of the pri | 629265.PMA4 | 1 | 0 | 0,000298541 | 3,16E-05 | 1 - | 0,000354731 - | 1 - | 0,000296564 - | 1 - | 0,000298545 - |
| 2632 | 1 | 3um_FIJJAFBF_667394 | Bacteria | Proteobacteri | Alphaproteob | Rhodobactera | Rhodobactera | Sulfitobacter | Sulfitobacter g | unclassified | unclassified | unclassified | unclassified | unclassified | 1,4 | 0,489897949 | 0,000297236 | 8,33E-05 | 1 - | 0,00025899 - | 1 - | 0,000216522 - | 1 - | 0,000217969 - |
| 2633 | 2 | 0.2um_NOKCPHJN_130260;3um_FIJJAF | Eukaryota | various | various | various | various | various | various | metabolism | Energy produc | COG1290 | Component of | 35128.Thapsd | 2,6 | 0,489897949 | 0,000769693 | 0,000196867 | 3 - | 0,001044393 - | 3 - | 0,000873138 - | 2 - | 0,000585981 - |
| 2634 | 1 | 3um_FIJJAFBF_537683 | Eukaryota | Bacillariophyt | Bacillariophyc | Naviculales | Naviculaceae | Fistulifera | Fistulifera sola | metabolism | Energy produc | 1DYB8 | PsaA and PsaE | 3055.DAA014 | 12,8 | 0,979795897 | 0,00106579 | 7,40E-05 | 11 - | 0,001094854 - | 14 - | 0,001164959 - | 13 - | 0,001088974 - |
| 2635 | 1 | 0.2um_NOKCPHJN_61811 | Bacteria | Bacteroidetes | Flavobacteriia | Flavobacterial | Flavobacteriac | Algibacter | Algibacter sp. | metabolism | Inorganic ion t | 05C27 | TonB-depende | 391587.KAOT | 16,8 | 2,638181192 | 0,001006278 | 9,45E-05 | 12 - | 0,000866967 - | 16 - | 0,000966408 - | 19 - | 0,001155276 - |
| 2636 | 1 | 0.2um_NOKCPHJN_858965 | Bacteria | Proteobacteri | Alphaproteob | Pelagibacteral | Pelagibacterac | Candidatus Pe | Candidatus Pe | poorly charact | Function unkn | 01QBG | Protein of unk | 684719.HIMB | 0,6 | 0,489897949 | 0,000206614 | 0,000170223 | 1 - | 0,000385815 - | 1 - | 0,000322551 - | 1 - | 0,000324706 - |
| 2637 | 2 | 3um_FIJJAFBF_56398;3um_FIJJAFBF_7 | Eukaryota | unclassified | Dinophyceae | Gonyaulacales | various | various | various | metabolism | Energy produc | 0FJ7X | This is one of t | various | 1 | 0 | 0,000179465 | 1,90E-05 | 1 - | 0,000213243 - | 1 - | 0,000178276 - | 1 - | 0,000179467 - |
| 2638 | 1 | 0.2um_NOKCPHJN_733220 | unclassified | unclassified | unclassified | unclassified | unclassified | unclassified | unclassified | unclassified | unclassified | unclassified | unclassified | unclassified | 1 | 0 | 9,37E-05 | 9,91E-06 | 1 - | 0,000111381 - | 1 - | 9,31E-05 - | 1 - | 9,37E-05 - |
| 2639 | 2 | 0.2um_NOKCPHJN_186789;3um_FIJJAF | various | various | various | various | various | various | various | various | various | various | various | various | 0,2 | 0,4 | 1,41E-05 | 2,83E-05 | 0 - | 0 - | 0 - | 0 - | 0 - | 0 - |
| 2640 | 1 | 0.2um_NOKCPHJN_401432 | Eukaryota | unclassified | Oomycetes | Albuginales | Albuginaceae | Albugo | Albugo laibach | unclassified | unclassified | unclassified | unclassified | unclassified | 0,6 | 0,489897949 | 4,88E-05 | 4,07E-05 | 1 - | 9,51E-05 - | 1 - | 7,95E-05 - | 0 - | 0 - |
| 2641 | 1 | 0.2um_NOKCPHJN_265489 | Eukaryota | unclassified | Pelagophycea | Pelagomonad | unclassified | Aureococcus | Aureococcus a | unclassified | unclassified | unclassified | unclassified | unclassified | 0,4 | 0,489897949 | 2,61E-05 | 3,22E-05 | 1 - | 7,09E-05 - | 0 - | 0 - | 1 - | 5,97E-05 - |
| 2642 | 1 | 3um_FIJJAFBF_517422 | Bacteria | Bacteroidetes | Flavobacteriia | Flavobacterial | Flavobacteriac | Polaribacter | Polaribacter a | information st | Translation, ri | 05C64 | Binds directly | 313594.PI23P | 1 | 1,095445115 | 0,000249833 | 0,000260995 | 0 - | 0 - | 1 - | 0,000273253 - | 0 - | 0 - |
| 2643 | 1 | 3um_FIJJAFBF_183089 | Eukaryota | Bacillariophyt | Mediophycea | Cymatosirales | Cymatosirace | Plagiogrammc | Plagiogrammc | cellular proces | Posttranslatio | 05C2Z | ATP-depender | 756067.Micva | 0,8 | 0,748331477 | 5,57E-05 | 4,86E-05 | 1 - | 8,47E-05 - | 1 - | 7,08E-05 - | 0 - | 0 - |
| 2644 | 1 | 3um_FIJJAFBF_254285 | Bacteria | Proteobacteri | Gammaprotec | Cellvibrionales | Spongiibacter | Oceanicoccus | Oceanicoccus | information st | Translation, ri | COG0087 | One of the pri | 247633.GP214 | 0,6 | 0,489897949 | 0,000166702 | 0,000136718 | 0 - | 0 - | 0 - | 0 - | 1 - | 0,000295741 - |
| 2645 | 1 | 0.2um_NOKCPHJN_205340 | unclassified | unclassified | unclassified | unclassified | unclassified | unclassified | unclassified | unclassified | unclassified | unclassified | unclassified | unclassified | 0,2 | 0,4 | 4,06E-05 | 8,13E-05 | 0 - | 0 - | 0 - | 0 - | 0 - | 0 - |
| 2646 | 1 | 3um_FIJJAFBF_311757 | Bacteria | Bacteroidetes | Flavobacteriia | Flavobacterial | Flavobacteriac | Polaribacter | Polaribacter a | information st | Translation, ri | COG0050 | This protein p | 313603.FB217 | 8,8 | 1,326649916 | 0,001388735 | 0,000147353 | 8 - | 0,001515913 - | 9 - | 0,001425758 - | 7 - | 0,001116331 - |
| 2647 | 1 | 3um_FIJJAFBF_550914 | Eukaryota | Bacillariophyt | Coscinodiscop | Thalassiosirale | Thalassiosirac | Thalassiosira | Thalassiosira c | unclassified | unclassified | unclassified | unclassified | unclassified | 3,8 | 0,4 | 0,002955187 | 0,000452743 | 4 - | 0,003696207 - | 4 - | 0,003090119 - | 3 - | 0,002333072 - |
| 2648 | 1 | 3um_FIJJAFBF_250489 | Eukaryota | unclassified | Oomycetes | Peronosporale | Peronosporac | Phytophthora | Phytophthora | metabolism | Amino acid tra | COG0137 | Citrulline--asp | 67593.Physo1 | 0,4 | 0,489897949 | 0,000159959 | 0,000195911 | 0 - | 0 - | 1 - | 0,000398566 - | 1 - | 0,000401229 - |
| 2649 | 1 | 3um_FIJJAFBF_44055 | Viruses | unclassified | unclassified | unclassified | various | various | various | unclassified | unclassified | unclassified | unclassified | unclassified | 1,8 | 0,4 | 0,000626441 | 0,00015747 | 2 - | 0,000827052 - | 2 - | 0,000691436 - | 1 - | 0,000348027 - |
| 2650 | 1 | 0.2um_NOKCPHJN_461932 | Eukaryota | Zoopagomyco | Basidiobolom | Basidiobolales | Basidiobolace | Basidiobolus | Basidiobolus r | metabolism | Secondary me | KOG0059 | ATP-binding c | 5786.XP_0032 | 0,8 | 0,4 | 2,75E-05 | 1,42E-05 | 1 - | 4,03E-05 - | 1 - | 3,37E-05 - | 1 - | 3,39E-05 - |
| 2651 | 1 | 0.2um_NOKCPHJN_70557 | Bacteria | Bacteroidetes | Flavobacteriia | Flavobacterial | Flavobacteriac | Polaribacter | Polaribacter s | poorly charact | Function unkn | 11PIQ | YCE I like fami | 313598.MED1 | 0,2 | 0,4 | 5,67E-05 | 0,00011335 | 0 - | 0 - | 0 - | 0 - | 0 - | 0 - |
| 2652 | 1 | 0.2um_NOKCPHJN_61814 | Bacteria | Bacteroidetes | Flavobacteriia | Flavobacterial | Flavobacteriac | Polaribacter | Polaribacter fi | poorly charact | Function unkn | 0XRCJ | NA | 391587.KAOT | 11,6 | 2,653299832 | 0,000677162 | 9,15E-05 | 8 - | 0,000567569 - | 12 - | 0,000711753 - | 10 - | 0,000597089 - |
| 2653 | 1 | 3um_FIJJAFBF_296153 | Bacteria | Bacteroidetes | unclassified | unclassified | unclassified | unclassified | Bacteroidetes | metabolism | Amino acid tra | 09AJS | Cleaves the N- | 1046627.BZAF | 0,2 | 0,4 | 2,66E-05 | 5,33E-05 | 0 - | 0 - | 0 - | 0 - | 0 - | 0 - |
| 2654 | 1 | 3um_FIJJAFBF_519562 | Bacteria | Proteobacteri | unclassified | unclassified | unclassified | unclassified | Proteobacteri | unclassified | unclassified | unclassified | unclassified | unclassified | 1 | 0,632455532 | 0,000126159 | 7,42E-05 | 0 - | 0 - | 1 - | 0,000133993 - | 1 - | 0,000134889 - |
| 2655 | 1 | 3um_FIJJAFBF_347366 | Eukaryota | unclassified | Bangiophycea | Bangiales | Bangiaceae | Porphyra | Porphyra umb | unclassified | unclassified | unclassified | unclassified | unclassified | 2 | 0,632455532 | 0,000187552 | 4,33E-05 | 1 - | 0,000115151 - | 2 - | 0,000192538 - | 2 - | 0,000193824 - |
| 2656 | 2 | 0.2um_NOKCPHJN_646842;3um_FIJJAF | Eukaryota | various | various | various | various | various | various | unclassified | unclassified | unclassified | unclassified | unclassified | 1 | 0 | 0,000154015 | 1,63E-05 | 1 - | 0,000183003 - | 1 - | 0,000152995 - | 1 - | 0,000154017 - |
| 2657 | 1 | 0.2um_NOKCPHJN_440521 | Bacteria | Proteobacteri | Gammaprotec | Chromatiales | Ectothiorhodc | Thioalkalivibri | Thioalkalivibri | unclassified | unclassified | unclassified | unclassified | unclassified | 0,2 | 0,4 | 7,64E-06 | 1,53E-05 | 0 - | 0 - | 1 - | 3,82E-05 - | 0 - | 0 - |
| 2658 | 1 | 3um_FIJJAFBF_189702 | Bacteria | Bacteroidetes | Flavobacteriia | Flavobacterial | Flavobacteriac | Polaribacter | Polaribacter fi | poorly charact | Function unkn | 09A4Z | rhodopsin | 313598.MED1 | 1 | 0 | 0,000262467 | 2,78E-05 | 1 - | 0,000311867 - | 1 - | 0,000260729 - | 1 - | 0,000262471 - |
| 2659 | 1 | 3um_FIJJAFBF_195324 | unclassified | unclassified | unclassified | unclassified | unclassified | unclassified | unclassified | unclassified | unclassified | unclassified | unclassified | unclassified | 12 | 2,19089023 | 0,001511933 | 0,000187868 | 10 - | 0,001518219 - | 13 - | 0,001650049 - | 9 - | 0,001149973 - |
| 2660 | 1 | 0.2um_NOKCPHJN_953814 | Eukaryota | unclassified | Oomycetes | Pythiales | Pythiaceae | Pythium | Pythium insidi | cellular proces | Signal transdu | 0XPQQ | CAMP-depenc | 6183.Smp_15 | 0,2 | 0,4 | 1,73E-05 | 3,46E-05 | 0 - | 0 - | 0 - | 0 - | 0 - | 0 - |
| 2661 | 1 | 3um_FIJJAFBF_48703 | Bacteria | Bacteroidetes | Flavobacteriia | Flavobacterial | Flavobacteriac | Tenacibaculur | Tenacibaculur | unclassified | unclassified | unclassified | unclassified | unclassified | 0,4 | 0,489897949 | 4,72E-05 | 5,80E-05 | 0 - | 0 - | 1 - | 0,000126159 - | 0 - | 0 - |
| 2662 | 1 | 3um_FIJJAFBF_554090 | Bacteria | Verrucomicrol | Opitutae | unclassified | unclassified | unclassified | Opitutae bact | unclassified | unclassified | unclassified | unclassified | unclassified | 2,2 | 0,4 | 0,001532109 | 0,000459075 | 3 - | 0,002440702 - | 2 - | 0,001360324 - | 2 - | 0,001369412 - |
| 2663 | 1 | 0.2um_NOKCPHJN_914416 | unclassified | unclassified | unclassified | unclassified | unclassified | unclassified | unclassified | unclassified | unclassified | unclassified | unclassified | unclassified | 1 | 0 | 0,000887214 | 9,38E-05 | 1 - | 0,0010542 - | 1 - | 0,000881337 - | 1 - | 0,000887224 - |
| 2664 | 1 | 0.2um_NOKCPHJN_249107 | unclassified | unclassified | unclassified | unclassified | unclassified | unclassified | unclassified | unclassified | unclassified | unclassified | unclassified | unclassified | 0,4 | 0,489897949 | 9,97E-05 | 0,000122104 | 0 - | 0 - | 0 - | 0 - | 1 - | 0,000255032 - |
| 2665 | 1 | 0.2um_NOKCPHJN_363535 | Eukaryota | unclassified | unclassified | unclassified | Apusomonadi | Thecamonas | Thecamonas t | cellular proces | Cell wall/mem | 05DAF | Involved in the | 869210.Marky | 0,4 | 0,489897949 | 4,78E-05 | 5,87E-05 | 0 - | 0 - | 0 - | 0 - | 0 - | 0 - |
| 2666 | 1 | 0.2um_NOKCPHJN_615874 | Eukaryota | unclassified | unclassified | Prymnesiales | Chrysochromu | Chrysochromu | Chrysochromu | unclassified | unclassified | unclassified | unclassified | unclassified | 0,6 | 0,489897949 | 0,000122019 | 0,000100072 | 0 - | 0 - | 0 - | 0 - | 1 - | 0,000216471 - |
| 2667 | 2 | 0.2um_NOKCPHJN_891700;0.2um_NO | various | various | various | various | various | various | various | unclassified | unclassified | unclassified | unclassified | unclassified | 0,2 | 0,4 | 1,82E-05 | 3,63E-05 | 0 - | 0 - | 0 - | 0 - | 1 - | 9,08E-05 - |
| 2668 | 1 | 0.2um_NOKCPHJN_480383 | Bacteria | Bacteroidetes | Flavobacteriia | Flavobacterial | Cryomorphac | Owenweeksia | Owenweeksia | metabolism | Amino acid tra | COG2171 | Catalyzes the | 926562.Oweh | 1 | 0 | 0,000335065 | 3,54E-05 | 1 - | 0,000398129 - | 1 - | 0,000332845 - | 1 - | 0,000335069 - |
| 2669 | 1 | 3um_FIJJAFBF_430146 | Eukaryota | Chlorophyta | Trebouxiophy | unclassified | Coccomyxace | Coccomyxa | Coccomyxa su | unclassified | unclassified | unclassified | unclassified | unclassified | 0,6 | 0,489897949 | 0,000292966 | 0,000244281 | 1 - | 0,00057136 - | 1 - | 0,000477671 - | 0 - | 0 - |
| 2670 | 1 | 3um_FIJJAFBF_815479 | Eukaryota | unclassified | Cryptophyta | Pyrenomonad | Geminigerace | Guillardia | Guillardia thet | metabolism | Lipid transpor | COG0439 | acetyl-CoA car | 2850.Phatr54 | 2,6 | 0,489897949 | 0,001196236 | 0,00016794 | 2 - | 0,001108862 - | 2 - | 0,000927036 - | 3 - | 0,001399843 - |
| 2671 | 1 | 0.2um_NOKCPHJN_587110 | Eukaryota | Basidiomycota | Pucciniomycet | Pucciniales | Pucciniaceae | Puccinia | Puccinia coron | unclassified | unclassified | unclassified | unclassified | unclassified | 0,4 | 0,489897949 | 0,000101261 | 0,000124561 | 0 - | 0 - | 0 - | 0 - | 1 - | 0,000271521 - |
| 2672 | 1 | 3um_FIJJAFBF_377213 | Bacteria | Bacteroidetes | Flavobacteriia | Flavobacterial | Flavobacteriac | Polaribacter | Polaribacter t | unclassified | unclassified | unclassified | unclassified | unclassified | 0,4 | 0,489897949 | 0,000269412 | 0,00033672 | 1 - | 0,000779669 - | 0 - | 0 - | 0 - | 0 - |
| 2673 | 1 | 0.2um_NOKCPHJN_216465 | Bacteria | Bacteroidetes | Cytophagia | Cytophagales | Hymenobacte | Hymenobacte | Hymenobacte | metabolism | Coenzyme tra | 09A4R | May play a ke | 388413.ALPR1 | 1 | 0 | 0,00014349 | 1,52E-05 | 1 - | 0,000170497 - | 1 - | 0,00014254 - | 1 - | 0,000143492 - |
| 2674 | 1 | 3um_FIJJAFBF_701411 | Eukaryota | Bacillariophyt | Mediophycea | Triceratiales | Triceratiaceae | Trieres | Trieres chinen | unclassified | unclassified | unclassified | unclassified | unclassified | 0,2 | 0,4 | 4,86E-05 | 9,73E-05 | 0 - | 0 - | 0 - | 0 - | 1 - | 0,000243216 - |
| 2675 | 1 | 3um_FIJJAFBF_365652 | Bacteria | Bacteroidetes | Flavobacteriia | Flavobacterial | Flavobacteriac | Aureitalea | Aureitalea ma | unclassified | unclassified | unclassified | unclassified | unclassified | 6,4 | 2,416609195 | 0,000439469 | 0,00012946 | 3 - | 0,000254297 - | 6 - | 0,000425198 - | 5 - | 0,000356698 - |
| 2676 | 1 | 0.2um_NOKCPHJN_358202 | Bacteria | Proteobacteri | Gammaprotec | Cellvibrionales | Microbulbifer | Microbulbifer | various | poorly charact | Function unkn | 05EC1 | AtP-binding pr | 313624.N9414 | 1 | 0 | 6,86E-05 | 7,26E-06 | 1 - | 8,15E-05 - | 1 - | 6,82E-05 - | 1 - | 6,86E-05 - |
| 2677 | 1 | 0.2um_NOKCPHJN_852447 | Bacteria | Bacteroidetes | Flavobacteriia | Flavobacterial | Flavobacteriac | Algibacter | Algibacter sp. | metabolism | Carbohydrate | 0XNQK | Transporter | 391587.KAOT | 0,6 | 0,489897949 | 7,80E-05 | 6,40E-05 | 0 - | 0 - | 0 - | 0 - | 1 - | 0,000138446 - |
| 2678 | 1 | 3um_FIJJAFBF_496029 | Eukaryota | Phaeophyceae | unclassified | Ectocarpales | Ectocarpaceae | Ectocarpus | Ectocarpus sili | unclassified | unclassified | unclassified | unclassified | unclassified | 2 | 0,632455532 | 0,000263623 | 8,31E-05 | 2 - | 0,000314488 - | 1 - | 0,00013146 - | 2 - | 0,000264676 - |
| 2679 | 1 | 3um_FIJJAFBF_390243 | Eukaryota | unclassified | Chrysophycea | Chromulinales | Chromulinace | Ochromonas | Ochromonas c | unclassified | unclassified | unclassified | unclassified | unclassified | 7,6 | 1,2 | 0,000753959 | 0,00011134 | 7 - | 0,000830328 - | 9 - | 0,000892511 - | 7 - | 0,000698812 - |
| 2680 | 1 | 0.2um_NOKCPHJN_335704 | Viruses | unclassified | unclassified | unclassified | Mimiviridae | Indivirus | Indivirus ILV1 | unclassified | unclassified | unclassified | unclassified | unclassified | 0,2 | 0,4 | 1,91E-05 | 3,82E-05 | 0 - | 0 - | 1 - | 9,55E-05 - | 0 - | 0 - |
| 2681 | 1 | 3um_FIJJAFBF_91117 | Eukaryota | Bacillariophyt | Coscinodiscop | Leptocylindral | Leptocylindra | Leptocylindru | Leptocylindru | unclassified | unclassified | unclassified | unclassified | unclassified | 1,8 | 0,4 | 0,001117346 | 0,000293322 | 2 - | 0,001467611 - | 2 - | 0,001226959 - | 2 - | 0,001235156 - |
| 2682 | 1 | 0.2um_NOKCPHJN_86871 | Bacteria | Proteobacteri | Gammaprotec | unclassified | unclassified | unclassified | Gammaprotec | information st | Transcription | COG0085 | DNA-depende | 247633.GP214 | 0,4 | 0,489897949 | 1,69E-05 | 2,07E-05 | 0 - | 0 - | 0 - | 0 - | 0 - | 0 - |
| 2683 | 1 | 3um_FIJJAFBF_211014 | Bacteria | Proteobacteri | Alphaproteob | Rhodobactera | Rhodobactera | Albidovulum | Albidovulum x | metabolism | Coenzyme tra | 01QD7 | Precorrin-2 C( | 391613.RTM1 | 0,6 | 0,489897949 | 0,000154538 | 0,000126852 | 0 - | 0 - | 1 - | 0,000268562 - | 1 - | 0,000270356 - |
| 2684 | 1 | 0.2um_NOKCPHJN_239282 | Eukaryota | unclassified | unclassified | Tritrichomona | Tritrichomona | Tritrichomona | Tritrichomona | unclassified | unclassified | unclassified | unclassified | unclassified | 0,2 | 0,4 | 6,75E-06 | 1,35E-05 | 0 - | 0 - | 0 - | 0 - | 0 - | 0 - |
| 2685 | 1 | 3um_FIJJAFBF_306499 | Bacteria | Proteobacteri | Betaproteoba | unclassified | unclassified | unclassified | Betaproteoba | unclassified | unclassified | unclassified | unclassified | unclassified | 3,2 | 0,748331477 | 0,000413128 | 9,06E-05 | 3 - | 0,000463935 - | 2 - | 0,000258574 - | 4 - | 0,000520603 - |
| 2686 | 1 | 3um_FIJJAFBF_140037 | Eukaryota | Bacillariophyt | Bacillariophyc | Naviculales | Phaeodactylac | Phaeodactylu | Phaeodactylu | unclassified | unclassified | unclassified | unclassified | unclassified | 0,2 | 0,4 | 4,90E-05 | 9,80E-05 | 0 - | 0 - | 0 - | 0 - | 1 - | 0,000245109 - |
| 2687 | 1 | 3um_FIJJAFBF_506614 | unclassified | unclassified | unclassified | unclassified | unclassified | unclassified | unclassified | unclassified | unclassified | unclassified | unclassified | unclassified | 1 | 0 | 0,000473625 | 5,01E-05 | 1 - | 0,000562768 - | 1 - | 0,000470488 - | 1 - | 0,000473631 - |
| 2688 | 1 | 0.2um_NOKCPHJN_567304 | Eukaryota | unclassified | Pelagophycea | Pelagomonad | unclassified | Aureococcus | Aureococcus a | unclassified | unclassified | unclassified | unclassified | unclassified | 9 | 0,632455532 | 0,002270052 | 0,000303067 | 9 - | 0,002694535 - | 9 - | 0,002252697 - | 10 - | 0,002519718 - |
| 2689 | 1 | 3um_FIJJAFBF_444691 | Eukaryota | unclassified | unclassified | Isochrysidales | Noelaerhabda | Emiliania | Emiliania huxl | metabolism | Carbohydrate | COG1850 | RuBisCO catal | 45157.CMV01 | 4,2 | 0,748331477 | 0,000532598 | 5,18E-05 | 3 - | 0,000460132 - | 4 - | 0,000512909 - | 4 - | 0,000516336 - |
| 2690 | 1 | 3um_FIJJAFBF_661387 | Bacteria | Bacteroidetes | Flavobacteriia | Flavobacterial | Flavobacteriac | Aquimarina | Aquimarina at | cellular proces | Cell wall/mem | 099US | Membrane bo | 391603.FBALC | 0,4 | 0,489897949 | 5,99E-05 | 7,37E-05 | 0 - | 0 - | 0 - | 0 - | 1 - | 0,000160696 - |
| 2691 | 1 | 0.2um_NOKCPHJN_807274 | Bacteria | Proteobacteri | Gammaprotec | Alteromonada | Pseudoalteror | Pseudoalteror | Pseudoalteror | cellular proces | Cell wall/mem | 0QTSX | Peptidoglycan | 722419.PH505 | 0,2 | 0,4 | 5,83E-05 | 0,000116512 | 0 - | 0 - | 0 - | 0 - | 0 - | 0 - |
| 2692 | 1 | 3um_FIJJAFBF_111317 | Viruses | unclassified | unclassified | unclassified | Mimiviridae | unclassified | Yellowstone la | unclassified | unclassified | unclassified | unclassified | unclassified | 1,6 | 0,489897949 | 0,0004969 | 0,000166646 | 2 - | 0,000737421 - | 1 - | 0,000308251 - | 1 - | 0,00031031 - |
| 2693 | 1 | 0.2um_NOKCPHJN_182716 | Eukaryota | Phaeophyceae | unclassified | Ectocarpales | Ectocarpaceae | Ectocarpus | Ectocarpus sili | cellular proces | Cytoskeleton | COG5245 | heavy chain | 653948.CCA1 | 0,2 | 0,4 | 2,34E-06 | 4,67E-06 | 0 - | 0 - | 0 - | 0 - | 0 - | 0 - |
| 2694 | 1 | 0.2um_NOKCPHJN_42076 | Eukaryota | unclassified | Oomycetes | Peronosporale | Peronosporac | Phytophthora | Phytophthora | unclassified | unclassified | unclassified | unclassified | unclassified | 0,2 | 0,4 | 6,40E-06 | 1,28E-05 | 1 - | 3,20E-05 - | 0 - | 0 - | 0 - | 0 - |
| 2695 | 1 | 3um_FIJJAFBF_95704 | Eukaryota | unclassified | Raphidophyce | Chattonellales | Chattonellace | Heterosigma | Heterosigma a | unclassified | unclassified | unclassified | unclassified | unclassified | 32 | 2,19089023 | 0,003785329 | 0,000180313 | 29 - | 0,004103209 - | 31 - | 0,003666961 - | 31 - | 0,003691458 - |
| 2696 | 1 | 0.2um_NOKCPHJN_611548 | Eukaryota | Phaeophyceae | unclassified | Ectocarpales | Ectocarpaceae | Ectocarpus | Ectocarpus sili | unclassified | unclassified | unclassified | unclassified | unclassified | 3,8 | 0,4 | 0,000411433 | 6,38E-05 | 4 - | 0,000514421 - | 3 - | 0,000322551 - | 4 - | 0,000432941 - |
| 2697 | 1 | 3um_FIJJAFBF_422288 | unclassified | unclassified | unclassified | unclassified | unclassified | unclassified | unclassified | unclassified | unclassified | unclassified | unclassified | unclassified | 0,4 | 0,489897949 | 0,000156911 | 0,000192559 | 0 - | 0 - | 0 - | 0 - | 0 - | 0 - |
| 2698 | 1 | 3um_FIJJAFBF_498429 | Bacteria | Proteobacteri | Gammaprotec | Arenicellales | Arenicellaceae | Arenicella | Arenicella xan | metabolism | Inorganic ion t | 177TK | phosphate-sel | 455436.GHTC | 3,2 | 0,979795897 | 0,000594124 | 0,000189073 | 3 - | 0,000662373 - | 5 - | 0,000922934 - | 2 - | 0,00037164 - |
| 2699 | 1 | 0.2um_NOKCPHJN_271556 | Eukaryota | Apicomplexa | Conoidasida | Eucoccidiorida | Eimeriidae | Eimeria | Eimeria praec | unclassified | unclassified | unclassified | unclassified | unclassified | 1,8 | 0,748331477 | 0,000248628 | 8,50E-05 | 1 - | 0,00016934 - | 2 - | 0,000283144 - | 2 - | 0,000285036 - |
| 2700 | 1 | 3um_FIJJAFBF_247107 | Eukaryota | Bacillariophyt | Bacillariophyc | Naviculales | Phaeodactylac | Phaeodactylu | Phaeodactylu | unclassified | unclassified | unclassified | unclassified | unclassified | 0,2 | 0,4 | 1,71E-05 | 3,42E-05 | 0 - | 0 - | 0 - | 0 - | 1 - | 8,55E-05 - |
| 2701 | 1 | 3um_FIJJAFBF_58623 | unclassified | unclassified | unclassified | unclassified | unclassified | unclassified | unclassified | unclassified | unclassified | unclassified | unclassified | unclassified | 3,8 | 0,4 | 0,00149971 | 0,000247475 | 4 - | 0,001871205 - | 4 - | 0,001564373 - | 4 - | 0,001574823 - |
| 2702 | 1 | 3um_FIJJAFBF_543287 | Bacteria | Bacteroidetes | Flavobacteriia | Flavobacterial | Flavobacteriac | Lacinutrix | Lacinutrix him | information st | Replication, re | 0PB0H | DNA-binding p | 983544.Lacal_ | 3,4 | 1,019803903 | 0,002332564 | 0,000620834 | 2 - | 0,001663293 - | 5 - | 0,003476384 - | 3 - | 0,002099765 - |
| 2703 | 1 | 3um_FIJJAFBF_376071 | Bacteria | Proteobacteri | Alphaproteob | Rhodobactera | Rhodobactera | Pseudorueger | Pseudorueger | unclassified | unclassified | unclassified | unclassified | unclassified | 1 | 0 | 0,000349957 | 3,70E-05 | 1 - | 0,000415823 - | 1 - | 0,000347638 - | 1 - | 0,000349961 - |
| 2704 | 1 | 0.2um_NOKCPHJN_401188 | Eukaryota | Apicomplexa | Aconoidasida | Haemosporida | Plasmodiidae | Plasmodium | Plasmodium fa | unclassified | unclassified | unclassified | unclassified | unclassified | 0,8 | 0,4 | 0,000311072 | 0,00016088 | 1 - | 0,000462026 - | 1 - | 0,000386265 - | 0 - | 0 - |
| 2705 | 1 | 0.2um_NOKCPHJN_744302 | Eukaryota | unclassified | unclassified | Isochrysidales | Noelaerhabda | Emiliania | Emiliania huxl | metabolism | Amino acid tra | 0HW4R | Glutaminase | 7217.FBpp011 | 0,4 | 0,489897949 | 2,95E-05 | 3,63E-05 | 0 - | 0 - | 1 - | 7,89E-05 - | 0 - | 0 - |
| 2706 | 1 | 0.2um_NOKCPHJN_702864 | Eukaryota | unclassified | Oomycetes | Pythiales | Pythiaceae | Pythium | Pythium insidi | unclassified | unclassified | unclassified | unclassified | unclassified | 2,2 | 0,4 | 0,000403771 | 4,94E-05 | 2 - | 0,000441582 - | 2 - | 0,000369174 - | 2 - | 0,00037164 - |
| 2707 | 1 | 0.2um_NOKCPHJN_521968 | Eukaryota | Chordata | Mammalia | Rodentia | Cricetidae | Mesocricetus | Mesocricetus | unclassified | unclassified | unclassified | unclassified | unclassified | 1 | 0 | 3,86E-05 | 4,09E-06 | 1 - | 4,59E-05 - | 1 - | 3,84E-05 - | 1 - | 3,86E-05 - |
| 2708 | 1 | 3um_FIJJAFBF_37426 | Bacteria | Proteobacteri | Alphaproteob | Rhodobactera | Rhodobactera | Loktanella | Loktanella sp. | metabolism | Energy produc | 0259B | 2-oxoacid deh | 991905.SL003 | 1 | 0 | 0,000196851 | 2,08E-05 | 1 - | 0,000233901 - | 1 - | 0,000195547 - | 1 - | 0,000196853 - |
| 2709 | 1 | 3um_FIJJAFBF_670384 | Eukaryota | unclassified | Dinophyceae | Peridiniales | Kryptoperidini | Kryptoperidini | Kryptoperidin | unclassified | unclassified | unclassified | unclassified | unclassified | 0,4 | 0,489897949 | 0,000173553 | 0,000212982 | 0 - | 0 - | 0 - | 0 - | 0 - | 0 - |
| 2710 | 1 | 0.2um_NOKCPHJN_364146 | Eukaryota | Ascomycota | Sordariomycet | Hypocreales | Ophiocordycip | Drechmeria | Drechmeria co | cellular proces | Nuclear struct | 19I79 | SRP40, C-term | 4932.YKR092C | 0,2 | 0,4 | 1,25E-05 | 2,49E-05 | 1 - | 6,24E-05 - | 0 - | 0 - | 0 - | 0 - |
| 2711 | 1 | 0.2um_NOKCPHJN_183124 | Viruses | unclassified | unclassified | unclassified | Phycodnavirid | unclassified | Aureococcus a | unclassified | unclassified | unclassified | unclassified | unclassified | 0,4 | 0,489897949 | 1,43E-05 | 1,76E-05 | 0 - | 0 - | 0 - | 0 - | 0 - | 0 - |
| 2712 | 1 | 3um_FIJJAFBF_95716 | Eukaryota | unclassified | Raphidophyce | Chattonellales | Chattonellace | Heterosigma | Heterosigma a | cellular proces | Posttranslatio | 0IEKR | One of the cor | 35128.Thapsd | 12,8 | 0,748331477 | 0,001755485 | 0,000161879 | 12 - | 0,001961088 - | 12 - | 0,001639518 - | 14 - | 0,001925548 - |
| 2713 | 1 | 3um_FIJJAFBF_369630 | Bacteria | Proteobacteri | Alphaproteob | Rhodobactera | Rhodobactera | Thalassobium | Thalassobium | unclassified | unclassified | unclassified | unclassified | unclassified | 0,6 | 0,489897949 | 6,04E-05 | 5,04E-05 | 1 - | 0,000117871 - | 1 - | 9,85E-05 - | 0 - | 0 - |
| 2714 | 1 | 0.2um_NOKCPHJN_404821 | Eukaryota | unclassified | Pelagophycea | Pelagomonad | unclassified | Aureococcus | Aureococcus a | cellular proces | Signal transdu | KOG0613 | fibronectin typ | 4787.PITG_01 | 0,2 | 0,4 | 4,94E-06 | 9,87E-06 | 0 - | 0 - | 0 - | 0 - | 1 - | 2,47E-05 - |
| 2715 | 1 | 3um_FIJJAFBF_116531 | Bacteria | Proteobacteri | Alphaproteob | unclassified | unclassified | Candidatus Pu | Candidatus Pu | metabolism | Carbohydrate | 0IVA1 | Photosystem I | 35128.Thapsd | 8,4 | 1,019803903 | 0,002722764 | 9,46E-05 | 7 - | 0,00272884 - | 8 - | 0,002607288 - | 8 - | 0,002624706 - |
| 2716 | 1 | 0.2um_NOKCPHJN_612933 | unclassified | unclassified | unclassified | unclassified | unclassified | unclassified | unclassified | unclassified | unclassified | unclassified | unclassified | unclassified | 0,2 | 0,4 | 9,63E-06 | 1,93E-05 | 0 - | 0 - | 0 - | 0 - | 0 - | 0 - |
| 2717 | 2 | 3um_FIJJAFBF_41903;3um_FIJJAFBF_5 | Bacteria | Bacteroidetes | Flavobacteriia | Flavobacterial | Flavobacteriac | Aquimarina | Aquimarina sp | poorly charact | Function unkn | 0PAPS | NA | 873517.HMPR | 1 | 0 | 0,000419948 | 4,44E-05 | 1 - | 0,000498988 - | 1 - | 0,000417166 - | 1 - | 0,000419953 - |
| 2718 | 1 | 3um_FIJJAFBF_567619 | Eukaryota | Bacillariophyt | Bacillariophyc | Naviculales | Naviculaceae | Fistulifera | Fistulifera sola | cellular proces | Signal transdu | COG5040 | Tyrosine 3-mc | 2850.Phatr56 | 1,8 | 0,4 | 0,001181008 | 0,000202951 | 1 - | 0,000796257 - | 2 - | 0,001331381 - | 2 - | 0,001340275 - |
| 2719 | 2 | 3um_FIJJAFBF_497643;3um_FIJJAFBF_ | Viruses | unclassified | unclassified | unclassified | Phycodnavirid | unclassified | Chrysochromu | cellular proces | Cell cycle cont | KOG0987 | DNA helicase | 44689.DDB02 | 0,8 | 0,4 | 0,000108841 | 5,63E-05 | 1 - | 0,000161659 - | 1 - | 0,000135151 - | 0 - | 0 - |
| 2720 | 1 | 3um_FIJJAFBF_411070 | Eukaryota | unclassified | Pelagophycea | Pelagomonad | unclassified | Aureococcus | Aureococcus a | unclassified | unclassified | unclassified | unclassified | unclassified | 1 | 0 | 0,000492126 | 5,20E-05 | 1 - | 0,000584751 - | 1 - | 0,000488867 - | 1 - | 0,000492132 - |
| 2721 | 1 | 0.2um_NOKCPHJN_331675 | Bacteria | Proteobacteri | Gammaprotec | Cellvibrionales | Halieaceae | Halioglobus | Halioglobus lu | poorly charact | Function unkn | 05EC1 | AtP-binding pr | 313624.N9414 | 0,2 | 0,4 | 9,40E-06 | 1,88E-05 | 0 - | 0 - | 0 - | 0 - | 0 - | 0 - |
| 2722 | 1 | 0.2um_NOKCPHJN_101547 | Bacteria | Bacteroidetes | Flavobacteriia | Flavobacterial | Flavobacteriac | Polaribacter | Polaribacter a | unclassified | unclassified | unclassified | unclassified | unclassified | 1 | 0 | 8,26E-05 | 8,73E-06 | 1 - | 9,81E-05 - | 1 - | 8,20E-05 - | 1 - | 8,26E-05 - |

| | | | | | | | | | | | | | | | | | | | | | | | | |
|---|---|---|---|---|---|---|---|---|---|---|---|---|---|---|---|---|---|---|---|---|---|---|---|---|
| 2723 | 1 | 3um_FIJJAFBF_501038 | Bacteria | Bacteroidetes | Flavobacteriia | Flavobacterial | Flavobacteriac | Flavobacteriu | Flavobacteriu | cellular proces | Cell wall/mem | 06B97 | phosphate-sel | 886377.Murru | 0,4 | 0,489897949 | 6,35E-05 | 7,79E-05 | 0 - | 0 - | 0 - | 0 - | 0 - | 0 - |
| 2724 | 1 | 3um_FIJJAFBF_56880 | Bacteria | Proteobacteri | Alphaproteob | Rhodobactera | Rhodobactera | Sulfitobacter | Sulfitobacter r | metabolism | Lipid transpor | 01SS3 | Dehydrogenas | 52598.EE36_1 | 1 | 0 | 0,000629922 | 6,66E-05 | 1 - | 0,000748482 - | 1 - | 0,000625749 - | 1 - | 0,000629929 - |
| 2725 | 1 | 0.2um_NOKCPHJN_552431 | Bacteria | Proteobacteri | Betaproteoba | Nitrosomonad | Sterolibacteri | Methyloversat | Methyloversat | unclassified | unclassified | unclassified | unclassified | unclassified | 0,4 | 0,489897949 | 4,70E-05 | 5,78E-05 | 0 - | 0 - | 1 - | 0,000125652 - | 0 - | 0 - |
| 2726 | 1 | 0.2um_NOKCPHJN_775240 | Eukaryota | unclassified | unclassified | Acytosteliales | Cavendeniacea | Cavendenia | Cavendenia fa | cellular proces | Cytoskeleton | 0S68Z | Inherit from a | 7460.GB11509 | 0,4 | 0,489897949 | 2,73E-05 | 3,36E-05 | 0 - | 0 - | 1 - | 7,30E-05 - | 0 - | 0 - |
| 2727 | 1 | 3um_FIJJAFBF_166373 | Eukaryota | Bacillariophyta | Bacillariophyc | Bacillariales | Bacillariaceae | Fragilariopsis | Fragilariopsis | cellular proces | Posttranslatio | 0XRV1 | UnCoupling Pr | 2850.Phatr359 | 0,2 | 0,4 | 3,78E-05 | 7,57E-05 | 0 - | 0 - | 0 - | 0 - | 1 - | 0,000189168 - |
| 2728 | 1 | 3um_FIJJAFBF_610024 | Eukaryota | Bacillariophyta | Coscinodiscop | Thalassiosirale | Thalassiosirac | Thalassiosira | Thalassiosira p | unclassified | unclassified | unclassified | unclassified | unclassified | 1 | 0 | 0,000133458 | 1,41E-05 | 1 - | 0,000158577 - | 1 - | 0,000132574 - | 1 - | 0,00013346 - |
| 2729 | 7 | 0.2um_NOKCPHJN_108163;0.2um_NO | Bacteria | Proteobacteri | Gammaprotec | Cellvibrionales | Porticoccacea | unclassified | various | information st | Replication, re | 05XQP | Phage integras | 314287.GB220 | 0,2 | 0,4 | 4,19E-05 | 8,37E-05 | 0 - | 0 - | 0 - | 0 - | 0 - | 0 - |
| 2730 | 1 | 0.2um_NOKCPHJN_205368 | Eukaryota | unclassified | Cryptophyta | Pyrenomonad | Geminigerace | Guillardia | Guillardia thet | poorly charact | Function unkn | 0IHFE | P4Hc | 486041.XP_00 | 0,4 | 0,489897949 | 4,66E-05 | 5,71E-05 | 0 - | 0 - | 0 - | 0 - | 0 - | 0 - |
| 2731 | 1 | 0.2um_NOKCPHJN_109945 | Eukaryota | unclassified | unclassified | Acytosteliales | Acytosteliacea | Acytostelium | Acytostelium s | unclassified | unclassified | unclassified | unclassified | unclassified | 0,8 | 0,4 | 3,33E-05 | 1,72E-05 | 1 - | 4,94E-05 - | 1 - | 4,13E-05 - | 0 - | 0 - |
| 2732 | 1 | 0.2um_NOKCPHJN_130339 | Eukaryota | Basidiomycota | Agaricomycete | Corticiales | Punctulariacea | Punctularia | Punctularia st | unclassified | unclassified | unclassified | unclassified | unclassified | 0,6 | 0,489897949 | 7,08E-05 | 5,83E-05 | 1 - | 0,000132241 - | 1 - | 0,000110556 - | 1 - | 0,000111295 - |
| 2733 | 1 | 3um_FIJJAFBF_358211 | Eukaryota | Chlorophyta | Mamiellophyc | Mamiellales | Mamiellaceae | Micromonas | Micromonas c | unclassified | unclassified | unclassified | unclassified | unclassified | 1 | 0 | 8,83E-05 | 9,34E-06 | 1 - | 0,000104976 - | 1 - | 8,78E-05 - | 1 - | 8,83E-05 - |
| 2734 | 1 | 0.2um_NOKCPHJN_1104834 | Eukaryota | Ascomycota | unclassified | unclassified | unclassified | Saitoella | Saitoella comp | unclassified | unclassified | unclassified | unclassified | unclassified | 0,2 | 0,4 | 3,25E-05 | 6,50E-05 | 0 - | 0 - | 0 - | 0 - | 0 - | 0 - |
| 2735 | 1 | 0.2um_NOKCPHJN_751162 | Eukaryota | Streptophyta | Marchantiopsi | Marchantiales | Marchantiacea | Marchantia | Marchantia po | cellular proces | Posttranslatio | COG0652 | peptidyl-proly | 4787.PITG_05 | 0,4 | 0,489897949 | 0,000279378 | 0,000342849 | 0 - | 0 - | 0 - | 0 - | 0 - | 0 - |
| 2736 | 1 | 0.2um_NOKCPHJN_335007 | Eukaryota | Ascomycota | Dothideomyce | Pleosporales | Corynesporas | Corynespora | Corynespora c | unclassified | unclassified | unclassified | unclassified | unclassified | 0,2 | 0,4 | 5,95E-06 | 1,19E-05 | 0 - | 0 - | 0 - | 0 - | 0 - | 0 - |
| 2737 | 1 | 3um_FIJJAFBF_53717 | Bacteria | Proteobacteri | Alphaproteob | Rhodobactera | Rhodobactera | Halocynthiibac | Halocynthiiba | metabolism | Inorganic ion t | 16PII | TRAP transpor | 391624.OIHEL | 0,6 | 0,489897949 | 0,000127968 | 0,000106879 | 1 - | 0,000252865 - | 0 - | 0 - | 0 - | 0 - |
| 2738 | 1 | 0.2um_NOKCPHJN_12461 | Bacteria | Bacteroidetes | Flavobacteriia | Flavobacterial | Flavobacteriac | Formosa | Formosa sp. H | unclassified | unclassified | unclassified | unclassified | unclassified | 2 | 0,632455532 | 0,001123757 | 0,000337232 | 1 - | 0,000680438 - | 2 - | 0,001137726 - | 3 - | 0,001717989 - |
| 2739 | 1 | 3um_FIJJAFBF_323987 | Bacteria | Proteobacteri | Gammaprotec | Alteromonada | Alteromonada | Aestuariibacte | Aestuariibacte | poorly charact | Function unkn | 0YV5D | NA | 203122.Sde_3 | 0,8 | 0,4 | 0,000309676 | 0,000160136 | 1 - | 0,000459191 - | 0 - | 0 - | 1 - | 0,00038646 - |
| 2740 | 2 | 3um_FIJJAFBF_553966;3um_FIJJAFBF_ | Eukaryota | Chlorophyta | Mamiellophyc | Monomastigal | Monomastiga | Monomastix | Monomastix s | metabolism | Energy produc | 0SSA5 | NADH dehydr | 7029.ACYPI00 | 1 | 0 | 0,000323037 | 3,42E-05 | 1 - | 0,000383837 - | 1 - | 0,000320897 - | 1 - | 0,000323041 - |
| 2741 | 1 | 0.2um_NOKCPHJN_361605 | Eukaryota | unclassified | unclassified | Isochrysidales | Noelaerhabda | Emiliania | Emiliania huxl | cellular proces | Posttranslatio | 0JF0E | P4Hc | 65071.PYU1_T | 1 | 0 | 0,000160285 | 1,70E-05 | 1 - | 0,000190453 - | 1 - | 0,000159224 - | 1 - | 0,000160287 - |
| 2742 | 1 | 0.2um_NOKCPHJN_461595 | Eukaryota | Arthropoda | Insecta | Hemiptera | Pentatomidae | Halyomorpha | Halyomorpha | poorly charact | Function unkn | 13MW6 | Family with se | 6085.XP_0021 | 0,6 | 0,489897949 | 6,87E-05 | 5,68E-05 | 1 - | 0,000129945 - | 0 - | 0 - | 1 - | 0,000109363 - |
| 2743 | 1 | 0.2um_NOKCPHJN_157840 | unclassified | unclassified | unclassified | unclassified | unclassified | unclassified | unclassified | unclassified | unclassified | unclassified | unclassified | unclassified | 0,8 | 0,4 | 6,74E-05 | 3,48E-05 | 1 - | 0,000100064 - | 1 - | 8,37E-05 - | 0 - | 0 - |
| 2744 | 1 | 0.2um_NOKCPHJN_990519 | Eukaryota | unclassified | Pelagophycea | Pelagomonada | unclassified | Aureococcus | Aureococcus a | poorly charact | Function unkn | 1DU18 | Inherit from e | 3055.EDP0909 | 0,4 | 0,489897949 | 3,93E-05 | 4,84E-05 | 0 - | 0 - | 1 - | 0,000105168 - | 0 - | 0 - |
| 2745 | 1 | 3um_FIJJAFBF_524441 | Bacteria | Bacteroidetes | Flavobacteriia | Flavobacterial | Flavobacteriac | Formosa | Formosa sp. H | unclassified | unclassified | unclassified | unclassified | unclassified | 0,4 | 0,489897949 | 0,000133765 | 0,000164481 | 0 - | 0 - | 1 - | 0,000357571 - | 0 - | 0 - |
| 2746 | 1 | 0.2um_NOKCPHJN_330840 | Eukaryota | unclassified | Cryptophyta | Pyrenomonad | Geminigerace | Guillardia | Guillardia thet | metabolism | Carbohydrate | COG1850 | RuBisCO catal | 35128.Thapsd | 2 | 0,632455532 | 0,000483578 | 0,000134237 | 2 - | 0,000582476 - | 2 - | 0,000486964 - | 1 - | 0,000245109 - |
| 2747 | 1 | 3um_FIJJAFBF_734139 | Eukaryota | Streptophyta | Liliopsida | Poales | Poaceae | Dichanthelium | Dichanthelium | unclassified | unclassified | unclassified | unclassified | unclassified | 1 | 0 | 0,000313394 | 3,31E-05 | 1 - | 0,000372379 - | 1 - | 0,000311318 - | 1 - | 0,000313398 - |
| 2748 | 1 | 3um_FIJJAFBF_278309 | Eukaryota | unclassified | Florideophyce | Batrachosperr | Batrachosperr | Kumanoa | Kumanoa ame | metabolism | Energy produc | COG0055 | Produces ATP | 7739.JGI12576 | 0,6 | 0,8 | 7,37E-05 | 0,000100315 | 0 - | 0 - | 0 - | 0 - | 0 - | 0 - |
| 2749 | 1 | 0.2um_NOKCPHJN_827194 | Eukaryota | unclassified | Oomycetes | Peronosporale | Peronosporac | Phytophthora | Phytophthora | cellular proces | Posttranslatio | KOG0019 | heat shock pro | 67593.Physo1 | 1 | 0 | 9,83E-05 | 1,04E-05 | 1 - | 0,000116768 - | 1 - | 9,76E-05 - | 1 - | 9,83E-05 - |
| 2750 | 1 | 3um_FIJJAFBF_284488 | Bacteria | Planctomycete | Planctomyceti | Planctomyceta | Planctomyceta | Fuerstia | Fuerstia maris | unclassified | unclassified | unclassified | unclassified | unclassified | 0,4 | 0,489897949 | 4,32E-05 | 5,28E-05 | 0 - | 0 - | 1 - | 0,000107517 - | 1 - | 0,000108235 - |
| 2751 | 1 | 3um_FIJJAFBF_87014 | Bacteria | Proteobacteri | Alphaproteob | Rhodospirillale | Rhodospirillac | Azospirillum | Azospirillum b | unclassified | unclassified | unclassified | unclassified | unclassified | 1,6 | 0,489897949 | 6,39E-05 | 1,62E-05 | 1 - | 4,86E-05 - | 2 - | 8,13E-05 - | 1 - | 4,09E-05 - |
| 2752 | 1 | 3um_FIJJAFBF_515657 | Bacteria | Proteobacteri | Alphaproteob | Rhodobactera | Rhodobactera | Ruegeria | Ruegeria sp. T | unclassified | unclassified | unclassified | unclassified | unclassified | 7,2 | 0,4 | 0,001451958 | 0,000237374 | 8 - | 0,00190696 - | 7 - | 0,001394982 - | 7 - | 0,001404301 - |
| 2753 | 1 | 0.2um_NOKCPHJN_39209 | Eukaryota | unclassified | Pelagophycea | Pelagomonada | unclassified | Aureoumbra | Aureoumbra l | metabolism | Energy produc | COG0055 | Produces ATP | 489825.LYNGI | 36,6 | 2,727636339 | 0,004806627 | 0,000464312 | 34 - | 0,005323929 - | 33 - | 0,004320026 - | 40 - | 0,005271376 - |
| 2754 | 2 | 3um_FIJJAFBF_151896;3um_FIJJAFBF_ | various | various | various | various | various | various | various | unclassified | unclassified | unclassified | unclassified | unclassified | 1 | 0 | 0,000252981 | 2,68E-05 | 1 - | 0,000300595 - | 1 - | 0,000251305 - | 1 - | 0,000252984 - |
| 2755 | 1 | 0.2um_NOKCPHJN_966897 | Eukaryota | unclassified | unclassified | Isochrysidales | Noelaerhabda | Emiliania | Emiliania huxl | unclassified | unclassified | unclassified | unclassified | unclassified | 1 | 0 | 0,000223377 | 2,36E-05 | 1 - | 0,000265419 - | 1 - | 0,000221897 - | 1 - | 0,000223379 - |
| 2756 | 1 | 0.2um_NOKCPHJN_380154 | Eukaryota | unclassified | Oomycetes | Pythiales | Pythiaceae | Pythium | Pythium insidi | information st | Translation, ri | KOG3184 | Ribosomal pro | 65071.PYU1_T | 1 | 0 | 0,000285032 | 3,01E-05 | 1 - | 0,00033868 - | 1 - | 0,000283144 - | 1 - | 0,000285036 - |
| 2757 | 1 | 0.2um_NOKCPHJN_177386 | Bacteria | Bacteroidetes | unclassified | unclassified | unclassified | unclassified | Bacteroidetes | information st | Transcription | COG0557 | 3'-5' exoribon | 709991.Odosp | 0,8 | 0,4 | 7,19E-05 | 3,71E-05 | 1 - | 0,000105569 - | 1 - | 8,83E-05 - | 1 - | 8,88E-05 - |
| 2758 | 2 | 0.2um_NOKCPHJN_308951;3um_FIJJAF | Eukaryota | various | various | various | various | various | various | unclassified | unclassified | unclassified | unclassified | unclassified | 0,2 | 0,4 | 1,22E-05 | 2,43E-05 | 0 - | 0 - | 0 - | 0 - | 0 - | 0 - |
| 2759 | 1 | 3um_FIJJAFBF_20711 | Eukaryota | unclassified | unclassified | Prymnesiales | Chrysochromu | Chrysochromu | Chrysochromu | unclassified | unclassified | unclassified | unclassified | unclassified | 1,8 | 0,748331477 | 0,000642652 | 0,000219833 | 1 - | 0,000437709 - | 2 - | 0,00073187 - | 2 - | 0,00073676 - |
| 2760 | 1 | 3um_FIJJAFBF_639895 | Bacteria | Proteobacteri | Gammaprotec | Oceanospirilla | various | various | various | poorly charact | Function unkn | COG3517 | type VI secreti | 83406.HDN1F | 0,2 | 0,4 | 2,27E-05 | 4,54E-05 | 0 - | 0 - | 0 - | 0 - | 0 - | 0 - |
| 2761 | 1 | 3um_FIJJAFBF_12518 | unclassified | unclassified | unclassified | unclassified | unclassified | unclassified | unclassified | unclassified | unclassified | unclassified | unclassified | unclassified | 0,4 | 0,489897949 | 0,000193462 | 0,000237886 | 0 - | 0 - | 1 - | 0,000517148 - | 0 - | 0 - |
| 2762 | 1 | 0.2um_NOKCPHJN_387795 | Eukaryota | Basidiomycota | Exobasidiomy | Microstromat | unclassified | Pseudomicros | Pseudomicros | unclassified | unclassified | unclassified | unclassified | unclassified | 0,2 | 0,4 | 4,72E-06 | 9,45E-06 | 0 - | 0 - | 1 - | 2,36E-05 - | 0 - | 0 - |
| 2763 | 1 | 0.2um_NOKCPHJN_157490 | Bacteria | Proteobacteri | Alphaproteob | Rhodobactera | Hyphomonada | Robiginitomac | Robiginitomac | unclassified | unclassified | unclassified | unclassified | unclassified | 0,4 | 0,489897949 | 2,85E-05 | 3,50E-05 | 0 - | 0 - | 0 - | 0 - | 0 - | 0 - |
| 2764 | 1 | 3um_FIJJAFBF_103096 | Eukaryota | unclassified | unclassified | Isochrysidales | Noelaerhabda | Emiliania | Emiliania huxl | unclassified | unclassified | unclassified | unclassified | unclassified | 5 | 0,632455532 | 0,001395844 | 0,000238788 | 5 - | 0,001655933 - | 6 - | 0,001661281 - | 5 - | 0,001393649 - |
| 2765 | 1 | 0.2um_NOKCPHJN_171095 | Eukaryota | Streptophyta | unclassified | Lamiales | Phrymaceae | Erythranthe | Erythranthe g | cellular proces | Cell cycle cont | 1BD3I | K06627 cyclin | 4577.GRMZM | 0,4 | 0,489897949 | 7,10E-05 | 8,78E-05 | 1 - | 0,000196969 - | 0 - | 0 - | 0 - | 0 - |
| 2766 | 1 | 3um_FIJJAFBF_331695 | Bacteria | Bacteroidetes | Flavobacteriia | Flavobacterial | Crocinitomica | unclassified | Crocinitomica | poorly charact | Function unkn | 09AR5 | Inherit from b | 313606.M231 | 1 | 0 | 0,000121372 | 1,28E-05 | 1 - | 0,000144216 - | 1 - | 0,000120568 - | 1 - | 0,000121374 - |
| 2767 | 1 | 3um_FIJJAFBF_715074 | Eukaryota | unclassified | Pelagophycea | Pelagomonada | unclassified | Pelagomonas | uncultured Pe | unclassified | unclassified | unclassified | unclassified | unclassified | 1,6 | 0,489897949 | 0,000807426 | 0,000239759 | 1 - | 0,000603614 - | 2 - | 0,001009273 - | 2 - | 0,001016015 - |
| 2768 | 1 | 3um_FIJJAFBF_177625 | Bacteria | Bacteroidetes | Flavobacteriia | Flavobacterial | Flavobacteriac | Tamlana | Tamlana sp. U | cellular proces | Cell wall/mem | 05C6Y | Udp-glucose 6 | 391598.FBBAL | 1 | 0 | 0,000136052 | 1,44E-05 | 1 - | 0,000161659 - | 1 - | 0,000135151 - | 1 - | 0,000136054 - |
| 2769 | 7 | 0.2um_NOKCPHJN_1041616;0.2um_N( | Bacteria | Bacteroidetes | Flavobacteriia | Flavobacterial | Flavobacteriac | various | various | unclassified | unclassified | unclassified | unclassified | unclassified | 0,8 | 0,748331477 | 0,000124735 | 0,000111999 | 0 - | 0 - | 0 - | 0 - | 1 - | 0,000169336 - |
| 2770 | 1 | 0.2um_NOKCPHJN_695286 | Bacteria | Proteobacteri | Alphaproteob | Rhodospirillale | Rhodospirillac | Candidatus En | Candidatus En | unclassified | unclassified | unclassified | unclassified | unclassified | 1 | 0 | 0,000258165 | 2,73E-05 | 1 - | 0,000306755 - | 1 - | 0,000256455 - | 1 - | 0,000258168 - |

| | | | | | | | | | | | | | | | | | | | | | | | | |
|---|---|---|---|---|---|---|---|---|---|---|---|---|---|---|---|---|---|---|---|---|---|---|---|---|
| 2723 | 1 | 3um_FIJJAFBF_501038 | Bacteria | Bacteroidetes | Flavobacteriia | Flavobacterial | Flavobacteriac | Flavobacteriu | Flavobacteriu | cellular proces | Cell wall/mem | 06B97 | phosphate-sel | 886377.Murru | 0,4 | 0,489897949 | 6,35E-05 | 7,79E-05 | 0 - | 0 - | 0 - | 0 - | 0 - | 0 - |
| 2724 | 1 | 3um_FIJJAFBF_56880 | Bacteria | Proteobacteri | Alphaproteob | Rhodobactera | Rhodobactera | Sulfitobacter | Sulfitobacter r | metabolism | Lipid transpor | 01SS3 | Dehydrogenas | 52598.EE36_1 | 1 | 0 | 0,000629922 | 6,66E-05 | 1 - | 0,000748482 - | 1 - | 0,000625749 - | 1 - | 0,000629929 - |
| 2725 | 1 | 0.2um_NOKCPHJN_552431 | Bacteria | Proteobacteri | Betaproteoba | Nitrosomonad | Sterolibacteri | Methyloversa | Methyloversa | unclassified | unclassified | unclassified | unclassified | unclassified | 0,4 | 0,489897949 | 4,70E-05 | 5,78E-05 | 0 - | 0 - | 1 - | 0,000125652 - | 0 - | 0 - |
| 2726 | 1 | 0.2um_NOKCPHJN_775240 | Eukaryota | unclassified | unclassified | Acytosteliales | Cavenderiacea | Cavenderia | Cavenderia fa | cellular proces | Cytoskeleton | 0S68Z | Inherit from a | 7460.GB1150 | 0,4 | 0,489897949 | 2,73E-05 | 3,36E-05 | 0 - | 0 - | 1 - | 7,30E-05 - | 0 - | 0 - |
| 2727 | 1 | 3um_FIJJAFBF_166373 | Eukaryota | Bacillariophyt | Bacillariophyc | Bacillariales | Bacillariaceae | Fragilariopsis | Fragilariopsis | cellular proces | Posttranslatio | 0XRV1 | UnCoupling Pr | 2850.Phatr35 | 0,2 | 0,4 | 3,78E-05 | 7,57E-05 | 0 - | 0 - | 0 - | 0 - | 1 - | 0,000189168 - |
| 2728 | 1 | 3um_FIJJAFBF_610024 | Eukaryota | Bacillariophyt | Coscinodiscop | Thalassiosirale | Thalassiosirac | Thalassiosira | Thalassiosira p | unclassified | unclassified | unclassified | unclassified | unclassified | 1 | 0 | 0,000133458 | 1,41E-05 | 1 - | 0,000158577 - | 1 - | 0,000132574 - | 1 - | 0,00013346 - |
| 2729 | 7 | 0.2um_NOKCPHJN_108163;0.2um_NO | Bacteria | Proteobacteri | Gammaprotec | Cellvibrionales | Porticoccacea | unclassified | various | information st | Replication, re | 05XQP | Phage integra | 314287.GB22 | 0,2 | 0,4 | 4,19E-05 | 8,37E-05 | 0 - | 0 - | 0 - | 0 - | 0 - | 0 - |
| 2730 | 1 | 0.2um_NOKCPHJN_205368 | Eukaryota | unclassified | Cryptophyta | Pyrenomonad | Geminigerace | Guillardia | Guillardia thet | poorly charact | Function unkn | 0IHFE | P4Hc | 486041.XP_00 | 0,4 | 0,489897949 | 4,66E-05 | 5,71E-05 | 0 - | 0 - | 0 - | 0 - | 0 - | 0 - |
| 2731 | 1 | 0.2um_NOKCPHJN_109945 | Eukaryota | unclassified | unclassified | Acytosteliales | Acytosteliacea | Acytostelium | Acytostelium | unclassified | unclassified | unclassified | unclassified | unclassified | 0,8 | 0,4 | 3,33E-05 | 1,72E-05 | 1 - | 4,94E-05 - | 1 - | 4,13E-05 - | 0 - | 0 - |
| 2732 | 1 | 0.2um_NOKCPHJN_130339 | Eukaryota | Basidiomycota | Agaricomycete | Corticiales | Punctulariacea | Punctularia | Punctularia st | unclassified | unclassified | unclassified | unclassified | unclassified | 0,6 | 0,489897949 | 7,08E-05 | 5,83E-05 | 1 - | 0,000132241 - | 1 - | 0,000110556 - | 1 - | 0,000111295 - |
| 2733 | 1 | 3um_FIJJAFBF_358211 | Eukaryota | Chlorophyta | Mamiellophyc | Mamiellales | Mamiellaceae | Micromonas | Micromonas c | unclassified | unclassified | unclassified | unclassified | unclassified | 1 | 0 | 8,83E-05 | 9,34E-06 | 1 - | 0,000104976 - | 1 - | 8,78E-05 - | 1 - | 8,83E-05 - |
| 2734 | 1 | 0.2um_NOKCPHJN_1104834 | Eukaryota | Ascomycota | unclassified | unclassified | unclassified | Saitoella | Saitoella comp | unclassified | unclassified | unclassified | unclassified | unclassified | 0,2 | 0,4 | 3,25E-05 | 6,50E-05 | 0 - | 0 - | 0 - | 0 - | 0 - | 0 - |
| 2735 | 1 | 0.2um_NOKCPHJN_751162 | Eukaryota | Streptophyta | Marchantiopsi | Marchantiales | Marchantiace | Marchantia | Marchantia pc | cellular proces | Posttranslatio | COG0652 | peptidyl-proly | 4787.PITG_05 | 0,4 | 0,489897949 | 0,000279378 | 0,000342849 | 0 - | 0 - | 0 - | 0 - | 0 - | 0 - |
| 2736 | 1 | 0.2um_NOKCPHJN_335007 | Eukaryota | Ascomycota | Dothideomyce | Pleosporales | Corynesporas | Corynespora | Corynespora c | unclassified | unclassified | unclassified | unclassified | unclassified | 0,2 | 0,4 | 5,95E-06 | 1,19E-05 | 0 - | 0 - | 0 - | 0 - | 0 - | 0 - |
| 2737 | 1 | 3um_FIJJAFBF_53717 | Bacteria | Proteobacteri | Alphaproteob | Rhodobactera | Rhodobactera | Halocynthiiba | Halocynthiiba | metabolism | Inorganic ion t | 16PII | TRAP transpor | 391624.OIHEL | 0,6 | 0,489897949 | 0,000127968 | 0,000106879 | 1 - | 0,000252865 - | 0 - | 0 - | 0 - | 0 - |
| 2738 | 1 | 0.2um_NOKCPHJN_12461 | Bacteria | Bacteroidetes | Flavobacteriia | Flavobacterial | Flavobacteriac | Formosa | Formosa sp. H | unclassified | unclassified | unclassified | unclassified | unclassified | 2 | 0,632455532 | 0,001123757 | 0,000337232 | 1 - | 0,000680438 - | 2 - | 0,001137726 - | 3 - | 0,001717989 - |
| 2739 | 1 | 3um_FIJJAFBF_323987 | Bacteria | Proteobacteri | Gammaprotec | Alteromonada | Alteromonada | Aestuariibacte | Aestuariibacte | poorly charact | Function unkn | 0YV5D | NA | 203122.Sde_3 | 0,8 | 0,4 | 0,000309676 | 0,000160136 | 1 - | 0,000459191 - | 0 - | 0 - | 1 - | 0,00038646 - |
| 2740 | 2 | 3um_FIJJAFBF_553966;3um_FIJJAFBF_ | Eukaryota | Chlorophyta | Mamiellophyc | Monomastigal | Monomastiga | Monomastix | Monomastix s | metabolism | Energy produc | 0SSA5 | NADH dehydr | 7029.ACYPI00 | 1 | 0 | 0,000323037 | 3,42E-05 | 1 - | 0,000383837 - | 1 - | 0,000320897 - | 1 - | 0,000323041 - |
| 2741 | 1 | 0.2um_NOKCPHJN_361605 | Eukaryota | unclassified | unclassified | Isochrysidales | Noelaerhabda | Emiliania | Emiliania huxl | cellular proces | Posttranslatio | 0JF0E | P4Hc | 65071.PYU1_T | 1 | 0 | 0,000160285 | 1,70E-05 | 1 - | 0,000190453 - | 1 - | 0,000159224 - | 1 - | 0,000160287 - |
| 2742 | 1 | 0.2um_NOKCPHJN_461595 | Eukaryota | Arthropoda | Insecta | Hemiptera | Pentatomidae | Halyomorpha | Halyomorpha | poorly charact | Function unkn | 13MW6 | Family with se | 6085.XP_0021 | 0,6 | 0,489897949 | 6,87E-05 | 5,68E-05 | 1 - | 0,000129945 - | 0 - | 0 - | 1 - | 0,000109363 - |
| 2743 | 1 | 0.2um_NOKCPHJN_157840 | unclassified | unclassified | unclassified | unclassified | unclassified | unclassified | unclassified | unclassified | unclassified | unclassified | unclassified | unclassified | 0,8 | 0,4 | 6,74E-05 | 3,48E-05 | 1 - | 0,000100064 - | 1 - | 8,37E-05 - | 0 - | 0 - |
| 2744 | 1 | 0.2um_NOKCPHJN_990519 | Eukaryota | unclassified | Pelagophycea | Pelagomonad | unclassified | Aureococcus | Aureococcus a | poorly charact | Function unkn | 1DU18 | Inherit from e | 3055.EDP0909 | 0,4 | 0,489897949 | 3,93E-05 | 4,84E-05 | 0 - | 0 - | 1 - | 0,000105168 - | 0 - | 0 - |
| 2745 | 1 | 3um_FIJJAFBF_524441 | Bacteria | Bacteroidetes | Flavobacteriia | Flavobacterial | Flavobacteriac | Formosa | Formosa sp. H | unclassified | unclassified | unclassified | unclassified | unclassified | 0,4 | 0,489897949 | 0,000133765 | 0,000164481 | 0 - | 0 - | 1 - | 0,000357571 - | 0 - | 0 - |
| 2746 | 1 | 0.2um_NOKCPHJN_330840 | Eukaryota | unclassified | Cryptophyta | Pyrenomonad | Geminigerace | Guillardia | Guillardia thet | metabolism | Carbohydrate | COG1850 | RuBisCO catal | 35128.Thapsd | 2 | 0,632455532 | 0,000483578 | 0,000134237 | 2 - | 0,000582476 - | 2 - | 0,000486964 - | 1 - | 0,000245109 - |
| 2747 | 1 | 3um_FIJJAFBF_734139 | Eukaryota | Streptophyta | Liliopsida | Poales | Poaceae | Dichanthelium | Dichanthelium | unclassified | unclassified | unclassified | unclassified | unclassified | 1 | 0 | 0,000313394 | 3,31E-05 | 1 - | 0,000372379 - | 1 - | 0,000311318 - | 1 - | 0,000313398 - |
| 2748 | 1 | 3um_FIJJAFBF_278309 | Eukaryota | unclassified | Florideophyce | Batrachosperr | Batrachosperr | Kumanoa | Kumanoa ame | metabolism | Energy produc | COG0055 | Produces ATP | 7739.JGI1257 | 0,6 | 0,8 | 7,37E-05 | 0,000100315 | 0 - | 0 - | 0 - | 0 - | 0 - | 0 - |
| 2749 | 1 | 0.2um_NOKCPHJN_827194 | Eukaryota | unclassified | Oomycetes | Peronosporale | Peronosporac | Phytophthora | Phytophthora | cellular proces | Posttranslatio | KOG0019 | heat shock pr | 67593.Physo1 | 1 | 0 | 9,83E-05 | 1,04E-05 | 1 - | 0,000116768 - | 1 - | 9,76E-05 - | 1 - | 9,83E-05 - |
| 2750 | 1 | 3um_FIJJAFBF_284488 | Bacteria | Planctomycete | Planctomyceti | Planctomycet | Planctomycet | Fuerstia | Fuerstia maris | unclassified | unclassified | unclassified | unclassified | unclassified | 0,4 | 0,489897949 | 4,32E-05 | 5,28E-05 | 0 - | 0 - | 1 - | 0,000107517 - | 1 - | 0,000108235 - |
| 2751 | 1 | 3um_FIJJAFBF_87014 | Bacteria | Proteobacteri | Alphaproteob | Rhodospirillal | Rhodospirillac | Azospirillum | Azospirillum b | unclassified | unclassified | unclassified | unclassified | unclassified | 1,6 | 0,489897949 | 6,39E-05 | 1,62E-05 | 1 - | 4,86E-05 - | 2 - | 8,13E-05 - | 1 - | 4,09E-05 - |
| 2752 | 1 | 3um_FIJJAFBF_515657 | Bacteria | Proteobacteri | Alphaproteob | Rhodobactera | Rhodobactera | Ruegeria | Ruegeria sp. T | unclassified | unclassified | unclassified | unclassified | unclassified | 7,2 | 0,4 | 0,001451958 | 0,000237374 | 8 - | 0,00190696 - | 7 - | 0,001394982 - | 7 - | 0,001404301 - |
| 2753 | 1 | 0.2um_NOKCPHJN_39209 | Eukaryota | unclassified | Pelagophycea | Pelagomonad | unclassified | Aureoumbra | Aureoumbra l | metabolism | Energy produc | COG0055 | Produces ATP | 489825.LYNG | 36,6 | 2,727636339 | 0,004806627 | 0,000464312 | 34 - | 0,005323929 - | 33 - | 0,004320026 - | 40 - | 0,005271376 - |
| 2754 | 2 | 3um_FIJJAFBF_151896;3um_FIJJAFBF_ | various | various | various | various | various | various | various | unclassified | unclassified | unclassified | unclassified | unclassified | 1 | 0 | 0,000252981 | 2,68E-05 | 1 - | 0,000300595 - | 1 - | 0,000251305 - | 1 - | 0,000252984 - |
| 2755 | 1 | 0.2um_NOKCPHJN_966897 | Eukaryota | unclassified | unclassified | Isochrysidales | Noelaerhabda | Emiliania | Emiliania huxl | unclassified | unclassified | unclassified | unclassified | unclassified | 1 | 0 | 0,000223377 | 2,36E-05 | 1 - | 0,000265419 - | 1 - | 0,000221897 - | 1 - | 0,000223379 - |
| 2756 | 1 | 0.2um_NOKCPHJN_380154 | Eukaryota | unclassified | Oomycetes | Pythiales | Pythiaceae | Pythium | Pythium insidi | information st | Translation, ri | KOG3184 | Ribosomal prc | 65071.PYU1_T | 1 | 0 | 0,000285032 | 3,01E-05 | 1 - | 0,00033868 - | 1 - | 0,000283144 - | 1 - | 0,000285036 - |
| 2757 | 1 | 0.2um_NOKCPHJN_177386 | Bacteria | Bacteroidetes | unclassified | unclassified | unclassified | unclassified | Bacteroidetes | information st | Transcription | COG0557 | 3'-5' exoribon | 709991.Odosp | 0,8 | 0,4 | 7,19E-05 | 3,71E-05 | 1 - | 0,000105569 - | 1 - | 8,83E-05 - | 1 - | 8,88E-05 - |
| 2758 | 2 | 0.2um_NOKCPHJN_308951;3um_FIJJAF | Eukaryota | various | various | various | various | various | various | unclassified | unclassified | unclassified | unclassified | unclassified | 0,2 | 0,4 | 1,22E-05 | 2,43E-05 | 0 - | 0 - | 0 - | 0 - | 0 - | 0 - |
| 2759 | 1 | 3um_FIJJAFBF_20711 | Eukaryota | unclassified | unclassified | Prymnesiales | Chrysochromu | Chrysochromu | Chrysochromu | unclassified | unclassified | unclassified | unclassified | unclassified | 1,8 | 0,748331477 | 0,000642652 | 0,000219833 | 1 - | 0,000437709 - | 2 - | 0,00073187 - | 2 - | 0,00073676 - |
| 2760 | 1 | 3um_FIJJAFBF_639895 | Bacteria | Proteobacteri | Gammaprotec | Oceanospirilla | various | various | various | poorly charact | Function unkn | COG3517 | type VI secreti | 83406.HDN1F | 0,2 | 0,4 | 2,27E-05 | 4,54E-05 | 0 - | 0 - | 0 - | 0 - | 0 - | 0 - |
| 2761 | 1 | 3um_FIJJAFBF_12518 | unclassified | unclassified | unclassified | unclassified | unclassified | unclassified | unclassified | unclassified | unclassified | unclassified | unclassified | unclassified | 0,4 | 0,489897949 | 0,000193462 | 0,000237886 | 0 - | 0 - | 1 - | 0,000517148 - | 0 - | 0 - |
| 2762 | 1 | 0.2um_NOKCPHJN_387795 | Eukaryota | Basidiomycota | Exobasidiomy | Microstromat | unclassified | Pseudomicros | Pseudomicros | unclassified | unclassified | unclassified | unclassified | unclassified | 0,2 | 0,4 | 4,72E-06 | 9,45E-06 | 0 - | 0 - | 1 - | 2,36E-05 - | 0 - | 0 - |
| 2763 | 1 | 0.2um_NOKCPHJN_157490 | Bacteria | Proteobacteri | Alphaproteob | Rhodobactera | Hyphomonada | Robiginitomac | Robiginitomac | unclassified | unclassified | unclassified | unclassified | unclassified | 0,4 | 0,489897949 | 2,85E-05 | 3,50E-05 | 0 - | 0 - | 0 - | 0 - | 0 - | 0 - |
| 2764 | 1 | 3um_FIJJAFBF_103096 | Eukaryota | unclassified | unclassified | Isochrysidales | Noelaerhabda | Emiliania | Emiliania huxl | unclassified | unclassified | unclassified | unclassified | unclassified | 5 | 0,632455532 | 0,001395844 | 0,000238788 | 5 - | 0,001655933 - | 6 - | 0,001661281 - | 5 - | 0,001393649 - |
| 2765 | 1 | 0.2um_NOKCPHJN_171095 | Eukaryota | Streptophyta | unclassified | Lamiales | Phrymaceae | Erythranthe | Erythranthe g | cellular proces | Cell cycle cont | 1BD3I | K06627 cyclin | 4577.GRMZM | 0,4 | 0,489897949 | 7,10E-05 | 8,78E-05 | 1 - | 0,000196969 - | 0 - | 0 - | 0 - | 0 - |
| 2766 | 1 | 3um_FIJJAFBF_331695 | Bacteria | Bacteroidetes | Flavobacteriia | Flavobacterial | Crocinitomica | unclassified | Crocinitomica | poorly charact | Function unkn | 09AR5 | Inherit from b | 313606.M231 | 1 | 0 | 0,000121372 | 1,28E-05 | 1 - | 0,000144216 - | 1 - | 0,000120568 - | 1 - | 0,000121374 - |
| 2767 | 1 | 3um_FIJJAFBF_715074 | Eukaryota | unclassified | Pelagophycea | Pelagomonad | unclassified | Pelagomonas | uncultured Pe | unclassified | unclassified | unclassified | unclassified | unclassified | 1,6 | 0,489897949 | 0,000807426 | 0,000239759 | 1 - | 0,000603614 - | 2 - | 0,001009273 - | 2 - | 0,001016015 - |
| 2768 | 1 | 3um_FIJJAFBF_177625 | Bacteria | Bacteroidetes | Flavobacteriia | Flavobacterial | Flavobacteriac | Tamlana | Tamlana sp. U | cellular proces | Cell wall/mem | 05C6Y | Udp-glucose 6 | 391598.FBBAL | 1 | 0 | 0,000136052 | 1,44E-05 | 1 - | 0,000161659 - | 1 - | 0,000135151 - | 1 - | 0,000136054 - |
| 2769 | 7 | 0.2um_NOKCPHJN_1041616;0.2um_N | Bacteria | Bacteroidetes | Flavobacteriia | Flavobacterial | Flavobacteriac | various | various | unclassified | unclassified | unclassified | unclassified | unclassified | 0,8 | 0,748331477 | 0,000124735 | 0,000111999 | 0 - | 0 - | 0 - | 0 - | 1 - | 0,000169336 - |
| 2770 | 1 | 0.2um_NOKCPHJN_695286 | Bacteria | Proteobacteri | Alphaproteob | Rhodospirillal | Rhodospirillac | Candidatus En | Candidatus En | unclassified | unclassified | unclassified | unclassified | unclassified | 1 | 0 | 0,000258165 | 2,73E-05 | 1 - | 0,000306755 - | 1 - | 0,000256455 - | 1 - | 0,000258168 - |

**Taxonomic assignment of all identified proteins on all filters (0.2, 3, 10 µm) using Prophane software tool**

**PROPHANE JOB INFORMATION**

**program versions**

prophane: 3.1.1
blast+: 2.7.1
diamond: 0.8.22
hmmer: 3.1b2
krona: 2.7

**tasks**

task: *taxonomic assignment*
task comment: tax_from_nr_20180808_qcover90
algorithm: diamond blastp
query: sequences without taxonomic classification (missing_taxa.faa)
database: NCBI NR database
database version: 2018-08-02
database comment: downloaded at 2018-08-02
database scope: tax
parameter:
- evalue 0.01
- query-cover 0.9
- max-target-seqs 1

| kingdom | Percentage on filter - kingdom level | | | phylum | Percentage on filter - phylum level | | | class | Percentage on filter - class level | | | order | Percentage on filter - order level | | | family | Percentage on filter - family level | | |
|---|---|---|---|---|---|---|---|---|---|---|---|---|---|---|---|---|---|---|---|
| | 0.2 µm | 3 µm | 10 µm | | 0.2 µm | 3 µm | 10 µm | | 0.2 µm | 3 µm | 10 µm | | 0.2 µm | 3 µm | 10 µm | | 0.2 µm | 3 µm | 10 µm |
| Bacteria | 85 | 45 | 31 | Proteobacteria | 55 | 41 | 39 | Gammaproteobacteria | 53 | 50 | 50 | Oceanospirillales | 43 | 11 | 12 | Saccharospirillaceae | 78 | 28 | 4 |
| | | | | | | | | | | | | | | | | Oceanospirillaceae | 19 | 48 | 65 |
| | | | | | | | | | | | | | | | | Halomonadaceae | 1 | 0 | 7 |
| | | | | | | | | | | | | | | | | unclassified | 1 | 3 | 1 |
| | | | | | | | | | | | | | | | | various | 0,5 | 2 | 0,3 |
| | | | | | | | | | | | | | | | | Alcanivoraceae | 0,3 | 0,7 | 1 |
| | | | | | | | | | | | | | | | | Oleiphilaceae | 0,3 | 0 | 2 |
| | | | | | | | | | | | | | | | | Kangiellaceae | 0,2 | 0 | 6 |
| | | | | | | | | | | | | | | | | Endozoicomonadaceae | 0,2 | 15 | 11 |
| | | | | | | | | | | | | | | | | Hahellaceae | 0,004 | 3 | 0,2 |
| | | | | | | | | | | | | Cellvibrionales | 20 | 33 | 37 | Porticoccaceae | 67 | 26 | 16 |
| | | | | | | | | | | | | | | | | Spongibacteraceae | 10 | 31 | 39 |
| | | | | | | | | | | | | | | | | Halieaceae | 9 | 18 | 26 |
| | | | | | | | | | | | | | | | | unclassified | 8 | 5 | 7 |
| | | | | | | | | | | | | | | | | Cellvibrionaceae | 4 | 10 | 7 |
| | | | | | | | | | | | | | | | | Microbulbiferaceae | 1 | 5 | 4 |
| | | | | | | | | | | | | | | | | various | 0,2 | 4 | 0,5 |
| | | | | | | | | | | | | Alteromonadales | 17 | 24 | 22 | Alteromonadaceae | 79 | 52 | 39 |
| | | | | | | | | | | | | | | | | Colwelliaceae | 9 | 19 | 24 |
| | | | | | | | | | | | | | | | | Pseudoalterononadaceae | 5 | 15 | 15 |
| | | | | | | | | | | | | | | | | Psychromonadaceae | 3 | 1 | 0,9 |
| | | | | | | | | | | | | | | | | unclassified | 2 | 2 | 9 |
| | | | | | | | | | | | | | | | | various | 1 | 6 | 8 |
| | | | | | | | | | | | | | | | | Ferrimonadaceae | 0,8 | 0,5 | 0 |
| | | | | | | | | | | | | | | | | Shewanellaceae | 0,6 | 0 | 0 |
| | | | | | | | | | | | | | | | | Idiomarinaceae | 0,3 | 4 | 4 |
| | | | | | | | | | | | | unclassified | 14 | 15 | 11 | unclassified | 100 | 94 | 94 |
| | | | | | | | | | | | | | | | | Competibacteraceae | 0,4 | 6 | 6 |
| | | | | | | | | | | | | Pseudomonadales | 2 | 10 | 7 | Pseudomonadaceae | 60 | 84 | 94 |
| | | | | | | | | | | | | | | | | Moraxellaceae | 40 | 16 | 6 |
| | | | | | | | | | | | | | | | | various | 0,6 | 0 | 0 |
| | | | | | | | | | | | | Arenicellales | 7 | 2 | 2 | Arenicellaceae | 100 | 100 | 100 |
| | | | | | | | | | | | | Chromatiales | 0,7 | 2 | 2 | unclassified | 42 | 0 | 0 |
| | | | | | | | | | | | | | | | | Ectothiorhodopiraceae | 27 | 53 | 45 |
| | | | | | | | | | | | | | | | | Chromatiaceae | 22 | 42 | 53 |
| | | | | | | | | | | | | | | | | Granulosicoccaceae | 10 | 5 | 2 |
| | | | | | | | | | | | | Vibrionales | 0,3 | 0,3 | 0,1 | Vibrionaceae | 100 | 100 | 100 |
| | | | | | | | | | | | | Thiotrichales | 0,3 | 0,2 | 0,1 | Thiotrichaceae | 48 | 0 | 14 |
| | | | | | | | | | | | | | | | | Piscirickettsiaceae | 43 | 100 | 77 |
| | | | | | | | | | | | | | | | | Francisellaceae | 0 | 0 | 9 |
| | | | | | | | | | | | | | | | | Thiolinaceae | 4 | 0 | 0 |
| | | | | | | | | | | | | | | | | Fastidiosibacteracea | 4 | 0 | 0 |
| | | | | | | | | | | | | various | 0,2 | 0,7 | 0,2 | various | 100 | 100 | 100 |
| | | | | | | | | | | | | Xanthomonadales | 0,1 | 0,6 | 2 | Rhodanobacteraceae | 64 | 0 | 79 |
| | | | | | | | | | | | | | | | | Xanthomonadaceae | 36 | 100 | 21 |
| | | | | | | | | | | | | Enterobacterales | 0,08 | 1 | 3 | Enterobacteriaceae | 84 | 49 | 70 |
| | | | | | | | | | | | | | | | | Yersiniaceae | 16 | 51 | 30 |
| | | | | | | | | | | | | Aeromonadales | 0,07 | 0,3 | 0 | Aeromonadaceae | 100 | 100 | 0 |
| | | | | | | | | | | | | Nevskiales | 0,07 | 0,3 | 0,03 | Sinobacteraceae | 97 | 0 | 0 |
| | | | | | | | | | | | | | | | | Algiphilaceae | 3 | 100 | 100 |
| | | | | | | | | | | | | Methylococcales | 0,03 | 0,2 | 0 | Metylococcaceae | 100 | 100 | 0 |
| | | | | | | | | | | | | Pasteurellales | 0,03 | 0,01 | 0 | Pasteurellaceae | 100 | 100 | 0 |
| | | | | | | | | | | | | Legionellales | 0,004 | 0 | 0 | Legionellaceae | 100 | 0 | 0 |
| | | | | | | | | Alphaproteobacteria | 40 | 42 | 41 | Rhodobacterales | 63 | 73 | 49 | Rhodobacteraceae | 77 | 78 | 82 |
| | | | | | | | | | | | | | | | | unclassified | 19 | 11 | 10 |
| | | | | | | | | | | | | | | | | various | 2 | 6 | 2 |
| | | | | | | | | | | | | | | | | Hyphomonadaceae | 2 | 4 | 5 |
| | | | | | | | | | | | | Pelagibacterales | 24 | 8 | 9 | Pelagibacteriaceae | 94 | 96 | 89 |
| | | | | | | | | | | | | | | | | unclassified | 3 | 4 | 11 |
| | | | | | | | | | | | | | | | | various | 2 | 0 | 0 |
| | | | | | | | | | | | | Rhodospirillales | 4 | 9 | 10 | Rhodospirilaceae | 91 | 92 | 92 |
| | | | | | | | | | | | | | | | | unclassified | 9 | 8 | 8 |
| | | | | | | | | | | | | unclassified | 3 | 19 | 29 | unclassified | 100 | 100 | 100 |
| | | | | | | | | | | | | Rhizobiales | 2 | 3 | 3 | unclassified | 72 | 71 | 41 |
| | | | | | | | | | | | | | | | | Hyphomicrobiaceae | 9 | 3 | 0 |
| | | | | | | | | | | | | | | | | Phyllobacteriaceae | 7 | 15 | 4 |
| | | | | | | | | | | | | | | | | Aurantimonadaceae | 4 | 0,6 | 0 |
| | | | | | | | | | | | | | | | | Bradyrhizobiaceae | 3 | 1 | 0 |
| | | | | | | | | | | | | | | | | Cohaesibacteraceae | 3 | 0 | 0 |
| | | | | | | | | | | | | | | | | Rhizobiaceae | 0,7 | 2 | 6 |
| | | | | | | | | | | | | | | | | Methylocystaceae | 0,1 | 0 | 0 |
| | | | | | | | | | | | | | | | | Methylobacteriaceae | 0,06 | 7 | 49 |
| | | | | | | | | | | | | various | 2 | 0,8 | 1 | various | 100 | 100 | 100 |
| | | | | | | | | | | | | Rickettsiales | 0,7 | 2 | 1 | unclassified | 97 | 91 | 84 |
| | | | | | | | | | | | | | | | | Canidatus Midichloriaceae | 2 | 6 | 13 |
| | | | | | | | | | | | | | | | | Anaplasmataceae | 0,8 | 4 | 3 |

| | | | | | | | | | | | | | | | |
|---|---|---|---|---|---|---|---|---|---|---|---|---|---|---|---|
| | | | | | | | | Sphingomonadales | 0,4 | 1 | 2 | Sphingomonadaceae | 86 | 33 | 16 |
| | | | | | | | | | | | | Erythrobacteraceae | 6 | 46 | 74 |
| | | | | | | | | | | | | unclassified | 6 | 0 | 3 |
| | | | | | | | | | | | | various | 3 | 21 | 7 |
| | | | | | | | | Kiloniellales | 0,1 | 0 | 0 | Kiloniellaceae | 100 | 0 | 0 |
| | | | | | | | | Kordiimonadales | 0,1 | 0 | 0 | unclassified | 100 | 0 | 0 |
| | | | | | | | | Caulobacterales | 0,06 | 0 | 0 | Caulobacteraceae | 81 | 0 | 0 |
| | | | | | | | | | | | | unclassified | 19 | 0 | 0 |
| | | | | | | | | Holosporales | 0,01 | 0 | 0 | Holosporaceae | 100 | 0 | |
| | | | | Betaproteobacteria | 7 | 5 | 5 | unclassified | 60 | 65 | 53 | unclassified | 100 | 100 | 100 |
| | | | | | | | | Burkholderiales | 34 | 14 | 37 | Comamondaceae | 78 | 80 | 58 |
| | | | | | | | | | | | | unclassified | 17 | 0 | 2 |
| | | | | | | | | | | | | various | 2 | 0 | 0 |
| | | | | | | | | | | | | Alcaligenaceae | 1 | 0 | 15 |
| | | | | | | | | | | | | Burkholderiaceae | 0,9 | 2 | 16 |
| | | | | | | | | | | | | Oxalobacteraceae | 0,4 | 17 | 9 |
| | | | | | | | | Nitrosomonadales | 4 | | 17 | unclassified | 68 | 65 | 78 |
| | | | | | | | | | | | | Thiobacillaceae | 11 | 0 | 5 |
| | | | | | | | | | | | | Methylophilaceae | 9 | 2 | 0 |
| | | | | | | | | | | | | Nitrosomonadaceae | 9 | 0 | 7 |
| | | | | | | | | | | | | Sterolobacteriaceae | 3 | 33 | 10 |
| | | | | | | | | | | | | Gallionellaceae | 0,2 | 0 | 0 |
| | | | | | | | | various | 2 | 4 | 0 | various | 100 | 100 | 100 |
| | | | | | | | | Ferrovales | 0 | 0 | 2 | Ferrovaceae | 0 | 0 | 100 |
| | | | | | | | | Rhodocyclales | 0,4 | 0 | 0 | Zoogloeaceae | 80 | 0 | 0 |
| | | | | | | | | | | | | Rhodocclaceae | 20 | 0 | 0 |
| | | | | | | | | Neisseriales | 0,4 | 0 | 0,3 | Neisseriaceae | 89 | 0 | 0 |
| | | | | | | | | | | | | Chromobacteriaceae | 11 | 0 | 0 |
| | | | | various | 0,3 | 0,2 | 0,2 | various | 100 | 100 | 100 | various | 100 | 0 | 0 |
| | | | | Deltaproteobacteria | 0,2 | 0,7 | 2 | Myxococcales | 47 | 82 | 75 | Nannocystaceae | 69 | 87 | 70 |
| | | | | | | | | | | | | Archangiaceae | 14 | 0 | 0 |
| | | | | | | | | | | | | Kofleriaceae | 10 | 0 | 0 |
| | | | | | | | | | | | | Anaeromyxobacteraceae | 4 | 0 | 1 |
| | | | | | | | | | | | | unclassified | 2 | 0 | 6 |
| | | | | | | | | | | | | Myxococcaceae | 0 | 13 | 0 |
| | | | | | | | | | | | | Polyangiaceae | 0 | 0 | 23 |
| | | | | | | | | unclassified | 31 | 13 | 23 | unclassified | 100 | 100 | 100 |
| | | | | | | | | Desulfobacterales | 13 | 0 | 2 | Desulfobulbaceae | 78 | 0 | 0 |
| | | | | | | | | | | | | Desulfobacteraceae | 22 | 0 | 100 |
| | | | | | | | | Desulfuromonadales | 6 | 0 | 0 | Geobacteraceae | 94 | 0 | 0 |
| | | | | | | | | | | | | Desulfuromonadaceae | 6 | 0 | 0 |
| | | | | | | | | Syntrophobacterales | 3 | 5 | 0 | Syntrophobacteraceae | 100 | 100 | 0 |
| | | | | unclassified | 0,1 | 2 | 1 | unclassified | 100 | 100 | 100 | unclassified | 100 | 100 | 100 |
| | | | | Epsilonproteobacteria | 0,05 | 0,05 | 0,7 | Campylobacterales | 55 | 72 | 100 | Campylobacteraceae | 65 | 84 | 89 |
| | | | | | | | | | | | | Helicobacteraceae | 35 | 16 | 11 |
| | | | | | | | | Nautiliales | 45 | 28 | 0 | Nautiliaceae | 100 | 100 | 0 |
| | | | | Oligoflexia | 0,006 | 0 | 0,01 | Bacteriovoracales | 100 | 0 | 0 | Bacteriovoracaceae | 100 | 0 | 0 |
| | | | | | | | | Oligoflexiales | 0 | 0 | 100 | Pseudobacteriovoracaceae | 0 | 0 | 100 |
| | | | | Zetaproteobacteria | 0,006 | 0 | 0 | unclassified | 100 | 0 | 0 | unclassified | 100 | 0 | 0 |
| | | | | Acidithiobacillia | 0,003 | 0 | 0 | Acidithiobacillales | 100 | 0 | 0 | unclassified | 100 | 0 | 0 |
| | | | | Canidatus Lambdaproteobact | 0 | 0 | 0,06 | unclassified | 0 | 0 | 100 | unclassified | 0 | 0 | 100 |
| Bacteroidetes | 40 | 48 | 47 | Flavobacteriia | 83 | 89 | 89 | Flavobacteriales | 90 | 98 | 99 | Flavobacteriaceae | 92 | 87 | 90 |
| | | | | | | | | | | | | Crocinitomicaceae | 6 | 10 | 7 |
| | | | | | | | | | | | | unclassified | 1 | 2 | 1 |
| | | | | | | | | | | | | Cryomorphaceae | 0,9 | 1 | 0,7 |
| | | | | | | | | | | | | various | 0,5 | 0,1 | 0 |
| | | | | | | | | | | | | Schleiferiaceae | 0,02 | 0 | 0 |
| | | | | | | | | unclassified | 6 | 0,9 | 0,2 | unclassified | 100 | 100 | 100 |
| | | | | | | | | various | 4 | 1 | 1 | various | 100 | 100 | 100 |
| | | | | unclassified | 9 | 4 | 4 | unclassified | 99 | 100 | 100 | unclassified | 100 | 100 | 100 |
| | | | | | | | | Bacteroidetes Order ll. Inc | 0,8 | 0 | 0 | Rhodothermaceae | 100 | 0 | 0 |
| | | | | Cytophagia | 7 | 4 | 4 | Cytophagales | 100 | 100 | 100 | Flammeovigaceae | 87 | 97 | 90 |
| | | | | | | | | | | | | Cytophagaceae | 9 | 2 | 5 |
| | | | | | | | | | | | | Cyclobacteriaceae | 2 | 0 | 0,3 |
| | | | | | | | | | | | | Catalimonadaceae | 0,7 | 0 | 0,5 |
| | | | | | | | | | | | | unclassified | 0,7 | 0 | 0 |
| | | | | | | | | | | | | Hymenobacteraceae | 0,4 | 0 | 3 |
| | | | | | | | | | | | | Microscillaceae | 0,2 | 0 | 0 |
| | | | | | | | | | | | | Bernardetiaceae | 0,03 | 0 | 0 |
| | | | | | | | | | | | | Amoebophilaceae | 0 | 0,5 | 0,7 |
| | | | | | | | | | | | | Persicobacteraceae | 0,02 | 0,2 | 1 |
| | | | | Saprospira | 0,5 | 2 | 2 | Saprospirales | 100 | 100 | 100 | Lewinellaceae | 64 | 16 | 14 |
| | | | | | | | | | | | | Saprospiraceae | 26 | 75 | 77 |
| | | | | | | | | | | | | Haliscomenobacteraceae | 10 | 9 | 10 |
| | | | | various | 0,3 | 0,7 | 0,2 | various | 100 | 100 | 100 | various | 100 | 100 | 100 |
| | | | | Bacteroidia | 0,1 | 0,3 | 0,4 | Marinilabiliales | 47 | 62 | 47 | unclassified | 63 | 8 | 0 |
| | | | | | | | | | | | | Prolixibacteraceae | 21 | 58 | 62 |
| | | | | | | | | | | | | Salinivirgaceae | 14 | 0 | 0 |

| | | | | | | | | | | | | | | | |
|---|---|---|---|---|---|---|---|---|---|---|---|---|---|---|---|
| | | | | | | | | | | | | Marinilabiliaceae | 2 | 34 | 38 |
| | | | | | | | | Bacteroidales | 38 | 38 | 53 | Porphyromonadaceae | 72 | 0 | 0 |
| | | | | | | | | | | | | Prevotellaceae | 24 | 100 | 100 |
| | | | | | | | | | | | | Bacteroidaceae | 4 | 0 | 0 |
| | | | | | | | | various | 15 | 0 | 0 | various | 100 | 0 | 0 |
| | | | | Sphingobacteriia | 0,07 | 0,04 | 0,3 | Sphingobacteriales | 100 | 0 | 100 | Sphingobacteriaceae | 100 | 0 | 100 |
| | | | | | | | | unclassified | 0 | 100 | 0 | unclassified | 0 | 100 | 0 |
| | | | | Chitinopagia | 0,01 | 0 | 0 | Chitinophagales | 100 | 0 | 0 | Chitinophagales | 100 | 0 | 0 |
| Actinobacteria | 2 | 1 | 2 | Actinobacteria | 96 | 69 | 64 | Micrococcales | 90 | 31 | 44 | Microbacteriaceae | 99 | 86 | 47 |
| | | | | | | | | | | | | Demequinaceae | 0,3 | 0 | 0 |
| | | | | | | | | | | | | Intrasporangiaceae | 0,3 | 14 | 53 |
| | | | | | | | | | | | | Cellulomonadaceae | 0,1 | 0 | 0 |
| | | | | | | | | Actinomycetales | 4 | 15 | 9 | unclassified | 100 | 100 | 100 |
| | | | | | | | | Streptosporagiales | 2 | 0 | 0 | Streptosporangiaceae | 100 | 0 | 0 |
| | | | | | | | | unclassified | 2 | 2 | 6 | unclassified | 100 | 100 | 100 |
| | | | | | | | | Bifidobacteriales | 1 | 4 | 6 | Bifidobacteriaceae | 100 | 100 | 100 |
| | | | | | | | | Corynebacteriales | 1 | 3 | 6 | Mycobacteriaceae | 50 | 100 | 100 |
| | | | | | | | | | | | | Corynebacteriaceae | 34 | 0 | 0 |
| | | | | | | | | | | | | Dietziaceae | 10 | 0 | 0 |
| | | | | | | | | | | | | Nocardiaceae | 6 | 0 | 0 |
| | | | | | | | | various | 0,9 | 0 | 0 | various | 100 | 0 | 0 |
| | | | | | | | | Streptomycetales | 0,3 | 43 | 28 | Streptomycetaceae | 100 | 100 | 100 |
| | | | | | | | | Jiangellales | 0,2 | 0 | 0 | Jiangellales | 100 | 0 | 0 |
| | | | | | | | | Nakamurellales | 0,1 | 0 | 0 | Nakamurellaceae | 100 | 0 | 0 |
| | | | | | | | | Pseudonocardiales | 0,007 | 0 | 0,7 | Pseudonocardiaceae | 100 | 0 | 100 |
| | | | | | | | | Micromonosporales | 0 | 2 | 0,5 | Micromonosporaceae | 0 | 100 | 100 |
| | | | | unclassified | 4 | 20 | 30 | unclassified | 100 | 100 | 100 | unclassified | 100 | 100 | 100 |
| | | | | various | 0,2 | 0 | 0 | various | 100 | 0 | 0 | various | 100 | 0 | 0 |
| | | | | Acidimicrobiia | 0,08 | 11 | 5 | Acidimicrobiales | 100 | 100 | 100 | Acidimicrobiaceae | 100 | 100 | 84 |
| | | | | | | | | | | | | Microthrixaceae | 0 | 0 | 9 |
| | | | | | | | | | | | | unclassified | 0 | 0 | 7 |
| | | | | Thermophilia | 0 | 0 | 0,6 | Solirubrobacterales | 0 | 0 | 100 | Patulibacteraceae | 0 | 0 | 100 |
| unclassified | 1 | 2 | 3 | unclassified | 100 | 100 | 100 | unclassified | 100 | 100 | 100 | unclassified | 100 | 100 | 100 |
| various | 0,7 | 1 | 1 | various | 100 | 100 | 100 | various | 100 | 100 | 100 | various | 100 | 100 | 100 |
| Verrucomicrobia | 0,5 | 3 | 2 | Opitutae | 75 | 67 | 67 | unclassified | 79 | 85 | 83 | unclassified | 100 | 100 | 100 |
| | | | | | | | | Opitutales | 12 | 8 | 13 | unclassified | 62 | 95 | 90 |
| | | | | | | | | | | | | Opitutaceae | 38 | 5 | 10 |
| | | | | | | | | Puniceicoccales | 9 | 8 | 4 | Puniceicoccaceae | 100 | 100 | 100 |
| | | | | unclassified | 13 | 6 | 9 | unclassified | 100 | 100 | 100 | unclassified | 100 | 100 | 100 |
| | | | | Verrucomicrobiae | 12 | 20 | 24 | Verrucomicrobiales | 70 | 89 | 83 | Rubritaleaceae | 58 | 18 | 41 |
| | | | | | | | | | | | | Verrucomicrobiaceae | 20 | 43 | 0 |
| | | | | | | | | | | | | Verrucomicrobia subdivision 3 | 12 | 26 | 45 |
| | | | | | | | | | | | | Akkermansiaceae | 10 | 7 | 13 |
| | | | | | | | | | | | | unclassified | 0 | 6 | 0 |
| | | | | | | | | unclassified | 28 | 11 | 17 | unclassified | 100 | 100 | 100 |
| | | | | | | | | various | 2 | 0 | 0 | various | 100 | 0 | 0 |
| Cyanobacteria | 0,3 | 1 | 3 | unclassified | 100 | 100 | 100 | Synechococcales | 72 | 52 | 54 | Synechococcaceae | 81 | 100 | 65 |
| | | | | | | | | | | | | Leptolyngbyaceae | 15 | 0 | 20 |
| | | | | | | | | | | | | Prochloraceae | 2 | 0 | 15 |
| | | | | | | | | | | | | Merismopediaceae | 2 | 0 | 0 |
| | | | | | | | | Oscillatoriales | 26 | 48 | 40 | Microcoleaceae | 71 | 67 | 56 |
| | | | | | | | | | | | | Cyanothecaceae | 20 | 31 | 21 |
| | | | | | | | | unclassified | 1 | 0 | 0 | unclassified | 8 | 0 | 21 |
| | | | | | | | | | | | | Oscillatoriaceae | 0,1 | 2 | 3 |
| | | | | | | | | | | | | unclassified | 100 | 0 | 0 |
| | | | | | | | | Nostocales | 0,3 | 1 | 0 | Nostocaceae | 100 | 100 | 0 |
| | | | | | | | | Chrococcales | 0 | 0 | 6 | Cyanobacteriaceae | 0 | 0 | 100 |
| Firmicutes | 0,09 | 0,5 | 0,6 | Bacilli | 57 | 56 | 57 | Bacillales | 58 | 97 | 82 | Bacillaceae | 86 | 100 | 91 |
| | | | | | | | | | | | | Paenibacillaceae | 7 | 0 | 4 |
| | | | | | | | | | | | | Staphylococcaceae | 4 | 0 | 5 |
| | | | | | | | | | | | | Thermoactinomycetaceae | 3 | 0 | 0 |
| | | | | | | | | Lactobacillales | 42 | 7 | 18 | Leuconostocaceae | 56 | 14 | 0 |
| | | | | | | | | | | | | Streptococcaceae | 42 | 0 | 0 |
| | | | | | | | | | | | | Enterococcaceae | 1 | 0 | 0 |
| | | | | | | | | | | | | Lactobacillaceae | 0,5 | 86 | 100 |
| | | | | Clostridia | 40 | 38 | 36 | Clostridiales | 100 | 100 | 100 | unclassified | 52 | 29 | 0 |
| | | | | | | | | | | | | Clostridiaceae | 38 | 71 | 100 |
| | | | | | | | | | | | | Eubacteriaceae | 6 | 0 | 0 |
| | | | | | | | | | | | | Ruminococcaceae | 4 | 0 | 0 |
| | | | | Negativicutes | 2 | 0 | 4 | Veillonellales | 100 | 0 | 100 | Veillonellaceae | 100 | 100 | 0 |
| | | | | Tissierellia | 0 | 7 | 3 | Tissierellales | 0 | 100 | 100 | Peptoniphilaceae | 0 | 100 | 100 |
| Plantomycetes | 0,09 | 1 | 0,9 | Planctomycetia | 70 | 39 | 61 | Planctomycetales | 98 | 100 | 100 | Planctomycetaceae | 95 | 100 | 100 |
| | | | | | | | | | | | | Gemmataceae | 5 | 0 | 0 |
| | | | | | | | | Canidatus Brocadiales | 2 | 0 | 0 | Canidatus Brocadiaceae | 100 | 0 | 0 |
| | | | | Phycisphaerae | 25 | 55 | 32 | Phycisphaerales | 100 | 100 | 100 | Phycisphaeraceae | 100 | 100 | 100 |
| | | | | unclassified | 5 | 7 | 7 | unclassified | 100 | 100 | 100 | unclassified | 100 | 100 | 100 |
| Acidobacteria | 0,07 | 4 | 0,4 | unclassified | 89 | 100 | 100 | unclassified | 100 | 100 | 100 | unclassified | 100 | 100 | 100 |
| | | | | Acidobacteriia | 11 | 0 | 0 | Acidobacteriales | 100 | 0 | 0 | Acidobacteriaceae | 100 | 0 | 0 |

| | | | | | | | | | | | | | | | |
|---|---|---|---|---|---|---|---|---|---|---|---|---|---|---|---|
| Nitrospinae | 0,03 | 0 | 0,009 | unclassified | 100 | 0 | 100 | unclassified | 100 | 0 | 100 | unclassified | 100 | 0 | 100 |
| Rhodothermaota | 0,02 | 0 | 0 | unclassified | 100 | 0 | 0 | unclassified | 100 | 0 | 0 | unclassified | 100 | 0 | 0 |
| Spirochaetes | 0,02 | 0,1 | 0,2 | Spirochaetia | 100 | 100 | 100 | unclassified | 97 | 100 | 97 | Leptospiraceae | 100 | 100 | 100 |
| | | | | | | | | Brachyspirales | 3 | 0 | 3 | Bachyspiraceae | 100 | 0 | 100 |
| Fusobacteria | 0,01 | 0,02 | 0,01 | Fusobacteriia | 100 | 100 | 100 | Fusobacteriales | 100 | 100 | 100 | Fusobacteriaceae | 100 | 100 | 100 |
| Ntrospirae | 0,01 | 0,04 | 0 | Nitrospira | 88 | 51 | 0 | Nitrospirales | 100 | 100 | 0 | Nitrospiraceae | 100 | 100 | 0 |
| | | | | unclassified | 12 | 49 | 0 | unclassified | 100 | 100 | 0 | unclassified | 100 | 100 | 0 |
| Chlamydiae | 0,007 | 0 | 0 | Chlamydiia | 100 | 0 | 0 | Parachlamydiales | 100 | 0 | 0 | Parachlamyciaceae | 100 | 0 | 0 |
| Canidatus Riflebact | 0,007 | 0 | 0 | unclassified | 100 | 0 | 0 | unclassified | 100 | 0 | 0 | unclassified | 100 | 0 | 0 |
| Gemmatimonadete | 0,007 | 0 | 0 | Gemmatimonadetes | 100 | 0 | 0 | Gemmatimonadales | 100 | 0 | 0 | Gemmatimonadaceae | 100 | 0 | 0 |
| Canidatus Marinim | 0,005 | 0 | 0,04 | unclassified | 100 | 0 | 100 | unclassified | 100 | 0 | 100 | unclassified | 100 | 0 | 100 |
| Ignavibacteria | 0,005 | 0 | 0 | Ignavibacteriae | 100 | 0 | 0 | Ignavibacteriales | 100 | 0 | 0 | unclassified | 100 | 0 | 0 |
| Canidatus Handelsr | 0,004 | 0 | 0,06 | unclassified | 100 | 0 | 100 | unclassified | 100 | 0 | 100 | unclassified | 100 | 0 | 100 |
| Lentisphaerae | 0,003 | 0 | 0,01 | unclassified | 100 | 0 | 0 | unclassified | 100 | 0 | 0 | unclassified | 100 | 0 | 0 |
| | | | | Lentisphaeria | 0 | 0 | 100 | Lentisphaerales | 0 | 100 | 0 | Lentisphaeraceae | 0 | 0 | 100 |
| Deferribactres | 0,003 | 0 | 0 | Deferribacteres | 100 | 0 | 0 | Deferribacterales | 100 | 0 | 0 | Deferribacteraceae | 100 | 0 | 0 |
| Chlorobi | 0,002 | 0 | 0,2 | Chlorobia | 100 | 0 | 100 | Chlorobiales | 100 | 0 | 100 | Chlorobiaceae | 100 | 0 | 100 |
| Canidatus Portonoy | 0,001 | 0 | 0 | unclassified | 100 | 0 | 0 | unclassified | 100 | 0 | 0 | unclassified | 100 | 0 | 0 |
| Canidatus Uhrbacte | 0,0009 | 0 | 0 | unclassified | 100 | 0 | 0 | unclassified | 100 | 0 | 0 | unclassified | 100 | 0 | 0 |
| Canidatus Sacchari | 0,0008 | 0 | 0 | unclassified | 100 | 0 | 0 | unclassified | 100 | 0 | 0 | unclassified | 100 | 0 | 0 |
| Tenericutes | 0,003 | 0,02 | 0,007 | Mollicutes | 100 | 100 | 100 | Mycoplasmatales | 100 | 100 | 100 | Mycoplasmataceae | 100 | 100 | 100 |
| Chloroflexi | 0,00007 | 0,03 | | unclassified | 100 | 85 | 0 | unclassified | 100 | 100 | 0 | unclassified | 100 | 100 | 0 |
| | | | | Chloroflexia | 0 | 5 | 0 | Herpetosiphonales | 0 | 100 | 0 | Herpetosiphonaceae | 0 | 100 | 0 |
| Canidatus Atribacte | 0 | 0,2 | 0,2 | unclassified | 0 | 100 | 100 | unclassified | 0 | 100 | 100 | unclassified | 0 | 100 | 100 |
| Canidatus Parcubac | 0 | 0,08 | 0,06 | unclassified | 0 | 100 | 100 | unclassified | 0 | 100 | 100 | unclassified | 0 | 100 | 100 |
| Fibrobacteres | 0 | 0,05 | 0,03 | Fibrobacteria | 0 | 100 | 100 | Fibrobacterales | 0 | 100 | 100 | Fibrobacteraceae | 0 | 100 | 100 |
| Deinococcus-Therm | 0 | 0,02 | 0,06 | Deinococci | 0 | 100 | 100 | Deinococcales | 0 | 100 | 100 | Deinococcaceae | 0 | 100 | 100 |
| Canidatus Wolfeba | 0 | 0,008 | 0,01 | unclassified | 0 | 100 | 100 | unclassified | 0 | 100 | 100 | unclassified | 0 | 100 | 100 |
| Canidatus Krefeldb | 0 | 0 | 0,1 | unclassified | 0 | 0 | 100 | unclassified | 0 | 0 | 100 | unclassified | 0 | 0 | 100 |

| | | | |
|---|---|---|---|
| Eukaryota | 11 | 43 | 54 |
| unclassified | 3 | 9 | 12 |
| Viruses | 1 | 3 | 3 |
| various | 0,04 | 0,2 | 0,3 |
| Archaea | 0,05 | 0,1 | 0,04 |
